# A humanized yeast phenomic model of deoxycytidine kinase to predict genetic buffering of nucleoside analog cytotoxicity

**DOI:** 10.1101/700153

**Authors:** Sean M. Santos, Mert Icyuz, Ilya Pound, Doreen William, Jingyu Guo, Brett A. McKinney, Michael Niederweis, John Rodgers, John L. Hartman

## Abstract

Knowledge about synthetic lethality can be applied to enhance the efficacy of anti-cancer therapies in individual patients harboring genetic alterations in their cancer that specifically render it vulnerable. We investigated the potential for high-resolution phenomic analysis in yeast to predict such genetic vulnerabilities by systematic, comprehensive, and quantitative assessment of drug-gene interaction for gemcitabine and cytarabine, substrates of deoxycytidine kinase that have similar molecular structures yet distinct anti-tumor efficacy. Human deoxycytidine kinase (dCK) was conditionally expressed in the *S. cerevisiae* genomic library of knockout and knockdown (YKO/KD) strains, to globally and quantitatively characterize differential drug-gene interaction for gemcitabine and cytarabine. Pathway enrichment analysis revealed that autophagy, histone modification, chromatin remodeling, and apoptosis-related processes influence gemcitabine specifically, while drug-gene interaction specific to cytarabine was less enriched in Gene Ontology. Processes having influence over both drugs were DNA repair and integrity checkpoints and vesicle transport and fusion. Non-GO-enriched genes were also informative. Yeast phenomic and cancer cell line pharmacogenomics data were integrated to identify yeast-human homologs with correlated differential gene expression and drug-efficacy, thus providing a unique resource to predict whether differential gene expression observed in cancer genetic profiles are causal in tumor-specific responses to cytotoxic agents.

## Introduction

Genomics has enabled targeted therapy aimed at cancer driver genes and oncogenic addiction [1], yet traditional cytotoxic chemotherapeutic agents remain among the most widely used and efficacious anti-cancer therapies [2]. Changes in the genetic network underlying cancer can produce vulnerabilities to cytotoxic chemotherapy that further influence the therapeutic window and provide additional insight into their mechanisms of action [3,4]. A potential advantage of so-called synthetic lethality-based treatment strategies is that they could have efficacy against passenger gene mutation or compensatory gene expression, while classic targeted therapies are directed primarily at driver genes (Fig. 1A). Quantitative high throughput cell array phenotyping of the yeast knockout and knockdown libraries provides a phenomic means for systems level, high-resolution modeling of gene interaction [5-9], which is applied here to predict cancer-relevant drug-gene interaction through integration with cancer pharmacogenomics resources (Fig. 1B).

**Figure 1.**
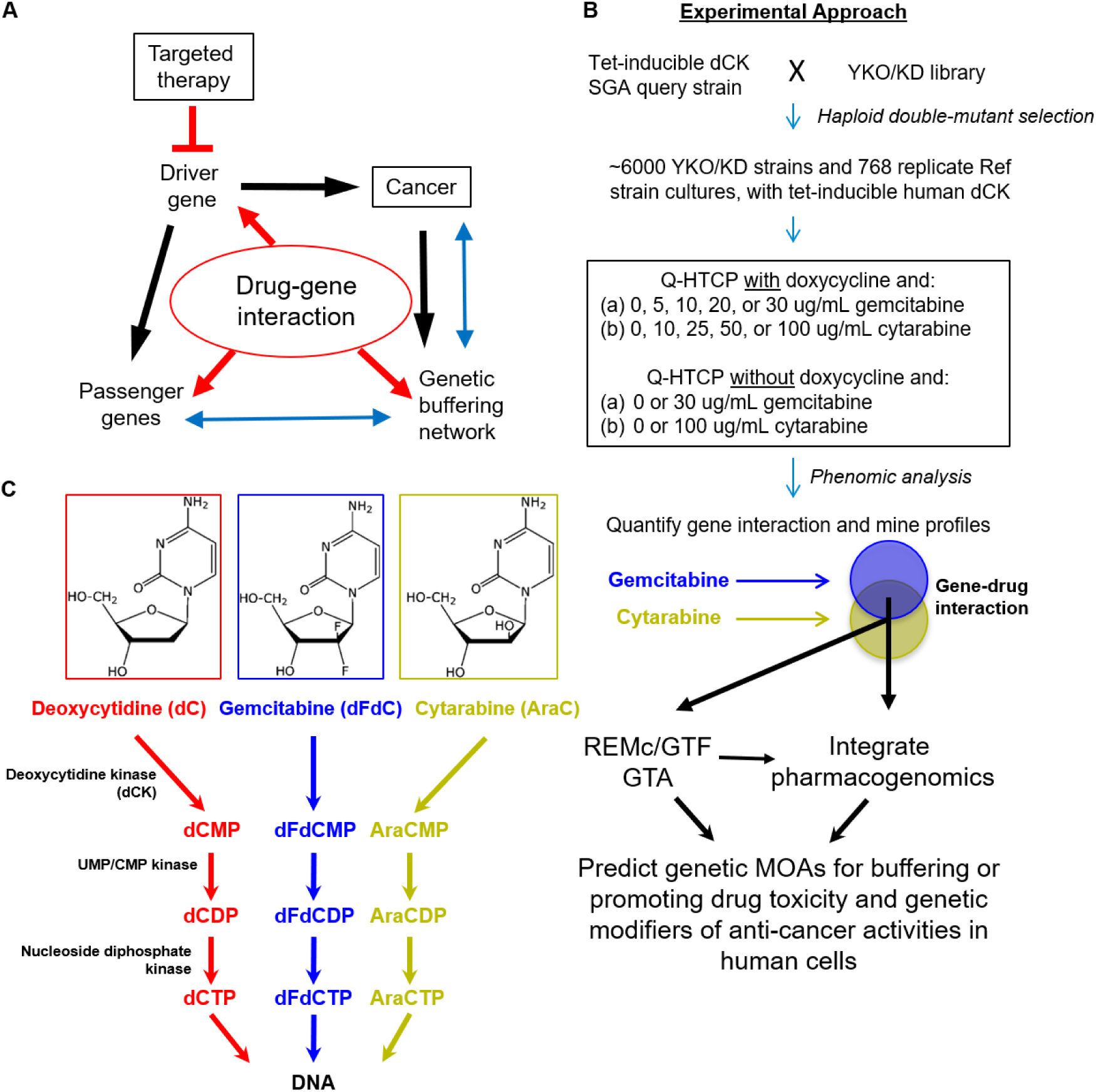
Experimental model of gemcitabine and cytarabine drug-gene interaction networks. (**A**) The strategy of cytotoxic anti-cancer drug-gene interaction is illustrated in the context of driver gene-mediated oncogenesis. Driver genes promote cancer and influence the expression of passenger genes (black arrows), which also leads to genomic instability and alterations in the genetic buffering network. The genetic buffering network (blue arrows) maintains cellular homeostasis, and is altered in cancer cells by genomic instability, thereby creating the potential for drug-gene interaction that increases the therapeutic window of anti-cancer agents (red arrows). Drug-gene interaction can either involve driver or passenger genes directly, or the compromised genetic buffering network, which are systematically characterized by the quantitative yeast phenomic model. (**B**) The synthetic genetic array (SGA) method was used to enable tet-inducible human dCK expression in the yeast knockout and knockdown (YKO/KD) collection. The phenomic model incorporates treatment of individually grown cultures of the YKO/KD collection, and 768 replicate Ref strain cultures, with increasing gemcitabine (0, 5, 10, 20, and 30 ug/mL) or cytarabine (0, 10, 25, 50, and 100 ug/mL) in HLD media, with dCK induced by addition of doxycycline. Drug-gene interaction profiles were subjected to REMc and GO term analysis to characterize phenomic modules with respect to drug-gene interaction for gemcitabine or cytarabine, and integrated with pharmacogenomics data to predict evolutionarily conserved drug-gene interactions relevant to precision oncology. (**C**) Structures and metabolism of deoxycytidine analogs.

Nucleoside analogs include a diverse group of compounds with anticancer, antiviral, and immunosuppressive efficacy [10]. The anti-cancer agents have tissue-specific efficacy ranging from solid tumors to leukemias, yet details about how these agents confer differential activity are unknown [10,11]. Gemcitabine (2’,2’-difluoro 2’-deoxycytidine, **dFdC**) and cytarabine (**Ara-C**) are deoxycytidine analogs that undergo the first step of conversion to their active triphosphate forms by deoxycytidine kinase (**dCK**) (Fig. 1C). The nucleoside triphosphate analogs can be incorporated into DNA and inhibit the functions of polymerases and other enzymes involved of DNA metabolism. For example, gemcitabine inhibits ribonucleotide reductase (**RNR**), which limits the production of deoxyribonucleotides (**dNTPs**) that are needed for DNA synthesis and repair [11]. Gemcitabine has been used as a single agent in the treatment of some cancers, such as pancreatic, and in combination with platinum-based drugs in non-small cell lung, breast, and ovarian cancers [12-15]. Cytarabine, on the other hand, has been an important agent in treatments for acute myeloid leukemia and acute lymphoblastic leukemia [16].

Deoxycytidine kinase (**dCK**) phosphorylates deoxycytidine to deoxycytidine monophosphate (**dCMP**), similarly phosphorylating gemcitabine and cytarabine to dFdCMP and AraCMP, respectively. UMP/CMP kinase and the nucleoside diphosphate kinase are subsequently involved in conversion to the triphosphate form (Fig. 1C). Reduced expression of dCK or high expression of RNR subunits *RRM1* and *RRM2* is associated with increased gemcitabine resistance [10,12,17-21]. Genomic analyses have suggested genetic influences on the efficacy of gemcitabine or cytarabine [22-26], which we model here at a systems level by surveying gene-drug interaction to elucidate biology underlying differential anti-cancer efficacies of the respective drugs, and thereby aid in predicting treatment outcomes based on individual patient cancer genetic profiles.

*Saccharomyces cerevisiae* does not have a dCK homolog and is thus naturally resistant to gemcitabine and cytarabine. To examine the gene-drug interaction networks for gemcitabine and cytarabine in yeast, we introduced human dCK into the yeast knockout and knockdown (**YKO/KD**) library by the synthetic genetic array (**SGA**) method [27-29], and conducted phenomic analysis on the resulting double mutant library by quantitative high throughput cell array phenotyping (**Q-HTCP**) [6-8,30], using multiple growth inhibitory concentrations of gemcitabine or cytarabine (Fig. 1B). Cell proliferation parameters (**CPPs**) obtained by Q-HTCP were used to quantify and compare drug-gene interaction for gemcitabine vs. cytarabine. The unbiased results provide a systems level resource of genetic and biological information about the cytotoxicity of these drugs, incorporating knowledge about genes that either buffer or promote their effects [3,5]

Recent advances in cancer pharmacogenomics have provided gene expression and drug sensitivity data from hundreds of cancer cell lines, establishing associations between gene expression and anti-cancer efficacy for many compounds, including gemcitabine and cytarabine [31-33]. We investigated the potential utility of a yeast phenomic model of chemotherapy sensitivity and resistance for predicting causality in correlations between differential gene expression and drug sensitivity by generating a network-level drug-gene interaction resource. The resource integrates cancer pharmacogenomic and yeast phenomic data, using the results to query the cancer genetics literature in order to obtain systems level biological insights about how yeast phenomic models help predict cytotoxic chemotherapy efficacy based on unique genetic alterations specific to each individual patient’s cancer (Fig. 1A).

## Materials and Methods

### Strains, media and drugs

We obtained the yeast gene knockout strain library (**YKO**) from Research Genetics (Huntsville, AL, USA) and the knockdown (**KD**) collection, also referred to as the Decreased Abundance of mRNA Production (**DAmP**) library, from Open Biosystems (Huntsville, AL, USA). The YKO library is in the genetic background of BY4741 (S288C MAT**a** *ura3-Δ0 his3-Δ1 leu2-Δ0 met17-Δ0*). Additional information and strains can be obtained at https://dharmacon.horizondiscovery.com/cdnas-and-orfs/non-mammalian-cdnas-and-orfs/yeast/#all. Some mutants appear multiple times in the library and they are treated independently in our analysis. HLD is a modified synthetic complete medium [8] and was used with 2% dextrose (**HLD**) as the carbon source. Doxycycline hydrochloride (BP26535) was obtained from Fisher Scientific. Gemcitabine (Gemzar) was obtained from Eli Lilly and Company (0002-7502-01). Cytarabine was obtained from Bedford Laboratories (55390-131-10).

A tet-inducible dCK query allele was constructed in the SGA background in the following way: An integrating plasmid for doxycycline-inducible gene expression was constructed by subcloning 3’UTR and 5’ORF targeting sequences from the LYP1 locus into pJH023 [34], creating pJH023_UO_lyp1, and the reverse VP16 transactivator (Tet-ON), obtained by PCR from pCM176 [35], was fused to the *ACT1* promoter by overlap PCR and subcloned into pJH023_UO_lyp1, replacing the VP16 transactivator (Tet-OFF) and creating the “Tet-ON” construct, pML1055 [36]. pML1055 was digested with NOT1 and transformed into strain 15578-1.2b_LYP1 (*MATα his3Δ1 leu2Δ0 ura3Δ0 can1Δ0::P_GAL1_-T_ADH1-_P_MFA1_-his5^+^_sp_ hmrΔ0::URA3ca*), which was derived by backcrossing 15578-1.2b (*MATα his3Δ1 leu2Δ0 ura3Δ0 can1Δ0::P_GAL1_-T_ADH1-_P_MFA1_-his5^+^_sp_ lyp1Δ0 hmrΔ0::URA3ca*) to restore the *LYP1* locus. The resulting chromosomal integration of pML1055 between the promoter and ORF at the *LYP1* locus was selected with nourseothricin, giving rise to yDW1 (*MATα his3Δ1 leu2Δ0 ura3Δ0 can1Δ0::P_GAL1_-T_ADH1-_ P_MFA1_-his5^+^_sp_ hmrΔ0::URA3ca Pact1-revTetR-VP16-*natMX-*PtetO7-LYP1*). Tet-inducible *LYP1* in yDW1 was verified phenotypically by doxycycline-dependent SAEC sensitivity [36]. Overlap PCR was performed to fuse deoxycytidine kinase (from a plasmid, gift of Bo Xu and William Parker, Southern Research) and the HPH gene (from pFA6a-HBH-hphMX4) [37], introducing flanking sequences for replacement of the *LYP1* ORF (see **Additional File 1, Table S1** for primers). The PCR product was transformed into yDW1 (**Additional File 2, Fig. S1**) and transformants selected on hygromycin were confirmed by doxycycline-induced sensitivity to gemcitabine and cytarabine, yielding yMI16.

The synthetic genetic array (**SGA**) method, a way to introduce an allele of interest into the YKO/KD library and recover haploid double mutants [28,29], was used to derive a haploid YKO/KD collection with doxycycline-inducible dCK expression.

### Quantitative high throughput cell array phenotyping (Q-HTCP)

Q-HTCP, an automated method of collecting growth curve phenotypes for the YKO/KD library arrayed onto agar media, was used to obtain phenomic data [38]. A Caliper Sciclone 3000 liquid handling robot was used for cell array printing, integrated with a custom imaging robot (Hartman laboratory) and Cytomat 6001 (Thermo Fisher Scientific, Asheville, NC, USA) incubator. Images of the 384-culture arrays were obtained approximately every 2-3 hours and analyzed as previously described [9,38]. To obtain CPPs, image analysis was performed in Matlab and data were fit to the logistic equation, G(t) = K/(1 + e^−r(t−l)^), assuming G(0) < K, where G(t) is the image intensity of a spotted culture vs. time, K is the carrying capacity, r is the maximum specific growth rate, and l is the moment of maximal absolute growth rate, occurring when G(t) = K/2 (the time to reach half of carrying capacity) [7]. The CPPs, primarily K and L, were used as phenotypes to measure drug-gene interaction.

### Quantification of drug-gene interaction

Gene interaction was defined by departure of the corresponding YKO/KD strain from its expected phenotypic response to gemcitabine or cytarabine. The expected phenotype was determined by cell proliferation phenotypes of the mutant without gemcitabine or cytarabine, and with 5ug/mL doxycycline, together with those of the reference strain with and without gemcitabine or cytarabine [5,6,9,34]. The concentrations of gemcitabine or cytarabine (ug/mL) were chosen based on phenotypic responses being functionally discriminating in the parental strain. Gemcitabine, cytarabine, or doxycycline, alone, did not alter cell proliferation (Fig. 2C-F; **Additional File 2, Fig. S2A-D**).

**Figure 2.**
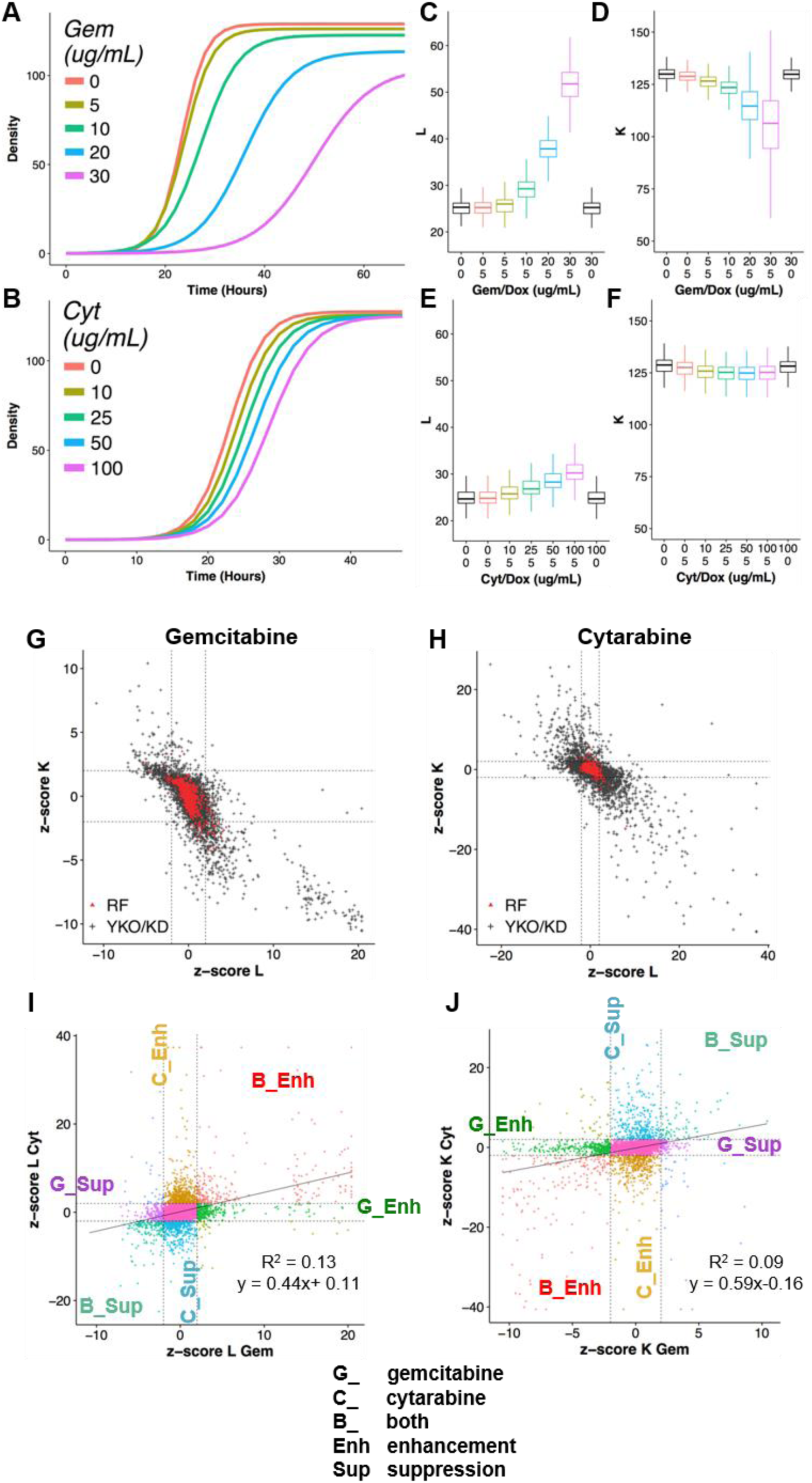
Phenomic analysis of drug-gene interaction for gemcitabine and cytarabine. Average growth curves (from fitting pixel intensity data of 768 replicate cultures to a logistic function) for the reference (RF) strain, treated with the indicated concentrations of (**A**) gemcitabine or (**B**) cytarabine. (**C-F**) CPP distributions from data depicted in panels A and B for (**C-D**) gemcitabine and (**E-F**) cytarabine for (**C, E**) L and (**D, F**) K. (**G, H**) Comparison of drug-gene interaction scores using either the L or K CPPs for (**G**) gemcitabine and (**H**) cytarabine. Score distributions of knockout/knockdown (YKO/KD, black) and non-mutant parental (Ref, red) strain cultures are indicated along with thresholds for deletion enhancement and suppression (dashed lines at +/- 2). (**I-J**) Differential drug-gene interaction using L (**I**) or K (**J**) as the CPP for gemcitabine *vs.* cytarabine, classified with respect to relative drug specificity of interactions. ‘G’, ‘C’, and ‘B’ indicate gemcitabine-, cytarabine-, or both drug-gene interactions, respectively. Deletion enhancement or suppression is indicated by ‘_Enh’ or ‘_Sup’, respectively.

Interaction scores were calculated as previously described [9,39], with slight modifications, as summarized below. All media conditions used for interaction score calculation had 5 ug/mL doxycycline to express dCK. Variables were defined as:

D_i_ = concentration (dose) of gemcitabine or cytarabine

R_i_ = observed mean growth parameter for parental Reference strain at D_i_

Y_i_ = observed growth parameter for the YKO/KD mutant strain at D_i_

K_i_ = Y_i_ – R_i_, the difference in growth parameter between the YKO/KD mutant (Y_i_) and Reference (R_i_) at D_i_

K_0_ = Y_0_ - R_0_, the effect of gene KO/KD on the observed phenotype in the absence of gemcitabine or cytarabine; this value is annotated as ‘shift’ and is subtracted from all K_i_ to obtain L_i_

L_i_ = K_i_ - K_0_, the interaction between (specific influence of) the KO/KD mutation on gemcitabine or cytarabine response, at D_i_

For cultures not generating a growth curve, Y_i_ = 0 for K and r, and the L parameter was assigned Y_i_ max, defined as the maximum observed Y_i_ among all cultures exhibiting a minimum carrying capacity (K) within 2 standard deviation (SD) of the parental reference strain mean at D_i_. Y_i_ max was also assigned to outlier values (*i.e.*, if Y_i_ > Y_i_ max).

Interaction was calculated by the following steps:

1. Compute the average value of the 768 reference cultures (R_i_) at each dose (D_i_):
2. Assign Y_i_ max (defined above) if growth curve is observed at D_0_, but not at D_i_, or if observed Y_i_ is greater than Y_i_ max.
3. Calculate K_i_ = Y_i_ - R_i_.
4. Calculate L_i_ = K_i_ – K_0_
5. Fit data by linear regression (least squares): L_i_ = A + B*D_i_
6. Compute the interaction value ‘INT’ at the max dose: INT = L_i_-max = A + B*D_max_
7. Calculate the mean and standard deviation of interaction scores for reference strains, mean(REF_INT_) and SD(REF_INT_); mean(REF_INT_) is expected to be approximately zero, with SD(REF_INT_) primarily useful for standardizing against variance (**Additional File 1, Tables S2-S5; Additional Files 3-4**).
8. Calculate interaction z-scores:

z-score(YKO_INT_) = (YKO_INT_ – mean(REF_INT_))/SD(REF_INT_)
z-score(YKO_INT_) > 2 for L or < −2 for K are referred to as gene deletion enhancers
 of gemcitabine or cytarabine cytotoxicity, and conversely, L interaction score < −2 or K interaction scores >2 are considered gene deletion suppressors. Because the CPP distributions for KD strains were different from the reference strain, we used the mean and standard deviation from the KD plates only as a conservative measure of variance where z-score(KD_INT_) = (KD_INT_ – mean(KD_INT_))/SD(KD_INT_).

### Recursive expectation-maximization clustering (**REMc**) and heatmap generation

REMc is a probability-based clustering method and was performed as previously described [40]. Clusters obtained by Weka 3.5, an EM-optimized Gaussian mixture-clustering module, were subjected to hierarchical clustering in R (http://www.r-project.org/) to further aid visualization with heatmaps. REMc was performed using L and K interaction z-scores (Fig. 3A). The effect of gene deletion on the CPP (in the absence of drug), termed ‘shift’ (K_0_), was not used for REMc, but was included for visualization in the final hierarchical clustering. **Additional File 5, Files A-B** contain REMc results in text files with associated data also displayed as heatmaps. In cases where a culture did not grow in the absence of drug, 0.0001 was assigned as the interaction score, so that associated data (‘NA’) could be easily indicated by red coloring in the shift columns of the heatmaps.

**Figure 3.**
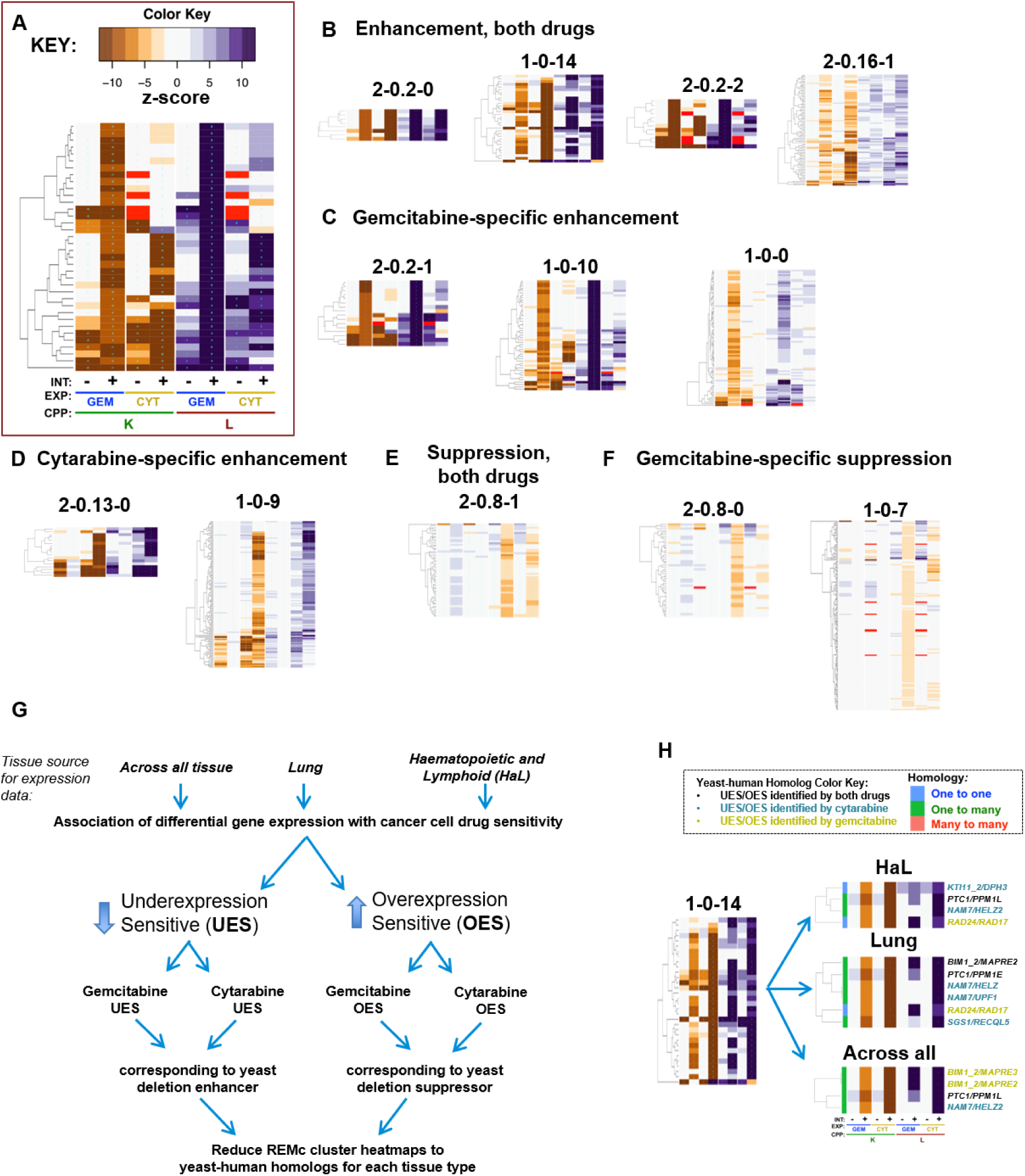
Prediction of drug-gene interaction in cancer cells by integration of yeast phenomic and human pharmacogenomic data. Recursive expectation-maximization clustering results were classified visually by their associated gene interaction profiles (see methods). (**A**) The data columns in all heatmaps are ordered from left to right, as shown in this example. K interactions for gemcitabine and cytarabine are in columns 2 and 4, respectively, with L interactions in columns 6 and 8. To the left of each interaction value (indicated by ‘+’), is the corresponding ‘shift’ value (indicated by ‘-‘), referring to the ΔCPP for the respective YKO/KD culture relative to the reference culture average in the absence of gemcitabine or cytarabine (see methods). (**B-F**) The relative strength of example clusters is ordered from left to right. **(B)** Enhancement, both drugs. (**C**) Gemcitabine-specific enhancement. (**D**) Cytarabine-specific enhancement. (**E**) Suppression, both drugs. (**F**) Gemcitabine-specific suppression. (**G**) Differential gene expression for cell lines from the GDSC database (either lung, hematopoietic and lymphoid, or across all tissues) was categorized in drug-sensitive cells as either underexpressed (**UES**) or overexpressed (**OES**), and filtered by correlation with yeast homologs being deletion enhancing or suppressing, respectively. (**H**) An example of yeast-human homologs identified as described in **G**. The category of homology assigned by *BiomaRt* is indicated in the left column of each heatmap (see homology color key). At right, the gene label indicates whether the human homolog was verified in PharmacoDB for both drugs (black), cytarabine (teal), or gemcitabine (gold). **Additional Files 5 (File B) and 8 (Files B-D)** contains all REMc heatmaps of the types indicated to the left and right, respectively, in panel H.

### Gene ontology term finder (**GTF**)

A python script was used to format REMc clusters for analysis with the command line version of the GO Term Finder (**GTF**) tool downloaded from http://search.cpan.org/dist/GO-TermFinder/ [41]. GTF reports on enrichment of Gene Ontology (**GO**) terms by comparing the ratio of genes assigned to a term within a cluster to the respective ratio involving all genes tested. **Additional File 5, File C** contains GTF analysis of all REMc clusters. GO-enriched terms from REMc were investigated with respect to genes representing the term and literature underlying their annotations [42].

### Gene ontology term averaging (**GTA**) analysis

In addition to using GTF to survey functional enrichment in REMc clusters, we developed GTA as a complementary workflow, using the GO information on SGD at https://downloads.yeastgenome.org/curation/literature/ to perform the following analysis:

1. Calculate the average and SD for interaction values of all genes in a GO term.
2. Filter results to obtain terms having GTA value greater than 2 or less than −2.
3. Obtain GTA scores defined as |GTA value| - gtaSD; filter for GTA score > 2.

The GTA analysis is contained in **Additional File 6** as tables and interactive plots created using the R *plotly* package https://CRAN.R-project.org/package=plotly. GTA results were analyzed using both the L and K interaction scores and are included in **Additional File 6 (Files A-C).**

### Prediction of human homologs that influence tumor response to gemcitabine or cytarabine

PharmacoDB holds pharmacogenomics data from cancer cell lines, including transcriptomics and drug sensitivity [33]. The *PharmacoGx* R/Bioconductor package [43] was used to analyze the GDSC1000 (https://pharmacodb.pmgenomics.ca/datasets/5) and gCSI (https://pharmacodb.pmgenomics.ca/datasets/4) datasets, which contained transcriptomic and drug sensitivity results. A p-value < 0.05 was used for differential gene expression and drug sensitivity. For gene expression, the sign of the standardized coefficient denotes increased (+) or decreased (-) expression. The *biomaRt* R package [44,45] was used with the Ensembl database [46] to match yeast and human homologs from the phenomic and transcriptomic data, classifying yeast-human homology as one to one, one to many, and many to many. The Princeton Protein Orthology Database (PPOD) was also used to manually review and further consider homology [47].

## Results

### Quantitative phenomic characterization of differential gene-drug interaction

The Q-HTCP workflow incorporates high-throughput kinetic imaging and analysis of proliferating 384-culture cell arrays plated on agar media to obtain CPPs for measuring gene-drug interaction, as previously described [7,9,38]. To apply it for analysis of dCK substrates, a tetracycline-inducible human dCK allele was introduced into the complete YKO/KD library by the synthetic genetic array method [29,48] (Figure 1B). The dependence of gemcitabine and cytarabine toxicity on dCK expression was demonstrated for the reference strain (Fig. 2A-F), as the two nucleosides exerted cytotoxicity only if dCK was induced by the addition of doxycycline. Induction of dCK had no effect on proliferation in the absence of gemcitabine or cytarabine (Fig. 2C-F).

Interaction scores were calculated by departure of the observed CPP for each YKO/KD strain from that expected based on the observed phenotypes for the reference strain treated and untreated with drug and the YKO/KD strain in the absence of drug, incorporating multiple drug concentrations, 768 replicate reference strain control cultures, and summarized by linear regression as z-scores [6-8,30,34,38]. Gene interaction scores with absolute value greater than two were selected for global analysis and termed deletion enhancers (z-score_L ≥ 2 or z-score_K ≤ −2) or deletion suppressors (z-score_L ≤ −2 or z-score_K ≥ 2) of drug cytotoxicity, revealing functions that buffer or promote drug cytotoxicity, respectively [39] (Fig. 2).

Growth inhibition was greater for gemcitabine than for cytarabine (Fig. 2A-F), however, the phenotypic variance was also less for cytarabine, such that interactions of smaller effect size were detectable and the range of scores was greater (**Additional File 1, Table S6**). The CPP, ‘L’, (the time at which half carrying capacity is reached), is most sensitive to growth inhibitory perturbation, while ‘K’ (carrying capacity) reports on more extreme growth differences (Fig. 2A-H). Low correlation between the gene-drug interaction profiles suggested differential buffering of these two drugs, consistent with their distinct anti-tumor efficacies (Fig. 2I-J).

### Functional analysis of gene-interaction modules

Recursive expectation-maximization clustering (**REMc**) was used to identify modules of genes that shared similar profiles of buffering or promoting nucleoside toxicity of gemcitabine or cytarabine [40] (see Fig. 3A-F; Table 1**; Additional File 5**). As described previously, REMc results were assessed with GO Term Finder for Gene Ontology functional enrichment [41] and heatmaps generated by first adding data regarding the main effect of the gene knockout or knockdown (*i.e.*, no drug) on cell proliferation, termed ‘shift’ (see methods), followed by hierarchical clustering [40,41]. GO Term Average (**GTA**) scores, which are based on the average and standard deviation of drug-gene interaction for all genes of each GO term [39], were used as a complement to REMc/GTF for identifying functions that buffer or promote drug effects (Table 2, Fig. 4, and **Additional File 6, Files A-C**). Yeast-human homologs were judged, regarding causality of differential gene expression associated with sensitivity to gemcitabine or cytarabine, by the correspondence of yeast phenomic and cancer pharmacogenomics results, thus establishing a model resource to test the utility of yeast phenomics to inform cancer genetic profiling for predicting drug-specific, anti-tumor efficacy (Fig. 3G-H). Heatmaps were also produced systematically to visualize drug-gene interaction profiles for all genes assigned to GO terms identified by REMc/GTF or GTA; these are referred to as term-specific heatmaps, and are grouped by GO term parent-child relationships (**Additional File 7**).

**Figure 4.**
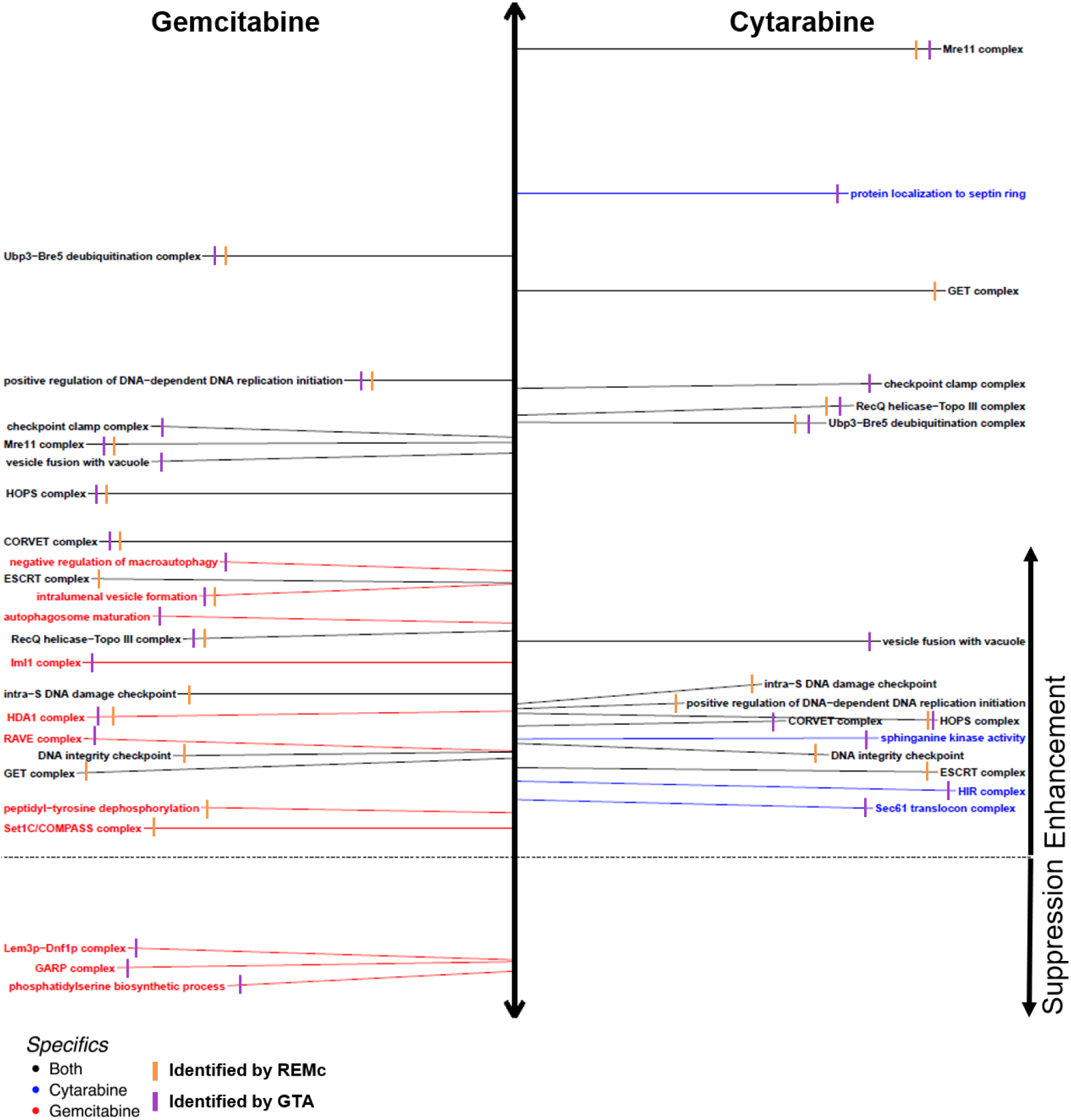
GO annotations associated with deletion enhancement or suppression of gemcitabine and/or cytarabine cytotoxicity. Representative GO terms are listed, which were identified by REMc/GTF (orange), GTA (purple), or both methods, for enhancement (above dashed line) or suppression (below dashed line) of gemcitabine (left, red), cytarabine (right, blue), or both media types (black). Term-specific heatmaps were manually reviewed for inclusion. Distance above or below the horizontal dashed line reflects the average interaction score for genes identified by REMc/GTF or the GTA score (see methods). See **Additional Files 5 and 6** for all REMc/GTF, and GTA results.

**Table 1.** GO terms enriched in REMc clusters.

| GO Term | Drug | INT | O | Cluster | Genes<br>in Term | p-value | Genes | Fig | GTA Gem L | GTA Cyt L |
| --- | --- | --- | --- | --- | --- | --- | --- | --- | --- | --- |
| Ubp3-Bre5 deubiquitination complex | Both | Enh | C | 2-0.2-0 | 2/2 | 2.57E-05 | UBP3:BRE5 | 5D | 19.8 | 14.32 |
| positive regulation of DNA-dependent DNA replication initiation | Both | Enh | P | 1-0-2 | 3/4 | 2.09E-04 | RFM1:FKH2:SUM1 | 5B | 15.7 | 4.9 |
| Mre11 complex | Both | Enh | C | 2-0.14-1 | 2/3 | 5.66E-04 | RAD50:XRS2 | 5B | 13.7 | 26.6 |
| HOPS complex | Both | Enh | C | 2-0.14-1 | 2/7 | 3.94E-03 | PEP3:VPS33 | 5D | 12.0 | 4.8 |
| CORVET complex | Both | Enh | C | 2-0.14-1 | 2/7 | 3.94E-03 | PEP3:VPS33 | 5D | 10.4 | 4.3 |
| RecQ helicase-Topo III complex | Both | Enh | C | 1-0-14 | 2/3 | 3.31E-03 | SGS1:RMI1 | 5B | 7.5 | 14.6 |
| GET complex | Both | Enh | C | 2-0.14-0 | 2/3 | 4.68E-04 | GET1:GET2 | 5D | 3.3 | 18.6 |
| DNA integrity checkpoint | Both | Enh | P | 1-0-14 | 4/40 | 3.85E-03 | DUN1:RAD17:RAD24:SGS1 | 5A | 4.8 | 4.8 |
| alpha-glucoside transmembrane transporter activity | Cyt | Enh | F | 2-0.17-3 | 2/2 | 5.98E-03 | MAL31:MAL11 | 7A | -0.7 | 2.2 |
| intraluminal vesicle formation | Gem | Enh | P | 1-0-10 | 3/7 | 2.90E-03 | DOA4:VPS24:BRO1 | 6A | 9.0 | 1.6 |
| HDA1 complex | Gem | Enh | C | 1-0-0 | 2/3 | 7.08E-02 | HDA1:HDA3 | 6B | 4.8 | 0.3 |
| Swr1 complex | Gem | Enh | C | 1-0-11 | 3/12 | 3.46E-02 | SWC3:VPS71:SWR1 | 6B | 2.9 | -1.6 |
| peptidyl-tyrosine dephosphorylation | Gem | Enh | P | 1-0-0 | 5/20 | 2.18E-03 | OCA2:SIW14:OCA1:OCA4:OCA6 | 6C | 1.5 | 0.5 |
| Set1C/COMPASS complex | Gem | Enh | C | 1-0-0 | 3/6 | 5.74E-03 | SDC1:SWD3:BRE2 | 6B | 1.0 | 0.6 |
| phospholipid-translocating ATPase activity | Gem | Sup | F | 1-0-8 | 3/7 | 9.70E-03 | DRS2:LEM3:DNF2 | 6D | -1.6 | -0.9 |

For each GO term, the table indicates which drugs interact with it, the interaction type (enhancing or suppressing), the ontology ('O') it derives from (cellular Process or Component, or molecular Function), the REMc cluster ID from which the term was most specific, the fraction of the genes in the term that were observed in the cluster and the p-value for enrichment of the genes. Relevant figures and associated GTA data are also given.

**Table 2.**
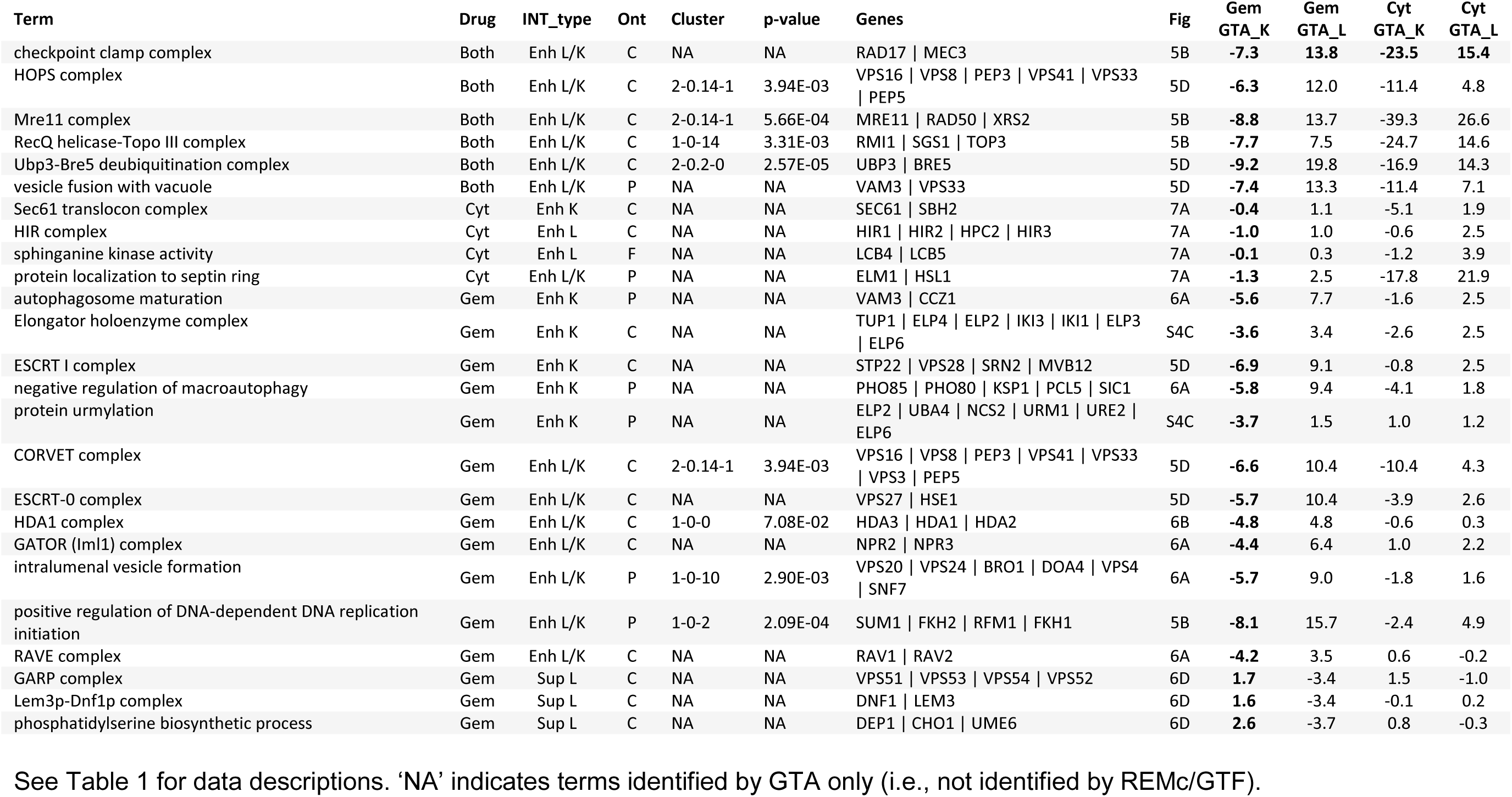
GO terms identified by GTA.

Cancer pharmacogenomics data in PharmacoDB were mined using *PharmacoGx* [43] and *biomaRt* [44,45] with the GDSC1000 [31,49] or gCSI [32,50] datasets to match yeast drug-gene interaction by homology to differential gene expression in gemcitabine or cytarabine sensitive cancer cell lines (Fig. 3G-H; **Additional File 8**). Yeast gene deletion enhancers identified human homologs underexpressed in gemcitabine or cytarabine sensitive cells, termed UES, while yeast gene deletion suppressors identified human homologs overexpressed in drug sensitive cells, termed OES (Fig. 3G).

The analysis was focused on the GDSC database, because it had expression data available for both gemcitabine and cytarabine; however, analysis of the gCSI data was also conducted for gemcitabine (**Additional File 8, File A**). Differential expression was analyzed: (1) across all tissue types, to consider interactions that might be applicable in novel treatment settings; (2) in hematopoietic & lymphoid tissue; and (3) in lung tissue, as cytarabine and gemcitabine are used to treat HaL and lung cancers, respectively. Gemcitabine is also used for pancreatic cancer; however, the number of cell lines tested (30) was lower than for lung (156) or HaL (152). Thus, yeast genes that were deletion enhancing or suppressing were catalogued with human homologs that were UES or OES in PharmacoDB (Figs. 3G-H, Tables 3-5, **and Additional File 8**).

**Table 3.** Yeast-human homologs predicted to similarly buffer or promote both gemcitabine and cytarabine toxicity.

| yGene | hGene | H | Drug | Cluster | Tissue | Gem K | Cyt K | Gem L | Cyt L | Ref | Description (Human) |
| --- | --- | --- | --- | --- | --- | --- | --- | --- | --- | --- | --- |
| NAM7 | HELZ | 2 | Cyt | 1-0-14 | L | -6.5 | -16.7 | 1.1 | 13.6 | [61-64] | helicase with zinc finger |
| NAM7 | HELZ2 | 2 | Cyt | 1-0-14 | A, H | -6.5 | -16.7 | 1.1 | 13.6 |  | helicase with zinc finger 2 |
| NAM7 | UPF1 | 2 | Cyt | 1-0-14 | L | -6.5 | -16.7 | 1.1 | 13.6 | [65-67] | UPF1, RNA helicase and ATPase |
| PTC1 | PPM1E | 2 | Both | 1-0-14 | L | -8.8 | -12.7 | 7.9 | 15.7 | [68] | protein phosphatase, Mg2+/Mn2+ dependent 1E |
| PTC1 | PPM1L | 2 | Both | 1-0-14 | A, H | -8.8 | -12.7 | 7.9 | 15.7 | [69] | protein phosphatase, Mg2+/Mn2+ dependent 1L |
| RAD24 | RAD17 | 1 | Gem | 1-0-14 | H, L | -7.4 | -27.6 | 14.2 | 8.3 | [70-74] | RAD17 checkpoint clamp loader component |
| SGS1 | RECQL5 | 2 | Cyt | 1-0-14 | L | -8.4 | -33.4 | 3.4 | 19.3 | [75,76] | RecQ like helicase 5 |
| KTI11_2 | DPH3 | 1 | Cyt | 1-0-14 | H | -7.7 | -10.3 | 6.5 | 9.1 | [77-79] | diphthamide biosynthesis 3 |
| BIM1_2 | MAPRE2 | 2 | Gem | 1-0-14 | A | -7.7 | -15.4 | 16.0 | 20.0 | [80] | microtubule associated protein RP/EB family member 2 |
| BIM1_2 | MAPRE2 | 2 | Both | 1-0-14 | L | -7.7 | -15.4 | 16.0 | 20.0 | [80] | microtubule associated protein RP/EB family member 2 |
| BIM1_2 | MAPRE3 | 2 | Gem | 1-0-14 | A | -7.7 | -15.4 | 16.0 | 20.0 | [81] | microtubule associated protein RP/EB family member 3 |
| ASF1 | ASF1B | 2 | Cyt | 2-0.16-1 | L | -6.1 | -9.5 | 4.1 | 8.3 | [82] | anti-silencing function 1B histone chaperone |
| AVL9 | AVL9 | 1 | Cyt | 2-0.16-1 | H | -4.3 | -2.5 | 0.2 | 2.9 | [83-85] | AVL9 cell migration associated |
| PMR1 | ATP1A1 | 2 | Cyt | 2-0.16-1 | A, H | -3.8 | -9.8 | 3.6 | 10.1 | [86] | ATPase Na+/K+ transporting subunit alpha 1 |
| PMR1 | ATP1A2 | 2 | Gem | 2-0.16-1 | A | -3.8 | -9.8 | 3.6 | 10.1 | [87] | ATPase Na+/K+ transporting subunit alpha 2 |
| PMR1 | ATP1A3 | 2 | Cyt | 2-0.16-1 | L | -3.8 | -9.8 | 3.6 | 10.1 |  | ATPase Na+/K+ transporting subunit alpha 3 |
| PMR1 | ATP1A4 | 2 | Gem | 2-0.16-1 | A | -3.8 | -9.8 | 3.6 | 10.1 |  | ATPase Na+/K+ transporting subunit alpha 4 |
| PMR1 | ATP1A4 | 2 | Both | 2-0.16-1 | H | -3.8 | -9.8 | 3.6 | 10.1 |  | ATPase Na+/K+ transporting subunit alpha 4 |
| PMR1 | ATP2C1 | 2 | Cyt | 2-0.16-1 | A | -3.8 | -9.8 | 3.6 | 10.1 | [88,89] | ATPase secretory pathway Ca2+ transporting 1 |
| PMR1 | ATP2C1 | 2 | Both | 2-0.16-1 | H | -3.8 | -9.8 | 3.6 | 10.1 | [88,89] | ATPase secretory pathway Ca2+ transporting 1 |
| TOP3 | TOP3A | 2 | Cyt | 2-0.16-1 | L | -5.2 | -4.0 | 3.3 | 3.4 | [90-92] | DNA topoisomerase III alpha |
| VPS21 | RAB21 | 3 | Cyt | 2-0.16-1 | A, H | -7.2 | -4.1 | -0.4 | 2.4 | [93,94] | RAB21, member RAS oncogene family |
| VPS21 | RAB22A | 3 | Gem | 2-0.16-1 | A | -7.2 | -4.1 | -0.4 | 2.4 | [95-97] | RAB22A, member RAS oncogene family |
| ACB1_2 | ACBD4 | 2 | Gem | 2-0.16-1 | H | -5.4 | -4.8 | 4.5 | 0.6 | [98,99] | acyl-CoA binding domain containing 4 |
| ACB1_2 | ACBD5 | 2 | Cyt | 2-0.16-1 | H | -5.4 | -4.8 | 4.5 | 0.6 | [100] | acyl-CoA binding domain containing 5 |
| ACB1_2 | DBI | 2 | Cyt | 2-0.16-1 | A, H | -5.4 | -4.8 | 4.5 | 0.6 | [101-103] | diazepam binding inhibitor, acyl-CoA binding protein |
| CPR3 | PPIA | 3 | Cyt | 2-0.8-1 | A, H | 2.1 | 1.6 | -4.1 | -2.8 | [104-106] | peptidylprolyl isomerase A |
| CPR3 | RGPD4 | 3 | Gem | 2-0.8-1 | A | 2.1 | 1.6 | -4.1 | -2.8 |  | RANBP2-like and GRIP domain containing 4 |
| ELO3 | ELOVL1 | 3 | Both | 2-0.8-1 | L | 2.2 | 1.3 | -3.4 | -4.0 | [107,108] | ELOVL fatty acid elongase 1 |
| ELO3 | ELOVL2 | 3 | Cyt | 2-0.8-1 | H | 2.2 | 1.3 | -3.4 | -4.0 | [109] | ELOVL fatty acid elongase 2 |
| ELO3 | ELOVL4 | 3 | Cyt | 2-0.8-1 | H | 2.2 | 1.3 | -3.4 | -4.0 |  | ELOVL fatty acid elongase 4 |
| ELO3 | ELOVL5 | 3 | Cyt | 2-0.8-1 | H | 2.2 | 1.3 | -3.4 | -4.0 |  | ELOVL fatty acid elongase 5 |
| ELO3 | ELOVL6 | 3 | Both | 2-0.8-1 | A, L | 2.2 | 1.3 | -3.4 | -4.0 | [110,111] | ELOVL fatty acid elongase 6 |
| MDL2 | ABCB10 | 3 | Gem | 2-0.8-1 | H | 2.5 | 1.5 | -3.0 | -3.0 | [112] | ATP binding cassette subfamily B member 10 |
| MDL2 | TAP1 | 3 | Cyt | 2-0.8-1 | L | 2.5 | 1.5 | -3.0 | -3.0 |  | transporter 1, ATP binding cassette subfamily B member |
| PIF1 | PIF1 | 2 | Gem | 2-0.8-1 | A | 2.2 | 1.5 | -4.5 | -3.4 | [113] | PIF1 5'-to-3' DNA helicase |
| RPS1B | RPS3A | 1 | Both | 2-0.8-1 | A | 2.3 | 0.9 | -3.9 | -2.3 | [114,115] | ribosomal protein S3A |
| SAC3 | MCM3AP | 2 | Gem | 2-0.8-1 | H | 2.2 | 1.5 | -5.2 | -3.8 | [116] | minichromosome maintenance complex component 3 associated protein |
| SAC3 | SAC3D1 | 2 | Cyt | 2-0.8-1 | H | 2.2 | 1.5 | -5.2 | -3.8 | [117,118] | SAC3 domain containing 1 |
| YTA7 | ATAD2 | 2 | Both | 2-0.8-1 | A, H | 1.8 | 1.0 | -6.0 | -3.6 | [119-125] | ATPase family, AAA domain containing 2 |
| YTA7 | ATAD2B | 2 | Both | 2-0.8-1 | H | 1.8 | 1.0 | -6.0 | -3.6 |  | ATPase family, AAA domain containing 2B |

**Table 4.**
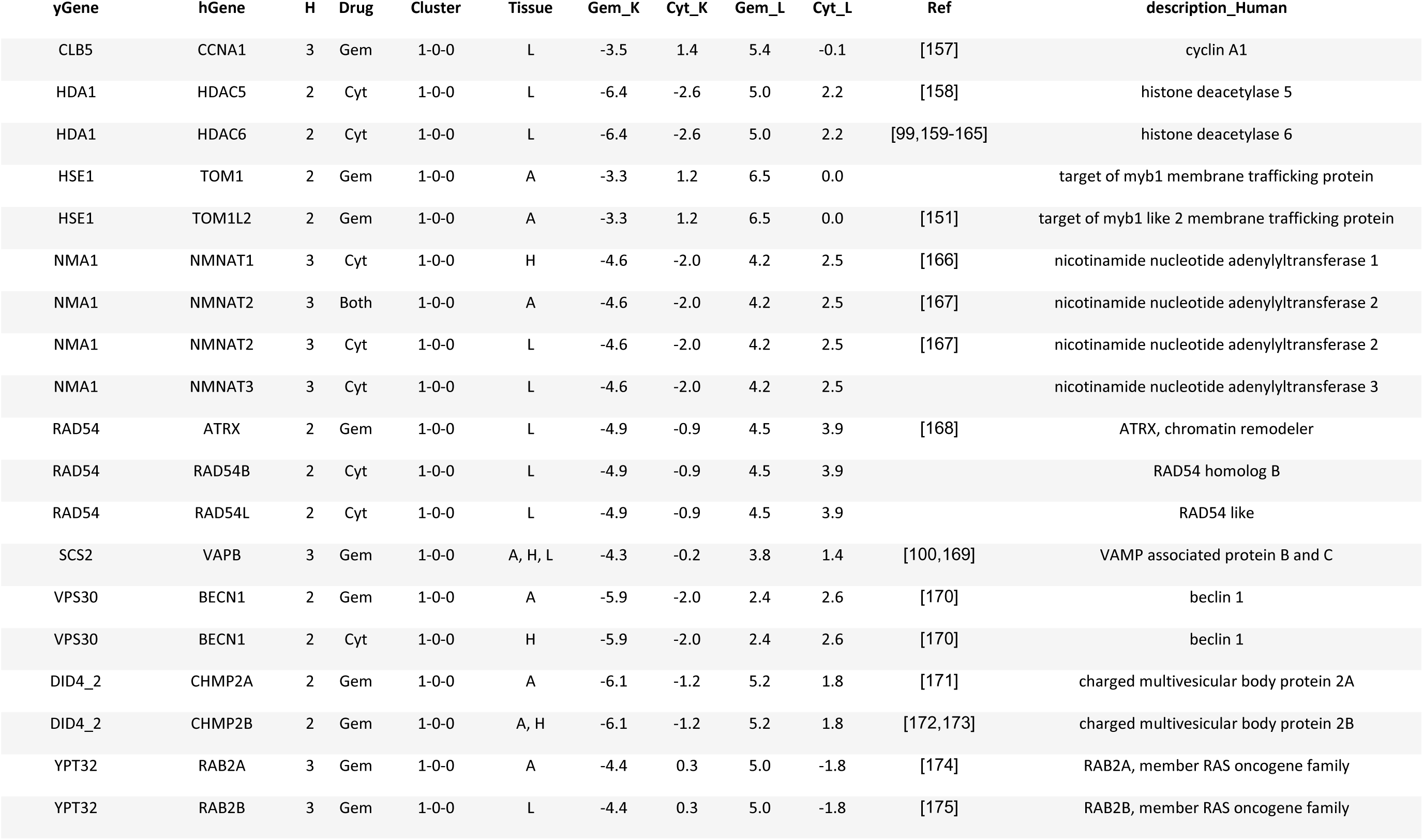
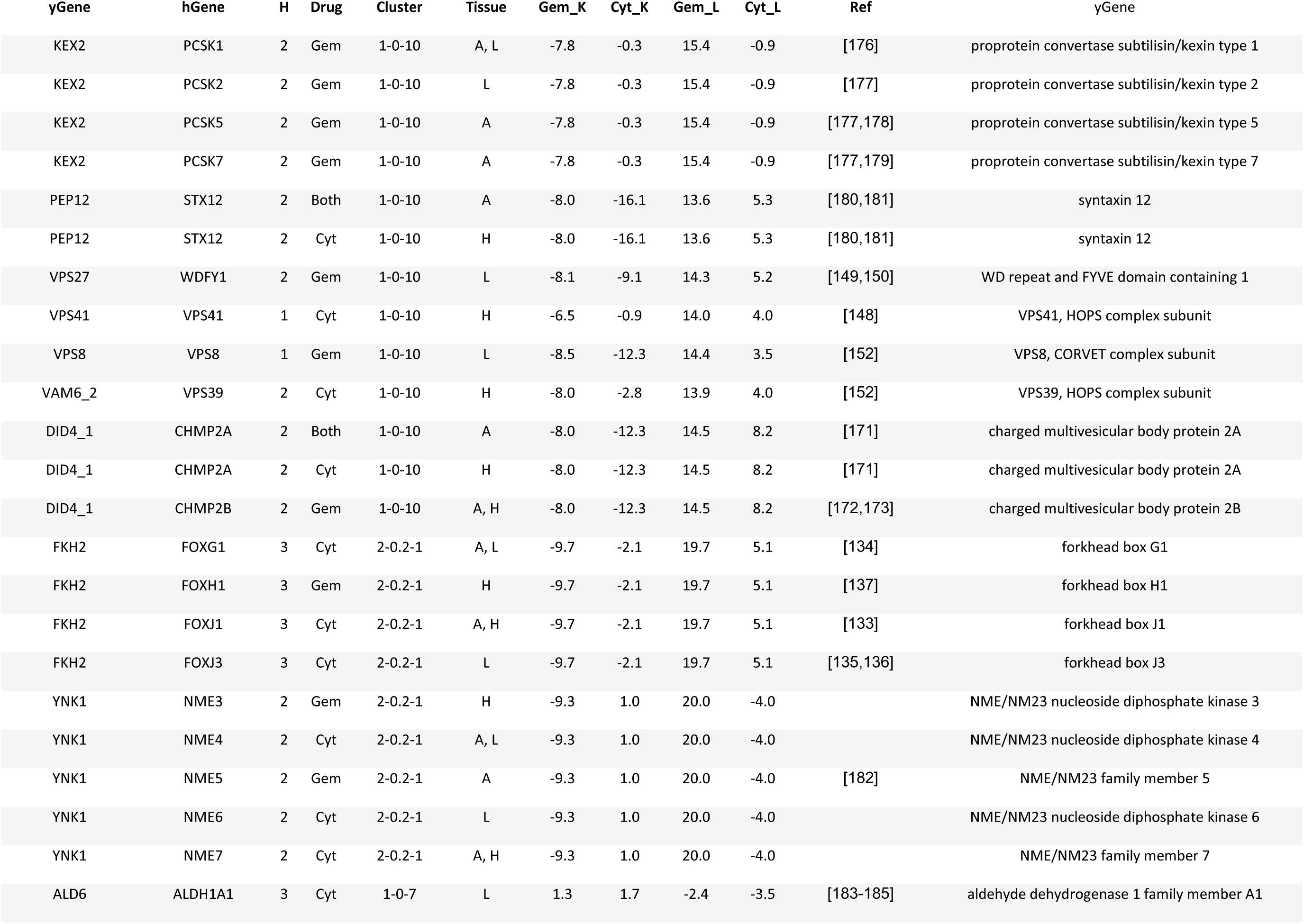
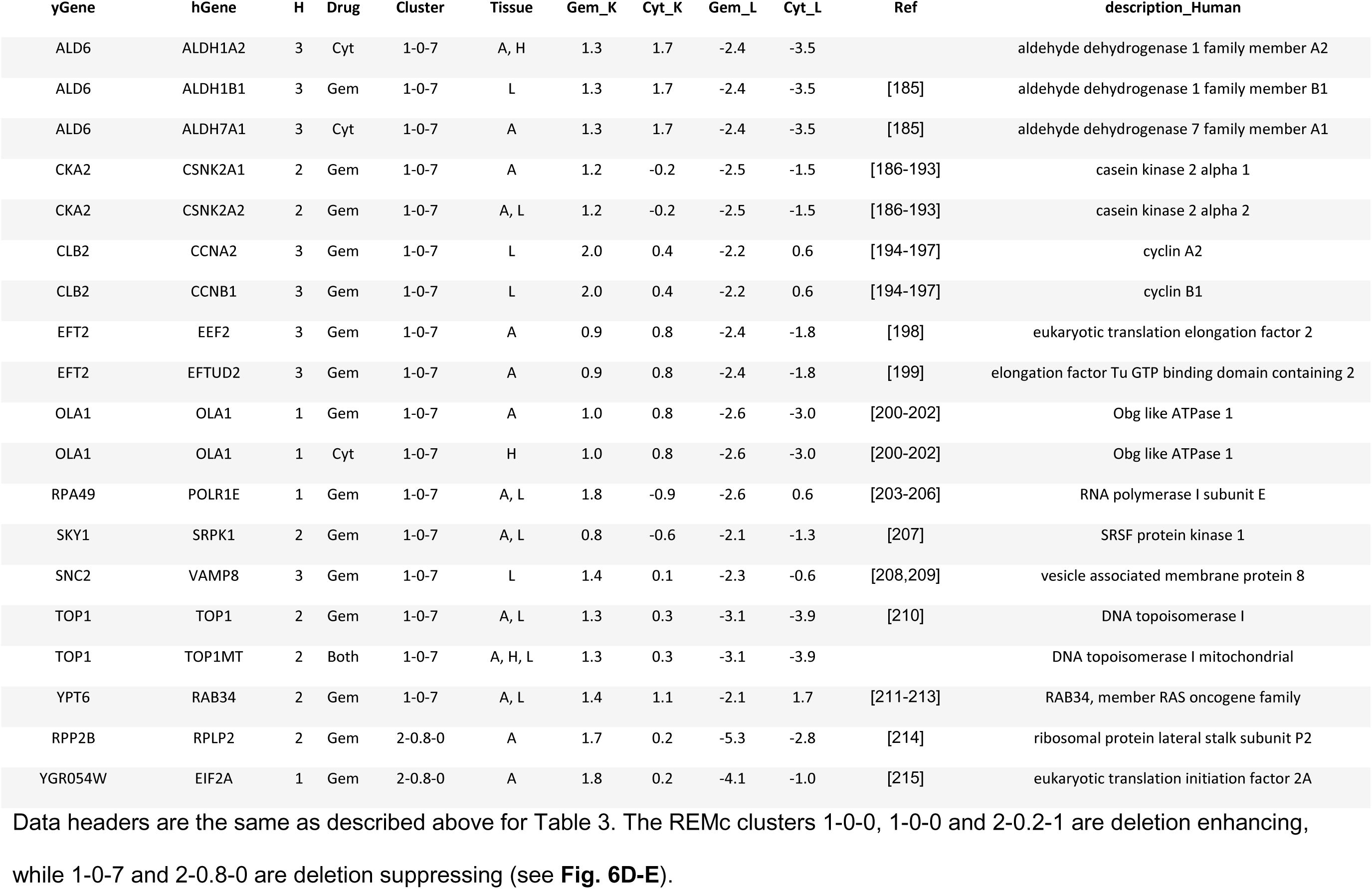
Yeast-human homologs predicted to buffer or promote gemcitabine to greater degree than cytarabine.

| yGene | hGene | H | Drug | Cluster | Tissue | Gem_K | Cyt_K | Gem_L | Cyt_L | Ref | description_Human |
| --- | --- | --- | --- | --- | --- | --- | --- | --- | --- | --- | --- |
| CLB5 | CCNA1 | 3 | Gem | 1-0-0 | L | -3.5 | 1.4 | 5.4 | -0.1 | [157] | cyclin A1 |
| HDA1 | HDAC5 | 2 | Cyt | 1-0-0 | L | -6.4 | -2.6 | 5.0 | 2.2 | [158] | histone deacetylase 5 |
| HDA1 | HDAC6 | 2 | Cyt | 1-0-0 | L | -6.4 | -2.6 | 5.0 | 2.2 | [99,159-165] | histone deacetylase 6 |
| HSE1 | TOM1 | 2 | Gem | 1-0-0 | A | -3.3 | 1.2 | 6.5 | 0.0 |  | target of myb1 membrane trafficking protein |
| HSE1 | TOM1L2 | 2 | Gem | 1-0-0 | A | -3.3 | 1.2 | 6.5 | 0.0 | [151] | target of myb1 like 2 membrane trafficking protein |
| NMA1 | NMNAT1 | 3 | Cyt | 1-0-0 | H | -4.6 | -2.0 | 4.2 | 2.5 | [166] | nicotinamide nucleotide adenyltransferase 1 |
| NMA1 | NMNAT2 | 3 | Both | 1-0-0 | A | -4.6 | -2.0 | 4.2 | 2.5 | [167] | nicotinamide nucleotide adenyltransferase 2 |
| NMA1 | NMNAT2 | 3 | Cyt | 1-0-0 | L | -4.6 | -2.0 | 4.2 | 2.5 | [167] | nicotinamide nucleotide adenyltransferase 2 |
| NMA1 | NMNAT3 | 3 | Cyt | 1-0-0 | L | -4.6 | -2.0 | 4.2 | 2.5 |  | nicotinamide nucleotide adenyltransferase 3 |
| RAD54 | ATRX | 2 | Gem | 1-0-0 | L | -4.9 | -0.9 | 4.5 | 3.9 | [168] | ATRX, chromatin remodeler |
| RAD54 | RAD54B | 2 | Cyt | 1-0-0 | L | -4.9 | -0.9 | 4.5 | 3.9 |  | RAD54 homolog B |
| RAD54 | RAD54L | 2 | Cyt | 1-0-0 | L | -4.9 | -0.9 | 4.5 | 3.9 |  | RAD54 like |
| SCS2 | VAPB | 3 | Gem | 1-0-0 | A, H, L | -4.3 | -0.2 | 3.8 | 1.4 | [100,169] | VAMP associated protein B and C |
| VPS30 | BECN1 | 2 | Gem | 1-0-0 | A | -5.9 | -2.0 | 2.4 | 2.6 | [170] | beclin 1 |
| VPS30 | BECN1 | 2 | Cyt | 1-0-0 | H | -5.9 | -2.0 | 2.4 | 2.6 | [170] | beclin 1 |
| DID4_2 | CHMP2A | 2 | Gem | 1-0-0 | A | -6.1 | -1.2 | 5.2 | 1.8 | [171] | charged multivesicular body protein 2A |
| DID4_2 | CHMP2B | 2 | Gem | 1-0-0 | A, H | -6.1 | -1.2 | 5.2 | 1.8 | [172,173] | charged multivesicular body protein 2B |
| YPT32 | RAB2A | 3 | Gem | 1-0-0 | A | -4.4 | 0.3 | 5.0 | -1.8 | [174] | RAB2A, member RAS oncogene family |
| YPT32 | RAB2B | 3 | Gem | 1-0-0 | L | -4.4 | 0.3 | 5.0 | -1.8 | [175] | RAB2B, member RAS oncogene family |

| yGene | hGene | H | Drug | Cluster | Tissue | Gem_K | Cyt_K | Gem_L | Cyt_L | Ref | yGene |
| --- | --- | --- | --- | --- | --- | --- | --- | --- | --- | --- | --- |
| KEX2 | PCSK1 | 2 | Gem | 1-0-10 | A, L | -7.8 | -0.3 | 15.4 | -0.9 | [176] | proprotein convertase subtilisin/kexin type 1 |
| KEX2 | PCSK2 | 2 | Gem | 1-0-10 | L | -7.8 | -0.3 | 15.4 | -0.9 | [177] | proprotein convertase subtilisin/kexin type 2 |
| KEX2 | PCSK5 | 2 | Gem | 1-0-10 | A | -7.8 | -0.3 | 15.4 | -0.9 | [177,178] | proprotein convertase subtilisin/kexin type 5 |
| KEX2 | PCSK7 | 2 | Gem | 1-0-10 | A | -7.8 | -0.3 | 15.4 | -0.9 | [177,179] | proprotein convertase subtilisin/kexin type 7 |
| PEP12 | STX12 | 2 | Both | 1-0-10 | A | -8.0 | -16.1 | 13.6 | 5.3 | [180,181] | syntaxin 12 |
| PEP12 | STX12 | 2 | Cyt | 1-0-10 | H | -8.0 | -16.1 | 13.6 | 5.3 | [180,181] | syntaxin 12 |
| VPS27 | WDFY1 | 2 | Gem | 1-0-10 | L | -8.1 | -9.1 | 14.3 | 5.2 | [149,150] | WD repeat and FYVE domain containing 1 |
| VPS41 | VPS41 | 1 | Cyt | 1-0-10 | H | -6.5 | -0.9 | 14.0 | 4.0 | [148] | VPS41, HOPS complex subunit |
| VPS8 | VPS8 | 1 | Gem | 1-0-10 | L | -8.5 | -12.3 | 14.4 | 3.5 | [152] | VPS8, CORVET complex subunit |
| VAM6_2 | VPS39 | 2 | Cyt | 1-0-10 | H | -8.0 | -2.8 | 13.9 | 4.0 | [152] | VPS39, HOPS complex subunit |
| DID4_1 | CHMP2A | 2 | Both | 1-0-10 | A | -8.0 | -12.3 | 14.5 | 8.2 | [171] | charged multivesicular body protein 2A |
| DID4_1 | CHMP2A | 2 | Cyt | 1-0-10 | H | -8.0 | -12.3 | 14.5 | 8.2 | [171] | charged multivesicular body protein 2A |
| DID4_1 | CHMP2B | 2 | Gem | 1-0-10 | A, H | -8.0 | -12.3 | 14.5 | 8.2 | [172,173] | charged multivesicular body protein 2B |
| FKH2 | FOXG1 | 3 | Cyt | 2-0.2-1 | A, L | -9.7 | -2.1 | 19.7 | 5.1 | [134] | forkhead box G1 |
| FKH2 | FOXH1 | 3 | Gem | 2-0.2-1 | H | -9.7 | -2.1 | 19.7 | 5.1 | [137] | forkhead box H1 |
| FKH2 | FOXJ1 | 3 | Cyt | 2-0.2-1 | A, H | -9.7 | -2.1 | 19.7 | 5.1 | [133] | forkhead box J1 |
| FKH2 | FOXJ3 | 3 | Cyt | 2-0.2-1 | L | -9.7 | -2.1 | 19.7 | 5.1 | [135,136] | forkhead box J3 |
| YNK1 | NME3 | 2 | Gem | 2-0.2-1 | H | -9.3 | 1.0 | 20.0 | -4.0 |  | NME/NM23 nucleoside diphosphate kinase 3 |
| YNK1 | NME4 | 2 | Cyt | 2-0.2-1 | A, L | -9.3 | 1.0 | 20.0 | -4.0 |  | NME/NM23 nucleoside diphosphate kinase 4 |
| YNK1 | NME5 | 2 | Gem | 2-0.2-1 | A | -9.3 | 1.0 | 20.0 | -4.0 | [182] | NME/NM23 family member 5 |
| YNK1 | NME6 | 2 | Cyt | 2-0.2-1 | L | -9.3 | 1.0 | 20.0 | -4.0 |  | NME/NM23 nucleoside diphosphate kinase 6 |
| YNK1 | NME7 | 2 | Cyt | 2-0.2-1 | A, H | -9.3 | 1.0 | 20.0 | -4.0 |  | NME/NM23 family member 7 |
| ALD6 | ALDH1A1 | 3 | Cyt | 1-0-7 | L | 1.3 | 1.7 | -2.4 | -3.5 | [183-185] | aldehyde dehydrogenase 1 family member A1 |
| ALD6 | ALDH1A2 | 3 | Cyt | 1-0-7 | A, H | 1.3 | 1.7 | -2.4 | -3.5 |  | aldehyde dehydrogenase 1 family member A2 |
| ALD6 | ALDH1B1 | 3 | Gem | 1-0-7 | L | 1.3 | 1.7 | -2.4 | -3.5 | [185] | aldehyde dehydrogenase 1 family member B1 |
| ALD6 | ALDH7A1 | 3 | Cyt | 1-0-7 | A | 1.3 | 1.7 | -2.4 | -3.5 | [185] | aldehyde dehydrogenase 7 family member A1 |
| CKA2 | CSNK2A1 | 2 | Gem | 1-0-7 | A | 1.2 | -0.2 | -2.5 | -1.5 | [186-193] | casein kinase 2 alpha 1 |
| CKA2 | CSNK2A2 | 2 | Gem | 1-0-7 | A, L | 1.2 | -0.2 | -2.5 | -1.5 | [186-193] | casein kinase 2 alpha 2 |
| CLB2 | CCNA2 | 3 | Gem | 1-0-7 | L | 2.0 | 0.4 | -2.2 | 0.6 | [194-197] | cyclin A2 |
| CLB2 | CCNB1 | 3 | Gem | 1-0-7 | L | 2.0 | 0.4 | -2.2 | 0.6 | [194-197] | cyclin B1 |
| EFT2 | EEF2 | 3 | Gem | 1-0-7 | A | 0.9 | 0.8 | -2.4 | -1.8 | [198] | eukaryotic translation elongation factor 2 |
| EFT2 | EFTUD2 | 3 | Gem | 1-0-7 | A | 0.9 | 0.8 | -2.4 | -1.8 | [199] | elongation factor Tu GTP binding domain containing 2 |
| OLA1 | OLA1 | 1 | Gem | 1-0-7 | A | 1.0 | 0.8 | -2.6 | -3.0 | [200-202] | Obg like ATPase 1 |
| OLA1 | OLA1 | 1 | Cyt | 1-0-7 | H | 1.0 | 0.8 | -2.6 | -3.0 | [200-202] | Obg like ATPase 1 |
| RPA49 | POLR1E | 1 | Gem | 1-0-7 | A, L | 1.8 | -0.9 | -2.6 | 0.6 | [203-206] | RNA polymerase I subunit E |
| SKY1 | SRPK1 | 2 | Gem | 1-0-7 | A, L | 0.8 | -0.6 | -2.1 | -1.3 | [207] | SRSF protein kinase 1 |
| SNC2 | VAMP8 | 3 | Gem | 1-0-7 | L | 1.4 | 0.1 | -2.3 | -0.6 | [208,209] | vesicle associated membrane protein 8 |
| TOP1 | TOP1 | 2 | Gem | 1-0-7 | A, L | 1.3 | 0.3 | -3.1 | -3.9 | [210] | DNA topoisomerase I |
| TOP1 | TOP1MT | 2 | Both | 1-0-7 | A, H, L | 1.3 | 0.3 | -3.1 | -3.9 |  | DNA topoisomerase I mitochondrial |
| YPT6 | RAB34 | 2 | Gem | 1-0-7 | A, L | 1.4 | 1.1 | -2.1 | 1.7 | [211-213] | RAB34, member RAS oncogene family |
| RPP2B | RPLP2 | 2 | Gem | 2-0.8-0 | A | 1.7 | 0.2 | -5.3 | -2.8 | [214] | ribosomal protein lateral stalk subunit P2 |
| YGR054W | EIF2A | 1 | Gem | 2-0.8-0 | A | 1.8 | 0.2 | -4.1 | -1.0 | [215] | eukaryotic translation initiation factor 2A |

**Table 5.** Yeast-human homologs predicted to buffer cytarabine to greater degree than gemcitabine.

| yGene | hGene | H | Drug | Cluster | Tissue | Gem_K | Cyt_K | Gem_L | Cyt_L | Ref | description_Human |
| --- | --- | --- | --- | --- | --- | --- | --- | --- | --- | --- | --- |
| CCH1 | CACNA1A | 2 | Cyt | 1-0-9 | A, L | 0.2 | -4.5 | 0.5 | 5.5 | [274] | calcium voltage-gated channel subunit alpha1 A |
| CCH1 | CACNA1B | 2 | Cyt | 1-0-9 | A, L | 0.2 | -4.5 | 0.5 | 5.5 | [274] | calcium voltage-gated channel subunit alpha1 B |
| CCH1 | CACNA1C | 2 | Cyt | 1-0-9 | A, H, L | 0.2 | -4.5 | 0.5 | 5.5 | [274] | calcium voltage-gated channel subunit alpha1 C |
| CCH1 | CACNA1E | 2 | Cyt | 1-0-9 | A, L | 0.2 | -4.5 | 0.5 | 5.5 | [274] | calcium voltage-gated channel subunit alpha1 E |
| CCH1 | CACNA1F | 2 | Cyt | 1-0-9 | A | 0.2 | -4.5 | 0.5 | 5.5 | [274] | calcium voltage-gated channel subunit alpha1 F |
| CCH1 | NALCN | 2 | Cyt | 1-0-9 | H | 0.2 | -4.5 | 0.5 | 5.5 |  | sodium leak channel, non-selective |
| FAT1 | SLC27A2 | 2 | Cyt | 1-0-9 | L | 0.7 | -8.5 | -0.9 | 8.9 | [275] | solute carrier family 27 member 2 |
| FAT1 | SLC27A3 | 2 | Cyt | 1-0-9 | L | 0.7 | -8.5 | -0.9 | 8.9 | [276] | solute carrier family 27 member 3 |
| FOL2 | GCH1 | 1 | Cyt | 1-0-9 | L | -0.9 | -9.4 | 0.7 | 7.1 | [277] | GTP cyclohydrolase 1 |
| HSL1 | BRSK1 | 3 | Cyt | 1-0-9 | A, L | 0.9 | -10.4 | 0.1 | 11.6 | [270,271] | BR serine/threonine kinase 1 |
| HSL1 | BRSK2 | 3 | Cyt | 1-0-9 | A, L | 0.9 | -10.4 | 0.1 | 11.6 | [272] | BR serine/threonine kinase 2 |
| IZH1 | ADIPOR1 | 3 | Cyt | 1-0-9 | H | 1.1 | -5.8 | -0.4 | 7.6 | [278,279] | adiponectin receptor 1 |
| IZH1 | PAQR4 | 3 | Cyt | 1-0-9 | A, L | 1.1 | -5.8 | -0.4 | 7.6 |  | progesterin and adipoQ receptor family member 4 |
| NAP1 | NAP1L3 | 2 | Cyt | 1-0-9 | A, L | 1.0 | -4.7 | -1.5 | 5.6 | [280] | nucleosome assembly protein 1 like 3 |
| NAP1 | NAP1L4 | 2 | Cyt | 1-0-9 | L | 1.0 | -4.7 | -1.5 | 5.6 |  | nucleosome assembly protein 1 like 4 |
| PTM1 | TMEM87B | 3 | Cyt | 1-0-9 | A, H | -0.7 | -3.8 | -0.2 | 5.7 | [281] | transmembrane protein 87B |

The data descriptions are the same as for Table 3.

In summary REMc, GTF, and GTA revealed functional genetic modules that alternatively buffer (deletion enhancing) or promote (deletion suppressing) drug cytotoxicity [5,40,51], and illustrated whether the effects were shared or differential between gemcitabine and cytarabine (Fig. 4). Yeast phenomic information was integrated with pharmacogenomics data results according to yeast-human gene homology to identify correlated differential gene expression associated with drug sensitivity in cancer cell lines (Figs. 5-7). This approach serves to generate hypotheses regarding whether differential expression of a particular gene is causal for increased drug sensitivity [52], and ultimately whether yeast phenomic models can improve the predictive value of cancer pharmacogenomics data in the context of precision oncology [53-58].

**Figure 5.**
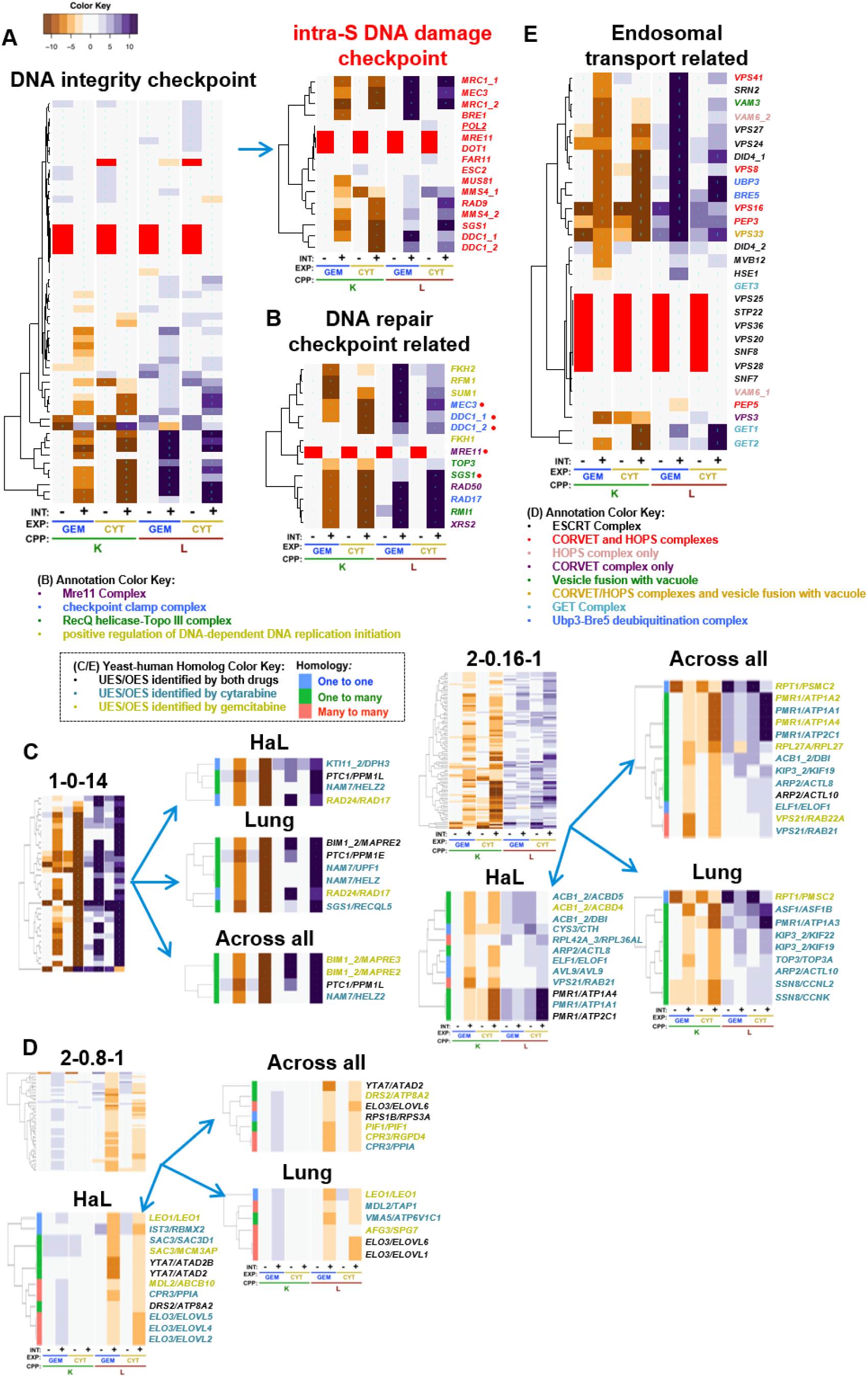
Drug-gene interaction common to gemcitabine and cytarabine. Genes that similarly influence the cytotoxicity of both gemcitabine and cytarabine suggest common pathways that buffer or promote toxicity, as illustrated by: (**A**) GO term-specific heatmaps for *DNA integrity checkpoint* and its child term *intra-S DNA damage checkpoint*, which buffer gemcitabine and cytarabine, along with (**B**) genes comprising other DNA checkpoint/repair related GO terms, such as *positive regulation of DNA-dependent DNA replication initiation*, and the Mre11, checkpoint clamp, and RecQ helicase-Topo III complexes; (**C, D**) REMc clusters filtered for PharmacoDB results for yeast-human homologs that exhibited (**C**) deletion enhancement and UES or (**D**) deletion suppression and OES; and (**E**) deletion enhancing endosomal-transport-related GO terms, including *vesicle fusion with vacuole*, and the CORVET/HOPS, ESCRT, GET, and Ubp3-Bre5 deubiquitination complexes. Gene labels are color-coordinated with legends in panels B and E, and as described in Fig. 3H for panels C and D.

**Figure 6.**
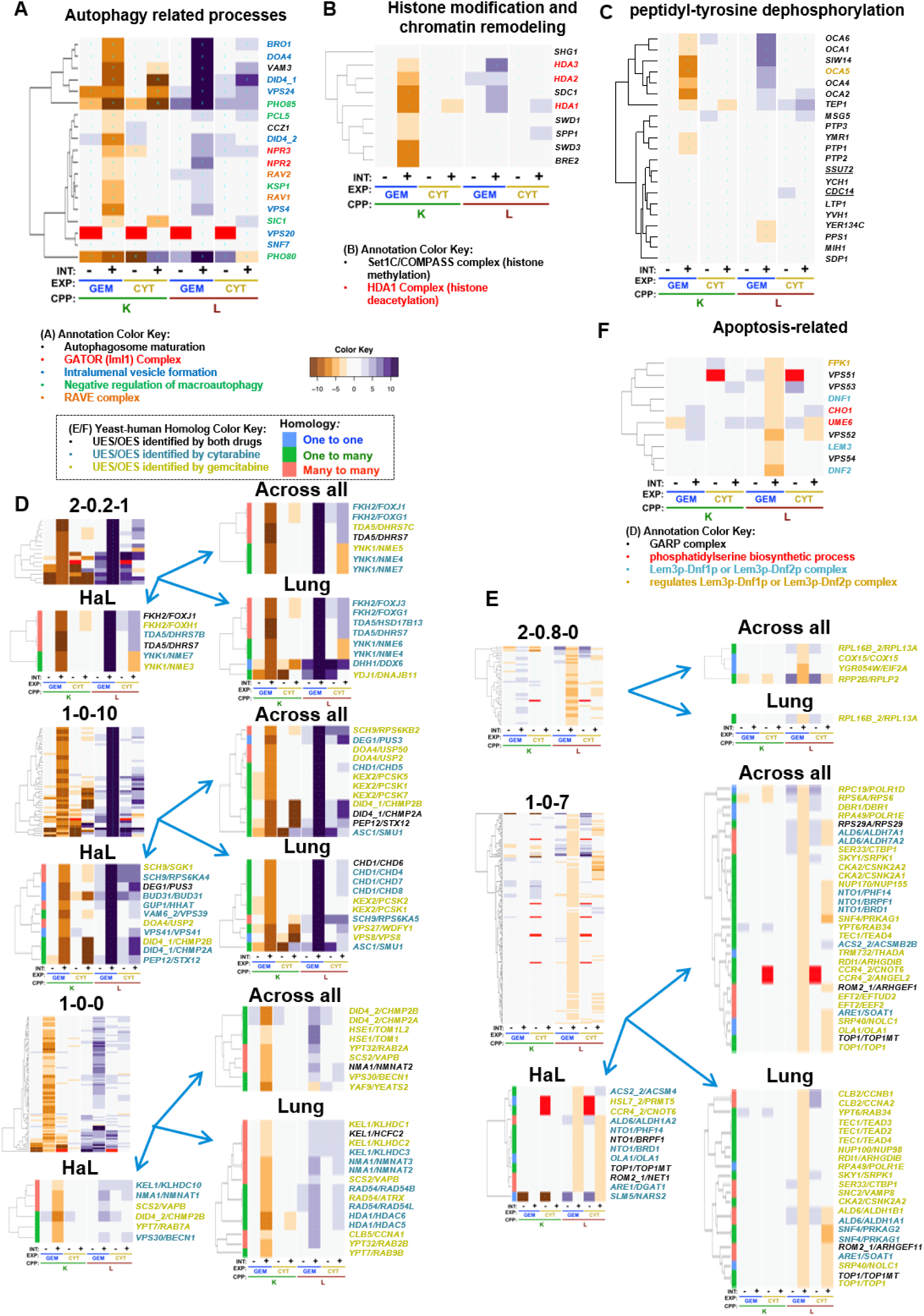
Gemcitabine-specific gene interaction. (**A-C**) Cellular processes that buffer gemcitabine to a greater extent than cytarabine included: (**A**) Autophagy-related processes; (**B**) Histone modification and chromatin remodeling (particularly for K interaction); and (**C**) Peptidyl-tyrosine dephosphorylation, representing the genes *OCA (1-6)* (*OCA5* was manually added to the panel (see text); *OCA3*/*SIW14* are aliases). (**D-E**) Human genes that are predicted to (**D**) buffer gemcitabine toxicity if they are UES and deletion enhancing, and to (**E**) promote gemcitabine toxicity if they are found to be OES and deletion suppressing, when comparing homolog interactions across yeast phenomic and cancer pharmacogenomic analyses. (**F**) Apoptosis-related genes and complexes were observed to promote toxicity of gemcitabine more than toxicity of cytarabine. Gene labels are color-coordinated with legends in panels A, B, and F, and as described in Fig. 3H for panels D and E.

**Figure 7.**
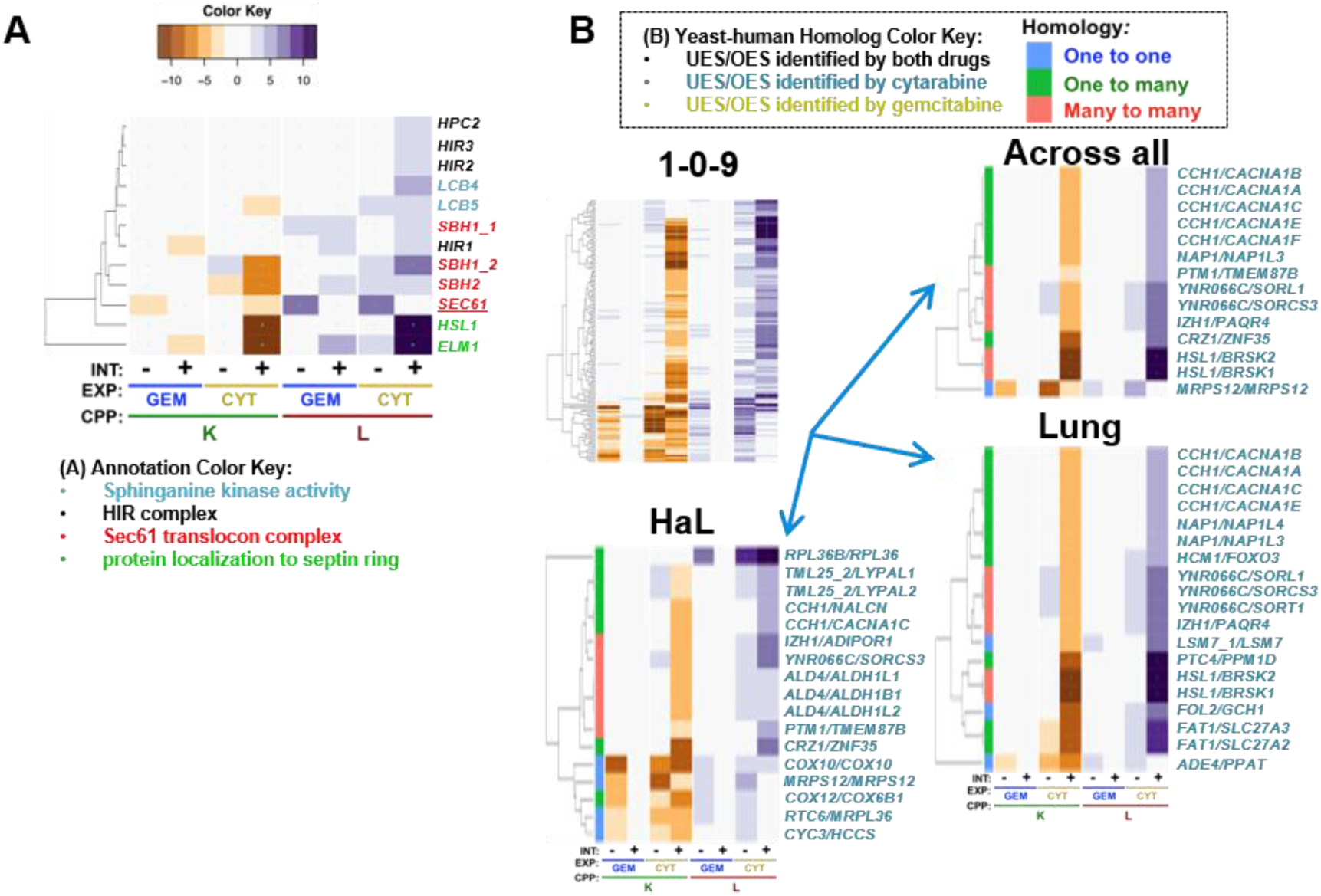
Cytarabine-specific gene interaction. (**A**) GO terms identified by GTA that revealed deletion enhancement to be greater for cytarabine than gemcitabine. (**B**) Human homologs of cytarabine-specific yeast gene deletion enhancers found to exhibit underexpression sensitivity for cytarabine in cancer cell lines.

### Functions that respond to gemcitabine and cytarabine similarly

#### Genetic modules that buffer cytotoxicity of both gemcitabine and cytarabine

To characterize gemcitabine and cytarabine, which have similar molecular structures and mechanisms of action, yet different spectra of anti-tumor efficacy, we first surveyed for buffering genes shared in common. Examples of genes with deletion enhancing interactions for both drugs are displayed in clusters 2-0.2-0, 1-0-14, 2-0.2-2 and 2-0.16-1 (Fig. 3B). GO enrichment was observed in these clusters for the *DNA integrity checkpoint, positive regulation of DNA replication,* and the Mre11, RecQ helicase-Topo III, CORVET, HOPS, GET, and Ubp3-Bre5 deubiquitination complexes (Fig. 4, Table 1). GTA identified many of the same functions and additionally revealed the terms *vesicle fusion with vacuole* and checkpoint clamp complex (Table 2). We mapped yeast gene-drug interactions to respective human homologs in PharmacoDB to find evidence for evolutionary conservation of gene-drug interaction (Fig. 5C-D, **Additional File 8, Files B-D**) and buffering mechanisms.

#### DNA integrity checkpoint and repair-related complexes

As gemcitabine and cytarabine triphosphate analogs are incorporated into DNA, we anticipated shared interactions with genes functioning in DNA metabolism and repair. Overlap was observed, however there were differential effects between genes assigned to the same gene ontology terms, such that GO TermFinder enrichment in REMc clusters was less than might have been expected. For example, deletion enhancing gene-drug interaction for the GO term, *DNA integrity checkpoint*, was enriched in cluster 1-0-14, which displayed deletion enhancement for both gemcitabine and cytarabine (Table 1, Figs. 3 **and** Fig. 5A). However, its child term, *intra-S DNA damage checkpoint*, was not GO-enriched because of differential clustering among drug-gene interactions associated with the term (**Additional File 5, File C**). Similarly, *intra-S DNA damage checkpoint* was not identified by GTA due to variation in interaction between genes assigned to the term, highlighting the utility in displaying the phenomic data for each GO term for manual review (Fig. 5A).

Enriched complexes functionally related to the DNA integrity checkpoint function included the RecQ helicase-Topo III, the checkpoint clamp, and the Mre11 complexes (Fig. 5B). Rmi1, Top3, and Sgs1 form the RecQ helicase Topo III complex, which is involved in Rad53 checkpoint activation and maintenance of genome integrity [59], and together with replication protein A functions in DNA decatenation and disentangling of chromosomes [60]. *RMI1* and *SGS1* deletion enhancement clustered together in 1-0-14, while *TOP3* had a similar, but slightly weaker interaction pattern in cluster 1-0-16 (**Additional File 5, File B**). The human homolog of *SGS1*, *RECQL5*, was UES for cytarabine in lung cancer cells (Fig. 5C; see **1-0-14** in 5C, all cluster heatmaps available in **Additional File 5, File B**). *RECQL5* preserves genome stability during transcription elongation, and deletion of *RECQL5* increases cancer susceptibility [75,76]. Human *TOP3A* was also UES for cytarabine in lung tissue (Fig 5C; **2-0.16-1**). *TOP3A* is underexpressed in ovarian cancer, and mutations in *TOP3A* are associated with increased risk for acute myeloid leukemia, myelodysplastic syndromes, suggesting potential cancer vulnerabilities if somatic, but can also occur in the germline, which would lead to enhanced host toxicity [90-92].

The checkpoint clamp in yeast is comprised of Rad17/hRad1, Ddc1, and Mec3, which function downstream of Rad24/hRad17 in the DNA damage checkpoint pathway [70-72] to recruit yDpb11/hTopB1 to stalled replication forks and activate the yMec1/hATR protein kinase activity, initiating the DNA damage response [73]. The human homolog of yeast *RAD24, RAD17*, was UES for gemcitabine in both lung and hematopoietic & lymphoid tissue (Fig. 5C; **1-0-14**), representing a synthetic lethal relationship of potential therapeutic relevance. Consistent with this finding in yeast, depletion of *hRAD17* can sensitize pancreatic cancer cells to gemcitabine [74].

Mre11, Xrs2, and Rad50 constitute the Mre11 complex, which participates in the formation and processing of double-strand DNA breaks involved in recombination and repair [126], and clustered together in 1-0-14 (Figs. 5B-C). Deficiency in the Mre1 complex is known to sensitize human cells to nucleoside analog toxicity [127], as also seen in cancer cell lines deficient for other checkpoint-signaling genes, such as Rad9, Chk1, or ATR, [128]. Single nucleotide polymorphisms in DNA damage response (*ATM* and *CHEK1*) have been associated with overall survival in pancreatic cancer patients treated with gemcitabine and radiation therapy [129]. Taken together, the results highlight evolutionarily conserved genes that function in DNA replication and recombination-based repair and are required to buffer the cytotoxic effects of both cytarabine and gemcitabine.

#### Positive regulation of DNA-dependent DNA replication initiation

The term, *positive regulation of DNA-dependent DNA replication initiation*, was identified by REMc/GTF and GTA for buffering interactions with both drugs, though stronger for gemcitabine (Tables 1 **and** 2). Genes representing this term were *FKH2, RFM1,* and *SUM1* (Fig. 5B). The origin binding protein, Sum1, is required for efficient replication initiation [130] and forms a complex with Rfm1 and the histone deacetylase, Hst1, which is recruited to replication origins to deacetylate H4K5 for initiation [131]. *HST1* was also a strong deletion enhancer but was observed only for the L parameter and clustered in 2-0.2-2. The forkhead box proteins, Fkh1 and Fkh2, contribute to proper replication origin timing and long range clustering of origins in G1 phase [132], and appear to buffer the cytotoxicity of gemcitabine more so than cytarabine, with *FKH2* deletion showing a stronger effect than its paralog (Fig. 5B). Multiple human forkhead box protein homologs (*yFKH2/hFOXJ1/FOXG1/FOXJ3/FOXH1*) (Fig. 6D) were observed as UES in PharmacoDB, of which *FOXJ1* underexpression is a marker of poor prognosis in gastric cancer [133], reduced expression of *FOXG1* is correlated with worse prognosis in breast cancer [134], *FOXJ3* is inhibited by miR-517a and associated with lung and colorectal cancer cell proliferation and invasion [135,136], and *FOXH1* is overexpressed in breast cancer, and *FOXH1* inhibition reduces proliferation in breast cancer cell lines [137]. Although not UES in PharmacoDB, inhibition of the *HST1* homolog, *SIRT1,* by Tenovin-6 inhibits the growth of acute lymphoblastic leukemia cells and enhances cytarabine cytotoxicity [138], enhances gemcitabine efficacy in pancreatic cancer cell lines, and improves survival in a pancreatic cancer mouse model [139]. Thus, loss of this gene module that positively regulates DNA replication initiation appears to be robustly involved in oncogenesis and is also synthetic lethal with gemcitabine and cytarabine.

#### Endosomal transport and related processes

GO annotated processes, enriched by REMc/GTF and GTA and having deletion enhancement profiles, related to endosome transport included *vesicle fusion with vacuole* (*VAM3* and *VPS33*), the CORVET/HOPS (V*PS41, VPS8, VPS16, PEP3, VPS33, VAM6,* and *VPS3*), ESCRT (*VPS27, VPS24, DID4, MVB12; HSE1* and *SRN2* were gemcitabine specific), GET complex (*GET1, GET2;* 2-0.14-0), and Ubp3-Bre5 deubiquitination (*UBP3* and *BRE5*) complexes (Tables 1-2, Fig. 5E). The CORVET and HOPS tethering complexes function in protein and lipid transport between endosomes and lysosomes/vacuoles, are required for vacuolar fusion, recognize SNARE complexes, help determine endomembrane identity, and interact with the ESCRT complex [140,141]. The ESCRT complex recognizes ubiquitinated endosomal proteins to mediate degradation through the multivesicular body pathway [142,143]. The Ubp3-Bre5 deubiquitination complex maintains Sec23, a subunit of COPII vesicles required for transport between the ER and Golgi, by cleaving its ubiquitinated form [144]. The GET complex (*GET1-3*) mediates insertion of tail-anchored proteins into the ER membrane, a critical process within the secretory pathway for vesicular trafficking [145-147]. Thus, these complexes, which function in processes related to endosomal transport, appear to be critical for buffering the toxicity of nucleoside analogs.

Several deletion enhancing, endosomal genes had human homologs associated with UES in cancer cell lines and/or reported roles in cancer biology (Figs. 5E, 6D), including: (1) *VPS41/VPS41*, in which a single nucleotide polymorphism is associated with familial melanoma [148]; (2) *VPS27/WDFY1*, which is regulated by *NPR2* to maintain the metastatic phenotype of cancer cells [149,150]; (3) human homologs of yeast *HSE1*, *TOM1* and *TOM1L2, TOM1L2* hypomorphic mice having increased tumor incidence associated with alterations in endosomal trafficking [151]; (4) *VPS8/VPS8* and *VAM6/VPS39,* which are predicted to be homologous members of the CORVET complex [152]; and (5) *VPS21/RAB21/RAB22A*, where *RAB21* promotes carcinoma-associated fibroblast invasion and knockdown inhibits glioma cell proliferation [93,94], and *RAB22A* promotes oncogenesis in lung, breast, and ovarian cancer [95-97]. Thus, it seems tumors arising in the context of deficiencies in certain endosomal trafficking genes could be vulnerable to gemcitabine and/or cytarabine.

#### ‘Non-GO-enriched’ homolog pairs with corresponding UES and deletion enhancement

We next explored yeast-human homologs exhibiting yeast deletion enhancement and underexpression sensitivity in cancer, systematically and regardless of whether their functions were enriched within Gene Ontology (Table 3, Fig. 5C). ‘Non-enriched’ interaction can be explained by a small total number of genes performing the function, only select genes annotated to a term impacting the phenotype, by genes contributing to a function without yet being annotated to it, by novel functions, and other possibilities.

With regard to the above, human homologs of the yeast type 2C protein phosphatase, *PTC1,* included *PPM1L* and *PPM1E* (Fig. 5C; **1-0-14**)*. PPM1L* has reduced expression in familial adenomatous polyposis [69], while *PPM1E* upregulation has been associated with cell proliferation in gastric cancer [68]. Such differential interactions of paralogs could result from tissue specific expression and functional differentiation of regulatory proteins. Previously, we reported *ptc1-Δ0* to buffer transcriptional repression of *RNR1* [34], which is upregulated as part of the DNA damage response to increase dNTP pools [153].

The microtubule binding proteins, y*BIM1/hMAPRE2/hMAPRE3*, were deletion enhancing in yeast and UES in cancer for gemcitabine (Fig. 5C; **1-0-14**), of which frameshift mutations were reported in *MAPRE3* for gastric and colorectal cancers [81], however, *MAPRE2* is upregulated in invasive pancreatic cancer cells [80], demonstrating that the yeast phenomic model could help distinguish causal influence in cases of paralogous gene expression having what appear to be opposing effects on phenotypic response of cancer cells to cytotoxic chemotherapy.

*NAM7* is a yeast RNA helicase that was deletion enhancing for both drugs, though slightly stronger for cytarabine, while its human homologs *HELZ, HELZ2*, and *UPF1*, were UES only with cytarabine (Fig. 5C; **1-0-14**). *HELZ* has differential influence in cancer, acting as a tumor suppressor or oncogene [61-64]. *UPF1* downregulation is associated with poor prognosis in gastric cancer and hepatocellular carcinoma, and mutations often occur in pancreatic adenosquamous carcinoma [65-67]. Thus, it is possible cytarabine could have efficacy for patients with mutational loss of function in members of this helicase family.

*ASF1/ASF1B* (Fig. 5C; **2-0.16-1**) functions in nucleosome assembly as an anti-silencing factor, and is one of the most overexpressed histone chaperones in cancer [82]. The yeast phenomic data suggest that anti-cancer approaches that target ASF1 as a driver [154] could be augmented by combination with gemcitabine or cytarabine.

*AVL9/AVL9* (Fig. 5C; **2-0.16-1**) functions in exocytic transport from the Golgi [83]. *AVL9* knockdown resulted in abnormal mitoses associated with defective protein trafficking, and increased cell migration with development of cysts [84], but also reduced cell proliferation and migration in other studies [85]. Regardless, the yeast phenomic model together with pharmacogenomics data would predict that functional loss of AVL9 renders cells vulnerable to cytarabine.

*PMR1* is a P-type ATPase that transports Mn++ and Ca++ into the Golgi. Several of its human homologs *ATP1A1, ATP1A2, ATP1A3, ATP1A4, ATP2C1* were UES, either for gemcitabine or cytarabine, in the PharmacoDB analysis (Fig. 5C; **2-0.16-1**). Reduced expression of *ATP1A1* can promote development of renal cell carcinoma [86], reduced expression of *ATP1A2* is associated with breast cancer [87], and mutations in *ATP2C1* impair the DNA damage response, and increase the incidence of squamous cell tumors in mice [88,89]. Like with *FKH2* (described above), *PMR1* deletion enhancement points to multiple human homologs that are both implicated in the cancer literature to promote cancer when underexpressed, yet are also UES in the pharmacogenomics data, suggesting a potentially clinically useful synthetic lethal vulnerability.

*KTI11/DPH3* (Fig. 5C; **1-0-14**), is a multi-functional protein involved in the biosynthesis of dipthamide and tRNA modifications important for regulation of translation, development and stress response [77,78], and has promoter mutations associated with skin cancer [79]. It was observed to be UES only for cytarabine and in hematopoietic and lymphoid cancer (the context cytarabine is used clinically).

*ACB1* binds acyl-CoA esters and transports them to acyl-CoA-consuming processes, which is upregulated in response to DNA replication stress [155]. Human homologs of *ACB1* exhibiting UES (Fig. 5C; **2-0.16-1**) included: (1) *DBI*, which is upregulated in hepatocellular carcinoma and lung cancer, and its expression is negatively associated with multidrug resistance in breast cancer [101-103]; (2) *ACBD4,* which promotes ER-peroxisome associations [98] and is upregulated by a histone deacetylase inhibitor, valproic acid, in a panel of cancer cell lines [99]; and (3) *ACBD5,* which also promotes ER-peroxisome associations, but its link to cancer is unclear [100].. Thus, it appears this gene family may influence epigenetic processes that buffer the cytotoxic effects of gemcitabine and cytarabine.

#### Deletion suppression of toxicity for both nucleosides

As opposed to deletion enhancing interactions, which represent functions that buffer the cytotoxic effects of the drugs, deletion suppression identifies genes that promote toxicity, thus predicting overexpression sensitivity (OES) in pharmacogenomics data that represent causal tumor vulnerabilities. REMc/GTF identified as deletion suppressing the GO terms *glutaminyl-tRNA(Gln) biosynthesis (*1-0*-*3), the nucleoplasmic THO (2-0.8-1), RNA cap binding (1-0-3), and the NuA3b histone acetyltransferase complexes (1-0-7) (**Additional File 5, File C**), while GTA identified *mitochondrial translational elongation* and the *nuclear cap binding complex* (**Additional File 6, File A**). However, the respective term-specific heatmaps revealed weak effects and high shift for many of the genes (**Additional File 2, Fig. S3**), highlighting the utility of this phenomic visualization tool for prioritizing findings, and leading us to shift our focus to individual yeast-human homologs identified in gene deletion suppressing clusters that were OES in the pharmacogenomics analysis, as detailed below.

In cluster 2-0.8-1, yeast-human homologs with correlated gene deletion suppression and OES for both gemcitabine and cytarabine (Fig. 5D; Table 3) included: (1) *YTA7/ATAD2/ATAD2B*, which localizes to chromatin and regulates histone gene expression. *ATAD2* overexpression portends poor prognosis in gastric, colorectal, cervical, hepatocellular carcinoma, lung, and breast cancer, and thus overexpression sensitivity could represent the potential to target a driver gene [119-125]; (2) *PIF1*/*PIF1*, a DNA helicase, which is involved in telomere regulation and is required during oncogenic stress [113]; (3) *RPS1B/RPS3A*, which is a small subunit ribosomal protein that is overexpressed in hepatitis B associated hepatocellular carcinoma and non-small cell lung cancer [114,115]; (4) *LEO1/LEO1*, which associates with the RNA polymerase II and acts as an oncogene in acute myelogenous leukemia [156]; (5) *ELO3/ELOVL1/ELOVL2/ ELOVL4/ELOVL6*, which constitutes a family of fatty acid elongases that function in sphingolipid biosynthesis, among which *ELOVL1* is overexpressed in breast and colorectal cancer tissue [107,108], *ELOVL2* is upregulated in hepatocellular carcinoma [109], and *ELOVL6* is overexpressed and associated with poor prognosis in liver and breast cancer [110,111]; (6) *MDL2/ABCB10*, which is a mitochondrial inner membrane ATP-binding cassette protein and is upregulated in breast cancer [112]; (7) *CPR3/PPIA*, which is a mitochondrial cyclophilin that is upregulated in lung cancer, esophageal, and pancreatic cancer [104-106]; and (8) *SAC3/MCM3AP/SAC3D1*, which is a nuclear pore-associated protein functioning in transcription and mRNA export, with *MCM3AP* being upregulated in glioma cells [116], while *SAC3D1* is upregulated in cervical cancer and hepatocellular carcinoma [117,118]. Yeast gene deletion suppression together with overexpression sensitivity of human homologs in cancer reveals potential therapeutic vulnerabilities that can be further explored in both systems.

### Gemcitabine-specific gene interaction modules

#### Gemcitabine-specific gene deletion enhancement

Gemcitabine-specific deletion enhancement indicates genes for which loss of function increases vulnerability to gemcitabine to a greater extent than cytarabine. Therefore, these genes provide insight into cytotoxic mechanisms that are unique between the two deoxycytidine analogs. Representative clusters were GO-enriched for *intralumenal vesicle formation (*1-0-10*), peptidyl-tyrosine dephosphorylation* (1-0-0), and the Set1C/COMPASS and HDA1 complexes (Fig. 3C, Fig. 6A-C, Table 1). GTA identified *negative regulation of macroautophagy, protein urmylation,* and the RAVE, GATOR (Iml1), and Elongator holoenzyme complexes (Fig. 6A, Table 2). Pharmacogenomics integration is highlighted for clusters 2-0.2-1, 1-0-10, and 1-0-0 (Fig. 6D; see also, **Additional File 8**). Taken together, the results suggest that autophagy-related processes and perhaps others less well characterized by GO buffer cytotoxicity of gemcitabine to a greater extent than cytarabine.

#### Autophagy related processes

Autophagy-related processes and complexes consisted of *intralumenal vesicle formation* (1-0-0; *BRO1, DOA4, DID4, VPS24, VPS4*), the GATOR/SEACIT/Iml1 complex (*NPR2, NPR3*), *autophagosome maturation* (*VAM3, CCZ1*), *negative regulation of macroautophagy* (*PHO85, PCL5, KSP1, SIC1, PHO80*), and the RAVE complex (*RAV1, RAV2*) (Fig. 6A).

Of the autophagy-related complexes, Npr2 and Npr3 form an evolutionarily conserved heterodimer involved in mediating induction of autophagy by inhibition of TORC1 signaling in response to amino acid starvation [216], and also promoting non-nitrogen starvation induced autophagy [217] (Fig. 6A). The RAVE complex (*RAV1/2*) promotes assembly of the vacuolar ATPase [218,219], which is required for vacuolar acidification and efficient autophagy [220]. Gene deletion strains in the term *negative regulation of macroautophagy* (*PHO85, PHO80,* and *SIC1*) [221], which seemed from the automated assessment to suggest an opposing effect, were less compelling following detailed visualization of the data, due to the associated high shift and cytarabine deletion enhancing interaction (Fig. 6A).

Regarding the term *intralumenal vesicle formation*, Vps24 and Did4 are components of the ESCRT-III complex (see Figs. 5E and 6A), which functions at endosomes, and the ATPase Vps4 is required for disassembly of the complex [222]. Doa4 interacts with Vps20 of ESCRT-III to promote intralumenal vesicle formation, which also requires *BRO1* [223]. Pharmacogenomics correlation revealed UES in cancer cell lines for *DID4/CHMP2A/CHMP2B (*Fig. 6D; **1-0-10**). During autophagy, *CHMP2A* translocates to the phagophore to regulate separation of the inner and outer autophagosomal membranes to form double-membrane autophagosomes [171]. *CHMP2B* is a member of the ESCRT-III complex required for efficient autophagy and has reduced expression in melanoma [172,173], raising the hypothesis that gemcitabine could have efficacy in that context.

Other genes involved in autophagy-related processes that had human homologs UES in cancer cell lines included: (1) *PEP12/STX12 (*Fig. 6D; **1-0-10**), a t-SNARE required for mitophagy [180], for which underexpression is associated with risk of recurrence [181]; and (2) *VPS30/BECN1,* knockdown of which enhances gemcitabine cytotoxicity in pancreatic cancer stem cells [170]. Furthermore, gemcitabine treatment has been found to upregulate autophagy in pancreatic or breast cancer, which buffers drug cytotoxicity as inferred by the combination of gemcitabine with autophagy inhibitors increased killing of cancer cells [224-226]. Thus, autophagy-related findings from the yeast model appear consistent with and to build upon previous cancer cell models.

#### Histone modification and chromatin remodeling

GTF/REMc identified the Hda1 and Set1C/COMPASS (1-0-0) complexes as gemcitabine-specific deletion enhancing, which was confirmed by term-specific heatmaps (Fig. 6B). The Set1C complex has been characterized to have a role in cell cycle coordination [227], which may be reflected by greater deletion enhancing interaction for the K than for the L CPP. The Set1C/COMPASS complex catalyzes mono-, di-, and tri-methylation of histone H3K4, which can differentially influence gene transcription depending on the number of methyl groups added [228-231], and was implicated by *BRE2*, *SWD1*, *SWD3*, *SDC1*, *SPP1,* and *SHG1* (Fig. 6B). The SWD1 ortholog, *RBBP5*, which was UES with gemcitabine in lung tissue (**Additional File 8, File C**; **1-0-4**), is upregulated in self-renewing cancer stem cells in glioblastoma and necessary for their self-renewal, is involved in the epithelial-mesenchymal transition in prostate cancer cells via its role in H3K4 trimethylation, and is upregulated in hepatocellular carcinoma [232-235]. Furthermore, gemcitabine sensitivity of pancreatic cancer cell lines was enhanced by H3K4me3 inhibition with verticillin A [233].

Histone deacetylases also influence cell cycle regulation [236], and the three genes that make up the yeast Hda1 deacetylase complex (homologous to mammalian class II Hda1-like proteins [237,238]) were gemcitabine deletion enhancers (Fig. 6B). Similar effects in cancer cells include *HDAC6* knockdown in pediatric acute myeloid leukemia cells, which enhances cytarabine-induced apoptosis [158-160] and the use of histone deacetylase inhibitors in combination with gemcitabine, which augments killing of pancreatic cancer cell lines [161-165] and HeLa cells [99].

#### Peptidyl-tyrosine dephosphorylation

REMc/GTF identified peptidyl-tyrosine dephosphorylation (1-0-0), for which the term specific heatmap (**Additional File 7**) revealed six genes previously characterized for their requirement in oxidant-induced cell cycle arrest and RNA virus replication [239,240], OCA1-6. Two additional tyrosine phosphatases, *YMR1,* and *PTP1*, had similar interaction profiles (Fig. 6C). *OCA1-3* deletions enhance growth defects associated with reactive oxygen species or caffeine treatment [239,240], and *OCA1-4* and *OCA6* are deletion suppressors of the cdc13-1 mutation [241]. Although it does not have a tyrosine phosphatase motif, Oca5 deletion also displayed gemcitabine-specific enhancement, consistent with the other genes annotated to this module (Fig. 6C). However, due to the regulatory nature and limited evolutionary conservation of tyrosine phosphorylation, it is not obvious how to predict functionally homologous genetic modules in cancer cells.

#### Elongator holoenzyme complex and protein urmylation

By GTA, K interactions revealed *protein urmylation* (*NCS6, NCS2, UBA4, ELP6, ELP2, URM1,* and *URE2*) and the Elongator holoenzyme complex (*IKI1, IKI3, ELP2, ELP3, ELP4*, and *ELP6*) (**Additional File 2, Fig. S4**). Protein urmylation involves the covalent modification of lysine residues with the ubiquitin-related modifier, Urm1 [242]. The Elongator holoenzyme complex has function in tRNA wobble position uridine thiolation (**Additional File 2, Fig. S4**), which occurs using Ure1 as a sulfur carrier [243-245]. The two processes share the *ELP2* and *ELP6* genes and may be distinct modules buffering gemcitabine cytotoxicity. However, several genes involved in tRNA wobble uridine modification have roles in cancer development and deficiency in this pathway enhances targeted therapy in melanoma [246,247], implicating this module as potentially important for personalized anti-cancer efficacy of gemcitabine.

#### Gemcitabine-buffering by non-GO-enriched yeast-human homologs

Homologs with correlated gemcitabine-specific yeast gene deletion enhancement and cancer cell UES (clusters 2-0.2-1, 1-0-10, and 1-0-0) included the family of nucleoside diphosphate kinases (NDKs) (Fig. 6D; Table 4). A single member of the NDK family, *YNK1*, exists in yeast, while the human genome encodes several paralogs (*NME* genes) (**Additional File 8, File A**). The NDKs transfer the gamma phosphate of ATP to nucleoside diphosphate as the final step of purine and pyrimidine nucleoside and deoxynucleoside triphosphate biosynthesis and salvage [248,249]. Thus, NDK appears to modulate gemcitabine toxicity by differential activity for endogenous substrates vs. nucleoside analog drugs. In yeast, deletion enhancement by YNK1 was selective for gemcitabine, however the effects in cancer cells are potentially more complex due to multiple NDK genes. In PharmacoDB, *NME3* and *5* were UES for gemcitabine, while *NME4*, *6*, and *7* were OES for cytarabine, implicating differential specificity of *NME* genes for natural and/or medicinal nucleosides as well as possibly influences of other kinases, which have, for example, been shown to act on gemcitabine diphosphate [250]. *NME5* overexpression was previously associated with gemcitabine-resistant cancer, and its knockdown can increase gemcitabine efficacy [182]. Thus, the anti-cancer efficacy of gemcitabine could be influenced by differential expression and activity of NDK isoforms across tissues [251], such that NME gene expression could be predictive of response to nucleoside analogs, or perhaps targeted for synergistic anti-tumor activity.

*KEX2* is the yeast member of the calcium-dependent proprotein convertase subtilisin/kexin type serine proteases, which functions in the secretory pathway. Four of the seven human homologs of KEX2 were UES in the pharmacogenomics analysis (Fig. 6D; **1-0-10**), including: (1) *PCSK1*, which can be downregulated by pancreatic cancer derived exosomes [176], (2) *PCSK2*, which has reduced expression in lung cancer [177], (3) *PCSK5*, which is also reduced in lung cancer and, furthermore, when reduced in triple negative breast cancer, leads to loss of the Gdf11 tumor suppressor [177,178], and (4) *PCSK7*, which has been reported both to have reduced expression in lung cancer and increased expression in gemcitabine resistant cells [177,179]. Thus, loss of this gene family may create cancer-specific vulnerabilities to gemcitabine cytotoxicity.

*NMA1* and its human homologs *NMNAT1, NMNAT2,* and *NMNAT3* are nicotinic acid mononucleotide adenylyltransferases involved in NAD biosynthesis and homeostasis, which were found to be UES for both gemcitabine and cytarabine (Fig. 6D, **1-0-0)**. Loss of function mutations and underexpression of *NMNAT1* are associated with increased rRNA expression and sensitivity to DNA damage in lung cancer cell lines [166], consistent with the hypothesis that they could have deletion enhancing therapeutic benefit in cancers treated with gemcitabine or cytarabine.

*RAD54* is a DNA-dependent ATPase that stimulates strand exchange in recombinational DNA repair, which is a known vulnerability of cancer [252]. The human homolog of *RAD54*, *ATRX*, was UES by PharmacoDB analysis (Fig. 6D, **1-0-0**), and loss of *ATRX* has been associated with improved response to gemcitabine plus radiation therapy in glioma patients with *IDH1* mutations [168].

*SCS2/VAPB* is an integral ER membrane protein that was deletion enhancing and UES for gemcitabine (Fig. 6D, **1-0-0**). *VAPB* regulates phospholipid metabolism and interacts with *ACBD5* (also described above) to promote ER-peroxisome tethering [100] and promotes proliferation in breast cancer via *AKT1* [169].

*YPT32/RAB2A/RAB2B* (Fig. 6D, **1-0-0**) is a Rab family GTPase involved in the trans-Golgi exocytic pathway, which accumulates during replication stress in yeast [155]. *RAB2A* overexpression promotes breast cancer stem cell expansion and tumorigenesis [174], and downregulation of *RAB2B* by miR-448 promotes cell cycle arrest and apoptosis in pancreatic cancer cells [175].

*CLB5*, a B-type cyclin, is involved in initiation of DNA replication and G1-S progression, for which promoter hypermethylation of the human homolog, *CCNA1*, is associated with multiple cancers [157], and which was found to be UES with gemcitabine (Fig. 6D, **1-0-0**).

#### Gemcitabine-specific gene deletion suppression

Representing this class of gene interaction, pharmacogenomics integration is highlighted for clusters 2-0.8-0 and 1-0-7 (Fig. 6E). Although there was limited Gene Ontology enrichment, the term *phosphatidylserine biosynthetic process* (*UME6* and *CHO1*), and the GARP (*VPS51*-*54*), and Lem3p-Dnf1p complexes were identified (Fig. 6F, Table 2). Ume6 is involved in both positive and negative regulation of the phosphatidylserine synthase, Cho1 [253,254]. Phosphatidylserine exposure to the plasma membrane is a marker of yeast and mammalian apoptosis [255], the latter of which is induced by gemcitabine [256]. In pancreatic cancer cells, addition of the sphingolipid, sphingomyelin, enhances gemcitabine cytotoxicity through increased apoptosis [256,257]. Moreover, GARP complex deficiency leads to reduction of sphingomyelin [258], and accumulation of sphingolipid intermediates, consistent with the hypothesis that reduced sphingolipid metabolism alleviates gemcitabine-mediated apoptosis. Lem3 complexes with Dnf1 or Dnf2 to form phospholipid flippases at the plasma and early endosome/trans-Golgi network membranes and regulate phosphatidylethanolamine and phosphatidylserine membrane content [259,260], potentially further influencing the apoptotic response. The Lem3-Dnf1 and Lem3-Dnf2 flippases are regulated by the serine/threonine kinase Fpk1 [261], which is also a gemcitabine-specific deletion suppressor (Fig. 6F).

#### Correlation of gemcitabine-specific gene deletion suppression with OES in cancer cells

Although yeast genes associated with GO-enriched terms from gemcitabine-specific deletion suppression (2-0.8-0 and 1-0-7) did not have human homologs that were OES in GDSC, several homologs of ‘non-GO-enriched’ genes were OES (Fig. 6E; Table 4). These included: (1) *YGR054W/EIF2A*, a eukaryotic initiation factor orthologous between yeast and human that has been implicated in translation of upstream ORFs as part of tumor initiation [215]. Thus, gemcitabine treatment in the context of EIF2A overexpression may increase efficacy; (2) *EFT2/EEF2/EFTUD2* (eukaryotic translation elongation factor 2), which further implicates translational regulation as a gemcitabine-targetable cancer driver. *EEF2* is overexpressed in numerous cancer types [198] and *EFTUD2* knockdown induces apoptosis in breast cancer cells [199]; (3) *RPP2B/RPLP2*, a component of the 60S ribosomal subunit stalk that is overexpressed in gynecologic cancer [214], again suggesting dysregulated translation promotes gemcitabine toxicity; (4) *RPA49/POLR1E*, a component of Pol1 [203-205] that has higher expression in bladder cancer and has been recently proposed as a novel target for anti-cancer therapy [206]; (5) *OLA1/OLA1* is a GTPase that is conserved from human to bacteria [200]. It is implicated in regulation of ribosomal translation [201] and has increased expression associated with poorer survival in lung cancer patients [202]. The interactions described above suggest gemcitabine may be more effective in the context of “oncogenic ribosomes” [262]. (6) *CKA2*, the alpha catalytic subunit of casein kinase 2, has two human homologs, *CSNK2A1* and CSNK2A2, which were OES with gemcitabine. They can be upregulated in cancer [186-191] and are considered targets for treatment [193]; (7) *CLB2/CCNA2/CCNB1,* a B-type cyclin involved in cell cycle progression, of which both *CCNA2* and *CCNB1* are overexpressed in breast and colorectal cancer [194-197]. Moreover, the observation that *CLB2* deletion (suppressing effect) opposes that of *CLB5* (deletion enhancing; see above Fig. 6D, 1-0-0) has been previously described in the context of loss of the S-phase checkpoint [263]; (8) *SKY1/SRPK1* (serine-arginine rich serine-threonine kinase), which is overexpressed in glioma, and prostate, breast, and lung cancer [207]; (9) *SNC2/VAMP8*, which functions in fusion of Golgi-derived vesicles with the plasma membrane and is overexpressed in glioma and breast cancer [208,209]; (10) *YPT6/RAB34*, which functions in fusion of endosome-derived vesicles with the late Golgi and is overexpressed in glioma, breast cancer, and hepatocellular carcinoma [211-213]; (11) *TOP1/TOP1/TOP1MT*, Topoisomerase I, which has increased copy number in pancreatic and bile duct cancer [210]; (12) *ALD6*, which encodes cytosolic aldehyde dehydrogenase, and was a deletion suppressor for both gemcitabine and cytarabine, having multiple homologs that were OES (*ALDH1A1*, *ALDH1A2*, *ALDH1B1*, and *ALDH7A1. ALDH1B1*). Overexpression of ALDH genes is observed in colorectal and pancreatic cancer [183,184] and is a prognostic marker of cancer stem cells [185].

### Cytarabine-specific gene interaction modules

#### Cytarabine-specific gene deletion enhancement

Cytarabine-specific deletion enhancement suggests functions that buffer cytotoxic effects of cytarabine to a greater extent than gemcitabine, potentially informing on differential activities of the drugs. There was no notable GO enrichment by REMc/GTF, but four functions of potential relevance were revealed by GTA (Fig. 7A, Table 2). Two of them, the *HIR complex* (*HIR1-3,* HPC2) and *sphinganine kinase activity* (*LCB4, LCB5*) were relatively weak, being deletion enhancing only for the L CPP (Fig. 7A). *LCB4/5* homologs that were UES in PharmacoDB included: (1) *CERKL* (**Additional File 8, Files B-C; 1-0-6**), a ceramide kinase like gene that regulates autophagy by stabilizing *SIRT1* [264], a gene mentioned above for its inhibition being synergistic with cytarabine against acute lymphoblastic leukemia cells [138], and (2) *AGK*, which is overexpressed in hepatocellular carcinoma, glioma, breast, and cervical squamous cell cancers [265-268]. Two stronger interaction modules, evidenced by deletion enhancement for both the K and L CPPs, were *protein localization to septin ring (HSL1* and *ELM1*) and the *Sec61 translocon complex* (*SBH1, SBH2*, and *SEC61*) (Fig. 7A, Table 2). In yeast, Hsl1 and Elm1 are annotated as “bud sensors” to recruit Hsl7 to the septin ring at the bud site to degrade the mitotic inhibitor, Swe1 [269]. The *HSL1* homologs, *BRSK1* and *BRSK2*, were UES in the cancer data. *BRSK1* is mutated in gastric and colorectal carcinoma [270] and its decreased expression is associated with breast cancer [271], but *BRSK2* is overexpressed in pancreatic cancer, where it is AKT-activating [272]. PharmacoDB also identified the *SEC61* homolog, *SEC61A1*, which is upregulated in colon adenocarcinoma tissue [273].

#### Human genes that have deletion enhancing yeast homologs and confer cytarabine UES

We identified human genes that were UES to cytarabine and homologous to yeast genes in REMc clusters (1-0-9 and 2-0.13-0) displaying a pattern of cytarabine-specific deletion enhancement (Fig. 7B; Table 5). Cancer-relevant examples include:

1. Ptm1, which is a protein of unknown function that copurifies with late Golgi vesicles containing the v-SNARE, Tlg2p, but interestingly, its human homologs, *TMEM87A* and *TMEM87B*, were UES for cytarabine and identified in a study focused on cytarabine efficacy in acute myelogenous leukemia [281].
2. *NAP1/NAP1L3/NAP1L4*, which is a nucleosome assembly protein involved in nuclear transport and exchange of histones H2A and H2B and also interacts with Clb2, is phosphorylated by CK2, and has protein abundance that increases in response to DNA replication stress [155]. *NAP1L3* is overexpressed in breast cancer [280].
3. *CCH1*, which is a voltage-gated high-affinity calcium channel with several homologs that were UES, including: *CACNA1A*, underexpressed in breast, colorectal, esophageal, gastric, and brain cancers; *CACNA1B*, underexpressed in breast and brain cancers; *CACNA1C*, underexpressed in brain, bladder, lung, lymphoma, prostate, and renal cancers; *CACNA1E*, underexpressed in breast, brain, gastric, leukemia, lung, and prostate cancers; and *CACNA1F*, underexpressed in lymphoma [274];
4. *IZH1*, a yeast membrane protein involved in zinc ion homeostasis, having a human homolog, *PAQR1/ADIPOR1* that encodes the adiponectin receptor protein 1, which is differentially regulated in breast cancers [278,279];
5. *FAT1*, a yeast fatty acid transporter and very long-chain fatty acyl-CoA synthetase that corresponds to *SLC27A2* (very long-chain acyl Co-A synthetase), which is underexpressed in lung cancer [275], and *SLC27A3* (long-chain fatty acid transport), which is hypermethylated in melanoma [276];
6. *FOL2/GCH1*, a GTP-cyclohydrolase that catalyzes the first step in folic acid biosynthesis. Downregulation of *GCH1* occurs in esophageal squamous cell carcinoma [277].

## Discussion

Informative phenomic models have been developed for multiple human diseases, including cystic fibrosis, neurodegenerative disorders, and cancer [30,282-284]. Molecular models include mutations in conserved residues of yeast homologs of a disease gene and introduction of human alleles into yeast. Complementation of gene functions by human homologs, and *vice versa*, has demonstrated evolutionary conservation of gene functions [285-287]. Like their basic functions, gene interactions are conserved [288,289] and yeast is unique in its capability to address complex genetic interactions experimentally [290]. Here, we model how yeast phenomic assessment of gene-drug interaction could be employed as part of a precision oncology paradigm to predict efficacy of cytotoxic chemotherapy based on the unique cancer genetic profiles of individual patients.

To model the networks that buffer deoxyribonucleoside analogs, we humanized yeast by introducing deoxycytidine kinase into the YKO/KD strain collection, as yeast do not encode dCK in their genomes, and thus cannot activate the unphosphorylated drugs. We hypothesized that gemcitabine and cytarabine would have different buffering profiles, despite their similar mechanisms of action, due to their distinct anti-cancer efficacies. Results of the unbiased yeast phenomic experiments confirmed this expectation, revealing distinct, though partially overlapping, gene interaction networks.

Differential interaction predominated despite the similarity of the molecules, illustrating that distinct mechanisms for buffering anti-cancer cytotoxic drug responses can be inferred from yeast phenomics and thus applied to predict how an individual’s cancer genome could influence responses to treatment [3,5]. Deletion enhancement of both gemcitabine and cytarabine suggested processes that function to buffer nucleoside analog cytotoxicity in common (Fig. 5), in contrast to buffering mechanisms that acted differentially in response to the drugs. Functionally enriched processes that buffered both drugs to a similar extent included the intra-S DNA damage checkpoint, positive regulation of DNA-dependent DNA replication initiation, vesicle fusion with vacuole, and the Mre11, checkpoint clamp, RecQ helicase-Topo III, CORVET, HOPS, ESCRT, GET, Ubp3-Bre5 deubiquitination complexes.

Among the drug-specific deletion enhancing interactions, autophagy, histone modification, chromatin remodeling, and peptidyl-tyrosine dephosphorylation buffered gemcitabine more so than cytarabine (Fig. 6). There were only a few cytarabine-specific deletion enhancing GO-enriched terms, but there were many individual genes with human homologs having cancer relevance that buffered cytarabine relatively specifically (Fig. 7). On the other hand, genes that preferentially promote cytotoxicity were observed primarily for gemcitabine, and enriched functions were related to apoptosis, including phosphatidylserine biosynthesis, and the GARP and Lem2/3 complexes (Fig. 6).

The model we constructed incorporates the powerful pharmacogenomics datasets and analysis tools from PharmacoDB, mining them by integration of yeast phenomic drug-gene interaction experiments. We integrated yeast phenomic and PharmacoDB data to identify, across the respective datasets, correlations between deletion enhancement and underexpression sensitivity or deletion suppression and overexpression sensitivity. Deletion enhancement indicates genes that are biomarkers and synergistic targets to augment drug efficacy and expand the therapeutic window, whereas deletion suppression identifies genes that promote drug cytotoxicity, and thus confer sensitivity when hyper-functional and resistance when deficient. A particularly attractive class of drug-gene interaction is overexpression sensitivity involving driver genes, however anti-cancer efficacy could be conferred by lethal drug-gene interactions involving passenger genes, tumor suppressor genes, or components of genetic buffering networks that become compromised due to genomic instability (Fig. 1A). The cancer literature revealed many deletion enhancing/UES and deletion suppressing/OES genes to have roles in cancer, suggesting that integration of yeast phenomic models and pharmacogenomics data could have clinical utility for choosing cytotoxic treatments based on gene expression profiles of individual cancers. While predictions sometimes involved GO-enriched processes, often the genes were identified individually.

The utility of phenomic data (i.e., Q-HTCP of the YKO/KD library) to help predict causal associations between gene expression changes and cell sensitivity in response to drugs derives from prior work demonstrating that genes differentially expressed after drug treatment do not significantly overlap with those that influence sensitivity and resistance [52]. As far as we know, this work represents the first application of this fundamental observation from yeast to systems level experimental data from human cells. Literature-based validations of the yeast phenomic model of nucleoside analogs in human cancer cell lines and other cancer models are exemplified in Table 6. These examples illustrate that integrative, systems level drug-gene interaction modeling employing the experimental power of *S. cerevisiae* phenomics could be applicable to cancer genomic profiling for systems level, precision oncology.

**Table 6.** Disease relevance of buffering interactions from the yeast phenomic model evidenced by the cancer biology literature.

| Gene<br>(yeast/human) | Process/Complex | Description (human) | Ref | Nucleoside analog relevance |
| --- | --- | --- | --- | --- |
| RAD24/RAD17 | DNA damage checkpoint | RAD17 checkpoint clamp loader component | [74] | Depletion of RAD17 sensitizes pancreatic cancer cells to gemcitabine |
| RAD50/RAD50 | Mre11 complex | RAD50 double strand break repair protein | [127] | Depletion of human Rad50 sensitizes Ataxia-telangiectasia (AT) fibroblasts to gemcitabine |
| HDA1/HDAC6 | Hda1 complex; histone deacetylation | histone deacetylase 6 | [160, 165] | HDAC inhibitors enhance sensitivity to gemcitabine in pancreatic cancer cells and are associated with reduction of HDAC6; HDAC6 inhibition induces apoptosis in cytarabine treated AML cells |
| RAD54/ATRX | Chromatin remodeling | ATRX, chromatin remodeler | [168] | Glioma patients with IDH1 mutations and loss of ATRX had improved response to gemcitabine plus radiation therapy |
| KEX2/PCSK7 | serine-type endopeptidase activity | proprotein convertase subtilisin/kexin type 7 | [179] | Overexpressed in gemcitabine resistant pancreatic cancer cell lines |
| YNK1/NME5 | Nucleoside diphosphate phosphorylation | NME/NM23 family member 5 | [182] | Depletion of NME5 sensitizes gemcitabine-resistant cancer cell lines to gemcitabine |
| VPS30/BECN1 | Autophagy | beclin 1 | [170] | Depletion of *BECN1* sensitizes pancreatic cancer stem cells to gemcitabine |
| LCB4/5/CERKL | sphinganine kinase activity | ceramide kinase like | [138, 264] | CERKL stabilizes SIRT1, SIRT1 chemical inhibition sensitizes acute myeloid leukemia cells to cytarabine |

Deletion-enhancing/UES drug-gene interactions are highlighted; most exemplify loss of buffering functions that lead to increased drug sensitivity; however, there is one instance (KEX2/PCSK7) of overexpression of the buffering gene that increases drug resistance.

In summary, the yeast phenomic model of nucleoside analog toxicity appears to serve as a valuable resource for interpreting cancer pharmacogenomics data regarding gene-drug interaction that could be predictive of patient-specific chemotherapeutic efficacy. Since it’s not possible to collect comparable phenomic information from human populations or cancerous tissue alone [5], systems level yeast phenomic models can help expand and integrate relevant (i.e., evolutionarily conserved) aspects of the extensive cancer literature with regard to cancer-specific vulnerabilities to cytotoxic therapies. A deeper understanding of how genomic instability influences the genetic network that buffers chemotherapeutic agents like nucleoside analogs could guide future research to personalize anti-cancer therapies based on cancer genomic profiles unique to individual patients. Thus, a future direction for this work should include development of algorithms that prospectively predict chemotherapy response in individual patient cancer cells, which could be tested as part of a prognostic evaluation. Most cytotoxic chemotherapeutic agents are used in combination, so another direction for yeast phenomic analysis of anti-cancer agents would be to characterize clinically relevant drug combinations.

## Conclusions

A humanized yeast phenomic model of deoxycytidine kinase was developed to map drug-gene interactions modulating anti-proliferative effects of nucleoside analogs in a eukaryotic cell and to investigate the relevance of the resulting networks for precision oncology by integration with cancer pharmacogenomics-derived associations between gene expression and cancer cell line drug sensitivity. The yeast phenomic model revealed gene-drug interaction for the two deoxycytidine analogs, gemcitabine and cytarabine, to be largely different, consistent with the distinct types of cancer for which they are used clinically. The model overall suggested evolutionary conservation of drug-gene interaction that could be used as a resource to predict anti-cancer therapeutic efficacy based on genetic information specific to individual patients’ tumors. Yeast phenomics affords a scalable, high-resolution approach to model, at a systems level, the genetic requirements for sensitivity and resistance to cytotoxic agents and thus the potential to resolve complex influences of genetic variation on drug response to more accurately. Global and quantitative models of the distinct genetic buffering networks required to maintain cellular homeostasis after exposure to chemotherapeutic agents could aid precision oncology paradigms aimed at identifying composite genomic derangements that create enhanced cancer cell-specific vulnerabilities to particular anti-cancer drugs.

## Supplementary Materials

**Additional File 1. Supplemental tables: Tables S1-S6. Table S1.** Primers used in strain construction (see **Fig. S1**). **Table S2.** YKO/KD strains with gemcitabine. **Table S3.** Reference cultures with gemcitabine. **Table S4.** YKO/KD strains with cytarabine. **Table S5.** Reference cultures with cytarabine. **Table S6**. Ranges of interaction z-scores for the YKO/YKD and Reference cultures from the phenomic analysis of gemcitabine and cytarabine drug-gene interaction.

**Additional File 2. Supplemental figures. Figure S1**. Construction of tet-inducible dCK allele. **Figure S2**. Reference r and AUC distributions with gemcitabine or cytarabine treatment. **Figure S3**. High shift or weak nucleoside analog gene deletion suppression modules. **Figure S4**. Elongator holoenzyme complex, protein urmylation, and tRNA wobble position uridine thiolation buffer gemcitabine cytotoxicity.

**Additional File 3. Interaction plots for gemcitabine.** Genome-wide analysis for **(A)** YKO, (**B**) KD, and **(C)** reference strains with gemcitabine. See also methods and **Additional File 2**.

**Additional File 4. Interaction plots for cytarabine.** Genome-wide analysis for **(A)** YKO, (**B**) KD, and **(C)** reference strains with cytarabine. See also methods and **Additional File 2**.

**Additional File 5. REMc results, plotted as drug-gene interaction profile heatmaps and assessed for Gene Ontology enrichment using GTF. File A** contains REMc results and associated gene interaction and shift data. **File B** is the heatmap representation of each REMc cluster after incorporating shift values and hierarchical clustering. **File C** contains the GTF results obtained for REMc clusters for the three ontologies – process, function, and component.

**Additional File 6. Gene Ontology Term Averaging (GTA) results and interactive plots. File A** contains all GTA values, cross-referenced with REMc-enriched terms. **File B** displays GTA L values associated with above-threshold GTA scores (see note below) plotted for gemcitabine *vs.* cytarabine. **File C** displays GTA K values associated with above-threshold GTA scores (see note below) plotted for gemcitabine *vs.* cytarabine. **Files B-C** should be opened in an Internet web browser so that embedded information from **Additional File 6A** can be viewed by scrolling over points on the graphs. Subsets in each of the plots can be toggled off and on by clicking on the respective legend label. In the embedded information, X1 represents gemcitabine and X2 represents cytarabine information.

**Additional File 7. GO term-specific heatmaps for REMc/GTF-enriched modules.** GO term-specific heatmaps for significant GO process terms were generated as described in methods and Figure 3. Any related child terms are presented in subsequent pages of the parent file name. GO terms with more than 100 children, with 2 or fewer genes annotated to the term, or a file size over 400KB are not shown. All heatmaps are generated with the same layout (see Fig. 3).

**Additional File 8. Application of yeast phenomic drug-gene interaction data to predict, from cancer cell line pharmacogenomic data (gene expression and drug sensitivity correlations), human genes that modify gemcitabine or cytarabine toxicity. (A)** Tables of UES and OES human genes and whether their yeast homologs were found to be deletion enhancing or deletion suppressing, respectively. **(B-D)** REMc heatmaps and tables of the yeast interaction scores corresponding to UES or OES human genes identified (**B**) across all tissue, (**C**) in lung, or (**D**) in hematopoietic and lymphoid tissue. See Fig. 3 for description of tables and the color keys (note: a teal color, which represents cytarabine-specific UES/OES in the heatmaps in the main manuscript figures, is represented as darker blue in the supplemental heatmaps, while gold, representing gemcitabine-specific UES/OES in the main manuscript, is represented as a brighter yellow in the supplemental heatmaps).

## List of abbreviations and glossary of terms

AraC: cytarabine; cytosine arabinoside
CPPs: Cell proliferation parameters: parameters of the logistic growth equation used to fit cell proliferation data obtained by Q-HTCP. The CPPs used to assess gene interaction in this study were K (carrying capacity) and L (time required to reach half of carrying capacity) [7-9,38].
DAmP: Decreased Abundance of mRNA Production: refers to a method of making YKD alleles, where the 3’ UTR of essential genes is disrupted, reducing mRNA stability and gene dosage [291].
dCK: deoxycytidine kinase
dCMP: deoxycytidine monophosphate
DE: Deletion enhancer: gene loss of function (knockout or knockdown) that results in enhancement / increase of drug sensitivity [9].
dFdC: 2’,2’-difluoro 2’-deoxycytidine, gemcitabine
dNTP: deoxyribonucleotide triphosphate
DS: Deletion suppressor: gene loss of function (knockout or knockdown) that results in suppression / reduction of drug sensitivity [9].
ESCRT: endosomal sorting complex required for transport
GARP complex: Golgi-associated retrograde protein complex.
gCSI: The Genentech Cell Line Screening Initiative: One of two pharmacogenomics datasets used in this study (https://pharmacodb.pmgenomics.ca/datasets/4).
GDSC1000: Genomics of Drug Sensitivity in Cancer: One of two pharmacogenomics datasets used in this study (https://pharmacodb.pmgenomics.ca/datasets/5)
GO: Gene ontology
GTF: Gene ontology term finder: an algorithm to assess GO term enrichment amongst a list of genes; applied to REMc (clustering) results [41].
GTA: Gene ontology term averaging: an assessment of GO term function obtained by averaging the gene interaction values for all genes of a GO term
GTA value: Gene ontology term average value
gtaSD: standard deviation of GTA value
GTA score: (GTA value - gtaSD)
HaL: hematopoietic & lymphoid tissue
HDAC: Histone deacetylase complex
HLD: Human-like media with dextrose [8]: the yeast media used in this study.
INT: Interaction score
NDK: nucleoside diphosphate kinase
OES: Overexpression sensitivity: refers to association of increased gene expression with drug sensitivity in pharmacogenomics data [33].
PharmacoDB: The resource used for cancer pharmacogenomics analysis [33].
PPOD: Princeton protein orthology database
Q-HTCP: Quantitative high throughput cell array phenotyping: a method of imaging, image analysis, and growth curve fitting to obtain cell proliferation parameters [7,38].
Ref: Reference: the genetic background from which the YKO/KD library was derived
REMc: Recursive expectation maximization clustering: a probabilistic clustering algorithm that determines a discrete number of clusters from a data matrix [40].
RNR: ribonucleotide reductase
SD: Standard deviation
SGA: Synthetic genetic array
SGD: Saccharomyces genome database
UES: Underexpression sensitivity: refers to association of reduced gene expression with drug sensitivity in pharmacogenomics data [33].
YKO: Yeast knockout
YKD: Yeast knockdown: DAmP alleles
YKO/KD: Yeast knockout or knockdown

## Acknowledgements

The authors thank the following funding agencies for their support: American Cancer Society (RSG-10-066-01-TBE), Howard Hughes Medical Institute (P/S ECA 57005927), NIH/NCI (P30 CA013148), NIH/NIA (R01 AG043076), and Cystic Fibrosis Foundation (HARTMA16G0). The authors also thank Amanda Stisher for assistance with constructing yDW1 and Bo Xu and William Parker for providing the deoxycytidine kinase cDNA cloned into a plasmid.

## Author Contributions

Conceptualization, J.L.H.; Methodology and Software, B.A.M., J.G., J.L.H., and S.M.S.; Formal Analysis, J.L.H and S.M.S.; Investigation, D.W., I.P., and M.I.; Resources, J.L.H. and M.N.; Data Curation, J.W.R.; Writing – Original Draft Preparation, J.L.H and S.M.S.; Writing, Review & Editing, J.L.H and S.M.S.

