## Additional File 2 for "A humanized yeast phenomic model of deoxycytidine kinase to predict genetic buffering of nucleoside analog cytotoxicity"

### Slide 1
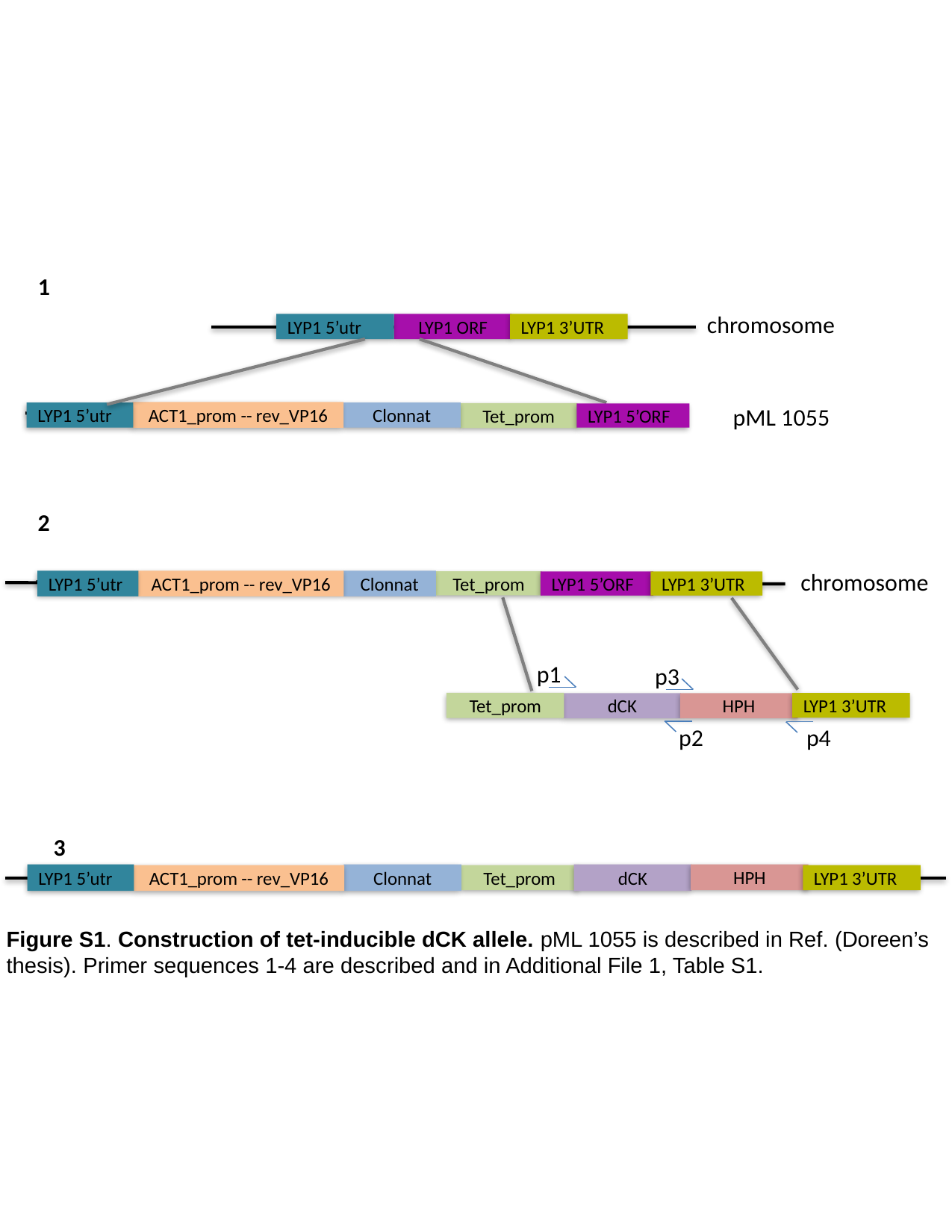

1
chromosome
LYP1 5’utr
LYP1 ORF
LYP1 3’UTR
pML 1055
ACT1_prom -- rev_VP16
LYP1 5’utr
Clonnat
Tet_prom
LYP1 5’ORF
2
chromosome
LYP1 5’utr
ACT1_prom -- rev_VP16
Clonnat
Tet_prom
LYP1 5’ORF
LYP1 3’UTR
p1
p3
Tet_prom
LYP1 3’UTR
dCK
HPH
p4
p2
3
dCK
HPH
LYP1 5’utr
Clonnat
LYP1 3’UTR
ACT1_prom -- rev_VP16
Tet_prom
Figure S1. Construction of tet-inducible dCK allele. pML 1055 is described in Ref. (Doreen’s thesis). Primer sequences 1-4 are described and in Additional File 1, Table S1.

### Slide 2
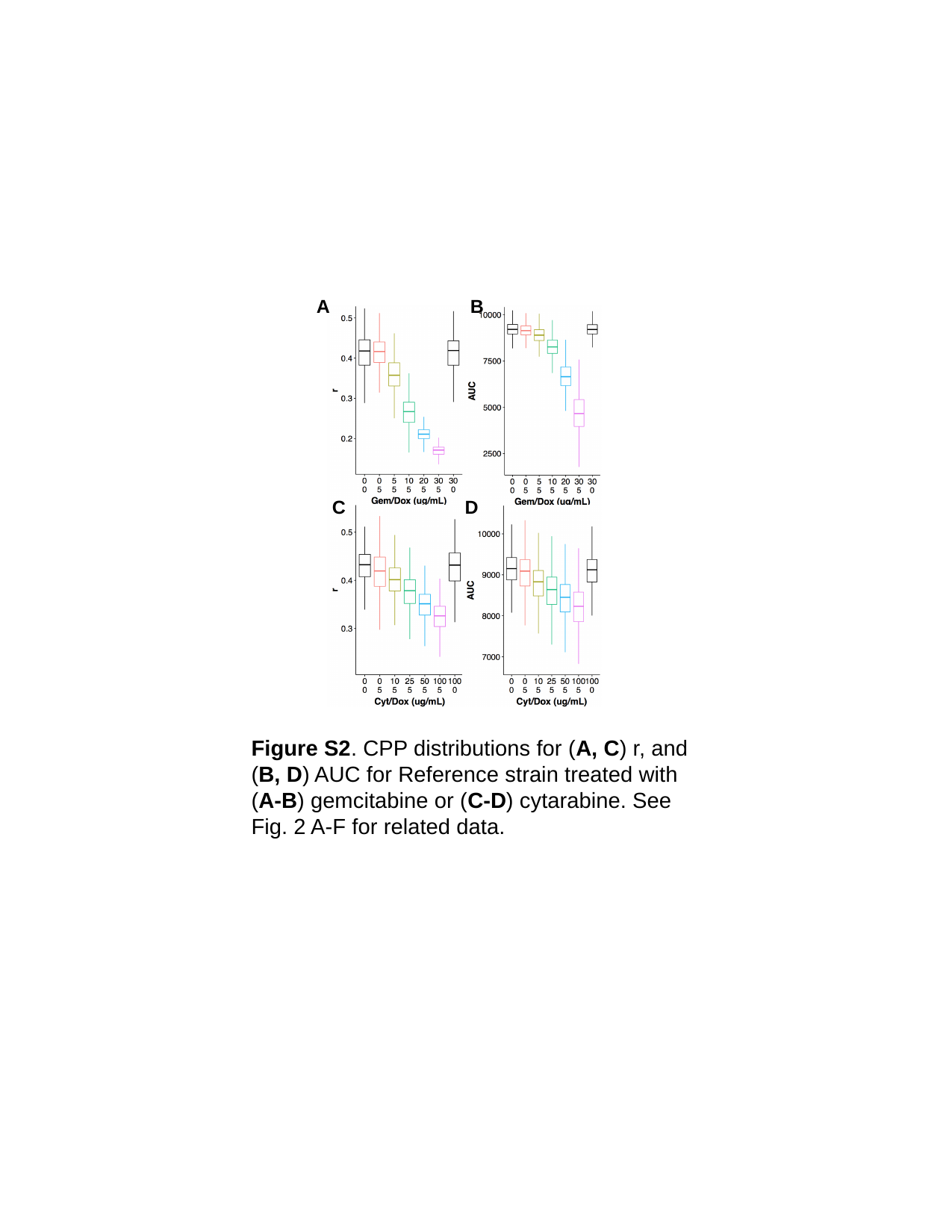

A
B
D
C
Figure S2. CPP distributions for (A, C) r, and (B, D) AUC for Reference strain treated with (A-B) gemcitabine or (C-D) cytarabine. See Fig. 2 A-F for related data.

### Slide 3
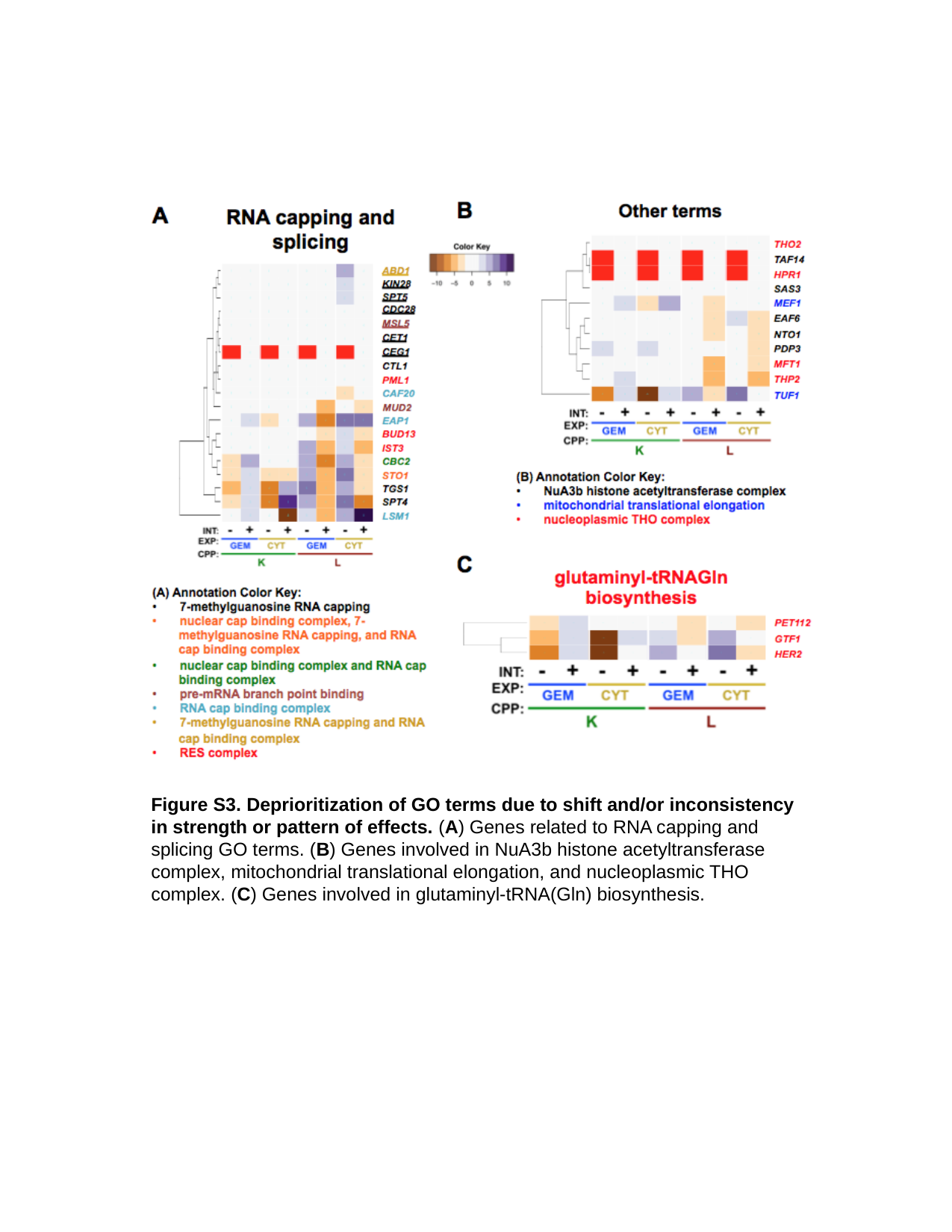

Figure S3. Deprioritization of GO terms due to shift and/or inconsistency in strength or pattern of effects. (A) Genes related to RNA capping and splicing GO terms. (B) Genes involved in NuA3b histone acetyltransferase complex, mitochondrial translational elongation, and nucleoplasmic THO complex. (C) Genes involved in glutaminyl-tRNA(Gln) biosynthesis.

### Slide 4
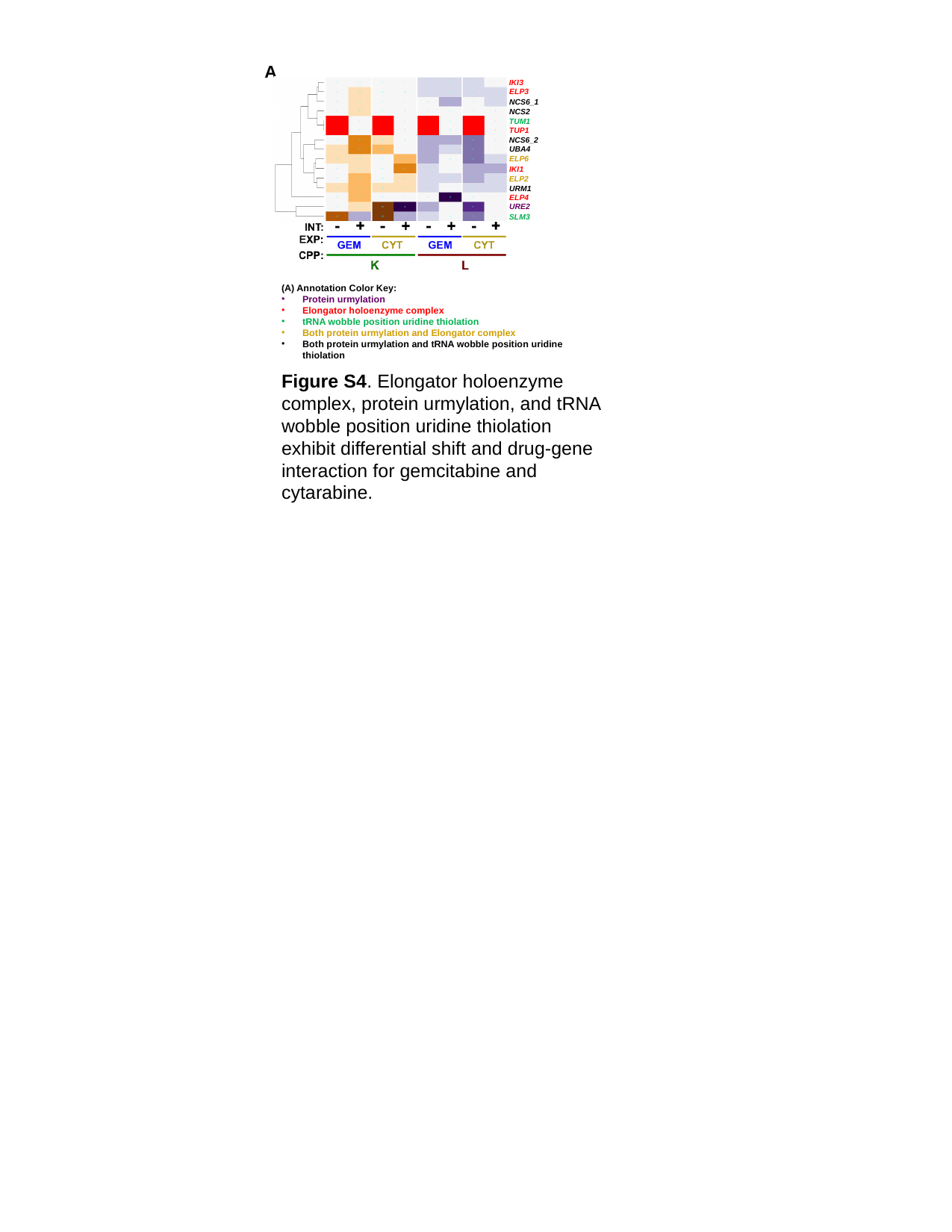

A
IKI3
ELP3
NCS6_1
NCS2
TUM1
TUP1
NCS6_2
UBA4
ELP6
IKI1
ELP2
URM1
ELP4
URE2
SLM3
(A) Annotation Color Key:
Protein urmylation
Elongator holoenzyme complex
tRNA wobble position uridine thiolation
Both protein urmylation and Elongator complex
Both protein urmylation and tRNA wobble position uridine thiolation
Figure S4. Elongator holoenzyme complex, protein urmylation, and tRNA wobble position uridine thiolation exhibit differential shift and drug-gene interaction for gemcitabine and cytarabine.
