## Additional File 3 for "A humanized yeast phenomic model of deoxycytidine kinase to predict genetic buffering of nucleoside analog cytotoxicity": A - InteractionPlots_Gemcitabine.pdf

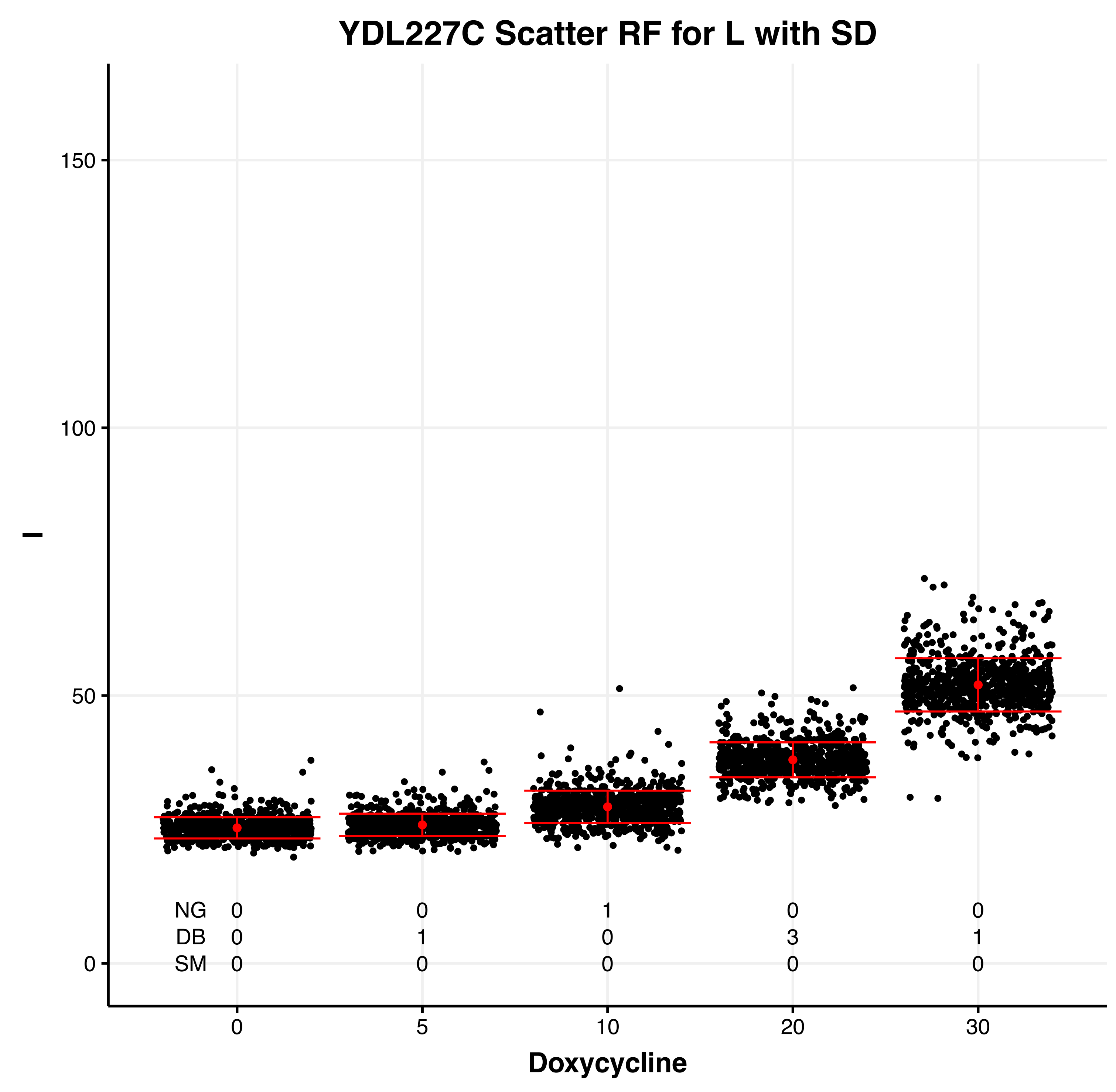
YDL227C Scatter RF for L with SD
l
150
100
50
0
NG 0 0 1 0 0
DB 0 1 0 3 1
SM 0 0 0 0 0
0 5 10 20 30
Doxycycline

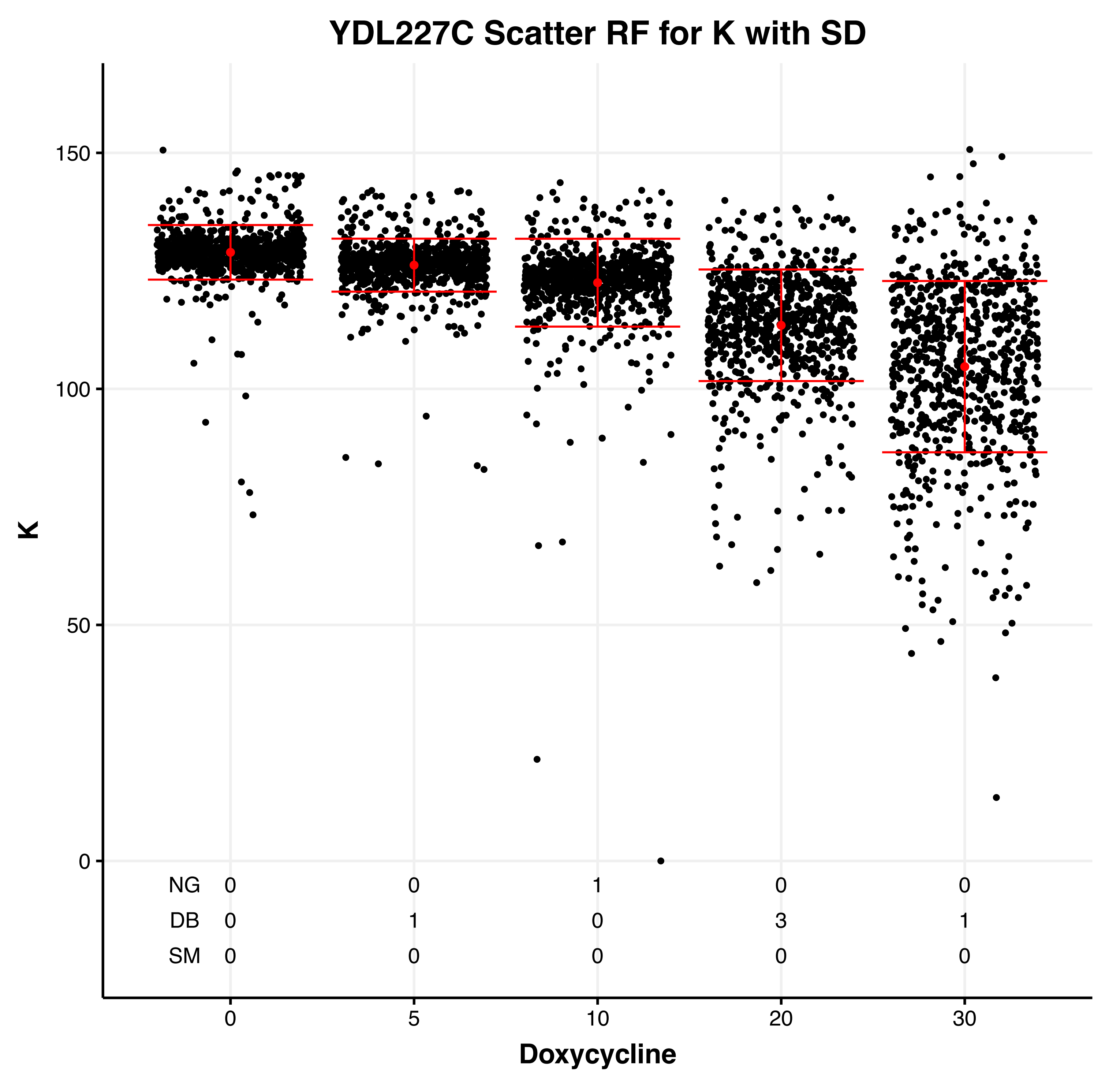
YDL227C Scatter RF for K with SD
K
150
100
50
0
NG 0 0 1 0 0
DB 0 1 0 3 1
SM 0 0 0 0 0
0 5 10 20 30
Doxycycline

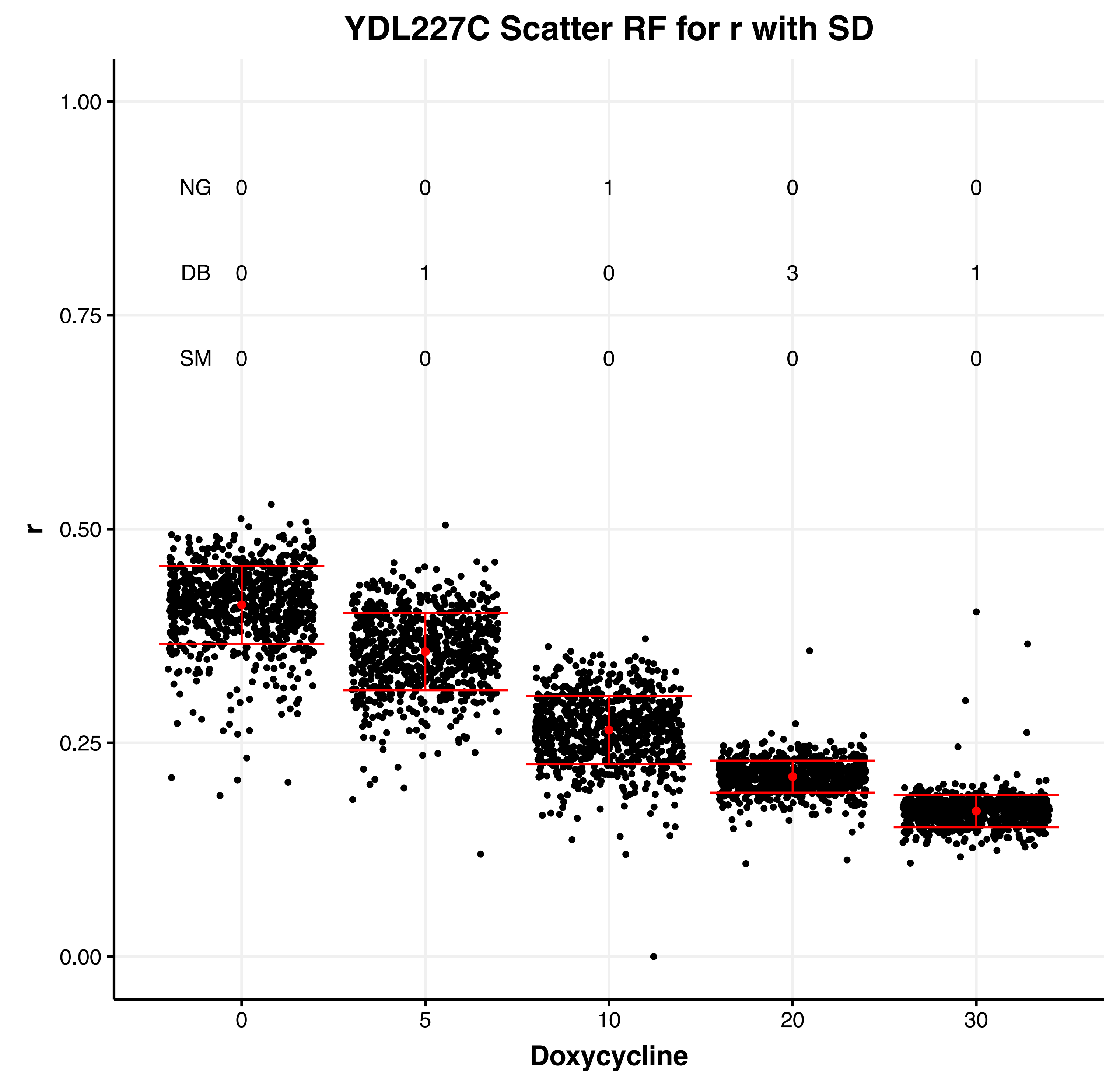
YDL227C Scatter RF for r with SD
r
1.00
0.75
0.50
0.25
0.00
NG 0 0 1 0 0
DB 0 1 0 3 1
SM 0 0 0 0 0
0 5 10 20 30
Doxycycline

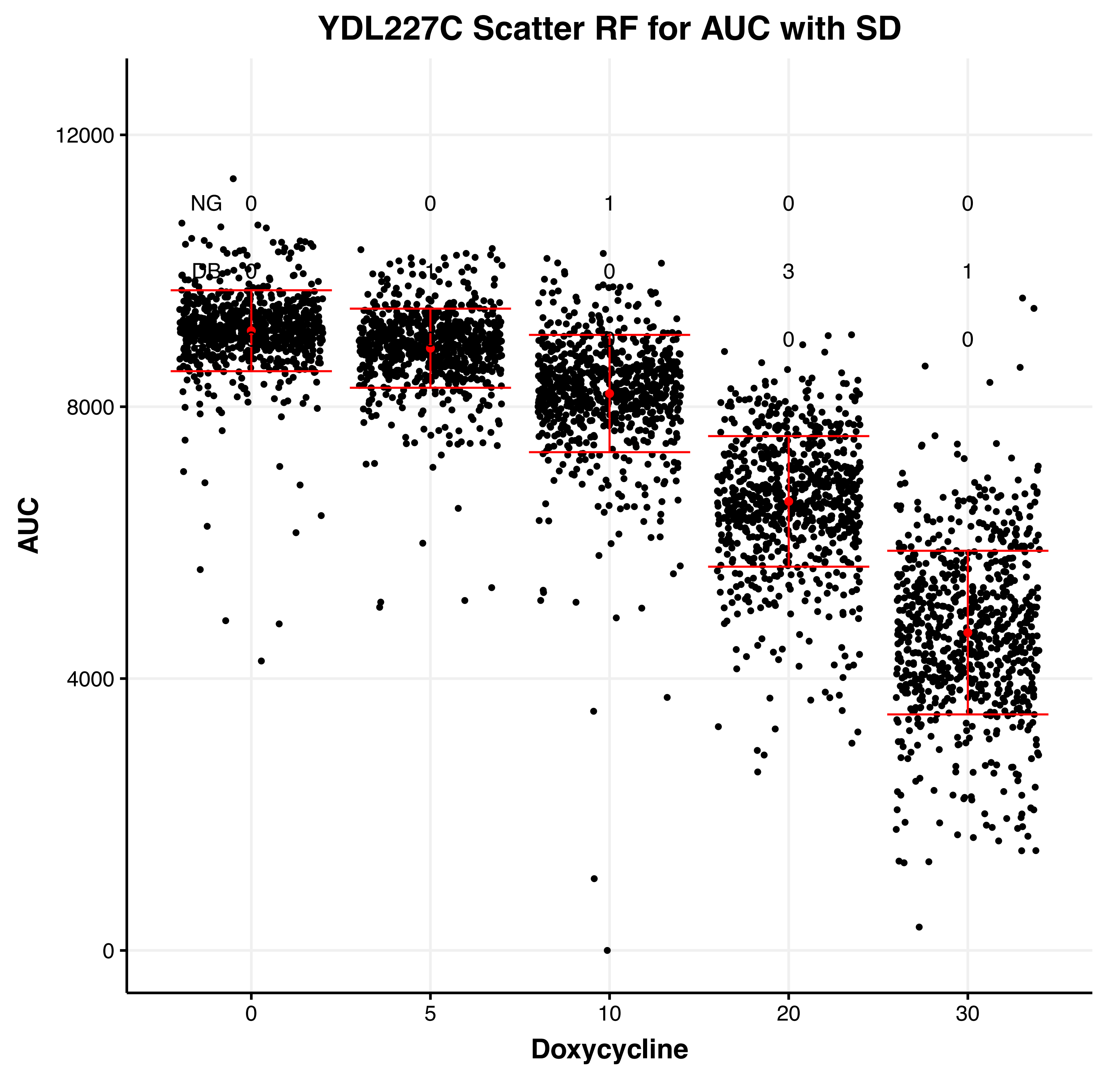
YDL227C Scatter RF for AUC with SD
AUC
12000
8000
4000
0
NG 0 0 1 0 0
DB 0 0 3 1
0 0
0 5 10 20 30
Doxycycline

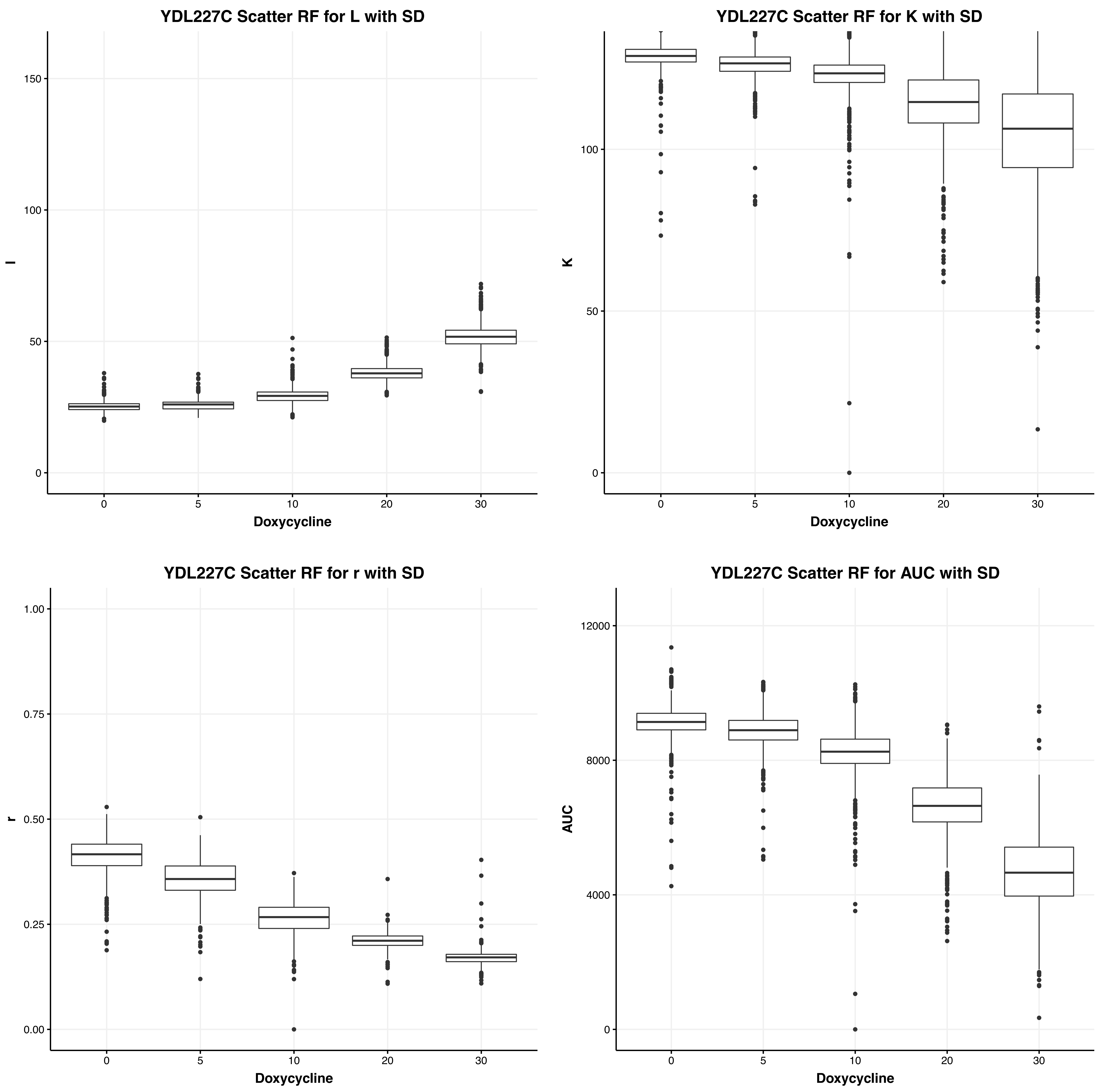
YDL227C Scatter RF for L with SD
l
Doxycycline
0
5
10
20
30
0
50
100
150
YDL227C Scatter RF for K with SD
K
Doxycycline
0
50
100
YDL227C Scatter RF for r with SD
r
Doxycycline
0.00
0.25
0.50
0.75
1.00
YDL227C Scatter RF for AUC with SD
AUC
Doxycycline
0
4000
8000
12000

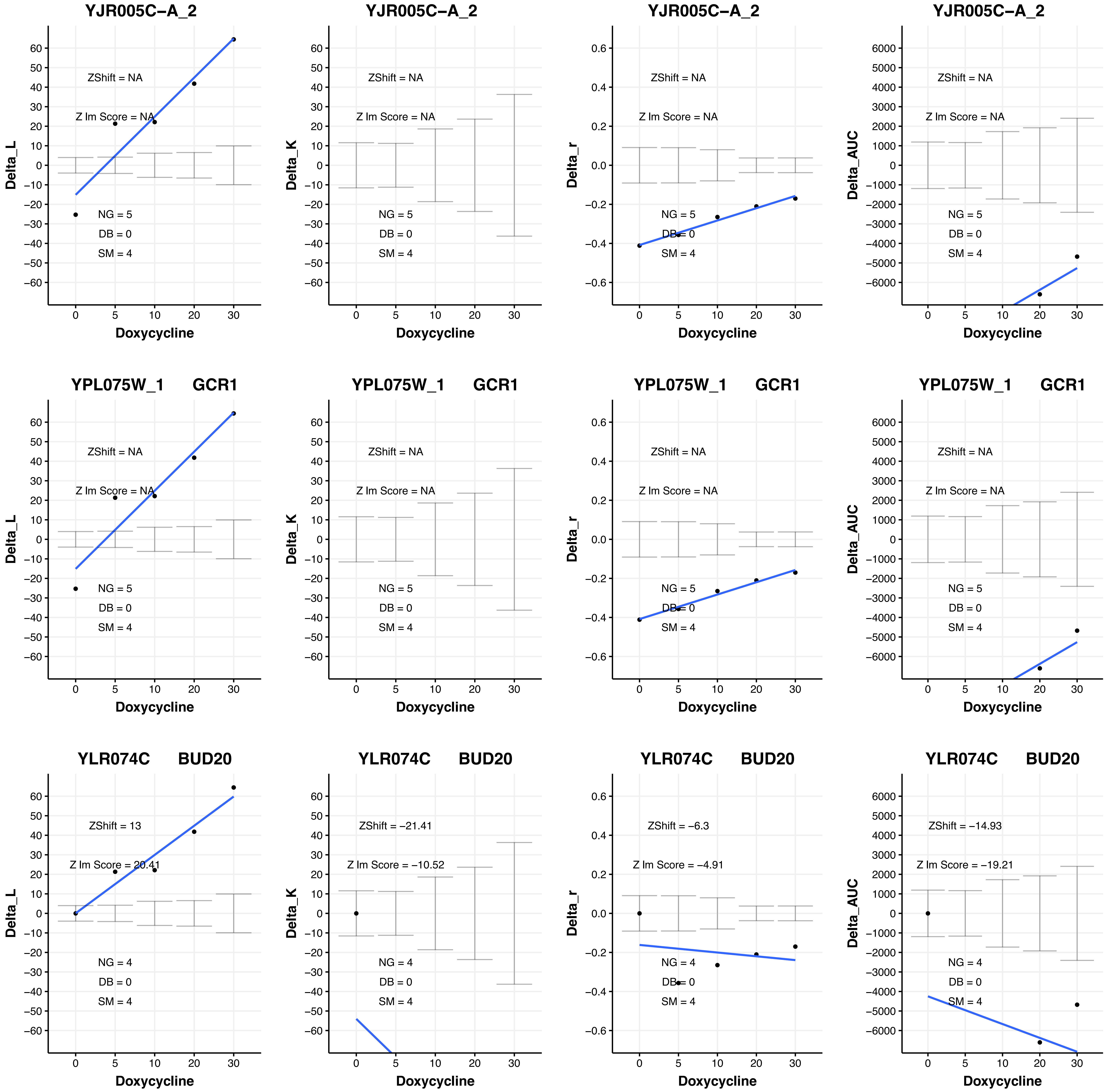

YJR005C−A_2
ZShift = NA
Z lm Score = NA
NG = 5
DB = 0
SM = 4
Delta_L
Doxycycline
YJR005C−A_2
ZShift = NA
Z lm Score = NA
NG = 5
DB = 0
SM = 4
Delta_K
Doxycycline
YJR005C−A_2
ZShift = NA
Z lm Score = NA
NG = 5
DB = 0
SM = 4
Delta_r
Doxycycline
YJR005C−A_2
ZShift = NA
Z lm Score = NA
NG = 5
DB = 0
SM = 4
Delta_AUC
Doxycycline
YPL075W_1 GCR1
ZShift = NA
Z lm Score = NA
NG = 5
DB = 0
SM = 4
Delta_L
Doxycycline
YPL075W_1 GCR1
ZShift = NA
Z lm Score = NA
NG = 5
DB = 0
SM = 4
Delta_K
Doxycycline
YPL075W_1 GCR1
ZShift = NA
Z lm Score = NA
NG = 5
DB = 0
SM = 4
Delta_r
Doxycycline
YPL075W_1 GCR1
ZShift = NA
Z lm Score = NA
NG = 5
DB = 0
SM = 4
Delta_AUC
Doxycycline
YLR074C BUD20
ZShift = 13
Z lm Score = 20.41
NG = 4
DB = 0
SM = 4
Delta_L
Doxycycline
YLR074C BUD20
ZShift = −21.41
Z lm Score = −10.52
NG = 4
DB = 0
SM = 4
Delta_K
Doxycycline
YLR074C BUD20
ZShift = −6.3
Z lm Score = −4.91
NG = 4
DB = 0
SM = 4
Delta_r
Doxycycline
YLR074C BUD20
ZShift = −14.93
Z lm Score = −19.21
NG = 4
DB = 0
SM = 4
Delta_AUC
Doxycycline

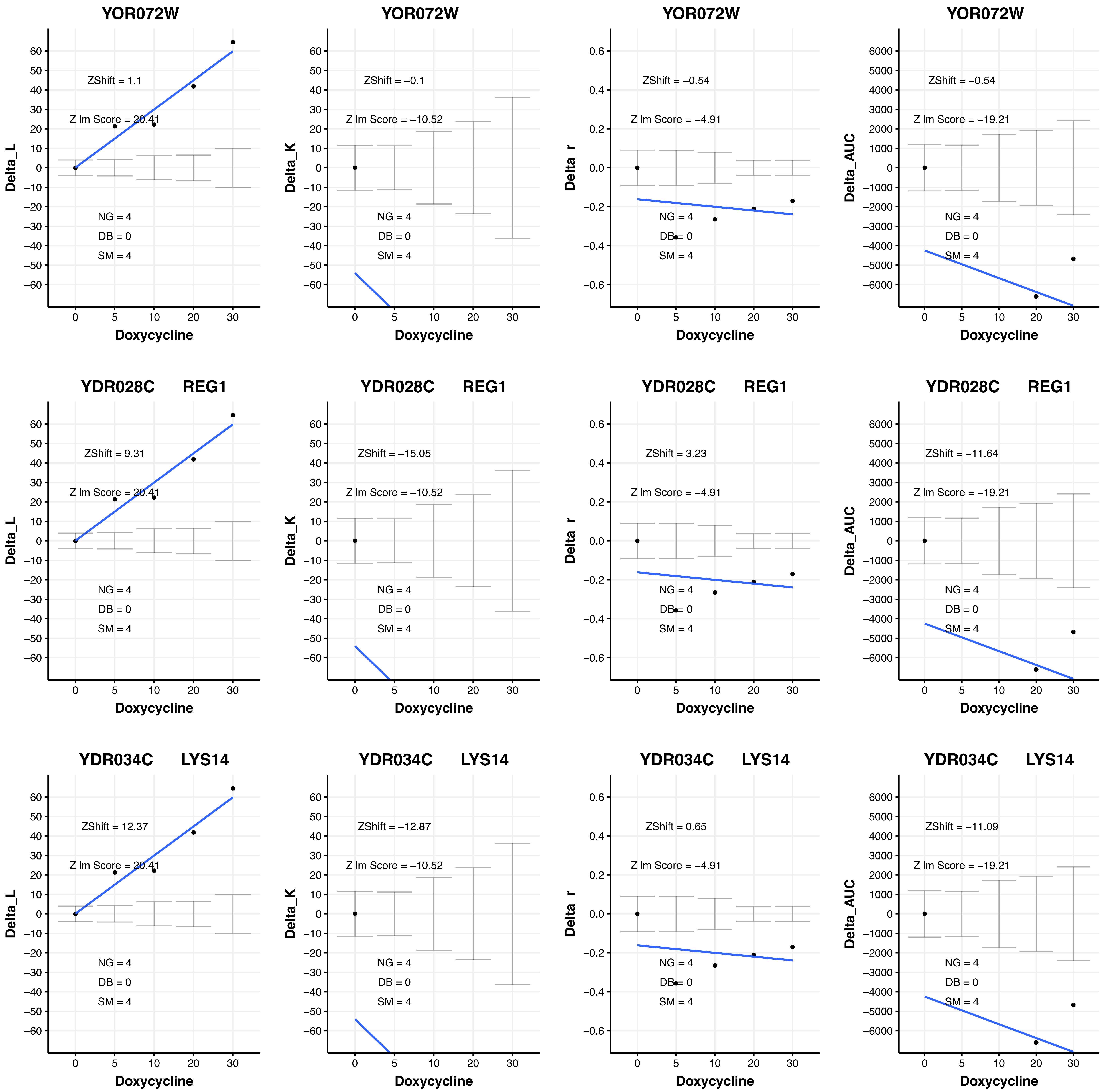
YOR072W
ZShift = 1.1
Z lm Score = 20.41
NG = 4
DB = 0
SM = 4
Delta_L
Doxycycline
YOR072W
ZShift = −0.1
Z lm Score = −10.52
NG = 4
DB = 0
SM = 4
Delta_K
Doxycycline
YOR072W
ZShift = −0.54
Z lm Score = −4.91
NG = 4
DB = 0
SM = 4
Delta_r
Doxycycline
YOR072W
ZShift = −0.54
Z lm Score = −19.21
NG = 4
DB = 0
SM = 4
Delta_AUC
Doxycycline
YDR028C REG1
ZShift = 9.31
Z lm Score = 20.41
NG = 4
DB = 0
SM = 4
Delta_L
Doxycycline
YDR028C REG1
ZShift = −15.05
Z lm Score = −10.52
NG = 4
DB = 0
SM = 4
Delta_K
Doxycycline
YDR028C REG1
ZShift = 3.23
Z lm Score = −4.91
NG = 4
DB = 0
SM = 4
Delta_r
Doxycycline
YDR028C REG1
ZShift = −11.64
Z lm Score = −19.21
NG = 4
DB = 0
SM = 4
Delta_AUC
Doxycycline
YDR034C LYS14
ZShift = 12.37
Z lm Score = 20.41
NG = 4
DB = 0
SM = 4
Delta_L
Doxycycline
YDR034C LYS14
ZShift = −12.87
Z lm Score = −10.52
NG = 4
DB = 0
SM = 4
Delta_K
Doxycycline
YDR034C LYS14
ZShift = 0.65
Z lm Score = −4.91
NG = 4
DB = 0
SM = 4
Delta_r
Doxycycline
YDR034C LYS14
ZShift = −11.09
Z lm Score = −19.21
NG = 4
DB = 0
SM = 4
Delta_AUC
Doxycycline

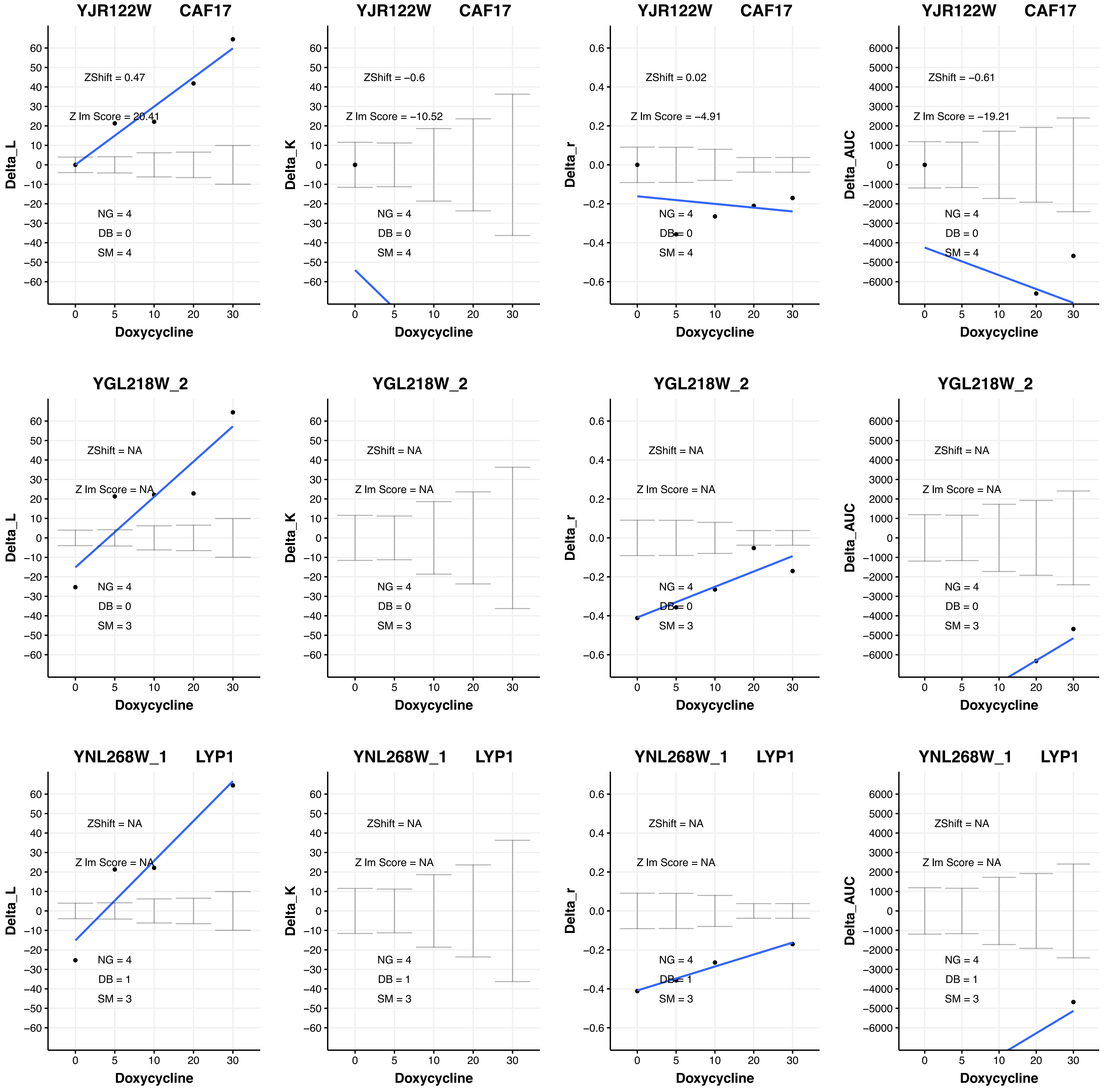
YJR122W CAF17
ZShift = 0.47
Z lm Score = 20.41
NG = 4
DB = 0
SM = 4
Delta_L
Doxycycline
YJR122W CAF17
ZShift = −0.6
Z lm Score = −10.52
NG = 4
DB = 0
SM = 4
Delta_K
Doxycycline
YJR122W CAF17
ZShift = 0.02
Z lm Score = −4.91
NG = 4
DB = 0
SM = 4
Delta_r
Doxycycline
YJR122W CAF17
ZShift = −0.61
Z lm Score = −19.21
NG = 4
DB = 0
SM = 4
Delta_AUC
Doxycycline
YGL218W_2
ZShift = NA
Z lm Score = NA
NG = 4
DB = 0
SM = 3
Delta_L
Doxycycline
YGL218W_2
ZShift = NA
Z lm Score = NA
NG = 4
DB = 0
SM = 3
Delta_K
Doxycycline
YGL218W_2
ZShift = NA
Z lm Score = NA
NG = 4
DB = 0
SM = 3
Delta_r
Doxycycline
YGL218W_2
ZShift = NA
Z lm Score = NA
NG = 4
DB = 0
SM = 3
Delta_AUC
Doxycycline
YNL268W_1 LYP1
ZShift = NA
Z lm Score = NA
NG = 4
DB = 1
SM = 3
Delta_L
Doxycycline
YNL268W_1 LYP1
ZShift = NA
Z lm Score = NA
NG = 4
DB = 1
SM = 3
Delta_K
Doxycycline
YNL268W_1 LYP1
ZShift = NA
Z lm Score = NA
NG = 4
DB = 1
SM = 3
Delta_r
Doxycycline
YNL268W_1 LYP1
ZShift = NA
Z lm Score = NA
NG = 4
DB = 1
SM = 3
Delta_AUC
Doxycycline

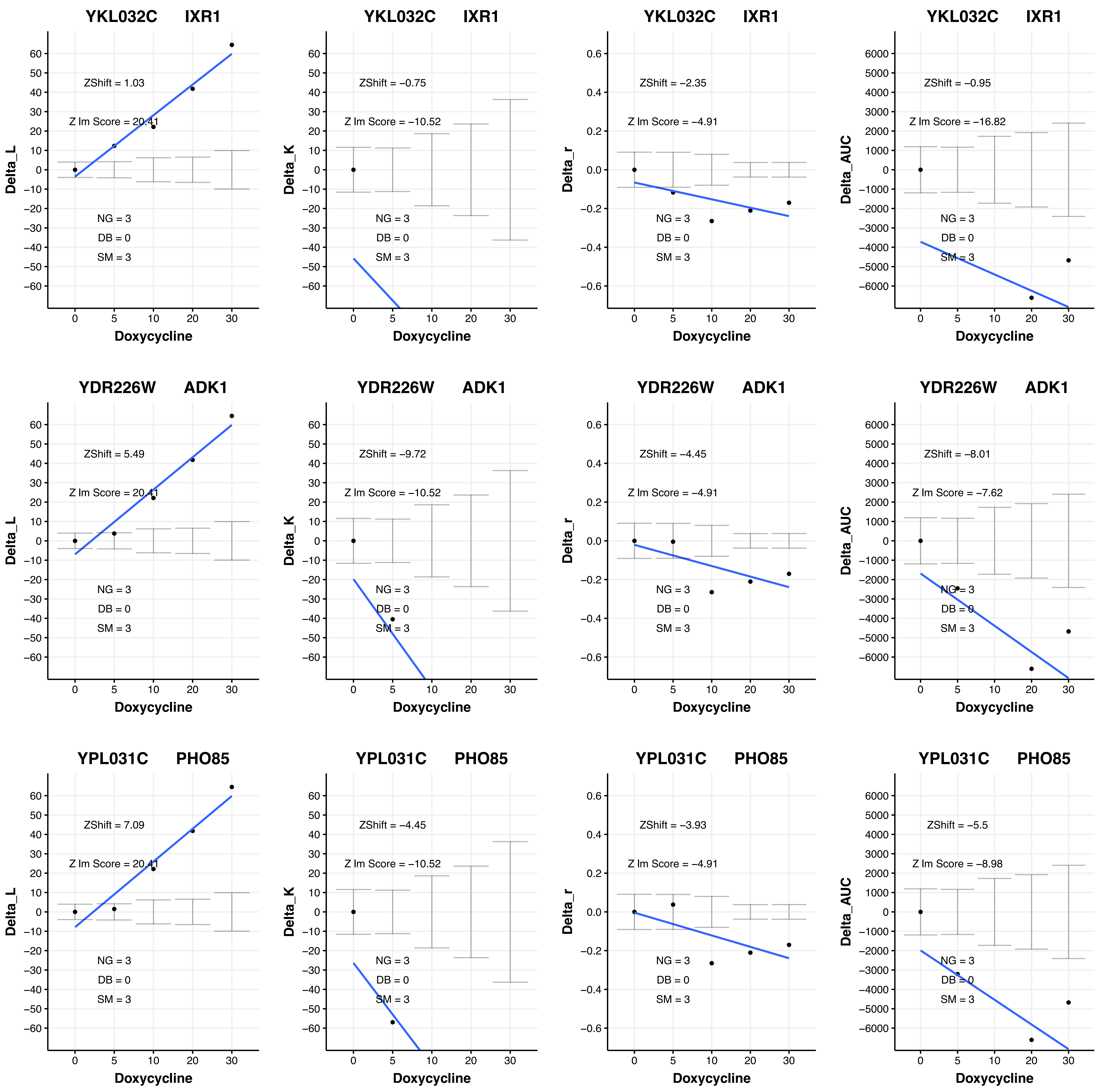

YKL032C IXR1
ZShift = 1.03
Z lm Score = 20.41
NG = 3
DB = 0
SM = 3
Delta_L
Doxycycline
YKL032C IXR1
ZShift = −0.75
Z lm Score = −10.52
NG = 3
DB = 0
SM = 3
Delta_K
Doxycycline
YKL032C IXR1
ZShift = −2.35
Z lm Score = −4.91
NG = 3
DB = 0
SM = 3
Delta_r
Doxycycline
YKL032C IXR1
ZShift = −0.95
Z lm Score = −16.82
NG = 3
DB = 0
SM = 3
Delta_AUC
Doxycycline
YDR226W ADK1
ZShift = 5.49
Z lm Score = 20.41
NG = 3
DB = 0
SM = 3
Delta_L
Doxycycline
YDR226W ADK1
ZShift = −9.72
Z lm Score = −10.52
NG = 3
DB = 0
SM = 3
Delta_K
Doxycycline
YDR226W ADK1
ZShift = −4.45
Z lm Score = −4.91
NG = 3
DB = 0
SM = 3
Delta_r
Doxycycline
YDR226W ADK1
ZShift = −8.01
Z lm Score = −7.62
NG = 3
DB = 0
SM = 3
Delta_AUC
Doxycycline
YPL031C PHO85
ZShift = 7.09
Z lm Score = 20.41
NG = 3
DB = 0
SM = 3
Delta_L
Doxycycline
YPL031C PHO85
ZShift = −4.45
Z lm Score = −10.52
NG = 3
DB = 0
SM = 3
Delta_K
Doxycycline
YPL031C PHO85
ZShift = −3.93
Z lm Score = −4.91
NG = 3
DB = 0
SM = 3
Delta_r
Doxycycline
YPL031C PHO85
ZShift = −5.5
Z lm Score = −8.98
NG = 3
DB = 0
SM = 3
Delta_AUC
Doxycycline

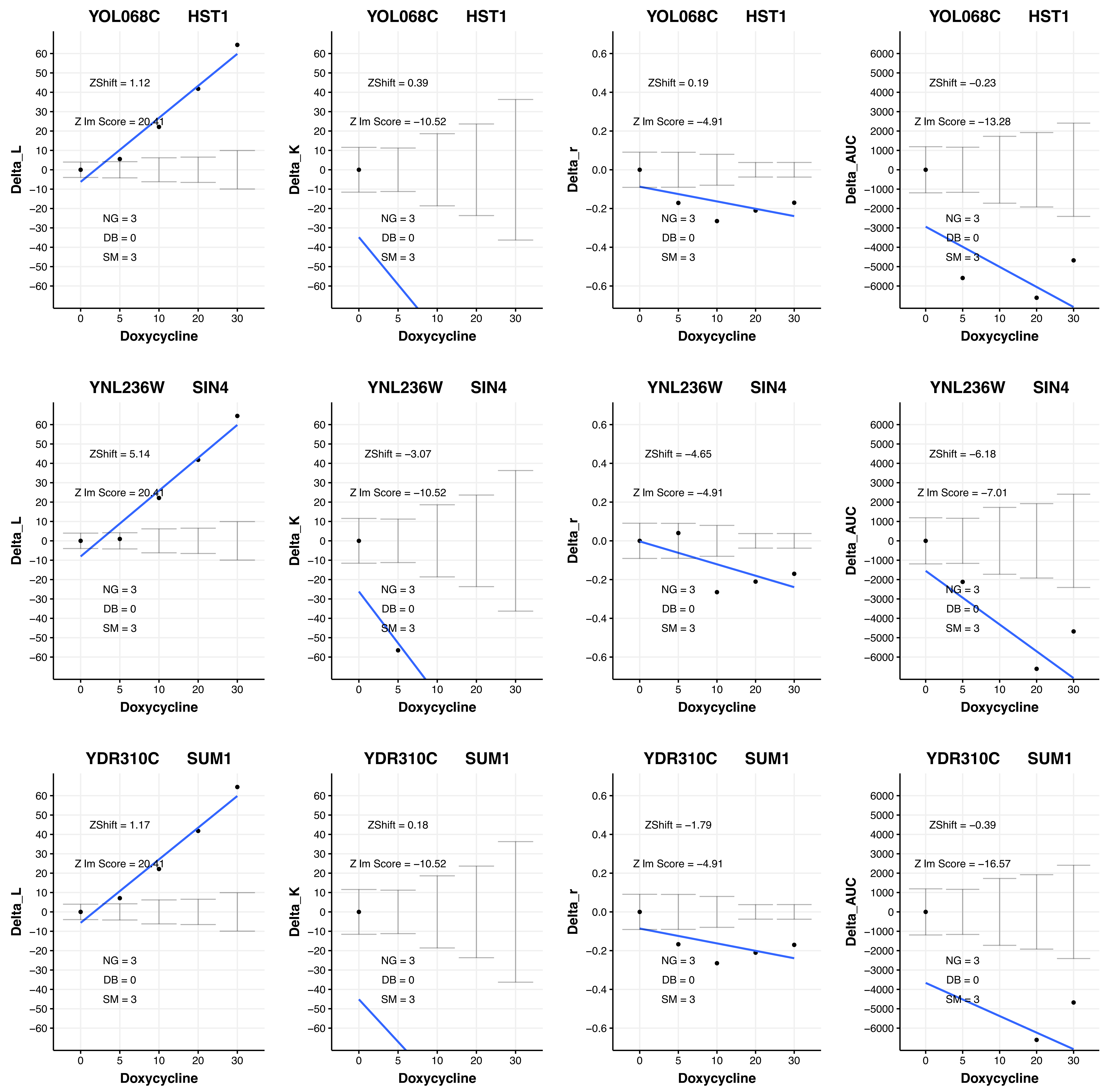
YOL068C HST1
ZShift = 1.12
Z lm Score = 20.41
NG = 3
DB = 0
SM = 3
Delta_L
Doxycycline
YOL068C HST1
ZShift = 0.39
Z lm Score = −10.52
NG = 3
DB = 0
SM = 3
Delta_K
Doxycycline
YOL068C HST1
ZShift = 0.19
Z lm Score = −4.91
NG = 3
DB = 0
SM = 3
Delta_r
Doxycycline
YOL068C HST1
ZShift = −0.23
Z lm Score = −13.28
NG = 3
DB = 0
SM = 3
Delta_AUC
Doxycycline
YNL236W SIN4
ZShift = 5.14
Z lm Score = 20.41
NG = 3
DB = 0
SM = 3
Delta_L
Doxycycline
YNL236W SIN4
ZShift = −3.07
Z lm Score = −10.52
NG = 3
DB = 0
SM = 3
Delta_K
Doxycycline
YNL236W SIN4
ZShift = −4.65
Z lm Score = −4.91
NG = 3
DB = 0
SM = 3
Delta_r
Doxycycline
YNL236W SIN4
ZShift = −6.18
Z lm Score = −7.01
NG = 3
DB = 0
SM = 3
Delta_AUC
Doxycycline
YDR310C SUM1
ZShift = 1.17
Z lm Score = 20.41
NG = 3
DB = 0
SM = 3
Delta_L
Doxycycline
YDR310C SUM1
ZShift = 0.18
Z lm Score = −10.52
NG = 3
DB = 0
SM = 3
Delta_K
Doxycycline
YDR310C SUM1
ZShift = −1.79
Z lm Score = −4.91
NG = 3
DB = 0
SM = 3
Delta_r
Doxycycline
YDR310C SUM1
ZShift = −0.39
Z lm Score = −16.57
NG = 3
DB = 0
SM = 3
Delta_AUC
Doxycycline

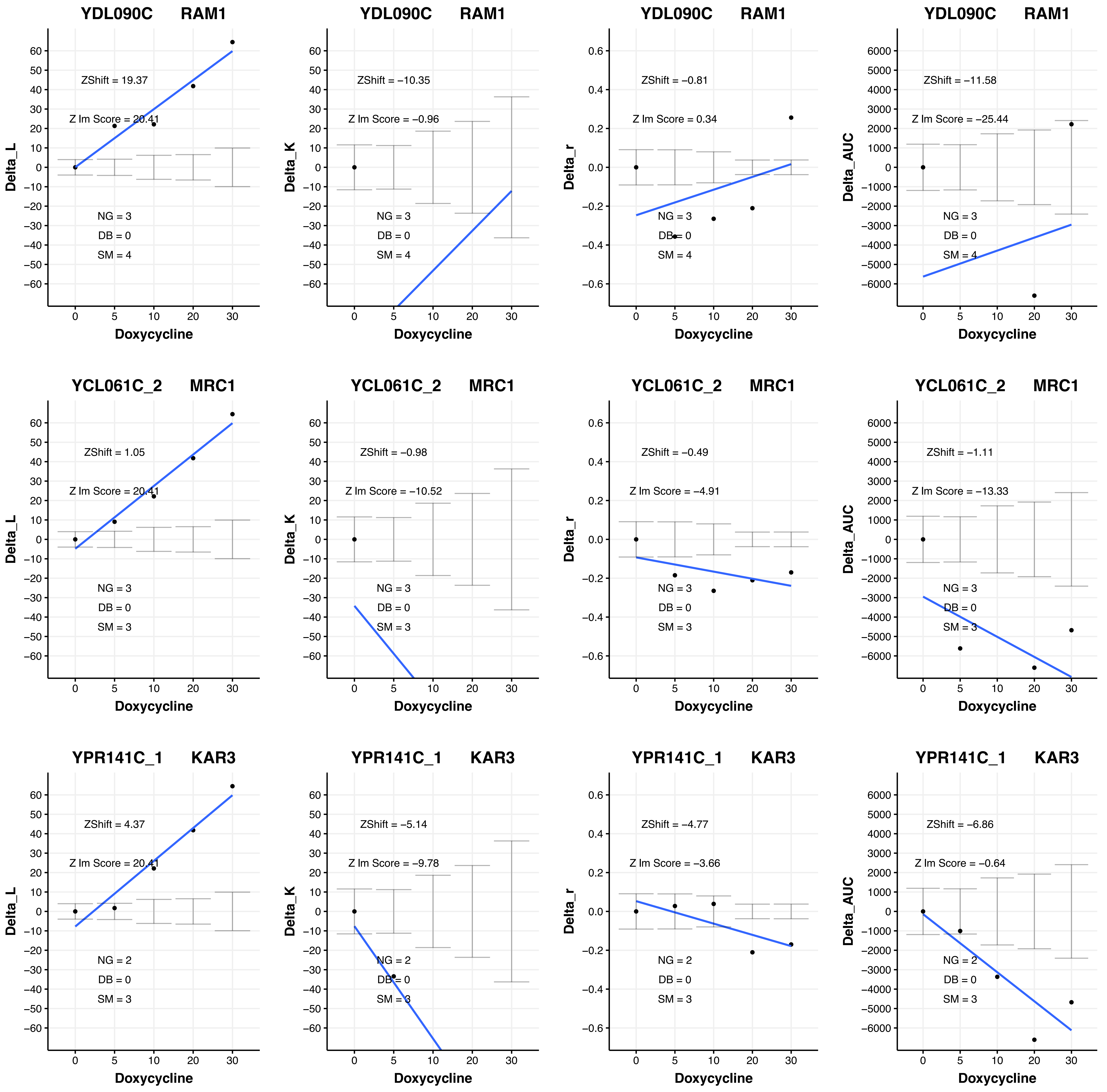
YDL090C RAM1
ZShift = 19.37
Z lm Score = 20.41
NG = 3
DB = 0
SM = 4
Delta_L
Doxycycline
YDL090C RAM1
ZShift = −10.35
Z lm Score = −0.96
NG = 3
DB = 0
SM = 4
Delta_K
Doxycycline
YDL090C RAM1
ZShift = −0.81
Z lm Score = 0.34
NG = 3
DB = 0
SM = 4
Delta_r
Doxycycline
YDL090C RAM1
ZShift = −11.58
Z lm Score = −25.44
NG = 3
DB = 0
SM = 4
Delta_AUC
Doxycycline
YCL061C_2 MRC1
ZShift = 1.05
Z lm Score = 20.41
NG = 3
DB = 0
SM = 3
Delta_L
Doxycycline
YCL061C_2 MRC1
ZShift = −0.98
Z lm Score = −10.52
NG = 3
DB = 0
SM = 3
Delta_K
Doxycycline
YCL061C_2 MRC1
ZShift = −0.49
Z lm Score = −4.91
NG = 3
DB = 0
SM = 3
Delta_r
Doxycycline
YCL061C_2 MRC1
ZShift = −1.11
Z lm Score = −13.33
NG = 3
DB = 0
SM = 3
Delta_AUC
Doxycycline
YPR141C_1 KAR3
ZShift = 4.37
Z lm Score = 20.41
NG = 2
DB = 0
SM = 3
Delta_L
Doxycycline
YPR141C_1 KAR3
ZShift = −5.14
Z lm Score = −9.78
NG = 2
DB = 0
SM = 3
Delta_K
Doxycycline
YPR141C_1 KAR3
ZShift = −4.77
Z lm Score = −3.66
NG = 2
DB = 0
SM = 3
Delta_r
Doxycycline
YPR141C_1 KAR3
ZShift = −6.86
Z lm Score = −0.64
NG = 2
DB = 0
SM = 3
Delta_AUC
Doxycycline

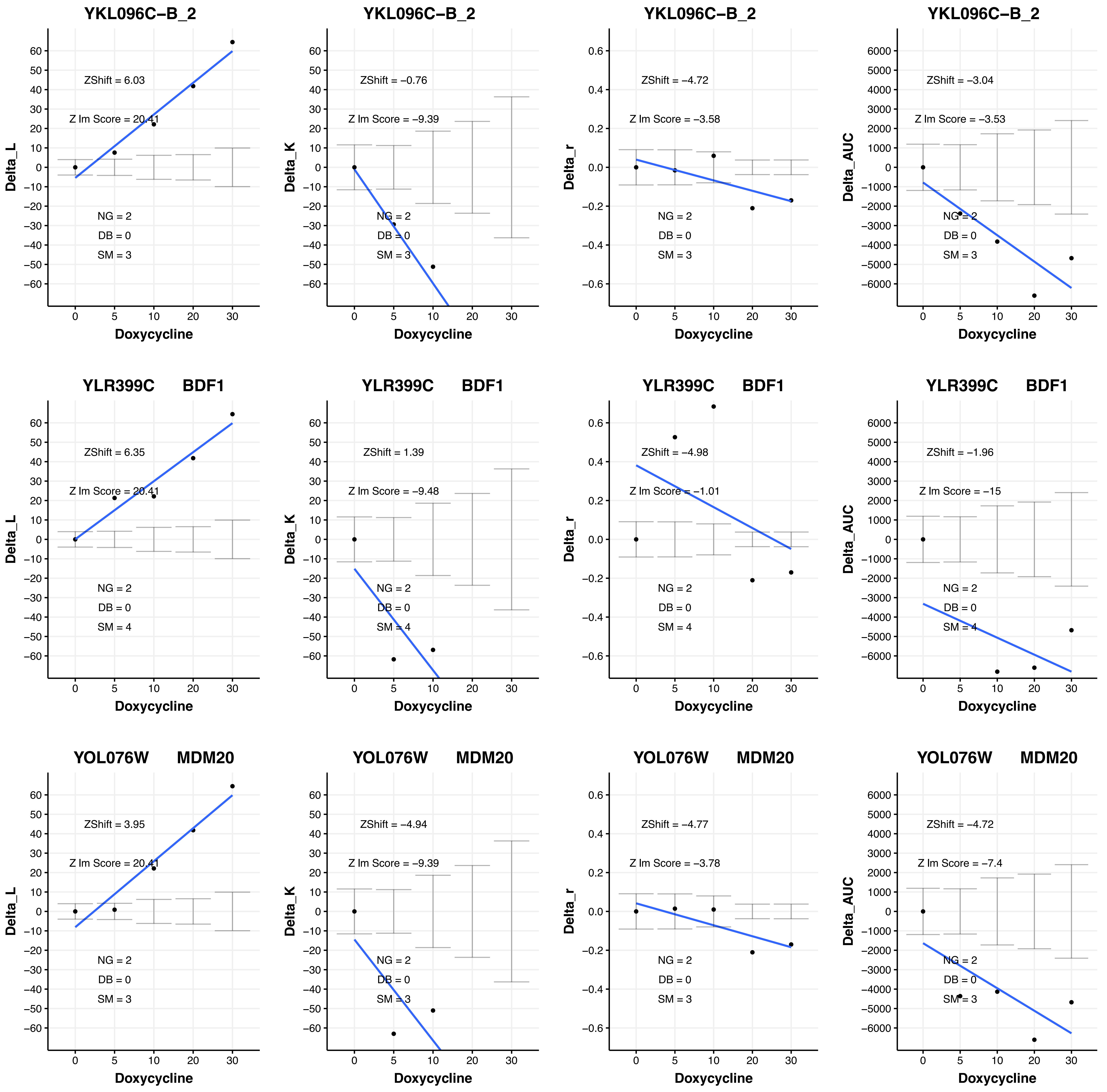

YKL096C−B_2
ZShift = 6.03
Z lm Score = 20.41
NG = 2
DB = 0
SM = 3
Delta_L
Doxycycline
YKL096C−B_2
ZShift = −0.76
Z lm Score = −9.39
NG = 2
DB = 0
SM = 3
Delta_K
Doxycycline
YKL096C−B_2
ZShift = −4.72
Z lm Score = −3.58
NG = 2
DB = 0
SM = 3
Delta_r
Doxycycline
YKL096C−B_2
ZShift = −3.04
Z lm Score = −3.53
NG = 2
DB = 0
SM = 3
Delta_AUC
Doxycycline
YLR399C BDF1
ZShift = 6.35
Z lm Score = 20.41
NG = 2
DB = 0
SM = 4
Delta_L
Doxycycline
YLR399C BDF1
ZShift = 1.39
Z lm Score = −9.48
NG = 2
DB = 0
SM = 4
Delta_K
Doxycycline
YLR399C BDF1
ZShift = −4.98
Z lm Score = −1.01
NG = 2
DB = 0
SM = 4
Delta_r
Doxycycline
YLR399C BDF1
ZShift = −1.96
Z lm Score = −15
NG = 2
DB = 0
SM = 4
Delta_AUC
Doxycycline
YOL076W MDM20
ZShift = 3.95
Z lm Score = 20.41
NG = 2
DB = 0
SM = 3
Delta_L
Doxycycline
YOL076W MDM20
ZShift = −4.94
Z lm Score = −9.39
NG = 2
DB = 0
SM = 3
Delta_K
Doxycycline
YOL076W MDM20
ZShift = −4.77
Z lm Score = −3.78
NG = 2
DB = 0
SM = 3
Delta_r
Doxycycline
YOL076W MDM20
ZShift = −4.72
Z lm Score = −7.4
NG = 2
DB = 0
SM = 3
Delta_AUC
Doxycycline

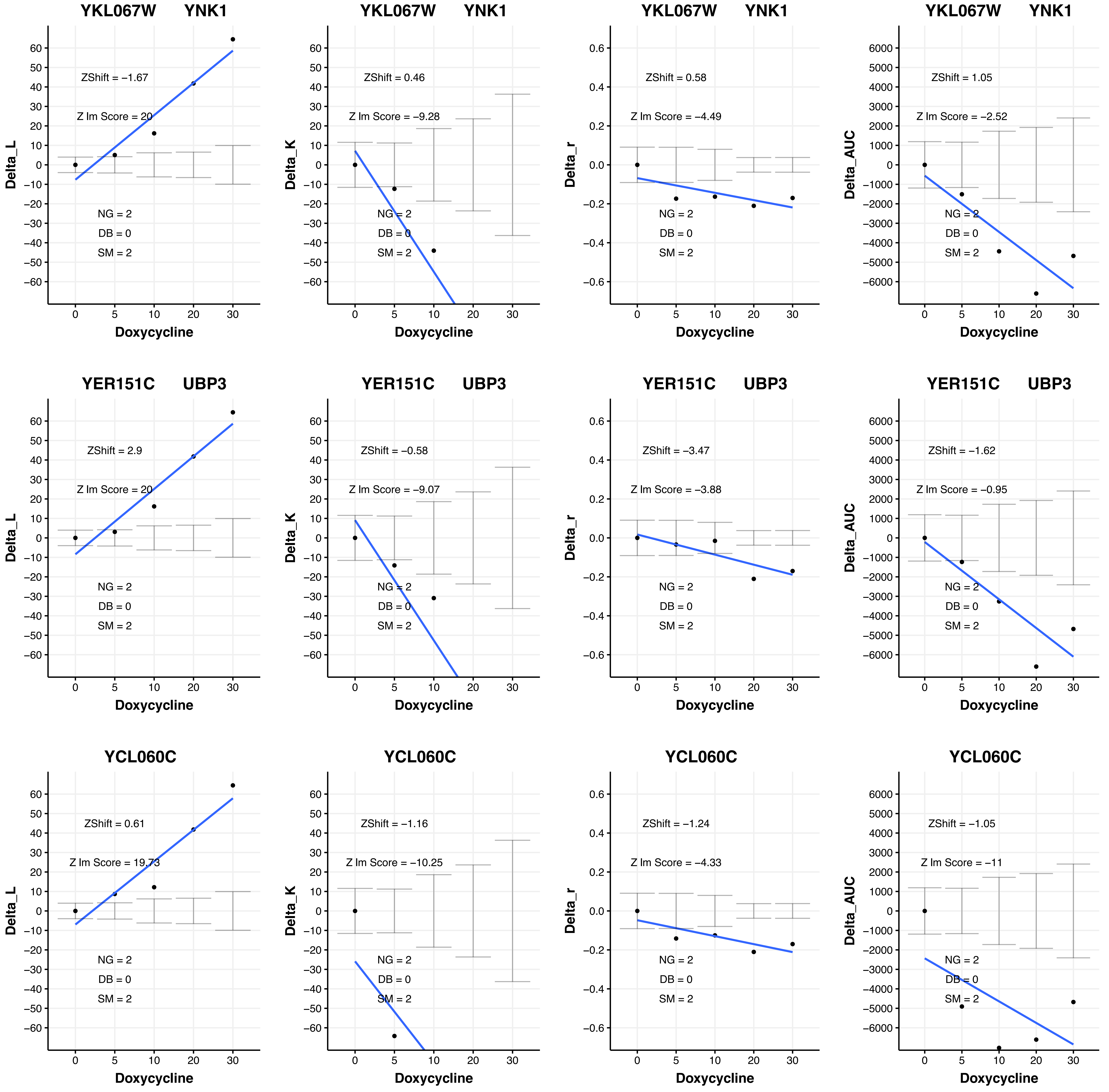
YKL067W YNK1
ZShift = −1.67
Z lm Score = 20
NG = 2
DB = 0
SM = 2
Delta_L
Doxycycline
YKL067W YNK1
ZShift = 0.46
Z lm Score = −9.28
NG = 2
DB = 0
SM = 2
Delta_K
Doxycycline
YKL067W YNK1
ZShift = 0.58
Z lm Score = −4.49
NG = 2
DB = 0
SM = 2
Delta_r
Doxycycline
YKL067W YNK1
ZShift = 1.05
Z lm Score = −2.52
NG = 2
DB = 0
SM = 2
Delta_AUC
Doxycycline
YER151C UBP3
ZShift = 2.9
Z lm Score = 20
NG = 2
DB = 0
SM = 2
Delta_L
Doxycycline
YER151C UBP3
ZShift = −0.58
Z lm Score = −9.07
NG = 2
DB = 0
SM = 2
Delta_K
Doxycycline
YER151C UBP3
ZShift = −3.47
Z lm Score = −3.88
NG = 2
DB = 0
SM = 2
Delta_r
Doxycycline
YER151C UBP3
ZShift = −1.62
Z lm Score = −0.95
NG = 2
DB = 0
SM = 2
Delta_AUC
Doxycycline
YCL060C
ZShift = 0.61
Z lm Score = 19.73
NG = 2
DB = 0
SM = 2
Delta_L
Doxycycline
YCL060C
ZShift = −1.16
Z lm Score = −10.25
NG = 2
DB = 0
SM = 2
Delta_K
Doxycycline
YCL060C
ZShift = −1.24
Z lm Score = −4.33
NG = 2
DB = 0
SM = 2
Delta_r
Doxycycline
YCL060C
ZShift = −1.05
Z lm Score = −11
NG = 2
DB = 0
SM = 2
Delta_AUC
Doxycycline

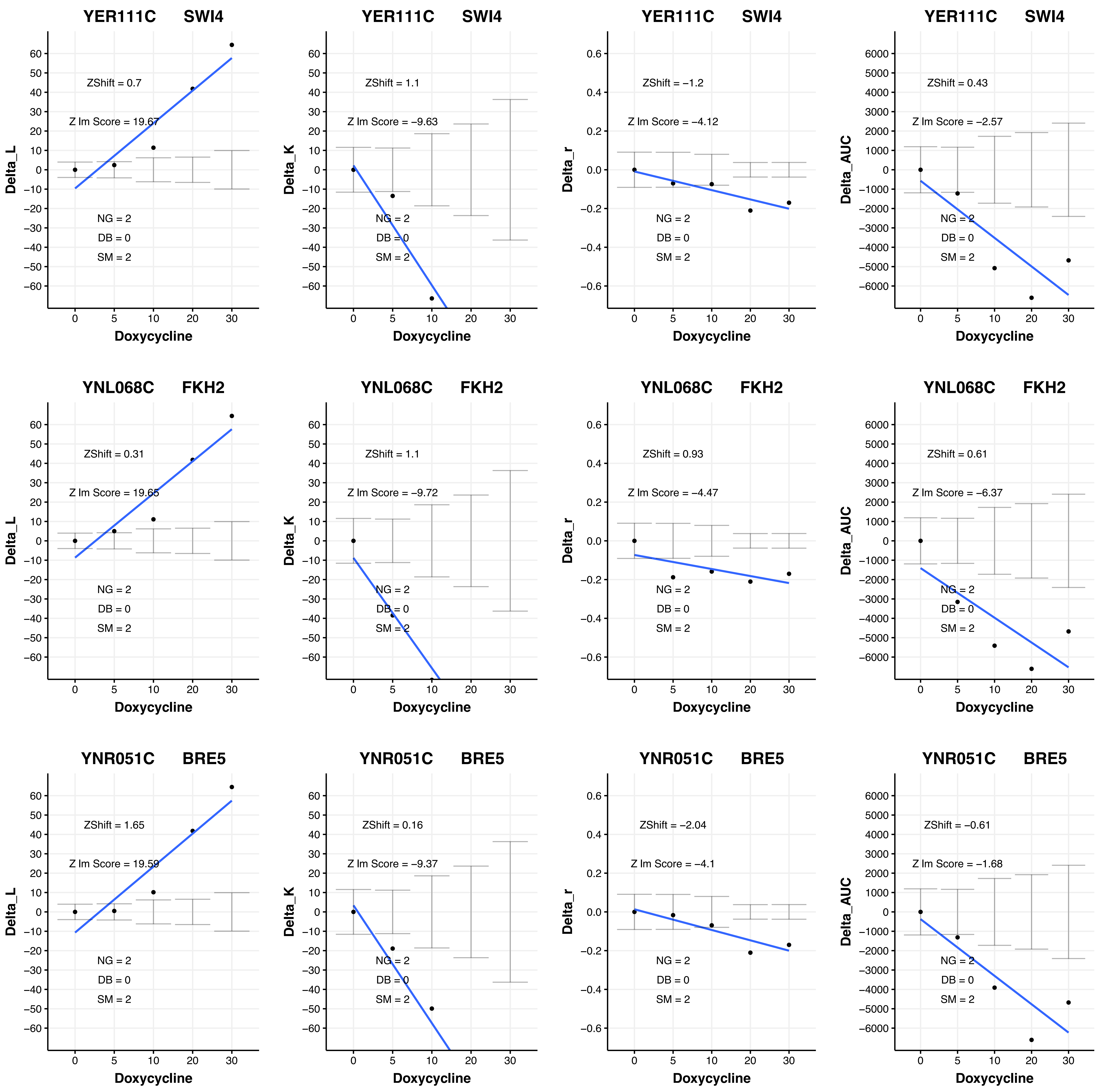
YER111C SWI4
ZShift = 0.7
Z lm Score = 19.67
NG = 2
DB = 0
SM = 2
Delta_L
Doxycycline
YER111C SWI4
ZShift = 1.1
Z lm Score = −9.63
NG = 2
DB = 0
SM = 2
Delta_K
Doxycycline
YER111C SWI4
ZShift = −1.2
Z lm Score = −4.12
NG = 2
DB = 0
SM = 2
Delta_r
Doxycycline
YER111C SWI4
ZShift = 0.43
Z lm Score = −2.57
NG = 2
DB = 0
SM = 2
Delta_AUC
Doxycycline
YNL068C FKH2
ZShift = 0.31
Z lm Score = 19.65
NG = 2
DB = 0
SM = 2
Delta_L
Doxycycline
YNL068C FKH2
ZShift = 1.1
Z lm Score = −9.72
NG = 2
DB = 0
SM = 2
Delta_K
Doxycycline
YNL068C FKH2
ZShift = 0.93
Z lm Score = −4.47
NG = 2
DB = 0
SM = 2
Delta_r
Doxycycline
YNL068C FKH2
ZShift = 0.61
Z lm Score = −6.37
NG = 2
DB = 0
SM = 2
Delta_AUC
Doxycycline
YNR051C BRE5
ZShift = 1.65
Z lm Score = 19.59
NG = 2
DB = 0
SM = 2
Delta_L
Doxycycline
YNR051C BRE5
ZShift = 0.16
Z lm Score = −9.37
NG = 2
DB = 0
SM = 2
Delta_K
Doxycycline
YNR051C BRE5
ZShift = −2.04
Z lm Score = −4.1
NG = 2
DB = 0
SM = 2
Delta_r
Doxycycline
YNR051C BRE5
ZShift = −0.61
Z lm Score = −1.68
NG = 2
DB = 0
SM = 2
Delta_AUC
Doxycycline

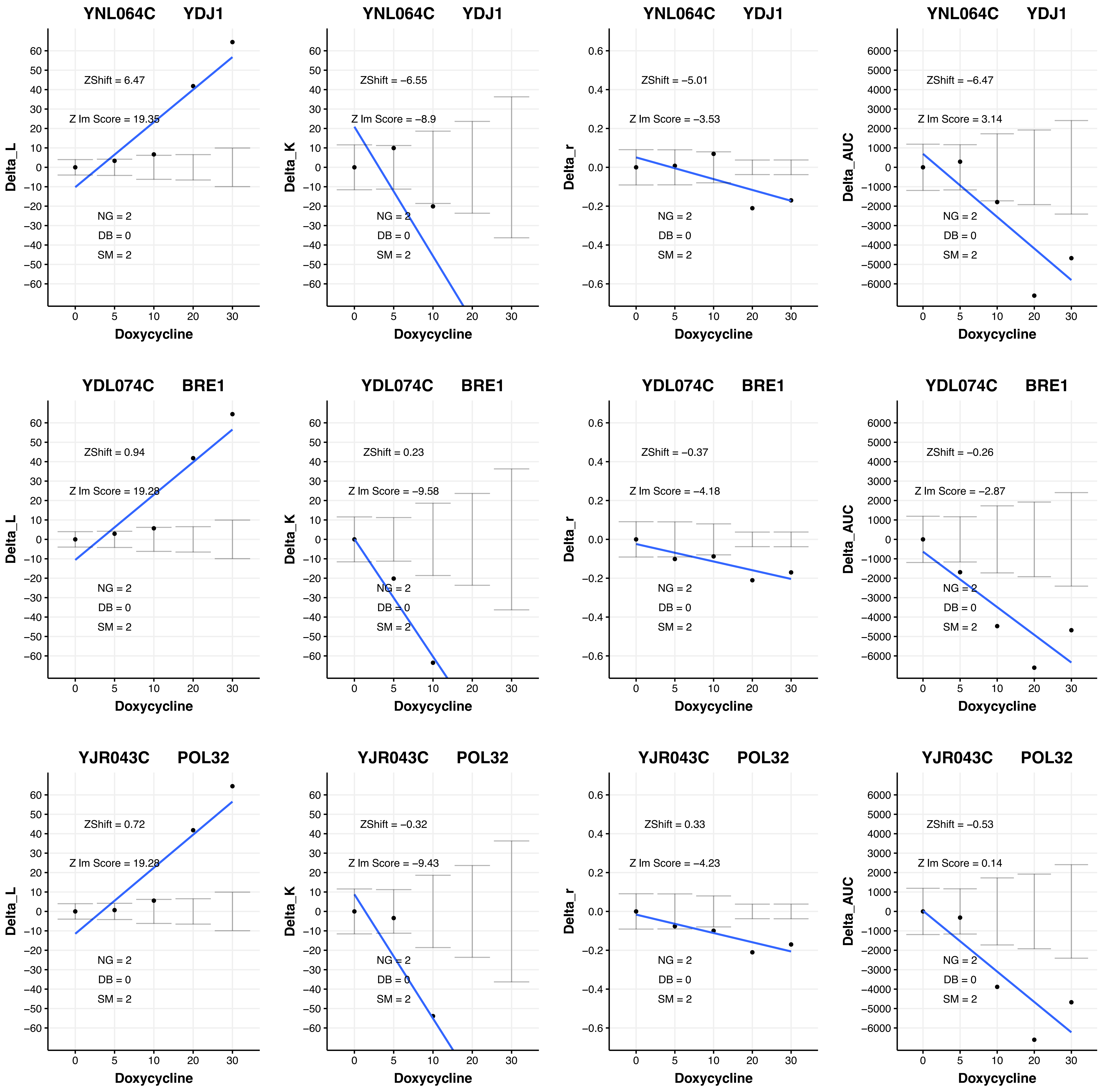
YNL064C YDJ1
ZShift = 6.47
Z lm Score = 19.35
NG = 2
DB = 0
SM = 2
Delta_L
Doxycycline
YNL064C YDJ1
ZShift = −6.55
Z lm Score = −8.9
NG = 2
DB = 0
SM = 2
Delta_K
Doxycycline
YNL064C YDJ1
ZShift = −5.01
Z lm Score = −3.53
NG = 2
DB = 0
SM = 2
Delta_r
Doxycycline
YNL064C YDJ1
ZShift = −6.47
Z lm Score = 3.14
NG = 2
DB = 0
SM = 2
Delta_AUC
Doxycycline
YDL074C BRE1
ZShift = 0.94
Z lm Score = 19.28
NG = 2
DB = 0
SM = 2
Delta_L
Doxycycline
YDL074C BRE1
ZShift = 0.23
Z lm Score = −9.58
NG = 2
DB = 0
SM = 2
Delta_K
Doxycycline
YDL074C BRE1
ZShift = −0.37
Z lm Score = −4.18
NG = 2
DB = 0
SM = 2
Delta_r
Doxycycline
YDL074C BRE1
ZShift = −0.26
Z lm Score = −2.87
NG = 2
DB = 0
SM = 2
Delta_AUC
Doxycycline
YJR043C POL32
ZShift = 0.72
Z lm Score = 19.28
NG = 2
DB = 0
SM = 2
Delta_L
Doxycycline
YJR043C POL32
ZShift = −0.32
Z lm Score = −9.43
NG = 2
DB = 0
SM = 2
Delta_K
Doxycycline
YJR043C POL32
ZShift = 0.33
Z lm Score = −4.23
NG = 2
DB = 0
SM = 2
Delta_r
Doxycycline
YJR043C POL32
ZShift = −0.53
Z lm Score = 0.14
NG = 2
DB = 0
SM = 2
Delta_AUC
Doxycycline

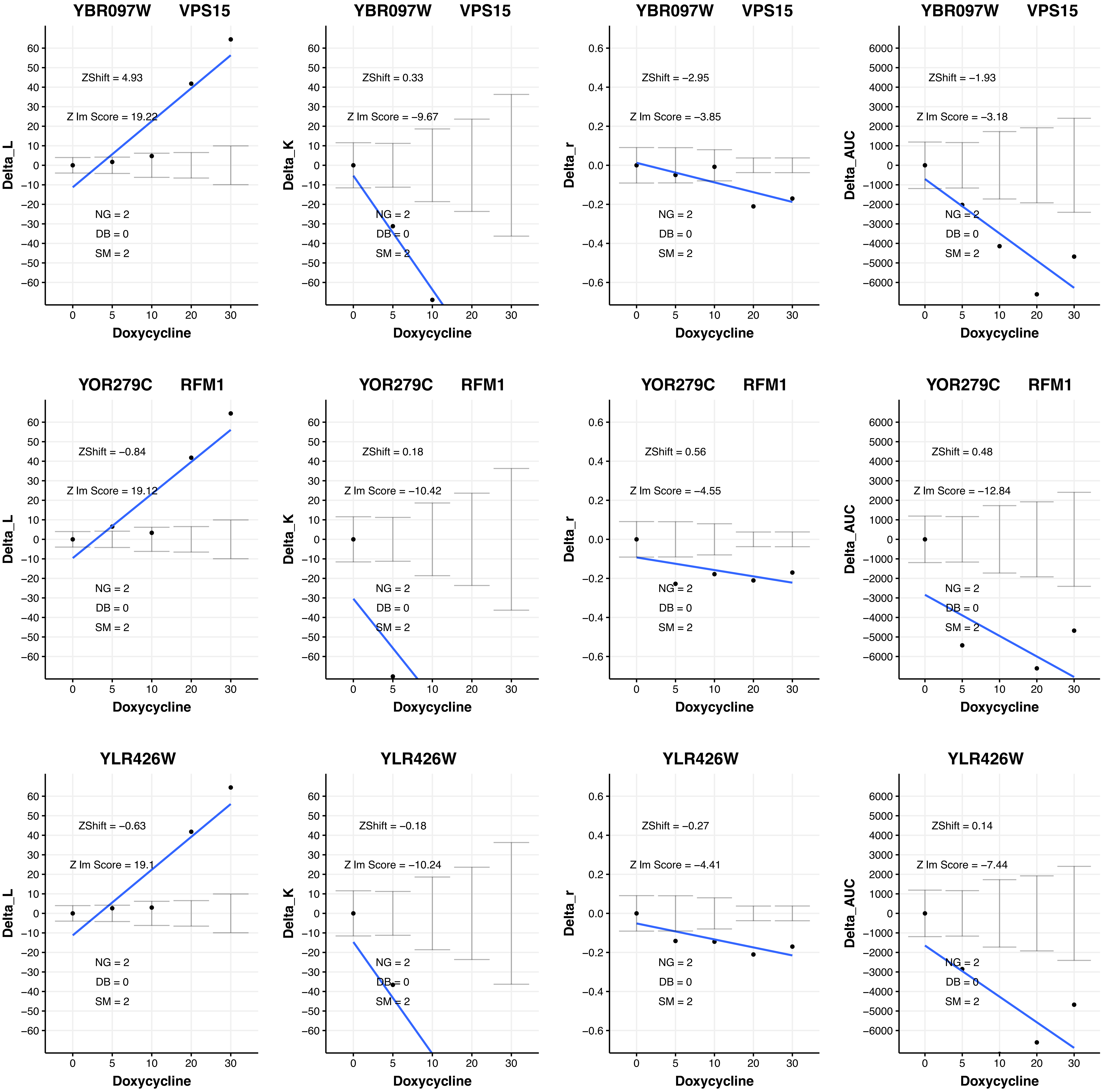

YBR097W VPS15
ZShift = 4.93
Z lm Score = 19.22
NG = 2
DB = 0
SM = 2
Delta_L
Doxycycline
YBR097W VPS15
ZShift = 0.33
Z lm Score = −9.67
NG = 2
DB = 0
SM = 2
Delta_K
Doxycycline
YBR097W VPS15
ZShift = −2.95
Z lm Score = −3.85
NG = 2
DB = 0
SM = 2
Delta_r
Doxycycline
YBR097W VPS15
ZShift = −1.93
Z lm Score = −3.18
NG = 2
DB = 0
SM = 2
Delta_AUC
Doxycycline
YOR279C RFM1
ZShift = −0.84
Z lm Score = 19.12
NG = 2
DB = 0
SM = 2
Delta_L
Doxycycline
YOR279C RFM1
ZShift = 0.18
Z lm Score = −10.42
NG = 2
DB = 0
SM = 2
Delta_K
Doxycycline
YOR279C RFM1
ZShift = 0.56
Z lm Score = −4.55
NG = 2
DB = 0
SM = 2
Delta_r
Doxycycline
YOR279C RFM1
ZShift = 0.48
Z lm Score = −12.84
NG = 2
DB = 0
SM = 2
Delta_AUC
Doxycycline
YLR426W
ZShift = −0.63
Z lm Score = 19.1
NG = 2
DB = 0
SM = 2
Delta_L
Doxycycline
YLR426W
ZShift = −0.18
Z lm Score = −10.24
NG = 2
DB = 0
SM = 2
Delta_K
Doxycycline
YLR426W
ZShift = −0.27
Z lm Score = −4.41
NG = 2
DB = 0
SM = 2
Delta_r
Doxycycline
YLR426W
ZShift = 0.14
Z lm Score = −7.44
NG = 2
DB = 0
SM = 2
Delta_AUC
Doxycycline

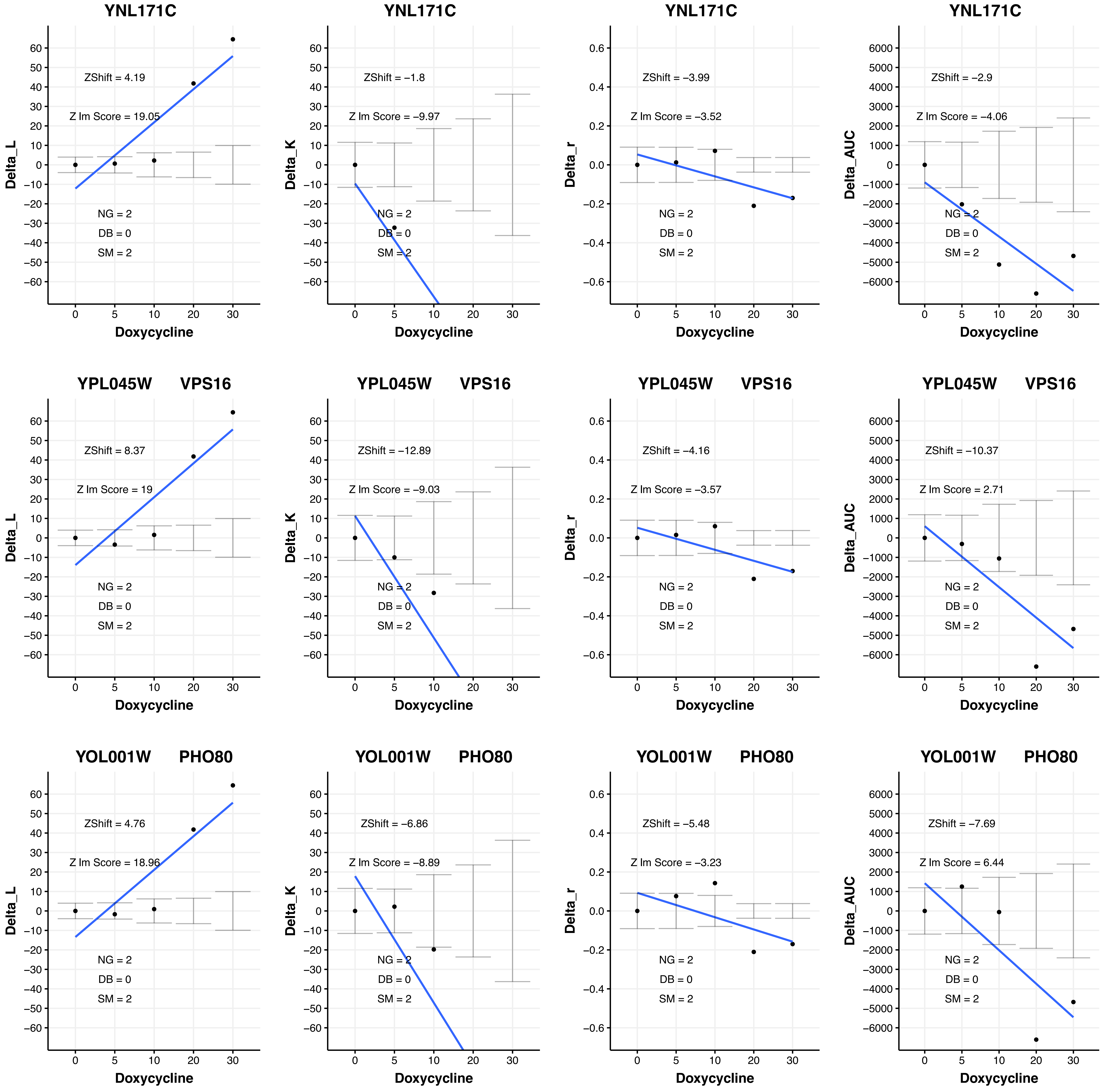
YNL171C
ZShift = 4.19
Z lm Score = 19.05
NG = 2
DB = 0
SM = 2
Delta_L
Doxycycline
YNL171C
ZShift = −1.8
Z lm Score = −9.97
NG = 2
DB = 0
SM = 2
Delta_K
Doxycycline
YNL171C
ZShift = −3.99
Z lm Score = −3.52
NG = 2
DB = 0
SM = 2
Delta_r
Doxycycline
YNL171C
ZShift = −2.9
Z lm Score = −4.06
NG = 2
DB = 0
SM = 2
Delta_AUC
Doxycycline
YPL045W VPS16
ZShift = 8.37
Z lm Score = 19
NG = 2
DB = 0
SM = 2
Delta_L
Doxycycline
YPL045W VPS16
ZShift = −12.89
Z lm Score = −9.03
NG = 2
DB = 0
SM = 2
Delta_K
Doxycycline
YPL045W VPS16
ZShift = −4.16
Z lm Score = −3.57
NG = 2
DB = 0
SM = 2
Delta_r
Doxycycline
YPL045W VPS16
ZShift = −10.37
Z lm Score = 2.71
NG = 2
DB = 0
SM = 2
Delta_AUC
Doxycycline
YOL001W PHO80
ZShift = 4.76
Z lm Score = 18.96
NG = 2
DB = 0
SM = 2
Delta_L
Doxycycline
YOL001W PHO80
ZShift = −6.86
Z lm Score = −8.89
NG = 2
DB = 0
SM = 2
Delta_K
Doxycycline
YOL001W PHO80
ZShift = −5.48
Z lm Score = −3.23
NG = 2
DB = 0
SM = 2
Delta_r
Doxycycline
YOL001W PHO80
ZShift = −7.69
Z lm Score = 6.44
NG = 2
DB = 0
SM = 2
Delta_AUC
Doxycycline

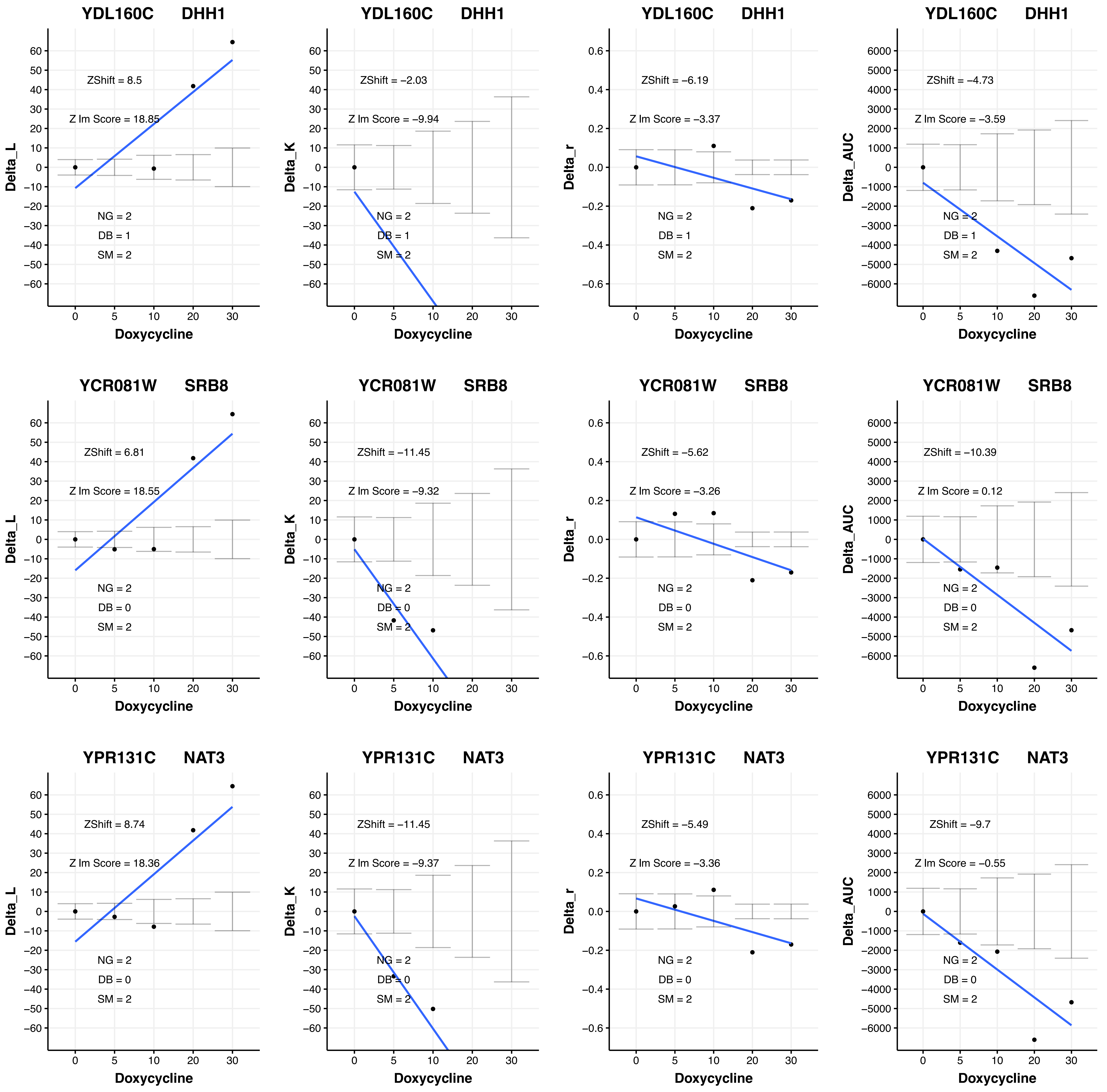
YDL160C DHH1
ZShift = 8.5
Z lm Score = 18.85
NG = 2
DB = 1
SM = 2
Delta_L
Doxycycline
YDL160C DHH1
ZShift = −2.03
Z lm Score = −9.94
NG = 2
DB = 1
SM = 2
Delta_K
Doxycycline
YDL160C DHH1
ZShift = −6.19
Z lm Score = −3.37
NG = 2
DB = 1
SM = 2
Delta_r
Doxycycline
YDL160C DHH1
ZShift = −4.73
Z lm Score = −3.59
NG = 2
DB = 1
SM = 2
Delta_AUC
Doxycycline
YCR081W SRB8
ZShift = 6.81
Z lm Score = 18.55
NG = 2
DB = 0
SM = 2
Delta_L
Doxycycline
YCR081W SRB8
ZShift = −11.45
Z lm Score = −9.32
NG = 2
DB = 0
SM = 2
Delta_K
Doxycycline
YCR081W SRB8
ZShift = −5.62
Z lm Score = −3.26
NG = 2
DB = 0
SM = 2
Delta_r
Doxycycline
YCR081W SRB8
ZShift = −10.39
Z lm Score = 0.12
NG = 2
DB = 0
SM = 2
Delta_AUC
Doxycycline
YPR131C NAT3
ZShift = 8.74
Z lm Score = 18.36
NG = 2
DB = 0
SM = 2
Delta_L
Doxycycline
YPR131C NAT3
ZShift = −11.45
Z lm Score = −9.37
NG = 2
DB = 0
SM = 2
Delta_K
Doxycycline
YPR131C NAT3
ZShift = −5.49
Z lm Score = −3.36
NG = 2
DB = 0
SM = 2
Delta_r
Doxycycline
YPR131C NAT3
ZShift = −9.7
Z lm Score = −0.55
NG = 2
DB = 0
SM = 2
Delta_AUC
Doxycycline

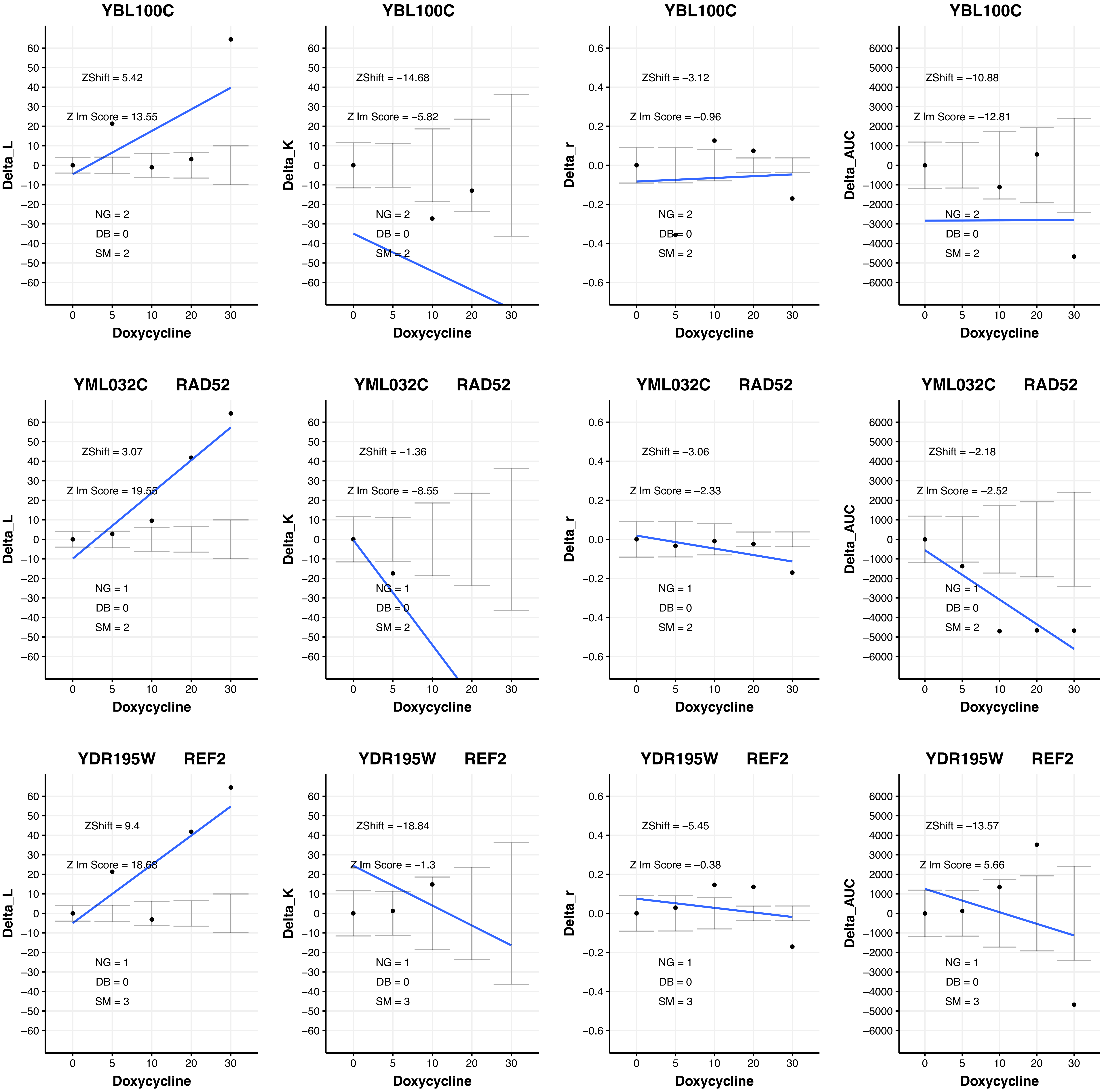

YBL100C
ZShift = 5.42
Z lm Score = 13.55
NG = 2
DB = 0
SM = 2
Delta_L
Doxycycline
YBL100C
ZShift = −14.68
Z lm Score = −5.82
NG = 2
DB = 0
SM = 2
Delta_K
Doxycycline
YBL100C
ZShift = −3.12
Z lm Score = −0.96
NG = 2
DB = 0
SM = 2
Delta_r
Doxycycline
YBL100C
ZShift = −10.88
Z lm Score = −12.81
NG = 2
DB = 0
SM = 2
Delta_AUC
Doxycycline
YML032C RAD52
ZShift = 3.07
Z lm Score = 19.55
NG = 1
DB = 0
SM = 2
Delta_L
Doxycycline
YML032C RAD52
ZShift = −1.36
Z lm Score = −8.55
NG = 1
DB = 0
SM = 2
Delta_K
Doxycycline
YML032C RAD52
ZShift = −3.06
Z lm Score = −2.33
NG = 1
DB = 0
SM = 2
Delta_r
Doxycycline
YML032C RAD52
ZShift = −2.18
Z lm Score = −2.52
NG = 1
DB = 0
SM = 2
Delta_AUC
Doxycycline
YDR195W REF2
ZShift = 9.4
Z lm Score = 18.68
NG = 1
DB = 0
SM = 3
Delta_L
Doxycycline
YDR195W REF2
ZShift = −18.84
Z lm Score = −1.3
NG = 1
DB = 0
SM = 3
Delta_K
Doxycycline
YDR195W REF2
ZShift = −5.45
Z lm Score = −0.38
NG = 1
DB = 0
SM = 3
Delta_r
Doxycycline
YDR195W REF2
ZShift = −13.57
Z lm Score = 5.66
NG = 1
DB = 0
SM = 3
Delta_AUC
Doxycycline

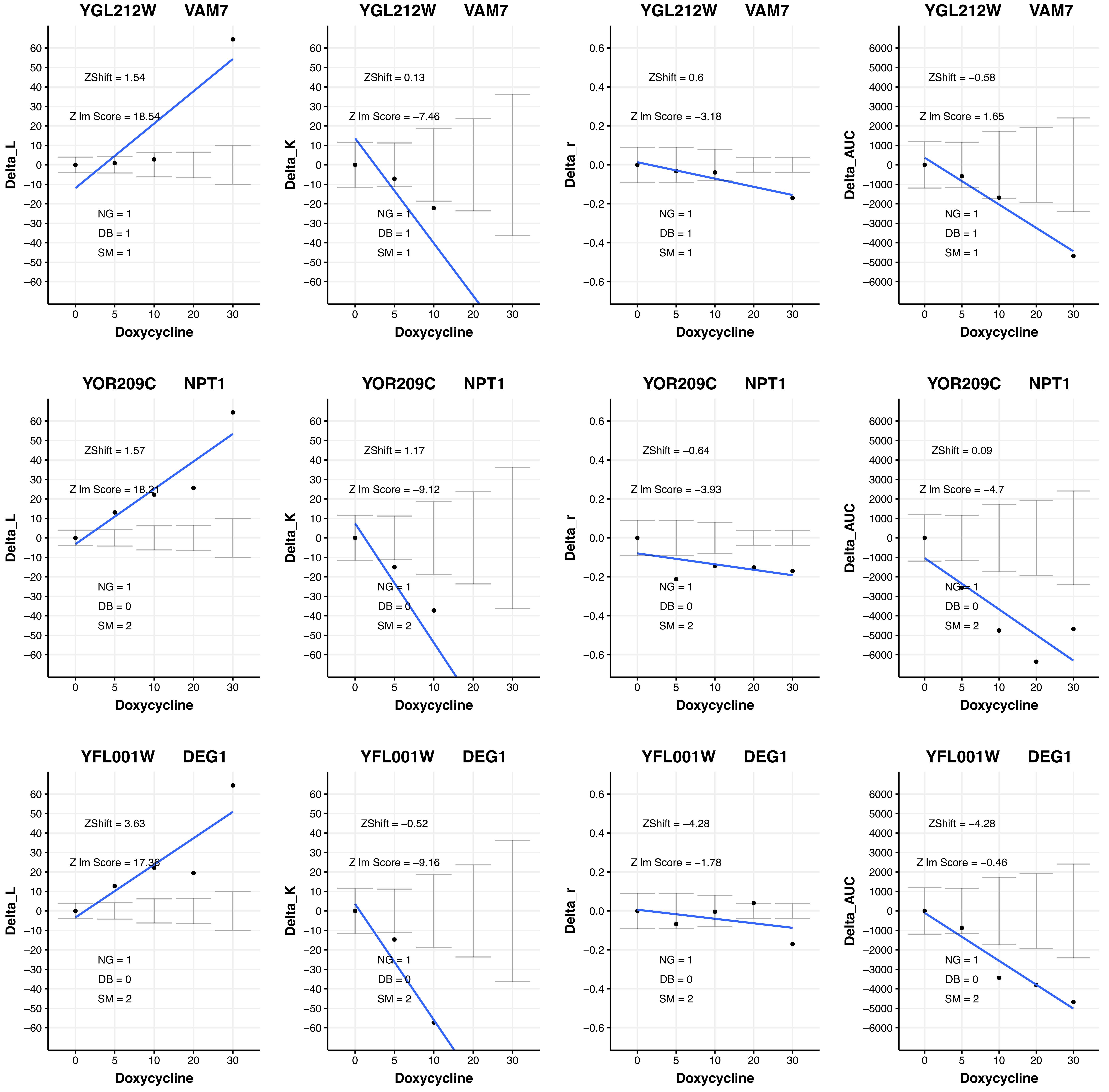
YGL212W VAM7
ZShift = 1.54
Z lm Score = 18.54
NG = 1
DB = 1
SM = 1
Delta_L
Doxycycline
YGL212W VAM7
ZShift = 0.13
Z lm Score = −7.46
NG = 1
DB = 1
SM = 1
Delta_K
Doxycycline
YGL212W VAM7
ZShift = 0.6
Z lm Score = −3.18
NG = 1
DB = 1
SM = 1
Delta_r
Doxycycline
YGL212W VAM7
ZShift = −0.58
Z lm Score = 1.65
NG = 1
DB = 1
SM = 1
Delta_AUC
Doxycycline
YOR209C NPT1
ZShift = 1.57
Z lm Score = 18.21
NG = 1
DB = 0
SM = 2
Delta_L
Doxycycline
YOR209C NPT1
ZShift = 1.17
Z lm Score = −9.12
NG = 1
DB = 0
SM = 2
Delta_K
Doxycycline
YOR209C NPT1
ZShift = −0.64
Z lm Score = −3.93
NG = 1
DB = 0
SM = 2
Delta_r
Doxycycline
YOR209C NPT1
ZShift = 0.09
Z lm Score = −4.7
NG = 1
DB = 0
SM = 2
Delta_AUC
Doxycycline
YFL001W DEG1
ZShift = 3.63
Z lm Score = 17.36
NG = 1
DB = 0
SM = 2
Delta_L
Doxycycline
YFL001W DEG1
ZShift = −0.52
Z lm Score = −9.16
NG = 1
DB = 0
SM = 2
Delta_K
Doxycycline
YFL001W DEG1
ZShift = −4.28
Z lm Score = −1.78
NG = 1
DB = 0
SM = 2
Delta_r
Doxycycline
YFL001W DEG1
ZShift = −4.28
Z lm Score = −0.46
NG = 1
DB = 0
SM = 2
Delta_AUC
Doxycycline

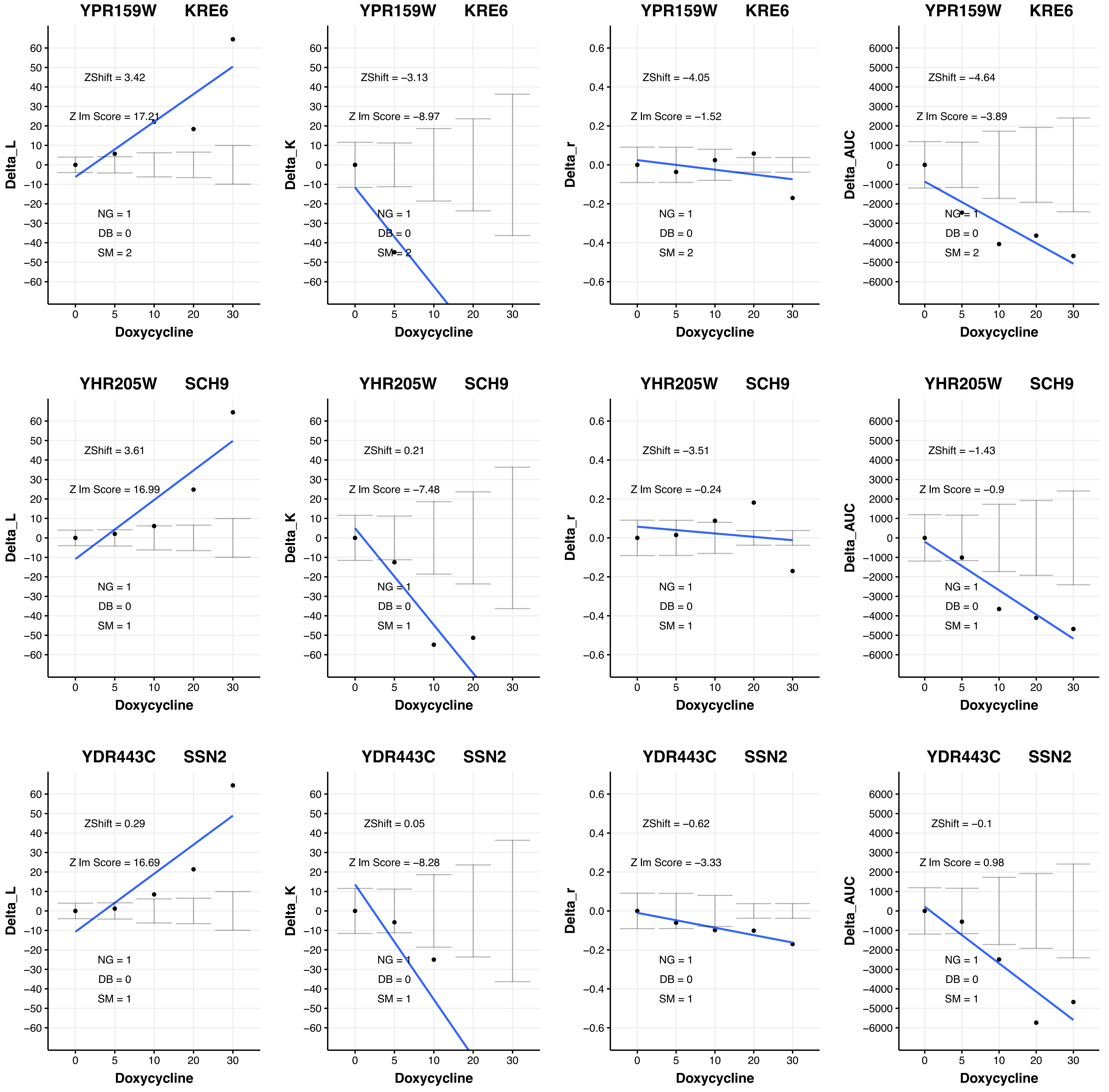

YPR159W KRE6
ZShift = 3.42
Z lm Score = 17.21
NG = 1
DB = 0
SM = 2
Delta_L
Doxycycline
YPR159W KRE6
ZShift = −3.13
Z lm Score = −8.97
NG = 1
DB = 0
SM = 2
Delta_K
Doxycycline
YPR159W KRE6
ZShift = −4.05
Z lm Score = −1.52
NG = 1
DB = 0
SM = 2
Delta_r
Doxycycline
YPR159W KRE6
ZShift = −4.64
Z lm Score = −3.89
NG = 1
DB = 0
SM = 2
Delta_AUC
Doxycycline
YHR205W SCH9
ZShift = 3.61
Z lm Score = 16.99
NG = 1
DB = 0
SM = 1
Delta_L
Doxycycline
YHR205W SCH9
ZShift = 0.21
Z lm Score = −7.48
NG = 1
DB = 0
SM = 1
Delta_K
Doxycycline
YHR205W SCH9
ZShift = −3.51
Z lm Score = −0.24
NG = 1
DB = 0
SM = 1
Delta_r
Doxycycline
YHR205W SCH9
ZShift = −1.43
Z lm Score = −0.9
NG = 1
DB = 0
SM = 1
Delta_AUC
Doxycycline
YDR443C SSN2
ZShift = 0.29
Z lm Score = 16.69
NG = 1
DB = 0
SM = 1
Delta_L
Doxycycline
YDR443C SSN2
ZShift = 0.05
Z lm Score = −8.28
NG = 1
DB = 0
SM = 1
Delta_K
Doxycycline
YDR443C SSN2
ZShift = −0.62
Z lm Score = −3.33
NG = 1
DB = 0
SM = 1
Delta_r
Doxycycline
YDR443C SSN2
ZShift = −0.1
Z lm Score = 0.98
NG = 1
DB = 0
SM = 1
Delta_AUC
Doxycycline

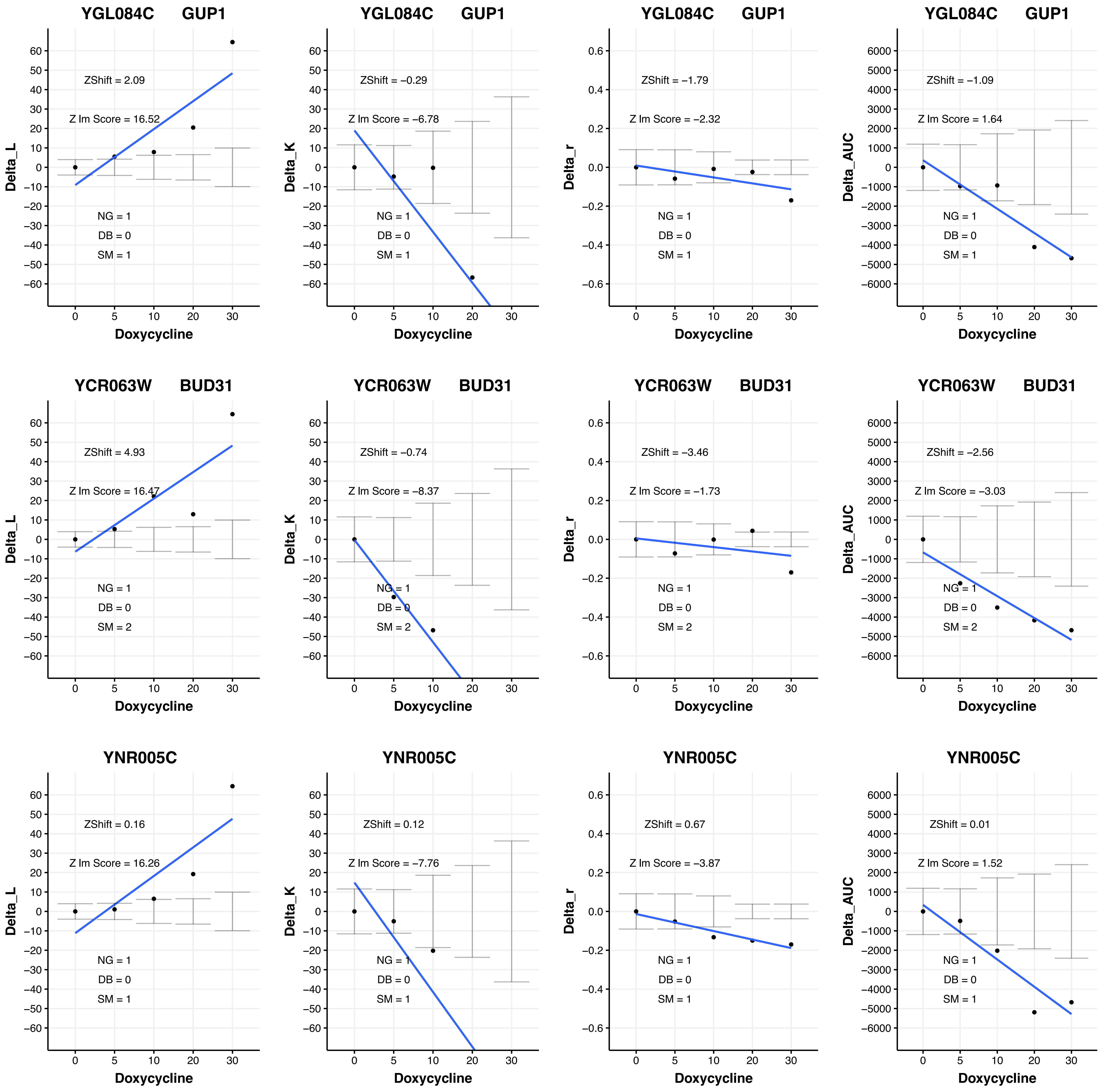

YGL084C GUP1
ZShift = 2.09
Z lm Score = 16.52
NG = 1
DB = 0
SM = 1
Delta_L
Doxycycline
YGL084C GUP1
ZShift = −0.29
Z lm Score = −6.78
NG = 1
DB = 0
SM = 1
Delta_K
Doxycycline
YGL084C GUP1
ZShift = −1.79
Z lm Score = −2.32
NG = 1
DB = 0
SM = 1
Delta_r
Doxycycline
YGL084C GUP1
ZShift = −1.09
Z lm Score = 1.64
NG = 1
DB = 0
SM = 1
Delta_AUC
Doxycycline
YCR063W BUD31
ZShift = 4.93
Z lm Score = 16.47
NG = 1
DB = 0
SM = 2
Delta_L
Doxycycline
YCR063W BUD31
ZShift = −0.74
Z lm Score = −8.37
NG = 1
DB = 0
SM = 2
Delta_K
Doxycycline
YCR063W BUD31
ZShift = −3.46
Z lm Score = −1.73
NG = 1
DB = 0
SM = 2
Delta_r
Doxycycline
YCR063W BUD31
ZShift = −2.56
Z lm Score = −3.03
NG = 1
DB = 0
SM = 2
Delta_AUC
Doxycycline
YNR005C
ZShift = 0.16
Z lm Score = 16.26
NG = 1
DB = 0
SM = 1
Delta_L
Doxycycline
YNR005C
ZShift = 0.12
Z lm Score = −7.76
NG = 1
DB = 0
SM = 1
Delta_K
Doxycycline
YNR005C
ZShift = 0.67
Z lm Score = −3.87
NG = 1
DB = 0
SM = 1
Delta_r
Doxycycline
YNR005C
ZShift = 0.01
Z lm Score = 1.52
NG = 1
DB = 0
SM = 1
Delta_AUC
Doxycycline

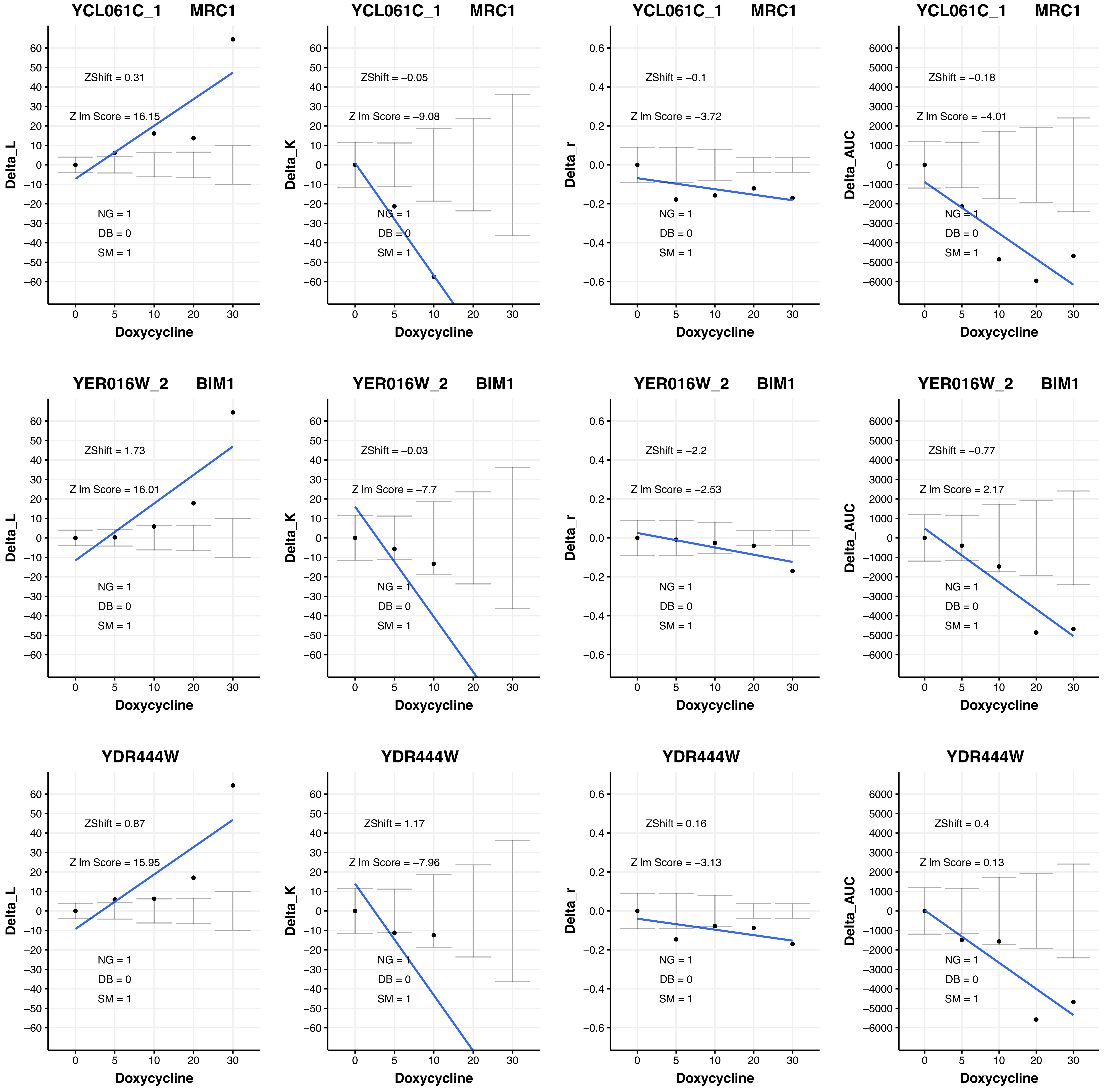

YCL061C_1 MRC1
ZShift = 0.31
Z lm Score = 16.15
NG = 1
DB = 0
SM = 1
Delta_L
Doxycycline
YCL061C_1 MRC1
ZShift = −0.05
Z lm Score = −9.08
NG = 1
DB = 0
SM = 1
Delta_K
Doxycycline
YCL061C_1 MRC1
ZShift = −0.1
Z lm Score = −3.72
NG = 1
DB = 0
SM = 1
Delta_r
Doxycycline
YCL061C_1 MRC1
ZShift = −0.18
Z lm Score = −4.01
NG = 1
DB = 0
SM = 1
Delta_AUC
Doxycycline
YER016W_2 BIM1
ZShift = 1.73
Z lm Score = 16.01
NG = 1
DB = 0
SM = 1
Delta_L
Doxycycline
YER016W_2 BIM1
ZShift = −0.03
Z lm Score = −7.7
NG = 1
DB = 0
SM = 1
Delta_K
Doxycycline
YER016W_2 BIM1
ZShift = −2.2
Z lm Score = −2.53
NG = 1
DB = 0
SM = 1
Delta_r
Doxycycline
YER016W_2 BIM1
ZShift = −0.77
Z lm Score = 2.17
NG = 1
DB = 0
SM = 1
Delta_AUC
Doxycycline
YDR444W
ZShift = 0.87
Z lm Score = 15.95
NG = 1
DB = 0
SM = 1
Delta_L
Doxycycline
YDR444W
ZShift = 1.17
Z lm Score = −7.96
NG = 1
DB = 0
SM = 1
Delta_K
Doxycycline
YDR444W
ZShift = 0.16
Z lm Score = −3.13
NG = 1
DB = 0
SM = 1
Delta_r
Doxycycline
YDR444W
ZShift = 0.4
Z lm Score = 0.13
NG = 1
DB = 0
SM = 1
Delta_AUC
Doxycycline

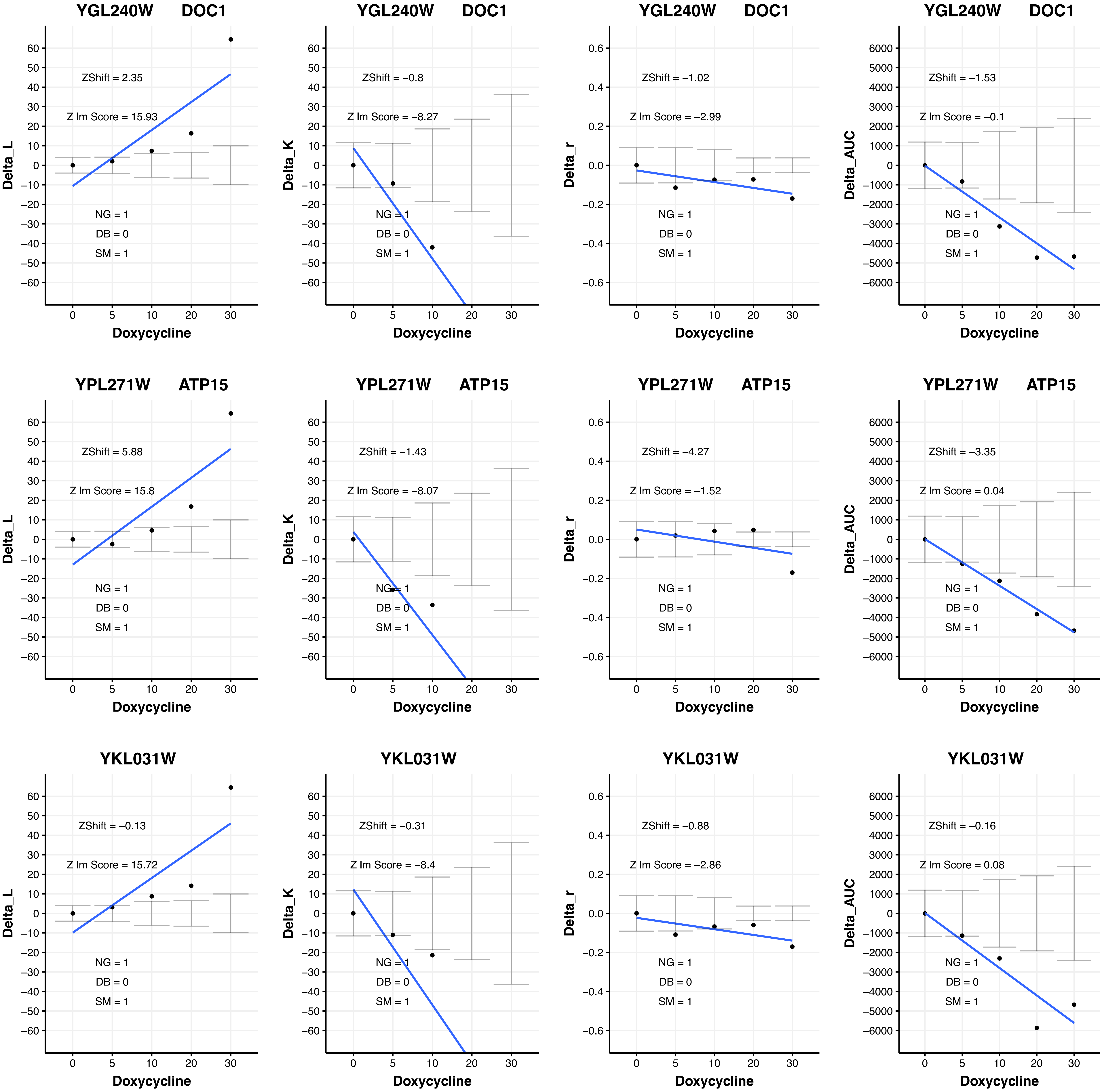
YGL240W DOC1
ZShift = 2.35
Z lm Score = 15.93
NG = 1
DB = 0
SM = 1
Delta_L
Doxycycline
YGL240W DOC1
ZShift = −0.8
Z lm Score = −8.27
NG = 1
DB = 0
SM = 1
Delta_K
Doxycycline
YGL240W DOC1
ZShift = −1.02
Z lm Score = −2.99
NG = 1
DB = 0
SM = 1
Delta_r
Doxycycline
YGL240W DOC1
ZShift = −1.53
Z lm Score = −0.1
NG = 1
DB = 0
SM = 1
Delta_AUC
Doxycycline
YPL271W ATP15
ZShift = 5.88
Z lm Score = 15.8
NG = 1
DB = 0
SM = 1
Delta_L
Doxycycline
YPL271W ATP15
ZShift = −1.43
Z lm Score = −8.07
NG = 1
DB = 0
SM = 1
Delta_K
Doxycycline
YPL271W ATP15
ZShift = −4.27
Z lm Score = −1.52
NG = 1
DB = 0
SM = 1
Delta_r
Doxycycline
YPL271W ATP15
ZShift = −3.35
Z lm Score = 0.04
NG = 1
DB = 0
SM = 1
Delta_AUC
Doxycycline
YKL031W
ZShift = −0.13
Z lm Score = 15.72
NG = 1
DB = 0
SM = 1
Delta_L
Doxycycline
YKL031W
ZShift = −0.31
Z lm Score = −8.4
NG = 1
DB = 0
SM = 1
Delta_K
Doxycycline
YKL031W
ZShift = −0.88
Z lm Score = −2.86
NG = 1
DB = 0
SM = 1
Delta_r
Doxycycline
YKL031W
ZShift = −0.16
Z lm Score = 0.08
NG = 1
DB = 0
SM = 1
Delta_AUC
Doxycycline

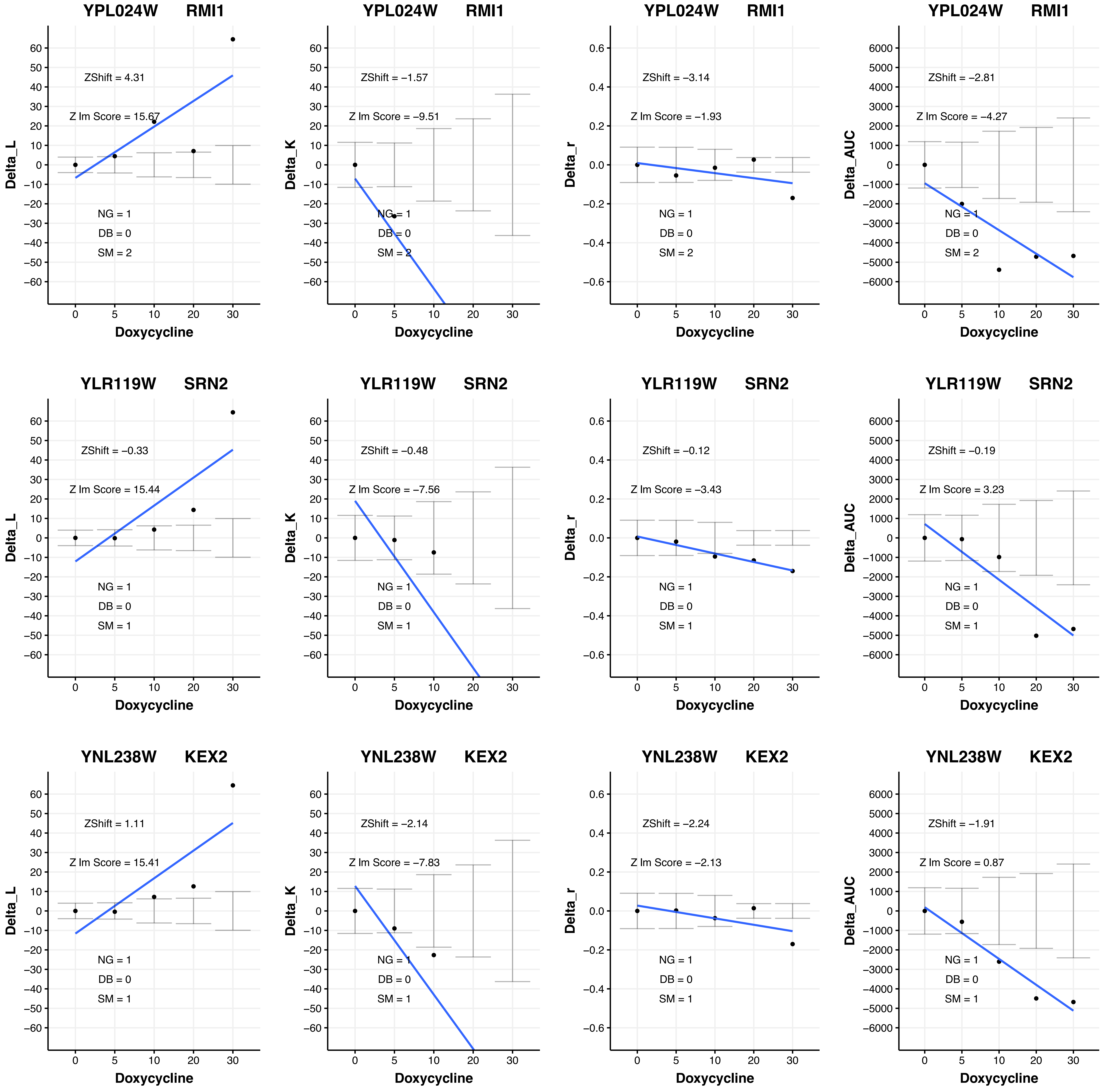
YPL024W RMI1
ZShift = 4.31
Z lm Score = 15.67
NG = 1
DB = 0
SM = 2
Delta_L
Doxycycline
YPL024W RMI1
ZShift = −1.57
Z lm Score = −9.51
NG = 1
DB = 0
SM = 2
Delta_K
Doxycycline
YPL024W RMI1
ZShift = −3.14
Z lm Score = −1.93
NG = 1
DB = 0
SM = 2
Delta_r
Doxycycline
YPL024W RMI1
ZShift = −2.81
Z lm Score = −4.27
NG = 1
DB = 0
SM = 2
Delta_AUC
Doxycycline
YLR119W SRN2
ZShift = −0.33
Z lm Score = 15.44
NG = 1
DB = 0
SM = 1
Delta_L
Doxycycline
YLR119W SRN2
ZShift = −0.48
Z lm Score = −7.56
NG = 1
DB = 0
SM = 1
Delta_K
Doxycycline
YLR119W SRN2
ZShift = −0.12
Z lm Score = −3.43
NG = 1
DB = 0
SM = 1
Delta_r
Doxycycline
YLR119W SRN2
ZShift = −0.19
Z lm Score = 3.23
NG = 1
DB = 0
SM = 1
Delta_AUC
Doxycycline
YNL238W KEX2
ZShift = 1.11
Z lm Score = 15.41
NG = 1
DB = 0
SM = 1
Delta_L
Doxycycline
YNL238W KEX2
ZShift = −2.14
Z lm Score = −7.83
NG = 1
DB = 0
SM = 1
Delta_K
Doxycycline
YNL238W KEX2
ZShift = −2.24
Z lm Score = −2.13
NG = 1
DB = 0
SM = 1
Delta_r
Doxycycline
YNL238W KEX2
ZShift = −1.91
Z lm Score = 0.87
NG = 1
DB = 0
SM = 1
Delta_AUC
Doxycycline

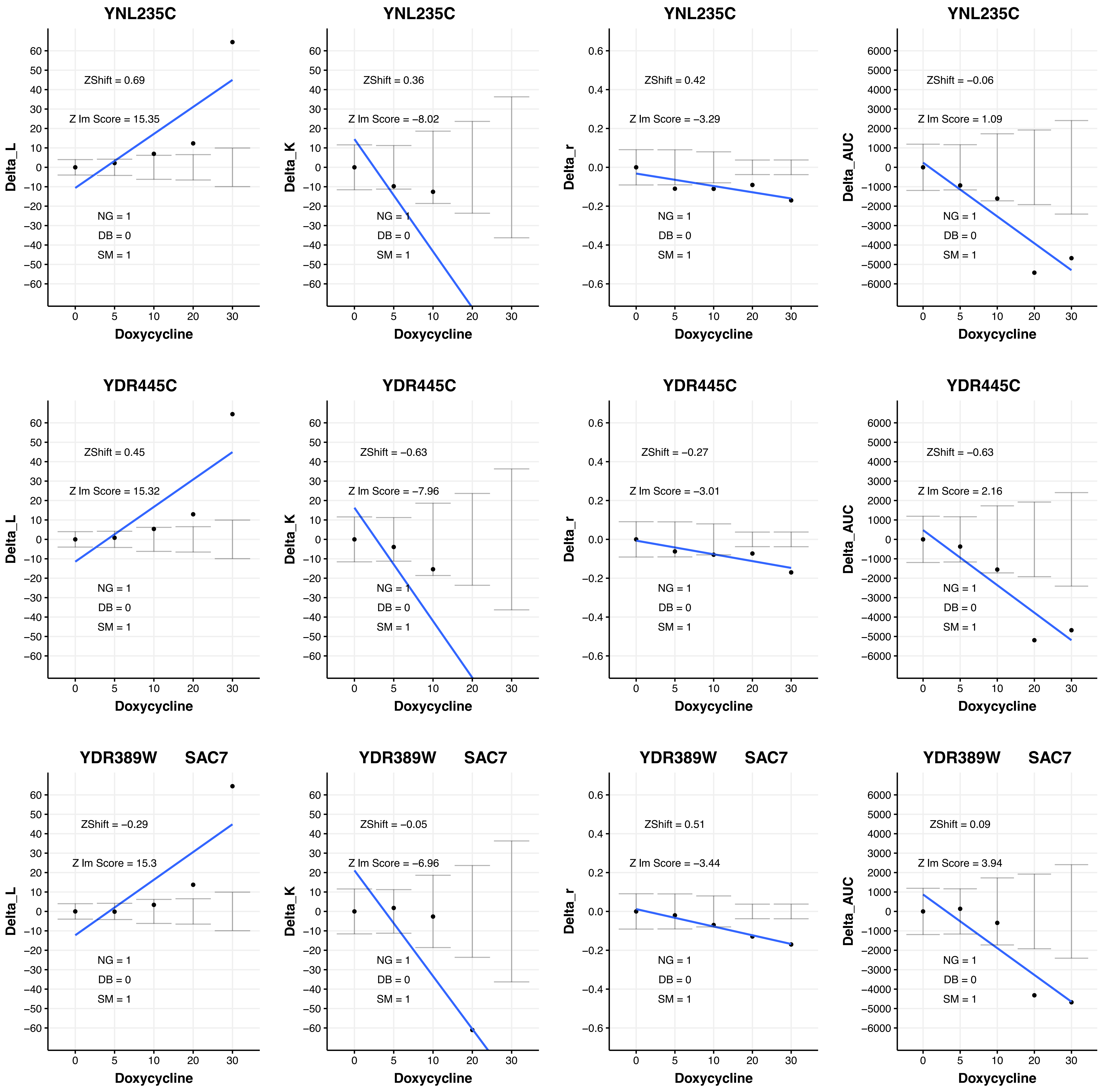
YNL235C
ZShift = 0.69
Z lm Score = 15.35
NG = 1
DB = 0
SM = 1
Delta_L
Doxycycline
YNL235C
ZShift = 0.36
Z lm Score = −8.02
NG = 1
DB = 0
SM = 1
Delta_K
Doxycycline
YNL235C
ZShift = 0.42
Z lm Score = −3.29
NG = 1
DB = 0
SM = 1
Delta_r
Doxycycline
YNL235C
ZShift = −0.06
Z lm Score = 1.09
NG = 1
DB = 0
SM = 1
Delta_AUC
Doxycycline
YDR445C
ZShift = 0.45
Z lm Score = 15.32
NG = 1
DB = 0
SM = 1
Delta_L
Doxycycline
YDR445C
ZShift = −0.63
Z lm Score = −7.96
NG = 1
DB = 0
SM = 1
Delta_K
Doxycycline
YDR445C
ZShift = −0.27
Z lm Score = −3.01
NG = 1
DB = 0
SM = 1
Delta_r
Doxycycline
YDR445C
ZShift = −0.63
Z lm Score = 2.16
NG = 1
DB = 0
SM = 1
Delta_AUC
Doxycycline
YDR389W SAC7
ZShift = −0.29
Z lm Score = 15.3
NG = 1
DB = 0
SM = 1
Delta_L
Doxycycline
YDR389W SAC7
ZShift = −0.05
Z lm Score = −6.96
NG = 1
DB = 0
SM = 1
Delta_K
Doxycycline
YDR389W SAC7
ZShift = 0.51
Z lm Score = −3.44
NG = 1
DB = 0
SM = 1
Delta_r
Doxycycline
YDR389W SAC7
ZShift = 0.09
Z lm Score = 3.94
NG = 1
DB = 0
SM = 1
Delta_AUC
Doxycycline

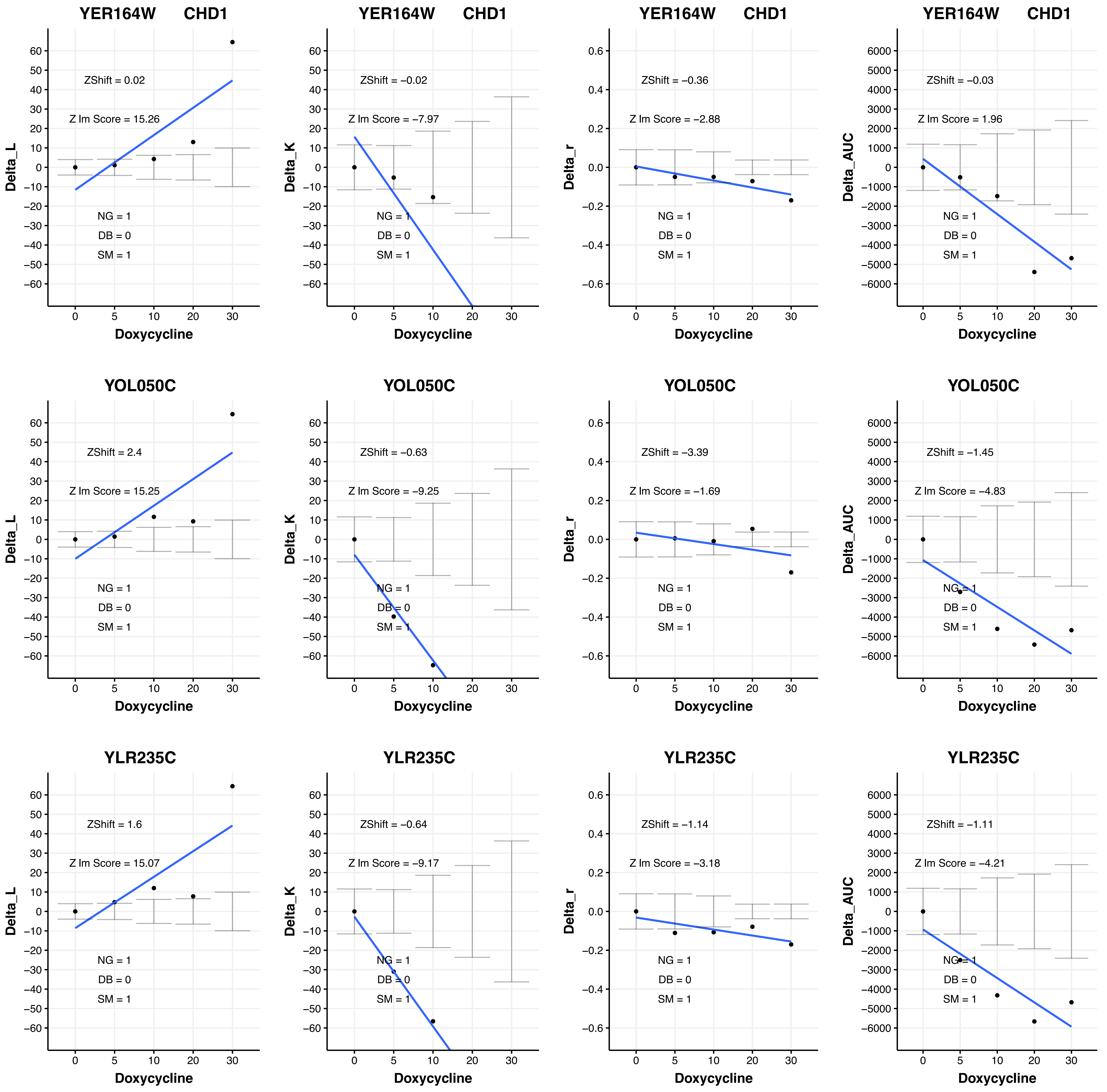
YER164W CHD1
ZShift = 0.02
Z lm Score = 15.26
NG = 1
DB = 0
SM = 1
Delta_L
Doxycycline
YER164W CHD1
ZShift = −0.02
Z lm Score = −7.97
NG = 1
DB = 0
SM = 1
Delta_K
Doxycycline
YER164W CHD1
ZShift = −0.36
Z lm Score = −2.88
NG = 1
DB = 0
SM = 1
Delta_r
Doxycycline
YER164W CHD1
ZShift = −0.03
Z lm Score = 1.96
NG = 1
DB = 0
SM = 1
Delta_AUC
Doxycycline
YOL050C
ZShift = 2.4
Z lm Score = 15.25
NG = 1
DB = 0
SM = 1
Delta_L
Doxycycline
YOL050C
ZShift = −0.63
Z lm Score = −9.25
NG = 1
DB = 0
SM = 1
Delta_K
Doxycycline
YOL050C
ZShift = −3.39
Z lm Score = −1.69
NG = 1
DB = 0
SM = 1
Delta_r
Doxycycline
YOL050C
ZShift = −1.45
Z lm Score = −4.83
NG = 1
DB = 0
SM = 1
Delta_AUC
Doxycycline
YLR235C
ZShift = 1.6
Z lm Score = 15.07
NG = 1
DB = 0
SM = 1
Delta_L
Doxycycline
YLR235C
ZShift = −0.64
Z lm Score = −9.17
NG = 1
DB = 0
SM = 1
Delta_K
Doxycycline
YLR235C
ZShift = −1.14
Z lm Score = −3.18
NG = 1
DB = 0
SM = 1
Delta_r
Doxycycline
YLR235C
ZShift = −1.11
Z lm Score = −4.21
NG = 1
DB = 0
SM = 1
Delta_AUC
Doxycycline

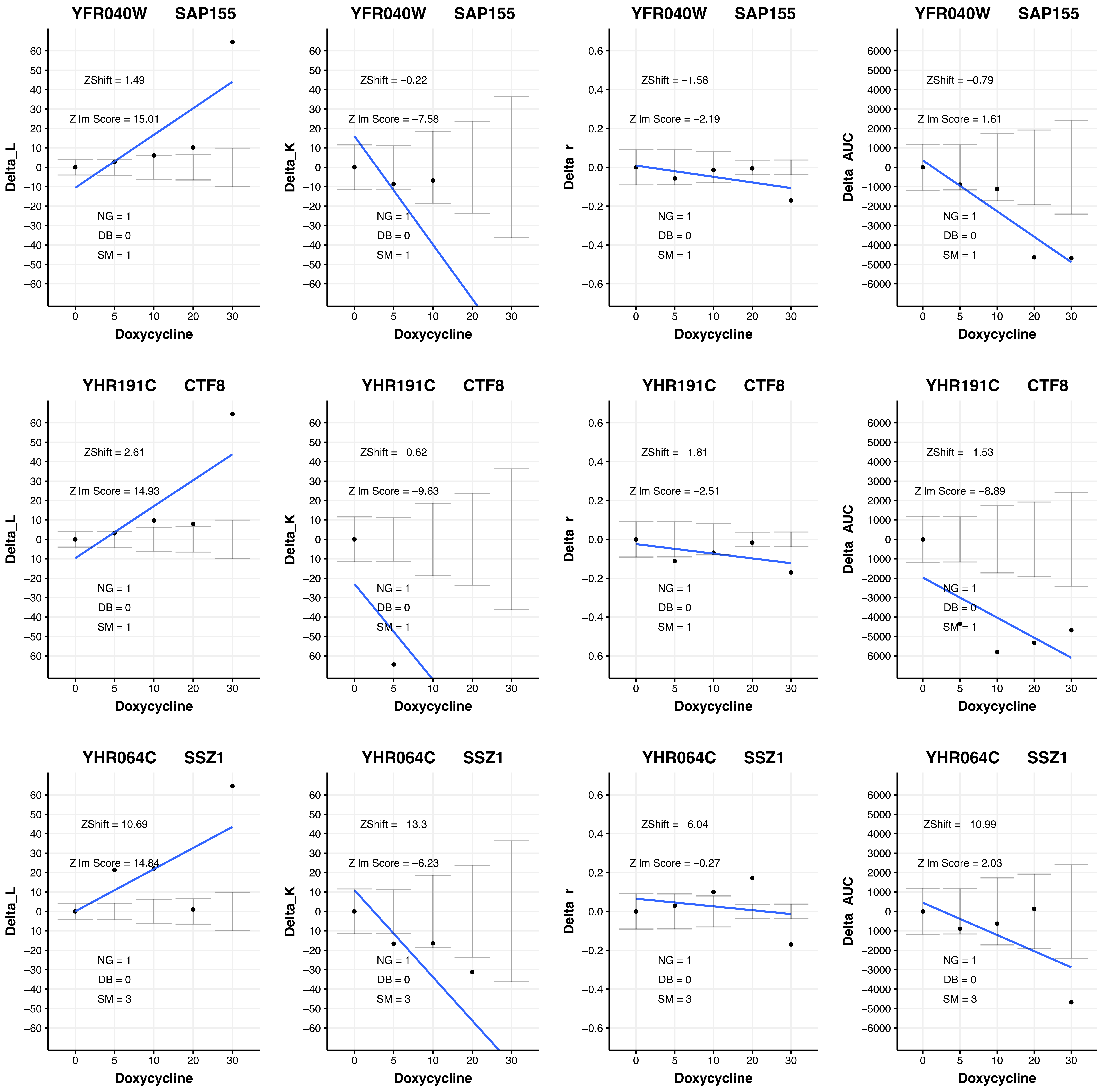

YFR040W SAP155
ZShift = 1.49
Z lm Score = 15.01
NG = 1
DB = 0
SM = 1
Delta_L
Doxycycline
YFR040W SAP155
ZShift = −0.22
Z lm Score = −7.58
NG = 1
DB = 0
SM = 1
Delta_K
Doxycycline
YFR040W SAP155
ZShift = −1.58
Z lm Score = −2.19
NG = 1
DB = 0
SM = 1
Delta_r
Doxycycline
YFR040W SAP155
ZShift = −0.79
Z lm Score = 1.61
NG = 1
DB = 0
SM = 1
Delta_AUC
Doxycycline
YHR191C CTF8
ZShift = 2.61
Z lm Score = 14.93
NG = 1
DB = 0
SM = 1
Delta_L
Doxycycline
YHR191C CTF8
ZShift = −0.62
Z lm Score = −9.63
NG = 1
DB = 0
SM = 1
Delta_K
Doxycycline
YHR191C CTF8
ZShift = −1.81
Z lm Score = −2.51
NG = 1
DB = 0
SM = 1
Delta_r
Doxycycline
YHR191C CTF8
ZShift = −1.53
Z lm Score = −8.89
NG = 1
DB = 0
SM = 1
Delta_AUC
Doxycycline
YHR064C SSZ1
ZShift = 10.69
Z lm Score = 14.84
NG = 1
DB = 0
SM = 3
Delta_L
Doxycycline
YHR064C SSZ1
ZShift = −13.3
Z lm Score = −6.23
NG = 1
DB = 0
SM = 3
Delta_K
Doxycycline
YHR064C SSZ1
ZShift = −6.04
Z lm Score = −0.27
NG = 1
DB = 0
SM = 3
Delta_r
Doxycycline
YHR064C SSZ1
ZShift = −10.99
Z lm Score = 2.03
NG = 1
DB = 0
SM = 3
Delta_AUC
Doxycycline

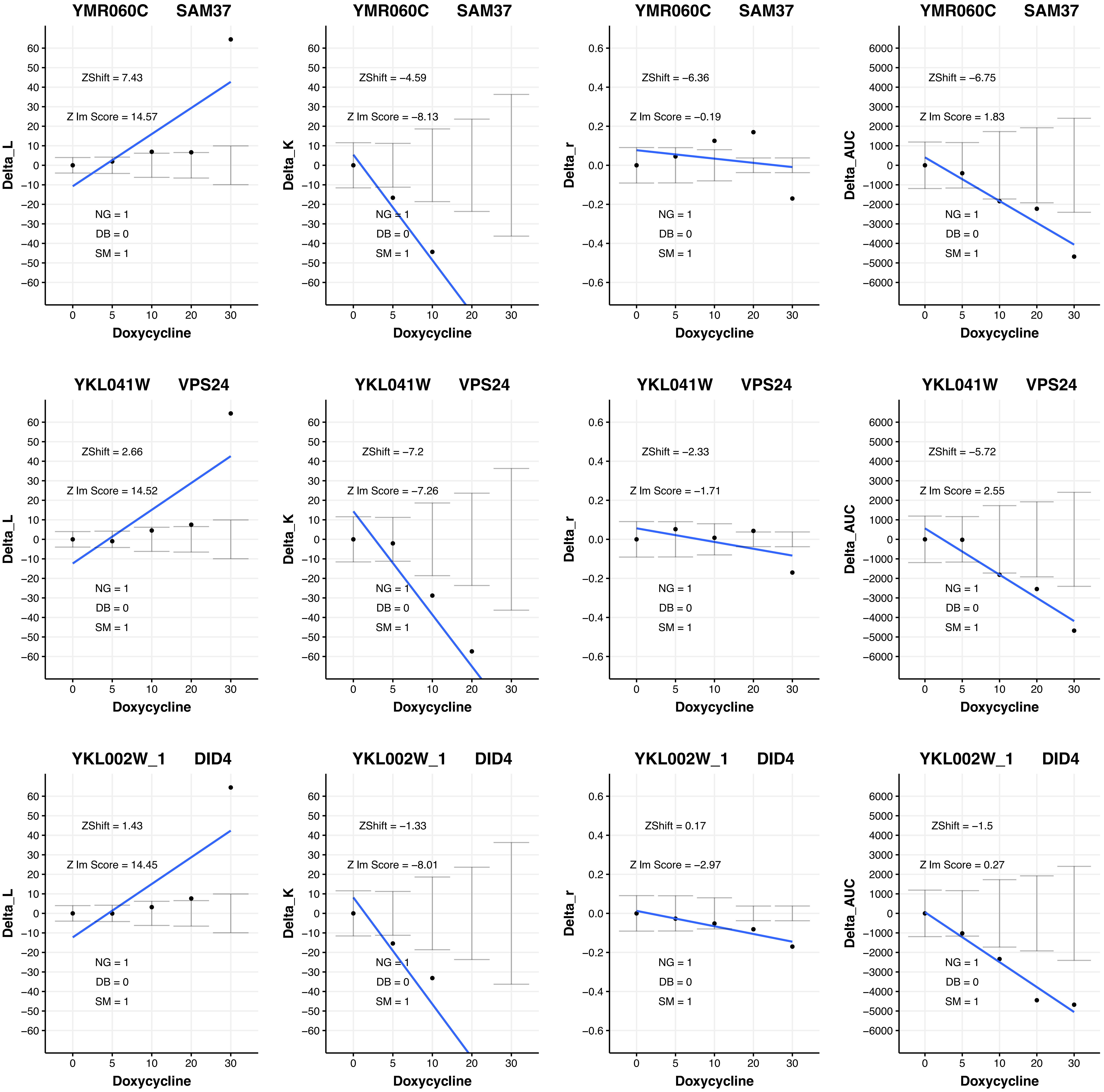

YMR060C SAM37
ZShift = 7.43
Z lm Score = 14.57
NG = 1
DB = 0
SM = 1
Delta_L
Doxycycline
YMR060C SAM37
ZShift = −4.59
Z lm Score = −8.13
NG = 1
DB = 0
SM = 1
Delta_K
Doxycycline
YMR060C SAM37
ZShift = −6.36
Z lm Score = −0.19
NG = 1
DB = 0
SM = 1
Delta_r
Doxycycline
YMR060C SAM37
ZShift = −6.75
Z lm Score = 1.83
NG = 1
DB = 0
SM = 1
Delta_AUC
Doxycycline
YKL041W VPS24
ZShift = 2.66
Z lm Score = 14.52
NG = 1
DB = 0
SM = 1
Delta_L
Doxycycline
YKL041W VPS24
ZShift = −7.2
Z lm Score = −7.26
NG = 1
DB = 0
SM = 1
Delta_K
Doxycycline
YKL041W VPS24
ZShift = −2.33
Z lm Score = −1.71
NG = 1
DB = 0
SM = 1
Delta_r
Doxycycline
YKL041W VPS24
ZShift = −5.72
Z lm Score = 2.55
NG = 1
DB = 0
SM = 1
Delta_AUC
Doxycycline
YKL002W_1 DID4
ZShift = 1.43
Z lm Score = 14.45
NG = 1
DB = 0
SM = 1
Delta_L
Doxycycline
YKL002W_1 DID4
ZShift = −1.33
Z lm Score = −8.01
NG = 1
DB = 0
SM = 1
Delta_K
Doxycycline
YKL002W_1 DID4
ZShift = 0.17
Z lm Score = −2.97
NG = 1
DB = 0
SM = 1
Delta_r
Doxycycline
YKL002W_1 DID4
ZShift = −1.5
Z lm Score = 0.27
NG = 1
DB = 0
SM = 1
Delta_AUC
Doxycycline

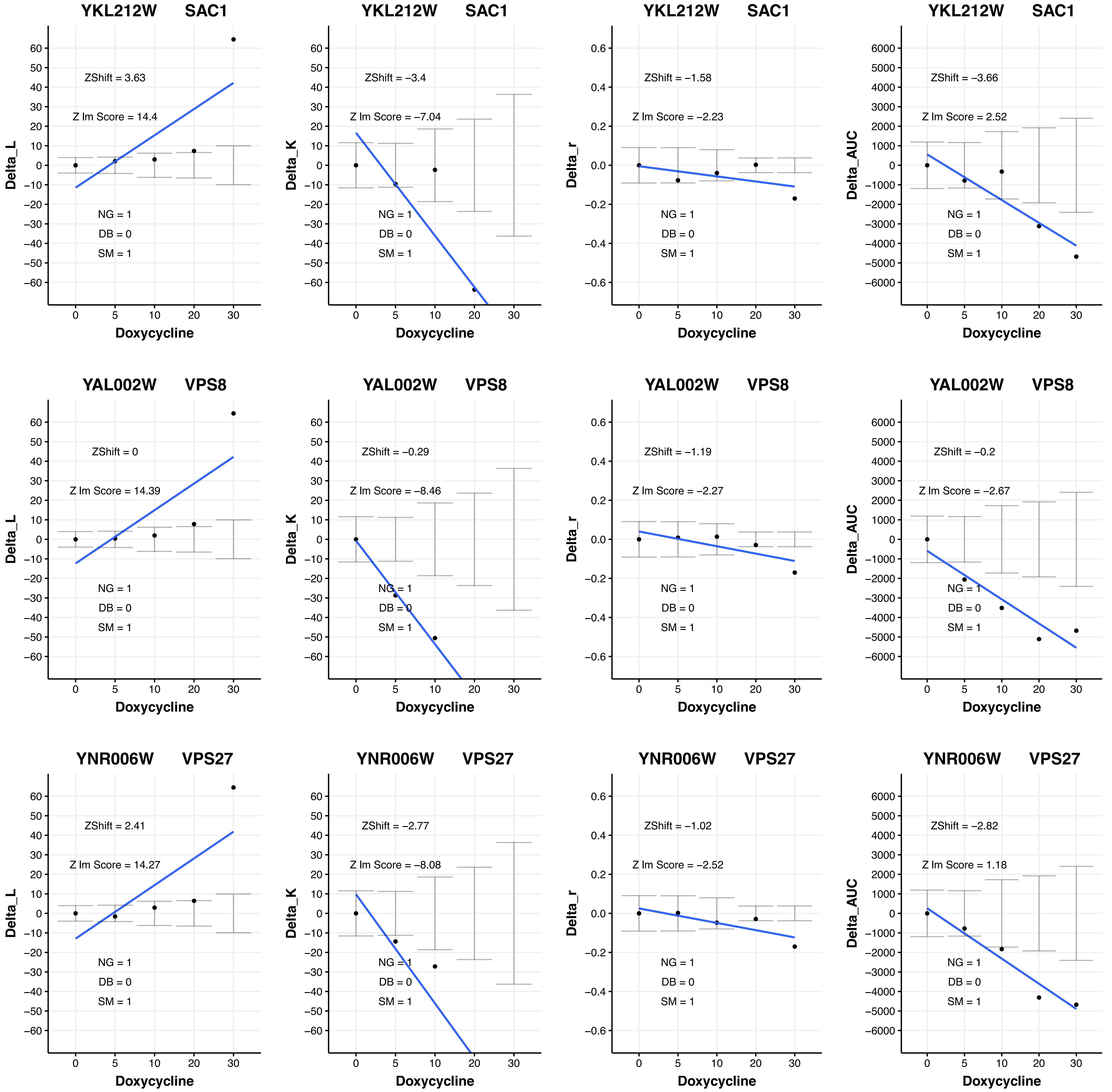

YKL212W SAC1
ZShift = 3.63
Z lm Score = 14.4
NG = 1
DB = 0
SM = 1
Delta_L
Doxycycline
YKL212W SAC1
ZShift = −3.4
Z lm Score = −7.04
NG = 1
DB = 0
SM = 1
Delta_K
Doxycycline
YKL212W SAC1
ZShift = −1.58
Z lm Score = −2.23
NG = 1
DB = 0
SM = 1
Delta_r
Doxycycline
YKL212W SAC1
ZShift = −3.66
Z lm Score = 2.52
NG = 1
DB = 0
SM = 1
Delta_AUC
Doxycycline
YAL002W VPS8
ZShift = 0
Z lm Score = 14.39
NG = 1
DB = 0
SM = 1
Delta_L
Doxycycline
YAL002W VPS8
ZShift = −0.29
Z lm Score = −8.46
NG = 1
DB = 0
SM = 1
Delta_K
Doxycycline
YAL002W VPS8
ZShift = −1.19
Z lm Score = −2.27
NG = 1
DB = 0
SM = 1
Delta_r
Doxycycline
YAL002W VPS8
ZShift = −0.2
Z lm Score = −2.67
NG = 1
DB = 0
SM = 1
Delta_AUC
Doxycycline
YNR006W VPS27
ZShift = 2.41
Z lm Score = 14.27
NG = 1
DB = 0
SM = 1
Delta_L
Doxycycline
YNR006W VPS27
ZShift = −2.77
Z lm Score = −8.08
NG = 1
DB = 0
SM = 1
Delta_K
Doxycycline
YNR006W VPS27
ZShift = −1.02
Z lm Score = −2.52
NG = 1
DB = 0
SM = 1
Delta_r
Doxycycline
YNR006W VPS27
ZShift = −2.82
Z lm Score = 1.18
NG = 1
DB = 0
SM = 1
Delta_AUC
Doxycycline

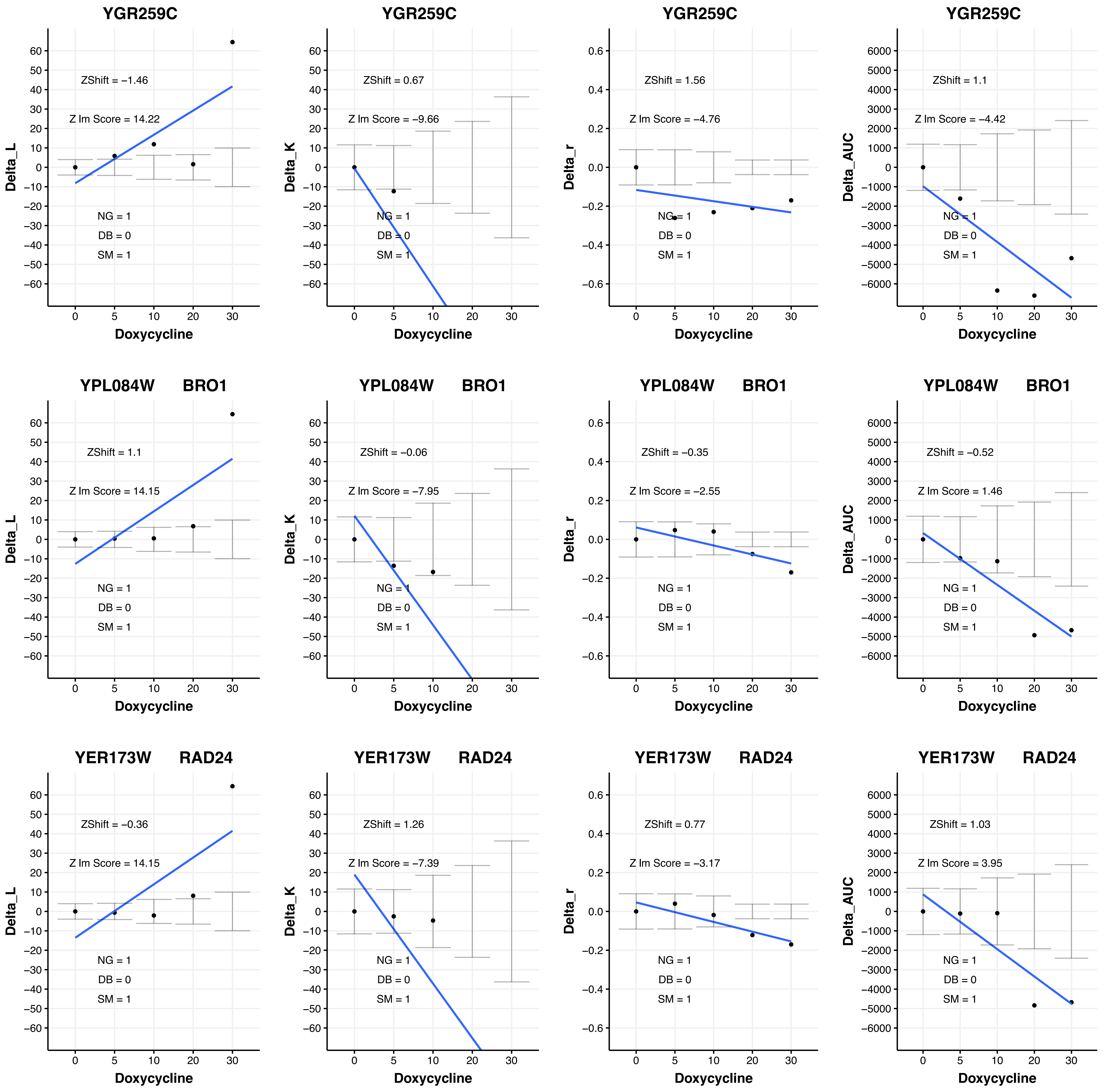

YGR259C
ZShift = −1.46
Z lm Score = 14.22
NG = 1
DB = 0
SM = 1
Delta_L
Doxycycline
YGR259C
ZShift = 0.67
Z lm Score = −9.66
NG = 1
DB = 0
SM = 1
Delta_K
Doxycycline
YGR259C
ZShift = 1.56
Z lm Score = −4.76
NG = 1
DB = 0
SM = 1
Delta_r
Doxycycline
YGR259C
ZShift = 1.1
Z lm Score = −4.42
NG = 1
DB = 0
SM = 1
Delta_AUC
Doxycycline
YPL084W BRO1
ZShift = 1.1
Z lm Score = 14.15
NG = 1
DB = 0
SM = 1
Delta_L
Doxycycline
YPL084W BRO1
ZShift = −0.06
Z lm Score = −7.95
NG = 1
DB = 0
SM = 1
Delta_K
Doxycycline
YPL084W BRO1
ZShift = −0.35
Z lm Score = −2.55
NG = 1
DB = 0
SM = 1
Delta_r
Doxycycline
YPL084W BRO1
ZShift = −0.52
Z lm Score = 1.46
NG = 1
DB = 0
SM = 1
Delta_AUC
Doxycycline
YER173W RAD24
ZShift = −0.36
Z lm Score = 14.15
NG = 1
DB = 0
SM = 1
Delta_L
Doxycycline
YER173W RAD24
ZShift = 1.26
Z lm Score = −7.39
NG = 1
DB = 0
SM = 1
Delta_K
Doxycycline
YER173W RAD24
ZShift = 0.77
Z lm Score = −3.17
NG = 1
DB = 0
SM = 1
Delta_r
Doxycycline
YER173W RAD24
ZShift = 1.03
Z lm Score = 3.95
NG = 1
DB = 0
SM = 1
Delta_AUC
Doxycycline

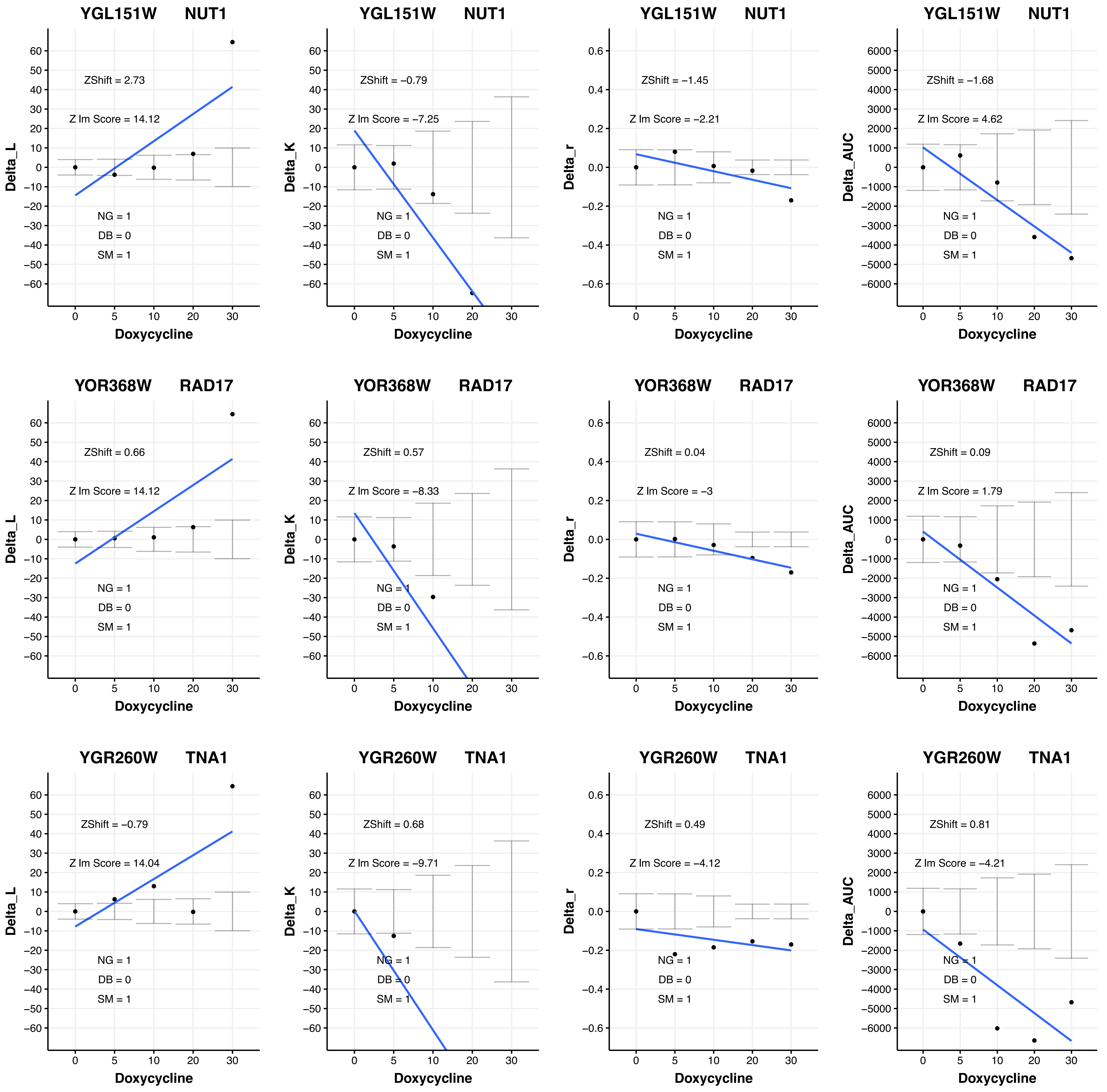

YGL151W NUT1
ZShift = 2.73
Z lm Score = 14.12
NG = 1
DB = 0
SM = 1
Delta_L
Doxycycline
YGL151W NUT1
ZShift = −0.79
Z lm Score = −7.25
NG = 1
DB = 0
SM = 1
Delta_K
Doxycycline
YGL151W NUT1
ZShift = −1.45
Z lm Score = −2.21
NG = 1
DB = 0
SM = 1
Delta_r
Doxycycline
YGL151W NUT1
ZShift = −1.68
Z lm Score = 4.62
NG = 1
DB = 0
SM = 1
Delta_AUC
Doxycycline
YOR368W RAD17
ZShift = 0.66
Z lm Score = 14.12
NG = 1
DB = 0
SM = 1
Delta_L
Doxycycline
YOR368W RAD17
ZShift = 0.57
Z lm Score = −8.33
NG = 1
DB = 0
SM = 1
Delta_K
Doxycycline
YOR368W RAD17
ZShift = 0.04
Z lm Score = −3
NG = 1
DB = 0
SM = 1
Delta_r
Doxycycline
YOR368W RAD17
ZShift = 0.09
Z lm Score = 1.79
NG = 1
DB = 0
SM = 1
Delta_AUC
Doxycycline
YGR260W TNA1
ZShift = −0.79
Z lm Score = 14.04
NG = 1
DB = 0
SM = 1
Delta_L
Doxycycline
YGR260W TNA1
ZShift = 0.68
Z lm Score = −9.71
NG = 1
DB = 0
SM = 1
Delta_K
Doxycycline
YGR260W TNA1
ZShift = 0.49
Z lm Score = −4.12
NG = 1
DB = 0
SM = 1
Delta_r
Doxycycline
YGR260W TNA1
ZShift = 0.81
Z lm Score = −4.21
NG = 1
DB = 0
SM = 1
Delta_AUC
Doxycycline

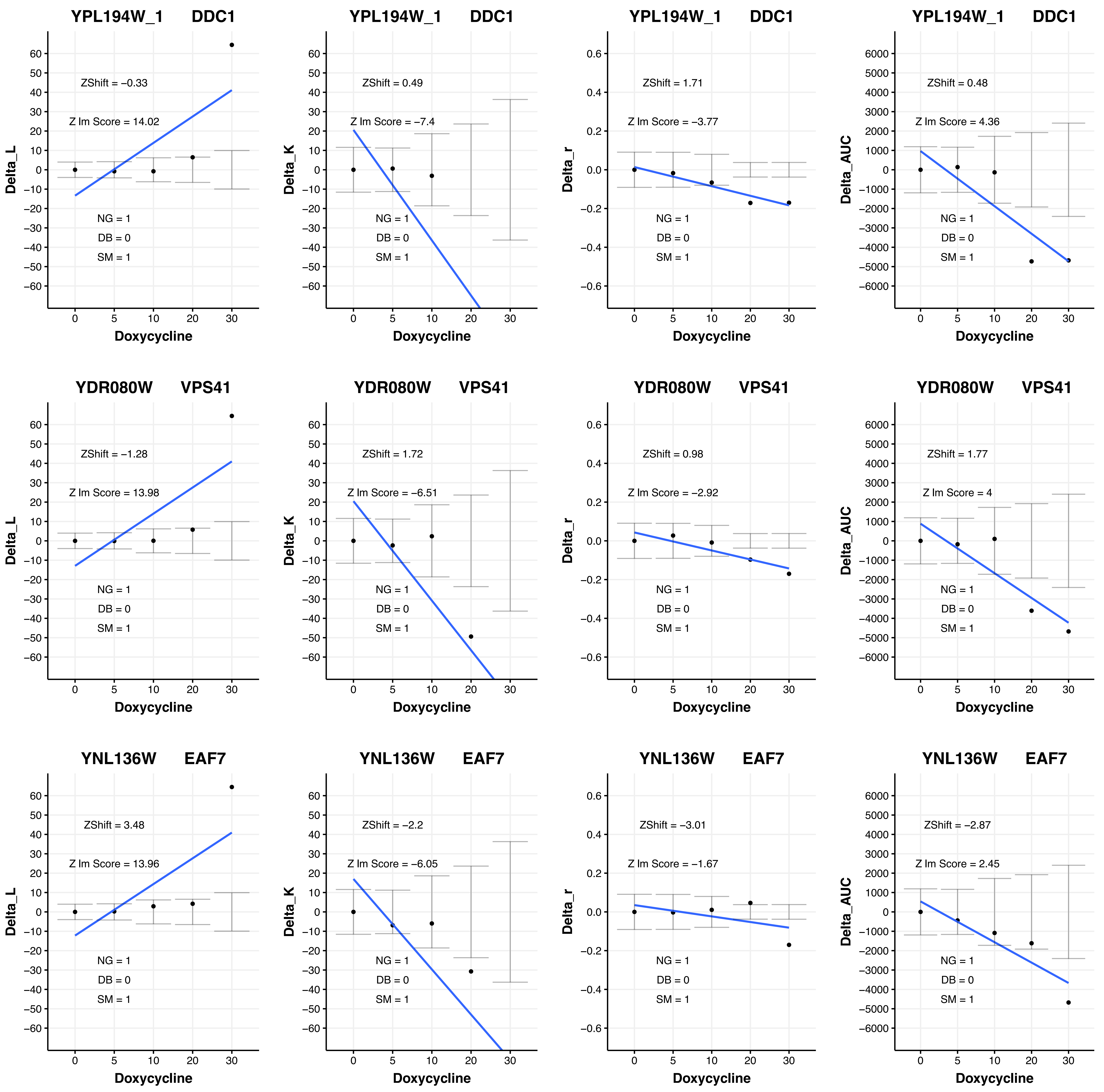
YPL194W_1 DDC1
ZShift = −0.33
Z lm Score = 14.02
NG = 1
DB = 0
SM = 1
Delta_L
Doxycycline
YPL194W_1 DDC1
ZShift = 0.49
Z lm Score = −7.4
NG = 1
DB = 0
SM = 1
Delta_K
Doxycycline
YPL194W_1 DDC1
ZShift = 1.71
Z lm Score = −3.77
NG = 1
DB = 0
SM = 1
Delta_r
Doxycycline
YPL194W_1 DDC1
ZShift = 0.48
Z lm Score = 4.36
NG = 1
DB = 0
SM = 1
Delta_AUC
Doxycycline
YDR080W VPS41
ZShift = −1.28
Z lm Score = 13.98
NG = 1
DB = 0
SM = 1
Delta_L
Doxycycline
YDR080W VPS41
ZShift = 1.72
Z lm Score = −6.51
NG = 1
DB = 0
SM = 1
Delta_K
Doxycycline
YDR080W VPS41
ZShift = 0.98
Z lm Score = −2.92
NG = 1
DB = 0
SM = 1
Delta_r
Doxycycline
YDR080W VPS41
ZShift = 1.77
Z lm Score = 4
NG = 1
DB = 0
SM = 1
Delta_AUC
Doxycycline
YNL136W EAF7
ZShift = 3.48
Z lm Score = 13.96
NG = 1
DB = 0
SM = 1
Delta_L
Doxycycline
YNL136W EAF7
ZShift = −2.2
Z lm Score = −6.05
NG = 1
DB = 0
SM = 1
Delta_K
Doxycycline
YNL136W EAF7
ZShift = −3.01
Z lm Score = −1.67
NG = 1
DB = 0
SM = 1
Delta_r
Doxycycline
YNL136W EAF7
ZShift = −2.87
Z lm Score = 2.45
NG = 1
DB = 0
SM = 1
Delta_AUC
Doxycycline

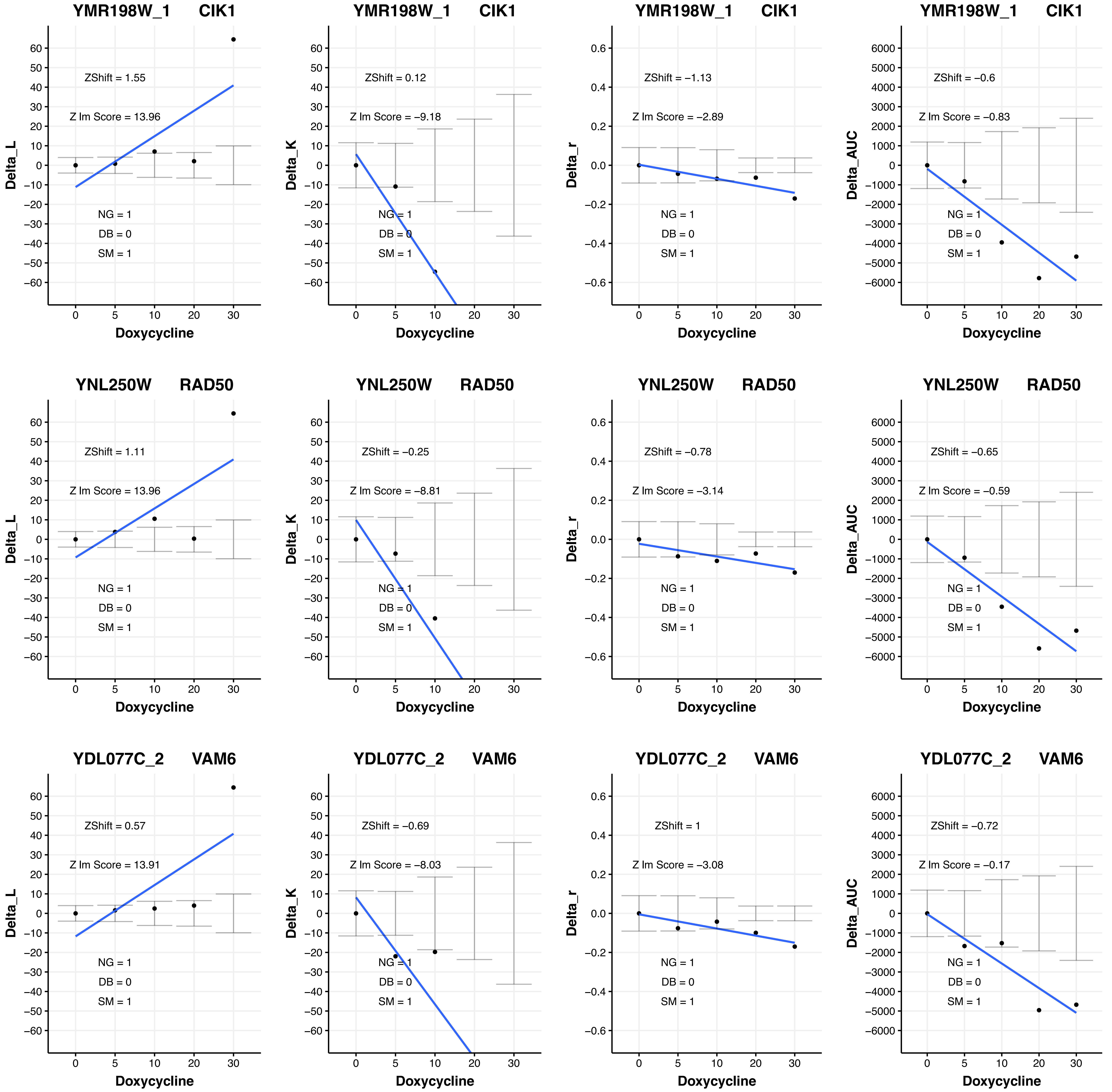

YMR198W_1 CIK1
ZShift = 1.55
Z lm Score = 13.96
NG = 1
DB = 0
SM = 1
Delta_L
Doxycycline
YMR198W_1 CIK1
ZShift = 0.12
Z lm Score = −9.18
NG = 1
DB = 0
SM = 1
Delta_K
Doxycycline
YMR198W_1 CIK1
ZShift = −1.13
Z lm Score = −2.89
NG = 1
DB = 0
SM = 1
Delta_r
Doxycycline
YMR198W_1 CIK1
ZShift = −0.6
Z lm Score = −0.83
NG = 1
DB = 0
SM = 1
Delta_AUC
Doxycycline
YNL250W RAD50
ZShift = 1.11
Z lm Score = 13.96
NG = 1
DB = 0
SM = 1
Delta_L
Doxycycline
YNL250W RAD50
ZShift = −0.25
Z lm Score = −8.81
NG = 1
DB = 0
SM = 1
Delta_K
Doxycycline
YNL250W RAD50
ZShift = −0.78
Z lm Score = −3.14
NG = 1
DB = 0
SM = 1
Delta_r
Doxycycline
YNL250W RAD50
ZShift = −0.65
Z lm Score = −0.59
NG = 1
DB = 0
SM = 1
Delta_AUC
Doxycycline
YDL077C_2 VAM6
ZShift = 0.57
Z lm Score = 13.91
NG = 1
DB = 0
SM = 1
Delta_L
Doxycycline
YDL077C_2 VAM6
ZShift = −0.69
Z lm Score = −8.03
NG = 1
DB = 0
SM = 1
Delta_K
Doxycycline
YDL077C_2 VAM6
ZShift = 1
Z lm Score = −3.08
NG = 1
DB = 0
SM = 1
Delta_r
Doxycycline
YDL077C_2 VAM6
ZShift = −0.72
Z lm Score = −0.17
NG = 1
DB = 0
SM = 1
Delta_AUC
Doxycycline

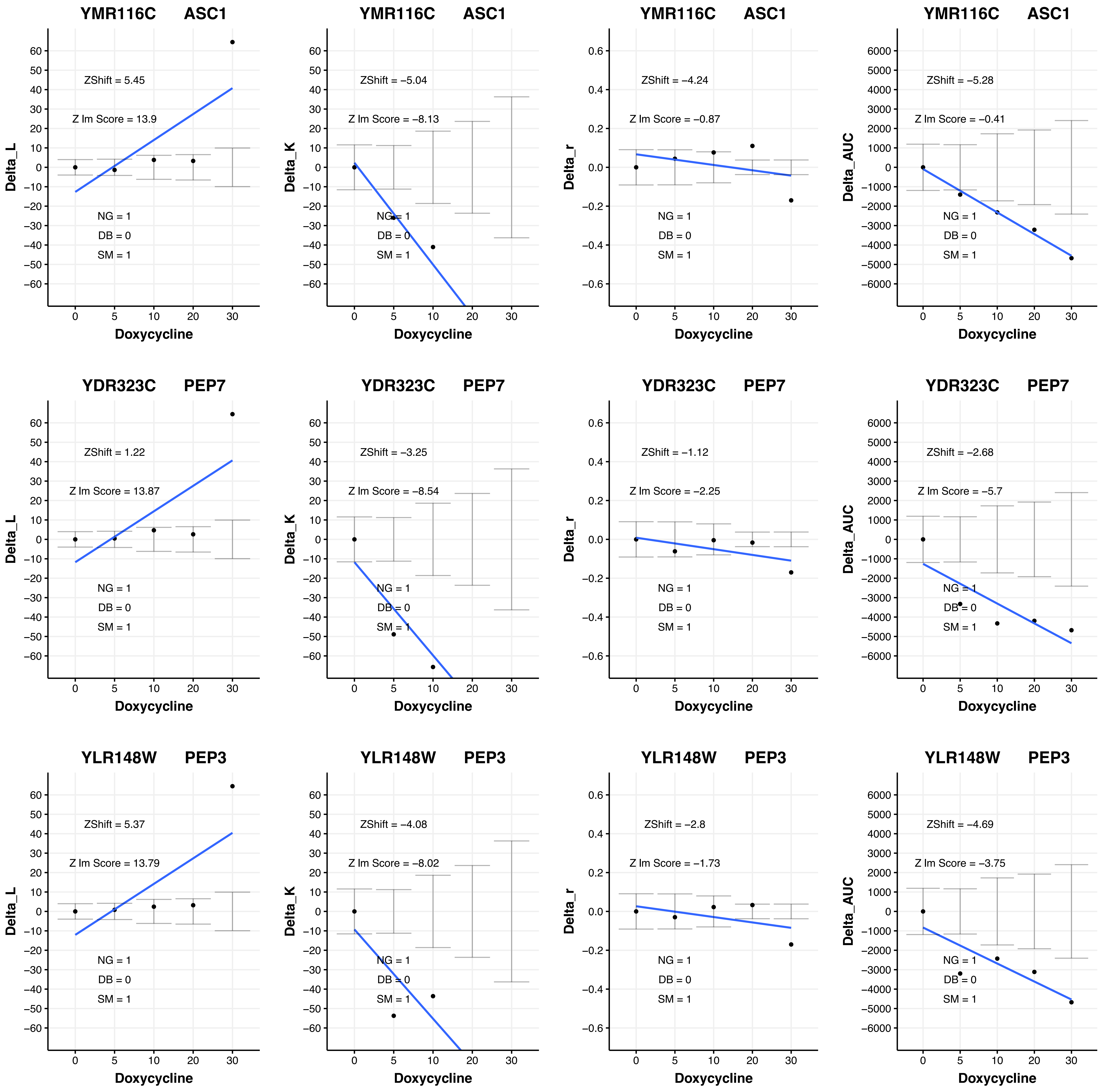
YMR116C ASC1
ZShift = 5.45
Z lm Score = 13.9
NG = 1
DB = 0
SM = 1
Delta_L
Doxycycline
YMR116C ASC1
ZShift = −5.04
Z lm Score = −8.13
NG = 1
DB = 0
SM = 1
Delta_K
Doxycycline
YMR116C ASC1
ZShift = −4.24
Z lm Score = −0.87
NG = 1
DB = 0
SM = 1
Delta_r
Doxycycline
YMR116C ASC1
ZShift = −5.28
Z lm Score = −0.41
NG = 1
DB = 0
SM = 1
Delta_AUC
Doxycycline
YDR323C PEP7
ZShift = 1.22
Z lm Score = 13.87
NG = 1
DB = 0
SM = 1
Delta_L
Doxycycline
YDR323C PEP7
ZShift = −3.25
Z lm Score = −8.54
NG = 1
DB = 0
SM = 1
Delta_K
Doxycycline
YDR323C PEP7
ZShift = −1.12
Z lm Score = −2.25
NG = 1
DB = 0
SM = 1
Delta_r
Doxycycline
YDR323C PEP7
ZShift = −2.68
Z lm Score = −5.7
NG = 1
DB = 0
SM = 1
Delta_AUC
Doxycycline
YLR148W PEP3
ZShift = 5.37
Z lm Score = 13.79
NG = 1
DB = 0
SM = 1
Delta_L
Doxycycline
YLR148W PEP3
ZShift = −4.08
Z lm Score = −8.02
NG = 1
DB = 0
SM = 1
Delta_K
Doxycycline
YLR148W PEP3
ZShift = −2.8
Z lm Score = −1.73
NG = 1
DB = 0
SM = 1
Delta_r
Doxycycline
YLR148W PEP3
ZShift = −4.69
Z lm Score = −3.75
NG = 1
DB = 0
SM = 1
Delta_AUC
Doxycycline

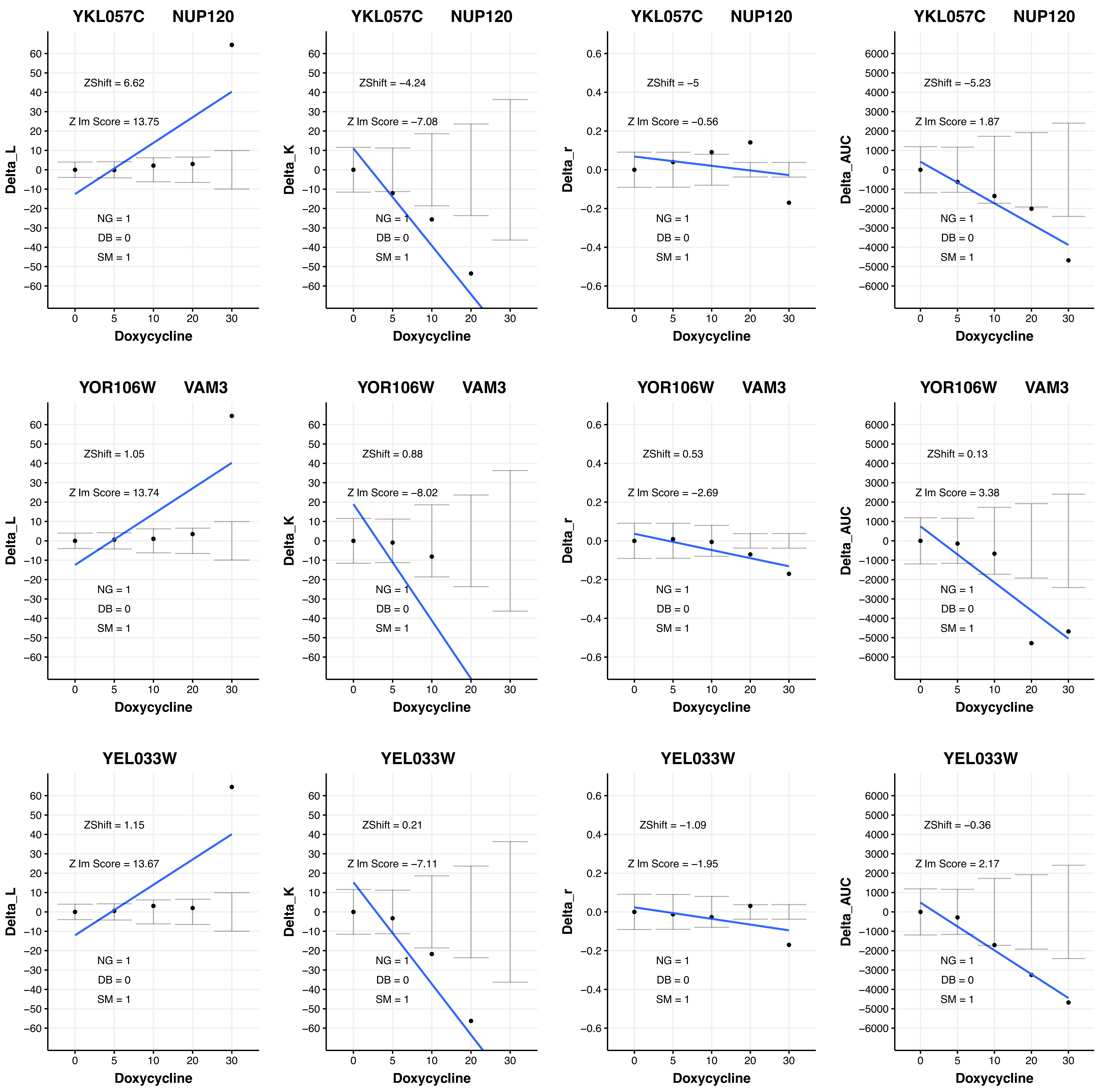

YKL057C NUP120
ZShift = 6.62
Z lm Score = 13.75
NG = 1
DB = 0
SM = 1
Delta_L
Doxycycline
YKL057C NUP120
ZShift = −4.24
Z lm Score = −7.08
NG = 1
DB = 0
SM = 1
Delta_K
Doxycycline
YKL057C NUP120
ZShift = −5
Z lm Score = −0.56
NG = 1
DB = 0
SM = 1
Delta_r
Doxycycline
YKL057C NUP120
ZShift = −5.23
Z lm Score = 1.87
NG = 1
DB = 0
SM = 1
Delta_AUC
Doxycycline
YOR106W VAM3
ZShift = 1.05
Z lm Score = 13.74
NG = 1
DB = 0
SM = 1
Delta_L
Doxycycline
YOR106W VAM3
ZShift = 0.88
Z lm Score = −8.02
NG = 1
DB = 0
SM = 1
Delta_K
Doxycycline
YOR106W VAM3
ZShift = 0.53
Z lm Score = −2.69
NG = 1
DB = 0
SM = 1
Delta_r
Doxycycline
YOR106W VAM3
ZShift = 0.13
Z lm Score = 3.38
NG = 1
DB = 0
SM = 1
Delta_AUC
Doxycycline
YEL033W
ZShift = 1.15
Z lm Score = 13.67
NG = 1
DB = 0
SM = 1
Delta_L
Doxycycline
YEL033W
ZShift = 0.21
Z lm Score = −7.11
NG = 1
DB = 0
SM = 1
Delta_K
Doxycycline
YEL033W
ZShift = −1.09
Z lm Score = −1.95
NG = 1
DB = 0
SM = 1
Delta_r
Doxycycline
YEL033W
ZShift = −0.36
Z lm Score = 2.17
NG = 1
DB = 0
SM = 1
Delta_AUC
Doxycycline

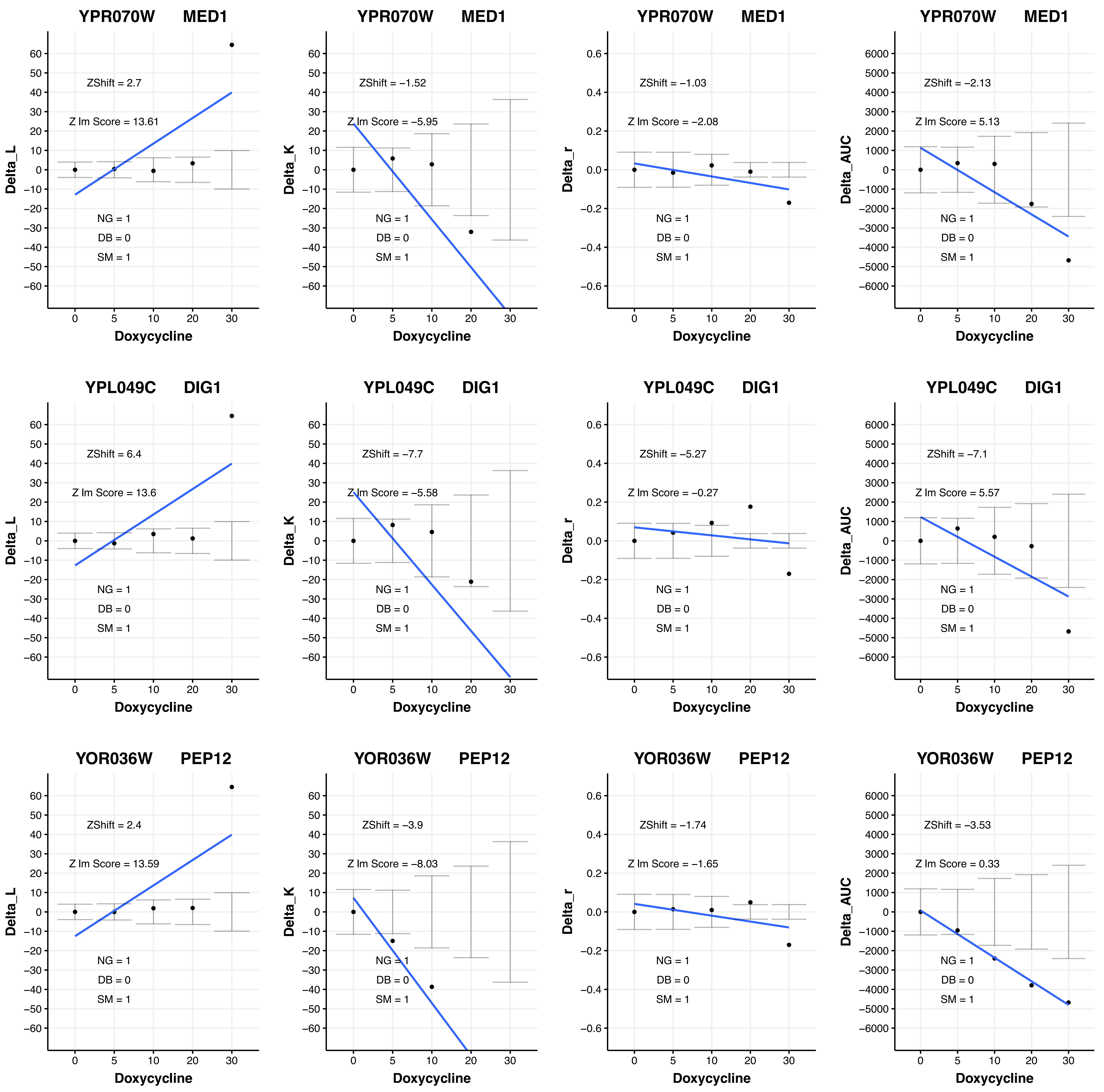
YPR070W MED1
ZShift = 2.7
Z lm Score = 13.61
NG = 1
DB = 0
SM = 1
Delta_L
Doxycycline
YPR070W MED1
ZShift = −1.52
Z lm Score = −5.95
NG = 1
DB = 0
SM = 1
Delta_K
Doxycycline
YPR070W MED1
ZShift = −1.03
Z lm Score = −2.08
NG = 1
DB = 0
SM = 1
Delta_r
Doxycycline
YPR070W MED1
ZShift = −2.13
Z lm Score = 5.13
NG = 1
DB = 0
SM = 1
Delta_AUC
Doxycycline
YPL049C DIG1
ZShift = 6.4
Z lm Score = 13.6
NG = 1
DB = 0
SM = 1
Delta_L
Doxycycline
YPL049C DIG1
ZShift = −7.7
Z lm Score = −5.58
NG = 1
DB = 0
SM = 1
Delta_K
Doxycycline
YPL049C DIG1
ZShift = −5.27
Z lm Score = −0.27
NG = 1
DB = 0
SM = 1
Delta_r
Doxycycline
YPL049C DIG1
ZShift = −7.1
Z lm Score = 5.57
NG = 1
DB = 0
SM = 1
Delta_AUC
Doxycycline
YOR036W PEP12
ZShift = 2.4
Z lm Score = 13.59
NG = 1
DB = 0
SM = 1
Delta_L
Doxycycline
YOR036W PEP12
ZShift = −3.9
Z lm Score = −8.03
NG = 1
DB = 0
SM = 1
Delta_K
Doxycycline
YOR036W PEP12
ZShift = −1.74
Z lm Score = −1.65
NG = 1
DB = 0
SM = 1
Delta_r
Doxycycline
YOR036W PEP12
ZShift = −3.53
Z lm Score = 0.33
NG = 1
DB = 0
SM = 1
Delta_AUC
Doxycycline

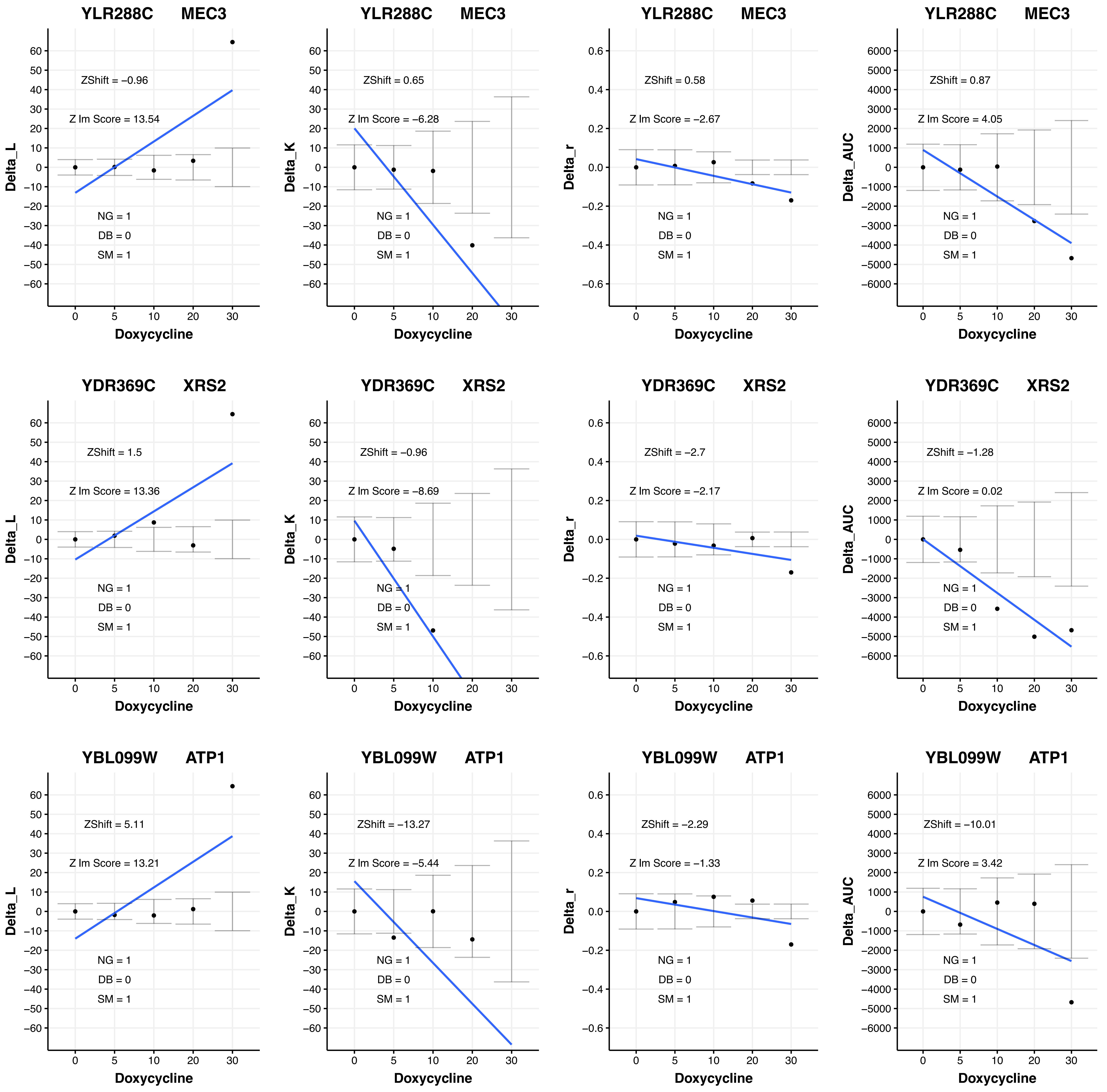

YLR288C MEC3
ZShift = −0.96
Z lm Score = 13.54
NG = 1
DB = 0
SM = 1
Delta_L
Doxycycline
YLR288C MEC3
ZShift = 0.65
Z lm Score = −6.28
NG = 1
DB = 0
SM = 1
Delta_K
Doxycycline
YLR288C MEC3
ZShift = 0.58
Z lm Score = −2.67
NG = 1
DB = 0
SM = 1
Delta_r
Doxycycline
YLR288C MEC3
ZShift = 0.87
Z lm Score = 4.05
NG = 1
DB = 0
SM = 1
Delta_AUC
Doxycycline
YDR369C XRS2
ZShift = 1.5
Z lm Score = 13.36
NG = 1
DB = 0
SM = 1
Delta_L
Doxycycline
YDR369C XRS2
ZShift = −0.96
Z lm Score = −8.69
NG = 1
DB = 0
SM = 1
Delta_K
Doxycycline
YDR369C XRS2
ZShift = −2.7
Z lm Score = −2.17
NG = 1
DB = 0
SM = 1
Delta_r
Doxycycline
YDR369C XRS2
ZShift = −1.28
Z lm Score = 0.02
NG = 1
DB = 0
SM = 1
Delta_AUC
Doxycycline
YBL099W ATP1
ZShift = 5.11
Z lm Score = 13.21
NG = 1
DB = 0
SM = 1
Delta_L
Doxycycline
YBL099W ATP1
ZShift = −13.27
Z lm Score = −5.44
NG = 1
DB = 0
SM = 1
Delta_K
Doxycycline
YBL099W ATP1
ZShift = −2.29
Z lm Score = −1.33
NG = 1
DB = 0
SM = 1
Delta_r
Doxycycline
YBL099W ATP1
ZShift = −10.01
Z lm Score = 3.42
NG = 1
DB = 0
SM = 1
Delta_AUC
Doxycycline

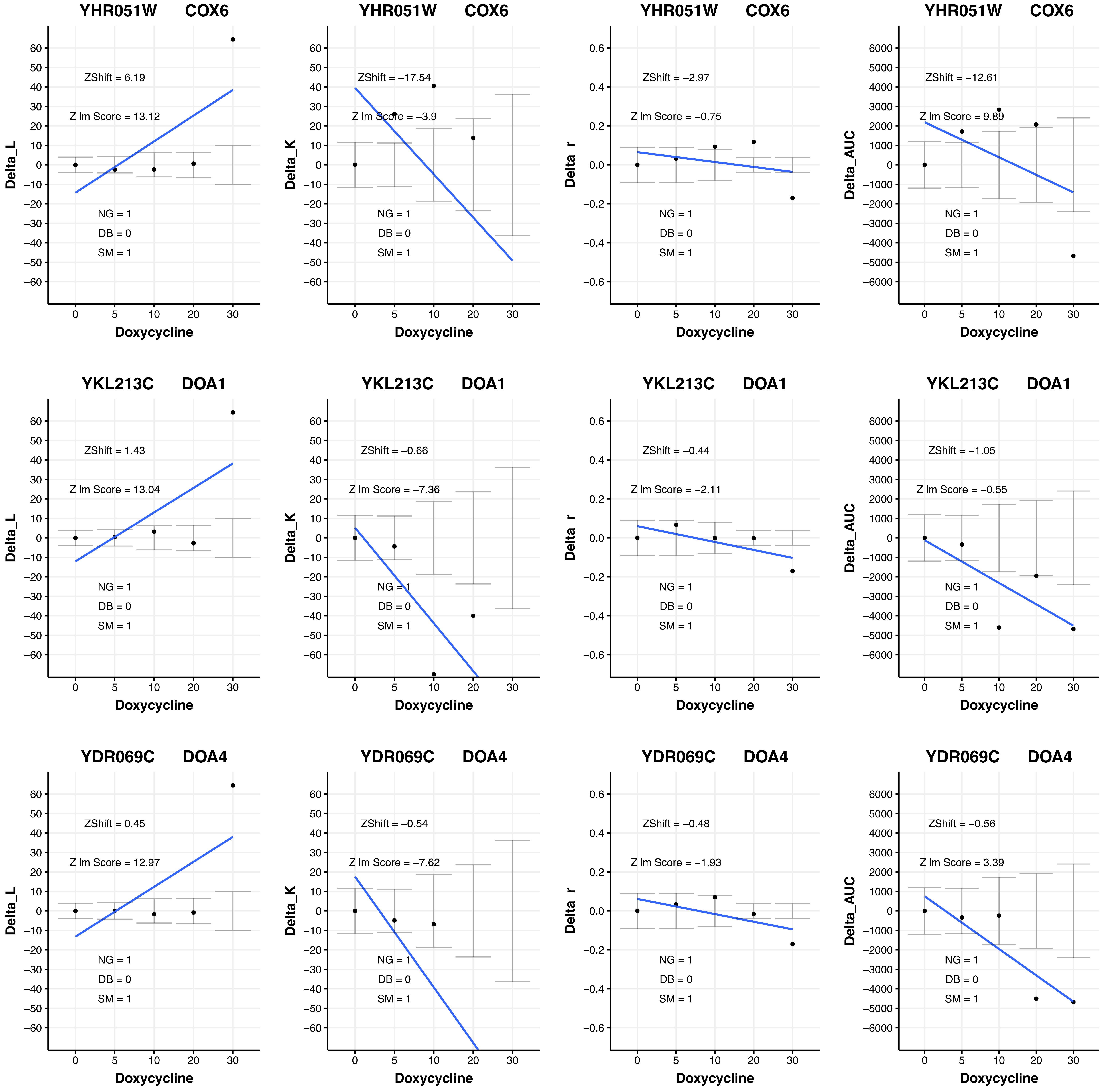

YHR051W COX6
ZShift = 6.19
Z lm Score = 13.12
NG = 1
DB = 0
SM = 1
Delta_L
Doxycycline
YHR051W COX6
ZShift = −17.54
Z lm Score = −3.9
NG = 1
DB = 0
SM = 1
Delta_K
Doxycycline
YHR051W COX6
ZShift = −2.97
Z lm Score = −0.75
NG = 1
DB = 0
SM = 1
Delta_r
Doxycycline
YHR051W COX6
ZShift = −12.61
Z lm Score = 9.89
NG = 1
DB = 0
SM = 1
Delta_AUC
Doxycycline
YKL213C DOA1
ZShift = 1.43
Z lm Score = 13.04
NG = 1
DB = 0
SM = 1
Delta_L
Doxycycline
YKL213C DOA1
ZShift = −0.66
Z lm Score = −7.36
NG = 1
DB = 0
SM = 1
Delta_K
Doxycycline
YKL213C DOA1
ZShift = −0.44
Z lm Score = −2.11
NG = 1
DB = 0
SM = 1
Delta_r
Doxycycline
YKL213C DOA1
ZShift = −1.05
Z lm Score = −0.55
NG = 1
DB = 0
SM = 1
Delta_AUC
Doxycycline
YDR069C DOA4
ZShift = 0.45
Z lm Score = 12.97
NG = 1
DB = 0
SM = 1
Delta_L
Doxycycline
YDR069C DOA4
ZShift = −0.54
Z lm Score = −7.62
NG = 1
DB = 0
SM = 1
Delta_K
Doxycycline
YDR069C DOA4
ZShift = −0.48
Z lm Score = −1.93
NG = 1
DB = 0
SM = 1
Delta_r
Doxycycline
YDR069C DOA4
ZShift = −0.56
Z lm Score = 3.39
NG = 1
DB = 0
SM = 1
Delta_AUC
Doxycycline

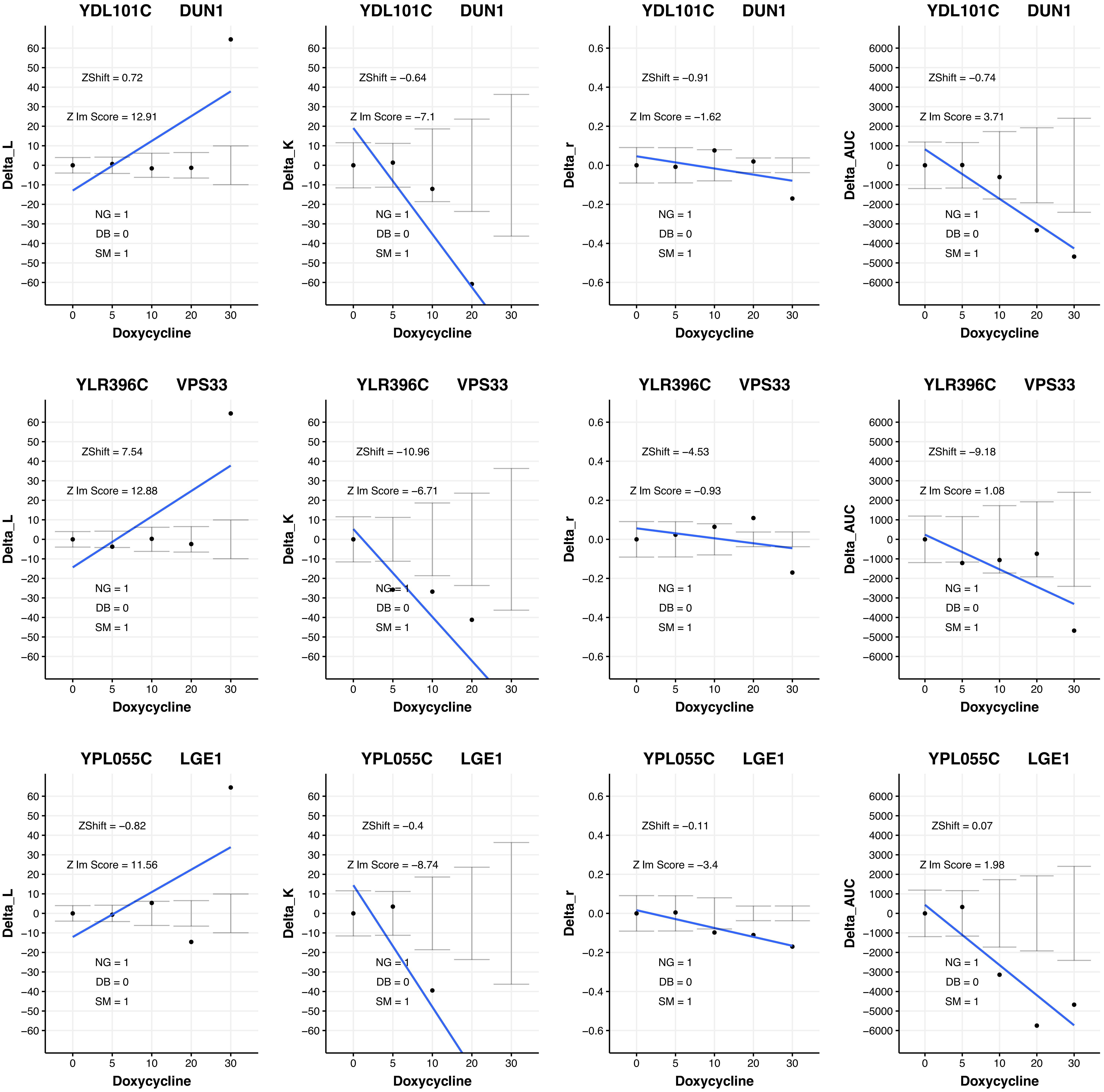

YDL101C DUN1
ZShift = 0.72
Z lm Score = 12.91
NG = 1
DB = 0
SM = 1
Delta_L
Doxycycline
YDL101C DUN1
ZShift = −0.64
Z lm Score = −7.1
NG = 1
DB = 0
SM = 1
Delta_K
Doxycycline
YDL101C DUN1
ZShift = −0.91
Z lm Score = −1.62
NG = 1
DB = 0
SM = 1
Delta_r
Doxycycline
YDL101C DUN1
ZShift = −0.74
Z lm Score = 3.71
NG = 1
DB = 0
SM = 1
Delta_AUC
Doxycycline
YLR396C VPS33
ZShift = 7.54
Z lm Score = 12.88
NG = 1
DB = 0
SM = 1
Delta_L
Doxycycline
YLR396C VPS33
ZShift = −10.96
Z lm Score = −6.71
NG = 1
DB = 0
SM = 1
Delta_K
Doxycycline
YLR396C VPS33
ZShift = −4.53
Z lm Score = −0.93
NG = 1
DB = 0
SM = 1
Delta_r
Doxycycline
YLR396C VPS33
ZShift = −9.18
Z lm Score = 1.08
NG = 1
DB = 0
SM = 1
Delta_AUC
Doxycycline
YPL055C LGE1
ZShift = −0.82
Z lm Score = 11.56
NG = 1
DB = 0
SM = 1
Delta_L
Doxycycline
YPL055C LGE1
ZShift = −0.4
Z lm Score = −8.74
NG = 1
DB = 0
SM = 1
Delta_K
Doxycycline
YPL055C LGE1
ZShift = −0.11
Z lm Score = −3.4
NG = 1
DB = 0
SM = 1
Delta_r
Doxycycline
YPL055C LGE1
ZShift = 0.07
Z lm Score = 1.98
NG = 1
DB = 0
SM = 1
Delta_AUC
Doxycycline

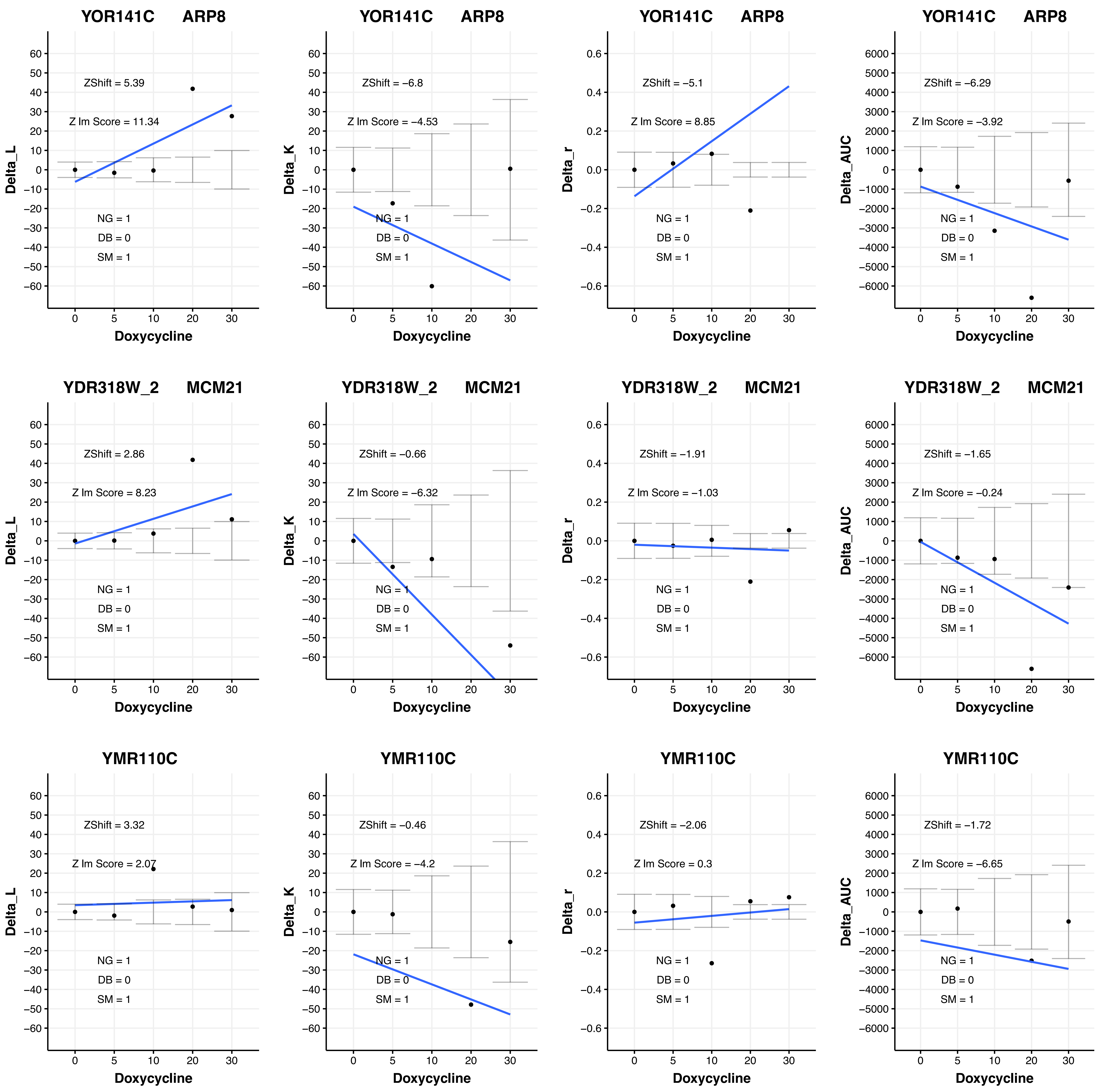

YOR141C ARP8
ZShift = 5.39
Z lm Score = 11.34
NG = 1
DB = 0
SM = 1
Delta_L
Doxycycline
YOR141C ARP8
ZShift = −6.8
Z lm Score = −4.53
NG = 1
DB = 0
SM = 1
Delta_K
Doxycycline
YOR141C ARP8
ZShift = −5.1
Z lm Score = 8.85
NG = 1
DB = 0
SM = 1
Delta_r
Doxycycline
YOR141C ARP8
ZShift = −6.29
Z lm Score = −3.92
NG = 1
DB = 0
SM = 1
Delta_AUC
Doxycycline
YDR318W_2 MCM21
ZShift = 2.86
Z lm Score = 8.23
NG = 1
DB = 0
SM = 1
Delta_L
Doxycycline
YDR318W_2 MCM21
ZShift = −0.66
Z lm Score = −6.32
NG = 1
DB = 0
SM = 1
Delta_K
Doxycycline
YDR318W_2 MCM21
ZShift = −1.91
Z lm Score = −1.03
NG = 1
DB = 0
SM = 1
Delta_r
Doxycycline
YDR318W_2 MCM21
ZShift = −1.65
Z lm Score = −0.24
NG = 1
DB = 0
SM = 1
Delta_AUC
Doxycycline
YMR110C
ZShift = 3.32
Z lm Score = 2.07
NG = 1
DB = 0
SM = 1
Delta_L
Doxycycline
YMR110C
ZShift = −0.46
Z lm Score = −4.2
NG = 1
DB = 0
SM = 1
Delta_K
Doxycycline
YMR110C
ZShift = −2.06
Z lm Score = 0.3
NG = 1
DB = 0
SM = 1
Delta_r
Doxycycline
YMR110C
ZShift = −1.72
Z lm Score = −6.65
NG = 1
DB = 0
SM = 1
Delta_AUC
Doxycycline

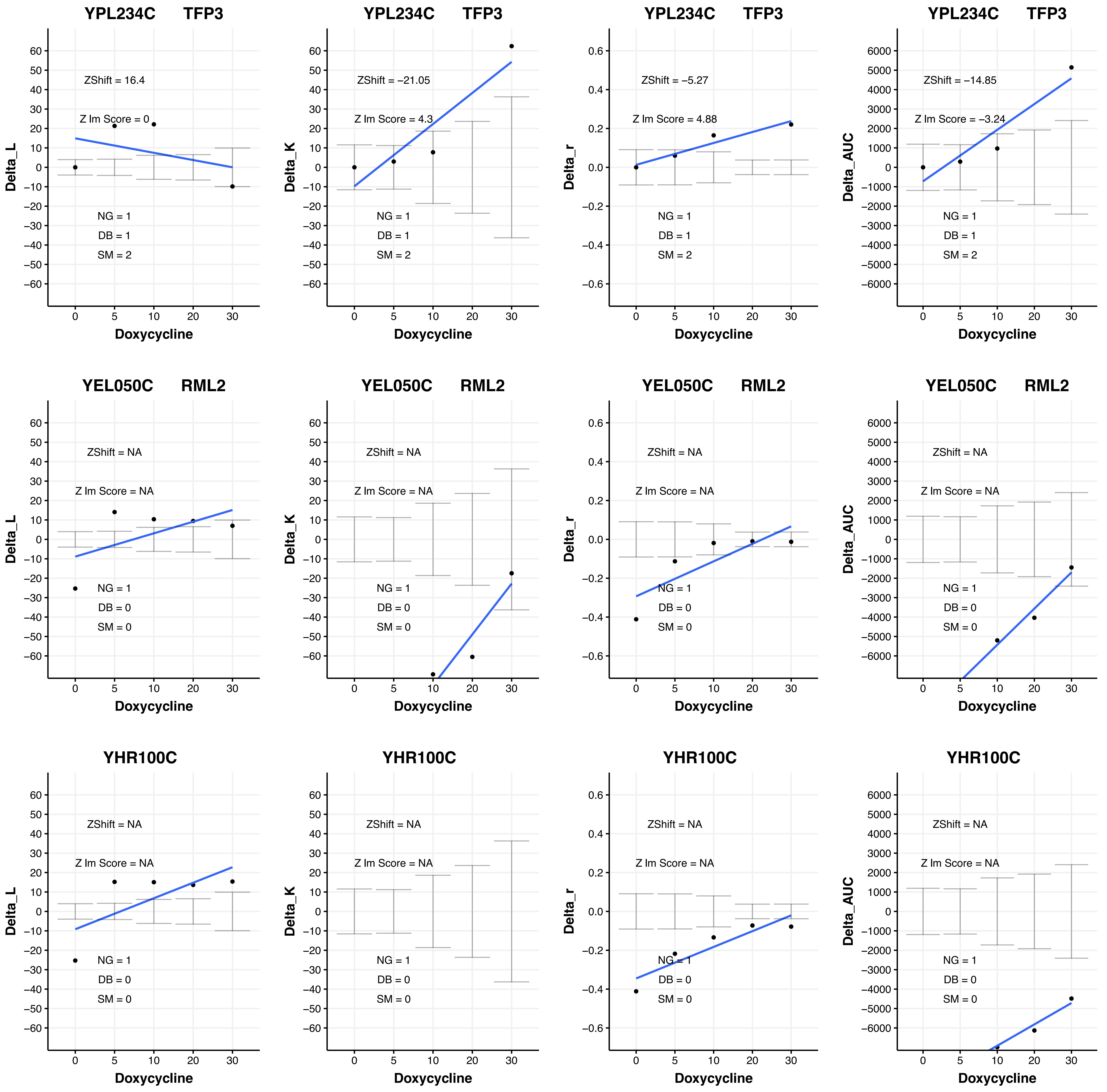
YPL234C TFP3
ZShift = 16.4
Z lm Score = 0
NG = 1
DB = 1
SM = 2
Delta_L
Doxycycline
YPL234C TFP3
ZShift = −21.05
Z lm Score = 4.3
NG = 1
DB = 1
SM = 2
Delta_K
Doxycycline
YPL234C TFP3
ZShift = −5.27
Z lm Score = 4.88
NG = 1
DB = 1
SM = 2
Delta_r
Doxycycline
YPL234C TFP3
ZShift = −14.85
Z lm Score = −3.24
NG = 1
DB = 1
SM = 2
Delta_AUC
Doxycycline
YEL050C RML2
ZShift = NA
Z lm Score = NA
NG = 1
DB = 0
SM = 0
Delta_L
Doxycycline
YEL050C RML2
ZShift = NA
Z lm Score = NA
NG = 1
DB = 0
SM = 0
Delta_K
Doxycycline
YEL050C RML2
ZShift = NA
Z lm Score = NA
NG = 1
DB = 0
SM = 0
Delta_r
Doxycycline
YEL050C RML2
ZShift = NA
Z lm Score = NA
NG = 1
DB = 0
SM = 0
Delta_AUC
Doxycycline
YHR100C
ZShift = NA
Z lm Score = NA
NG = 1
DB = 0
SM = 0
Delta_L
Doxycycline
YHR100C
ZShift = NA
Z lm Score = NA
NG = 1
DB = 0
SM = 0
Delta_K
Doxycycline
YHR100C
ZShift = NA
Z lm Score = NA
NG = 1
DB = 0
SM = 0
Delta_r
Doxycycline
YHR100C
ZShift = NA
Z lm Score = NA
NG = 1
DB = 0
SM = 0
Delta_AUC
Doxycycline

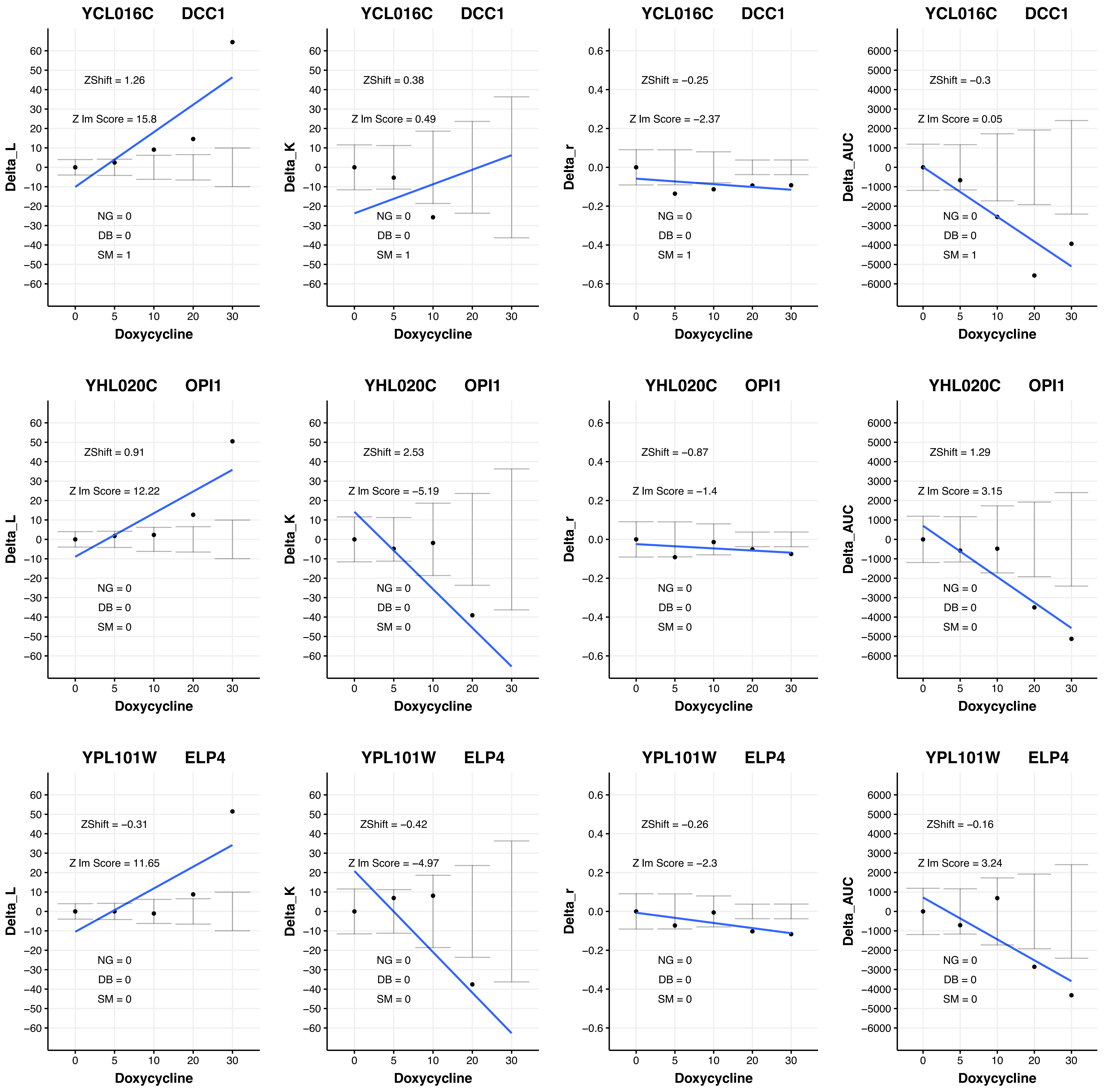

YCL016C DCC1
ZShift = 1.26
Z lm Score = 15.8
NG = 0
DB = 0
SM = 1
Delta_L
Doxycycline
YCL016C DCC1
ZShift = 0.38
Z lm Score = 0.49
NG = 0
DB = 0
SM = 1
Delta_K
Doxycycline
YCL016C DCC1
ZShift = −0.25
Z lm Score = −2.37
NG = 0
DB = 0
SM = 1
Delta_r
Doxycycline
YCL016C DCC1
ZShift = −0.3
Z lm Score = 0.05
NG = 0
DB = 0
SM = 1
Delta_AUC
Doxycycline
YHL020C OPI1
ZShift = 0.91
Z lm Score = 12.22
NG = 0
DB = 0
SM = 0
Delta_L
Doxycycline
YHL020C OPI1
ZShift = 2.53
Z lm Score = −5.19
NG = 0
DB = 0
SM = 0
Delta_K
Doxycycline
YHL020C OPI1
ZShift = −0.87
Z lm Score = −1.4
NG = 0
DB = 0
SM = 0
Delta_r
Doxycycline
YHL020C OPI1
ZShift = 1.29
Z lm Score = 3.15
NG = 0
DB = 0
SM = 0
Delta_AUC
Doxycycline
YPL101W ELP4
ZShift = −0.31
Z lm Score = 11.65
NG = 0
DB = 0
SM = 0
Delta_L
Doxycycline
YPL101W ELP4
ZShift = −0.42
Z lm Score = −4.97
NG = 0
DB = 0
SM = 0
Delta_K
Doxycycline
YPL101W ELP4
ZShift = −0.26
Z lm Score = −2.3
NG = 0
DB = 0
SM = 0
Delta_r
Doxycycline
YPL101W ELP4
ZShift = −0.16
Z lm Score = 3.24
NG = 0
DB = 0
SM = 0
Delta_AUC
Doxycycline

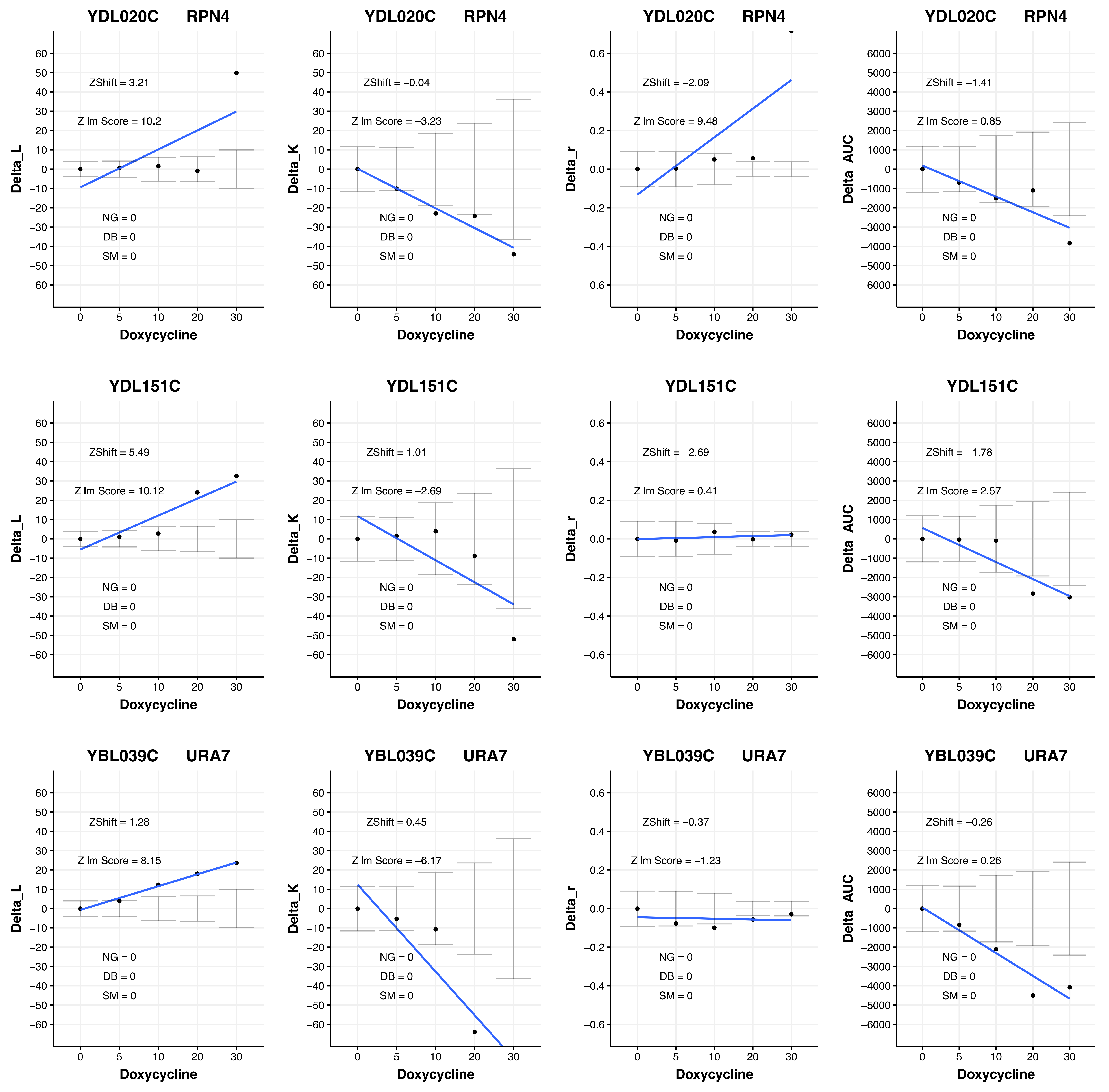
YDL020C RPN4
ZShift = 3.21
Z lm Score = 10.2
NG = 0
DB = 0
SM = 0
Delta_L
Doxycycline
YDL020C RPN4
ZShift = −0.04
Z lm Score = −3.23
NG = 0
DB = 0
SM = 0
Delta_K
Doxycycline
YDL020C RPN4
ZShift = −2.09
Z lm Score = 9.48
NG = 0
DB = 0
SM = 0
Delta_r
Doxycycline
YDL020C RPN4
ZShift = −1.41
Z lm Score = 0.85
NG = 0
DB = 0
SM = 0
Delta_AUC
Doxycycline
YDL151C
ZShift = 5.49
Z lm Score = 10.12
NG = 0
DB = 0
SM = 0
Delta_L
Doxycycline
YDL151C
ZShift = 1.01
Z lm Score = −2.69
NG = 0
DB = 0
SM = 0
Delta_K
Doxycycline
YDL151C
ZShift = −2.69
Z lm Score = 0.41
NG = 0
DB = 0
SM = 0
Delta_r
Doxycycline
YDL151C
ZShift = −1.78
Z lm Score = 2.57
NG = 0
DB = 0
SM = 0
Delta_AUC
Doxycycline
YBL039C URA7
ZShift = 1.28
Z lm Score = 8.15
NG = 0
DB = 0
SM = 0
Delta_L
Doxycycline
YBL039C URA7
ZShift = 0.45
Z lm Score = −6.17
NG = 0
DB = 0
SM = 0
Delta_K
Doxycycline
YBL039C URA7
ZShift = −0.37
Z lm Score = −1.23
NG = 0
DB = 0
SM = 0
Delta_r
Doxycycline
YBL039C URA7
ZShift = −0.26
Z lm Score = 0.26
NG = 0
DB = 0
SM = 0
Delta_AUC
Doxycycline

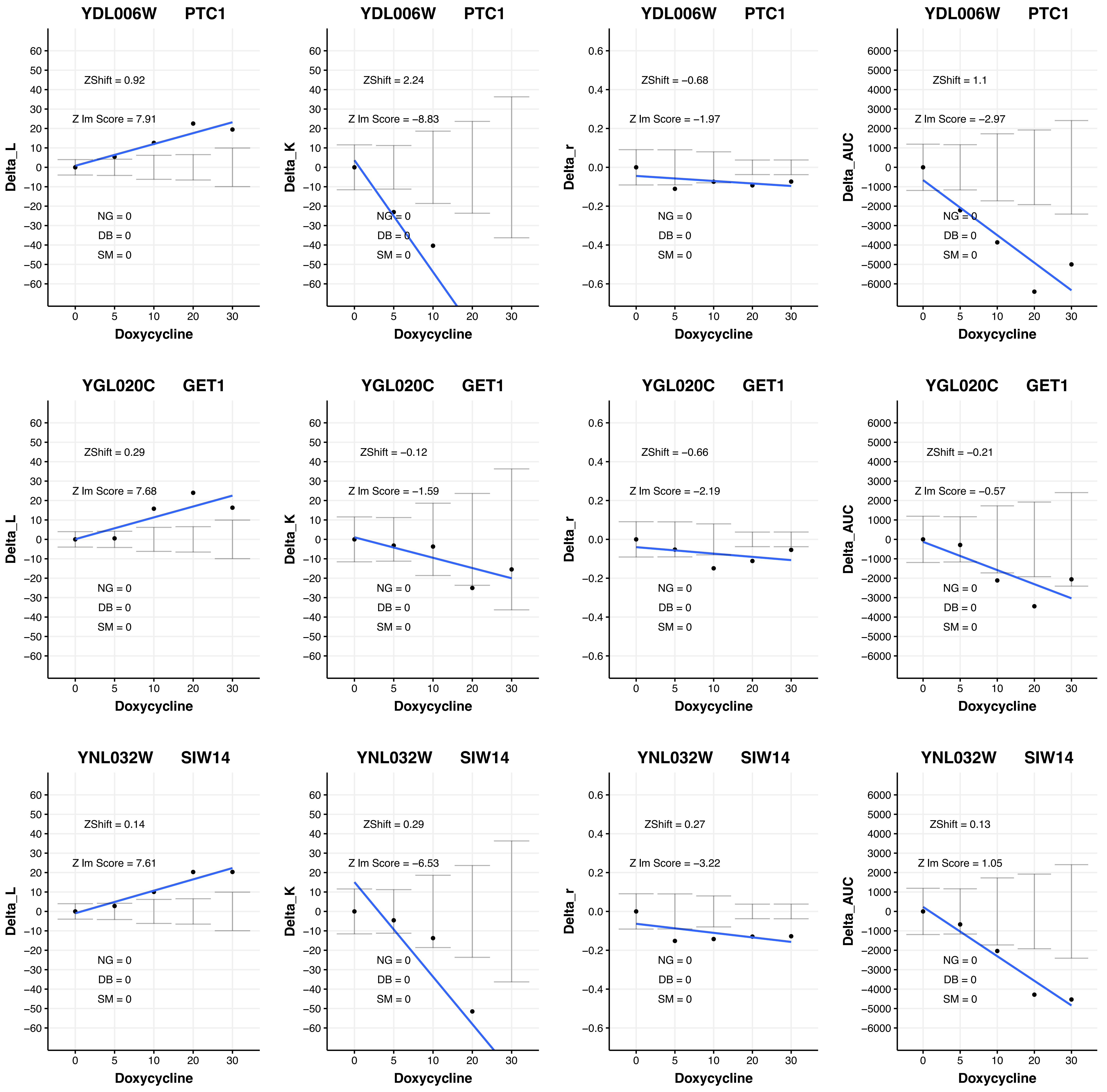

YDL006W PTC1
Delta_L
ZShift = 0.92
Z lm Score = 7.91
NG = 0
DB = 0
SM = 0
Doxycycline
YDL006W PTC1
Delta_K
ZShift = 2.24
Z lm Score = −8.83
NG = 0
DB = 0
SM = 0
Doxycycline
YDL006W PTC1
Delta_r
ZShift = −0.68
Z lm Score = −1.97
NG = 0
DB = 0
SM = 0
Doxycycline
YDL006W PTC1
Delta_AUC
ZShift = 1.1
Z lm Score = −2.97
NG = 0
DB = 0
SM = 0
Doxycycline
YGL020C GET1
Delta_L
ZShift = 0.29
Z lm Score = 7.68
NG = 0
DB = 0
SM = 0
Doxycycline
YGL020C GET1
Delta_K
ZShift = −0.12
Z lm Score = −1.59
NG = 0
DB = 0
SM = 0
Doxycycline
YGL020C GET1
Delta_r
ZShift = −0.66
Z lm Score = −2.19
NG = 0
DB = 0
SM = 0
Doxycycline
YGL020C GET1
Delta_AUC
ZShift = −0.21
Z lm Score = −0.57
NG = 0
DB = 0
SM = 0
Doxycycline
YNL032W SIW14
Delta_L
ZShift = 0.14
Z lm Score = 7.61
NG = 0
DB = 0
SM = 0
Doxycycline
YNL032W SIW14
Delta_K
ZShift = 0.29
Z lm Score = −6.53
NG = 0
DB = 0
SM = 0
Doxycycline
YNL032W SIW14
Delta_r
ZShift = 0.27
Z lm Score = −3.22
NG = 0
DB = 0
SM = 0
Doxycycline
YNL032W SIW14
Delta_AUC
ZShift = 0.13
Z lm Score = 1.05
NG = 0
DB = 0
SM = 0
Doxycycline

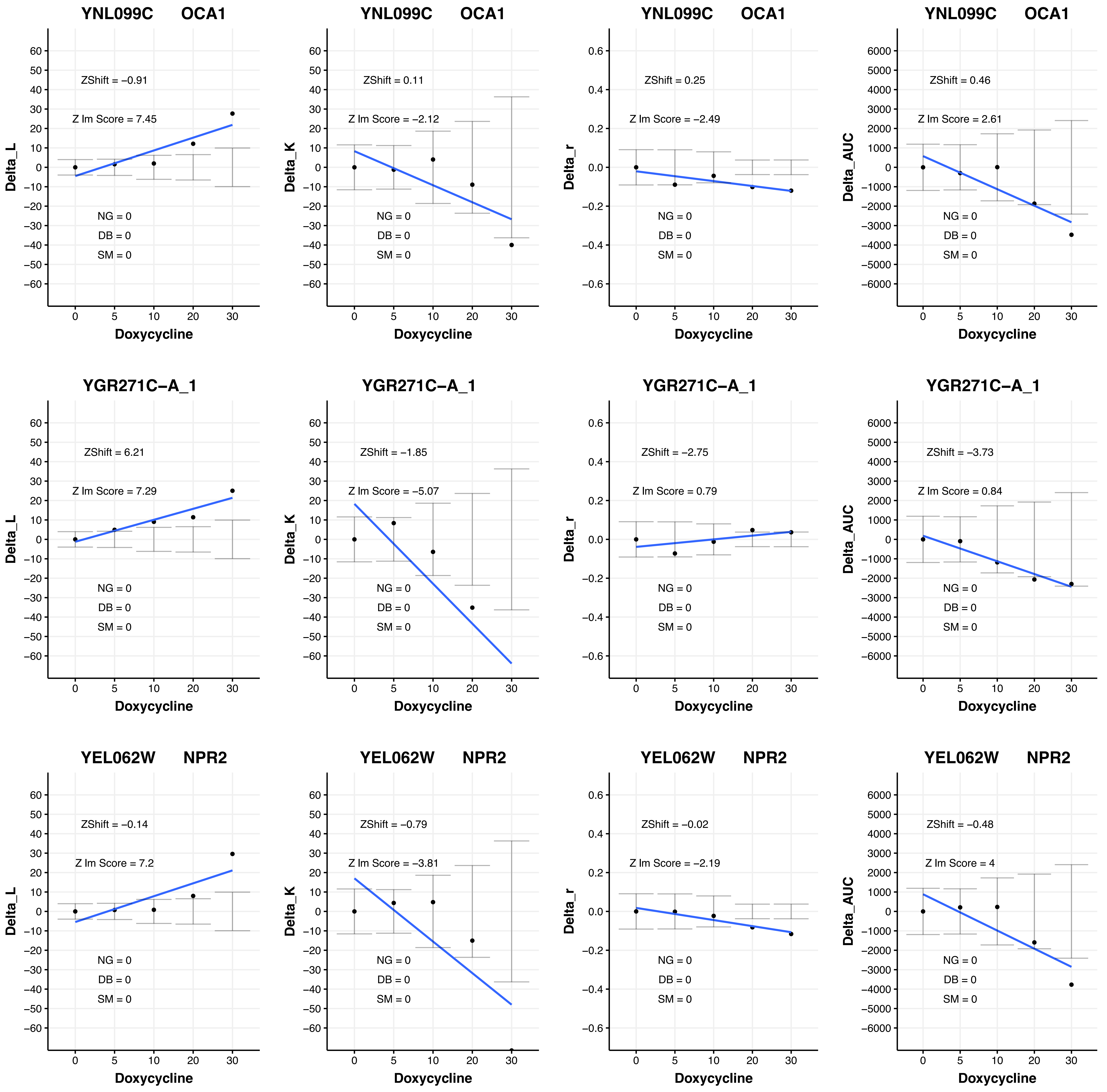
YNL099C OCA1
ZShift = −0.91
Z lm Score = 7.45
NG = 0
DB = 0
SM = 0
Delta_L
Doxycycline
YNL099C OCA1
ZShift = 0.11
Z lm Score = −2.12
NG = 0
DB = 0
SM = 0
Delta_K
Doxycycline
YNL099C OCA1
ZShift = 0.25
Z lm Score = −2.49
NG = 0
DB = 0
SM = 0
Delta_r
Doxycycline
YNL099C OCA1
ZShift = 0.46
Z lm Score = 2.61
NG = 0
DB = 0
SM = 0
Delta_AUC
Doxycycline
YGR271C−A_1
ZShift = 6.21
Z lm Score = 7.29
NG = 0
DB = 0
SM = 0
Delta_L
Doxycycline
YGR271C−A_1
ZShift = −1.85
Z lm Score = −5.07
NG = 0
DB = 0
SM = 0
Delta_K
Doxycycline
YGR271C−A_1
ZShift = −2.75
Z lm Score = 0.79
NG = 0
DB = 0
SM = 0
Delta_r
Doxycycline
YGR271C−A_1
ZShift = −3.73
Z lm Score = 0.84
NG = 0
DB = 0
SM = 0
Delta_AUC
Doxycycline
YEL062W NPR2
ZShift = −0.14
Z lm Score = 7.2
NG = 0
DB = 0
SM = 0
Delta_L
Doxycycline
YEL062W NPR2
ZShift = −0.79
Z lm Score = −3.81
NG = 0
DB = 0
SM = 0
Delta_K
Doxycycline
YEL062W NPR2
ZShift = −0.02
Z lm Score = −2.19
NG = 0
DB = 0
SM = 0
Delta_r
Doxycycline
YEL062W NPR2
ZShift = −0.48
Z lm Score = 4
NG = 0
DB = 0
SM = 0
Delta_AUC
Doxycycline

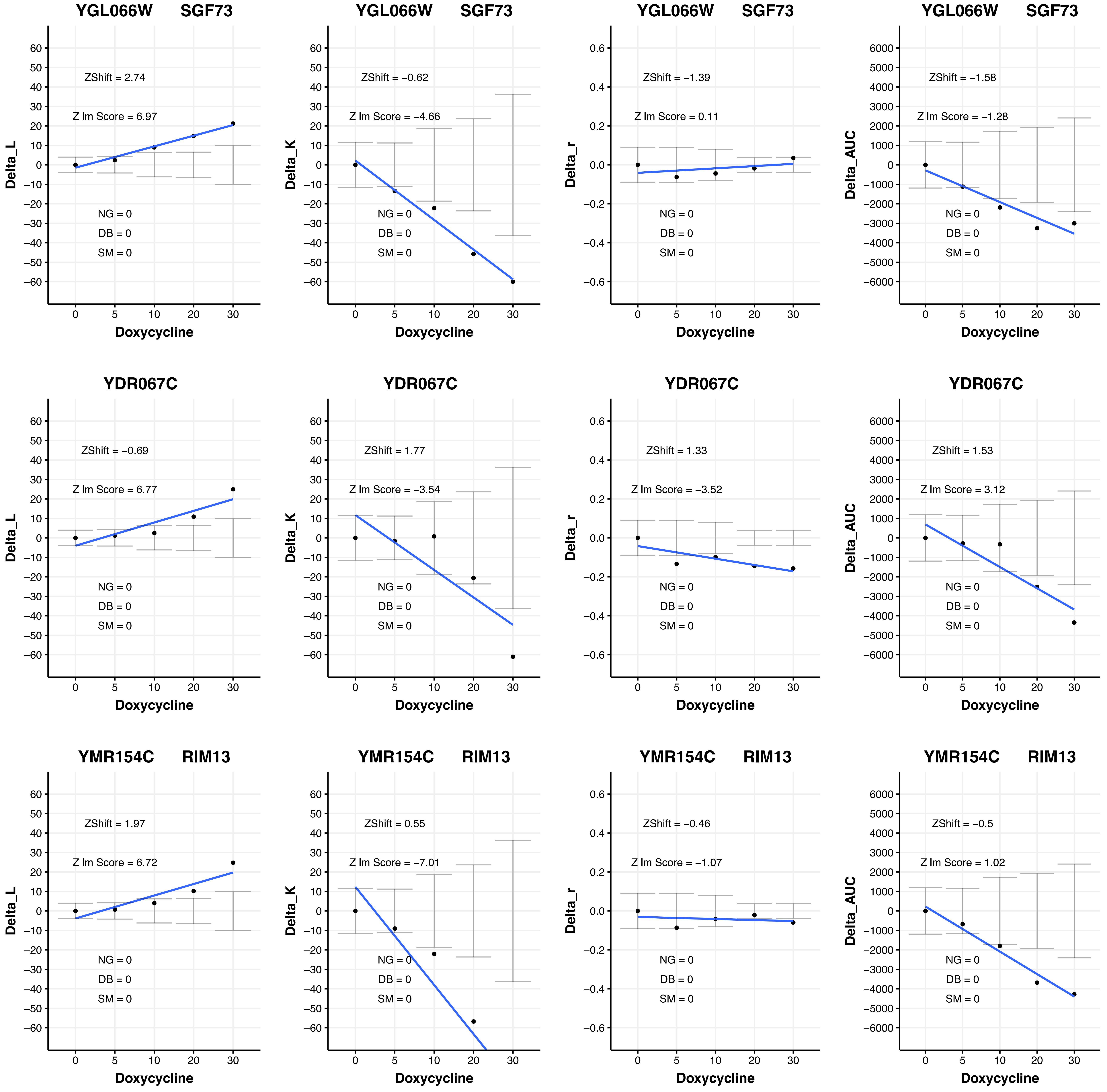
YGL066W SGF73
ZShift = 2.74
Z lm Score = 6.97
NG = 0
DB = 0
SM = 0
Delta_L
Doxycycline
YGL066W SGF73
ZShift = −0.62
Z lm Score = −4.66
NG = 0
DB = 0
SM = 0
Delta_K
Doxycycline
YGL066W SGF73
ZShift = −1.39
Z lm Score = 0.11
NG = 0
DB = 0
SM = 0
Delta_r
Doxycycline
YGL066W SGF73
ZShift = −1.58
Z lm Score = −1.28
NG = 0
DB = 0
SM = 0
Delta_AUC
Doxycycline
YDR067C
ZShift = −0.69
Z lm Score = 6.77
NG = 0
DB = 0
SM = 0
Delta_L
Doxycycline
YDR067C
ZShift = 1.77
Z lm Score = −3.54
NG = 0
DB = 0
SM = 0
Delta_K
Doxycycline
YDR067C
ZShift = 1.33
Z lm Score = −3.52
NG = 0
DB = 0
SM = 0
Delta_r
Doxycycline
YDR067C
ZShift = 1.53
Z lm Score = 3.12
NG = 0
DB = 0
SM = 0
Delta_AUC
Doxycycline
YMR154C RIM13
ZShift = 1.97
Z lm Score = 6.72
NG = 0
DB = 0
SM = 0
Delta_L
Doxycycline
YMR154C RIM13
ZShift = 0.55
Z lm Score = −7.01
NG = 0
DB = 0
SM = 0
Delta_K
Doxycycline
YMR154C RIM13
ZShift = −0.46
Z lm Score = −1.07
NG = 0
DB = 0
SM = 0
Delta_r
Doxycycline
YMR154C RIM13
ZShift = −0.5
Z lm Score = 1.02
NG = 0
DB = 0
SM = 0
Delta_AUC
Doxycycline

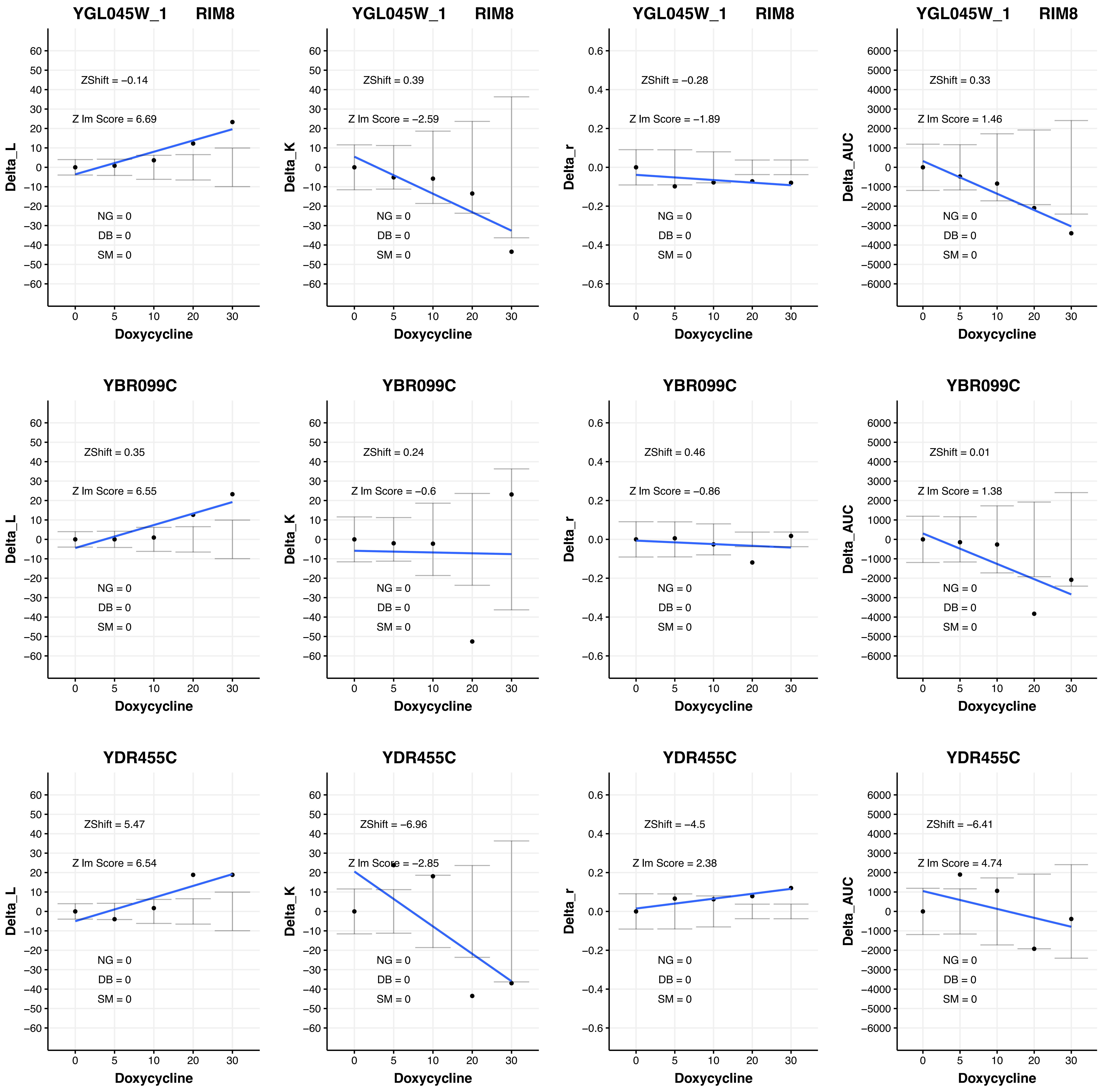
YGL045W_1 RIM8
ZShift = −0.14
Z lm Score = 6.69
NG = 0
DB = 0
SM = 0
Delta_L
Doxycycline
YGL045W_1 RIM8
ZShift = 0.39
Z lm Score = −2.59
NG = 0
DB = 0
SM = 0
Delta_K
Doxycycline
YGL045W_1 RIM8
ZShift = −0.28
Z lm Score = −1.89
NG = 0
DB = 0
SM = 0
Delta_r
Doxycycline
YGL045W_1 RIM8
ZShift = 0.33
Z lm Score = 1.46
NG = 0
DB = 0
SM = 0
Delta_AUC
Doxycycline
YBR099C
ZShift = 0.35
Z lm Score = 6.55
NG = 0
DB = 0
SM = 0
Delta_L
Doxycycline
YBR099C
ZShift = 0.24
Z lm Score = −0.6
NG = 0
DB = 0
SM = 0
Delta_K
Doxycycline
YBR099C
ZShift = 0.46
Z lm Score = −0.86
NG = 0
DB = 0
SM = 0
Delta_r
Doxycycline
YBR099C
ZShift = 0.01
Z lm Score = 1.38
NG = 0
DB = 0
SM = 0
Delta_AUC
Doxycycline
YDR455C
ZShift = 5.47
Z lm Score = 6.54
NG = 0
DB = 0
SM = 0
Delta_L
Doxycycline
YDR455C
ZShift = −6.96
Z lm Score = −2.85
NG = 0
DB = 0
SM = 0
Delta_K
Doxycycline
YDR455C
ZShift = −4.5
Z lm Score = 2.38
NG = 0
DB = 0
SM = 0
Delta_r
Doxycycline
YDR455C
ZShift = −6.41
Z lm Score = 4.74
NG = 0
DB = 0
SM = 0
Delta_AUC
Doxycycline

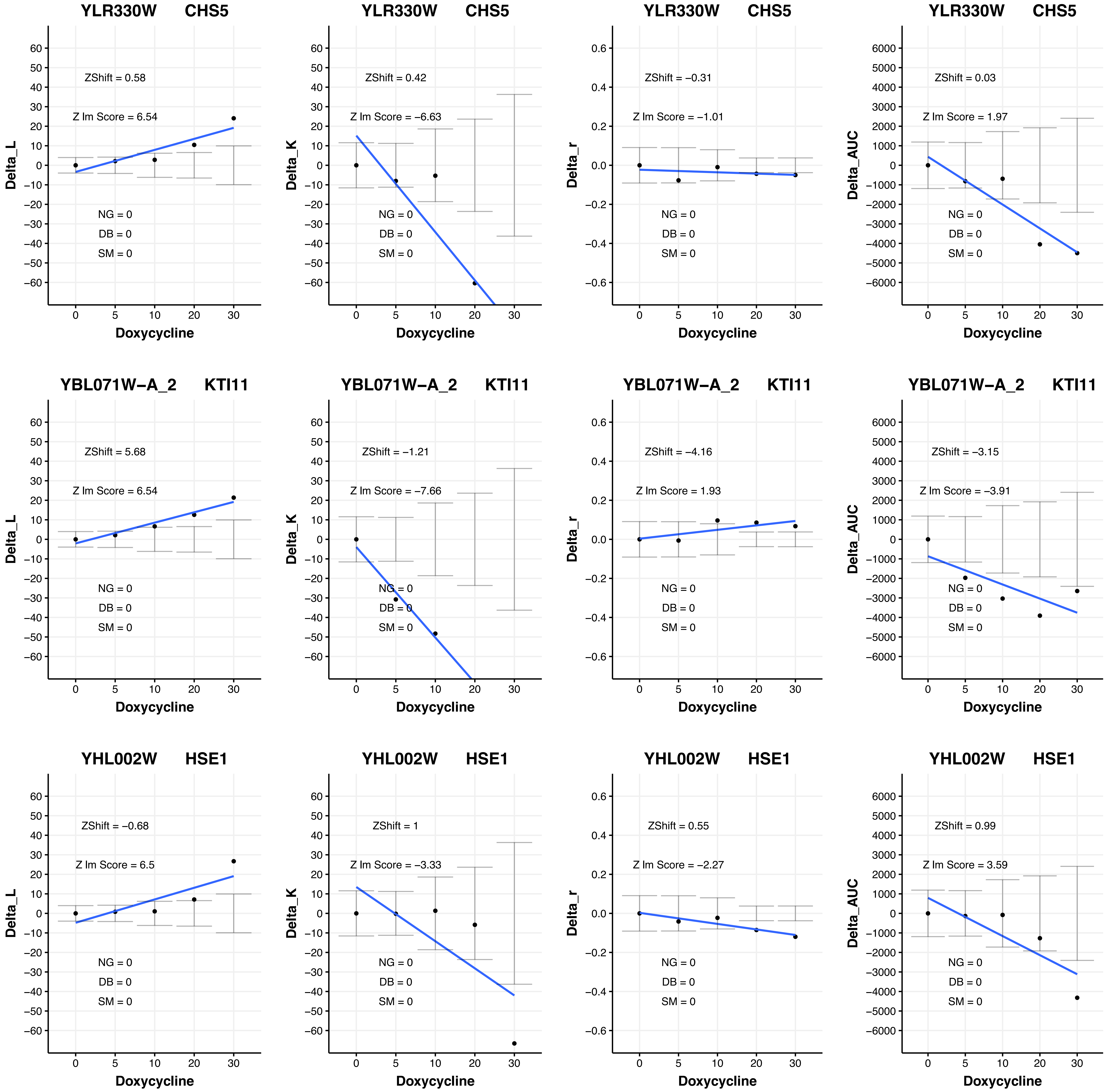
YLR330W CHS5
ZShift = 0.58
Z lm Score = 6.54
NG = 0
DB = 0
SM = 0
Delta_L
Doxycycline
YLR330W CHS5
ZShift = 0.42
Z lm Score = −6.63
NG = 0
DB = 0
SM = 0
Delta_K
Doxycycline
YLR330W CHS5
ZShift = −0.31
Z lm Score = −1.01
NG = 0
DB = 0
SM = 0
Delta_r
Doxycycline
YLR330W CHS5
ZShift = 0.03
Z lm Score = 1.97
NG = 0
DB = 0
SM = 0
Delta_AUC
Doxycycline
YBL071W−A_2 KTI11
ZShift = 5.68
Z lm Score = 6.54
NG = 0
DB = 0
SM = 0
Delta_L
Doxycycline
YBL071W−A_2 KTI11
ZShift = −1.21
Z lm Score = −7.66
NG = 0
DB = 0
SM = 0
Delta_K
Doxycycline
YBL071W−A_2 KTI11
ZShift = −4.16
Z lm Score = 1.93
NG = 0
DB = 0
SM = 0
Delta_r
Doxycycline
YBL071W−A_2 KTI11
ZShift = −3.15
Z lm Score = −3.91
NG = 0
DB = 0
SM = 0
Delta_AUC
Doxycycline
YHL002W HSE1
ZShift = −0.68
Z lm Score = 6.5
NG = 0
DB = 0
SM = 0
Delta_L
Doxycycline
YHL002W HSE1
ZShift = 1
Z lm Score = −3.33
NG = 0
DB = 0
SM = 0
Delta_K
Doxycycline
YHL002W HSE1
ZShift = 0.55
Z lm Score = −2.27
NG = 0
DB = 0
SM = 0
Delta_r
Doxycycline
YHL002W HSE1
ZShift = 0.99
Z lm Score = 3.59
NG = 0
DB = 0
SM = 0
Delta_AUC
Doxycycline

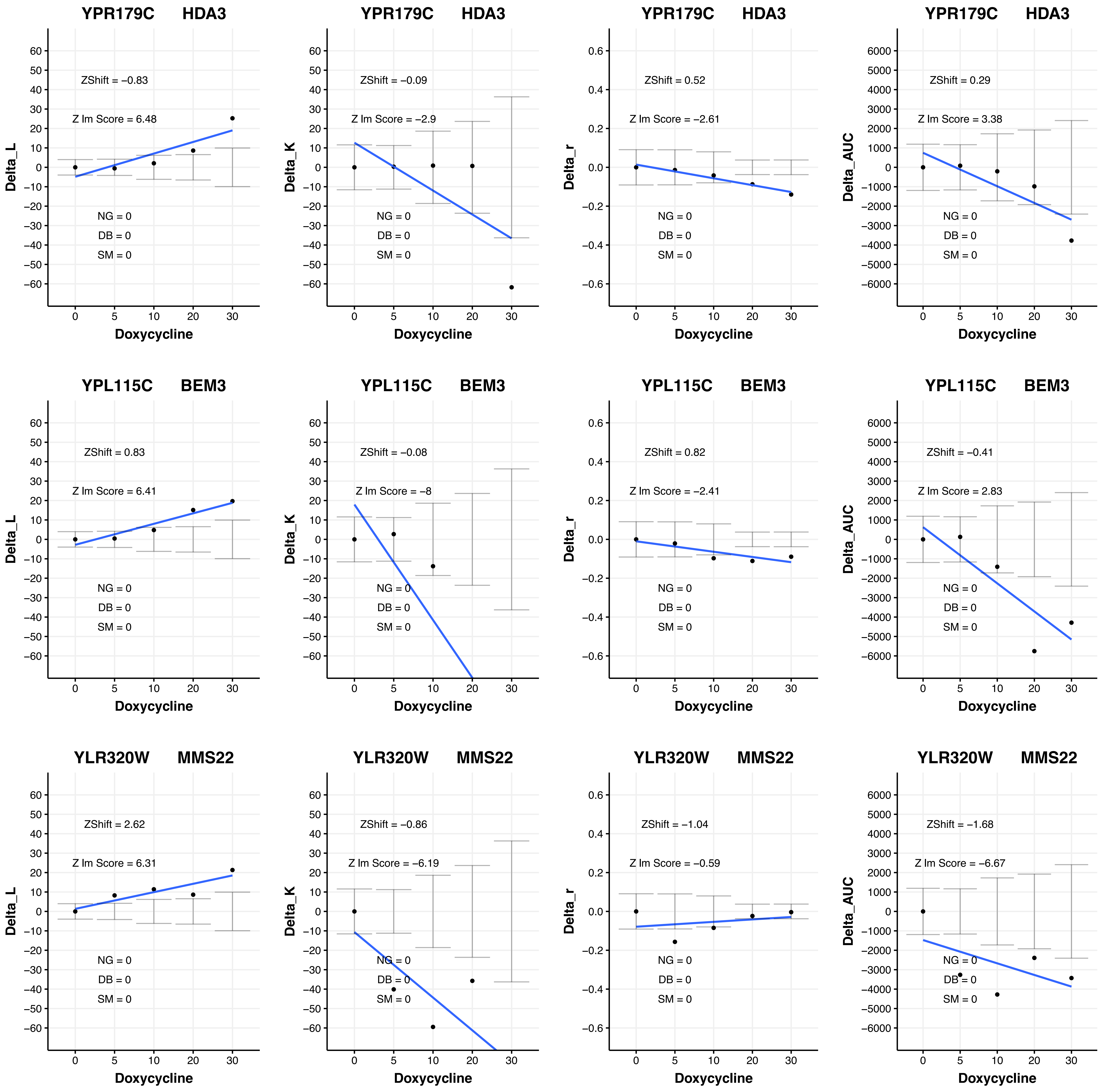

YPR179C HDA3
ZShift = −0.83
Z lm Score = 6.48
NG = 0
DB = 0
SM = 0
Delta_L
Doxycycline
YPR179C HDA3
ZShift = −0.09
Z lm Score = −2.9
NG = 0
DB = 0
SM = 0
Delta_K
Doxycycline
YPR179C HDA3
ZShift = 0.52
Z lm Score = −2.61
NG = 0
DB = 0
SM = 0
Delta_r
Doxycycline
YPR179C HDA3
ZShift = 0.29
Z lm Score = 3.38
NG = 0
DB = 0
SM = 0
Delta_AUC
Doxycycline
YPL115C BEM3
ZShift = 0.83
Z lm Score = 6.41
NG = 0
DB = 0
SM = 0
Delta_L
Doxycycline
YPL115C BEM3
ZShift = −0.08
Z lm Score = −8
NG = 0
DB = 0
SM = 0
Delta_K
Doxycycline
YPL115C BEM3
ZShift = 0.82
Z lm Score = −2.41
NG = 0
DB = 0
SM = 0
Delta_r
Doxycycline
YPL115C BEM3
ZShift = −0.41
Z lm Score = 2.83
NG = 0
DB = 0
SM = 0
Delta_AUC
Doxycycline
YLR320W MMS22
ZShift = 2.62
Z lm Score = 6.31
NG = 0
DB = 0
SM = 0
Delta_L
Doxycycline
YLR320W MMS22
ZShift = −0.86
Z lm Score = −6.19
NG = 0
DB = 0
SM = 0
Delta_K
Doxycycline
YLR320W MMS22
ZShift = −1.04
Z lm Score = −0.59
NG = 0
DB = 0
SM = 0
Delta_r
Doxycycline
YLR320W MMS22
ZShift = −1.68
Z lm Score = −6.67
NG = 0
DB = 0
SM = 0
Delta_AUC
Doxycycline

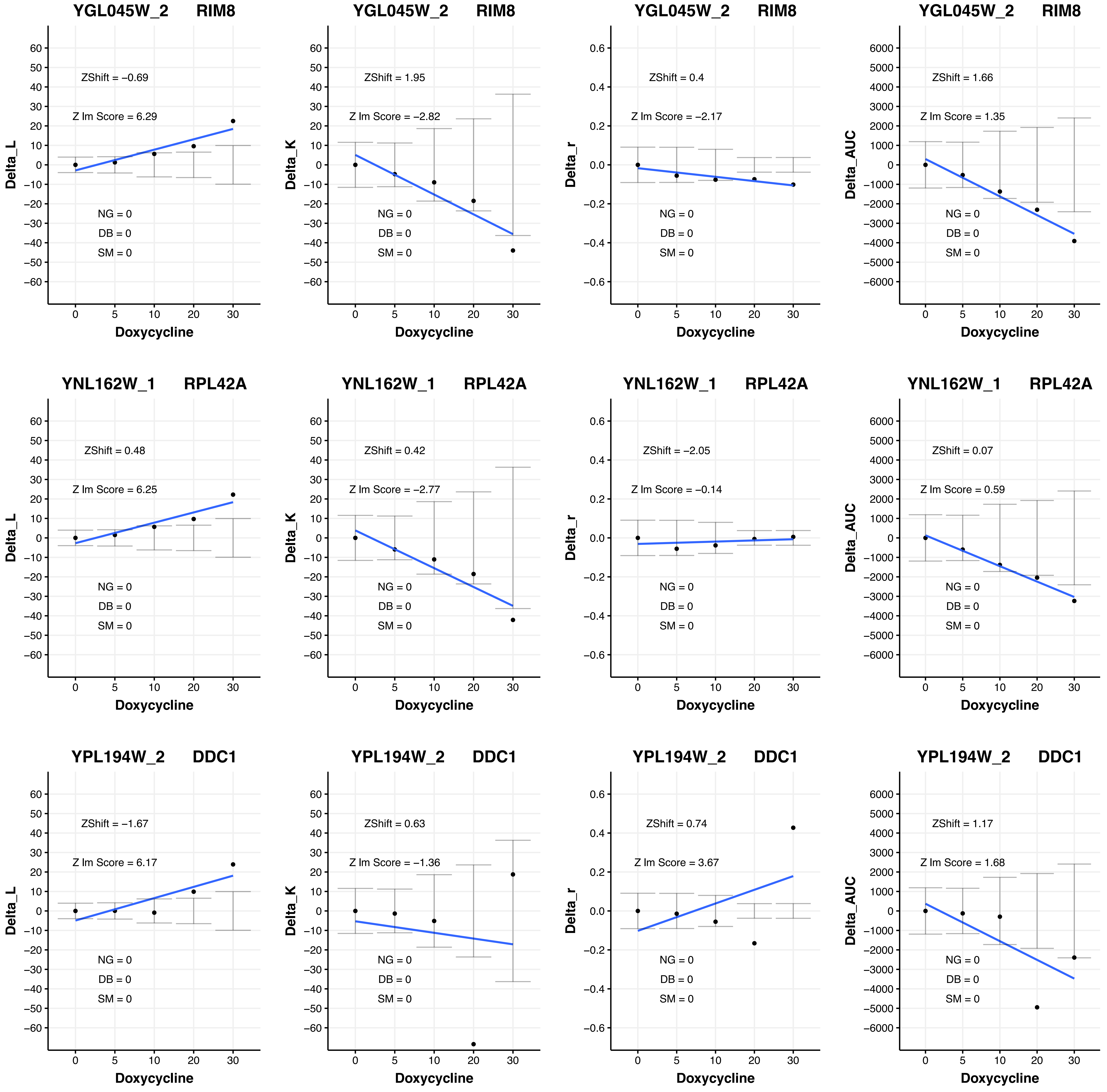
YGL045W_2 RIM8
ZShift = −0.69
Z lm Score = 6.29
NG = 0
DB = 0
SM = 0
Delta_L
Doxycycline
YGL045W_2 RIM8
ZShift = 1.95
Z lm Score = −2.82
NG = 0
DB = 0
SM = 0
Delta_K
Doxycycline
YGL045W_2 RIM8
ZShift = 0.4
Z lm Score = −2.17
NG = 0
DB = 0
SM = 0
Delta_r
Doxycycline
YGL045W_2 RIM8
ZShift = 1.66
Z lm Score = 1.35
NG = 0
DB = 0
SM = 0
Delta_AUC
Doxycycline
YNL162W_1 RPL42A
ZShift = 0.48
Z lm Score = 6.25
NG = 0
DB = 0
SM = 0
Delta_L
Doxycycline
YNL162W_1 RPL42A
ZShift = 0.42
Z lm Score = −2.77
NG = 0
DB = 0
SM = 0
Delta_K
Doxycycline
YNL162W_1 RPL42A
ZShift = −2.05
Z lm Score = −0.14
NG = 0
DB = 0
SM = 0
Delta_r
Doxycycline
YNL162W_1 RPL42A
ZShift = 0.07
Z lm Score = 0.59
NG = 0
DB = 0
SM = 0
Delta_AUC
Doxycycline
YPL194W_2 DDC1
ZShift = −1.67
Z lm Score = 6.17
NG = 0
DB = 0
SM = 0
Delta_L
Doxycycline
YPL194W_2 DDC1
ZShift = 0.63
Z lm Score = −1.36
NG = 0
DB = 0
SM = 0
Delta_K
Doxycycline
YPL194W_2 DDC1
ZShift = 0.74
Z lm Score = 3.67
NG = 0
DB = 0
SM = 0
Delta_r
Doxycycline
YPL194W_2 DDC1
ZShift = 1.17
Z lm Score = 1.68
NG = 0
DB = 0
SM = 0
Delta_AUC
Doxycycline

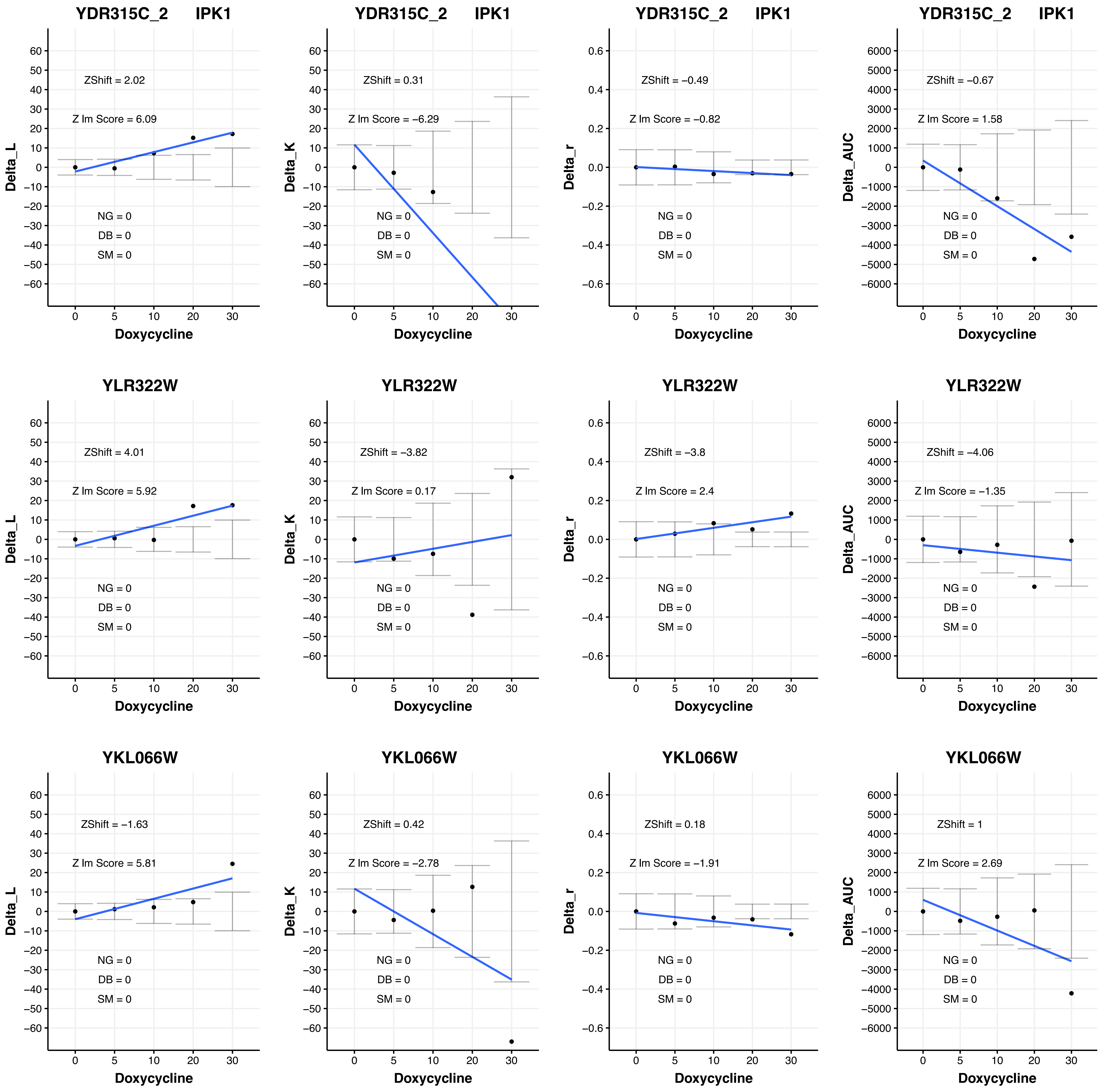
YDR315C_2 IPK1
ZShift = 2.02
Z lm Score = 6.09
NG = 0
DB = 0
SM = 0
Delta_L
Doxycycline
YDR315C_2 IPK1
ZShift = 0.31
Z lm Score = −6.29
NG = 0
DB = 0
SM = 0
Delta_K
Doxycycline
YDR315C_2 IPK1
ZShift = −0.49
Z lm Score = −0.82
NG = 0
DB = 0
SM = 0
Delta_r
Doxycycline
YDR315C_2 IPK1
ZShift = −0.67
Z lm Score = 1.58
NG = 0
DB = 0
SM = 0
Delta_AUC
Doxycycline
YLR322W
ZShift = 4.01
Z lm Score = 5.92
NG = 0
DB = 0
SM = 0
Delta_L
Doxycycline
YLR322W
ZShift = −3.82
Z lm Score = 0.17
NG = 0
DB = 0
SM = 0
Delta_K
Doxycycline
YLR322W
ZShift = −3.8
Z lm Score = 2.4
NG = 0
DB = 0
SM = 0
Delta_r
Doxycycline
YLR322W
ZShift = −4.06
Z lm Score = −1.35
NG = 0
DB = 0
SM = 0
Delta_AUC
Doxycycline
YKL066W
ZShift = −1.63
Z lm Score = 5.81
NG = 0
DB = 0
SM = 0
Delta_L
Doxycycline
YKL066W
ZShift = 0.42
Z lm Score = −2.78
NG = 0
DB = 0
SM = 0
Delta_K
Doxycycline
YKL066W
ZShift = 0.18
Z lm Score = −1.91
NG = 0
DB = 0
SM = 0
Delta_r
Doxycycline
YKL066W
ZShift = 1
Z lm Score = 2.69
NG = 0
DB = 0
SM = 0
Delta_AUC
Doxycycline

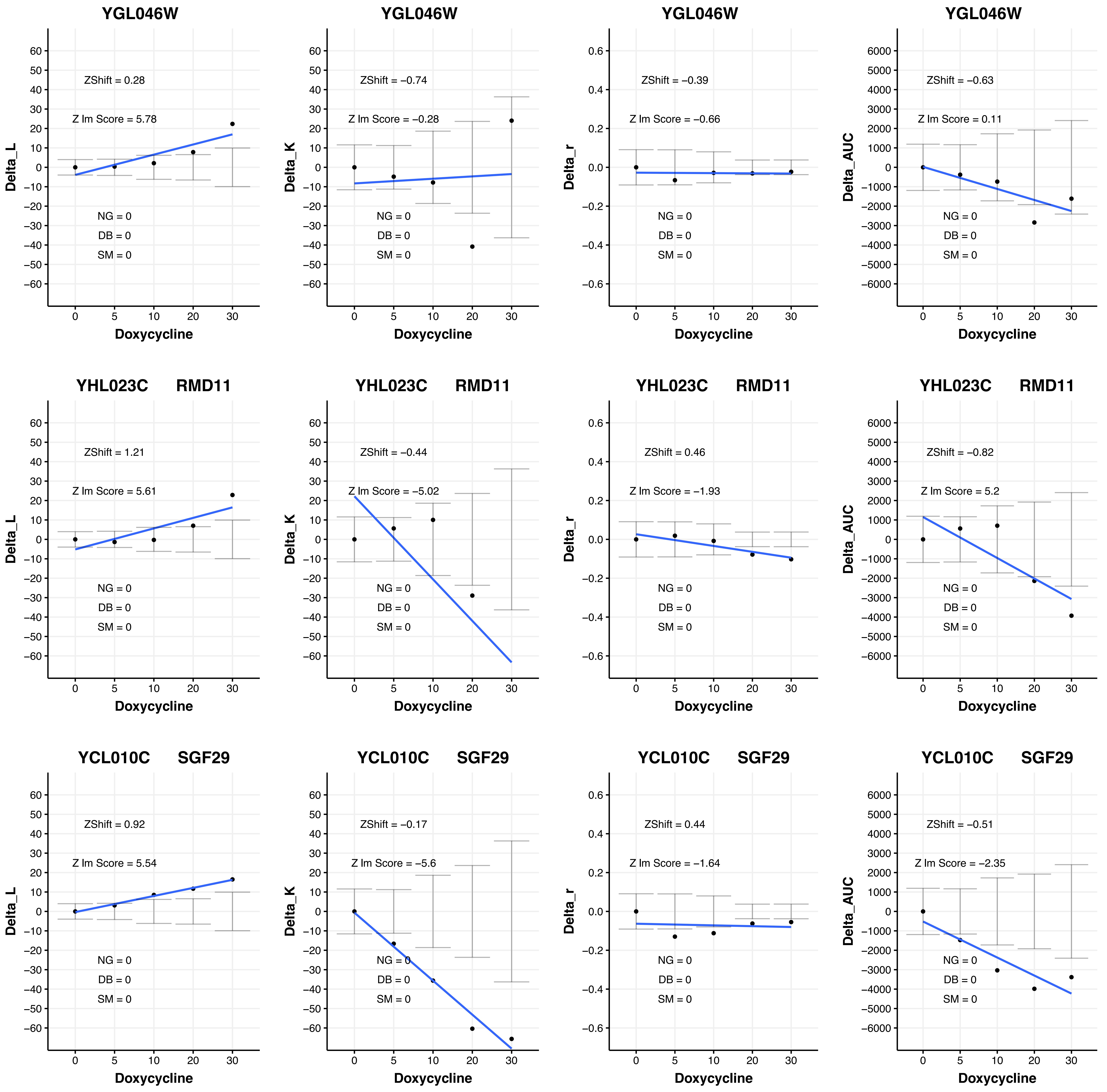
YGL046W
ZShift = 0.28
Z lm Score = 5.78
NG = 0
DB = 0
SM = 0
Delta_L
Doxycycline
YGL046W
ZShift = −0.74
Z lm Score = −0.28
NG = 0
DB = 0
SM = 0
Delta_K
Doxycycline
YGL046W
ZShift = −0.39
Z lm Score = −0.66
NG = 0
DB = 0
SM = 0
Delta_r
Doxycycline
YGL046W
ZShift = −0.63
Z lm Score = 0.11
NG = 0
DB = 0
SM = 0
Delta_AUC
Doxycycline
YHL023C RMD11
ZShift = 1.21
Z lm Score = 5.61
NG = 0
DB = 0
SM = 0
Delta_L
Doxycycline
YHL023C RMD11
ZShift = −0.44
Z lm Score = −5.02
NG = 0
DB = 0
SM = 0
Delta_K
Doxycycline
YHL023C RMD11
ZShift = 0.46
Z lm Score = −1.93
NG = 0
DB = 0
SM = 0
Delta_r
Doxycycline
YHL023C RMD11
ZShift = −0.82
Z lm Score = 5.2
NG = 0
DB = 0
SM = 0
Delta_AUC
Doxycycline
YCL010C SGF29
ZShift = 0.92
Z lm Score = 5.54
NG = 0
DB = 0
SM = 0
Delta_L
Doxycycline
YCL010C SGF29
ZShift = −0.17
Z lm Score = −5.6
NG = 0
DB = 0
SM = 0
Delta_K
Doxycycline
YCL010C SGF29
ZShift = 0.44
Z lm Score = −1.64
NG = 0
DB = 0
SM = 0
Delta_r
Doxycycline
YCL010C SGF29
ZShift = −0.51
Z lm Score = −2.35
NG = 0
DB = 0
SM = 0
Delta_AUC
Doxycycline

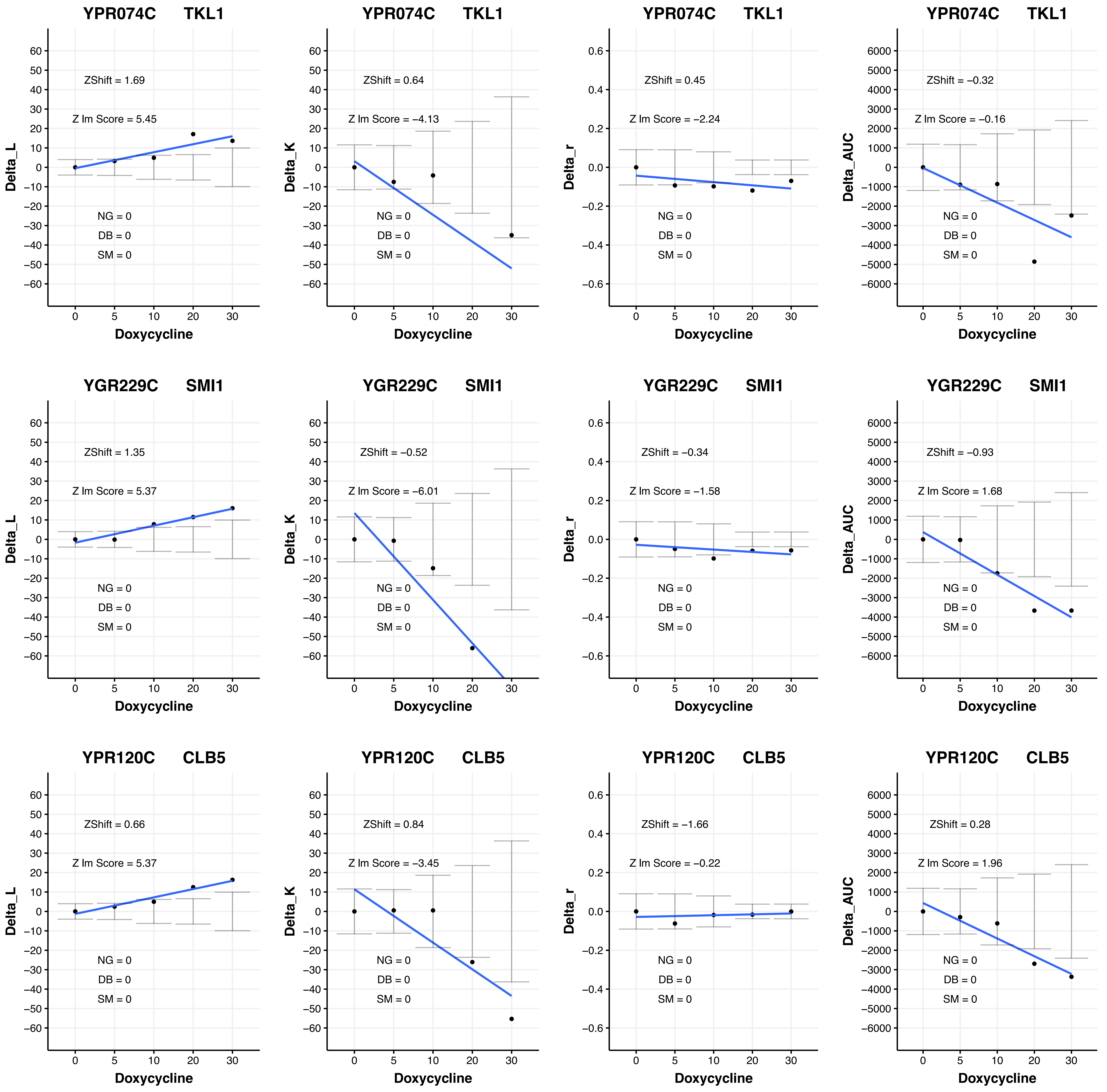

YPR074C TKL1
ZShift = 1.69
Z lm Score = 5.45
NG = 0
DB = 0
SM = 0
Delta_L
Doxycycline
YPR074C TKL1
ZShift = 0.64
Z lm Score = −4.13
NG = 0
DB = 0
SM = 0
Delta_K
Doxycycline
YPR074C TKL1
ZShift = 0.45
Z lm Score = −2.24
NG = 0
DB = 0
SM = 0
Delta_r
Doxycycline
YPR074C TKL1
ZShift = −0.32
Z lm Score = −0.16
NG = 0
DB = 0
SM = 0
Delta_AUC
Doxycycline
YGR229C SMI1
ZShift = 1.35
Z lm Score = 5.37
NG = 0
DB = 0
SM = 0
Delta_L
Doxycycline
YGR229C SMI1
ZShift = −0.52
Z lm Score = −6.01
NG = 0
DB = 0
SM = 0
Delta_K
Doxycycline
YGR229C SMI1
ZShift = −0.34
Z lm Score = −1.58
NG = 0
DB = 0
SM = 0
Delta_r
Doxycycline
YGR229C SMI1
ZShift = −0.93
Z lm Score = 1.68
NG = 0
DB = 0
SM = 0
Delta_AUC
Doxycycline
YPR120C CLB5
ZShift = 0.66
Z lm Score = 5.37
NG = 0
DB = 0
SM = 0
Delta_L
Doxycycline
YPR120C CLB5
ZShift = 0.84
Z lm Score = −3.45
NG = 0
DB = 0
SM = 0
Delta_K
Doxycycline
YPR120C CLB5
ZShift = −1.66
Z lm Score = −0.22
NG = 0
DB = 0
SM = 0
Delta_r
Doxycycline
YPR120C CLB5
ZShift = 0.28
Z lm Score = 1.96
NG = 0
DB = 0
SM = 0
Delta_AUC
Doxycycline

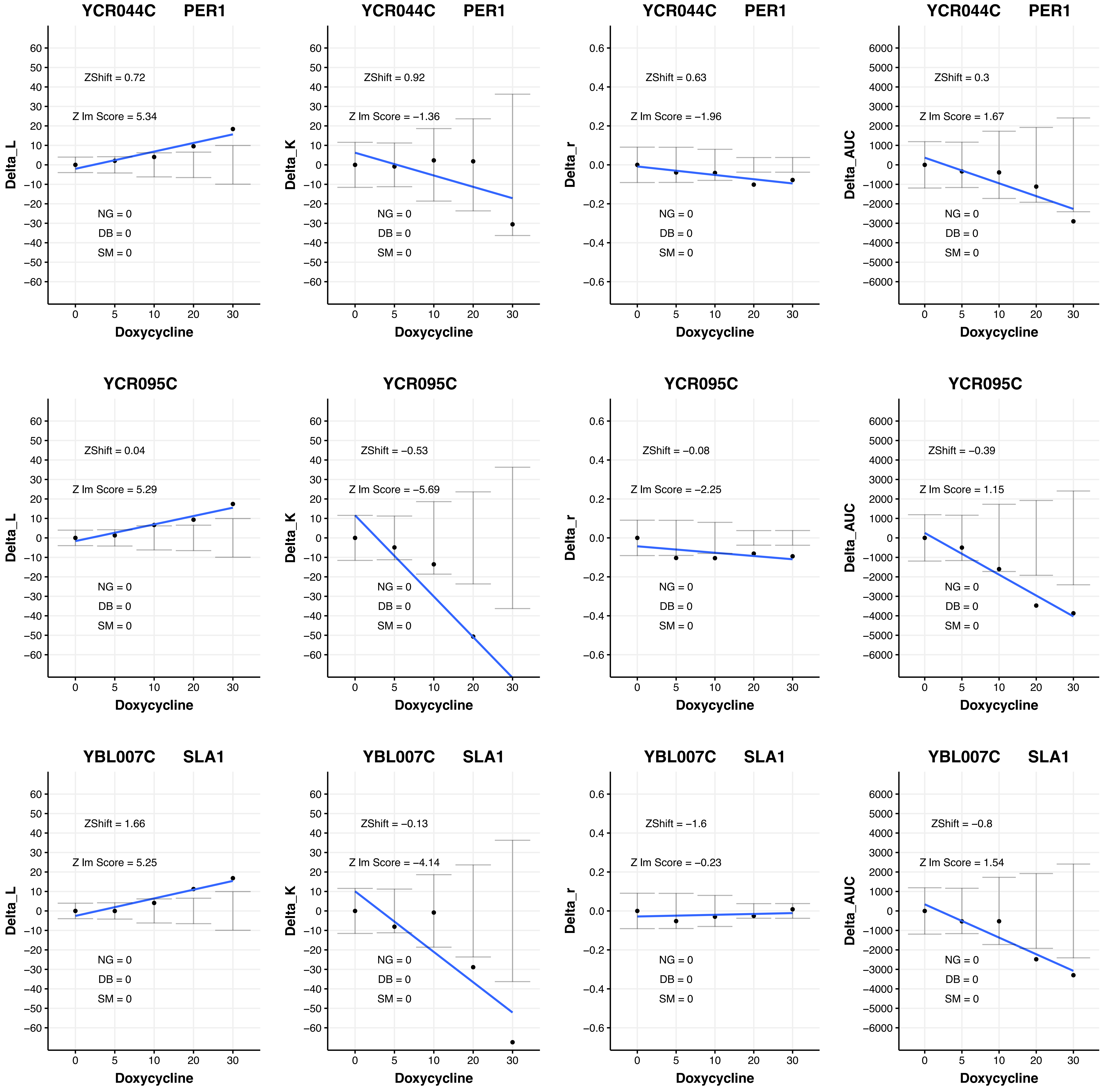

YCR044C PER1
ZShift = 0.72
Z lm Score = 5.34
NG = 0
DB = 0
SM = 0
Delta_L
Doxycycline
YCR044C PER1
ZShift = 0.92
Z lm Score = −1.36
NG = 0
DB = 0
SM = 0
Delta_K
Doxycycline
YCR044C PER1
ZShift = 0.63
Z lm Score = −1.96
NG = 0
DB = 0
SM = 0
Delta_r
Doxycycline
YCR044C PER1
ZShift = 0.3
Z lm Score = 1.67
NG = 0
DB = 0
SM = 0
Delta_AUC
Doxycycline
YCR095C
ZShift = 0.04
Z lm Score = 5.29
NG = 0
DB = 0
SM = 0
Delta_L
Doxycycline
YCR095C
ZShift = −0.53
Z lm Score = −5.69
NG = 0
DB = 0
SM = 0
Delta_K
Doxycycline
YCR095C
ZShift = −0.08
Z lm Score = −2.25
NG = 0
DB = 0
SM = 0
Delta_r
Doxycycline
YCR095C
ZShift = −0.39
Z lm Score = 1.15
NG = 0
DB = 0
SM = 0
Delta_AUC
Doxycycline
YBL007C SLA1
ZShift = 1.66
Z lm Score = 5.25
NG = 0
DB = 0
SM = 0
Delta_L
Doxycycline
YBL007C SLA1
ZShift = −0.13
Z lm Score = −4.14
NG = 0
DB = 0
SM = 0
Delta_K
Doxycycline
YBL007C SLA1
ZShift = −1.6
Z lm Score = −0.23
NG = 0
DB = 0
SM = 0
Delta_r
Doxycycline
YBL007C SLA1
ZShift = −0.8
Z lm Score = 1.54
NG = 0
DB = 0
SM = 0
Delta_AUC
Doxycycline

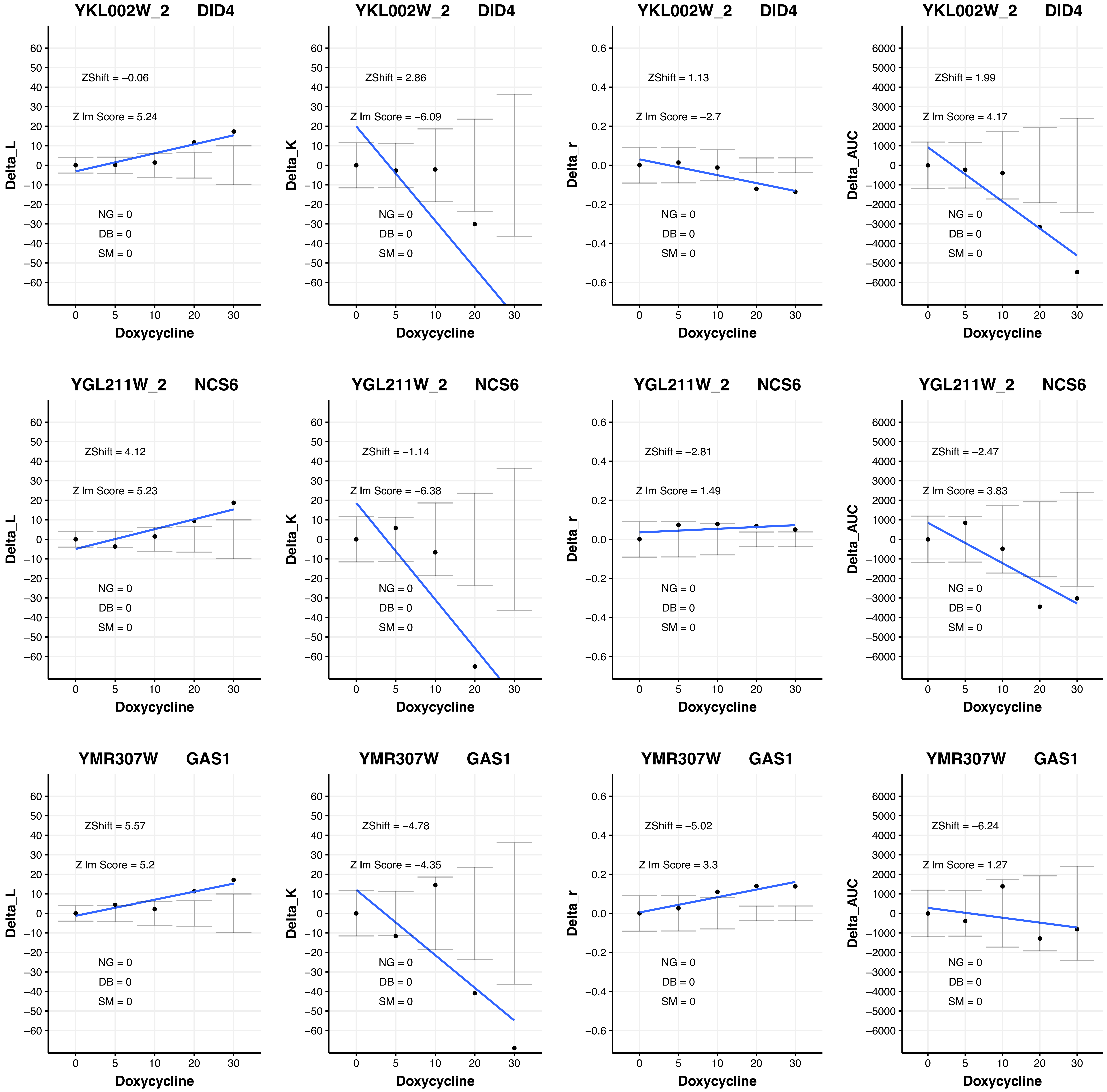
YKL002W_2 DID4
ZShift = −0.06
Z lm Score = 5.24
NG = 0
DB = 0
SM = 0
Delta_L
Doxycycline
YKL002W_2 DID4
ZShift = 2.86
Z lm Score = −6.09
NG = 0
DB = 0
SM = 0
Delta_K
Doxycycline
YKL002W_2 DID4
ZShift = 1.13
Z lm Score = −2.7
NG = 0
DB = 0
SM = 0
Delta_r
Doxycycline
YKL002W_2 DID4
ZShift = 1.99
Z lm Score = 4.17
NG = 0
DB = 0
SM = 0
Delta_AUC
Doxycycline
YGL211W_2 NCS6
ZShift = 4.12
Z lm Score = 5.23
NG = 0
DB = 0
SM = 0
Delta_L
Doxycycline
YGL211W_2 NCS6
ZShift = −1.14
Z lm Score = −6.38
NG = 0
DB = 0
SM = 0
Delta_K
Doxycycline
YGL211W_2 NCS6
ZShift = −2.81
Z lm Score = 1.49
NG = 0
DB = 0
SM = 0
Delta_r
Doxycycline
YGL211W_2 NCS6
ZShift = −2.47
Z lm Score = 3.83
NG = 0
DB = 0
SM = 0
Delta_AUC
Doxycycline
YMR307W GAS1
ZShift = 5.57
Z lm Score = 5.2
NG = 0
DB = 0
SM = 0
Delta_L
Doxycycline
YMR307W GAS1
ZShift = −4.78
Z lm Score = −4.35
NG = 0
DB = 0
SM = 0
Delta_K
Doxycycline
YMR307W GAS1
ZShift = −5.02
Z lm Score = 3.3
NG = 0
DB = 0
SM = 0
Delta_r
Doxycycline
YMR307W GAS1
ZShift = −6.24
Z lm Score = 1.27
NG = 0
DB = 0
SM = 0
Delta_AUC
Doxycycline

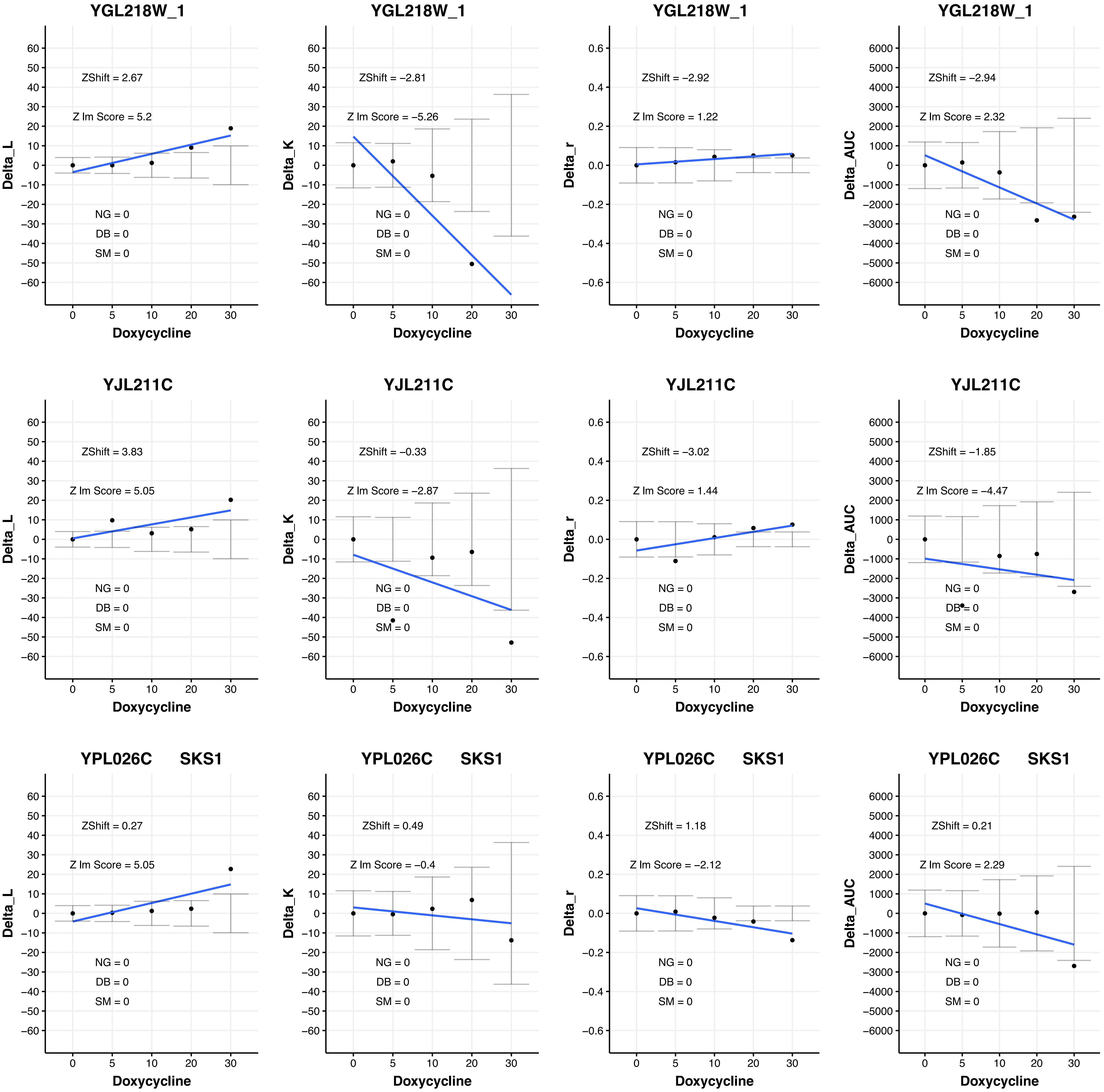
YGL218W_1
ZShift = 2.67
Z lm Score = 5.2
NG = 0
DB = 0
SM = 0
Delta_L
Doxycycline
YGL218W_1
ZShift = −2.81
Z lm Score = −5.26
NG = 0
DB = 0
SM = 0
Delta_K
Doxycycline
YGL218W_1
ZShift = −2.92
Z lm Score = 1.22
NG = 0
DB = 0
SM = 0
Delta_r
Doxycycline
YGL218W_1
ZShift = −2.94
Z lm Score = 2.32
NG = 0
DB = 0
SM = 0
Delta_AUC
Doxycycline
YJL211C
ZShift = 3.83
Z lm Score = 5.05
NG = 0
DB = 0
SM = 0
Delta_L
Doxycycline
YJL211C
ZShift = −0.33
Z lm Score = −2.87
NG = 0
DB = 0
SM = 0
Delta_K
Doxycycline
YJL211C
ZShift = −3.02
Z lm Score = 1.44
NG = 0
DB = 0
SM = 0
Delta_r
Doxycycline
YJL211C
ZShift = −1.85
Z lm Score = −4.47
NG = 0
DB = 0
SM = 0
Delta_AUC
Doxycycline
YPL026C SKS1
ZShift = 0.27
Z lm Score = 5.05
NG = 0
DB = 0
SM = 0
Delta_L
Doxycycline
YPL026C SKS1
ZShift = 0.49
Z lm Score = −0.4
NG = 0
DB = 0
SM = 0
Delta_K
Doxycycline
YPL026C SKS1
ZShift = 1.18
Z lm Score = −2.12
NG = 0
DB = 0
SM = 0
Delta_r
Doxycycline
YPL026C SKS1
ZShift = 0.21
Z lm Score = 2.29
NG = 0
DB = 0
SM = 0
Delta_AUC
Doxycycline

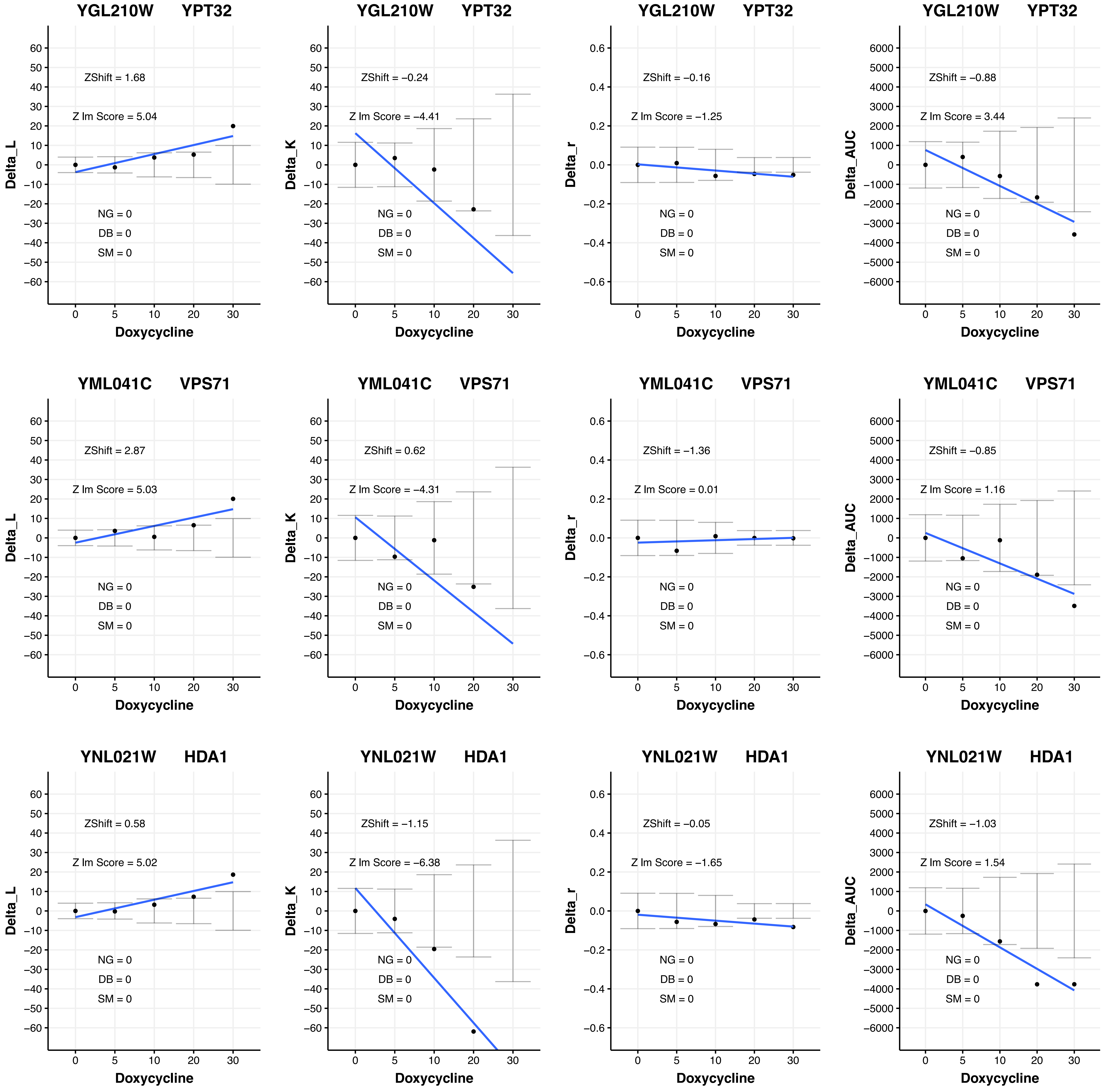

YGL210W YPT32
ZShift = 1.68
Z lm Score = 5.04
NG = 0
DB = 0
SM = 0
Delta_L
Doxycycline
YGL210W YPT32
ZShift = −0.24
Z lm Score = −4.41
NG = 0
DB = 0
SM = 0
Delta_K
Doxycycline
YGL210W YPT32
ZShift = −0.16
Z lm Score = −1.25
NG = 0
DB = 0
SM = 0
Delta_r
Doxycycline
YGL210W YPT32
ZShift = −0.88
Z lm Score = 3.44
NG = 0
DB = 0
SM = 0
Delta_AUC
Doxycycline
YML041C VPS71
ZShift = 2.87
Z lm Score = 5.03
NG = 0
DB = 0
SM = 0
Delta_L
Doxycycline
YML041C VPS71
ZShift = 0.62
Z lm Score = −4.31
NG = 0
DB = 0
SM = 0
Delta_K
Doxycycline
YML041C VPS71
ZShift = −1.36
Z lm Score = 0.01
NG = 0
DB = 0
SM = 0
Delta_r
Doxycycline
YML041C VPS71
ZShift = −0.85
Z lm Score = 1.16
NG = 0
DB = 0
SM = 0
Delta_AUC
Doxycycline
YNL021W HDA1
ZShift = 0.58
Z lm Score = 5.02
NG = 0
DB = 0
SM = 0
Delta_L
Doxycycline
YNL021W HDA1
ZShift = −1.15
Z lm Score = −6.38
NG = 0
DB = 0
SM = 0
Delta_K
Doxycycline
YNL021W HDA1
ZShift = −0.05
Z lm Score = −1.65
NG = 0
DB = 0
SM = 0
Delta_r
Doxycycline
YNL021W HDA1
ZShift = −1.03
Z lm Score = 1.54
NG = 0
DB = 0
SM = 0
Delta_AUC
Doxycycline

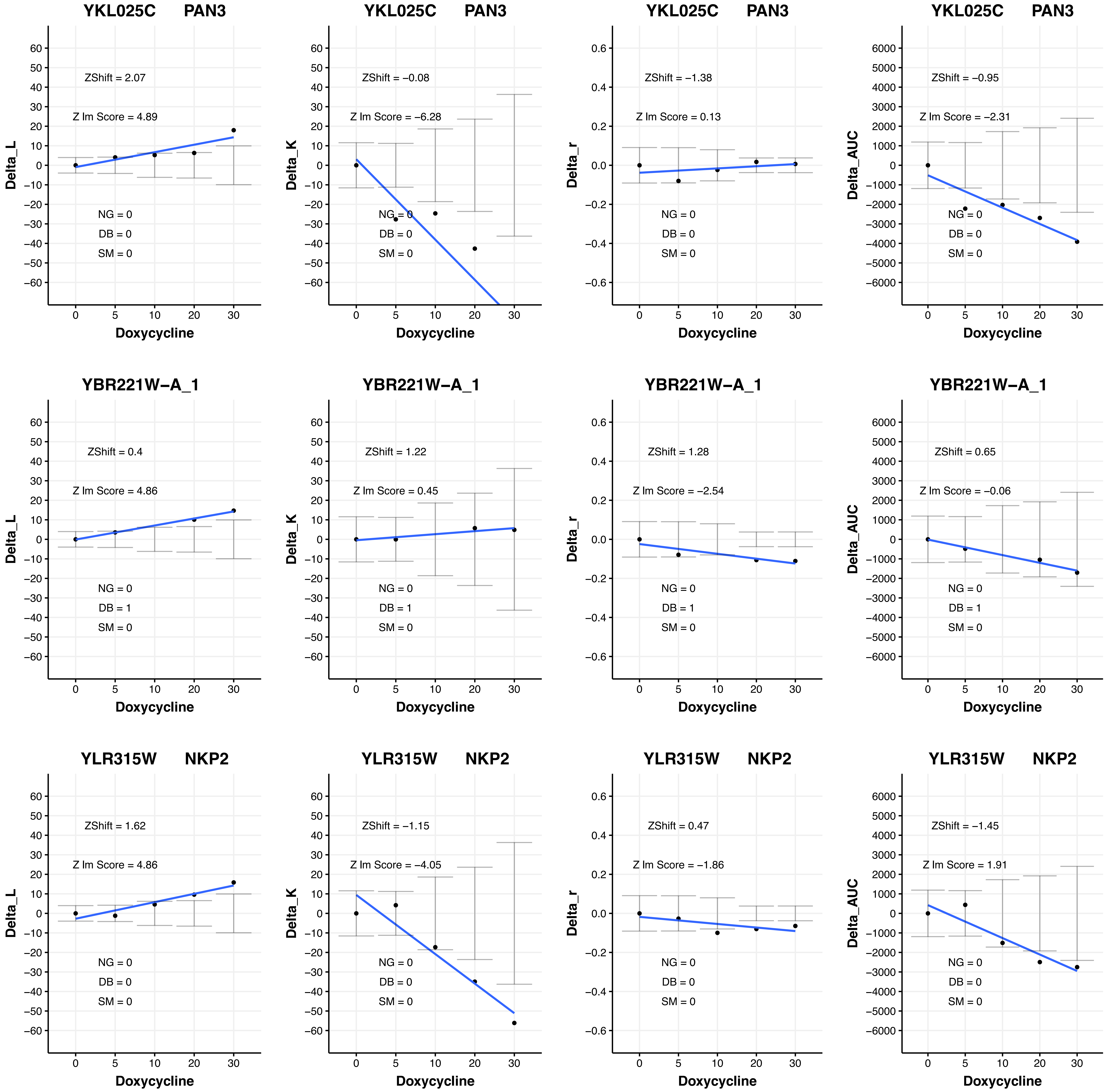

YKL025C PAN3
ZShift = 2.07
Z lm Score = 4.89
NG = 0
DB = 0
SM = 0
Delta_L
Doxycycline
YKL025C PAN3
ZShift = −0.08
Z lm Score = −6.28
NG = 0
DB = 0
SM = 0
Delta_K
Doxycycline
YKL025C PAN3
ZShift = −1.38
Z lm Score = 0.13
NG = 0
DB = 0
SM = 0
Delta_r
Doxycycline
YKL025C PAN3
ZShift = −0.95
Z lm Score = −2.31
NG = 0
DB = 0
SM = 0
Delta_AUC
Doxycycline
YBR221W−A_1
ZShift = 0.4
Z lm Score = 4.86
NG = 0
DB = 1
SM = 0
Delta_L
Doxycycline
YBR221W−A_1
ZShift = 1.22
Z lm Score = 0.45
NG = 0
DB = 1
SM = 0
Delta_K
Doxycycline
YBR221W−A_1
ZShift = 1.28
Z lm Score = −2.54
NG = 0
DB = 1
SM = 0
Delta_r
Doxycycline
YBR221W−A_1
ZShift = 0.65
Z lm Score = −0.06
NG = 0
DB = 1
SM = 0
Delta_AUC
Doxycycline
YLR315W NKP2
ZShift = 1.62
Z lm Score = 4.86
NG = 0
DB = 0
SM = 0
Delta_L
Doxycycline
YLR315W NKP2
ZShift = −1.15
Z lm Score = −4.05
NG = 0
DB = 0
SM = 0
Delta_K
Doxycycline
YLR315W NKP2
ZShift = 0.47
Z lm Score = −1.86
NG = 0
DB = 0
SM = 0
Delta_r
Doxycycline
YLR315W NKP2
ZShift = −1.45
Z lm Score = 1.91
NG = 0
DB = 0
SM = 0
Delta_AUC
Doxycycline

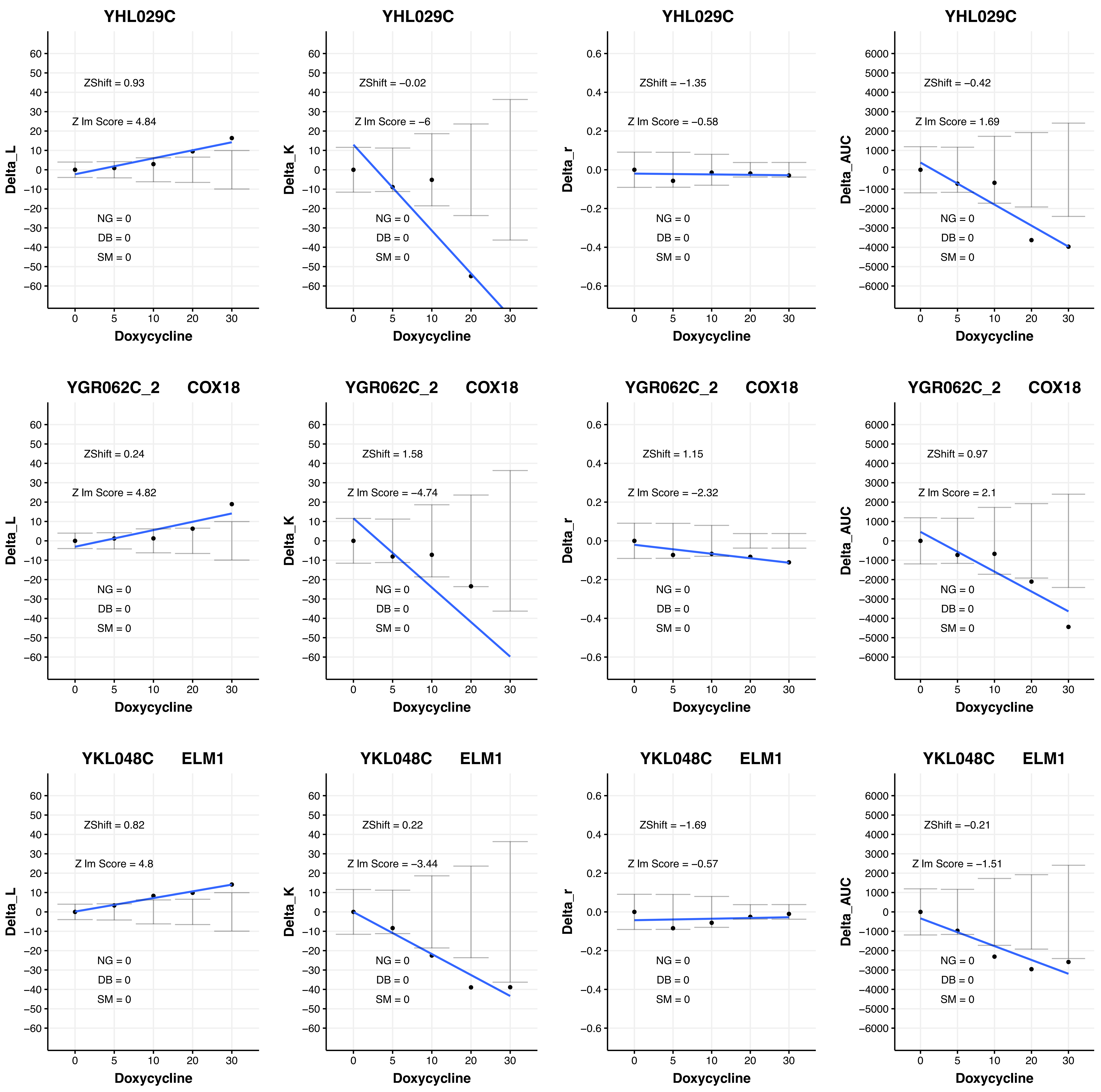
YHL029C
Delta_L
ZShift = 0.93
Z lm Score = 4.84
NG = 0
DB = 0
SM = 0
Doxycycline
YHL029C
Delta_K
ZShift = −0.02
Z lm Score = −6
NG = 0
DB = 0
SM = 0
Doxycycline
YHL029C
Delta_r
ZShift = −1.35
Z lm Score = −0.58
NG = 0
DB = 0
SM = 0
Doxycycline
YHL029C
Delta_AUC
ZShift = −0.42
Z lm Score = 1.69
NG = 0
DB = 0
SM = 0
Doxycycline
YGR062C_2 COX18
Delta_L
ZShift = 0.24
Z lm Score = 4.82
NG = 0
DB = 0
SM = 0
Doxycycline
YGR062C_2 COX18
Delta_K
ZShift = 1.58
Z lm Score = −4.74
NG = 0
DB = 0
SM = 0
Doxycycline
YGR062C_2 COX18
Delta_r
ZShift = 1.15
Z lm Score = −2.32
NG = 0
DB = 0
SM = 0
Doxycycline
YGR062C_2 COX18
Delta_AUC
ZShift = 0.97
Z lm Score = 2.1
NG = 0
DB = 0
SM = 0
Doxycycline
YKL048C ELM1
Delta_L
ZShift = 0.82
Z lm Score = 4.8
NG = 0
DB = 0
SM = 0
Doxycycline
YKL048C ELM1
Delta_K
ZShift = 0.22
Z lm Score = −3.44
NG = 0
DB = 0
SM = 0
Doxycycline
YKL048C ELM1
Delta_r
ZShift = −1.69
Z lm Score = −0.57
NG = 0
DB = 0
SM = 0
Doxycycline
YKL048C ELM1
Delta_AUC
ZShift = −0.21
Z lm Score = −1.51
NG = 0
DB = 0
SM = 0
Doxycycline

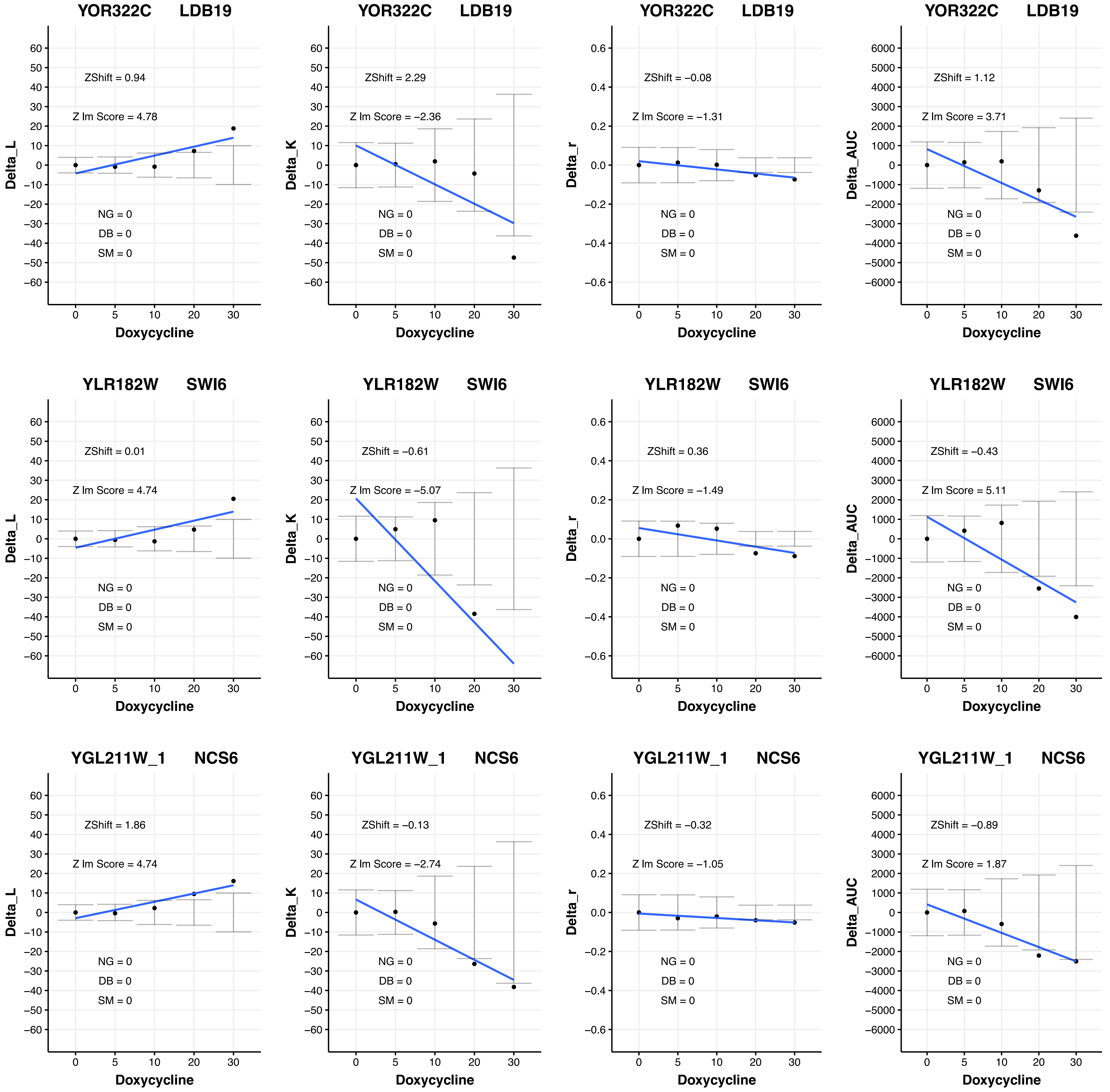
YOR322C LDB19
ZShift = 0.94
Z lm Score = 4.78
NG = 0
DB = 0
SM = 0
Delta_L
Doxycycline
YOR322C LDB19
ZShift = 2.29
Z lm Score = −2.36
NG = 0
DB = 0
SM = 0
Delta_K
Doxycycline
YOR322C LDB19
ZShift = −0.08
Z lm Score = −1.31
NG = 0
DB = 0
SM = 0
Delta_r
Doxycycline
YOR322C LDB19
ZShift = 1.12
Z lm Score = 3.71
NG = 0
DB = 0
SM = 0
Delta_AUC
Doxycycline
YLR182W SWI6
ZShift = 0.01
Z lm Score = 4.74
NG = 0
DB = 0
SM = 0
Delta_L
Doxycycline
YLR182W SWI6
ZShift = −0.61
Z lm Score = −5.07
NG = 0
DB = 0
SM = 0
Delta_K
Doxycycline
YLR182W SWI6
ZShift = 0.36
Z lm Score = −1.49
NG = 0
DB = 0
SM = 0
Delta_r
Doxycycline
YLR182W SWI6
ZShift = −0.43
Z lm Score = 5.11
NG = 0
DB = 0
SM = 0
Delta_AUC
Doxycycline
YGL211W_1 NCS6
ZShift = 1.86
Z lm Score = 4.74
NG = 0
DB = 0
SM = 0
Delta_L
Doxycycline
YGL211W_1 NCS6
ZShift = −0.13
Z lm Score = −2.74
NG = 0
DB = 0
SM = 0
Delta_K
Doxycycline
YGL211W_1 NCS6
ZShift = −0.32
Z lm Score = −1.05
NG = 0
DB = 0
SM = 0
Delta_r
Doxycycline
YGL211W_1 NCS6
ZShift = −0.89
Z lm Score = 1.87
NG = 0
DB = 0
SM = 0
Delta_AUC
Doxycycline

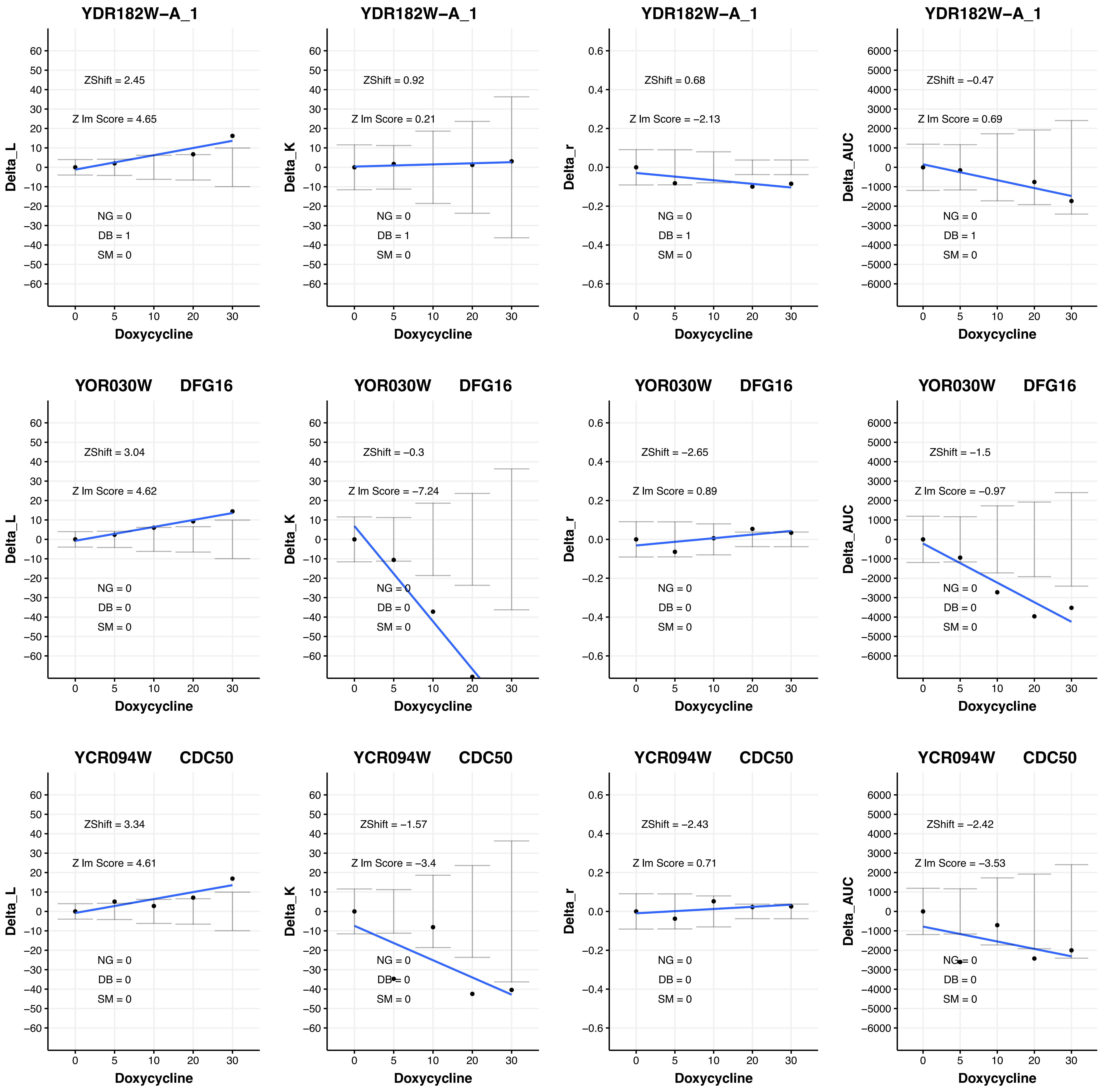

YDR182W−A_1
ZShift = 2.45
Z lm Score = 4.65
NG = 0
DB = 1
SM = 0
Delta_L
Doxycycline
YDR182W−A_1
ZShift = 0.92
Z lm Score = 0.21
NG = 0
DB = 1
SM = 0
Delta_K
Doxycycline
YDR182W−A_1
ZShift = 0.68
Z lm Score = −2.13
NG = 0
DB = 1
SM = 0
Delta_r
Doxycycline
YDR182W−A_1
ZShift = −0.47
Z lm Score = 0.69
NG = 0
DB = 1
SM = 0
Delta_AUC
Doxycycline
YOR030W DFG16
ZShift = 3.04
Z lm Score = 4.62
NG = 0
DB = 0
SM = 0
Delta_L
Doxycycline
YOR030W DFG16
ZShift = −0.3
Z lm Score = −7.24
NG = 0
DB = 0
SM = 0
Delta_K
Doxycycline
YOR030W DFG16
ZShift = −2.65
Z lm Score = 0.89
NG = 0
DB = 0
SM = 0
Delta_r
Doxycycline
YOR030W DFG16
ZShift = −1.5
Z lm Score = −0.97
NG = 0
DB = 0
SM = 0
Delta_AUC
Doxycycline
YCR094W CDC50
ZShift = 3.34
Z lm Score = 4.61
NG = 0
DB = 0
SM = 0
Delta_L
Doxycycline
YCR094W CDC50
ZShift = −1.57
Z lm Score = −3.4
NG = 0
DB = 0
SM = 0
Delta_K
Doxycycline
YCR094W CDC50
ZShift = −2.43
Z lm Score = 0.71
NG = 0
DB = 0
SM = 0
Delta_r
Doxycycline
YCR094W CDC50
ZShift = −2.42
Z lm Score = −3.53
NG = 0
DB = 0
SM = 0
Delta_AUC
Doxycycline

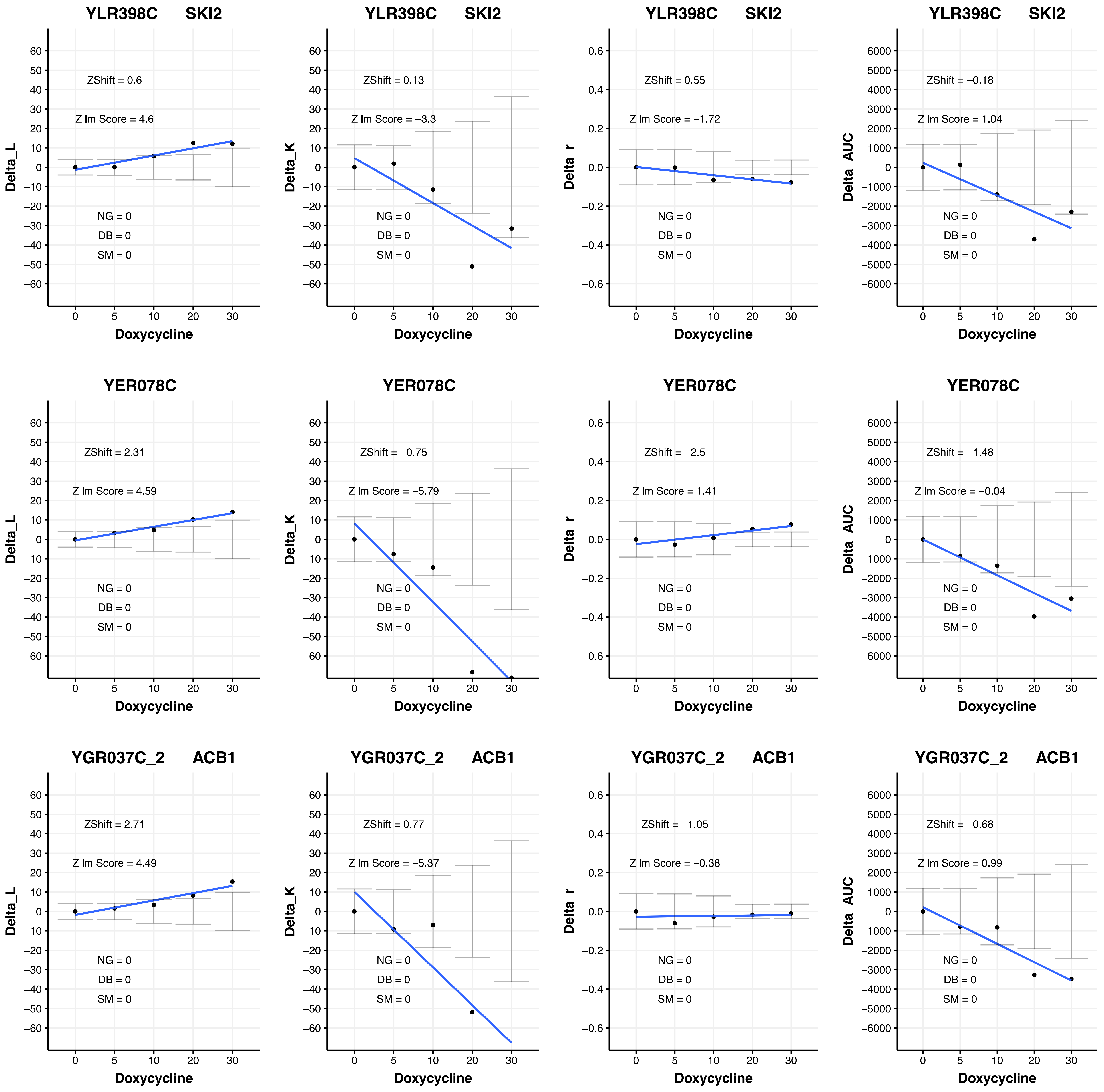
YLR398C SKI2
ZShift = 0.6
Z lm Score = 4.6
NG = 0
DB = 0
SM = 0
Delta_L
Doxycycline
YLR398C SKI2
ZShift = 0.13
Z lm Score = −3.3
NG = 0
DB = 0
SM = 0
Delta_K
Doxycycline
YLR398C SKI2
ZShift = 0.55
Z lm Score = −1.72
NG = 0
DB = 0
SM = 0
Delta_r
Doxycycline
YLR398C SKI2
ZShift = −0.18
Z lm Score = 1.04
NG = 0
DB = 0
SM = 0
Delta_AUC
Doxycycline
YER078C
ZShift = 2.31
Z lm Score = 4.59
NG = 0
DB = 0
SM = 0
Delta_L
Doxycycline
YER078C
ZShift = −0.75
Z lm Score = −5.79
NG = 0
DB = 0
SM = 0
Delta_K
Doxycycline
YER078C
ZShift = −2.5
Z lm Score = 1.41
NG = 0
DB = 0
SM = 0
Delta_r
Doxycycline
YER078C
ZShift = −1.48
Z lm Score = −0.04
NG = 0
DB = 0
SM = 0
Delta_AUC
Doxycycline
YGR037C_2 ACB1
ZShift = 2.71
Z lm Score = 4.49
NG = 0
DB = 0
SM = 0
Delta_L
Doxycycline
YGR037C_2 ACB1
ZShift = 0.77
Z lm Score = −5.37
NG = 0
DB = 0
SM = 0
Delta_K
Doxycycline
YGR037C_2 ACB1
ZShift = −1.05
Z lm Score = −0.38
NG = 0
DB = 0
SM = 0
Delta_r
Doxycycline
YGR037C_2 ACB1
ZShift = −0.68
Z lm Score = 0.99
NG = 0
DB = 0
SM = 0
Delta_AUC
Doxycycline

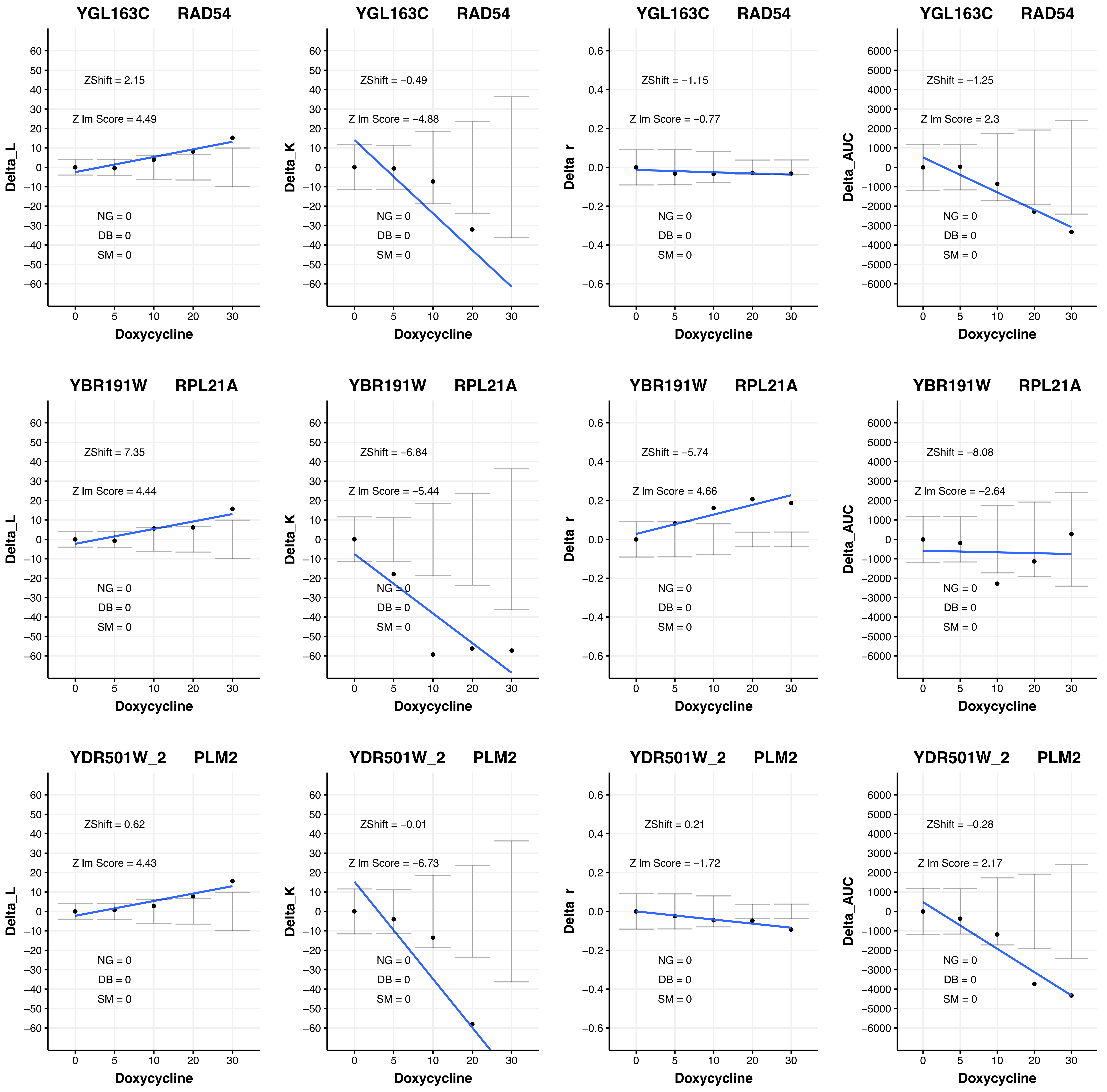
YGL163C RAD54
ZShift = 2.15
Z lm Score = 4.49
NG = 0
DB = 0
SM = 0
Delta_L
Doxycycline
YGL163C RAD54
ZShift = −0.49
Z lm Score = −4.88
NG = 0
DB = 0
SM = 0
Delta_K
Doxycycline
YGL163C RAD54
ZShift = −1.15
Z lm Score = −0.77
NG = 0
DB = 0
SM = 0
Delta_r
Doxycycline
YGL163C RAD54
ZShift = −1.25
Z lm Score = 2.3
NG = 0
DB = 0
SM = 0
Delta_AUC
Doxycycline
YBR191W RPL21A
ZShift = 7.35
Z lm Score = 4.44
NG = 0
DB = 0
SM = 0
Delta_L
Doxycycline
YBR191W RPL21A
ZShift = −6.84
Z lm Score = −5.44
NG = 0
DB = 0
SM = 0
Delta_K
Doxycycline
YBR191W RPL21A
ZShift = −5.74
Z lm Score = 4.66
NG = 0
DB = 0
SM = 0
Delta_r
Doxycycline
YBR191W RPL21A
ZShift = −8.08
Z lm Score = −2.64
NG = 0
DB = 0
SM = 0
Delta_AUC
Doxycycline
YDR501W_2 PLM2
ZShift = 0.62
Z lm Score = 4.43
NG = 0
DB = 0
SM = 0
Delta_L
Doxycycline
YDR501W_2 PLM2
ZShift = −0.01
Z lm Score = −6.73
NG = 0
DB = 0
SM = 0
Delta_K
Doxycycline
YDR501W_2 PLM2
ZShift = 0.21
Z lm Score = −1.72
NG = 0
DB = 0
SM = 0
Delta_r
Doxycycline
YDR501W_2 PLM2
ZShift = −0.28
Z lm Score = 2.17
NG = 0
DB = 0
SM = 0
Delta_AUC
Doxycycline

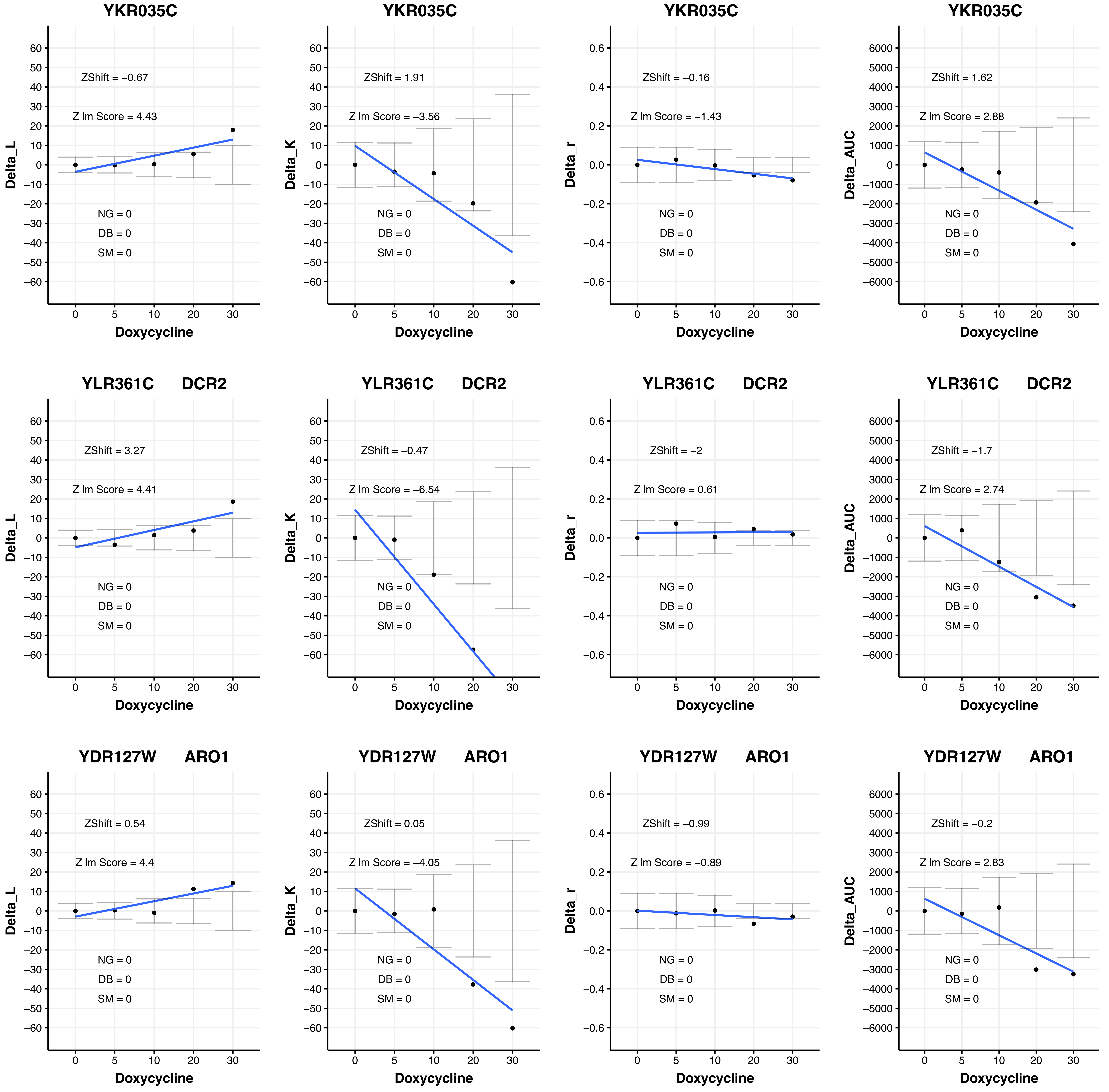

YKR035C
ZShift = −0.67
Z lm Score = 4.43
NG = 0
DB = 0
SM = 0
Delta_L
Doxycycline
YKR035C
ZShift = 1.91
Z lm Score = −3.56
NG = 0
DB = 0
SM = 0
Delta_K
Doxycycline
YKR035C
ZShift = −0.16
Z lm Score = −1.43
NG = 0
DB = 0
SM = 0
Delta_r
Doxycycline
YKR035C
ZShift = 1.62
Z lm Score = 2.88
NG = 0
DB = 0
SM = 0
Delta_AUC
Doxycycline
YLR361C DCR2
ZShift = 3.27
Z lm Score = 4.41
NG = 0
DB = 0
SM = 0
Delta_L
Doxycycline
YLR361C DCR2
ZShift = −0.47
Z lm Score = −6.54
NG = 0
DB = 0
SM = 0
Delta_K
Doxycycline
YLR361C DCR2
ZShift = −2
Z lm Score = 0.61
NG = 0
DB = 0
SM = 0
Delta_r
Doxycycline
YLR361C DCR2
ZShift = −1.7
Z lm Score = 2.74
NG = 0
DB = 0
SM = 0
Delta_AUC
Doxycycline
YDR127W ARO1
ZShift = 0.54
Z lm Score = 4.4
NG = 0
DB = 0
SM = 0
Delta_L
Doxycycline
YDR127W ARO1
ZShift = 0.05
Z lm Score = −4.05
NG = 0
DB = 0
SM = 0
Delta_K
Doxycycline
YDR127W ARO1
ZShift = −0.99
Z lm Score = −0.89
NG = 0
DB = 0
SM = 0
Delta_r
Doxycycline
YDR127W ARO1
ZShift = −0.2
Z lm Score = 2.83
NG = 0
DB = 0
SM = 0
Delta_AUC
Doxycycline

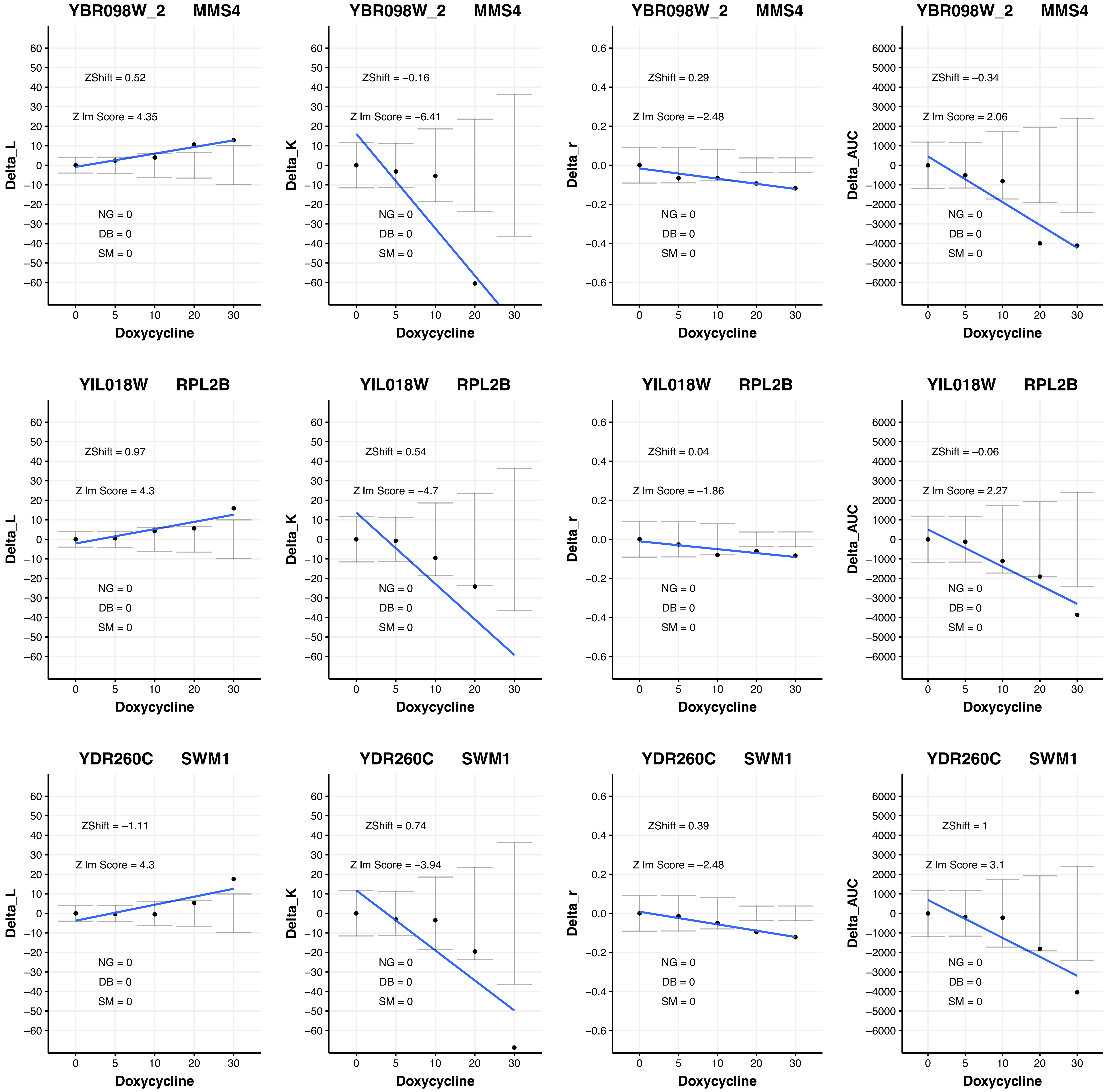

YBR098W_2 MMS4
ZShift = 0.52
Z lm Score = 4.35
ZShift = −0.16
Z lm Score = −6.41
ZShift = 0.29
Z lm Score = −2.48
ZShift = −0.34
Z lm Score = 2.06
YIL018W RPL2B
ZShift = 0.97
Z lm Score = 4.3
ZShift = 0.54
Z lm Score = −4.7
ZShift = 0.04
Z lm Score = −1.86
ZShift = −0.06
Z lm Score = 2.27
YDR260C SWM1
ZShift = −1.11
Z lm Score = 4.3
ZShift = 0.74
Z lm Score = −3.94
ZShift = 0.39
Z lm Score = −2.48
ZShift = 1
Z lm Score = 3.1
NG = 0
DB = 0
SM = 0
Delta_L
Delta_K
Delta_r
Delta_AUC
Doxycycline

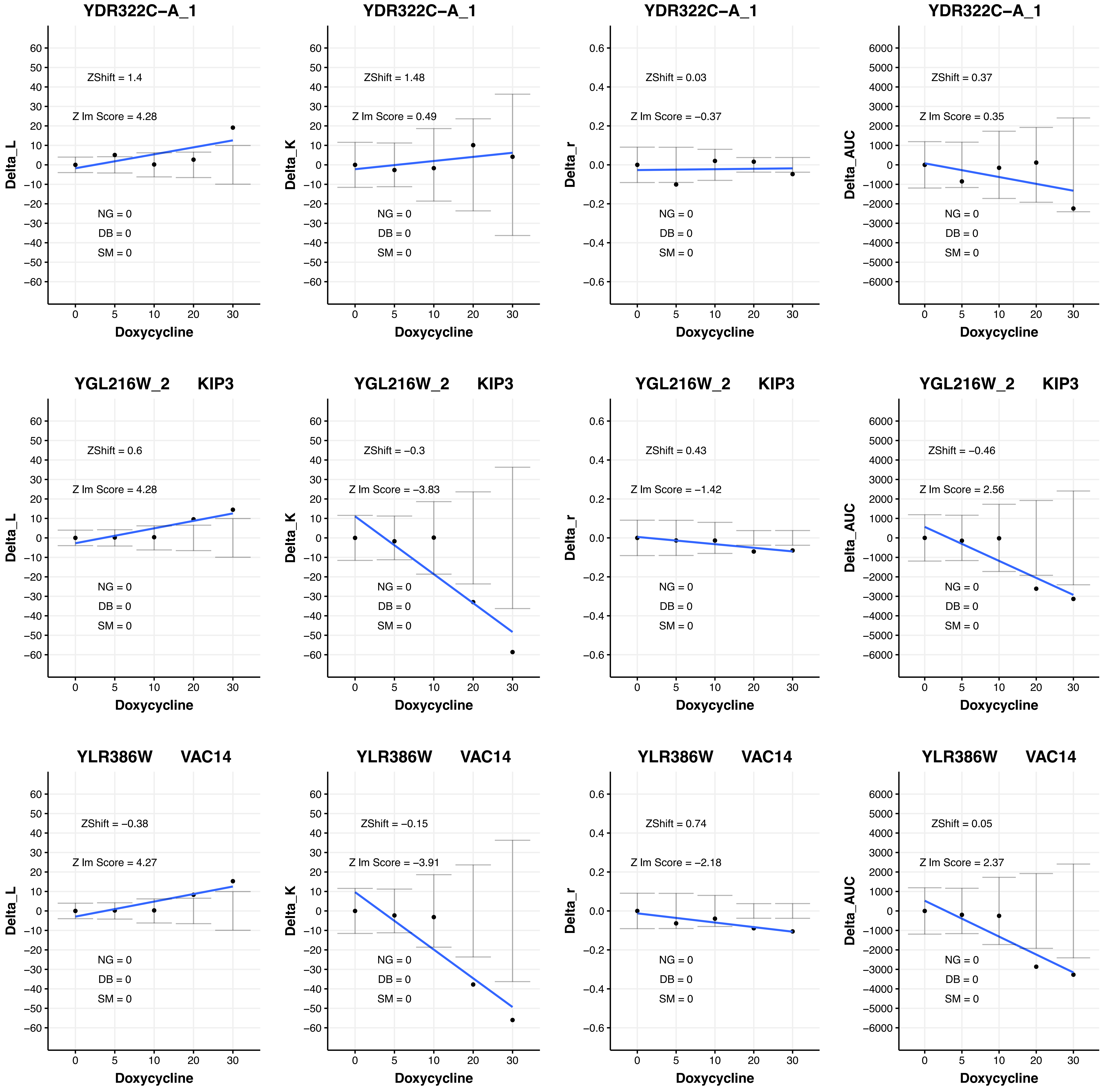

YDR322C−A_1
ZShift = 1.4
Z lm Score = 4.28
NG = 0
DB = 0
SM = 0
Delta_L
Doxycycline
YDR322C−A_1
ZShift = 1.48
Z lm Score = 0.49
NG = 0
DB = 0
SM = 0
Delta_K
Doxycycline
YDR322C−A_1
ZShift = 0.03
Z lm Score = −0.37
NG = 0
DB = 0
SM = 0
Delta_r
Doxycycline
YDR322C−A_1
ZShift = 0.37
Z lm Score = 0.35
NG = 0
DB = 0
SM = 0
Delta_AUC
Doxycycline
YGL216W_2 KIP3
ZShift = 0.6
Z lm Score = 4.28
NG = 0
DB = 0
SM = 0
Delta_L
Doxycycline
YGL216W_2 KIP3
ZShift = −0.3
Z lm Score = −3.83
NG = 0
DB = 0
SM = 0
Delta_K
Doxycycline
YGL216W_2 KIP3
ZShift = 0.43
Z lm Score = −1.42
NG = 0
DB = 0
SM = 0
Delta_r
Doxycycline
YGL216W_2 KIP3
ZShift = −0.46
Z lm Score = 2.56
NG = 0
DB = 0
SM = 0
Delta_AUC
Doxycycline
YLR386W VAC14
ZShift = −0.38
Z lm Score = 4.27
NG = 0
DB = 0
SM = 0
Delta_L
Doxycycline
YLR386W VAC14
ZShift = −0.15
Z lm Score = −3.91
NG = 0
DB = 0
SM = 0
Delta_K
Doxycycline
YLR386W VAC14
ZShift = 0.74
Z lm Score = −2.18
NG = 0
DB = 0
SM = 0
Delta_r
Doxycycline
YLR386W VAC14
ZShift = 0.05
Z lm Score = 2.37
NG = 0
DB = 0
SM = 0
Delta_AUC
Doxycycline

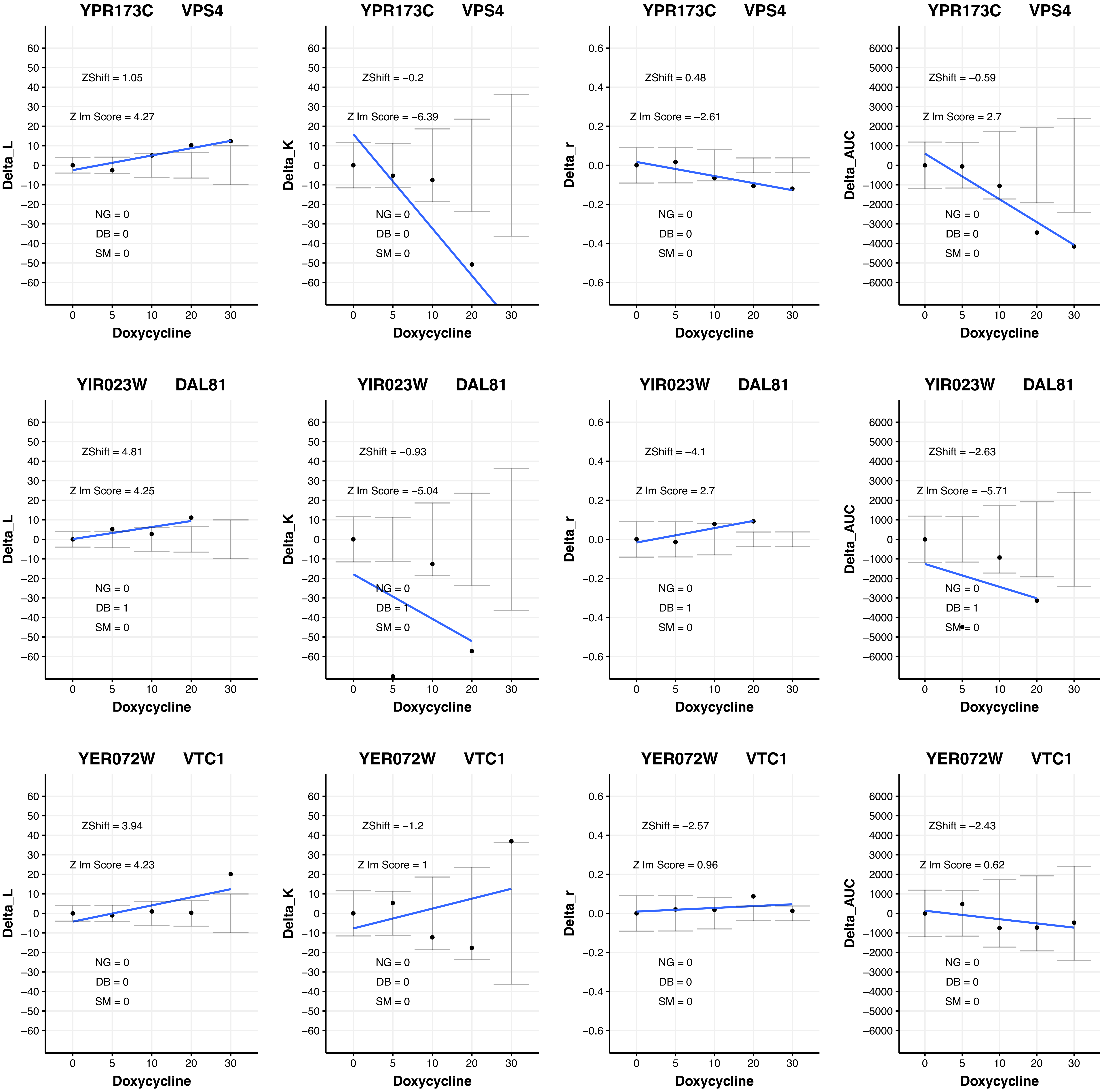
YPR173C VPS4
Delta_L
ZShift = 1.05
Z lm Score = 4.27
NG = 0
DB = 0
SM = 0
YPR173C VPS4
Delta_K
ZShift = −0.2
Z lm Score = −6.39
NG = 0
DB = 0
SM = 0
YPR173C VPS4
Delta_r
ZShift = 0.48
Z lm Score = −2.61
NG = 0
DB = 0
SM = 0
YPR173C VPS4
Delta_AUC
ZShift = −0.59
Z lm Score = 2.7
NG = 0
DB = 0
SM = 0
YIR023W DAL81
Delta_L
ZShift = 4.81
Z lm Score = 4.25
NG = 0
DB = 1
SM = 0
YIR023W DAL81
Delta_K
ZShift = −0.93
Z lm Score = −5.04
NG = 0
DB = 1
SM = 0
YIR023W DAL81
Delta_r
ZShift = −4.1
Z lm Score = 2.7
NG = 0
DB = 1
SM = 0
YIR023W DAL81
Delta_AUC
ZShift = −2.63
Z lm Score = −5.71
NG = 0
DB = 1
SM = 0
YER072W VTC1
Delta_L
ZShift = 3.94
Z lm Score = 4.23
NG = 0
DB = 0
SM = 0
YER072W VTC1
Delta_K
ZShift = −1.2
Z lm Score = 1
NG = 0
DB = 0
SM = 0
YER072W VTC1
Delta_r
ZShift = −2.57
Z lm Score = 0.96
NG = 0
DB = 0
SM = 0
YER072W VTC1
Delta_AUC
ZShift = −2.43
Z lm Score = 0.62
NG = 0
DB = 0
SM = 0
Doxycycline
0 5 10 20 30

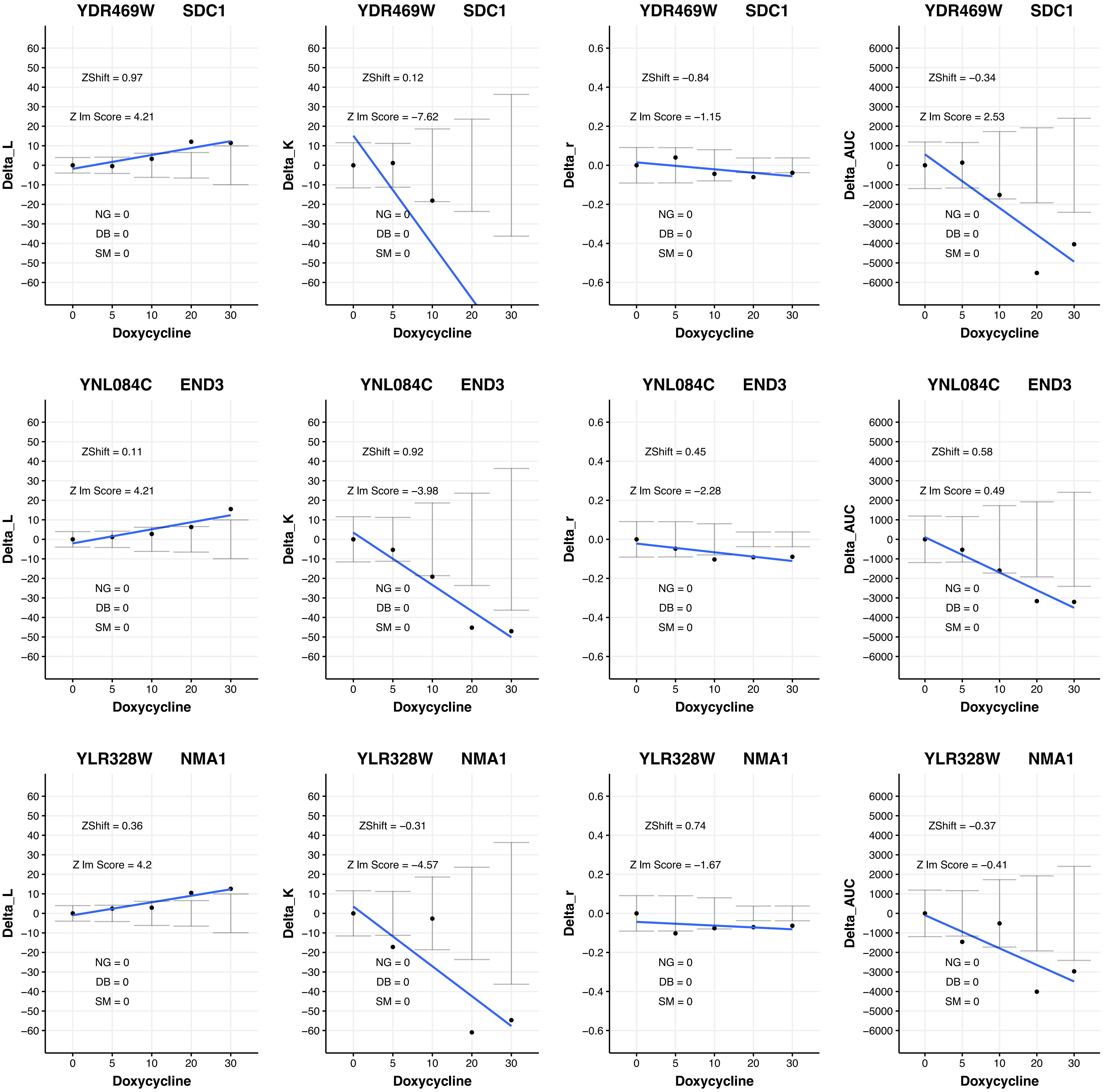
YDR469W SDC1
ZShift = 0.97
Z lm Score = 4.21
NG = 0
DB = 0
SM = 0
Delta_L
Doxycycline
YDR469W SDC1
ZShift = 0.12
Z lm Score = −7.62
NG = 0
DB = 0
SM = 0
Delta_K
Doxycycline
YDR469W SDC1
ZShift = −0.84
Z lm Score = −1.15
NG = 0
DB = 0
SM = 0
Delta_r
Doxycycline
YDR469W SDC1
ZShift = −0.34
Z lm Score = 2.53
NG = 0
DB = 0
SM = 0
Delta_AUC
Doxycycline
YNL084C END3
ZShift = 0.11
Z lm Score = 4.21
NG = 0
DB = 0
SM = 0
Delta_L
Doxycycline
YNL084C END3
ZShift = 0.92
Z lm Score = −3.98
NG = 0
DB = 0
SM = 0
Delta_K
Doxycycline
YNL084C END3
ZShift = 0.45
Z lm Score = −2.28
NG = 0
DB = 0
SM = 0
Delta_r
Doxycycline
YNL084C END3
ZShift = 0.58
Z lm Score = 0.49
NG = 0
DB = 0
SM = 0
Delta_AUC
Doxycycline
YLR328W NMA1
ZShift = 0.36
Z lm Score = 4.2
NG = 0
DB = 0
SM = 0
Delta_L
Doxycycline
YLR328W NMA1
ZShift = −0.31
Z lm Score = −4.57
NG = 0
DB = 0
SM = 0
Delta_K
Doxycycline
YLR328W NMA1
ZShift = 0.74
Z lm Score = −1.67
NG = 0
DB = 0
SM = 0
Delta_r
Doxycycline
YLR328W NMA1
ZShift = −0.37
Z lm Score = −0.41
NG = 0
DB = 0
SM = 0
Delta_AUC
Doxycycline

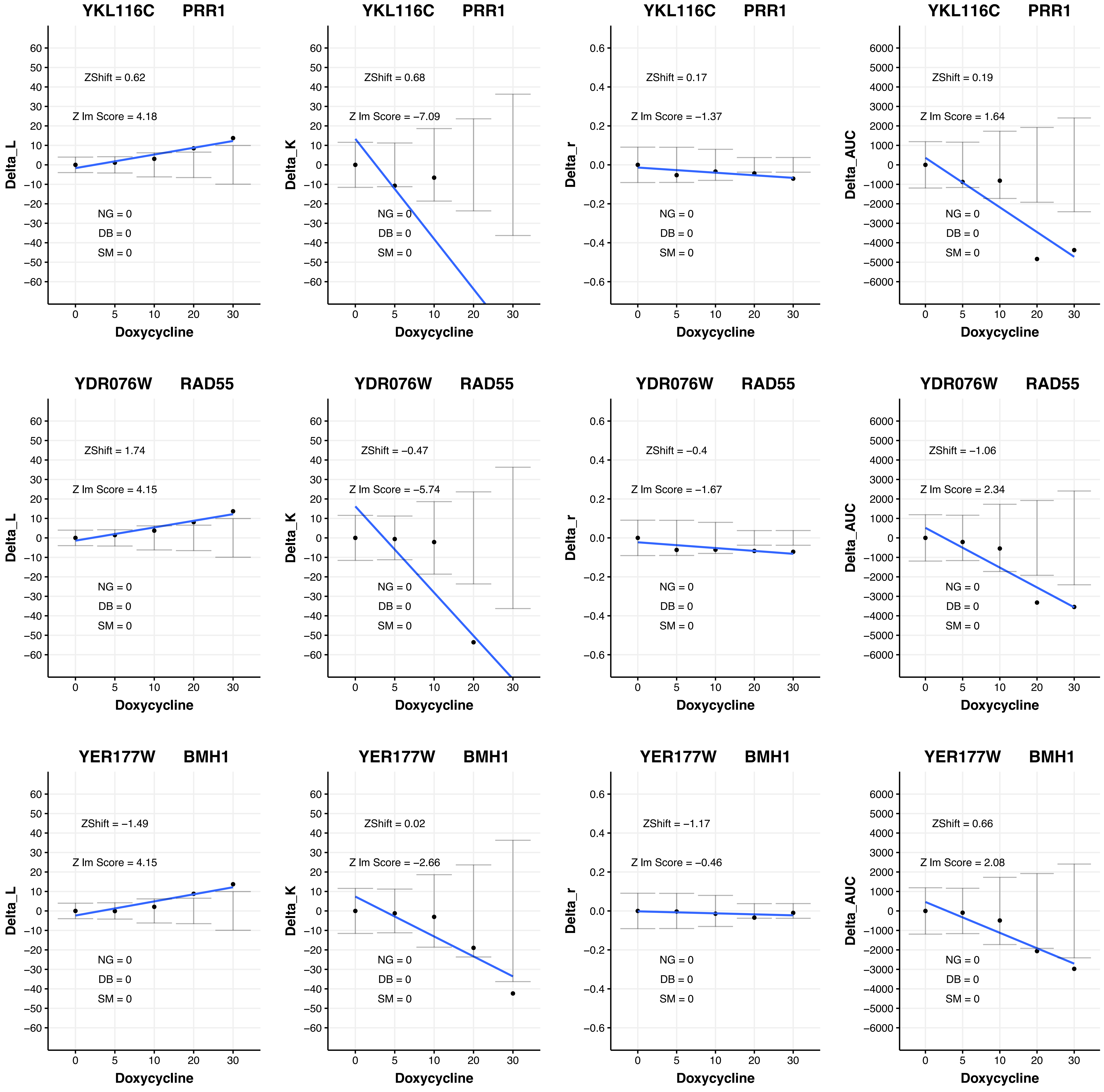

YKL116C PRR1
Delta_L
ZShift = 0.62
Z lm Score = 4.18
NG = 0
DB = 0
SM = 0
Doxycycline
YKL116C PRR1
Delta_K
ZShift = 0.68
Z lm Score = −7.09
NG = 0
DB = 0
SM = 0
Doxycycline
YKL116C PRR1
Delta_r
ZShift = 0.17
Z lm Score = −1.37
NG = 0
DB = 0
SM = 0
Doxycycline
YKL116C PRR1
Delta_AUC
ZShift = 0.19
Z lm Score = 1.64
NG = 0
DB = 0
SM = 0
Doxycycline
YDR076W RAD55
Delta_L
ZShift = 1.74
Z lm Score = 4.15
NG = 0
DB = 0
SM = 0
Doxycycline
YDR076W RAD55
Delta_K
ZShift = −0.47
Z lm Score = −5.74
NG = 0
DB = 0
SM = 0
Doxycycline
YDR076W RAD55
Delta_r
ZShift = −0.4
Z lm Score = −1.67
NG = 0
DB = 0
SM = 0
Doxycycline
YDR076W RAD55
Delta_AUC
ZShift = −1.06
Z lm Score = 2.34
NG = 0
DB = 0
SM = 0
Doxycycline
YER177W BMH1
Delta_L
ZShift = −1.49
Z lm Score = 4.15
NG = 0
DB = 0
SM = 0
Doxycycline
YER177W BMH1
Delta_K
ZShift = 0.02
Z lm Score = −2.66
NG = 0
DB = 0
SM = 0
Doxycycline
YER177W BMH1
Delta_r
ZShift = −1.17
Z lm Score = −0.46
NG = 0
DB = 0
SM = 0
Doxycycline
YER177W BMH1
Delta_AUC
ZShift = 0.66
Z lm Score = 2.08
NG = 0
DB = 0
SM = 0
Doxycycline

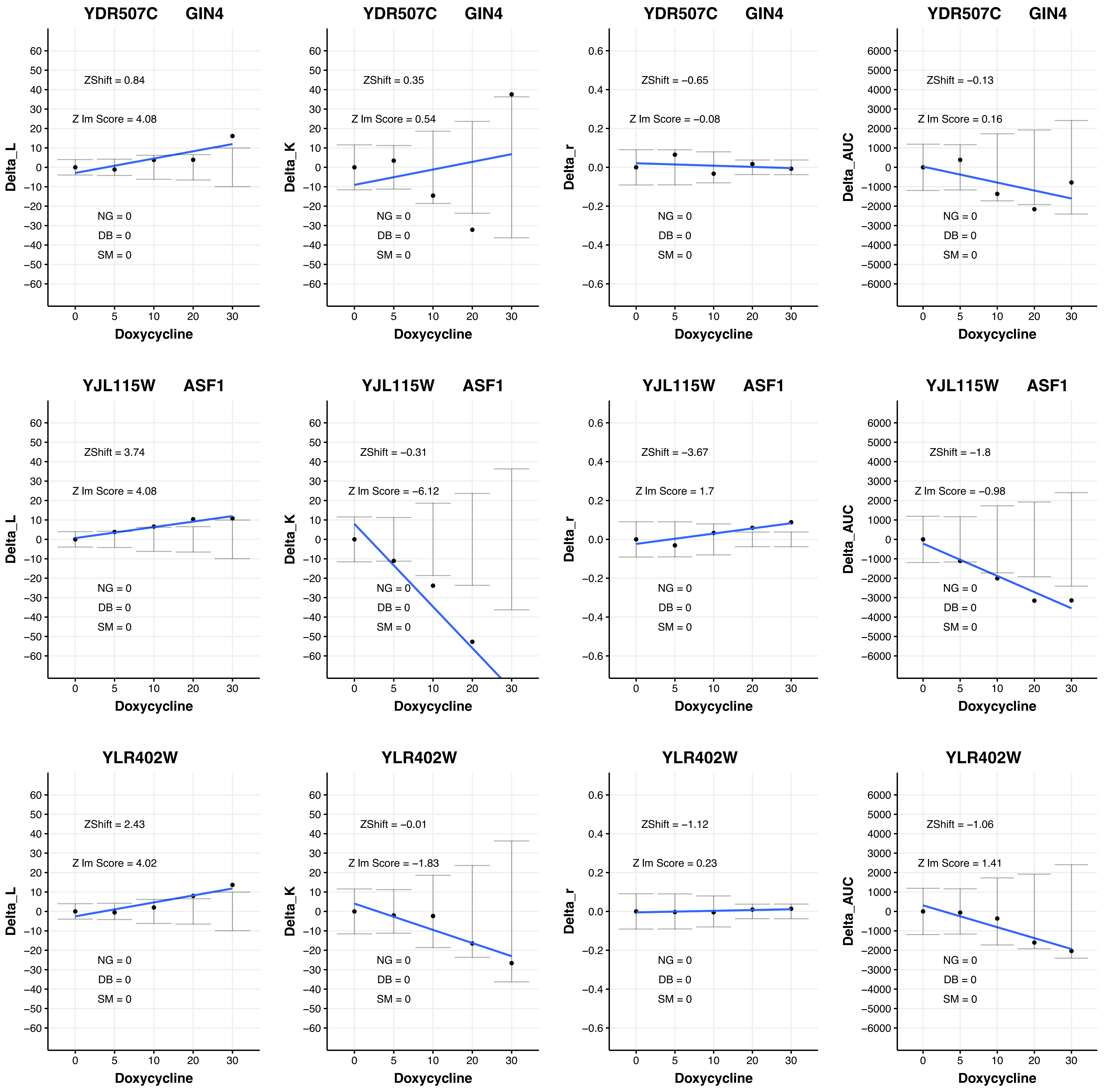

YDR507C GIN4
ZShift = 0.84
Z lm Score = 4.08
NG = 0
DB = 0
SM = 0
Delta_L
Doxycycline
YDR507C GIN4
ZShift = 0.35
Z lm Score = 0.54
NG = 0
DB = 0
SM = 0
Delta_K
Doxycycline
YDR507C GIN4
ZShift = −0.65
Z lm Score = −0.08
NG = 0
DB = 0
SM = 0
Delta_r
Doxycycline
YDR507C GIN4
ZShift = −0.13
Z lm Score = 0.16
NG = 0
DB = 0
SM = 0
Delta_AUC
Doxycycline
YJL115W ASF1
ZShift = 3.74
Z lm Score = 4.08
NG = 0
DB = 0
SM = 0
Delta_L
Doxycycline
YJL115W ASF1
ZShift = −0.31
Z lm Score = −6.12
NG = 0
DB = 0
SM = 0
Delta_K
Doxycycline
YJL115W ASF1
ZShift = −3.67
Z lm Score = 1.7
NG = 0
DB = 0
SM = 0
Delta_r
Doxycycline
YJL115W ASF1
ZShift = −1.8
Z lm Score = −0.98
NG = 0
DB = 0
SM = 0
Delta_AUC
Doxycycline
YLR402W
ZShift = 2.43
Z lm Score = 4.02
NG = 0
DB = 0
SM = 0
Delta_L
Doxycycline
YLR402W
ZShift = −0.01
Z lm Score = −1.83
NG = 0
DB = 0
SM = 0
Delta_K
Doxycycline
YLR402W
ZShift = −1.12
Z lm Score = 0.23
NG = 0
DB = 0
SM = 0
Delta_r
Doxycycline
YLR402W
ZShift = −1.06
Z lm Score = 1.41
NG = 0
DB = 0
SM = 0
Delta_AUC
Doxycycline

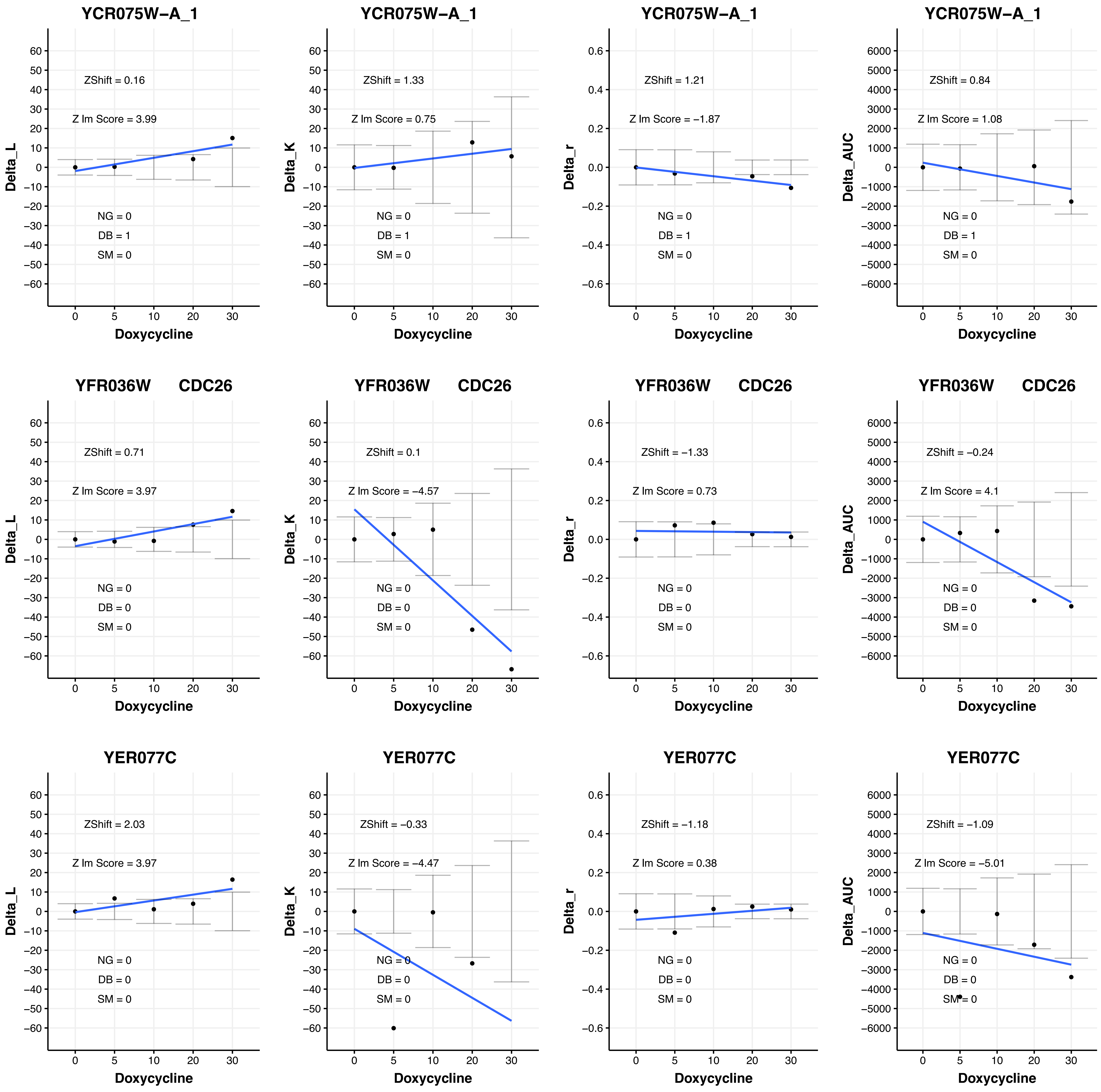

YCR075W−A_1
ZShift = 0.16
Z lm Score = 3.99
NG = 0
DB = 1
SM = 0
Delta_L
Doxycycline
YCR075W−A_1
ZShift = 1.33
Z lm Score = 0.75
NG = 0
DB = 1
SM = 0
Delta_K
Doxycycline
YCR075W−A_1
ZShift = 1.21
Z lm Score = −1.87
NG = 0
DB = 1
SM = 0
Delta_r
Doxycycline
YCR075W−A_1
ZShift = 0.84
Z lm Score = 1.08
NG = 0
DB = 1
SM = 0
Delta_AUC
Doxycycline
YFR036W CDC26
ZShift = 0.71
Z lm Score = 3.97
NG = 0
DB = 0
SM = 0
Delta_L
Doxycycline
YFR036W CDC26
ZShift = 0.1
Z lm Score = −4.57
NG = 0
DB = 0
SM = 0
Delta_K
Doxycycline
YFR036W CDC26
ZShift = −1.33
Z lm Score = 0.73
NG = 0
DB = 0
SM = 0
Delta_r
Doxycycline
YFR036W CDC26
ZShift = −0.24
Z lm Score = 4.1
NG = 0
DB = 0
SM = 0
Delta_AUC
Doxycycline
YER077C
ZShift = 2.03
Z lm Score = 3.97
NG = 0
DB = 0
SM = 0
Delta_L
Doxycycline
YER077C
ZShift = −0.33
Z lm Score = −4.47
NG = 0
DB = 0
SM = 0
Delta_K
Doxycycline
YER077C
ZShift = −1.18
Z lm Score = 0.38
NG = 0
DB = 0
SM = 0
Delta_r
Doxycycline
YER077C
ZShift = −1.09
Z lm Score = −5.01
NG = 0
DB = 0
SM = 0
Delta_AUC
Doxycycline

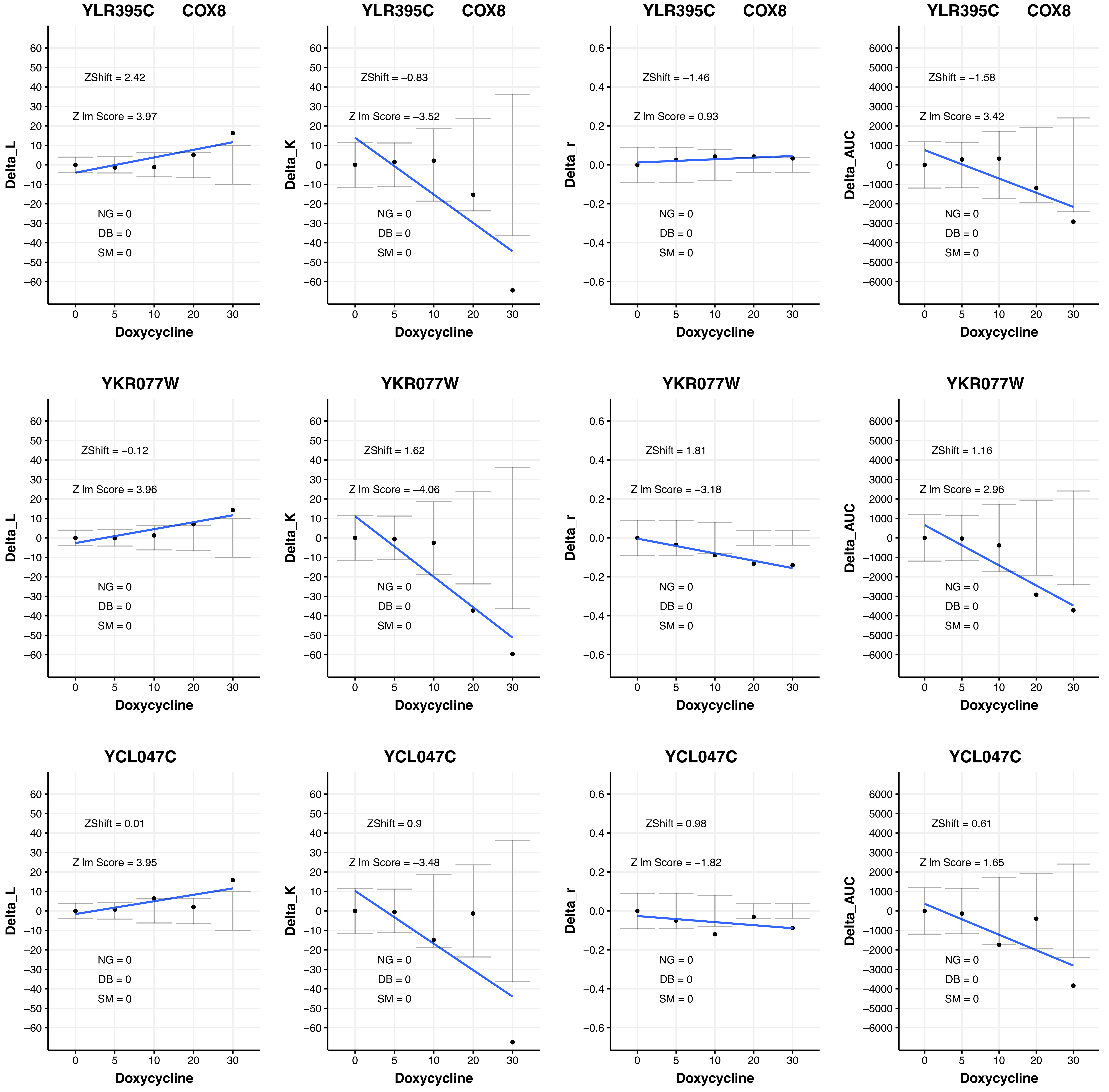

YLR395C COX8
ZShift = 2.42
Z lm Score = 3.97
NG = 0
DB = 0
SM = 0
Delta_L
Doxycycline
YLR395C COX8
ZShift = −0.83
Z lm Score = −3.52
NG = 0
DB = 0
SM = 0
Delta_K
Doxycycline
YLR395C COX8
ZShift = −1.46
Z lm Score = 0.93
NG = 0
DB = 0
SM = 0
Delta_r
Doxycycline
YLR395C COX8
ZShift = −1.58
Z lm Score = 3.42
NG = 0
DB = 0
SM = 0
Delta_AUC
Doxycycline
YKR077W
ZShift = −0.12
Z lm Score = 3.96
NG = 0
DB = 0
SM = 0
Delta_L
Doxycycline
YKR077W
ZShift = 1.62
Z lm Score = −4.06
NG = 0
DB = 0
SM = 0
Delta_K
Doxycycline
YKR077W
ZShift = 1.81
Z lm Score = −3.18
NG = 0
DB = 0
SM = 0
Delta_r
Doxycycline
YKR077W
ZShift = 1.16
Z lm Score = 2.96
NG = 0
DB = 0
SM = 0
Delta_AUC
Doxycycline
YCL047C
ZShift = 0.01
Z lm Score = 3.95
NG = 0
DB = 0
SM = 0
Delta_L
Doxycycline
YCL047C
ZShift = 0.9
Z lm Score = −3.48
NG = 0
DB = 0
SM = 0
Delta_K
Doxycycline
YCL047C
ZShift = 0.98
Z lm Score = −1.82
NG = 0
DB = 0
SM = 0
Delta_r
Doxycycline
YCL047C
ZShift = 0.61
Z lm Score = 1.65
NG = 0
DB = 0
SM = 0
Delta_AUC
Doxycycline

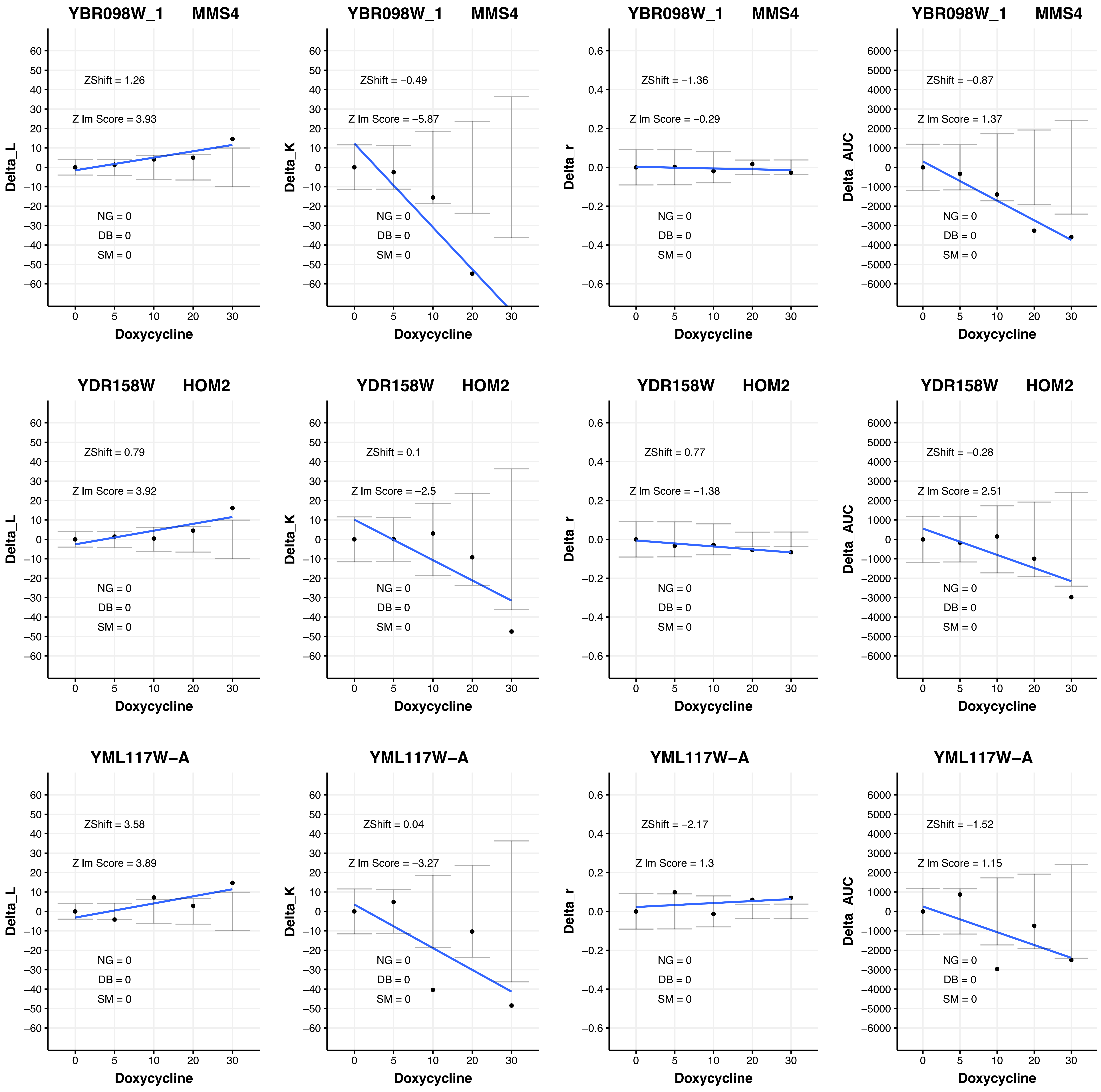
YBR098W_1 MMS4
ZShift = 1.26
Z lm Score = 3.93
NG = 0
DB = 0
SM = 0
Delta_L
Doxycycline
YBR098W_1 MMS4
ZShift = −0.49
Z lm Score = −5.87
NG = 0
DB = 0
SM = 0
Delta_K
Doxycycline
YBR098W_1 MMS4
ZShift = −1.36
Z lm Score = −0.29
NG = 0
DB = 0
SM = 0
Delta_r
Doxycycline
YBR098W_1 MMS4
ZShift = −0.87
Z lm Score = 1.37
NG = 0
DB = 0
SM = 0
Delta_AUC
Doxycycline
YDR158W HOM2
ZShift = 0.79
Z lm Score = 3.92
NG = 0
DB = 0
SM = 0
Delta_L
Doxycycline
YDR158W HOM2
ZShift = 0.1
Z lm Score = −2.5
NG = 0
DB = 0
SM = 0
Delta_K
Doxycycline
YDR158W HOM2
ZShift = 0.77
Z lm Score = −1.38
NG = 0
DB = 0
SM = 0
Delta_r
Doxycycline
YDR158W HOM2
ZShift = −0.28
Z lm Score = 2.51
NG = 0
DB = 0
SM = 0
Delta_AUC
Doxycycline
YML117W−A
ZShift = 3.58
Z lm Score = 3.89
NG = 0
DB = 0
SM = 0
Delta_L
Doxycycline
YML117W−A
ZShift = 0.04
Z lm Score = −3.27
NG = 0
DB = 0
SM = 0
Delta_K
Doxycycline
YML117W−A
ZShift = −2.17
Z lm Score = 1.3
NG = 0
DB = 0
SM = 0
Delta_r
Doxycycline
YML117W−A
ZShift = −1.52
Z lm Score = 1.15
NG = 0
DB = 0
SM = 0
Delta_AUC
Doxycycline

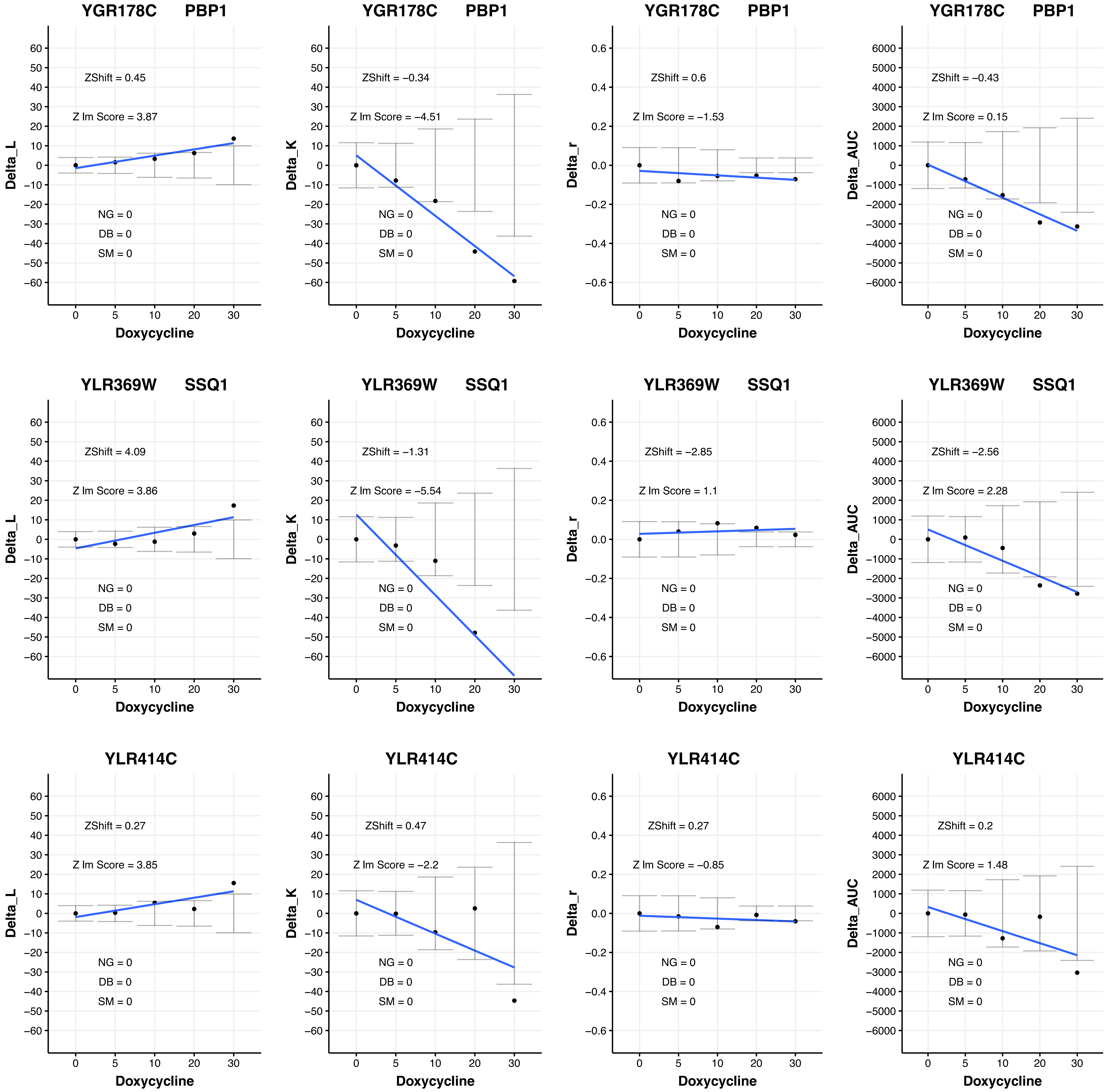
YGR178C PBP1
ZShift = 0.45
Z lm Score = 3.87
NG = 0
DB = 0
SM = 0
Delta_L
Doxycycline
YGR178C PBP1
ZShift = −0.34
Z lm Score = −4.51
NG = 0
DB = 0
SM = 0
Delta_K
Doxycycline
YGR178C PBP1
ZShift = 0.6
Z lm Score = −1.53
NG = 0
DB = 0
SM = 0
Delta_r
Doxycycline
YGR178C PBP1
ZShift = −0.43
Z lm Score = 0.15
NG = 0
DB = 0
SM = 0
Delta_AUC
Doxycycline
YLR369W SSQ1
ZShift = 4.09
Z lm Score = 3.86
NG = 0
DB = 0
SM = 0
Delta_L
Doxycycline
YLR369W SSQ1
ZShift = −1.31
Z lm Score = −5.54
NG = 0
DB = 0
SM = 0
Delta_K
Doxycycline
YLR369W SSQ1
ZShift = −2.85
Z lm Score = 1.1
NG = 0
DB = 0
SM = 0
Delta_r
Doxycycline
YLR369W SSQ1
ZShift = −2.56
Z lm Score = 2.28
NG = 0
DB = 0
SM = 0
Delta_AUC
Doxycycline
YLR414C
ZShift = 0.27
Z lm Score = 3.85
NG = 0
DB = 0
SM = 0
Delta_L
Doxycycline
YLR414C
ZShift = 0.47
Z lm Score = −2.2
NG = 0
DB = 0
SM = 0
Delta_K
Doxycycline
YLR414C
ZShift = 0.27
Z lm Score = −0.85
NG = 0
DB = 0
SM = 0
Delta_r
Doxycycline
YLR414C
ZShift = 0.2
Z lm Score = 1.48
NG = 0
DB = 0
SM = 0
Delta_AUC
Doxycycline

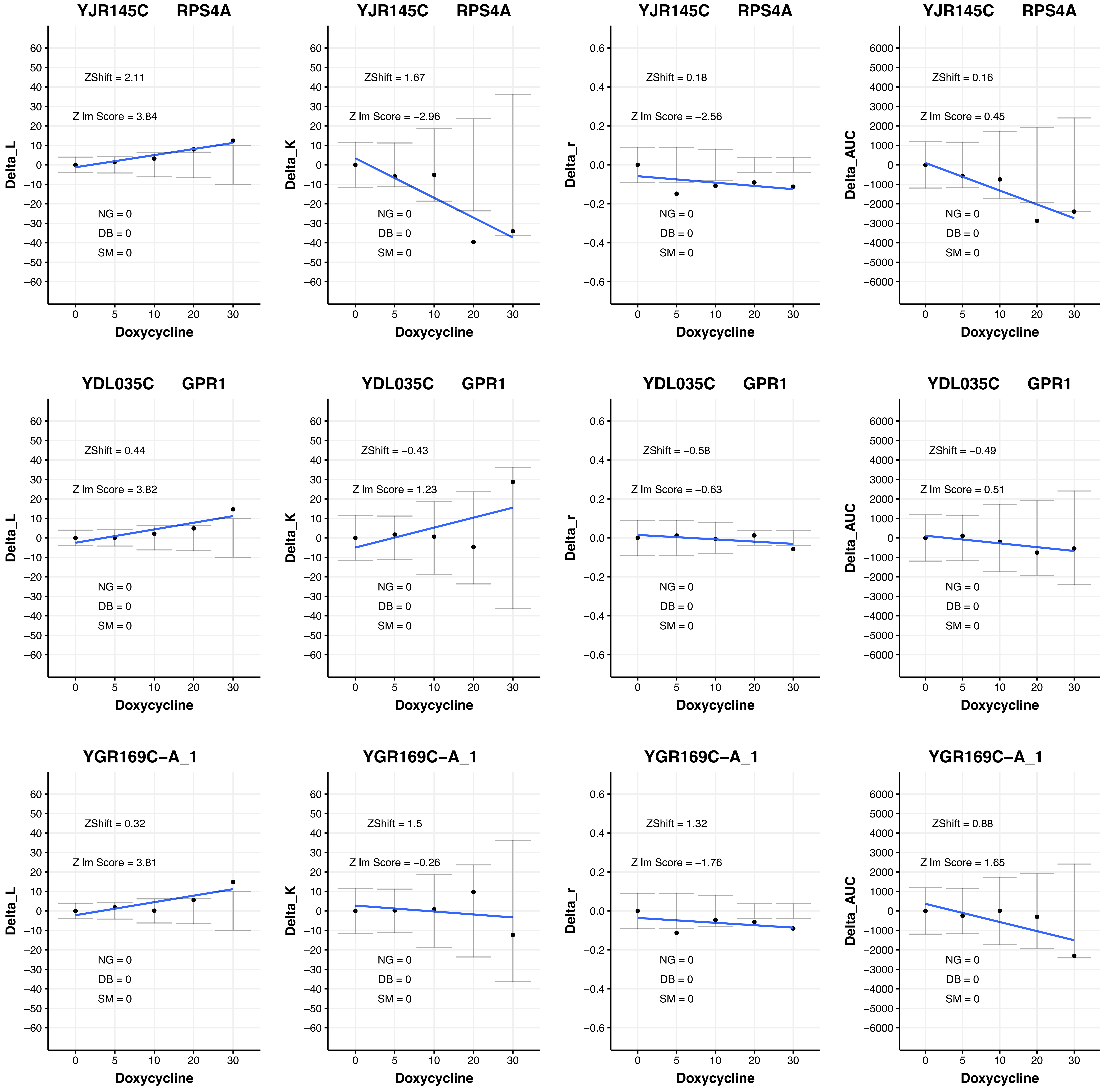

YJR145C RPS4A
ZShift = 2.11
Z lm Score = 3.84
NG = 0
DB = 0
SM = 0
Delta_L
Doxycycline
YJR145C RPS4A
ZShift = 1.67
Z lm Score = −2.96
NG = 0
DB = 0
SM = 0
Delta_K
Doxycycline
YJR145C RPS4A
ZShift = 0.18
Z lm Score = −2.56
NG = 0
DB = 0
SM = 0
Delta_r
Doxycycline
YJR145C RPS4A
ZShift = 0.16
Z lm Score = 0.45
NG = 0
DB = 0
SM = 0
Delta_AUC
Doxycycline
YDL035C GPR1
ZShift = 0.44
Z lm Score = 3.82
NG = 0
DB = 0
SM = 0
Delta_L
Doxycycline
YDL035C GPR1
ZShift = −0.43
Z lm Score = 1.23
NG = 0
DB = 0
SM = 0
Delta_K
Doxycycline
YDL035C GPR1
ZShift = −0.58
Z lm Score = −0.63
NG = 0
DB = 0
SM = 0
Delta_r
Doxycycline
YDL035C GPR1
ZShift = −0.49
Z lm Score = 0.51
NG = 0
DB = 0
SM = 0
Delta_AUC
Doxycycline
YGR169C−A_1
ZShift = 0.32
Z lm Score = 3.81
NG = 0
DB = 0
SM = 0
Delta_L
Doxycycline
YGR169C−A_1
ZShift = 1.5
Z lm Score = −0.26
NG = 0
DB = 0
SM = 0
Delta_K
Doxycycline
YGR169C−A_1
ZShift = 1.32
Z lm Score = −1.76
NG = 0
DB = 0
SM = 0
Delta_r
Doxycycline
YGR169C−A_1
ZShift = 0.88
Z lm Score = 1.65
NG = 0
DB = 0
SM = 0
Delta_AUC
Doxycycline

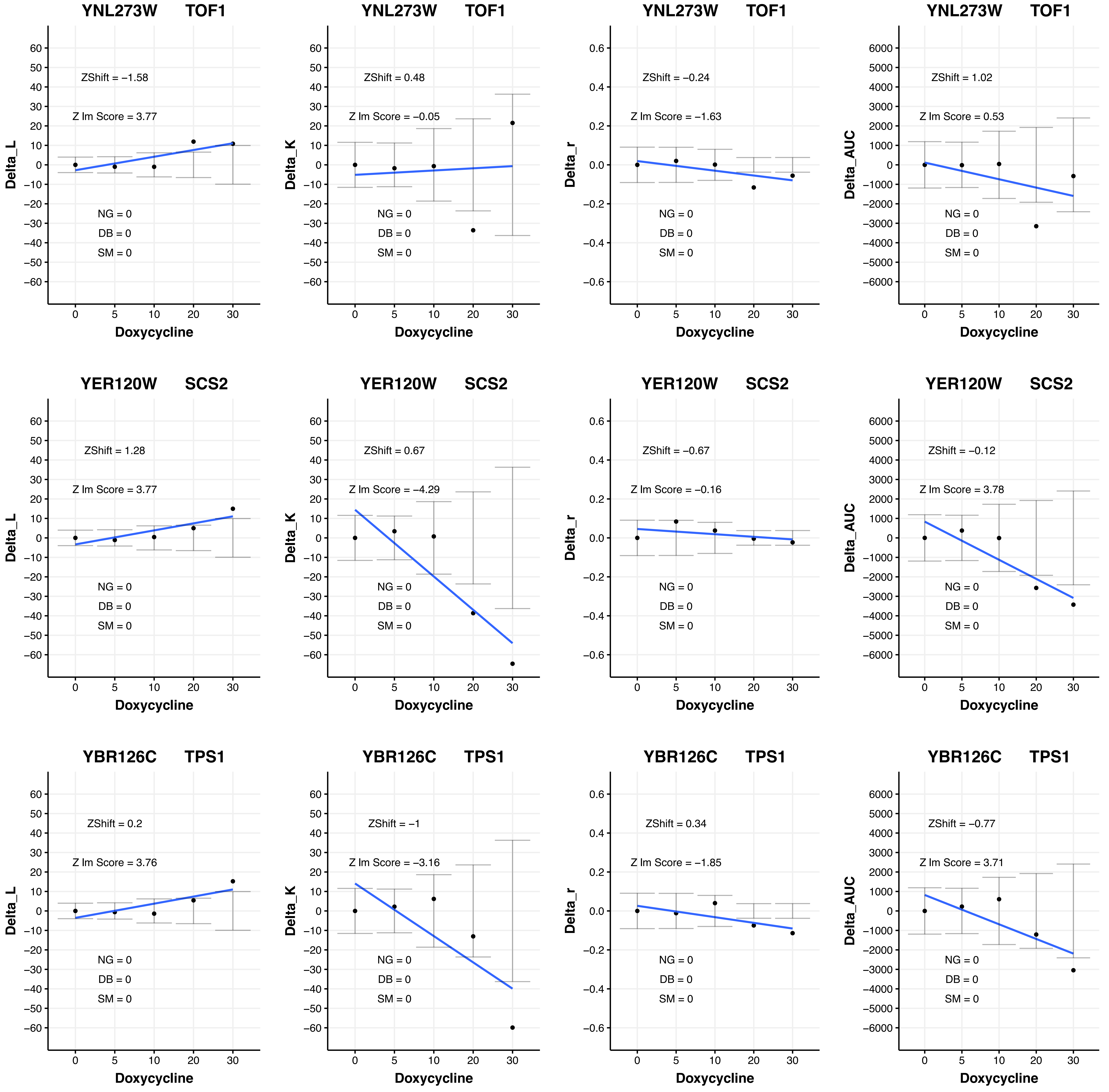
YNL273W TOF1
ZShift = −1.58
Z lm Score = 3.77
NG = 0
DB = 0
SM = 0
Delta_L
Doxycycline
YNL273W TOF1
ZShift = 0.48
Z lm Score = −0.05
NG = 0
DB = 0
SM = 0
Delta_K
Doxycycline
YNL273W TOF1
ZShift = −0.24
Z lm Score = −1.63
NG = 0
DB = 0
SM = 0
Delta_r
Doxycycline
YNL273W TOF1
ZShift = 1.02
Z lm Score = 0.53
NG = 0
DB = 0
SM = 0
Delta_AUC
Doxycycline
YER120W SCS2
ZShift = 1.28
Z lm Score = 3.77
NG = 0
DB = 0
SM = 0
Delta_L
Doxycycline
YER120W SCS2
ZShift = 0.67
Z lm Score = −4.29
NG = 0
DB = 0
SM = 0
Delta_K
Doxycycline
YER120W SCS2
ZShift = −0.67
Z lm Score = −0.16
NG = 0
DB = 0
SM = 0
Delta_r
Doxycycline
YER120W SCS2
ZShift = −0.12
Z lm Score = 3.78
NG = 0
DB = 0
SM = 0
Delta_AUC
Doxycycline
YBR126C TPS1
ZShift = 0.2
Z lm Score = 3.76
NG = 0
DB = 0
SM = 0
Delta_L
Doxycycline
YBR126C TPS1
ZShift = −1
Z lm Score = −3.16
NG = 0
DB = 0
SM = 0
Delta_K
Doxycycline
YBR126C TPS1
ZShift = 0.34
Z lm Score = −1.85
NG = 0
DB = 0
SM = 0
Delta_r
Doxycycline
YBR126C TPS1
ZShift = −0.77
Z lm Score = 3.71
NG = 0
DB = 0
SM = 0
Delta_AUC
Doxycycline

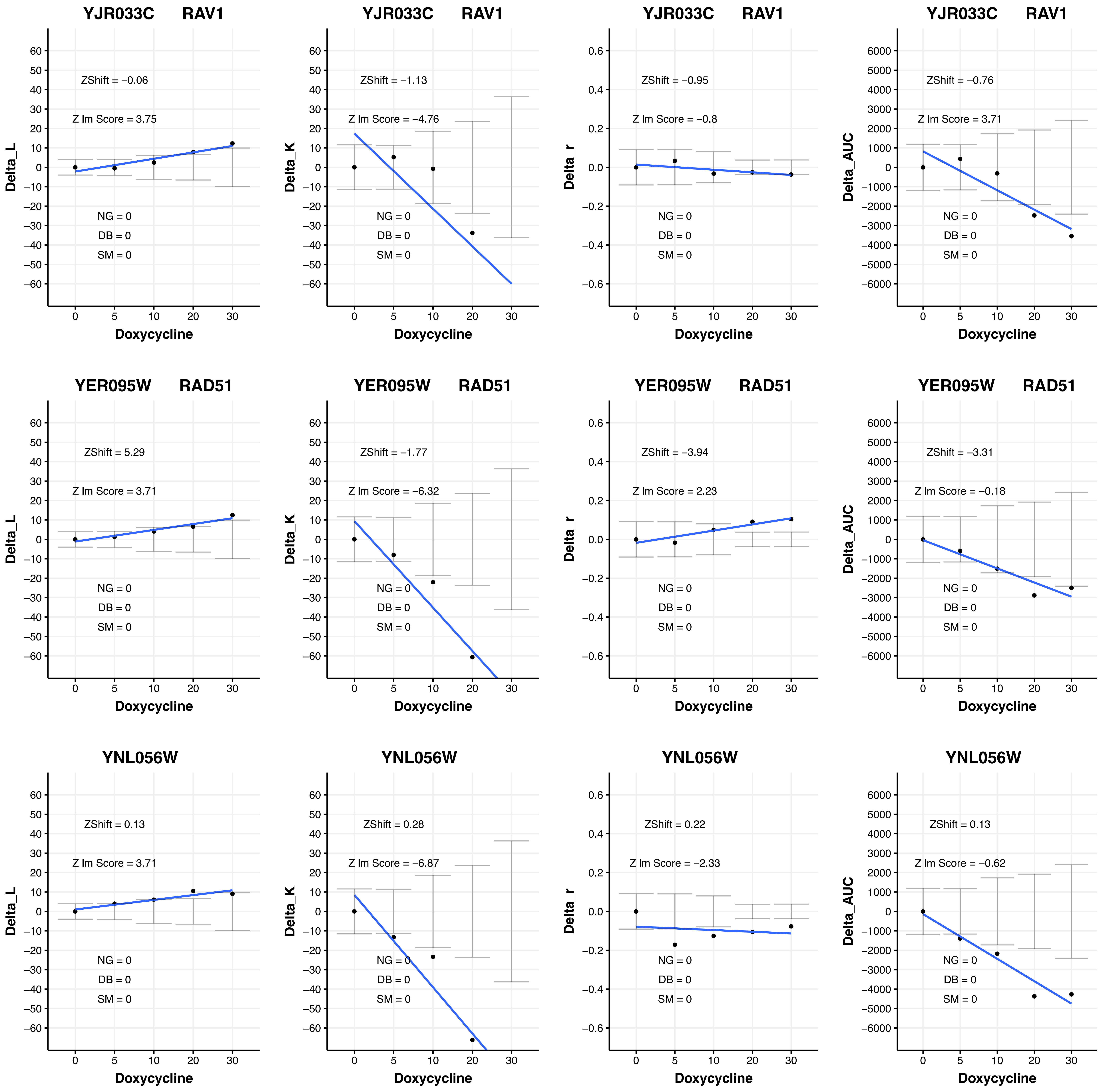

YJR033C RAV1
ZShift = −0.06
Z lm Score = 3.75
NG = 0
DB = 0
SM = 0
Delta_L
Doxycycline
YJR033C RAV1
ZShift = −1.13
Z lm Score = −4.76
NG = 0
DB = 0
SM = 0
Delta_K
Doxycycline
YJR033C RAV1
ZShift = −0.95
Z lm Score = −0.8
NG = 0
DB = 0
SM = 0
Delta_r
Doxycycline
YJR033C RAV1
ZShift = −0.76
Z lm Score = 3.71
NG = 0
DB = 0
SM = 0
Delta_AUC
Doxycycline
YER095W RAD51
ZShift = 5.29
Z lm Score = 3.71
NG = 0
DB = 0
SM = 0
Delta_L
Doxycycline
YER095W RAD51
ZShift = −1.77
Z lm Score = −6.32
NG = 0
DB = 0
SM = 0
Delta_K
Doxycycline
YER095W RAD51
ZShift = −3.94
Z lm Score = 2.23
NG = 0
DB = 0
SM = 0
Delta_r
Doxycycline
YER095W RAD51
ZShift = −3.31
Z lm Score = −0.18
NG = 0
DB = 0
SM = 0
Delta_AUC
Doxycycline
YNL056W
ZShift = 0.13
Z lm Score = 3.71
NG = 0
DB = 0
SM = 0
Delta_L
Doxycycline
YNL056W
ZShift = 0.28
Z lm Score = −6.87
NG = 0
DB = 0
SM = 0
Delta_K
Doxycycline
YNL056W
ZShift = 0.22
Z lm Score = −2.33
NG = 0
DB = 0
SM = 0
Delta_r
Doxycycline
YNL056W
ZShift = 0.13
Z lm Score = −0.62
NG = 0
DB = 0
SM = 0
Delta_AUC
Doxycycline

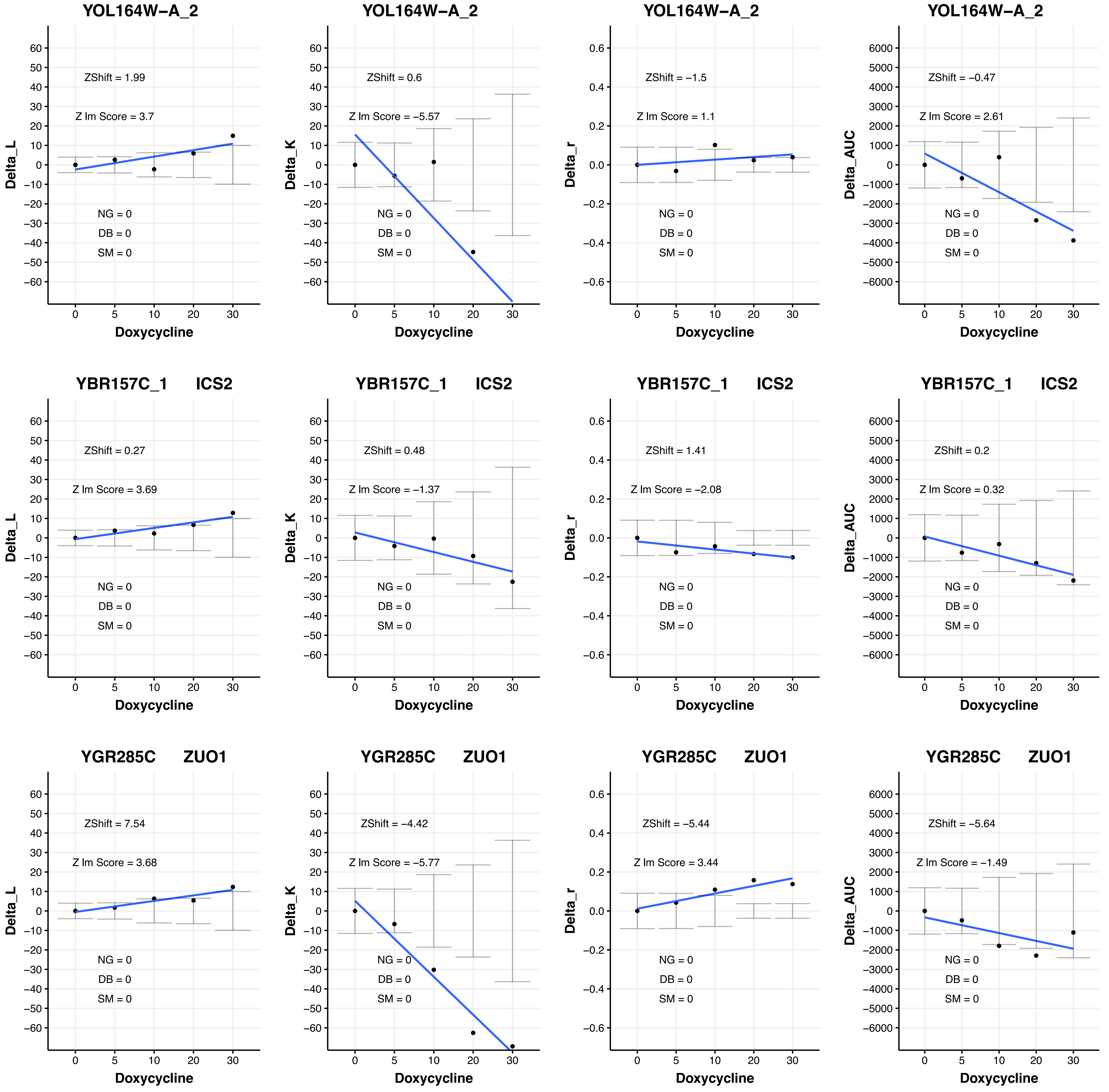

YOL164W−A_2
ZShift = 1.99
Z lm Score = 3.7
NG = 0
DB = 0
SM = 0
Delta_L
Doxycycline
YOL164W−A_2
ZShift = 0.6
Z lm Score = −5.57
NG = 0
DB = 0
SM = 0
Delta_K
Doxycycline
YOL164W−A_2
ZShift = −1.5
Z lm Score = 1.1
NG = 0
DB = 0
SM = 0
Delta_r
Doxycycline
YOL164W−A_2
ZShift = −0.47
Z lm Score = 2.61
NG = 0
DB = 0
SM = 0
Delta_AUC
Doxycycline
YBR157C_1 ICS2
ZShift = 0.27
Z lm Score = 3.69
NG = 0
DB = 0
SM = 0
Delta_L
Doxycycline
YBR157C_1 ICS2
ZShift = 0.48
Z lm Score = −1.37
NG = 0
DB = 0
SM = 0
Delta_K
Doxycycline
YBR157C_1 ICS2
ZShift = 1.41
Z lm Score = −2.08
NG = 0
DB = 0
SM = 0
Delta_r
Doxycycline
YBR157C_1 ICS2
ZShift = 0.2
Z lm Score = 0.32
NG = 0
DB = 0
SM = 0
Delta_AUC
Doxycycline
YGR285C ZUO1
ZShift = 7.54
Z lm Score = 3.68
NG = 0
DB = 0
SM = 0
Delta_L
Doxycycline
YGR285C ZUO1
ZShift = −4.42
Z lm Score = −5.77
NG = 0
DB = 0
SM = 0
Delta_K
Doxycycline
YGR285C ZUO1
ZShift = −5.44
Z lm Score = 3.44
NG = 0
DB = 0
SM = 0
Delta_r
Doxycycline
YGR285C ZUO1
ZShift = −5.64
Z lm Score = −1.49
NG = 0
DB = 0
SM = 0
Delta_AUC
Doxycycline

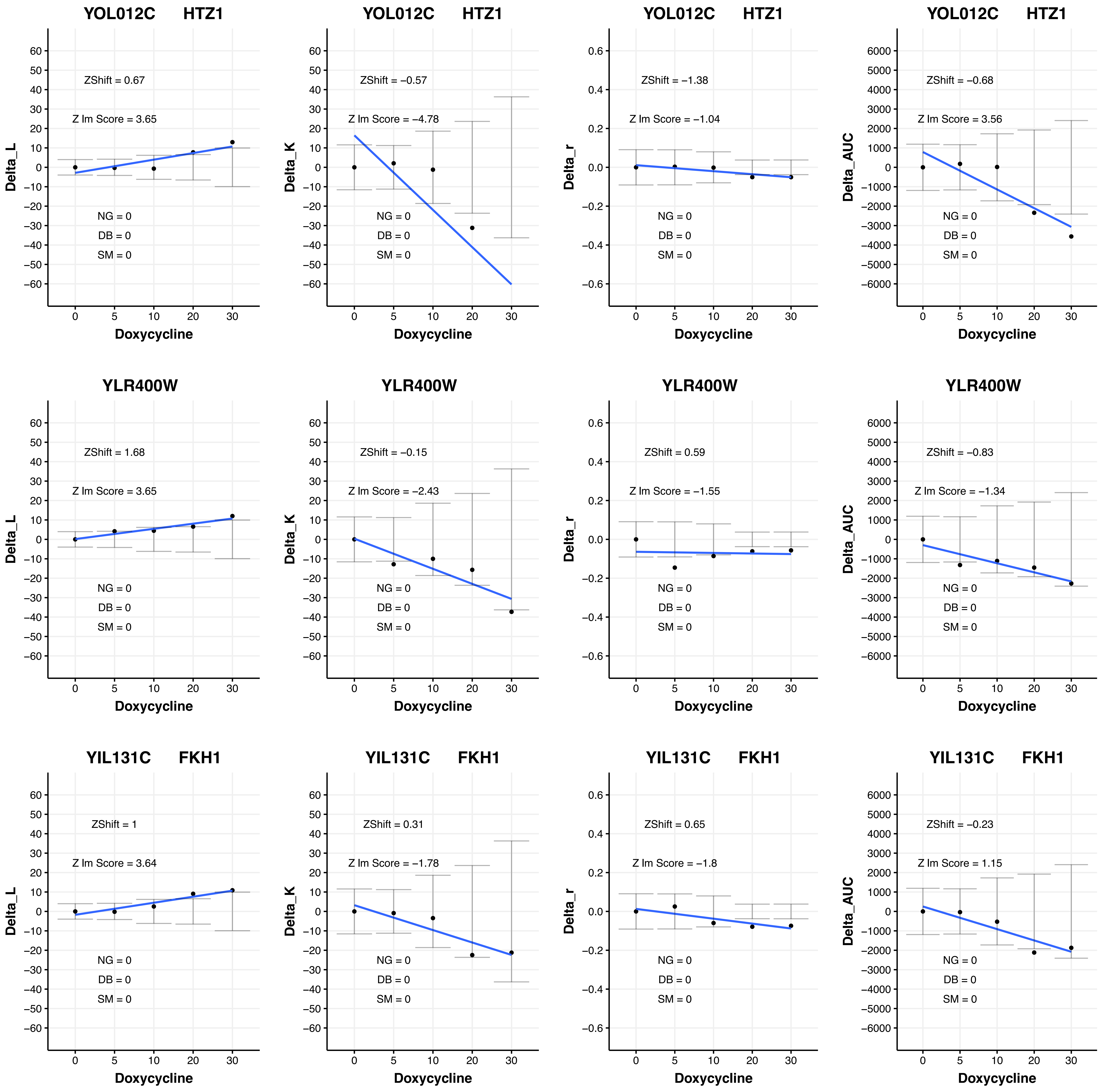
YOL012C HTZ1
ZShift = 0.67
Z lm Score = 3.65
NG = 0
DB = 0
SM = 0
Delta_L
Doxycycline
YOL012C HTZ1
ZShift = −0.57
Z lm Score = −4.78
NG = 0
DB = 0
SM = 0
Delta_K
Doxycycline
YOL012C HTZ1
ZShift = −1.38
Z lm Score = −1.04
NG = 0
DB = 0
SM = 0
Delta_r
Doxycycline
YOL012C HTZ1
ZShift = −0.68
Z lm Score = 3.56
NG = 0
DB = 0
SM = 0
Delta_AUC
Doxycycline
YLR400W
ZShift = 1.68
Z lm Score = 3.65
NG = 0
DB = 0
SM = 0
Delta_L
Doxycycline
YLR400W
ZShift = −0.15
Z lm Score = −2.43
NG = 0
DB = 0
SM = 0
Delta_K
Doxycycline
YLR400W
ZShift = 0.59
Z lm Score = −1.55
NG = 0
DB = 0
SM = 0
Delta_r
Doxycycline
YLR400W
ZShift = −0.83
Z lm Score = −1.34
NG = 0
DB = 0
SM = 0
Delta_AUC
Doxycycline
YIL131C FKH1
ZShift = 1
Z lm Score = 3.64
NG = 0
DB = 0
SM = 0
Delta_L
Doxycycline
YIL131C FKH1
ZShift = 0.31
Z lm Score = −1.78
NG = 0
DB = 0
SM = 0
Delta_K
Doxycycline
YIL131C FKH1
ZShift = 0.65
Z lm Score = −1.8
NG = 0
DB = 0
SM = 0
Delta_r
Doxycycline
YIL131C FKH1
ZShift = −0.23
Z lm Score = 1.15
NG = 0
DB = 0
SM = 0
Delta_AUC
Doxycycline

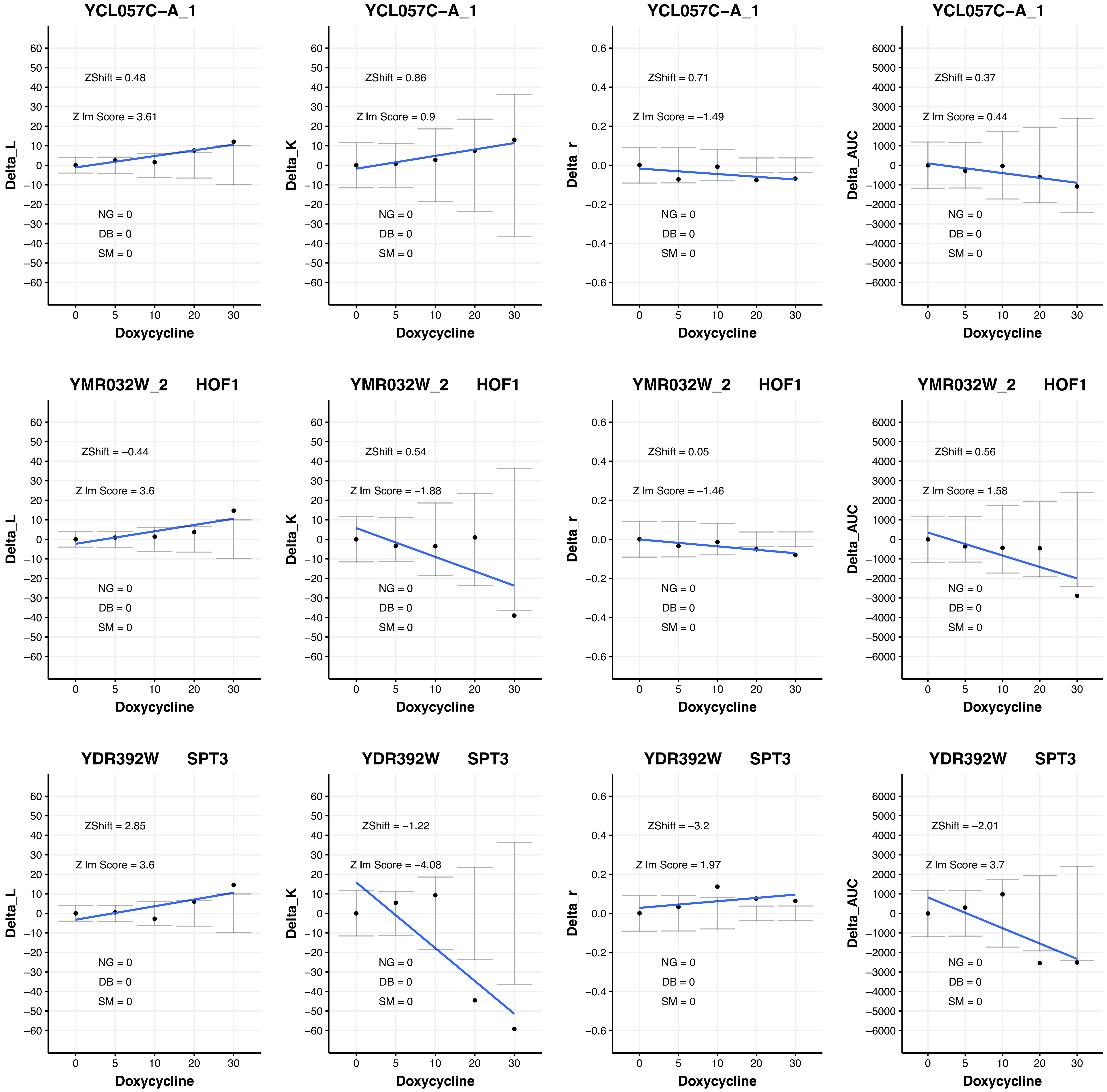
YCL057C−A_1
ZShift = 0.48
Z lm Score = 3.61
NG = 0
DB = 0
SM = 0
Delta_L
Doxycycline
YCL057C−A_1
ZShift = 0.86
Z lm Score = 0.9
Delta_K
YCL057C−A_1
ZShift = 0.71
Z lm Score = −1.49
Delta_r
YCL057C−A_1
ZShift = 0.37
Z lm Score = 0.44
Delta_AUC
YMR032W_2 HOF1
ZShift = −0.44
Z lm Score = 3.6
ZShift = 0.54
Z lm Score = −1.88
ZShift = 0.05
Z lm Score = −1.46
ZShift = 0.56
Z lm Score = 1.58
YDR392W SPT3
ZShift = 2.85
Z lm Score = 3.6
ZShift = −1.22
Z lm Score = −4.08
ZShift = −3.2
Z lm Score = 1.97
ZShift = −2.01
Z lm Score = 3.7

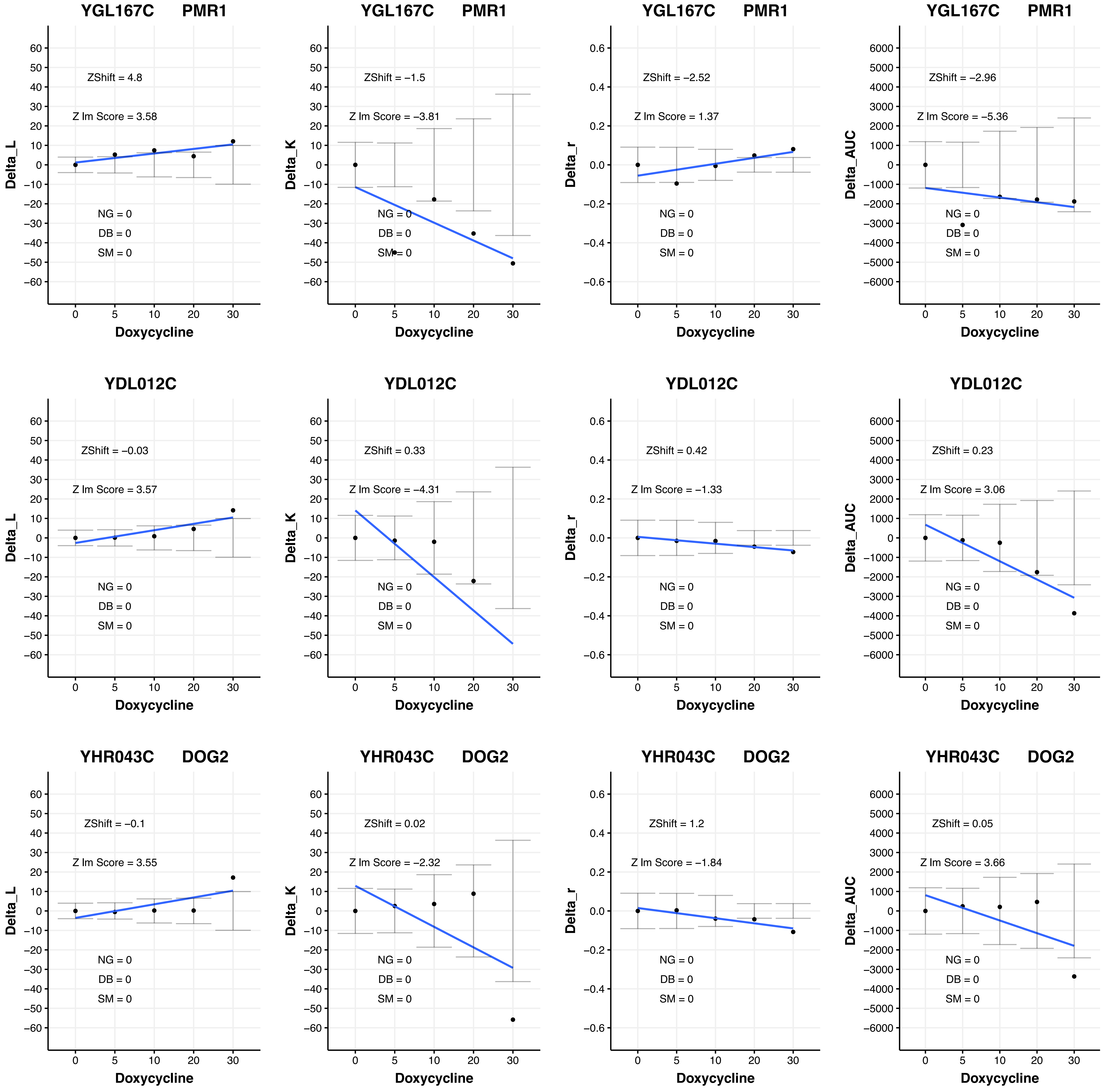

YGL167C PMR1
ZShift = 4.8
Z lm Score = 3.58
NG = 0
DB = 0
SM = 0
Delta_L
Doxycycline
YGL167C PMR1
ZShift = −1.5
Z lm Score = −3.81
NG = 0
DB = 0
SM = 0
Delta_K
Doxycycline
YGL167C PMR1
ZShift = −2.52
Z lm Score = 1.37
NG = 0
DB = 0
SM = 0
Delta_r
Doxycycline
YGL167C PMR1
ZShift = −2.96
Z lm Score = −5.36
NG = 0
DB = 0
SM = 0
Delta_AUC
Doxycycline
YDL012C
ZShift = −0.03
Z lm Score = 3.57
NG = 0
DB = 0
SM = 0
Delta_L
Doxycycline
YDL012C
ZShift = 0.33
Z lm Score = −4.31
NG = 0
DB = 0
SM = 0
Delta_K
Doxycycline
YDL012C
ZShift = 0.42
Z lm Score = −1.33
NG = 0
DB = 0
SM = 0
Delta_r
Doxycycline
YDL012C
ZShift = 0.23
Z lm Score = 3.06
NG = 0
DB = 0
SM = 0
Delta_AUC
Doxycycline
YHR043C DOG2
ZShift = −0.1
Z lm Score = 3.55
NG = 0
DB = 0
SM = 0
Delta_L
Doxycycline
YHR043C DOG2
ZShift = 0.02
Z lm Score = −2.32
NG = 0
DB = 0
SM = 0
Delta_K
Doxycycline
YHR043C DOG2
ZShift = 1.2
Z lm Score = −1.84
NG = 0
DB = 0
SM = 0
Delta_r
Doxycycline
YHR043C DOG2
ZShift = 0.05
Z lm Score = 3.66
NG = 0
DB = 0
SM = 0
Delta_AUC
Doxycycline

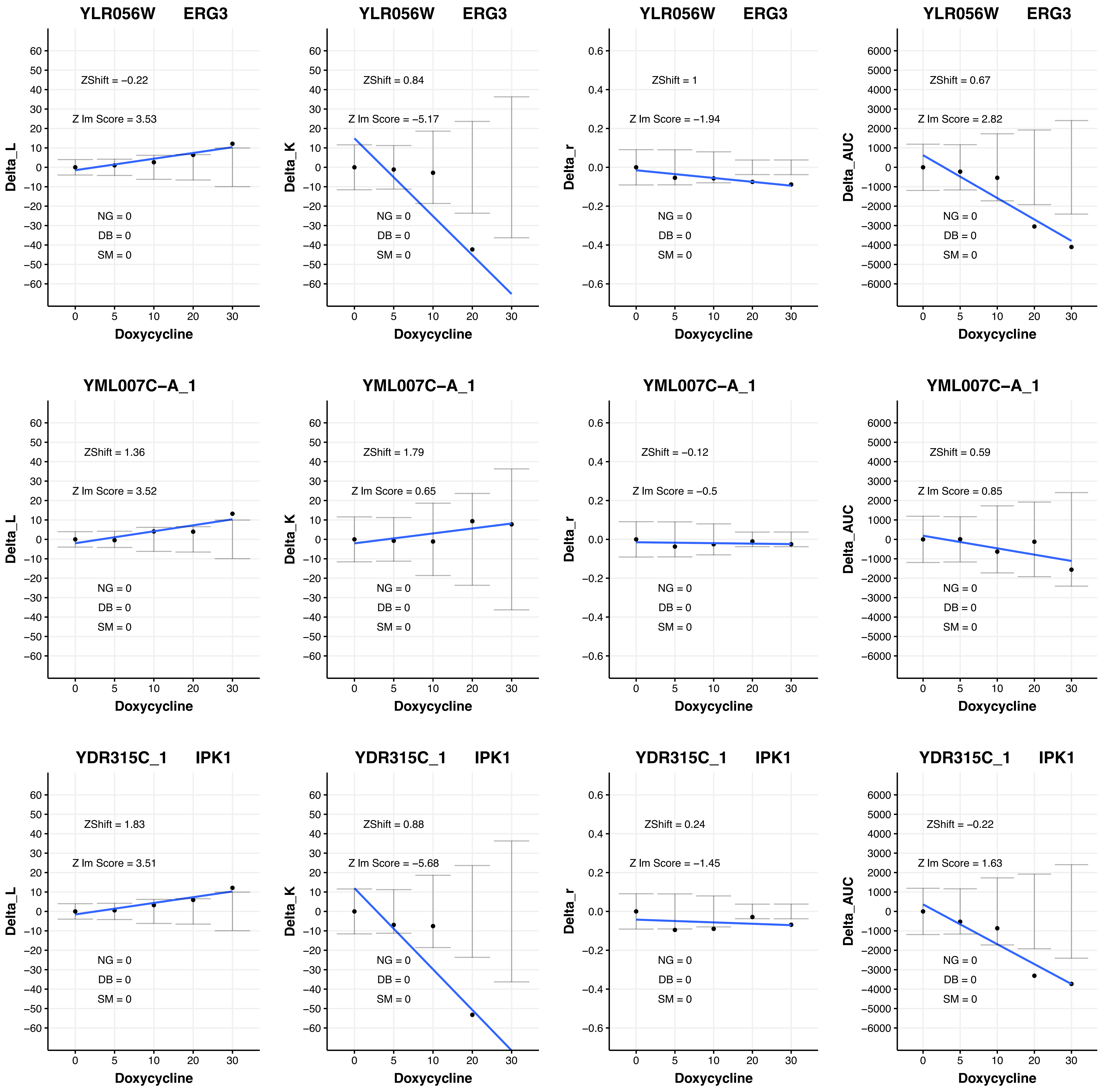
YLR056W ERG3
ZShift = −0.22
Z lm Score = 3.53
NG = 0
DB = 0
SM = 0
Delta_L
Doxycycline
YLR056W ERG3
ZShift = 0.84
Z lm Score = −5.17
NG = 0
DB = 0
SM = 0
Delta_K
Doxycycline
YLR056W ERG3
ZShift = 1
Z lm Score = −1.94
NG = 0
DB = 0
SM = 0
Delta_r
Doxycycline
YLR056W ERG3
ZShift = 0.67
Z lm Score = 2.82
NG = 0
DB = 0
SM = 0
Delta_AUC
Doxycycline
YML007C−A_1
ZShift = 1.36
Z lm Score = 3.52
NG = 0
DB = 0
SM = 0
Delta_L
Doxycycline
YML007C−A_1
ZShift = 1.79
Z lm Score = 0.65
NG = 0
DB = 0
SM = 0
Delta_K
Doxycycline
YML007C−A_1
ZShift = −0.12
Z lm Score = −0.5
NG = 0
DB = 0
SM = 0
Delta_r
Doxycycline
YML007C−A_1
ZShift = 0.59
Z lm Score = 0.85
NG = 0
DB = 0
SM = 0
Delta_AUC
Doxycycline
YDR315C_1 IPK1
ZShift = 1.83
Z lm Score = 3.51
NG = 0
DB = 0
SM = 0
Delta_L
Doxycycline
YDR315C_1 IPK1
ZShift = 0.88
Z lm Score = −5.68
NG = 0
DB = 0
SM = 0
Delta_K
Doxycycline
YDR315C_1 IPK1
ZShift = 0.24
Z lm Score = −1.45
NG = 0
DB = 0
SM = 0
Delta_r
Doxycycline
YDR315C_1 IPK1
ZShift = −0.22
Z lm Score = 1.63
NG = 0
DB = 0
SM = 0
Delta_AUC
Doxycycline

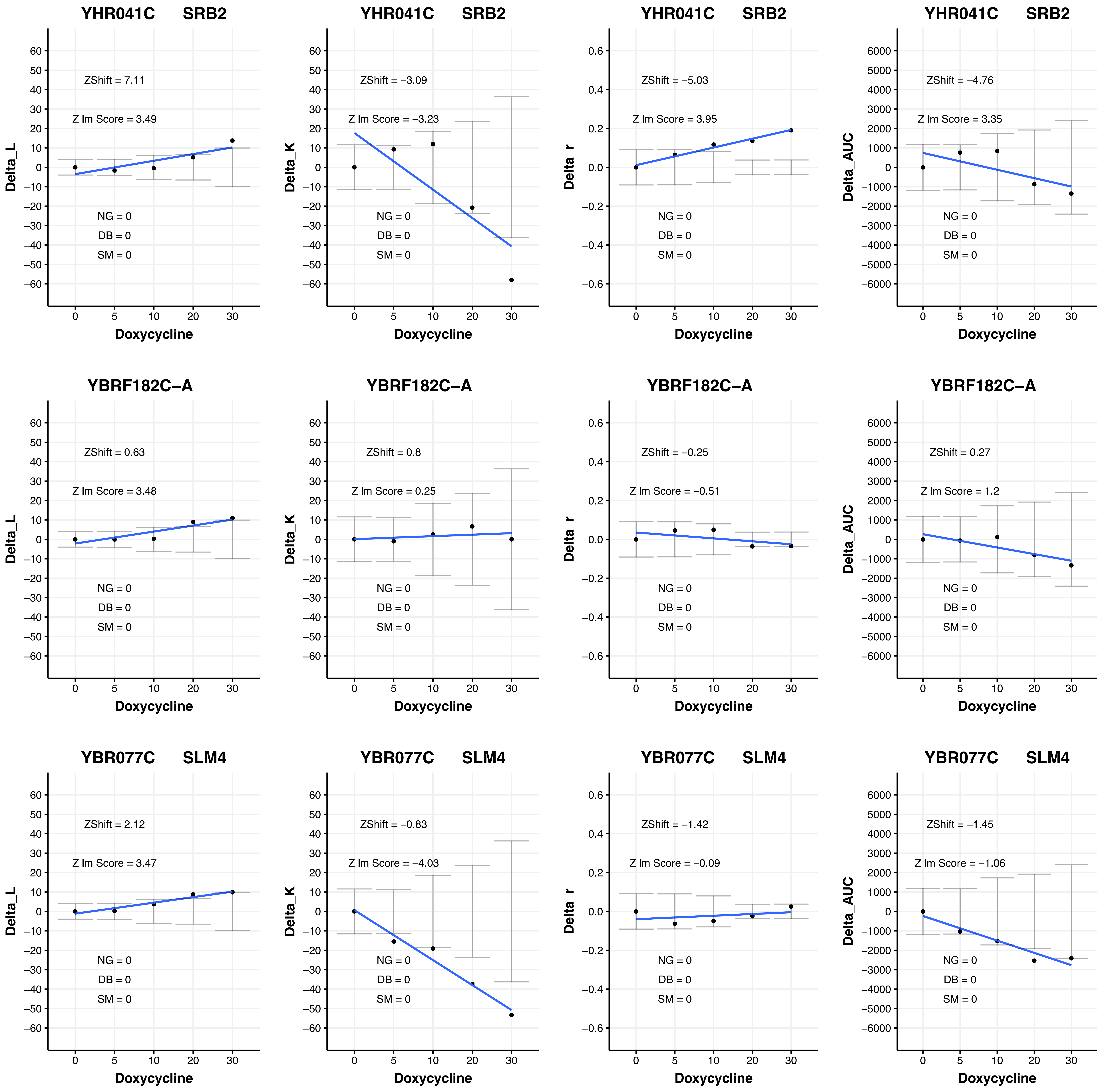

YHR041C SRB2
ZShift = 7.11
Z lm Score = 3.49
Delta_L
NG = 0
DB = 0
SM = 0
Doxycycline
YHR041C SRB2
ZShift = −3.09
Z lm Score = −3.23
Delta_K
NG = 0
DB = 0
SM = 0
Doxycycline
YHR041C SRB2
ZShift = −5.03
Z lm Score = 3.95
Delta_r
NG = 0
DB = 0
SM = 0
Doxycycline
YHR041C SRB2
ZShift = −4.76
Z lm Score = 3.35
Delta_AUC
NG = 0
DB = 0
SM = 0
Doxycycline
YBRF182C−A
ZShift = 0.63
Z lm Score = 3.48
Delta_L
NG = 0
DB = 0
SM = 0
Doxycycline
YBRF182C−A
ZShift = 0.8
Z lm Score = 0.25
Delta_K
NG = 0
DB = 0
SM = 0
Doxycycline
YBRF182C−A
ZShift = −0.25
Z lm Score = −0.51
Delta_r
NG = 0
DB = 0
SM = 0
Doxycycline
YBRF182C−A
ZShift = 0.27
Z lm Score = 1.2
Delta_AUC
NG = 0
DB = 0
SM = 0
Doxycycline
YBR077C SLM4
ZShift = 2.12
Z lm Score = 3.47
Delta_L
NG = 0
DB = 0
SM = 0
Doxycycline
YBR077C SLM4
ZShift = −0.83
Z lm Score = −4.03
Delta_K
NG = 0
DB = 0
SM = 0
Doxycycline
YBR077C SLM4
ZShift = −1.42
Z lm Score = −0.09
Delta_r
NG = 0
DB = 0
SM = 0
Doxycycline
YBR077C SLM4
ZShift = −1.45
Z lm Score = −1.06
Delta_AUC
NG = 0
DB = 0
SM = 0
Doxycycline

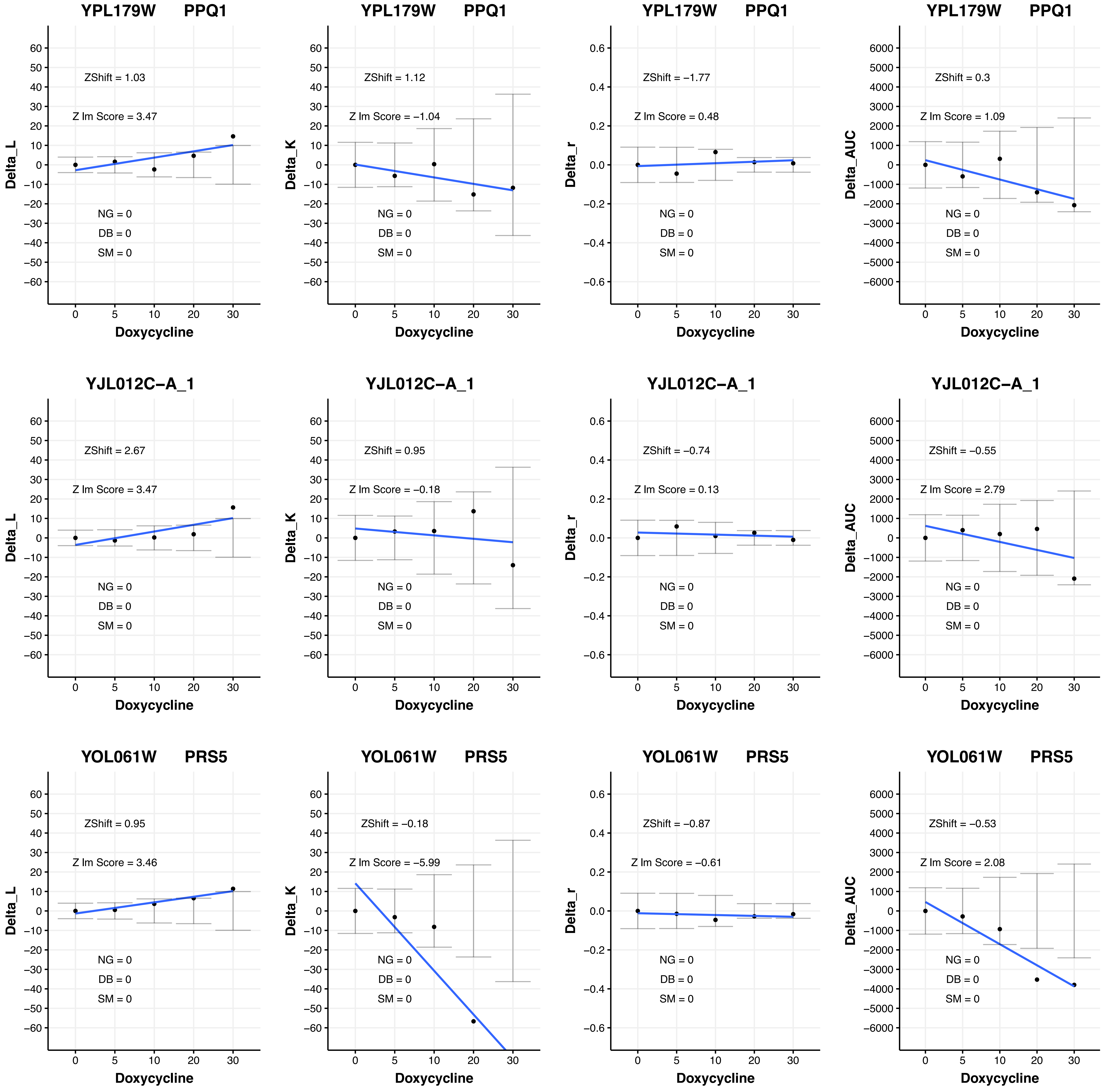

YPL179W PPQ1
ZShift = 1.03
Z lm Score = 3.47
NG = 0
DB = 0
SM = 0
Delta_L
Doxycycline
YPL179W PPQ1
ZShift = 1.12
Z lm Score = −1.04
NG = 0
DB = 0
SM = 0
Delta_K
Doxycycline
YPL179W PPQ1
ZShift = −1.77
Z lm Score = 0.48
NG = 0
DB = 0
SM = 0
Delta_r
Doxycycline
YPL179W PPQ1
ZShift = 0.3
Z lm Score = 1.09
NG = 0
DB = 0
SM = 0
Delta_AUC
Doxycycline
YJL012C−A_1
ZShift = 2.67
Z lm Score = 3.47
NG = 0
DB = 0
SM = 0
Delta_L
Doxycycline
YJL012C−A_1
ZShift = 0.95
Z lm Score = −0.18
NG = 0
DB = 0
SM = 0
Delta_K
Doxycycline
YJL012C−A_1
ZShift = −0.74
Z lm Score = 0.13
NG = 0
DB = 0
SM = 0
Delta_r
Doxycycline
YJL012C−A_1
ZShift = −0.55
Z lm Score = 2.79
NG = 0
DB = 0
SM = 0
Delta_AUC
Doxycycline
YOL061W PRS5
ZShift = 0.95
Z lm Score = 3.46
NG = 0
DB = 0
SM = 0
Delta_L
Doxycycline
YOL061W PRS5
ZShift = −0.18
Z lm Score = −5.99
NG = 0
DB = 0
SM = 0
Delta_K
Doxycycline
YOL061W PRS5
ZShift = −0.87
Z lm Score = −0.61
NG = 0
DB = 0
SM = 0
Delta_r
Doxycycline
YOL061W PRS5
ZShift = −0.53
Z lm Score = 2.08
NG = 0
DB = 0
SM = 0
Delta_AUC
Doxycycline

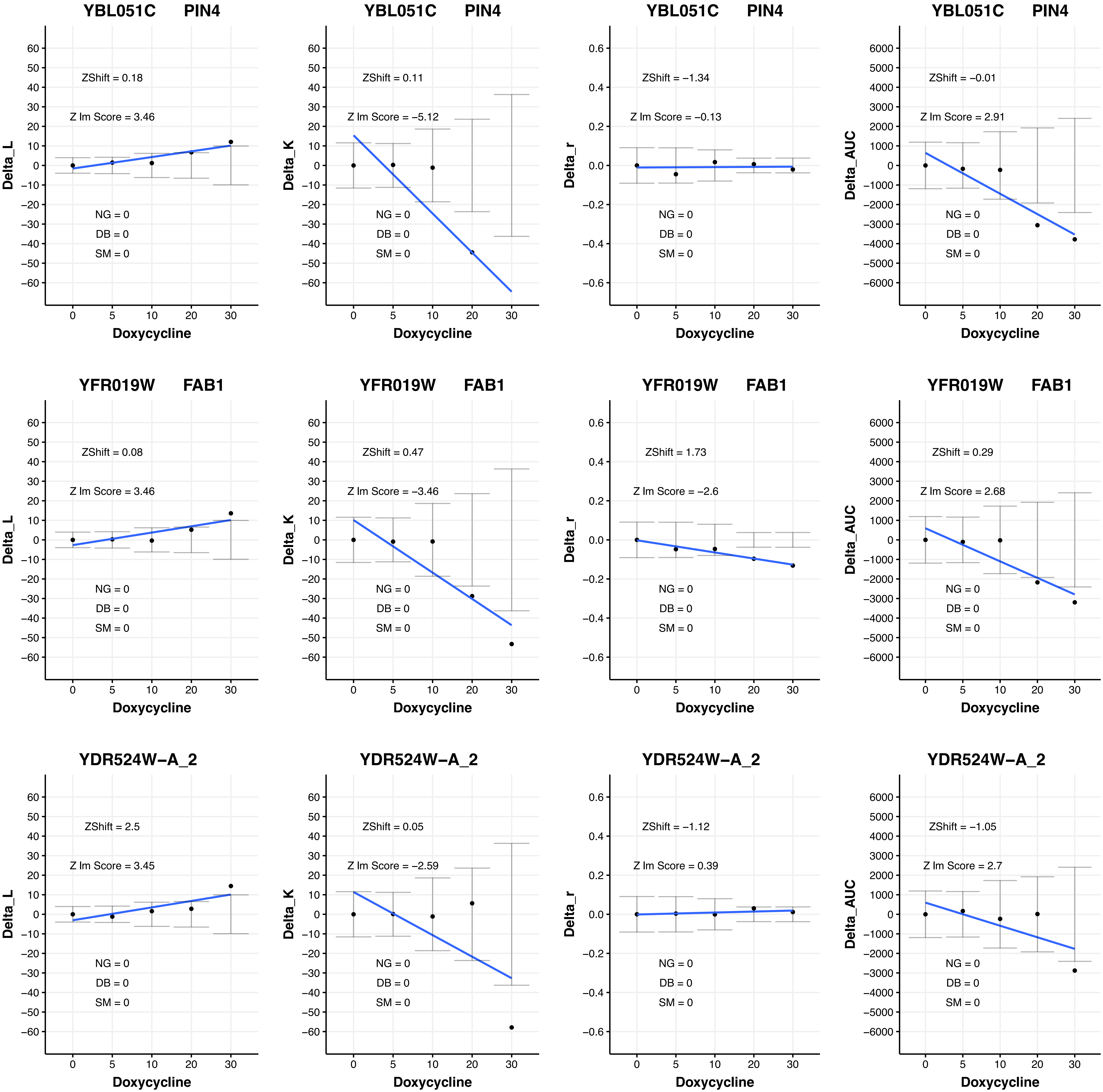
YBL051C PIN4
Delta_L
Doxycycline
ZShift = 0.18
Z lm Score = 3.46
NG = 0
DB = 0
SM = 0
YBL051C PIN4
Delta_K
Doxycycline
ZShift = 0.11
Z lm Score = −5.12
NG = 0
DB = 0
SM = 0
YBL051C PIN4
Delta_r
Doxycycline
ZShift = −1.34
Z lm Score = −0.13
NG = 0
DB = 0
SM = 0
YBL051C PIN4
Delta_AUC
Doxycycline
ZShift = −0.01
Z lm Score = 2.91
NG = 0
DB = 0
SM = 0
YFR019W FAB1
Delta_L
Doxycycline
ZShift = 0.08
Z lm Score = 3.46
NG = 0
DB = 0
SM = 0
YFR019W FAB1
Delta_K
Doxycycline
ZShift = 0.47
Z lm Score = −3.46
NG = 0
DB = 0
SM = 0
YFR019W FAB1
Delta_r
Doxycycline
ZShift = 1.73
Z lm Score = −2.6
NG = 0
DB = 0
SM = 0
YFR019W FAB1
Delta_AUC
Doxycycline
ZShift = 0.29
Z lm Score = 2.68
NG = 0
DB = 0
SM = 0
YDR524W−A_2
Delta_L
Doxycycline
ZShift = 2.5
Z lm Score = 3.45
NG = 0
DB = 0
SM = 0
YDR524W−A_2
Delta_K
Doxycycline
ZShift = 0.05
Z lm Score = −2.59
NG = 0
DB = 0
SM = 0
YDR524W−A_2
Delta_r
Doxycycline
ZShift = −1.12
Z lm Score = 0.39
NG = 0
DB = 0
SM = 0
YDR524W−A_2
Delta_AUC
Doxycycline
ZShift = −1.05
Z lm Score = 2.7
NG = 0
DB = 0
SM = 0

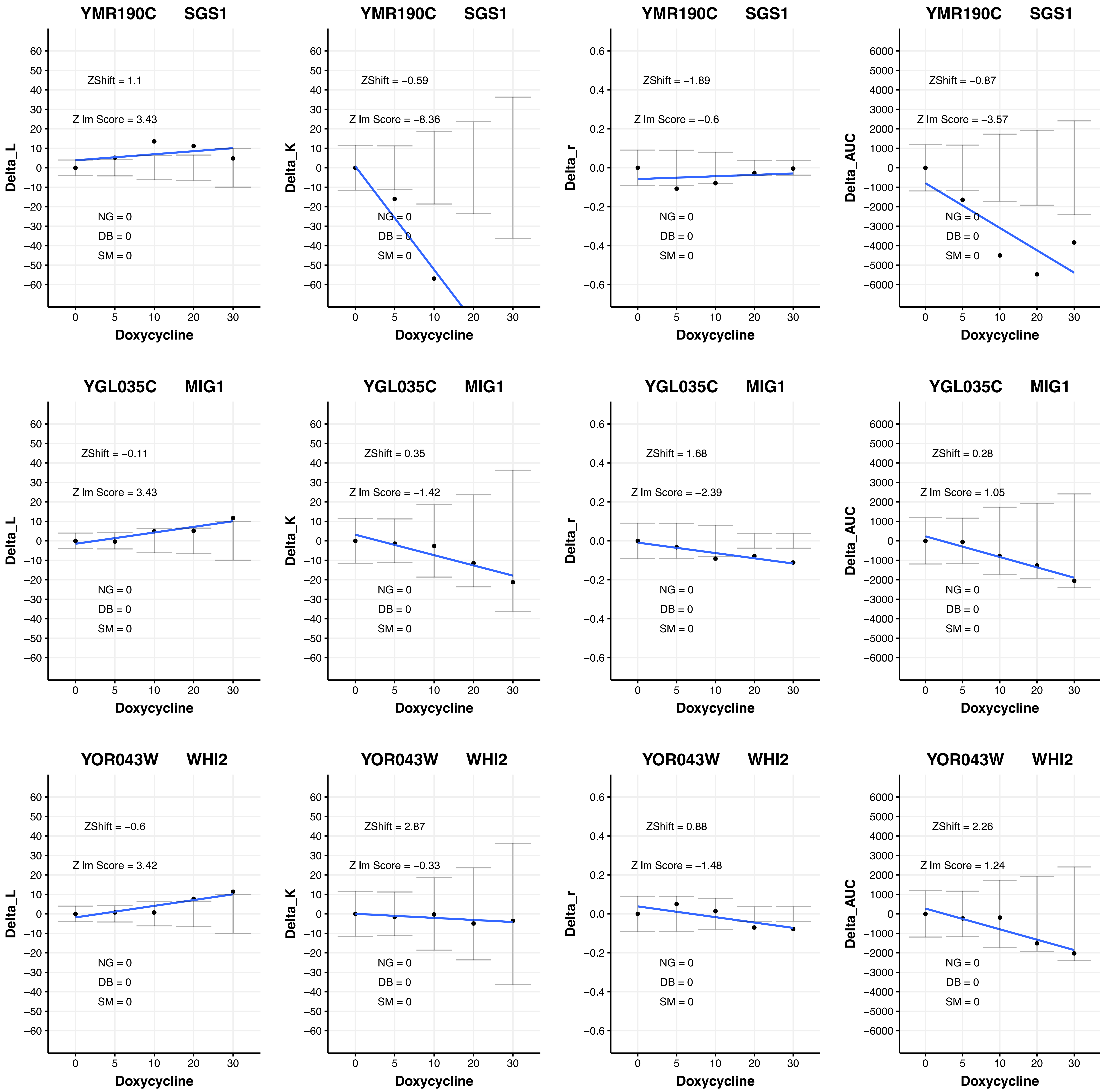

YMR190C SGS1
ZShift = 1.1
Z lm Score = 3.43
NG = 0
DB = 0
SM = 0
Delta_L
Doxycycline
YMR190C SGS1
ZShift = −0.59
Z lm Score = −8.36
NG = 0
DB = 0
SM = 0
Delta_K
Doxycycline
YMR190C SGS1
ZShift = −1.89
Z lm Score = −0.6
NG = 0
DB = 0
SM = 0
Delta_r
Doxycycline
YMR190C SGS1
ZShift = −0.87
Z lm Score = −3.57
NG = 0
DB = 0
SM = 0
Delta_AUC
Doxycycline
YGL035C MIG1
ZShift = −0.11
Z lm Score = 3.43
NG = 0
DB = 0
SM = 0
Delta_L
Doxycycline
YGL035C MIG1
ZShift = 0.35
Z lm Score = −1.42
NG = 0
DB = 0
SM = 0
Delta_K
Doxycycline
YGL035C MIG1
ZShift = 1.68
Z lm Score = −2.39
NG = 0
DB = 0
SM = 0
Delta_r
Doxycycline
YGL035C MIG1
ZShift = 0.28
Z lm Score = 1.05
NG = 0
DB = 0
SM = 0
Delta_AUC
Doxycycline
YOR043W WHI2
ZShift = −0.6
Z lm Score = 3.42
NG = 0
DB = 0
SM = 0
Delta_L
Doxycycline
YOR043W WHI2
ZShift = 2.87
Z lm Score = −0.33
NG = 0
DB = 0
SM = 0
Delta_K
Doxycycline
YOR043W WHI2
ZShift = 0.88
Z lm Score = −1.48
NG = 0
DB = 0
SM = 0
Delta_r
Doxycycline
YOR043W WHI2
ZShift = 2.26
Z lm Score = 1.24
NG = 0
DB = 0
SM = 0
Delta_AUC
Doxycycline

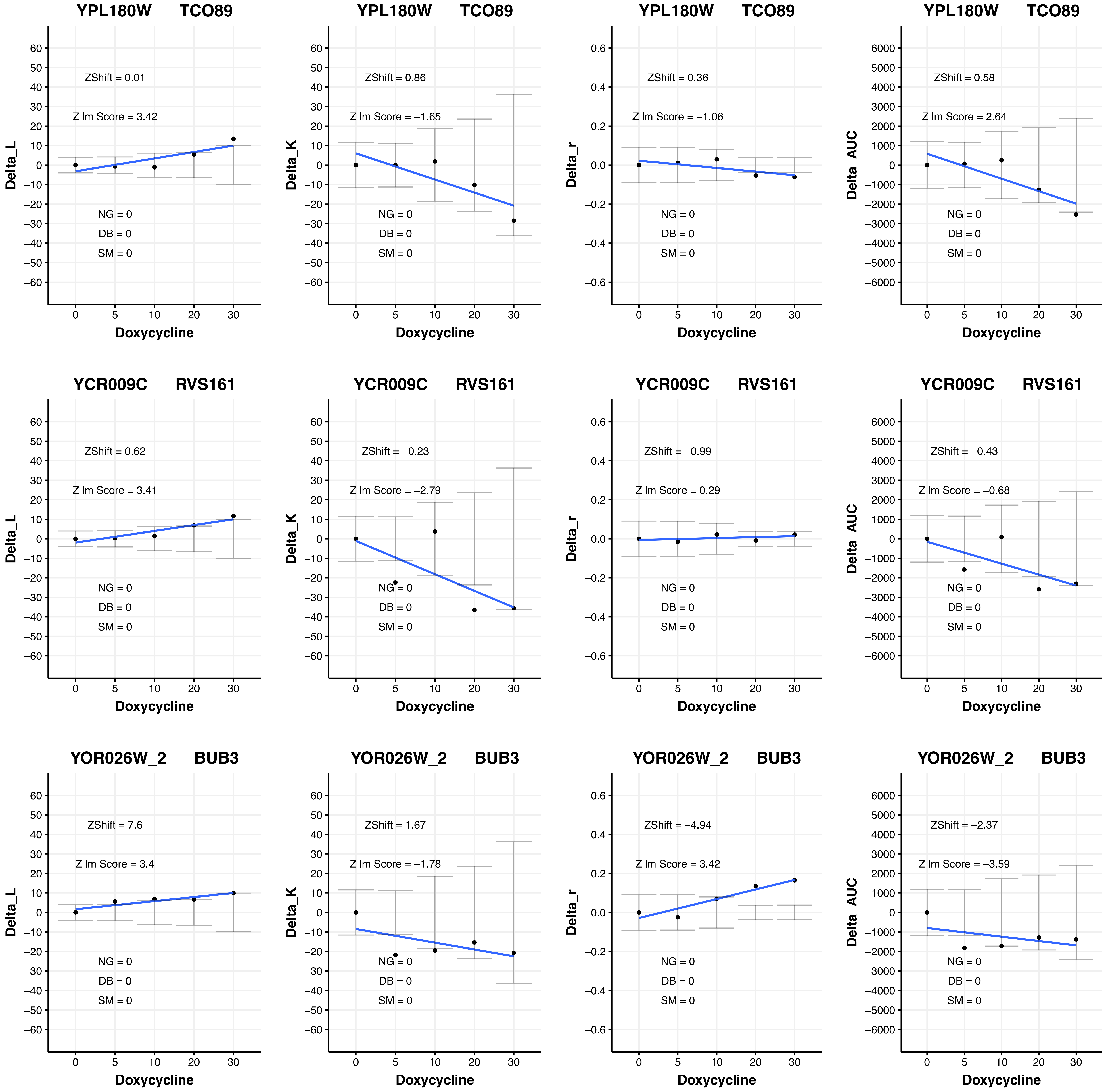
YPL180W TCO89
ZShift = 0.01
Z lm Score = 3.42
NG = 0
DB = 0
SM = 0
Delta_L
Doxycycline
YPL180W TCO89
ZShift = 0.86
Z lm Score = −1.65
NG = 0
DB = 0
SM = 0
Delta_K
Doxycycline
YPL180W TCO89
ZShift = 0.36
Z lm Score = −1.06
NG = 0
DB = 0
SM = 0
Delta_r
Doxycycline
YPL180W TCO89
ZShift = 0.58
Z lm Score = 2.64
NG = 0
DB = 0
SM = 0
Delta_AUC
Doxycycline
YCR009C RVS161
ZShift = 0.62
Z lm Score = 3.41
NG = 0
DB = 0
SM = 0
Delta_L
Doxycycline
YCR009C RVS161
ZShift = −0.23
Z lm Score = −2.79
NG = 0
DB = 0
SM = 0
Delta_K
Doxycycline
YCR009C RVS161
ZShift = −0.99
Z lm Score = 0.29
NG = 0
DB = 0
SM = 0
Delta_r
Doxycycline
YCR009C RVS161
ZShift = −0.43
Z lm Score = −0.68
NG = 0
DB = 0
SM = 0
Delta_AUC
Doxycycline
YOR026W_2 BUB3
ZShift = 7.6
Z lm Score = 3.4
NG = 0
DB = 0
SM = 0
Delta_L
Doxycycline
YOR026W_2 BUB3
ZShift = 1.67
Z lm Score = −1.78
NG = 0
DB = 0
SM = 0
Delta_K
Doxycycline
YOR026W_2 BUB3
ZShift = −4.94
Z lm Score = 3.42
NG = 0
DB = 0
SM = 0
Delta_r
Doxycycline
YOR026W_2 BUB3
ZShift = −2.37
Z lm Score = −3.59
NG = 0
DB = 0
SM = 0
Delta_AUC
Doxycycline

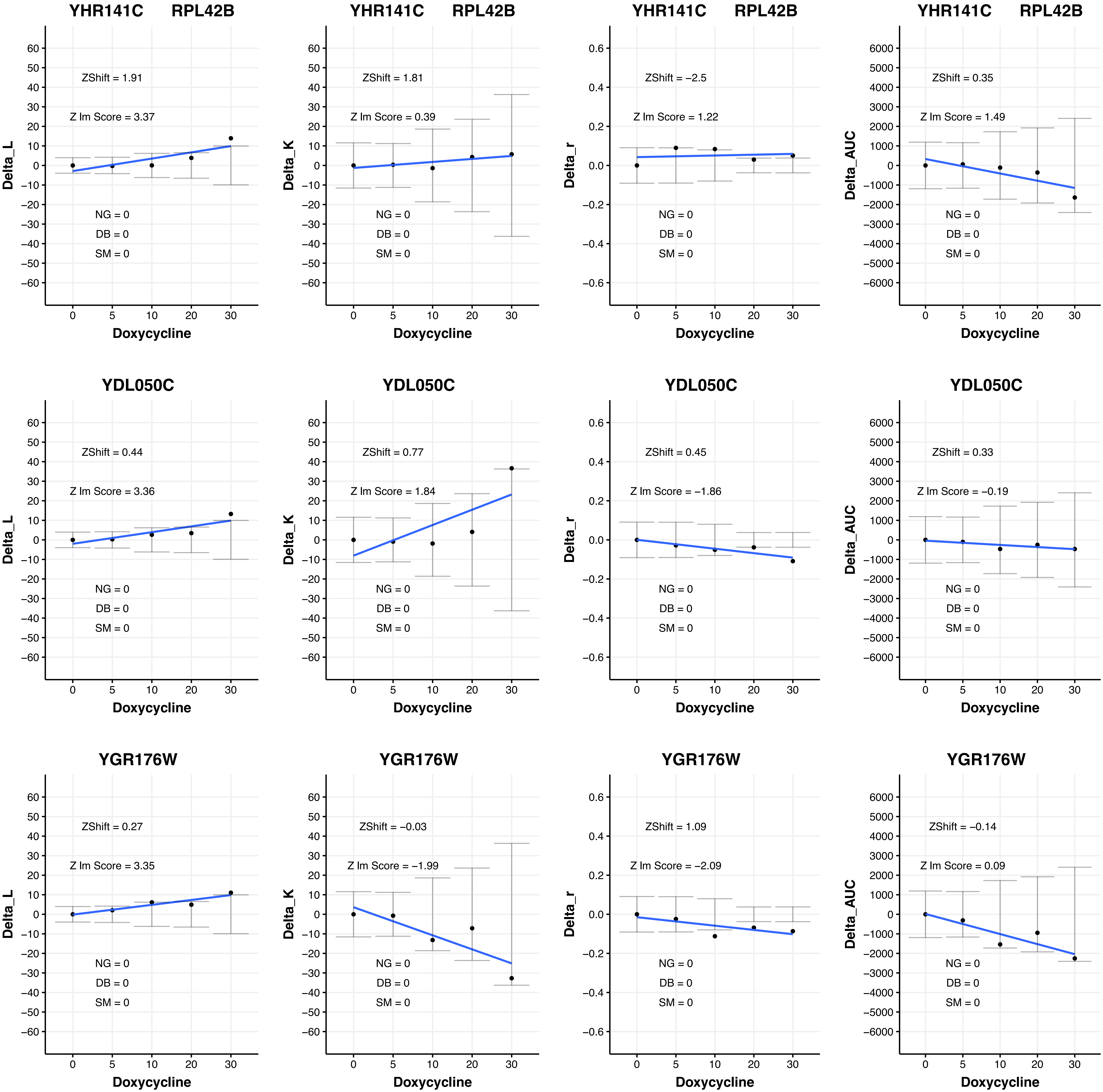

YHR141C RPL42B
ZShift = 1.91
Z lm Score = 3.37
NG = 0
DB = 0
SM = 0
Delta_L
Doxycycline
YHR141C RPL42B
ZShift = 1.81
Z lm Score = 0.39
NG = 0
DB = 0
SM = 0
Delta_K
Doxycycline
YHR141C RPL42B
ZShift = −2.5
Z lm Score = 1.22
NG = 0
DB = 0
SM = 0
Delta_r
Doxycycline
YHR141C RPL42B
ZShift = 0.35
Z lm Score = 1.49
NG = 0
DB = 0
SM = 0
Delta_AUC
Doxycycline
YDL050C
ZShift = 0.44
Z lm Score = 3.36
NG = 0
DB = 0
SM = 0
Delta_L
Doxycycline
YDL050C
ZShift = 0.77
Z lm Score = 1.84
NG = 0
DB = 0
SM = 0
Delta_K
Doxycycline
YDL050C
ZShift = 0.45
Z lm Score = −1.86
NG = 0
DB = 0
SM = 0
Delta_r
Doxycycline
YDL050C
ZShift = 0.33
Z lm Score = −0.19
NG = 0
DB = 0
SM = 0
Delta_AUC
Doxycycline
YGR176W
ZShift = 0.27
Z lm Score = 3.35
NG = 0
DB = 0
SM = 0
Delta_L
Doxycycline
YGR176W
ZShift = −0.03
Z lm Score = −1.99
NG = 0
DB = 0
SM = 0
Delta_K
Doxycycline
YGR176W
ZShift = 1.09
Z lm Score = −2.09
NG = 0
DB = 0
SM = 0
Delta_r
Doxycycline
YGR176W
ZShift = −0.14
Z lm Score = 0.09
NG = 0
DB = 0
SM = 0
Delta_AUC
Doxycycline

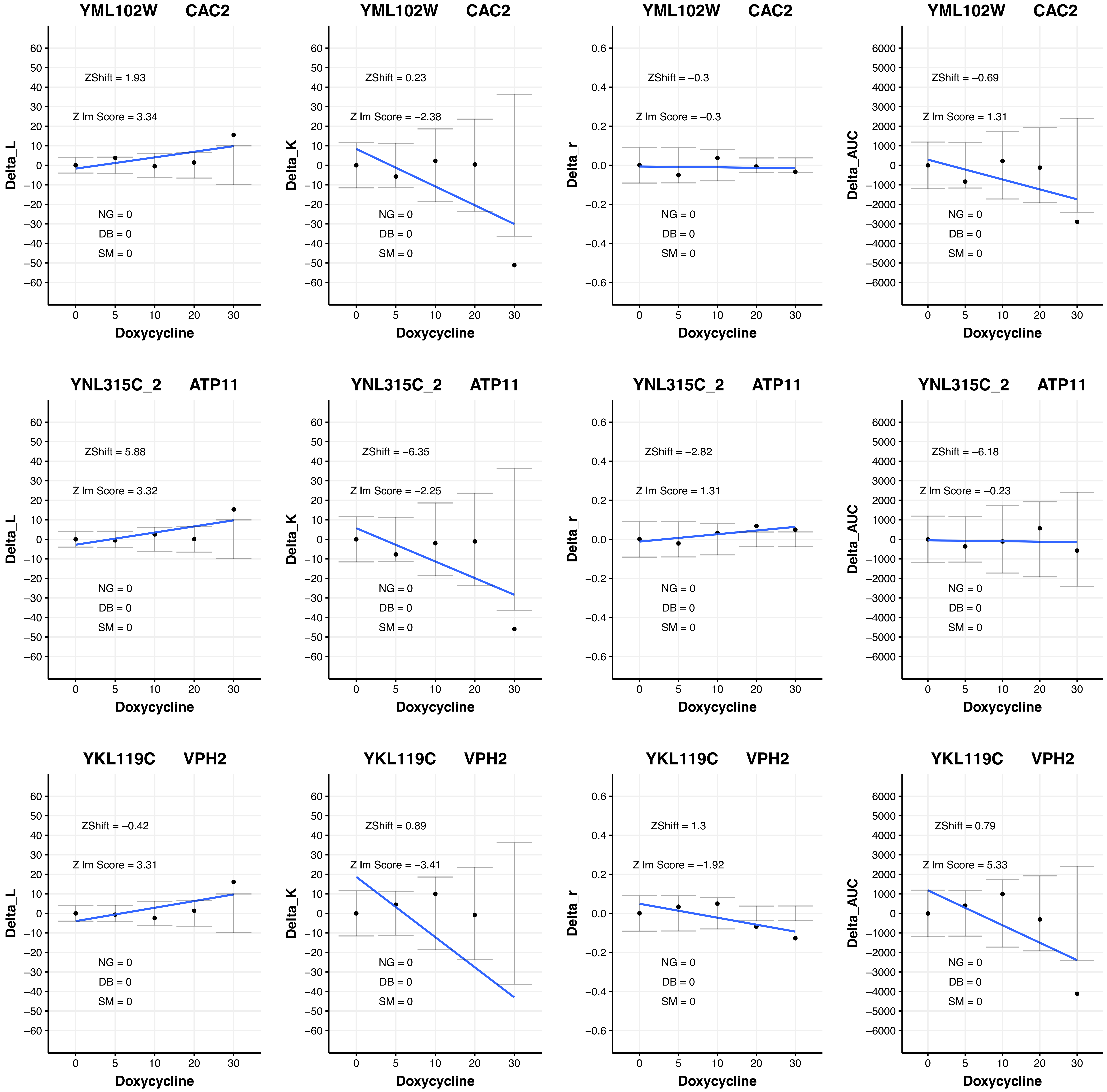
YML102W CAC2
ZShift = 1.93
Z lm Score = 3.34
NG = 0
DB = 0
SM = 0
Delta_L
Doxycycline
YML102W CAC2
ZShift = 0.23
Z lm Score = −2.38
NG = 0
DB = 0
SM = 0
Delta_K
Doxycycline
YML102W CAC2
ZShift = −0.3
Z lm Score = −0.3
NG = 0
DB = 0
SM = 0
Delta_r
Doxycycline
YML102W CAC2
ZShift = −0.69
Z lm Score = 1.31
NG = 0
DB = 0
SM = 0
Delta_AUC
Doxycycline
YNL315C_2 ATP11
ZShift = 5.88
Z lm Score = 3.32
NG = 0
DB = 0
SM = 0
Delta_L
Doxycycline
YNL315C_2 ATP11
ZShift = −6.35
Z lm Score = −2.25
NG = 0
DB = 0
SM = 0
Delta_K
Doxycycline
YNL315C_2 ATP11
ZShift = −2.82
Z lm Score = 1.31
NG = 0
DB = 0
SM = 0
Delta_r
Doxycycline
YNL315C_2 ATP11
ZShift = −6.18
Z lm Score = −0.23
NG = 0
DB = 0
SM = 0
Delta_AUC
Doxycycline
YKL119C VPH2
ZShift = −0.42
Z lm Score = 3.31
NG = 0
DB = 0
SM = 0
Delta_L
Doxycycline
YKL119C VPH2
ZShift = 0.89
Z lm Score = −3.41
NG = 0
DB = 0
SM = 0
Delta_K
Doxycycline
YKL119C VPH2
ZShift = 1.3
Z lm Score = −1.92
NG = 0
DB = 0
SM = 0
Delta_r
Doxycycline
YKL119C VPH2
ZShift = 0.79
Z lm Score = 5.33
NG = 0
DB = 0
SM = 0
Delta_AUC
Doxycycline

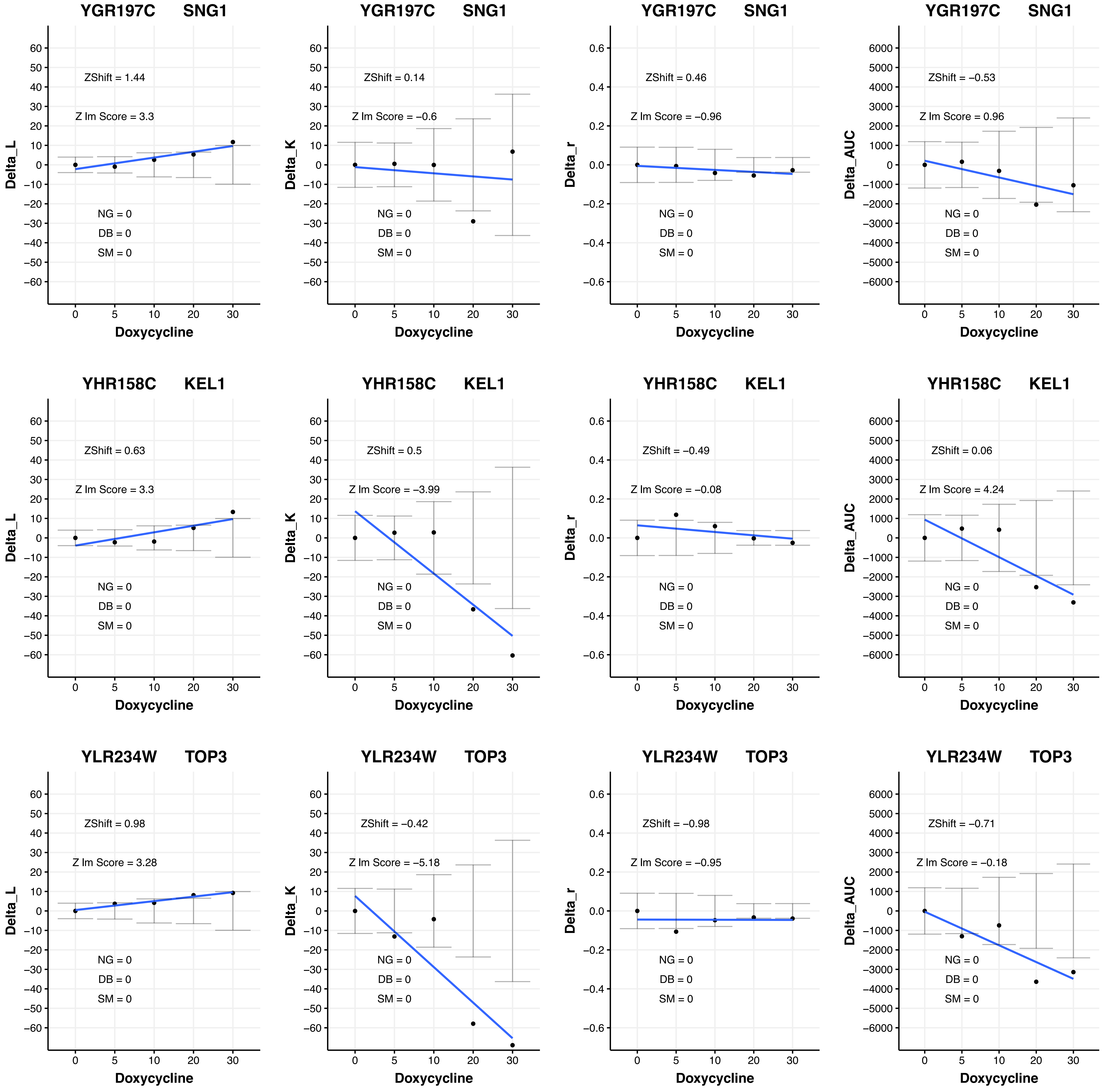

YGR197C SNG1
ZShift = 1.44
Z lm Score = 3.3
NG = 0
DB = 0
SM = 0
Delta_L
Doxycycline
YGR197C SNG1
ZShift = 0.14
Z lm Score = −0.6
NG = 0
DB = 0
SM = 0
Delta_K
Doxycycline
YGR197C SNG1
ZShift = 0.46
Z lm Score = −0.96
NG = 0
DB = 0
SM = 0
Delta_r
Doxycycline
YGR197C SNG1
ZShift = −0.53
Z lm Score = 0.96
NG = 0
DB = 0
SM = 0
Delta_AUC
Doxycycline
YHR158C KEL1
ZShift = 0.63
Z lm Score = 3.3
NG = 0
DB = 0
SM = 0
Delta_L
Doxycycline
YHR158C KEL1
ZShift = 0.5
Z lm Score = −3.99
NG = 0
DB = 0
SM = 0
Delta_K
Doxycycline
YHR158C KEL1
ZShift = −0.49
Z lm Score = −0.08
NG = 0
DB = 0
SM = 0
Delta_r
Doxycycline
YHR158C KEL1
ZShift = 0.06
Z lm Score = 4.24
NG = 0
DB = 0
SM = 0
Delta_AUC
Doxycycline
YLR234W TOP3
ZShift = 0.98
Z lm Score = 3.28
NG = 0
DB = 0
SM = 0
Delta_L
Doxycycline
YLR234W TOP3
ZShift = −0.42
Z lm Score = −5.18
NG = 0
DB = 0
SM = 0
Delta_K
Doxycycline
YLR234W TOP3
ZShift = −0.98
Z lm Score = −0.95
NG = 0
DB = 0
SM = 0
Delta_r
Doxycycline
YLR234W TOP3
ZShift = −0.71
Z lm Score = −0.18
NG = 0
DB = 0
SM = 0
Delta_AUC
Doxycycline

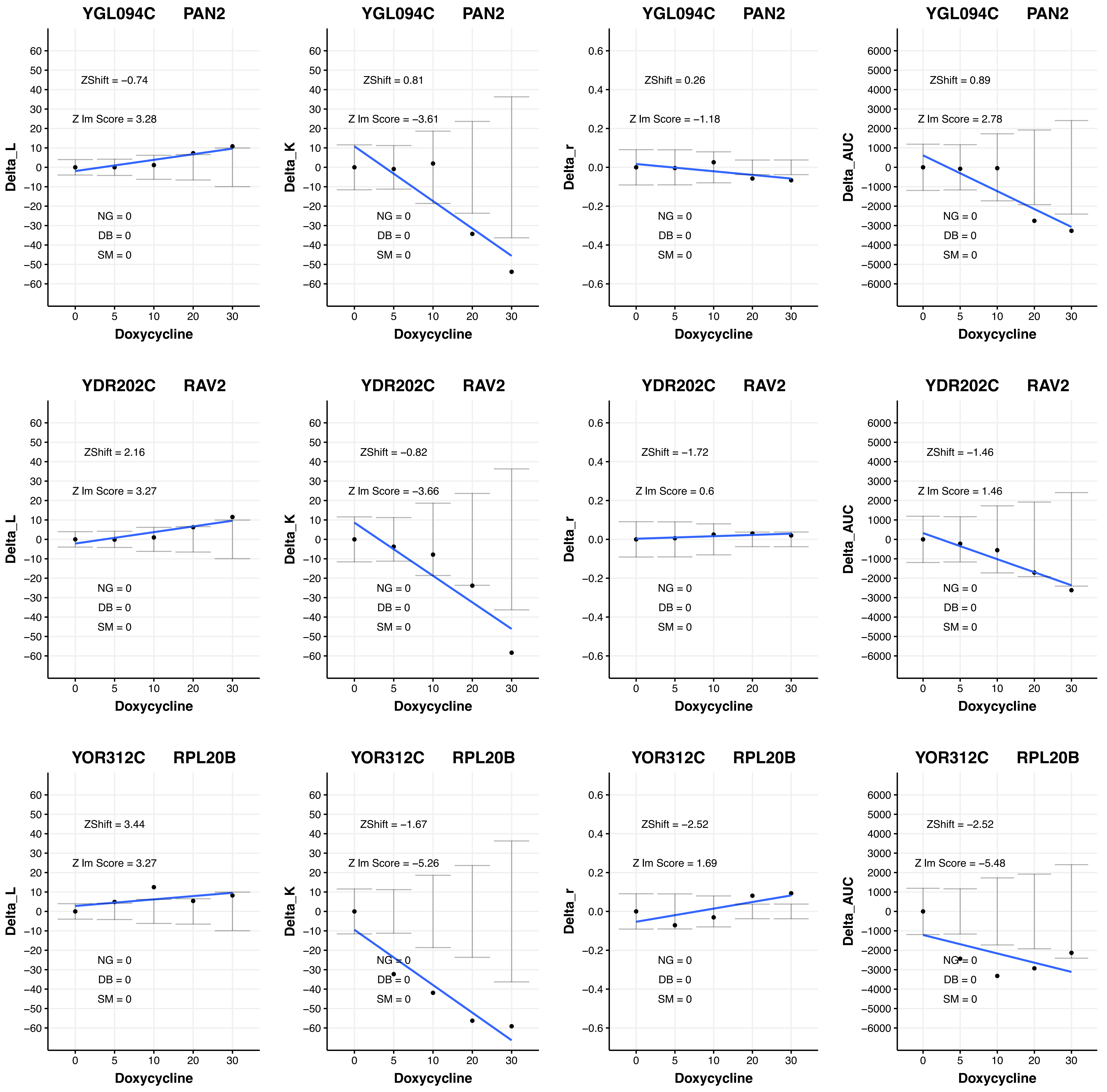
YGL094C PAN2
ZShift = −0.74
Z lm Score = 3.28
NG = 0
DB = 0
SM = 0
Delta_L
Doxycycline
YGL094C PAN2
ZShift = 0.81
Z lm Score = −3.61
NG = 0
DB = 0
SM = 0
Delta_K
Doxycycline
YGL094C PAN2
ZShift = 0.26
Z lm Score = −1.18
NG = 0
DB = 0
SM = 0
Delta_r
Doxycycline
YGL094C PAN2
ZShift = 0.89
Z lm Score = 2.78
NG = 0
DB = 0
SM = 0
Delta_AUC
Doxycycline
YDR202C RAV2
ZShift = 2.16
Z lm Score = 3.27
NG = 0
DB = 0
SM = 0
Delta_L
Doxycycline
YDR202C RAV2
ZShift = −0.82
Z lm Score = −3.66
NG = 0
DB = 0
SM = 0
Delta_K
Doxycycline
YDR202C RAV2
ZShift = −1.72
Z lm Score = 0.6
NG = 0
DB = 0
SM = 0
Delta_r
Doxycycline
YDR202C RAV2
ZShift = −1.46
Z lm Score = 1.46
NG = 0
DB = 0
SM = 0
Delta_AUC
Doxycycline
YOR312C RPL20B
ZShift = 3.44
Z lm Score = 3.27
NG = 0
DB = 0
SM = 0
Delta_L
Doxycycline
YOR312C RPL20B
ZShift = −1.67
Z lm Score = −5.26
NG = 0
DB = 0
SM = 0
Delta_K
Doxycycline
YOR312C RPL20B
ZShift = −2.52
Z lm Score = 1.69
NG = 0
DB = 0
SM = 0
Delta_r
Doxycycline
YOR312C RPL20B
ZShift = −2.52
Z lm Score = −5.48
NG = 0
DB = 0
SM = 0
Delta_AUC
Doxycycline

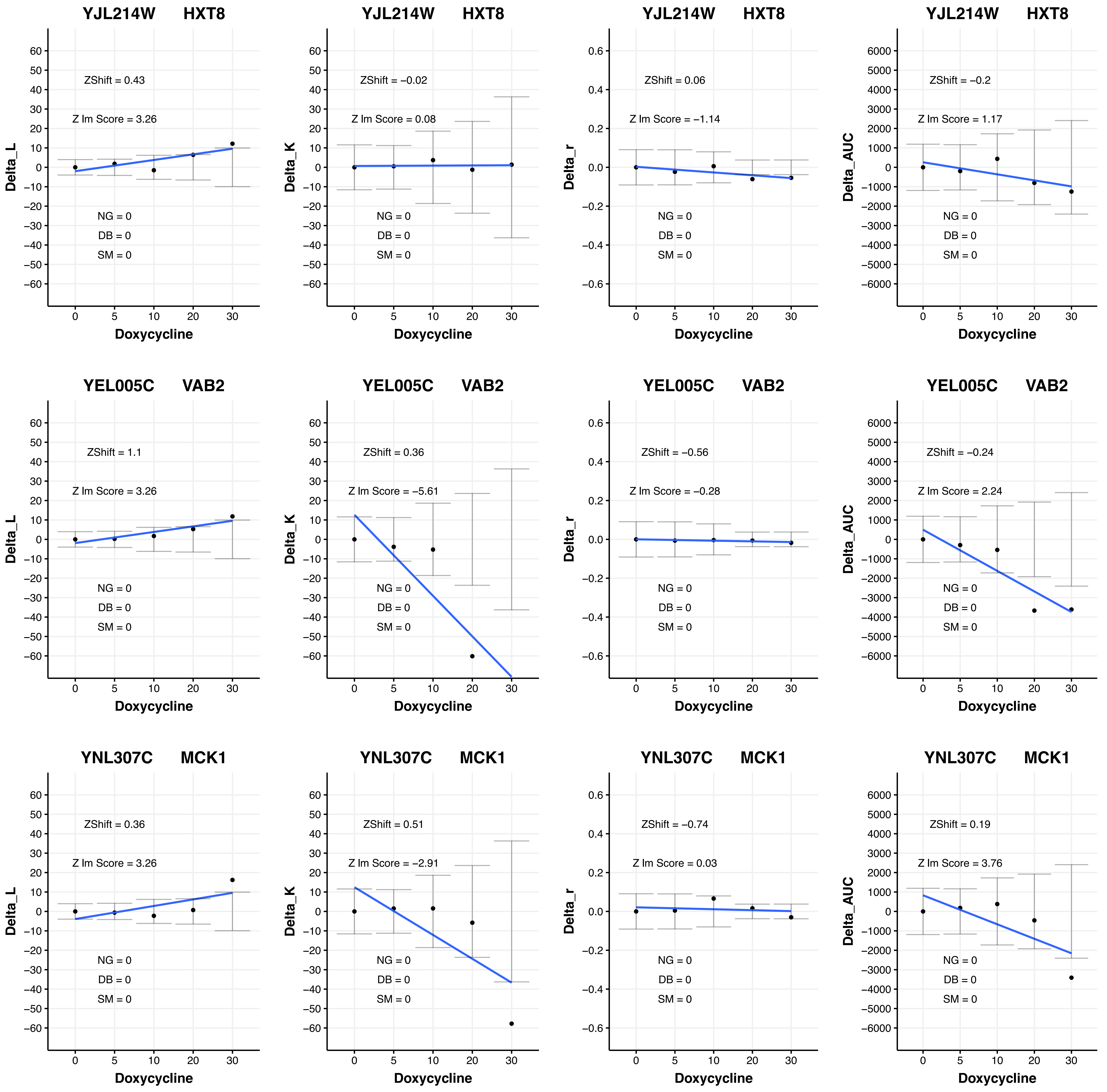

YJL214W HXT8
ZShift = 0.43
Z lm Score = 3.26
NG = 0
DB = 0
SM = 0
Delta_L
Doxycycline
YJL214W HXT8
ZShift = −0.02
Z lm Score = 0.08
NG = 0
DB = 0
SM = 0
Delta_K
Doxycycline
YJL214W HXT8
ZShift = 0.06
Z lm Score = −1.14
NG = 0
DB = 0
SM = 0
Delta_r
Doxycycline
YJL214W HXT8
ZShift = −0.2
Z lm Score = 1.17
NG = 0
DB = 0
SM = 0
Delta_AUC
Doxycycline
YEL005C VAB2
ZShift = 1.1
Z lm Score = 3.26
NG = 0
DB = 0
SM = 0
Delta_L
Doxycycline
YEL005C VAB2
ZShift = 0.36
Z lm Score = −5.61
NG = 0
DB = 0
SM = 0
Delta_K
Doxycycline
YEL005C VAB2
ZShift = −0.56
Z lm Score = −0.28
NG = 0
DB = 0
SM = 0
Delta_r
Doxycycline
YEL005C VAB2
ZShift = −0.24
Z lm Score = 2.24
NG = 0
DB = 0
SM = 0
Delta_AUC
Doxycycline
YNL307C MCK1
ZShift = 0.36
Z lm Score = 3.26
NG = 0
DB = 0
SM = 0
Delta_L
Doxycycline
YNL307C MCK1
ZShift = 0.51
Z lm Score = −2.91
NG = 0
DB = 0
SM = 0
Delta_K
Doxycycline
YNL307C MCK1
ZShift = −0.74
Z lm Score = 0.03
NG = 0
DB = 0
SM = 0
Delta_r
Doxycycline
YNL307C MCK1
ZShift = 0.19
Z lm Score = 3.76
NG = 0
DB = 0
SM = 0
Delta_AUC
Doxycycline

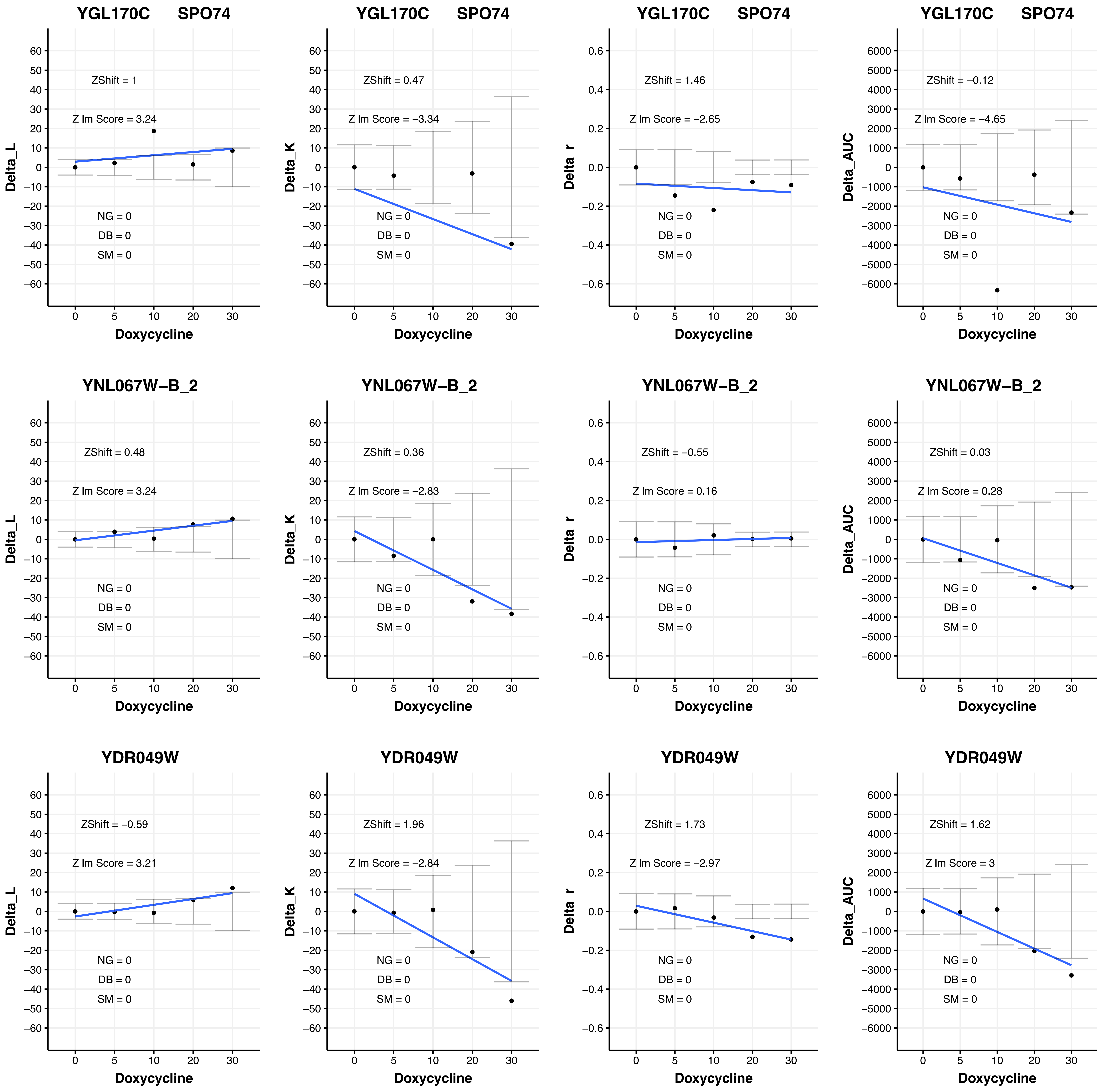
YGL170C SPO74
ZShift = 1
Z lm Score = 3.24
NG = 0
DB = 0
SM = 0
Delta_L
Doxycycline
YGL170C SPO74
ZShift = 0.47
Z lm Score = −3.34
NG = 0
DB = 0
SM = 0
Delta_K
Doxycycline
YGL170C SPO74
ZShift = 1.46
Z lm Score = −2.65
NG = 0
DB = 0
SM = 0
Delta_r
Doxycycline
YGL170C SPO74
ZShift = −0.12
Z lm Score = −4.65
NG = 0
DB = 0
SM = 0
Delta_AUC
Doxycycline
YNL067W−B_2
ZShift = 0.48
Z lm Score = 3.24
NG = 0
DB = 0
SM = 0
Delta_L
Doxycycline
YNL067W−B_2
ZShift = 0.36
Z lm Score = −2.83
NG = 0
DB = 0
SM = 0
Delta_K
Doxycycline
YNL067W−B_2
ZShift = −0.55
Z lm Score = 0.16
NG = 0
DB = 0
SM = 0
Delta_r
Doxycycline
YNL067W−B_2
ZShift = 0.03
Z lm Score = 0.28
NG = 0
DB = 0
SM = 0
Delta_AUC
Doxycycline
YDR049W
ZShift = −0.59
Z lm Score = 3.21
NG = 0
DB = 0
SM = 0
Delta_L
Doxycycline
YDR049W
ZShift = 1.96
Z lm Score = −2.84
NG = 0
DB = 0
SM = 0
Delta_K
Doxycycline
YDR049W
ZShift = 1.73
Z lm Score = −2.97
NG = 0
DB = 0
SM = 0
Delta_r
Doxycycline
YDR049W
ZShift = 1.62
Z lm Score = 3
NG = 0
DB = 0
SM = 0
Delta_AUC
Doxycycline

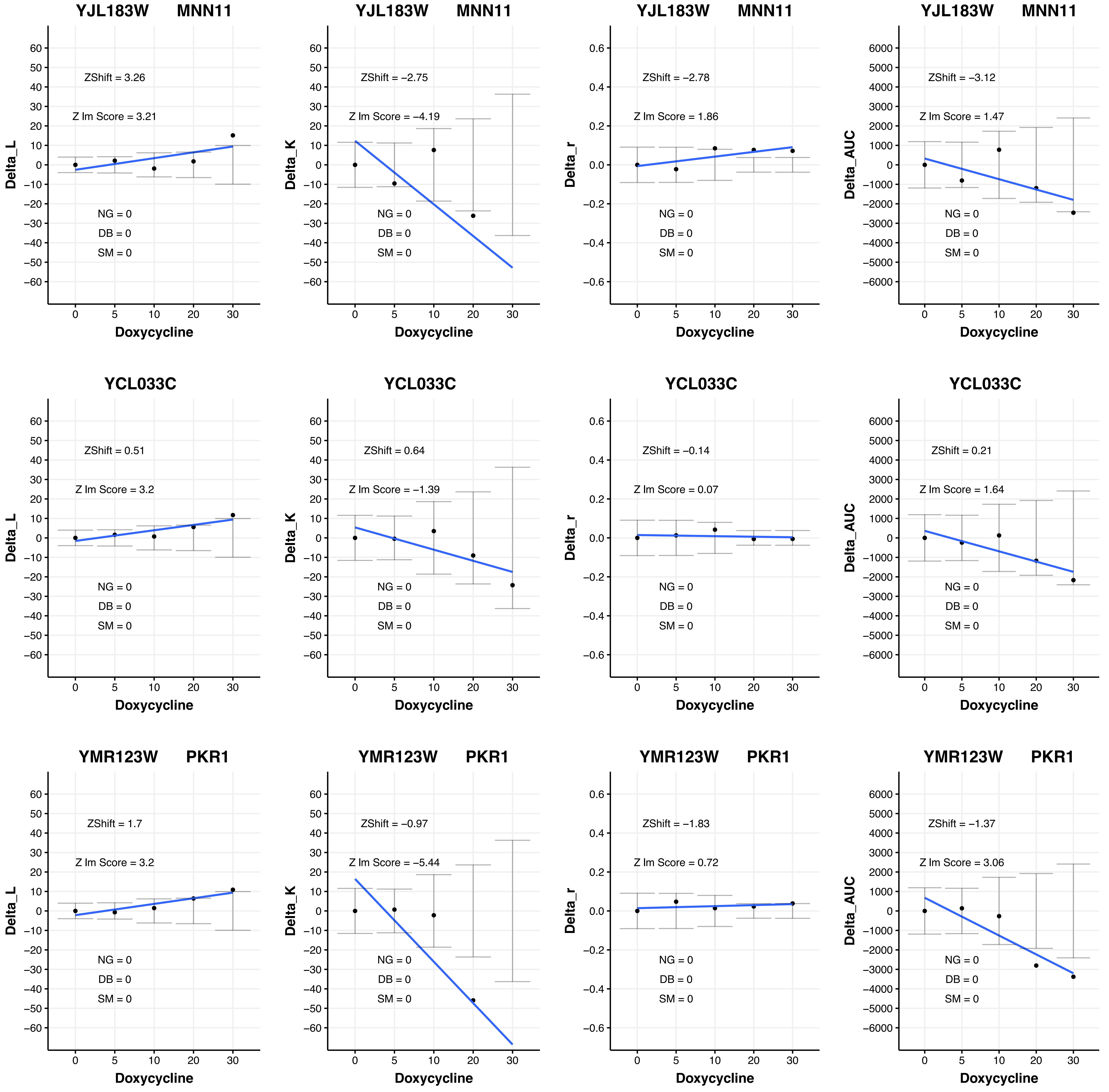
YJL183W MNN11
ZShift = 3.26
Z lm Score = 3.21
NG = 0
DB = 0
SM = 0
Delta_L
Doxycycline
YJL183W MNN11
ZShift = −2.75
Z lm Score = −4.19
NG = 0
DB = 0
SM = 0
Delta_K
Doxycycline
YJL183W MNN11
ZShift = −2.78
Z lm Score = 1.86
NG = 0
DB = 0
SM = 0
Delta_r
Doxycycline
YJL183W MNN11
ZShift = −3.12
Z lm Score = 1.47
NG = 0
DB = 0
SM = 0
Delta_AUC
Doxycycline
YCL033C
ZShift = 0.51
Z lm Score = 3.2
NG = 0
DB = 0
SM = 0
Delta_L
Doxycycline
YCL033C
ZShift = 0.64
Z lm Score = −1.39
NG = 0
DB = 0
SM = 0
Delta_K
Doxycycline
YCL033C
ZShift = −0.14
Z lm Score = 0.07
NG = 0
DB = 0
SM = 0
Delta_r
Doxycycline
YCL033C
ZShift = 0.21
Z lm Score = 1.64
NG = 0
DB = 0
SM = 0
Delta_AUC
Doxycycline
YMR123W PKR1
ZShift = 1.7
Z lm Score = 3.2
NG = 0
DB = 0
SM = 0
Delta_L
Doxycycline
YMR123W PKR1
ZShift = −0.97
Z lm Score = −5.44
NG = 0
DB = 0
SM = 0
Delta_K
Doxycycline
YMR123W PKR1
ZShift = −1.83
Z lm Score = 0.72
NG = 0
DB = 0
SM = 0
Delta_r
Doxycycline
YMR123W PKR1
ZShift = −1.37
Z lm Score = 3.06
NG = 0
DB = 0
SM = 0
Delta_AUC
Doxycycline

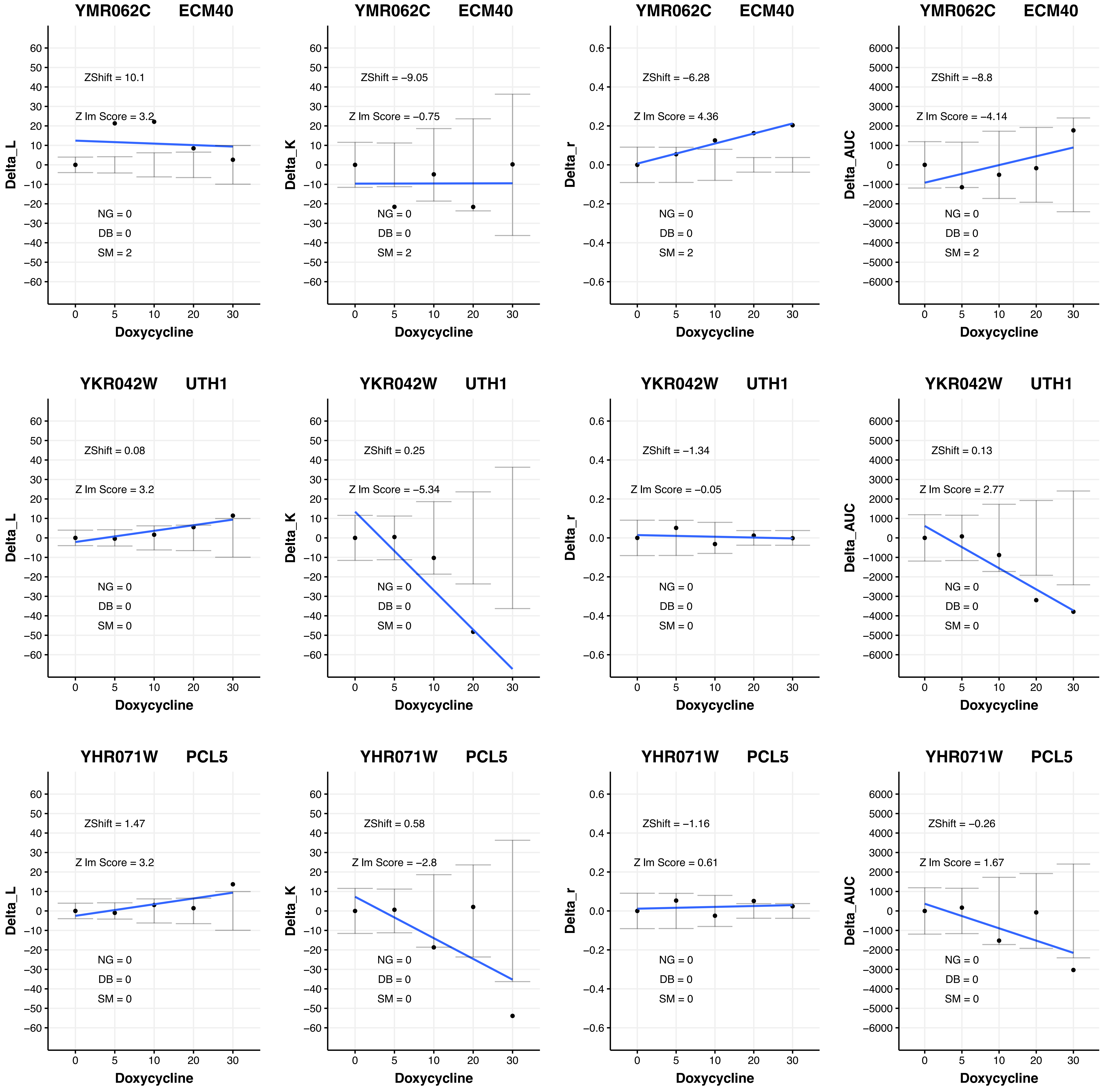
YMR062C ECM40
ZShift = 10.1
Z lm Score = 3.2
NG = 0
DB = 0
SM = 2
Delta_L
Doxycycline
YMR062C ECM40
ZShift = −9.05
Z lm Score = −0.75
NG = 0
DB = 0
SM = 2
Delta_K
Doxycycline
YMR062C ECM40
ZShift = −6.28
Z lm Score = 4.36
NG = 0
DB = 0
SM = 2
Delta_r
Doxycycline
YMR062C ECM40
ZShift = −8.8
Z lm Score = −4.14
NG = 0
DB = 0
SM = 2
Delta_AUC
Doxycycline
YKR042W UTH1
ZShift = 0.08
Z lm Score = 3.2
NG = 0
DB = 0
SM = 0
Delta_L
Doxycycline
YKR042W UTH1
ZShift = 0.25
Z lm Score = −5.34
NG = 0
DB = 0
SM = 0
Delta_K
Doxycycline
YKR042W UTH1
ZShift = −1.34
Z lm Score = −0.05
NG = 0
DB = 0
SM = 0
Delta_r
Doxycycline
YKR042W UTH1
ZShift = 0.13
Z lm Score = 2.77
NG = 0
DB = 0
SM = 0
Delta_AUC
Doxycycline
YHR071W PCL5
ZShift = 1.47
Z lm Score = 3.2
NG = 0
DB = 0
SM = 0
Delta_L
Doxycycline
YHR071W PCL5
ZShift = 0.58
Z lm Score = −2.8
NG = 0
DB = 0
SM = 0
Delta_K
Doxycycline
YHR071W PCL5
ZShift = −1.16
Z lm Score = 0.61
NG = 0
DB = 0
SM = 0
Delta_r
Doxycycline
YHR071W PCL5
ZShift = −0.26
Z lm Score = 1.67
NG = 0
DB = 0
SM = 0
Delta_AUC
Doxycycline

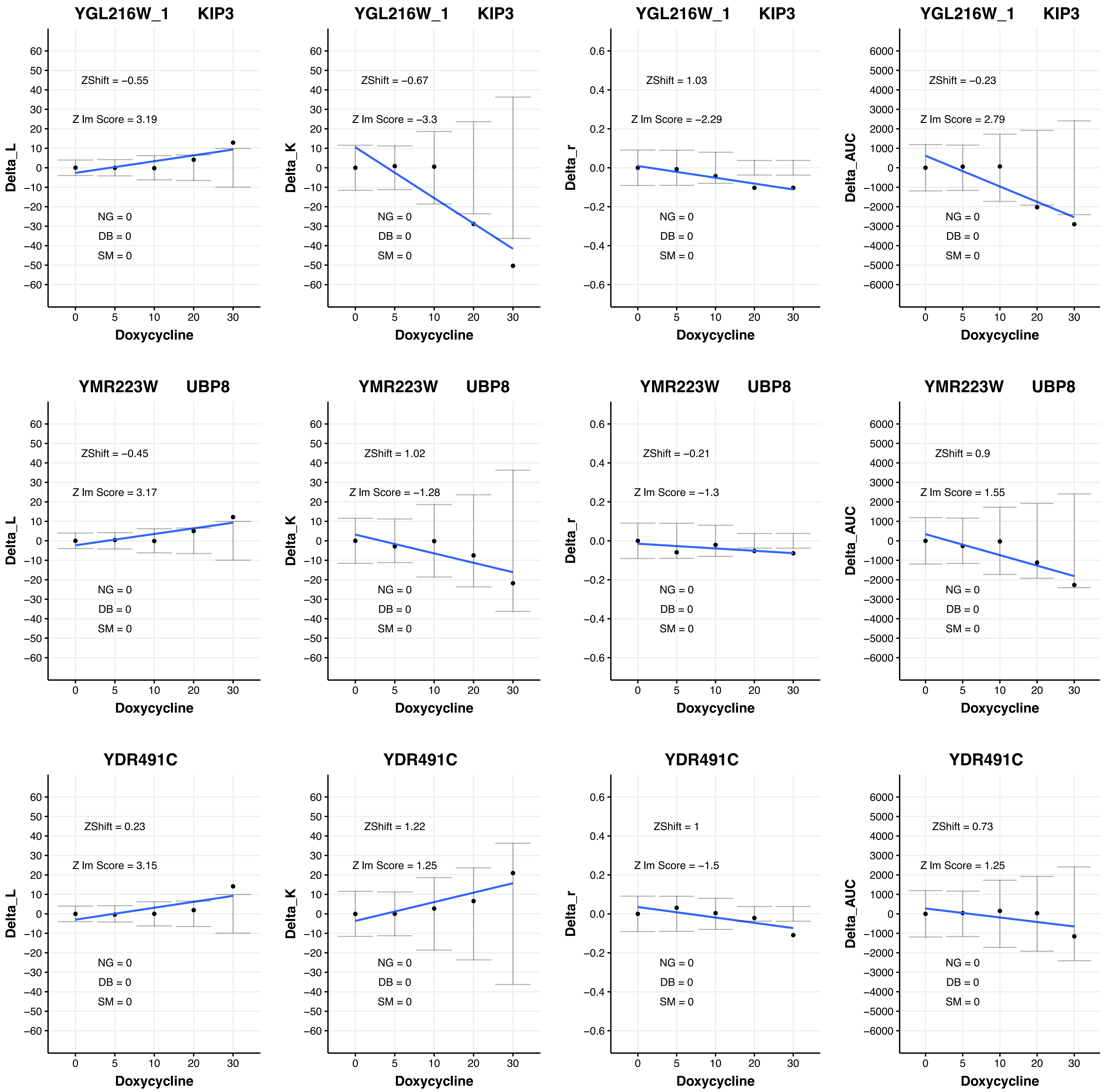

YGL216W_1 KIP3
ZShift = −0.55
Z lm Score = 3.19
NG = 0
DB = 0
SM = 0
Delta_L
Doxycycline
YGL216W_1 KIP3
ZShift = −0.67
Z lm Score = −3.3
NG = 0
DB = 0
SM = 0
Delta_K
Doxycycline
YGL216W_1 KIP3
ZShift = 1.03
Z lm Score = −2.29
NG = 0
DB = 0
SM = 0
Delta_r
Doxycycline
YGL216W_1 KIP3
ZShift = −0.23
Z lm Score = 2.79
NG = 0
DB = 0
SM = 0
Delta_AUC
Doxycycline
YMR223W UBP8
ZShift = −0.45
Z lm Score = 3.17
NG = 0
DB = 0
SM = 0
Delta_L
Doxycycline
YMR223W UBP8
ZShift = 1.02
Z lm Score = −1.28
NG = 0
DB = 0
SM = 0
Delta_K
Doxycycline
YMR223W UBP8
ZShift = −0.21
Z lm Score = −1.3
NG = 0
DB = 0
SM = 0
Delta_r
Doxycycline
YMR223W UBP8
ZShift = 0.9
Z lm Score = 1.55
NG = 0
DB = 0
SM = 0
Delta_AUC
Doxycycline
YDR491C
ZShift = 0.23
Z lm Score = 3.15
NG = 0
DB = 0
SM = 0
Delta_L
Doxycycline
YDR491C
ZShift = 1.22
Z lm Score = 1.25
NG = 0
DB = 0
SM = 0
Delta_K
Doxycycline
YDR491C
ZShift = 1
Z lm Score = −1.5
NG = 0
DB = 0
SM = 0
Delta_r
Doxycycline
YDR491C
ZShift = 0.73
Z lm Score = 1.25
NG = 0
DB = 0
SM = 0
Delta_AUC
Doxycycline

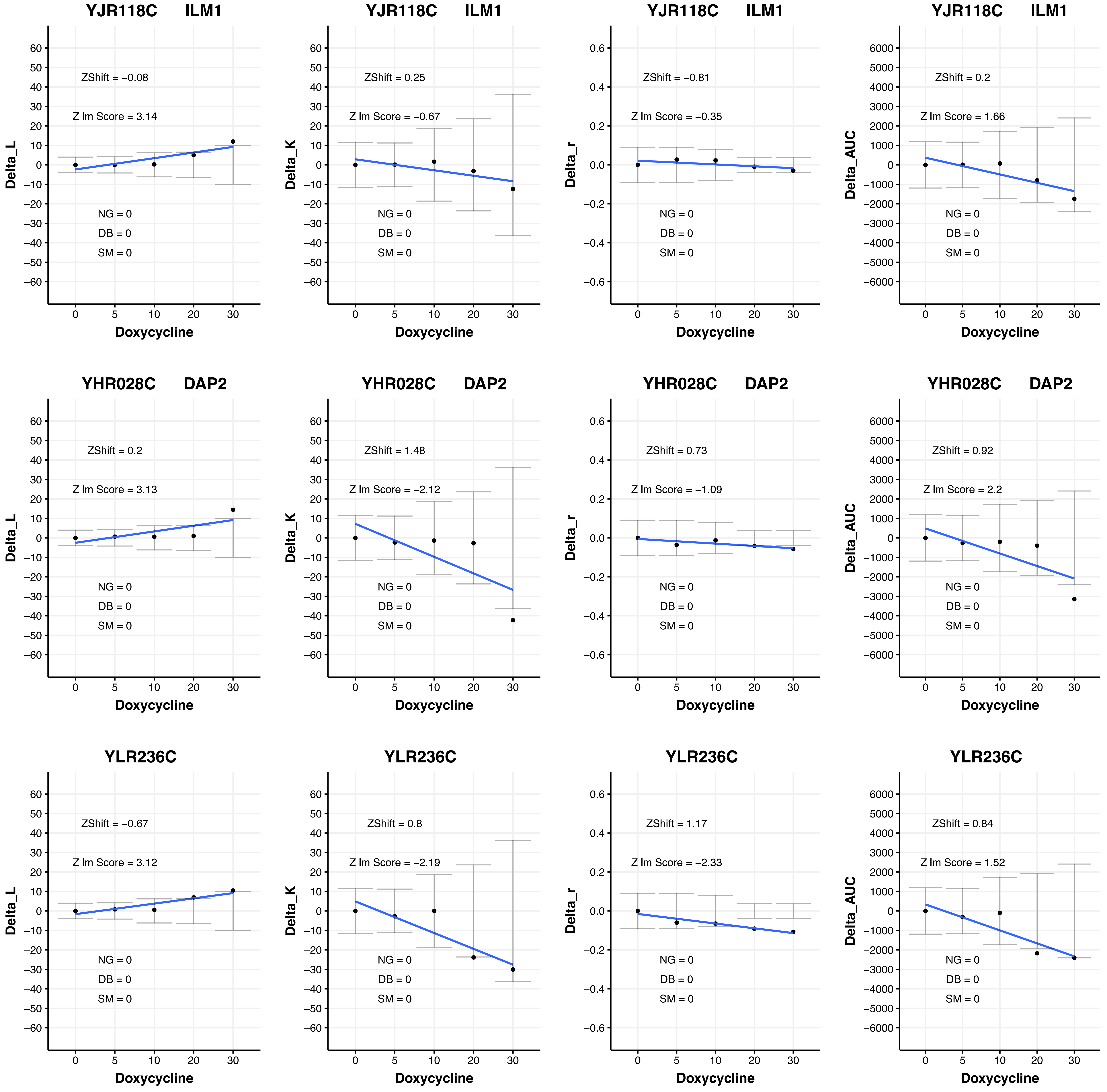

YJR118C ILM1
ZShift = −0.08
Z lm Score = 3.14
NG = 0
DB = 0
SM = 0
Delta_L
Doxycycline
YJR118C ILM1
ZShift = 0.25
Z lm Score = −0.67
NG = 0
DB = 0
SM = 0
Delta_K
Doxycycline
YJR118C ILM1
ZShift = −0.81
Z lm Score = −0.35
NG = 0
DB = 0
SM = 0
Delta_r
Doxycycline
YJR118C ILM1
ZShift = 0.2
Z lm Score = 1.66
NG = 0
DB = 0
SM = 0
Delta_AUC
Doxycycline
YHR028C DAP2
ZShift = 0.2
Z lm Score = 3.13
NG = 0
DB = 0
SM = 0
Delta_L
Doxycycline
YHR028C DAP2
ZShift = 1.48
Z lm Score = −2.12
NG = 0
DB = 0
SM = 0
Delta_K
Doxycycline
YHR028C DAP2
ZShift = 0.73
Z lm Score = −1.09
NG = 0
DB = 0
SM = 0
Delta_r
Doxycycline
YHR028C DAP2
ZShift = 0.92
Z lm Score = 2.2
NG = 0
DB = 0
SM = 0
Delta_AUC
Doxycycline
YLR236C
ZShift = −0.67
Z lm Score = 3.12
NG = 0
DB = 0
SM = 0
Delta_L
Doxycycline
YLR236C
ZShift = 0.8
Z lm Score = −2.19
NG = 0
DB = 0
SM = 0
Delta_K
Doxycycline
YLR236C
ZShift = 1.17
Z lm Score = −2.33
NG = 0
DB = 0
SM = 0
Delta_r
Doxycycline
YLR236C
ZShift = 0.84
Z lm Score = 1.52
NG = 0
DB = 0
SM = 0
Delta_AUC
Doxycycline

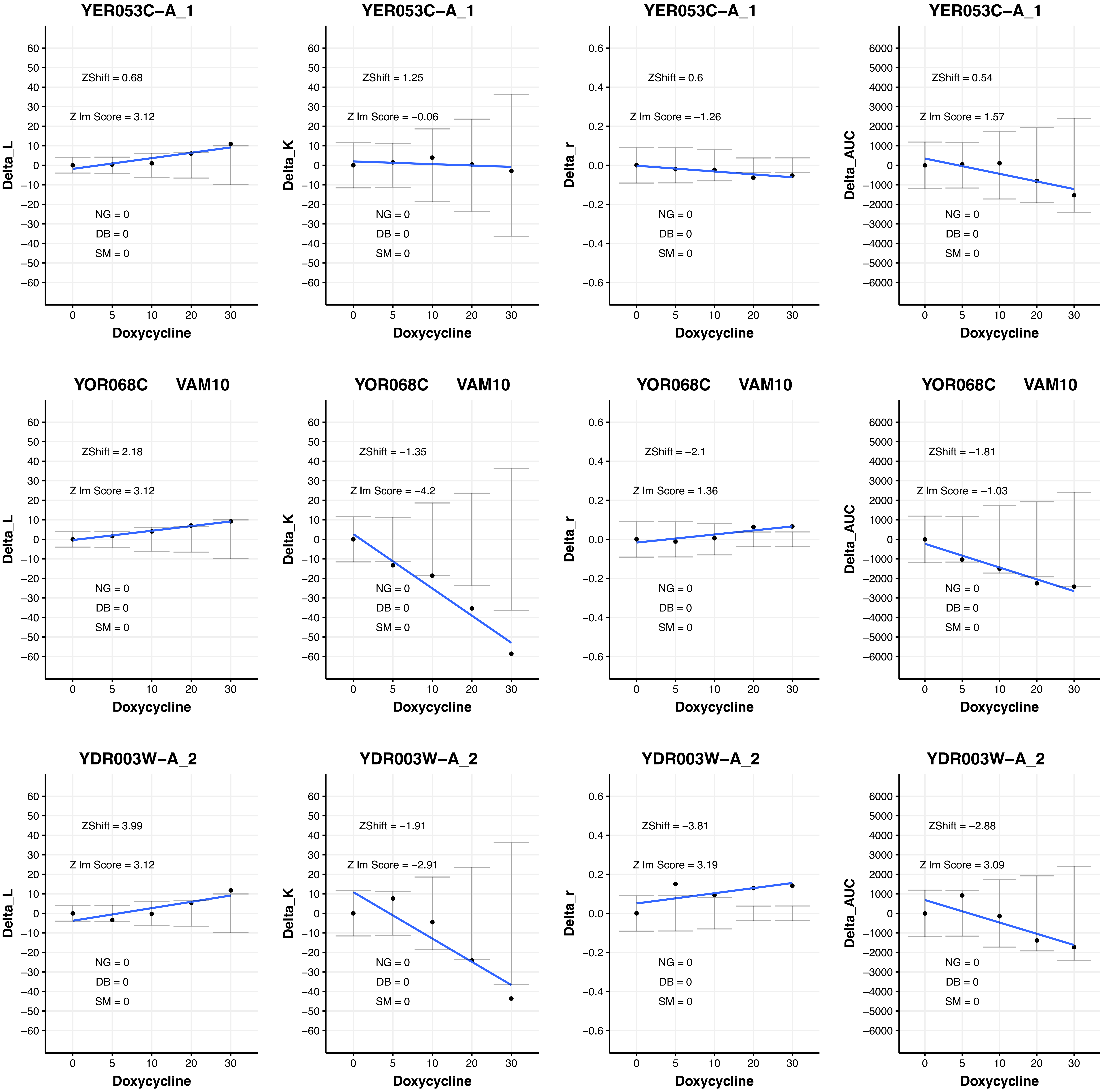
YER053C−A_1
ZShift = 0.68
Z lm Score = 3.12
NG = 0
DB = 0
SM = 0
Delta_L
Doxycycline
YER053C−A_1
ZShift = 1.25
Z lm Score = −0.06
NG = 0
DB = 0
SM = 0
Delta_K
Doxycycline
YER053C−A_1
ZShift = 0.6
Z lm Score = −1.26
NG = 0
DB = 0
SM = 0
Delta_r
Doxycycline
YER053C−A_1
ZShift = 0.54
Z lm Score = 1.57
NG = 0
DB = 0
SM = 0
Delta_AUC
Doxycycline
YOR068C VAM10
ZShift = 2.18
Z lm Score = 3.12
NG = 0
DB = 0
SM = 0
Delta_L
Doxycycline
YOR068C VAM10
ZShift = −1.35
Z lm Score = −4.2
NG = 0
DB = 0
SM = 0
Delta_K
Doxycycline
YOR068C VAM10
ZShift = −2.1
Z lm Score = 1.36
NG = 0
DB = 0
SM = 0
Delta_r
Doxycycline
YOR068C VAM10
ZShift = −1.81
Z lm Score = −1.03
NG = 0
DB = 0
SM = 0
Delta_AUC
Doxycycline
YDR003W−A_2
ZShift = 3.99
Z lm Score = 3.12
NG = 0
DB = 0
SM = 0
Delta_L
Doxycycline
YDR003W−A_2
ZShift = −1.91
Z lm Score = −2.91
NG = 0
DB = 0
SM = 0
Delta_K
Doxycycline
YDR003W−A_2
ZShift = −3.81
Z lm Score = 3.19
NG = 0
DB = 0
SM = 0
Delta_r
Doxycycline
YDR003W−A_2
ZShift = −2.88
Z lm Score = 3.09
NG = 0
DB = 0
SM = 0
Delta_AUC
Doxycycline

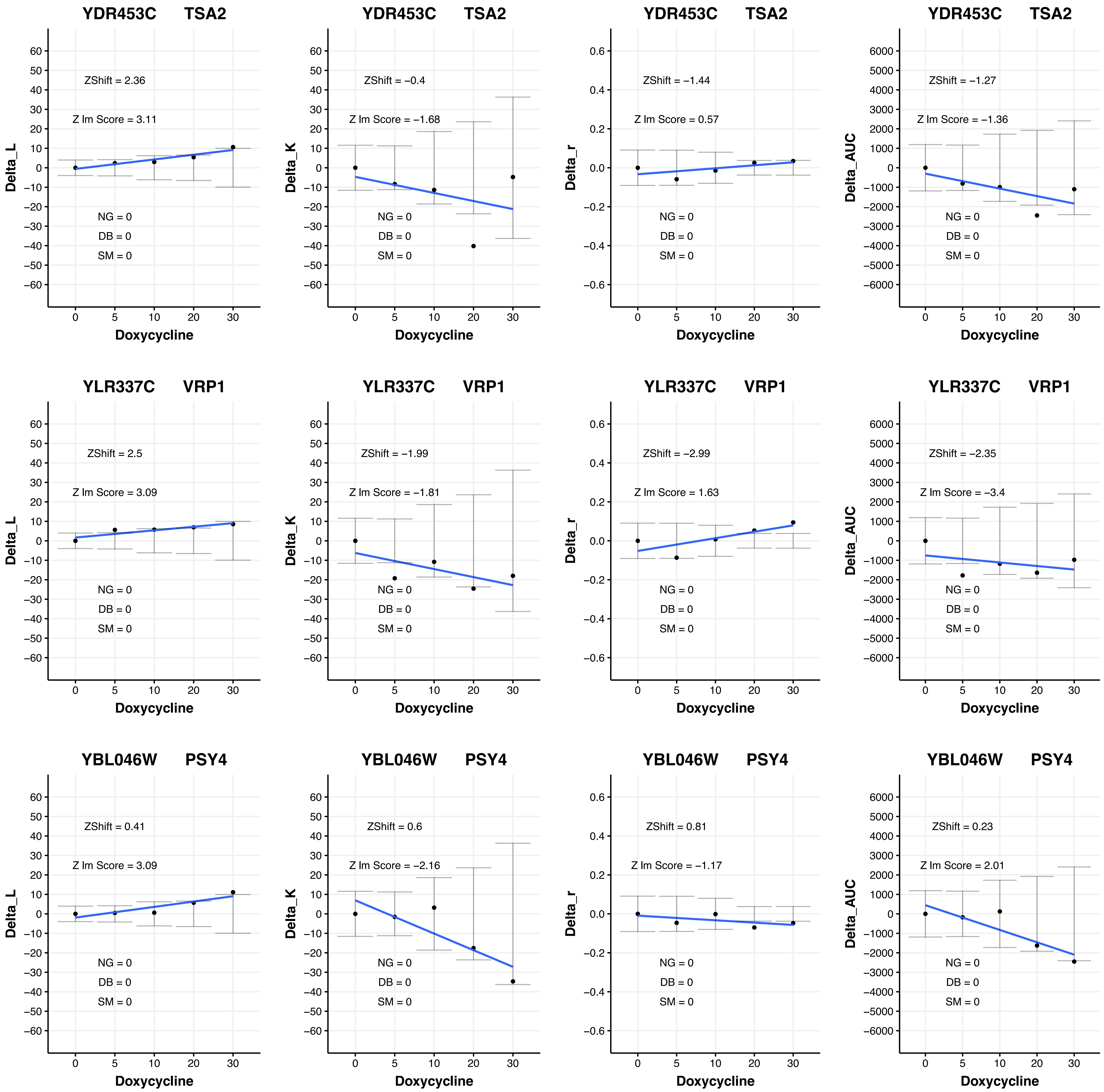

YDR453C TSA2
ZShift = 2.36
Z lm Score = 3.11
NG = 0
DB = 0
SM = 0
Delta_L
Doxycycline
YDR453C TSA2
ZShift = −0.4
Z lm Score = −1.68
NG = 0
DB = 0
SM = 0
Delta_K
Doxycycline
YDR453C TSA2
ZShift = −1.44
Z lm Score = 0.57
NG = 0
DB = 0
SM = 0
Delta_r
Doxycycline
YDR453C TSA2
ZShift = −1.27
Z lm Score = −1.36
NG = 0
DB = 0
SM = 0
Delta_AUC
Doxycycline
YLR337C VRP1
ZShift = 2.5
Z lm Score = 3.09
NG = 0
DB = 0
SM = 0
Delta_L
Doxycycline
YLR337C VRP1
ZShift = −1.99
Z lm Score = −1.81
NG = 0
DB = 0
SM = 0
Delta_K
Doxycycline
YLR337C VRP1
ZShift = −2.99
Z lm Score = 1.63
NG = 0
DB = 0
SM = 0
Delta_r
Doxycycline
YLR337C VRP1
ZShift = −2.35
Z lm Score = −3.4
NG = 0
DB = 0
SM = 0
Delta_AUC
Doxycycline
YBL046W PSY4
ZShift = 0.41
Z lm Score = 3.09
NG = 0
DB = 0
SM = 0
Delta_L
Doxycycline
YBL046W PSY4
ZShift = 0.6
Z lm Score = −2.16
NG = 0
DB = 0
SM = 0
Delta_K
Doxycycline
YBL046W PSY4
ZShift = 0.81
Z lm Score = −1.17
NG = 0
DB = 0
SM = 0
Delta_r
Doxycycline
YBL046W PSY4
ZShift = 0.23
Z lm Score = 2.01
NG = 0
DB = 0
SM = 0
Delta_AUC
Doxycycline

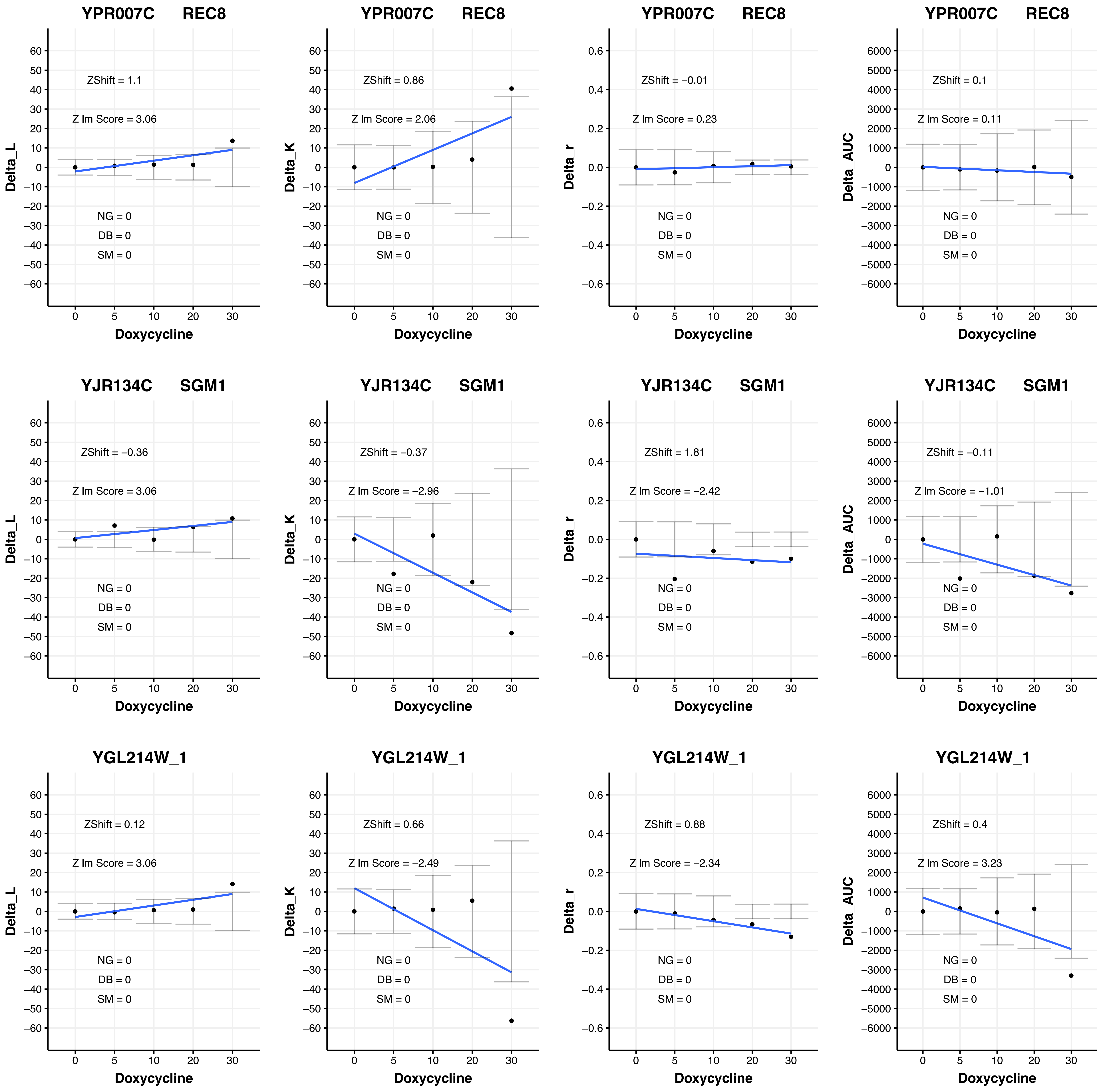

YPR007C REC8
ZShift = 1.1
Z lm Score = 3.06
NG = 0
DB = 0
SM = 0
Delta_L
Doxycycline
YPR007C REC8
ZShift = 0.86
Z lm Score = 2.06
NG = 0
DB = 0
SM = 0
Delta_K
Doxycycline
YPR007C REC8
ZShift = −0.01
Z lm Score = 0.23
NG = 0
DB = 0
SM = 0
Delta_r
Doxycycline
YPR007C REC8
ZShift = 0.1
Z lm Score = 0.11
NG = 0
DB = 0
SM = 0
Delta_AUC
Doxycycline
YJR134C SGM1
ZShift = −0.36
Z lm Score = 3.06
NG = 0
DB = 0
SM = 0
Delta_L
Doxycycline
YJR134C SGM1
ZShift = −0.37
Z lm Score = −2.96
NG = 0
DB = 0
SM = 0
Delta_K
Doxycycline
YJR134C SGM1
ZShift = 1.81
Z lm Score = −2.42
NG = 0
DB = 0
SM = 0
Delta_r
Doxycycline
YJR134C SGM1
ZShift = −0.11
Z lm Score = −1.01
NG = 0
DB = 0
SM = 0
Delta_AUC
Doxycycline
YGL214W_1
ZShift = 0.12
Z lm Score = 3.06
NG = 0
DB = 0
SM = 0
Delta_L
Doxycycline
YGL214W_1
ZShift = 0.66
Z lm Score = −2.49
NG = 0
DB = 0
SM = 0
Delta_K
Doxycycline
YGL214W_1
ZShift = 0.88
Z lm Score = −2.34
NG = 0
DB = 0
SM = 0
Delta_r
Doxycycline
YGL214W_1
ZShift = 0.4
Z lm Score = 3.23
NG = 0
DB = 0
SM = 0
Delta_AUC
Doxycycline

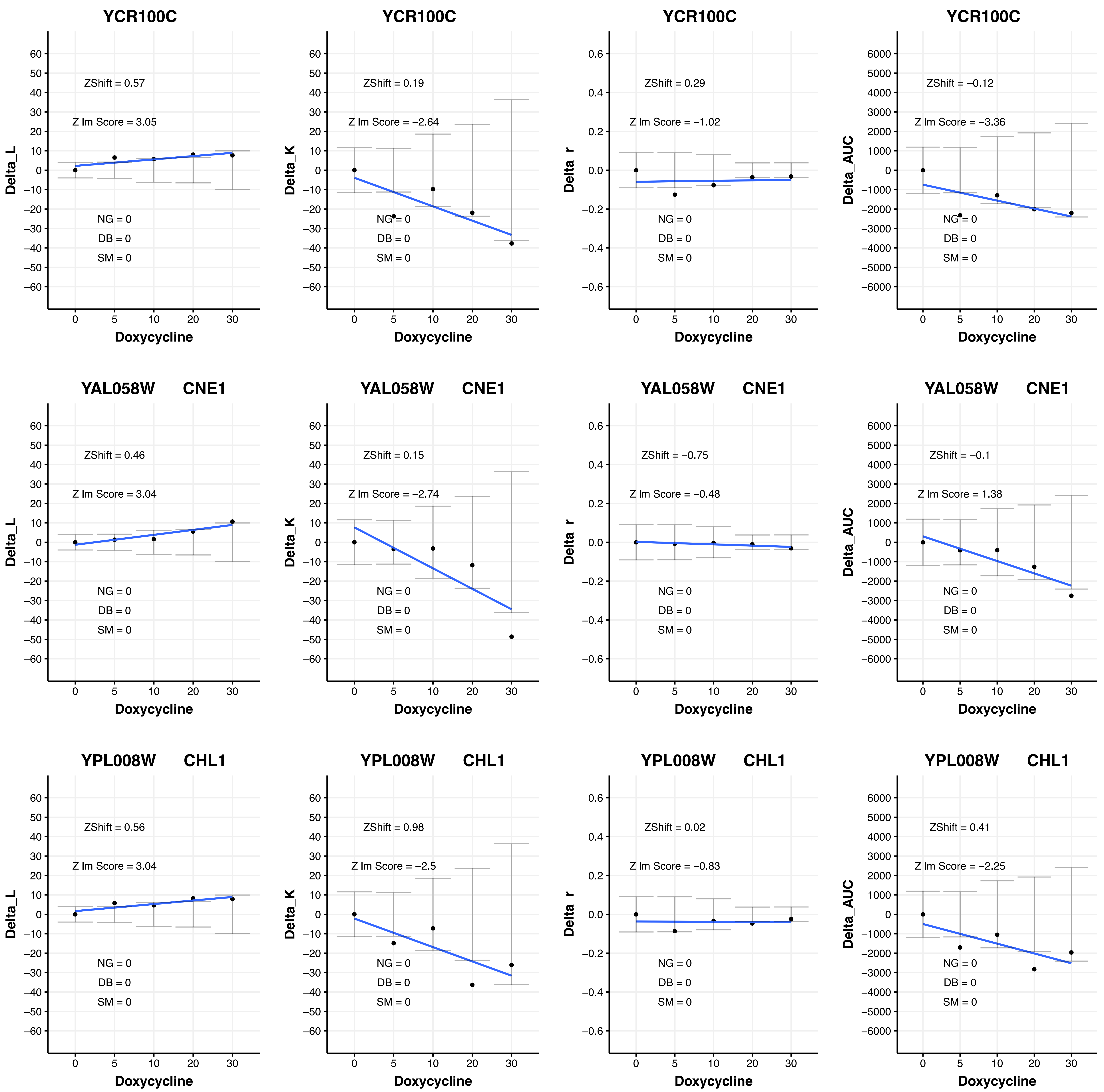

YCR100C
ZShift = 0.57
Z lm Score = 3.05
NG = 0
DB = 0
SM = 0
Delta_L
Doxycycline
YCR100C
ZShift = 0.19
Z lm Score = −2.64
NG = 0
DB = 0
SM = 0
Delta_K
Doxycycline
YCR100C
ZShift = 0.29
Z lm Score = −1.02
NG = 0
DB = 0
SM = 0
Delta_r
Doxycycline
YCR100C
ZShift = −0.12
Z lm Score = −3.36
NG = 0
DB = 0
SM = 0
Delta_AUC
Doxycycline
YAL058W CNE1
ZShift = 0.46
Z lm Score = 3.04
NG = 0
DB = 0
SM = 0
Delta_L
Doxycycline
YAL058W CNE1
ZShift = 0.15
Z lm Score = −2.74
NG = 0
DB = 0
SM = 0
Delta_K
Doxycycline
YAL058W CNE1
ZShift = −0.75
Z lm Score = −0.48
NG = 0
DB = 0
SM = 0
Delta_r
Doxycycline
YAL058W CNE1
ZShift = −0.1
Z lm Score = 1.38
NG = 0
DB = 0
SM = 0
Delta_AUC
Doxycycline
YPL008W CHL1
ZShift = 0.56
Z lm Score = 3.04
NG = 0
DB = 0
SM = 0
Delta_L
Doxycycline
YPL008W CHL1
ZShift = 0.98
Z lm Score = −2.5
NG = 0
DB = 0
SM = 0
Delta_K
Doxycycline
YPL008W CHL1
ZShift = 0.02
Z lm Score = −0.83
NG = 0
DB = 0
SM = 0
Delta_r
Doxycycline
YPL008W CHL1
ZShift = 0.41
Z lm Score = −2.25
NG = 0
DB = 0
SM = 0
Delta_AUC
Doxycycline

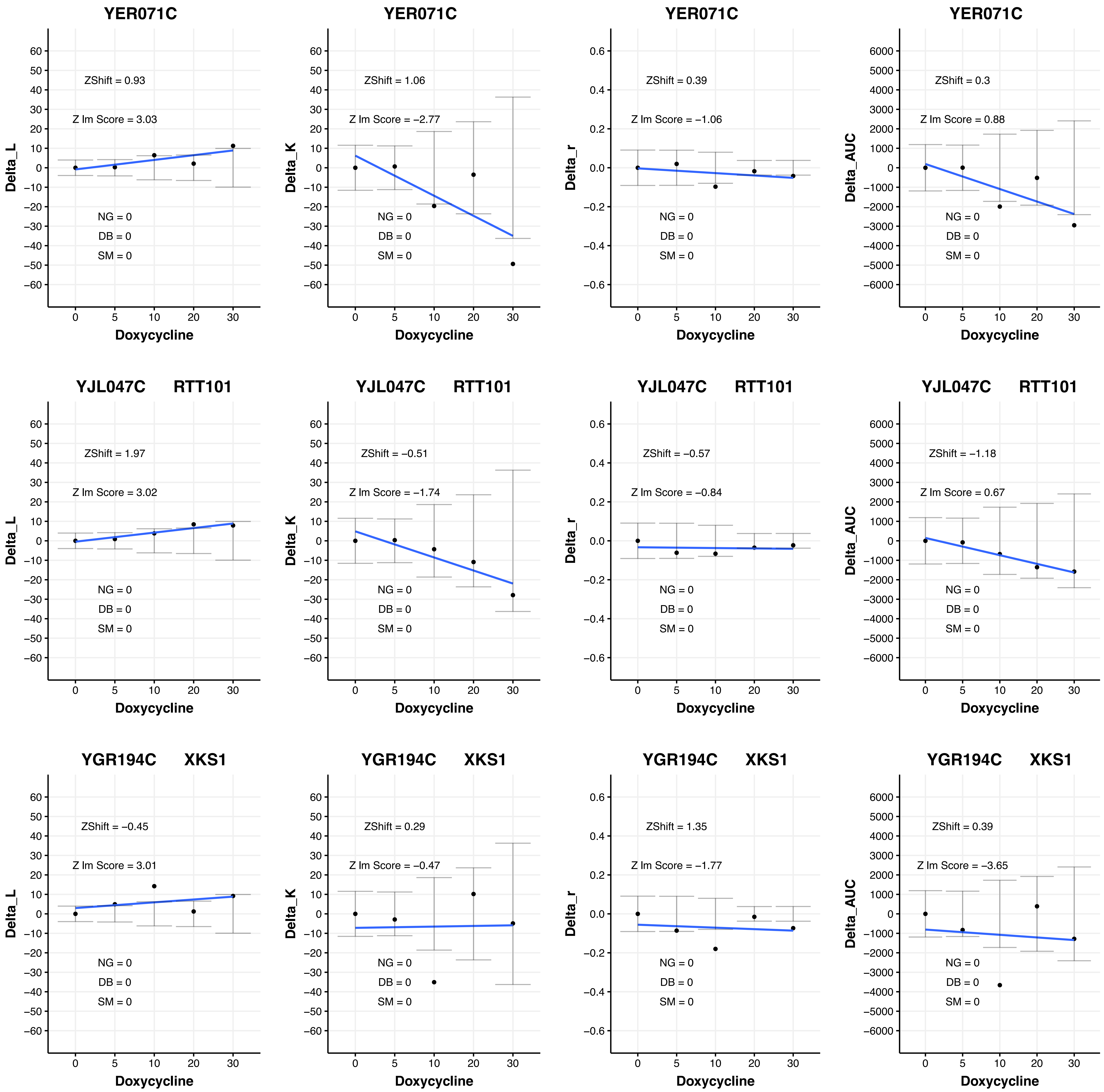
YER071C
ZShift = 0.93
Z lm Score = 3.03
NG = 0
DB = 0
SM = 0
Delta_L
Doxycycline
YER071C
ZShift = 1.06
Z lm Score = −2.77
NG = 0
DB = 0
SM = 0
Delta_K
Doxycycline
YER071C
ZShift = 0.39
Z lm Score = −1.06
NG = 0
DB = 0
SM = 0
Delta_r
Doxycycline
YER071C
ZShift = 0.3
Z lm Score = 0.88
NG = 0
DB = 0
SM = 0
Delta_AUC
Doxycycline
YJL047C RTT101
ZShift = 1.97
Z lm Score = 3.02
NG = 0
DB = 0
SM = 0
Delta_L
Doxycycline
YJL047C RTT101
ZShift = −0.51
Z lm Score = −1.74
NG = 0
DB = 0
SM = 0
Delta_K
Doxycycline
YJL047C RTT101
ZShift = −0.57
Z lm Score = −0.84
NG = 0
DB = 0
SM = 0
Delta_r
Doxycycline
YJL047C RTT101
ZShift = −1.18
Z lm Score = 0.67
NG = 0
DB = 0
SM = 0
Delta_AUC
Doxycycline
YGR194C XKS1
ZShift = −0.45
Z lm Score = 3.01
NG = 0
DB = 0
SM = 0
Delta_L
Doxycycline
YGR194C XKS1
ZShift = 0.29
Z lm Score = −0.47
NG = 0
DB = 0
SM = 0
Delta_K
Doxycycline
YGR194C XKS1
ZShift = 1.35
Z lm Score = −1.77
NG = 0
DB = 0
SM = 0
Delta_r
Doxycycline
YGR194C XKS1
ZShift = 0.39
Z lm Score = −3.65
NG = 0
DB = 0
SM = 0
Delta_AUC
Doxycycline

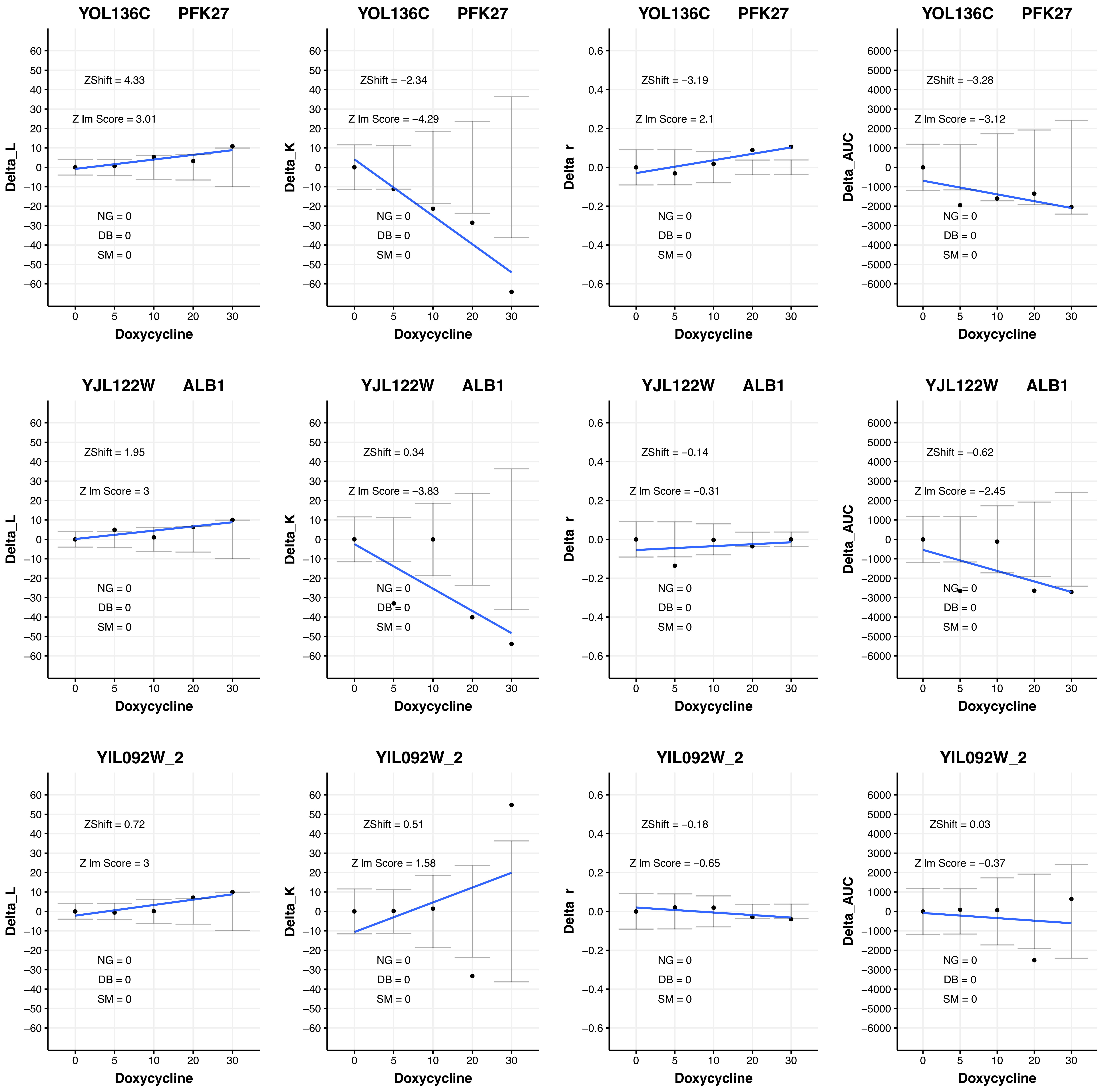
YOL136C PFK27
ZShift = 4.33
Z lm Score = 3.01
NG = 0
DB = 0
SM = 0
Delta_L
Doxycycline
YOL136C PFK27
ZShift = −2.34
Z lm Score = −4.29
NG = 0
DB = 0
SM = 0
Delta_K
Doxycycline
YOL136C PFK27
ZShift = −3.19
Z lm Score = 2.1
NG = 0
DB = 0
SM = 0
Delta_r
Doxycycline
YOL136C PFK27
ZShift = −3.28
Z lm Score = −3.12
NG = 0
DB = 0
SM = 0
Delta_AUC
Doxycycline
YJL122W ALB1
ZShift = 1.95
Z lm Score = 3
NG = 0
DB = 0
SM = 0
Delta_L
Doxycycline
YJL122W ALB1
ZShift = 0.34
Z lm Score = −3.83
NG = 0
DB = 0
SM = 0
Delta_K
Doxycycline
YJL122W ALB1
ZShift = −0.14
Z lm Score = −0.31
NG = 0
DB = 0
SM = 0
Delta_r
Doxycycline
YJL122W ALB1
ZShift = −0.62
Z lm Score = −2.45
NG = 0
DB = 0
SM = 0
Delta_AUC
Doxycycline
YIL092W_2
ZShift = 0.72
Z lm Score = 3
NG = 0
DB = 0
SM = 0
Delta_L
Doxycycline
YIL092W_2
ZShift = 0.51
Z lm Score = 1.58
NG = 0
DB = 0
SM = 0
Delta_K
Doxycycline
YIL092W_2
ZShift = −0.18
Z lm Score = −0.65
NG = 0
DB = 0
SM = 0
Delta_r
Doxycycline
YIL092W_2
ZShift = 0.03
Z lm Score = −0.37
NG = 0
DB = 0
SM = 0
Delta_AUC
Doxycycline

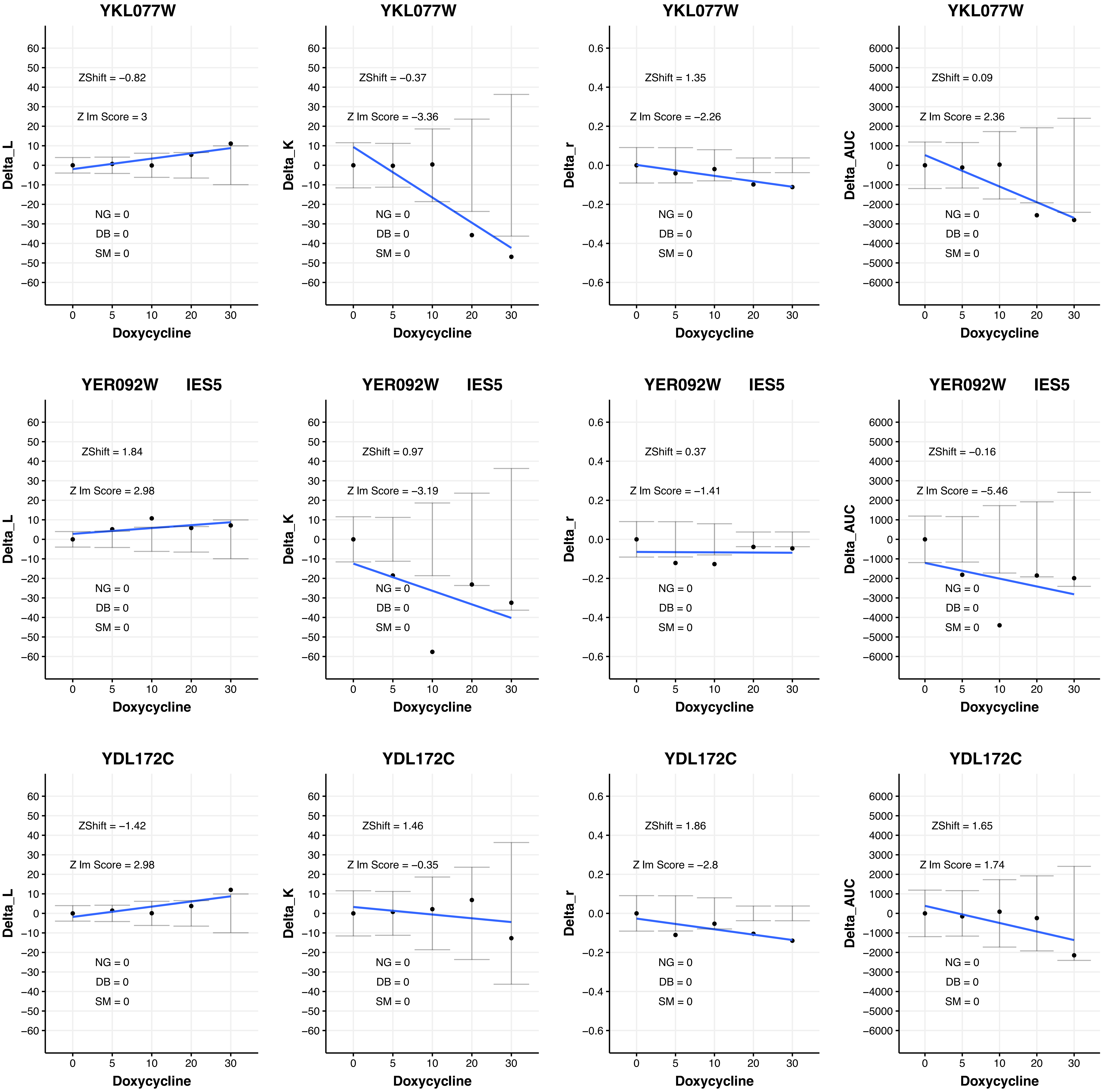

YKL077W
ZShift = −0.82
Z lm Score = 3
NG = 0
DB = 0
SM = 0
Delta_L
Doxycycline
YKL077W
ZShift = −0.37
Z lm Score = −3.36
NG = 0
DB = 0
SM = 0
Delta_K
Doxycycline
YKL077W
ZShift = 1.35
Z lm Score = −2.26
NG = 0
DB = 0
SM = 0
Delta_r
Doxycycline
YKL077W
ZShift = 0.09
Z lm Score = 2.36
NG = 0
DB = 0
SM = 0
Delta_AUC
Doxycycline
YER092W IES5
ZShift = 1.84
Z lm Score = 2.98
NG = 0
DB = 0
SM = 0
Delta_L
Doxycycline
YER092W IES5
ZShift = 0.97
Z lm Score = −3.19
NG = 0
DB = 0
SM = 0
Delta_K
Doxycycline
YER092W IES5
ZShift = 0.37
Z lm Score = −1.41
NG = 0
DB = 0
SM = 0
Delta_r
Doxycycline
YER092W IES5
ZShift = −0.16
Z lm Score = −5.46
NG = 0
DB = 0
SM = 0
Delta_AUC
Doxycycline
YDL172C
ZShift = −1.42
Z lm Score = 2.98
NG = 0
DB = 0
SM = 0
Delta_L
Doxycycline
YDL172C
ZShift = 1.46
Z lm Score = −0.35
NG = 0
DB = 0
SM = 0
Delta_K
Doxycycline
YDL172C
ZShift = 1.86
Z lm Score = −2.8
NG = 0
DB = 0
SM = 0
Delta_r
Doxycycline
YDL172C
ZShift = 1.65
Z lm Score = 1.74
NG = 0
DB = 0
SM = 0
Delta_AUC
Doxycycline

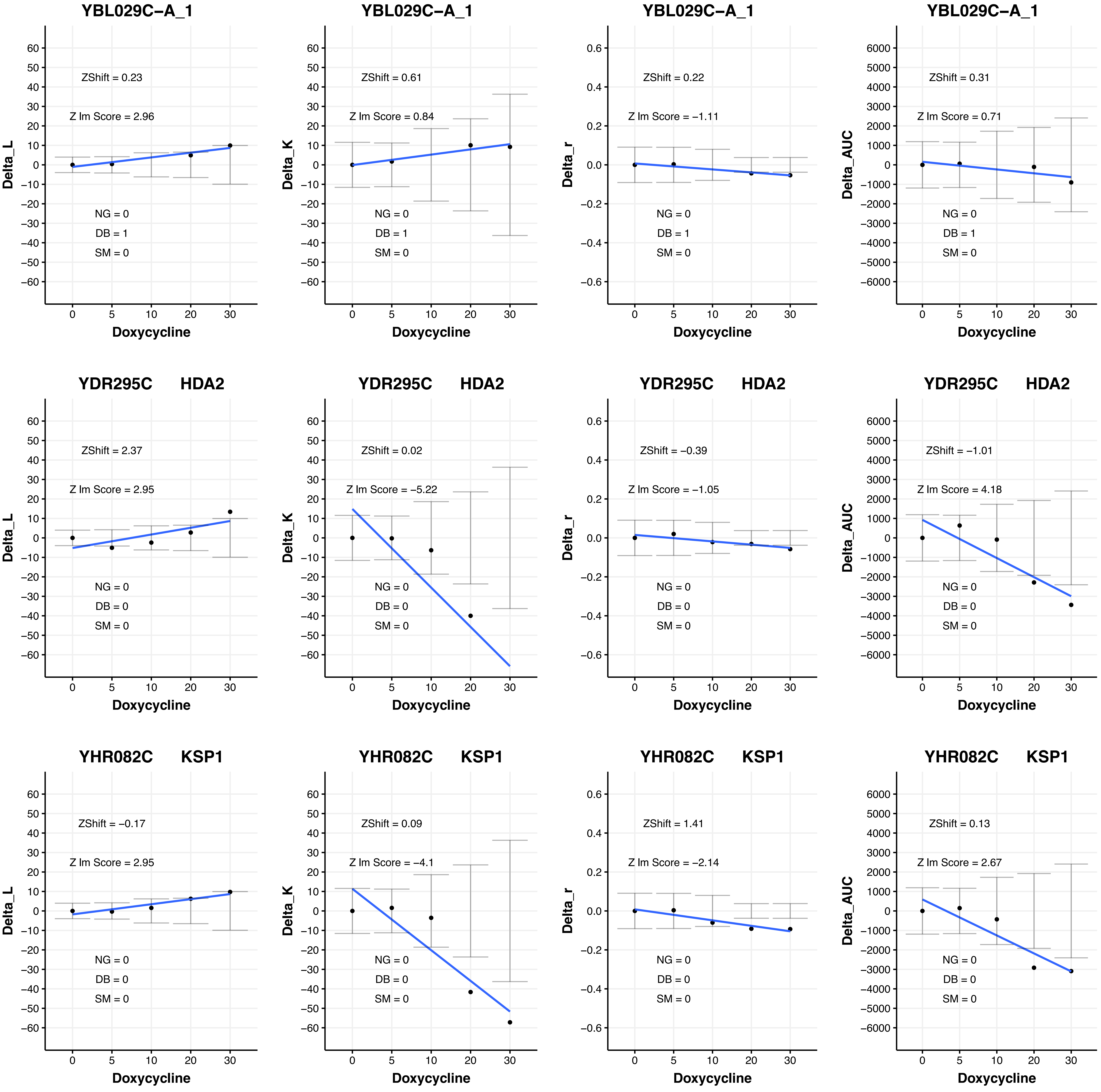

YBL029C−A_1
ZShift = 0.23
Z lm Score = 2.96
NG = 0
DB = 1
SM = 0
Delta_L
YBL029C−A_1
ZShift = 0.61
Z lm Score = 0.84
NG = 0
DB = 1
SM = 0
Delta_K
YBL029C−A_1
ZShift = 0.22
Z lm Score = −1.11
NG = 0
DB = 1
SM = 0
Delta_r
YBL029C−A_1
ZShift = 0.31
Z lm Score = 0.71
NG = 0
DB = 1
SM = 0
Delta_AUC
YDR295C HDA2
ZShift = 2.37
Z lm Score = 2.95
NG = 0
DB = 0
SM = 0
Delta_L
YDR295C HDA2
ZShift = 0.02
Z lm Score = −5.22
NG = 0
DB = 0
SM = 0
Delta_K
YDR295C HDA2
ZShift = −0.39
Z lm Score = −1.05
NG = 0
DB = 0
SM = 0
Delta_r
YDR295C HDA2
ZShift = −1.01
Z lm Score = 4.18
NG = 0
DB = 0
SM = 0
Delta_AUC
YHR082C KSP1
ZShift = −0.17
Z lm Score = 2.95
NG = 0
DB = 0
SM = 0
Delta_L
YHR082C KSP1
ZShift = 0.09
Z lm Score = −4.1
NG = 0
DB = 0
SM = 0
Delta_K
YHR082C KSP1
ZShift = 1.41
Z lm Score = −2.14
NG = 0
DB = 0
SM = 0
Delta_r
YHR082C KSP1
ZShift = 0.13
Z lm Score = 2.67
NG = 0
DB = 0
SM = 0
Delta_AUC
Doxycycline
0 5 10 20 30

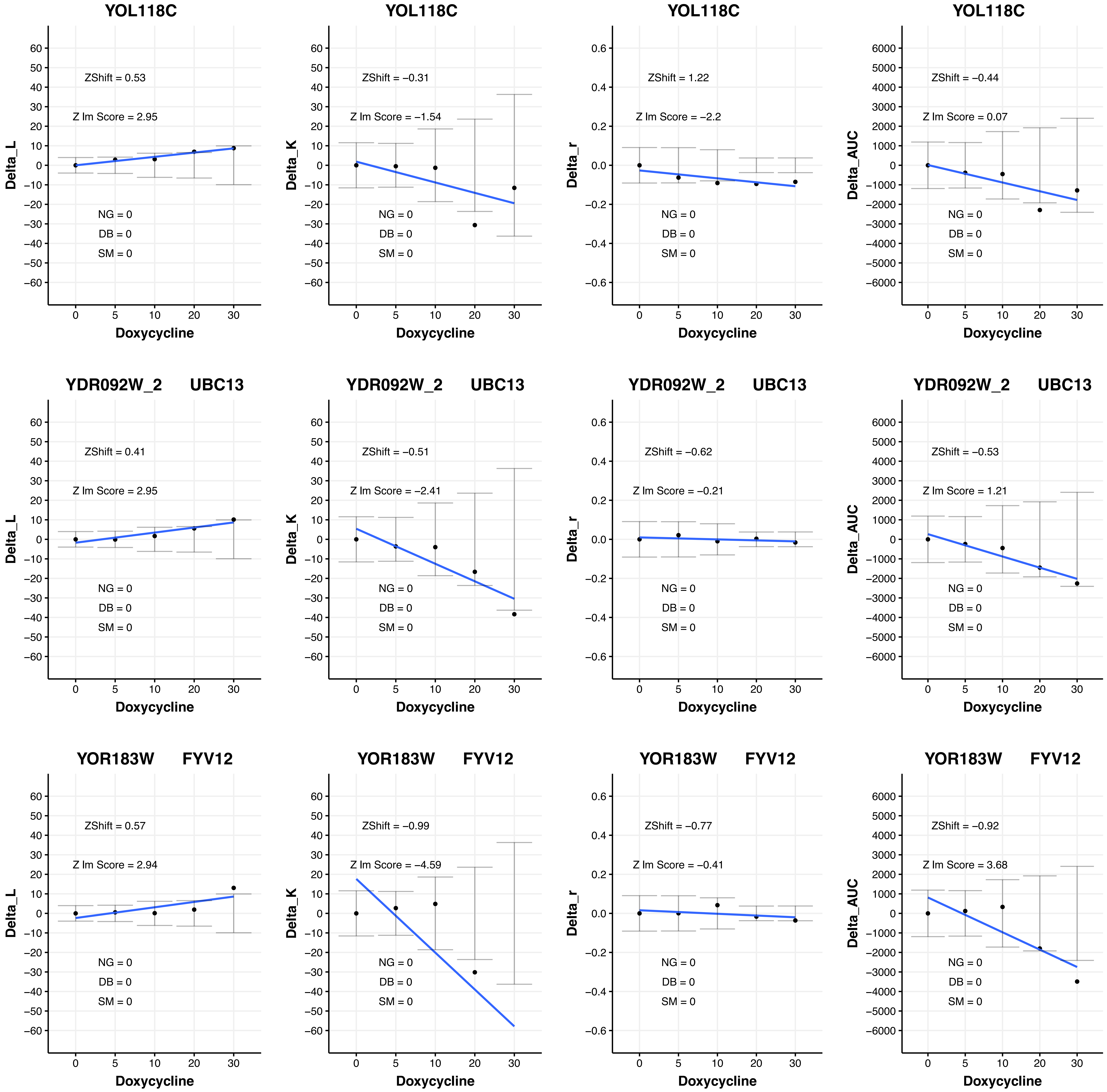
YOL118C
ZShift = 0.53
Z lm Score = 2.95
NG = 0
DB = 0
SM = 0
Delta_L
Doxycycline
YOL118C
ZShift = −0.31
Z lm Score = −1.54
NG = 0
DB = 0
SM = 0
Delta_K
Doxycycline
YOL118C
ZShift = 1.22
Z lm Score = −2.2
NG = 0
DB = 0
SM = 0
Delta_r
Doxycycline
YOL118C
ZShift = −0.44
Z lm Score = 0.07
NG = 0
DB = 0
SM = 0
Delta_AUC
Doxycycline
YDR092W_2 UBC13
ZShift = 0.41
Z lm Score = 2.95
NG = 0
DB = 0
SM = 0
Delta_L
Doxycycline
YDR092W_2 UBC13
ZShift = −0.51
Z lm Score = −2.41
NG = 0
DB = 0
SM = 0
Delta_K
Doxycycline
YDR092W_2 UBC13
ZShift = −0.62
Z lm Score = −0.21
NG = 0
DB = 0
SM = 0
Delta_r
Doxycycline
YDR092W_2 UBC13
ZShift = −0.53
Z lm Score = 1.21
NG = 0
DB = 0
SM = 0
Delta_AUC
Doxycycline
YOR183W FYV12
ZShift = 0.57
Z lm Score = 2.94
NG = 0
DB = 0
SM = 0
Delta_L
Doxycycline
YOR183W FYV12
ZShift = −0.99
Z lm Score = −4.59
NG = 0
DB = 0
SM = 0
Delta_K
Doxycycline
YOR183W FYV12
ZShift = −0.77
Z lm Score = −0.41
NG = 0
DB = 0
SM = 0
Delta_r
Doxycycline
YOR183W FYV12
ZShift = −0.92
Z lm Score = 3.68
NG = 0
DB = 0
SM = 0
Delta_AUC
Doxycycline

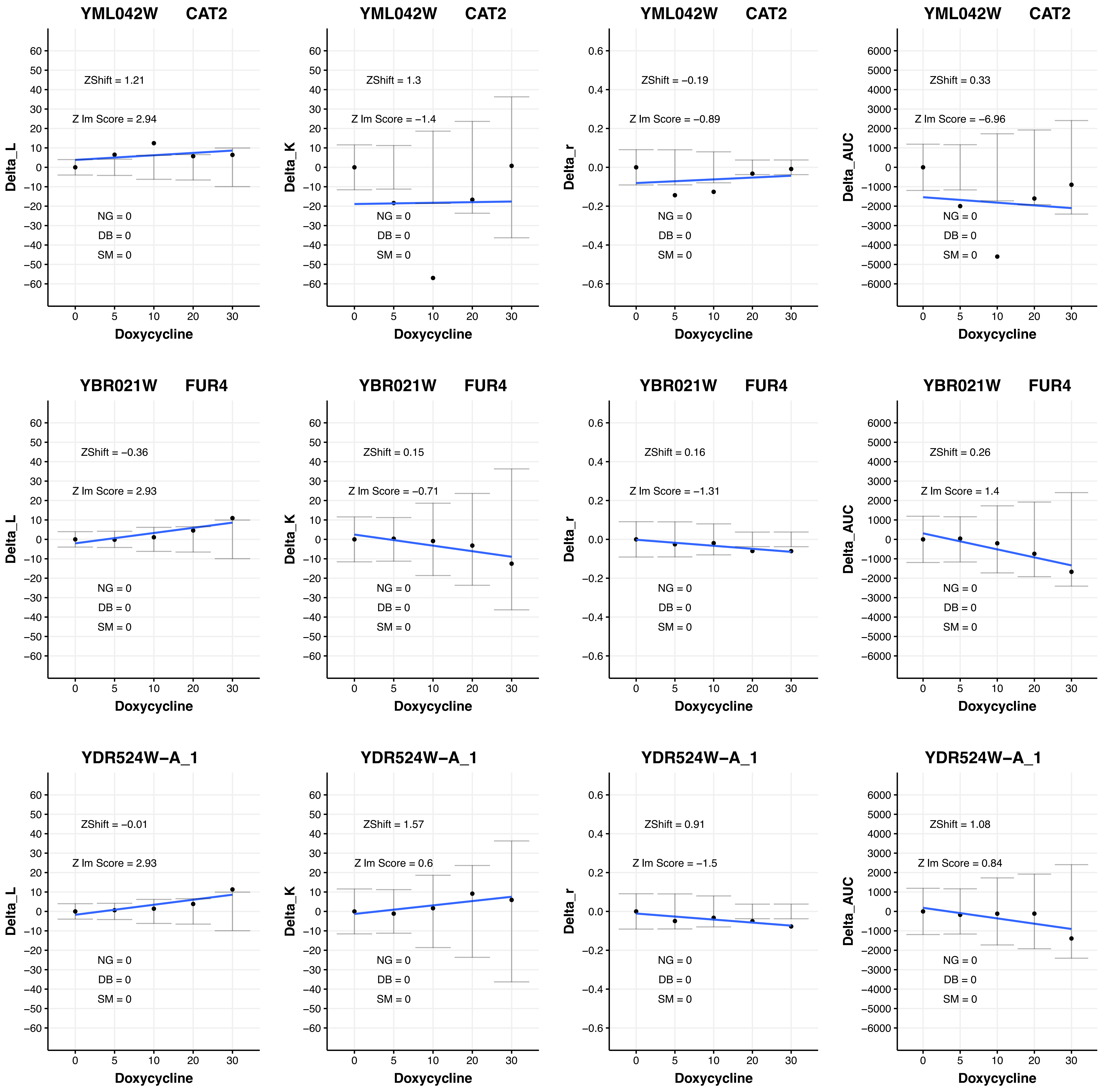

YML042W CAT2
ZShift = 1.21
Z lm Score = 2.94
NG = 0
DB = 0
SM = 0
Delta_L
Doxycycline
YML042W CAT2
ZShift = 1.3
Z lm Score = −1.4
NG = 0
DB = 0
SM = 0
Delta_K
Doxycycline
YML042W CAT2
ZShift = −0.19
Z lm Score = −0.89
NG = 0
DB = 0
SM = 0
Delta_r
Doxycycline
YML042W CAT2
ZShift = 0.33
Z lm Score = −6.96
NG = 0
DB = 0
SM = 0
Delta_AUC
Doxycycline
YBR021W FUR4
ZShift = −0.36
Z lm Score = 2.93
NG = 0
DB = 0
SM = 0
Delta_L
Doxycycline
YBR021W FUR4
ZShift = 0.15
Z lm Score = −0.71
NG = 0
DB = 0
SM = 0
Delta_K
Doxycycline
YBR021W FUR4
ZShift = 0.16
Z lm Score = −1.31
NG = 0
DB = 0
SM = 0
Delta_r
Doxycycline
YBR021W FUR4
ZShift = 0.26
Z lm Score = 1.4
NG = 0
DB = 0
SM = 0
Delta_AUC
Doxycycline
YDR524W−A_1
ZShift = −0.01
Z lm Score = 2.93
NG = 0
DB = 0
SM = 0
Delta_L
Doxycycline
YDR524W−A_1
ZShift = 1.57
Z lm Score = 0.6
NG = 0
DB = 0
SM = 0
Delta_K
Doxycycline
YDR524W−A_1
ZShift = 0.91
Z lm Score = −1.5
NG = 0
DB = 0
SM = 0
Delta_r
Doxycycline
YDR524W−A_1
ZShift = 1.08
Z lm Score = 0.84
NG = 0
DB = 0
SM = 0
Delta_AUC
Doxycycline

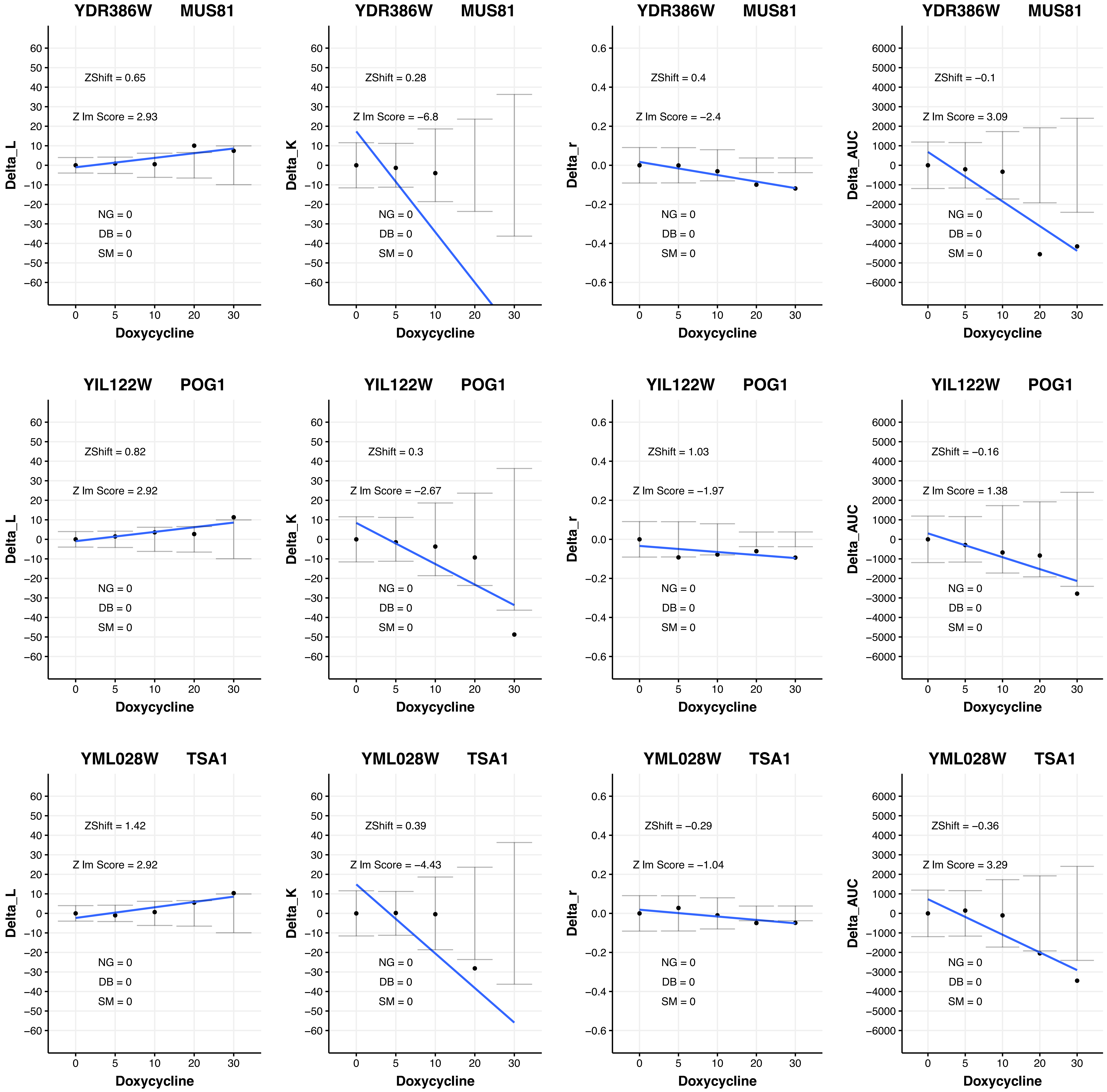
YDR386W MUS81
ZShift = 0.65
Z lm Score = 2.93
NG = 0
DB = 0
SM = 0
Delta_L
Doxycycline
YDR386W MUS81
ZShift = 0.28
Z lm Score = −6.8
NG = 0
DB = 0
SM = 0
Delta_K
Doxycycline
YDR386W MUS81
ZShift = 0.4
Z lm Score = −2.4
NG = 0
DB = 0
SM = 0
Delta_r
Doxycycline
YDR386W MUS81
ZShift = −0.1
Z lm Score = 3.09
NG = 0
DB = 0
SM = 0
Delta_AUC
Doxycycline
YIL122W POG1
ZShift = 0.82
Z lm Score = 2.92
NG = 0
DB = 0
SM = 0
Delta_L
Doxycycline
YIL122W POG1
ZShift = 0.3
Z lm Score = −2.67
NG = 0
DB = 0
SM = 0
Delta_K
Doxycycline
YIL122W POG1
ZShift = 1.03
Z lm Score = −1.97
NG = 0
DB = 0
SM = 0
Delta_r
Doxycycline
YIL122W POG1
ZShift = −0.16
Z lm Score = 1.38
NG = 0
DB = 0
SM = 0
Delta_AUC
Doxycycline
YML028W TSA1
ZShift = 1.42
Z lm Score = 2.92
NG = 0
DB = 0
SM = 0
Delta_L
Doxycycline
YML028W TSA1
ZShift = 0.39
Z lm Score = −4.43
NG = 0
DB = 0
SM = 0
Delta_K
Doxycycline
YML028W TSA1
ZShift = −0.29
Z lm Score = −1.04
NG = 0
DB = 0
SM = 0
Delta_r
Doxycycline
YML028W TSA1
ZShift = −0.36
Z lm Score = 3.29
NG = 0
DB = 0
SM = 0
Delta_AUC
Doxycycline

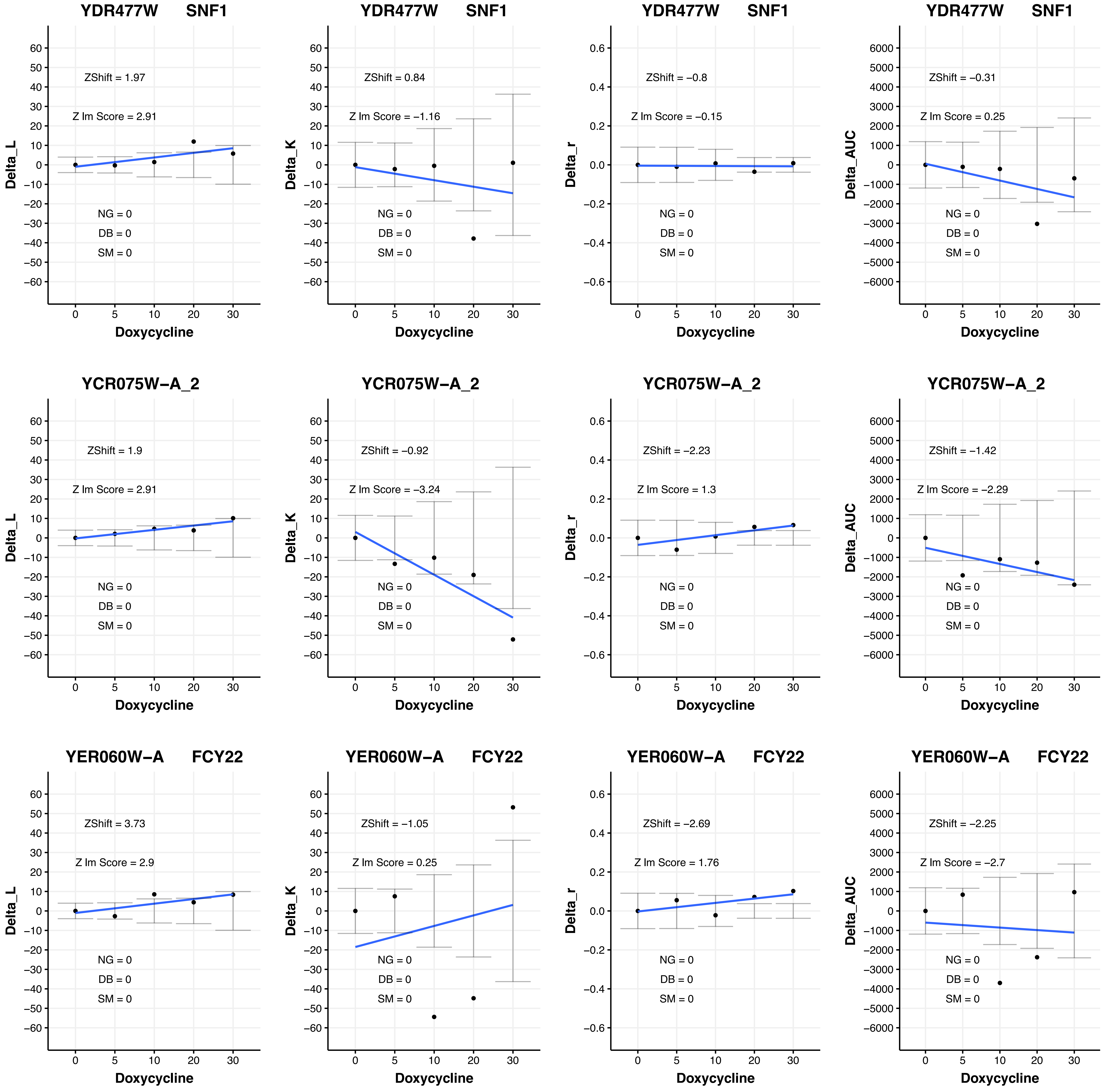
YDR477W SNF1
ZShift = 1.97
Z lm Score = 2.91
NG = 0
DB = 0
SM = 0
Delta_L
Doxycycline
YDR477W SNF1
ZShift = 0.84
Z lm Score = −1.16
NG = 0
DB = 0
SM = 0
Delta_K
Doxycycline
YDR477W SNF1
ZShift = −0.8
Z lm Score = −0.15
NG = 0
DB = 0
SM = 0
Delta_r
Doxycycline
YDR477W SNF1
ZShift = −0.31
Z lm Score = 0.25
NG = 0
DB = 0
SM = 0
Delta_AUC
Doxycycline
YCR075W−A_2
ZShift = 1.9
Z lm Score = 2.91
NG = 0
DB = 0
SM = 0
Delta_L
Doxycycline
YCR075W−A_2
ZShift = −0.92
Z lm Score = −3.24
NG = 0
DB = 0
SM = 0
Delta_K
Doxycycline
YCR075W−A_2
ZShift = −2.23
Z lm Score = 1.3
NG = 0
DB = 0
SM = 0
Delta_r
Doxycycline
YCR075W−A_2
ZShift = −1.42
Z lm Score = −2.29
NG = 0
DB = 0
SM = 0
Delta_AUC
Doxycycline
YER060W−A FCY22
ZShift = 3.73
Z lm Score = 2.9
NG = 0
DB = 0
SM = 0
Delta_L
Doxycycline
YER060W−A FCY22
ZShift = −1.05
Z lm Score = 0.25
NG = 0
DB = 0
SM = 0
Delta_K
Doxycycline
YER060W−A FCY22
ZShift = −2.69
Z lm Score = 1.76
NG = 0
DB = 0
SM = 0
Delta_r
Doxycycline
YER060W−A FCY22
ZShift = −2.25
Z lm Score = −2.7
NG = 0
DB = 0
SM = 0
Delta_AUC
Doxycycline

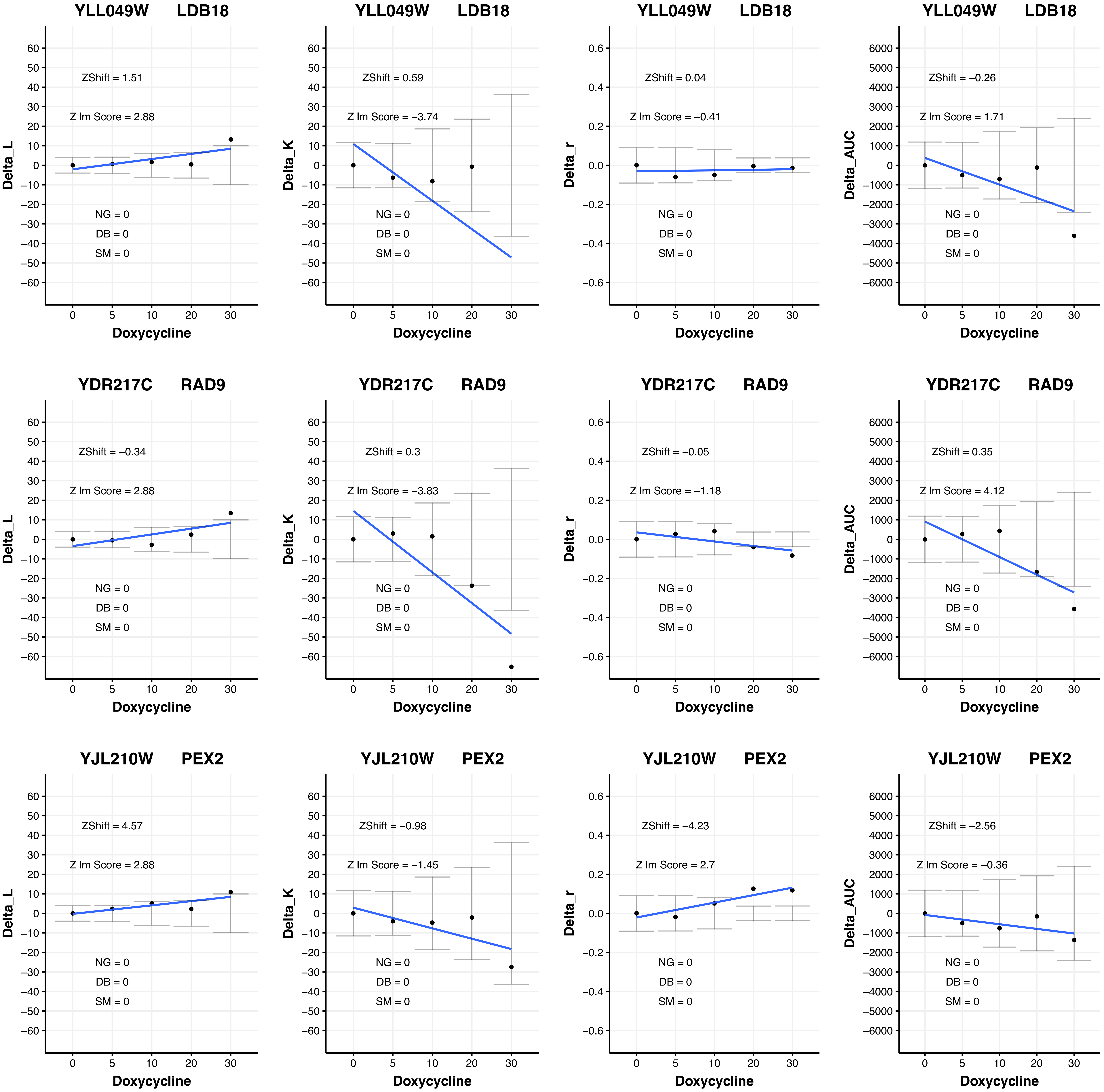
YLL049W LDB18
ZShift = 1.51
Z lm Score = 2.88
NG = 0
DB = 0
SM = 0
Delta_L
Doxycycline
YLL049W LDB18
ZShift = 0.59
Z lm Score = −3.74
NG = 0
DB = 0
SM = 0
Delta_K
Doxycycline
YLL049W LDB18
ZShift = 0.04
Z lm Score = −0.41
NG = 0
DB = 0
SM = 0
Delta_r
Doxycycline
YLL049W LDB18
ZShift = −0.26
Z lm Score = 1.71
NG = 0
DB = 0
SM = 0
Delta_AUC
Doxycycline
YDR217C RAD9
ZShift = −0.34
Z lm Score = 2.88
NG = 0
DB = 0
SM = 0
Delta_L
Doxycycline
YDR217C RAD9
ZShift = 0.3
Z lm Score = −3.83
NG = 0
DB = 0
SM = 0
Delta_K
Doxycycline
YDR217C RAD9
ZShift = −0.05
Z lm Score = −1.18
NG = 0
DB = 0
SM = 0
Delta_r
Doxycycline
YDR217C RAD9
ZShift = 0.35
Z lm Score = 4.12
NG = 0
DB = 0
SM = 0
Delta_AUC
Doxycycline
YJL210W PEX2
ZShift = 4.57
Z lm Score = 2.88
NG = 0
DB = 0
SM = 0
Delta_L
Doxycycline
YJL210W PEX2
ZShift = −0.98
Z lm Score = −1.45
NG = 0
DB = 0
SM = 0
Delta_K
Doxycycline
YJL210W PEX2
ZShift = −4.23
Z lm Score = 2.7
NG = 0
DB = 0
SM = 0
Delta_r
Doxycycline
YJL210W PEX2
ZShift = −2.56
Z lm Score = −0.36
NG = 0
DB = 0
SM = 0
Delta_AUC
Doxycycline

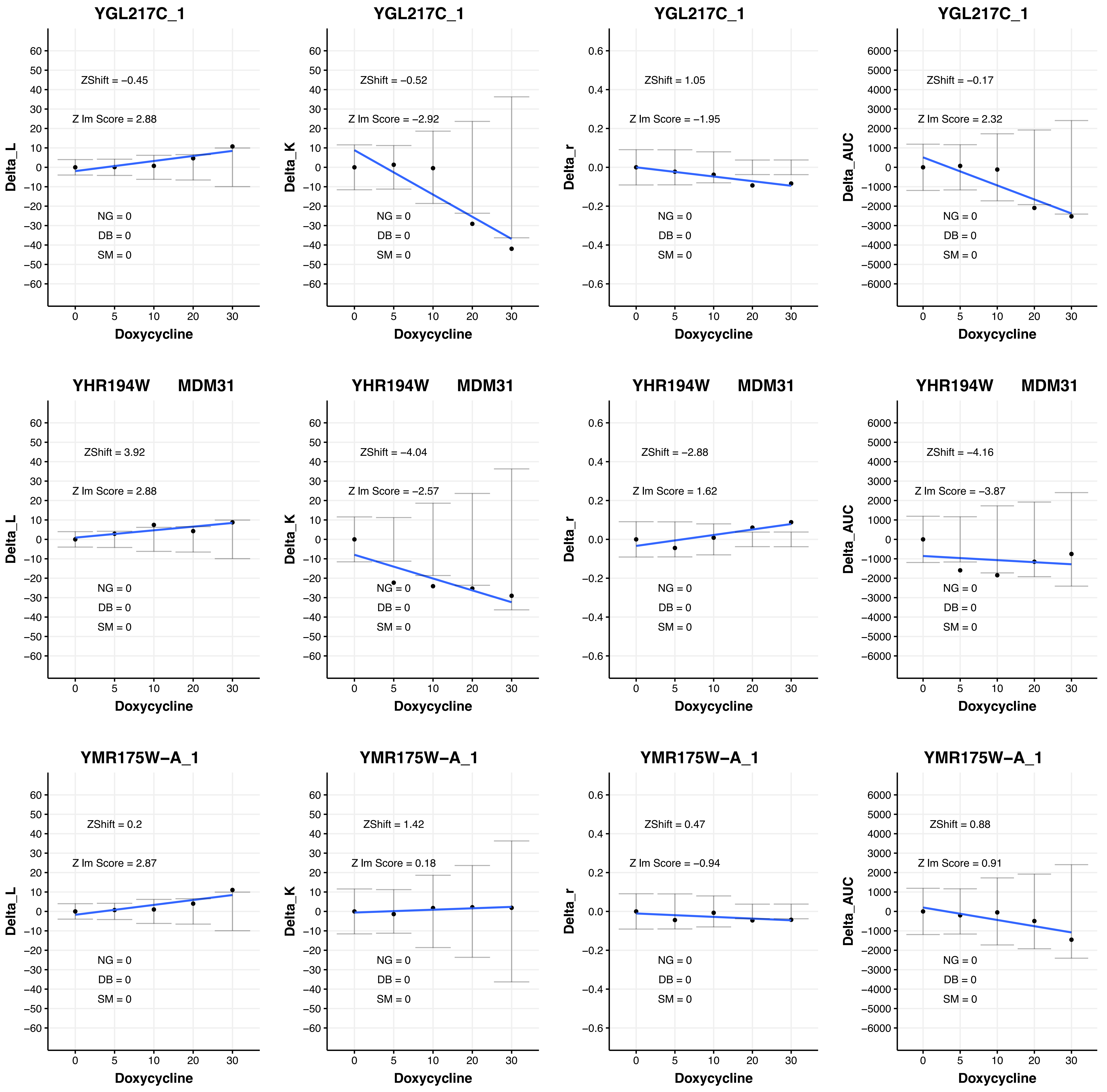

YGL217C_1
ZShift = −0.45
Z lm Score = 2.88
NG = 0
DB = 0
SM = 0
Delta_L
Doxycycline
YGL217C_1
ZShift = −0.52
Z lm Score = −2.92
NG = 0
DB = 0
SM = 0
Delta_K
Doxycycline
YGL217C_1
ZShift = 1.05
Z lm Score = −1.95
NG = 0
DB = 0
SM = 0
Delta_r
Doxycycline
YGL217C_1
ZShift = −0.17
Z lm Score = 2.32
NG = 0
DB = 0
SM = 0
Delta_AUC
Doxycycline
YHR194W MDM31
ZShift = 3.92
Z lm Score = 2.88
NG = 0
DB = 0
SM = 0
Delta_L
Doxycycline
YHR194W MDM31
ZShift = −4.04
Z lm Score = −2.57
NG = 0
DB = 0
SM = 0
Delta_K
Doxycycline
YHR194W MDM31
ZShift = −2.88
Z lm Score = 1.62
NG = 0
DB = 0
SM = 0
Delta_r
Doxycycline
YHR194W MDM31
ZShift = −4.16
Z lm Score = −3.87
NG = 0
DB = 0
SM = 0
Delta_AUC
Doxycycline
YMR175W−A_1
ZShift = 0.2
Z lm Score = 2.87
NG = 0
DB = 0
SM = 0
Delta_L
Doxycycline
YMR175W−A_1
ZShift = 1.42
Z lm Score = 0.18
NG = 0
DB = 0
SM = 0
Delta_K
Doxycycline
YMR175W−A_1
ZShift = 0.47
Z lm Score = −0.94
NG = 0
DB = 0
SM = 0
Delta_r
Doxycycline
YMR175W−A_1
ZShift = 0.88
Z lm Score = 0.91
NG = 0
DB = 0
SM = 0
Delta_AUC
Doxycycline

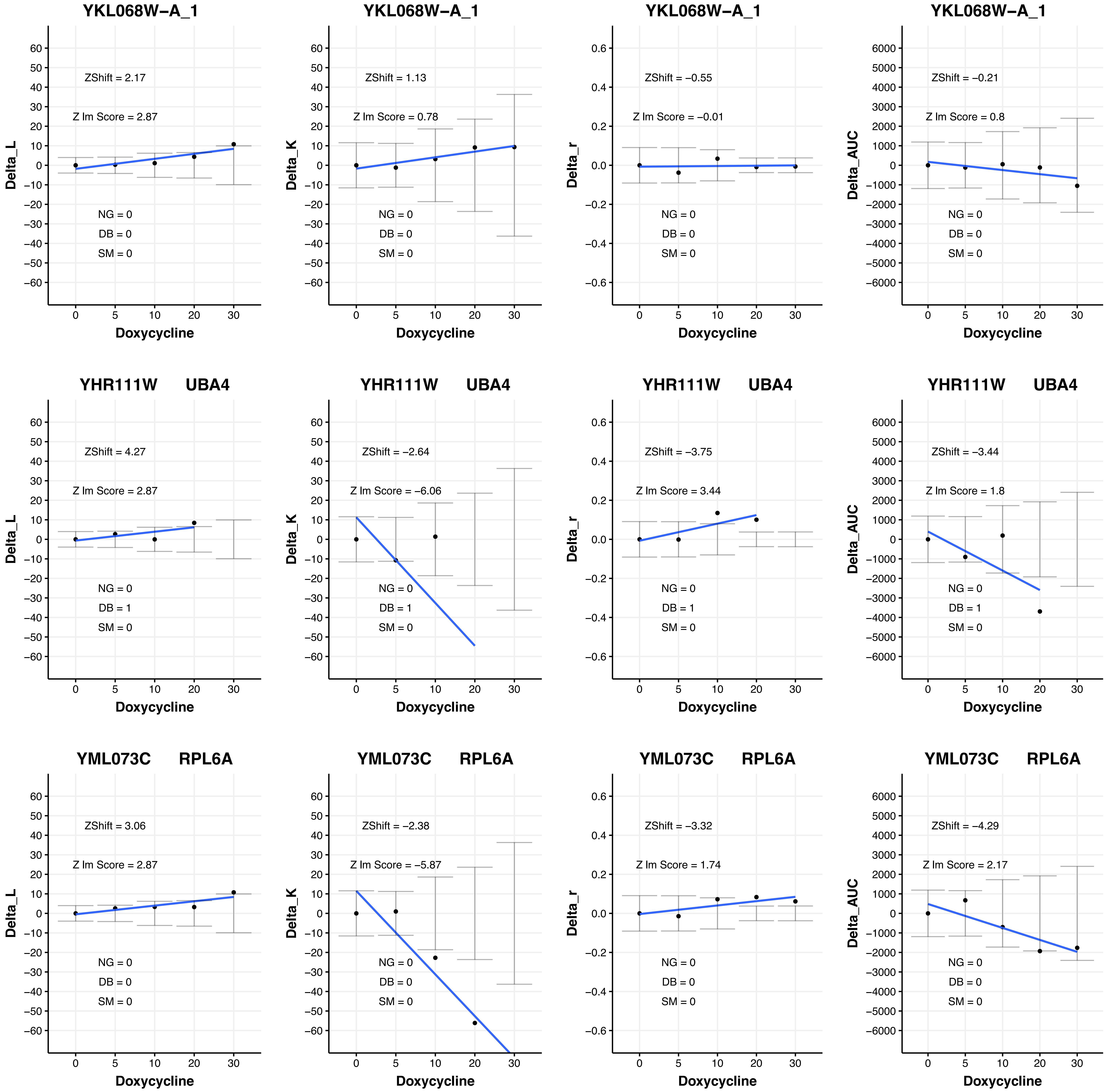
YKL068W−A_1
ZShift = 2.17
Z lm Score = 2.87
NG = 0
DB = 0
SM = 0
Delta_L
Doxycycline
YKL068W−A_1
ZShift = 1.13
Z lm Score = 0.78
NG = 0
DB = 0
SM = 0
Delta_K
YKL068W−A_1
ZShift = −0.55
Z lm Score = −0.01
NG = 0
DB = 0
SM = 0
Delta_r
YKL068W−A_1
ZShift = −0.21
Z lm Score = 0.8
NG = 0
DB = 0
SM = 0
Delta_AUC
YHR111W UBA4
ZShift = 4.27
Z lm Score = 2.87
NG = 0
DB = 1
SM = 0
YHR111W UBA4
ZShift = −2.64
Z lm Score = −6.06
NG = 0
DB = 1
SM = 0
YHR111W UBA4
ZShift = −3.75
Z lm Score = 3.44
NG = 0
DB = 1
SM = 0
YHR111W UBA4
ZShift = −3.44
Z lm Score = 1.8
NG = 0
DB = 1
SM = 0
YML073C RPL6A
ZShift = 3.06
Z lm Score = 2.87
NG = 0
DB = 0
SM = 0
YML073C RPL6A
ZShift = −2.38
Z lm Score = −5.87
NG = 0
DB = 0
SM = 0
YML073C RPL6A
ZShift = −3.32
Z lm Score = 1.74
NG = 0
DB = 0
SM = 0
YML073C RPL6A
ZShift = −4.29
Z lm Score = 2.17
NG = 0
DB = 0
SM = 0

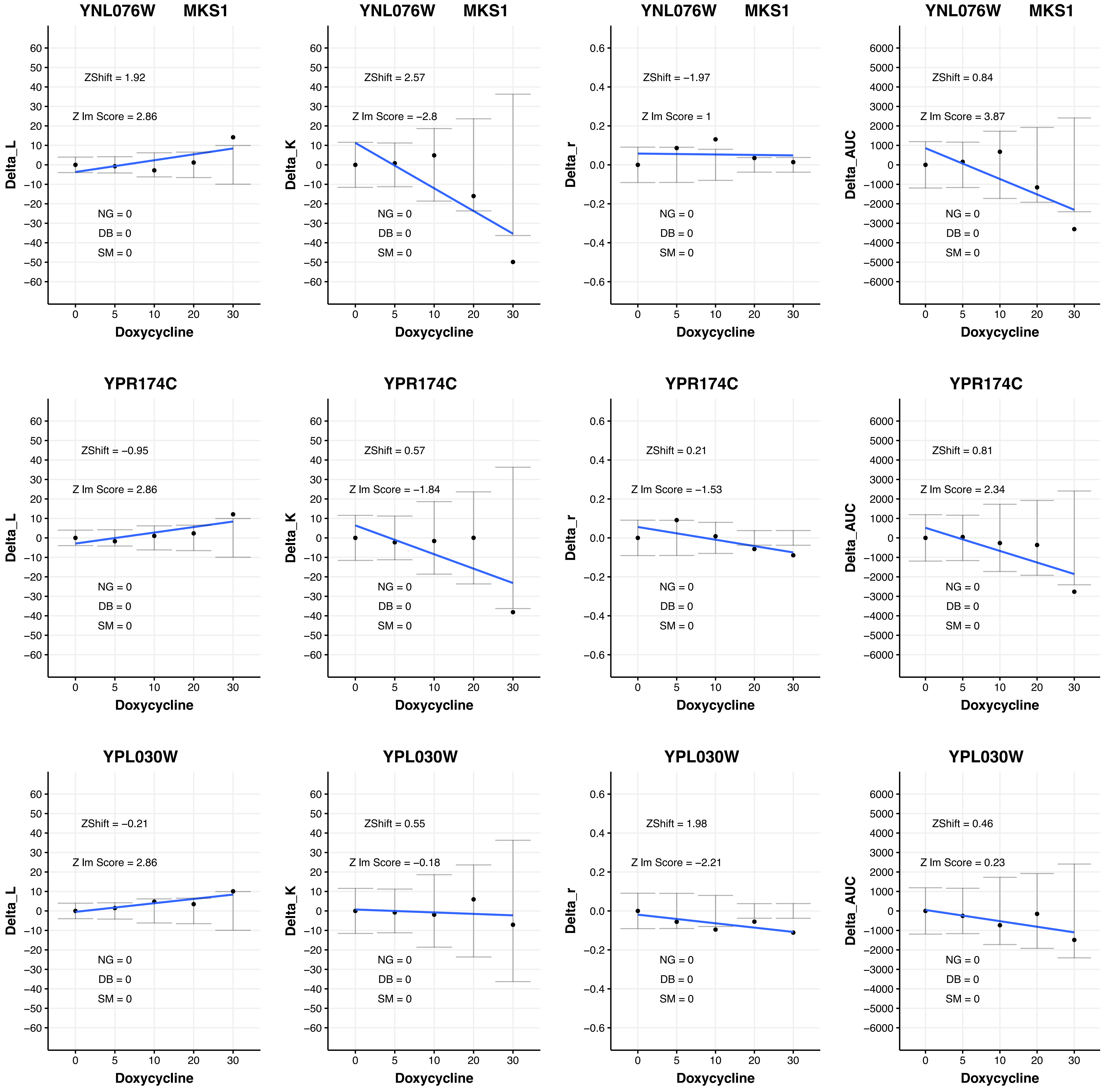

YNL076W MKS1
ZShift = 1.92
Z lm Score = 2.86
NG = 0
DB = 0
SM = 0
Delta_L
Doxycycline
YNL076W MKS1
ZShift = 2.57
Z lm Score = −2.8
NG = 0
DB = 0
SM = 0
Delta_K
Doxycycline
YNL076W MKS1
ZShift = −1.97
Z lm Score = 1
NG = 0
DB = 0
SM = 0
Delta_r
Doxycycline
YNL076W MKS1
ZShift = 0.84
Z lm Score = 3.87
NG = 0
DB = 0
SM = 0
Delta_AUC
Doxycycline
YPR174C
ZShift = −0.95
Z lm Score = 2.86
NG = 0
DB = 0
SM = 0
Delta_L
Doxycycline
YPR174C
ZShift = 0.57
Z lm Score = −1.84
NG = 0
DB = 0
SM = 0
Delta_K
Doxycycline
YPR174C
ZShift = 0.21
Z lm Score = −1.53
NG = 0
DB = 0
SM = 0
Delta_r
Doxycycline
YPR174C
ZShift = 0.81
Z lm Score = 2.34
NG = 0
DB = 0
SM = 0
Delta_AUC
Doxycycline
YPL030W
ZShift = −0.21
Z lm Score = 2.86
NG = 0
DB = 0
SM = 0
Delta_L
Doxycycline
YPL030W
ZShift = 0.55
Z lm Score = −0.18
NG = 0
DB = 0
SM = 0
Delta_K
Doxycycline
YPL030W
ZShift = 1.98
Z lm Score = −2.21
NG = 0
DB = 0
SM = 0
Delta_r
Doxycycline
YPL030W
ZShift = 0.46
Z lm Score = 0.23
NG = 0
DB = 0
SM = 0
Delta_AUC
Doxycycline

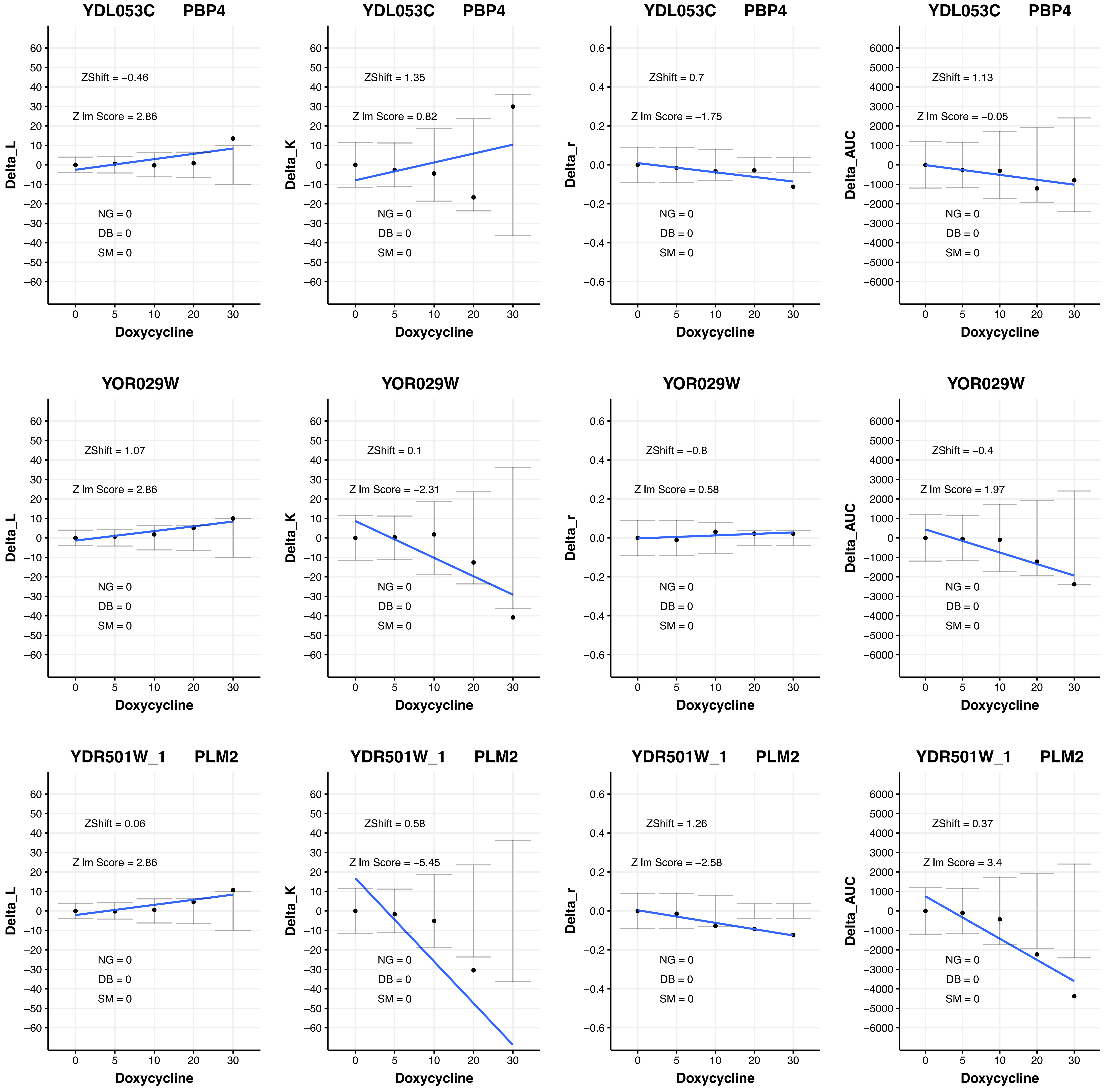
YDL053C PBP4
ZShift = −0.46
Z lm Score = 2.86
NG = 0
DB = 0
SM = 0
Delta_L
Doxycycline
YDL053C PBP4
ZShift = 1.35
Z lm Score = 0.82
NG = 0
DB = 0
SM = 0
Delta_K
Doxycycline
YDL053C PBP4
ZShift = 0.7
Z lm Score = −1.75
NG = 0
DB = 0
SM = 0
Delta_r
Doxycycline
YDL053C PBP4
ZShift = 1.13
Z lm Score = −0.05
NG = 0
DB = 0
SM = 0
Delta_AUC
Doxycycline
YOR029W
ZShift = 1.07
Z lm Score = 2.86
NG = 0
DB = 0
SM = 0
Delta_L
Doxycycline
YOR029W
ZShift = 0.1
Z lm Score = −2.31
NG = 0
DB = 0
SM = 0
Delta_K
Doxycycline
YOR029W
ZShift = −0.8
Z lm Score = 0.58
NG = 0
DB = 0
SM = 0
Delta_r
Doxycycline
YOR029W
ZShift = −0.4
Z lm Score = 1.97
NG = 0
DB = 0
SM = 0
Delta_AUC
Doxycycline
YDR501W_1 PLM2
ZShift = 0.06
Z lm Score = 2.86
NG = 0
DB = 0
SM = 0
Delta_L
Doxycycline
YDR501W_1 PLM2
ZShift = 0.58
Z lm Score = −5.45
NG = 0
DB = 0
SM = 0
Delta_K
Doxycycline
YDR501W_1 PLM2
ZShift = 1.26
Z lm Score = −2.58
NG = 0
DB = 0
SM = 0
Delta_r
Doxycycline
YDR501W_1 PLM2
ZShift = 0.37
Z lm Score = 3.4
NG = 0
DB = 0
SM = 0
Delta_AUC
Doxycycline

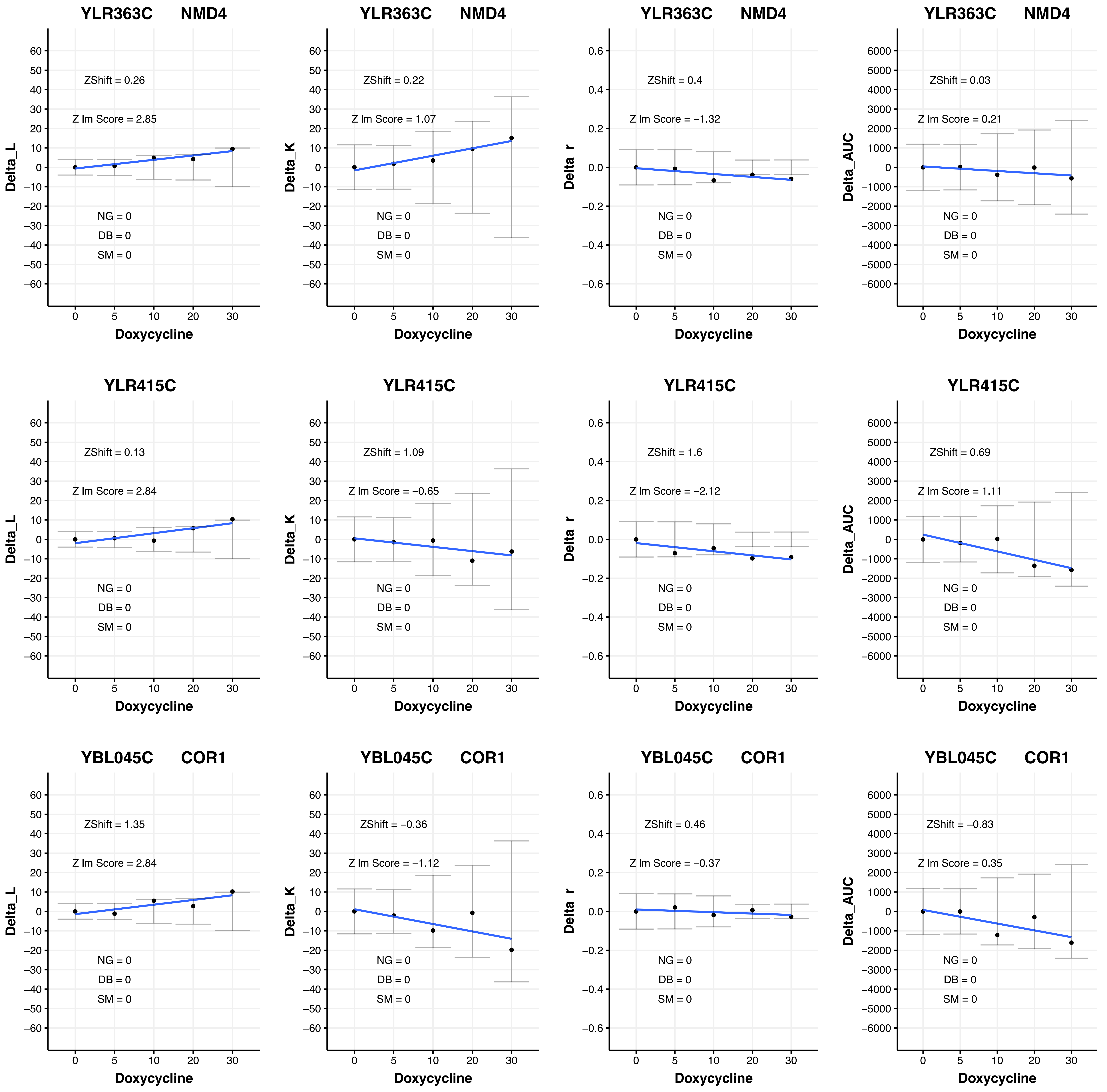
YLR363C NMD4
ZShift = 0.26
Z lm Score = 2.85
NG = 0
DB = 0
SM = 0
Delta_L
Doxycycline
YLR363C NMD4
ZShift = 0.22
Z lm Score = 1.07
NG = 0
DB = 0
SM = 0
Delta_K
Doxycycline
YLR363C NMD4
ZShift = 0.4
Z lm Score = −1.32
NG = 0
DB = 0
SM = 0
Delta_r
Doxycycline
YLR363C NMD4
ZShift = 0.03
Z lm Score = 0.21
NG = 0
DB = 0
SM = 0
Delta_AUC
Doxycycline
YLR415C
ZShift = 0.13
Z lm Score = 2.84
NG = 0
DB = 0
SM = 0
Delta_L
Doxycycline
YLR415C
ZShift = 1.09
Z lm Score = −0.65
NG = 0
DB = 0
SM = 0
Delta_K
Doxycycline
YLR415C
ZShift = 1.6
Z lm Score = −2.12
NG = 0
DB = 0
SM = 0
Delta_r
Doxycycline
YLR415C
ZShift = 0.69
Z lm Score = 1.11
NG = 0
DB = 0
SM = 0
Delta_AUC
Doxycycline
YBL045C COR1
ZShift = 1.35
Z lm Score = 2.84
NG = 0
DB = 0
SM = 0
Delta_L
Doxycycline
YBL045C COR1
ZShift = −0.36
Z lm Score = −1.12
NG = 0
DB = 0
SM = 0
Delta_K
Doxycycline
YBL045C COR1
ZShift = 0.46
Z lm Score = −0.37
NG = 0
DB = 0
SM = 0
Delta_r
Doxycycline
YBL045C COR1
ZShift = −0.83
Z lm Score = 0.35
NG = 0
DB = 0
SM = 0
Delta_AUC
Doxycycline

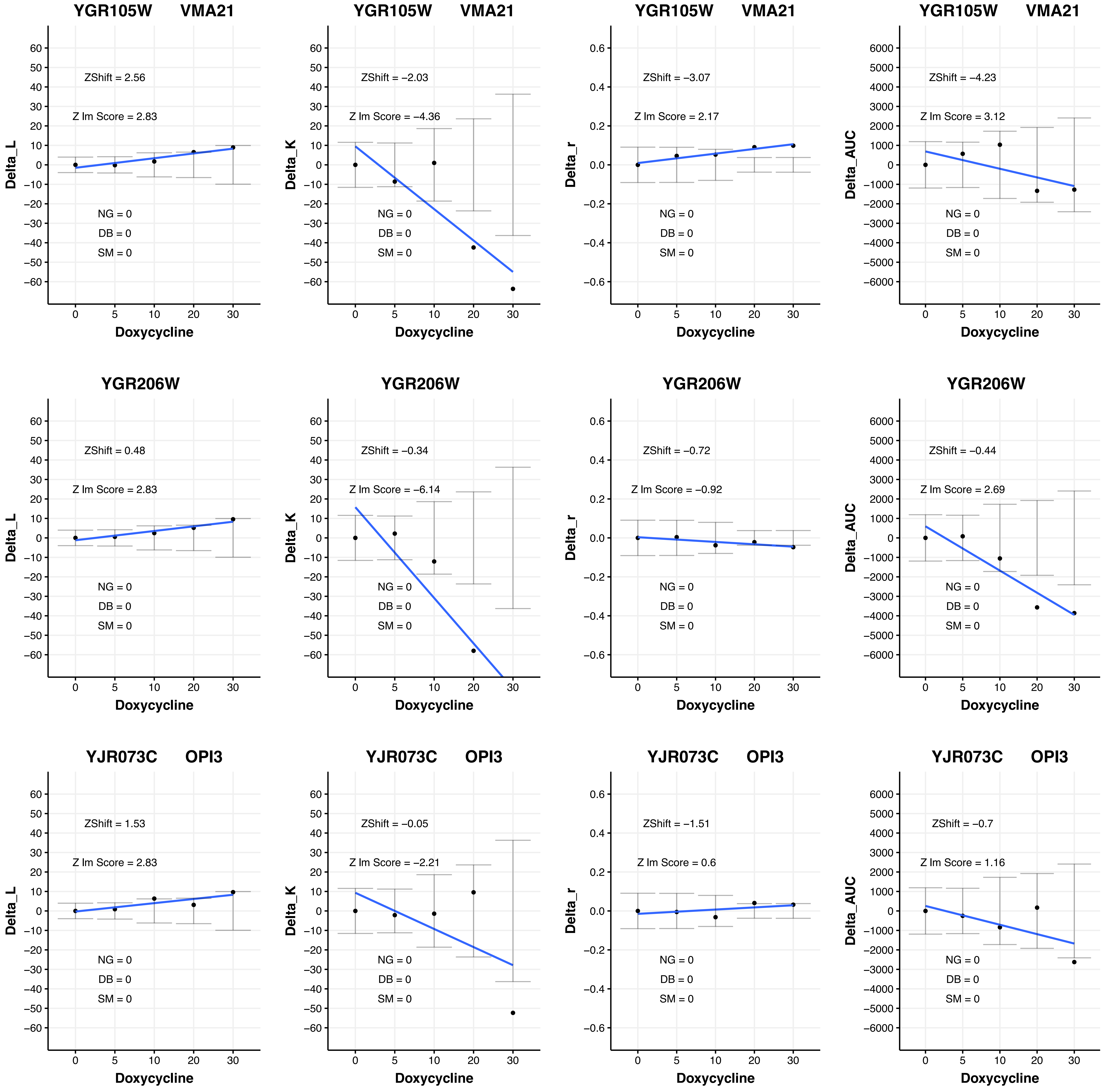
YGR105W VMA21
ZShift = 2.56
Z lm Score = 2.83
NG = 0
DB = 0
SM = 0
Delta_L
Doxycycline
YGR105W VMA21
ZShift = −2.03
Z lm Score = −4.36
Delta_K
YGR105W VMA21
ZShift = −3.07
Z lm Score = 2.17
Delta_r
YGR105W VMA21
ZShift = −4.23
Z lm Score = 3.12
Delta_AUC
YGR206W
ZShift = 0.48
Z lm Score = 2.83
YGR206W
ZShift = −0.34
Z lm Score = −6.14
YGR206W
ZShift = −0.72
Z lm Score = −0.92
YGR206W
ZShift = −0.44
Z lm Score = 2.69
YJR073C OPI3
ZShift = 1.53
Z lm Score = 2.83
YJR073C OPI3
ZShift = −0.05
Z lm Score = −2.21
YJR073C OPI3
ZShift = −1.51
Z lm Score = 0.6
YJR073C OPI3
ZShift = −0.7
Z lm Score = 1.16

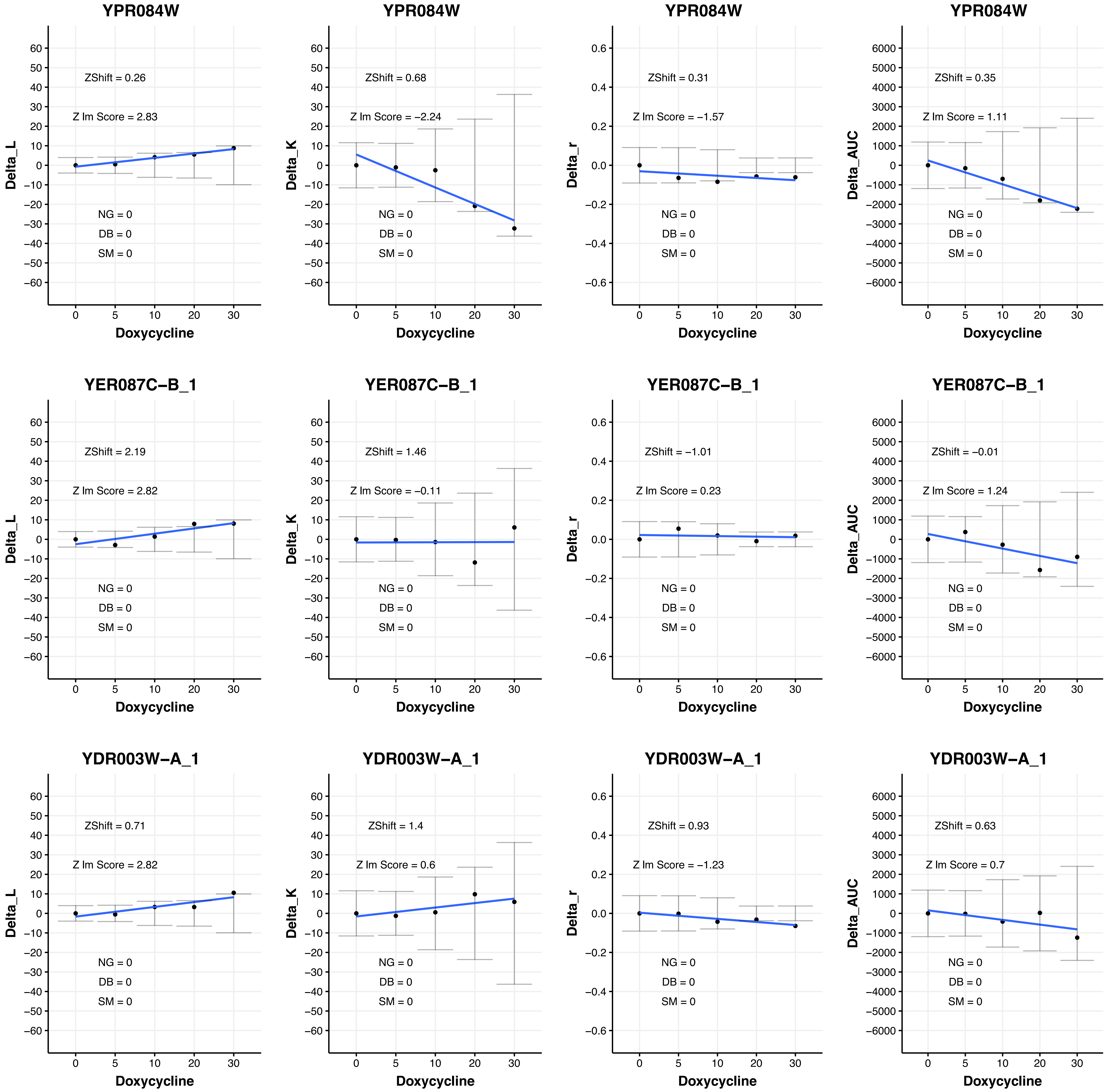
YPR084W
ZShift = 0.26
Z lm Score = 2.83
NG = 0
DB = 0
SM = 0
Delta_L
Doxycycline
YPR084W
ZShift = 0.68
Z lm Score = −2.24
NG = 0
DB = 0
SM = 0
Delta_K
Doxycycline
YPR084W
ZShift = 0.31
Z lm Score = −1.57
NG = 0
DB = 0
SM = 0
Delta_r
Doxycycline
YPR084W
ZShift = 0.35
Z lm Score = 1.11
NG = 0
DB = 0
SM = 0
Delta_AUC
Doxycycline
YER087C−B_1
ZShift = 2.19
Z lm Score = 2.82
NG = 0
DB = 0
SM = 0
Delta_L
Doxycycline
YER087C−B_1
ZShift = 1.46
Z lm Score = −0.11
NG = 0
DB = 0
SM = 0
Delta_K
Doxycycline
YER087C−B_1
ZShift = −1.01
Z lm Score = 0.23
NG = 0
DB = 0
SM = 0
Delta_r
Doxycycline
YER087C−B_1
ZShift = −0.01
Z lm Score = 1.24
NG = 0
DB = 0
SM = 0
Delta_AUC
Doxycycline
YDR003W−A_1
ZShift = 0.71
Z lm Score = 2.82
NG = 0
DB = 0
SM = 0
Delta_L
Doxycycline
YDR003W−A_1
ZShift = 1.4
Z lm Score = 0.6
NG = 0
DB = 0
SM = 0
Delta_K
Doxycycline
YDR003W−A_1
ZShift = 0.93
Z lm Score = −1.23
NG = 0
DB = 0
SM = 0
Delta_r
Doxycycline
YDR003W−A_1
ZShift = 0.63
Z lm Score = 0.7
NG = 0
DB = 0
SM = 0
Delta_AUC
Doxycycline

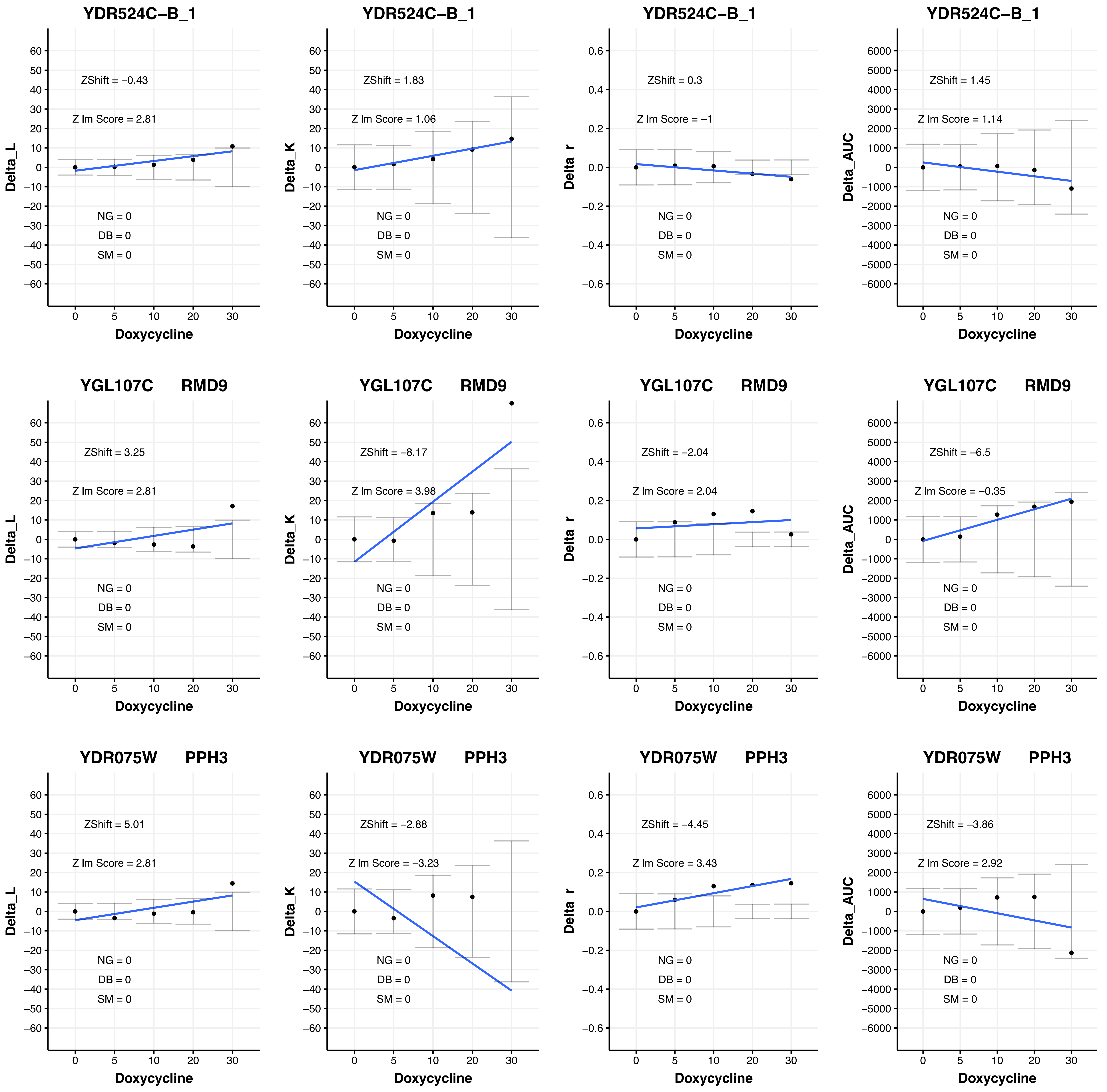

YDR524C–B_1
ZShift = –0.43
Z lm Score = 2.81
NG = 0
DB = 0
SM = 0
Delta_L
Doxycycline
YDR524C–B_1
ZShift = 1.83
Z lm Score = 1.06
NG = 0
DB = 0
SM = 0
Delta_K
Doxycycline
YDR524C–B_1
ZShift = 0.3
Z lm Score = –1
NG = 0
DB = 0
SM = 0
Delta_r
Doxycycline
YDR524C–B_1
ZShift = 1.45
Z lm Score = 1.14
NG = 0
DB = 0
SM = 0
Delta_AUC
Doxycycline
YGL107C RMD9
ZShift = 3.25
Z lm Score = 2.81
NG = 0
DB = 0
SM = 0
Delta_L
Doxycycline
YGL107C RMD9
ZShift = –8.17
Z lm Score = 3.98
NG = 0
DB = 0
SM = 0
Delta_K
Doxycycline
YGL107C RMD9
ZShift = –2.04
Z lm Score = 2.04
NG = 0
DB = 0
SM = 0
Delta_r
Doxycycline
YGL107C RMD9
ZShift = –6.5
Z lm Score = –0.35
NG = 0
DB = 0
SM = 0
Delta_AUC
Doxycycline
YDR075W PPH3
ZShift = 5.01
Z lm Score = 2.81
NG = 0
DB = 0
SM = 0
Delta_L
Doxycycline
YDR075W PPH3
ZShift = –2.88
Z lm Score = –3.23
NG = 0
DB = 0
SM = 0
Delta_K
Doxycycline
YDR075W PPH3
ZShift = –4.45
Z lm Score = 3.43
NG = 0
DB = 0
SM = 0
Delta_r
Doxycycline
YDR075W PPH3
ZShift = –3.86
Z lm Score = 2.92
NG = 0
DB = 0
SM = 0
Delta_AUC
Doxycycline

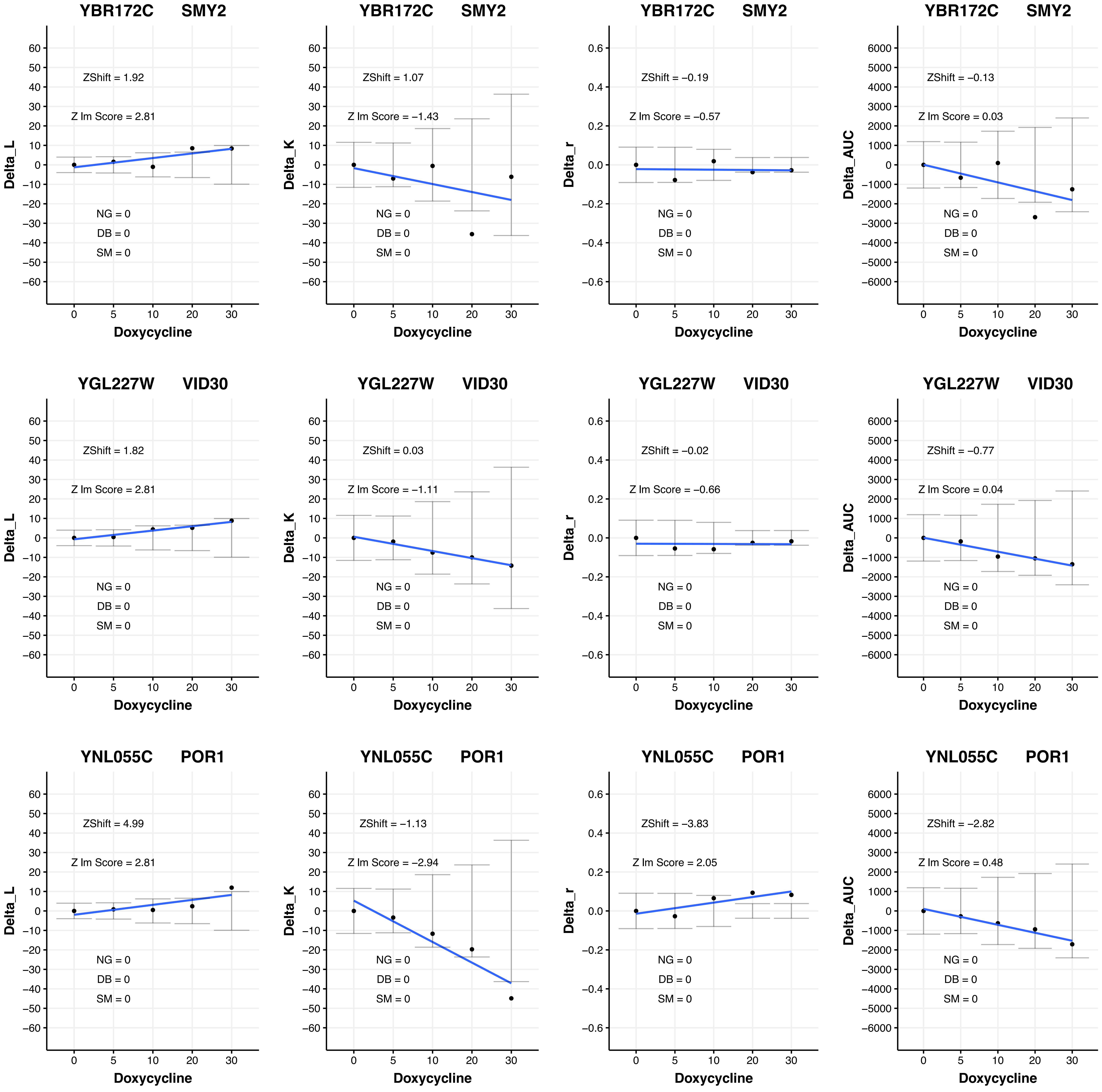

YBR172C SMY2
ZShift = 1.92
Z lm Score = 2.81
NG = 0
DB = 0
SM = 0
Delta_L
Doxycycline
YBR172C SMY2
ZShift = 1.07
Z lm Score = −1.43
NG = 0
DB = 0
SM = 0
Delta_K
Doxycycline
YBR172C SMY2
ZShift = −0.19
Z lm Score = −0.57
NG = 0
DB = 0
SM = 0
Delta_r
Doxycycline
YBR172C SMY2
ZShift = −0.13
Z lm Score = 0.03
NG = 0
DB = 0
SM = 0
Delta_AUC
Doxycycline
YGL227W VID30
ZShift = 1.82
Z lm Score = 2.81
NG = 0
DB = 0
SM = 0
Delta_L
Doxycycline
YGL227W VID30
ZShift = 0.03
Z lm Score = −1.11
NG = 0
DB = 0
SM = 0
Delta_K
Doxycycline
YGL227W VID30
ZShift = −0.02
Z lm Score = −0.66
NG = 0
DB = 0
SM = 0
Delta_r
Doxycycline
YGL227W VID30
ZShift = −0.77
Z lm Score = 0.04
NG = 0
DB = 0
SM = 0
Delta_AUC
Doxycycline
YNL055C POR1
ZShift = 4.99
Z lm Score = 2.81
NG = 0
DB = 0
SM = 0
Delta_L
Doxycycline
YNL055C POR1
ZShift = −1.13
Z lm Score = −2.94
NG = 0
DB = 0
SM = 0
Delta_K
Doxycycline
YNL055C POR1
ZShift = −3.83
Z lm Score = 2.05
NG = 0
DB = 0
SM = 0
Delta_r
Doxycycline
YNL055C POR1
ZShift = −2.82
Z lm Score = 0.48
NG = 0
DB = 0
SM = 0
Delta_AUC
Doxycycline

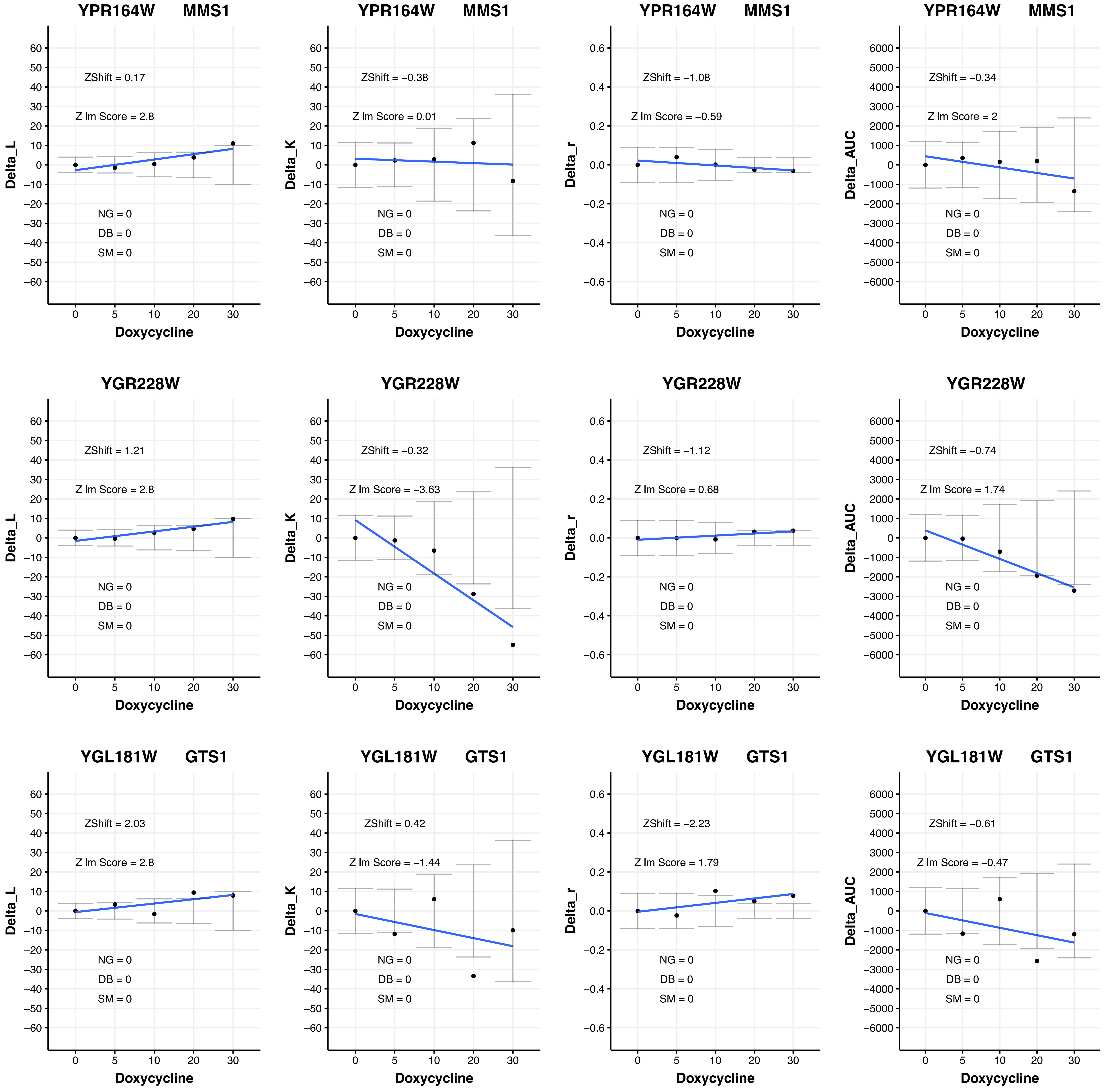

YPR164W MMS1
Delta_L
ZShift = 0.17
Z lm Score = 2.8
NG = 0
DB = 0
SM = 0
Doxycycline
YPR164W MMS1
Delta_K
ZShift = −0.38
Z lm Score = 0.01
NG = 0
DB = 0
SM = 0
Doxycycline
YPR164W MMS1
Delta_r
ZShift = −1.08
Z lm Score = −0.59
NG = 0
DB = 0
SM = 0
Doxycycline
YPR164W MMS1
Delta_AUC
ZShift = −0.34
Z lm Score = 2
NG = 0
DB = 0
SM = 0
Doxycycline
YGR228W
Delta_L
ZShift = 1.21
Z lm Score = 2.8
NG = 0
DB = 0
SM = 0
Doxycycline
YGR228W
Delta_K
ZShift = −0.32
Z lm Score = −3.63
NG = 0
DB = 0
SM = 0
Doxycycline
YGR228W
Delta_r
ZShift = −1.12
Z lm Score = 0.68
NG = 0
DB = 0
SM = 0
Doxycycline
YGR228W
Delta_AUC
ZShift = −0.74
Z lm Score = 1.74
NG = 0
DB = 0
SM = 0
Doxycycline
YGL181W GTS1
Delta_L
ZShift = 2.03
Z lm Score = 2.8
NG = 0
DB = 0
SM = 0
Doxycycline
YGL181W GTS1
Delta_K
ZShift = 0.42
Z lm Score = −1.44
NG = 0
DB = 0
SM = 0
Doxycycline
YGL181W GTS1
Delta_r
ZShift = −2.23
Z lm Score = 1.79
NG = 0
DB = 0
SM = 0
Doxycycline
YGL181W GTS1
Delta_AUC
ZShift = −0.61
Z lm Score = −0.47
NG = 0
DB = 0
SM = 0
Doxycycline

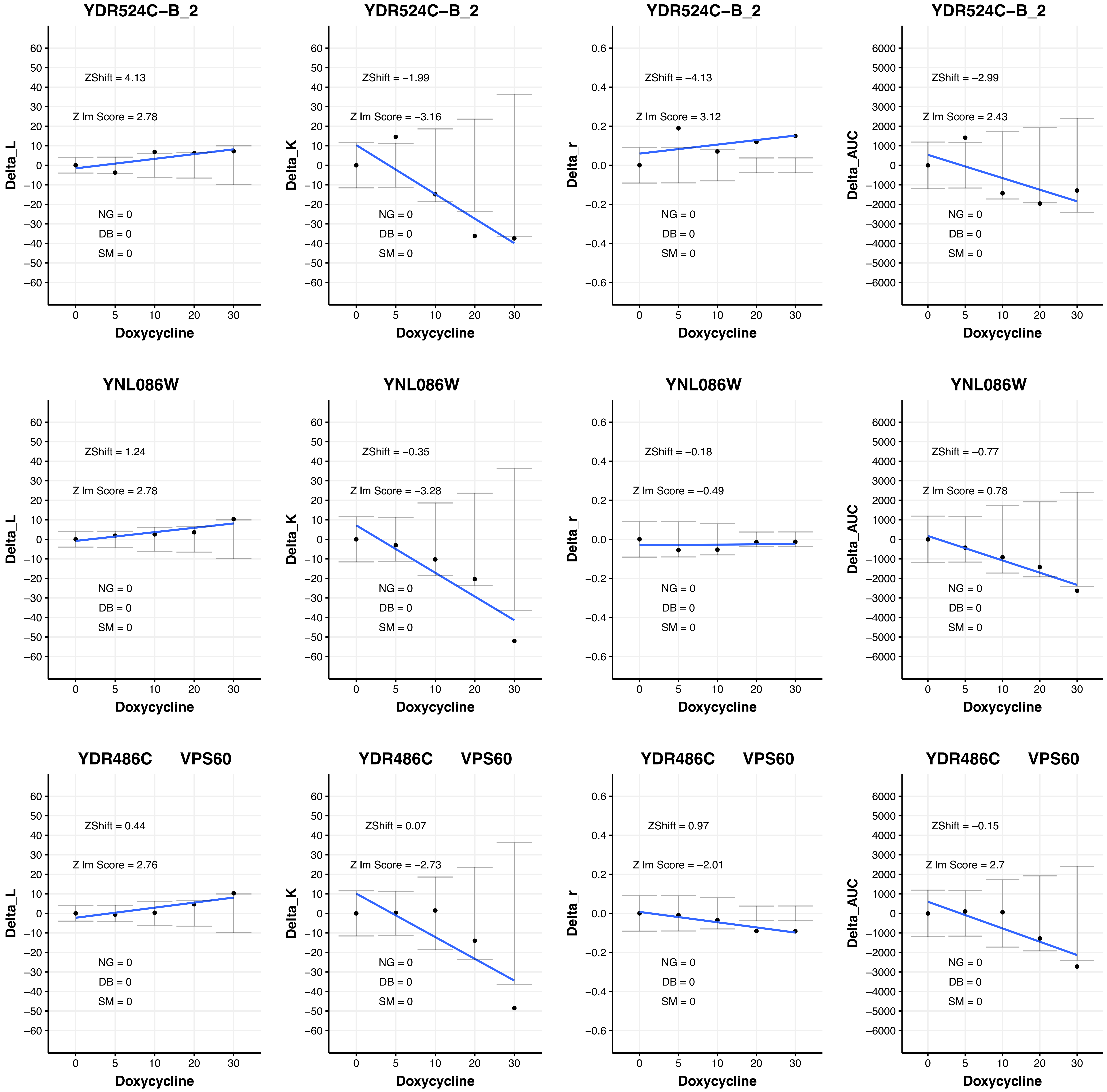

YDR524C−B_2
ZShift = 4.13
Z lm Score = 2.78
NG = 0
DB = 0
SM = 0
Delta_L
Doxycycline
YDR524C−B_2
ZShift = −1.99
Z lm Score = −3.16
NG = 0
DB = 0
SM = 0
Delta_K
Doxycycline
YDR524C−B_2
ZShift = −4.13
Z lm Score = 3.12
NG = 0
DB = 0
SM = 0
Delta_r
Doxycycline
YDR524C−B_2
ZShift = −2.99
Z lm Score = 2.43
NG = 0
DB = 0
SM = 0
Delta_AUC
Doxycycline
YNL086W
ZShift = 1.24
Z lm Score = 2.78
NG = 0
DB = 0
SM = 0
Delta_L
Doxycycline
YNL086W
ZShift = −0.35
Z lm Score = −3.28
NG = 0
DB = 0
SM = 0
Delta_K
Doxycycline
YNL086W
ZShift = −0.18
Z lm Score = −0.49
NG = 0
DB = 0
SM = 0
Delta_r
Doxycycline
YNL086W
ZShift = −0.77
Z lm Score = 0.78
NG = 0
DB = 0
SM = 0
Delta_AUC
Doxycycline
YDR486C VPS60
ZShift = 0.44
Z lm Score = 2.76
NG = 0
DB = 0
SM = 0
Delta_L
Doxycycline
YDR486C VPS60
ZShift = 0.07
Z lm Score = −2.73
NG = 0
DB = 0
SM = 0
Delta_K
Doxycycline
YDR486C VPS60
ZShift = 0.97
Z lm Score = −2.01
NG = 0
DB = 0
SM = 0
Delta_r
Doxycycline
YDR486C VPS60
ZShift = −0.15
Z lm Score = 2.7
NG = 0
DB = 0
SM = 0
Delta_AUC
Doxycycline

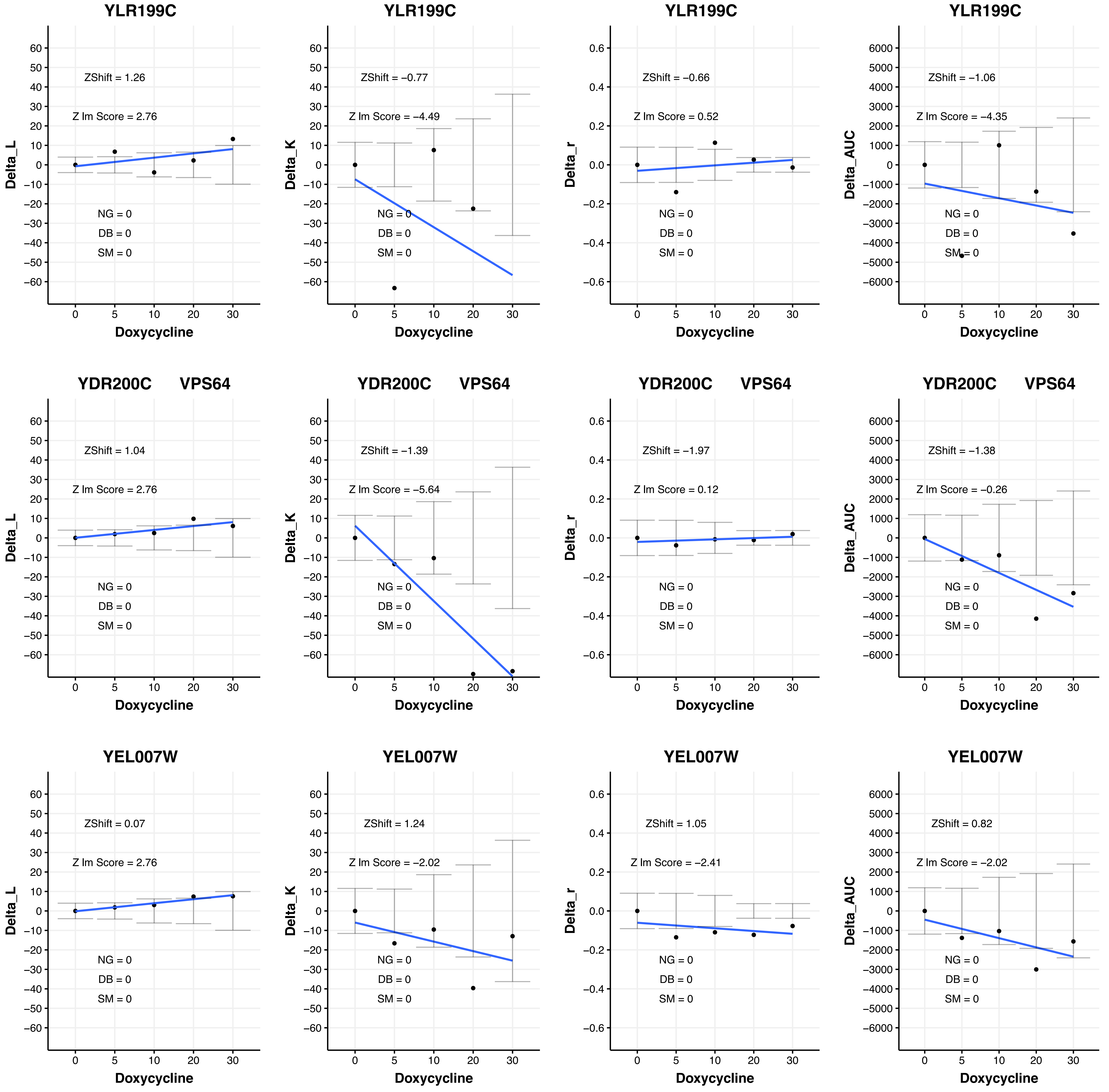
YLR199C
ZShift = 1.26
Z lm Score = 2.76
NG = 0
DB = 0
SM = 0
Delta_L
Doxycycline
YLR199C
ZShift = −0.77
Z lm Score = −4.49
NG = 0
DB = 0
SM = 0
Delta_K
Doxycycline
YLR199C
ZShift = −0.66
Z lm Score = 0.52
NG = 0
DB = 0
SM = 0
Delta_r
Doxycycline
YLR199C
ZShift = −1.06
Z lm Score = −4.35
NG = 0
DB = 0
SM = 0
Delta_AUC
Doxycycline
YDR200C VPS64
ZShift = 1.04
Z lm Score = 2.76
NG = 0
DB = 0
SM = 0
Delta_L
Doxycycline
YDR200C VPS64
ZShift = −1.39
Z lm Score = −5.64
NG = 0
DB = 0
SM = 0
Delta_K
Doxycycline
YDR200C VPS64
ZShift = −1.97
Z lm Score = 0.12
NG = 0
DB = 0
SM = 0
Delta_r
Doxycycline
YDR200C VPS64
ZShift = −1.38
Z lm Score = −0.26
NG = 0
DB = 0
SM = 0
Delta_AUC
Doxycycline
YEL007W
ZShift = 0.07
Z lm Score = 2.76
NG = 0
DB = 0
SM = 0
Delta_L
Doxycycline
YEL007W
ZShift = 1.24
Z lm Score = −2.02
NG = 0
DB = 0
SM = 0
Delta_K
Doxycycline
YEL007W
ZShift = 1.05
Z lm Score = −2.41
NG = 0
DB = 0
SM = 0
Delta_r
Doxycycline
YEL007W
ZShift = 0.82
Z lm Score = −2.02
NG = 0
DB = 0
SM = 0
Delta_AUC
Doxycycline

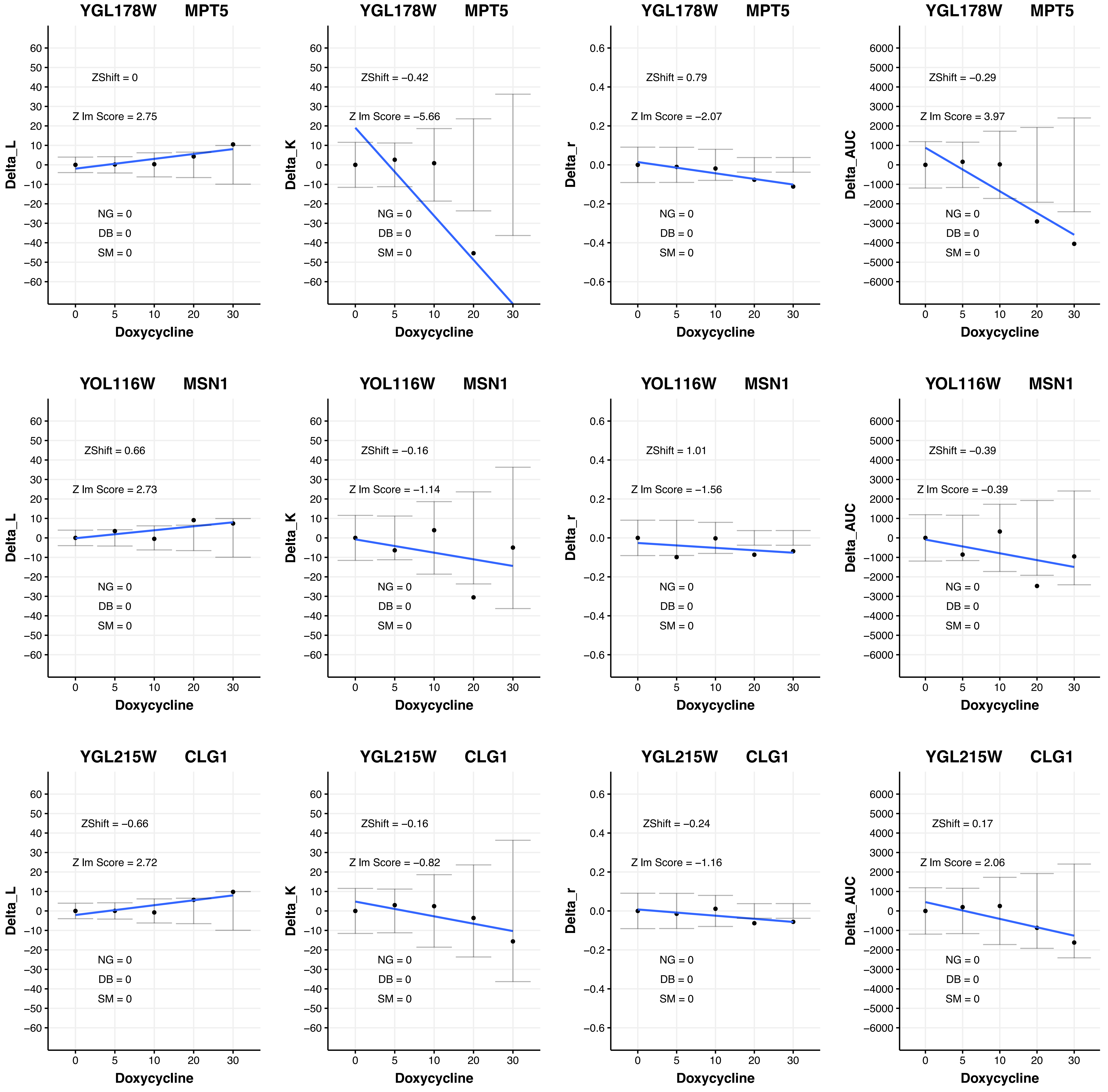

YGL178W MPT5
ZShift = 0
Z lm Score = 2.75
NG = 0
DB = 0
SM = 0
Delta_L
Doxycycline
YGL178W MPT5
ZShift = −0.42
Z lm Score = −5.66
NG = 0
DB = 0
SM = 0
Delta_K
Doxycycline
YGL178W MPT5
ZShift = 0.79
Z lm Score = −2.07
NG = 0
DB = 0
SM = 0
Delta_r
Doxycycline
YGL178W MPT5
ZShift = −0.29
Z lm Score = 3.97
NG = 0
DB = 0
SM = 0
Delta_AUC
Doxycycline
YOL116W MSN1
ZShift = 0.66
Z lm Score = 2.73
NG = 0
DB = 0
SM = 0
Delta_L
Doxycycline
YOL116W MSN1
ZShift = −0.16
Z lm Score = −1.14
NG = 0
DB = 0
SM = 0
Delta_K
Doxycycline
YOL116W MSN1
ZShift = 1.01
Z lm Score = −1.56
NG = 0
DB = 0
SM = 0
Delta_r
Doxycycline
YOL116W MSN1
ZShift = −0.39
Z lm Score = −0.39
NG = 0
DB = 0
SM = 0
Delta_AUC
Doxycycline
YGL215W CLG1
ZShift = −0.66
Z lm Score = 2.72
NG = 0
DB = 0
SM = 0
Delta_L
Doxycycline
YGL215W CLG1
ZShift = −0.16
Z lm Score = −0.82
NG = 0
DB = 0
SM = 0
Delta_K
Doxycycline
YGL215W CLG1
ZShift = −0.24
Z lm Score = −1.16
NG = 0
DB = 0
SM = 0
Delta_r
Doxycycline
YGL215W CLG1
ZShift = 0.17
Z lm Score = 2.06
NG = 0
DB = 0
SM = 0
Delta_AUC
Doxycycline

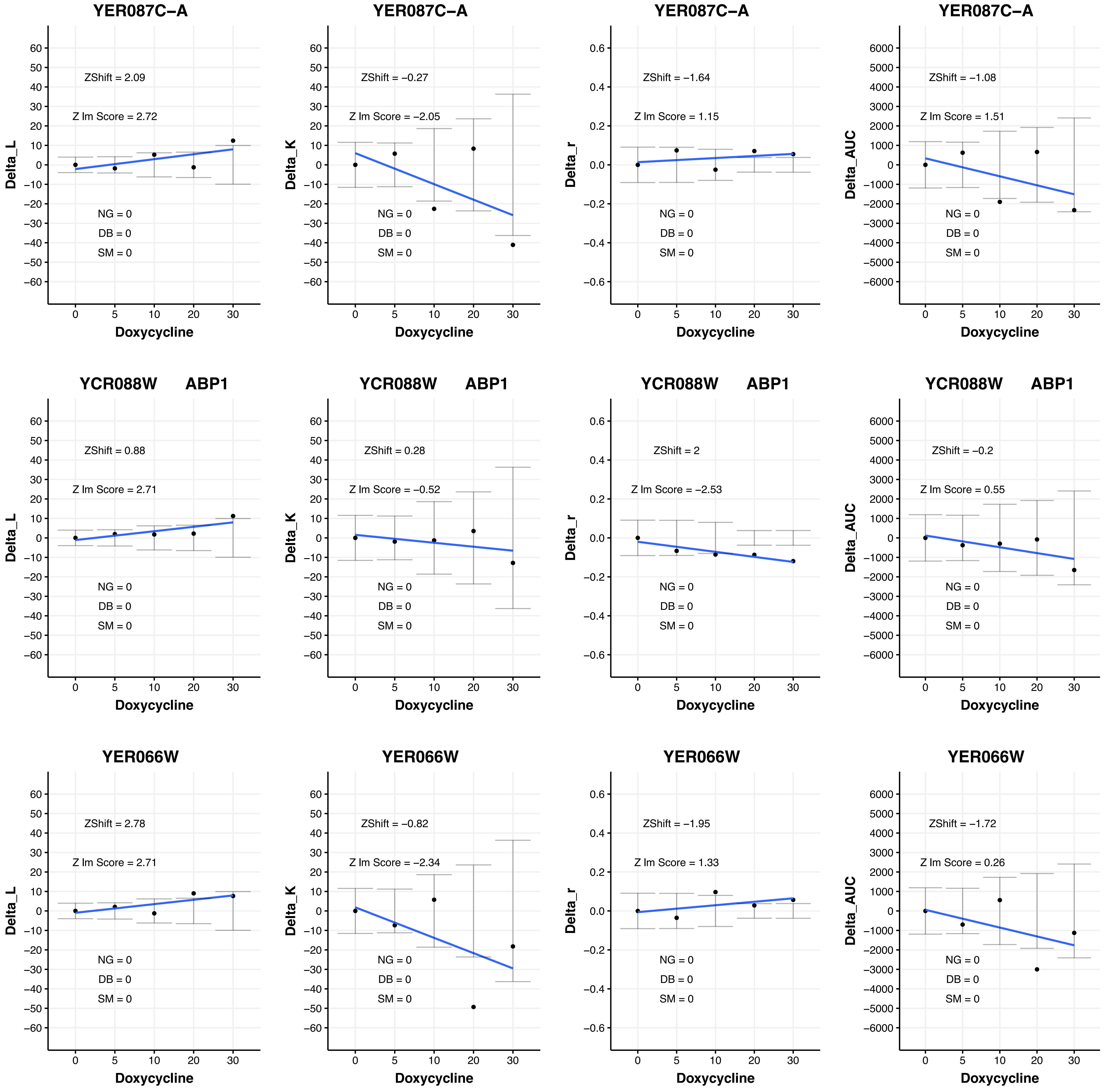
YER087C−A
ZShift = 2.09
Z lm Score = 2.72
NG = 0
DB = 0
SM = 0
Delta_L
Doxycycline
YER087C−A
ZShift = −0.27
Z lm Score = −2.05
NG = 0
DB = 0
SM = 0
Delta_K
Doxycycline
YER087C−A
ZShift = −1.64
Z lm Score = 1.15
NG = 0
DB = 0
SM = 0
Delta_r
Doxycycline
YER087C−A
ZShift = −1.08
Z lm Score = 1.51
NG = 0
DB = 0
SM = 0
Delta_AUC
Doxycycline
YCR088W ABP1
ZShift = 0.88
Z lm Score = 2.71
NG = 0
DB = 0
SM = 0
Delta_L
Doxycycline
YCR088W ABP1
ZShift = 0.28
Z lm Score = −0.52
NG = 0
DB = 0
SM = 0
Delta_K
Doxycycline
YCR088W ABP1
ZShift = 2
Z lm Score = −2.53
NG = 0
DB = 0
SM = 0
Delta_r
Doxycycline
YCR088W ABP1
ZShift = −0.2
Z lm Score = 0.55
NG = 0
DB = 0
SM = 0
Delta_AUC
Doxycycline
YER066W
ZShift = 2.78
Z lm Score = 2.71
NG = 0
DB = 0
SM = 0
Delta_L
Doxycycline
YER066W
ZShift = −0.82
Z lm Score = −2.34
NG = 0
DB = 0
SM = 0
Delta_K
Doxycycline
YER066W
ZShift = −1.95
Z lm Score = 1.33
NG = 0
DB = 0
SM = 0
Delta_r
Doxycycline
YER066W
ZShift = −1.72
Z lm Score = 0.26
NG = 0
DB = 0
SM = 0
Delta_AUC
Doxycycline

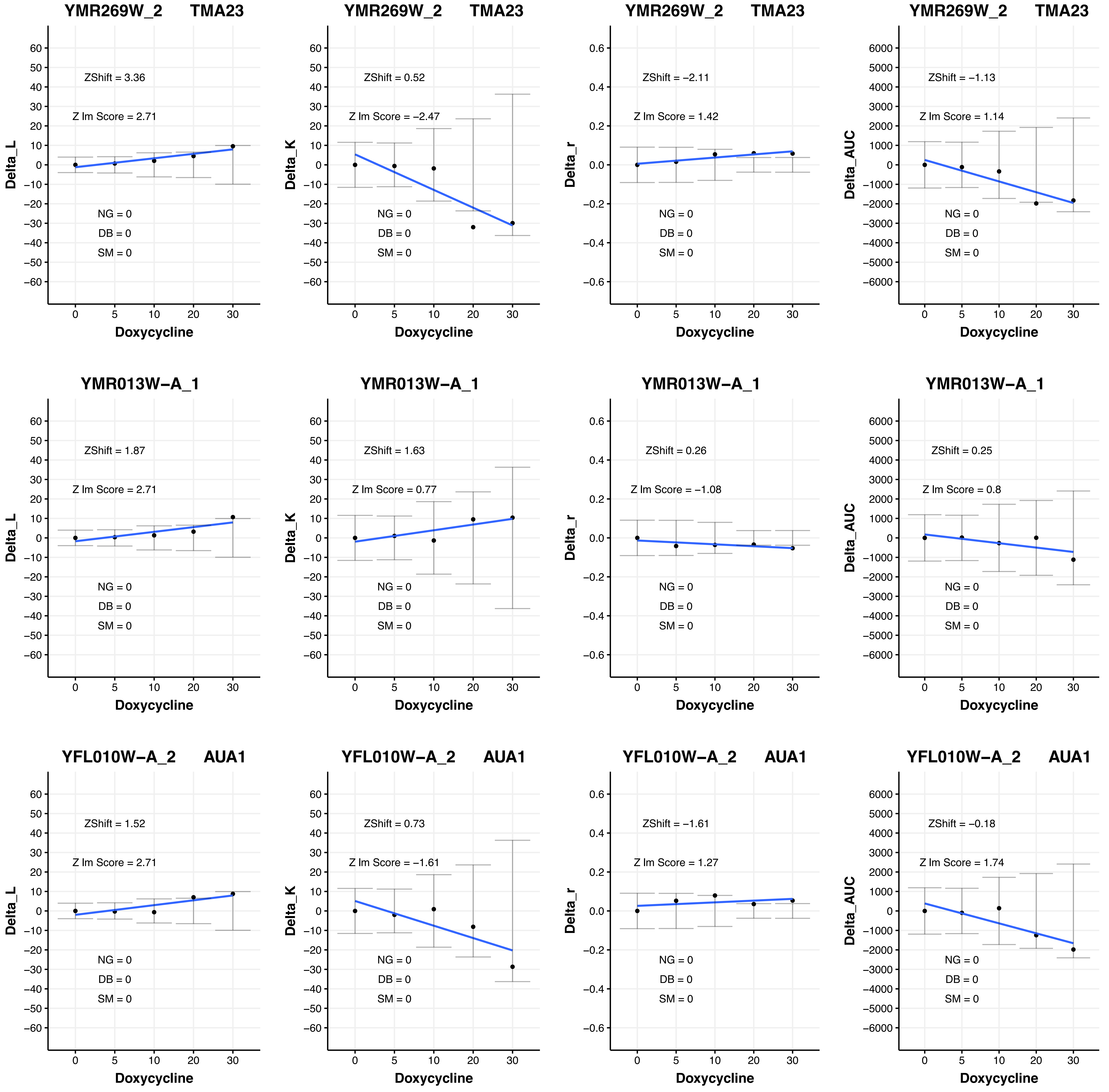
YMR269W_2 TMA23
ZShift = 3.36
Z lm Score = 2.71
NG = 0
DB = 0
SM = 0
Delta_L
Doxycycline
YMR269W_2 TMA23
ZShift = 0.52
Z lm Score = −2.47
NG = 0
DB = 0
SM = 0
Delta_K
Doxycycline
YMR269W_2 TMA23
ZShift = −2.11
Z lm Score = 1.42
NG = 0
DB = 0
SM = 0
Delta_r
Doxycycline
YMR269W_2 TMA23
ZShift = −1.13
Z lm Score = 1.14
NG = 0
DB = 0
SM = 0
Delta_AUC
Doxycycline
YMR013W−A_1
ZShift = 1.87
Z lm Score = 2.71
NG = 0
DB = 0
SM = 0
Delta_L
Doxycycline
YMR013W−A_1
ZShift = 1.63
Z lm Score = 0.77
NG = 0
DB = 0
SM = 0
Delta_K
Doxycycline
YMR013W−A_1
ZShift = 0.26
Z lm Score = −1.08
NG = 0
DB = 0
SM = 0
Delta_r
Doxycycline
YMR013W−A_1
ZShift = 0.25
Z lm Score = 0.8
NG = 0
DB = 0
SM = 0
Delta_AUC
Doxycycline
YFL010W−A_2 AUA1
ZShift = 1.52
Z lm Score = 2.71
NG = 0
DB = 0
SM = 0
Delta_L
Doxycycline
YFL010W−A_2 AUA1
ZShift = 0.73
Z lm Score = −1.61
NG = 0
DB = 0
SM = 0
Delta_K
Doxycycline
YFL010W−A_2 AUA1
ZShift = −1.61
Z lm Score = 1.27
NG = 0
DB = 0
SM = 0
Delta_r
Doxycycline
YFL010W−A_2 AUA1
ZShift = −0.18
Z lm Score = 1.74
NG = 0
DB = 0
SM = 0
Delta_AUC
Doxycycline

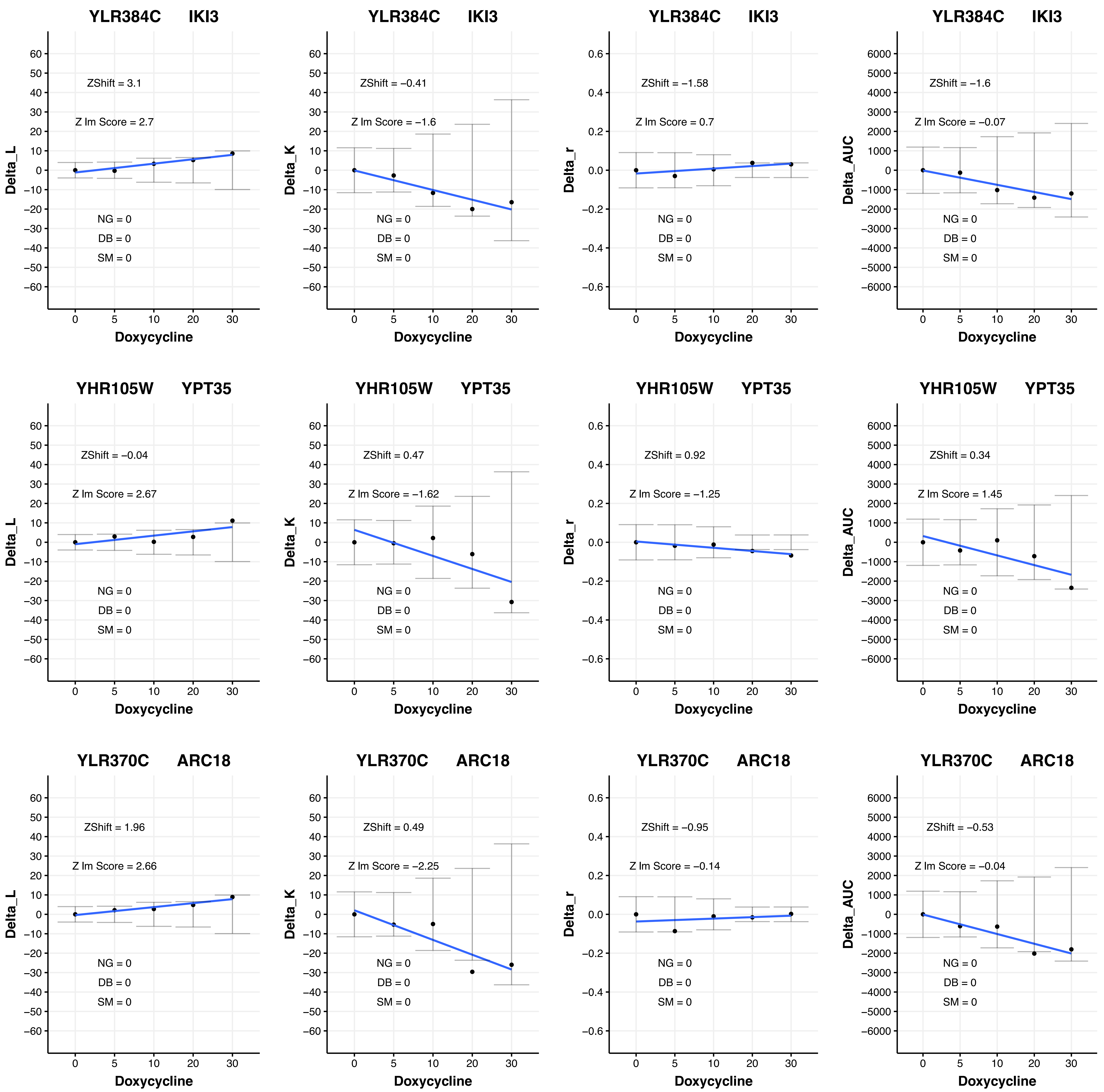
YLR384C IKI3
ZShift = 3.1
Z lm Score = 2.7
NG = 0
DB = 0
SM = 0
Delta_L
Doxycycline
YLR384C IKI3
ZShift = −0.41
Z lm Score = −1.6
NG = 0
DB = 0
SM = 0
Delta_K
Doxycycline
YLR384C IKI3
ZShift = −1.58
Z lm Score = 0.7
NG = 0
DB = 0
SM = 0
Delta_r
Doxycycline
YLR384C IKI3
ZShift = −1.6
Z lm Score = −0.07
NG = 0
DB = 0
SM = 0
Delta_AUC
Doxycycline
YHR105W YPT35
ZShift = −0.04
Z lm Score = 2.67
NG = 0
DB = 0
SM = 0
Delta_L
Doxycycline
YHR105W YPT35
ZShift = 0.47
Z lm Score = −1.62
NG = 0
DB = 0
SM = 0
Delta_K
Doxycycline
YHR105W YPT35
ZShift = 0.92
Z lm Score = −1.25
NG = 0
DB = 0
SM = 0
Delta_r
Doxycycline
YHR105W YPT35
ZShift = 0.34
Z lm Score = 1.45
NG = 0
DB = 0
SM = 0
Delta_AUC
Doxycycline
YLR370C ARC18
ZShift = 1.96
Z lm Score = 2.66
NG = 0
DB = 0
SM = 0
Delta_L
Doxycycline
YLR370C ARC18
ZShift = 0.49
Z lm Score = −2.25
NG = 0
DB = 0
SM = 0
Delta_K
Doxycycline
YLR370C ARC18
ZShift = −0.95
Z lm Score = −0.14
NG = 0
DB = 0
SM = 0
Delta_r
Doxycycline
YLR370C ARC18
ZShift = −0.53
Z lm Score = −0.04
NG = 0
DB = 0
SM = 0
Delta_AUC
Doxycycline

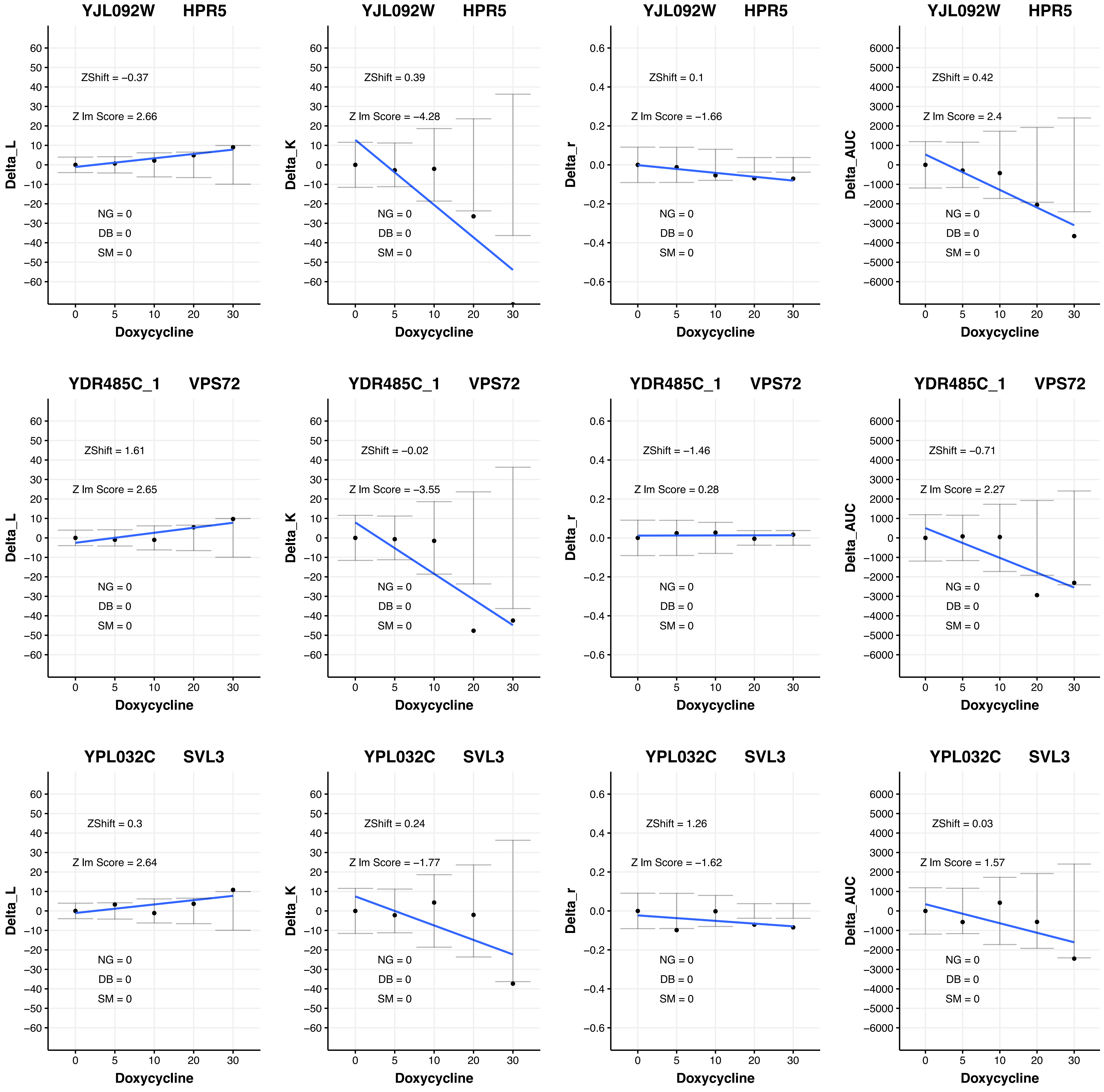

YJL092W HPR5
ZShift = −0.37
Z lm Score = 2.66
NG = 0
DB = 0
SM = 0
Delta_L
Doxycycline
YJL092W HPR5
ZShift = 0.39
Z lm Score = −4.28
NG = 0
DB = 0
SM = 0
Delta_K
Doxycycline
YJL092W HPR5
ZShift = 0.1
Z lm Score = −1.66
NG = 0
DB = 0
SM = 0
Delta_r
Doxycycline
YJL092W HPR5
ZShift = 0.42
Z lm Score = 2.4
NG = 0
DB = 0
SM = 0
Delta_AUC
Doxycycline
YDR485C_1 VPS72
ZShift = 1.61
Z lm Score = 2.65
NG = 0
DB = 0
SM = 0
Delta_L
Doxycycline
YDR485C_1 VPS72
ZShift = −0.02
Z lm Score = −3.55
NG = 0
DB = 0
SM = 0
Delta_K
Doxycycline
YDR485C_1 VPS72
ZShift = −1.46
Z lm Score = 0.28
NG = 0
DB = 0
SM = 0
Delta_r
Doxycycline
YDR485C_1 VPS72
ZShift = −0.71
Z lm Score = 2.27
NG = 0
DB = 0
SM = 0
Delta_AUC
Doxycycline
YPL032C SVL3
ZShift = 0.3
Z lm Score = 2.64
NG = 0
DB = 0
SM = 0
Delta_L
Doxycycline
YPL032C SVL3
ZShift = 0.24
Z lm Score = −1.77
NG = 0
DB = 0
SM = 0
Delta_K
Doxycycline
YPL032C SVL3
ZShift = 1.26
Z lm Score = −1.62
NG = 0
DB = 0
SM = 0
Delta_r
Doxycycline
YPL032C SVL3
ZShift = 0.03
Z lm Score = 1.57
NG = 0
DB = 0
SM = 0
Delta_AUC
Doxycycline

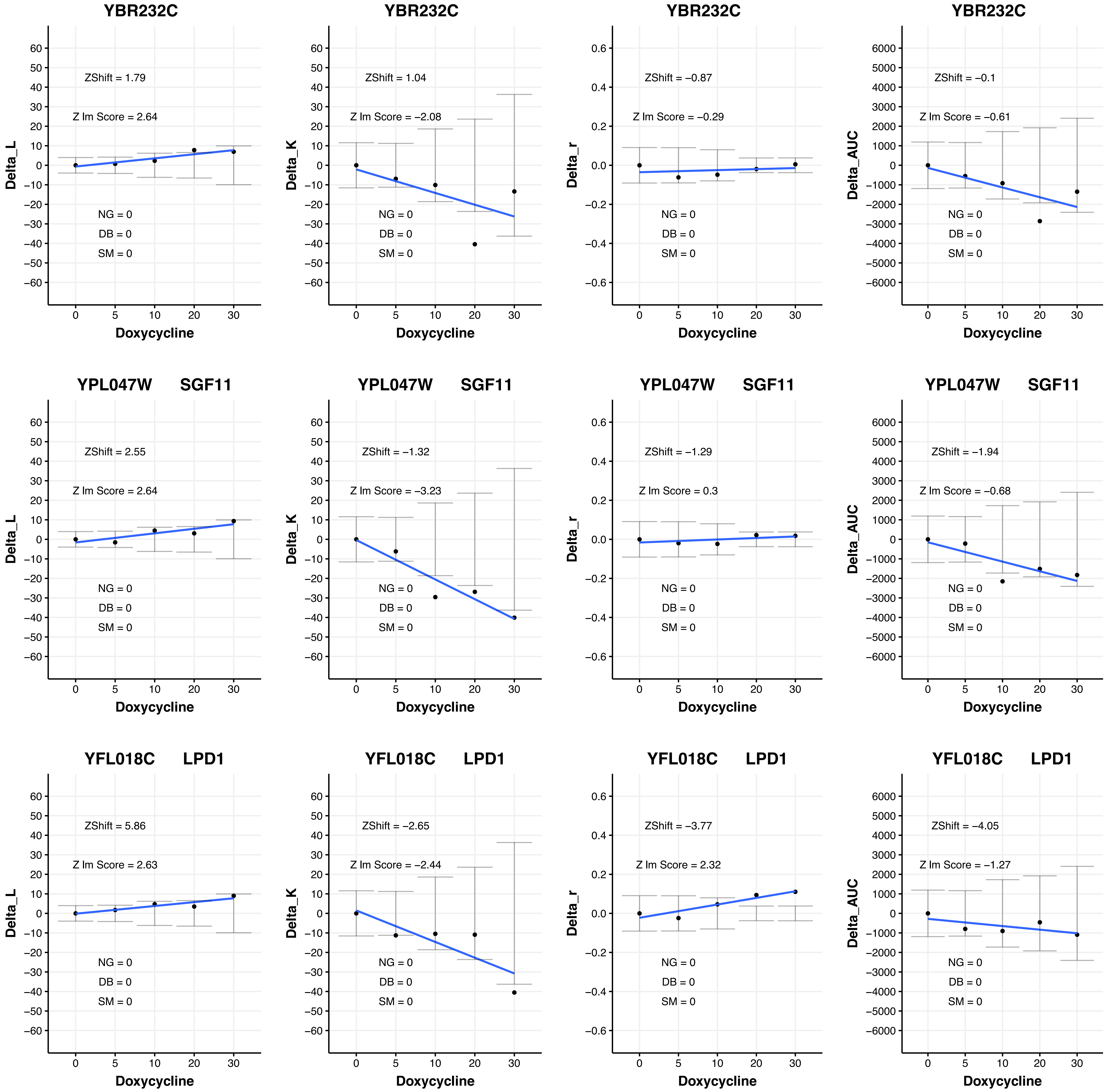
YBR232C
ZShift = 1.79
Z lm Score = 2.64
NG = 0
DB = 0
SM = 0
Delta_L
Doxycycline
YBR232C
ZShift = 1.04
Z lm Score = −2.08
NG = 0
DB = 0
SM = 0
Delta_K
Doxycycline
YBR232C
ZShift = −0.87
Z lm Score = −0.29
NG = 0
DB = 0
SM = 0
Delta_r
Doxycycline
YBR232C
ZShift = −0.1
Z lm Score = −0.61
NG = 0
DB = 0
SM = 0
Delta_AUC
Doxycycline
YPL047W SGF11
ZShift = 2.55
Z lm Score = 2.64
NG = 0
DB = 0
SM = 0
Delta_L
Doxycycline
YPL047W SGF11
ZShift = −1.32
Z lm Score = −3.23
NG = 0
DB = 0
SM = 0
Delta_K
Doxycycline
YPL047W SGF11
ZShift = −1.29
Z lm Score = 0.3
NG = 0
DB = 0
SM = 0
Delta_r
Doxycycline
YPL047W SGF11
ZShift = −1.94
Z lm Score = −0.68
NG = 0
DB = 0
SM = 0
Delta_AUC
Doxycycline
YFL018C LPD1
ZShift = 5.86
Z lm Score = 2.63
NG = 0
DB = 0
SM = 0
Delta_L
Doxycycline
YFL018C LPD1
ZShift = −2.65
Z lm Score = −2.44
NG = 0
DB = 0
SM = 0
Delta_K
Doxycycline
YFL018C LPD1
ZShift = −3.77
Z lm Score = 2.32
NG = 0
DB = 0
SM = 0
Delta_r
Doxycycline
YFL018C LPD1
ZShift = −4.05
Z lm Score = −1.27
NG = 0
DB = 0
SM = 0
Delta_AUC
Doxycycline

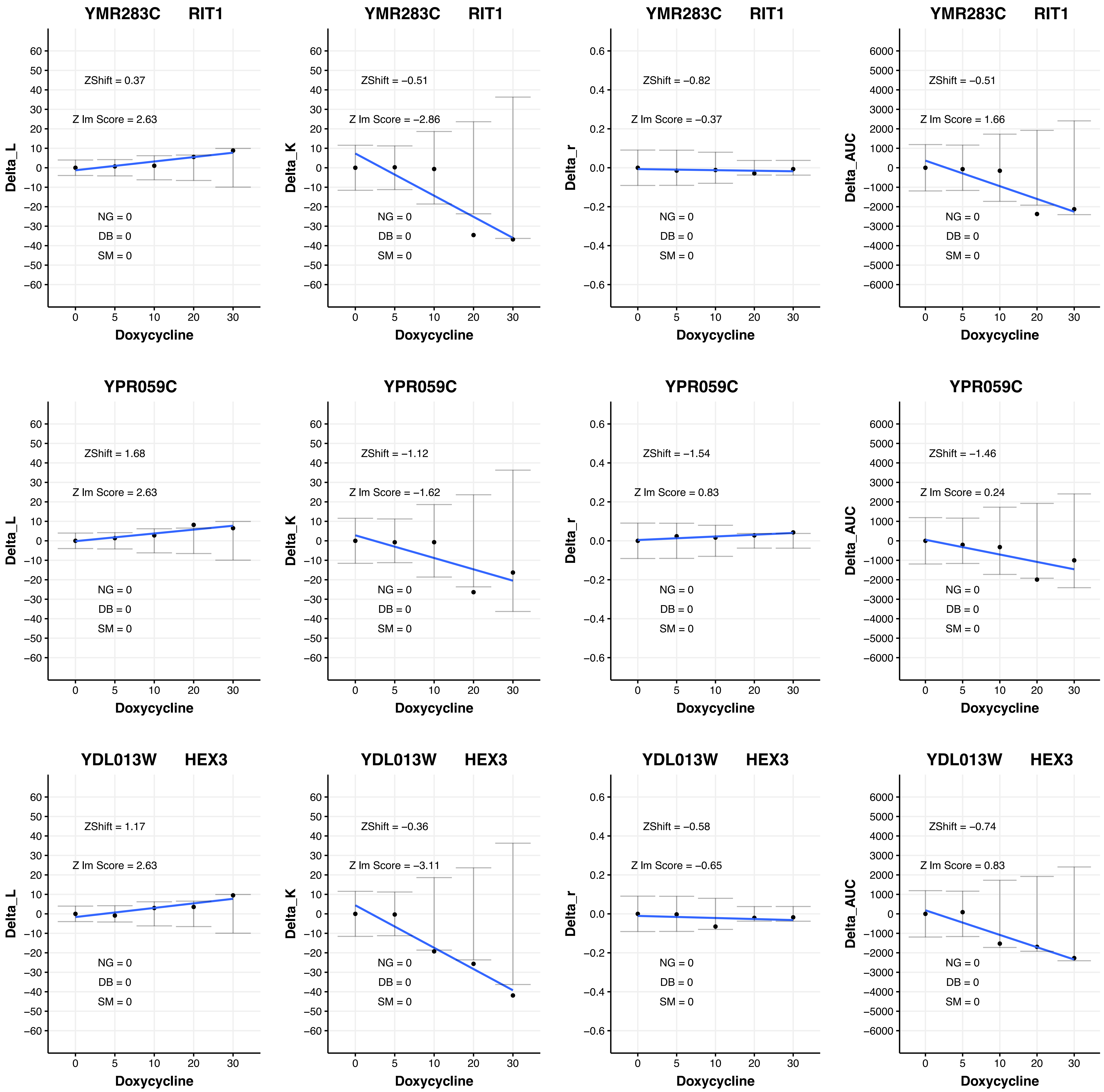
YMR283C RIT1
ZShift = 0.37
Z lm Score = 2.63
NG = 0
DB = 0
SM = 0
Delta_L
Doxycycline
YMR283C RIT1
ZShift = −0.51
Z lm Score = −2.86
NG = 0
DB = 0
SM = 0
Delta_K
Doxycycline
YMR283C RIT1
ZShift = −0.82
Z lm Score = −0.37
NG = 0
DB = 0
SM = 0
Delta_r
Doxycycline
YMR283C RIT1
ZShift = −0.51
Z lm Score = 1.66
NG = 0
DB = 0
SM = 0
Delta_AUC
Doxycycline
YPR059C
ZShift = 1.68
Z lm Score = 2.63
NG = 0
DB = 0
SM = 0
Delta_L
Doxycycline
YPR059C
ZShift = −1.12
Z lm Score = −1.62
NG = 0
DB = 0
SM = 0
Delta_K
Doxycycline
YPR059C
ZShift = −1.54
Z lm Score = 0.83
NG = 0
DB = 0
SM = 0
Delta_r
Doxycycline
YPR059C
ZShift = −1.46
Z lm Score = 0.24
NG = 0
DB = 0
SM = 0
Delta_AUC
Doxycycline
YDL013W HEX3
ZShift = 1.17
Z lm Score = 2.63
NG = 0
DB = 0
SM = 0
Delta_L
Doxycycline
YDL013W HEX3
ZShift = −0.36
Z lm Score = −3.11
NG = 0
DB = 0
SM = 0
Delta_K
Doxycycline
YDL013W HEX3
ZShift = −0.58
Z lm Score = −0.65
NG = 0
DB = 0
SM = 0
Delta_r
Doxycycline
YDL013W HEX3
ZShift = −0.74
Z lm Score = 0.83
NG = 0
DB = 0
SM = 0
Delta_AUC
Doxycycline

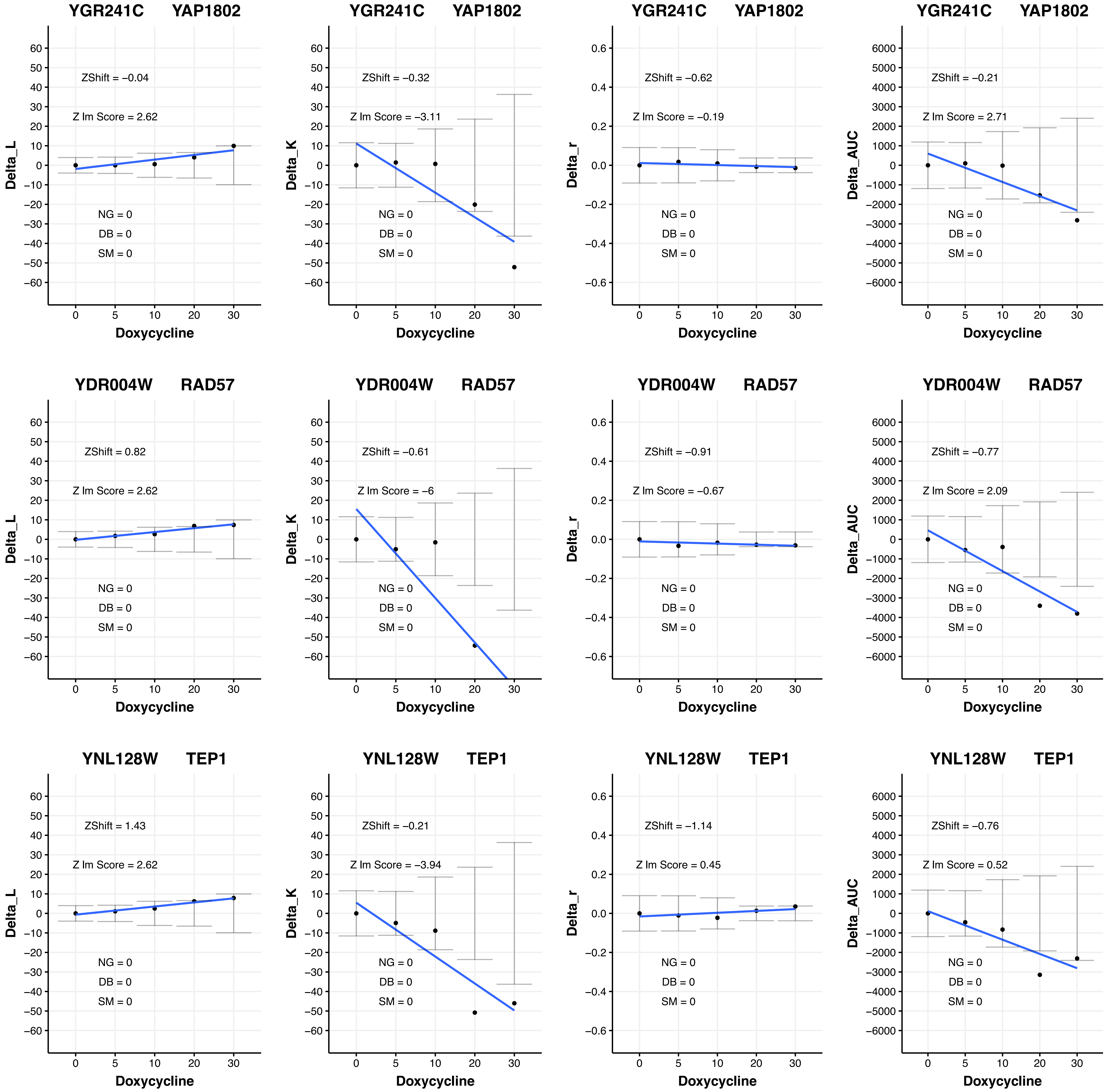
YGR241C YAP1802
ZShift = −0.04
Z lm Score = 2.62
NG = 0
DB = 0
SM = 0
Delta_L
Doxycycline
YGR241C YAP1802
ZShift = −0.32
Z lm Score = −3.11
NG = 0
DB = 0
SM = 0
Delta_K
Doxycycline
YGR241C YAP1802
ZShift = −0.62
Z lm Score = −0.19
NG = 0
DB = 0
SM = 0
Delta_r
Doxycycline
YGR241C YAP1802
ZShift = −0.21
Z lm Score = 2.71
NG = 0
DB = 0
SM = 0
Delta_AUC
Doxycycline
YDR004W RAD57
ZShift = 0.82
Z lm Score = 2.62
NG = 0
DB = 0
SM = 0
Delta_L
Doxycycline
YDR004W RAD57
ZShift = −0.61
Z lm Score = −6
NG = 0
DB = 0
SM = 0
Delta_K
Doxycycline
YDR004W RAD57
ZShift = −0.91
Z lm Score = −0.67
NG = 0
DB = 0
SM = 0
Delta_r
Doxycycline
YDR004W RAD57
ZShift = −0.77
Z lm Score = 2.09
NG = 0
DB = 0
SM = 0
Delta_AUC
Doxycycline
YNL128W TEP1
ZShift = 1.43
Z lm Score = 2.62
NG = 0
DB = 0
SM = 0
Delta_L
Doxycycline
YNL128W TEP1
ZShift = −0.21
Z lm Score = −3.94
NG = 0
DB = 0
SM = 0
Delta_K
Doxycycline
YNL128W TEP1
ZShift = −1.14
Z lm Score = 0.45
NG = 0
DB = 0
SM = 0
Delta_r
Doxycycline
YNL128W TEP1
ZShift = −0.76
Z lm Score = 0.52
NG = 0
DB = 0
SM = 0
Delta_AUC
Doxycycline

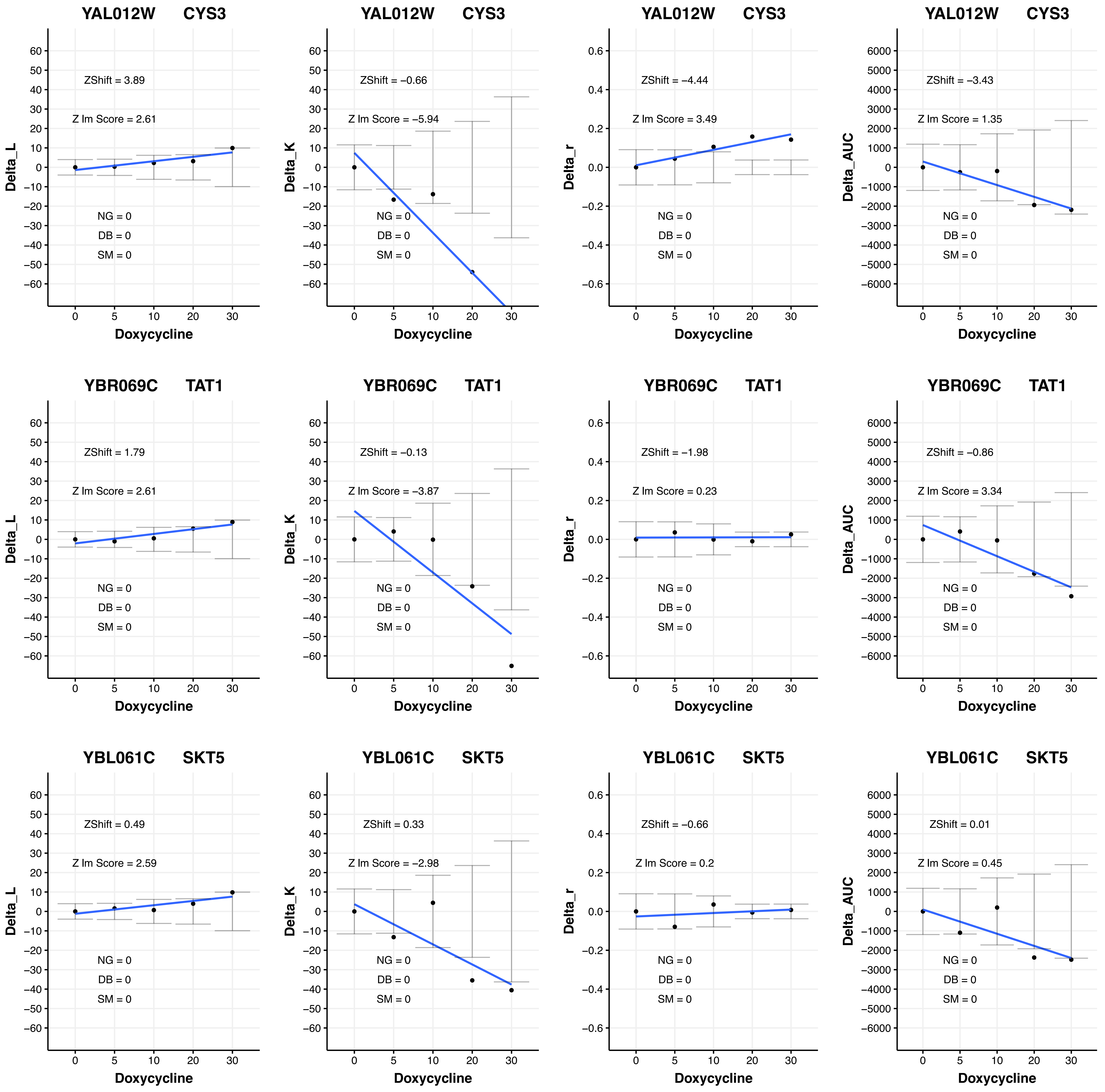

YAL012W CYS3
ZShift = 3.89
Z lm Score = 2.61
NG = 0
DB = 0
SM = 0
Delta_L
Doxycycline
YAL012W CYS3
ZShift = −0.66
Z lm Score = −5.94
NG = 0
DB = 0
SM = 0
Delta_K
Doxycycline
YAL012W CYS3
ZShift = −4.44
Z lm Score = 3.49
NG = 0
DB = 0
SM = 0
Delta_r
Doxycycline
YAL012W CYS3
ZShift = −3.43
Z lm Score = 1.35
NG = 0
DB = 0
SM = 0
Delta_AUC
Doxycycline
YBR069C TAT1
ZShift = 1.79
Z lm Score = 2.61
NG = 0
DB = 0
SM = 0
Delta_L
Doxycycline
YBR069C TAT1
ZShift = −0.13
Z lm Score = −3.87
NG = 0
DB = 0
SM = 0
Delta_K
Doxycycline
YBR069C TAT1
ZShift = −1.98
Z lm Score = 0.23
NG = 0
DB = 0
SM = 0
Delta_r
Doxycycline
YBR069C TAT1
ZShift = −0.86
Z lm Score = 3.34
NG = 0
DB = 0
SM = 0
Delta_AUC
Doxycycline
YBL061C SKT5
ZShift = 0.49
Z lm Score = 2.59
NG = 0
DB = 0
SM = 0
Delta_L
Doxycycline
YBL061C SKT5
ZShift = 0.33
Z lm Score = −2.98
NG = 0
DB = 0
SM = 0
Delta_K
Doxycycline
YBL061C SKT5
ZShift = −0.66
Z lm Score = 0.2
NG = 0
DB = 0
SM = 0
Delta_r
Doxycycline
YBL061C SKT5
ZShift = 0.01
Z lm Score = 0.45
NG = 0
DB = 0
SM = 0
Delta_AUC
Doxycycline

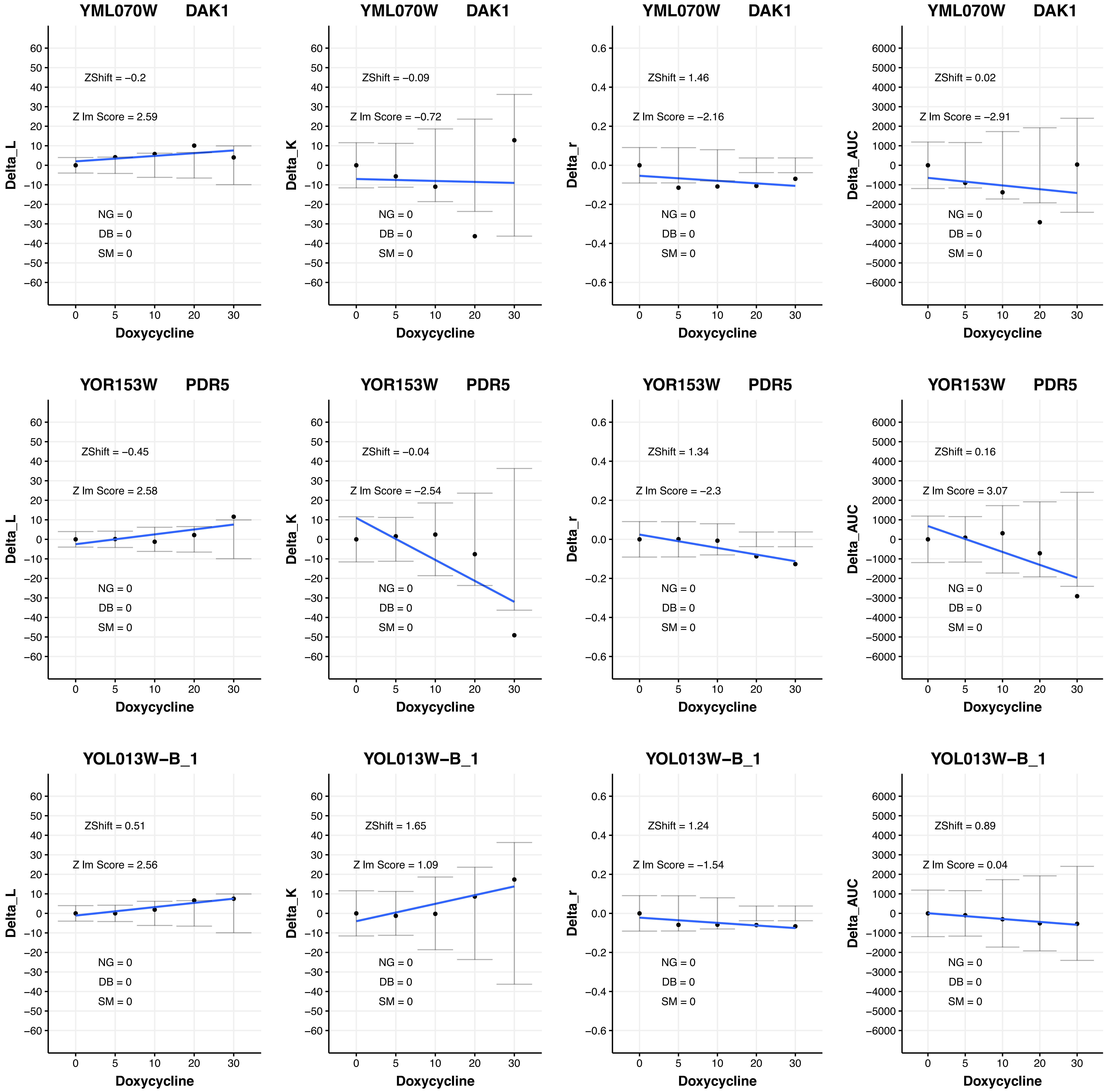
YML070W DAK1
ZShift = −0.2
Z lm Score = 2.59
NG = 0
DB = 0
SM = 0
Delta_L
Doxycycline
YML070W DAK1
ZShift = −0.09
Z lm Score = −0.72
NG = 0
DB = 0
SM = 0
Delta_K
Doxycycline
YML070W DAK1
ZShift = 1.46
Z lm Score = −2.16
NG = 0
DB = 0
SM = 0
Delta_r
Doxycycline
YML070W DAK1
ZShift = 0.02
Z lm Score = −2.91
NG = 0
DB = 0
SM = 0
Delta_AUC
Doxycycline
YOR153W PDR5
ZShift = −0.45
Z lm Score = 2.58
NG = 0
DB = 0
SM = 0
Delta_L
Doxycycline
YOR153W PDR5
ZShift = −0.04
Z lm Score = −2.54
NG = 0
DB = 0
SM = 0
Delta_K
Doxycycline
YOR153W PDR5
ZShift = 1.34
Z lm Score = −2.3
NG = 0
DB = 0
SM = 0
Delta_r
Doxycycline
YOR153W PDR5
ZShift = 0.16
Z lm Score = 3.07
NG = 0
DB = 0
SM = 0
Delta_AUC
Doxycycline
YOL013W−B_1
ZShift = 0.51
Z lm Score = 2.56
NG = 0
DB = 0
SM = 0
Delta_L
Doxycycline
YOL013W−B_1
ZShift = 1.65
Z lm Score = 1.09
NG = 0
DB = 0
SM = 0
Delta_K
Doxycycline
YOL013W−B_1
ZShift = 1.24
Z lm Score = −1.54
NG = 0
DB = 0
SM = 0
Delta_r
Doxycycline
YOL013W−B_1
ZShift = 0.89
Z lm Score = 0.04
NG = 0
DB = 0
SM = 0
Delta_AUC
Doxycycline

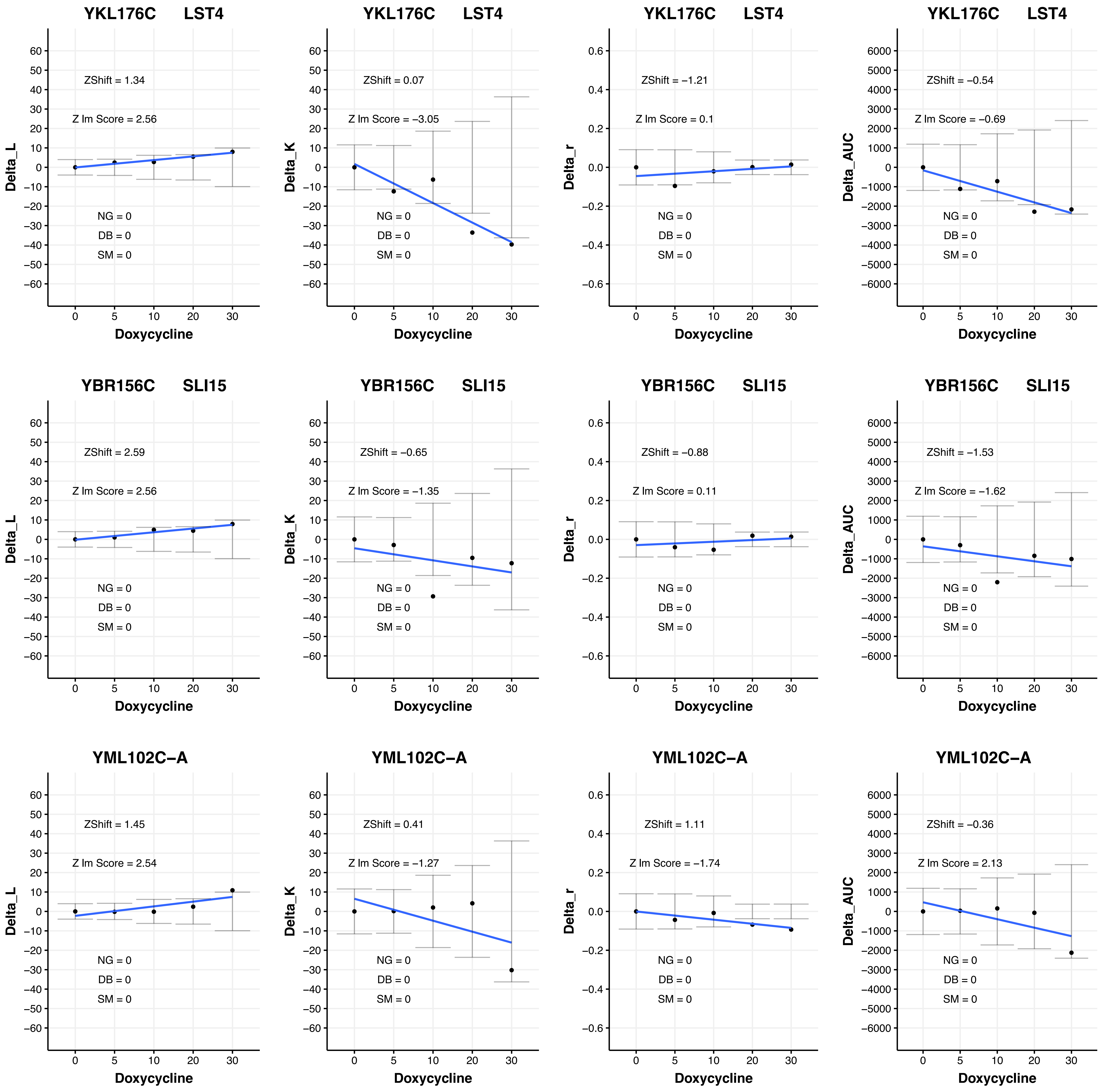
YKL176C LST4
ZShift = 1.34
Z lm Score = 2.56
NG = 0
DB = 0
SM = 0
Delta_L
Doxycycline
YKL176C LST4
ZShift = 0.07
Z lm Score = −3.05
NG = 0
DB = 0
SM = 0
Delta_K
Doxycycline
YKL176C LST4
ZShift = −1.21
Z lm Score = 0.1
NG = 0
DB = 0
SM = 0
Delta_r
Doxycycline
YKL176C LST4
ZShift = −0.54
Z lm Score = −0.69
NG = 0
DB = 0
SM = 0
Delta_AUC
Doxycycline
YBR156C SLI15
ZShift = 2.59
Z lm Score = 2.56
NG = 0
DB = 0
SM = 0
Delta_L
Doxycycline
YBR156C SLI15
ZShift = −0.65
Z lm Score = −1.35
NG = 0
DB = 0
SM = 0
Delta_K
Doxycycline
YBR156C SLI15
ZShift = −0.88
Z lm Score = 0.11
NG = 0
DB = 0
SM = 0
Delta_r
Doxycycline
YBR156C SLI15
ZShift = −1.53
Z lm Score = −1.62
NG = 0
DB = 0
SM = 0
Delta_AUC
Doxycycline
YML102C−A
ZShift = 1.45
Z lm Score = 2.54
NG = 0
DB = 0
SM = 0
Delta_L
Doxycycline
YML102C−A
ZShift = 0.41
Z lm Score = −1.27
NG = 0
DB = 0
SM = 0
Delta_K
Doxycycline
YML102C−A
ZShift = 1.11
Z lm Score = −1.74
NG = 0
DB = 0
SM = 0
Delta_r
Doxycycline
YML102C−A
ZShift = −0.36
Z lm Score = 2.13
NG = 0
DB = 0
SM = 0
Delta_AUC
Doxycycline

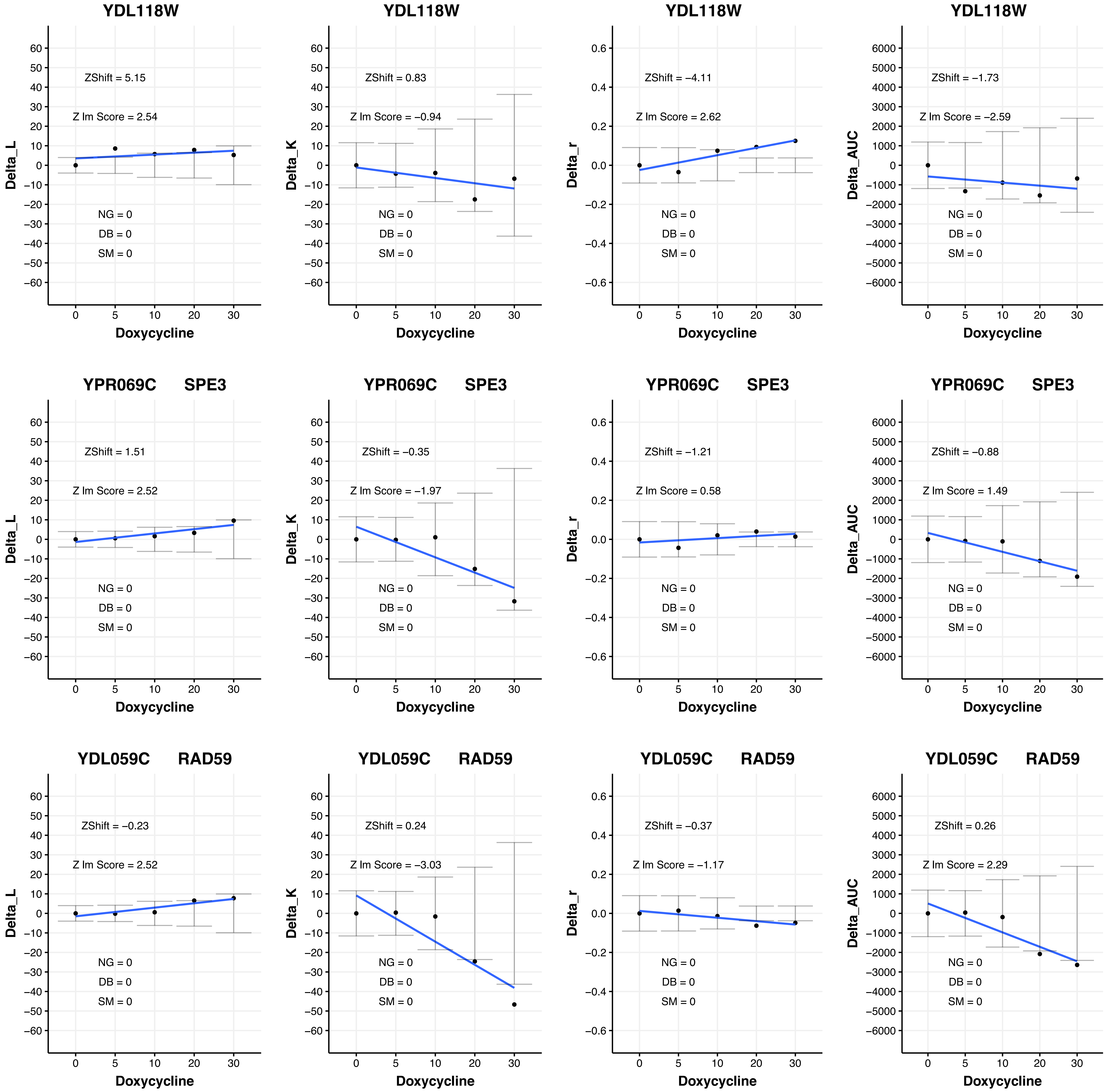

YDL118W
Delta_L
ZShift = 5.15
Z lm Score = 2.54
NG = 0
DB = 0
SM = 0
Doxycycline
YDL118W
Delta_K
ZShift = 0.83
Z lm Score = −0.94
NG = 0
DB = 0
SM = 0
Doxycycline
YDL118W
Delta_r
ZShift = −4.11
Z lm Score = 2.62
NG = 0
DB = 0
SM = 0
Doxycycline
YDL118W
Delta_AUC
ZShift = −1.73
Z lm Score = −2.59
NG = 0
DB = 0
SM = 0
Doxycycline
YPR069C SPE3
Delta_L
ZShift = 1.51
Z lm Score = 2.52
NG = 0
DB = 0
SM = 0
Doxycycline
YPR069C SPE3
Delta_K
ZShift = −0.35
Z lm Score = −1.97
NG = 0
DB = 0
SM = 0
Doxycycline
YPR069C SPE3
Delta_r
ZShift = −1.21
Z lm Score = 0.58
NG = 0
DB = 0
SM = 0
Doxycycline
YPR069C SPE3
Delta_AUC
ZShift = −0.88
Z lm Score = 1.49
NG = 0
DB = 0
SM = 0
Doxycycline
YDL059C RAD59
Delta_L
ZShift = −0.23
Z lm Score = 2.52
NG = 0
DB = 0
SM = 0
Doxycycline
YDL059C RAD59
Delta_K
ZShift = 0.24
Z lm Score = −3.03
NG = 0
DB = 0
SM = 0
Doxycycline
YDL059C RAD59
Delta_r
ZShift = −0.37
Z lm Score = −1.17
NG = 0
DB = 0
SM = 0
Doxycycline
YDL059C RAD59
Delta_AUC
ZShift = 0.26
Z lm Score = 2.29
NG = 0
DB = 0
SM = 0
Doxycycline

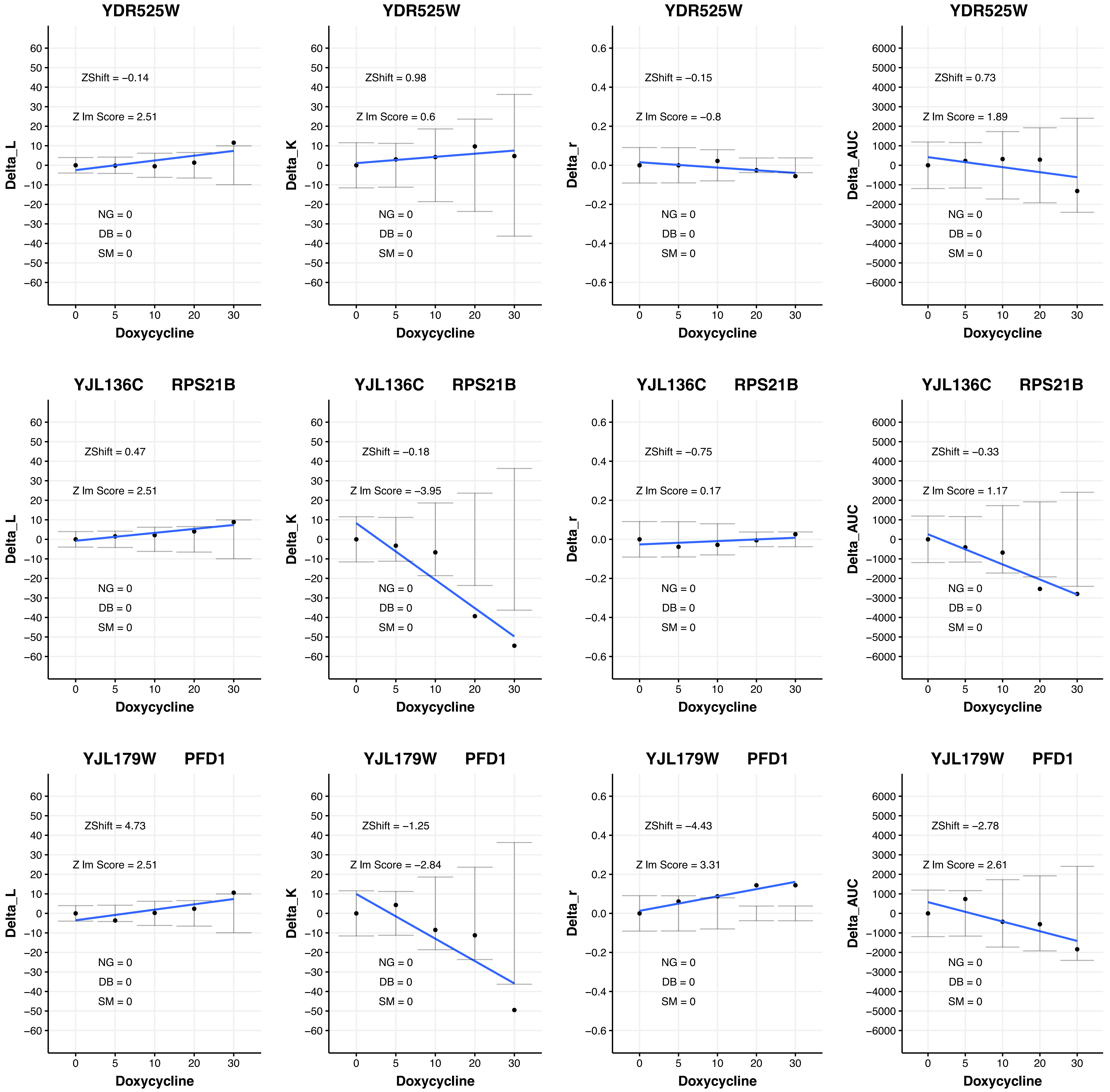
YDR525W
ZShift = −0.14
Z lm Score = 2.51
NG = 0
DB = 0
SM = 0
Delta_L
Doxycycline
YDR525W
ZShift = 0.98
Z lm Score = 0.6
NG = 0
DB = 0
SM = 0
Delta_K
Doxycycline
YDR525W
ZShift = −0.15
Z lm Score = −0.8
NG = 0
DB = 0
SM = 0
Delta_r
Doxycycline
YDR525W
ZShift = 0.73
Z lm Score = 1.89
NG = 0
DB = 0
SM = 0
Delta_AUC
Doxycycline
YJL136C RPS21B
ZShift = 0.47
Z lm Score = 2.51
NG = 0
DB = 0
SM = 0
Delta_L
Doxycycline
YJL136C RPS21B
ZShift = −0.18
Z lm Score = −3.95
NG = 0
DB = 0
SM = 0
Delta_K
Doxycycline
YJL136C RPS21B
ZShift = −0.75
Z lm Score = 0.17
NG = 0
DB = 0
SM = 0
Delta_r
Doxycycline
YJL136C RPS21B
ZShift = −0.33
Z lm Score = 1.17
NG = 0
DB = 0
SM = 0
Delta_AUC
Doxycycline
YJL179W PFD1
ZShift = 4.73
Z lm Score = 2.51
NG = 0
DB = 0
SM = 0
Delta_L
Doxycycline
YJL179W PFD1
ZShift = −1.25
Z lm Score = −2.84
NG = 0
DB = 0
SM = 0
Delta_K
Doxycycline
YJL179W PFD1
ZShift = −4.43
Z lm Score = 3.31
NG = 0
DB = 0
SM = 0
Delta_r
Doxycycline
YJL179W PFD1
ZShift = −2.78
Z lm Score = 2.61
NG = 0
DB = 0
SM = 0
Delta_AUC
Doxycycline

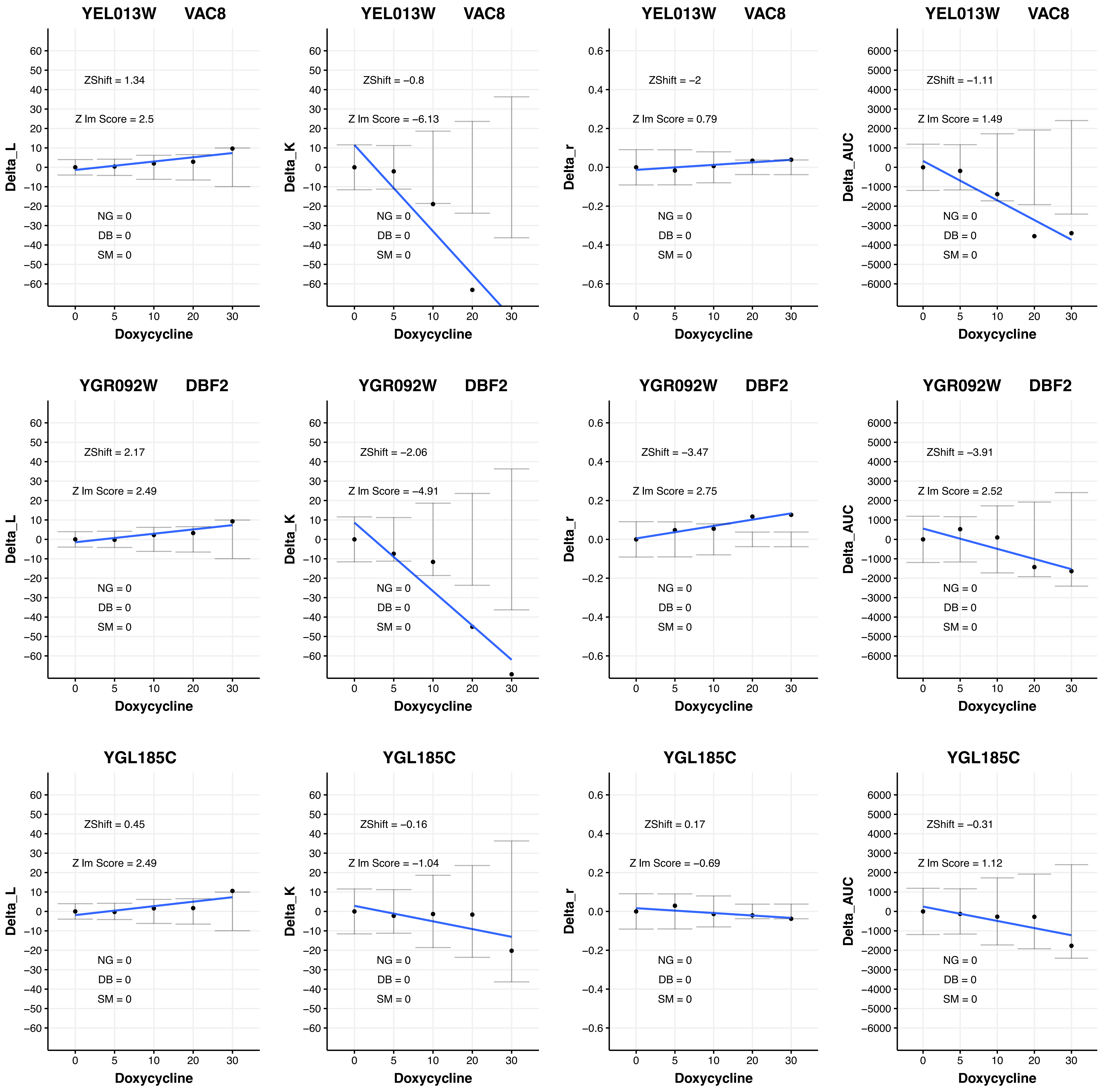
YEL013W VAC8
ZShift = 1.34
Z lm Score = 2.5
NG = 0
DB = 0
SM = 0
Delta_L
Doxycycline
YEL013W VAC8
ZShift = −0.8
Z lm Score = −6.13
NG = 0
DB = 0
SM = 0
Delta_K
Doxycycline
YEL013W VAC8
ZShift = −2
Z lm Score = 0.79
NG = 0
DB = 0
SM = 0
Delta_r
Doxycycline
YEL013W VAC8
ZShift = −1.11
Z lm Score = 1.49
NG = 0
DB = 0
SM = 0
Delta_AUC
Doxycycline
YGR092W DBF2
ZShift = 2.17
Z lm Score = 2.49
NG = 0
DB = 0
SM = 0
Delta_L
Doxycycline
YGR092W DBF2
ZShift = −2.06
Z lm Score = −4.91
NG = 0
DB = 0
SM = 0
Delta_K
Doxycycline
YGR092W DBF2
ZShift = −3.47
Z lm Score = 2.75
NG = 0
DB = 0
SM = 0
Delta_r
Doxycycline
YGR092W DBF2
ZShift = −3.91
Z lm Score = 2.52
NG = 0
DB = 0
SM = 0
Delta_AUC
Doxycycline
YGL185C
ZShift = 0.45
Z lm Score = 2.49
NG = 0
DB = 0
SM = 0
Delta_L
Doxycycline
YGL185C
ZShift = −0.16
Z lm Score = −1.04
NG = 0
DB = 0
SM = 0
Delta_K
Doxycycline
YGL185C
ZShift = 0.17
Z lm Score = −0.69
NG = 0
DB = 0
SM = 0
Delta_r
Doxycycline
YGL185C
ZShift = −0.31
Z lm Score = 1.12
NG = 0
DB = 0
SM = 0
Delta_AUC
Doxycycline

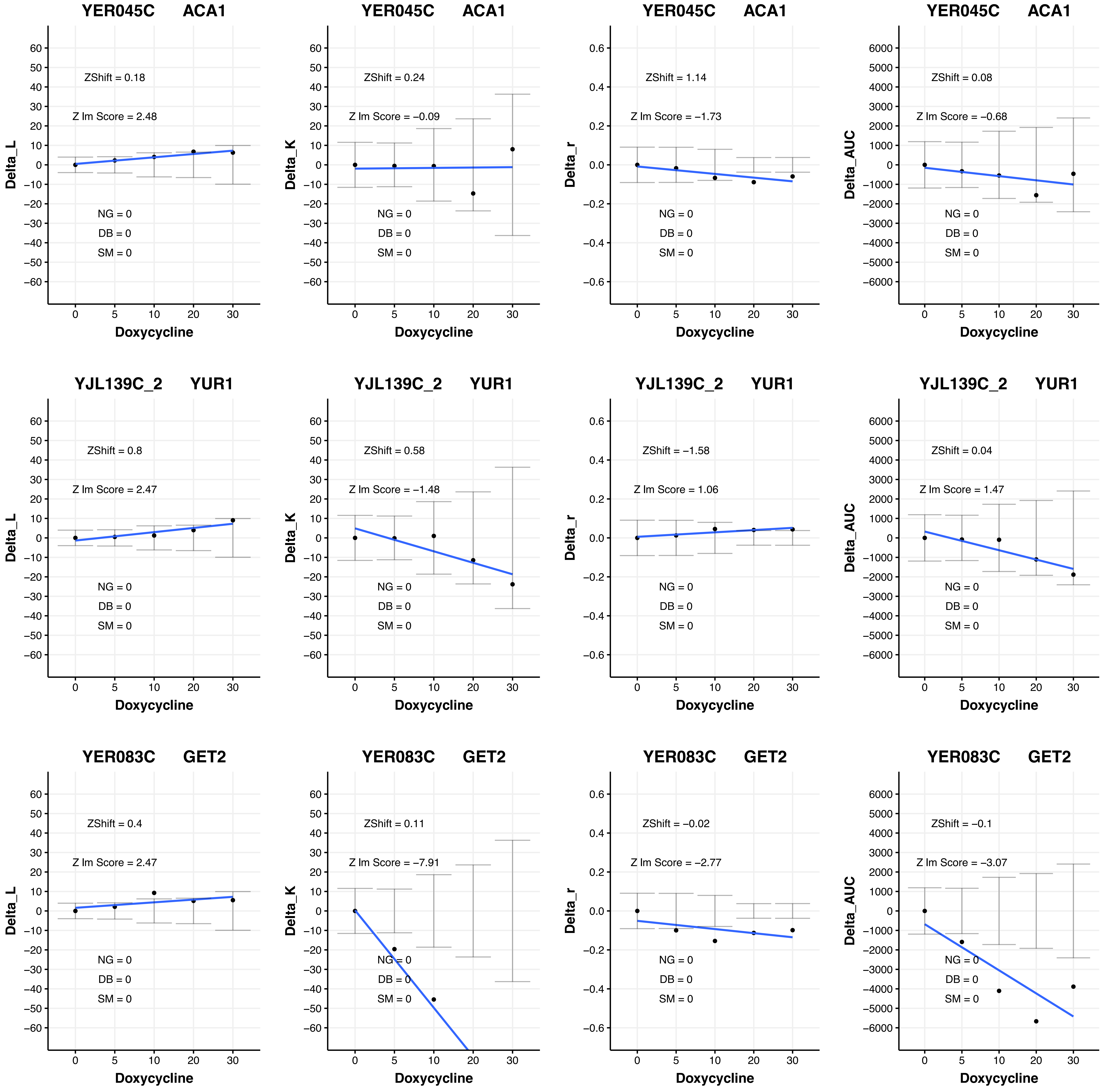
YER045C ACA1
ZShift = 0.18
Z lm Score = 2.48
NG = 0
DB = 0
SM = 0
Delta_L
Doxycycline
YER045C ACA1
ZShift = 0.24
Z lm Score = −0.09
NG = 0
DB = 0
SM = 0
Delta_K
Doxycycline
YER045C ACA1
ZShift = 1.14
Z lm Score = −1.73
NG = 0
DB = 0
SM = 0
Delta_r
Doxycycline
YER045C ACA1
ZShift = 0.08
Z lm Score = −0.68
NG = 0
DB = 0
SM = 0
Delta_AUC
Doxycycline
YJL139C_2 YUR1
ZShift = 0.8
Z lm Score = 2.47
NG = 0
DB = 0
SM = 0
Delta_L
Doxycycline
YJL139C_2 YUR1
ZShift = 0.58
Z lm Score = −1.48
NG = 0
DB = 0
SM = 0
Delta_K
Doxycycline
YJL139C_2 YUR1
ZShift = −1.58
Z lm Score = 1.06
NG = 0
DB = 0
SM = 0
Delta_r
Doxycycline
YJL139C_2 YUR1
ZShift = 0.04
Z lm Score = 1.47
NG = 0
DB = 0
SM = 0
Delta_AUC
Doxycycline
YER083C GET2
ZShift = 0.4
Z lm Score = 2.47
NG = 0
DB = 0
SM = 0
Delta_L
Doxycycline
YER083C GET2
ZShift = 0.11
Z lm Score = −7.91
NG = 0
DB = 0
SM = 0
Delta_K
Doxycycline
YER083C GET2
ZShift = −0.02
Z lm Score = −2.77
NG = 0
DB = 0
SM = 0
Delta_r
Doxycycline
YER083C GET2
ZShift = −0.1
Z lm Score = −3.07
NG = 0
DB = 0
SM = 0
Delta_AUC
Doxycycline

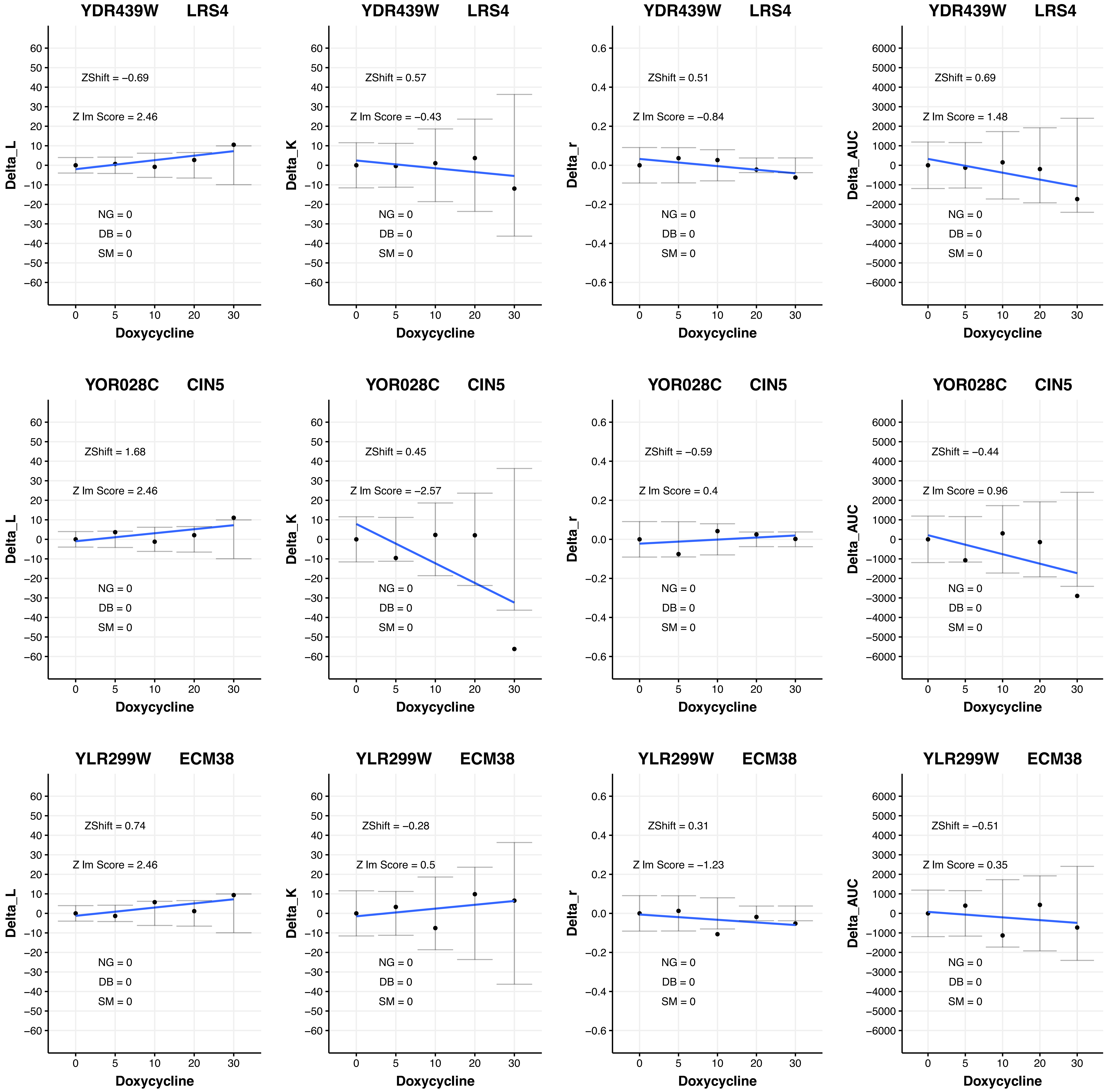
YDR439W LRS4
ZShift = −0.69
Z lm Score = 2.46
NG = 0
DB = 0
SM = 0
Delta_L
Doxycycline
YDR439W LRS4
ZShift = 0.57
Z lm Score = −0.43
NG = 0
DB = 0
SM = 0
Delta_K
Doxycycline
YDR439W LRS4
ZShift = 0.51
Z lm Score = −0.84
NG = 0
DB = 0
SM = 0
Delta_r
Doxycycline
YDR439W LRS4
ZShift = 0.69
Z lm Score = 1.48
NG = 0
DB = 0
SM = 0
Delta_AUC
Doxycycline
YOR028C CIN5
ZShift = 1.68
Z lm Score = 2.46
NG = 0
DB = 0
SM = 0
Delta_L
Doxycycline
YOR028C CIN5
ZShift = 0.45
Z lm Score = −2.57
NG = 0
DB = 0
SM = 0
Delta_K
Doxycycline
YOR028C CIN5
ZShift = −0.59
Z lm Score = 0.4
NG = 0
DB = 0
SM = 0
Delta_r
Doxycycline
YOR028C CIN5
ZShift = −0.44
Z lm Score = 0.96
NG = 0
DB = 0
SM = 0
Delta_AUC
Doxycycline
YLR299W ECM38
ZShift = 0.74
Z lm Score = 2.46
NG = 0
DB = 0
SM = 0
Delta_L
Doxycycline
YLR299W ECM38
ZShift = −0.28
Z lm Score = 0.5
NG = 0
DB = 0
SM = 0
Delta_K
Doxycycline
YLR299W ECM38
ZShift = 0.31
Z lm Score = −1.23
NG = 0
DB = 0
SM = 0
Delta_r
Doxycycline
YLR299W ECM38
ZShift = −0.51
Z lm Score = 0.35
NG = 0
DB = 0
SM = 0
Delta_AUC
Doxycycline

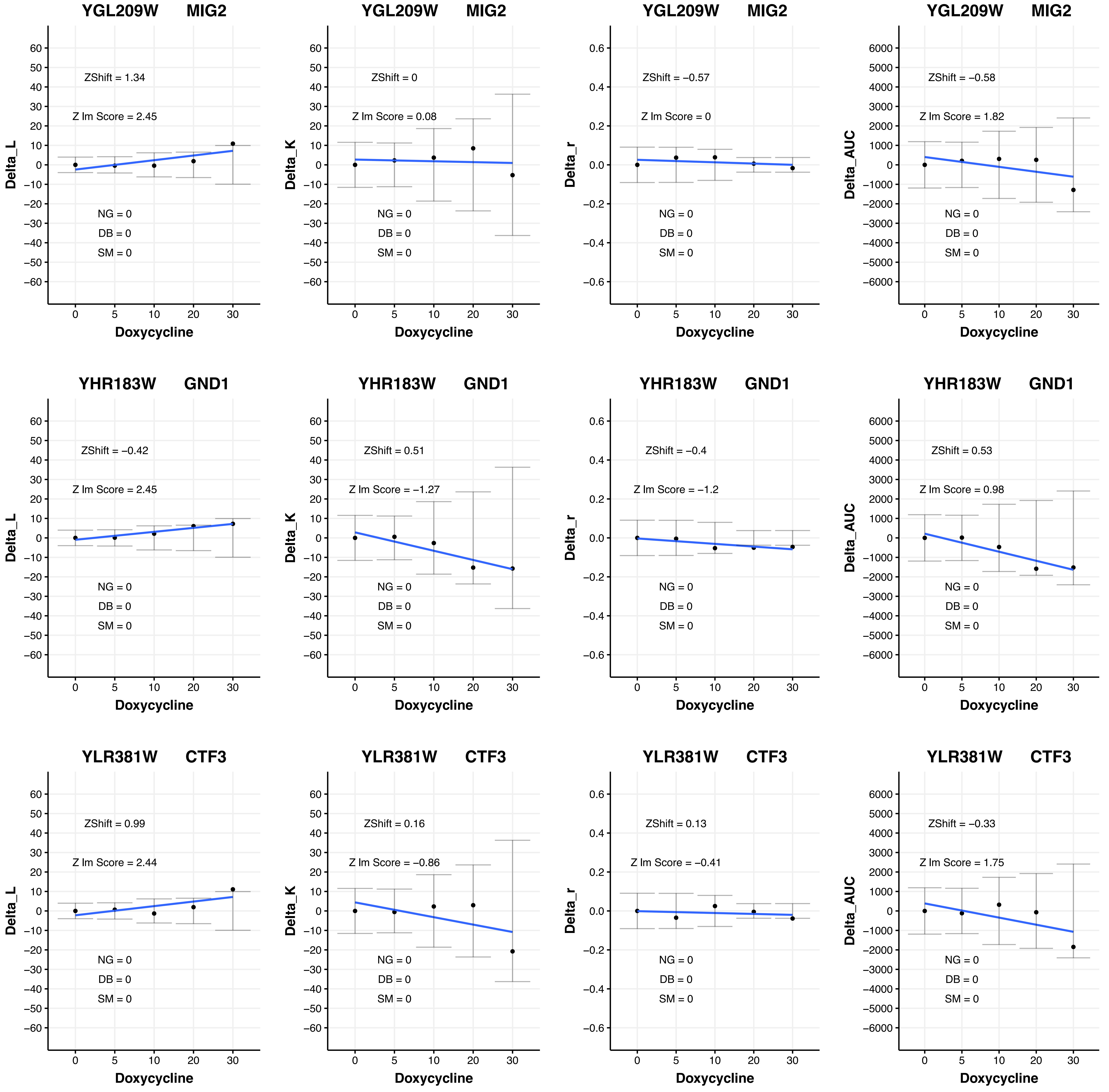
YGL209W MIG2
ZShift = 1.34
Z lm Score = 2.45
NG = 0
DB = 0
SM = 0
Delta_L
Doxycycline
YGL209W MIG2
ZShift = 0
Z lm Score = 0.08
NG = 0
DB = 0
SM = 0
Delta_K
Doxycycline
YGL209W MIG2
ZShift = −0.57
Z lm Score = 0
NG = 0
DB = 0
SM = 0
Delta_r
Doxycycline
YGL209W MIG2
ZShift = −0.58
Z lm Score = 1.82
NG = 0
DB = 0
SM = 0
Delta_AUC
Doxycycline
YHR183W GND1
ZShift = −0.42
Z lm Score = 2.45
NG = 0
DB = 0
SM = 0
Delta_L
Doxycycline
YHR183W GND1
ZShift = 0.51
Z lm Score = −1.27
NG = 0
DB = 0
SM = 0
Delta_K
Doxycycline
YHR183W GND1
ZShift = −0.4
Z lm Score = −1.2
NG = 0
DB = 0
SM = 0
Delta_r
Doxycycline
YHR183W GND1
ZShift = 0.53
Z lm Score = 0.98
NG = 0
DB = 0
SM = 0
Delta_AUC
Doxycycline
YLR381W CTF3
ZShift = 0.99
Z lm Score = 2.44
NG = 0
DB = 0
SM = 0
Delta_L
Doxycycline
YLR381W CTF3
ZShift = 0.16
Z lm Score = −0.86
NG = 0
DB = 0
SM = 0
Delta_K
Doxycycline
YLR381W CTF3
ZShift = 0.13
Z lm Score = −0.41
NG = 0
DB = 0
SM = 0
Delta_r
Doxycycline
YLR381W CTF3
ZShift = −0.33
Z lm Score = 1.75
NG = 0
DB = 0
SM = 0
Delta_AUC
Doxycycline

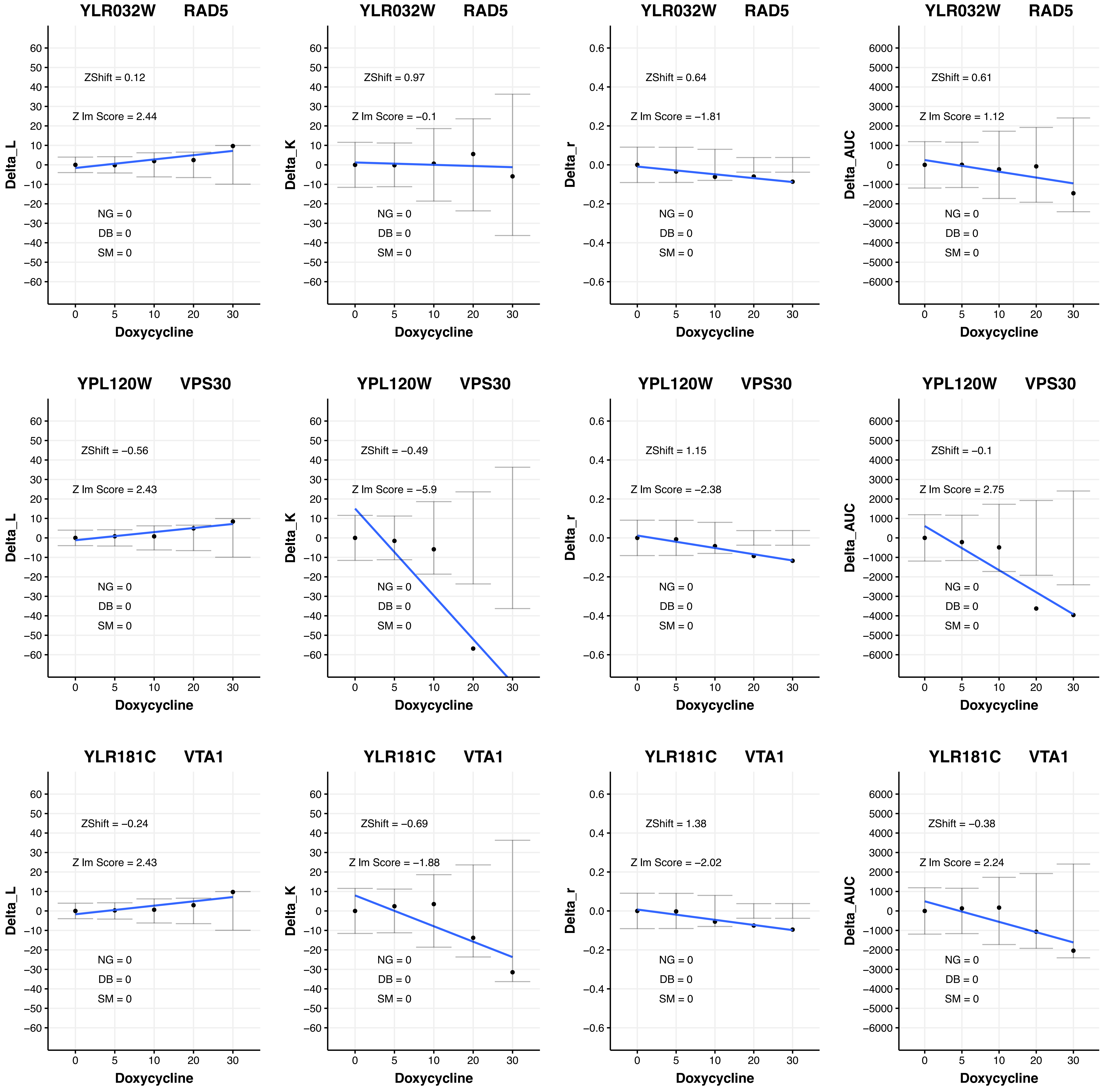
YLR032W RAD5
ZShift = 0.12
Z lm Score = 2.44
NG = 0
DB = 0
SM = 0
Delta_L
Doxycycline
YLR032W RAD5
ZShift = 0.97
Z lm Score = −0.1
NG = 0
DB = 0
SM = 0
Delta_K
Doxycycline
YLR032W RAD5
ZShift = 0.64
Z lm Score = −1.81
NG = 0
DB = 0
SM = 0
Delta_r
Doxycycline
YLR032W RAD5
ZShift = 0.61
Z lm Score = 1.12
NG = 0
DB = 0
SM = 0
Delta_AUC
Doxycycline
YPL120W VPS30
ZShift = −0.56
Z lm Score = 2.43
NG = 0
DB = 0
SM = 0
Delta_L
Doxycycline
YPL120W VPS30
ZShift = −0.49
Z lm Score = −5.9
NG = 0
DB = 0
SM = 0
Delta_K
Doxycycline
YPL120W VPS30
ZShift = 1.15
Z lm Score = −2.38
NG = 0
DB = 0
SM = 0
Delta_r
Doxycycline
YPL120W VPS30
ZShift = −0.1
Z lm Score = 2.75
NG = 0
DB = 0
SM = 0
Delta_AUC
Doxycycline
YLR181C VTA1
ZShift = −0.24
Z lm Score = 2.43
NG = 0
DB = 0
SM = 0
Delta_L
Doxycycline
YLR181C VTA1
ZShift = −0.69
Z lm Score = −1.88
NG = 0
DB = 0
SM = 0
Delta_K
Doxycycline
YLR181C VTA1
ZShift = 1.38
Z lm Score = −2.02
NG = 0
DB = 0
SM = 0
Delta_r
Doxycycline
YLR181C VTA1
ZShift = −0.38
Z lm Score = 2.24
NG = 0
DB = 0
SM = 0
Delta_AUC
Doxycycline

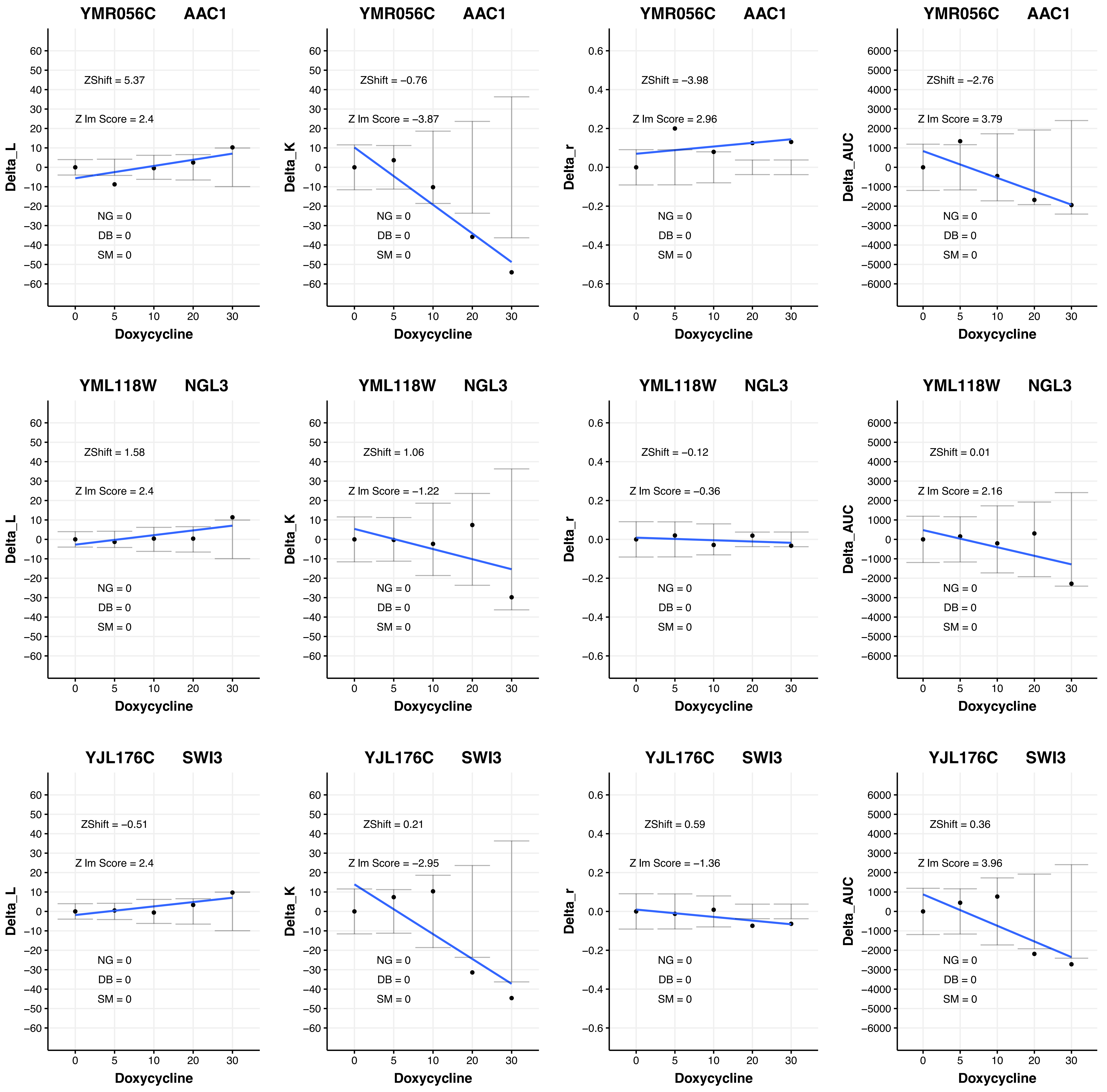
YMR056C AAC1
ZShift = 5.37
Z lm Score = 2.4
NG = 0
DB = 0
SM = 0
Delta_L
Doxycycline
YMR056C AAC1
ZShift = −0.76
Z lm Score = −3.87
NG = 0
DB = 0
SM = 0
Delta_K
Doxycycline
YMR056C AAC1
ZShift = −3.98
Z lm Score = 2.96
NG = 0
DB = 0
SM = 0
Delta_r
Doxycycline
YMR056C AAC1
ZShift = −2.76
Z lm Score = 3.79
NG = 0
DB = 0
SM = 0
Delta_AUC
Doxycycline
YML118W NGL3
ZShift = 1.58
Z lm Score = 2.4
NG = 0
DB = 0
SM = 0
Delta_L
Doxycycline
YML118W NGL3
ZShift = 1.06
Z lm Score = −1.22
NG = 0
DB = 0
SM = 0
Delta_K
Doxycycline
YML118W NGL3
ZShift = −0.12
Z lm Score = −0.36
NG = 0
DB = 0
SM = 0
Delta_r
Doxycycline
YML118W NGL3
ZShift = 0.01
Z lm Score = 2.16
NG = 0
DB = 0
SM = 0
Delta_AUC
Doxycycline
YJL176C SWI3
ZShift = −0.51
Z lm Score = 2.4
NG = 0
DB = 0
SM = 0
Delta_L
Doxycycline
YJL176C SWI3
ZShift = 0.21
Z lm Score = −2.95
NG = 0
DB = 0
SM = 0
Delta_K
Doxycycline
YJL176C SWI3
ZShift = 0.59
Z lm Score = −1.36
NG = 0
DB = 0
SM = 0
Delta_r
Doxycycline
YJL176C SWI3
ZShift = 0.36
Z lm Score = 3.96
NG = 0
DB = 0
SM = 0
Delta_AUC
Doxycycline

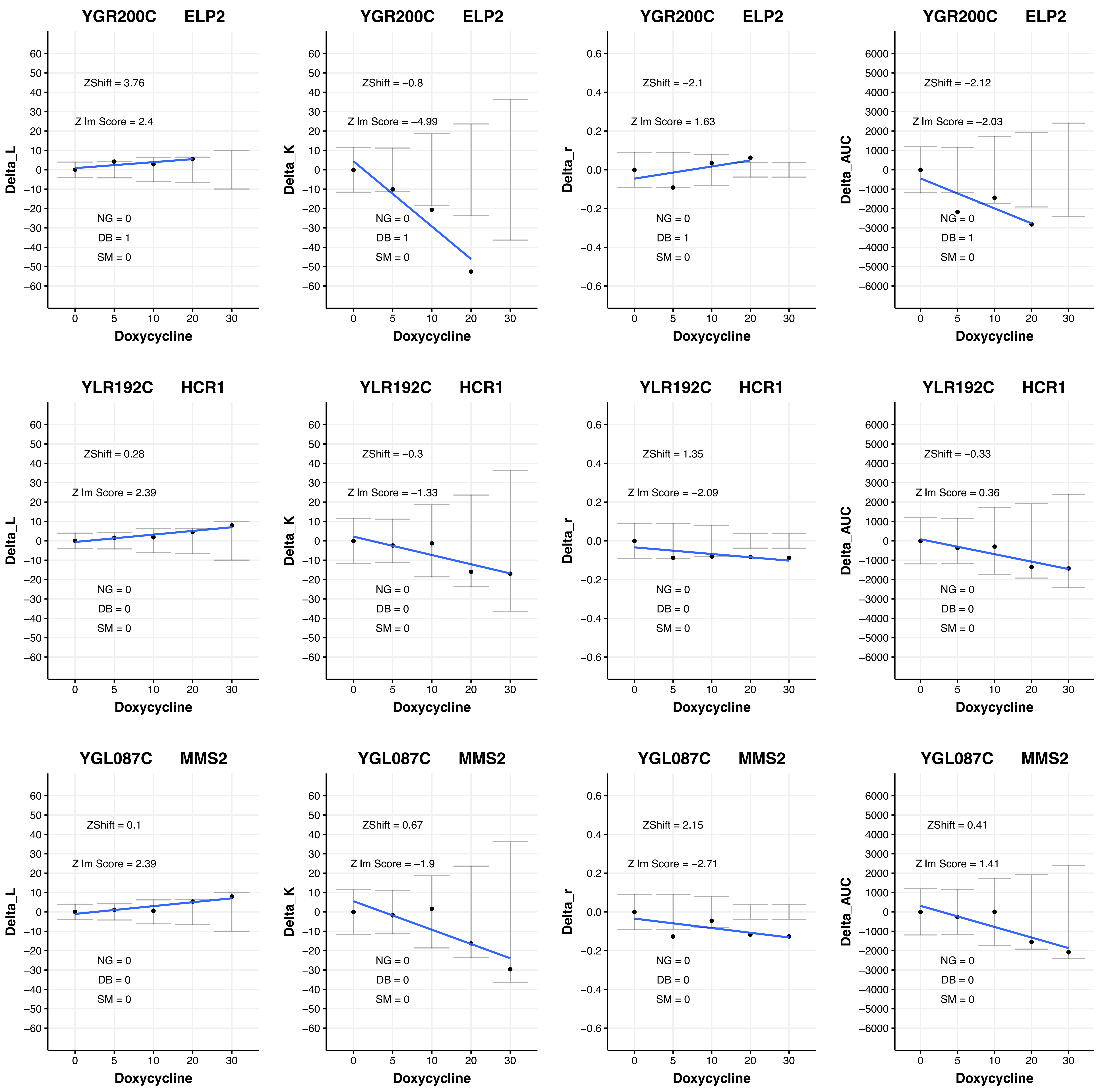
YGR200C ELP2
ZShift = 3.76
Z lm Score = 2.4
NG = 0
DB = 1
SM = 0
Delta_L
Doxycycline
YGR200C ELP2
ZShift = −0.8
Z lm Score = −4.99
NG = 0
DB = 1
SM = 0
Delta_K
Doxycycline
YGR200C ELP2
ZShift = −2.1
Z lm Score = 1.63
NG = 0
DB = 1
SM = 0
Delta_r
Doxycycline
YGR200C ELP2
ZShift = −2.12
Z lm Score = −2.03
NG = 0
DB = 1
SM = 0
Delta_AUC
Doxycycline
YLR192C HCR1
ZShift = 0.28
Z lm Score = 2.39
NG = 0
DB = 0
SM = 0
Delta_L
Doxycycline
YLR192C HCR1
ZShift = −0.3
Z lm Score = −1.33
NG = 0
DB = 0
SM = 0
Delta_K
Doxycycline
YLR192C HCR1
ZShift = 1.35
Z lm Score = −2.09
NG = 0
DB = 0
SM = 0
Delta_r
Doxycycline
YLR192C HCR1
ZShift = −0.33
Z lm Score = 0.36
NG = 0
DB = 0
SM = 0
Delta_AUC
Doxycycline
YGL087C MMS2
ZShift = 0.1
Z lm Score = 2.39
NG = 0
DB = 0
SM = 0
Delta_L
Doxycycline
YGL087C MMS2
ZShift = 0.67
Z lm Score = −1.9
NG = 0
DB = 0
SM = 0
Delta_K
Doxycycline
YGL087C MMS2
ZShift = 2.15
Z lm Score = −2.71
NG = 0
DB = 0
SM = 0
Delta_r
Doxycycline
YGL087C MMS2
ZShift = 0.41
Z lm Score = 1.41
NG = 0
DB = 0
SM = 0
Delta_AUC
Doxycycline

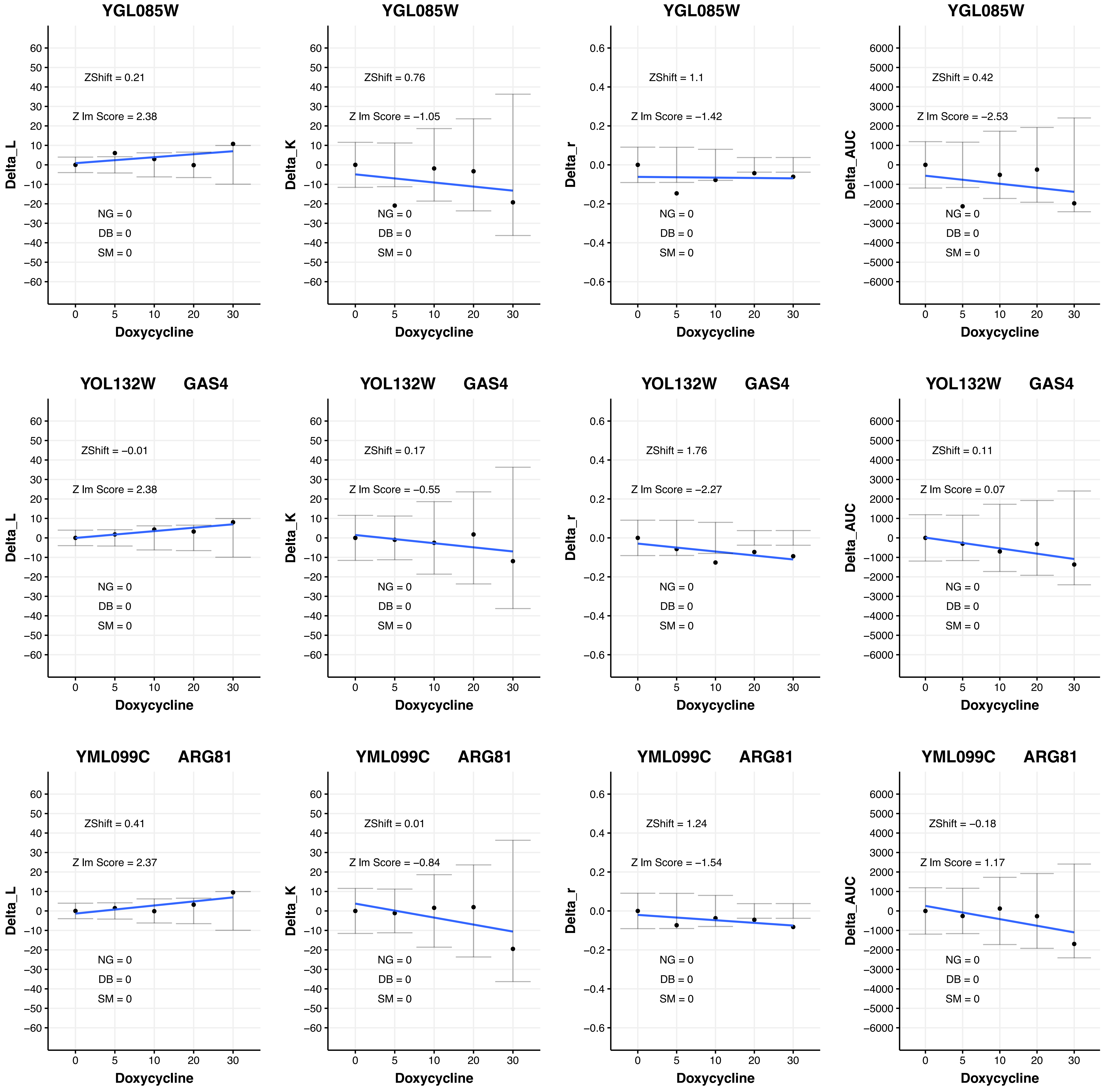
YGL085W
ZShift = 0.21
Z lm Score = 2.38
NG = 0
DB = 0
SM = 0
Delta_L
Doxycycline
YGL085W
ZShift = 0.76
Z lm Score = −1.05
NG = 0
DB = 0
SM = 0
Delta_K
Doxycycline
YGL085W
ZShift = 1.1
Z lm Score = −1.42
NG = 0
DB = 0
SM = 0
Delta_r
Doxycycline
YGL085W
ZShift = 0.42
Z lm Score = −2.53
NG = 0
DB = 0
SM = 0
Delta_AUC
Doxycycline
YOL132W GAS4
ZShift = −0.01
Z lm Score = 2.38
NG = 0
DB = 0
SM = 0
Delta_L
Doxycycline
YOL132W GAS4
ZShift = 0.17
Z lm Score = −0.55
NG = 0
DB = 0
SM = 0
Delta_K
Doxycycline
YOL132W GAS4
ZShift = 1.76
Z lm Score = −2.27
NG = 0
DB = 0
SM = 0
Delta_r
Doxycycline
YOL132W GAS4
ZShift = 0.11
Z lm Score = 0.07
NG = 0
DB = 0
SM = 0
Delta_AUC
Doxycycline
YML099C ARG81
ZShift = 0.41
Z lm Score = 2.37
NG = 0
DB = 0
SM = 0
Delta_L
Doxycycline
YML099C ARG81
ZShift = 0.01
Z lm Score = −0.84
NG = 0
DB = 0
SM = 0
Delta_K
Doxycycline
YML099C ARG81
ZShift = 1.24
Z lm Score = −1.54
NG = 0
DB = 0
SM = 0
Delta_r
Doxycycline
YML099C ARG81
ZShift = −0.18
Z lm Score = 1.17
NG = 0
DB = 0
SM = 0
Delta_AUC
Doxycycline

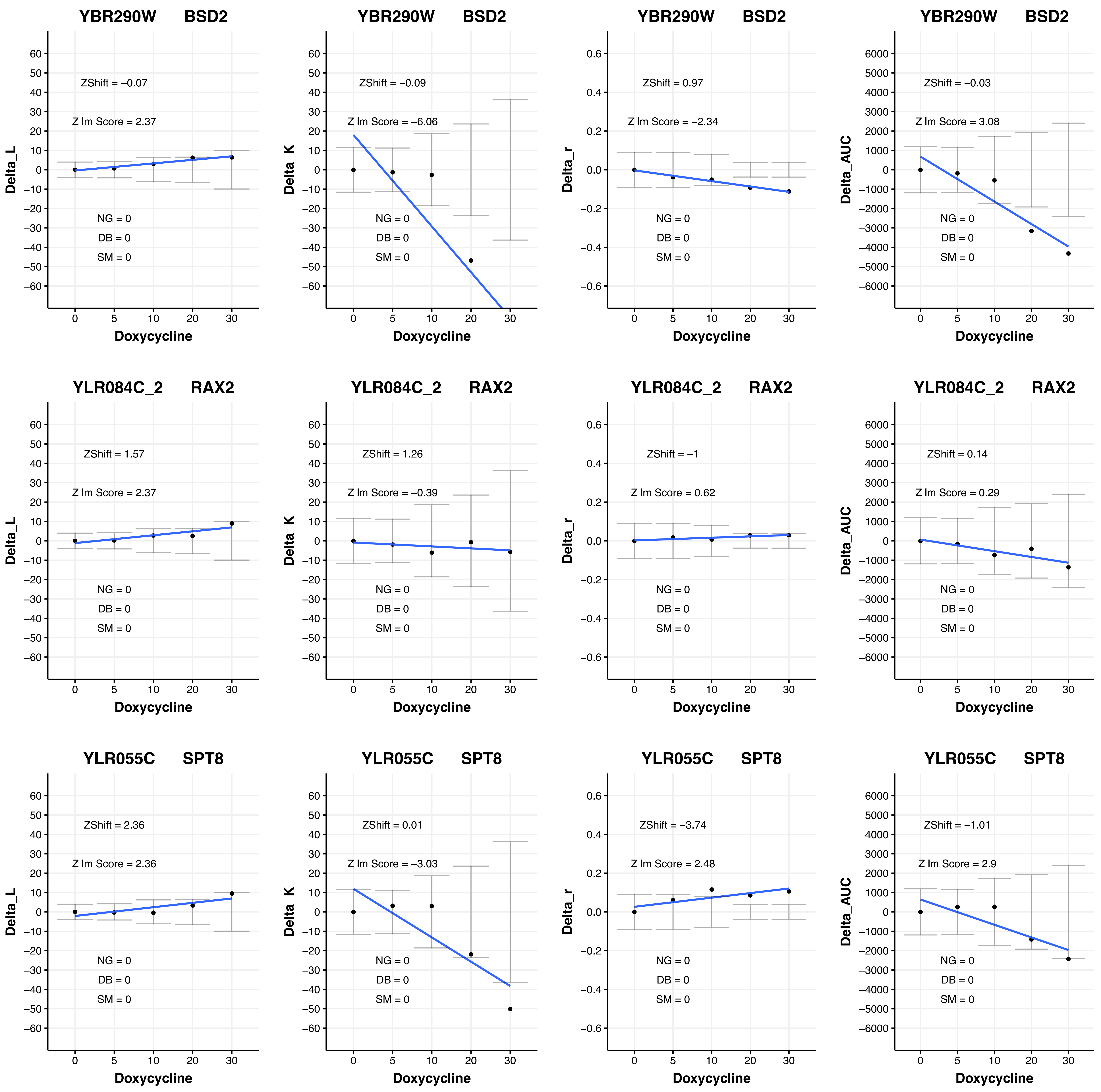
YBR290W BSD2
ZShift = −0.07
Z lm Score = 2.37
NG = 0
DB = 0
SM = 0
Delta_L
Doxycycline
YBR290W BSD2
ZShift = −0.09
Z lm Score = −6.06
NG = 0
DB = 0
SM = 0
Delta_K
Doxycycline
YBR290W BSD2
ZShift = 0.97
Z lm Score = −2.34
NG = 0
DB = 0
SM = 0
Delta_r
Doxycycline
YBR290W BSD2
ZShift = −0.03
Z lm Score = 3.08
NG = 0
DB = 0
SM = 0
Delta_AUC
Doxycycline
YLR084C_2 RAX2
ZShift = 1.57
Z lm Score = 2.37
NG = 0
DB = 0
SM = 0
Delta_L
Doxycycline
YLR084C_2 RAX2
ZShift = 1.26
Z lm Score = −0.39
NG = 0
DB = 0
SM = 0
Delta_K
Doxycycline
YLR084C_2 RAX2
ZShift = −1
Z lm Score = 0.62
NG = 0
DB = 0
SM = 0
Delta_r
Doxycycline
YLR084C_2 RAX2
ZShift = 0.14
Z lm Score = 0.29
NG = 0
DB = 0
SM = 0
Delta_AUC
Doxycycline
YLR055C SPT8
ZShift = 2.36
Z lm Score = 2.36
NG = 0
DB = 0
SM = 0
Delta_L
Doxycycline
YLR055C SPT8
ZShift = 0.01
Z lm Score = −3.03
NG = 0
DB = 0
SM = 0
Delta_K
Doxycycline
YLR055C SPT8
ZShift = −3.74
Z lm Score = 2.48
NG = 0
DB = 0
SM = 0
Delta_r
Doxycycline
YLR055C SPT8
ZShift = −1.01
Z lm Score = 2.9
NG = 0
DB = 0
SM = 0
Delta_AUC
Doxycycline

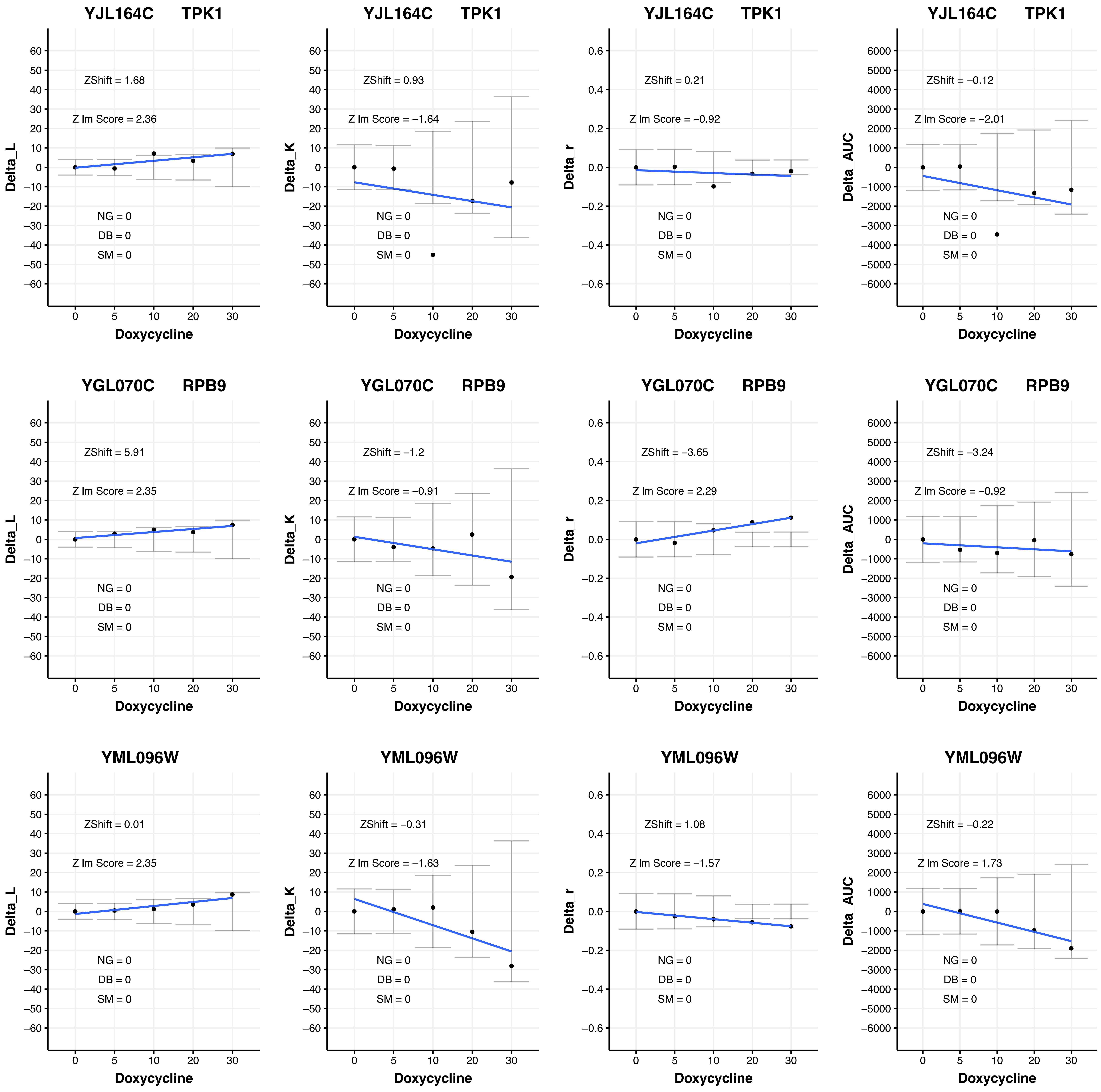
YJL164C TPK1
ZShift = 1.68
Z lm Score = 2.36
NG = 0
DB = 0
SM = 0
Delta_L
Doxycycline
YJL164C TPK1
ZShift = 0.93
Z lm Score = −1.64
NG = 0
DB = 0
SM = 0
Delta_K
Doxycycline
YJL164C TPK1
ZShift = 0.21
Z lm Score = −0.92
NG = 0
DB = 0
SM = 0
Delta_r
Doxycycline
YJL164C TPK1
ZShift = −0.12
Z lm Score = −2.01
NG = 0
DB = 0
SM = 0
Delta_AUC
Doxycycline
YGL070C RPB9
ZShift = 5.91
Z lm Score = 2.35
NG = 0
DB = 0
SM = 0
Delta_L
Doxycycline
YGL070C RPB9
ZShift = −1.2
Z lm Score = −0.91
NG = 0
DB = 0
SM = 0
Delta_K
Doxycycline
YGL070C RPB9
ZShift = −3.65
Z lm Score = 2.29
NG = 0
DB = 0
SM = 0
Delta_r
Doxycycline
YGL070C RPB9
ZShift = −3.24
Z lm Score = −0.92
NG = 0
DB = 0
SM = 0
Delta_AUC
Doxycycline
YML096W
ZShift = 0.01
Z lm Score = 2.35
NG = 0
DB = 0
SM = 0
Delta_L
Doxycycline
YML096W
ZShift = −0.31
Z lm Score = −1.63
NG = 0
DB = 0
SM = 0
Delta_K
Doxycycline
YML096W
ZShift = 1.08
Z lm Score = −1.57
NG = 0
DB = 0
SM = 0
Delta_r
Doxycycline
YML096W
ZShift = −0.22
Z lm Score = 1.73
NG = 0
DB = 0
SM = 0
Delta_AUC
Doxycycline

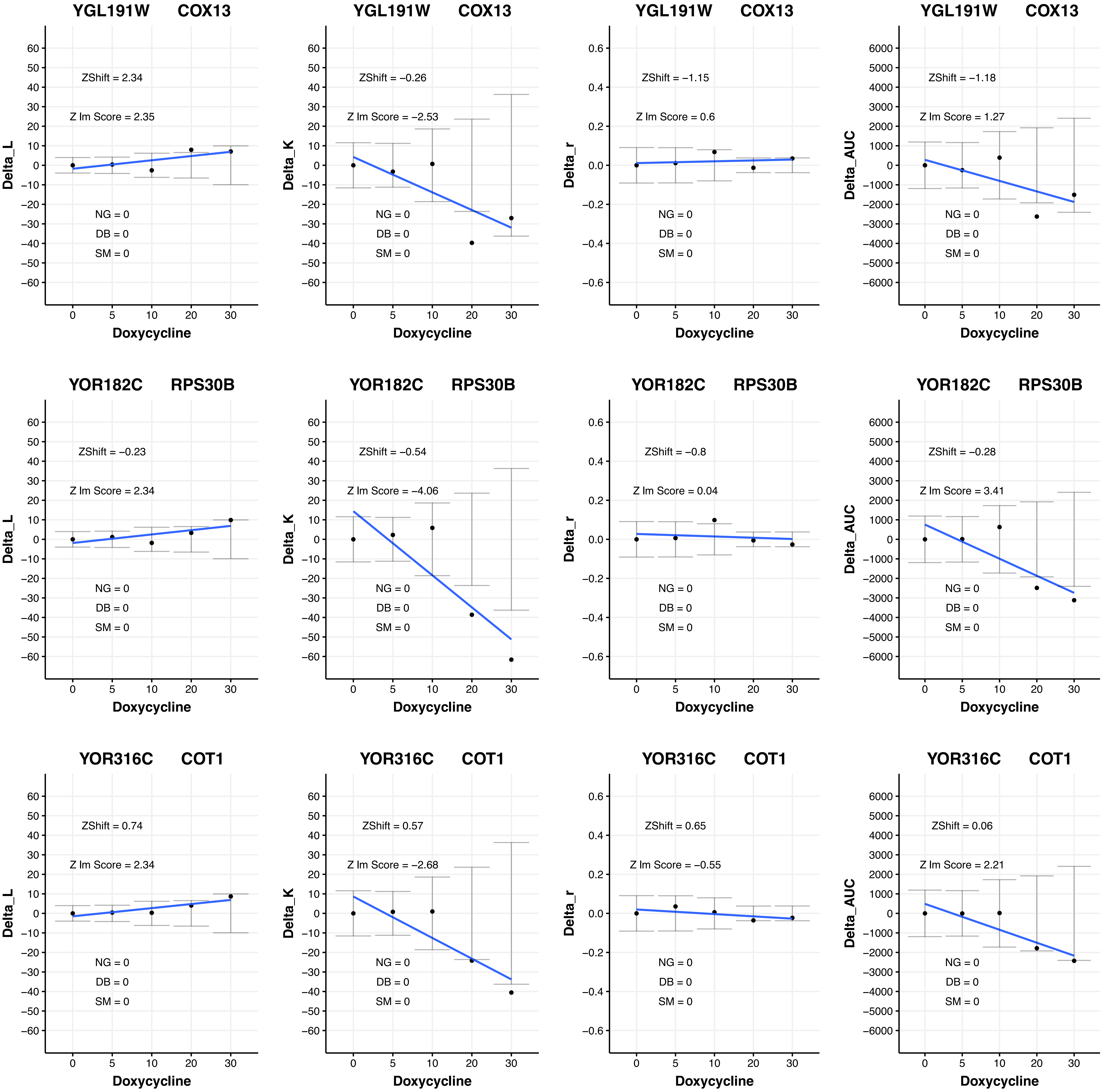

YGL191W COX13
ZShift = 2.34
Z lm Score = 2.35
NG = 0
DB = 0
SM = 0
Delta_L
Doxycycline
YGL191W COX13
ZShift = −0.26
Z lm Score = −2.53
NG = 0
DB = 0
SM = 0
Delta_K
Doxycycline
YGL191W COX13
ZShift = −1.15
Z lm Score = 0.6
NG = 0
DB = 0
SM = 0
Delta_r
Doxycycline
YGL191W COX13
ZShift = −1.18
Z lm Score = 1.27
NG = 0
DB = 0
SM = 0
Delta_AUC
Doxycycline
YOR182C RPS30B
ZShift = −0.23
Z lm Score = 2.34
NG = 0
DB = 0
SM = 0
Delta_L
Doxycycline
YOR182C RPS30B
ZShift = −0.54
Z lm Score = −4.06
NG = 0
DB = 0
SM = 0
Delta_K
Doxycycline
YOR182C RPS30B
ZShift = −0.8
Z lm Score = 0.04
NG = 0
DB = 0
SM = 0
Delta_r
Doxycycline
YOR182C RPS30B
ZShift = −0.28
Z lm Score = 3.41
NG = 0
DB = 0
SM = 0
Delta_AUC
Doxycycline
YOR316C COT1
ZShift = 0.74
Z lm Score = 2.34
NG = 0
DB = 0
SM = 0
Delta_L
Doxycycline
YOR316C COT1
ZShift = 0.57
Z lm Score = −2.68
NG = 0
DB = 0
SM = 0
Delta_K
Doxycycline
YOR316C COT1
ZShift = 0.65
Z lm Score = −0.55
NG = 0
DB = 0
SM = 0
Delta_r
Doxycycline
YOR316C COT1
ZShift = 0.06
Z lm Score = 2.21
NG = 0
DB = 0
SM = 0
Delta_AUC
Doxycycline

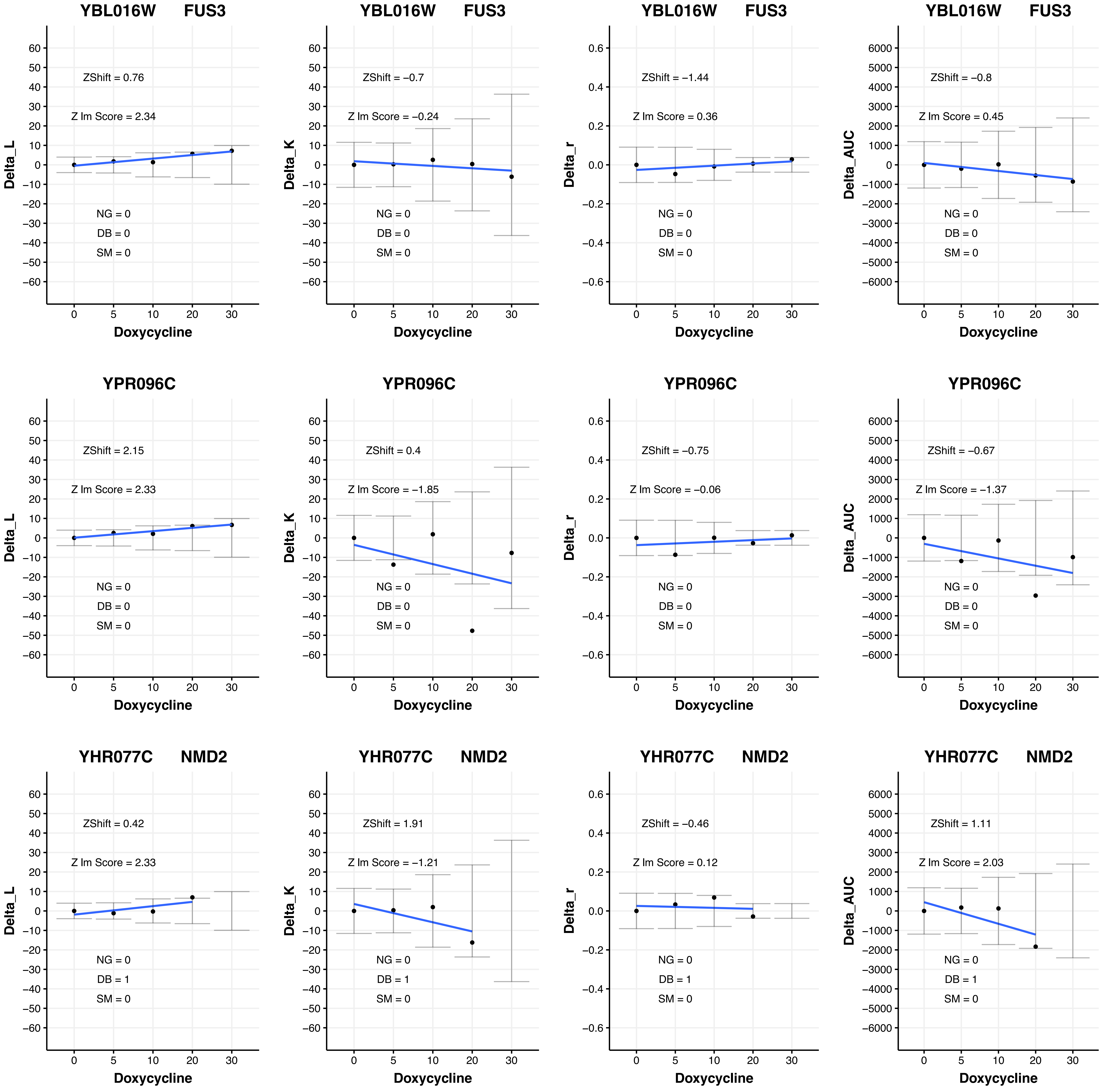
YBL016W FUS3
ZShift = 0.76
Z lm Score = 2.34
NG = 0
DB = 0
SM = 0
Delta_L
Doxycycline
YBL016W FUS3
ZShift = −0.7
Z lm Score = −0.24
NG = 0
DB = 0
SM = 0
Delta_K
Doxycycline
YBL016W FUS3
ZShift = −1.44
Z lm Score = 0.36
NG = 0
DB = 0
SM = 0
Delta_r
Doxycycline
YBL016W FUS3
ZShift = −0.8
Z lm Score = 0.45
NG = 0
DB = 0
SM = 0
Delta_AUC
Doxycycline
YPR096C
ZShift = 2.15
Z lm Score = 2.33
NG = 0
DB = 0
SM = 0
Delta_L
Doxycycline
YPR096C
ZShift = 0.4
Z lm Score = −1.85
NG = 0
DB = 0
SM = 0
Delta_K
Doxycycline
YPR096C
ZShift = −0.75
Z lm Score = −0.06
NG = 0
DB = 0
SM = 0
Delta_r
Doxycycline
YPR096C
ZShift = −0.67
Z lm Score = −1.37
NG = 0
DB = 0
SM = 0
Delta_AUC
Doxycycline
YHR077C NMD2
ZShift = 0.42
Z lm Score = 2.33
NG = 0
DB = 1
SM = 0
Delta_L
Doxycycline
YHR077C NMD2
ZShift = 1.91
Z lm Score = −1.21
NG = 0
DB = 1
SM = 0
Delta_K
Doxycycline
YHR077C NMD2
ZShift = −0.46
Z lm Score = 0.12
NG = 0
DB = 1
SM = 0
Delta_r
Doxycycline
YHR077C NMD2
ZShift = 1.11
Z lm Score = 2.03
NG = 0
DB = 1
SM = 0
Delta_AUC
Doxycycline

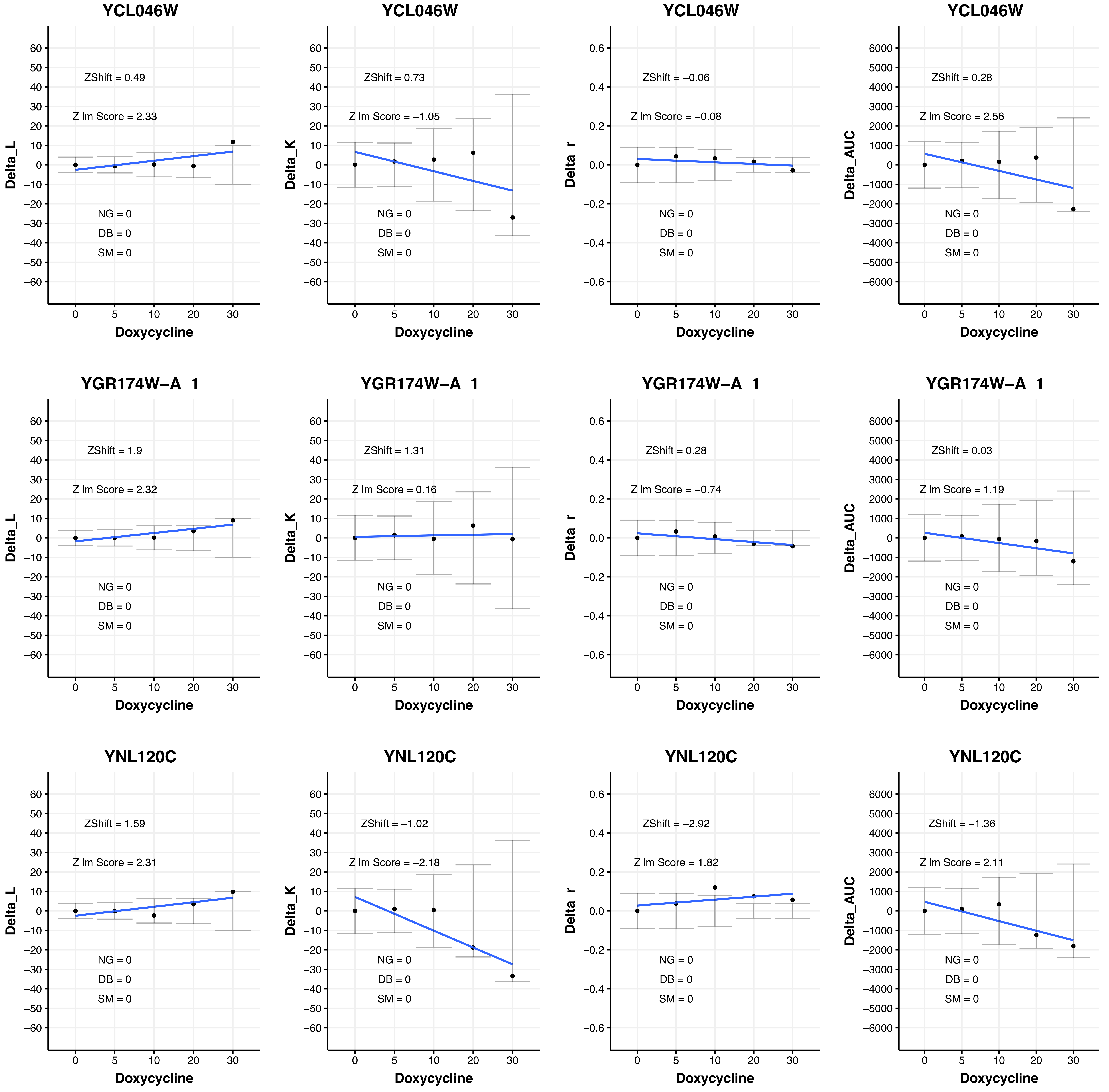

YCL046W
ZShift = 0.49
Z lm Score = 2.33
NG = 0
DB = 0
SM = 0
Delta_L
Doxycycline
YCL046W
ZShift = 0.73
Z lm Score = −1.05
NG = 0
DB = 0
SM = 0
Delta_K
Doxycycline
YCL046W
ZShift = −0.06
Z lm Score = −0.08
NG = 0
DB = 0
SM = 0
Delta_r
Doxycycline
YCL046W
ZShift = 0.28
Z lm Score = 2.56
NG = 0
DB = 0
SM = 0
Delta_AUC
Doxycycline
YGR174W−A_1
ZShift = 1.9
Z lm Score = 2.32
NG = 0
DB = 0
SM = 0
Delta_L
Doxycycline
YGR174W−A_1
ZShift = 1.31
Z lm Score = 0.16
NG = 0
DB = 0
SM = 0
Delta_K
Doxycycline
YGR174W−A_1
ZShift = 0.28
Z lm Score = −0.74
NG = 0
DB = 0
SM = 0
Delta_r
Doxycycline
YGR174W−A_1
ZShift = 0.03
Z lm Score = 1.19
NG = 0
DB = 0
SM = 0
Delta_AUC
Doxycycline
YNL120C
ZShift = 1.59
Z lm Score = 2.31
NG = 0
DB = 0
SM = 0
Delta_L
Doxycycline
YNL120C
ZShift = −1.02
Z lm Score = −2.18
NG = 0
DB = 0
SM = 0
Delta_K
Doxycycline
YNL120C
ZShift = −2.92
Z lm Score = 1.82
NG = 0
DB = 0
SM = 0
Delta_r
Doxycycline
YNL120C
ZShift = −1.36
Z lm Score = 2.11
NG = 0
DB = 0
SM = 0
Delta_AUC
Doxycycline

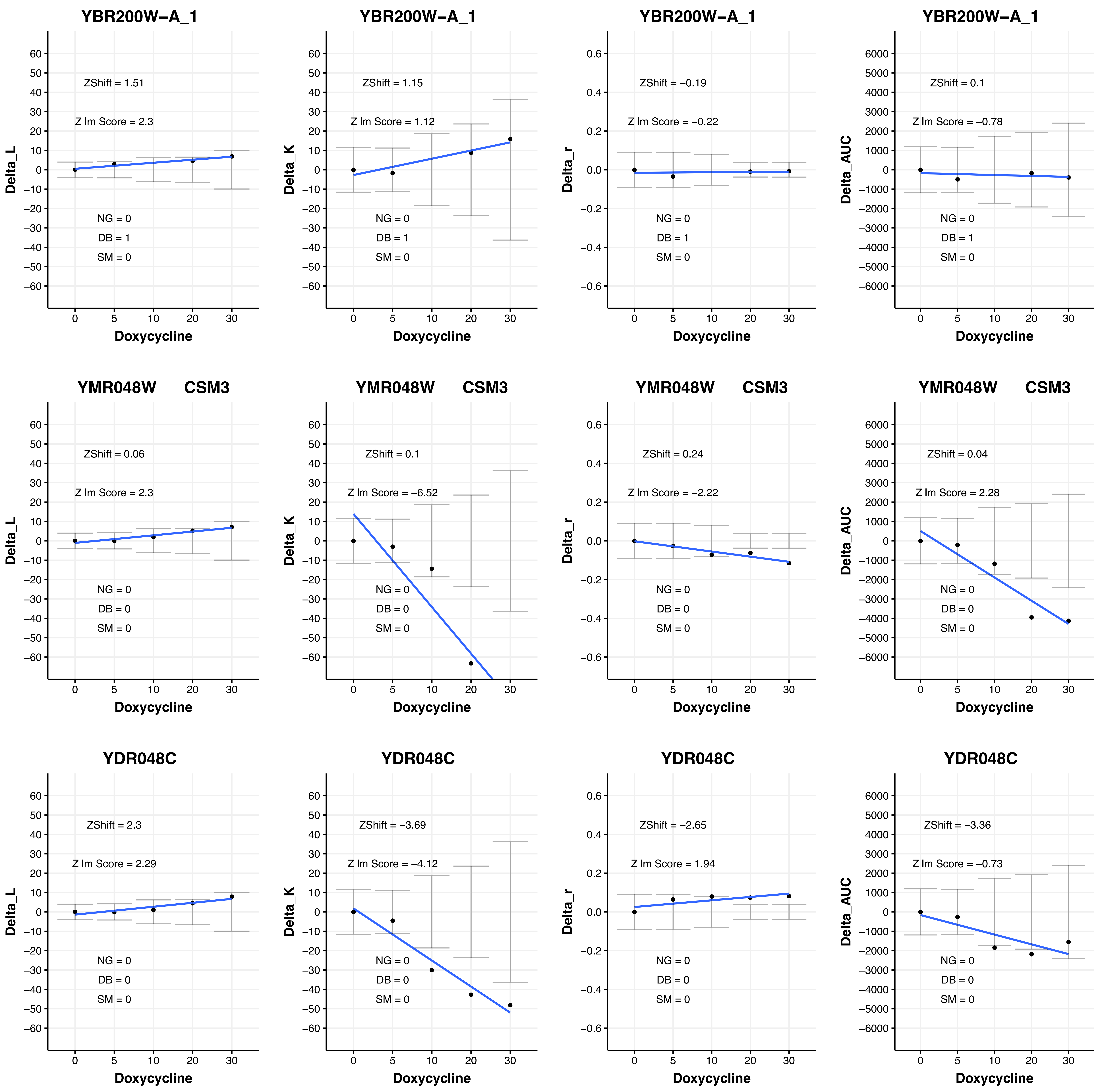

YBR200W-A_1
ZShift = 1.51
Z lm Score = 2.3
NG = 0
DB = 1
SM = 0
Delta_L
Doxycycline
YBR200W-A_1
ZShift = 1.15
Z lm Score = 1.12
NG = 0
DB = 1
SM = 0
Delta_K
Doxycycline
YBR200W-A_1
ZShift = −0.19
Z lm Score = −0.22
NG = 0
DB = 1
SM = 0
Delta_r
Doxycycline
YBR200W-A_1
ZShift = 0.1
Z lm Score = −0.78
NG = 0
DB = 1
SM = 0
Delta_AUC
Doxycycline
YMR048W CSM3
ZShift = 0.06
Z lm Score = 2.3
NG = 0
DB = 0
SM = 0
Delta_L
Doxycycline
YMR048W CSM3
ZShift = 0.1
Z lm Score = −6.52
NG = 0
DB = 0
SM = 0
Delta_K
Doxycycline
YMR048W CSM3
ZShift = 0.24
Z lm Score = −2.22
NG = 0
DB = 0
SM = 0
Delta_r
Doxycycline
YMR048W CSM3
ZShift = 0.04
Z lm Score = 2.28
NG = 0
DB = 0
SM = 0
Delta_AUC
Doxycycline
YDR048C
ZShift = 2.3
Z lm Score = 2.29
NG = 0
DB = 0
SM = 0
Delta_L
Doxycycline
YDR048C
ZShift = −3.69
Z lm Score = −4.12
NG = 0
DB = 0
SM = 0
Delta_K
Doxycycline
YDR048C
ZShift = −2.65
Z lm Score = 1.94
NG = 0
DB = 0
SM = 0
Delta_r
Doxycycline
YDR048C
ZShift = −3.36
Z lm Score = −0.73
NG = 0
DB = 0
SM = 0
Delta_AUC
Doxycycline

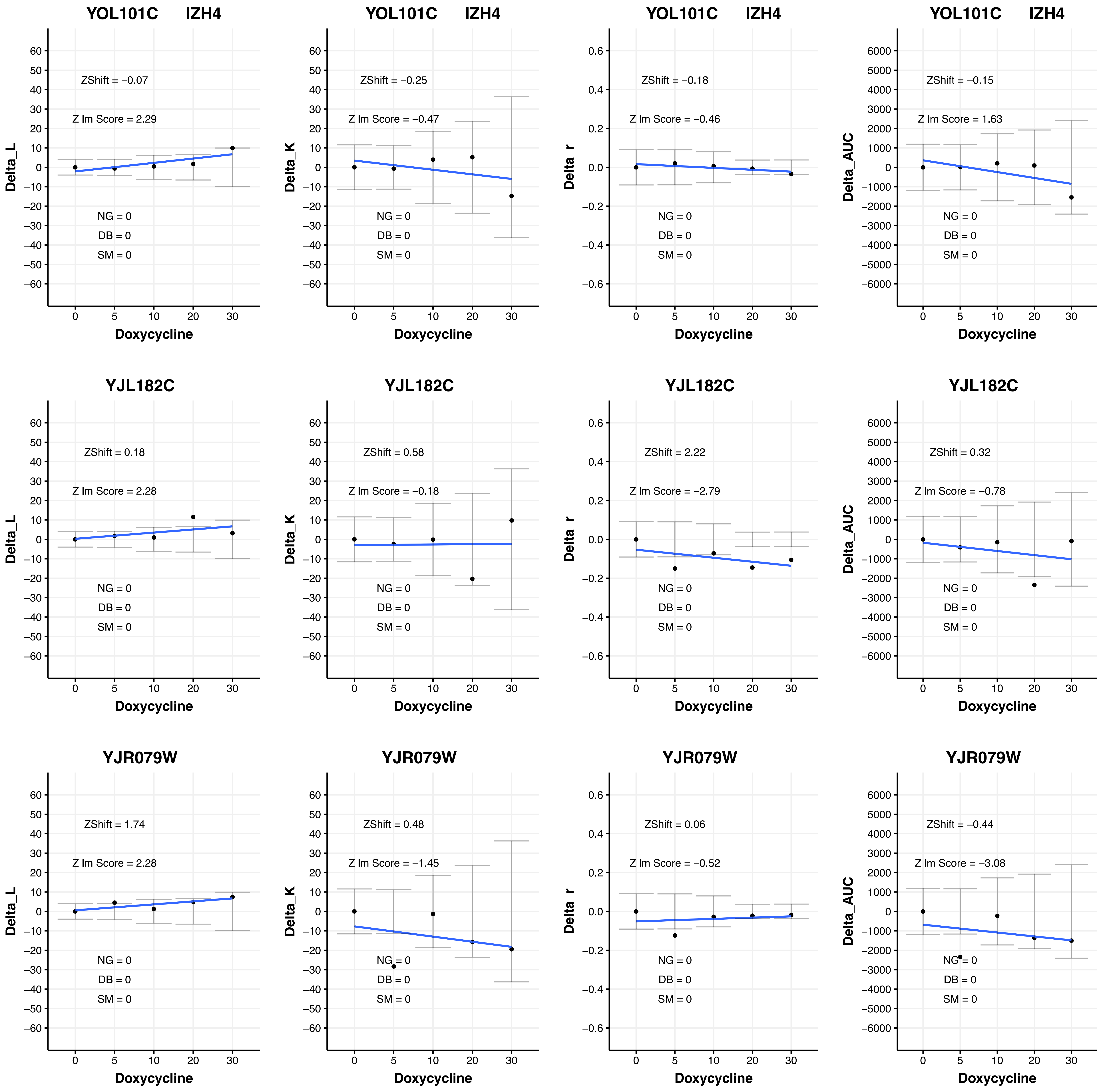

YOL101C IZH4
ZShift = −0.07
Z lm Score = 2.29
NG = 0
DB = 0
SM = 0
Delta_L
Doxycycline
YOL101C IZH4
ZShift = −0.25
Z lm Score = −0.47
NG = 0
DB = 0
SM = 0
Delta_K
Doxycycline
YOL101C IZH4
ZShift = −0.18
Z lm Score = −0.46
NG = 0
DB = 0
SM = 0
Delta_r
Doxycycline
YOL101C IZH4
ZShift = −0.15
Z lm Score = 1.63
NG = 0
DB = 0
SM = 0
Delta_AUC
Doxycycline
YJL182C
ZShift = 0.18
Z lm Score = 2.28
NG = 0
DB = 0
SM = 0
Delta_L
Doxycycline
YJL182C
ZShift = 0.58
Z lm Score = −0.18
NG = 0
DB = 0
SM = 0
Delta_K
Doxycycline
YJL182C
ZShift = 2.22
Z lm Score = −2.79
NG = 0
DB = 0
SM = 0
Delta_r
Doxycycline
YJL182C
ZShift = 0.32
Z lm Score = −0.78
NG = 0
DB = 0
SM = 0
Delta_AUC
Doxycycline
YJR079W
ZShift = 1.74
Z lm Score = 2.28
NG = 0
DB = 0
SM = 0
Delta_L
Doxycycline
YJR079W
ZShift = 0.48
Z lm Score = −1.45
NG = 0
DB = 0
SM = 0
Delta_K
Doxycycline
YJR079W
ZShift = 0.06
Z lm Score = −0.52
NG = 0
DB = 0
SM = 0
Delta_r
Doxycycline
YJR079W
ZShift = −0.44
Z lm Score = −3.08
NG = 0
DB = 0
SM = 0
Delta_AUC
Doxycycline
0 5 10 20 30

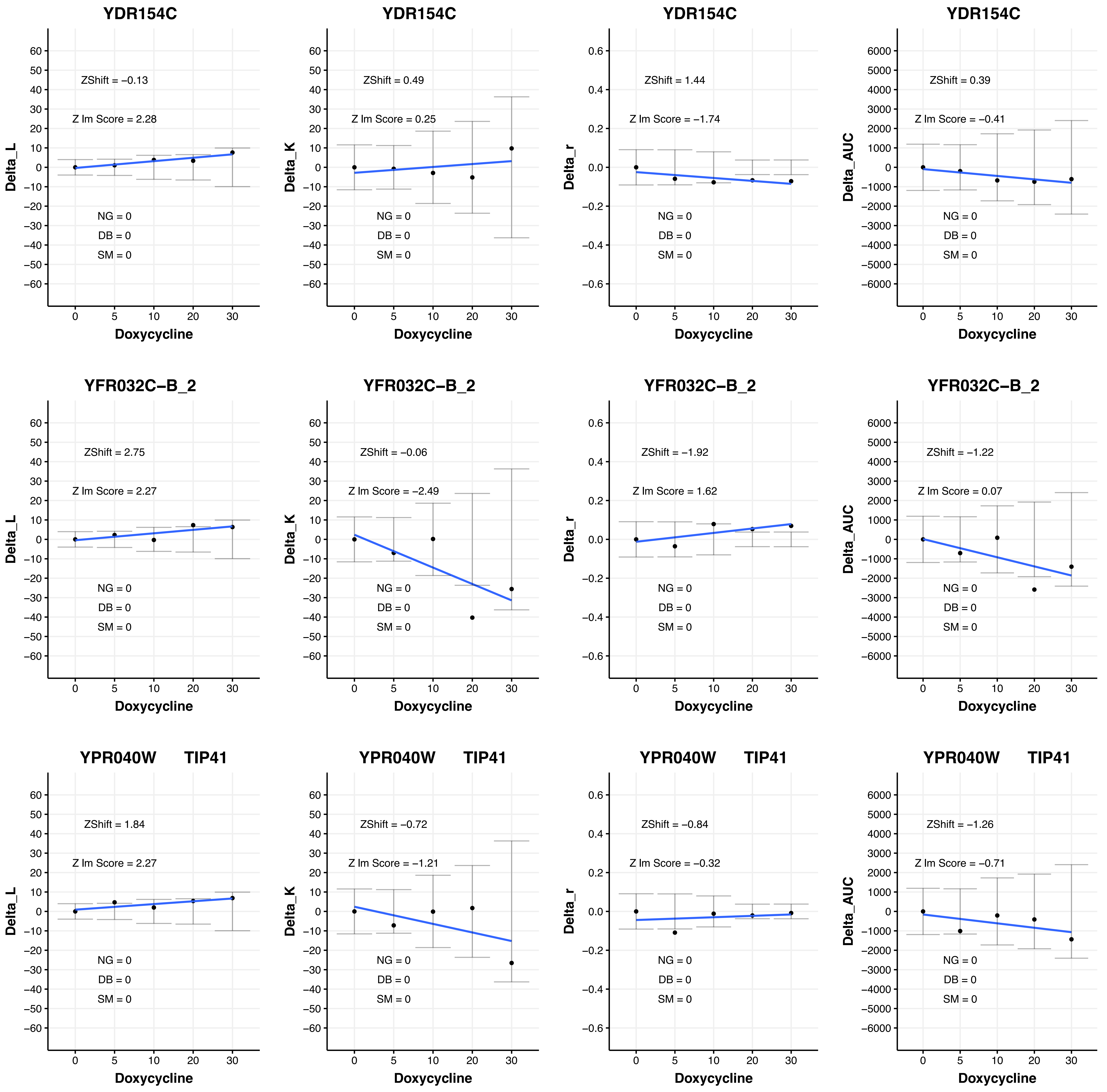

YDR154C
Delta_L
ZShift = −0.13
Z lm Score = 2.28
NG = 0
DB = 0
SM = 0
Doxycycline
YDR154C
Delta_K
ZShift = 0.49
Z lm Score = 0.25
NG = 0
DB = 0
SM = 0
Doxycycline
YDR154C
Delta_r
ZShift = 1.44
Z lm Score = −1.74
NG = 0
DB = 0
SM = 0
Doxycycline
YDR154C
Delta_AUC
ZShift = 0.39
Z lm Score = −0.41
NG = 0
DB = 0
SM = 0
Doxycycline
YFR032C−B_2
Delta_L
ZShift = 2.75
Z lm Score = 2.27
NG = 0
DB = 0
SM = 0
Doxycycline
YFR032C−B_2
Delta_K
ZShift = −0.06
Z lm Score = −2.49
NG = 0
DB = 0
SM = 0
Doxycycline
YFR032C−B_2
Delta_r
ZShift = −1.92
Z lm Score = 1.62
NG = 0
DB = 0
SM = 0
Doxycycline
YFR032C−B_2
Delta_AUC
ZShift = −1.22
Z lm Score = 0.07
NG = 0
DB = 0
SM = 0
Doxycycline
YPR040W TIP41
Delta_L
ZShift = 1.84
Z lm Score = 2.27
NG = 0
DB = 0
SM = 0
Doxycycline
YPR040W TIP41
Delta_K
ZShift = −0.72
Z lm Score = −1.21
NG = 0
DB = 0
SM = 0
Doxycycline
YPR040W TIP41
Delta_r
ZShift = −0.84
Z lm Score = −0.32
NG = 0
DB = 0
SM = 0
Doxycycline
YPR040W TIP41
Delta_AUC
ZShift = −1.26
Z lm Score = −0.71
NG = 0
DB = 0
SM = 0
Doxycycline

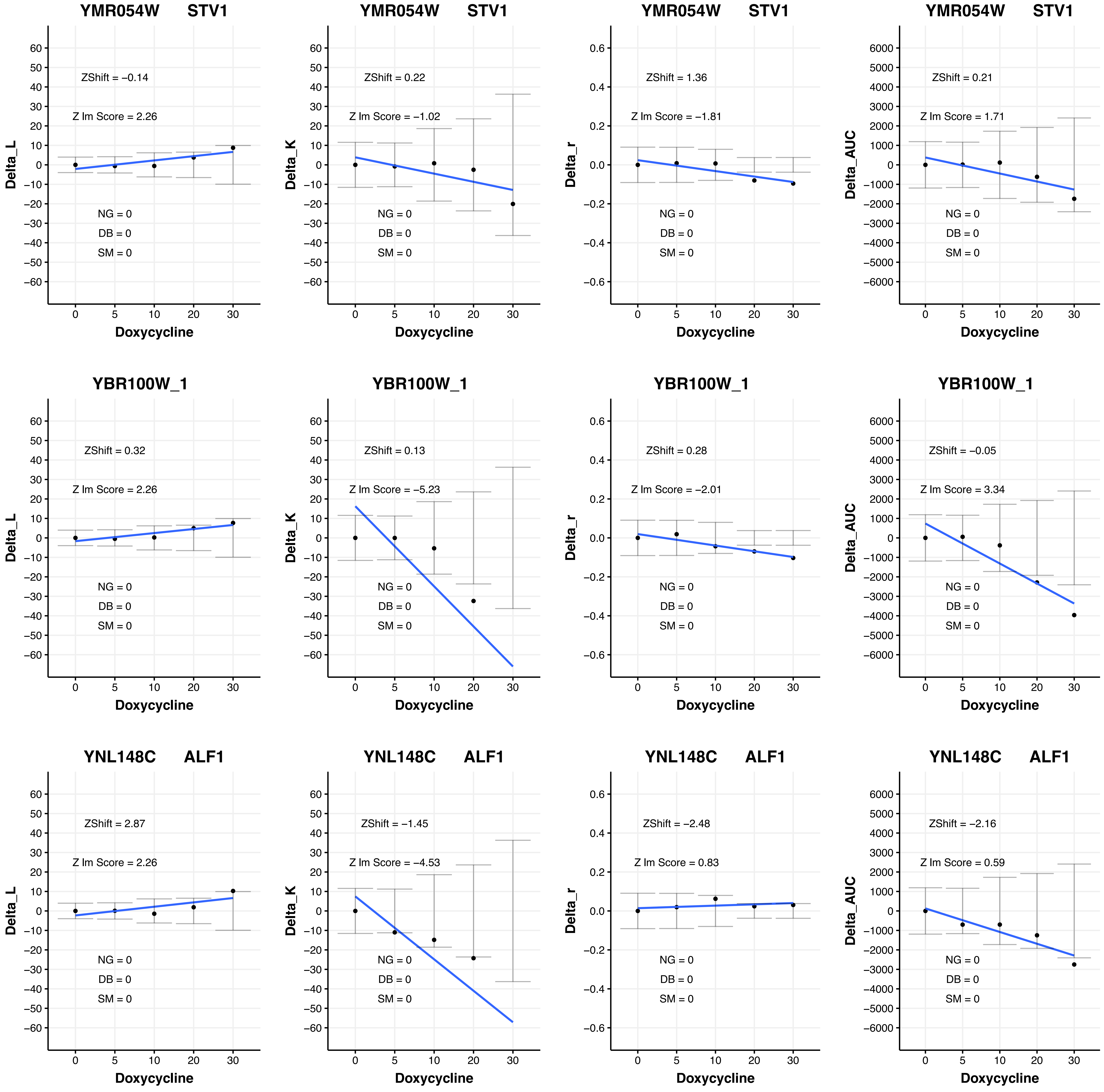
YMR054W STV1
ZShift = −0.14
Z lm Score = 2.26
NG = 0
DB = 0
SM = 0
Delta_L
Doxycycline
YMR054W STV1
ZShift = 0.22
Z lm Score = −1.02
NG = 0
DB = 0
SM = 0
Delta_K
Doxycycline
YMR054W STV1
ZShift = 1.36
Z lm Score = −1.81
NG = 0
DB = 0
SM = 0
Delta_r
Doxycycline
YMR054W STV1
ZShift = 0.21
Z lm Score = 1.71
NG = 0
DB = 0
SM = 0
Delta_AUC
Doxycycline
YBR100W_1
ZShift = 0.32
Z lm Score = 2.26
NG = 0
DB = 0
SM = 0
Delta_L
Doxycycline
YBR100W_1
ZShift = 0.13
Z lm Score = −5.23
NG = 0
DB = 0
SM = 0
Delta_K
Doxycycline
YBR100W_1
ZShift = 0.28
Z lm Score = −2.01
NG = 0
DB = 0
SM = 0
Delta_r
Doxycycline
YBR100W_1
ZShift = −0.05
Z lm Score = 3.34
NG = 0
DB = 0
SM = 0
Delta_AUC
Doxycycline
YNL148C ALF1
ZShift = 2.87
Z lm Score = 2.26
NG = 0
DB = 0
SM = 0
Delta_L
Doxycycline
YNL148C ALF1
ZShift = −1.45
Z lm Score = −4.53
NG = 0
DB = 0
SM = 0
Delta_K
Doxycycline
YNL148C ALF1
ZShift = −2.48
Z lm Score = 0.83
NG = 0
DB = 0
SM = 0
Delta_r
Doxycycline
YNL148C ALF1
ZShift = −2.16
Z lm Score = 0.59
NG = 0
DB = 0
SM = 0
Delta_AUC
Doxycycline

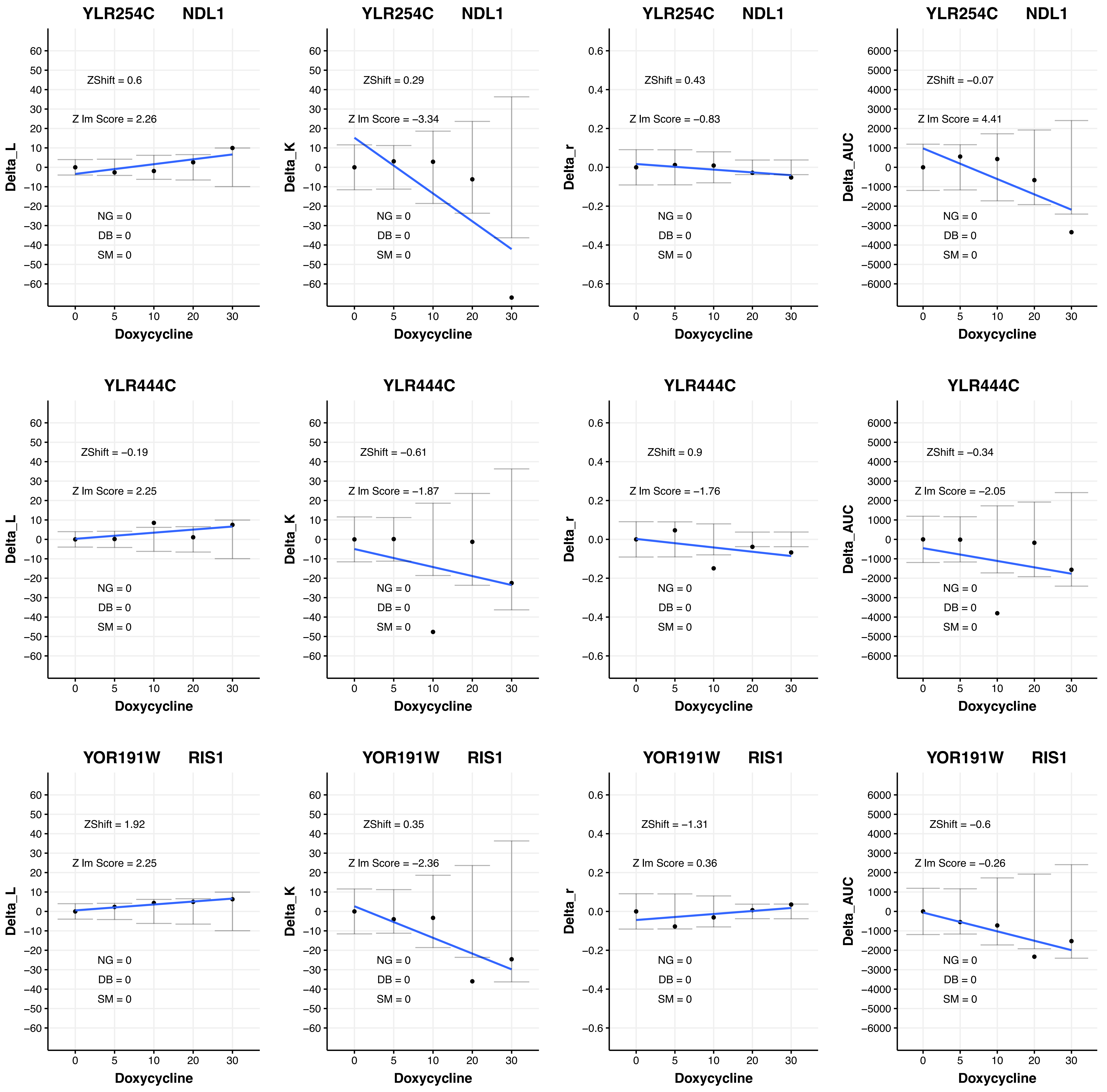
YLR254C NDL1
ZShift = 0.6
Z lm Score = 2.26
NG = 0
DB = 0
SM = 0
Delta_L
Doxycycline
YLR254C NDL1
ZShift = 0.29
Z lm Score = −3.34
NG = 0
DB = 0
SM = 0
Delta_K
Doxycycline
YLR254C NDL1
ZShift = 0.43
Z lm Score = −0.83
NG = 0
DB = 0
SM = 0
Delta_r
Doxycycline
YLR254C NDL1
ZShift = −0.07
Z lm Score = 4.41
NG = 0
DB = 0
SM = 0
Delta_AUC
Doxycycline
YLR444C
ZShift = −0.19
Z lm Score = 2.25
NG = 0
DB = 0
SM = 0
Delta_L
Doxycycline
YLR444C
ZShift = −0.61
Z lm Score = −1.87
NG = 0
DB = 0
SM = 0
Delta_K
Doxycycline
YLR444C
ZShift = 0.9
Z lm Score = −1.76
NG = 0
DB = 0
SM = 0
Delta_r
Doxycycline
YLR444C
ZShift = −0.34
Z lm Score = −2.05
NG = 0
DB = 0
SM = 0
Delta_AUC
Doxycycline
YOR191W RIS1
ZShift = 1.92
Z lm Score = 2.25
NG = 0
DB = 0
SM = 0
Delta_L
Doxycycline
YOR191W RIS1
ZShift = 0.35
Z lm Score = −2.36
NG = 0
DB = 0
SM = 0
Delta_K
Doxycycline
YOR191W RIS1
ZShift = −1.31
Z lm Score = 0.36
NG = 0
DB = 0
SM = 0
Delta_r
Doxycycline
YOR191W RIS1
ZShift = −0.6
Z lm Score = −0.26
NG = 0
DB = 0
SM = 0
Delta_AUC
Doxycycline

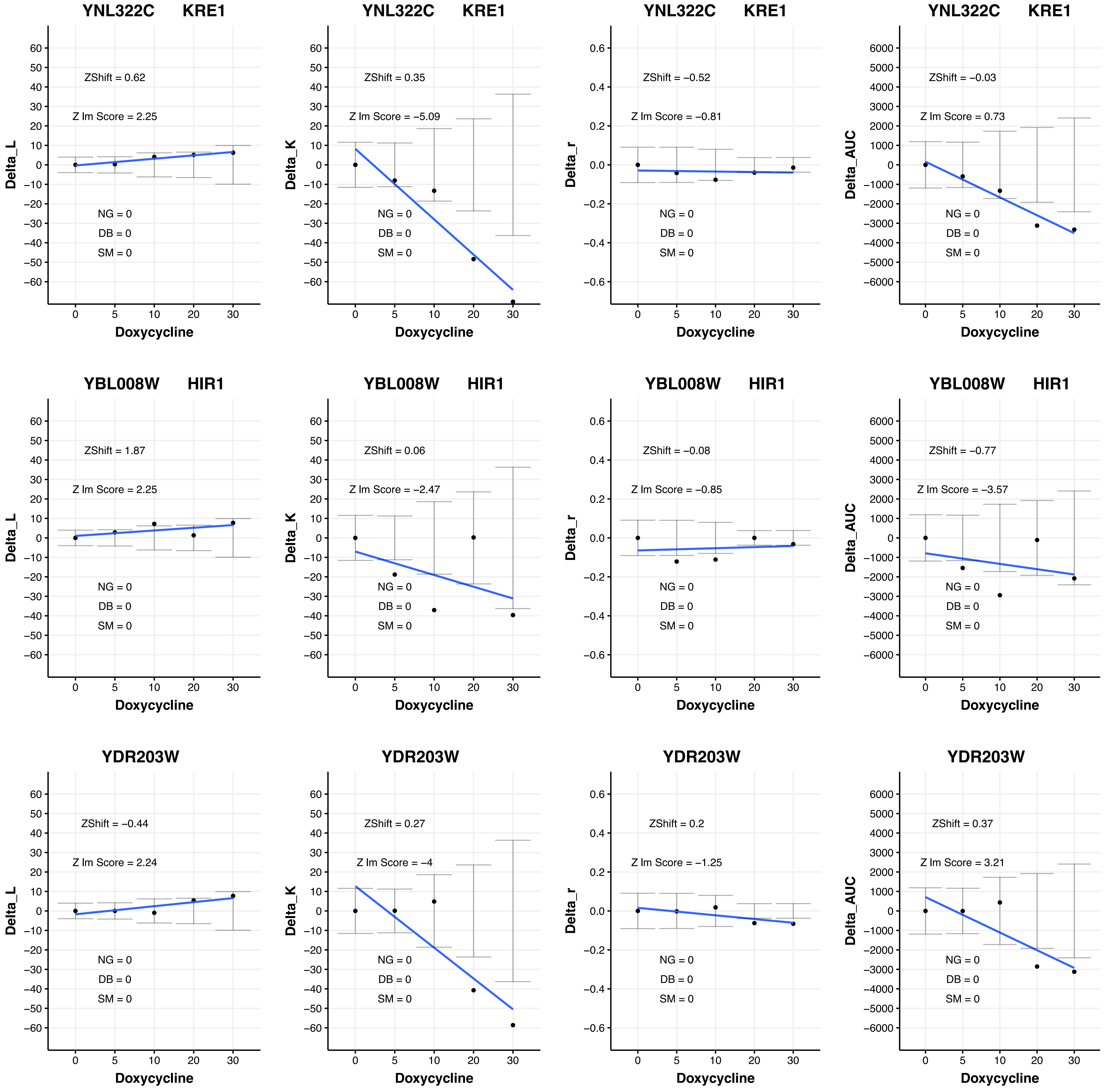

YNL322C KRE1
ZShift = 0.62
Z lm Score = 2.25
NG = 0
DB = 0
SM = 0
Delta_L
Doxycycline
YNL322C KRE1
ZShift = 0.35
Z lm Score = −5.09
NG = 0
DB = 0
SM = 0
Delta_K
Doxycycline
YNL322C KRE1
ZShift = −0.52
Z lm Score = −0.81
NG = 0
DB = 0
SM = 0
Delta_r
Doxycycline
YNL322C KRE1
ZShift = −0.03
Z lm Score = 0.73
NG = 0
DB = 0
SM = 0
Delta_AUC
Doxycycline
YBL008W HIR1
ZShift = 1.87
Z lm Score = 2.25
NG = 0
DB = 0
SM = 0
Delta_L
Doxycycline
YBL008W HIR1
ZShift = 0.06
Z lm Score = −2.47
NG = 0
DB = 0
SM = 0
Delta_K
Doxycycline
YBL008W HIR1
ZShift = −0.08
Z lm Score = −0.85
NG = 0
DB = 0
SM = 0
Delta_r
Doxycycline
YBL008W HIR1
ZShift = −0.77
Z lm Score = −3.57
NG = 0
DB = 0
SM = 0
Delta_AUC
Doxycycline
YDR203W
ZShift = −0.44
Z lm Score = 2.24
NG = 0
DB = 0
SM = 0
Delta_L
Doxycycline
YDR203W
ZShift = 0.27
Z lm Score = −4
NG = 0
DB = 0
SM = 0
Delta_K
Doxycycline
YDR203W
ZShift = 0.2
Z lm Score = −1.25
NG = 0
DB = 0
SM = 0
Delta_r
Doxycycline
YDR203W
ZShift = 0.37
Z lm Score = 3.21
NG = 0
DB = 0
SM = 0
Delta_AUC
Doxycycline

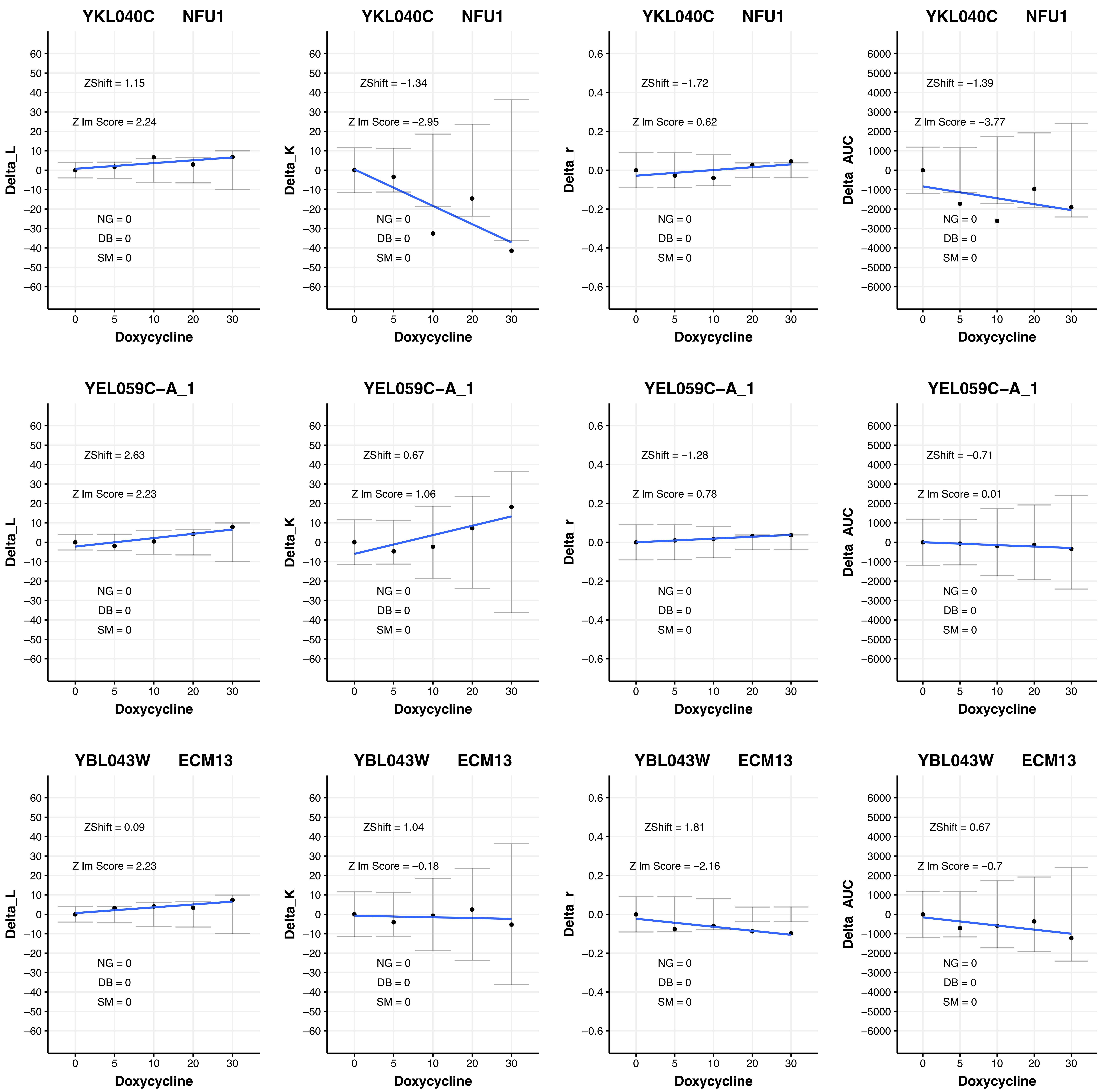

YKL040C NFU1
ZShift = 1.15
Z lm Score = 2.24
NG = 0
DB = 0
SM = 0
Delta_L
Doxycycline
YKL040C NFU1
ZShift = −1.34
Z lm Score = −2.95
NG = 0
DB = 0
SM = 0
Delta_K
Doxycycline
YKL040C NFU1
ZShift = −1.72
Z lm Score = 0.62
NG = 0
DB = 0
SM = 0
Delta_r
Doxycycline
YKL040C NFU1
ZShift = −1.39
Z lm Score = −3.77
NG = 0
DB = 0
SM = 0
Delta_AUC
Doxycycline
YEL059C−A_1
ZShift = 2.63
Z lm Score = 2.23
NG = 0
DB = 0
SM = 0
Delta_L
Doxycycline
YEL059C−A_1
ZShift = 0.67
Z lm Score = 1.06
NG = 0
DB = 0
SM = 0
Delta_K
Doxycycline
YEL059C−A_1
ZShift = −1.28
Z lm Score = 0.78
NG = 0
DB = 0
SM = 0
Delta_r
Doxycycline
YEL059C−A_1
ZShift = −0.71
Z lm Score = 0.01
NG = 0
DB = 0
SM = 0
Delta_AUC
Doxycycline
YBL043W ECM13
ZShift = 0.09
Z lm Score = 2.23
NG = 0
DB = 0
SM = 0
Delta_L
Doxycycline
YBL043W ECM13
ZShift = 1.04
Z lm Score = −0.18
NG = 0
DB = 0
SM = 0
Delta_K
Doxycycline
YBL043W ECM13
ZShift = 1.81
Z lm Score = −2.16
NG = 0
DB = 0
SM = 0
Delta_r
Doxycycline
YBL043W ECM13
ZShift = 0.67
Z lm Score = −0.7
NG = 0
DB = 0
SM = 0
Delta_AUC
Doxycycline

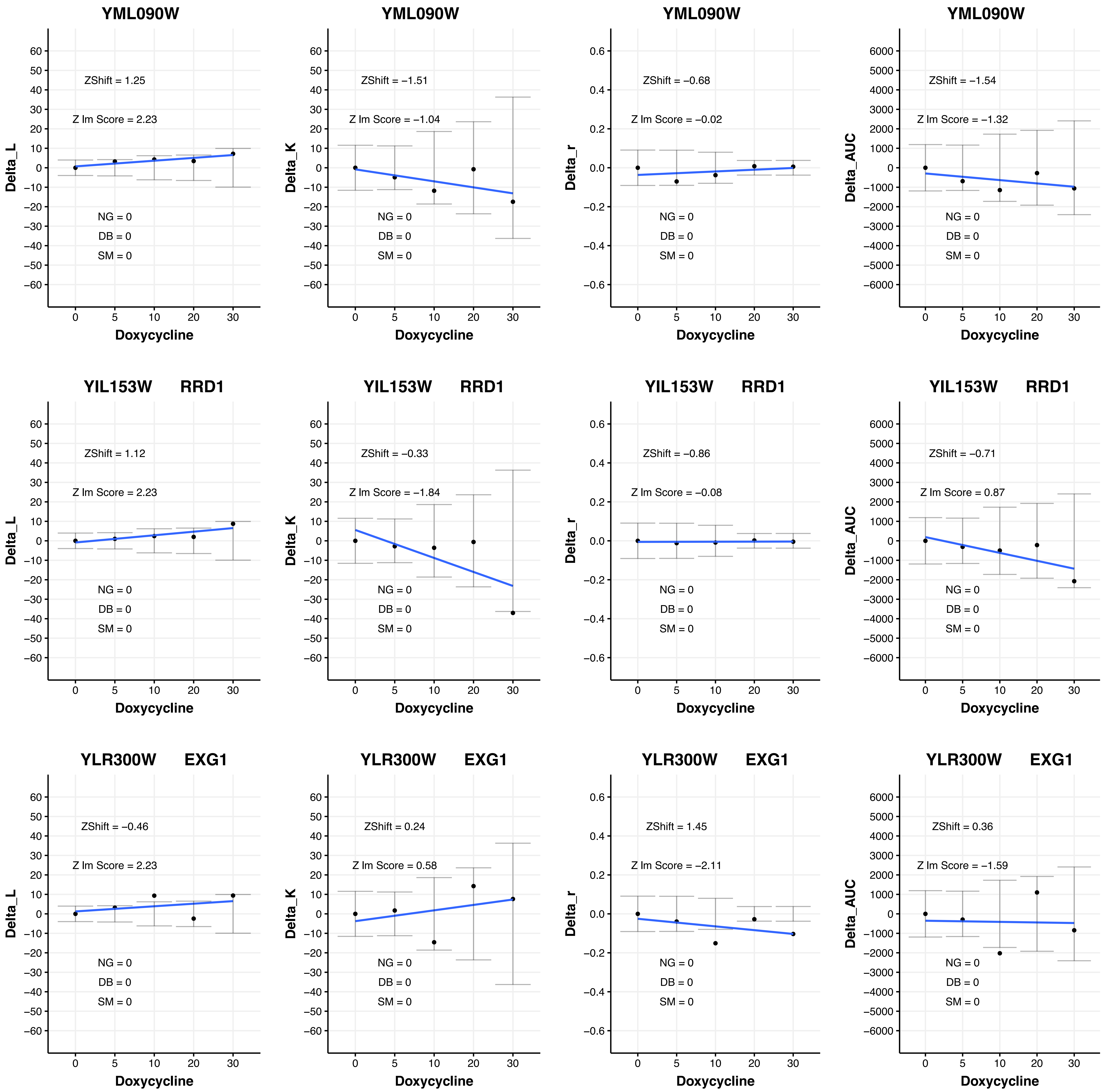
YML090W
ZShift = 1.25
Z lm Score = 2.23
NG = 0
DB = 0
SM = 0
Delta_L
Doxycycline
YML090W
ZShift = −1.51
Z lm Score = −1.04
NG = 0
DB = 0
SM = 0
Delta_K
Doxycycline
YML090W
ZShift = −0.68
Z lm Score = −0.02
NG = 0
DB = 0
SM = 0
Delta_r
Doxycycline
YML090W
ZShift = −1.54
Z lm Score = −1.32
NG = 0
DB = 0
SM = 0
Delta_AUC
Doxycycline
YIL153W RRD1
ZShift = 1.12
Z lm Score = 2.23
NG = 0
DB = 0
SM = 0
Delta_L
Doxycycline
YIL153W RRD1
ZShift = −0.33
Z lm Score = −1.84
NG = 0
DB = 0
SM = 0
Delta_K
Doxycycline
YIL153W RRD1
ZShift = −0.86
Z lm Score = −0.08
NG = 0
DB = 0
SM = 0
Delta_r
Doxycycline
YIL153W RRD1
ZShift = −0.71
Z lm Score = 0.87
NG = 0
DB = 0
SM = 0
Delta_AUC
Doxycycline
YLR300W EXG1
ZShift = −0.46
Z lm Score = 2.23
NG = 0
DB = 0
SM = 0
Delta_L
Doxycycline
YLR300W EXG1
ZShift = 0.24
Z lm Score = 0.58
NG = 0
DB = 0
SM = 0
Delta_K
Doxycycline
YLR300W EXG1
ZShift = 1.45
Z lm Score = −2.11
NG = 0
DB = 0
SM = 0
Delta_r
Doxycycline
YLR300W EXG1
ZShift = 0.36
Z lm Score = −1.59
NG = 0
DB = 0
SM = 0
Delta_AUC
Doxycycline

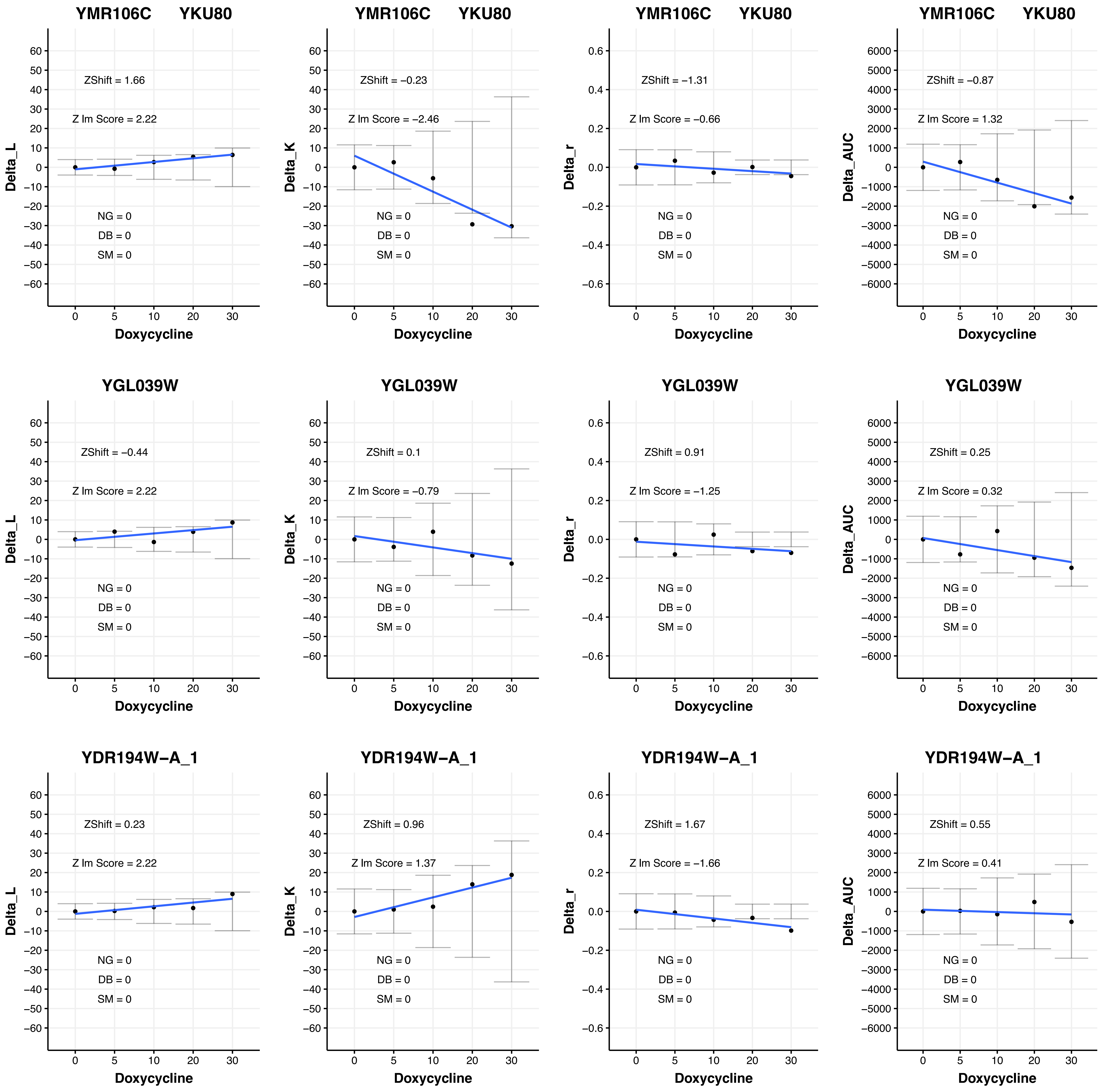

YMR106C YKU80
ZShift = 1.66
Z lm Score = 2.22
NG = 0
DB = 0
SM = 0
Delta_L
Doxycycline
YMR106C YKU80
ZShift = −0.23
Z lm Score = −2.46
NG = 0
DB = 0
SM = 0
Delta_K
Doxycycline
YMR106C YKU80
ZShift = −1.31
Z lm Score = −0.66
NG = 0
DB = 0
SM = 0
Delta_r
Doxycycline
YMR106C YKU80
ZShift = −0.87
Z lm Score = 1.32
NG = 0
DB = 0
SM = 0
Delta_AUC
Doxycycline
YGL039W
ZShift = −0.44
Z lm Score = 2.22
NG = 0
DB = 0
SM = 0
Delta_L
Doxycycline
YGL039W
ZShift = 0.1
Z lm Score = −0.79
NG = 0
DB = 0
SM = 0
Delta_K
Doxycycline
YGL039W
ZShift = 0.91
Z lm Score = −1.25
NG = 0
DB = 0
SM = 0
Delta_r
Doxycycline
YGL039W
ZShift = 0.25
Z lm Score = 0.32
NG = 0
DB = 0
SM = 0
Delta_AUC
Doxycycline
YDR194W−A_1
ZShift = 0.23
Z lm Score = 2.22
NG = 0
DB = 0
SM = 0
Delta_L
Doxycycline
YDR194W−A_1
ZShift = 0.96
Z lm Score = 1.37
NG = 0
DB = 0
SM = 0
Delta_K
Doxycycline
YDR194W−A_1
ZShift = 1.67
Z lm Score = −1.66
NG = 0
DB = 0
SM = 0
Delta_r
Doxycycline
YDR194W−A_1
ZShift = 0.55
Z lm Score = 0.41
NG = 0
DB = 0
SM = 0
Delta_AUC
Doxycycline

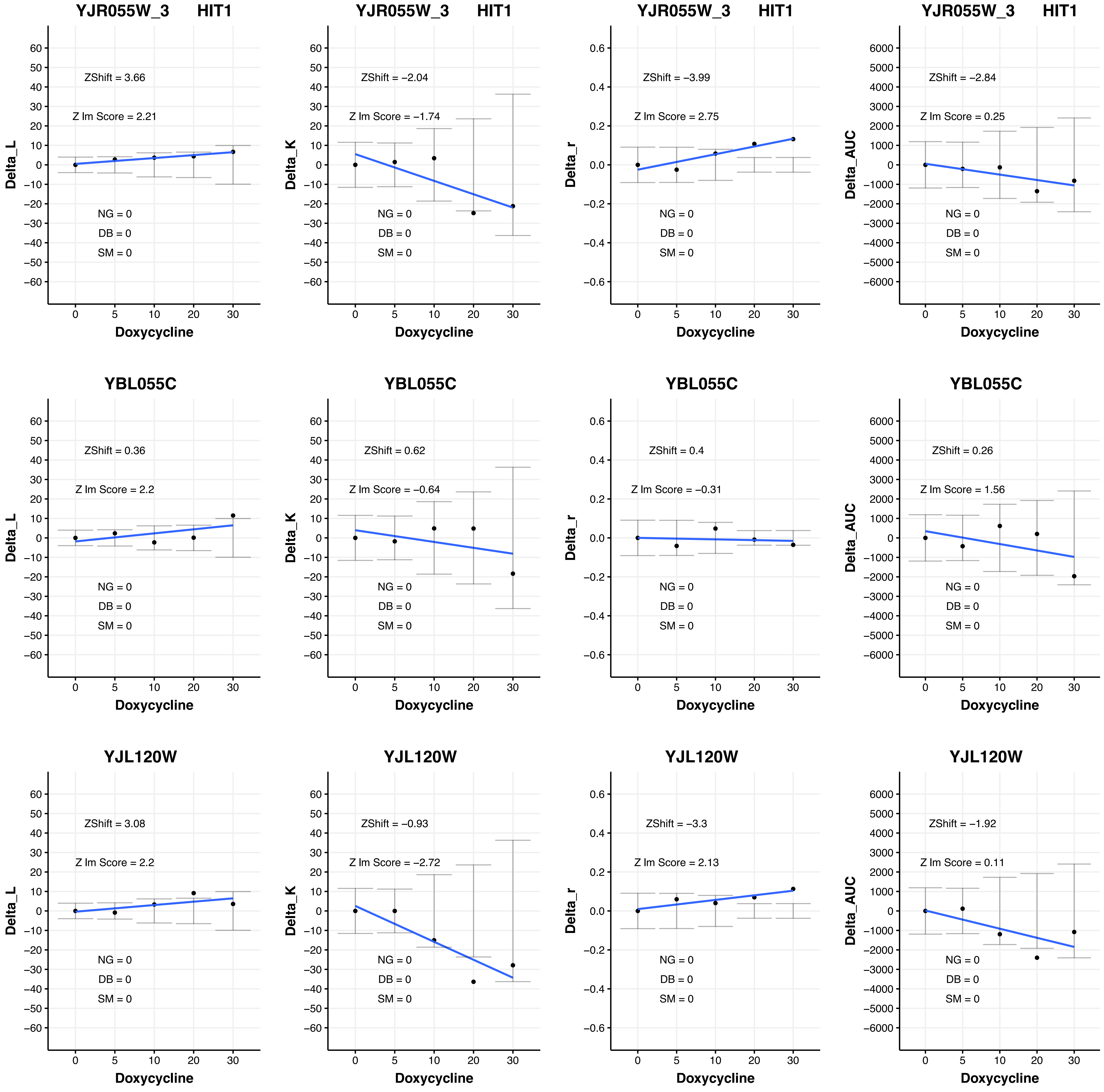
YJR055W_3 HIT1
ZShift = 3.66
Z lm Score = 2.21
NG = 0
DB = 0
SM = 0
Delta_L
Doxycycline
YJR055W_3 HIT1
ZShift = −2.04
Z lm Score = −1.74
NG = 0
DB = 0
SM = 0
Delta_K
Doxycycline
YJR055W_3 HIT1
ZShift = −3.99
Z lm Score = 2.75
NG = 0
DB = 0
SM = 0
Delta_r
Doxycycline
YJR055W_3 HIT1
ZShift = −2.84
Z lm Score = 0.25
NG = 0
DB = 0
SM = 0
Delta_AUC
Doxycycline
YBL055C
ZShift = 0.36
Z lm Score = 2.2
NG = 0
DB = 0
SM = 0
Delta_L
Doxycycline
YBL055C
ZShift = 0.62
Z lm Score = −0.64
NG = 0
DB = 0
SM = 0
Delta_K
Doxycycline
YBL055C
ZShift = 0.4
Z lm Score = −0.31
NG = 0
DB = 0
SM = 0
Delta_r
Doxycycline
YBL055C
ZShift = 0.26
Z lm Score = 1.56
NG = 0
DB = 0
SM = 0
Delta_AUC
Doxycycline
YJL120W
ZShift = 3.08
Z lm Score = 2.2
NG = 0
DB = 0
SM = 0
Delta_L
Doxycycline
YJL120W
ZShift = −0.93
Z lm Score = −2.72
NG = 0
DB = 0
SM = 0
Delta_K
Doxycycline
YJL120W
ZShift = −3.3
Z lm Score = 2.13
NG = 0
DB = 0
SM = 0
Delta_r
Doxycycline
YJL120W
ZShift = −1.92
Z lm Score = 0.11
NG = 0
DB = 0
SM = 0
Delta_AUC
Doxycycline

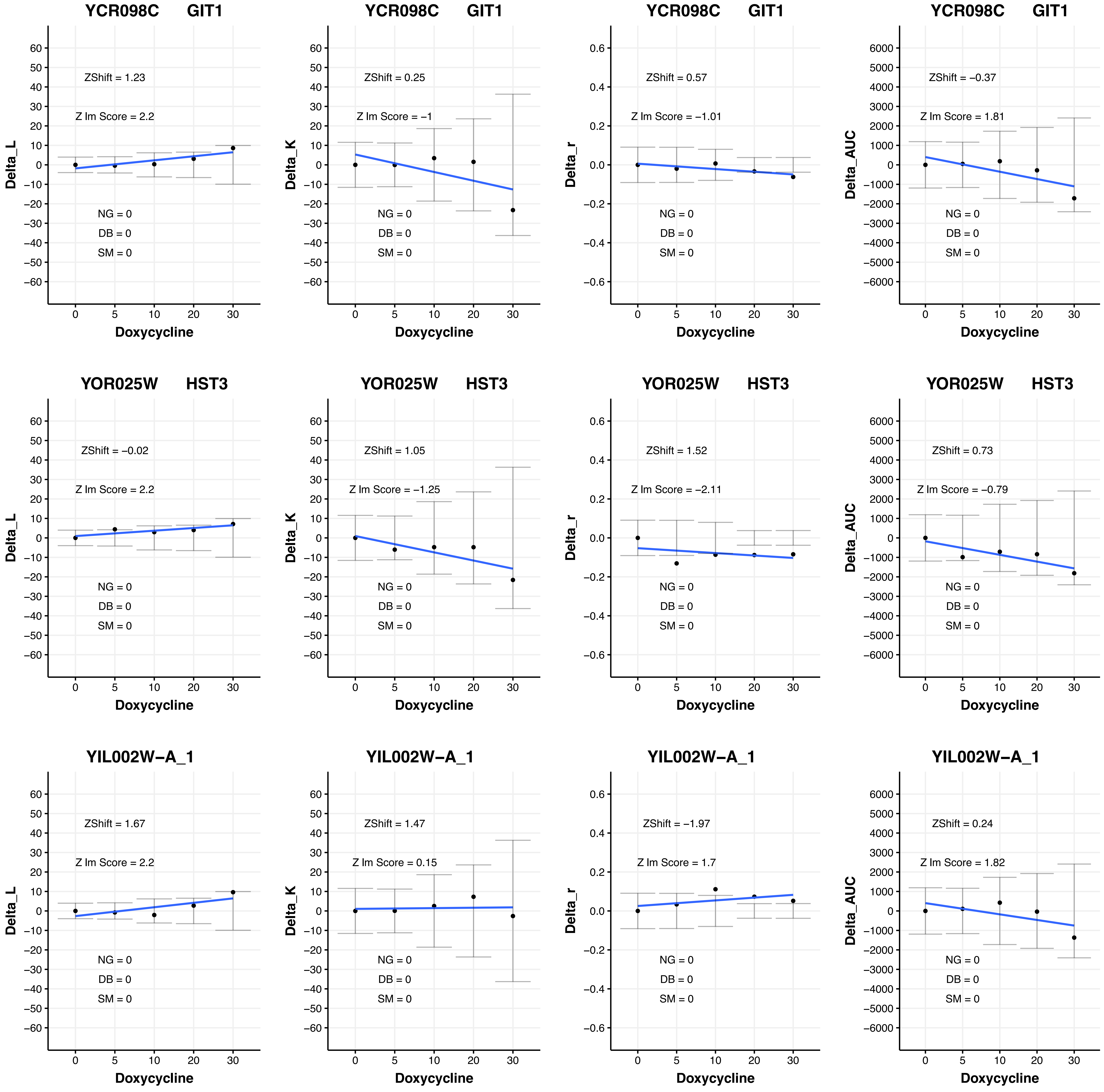
YCR098C GIT1
ZShift = 1.23
Z lm Score = 2.2
NG = 0
DB = 0
SM = 0
Delta_L
Doxycycline
YCR098C GIT1
ZShift = 0.25
Z lm Score = −1
NG = 0
DB = 0
SM = 0
Delta_K
Doxycycline
YCR098C GIT1
ZShift = 0.57
Z lm Score = −1.01
NG = 0
DB = 0
SM = 0
Delta_r
Doxycycline
YCR098C GIT1
ZShift = −0.37
Z lm Score = 1.81
NG = 0
DB = 0
SM = 0
Delta_AUC
Doxycycline
YOR025W HST3
ZShift = −0.02
Z lm Score = 2.2
NG = 0
DB = 0
SM = 0
Delta_L
Doxycycline
YOR025W HST3
ZShift = 1.05
Z lm Score = −1.25
NG = 0
DB = 0
SM = 0
Delta_K
Doxycycline
YOR025W HST3
ZShift = 1.52
Z lm Score = −2.11
NG = 0
DB = 0
SM = 0
Delta_r
Doxycycline
YOR025W HST3
ZShift = 0.73
Z lm Score = −0.79
NG = 0
DB = 0
SM = 0
Delta_AUC
Doxycycline
YIL002W−A_1
ZShift = 1.67
Z lm Score = 2.2
NG = 0
DB = 0
SM = 0
Delta_L
Doxycycline
YIL002W−A_1
ZShift = 1.47
Z lm Score = 0.15
NG = 0
DB = 0
SM = 0
Delta_K
Doxycycline
YIL002W−A_1
ZShift = −1.97
Z lm Score = 1.7
NG = 0
DB = 0
SM = 0
Delta_r
Doxycycline
YIL002W−A_1
ZShift = 0.24
Z lm Score = 1.82
NG = 0
DB = 0
SM = 0
Delta_AUC
Doxycycline

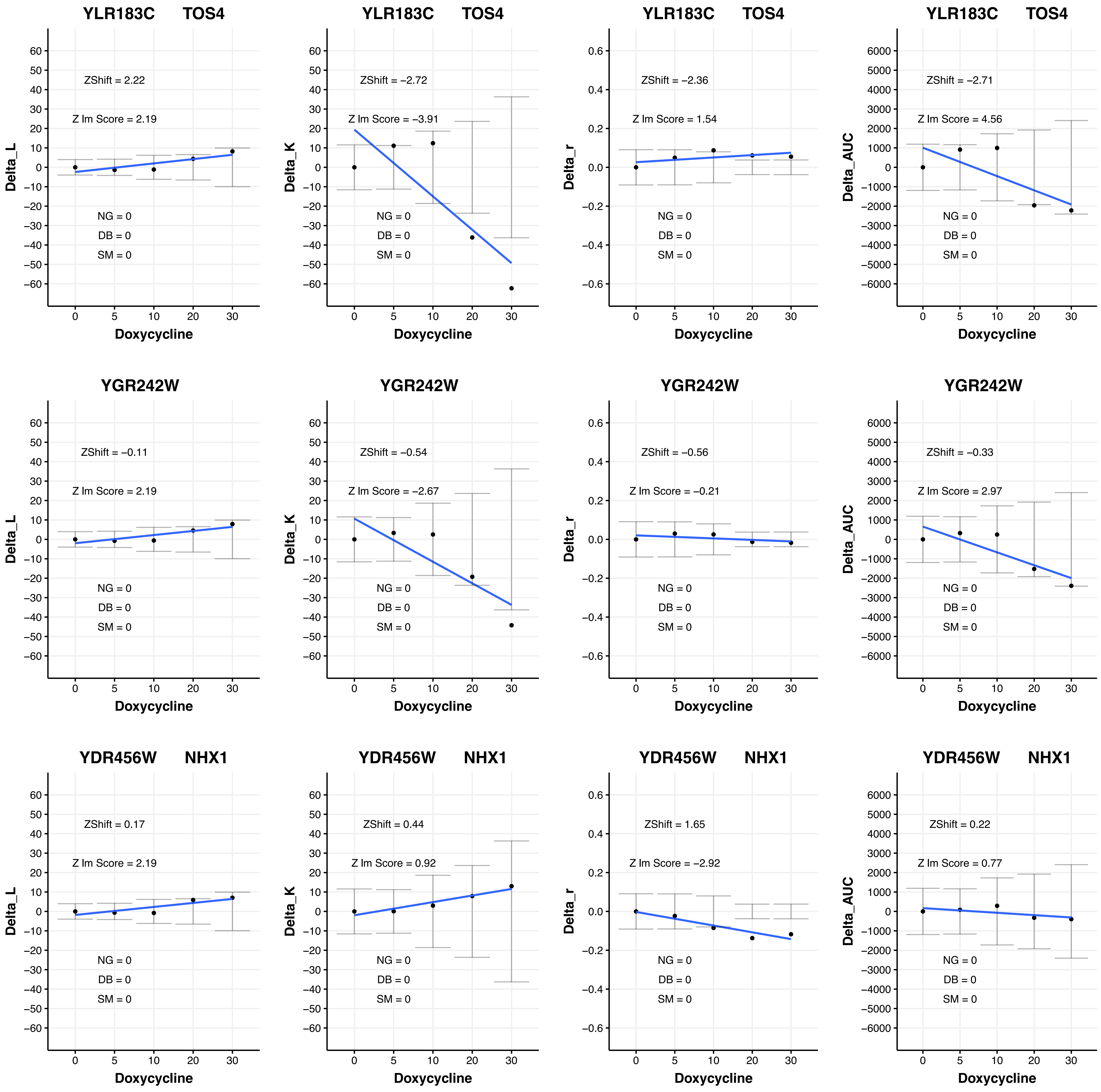
YLR183C TOS4
ZShift = 2.22
Z lm Score = 2.19
NG = 0
DB = 0
SM = 0
Delta_L
Doxycycline
YLR183C TOS4
ZShift = −2.72
Z lm Score = −3.91
NG = 0
DB = 0
SM = 0
Delta_K
Doxycycline
YLR183C TOS4
ZShift = −2.36
Z lm Score = 1.54
NG = 0
DB = 0
SM = 0
Delta_r
Doxycycline
YLR183C TOS4
ZShift = −2.71
Z lm Score = 4.56
NG = 0
DB = 0
SM = 0
Delta_AUC
Doxycycline
YGR242W
ZShift = −0.11
Z lm Score = 2.19
NG = 0
DB = 0
SM = 0
Delta_L
Doxycycline
YGR242W
ZShift = −0.54
Z lm Score = −2.67
NG = 0
DB = 0
SM = 0
Delta_K
Doxycycline
YGR242W
ZShift = −0.56
Z lm Score = −0.21
NG = 0
DB = 0
SM = 0
Delta_r
Doxycycline
YGR242W
ZShift = −0.33
Z lm Score = 2.97
NG = 0
DB = 0
SM = 0
Delta_AUC
Doxycycline
YDR456W NHX1
ZShift = 0.17
Z lm Score = 2.19
NG = 0
DB = 0
SM = 0
Delta_L
Doxycycline
YDR456W NHX1
ZShift = 0.44
Z lm Score = 0.92
NG = 0
DB = 0
SM = 0
Delta_K
Doxycycline
YDR456W NHX1
ZShift = 1.65
Z lm Score = −2.92
NG = 0
DB = 0
SM = 0
Delta_r
Doxycycline
YDR456W NHX1
ZShift = 0.22
Z lm Score = 0.77
NG = 0
DB = 0
SM = 0
Delta_AUC
Doxycycline

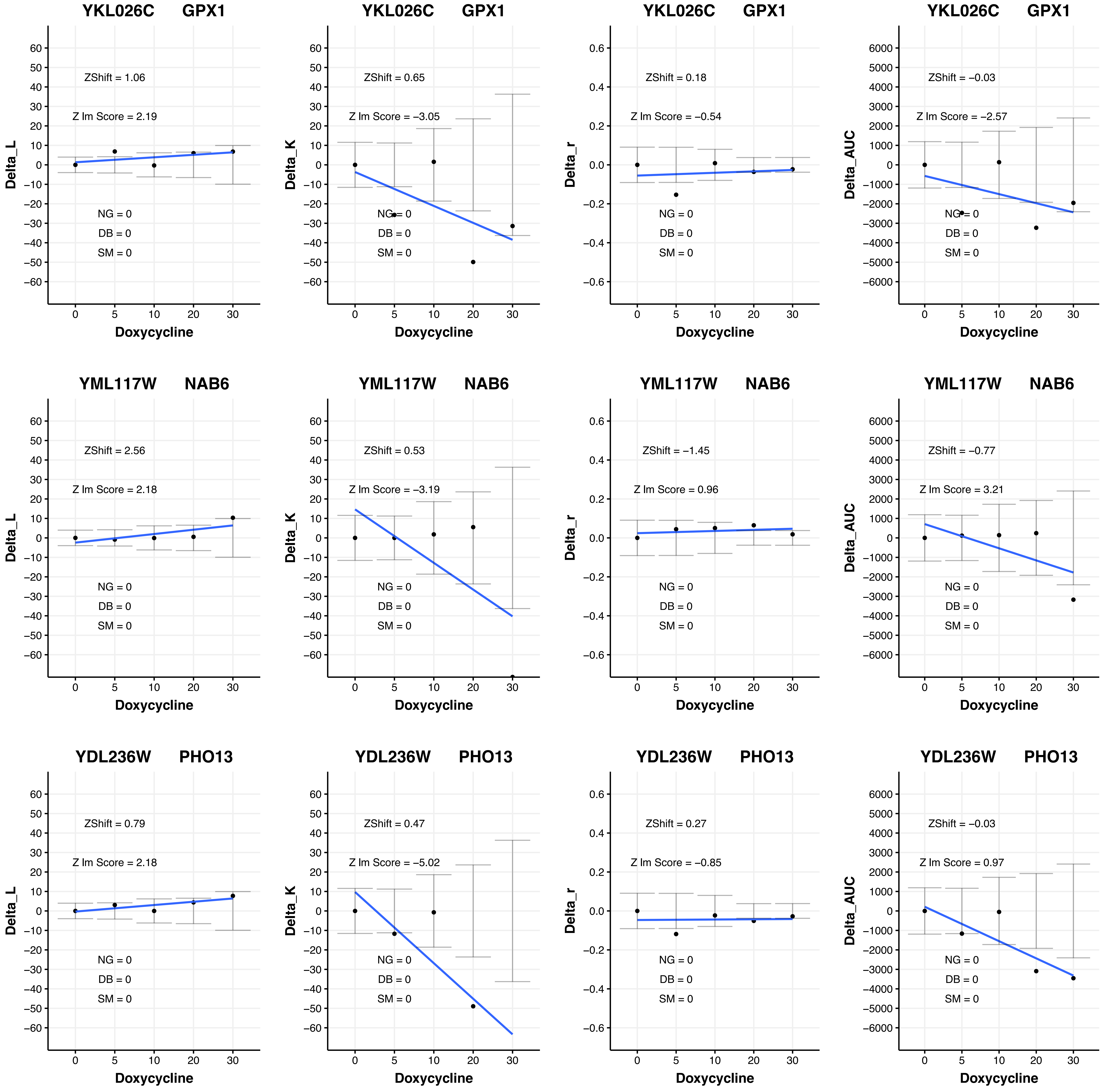

YKL026C GPX1
ZShift = 1.06
Z lm Score = 2.19
NG = 0
DB = 0
SM = 0
Delta_L
Doxycycline
YKL026C GPX1
ZShift = 0.65
Z lm Score = −3.05
NG = 0
DB = 0
SM = 0
Delta_K
Doxycycline
YKL026C GPX1
ZShift = 0.18
Z lm Score = −0.54
NG = 0
DB = 0
SM = 0
Delta_r
Doxycycline
YKL026C GPX1
ZShift = −0.03
Z lm Score = −2.57
NG = 0
DB = 0
SM = 0
Delta_AUC
Doxycycline
YML117W NAB6
ZShift = 2.56
Z lm Score = 2.18
NG = 0
DB = 0
SM = 0
Delta_L
Doxycycline
YML117W NAB6
ZShift = 0.53
Z lm Score = −3.19
NG = 0
DB = 0
SM = 0
Delta_K
Doxycycline
YML117W NAB6
ZShift = −1.45
Z lm Score = 0.96
NG = 0
DB = 0
SM = 0
Delta_r
Doxycycline
YML117W NAB6
ZShift = −0.77
Z lm Score = 3.21
NG = 0
DB = 0
SM = 0
Delta_AUC
Doxycycline
YDL236W PHO13
ZShift = 0.79
Z lm Score = 2.18
NG = 0
DB = 0
SM = 0
Delta_L
Doxycycline
YDL236W PHO13
ZShift = 0.47
Z lm Score = −5.02
NG = 0
DB = 0
SM = 0
Delta_K
Doxycycline
YDL236W PHO13
ZShift = 0.27
Z lm Score = −0.85
NG = 0
DB = 0
SM = 0
Delta_r
Doxycycline
YDL236W PHO13
ZShift = −0.03
Z lm Score = 0.97
NG = 0
DB = 0
SM = 0
Delta_AUC
Doxycycline

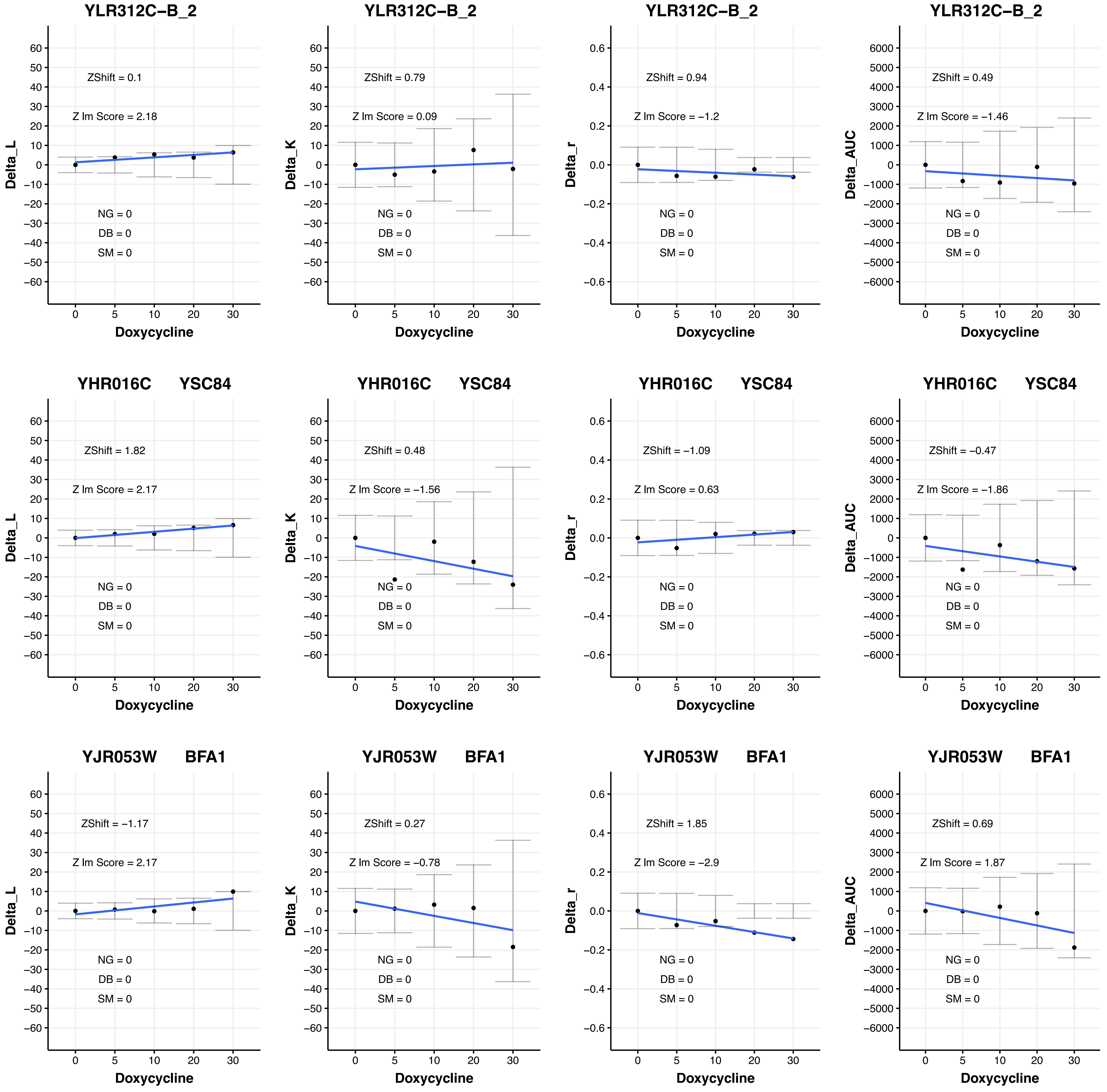

YLR312C−B_2
ZShift = 0.1
Z lm Score = 2.18
NG = 0
DB = 0
SM = 0
Delta_L
Doxycycline
YLR312C−B_2
ZShift = 0.79
Z lm Score = 0.09
NG = 0
DB = 0
SM = 0
Delta_K
Doxycycline
YLR312C−B_2
ZShift = 0.94
Z lm Score = −1.2
NG = 0
DB = 0
SM = 0
Delta_r
Doxycycline
YLR312C−B_2
ZShift = 0.49
Z lm Score = −1.46
NG = 0
DB = 0
SM = 0
Delta_AUC
Doxycycline
YHR016C YSC84
ZShift = 1.82
Z lm Score = 2.17
NG = 0
DB = 0
SM = 0
Delta_L
Doxycycline
YHR016C YSC84
ZShift = 0.48
Z lm Score = −1.56
NG = 0
DB = 0
SM = 0
Delta_K
Doxycycline
YHR016C YSC84
ZShift = −1.09
Z lm Score = 0.63
NG = 0
DB = 0
SM = 0
Delta_r
Doxycycline
YHR016C YSC84
ZShift = −0.47
Z lm Score = −1.86
NG = 0
DB = 0
SM = 0
Delta_AUC
Doxycycline
YJR053W BFA1
ZShift = −1.17
Z lm Score = 2.17
NG = 0
DB = 0
SM = 0
Delta_L
Doxycycline
YJR053W BFA1
ZShift = 0.27
Z lm Score = −0.78
NG = 0
DB = 0
SM = 0
Delta_K
Doxycycline
YJR053W BFA1
ZShift = 1.85
Z lm Score = −2.9
NG = 0
DB = 0
SM = 0
Delta_r
Doxycycline
YJR053W BFA1
ZShift = 0.69
Z lm Score = 1.87
NG = 0
DB = 0
SM = 0
Delta_AUC
Doxycycline

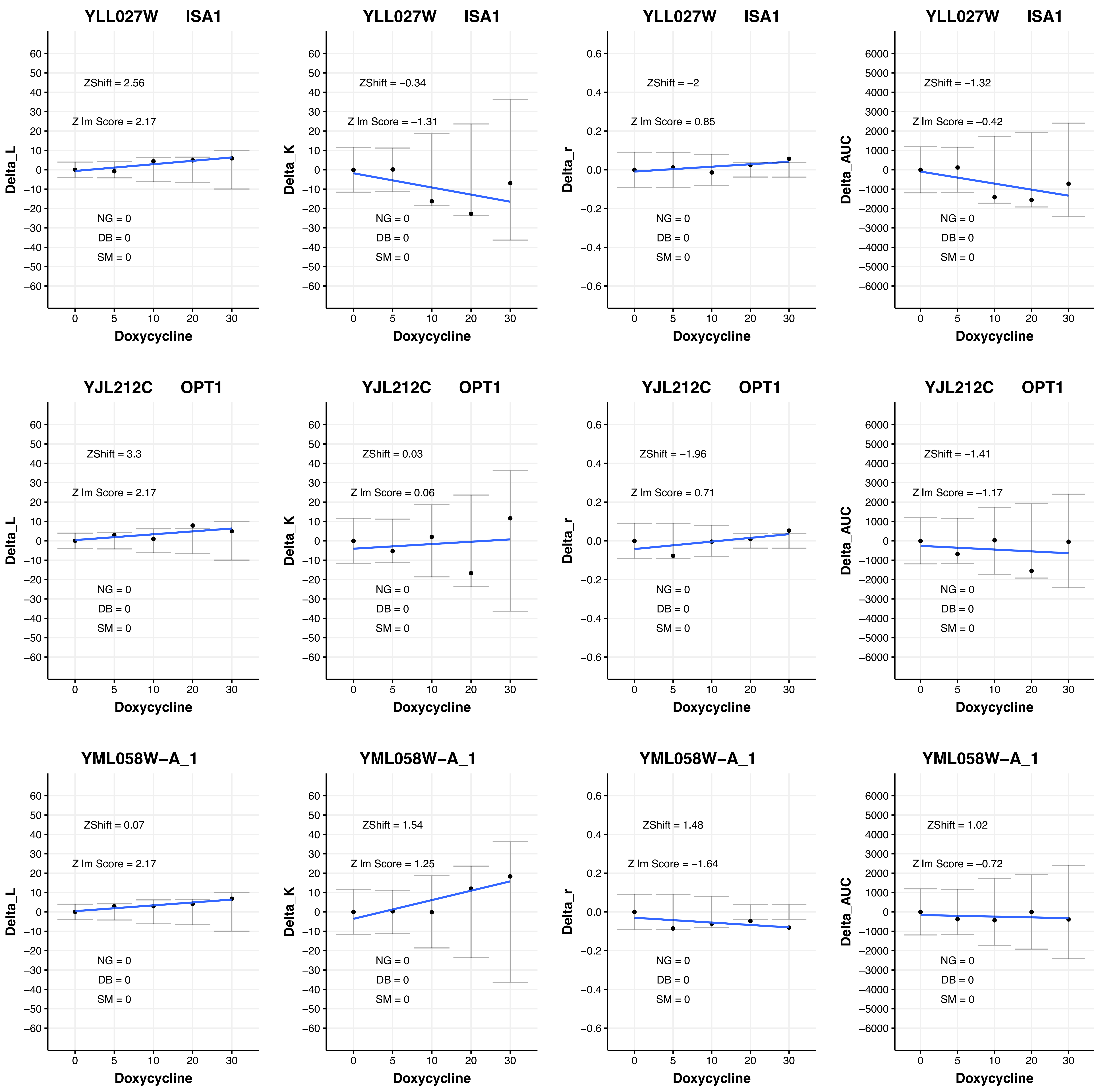

YLL027W ISA1
ZShift = 2.56
Z lm Score = 2.17
NG = 0
DB = 0
SM = 0
Delta_L
Doxycycline
YLL027W ISA1
ZShift = −0.34
Z lm Score = −1.31
NG = 0
DB = 0
SM = 0
Delta_K
Doxycycline
YLL027W ISA1
ZShift = −2
Z lm Score = 0.85
NG = 0
DB = 0
SM = 0
Delta_r
Doxycycline
YLL027W ISA1
ZShift = −1.32
Z lm Score = −0.42
NG = 0
DB = 0
SM = 0
Delta_AUC
Doxycycline
YJL212C OPT1
ZShift = 3.3
Z lm Score = 2.17
NG = 0
DB = 0
SM = 0
Delta_L
Doxycycline
YJL212C OPT1
ZShift = 0.03
Z lm Score = 0.06
NG = 0
DB = 0
SM = 0
Delta_K
Doxycycline
YJL212C OPT1
ZShift = −1.96
Z lm Score = 0.71
NG = 0
DB = 0
SM = 0
Delta_r
Doxycycline
YJL212C OPT1
ZShift = −1.41
Z lm Score = −1.17
NG = 0
DB = 0
SM = 0
Delta_AUC
Doxycycline
YML058W−A_1
ZShift = 0.07
Z lm Score = 2.17
NG = 0
DB = 0
SM = 0
Delta_L
Doxycycline
YML058W−A_1
ZShift = 1.54
Z lm Score = 1.25
NG = 0
DB = 0
SM = 0
Delta_K
Doxycycline
YML058W−A_1
ZShift = 1.48
Z lm Score = −1.64
NG = 0
DB = 0
SM = 0
Delta_r
Doxycycline
YML058W−A_1
ZShift = 1.02
Z lm Score = −0.72
NG = 0
DB = 0
SM = 0
Delta_AUC
Doxycycline

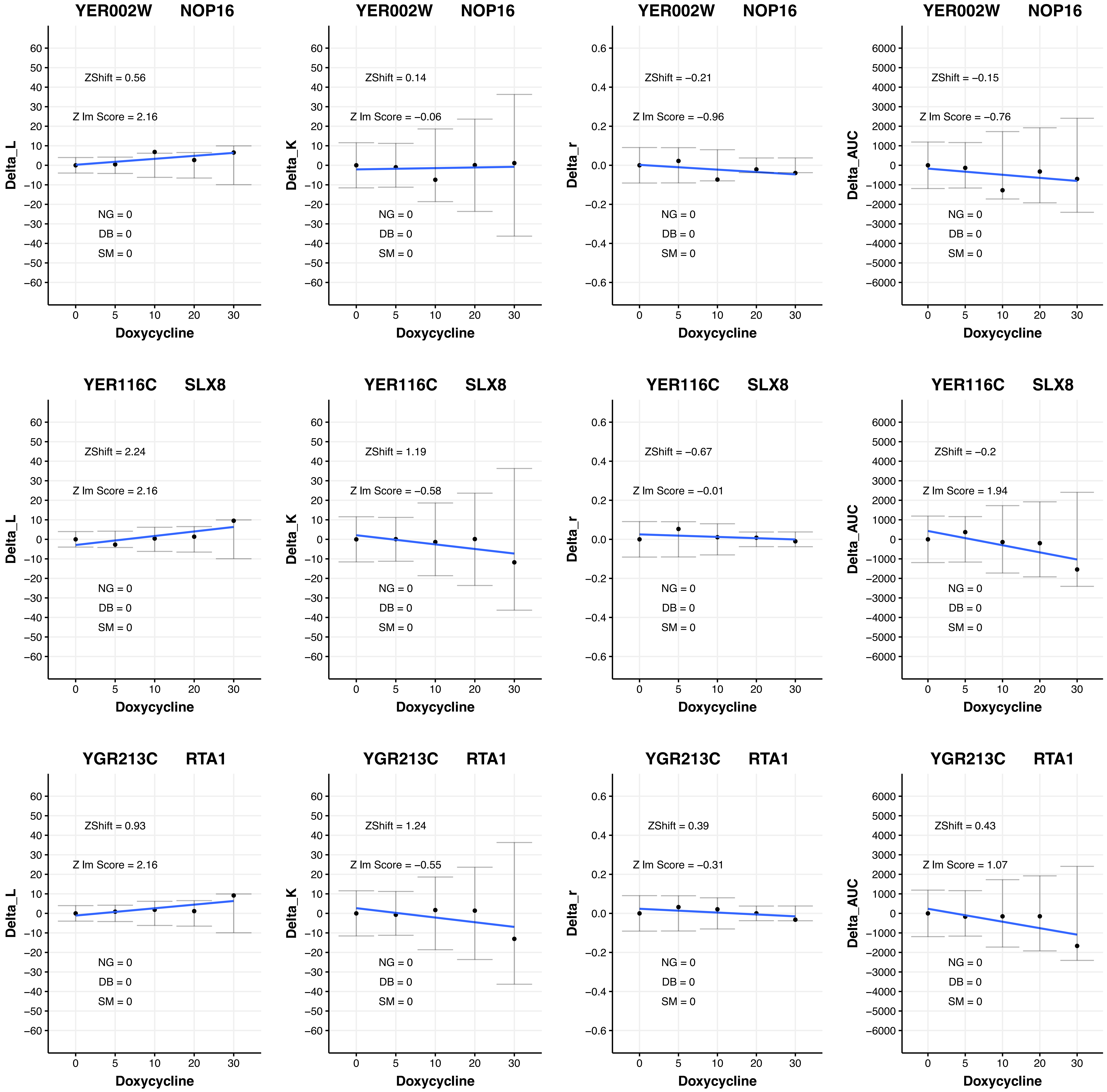

YER002W NOP16
ZShift = 0.56
Z lm Score = 2.16
NG = 0
DB = 0
SM = 0
Delta_L
Doxycycline
YER002W NOP16
ZShift = 0.14
Z lm Score = −0.06
NG = 0
DB = 0
SM = 0
Delta_K
Doxycycline
YER002W NOP16
ZShift = −0.21
Z lm Score = −0.96
NG = 0
DB = 0
SM = 0
Delta_r
Doxycycline
YER002W NOP16
ZShift = −0.15
Z lm Score = −0.76
NG = 0
DB = 0
SM = 0
Delta_AUC
Doxycycline
YER116C SLX8
ZShift = 2.24
Z lm Score = 2.16
NG = 0
DB = 0
SM = 0
Delta_L
Doxycycline
YER116C SLX8
ZShift = 1.19
Z lm Score = −0.58
NG = 0
DB = 0
SM = 0
Delta_K
Doxycycline
YER116C SLX8
ZShift = −0.67
Z lm Score = −0.01
NG = 0
DB = 0
SM = 0
Delta_r
Doxycycline
YER116C SLX8
ZShift = −0.2
Z lm Score = 1.94
NG = 0
DB = 0
SM = 0
Delta_AUC
Doxycycline
YGR213C RTA1
ZShift = 0.93
Z lm Score = 2.16
NG = 0
DB = 0
SM = 0
Delta_L
Doxycycline
YGR213C RTA1
ZShift = 1.24
Z lm Score = −0.55
NG = 0
DB = 0
SM = 0
Delta_K
Doxycycline
YGR213C RTA1
ZShift = 0.39
Z lm Score = −0.31
NG = 0
DB = 0
SM = 0
Delta_r
Doxycycline
YGR213C RTA1
ZShift = 0.43
Z lm Score = 1.07
NG = 0
DB = 0
SM = 0
Delta_AUC
Doxycycline

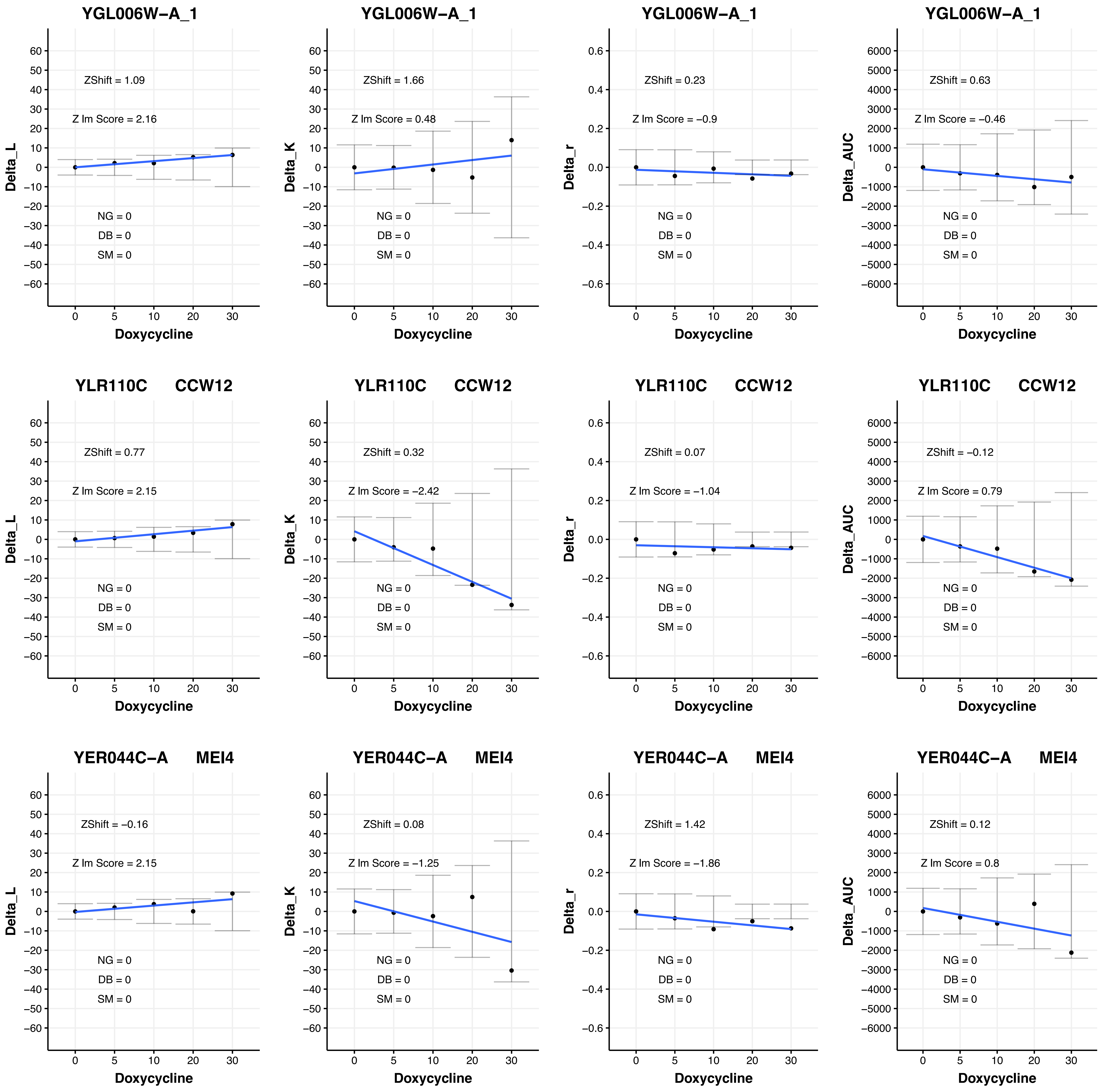

YGL006W-A_1
ZShift = 1.09
Z lm Score = 2.16
NG = 0
DB = 0
SM = 0
Delta_L
Doxycycline
YGL006W-A_1
ZShift = 1.66
Z lm Score = 0.48
NG = 0
DB = 0
SM = 0
Delta_K
Doxycycline
YGL006W-A_1
ZShift = 0.23
Z lm Score = −0.9
NG = 0
DB = 0
SM = 0
Delta_r
Doxycycline
YGL006W-A_1
ZShift = 0.63
Z lm Score = −0.46
NG = 0
DB = 0
SM = 0
Delta_AUC
Doxycycline
YLR110C CCW12
ZShift = 0.77
Z lm Score = 2.15
NG = 0
DB = 0
SM = 0
Delta_L
Doxycycline
YLR110C CCW12
ZShift = 0.32
Z lm Score = −2.42
NG = 0
DB = 0
SM = 0
Delta_K
Doxycycline
YLR110C CCW12
ZShift = 0.07
Z lm Score = −1.04
NG = 0
DB = 0
SM = 0
Delta_r
Doxycycline
YLR110C CCW12
ZShift = −0.12
Z lm Score = 0.79
NG = 0
DB = 0
SM = 0
Delta_AUC
Doxycycline
YER044C-A MEI4
ZShift = −0.16
Z lm Score = 2.15
NG = 0
DB = 0
SM = 0
Delta_L
Doxycycline
YER044C-A MEI4
ZShift = 0.08
Z lm Score = −1.25
NG = 0
DB = 0
SM = 0
Delta_K
Doxycycline
YER044C-A MEI4
ZShift = 1.42
Z lm Score = −1.86
NG = 0
DB = 0
SM = 0
Delta_r
Doxycycline
YER044C-A MEI4
ZShift = 0.12
Z lm Score = 0.8
NG = 0
DB = 0
SM = 0
Delta_AUC
Doxycycline

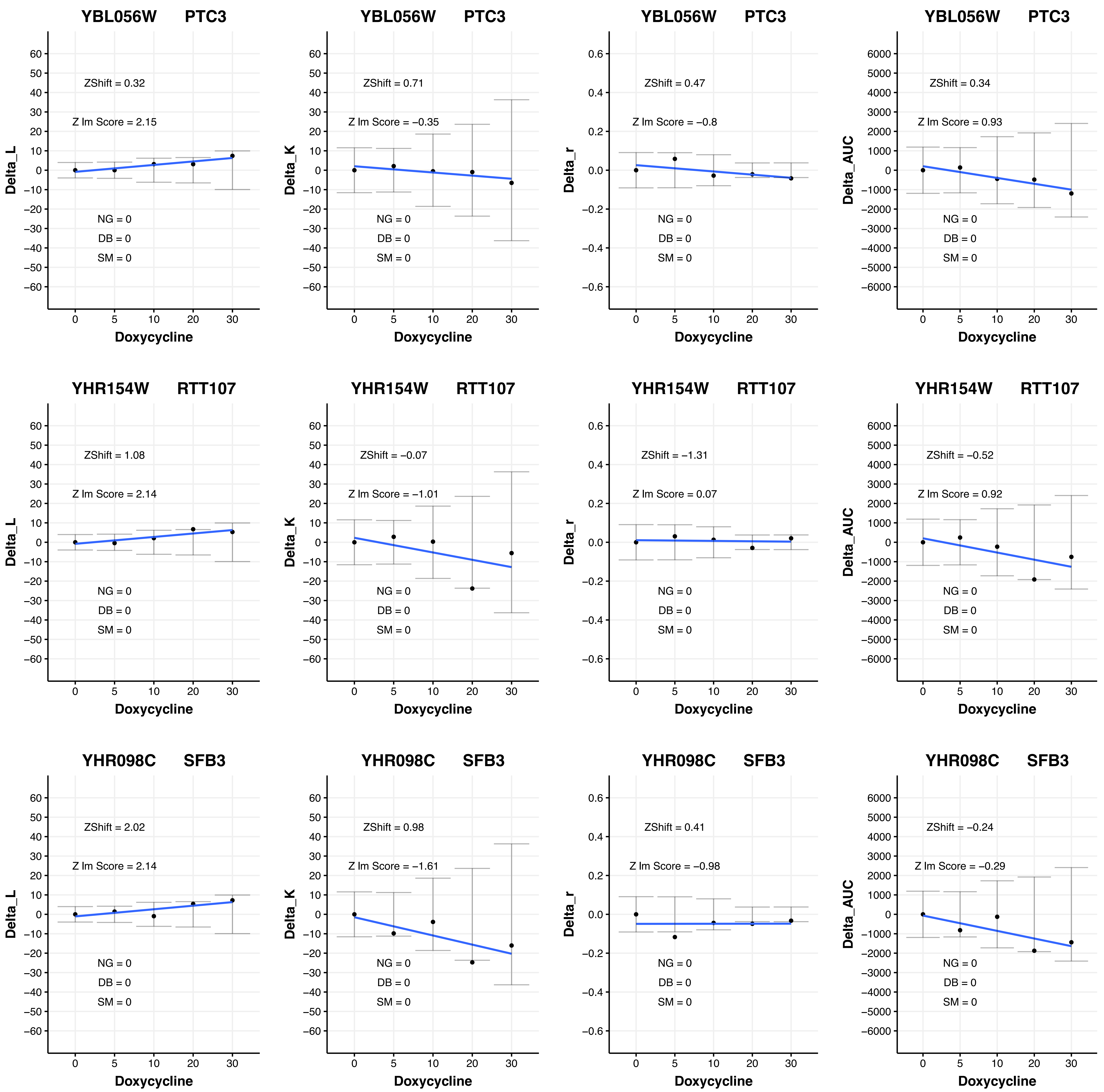
YBL056W PTC3
ZShift = 0.32
Z lm Score = 2.15
NG = 0
DB = 0
SM = 0
Delta_L
Doxycycline
YBL056W PTC3
ZShift = 0.71
Z lm Score = −0.35
Delta_K
YBL056W PTC3
ZShift = 0.47
Z lm Score = −0.8
Delta_r
YBL056W PTC3
ZShift = 0.34
Z lm Score = 0.93
Delta_AUC
YHR154W RTT107
ZShift = 1.08
Z lm Score = 2.14
YHR154W RTT107
ZShift = −0.07
Z lm Score = −1.01
YHR154W RTT107
ZShift = −1.31
Z lm Score = 0.07
YHR154W RTT107
ZShift = −0.52
Z lm Score = 0.92
YHR098C SFB3
ZShift = 2.02
Z lm Score = 2.14
YHR098C SFB3
ZShift = 0.98
Z lm Score = −1.61
YHR098C SFB3
ZShift = 0.41
Z lm Score = −0.98
YHR098C SFB3
ZShift = −0.24
Z lm Score = −0.29

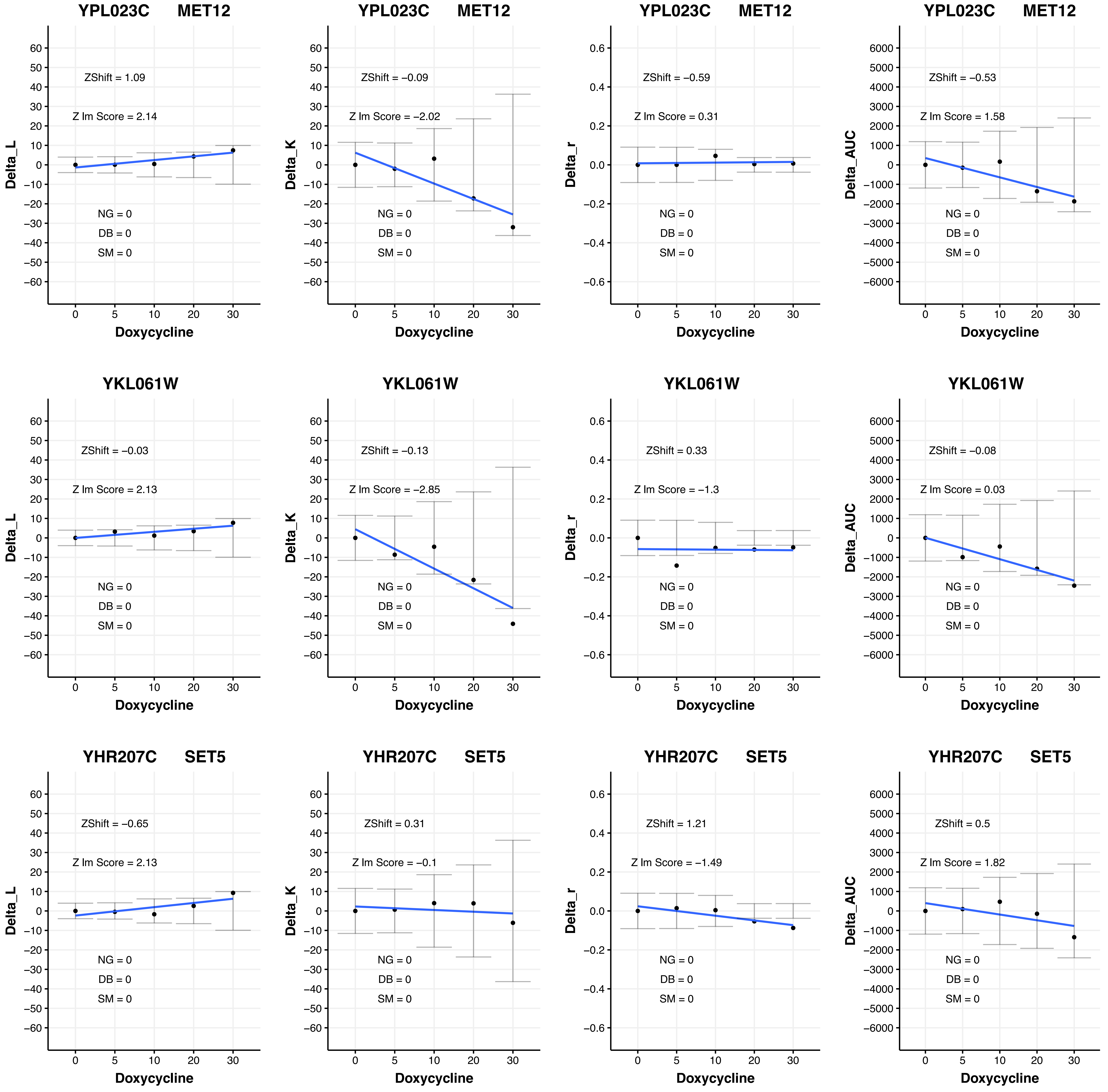

YPL023C MET12
ZShift = 1.09
Z lm Score = 2.14
NG = 0
DB = 0
SM = 0
Delta_L
Doxycycline
YPL023C MET12
ZShift = −0.09
Z lm Score = −2.02
NG = 0
DB = 0
SM = 0
Delta_K
Doxycycline
YPL023C MET12
ZShift = −0.59
Z lm Score = 0.31
NG = 0
DB = 0
SM = 0
Delta_r
Doxycycline
YPL023C MET12
ZShift = −0.53
Z lm Score = 1.58
NG = 0
DB = 0
SM = 0
Delta_AUC
Doxycycline
YKL061W
ZShift = −0.03
Z lm Score = 2.13
NG = 0
DB = 0
SM = 0
Delta_L
Doxycycline
YKL061W
ZShift = −0.13
Z lm Score = −2.85
NG = 0
DB = 0
SM = 0
Delta_K
Doxycycline
YKL061W
ZShift = 0.33
Z lm Score = −1.3
NG = 0
DB = 0
SM = 0
Delta_r
Doxycycline
YKL061W
ZShift = −0.08
Z lm Score = 0.03
NG = 0
DB = 0
SM = 0
Delta_AUC
Doxycycline
YHR207C SET5
ZShift = −0.65
Z lm Score = 2.13
NG = 0
DB = 0
SM = 0
Delta_L
Doxycycline
YHR207C SET5
ZShift = 0.31
Z lm Score = −0.1
NG = 0
DB = 0
SM = 0
Delta_K
Doxycycline
YHR207C SET5
ZShift = 1.21
Z lm Score = −1.49
NG = 0
DB = 0
SM = 0
Delta_r
Doxycycline
YHR207C SET5
ZShift = 0.5
Z lm Score = 1.82
NG = 0
DB = 0
SM = 0
Delta_AUC
Doxycycline

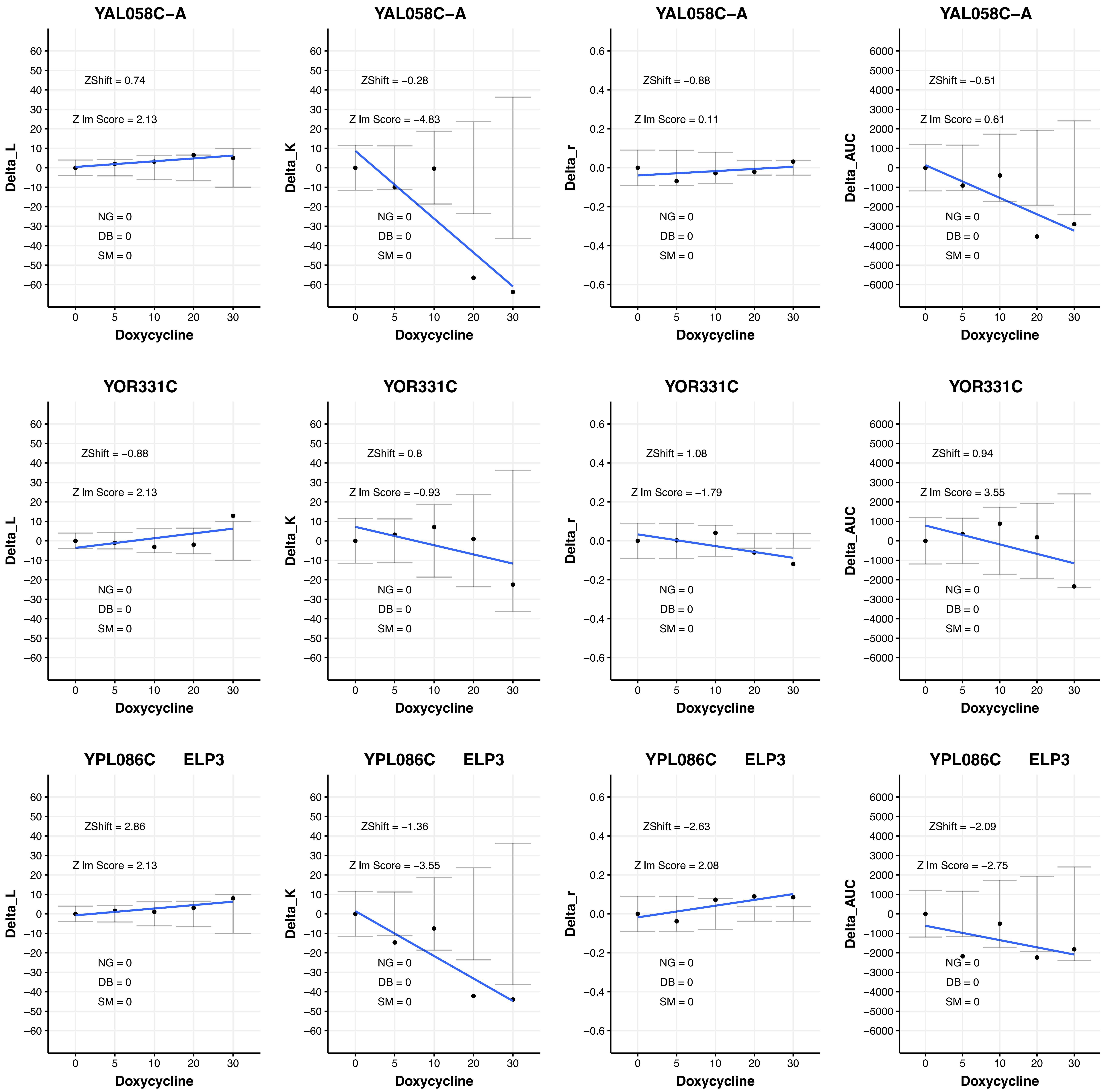
YAL058C–A
ZShift = 0.74
Z lm Score = 2.13
NG = 0
DB = 0
SM = 0
Delta_L
Doxycycline
YAL058C–A
ZShift = –0.28
Z lm Score = –4.83
NG = 0
DB = 0
SM = 0
Delta_K
Doxycycline
YAL058C–A
ZShift = –0.88
Z lm Score = 0.11
NG = 0
DB = 0
SM = 0
Delta_r
Doxycycline
YAL058C–A
ZShift = –0.51
Z lm Score = 0.61
NG = 0
DB = 0
SM = 0
Delta_AUC
Doxycycline
YOR331C
ZShift = –0.88
Z lm Score = 2.13
NG = 0
DB = 0
SM = 0
Delta_L
Doxycycline
YOR331C
ZShift = 0.8
Z lm Score = –0.93
NG = 0
DB = 0
SM = 0
Delta_K
Doxycycline
YOR331C
ZShift = 1.08
Z lm Score = –1.79
NG = 0
DB = 0
SM = 0
Delta_r
Doxycycline
YOR331C
ZShift = 0.94
Z lm Score = 3.55
NG = 0
DB = 0
SM = 0
Delta_AUC
Doxycycline
YPL086C ELP3
ZShift = 2.86
Z lm Score = 2.13
NG = 0
DB = 0
SM = 0
Delta_L
Doxycycline
YPL086C ELP3
ZShift = –1.36
Z lm Score = –3.55
NG = 0
DB = 0
SM = 0
Delta_K
Doxycycline
YPL086C ELP3
ZShift = –2.63
Z lm Score = 2.08
NG = 0
DB = 0
SM = 0
Delta_r
Doxycycline
YPL086C ELP3
ZShift = –2.09
Z lm Score = –2.75
NG = 0
DB = 0
SM = 0
Delta_AUC
Doxycycline

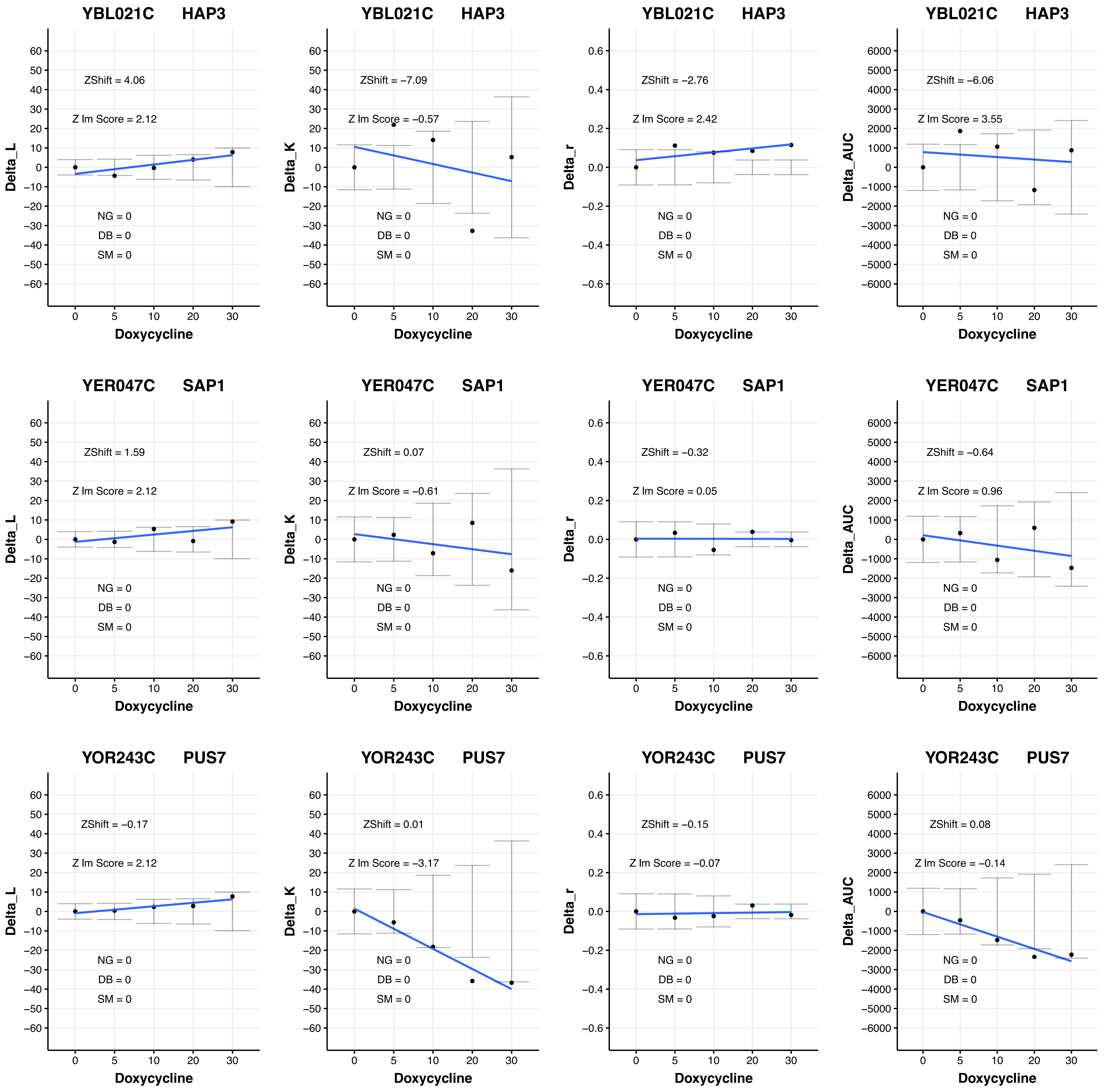
YBL021C HAP3
ZShift = 4.06
Z lm Score = 2.12
NG = 0
DB = 0
SM = 0
Delta_L
Doxycycline
YBL021C HAP3
ZShift = −7.09
Z lm Score = −0.57
NG = 0
DB = 0
SM = 0
Delta_K
Doxycycline
YBL021C HAP3
ZShift = −2.76
Z lm Score = 2.42
NG = 0
DB = 0
SM = 0
Delta_r
Doxycycline
YBL021C HAP3
ZShift = −6.06
Z lm Score = 3.55
NG = 0
DB = 0
SM = 0
Delta_AUC
Doxycycline
YER047C SAP1
ZShift = 1.59
Z lm Score = 2.12
NG = 0
DB = 0
SM = 0
Delta_L
Doxycycline
YER047C SAP1
ZShift = 0.07
Z lm Score = −0.61
NG = 0
DB = 0
SM = 0
Delta_K
Doxycycline
YER047C SAP1
ZShift = −0.32
Z lm Score = 0.05
NG = 0
DB = 0
SM = 0
Delta_r
Doxycycline
YER047C SAP1
ZShift = −0.64
Z lm Score = 0.96
NG = 0
DB = 0
SM = 0
Delta_AUC
Doxycycline
YOR243C PUS7
ZShift = −0.17
Z lm Score = 2.12
NG = 0
DB = 0
SM = 0
Delta_L
Doxycycline
YOR243C PUS7
ZShift = 0.01
Z lm Score = −3.17
NG = 0
DB = 0
SM = 0
Delta_K
Doxycycline
YOR243C PUS7
ZShift = −0.15
Z lm Score = −0.07
NG = 0
DB = 0
SM = 0
Delta_r
Doxycycline
YOR243C PUS7
ZShift = 0.08
Z lm Score = −0.14
NG = 0
DB = 0
SM = 0
Delta_AUC
Doxycycline

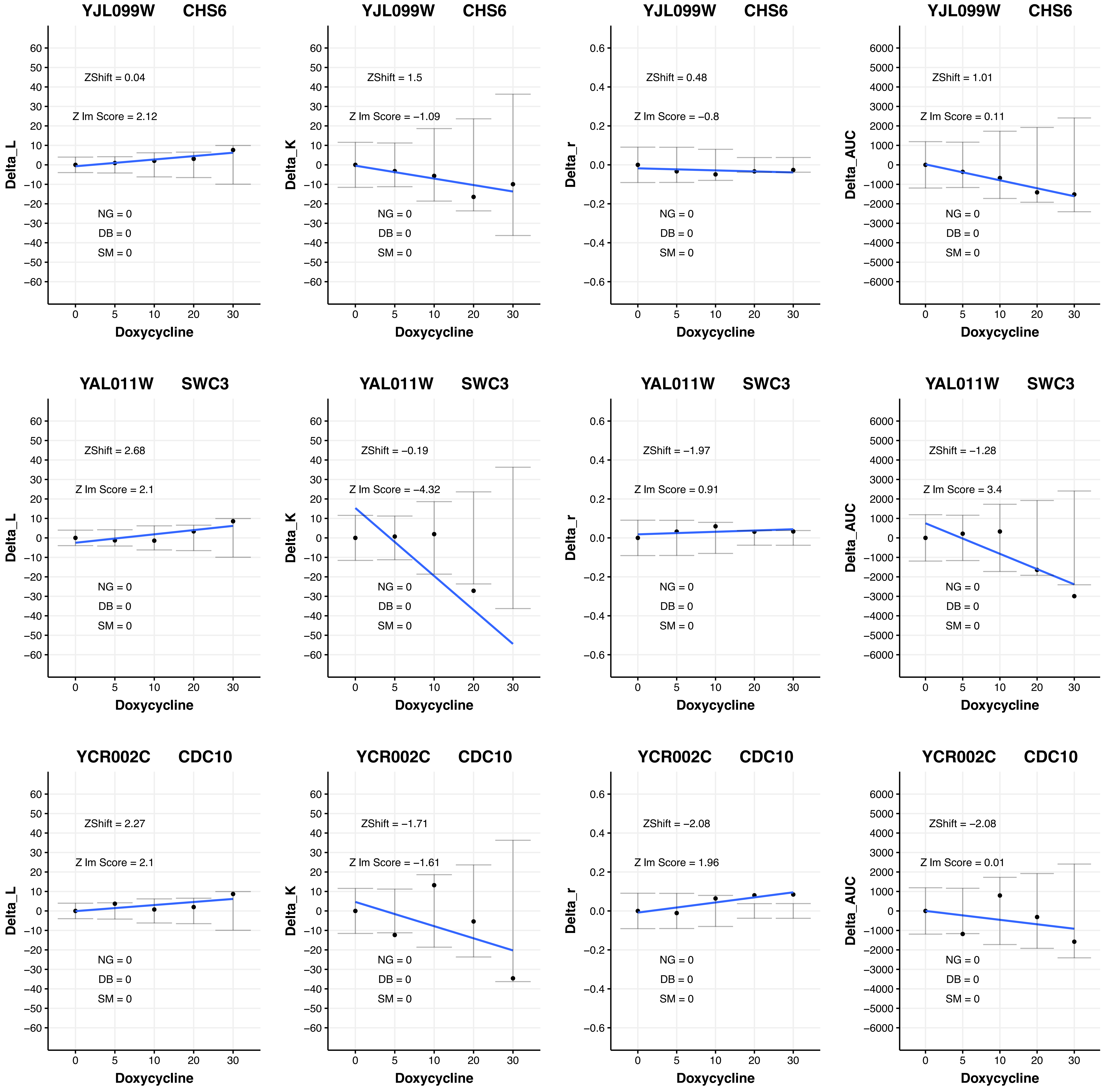
YJL099W CHS6
ZShift = 0.04
Z lm Score = 2.12
NG = 0
DB = 0
SM = 0
Delta_L
Doxycycline
YJL099W CHS6
ZShift = 1.5
Z lm Score = −1.09
NG = 0
DB = 0
SM = 0
Delta_K
Doxycycline
YJL099W CHS6
ZShift = 0.48
Z lm Score = −0.8
NG = 0
DB = 0
SM = 0
Delta_r
Doxycycline
YJL099W CHS6
ZShift = 1.01
Z lm Score = 0.11
NG = 0
DB = 0
SM = 0
Delta_AUC
Doxycycline
YAL011W SWC3
ZShift = 2.68
Z lm Score = 2.1
NG = 0
DB = 0
SM = 0
Delta_L
Doxycycline
YAL011W SWC3
ZShift = −0.19
Z lm Score = −4.32
NG = 0
DB = 0
SM = 0
Delta_K
Doxycycline
YAL011W SWC3
ZShift = −1.97
Z lm Score = 0.91
NG = 0
DB = 0
SM = 0
Delta_r
Doxycycline
YAL011W SWC3
ZShift = −1.28
Z lm Score = 3.4
NG = 0
DB = 0
SM = 0
Delta_AUC
Doxycycline
YCR002C CDC10
ZShift = 2.27
Z lm Score = 2.1
NG = 0
DB = 0
SM = 0
Delta_L
Doxycycline
YCR002C CDC10
ZShift = −1.71
Z lm Score = −1.61
NG = 0
DB = 0
SM = 0
Delta_K
Doxycycline
YCR002C CDC10
ZShift = −2.08
Z lm Score = 1.96
NG = 0
DB = 0
SM = 0
Delta_r
Doxycycline
YCR002C CDC10
ZShift = −2.08
Z lm Score = 0.01
NG = 0
DB = 0
SM = 0
Delta_AUC
Doxycycline

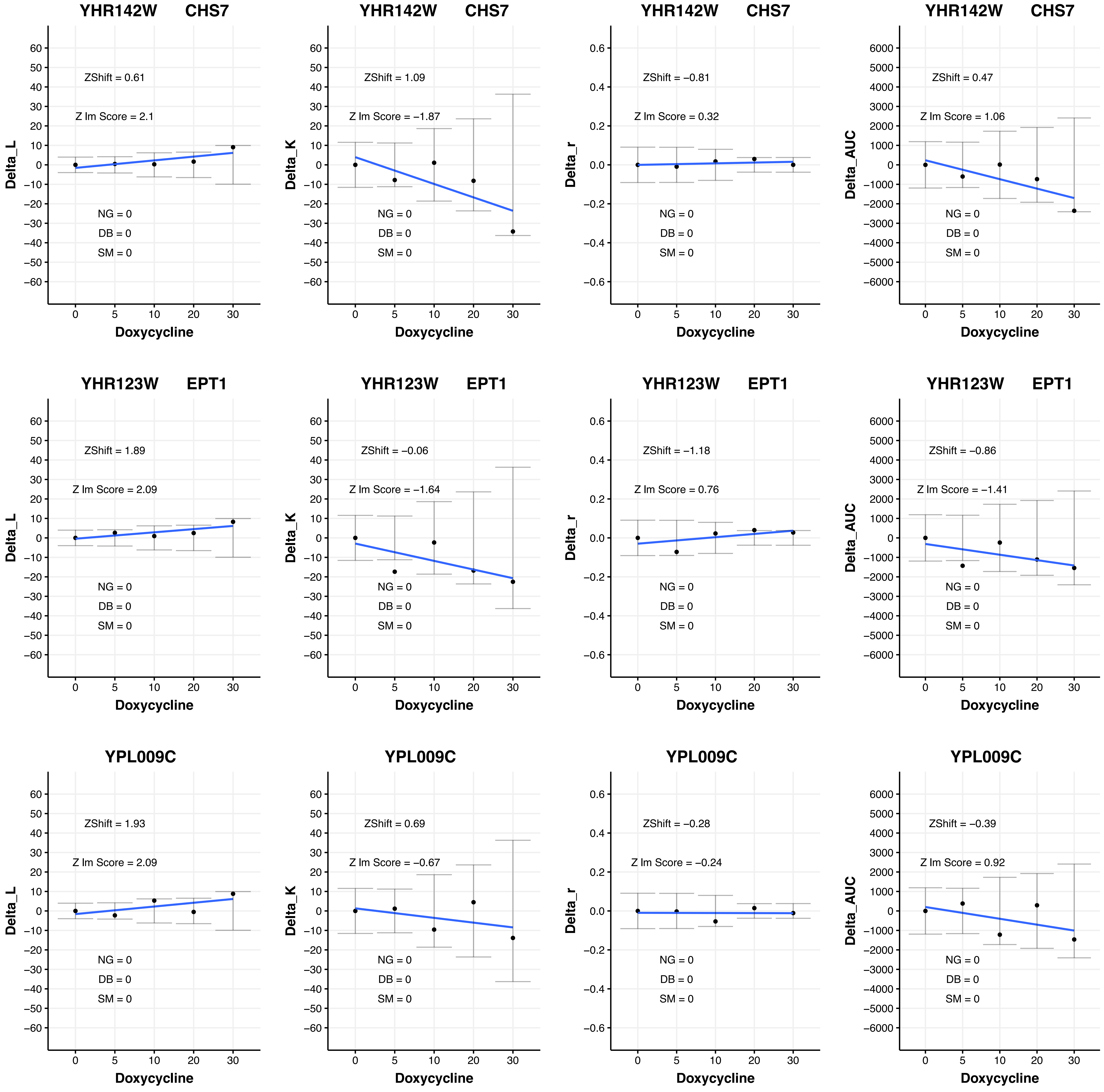
YHR142W CHS7
ZShift = 0.61
Z lm Score = 2.1
NG = 0
DB = 0
SM = 0
Delta_L
Doxycycline
YHR142W CHS7
ZShift = 1.09
Z lm Score = −1.87
NG = 0
DB = 0
SM = 0
Delta_K
Doxycycline
YHR142W CHS7
ZShift = −0.81
Z lm Score = 0.32
NG = 0
DB = 0
SM = 0
Delta_r
Doxycycline
YHR142W CHS7
ZShift = 0.47
Z lm Score = 1.06
NG = 0
DB = 0
SM = 0
Delta_AUC
Doxycycline
YHR123W EPT1
ZShift = 1.89
Z lm Score = 2.09
NG = 0
DB = 0
SM = 0
Delta_L
Doxycycline
YHR123W EPT1
ZShift = −0.06
Z lm Score = −1.64
NG = 0
DB = 0
SM = 0
Delta_K
Doxycycline
YHR123W EPT1
ZShift = −1.18
Z lm Score = 0.76
NG = 0
DB = 0
SM = 0
Delta_r
Doxycycline
YHR123W EPT1
ZShift = −0.86
Z lm Score = −1.41
NG = 0
DB = 0
SM = 0
Delta_AUC
Doxycycline
YPL009C
ZShift = 1.93
Z lm Score = 2.09
NG = 0
DB = 0
SM = 0
Delta_L
Doxycycline
YPL009C
ZShift = 0.69
Z lm Score = −0.67
NG = 0
DB = 0
SM = 0
Delta_K
Doxycycline
YPL009C
ZShift = −0.28
Z lm Score = −0.24
NG = 0
DB = 0
SM = 0
Delta_r
Doxycycline
YPL009C
ZShift = −0.39
Z lm Score = 0.92
NG = 0
DB = 0
SM = 0
Delta_AUC
Doxycycline

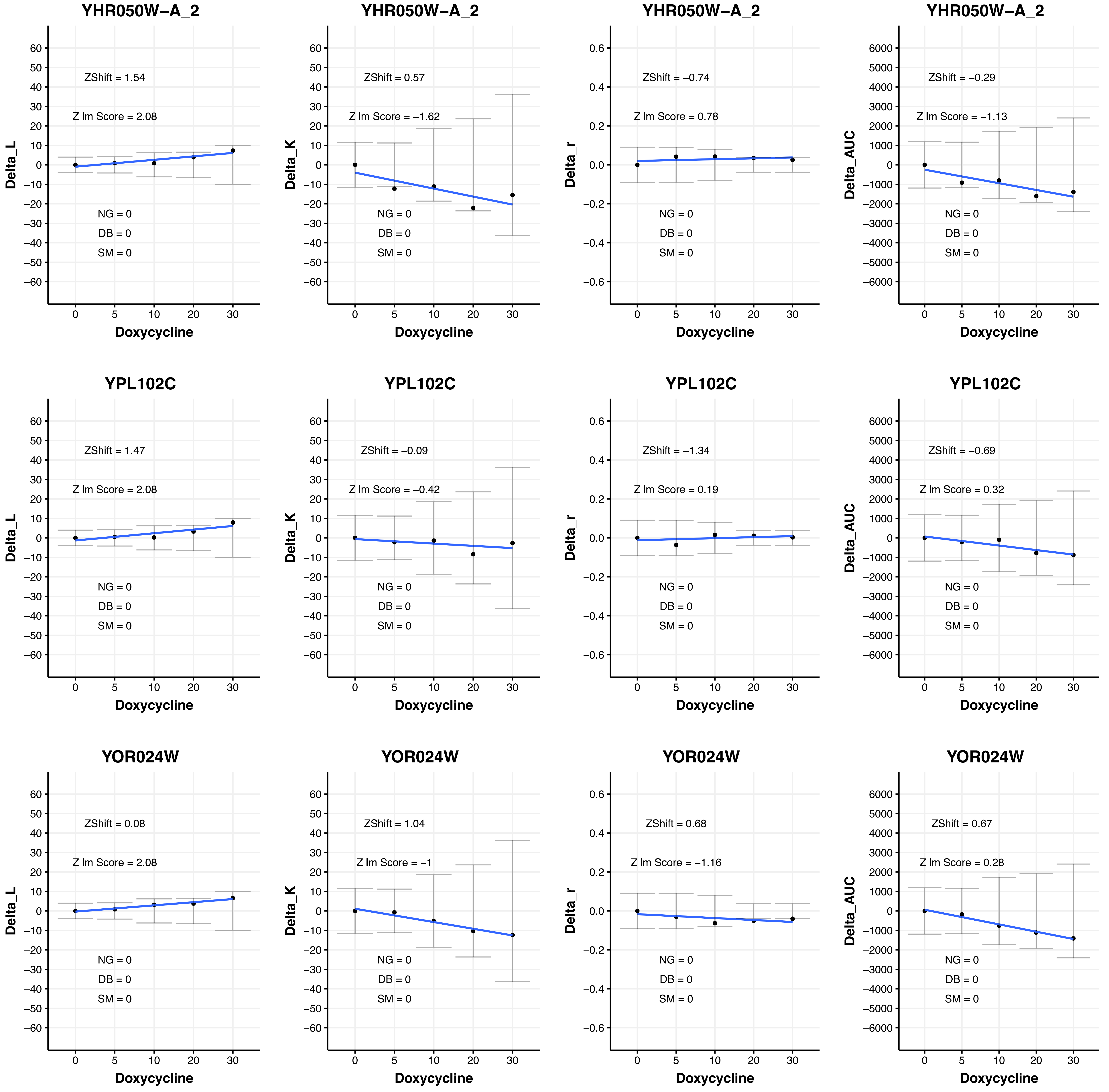
YHR050W−A_2
ZShift = 1.54
Z lm Score = 2.08
NG = 0
DB = 0
SM = 0
Delta_L
Doxycycline
YHR050W−A_2
ZShift = 0.57
Z lm Score = −1.62
NG = 0
DB = 0
SM = 0
Delta_K
Doxycycline
YHR050W−A_2
ZShift = −0.74
Z lm Score = 0.78
NG = 0
DB = 0
SM = 0
Delta_r
Doxycycline
YHR050W−A_2
ZShift = −0.29
Z lm Score = −1.13
NG = 0
DB = 0
SM = 0
Delta_AUC
Doxycycline
YPL102C
ZShift = 1.47
Z lm Score = 2.08
NG = 0
DB = 0
SM = 0
Delta_L
Doxycycline
YPL102C
ZShift = −0.09
Z lm Score = −0.42
NG = 0
DB = 0
SM = 0
Delta_K
Doxycycline
YPL102C
ZShift = −1.34
Z lm Score = 0.19
NG = 0
DB = 0
SM = 0
Delta_r
Doxycycline
YPL102C
ZShift = −0.69
Z lm Score = 0.32
NG = 0
DB = 0
SM = 0
Delta_AUC
Doxycycline
YOR024W
ZShift = 0.08
Z lm Score = 2.08
NG = 0
DB = 0
SM = 0
Delta_L
Doxycycline
YOR024W
ZShift = 1.04
Z lm Score = −1
NG = 0
DB = 0
SM = 0
Delta_K
Doxycycline
YOR024W
ZShift = 0.68
Z lm Score = −1.16
NG = 0
DB = 0
SM = 0
Delta_r
Doxycycline
YOR024W
ZShift = 0.67
Z lm Score = 0.28
NG = 0
DB = 0
SM = 0
Delta_AUC
Doxycycline

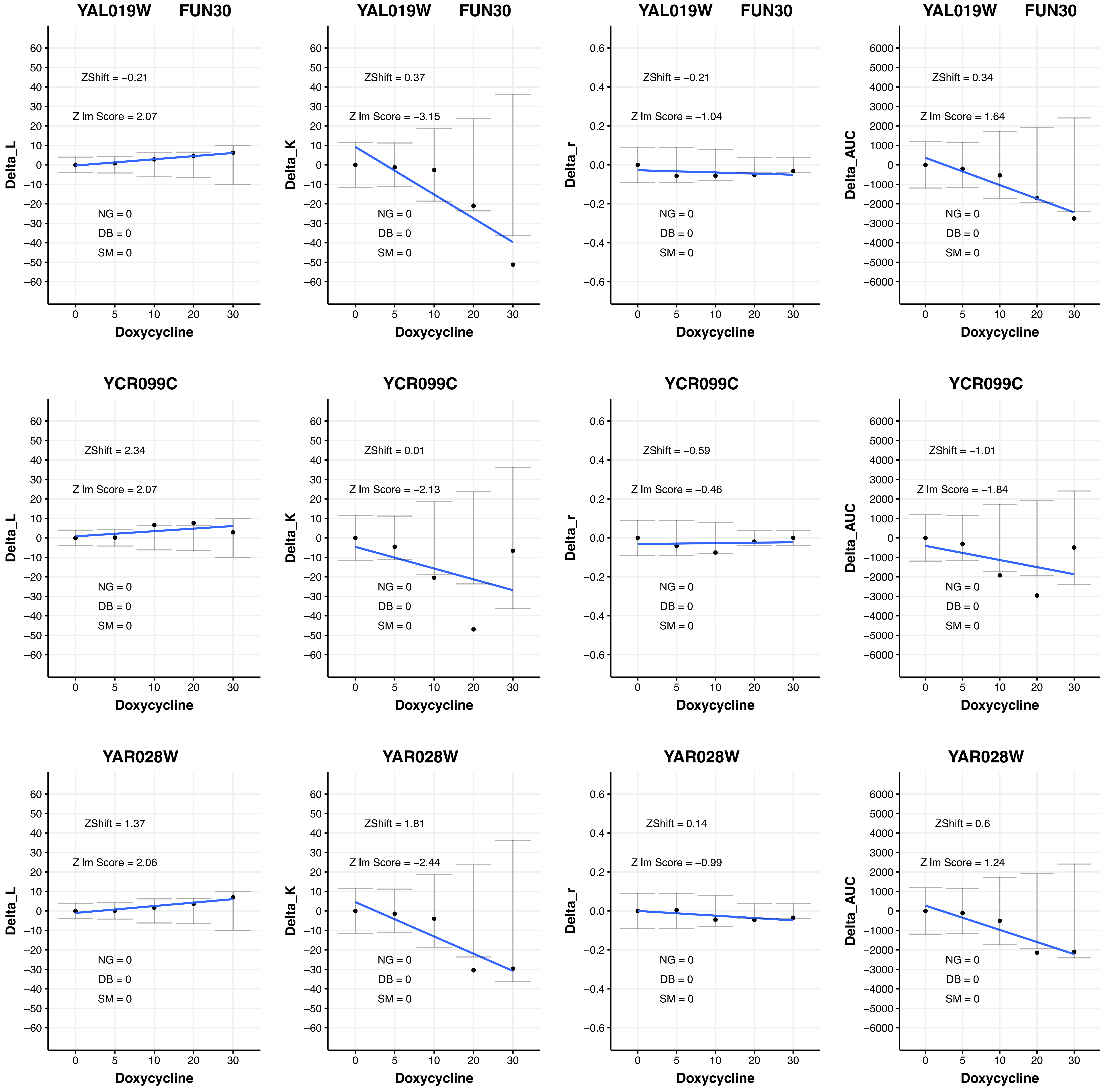

YAL019W FUN30
ZShift = −0.21
Z lm Score = 2.07
NG = 0
DB = 0
SM = 0
Delta_L
Doxycycline
YAL019W FUN30
ZShift = 0.37
Z lm Score = −3.15
NG = 0
DB = 0
SM = 0
Delta_K
Doxycycline
YAL019W FUN30
ZShift = −0.21
Z lm Score = −1.04
NG = 0
DB = 0
SM = 0
Delta_r
Doxycycline
YAL019W FUN30
ZShift = 0.34
Z lm Score = 1.64
NG = 0
DB = 0
SM = 0
Delta_AUC
Doxycycline
YCR099C
ZShift = 2.34
Z lm Score = 2.07
NG = 0
DB = 0
SM = 0
Delta_L
Doxycycline
YCR099C
ZShift = 0.01
Z lm Score = −2.13
NG = 0
DB = 0
SM = 0
Delta_K
Doxycycline
YCR099C
ZShift = −0.59
Z lm Score = −0.46
NG = 0
DB = 0
SM = 0
Delta_r
Doxycycline
YCR099C
ZShift = −1.01
Z lm Score = −1.84
NG = 0
DB = 0
SM = 0
Delta_AUC
Doxycycline
YAR028W
ZShift = 1.37
Z lm Score = 2.06
NG = 0
DB = 0
SM = 0
Delta_L
Doxycycline
YAR028W
ZShift = 1.81
Z lm Score = −2.44
NG = 0
DB = 0
SM = 0
Delta_K
Doxycycline
YAR028W
ZShift = 0.14
Z lm Score = −0.99
NG = 0
DB = 0
SM = 0
Delta_r
Doxycycline
YAR028W
ZShift = 0.6
Z lm Score = 1.24
NG = 0
DB = 0
SM = 0
Delta_AUC
Doxycycline

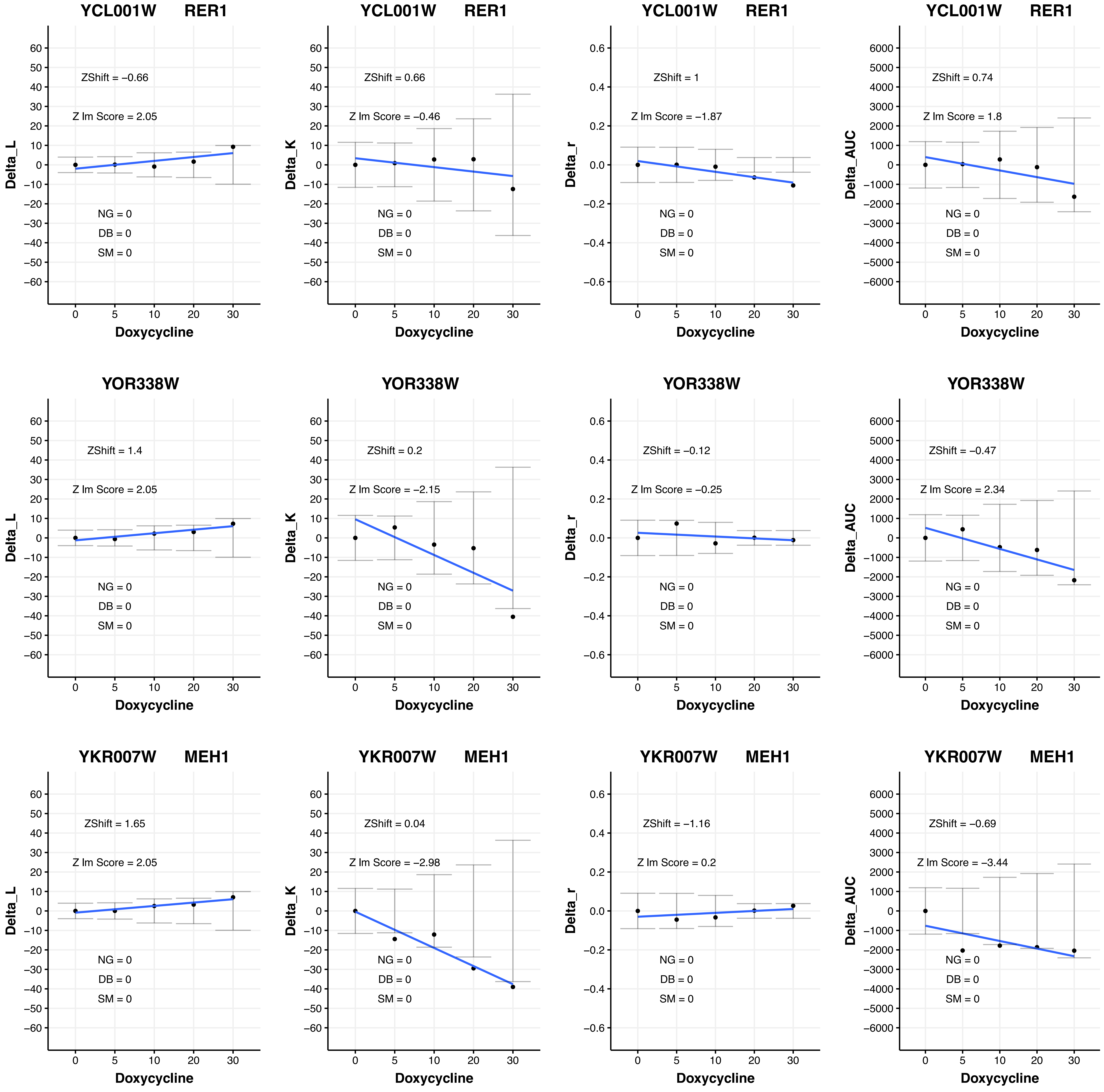
YCL001W RER1
ZShift = −0.66
Z lm Score = 2.05
NG = 0
DB = 0
SM = 0
Delta_L
Doxycycline
YCL001W RER1
ZShift = 0.66
Z lm Score = −0.46
NG = 0
DB = 0
SM = 0
Delta_K
Doxycycline
YCL001W RER1
ZShift = 1
Z lm Score = −1.87
NG = 0
DB = 0
SM = 0
Delta_r
Doxycycline
YCL001W RER1
ZShift = 0.74
Z lm Score = 1.8
NG = 0
DB = 0
SM = 0
Delta_AUC
Doxycycline
YOR338W
ZShift = 1.4
Z lm Score = 2.05
NG = 0
DB = 0
SM = 0
Delta_L
Doxycycline
YOR338W
ZShift = 0.2
Z lm Score = −2.15
NG = 0
DB = 0
SM = 0
Delta_K
Doxycycline
YOR338W
ZShift = −0.12
Z lm Score = −0.25
NG = 0
DB = 0
SM = 0
Delta_r
Doxycycline
YOR338W
ZShift = −0.47
Z lm Score = 2.34
NG = 0
DB = 0
SM = 0
Delta_AUC
Doxycycline
YKR007W MEH1
ZShift = 1.65
Z lm Score = 2.05
NG = 0
DB = 0
SM = 0
Delta_L
Doxycycline
YKR007W MEH1
ZShift = 0.04
Z lm Score = −2.98
NG = 0
DB = 0
SM = 0
Delta_K
Doxycycline
YKR007W MEH1
ZShift = −1.16
Z lm Score = 0.2
NG = 0
DB = 0
SM = 0
Delta_r
Doxycycline
YKR007W MEH1
ZShift = −0.69
Z lm Score = −3.44
NG = 0
DB = 0
SM = 0
Delta_AUC
Doxycycline

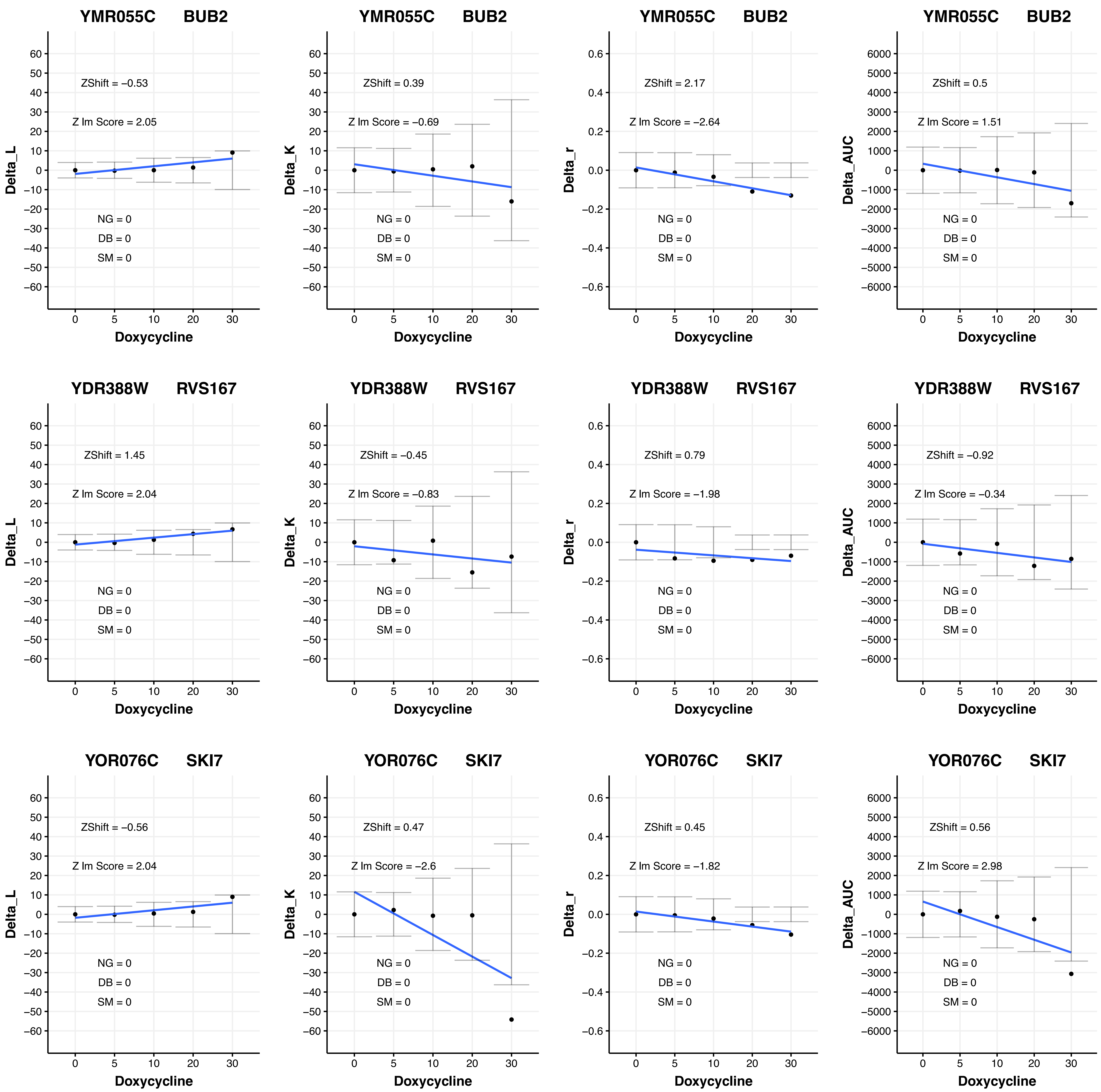
YMR055C BUB2
ZShift = −0.53
Z lm Score = 2.05
NG = 0
DB = 0
SM = 0
Delta_L
Doxycycline
YMR055C BUB2
ZShift = 0.39
Z lm Score = −0.69
NG = 0
DB = 0
SM = 0
Delta_K
Doxycycline
YMR055C BUB2
ZShift = 2.17
Z lm Score = −2.64
NG = 0
DB = 0
SM = 0
Delta_r
Doxycycline
YMR055C BUB2
ZShift = 0.5
Z lm Score = 1.51
NG = 0
DB = 0
SM = 0
Delta_AUC
Doxycycline
YDR388W RVS167
ZShift = 1.45
Z lm Score = 2.04
NG = 0
DB = 0
SM = 0
Delta_L
Doxycycline
YDR388W RVS167
ZShift = −0.45
Z lm Score = −0.83
NG = 0
DB = 0
SM = 0
Delta_K
Doxycycline
YDR388W RVS167
ZShift = 0.79
Z lm Score = −1.98
NG = 0
DB = 0
SM = 0
Delta_r
Doxycycline
YDR388W RVS167
ZShift = −0.92
Z lm Score = −0.34
NG = 0
DB = 0
SM = 0
Delta_AUC
Doxycycline
YOR076C SKI7
ZShift = −0.56
Z lm Score = 2.04
NG = 0
DB = 0
SM = 0
Delta_L
Doxycycline
YOR076C SKI7
ZShift = 0.47
Z lm Score = −2.6
NG = 0
DB = 0
SM = 0
Delta_K
Doxycycline
YOR076C SKI7
ZShift = 0.45
Z lm Score = −1.82
NG = 0
DB = 0
SM = 0
Delta_r
Doxycycline
YOR076C SKI7
ZShift = 0.56
Z lm Score = 2.98
NG = 0
DB = 0
SM = 0
Delta_AUC
Doxycycline

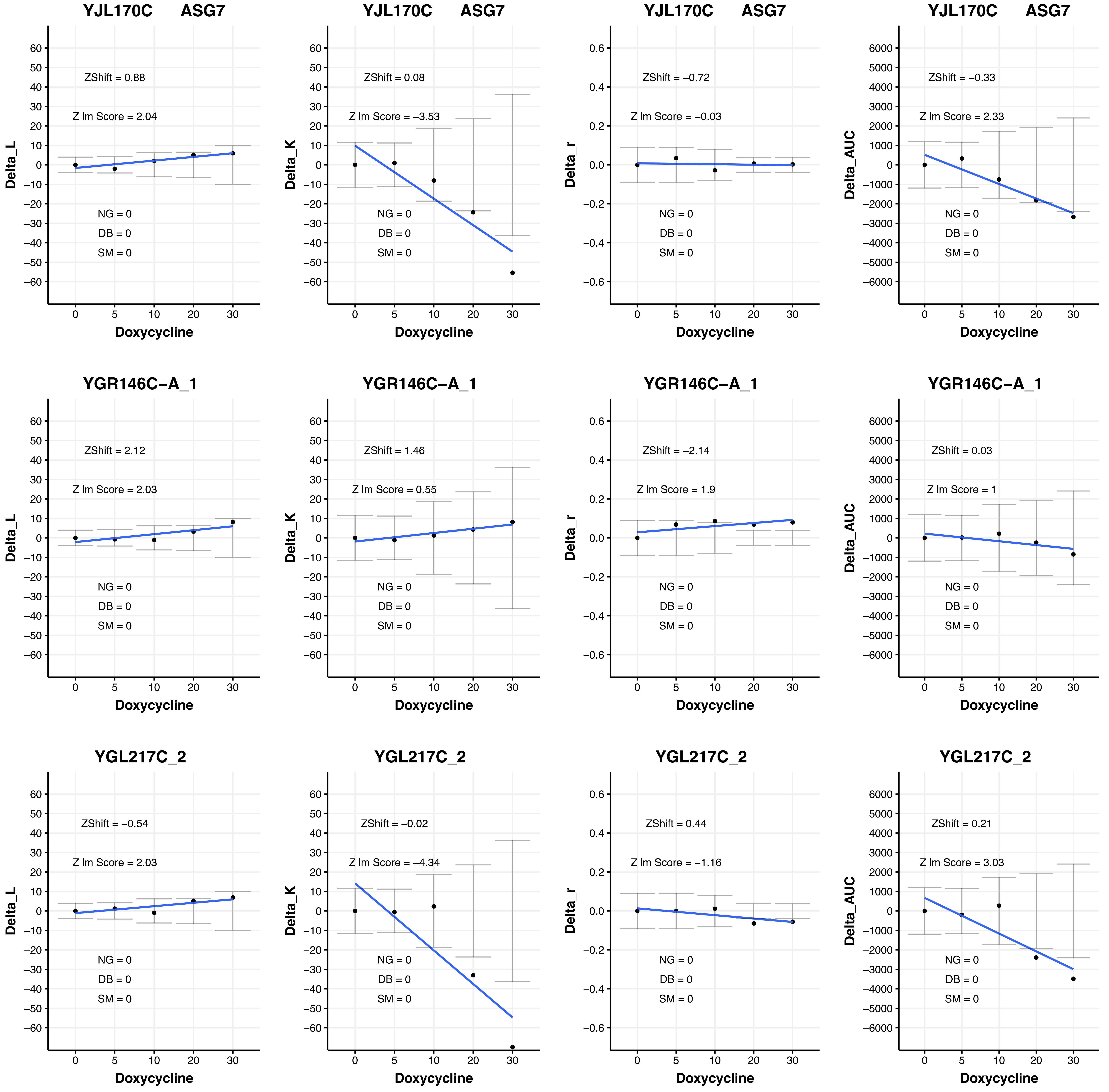

YJL170C ASG7
ZShift = 0.88
Z lm Score = 2.04
NG = 0
DB = 0
SM = 0
Delta_L
Doxycycline
YJL170C ASG7
ZShift = 0.08
Z lm Score = −3.53
NG = 0
DB = 0
SM = 0
Delta_K
Doxycycline
YJL170C ASG7
ZShift = −0.72
Z lm Score = −0.03
NG = 0
DB = 0
SM = 0
Delta_r
Doxycycline
YJL170C ASG7
ZShift = −0.33
Z lm Score = 2.33
NG = 0
DB = 0
SM = 0
Delta_AUC
Doxycycline
YGR146C−A_1
ZShift = 2.12
Z lm Score = 2.03
NG = 0
DB = 0
SM = 0
Delta_L
Doxycycline
YGR146C−A_1
ZShift = 1.46
Z lm Score = 0.55
NG = 0
DB = 0
SM = 0
Delta_K
Doxycycline
YGR146C−A_1
ZShift = −2.14
Z lm Score = 1.9
NG = 0
DB = 0
SM = 0
Delta_r
Doxycycline
YGR146C−A_1
ZShift = 0.03
Z lm Score = 1
NG = 0
DB = 0
SM = 0
Delta_AUC
Doxycycline
YGL217C_2
ZShift = −0.54
Z lm Score = 2.03
NG = 0
DB = 0
SM = 0
Delta_L
Doxycycline
YGL217C_2
ZShift = −0.02
Z lm Score = −4.34
NG = 0
DB = 0
SM = 0
Delta_K
Doxycycline
YGL217C_2
ZShift = 0.44
Z lm Score = −1.16
NG = 0
DB = 0
SM = 0
Delta_r
Doxycycline
YGL217C_2
ZShift = 0.21
Z lm Score = 3.03
NG = 0
DB = 0
SM = 0
Delta_AUC
Doxycycline

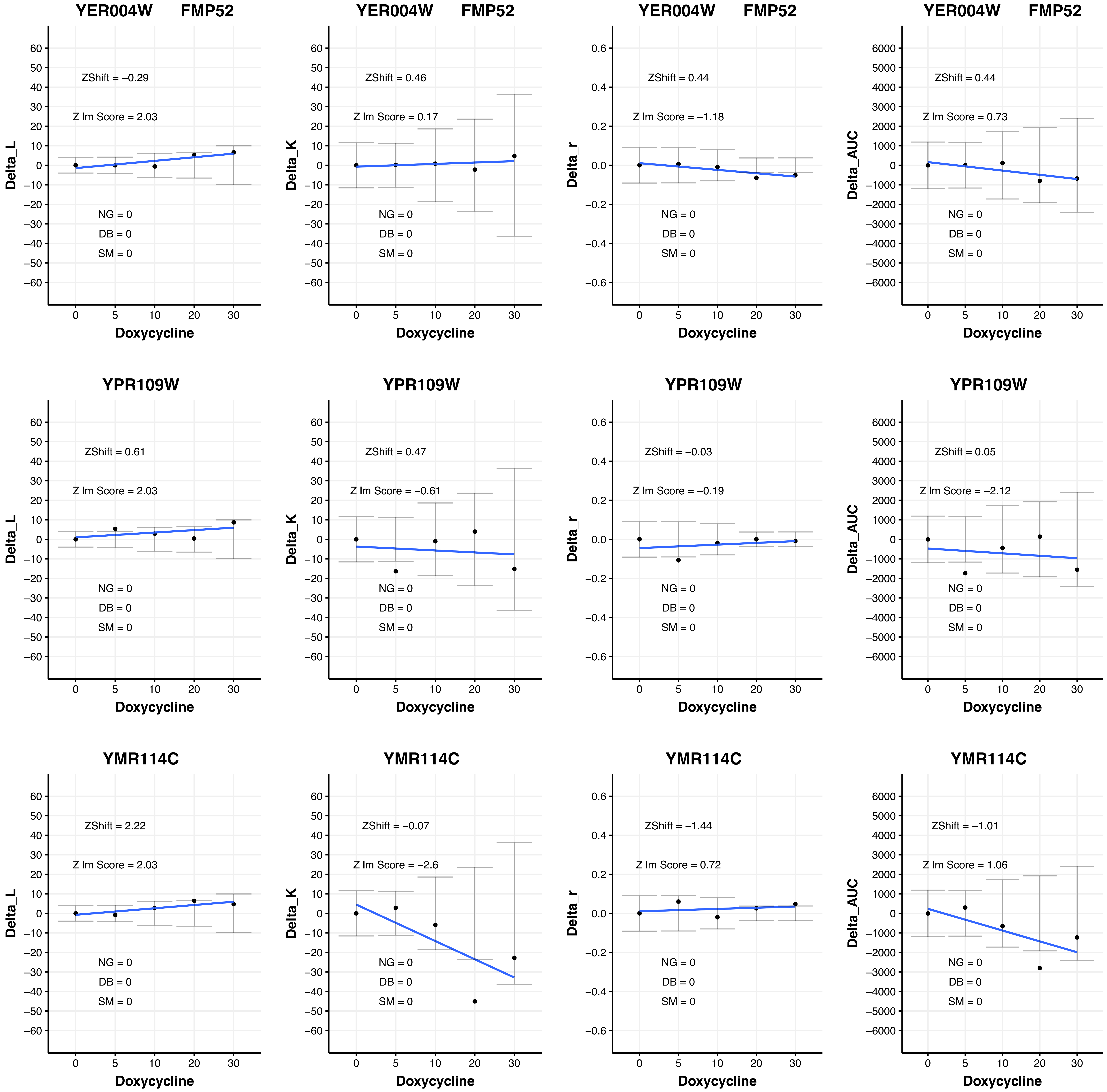

YER004W FMP52
ZShift = −0.29
Z lm Score = 2.03
NG = 0
DB = 0
SM = 0
Delta_L
Doxycycline
YER004W FMP52
ZShift = 0.46
Z lm Score = 0.17
NG = 0
DB = 0
SM = 0
Delta_K
Doxycycline
YER004W FMP52
ZShift = 0.44
Z lm Score = −1.18
NG = 0
DB = 0
SM = 0
Delta_r
Doxycycline
YER004W FMP52
ZShift = 0.44
Z lm Score = 0.73
NG = 0
DB = 0
SM = 0
Delta_AUC
Doxycycline
YPR109W
ZShift = 0.61
Z lm Score = 2.03
NG = 0
DB = 0
SM = 0
Delta_L
Doxycycline
YPR109W
ZShift = 0.47
Z lm Score = −0.61
NG = 0
DB = 0
SM = 0
Delta_K
Doxycycline
YPR109W
ZShift = −0.03
Z lm Score = −0.19
NG = 0
DB = 0
SM = 0
Delta_r
Doxycycline
YPR109W
ZShift = 0.05
Z lm Score = −2.12
NG = 0
DB = 0
SM = 0
Delta_AUC
Doxycycline
YMR114C
ZShift = 2.22
Z lm Score = 2.03
NG = 0
DB = 0
SM = 0
Delta_L
Doxycycline
YMR114C
ZShift = −0.07
Z lm Score = −2.6
NG = 0
DB = 0
SM = 0
Delta_K
Doxycycline
YMR114C
ZShift = −1.44
Z lm Score = 0.72
NG = 0
DB = 0
SM = 0
Delta_r
Doxycycline
YMR114C
ZShift = −1.01
Z lm Score = 1.06
NG = 0
DB = 0
SM = 0
Delta_AUC
Doxycycline

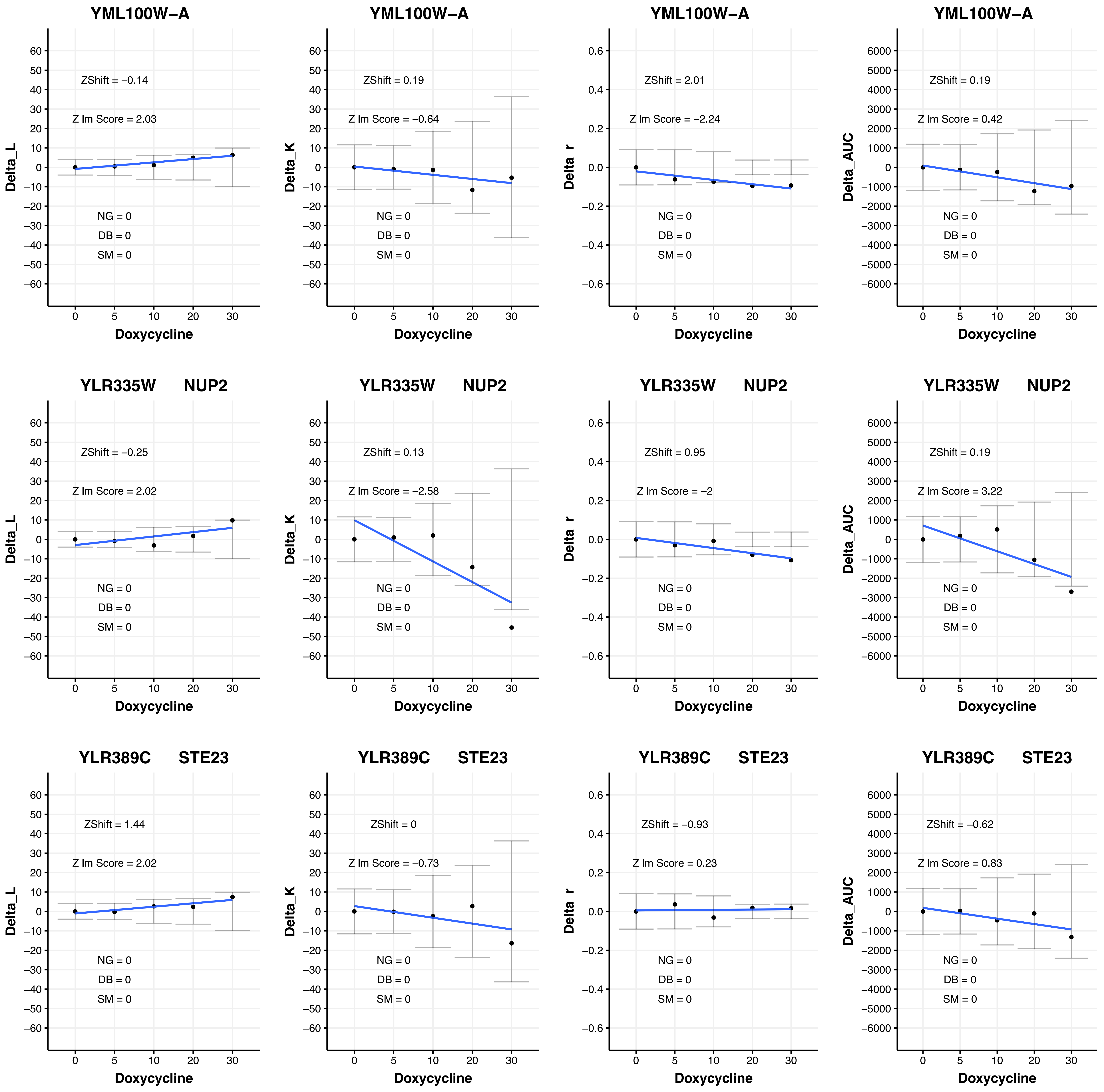
YML100W−A
ZShift = −0.14
Z lm Score = 2.03
NG = 0
DB = 0
SM = 0
Delta_L
Doxycycline
YML100W−A
ZShift = 0.19
Z lm Score = −0.64
NG = 0
DB = 0
SM = 0
Delta_K
Doxycycline
YML100W−A
ZShift = 2.01
Z lm Score = −2.24
NG = 0
DB = 0
SM = 0
Delta_r
Doxycycline
YML100W−A
ZShift = 0.19
Z lm Score = 0.42
NG = 0
DB = 0
SM = 0
Delta_AUC
Doxycycline
YLR335W NUP2
ZShift = −0.25
Z lm Score = 2.02
NG = 0
DB = 0
SM = 0
Delta_L
Doxycycline
YLR335W NUP2
ZShift = 0.13
Z lm Score = −2.58
NG = 0
DB = 0
SM = 0
Delta_K
Doxycycline
YLR335W NUP2
ZShift = 0.95
Z lm Score = −2
NG = 0
DB = 0
SM = 0
Delta_r
Doxycycline
YLR335W NUP2
ZShift = 0.19
Z lm Score = 3.22
NG = 0
DB = 0
SM = 0
Delta_AUC
Doxycycline
YLR389C STE23
ZShift = 1.44
Z lm Score = 2.02
NG = 0
DB = 0
SM = 0
Delta_L
Doxycycline
YLR389C STE23
ZShift = 0
Z lm Score = −0.73
NG = 0
DB = 0
SM = 0
Delta_K
Doxycycline
YLR389C STE23
ZShift = −0.93
Z lm Score = 0.23
NG = 0
DB = 0
SM = 0
Delta_r
Doxycycline
YLR389C STE23
ZShift = −0.62
Z lm Score = 0.83
NG = 0
DB = 0
SM = 0
Delta_AUC
Doxycycline

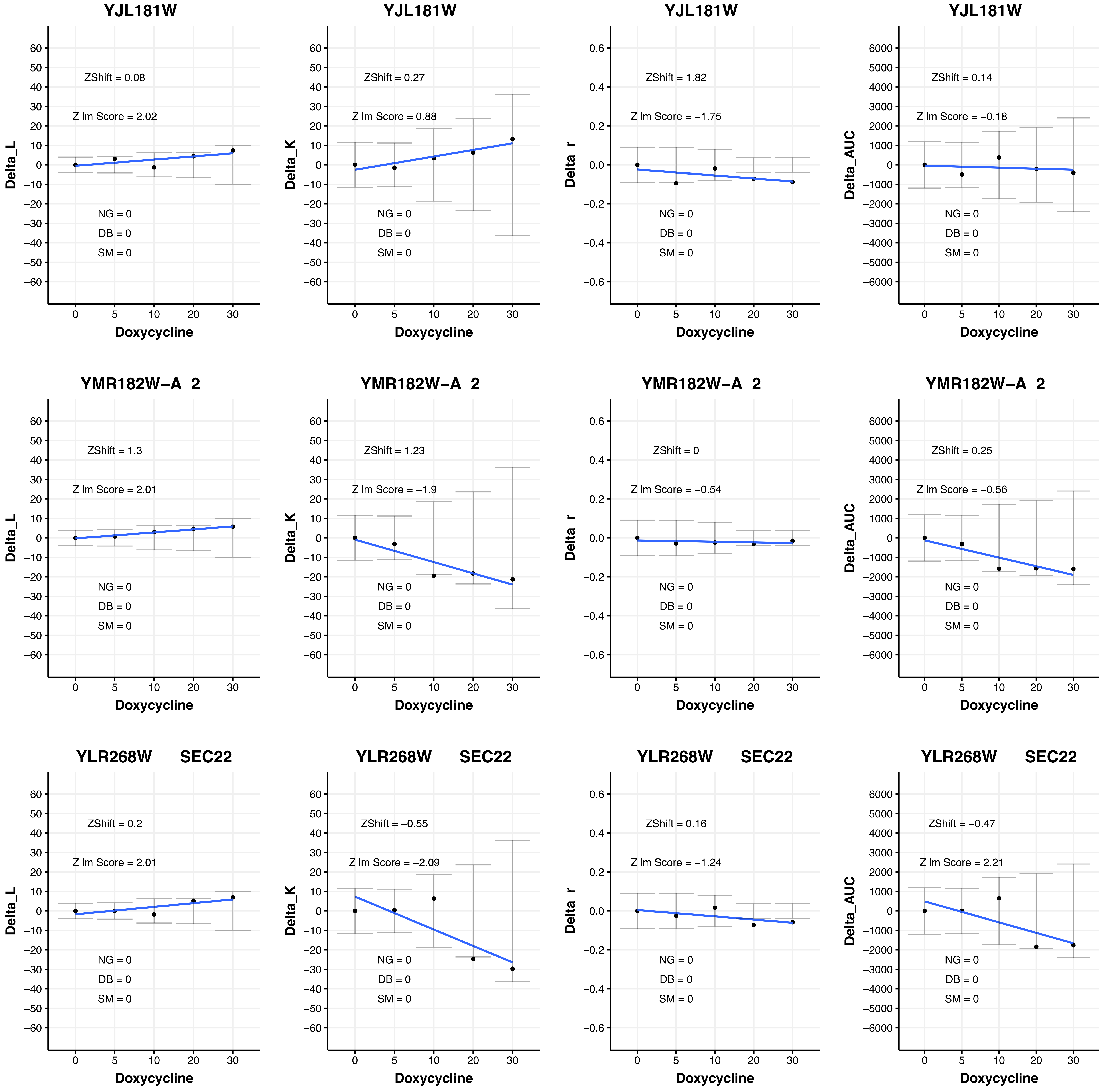
YJL181W
ZShift = 0.08
Z lm Score = 2.02
NG = 0
DB = 0
SM = 0
Delta_L
Doxycycline
YJL181W
ZShift = 0.27
Z lm Score = 0.88
NG = 0
DB = 0
SM = 0
Delta_K
Doxycycline
YJL181W
ZShift = 1.82
Z lm Score = −1.75
NG = 0
DB = 0
SM = 0
Delta_r
Doxycycline
YJL181W
ZShift = 0.14
Z lm Score = −0.18
NG = 0
DB = 0
SM = 0
Delta_AUC
Doxycycline
YMR182W−A_2
ZShift = 1.3
Z lm Score = 2.01
NG = 0
DB = 0
SM = 0
Delta_L
Doxycycline
YMR182W−A_2
ZShift = 1.23
Z lm Score = −1.9
NG = 0
DB = 0
SM = 0
Delta_K
Doxycycline
YMR182W−A_2
ZShift = 0
Z lm Score = −0.54
NG = 0
DB = 0
SM = 0
Delta_r
Doxycycline
YMR182W−A_2
ZShift = 0.25
Z lm Score = −0.56
NG = 0
DB = 0
SM = 0
Delta_AUC
Doxycycline
YLR268W SEC22
ZShift = 0.2
Z lm Score = 2.01
NG = 0
DB = 0
SM = 0
Delta_L
Doxycycline
YLR268W SEC22
ZShift = −0.55
Z lm Score = −2.09
NG = 0
DB = 0
SM = 0
Delta_K
Doxycycline
YLR268W SEC22
ZShift = 0.16
Z lm Score = −1.24
NG = 0
DB = 0
SM = 0
Delta_r
Doxycycline
YLR268W SEC22
ZShift = −0.47
Z lm Score = 2.21
NG = 0
DB = 0
SM = 0
Delta_AUC
Doxycycline

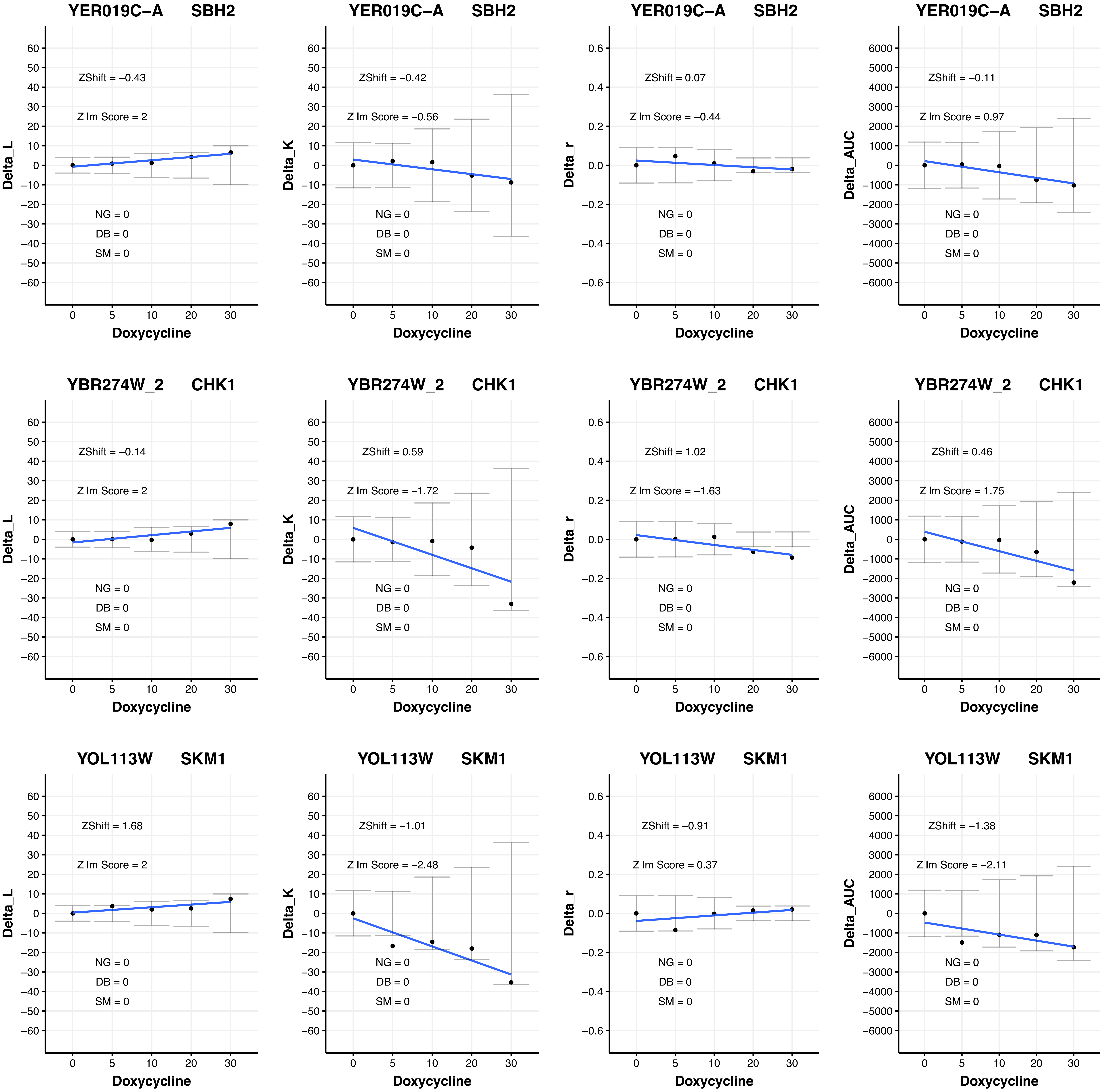
YER019C−A SBH2
ZShift = −0.43
Z lm Score = 2
NG = 0
DB = 0
SM = 0
Delta_L
Doxycycline
YER019C−A SBH2
ZShift = −0.42
Z lm Score = −0.56
NG = 0
DB = 0
SM = 0
Delta_K
Doxycycline
YER019C−A SBH2
ZShift = 0.07
Z lm Score = −0.44
NG = 0
DB = 0
SM = 0
Delta_r
Doxycycline
YER019C−A SBH2
ZShift = −0.11
Z lm Score = 0.97
NG = 0
DB = 0
SM = 0
Delta_AUC
Doxycycline
YBR274W_2 CHK1
ZShift = −0.14
Z lm Score = 2
NG = 0
DB = 0
SM = 0
Delta_L
Doxycycline
YBR274W_2 CHK1
ZShift = 0.59
Z lm Score = −1.72
NG = 0
DB = 0
SM = 0
Delta_K
Doxycycline
YBR274W_2 CHK1
ZShift = 1.02
Z lm Score = −1.63
NG = 0
DB = 0
SM = 0
Delta_r
Doxycycline
YBR274W_2 CHK1
ZShift = 0.46
Z lm Score = 1.75
NG = 0
DB = 0
SM = 0
Delta_AUC
Doxycycline
YOL113W SKM1
ZShift = 1.68
Z lm Score = 2
NG = 0
DB = 0
SM = 0
Delta_L
Doxycycline
YOL113W SKM1
ZShift = −1.01
Z lm Score = −2.48
NG = 0
DB = 0
SM = 0
Delta_K
Doxycycline
YOL113W SKM1
ZShift = −0.91
Z lm Score = 0.37
NG = 0
DB = 0
SM = 0
Delta_r
Doxycycline
YOL113W SKM1
ZShift = −1.38
Z lm Score = −2.11
NG = 0
DB = 0
SM = 0
Delta_AUC
Doxycycline

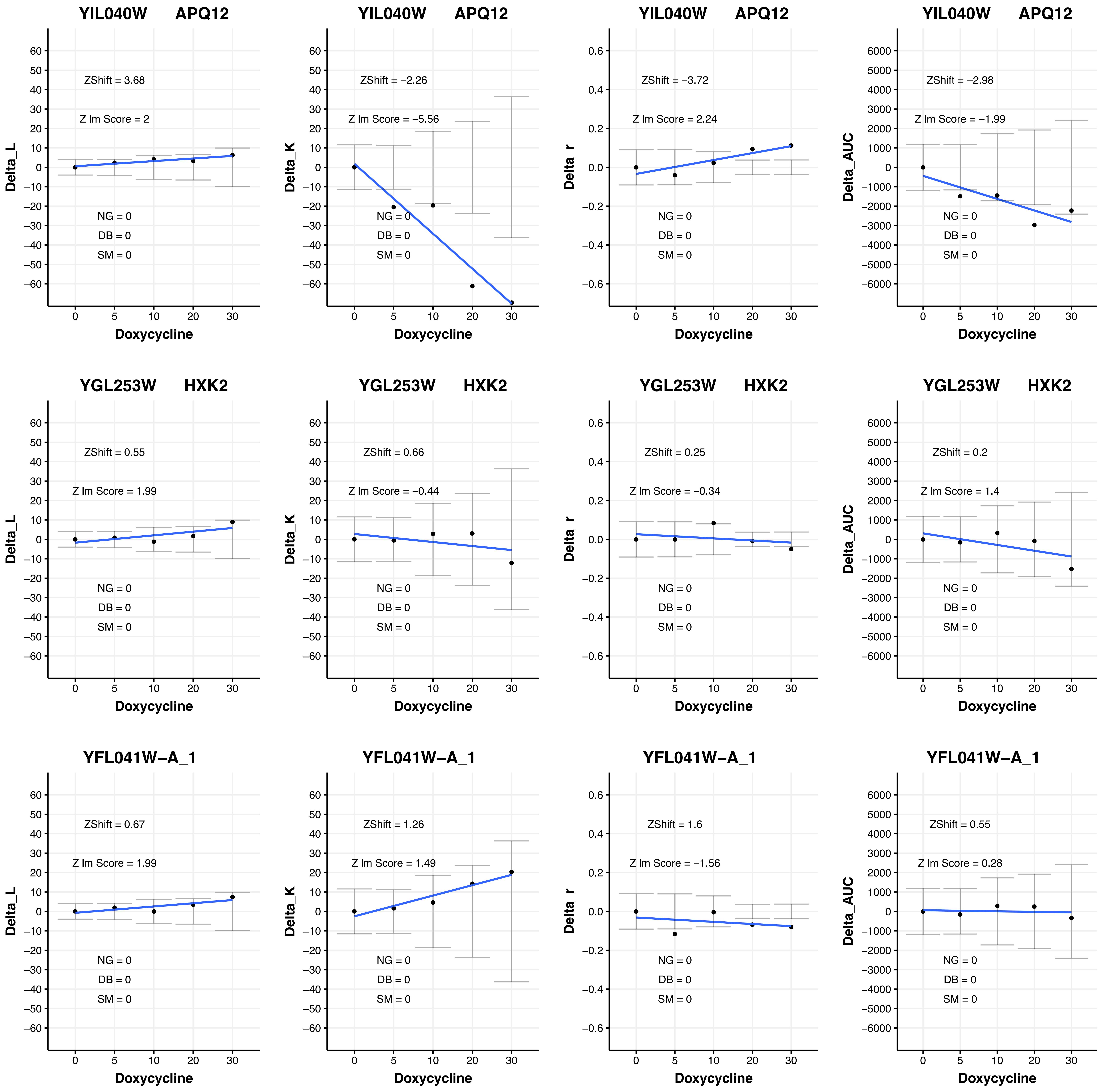
YIL040W APQ12
ZShift = 3.68
Z lm Score = 2
NG = 0
DB = 0
SM = 0
Delta_L
Doxycycline
YIL040W APQ12
ZShift = −2.26
Z lm Score = −5.56
NG = 0
DB = 0
SM = 0
Delta_K
Doxycycline
YIL040W APQ12
ZShift = −3.72
Z lm Score = 2.24
NG = 0
DB = 0
SM = 0
Delta_r
Doxycycline
YIL040W APQ12
ZShift = −2.98
Z lm Score = −1.99
NG = 0
DB = 0
SM = 0
Delta_AUC
Doxycycline
YGL253W HXK2
ZShift = 0.55
Z lm Score = 1.99
NG = 0
DB = 0
SM = 0
Delta_L
Doxycycline
YGL253W HXK2
ZShift = 0.66
Z lm Score = −0.44
NG = 0
DB = 0
SM = 0
Delta_K
Doxycycline
YGL253W HXK2
ZShift = 0.25
Z lm Score = −0.34
NG = 0
DB = 0
SM = 0
Delta_r
Doxycycline
YGL253W HXK2
ZShift = 0.2
Z lm Score = 1.4
NG = 0
DB = 0
SM = 0
Delta_AUC
Doxycycline
YFL041W−A_1
ZShift = 0.67
Z lm Score = 1.99
NG = 0
DB = 0
SM = 0
Delta_L
Doxycycline
YFL041W−A_1
ZShift = 1.26
Z lm Score = 1.49
NG = 0
DB = 0
SM = 0
Delta_K
Doxycycline
YFL041W−A_1
ZShift = 1.6
Z lm Score = −1.56
NG = 0
DB = 0
SM = 0
Delta_r
Doxycycline
YFL041W−A_1
ZShift = 0.55
Z lm Score = 0.28
NG = 0
DB = 0
SM = 0
Delta_AUC
Doxycycline

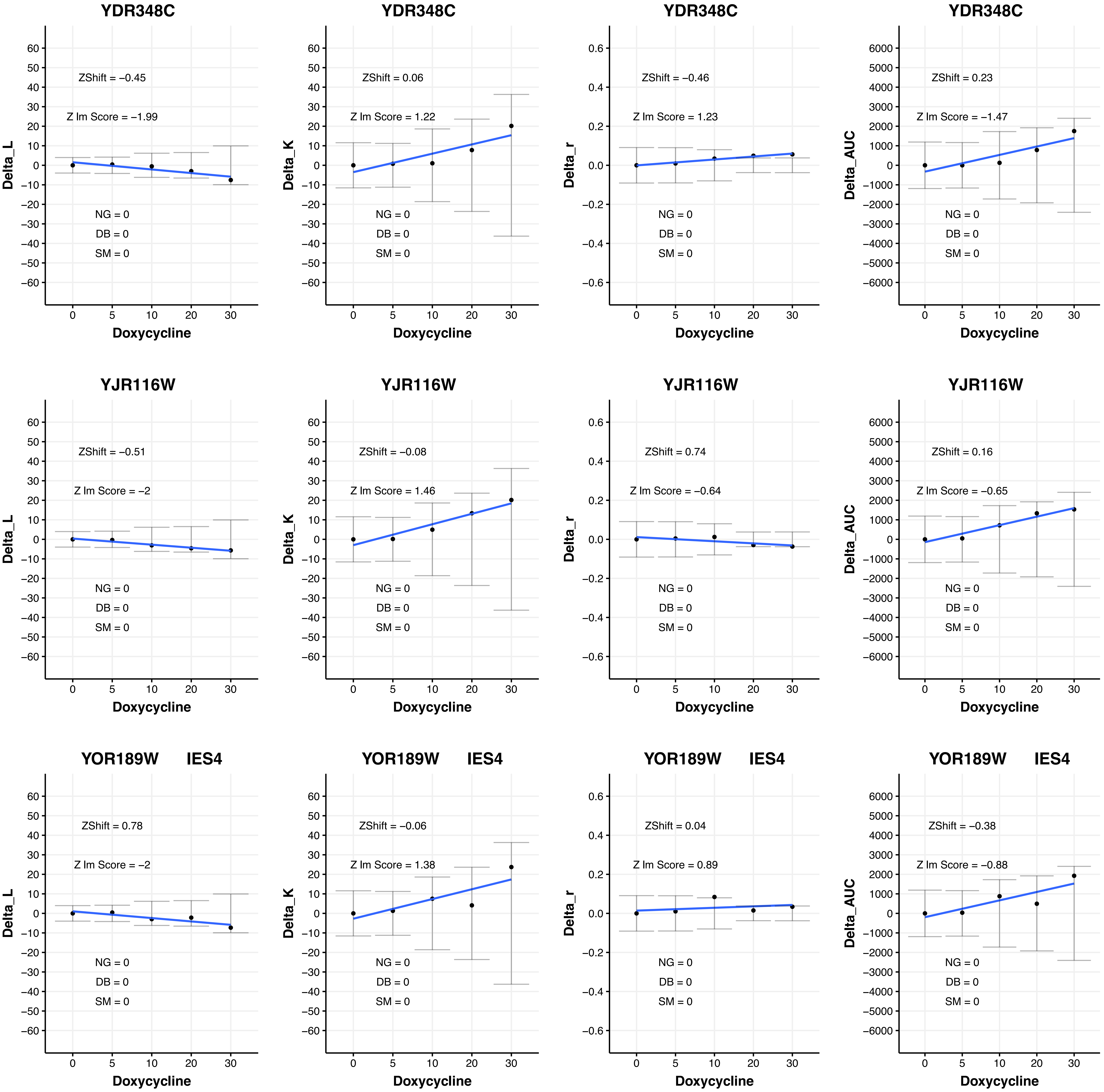
YDR348C
ZShift = −0.45
Z lm Score = −1.99
NG = 0
DB = 0
SM = 0
Delta_L
Doxycycline
YDR348C
ZShift = 0.06
Z lm Score = 1.22
NG = 0
DB = 0
SM = 0
Delta_K
Doxycycline
YDR348C
ZShift = −0.46
Z lm Score = 1.23
NG = 0
DB = 0
SM = 0
Delta_r
Doxycycline
YDR348C
ZShift = 0.23
Z lm Score = −1.47
NG = 0
DB = 0
SM = 0
Delta_AUC
Doxycycline
YJR116W
ZShift = −0.51
Z lm Score = −2
NG = 0
DB = 0
SM = 0
Delta_L
Doxycycline
YJR116W
ZShift = −0.08
Z lm Score = 1.46
NG = 0
DB = 0
SM = 0
Delta_K
Doxycycline
YJR116W
ZShift = 0.74
Z lm Score = −0.64
NG = 0
DB = 0
SM = 0
Delta_r
Doxycycline
YJR116W
ZShift = 0.16
Z lm Score = −0.65
NG = 0
DB = 0
SM = 0
Delta_AUC
Doxycycline
YOR189W IES4
ZShift = 0.78
Z lm Score = −2
NG = 0
DB = 0
SM = 0
Delta_L
Doxycycline
YOR189W IES4
ZShift = −0.06
Z lm Score = 1.38
NG = 0
DB = 0
SM = 0
Delta_K
Doxycycline
YOR189W IES4
ZShift = 0.04
Z lm Score = 0.89
NG = 0
DB = 0
SM = 0
Delta_r
Doxycycline
YOR189W IES4
ZShift = −0.38
Z lm Score = −0.88
NG = 0
DB = 0
SM = 0
Delta_AUC
Doxycycline

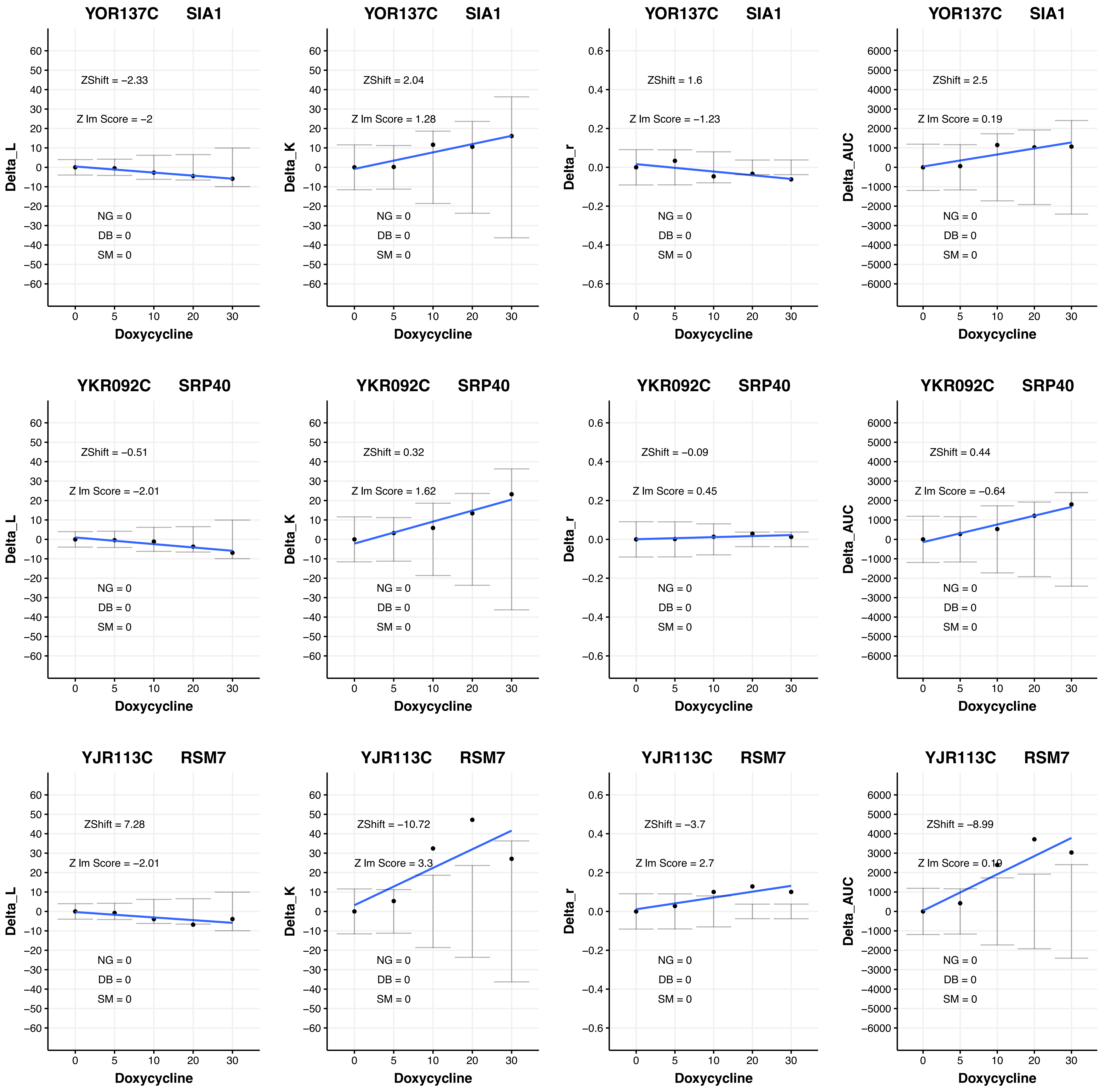

YOR137C SIA1
ZShift = −2.33
Z lm Score = −2
NG = 0
DB = 0
SM = 0
Delta_L
Doxycycline
YOR137C SIA1
ZShift = 2.04
Z lm Score = 1.28
NG = 0
DB = 0
SM = 0
Delta_K
Doxycycline
YOR137C SIA1
ZShift = 1.6
Z lm Score = −1.23
NG = 0
DB = 0
SM = 0
Delta_r
Doxycycline
YOR137C SIA1
ZShift = 2.5
Z lm Score = 0.19
NG = 0
DB = 0
SM = 0
Delta_AUC
Doxycycline
YKR092C SRP40
ZShift = −0.51
Z lm Score = −2.01
NG = 0
DB = 0
SM = 0
Delta_L
Doxycycline
YKR092C SRP40
ZShift = 0.32
Z lm Score = 1.62
NG = 0
DB = 0
SM = 0
Delta_K
Doxycycline
YKR092C SRP40
ZShift = −0.09
Z lm Score = 0.45
NG = 0
DB = 0
SM = 0
Delta_r
Doxycycline
YKR092C SRP40
ZShift = 0.44
Z lm Score = −0.64
NG = 0
DB = 0
SM = 0
Delta_AUC
Doxycycline
YJR113C RSM7
ZShift = 7.28
Z lm Score = −2.01
NG = 0
DB = 0
SM = 0
Delta_L
Doxycycline
YJR113C RSM7
ZShift = −10.72
Z lm Score = 3.3
NG = 0
DB = 0
SM = 0
Delta_K
Doxycycline
YJR113C RSM7
ZShift = −3.7
Z lm Score = 2.7
NG = 0
DB = 0
SM = 0
Delta_r
Doxycycline
YJR113C RSM7
ZShift = −8.99
Z lm Score = 0.19
NG = 0
DB = 0
SM = 0
Delta_AUC
Doxycycline

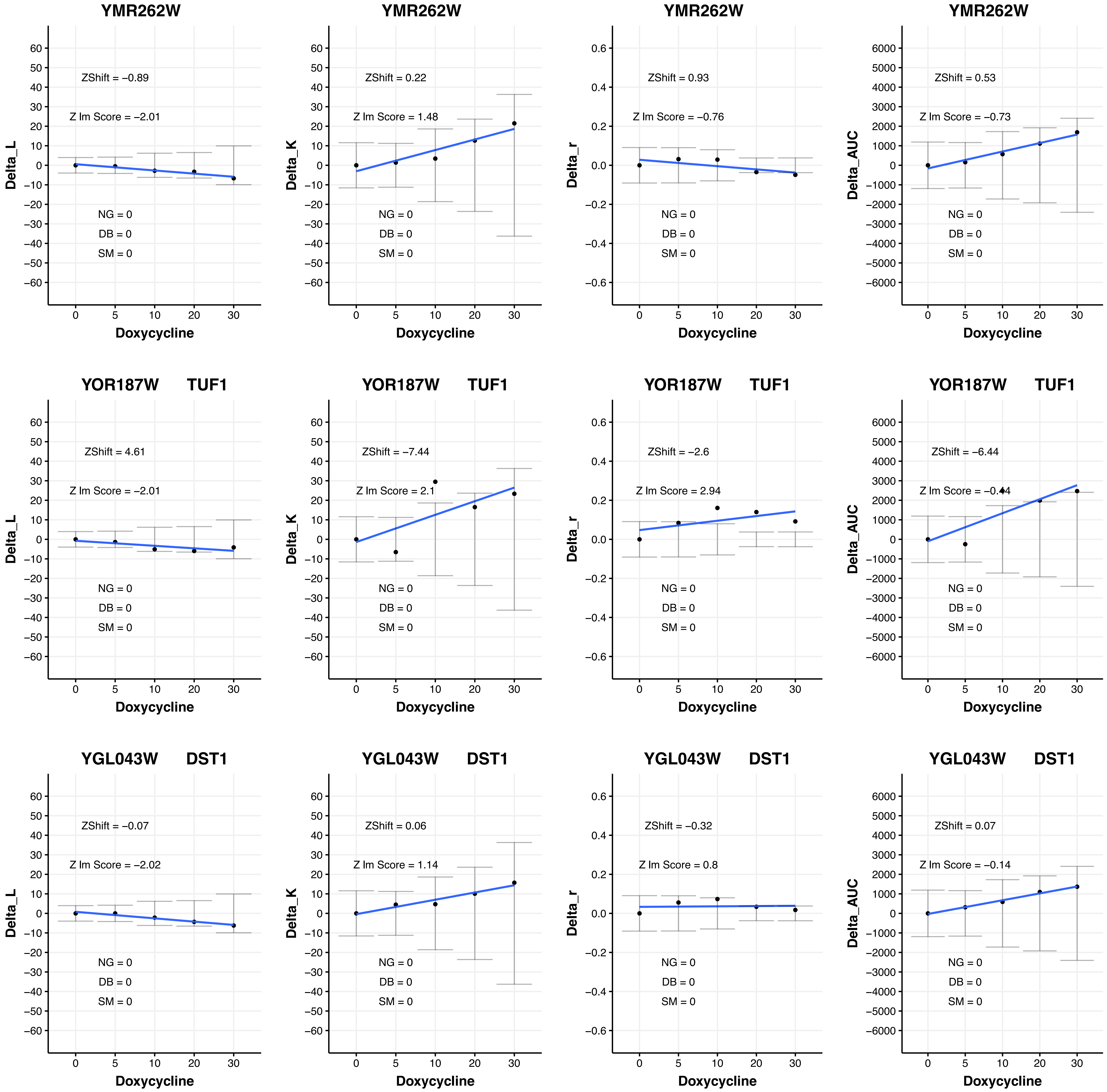

YMR262W
ZShift = −0.89
Z lm Score = −2.01
NG = 0
DB = 0
SM = 0
Delta_L
Doxycycline
YMR262W
ZShift = 0.22
Z lm Score = 1.48
NG = 0
DB = 0
SM = 0
Delta_K
Doxycycline
YMR262W
ZShift = 0.93
Z lm Score = −0.76
NG = 0
DB = 0
SM = 0
Delta_r
Doxycycline
YMR262W
ZShift = 0.53
Z lm Score = −0.73
NG = 0
DB = 0
SM = 0
Delta_AUC
Doxycycline
YOR187W TUF1
ZShift = 4.61
Z lm Score = −2.01
NG = 0
DB = 0
SM = 0
Delta_L
Doxycycline
YOR187W TUF1
ZShift = −7.44
Z lm Score = 2.1
NG = 0
DB = 0
SM = 0
Delta_K
Doxycycline
YOR187W TUF1
ZShift = −2.6
Z lm Score = 2.94
NG = 0
DB = 0
SM = 0
Delta_r
Doxycycline
YOR187W TUF1
ZShift = −6.44
Z lm Score = −0.44
NG = 0
DB = 0
SM = 0
Delta_AUC
Doxycycline
YGL043W DST1
ZShift = −0.07
Z lm Score = −2.02
NG = 0
DB = 0
SM = 0
Delta_L
Doxycycline
YGL043W DST1
ZShift = 0.06
Z lm Score = 1.14
NG = 0
DB = 0
SM = 0
Delta_K
Doxycycline
YGL043W DST1
ZShift = −0.32
Z lm Score = 0.8
NG = 0
DB = 0
SM = 0
Delta_r
Doxycycline
YGL043W DST1
ZShift = 0.07
Z lm Score = −0.14
NG = 0
DB = 0
SM = 0
Delta_AUC
Doxycycline

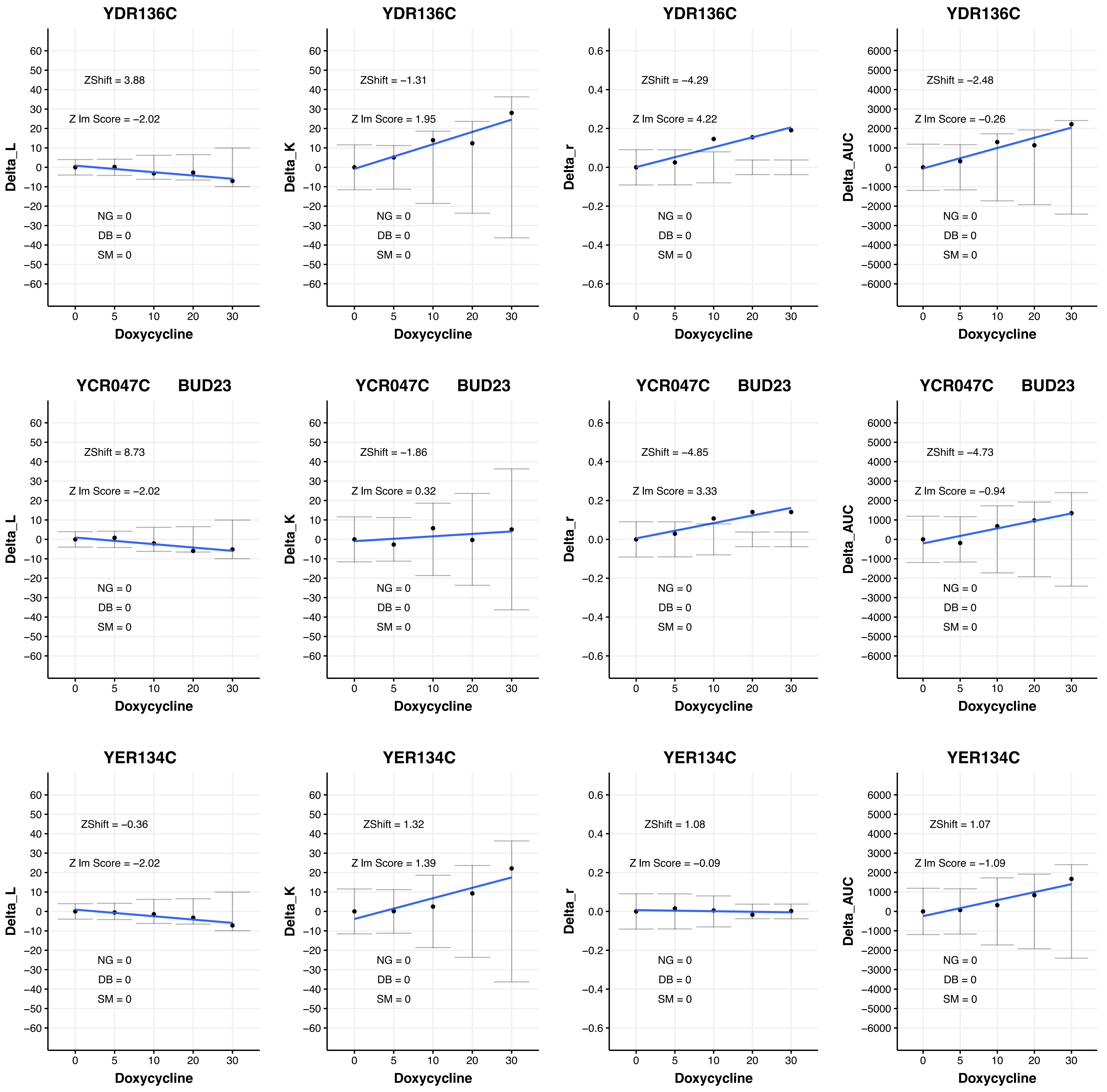

YDR136C
ZShift = 3.88
Z lm Score = −2.02
NG = 0
DB = 0
SM = 0
Delta_L
Doxycycline
YDR136C
ZShift = −1.31
Z lm Score = 1.95
NG = 0
DB = 0
SM = 0
Delta_K
Doxycycline
YDR136C
ZShift = −4.29
Z lm Score = 4.22
NG = 0
DB = 0
SM = 0
Delta_r
Doxycycline
YDR136C
ZShift = −2.48
Z lm Score = −0.26
NG = 0
DB = 0
SM = 0
Delta_AUC
Doxycycline
YCR047C BUD23
ZShift = 8.73
Z lm Score = −2.02
NG = 0
DB = 0
SM = 0
Delta_L
Doxycycline
YCR047C BUD23
ZShift = −1.86
Z lm Score = 0.32
NG = 0
DB = 0
SM = 0
Delta_K
Doxycycline
YCR047C BUD23
ZShift = −4.85
Z lm Score = 3.33
NG = 0
DB = 0
SM = 0
Delta_r
Doxycycline
YCR047C BUD23
ZShift = −4.73
Z lm Score = −0.94
NG = 0
DB = 0
SM = 0
Delta_AUC
Doxycycline
YER134C
ZShift = −0.36
Z lm Score = −2.02
NG = 0
DB = 0
SM = 0
Delta_L
Doxycycline
YER134C
ZShift = 1.32
Z lm Score = 1.39
NG = 0
DB = 0
SM = 0
Delta_K
Doxycycline
YER134C
ZShift = 1.08
Z lm Score = −0.09
NG = 0
DB = 0
SM = 0
Delta_r
Doxycycline
YER134C
ZShift = 1.07
Z lm Score = −1.09
NG = 0
DB = 0
SM = 0
Delta_AUC
Doxycycline

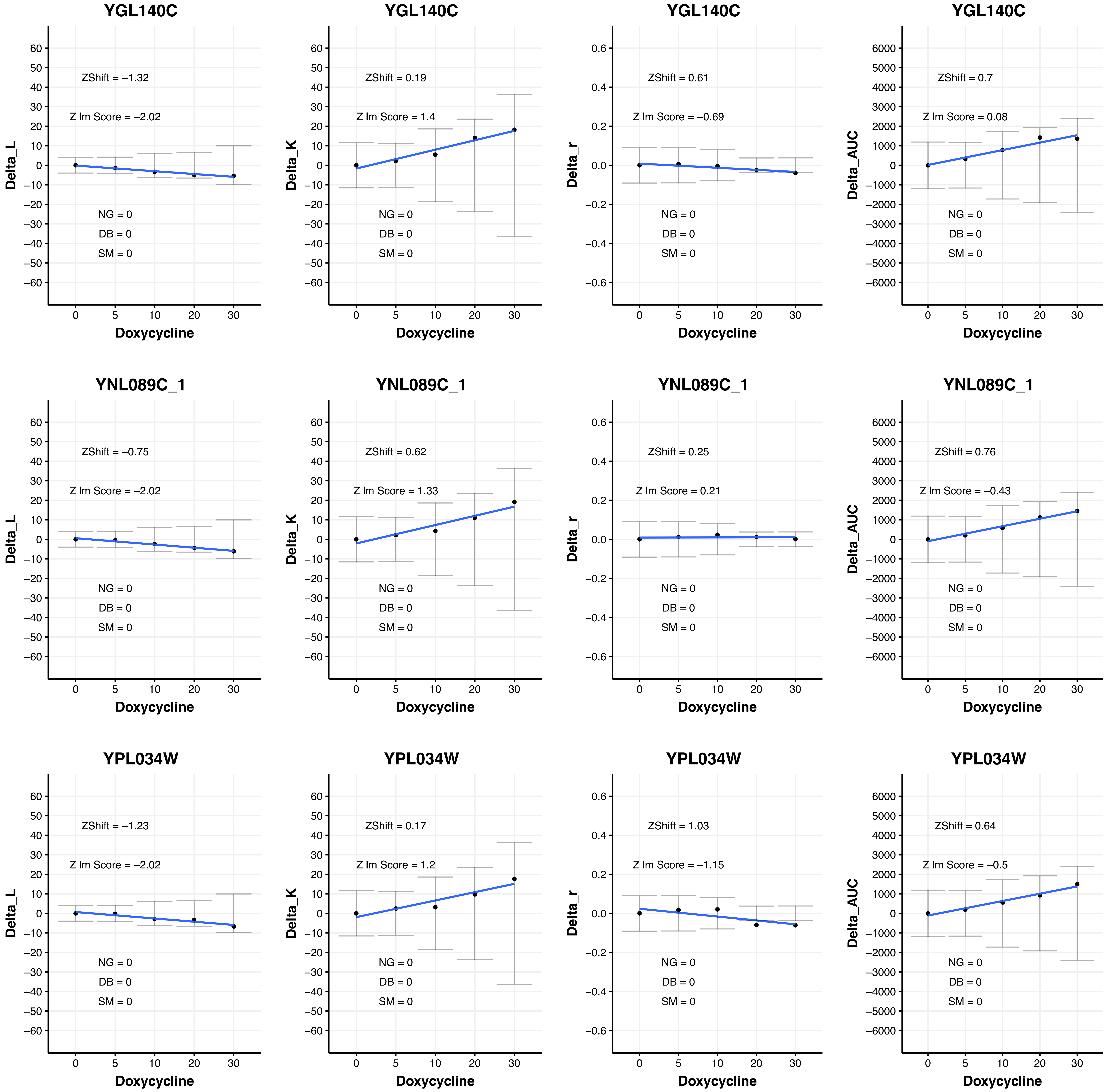
YGL140C
Delta_L
ZShift = −1.32
Z lm Score = −2.02
NG = 0
DB = 0
SM = 0
Doxycycline
YGL140C
Delta_K
ZShift = 0.19
Z lm Score = 1.4
NG = 0
DB = 0
SM = 0
Doxycycline
YGL140C
Delta_r
ZShift = 0.61
Z lm Score = −0.69
NG = 0
DB = 0
SM = 0
Doxycycline
YGL140C
Delta_AUC
ZShift = 0.7
Z lm Score = 0.08
NG = 0
DB = 0
SM = 0
Doxycycline
YNL089C_1
Delta_L
ZShift = −0.75
Z lm Score = −2.02
NG = 0
DB = 0
SM = 0
Doxycycline
YNL089C_1
Delta_K
ZShift = 0.62
Z lm Score = 1.33
NG = 0
DB = 0
SM = 0
Doxycycline
YNL089C_1
Delta_r
ZShift = 0.25
Z lm Score = 0.21
NG = 0
DB = 0
SM = 0
Doxycycline
YNL089C_1
Delta_AUC
ZShift = 0.76
Z lm Score = −0.43
NG = 0
DB = 0
SM = 0
Doxycycline
YPL034W
Delta_L
ZShift = −1.23
Z lm Score = −2.02
NG = 0
DB = 0
SM = 0
Doxycycline
YPL034W
Delta_K
ZShift = 0.17
Z lm Score = 1.2
NG = 0
DB = 0
SM = 0
Doxycycline
YPL034W
Delta_r
ZShift = 1.03
Z lm Score = −1.15
NG = 0
DB = 0
SM = 0
Doxycycline
YPL034W
Delta_AUC
ZShift = 0.64
Z lm Score = −0.5
NG = 0
DB = 0
SM = 0
Doxycycline

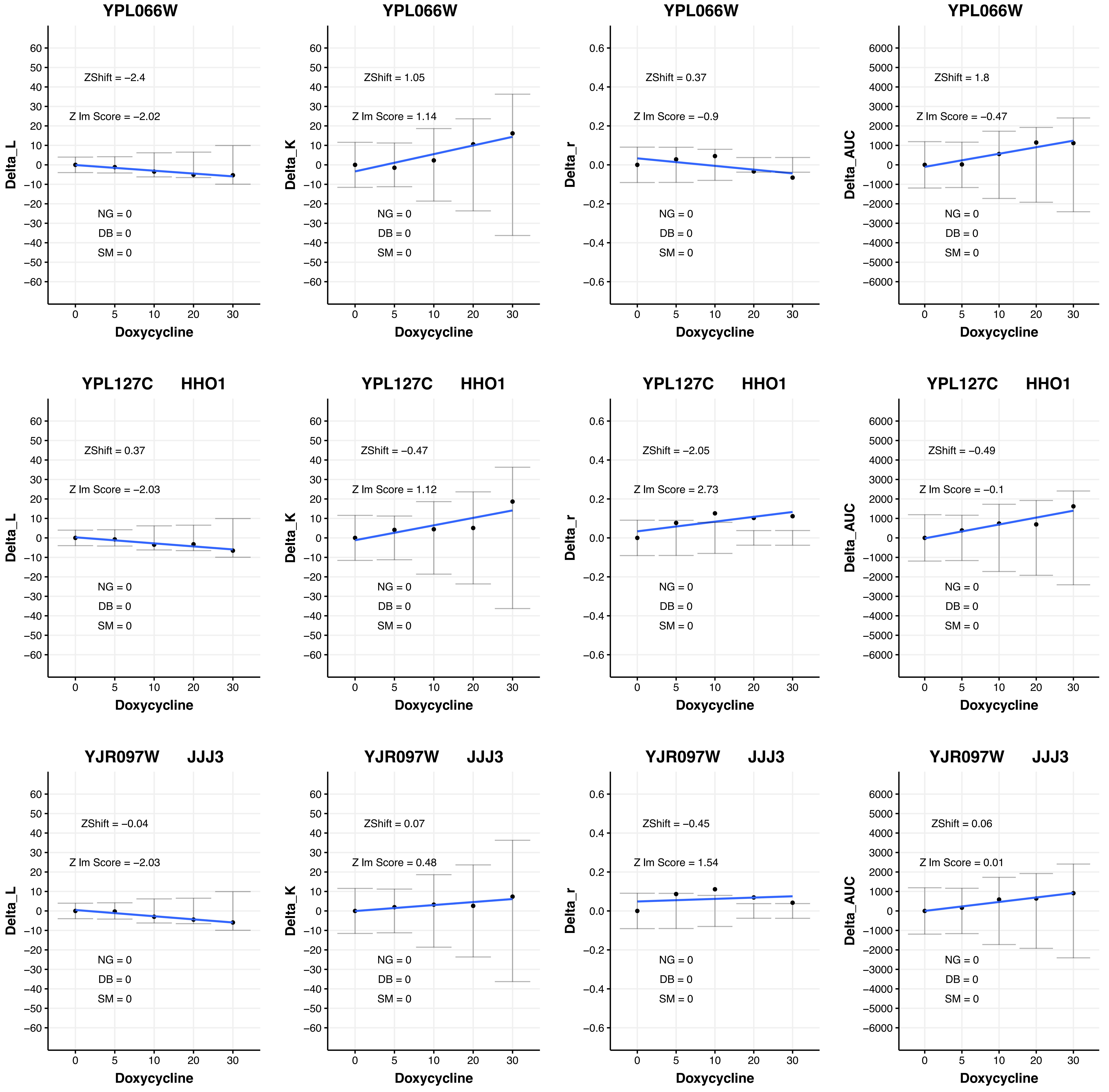
YPL066W
ZShift = −2.4
Z lm Score = −2.02
NG = 0
DB = 0
SM = 0
Delta_L
Doxycycline
YPL066W
ZShift = 1.05
Z lm Score = 1.14
NG = 0
DB = 0
SM = 0
Delta_K
Doxycycline
YPL066W
ZShift = 0.37
Z lm Score = −0.9
NG = 0
DB = 0
SM = 0
Delta_r
Doxycycline
YPL066W
ZShift = 1.8
Z lm Score = −0.47
NG = 0
DB = 0
SM = 0
Delta_AUC
Doxycycline
YPL127C HHO1
ZShift = 0.37
Z lm Score = −2.03
NG = 0
DB = 0
SM = 0
Delta_L
Doxycycline
YPL127C HHO1
ZShift = −0.47
Z lm Score = 1.12
NG = 0
DB = 0
SM = 0
Delta_K
Doxycycline
YPL127C HHO1
ZShift = −2.05
Z lm Score = 2.73
NG = 0
DB = 0
SM = 0
Delta_r
Doxycycline
YPL127C HHO1
ZShift = −0.49
Z lm Score = −0.1
NG = 0
DB = 0
SM = 0
Delta_AUC
Doxycycline
YJR097W JJJ3
ZShift = −0.04
Z lm Score = −2.03
NG = 0
DB = 0
SM = 0
Delta_L
Doxycycline
YJR097W JJJ3
ZShift = 0.07
Z lm Score = 0.48
NG = 0
DB = 0
SM = 0
Delta_K
Doxycycline
YJR097W JJJ3
ZShift = −0.45
Z lm Score = 1.54
NG = 0
DB = 0
SM = 0
Delta_r
Doxycycline
YJR097W JJJ3
ZShift = 0.06
Z lm Score = 0.01
NG = 0
DB = 0
SM = 0
Delta_AUC
Doxycycline

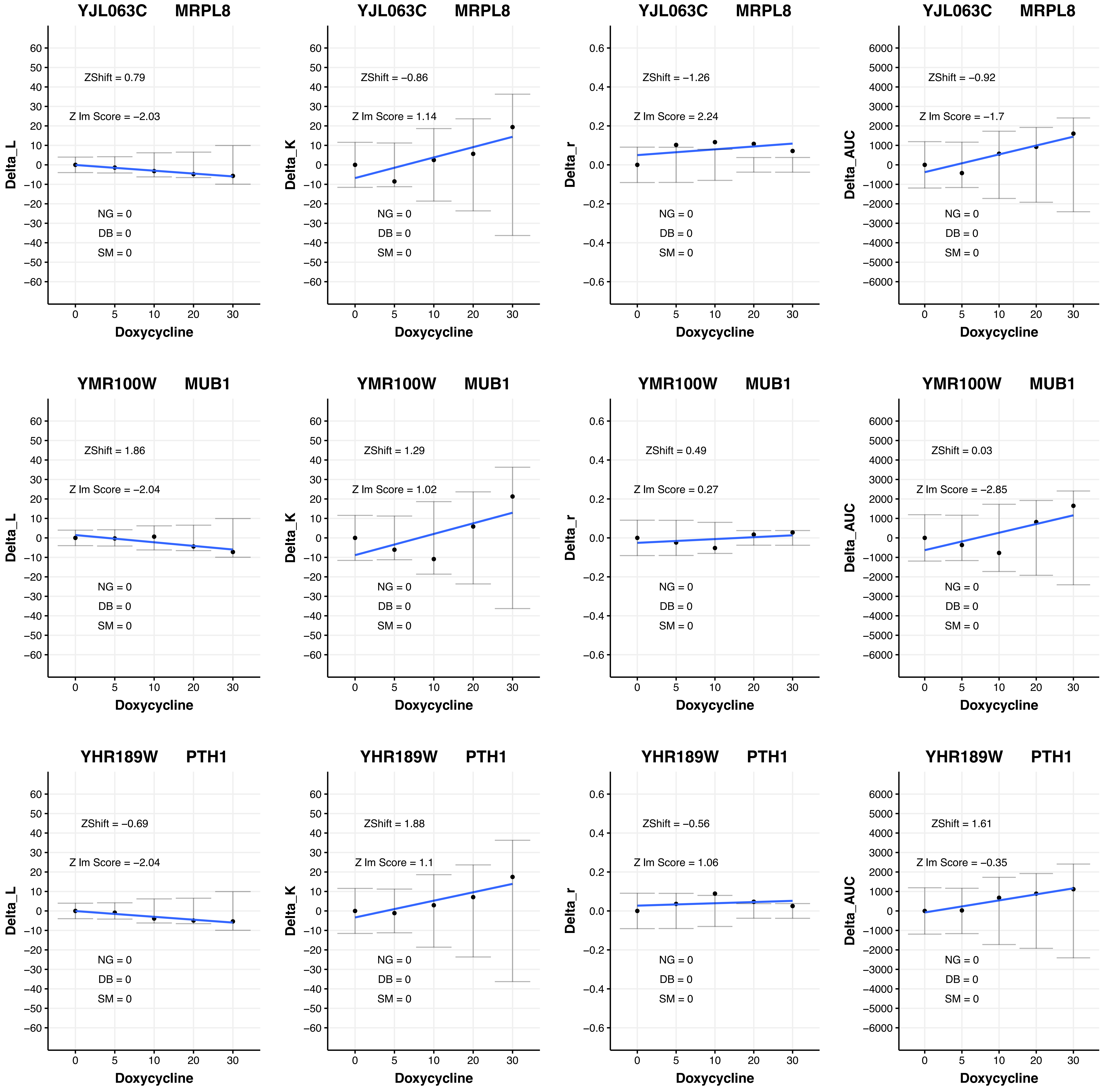
YJL063C MRPL8
ZShift = 0.79
Z lm Score = −2.03
NG = 0
DB = 0
SM = 0
Delta_L
Doxycycline
YJL063C MRPL8
ZShift = −0.86
Z lm Score = 1.14
NG = 0
DB = 0
SM = 0
Delta_K
Doxycycline
YJL063C MRPL8
ZShift = −1.26
Z lm Score = 2.24
NG = 0
DB = 0
SM = 0
Delta_r
Doxycycline
YJL063C MRPL8
ZShift = −0.92
Z lm Score = −1.7
NG = 0
DB = 0
SM = 0
Delta_AUC
Doxycycline
YMR100W MUB1
ZShift = 1.86
Z lm Score = −2.04
NG = 0
DB = 0
SM = 0
Delta_L
Doxycycline
YMR100W MUB1
ZShift = 1.29
Z lm Score = 1.02
NG = 0
DB = 0
SM = 0
Delta_K
Doxycycline
YMR100W MUB1
ZShift = 0.49
Z lm Score = 0.27
NG = 0
DB = 0
SM = 0
Delta_r
Doxycycline
YMR100W MUB1
ZShift = 0.03
Z lm Score = −2.85
NG = 0
DB = 0
SM = 0
Delta_AUC
Doxycycline
YHR189W PTH1
ZShift = −0.69
Z lm Score = −2.04
NG = 0
DB = 0
SM = 0
Delta_L
Doxycycline
YHR189W PTH1
ZShift = 1.88
Z lm Score = 1.1
NG = 0
DB = 0
SM = 0
Delta_K
Doxycycline
YHR189W PTH1
ZShift = −0.56
Z lm Score = 1.06
NG = 0
DB = 0
SM = 0
Delta_r
Doxycycline
YHR189W PTH1
ZShift = 1.61
Z lm Score = −0.35
NG = 0
DB = 0
SM = 0
Delta_AUC
Doxycycline

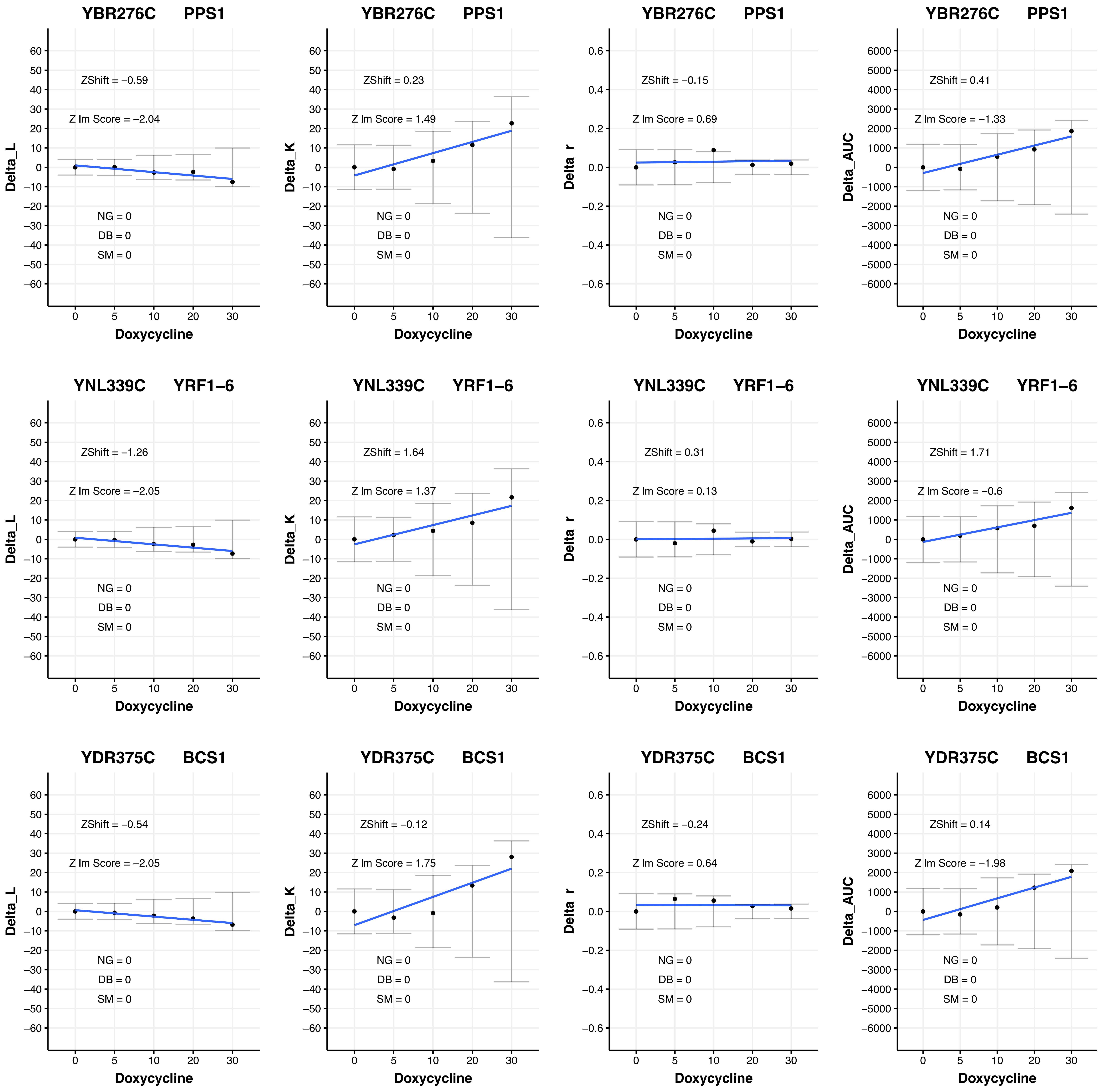

YBR276C PPS1
ZShift = −0.59
Z lm Score = −2.04
NG = 0
DB = 0
SM = 0
Delta_L
Doxycycline
YBR276C PPS1
ZShift = 0.23
Z lm Score = 1.49
NG = 0
DB = 0
SM = 0
Delta_K
Doxycycline
YBR276C PPS1
ZShift = −0.15
Z lm Score = 0.69
NG = 0
DB = 0
SM = 0
Delta_r
Doxycycline
YBR276C PPS1
ZShift = 0.41
Z lm Score = −1.33
NG = 0
DB = 0
SM = 0
Delta_AUC
Doxycycline
YNL339C YRF1−6
ZShift = −1.26
Z lm Score = −2.05
NG = 0
DB = 0
SM = 0
Delta_L
Doxycycline
YNL339C YRF1−6
ZShift = 1.64
Z lm Score = 1.37
NG = 0
DB = 0
SM = 0
Delta_K
Doxycycline
YNL339C YRF1−6
ZShift = 0.31
Z lm Score = 0.13
NG = 0
DB = 0
SM = 0
Delta_r
Doxycycline
YNL339C YRF1−6
ZShift = 1.71
Z lm Score = −0.6
NG = 0
DB = 0
SM = 0
Delta_AUC
Doxycycline
YDR375C BCS1
ZShift = −0.54
Z lm Score = −2.05
NG = 0
DB = 0
SM = 0
Delta_L
Doxycycline
YDR375C BCS1
ZShift = −0.12
Z lm Score = 1.75
NG = 0
DB = 0
SM = 0
Delta_K
Doxycycline
YDR375C BCS1
ZShift = −0.24
Z lm Score = 0.64
NG = 0
DB = 0
SM = 0
Delta_r
Doxycycline
YDR375C BCS1
ZShift = 0.14
Z lm Score = −1.98
NG = 0
DB = 0
SM = 0
Delta_AUC
Doxycycline

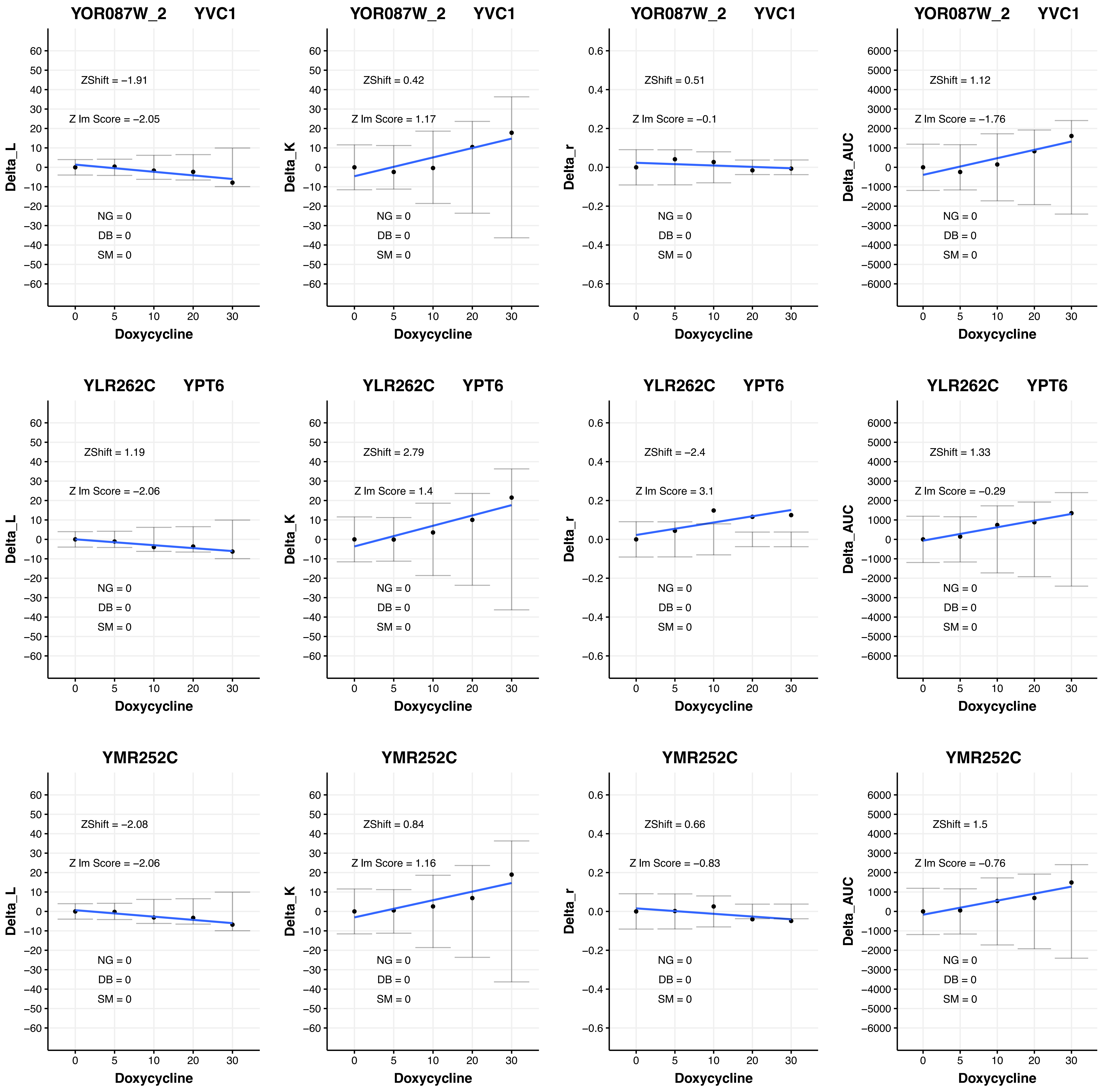
YOR087W_2 YVC1
ZShift = −1.91
Z lm Score = −2.05
NG = 0
DB = 0
SM = 0
Delta_L
Doxycycline
YOR087W_2 YVC1
ZShift = 0.42
Z lm Score = 1.17
NG = 0
DB = 0
SM = 0
Delta_K
Doxycycline
YOR087W_2 YVC1
ZShift = 0.51
Z lm Score = −0.1
NG = 0
DB = 0
SM = 0
Delta_r
Doxycycline
YOR087W_2 YVC1
ZShift = 1.12
Z lm Score = −1.76
NG = 0
DB = 0
SM = 0
Delta_AUC
Doxycycline
YLR262C YPT6
ZShift = 1.19
Z lm Score = −2.06
NG = 0
DB = 0
SM = 0
Delta_L
Doxycycline
YLR262C YPT6
ZShift = 2.79
Z lm Score = 1.4
NG = 0
DB = 0
SM = 0
Delta_K
Doxycycline
YLR262C YPT6
ZShift = −2.4
Z lm Score = 3.1
NG = 0
DB = 0
SM = 0
Delta_r
Doxycycline
YLR262C YPT6
ZShift = 1.33
Z lm Score = −0.29
NG = 0
DB = 0
SM = 0
Delta_AUC
Doxycycline
YMR252C
ZShift = −2.08
Z lm Score = −2.06
NG = 0
DB = 0
SM = 0
Delta_L
Doxycycline
YMR252C
ZShift = 0.84
Z lm Score = 1.16
NG = 0
DB = 0
SM = 0
Delta_K
Doxycycline
YMR252C
ZShift = 0.66
Z lm Score = −0.83
NG = 0
DB = 0
SM = 0
Delta_r
Doxycycline
YMR252C
ZShift = 1.5
Z lm Score = −0.76
NG = 0
DB = 0
SM = 0
Delta_AUC
Doxycycline

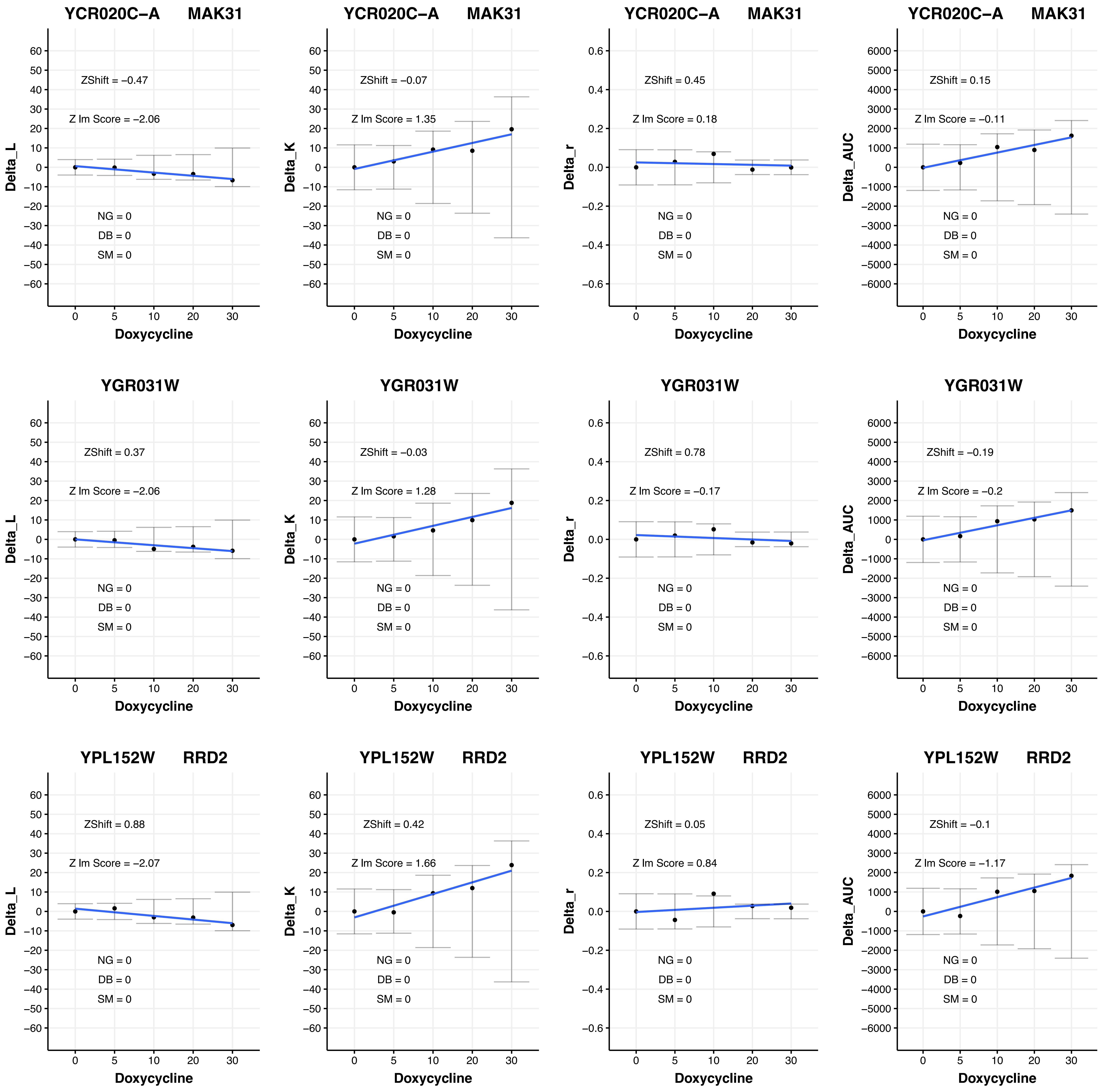

YCR020C−A MAK31
ZShift = −0.47
Z lm Score = −2.06
NG = 0
DB = 0
SM = 0
Delta_L
Doxycycline
YCR020C−A MAK31
ZShift = −0.07
Z lm Score = 1.35
NG = 0
DB = 0
SM = 0
Delta_K
Doxycycline
YCR020C−A MAK31
ZShift = 0.45
Z lm Score = 0.18
NG = 0
DB = 0
SM = 0
Delta_r
Doxycycline
YCR020C−A MAK31
ZShift = 0.15
Z lm Score = −0.11
NG = 0
DB = 0
SM = 0
Delta_AUC
Doxycycline
YGR031W
ZShift = 0.37
Z lm Score = −2.06
NG = 0
DB = 0
SM = 0
Delta_L
Doxycycline
YGR031W
ZShift = −0.03
Z lm Score = 1.28
NG = 0
DB = 0
SM = 0
Delta_K
Doxycycline
YGR031W
ZShift = 0.78
Z lm Score = −0.17
NG = 0
DB = 0
SM = 0
Delta_r
Doxycycline
YGR031W
ZShift = −0.19
Z lm Score = −0.2
NG = 0
DB = 0
SM = 0
Delta_AUC
Doxycycline
YPL152W RRD2
ZShift = 0.88
Z lm Score = −2.07
NG = 0
DB = 0
SM = 0
Delta_L
Doxycycline
YPL152W RRD2
ZShift = 0.42
Z lm Score = 1.66
NG = 0
DB = 0
SM = 0
Delta_K
Doxycycline
YPL152W RRD2
ZShift = 0.05
Z lm Score = 0.84
NG = 0
DB = 0
SM = 0
Delta_r
Doxycycline
YPL152W RRD2
ZShift = −0.1
Z lm Score = −1.17
NG = 0
DB = 0
SM = 0
Delta_AUC
Doxycycline

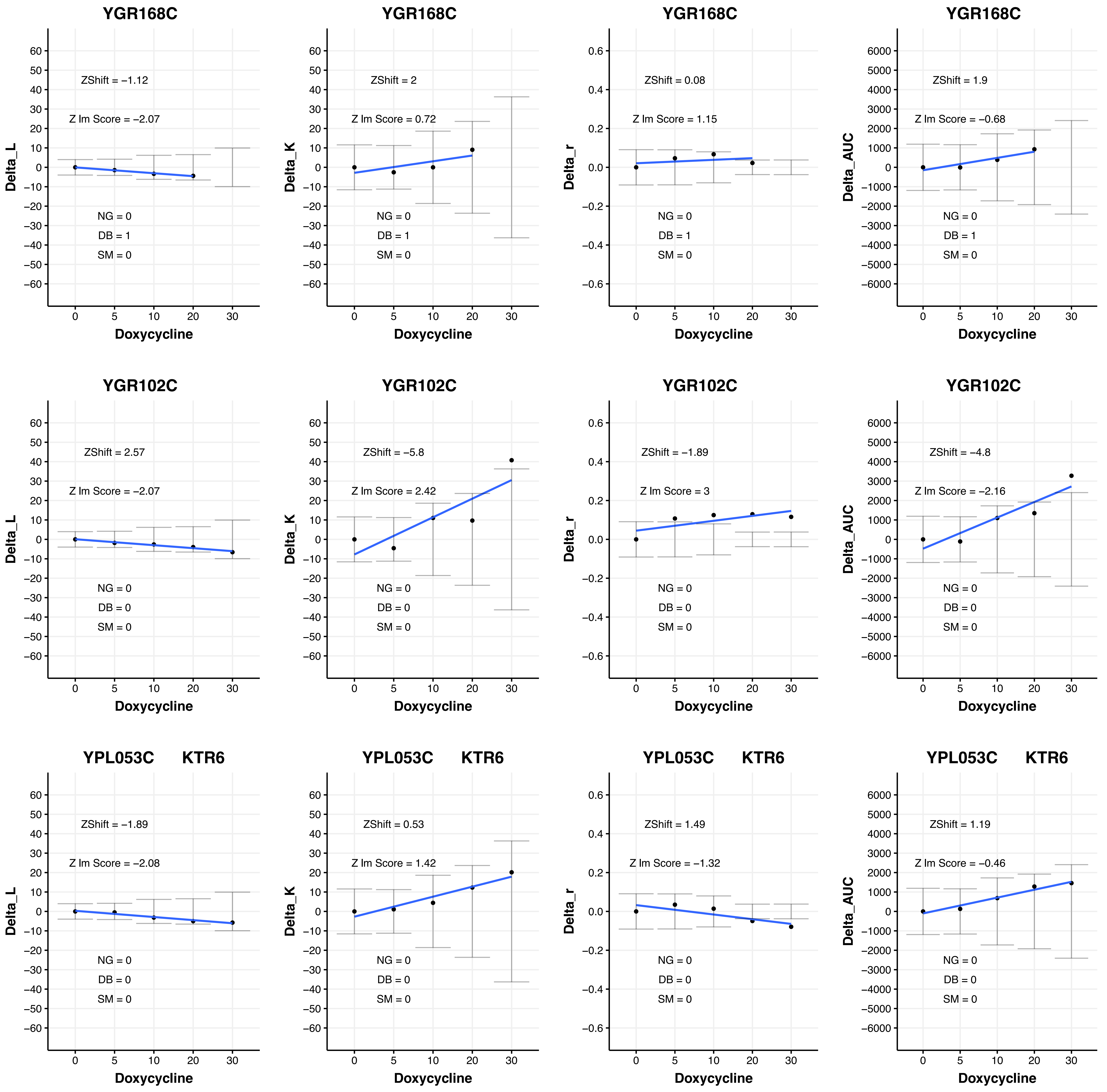

YGR168C
ZShift = −1.12
Z lm Score = −2.07
NG = 0
DB = 1
SM = 0
Delta_L
Doxycycline
YGR168C
ZShift = 2
Z lm Score = 0.72
NG = 0
DB = 1
SM = 0
Delta_K
Doxycycline
YGR168C
ZShift = 0.08
Z lm Score = 1.15
NG = 0
DB = 1
SM = 0
Delta_r
Doxycycline
YGR168C
ZShift = 1.9
Z lm Score = −0.68
NG = 0
DB = 1
SM = 0
Delta_AUC
Doxycycline
YGR102C
ZShift = 2.57
Z lm Score = −2.07
NG = 0
DB = 0
SM = 0
Delta_L
Doxycycline
YGR102C
ZShift = −5.8
Z lm Score = 2.42
NG = 0
DB = 0
SM = 0
Delta_K
Doxycycline
YGR102C
ZShift = −1.89
Z lm Score = 3
NG = 0
DB = 0
SM = 0
Delta_r
Doxycycline
YGR102C
ZShift = −4.8
Z lm Score = −2.16
NG = 0
DB = 0
SM = 0
Delta_AUC
Doxycycline
YPL053C KTR6
ZShift = −1.89
Z lm Score = −2.08
NG = 0
DB = 0
SM = 0
Delta_L
Doxycycline
YPL053C KTR6
ZShift = 0.53
Z lm Score = 1.42
NG = 0
DB = 0
SM = 0
Delta_K
Doxycycline
YPL053C KTR6
ZShift = 1.49
Z lm Score = −1.32
NG = 0
DB = 0
SM = 0
Delta_r
Doxycycline
YPL053C KTR6
ZShift = 1.19
Z lm Score = −0.46
NG = 0
DB = 0
SM = 0
Delta_AUC
Doxycycline

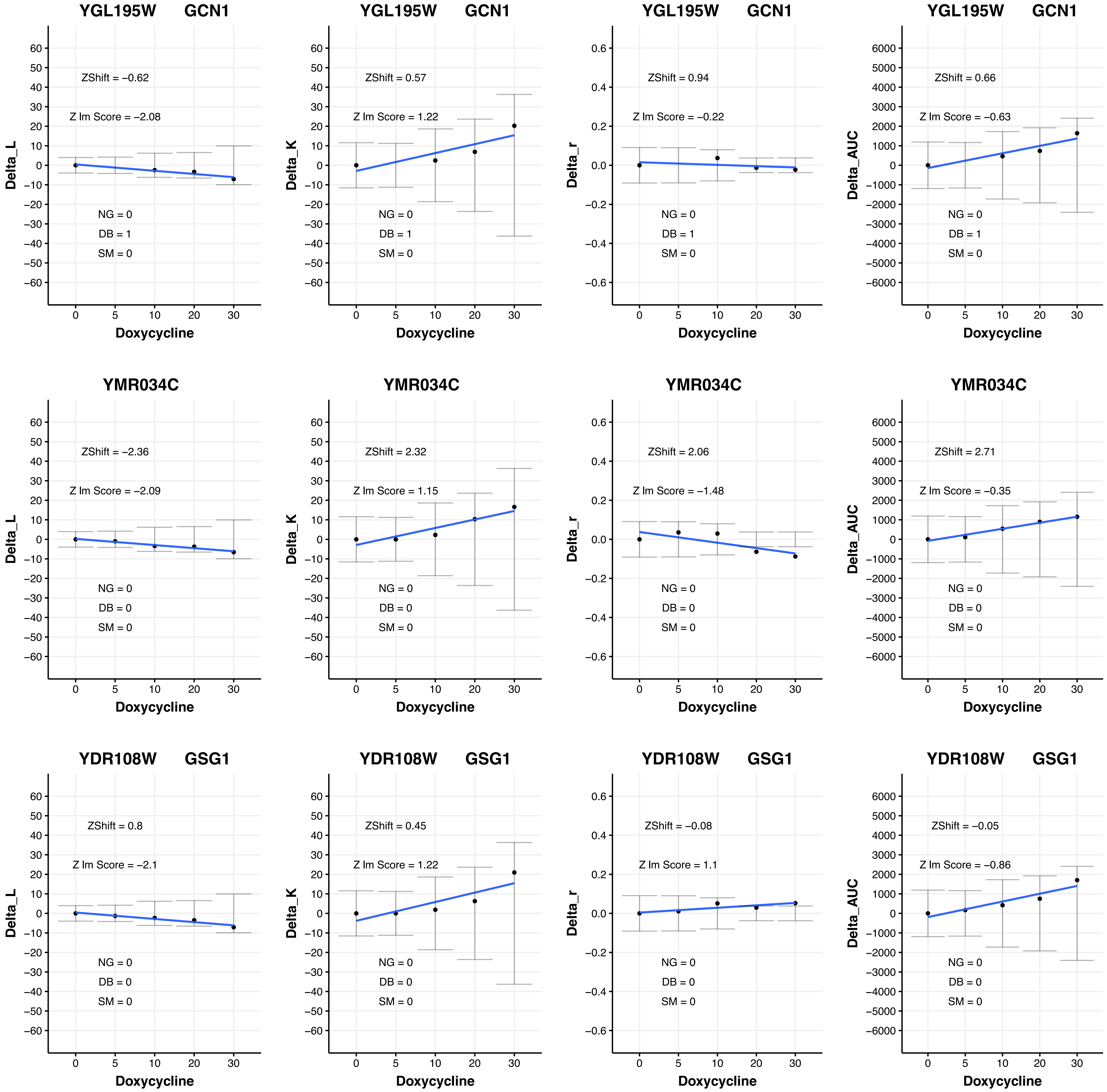

YGL195W GCN1
ZShift = −0.62
Z lm Score = −2.08
NG = 0
DB = 1
SM = 0
Delta_L
Doxycycline
YGL195W GCN1
ZShift = 0.57
Z lm Score = 1.22
NG = 0
DB = 1
SM = 0
Delta_K
Doxycycline
YGL195W GCN1
ZShift = 0.94
Z lm Score = −0.22
NG = 0
DB = 1
SM = 0
Delta_r
Doxycycline
YGL195W GCN1
ZShift = 0.66
Z lm Score = −0.63
NG = 0
DB = 1
SM = 0
Delta_AUC
Doxycycline
YMR034C
ZShift = −2.36
Z lm Score = −2.09
NG = 0
DB = 0
SM = 0
Delta_L
Doxycycline
YMR034C
ZShift = 2.32
Z lm Score = 1.15
NG = 0
DB = 0
SM = 0
Delta_K
Doxycycline
YMR034C
ZShift = 2.06
Z lm Score = −1.48
NG = 0
DB = 0
SM = 0
Delta_r
Doxycycline
YMR034C
ZShift = 2.71
Z lm Score = −0.35
NG = 0
DB = 0
SM = 0
Delta_AUC
Doxycycline
YDR108W GSG1
ZShift = 0.8
Z lm Score = −2.1
NG = 0
DB = 0
SM = 0
Delta_L
Doxycycline
YDR108W GSG1
ZShift = 0.45
Z lm Score = 1.22
NG = 0
DB = 0
SM = 0
Delta_K
Doxycycline
YDR108W GSG1
ZShift = −0.08
Z lm Score = 1.1
NG = 0
DB = 0
SM = 0
Delta_r
Doxycycline
YDR108W GSG1
ZShift = −0.05
Z lm Score = −0.86
NG = 0
DB = 0
SM = 0
Delta_AUC
Doxycycline

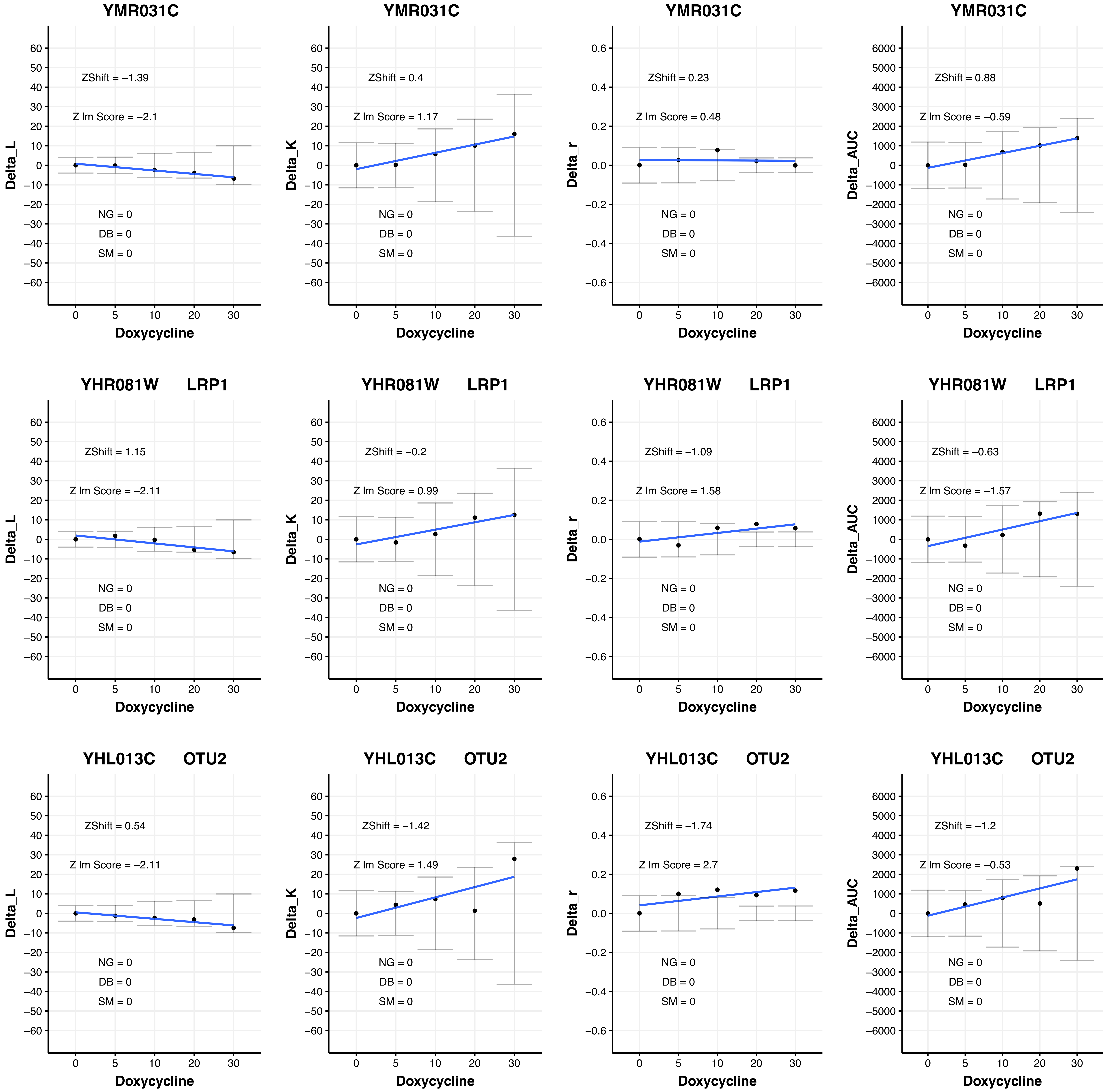

YMR031C
ZShift = −1.39
Z lm Score = −2.1
NG = 0
DB = 0
SM = 0
Delta_L
Doxycycline
YMR031C
ZShift = 0.4
Z lm Score = 1.17
NG = 0
DB = 0
SM = 0
Delta_K
Doxycycline
YMR031C
ZShift = 0.23
Z lm Score = 0.48
NG = 0
DB = 0
SM = 0
Delta_r
Doxycycline
YMR031C
ZShift = 0.88
Z lm Score = −0.59
NG = 0
DB = 0
SM = 0
Delta_AUC
Doxycycline
YHR081W LRP1
ZShift = 1.15
Z lm Score = −2.11
NG = 0
DB = 0
SM = 0
Delta_L
Doxycycline
YHR081W LRP1
ZShift = −0.2
Z lm Score = 0.99
NG = 0
DB = 0
SM = 0
Delta_K
Doxycycline
YHR081W LRP1
ZShift = −1.09
Z lm Score = 1.58
NG = 0
DB = 0
SM = 0
Delta_r
Doxycycline
YHR081W LRP1
ZShift = −0.63
Z lm Score = −1.57
NG = 0
DB = 0
SM = 0
Delta_AUC
Doxycycline
YHL013C OTU2
ZShift = 0.54
Z lm Score = −2.11
NG = 0
DB = 0
SM = 0
Delta_L
Doxycycline
YHL013C OTU2
ZShift = −1.42
Z lm Score = 1.49
NG = 0
DB = 0
SM = 0
Delta_K
Doxycycline
YHL013C OTU2
ZShift = −1.74
Z lm Score = 2.7
NG = 0
DB = 0
SM = 0
Delta_r
Doxycycline
YHL013C OTU2
ZShift = −1.2
Z lm Score = −0.53
NG = 0
DB = 0
SM = 0
Delta_AUC
Doxycycline

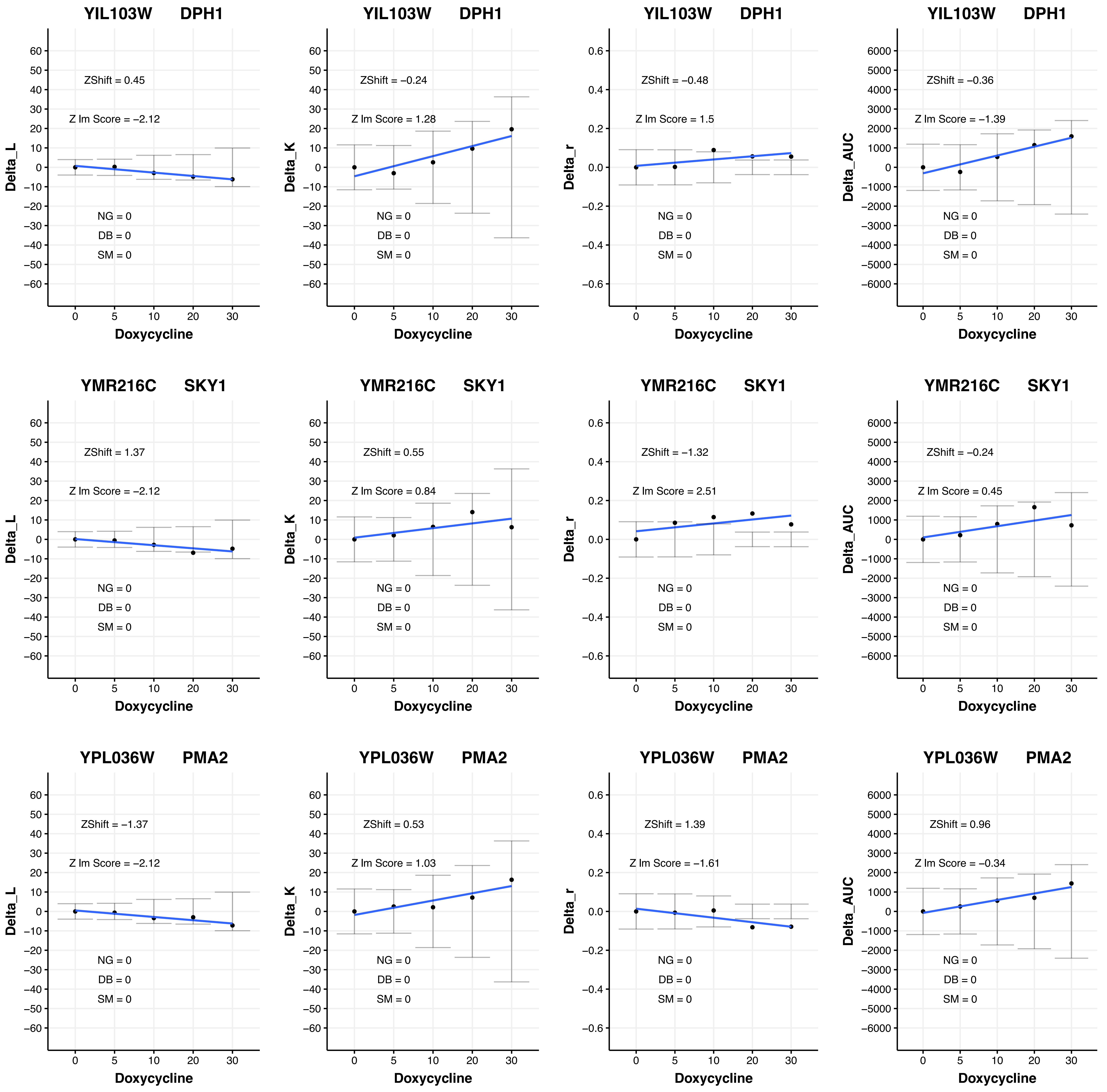

YIL103W DPH1
ZShift = 0.45
Z lm Score = −2.12
NG = 0
DB = 0
SM = 0
Delta_L
Doxycycline
YIL103W DPH1
ZShift = −0.24
Z lm Score = 1.28
NG = 0
DB = 0
SM = 0
Delta_K
Doxycycline
YIL103W DPH1
ZShift = −0.48
Z lm Score = 1.5
NG = 0
DB = 0
SM = 0
Delta_r
Doxycycline
YIL103W DPH1
ZShift = −0.36
Z lm Score = −1.39
NG = 0
DB = 0
SM = 0
Delta_AUC
Doxycycline
YMR216C SKY1
ZShift = 1.37
Z lm Score = −2.12
NG = 0
DB = 0
SM = 0
Delta_L
Doxycycline
YMR216C SKY1
ZShift = 0.55
Z lm Score = 0.84
NG = 0
DB = 0
SM = 0
Delta_K
Doxycycline
YMR216C SKY1
ZShift = −1.32
Z lm Score = 2.51
NG = 0
DB = 0
SM = 0
Delta_r
Doxycycline
YMR216C SKY1
ZShift = −0.24
Z lm Score = 0.45
NG = 0
DB = 0
SM = 0
Delta_AUC
Doxycycline
YPL036W PMA2
ZShift = −1.37
Z lm Score = −2.12
NG = 0
DB = 0
SM = 0
Delta_L
Doxycycline
YPL036W PMA2
ZShift = 0.53
Z lm Score = 1.03
NG = 0
DB = 0
SM = 0
Delta_K
Doxycycline
YPL036W PMA2
ZShift = 1.39
Z lm Score = −1.61
NG = 0
DB = 0
SM = 0
Delta_r
Doxycycline
YPL036W PMA2
ZShift = 0.96
Z lm Score = −0.34
NG = 0
DB = 0
SM = 0
Delta_AUC
Doxycycline

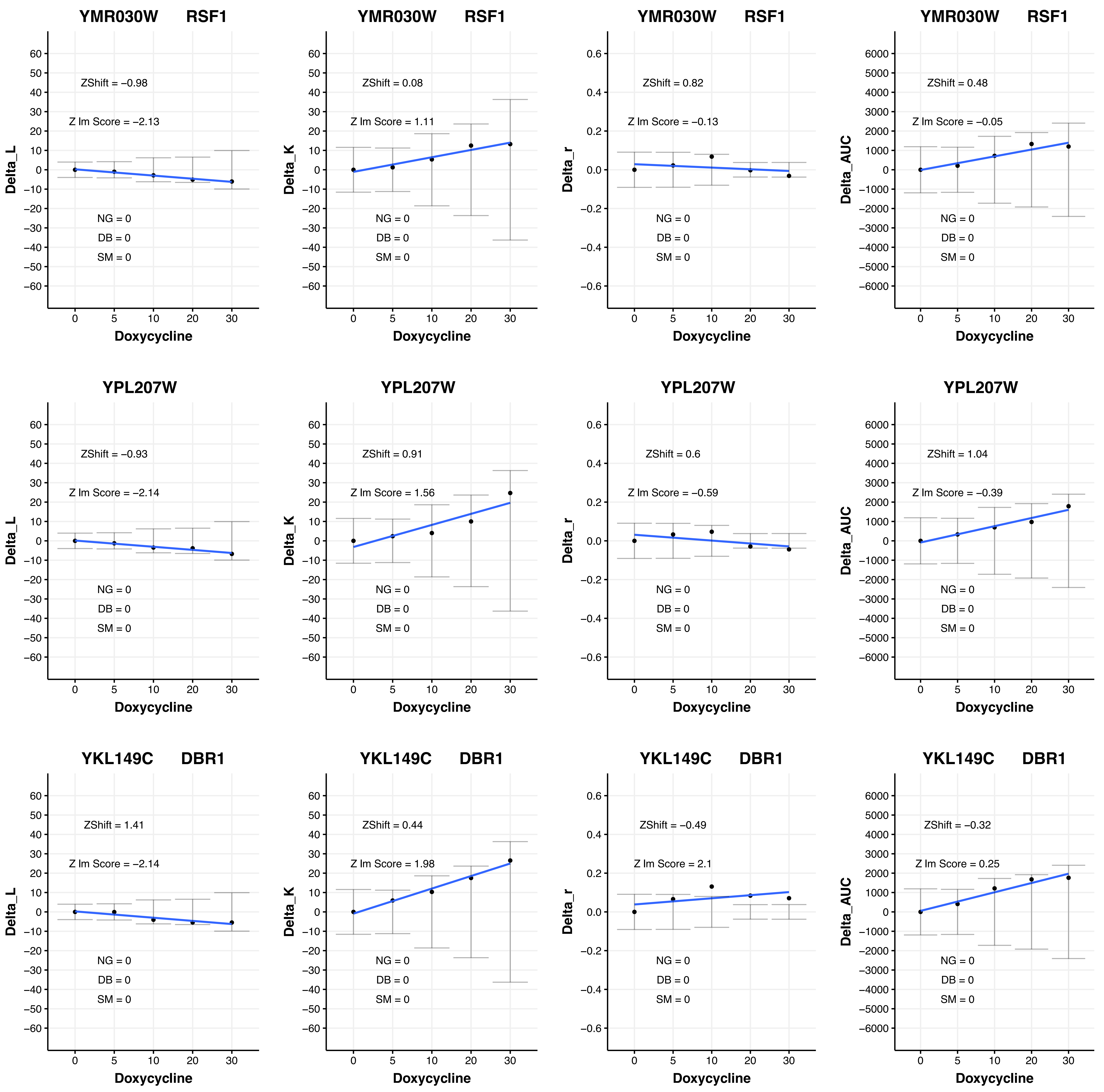

YMR030W RSF1
ZShift = −0.98
Z lm Score = −2.13
NG = 0
DB = 0
SM = 0
Delta_L
Doxycycline
YMR030W RSF1
ZShift = 0.08
Z lm Score = 1.11
NG = 0
DB = 0
SM = 0
Delta_K
Doxycycline
YMR030W RSF1
ZShift = 0.82
Z lm Score = −0.13
NG = 0
DB = 0
SM = 0
Delta_r
Doxycycline
YMR030W RSF1
ZShift = 0.48
Z lm Score = −0.05
NG = 0
DB = 0
SM = 0
Delta_AUC
Doxycycline
YPL207W
ZShift = −0.93
Z lm Score = −2.14
NG = 0
DB = 0
SM = 0
Delta_L
Doxycycline
YPL207W
ZShift = 0.91
Z lm Score = 1.56
NG = 0
DB = 0
SM = 0
Delta_K
Doxycycline
YPL207W
ZShift = 0.6
Z lm Score = −0.59
NG = 0
DB = 0
SM = 0
Delta_r
Doxycycline
YPL207W
ZShift = 1.04
Z lm Score = −0.39
NG = 0
DB = 0
SM = 0
Delta_AUC
Doxycycline
YKL149C DBR1
ZShift = 1.41
Z lm Score = −2.14
NG = 0
DB = 0
SM = 0
Delta_L
Doxycycline
YKL149C DBR1
ZShift = 0.44
Z lm Score = 1.98
NG = 0
DB = 0
SM = 0
Delta_K
Doxycycline
YKL149C DBR1
ZShift = −0.49
Z lm Score = 2.1
NG = 0
DB = 0
SM = 0
Delta_r
Doxycycline
YKL149C DBR1
ZShift = −0.32
Z lm Score = 0.25
NG = 0
DB = 0
SM = 0
Delta_AUC
Doxycycline

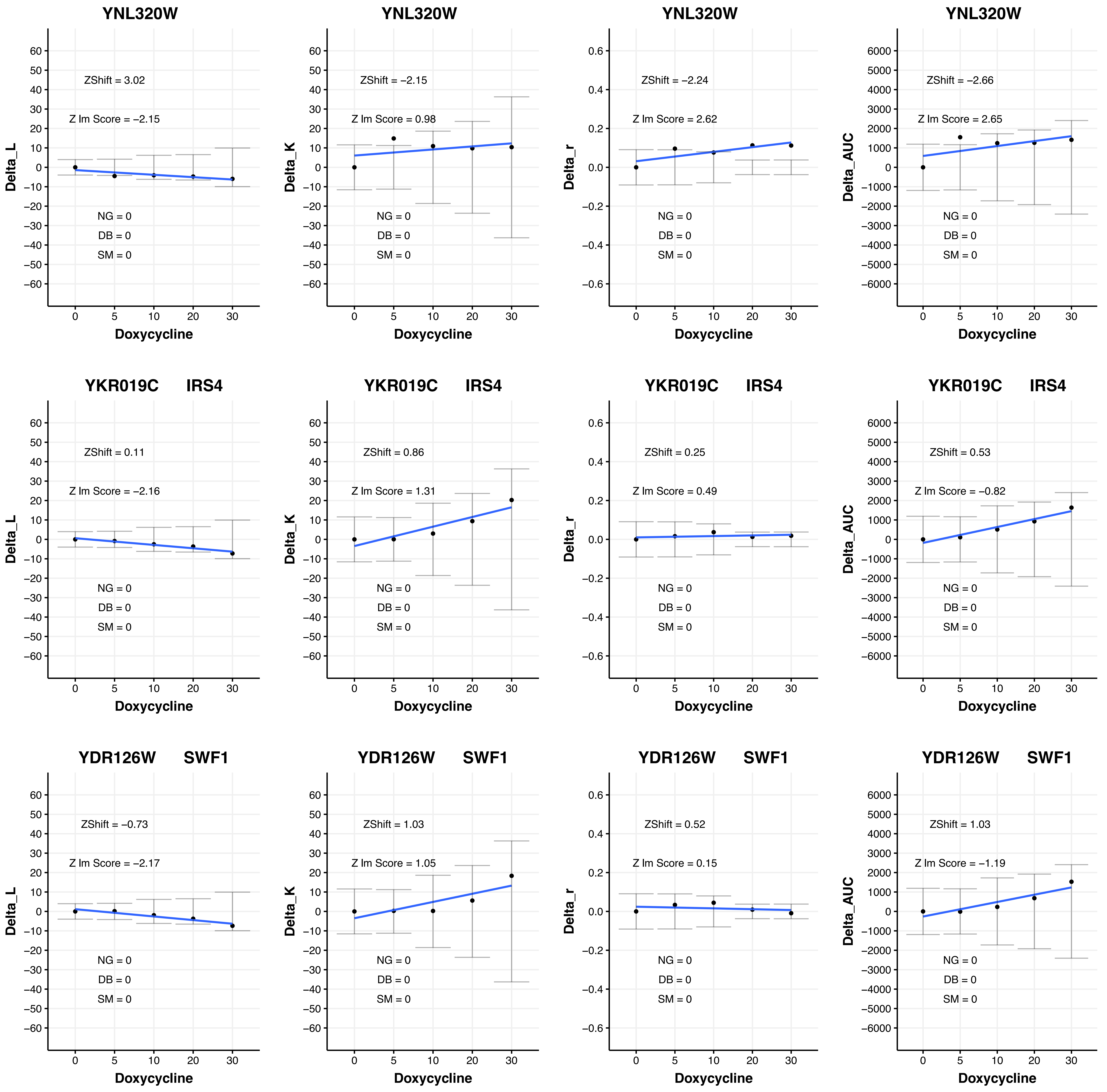
YNL320W
ZShift = 3.02
Z lm Score = −2.15
NG = 0
DB = 0
SM = 0
Delta_L
Doxycycline
YNL320W
ZShift = −2.15
Z lm Score = 0.98
NG = 0
DB = 0
SM = 0
Delta_K
Doxycycline
YNL320W
ZShift = −2.24
Z lm Score = 2.62
NG = 0
DB = 0
SM = 0
Delta_r
Doxycycline
YNL320W
ZShift = −2.66
Z lm Score = 2.65
NG = 0
DB = 0
SM = 0
Delta_AUC
Doxycycline
YKR019C IRS4
ZShift = 0.11
Z lm Score = −2.16
NG = 0
DB = 0
SM = 0
Delta_L
Doxycycline
YKR019C IRS4
ZShift = 0.86
Z lm Score = 1.31
NG = 0
DB = 0
SM = 0
Delta_K
Doxycycline
YKR019C IRS4
ZShift = 0.25
Z lm Score = 0.49
NG = 0
DB = 0
SM = 0
Delta_r
Doxycycline
YKR019C IRS4
ZShift = 0.53
Z lm Score = −0.82
NG = 0
DB = 0
SM = 0
Delta_AUC
Doxycycline
YDR126W SWF1
ZShift = −0.73
Z lm Score = −2.17
NG = 0
DB = 0
SM = 0
Delta_L
Doxycycline
YDR126W SWF1
ZShift = 1.03
Z lm Score = 1.05
NG = 0
DB = 0
SM = 0
Delta_K
Doxycycline
YDR126W SWF1
ZShift = 0.52
Z lm Score = 0.15
NG = 0
DB = 0
SM = 0
Delta_r
Doxycycline
YDR126W SWF1
ZShift = 1.03
Z lm Score = −1.19
NG = 0
DB = 0
SM = 0
Delta_AUC
Doxycycline

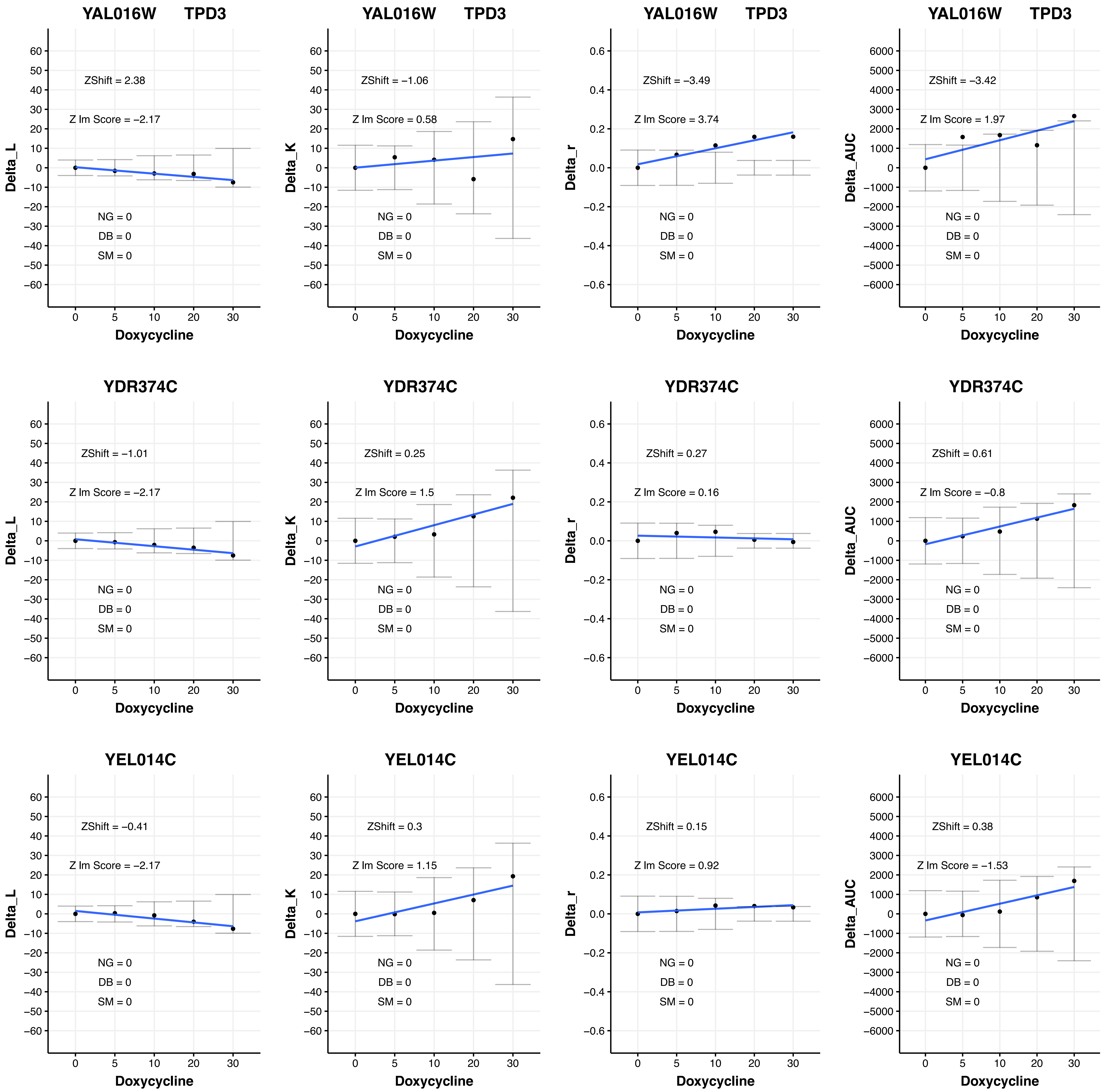

YAL016W TPD3
ZShift = 2.38
Z lm Score = −2.17
NG = 0
DB = 0
SM = 0
Delta_L
YAL016W TPD3
ZShift = −1.06
Z lm Score = 0.58
NG = 0
DB = 0
SM = 0
Delta_K
YAL016W TPD3
ZShift = −3.49
Z lm Score = 3.74
NG = 0
DB = 0
SM = 0
Delta_r
YAL016W TPD3
ZShift = −3.42
Z lm Score = 1.97
NG = 0
DB = 0
SM = 0
Delta_AUC
YDR374C
ZShift = −1.01
Z lm Score = −2.17
NG = 0
DB = 0
SM = 0
Delta_L
YDR374C
ZShift = 0.25
Z lm Score = 1.5
NG = 0
DB = 0
SM = 0
Delta_K
YDR374C
ZShift = 0.27
Z lm Score = 0.16
NG = 0
DB = 0
SM = 0
Delta_r
YDR374C
ZShift = 0.61
Z lm Score = −0.8
NG = 0
DB = 0
SM = 0
Delta_AUC
YEL014C
ZShift = −0.41
Z lm Score = −2.17
NG = 0
DB = 0
SM = 0
Delta_L
YEL014C
ZShift = 0.3
Z lm Score = 1.15
NG = 0
DB = 0
SM = 0
Delta_K
YEL014C
ZShift = 0.15
Z lm Score = 0.92
NG = 0
DB = 0
SM = 0
Delta_r
YEL014C
ZShift = 0.38
Z lm Score = −1.53
NG = 0
DB = 0
SM = 0
Delta_AUC
Doxycycline
0
5
10
20
30

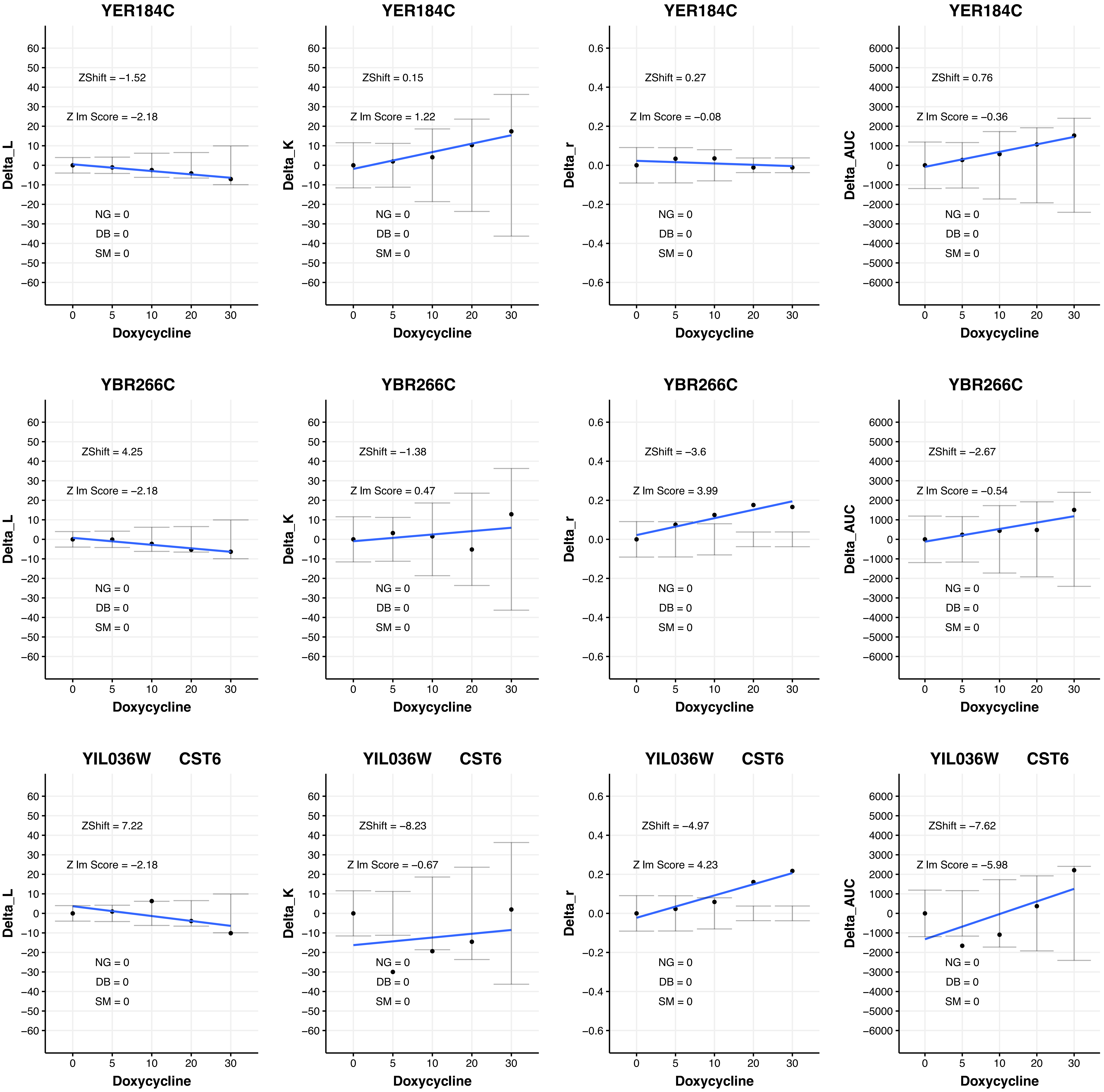

YER184C
ZShift = −1.52
Z lm Score = −2.18
NG = 0
DB = 0
SM = 0
Delta_L
Doxycycline
YER184C
ZShift = 0.15
Z lm Score = 1.22
NG = 0
DB = 0
SM = 0
Delta_K
Doxycycline
YER184C
ZShift = 0.27
Z lm Score = −0.08
NG = 0
DB = 0
SM = 0
Delta_r
Doxycycline
YER184C
ZShift = 0.76
Z lm Score = −0.36
NG = 0
DB = 0
SM = 0
Delta_AUC
Doxycycline
YBR266C
ZShift = 4.25
Z lm Score = −2.18
NG = 0
DB = 0
SM = 0
Delta_L
Doxycycline
YBR266C
ZShift = −1.38
Z lm Score = 0.47
NG = 0
DB = 0
SM = 0
Delta_K
Doxycycline
YBR266C
ZShift = −3.6
Z lm Score = 3.99
NG = 0
DB = 0
SM = 0
Delta_r
Doxycycline
YBR266C
ZShift = −2.67
Z lm Score = −0.54
NG = 0
DB = 0
SM = 0
Delta_AUC
Doxycycline
YIL036W CST6
ZShift = 7.22
Z lm Score = −2.18
NG = 0
DB = 0
SM = 0
Delta_L
Doxycycline
YIL036W CST6
ZShift = −8.23
Z lm Score = −0.67
NG = 0
DB = 0
SM = 0
Delta_K
Doxycycline
YIL036W CST6
ZShift = −4.97
Z lm Score = 4.23
NG = 0
DB = 0
SM = 0
Delta_r
Doxycycline
YIL036W CST6
ZShift = −7.62
Z lm Score = −5.98
NG = 0
DB = 0
SM = 0
Delta_AUC
Doxycycline

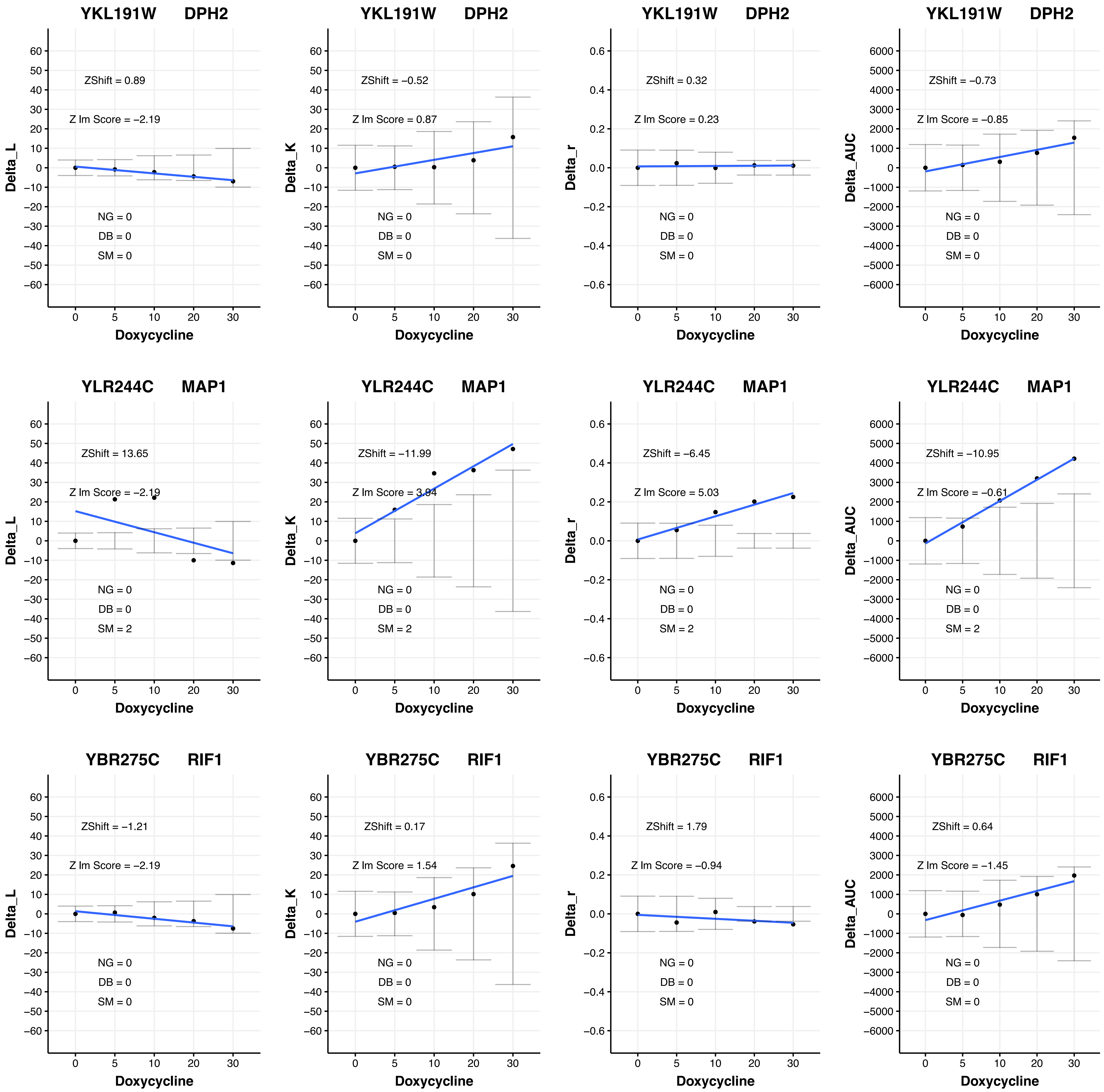
YKL191W DPH2
ZShift = 0.89
Z lm Score = −2.19
NG = 0
DB = 0
SM = 0
Delta_L
Doxycycline
YKL191W DPH2
ZShift = −0.52
Z lm Score = 0.87
NG = 0
DB = 0
SM = 0
Delta_K
Doxycycline
YKL191W DPH2
ZShift = 0.32
Z lm Score = 0.23
NG = 0
DB = 0
SM = 0
Delta_r
Doxycycline
YKL191W DPH2
ZShift = −0.73
Z lm Score = −0.85
NG = 0
DB = 0
SM = 0
Delta_AUC
Doxycycline
YLR244C MAP1
ZShift = 13.65
Z lm Score = −2.19
NG = 0
DB = 0
SM = 2
Delta_L
Doxycycline
YLR244C MAP1
ZShift = −11.99
Z lm Score = 3.94
NG = 0
DB = 0
SM = 2
Delta_K
Doxycycline
YLR244C MAP1
ZShift = −6.45
Z lm Score = 5.03
NG = 0
DB = 0
SM = 2
Delta_r
Doxycycline
YLR244C MAP1
ZShift = −10.95
Z lm Score = −0.61
NG = 0
DB = 0
SM = 2
Delta_AUC
Doxycycline
YBR275C RIF1
ZShift = −1.21
Z lm Score = −2.19
NG = 0
DB = 0
SM = 0
Delta_L
Doxycycline
YBR275C RIF1
ZShift = 0.17
Z lm Score = 1.54
NG = 0
DB = 0
SM = 0
Delta_K
Doxycycline
YBR275C RIF1
ZShift = 1.79
Z lm Score = −0.94
NG = 0
DB = 0
SM = 0
Delta_r
Doxycycline
YBR275C RIF1
ZShift = 0.64
Z lm Score = −1.45
NG = 0
DB = 0
SM = 0
Delta_AUC
Doxycycline

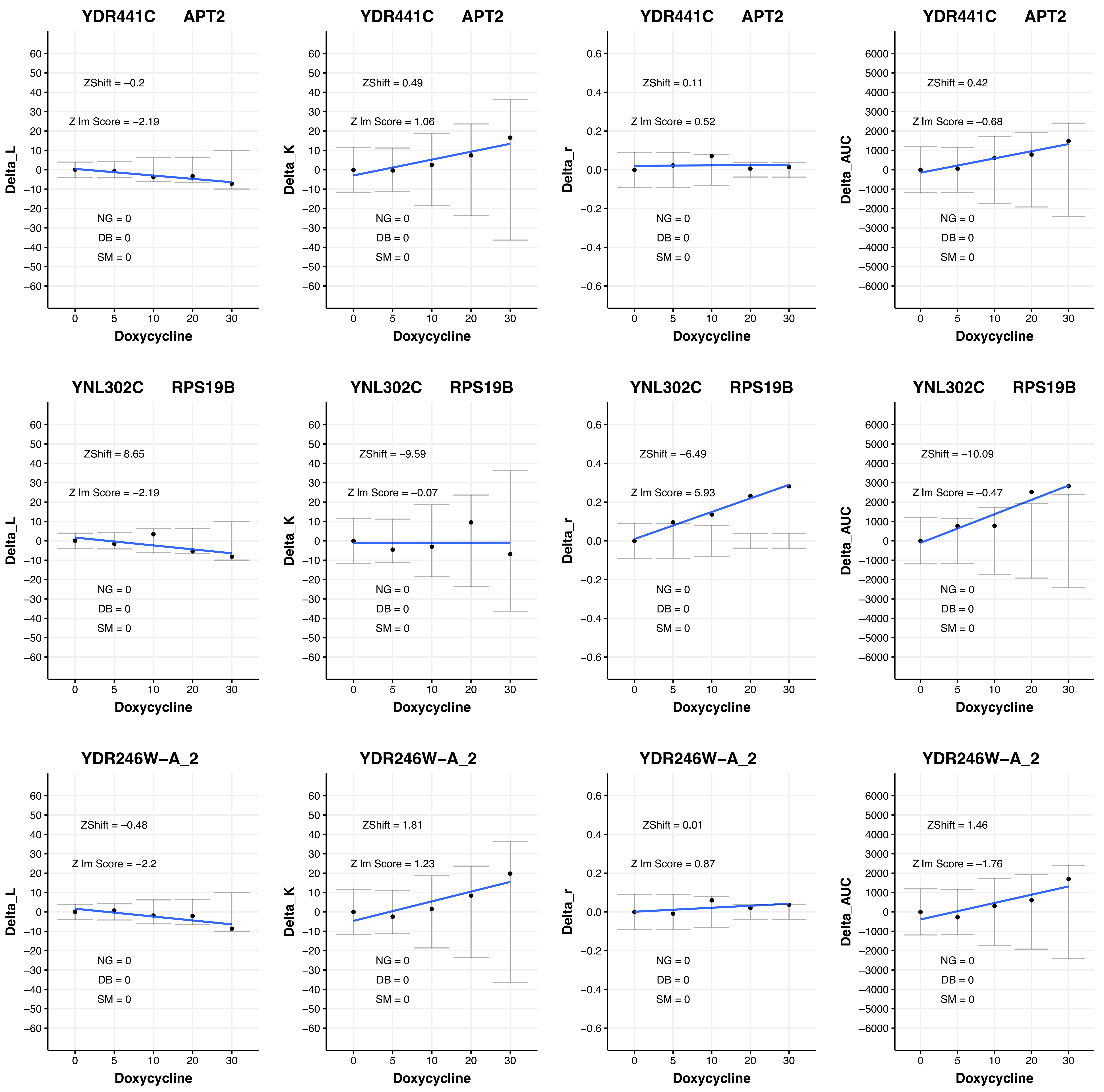
YDR441C APT2
ZShift = −0.2
Z lm Score = −2.19
NG = 0
DB = 0
SM = 0
Delta_L
Doxycycline
YDR441C APT2
ZShift = 0.49
Z lm Score = 1.06
NG = 0
DB = 0
SM = 0
Delta_K
Doxycycline
YDR441C APT2
ZShift = 0.11
Z lm Score = 0.52
NG = 0
DB = 0
SM = 0
Delta_r
Doxycycline
YDR441C APT2
ZShift = 0.42
Z lm Score = −0.68
NG = 0
DB = 0
SM = 0
Delta_AUC
Doxycycline
YNL302C RPS19B
ZShift = 8.65
Z lm Score = −2.19
NG = 0
DB = 0
SM = 0
Delta_L
Doxycycline
YNL302C RPS19B
ZShift = −9.59
Z lm Score = −0.07
NG = 0
DB = 0
SM = 0
Delta_K
Doxycycline
YNL302C RPS19B
ZShift = −6.49
Z lm Score = 5.93
NG = 0
DB = 0
SM = 0
Delta_r
Doxycycline
YNL302C RPS19B
ZShift = −10.09
Z lm Score = −0.47
NG = 0
DB = 0
SM = 0
Delta_AUC
Doxycycline
YDR246W−A_2
ZShift = −0.48
Z lm Score = −2.2
NG = 0
DB = 0
SM = 0
Delta_L
Doxycycline
YDR246W−A_2
ZShift = 1.81
Z lm Score = 1.23
NG = 0
DB = 0
SM = 0
Delta_K
Doxycycline
YDR246W−A_2
ZShift = 0.01
Z lm Score = 0.87
NG = 0
DB = 0
SM = 0
Delta_r
Doxycycline
YDR246W−A_2
ZShift = 1.46
Z lm Score = −1.76
NG = 0
DB = 0
SM = 0
Delta_AUC
Doxycycline

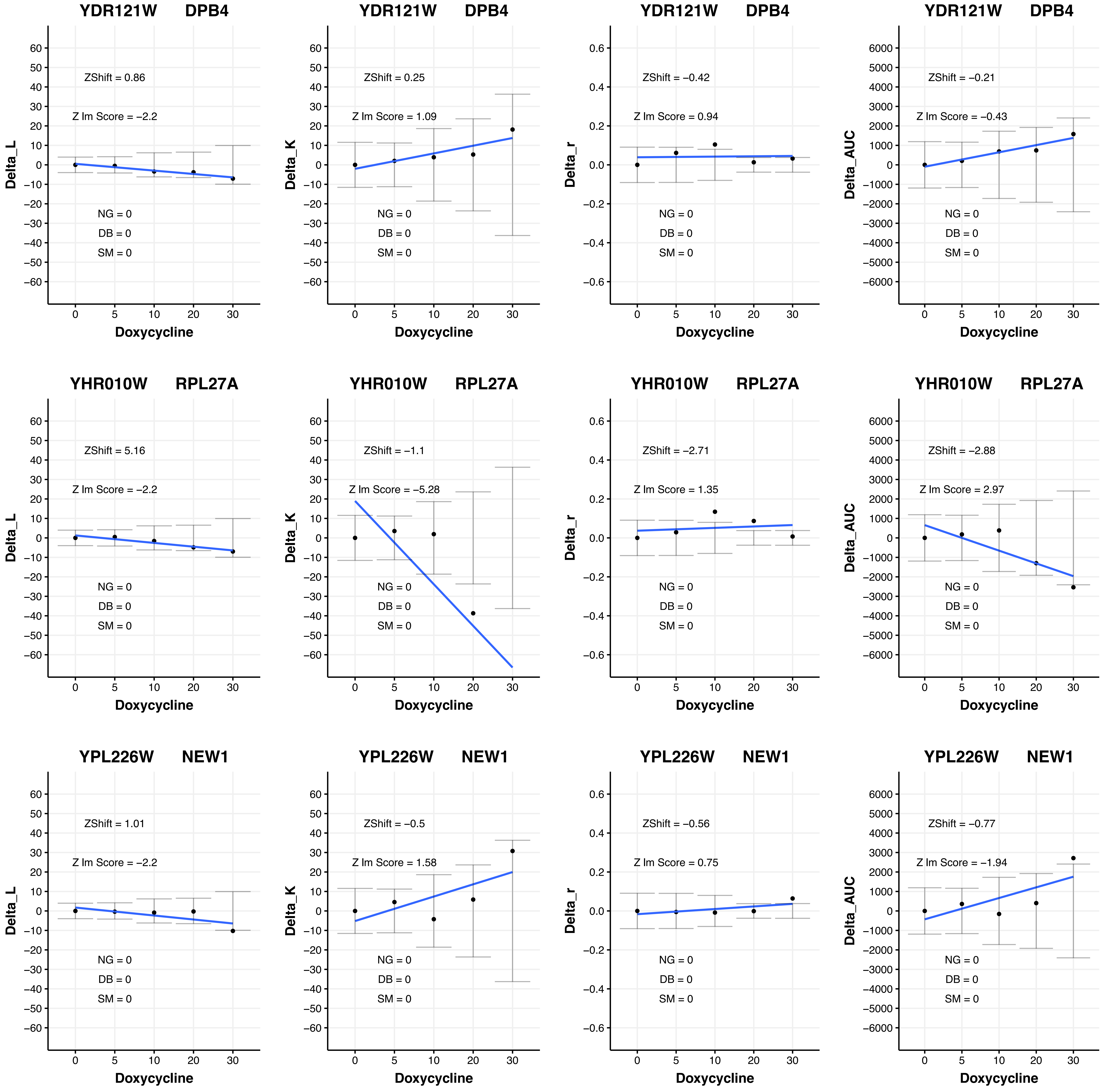

YDR121W DPB4
ZShift = 0.86
Z lm Score = −2.2
NG = 0
DB = 0
SM = 0
Delta_L
Doxycycline
YDR121W DPB4
ZShift = 0.25
Z lm Score = 1.09
NG = 0
DB = 0
SM = 0
Delta_K
Doxycycline
YDR121W DPB4
ZShift = −0.42
Z lm Score = 0.94
NG = 0
DB = 0
SM = 0
Delta_r
Doxycycline
YDR121W DPB4
ZShift = −0.21
Z lm Score = −0.43
NG = 0
DB = 0
SM = 0
Delta_AUC
Doxycycline
YHR010W RPL27A
ZShift = 5.16
Z lm Score = −2.2
NG = 0
DB = 0
SM = 0
Delta_L
Doxycycline
YHR010W RPL27A
ZShift = −1.1
Z lm Score = −5.28
NG = 0
DB = 0
SM = 0
Delta_K
Doxycycline
YHR010W RPL27A
ZShift = −2.71
Z lm Score = 1.35
NG = 0
DB = 0
SM = 0
Delta_r
Doxycycline
YHR010W RPL27A
ZShift = −2.88
Z lm Score = 2.97
NG = 0
DB = 0
SM = 0
Delta_AUC
Doxycycline
YPL226W NEW1
ZShift = 1.01
Z lm Score = −2.2
NG = 0
DB = 0
SM = 0
Delta_L
Doxycycline
YPL226W NEW1
ZShift = −0.5
Z lm Score = 1.58
NG = 0
DB = 0
SM = 0
Delta_K
Doxycycline
YPL226W NEW1
ZShift = −0.56
Z lm Score = 0.75
NG = 0
DB = 0
SM = 0
Delta_r
Doxycycline
YPL226W NEW1
ZShift = −0.77
Z lm Score = −1.94
NG = 0
DB = 0
SM = 0
Delta_AUC
Doxycycline

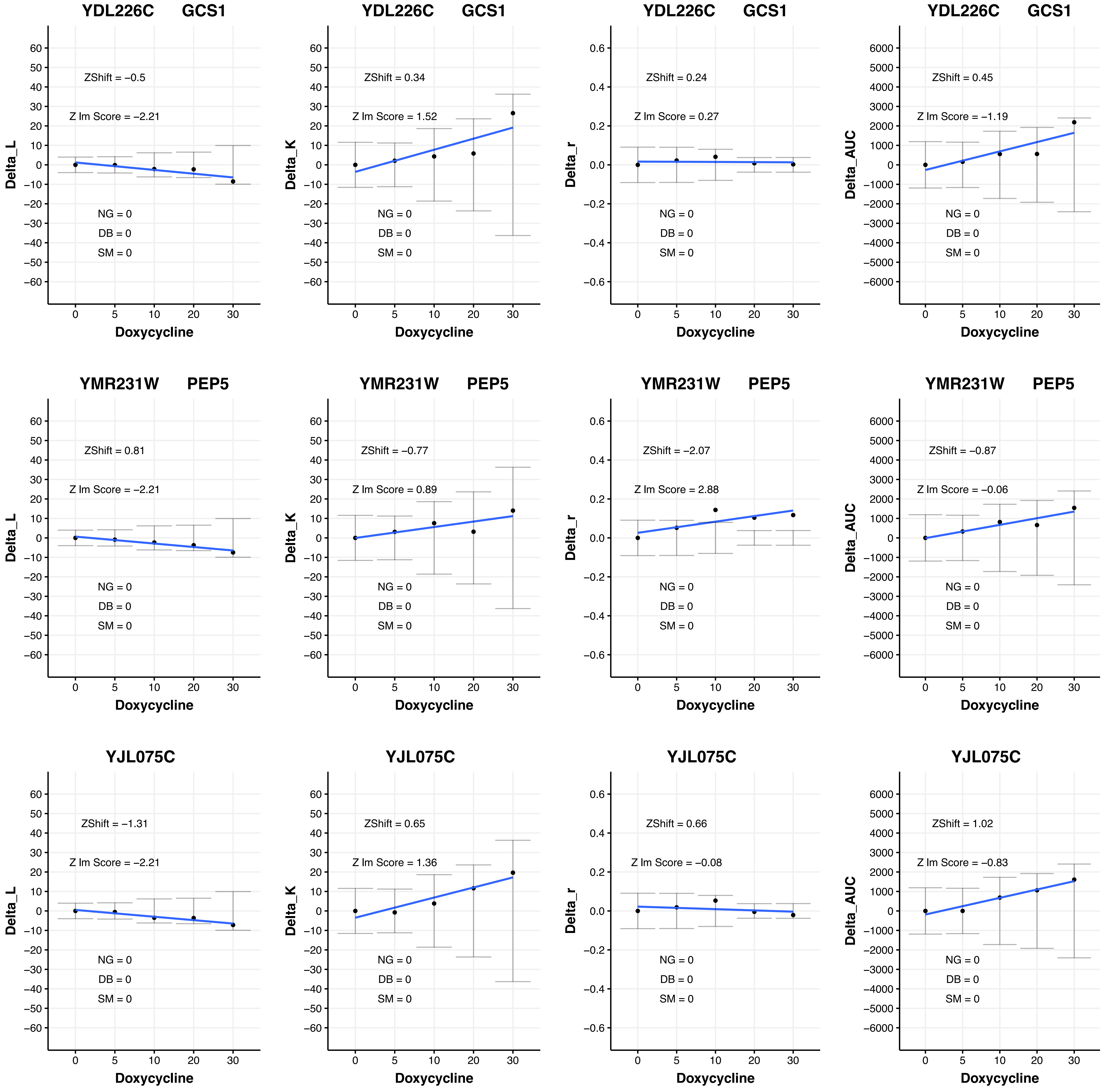
YDL226C GCS1
ZShift = −0.5
Z lm Score = −2.21
NG = 0
DB = 0
SM = 0
Delta_L
Doxycycline
YDL226C GCS1
ZShift = 0.34
Z lm Score = 1.52
NG = 0
DB = 0
SM = 0
Delta_K
Doxycycline
YDL226C GCS1
ZShift = 0.24
Z lm Score = 0.27
NG = 0
DB = 0
SM = 0
Delta_r
Doxycycline
YDL226C GCS1
ZShift = 0.45
Z lm Score = −1.19
NG = 0
DB = 0
SM = 0
Delta_AUC
Doxycycline
YMR231W PEP5
ZShift = 0.81
Z lm Score = −2.21
NG = 0
DB = 0
SM = 0
Delta_L
Doxycycline
YMR231W PEP5
ZShift = −0.77
Z lm Score = 0.89
NG = 0
DB = 0
SM = 0
Delta_K
Doxycycline
YMR231W PEP5
ZShift = −2.07
Z lm Score = 2.88
NG = 0
DB = 0
SM = 0
Delta_r
Doxycycline
YMR231W PEP5
ZShift = −0.87
Z lm Score = −0.06
NG = 0
DB = 0
SM = 0
Delta_AUC
Doxycycline
YJL075C
ZShift = −1.31
Z lm Score = −2.21
NG = 0
DB = 0
SM = 0
Delta_L
Doxycycline
YJL075C
ZShift = 0.65
Z lm Score = 1.36
NG = 0
DB = 0
SM = 0
Delta_K
Doxycycline
YJL075C
ZShift = 0.66
Z lm Score = −0.08
NG = 0
DB = 0
SM = 0
Delta_r
Doxycycline
YJL075C
ZShift = 1.02
Z lm Score = −0.83
NG = 0
DB = 0
SM = 0
Delta_AUC
Doxycycline

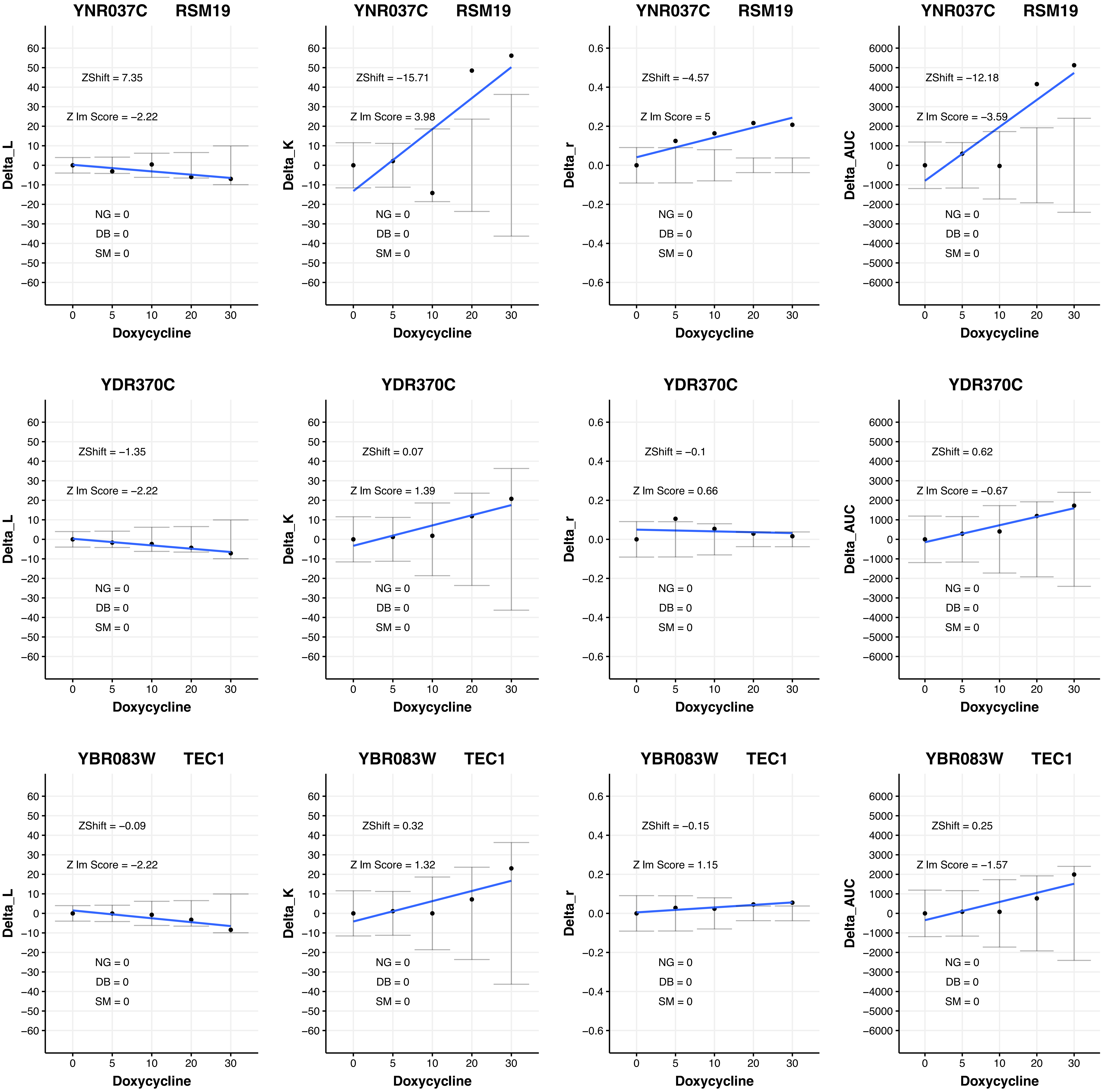
YNR037C RSM19
ZShift = 7.35
Z lm Score = −2.22
NG = 0
DB = 0
SM = 0
Delta_L
Doxycycline
YNR037C RSM19
ZShift = −15.71
Z lm Score = 3.98
NG = 0
DB = 0
SM = 0
Delta_K
Doxycycline
YNR037C RSM19
ZShift = −4.57
Z lm Score = 5
NG = 0
DB = 0
SM = 0
Delta_r
Doxycycline
YNR037C RSM19
ZShift = −12.18
Z lm Score = −3.59
NG = 0
DB = 0
SM = 0
Delta_AUC
Doxycycline
YDR370C
ZShift = −1.35
Z lm Score = −2.22
NG = 0
DB = 0
SM = 0
Delta_L
Doxycycline
YDR370C
ZShift = 0.07
Z lm Score = 1.39
NG = 0
DB = 0
SM = 0
Delta_K
Doxycycline
YDR370C
ZShift = −0.1
Z lm Score = 0.66
NG = 0
DB = 0
SM = 0
Delta_r
Doxycycline
YDR370C
ZShift = 0.62
Z lm Score = −0.67
NG = 0
DB = 0
SM = 0
Delta_AUC
Doxycycline
YBR083W TEC1
ZShift = −0.09
Z lm Score = −2.22
NG = 0
DB = 0
SM = 0
Delta_L
Doxycycline
YBR083W TEC1
ZShift = 0.32
Z lm Score = 1.32
NG = 0
DB = 0
SM = 0
Delta_K
Doxycycline
YBR083W TEC1
ZShift = −0.15
Z lm Score = 1.15
NG = 0
DB = 0
SM = 0
Delta_r
Doxycycline
YBR083W TEC1
ZShift = 0.25
Z lm Score = −1.57
NG = 0
DB = 0
SM = 0
Delta_AUC
Doxycycline

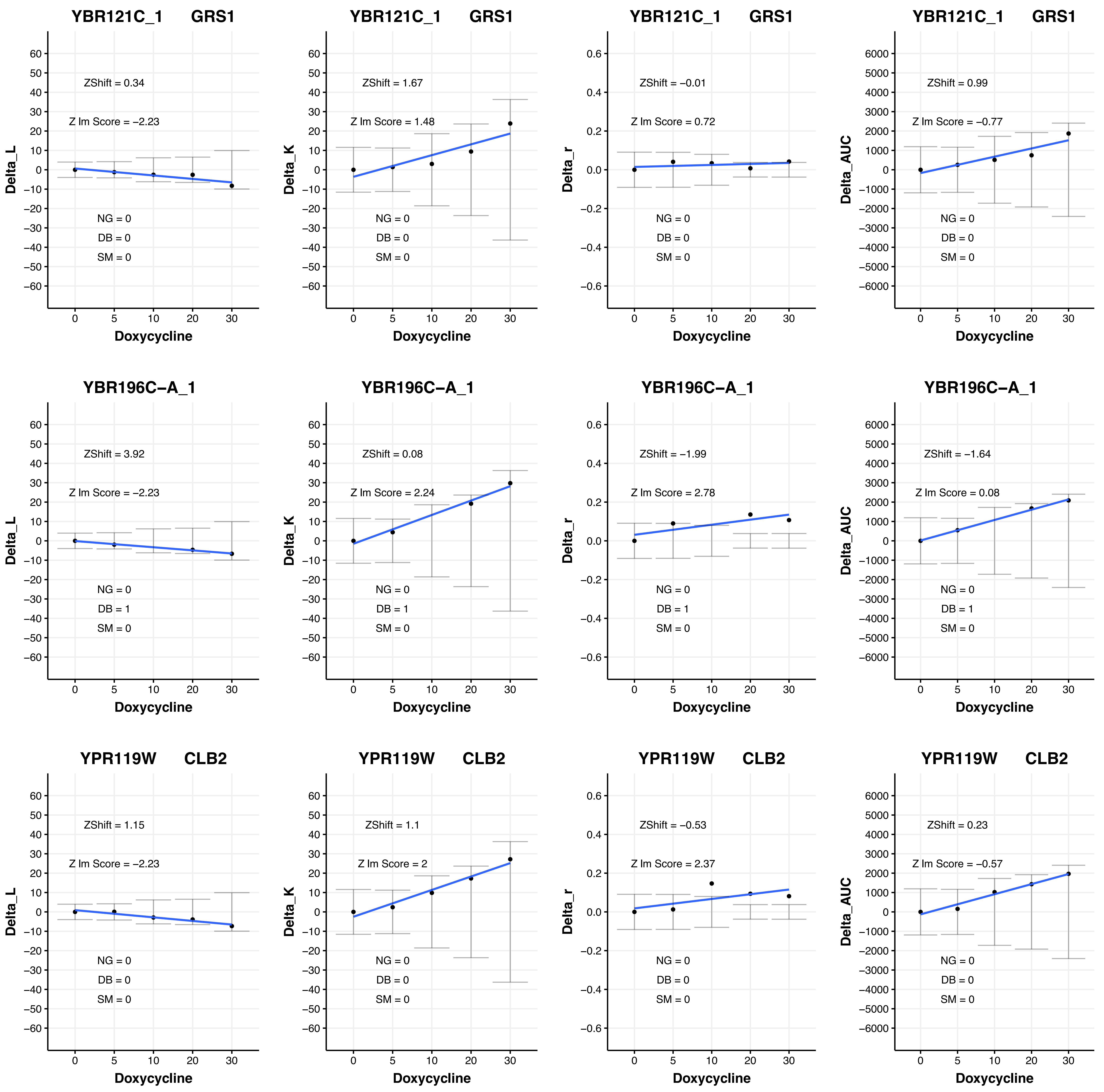

YBR121C_1 GRS1
ZShift = 0.34
Z lm Score = −2.23
NG = 0
DB = 0
SM = 0
Delta_L
Doxycycline
YBR121C_1 GRS1
ZShift = 1.67
Z lm Score = 1.48
NG = 0
DB = 0
SM = 0
Delta_K
Doxycycline
YBR121C_1 GRS1
ZShift = −0.01
Z lm Score = 0.72
NG = 0
DB = 0
SM = 0
Delta_r
Doxycycline
YBR121C_1 GRS1
ZShift = 0.99
Z lm Score = −0.77
NG = 0
DB = 0
SM = 0
Delta_AUC
Doxycycline
YBR196C−A_1
ZShift = 3.92
Z lm Score = −2.23
NG = 0
DB = 1
SM = 0
Delta_L
Doxycycline
YBR196C−A_1
ZShift = 0.08
Z lm Score = 2.24
NG = 0
DB = 1
SM = 0
Delta_K
Doxycycline
YBR196C−A_1
ZShift = −1.99
Z lm Score = 2.78
NG = 0
DB = 1
SM = 0
Delta_r
Doxycycline
YBR196C−A_1
ZShift = −1.64
Z lm Score = 0.08
NG = 0
DB = 1
SM = 0
Delta_AUC
Doxycycline
YPR119W CLB2
ZShift = 1.15
Z lm Score = −2.23
NG = 0
DB = 0
SM = 0
Delta_L
Doxycycline
YPR119W CLB2
ZShift = 1.1
Z lm Score = 2
NG = 0
DB = 0
SM = 0
Delta_K
Doxycycline
YPR119W CLB2
ZShift = −0.53
Z lm Score = 2.37
NG = 0
DB = 0
SM = 0
Delta_r
Doxycycline
YPR119W CLB2
ZShift = 0.23
Z lm Score = −0.57
NG = 0
DB = 0
SM = 0
Delta_AUC
Doxycycline

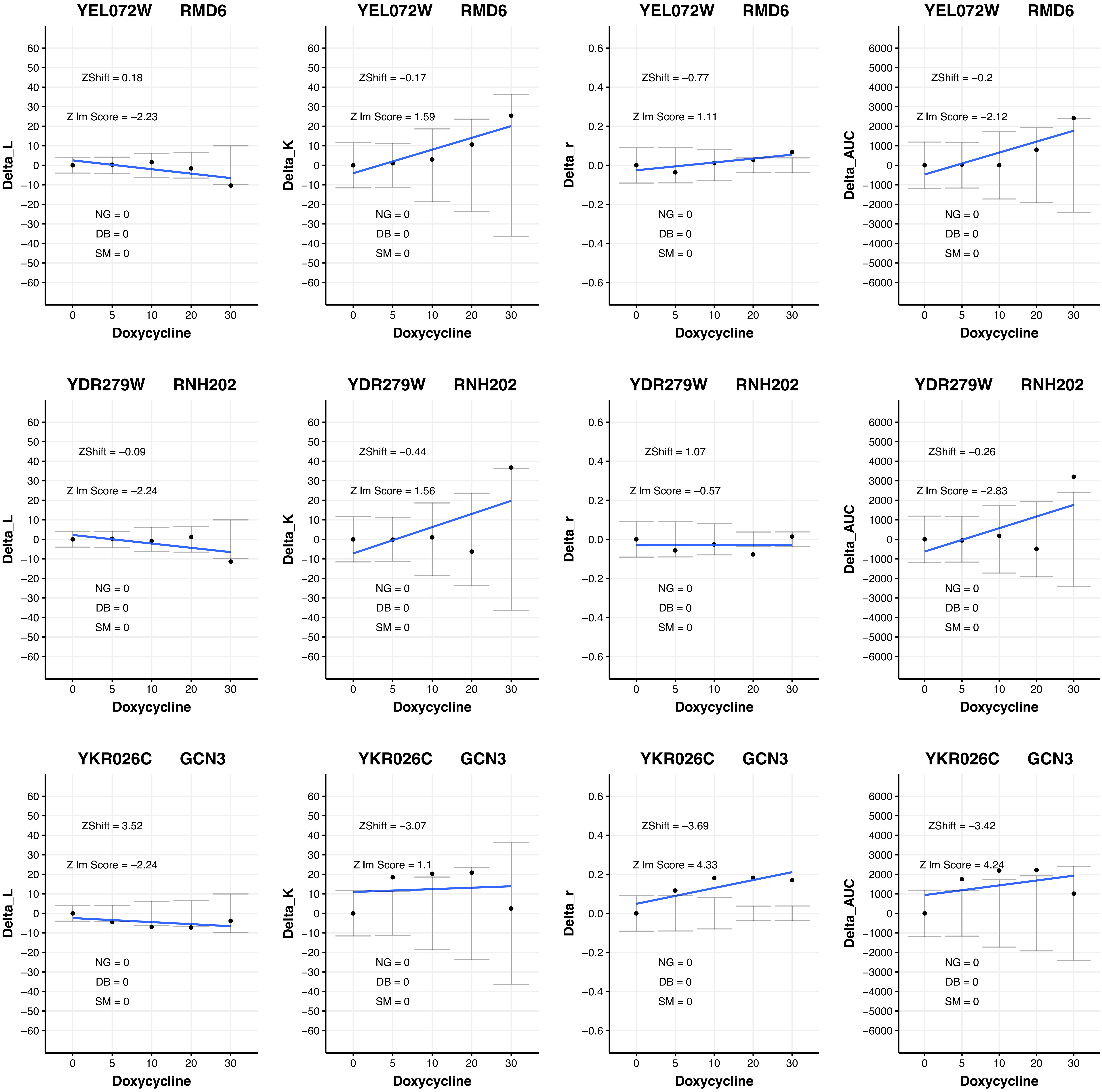

YEL072W RMD6
ZShift = 0.18
Z lm Score = −2.23
NG = 0
DB = 0
SM = 0
Delta_L
Doxycycline
YEL072W RMD6
ZShift = −0.17
Z lm Score = 1.59
NG = 0
DB = 0
SM = 0
Delta_K
Doxycycline
YEL072W RMD6
ZShift = −0.77
Z lm Score = 1.11
NG = 0
DB = 0
SM = 0
Delta_r
Doxycycline
YEL072W RMD6
ZShift = −0.2
Z lm Score = −2.12
NG = 0
DB = 0
SM = 0
Delta_AUC
Doxycycline
YDR279W RNH202
ZShift = −0.09
Z lm Score = −2.24
NG = 0
DB = 0
SM = 0
Delta_L
Doxycycline
YDR279W RNH202
ZShift = −0.44
Z lm Score = 1.56
NG = 0
DB = 0
SM = 0
Delta_K
Doxycycline
YDR279W RNH202
ZShift = 1.07
Z lm Score = −0.57
NG = 0
DB = 0
SM = 0
Delta_r
Doxycycline
YDR279W RNH202
ZShift = −0.26
Z lm Score = −2.83
NG = 0
DB = 0
SM = 0
Delta_AUC
Doxycycline
YKR026C GCN3
ZShift = 3.52
Z lm Score = −2.24
NG = 0
DB = 0
SM = 0
Delta_L
Doxycycline
YKR026C GCN3
ZShift = −3.07
Z lm Score = 1.1
NG = 0
DB = 0
SM = 0
Delta_K
Doxycycline
YKR026C GCN3
ZShift = −3.69
Z lm Score = 4.33
NG = 0
DB = 0
SM = 0
Delta_r
Doxycycline
YKR026C GCN3
ZShift = −3.42
Z lm Score = 4.24
NG = 0
DB = 0
SM = 0
Delta_AUC
Doxycycline

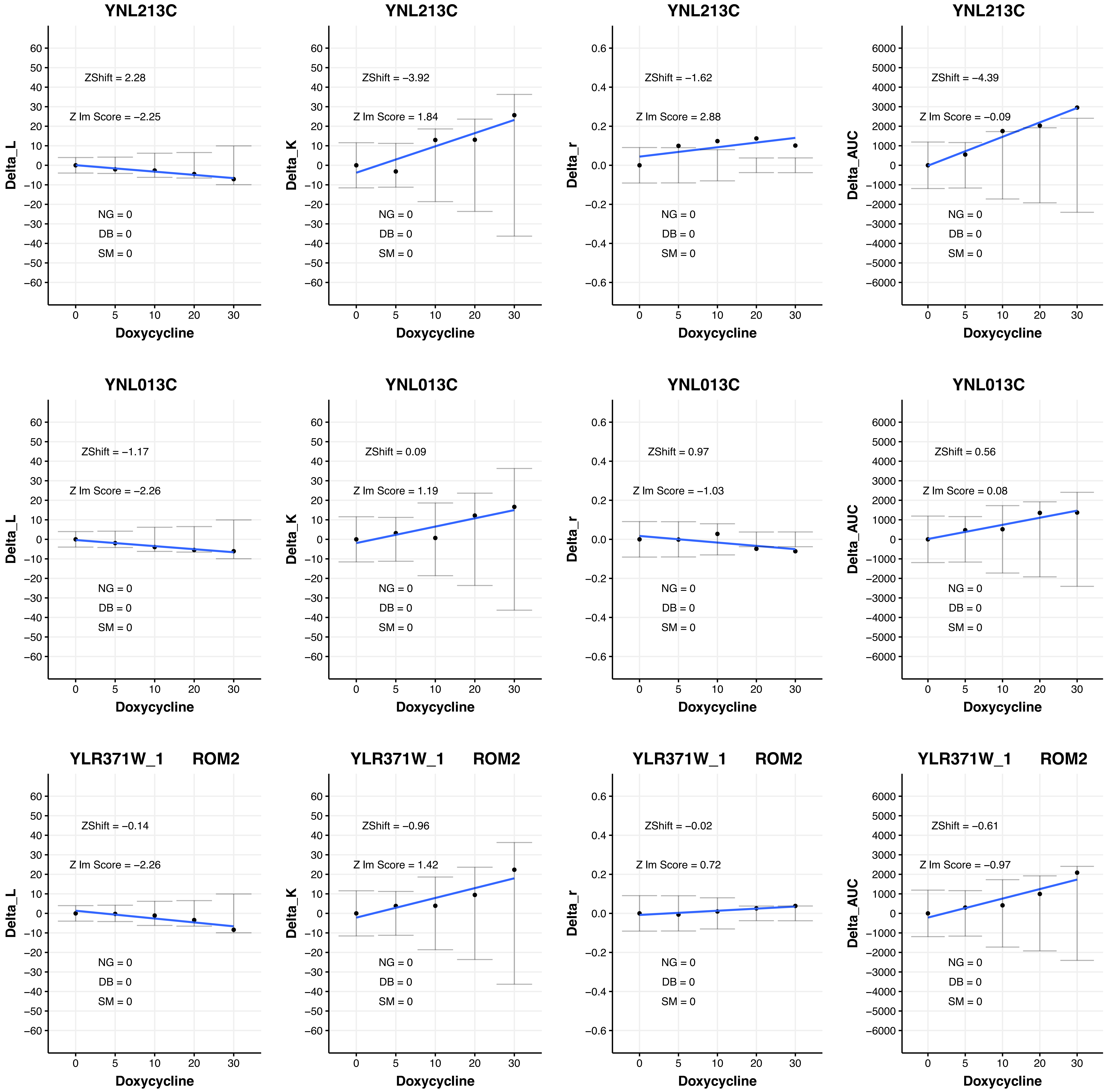
YNL213C
ZShift = 2.28
Z lm Score = −2.25
NG = 0
DB = 0
SM = 0
Delta_L
Doxycycline
YNL213C
ZShift = −3.92
Z lm Score = 1.84
NG = 0
DB = 0
SM = 0
Delta_K
Doxycycline
YNL213C
ZShift = −1.62
Z lm Score = 2.88
NG = 0
DB = 0
SM = 0
Delta_r
Doxycycline
YNL213C
ZShift = −4.39
Z lm Score = −0.09
NG = 0
DB = 0
SM = 0
Delta_AUC
Doxycycline
YNL013C
ZShift = −1.17
Z lm Score = −2.26
NG = 0
DB = 0
SM = 0
Delta_L
Doxycycline
YNL013C
ZShift = 0.09
Z lm Score = 1.19
NG = 0
DB = 0
SM = 0
Delta_K
Doxycycline
YNL013C
ZShift = 0.97
Z lm Score = −1.03
NG = 0
DB = 0
SM = 0
Delta_r
Doxycycline
YNL013C
ZShift = 0.56
Z lm Score = 0.08
NG = 0
DB = 0
SM = 0
Delta_AUC
Doxycycline
YLR371W_1 ROM2
ZShift = −0.14
Z lm Score = −2.26
NG = 0
DB = 0
SM = 0
Delta_L
Doxycycline
YLR371W_1 ROM2
ZShift = −0.96
Z lm Score = 1.42
NG = 0
DB = 0
SM = 0
Delta_K
Doxycycline
YLR371W_1 ROM2
ZShift = −0.02
Z lm Score = 0.72
NG = 0
DB = 0
SM = 0
Delta_r
Doxycycline
YLR371W_1 ROM2
ZShift = −0.61
Z lm Score = −0.97
NG = 0
DB = 0
SM = 0
Delta_AUC
Doxycycline

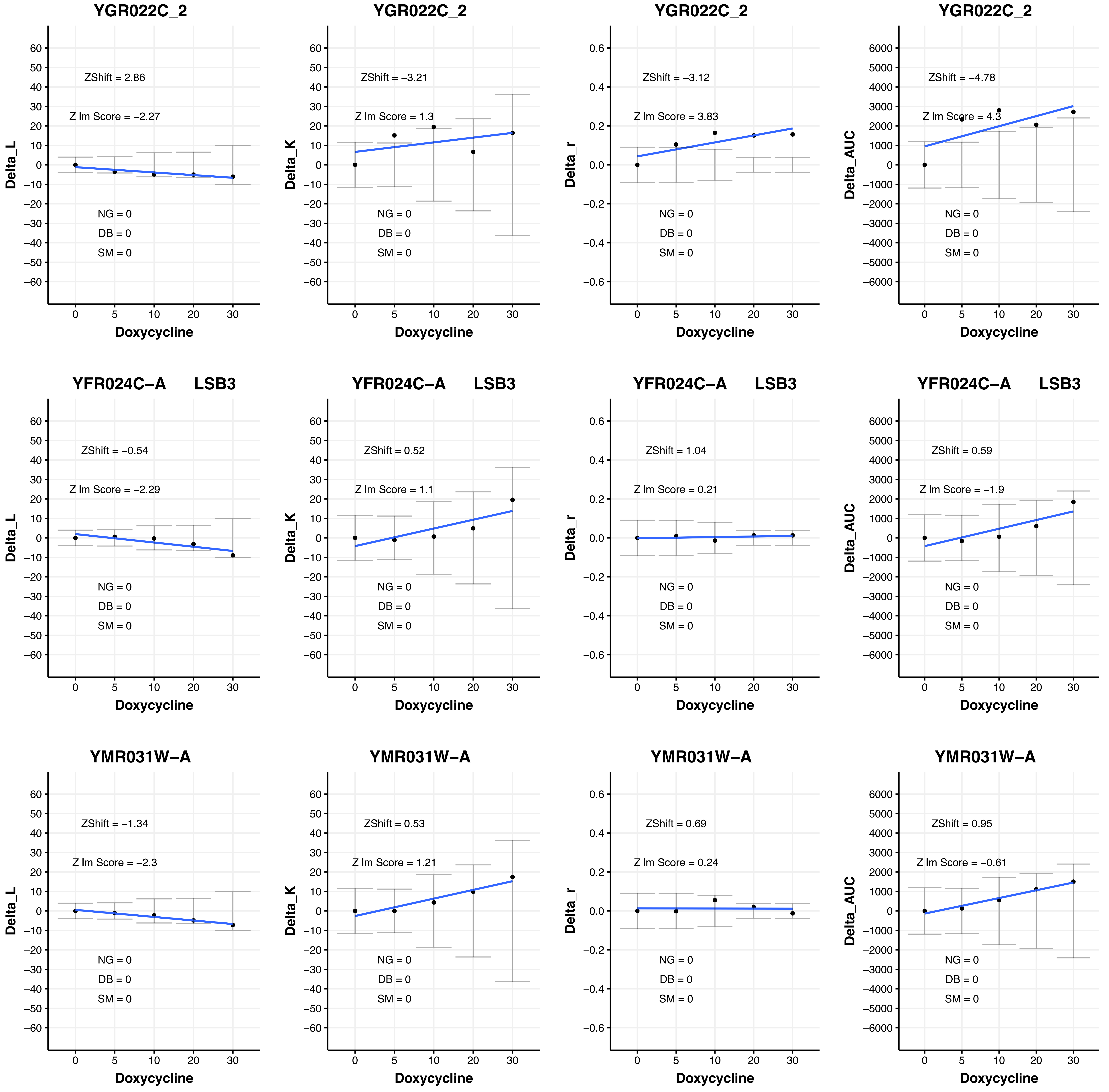

YGR022C_2
Delta_L
ZShift = 2.86
Z lm Score = −2.27
NG = 0
DB = 0
SM = 0
Doxycycline
YGR022C_2
Delta_K
ZShift = −3.21
Z lm Score = 1.3
NG = 0
DB = 0
SM = 0
Doxycycline
YGR022C_2
Delta_r
ZShift = −3.12
Z lm Score = 3.83
NG = 0
DB = 0
SM = 0
Doxycycline
YGR022C_2
Delta_AUC
ZShift = −4.78
Z lm Score = 4.3
NG = 0
DB = 0
SM = 0
Doxycycline
YFR024C−A LSB3
Delta_L
ZShift = −0.54
Z lm Score = −2.29
NG = 0
DB = 0
SM = 0
Doxycycline
YFR024C−A LSB3
Delta_K
ZShift = 0.52
Z lm Score = 1.1
NG = 0
DB = 0
SM = 0
Doxycycline
YFR024C−A LSB3
Delta_r
ZShift = 1.04
Z lm Score = 0.21
NG = 0
DB = 0
SM = 0
Doxycycline
YFR024C−A LSB3
Delta_AUC
ZShift = 0.59
Z lm Score = −1.9
NG = 0
DB = 0
SM = 0
Doxycycline
YMR031W−A
Delta_L
ZShift = −1.34
Z lm Score = −2.3
NG = 0
DB = 0
SM = 0
Doxycycline
YMR031W−A
Delta_K
ZShift = 0.53
Z lm Score = 1.21
NG = 0
DB = 0
SM = 0
Doxycycline
YMR031W−A
Delta_r
ZShift = 0.69
Z lm Score = 0.24
NG = 0
DB = 0
SM = 0
Doxycycline
YMR031W−A
Delta_AUC
ZShift = 0.95
Z lm Score = −0.61
NG = 0
DB = 0
SM = 0
Doxycycline

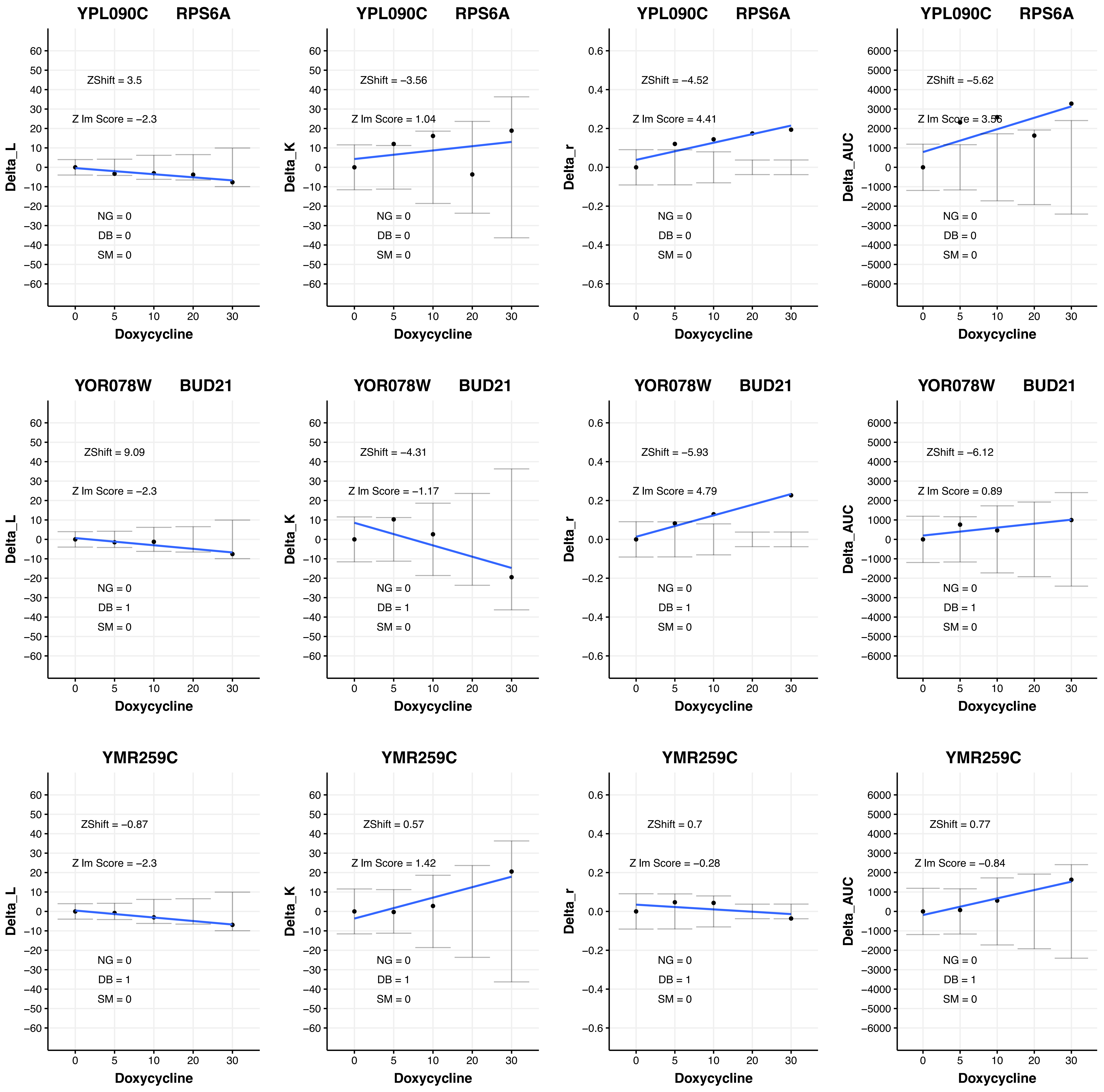

YPL090C RPS6A
ZShift = 3.5
Z lm Score = −2.3
NG = 0
DB = 0
SM = 0
Delta_L
Doxycycline
YPL090C RPS6A
ZShift = −3.56
Z lm Score = 1.04
NG = 0
DB = 0
SM = 0
Delta_K
Doxycycline
YPL090C RPS6A
ZShift = −4.52
Z lm Score = 4.41
NG = 0
DB = 0
SM = 0
Delta_r
Doxycycline
YPL090C RPS6A
ZShift = −5.62
Z lm Score = 3.56
NG = 0
DB = 0
SM = 0
Delta_AUC
Doxycycline
YOR078W BUD21
ZShift = 9.09
Z lm Score = −2.3
NG = 0
DB = 1
SM = 0
Delta_L
Doxycycline
YOR078W BUD21
ZShift = −4.31
Z lm Score = −1.17
NG = 0
DB = 1
SM = 0
Delta_K
Doxycycline
YOR078W BUD21
ZShift = −5.93
Z lm Score = 4.79
NG = 0
DB = 1
SM = 0
Delta_r
Doxycycline
YOR078W BUD21
ZShift = −6.12
Z lm Score = 0.89
NG = 0
DB = 1
SM = 0
Delta_AUC
Doxycycline
YMR259C
ZShift = −0.87
Z lm Score = −2.3
NG = 0
DB = 1
SM = 0
Delta_L
Doxycycline
YMR259C
ZShift = 0.57
Z lm Score = 1.42
NG = 0
DB = 1
SM = 0
Delta_K
Doxycycline
YMR259C
ZShift = 0.7
Z lm Score = −0.28
NG = 0
DB = 1
SM = 0
Delta_r
Doxycycline
YMR259C
ZShift = 0.77
Z lm Score = −0.84
NG = 0
DB = 1
SM = 0
Delta_AUC
Doxycycline

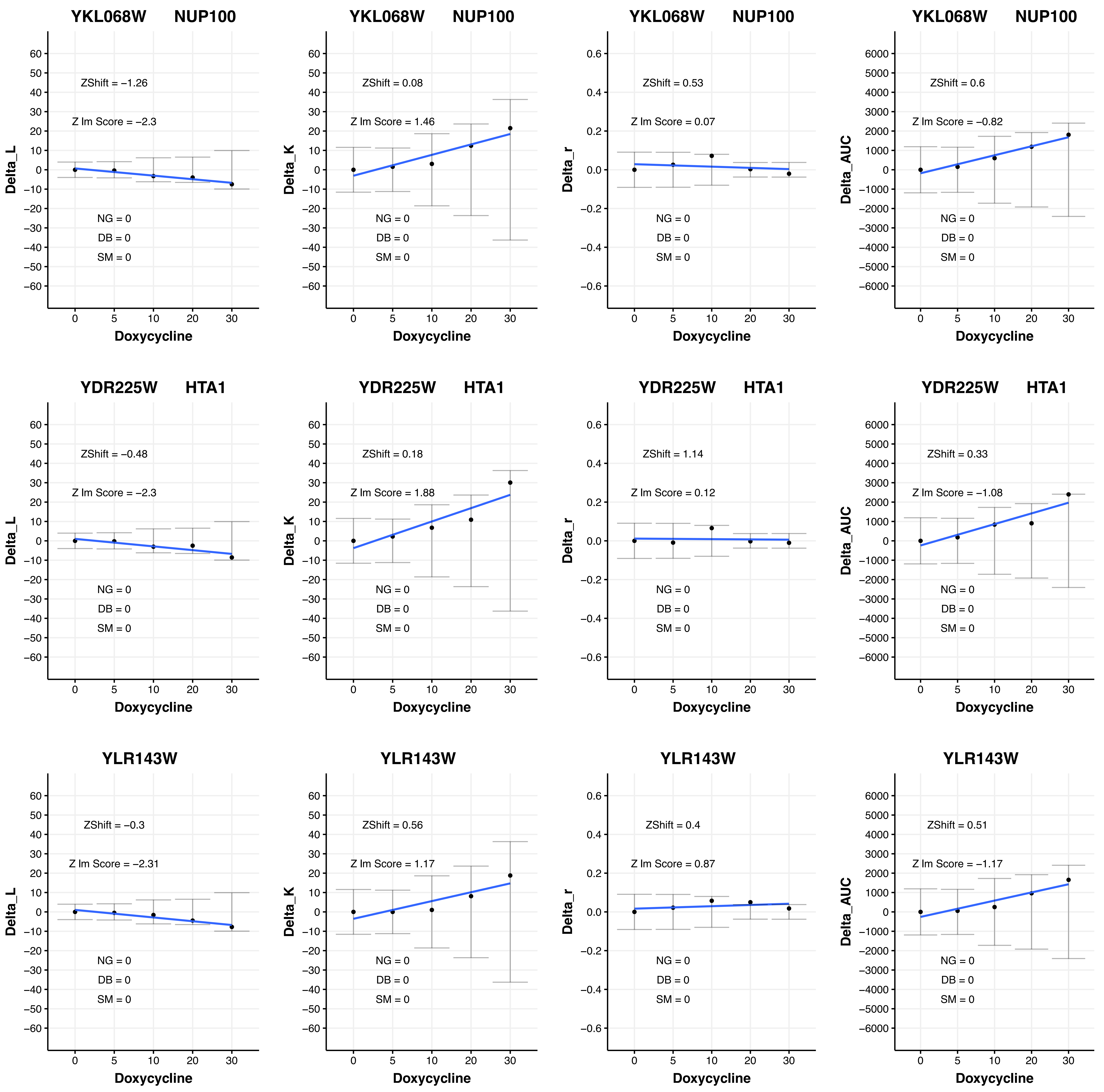

YKL068W NUP100
ZShift = −1.26
Z lm Score = −2.3
NG = 0
DB = 0
SM = 0
Delta_L
Doxycycline
YKL068W NUP100
ZShift = 0.08
Z lm Score = 1.46
NG = 0
DB = 0
SM = 0
Delta_K
Doxycycline
YKL068W NUP100
ZShift = 0.53
Z lm Score = 0.07
NG = 0
DB = 0
SM = 0
Delta_r
Doxycycline
YKL068W NUP100
ZShift = 0.6
Z lm Score = −0.82
NG = 0
DB = 0
SM = 0
Delta_AUC
Doxycycline
YDR225W HTA1
ZShift = −0.48
Z lm Score = −2.3
NG = 0
DB = 0
SM = 0
Delta_L
Doxycycline
YDR225W HTA1
ZShift = 0.18
Z lm Score = 1.88
NG = 0
DB = 0
SM = 0
Delta_K
Doxycycline
YDR225W HTA1
ZShift = 1.14
Z lm Score = 0.12
NG = 0
DB = 0
SM = 0
Delta_r
Doxycycline
YDR225W HTA1
ZShift = 0.33
Z lm Score = −1.08
NG = 0
DB = 0
SM = 0
Delta_AUC
Doxycycline
YLR143W
ZShift = −0.3
Z lm Score = −2.31
NG = 0
DB = 0
SM = 0
Delta_L
Doxycycline
YLR143W
ZShift = 0.56
Z lm Score = 1.17
NG = 0
DB = 0
SM = 0
Delta_K
Doxycycline
YLR143W
ZShift = 0.4
Z lm Score = 0.87
NG = 0
DB = 0
SM = 0
Delta_r
Doxycycline
YLR143W
ZShift = 0.51
Z lm Score = −1.17
NG = 0
DB = 0
SM = 0
Delta_AUC
Doxycycline

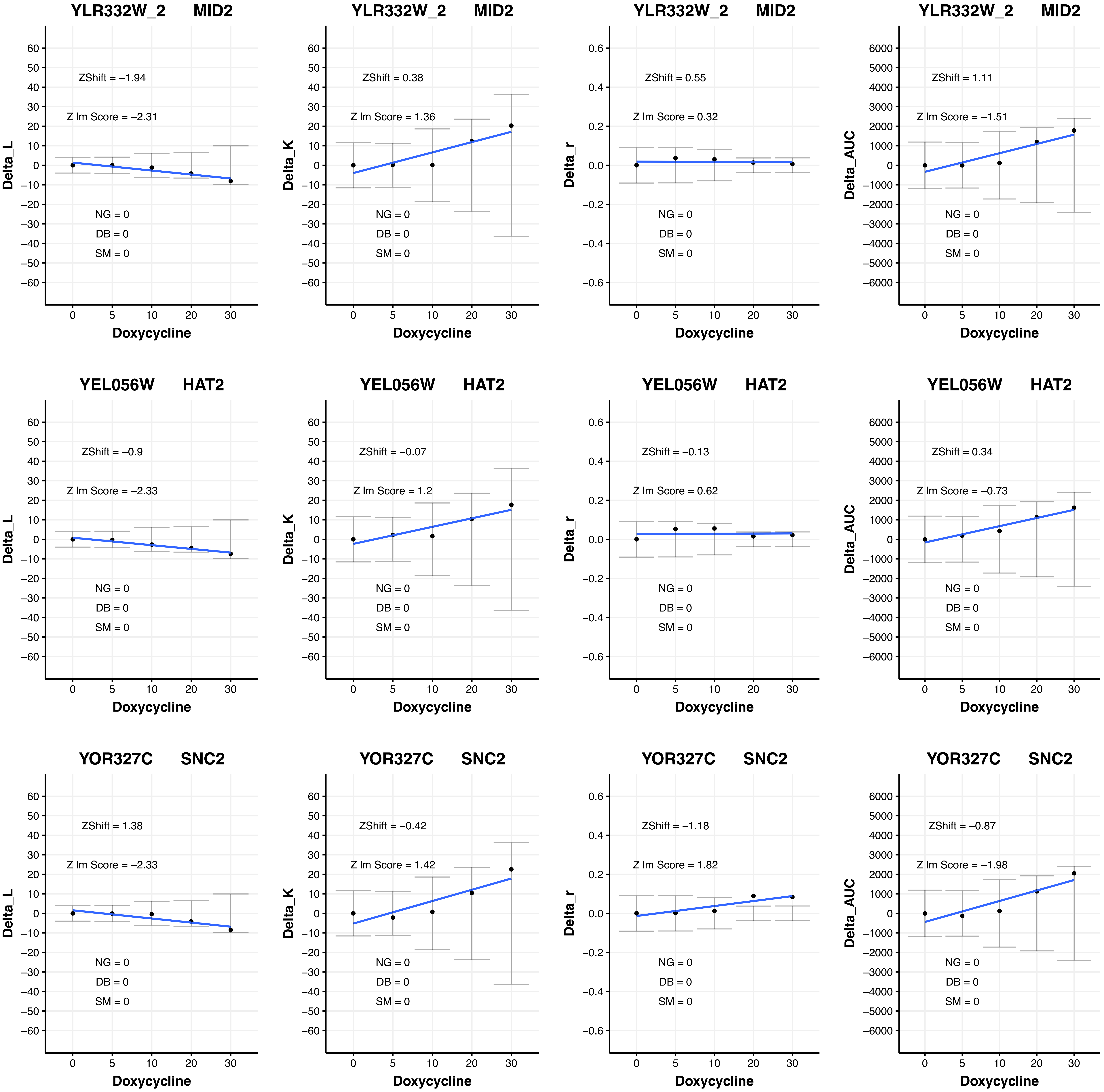

YLR332W_2 MID2
ZShift = −1.94
Z lm Score = −2.31
NG = 0
DB = 0
SM = 0
Delta_L
Doxycycline
YLR332W_2 MID2
ZShift = 0.38
Z lm Score = 1.36
NG = 0
DB = 0
SM = 0
Delta_K
Doxycycline
YLR332W_2 MID2
ZShift = 0.55
Z lm Score = 0.32
NG = 0
DB = 0
SM = 0
Delta_r
Doxycycline
YLR332W_2 MID2
ZShift = 1.11
Z lm Score = −1.51
NG = 0
DB = 0
SM = 0
Delta_AUC
Doxycycline
YEL056W HAT2
ZShift = −0.9
Z lm Score = −2.33
NG = 0
DB = 0
SM = 0
Delta_L
Doxycycline
YEL056W HAT2
ZShift = −0.07
Z lm Score = 1.2
NG = 0
DB = 0
SM = 0
Delta_K
Doxycycline
YEL056W HAT2
ZShift = −0.13
Z lm Score = 0.62
NG = 0
DB = 0
SM = 0
Delta_r
Doxycycline
YEL056W HAT2
ZShift = 0.34
Z lm Score = −0.73
NG = 0
DB = 0
SM = 0
Delta_AUC
Doxycycline
YOR327C SNC2
ZShift = 1.38
Z lm Score = −2.33
NG = 0
DB = 0
SM = 0
Delta_L
Doxycycline
YOR327C SNC2
ZShift = −0.42
Z lm Score = 1.42
NG = 0
DB = 0
SM = 0
Delta_K
Doxycycline
YOR327C SNC2
ZShift = −1.18
Z lm Score = 1.82
NG = 0
DB = 0
SM = 0
Delta_r
Doxycycline
YOR327C SNC2
ZShift = −0.87
Z lm Score = −1.98
NG = 0
DB = 0
SM = 0
Delta_AUC
Doxycycline

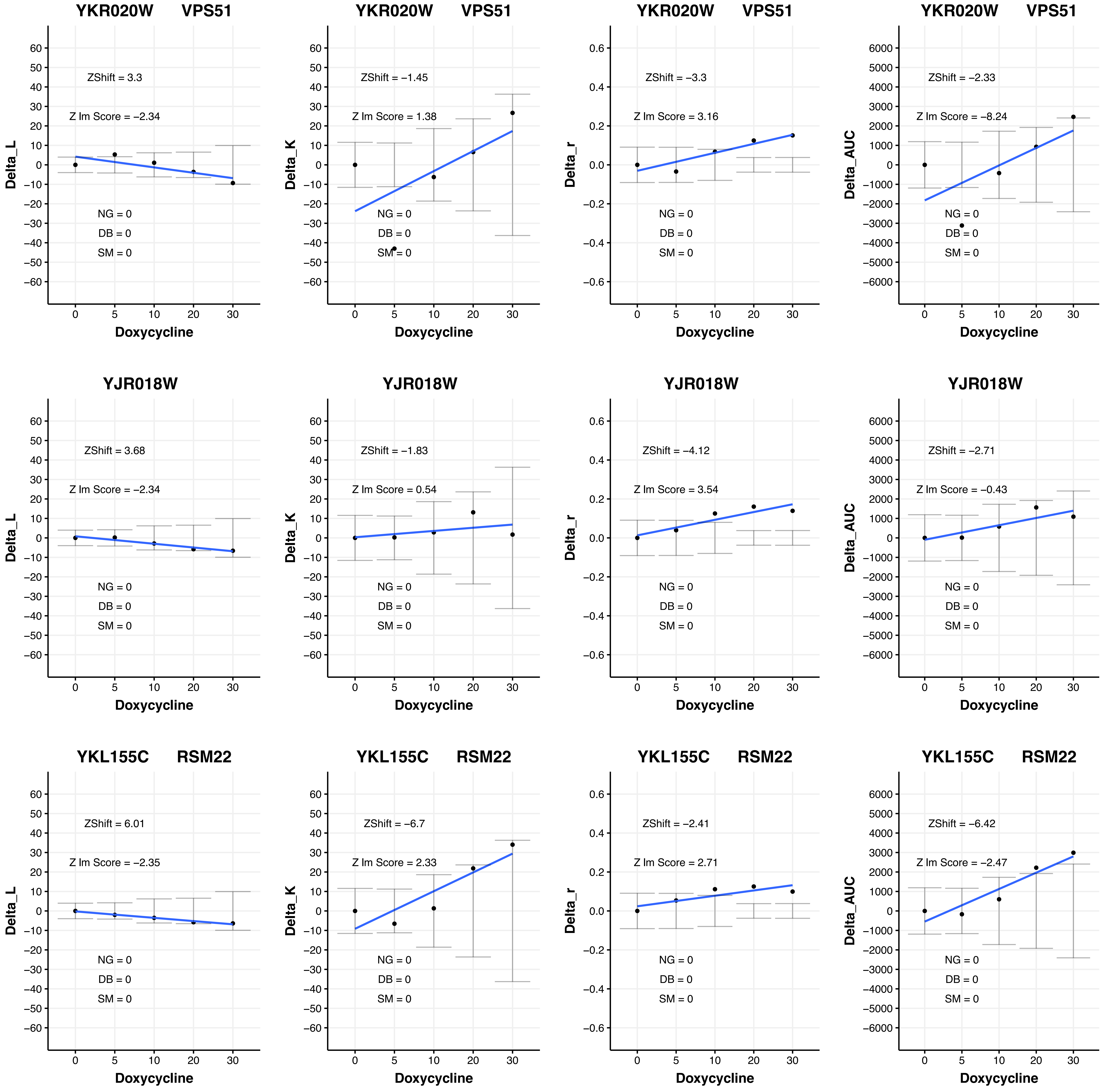
YKR020W VPS51
ZShift = 3.3
Z lm Score = −2.34
NG = 0
DB = 0
SM = 0
Delta_L
Doxycycline
YKR020W VPS51
ZShift = −1.45
Z lm Score = 1.38
NG = 0
DB = 0
SM = 0
Delta_K
Doxycycline
YKR020W VPS51
ZShift = −3.3
Z lm Score = 3.16
NG = 0
DB = 0
SM = 0
Delta_r
Doxycycline
YKR020W VPS51
ZShift = −2.33
Z lm Score = −8.24
NG = 0
DB = 0
SM = 0
Delta_AUC
Doxycycline
YJR018W
ZShift = 3.68
Z lm Score = −2.34
NG = 0
DB = 0
SM = 0
Delta_L
Doxycycline
YJR018W
ZShift = −1.83
Z lm Score = 0.54
NG = 0
DB = 0
SM = 0
Delta_K
Doxycycline
YJR018W
ZShift = −4.12
Z lm Score = 3.54
NG = 0
DB = 0
SM = 0
Delta_r
Doxycycline
YJR018W
ZShift = −2.71
Z lm Score = −0.43
NG = 0
DB = 0
SM = 0
Delta_AUC
Doxycycline
YKL155C RSM22
ZShift = 6.01
Z lm Score = −2.35
NG = 0
DB = 0
SM = 0
Delta_L
Doxycycline
YKL155C RSM22
ZShift = −6.7
Z lm Score = 2.33
NG = 0
DB = 0
SM = 0
Delta_K
Doxycycline
YKL155C RSM22
ZShift = −2.41
Z lm Score = 2.71
NG = 0
DB = 0
SM = 0
Delta_r
Doxycycline
YKL155C RSM22
ZShift = −6.42
Z lm Score = −2.47
NG = 0
DB = 0
SM = 0
Delta_AUC
Doxycycline

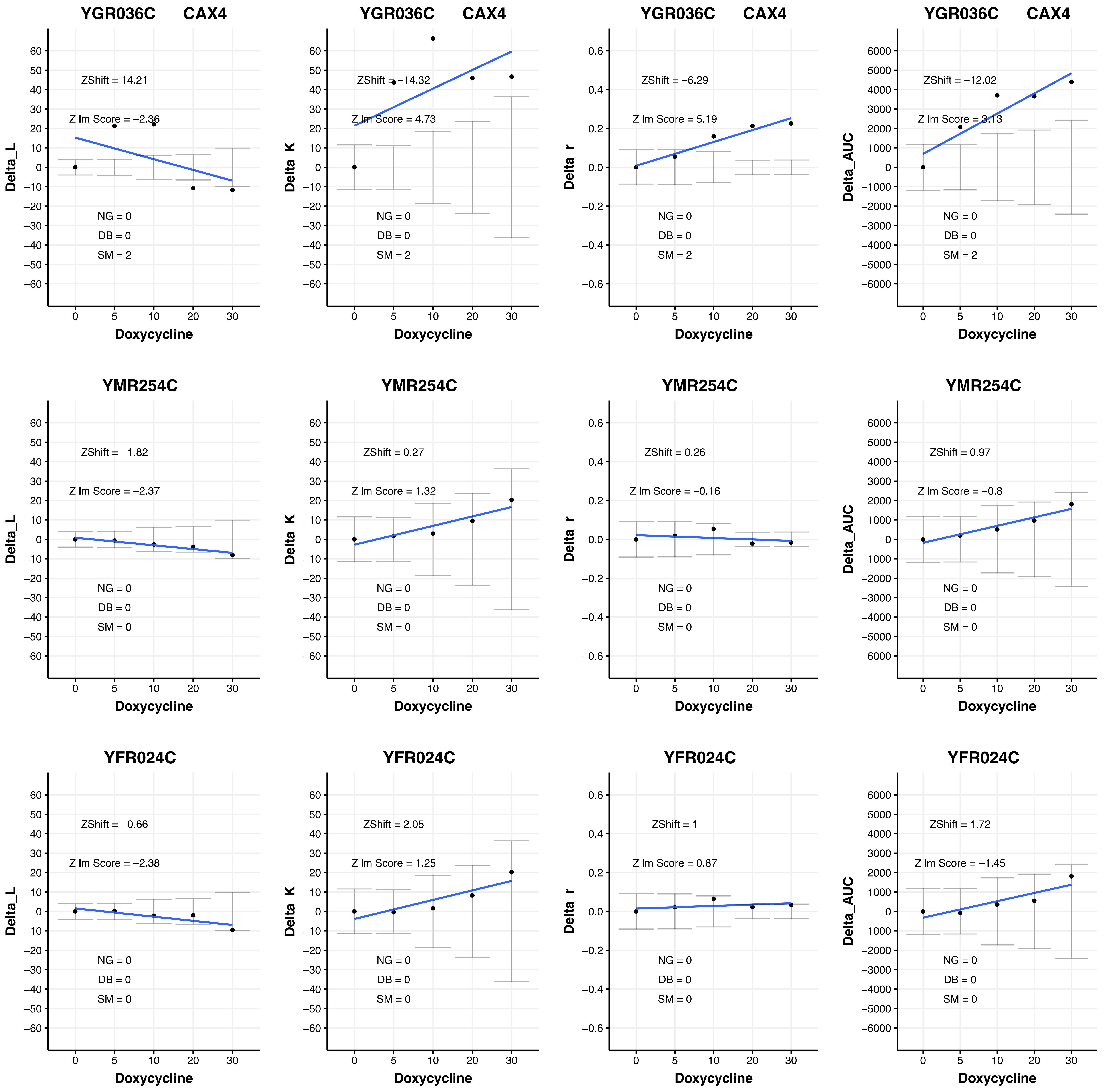
YGR036C CAX4
Delta_L
ZShift = 14.21
Z lm Score = −2.36
NG = 0
DB = 0
SM = 2
Doxycycline
YGR036C CAX4
Delta_K
ZShift = −14.32
Z lm Score = 4.73
NG = 0
DB = 0
SM = 2
Doxycycline
YGR036C CAX4
Delta_r
ZShift = −6.29
Z lm Score = 5.19
NG = 0
DB = 0
SM = 2
Doxycycline
YGR036C CAX4
Delta_AUC
ZShift = −12.02
Z lm Score = 3.13
NG = 0
DB = 0
SM = 2
Doxycycline
YMR254C
Delta_L
ZShift = −1.82
Z lm Score = −2.37
NG = 0
DB = 0
SM = 0
Doxycycline
YMR254C
Delta_K
ZShift = 0.27
Z lm Score = 1.32
NG = 0
DB = 0
SM = 0
Doxycycline
YMR254C
Delta_r
ZShift = 0.26
Z lm Score = −0.16
NG = 0
DB = 0
SM = 0
Doxycycline
YMR254C
Delta_AUC
ZShift = 0.97
Z lm Score = −0.8
NG = 0
DB = 0
SM = 0
Doxycycline
YFR024C
Delta_L
ZShift = −0.66
Z lm Score = −2.38
NG = 0
DB = 0
SM = 0
Doxycycline
YFR024C
Delta_K
ZShift = 2.05
Z lm Score = 1.25
NG = 0
DB = 0
SM = 0
Doxycycline
YFR024C
Delta_r
ZShift = 1
Z lm Score = 0.87
NG = 0
DB = 0
SM = 0
Doxycycline
YFR024C
Delta_AUC
ZShift = 1.72
Z lm Score = −1.45
NG = 0
DB = 0
SM = 0
Doxycycline

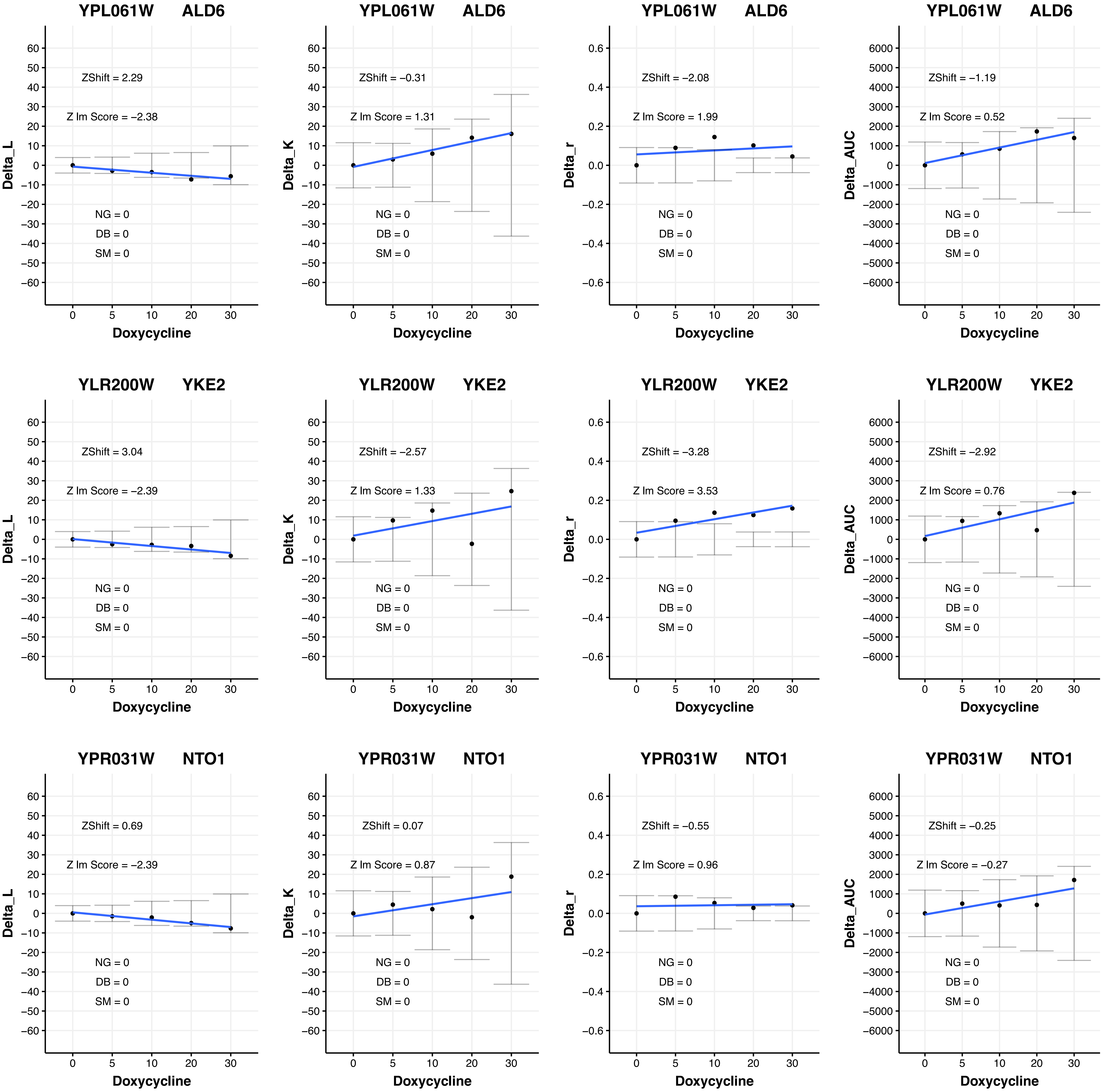
YPL061W ALD6
ZShift = 2.29
Z lm Score = −2.38
NG = 0
DB = 0
SM = 0
Delta_L
Doxycycline
YPL061W ALD6
ZShift = −0.31
Z lm Score = 1.31
NG = 0
DB = 0
SM = 0
Delta_K
Doxycycline
YPL061W ALD6
ZShift = −2.08
Z lm Score = 1.99
NG = 0
DB = 0
SM = 0
Delta_r
Doxycycline
YPL061W ALD6
ZShift = −1.19
Z lm Score = 0.52
NG = 0
DB = 0
SM = 0
Delta_AUC
Doxycycline
YLR200W YKE2
ZShift = 3.04
Z lm Score = −2.39
NG = 0
DB = 0
SM = 0
Delta_L
Doxycycline
YLR200W YKE2
ZShift = −2.57
Z lm Score = 1.33
NG = 0
DB = 0
SM = 0
Delta_K
Doxycycline
YLR200W YKE2
ZShift = −3.28
Z lm Score = 3.53
NG = 0
DB = 0
SM = 0
Delta_r
Doxycycline
YLR200W YKE2
ZShift = −2.92
Z lm Score = 0.76
NG = 0
DB = 0
SM = 0
Delta_AUC
Doxycycline
YPR031W NTO1
ZShift = 0.69
Z lm Score = −2.39
NG = 0
DB = 0
SM = 0
Delta_L
Doxycycline
YPR031W NTO1
ZShift = 0.07
Z lm Score = 0.87
NG = 0
DB = 0
SM = 0
Delta_K
Doxycycline
YPR031W NTO1
ZShift = −0.55
Z lm Score = 0.96
NG = 0
DB = 0
SM = 0
Delta_r
Doxycycline
YPR031W NTO1
ZShift = −0.25
Z lm Score = −0.27
NG = 0
DB = 0
SM = 0
Delta_AUC
Doxycycline

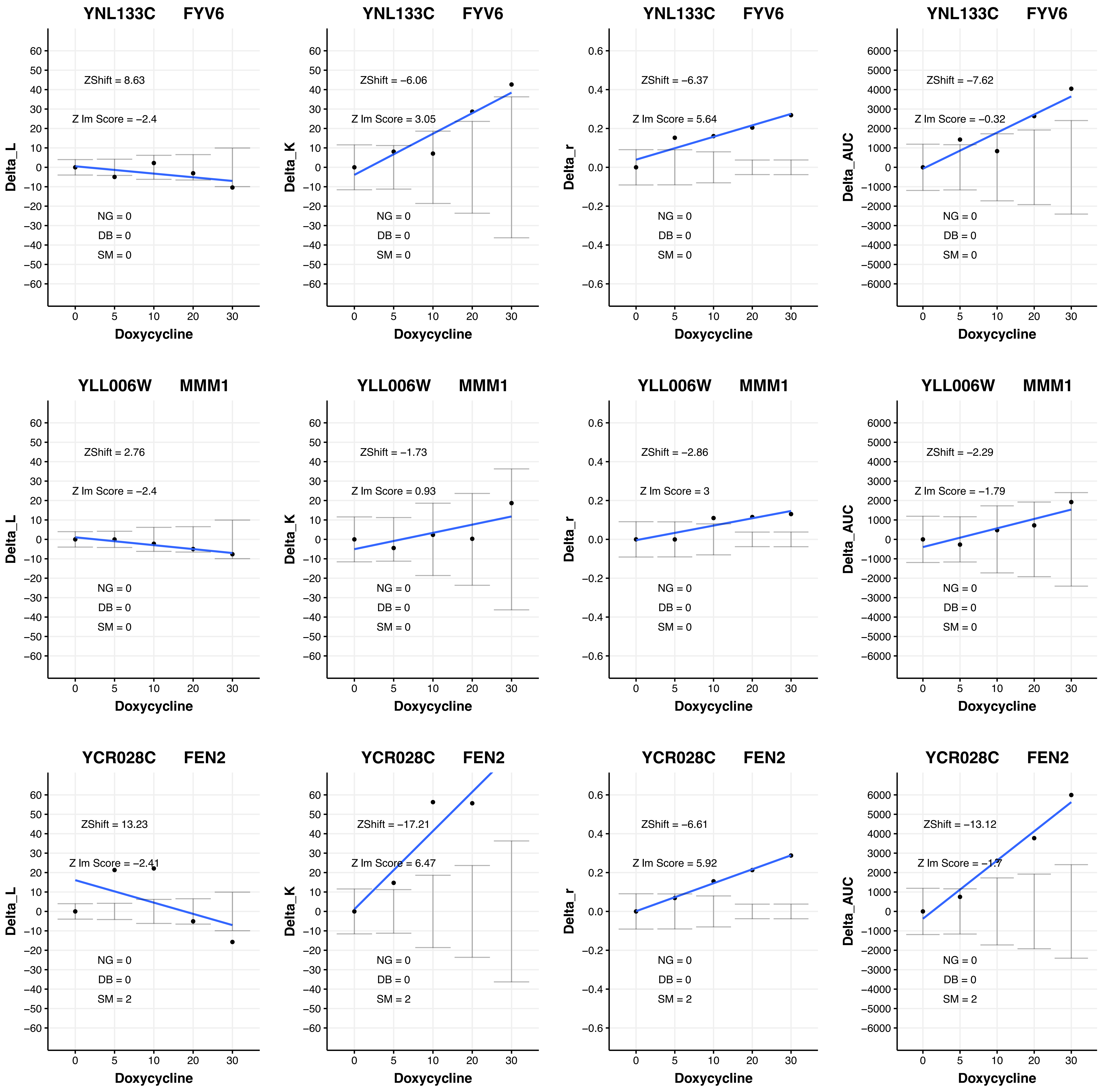

YNL133C FYV6
ZShift = 8.63
Z lm Score = −2.4
NG = 0
DB = 0
SM = 0
Delta_L
Doxycycline
YNL133C FYV6
ZShift = −6.06
Z lm Score = 3.05
NG = 0
DB = 0
SM = 0
Delta_K
Doxycycline
YNL133C FYV6
ZShift = −6.37
Z lm Score = 5.64
NG = 0
DB = 0
SM = 0
Delta_r
Doxycycline
YNL133C FYV6
ZShift = −7.62
Z lm Score = −0.32
NG = 0
DB = 0
SM = 0
Delta_AUC
Doxycycline
YLL006W MMM1
ZShift = 2.76
Z lm Score = −2.4
NG = 0
DB = 0
SM = 0
Delta_L
Doxycycline
YLL006W MMM1
ZShift = −1.73
Z lm Score = 0.93
NG = 0
DB = 0
SM = 0
Delta_K
Doxycycline
YLL006W MMM1
ZShift = −2.86
Z lm Score = 3
NG = 0
DB = 0
SM = 0
Delta_r
Doxycycline
YLL006W MMM1
ZShift = −2.29
Z lm Score = −1.79
NG = 0
DB = 0
SM = 0
Delta_AUC
Doxycycline
YCR028C FEN2
ZShift = 13.23
Z lm Score = −2.41
NG = 0
DB = 0
SM = 2
Delta_L
Doxycycline
YCR028C FEN2
ZShift = −17.21
Z lm Score = 6.47
NG = 0
DB = 0
SM = 2
Delta_K
Doxycycline
YCR028C FEN2
ZShift = −6.61
Z lm Score = 5.92
NG = 0
DB = 0
SM = 2
Delta_r
Doxycycline
YCR028C FEN2
ZShift = −13.12
Z lm Score = −1.7
NG = 0
DB = 0
SM = 2
Delta_AUC
Doxycycline

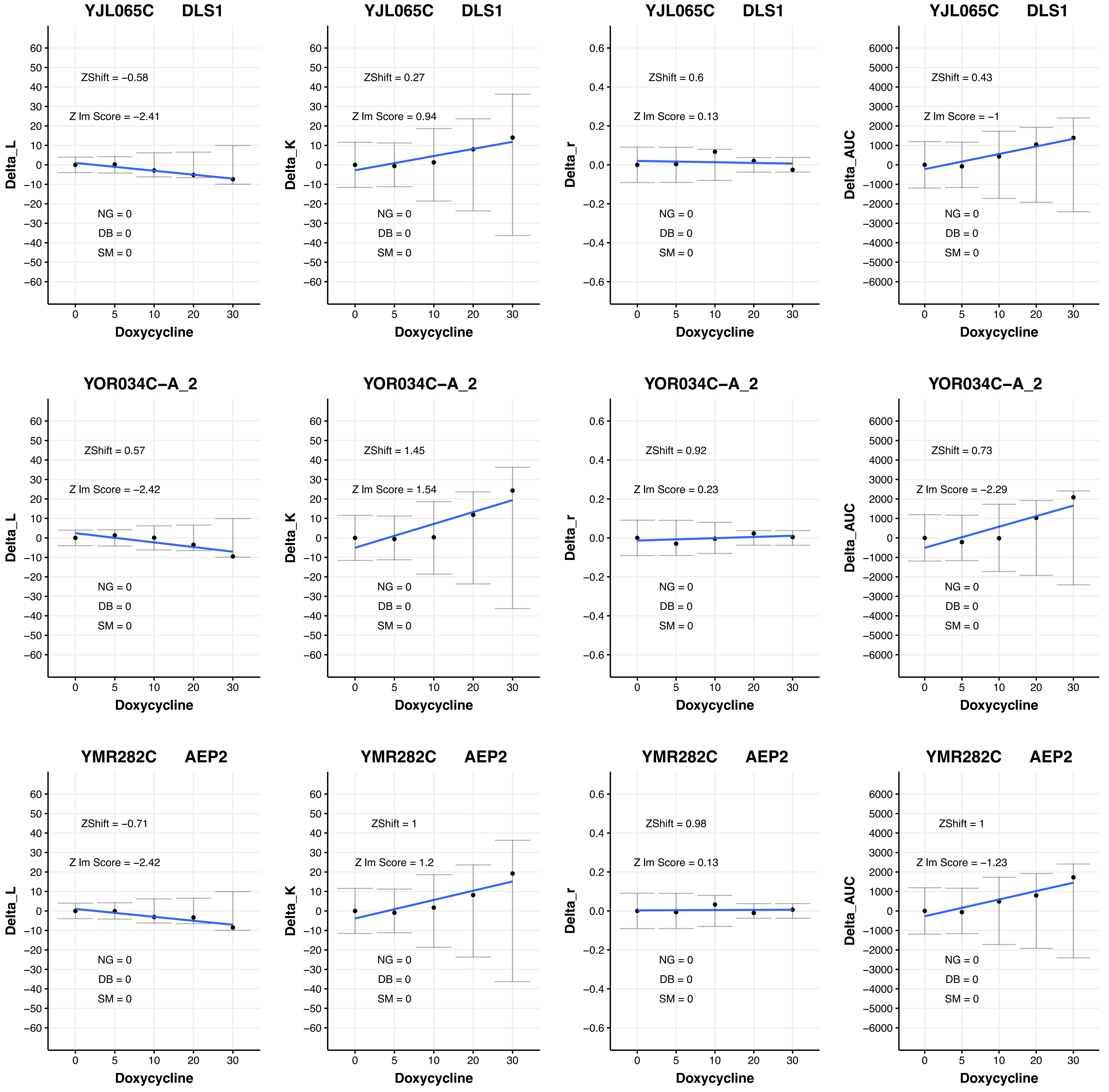

YJL065C DLS1
ZShift = −0.58
Z Im Score = −2.41
NG = 0
DB = 0
SM = 0
Delta_L
Doxycycline
YJL065C DLS1
ZShift = 0.27
Z Im Score = 0.94
NG = 0
DB = 0
SM = 0
Delta_K
Doxycycline
YJL065C DLS1
ZShift = 0.6
Z Im Score = 0.13
NG = 0
DB = 0
SM = 0
Delta_r
Doxycycline
YJL065C DLS1
ZShift = 0.43
Z Im Score = −1
NG = 0
DB = 0
SM = 0
Delta_AUC
Doxycycline
YOR034C−A_2
ZShift = 0.57
Z Im Score = −2.42
NG = 0
DB = 0
SM = 0
Delta_L
Doxycycline
YOR034C−A_2
ZShift = 1.45
Z Im Score = 1.54
NG = 0
DB = 0
SM = 0
Delta_K
Doxycycline
YOR034C−A_2
ZShift = 0.92
Z Im Score = 0.23
NG = 0
DB = 0
SM = 0
Delta_r
Doxycycline
YOR034C−A_2
ZShift = 0.73
Z Im Score = −2.29
NG = 0
DB = 0
SM = 0
Delta_AUC
Doxycycline
YMR282C AEP2
ZShift = −0.71
Z Im Score = −2.42
NG = 0
DB = 0
SM = 0
Delta_L
Doxycycline
YMR282C AEP2
ZShift = 1
Z Im Score = 1.2
NG = 0
DB = 0
SM = 0
Delta_K
Doxycycline
YMR282C AEP2
ZShift = 0.98
Z Im Score = 0.13
NG = 0
DB = 0
SM = 0
Delta_r
Doxycycline
YMR282C AEP2
ZShift = 1
Z Im Score = −1.23
NG = 0
DB = 0
SM = 0
Delta_AUC
Doxycycline

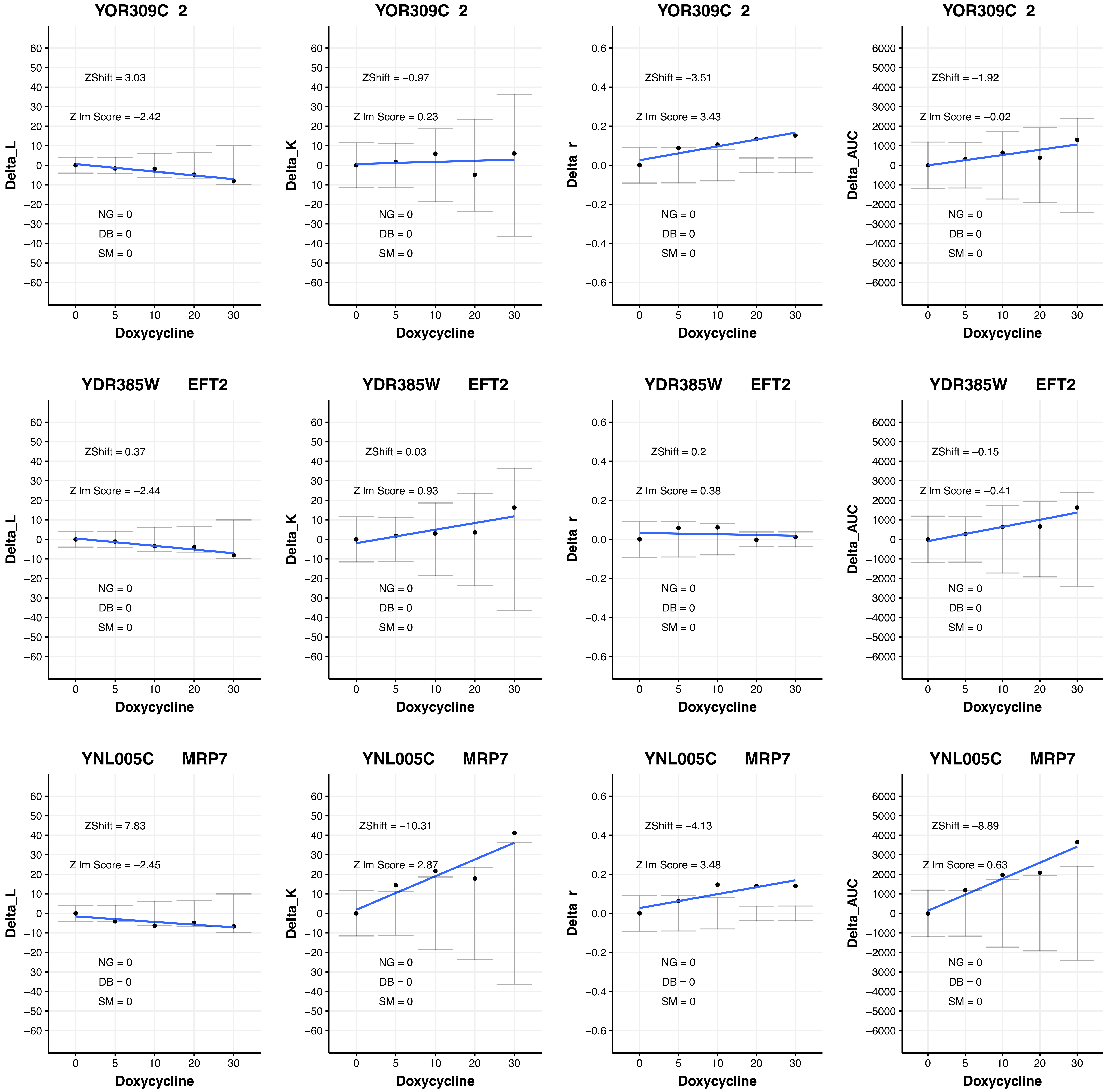

YOR309C_2
ZShift = 3.03
Z lm Score = −2.42
NG = 0
DB = 0
SM = 0
Delta_L
Doxycycline
YOR309C_2
ZShift = −0.97
Z lm Score = 0.23
NG = 0
DB = 0
SM = 0
Delta_K
Doxycycline
YOR309C_2
ZShift = −3.51
Z lm Score = 3.43
NG = 0
DB = 0
SM = 0
Delta_r
Doxycycline
YOR309C_2
ZShift = −1.92
Z lm Score = −0.02
NG = 0
DB = 0
SM = 0
Delta_AUC
Doxycycline
YDR385W EFT2
ZShift = 0.37
Z lm Score = −2.44
NG = 0
DB = 0
SM = 0
Delta_L
Doxycycline
YDR385W EFT2
ZShift = 0.03
Z lm Score = 0.93
NG = 0
DB = 0
SM = 0
Delta_K
Doxycycline
YDR385W EFT2
ZShift = 0.2
Z lm Score = 0.38
NG = 0
DB = 0
SM = 0
Delta_r
Doxycycline
YDR385W EFT2
ZShift = −0.15
Z lm Score = −0.41
NG = 0
DB = 0
SM = 0
Delta_AUC
Doxycycline
YNL005C MRP7
ZShift = 7.83
Z lm Score = −2.45
NG = 0
DB = 0
SM = 0
Delta_L
Doxycycline
YNL005C MRP7
ZShift = −10.31
Z lm Score = 2.87
NG = 0
DB = 0
SM = 0
Delta_K
Doxycycline
YNL005C MRP7
ZShift = −4.13
Z lm Score = 3.48
NG = 0
DB = 0
SM = 0
Delta_r
Doxycycline
YNL005C MRP7
ZShift = −8.89
Z lm Score = 0.63
NG = 0
DB = 0
SM = 0
Delta_AUC
Doxycycline

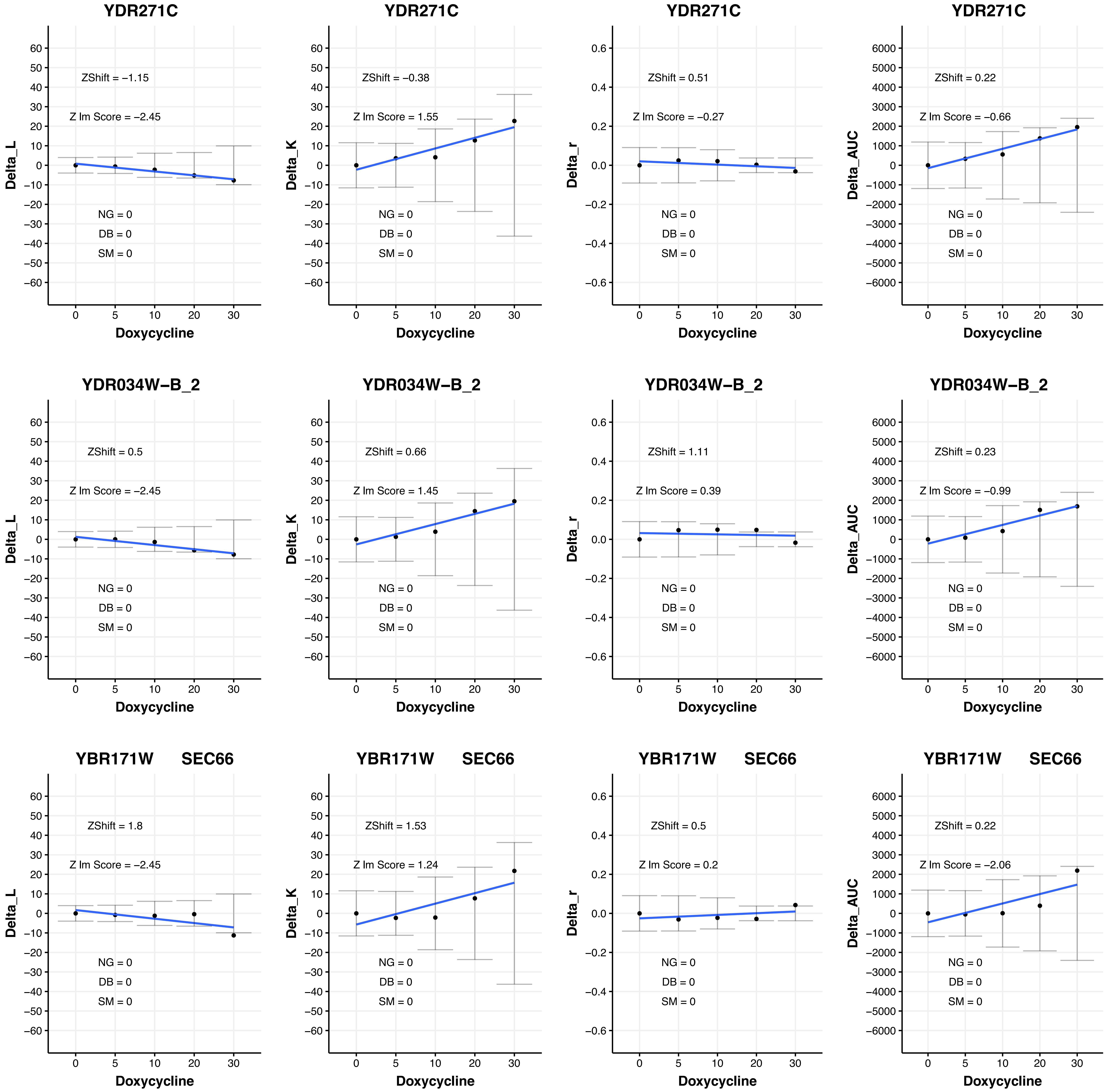

YDR271C
ZShift = −1.15
Z lm Score = −2.45
NG = 0
DB = 0
SM = 0
Delta_L
Doxycycline
YDR271C
ZShift = −0.38
Z lm Score = 1.55
NG = 0
DB = 0
SM = 0
Delta_K
Doxycycline
YDR271C
ZShift = 0.51
Z lm Score = −0.27
NG = 0
DB = 0
SM = 0
Delta_r
Doxycycline
YDR271C
ZShift = 0.22
Z lm Score = −0.66
NG = 0
DB = 0
SM = 0
Delta_AUC
Doxycycline
YDR034W−B_2
ZShift = 0.5
Z lm Score = −2.45
NG = 0
DB = 0
SM = 0
Delta_L
Doxycycline
YDR034W−B_2
ZShift = 0.66
Z lm Score = 1.45
NG = 0
DB = 0
SM = 0
Delta_K
Doxycycline
YDR034W−B_2
ZShift = 1.11
Z lm Score = 0.39
NG = 0
DB = 0
SM = 0
Delta_r
Doxycycline
YDR034W−B_2
ZShift = 0.23
Z lm Score = −0.99
NG = 0
DB = 0
SM = 0
Delta_AUC
Doxycycline
YBR171W SEC66
ZShift = 1.8
Z lm Score = −2.45
NG = 0
DB = 0
SM = 0
Delta_L
Doxycycline
YBR171W SEC66
ZShift = 1.53
Z lm Score = 1.24
NG = 0
DB = 0
SM = 0
Delta_K
Doxycycline
YBR171W SEC66
ZShift = 0.5
Z lm Score = 0.2
NG = 0
DB = 0
SM = 0
Delta_r
Doxycycline
YBR171W SEC66
ZShift = 0.22
Z lm Score = −2.06
NG = 0
DB = 0
SM = 0
Delta_AUC
Doxycycline

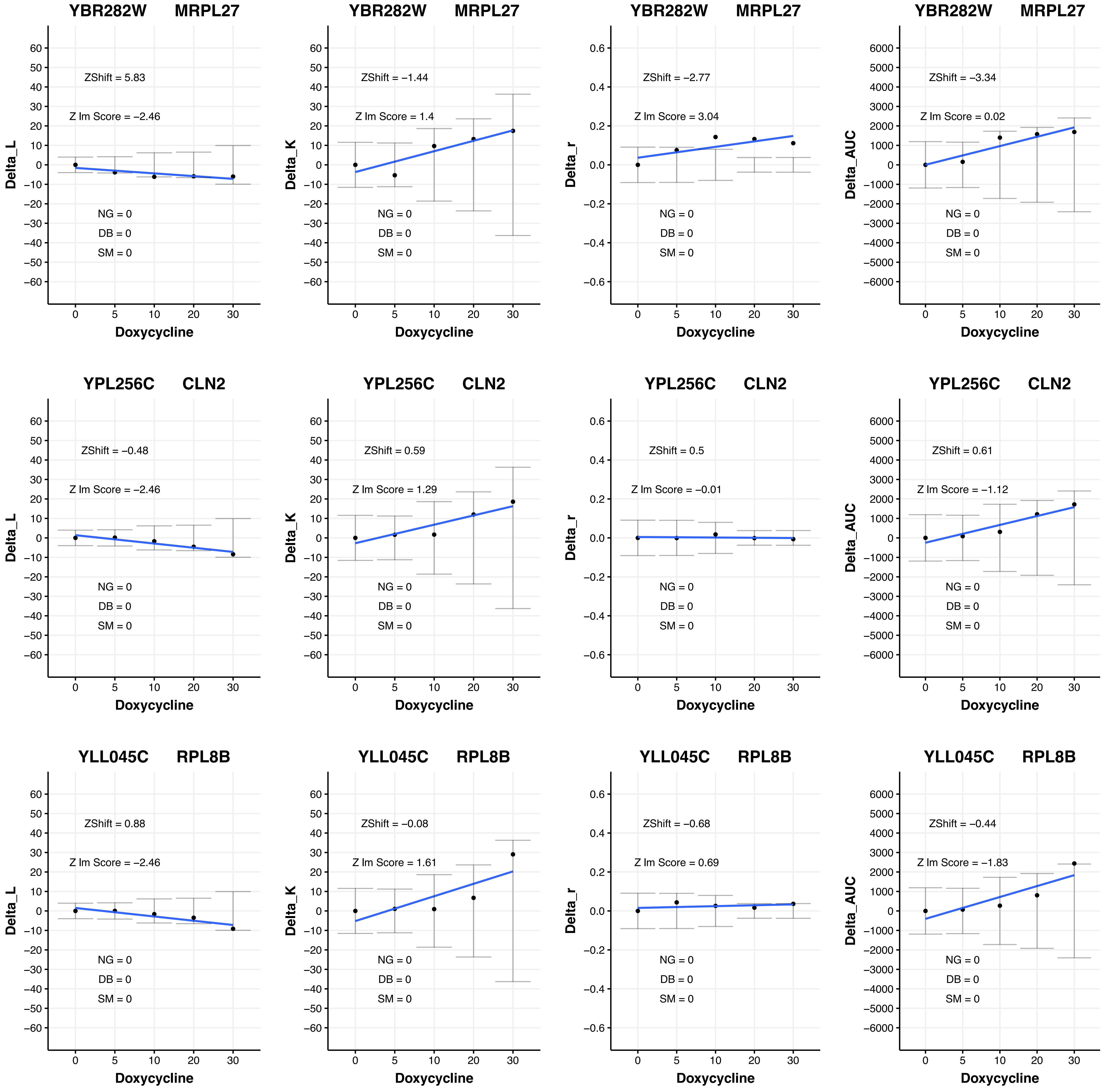

YBR282W MRPL27
ZShift = 5.83
Z lm Score = −2.46
NG = 0
DB = 0
SM = 0
Delta_L
Doxycycline
YBR282W MRPL27
ZShift = −1.44
Z lm Score = 1.4
NG = 0
DB = 0
SM = 0
Delta_K
Doxycycline
YBR282W MRPL27
ZShift = −2.77
Z lm Score = 3.04
NG = 0
DB = 0
SM = 0
Delta_r
Doxycycline
YBR282W MRPL27
ZShift = −3.34
Z lm Score = 0.02
NG = 0
DB = 0
SM = 0
Delta_AUC
Doxycycline
YPL256C CLN2
ZShift = −0.48
Z lm Score = −2.46
NG = 0
DB = 0
SM = 0
Delta_L
Doxycycline
YPL256C CLN2
ZShift = 0.59
Z lm Score = 1.29
NG = 0
DB = 0
SM = 0
Delta_K
Doxycycline
YPL256C CLN2
ZShift = 0.5
Z lm Score = −0.01
NG = 0
DB = 0
SM = 0
Delta_r
Doxycycline
YPL256C CLN2
ZShift = 0.61
Z lm Score = −1.12
NG = 0
DB = 0
SM = 0
Delta_AUC
Doxycycline
YLL045C RPL8B
ZShift = 0.88
Z lm Score = −2.46
NG = 0
DB = 0
SM = 0
Delta_L
Doxycycline
YLL045C RPL8B
ZShift = −0.08
Z lm Score = 1.61
NG = 0
DB = 0
SM = 0
Delta_K
Doxycycline
YLL045C RPL8B
ZShift = −0.68
Z lm Score = 0.69
NG = 0
DB = 0
SM = 0
Delta_r
Doxycycline
YLL045C RPL8B
ZShift = −0.44
Z lm Score = −1.83
NG = 0
DB = 0
SM = 0
Delta_AUC
Doxycycline

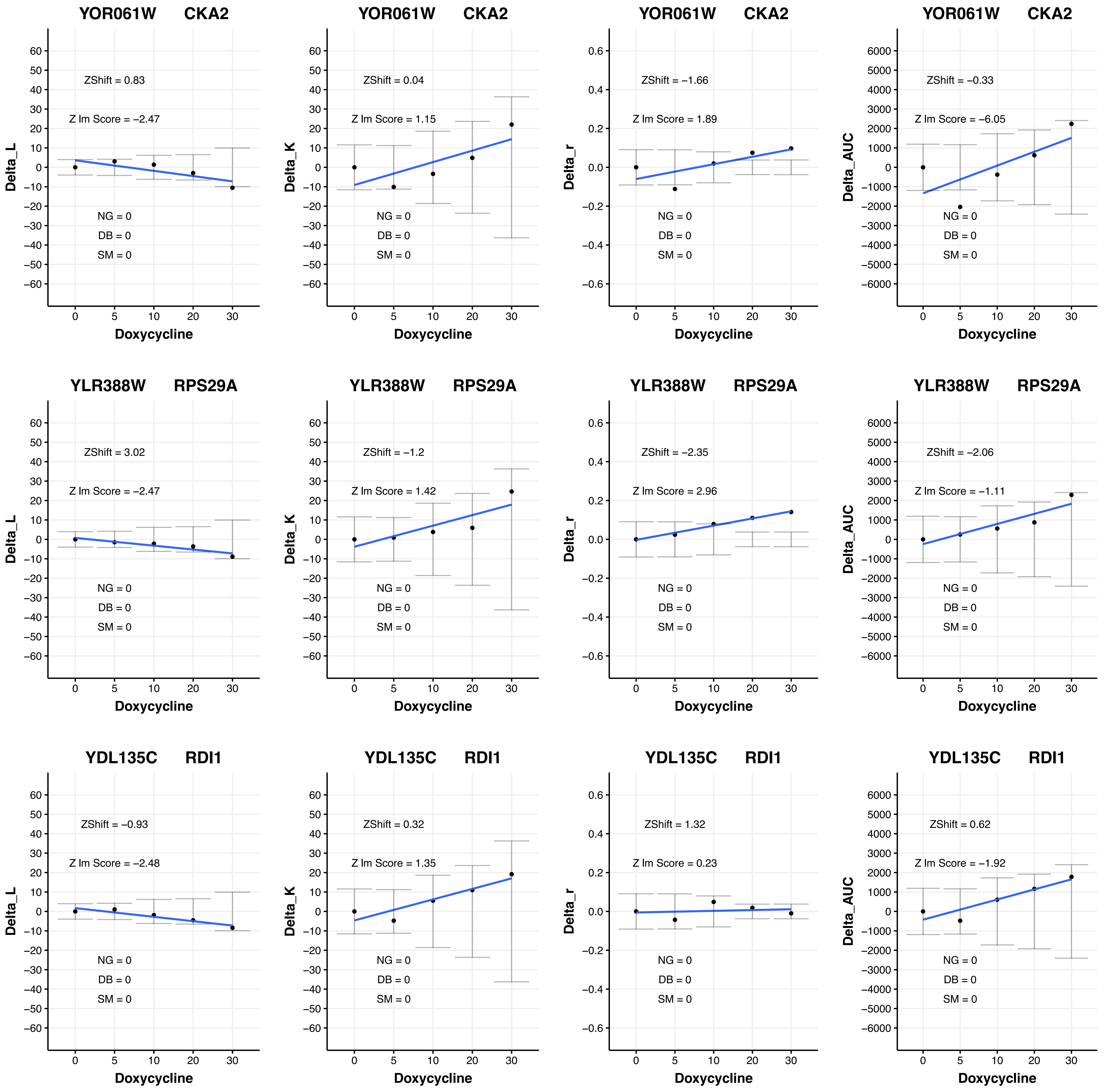
YOR061W CKA2
ZShift = 0.83
Z lm Score = −2.47
NG = 0
DB = 0
SM = 0
Delta_L
Doxycycline
YOR061W CKA2
ZShift = 0.04
Z lm Score = 1.15
NG = 0
DB = 0
SM = 0
Delta_K
Doxycycline
YOR061W CKA2
ZShift = −1.66
Z lm Score = 1.89
NG = 0
DB = 0
SM = 0
Delta_r
Doxycycline
YOR061W CKA2
ZShift = −0.33
Z lm Score = −6.05
NG = 0
DB = 0
SM = 0
Delta_AUC
Doxycycline
YLR388W RPS29A
ZShift = 3.02
Z lm Score = −2.47
NG = 0
DB = 0
SM = 0
Delta_L
Doxycycline
YLR388W RPS29A
ZShift = −1.2
Z lm Score = 1.42
NG = 0
DB = 0
SM = 0
Delta_K
Doxycycline
YLR388W RPS29A
ZShift = −2.35
Z lm Score = 2.96
NG = 0
DB = 0
SM = 0
Delta_r
Doxycycline
YLR388W RPS29A
ZShift = −2.06
Z lm Score = −1.11
NG = 0
DB = 0
SM = 0
Delta_AUC
Doxycycline
YDL135C RDI1
ZShift = −0.93
Z lm Score = −2.48
NG = 0
DB = 0
SM = 0
Delta_L
Doxycycline
YDL135C RDI1
ZShift = 0.32
Z lm Score = 1.35
NG = 0
DB = 0
SM = 0
Delta_K
Doxycycline
YDL135C RDI1
ZShift = 1.32
Z lm Score = 0.23
NG = 0
DB = 0
SM = 0
Delta_r
Doxycycline
YDL135C RDI1
ZShift = 0.62
Z lm Score = −1.92
NG = 0
DB = 0
SM = 0
Delta_AUC
Doxycycline

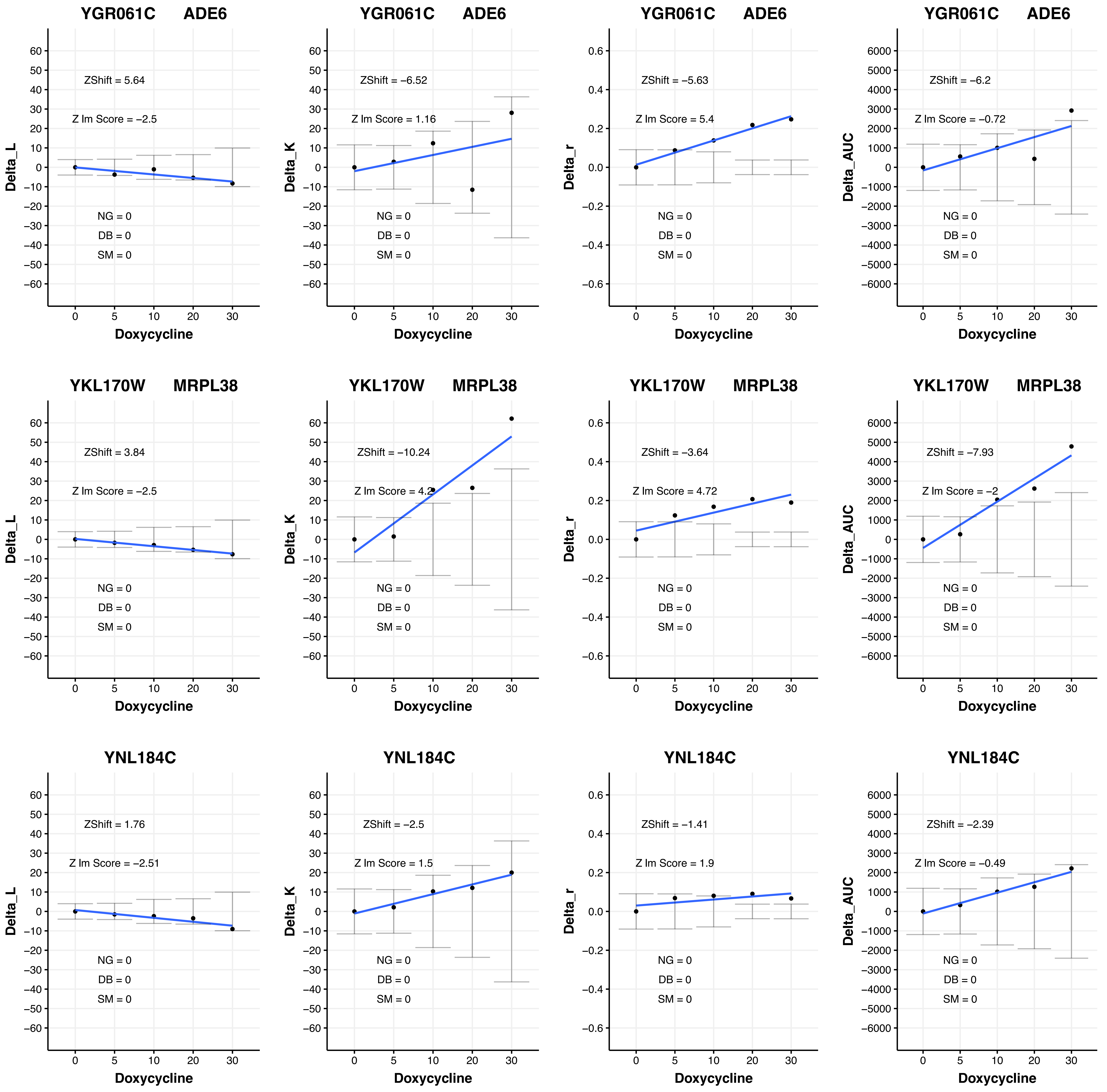
YGR061C ADE6
ZShift = 5.64
Z lm Score = −2.5
NG = 0
DB = 0
SM = 0
Delta_L
Doxycycline
YGR061C ADE6
ZShift = −6.52
Z lm Score = 1.16
NG = 0
DB = 0
SM = 0
Delta_K
Doxycycline
YGR061C ADE6
ZShift = −5.63
Z lm Score = 5.4
NG = 0
DB = 0
SM = 0
Delta_r
Doxycycline
YGR061C ADE6
ZShift = −6.2
Z lm Score = −0.72
NG = 0
DB = 0
SM = 0
Delta_AUC
Doxycycline
YKL170W MRPL38
ZShift = 3.84
Z lm Score = −2.5
NG = 0
DB = 0
SM = 0
Delta_L
Doxycycline
YKL170W MRPL38
ZShift = −10.24
Z lm Score = 4.2
NG = 0
DB = 0
SM = 0
Delta_K
Doxycycline
YKL170W MRPL38
ZShift = −3.64
Z lm Score = 4.72
NG = 0
DB = 0
SM = 0
Delta_r
Doxycycline
YKL170W MRPL38
ZShift = −7.93
Z lm Score = −2
NG = 0
DB = 0
SM = 0
Delta_AUC
Doxycycline
YNL184C
ZShift = 1.76
Z lm Score = −2.51
NG = 0
DB = 0
SM = 0
Delta_L
Doxycycline
YNL184C
ZShift = −2.5
Z lm Score = 1.5
NG = 0
DB = 0
SM = 0
Delta_K
Doxycycline
YNL184C
ZShift = −1.41
Z lm Score = 1.9
NG = 0
DB = 0
SM = 0
Delta_r
Doxycycline
YNL184C
ZShift = −2.39
Z lm Score = −0.49
NG = 0
DB = 0
SM = 0
Delta_AUC
Doxycycline

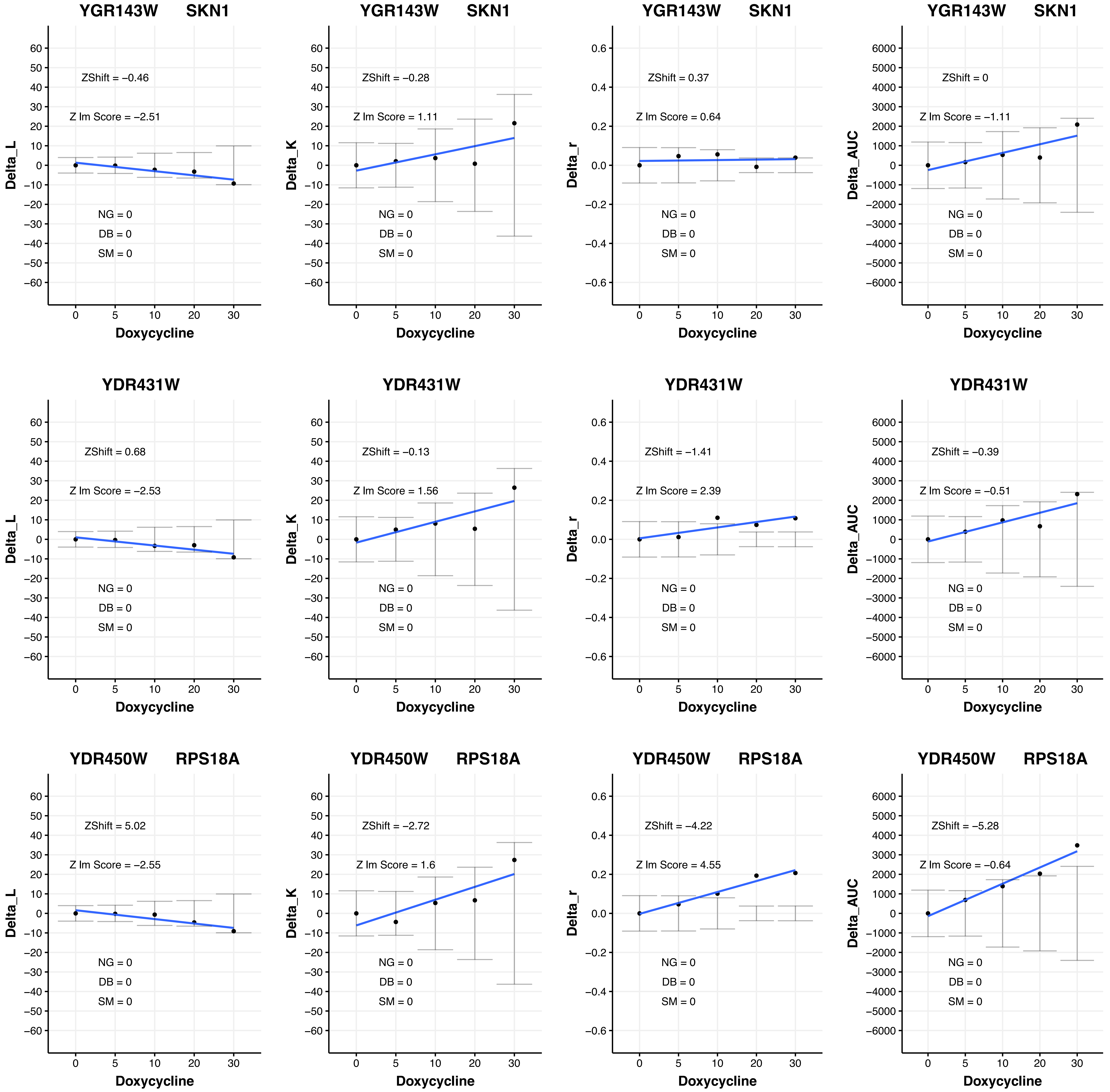

YGR143W SKN1
ZShift = −0.46
Z lm Score = −2.51
NG = 0
DB = 0
SM = 0
Delta_L
Doxycycline
YGR143W SKN1
ZShift = −0.28
Z lm Score = 1.11
NG = 0
DB = 0
SM = 0
Delta_K
Doxycycline
YGR143W SKN1
ZShift = 0.37
Z lm Score = 0.64
NG = 0
DB = 0
SM = 0
Delta_r
Doxycycline
YGR143W SKN1
ZShift = 0
Z lm Score = −1.11
NG = 0
DB = 0
SM = 0
Delta_AUC
Doxycycline
YDR431W
ZShift = 0.68
Z lm Score = −2.53
NG = 0
DB = 0
SM = 0
Delta_L
Doxycycline
YDR431W
ZShift = −0.13
Z lm Score = 1.56
NG = 0
DB = 0
SM = 0
Delta_K
Doxycycline
YDR431W
ZShift = −1.41
Z lm Score = 2.39
NG = 0
DB = 0
SM = 0
Delta_r
Doxycycline
YDR431W
ZShift = −0.39
Z lm Score = −0.51
NG = 0
DB = 0
SM = 0
Delta_AUC
Doxycycline
YDR450W RPS18A
ZShift = 5.02
Z lm Score = −2.55
NG = 0
DB = 0
SM = 0
Delta_L
Doxycycline
YDR450W RPS18A
ZShift = −2.72
Z lm Score = 1.6
NG = 0
DB = 0
SM = 0
Delta_K
Doxycycline
YDR450W RPS18A
ZShift = −4.22
Z lm Score = 4.55
NG = 0
DB = 0
SM = 0
Delta_r
Doxycycline
YDR450W RPS18A
ZShift = −5.28
Z lm Score = −0.64
NG = 0
DB = 0
SM = 0
Delta_AUC
Doxycycline

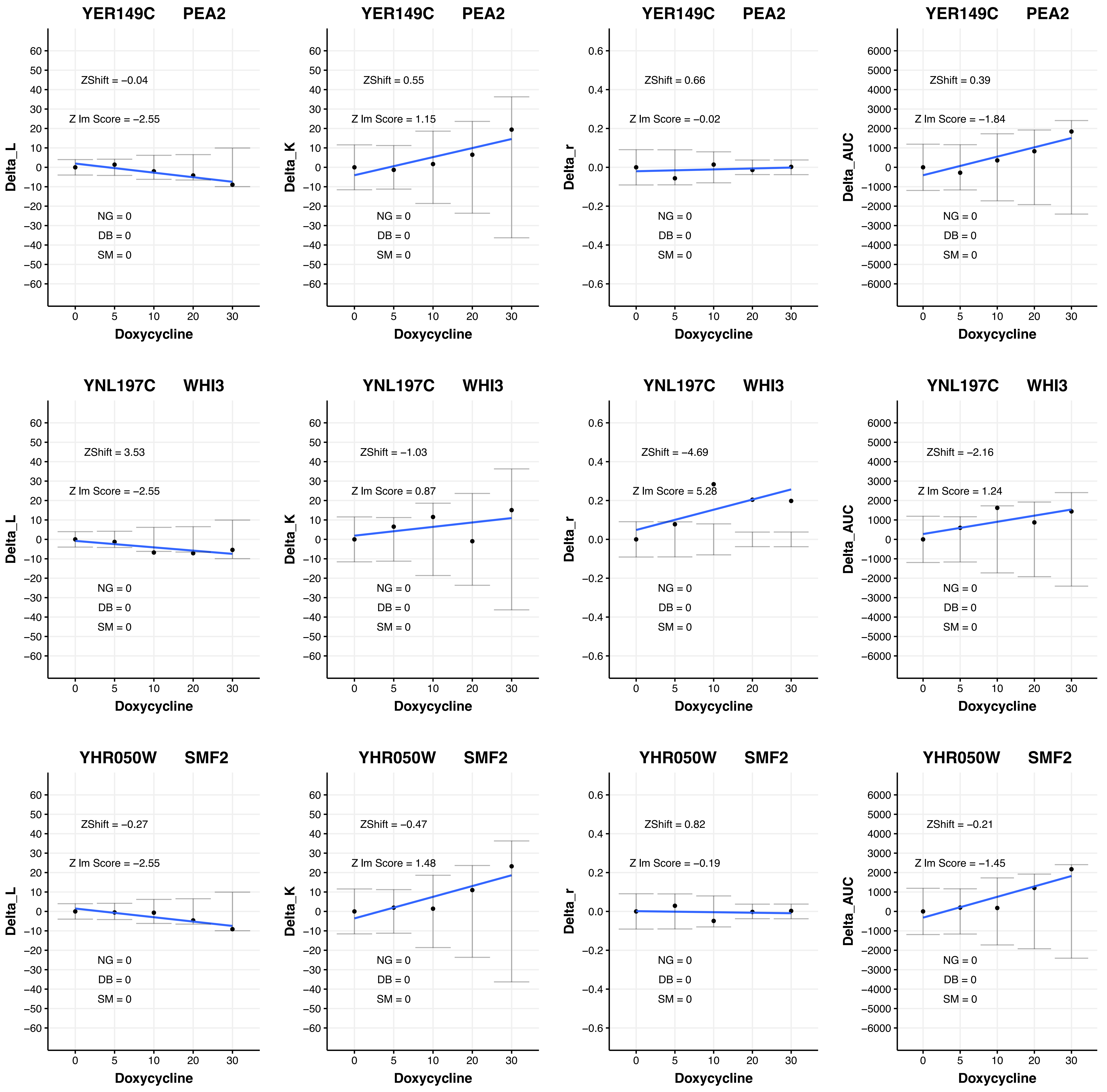
YER149C PEA2
ZShift = −0.04
Z lm Score = −2.55
NG = 0
DB = 0
SM = 0
Delta_L
Doxycycline
YER149C PEA2
ZShift = 0.55
Z lm Score = 1.15
NG = 0
DB = 0
SM = 0
Delta_K
Doxycycline
YER149C PEA2
ZShift = 0.66
Z lm Score = −0.02
NG = 0
DB = 0
SM = 0
Delta_r
Doxycycline
YER149C PEA2
ZShift = 0.39
Z lm Score = −1.84
NG = 0
DB = 0
SM = 0
Delta_AUC
Doxycycline
YNL197C WHI3
ZShift = 3.53
Z lm Score = −2.55
NG = 0
DB = 0
SM = 0
Delta_L
Doxycycline
YNL197C WHI3
ZShift = −1.03
Z lm Score = 0.87
NG = 0
DB = 0
SM = 0
Delta_K
Doxycycline
YNL197C WHI3
ZShift = −4.69
Z lm Score = 5.28
NG = 0
DB = 0
SM = 0
Delta_r
Doxycycline
YNL197C WHI3
ZShift = −2.16
Z lm Score = 1.24
NG = 0
DB = 0
SM = 0
Delta_AUC
Doxycycline
YHR050W SMF2
ZShift = −0.27
Z lm Score = −2.55
NG = 0
DB = 0
SM = 0
Delta_L
Doxycycline
YHR050W SMF2
ZShift = −0.47
Z lm Score = 1.48
NG = 0
DB = 0
SM = 0
Delta_K
Doxycycline
YHR050W SMF2
ZShift = 0.82
Z lm Score = −0.19
NG = 0
DB = 0
SM = 0
Delta_r
Doxycycline
YHR050W SMF2
ZShift = −0.21
Z lm Score = −1.45
NG = 0
DB = 0
SM = 0
Delta_AUC
Doxycycline

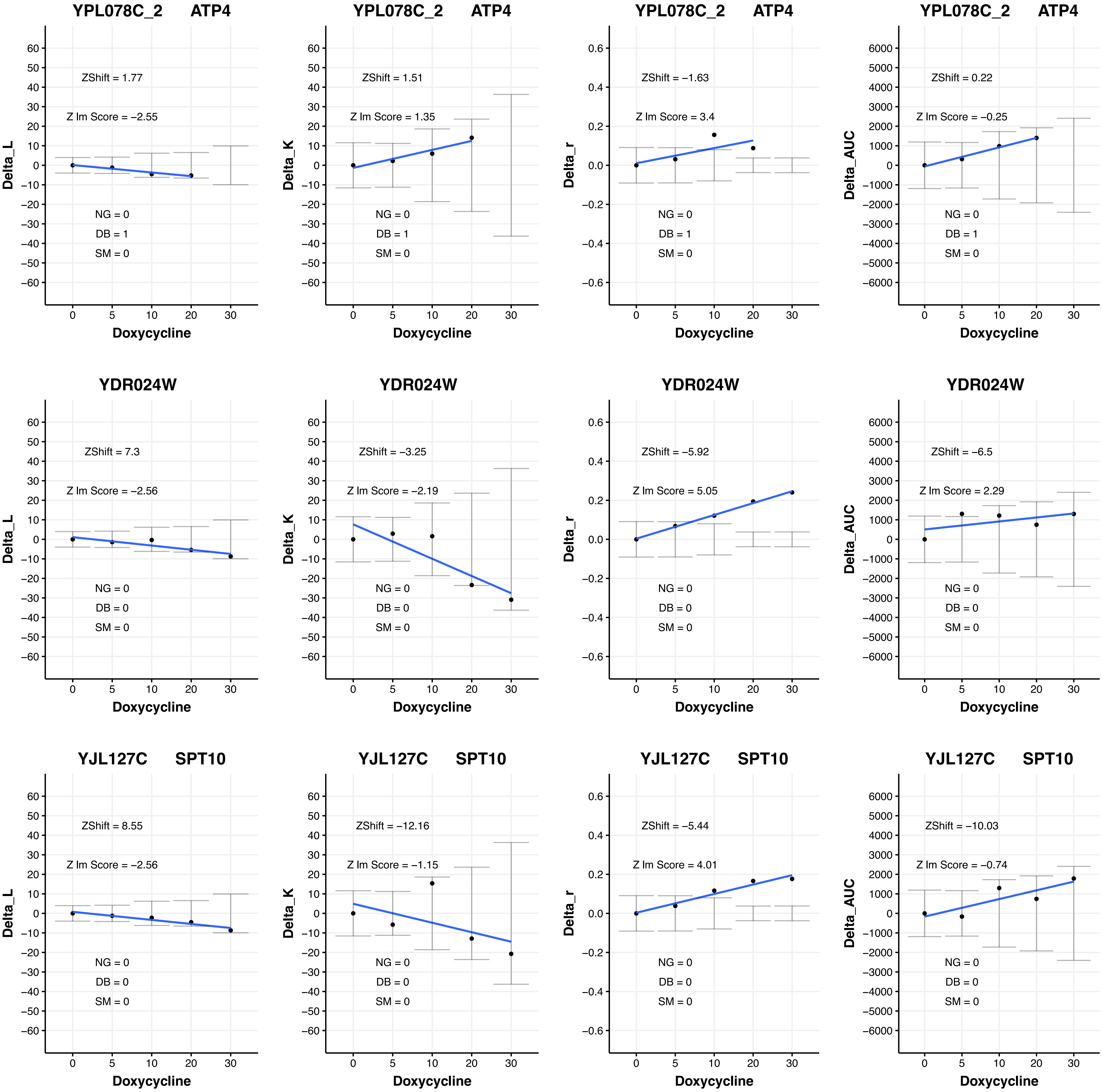

YPL078C_2 ATP4
ZShift = 1.77
Z lm Score = −2.55
NG = 0
DB = 1
SM = 0
Delta_L
Doxycycline
YPL078C_2 ATP4
ZShift = 1.51
Z lm Score = 1.35
NG = 0
DB = 1
SM = 0
Delta_K
Doxycycline
YPL078C_2 ATP4
ZShift = −1.63
Z lm Score = 3.4
NG = 0
DB = 1
SM = 0
Delta_r
Doxycycline
YPL078C_2 ATP4
ZShift = 0.22
Z lm Score = −0.25
NG = 0
DB = 1
SM = 0
Delta_AUC
Doxycycline
YDR024W
ZShift = 7.3
Z lm Score = −2.56
NG = 0
DB = 0
SM = 0
Delta_L
Doxycycline
YDR024W
ZShift = −3.25
Z lm Score = −2.19
NG = 0
DB = 0
SM = 0
Delta_K
Doxycycline
YDR024W
ZShift = −5.92
Z lm Score = 5.05
NG = 0
DB = 0
SM = 0
Delta_r
Doxycycline
YDR024W
ZShift = −6.5
Z lm Score = 2.29
NG = 0
DB = 0
SM = 0
Delta_AUC
Doxycycline
YJL127C SPT10
ZShift = 8.55
Z lm Score = −2.56
NG = 0
DB = 0
SM = 0
Delta_L
Doxycycline
YJL127C SPT10
ZShift = −12.16
Z lm Score = −1.15
NG = 0
DB = 0
SM = 0
Delta_K
Doxycycline
YJL127C SPT10
ZShift = −5.44
Z lm Score = 4.01
NG = 0
DB = 0
SM = 0
Delta_r
Doxycycline
YJL127C SPT10
ZShift = −10.03
Z lm Score = −0.74
NG = 0
DB = 0
SM = 0
Delta_AUC
Doxycycline

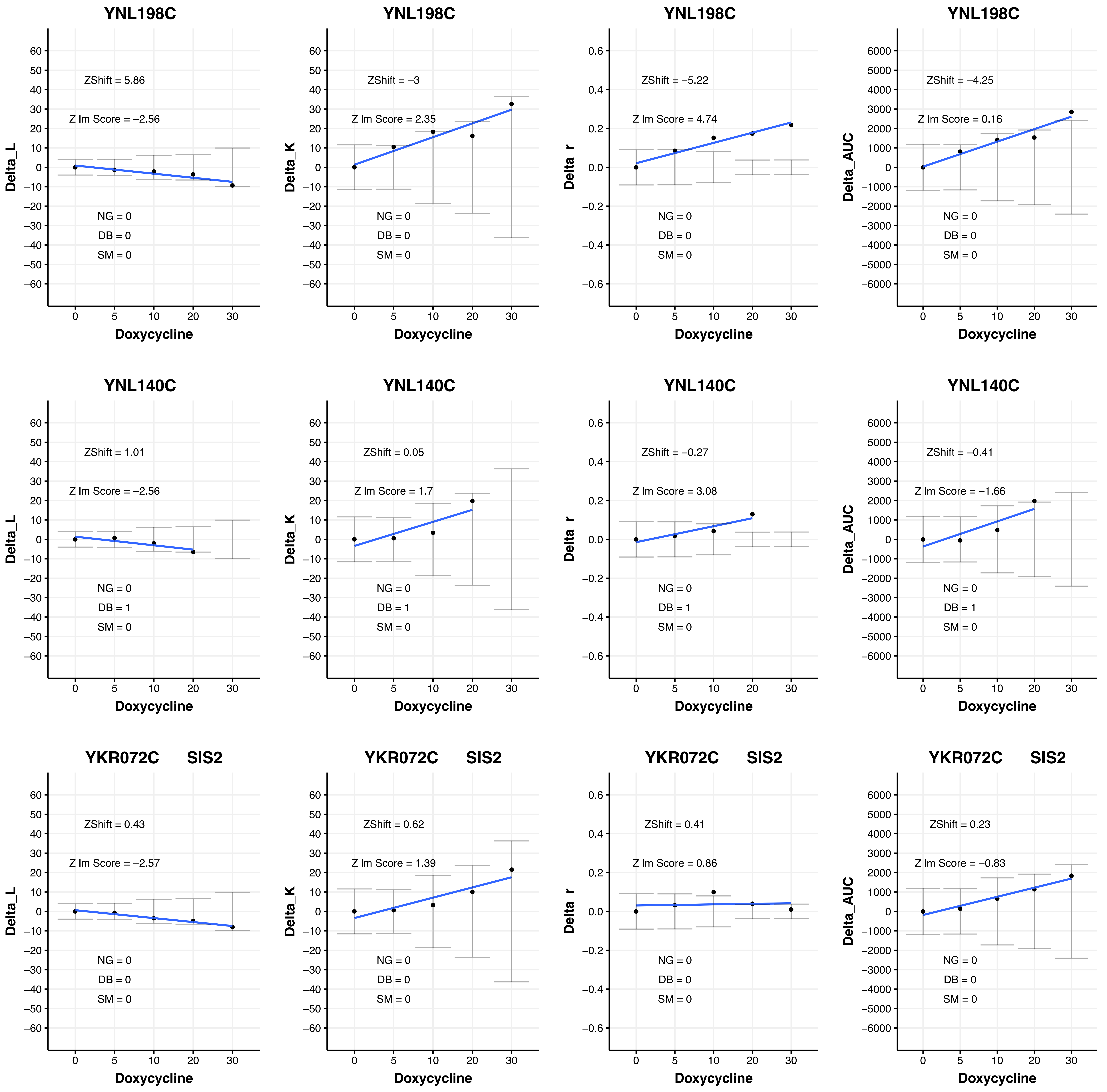
YNL198C
ZShift = 5.86
Z lm Score = −2.56
NG = 0
DB = 0
SM = 0
Delta_L
Doxycycline
YNL198C
ZShift = −3
Z lm Score = 2.35
NG = 0
DB = 0
SM = 0
Delta_K
Doxycycline
YNL198C
ZShift = −5.22
Z lm Score = 4.74
NG = 0
DB = 0
SM = 0
Delta_r
Doxycycline
YNL198C
ZShift = −4.25
Z lm Score = 0.16
NG = 0
DB = 0
SM = 0
Delta_AUC
Doxycycline
YNL140C
ZShift = 1.01
Z lm Score = −2.56
NG = 0
DB = 1
SM = 0
Delta_L
Doxycycline
YNL140C
ZShift = 0.05
Z lm Score = 1.7
NG = 0
DB = 1
SM = 0
Delta_K
Doxycycline
YNL140C
ZShift = −0.27
Z lm Score = 3.08
NG = 0
DB = 1
SM = 0
Delta_r
Doxycycline
YNL140C
ZShift = −0.41
Z lm Score = −1.66
NG = 0
DB = 1
SM = 0
Delta_AUC
Doxycycline
YKR072C SIS2
ZShift = 0.43
Z lm Score = −2.57
NG = 0
DB = 0
SM = 0
Delta_L
Doxycycline
YKR072C SIS2
ZShift = 0.62
Z lm Score = 1.39
NG = 0
DB = 0
SM = 0
Delta_K
Doxycycline
YKR072C SIS2
ZShift = 0.41
Z lm Score = 0.86
NG = 0
DB = 0
SM = 0
Delta_r
Doxycycline
YKR072C SIS2
ZShift = 0.23
Z lm Score = −0.83
NG = 0
DB = 0
SM = 0
Delta_AUC
Doxycycline

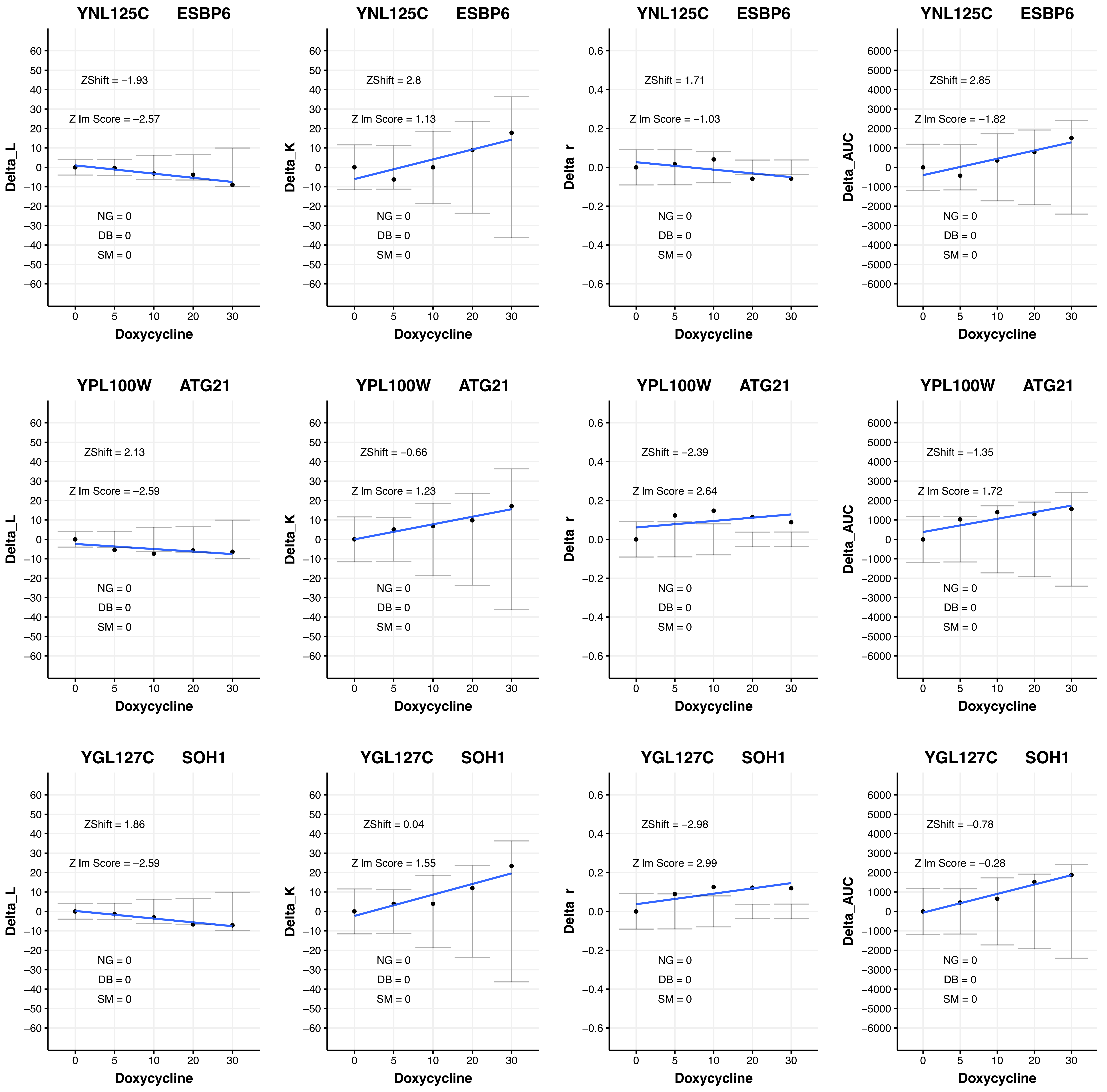
YNL125C ESBP6
ZShift = −1.93
Z lm Score = −2.57
NG = 0
DB = 0
SM = 0
Delta_L
Doxycycline
YNL125C ESBP6
ZShift = 2.8
Z lm Score = 1.13
NG = 0
DB = 0
SM = 0
Delta_K
Doxycycline
YNL125C ESBP6
ZShift = 1.71
Z lm Score = −1.03
NG = 0
DB = 0
SM = 0
Delta_r
Doxycycline
YNL125C ESBP6
ZShift = 2.85
Z lm Score = −1.82
NG = 0
DB = 0
SM = 0
Delta_AUC
Doxycycline
YPL100W ATG21
ZShift = 2.13
Z lm Score = −2.59
NG = 0
DB = 0
SM = 0
Delta_L
Doxycycline
YPL100W ATG21
ZShift = −0.66
Z lm Score = 1.23
NG = 0
DB = 0
SM = 0
Delta_K
Doxycycline
YPL100W ATG21
ZShift = −2.39
Z lm Score = 2.64
NG = 0
DB = 0
SM = 0
Delta_r
Doxycycline
YPL100W ATG21
ZShift = −1.35
Z lm Score = 1.72
NG = 0
DB = 0
SM = 0
Delta_AUC
Doxycycline
YGL127C SOH1
ZShift = 1.86
Z lm Score = −2.59
NG = 0
DB = 0
SM = 0
Delta_L
Doxycycline
YGL127C SOH1
ZShift = 0.04
Z lm Score = 1.55
NG = 0
DB = 0
SM = 0
Delta_K
Doxycycline
YGL127C SOH1
ZShift = −2.98
Z lm Score = 2.99
NG = 0
DB = 0
SM = 0
Delta_r
Doxycycline
YGL127C SOH1
ZShift = −0.78
Z lm Score = −0.28
NG = 0
DB = 0
SM = 0
Delta_AUC
Doxycycline

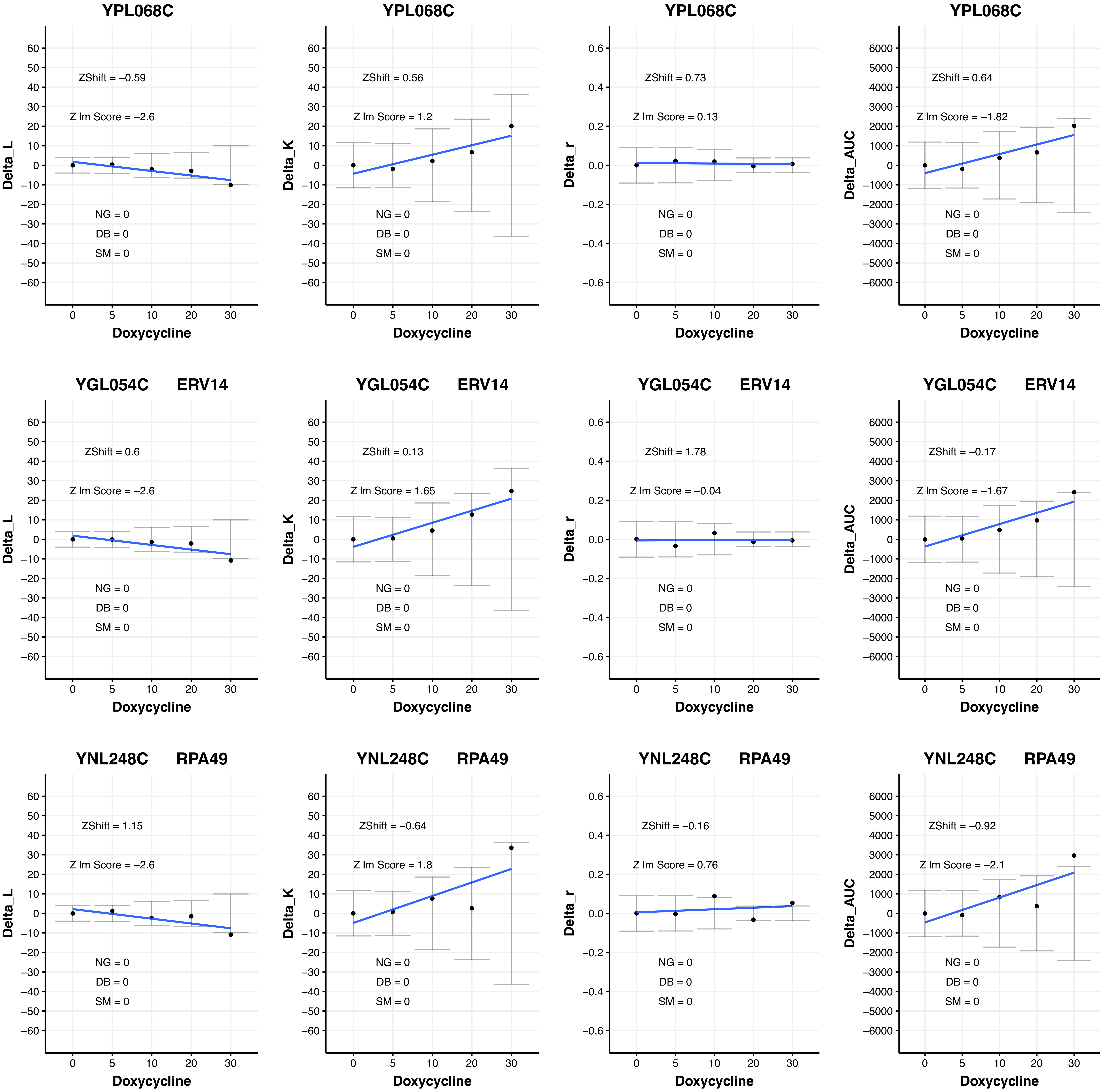
YPL068C
ZShift = −0.59
Z lm Score = −2.6
NG = 0
DB = 0
SM = 0
Delta_L
Doxycycline
YPL068C
ZShift = 0.56
Z lm Score = 1.2
NG = 0
DB = 0
SM = 0
Delta_K
Doxycycline
YPL068C
ZShift = 0.73
Z lm Score = 0.13
NG = 0
DB = 0
SM = 0
Delta_r
Doxycycline
YPL068C
ZShift = 0.64
Z lm Score = −1.82
NG = 0
DB = 0
SM = 0
Delta_AUC
Doxycycline
YGL054C ERV14
ZShift = 0.6
Z lm Score = −2.6
NG = 0
DB = 0
SM = 0
Delta_L
Doxycycline
YGL054C ERV14
ZShift = 0.13
Z lm Score = 1.65
NG = 0
DB = 0
SM = 0
Delta_K
Doxycycline
YGL054C ERV14
ZShift = 1.78
Z lm Score = −0.04
NG = 0
DB = 0
SM = 0
Delta_r
Doxycycline
YGL054C ERV14
ZShift = −0.17
Z lm Score = −1.67
NG = 0
DB = 0
SM = 0
Delta_AUC
Doxycycline
YNL248C RPA49
ZShift = 1.15
Z lm Score = −2.6
NG = 0
DB = 0
SM = 0
Delta_L
Doxycycline
YNL248C RPA49
ZShift = −0.64
Z lm Score = 1.8
NG = 0
DB = 0
SM = 0
Delta_K
Doxycycline
YNL248C RPA49
ZShift = −0.16
Z lm Score = 0.76
NG = 0
DB = 0
SM = 0
Delta_r
Doxycycline
YNL248C RPA49
ZShift = −0.92
Z lm Score = −2.1
NG = 0
DB = 0
SM = 0
Delta_AUC
Doxycycline

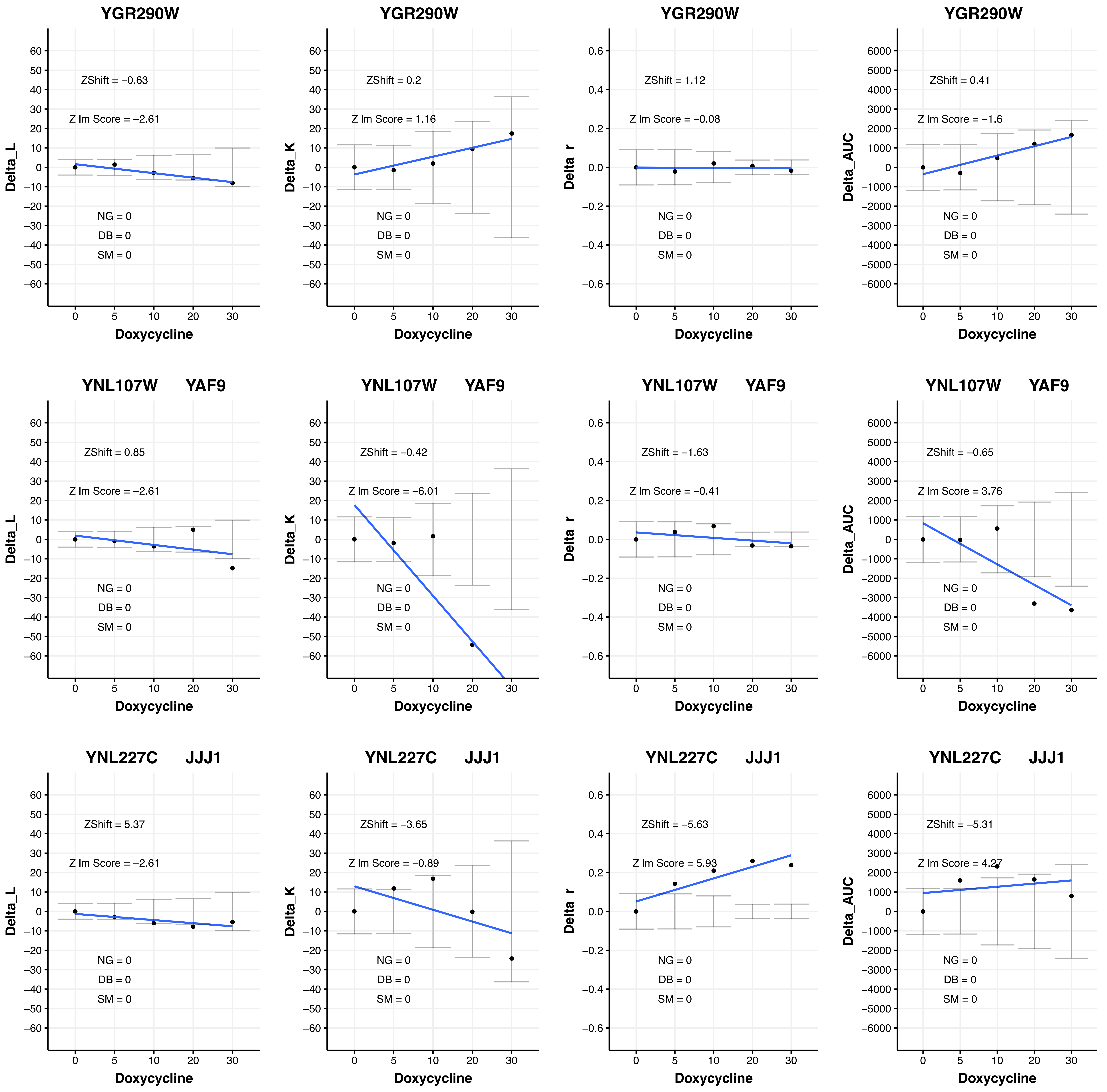

YGR290W
ZShift = −0.63
Z lm Score = −2.61
NG = 0
DB = 0
SM = 0
Delta_L
Doxycycline
YGR290W
ZShift = 0.2
Z lm Score = 1.16
NG = 0
DB = 0
SM = 0
Delta_K
Doxycycline
YGR290W
ZShift = 1.12
Z lm Score = −0.08
NG = 0
DB = 0
SM = 0
Delta_r
Doxycycline
YGR290W
ZShift = 0.41
Z lm Score = −1.6
NG = 0
DB = 0
SM = 0
Delta_AUC
Doxycycline
YNL107W YAF9
ZShift = 0.85
Z lm Score = −2.61
NG = 0
DB = 0
SM = 0
Delta_L
Doxycycline
YNL107W YAF9
ZShift = −0.42
Z lm Score = −6.01
NG = 0
DB = 0
SM = 0
Delta_K
Doxycycline
YNL107W YAF9
ZShift = −1.63
Z lm Score = −0.41
NG = 0
DB = 0
SM = 0
Delta_r
Doxycycline
YNL107W YAF9
ZShift = −0.65
Z lm Score = 3.76
NG = 0
DB = 0
SM = 0
Delta_AUC
Doxycycline
YNL227C JJJ1
ZShift = 5.37
Z lm Score = −2.61
NG = 0
DB = 0
SM = 0
Delta_L
Doxycycline
YNL227C JJJ1
ZShift = −3.65
Z lm Score = −0.89
NG = 0
DB = 0
SM = 0
Delta_K
Doxycycline
YNL227C JJJ1
ZShift = −5.63
Z lm Score = 5.93
NG = 0
DB = 0
SM = 0
Delta_r
Doxycycline
YNL227C JJJ1
ZShift = −5.31
Z lm Score = 4.27
NG = 0
DB = 0
SM = 0
Delta_AUC
Doxycycline

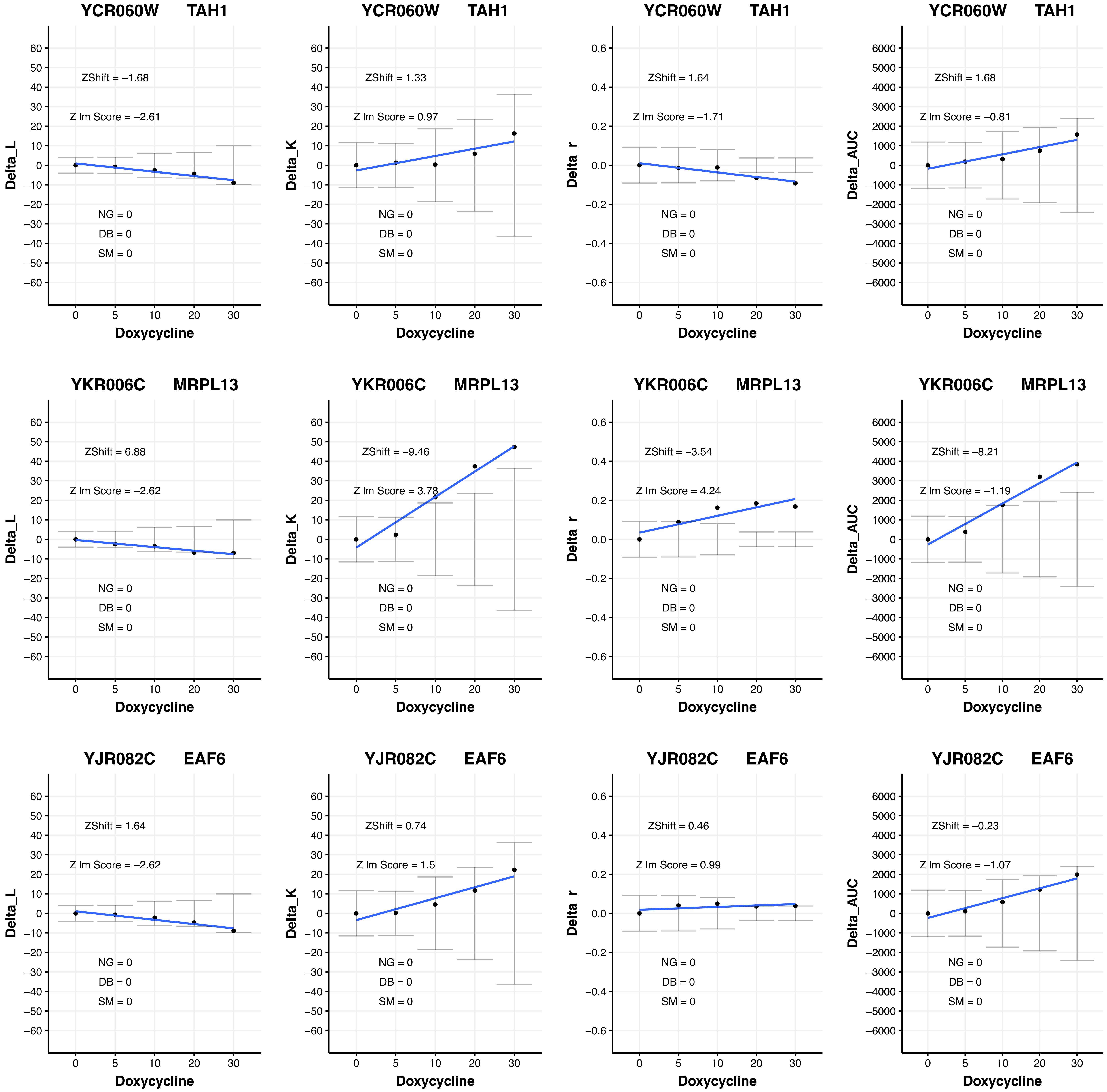
YCR060W TAH1
ZShift = −1.68
Z lm Score = −2.61
NG = 0
DB = 0
SM = 0
Delta_L
Doxycycline
YCR060W TAH1
ZShift = 1.33
Z lm Score = 0.97
NG = 0
DB = 0
SM = 0
Delta_K
Doxycycline
YCR060W TAH1
ZShift = 1.64
Z lm Score = −1.71
NG = 0
DB = 0
SM = 0
Delta_r
Doxycycline
YCR060W TAH1
ZShift = 1.68
Z lm Score = −0.81
NG = 0
DB = 0
SM = 0
Delta_AUC
Doxycycline
YKR006C MRPL13
ZShift = 6.88
Z lm Score = −2.62
NG = 0
DB = 0
SM = 0
Delta_L
Doxycycline
YKR006C MRPL13
ZShift = −9.46
Z lm Score = 3.78
NG = 0
DB = 0
SM = 0
Delta_K
Doxycycline
YKR006C MRPL13
ZShift = −3.54
Z lm Score = 4.24
NG = 0
DB = 0
SM = 0
Delta_r
Doxycycline
YKR006C MRPL13
ZShift = −8.21
Z lm Score = −1.19
NG = 0
DB = 0
SM = 0
Delta_AUC
Doxycycline
YJR082C EAF6
ZShift = 1.64
Z lm Score = −2.62
NG = 0
DB = 0
SM = 0
Delta_L
Doxycycline
YJR082C EAF6
ZShift = 0.74
Z lm Score = 1.5
NG = 0
DB = 0
SM = 0
Delta_K
Doxycycline
YJR082C EAF6
ZShift = 0.46
Z lm Score = 0.99
NG = 0
DB = 0
SM = 0
Delta_r
Doxycycline
YJR082C EAF6
ZShift = −0.23
Z lm Score = −1.07
NG = 0
DB = 0
SM = 0
Delta_AUC
Doxycycline

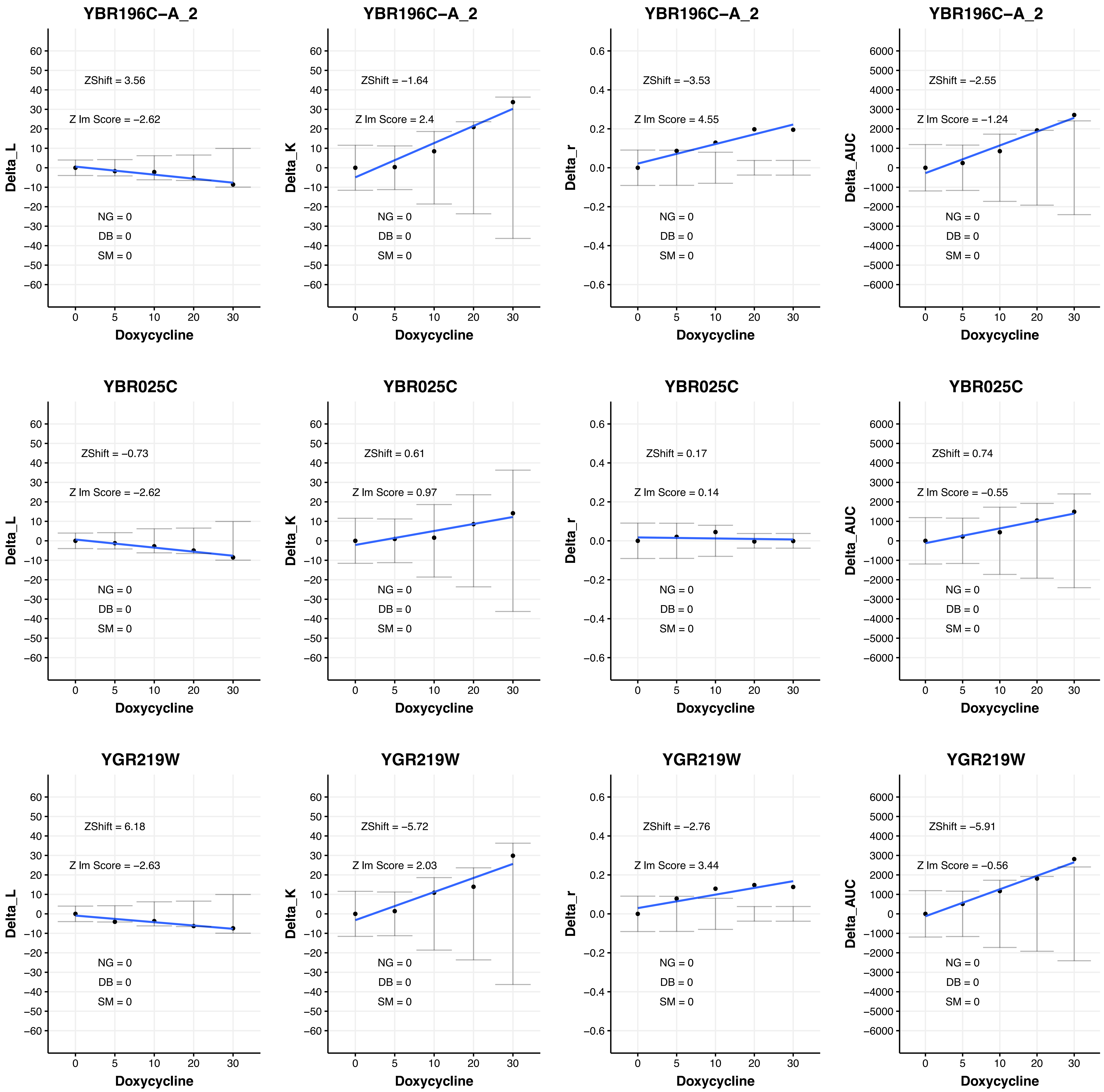

YBR196C−A_2
ZShift = 3.56
Z lm Score = −2.62
NG = 0
DB = 0
SM = 0
Delta_L
Doxycycline
YBR196C−A_2
ZShift = −1.64
Z lm Score = 2.4
NG = 0
DB = 0
SM = 0
Delta_K
Doxycycline
YBR196C−A_2
ZShift = −3.53
Z lm Score = 4.55
NG = 0
DB = 0
SM = 0
Delta_r
Doxycycline
YBR196C−A_2
ZShift = −2.55
Z lm Score = −1.24
NG = 0
DB = 0
SM = 0
Delta_AUC
Doxycycline
YBR025C
ZShift = −0.73
Z lm Score = −2.62
NG = 0
DB = 0
SM = 0
Delta_L
Doxycycline
YBR025C
ZShift = 0.61
Z lm Score = 0.97
NG = 0
DB = 0
SM = 0
Delta_K
Doxycycline
YBR025C
ZShift = 0.17
Z lm Score = 0.14
NG = 0
DB = 0
SM = 0
Delta_r
Doxycycline
YBR025C
ZShift = 0.74
Z lm Score = −0.55
NG = 0
DB = 0
SM = 0
Delta_AUC
Doxycycline
YGR219W
ZShift = 6.18
Z lm Score = −2.63
NG = 0
DB = 0
SM = 0
Delta_L
Doxycycline
YGR219W
ZShift = −5.72
Z lm Score = 2.03
NG = 0
DB = 0
SM = 0
Delta_K
Doxycycline
YGR219W
ZShift = −2.76
Z lm Score = 3.44
NG = 0
DB = 0
SM = 0
Delta_r
Doxycycline
YGR219W
ZShift = −5.91
Z lm Score = −0.56
NG = 0
DB = 0
SM = 0
Delta_AUC
Doxycycline

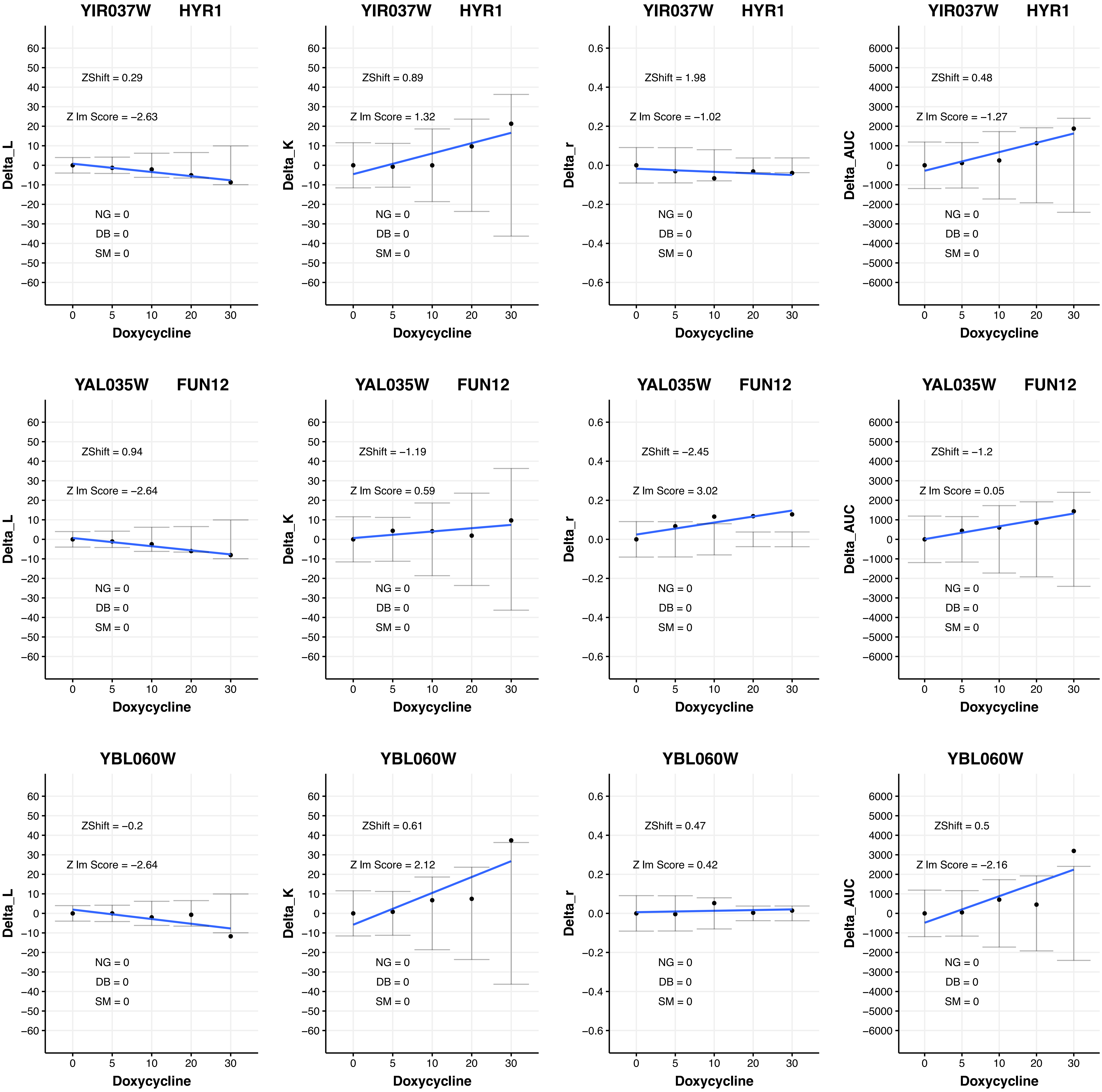
YIR037W HYR1
ZShift = 0.29
Z lm Score = −2.63
NG = 0
DB = 0
SM = 0
Delta_L
Doxycycline
YIR037W HYR1
ZShift = 0.89
Z lm Score = 1.32
NG = 0
DB = 0
SM = 0
Delta_K
Doxycycline
YIR037W HYR1
ZShift = 1.98
Z lm Score = −1.02
NG = 0
DB = 0
SM = 0
Delta_r
Doxycycline
YIR037W HYR1
ZShift = 0.48
Z lm Score = −1.27
NG = 0
DB = 0
SM = 0
Delta_AUC
Doxycycline
YAL035W FUN12
ZShift = 0.94
Z lm Score = −2.64
NG = 0
DB = 0
SM = 0
Delta_L
Doxycycline
YAL035W FUN12
ZShift = −1.19
Z lm Score = 0.59
NG = 0
DB = 0
SM = 0
Delta_K
Doxycycline
YAL035W FUN12
ZShift = −2.45
Z lm Score = 3.02
NG = 0
DB = 0
SM = 0
Delta_r
Doxycycline
YAL035W FUN12
ZShift = −1.2
Z lm Score = 0.05
NG = 0
DB = 0
SM = 0
Delta_AUC
Doxycycline
YBL060W
ZShift = −0.2
Z lm Score = −2.64
NG = 0
DB = 0
SM = 0
Delta_L
Doxycycline
YBL060W
ZShift = 0.61
Z lm Score = 2.12
NG = 0
DB = 0
SM = 0
Delta_K
Doxycycline
YBL060W
ZShift = 0.47
Z lm Score = 0.42
NG = 0
DB = 0
SM = 0
Delta_r
Doxycycline
YBL060W
ZShift = 0.5
Z lm Score = −2.16
NG = 0
DB = 0
SM = 0
Delta_AUC
Doxycycline

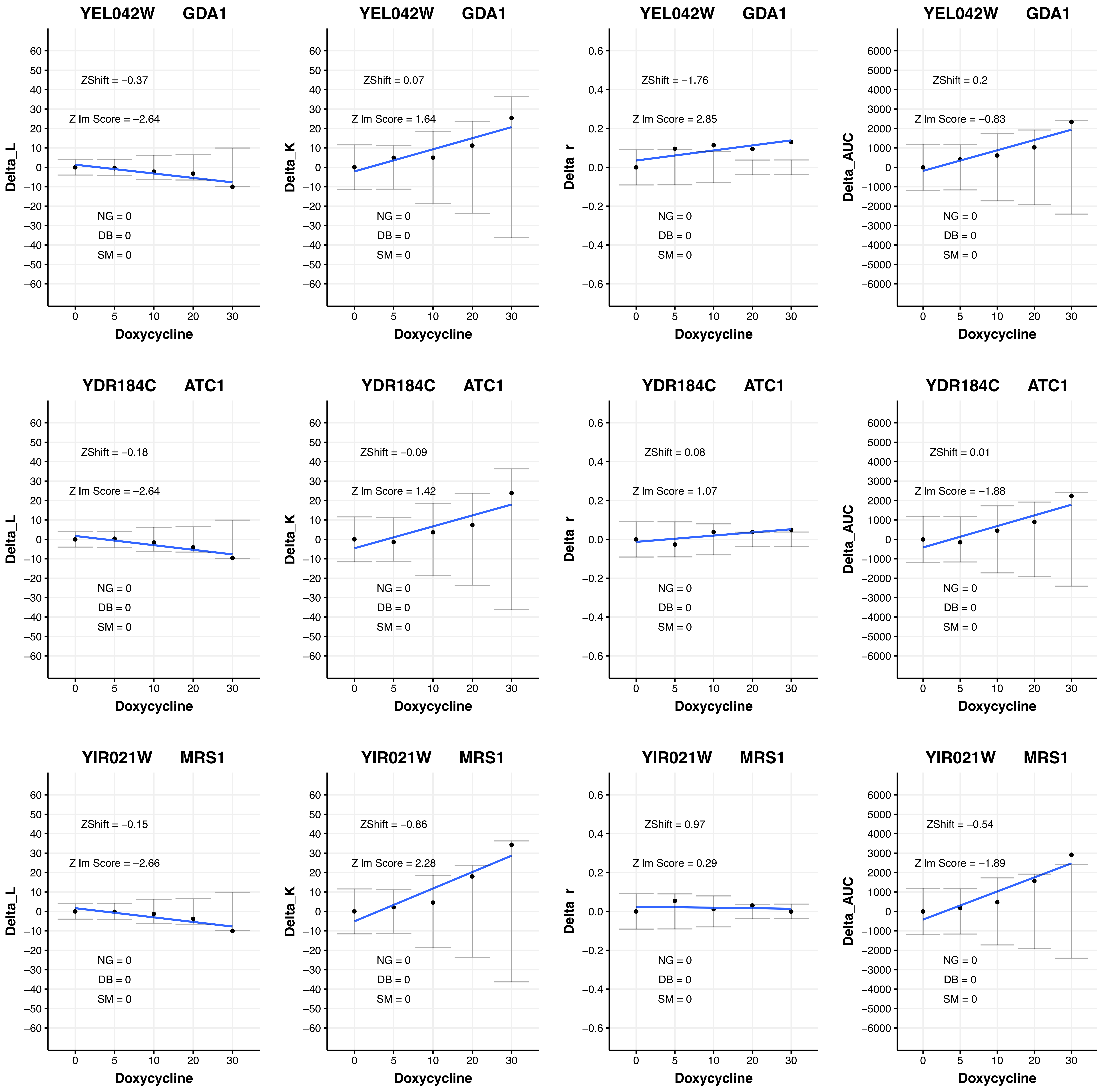
YEL042W GDA1
ZShift = −0.37
Z lm Score = −2.64
NG = 0
DB = 0
SM = 0
Delta_L
Doxycycline
YEL042W GDA1
ZShift = 0.07
Z lm Score = 1.64
NG = 0
DB = 0
SM = 0
Delta_K
Doxycycline
YEL042W GDA1
ZShift = −1.76
Z lm Score = 2.85
NG = 0
DB = 0
SM = 0
Delta_r
Doxycycline
YEL042W GDA1
ZShift = 0.2
Z lm Score = −0.83
NG = 0
DB = 0
SM = 0
Delta_AUC
Doxycycline
YDR184C ATC1
ZShift = −0.18
Z lm Score = −2.64
NG = 0
DB = 0
SM = 0
Delta_L
Doxycycline
YDR184C ATC1
ZShift = −0.09
Z lm Score = 1.42
NG = 0
DB = 0
SM = 0
Delta_K
Doxycycline
YDR184C ATC1
ZShift = 0.08
Z lm Score = 1.07
NG = 0
DB = 0
SM = 0
Delta_r
Doxycycline
YDR184C ATC1
ZShift = 0.01
Z lm Score = −1.88
NG = 0
DB = 0
SM = 0
Delta_AUC
Doxycycline
YIR021W MRS1
ZShift = −0.15
Z lm Score = −2.66
NG = 0
DB = 0
SM = 0
Delta_L
Doxycycline
YIR021W MRS1
ZShift = −0.86
Z lm Score = 2.28
NG = 0
DB = 0
SM = 0
Delta_K
Doxycycline
YIR021W MRS1
ZShift = 0.97
Z lm Score = 0.29
NG = 0
DB = 0
SM = 0
Delta_r
Doxycycline
YIR021W MRS1
ZShift = −0.54
Z lm Score = −1.89
NG = 0
DB = 0
SM = 0
Delta_AUC
Doxycycline

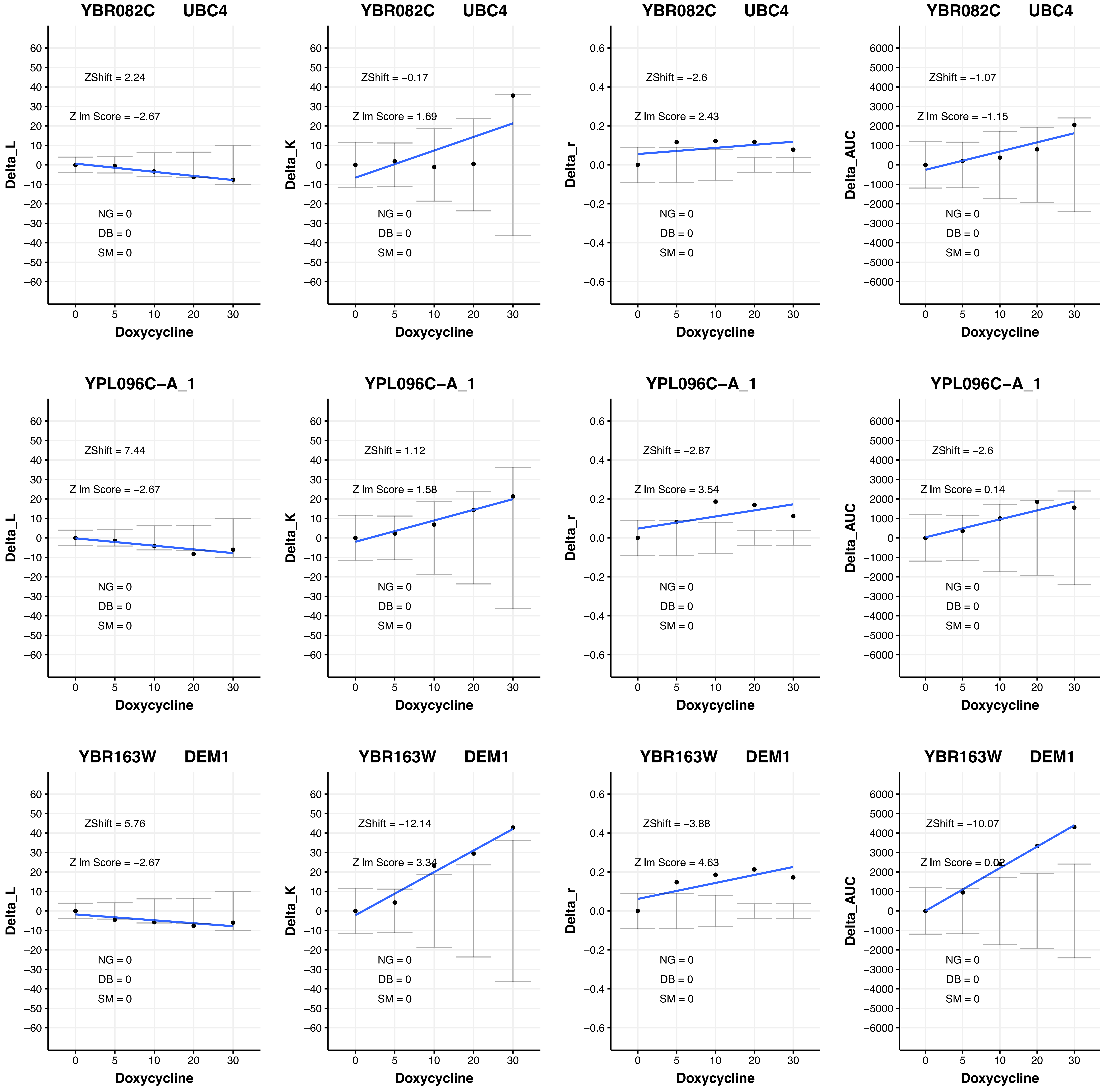
YBR082C UBC4
ZShift = 2.24
Z lm Score = −2.67
NG = 0
DB = 0
SM = 0
Delta_L
Doxycycline
YBR082C UBC4
ZShift = −0.17
Z lm Score = 1.69
NG = 0
DB = 0
SM = 0
Delta_K
Doxycycline
YBR082C UBC4
ZShift = −2.6
Z lm Score = 2.43
NG = 0
DB = 0
SM = 0
Delta_r
Doxycycline
YBR082C UBC4
ZShift = −1.07
Z lm Score = −1.15
NG = 0
DB = 0
SM = 0
Delta_AUC
Doxycycline
YPL096C−A_1
ZShift = 7.44
Z lm Score = −2.67
NG = 0
DB = 0
SM = 0
Delta_L
Doxycycline
YPL096C−A_1
ZShift = 1.12
Z lm Score = 1.58
NG = 0
DB = 0
SM = 0
Delta_K
Doxycycline
YPL096C−A_1
ZShift = −2.87
Z lm Score = 3.54
NG = 0
DB = 0
SM = 0
Delta_r
Doxycycline
YPL096C−A_1
ZShift = −2.6
Z lm Score = 0.14
NG = 0
DB = 0
SM = 0
Delta_AUC
Doxycycline
YBR163W DEM1
ZShift = 5.76
Z lm Score = −2.67
NG = 0
DB = 0
SM = 0
Delta_L
Doxycycline
YBR163W DEM1
ZShift = −12.14
Z lm Score = 3.34
NG = 0
DB = 0
SM = 0
Delta_K
Doxycycline
YBR163W DEM1
ZShift = −3.88
Z lm Score = 4.63
NG = 0
DB = 0
SM = 0
Delta_r
Doxycycline
YBR163W DEM1
ZShift = −10.07
Z lm Score = 0.02
NG = 0
DB = 0
SM = 0
Delta_AUC
Doxycycline

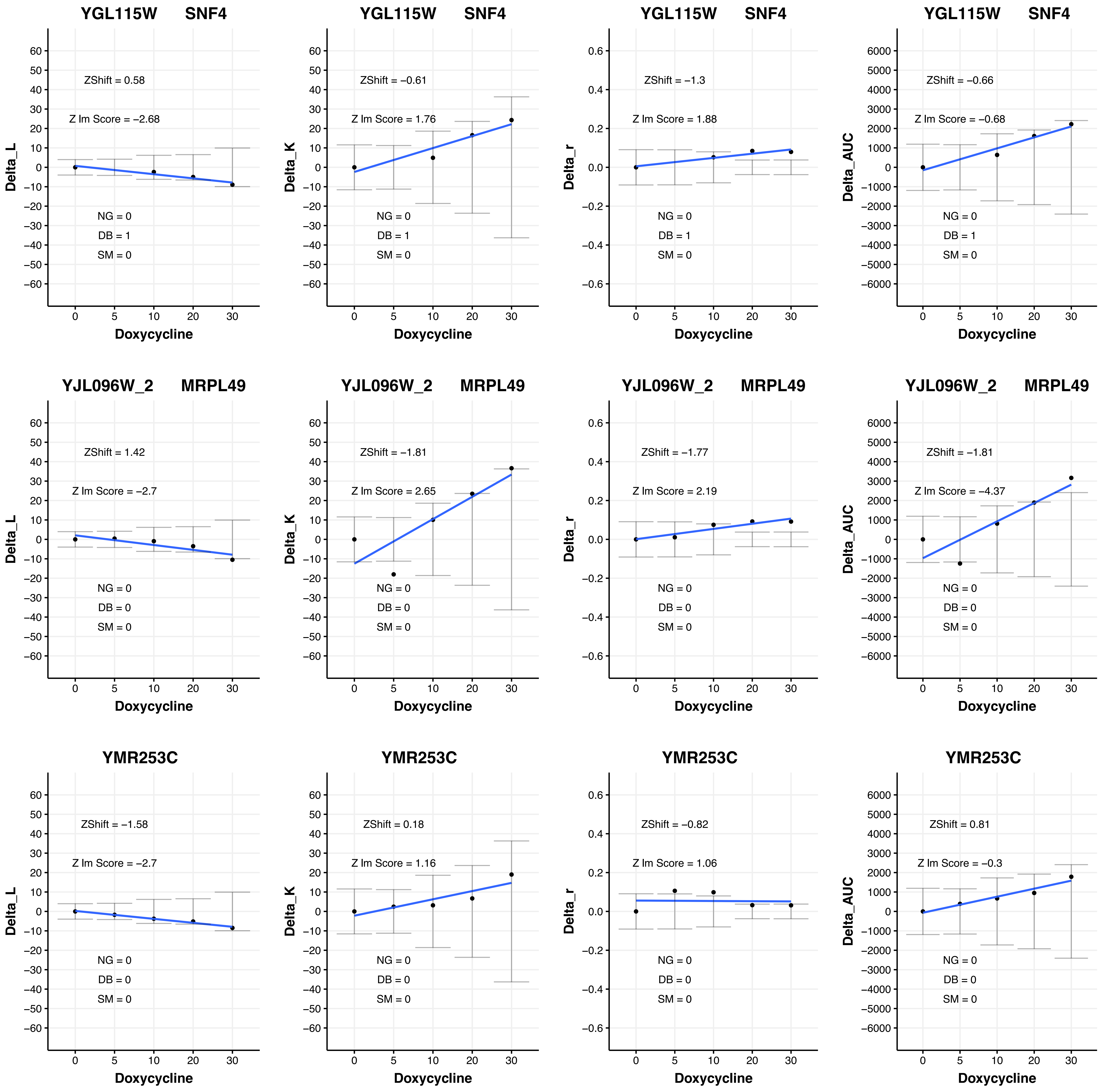
YGL115W SNF4
ZShift = 0.58
Z lm Score = −2.68
NG = 0
DB = 1
SM = 0
Delta_L
Doxycycline
YGL115W SNF4
ZShift = −0.61
Z lm Score = 1.76
NG = 0
DB = 1
SM = 0
Delta_K
Doxycycline
YGL115W SNF4
ZShift = −1.3
Z lm Score = 1.88
NG = 0
DB = 1
SM = 0
Delta_r
Doxycycline
YGL115W SNF4
ZShift = −0.66
Z lm Score = −0.68
NG = 0
DB = 1
SM = 0
Delta_AUC
Doxycycline
YJL096W_2 MRPL49
ZShift = 1.42
Z lm Score = −2.7
NG = 0
DB = 0
SM = 0
Delta_L
Doxycycline
YJL096W_2 MRPL49
ZShift = −1.81
Z lm Score = 2.65
NG = 0
DB = 0
SM = 0
Delta_K
Doxycycline
YJL096W_2 MRPL49
ZShift = −1.77
Z lm Score = 2.19
NG = 0
DB = 0
SM = 0
Delta_r
Doxycycline
YJL096W_2 MRPL49
ZShift = −1.81
Z lm Score = −4.37
NG = 0
DB = 0
SM = 0
Delta_AUC
Doxycycline
YMR253C
ZShift = −1.58
Z lm Score = −2.7
NG = 0
DB = 0
SM = 0
Delta_L
Doxycycline
YMR253C
ZShift = 0.18
Z lm Score = 1.16
NG = 0
DB = 0
SM = 0
Delta_K
Doxycycline
YMR253C
ZShift = −0.82
Z lm Score = 1.06
NG = 0
DB = 0
SM = 0
Delta_r
Doxycycline
YMR253C
ZShift = 0.81
Z lm Score = −0.3
NG = 0
DB = 0
SM = 0
Delta_AUC
Doxycycline

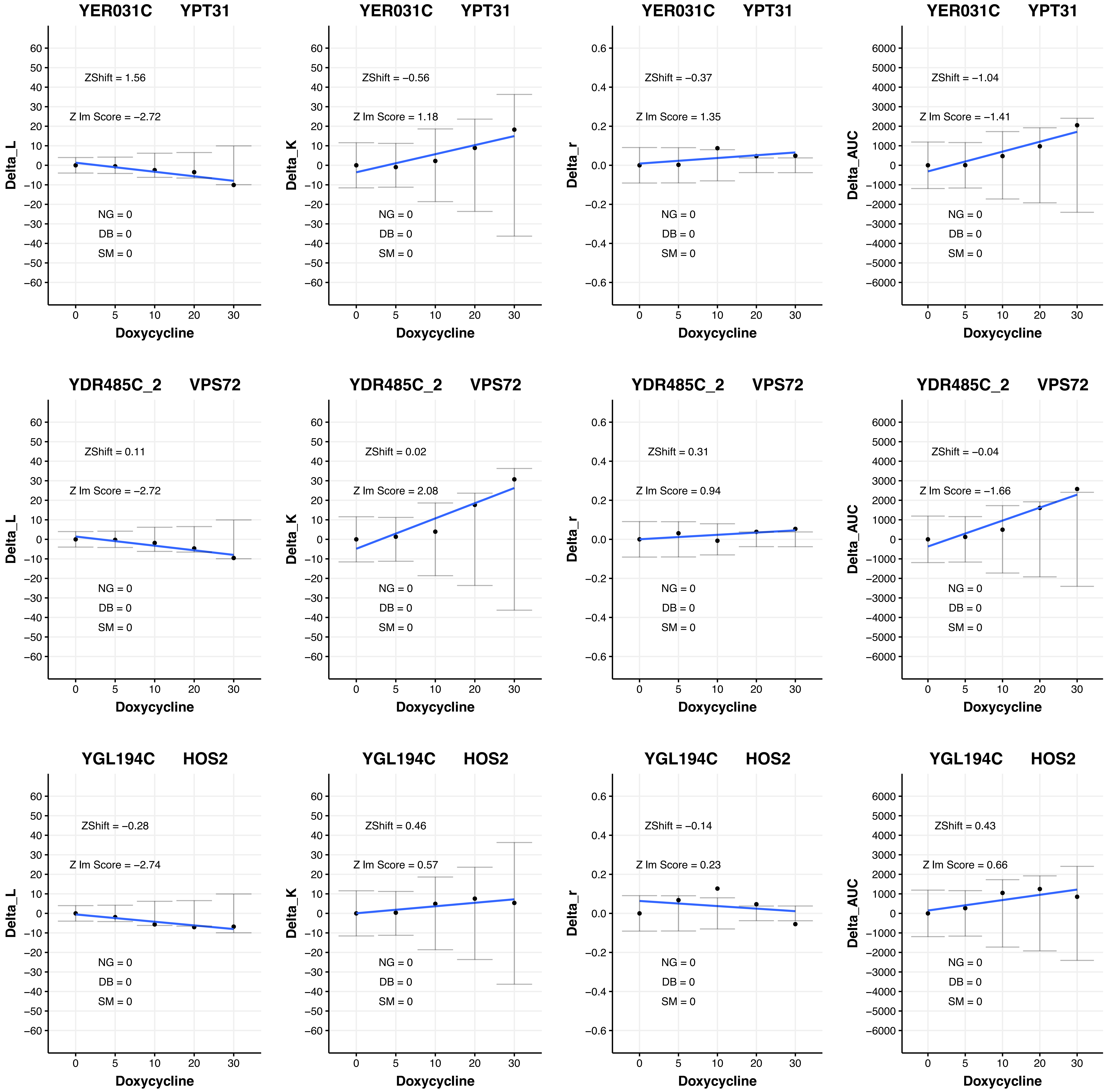
YER031C YPT31
ZShift = 1.56
Z lm Score = −2.72
NG = 0
DB = 0
SM = 0
Delta_L
Doxycycline
YER031C YPT31
ZShift = −0.56
Z lm Score = 1.18
NG = 0
DB = 0
SM = 0
Delta_K
Doxycycline
YER031C YPT31
ZShift = −0.37
Z lm Score = 1.35
NG = 0
DB = 0
SM = 0
Delta_r
Doxycycline
YER031C YPT31
ZShift = −1.04
Z lm Score = −1.41
NG = 0
DB = 0
SM = 0
Delta_AUC
Doxycycline
YDR485C_2 VPS72
ZShift = 0.11
Z lm Score = −2.72
NG = 0
DB = 0
SM = 0
Delta_L
Doxycycline
YDR485C_2 VPS72
ZShift = 0.02
Z lm Score = 2.08
NG = 0
DB = 0
SM = 0
Delta_K
Doxycycline
YDR485C_2 VPS72
ZShift = 0.31
Z lm Score = 0.94
NG = 0
DB = 0
SM = 0
Delta_r
Doxycycline
YDR485C_2 VPS72
ZShift = −0.04
Z lm Score = −1.66
NG = 0
DB = 0
SM = 0
Delta_AUC
Doxycycline
YGL194C HOS2
ZShift = −0.28
Z lm Score = −2.74
NG = 0
DB = 0
SM = 0
Delta_L
Doxycycline
YGL194C HOS2
ZShift = 0.46
Z lm Score = 0.57
NG = 0
DB = 0
SM = 0
Delta_K
Doxycycline
YGL194C HOS2
ZShift = −0.14
Z lm Score = 0.23
NG = 0
DB = 0
SM = 0
Delta_r
Doxycycline
YGL194C HOS2
ZShift = 0.43
Z lm Score = 0.66
NG = 0
DB = 0
SM = 0
Delta_AUC
Doxycycline

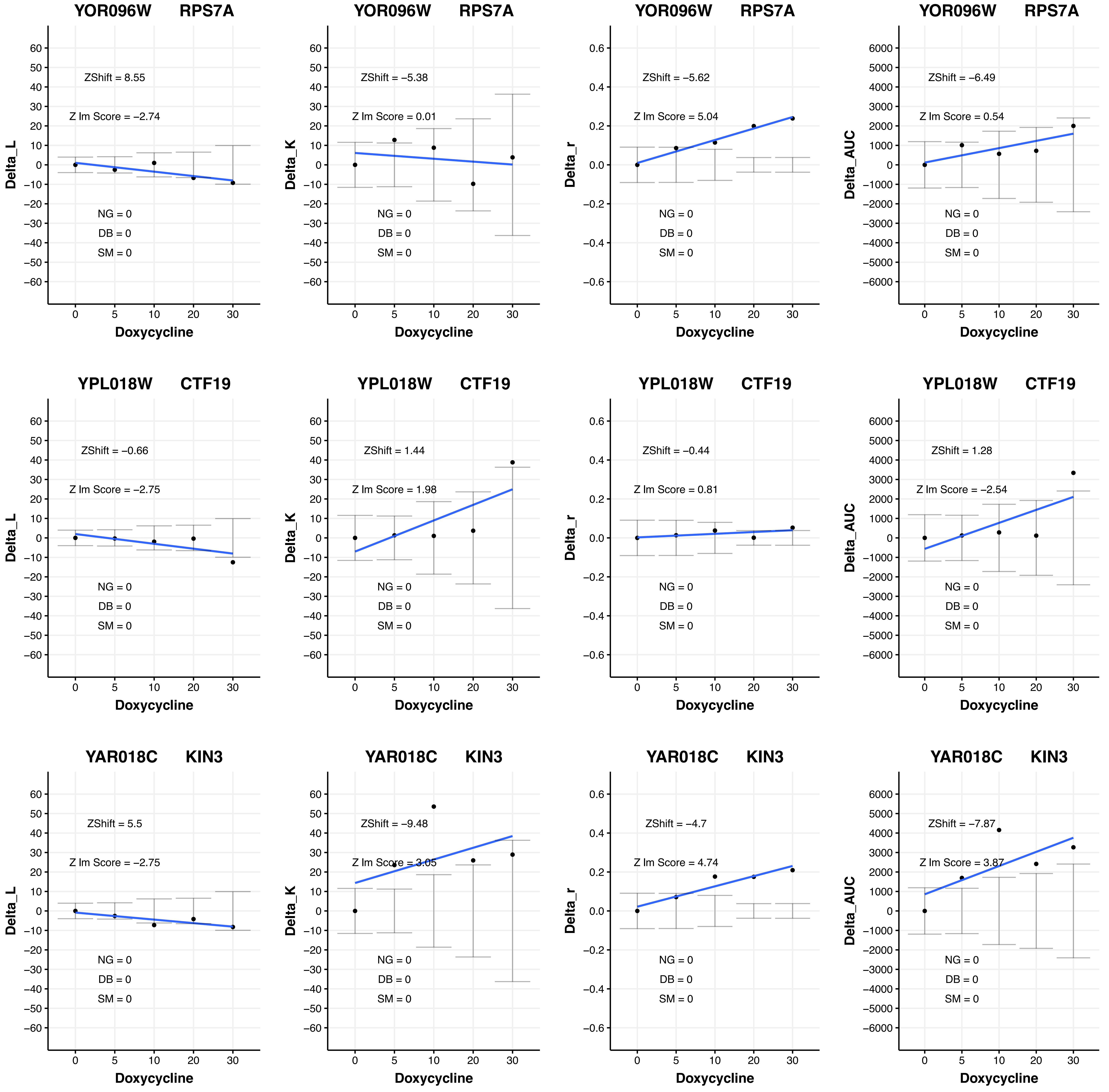
YOR096W RPS7A
ZShift = 8.55
Z lm Score = −2.74
NG = 0
DB = 0
SM = 0
Delta_L
Doxycycline
YOR096W RPS7A
ZShift = −5.38
Z lm Score = 0.01
NG = 0
DB = 0
SM = 0
Delta_K
Doxycycline
YOR096W RPS7A
ZShift = −5.62
Z lm Score = 5.04
NG = 0
DB = 0
SM = 0
Delta_r
Doxycycline
YOR096W RPS7A
ZShift = −6.49
Z lm Score = 0.54
NG = 0
DB = 0
SM = 0
Delta_AUC
Doxycycline
YPL018W CTF19
ZShift = −0.66
Z lm Score = −2.75
NG = 0
DB = 0
SM = 0
Delta_L
Doxycycline
YPL018W CTF19
ZShift = 1.44
Z lm Score = 1.98
NG = 0
DB = 0
SM = 0
Delta_K
Doxycycline
YPL018W CTF19
ZShift = −0.44
Z lm Score = 0.81
NG = 0
DB = 0
SM = 0
Delta_r
Doxycycline
YPL018W CTF19
ZShift = 1.28
Z lm Score = −2.54
NG = 0
DB = 0
SM = 0
Delta_AUC
Doxycycline
YAR018C KIN3
ZShift = 5.5
Z lm Score = −2.75
NG = 0
DB = 0
SM = 0
Delta_L
Doxycycline
YAR018C KIN3
ZShift = −9.48
Z lm Score = 3.05
NG = 0
DB = 0
SM = 0
Delta_K
Doxycycline
YAR018C KIN3
ZShift = −4.7
Z lm Score = 4.74
NG = 0
DB = 0
SM = 0
Delta_r
Doxycycline
YAR018C KIN3
ZShift = −7.87
Z lm Score = 3.87
NG = 0
DB = 0
SM = 0
Delta_AUC
Doxycycline

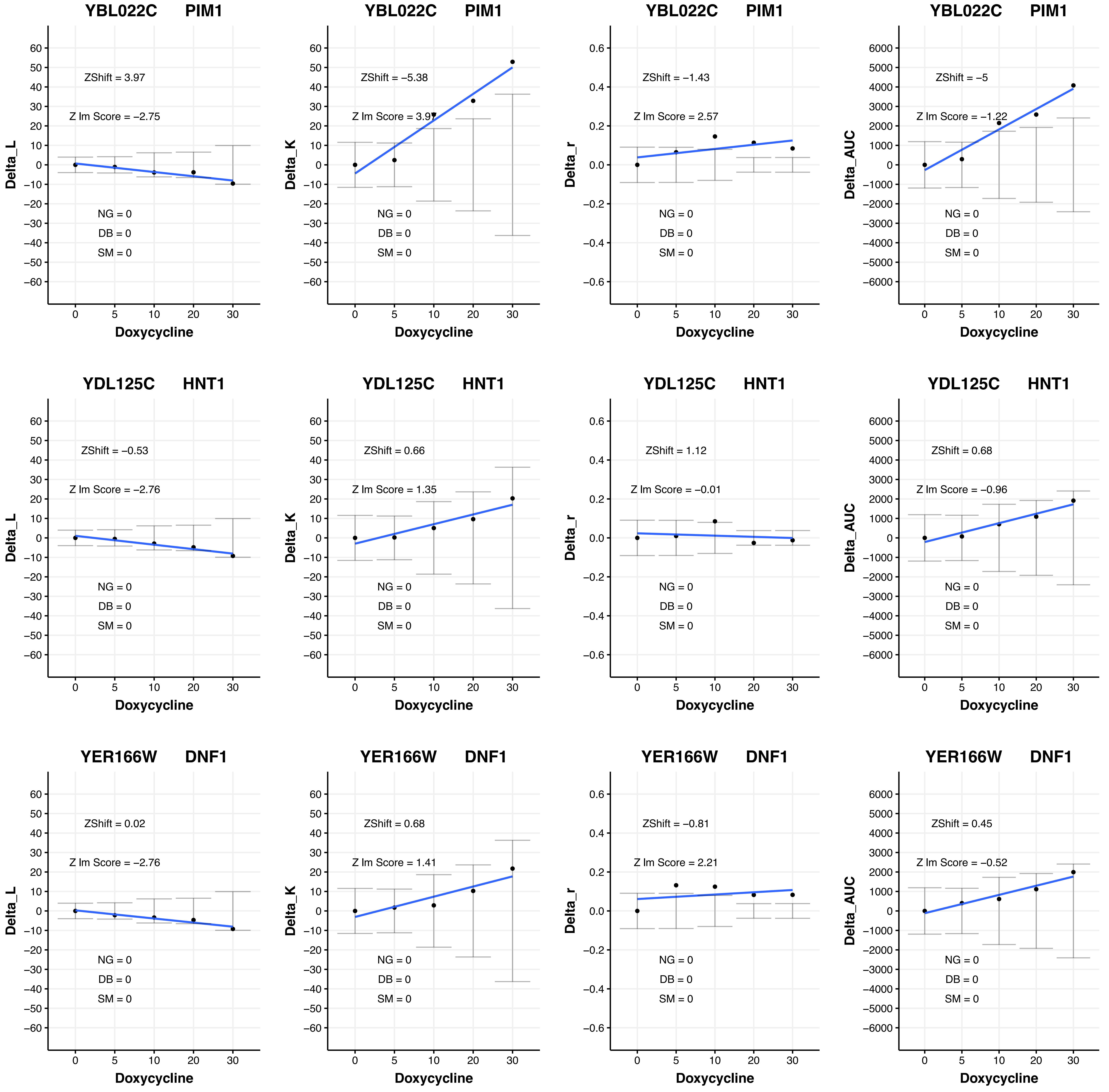
YBL022C PIM1
ZShift = 3.97
Z lm Score = −2.75
NG = 0
DB = 0
SM = 0
Delta_L
Doxycycline
YBL022C PIM1
ZShift = −5.38
Z lm Score = 3.97
Delta_K
YBL022C PIM1
ZShift = −1.43
Z lm Score = 2.57
Delta_r
YBL022C PIM1
ZShift = −5
Z lm Score = −1.22
Delta_AUC
YDL125C HNT1
ZShift = −0.53
Z lm Score = −2.76
YDL125C HNT1
ZShift = 0.66
Z lm Score = 1.35
YDL125C HNT1
ZShift = 1.12
Z lm Score = −0.01
YDL125C HNT1
ZShift = 0.68
Z lm Score = −0.96
YER166W DNF1
ZShift = 0.02
Z lm Score = −2.76
YER166W DNF1
ZShift = 0.68
Z lm Score = 1.41
YER166W DNF1
ZShift = −0.81
Z lm Score = 2.21
YER166W DNF1
ZShift = 0.45
Z lm Score = −0.52

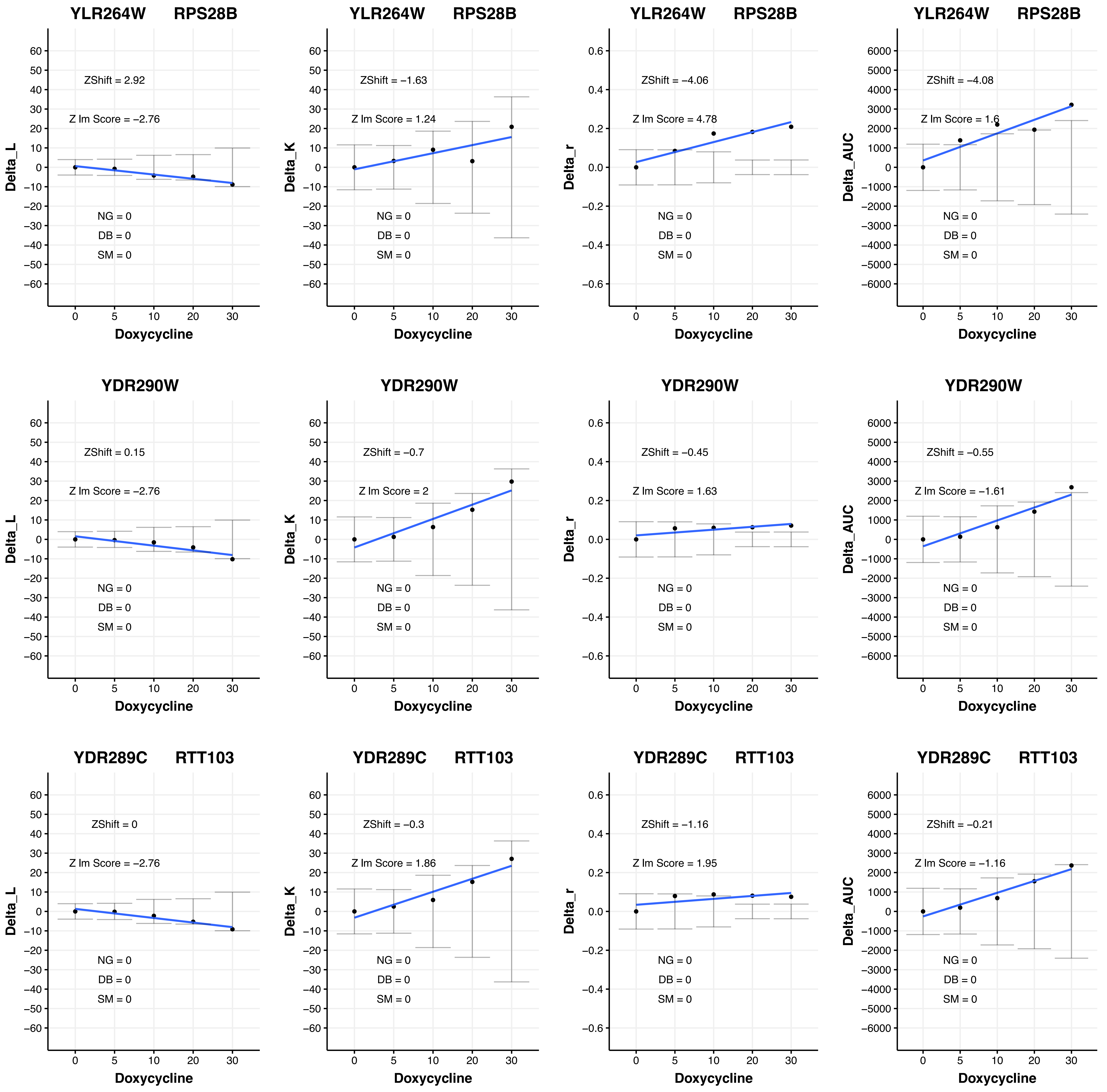
YLR264W RPS28B
ZShift = 2.92
Z lm Score = −2.76
NG = 0
DB = 0
SM = 0
Delta_L
Doxycycline
YLR264W RPS28B
ZShift = −1.63
Z lm Score = 1.24
NG = 0
DB = 0
SM = 0
Delta_K
Doxycycline
YLR264W RPS28B
ZShift = −4.06
Z lm Score = 4.78
NG = 0
DB = 0
SM = 0
Delta_r
Doxycycline
YLR264W RPS28B
ZShift = −4.08
Z lm Score = 1.6
NG = 0
DB = 0
SM = 0
Delta_AUC
Doxycycline
YDR290W
ZShift = 0.15
Z lm Score = −2.76
NG = 0
DB = 0
SM = 0
Delta_L
Doxycycline
YDR290W
ZShift = −0.7
Z lm Score = 2
NG = 0
DB = 0
SM = 0
Delta_K
Doxycycline
YDR290W
ZShift = −0.45
Z lm Score = 1.63
NG = 0
DB = 0
SM = 0
Delta_r
Doxycycline
YDR290W
ZShift = −0.55
Z lm Score = −1.61
NG = 0
DB = 0
SM = 0
Delta_AUC
Doxycycline
YDR289C RTT103
ZShift = 0
Z lm Score = −2.76
NG = 0
DB = 0
SM = 0
Delta_L
Doxycycline
YDR289C RTT103
ZShift = −0.3
Z lm Score = 1.86
NG = 0
DB = 0
SM = 0
Delta_K
Doxycycline
YDR289C RTT103
ZShift = −1.16
Z lm Score = 1.95
NG = 0
DB = 0
SM = 0
Delta_r
Doxycycline
YDR289C RTT103
ZShift = −0.21
Z lm Score = −1.16
NG = 0
DB = 0
SM = 0
Delta_AUC
Doxycycline

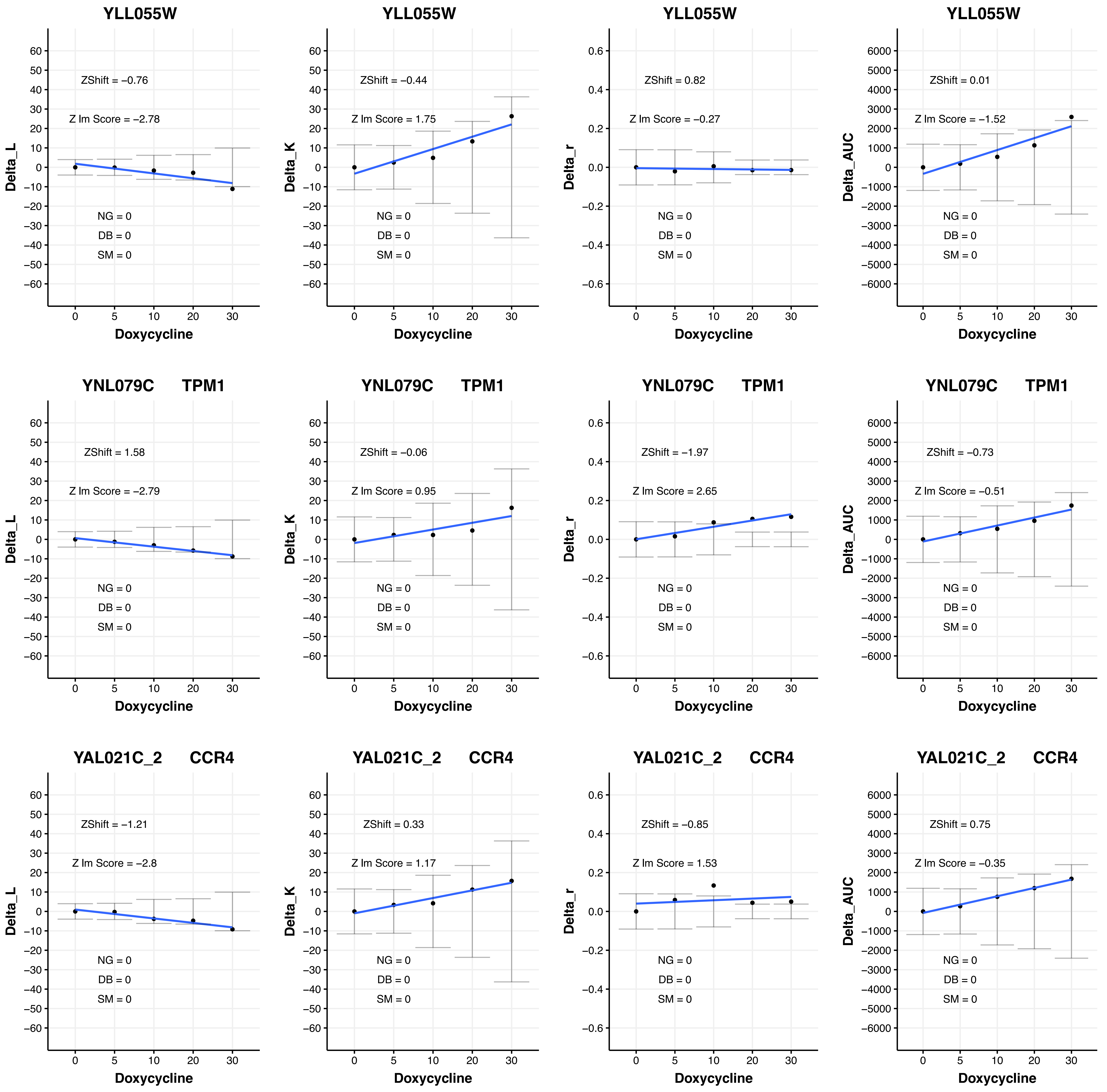
YLL055W
ZShift = −0.76
Z lm Score = −2.78
NG = 0
DB = 0
SM = 0
Delta_L
Doxycycline
YLL055W
ZShift = −0.44
Z lm Score = 1.75
NG = 0
DB = 0
SM = 0
Delta_K
Doxycycline
YLL055W
ZShift = 0.82
Z lm Score = −0.27
NG = 0
DB = 0
SM = 0
Delta_r
Doxycycline
YLL055W
ZShift = 0.01
Z lm Score = −1.52
NG = 0
DB = 0
SM = 0
Delta_AUC
Doxycycline
YNL079C TPM1
ZShift = 1.58
Z lm Score = −2.79
NG = 0
DB = 0
SM = 0
Delta_L
Doxycycline
YNL079C TPM1
ZShift = −0.06
Z lm Score = 0.95
NG = 0
DB = 0
SM = 0
Delta_K
Doxycycline
YNL079C TPM1
ZShift = −1.97
Z lm Score = 2.65
NG = 0
DB = 0
SM = 0
Delta_r
Doxycycline
YNL079C TPM1
ZShift = −0.73
Z lm Score = −0.51
NG = 0
DB = 0
SM = 0
Delta_AUC
Doxycycline
YAL021C_2 CCR4
ZShift = −1.21
Z lm Score = −2.8
NG = 0
DB = 0
SM = 0
Delta_L
Doxycycline
YAL021C_2 CCR4
ZShift = 0.33
Z lm Score = 1.17
NG = 0
DB = 0
SM = 0
Delta_K
Doxycycline
YAL021C_2 CCR4
ZShift = −0.85
Z lm Score = 1.53
NG = 0
DB = 0
SM = 0
Delta_r
Doxycycline
YAL021C_2 CCR4
ZShift = 0.75
Z lm Score = −0.35
NG = 0
DB = 0
SM = 0
Delta_AUC
Doxycycline

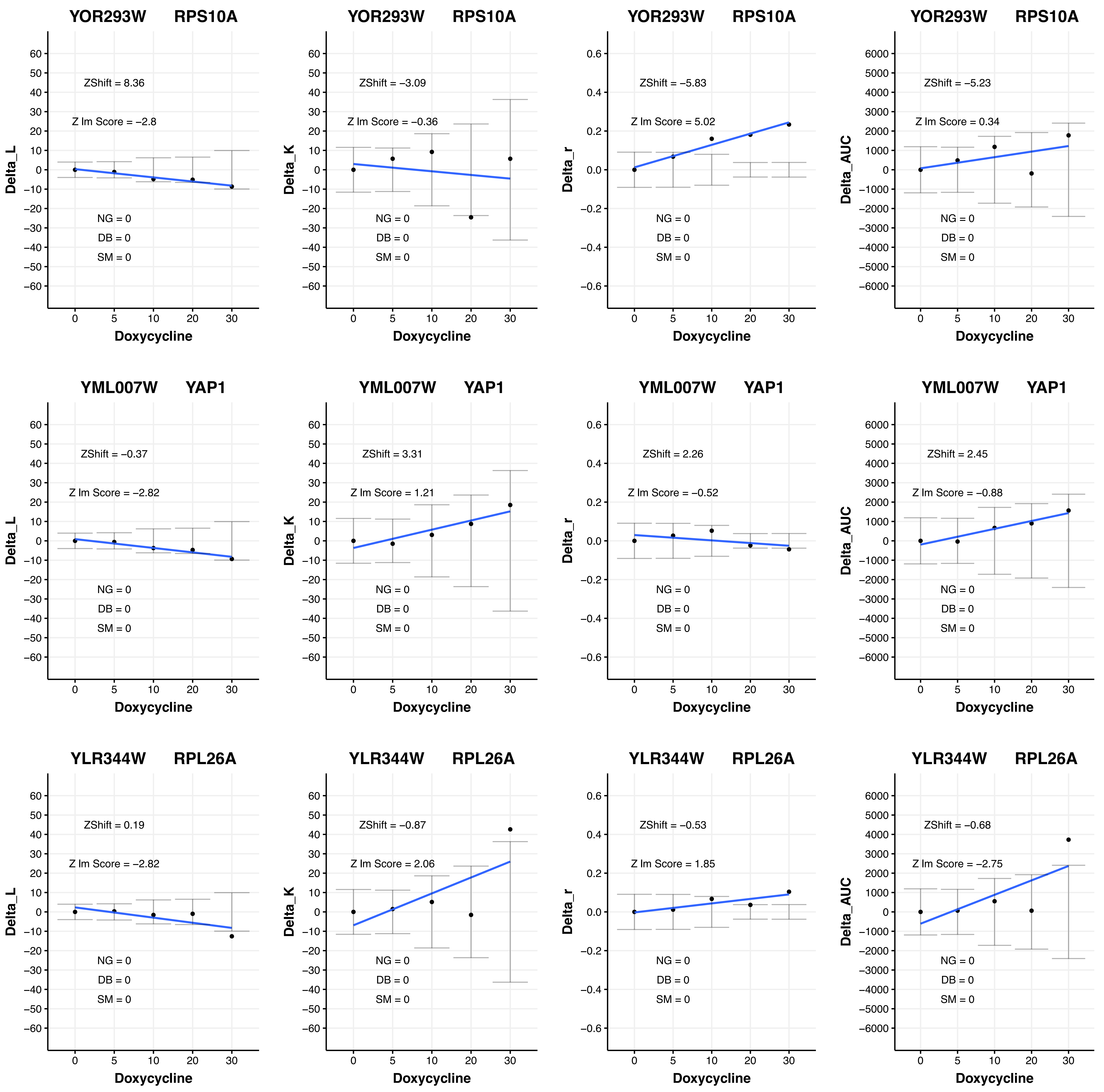

YOR293W RPS10A
ZShift = 8.36
Z lm Score = −2.8
NG = 0
DB = 0
SM = 0
Delta_L
Doxycycline
YOR293W RPS10A
ZShift = −3.09
Z lm Score = −0.36
NG = 0
DB = 0
SM = 0
Delta_K
Doxycycline
YOR293W RPS10A
ZShift = −5.83
Z lm Score = 5.02
NG = 0
DB = 0
SM = 0
Delta_r
Doxycycline
YOR293W RPS10A
ZShift = −5.23
Z lm Score = 0.34
NG = 0
DB = 0
SM = 0
Delta_AUC
Doxycycline
YML007W YAP1
ZShift = −0.37
Z lm Score = −2.82
NG = 0
DB = 0
SM = 0
Delta_L
Doxycycline
YML007W YAP1
ZShift = 3.31
Z lm Score = 1.21
NG = 0
DB = 0
SM = 0
Delta_K
Doxycycline
YML007W YAP1
ZShift = 2.26
Z lm Score = −0.52
NG = 0
DB = 0
SM = 0
Delta_r
Doxycycline
YML007W YAP1
ZShift = 2.45
Z lm Score = −0.88
NG = 0
DB = 0
SM = 0
Delta_AUC
Doxycycline
YLR344W RPL26A
ZShift = 0.19
Z lm Score = −2.82
NG = 0
DB = 0
SM = 0
Delta_L
Doxycycline
YLR344W RPL26A
ZShift = −0.87
Z lm Score = 2.06
NG = 0
DB = 0
SM = 0
Delta_K
Doxycycline
YLR344W RPL26A
ZShift = −0.53
Z lm Score = 1.85
NG = 0
DB = 0
SM = 0
Delta_r
Doxycycline
YLR344W RPL26A
ZShift = −0.68
Z lm Score = −2.75
NG = 0
DB = 0
SM = 0
Delta_AUC
Doxycycline

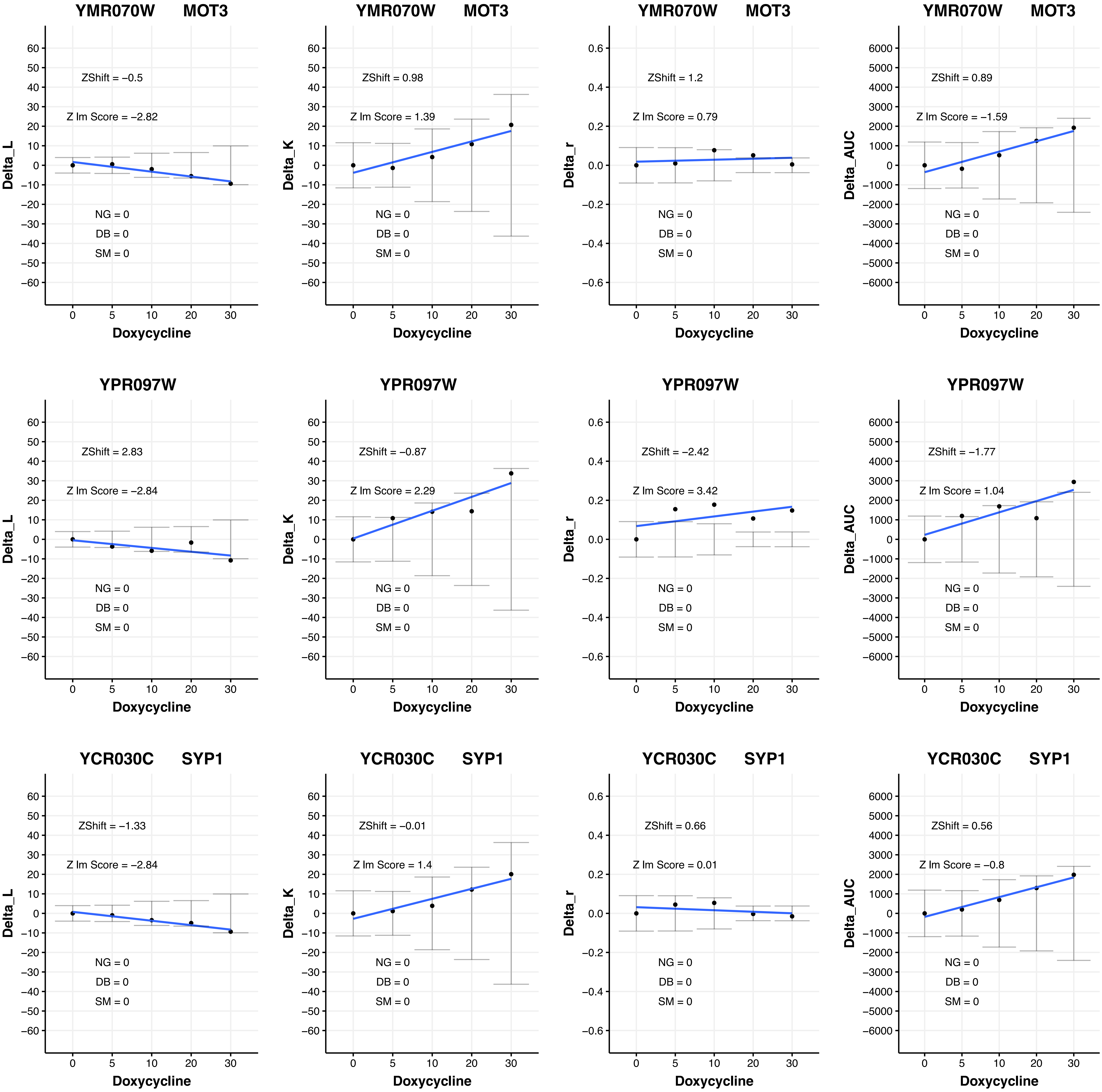

YMR070W MOT3
ZShift = −0.5
Z lm Score = −2.82
NG = 0
DB = 0
SM = 0
Delta_L
Doxycycline
YMR070W MOT3
ZShift = 0.98
Z lm Score = 1.39
NG = 0
DB = 0
SM = 0
Delta_K
Doxycycline
YMR070W MOT3
ZShift = 1.2
Z lm Score = 0.79
NG = 0
DB = 0
SM = 0
Delta_r
Doxycycline
YMR070W MOT3
ZShift = 0.89
Z lm Score = −1.59
NG = 0
DB = 0
SM = 0
Delta_AUC
Doxycycline
YPR097W
ZShift = 2.83
Z lm Score = −2.84
NG = 0
DB = 0
SM = 0
Delta_L
Doxycycline
YPR097W
ZShift = −0.87
Z lm Score = 2.29
NG = 0
DB = 0
SM = 0
Delta_K
Doxycycline
YPR097W
ZShift = −2.42
Z lm Score = 3.42
NG = 0
DB = 0
SM = 0
Delta_r
Doxycycline
YPR097W
ZShift = −1.77
Z lm Score = 1.04
NG = 0
DB = 0
SM = 0
Delta_AUC
Doxycycline
YCR030C SYP1
ZShift = −1.33
Z lm Score = −2.84
NG = 0
DB = 0
SM = 0
Delta_L
Doxycycline
YCR030C SYP1
ZShift = −0.01
Z lm Score = 1.4
NG = 0
DB = 0
SM = 0
Delta_K
Doxycycline
YCR030C SYP1
ZShift = 0.66
Z lm Score = 0.01
NG = 0
DB = 0
SM = 0
Delta_r
Doxycycline
YCR030C SYP1
ZShift = 0.56
Z lm Score = −0.8
NG = 0
DB = 0
SM = 0
Delta_AUC
Doxycycline

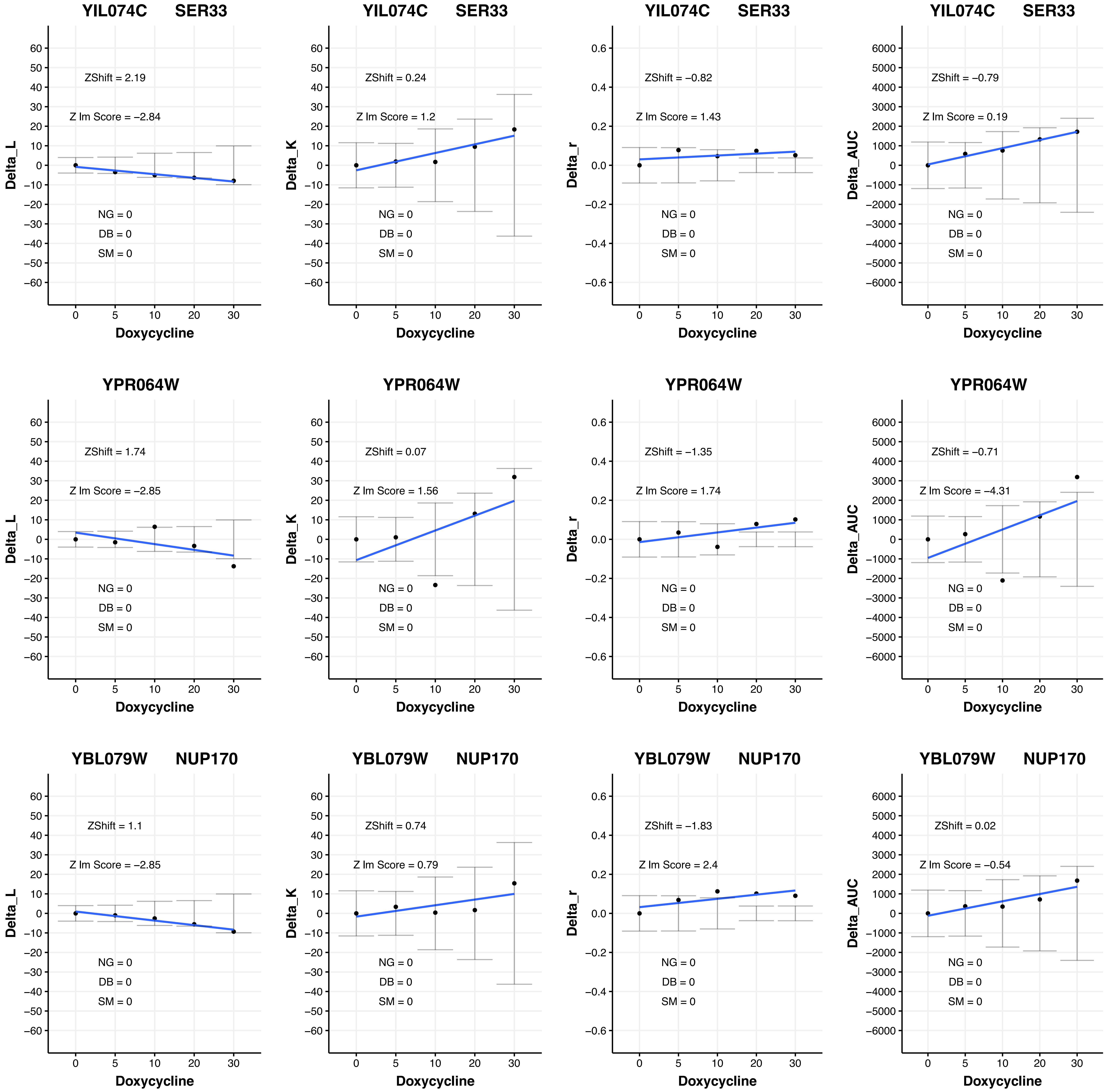
YIL074C SER33
ZShift = 2.19
Z lm Score = −2.84
NG = 0
DB = 0
SM = 0
Delta_L
Doxycycline
YIL074C SER33
ZShift = 0.24
Z lm Score = 1.2
NG = 0
DB = 0
SM = 0
Delta_K
Doxycycline
YIL074C SER33
ZShift = −0.82
Z lm Score = 1.43
NG = 0
DB = 0
SM = 0
Delta_r
Doxycycline
YIL074C SER33
ZShift = −0.79
Z lm Score = 0.19
NG = 0
DB = 0
SM = 0
Delta_AUC
Doxycycline
YPR064W
ZShift = 1.74
Z lm Score = −2.85
NG = 0
DB = 0
SM = 0
Delta_L
Doxycycline
YPR064W
ZShift = 0.07
Z lm Score = 1.56
NG = 0
DB = 0
SM = 0
Delta_K
Doxycycline
YPR064W
ZShift = −1.35
Z lm Score = 1.74
NG = 0
DB = 0
SM = 0
Delta_r
Doxycycline
YPR064W
ZShift = −0.71
Z lm Score = −4.31
NG = 0
DB = 0
SM = 0
Delta_AUC
Doxycycline
YBL079W NUP170
ZShift = 1.1
Z lm Score = −2.85
NG = 0
DB = 0
SM = 0
Delta_L
Doxycycline
YBL079W NUP170
ZShift = 0.74
Z lm Score = 0.79
NG = 0
DB = 0
SM = 0
Delta_K
Doxycycline
YBL079W NUP170
ZShift = −1.83
Z lm Score = 2.4
NG = 0
DB = 0
SM = 0
Delta_r
Doxycycline
YBL079W NUP170
ZShift = 0.02
Z lm Score = −0.54
NG = 0
DB = 0
SM = 0
Delta_AUC
Doxycycline

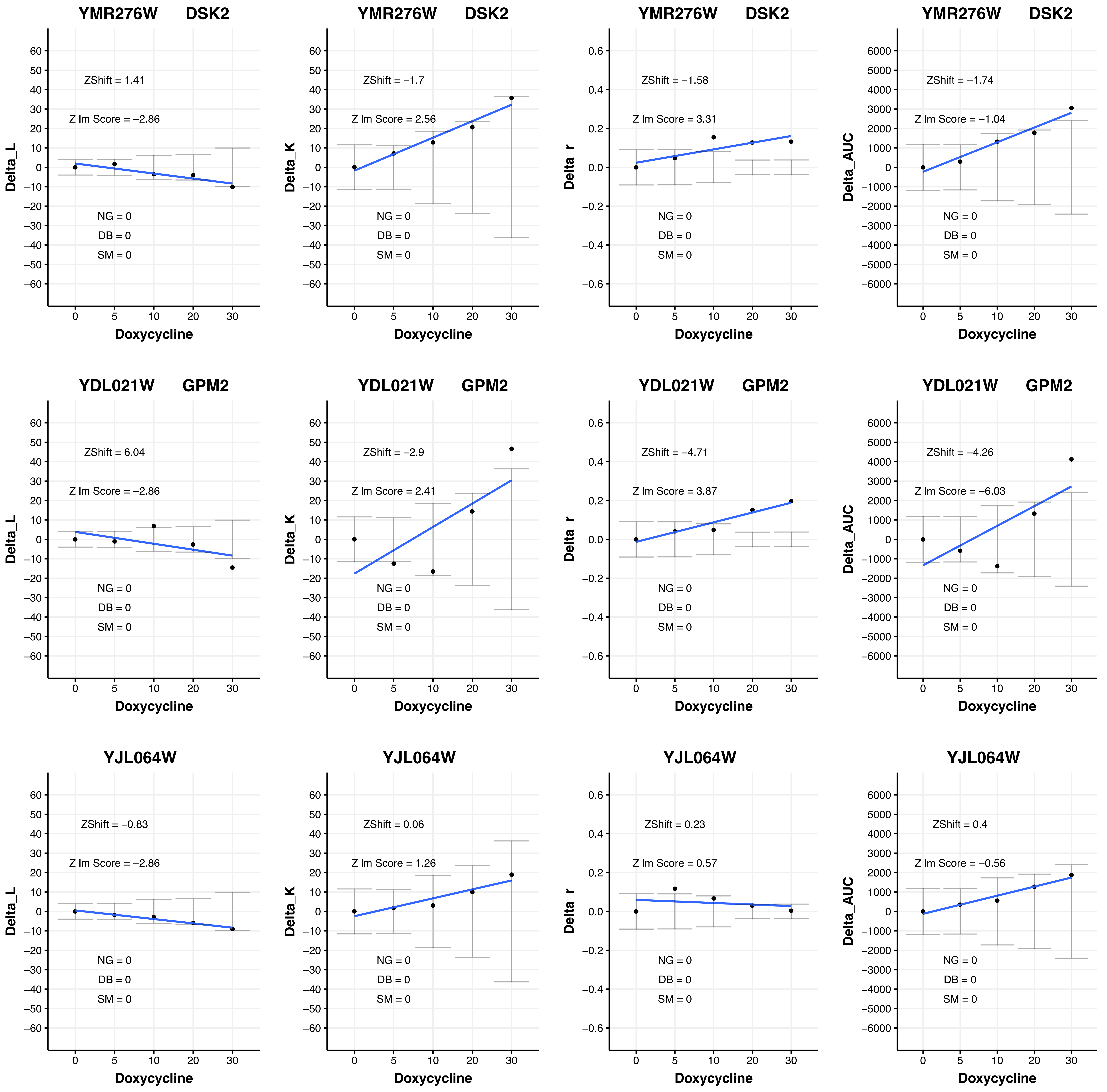
YMR276W DSK2
ZShift = 1.41
Z lm Score = −2.86
NG = 0
DB = 0
SM = 0
Delta_L
Doxycycline
YMR276W DSK2
ZShift = −1.7
Z lm Score = 2.56
NG = 0
DB = 0
SM = 0
Delta_K
Doxycycline
YMR276W DSK2
ZShift = −1.58
Z lm Score = 3.31
NG = 0
DB = 0
SM = 0
Delta_r
Doxycycline
YMR276W DSK2
ZShift = −1.74
Z lm Score = −1.04
NG = 0
DB = 0
SM = 0
Delta_AUC
Doxycycline
YDL021W GPM2
ZShift = 6.04
Z lm Score = −2.86
NG = 0
DB = 0
SM = 0
Delta_L
Doxycycline
YDL021W GPM2
ZShift = −2.9
Z lm Score = 2.41
NG = 0
DB = 0
SM = 0
Delta_K
Doxycycline
YDL021W GPM2
ZShift = −4.71
Z lm Score = 3.87
NG = 0
DB = 0
SM = 0
Delta_r
Doxycycline
YDL021W GPM2
ZShift = −4.26
Z lm Score = −6.03
NG = 0
DB = 0
SM = 0
Delta_AUC
Doxycycline
YJL064W
ZShift = −0.83
Z lm Score = −2.86
NG = 0
DB = 0
SM = 0
Delta_L
Doxycycline
YJL064W
ZShift = 0.06
Z lm Score = 1.26
NG = 0
DB = 0
SM = 0
Delta_K
Doxycycline
YJL064W
ZShift = 0.23
Z lm Score = 0.57
NG = 0
DB = 0
SM = 0
Delta_r
Doxycycline
YJL064W
ZShift = 0.4
Z lm Score = −0.56
NG = 0
DB = 0
SM = 0
Delta_AUC
Doxycycline

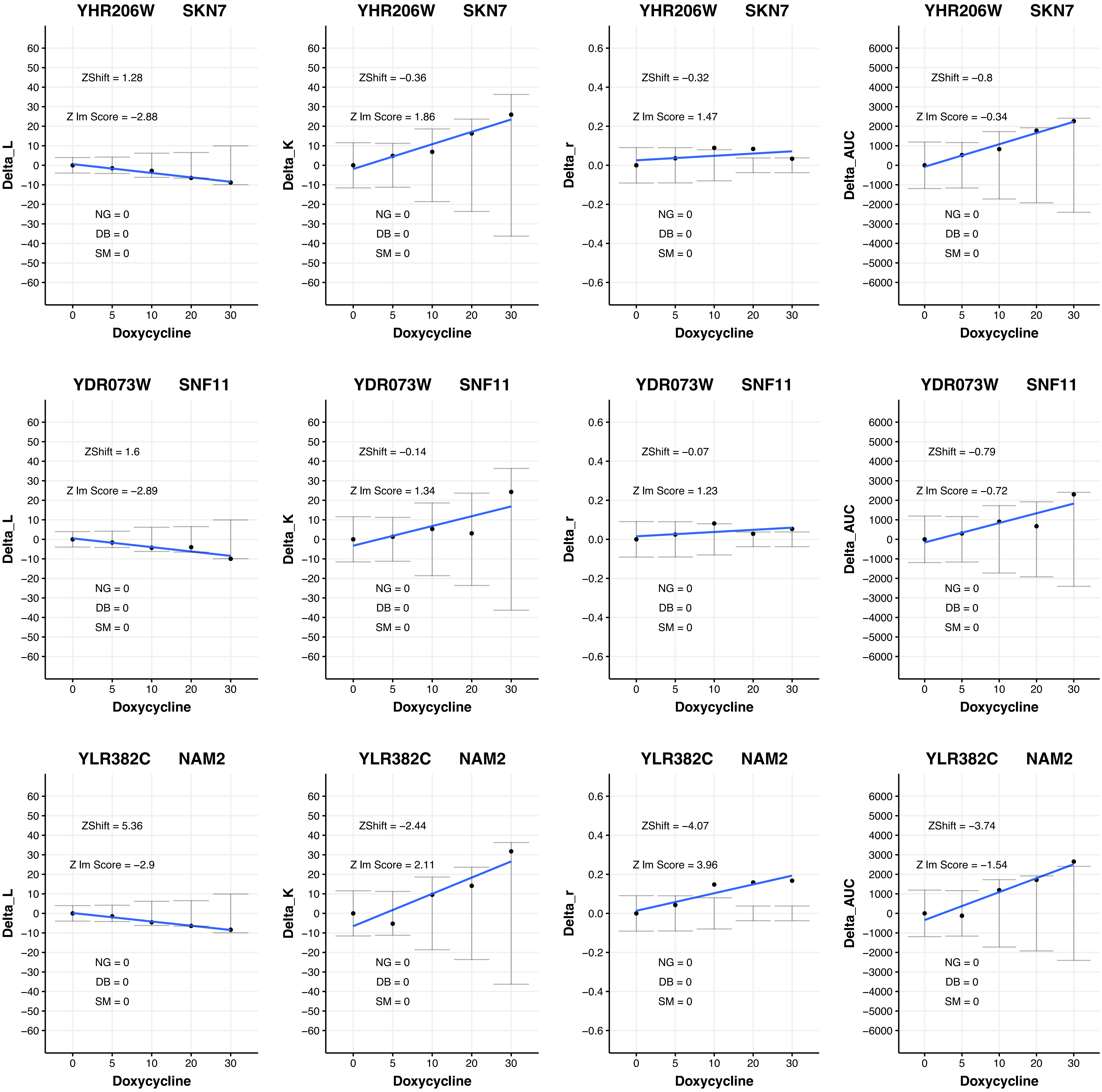

YHR206W SKN7
ZShift = 1.28
Z lm Score = −2.88
NG = 0
DB = 0
SM = 0
Delta_L
Doxycycline
YHR206W SKN7
ZShift = −0.36
Z lm Score = 1.86
NG = 0
DB = 0
SM = 0
Delta_K
Doxycycline
YHR206W SKN7
ZShift = −0.32
Z lm Score = 1.47
NG = 0
DB = 0
SM = 0
Delta_r
Doxycycline
YHR206W SKN7
ZShift = −0.8
Z lm Score = −0.34
NG = 0
DB = 0
SM = 0
Delta_AUC
Doxycycline
YDR073W SNF11
ZShift = 1.6
Z lm Score = −2.89
NG = 0
DB = 0
SM = 0
Delta_L
Doxycycline
YDR073W SNF11
ZShift = −0.14
Z lm Score = 1.34
NG = 0
DB = 0
SM = 0
Delta_K
Doxycycline
YDR073W SNF11
ZShift = −0.07
Z lm Score = 1.23
NG = 0
DB = 0
SM = 0
Delta_r
Doxycycline
YDR073W SNF11
ZShift = −0.79
Z lm Score = −0.72
NG = 0
DB = 0
SM = 0
Delta_AUC
Doxycycline
YLR382C NAM2
ZShift = 5.36
Z lm Score = −2.9
NG = 0
DB = 0
SM = 0
Delta_L
Doxycycline
YLR382C NAM2
ZShift = −2.44
Z lm Score = 2.11
NG = 0
DB = 0
SM = 0
Delta_K
Doxycycline
YLR382C NAM2
ZShift = −4.07
Z lm Score = 3.96
NG = 0
DB = 0
SM = 0
Delta_r
Doxycycline
YLR382C NAM2
ZShift = −3.74
Z lm Score = −1.54
NG = 0
DB = 0
SM = 0
Delta_AUC
Doxycycline

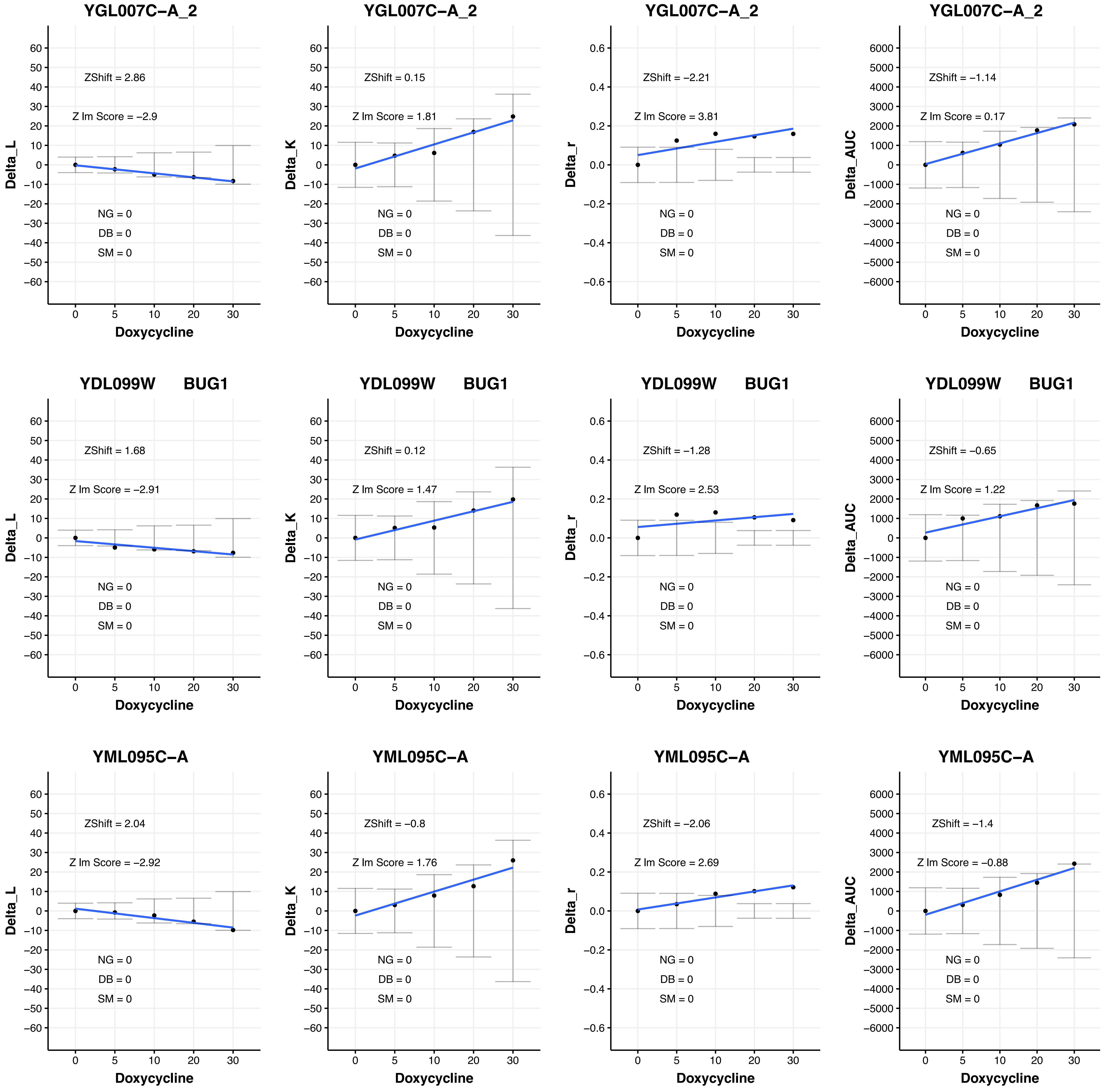
YGL007C−A_2
ZShift = 2.86
Z lm Score = −2.9
NG = 0
DB = 0
SM = 0
Delta_L
Doxycycline
YGL007C−A_2
ZShift = 0.15
Z lm Score = 1.81
NG = 0
DB = 0
SM = 0
Delta_K
Doxycycline
YGL007C−A_2
ZShift = −2.21
Z lm Score = 3.81
NG = 0
DB = 0
SM = 0
Delta_r
Doxycycline
YGL007C−A_2
ZShift = −1.14
Z lm Score = 0.17
NG = 0
DB = 0
SM = 0
Delta_AUC
Doxycycline
YDL099W BUG1
ZShift = 1.68
Z lm Score = −2.91
NG = 0
DB = 0
SM = 0
Delta_L
Doxycycline
YDL099W BUG1
ZShift = 0.12
Z lm Score = 1.47
NG = 0
DB = 0
SM = 0
Delta_K
Doxycycline
YDL099W BUG1
ZShift = −1.28
Z lm Score = 2.53
NG = 0
DB = 0
SM = 0
Delta_r
Doxycycline
YDL099W BUG1
ZShift = −0.65
Z lm Score = 1.22
NG = 0
DB = 0
SM = 0
Delta_AUC
Doxycycline
YML095C−A
ZShift = 2.04
Z lm Score = −2.92
NG = 0
DB = 0
SM = 0
Delta_L
Doxycycline
YML095C−A
ZShift = −0.8
Z lm Score = 1.76
NG = 0
DB = 0
SM = 0
Delta_K
Doxycycline
YML095C−A
ZShift = −2.06
Z lm Score = 2.69
NG = 0
DB = 0
SM = 0
Delta_r
Doxycycline
YML095C−A
ZShift = −1.4
Z lm Score = −0.88
NG = 0
DB = 0
SM = 0
Delta_AUC
Doxycycline

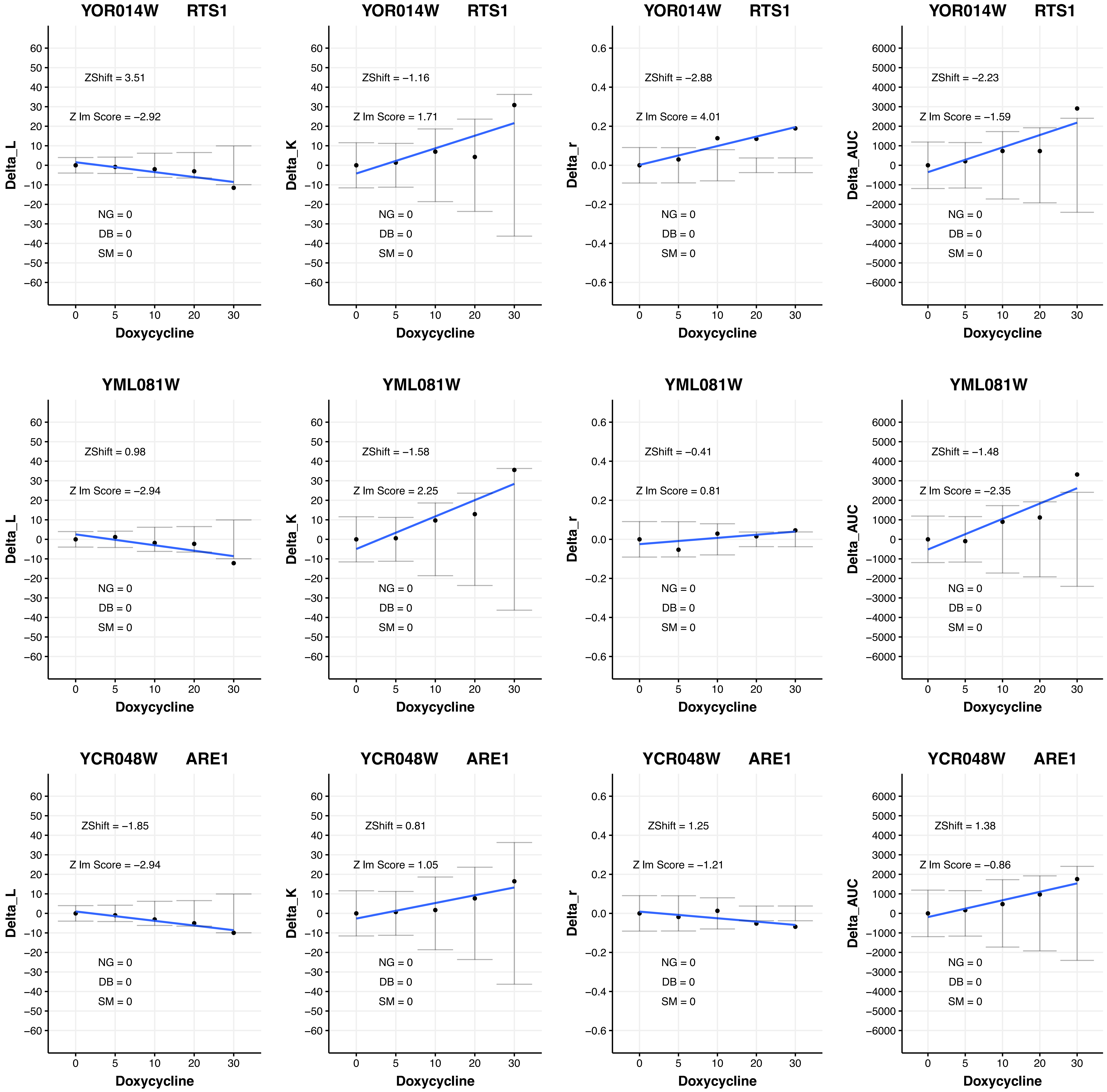

YOR014W RTS1
ZShift = 3.51
Z lm Score = −2.92
NG = 0
DB = 0
SM = 0
Delta_L
Doxycycline
YOR014W RTS1
ZShift = −1.16
Z lm Score = 1.71
NG = 0
DB = 0
SM = 0
Delta_K
Doxycycline
YOR014W RTS1
ZShift = −2.88
Z lm Score = 4.01
NG = 0
DB = 0
SM = 0
Delta_r
Doxycycline
YOR014W RTS1
ZShift = −2.23
Z lm Score = −1.59
NG = 0
DB = 0
SM = 0
Delta_AUC
Doxycycline
YML081W
ZShift = 0.98
Z lm Score = −2.94
NG = 0
DB = 0
SM = 0
Delta_L
Doxycycline
YML081W
ZShift = −1.58
Z lm Score = 2.25
NG = 0
DB = 0
SM = 0
Delta_K
Doxycycline
YML081W
ZShift = −0.41
Z lm Score = 0.81
NG = 0
DB = 0
SM = 0
Delta_r
Doxycycline
YML081W
ZShift = −1.48
Z lm Score = −2.35
NG = 0
DB = 0
SM = 0
Delta_AUC
Doxycycline
YCR048W ARE1
ZShift = −1.85
Z lm Score = −2.94
NG = 0
DB = 0
SM = 0
Delta_L
Doxycycline
YCR048W ARE1
ZShift = 0.81
Z lm Score = 1.05
NG = 0
DB = 0
SM = 0
Delta_K
Doxycycline
YCR048W ARE1
ZShift = 1.25
Z lm Score = −1.21
NG = 0
DB = 0
SM = 0
Delta_r
Doxycycline
YCR048W ARE1
ZShift = 1.38
Z lm Score = −0.86
NG = 0
DB = 0
SM = 0
Delta_AUC
Doxycycline

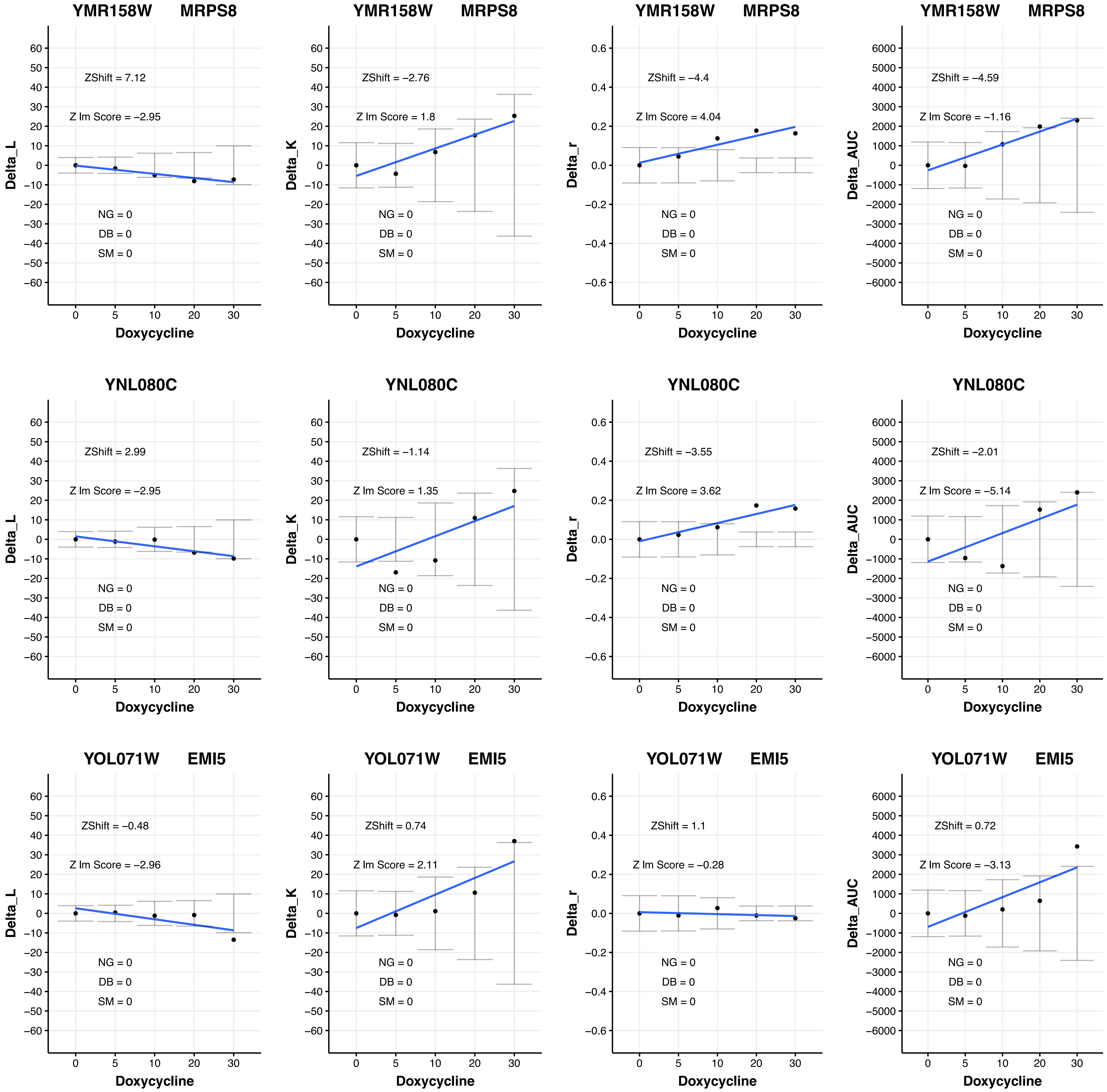
YMR158W MRPS8
ZShift = 7.12
Z lm Score = −2.95
NG = 0
DB = 0
SM = 0
Delta_L
Doxycycline
ZShift = −2.76
Z lm Score = 1.8
Delta_K
ZShift = −4.4
Z lm Score = 4.04
Delta_r
ZShift = −4.59
Z lm Score = −1.16
Delta_AUC
YNL080C
ZShift = 2.99
Z lm Score = −2.95
ZShift = −1.14
Z lm Score = 1.35
ZShift = −3.55
Z lm Score = 3.62
ZShift = −2.01
Z lm Score = −5.14
YOL071W EMI5
ZShift = −0.48
Z lm Score = −2.96
ZShift = 0.74
Z lm Score = 2.11
ZShift = 1.1
Z lm Score = −0.28
ZShift = 0.72
Z lm Score = −3.13

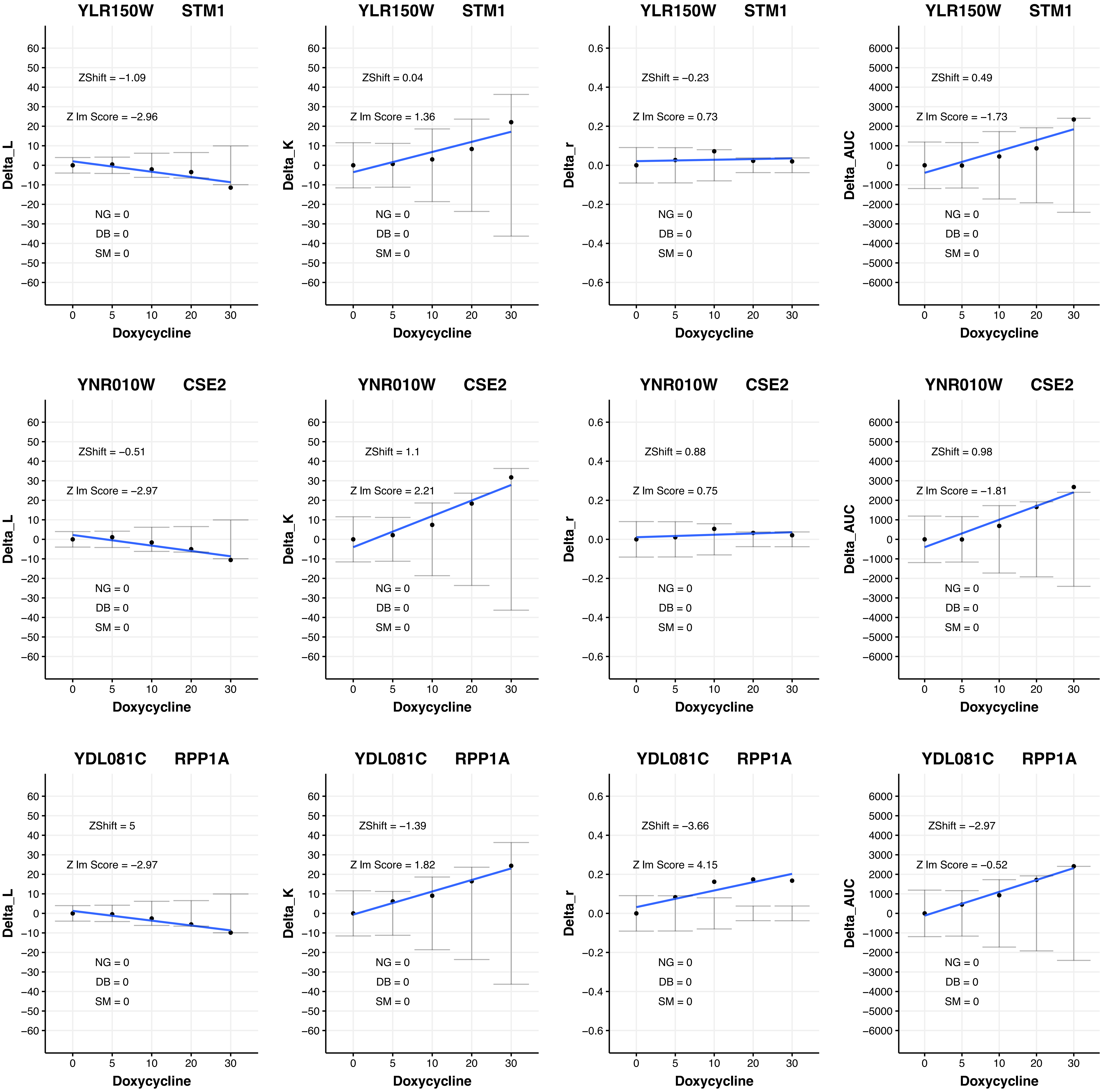

YLR150W STM1
ZShift = −1.09
Z lm Score = −2.96
NG = 0
DB = 0
SM = 0
Delta_L
Doxycycline
YLR150W STM1
ZShift = 0.04
Z lm Score = 1.36
NG = 0
DB = 0
SM = 0
Delta_K
Doxycycline
YLR150W STM1
ZShift = −0.23
Z lm Score = 0.73
NG = 0
DB = 0
SM = 0
Delta_r
Doxycycline
YLR150W STM1
ZShift = 0.49
Z lm Score = −1.73
NG = 0
DB = 0
SM = 0
Delta_AUC
Doxycycline
YNR010W CSE2
ZShift = −0.51
Z lm Score = −2.97
NG = 0
DB = 0
SM = 0
Delta_L
Doxycycline
YNR010W CSE2
ZShift = 1.1
Z lm Score = 2.21
NG = 0
DB = 0
SM = 0
Delta_K
Doxycycline
YNR010W CSE2
ZShift = 0.88
Z lm Score = 0.75
NG = 0
DB = 0
SM = 0
Delta_r
Doxycycline
YNR010W CSE2
ZShift = 0.98
Z lm Score = −1.81
NG = 0
DB = 0
SM = 0
Delta_AUC
Doxycycline
YDL081C RPP1A
ZShift = 5
Z lm Score = −2.97
NG = 0
DB = 0
SM = 0
Delta_L
Doxycycline
YDL081C RPP1A
ZShift = −1.39
Z lm Score = 1.82
NG = 0
DB = 0
SM = 0
Delta_K
Doxycycline
YDL081C RPP1A
ZShift = −3.66
Z lm Score = 4.15
NG = 0
DB = 0
SM = 0
Delta_r
Doxycycline
YDL081C RPP1A
ZShift = −2.97
Z lm Score = −0.52
NG = 0
DB = 0
SM = 0
Delta_AUC
Doxycycline

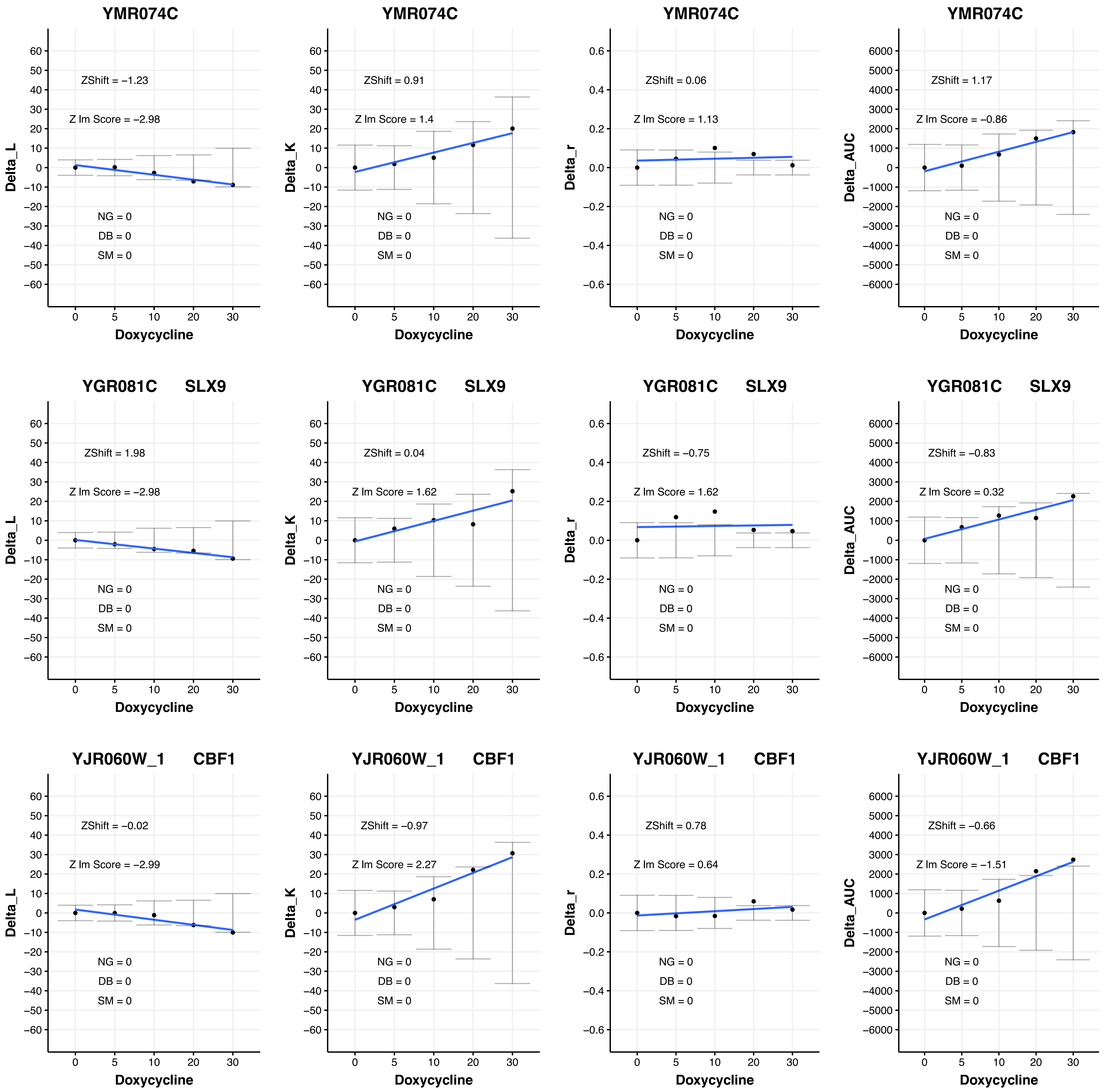
YMR074C
ZShift = −1.23
Z lm Score = −2.98
ZShift = 0.91
Z lm Score = 1.4
ZShift = 0.06
Z lm Score = 1.13
ZShift = 1.17
Z lm Score = −0.86
YGR081C SLX9
ZShift = 1.98
Z lm Score = −2.98
ZShift = 0.04
Z lm Score = 1.62
ZShift = −0.75
Z lm Score = 1.62
ZShift = −0.83
Z lm Score = 0.32
YJR060W_1 CBF1
ZShift = −0.02
Z lm Score = −2.99
ZShift = −0.97
Z lm Score = 2.27
ZShift = 0.78
Z lm Score = 0.64
ZShift = −0.66
Z lm Score = −1.51
NG = 0
DB = 0
SM = 0
Delta_L
Delta_K
Delta_r
Delta_AUC
Doxycycline

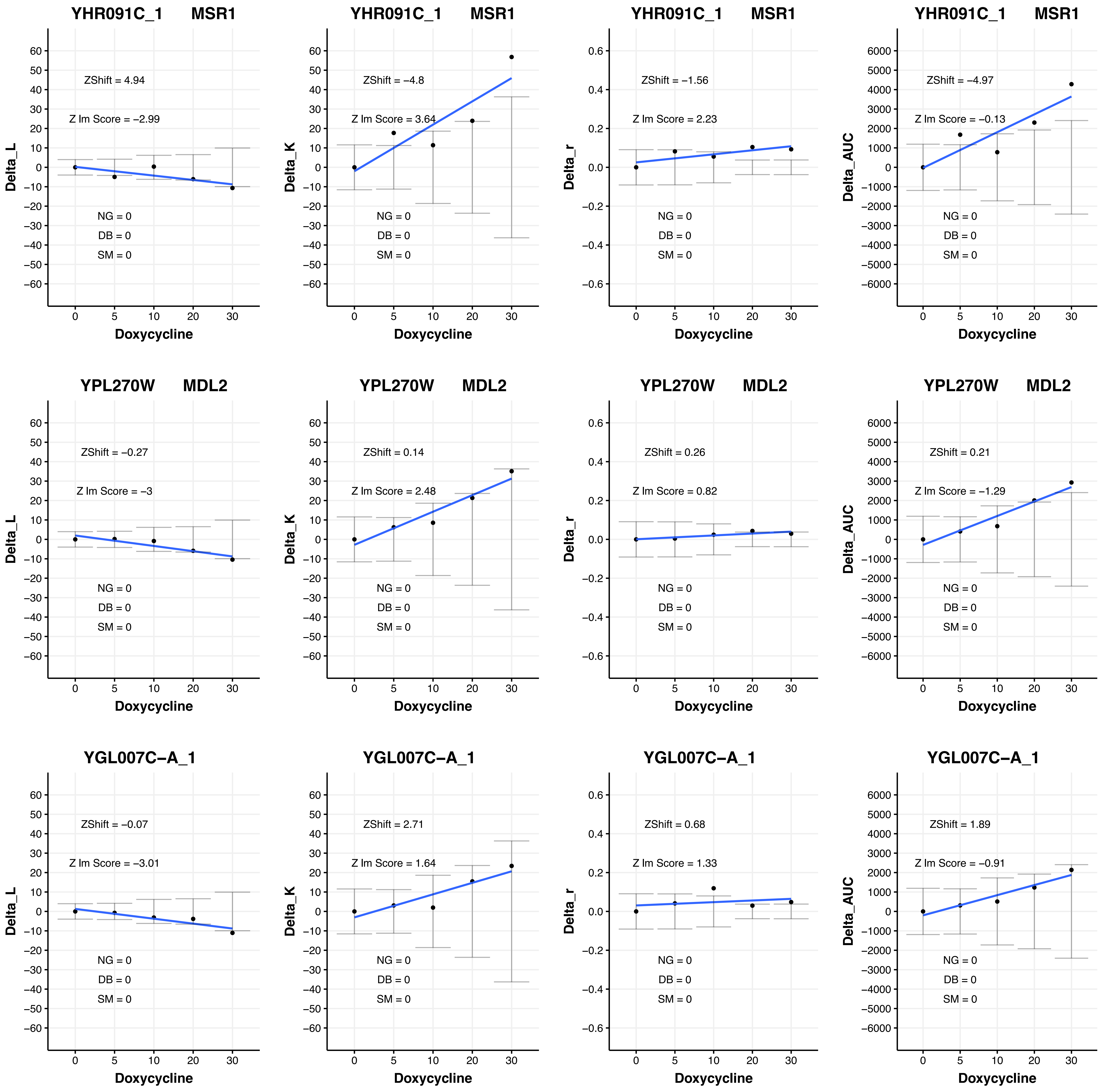
YHR091C_1 MSR1
ZShift = 4.94
Z lm Score = −2.99
NG = 0
DB = 0
SM = 0
Delta_L
Doxycycline
YHR091C_1 MSR1
ZShift = −4.8
Z lm Score = 3.64
NG = 0
DB = 0
SM = 0
Delta_K
Doxycycline
YHR091C_1 MSR1
ZShift = −1.56
Z lm Score = 2.23
NG = 0
DB = 0
SM = 0
Delta_r
Doxycycline
YHR091C_1 MSR1
ZShift = −4.97
Z lm Score = −0.13
NG = 0
DB = 0
SM = 0
Delta_AUC
Doxycycline
YPL270W MDL2
ZShift = −0.27
Z lm Score = −3
NG = 0
DB = 0
SM = 0
Delta_L
Doxycycline
YPL270W MDL2
ZShift = 0.14
Z lm Score = 2.48
NG = 0
DB = 0
SM = 0
Delta_K
Doxycycline
YPL270W MDL2
ZShift = 0.26
Z lm Score = 0.82
NG = 0
DB = 0
SM = 0
Delta_r
Doxycycline
YPL270W MDL2
ZShift = 0.21
Z lm Score = −1.29
NG = 0
DB = 0
SM = 0
Delta_AUC
Doxycycline
YGL007C−A_1
ZShift = −0.07
Z lm Score = −3.01
NG = 0
DB = 0
SM = 0
Delta_L
Doxycycline
YGL007C−A_1
ZShift = 2.71
Z lm Score = 1.64
NG = 0
DB = 0
SM = 0
Delta_K
Doxycycline
YGL007C−A_1
ZShift = 0.68
Z lm Score = 1.33
NG = 0
DB = 0
SM = 0
Delta_r
Doxycycline
YGL007C−A_1
ZShift = 1.89
Z lm Score = −0.91
NG = 0
DB = 0
SM = 0
Delta_AUC
Doxycycline

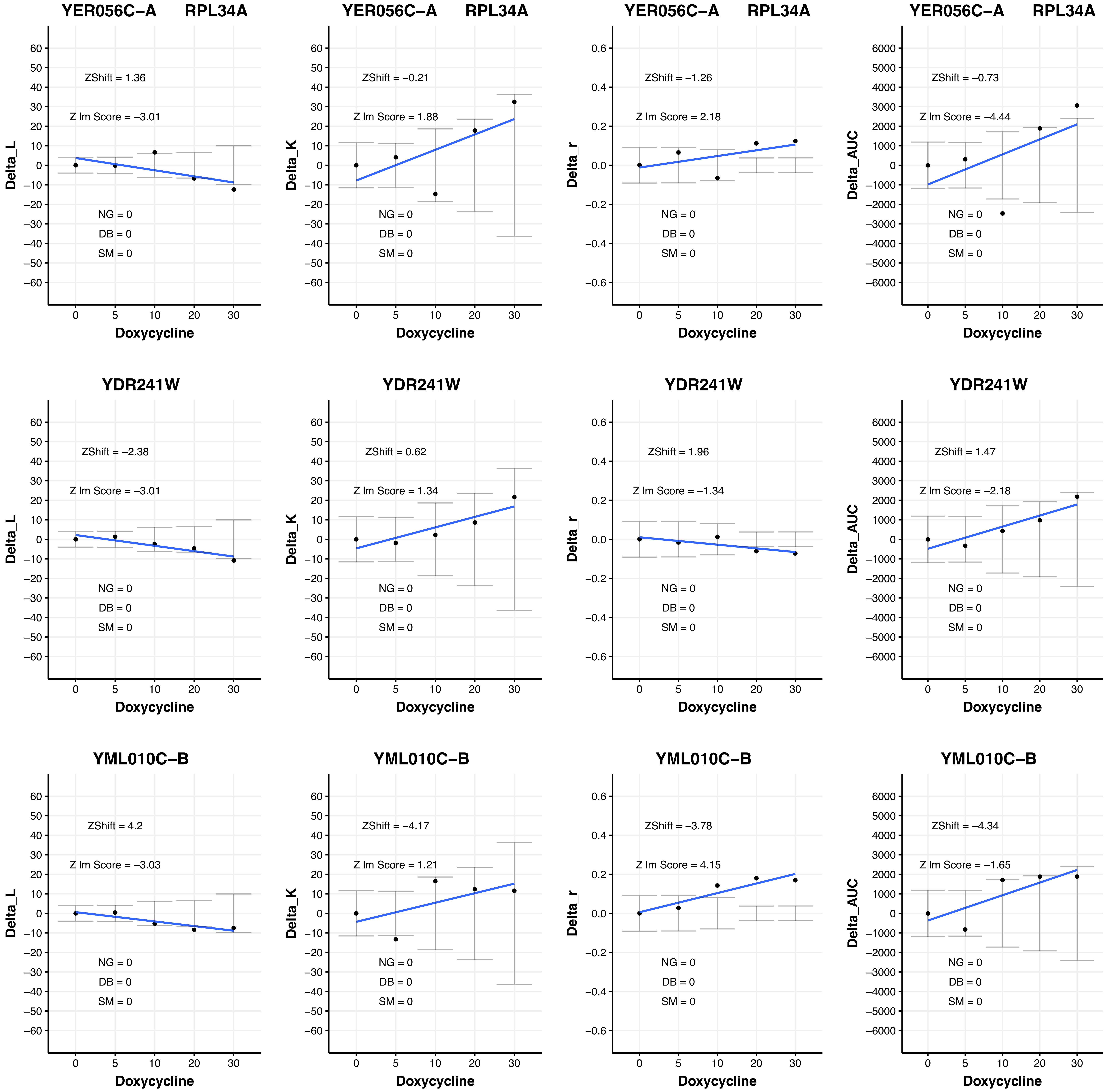
YER056C−A RPL34A
ZShift = 1.36
Z lm Score = −3.01
NG = 0
DB = 0
SM = 0
Delta_L
Doxycycline
YER056C−A RPL34A
ZShift = −0.21
Z lm Score = 1.88
NG = 0
DB = 0
SM = 0
Delta_K
Doxycycline
YER056C−A RPL34A
ZShift = −1.26
Z lm Score = 2.18
NG = 0
DB = 0
SM = 0
Delta_r
Doxycycline
YER056C−A RPL34A
ZShift = −0.73
Z lm Score = −4.44
NG = 0
DB = 0
SM = 0
Delta_AUC
Doxycycline
YDR241W
ZShift = −2.38
Z lm Score = −3.01
NG = 0
DB = 0
SM = 0
Delta_L
Doxycycline
YDR241W
ZShift = 0.62
Z lm Score = 1.34
NG = 0
DB = 0
SM = 0
Delta_K
Doxycycline
YDR241W
ZShift = 1.96
Z lm Score = −1.34
NG = 0
DB = 0
SM = 0
Delta_r
Doxycycline
YDR241W
ZShift = 1.47
Z lm Score = −2.18
NG = 0
DB = 0
SM = 0
Delta_AUC
Doxycycline
YML010C−B
ZShift = 4.2
Z lm Score = −3.03
NG = 0
DB = 0
SM = 0
Delta_L
Doxycycline
YML010C−B
ZShift = −4.17
Z lm Score = 1.21
NG = 0
DB = 0
SM = 0
Delta_K
Doxycycline
YML010C−B
ZShift = −3.78
Z lm Score = 4.15
NG = 0
DB = 0
SM = 0
Delta_r
Doxycycline
YML010C−B
ZShift = −4.34
Z lm Score = −1.65
NG = 0
DB = 0
SM = 0
Delta_AUC
Doxycycline

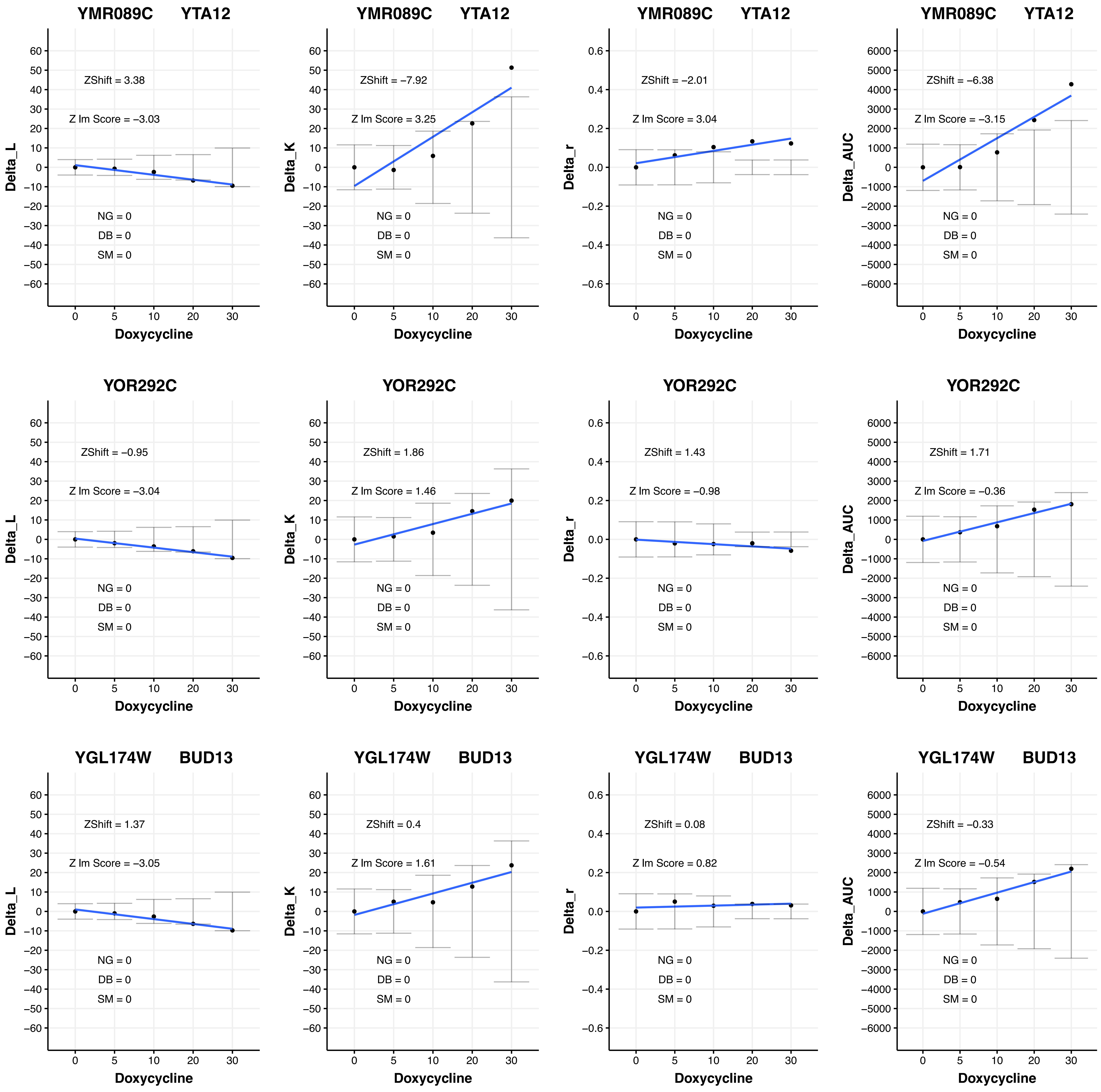

YMR089C YTA12
ZShift = 3.38
Z lm Score = −3.03
NG = 0
DB = 0
SM = 0
Delta_L
Doxycycline
YMR089C YTA12
ZShift = −7.92
Z lm Score = 3.25
NG = 0
DB = 0
SM = 0
Delta_K
Doxycycline
YMR089C YTA12
ZShift = −2.01
Z lm Score = 3.04
NG = 0
DB = 0
SM = 0
Delta_r
Doxycycline
YMR089C YTA12
ZShift = −6.38
Z lm Score = −3.15
NG = 0
DB = 0
SM = 0
Delta_AUC
Doxycycline
YOR292C
ZShift = −0.95
Z lm Score = −3.04
NG = 0
DB = 0
SM = 0
Delta_L
Doxycycline
YOR292C
ZShift = 1.86
Z lm Score = 1.46
NG = 0
DB = 0
SM = 0
Delta_K
Doxycycline
YOR292C
ZShift = 1.43
Z lm Score = −0.98
NG = 0
DB = 0
SM = 0
Delta_r
Doxycycline
YOR292C
ZShift = 1.71
Z lm Score = −0.36
NG = 0
DB = 0
SM = 0
Delta_AUC
Doxycycline
YGL174W BUD13
ZShift = 1.37
Z lm Score = −3.05
NG = 0
DB = 0
SM = 0
Delta_L
Doxycycline
YGL174W BUD13
ZShift = 0.4
Z lm Score = 1.61
NG = 0
DB = 0
SM = 0
Delta_K
Doxycycline
YGL174W BUD13
ZShift = 0.08
Z lm Score = 0.82
NG = 0
DB = 0
SM = 0
Delta_r
Doxycycline
YGL174W BUD13
ZShift = −0.33
Z lm Score = −0.54
NG = 0
DB = 0
SM = 0
Delta_AUC
Doxycycline

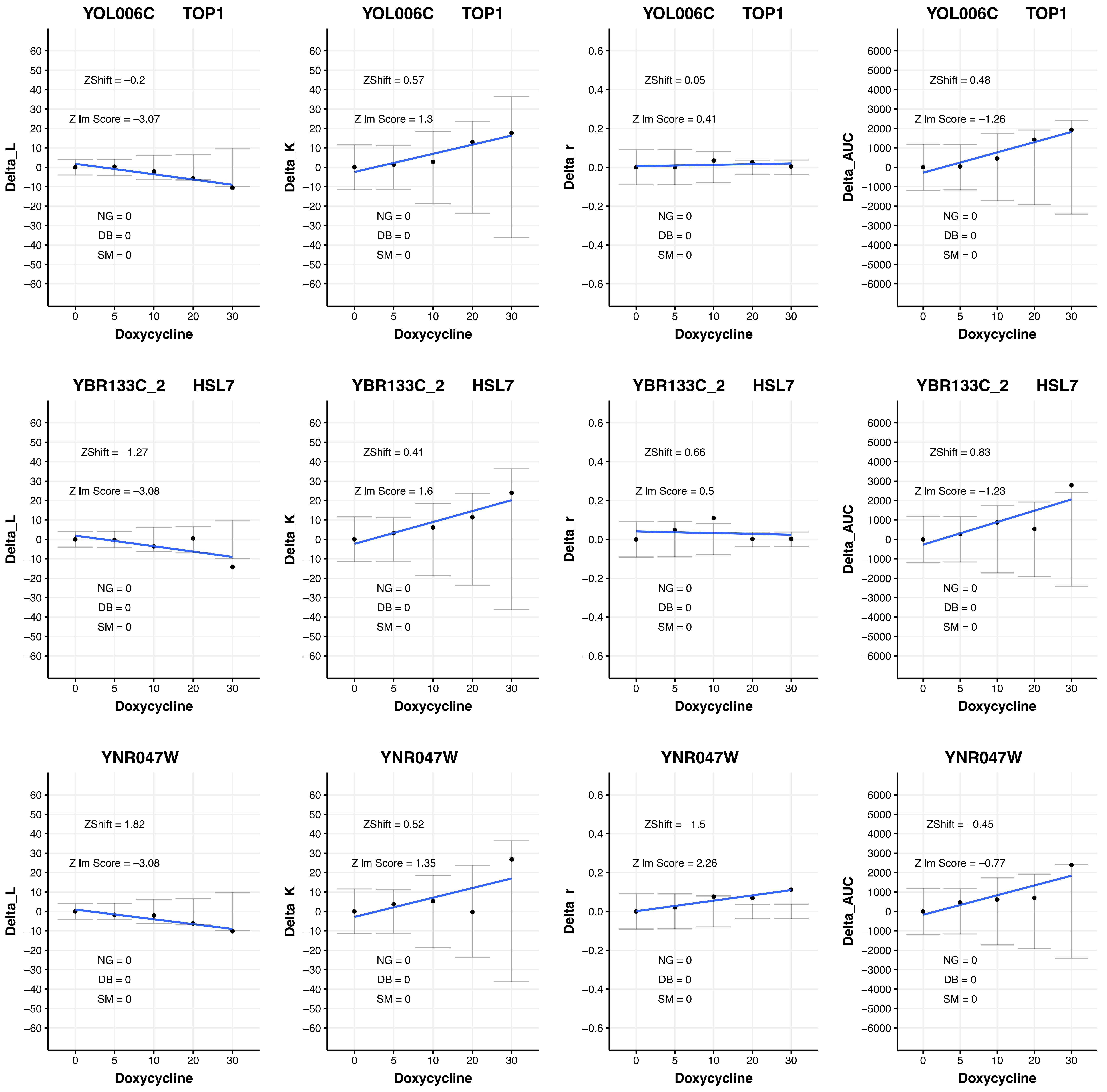

YOL006C TOP1
ZShift = −0.2
Z lm Score = −3.07
NG = 0
DB = 0
SM = 0
Delta_L
Doxycycline
YOL006C TOP1
ZShift = 0.57
Z lm Score = 1.3
NG = 0
DB = 0
SM = 0
Delta_K
Doxycycline
YOL006C TOP1
ZShift = 0.05
Z lm Score = 0.41
NG = 0
DB = 0
SM = 0
Delta_r
Doxycycline
YOL006C TOP1
ZShift = 0.48
Z lm Score = −1.26
NG = 0
DB = 0
SM = 0
Delta_AUC
Doxycycline
YBR133C_2 HSL7
ZShift = −1.27
Z lm Score = −3.08
NG = 0
DB = 0
SM = 0
Delta_L
Doxycycline
YBR133C_2 HSL7
ZShift = 0.41
Z lm Score = 1.6
NG = 0
DB = 0
SM = 0
Delta_K
Doxycycline
YBR133C_2 HSL7
ZShift = 0.66
Z lm Score = 0.5
NG = 0
DB = 0
SM = 0
Delta_r
Doxycycline
YBR133C_2 HSL7
ZShift = 0.83
Z lm Score = −1.23
NG = 0
DB = 0
SM = 0
Delta_AUC
Doxycycline
YNR047W
ZShift = 1.82
Z lm Score = −3.08
NG = 0
DB = 0
SM = 0
Delta_L
Doxycycline
YNR047W
ZShift = 0.52
Z lm Score = 1.35
NG = 0
DB = 0
SM = 0
Delta_K
Doxycycline
YNR047W
ZShift = −1.5
Z lm Score = 2.26
NG = 0
DB = 0
SM = 0
Delta_r
Doxycycline
YNR047W
ZShift = −0.45
Z lm Score = −0.77
NG = 0
DB = 0
SM = 0
Delta_AUC
Doxycycline

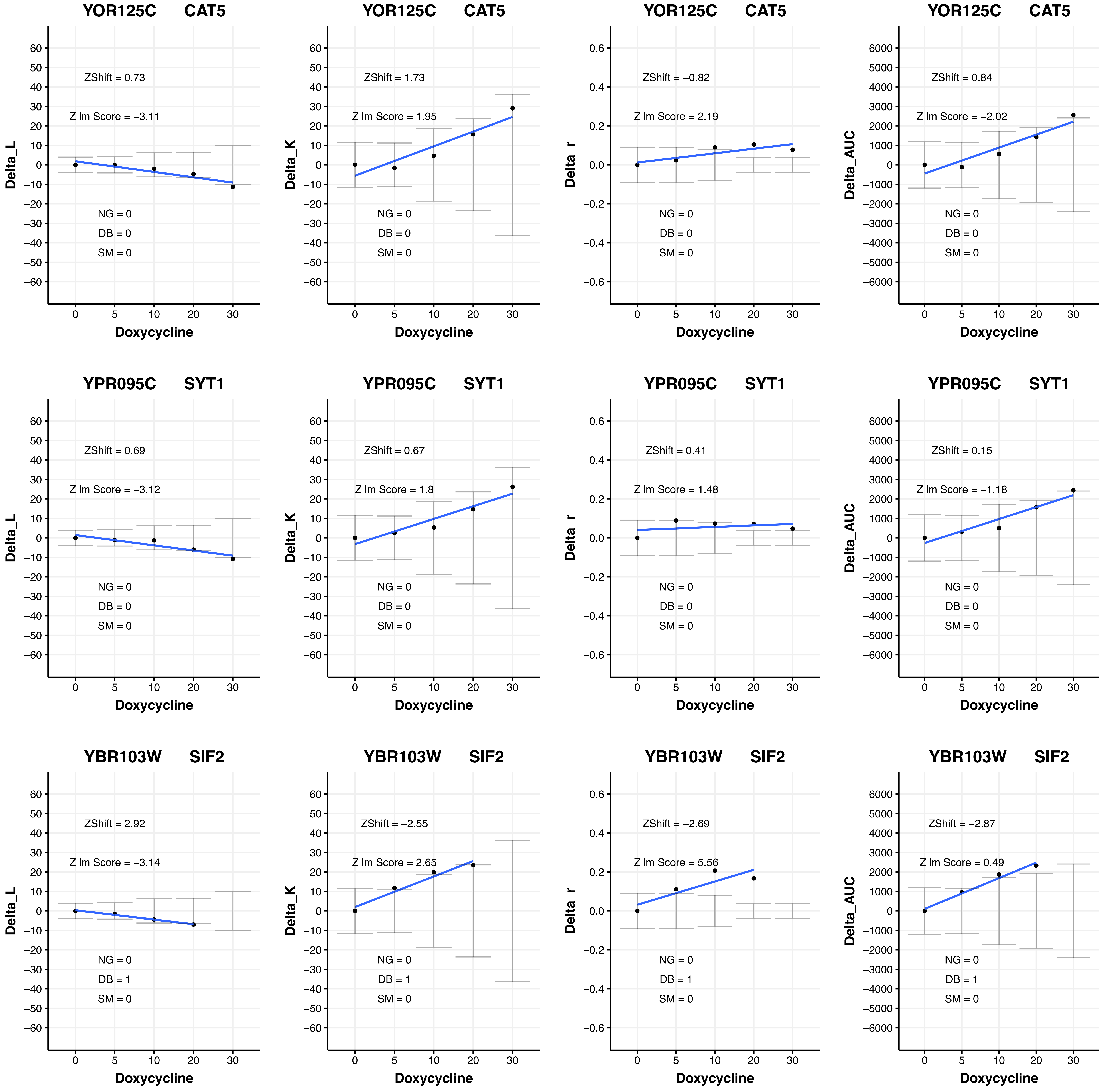
YOR125C CAT5
ZShift = 0.73
Z lm Score = −3.11
NG = 0
DB = 0
SM = 0
Delta_L
Doxycycline
YOR125C CAT5
ZShift = 1.73
Z lm Score = 1.95
NG = 0
DB = 0
SM = 0
Delta_K
Doxycycline
YOR125C CAT5
ZShift = −0.82
Z lm Score = 2.19
NG = 0
DB = 0
SM = 0
Delta_r
Doxycycline
YOR125C CAT5
ZShift = 0.84
Z lm Score = −2.02
NG = 0
DB = 0
SM = 0
Delta_AUC
Doxycycline
YPR095C SYT1
ZShift = 0.69
Z lm Score = −3.12
NG = 0
DB = 0
SM = 0
Delta_L
Doxycycline
YPR095C SYT1
ZShift = 0.67
Z lm Score = 1.8
NG = 0
DB = 0
SM = 0
Delta_K
Doxycycline
YPR095C SYT1
ZShift = 0.41
Z lm Score = 1.48
NG = 0
DB = 0
SM = 0
Delta_r
Doxycycline
YPR095C SYT1
ZShift = 0.15
Z lm Score = −1.18
NG = 0
DB = 0
SM = 0
Delta_AUC
Doxycycline
YBR103W SIF2
ZShift = 2.92
Z lm Score = −3.14
NG = 0
DB = 1
SM = 0
Delta_L
Doxycycline
YBR103W SIF2
ZShift = −2.55
Z lm Score = 2.65
NG = 0
DB = 1
SM = 0
Delta_K
Doxycycline
YBR103W SIF2
ZShift = −2.69
Z lm Score = 5.56
NG = 0
DB = 1
SM = 0
Delta_r
Doxycycline
YBR103W SIF2
ZShift = −2.87
Z lm Score = 0.49
NG = 0
DB = 1
SM = 0
Delta_AUC
Doxycycline

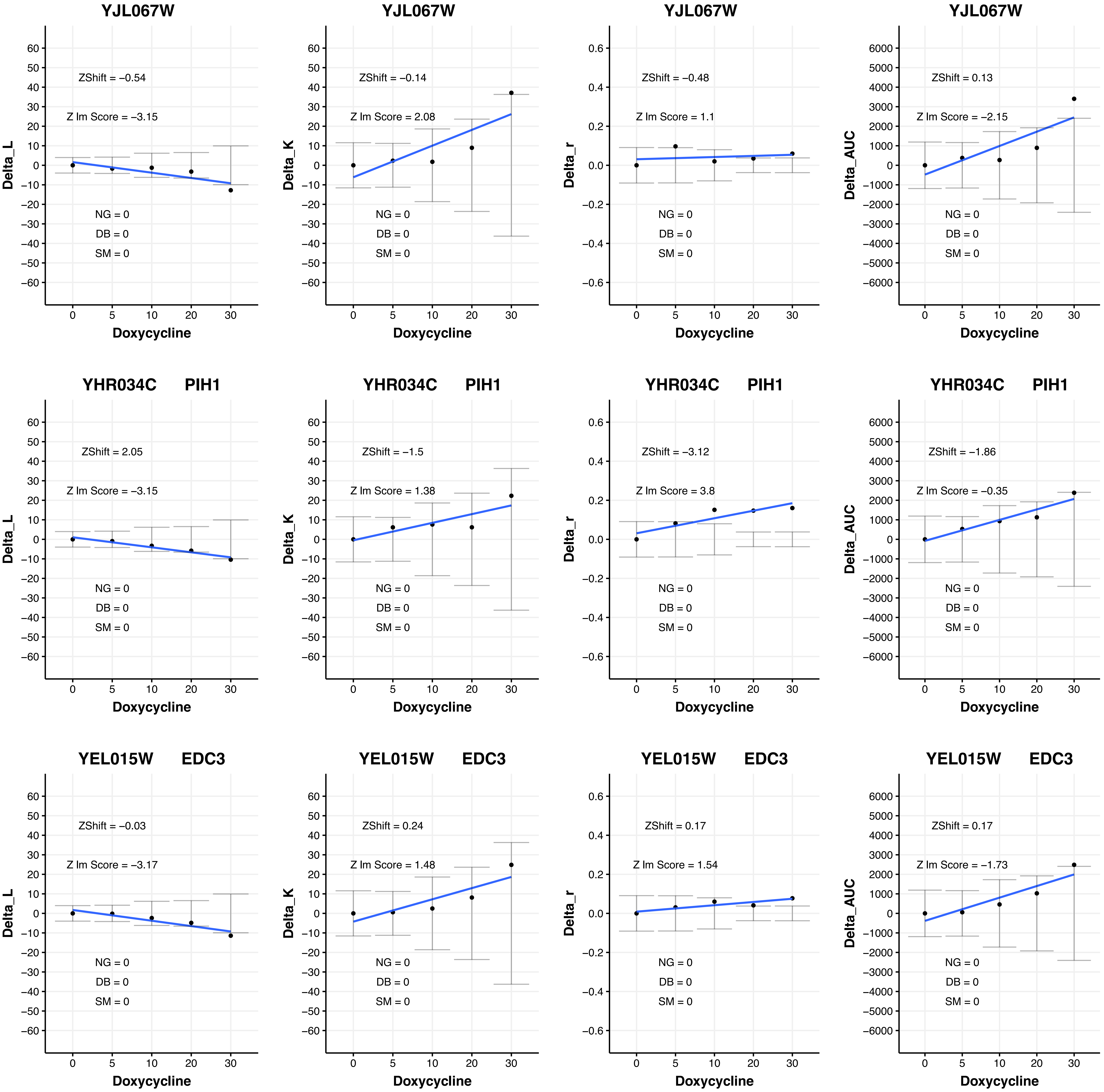

YJL067W
ZShift = −0.54
Z lm Score = −3.15
NG = 0
DB = 0
SM = 0
Delta_L
Doxycycline
YJL067W
ZShift = −0.14
Z lm Score = 2.08
NG = 0
DB = 0
SM = 0
Delta_K
Doxycycline
YJL067W
ZShift = −0.48
Z lm Score = 1.1
NG = 0
DB = 0
SM = 0
Delta_r
Doxycycline
YJL067W
ZShift = 0.13
Z lm Score = −2.15
NG = 0
DB = 0
SM = 0
Delta_AUC
Doxycycline
YHR034C PIH1
ZShift = 2.05
Z lm Score = −3.15
NG = 0
DB = 0
SM = 0
Delta_L
Doxycycline
YHR034C PIH1
ZShift = −1.5
Z lm Score = 1.38
NG = 0
DB = 0
SM = 0
Delta_K
Doxycycline
YHR034C PIH1
ZShift = −3.12
Z lm Score = 3.8
NG = 0
DB = 0
SM = 0
Delta_r
Doxycycline
YHR034C PIH1
ZShift = −1.86
Z lm Score = −0.35
NG = 0
DB = 0
SM = 0
Delta_AUC
Doxycycline
YEL015W EDC3
ZShift = −0.03
Z lm Score = −3.17
NG = 0
DB = 0
SM = 0
Delta_L
Doxycycline
YEL015W EDC3
ZShift = 0.24
Z lm Score = 1.48
NG = 0
DB = 0
SM = 0
Delta_K
Doxycycline
YEL015W EDC3
ZShift = 0.17
Z lm Score = 1.54
NG = 0
DB = 0
SM = 0
Delta_r
Doxycycline
YEL015W EDC3
ZShift = 0.17
Z lm Score = −1.73
NG = 0
DB = 0
SM = 0
Delta_AUC
Doxycycline

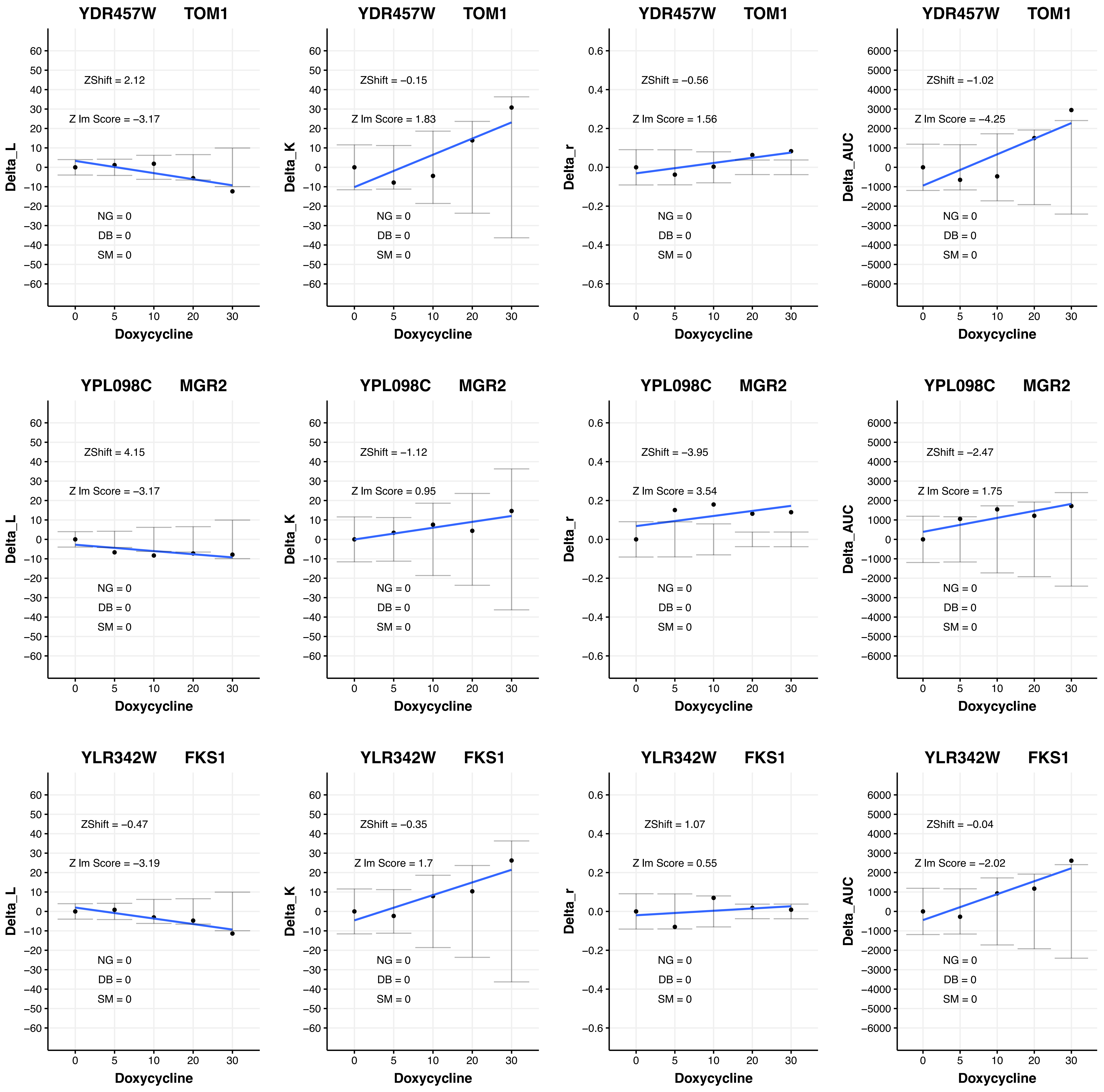

YDR457W TOM1
ZShift = 2.12
Z lm Score = −3.17
NG = 0
DB = 0
SM = 0
Delta_L
Doxycycline
YDR457W TOM1
ZShift = −0.15
Z lm Score = 1.83
NG = 0
DB = 0
SM = 0
Delta_K
Doxycycline
YDR457W TOM1
ZShift = −0.56
Z lm Score = 1.56
NG = 0
DB = 0
SM = 0
Delta_r
Doxycycline
YDR457W TOM1
ZShift = −1.02
Z lm Score = −4.25
NG = 0
DB = 0
SM = 0
Delta_AUC
Doxycycline
YPL098C MGR2
ZShift = 4.15
Z lm Score = −3.17
NG = 0
DB = 0
SM = 0
Delta_L
Doxycycline
YPL098C MGR2
ZShift = −1.12
Z lm Score = 0.95
NG = 0
DB = 0
SM = 0
Delta_K
Doxycycline
YPL098C MGR2
ZShift = −3.95
Z lm Score = 3.54
NG = 0
DB = 0
SM = 0
Delta_r
Doxycycline
YPL098C MGR2
ZShift = −2.47
Z lm Score = 1.75
NG = 0
DB = 0
SM = 0
Delta_AUC
Doxycycline
YLR342W FKS1
ZShift = −0.47
Z lm Score = −3.19
NG = 0
DB = 0
SM = 0
Delta_L
Doxycycline
YLR342W FKS1
ZShift = −0.35
Z lm Score = 1.7
NG = 0
DB = 0
SM = 0
Delta_K
Doxycycline
YLR342W FKS1
ZShift = 1.07
Z lm Score = 0.55
NG = 0
DB = 0
SM = 0
Delta_r
Doxycycline
YLR342W FKS1
ZShift = −0.04
Z lm Score = −2.02
NG = 0
DB = 0
SM = 0
Delta_AUC
Doxycycline

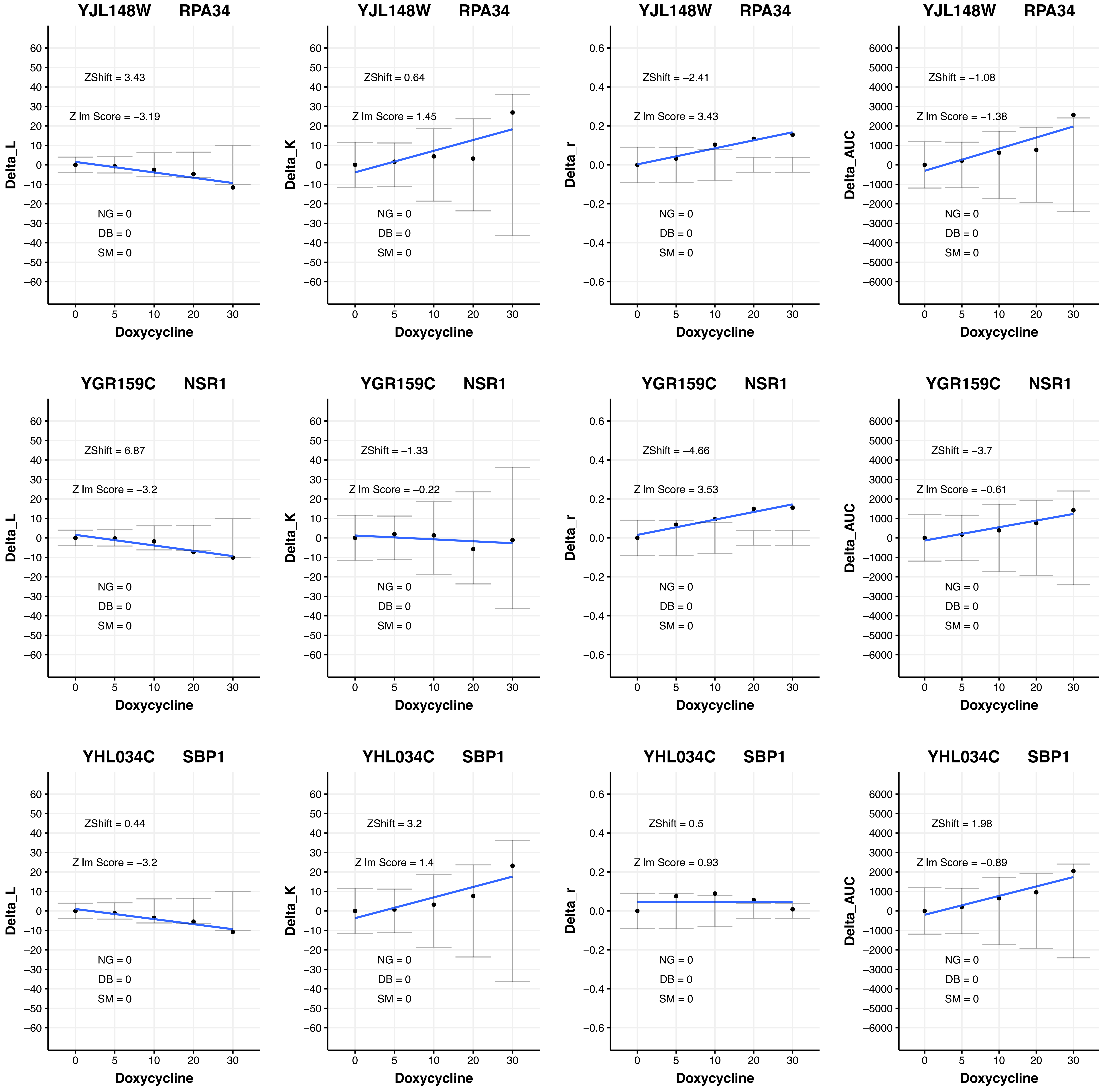
YJL148W RPA34
ZShift = 3.43
Z lm Score = −3.19
NG = 0
DB = 0
SM = 0
Delta_L
Doxycycline
YJL148W RPA34
ZShift = 0.64
Z lm Score = 1.45
NG = 0
DB = 0
SM = 0
Delta_K
Doxycycline
YJL148W RPA34
ZShift = −2.41
Z lm Score = 3.43
NG = 0
DB = 0
SM = 0
Delta_r
Doxycycline
YJL148W RPA34
ZShift = −1.08
Z lm Score = −1.38
NG = 0
DB = 0
SM = 0
Delta_AUC
Doxycycline
YGR159C NSR1
ZShift = 6.87
Z lm Score = −3.2
NG = 0
DB = 0
SM = 0
Delta_L
Doxycycline
YGR159C NSR1
ZShift = −1.33
Z lm Score = −0.22
NG = 0
DB = 0
SM = 0
Delta_K
Doxycycline
YGR159C NSR1
ZShift = −4.66
Z lm Score = 3.53
NG = 0
DB = 0
SM = 0
Delta_r
Doxycycline
YGR159C NSR1
ZShift = −3.7
Z lm Score = −0.61
NG = 0
DB = 0
SM = 0
Delta_AUC
Doxycycline
YHL034C SBP1
ZShift = 0.44
Z lm Score = −3.2
NG = 0
DB = 0
SM = 0
Delta_L
Doxycycline
YHL034C SBP1
ZShift = 3.2
Z lm Score = 1.4
NG = 0
DB = 0
SM = 0
Delta_K
Doxycycline
YHL034C SBP1
ZShift = 0.5
Z lm Score = 0.93
NG = 0
DB = 0
SM = 0
Delta_r
Doxycycline
YHL034C SBP1
ZShift = 1.98
Z lm Score = −0.89
NG = 0
DB = 0
SM = 0
Delta_AUC
Doxycycline

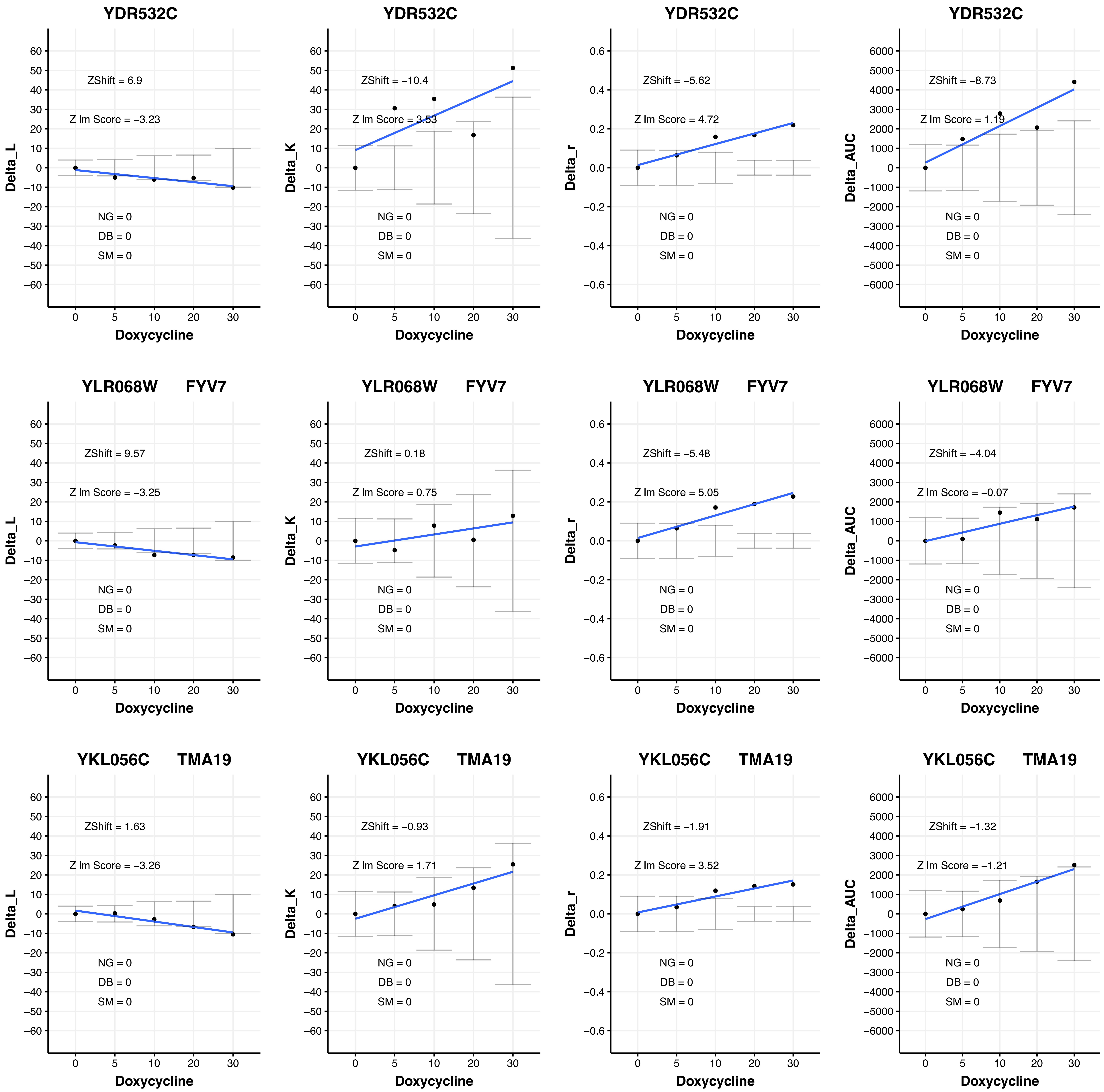

YDR532C
ZShift = 6.9
Z lm Score = −3.23
NG = 0
DB = 0
SM = 0
Delta_L
Doxycycline
YDR532C
ZShift = −10.4
Z lm Score = 3.53
NG = 0
DB = 0
SM = 0
Delta_K
Doxycycline
YDR532C
ZShift = −5.62
Z lm Score = 4.72
NG = 0
DB = 0
SM = 0
Delta_r
Doxycycline
YDR532C
ZShift = −8.73
Z lm Score = 1.19
NG = 0
DB = 0
SM = 0
Delta_AUC
Doxycycline
YLR068W FYV7
ZShift = 9.57
Z lm Score = −3.25
NG = 0
DB = 0
SM = 0
Delta_L
Doxycycline
YLR068W FYV7
ZShift = 0.18
Z lm Score = 0.75
NG = 0
DB = 0
SM = 0
Delta_K
Doxycycline
YLR068W FYV7
ZShift = −5.48
Z lm Score = 5.05
NG = 0
DB = 0
SM = 0
Delta_r
Doxycycline
YLR068W FYV7
ZShift = −4.04
Z lm Score = −0.07
NG = 0
DB = 0
SM = 0
Delta_AUC
Doxycycline
YKL056C TMA19
ZShift = 1.63
Z lm Score = −3.26
NG = 0
DB = 0
SM = 0
Delta_L
Doxycycline
YKL056C TMA19
ZShift = −0.93
Z lm Score = 1.71
NG = 0
DB = 0
SM = 0
Delta_K
Doxycycline
YKL056C TMA19
ZShift = −1.91
Z lm Score = 3.52
NG = 0
DB = 0
SM = 0
Delta_r
Doxycycline
YKL056C TMA19
ZShift = −1.32
Z lm Score = −1.21
NG = 0
DB = 0
SM = 0
Delta_AUC
Doxycycline

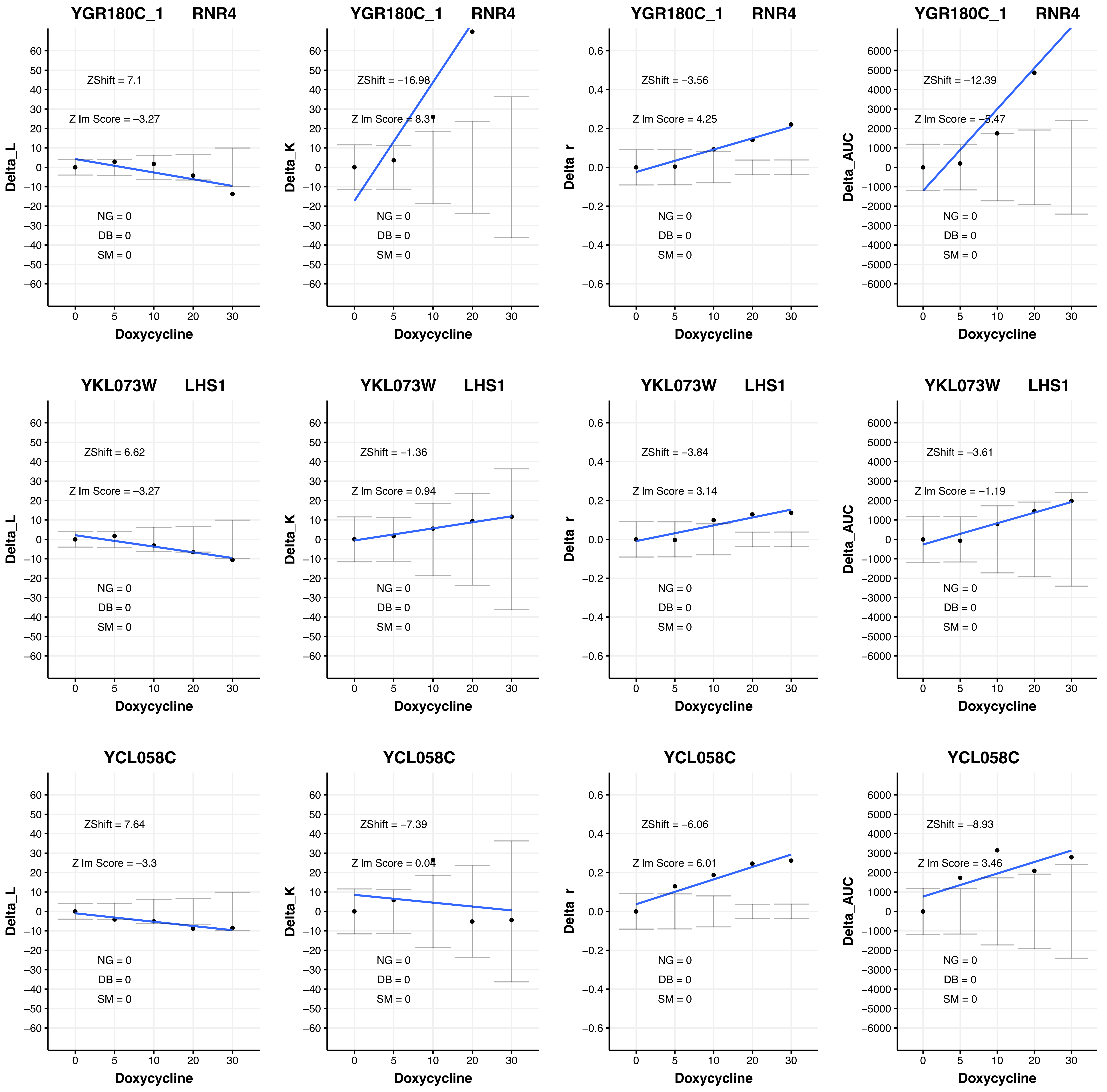
YGR180C_1 RNR4
ZShift = 7.1
Z lm Score = −3.27
NG = 0
DB = 0
SM = 0
Delta_L
Doxycycline
YGR180C_1 RNR4
ZShift = −16.98
Z lm Score = 8.31
NG = 0
DB = 0
SM = 0
Delta_K
Doxycycline
YGR180C_1 RNR4
ZShift = −3.56
Z lm Score = 4.25
NG = 0
DB = 0
SM = 0
Delta_r
Doxycycline
YGR180C_1 RNR4
ZShift = −12.39
Z lm Score = −5.47
NG = 0
DB = 0
SM = 0
Delta_AUC
Doxycycline
YKL073W LHS1
ZShift = 6.62
Z lm Score = −3.27
NG = 0
DB = 0
SM = 0
Delta_L
Doxycycline
YKL073W LHS1
ZShift = −1.36
Z lm Score = 0.94
NG = 0
DB = 0
SM = 0
Delta_K
Doxycycline
YKL073W LHS1
ZShift = −3.84
Z lm Score = 3.14
NG = 0
DB = 0
SM = 0
Delta_r
Doxycycline
YKL073W LHS1
ZShift = −3.61
Z lm Score = −1.19
NG = 0
DB = 0
SM = 0
Delta_AUC
Doxycycline
YCL058C
ZShift = 7.64
Z lm Score = −3.3
NG = 0
DB = 0
SM = 0
Delta_L
Doxycycline
YCL058C
ZShift = −7.39
Z lm Score = 0.04
NG = 0
DB = 0
SM = 0
Delta_K
Doxycycline
YCL058C
ZShift = −6.06
Z lm Score = 6.01
NG = 0
DB = 0
SM = 0
Delta_r
Doxycycline
YCL058C
ZShift = −8.93
Z lm Score = 3.46
NG = 0
DB = 0
SM = 0
Delta_AUC
Doxycycline

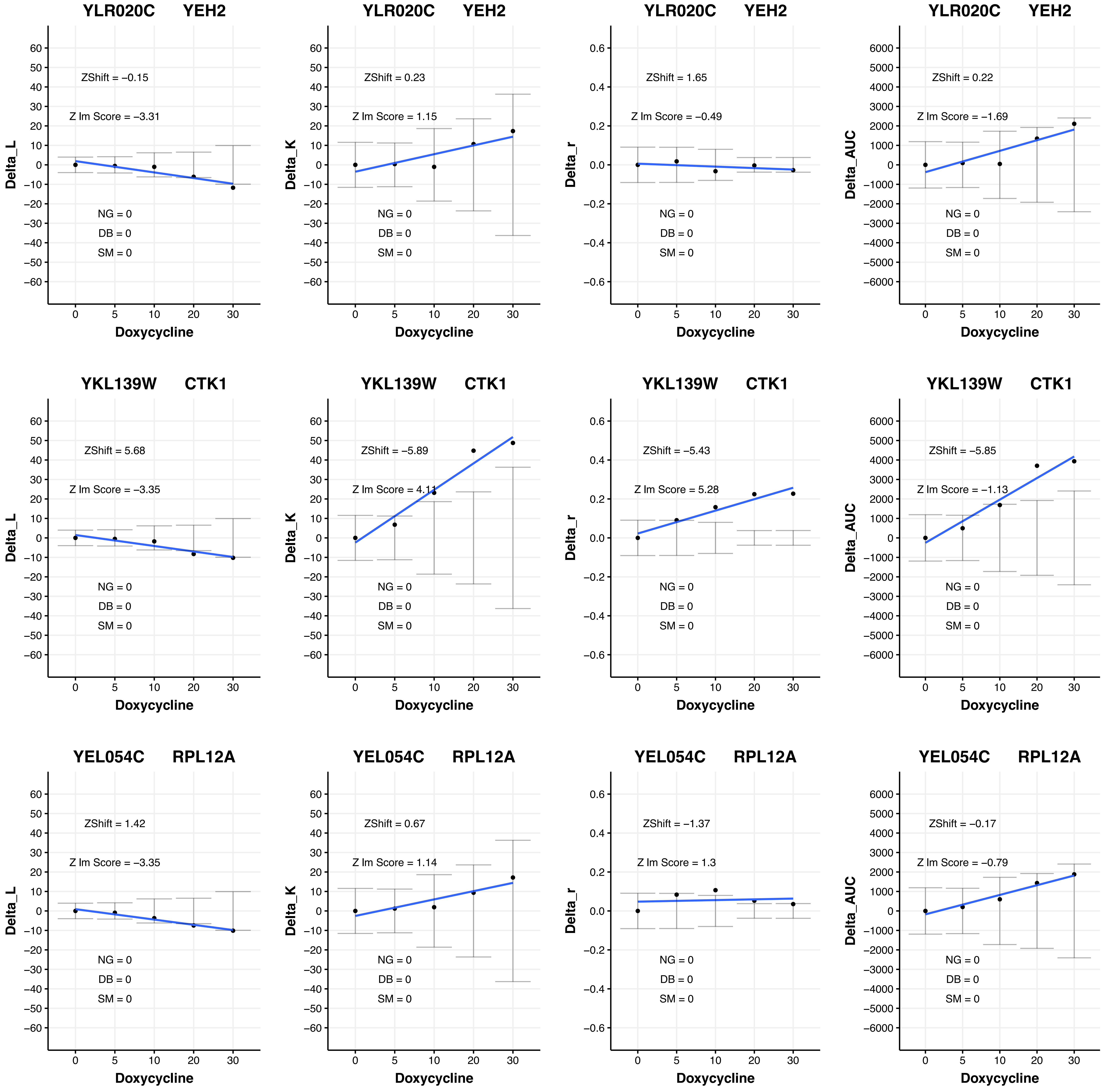

YLR020C YEH2
ZShift = −0.15
Z lm Score = −3.31
NG = 0
DB = 0
SM = 0
Delta_L
Doxycycline
YLR020C YEH2
ZShift = 0.23
Z lm Score = 1.15
NG = 0
DB = 0
SM = 0
Delta_K
Doxycycline
YLR020C YEH2
ZShift = 1.65
Z lm Score = −0.49
NG = 0
DB = 0
SM = 0
Delta_r
Doxycycline
YLR020C YEH2
ZShift = 0.22
Z lm Score = −1.69
NG = 0
DB = 0
SM = 0
Delta_AUC
Doxycycline
YKL139W CTK1
ZShift = 5.68
Z lm Score = −3.35
NG = 0
DB = 0
SM = 0
Delta_L
Doxycycline
YKL139W CTK1
ZShift = −5.89
Z lm Score = 4.11
NG = 0
DB = 0
SM = 0
Delta_K
Doxycycline
YKL139W CTK1
ZShift = −5.43
Z lm Score = 5.28
NG = 0
DB = 0
SM = 0
Delta_r
Doxycycline
YKL139W CTK1
ZShift = −5.85
Z lm Score = −1.13
NG = 0
DB = 0
SM = 0
Delta_AUC
Doxycycline
YEL054C RPL12A
ZShift = 1.42
Z lm Score = −3.35
NG = 0
DB = 0
SM = 0
Delta_L
Doxycycline
YEL054C RPL12A
ZShift = 0.67
Z lm Score = 1.14
NG = 0
DB = 0
SM = 0
Delta_K
Doxycycline
YEL054C RPL12A
ZShift = −1.37
Z lm Score = 1.3
NG = 0
DB = 0
SM = 0
Delta_r
Doxycycline
YEL054C RPL12A
ZShift = −0.17
Z lm Score = −0.79
NG = 0
DB = 0
SM = 0
Delta_AUC
Doxycycline

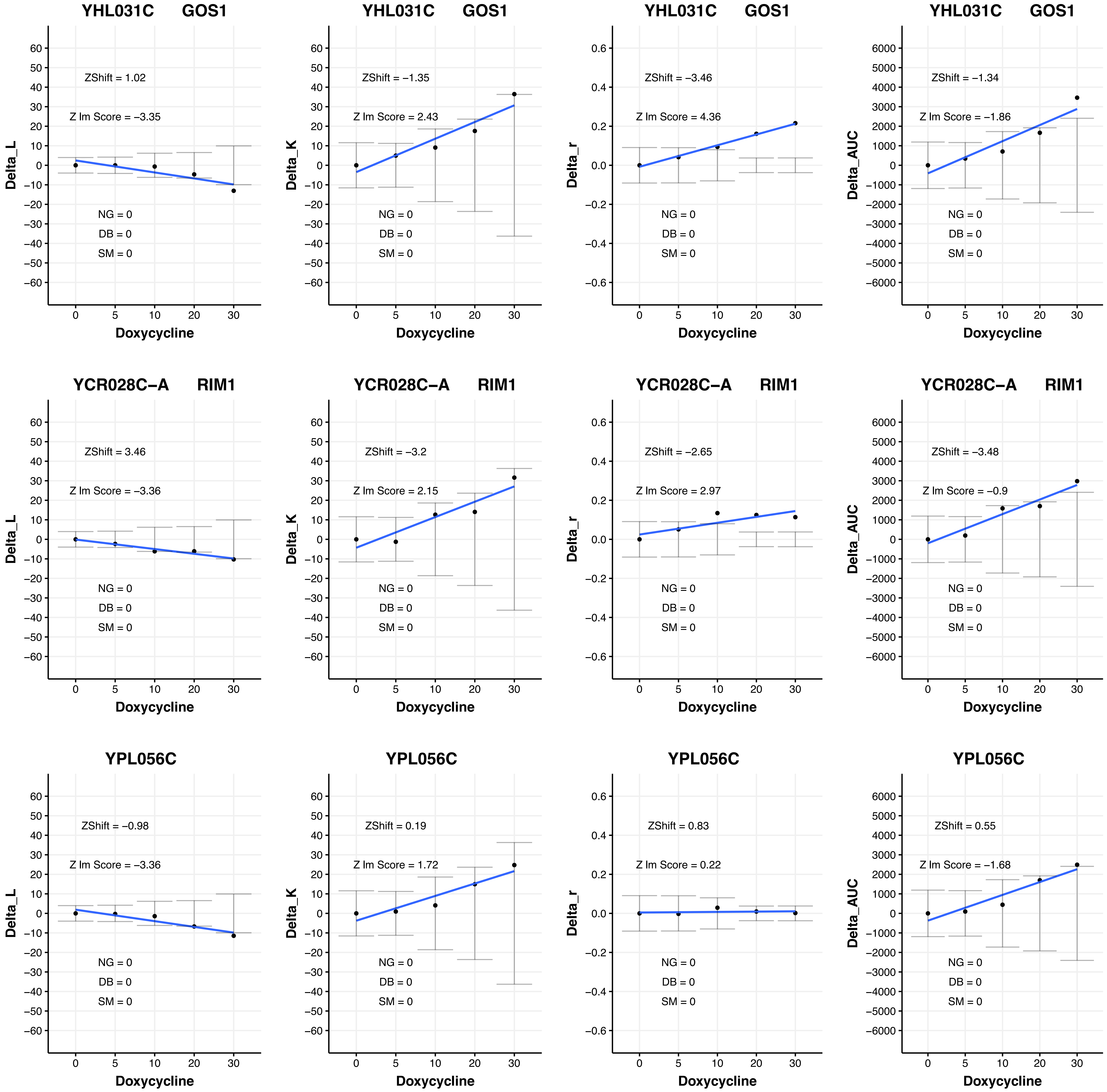

YHL031C GOS1
ZShift = 1.02
Z lm Score = −3.35
NG = 0
DB = 0
SM = 0
Delta_L
Doxycycline
YHL031C GOS1
ZShift = −1.35
Z lm Score = 2.43
NG = 0
DB = 0
SM = 0
Delta_K
Doxycycline
YHL031C GOS1
ZShift = −3.46
Z lm Score = 4.36
NG = 0
DB = 0
SM = 0
Delta_r
Doxycycline
YHL031C GOS1
ZShift = −1.34
Z lm Score = −1.86
NG = 0
DB = 0
SM = 0
Delta_AUC
Doxycycline
YCR028C−A RIM1
ZShift = 3.46
Z lm Score = −3.36
NG = 0
DB = 0
SM = 0
Delta_L
Doxycycline
YCR028C−A RIM1
ZShift = −3.2
Z lm Score = 2.15
NG = 0
DB = 0
SM = 0
Delta_K
Doxycycline
YCR028C−A RIM1
ZShift = −2.65
Z lm Score = 2.97
NG = 0
DB = 0
SM = 0
Delta_r
Doxycycline
YCR028C−A RIM1
ZShift = −3.48
Z lm Score = −0.9
NG = 0
DB = 0
SM = 0
Delta_AUC
Doxycycline
YPL056C
ZShift = −0.98
Z lm Score = −3.36
NG = 0
DB = 0
SM = 0
Delta_L
Doxycycline
YPL056C
ZShift = 0.19
Z lm Score = 1.72
NG = 0
DB = 0
SM = 0
Delta_K
Doxycycline
YPL056C
ZShift = 0.83
Z lm Score = 0.22
NG = 0
DB = 0
SM = 0
Delta_r
Doxycycline
YPL056C
ZShift = 0.55
Z lm Score = −1.68
NG = 0
DB = 0
SM = 0
Delta_AUC
Doxycycline

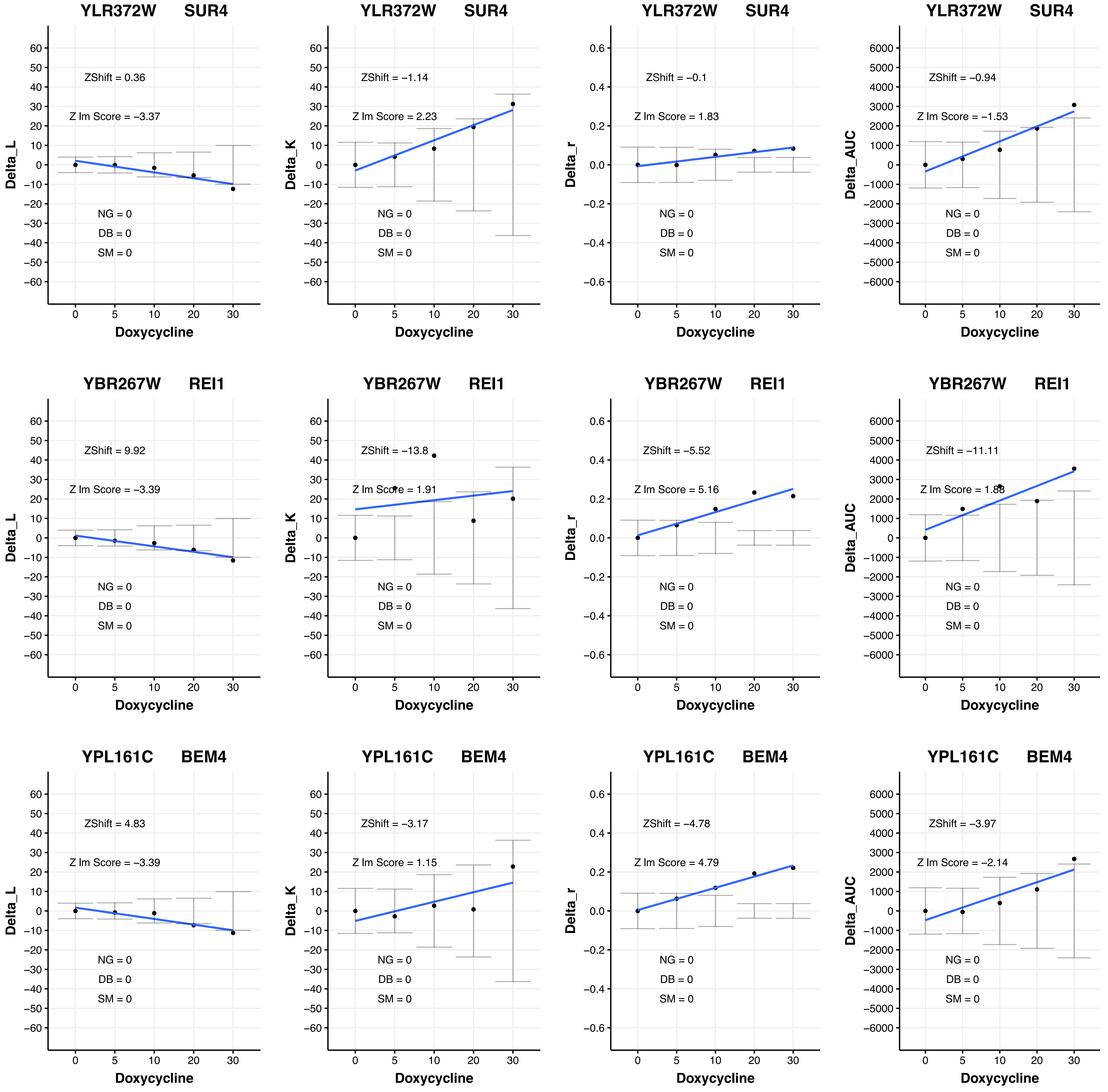

YLR372W SUR4
ZShift = 0.36
Z lm Score = −3.37
NG = 0
DB = 0
SM = 0
Delta_L
Doxycycline
YLR372W SUR4
ZShift = −1.14
Z lm Score = 2.23
NG = 0
DB = 0
SM = 0
Delta_K
Doxycycline
YLR372W SUR4
ZShift = −0.1
Z lm Score = 1.83
NG = 0
DB = 0
SM = 0
Delta_r
Doxycycline
YLR372W SUR4
ZShift = −0.94
Z lm Score = −1.53
NG = 0
DB = 0
SM = 0
Delta_AUC
Doxycycline
YBR267W REI1
ZShift = 9.92
Z lm Score = −3.39
NG = 0
DB = 0
SM = 0
Delta_L
Doxycycline
YBR267W REI1
ZShift = −13.8
Z lm Score = 1.91
NG = 0
DB = 0
SM = 0
Delta_K
Doxycycline
YBR267W REI1
ZShift = −5.52
Z lm Score = 5.16
NG = 0
DB = 0
SM = 0
Delta_r
Doxycycline
YBR267W REI1
ZShift = −11.11
Z lm Score = 1.88
NG = 0
DB = 0
SM = 0
Delta_AUC
Doxycycline
YPL161C BEM4
ZShift = 4.83
Z lm Score = −3.39
NG = 0
DB = 0
SM = 0
Delta_L
Doxycycline
YPL161C BEM4
ZShift = −3.17
Z lm Score = 1.15
NG = 0
DB = 0
SM = 0
Delta_K
Doxycycline
YPL161C BEM4
ZShift = −4.78
Z lm Score = 4.79
NG = 0
DB = 0
SM = 0
Delta_r
Doxycycline
YPL161C BEM4
ZShift = −3.97
Z lm Score = −2.14
NG = 0
DB = 0
SM = 0
Delta_AUC
Doxycycline

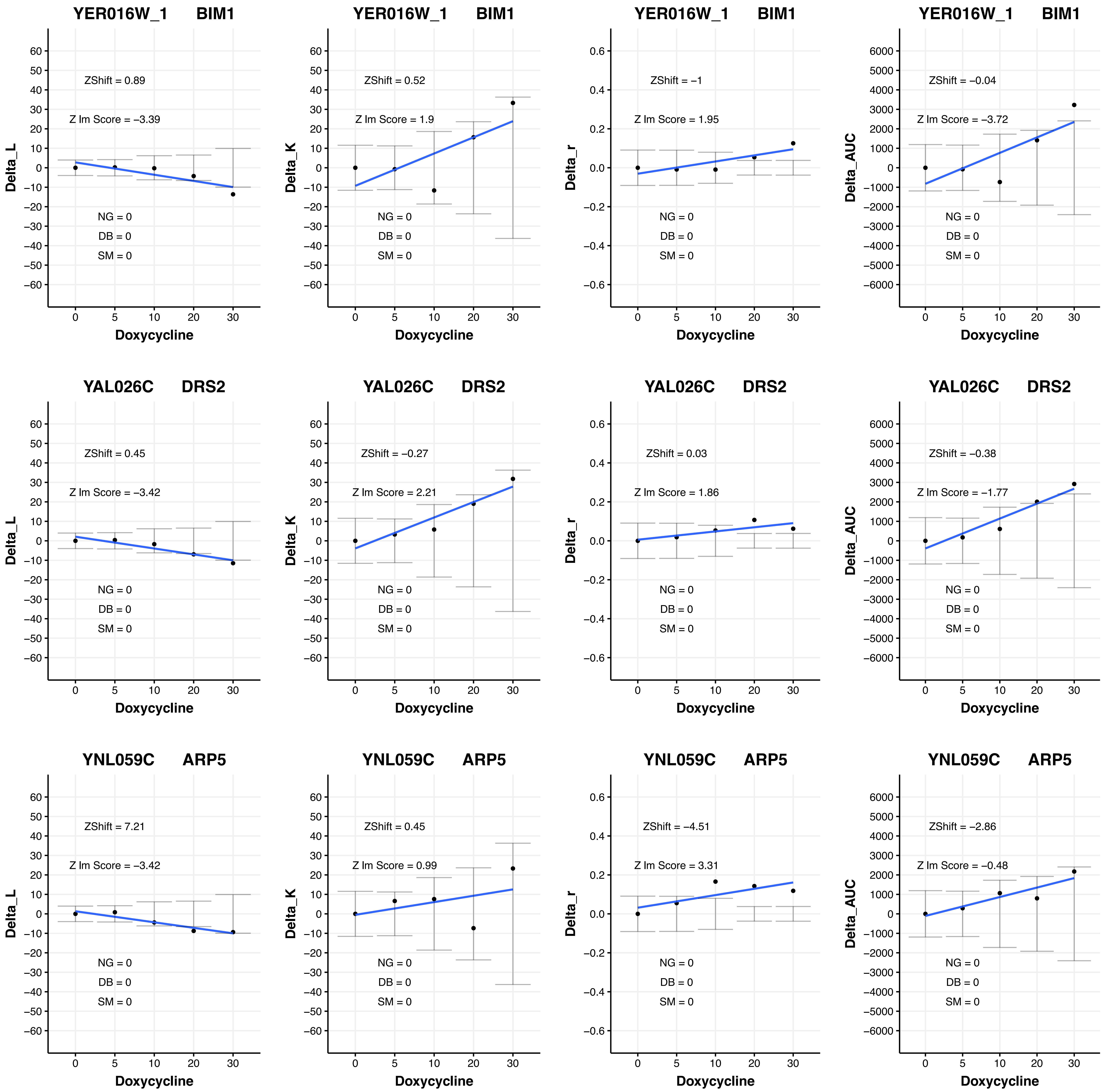
YER016W_1 BIM1
ZShift = 0.89
Z lm Score = −3.39
NG = 0
DB = 0
SM = 0
Delta_L
Doxycycline
YER016W_1 BIM1
ZShift = 0.52
Z lm Score = 1.9
NG = 0
DB = 0
SM = 0
Delta_K
Doxycycline
YER016W_1 BIM1
ZShift = −1
Z lm Score = 1.95
NG = 0
DB = 0
SM = 0
Delta_r
Doxycycline
YER016W_1 BIM1
ZShift = −0.04
Z lm Score = −3.72
NG = 0
DB = 0
SM = 0
Delta_AUC
Doxycycline
YAL026C DRS2
ZShift = 0.45
Z lm Score = −3.42
NG = 0
DB = 0
SM = 0
Delta_L
Doxycycline
YAL026C DRS2
ZShift = −0.27
Z lm Score = 2.21
NG = 0
DB = 0
SM = 0
Delta_K
Doxycycline
YAL026C DRS2
ZShift = 0.03
Z lm Score = 1.86
NG = 0
DB = 0
SM = 0
Delta_r
Doxycycline
YAL026C DRS2
ZShift = −0.38
Z lm Score = −1.77
NG = 0
DB = 0
SM = 0
Delta_AUC
Doxycycline
YNL059C ARP5
ZShift = 7.21
Z lm Score = −3.42
NG = 0
DB = 0
SM = 0
Delta_L
Doxycycline
YNL059C ARP5
ZShift = 0.45
Z lm Score = 0.99
NG = 0
DB = 0
SM = 0
Delta_K
Doxycycline
YNL059C ARP5
ZShift = −4.51
Z lm Score = 3.31
NG = 0
DB = 0
SM = 0
Delta_r
Doxycycline
YNL059C ARP5
ZShift = −2.86
Z lm Score = −0.48
NG = 0
DB = 0
SM = 0
Delta_AUC
Doxycycline

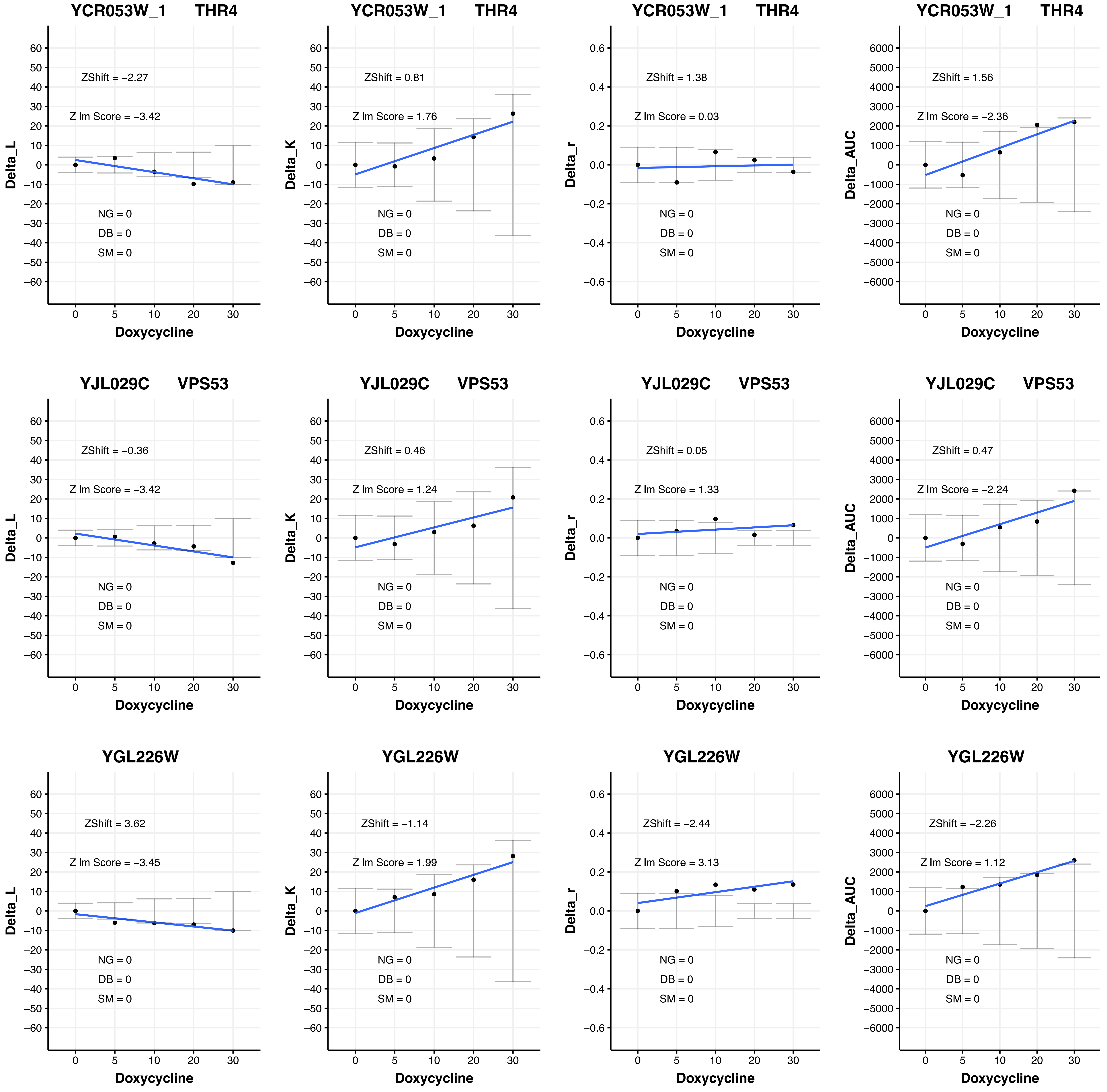

YCR053W_1 THR4
ZShift = −2.27
Z lm Score = −3.42
NG = 0
DB = 0
SM = 0
Delta_L
Doxycycline
YCR053W_1 THR4
ZShift = 0.81
Z lm Score = 1.76
NG = 0
DB = 0
SM = 0
Delta_K
Doxycycline
YCR053W_1 THR4
ZShift = 1.38
Z lm Score = 0.03
NG = 0
DB = 0
SM = 0
Delta_r
Doxycycline
YCR053W_1 THR4
ZShift = 1.56
Z lm Score = −2.36
NG = 0
DB = 0
SM = 0
Delta_AUC
Doxycycline
YJL029C VPS53
ZShift = −0.36
Z lm Score = −3.42
NG = 0
DB = 0
SM = 0
Delta_L
Doxycycline
YJL029C VPS53
ZShift = 0.46
Z lm Score = 1.24
NG = 0
DB = 0
SM = 0
Delta_K
Doxycycline
YJL029C VPS53
ZShift = 0.05
Z lm Score = 1.33
NG = 0
DB = 0
SM = 0
Delta_r
Doxycycline
YJL029C VPS53
ZShift = 0.47
Z lm Score = −2.24
NG = 0
DB = 0
SM = 0
Delta_AUC
Doxycycline
YGL226W
ZShift = 3.62
Z lm Score = −3.45
NG = 0
DB = 0
SM = 0
Delta_L
Doxycycline
YGL226W
ZShift = −1.14
Z lm Score = 1.99
NG = 0
DB = 0
SM = 0
Delta_K
Doxycycline
YGL226W
ZShift = −2.44
Z lm Score = 3.13
NG = 0
DB = 0
SM = 0
Delta_r
Doxycycline
YGL226W
ZShift = −2.26
Z lm Score = 1.12
NG = 0
DB = 0
SM = 0
Delta_AUC
Doxycycline

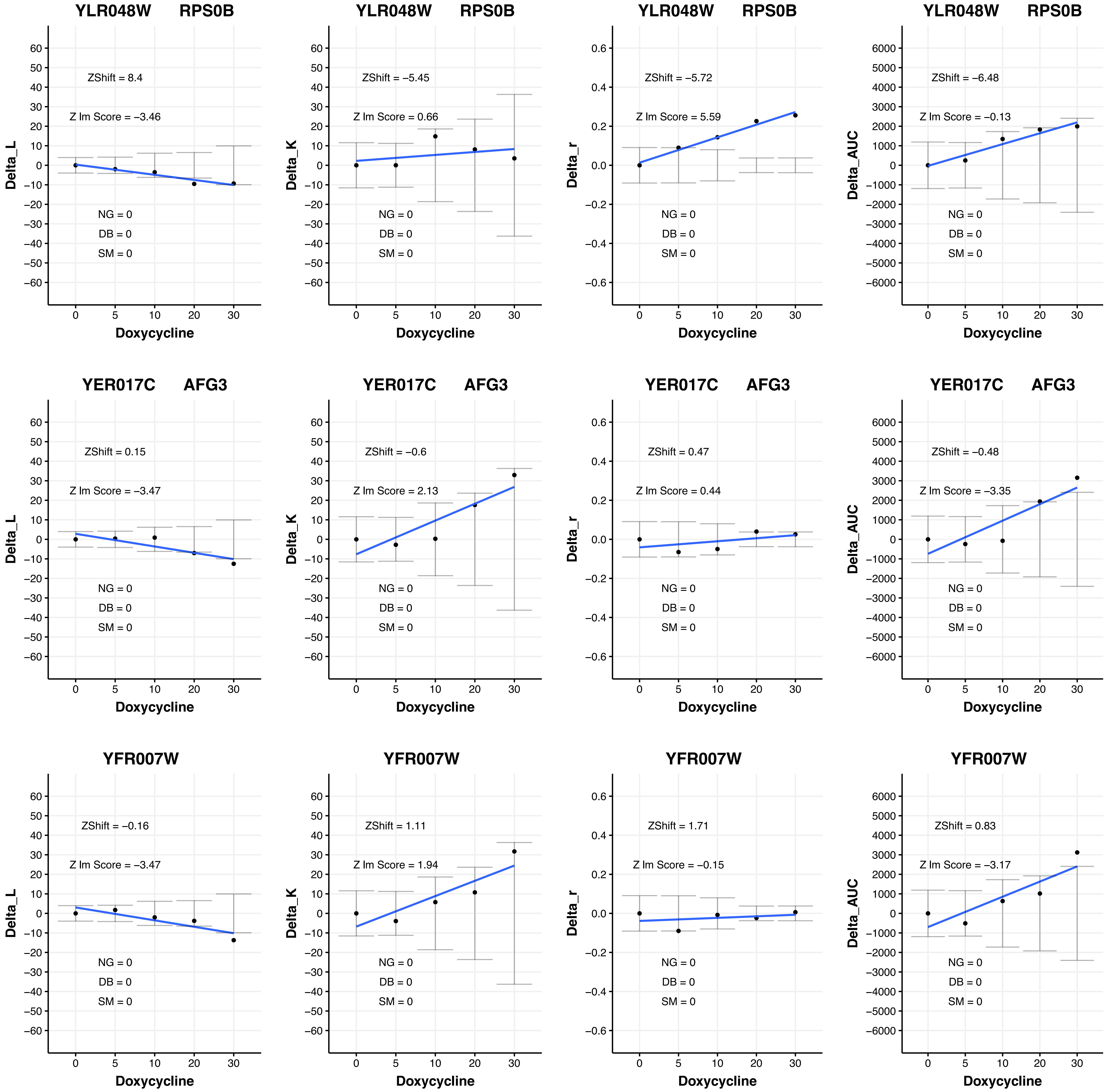

YLR048W RPS0B
ZShift = 8.4
Z lm Score = −3.46
NG = 0
DB = 0
SM = 0
Delta_L
Doxycycline
YLR048W RPS0B
ZShift = −5.45
Z lm Score = 0.66
NG = 0
DB = 0
SM = 0
Delta_K
Doxycycline
YLR048W RPS0B
ZShift = −5.72
Z lm Score = 5.59
NG = 0
DB = 0
SM = 0
Delta_r
Doxycycline
YLR048W RPS0B
ZShift = −6.48
Z lm Score = −0.13
NG = 0
DB = 0
SM = 0
Delta_AUC
Doxycycline
YER017C AFG3
ZShift = 0.15
Z lm Score = −3.47
NG = 0
DB = 0
SM = 0
Delta_L
Doxycycline
YER017C AFG3
ZShift = −0.6
Z lm Score = 2.13
NG = 0
DB = 0
SM = 0
Delta_K
Doxycycline
YER017C AFG3
ZShift = 0.47
Z lm Score = 0.44
NG = 0
DB = 0
SM = 0
Delta_r
Doxycycline
YER017C AFG3
ZShift = −0.48
Z lm Score = −3.35
NG = 0
DB = 0
SM = 0
Delta_AUC
Doxycycline
YFR007W
ZShift = −0.16
Z lm Score = −3.47
NG = 0
DB = 0
SM = 0
Delta_L
Doxycycline
YFR007W
ZShift = 1.11
Z lm Score = 1.94
NG = 0
DB = 0
SM = 0
Delta_K
Doxycycline
YFR007W
ZShift = 1.71
Z lm Score = −0.15
NG = 0
DB = 0
SM = 0
Delta_r
Doxycycline
YFR007W
ZShift = 0.83
Z lm Score = −3.17
NG = 0
DB = 0
SM = 0
Delta_AUC
Doxycycline

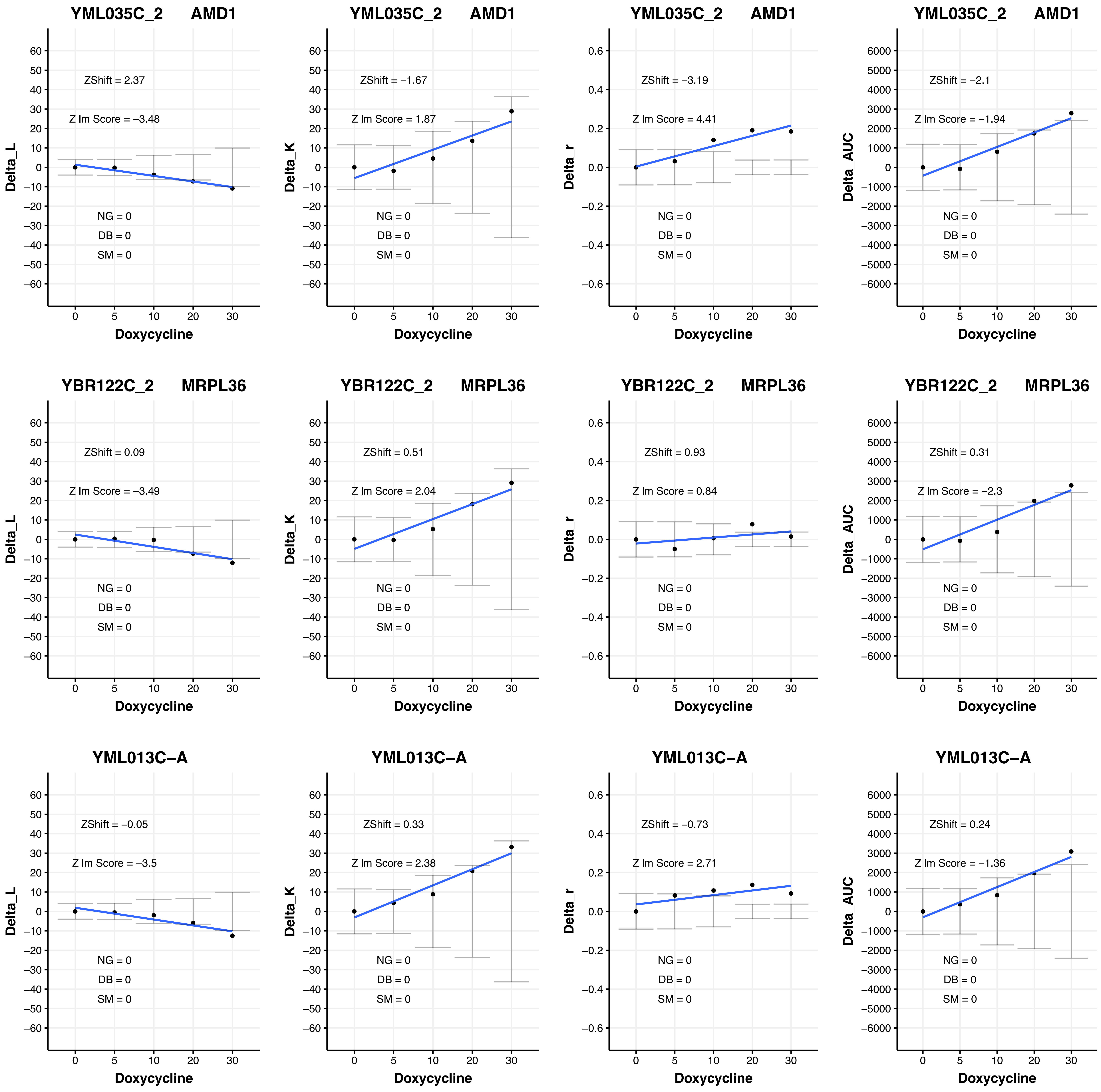
YML035C_2 AMD1
ZShift = 2.37
Z lm Score = −3.48
NG = 0
DB = 0
SM = 0
Delta_L
Doxycycline
YML035C_2 AMD1
ZShift = −1.67
Z lm Score = 1.87
NG = 0
DB = 0
SM = 0
Delta_K
Doxycycline
YML035C_2 AMD1
ZShift = −3.19
Z lm Score = 4.41
NG = 0
DB = 0
SM = 0
Delta_r
Doxycycline
YML035C_2 AMD1
ZShift = −2.1
Z lm Score = −1.94
NG = 0
DB = 0
SM = 0
Delta_AUC
Doxycycline
YBR122C_2 MRPL36
ZShift = 0.09
Z lm Score = −3.49
NG = 0
DB = 0
SM = 0
Delta_L
Doxycycline
YBR122C_2 MRPL36
ZShift = 0.51
Z lm Score = 2.04
NG = 0
DB = 0
SM = 0
Delta_K
Doxycycline
YBR122C_2 MRPL36
ZShift = 0.93
Z lm Score = 0.84
NG = 0
DB = 0
SM = 0
Delta_r
Doxycycline
YBR122C_2 MRPL36
ZShift = 0.31
Z lm Score = −2.3
NG = 0
DB = 0
SM = 0
Delta_AUC
Doxycycline
YML013C−A
ZShift = −0.05
Z lm Score = −3.5
NG = 0
DB = 0
SM = 0
Delta_L
Doxycycline
YML013C−A
ZShift = 0.33
Z lm Score = 2.38
NG = 0
DB = 0
SM = 0
Delta_K
Doxycycline
YML013C−A
ZShift = −0.73
Z lm Score = 2.71
NG = 0
DB = 0
SM = 0
Delta_r
Doxycycline
YML013C−A
ZShift = 0.24
Z lm Score = −1.36
NG = 0
DB = 0
SM = 0
Delta_AUC
Doxycycline

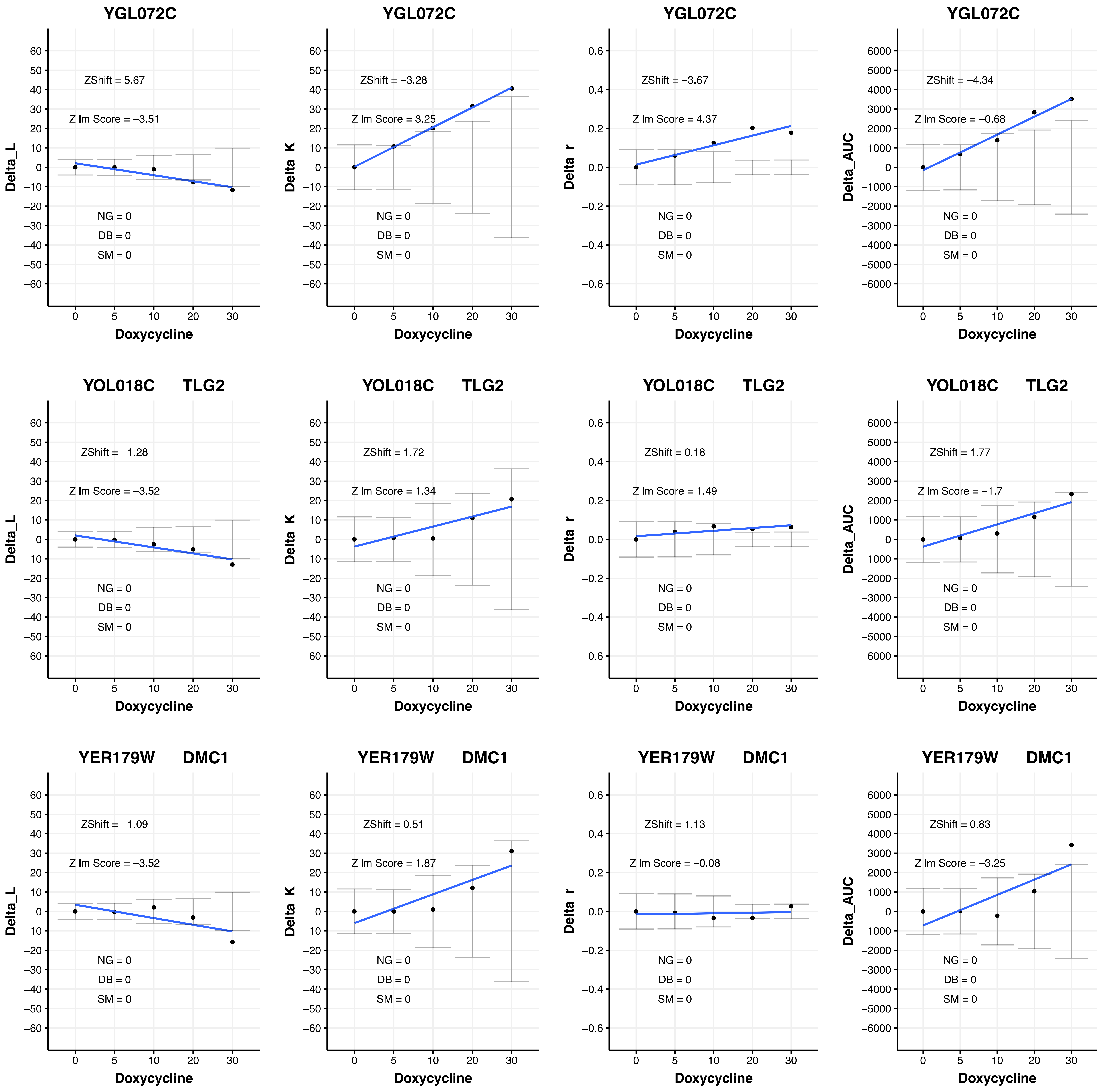
YGL072C
ZShift = 5.67
Z lm Score = −3.51
NG = 0
DB = 0
SM = 0
Delta_L
Doxycycline
YGL072C
ZShift = −3.28
Z lm Score = 3.25
NG = 0
DB = 0
SM = 0
Delta_K
Doxycycline
YGL072C
ZShift = −3.67
Z lm Score = 4.37
NG = 0
DB = 0
SM = 0
Delta_r
Doxycycline
YGL072C
ZShift = −4.34
Z lm Score = −0.68
NG = 0
DB = 0
SM = 0
Delta_AUC
Doxycycline
YOL018C TLG2
ZShift = −1.28
Z lm Score = −3.52
NG = 0
DB = 0
SM = 0
Delta_L
Doxycycline
YOL018C TLG2
ZShift = 1.72
Z lm Score = 1.34
NG = 0
DB = 0
SM = 0
Delta_K
Doxycycline
YOL018C TLG2
ZShift = 0.18
Z lm Score = 1.49
NG = 0
DB = 0
SM = 0
Delta_r
Doxycycline
YOL018C TLG2
ZShift = 1.77
Z lm Score = −1.7
NG = 0
DB = 0
SM = 0
Delta_AUC
Doxycycline
YER179W DMC1
ZShift = −1.09
Z lm Score = −3.52
NG = 0
DB = 0
SM = 0
Delta_L
Doxycycline
YER179W DMC1
ZShift = 0.51
Z lm Score = 1.87
NG = 0
DB = 0
SM = 0
Delta_K
Doxycycline
YER179W DMC1
ZShift = 1.13
Z lm Score = −0.08
NG = 0
DB = 0
SM = 0
Delta_r
Doxycycline
YER179W DMC1
ZShift = 0.83
Z lm Score = −3.25
NG = 0
DB = 0
SM = 0
Delta_AUC
Doxycycline

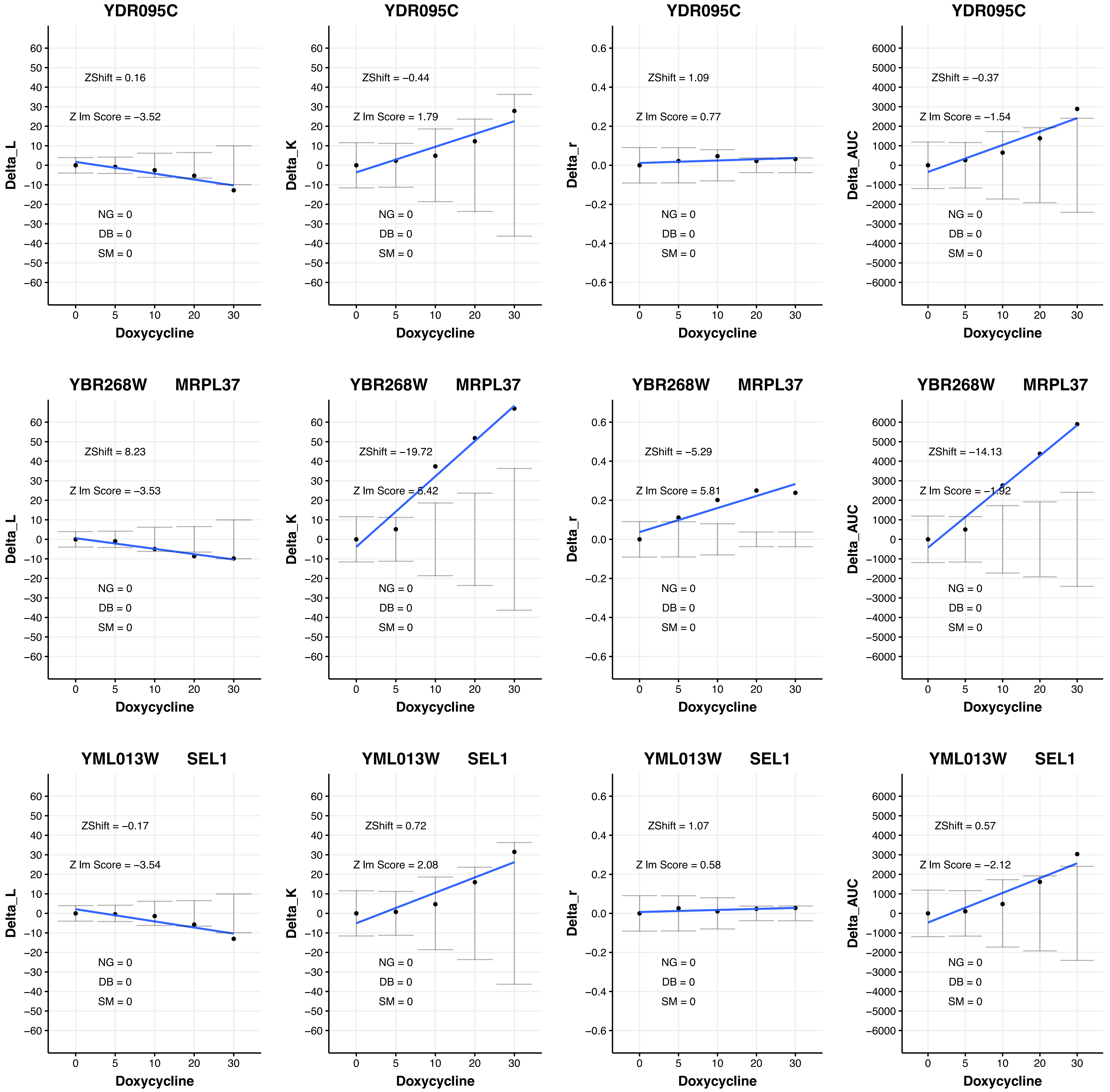

YDR095C
ZShift = 0.16
Z lm Score = −3.52
NG = 0
DB = 0
SM = 0
Delta_L
Doxycycline
YDR095C
ZShift = −0.44
Z lm Score = 1.79
NG = 0
DB = 0
SM = 0
Delta_K
Doxycycline
YDR095C
ZShift = 1.09
Z lm Score = 0.77
NG = 0
DB = 0
SM = 0
Delta_r
Doxycycline
YDR095C
ZShift = −0.37
Z lm Score = −1.54
NG = 0
DB = 0
SM = 0
Delta_AUC
Doxycycline
YBR268W MRPL37
ZShift = 8.23
Z lm Score = −3.53
NG = 0
DB = 0
SM = 0
Delta_L
Doxycycline
YBR268W MRPL37
ZShift = −19.72
Z lm Score = 5.42
NG = 0
DB = 0
SM = 0
Delta_K
Doxycycline
YBR268W MRPL37
ZShift = −5.29
Z lm Score = 5.81
NG = 0
DB = 0
SM = 0
Delta_r
Doxycycline
YBR268W MRPL37
ZShift = −14.13
Z lm Score = −1.92
NG = 0
DB = 0
SM = 0
Delta_AUC
Doxycycline
YML013W SEL1
ZShift = −0.17
Z lm Score = −3.54
NG = 0
DB = 0
SM = 0
Delta_L
Doxycycline
YML013W SEL1
ZShift = 0.72
Z lm Score = 2.08
NG = 0
DB = 0
SM = 0
Delta_K
Doxycycline
YML013W SEL1
ZShift = 1.07
Z lm Score = 0.58
NG = 0
DB = 0
SM = 0
Delta_r
Doxycycline
YML013W SEL1
ZShift = 0.57
Z lm Score = −2.12
NG = 0
DB = 0
SM = 0
Delta_AUC
Doxycycline

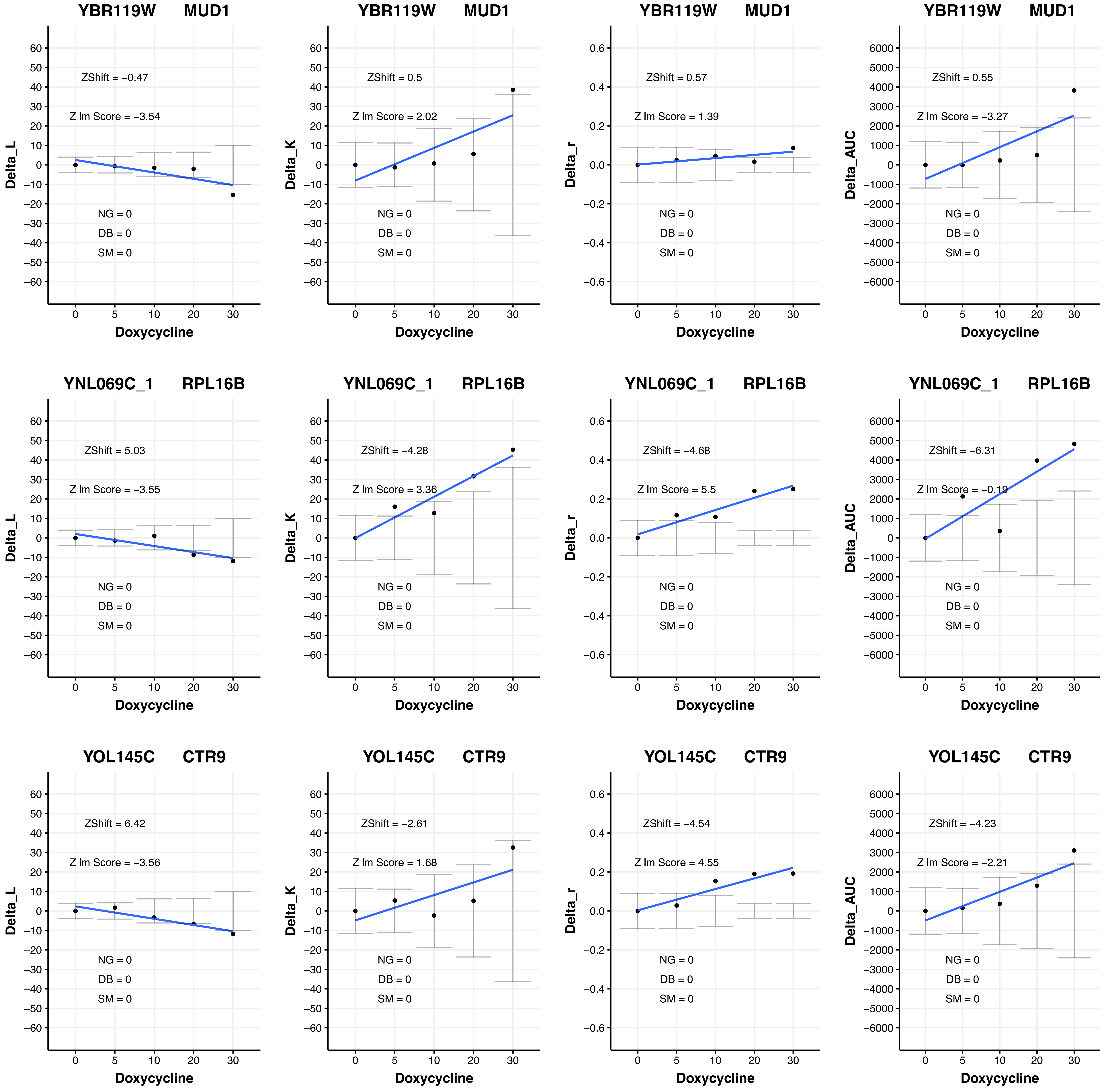
YBR119W MUD1
ZShift = −0.47
Z lm Score = −3.54
NG = 0
DB = 0
SM = 0
Delta_L
Doxycycline
YBR119W MUD1
ZShift = 0.5
Z lm Score = 2.02
NG = 0
DB = 0
SM = 0
Delta_K
Doxycycline
YBR119W MUD1
ZShift = 0.57
Z lm Score = 1.39
NG = 0
DB = 0
SM = 0
Delta_r
Doxycycline
YBR119W MUD1
ZShift = 0.55
Z lm Score = −3.27
NG = 0
DB = 0
SM = 0
Delta_AUC
Doxycycline
YNL069C_1 RPL16B
ZShift = 5.03
Z lm Score = −3.55
NG = 0
DB = 0
SM = 0
Delta_L
Doxycycline
YNL069C_1 RPL16B
ZShift = −4.28
Z lm Score = 3.36
NG = 0
DB = 0
SM = 0
Delta_K
Doxycycline
YNL069C_1 RPL16B
ZShift = −4.68
Z lm Score = 5.5
NG = 0
DB = 0
SM = 0
Delta_r
Doxycycline
YNL069C_1 RPL16B
ZShift = −6.31
Z lm Score = −0.19
NG = 0
DB = 0
SM = 0
Delta_AUC
Doxycycline
YOL145C CTR9
ZShift = 6.42
Z lm Score = −3.56
NG = 0
DB = 0
SM = 0
Delta_L
Doxycycline
YOL145C CTR9
ZShift = −2.61
Z lm Score = 1.68
NG = 0
DB = 0
SM = 0
Delta_K
Doxycycline
YOL145C CTR9
ZShift = −4.54
Z lm Score = 4.55
NG = 0
DB = 0
SM = 0
Delta_r
Doxycycline
YOL145C CTR9
ZShift = −4.23
Z lm Score = −2.21
NG = 0
DB = 0
SM = 0
Delta_AUC
Doxycycline

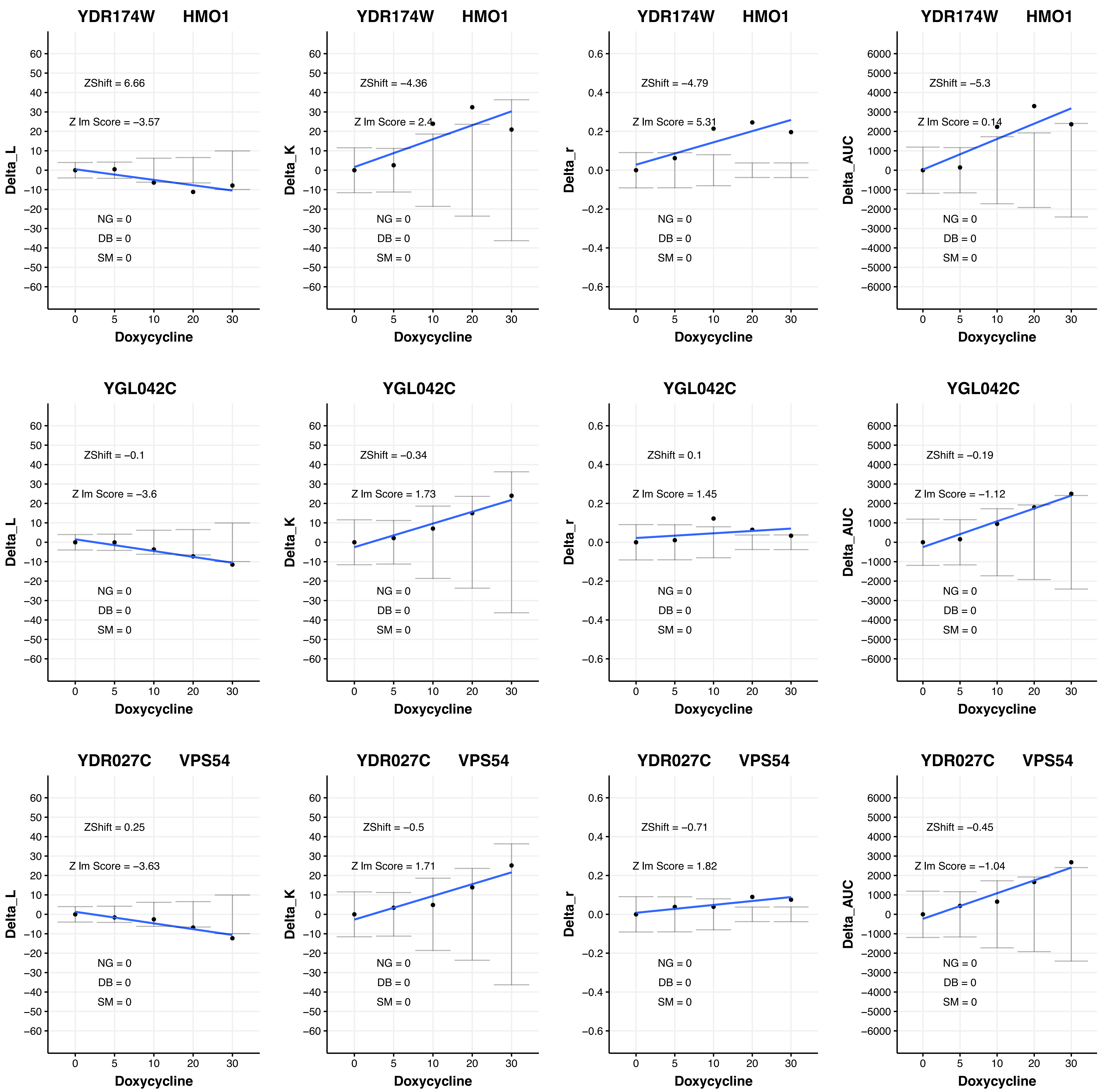
YDR174W HMO1
ZShift = 6.66
Z lm Score = −3.57
ZShift = −4.36
Z lm Score = 2.4
ZShift = −4.79
Z lm Score = 5.31
ZShift = −5.3
Z lm Score = 0.14
YGL042C
ZShift = −0.1
Z lm Score = −3.6
ZShift = −0.34
Z lm Score = 1.73
ZShift = 0.1
Z lm Score = 1.45
ZShift = −0.19
Z lm Score = −1.12
YDR027C VPS54
ZShift = 0.25
Z lm Score = −3.63
ZShift = −0.5
Z lm Score = 1.71
ZShift = −0.71
Z lm Score = 1.82
ZShift = −0.45
Z lm Score = −1.04
NG = 0
DB = 0
SM = 0
Delta_L
Delta_K
Delta_r
Delta_AUC
Doxycycline

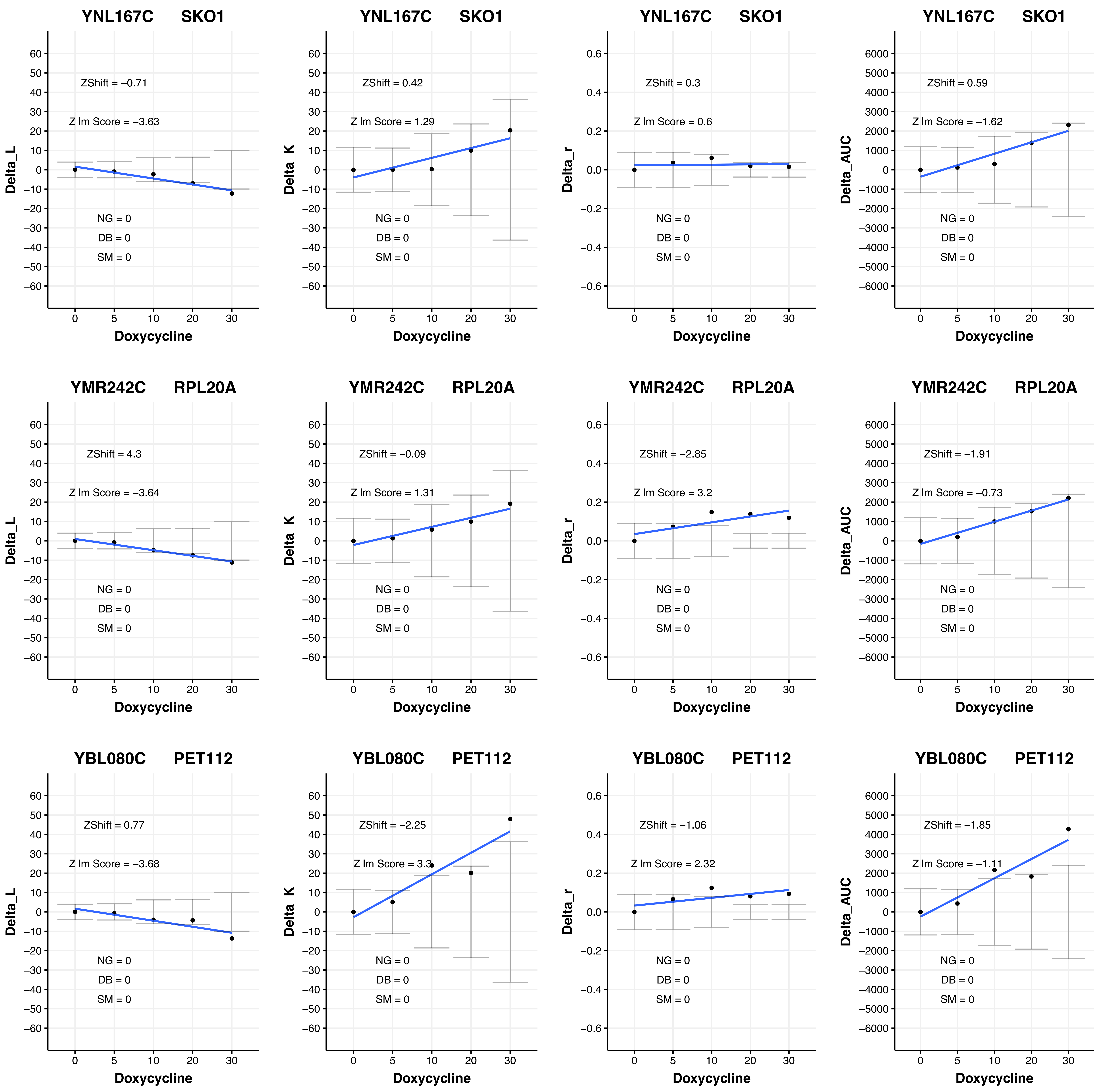
YNL167C SKO1
ZShift = −0.71
Z lm Score = −3.63
NG = 0
DB = 0
SM = 0
Delta_L
Doxycycline
YNL167C SKO1
ZShift = 0.42
Z lm Score = 1.29
NG = 0
DB = 0
SM = 0
Delta_K
Doxycycline
YNL167C SKO1
ZShift = 0.3
Z lm Score = 0.6
NG = 0
DB = 0
SM = 0
Delta_r
Doxycycline
YNL167C SKO1
ZShift = 0.59
Z lm Score = −1.62
NG = 0
DB = 0
SM = 0
Delta_AUC
Doxycycline
YMR242C RPL20A
ZShift = 4.3
Z lm Score = −3.64
NG = 0
DB = 0
SM = 0
Delta_L
Doxycycline
YMR242C RPL20A
ZShift = −0.09
Z lm Score = 1.31
NG = 0
DB = 0
SM = 0
Delta_K
Doxycycline
YMR242C RPL20A
ZShift = −2.85
Z lm Score = 3.2
NG = 0
DB = 0
SM = 0
Delta_r
Doxycycline
YMR242C RPL20A
ZShift = −1.91
Z lm Score = −0.73
NG = 0
DB = 0
SM = 0
Delta_AUC
Doxycycline
YBL080C PET112
ZShift = 0.77
Z lm Score = −3.68
NG = 0
DB = 0
SM = 0
Delta_L
Doxycycline
YBL080C PET112
ZShift = −2.25
Z lm Score = 3.3
NG = 0
DB = 0
SM = 0
Delta_K
Doxycycline
YBL080C PET112
ZShift = −1.06
Z lm Score = 2.32
NG = 0
DB = 0
SM = 0
Delta_r
Doxycycline
YBL080C PET112
ZShift = −1.85
Z lm Score = −1.11
NG = 0
DB = 0
SM = 0
Delta_AUC
Doxycycline

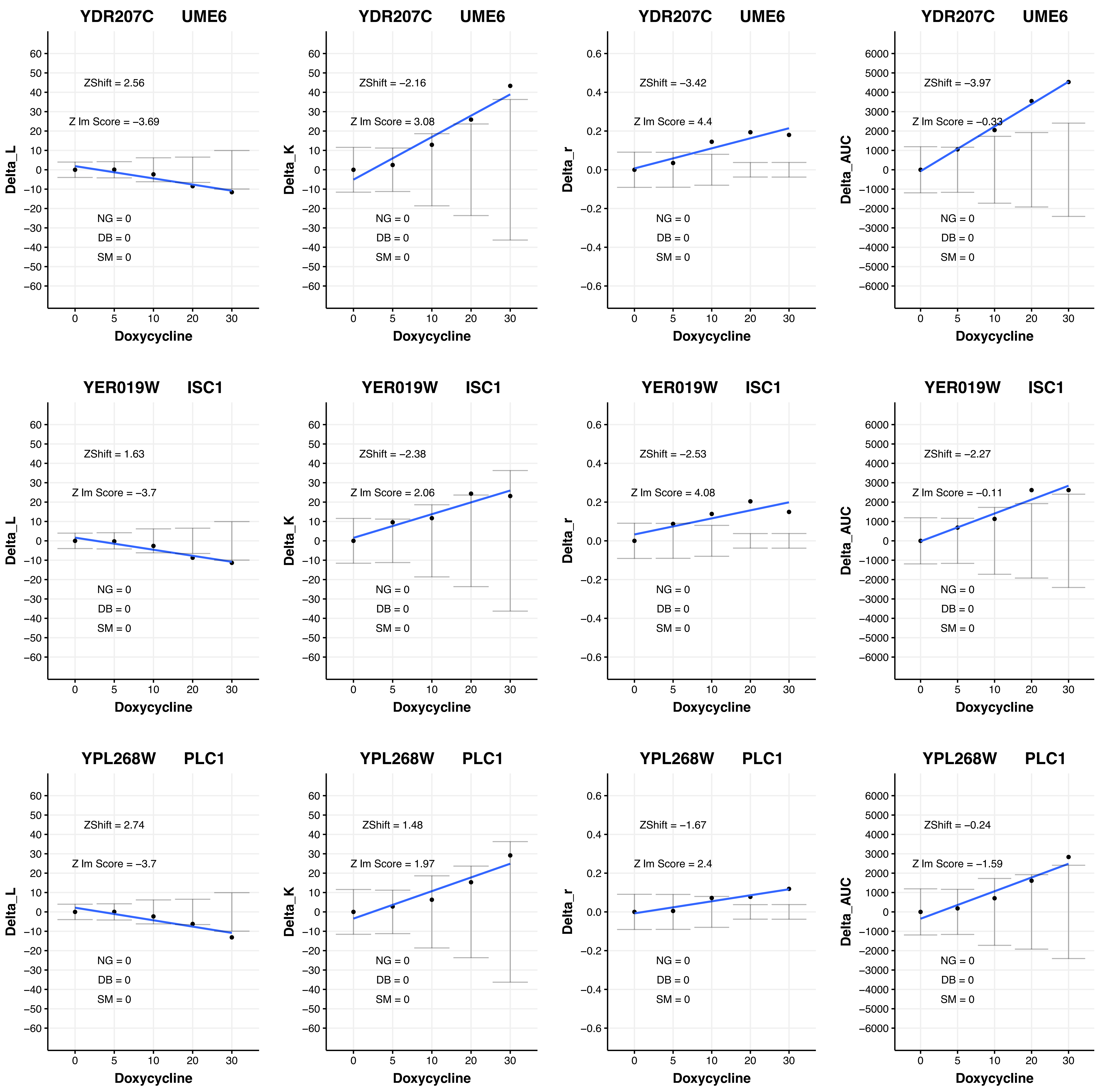
YDR207C UME6
ZShift = 2.56
Z lm Score = −3.69
NG = 0
DB = 0
SM = 0
Delta_L
Doxycycline
YDR207C UME6
ZShift = −2.16
Z lm Score = 3.08
NG = 0
DB = 0
SM = 0
Delta_K
Doxycycline
YDR207C UME6
ZShift = −3.42
Z lm Score = 4.4
NG = 0
DB = 0
SM = 0
Delta_r
Doxycycline
YDR207C UME6
ZShift = −3.97
Z lm Score = −0.33
NG = 0
DB = 0
SM = 0
Delta_AUC
Doxycycline
YER019W ISC1
ZShift = 1.63
Z lm Score = −3.7
NG = 0
DB = 0
SM = 0
Delta_L
Doxycycline
YER019W ISC1
ZShift = −2.38
Z lm Score = 2.06
NG = 0
DB = 0
SM = 0
Delta_K
Doxycycline
YER019W ISC1
ZShift = −2.53
Z lm Score = 4.08
NG = 0
DB = 0
SM = 0
Delta_r
Doxycycline
YER019W ISC1
ZShift = −2.27
Z lm Score = −0.11
NG = 0
DB = 0
SM = 0
Delta_AUC
Doxycycline
YPL268W PLC1
ZShift = 2.74
Z lm Score = −3.7
NG = 0
DB = 0
SM = 0
Delta_L
Doxycycline
YPL268W PLC1
ZShift = 1.48
Z lm Score = 1.97
NG = 0
DB = 0
SM = 0
Delta_K
Doxycycline
YPL268W PLC1
ZShift = −1.67
Z lm Score = 2.4
NG = 0
DB = 0
SM = 0
Delta_r
Doxycycline
YPL268W PLC1
ZShift = −0.24
Z lm Score = −1.59
NG = 0
DB = 0
SM = 0
Delta_AUC
Doxycycline

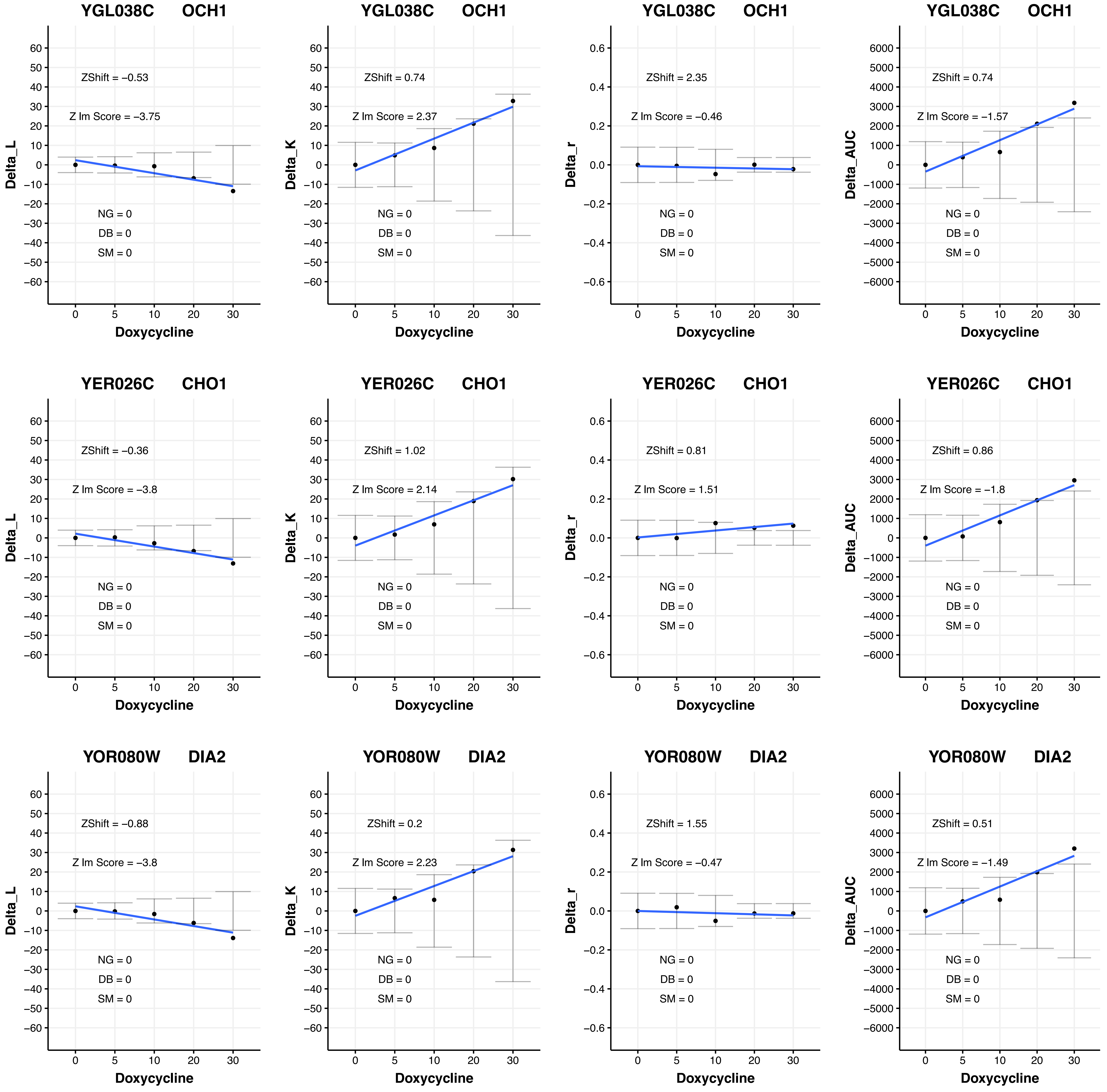

YGL038C OCH1
ZShift = −0.53
Z lm Score = −3.75
NG = 0
DB = 0
SM = 0
Delta_L
Doxycycline
YGL038C OCH1
ZShift = 0.74
Z lm Score = 2.37
NG = 0
DB = 0
SM = 0
Delta_K
Doxycycline
YGL038C OCH1
ZShift = 2.35
Z lm Score = −0.46
NG = 0
DB = 0
SM = 0
Delta_r
Doxycycline
YGL038C OCH1
ZShift = 0.74
Z lm Score = −1.57
NG = 0
DB = 0
SM = 0
Delta_AUC
Doxycycline
YER026C CHO1
ZShift = −0.36
Z lm Score = −3.8
NG = 0
DB = 0
SM = 0
Delta_L
Doxycycline
YER026C CHO1
ZShift = 1.02
Z lm Score = 2.14
NG = 0
DB = 0
SM = 0
Delta_K
Doxycycline
YER026C CHO1
ZShift = 0.81
Z lm Score = 1.51
NG = 0
DB = 0
SM = 0
Delta_r
Doxycycline
YER026C CHO1
ZShift = 0.86
Z lm Score = −1.8
NG = 0
DB = 0
SM = 0
Delta_AUC
Doxycycline
YOR080W DIA2
ZShift = −0.88
Z lm Score = −3.8
NG = 0
DB = 0
SM = 0
Delta_L
Doxycycline
YOR080W DIA2
ZShift = 0.2
Z lm Score = 2.23
NG = 0
DB = 0
SM = 0
Delta_K
Doxycycline
YOR080W DIA2
ZShift = 1.55
Z lm Score = −0.47
NG = 0
DB = 0
SM = 0
Delta_r
Doxycycline
YOR080W DIA2
ZShift = 0.51
Z lm Score = −1.49
NG = 0
DB = 0
SM = 0
Delta_AUC
Doxycycline

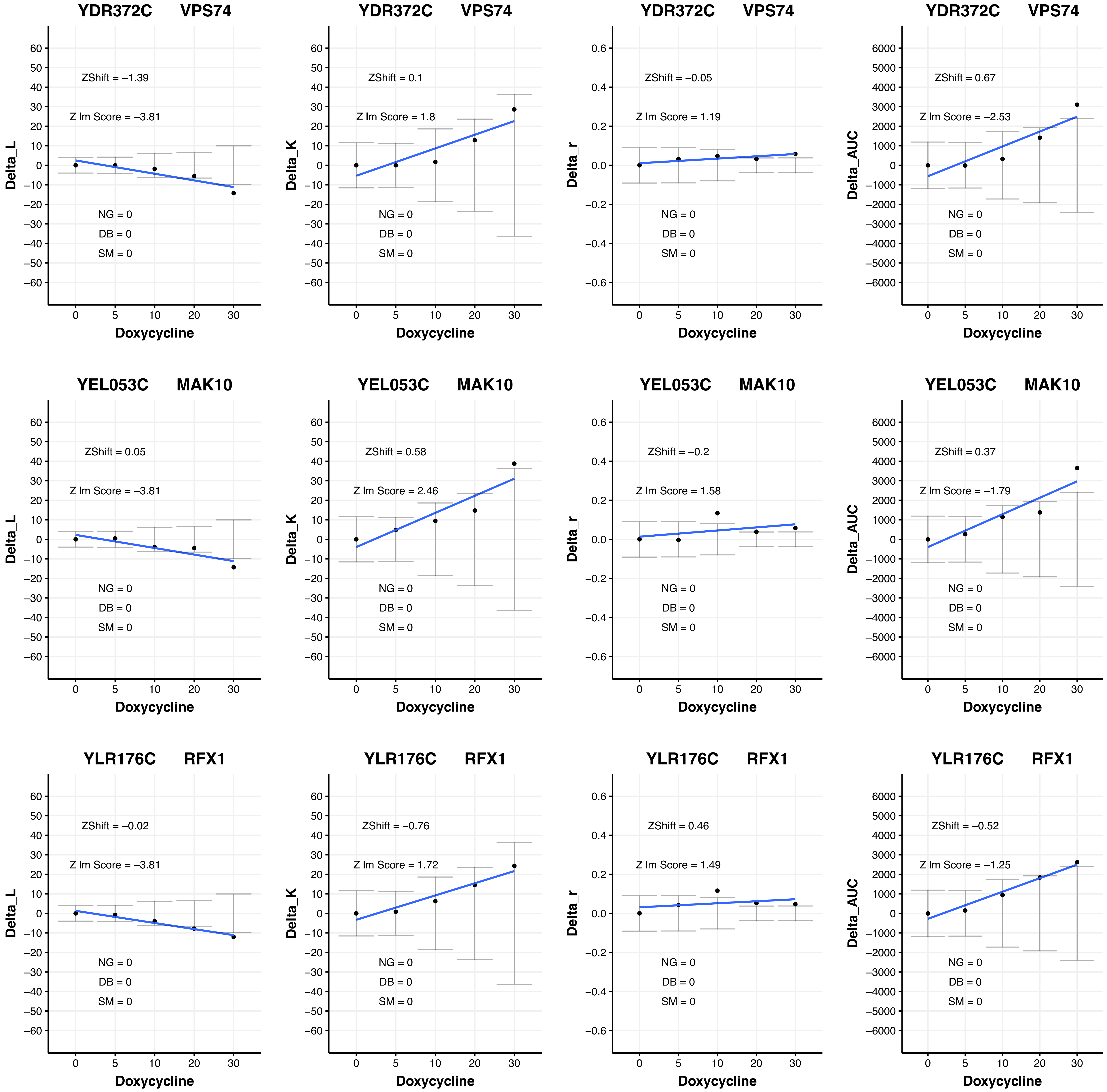

YDR372C VPS74
ZShift = –1.39
Z lm Score = –3.81
NG = 0
DB = 0
SM = 0
Delta_L
Doxycycline
YDR372C VPS74
ZShift = 0.1
Z lm Score = 1.8
NG = 0
DB = 0
SM = 0
Delta_K
Doxycycline
YDR372C VPS74
ZShift = –0.05
Z lm Score = 1.19
NG = 0
DB = 0
SM = 0
Delta_r
Doxycycline
YDR372C VPS74
ZShift = 0.67
Z lm Score = –2.53
NG = 0
DB = 0
SM = 0
Delta_AUC
Doxycycline
YEL053C MAK10
ZShift = 0.05
Z lm Score = –3.81
NG = 0
DB = 0
SM = 0
Delta_L
Doxycycline
YEL053C MAK10
ZShift = 0.58
Z lm Score = 2.46
NG = 0
DB = 0
SM = 0
Delta_K
Doxycycline
YEL053C MAK10
ZShift = –0.2
Z lm Score = 1.58
NG = 0
DB = 0
SM = 0
Delta_r
Doxycycline
YEL053C MAK10
ZShift = 0.37
Z lm Score = –1.79
NG = 0
DB = 0
SM = 0
Delta_AUC
Doxycycline
YLR176C RFX1
ZShift = –0.02
Z lm Score = –3.81
NG = 0
DB = 0
SM = 0
Delta_L
Doxycycline
YLR176C RFX1
ZShift = –0.76
Z lm Score = 1.72
NG = 0
DB = 0
SM = 0
Delta_K
Doxycycline
YLR176C RFX1
ZShift = 0.46
Z lm Score = 1.49
NG = 0
DB = 0
SM = 0
Delta_r
Doxycycline
YLR176C RFX1
ZShift = –0.52
Z lm Score = –1.25
NG = 0
DB = 0
SM = 0
Delta_AUC
Doxycycline

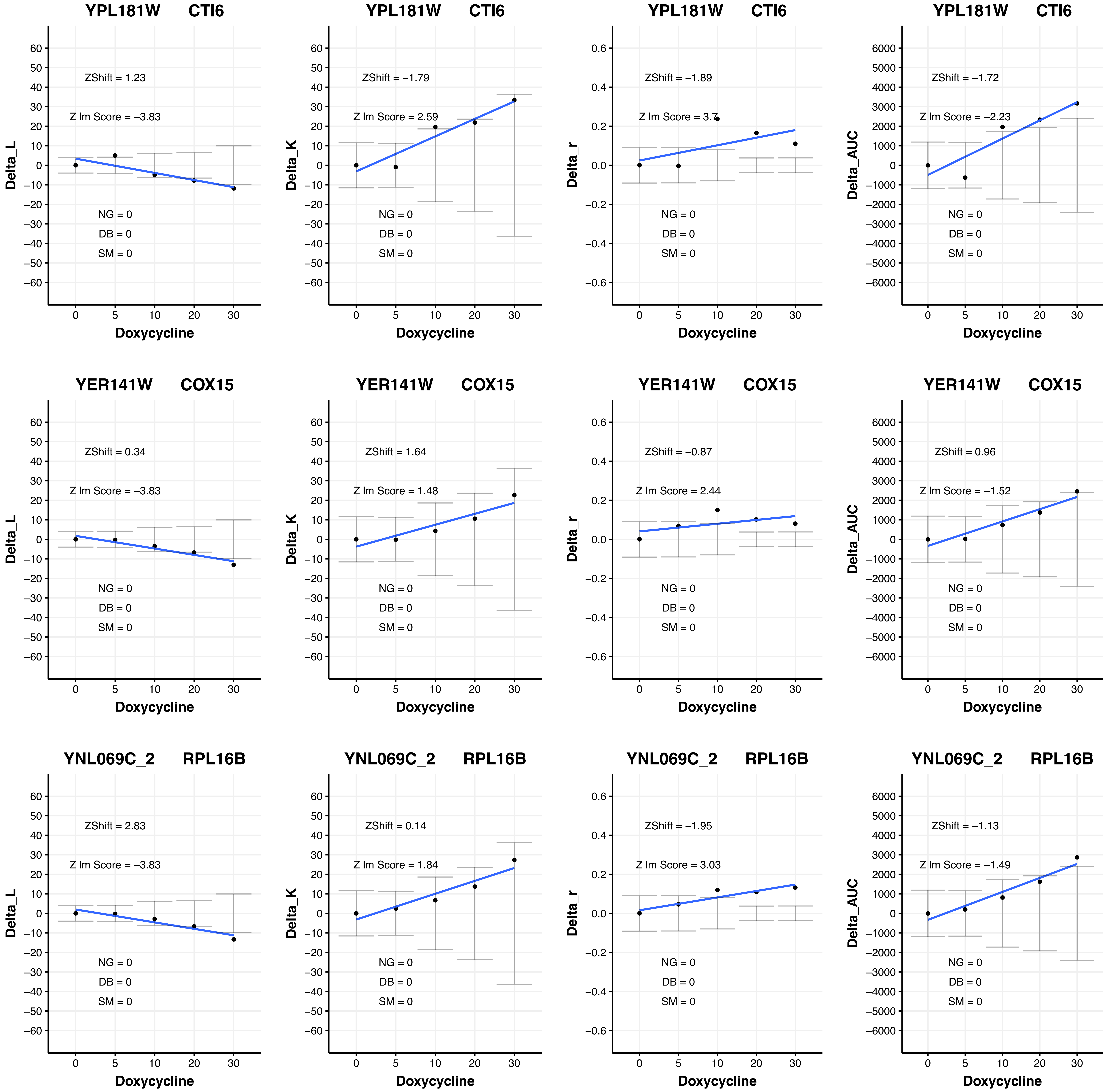

YPL181W CTI6
ZShift = 1.23
Z lm Score = −3.83
NG = 0
DB = 0
SM = 0
Delta_L
Doxycycline
YPL181W CTI6
ZShift = −1.79
Z lm Score = 2.59
NG = 0
DB = 0
SM = 0
Delta_K
Doxycycline
YPL181W CTI6
ZShift = −1.89
Z lm Score = 3.7
NG = 0
DB = 0
SM = 0
Delta_r
Doxycycline
YPL181W CTI6
ZShift = −1.72
Z lm Score = −2.23
NG = 0
DB = 0
SM = 0
Delta_AUC
Doxycycline
YER141W COX15
ZShift = 0.34
Z lm Score = −3.83
NG = 0
DB = 0
SM = 0
Delta_L
Doxycycline
YER141W COX15
ZShift = 1.64
Z lm Score = 1.48
NG = 0
DB = 0
SM = 0
Delta_K
Doxycycline
YER141W COX15
ZShift = −0.87
Z lm Score = 2.44
NG = 0
DB = 0
SM = 0
Delta_r
Doxycycline
YER141W COX15
ZShift = 0.96
Z lm Score = −1.52
NG = 0
DB = 0
SM = 0
Delta_AUC
Doxycycline
YNL069C_2 RPL16B
ZShift = 2.83
Z lm Score = −3.83
NG = 0
DB = 0
SM = 0
Delta_L
Doxycycline
YNL069C_2 RPL16B
ZShift = 0.14
Z lm Score = 1.84
NG = 0
DB = 0
SM = 0
Delta_K
Doxycycline
YNL069C_2 RPL16B
ZShift = −1.95
Z lm Score = 3.03
NG = 0
DB = 0
SM = 0
Delta_r
Doxycycline
YNL069C_2 RPL16B
ZShift = −1.13
Z lm Score = −1.49
NG = 0
DB = 0
SM = 0
Delta_AUC
Doxycycline

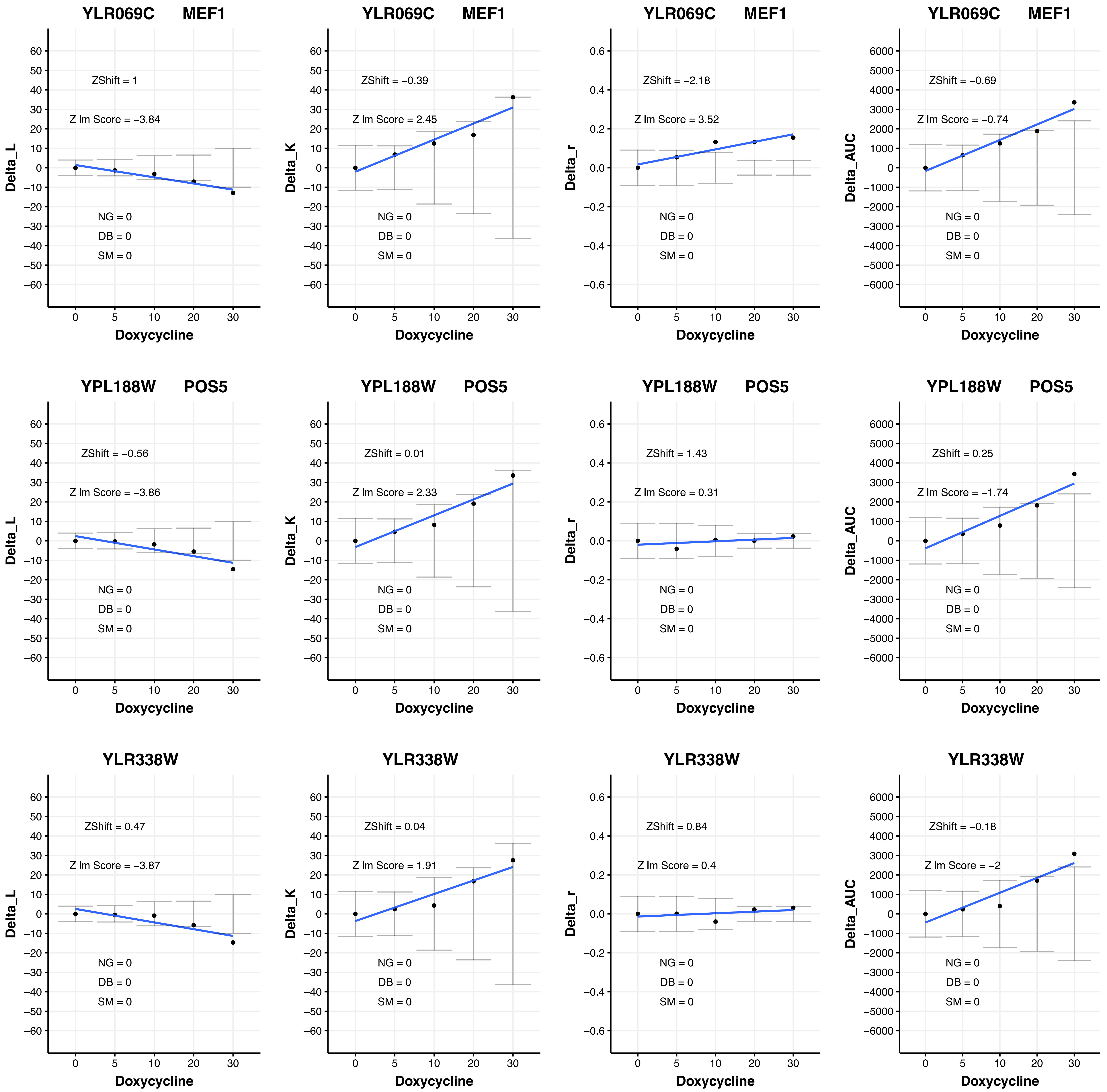

YLR069C MEF1
ZShift = 1
Z lm Score = −3.84
NG = 0
DB = 0
SM = 0
Delta_L
Doxycycline
YLR069C MEF1
ZShift = −0.39
Z lm Score = 2.45
NG = 0
DB = 0
SM = 0
Delta_K
Doxycycline
YLR069C MEF1
ZShift = −2.18
Z lm Score = 3.52
NG = 0
DB = 0
SM = 0
Delta_r
Doxycycline
YLR069C MEF1
ZShift = −0.69
Z lm Score = −0.74
NG = 0
DB = 0
SM = 0
Delta_AUC
Doxycycline
YPL188W POS5
ZShift = −0.56
Z lm Score = −3.86
NG = 0
DB = 0
SM = 0
Delta_L
Doxycycline
YPL188W POS5
ZShift = 0.01
Z lm Score = 2.33
NG = 0
DB = 0
SM = 0
Delta_K
Doxycycline
YPL188W POS5
ZShift = 1.43
Z lm Score = 0.31
NG = 0
DB = 0
SM = 0
Delta_r
Doxycycline
YPL188W POS5
ZShift = 0.25
Z lm Score = −1.74
NG = 0
DB = 0
SM = 0
Delta_AUC
Doxycycline
YLR338W
ZShift = 0.47
Z lm Score = −3.87
NG = 0
DB = 0
SM = 0
Delta_L
Doxycycline
YLR338W
ZShift = 0.04
Z lm Score = 1.91
NG = 0
DB = 0
SM = 0
Delta_K
Doxycycline
YLR338W
ZShift = 0.84
Z lm Score = 0.4
NG = 0
DB = 0
SM = 0
Delta_r
Doxycycline
YLR338W
ZShift = −0.18
Z lm Score = −2
NG = 0
DB = 0
SM = 0
Delta_AUC
Doxycycline

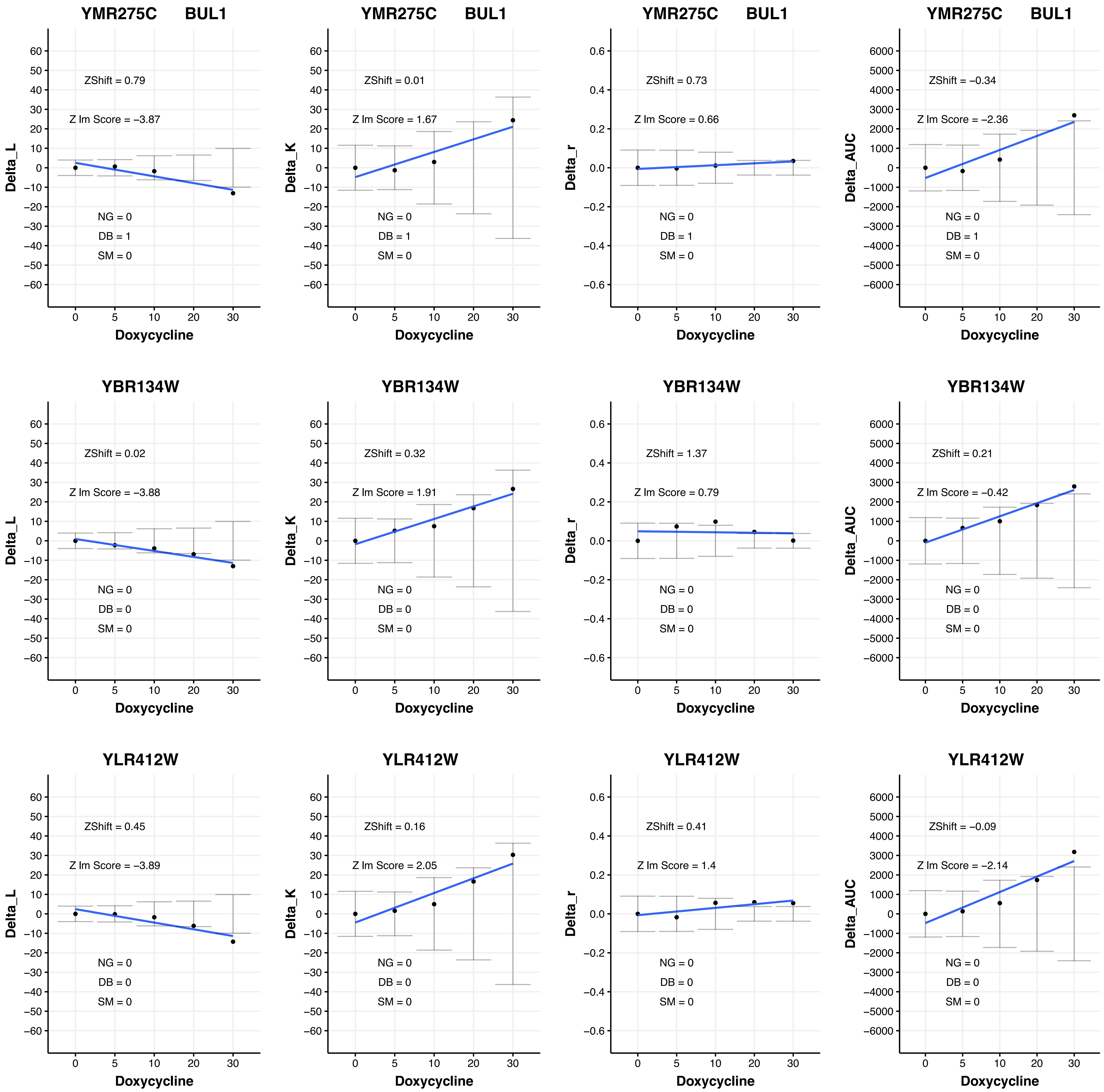
YMR275C BUL1
ZShift = 0.79
Z lm Score = −3.87
NG = 0
DB = 1
SM = 0
Delta_L
Doxycycline
YMR275C BUL1
ZShift = 0.01
Z lm Score = 1.67
NG = 0
DB = 1
SM = 0
Delta_K
Doxycycline
YMR275C BUL1
ZShift = 0.73
Z lm Score = 0.66
NG = 0
DB = 1
SM = 0
Delta_r
Doxycycline
YMR275C BUL1
ZShift = −0.34
Z lm Score = −2.36
NG = 0
DB = 1
SM = 0
Delta_AUC
Doxycycline
YBR134W
ZShift = 0.02
Z lm Score = −3.88
NG = 0
DB = 0
SM = 0
Delta_L
Doxycycline
YBR134W
ZShift = 0.32
Z lm Score = 1.91
NG = 0
DB = 0
SM = 0
Delta_K
Doxycycline
YBR134W
ZShift = 1.37
Z lm Score = 0.79
NG = 0
DB = 0
SM = 0
Delta_r
Doxycycline
YBR134W
ZShift = 0.21
Z lm Score = −0.42
NG = 0
DB = 0
SM = 0
Delta_AUC
Doxycycline
YLR412W
ZShift = 0.45
Z lm Score = −3.89
NG = 0
DB = 0
SM = 0
Delta_L
Doxycycline
YLR412W
ZShift = 0.16
Z lm Score = 2.05
NG = 0
DB = 0
SM = 0
Delta_K
Doxycycline
YLR412W
ZShift = 0.41
Z lm Score = 1.4
NG = 0
DB = 0
SM = 0
Delta_r
Doxycycline
YLR412W
ZShift = −0.09
Z lm Score = −2.14
NG = 0
DB = 0
SM = 0
Delta_AUC
Doxycycline

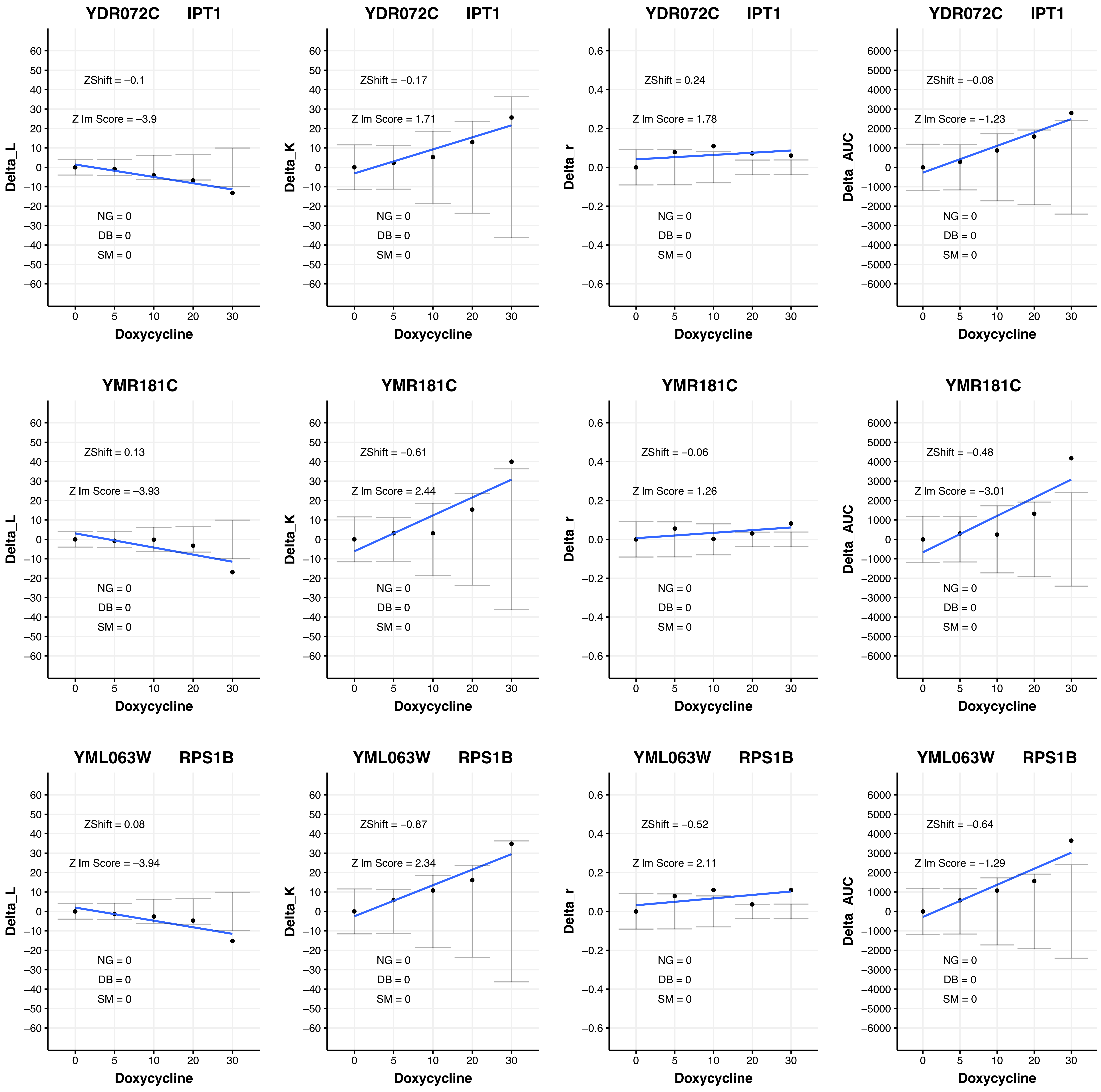

YDR072C IPT1
ZShift = −0.1
Z lm Score = −3.9
NG = 0
DB = 0
SM = 0
Delta_L
Doxycycline
YDR072C IPT1
ZShift = −0.17
Z lm Score = 1.71
NG = 0
DB = 0
SM = 0
Delta_K
Doxycycline
YDR072C IPT1
ZShift = 0.24
Z lm Score = 1.78
NG = 0
DB = 0
SM = 0
Delta_r
Doxycycline
YDR072C IPT1
ZShift = −0.08
Z lm Score = −1.23
NG = 0
DB = 0
SM = 0
Delta_AUC
Doxycycline
YMR181C
ZShift = 0.13
Z lm Score = −3.93
NG = 0
DB = 0
SM = 0
Delta_L
Doxycycline
YMR181C
ZShift = −0.61
Z lm Score = 2.44
NG = 0
DB = 0
SM = 0
Delta_K
Doxycycline
YMR181C
ZShift = −0.06
Z lm Score = 1.26
NG = 0
DB = 0
SM = 0
Delta_r
Doxycycline
YMR181C
ZShift = −0.48
Z lm Score = −3.01
NG = 0
DB = 0
SM = 0
Delta_AUC
Doxycycline
YML063W RPS1B
ZShift = 0.08
Z lm Score = −3.94
NG = 0
DB = 0
SM = 0
Delta_L
Doxycycline
YML063W RPS1B
ZShift = −0.87
Z lm Score = 2.34
NG = 0
DB = 0
SM = 0
Delta_K
Doxycycline
YML063W RPS1B
ZShift = −0.52
Z lm Score = 2.11
NG = 0
DB = 0
SM = 0
Delta_r
Doxycycline
YML063W RPS1B
ZShift = −0.64
Z lm Score = −1.29
NG = 0
DB = 0
SM = 0
Delta_AUC
Doxycycline

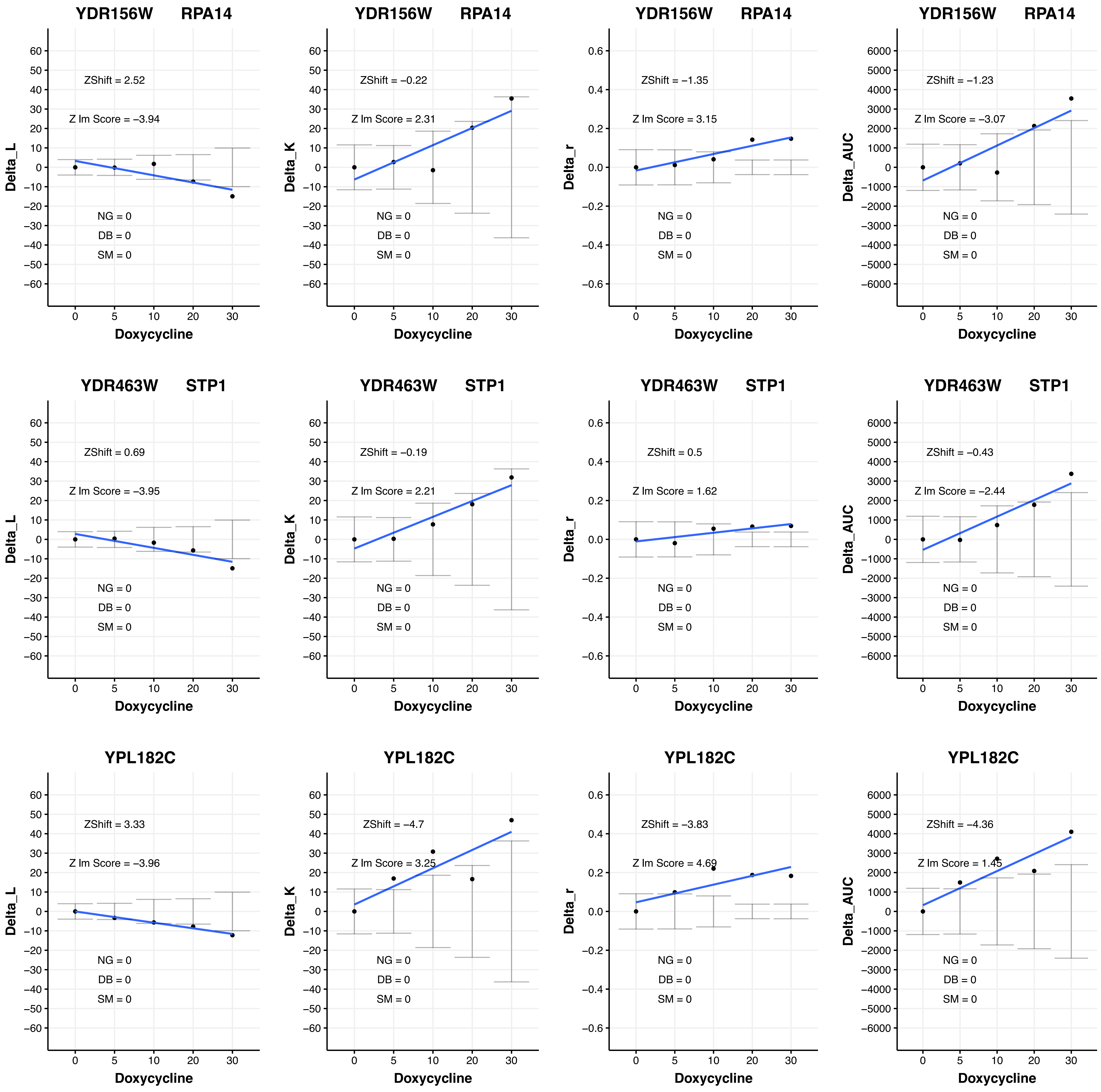

YDR156W RPA14
ZShift = 2.52
Z lm Score = −3.94
NG = 0
DB = 0
SM = 0
Delta_L
Doxycycline
YDR156W RPA14
ZShift = −0.22
Z lm Score = 2.31
NG = 0
DB = 0
SM = 0
Delta_K
Doxycycline
YDR156W RPA14
ZShift = −1.35
Z lm Score = 3.15
NG = 0
DB = 0
SM = 0
Delta_r
Doxycycline
YDR156W RPA14
ZShift = −1.23
Z lm Score = −3.07
NG = 0
DB = 0
SM = 0
Delta_AUC
Doxycycline
YDR463W STP1
ZShift = 0.69
Z lm Score = −3.95
NG = 0
DB = 0
SM = 0
Delta_L
Doxycycline
YDR463W STP1
ZShift = −0.19
Z lm Score = 2.21
NG = 0
DB = 0
SM = 0
Delta_K
Doxycycline
YDR463W STP1
ZShift = 0.5
Z lm Score = 1.62
NG = 0
DB = 0
SM = 0
Delta_r
Doxycycline
YDR463W STP1
ZShift = −0.43
Z lm Score = −2.44
NG = 0
DB = 0
SM = 0
Delta_AUC
Doxycycline
YPL182C
ZShift = 3.33
Z lm Score = −3.96
NG = 0
DB = 0
SM = 0
Delta_L
Doxycycline
YPL182C
ZShift = −4.7
Z lm Score = 3.25
NG = 0
DB = 0
SM = 0
Delta_K
Doxycycline
YPL182C
ZShift = −3.83
Z lm Score = 4.69
NG = 0
DB = 0
SM = 0
Delta_r
Doxycycline
YPL182C
ZShift = −4.36
Z lm Score = 1.45
NG = 0
DB = 0
SM = 0
Delta_AUC
Doxycycline

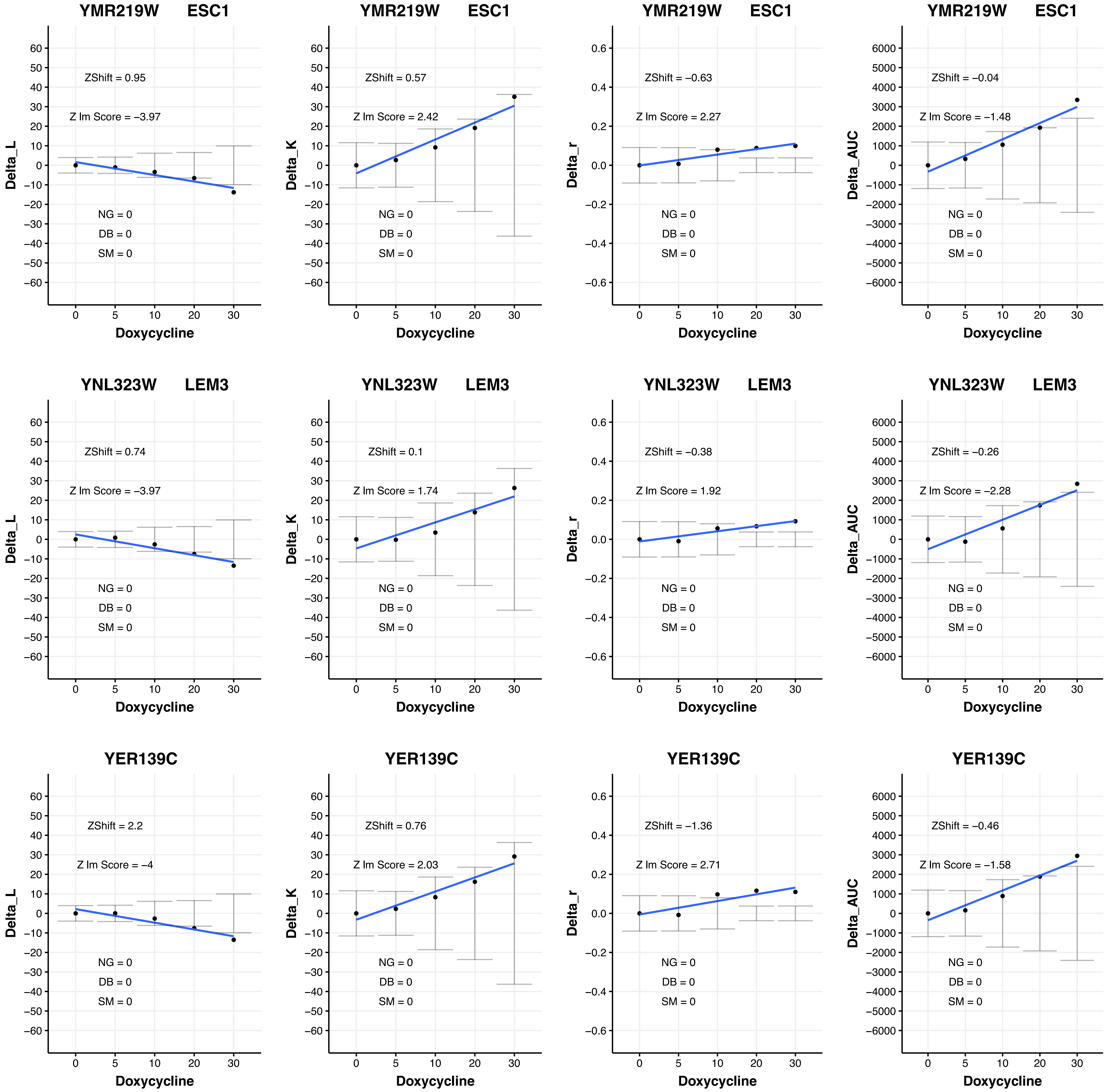

YMR219W ESC1
ZShift = 0.95
Z lm Score = −3.97
ZShift = 0.57
Z lm Score = 2.42
ZShift = −0.63
Z lm Score = 2.27
ZShift = −0.04
Z lm Score = −1.48
YNL323W LEM3
ZShift = 0.74
Z lm Score = −3.97
ZShift = 0.1
Z lm Score = 1.74
ZShift = −0.38
Z lm Score = 1.92
ZShift = −0.26
Z lm Score = −2.28
YER139C
ZShift = 2.2
Z lm Score = −4
ZShift = 0.76
Z lm Score = 2.03
ZShift = −1.36
Z lm Score = 2.71
ZShift = −0.46
Z lm Score = −1.58
NG = 0
DB = 0
SM = 0
Delta_L
Delta_K
Delta_r
Delta_AUC
Doxycycline

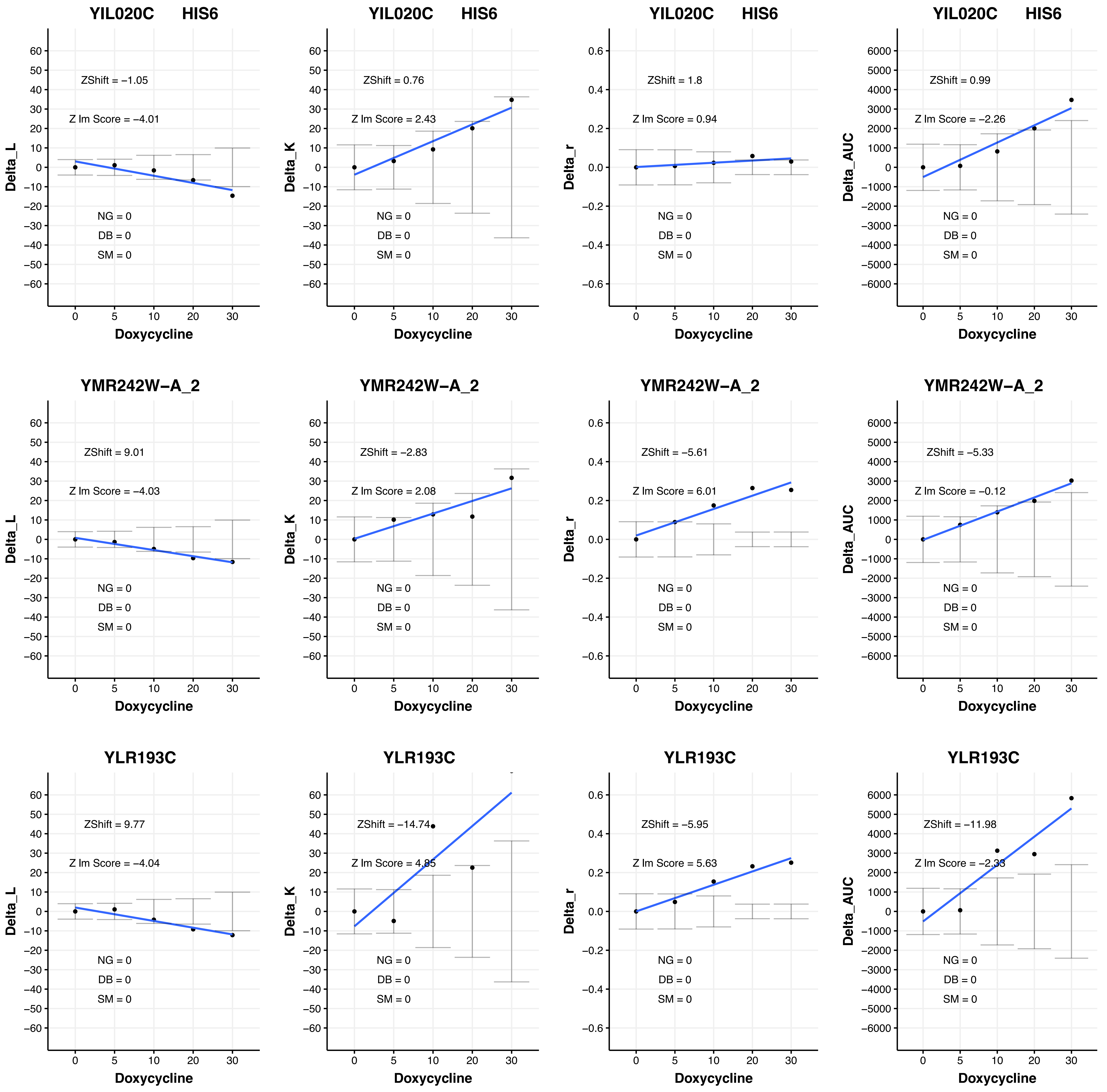
YIL020C HIS6
ZShift = −1.05
Z lm Score = −4.01
NG = 0
DB = 0
SM = 0
Delta_L
Doxycycline
YIL020C HIS6
ZShift = 0.76
Z lm Score = 2.43
NG = 0
DB = 0
SM = 0
Delta_K
Doxycycline
YIL020C HIS6
ZShift = 1.8
Z lm Score = 0.94
NG = 0
DB = 0
SM = 0
Delta_r
Doxycycline
YIL020C HIS6
ZShift = 0.99
Z lm Score = −2.26
NG = 0
DB = 0
SM = 0
Delta_AUC
Doxycycline
YMR242W−A_2
ZShift = 9.01
Z lm Score = −4.03
NG = 0
DB = 0
SM = 0
Delta_L
Doxycycline
YMR242W−A_2
ZShift = −2.83
Z lm Score = 2.08
NG = 0
DB = 0
SM = 0
Delta_K
Doxycycline
YMR242W−A_2
ZShift = −5.61
Z lm Score = 6.01
NG = 0
DB = 0
SM = 0
Delta_r
Doxycycline
YMR242W−A_2
ZShift = −5.33
Z lm Score = −0.12
NG = 0
DB = 0
SM = 0
Delta_AUC
Doxycycline
YLR193C
ZShift = 9.77
Z lm Score = −4.04
NG = 0
DB = 0
SM = 0
Delta_L
Doxycycline
YLR193C
ZShift = −14.74
Z lm Score = 4.85
NG = 0
DB = 0
SM = 0
Delta_K
Doxycycline
YLR193C
ZShift = −5.95
Z lm Score = 5.63
NG = 0
DB = 0
SM = 0
Delta_r
Doxycycline
YLR193C
ZShift = −11.98
Z lm Score = −2.33
NG = 0
DB = 0
SM = 0
Delta_AUC
Doxycycline

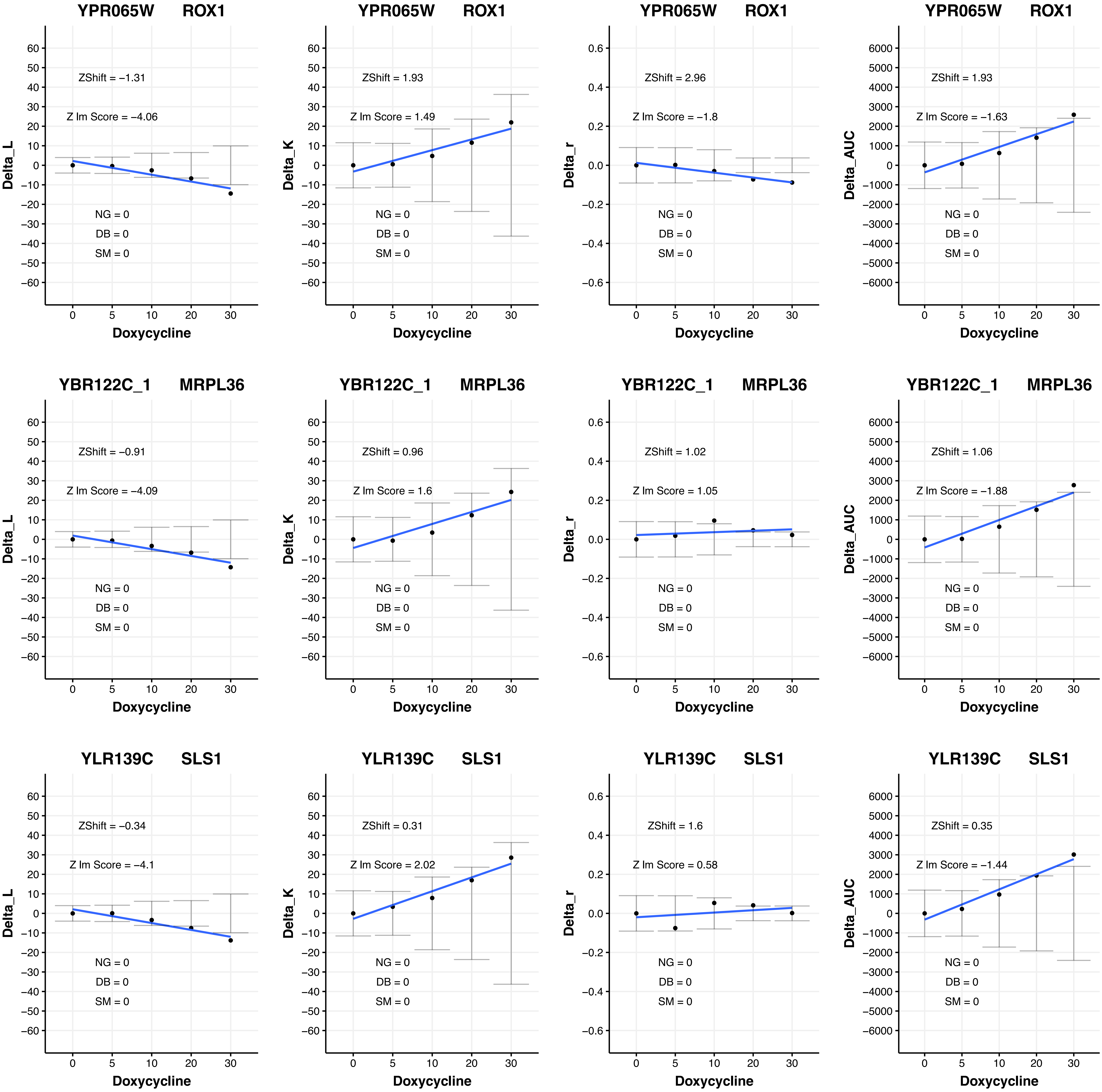
YPR065W ROX1
ZShift = −1.31
Z lm Score = −4.06
NG = 0
DB = 0
SM = 0
Delta_L
Doxycycline
YPR065W ROX1
ZShift = 1.93
Z lm Score = 1.49
NG = 0
DB = 0
SM = 0
Delta_K
Doxycycline
YPR065W ROX1
ZShift = 2.96
Z lm Score = −1.8
NG = 0
DB = 0
SM = 0
Delta_r
Doxycycline
YPR065W ROX1
ZShift = 1.93
Z lm Score = −1.63
NG = 0
DB = 0
SM = 0
Delta_AUC
Doxycycline
YBR122C_1 MRPL36
ZShift = −0.91
Z lm Score = −4.09
NG = 0
DB = 0
SM = 0
Delta_L
Doxycycline
YBR122C_1 MRPL36
ZShift = 0.96
Z lm Score = 1.6
NG = 0
DB = 0
SM = 0
Delta_K
Doxycycline
YBR122C_1 MRPL36
ZShift = 1.02
Z lm Score = 1.05
NG = 0
DB = 0
SM = 0
Delta_r
Doxycycline
YBR122C_1 MRPL36
ZShift = 1.06
Z lm Score = −1.88
NG = 0
DB = 0
SM = 0
Delta_AUC
Doxycycline
YLR139C SLS1
ZShift = −0.34
Z lm Score = −4.1
NG = 0
DB = 0
SM = 0
Delta_L
Doxycycline
YLR139C SLS1
ZShift = 0.31
Z lm Score = 2.02
NG = 0
DB = 0
SM = 0
Delta_K
Doxycycline
YLR139C SLS1
ZShift = 1.6
Z lm Score = 0.58
NG = 0
DB = 0
SM = 0
Delta_r
Doxycycline
YLR139C SLS1
ZShift = 0.35
Z lm Score = −1.44
NG = 0
DB = 0
SM = 0
Delta_AUC
Doxycycline

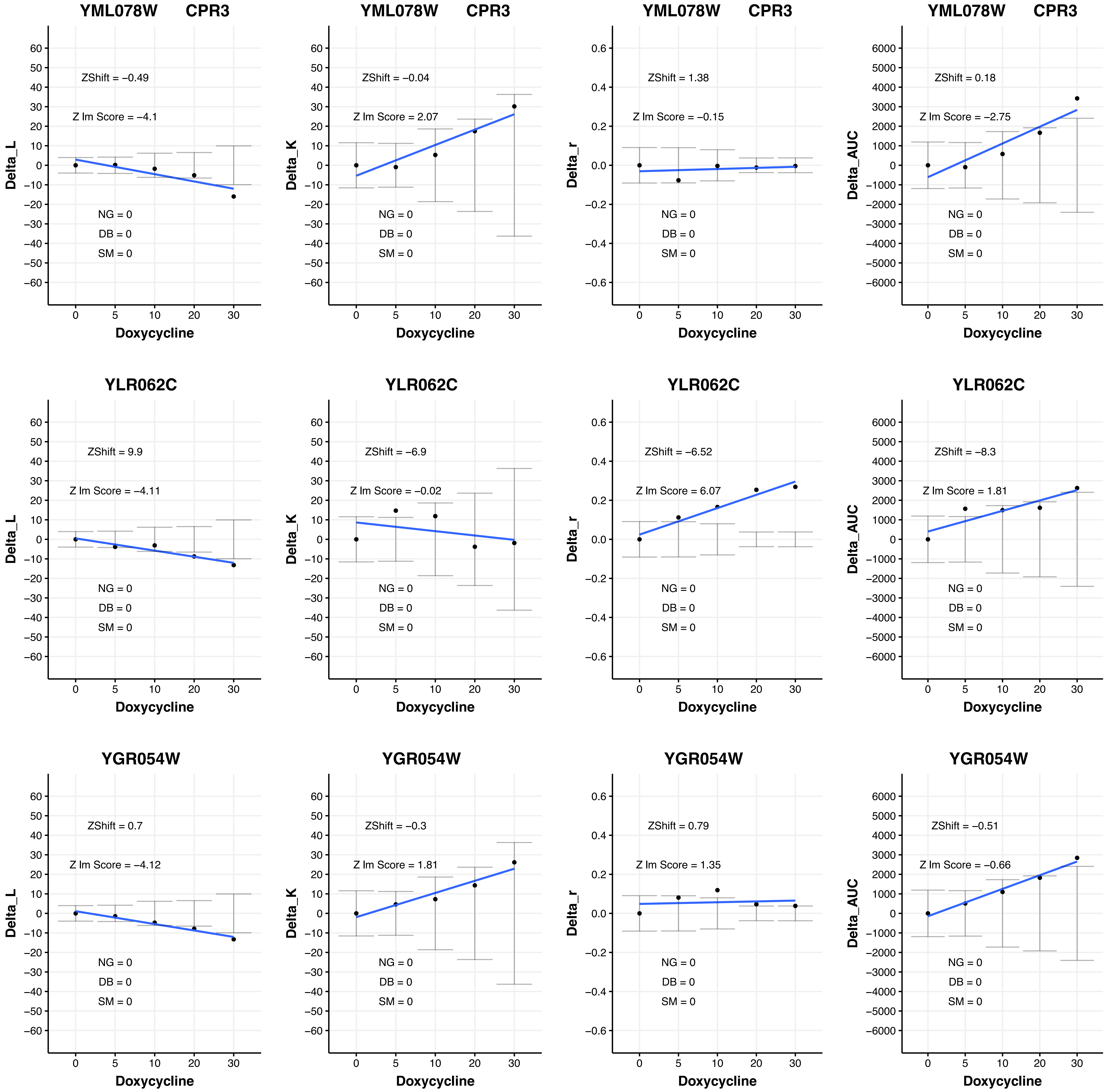

YML078W CPR3
ZShift = −0.49
Z lm Score = −4.1
NG = 0
DB = 0
SM = 0
Delta_L
Doxycycline
YML078W CPR3
ZShift = −0.04
Z lm Score = 2.07
NG = 0
DB = 0
SM = 0
Delta_K
Doxycycline
YML078W CPR3
ZShift = 1.38
Z lm Score = −0.15
NG = 0
DB = 0
SM = 0
Delta_r
Doxycycline
YML078W CPR3
ZShift = 0.18
Z lm Score = −2.75
NG = 0
DB = 0
SM = 0
Delta_AUC
Doxycycline
YLR062C
ZShift = 9.9
Z lm Score = −4.11
NG = 0
DB = 0
SM = 0
Delta_L
Doxycycline
YLR062C
ZShift = −6.9
Z lm Score = −0.02
NG = 0
DB = 0
SM = 0
Delta_K
Doxycycline
YLR062C
ZShift = −6.52
Z lm Score = 6.07
NG = 0
DB = 0
SM = 0
Delta_r
Doxycycline
YLR062C
ZShift = −8.3
Z lm Score = 1.81
NG = 0
DB = 0
SM = 0
Delta_AUC
Doxycycline
YGR054W
ZShift = 0.7
Z lm Score = −4.12
NG = 0
DB = 0
SM = 0
Delta_L
Doxycycline
YGR054W
ZShift = −0.3
Z lm Score = 1.81
NG = 0
DB = 0
SM = 0
Delta_K
Doxycycline
YGR054W
ZShift = 0.79
Z lm Score = 1.35
NG = 0
DB = 0
SM = 0
Delta_r
Doxycycline
YGR054W
ZShift = −0.51
Z lm Score = −0.66
NG = 0
DB = 0
SM = 0
Delta_AUC
Doxycycline

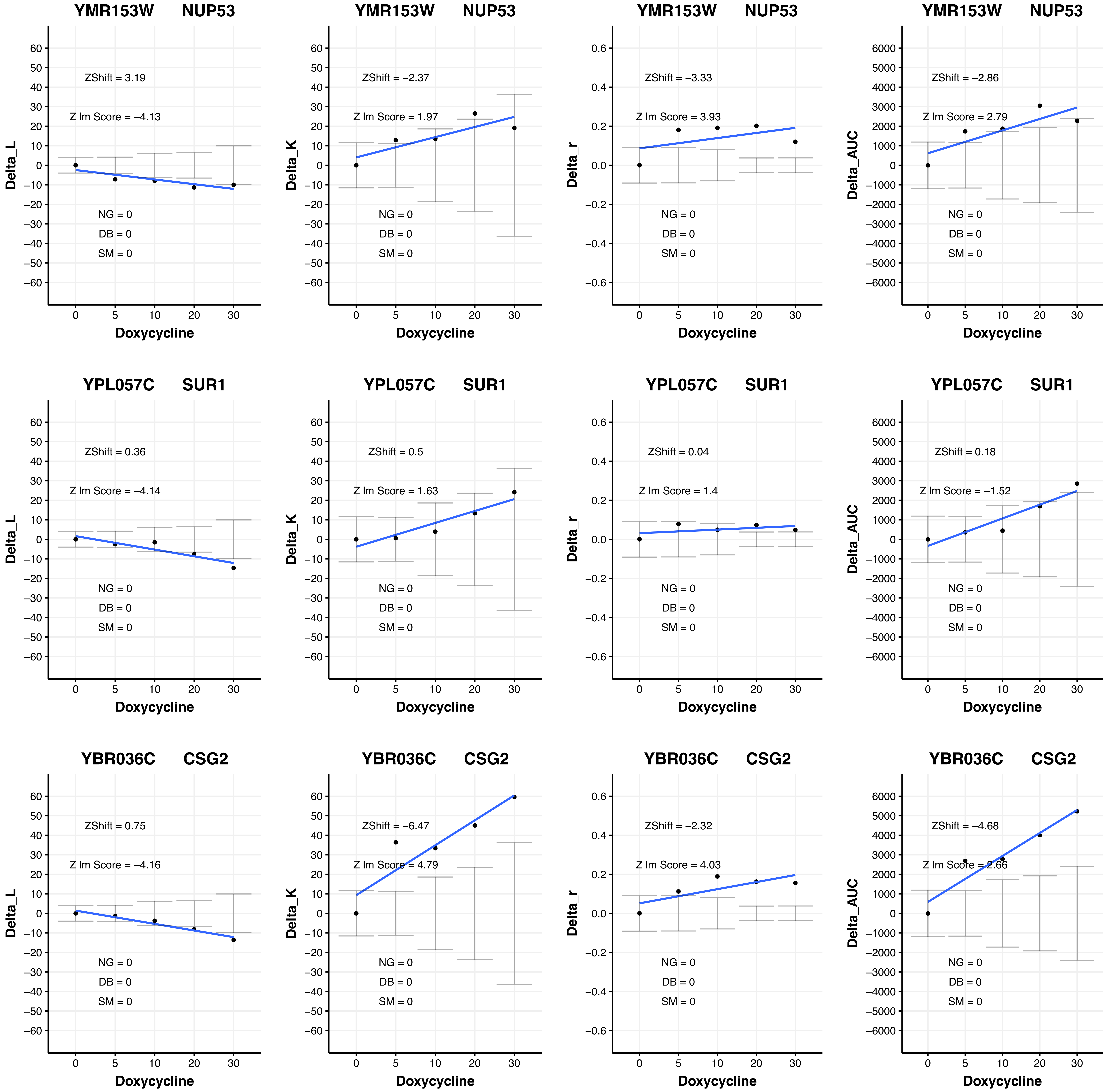

YMR153W NUP53
ZShift = 3.19
Z lm Score = −4.13
NG = 0
DB = 0
SM = 0
Delta_L
Doxycycline
YMR153W NUP53
ZShift = −2.37
Z lm Score = 1.97
NG = 0
DB = 0
SM = 0
Delta_K
Doxycycline
YMR153W NUP53
ZShift = −3.33
Z lm Score = 3.93
NG = 0
DB = 0
SM = 0
Delta_r
Doxycycline
YMR153W NUP53
ZShift = −2.86
Z lm Score = 2.79
NG = 0
DB = 0
SM = 0
Delta_AUC
Doxycycline
YPL057C SUR1
ZShift = 0.36
Z lm Score = −4.14
NG = 0
DB = 0
SM = 0
Delta_L
Doxycycline
YPL057C SUR1
ZShift = 0.5
Z lm Score = 1.63
NG = 0
DB = 0
SM = 0
Delta_K
Doxycycline
YPL057C SUR1
ZShift = 0.04
Z lm Score = 1.4
NG = 0
DB = 0
SM = 0
Delta_r
Doxycycline
YPL057C SUR1
ZShift = 0.18
Z lm Score = −1.52
NG = 0
DB = 0
SM = 0
Delta_AUC
Doxycycline
YBR036C CSG2
ZShift = 0.75
Z lm Score = −4.16
NG = 0
DB = 0
SM = 0
Delta_L
Doxycycline
YBR036C CSG2
ZShift = −6.47
Z lm Score = 4.79
NG = 0
DB = 0
SM = 0
Delta_K
Doxycycline
YBR036C CSG2
ZShift = −2.32
Z lm Score = 4.03
NG = 0
DB = 0
SM = 0
Delta_r
Doxycycline
YBR036C CSG2
ZShift = −4.68
Z lm Score = 2.66
NG = 0
DB = 0
SM = 0
Delta_AUC
Doxycycline

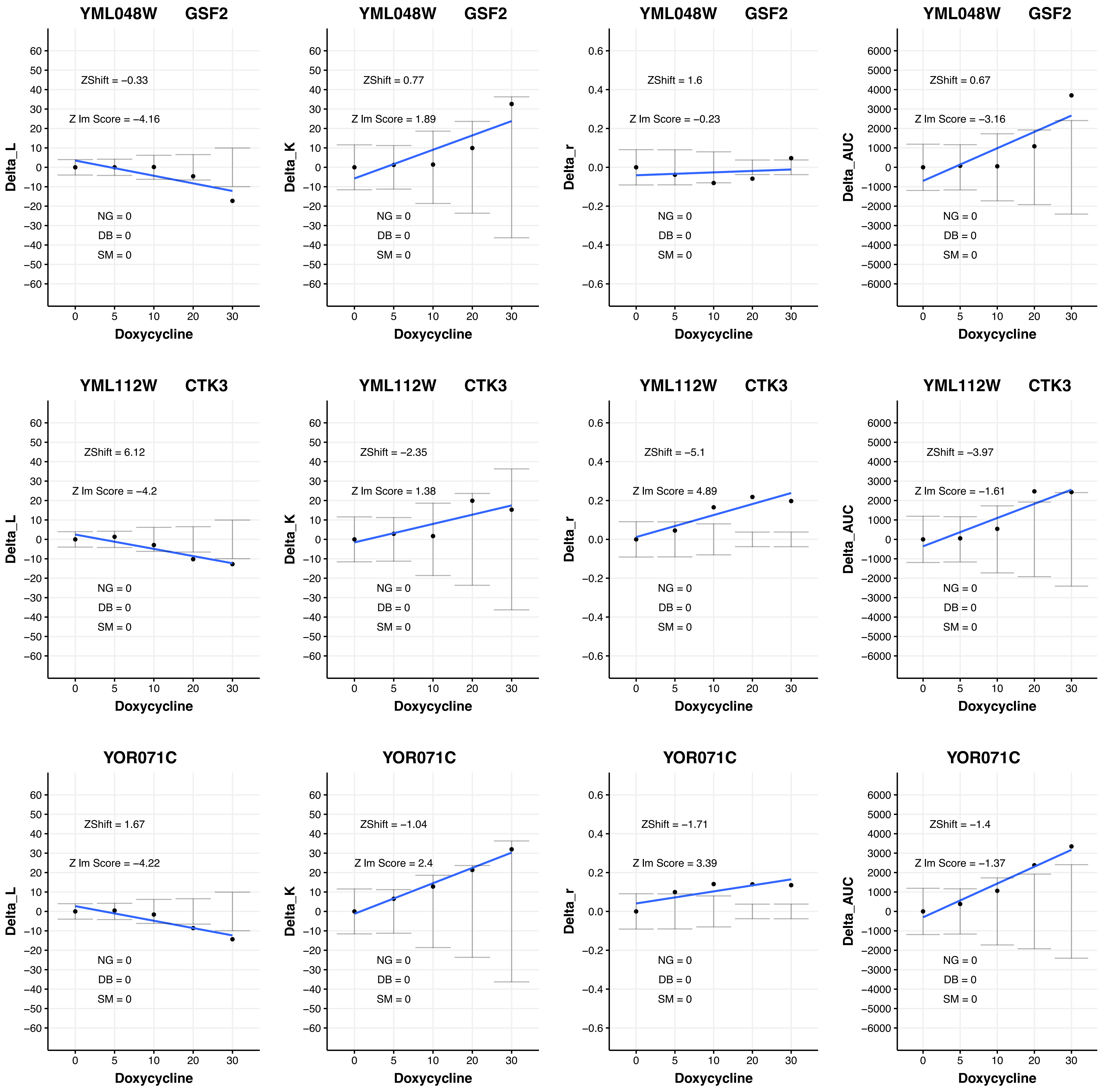
YML048W GSF2
ZShift = −0.33
Z lm Score = −4.16
NG = 0
DB = 0
SM = 0
Delta_L
Doxycycline
YML048W GSF2
ZShift = 0.77
Z lm Score = 1.89
NG = 0
DB = 0
SM = 0
Delta_K
Doxycycline
YML048W GSF2
ZShift = 1.6
Z lm Score = −0.23
NG = 0
DB = 0
SM = 0
Delta_r
Doxycycline
YML048W GSF2
ZShift = 0.67
Z lm Score = −3.16
NG = 0
DB = 0
SM = 0
Delta_AUC
Doxycycline
YML112W CTK3
ZShift = 6.12
Z lm Score = −4.2
NG = 0
DB = 0
SM = 0
Delta_L
Doxycycline
YML112W CTK3
ZShift = −2.35
Z lm Score = 1.38
NG = 0
DB = 0
SM = 0
Delta_K
Doxycycline
YML112W CTK3
ZShift = −5.1
Z lm Score = 4.89
NG = 0
DB = 0
SM = 0
Delta_r
Doxycycline
YML112W CTK3
ZShift = −3.97
Z lm Score = −1.61
NG = 0
DB = 0
SM = 0
Delta_AUC
Doxycycline
YOR071C
ZShift = 1.67
Z lm Score = −4.22
NG = 0
DB = 0
SM = 0
Delta_L
Doxycycline
YOR071C
ZShift = −1.04
Z lm Score = 2.4
NG = 0
DB = 0
SM = 0
Delta_K
Doxycycline
YOR071C
ZShift = −1.71
Z lm Score = 3.39
NG = 0
DB = 0
SM = 0
Delta_r
Doxycycline
YOR071C
ZShift = −1.4
Z lm Score = −1.37
NG = 0
DB = 0
SM = 0
Delta_AUC
Doxycycline

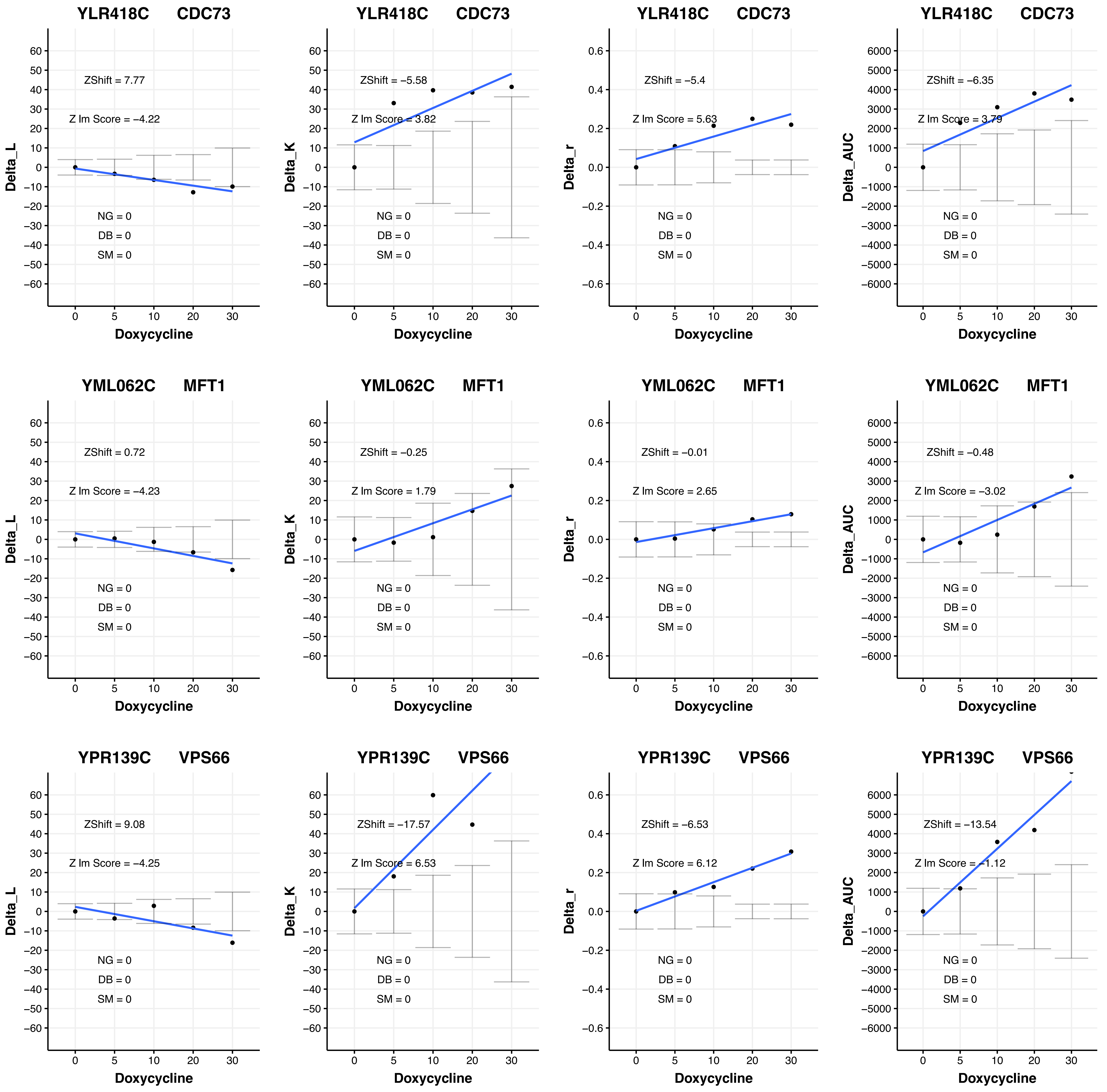

YLR418C CDC73
ZShift = 7.77
Z Im Score = −4.22
NG = 0
DB = 0
SM = 0
Delta_L
Doxycycline
YLR418C CDC73
ZShift = −5.58
Z Im Score = 3.82
NG = 0
DB = 0
SM = 0
Delta_K
Doxycycline
YLR418C CDC73
ZShift = −5.4
Z Im Score = 5.63
NG = 0
DB = 0
SM = 0
Delta_r
Doxycycline
YLR418C CDC73
ZShift = −6.35
Z Im Score = 3.79
NG = 0
DB = 0
SM = 0
Delta_AUC
Doxycycline
YML062C MFT1
ZShift = 0.72
Z Im Score = −4.23
NG = 0
DB = 0
SM = 0
Delta_L
Doxycycline
YML062C MFT1
ZShift = −0.25
Z Im Score = 1.79
NG = 0
DB = 0
SM = 0
Delta_K
Doxycycline
YML062C MFT1
ZShift = −0.01
Z Im Score = 2.65
NG = 0
DB = 0
SM = 0
Delta_r
Doxycycline
YML062C MFT1
ZShift = −0.48
Z Im Score = −3.02
NG = 0
DB = 0
SM = 0
Delta_AUC
Doxycycline
YPR139C VPS66
ZShift = 9.08
Z Im Score = −4.25
NG = 0
DB = 0
SM = 0
Delta_L
Doxycycline
YPR139C VPS66
ZShift = −17.57
Z Im Score = 6.53
NG = 0
DB = 0
SM = 0
Delta_K
Doxycycline
YPR139C VPS66
ZShift = −6.53
Z Im Score = 6.12
NG = 0
DB = 0
SM = 0
Delta_r
Doxycycline
YPR139C VPS66
ZShift = −13.54
Z Im Score = −1.12
NG = 0
DB = 0
SM = 0
Delta_AUC
Doxycycline

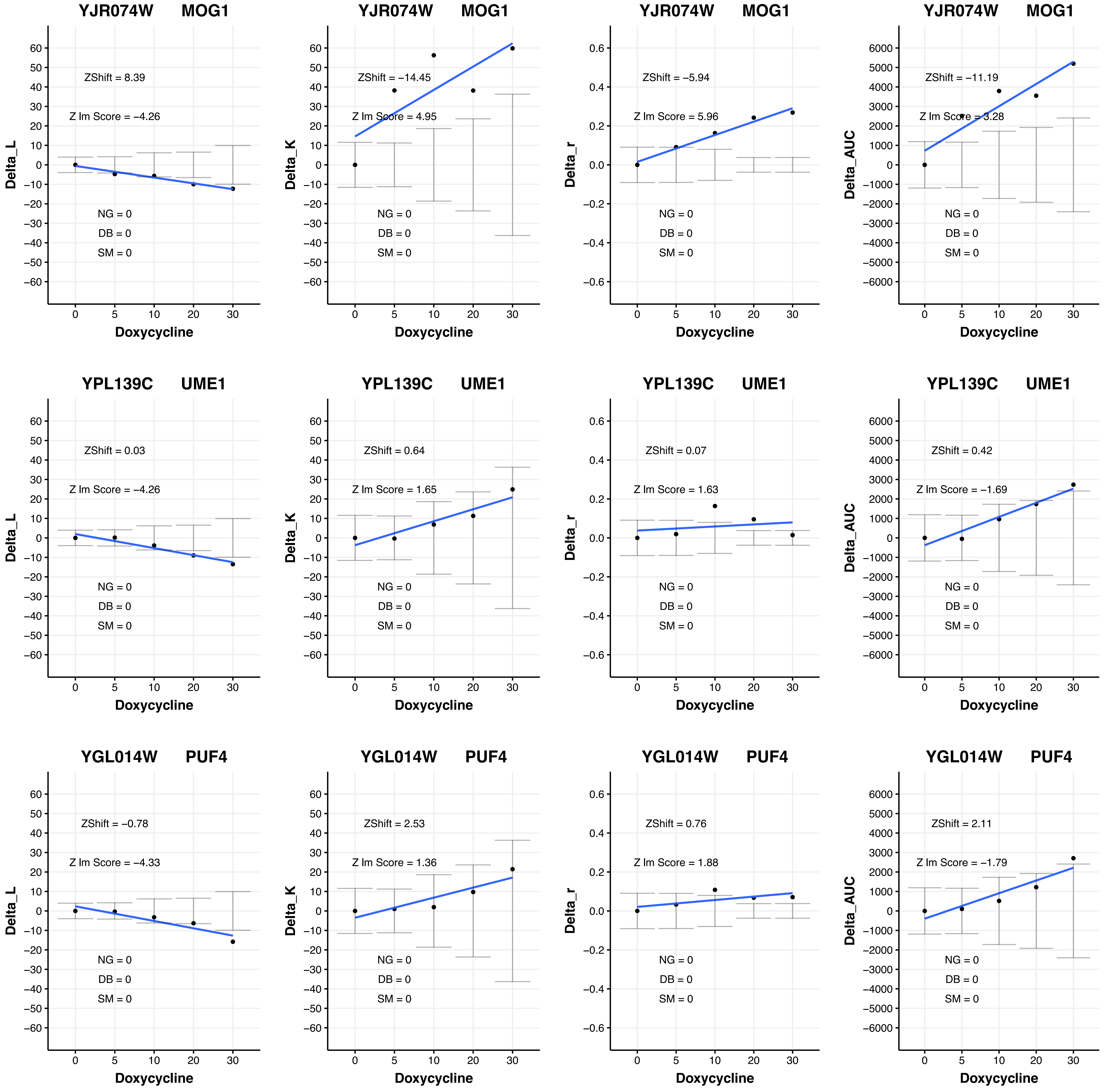
YJR074W MOG1
ZShift = 8.39
Z lm Score = −4.26
NG = 0
DB = 0
SM = 0
Delta_L
Doxycycline
YJR074W MOG1
ZShift = −14.45
Z lm Score = 4.95
NG = 0
DB = 0
SM = 0
Delta_K
Doxycycline
YJR074W MOG1
ZShift = −5.94
Z lm Score = 5.96
NG = 0
DB = 0
SM = 0
Delta_r
Doxycycline
YJR074W MOG1
ZShift = −11.19
Z lm Score = 3.28
NG = 0
DB = 0
SM = 0
Delta_AUC
Doxycycline
YPL139C UME1
ZShift = 0.03
Z lm Score = −4.26
NG = 0
DB = 0
SM = 0
Delta_L
Doxycycline
YPL139C UME1
ZShift = 0.64
Z lm Score = 1.65
NG = 0
DB = 0
SM = 0
Delta_K
Doxycycline
YPL139C UME1
ZShift = 0.07
Z lm Score = 1.63
NG = 0
DB = 0
SM = 0
Delta_r
Doxycycline
YPL139C UME1
ZShift = 0.42
Z lm Score = −1.69
NG = 0
DB = 0
SM = 0
Delta_AUC
Doxycycline
YGL014W PUF4
ZShift = −0.78
Z lm Score = −4.33
NG = 0
DB = 0
SM = 0
Delta_L
Doxycycline
YGL014W PUF4
ZShift = 2.53
Z lm Score = 1.36
NG = 0
DB = 0
SM = 0
Delta_K
Doxycycline
YGL014W PUF4
ZShift = 0.76
Z lm Score = 1.88
NG = 0
DB = 0
SM = 0
Delta_r
Doxycycline
YGL014W PUF4
ZShift = 2.11
Z lm Score = −1.79
NG = 0
DB = 0
SM = 0
Delta_AUC
Doxycycline

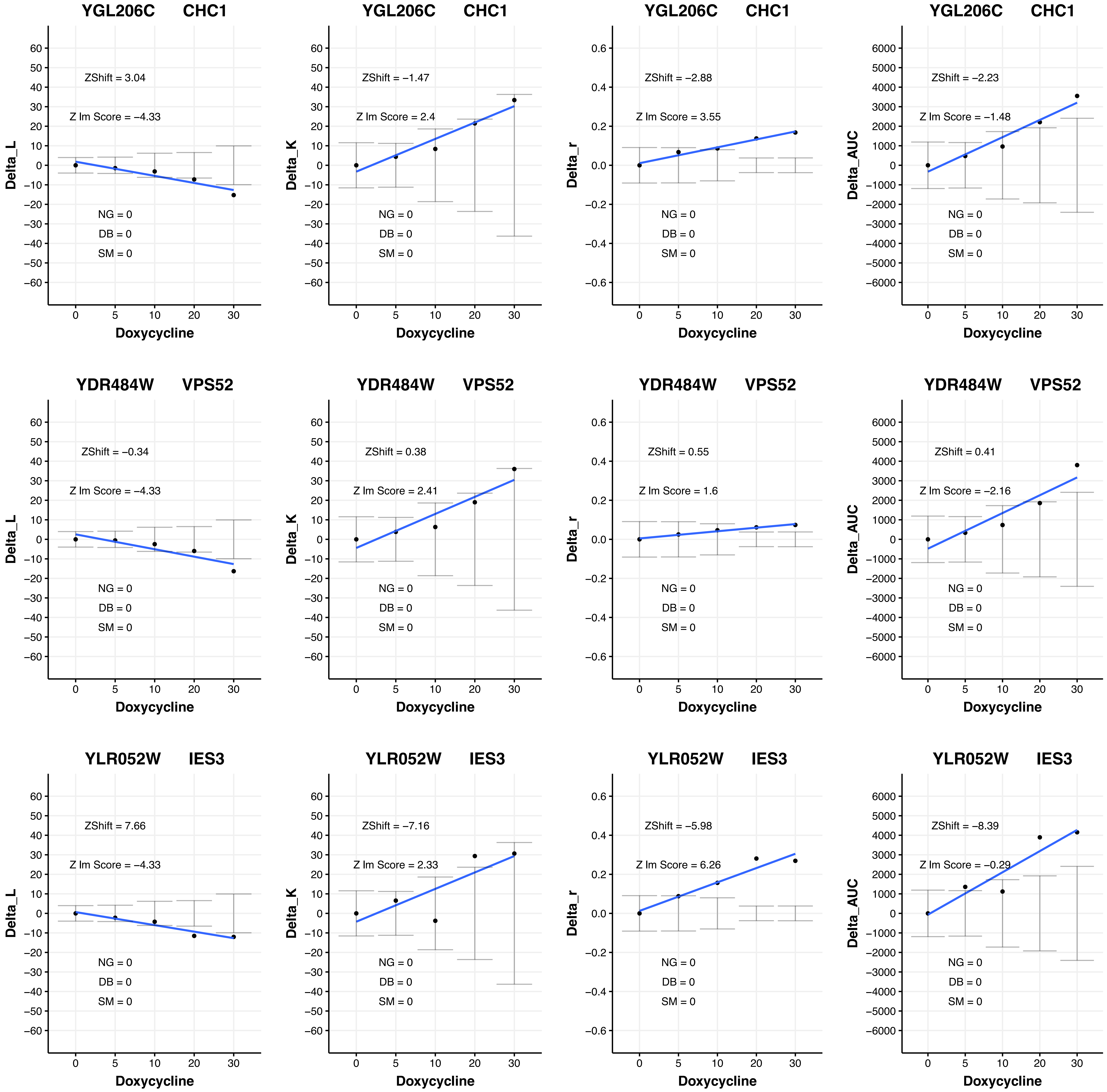

YGL206C CHC1
ZShift = 3.04
Z lm Score = −4.33
NG = 0
DB = 0
SM = 0
Delta_L
Doxycycline
YGL206C CHC1
ZShift = −1.47
Z lm Score = 2.4
NG = 0
DB = 0
SM = 0
Delta_K
Doxycycline
YGL206C CHC1
ZShift = −2.88
Z lm Score = 3.55
NG = 0
DB = 0
SM = 0
Delta_r
Doxycycline
YGL206C CHC1
ZShift = −2.23
Z lm Score = −1.48
NG = 0
DB = 0
SM = 0
Delta_AUC
Doxycycline
YDR484W VPS52
ZShift = −0.34
Z lm Score = −4.33
NG = 0
DB = 0
SM = 0
Delta_L
Doxycycline
YDR484W VPS52
ZShift = 0.38
Z lm Score = 2.41
NG = 0
DB = 0
SM = 0
Delta_K
Doxycycline
YDR484W VPS52
ZShift = 0.55
Z lm Score = 1.6
NG = 0
DB = 0
SM = 0
Delta_r
Doxycycline
YDR484W VPS52
ZShift = 0.41
Z lm Score = −2.16
NG = 0
DB = 0
SM = 0
Delta_AUC
Doxycycline
YLR052W IES3
ZShift = 7.66
Z lm Score = −4.33
NG = 0
DB = 0
SM = 0
Delta_L
Doxycycline
YLR052W IES3
ZShift = −7.16
Z lm Score = 2.33
NG = 0
DB = 0
SM = 0
Delta_K
Doxycycline
YLR052W IES3
ZShift = −5.98
Z lm Score = 6.26
NG = 0
DB = 0
SM = 0
Delta_r
Doxycycline
YLR052W IES3
ZShift = −8.39
Z lm Score = −0.29
NG = 0
DB = 0
SM = 0
Delta_AUC
Doxycycline

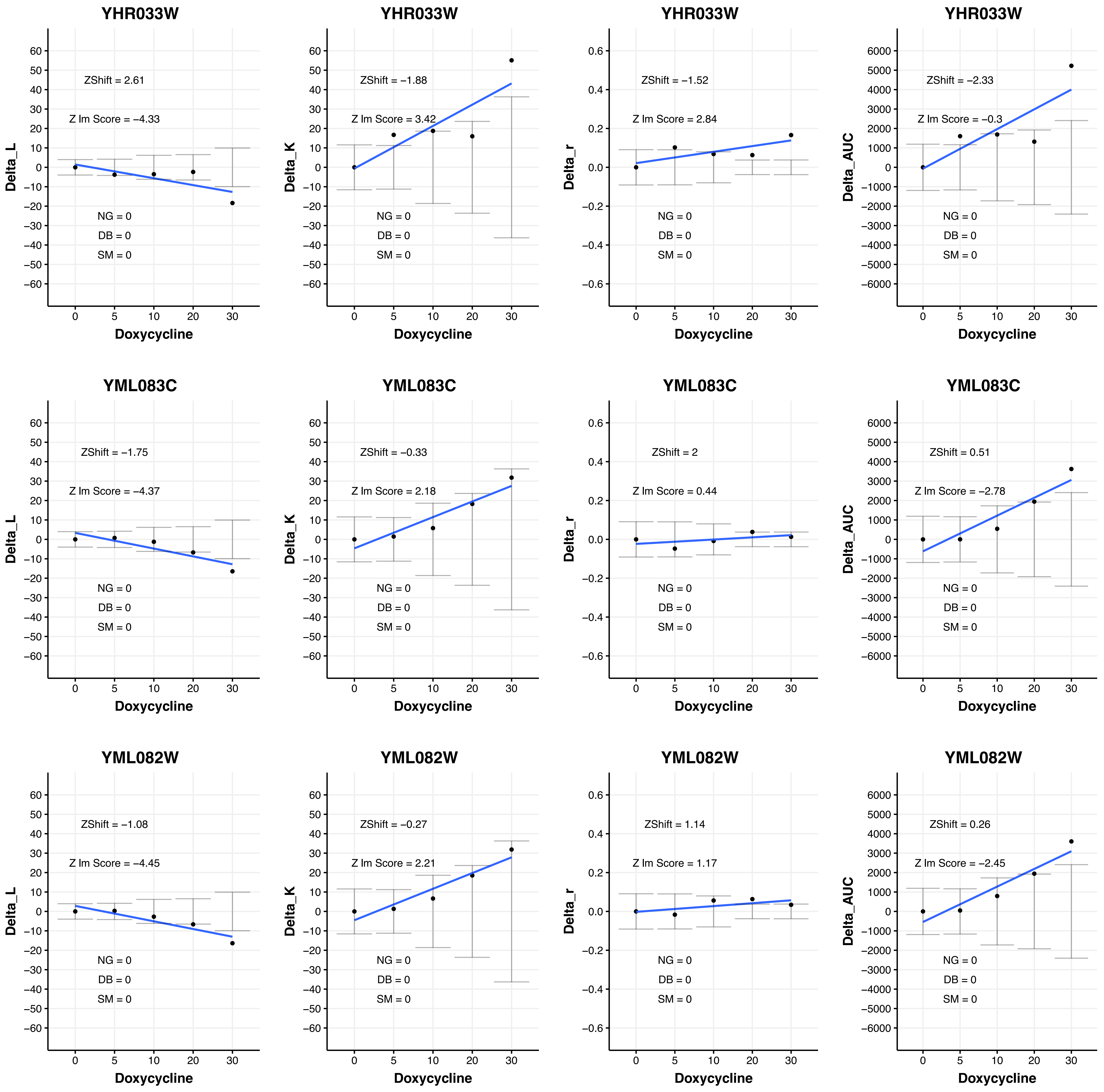
YHR033W
ZShift = 2.61
Z lm Score = −4.33
NG = 0
DB = 0
SM = 0
Delta_L
Doxycycline
YHR033W
ZShift = −1.88
Z lm Score = 3.42
NG = 0
DB = 0
SM = 0
Delta_K
Doxycycline
YHR033W
ZShift = −1.52
Z lm Score = 2.84
NG = 0
DB = 0
SM = 0
Delta_r
Doxycycline
YHR033W
ZShift = −2.33
Z lm Score = −0.3
NG = 0
DB = 0
SM = 0
Delta_AUC
Doxycycline
YML083C
ZShift = −1.75
Z lm Score = −4.37
NG = 0
DB = 0
SM = 0
Delta_L
Doxycycline
YML083C
ZShift = −0.33
Z lm Score = 2.18
NG = 0
DB = 0
SM = 0
Delta_K
Doxycycline
YML083C
ZShift = 2
Z lm Score = 0.44
NG = 0
DB = 0
SM = 0
Delta_r
Doxycycline
YML083C
ZShift = 0.51
Z lm Score = −2.78
NG = 0
DB = 0
SM = 0
Delta_AUC
Doxycycline
YML082W
ZShift = −1.08
Z lm Score = −4.45
NG = 0
DB = 0
SM = 0
Delta_L
Doxycycline
YML082W
ZShift = −0.27
Z lm Score = 2.21
NG = 0
DB = 0
SM = 0
Delta_K
Doxycycline
YML082W
ZShift = 1.14
Z lm Score = 1.17
NG = 0
DB = 0
SM = 0
Delta_r
Doxycycline
YML082W
ZShift = 0.26
Z lm Score = −2.45
NG = 0
DB = 0
SM = 0
Delta_AUC
Doxycycline

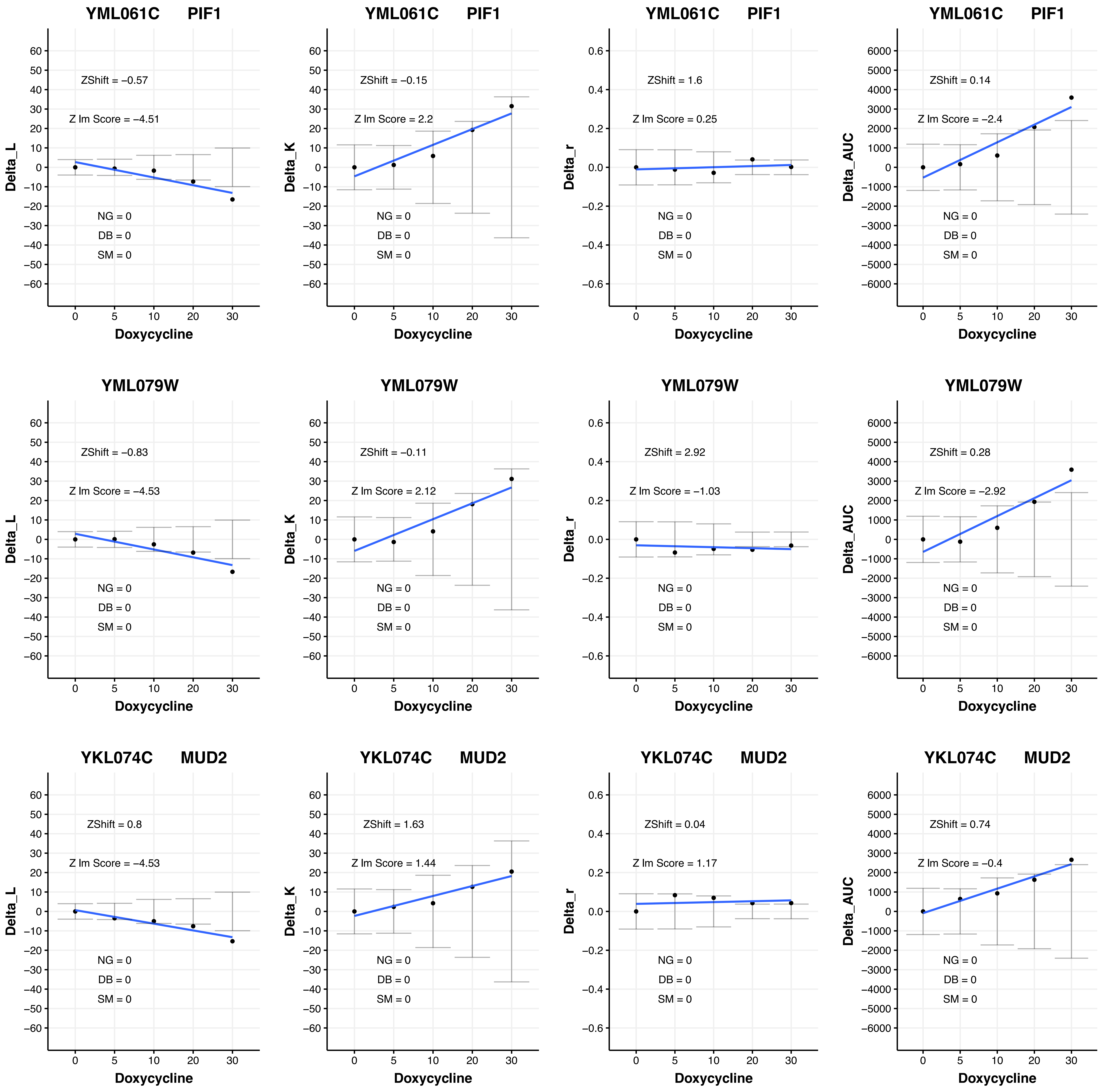

YML061C PIF1
ZShift = −0.57
Z lm Score = −4.51
NG = 0
DB = 0
SM = 0
Delta_L
Doxycycline
YML061C PIF1
ZShift = −0.15
Z lm Score = 2.2
NG = 0
DB = 0
SM = 0
Delta_K
Doxycycline
YML061C PIF1
ZShift = 1.6
Z lm Score = 0.25
NG = 0
DB = 0
SM = 0
Delta_r
Doxycycline
YML061C PIF1
ZShift = 0.14
Z lm Score = −2.4
NG = 0
DB = 0
SM = 0
Delta_AUC
Doxycycline
YML079W
ZShift = −0.83
Z lm Score = −4.53
NG = 0
DB = 0
SM = 0
Delta_L
Doxycycline
YML079W
ZShift = −0.11
Z lm Score = 2.12
NG = 0
DB = 0
SM = 0
Delta_K
Doxycycline
YML079W
ZShift = 2.92
Z lm Score = −1.03
NG = 0
DB = 0
SM = 0
Delta_r
Doxycycline
YML079W
ZShift = 0.28
Z lm Score = −2.92
NG = 0
DB = 0
SM = 0
Delta_AUC
Doxycycline
YKL074C MUD2
ZShift = 0.8
Z lm Score = −4.53
NG = 0
DB = 0
SM = 0
Delta_L
Doxycycline
YKL074C MUD2
ZShift = 1.63
Z lm Score = 1.44
NG = 0
DB = 0
SM = 0
Delta_K
Doxycycline
YKL074C MUD2
ZShift = 0.04
Z lm Score = 1.17
NG = 0
DB = 0
SM = 0
Delta_r
Doxycycline
YKL074C MUD2
ZShift = 0.74
Z lm Score = −0.4
NG = 0
DB = 0
SM = 0
Delta_AUC
Doxycycline

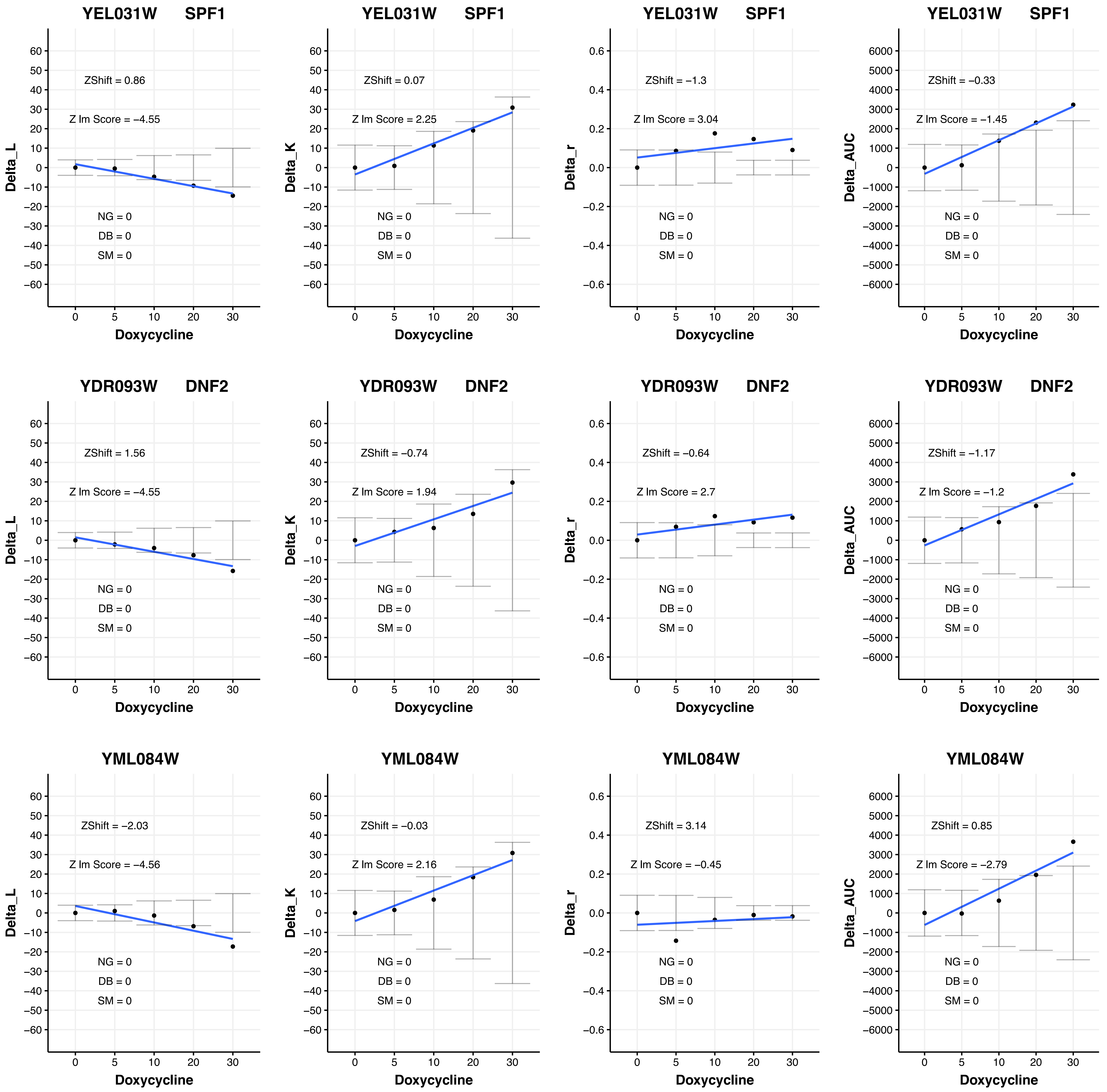
YEL031W SPF1
Delta_L
ZShift = 0.86
Z lm Score = −4.55
NG = 0
DB = 0
SM = 0
Doxycycline
YEL031W SPF1
Delta_K
ZShift = 0.07
Z lm Score = 2.25
NG = 0
DB = 0
SM = 0
Doxycycline
YEL031W SPF1
Delta_r
ZShift = −1.3
Z lm Score = 3.04
NG = 0
DB = 0
SM = 0
Doxycycline
YEL031W SPF1
Delta_AUC
ZShift = −0.33
Z lm Score = −1.45
NG = 0
DB = 0
SM = 0
Doxycycline
YDR093W DNF2
Delta_L
ZShift = 1.56
Z lm Score = −4.55
NG = 0
DB = 0
SM = 0
Doxycycline
YDR093W DNF2
Delta_K
ZShift = −0.74
Z lm Score = 1.94
NG = 0
DB = 0
SM = 0
Doxycycline
YDR093W DNF2
Delta_r
ZShift = −0.64
Z lm Score = 2.7
NG = 0
DB = 0
SM = 0
Doxycycline
YDR093W DNF2
Delta_AUC
ZShift = −1.17
Z lm Score = −1.2
NG = 0
DB = 0
SM = 0
Doxycycline
YML084W
Delta_L
ZShift = −2.03
Z lm Score = −4.56
NG = 0
DB = 0
SM = 0
Doxycycline
YML084W
Delta_K
ZShift = −0.03
Z lm Score = 2.16
NG = 0
DB = 0
SM = 0
Doxycycline
YML084W
Delta_r
ZShift = 3.14
Z lm Score = −0.45
NG = 0
DB = 0
SM = 0
Doxycycline
YML084W
Delta_AUC
ZShift = 0.85
Z lm Score = −2.79
NG = 0
DB = 0
SM = 0
Doxycycline

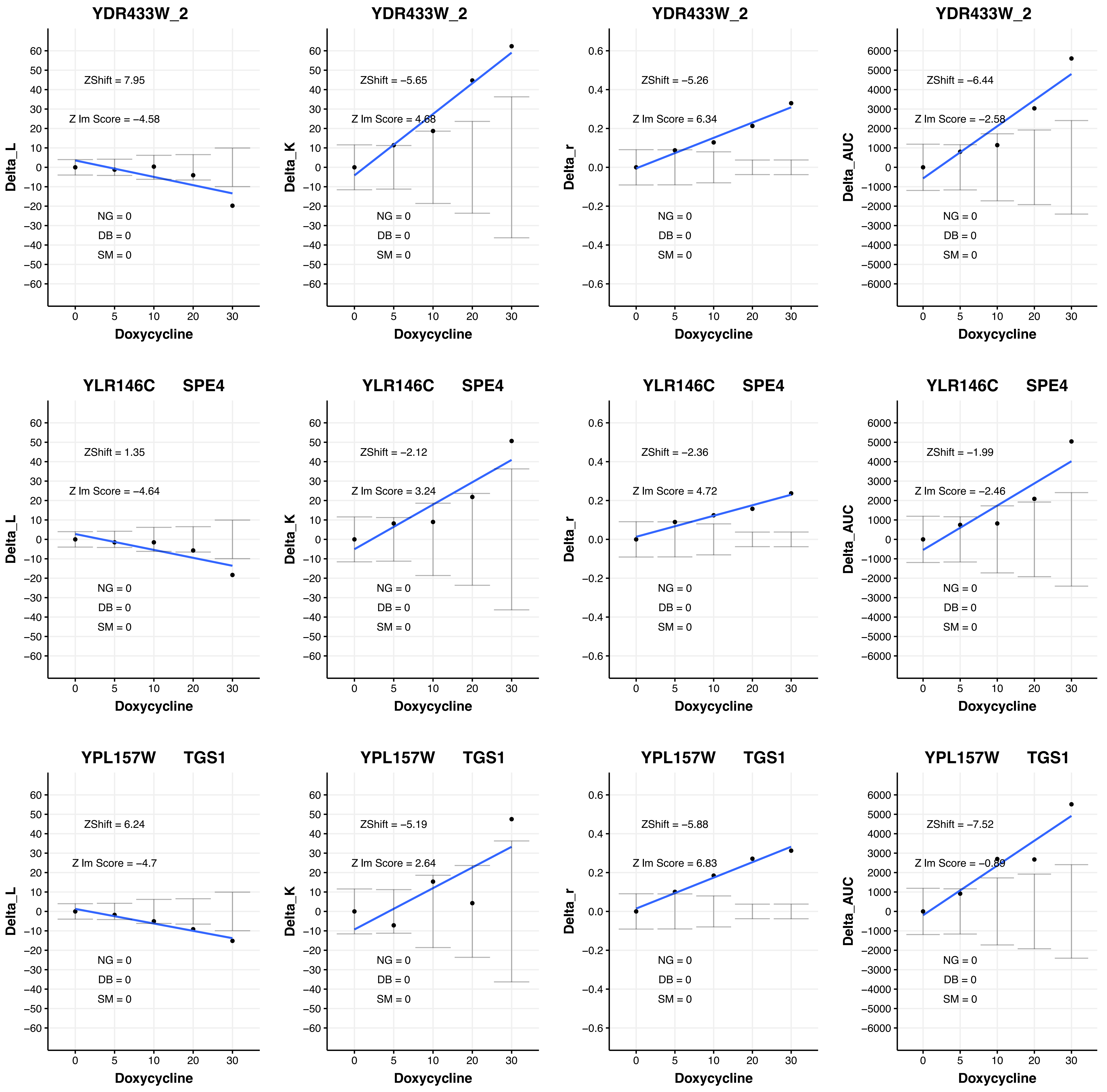

YDR433W_2
ZShift = 7.95
Z lm Score = −4.58
NG = 0
DB = 0
SM = 0
Delta_L
Doxycycline
YDR433W_2
ZShift = −5.65
Z lm Score = 4.68
NG = 0
DB = 0
SM = 0
Delta_K
Doxycycline
YDR433W_2
ZShift = −5.26
Z lm Score = 6.34
NG = 0
DB = 0
SM = 0
Delta_r
Doxycycline
YDR433W_2
ZShift = −6.44
Z lm Score = −2.58
NG = 0
DB = 0
SM = 0
Delta_AUC
Doxycycline
YLR146C SPE4
ZShift = 1.35
Z lm Score = −4.64
NG = 0
DB = 0
SM = 0
Delta_L
Doxycycline
YLR146C SPE4
ZShift = −2.12
Z lm Score = 3.24
NG = 0
DB = 0
SM = 0
Delta_K
Doxycycline
YLR146C SPE4
ZShift = −2.36
Z lm Score = 4.72
NG = 0
DB = 0
SM = 0
Delta_r
Doxycycline
YLR146C SPE4
ZShift = −1.99
Z lm Score = −2.46
NG = 0
DB = 0
SM = 0
Delta_AUC
Doxycycline
YPL157W TGS1
ZShift = 6.24
Z lm Score = −4.7
NG = 0
DB = 0
SM = 0
Delta_L
Doxycycline
YPL157W TGS1
ZShift = −5.19
Z lm Score = 2.64
NG = 0
DB = 0
SM = 0
Delta_K
Doxycycline
YPL157W TGS1
ZShift = −5.88
Z lm Score = 6.83
NG = 0
DB = 0
SM = 0
Delta_r
Doxycycline
YPL157W TGS1
ZShift = −7.52
Z lm Score = −0.89
NG = 0
DB = 0
SM = 0
Delta_AUC
Doxycycline

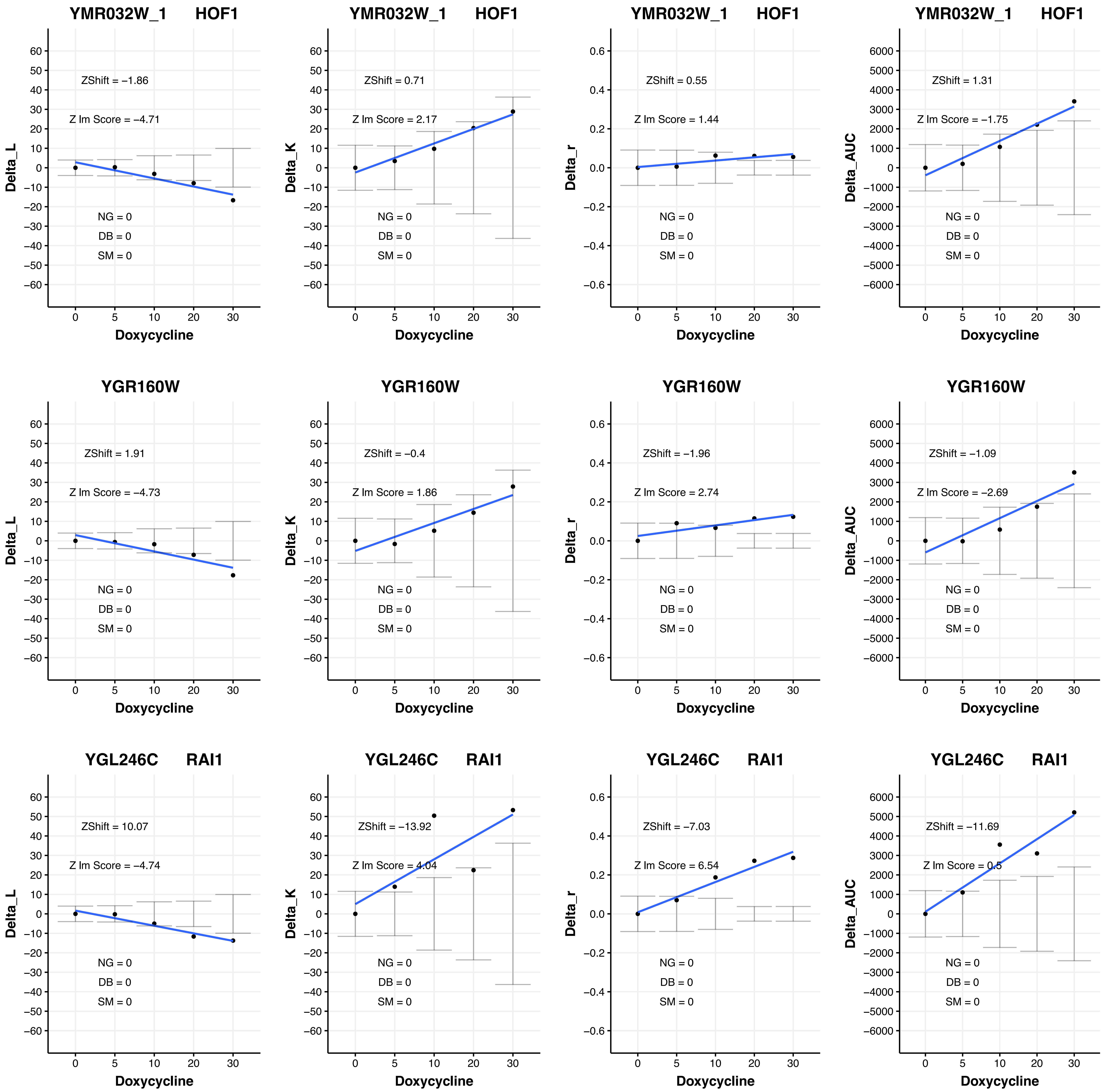
YMR032W_1 HOF1
ZShift = −1.86
Z lm Score = −4.71
NG = 0
DB = 0
SM = 0
Delta_L
Doxycycline
YMR032W_1 HOF1
ZShift = 0.71
Z lm Score = 2.17
NG = 0
DB = 0
SM = 0
Delta_K
Doxycycline
YMR032W_1 HOF1
ZShift = 0.55
Z lm Score = 1.44
NG = 0
DB = 0
SM = 0
Delta_r
Doxycycline
YMR032W_1 HOF1
ZShift = 1.31
Z lm Score = −1.75
NG = 0
DB = 0
SM = 0
Delta_AUC
Doxycycline
YGR160W
ZShift = 1.91
Z lm Score = −4.73
NG = 0
DB = 0
SM = 0
Delta_L
Doxycycline
YGR160W
ZShift = −0.4
Z lm Score = 1.86
NG = 0
DB = 0
SM = 0
Delta_K
Doxycycline
YGR160W
ZShift = −1.96
Z lm Score = 2.74
NG = 0
DB = 0
SM = 0
Delta_r
Doxycycline
YGR160W
ZShift = −1.09
Z lm Score = −2.69
NG = 0
DB = 0
SM = 0
Delta_AUC
Doxycycline
YGL246C RAI1
ZShift = 10.07
Z lm Score = −4.74
NG = 0
DB = 0
SM = 0
Delta_L
Doxycycline
YGL246C RAI1
ZShift = −13.92
Z lm Score = 4.04
NG = 0
DB = 0
SM = 0
Delta_K
Doxycycline
YGL246C RAI1
ZShift = −7.03
Z lm Score = 6.54
NG = 0
DB = 0
SM = 0
Delta_r
Doxycycline
YGL246C RAI1
ZShift = −11.69
Z lm Score = 0.5
NG = 0
DB = 0
SM = 0
Delta_AUC
Doxycycline

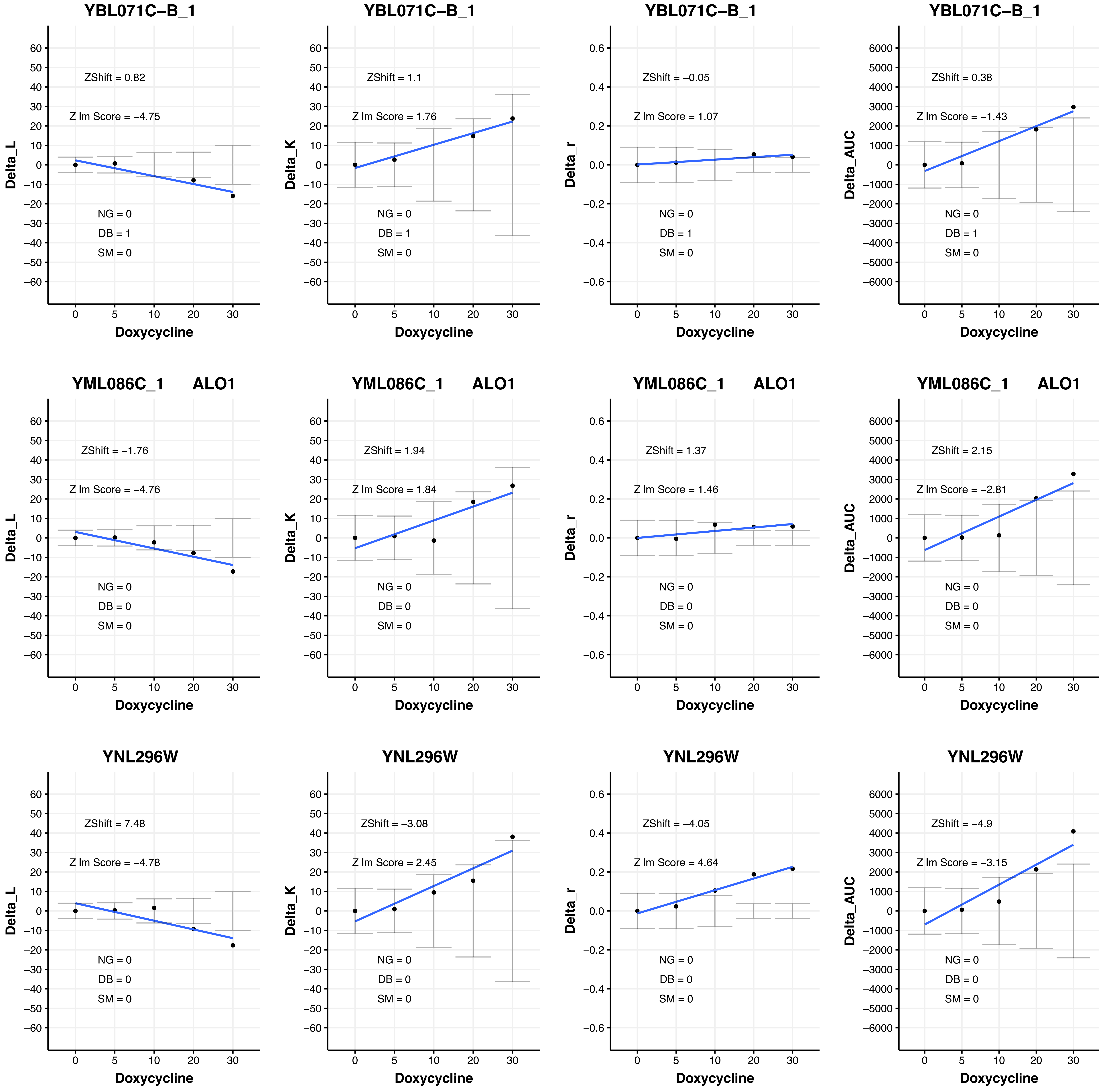
YBL071C–B_1
ZShift = 0.82
Z lm Score = –4.75
NG = 0
DB = 1
SM = 0
Delta_L
Doxycycline
YBL071C–B_1
ZShift = 1.1
Z lm Score = 1.76
NG = 0
DB = 1
SM = 0
Delta_K
Doxycycline
YBL071C–B_1
ZShift = –0.05
Z lm Score = 1.07
NG = 0
DB = 1
SM = 0
Delta_r
Doxycycline
YBL071C–B_1
ZShift = 0.38
Z lm Score = –1.43
NG = 0
DB = 1
SM = 0
Delta_AUC
Doxycycline
YML086C_1 ALO1
ZShift = –1.76
Z lm Score = –4.76
NG = 0
DB = 0
SM = 0
Delta_L
Doxycycline
YML086C_1 ALO1
ZShift = 1.94
Z lm Score = 1.84
NG = 0
DB = 0
SM = 0
Delta_K
Doxycycline
YML086C_1 ALO1
ZShift = 1.37
Z lm Score = 1.46
NG = 0
DB = 0
SM = 0
Delta_r
Doxycycline
YML086C_1 ALO1
ZShift = 2.15
Z lm Score = –2.81
NG = 0
DB = 0
SM = 0
Delta_AUC
Doxycycline
YNL296W
ZShift = 7.48
Z lm Score = –4.78
NG = 0
DB = 0
SM = 0
Delta_L
Doxycycline
YNL296W
ZShift = –3.08
Z lm Score = 2.45
NG = 0
DB = 0
SM = 0
Delta_K
Doxycycline
YNL296W
ZShift = –4.05
Z lm Score = 4.64
NG = 0
DB = 0
SM = 0
Delta_r
Doxycycline
YNL296W
ZShift = –4.9
Z lm Score = –3.15
NG = 0
DB = 0
SM = 0
Delta_AUC
Doxycycline

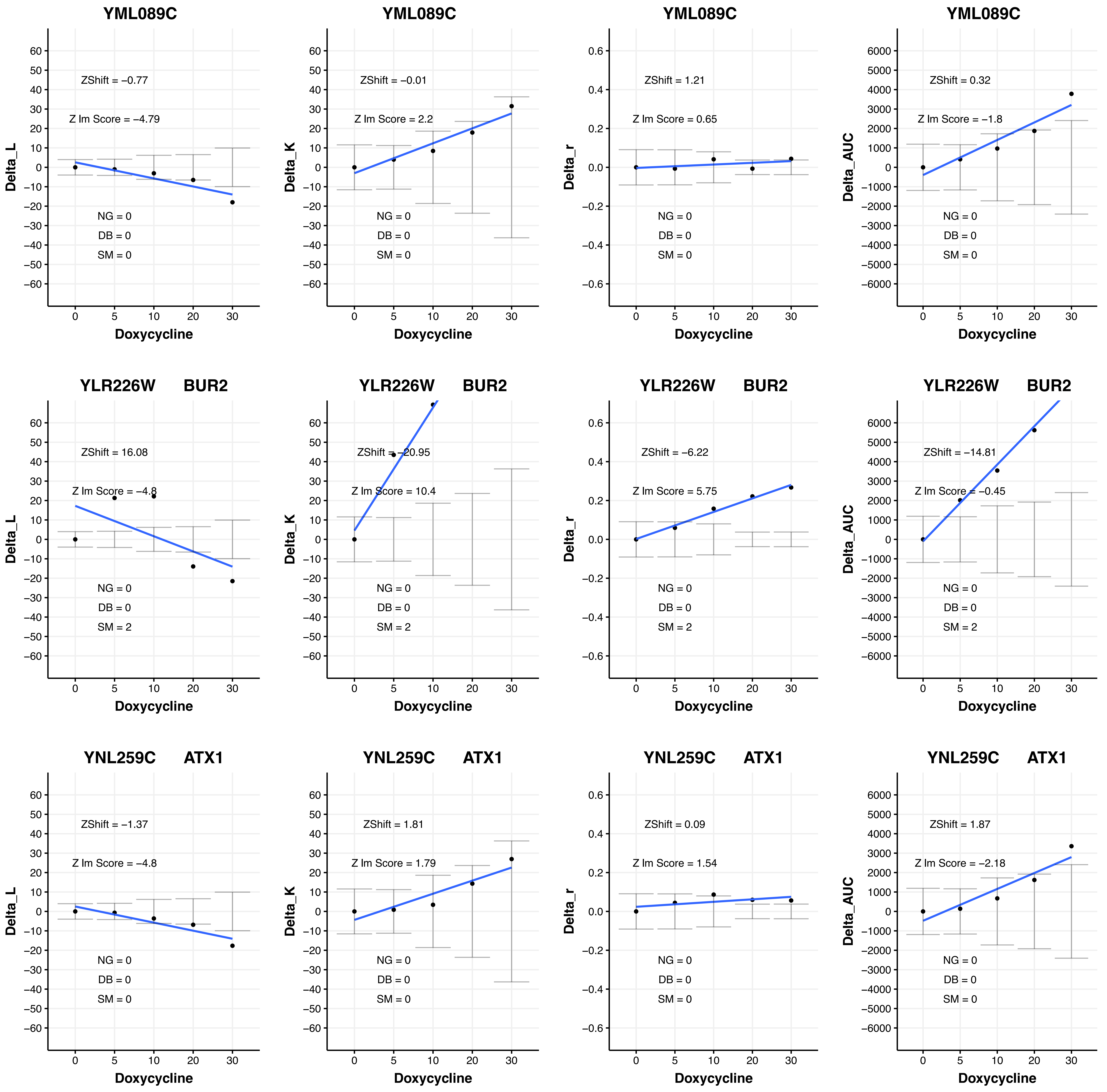
YML089C
ZShift = −0.77
Z lm Score = −4.79
NG = 0
DB = 0
SM = 0
Delta_L
Doxycycline
YML089C
ZShift = −0.01
Z lm Score = 2.2
NG = 0
DB = 0
SM = 0
Delta_K
Doxycycline
YML089C
ZShift = 1.21
Z lm Score = 0.65
NG = 0
DB = 0
SM = 0
Delta_r
Doxycycline
YML089C
ZShift = 0.32
Z lm Score = −1.8
NG = 0
DB = 0
SM = 0
Delta_AUC
Doxycycline
YLR226W BUR2
ZShift = 16.08
Z lm Score = −4.8
NG = 0
DB = 0
SM = 2
Delta_L
Doxycycline
YLR226W BUR2
ZShift = −20.95
Z lm Score = 10.4
NG = 0
DB = 0
SM = 2
Delta_K
Doxycycline
YLR226W BUR2
ZShift = −6.22
Z lm Score = 5.75
NG = 0
DB = 0
SM = 2
Delta_r
Doxycycline
YLR226W BUR2
ZShift = −14.81
Z lm Score = −0.45
NG = 0
DB = 0
SM = 2
Delta_AUC
Doxycycline
YNL259C ATX1
ZShift = −1.37
Z lm Score = −4.8
NG = 0
DB = 0
SM = 0
Delta_L
Doxycycline
YNL259C ATX1
ZShift = 1.81
Z lm Score = 1.79
NG = 0
DB = 0
SM = 0
Delta_K
Doxycycline
YNL259C ATX1
ZShift = 0.09
Z lm Score = 1.54
NG = 0
DB = 0
SM = 0
Delta_r
Doxycycline
YNL259C ATX1
ZShift = 1.87
Z lm Score = −2.18
NG = 0
DB = 0
SM = 0
Delta_AUC
Doxycycline

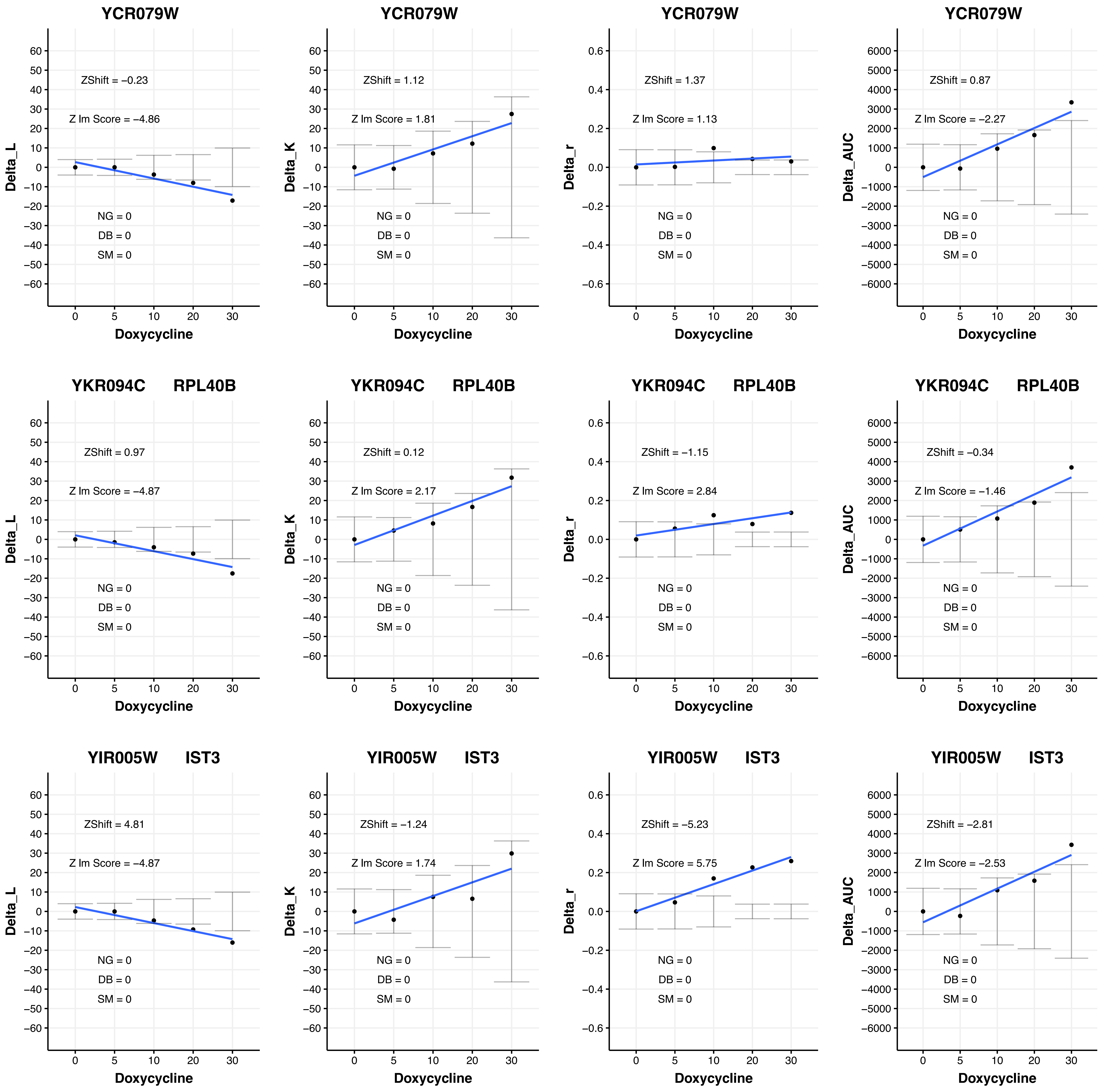

YCR079W
ZShift = −0.23
Z lm Score = −4.86
NG = 0
DB = 0
SM = 0
Delta_L
Doxycycline
YCR079W
ZShift = 1.12
Z lm Score = 1.81
NG = 0
DB = 0
SM = 0
Delta_K
Doxycycline
YCR079W
ZShift = 1.37
Z lm Score = 1.13
NG = 0
DB = 0
SM = 0
Delta_r
Doxycycline
YCR079W
ZShift = 0.87
Z lm Score = −2.27
NG = 0
DB = 0
SM = 0
Delta_AUC
Doxycycline
YKR094C RPL40B
ZShift = 0.97
Z lm Score = −4.87
NG = 0
DB = 0
SM = 0
Delta_L
Doxycycline
YKR094C RPL40B
ZShift = 0.12
Z lm Score = 2.17
NG = 0
DB = 0
SM = 0
Delta_K
Doxycycline
YKR094C RPL40B
ZShift = −1.15
Z lm Score = 2.84
NG = 0
DB = 0
SM = 0
Delta_r
Doxycycline
YKR094C RPL40B
ZShift = −0.34
Z lm Score = −1.46
NG = 0
DB = 0
SM = 0
Delta_AUC
Doxycycline
YIR005W IST3
ZShift = 4.81
Z lm Score = −4.87
NG = 0
DB = 0
SM = 0
Delta_L
Doxycycline
YIR005W IST3
ZShift = −1.24
Z lm Score = 1.74
NG = 0
DB = 0
SM = 0
Delta_K
Doxycycline
YIR005W IST3
ZShift = −5.23
Z lm Score = 5.75
NG = 0
DB = 0
SM = 0
Delta_r
Doxycycline
YIR005W IST3
ZShift = −2.81
Z lm Score = −2.53
NG = 0
DB = 0
SM = 0
Delta_AUC
Doxycycline

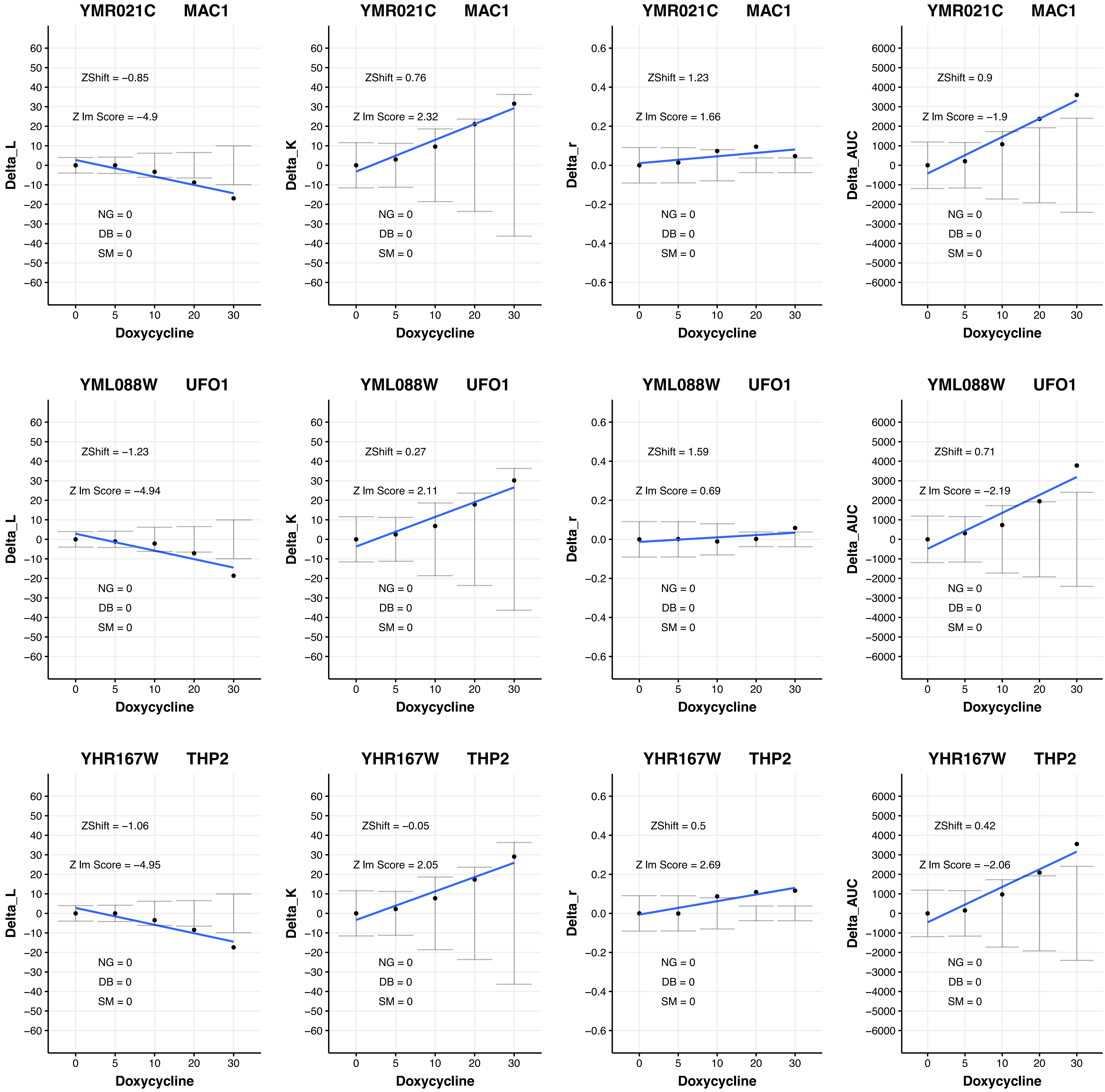
YMR021C MAC1
ZShift = −0.85
Z lm Score = −4.9
NG = 0
DB = 0
SM = 0
Delta_L
Doxycycline
YMR021C MAC1
ZShift = 0.76
Z lm Score = 2.32
NG = 0
DB = 0
SM = 0
Delta_K
Doxycycline
YMR021C MAC1
ZShift = 1.23
Z lm Score = 1.66
NG = 0
DB = 0
SM = 0
Delta_r
Doxycycline
YMR021C MAC1
ZShift = 0.9
Z lm Score = −1.9
NG = 0
DB = 0
SM = 0
Delta_AUC
Doxycycline
YML088W UFO1
ZShift = −1.23
Z lm Score = −4.94
NG = 0
DB = 0
SM = 0
Delta_L
Doxycycline
YML088W UFO1
ZShift = 0.27
Z lm Score = 2.11
NG = 0
DB = 0
SM = 0
Delta_K
Doxycycline
YML088W UFO1
ZShift = 1.59
Z lm Score = 0.69
NG = 0
DB = 0
SM = 0
Delta_r
Doxycycline
YML088W UFO1
ZShift = 0.71
Z lm Score = −2.19
NG = 0
DB = 0
SM = 0
Delta_AUC
Doxycycline
YHR167W THP2
ZShift = −1.06
Z lm Score = −4.95
NG = 0
DB = 0
SM = 0
Delta_L
Doxycycline
YHR167W THP2
ZShift = −0.05
Z lm Score = 2.05
NG = 0
DB = 0
SM = 0
Delta_K
Doxycycline
YHR167W THP2
ZShift = 0.5
Z lm Score = 2.69
NG = 0
DB = 0
SM = 0
Delta_r
Doxycycline
YHR167W THP2
ZShift = 0.42
Z lm Score = −2.06
NG = 0
DB = 0
SM = 0
Delta_AUC
Doxycycline

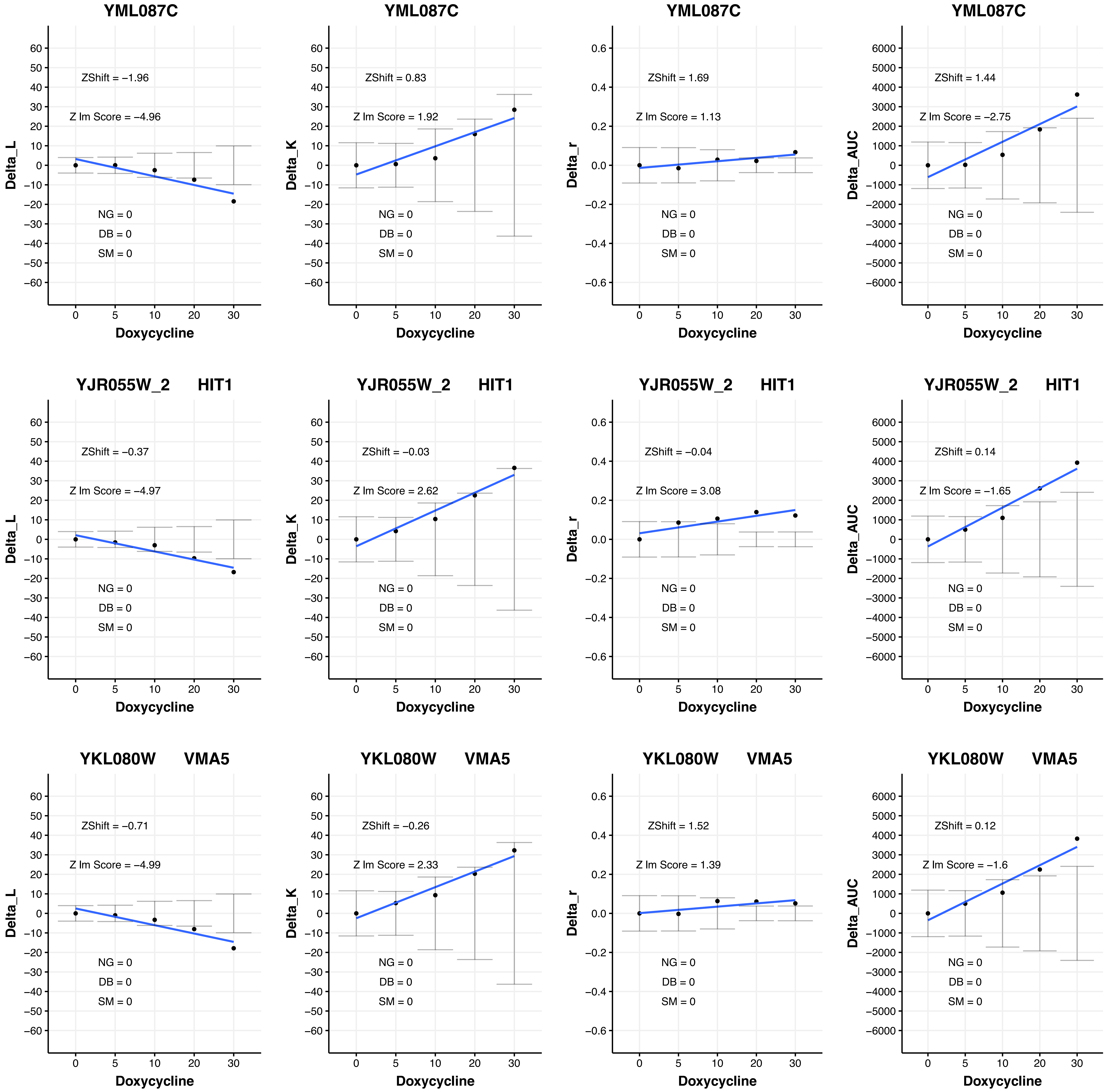

YML087C
ZShift = −1.96
Z lm Score = −4.96
NG = 0
DB = 0
SM = 0
Delta_L
Doxycycline
YML087C
ZShift = 0.83
Z lm Score = 1.92
NG = 0
DB = 0
SM = 0
Delta_K
Doxycycline
YML087C
ZShift = 1.69
Z lm Score = 1.13
NG = 0
DB = 0
SM = 0
Delta_r
Doxycycline
YML087C
ZShift = 1.44
Z lm Score = −2.75
NG = 0
DB = 0
SM = 0
Delta_AUC
Doxycycline
YJR055W_2 HIT1
ZShift = −0.37
Z lm Score = −4.97
NG = 0
DB = 0
SM = 0
Delta_L
Doxycycline
YJR055W_2 HIT1
ZShift = −0.03
Z lm Score = 2.62
NG = 0
DB = 0
SM = 0
Delta_K
Doxycycline
YJR055W_2 HIT1
ZShift = −0.04
Z lm Score = 3.08
NG = 0
DB = 0
SM = 0
Delta_r
Doxycycline
YJR055W_2 HIT1
ZShift = 0.14
Z lm Score = −1.65
NG = 0
DB = 0
SM = 0
Delta_AUC
Doxycycline
YKL080W VMA5
ZShift = −0.71
Z lm Score = −4.99
NG = 0
DB = 0
SM = 0
Delta_L
Doxycycline
YKL080W VMA5
ZShift = −0.26
Z lm Score = 2.33
NG = 0
DB = 0
SM = 0
Delta_K
Doxycycline
YKL080W VMA5
ZShift = 1.52
Z lm Score = 1.39
NG = 0
DB = 0
SM = 0
Delta_r
Doxycycline
YKL080W VMA5
ZShift = 0.12
Z lm Score = −1.6
NG = 0
DB = 0
SM = 0
Delta_AUC
Doxycycline

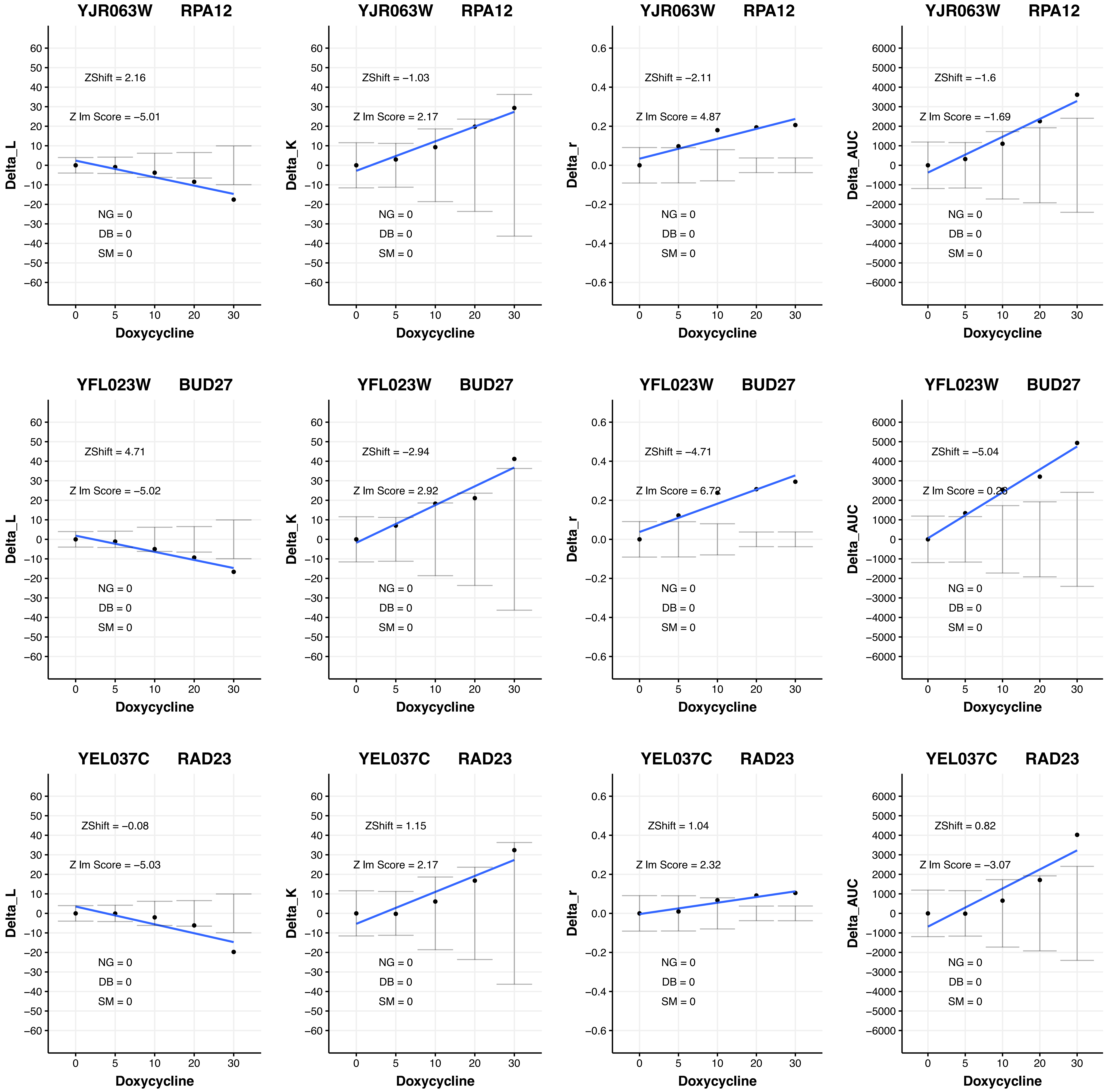

YJR063W RPA12
ZShift = 2.16
Z lm Score = −5.01
NG = 0
DB = 0
SM = 0
Delta_L
Doxycycline
YJR063W RPA12
ZShift = −1.03
Z lm Score = 2.17
NG = 0
DB = 0
SM = 0
Delta_K
Doxycycline
YJR063W RPA12
ZShift = −2.11
Z lm Score = 4.87
NG = 0
DB = 0
SM = 0
Delta_r
Doxycycline
YJR063W RPA12
ZShift = −1.6
Z lm Score = −1.69
NG = 0
DB = 0
SM = 0
Delta_AUC
Doxycycline
YFL023W BUD27
ZShift = 4.71
Z lm Score = −5.02
NG = 0
DB = 0
SM = 0
Delta_L
Doxycycline
YFL023W BUD27
ZShift = −2.94
Z lm Score = 2.92
NG = 0
DB = 0
SM = 0
Delta_K
Doxycycline
YFL023W BUD27
ZShift = −4.71
Z lm Score = 6.72
NG = 0
DB = 0
SM = 0
Delta_r
Doxycycline
YFL023W BUD27
ZShift = −5.04
Z lm Score = 0.28
NG = 0
DB = 0
SM = 0
Delta_AUC
Doxycycline
YEL037C RAD23
ZShift = −0.08
Z lm Score = −5.03
NG = 0
DB = 0
SM = 0
Delta_L
Doxycycline
YEL037C RAD23
ZShift = 1.15
Z lm Score = 2.17
NG = 0
DB = 0
SM = 0
Delta_K
Doxycycline
YEL037C RAD23
ZShift = 1.04
Z lm Score = 2.32
NG = 0
DB = 0
SM = 0
Delta_r
Doxycycline
YEL037C RAD23
ZShift = 0.82
Z lm Score = −3.07
NG = 0
DB = 0
SM = 0
Delta_AUC
Doxycycline

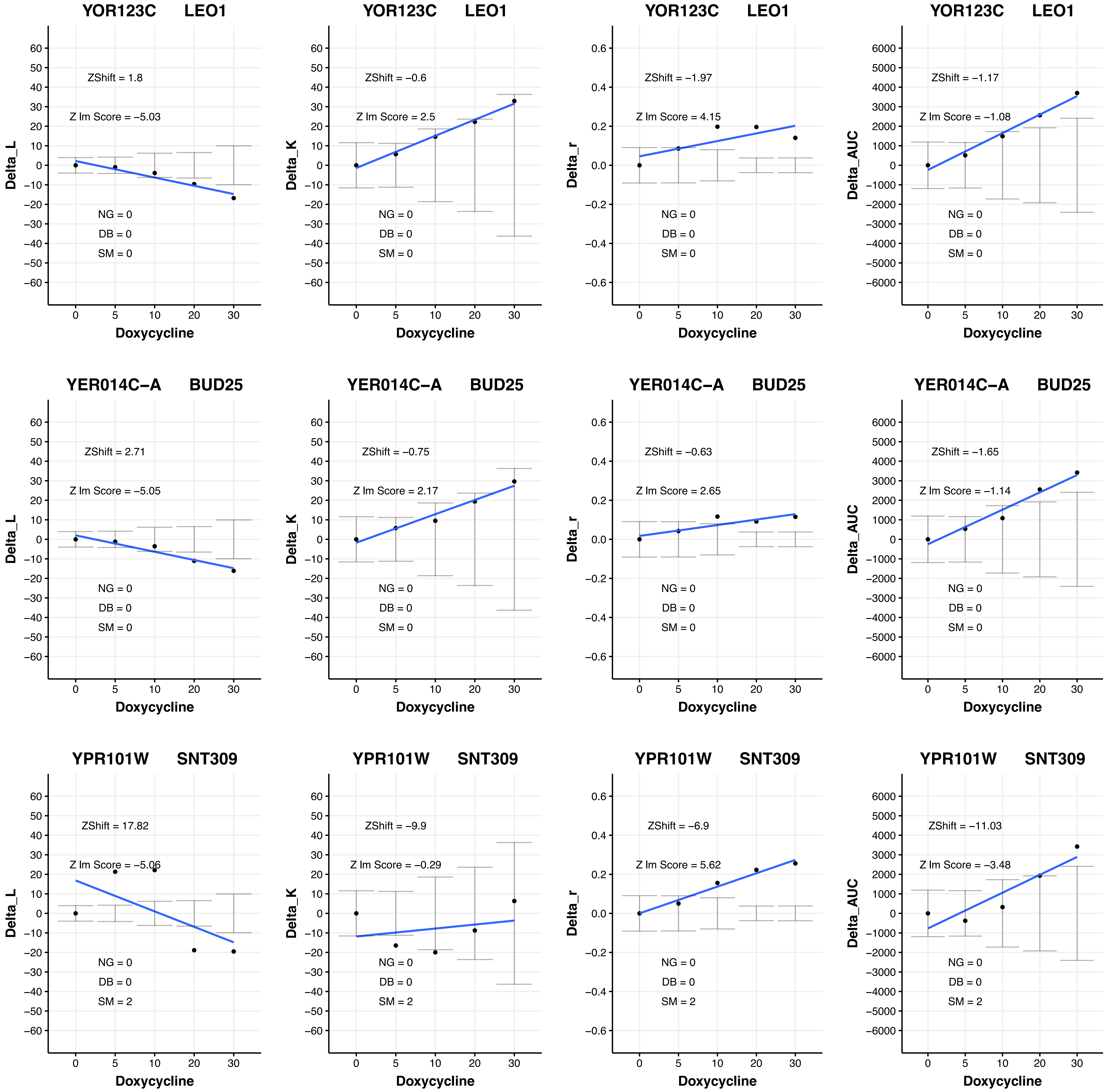

YOR123C LEO1
ZShift = 1.8
Z Im Score = −5.03
NG = 0
DB = 0
SM = 0
Delta_L
Doxycycline
YOR123C LEO1
ZShift = −0.6
Z Im Score = 2.5
NG = 0
DB = 0
SM = 0
Delta_K
Doxycycline
YOR123C LEO1
ZShift = −1.97
Z Im Score = 4.15
NG = 0
DB = 0
SM = 0
Delta_r
Doxycycline
YOR123C LEO1
ZShift = −1.17
Z Im Score = −1.08
NG = 0
DB = 0
SM = 0
Delta_AUC
Doxycycline
YER014C−A BUD25
ZShift = 2.71
Z Im Score = −5.05
NG = 0
DB = 0
SM = 0
Delta_L
Doxycycline
YER014C−A BUD25
ZShift = −0.75
Z Im Score = 2.17
NG = 0
DB = 0
SM = 0
Delta_K
Doxycycline
YER014C−A BUD25
ZShift = −0.63
Z Im Score = 2.65
NG = 0
DB = 0
SM = 0
Delta_r
Doxycycline
YER014C−A BUD25
ZShift = −1.65
Z Im Score = −1.14
NG = 0
DB = 0
SM = 0
Delta_AUC
Doxycycline
YPR101W SNT309
ZShift = 17.82
Z Im Score = −5.06
NG = 0
DB = 0
SM = 2
Delta_L
Doxycycline
YPR101W SNT309
ZShift = −9.9
Z Im Score = −0.29
NG = 0
DB = 0
SM = 2
Delta_K
Doxycycline
YPR101W SNT309
ZShift = −6.9
Z Im Score = 5.62
NG = 0
DB = 0
SM = 2
Delta_r
Doxycycline
YPR101W SNT309
ZShift = −11.03
Z Im Score = −3.48
NG = 0
DB = 0
SM = 2
Delta_AUC
Doxycycline

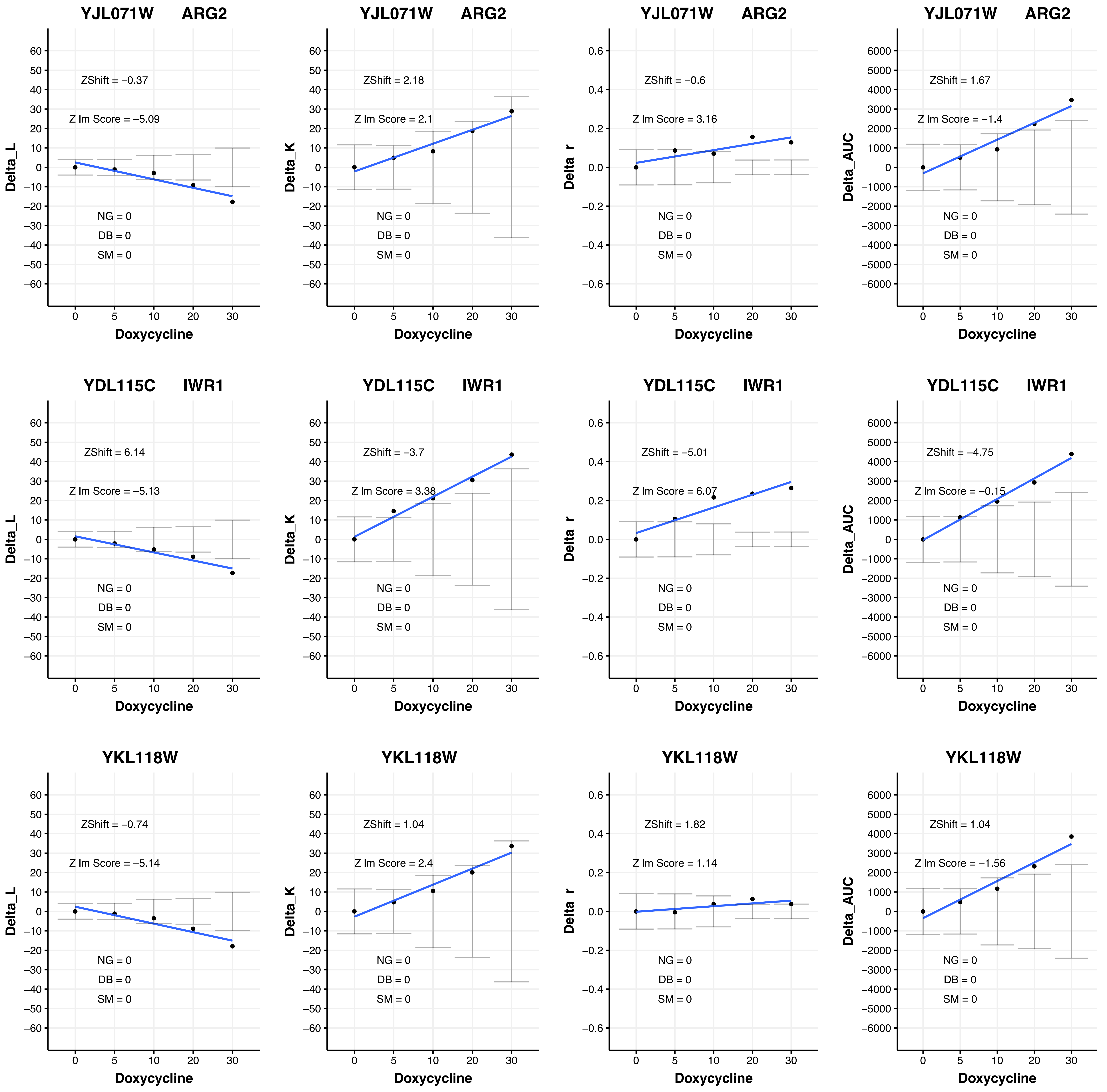

YJL071W ARG2
ZShift = −0.37
Z lm Score = −5.09
NG = 0
DB = 0
SM = 0
Delta_L
Doxycycline
YJL071W ARG2
ZShift = 2.18
Z lm Score = 2.1
NG = 0
DB = 0
SM = 0
Delta_K
Doxycycline
YJL071W ARG2
ZShift = −0.6
Z lm Score = 3.16
NG = 0
DB = 0
SM = 0
Delta_r
Doxycycline
YJL071W ARG2
ZShift = 1.67
Z lm Score = −1.4
NG = 0
DB = 0
SM = 0
Delta_AUC
Doxycycline
YDL115C IWR1
ZShift = 6.14
Z lm Score = −5.13
NG = 0
DB = 0
SM = 0
Delta_L
Doxycycline
YDL115C IWR1
ZShift = −3.7
Z lm Score = 3.38
NG = 0
DB = 0
SM = 0
Delta_K
Doxycycline
YDL115C IWR1
ZShift = −5.01
Z lm Score = 6.07
NG = 0
DB = 0
SM = 0
Delta_r
Doxycycline
YDL115C IWR1
ZShift = −4.75
Z lm Score = −0.15
NG = 0
DB = 0
SM = 0
Delta_AUC
Doxycycline
YKL118W
ZShift = −0.74
Z lm Score = −5.14
NG = 0
DB = 0
SM = 0
Delta_L
Doxycycline
YKL118W
ZShift = 1.04
Z lm Score = 2.4
NG = 0
DB = 0
SM = 0
Delta_K
Doxycycline
YKL118W
ZShift = 1.82
Z lm Score = 1.14
NG = 0
DB = 0
SM = 0
Delta_r
Doxycycline
YKL118W
ZShift = 1.04
Z lm Score = −1.56
NG = 0
DB = 0
SM = 0
Delta_AUC
Doxycycline

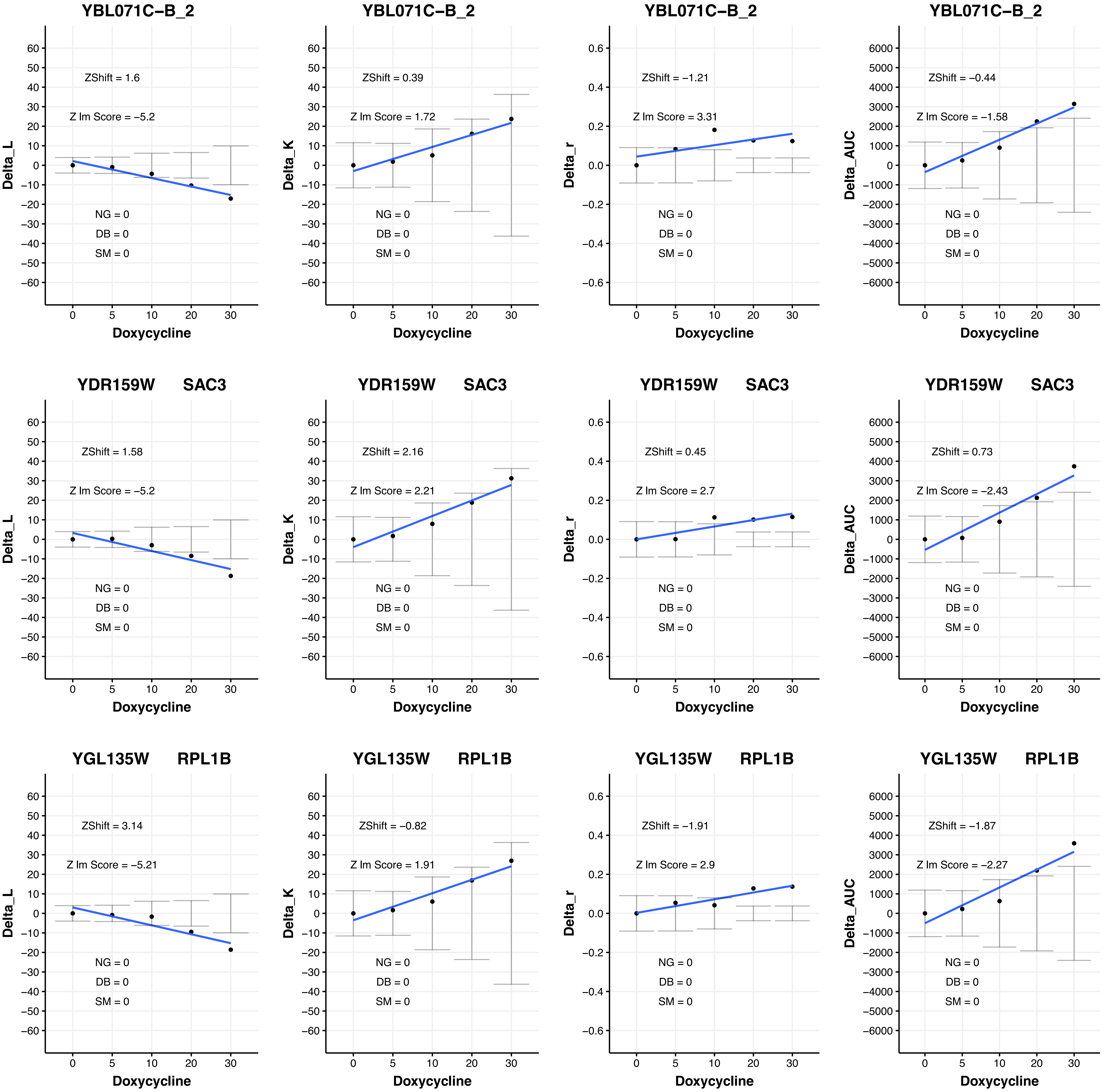
YBL071C–B_2
ZShift = 1.6
Z lm Score = −5.2
NG = 0
DB = 0
SM = 0
Delta_L
Doxycycline
YBL071C–B_2
ZShift = 0.39
Z lm Score = 1.72
NG = 0
DB = 0
SM = 0
Delta_K
Doxycycline
YBL071C–B_2
ZShift = −1.21
Z lm Score = 3.31
NG = 0
DB = 0
SM = 0
Delta_r
Doxycycline
YBL071C–B_2
ZShift = −0.44
Z lm Score = −1.58
NG = 0
DB = 0
SM = 0
Delta_AUC
Doxycycline
YDR159W SAC3
ZShift = 1.58
Z lm Score = −5.2
NG = 0
DB = 0
SM = 0
Delta_L
Doxycycline
YDR159W SAC3
ZShift = 2.16
Z lm Score = 2.21
NG = 0
DB = 0
SM = 0
Delta_K
Doxycycline
YDR159W SAC3
ZShift = 0.45
Z lm Score = 2.7
NG = 0
DB = 0
SM = 0
Delta_r
Doxycycline
YDR159W SAC3
ZShift = 0.73
Z lm Score = −2.43
NG = 0
DB = 0
SM = 0
Delta_AUC
Doxycycline
YGL135W RPL1B
ZShift = 3.14
Z lm Score = −5.21
NG = 0
DB = 0
SM = 0
Delta_L
Doxycycline
YGL135W RPL1B
ZShift = −0.82
Z lm Score = 1.91
NG = 0
DB = 0
SM = 0
Delta_K
Doxycycline
YGL135W RPL1B
ZShift = −1.91
Z lm Score = 2.9
NG = 0
DB = 0
SM = 0
Delta_r
Doxycycline
YGL135W RPL1B
ZShift = −1.87
Z lm Score = −2.27
NG = 0
DB = 0
SM = 0
Delta_AUC
Doxycycline

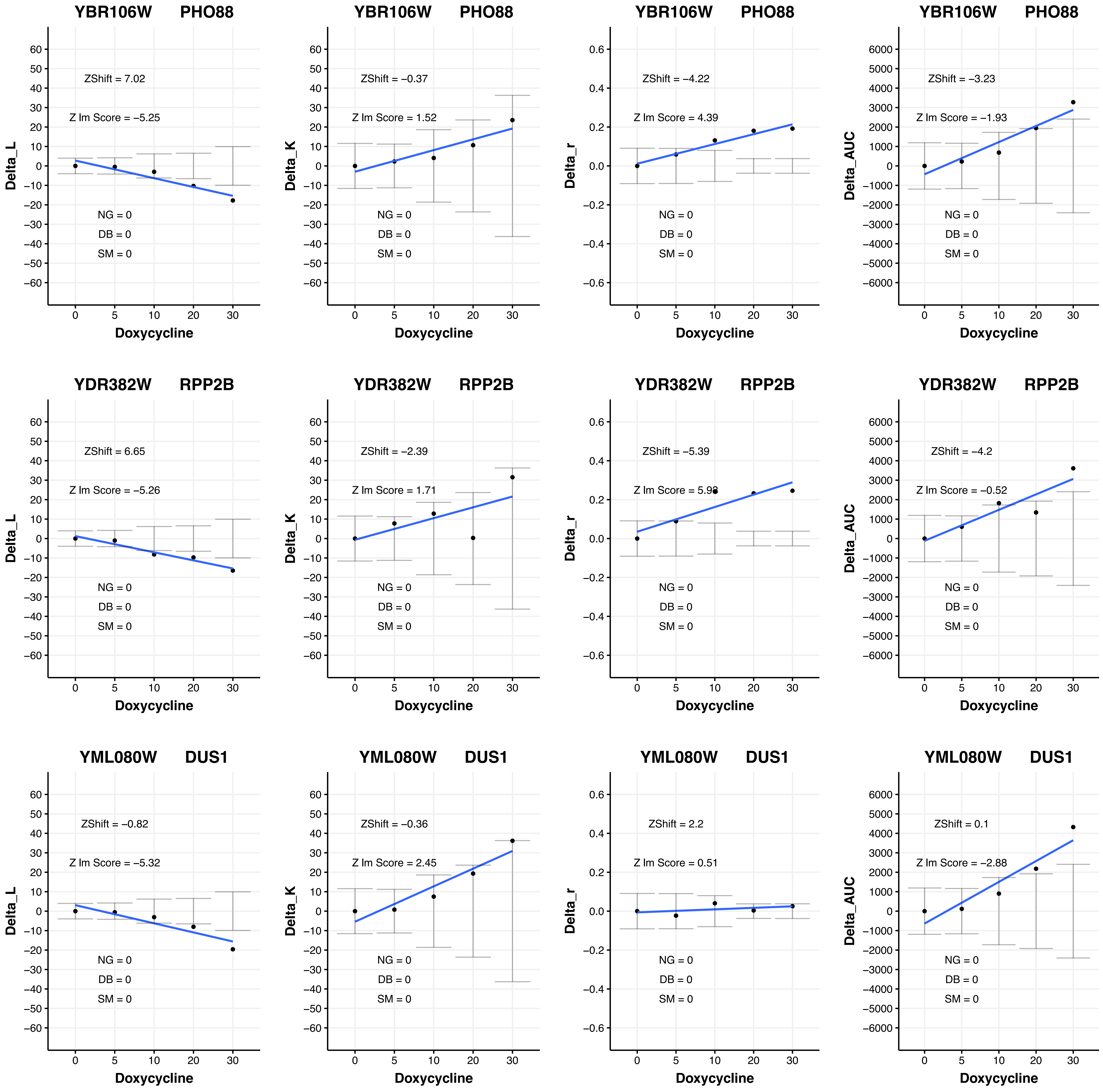
YBR106W PHO88
Delta_L
ZShift = 7.02
Z lm Score = −5.25
NG = 0
DB = 0
SM = 0
Doxycycline
YBR106W PHO88
Delta_K
ZShift = −0.37
Z lm Score = 1.52
NG = 0
DB = 0
SM = 0
Doxycycline
YBR106W PHO88
Delta_r
ZShift = −4.22
Z lm Score = 4.39
NG = 0
DB = 0
SM = 0
Doxycycline
YBR106W PHO88
Delta_AUC
ZShift = −3.23
Z lm Score = −1.93
NG = 0
DB = 0
SM = 0
Doxycycline
YDR382W RPP2B
Delta_L
ZShift = 6.65
Z lm Score = −5.26
NG = 0
DB = 0
SM = 0
Doxycycline
YDR382W RPP2B
Delta_K
ZShift = −2.39
Z lm Score = 1.71
NG = 0
DB = 0
SM = 0
Doxycycline
YDR382W RPP2B
Delta_r
ZShift = −5.39
NG = 0
DB = 0
SM = 0
Doxycycline
YDR382W RPP2B
Delta_AUC
ZShift = −4.2
Z lm Score = −0.52
NG = 0
DB = 0
SM = 0
Doxycycline
YML080W DUS1
Delta_L
ZShift = −0.82
Z lm Score = −5.32
NG = 0
DB = 0
SM = 0
Doxycycline
YML080W DUS1
Delta_K
ZShift = −0.36
Z lm Score = 2.45
NG = 0
DB = 0
SM = 0
Doxycycline
YML080W DUS1
Delta_r
ZShift = 2.2
Z lm Score = 0.51
NG = 0
DB = 0
SM = 0
Doxycycline
YML080W DUS1
Delta_AUC
ZShift = 0.1
Z lm Score = −2.88
NG = 0
DB = 0
SM = 0
Doxycycline

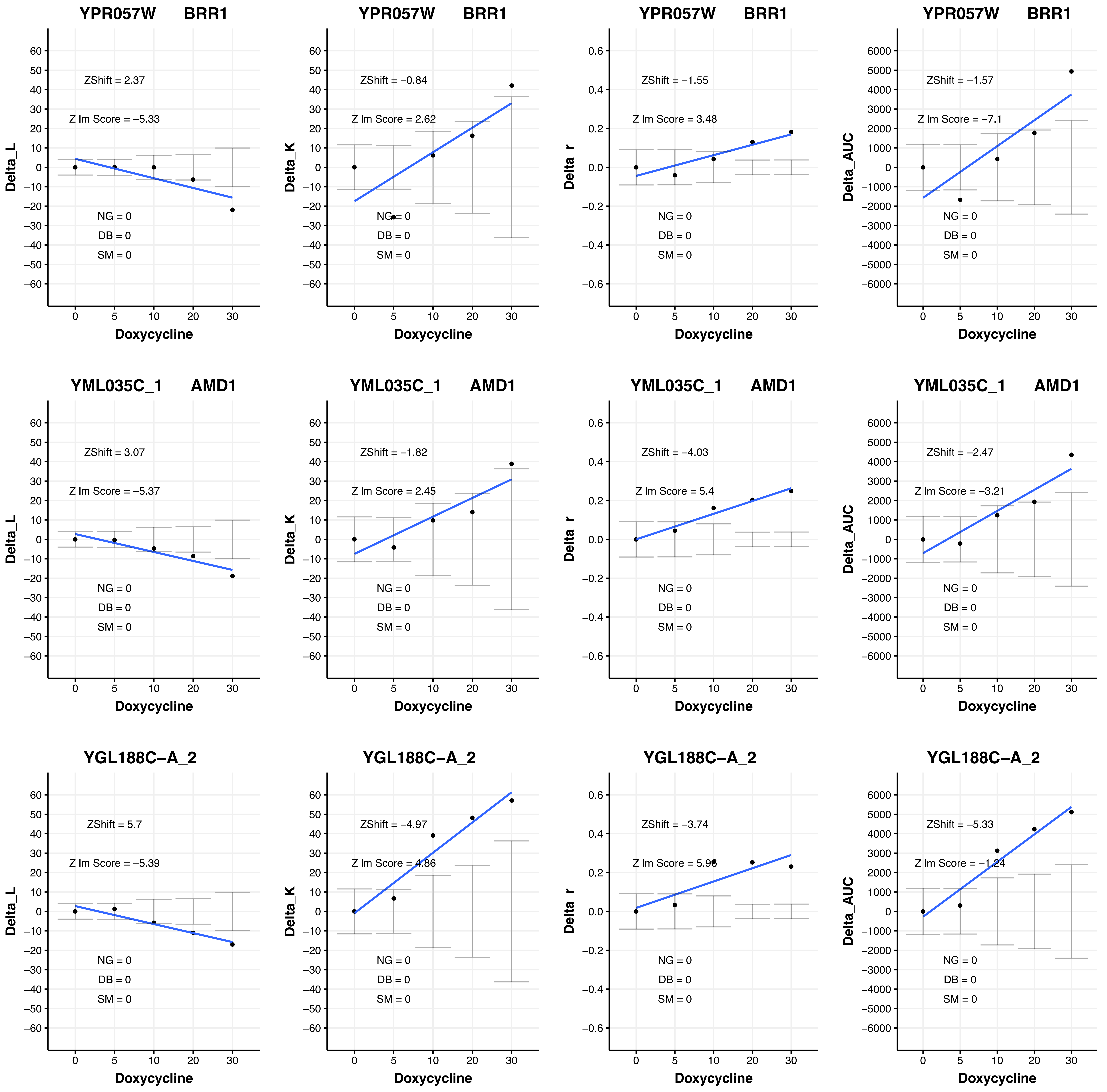

YPR057W BRR1
ZShift = 2.37
Z lm Score = −5.33
NG = 0
DB = 0
SM = 0
Delta_L
Doxycycline
YPR057W BRR1
ZShift = −0.84
Z lm Score = 2.62
NG = 0
DB = 0
SM = 0
Delta_K
Doxycycline
YPR057W BRR1
ZShift = −1.55
Z lm Score = 3.48
NG = 0
DB = 0
SM = 0
Delta_r
Doxycycline
YPR057W BRR1
ZShift = −1.57
Z lm Score = −7.1
NG = 0
DB = 0
SM = 0
Delta_AUC
Doxycycline
YML035C_1 AMD1
ZShift = 3.07
Z lm Score = −5.37
NG = 0
DB = 0
SM = 0
Delta_L
Doxycycline
YML035C_1 AMD1
ZShift = −1.82
Z lm Score = 2.45
NG = 0
DB = 0
SM = 0
Delta_K
Doxycycline
YML035C_1 AMD1
ZShift = −4.03
Z lm Score = 5.4
NG = 0
DB = 0
SM = 0
Delta_r
Doxycycline
YML035C_1 AMD1
ZShift = −2.47
Z lm Score = −3.21
NG = 0
DB = 0
SM = 0
Delta_AUC
Doxycycline
YGL188C−A_2
ZShift = 5.7
Z lm Score = −5.39
NG = 0
DB = 0
SM = 0
Delta_L
Doxycycline
YGL188C−A_2
ZShift = −4.97
Z lm Score = 4.86
NG = 0
DB = 0
SM = 0
Delta_K
Doxycycline
YGL188C−A_2
ZShift = −3.74
Z lm Score = 5.96
NG = 0
DB = 0
SM = 0
Delta_r
Doxycycline
YGL188C−A_2
ZShift = −5.33
Z lm Score = −1.24
NG = 0
DB = 0
SM = 0
Delta_AUC
Doxycycline

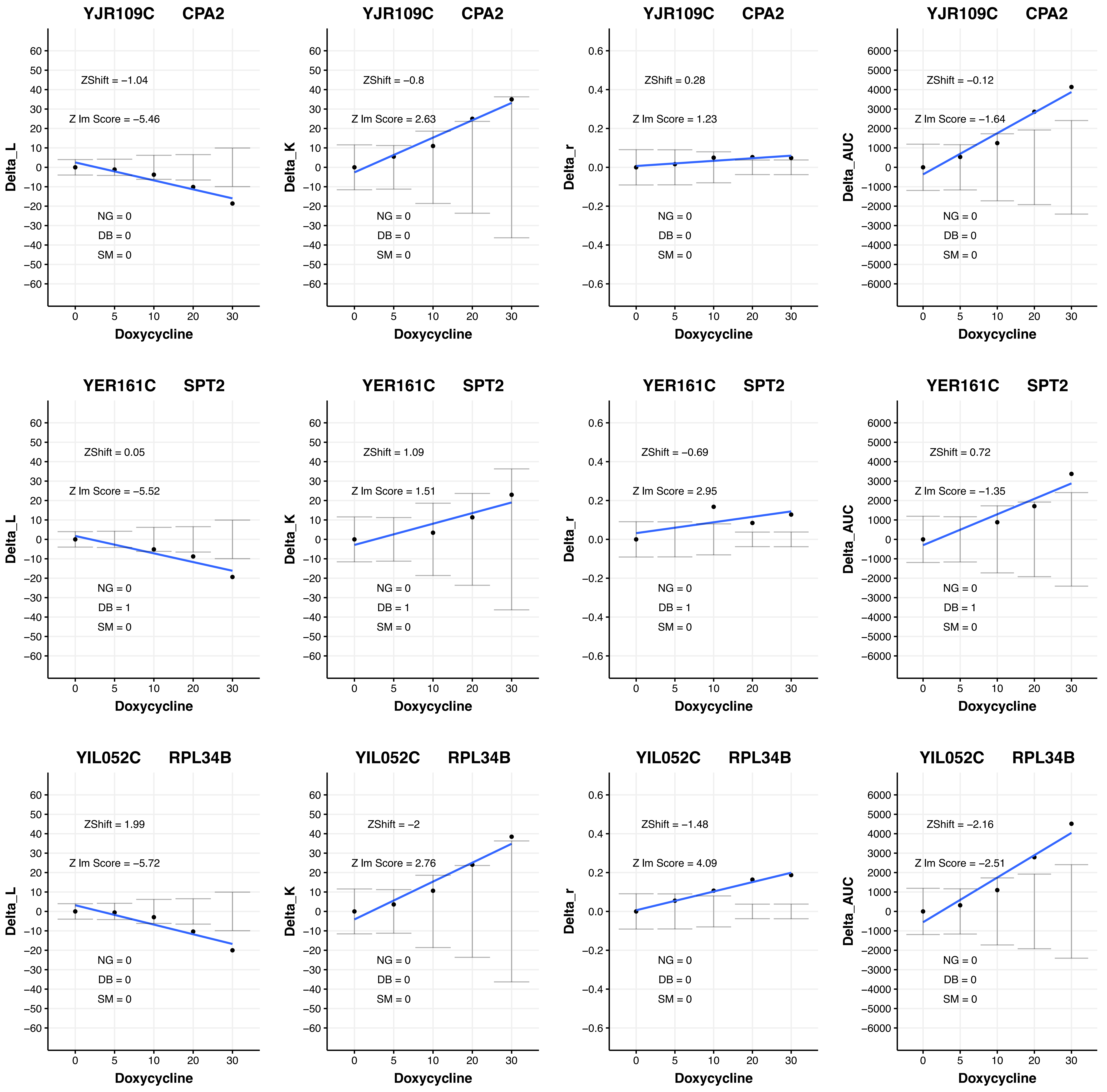
YJR109C CPA2
ZShift = −1.04
Z lm Score = −5.46
NG = 0
DB = 0
SM = 0
Delta_L
Doxycycline
YJR109C CPA2
ZShift = −0.8
Z lm Score = 2.63
NG = 0
DB = 0
SM = 0
Delta_K
Doxycycline
YJR109C CPA2
ZShift = 0.28
Z lm Score = 1.23
NG = 0
DB = 0
SM = 0
Delta_r
Doxycycline
YJR109C CPA2
ZShift = −0.12
Z lm Score = −1.64
NG = 0
DB = 0
SM = 0
Delta_AUC
Doxycycline
YER161C SPT2
ZShift = 0.05
Z lm Score = −5.52
NG = 0
DB = 1
SM = 0
Delta_L
Doxycycline
YER161C SPT2
ZShift = 1.09
Z lm Score = 1.51
NG = 0
DB = 1
SM = 0
Delta_K
Doxycycline
YER161C SPT2
ZShift = −0.69
Z lm Score = 2.95
NG = 0
DB = 1
SM = 0
Delta_r
Doxycycline
YER161C SPT2
ZShift = 0.72
Z lm Score = −1.35
NG = 0
DB = 1
SM = 0
Delta_AUC
Doxycycline
YIL052C RPL34B
ZShift = 1.99
Z lm Score = −5.72
NG = 0
DB = 0
SM = 0
Delta_L
Doxycycline
YIL052C RPL34B
ZShift = −2
Z lm Score = 2.76
NG = 0
DB = 0
SM = 0
Delta_K
Doxycycline
YIL052C RPL34B
ZShift = −1.48
Z lm Score = 4.09
NG = 0
DB = 0
SM = 0
Delta_r
Doxycycline
YIL052C RPL34B
ZShift = −2.16
Z lm Score = −2.51
NG = 0
DB = 0
SM = 0
Delta_AUC
Doxycycline

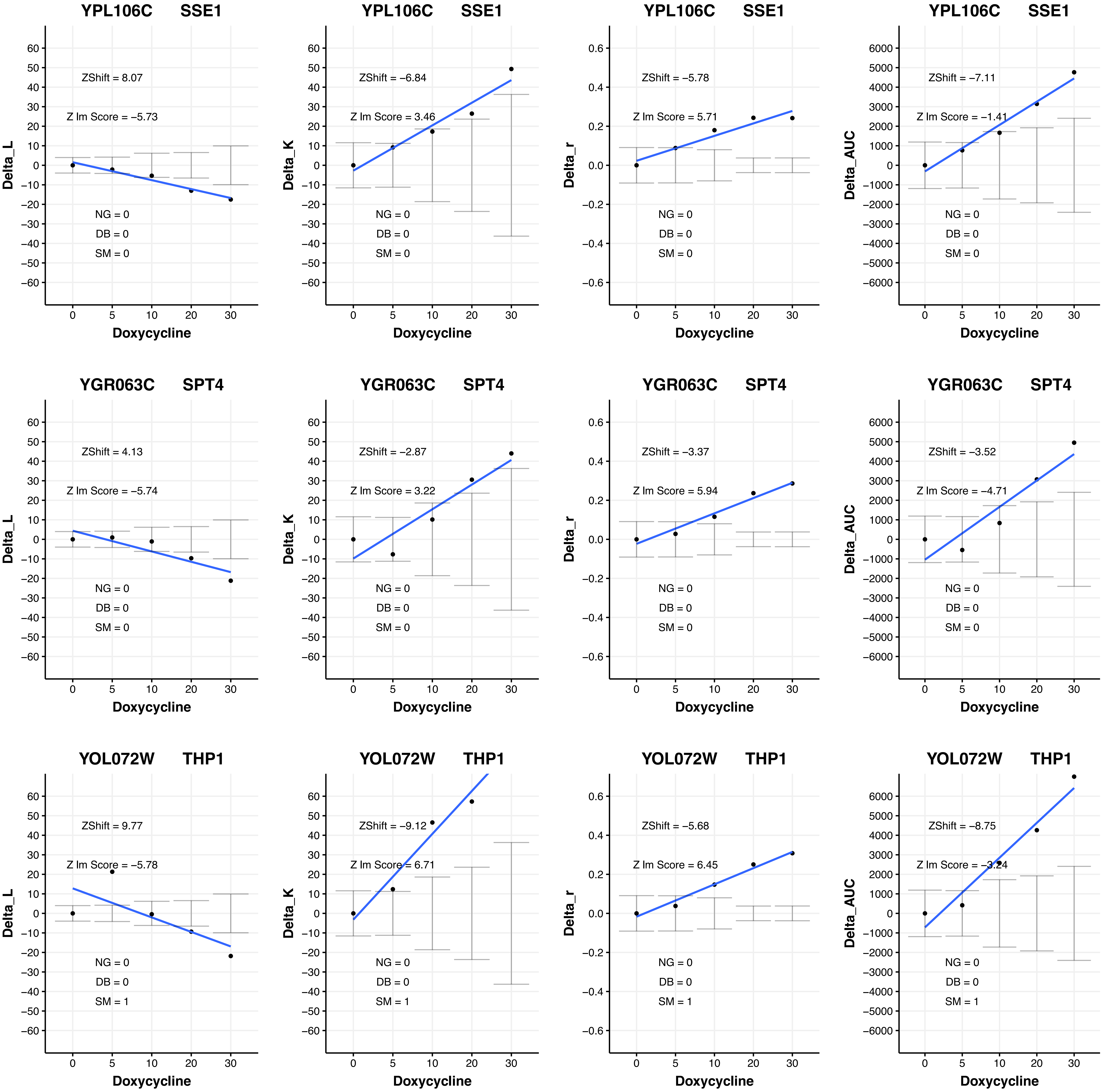

YPL106C SSE1
ZShift = 8.07
Z lm Score = −5.73
NG = 0
DB = 0
SM = 0
Delta_L
Doxycycline
YPL106C SSE1
ZShift = −6.84
Z lm Score = 3.46
NG = 0
DB = 0
SM = 0
Delta_K
Doxycycline
YPL106C SSE1
ZShift = −5.78
Z lm Score = 5.71
NG = 0
DB = 0
SM = 0
Delta_r
Doxycycline
YPL106C SSE1
ZShift = −7.11
Z lm Score = −1.41
NG = 0
DB = 0
SM = 0
Delta_AUC
Doxycycline
YGR063C SPT4
ZShift = 4.13
Z lm Score = −5.74
NG = 0
DB = 0
SM = 0
Delta_L
Doxycycline
YGR063C SPT4
ZShift = −2.87
Z lm Score = 3.22
NG = 0
DB = 0
SM = 0
Delta_K
Doxycycline
YGR063C SPT4
ZShift = −3.37
Z lm Score = 5.94
NG = 0
DB = 0
SM = 0
Delta_r
Doxycycline
YGR063C SPT4
ZShift = −3.52
Z lm Score = −4.71
NG = 0
DB = 0
SM = 0
Delta_AUC
Doxycycline
YOL072W THP1
ZShift = 9.77
Z lm Score = −5.78
NG = 0
DB = 0
SM = 1
Delta_L
Doxycycline
YOL072W THP1
ZShift = −9.12
Z lm Score = 6.71
NG = 0
DB = 0
SM = 1
Delta_K
Doxycycline
YOL072W THP1
ZShift = −5.68
Z lm Score = 6.45
NG = 0
DB = 0
SM = 1
Delta_r
Doxycycline
YOL072W THP1
ZShift = −8.75
Z lm Score = −3.24
NG = 0
DB = 0
SM = 1
Delta_AUC
Doxycycline

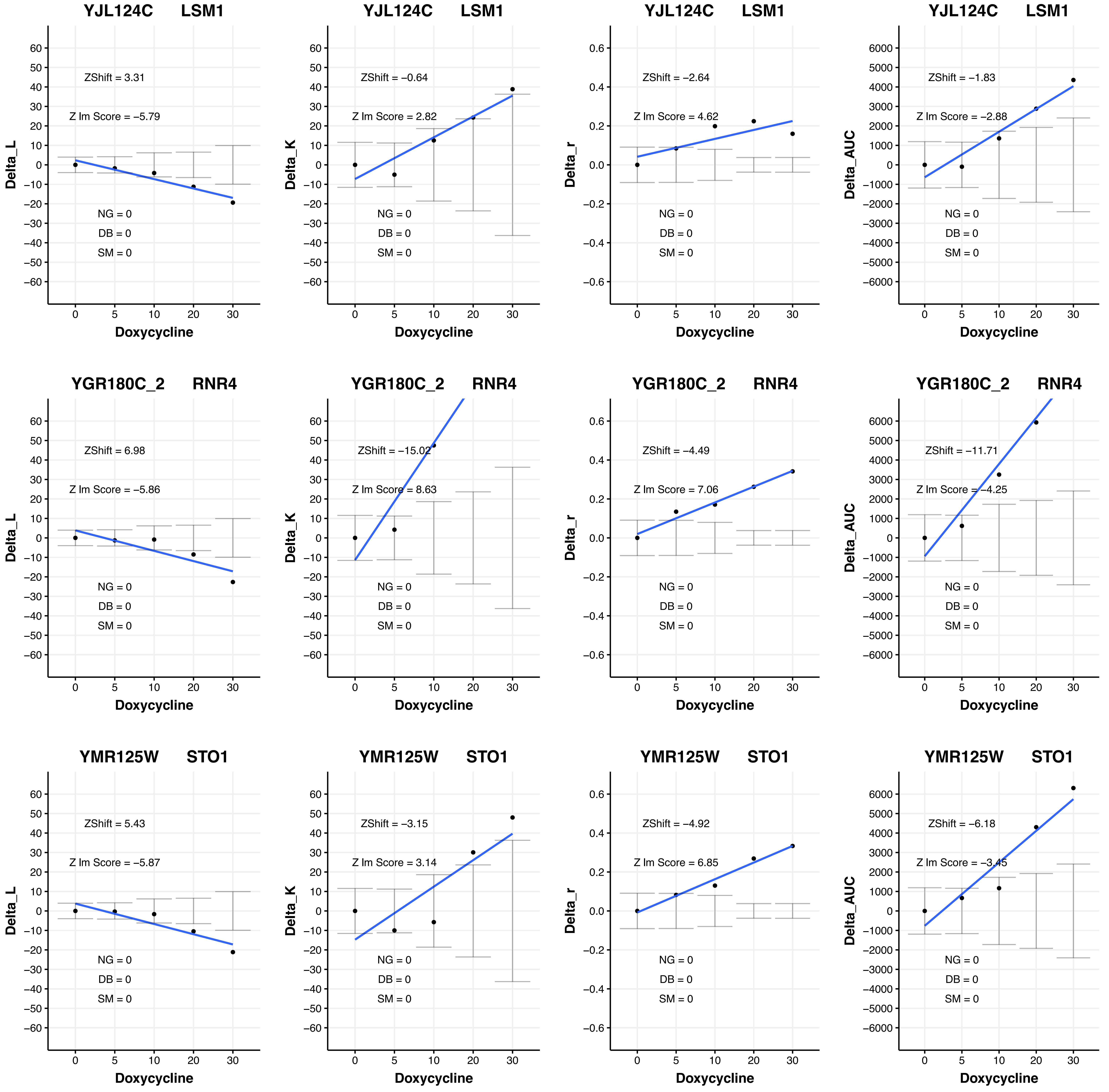

YJL124C LSM1
ZShift = 3.31
Z lm Score = −5.79
NG = 0
DB = 0
SM = 0
Delta_L
Doxycycline
YJL124C LSM1
ZShift = −0.64
Z lm Score = 2.82
NG = 0
DB = 0
SM = 0
Delta_K
Doxycycline
YJL124C LSM1
ZShift = −2.64
Z lm Score = 4.62
NG = 0
DB = 0
SM = 0
Delta_r
Doxycycline
YJL124C LSM1
ZShift = −1.83
Z lm Score = −2.88
NG = 0
DB = 0
SM = 0
Delta_AUC
Doxycycline
YGR180C_2 RNR4
ZShift = 6.98
Z lm Score = −5.86
NG = 0
DB = 0
SM = 0
Delta_L
Doxycycline
YGR180C_2 RNR4
ZShift = −15.02
Z lm Score = 8.63
NG = 0
DB = 0
SM = 0
Delta_K
Doxycycline
YGR180C_2 RNR4
ZShift = −4.49
Z lm Score = 7.06
NG = 0
DB = 0
SM = 0
Delta_r
Doxycycline
YGR180C_2 RNR4
ZShift = −11.71
Z lm Score = −4.25
NG = 0
DB = 0
SM = 0
Delta_AUC
Doxycycline
YMR125W STO1
ZShift = 5.43
Z lm Score = −5.87
NG = 0
DB = 0
SM = 0
Delta_L
Doxycycline
YMR125W STO1
ZShift = −3.15
Z lm Score = 3.14
NG = 0
DB = 0
SM = 0
Delta_K
Doxycycline
YMR125W STO1
ZShift = −4.92
Z lm Score = 6.85
NG = 0
DB = 0
SM = 0
Delta_r
Doxycycline
YMR125W STO1
ZShift = −6.18
Z lm Score = −3.45
NG = 0
DB = 0
SM = 0
Delta_AUC
Doxycycline

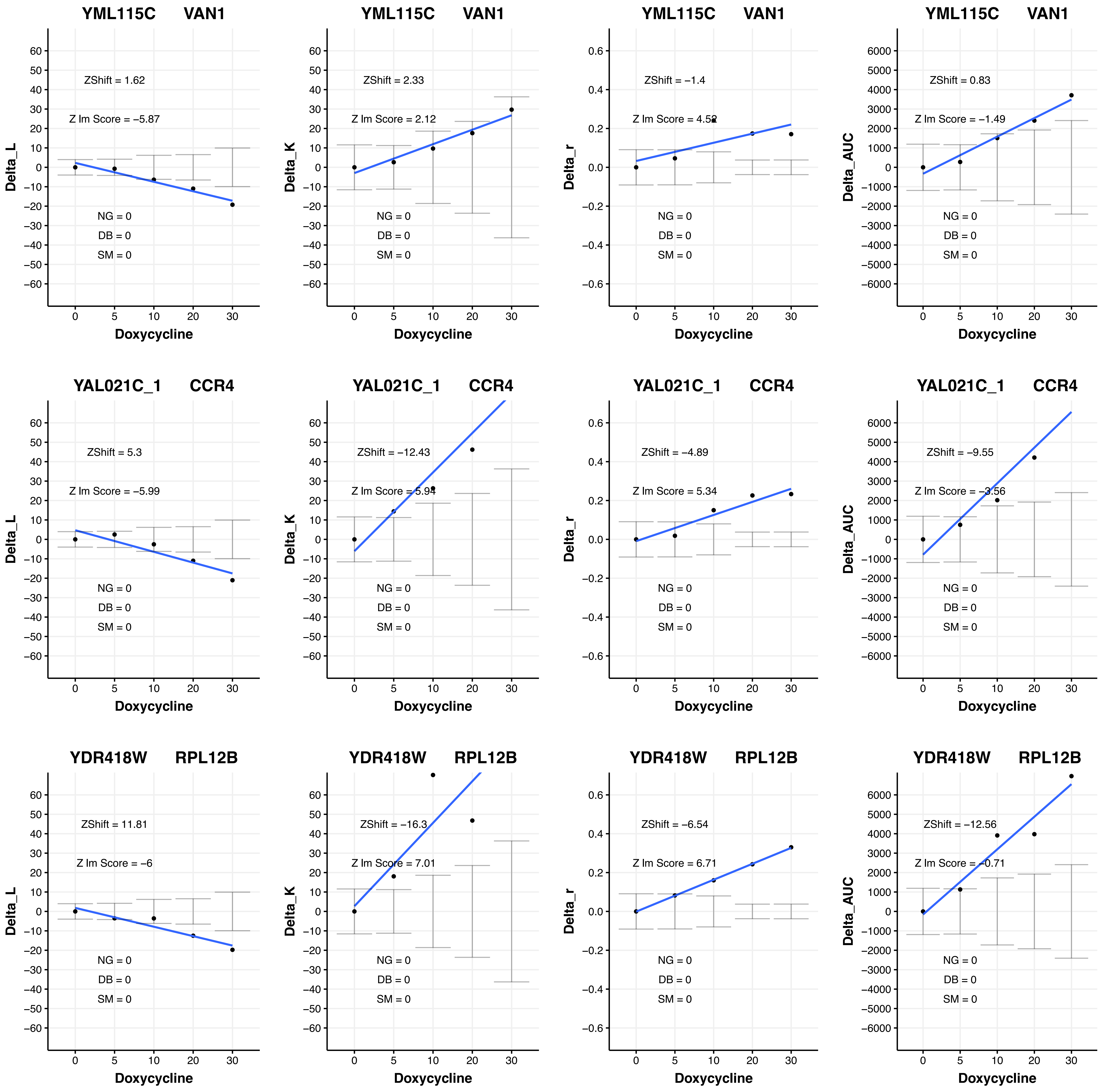

YML115C VAN1
ZShift = 1.62
Z lm Score = −5.87
NG = 0
DB = 0
SM = 0
Delta_L
Doxycycline
YML115C VAN1
ZShift = 2.33
Z lm Score = 2.12
NG = 0
DB = 0
SM = 0
Delta_K
Doxycycline
YML115C VAN1
ZShift = −1.4
Z lm Score = 4.52
NG = 0
DB = 0
SM = 0
Delta_r
Doxycycline
YML115C VAN1
ZShift = 0.83
Z lm Score = −1.49
NG = 0
DB = 0
SM = 0
Delta_AUC
Doxycycline
YAL021C_1 CCR4
ZShift = 5.3
Z lm Score = −5.99
NG = 0
DB = 0
SM = 0
Delta_L
Doxycycline
YAL021C_1 CCR4
ZShift = −12.43
Z lm Score = 5.94
NG = 0
DB = 0
SM = 0
Delta_K
Doxycycline
YAL021C_1 CCR4
ZShift = −4.89
Z lm Score = 5.34
NG = 0
DB = 0
SM = 0
Delta_r
Doxycycline
YAL021C_1 CCR4
ZShift = −9.55
Z lm Score = −3.56
NG = 0
DB = 0
SM = 0
Delta_AUC
Doxycycline
YDR418W RPL12B
ZShift = 11.81
Z lm Score = −6
NG = 0
DB = 0
SM = 0
Delta_L
Doxycycline
YDR418W RPL12B
ZShift = −16.3
Z lm Score = 7.01
NG = 0
DB = 0
SM = 0
Delta_K
Doxycycline
YDR418W RPL12B
ZShift = −6.54
Z lm Score = 6.71
NG = 0
DB = 0
SM = 0
Delta_r
Doxycycline
YDR418W RPL12B
ZShift = −12.56
Z lm Score = −0.71
NG = 0
DB = 0
SM = 0
Delta_AUC
Doxycycline

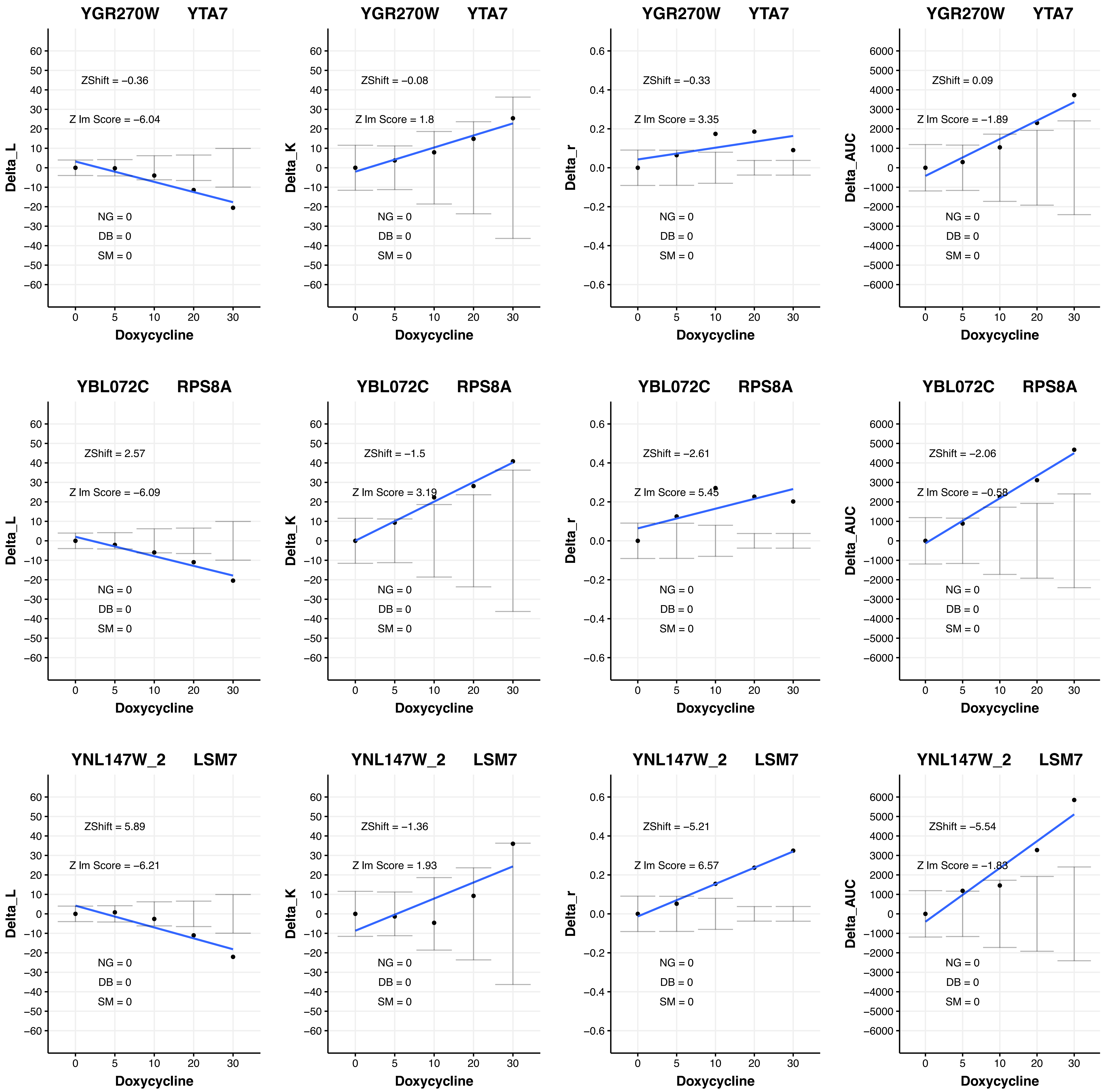
YGR270W YTA7
ZShift = −0.36
Z lm Score = −6.04
NG = 0
DB = 0
SM = 0
Delta_L
Doxycycline
YGR270W YTA7
ZShift = −0.08
Z lm Score = 1.8
NG = 0
DB = 0
SM = 0
Delta_K
Doxycycline
YGR270W YTA7
ZShift = −0.33
Z lm Score = 3.35
NG = 0
DB = 0
SM = 0
Delta_r
Doxycycline
YGR270W YTA7
ZShift = 0.09
Z lm Score = −1.89
NG = 0
DB = 0
SM = 0
Delta_AUC
Doxycycline
YBL072C RPS8A
ZShift = 2.57
Z lm Score = −6.09
NG = 0
DB = 0
SM = 0
Delta_L
Doxycycline
YBL072C RPS8A
ZShift = −1.5
Z lm Score = 3.19
NG = 0
DB = 0
SM = 0
Delta_K
Doxycycline
YBL072C RPS8A
ZShift = −2.61
Z lm Score = 5.45
NG = 0
DB = 0
SM = 0
Delta_r
Doxycycline
YBL072C RPS8A
ZShift = −2.06
Z lm Score = −0.58
NG = 0
DB = 0
SM = 0
Delta_AUC
Doxycycline
YNL147W_2 LSM7
ZShift = 5.89
Z lm Score = −6.21
NG = 0
DB = 0
SM = 0
Delta_L
Doxycycline
YNL147W_2 LSM7
ZShift = −1.36
Z lm Score = 1.93
NG = 0
DB = 0
SM = 0
Delta_K
Doxycycline
YNL147W_2 LSM7
ZShift = −5.21
Z lm Score = 6.57
NG = 0
DB = 0
SM = 0
Delta_r
Doxycycline
YNL147W_2 LSM7
ZShift = −5.54
Z lm Score = −1.83
NG = 0
DB = 0
SM = 0
Delta_AUC
Doxycycline

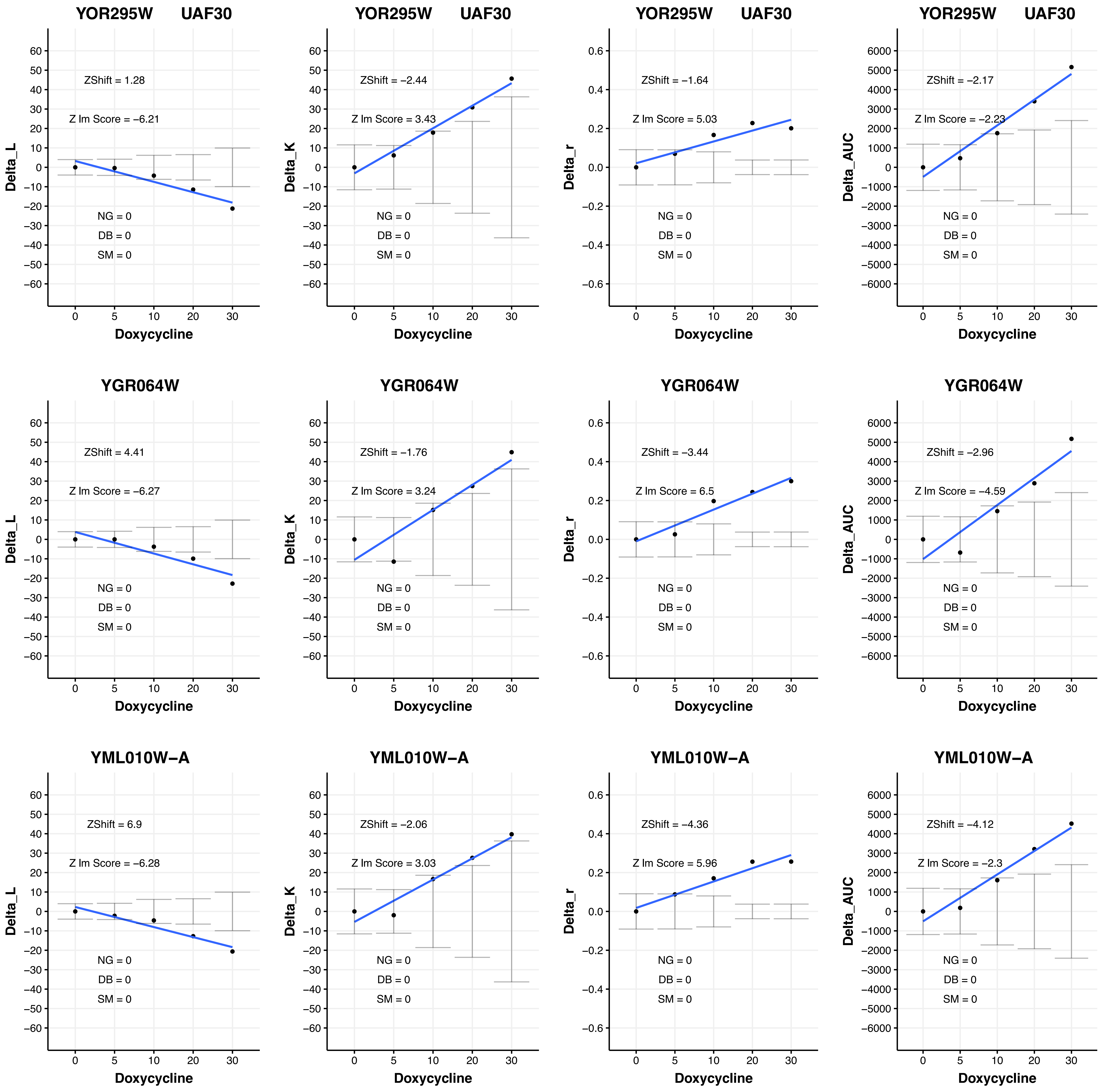

YOR295W UAF30
ZShift = 1.28
Z lm Score = −6.21
NG = 0
DB = 0
SM = 0
Delta_L
Doxycycline
YOR295W UAF30
ZShift = −2.44
Z lm Score = 3.43
NG = 0
DB = 0
SM = 0
Delta_K
Doxycycline
YOR295W UAF30
ZShift = −1.64
Z lm Score = 5.03
NG = 0
DB = 0
SM = 0
Delta_r
Doxycycline
YOR295W UAF30
ZShift = −2.17
Z lm Score = −2.23
NG = 0
DB = 0
SM = 0
Delta_AUC
Doxycycline
YGR064W
ZShift = 4.41
Z lm Score = −6.27
NG = 0
DB = 0
SM = 0
Delta_L
Doxycycline
YGR064W
ZShift = −1.76
Z lm Score = 3.24
NG = 0
DB = 0
SM = 0
Delta_K
Doxycycline
YGR064W
ZShift = −3.44
Z lm Score = 6.5
NG = 0
DB = 0
SM = 0
Delta_r
Doxycycline
YGR064W
ZShift = −2.96
Z lm Score = −4.59
NG = 0
DB = 0
SM = 0
Delta_AUC
Doxycycline
YML010W−A
ZShift = 6.9
Z lm Score = −6.28
NG = 0
DB = 0
SM = 0
Delta_L
Doxycycline
YML010W−A
ZShift = −2.06
Z lm Score = 3.03
NG = 0
DB = 0
SM = 0
Delta_K
Doxycycline
YML010W−A
ZShift = −4.36
Z lm Score = 5.96
NG = 0
DB = 0
SM = 0
Delta_r
Doxycycline
YML010W−A
ZShift = −4.12
Z lm Score = −2.3
NG = 0
DB = 0
SM = 0
Delta_AUC
Doxycycline

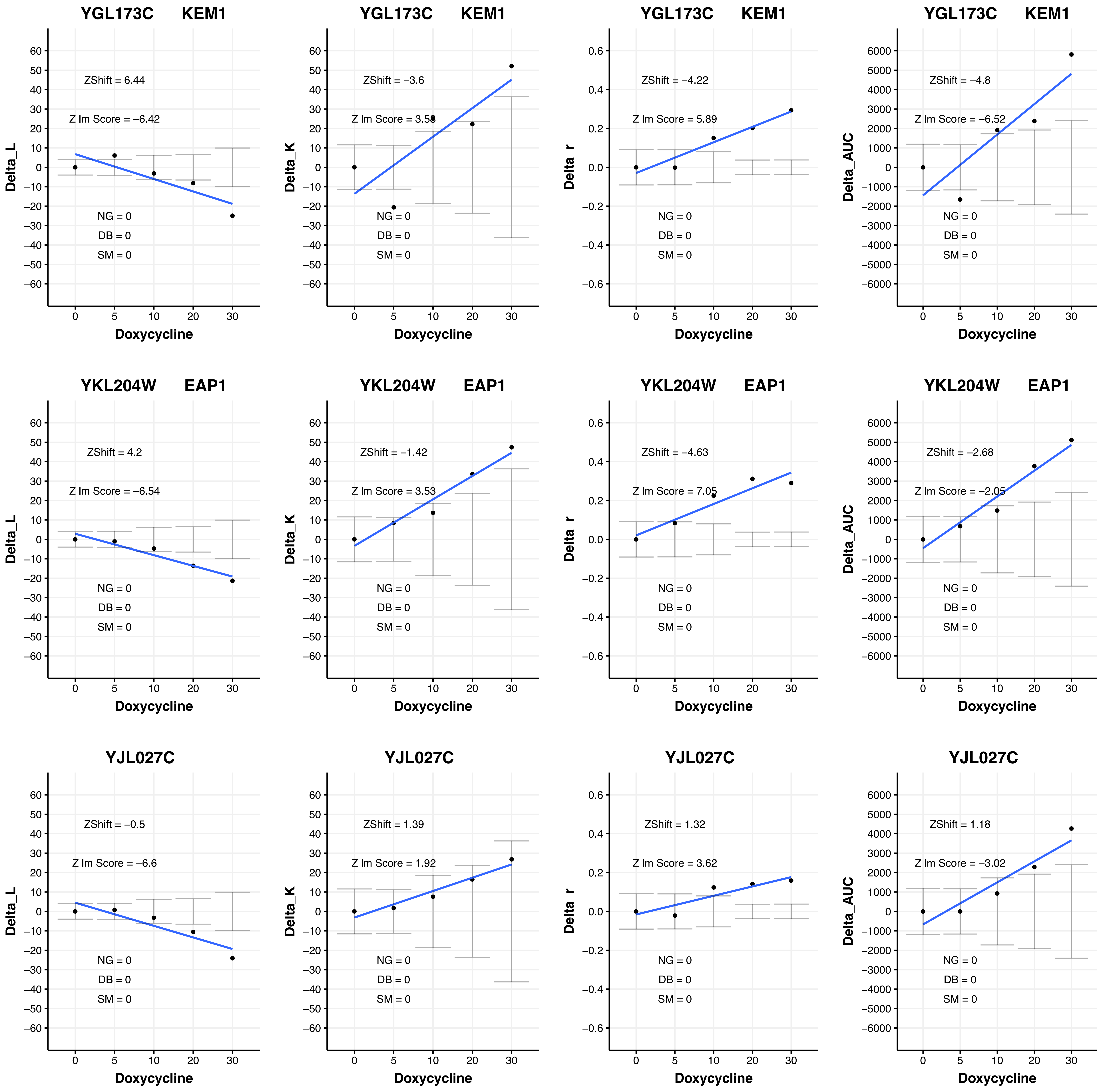

YGL173C KEM1
ZShift = 6.44
Z lm Score = −6.42
NG = 0
DB = 0
SM = 0
Delta_L
Doxycycline
YGL173C KEM1
ZShift = −3.6
Z lm Score = 3.58
NG = 0
DB = 0
SM = 0
Delta_K
Doxycycline
YGL173C KEM1
ZShift = −4.22
Z lm Score = 5.89
NG = 0
DB = 0
SM = 0
Delta_r
Doxycycline
YGL173C KEM1
ZShift = −4.8
Z lm Score = −6.52
NG = 0
DB = 0
SM = 0
Delta_AUC
Doxycycline
YKL204W EAP1
ZShift = 4.2
Z lm Score = −6.54
NG = 0
DB = 0
SM = 0
Delta_L
Doxycycline
YKL204W EAP1
ZShift = −1.42
Z lm Score = 3.53
NG = 0
DB = 0
SM = 0
Delta_K
Doxycycline
YKL204W EAP1
ZShift = −4.63
Z lm Score = 7.05
NG = 0
DB = 0
SM = 0
Delta_r
Doxycycline
YKL204W EAP1
ZShift = −2.68
Z lm Score = −2.05
NG = 0
DB = 0
SM = 0
Delta_AUC
Doxycycline
YJL027C
ZShift = −0.5
Z lm Score = −6.6
NG = 0
DB = 0
SM = 0
Delta_L
Doxycycline
YJL027C
ZShift = 1.39
Z lm Score = 1.92
NG = 0
DB = 0
SM = 0
Delta_K
Doxycycline
YJL027C
ZShift = 1.32
Z lm Score = 3.62
NG = 0
DB = 0
SM = 0
Delta_r
Doxycycline
YJL027C
ZShift = 1.18
Z lm Score = −3.02
NG = 0
DB = 0
SM = 0
Delta_AUC
Doxycycline

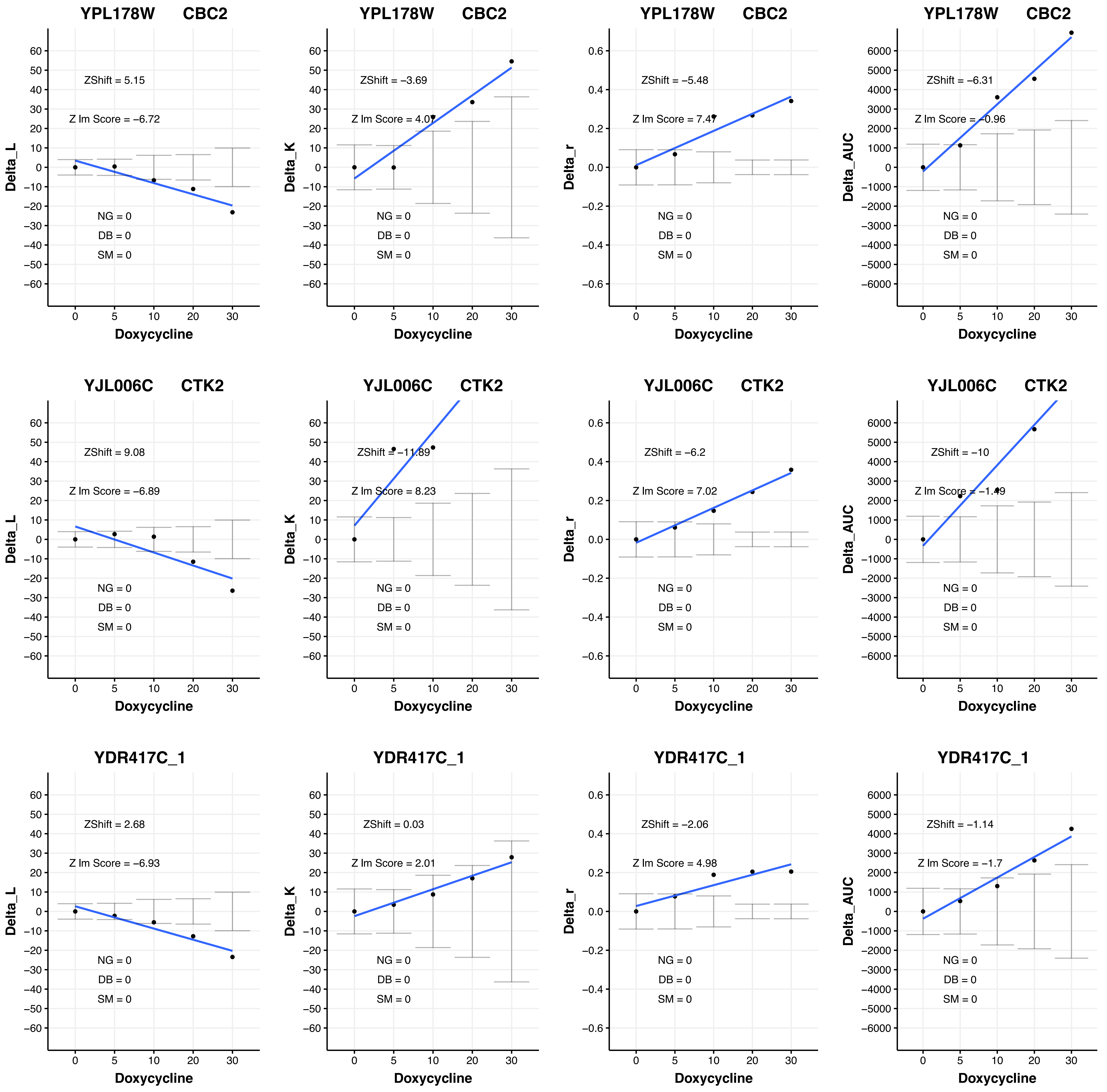
YPL178W CBC2
ZShift = 5.15
Z lm Score = −6.72
Delta_L
NG = 0
DB = 0
SM = 0
Doxycycline
YPL178W CBC2
ZShift = −3.69
Z lm Score = 4.07
Delta_K
NG = 0
DB = 0
SM = 0
Doxycycline
YPL178W CBC2
ZShift = −5.48
Z lm Score = 7.47
Delta_r
NG = 0
DB = 0
SM = 0
Doxycycline
YPL178W CBC2
ZShift = −6.31
Z lm Score = −0.96
Delta_AUC
NG = 0
DB = 0
SM = 0
Doxycycline
YJL006C CTK2
ZShift = 9.08
Z lm Score = −6.89
Delta_L
NG = 0
DB = 0
SM = 0
Doxycycline
YJL006C CTK2
ZShift = −11.89
Z lm Score = 8.23
Delta_K
NG = 0
DB = 0
SM = 0
Doxycycline
YJL006C CTK2
ZShift = −6.2
Z lm Score = 7.02
Delta_r
NG = 0
DB = 0
SM = 0
Doxycycline
YJL006C CTK2
ZShift = −10
Z lm Score = −1.49
Delta_AUC
NG = 0
DB = 0
SM = 0
Doxycycline
YDR417C_1
ZShift = 2.68
Z lm Score = −6.93
Delta_L
NG = 0
DB = 0
SM = 0
Doxycycline
YDR417C_1
ZShift = 0.03
Z lm Score = 2.01
Delta_K
NG = 0
DB = 0
SM = 0
Doxycycline
YDR417C_1
ZShift = −2.06
Z lm Score = 4.98
Delta_r
NG = 0
DB = 0
SM = 0
Doxycycline
YDR417C_1
ZShift = −1.14
Z lm Score = −1.7
Delta_AUC
NG = 0
DB = 0
SM = 0
Doxycycline

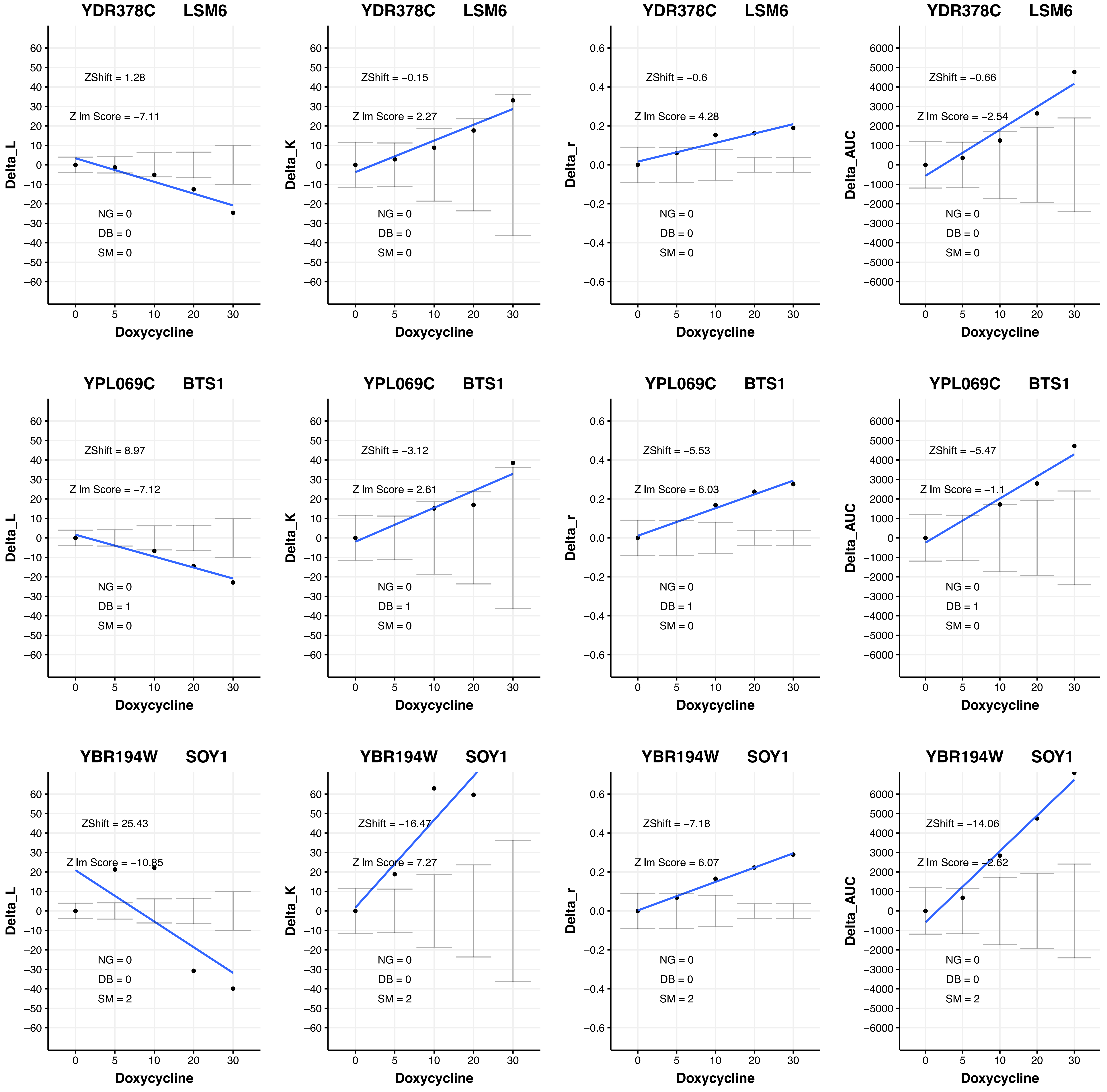
YDR378C LSM6
ZShift = 1.28
Z lm Score = −7.11
NG = 0
DB = 0
SM = 0
Delta_L
Doxycycline
YDR378C LSM6
ZShift = −0.15
Z lm Score = 2.27
NG = 0
DB = 0
SM = 0
Delta_K
Doxycycline
YDR378C LSM6
ZShift = −0.6
Z lm Score = 4.28
NG = 0
DB = 0
SM = 0
Delta_r
Doxycycline
YDR378C LSM6
ZShift = −0.66
Z lm Score = −2.54
NG = 0
DB = 0
SM = 0
Delta_AUC
Doxycycline
YPL069C BTS1
ZShift = 8.97
Z lm Score = −7.12
NG = 0
DB = 1
SM = 0
Delta_L
Doxycycline
YPL069C BTS1
ZShift = −3.12
Z lm Score = 2.61
NG = 0
DB = 1
SM = 0
Delta_K
Doxycycline
YPL069C BTS1
ZShift = −5.53
Z lm Score = 6.03
NG = 0
DB = 1
SM = 0
Delta_r
Doxycycline
YPL069C BTS1
ZShift = −5.47
Z lm Score = −1.1
NG = 0
DB = 1
SM = 0
Delta_AUC
Doxycycline
YBR194W SOY1
ZShift = 25.43
Z lm Score = −10.85
NG = 0
DB = 0
SM = 2
Delta_L
Doxycycline
YBR194W SOY1
ZShift = −16.47
Z lm Score = 7.27
NG = 0
DB = 0
SM = 2
Delta_K
Doxycycline
YBR194W SOY1
ZShift = −7.18
Z lm Score = 6.07
NG = 0
DB = 0
SM = 2
Delta_r
Doxycycline
YBR194W SOY1
ZShift = −14.06
Z lm Score = −2.62
NG = 0
DB = 0
SM = 2
Delta_AUC
Doxycycline
