## Additional File 3 for "A humanized yeast phenomic model of deoxycytidine kinase to predict genetic buffering of nucleoside analog cytotoxicity": B - InteractionPlots_DAmPs_Gemcitabine.pdf

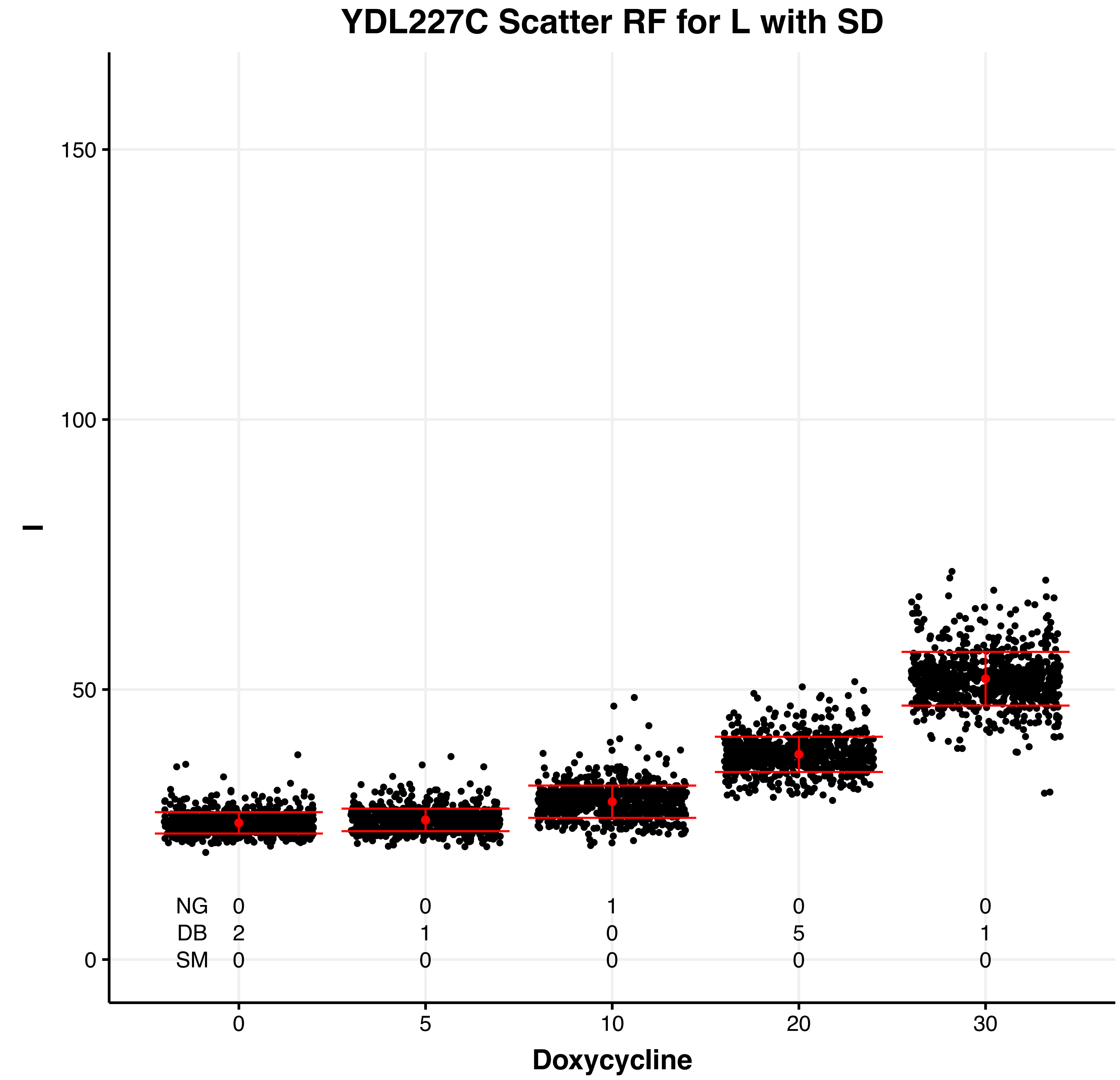
YDL227C Scatter RF for L with SD
l
Doxycycline
0
5
10
20
30
0
50
100
150
NG 0 0 1 0 0
DB 2 1 0 5 1
SM 0 0 0 0 0
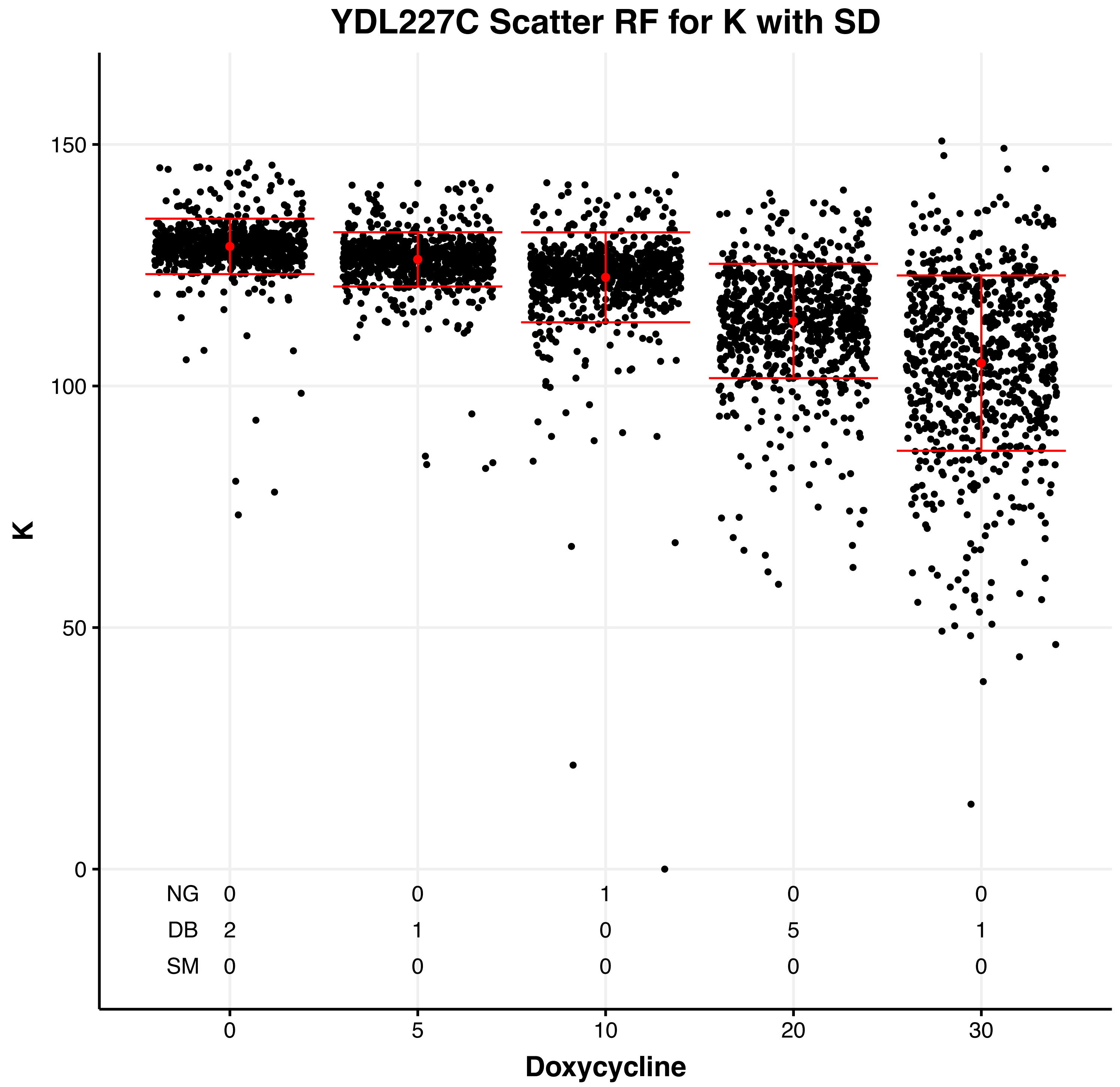
YDL227C Scatter RF for K with SD
K
Doxycycline
0
5
10
20
30
0
50
100
150
NG 0 0 1 0 0
DB 2 1 0 5 1
SM 0 0 0 0 0
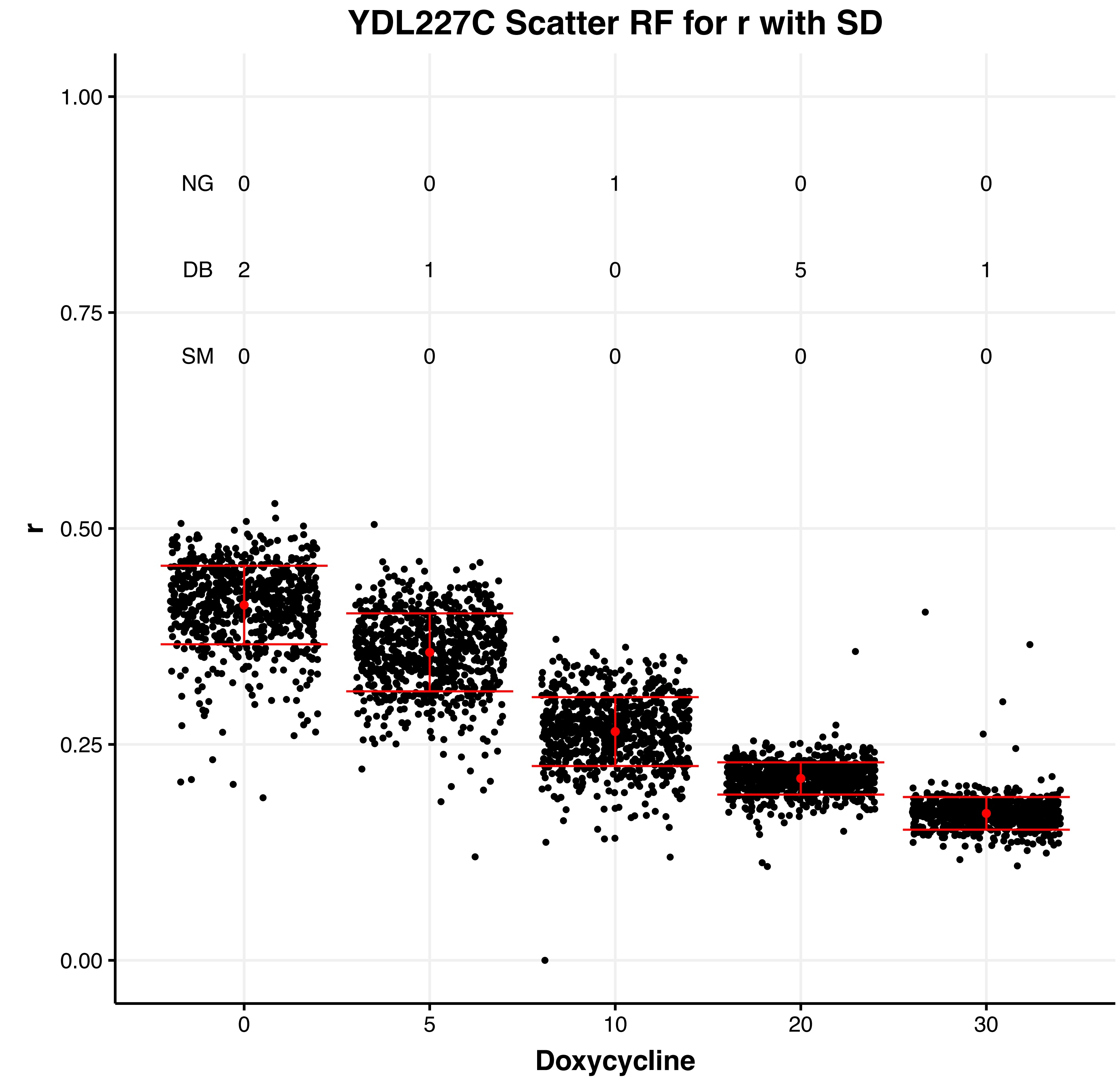
YDL227C Scatter RF for r with SD
r
Doxycycline
0
5
10
20
30
0.00
0.25
0.50
0.75
1.00
NG 0 0 1 0 0
DB 2 1 0 5 1
SM 0 0 0 0 0
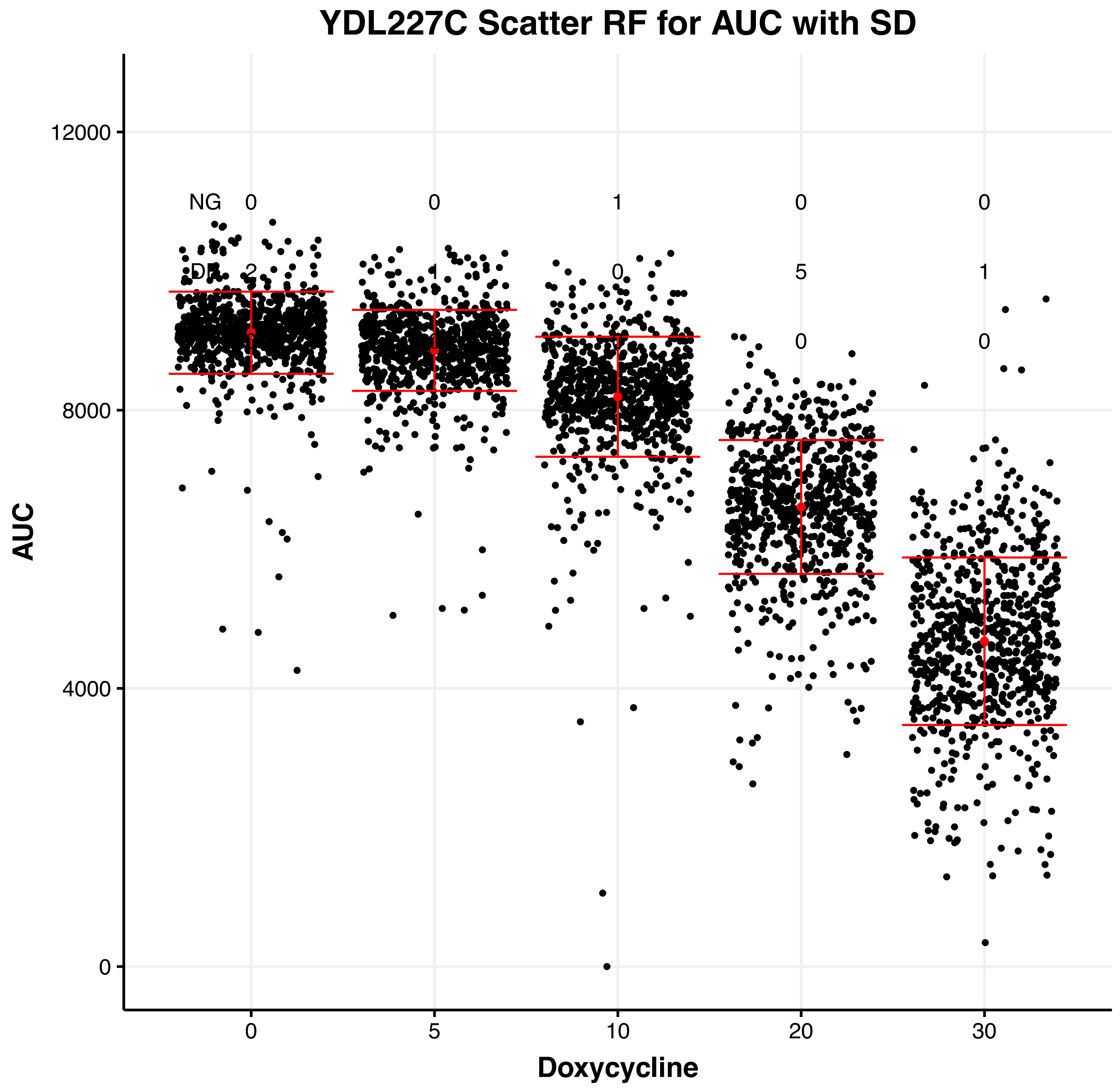
YDL227C Scatter RF for AUC with SD
AUC
Doxycycline
0
5
10
20
30
0
4000
8000
12000
NG 0 0 1 0 0
DB 2 1 0 5 1
SM 0 0 0 0 0

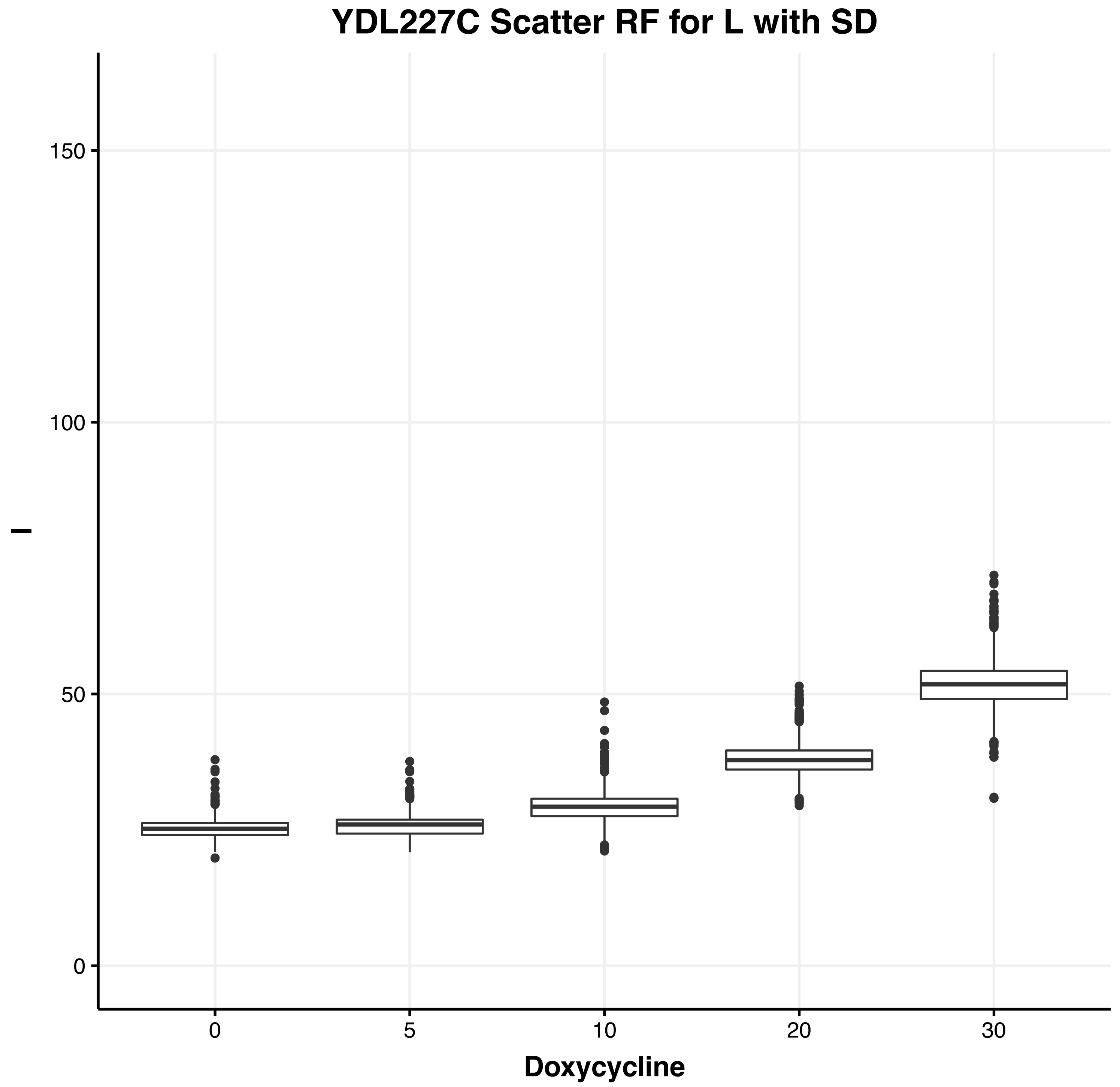
YDL227C Scatter RF for L with SD
l
Doxycycline
0
50
100
150
0
5
10
20
30

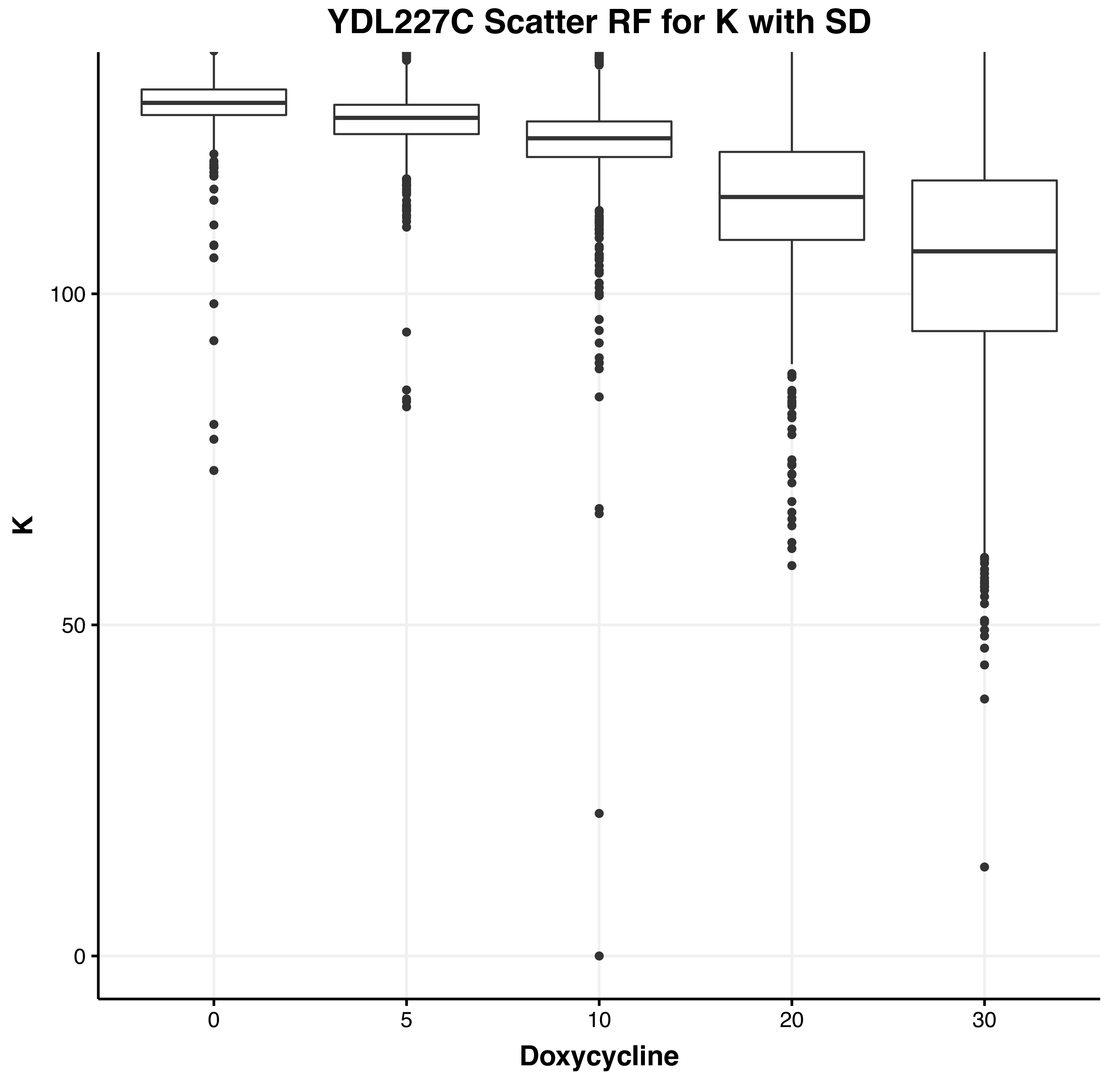
YDL227C Scatter RF for K with SD
K
Doxycycline
0
50
100
0
5
10
20
30

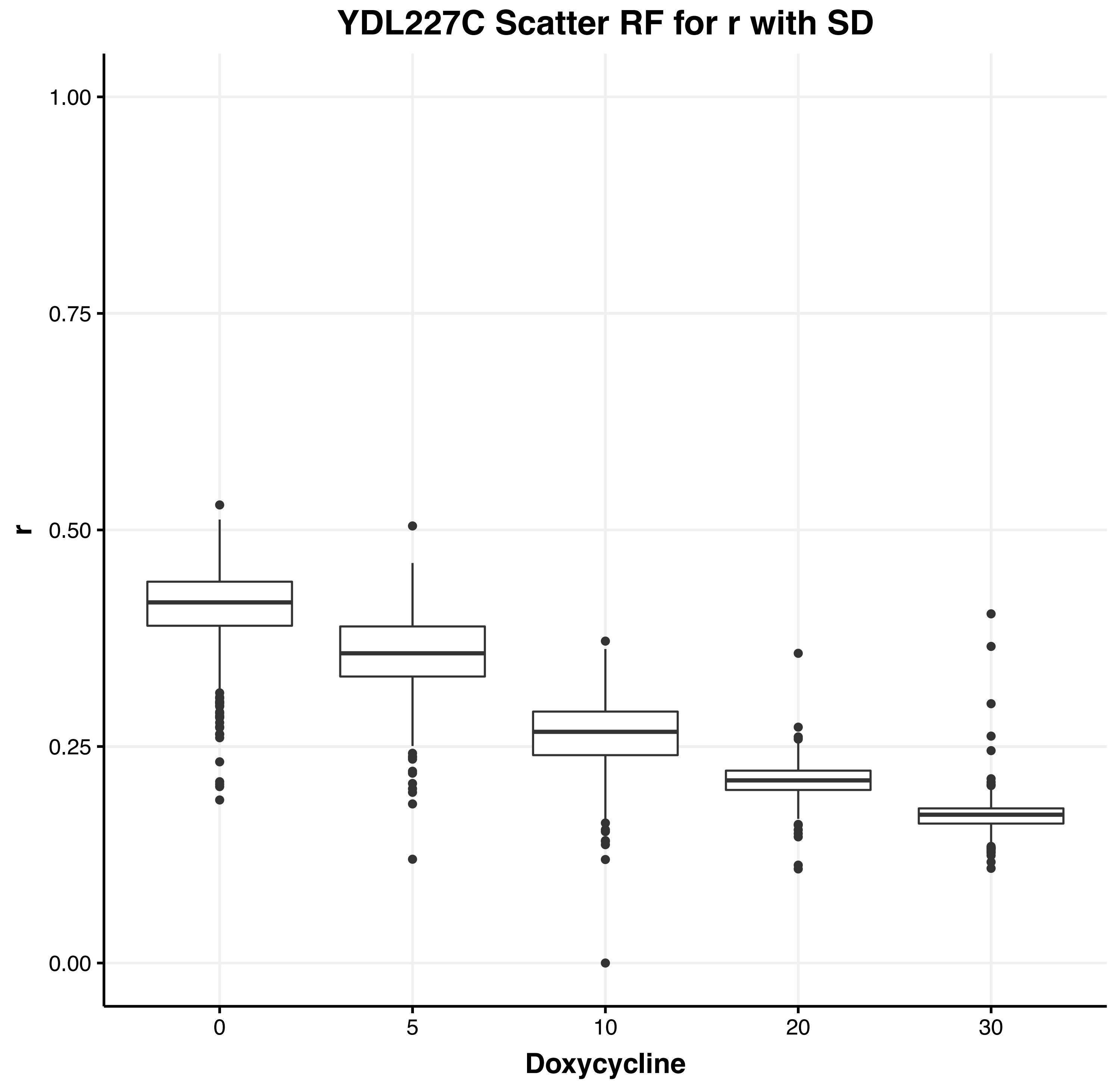
YDL227C Scatter RF for r with SD
r
Doxycycline
0.00
0.25
0.50
0.75
1.00
0
5
10
20
30

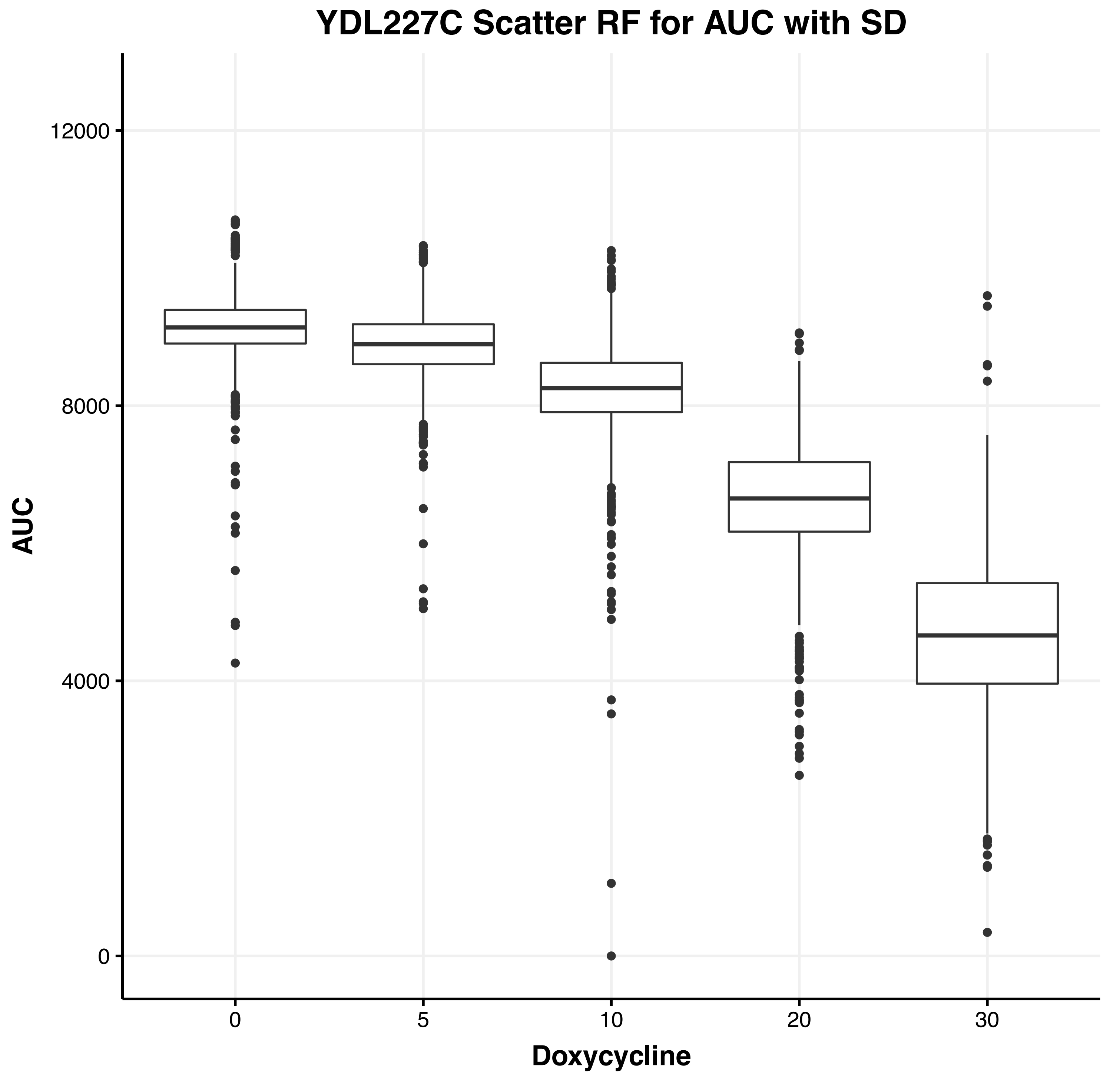
YDL227C Scatter RF for AUC with SD
AUC
Doxycycline
0
4000
8000
12000
0
5
10
20
30

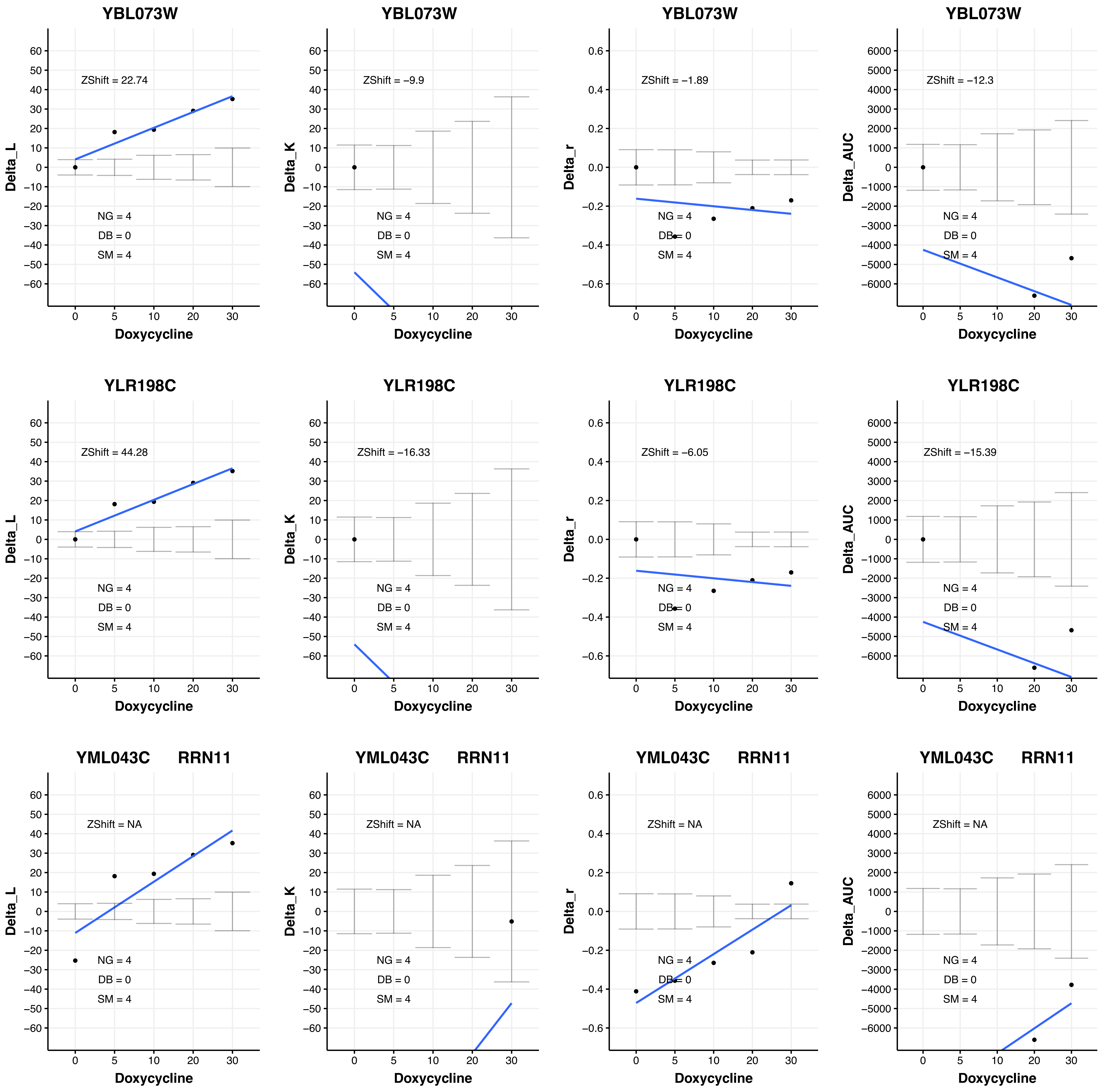
YBL073W
ZShift = 22.74
NG = 4
DB = 0
SM = 4
Delta_L
Doxycycline
YBL073W
ZShift = −9.9
NG = 4
DB = 0
SM = 4
Delta_K
Doxycycline
YBL073W
ZShift = −1.89
NG = 4
DB = 0
SM = 4
Delta_r
Doxycycline
YBL073W
ZShift = −12.3
NG = 4
DB = 0
SM = 4
Delta_AUC
Doxycycline
YLR198C
ZShift = 44.28
NG = 4
DB = 0
SM = 4
Delta_L
Doxycycline
YLR198C
ZShift = −16.33
NG = 4
DB = 0
SM = 4
Delta_K
Doxycycline
YLR198C
ZShift = −6.05
NG = 4
DB = 0
SM = 4
Delta_r
Doxycycline
YLR198C
ZShift = −15.39
NG = 4
DB = 0
SM = 4
Delta_AUC
Doxycycline
YML043C RRN11
ZShift = NA
NG = 4
DB = 0
SM = 4
Delta_L
Doxycycline
YML043C RRN11
ZShift = NA
NG = 4
DB = 0
SM = 4
Delta_K
Doxycycline
YML043C RRN11
ZShift = NA
NG = 4
DB = 0
SM = 4
Delta_r
Doxycycline
YML043C RRN11
ZShift = NA
NG = 4
DB = 0
SM = 4
Delta_AUC
Doxycycline

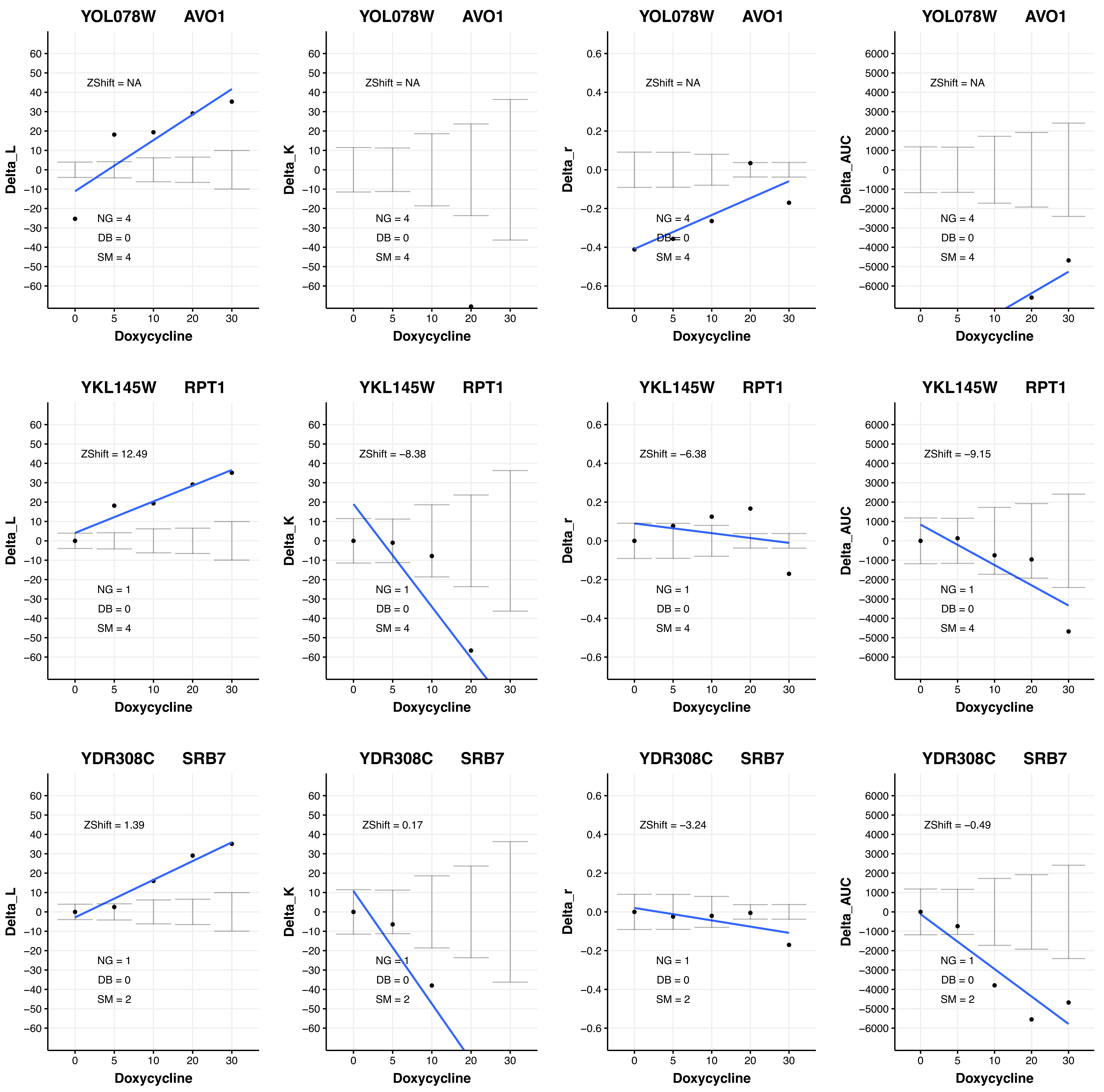

YOL078W AVO1
ZShift = NA
NG = 4
DB = 0
SM = 4
Delta_L
Delta_K
Delta_r
Delta_AUC
Doxycycline
YKL145W RPT1
ZShift = 12.49
ZShift = −8.38
ZShift = −6.38
ZShift = −9.15
NG = 1
DB = 0
SM = 4
YDR308C SRB7
ZShift = 1.39
ZShift = 0.17
ZShift = −3.24
ZShift = −0.49
NG = 1
DB = 0
SM = 2

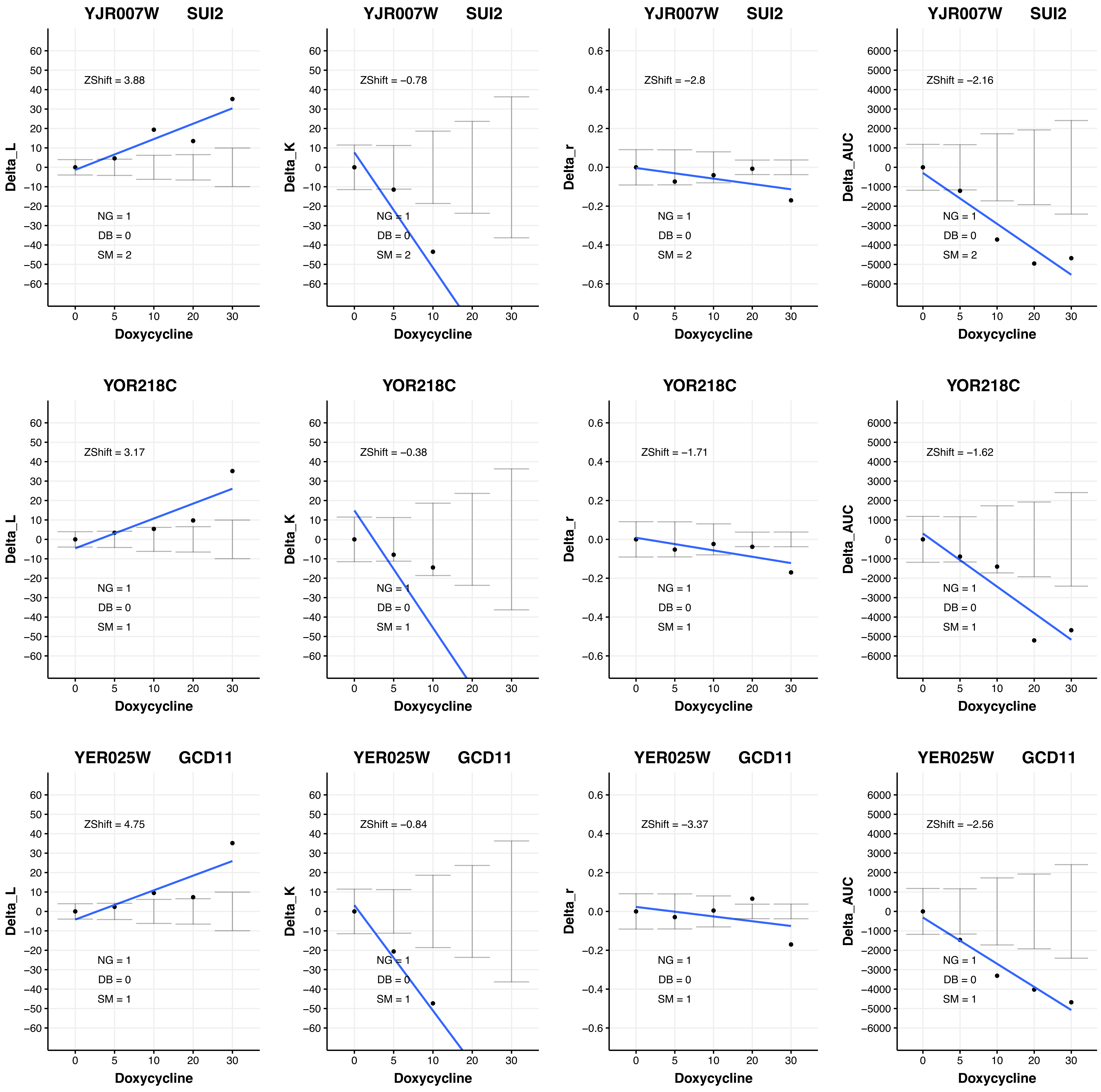
YJR007W SUI2
ZShift = 3.88
NG = 1
DB = 0
SM = 2
Delta_L
Doxycycline
YJR007W SUI2
ZShift = −0.78
NG = 1
DB = 0
SM = 2
Delta_K
Doxycycline
YJR007W SUI2
ZShift = −2.8
NG = 1
DB = 0
SM = 2
Delta_r
Doxycycline
YJR007W SUI2
ZShift = −2.16
NG = 1
DB = 0
SM = 2
Delta_AUC
Doxycycline
YOR218C
ZShift = 3.17
NG = 1
DB = 0
SM = 1
Delta_L
Doxycycline
YOR218C
ZShift = −0.38
NG = 1
DB = 0
SM = 1
Delta_K
Doxycycline
YOR218C
ZShift = −1.71
NG = 1
DB = 0
SM = 1
Delta_r
Doxycycline
YOR218C
ZShift = −1.62
NG = 1
DB = 0
SM = 1
Delta_AUC
Doxycycline
YER025W GCD11
ZShift = 4.75
NG = 1
DB = 0
SM = 1
Delta_L
Doxycycline
YER025W GCD11
ZShift = −0.84
NG = 1
DB = 0
SM = 1
Delta_K
Doxycycline
YER025W GCD11
ZShift = −3.37
NG = 1
DB = 0
SM = 1
Delta_r
Doxycycline
YER025W GCD11
ZShift = −2.56
NG = 1
DB = 0
SM = 1
Delta_AUC
Doxycycline

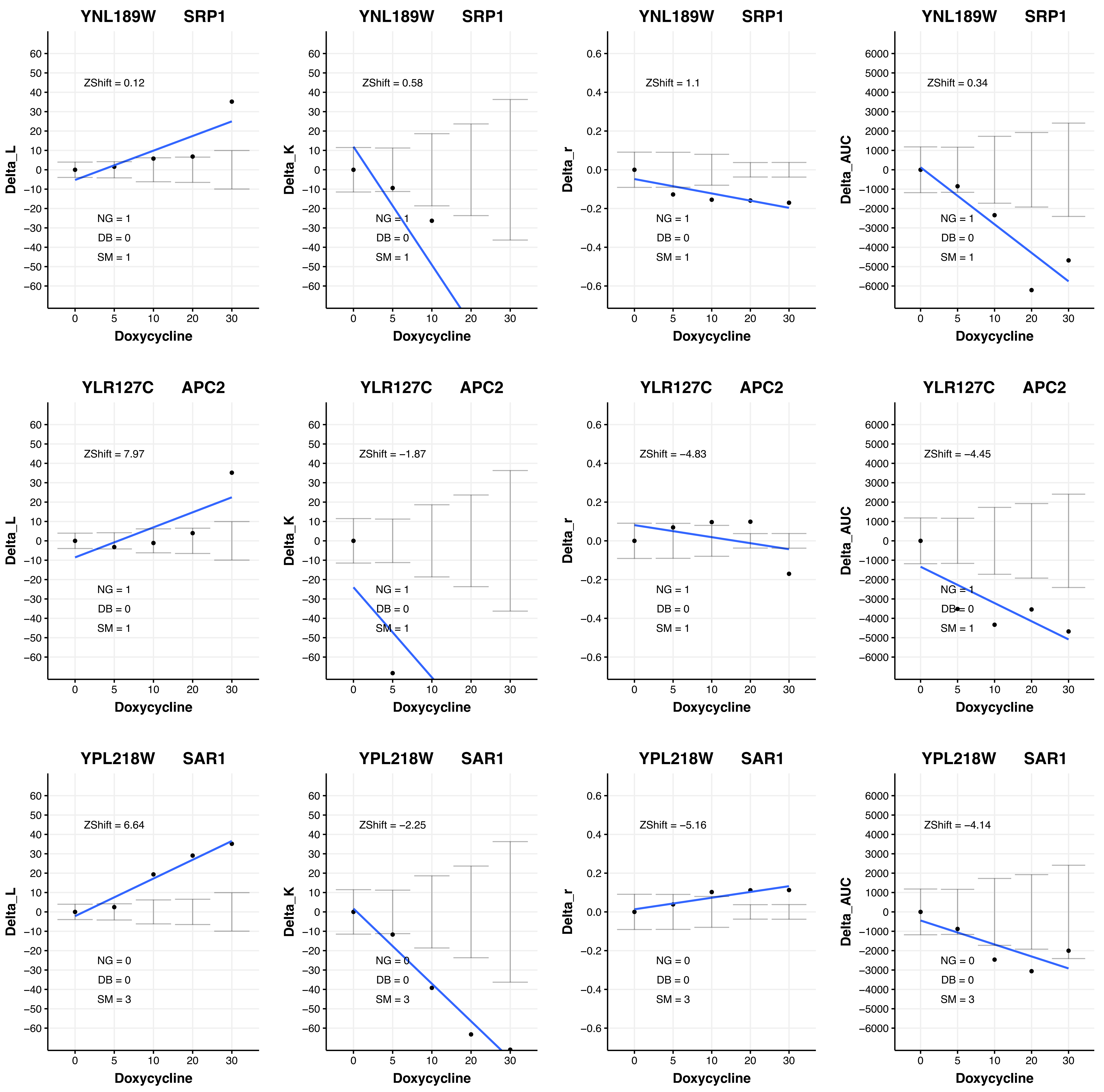
YNL189W SRP1
ZShift = 0.12
NG = 1
DB = 0
SM = 1
Delta_L
Doxycycline
YNL189W SRP1
ZShift = 0.58
NG = 1
DB = 0
SM = 1
Delta_K
Doxycycline
YNL189W SRP1
ZShift = 1.1
NG = 1
DB = 0
SM = 1
Delta_r
Doxycycline
YNL189W SRP1
ZShift = 0.34
NG = 1
DB = 0
SM = 1
Delta_AUC
Doxycycline
YLR127C APC2
ZShift = 7.97
NG = 1
DB = 0
SM = 1
Delta_L
Doxycycline
YLR127C APC2
ZShift = −1.87
NG = 1
DB = 0
SM = 1
Delta_K
Doxycycline
YLR127C APC2
ZShift = −4.83
NG = 1
DB = 0
SM = 1
Delta_r
Doxycycline
YLR127C APC2
ZShift = −4.45
NG = 1
DB = 0
SM = 1
Delta_AUC
Doxycycline
YPL218W SAR1
ZShift = 6.64
NG = 0
DB = 0
SM = 3
Delta_L
Doxycycline
YPL218W SAR1
ZShift = −2.25
NG = 0
DB = 0
SM = 3
Delta_K
Doxycycline
YPL218W SAR1
ZShift = −5.16
NG = 0
DB = 0
SM = 3
Delta_r
Doxycycline
YPL218W SAR1
ZShift = −4.14
NG = 0
DB = 0
SM = 3
Delta_AUC
Doxycycline

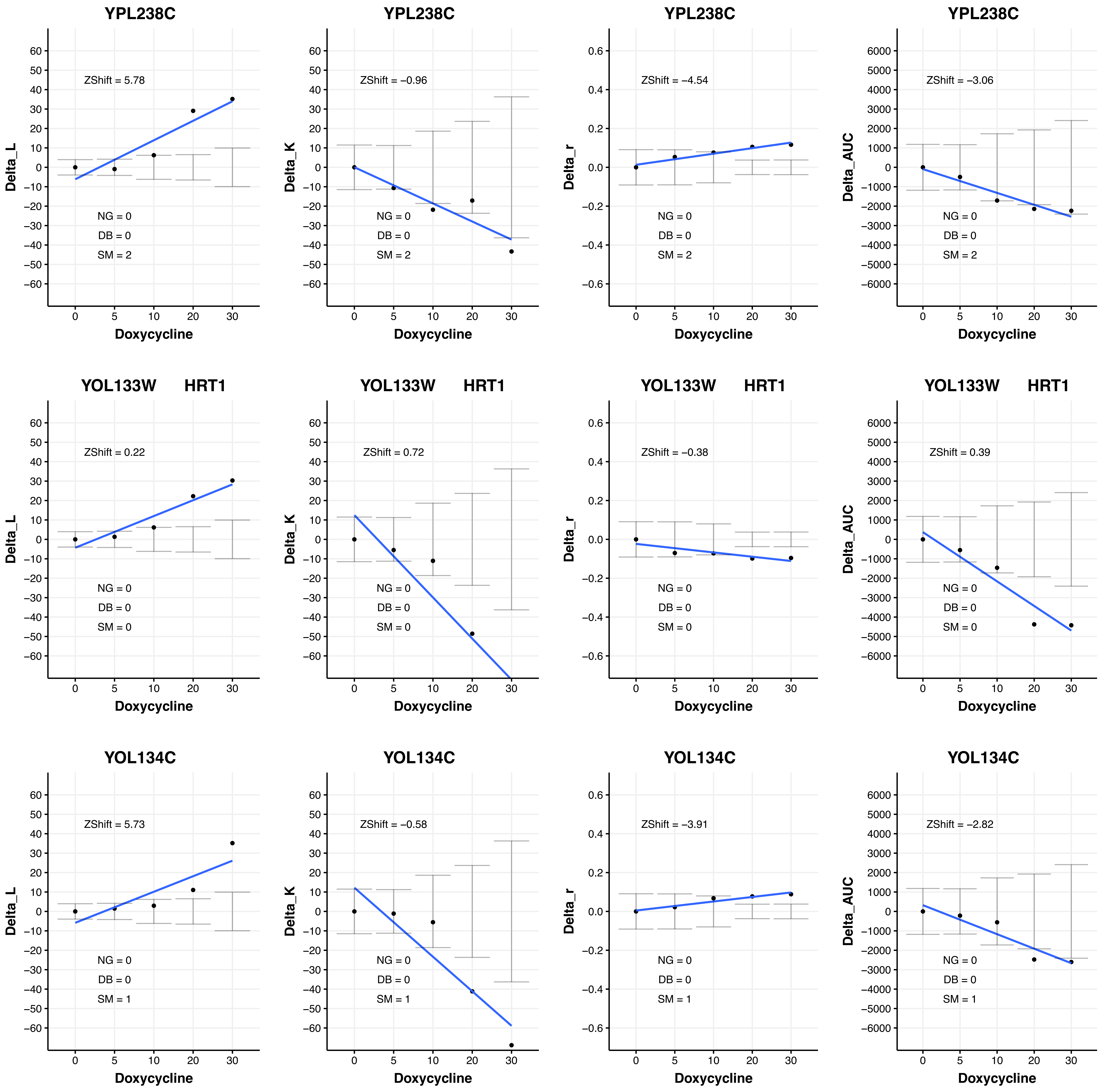

YPL238C
ZShift = 5.78
NG = 0
DB = 0
SM = 2
Delta_L
Doxycycline
YPL238C
ZShift = −0.96
NG = 0
DB = 0
SM = 2
Delta_K
Doxycycline
YPL238C
ZShift = −4.54
NG = 0
DB = 0
SM = 2
Delta_r
Doxycycline
YPL238C
ZShift = −3.06
NG = 0
DB = 0
SM = 2
Delta_AUC
Doxycycline
YOL133W HRT1
ZShift = 0.22
NG = 0
DB = 0
SM = 0
Delta_L
Doxycycline
YOL133W HRT1
ZShift = 0.72
NG = 0
DB = 0
SM = 0
Delta_K
Doxycycline
YOL133W HRT1
ZShift = −0.38
NG = 0
DB = 0
SM = 0
Delta_r
Doxycycline
YOL133W HRT1
ZShift = 0.39
NG = 0
DB = 0
SM = 0
Delta_AUC
Doxycycline
YOL134C
ZShift = 5.73
NG = 0
DB = 0
SM = 1
Delta_L
Doxycycline
YOL134C
ZShift = −0.58
NG = 0
DB = 0
SM = 1
Delta_K
Doxycycline
YOL134C
ZShift = −3.91
NG = 0
DB = 0
SM = 1
Delta_r
Doxycycline
YOL134C
ZShift = −2.82
NG = 0
DB = 0
SM = 1
Delta_AUC
Doxycycline

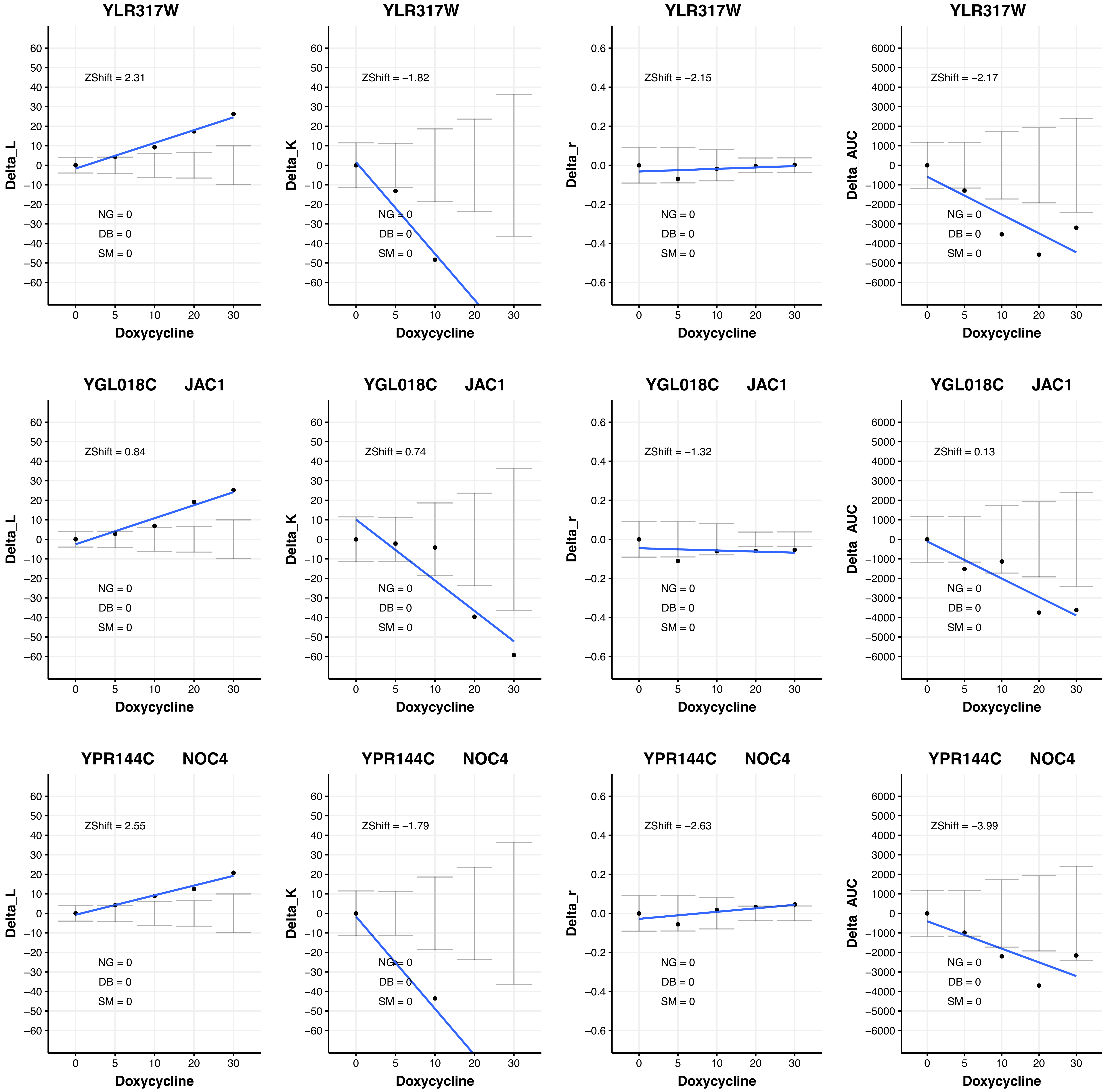

YLR317W
ZShift = 2.31
Delta_L
NG = 0
DB = 0
SM = 0
Doxycycline
YLR317W
ZShift = −1.82
Delta_K
NG = 0
DB = 0
SM = 0
Doxycycline
YLR317W
ZShift = −2.15
Delta_r
NG = 0
DB = 0
SM = 0
Doxycycline
YLR317W
ZShift = −2.17
Delta_AUC
NG = 0
DB = 0
SM = 0
Doxycycline
YGL018C JAC1
ZShift = 0.84
Delta_L
NG = 0
DB = 0
SM = 0
Doxycycline
YGL018C JAC1
ZShift = 0.74
Delta_K
NG = 0
DB = 0
SM = 0
Doxycycline
YGL018C JAC1
ZShift = −1.32
Delta_r
NG = 0
DB = 0
SM = 0
Doxycycline
YGL018C JAC1
ZShift = 0.13
Delta_AUC
NG = 0
DB = 0
SM = 0
Doxycycline
YPR144C NOC4
ZShift = 2.55
Delta_L
NG = 0
DB = 0
SM = 0
Doxycycline
YPR144C NOC4
ZShift = −1.79
Delta_K
NG = 0
DB = 0
SM = 0
Doxycycline
YPR144C NOC4
ZShift = −2.63
Delta_r
NG = 0
DB = 0
SM = 0
Doxycycline
YPR144C NOC4
ZShift = −3.99
Delta_AUC
NG = 0
DB = 0
SM = 0
Doxycycline

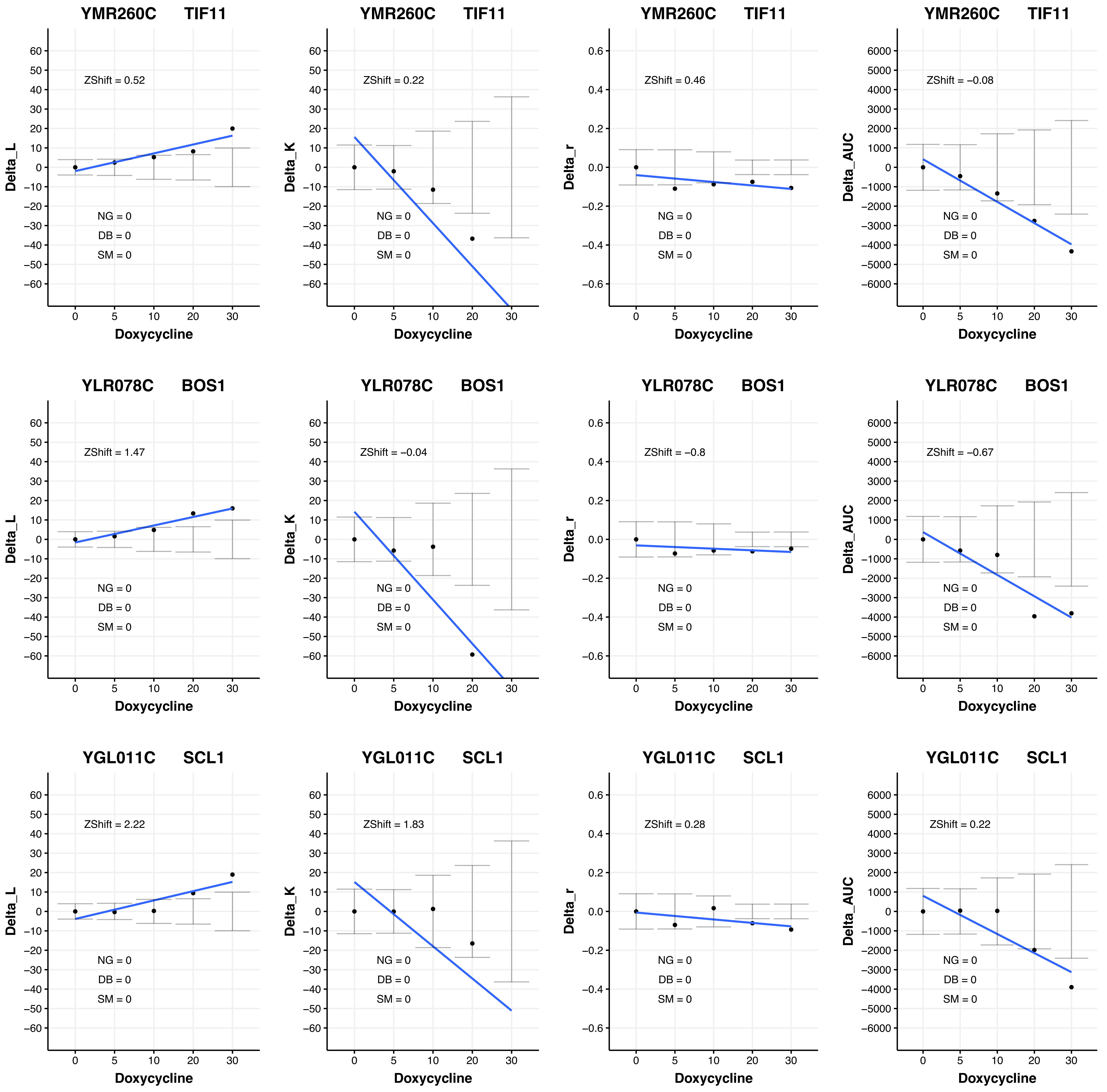

YMR260C TIF11
ZShift = 0.52
NG = 0
DB = 0
SM = 0
Delta_L
Doxycycline
YMR260C TIF11
ZShift = 0.22
NG = 0
DB = 0
SM = 0
Delta_K
Doxycycline
YMR260C TIF11
ZShift = 0.46
NG = 0
DB = 0
SM = 0
Delta_r
Doxycycline
YMR260C TIF11
ZShift = −0.08
NG = 0
DB = 0
SM = 0
Delta_AUC
Doxycycline
YLR078C BOS1
ZShift = 1.47
NG = 0
DB = 0
SM = 0
Delta_L
Doxycycline
YLR078C BOS1
ZShift = −0.04
NG = 0
DB = 0
SM = 0
Delta_K
Doxycycline
YLR078C BOS1
ZShift = −0.8
NG = 0
DB = 0
SM = 0
Delta_r
Doxycycline
YLR078C BOS1
ZShift = −0.67
NG = 0
DB = 0
SM = 0
Delta_AUC
Doxycycline
YGL011C SCL1
ZShift = 2.22
NG = 0
DB = 0
SM = 0
Delta_L
Doxycycline
YGL011C SCL1
ZShift = 1.83
NG = 0
DB = 0
SM = 0
Delta_K
Doxycycline
YGL011C SCL1
ZShift = 0.28
NG = 0
DB = 0
SM = 0
Delta_r
Doxycycline
YGL011C SCL1
ZShift = 0.22
NG = 0
DB = 0
SM = 0
Delta_AUC
Doxycycline

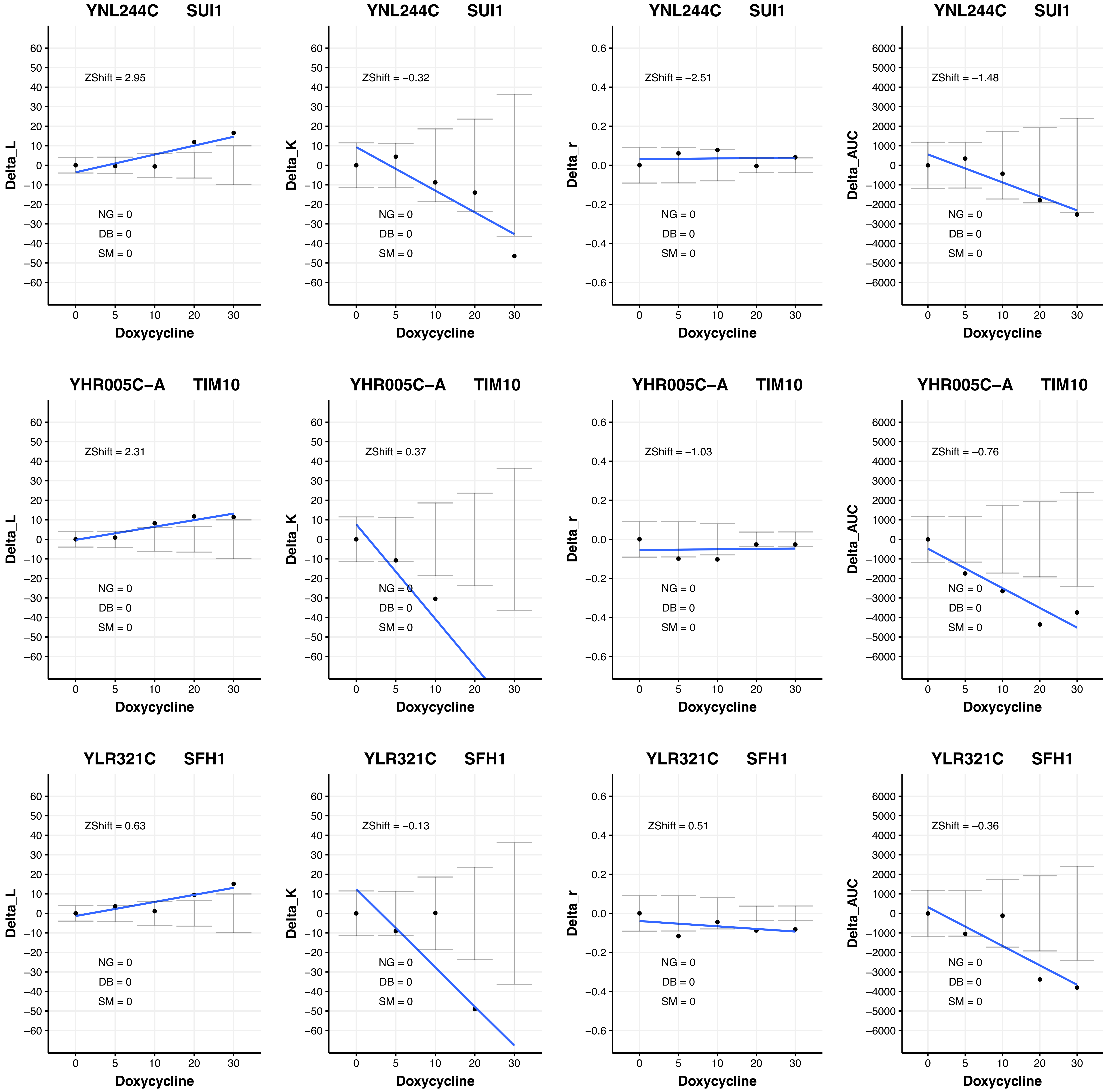

YNL244C SUI1
ZShift = 2.95
Delta_L
NG = 0
DB = 0
SM = 0
Doxycycline
YNL244C SUI1
ZShift = −0.32
Delta_K
NG = 0
DB = 0
SM = 0
Doxycycline
YNL244C SUI1
ZShift = −2.51
Delta_r
NG = 0
DB = 0
SM = 0
Doxycycline
YNL244C SUI1
ZShift = −1.48
Delta_AUC
NG = 0
DB = 0
SM = 0
Doxycycline
YHR005C−A TIM10
ZShift = 2.31
Delta_L
NG = 0
DB = 0
SM = 0
Doxycycline
YHR005C−A TIM10
ZShift = 0.37
Delta_K
NG = 0
DB = 0
SM = 0
Doxycycline
YHR005C−A TIM10
ZShift = −1.03
Delta_r
NG = 0
DB = 0
SM = 0
Doxycycline
YHR005C−A TIM10
ZShift = −0.76
Delta_AUC
NG = 0
DB = 0
SM = 0
Doxycycline
YLR321C SFH1
ZShift = 0.63
Delta_L
NG = 0
DB = 0
SM = 0
Doxycycline
YLR321C SFH1
ZShift = −0.13
Delta_K
NG = 0
DB = 0
SM = 0
Doxycycline
YLR321C SFH1
ZShift = 0.51
Delta_r
NG = 0
DB = 0
SM = 0
Doxycycline
YLR321C SFH1
ZShift = −0.36
Delta_AUC
NG = 0
DB = 0
SM = 0
Doxycycline

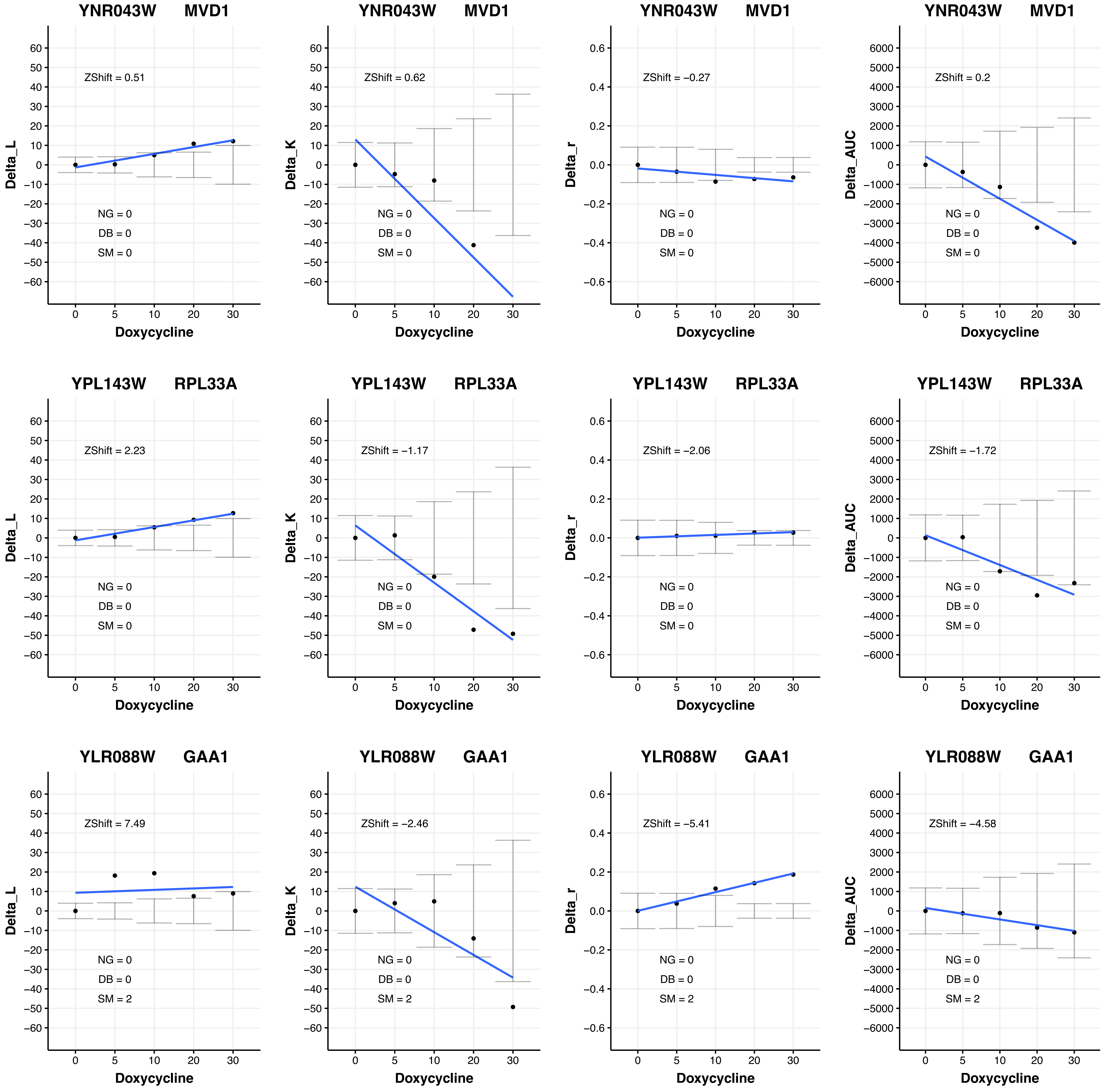

YNR043W MVD1
ZShift = 0.51
NG = 0
DB = 0
SM = 0
Delta_L
Doxycycline
YNR043W MVD1
ZShift = 0.62
NG = 0
DB = 0
SM = 0
Delta_K
Doxycycline
YNR043W MVD1
ZShift = −0.27
NG = 0
DB = 0
SM = 0
Delta_r
Doxycycline
YNR043W MVD1
ZShift = 0.2
NG = 0
DB = 0
SM = 0
Delta_AUC
Doxycycline
YPL143W RPL33A
ZShift = 2.23
NG = 0
DB = 0
SM = 0
Delta_L
Doxycycline
YPL143W RPL33A
ZShift = −1.17
NG = 0
DB = 0
SM = 0
Delta_K
Doxycycline
YPL143W RPL33A
ZShift = −2.06
NG = 0
DB = 0
SM = 0
Delta_r
Doxycycline
YPL143W RPL33A
ZShift = −1.72
NG = 0
DB = 0
SM = 0
Delta_AUC
Doxycycline
YLR088W GAA1
ZShift = 7.49
NG = 0
DB = 0
SM = 2
Delta_L
Doxycycline
YLR088W GAA1
ZShift = −2.46
NG = 0
DB = 0
SM = 2
Delta_K
Doxycycline
YLR088W GAA1
ZShift = −5.41
NG = 0
DB = 0
SM = 2
Delta_r
Doxycycline
YLR088W GAA1
ZShift = −4.58
NG = 0
DB = 0
SM = 2
Delta_AUC
Doxycycline

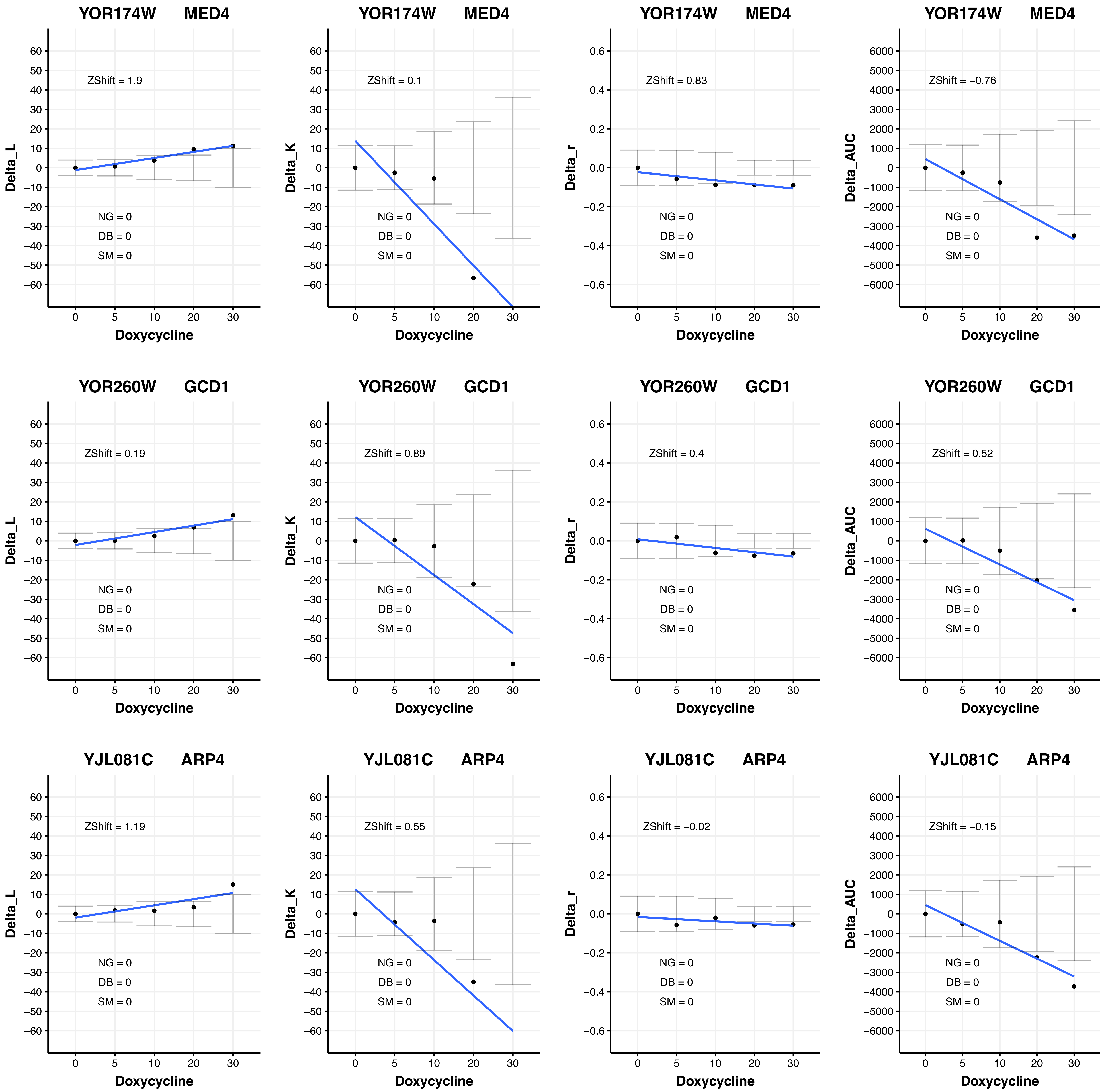
YOR174W MED4
ZShift = 1.9
NG = 0
DB = 0
SM = 0
Delta_L
Doxycycline
YOR174W MED4
ZShift = 0.1
NG = 0
DB = 0
SM = 0
Delta_K
Doxycycline
YOR174W MED4
ZShift = 0.83
NG = 0
DB = 0
SM = 0
Delta_r
Doxycycline
YOR174W MED4
ZShift = −0.76
NG = 0
DB = 0
SM = 0
Delta_AUC
Doxycycline
YOR260W GCD1
ZShift = 0.19
NG = 0
DB = 0
SM = 0
Delta_L
Doxycycline
YOR260W GCD1
ZShift = 0.89
NG = 0
DB = 0
SM = 0
Delta_K
Doxycycline
YOR260W GCD1
ZShift = 0.4
NG = 0
DB = 0
SM = 0
Delta_r
Doxycycline
YOR260W GCD1
ZShift = 0.52
NG = 0
DB = 0
SM = 0
Delta_AUC
Doxycycline
YJL081C ARP4
ZShift = 1.19
NG = 0
DB = 0
SM = 0
Delta_L
Doxycycline
YJL081C ARP4
ZShift = 0.55
NG = 0
DB = 0
SM = 0
Delta_K
Doxycycline
YJL081C ARP4
ZShift = −0.02
NG = 0
DB = 0
SM = 0
Delta_r
Doxycycline
YJL081C ARP4
ZShift = −0.15
NG = 0
DB = 0
SM = 0
Delta_AUC
Doxycycline
0
5
10
20
30

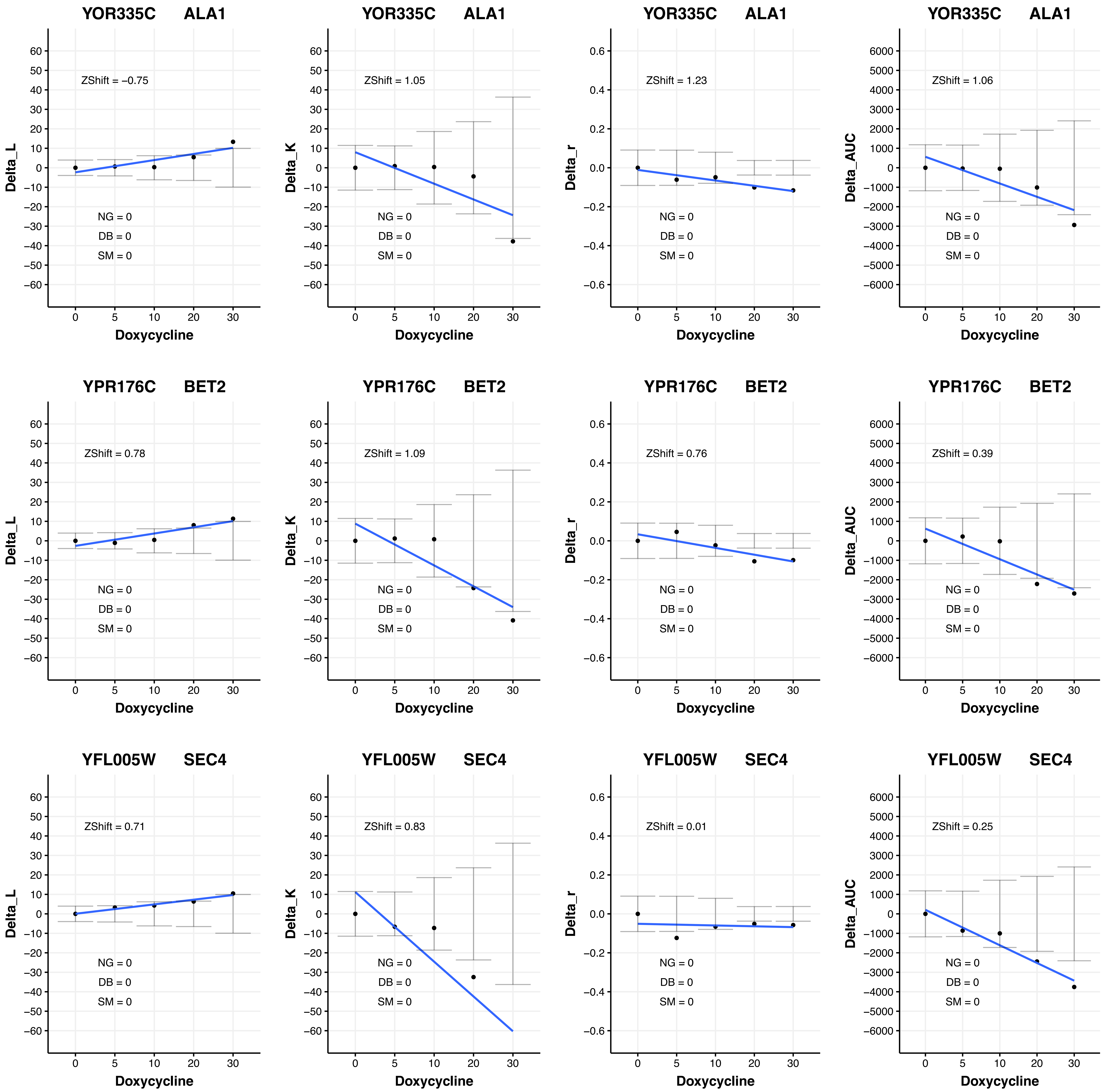
YOR335C ALA1
ZShift = −0.75
NG = 0
DB = 0
SM = 0
Delta_L
Doxycycline
YOR335C ALA1
ZShift = 1.05
NG = 0
DB = 0
SM = 0
Delta_K
Doxycycline
YOR335C ALA1
ZShift = 1.23
NG = 0
DB = 0
SM = 0
Delta_r
Doxycycline
YOR335C ALA1
ZShift = 1.06
NG = 0
DB = 0
SM = 0
Delta_AUC
Doxycycline
YPR176C BET2
ZShift = 0.78
NG = 0
DB = 0
SM = 0
Delta_L
Doxycycline
YPR176C BET2
ZShift = 1.09
NG = 0
DB = 0
SM = 0
Delta_K
Doxycycline
YPR176C BET2
ZShift = 0.76
NG = 0
DB = 0
SM = 0
Delta_r
Doxycycline
YPR176C BET2
ZShift = 0.39
NG = 0
DB = 0
SM = 0
Delta_AUC
Doxycycline
YFL005W SEC4
ZShift = 0.71
NG = 0
DB = 0
SM = 0
Delta_L
Doxycycline
YFL005W SEC4
ZShift = 0.83
NG = 0
DB = 0
SM = 0
Delta_K
Doxycycline
YFL005W SEC4
ZShift = 0.01
NG = 0
DB = 0
SM = 0
Delta_r
Doxycycline
YFL005W SEC4
ZShift = 0.25
NG = 0
DB = 0
SM = 0
Delta_AUC
Doxycycline

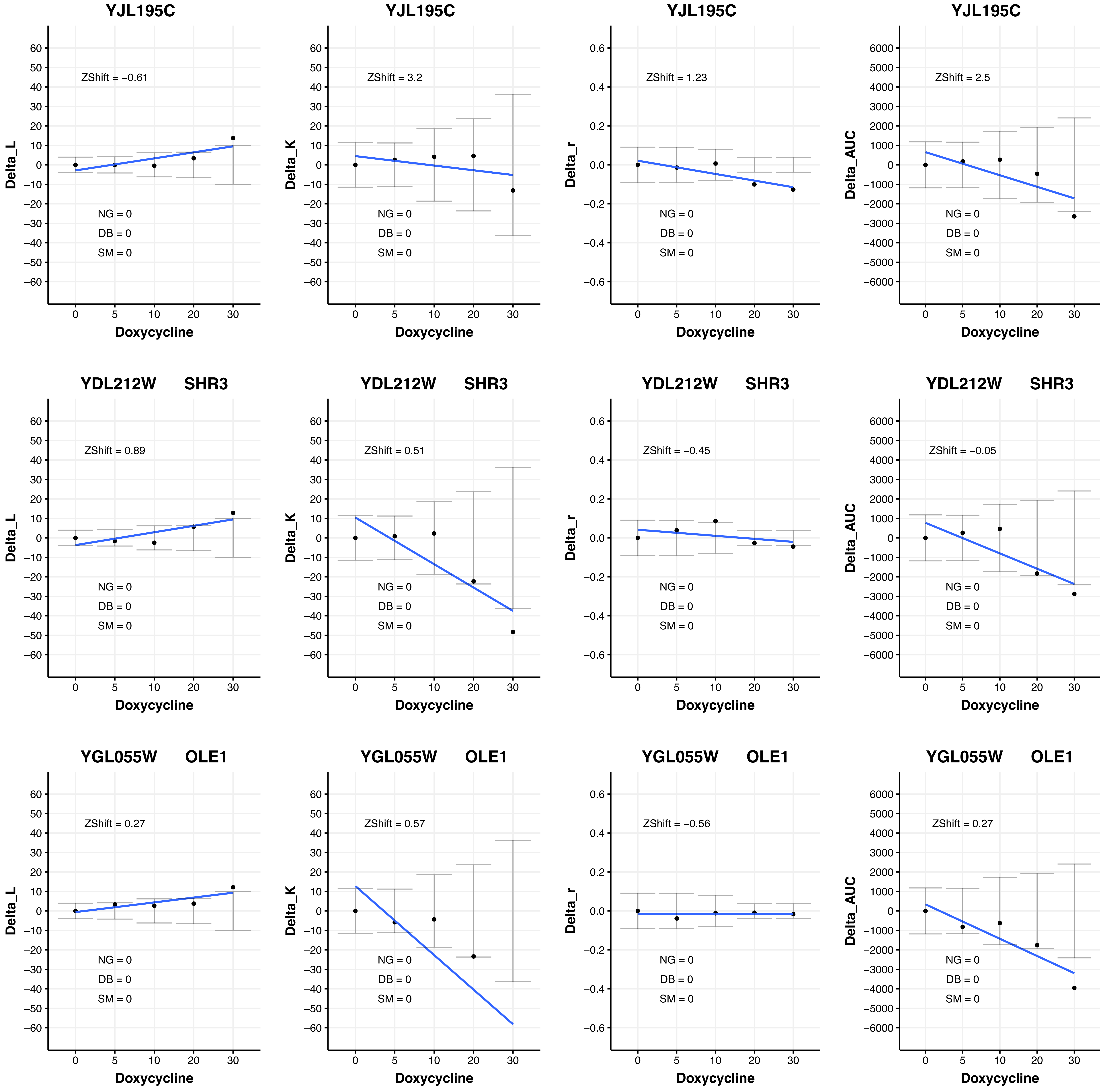

YJL195C
ZShift = −0.61
NG = 0
DB = 0
SM = 0
Delta_L
Doxycycline
YJL195C
ZShift = 3.2
NG = 0
DB = 0
SM = 0
Delta_K
Doxycycline
YJL195C
ZShift = 1.23
NG = 0
DB = 0
SM = 0
Delta_r
Doxycycline
YJL195C
ZShift = 2.5
NG = 0
DB = 0
SM = 0
Delta_AUC
Doxycycline
YDL212W SHR3
ZShift = 0.89
NG = 0
DB = 0
SM = 0
Delta_L
Doxycycline
YDL212W SHR3
ZShift = 0.51
NG = 0
DB = 0
SM = 0
Delta_K
Doxycycline
YDL212W SHR3
ZShift = −0.45
NG = 0
DB = 0
SM = 0
Delta_r
Doxycycline
YDL212W SHR3
ZShift = −0.05
NG = 0
DB = 0
SM = 0
Delta_AUC
Doxycycline
YGL055W OLE1
ZShift = 0.27
NG = 0
DB = 0
SM = 0
Delta_L
Doxycycline
YGL055W OLE1
ZShift = 0.57
NG = 0
DB = 0
SM = 0
Delta_K
Doxycycline
YGL055W OLE1
ZShift = −0.56
NG = 0
DB = 0
SM = 0
Delta_r
Doxycycline
YGL055W OLE1
ZShift = 0.27
NG = 0
DB = 0
SM = 0
Delta_AUC
Doxycycline

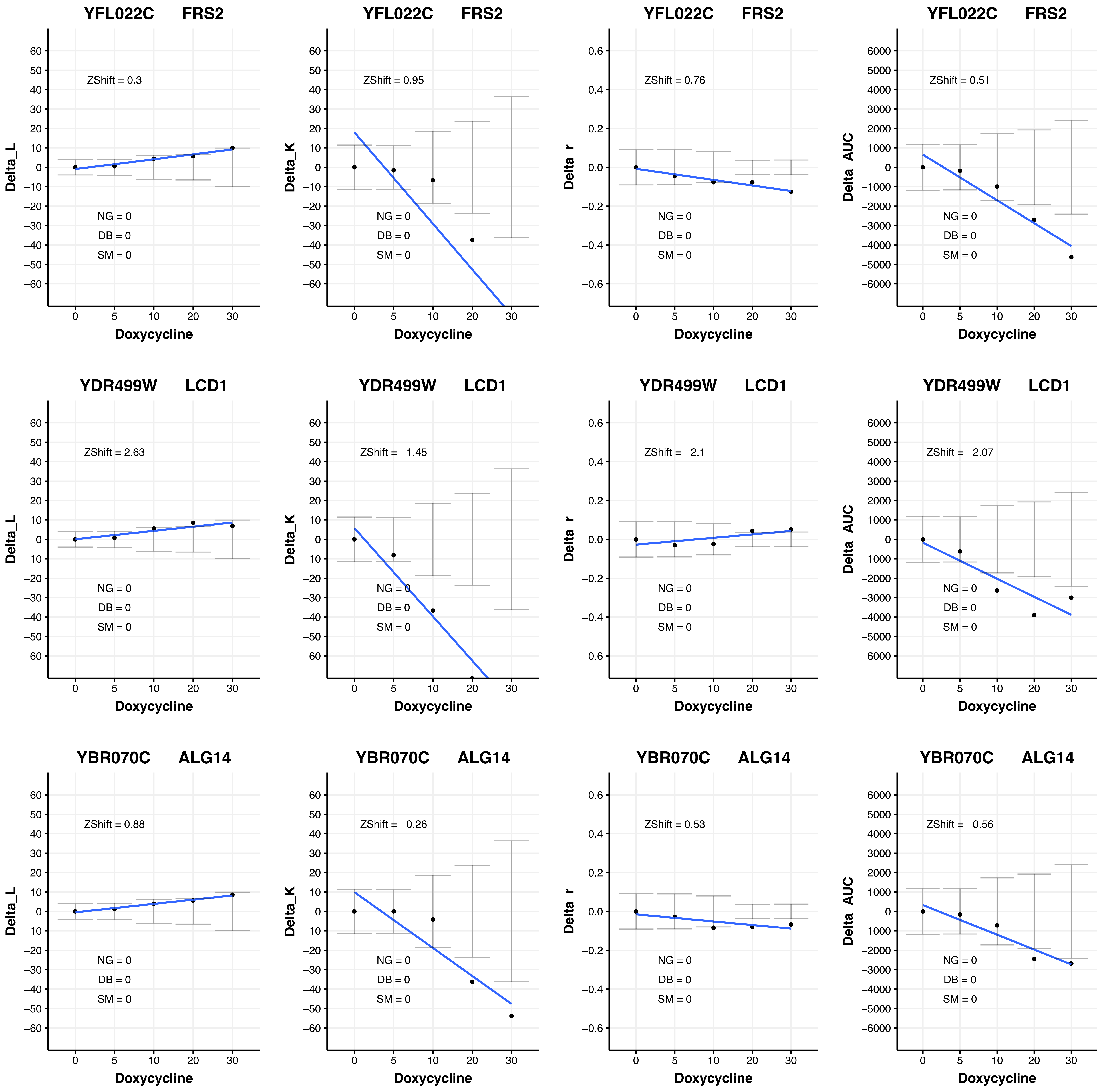
YFL022C FRS2
ZShift = 0.3
NG = 0
DB = 0
SM = 0
Delta_L
Doxycycline
YFL022C FRS2
ZShift = 0.95
NG = 0
DB = 0
SM = 0
Delta_K
Doxycycline
YFL022C FRS2
ZShift = 0.76
NG = 0
DB = 0
SM = 0
Delta_r
Doxycycline
YFL022C FRS2
ZShift = 0.51
NG = 0
DB = 0
SM = 0
Delta_AUC
Doxycycline
YDR499W LCD1
ZShift = 2.63
NG = 0
DB = 0
SM = 0
Delta_L
Doxycycline
YDR499W LCD1
ZShift = −1.45
NG = 0
DB = 0
SM = 0
Delta_K
Doxycycline
YDR499W LCD1
ZShift = −2.1
NG = 0
DB = 0
SM = 0
Delta_r
Doxycycline
YDR499W LCD1
ZShift = −2.07
NG = 0
DB = 0
SM = 0
Delta_AUC
Doxycycline
YBR070C ALG14
ZShift = 0.88
NG = 0
DB = 0
SM = 0
Delta_L
Doxycycline
YBR070C ALG14
ZShift = −0.26
NG = 0
DB = 0
SM = 0
Delta_K
Doxycycline
YBR070C ALG14
ZShift = 0.53
NG = 0
DB = 0
SM = 0
Delta_r
Doxycycline
YBR070C ALG14
ZShift = −0.56
NG = 0
DB = 0
SM = 0
Delta_AUC
Doxycycline

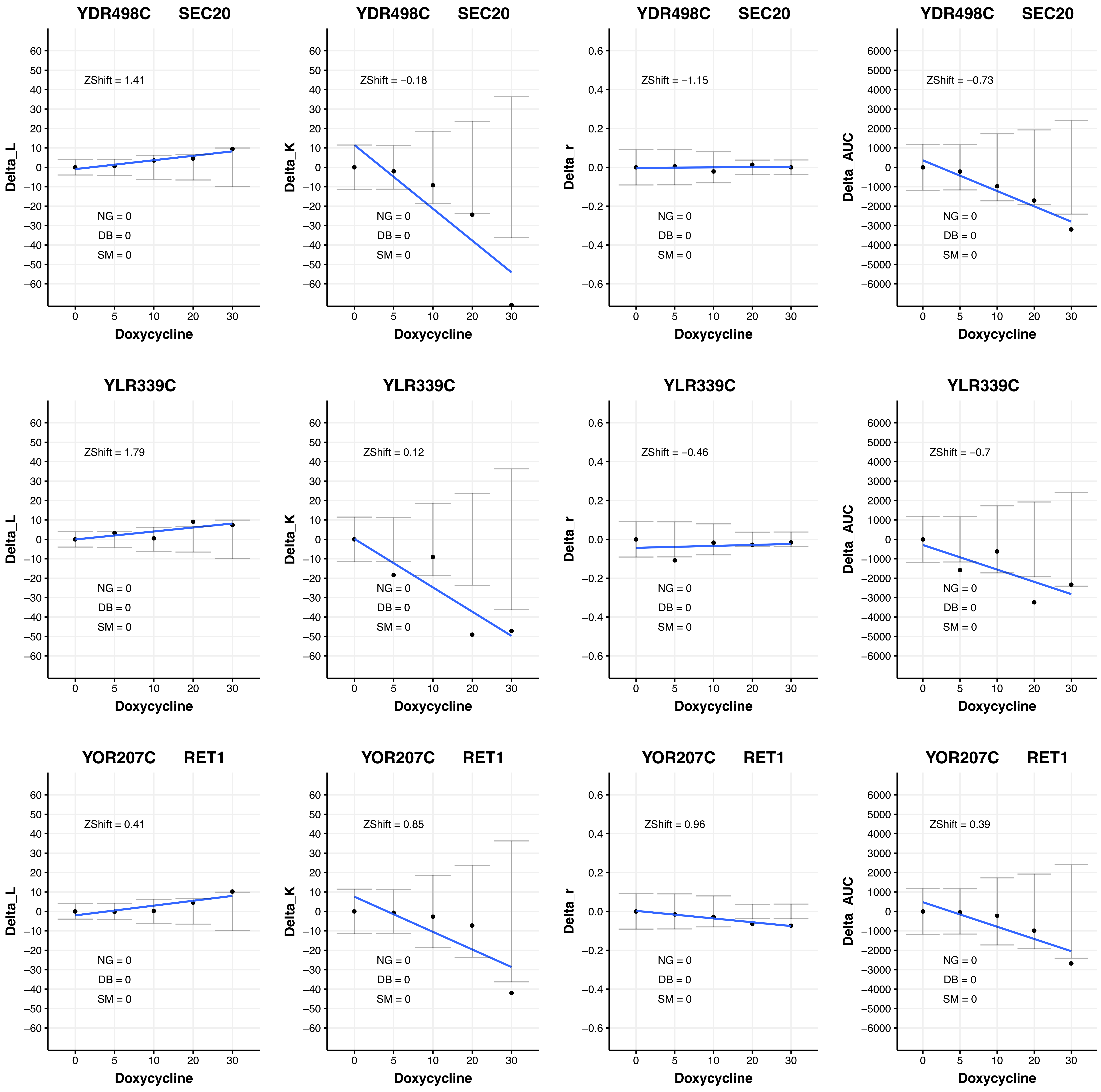
YDR498C SEC20
ZShift = 1.41
NG = 0
DB = 0
SM = 0
Delta_L
Doxycycline
YDR498C SEC20
ZShift = −0.18
NG = 0
DB = 0
SM = 0
Delta_K
Doxycycline
YDR498C SEC20
ZShift = −1.15
NG = 0
DB = 0
SM = 0
Delta_r
Doxycycline
YDR498C SEC20
ZShift = −0.73
NG = 0
DB = 0
SM = 0
Delta_AUC
Doxycycline
YLR339C
ZShift = 1.79
NG = 0
DB = 0
SM = 0
Delta_L
Doxycycline
YLR339C
ZShift = 0.12
NG = 0
DB = 0
SM = 0
Delta_K
Doxycycline
YLR339C
ZShift = −0.46
NG = 0
DB = 0
SM = 0
Delta_r
Doxycycline
YLR339C
ZShift = −0.7
NG = 0
DB = 0
SM = 0
Delta_AUC
Doxycycline
YOR207C RET1
ZShift = 0.41
NG = 0
DB = 0
SM = 0
Delta_L
Doxycycline
YOR207C RET1
ZShift = 0.85
NG = 0
DB = 0
SM = 0
Delta_K
Doxycycline
YOR207C RET1
ZShift = 0.96
NG = 0
DB = 0
SM = 0
Delta_r
Doxycycline
YOR207C RET1
ZShift = 0.39
NG = 0
DB = 0
SM = 0
Delta_AUC
Doxycycline

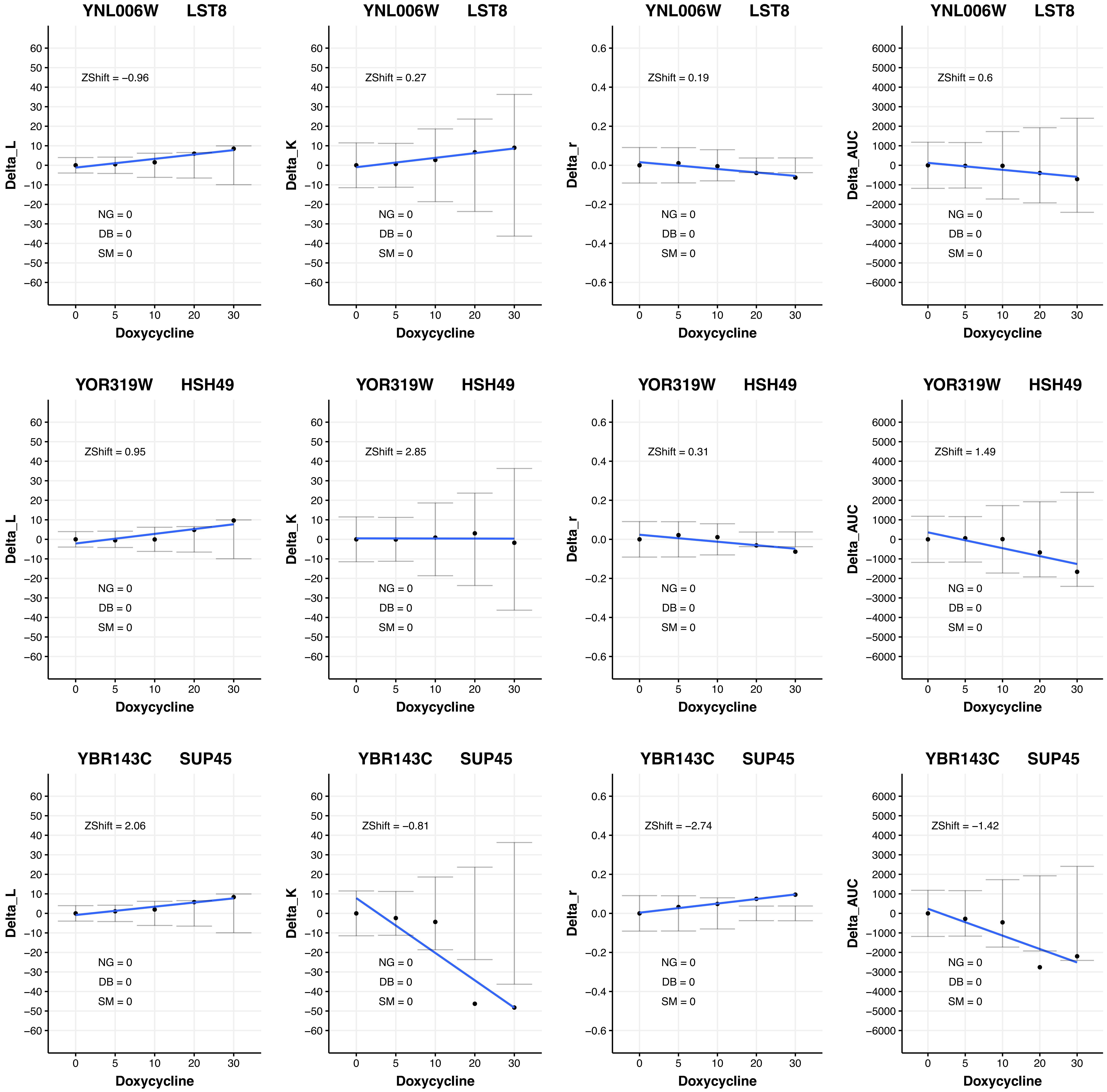

YNL006W LST8
ZShift = −0.96
NG = 0
DB = 0
SM = 0
Delta_L
Doxycycline
YNL006W LST8
ZShift = 0.27
NG = 0
DB = 0
SM = 0
Delta_K
Doxycycline
YNL006W LST8
ZShift = 0.19
NG = 0
DB = 0
SM = 0
Delta_r
Doxycycline
YNL006W LST8
ZShift = 0.6
NG = 0
DB = 0
SM = 0
Delta_AUC
Doxycycline
YOR319W HSH49
ZShift = 0.95
NG = 0
DB = 0
SM = 0
Delta_L
Doxycycline
YOR319W HSH49
ZShift = 2.85
NG = 0
DB = 0
SM = 0
Delta_K
Doxycycline
YOR319W HSH49
ZShift = 0.31
NG = 0
DB = 0
SM = 0
Delta_r
Doxycycline
YOR319W HSH49
ZShift = 1.49
NG = 0
DB = 0
SM = 0
Delta_AUC
Doxycycline
YBR143C SUP45
ZShift = 2.06
NG = 0
DB = 0
SM = 0
Delta_L
Doxycycline
YBR143C SUP45
ZShift = −0.81
NG = 0
DB = 0
SM = 0
Delta_K
Doxycycline
YBR143C SUP45
ZShift = −2.74
NG = 0
DB = 0
SM = 0
Delta_r
Doxycycline
YBR143C SUP45
ZShift = −1.42
NG = 0
DB = 0
SM = 0
Delta_AUC
Doxycycline

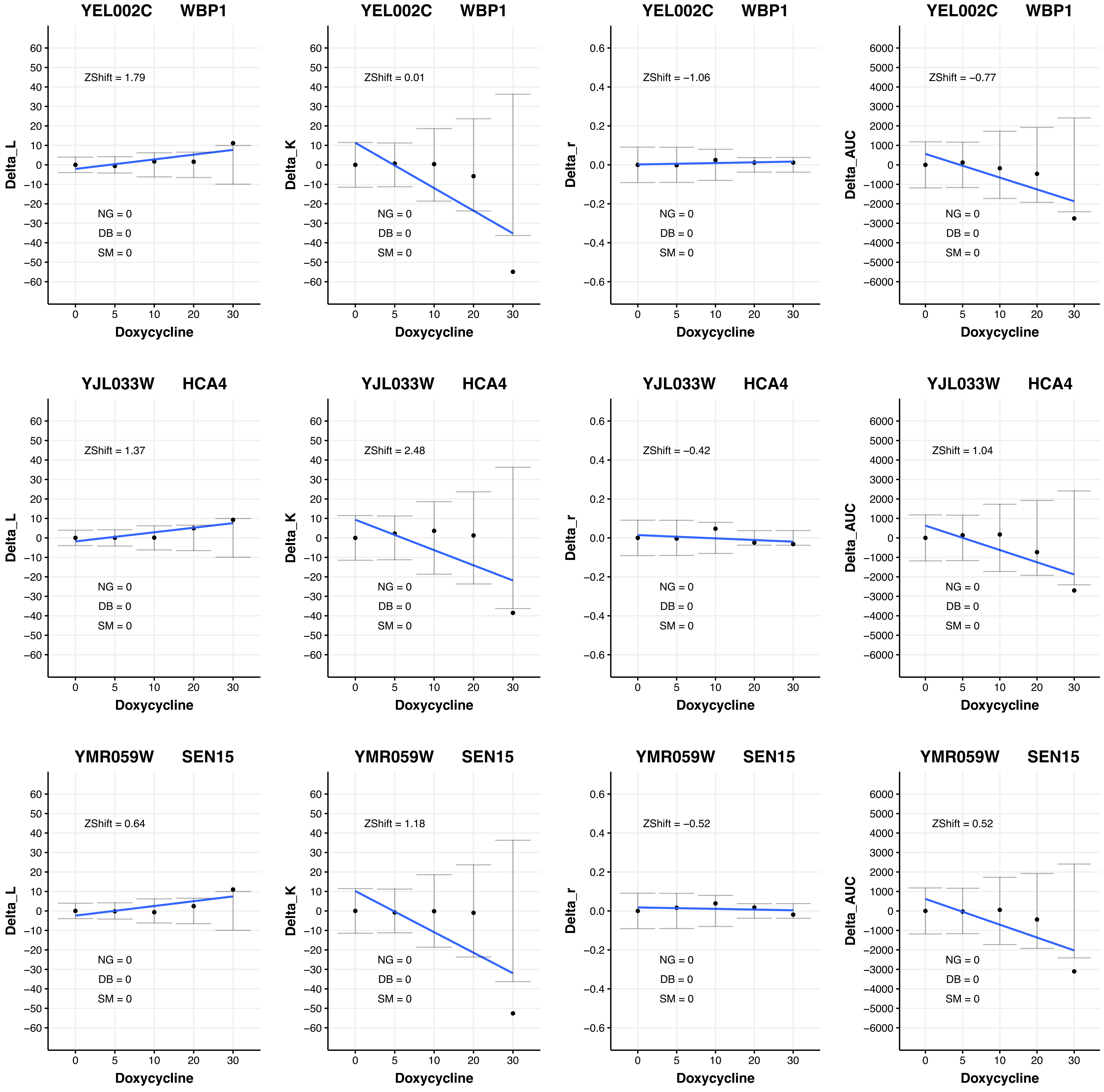

YEL002C WBP1
ZShift = 1.79
NG = 0
DB = 0
SM = 0
Delta_L
Doxycycline
YEL002C WBP1
ZShift = 0.01
NG = 0
DB = 0
SM = 0
Delta_K
Doxycycline
YEL002C WBP1
ZShift = −1.06
NG = 0
DB = 0
SM = 0
Delta_r
Doxycycline
YEL002C WBP1
ZShift = −0.77
NG = 0
DB = 0
SM = 0
Delta_AUC
Doxycycline
YJL033W HCA4
ZShift = 1.37
NG = 0
DB = 0
SM = 0
Delta_L
Doxycycline
YJL033W HCA4
ZShift = 2.48
NG = 0
DB = 0
SM = 0
Delta_K
Doxycycline
YJL033W HCA4
ZShift = −0.42
NG = 0
DB = 0
SM = 0
Delta_r
Doxycycline
YJL033W HCA4
ZShift = 1.04
NG = 0
DB = 0
SM = 0
Delta_AUC
Doxycycline
YMR059W SEN15
ZShift = 0.64
NG = 0
DB = 0
SM = 0
Delta_L
Doxycycline
YMR059W SEN15
ZShift = 1.18
NG = 0
DB = 0
SM = 0
Delta_K
Doxycycline
YMR059W SEN15
ZShift = −0.52
NG = 0
DB = 0
SM = 0
Delta_r
Doxycycline
YMR059W SEN15
ZShift = 0.52
NG = 0
DB = 0
SM = 0
Delta_AUC
Doxycycline

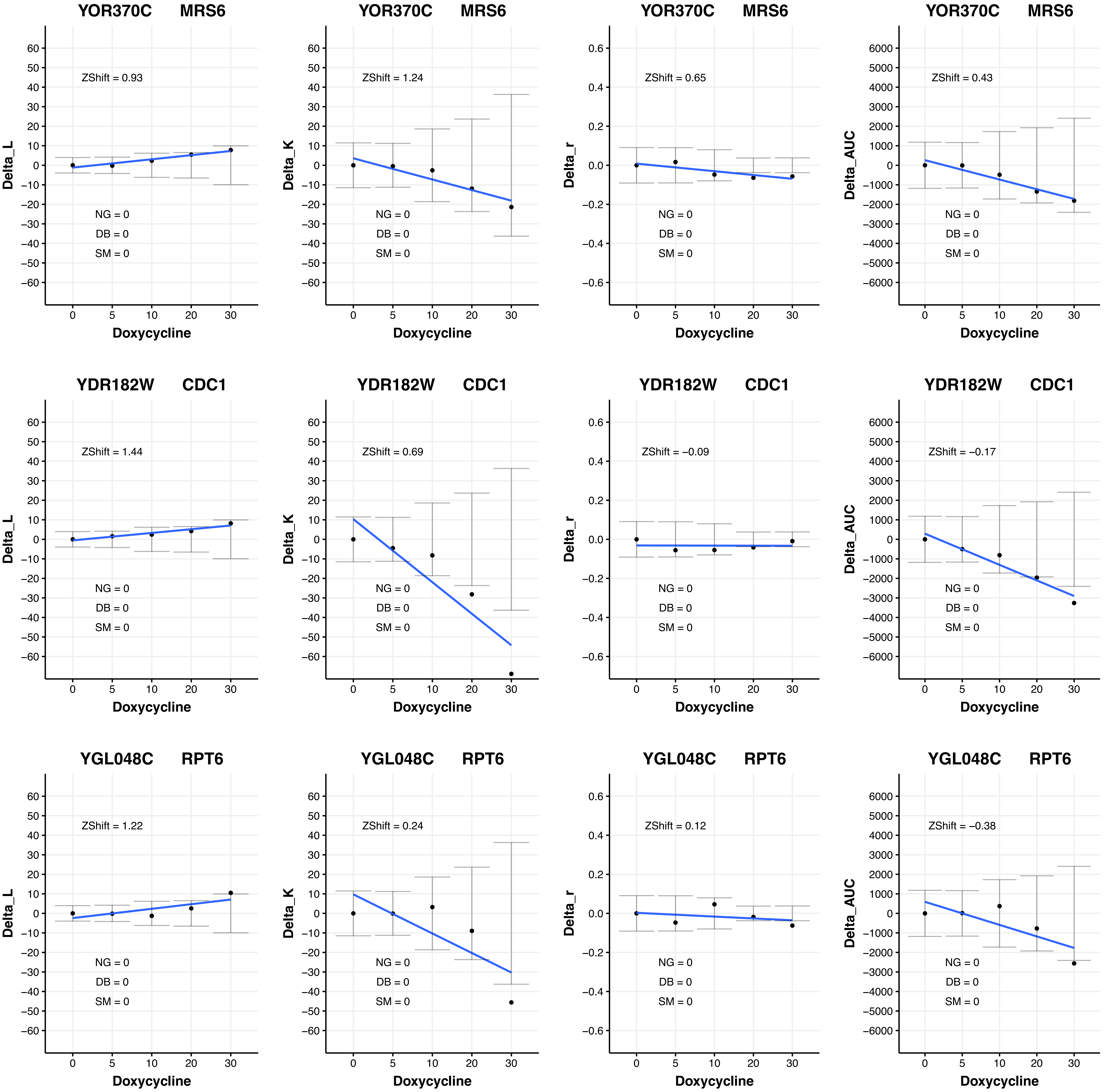
YOR370C MRS6
ZShift = 0.93
ZShift = 1.24
ZShift = 0.65
ZShift = 0.43
YDR182W CDC1
ZShift = 1.44
ZShift = 0.69
ZShift = −0.09
ZShift = −0.17
YGL048C RPT6
ZShift = 1.22
ZShift = 0.24
ZShift = 0.12
ZShift = −0.38
NG = 0
DB = 0
SM = 0
Delta_L
Delta_K
Delta_r
Delta_AUC
Doxycycline
0 5 10 20 30

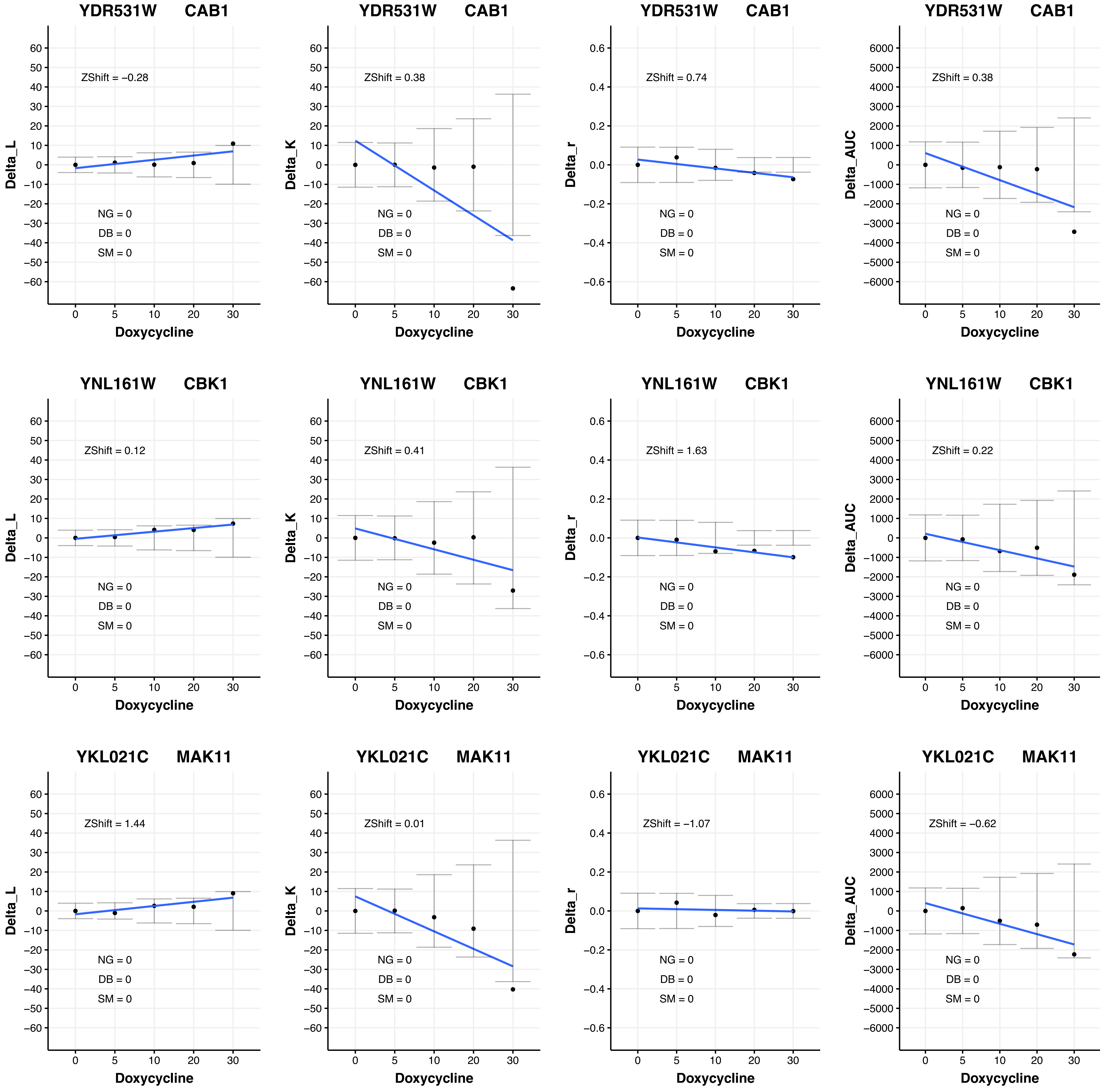
YDR531W CAB1
ZShift = −0.28
NG = 0
DB = 0
SM = 0
Delta_L
Doxycycline
YDR531W CAB1
ZShift = 0.38
NG = 0
DB = 0
SM = 0
Delta_K
Doxycycline
YDR531W CAB1
ZShift = 0.74
NG = 0
DB = 0
SM = 0
Delta_r
Doxycycline
YDR531W CAB1
ZShift = 0.38
NG = 0
DB = 0
SM = 0
Delta_AUC
Doxycycline
YNL161W CBK1
ZShift = 0.12
NG = 0
DB = 0
SM = 0
Delta_L
Doxycycline
YNL161W CBK1
ZShift = 0.41
NG = 0
DB = 0
SM = 0
Delta_K
Doxycycline
YNL161W CBK1
ZShift = 1.63
NG = 0
DB = 0
SM = 0
Delta_r
Doxycycline
YNL161W CBK1
ZShift = 0.22
NG = 0
DB = 0
SM = 0
Delta_AUC
Doxycycline
YKL021C MAK11
ZShift = 1.44
NG = 0
DB = 0
SM = 0
Delta_L
Doxycycline
YKL021C MAK11
ZShift = 0.01
NG = 0
DB = 0
SM = 0
Delta_K
Doxycycline
YKL021C MAK11
ZShift = −1.07
NG = 0
DB = 0
SM = 0
Delta_r
Doxycycline
YKL021C MAK11
ZShift = −0.62
NG = 0
DB = 0
SM = 0
Delta_AUC
Doxycycline

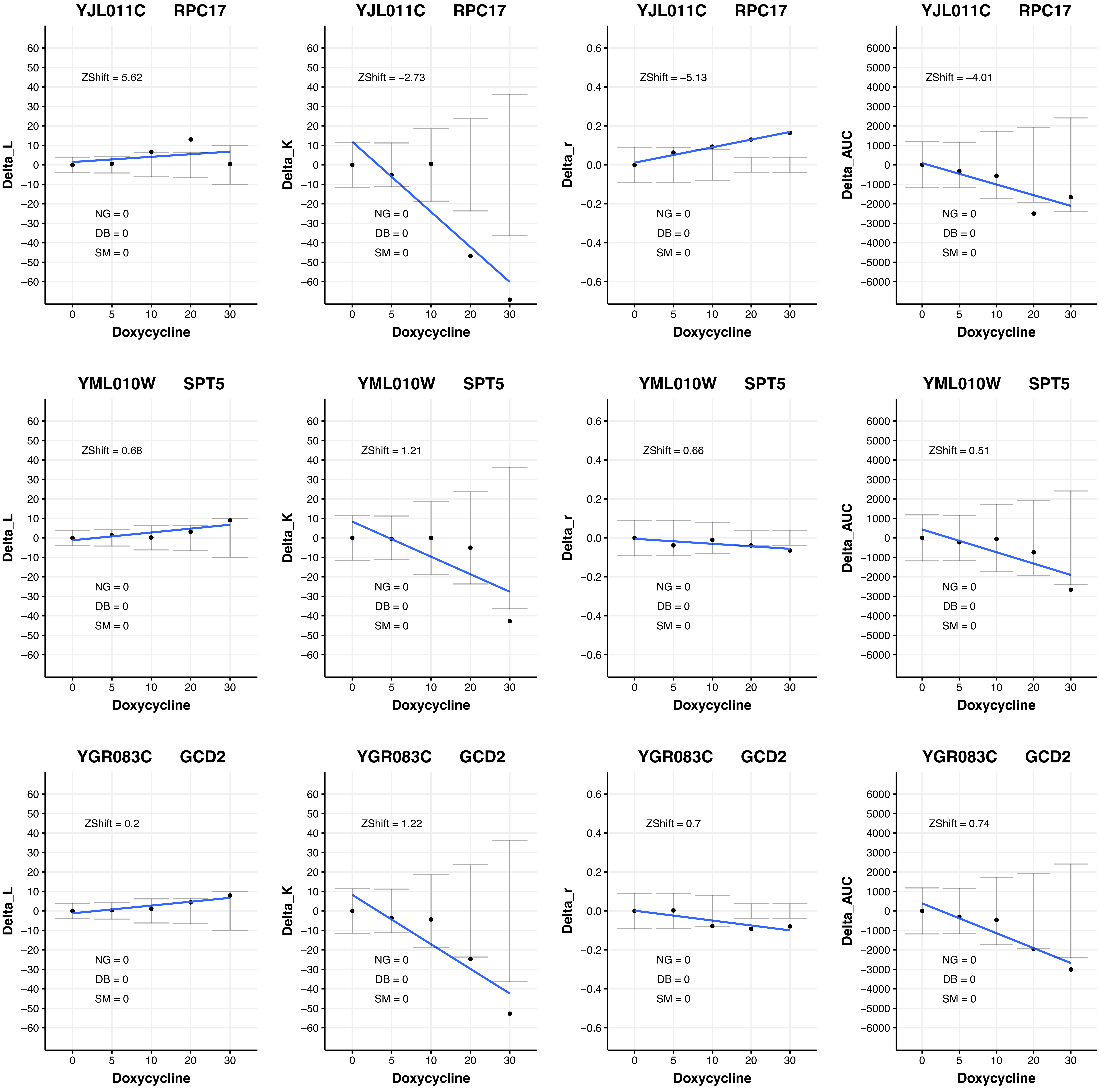
YJL011C RPC17
ZShift = 5.62
NG = 0
DB = 0
SM = 0
Delta_L
Doxycycline
YJL011C RPC17
ZShift = −2.73
NG = 0
DB = 0
SM = 0
Delta_K
Doxycycline
YJL011C RPC17
ZShift = −5.13
NG = 0
DB = 0
SM = 0
Delta_r
Doxycycline
YJL011C RPC17
ZShift = −4.01
NG = 0
DB = 0
SM = 0
Delta_AUC
Doxycycline
YML010W SPT5
ZShift = 0.68
NG = 0
DB = 0
SM = 0
Delta_L
Doxycycline
YML010W SPT5
ZShift = 1.21
NG = 0
DB = 0
SM = 0
Delta_K
Doxycycline
YML010W SPT5
ZShift = 0.66
NG = 0
DB = 0
SM = 0
Delta_r
Doxycycline
YML010W SPT5
ZShift = 0.51
NG = 0
DB = 0
SM = 0
Delta_AUC
Doxycycline
YGR083C GCD2
ZShift = 0.2
NG = 0
DB = 0
SM = 0
Delta_L
Doxycycline
YGR083C GCD2
ZShift = 1.22
NG = 0
DB = 0
SM = 0
Delta_K
Doxycycline
YGR083C GCD2
ZShift = 0.7
NG = 0
DB = 0
SM = 0
Delta_r
Doxycycline
YGR083C GCD2
ZShift = 0.74
NG = 0
DB = 0
SM = 0
Delta_AUC
Doxycycline

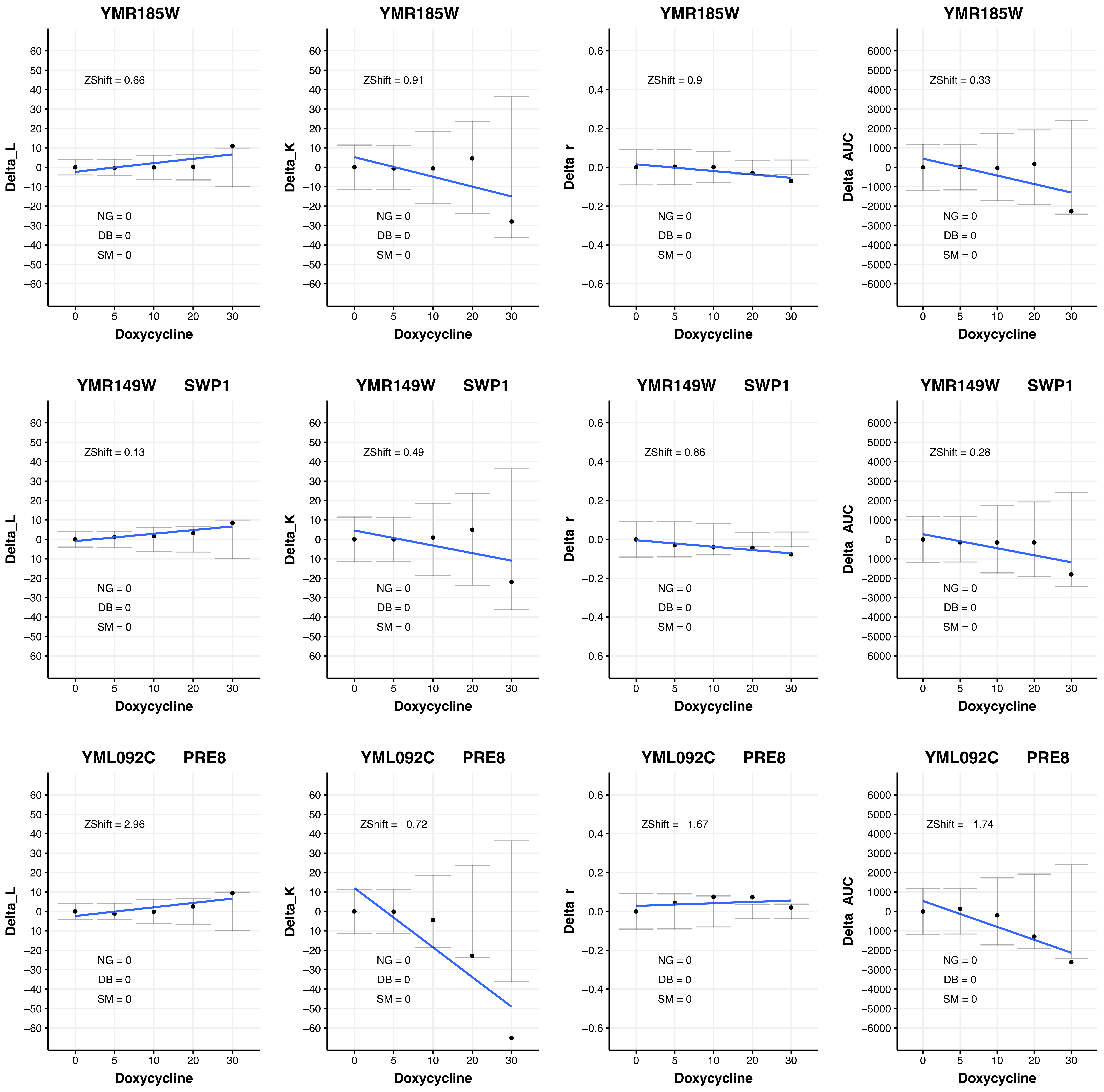
YMR185W
ZShift = 0.66
NG = 0
DB = 0
SM = 0
Delta_L
Doxycycline
YMR185W
ZShift = 0.91
NG = 0
DB = 0
SM = 0
Delta_K
Doxycycline
YMR185W
ZShift = 0.9
NG = 0
DB = 0
SM = 0
Delta_r
Doxycycline
YMR185W
ZShift = 0.33
NG = 0
DB = 0
SM = 0
Delta_AUC
Doxycycline
YMR149W SWP1
ZShift = 0.13
NG = 0
DB = 0
SM = 0
Delta_L
Doxycycline
YMR149W SWP1
ZShift = 0.49
NG = 0
DB = 0
SM = 0
Delta_K
Doxycycline
YMR149W SWP1
ZShift = 0.86
NG = 0
DB = 0
SM = 0
Delta_r
Doxycycline
YMR149W SWP1
ZShift = 0.28
NG = 0
DB = 0
SM = 0
Delta_AUC
Doxycycline
YML092C PRE8
ZShift = 2.96
NG = 0
DB = 0
SM = 0
Delta_L
Doxycycline
YML092C PRE8
ZShift = −0.72
NG = 0
DB = 0
SM = 0
Delta_K
Doxycycline
YML092C PRE8
ZShift = −1.67
NG = 0
DB = 0
SM = 0
Delta_r
Doxycycline
YML092C PRE8
ZShift = −1.74
NG = 0
DB = 0
SM = 0
Delta_AUC
Doxycycline

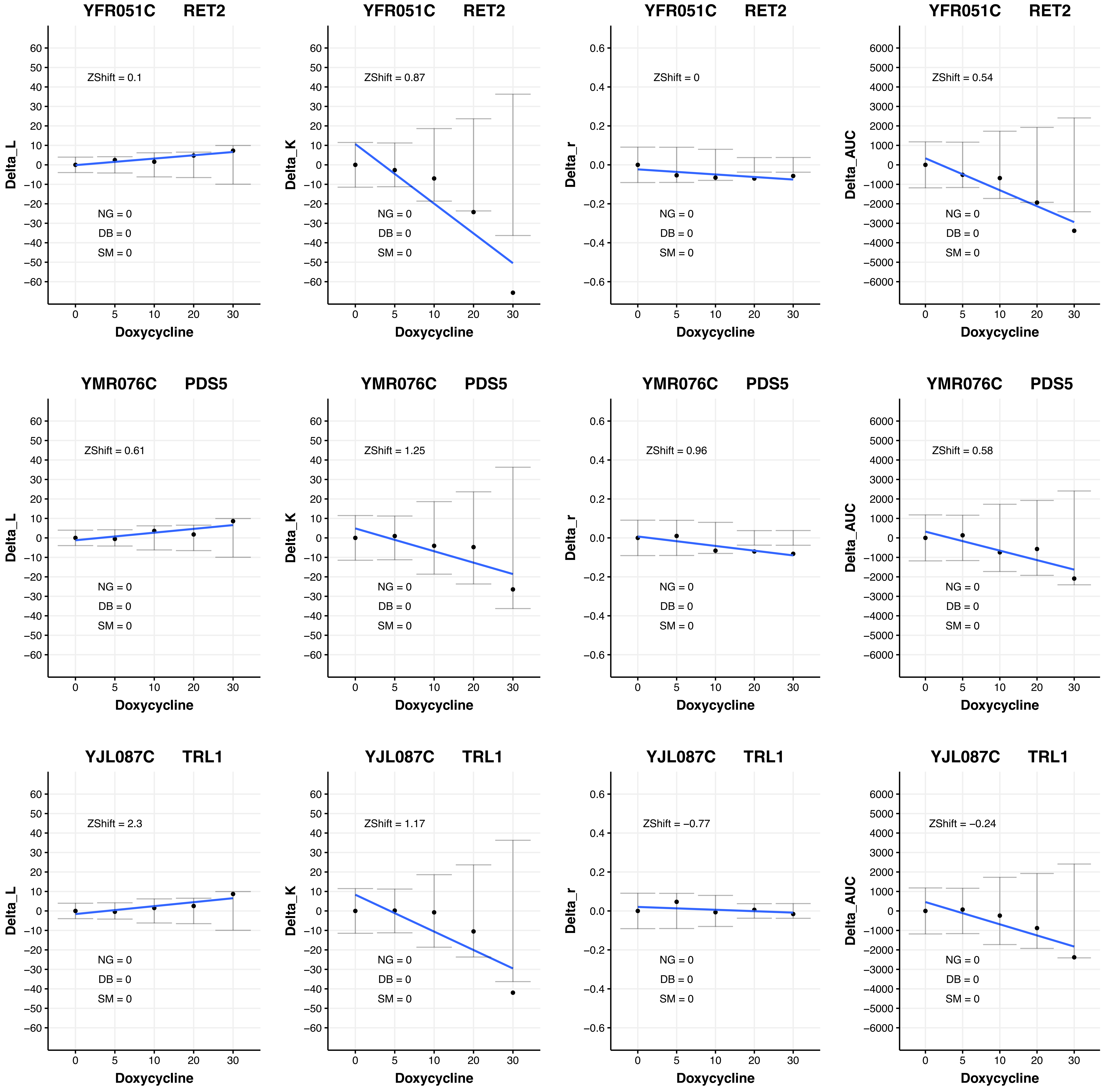
YFR051C RET2
ZShift = 0.1
NG = 0
DB = 0
SM = 0
Delta_L
Doxycycline
YFR051C RET2
ZShift = 0.87
NG = 0
DB = 0
SM = 0
Delta_K
Doxycycline
YFR051C RET2
ZShift = 0
NG = 0
DB = 0
SM = 0
Delta_r
Doxycycline
YFR051C RET2
ZShift = 0.54
NG = 0
DB = 0
SM = 0
Delta_AUC
Doxycycline
YMR076C PDS5
ZShift = 0.61
NG = 0
DB = 0
SM = 0
Delta_L
Doxycycline
YMR076C PDS5
ZShift = 1.25
NG = 0
DB = 0
SM = 0
Delta_K
Doxycycline
YMR076C PDS5
ZShift = 0.96
NG = 0
DB = 0
SM = 0
Delta_r
Doxycycline
YMR076C PDS5
ZShift = 0.58
NG = 0
DB = 0
SM = 0
Delta_AUC
Doxycycline
YJL087C TRL1
ZShift = 2.3
NG = 0
DB = 0
SM = 0
Delta_L
Doxycycline
YJL087C TRL1
ZShift = 1.17
NG = 0
DB = 0
SM = 0
Delta_K
Doxycycline
YJL087C TRL1
ZShift = −0.77
NG = 0
DB = 0
SM = 0
Delta_r
Doxycycline
YJL087C TRL1
ZShift = −0.24
NG = 0
DB = 0
SM = 0
Delta_AUC
Doxycycline

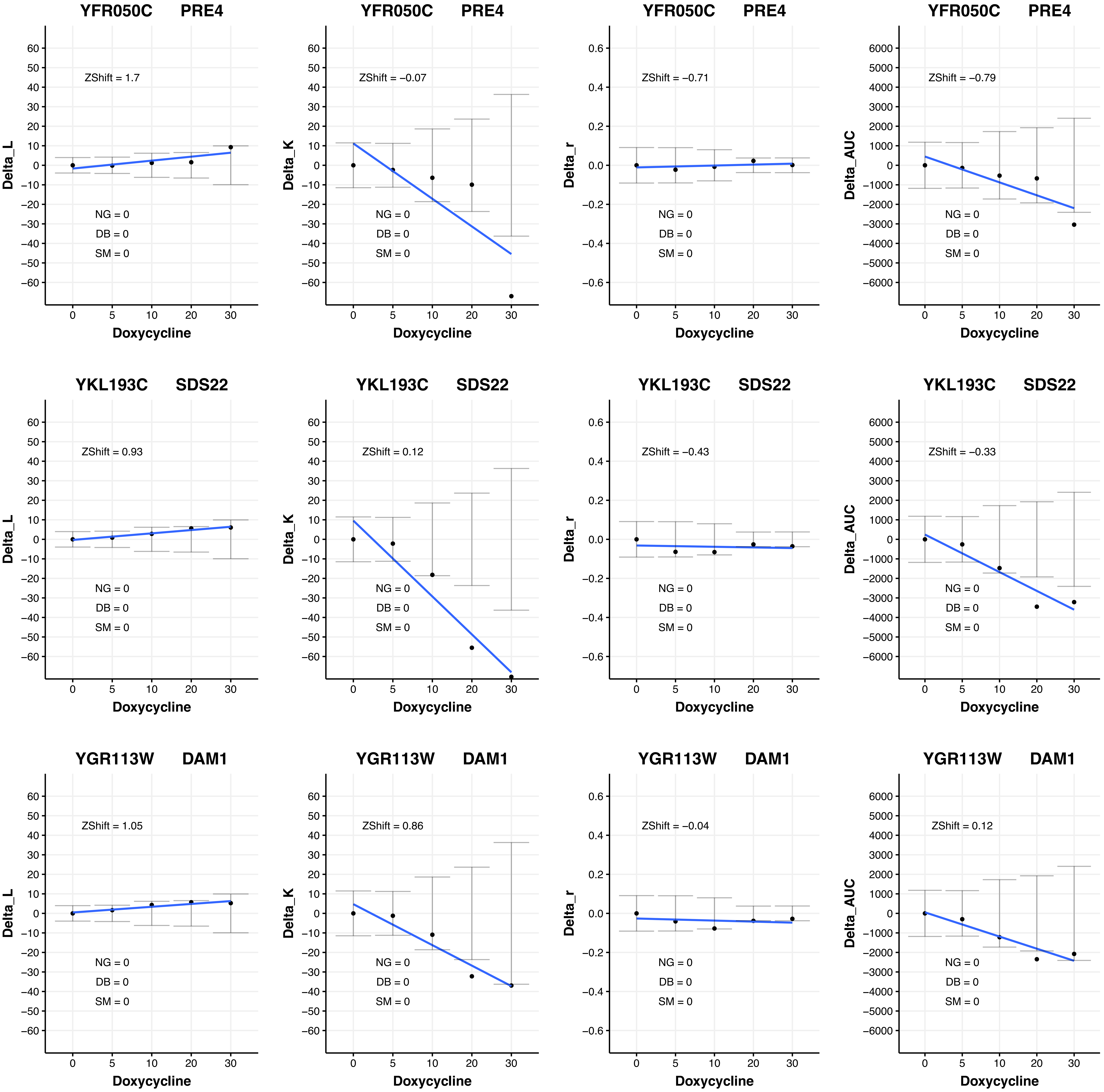
YFR050C PRE4
ZShift = 1.7
NG = 0
DB = 0
SM = 0
Delta_L
Doxycycline
YFR050C PRE4
ZShift = −0.07
NG = 0
DB = 0
SM = 0
Delta_K
Doxycycline
YFR050C PRE4
ZShift = −0.71
NG = 0
DB = 0
SM = 0
Delta_r
Doxycycline
YFR050C PRE4
ZShift = −0.79
NG = 0
DB = 0
SM = 0
Delta_AUC
Doxycycline
YKL193C SDS22
ZShift = 0.93
NG = 0
DB = 0
SM = 0
Delta_L
Doxycycline
YKL193C SDS22
ZShift = 0.12
NG = 0
DB = 0
SM = 0
Delta_K
Doxycycline
YKL193C SDS22
ZShift = −0.43
NG = 0
DB = 0
SM = 0
Delta_r
Doxycycline
YKL193C SDS22
ZShift = −0.33
NG = 0
DB = 0
SM = 0
Delta_AUC
Doxycycline
YGR113W DAM1
ZShift = 1.05
NG = 0
DB = 0
SM = 0
Delta_L
Doxycycline
YGR113W DAM1
ZShift = 0.86
NG = 0
DB = 0
SM = 0
Delta_K
Doxycycline
YGR113W DAM1
ZShift = −0.04
NG = 0
DB = 0
SM = 0
Delta_r
Doxycycline
YGR113W DAM1
ZShift = 0.12
NG = 0
DB = 0
SM = 0
Delta_AUC
Doxycycline

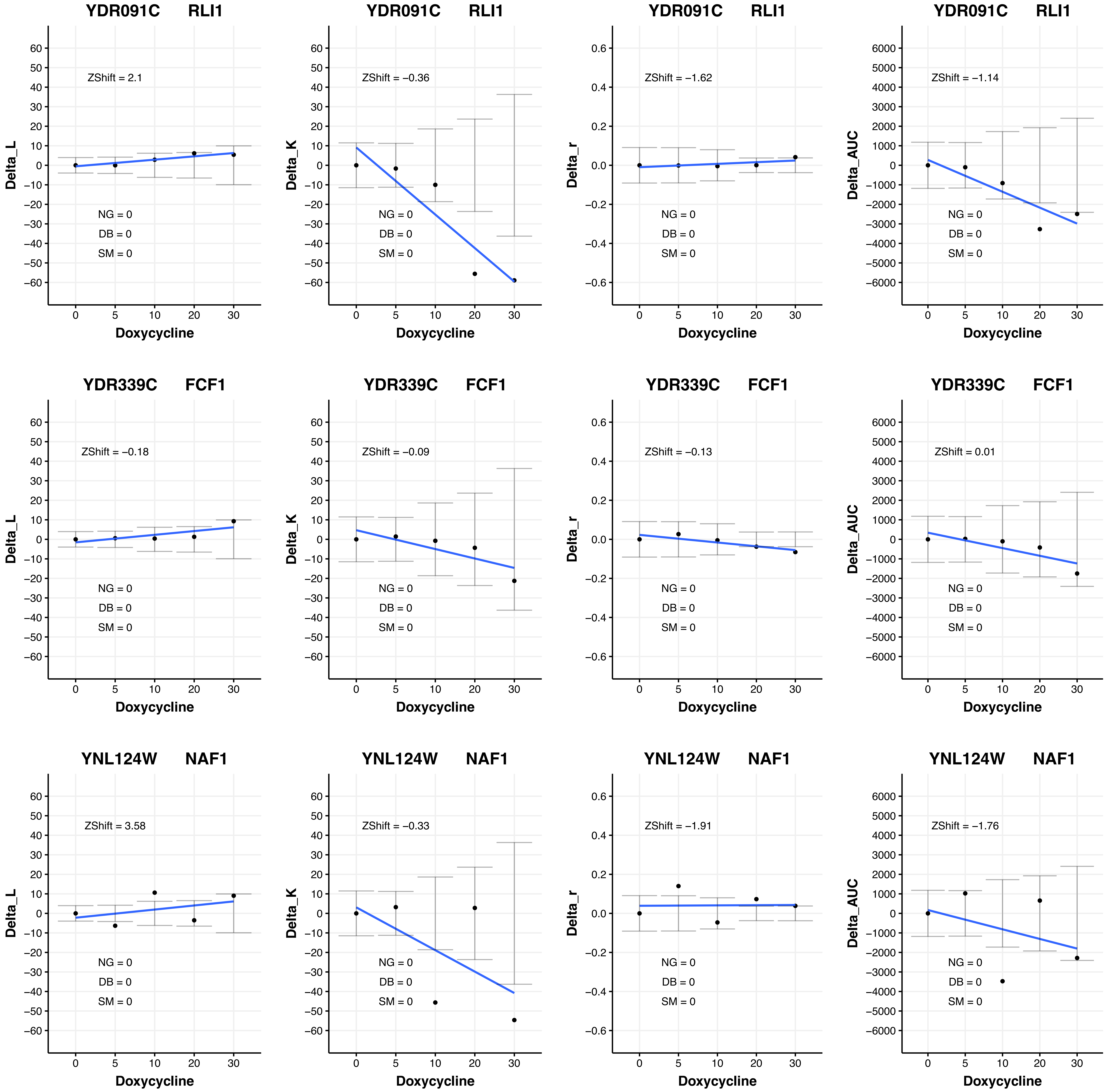

YDR091C RLI1
ZShift = 2.1
NG = 0
DB = 0
SM = 0
Delta_L
Doxycycline
YDR091C RLI1
ZShift = −0.36
NG = 0
DB = 0
SM = 0
Delta_K
Doxycycline
YDR091C RLI1
ZShift = −1.62
NG = 0
DB = 0
SM = 0
Delta_r
Doxycycline
YDR091C RLI1
ZShift = −1.14
NG = 0
DB = 0
SM = 0
Delta_AUC
Doxycycline
YDR339C FCF1
ZShift = −0.18
NG = 0
DB = 0
SM = 0
Delta_L
Doxycycline
YDR339C FCF1
ZShift = −0.09
NG = 0
DB = 0
SM = 0
Delta_K
Doxycycline
YDR339C FCF1
ZShift = −0.13
NG = 0
DB = 0
SM = 0
Delta_r
Doxycycline
YDR339C FCF1
ZShift = 0.01
NG = 0
DB = 0
SM = 0
Delta_AUC
Doxycycline
YNL124W NAF1
ZShift = 3.58
NG = 0
DB = 0
SM = 0
Delta_L
Doxycycline
YNL124W NAF1
ZShift = −0.33
NG = 0
DB = 0
SM = 0
Delta_K
Doxycycline
YNL124W NAF1
ZShift = −1.91
NG = 0
DB = 0
SM = 0
Delta_r
Doxycycline
YNL124W NAF1
ZShift = −1.76
NG = 0
DB = 0
SM = 0
Delta_AUC
Doxycycline

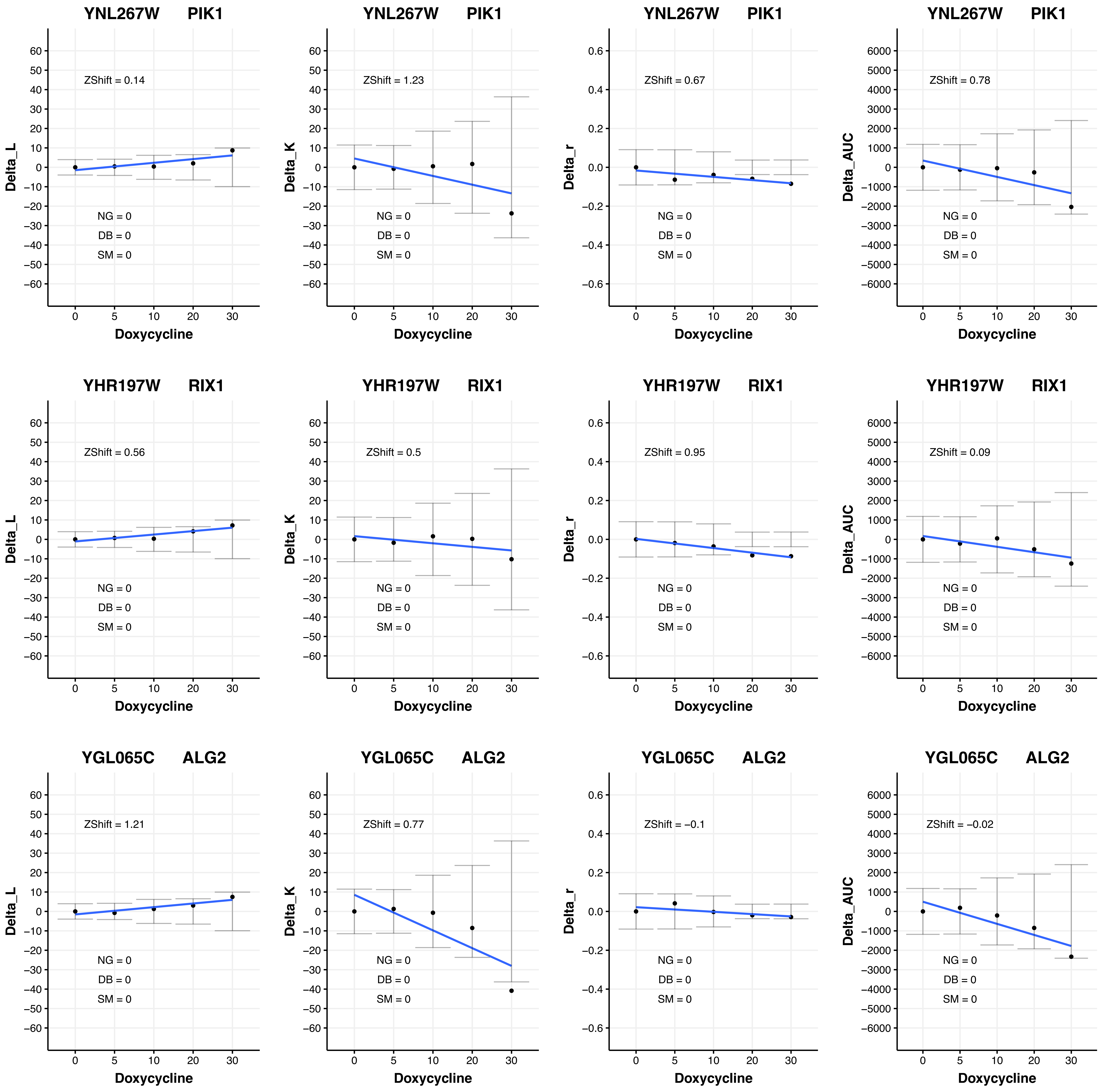

YNL267W PIK1
ZShift = 0.14
NG = 0
DB = 0
SM = 0
Delta_L
Doxycycline
YNL267W PIK1
ZShift = 1.23
NG = 0
DB = 0
SM = 0
Delta_K
Doxycycline
YNL267W PIK1
ZShift = 0.67
NG = 0
DB = 0
SM = 0
Delta_r
Doxycycline
YNL267W PIK1
ZShift = 0.78
NG = 0
DB = 0
SM = 0
Delta_AUC
Doxycycline
YHR197W RIX1
ZShift = 0.56
NG = 0
DB = 0
SM = 0
Delta_L
Doxycycline
YHR197W RIX1
ZShift = 0.5
NG = 0
DB = 0
SM = 0
Delta_K
Doxycycline
YHR197W RIX1
ZShift = 0.95
NG = 0
DB = 0
SM = 0
Delta_r
Doxycycline
YHR197W RIX1
ZShift = 0.09
NG = 0
DB = 0
SM = 0
Delta_AUC
Doxycycline
YGL065C ALG2
ZShift = 1.21
NG = 0
DB = 0
SM = 0
Delta_L
Doxycycline
YGL065C ALG2
ZShift = 0.77
NG = 0
DB = 0
SM = 0
Delta_K
Doxycycline
YGL065C ALG2
ZShift = −0.1
NG = 0
DB = 0
SM = 0
Delta_r
Doxycycline
YGL065C ALG2
ZShift = −0.02
NG = 0
DB = 0
SM = 0
Delta_AUC
Doxycycline

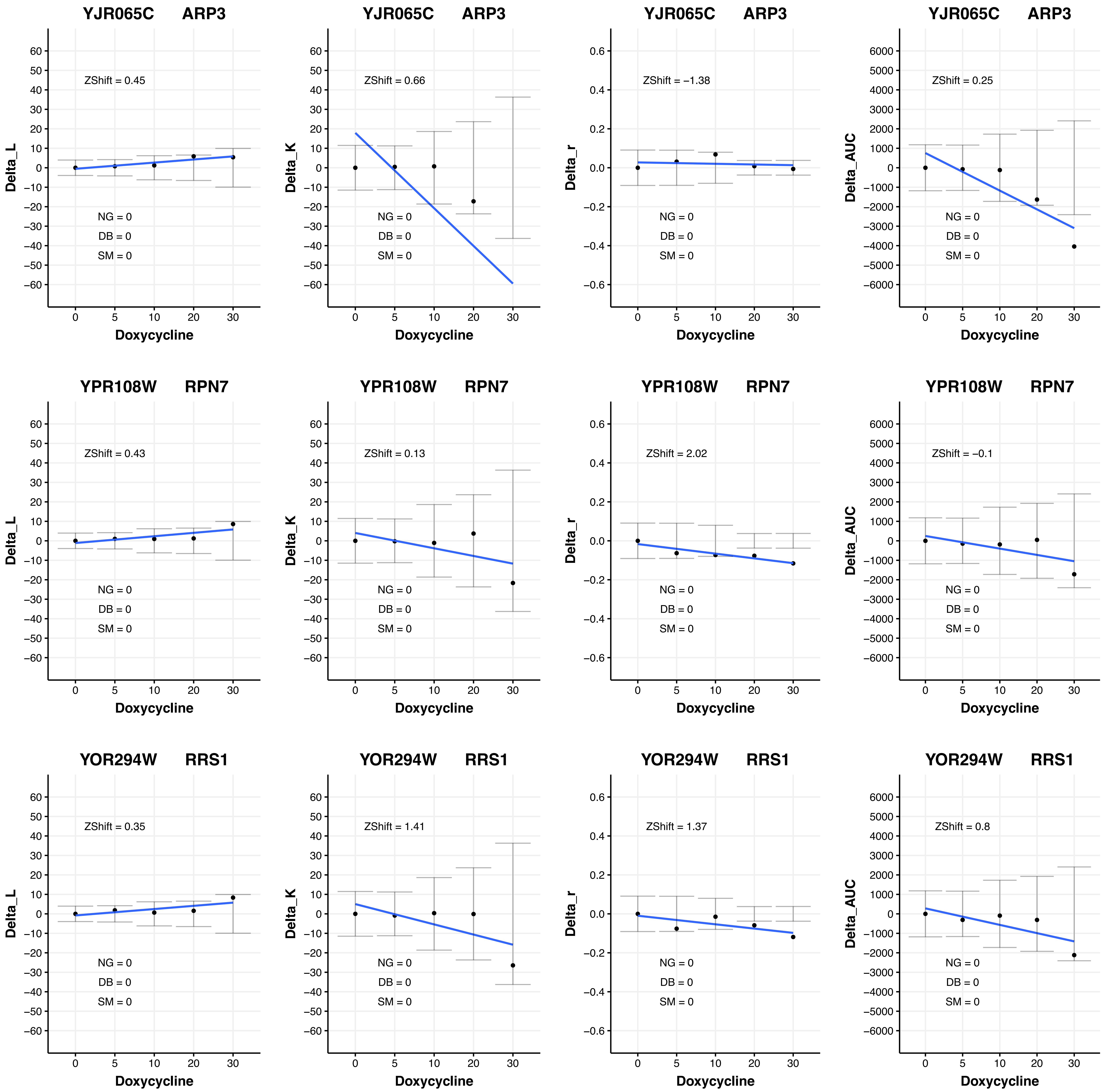
YJR065C ARP3
ZShift = 0.45
NG = 0
DB = 0
SM = 0
Delta_L
Doxycycline
YJR065C ARP3
ZShift = 0.66
NG = 0
DB = 0
SM = 0
Delta_K
Doxycycline
YJR065C ARP3
ZShift = −1.38
NG = 0
DB = 0
SM = 0
Delta_r
Doxycycline
YJR065C ARP3
ZShift = 0.25
NG = 0
DB = 0
SM = 0
Delta_AUC
Doxycycline
YPR108W RPN7
ZShift = 0.43
NG = 0
DB = 0
SM = 0
Delta_L
Doxycycline
YPR108W RPN7
ZShift = 0.13
NG = 0
DB = 0
SM = 0
Delta_K
Doxycycline
YPR108W RPN7
ZShift = 2.02
NG = 0
DB = 0
SM = 0
Delta_r
Doxycycline
YPR108W RPN7
ZShift = −0.1
NG = 0
DB = 0
SM = 0
Delta_AUC
Doxycycline
YOR294W RRS1
ZShift = 0.35
NG = 0
DB = 0
SM = 0
Delta_L
Doxycycline
YOR294W RRS1
ZShift = 1.41
NG = 0
DB = 0
SM = 0
Delta_K
Doxycycline
YOR294W RRS1
ZShift = 1.37
NG = 0
DB = 0
SM = 0
Delta_r
Doxycycline
YOR294W RRS1
ZShift = 0.8
NG = 0
DB = 0
SM = 0
Delta_AUC
Doxycycline

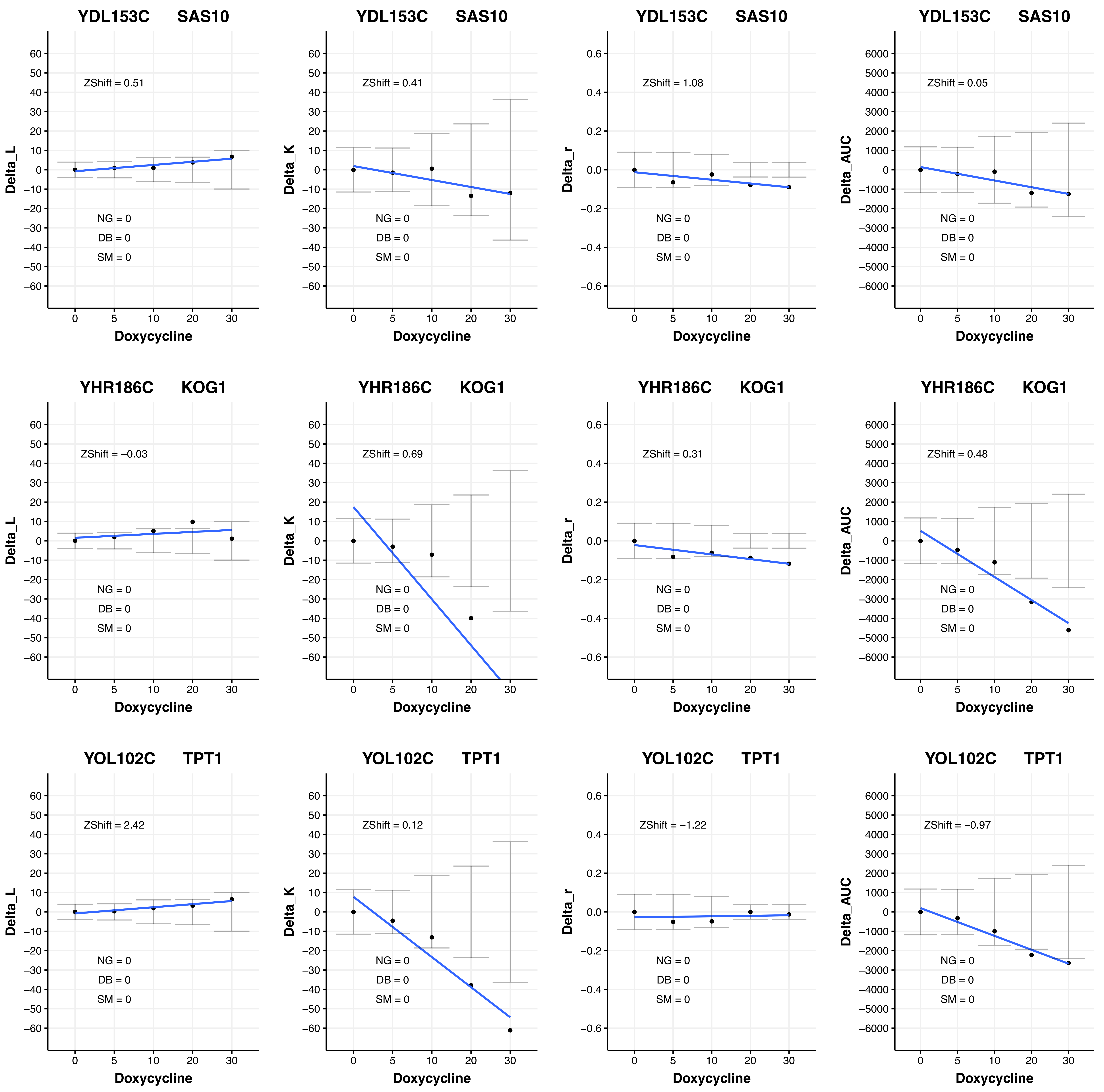

YDL153C SAS10
ZShift = 0.51
NG = 0
DB = 0
SM = 0
Delta_L
Doxycycline
YDL153C SAS10
ZShift = 0.41
NG = 0
DB = 0
SM = 0
Delta_K
Doxycycline
YDL153C SAS10
ZShift = 1.08
NG = 0
DB = 0
SM = 0
Delta_r
Doxycycline
YDL153C SAS10
ZShift = 0.05
NG = 0
DB = 0
SM = 0
Delta_AUC
Doxycycline
YHR186C KOG1
ZShift = −0.03
NG = 0
DB = 0
SM = 0
Delta_L
Doxycycline
YHR186C KOG1
ZShift = 0.69
NG = 0
DB = 0
SM = 0
Delta_K
Doxycycline
YHR186C KOG1
ZShift = 0.31
NG = 0
DB = 0
SM = 0
Delta_r
Doxycycline
YHR186C KOG1
ZShift = 0.48
NG = 0
DB = 0
SM = 0
Delta_AUC
Doxycycline
YOL102C TPT1
ZShift = 2.42
NG = 0
DB = 0
SM = 0
Delta_L
Doxycycline
YOL102C TPT1
ZShift = 0.12
NG = 0
DB = 0
SM = 0
Delta_K
Doxycycline
YOL102C TPT1
ZShift = −1.22
NG = 0
DB = 0
SM = 0
Delta_r
Doxycycline
YOL102C TPT1
ZShift = −0.97
NG = 0
DB = 0
SM = 0
Delta_AUC
Doxycycline

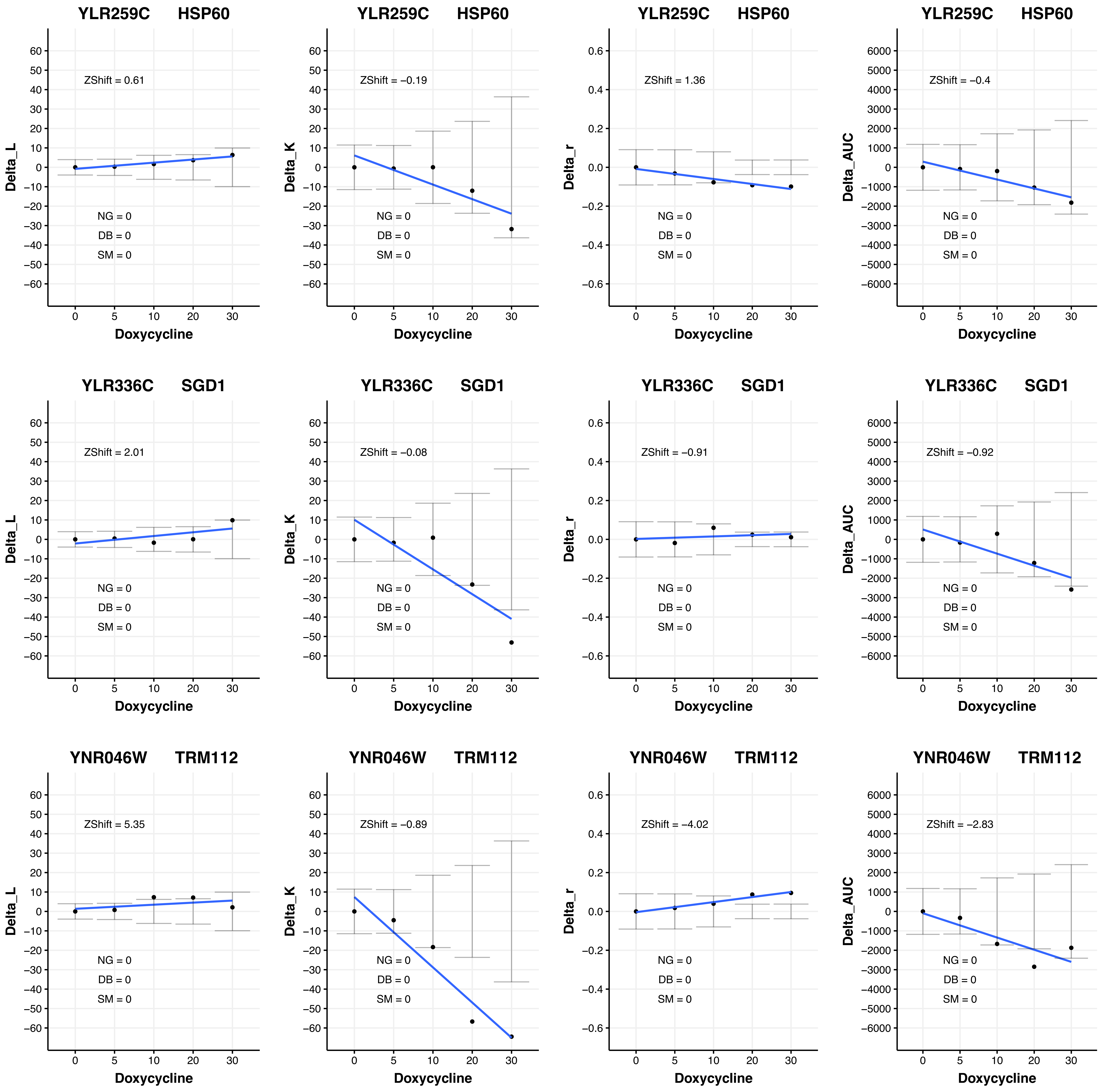

YLR259C HSP60
ZShift = 0.61
Delta_L
NG = 0
DB = 0
SM = 0
Doxycycline
YLR259C HSP60
ZShift = −0.19
Delta_K
NG = 0
DB = 0
SM = 0
Doxycycline
YLR259C HSP60
ZShift = 1.36
Delta_r
NG = 0
DB = 0
SM = 0
Doxycycline
YLR259C HSP60
ZShift = −0.4
Delta_AUC
NG = 0
DB = 0
SM = 0
Doxycycline
YLR336C SGD1
ZShift = 2.01
Delta_L
NG = 0
DB = 0
SM = 0
Doxycycline
YLR336C SGD1
ZShift = −0.08
Delta_K
NG = 0
DB = 0
SM = 0
Doxycycline
YLR336C SGD1
ZShift = −0.91
Delta_r
NG = 0
DB = 0
SM = 0
Doxycycline
YLR336C SGD1
ZShift = −0.92
Delta_AUC
NG = 0
DB = 0
SM = 0
Doxycycline
YNR046W TRM112
ZShift = 5.35
Delta_L
NG = 0
DB = 0
SM = 0
Doxycycline
YNR046W TRM112
ZShift = −0.89
Delta_K
NG = 0
DB = 0
SM = 0
Doxycycline
YNR046W TRM112
ZShift = −4.02
Delta_r
NG = 0
DB = 0
SM = 0
Doxycycline
YNR046W TRM112
ZShift = −2.83
Delta_AUC
NG = 0
DB = 0
SM = 0
Doxycycline

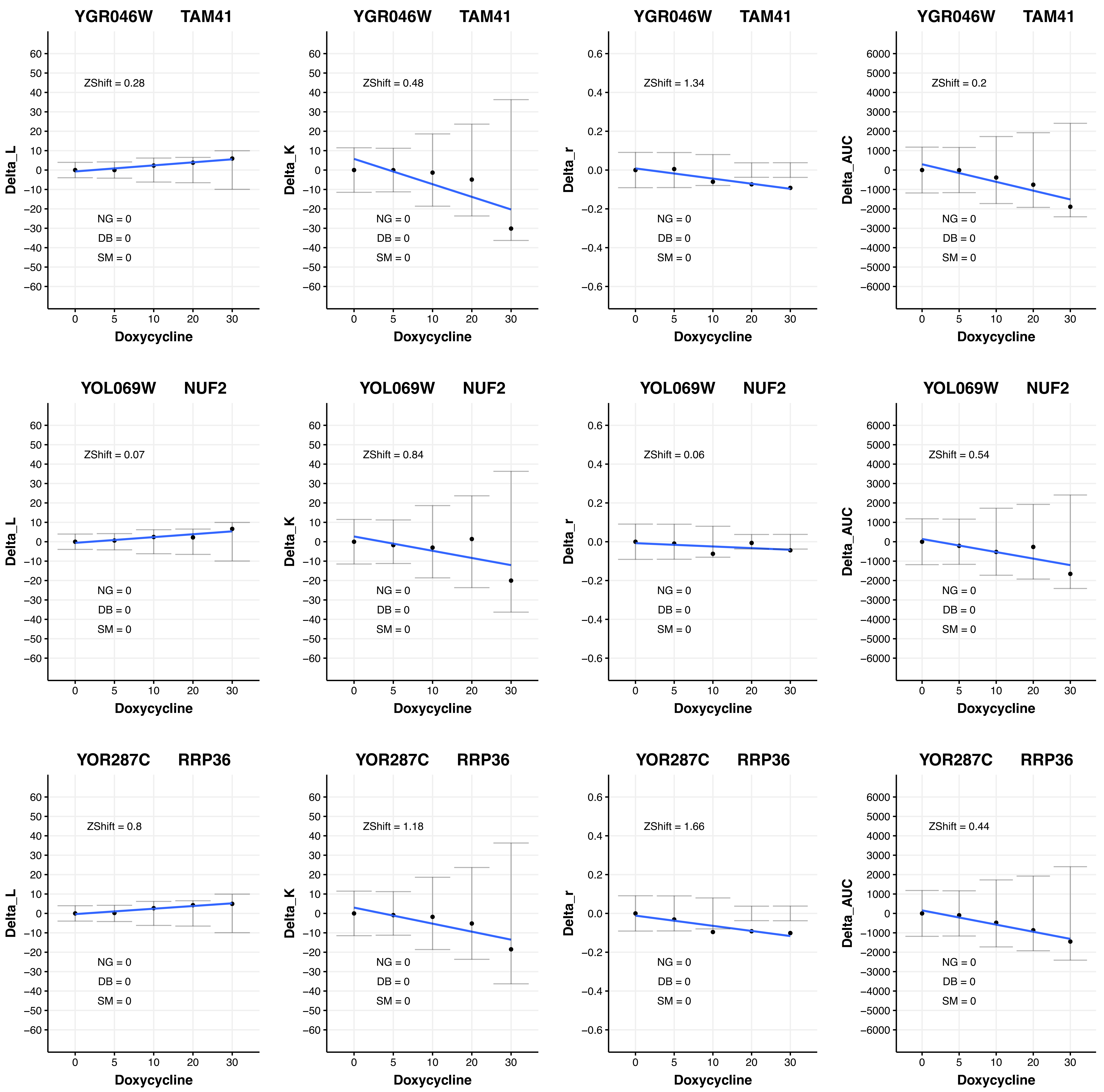

YGR046W TAM41
Delta_L
ZShift = 0.28
NG = 0
DB = 0
SM = 0
Doxycycline
YGR046W TAM41
Delta_K
ZShift = 0.48
NG = 0
DB = 0
SM = 0
Doxycycline
YGR046W TAM41
Delta_r
ZShift = 1.34
NG = 0
DB = 0
SM = 0
Doxycycline
YGR046W TAM41
Delta_AUC
ZShift = 0.2
NG = 0
DB = 0
SM = 0
Doxycycline
YOL069W NUF2
Delta_L
ZShift = 0.07
NG = 0
DB = 0
SM = 0
Doxycycline
YOL069W NUF2
Delta_K
ZShift = 0.84
NG = 0
DB = 0
SM = 0
Doxycycline
YOL069W NUF2
Delta_r
ZShift = 0.06
NG = 0
DB = 0
SM = 0
Doxycycline
YOL069W NUF2
Delta_AUC
ZShift = 0.54
NG = 0
DB = 0
SM = 0
Doxycycline
YOR287C RRP36
Delta_L
ZShift = 0.8
NG = 0
DB = 0
SM = 0
Doxycycline
YOR287C RRP36
Delta_K
ZShift = 1.18
NG = 0
DB = 0
SM = 0
Doxycycline
YOR287C RRP36
Delta_r
ZShift = 1.66
NG = 0
DB = 0
SM = 0
Doxycycline
YOR287C RRP36
Delta_AUC
ZShift = 0.44
NG = 0
DB = 0
SM = 0
Doxycycline

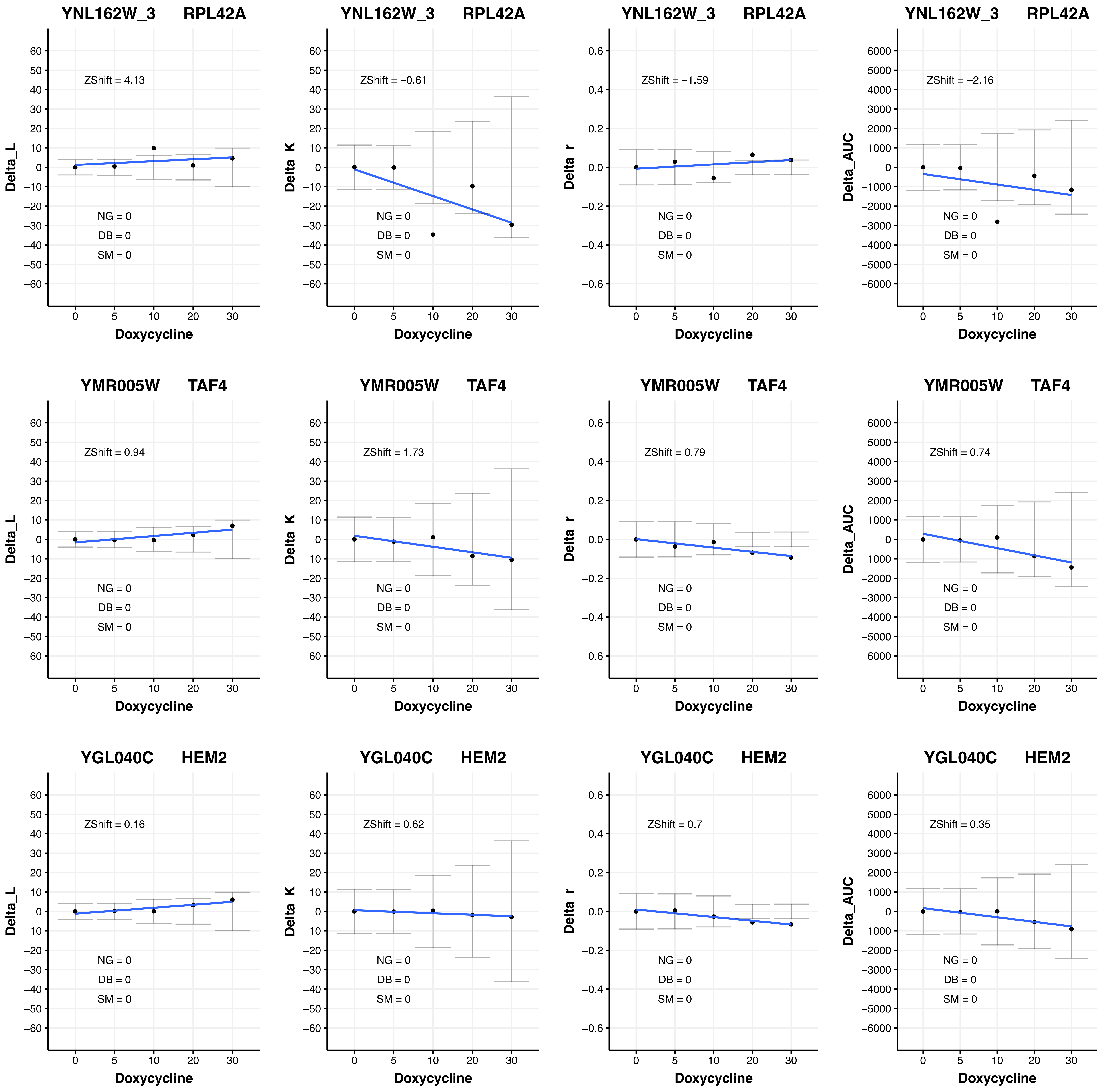

YNL162W_3 RPL42A
ZShift = 4.13
NG = 0
DB = 0
SM = 0
Delta_L
Doxycycline
YNL162W_3 RPL42A
ZShift = −0.61
NG = 0
DB = 0
SM = 0
Delta_K
Doxycycline
YNL162W_3 RPL42A
ZShift = −1.59
NG = 0
DB = 0
SM = 0
Delta_r
Doxycycline
YNL162W_3 RPL42A
ZShift = −2.16
NG = 0
DB = 0
SM = 0
Delta_AUC
Doxycycline
YMR005W TAF4
ZShift = 0.94
NG = 0
DB = 0
SM = 0
Delta_L
Doxycycline
YMR005W TAF4
ZShift = 1.73
NG = 0
DB = 0
SM = 0
Delta_K
Doxycycline
YMR005W TAF4
ZShift = 0.79
NG = 0
DB = 0
SM = 0
Delta_r
Doxycycline
YMR005W TAF4
ZShift = 0.74
NG = 0
DB = 0
SM = 0
Delta_AUC
Doxycycline
YGL040C HEM2
ZShift = 0.16
NG = 0
DB = 0
SM = 0
Delta_L
Doxycycline
YGL040C HEM2
ZShift = 0.62
NG = 0
DB = 0
SM = 0
Delta_K
Doxycycline
YGL040C HEM2
ZShift = 0.7
NG = 0
DB = 0
SM = 0
Delta_r
Doxycycline
YGL040C HEM2
ZShift = 0.35
NG = 0
DB = 0
SM = 0
Delta_AUC
Doxycycline

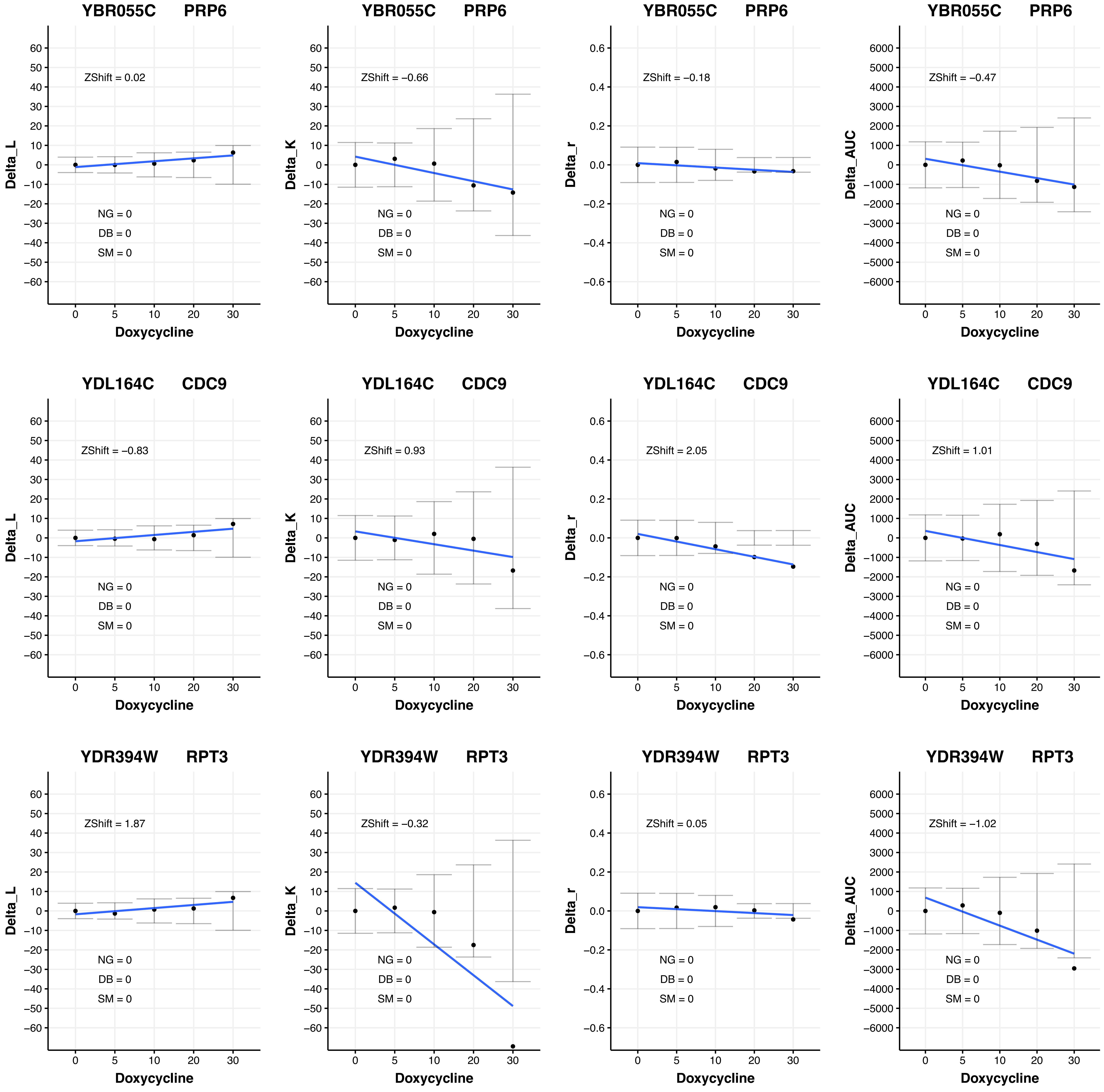
YBR055C PRP6
ZShift = 0.02
NG = 0
DB = 0
SM = 0
Delta_L
Doxycycline
YBR055C PRP6
ZShift = −0.66
NG = 0
DB = 0
SM = 0
Delta_K
Doxycycline
YBR055C PRP6
ZShift = −0.18
NG = 0
DB = 0
SM = 0
Delta_r
Doxycycline
YBR055C PRP6
ZShift = −0.47
NG = 0
DB = 0
SM = 0
Delta_AUC
Doxycycline
YDL164C CDC9
ZShift = −0.83
NG = 0
DB = 0
SM = 0
Delta_L
Doxycycline
YDL164C CDC9
ZShift = 0.93
NG = 0
DB = 0
SM = 0
Delta_K
Doxycycline
YDL164C CDC9
ZShift = 2.05
NG = 0
DB = 0
SM = 0
Delta_r
Doxycycline
YDL164C CDC9
ZShift = 1.01
NG = 0
DB = 0
SM = 0
Delta_AUC
Doxycycline
YDR394W RPT3
ZShift = 1.87
NG = 0
DB = 0
SM = 0
Delta_L
Doxycycline
YDR394W RPT3
ZShift = −0.32
NG = 0
DB = 0
SM = 0
Delta_K
Doxycycline
YDR394W RPT3
ZShift = 0.05
NG = 0
DB = 0
SM = 0
Delta_r
Doxycycline
YDR394W RPT3
ZShift = −1.02
NG = 0
DB = 0
SM = 0
Delta_AUC
Doxycycline

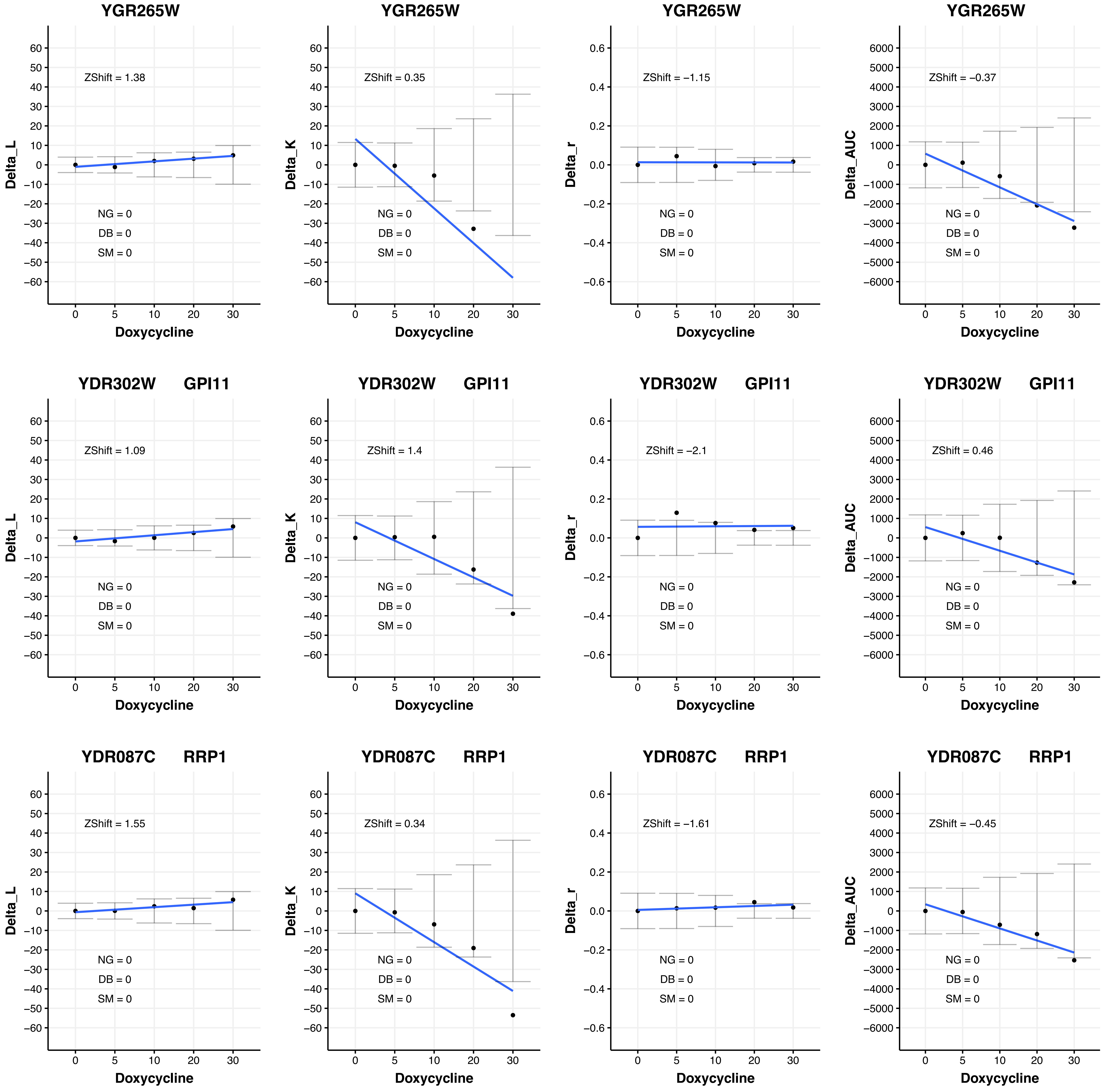
YGR265W
ZShift = 1.38
NG = 0
DB = 0
SM = 0
Delta_L
Doxycycline
YGR265W
ZShift = 0.35
NG = 0
DB = 0
SM = 0
Delta_K
Doxycycline
YGR265W
ZShift = −1.15
NG = 0
DB = 0
SM = 0
Delta_r
Doxycycline
YGR265W
ZShift = −0.37
NG = 0
DB = 0
SM = 0
Delta_AUC
Doxycycline
YDR302W GPI11
ZShift = 1.09
NG = 0
DB = 0
SM = 0
Delta_L
Doxycycline
YDR302W GPI11
ZShift = 1.4
NG = 0
DB = 0
SM = 0
Delta_K
Doxycycline
YDR302W GPI11
ZShift = −2.1
NG = 0
DB = 0
SM = 0
Delta_r
Doxycycline
YDR302W GPI11
ZShift = 0.46
NG = 0
DB = 0
SM = 0
Delta_AUC
Doxycycline
YDR087C RRP1
ZShift = 1.55
NG = 0
DB = 0
SM = 0
Delta_L
Doxycycline
YDR087C RRP1
ZShift = 0.34
NG = 0
DB = 0
SM = 0
Delta_K
Doxycycline
YDR087C RRP1
ZShift = −1.61
NG = 0
DB = 0
SM = 0
Delta_r
Doxycycline
YDR087C RRP1
ZShift = −0.45
NG = 0
DB = 0
SM = 0
Delta_AUC
Doxycycline

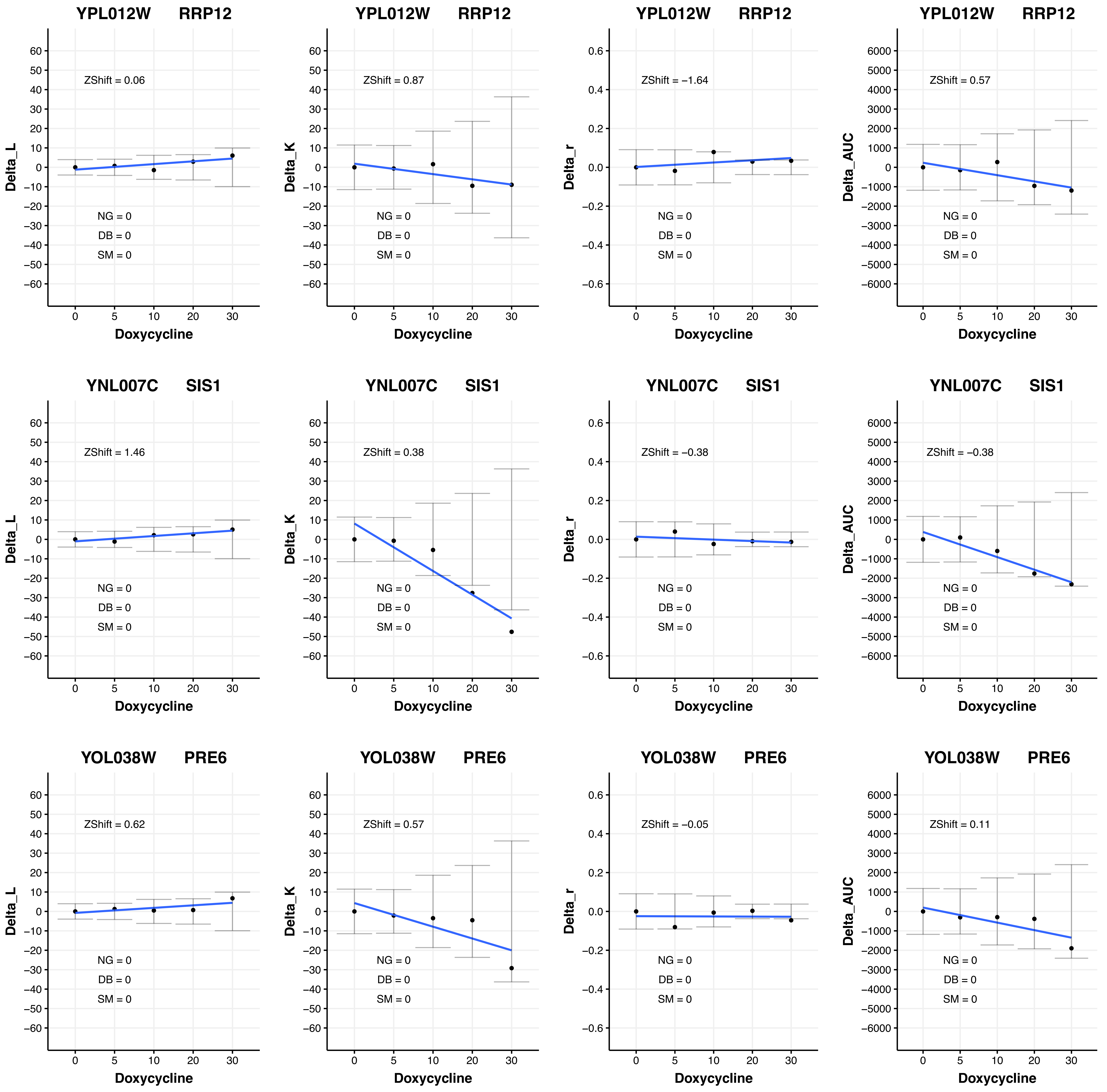
YPL012W RRP12
ZShift = 0.06
Delta_L
NG = 0
DB = 0
SM = 0
Doxycycline
YPL012W RRP12
ZShift = 0.87
Delta_K
NG = 0
DB = 0
SM = 0
Doxycycline
YPL012W RRP12
ZShift = −1.64
Delta_r
NG = 0
DB = 0
SM = 0
Doxycycline
YPL012W RRP12
ZShift = 0.57
Delta_AUC
NG = 0
DB = 0
SM = 0
Doxycycline
YNL007C SIS1
ZShift = 1.46
Delta_L
NG = 0
DB = 0
SM = 0
Doxycycline
YNL007C SIS1
ZShift = 0.38
Delta_K
NG = 0
DB = 0
SM = 0
Doxycycline
YNL007C SIS1
ZShift = −0.38
Delta_r
NG = 0
DB = 0
SM = 0
Doxycycline
YNL007C SIS1
ZShift = −0.38
Delta_AUC
NG = 0
DB = 0
SM = 0
Doxycycline
YOL038W PRE6
ZShift = 0.62
Delta_L
NG = 0
DB = 0
SM = 0
Doxycycline
YOL038W PRE6
ZShift = 0.57
Delta_K
NG = 0
DB = 0
SM = 0
Doxycycline
YOL038W PRE6
ZShift = −0.05
Delta_r
NG = 0
DB = 0
SM = 0
Doxycycline
YOL038W PRE6
ZShift = 0.11
Delta_AUC
NG = 0
DB = 0
SM = 0
Doxycycline

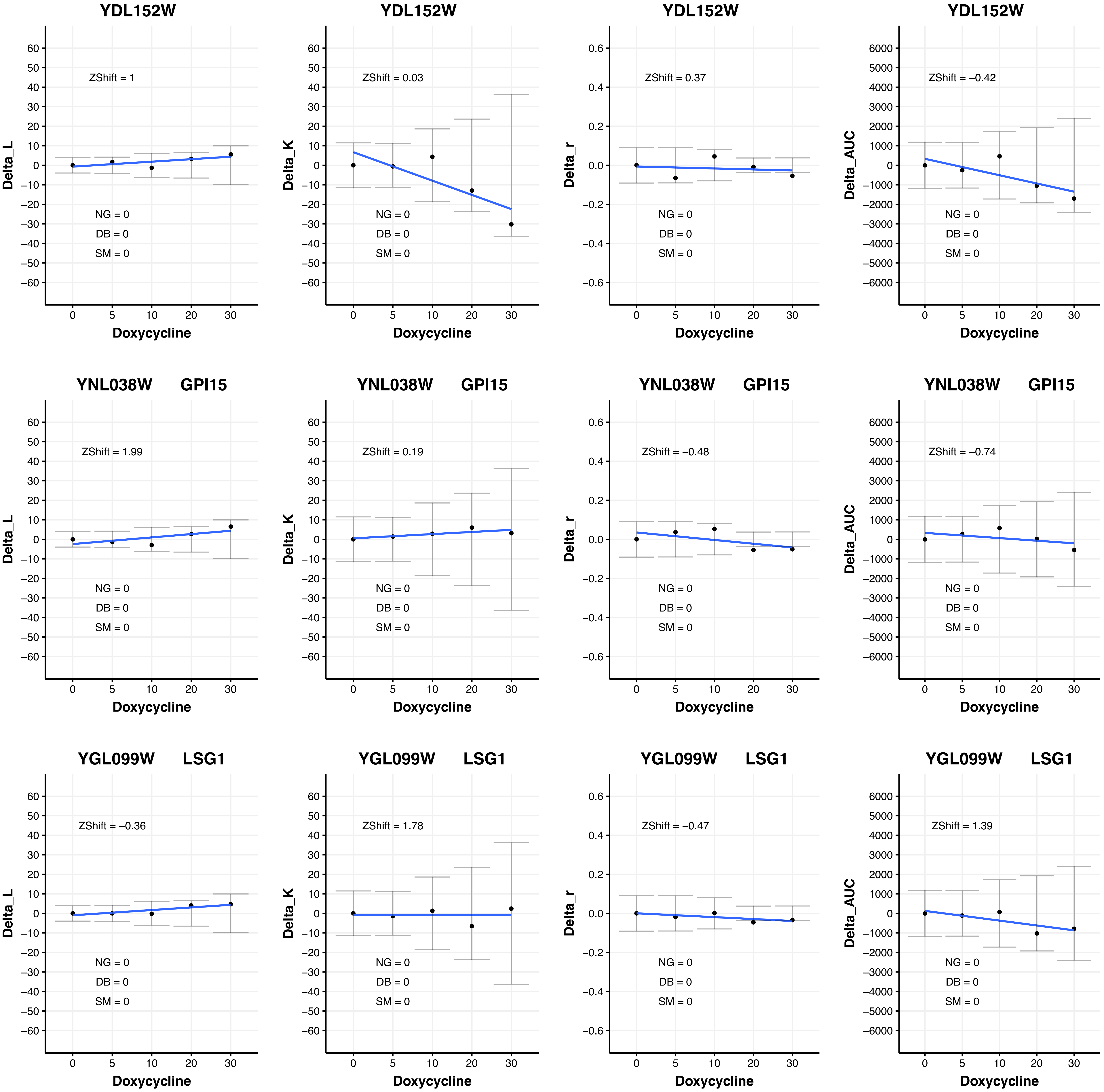
YDL152W
ZShift = 1
NG = 0
DB = 0
SM = 0
Delta_L
Doxycycline
YDL152W
ZShift = 0.03
NG = 0
DB = 0
SM = 0
Delta_K
Doxycycline
YDL152W
ZShift = 0.37
NG = 0
DB = 0
SM = 0
Delta_r
Doxycycline
YDL152W
ZShift = −0.42
NG = 0
DB = 0
SM = 0
Delta_AUC
Doxycycline
YNL038W GPI15
ZShift = 1.99
NG = 0
DB = 0
SM = 0
Delta_L
Doxycycline
YNL038W GPI15
ZShift = 0.19
NG = 0
DB = 0
SM = 0
Delta_K
Doxycycline
YNL038W GPI15
ZShift = −0.48
NG = 0
DB = 0
SM = 0
Delta_r
Doxycycline
YNL038W GPI15
ZShift = −0.74
NG = 0
DB = 0
SM = 0
Delta_AUC
Doxycycline
YGL099W LSG1
ZShift = −0.36
NG = 0
DB = 0
SM = 0
Delta_L
Doxycycline
YGL099W LSG1
ZShift = 1.78
NG = 0
DB = 0
SM = 0
Delta_K
Doxycycline
YGL099W LSG1
ZShift = −0.47
NG = 0
DB = 0
SM = 0
Delta_r
Doxycycline
YGL099W LSG1
ZShift = 1.39
NG = 0
DB = 0
SM = 0
Delta_AUC
Doxycycline

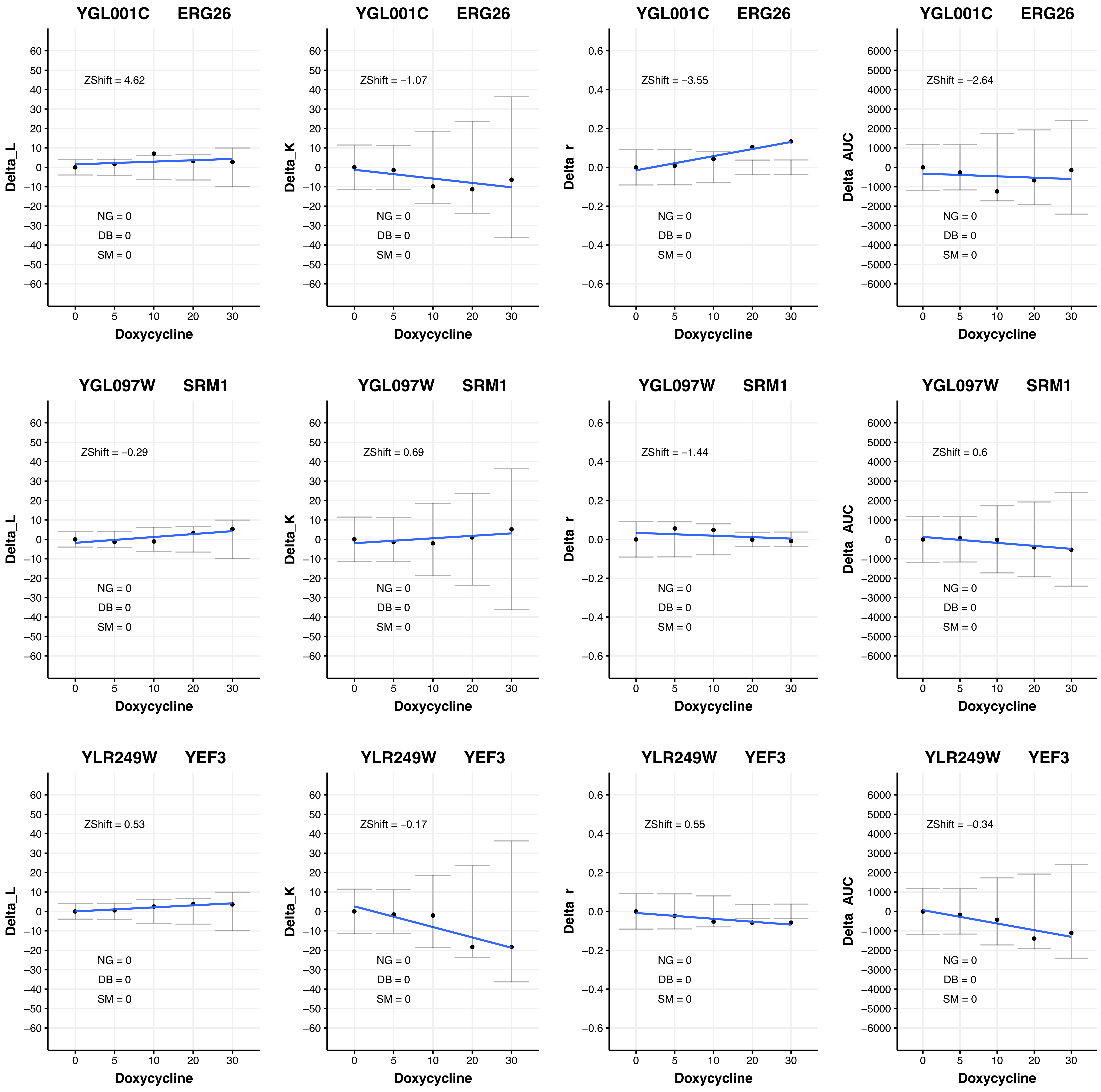
YGL001C ERG26
ZShift = 4.62
NG = 0
DB = 0
SM = 0
Delta_L
Doxycycline
YGL001C ERG26
ZShift = −1.07
NG = 0
DB = 0
SM = 0
Delta_K
Doxycycline
YGL001C ERG26
ZShift = −3.55
NG = 0
DB = 0
SM = 0
Delta_r
Doxycycline
YGL001C ERG26
ZShift = −2.64
NG = 0
DB = 0
SM = 0
Delta_AUC
Doxycycline
YGL097W SRM1
ZShift = −0.29
NG = 0
DB = 0
SM = 0
Delta_L
Doxycycline
YGL097W SRM1
ZShift = 0.69
NG = 0
DB = 0
SM = 0
Delta_K
Doxycycline
YGL097W SRM1
ZShift = −1.44
NG = 0
DB = 0
SM = 0
Delta_r
Doxycycline
YGL097W SRM1
ZShift = 0.6
NG = 0
DB = 0
SM = 0
Delta_AUC
Doxycycline
YLR249W YEF3
ZShift = 0.53
NG = 0
DB = 0
SM = 0
Delta_L
Doxycycline
YLR249W YEF3
ZShift = −0.17
NG = 0
DB = 0
SM = 0
Delta_K
Doxycycline
YLR249W YEF3
ZShift = 0.55
NG = 0
DB = 0
SM = 0
Delta_r
Doxycycline
YLR249W YEF3
ZShift = −0.34
NG = 0
DB = 0
SM = 0
Delta_AUC
Doxycycline

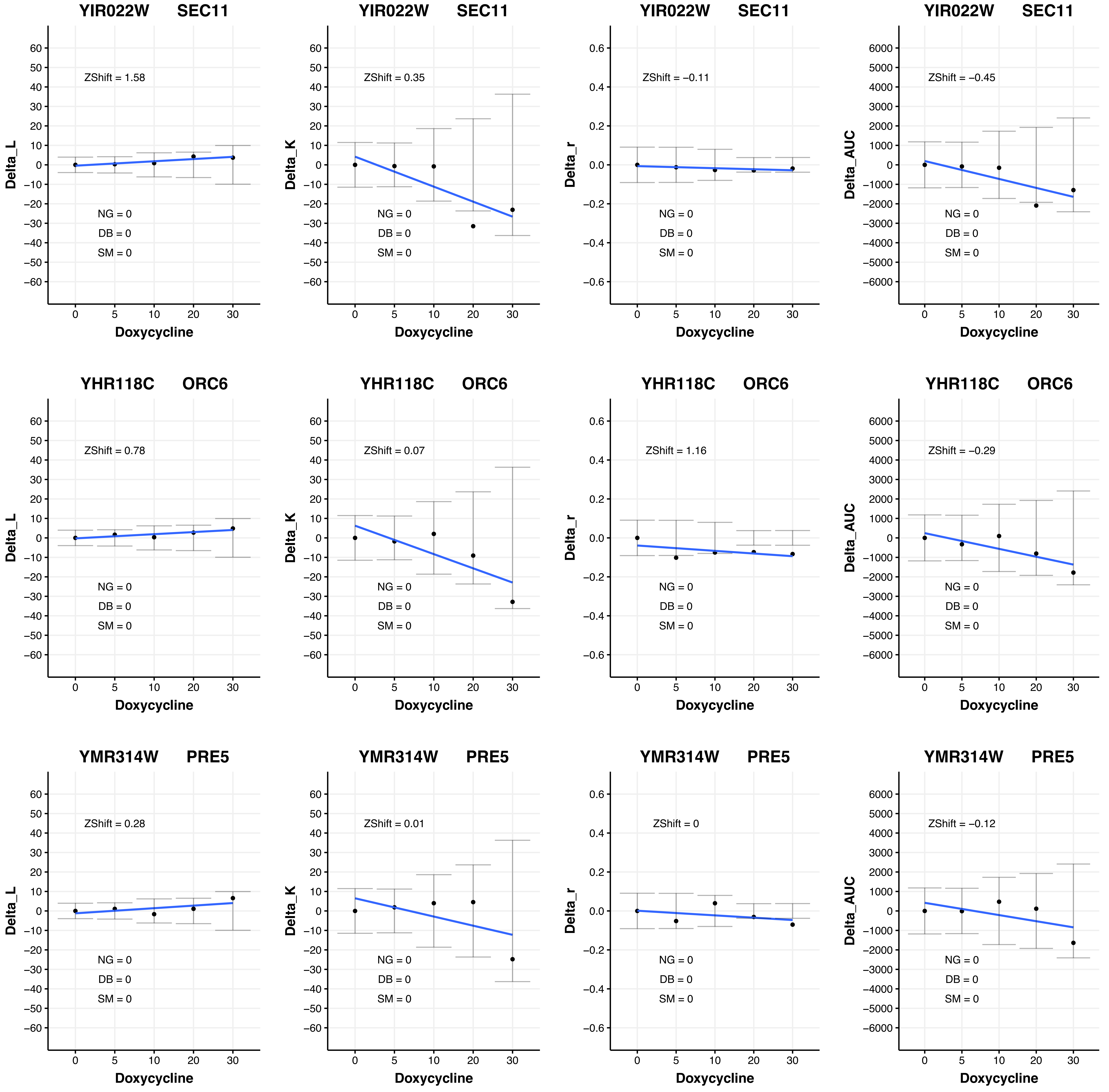
YIR022W SEC11
ZShift = 1.58
NG = 0
DB = 0
SM = 0
Delta_L
Doxycycline
YIR022W SEC11
ZShift = 0.35
NG = 0
DB = 0
SM = 0
Delta_K
Doxycycline
YIR022W SEC11
ZShift = −0.11
NG = 0
DB = 0
SM = 0
Delta_r
Doxycycline
YIR022W SEC11
ZShift = −0.45
NG = 0
DB = 0
SM = 0
Delta_AUC
Doxycycline
YHR118C ORC6
ZShift = 0.78
NG = 0
DB = 0
SM = 0
Delta_L
Doxycycline
YHR118C ORC6
ZShift = 0.07
NG = 0
DB = 0
SM = 0
Delta_K
Doxycycline
YHR118C ORC6
ZShift = 1.16
NG = 0
DB = 0
SM = 0
Delta_r
Doxycycline
YHR118C ORC6
ZShift = −0.29
NG = 0
DB = 0
SM = 0
Delta_AUC
Doxycycline
YMR314W PRE5
ZShift = 0.28
NG = 0
DB = 0
SM = 0
Delta_L
Doxycycline
YMR314W PRE5
ZShift = 0.01
NG = 0
DB = 0
SM = 0
Delta_K
Doxycycline
YMR314W PRE5
ZShift = 0
NG = 0
DB = 0
SM = 0
Delta_r
Doxycycline
YMR314W PRE5
ZShift = −0.12
NG = 0
DB = 0
SM = 0
Delta_AUC
Doxycycline

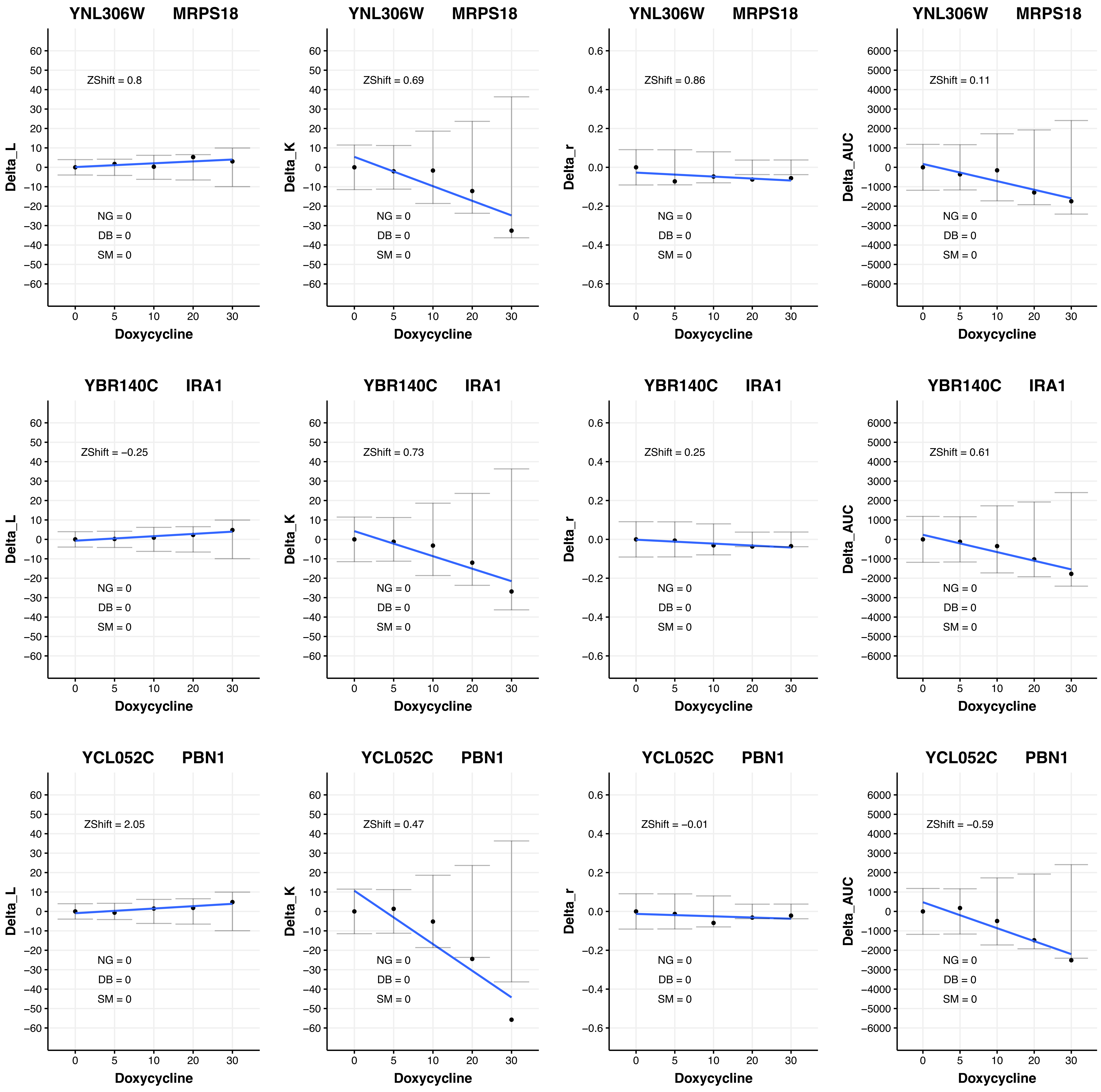

YNL306W MRPS18
ZShift = 0.8
ZShift = 0.69
ZShift = 0.86
ZShift = 0.11
YBR140C IRA1
ZShift = −0.25
ZShift = 0.73
ZShift = 0.25
ZShift = 0.61
YCL052C PBN1
ZShift = 2.05
ZShift = 0.47
ZShift = −0.01
ZShift = −0.59
NG = 0
DB = 0
SM = 0
Delta_L
Delta_K
Delta_r
Delta_AUC
Doxycycline

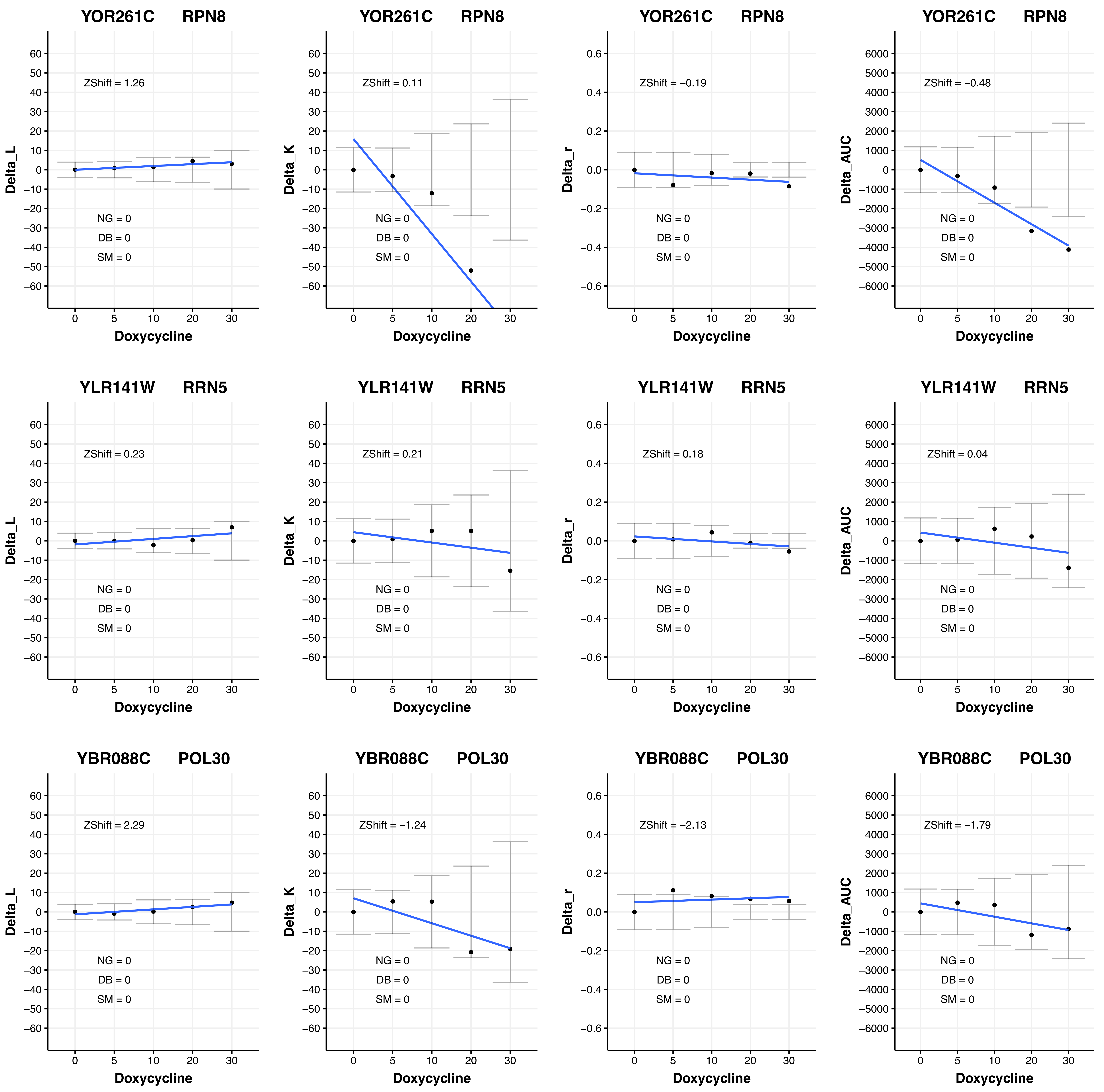

YOR261C RPN8
ZShift = 1.26
NG = 0
DB = 0
SM = 0
Delta_L
Doxycycline
YOR261C RPN8
ZShift = 0.11
NG = 0
DB = 0
SM = 0
Delta_K
Doxycycline
YOR261C RPN8
ZShift = −0.19
NG = 0
DB = 0
SM = 0
Delta_r
Doxycycline
YOR261C RPN8
ZShift = −0.48
NG = 0
DB = 0
SM = 0
Delta_AUC
Doxycycline
YLR141W RRN5
ZShift = 0.23
NG = 0
DB = 0
SM = 0
Delta_L
Doxycycline
YLR141W RRN5
ZShift = 0.21
NG = 0
DB = 0
SM = 0
Delta_K
Doxycycline
YLR141W RRN5
ZShift = 0.18
NG = 0
DB = 0
SM = 0
Delta_r
Doxycycline
YLR141W RRN5
ZShift = 0.04
NG = 0
DB = 0
SM = 0
Delta_AUC
Doxycycline
YBR088C POL30
ZShift = 2.29
NG = 0
DB = 0
SM = 0
Delta_L
Doxycycline
YBR088C POL30
ZShift = −1.24
NG = 0
DB = 0
SM = 0
Delta_K
Doxycycline
YBR088C POL30
ZShift = −2.13
NG = 0
DB = 0
SM = 0
Delta_r
Doxycycline
YBR088C POL30
ZShift = −1.79
NG = 0
DB = 0
SM = 0
Delta_AUC
Doxycycline

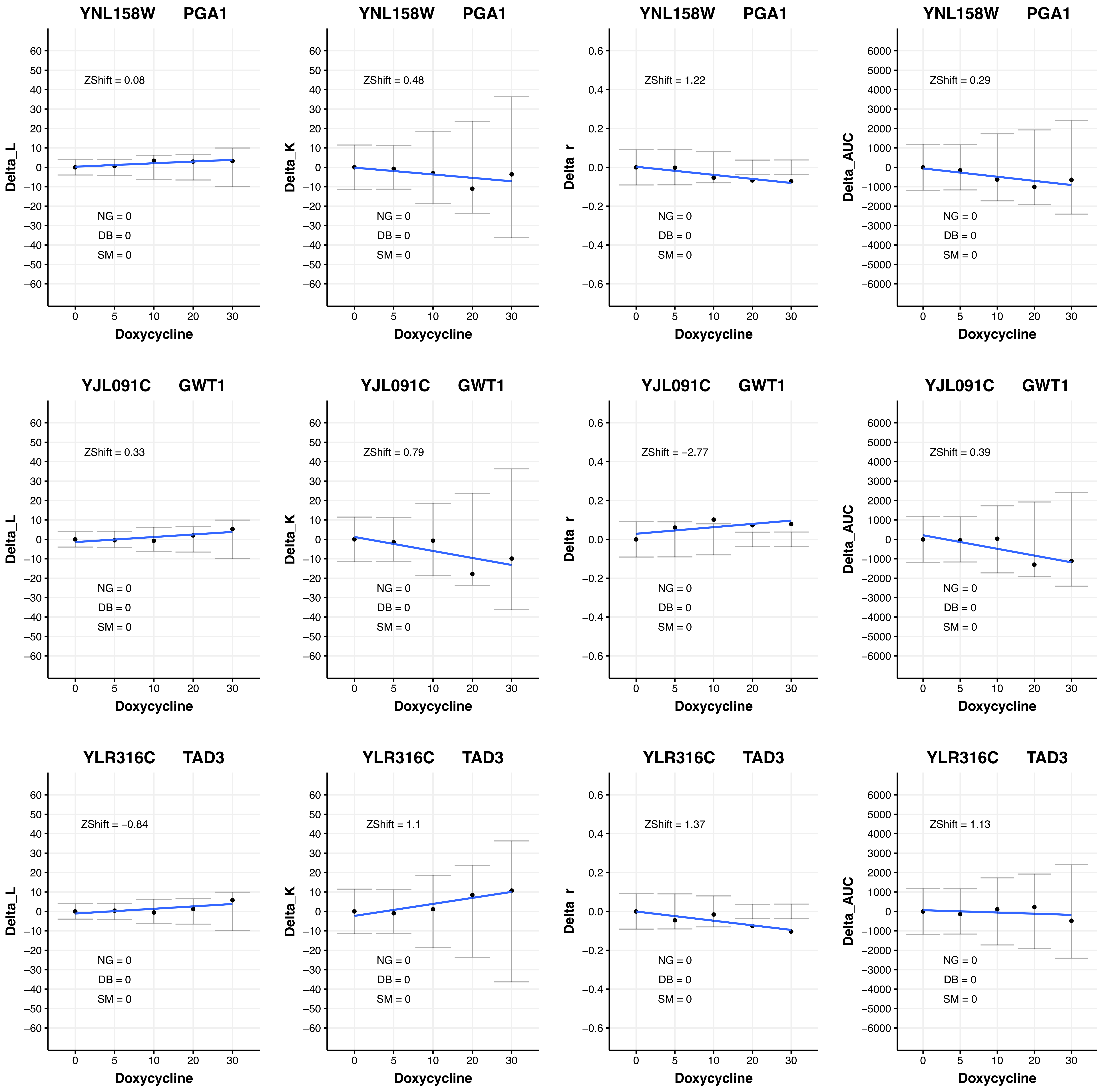
YNL158W PGA1
ZShift = 0.08
Delta_L
NG = 0
DB = 0
SM = 0
Doxycycline
YNL158W PGA1
ZShift = 0.48
Delta_K
NG = 0
DB = 0
SM = 0
Doxycycline
YNL158W PGA1
ZShift = 1.22
Delta_r
NG = 0
DB = 0
SM = 0
Doxycycline
YNL158W PGA1
ZShift = 0.29
Delta_AUC
NG = 0
DB = 0
SM = 0
Doxycycline
YJL091C GWT1
ZShift = 0.33
Delta_L
NG = 0
DB = 0
SM = 0
Doxycycline
YJL091C GWT1
ZShift = 0.79
Delta_K
NG = 0
DB = 0
SM = 0
Doxycycline
YJL091C GWT1
ZShift = −2.77
Delta_r
NG = 0
DB = 0
SM = 0
Doxycycline
YJL091C GWT1
ZShift = 0.39
Delta_AUC
NG = 0
DB = 0
SM = 0
Doxycycline
YLR316C TAD3
ZShift = −0.84
Delta_L
NG = 0
DB = 0
SM = 0
Doxycycline
YLR316C TAD3
ZShift = 1.1
Delta_K
NG = 0
DB = 0
SM = 0
Doxycycline
YLR316C TAD3
ZShift = 1.37
Delta_r
NG = 0
DB = 0
SM = 0
Doxycycline
YLR316C TAD3
ZShift = 1.13
Delta_AUC
NG = 0
DB = 0
SM = 0
Doxycycline

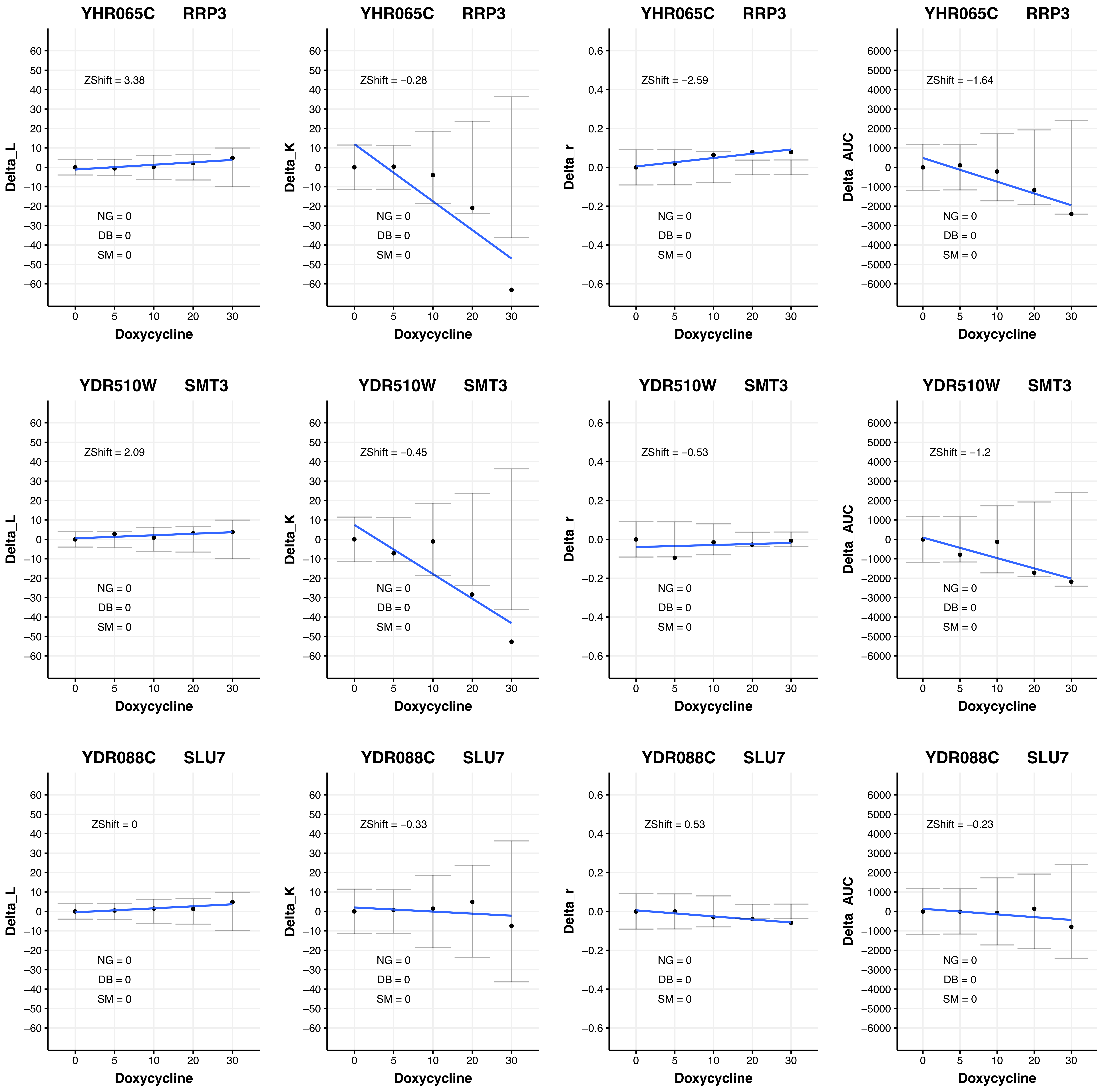

YHR065C RRP3
ZShift = 3.38
NG = 0
DB = 0
SM = 0
Delta_L
Doxycycline
YHR065C RRP3
ZShift = −0.28
NG = 0
DB = 0
SM = 0
Delta_K
Doxycycline
YHR065C RRP3
ZShift = −2.59
NG = 0
DB = 0
SM = 0
Delta_r
Doxycycline
YHR065C RRP3
ZShift = −1.64
NG = 0
DB = 0
SM = 0
Delta_AUC
Doxycycline
YDR510W SMT3
ZShift = 2.09
NG = 0
DB = 0
SM = 0
Delta_L
Doxycycline
YDR510W SMT3
ZShift = −0.45
NG = 0
DB = 0
SM = 0
Delta_K
Doxycycline
YDR510W SMT3
ZShift = −0.53
NG = 0
DB = 0
SM = 0
Delta_r
Doxycycline
YDR510W SMT3
ZShift = −1.2
NG = 0
DB = 0
SM = 0
Delta_AUC
Doxycycline
YDR088C SLU7
ZShift = 0
NG = 0
DB = 0
SM = 0
Delta_L
Doxycycline
YDR088C SLU7
ZShift = −0.33
NG = 0
DB = 0
SM = 0
Delta_K
Doxycycline
YDR088C SLU7
ZShift = 0.53
NG = 0
DB = 0
SM = 0
Delta_r
Doxycycline
YDR088C SLU7
ZShift = −0.23
NG = 0
DB = 0
SM = 0
Delta_AUC
Doxycycline

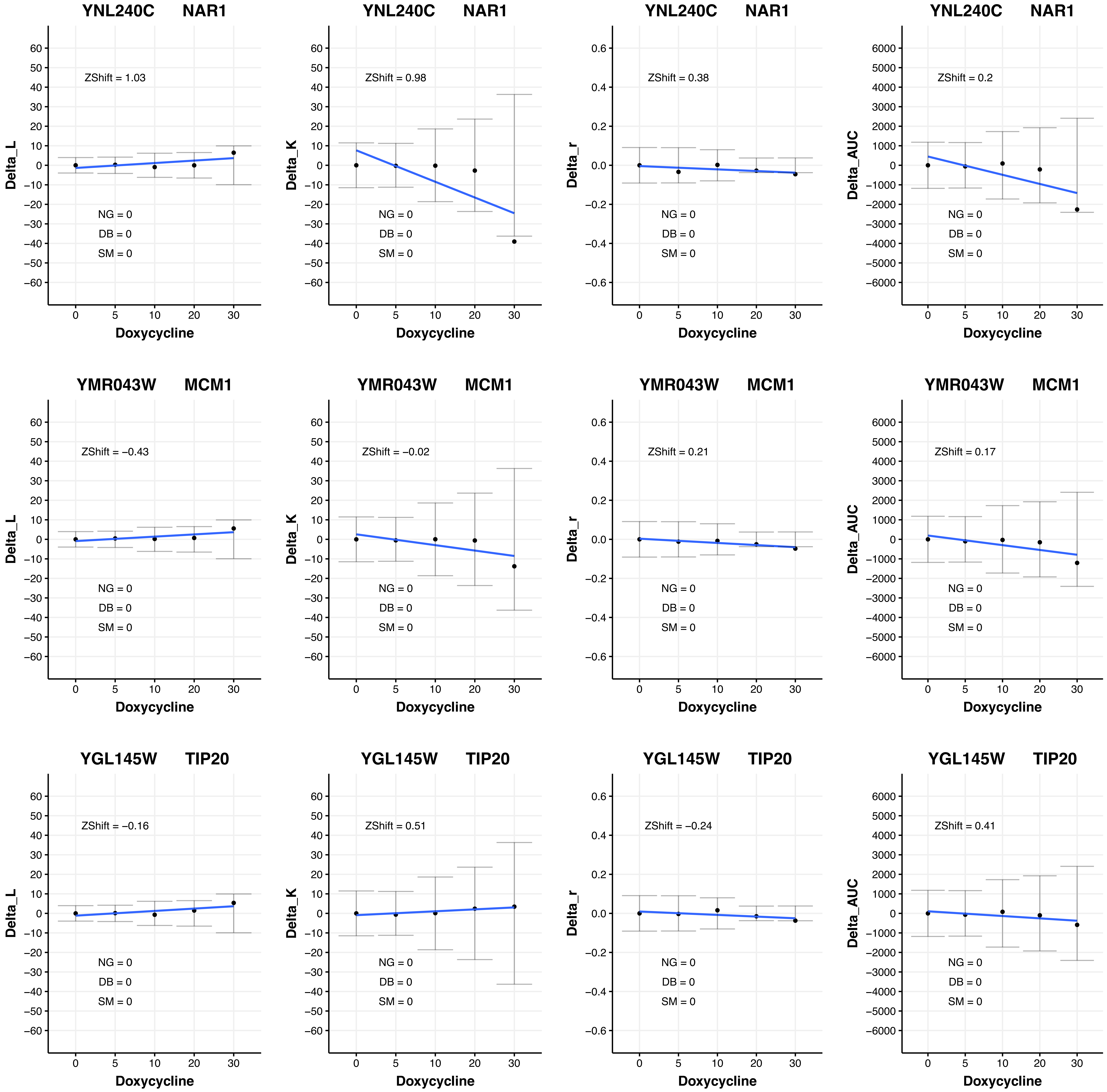
YNL240C NAR1
ZShift = 1.03
NG = 0
DB = 0
SM = 0
Delta_L
Doxycycline
YNL240C NAR1
ZShift = 0.98
NG = 0
DB = 0
SM = 0
Delta_K
Doxycycline
YNL240C NAR1
ZShift = 0.38
NG = 0
DB = 0
SM = 0
Delta_r
Doxycycline
YNL240C NAR1
ZShift = 0.2
NG = 0
DB = 0
SM = 0
Delta_AUC
Doxycycline
YMR043W MCM1
ZShift = −0.43
NG = 0
DB = 0
SM = 0
Delta_L
Doxycycline
YMR043W MCM1
ZShift = −0.02
NG = 0
DB = 0
SM = 0
Delta_K
Doxycycline
YMR043W MCM1
ZShift = 0.21
NG = 0
DB = 0
SM = 0
Delta_r
Doxycycline
YMR043W MCM1
ZShift = 0.17
NG = 0
DB = 0
SM = 0
Delta_AUC
Doxycycline
YGL145W TIP20
ZShift = −0.16
NG = 0
DB = 0
SM = 0
Delta_L
Doxycycline
YGL145W TIP20
ZShift = 0.51
NG = 0
DB = 0
SM = 0
Delta_K
Doxycycline
YGL145W TIP20
ZShift = −0.24
NG = 0
DB = 0
SM = 0
Delta_r
Doxycycline
YGL145W TIP20
ZShift = 0.41
NG = 0
DB = 0
SM = 0
Delta_AUC
Doxycycline

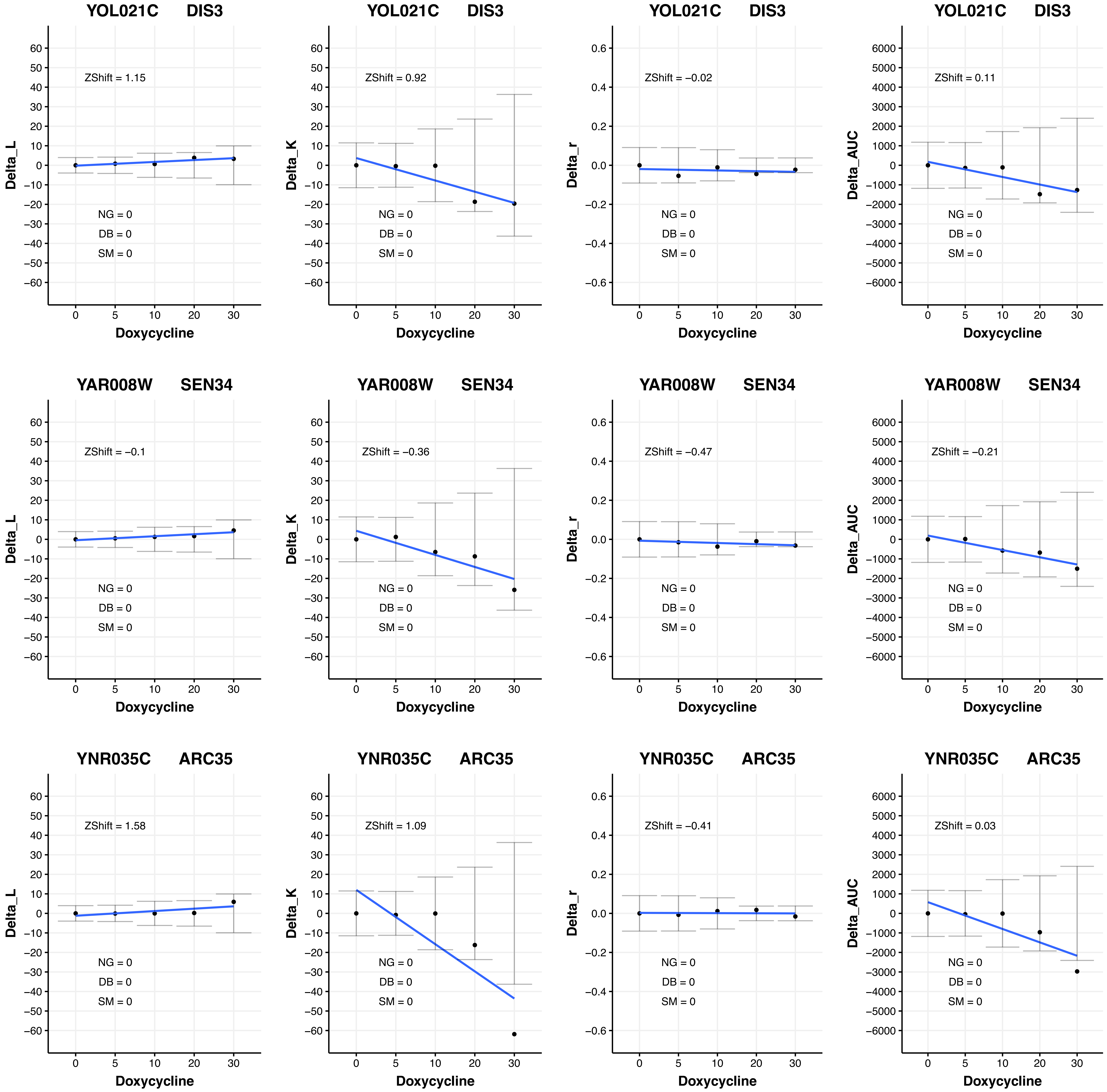
YOL021C DIS3
ZShift = 1.15
NG = 0
DB = 0
SM = 0
Delta_L
Doxycycline
YOL021C DIS3
ZShift = 0.92
NG = 0
DB = 0
SM = 0
Delta_K
Doxycycline
YOL021C DIS3
ZShift = −0.02
NG = 0
DB = 0
SM = 0
Delta_r
Doxycycline
YOL021C DIS3
ZShift = 0.11
NG = 0
DB = 0
SM = 0
Delta_AUC
Doxycycline
YAR008W SEN34
ZShift = −0.1
NG = 0
DB = 0
SM = 0
Delta_L
Doxycycline
YAR008W SEN34
ZShift = −0.36
NG = 0
DB = 0
SM = 0
Delta_K
Doxycycline
YAR008W SEN34
ZShift = −0.47
NG = 0
DB = 0
SM = 0
Delta_r
Doxycycline
YAR008W SEN34
ZShift = −0.21
NG = 0
DB = 0
SM = 0
Delta_AUC
Doxycycline
YNR035C ARC35
ZShift = 1.58
NG = 0
DB = 0
SM = 0
Delta_L
Doxycycline
YNR035C ARC35
ZShift = 1.09
NG = 0
DB = 0
SM = 0
Delta_K
Doxycycline
YNR035C ARC35
ZShift = −0.41
NG = 0
DB = 0
SM = 0
Delta_r
Doxycycline
YNR035C ARC35
ZShift = 0.03
NG = 0
DB = 0
SM = 0
Delta_AUC
Doxycycline

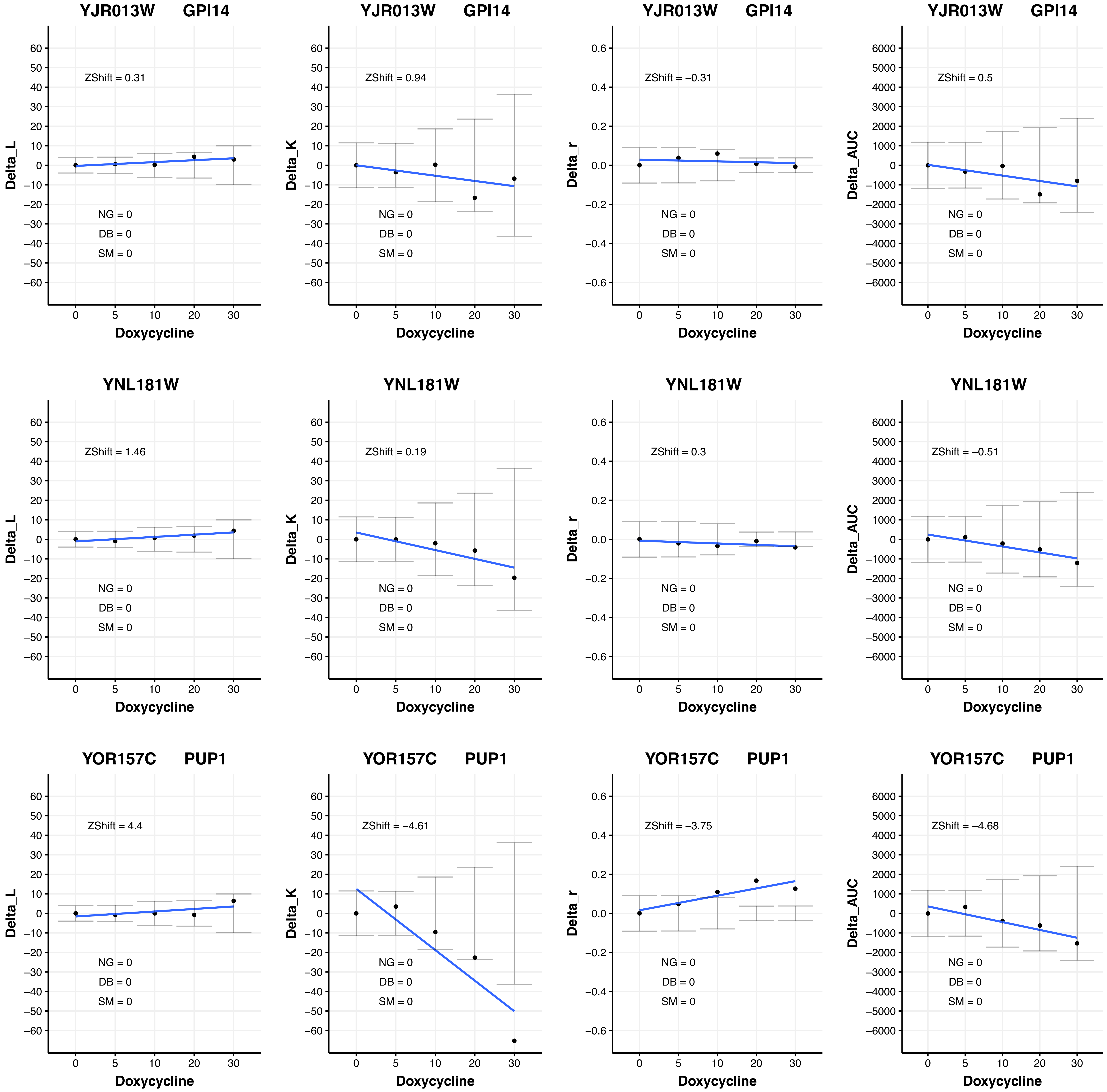

YJR013W GPI14
ZShift = 0.31
NG = 0
DB = 0
SM = 0
Delta_L
Doxycycline
YJR013W GPI14
ZShift = 0.94
NG = 0
DB = 0
SM = 0
Delta_K
Doxycycline
YJR013W GPI14
ZShift = −0.31
NG = 0
DB = 0
SM = 0
Delta_r
Doxycycline
YJR013W GPI14
ZShift = 0.5
NG = 0
DB = 0
SM = 0
Delta_AUC
Doxycycline
YNL181W
ZShift = 1.46
NG = 0
DB = 0
SM = 0
Delta_L
Doxycycline
YNL181W
ZShift = 0.19
NG = 0
DB = 0
SM = 0
Delta_K
Doxycycline
YNL181W
ZShift = 0.3
NG = 0
DB = 0
SM = 0
Delta_r
Doxycycline
YNL181W
ZShift = −0.51
NG = 0
DB = 0
SM = 0
Delta_AUC
Doxycycline
YOR157C PUP1
ZShift = 4.4
NG = 0
DB = 0
SM = 0
Delta_L
Doxycycline
YOR157C PUP1
ZShift = −4.61
NG = 0
DB = 0
SM = 0
Delta_K
Doxycycline
YOR157C PUP1
ZShift = −3.75
NG = 0
DB = 0
SM = 0
Delta_r
Doxycycline
YOR157C PUP1
ZShift = −4.68
NG = 0
DB = 0
SM = 0
Delta_AUC
Doxycycline

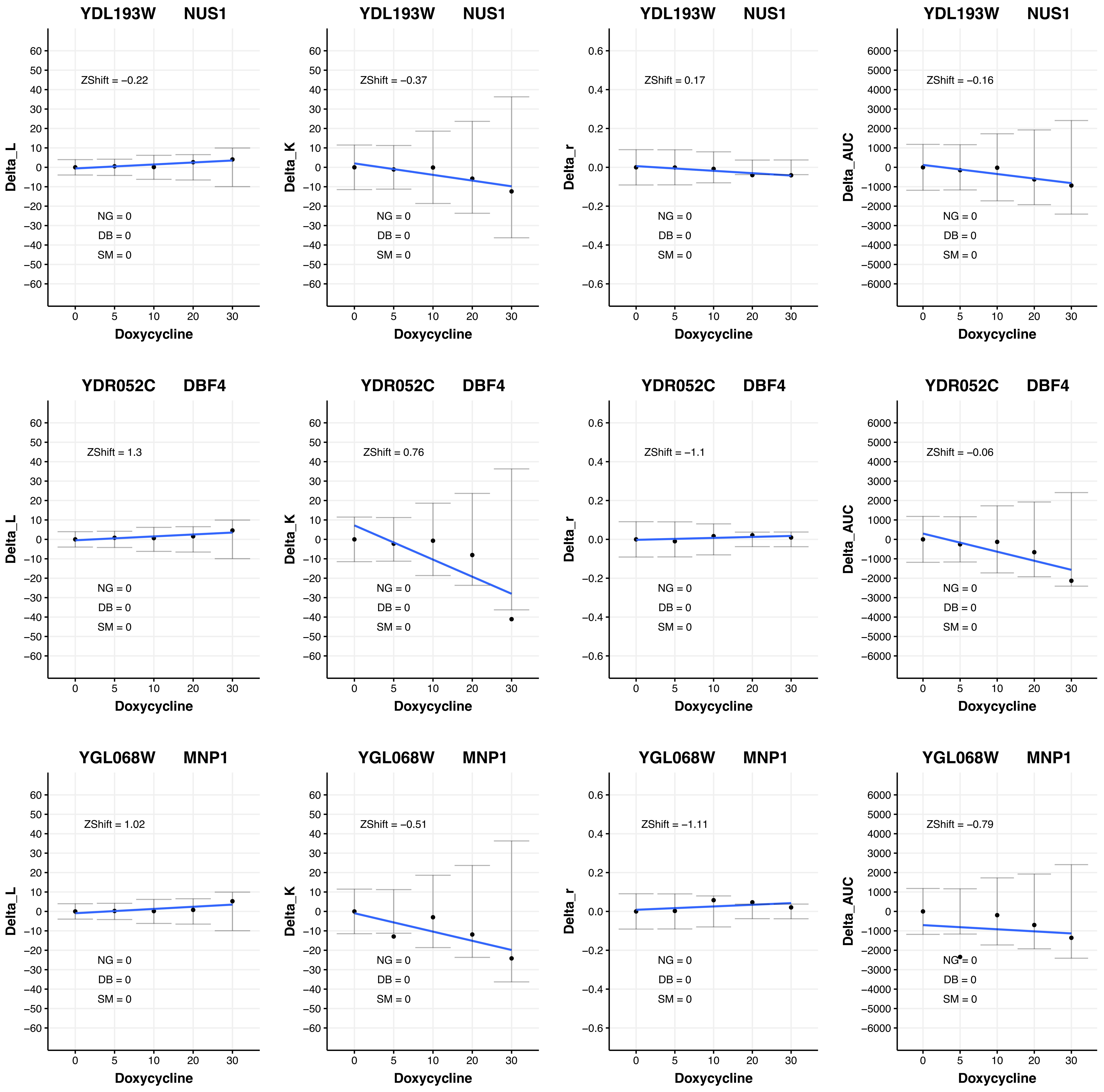
YDL193W NUS1
Delta_L
ZShift = −0.22
NG = 0
DB = 0
SM = 0
Doxycycline
YDL193W NUS1
Delta_K
ZShift = −0.37
NG = 0
DB = 0
SM = 0
Doxycycline
YDL193W NUS1
Delta_r
ZShift = 0.17
NG = 0
DB = 0
SM = 0
Doxycycline
YDL193W NUS1
Delta_AUC
ZShift = −0.16
NG = 0
DB = 0
SM = 0
Doxycycline
YDR052C DBF4
Delta_L
ZShift = 1.3
NG = 0
DB = 0
SM = 0
Doxycycline
YDR052C DBF4
Delta_K
ZShift = 0.76
NG = 0
DB = 0
SM = 0
Doxycycline
YDR052C DBF4
Delta_r
ZShift = −1.1
NG = 0
DB = 0
SM = 0
Doxycycline
YDR052C DBF4
Delta_AUC
ZShift = −0.06
NG = 0
DB = 0
SM = 0
Doxycycline
YGL068W MNP1
Delta_L
ZShift = 1.02
NG = 0
DB = 0
SM = 0
Doxycycline
YGL068W MNP1
Delta_K
ZShift = −0.51
NG = 0
DB = 0
SM = 0
Doxycycline
YGL068W MNP1
Delta_r
ZShift = −1.11
NG = 0
DB = 0
SM = 0
Doxycycline
YGL068W MNP1
Delta_AUC
ZShift = −0.79
NG = 0
DB = 0
SM = 0
Doxycycline

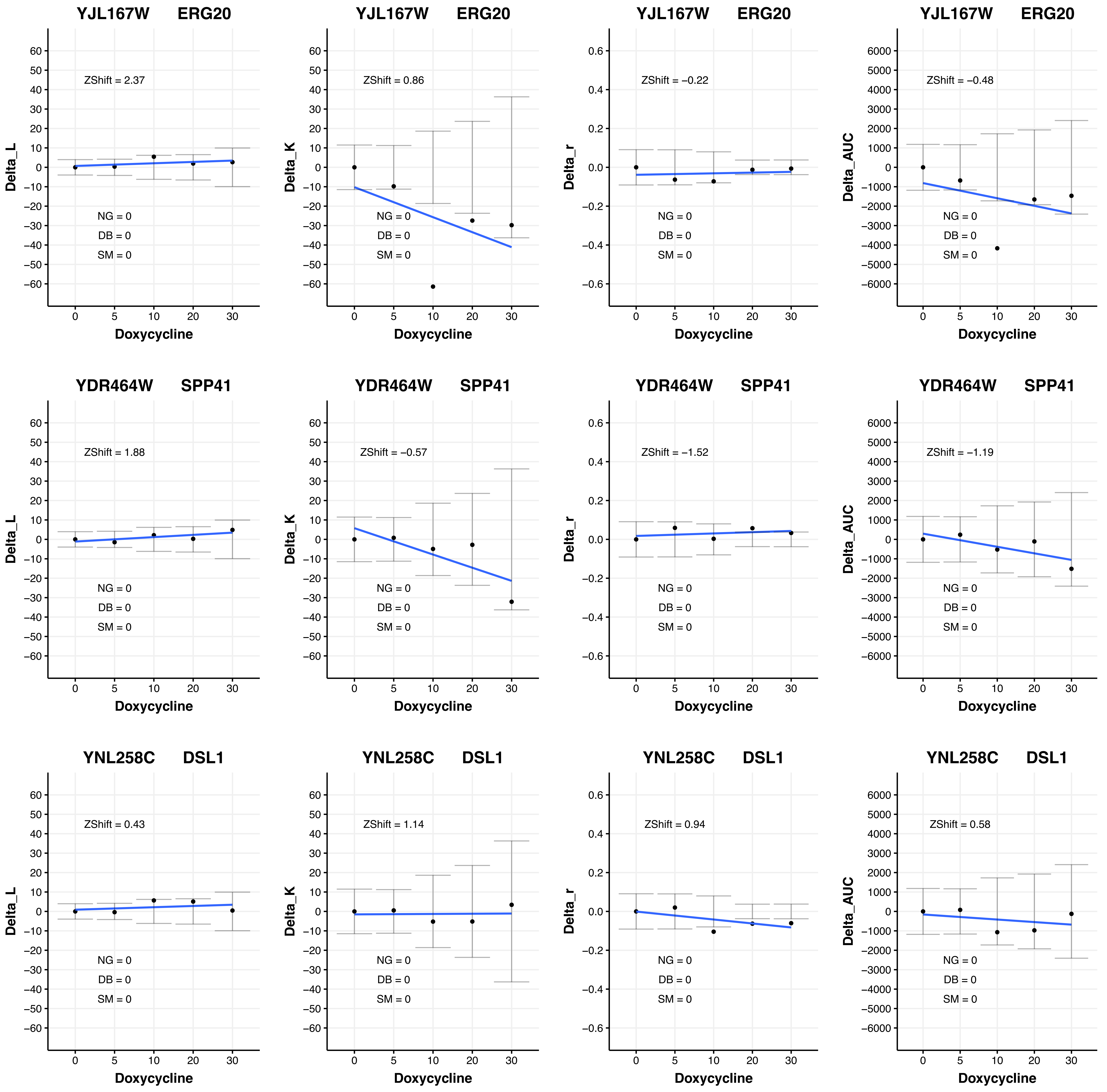
YJL167W ERG20
Delta_L
ZShift = 2.37
NG = 0
DB = 0
SM = 0
Doxycycline
YJL167W ERG20
Delta_K
ZShift = 0.86
NG = 0
DB = 0
SM = 0
Doxycycline
YJL167W ERG20
Delta_r
ZShift = −0.22
NG = 0
DB = 0
SM = 0
Doxycycline
YJL167W ERG20
Delta_AUC
ZShift = −0.48
NG = 0
DB = 0
SM = 0
Doxycycline
YDR464W SPP41
Delta_L
ZShift = 1.88
NG = 0
DB = 0
SM = 0
Doxycycline
YDR464W SPP41
Delta_K
ZShift = −0.57
NG = 0
DB = 0
SM = 0
Doxycycline
YDR464W SPP41
Delta_r
ZShift = −1.52
NG = 0
DB = 0
SM = 0
Doxycycline
YDR464W SPP41
Delta_AUC
ZShift = −1.19
NG = 0
DB = 0
SM = 0
Doxycycline
YNL258C DSL1
Delta_L
ZShift = 0.43
NG = 0
DB = 0
SM = 0
Doxycycline
YNL258C DSL1
Delta_K
ZShift = 1.14
NG = 0
DB = 0
SM = 0
Doxycycline
YNL258C DSL1
Delta_r
ZShift = 0.94
NG = 0
DB = 0
SM = 0
Doxycycline
YNL258C DSL1
Delta_AUC
ZShift = 0.58
NG = 0
DB = 0
SM = 0
Doxycycline

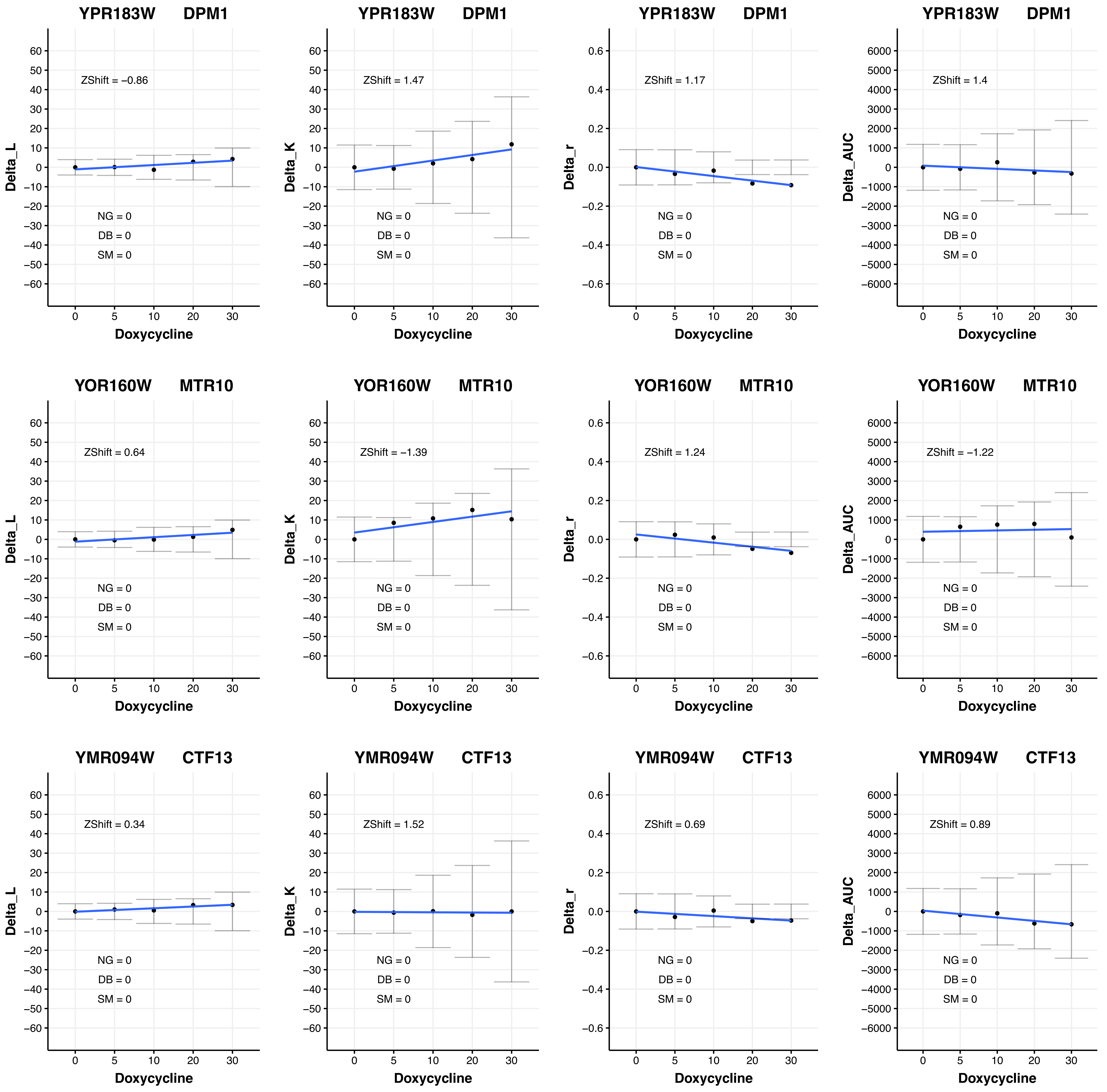

YPR183W DPM1
Delta_L
ZShift = −0.86
NG = 0
DB = 0
SM = 0
Doxycycline
YPR183W DPM1
Delta_K
ZShift = 1.47
NG = 0
DB = 0
SM = 0
Doxycycline
YPR183W DPM1
Delta_r
ZShift = 1.17
NG = 0
DB = 0
SM = 0
Doxycycline
YPR183W DPM1
Delta_AUC
ZShift = 1.4
NG = 0
DB = 0
SM = 0
Doxycycline
YOR160W MTR10
Delta_L
ZShift = 0.64
NG = 0
DB = 0
SM = 0
Doxycycline
YOR160W MTR10
Delta_K
ZShift = −1.39
NG = 0
DB = 0
SM = 0
Doxycycline
YOR160W MTR10
Delta_r
ZShift = 1.24
NG = 0
DB = 0
SM = 0
Doxycycline
YOR160W MTR10
Delta_AUC
ZShift = −1.22
NG = 0
DB = 0
SM = 0
Doxycycline
YMR094W CTF13
Delta_L
ZShift = 0.34
NG = 0
DB = 0
SM = 0
Doxycycline
YMR094W CTF13
Delta_K
ZShift = 1.52
NG = 0
DB = 0
SM = 0
Doxycycline
YMR094W CTF13
Delta_r
ZShift = 0.69
NG = 0
DB = 0
SM = 0
Doxycycline
YMR094W CTF13
Delta_AUC
ZShift = 0.89
NG = 0
DB = 0
SM = 0
Doxycycline

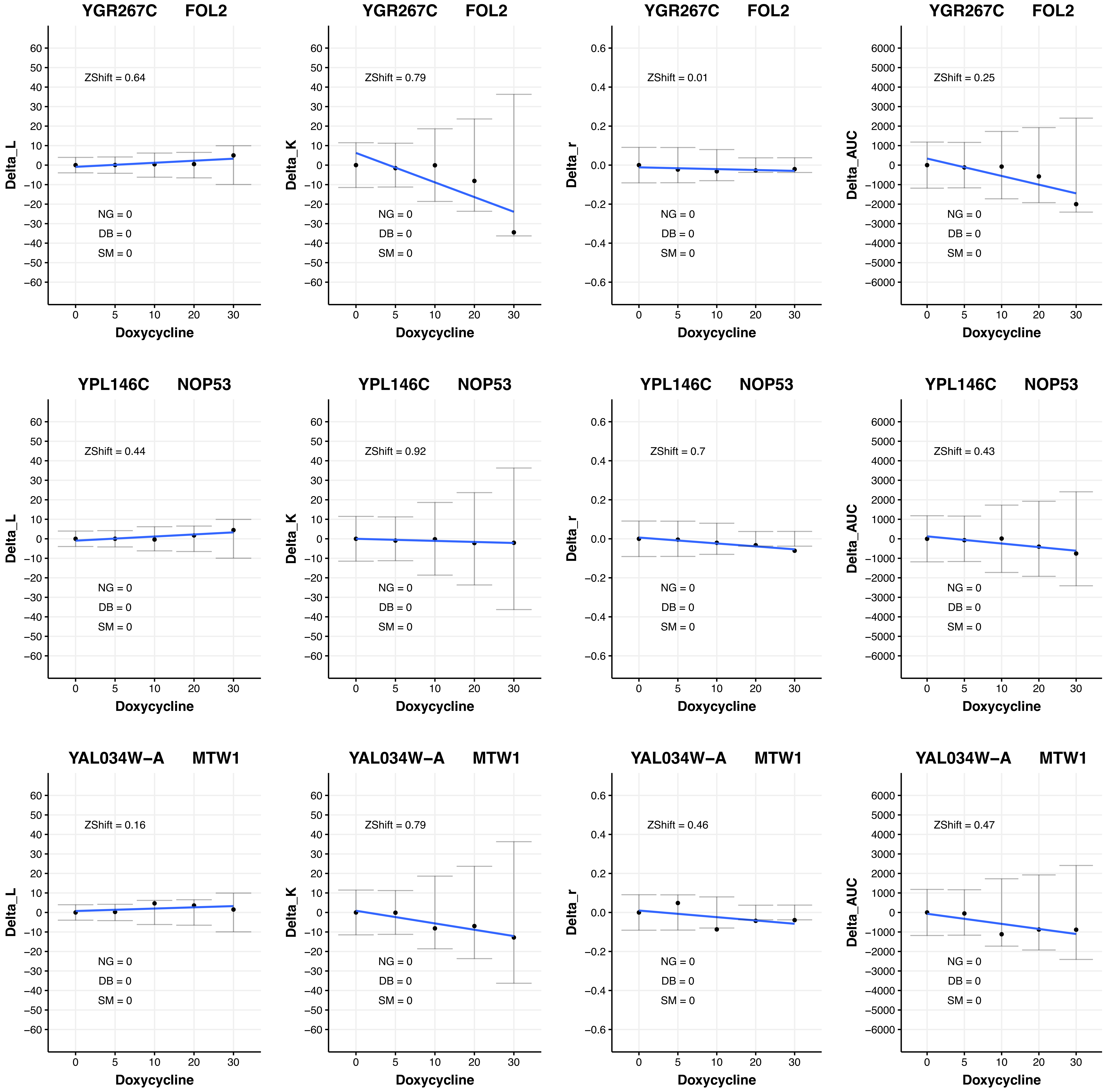
YGR267C FOL2
ZShift = 0.64
NG = 0
DB = 0
SM = 0
Delta_L
Doxycycline
YGR267C FOL2
ZShift = 0.79
NG = 0
DB = 0
SM = 0
Delta_K
Doxycycline
YGR267C FOL2
ZShift = 0.01
NG = 0
DB = 0
SM = 0
Delta_r
Doxycycline
YGR267C FOL2
ZShift = 0.25
NG = 0
DB = 0
SM = 0
Delta_AUC
Doxycycline
YPL146C NOP53
ZShift = 0.44
NG = 0
DB = 0
SM = 0
Delta_L
Doxycycline
YPL146C NOP53
ZShift = 0.92
NG = 0
DB = 0
SM = 0
Delta_K
Doxycycline
YPL146C NOP53
ZShift = 0.7
NG = 0
DB = 0
SM = 0
Delta_r
Doxycycline
YPL146C NOP53
ZShift = 0.43
NG = 0
DB = 0
SM = 0
Delta_AUC
Doxycycline
YAL034W−A MTW1
ZShift = 0.16
NG = 0
DB = 0
SM = 0
Delta_L
Doxycycline
YAL034W−A MTW1
ZShift = 0.79
NG = 0
DB = 0
SM = 0
Delta_K
Doxycycline
YAL034W−A MTW1
ZShift = 0.46
NG = 0
DB = 0
SM = 0
Delta_r
Doxycycline
YAL034W−A MTW1
ZShift = 0.47
NG = 0
DB = 0
SM = 0
Delta_AUC
Doxycycline

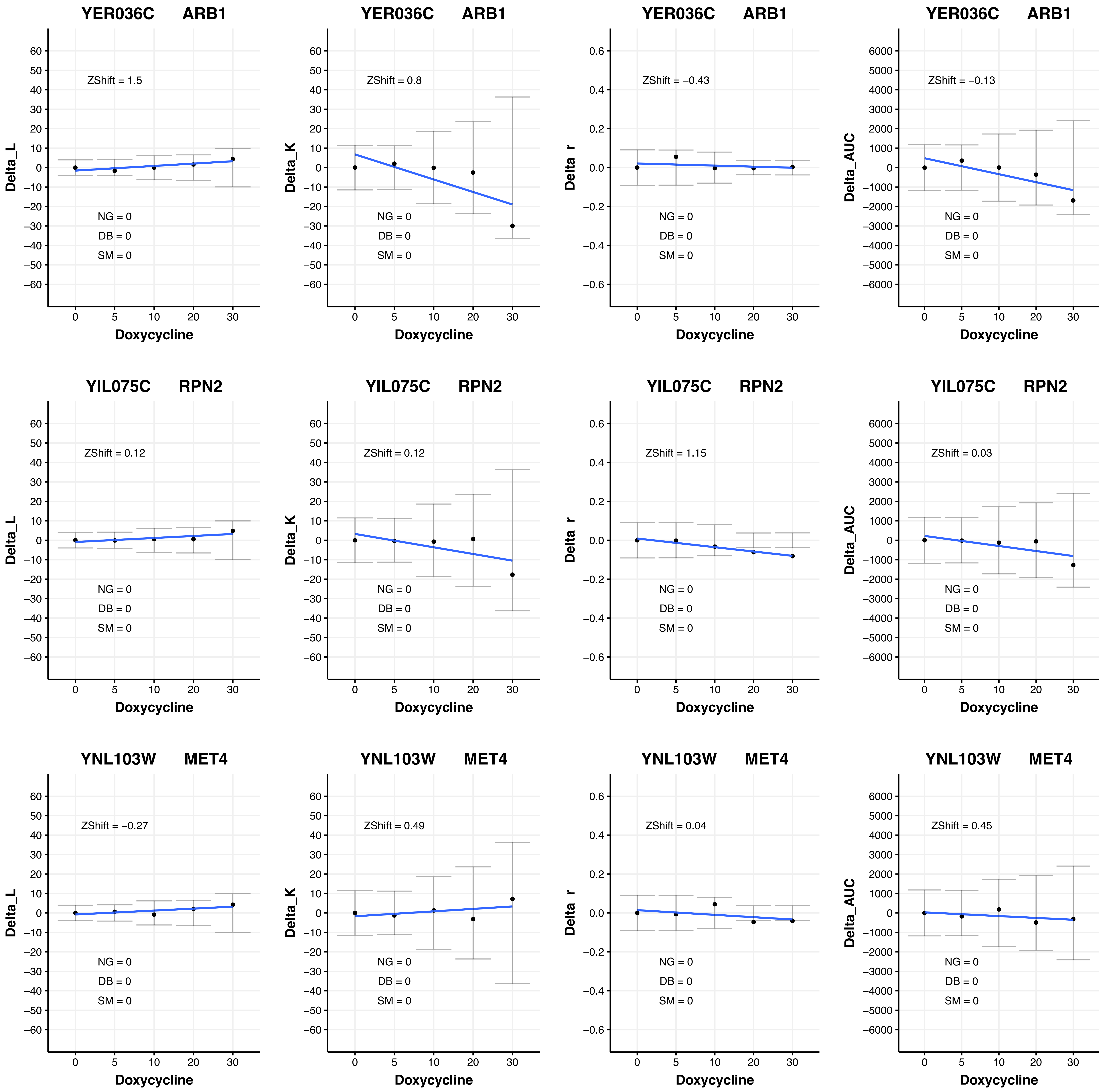

YER036C ARB1
ZShift = 1.5
ZShift = 0.8
ZShift = −0.43
ZShift = −0.13
YIL075C RPN2
ZShift = 0.12
ZShift = 0.12
ZShift = 1.15
ZShift = 0.03
YNL103W MET4
ZShift = −0.27
ZShift = 0.49
ZShift = 0.04
ZShift = 0.45
NG = 0
DB = 0
SM = 0
Delta_L
Delta_K
Delta_r
Delta_AUC
Doxycycline

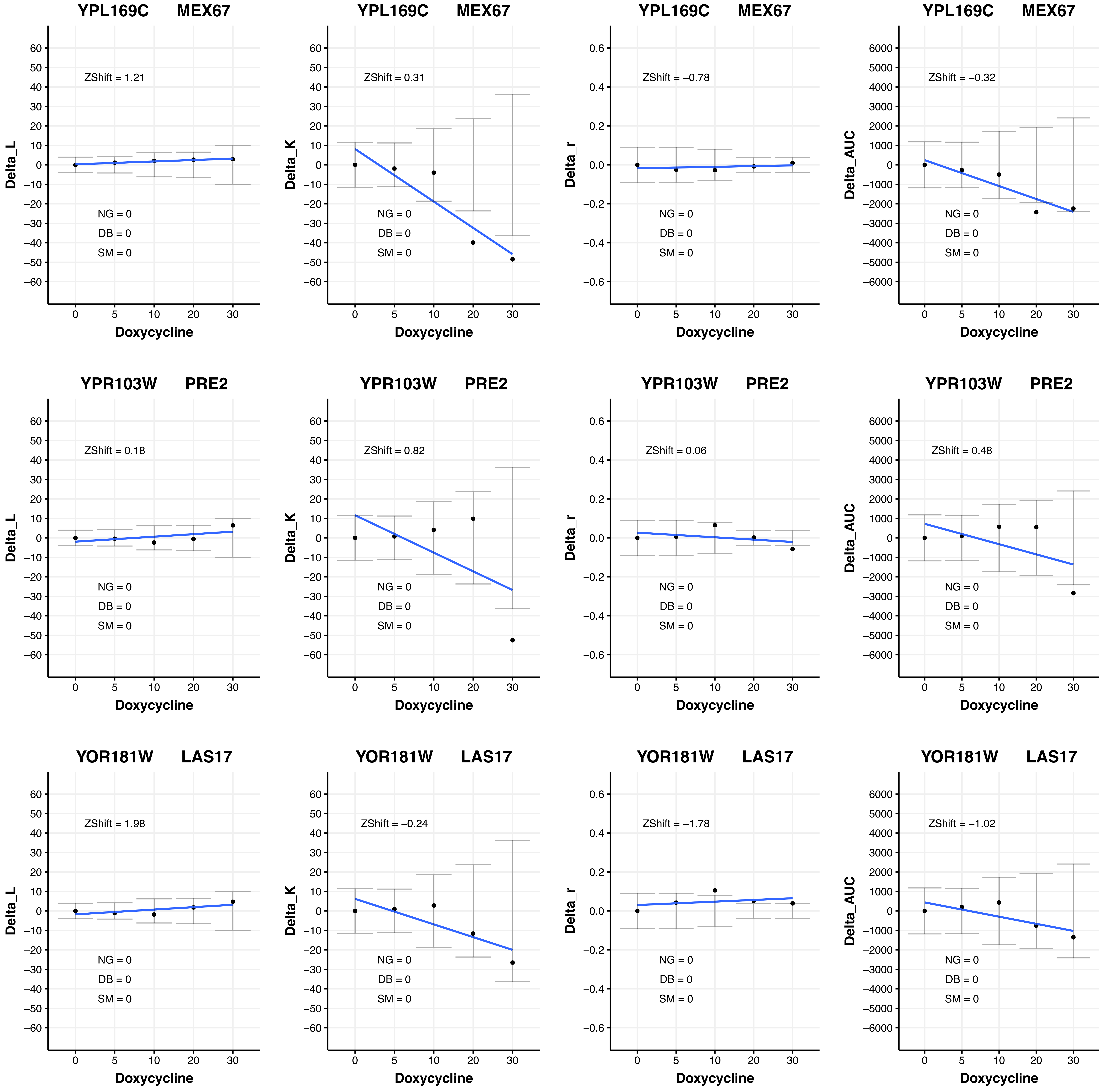

YPL169C MEX67
ZShift = 1.21
Delta_L
NG = 0
DB = 0
SM = 0
Doxycycline
YPL169C MEX67
ZShift = 0.31
Delta_K
NG = 0
DB = 0
SM = 0
Doxycycline
YPL169C MEX67
ZShift = −0.78
Delta_r
NG = 0
DB = 0
SM = 0
Doxycycline
YPL169C MEX67
ZShift = −0.32
Delta_AUC
NG = 0
DB = 0
SM = 0
Doxycycline
YPR103W PRE2
ZShift = 0.18
Delta_L
NG = 0
DB = 0
SM = 0
Doxycycline
YPR103W PRE2
ZShift = 0.82
Delta_K
NG = 0
DB = 0
SM = 0
Doxycycline
YPR103W PRE2
ZShift = 0.06
Delta_r
NG = 0
DB = 0
SM = 0
Doxycycline
YPR103W PRE2
ZShift = 0.48
Delta_AUC
NG = 0
DB = 0
SM = 0
Doxycycline
YOR181W LAS17
ZShift = 1.98
Delta_L
NG = 0
DB = 0
SM = 0
Doxycycline
YOR181W LAS17
ZShift = −0.24
Delta_K
NG = 0
DB = 0
SM = 0
Doxycycline
YOR181W LAS17
ZShift = −1.78
Delta_r
NG = 0
DB = 0
SM = 0
Doxycycline
YOR181W LAS17
ZShift = −1.02
Delta_AUC
NG = 0
DB = 0
SM = 0
Doxycycline

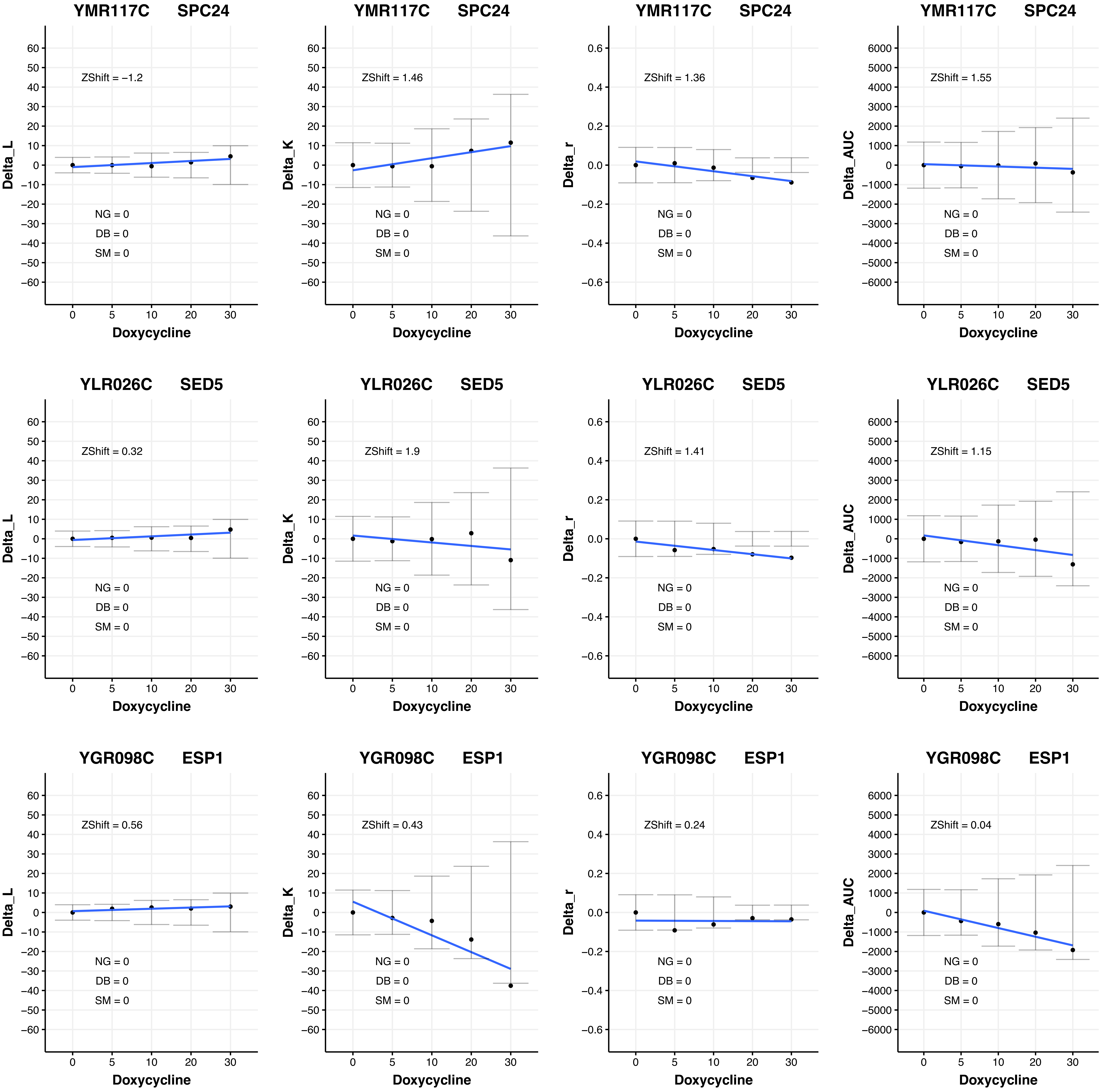

YMR117C SPC24
ZShift = −1.2
Delta_L
NG = 0
DB = 0
SM = 0
Doxycycline
YMR117C SPC24
ZShift = 1.46
Delta_K
NG = 0
DB = 0
SM = 0
Doxycycline
YMR117C SPC24
ZShift = 1.36
Delta_r
NG = 0
DB = 0
SM = 0
Doxycycline
YMR117C SPC24
ZShift = 1.55
Delta_AUC
NG = 0
DB = 0
SM = 0
Doxycycline
YLR026C SED5
ZShift = 0.32
Delta_L
NG = 0
DB = 0
SM = 0
Doxycycline
YLR026C SED5
ZShift = 1.9
Delta_K
NG = 0
DB = 0
SM = 0
Doxycycline
YLR026C SED5
ZShift = 1.41
Delta_r
NG = 0
DB = 0
SM = 0
Doxycycline
YLR026C SED5
ZShift = 1.15
Delta_AUC
NG = 0
DB = 0
SM = 0
Doxycycline
YGR098C ESP1
ZShift = 0.56
Delta_L
NG = 0
DB = 0
SM = 0
Doxycycline
YGR098C ESP1
ZShift = 0.43
Delta_K
NG = 0
DB = 0
SM = 0
Doxycycline
YGR098C ESP1
ZShift = 0.24
Delta_r
NG = 0
DB = 0
SM = 0
Doxycycline
YGR098C ESP1
ZShift = 0.04
Delta_AUC
NG = 0
DB = 0
SM = 0
Doxycycline

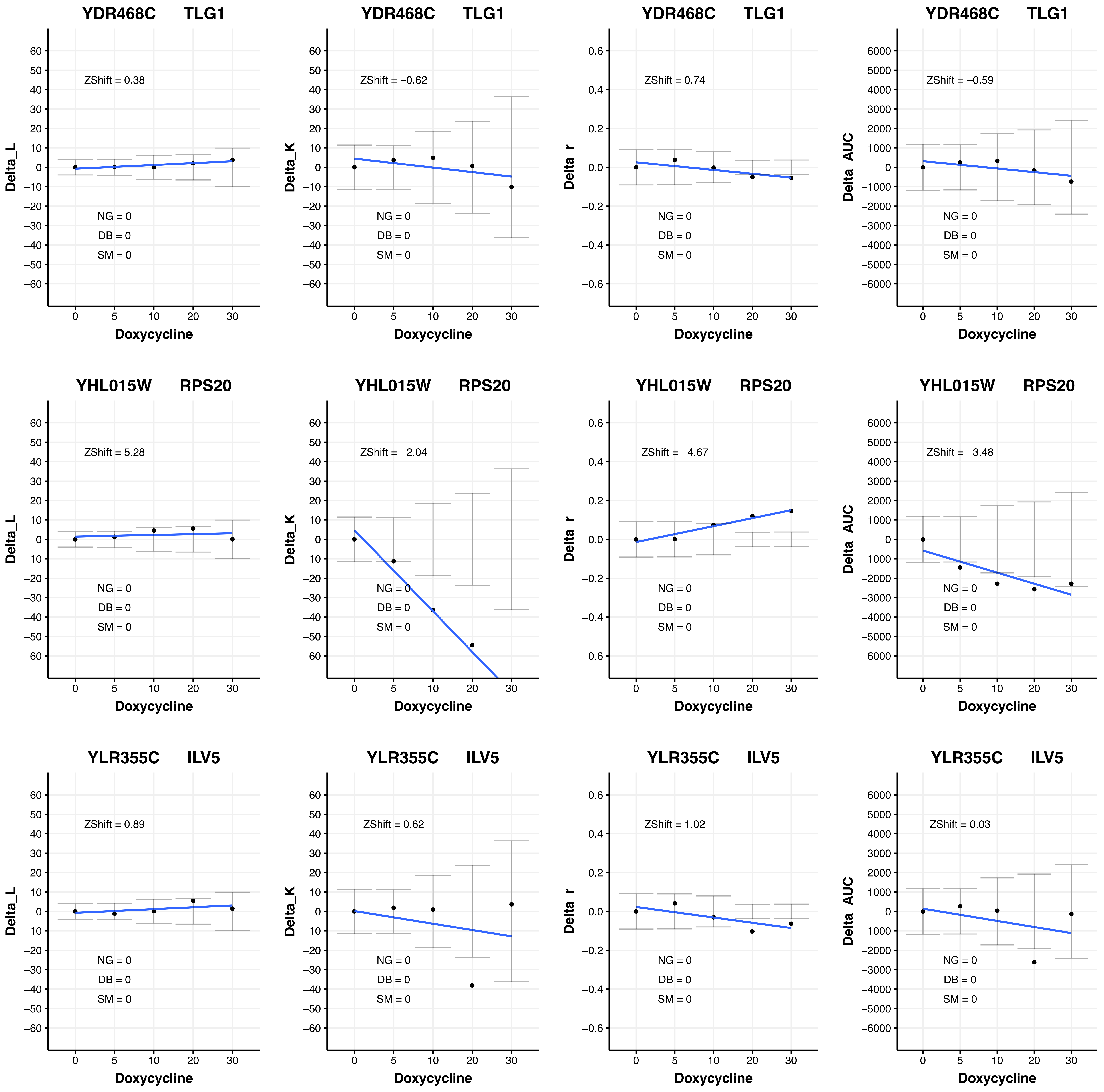
YDR468C TLG1
ZShift = 0.38
NG = 0
DB = 0
SM = 0
Delta_L
Doxycycline
YDR468C TLG1
ZShift = −0.62
NG = 0
DB = 0
SM = 0
Delta_K
Doxycycline
YDR468C TLG1
ZShift = 0.74
NG = 0
DB = 0
SM = 0
Delta_r
Doxycycline
YDR468C TLG1
ZShift = −0.59
NG = 0
DB = 0
SM = 0
Delta_AUC
Doxycycline
YHL015W RPS20
ZShift = 5.28
NG = 0
DB = 0
SM = 0
Delta_L
Doxycycline
YHL015W RPS20
ZShift = −2.04
NG = 0
DB = 0
SM = 0
Delta_K
Doxycycline
YHL015W RPS20
ZShift = −4.67
NG = 0
DB = 0
SM = 0
Delta_r
Doxycycline
YHL015W RPS20
ZShift = −3.48
NG = 0
DB = 0
SM = 0
Delta_AUC
Doxycycline
YLR355C ILV5
ZShift = 0.89
NG = 0
DB = 0
SM = 0
Delta_L
Doxycycline
YLR355C ILV5
ZShift = 0.62
NG = 0
DB = 0
SM = 0
Delta_K
Doxycycline
YLR355C ILV5
ZShift = 1.02
NG = 0
DB = 0
SM = 0
Delta_r
Doxycycline
YLR355C ILV5
ZShift = 0.03
NG = 0
DB = 0
SM = 0
Delta_AUC
Doxycycline

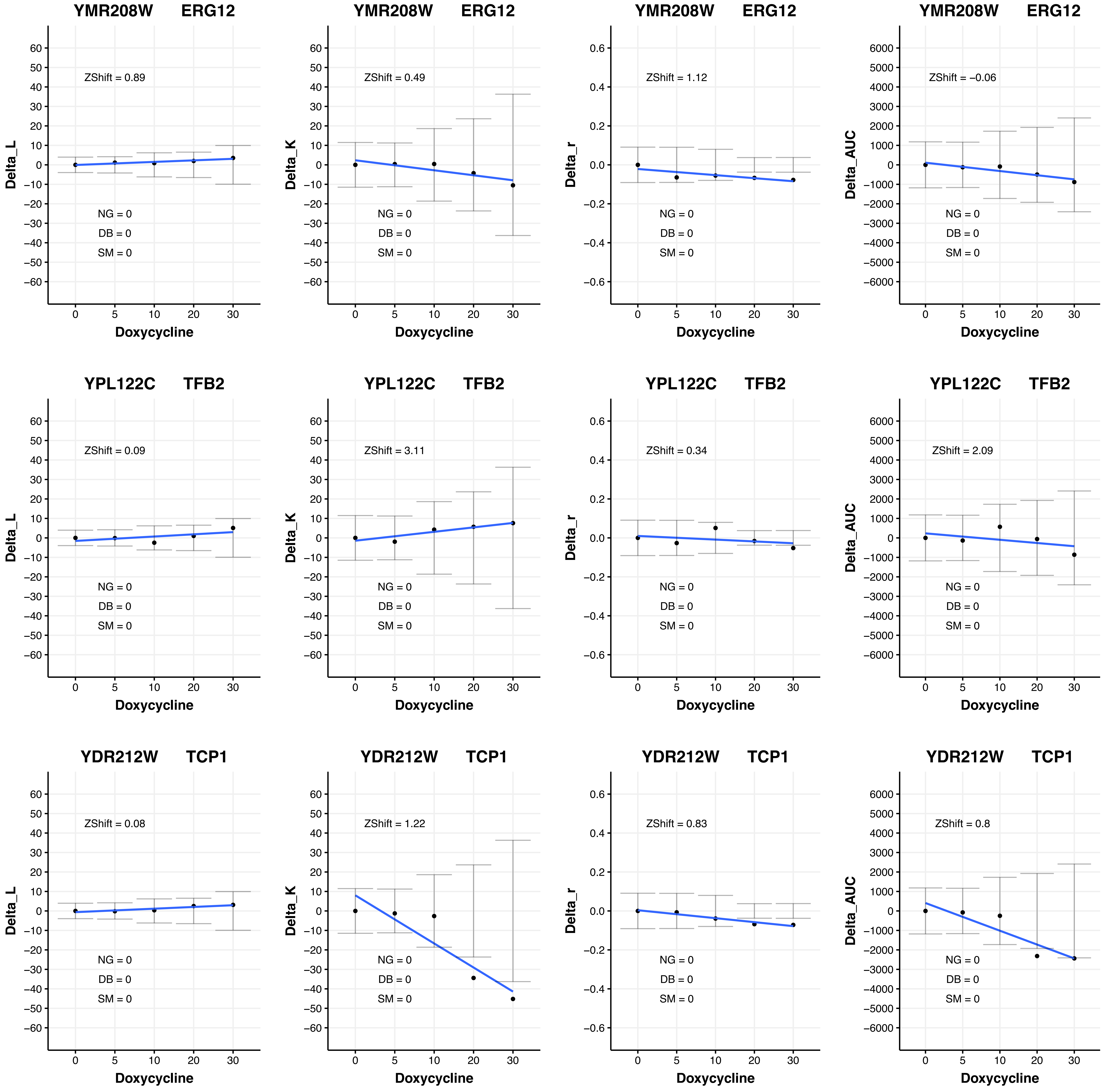
YMR208W ERG12
ZShift = 0.89
NG = 0
DB = 0
SM = 0
Delta_L
Doxycycline
YMR208W ERG12
ZShift = 0.49
NG = 0
DB = 0
SM = 0
Delta_K
Doxycycline
YMR208W ERG12
ZShift = 1.12
NG = 0
DB = 0
SM = 0
Delta_r
Doxycycline
YMR208W ERG12
ZShift = −0.06
NG = 0
DB = 0
SM = 0
Delta_AUC
Doxycycline
YPL122C TFB2
ZShift = 0.09
NG = 0
DB = 0
SM = 0
Delta_L
Doxycycline
YPL122C TFB2
ZShift = 3.11
NG = 0
DB = 0
SM = 0
Delta_K
Doxycycline
YPL122C TFB2
ZShift = 0.34
NG = 0
DB = 0
SM = 0
Delta_r
Doxycycline
YPL122C TFB2
ZShift = 2.09
NG = 0
DB = 0
SM = 0
Delta_AUC
Doxycycline
YDR212W TCP1
ZShift = 0.08
NG = 0
DB = 0
SM = 0
Delta_L
Doxycycline
YDR212W TCP1
ZShift = 1.22
NG = 0
DB = 0
SM = 0
Delta_K
Doxycycline
YDR212W TCP1
ZShift = 0.83
NG = 0
DB = 0
SM = 0
Delta_r
Doxycycline
YDR212W TCP1
ZShift = 0.8
NG = 0
DB = 0
SM = 0
Delta_AUC
Doxycycline

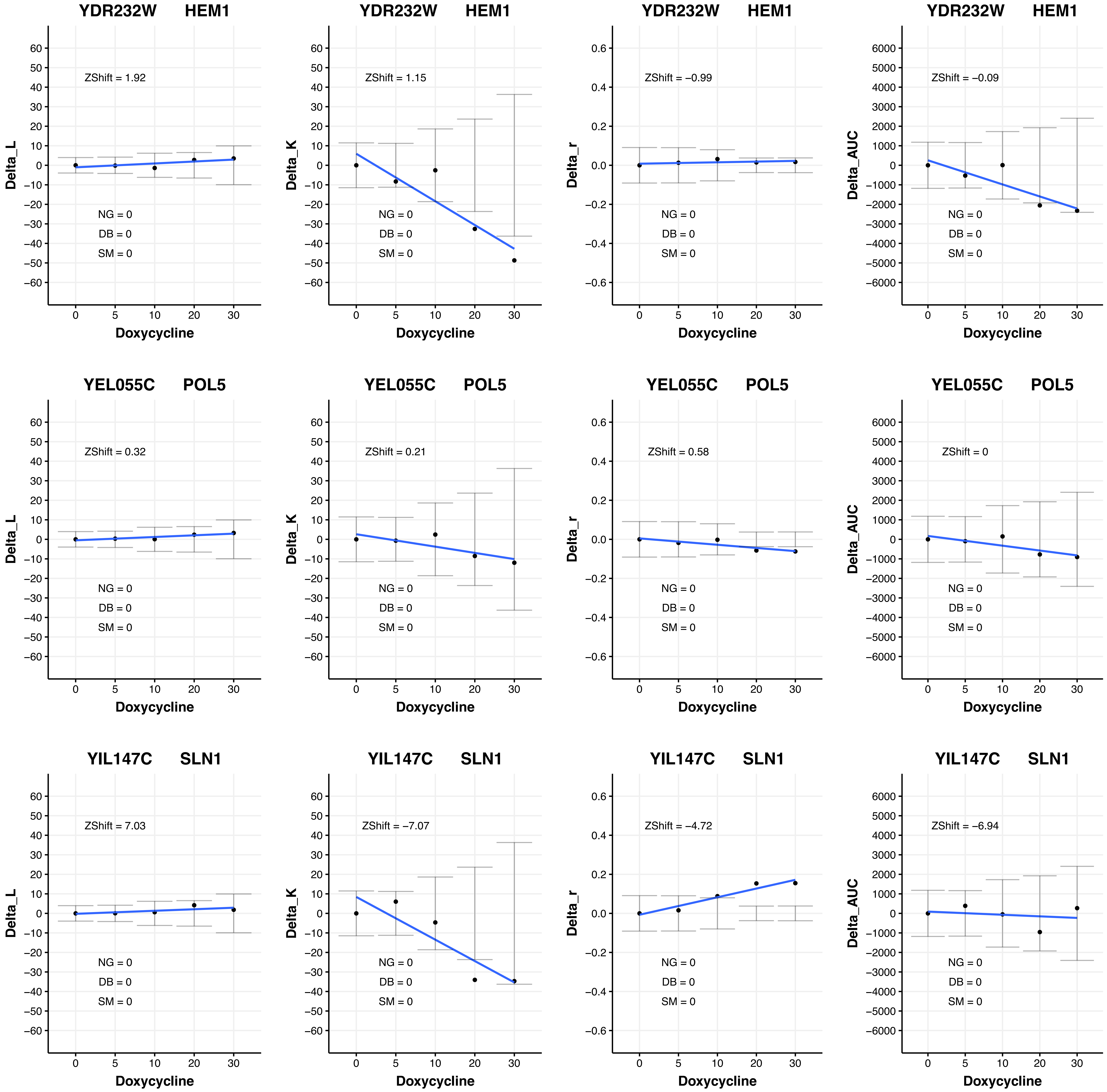
YDR232W HEM1
ZShift = 1.92
NG = 0
DB = 0
SM = 0
Delta_L
Doxycycline
YDR232W HEM1
ZShift = 1.15
NG = 0
DB = 0
SM = 0
Delta_K
Doxycycline
YDR232W HEM1
ZShift = −0.99
NG = 0
DB = 0
SM = 0
Delta_r
Doxycycline
YDR232W HEM1
ZShift = −0.09
NG = 0
DB = 0
SM = 0
Delta_AUC
Doxycycline
YEL055C POL5
ZShift = 0.32
NG = 0
DB = 0
SM = 0
Delta_L
Doxycycline
YEL055C POL5
ZShift = 0.21
NG = 0
DB = 0
SM = 0
Delta_K
Doxycycline
YEL055C POL5
ZShift = 0.58
NG = 0
DB = 0
SM = 0
Delta_r
Doxycycline
YEL055C POL5
ZShift = 0
NG = 0
DB = 0
SM = 0
Delta_AUC
Doxycycline
YIL147C SLN1
ZShift = 7.03
NG = 0
DB = 0
SM = 0
Delta_L
Doxycycline
YIL147C SLN1
ZShift = −7.07
NG = 0
DB = 0
SM = 0
Delta_K
Doxycycline
YIL147C SLN1
ZShift = −4.72
NG = 0
DB = 0
SM = 0
Delta_r
Doxycycline
YIL147C SLN1
ZShift = −6.94
NG = 0
DB = 0
SM = 0
Delta_AUC
Doxycycline

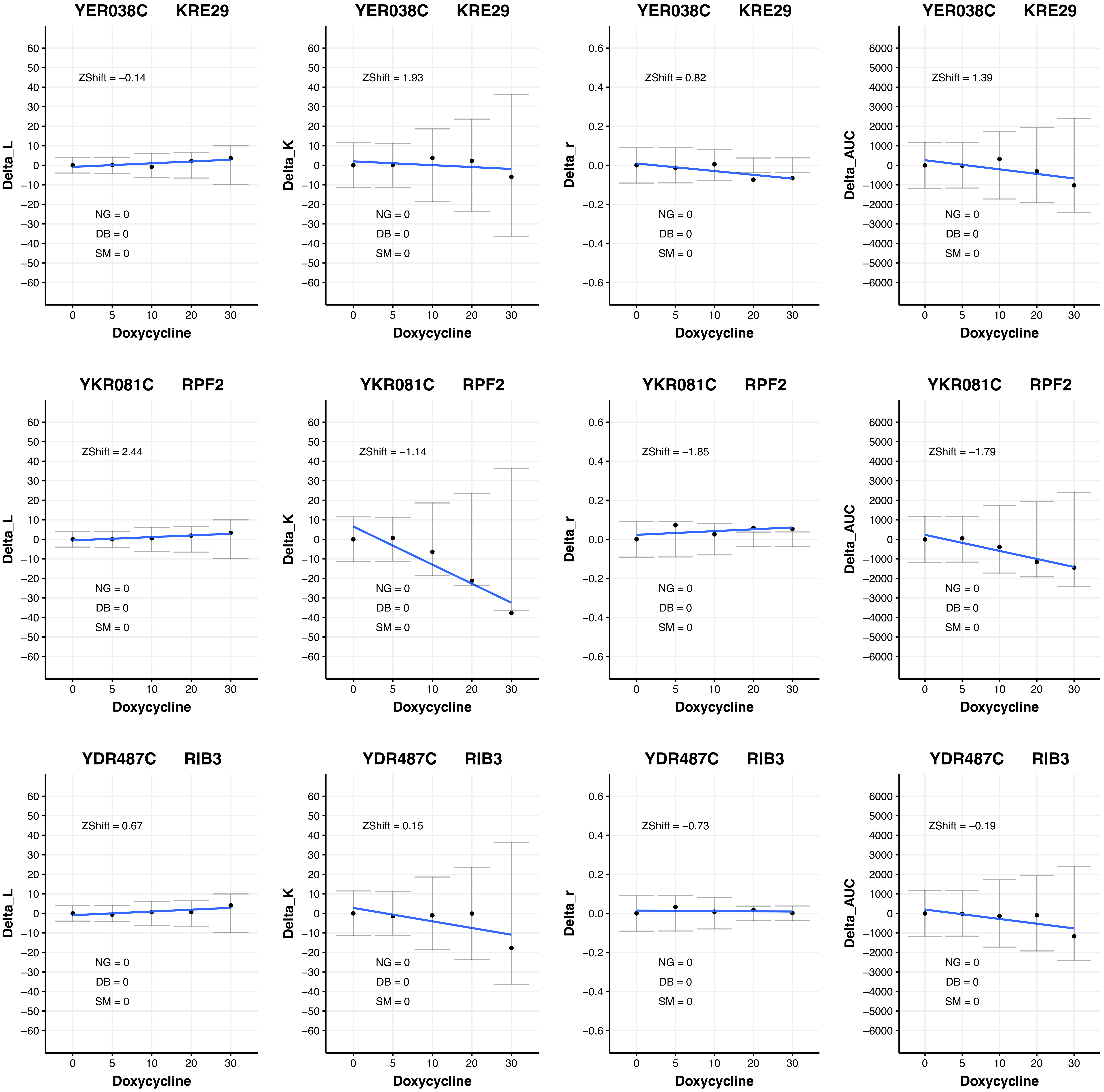
YER038C KRE29
ZShift = −0.14
NG = 0
DB = 0
SM = 0
Delta_L
60 50 40 30 20 10 0 −10 −20 −30 −40 −50 −60
0 5 10 20 30
Doxycycline
YER038C KRE29
ZShift = 1.93
NG = 0
DB = 0
SM = 0
Delta_K
60 50 40 30 20 10 0 −10 −20 −30 −40 −50 −60
0 5 10 20 30
Doxycycline
YER038C KRE29
ZShift = 0.82
NG = 0
DB = 0
SM = 0
Delta_r
0.6 0.4 0.2 0.0 −0.2 −0.4 −0.6
0 5 10 20 30
Doxycycline
YER038C KRE29
ZShift = 1.39
NG = 0
DB = 0
SM = 0
Delta_AUC
6000 5000 4000 3000 2000 1000 0 −1000 −2000 −3000 −4000 −5000 −6000
0 5 10 20 30
Doxycycline
YKR081C RPF2
ZShift = 2.44
NG = 0
DB = 0
SM = 0
Delta_L
60 50 40 30 20 10 0 −10 −20 −30 −40 −50 −60
0 5 10 20 30
Doxycycline
YKR081C RPF2
ZShift = −1.14
NG = 0
DB = 0
SM = 0
Delta_K
60 50 40 30 20 10 0 −10 −20 −30 −40 −50 −60
0 5 10 20 30
Doxycycline
YKR081C RPF2
ZShift = −1.85
NG = 0
DB = 0
SM = 0
Delta_r
0.6 0.4 0.2 0.0 −0.2 −0.4 −0.6
0 5 10 20 30
Doxycycline
YKR081C RPF2
ZShift = −1.79
NG = 0
DB = 0
SM = 0
Delta_AUC
6000 5000 4000 3000 2000 1000 0 −1000 −2000 −3000 −4000 −5000 −6000
0 5 10 20 30
Doxycycline
YDR487C RIB3
ZShift = 0.67
NG = 0
DB = 0
SM = 0
Delta_L
60 50 40 30 20 10 0 −10 −20 −30 −40 −50 −60
0 5 10 20 30
Doxycycline
YDR487C RIB3
ZShift = 0.15
NG = 0
DB = 0
SM = 0
Delta_K
60 50 40 30 20 10 0 −10 −20 −30 −40 −50 −60
0 5 10 20 30
Doxycycline
YDR487C RIB3
ZShift = −0.73
NG = 0
DB = 0
SM = 0
Delta_r
0.6 0.4 0.2 0.0 −0.2 −0.4 −0.6
0 5 10 20 30
Doxycycline
YDR487C RIB3
ZShift = −0.19
NG = 0
DB = 0
SM = 0
Delta_AUC
6000 5000 4000 3000 2000 1000 0 −1000 −2000 −3000 −4000 −5000 −6000
0 5 10 20 30
Doxycycline

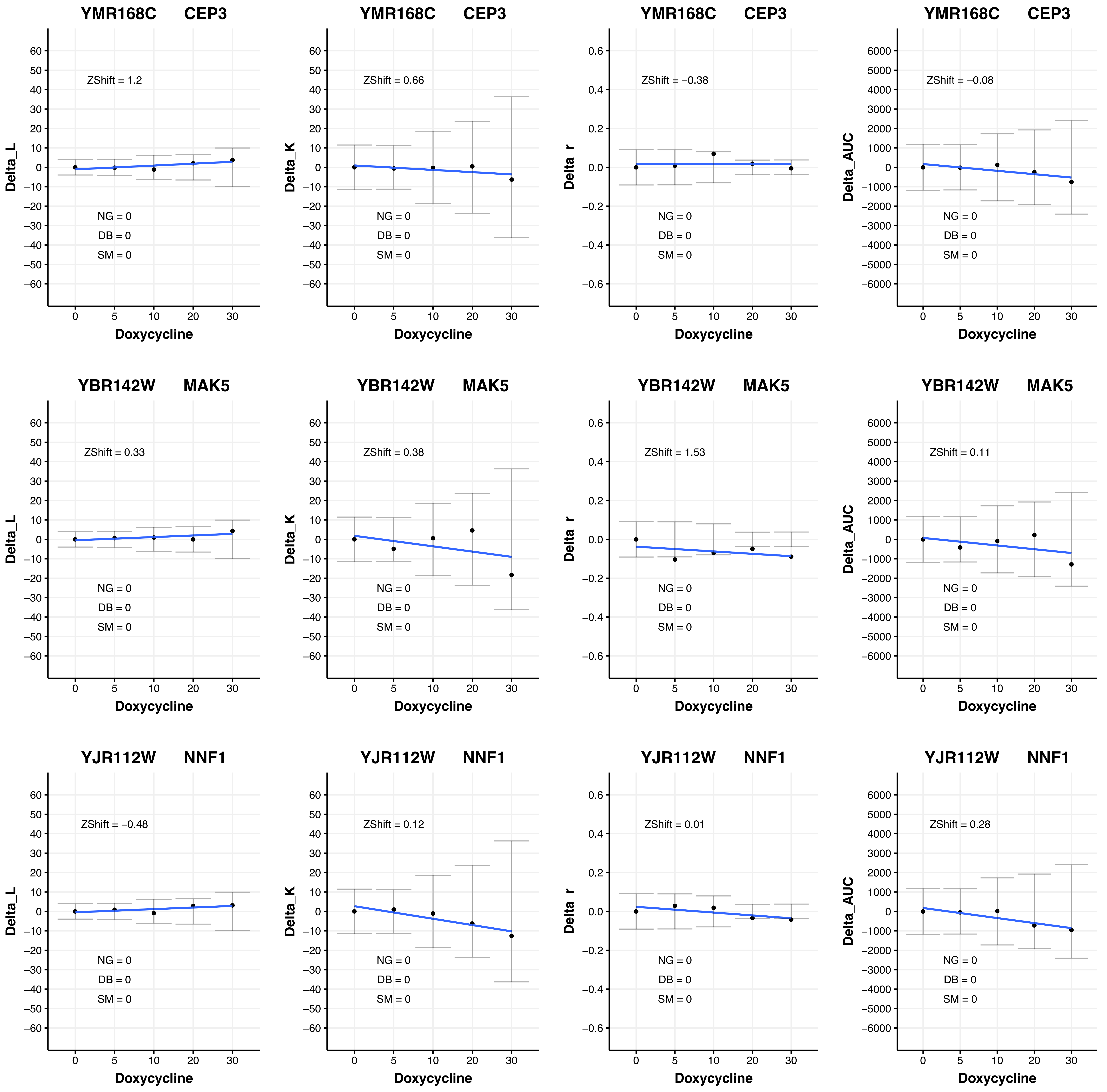

YMR168C CEP3
ZShift = 1.2
Delta_L
NG = 0
DB = 0
SM = 0
Doxycycline
YMR168C CEP3
ZShift = 0.66
Delta_K
NG = 0
DB = 0
SM = 0
Doxycycline
YMR168C CEP3
ZShift = −0.38
Delta_r
NG = 0
DB = 0
SM = 0
Doxycycline
YMR168C CEP3
ZShift = −0.08
Delta_AUC
NG = 0
DB = 0
SM = 0
Doxycycline
YBR142W MAK5
ZShift = 0.33
Delta_L
NG = 0
DB = 0
SM = 0
Doxycycline
YBR142W MAK5
ZShift = 0.38
Delta_K
NG = 0
DB = 0
SM = 0
Doxycycline
YBR142W MAK5
ZShift = 1.53
Delta_r
NG = 0
DB = 0
SM = 0
Doxycycline
YBR142W MAK5
ZShift = 0.11
Delta_AUC
NG = 0
DB = 0
SM = 0
Doxycycline
YJR112W NNF1
ZShift = −0.48
Delta_L
NG = 0
DB = 0
SM = 0
Doxycycline
YJR112W NNF1
ZShift = 0.12
Delta_K
NG = 0
DB = 0
SM = 0
Doxycycline
YJR112W NNF1
ZShift = 0.01
Delta_r
NG = 0
DB = 0
SM = 0
Doxycycline
YJR112W NNF1
ZShift = 0.28
Delta_AUC
NG = 0
DB = 0
SM = 0
Doxycycline

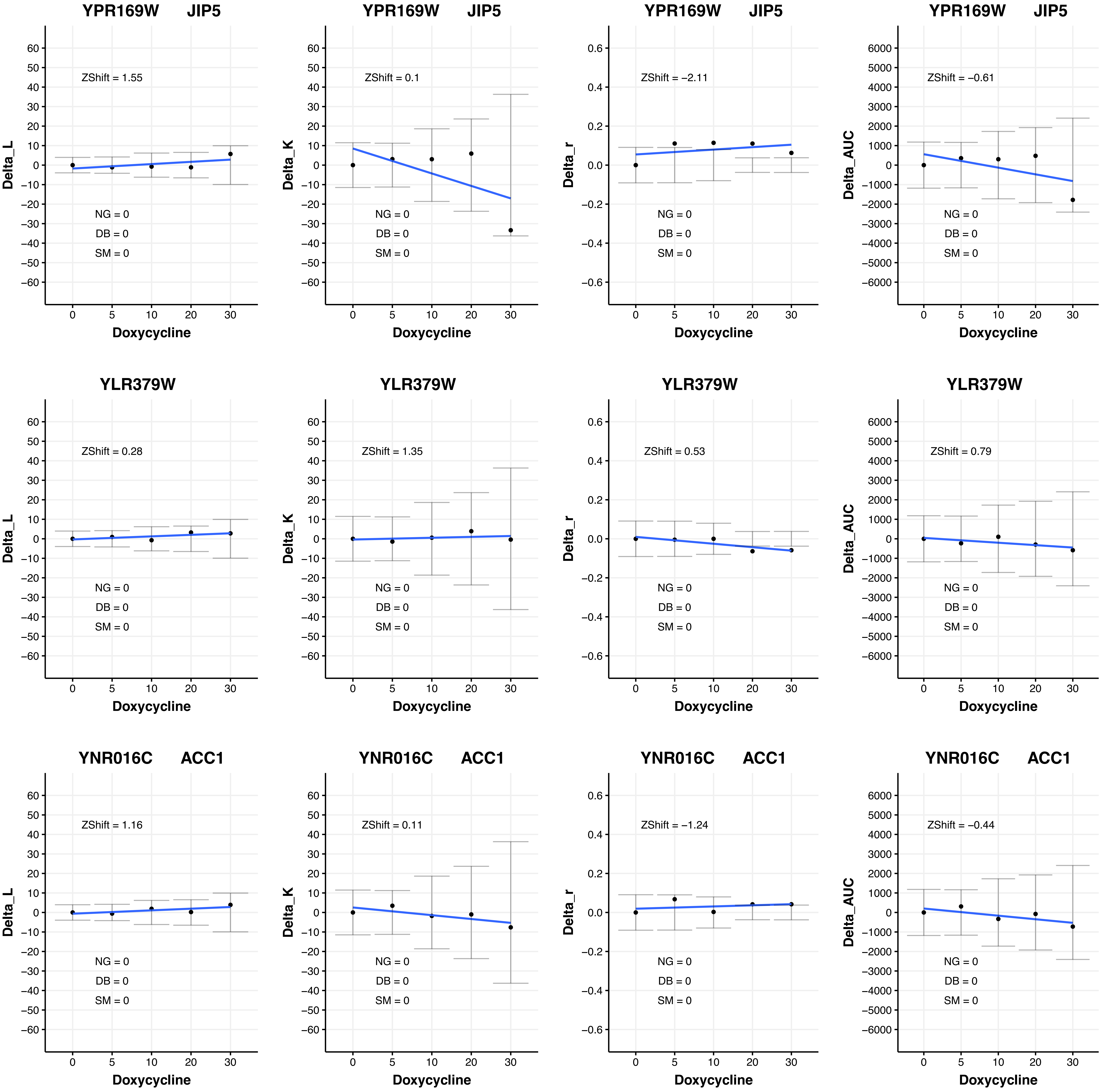
YPR169W JIP5
ZShift = 1.55
NG = 0
DB = 0
SM = 0
Delta_L
Doxycycline
YPR169W JIP5
ZShift = 0.1
NG = 0
DB = 0
SM = 0
Delta_K
Doxycycline
YPR169W JIP5
ZShift = −2.11
NG = 0
DB = 0
SM = 0
Delta_r
Doxycycline
YPR169W JIP5
ZShift = −0.61
NG = 0
DB = 0
SM = 0
Delta_AUC
Doxycycline
YLR379W
ZShift = 0.28
NG = 0
DB = 0
SM = 0
Delta_L
Doxycycline
YLR379W
ZShift = 1.35
NG = 0
DB = 0
SM = 0
Delta_K
Doxycycline
YLR379W
ZShift = 0.53
NG = 0
DB = 0
SM = 0
Delta_r
Doxycycline
YLR379W
ZShift = 0.79
NG = 0
DB = 0
SM = 0
Delta_AUC
Doxycycline
YNR016C ACC1
ZShift = 1.16
NG = 0
DB = 0
SM = 0
Delta_L
Doxycycline
YNR016C ACC1
ZShift = 0.11
NG = 0
DB = 0
SM = 0
Delta_K
Doxycycline
YNR016C ACC1
ZShift = −1.24
NG = 0
DB = 0
SM = 0
Delta_r
Doxycycline
YNR016C ACC1
ZShift = −0.44
NG = 0
DB = 0
SM = 0
Delta_AUC
Doxycycline

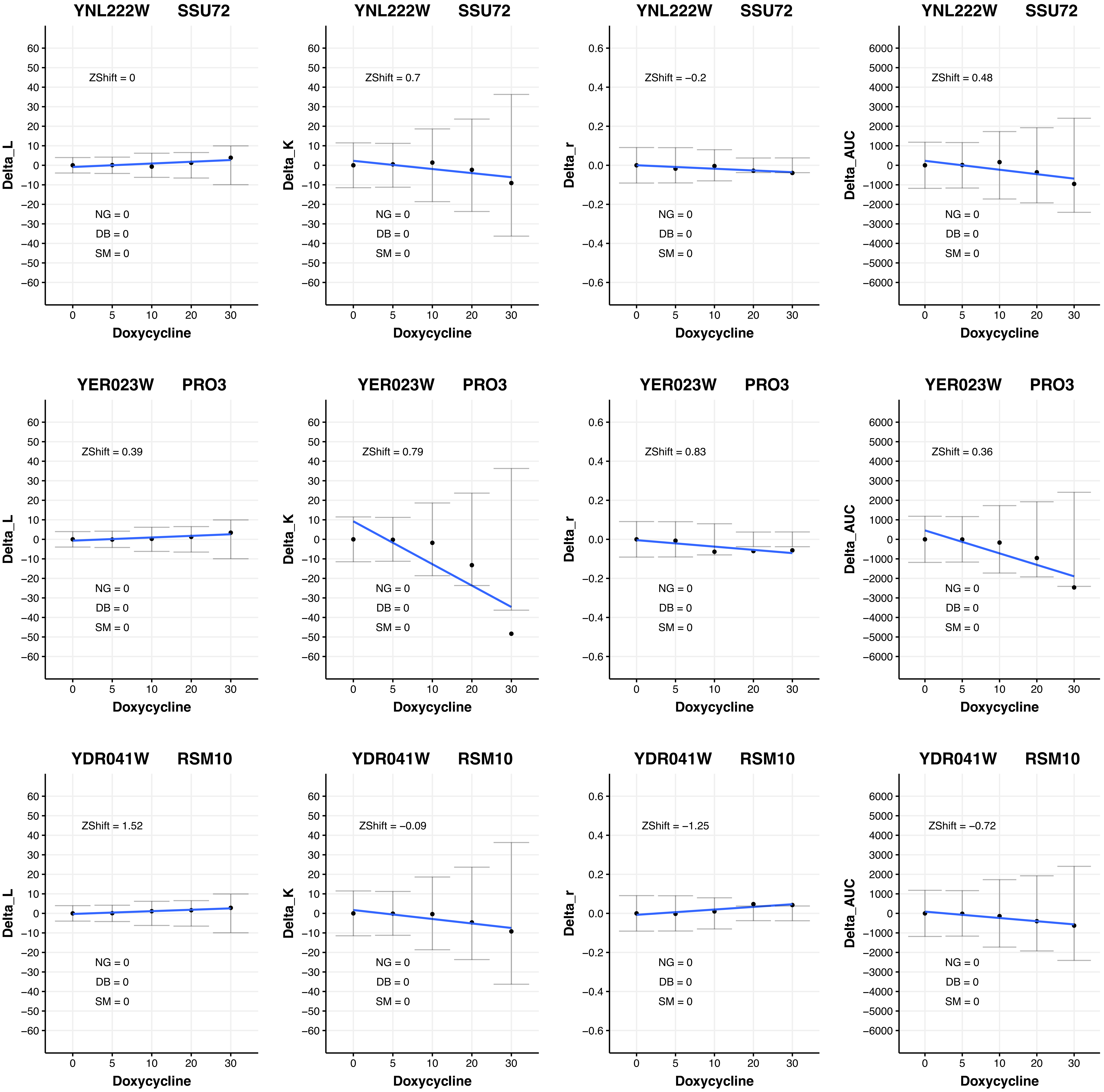

YNL222W SSU72
ZShift = 0
Delta_L
NG = 0
DB = 0
SM = 0
Doxycycline
YNL222W SSU72
ZShift = 0.7
Delta_K
NG = 0
DB = 0
SM = 0
Doxycycline
YNL222W SSU72
ZShift = −0.2
Delta_r
NG = 0
DB = 0
SM = 0
Doxycycline
YNL222W SSU72
ZShift = 0.48
Delta_AUC
NG = 0
DB = 0
SM = 0
Doxycycline
YER023W PRO3
ZShift = 0.39
Delta_L
NG = 0
DB = 0
SM = 0
Doxycycline
YER023W PRO3
ZShift = 0.79
Delta_K
NG = 0
DB = 0
SM = 0
Doxycycline
YER023W PRO3
ZShift = 0.83
Delta_r
NG = 0
DB = 0
SM = 0
Doxycycline
YER023W PRO3
ZShift = 0.36
Delta_AUC
NG = 0
DB = 0
SM = 0
Doxycycline
YDR041W RSM10
ZShift = 1.52
Delta_L
NG = 0
DB = 0
SM = 0
Doxycycline
YDR041W RSM10
ZShift = −0.09
Delta_K
NG = 0
DB = 0
SM = 0
Doxycycline
YDR041W RSM10
ZShift = −1.25
Delta_r
NG = 0
DB = 0
SM = 0
Doxycycline
YDR041W RSM10
ZShift = −0.72
Delta_AUC
NG = 0
DB = 0
SM = 0
Doxycycline

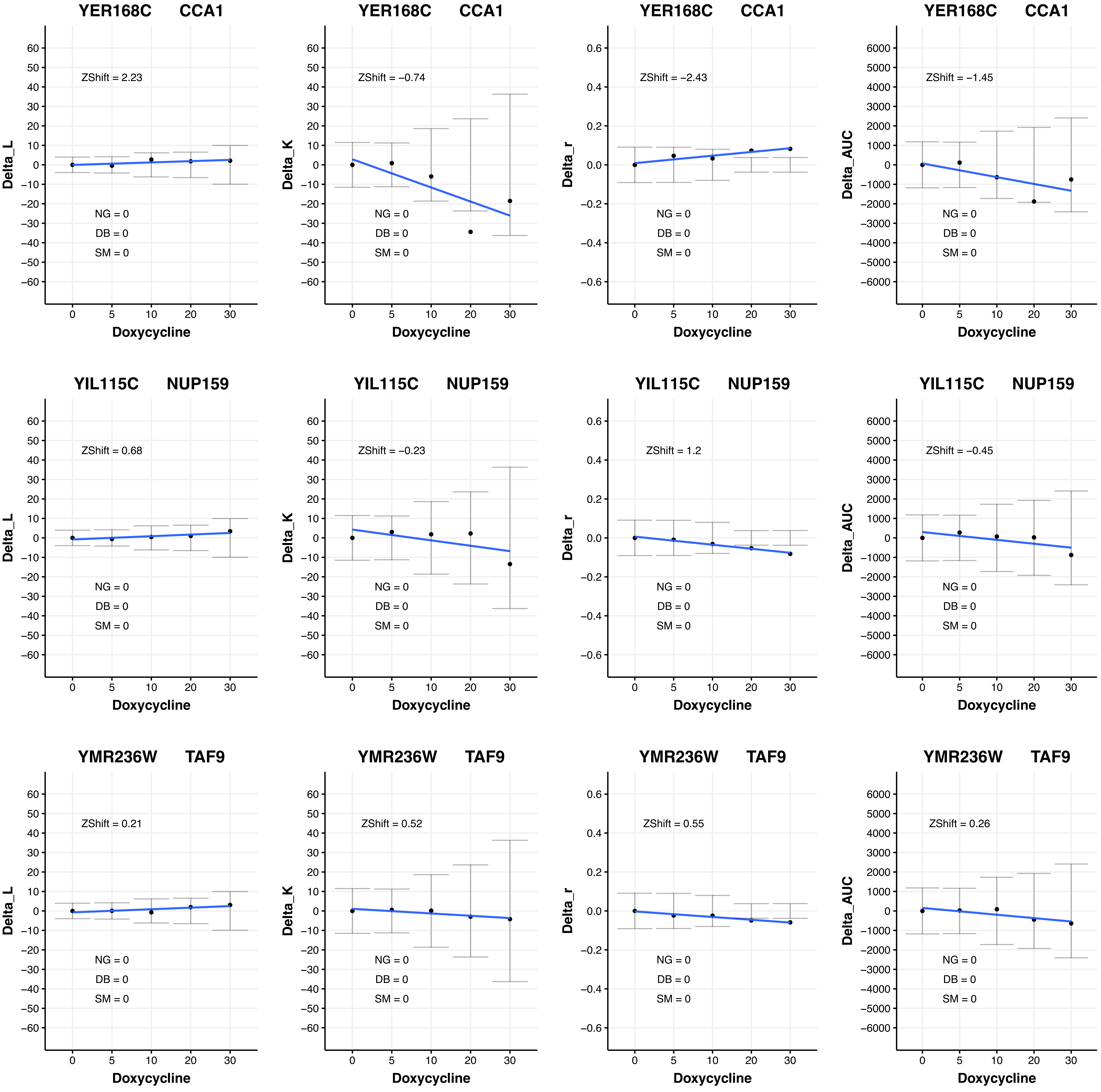
YER168C CCA1
ZShift = 2.23
NG = 0
DB = 0
SM = 0
Delta_L
Doxycycline
YER168C CCA1
ZShift = −0.74
NG = 0
DB = 0
SM = 0
Delta_K
Doxycycline
YER168C CCA1
ZShift = −2.43
NG = 0
DB = 0
SM = 0
Delta_r
Doxycycline
YER168C CCA1
ZShift = −1.45
NG = 0
DB = 0
SM = 0
Delta_AUC
Doxycycline
YIL115C NUP159
ZShift = 0.68
NG = 0
DB = 0
SM = 0
Delta_L
Doxycycline
YIL115C NUP159
ZShift = −0.23
NG = 0
DB = 0
SM = 0
Delta_K
Doxycycline
YIL115C NUP159
ZShift = 1.2
NG = 0
DB = 0
SM = 0
Delta_r
Doxycycline
YIL115C NUP159
ZShift = −0.45
NG = 0
DB = 0
SM = 0
Delta_AUC
Doxycycline
YMR236W TAF9
ZShift = 0.21
NG = 0
DB = 0
SM = 0
Delta_L
Doxycycline
YMR236W TAF9
ZShift = 0.52
NG = 0
DB = 0
SM = 0
Delta_K
Doxycycline
YMR236W TAF9
ZShift = 0.55
NG = 0
DB = 0
SM = 0
Delta_r
Doxycycline
YMR236W TAF9
ZShift = 0.26
NG = 0
DB = 0
SM = 0
Delta_AUC
Doxycycline

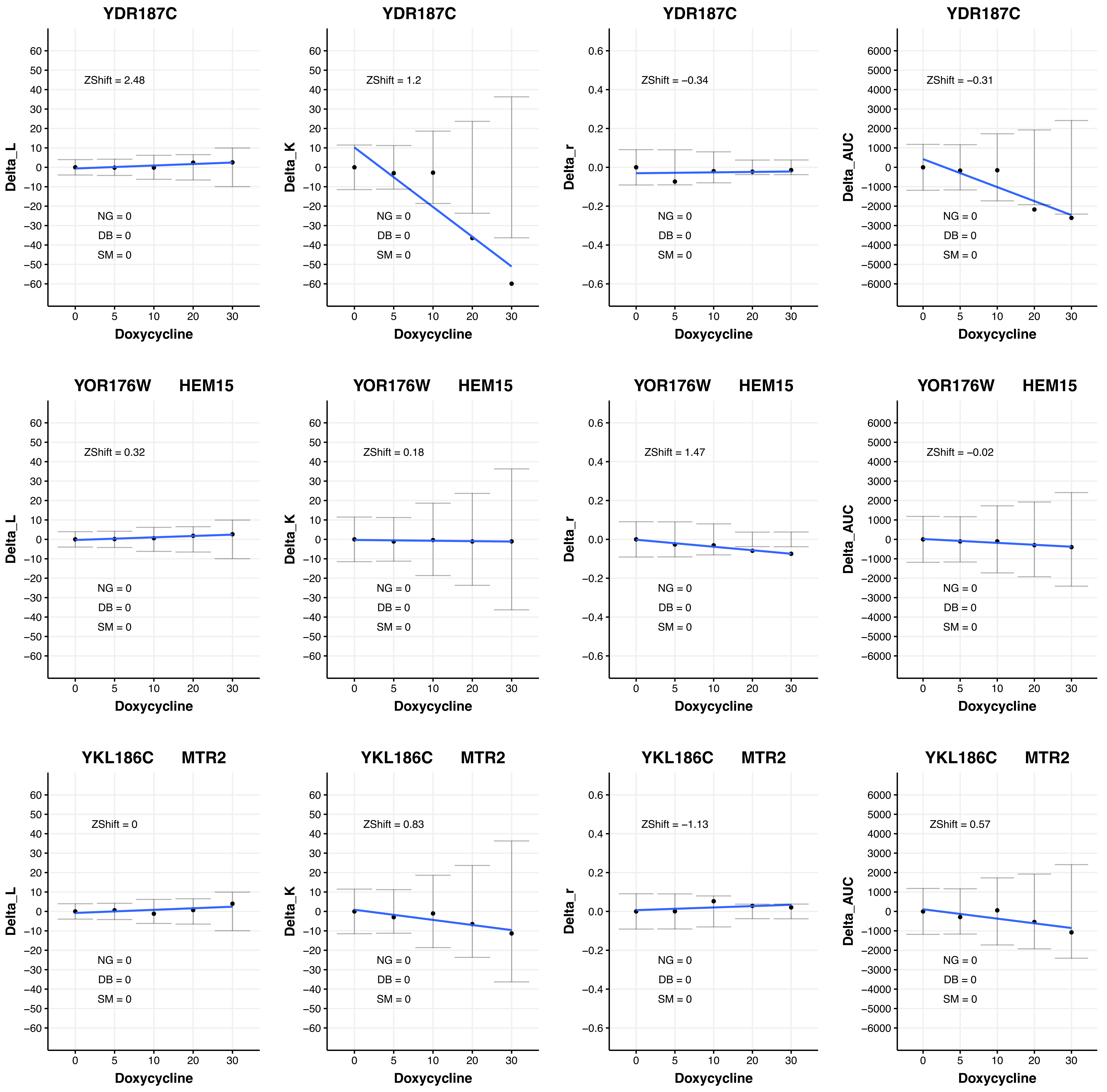
YDR187C
ZShift = 2.48
NG = 0
DB = 0
SM = 0
Delta_L
Doxycycline
YDR187C
ZShift = 1.2
NG = 0
DB = 0
SM = 0
Delta_K
Doxycycline
YDR187C
ZShift = −0.34
NG = 0
DB = 0
SM = 0
Delta_r
Doxycycline
YDR187C
ZShift = −0.31
NG = 0
DB = 0
SM = 0
Delta_AUC
Doxycycline
YOR176W HEM15
ZShift = 0.32
NG = 0
DB = 0
SM = 0
Delta_L
Doxycycline
YOR176W HEM15
ZShift = 0.18
NG = 0
DB = 0
SM = 0
Delta_K
Doxycycline
YOR176W HEM15
ZShift = 1.47
NG = 0
DB = 0
SM = 0
Delta_r
Doxycycline
YOR176W HEM15
ZShift = −0.02
NG = 0
DB = 0
SM = 0
Delta_AUC
Doxycycline
YKL186C MTR2
ZShift = 0
NG = 0
DB = 0
SM = 0
Delta_L
Doxycycline
YKL186C MTR2
ZShift = 0.83
NG = 0
DB = 0
SM = 0
Delta_K
Doxycycline
YKL186C MTR2
ZShift = −1.13
NG = 0
DB = 0
SM = 0
Delta_r
Doxycycline
YKL186C MTR2
ZShift = 0.57
NG = 0
DB = 0
SM = 0
Delta_AUC
Doxycycline

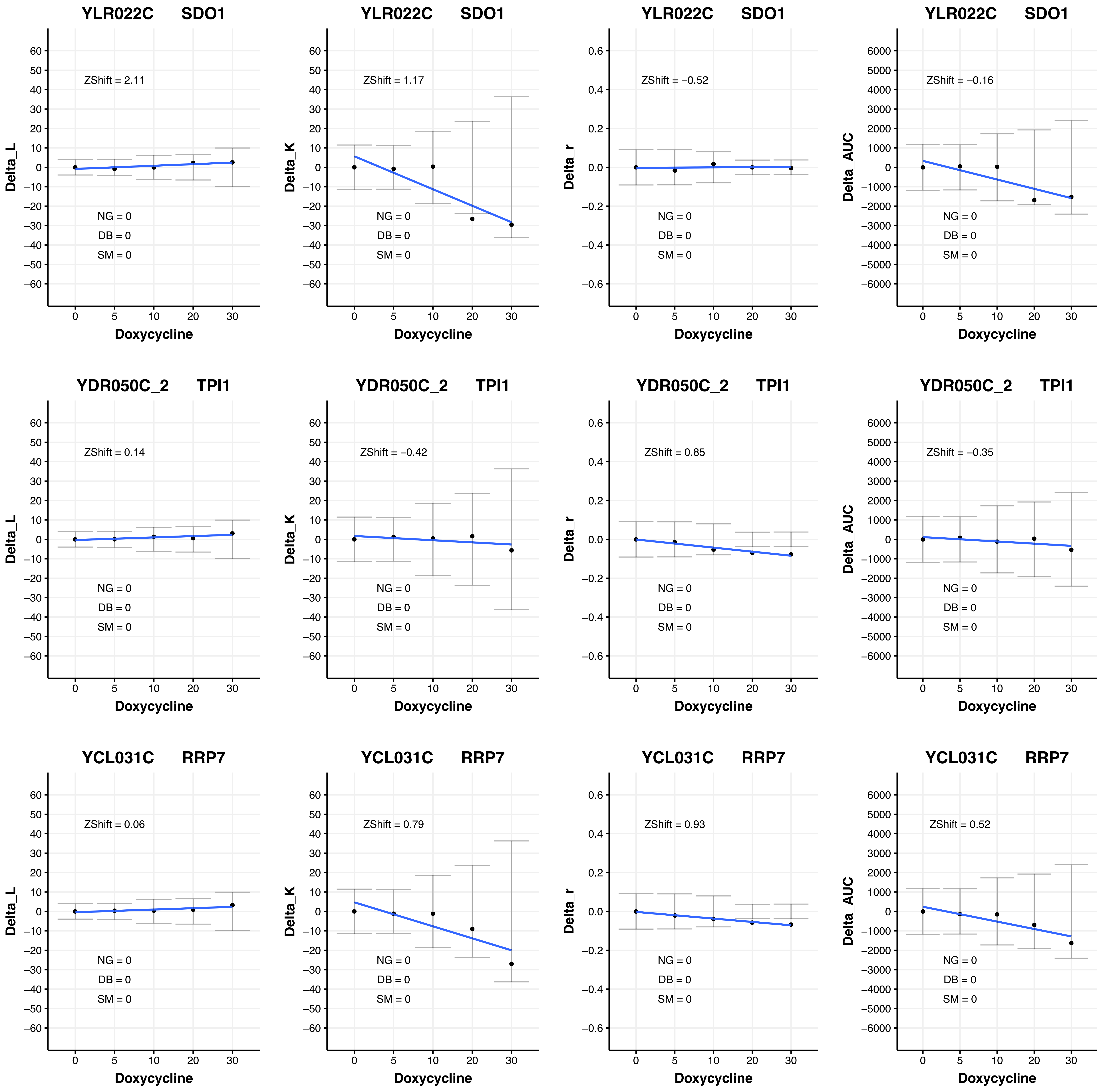

YLR022C SDO1
ZShift = 2.11
NG = 0
DB = 0
SM = 0
Delta_L
Doxycycline
YLR022C SDO1
ZShift = 1.17
NG = 0
DB = 0
SM = 0
Delta_K
Doxycycline
YLR022C SDO1
ZShift = −0.52
NG = 0
DB = 0
SM = 0
Delta_r
Doxycycline
YLR022C SDO1
ZShift = −0.16
NG = 0
DB = 0
SM = 0
Delta_AUC
Doxycycline
YDR050C_2 TPI1
ZShift = 0.14
NG = 0
DB = 0
SM = 0
Delta_L
Doxycycline
YDR050C_2 TPI1
ZShift = −0.42
NG = 0
DB = 0
SM = 0
Delta_K
Doxycycline
YDR050C_2 TPI1
ZShift = 0.85
NG = 0
DB = 0
SM = 0
Delta_r
Doxycycline
YDR050C_2 TPI1
ZShift = −0.35
NG = 0
DB = 0
SM = 0
Delta_AUC
Doxycycline
YCL031C RRP7
ZShift = 0.06
NG = 0
DB = 0
SM = 0
Delta_L
Doxycycline
YCL031C RRP7
ZShift = 0.79
NG = 0
DB = 0
SM = 0
Delta_K
Doxycycline
YCL031C RRP7
ZShift = 0.93
NG = 0
DB = 0
SM = 0
Delta_r
Doxycycline
YCL031C RRP7
ZShift = 0.52
NG = 0
DB = 0
SM = 0
Delta_AUC
Doxycycline

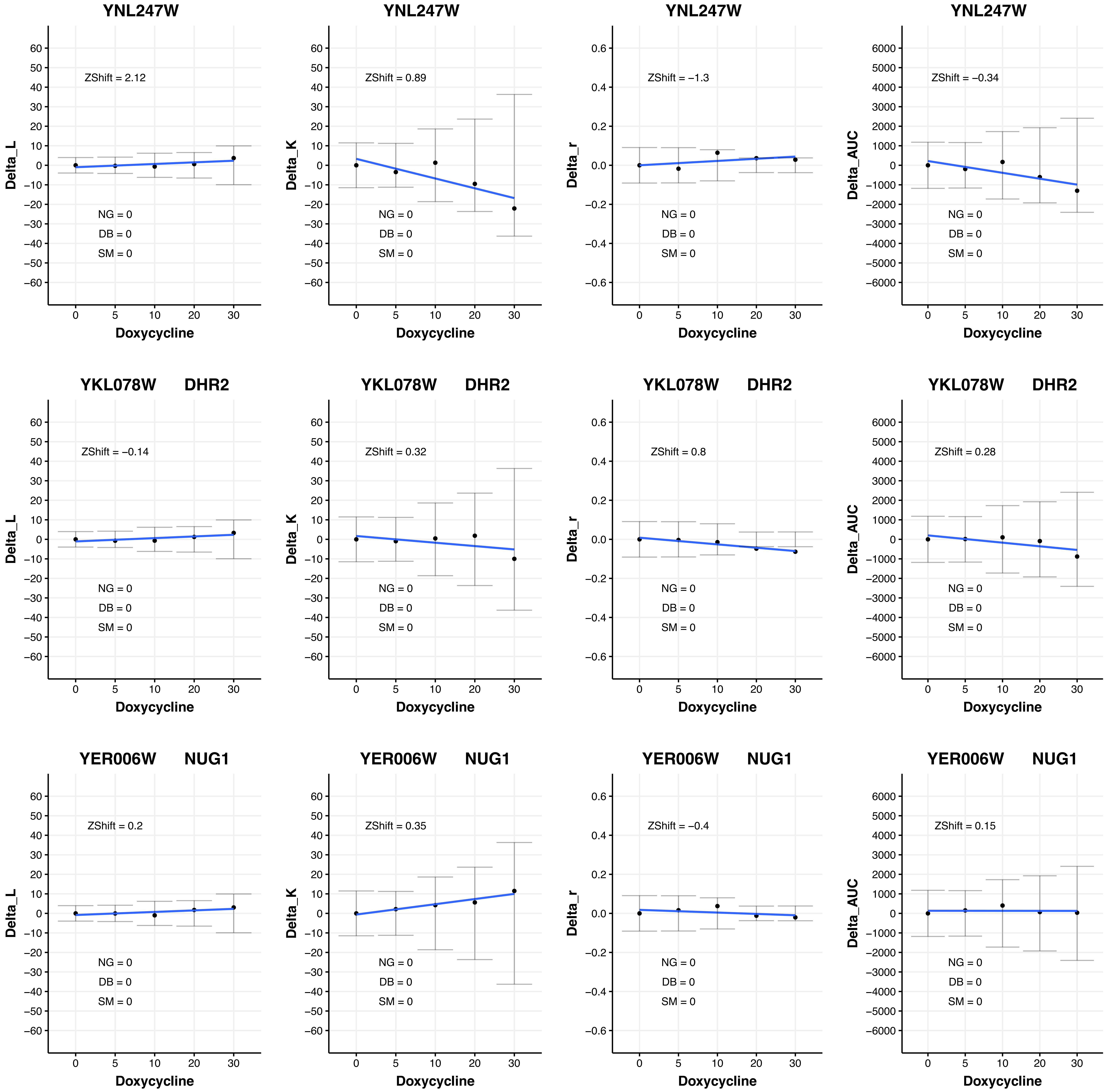

YNL247W
ZShift = 2.12
NG = 0
DB = 0
SM = 0
Delta_L
Doxycycline
YNL247W
ZShift = 0.89
NG = 0
DB = 0
SM = 0
Delta_K
Doxycycline
YNL247W
ZShift = −1.3
NG = 0
DB = 0
SM = 0
Delta_r
Doxycycline
YNL247W
ZShift = −0.34
NG = 0
DB = 0
SM = 0
Delta_AUC
Doxycycline
YKL078W DHR2
ZShift = −0.14
NG = 0
DB = 0
SM = 0
Delta_L
Doxycycline
YKL078W DHR2
ZShift = 0.32
NG = 0
DB = 0
SM = 0
Delta_K
Doxycycline
YKL078W DHR2
ZShift = 0.8
NG = 0
DB = 0
SM = 0
Delta_r
Doxycycline
YKL078W DHR2
ZShift = 0.28
NG = 0
DB = 0
SM = 0
Delta_AUC
Doxycycline
YER006W NUG1
ZShift = 0.2
NG = 0
DB = 0
SM = 0
Delta_L
Doxycycline
YER006W NUG1
ZShift = 0.35
NG = 0
DB = 0
SM = 0
Delta_K
Doxycycline
YER006W NUG1
ZShift = −0.4
NG = 0
DB = 0
SM = 0
Delta_r
Doxycycline
YER006W NUG1
ZShift = 0.15
NG = 0
DB = 0
SM = 0
Delta_AUC
Doxycycline

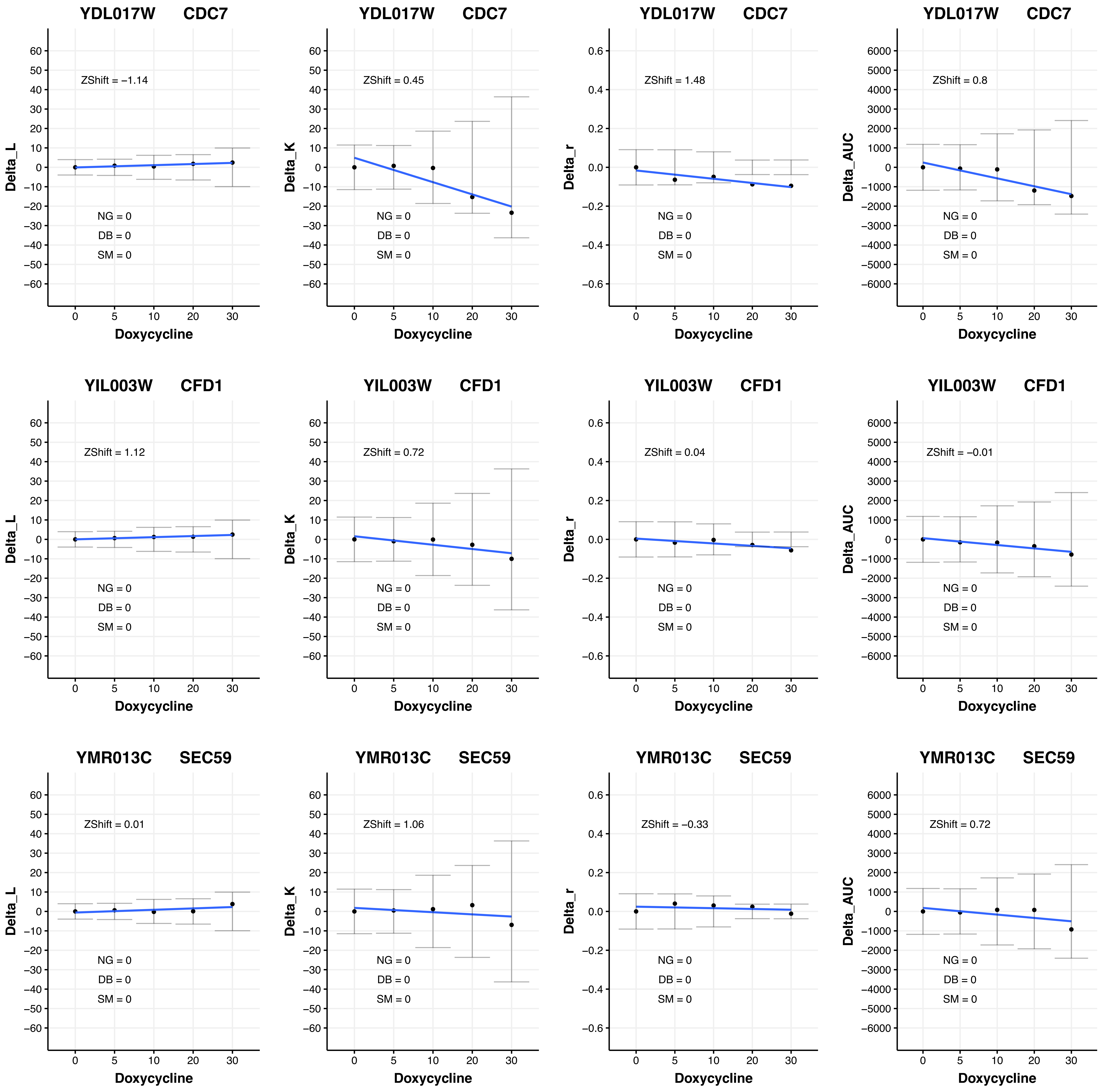
YDL017W CDC7
ZShift = −1.14
NG = 0
DB = 0
SM = 0
Delta_L
Doxycycline
YDL017W CDC7
ZShift = 0.45
NG = 0
DB = 0
SM = 0
Delta_K
Doxycycline
YDL017W CDC7
ZShift = 1.48
NG = 0
DB = 0
SM = 0
Delta_r
Doxycycline
YDL017W CDC7
ZShift = 0.8
NG = 0
DB = 0
SM = 0
Delta_AUC
Doxycycline
YIL003W CFD1
ZShift = 1.12
NG = 0
DB = 0
SM = 0
Delta_L
Doxycycline
YIL003W CFD1
ZShift = 0.72
NG = 0
DB = 0
SM = 0
Delta_K
Doxycycline
YIL003W CFD1
ZShift = 0.04
NG = 0
DB = 0
SM = 0
Delta_r
Doxycycline
YIL003W CFD1
ZShift = −0.01
NG = 0
DB = 0
SM = 0
Delta_AUC
Doxycycline
YMR013C SEC59
ZShift = 0.01
NG = 0
DB = 0
SM = 0
Delta_L
Doxycycline
YMR013C SEC59
ZShift = 1.06
NG = 0
DB = 0
SM = 0
Delta_K
Doxycycline
YMR013C SEC59
ZShift = −0.33
NG = 0
DB = 0
SM = 0
Delta_r
Doxycycline
YMR013C SEC59
ZShift = 0.72
NG = 0
DB = 0
SM = 0
Delta_AUC
Doxycycline

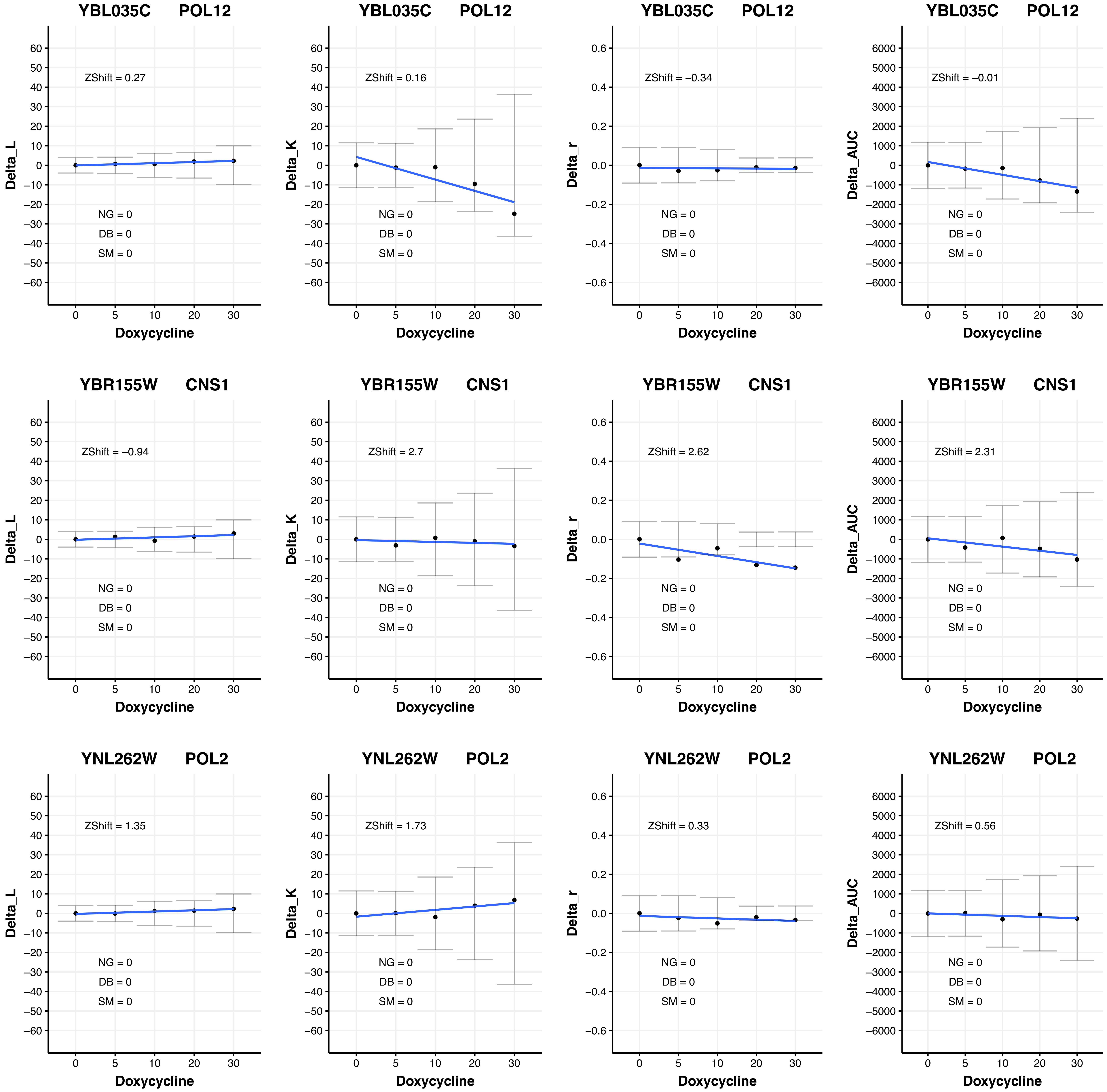
YBL035C POL12
ZShift = 0.27
NG = 0
DB = 0
SM = 0
Delta_L
YBL035C POL12
ZShift = 0.16
NG = 0
DB = 0
SM = 0
Delta_K
YBL035C POL12
ZShift = −0.34
NG = 0
DB = 0
SM = 0
Delta_r
YBL035C POL12
ZShift = −0.01
NG = 0
DB = 0
SM = 0
Delta_AUC
YBR155W CNS1
ZShift = −0.94
NG = 0
DB = 0
SM = 0
Delta_L
YBR155W CNS1
ZShift = 2.7
NG = 0
DB = 0
SM = 0
Delta_K
YBR155W CNS1
ZShift = 2.62
NG = 0
DB = 0
SM = 0
Delta_r
YBR155W CNS1
ZShift = 2.31
NG = 0
DB = 0
SM = 0
Delta_AUC
YNL262W POL2
ZShift = 1.35
NG = 0
DB = 0
SM = 0
Delta_L
YNL262W POL2
ZShift = 1.73
NG = 0
DB = 0
SM = 0
Delta_K
YNL262W POL2
ZShift = 0.33
NG = 0
DB = 0
SM = 0
Delta_r
YNL262W POL2
ZShift = 0.56
NG = 0
DB = 0
SM = 0
Delta_AUC
Doxycycline
0
5
10
20
30

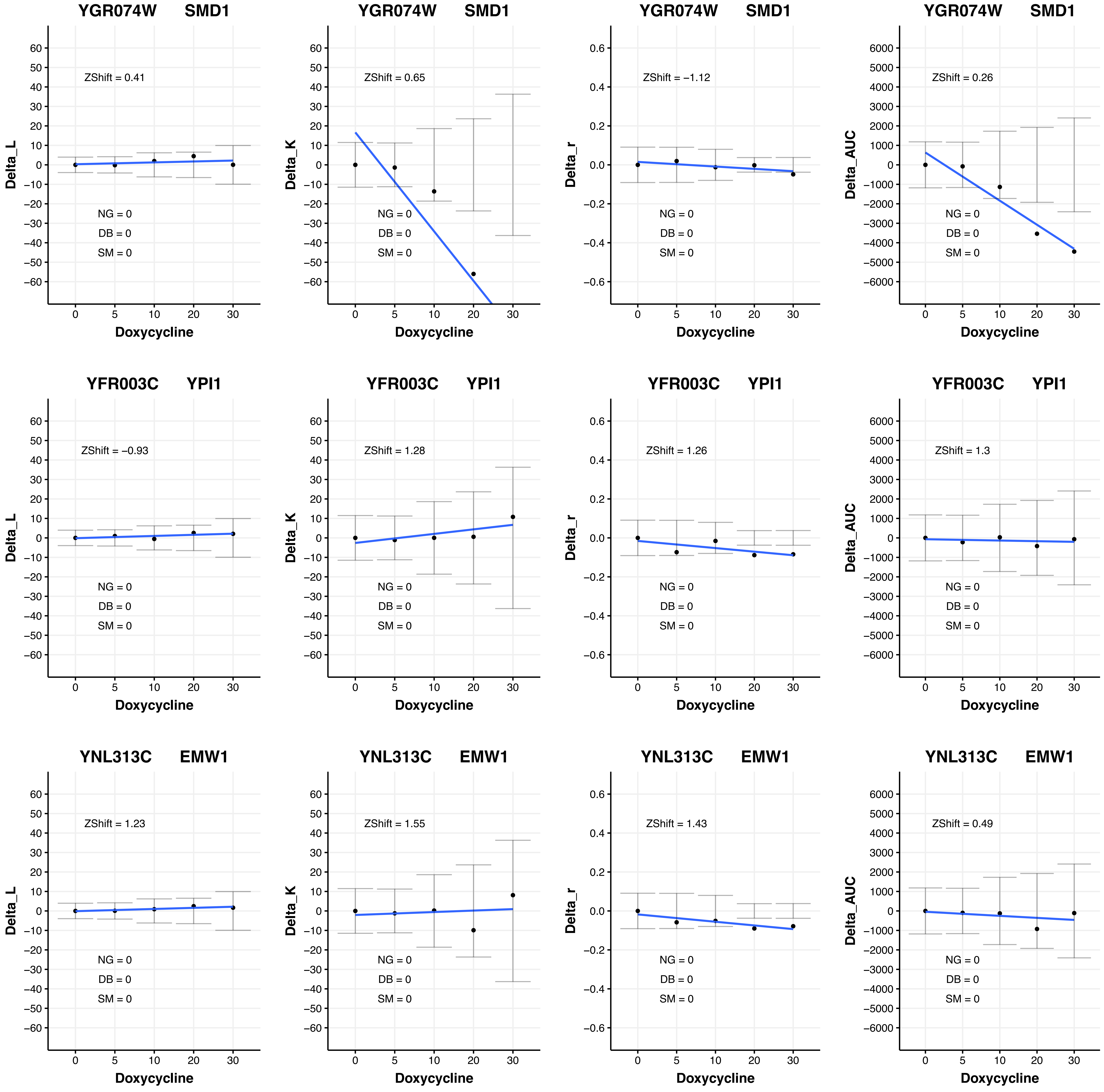
YGR074W SMD1
ZShift = 0.41
NG = 0
DB = 0
SM = 0
Delta_L
Doxycycline
YGR074W SMD1
ZShift = 0.65
NG = 0
DB = 0
SM = 0
Delta_K
Doxycycline
YGR074W SMD1
ZShift = −1.12
NG = 0
DB = 0
SM = 0
Delta_r
Doxycycline
YGR074W SMD1
ZShift = 0.26
NG = 0
DB = 0
SM = 0
Delta_AUC
Doxycycline
YFR003C YPI1
ZShift = −0.93
NG = 0
DB = 0
SM = 0
Delta_L
Doxycycline
YFR003C YPI1
ZShift = 1.28
NG = 0
DB = 0
SM = 0
Delta_K
Doxycycline
YFR003C YPI1
ZShift = 1.26
NG = 0
DB = 0
SM = 0
Delta_r
Doxycycline
YFR003C YPI1
ZShift = 1.3
NG = 0
DB = 0
SM = 0
Delta_AUC
Doxycycline
YNL313C EMW1
ZShift = 1.23
NG = 0
DB = 0
SM = 0
Delta_L
Doxycycline
YNL313C EMW1
ZShift = 1.55
NG = 0
DB = 0
SM = 0
Delta_K
Doxycycline
YNL313C EMW1
ZShift = 1.43
NG = 0
DB = 0
SM = 0
Delta_r
Doxycycline
YNL313C EMW1
ZShift = 0.49
NG = 0
DB = 0
SM = 0
Delta_AUC
Doxycycline

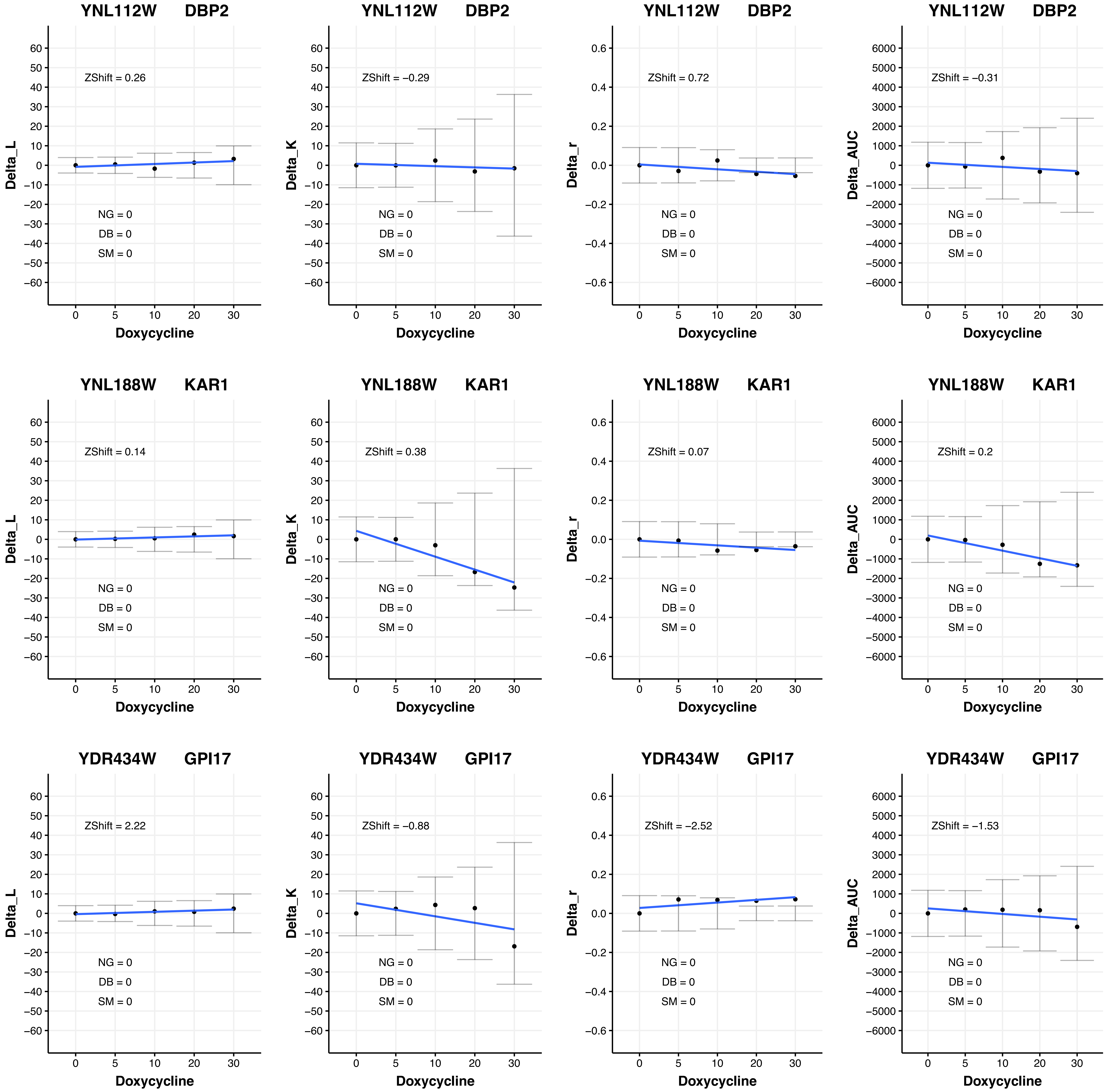
YNL112W DBP2
ZShift = 0.26
NG = 0
DB = 0
SM = 0
Delta_L
Doxycycline
YNL112W DBP2
ZShift = −0.29
NG = 0
DB = 0
SM = 0
Delta_K
Doxycycline
YNL112W DBP2
ZShift = 0.72
NG = 0
DB = 0
SM = 0
Delta_r
Doxycycline
YNL112W DBP2
ZShift = −0.31
NG = 0
DB = 0
SM = 0
Delta_AUC
Doxycycline
YNL188W KAR1
ZShift = 0.14
NG = 0
DB = 0
SM = 0
Delta_L
Doxycycline
YNL188W KAR1
ZShift = 0.38
NG = 0
DB = 0
SM = 0
Delta_K
Doxycycline
YNL188W KAR1
ZShift = 0.07
NG = 0
DB = 0
SM = 0
Delta_r
Doxycycline
YNL188W KAR1
ZShift = 0.2
NG = 0
DB = 0
SM = 0
Delta_AUC
Doxycycline
YDR434W GPI17
ZShift = 2.22
NG = 0
DB = 0
SM = 0
Delta_L
Doxycycline
YDR434W GPI17
ZShift = −0.88
NG = 0
DB = 0
SM = 0
Delta_K
Doxycycline
YDR434W GPI17
ZShift = −2.52
NG = 0
DB = 0
SM = 0
Delta_r
Doxycycline
YDR434W GPI17
ZShift = −1.53
NG = 0
DB = 0
SM = 0
Delta_AUC
Doxycycline

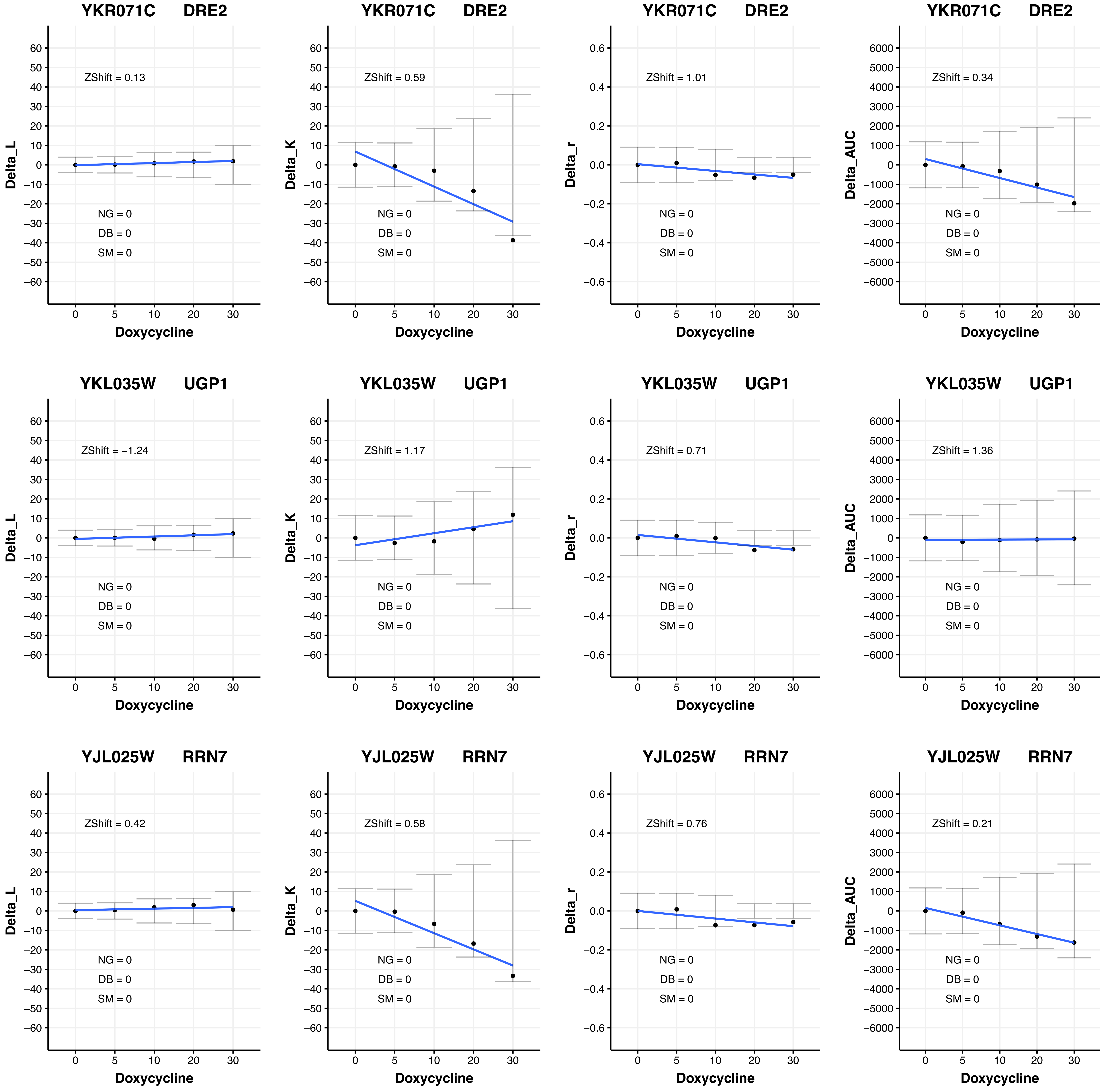
YKR071C DRE2
ZShift = 0.13
NG = 0
DB = 0
SM = 0
Delta_L
Doxycycline
YKR071C DRE2
ZShift = 0.59
NG = 0
DB = 0
SM = 0
Delta_K
Doxycycline
YKR071C DRE2
ZShift = 1.01
NG = 0
DB = 0
SM = 0
Delta_r
Doxycycline
YKR071C DRE2
ZShift = 0.34
NG = 0
DB = 0
SM = 0
Delta_AUC
Doxycycline
YKL035W UGP1
ZShift = −1.24
NG = 0
DB = 0
SM = 0
Delta_L
Doxycycline
YKL035W UGP1
ZShift = 1.17
NG = 0
DB = 0
SM = 0
Delta_K
Doxycycline
YKL035W UGP1
ZShift = 0.71
NG = 0
DB = 0
SM = 0
Delta_r
Doxycycline
YKL035W UGP1
ZShift = 1.36
NG = 0
DB = 0
SM = 0
Delta_AUC
Doxycycline
YJL025W RRN7
ZShift = 0.42
NG = 0
DB = 0
SM = 0
Delta_L
Doxycycline
YJL025W RRN7
ZShift = 0.58
NG = 0
DB = 0
SM = 0
Delta_K
Doxycycline
YJL025W RRN7
ZShift = 0.76
NG = 0
DB = 0
SM = 0
Delta_r
Doxycycline
YJL025W RRN7
ZShift = 0.21
NG = 0
DB = 0
SM = 0
Delta_AUC
Doxycycline

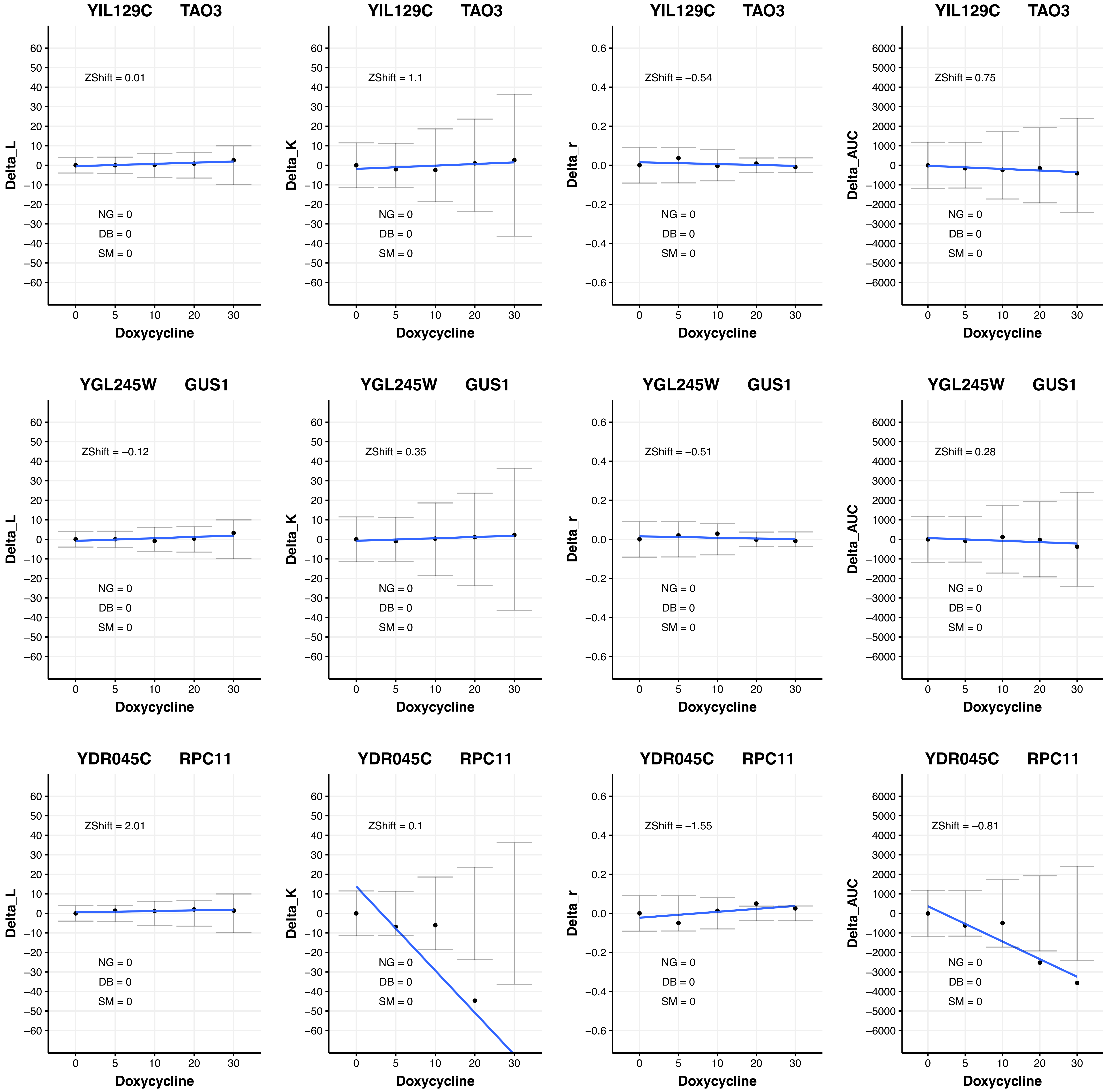
YIL129C TAO3
ZShift = 0.01
NG = 0
DB = 0
SM = 0
Delta_L
Doxycycline
YIL129C TAO3
ZShift = 1.1
NG = 0
DB = 0
SM = 0
Delta_K
Doxycycline
YIL129C TAO3
ZShift = −0.54
NG = 0
DB = 0
SM = 0
Delta_r
Doxycycline
YIL129C TAO3
ZShift = 0.75
NG = 0
DB = 0
SM = 0
Delta_AUC
Doxycycline
YGL245W GUS1
ZShift = −0.12
NG = 0
DB = 0
SM = 0
Delta_L
Doxycycline
YGL245W GUS1
ZShift = 0.35
NG = 0
DB = 0
SM = 0
Delta_K
Doxycycline
YGL245W GUS1
ZShift = −0.51
NG = 0
DB = 0
SM = 0
Delta_r
Doxycycline
YGL245W GUS1
ZShift = 0.28
NG = 0
DB = 0
SM = 0
Delta_AUC
Doxycycline
YDR045C RPC11
ZShift = 2.01
NG = 0
DB = 0
SM = 0
Delta_L
Doxycycline
YDR045C RPC11
ZShift = 0.1
NG = 0
DB = 0
SM = 0
Delta_K
Doxycycline
YDR045C RPC11
ZShift = −1.55
NG = 0
DB = 0
SM = 0
Delta_r
Doxycycline
YDR045C RPC11
ZShift = −0.81
NG = 0
DB = 0
SM = 0
Delta_AUC
Doxycycline

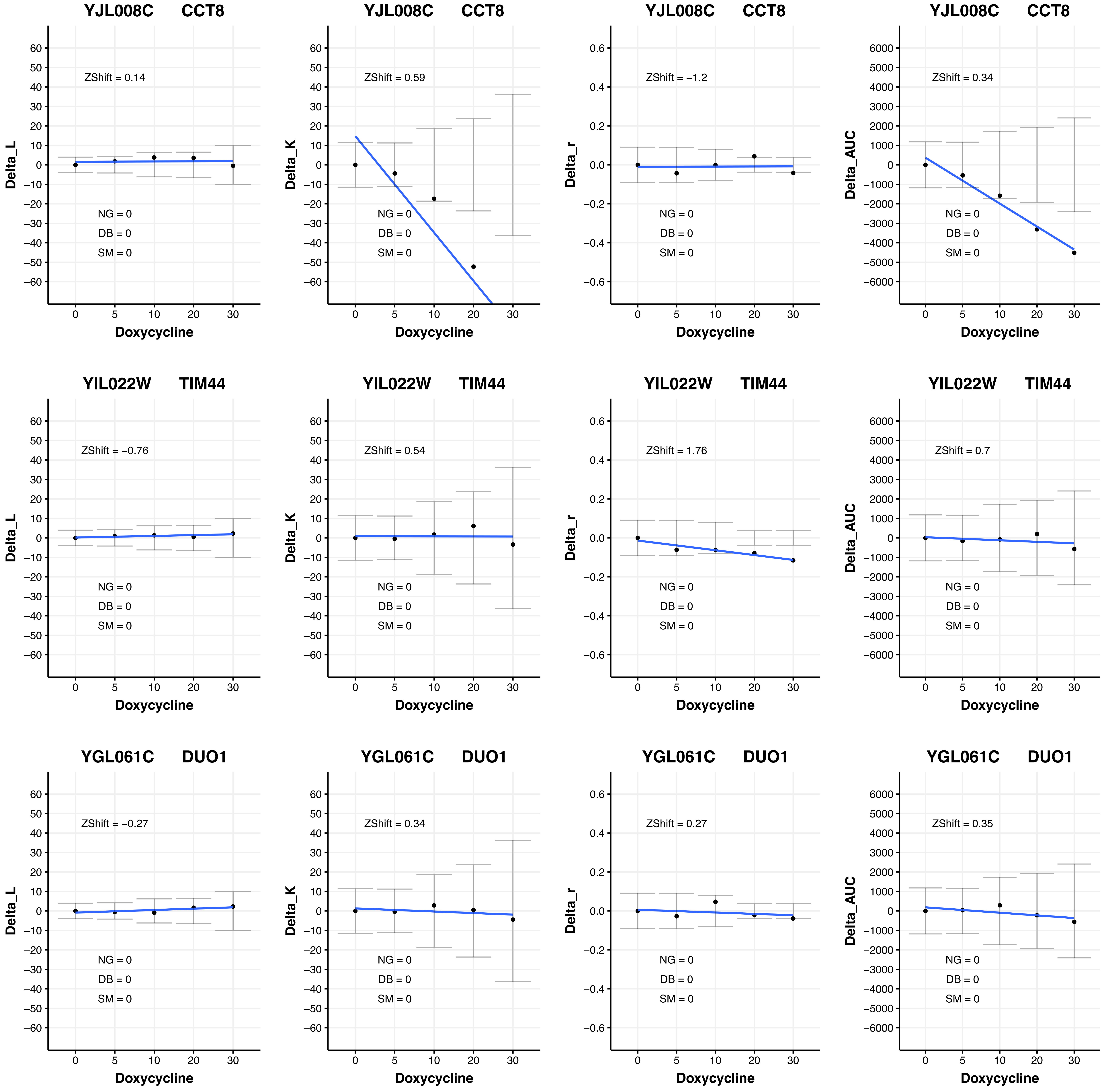
YJL008C CCT8
ZShift = 0.14
NG = 0
DB = 0
SM = 0
Delta_L
Doxycycline
YJL008C CCT8
ZShift = 0.59
NG = 0
DB = 0
SM = 0
Delta_K
Doxycycline
YJL008C CCT8
ZShift = −1.2
NG = 0
DB = 0
SM = 0
Delta_r
Doxycycline
YJL008C CCT8
ZShift = 0.34
NG = 0
DB = 0
SM = 0
Delta_AUC
Doxycycline
YIL022W TIM44
ZShift = −0.76
NG = 0
DB = 0
SM = 0
Delta_L
Doxycycline
YIL022W TIM44
ZShift = 0.54
NG = 0
DB = 0
SM = 0
Delta_K
Doxycycline
YIL022W TIM44
ZShift = 1.76
NG = 0
DB = 0
SM = 0
Delta_r
Doxycycline
YIL022W TIM44
ZShift = 0.7
NG = 0
DB = 0
SM = 0
Delta_AUC
Doxycycline
YGL061C DUO1
ZShift = −0.27
NG = 0
DB = 0
SM = 0
Delta_L
Doxycycline
YGL061C DUO1
ZShift = 0.34
NG = 0
DB = 0
SM = 0
Delta_K
Doxycycline
YGL061C DUO1
ZShift = 0.27
NG = 0
DB = 0
SM = 0
Delta_r
Doxycycline
YGL061C DUO1
ZShift = 0.35
NG = 0
DB = 0
SM = 0
Delta_AUC
Doxycycline

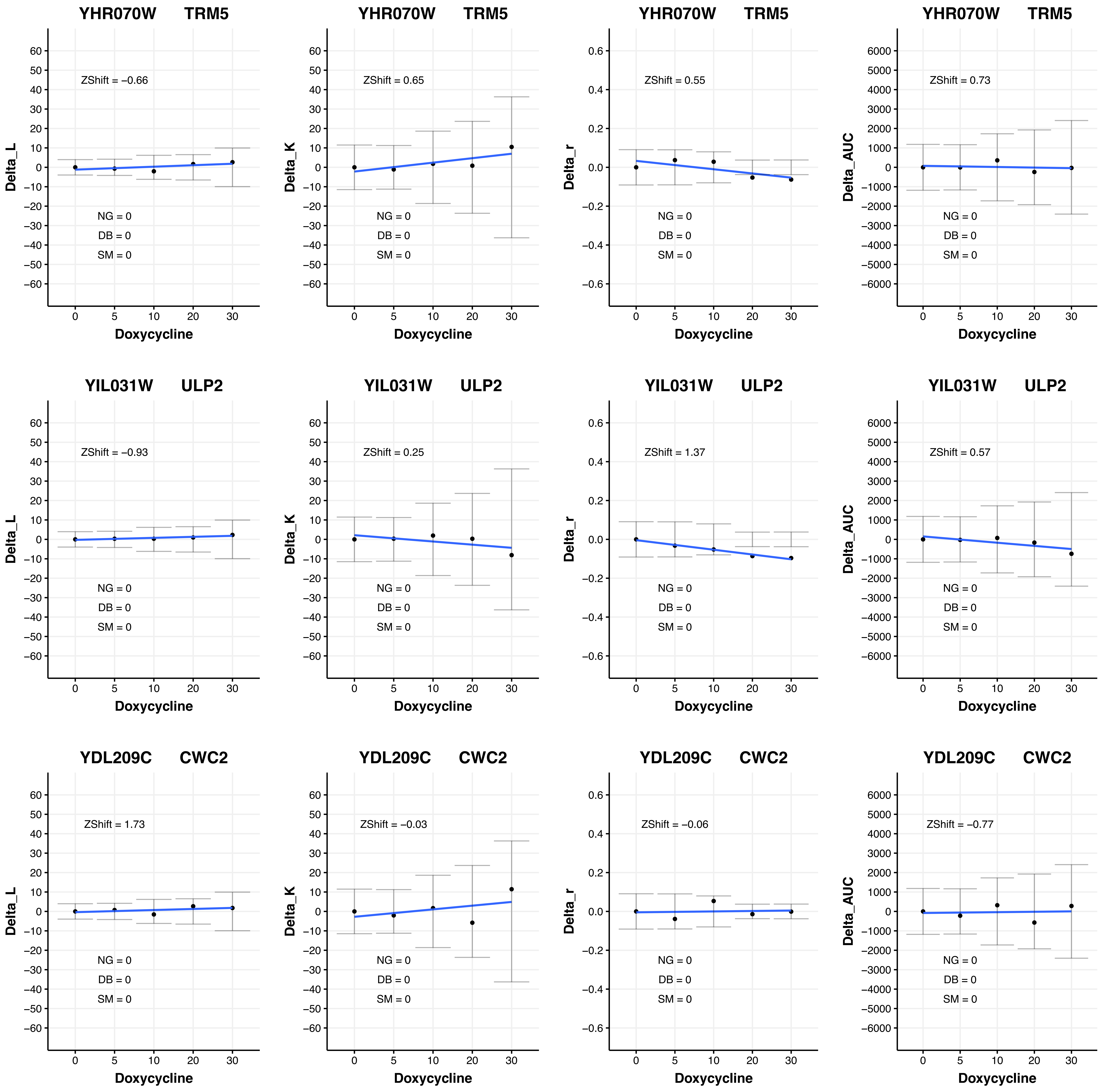
YHR070W TRM5
ZShift = −0.66
NG = 0
DB = 0
SM = 0
Delta_L
Doxycycline
YHR070W TRM5
ZShift = 0.65
NG = 0
DB = 0
SM = 0
Delta_K
Doxycycline
YHR070W TRM5
ZShift = 0.55
NG = 0
DB = 0
SM = 0
Delta_r
Doxycycline
YHR070W TRM5
ZShift = 0.73
NG = 0
DB = 0
SM = 0
Delta_AUC
Doxycycline
YIL031W ULP2
ZShift = −0.93
NG = 0
DB = 0
SM = 0
Delta_L
Doxycycline
YIL031W ULP2
ZShift = 0.25
NG = 0
DB = 0
SM = 0
Delta_K
Doxycycline
YIL031W ULP2
ZShift = 1.37
NG = 0
DB = 0
SM = 0
Delta_r
Doxycycline
YIL031W ULP2
ZShift = 0.57
NG = 0
DB = 0
SM = 0
Delta_AUC
Doxycycline
YDL209C CWC2
ZShift = 1.73
NG = 0
DB = 0
SM = 0
Delta_L
Doxycycline
YDL209C CWC2
ZShift = −0.03
NG = 0
DB = 0
SM = 0
Delta_K
Doxycycline
YDL209C CWC2
ZShift = −0.06
NG = 0
DB = 0
SM = 0
Delta_r
Doxycycline
YDL209C CWC2
ZShift = −0.77
NG = 0
DB = 0
SM = 0
Delta_AUC
Doxycycline

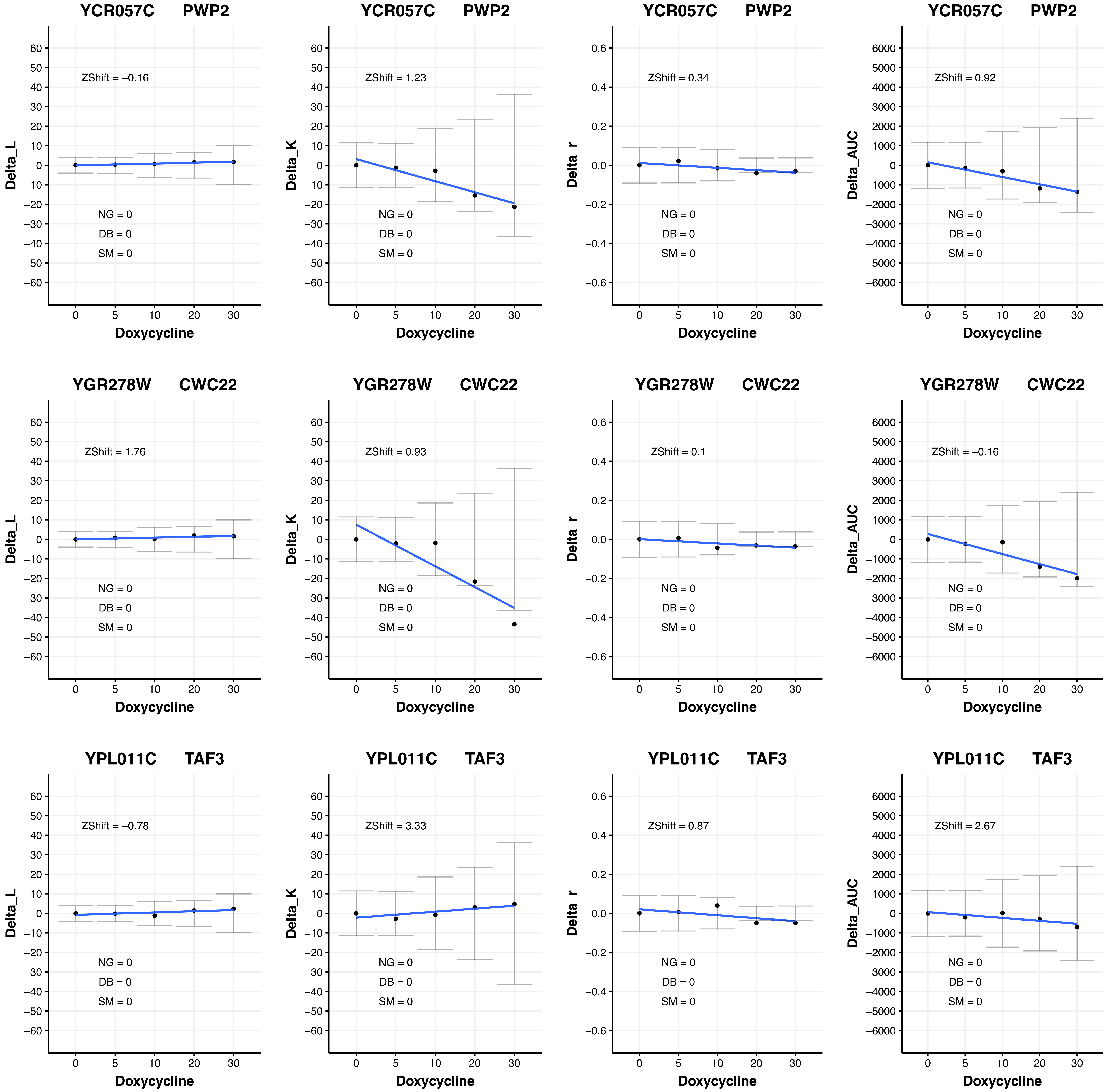

YCR057C PWP2
Delta_L
ZShift = −0.16
NG = 0
DB = 0
SM = 0
Doxycycline
YCR057C PWP2
Delta_K
ZShift = 1.23
NG = 0
DB = 0
SM = 0
Doxycycline
YCR057C PWP2
Delta_r
ZShift = 0.34
NG = 0
DB = 0
SM = 0
Doxycycline
YCR057C PWP2
Delta_AUC
ZShift = 0.92
NG = 0
DB = 0
SM = 0
Doxycycline
YGR278W CWC22
Delta_L
ZShift = 1.76
NG = 0
DB = 0
SM = 0
Doxycycline
YGR278W CWC22
Delta_K
ZShift = 0.93
NG = 0
DB = 0
SM = 0
Doxycycline
YGR278W CWC22
Delta_r
ZShift = 0.1
NG = 0
DB = 0
SM = 0
Doxycycline
YGR278W CWC22
Delta_AUC
ZShift = −0.16
NG = 0
DB = 0
SM = 0
Doxycycline
YPL011C TAF3
Delta_L
ZShift = −0.78
NG = 0
DB = 0
SM = 0
Doxycycline
YPL011C TAF3
Delta_K
ZShift = 3.33
NG = 0
DB = 0
SM = 0
Doxycycline
YPL011C TAF3
Delta_r
ZShift = 0.87
NG = 0
DB = 0
SM = 0
Doxycycline
YPL011C TAF3
Delta_AUC
ZShift = 2.67
NG = 0
DB = 0
SM = 0
Doxycycline

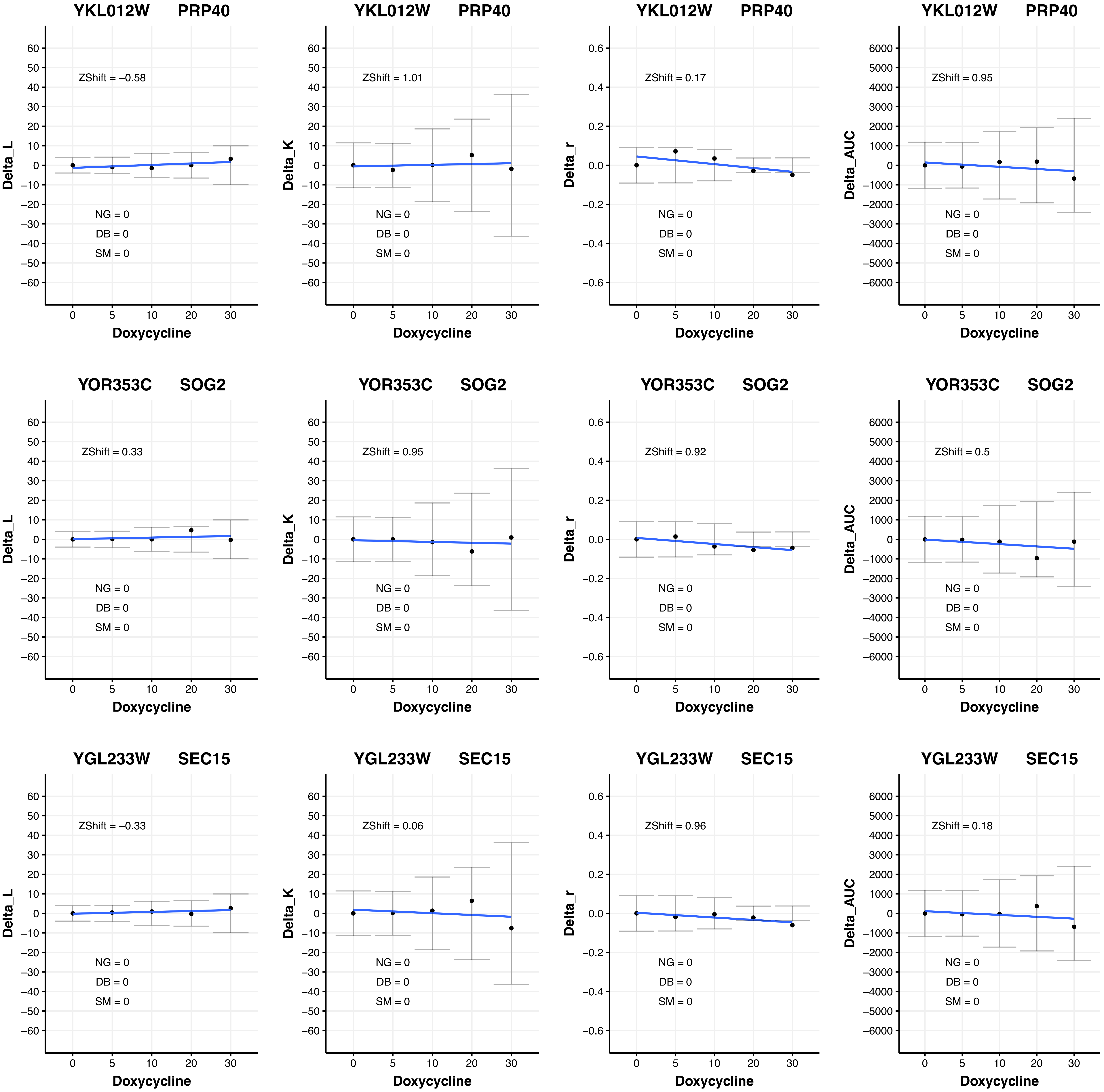
YKL012W PRP40
ZShift = −0.58
NG = 0
DB = 0
SM = 0
Delta_L
Doxycycline
YKL012W PRP40
ZShift = 1.01
NG = 0
DB = 0
SM = 0
Delta_K
Doxycycline
YKL012W PRP40
ZShift = 0.17
NG = 0
DB = 0
SM = 0
Delta_r
Doxycycline
YKL012W PRP40
ZShift = 0.95
NG = 0
DB = 0
SM = 0
Delta_AUC
Doxycycline
YOR353C SOG2
ZShift = 0.33
NG = 0
DB = 0
SM = 0
Delta_L
Doxycycline
YOR353C SOG2
ZShift = 0.95
NG = 0
DB = 0
SM = 0
Delta_K
Doxycycline
YOR353C SOG2
ZShift = 0.92
NG = 0
DB = 0
SM = 0
Delta_r
Doxycycline
YOR353C SOG2
ZShift = 0.5
NG = 0
DB = 0
SM = 0
Delta_AUC
Doxycycline
YGL233W SEC15
ZShift = −0.33
NG = 0
DB = 0
SM = 0
Delta_L
Doxycycline
YGL233W SEC15
ZShift = 0.06
NG = 0
DB = 0
SM = 0
Delta_K
Doxycycline
YGL233W SEC15
ZShift = 0.96
NG = 0
DB = 0
SM = 0
Delta_r
Doxycycline
YGL233W SEC15
ZShift = 0.18
NG = 0
DB = 0
SM = 0
Delta_AUC
Doxycycline

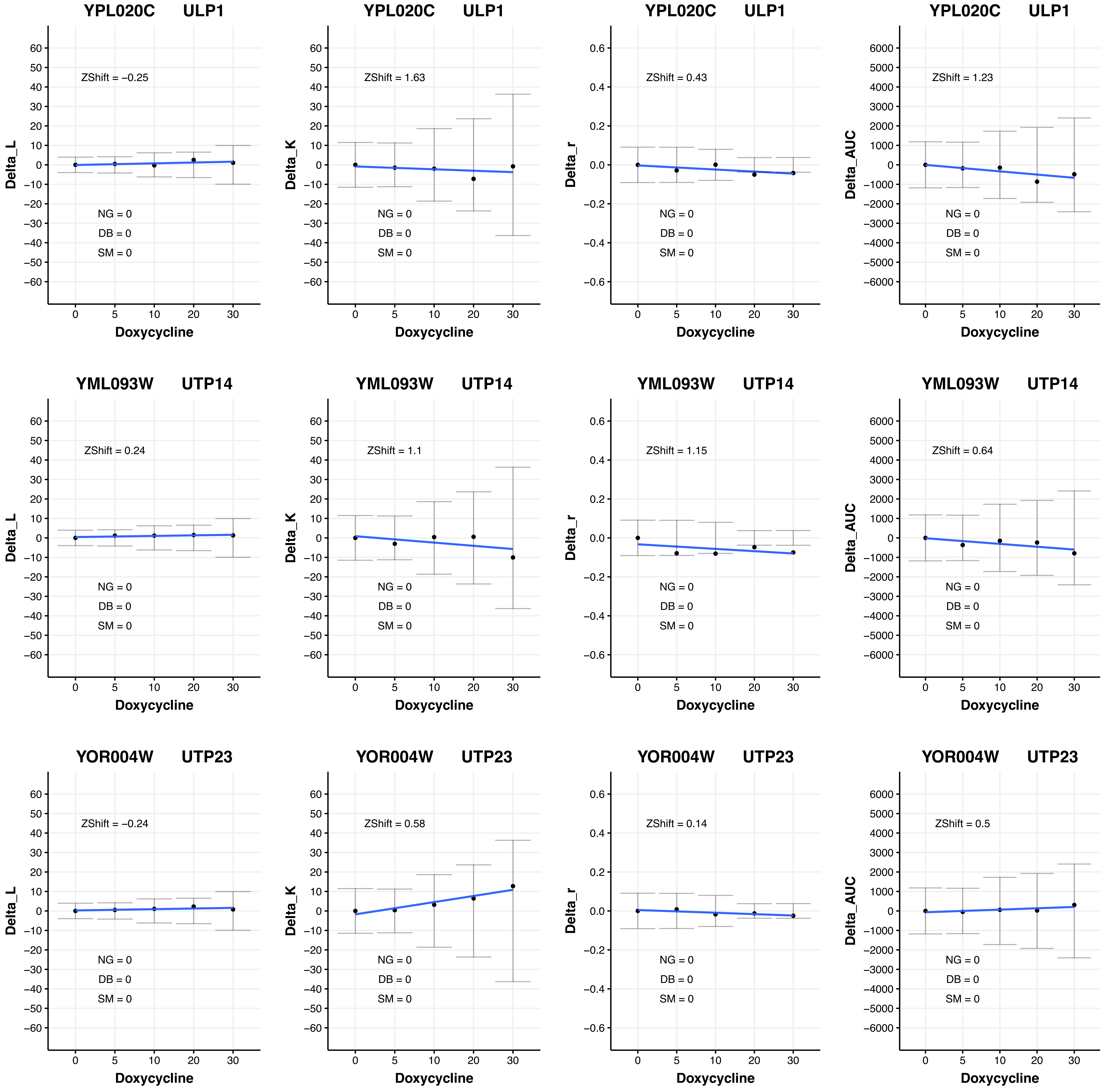
YPL020C ULP1
ZShift = −0.25
NG = 0
DB = 0
SM = 0
Delta_L
Doxycycline
YPL020C ULP1
ZShift = 1.63
NG = 0
DB = 0
SM = 0
Delta_K
Doxycycline
YPL020C ULP1
ZShift = 0.43
NG = 0
DB = 0
SM = 0
Delta_r
Doxycycline
YPL020C ULP1
ZShift = 1.23
NG = 0
DB = 0
SM = 0
Delta_AUC
Doxycycline
YML093W UTP14
ZShift = 0.24
NG = 0
DB = 0
SM = 0
Delta_L
Doxycycline
YML093W UTP14
ZShift = 1.1
NG = 0
DB = 0
SM = 0
Delta_K
Doxycycline
YML093W UTP14
ZShift = 1.15
NG = 0
DB = 0
SM = 0
Delta_r
Doxycycline
YML093W UTP14
ZShift = 0.64
NG = 0
DB = 0
SM = 0
Delta_AUC
Doxycycline
YOR004W UTP23
ZShift = −0.24
NG = 0
DB = 0
SM = 0
Delta_L
Doxycycline
YOR004W UTP23
ZShift = 0.58
NG = 0
DB = 0
SM = 0
Delta_K
Doxycycline
YOR004W UTP23
ZShift = 0.14
NG = 0
DB = 0
SM = 0
Delta_r
Doxycycline
YOR004W UTP23
ZShift = 0.5
NG = 0
DB = 0
SM = 0
Delta_AUC
Doxycycline

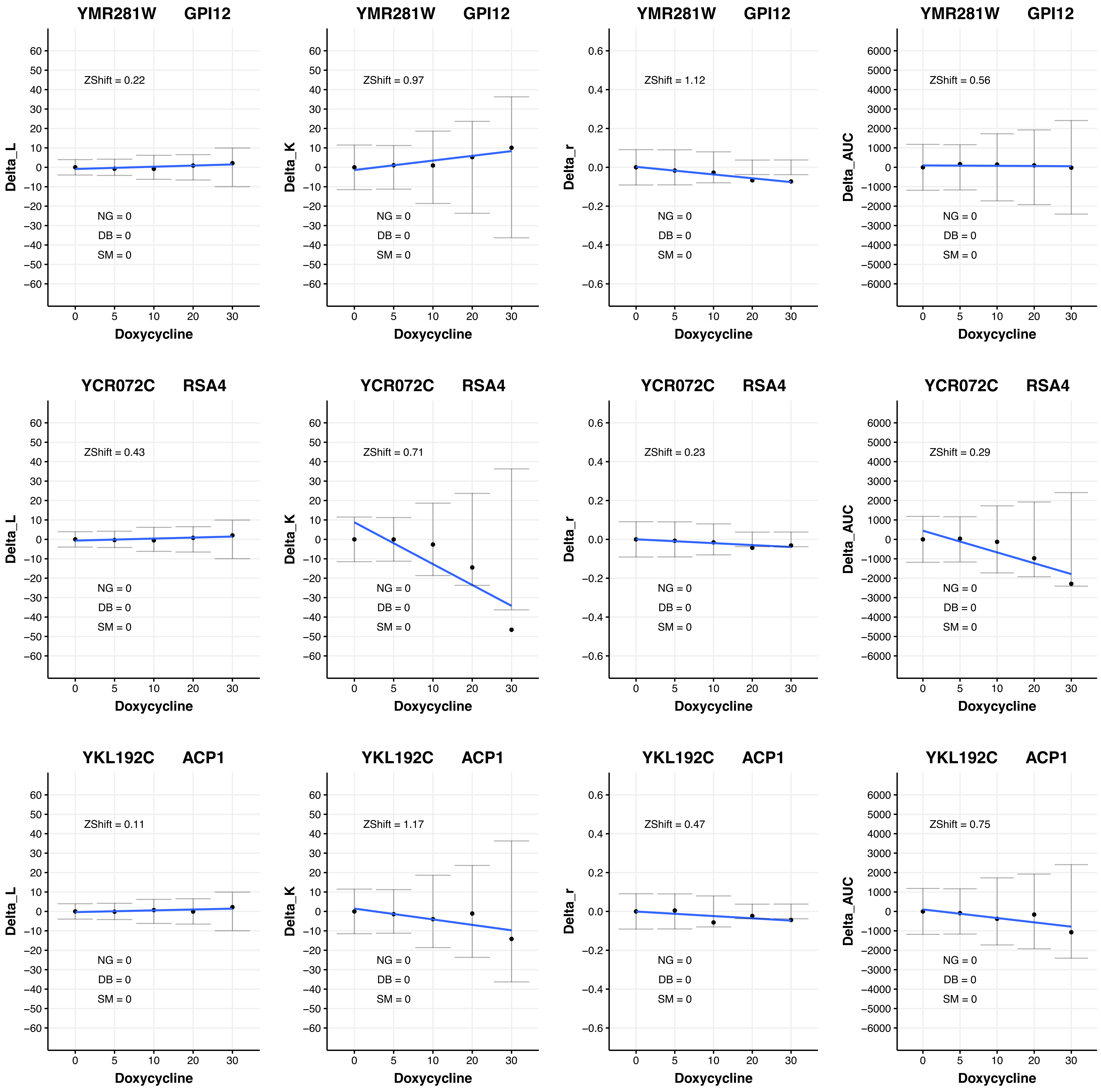
YMR281W GPI12
ZShift = 0.22
NG = 0
DB = 0
SM = 0
Delta_L
Doxycycline
YMR281W GPI12
ZShift = 0.97
NG = 0
DB = 0
SM = 0
Delta_K
Doxycycline
YMR281W GPI12
ZShift = 1.12
NG = 0
DB = 0
SM = 0
Delta_r
Doxycycline
YMR281W GPI12
ZShift = 0.56
NG = 0
DB = 0
SM = 0
Delta_AUC
Doxycycline
YCR072C RSA4
ZShift = 0.43
NG = 0
DB = 0
SM = 0
Delta_L
Doxycycline
YCR072C RSA4
ZShift = 0.71
NG = 0
DB = 0
SM = 0
Delta_K
Doxycycline
YCR072C RSA4
ZShift = 0.23
NG = 0
DB = 0
SM = 0
Delta_r
Doxycycline
YCR072C RSA4
ZShift = 0.29
NG = 0
DB = 0
SM = 0
Delta_AUC
Doxycycline
YKL192C ACP1
ZShift = 0.11
NG = 0
DB = 0
SM = 0
Delta_L
Doxycycline
YKL192C ACP1
ZShift = 1.17
NG = 0
DB = 0
SM = 0
Delta_K
Doxycycline
YKL192C ACP1
ZShift = 0.47
NG = 0
DB = 0
SM = 0
Delta_r
Doxycycline
YKL192C ACP1
ZShift = 0.75
NG = 0
DB = 0
SM = 0
Delta_AUC
Doxycycline
0 5 10 20 30
60 50 40 30 20 10 0 −10 −20 −30 −40 −50 −60
0.6 0.4 0.2 0.0 −0.2 −0.4 −0.6
6000 5000 4000 3000 2000 1000 0 −1000 −2000 −3000 −4000 −5000 −6000

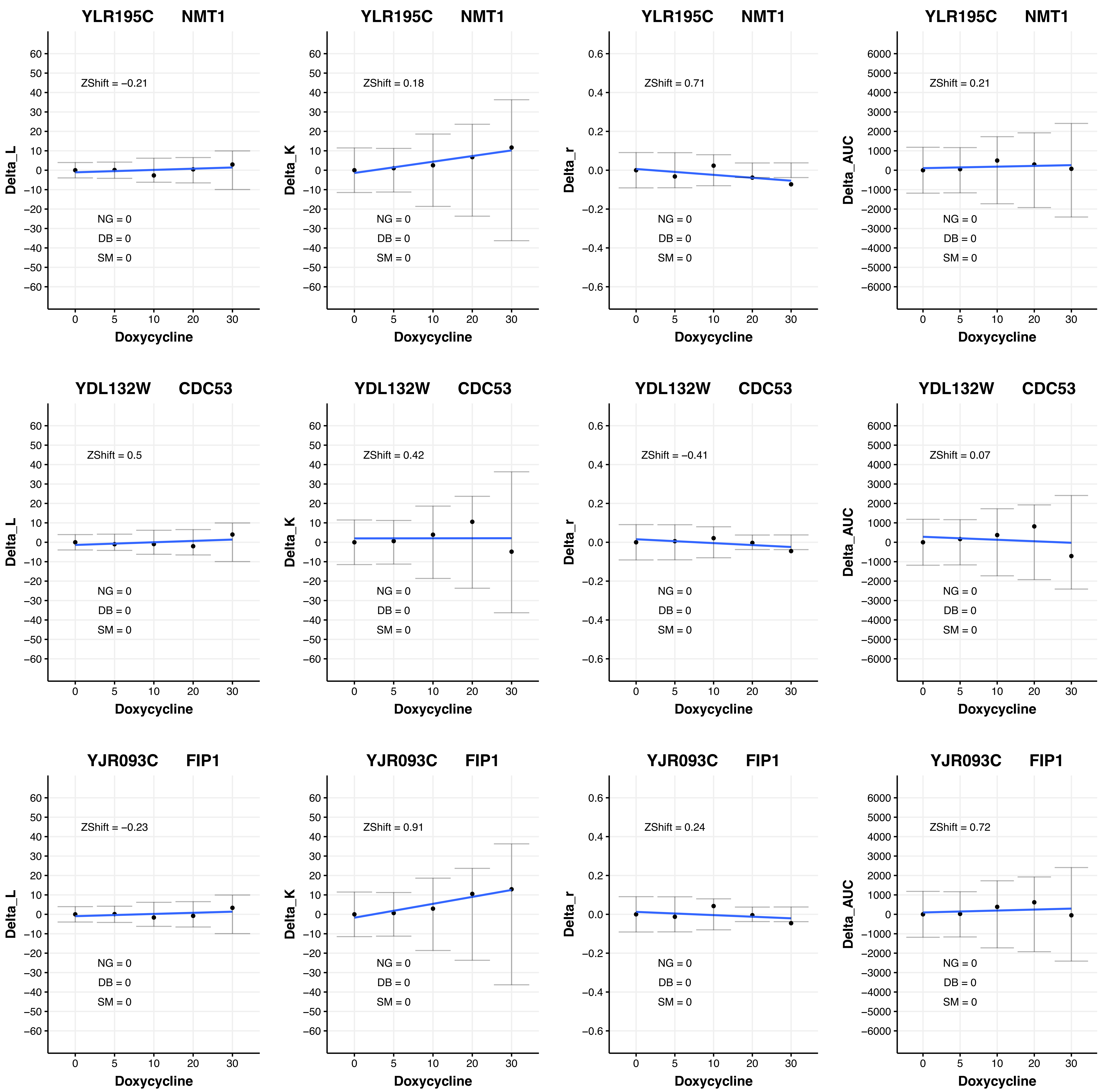
YLR195C NMT1
ZShift = −0.21
NG = 0
DB = 0
SM = 0
Delta_L
Doxycycline
YLR195C NMT1
ZShift = 0.18
NG = 0
DB = 0
SM = 0
Delta_K
Doxycycline
YLR195C NMT1
ZShift = 0.71
NG = 0
DB = 0
SM = 0
Delta_r
Doxycycline
YLR195C NMT1
ZShift = 0.21
NG = 0
DB = 0
SM = 0
Delta_AUC
Doxycycline
YDL132W CDC53
ZShift = 0.5
NG = 0
DB = 0
SM = 0
Delta_L
Doxycycline
YDL132W CDC53
ZShift = 0.42
NG = 0
DB = 0
SM = 0
Delta_K
Doxycycline
YDL132W CDC53
ZShift = −0.41
NG = 0
DB = 0
SM = 0
Delta_r
Doxycycline
YDL132W CDC53
ZShift = 0.07
NG = 0
DB = 0
SM = 0
Delta_AUC
Doxycycline
YJR093C FIP1
ZShift = −0.23
NG = 0
DB = 0
SM = 0
Delta_L
Doxycycline
YJR093C FIP1
ZShift = 0.91
NG = 0
DB = 0
SM = 0
Delta_K
Doxycycline
YJR093C FIP1
ZShift = 0.24
NG = 0
DB = 0
SM = 0
Delta_r
Doxycycline
YJR093C FIP1
ZShift = 0.72
NG = 0
DB = 0
SM = 0
Delta_AUC
Doxycycline

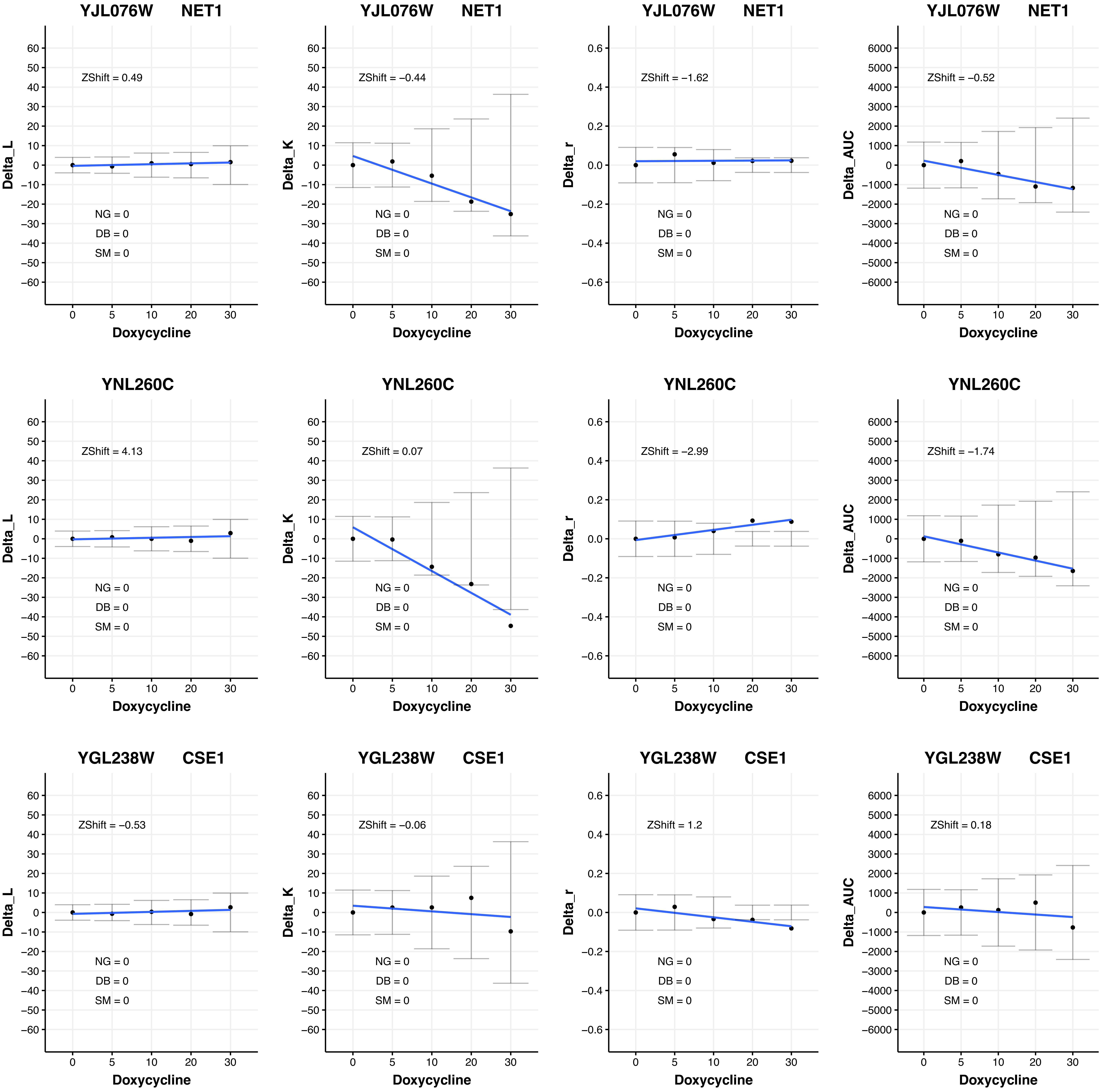
YJL076W NET1
ZShift = 0.49
NG = 0
DB = 0
SM = 0
Delta_L
Doxycycline
YJL076W NET1
ZShift = −0.44
NG = 0
DB = 0
SM = 0
Delta_K
Doxycycline
YJL076W NET1
ZShift = −1.62
NG = 0
DB = 0
SM = 0
Delta_r
Doxycycline
YJL076W NET1
ZShift = −0.52
NG = 0
DB = 0
SM = 0
Delta_AUC
Doxycycline
YNL260C
ZShift = 4.13
NG = 0
DB = 0
SM = 0
Delta_L
Doxycycline
YNL260C
ZShift = 0.07
NG = 0
DB = 0
SM = 0
Delta_K
Doxycycline
YNL260C
ZShift = −2.99
NG = 0
DB = 0
SM = 0
Delta_r
Doxycycline
YNL260C
ZShift = −1.74
NG = 0
DB = 0
SM = 0
Delta_AUC
Doxycycline
YGL238W CSE1
ZShift = −0.53
NG = 0
DB = 0
SM = 0
Delta_L
Doxycycline
YGL238W CSE1
ZShift = −0.06
NG = 0
DB = 0
SM = 0
Delta_K
Doxycycline
YGL238W CSE1
ZShift = 1.2
NG = 0
DB = 0
SM = 0
Delta_r
Doxycycline
YGL238W CSE1
ZShift = 0.18
NG = 0
DB = 0
SM = 0
Delta_AUC
Doxycycline

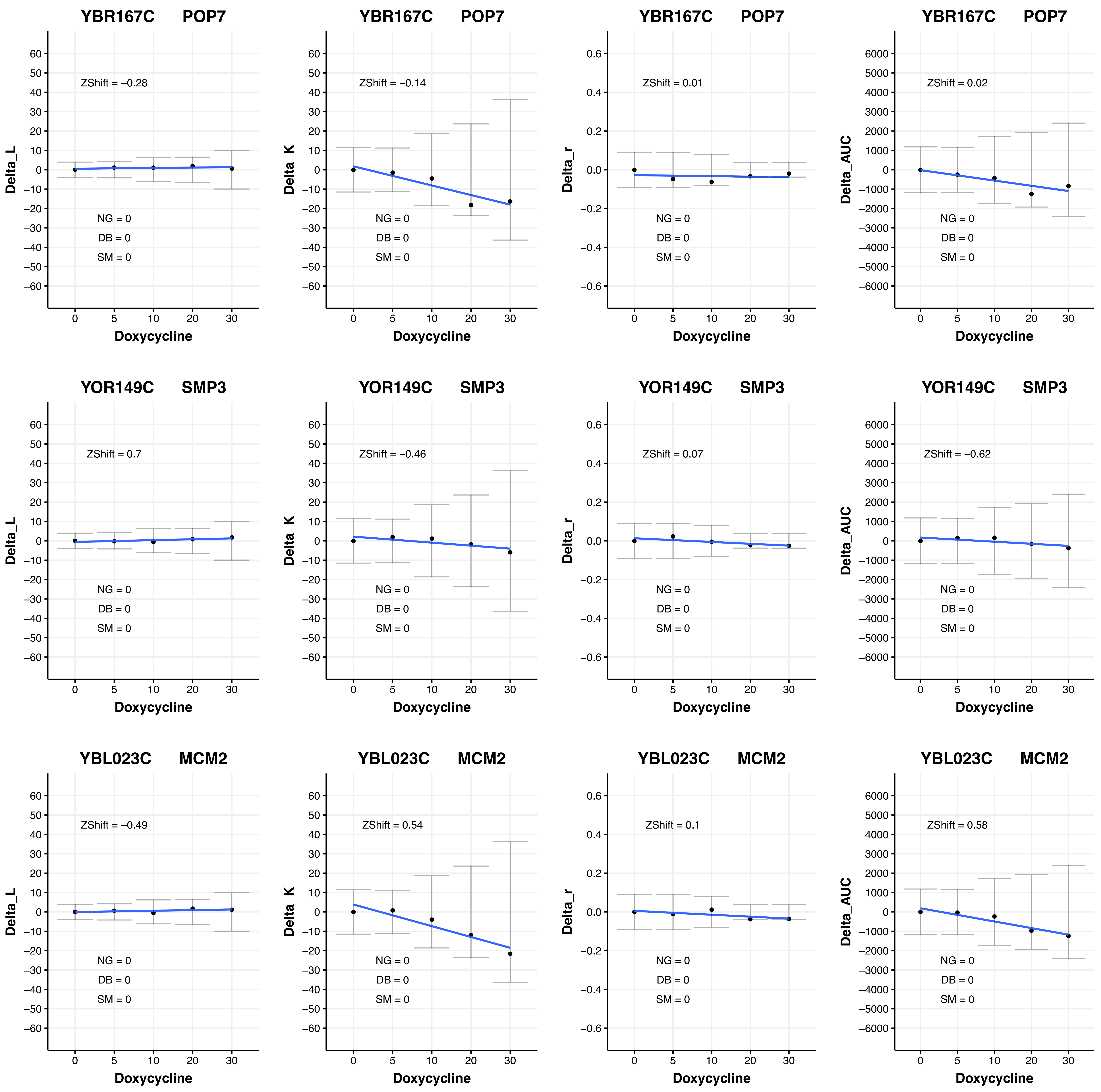
YBR167C POP7
ZShift = −0.28
NG = 0
DB = 0
SM = 0
Delta_L
Doxycycline
YBR167C POP7
ZShift = −0.14
NG = 0
DB = 0
SM = 0
Delta_K
Doxycycline
YBR167C POP7
ZShift = 0.01
NG = 0
DB = 0
SM = 0
Delta_r
Doxycycline
YBR167C POP7
ZShift = 0.02
NG = 0
DB = 0
SM = 0
Delta_AUC
Doxycycline
YOR149C SMP3
ZShift = 0.7
NG = 0
DB = 0
SM = 0
Delta_L
Doxycycline
YOR149C SMP3
ZShift = −0.46
NG = 0
DB = 0
SM = 0
Delta_K
Doxycycline
YOR149C SMP3
ZShift = 0.07
NG = 0
DB = 0
SM = 0
Delta_r
Doxycycline
YOR149C SMP3
ZShift = −0.62
NG = 0
DB = 0
SM = 0
Delta_AUC
Doxycycline
YBL023C MCM2
ZShift = −0.49
NG = 0
DB = 0
SM = 0
Delta_L
Doxycycline
YBL023C MCM2
ZShift = 0.54
NG = 0
DB = 0
SM = 0
Delta_K
Doxycycline
YBL023C MCM2
ZShift = 0.1
NG = 0
DB = 0
SM = 0
Delta_r
Doxycycline
YBL023C MCM2
ZShift = 0.58
NG = 0
DB = 0
SM = 0
Delta_AUC
Doxycycline

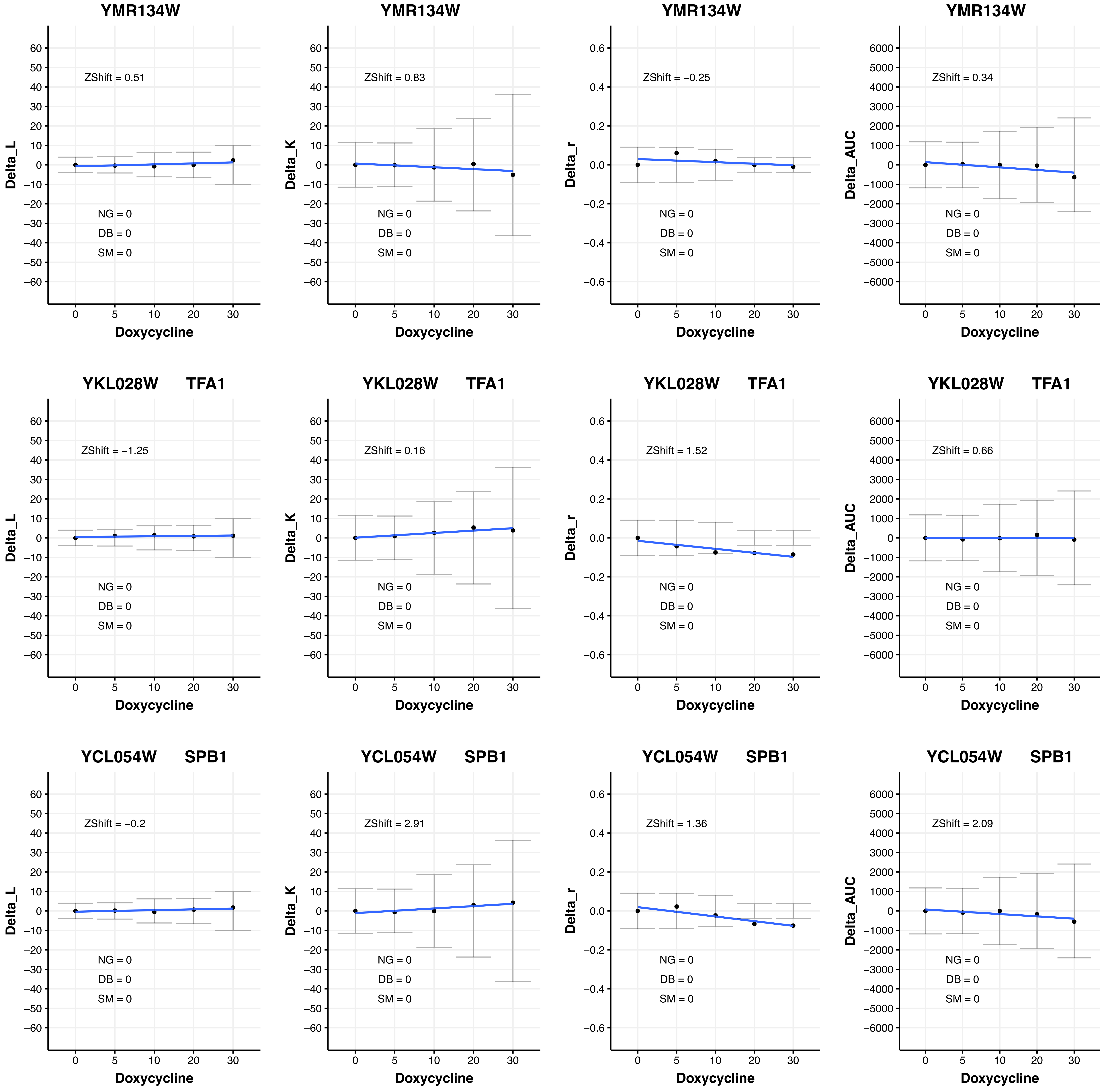

YMR134W
ZShift = 0.51
Delta_L
NG = 0
DB = 0
SM = 0
Doxycycline
YMR134W
ZShift = 0.83
Delta_K
NG = 0
DB = 0
SM = 0
Doxycycline
YMR134W
ZShift = −0.25
Delta_r
NG = 0
DB = 0
SM = 0
Doxycycline
YMR134W
ZShift = 0.34
Delta_AUC
NG = 0
DB = 0
SM = 0
Doxycycline
YKL028W TFA1
ZShift = −1.25
Delta_L
NG = 0
DB = 0
SM = 0
Doxycycline
YKL028W TFA1
ZShift = 0.16
Delta_K
NG = 0
DB = 0
SM = 0
Doxycycline
YKL028W TFA1
ZShift = 1.52
Delta_r
NG = 0
DB = 0
SM = 0
Doxycycline
YKL028W TFA1
ZShift = 0.66
Delta_AUC
NG = 0
DB = 0
SM = 0
Doxycycline
YCL054W SPB1
ZShift = −0.2
Delta_L
NG = 0
DB = 0
SM = 0
Doxycycline
YCL054W SPB1
ZShift = 2.91
Delta_K
NG = 0
DB = 0
SM = 0
Doxycycline
YCL054W SPB1
ZShift = 1.36
Delta_r
NG = 0
DB = 0
SM = 0
Doxycycline
YCL054W SPB1
ZShift = 2.09
Delta_AUC
NG = 0
DB = 0
SM = 0
Doxycycline

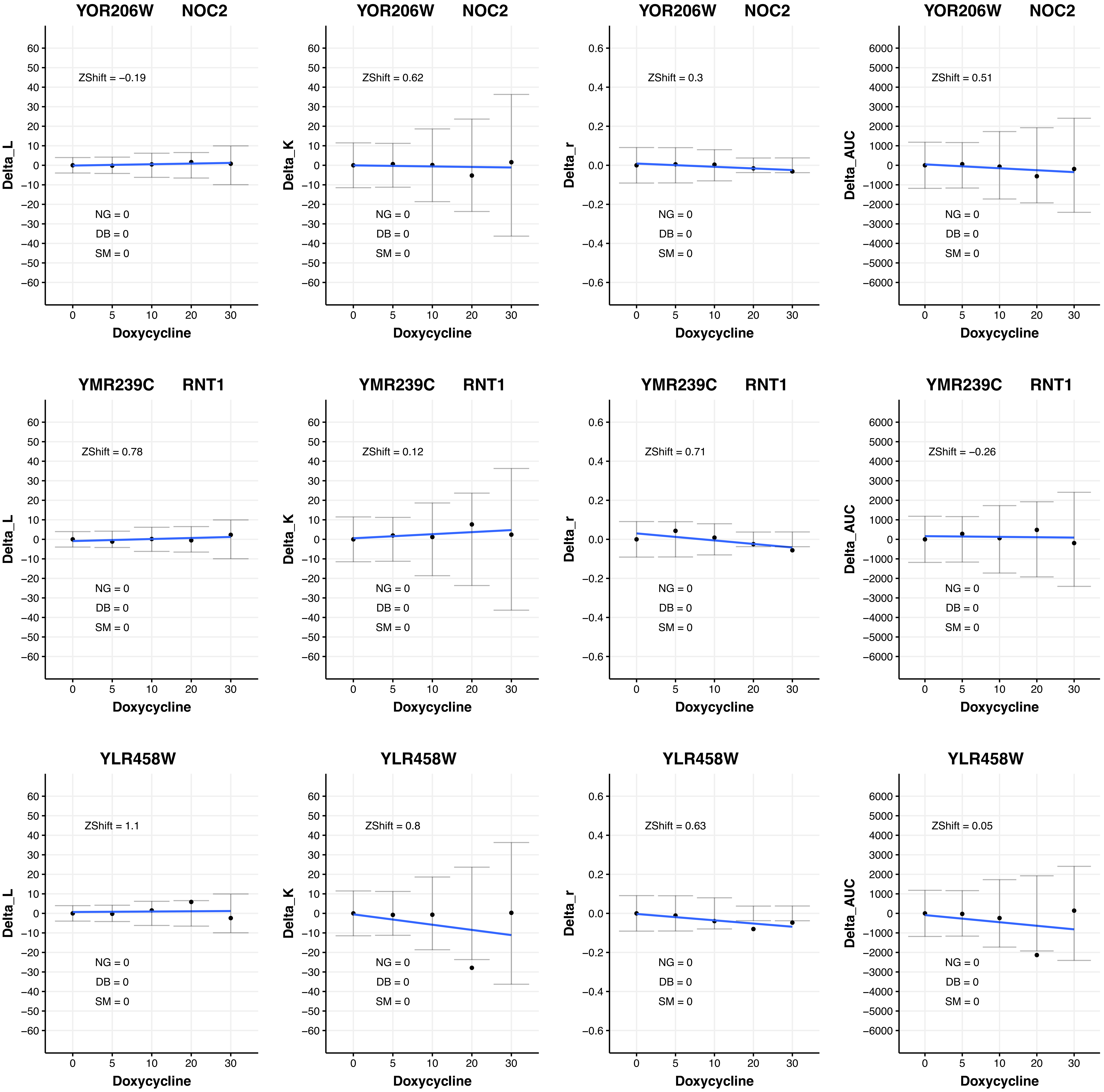
YOR206W NOC2
ZShift = −0.19
NG = 0
DB = 0
SM = 0
Delta_L
Doxycycline
0 5 10 20 30
60 50 40 30 20 10 0 −10 −20 −30 −40 −50 −60
YOR206W NOC2
ZShift = 0.62
NG = 0
DB = 0
SM = 0
Delta_K
Doxycycline
YOR206W NOC2
ZShift = 0.3
NG = 0
DB = 0
SM = 0
Delta_r
Doxycycline
0.6 0.4 0.2 0.0 −0.2 −0.4 −0.6
YOR206W NOC2
ZShift = 0.51
NG = 0
DB = 0
SM = 0
Delta_AUC
Doxycycline
6000 5000 4000 3000 2000 1000 0 −1000 −2000 −3000 −4000 −5000 −6000
YMR239C RNT1
ZShift = 0.78
NG = 0
DB = 0
SM = 0
Delta_L
Doxycycline
YMR239C RNT1
ZShift = 0.12
NG = 0
DB = 0
SM = 0
Delta_K
Doxycycline
YMR239C RNT1
ZShift = 0.71
NG = 0
DB = 0
SM = 0
Delta_r
Doxycycline
YMR239C RNT1
ZShift = −0.26
NG = 0
DB = 0
SM = 0
Delta_AUC
Doxycycline
YLR458W
ZShift = 1.1
NG = 0
DB = 0
SM = 0
Delta_L
Doxycycline
YLR458W
ZShift = 0.8
NG = 0
DB = 0
SM = 0
Delta_K
Doxycycline
YLR458W
ZShift = 0.63
NG = 0
DB = 0
SM = 0
Delta_r
Doxycycline
YLR458W
ZShift = 0.05
NG = 0
DB = 0
SM = 0
Delta_AUC
Doxycycline

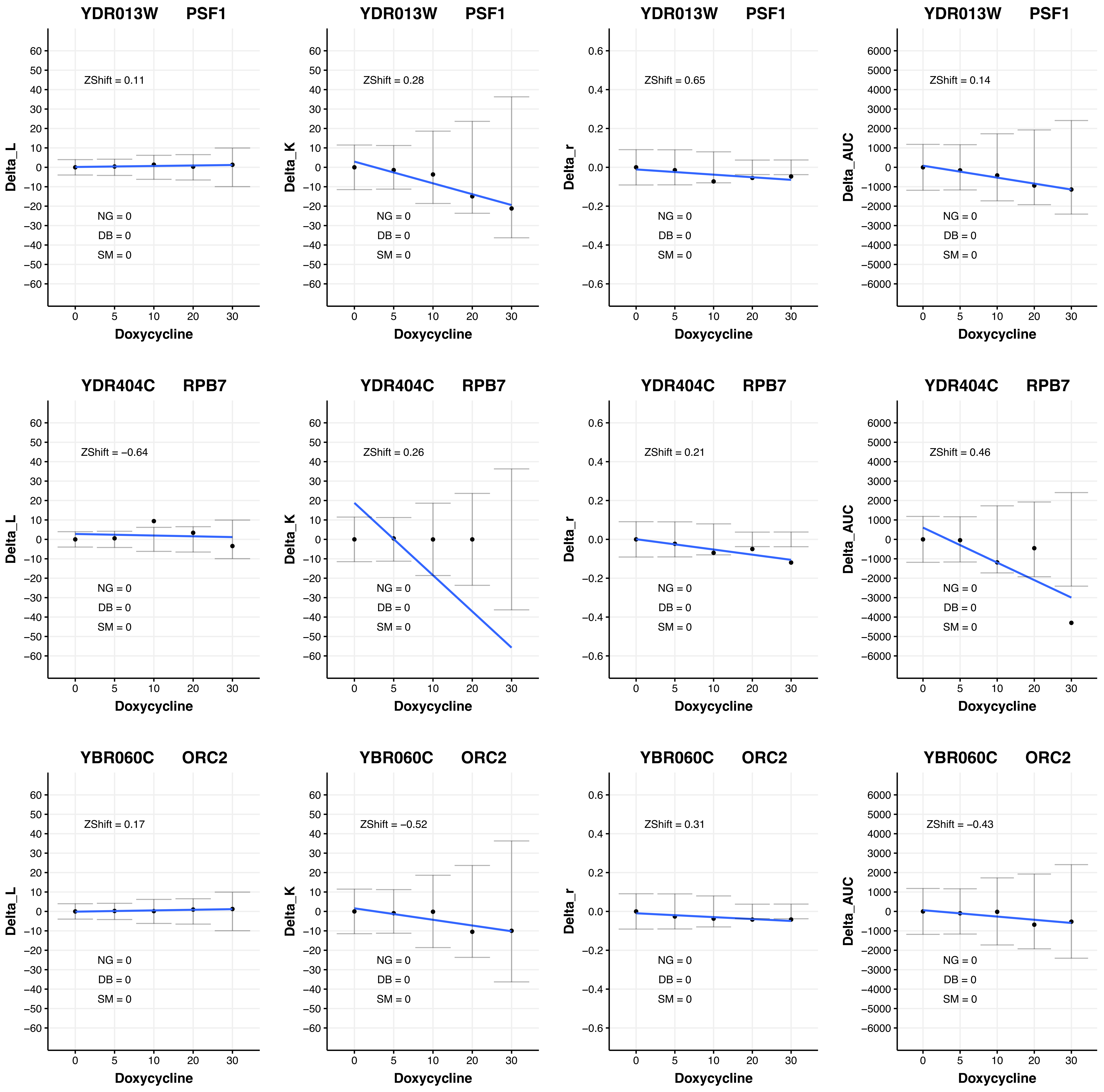

YDR013W PSF1
ZShift = 0.11
NG = 0
DB = 0
SM = 0
Delta_L
Doxycycline
YDR013W PSF1
ZShift = 0.28
NG = 0
DB = 0
SM = 0
Delta_K
Doxycycline
YDR013W PSF1
ZShift = 0.65
NG = 0
DB = 0
SM = 0
Delta_r
Doxycycline
YDR013W PSF1
ZShift = 0.14
NG = 0
DB = 0
SM = 0
Delta_AUC
Doxycycline
YDR404C RPB7
ZShift = −0.64
NG = 0
DB = 0
SM = 0
Delta_L
Doxycycline
YDR404C RPB7
ZShift = 0.26
NG = 0
DB = 0
SM = 0
Delta_K
Doxycycline
YDR404C RPB7
ZShift = 0.21
NG = 0
DB = 0
SM = 0
Delta_r
Doxycycline
YDR404C RPB7
ZShift = 0.46
NG = 0
DB = 0
SM = 0
Delta_AUC
Doxycycline
YBR060C ORC2
ZShift = 0.17
NG = 0
DB = 0
SM = 0
Delta_L
Doxycycline
YBR060C ORC2
ZShift = −0.52
NG = 0
DB = 0
SM = 0
Delta_K
Doxycycline
YBR060C ORC2
ZShift = 0.31
NG = 0
DB = 0
SM = 0
Delta_r
Doxycycline
YBR060C ORC2
ZShift = −0.43
NG = 0
DB = 0
SM = 0
Delta_AUC
Doxycycline

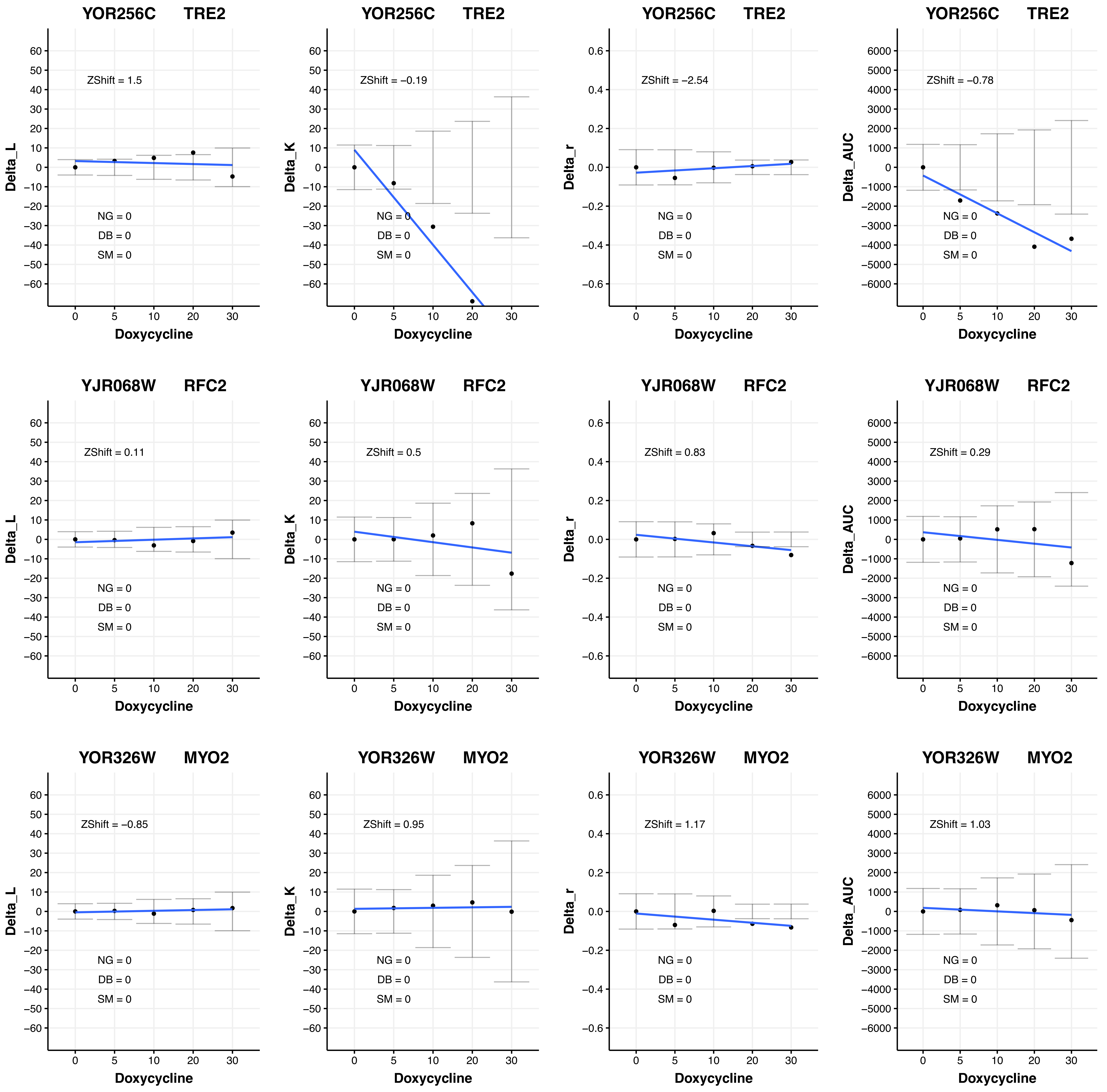
YOR256C TRE2
ZShift = 1.5
NG = 0
DB = 0
SM = 0
Delta_L
Doxycycline
YOR256C TRE2
ZShift = −0.19
NG = 0
DB = 0
SM = 0
Delta_K
Doxycycline
YOR256C TRE2
ZShift = −2.54
NG = 0
DB = 0
SM = 0
Delta_r
Doxycycline
YOR256C TRE2
ZShift = −0.78
NG = 0
DB = 0
SM = 0
Delta_AUC
Doxycycline
YJR068W RFC2
ZShift = 0.11
NG = 0
DB = 0
SM = 0
Delta_L
Doxycycline
YJR068W RFC2
ZShift = 0.5
NG = 0
DB = 0
SM = 0
Delta_K
Doxycycline
YJR068W RFC2
ZShift = 0.83
NG = 0
DB = 0
SM = 0
Delta_r
Doxycycline
YJR068W RFC2
ZShift = 0.29
NG = 0
DB = 0
SM = 0
Delta_AUC
Doxycycline
YOR326W MYO2
ZShift = −0.85
NG = 0
DB = 0
SM = 0
Delta_L
Doxycycline
YOR326W MYO2
ZShift = 0.95
NG = 0
DB = 0
SM = 0
Delta_K
Doxycycline
YOR326W MYO2
ZShift = 1.17
NG = 0
DB = 0
SM = 0
Delta_r
Doxycycline
YOR326W MYO2
ZShift = 1.03
NG = 0
DB = 0
SM = 0
Delta_AUC
Doxycycline

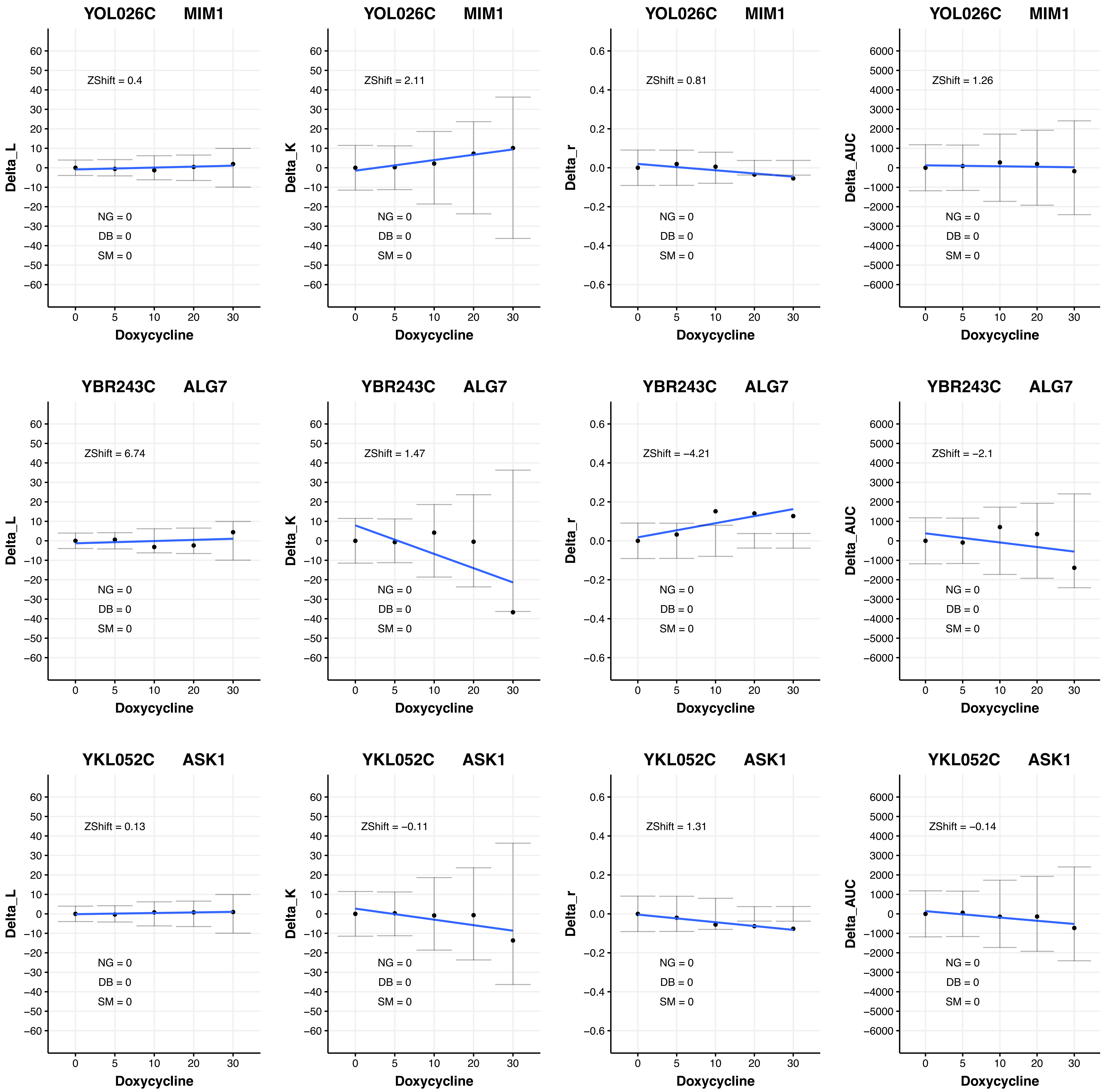
YOL026C MIM1
ZShift = 0.4
NG = 0
DB = 0
SM = 0
Delta_L
Doxycycline
YOL026C MIM1
ZShift = 2.11
NG = 0
DB = 0
SM = 0
Delta_K
Doxycycline
YOL026C MIM1
ZShift = 0.81
NG = 0
DB = 0
SM = 0
Delta_r
Doxycycline
YOL026C MIM1
ZShift = 1.26
NG = 0
DB = 0
SM = 0
Delta_AUC
Doxycycline
YBR243C ALG7
ZShift = 6.74
NG = 0
DB = 0
SM = 0
Delta_L
Doxycycline
YBR243C ALG7
ZShift = 1.47
NG = 0
DB = 0
SM = 0
Delta_K
Doxycycline
YBR243C ALG7
ZShift = −4.21
NG = 0
DB = 0
SM = 0
Delta_r
Doxycycline
YBR243C ALG7
ZShift = −2.1
NG = 0
DB = 0
SM = 0
Delta_AUC
Doxycycline
YKL052C ASK1
ZShift = 0.13
NG = 0
DB = 0
SM = 0
Delta_L
Doxycycline
YKL052C ASK1
ZShift = −0.11
NG = 0
DB = 0
SM = 0
Delta_K
Doxycycline
YKL052C ASK1
ZShift = 1.31
NG = 0
DB = 0
SM = 0
Delta_r
Doxycycline
YKL052C ASK1
ZShift = −0.14
NG = 0
DB = 0
SM = 0
Delta_AUC
Doxycycline

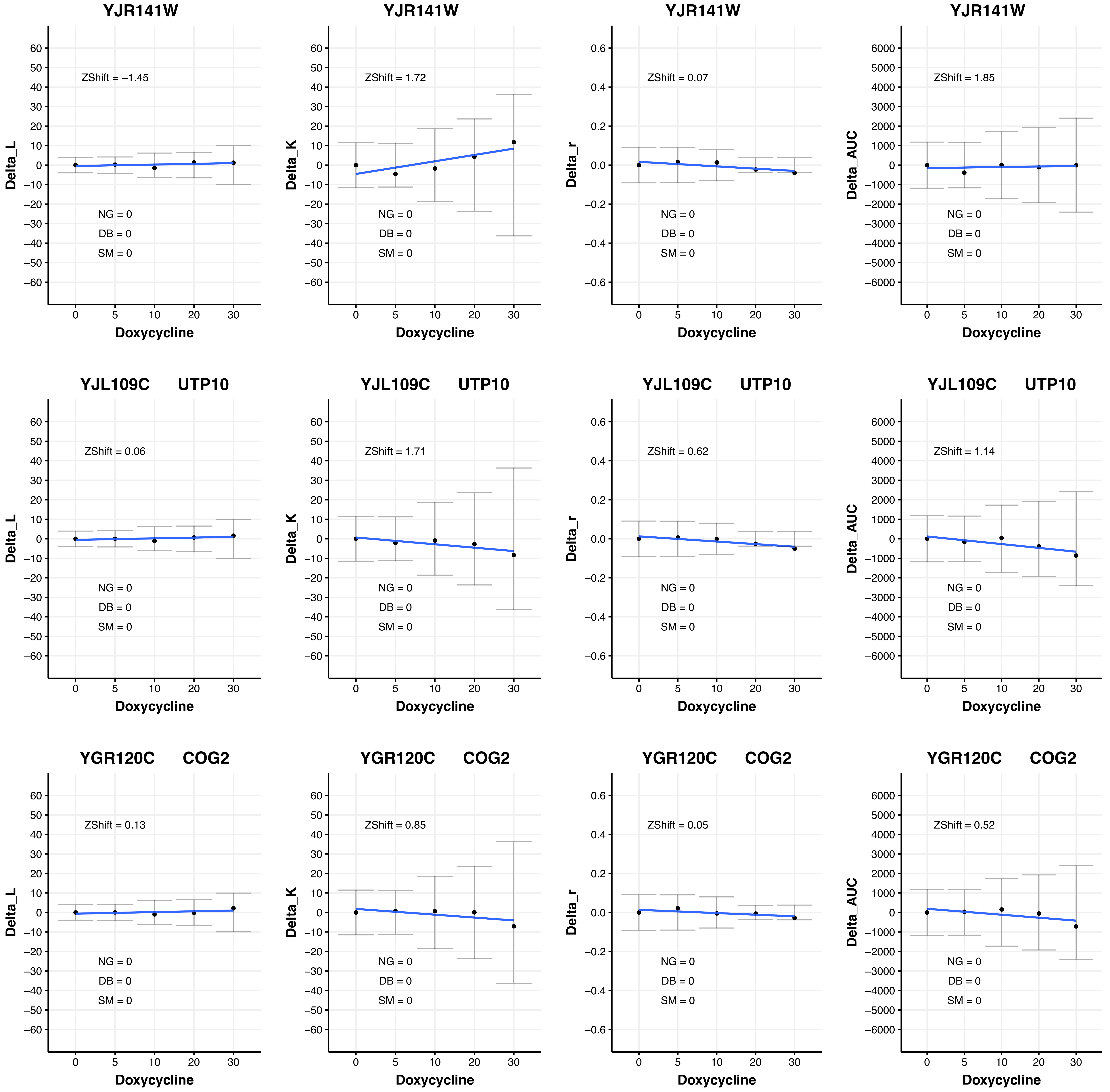
YJR141W
ZShift = −1.45
NG = 0
DB = 0
SM = 0
Delta_L
Doxycycline
YJR141W
ZShift = 1.72
NG = 0
DB = 0
SM = 0
Delta_K
Doxycycline
YJR141W
ZShift = 0.07
NG = 0
DB = 0
SM = 0
Delta_r
Doxycycline
YJR141W
ZShift = 1.85
NG = 0
DB = 0
SM = 0
Delta_AUC
Doxycycline
YJL109C UTP10
ZShift = 0.06
NG = 0
DB = 0
SM = 0
Delta_L
Doxycycline
YJL109C UTP10
ZShift = 1.71
NG = 0
DB = 0
SM = 0
Delta_K
Doxycycline
YJL109C UTP10
ZShift = 0.62
NG = 0
DB = 0
SM = 0
Delta_r
Doxycycline
YJL109C UTP10
ZShift = 1.14
NG = 0
DB = 0
SM = 0
Delta_AUC
Doxycycline
YGR120C COG2
ZShift = 0.13
NG = 0
DB = 0
SM = 0
Delta_L
Doxycycline
YGR120C COG2
ZShift = 0.85
NG = 0
DB = 0
SM = 0
Delta_K
Doxycycline
YGR120C COG2
ZShift = 0.05
NG = 0
DB = 0
SM = 0
Delta_r
Doxycycline
YGR120C COG2
ZShift = 0.52
NG = 0
DB = 0
SM = 0
Delta_AUC
Doxycycline

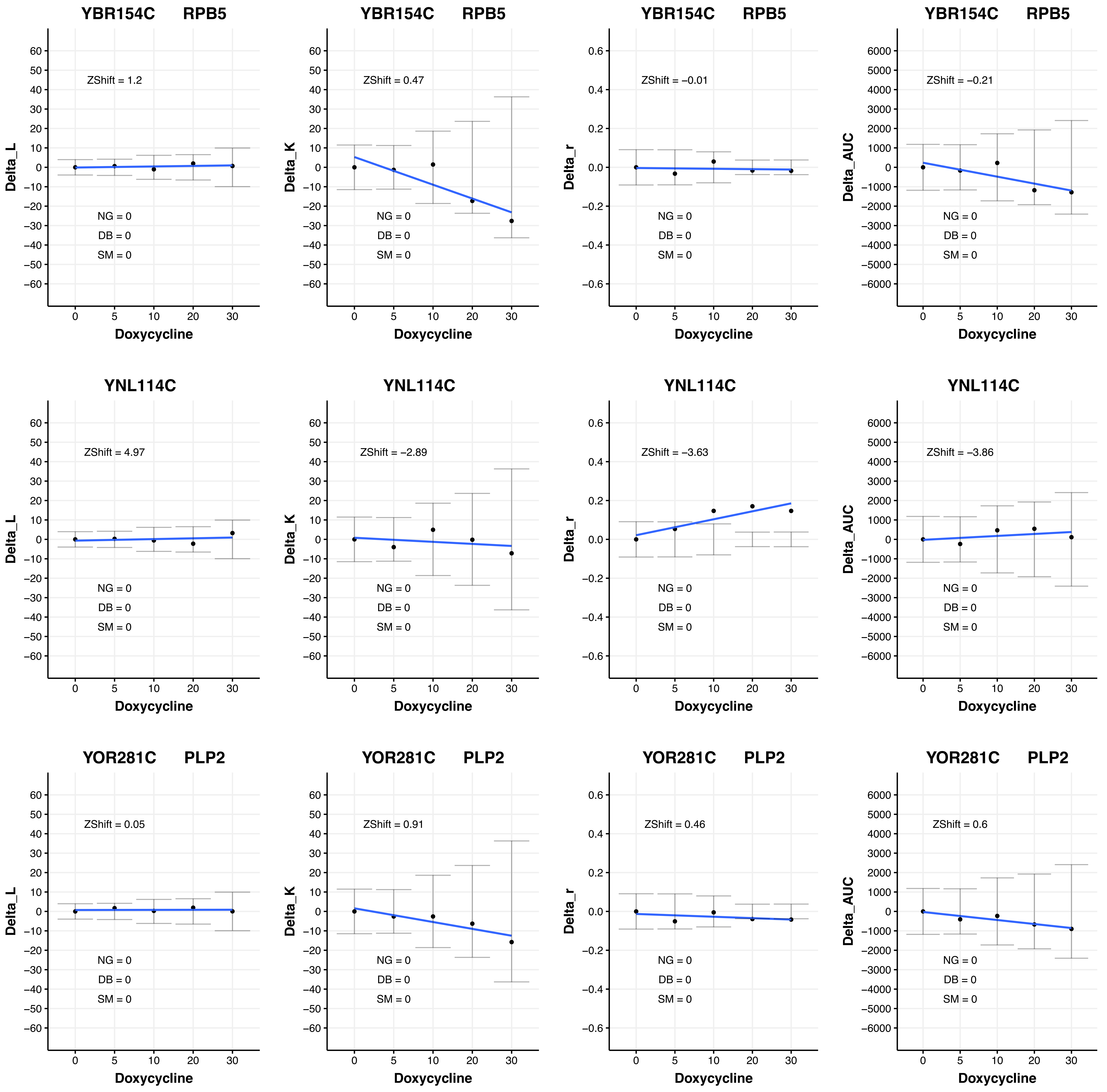
YBR154C RPB5
ZShift = 1.2
NG = 0
DB = 0
SM = 0
Delta_L
Doxycycline
YBR154C RPB5
ZShift = 0.47
NG = 0
DB = 0
SM = 0
Delta_K
Doxycycline
YBR154C RPB5
ZShift = −0.01
NG = 0
DB = 0
SM = 0
Delta_r
Doxycycline
YBR154C RPB5
ZShift = −0.21
NG = 0
DB = 0
SM = 0
Delta_AUC
Doxycycline
YNL114C
ZShift = 4.97
NG = 0
DB = 0
SM = 0
Delta_L
Doxycycline
YNL114C
ZShift = −2.89
NG = 0
DB = 0
SM = 0
Delta_K
Doxycycline
YNL114C
ZShift = −3.63
NG = 0
DB = 0
SM = 0
Delta_r
Doxycycline
YNL114C
ZShift = −3.86
NG = 0
DB = 0
SM = 0
Delta_AUC
Doxycycline
YOR281C PLP2
ZShift = 0.05
NG = 0
DB = 0
SM = 0
Delta_L
Doxycycline
YOR281C PLP2
ZShift = 0.91
NG = 0
DB = 0
SM = 0
Delta_K
Doxycycline
YOR281C PLP2
ZShift = 0.46
NG = 0
DB = 0
SM = 0
Delta_r
Doxycycline
YOR281C PLP2
ZShift = 0.6
NG = 0
DB = 0
SM = 0
Delta_AUC
Doxycycline

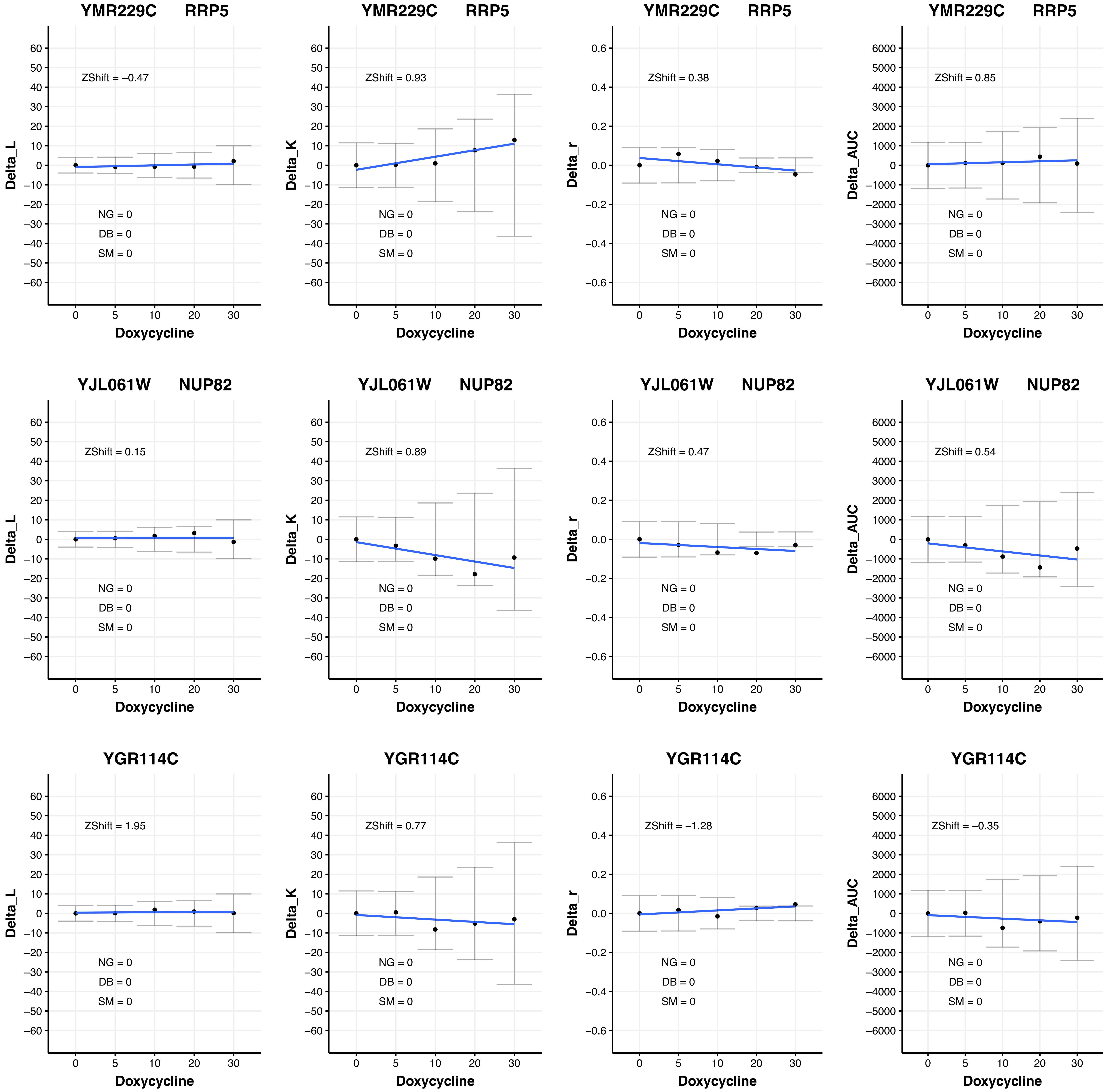

YMR229C RRP5
ZShift = −0.47
Delta_L
NG = 0
DB = 0
SM = 0
Doxycycline
YMR229C RRP5
ZShift = 0.93
Delta_K
NG = 0
DB = 0
SM = 0
Doxycycline
YMR229C RRP5
ZShift = 0.38
Delta_r
NG = 0
DB = 0
SM = 0
Doxycycline
YMR229C RRP5
ZShift = 0.85
Delta_AUC
NG = 0
DB = 0
SM = 0
Doxycycline
YJL061W NUP82
ZShift = 0.15
Delta_L
NG = 0
DB = 0
SM = 0
Doxycycline
YJL061W NUP82
ZShift = 0.89
Delta_K
NG = 0
DB = 0
SM = 0
Doxycycline
YJL061W NUP82
ZShift = 0.47
Delta_r
NG = 0
DB = 0
SM = 0
Doxycycline
YJL061W NUP82
ZShift = 0.54
Delta_AUC
NG = 0
DB = 0
SM = 0
Doxycycline
YGR114C
ZShift = 1.95
Delta_L
NG = 0
DB = 0
SM = 0
Doxycycline
YGR114C
ZShift = 0.77
Delta_K
NG = 0
DB = 0
SM = 0
Doxycycline
YGR114C
ZShift = −1.28
Delta_r
NG = 0
DB = 0
SM = 0
Doxycycline
YGR114C
ZShift = −0.35
Delta_AUC
NG = 0
DB = 0
SM = 0
Doxycycline

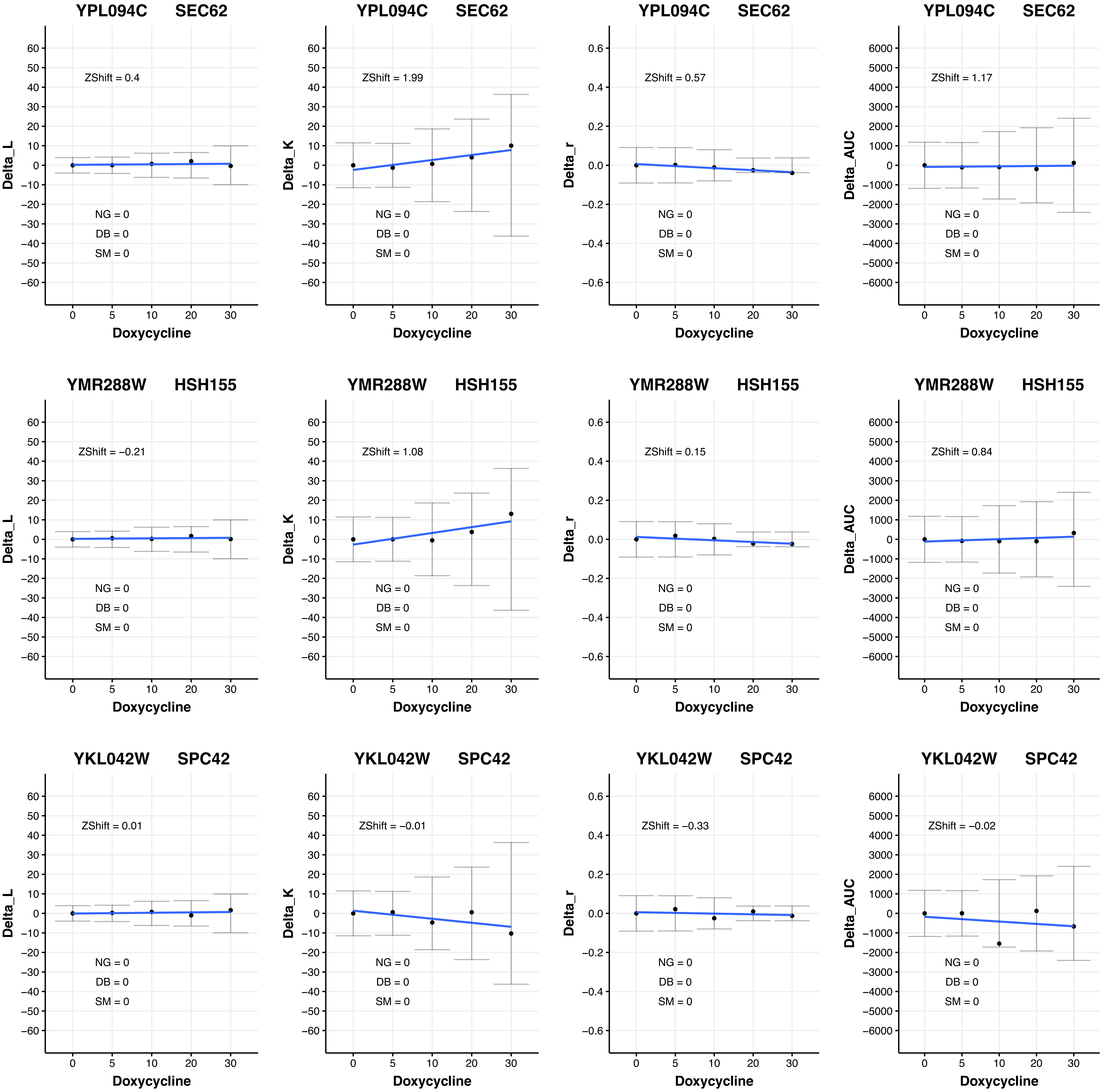
YPL094C SEC62
ZShift = 0.4
NG = 0
DB = 0
SM = 0
Delta_L
Doxycycline
YPL094C SEC62
ZShift = 1.99
NG = 0
DB = 0
SM = 0
Delta_K
Doxycycline
YPL094C SEC62
ZShift = 0.57
NG = 0
DB = 0
SM = 0
Delta_r
Doxycycline
YPL094C SEC62
ZShift = 1.17
NG = 0
DB = 0
SM = 0
Delta_AUC
Doxycycline
YMR288W HSH155
ZShift = −0.21
NG = 0
DB = 0
SM = 0
Delta_L
Doxycycline
YMR288W HSH155
ZShift = 1.08
NG = 0
DB = 0
SM = 0
Delta_K
Doxycycline
YMR288W HSH155
ZShift = 0.15
NG = 0
DB = 0
SM = 0
Delta_r
Doxycycline
YMR288W HSH155
ZShift = 0.84
NG = 0
DB = 0
SM = 0
Delta_AUC
Doxycycline
YKL042W SPC42
ZShift = 0.01
NG = 0
DB = 0
SM = 0
Delta_L
Doxycycline
YKL042W SPC42
ZShift = −0.01
NG = 0
DB = 0
SM = 0
Delta_K
Doxycycline
YKL042W SPC42
ZShift = −0.33
NG = 0
DB = 0
SM = 0
Delta_r
Doxycycline
YKL042W SPC42
ZShift = −0.02
NG = 0
DB = 0
SM = 0
Delta_AUC
Doxycycline

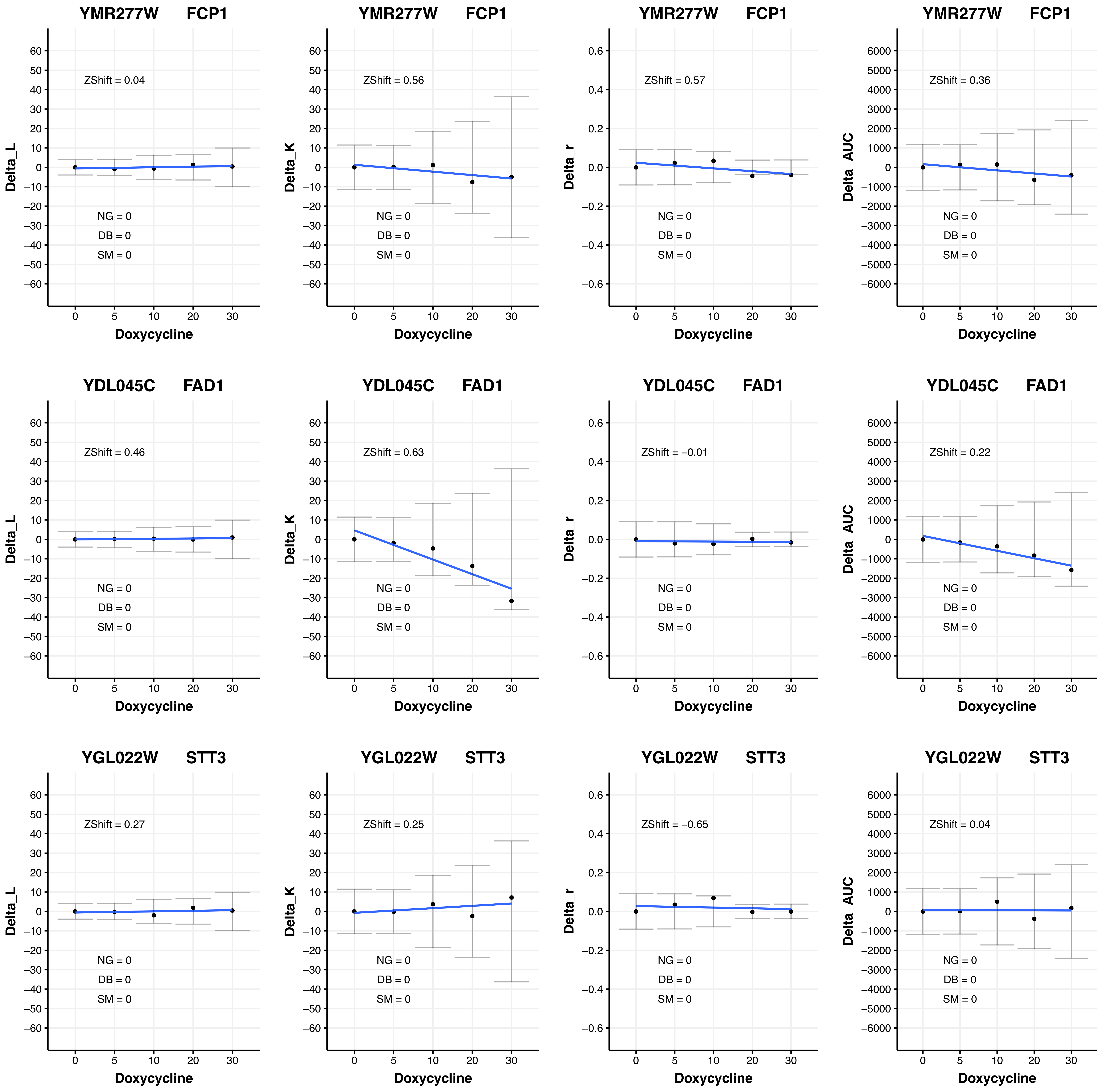
YMR277W FCP1
ZShift = 0.04
NG = 0
DB = 0
SM = 0
Delta_L
Doxycycline
YMR277W FCP1
ZShift = 0.56
NG = 0
DB = 0
SM = 0
Delta_K
Doxycycline
YMR277W FCP1
ZShift = 0.57
NG = 0
DB = 0
SM = 0
Delta_r
Doxycycline
YMR277W FCP1
ZShift = 0.36
NG = 0
DB = 0
SM = 0
Delta_AUC
Doxycycline
YDL045C FAD1
ZShift = 0.46
NG = 0
DB = 0
SM = 0
Delta_L
Doxycycline
YDL045C FAD1
ZShift = 0.63
NG = 0
DB = 0
SM = 0
Delta_K
Doxycycline
YDL045C FAD1
ZShift = −0.01
NG = 0
DB = 0
SM = 0
Delta_r
Doxycycline
YDL045C FAD1
ZShift = 0.22
NG = 0
DB = 0
SM = 0
Delta_AUC
Doxycycline
YGL022W STT3
ZShift = 0.27
NG = 0
DB = 0
SM = 0
Delta_L
Doxycycline
YGL022W STT3
ZShift = 0.25
NG = 0
DB = 0
SM = 0
Delta_K
Doxycycline
YGL022W STT3
ZShift = −0.65
NG = 0
DB = 0
SM = 0
Delta_r
Doxycycline
YGL022W STT3
ZShift = 0.04
NG = 0
DB = 0
SM = 0
Delta_AUC
Doxycycline

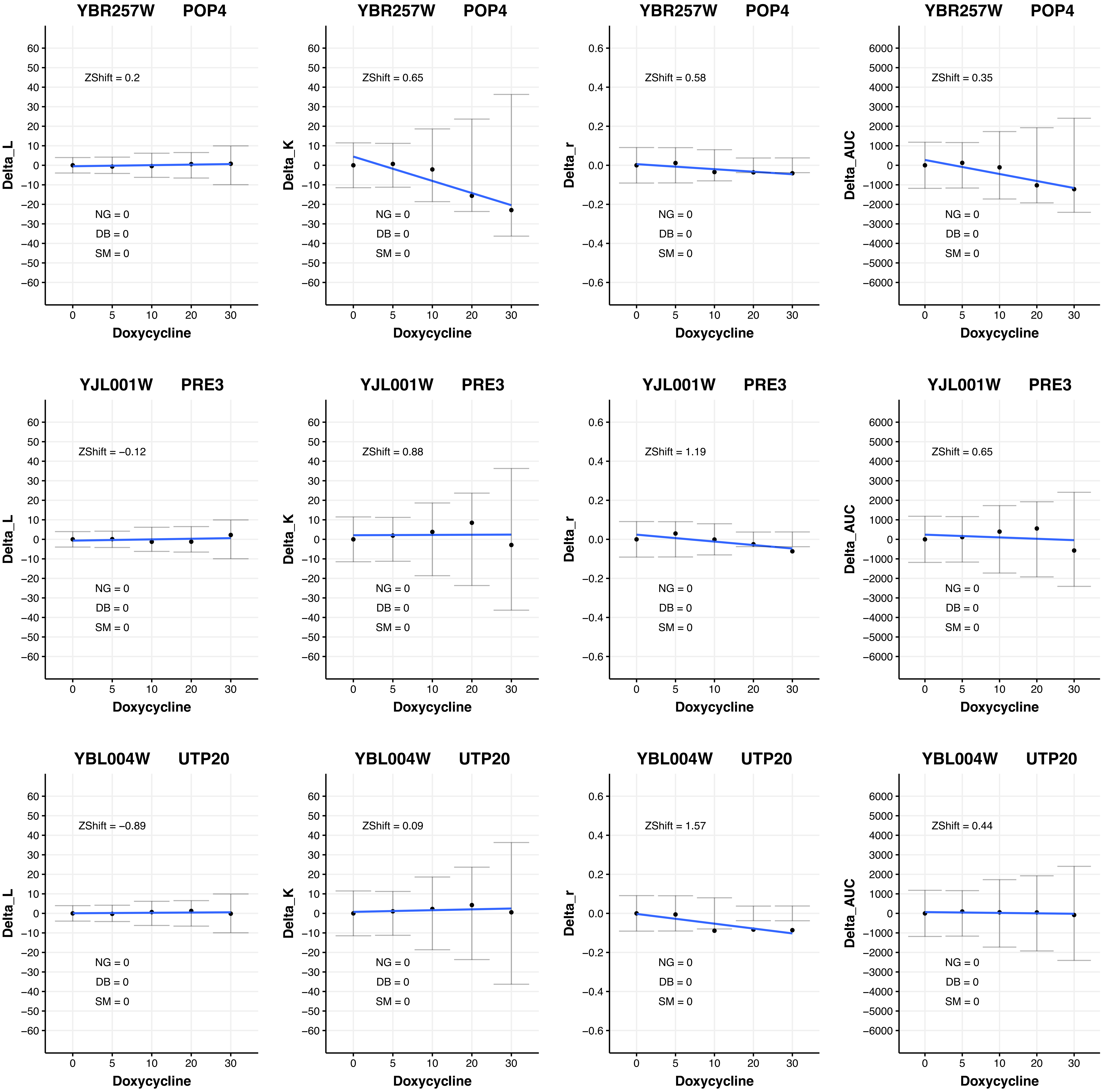

YBR257W POP4
ZShift = 0.2
Delta_L
NG = 0
DB = 0
SM = 0
Doxycycline
YBR257W POP4
ZShift = 0.65
Delta_K
NG = 0
DB = 0
SM = 0
Doxycycline
YBR257W POP4
ZShift = 0.58
Delta_r
NG = 0
DB = 0
SM = 0
Doxycycline
YBR257W POP4
ZShift = 0.35
Delta_AUC
NG = 0
DB = 0
SM = 0
Doxycycline
YJL001W PRE3
ZShift = −0.12
Delta_L
NG = 0
DB = 0
SM = 0
Doxycycline
YJL001W PRE3
ZShift = 0.88
Delta_K
NG = 0
DB = 0
SM = 0
Doxycycline
YJL001W PRE3
ZShift = 1.19
Delta_r
NG = 0
DB = 0
SM = 0
Doxycycline
YJL001W PRE3
ZShift = 0.65
Delta_AUC
NG = 0
DB = 0
SM = 0
Doxycycline
YBL004W UTP20
ZShift = −0.89
Delta_L
NG = 0
DB = 0
SM = 0
Doxycycline
YBL004W UTP20
ZShift = 0.09
Delta_K
NG = 0
DB = 0
SM = 0
Doxycycline
YBL004W UTP20
ZShift = 1.57
Delta_r
NG = 0
DB = 0
SM = 0
Doxycycline
YBL004W UTP20
ZShift = 0.44
Delta_AUC
NG = 0
DB = 0
SM = 0
Doxycycline

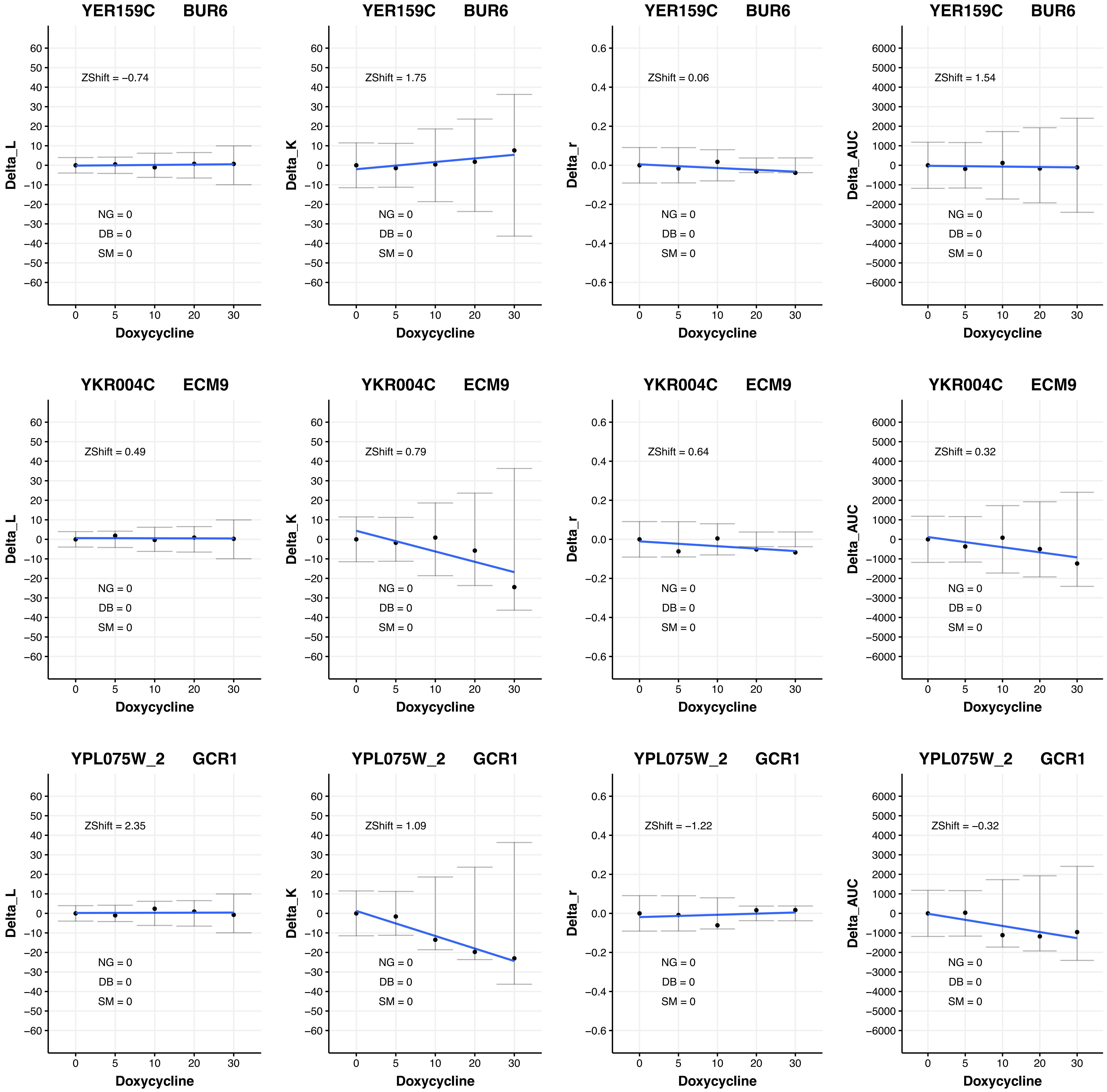

YER159C BUR6
Delta_L
ZShift = −0.74
NG = 0
DB = 0
SM = 0
Doxycycline
YER159C BUR6
Delta_K
ZShift = 1.75
NG = 0
DB = 0
SM = 0
Doxycycline
YER159C BUR6
Delta_r
ZShift = 0.06
NG = 0
DB = 0
SM = 0
Doxycycline
YER159C BUR6
Delta_AUC
ZShift = 1.54
NG = 0
DB = 0
SM = 0
Doxycycline
YKR004C ECM9
Delta_L
ZShift = 0.49
NG = 0
DB = 0
SM = 0
Doxycycline
YKR004C ECM9
Delta_K
ZShift = 0.79
NG = 0
DB = 0
SM = 0
Doxycycline
YKR004C ECM9
Delta_r
ZShift = 0.64
NG = 0
DB = 0
SM = 0
Doxycycline
YKR004C ECM9
Delta_AUC
ZShift = 0.32
NG = 0
DB = 0
SM = 0
Doxycycline
YPL075W_2 GCR1
Delta_L
ZShift = 2.35
NG = 0
DB = 0
SM = 0
Doxycycline
YPL075W_2 GCR1
Delta_K
ZShift = 1.09
NG = 0
DB = 0
SM = 0
Doxycycline
YPL075W_2 GCR1
Delta_r
ZShift = −1.22
NG = 0
DB = 0
SM = 0
Doxycycline
YPL075W_2 GCR1
Delta_AUC
ZShift = −0.32
NG = 0
DB = 0
SM = 0
Doxycycline

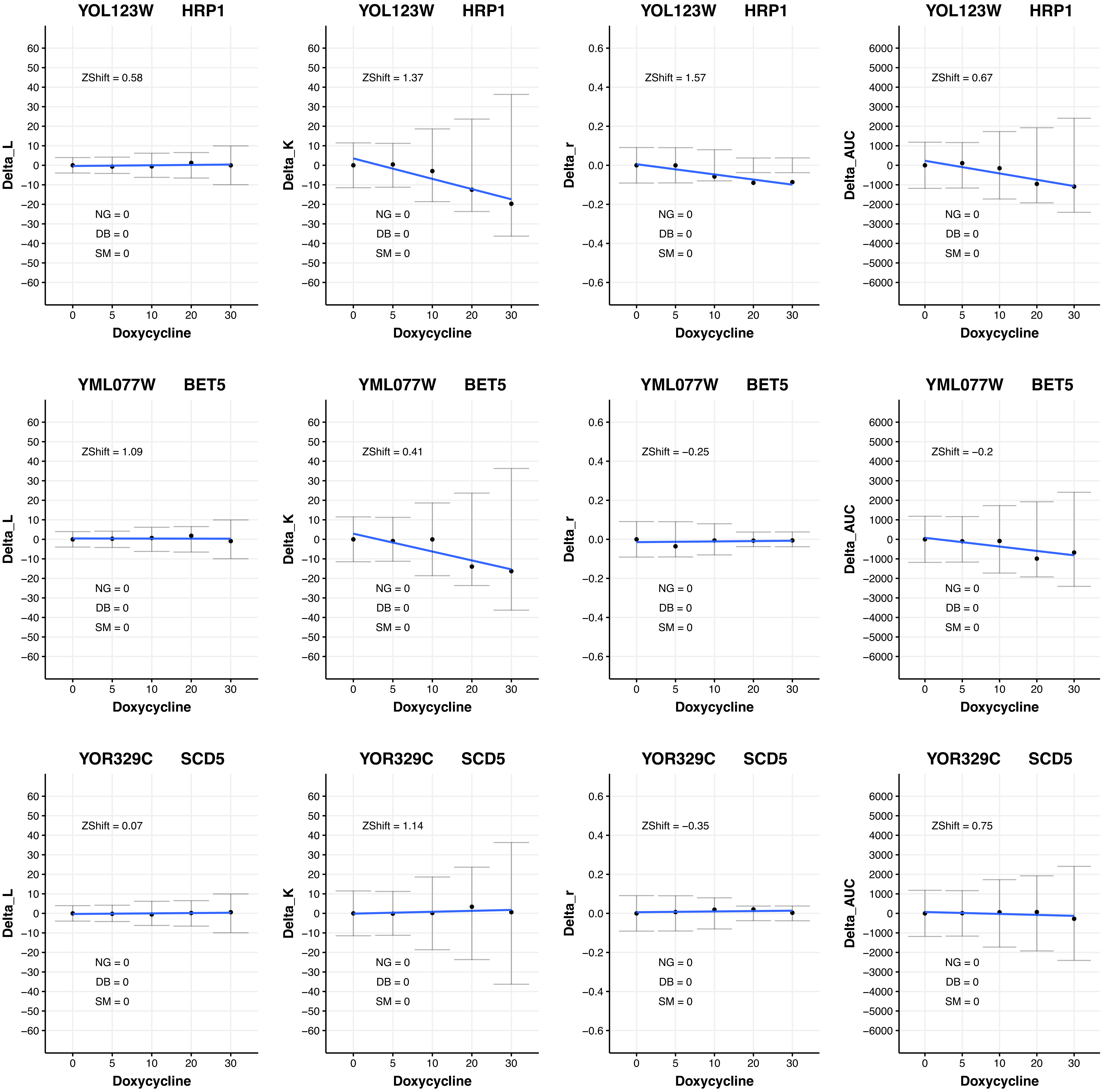

YOL123W HRP1
ZShift = 0.58
ZShift = 1.37
ZShift = 1.57
ZShift = 0.67
YML077W BET5
ZShift = 1.09
ZShift = 0.41
ZShift = −0.25
ZShift = −0.2
YOR329C SCD5
ZShift = 0.07
ZShift = 1.14
ZShift = −0.35
ZShift = 0.75
NG = 0
DB = 0
SM = 0
Delta_L
Delta_K
Delta_r
Delta_AUC
Doxycycline

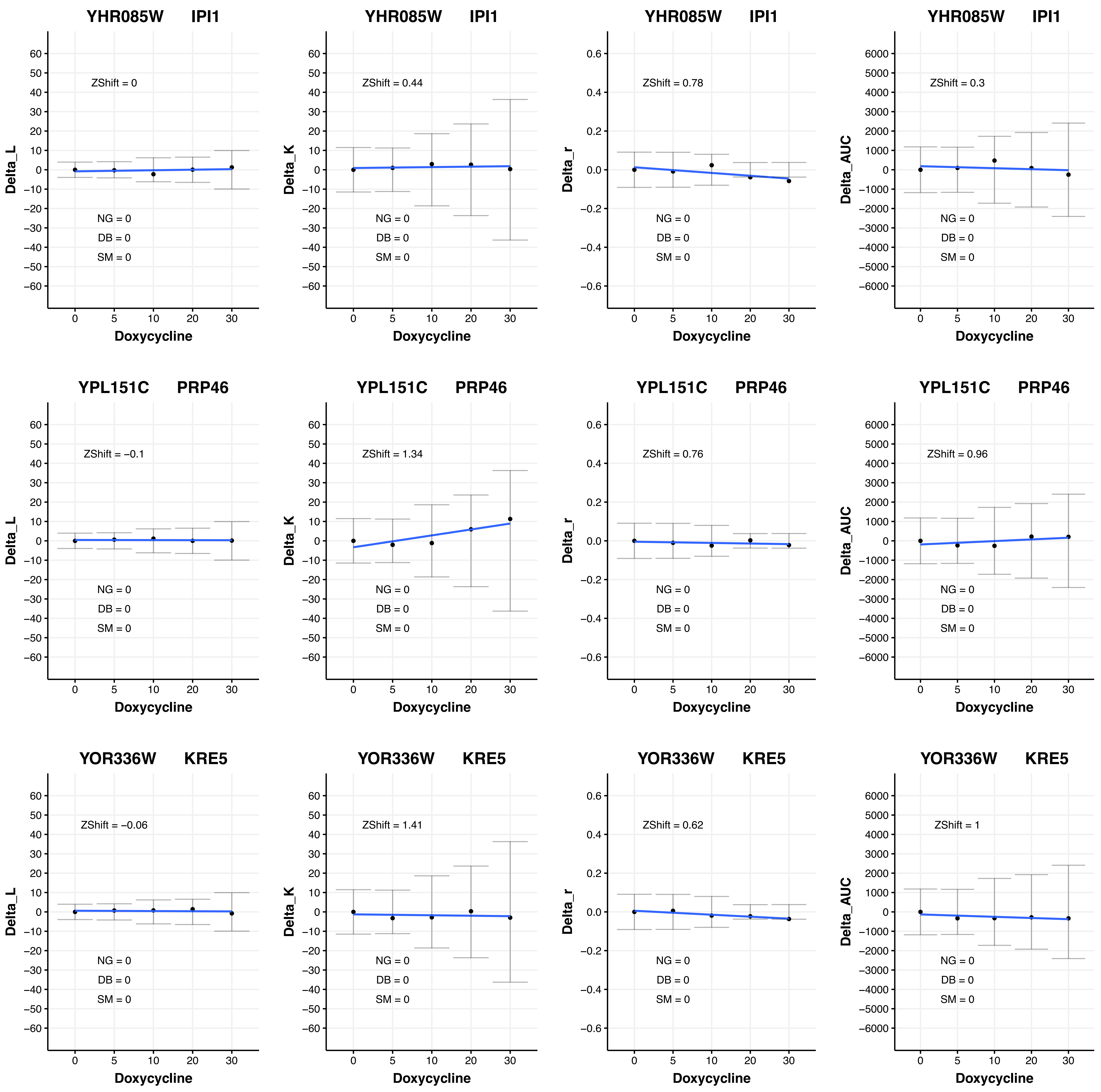
YHR085W IPI1
ZShift = 0
Delta_L
NG = 0
DB = 0
SM = 0
Doxycycline
YHR085W IPI1
ZShift = 0.44
Delta_K
NG = 0
DB = 0
SM = 0
Doxycycline
YHR085W IPI1
ZShift = 0.78
Delta_r
NG = 0
DB = 0
SM = 0
Doxycycline
YHR085W IPI1
ZShift = 0.3
Delta_AUC
NG = 0
DB = 0
SM = 0
Doxycycline
YPL151C PRP46
ZShift = −0.1
Delta_L
NG = 0
DB = 0
SM = 0
Doxycycline
YPL151C PRP46
ZShift = 1.34
Delta_K
NG = 0
DB = 0
SM = 0
Doxycycline
YPL151C PRP46
ZShift = 0.76
Delta_r
NG = 0
DB = 0
SM = 0
Doxycycline
YPL151C PRP46
ZShift = 0.96
Delta_AUC
NG = 0
DB = 0
SM = 0
Doxycycline
YOR336W KRE5
ZShift = −0.06
Delta_L
NG = 0
DB = 0
SM = 0
Doxycycline
YOR336W KRE5
ZShift = 1.41
Delta_K
NG = 0
DB = 0
SM = 0
Doxycycline
YOR336W KRE5
ZShift = 0.62
Delta_r
NG = 0
DB = 0
SM = 0
Doxycycline
YOR336W KRE5
ZShift = 1
Delta_AUC
NG = 0
DB = 0
SM = 0
Doxycycline

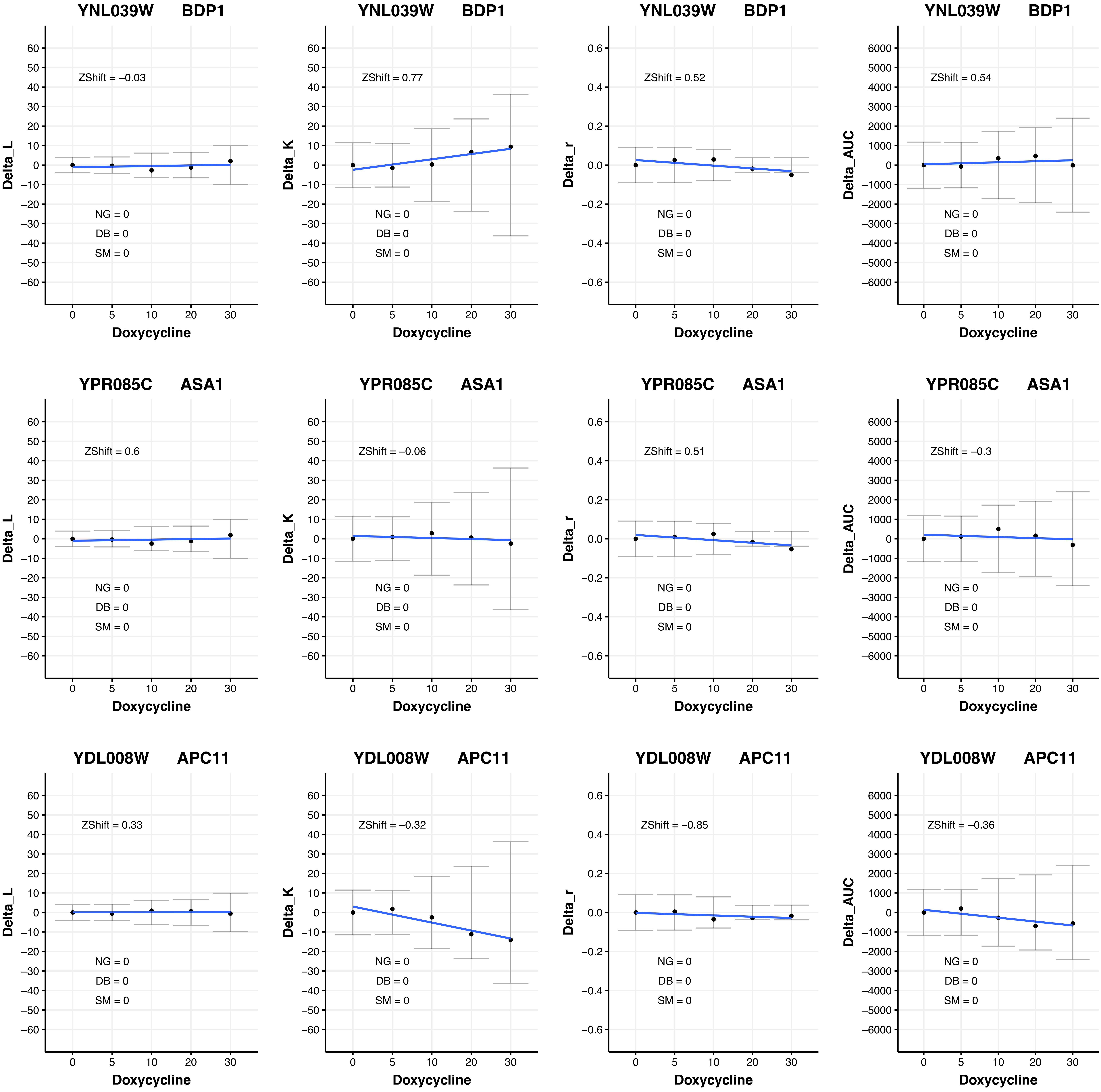
YNL039W BDP1
ZShift = −0.03
NG = 0
DB = 0
SM = 0
Delta_L
Doxycycline
YNL039W BDP1
ZShift = 0.77
NG = 0
DB = 0
SM = 0
Delta_K
Doxycycline
YNL039W BDP1
ZShift = 0.52
NG = 0
DB = 0
SM = 0
Delta_r
Doxycycline
YNL039W BDP1
ZShift = 0.54
NG = 0
DB = 0
SM = 0
Delta_AUC
Doxycycline
YPR085C ASA1
ZShift = 0.6
NG = 0
DB = 0
SM = 0
Delta_L
Doxycycline
YPR085C ASA1
ZShift = −0.06
NG = 0
DB = 0
SM = 0
Delta_K
Doxycycline
YPR085C ASA1
ZShift = 0.51
NG = 0
DB = 0
SM = 0
Delta_r
Doxycycline
YPR085C ASA1
ZShift = −0.3
NG = 0
DB = 0
SM = 0
Delta_AUC
Doxycycline
YDL008W APC11
ZShift = 0.33
NG = 0
DB = 0
SM = 0
Delta_L
Doxycycline
YDL008W APC11
ZShift = −0.32
NG = 0
DB = 0
SM = 0
Delta_K
Doxycycline
YDL008W APC11
ZShift = −0.85
NG = 0
DB = 0
SM = 0
Delta_r
Doxycycline
YDL008W APC11
ZShift = −0.36
NG = 0
DB = 0
SM = 0
Delta_AUC
Doxycycline

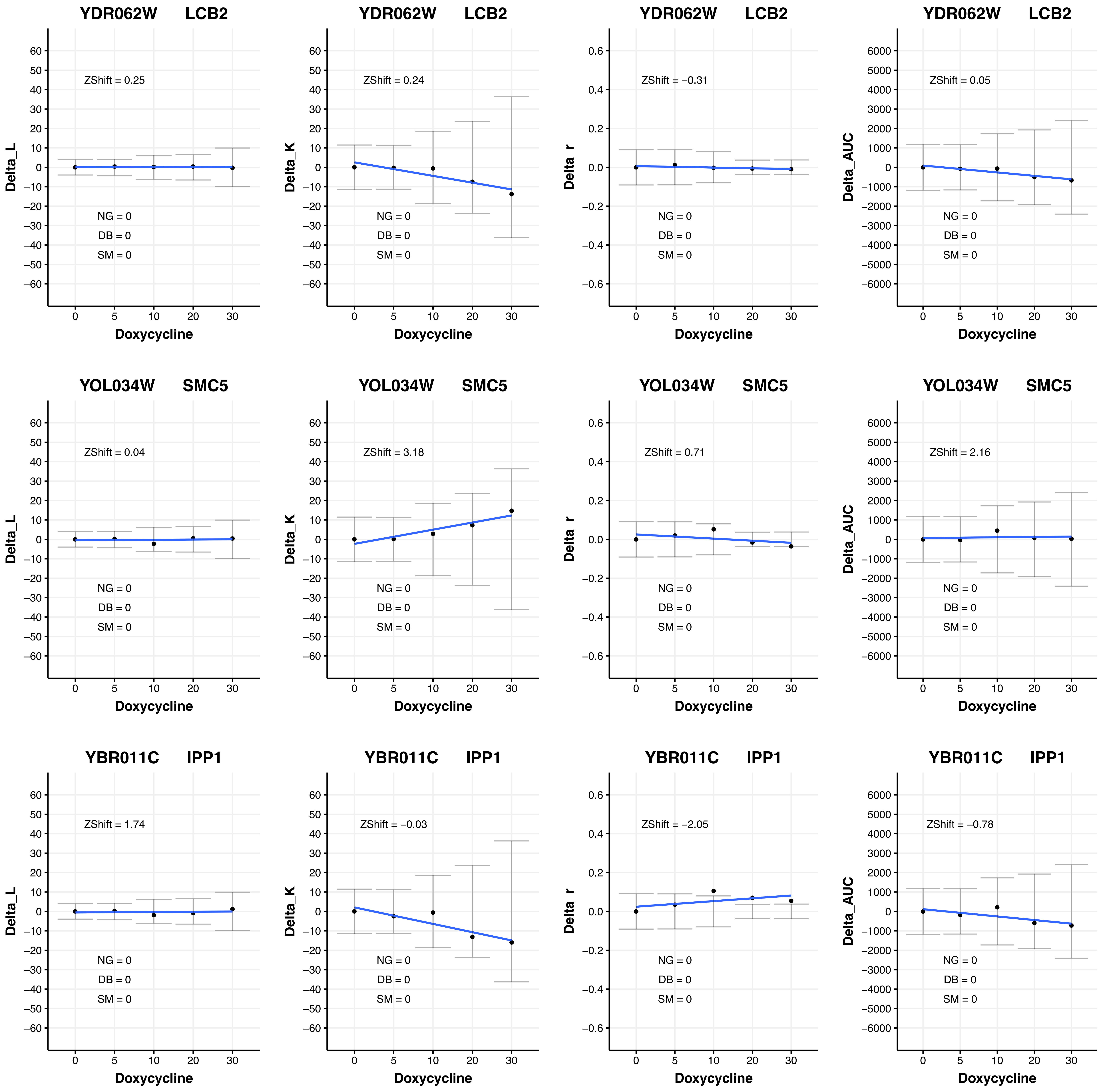

YDR062W LCB2
ZShift = 0.25
Delta_L
NG = 0
DB = 0
SM = 0
Doxycycline
YDR062W LCB2
ZShift = 0.24
Delta_K
NG = 0
DB = 0
SM = 0
Doxycycline
YDR062W LCB2
ZShift = −0.31
Delta_r
NG = 0
DB = 0
SM = 0
Doxycycline
YDR062W LCB2
ZShift = 0.05
Delta_AUC
NG = 0
DB = 0
SM = 0
Doxycycline
YOL034W SMC5
ZShift = 0.04
Delta_L
NG = 0
DB = 0
SM = 0
Doxycycline
YOL034W SMC5
ZShift = 3.18
Delta_K
NG = 0
DB = 0
SM = 0
Doxycycline
YOL034W SMC5
ZShift = 0.71
Delta_r
NG = 0
DB = 0
SM = 0
Doxycycline
YOL034W SMC5
ZShift = 2.16
Delta_AUC
NG = 0
DB = 0
SM = 0
Doxycycline
YBR011C IPP1
ZShift = 1.74
Delta_L
NG = 0
DB = 0
SM = 0
Doxycycline
YBR011C IPP1
ZShift = −0.03
Delta_K
NG = 0
DB = 0
SM = 0
Doxycycline
YBR011C IPP1
ZShift = −2.05
Delta_r
NG = 0
DB = 0
SM = 0
Doxycycline
YBR011C IPP1
ZShift = −0.78
Delta_AUC
NG = 0
DB = 0
SM = 0
Doxycycline

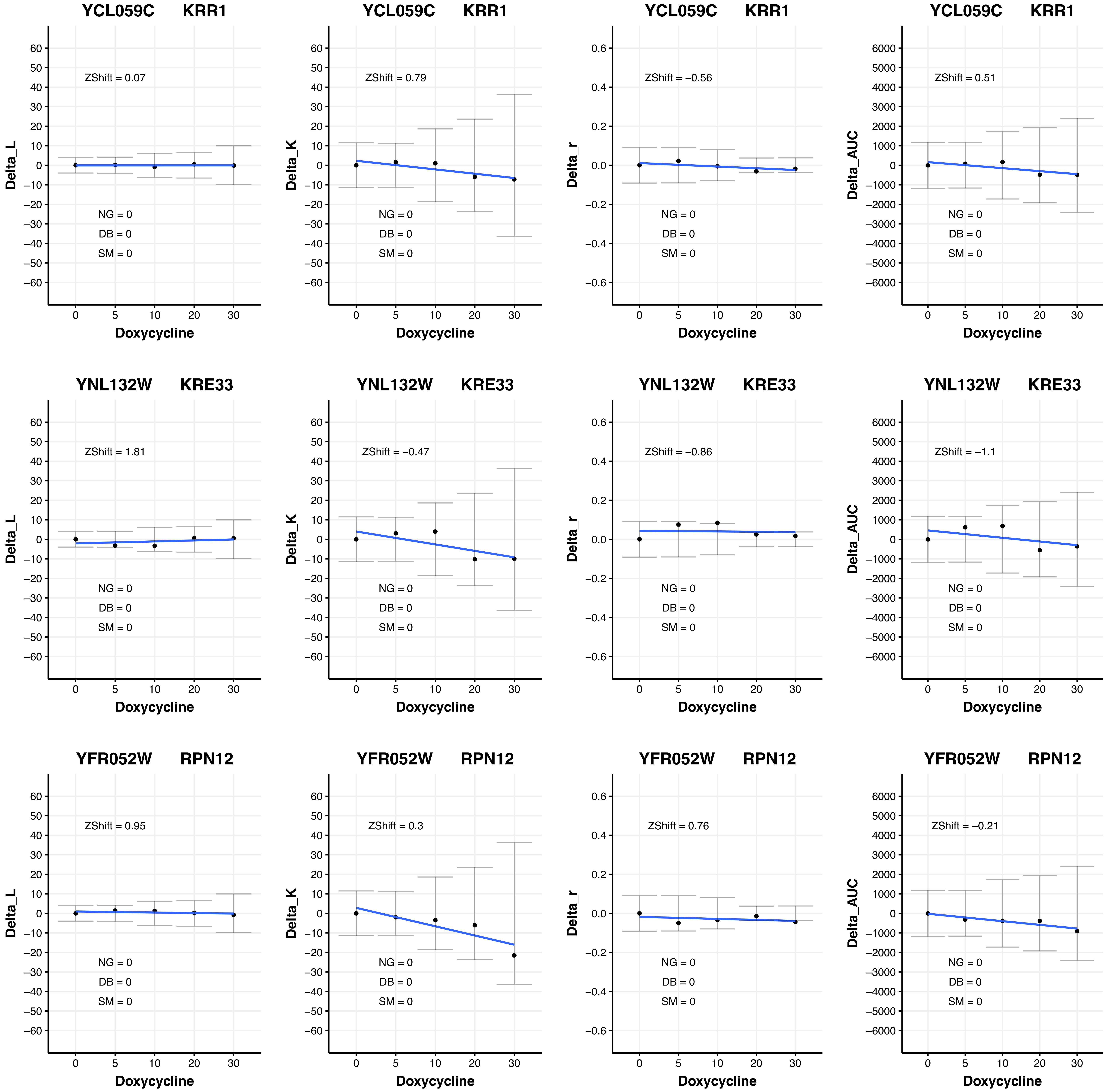

YCL059C KRR1
ZShift = 0.07
Delta_L
NG = 0
DB = 0
SM = 0
Doxycycline
YCL059C KRR1
ZShift = 0.79
Delta_K
NG = 0
DB = 0
SM = 0
Doxycycline
YCL059C KRR1
ZShift = −0.56
Delta_r
NG = 0
DB = 0
SM = 0
Doxycycline
YCL059C KRR1
ZShift = 0.51
Delta_AUC
NG = 0
DB = 0
SM = 0
Doxycycline
YNL132W KRE33
ZShift = 1.81
Delta_L
NG = 0
DB = 0
SM = 0
Doxycycline
YNL132W KRE33
ZShift = −0.47
Delta_K
NG = 0
DB = 0
SM = 0
Doxycycline
YNL132W KRE33
ZShift = −0.86
Delta_r
NG = 0
DB = 0
SM = 0
Doxycycline
YNL132W KRE33
ZShift = −1.1
Delta_AUC
NG = 0
DB = 0
SM = 0
Doxycycline
YFR052W RPN12
ZShift = 0.95
Delta_L
NG = 0
DB = 0
SM = 0
Doxycycline
YFR052W RPN12
ZShift = 0.3
Delta_K
NG = 0
DB = 0
SM = 0
Doxycycline
YFR052W RPN12
ZShift = 0.76
Delta_r
NG = 0
DB = 0
SM = 0
Doxycycline
YFR052W RPN12
ZShift = −0.21
Delta_AUC
NG = 0
DB = 0
SM = 0
Doxycycline

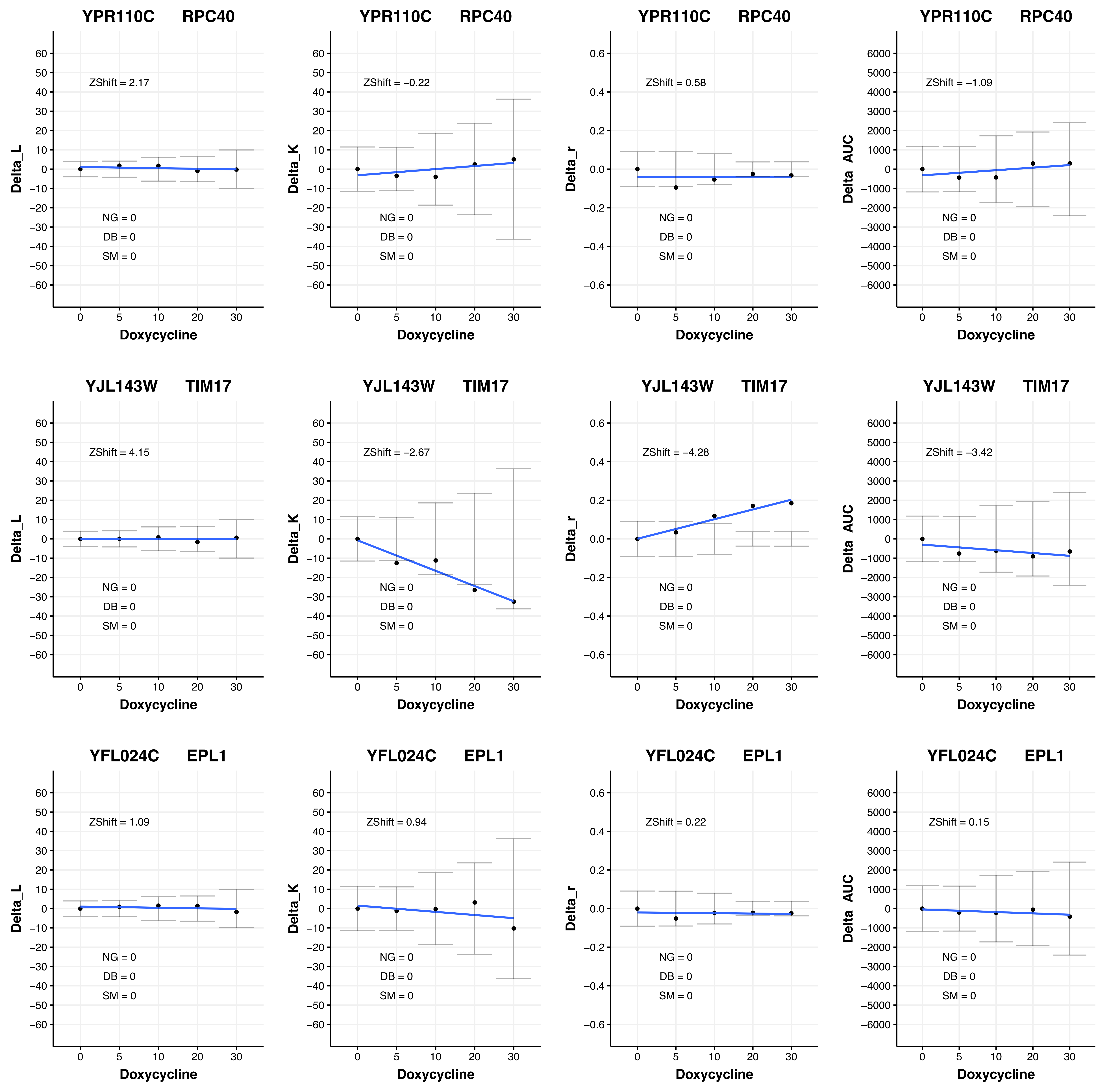

YPR110C RPC40
ZShift = 2.17
ZShift = −0.22
ZShift = 0.58
ZShift = −1.09
YJL143W TIM17
ZShift = 4.15
ZShift = −2.67
ZShift = −4.28
ZShift = −3.42
YFL024C EPL1
ZShift = 1.09
ZShift = 0.94
ZShift = 0.22
ZShift = 0.15
NG = 0
DB = 0
SM = 0
Delta_L
Delta_K
Delta_r
Delta_AUC
Doxycycline

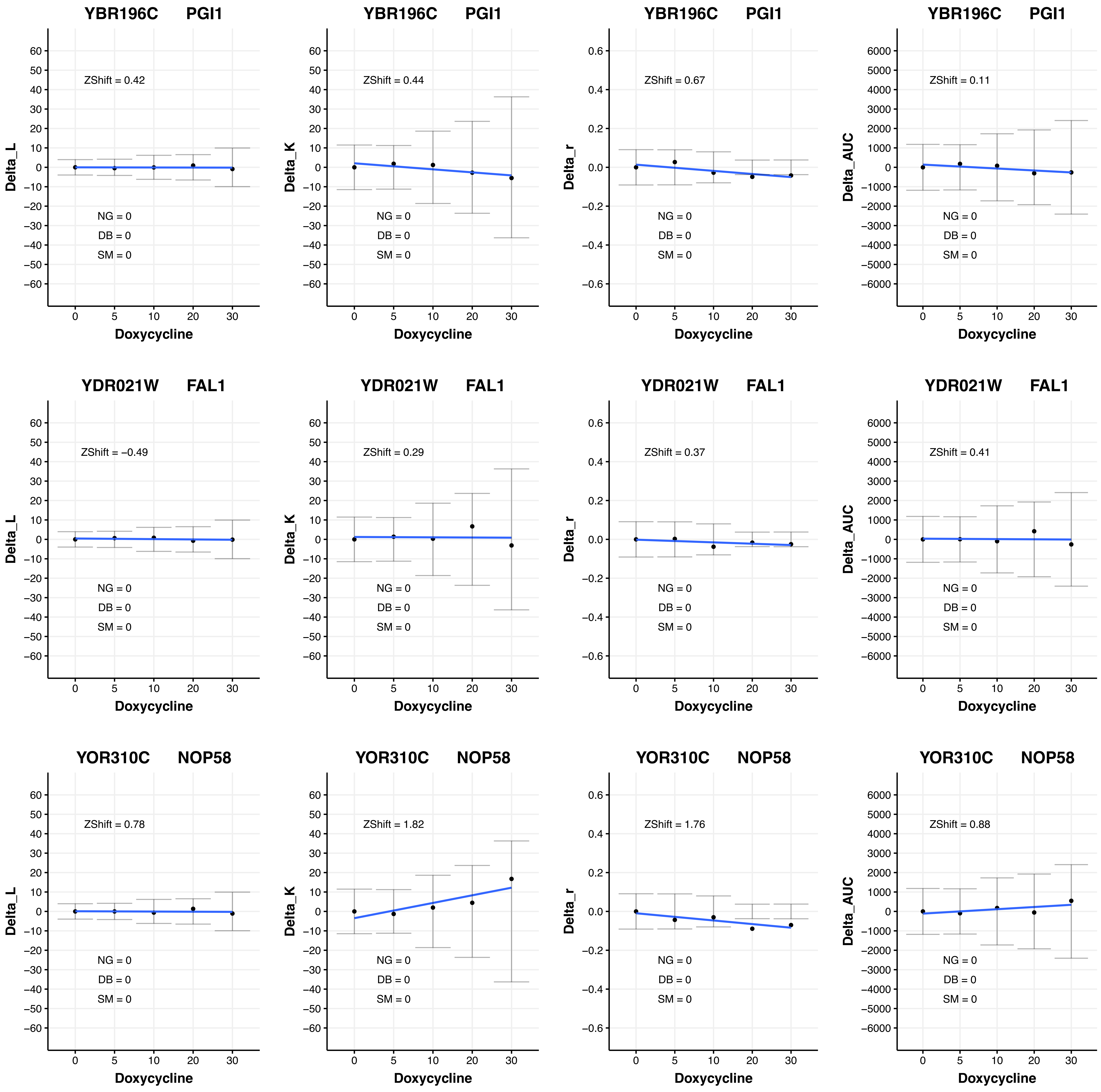

YBR196C PGI1
ZShift = 0.42
Delta_L
NG = 0
DB = 0
SM = 0
Doxycycline
YBR196C PGI1
ZShift = 0.44
Delta_K
NG = 0
DB = 0
SM = 0
Doxycycline
YBR196C PGI1
ZShift = 0.67
Delta_r
NG = 0
DB = 0
SM = 0
Doxycycline
YBR196C PGI1
ZShift = 0.11
Delta_AUC
NG = 0
DB = 0
SM = 0
Doxycycline
YDR021W FAL1
ZShift = −0.49
Delta_L
NG = 0
DB = 0
SM = 0
Doxycycline
YDR021W FAL1
ZShift = 0.29
Delta_K
NG = 0
DB = 0
SM = 0
Doxycycline
YDR021W FAL1
ZShift = 0.37
Delta_r
NG = 0
DB = 0
SM = 0
Doxycycline
YDR021W FAL1
ZShift = 0.41
Delta_AUC
NG = 0
DB = 0
SM = 0
Doxycycline
YOR310C NOP58
ZShift = 0.78
Delta_L
NG = 0
DB = 0
SM = 0
Doxycycline
YOR310C NOP58
ZShift = 1.82
Delta_K
NG = 0
DB = 0
SM = 0
Doxycycline
YOR310C NOP58
ZShift = 1.76
Delta_r
NG = 0
DB = 0
SM = 0
Doxycycline
YOR310C NOP58
ZShift = 0.88
Delta_AUC
NG = 0
DB = 0
SM = 0
Doxycycline

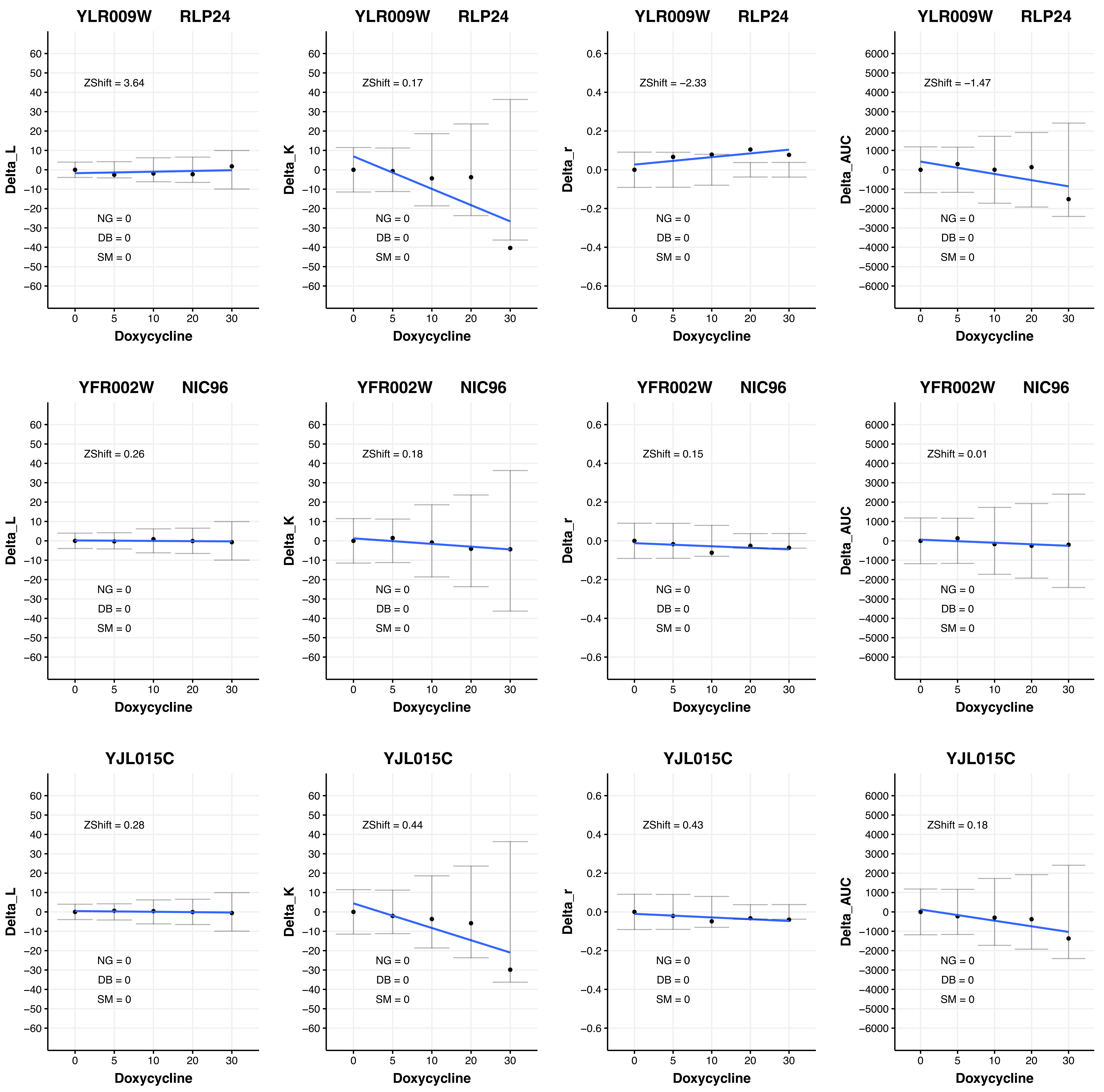

YLR009W RLP24
ZShift = 3.64
Delta_L
NG = 0
DB = 0
SM = 0
Doxycycline
YLR009W RLP24
ZShift = 0.17
Delta_K
NG = 0
DB = 0
SM = 0
Doxycycline
YLR009W RLP24
ZShift = −2.33
Delta_r
NG = 0
DB = 0
SM = 0
Doxycycline
YLR009W RLP24
ZShift = −1.47
Delta_AUC
NG = 0
DB = 0
SM = 0
Doxycycline
YFR002W NIC96
ZShift = 0.26
Delta_L
NG = 0
DB = 0
SM = 0
Doxycycline
YFR002W NIC96
ZShift = 0.18
Delta_K
NG = 0
DB = 0
SM = 0
Doxycycline
YFR002W NIC96
ZShift = 0.15
Delta_r
NG = 0
DB = 0
SM = 0
Doxycycline
YFR002W NIC96
ZShift = 0.01
Delta_AUC
NG = 0
DB = 0
SM = 0
Doxycycline
YJL015C
ZShift = 0.28
Delta_L
NG = 0
DB = 0
SM = 0
Doxycycline
YJL015C
ZShift = 0.44
Delta_K
NG = 0
DB = 0
SM = 0
Doxycycline
YJL015C
ZShift = 0.43
Delta_r
NG = 0
DB = 0
SM = 0
Doxycycline
YJL015C
ZShift = 0.18
Delta_AUC
NG = 0
DB = 0
SM = 0
Doxycycline

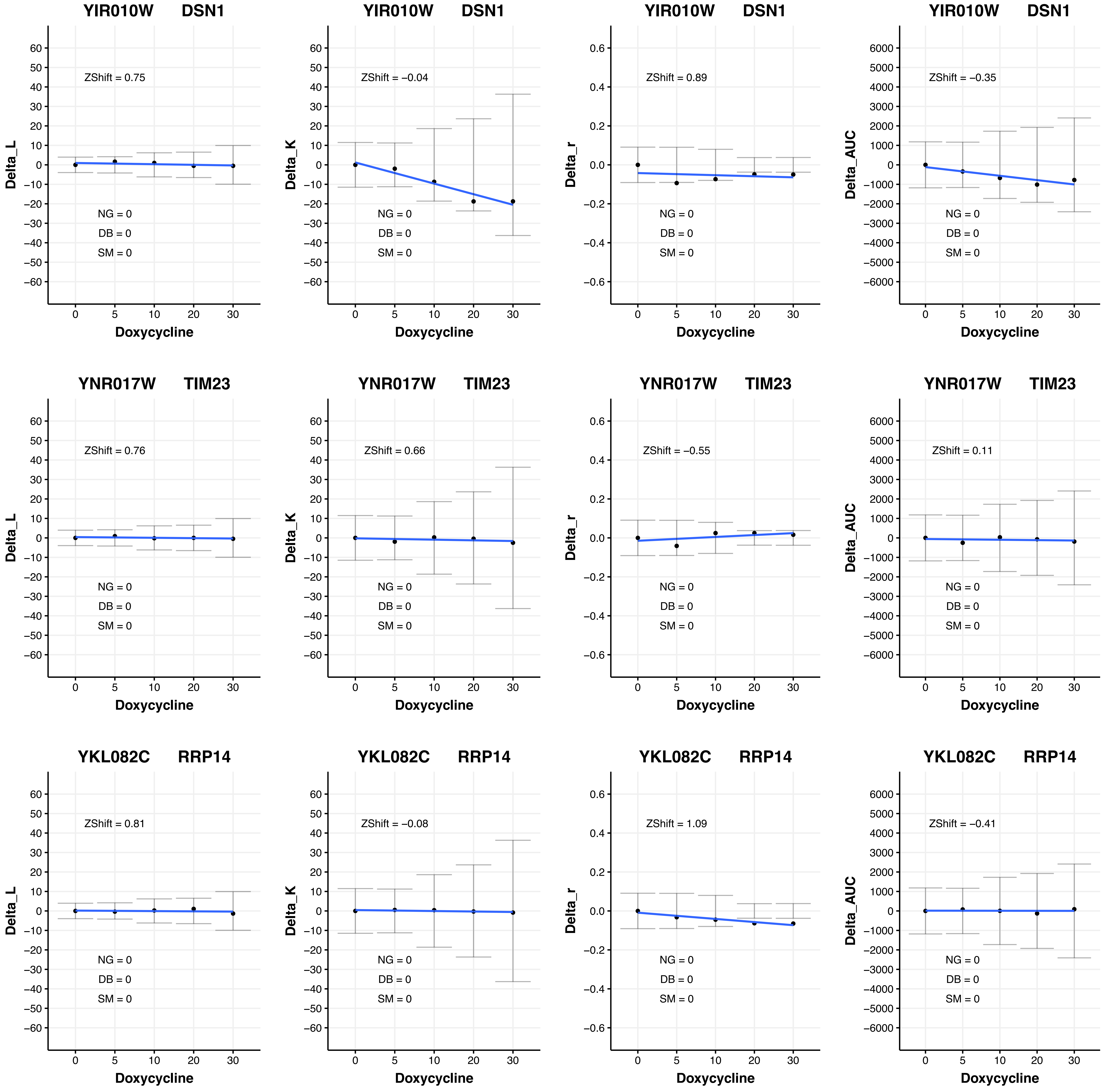

YIR010W DSN1
ZShift = 0.75
NG = 0
DB = 0
SM = 0
Delta_L
Doxycycline
YIR010W DSN1
ZShift = −0.04
NG = 0
DB = 0
SM = 0
Delta_K
Doxycycline
YIR010W DSN1
ZShift = 0.89
NG = 0
DB = 0
SM = 0
Delta_r
Doxycycline
YIR010W DSN1
ZShift = −0.35
NG = 0
DB = 0
SM = 0
Delta_AUC
Doxycycline
YNR017W TIM23
ZShift = 0.76
NG = 0
DB = 0
SM = 0
Delta_L
Doxycycline
YNR017W TIM23
ZShift = 0.66
NG = 0
DB = 0
SM = 0
Delta_K
Doxycycline
YNR017W TIM23
ZShift = −0.55
NG = 0
DB = 0
SM = 0
Delta_r
Doxycycline
YNR017W TIM23
ZShift = 0.11
NG = 0
DB = 0
SM = 0
Delta_AUC
Doxycycline
YKL082C RRP14
ZShift = 0.81
NG = 0
DB = 0
SM = 0
Delta_L
Doxycycline
YKL082C RRP14
ZShift = −0.08
NG = 0
DB = 0
SM = 0
Delta_K
Doxycycline
YKL082C RRP14
ZShift = 1.09
NG = 0
DB = 0
SM = 0
Delta_r
Doxycycline
YKL082C RRP14
ZShift = −0.41
NG = 0
DB = 0
SM = 0
Delta_AUC
Doxycycline

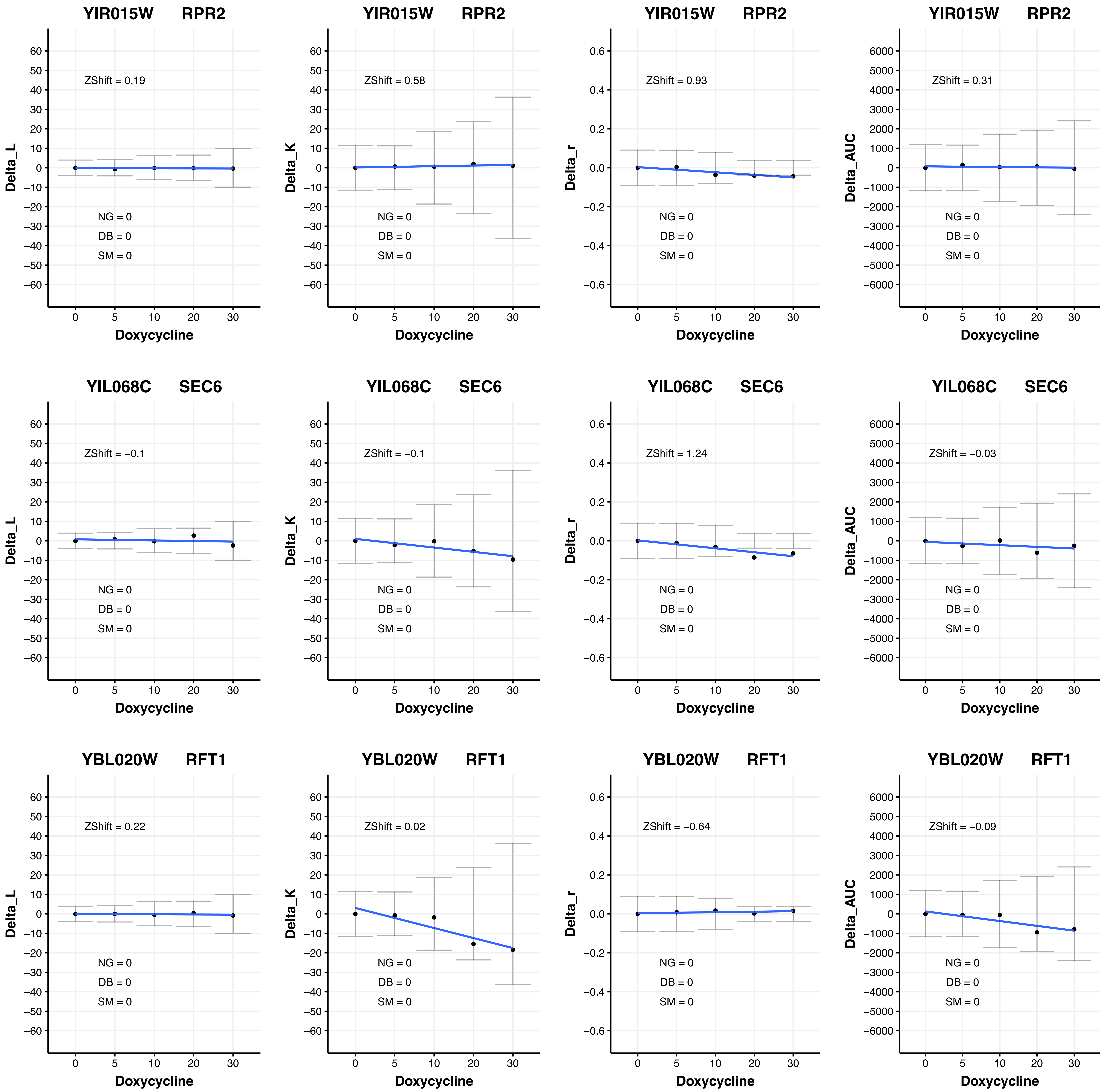

YIR015W RPR2
ZShift = 0.19
NG = 0
DB = 0
SM = 0
Delta_L
Doxycycline
YIR015W RPR2
ZShift = 0.58
NG = 0
DB = 0
SM = 0
Delta_K
Doxycycline
YIR015W RPR2
ZShift = 0.93
NG = 0
DB = 0
SM = 0
Delta_r
Doxycycline
YIR015W RPR2
ZShift = 0.31
NG = 0
DB = 0
SM = 0
Delta_AUC
Doxycycline
YIL068C SEC6
ZShift = −0.1
NG = 0
DB = 0
SM = 0
Delta_L
Doxycycline
YIL068C SEC6
ZShift = −0.1
NG = 0
DB = 0
SM = 0
Delta_K
Doxycycline
YIL068C SEC6
ZShift = 1.24
NG = 0
DB = 0
SM = 0
Delta_r
Doxycycline
YIL068C SEC6
ZShift = −0.03
NG = 0
DB = 0
SM = 0
Delta_AUC
Doxycycline
YBL020W RFT1
ZShift = 0.22
NG = 0
DB = 0
SM = 0
Delta_L
Doxycycline
YBL020W RFT1
ZShift = 0.02
NG = 0
DB = 0
SM = 0
Delta_K
Doxycycline
YBL020W RFT1
ZShift = −0.64
NG = 0
DB = 0
SM = 0
Delta_r
Doxycycline
YBL020W RFT1
ZShift = −0.09
NG = 0
DB = 0
SM = 0
Delta_AUC
Doxycycline

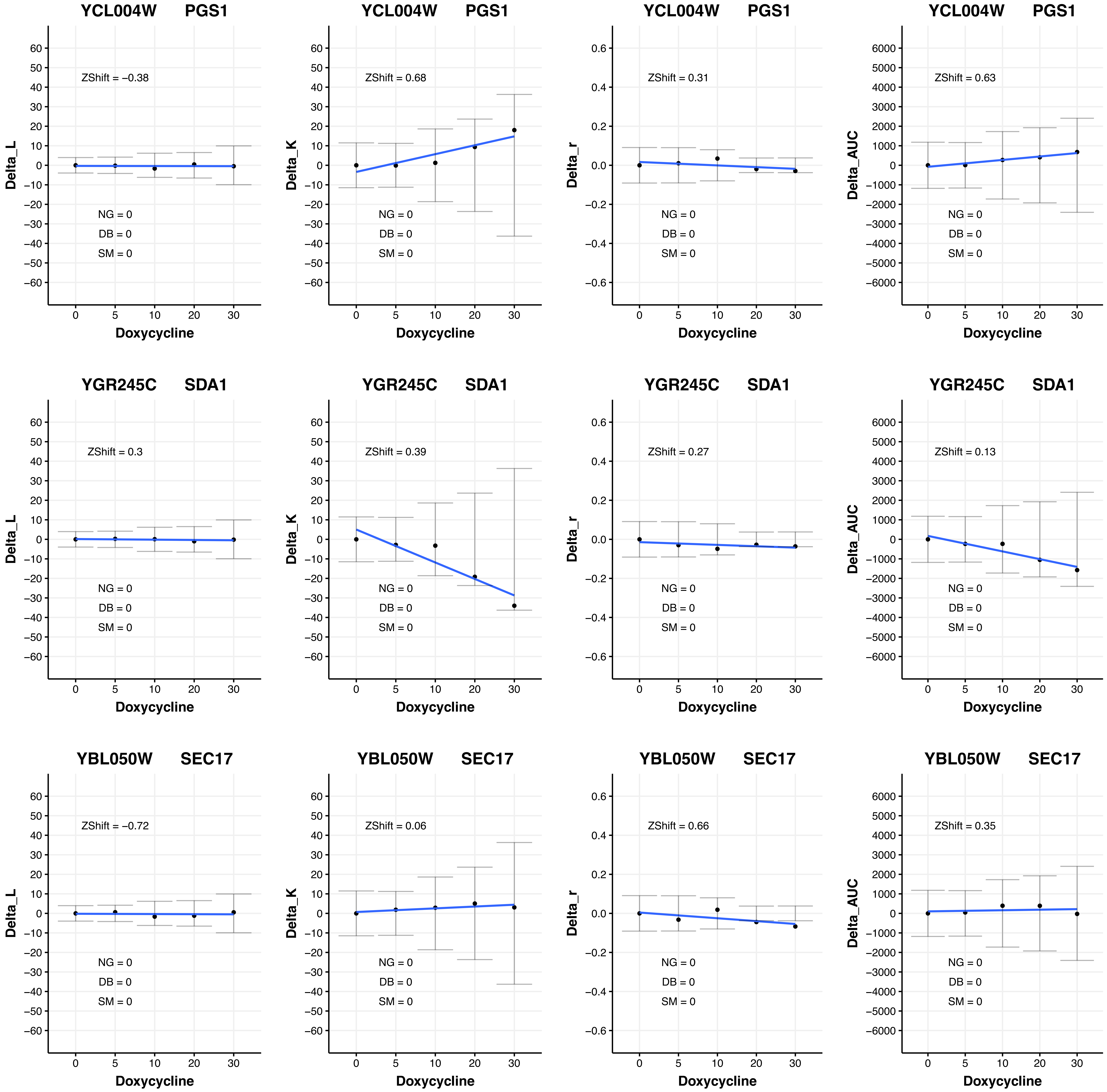
YCL004W PGS1
ZShift = −0.38
NG = 0
DB = 0
SM = 0
Delta_L
Doxycycline
YCL004W PGS1
ZShift = 0.68
NG = 0
DB = 0
SM = 0
Delta_K
Doxycycline
YCL004W PGS1
ZShift = 0.31
NG = 0
DB = 0
SM = 0
Delta_r
Doxycycline
YCL004W PGS1
ZShift = 0.63
NG = 0
DB = 0
SM = 0
Delta_AUC
Doxycycline
YGR245C SDA1
ZShift = 0.3
NG = 0
DB = 0
SM = 0
Delta_L
Doxycycline
YGR245C SDA1
ZShift = 0.39
NG = 0
DB = 0
SM = 0
Delta_K
Doxycycline
YGR245C SDA1
ZShift = 0.27
NG = 0
DB = 0
SM = 0
Delta_r
Doxycycline
YGR245C SDA1
ZShift = 0.13
NG = 0
DB = 0
SM = 0
Delta_AUC
Doxycycline
YBL050W SEC17
ZShift = −0.72
NG = 0
DB = 0
SM = 0
Delta_L
Doxycycline
YBL050W SEC17
ZShift = 0.06
NG = 0
DB = 0
SM = 0
Delta_K
Doxycycline
YBL050W SEC17
ZShift = 0.66
NG = 0
DB = 0
SM = 0
Delta_r
Doxycycline
YBL050W SEC17
ZShift = 0.35
NG = 0
DB = 0
SM = 0
Delta_AUC
Doxycycline

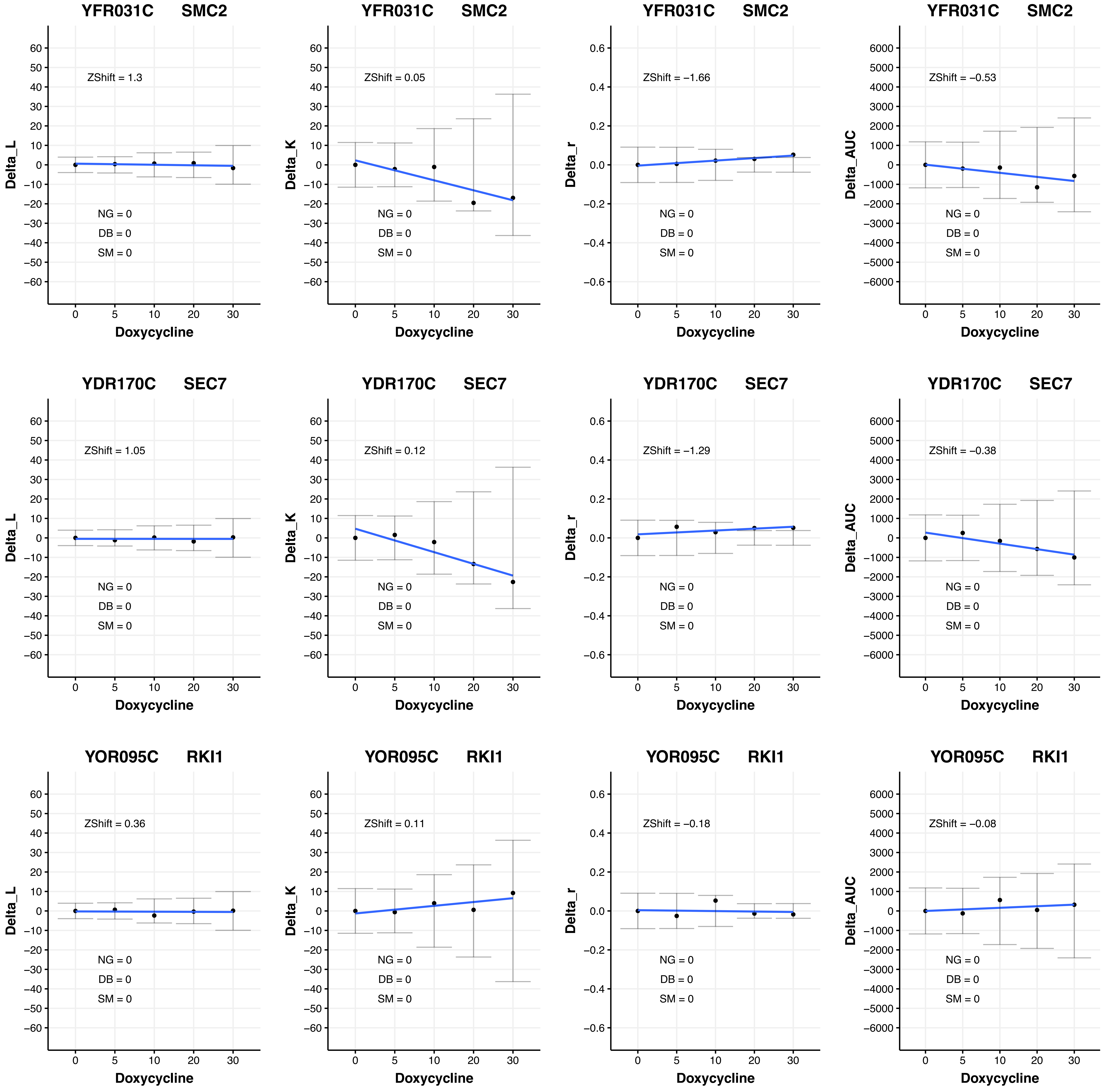

YFR031C SMC2
ZShift = 1.3
ZShift = 0.05
ZShift = −1.66
ZShift = −0.53
YDR170C SEC7
ZShift = 1.05
ZShift = 0.12
ZShift = −1.29
ZShift = −0.38
YOR095C RKI1
ZShift = 0.36
ZShift = 0.11
ZShift = −0.18
ZShift = −0.08
NG = 0
DB = 0
SM = 0
Delta_L
Delta_K
Delta_r
Delta_AUC
Doxycycline

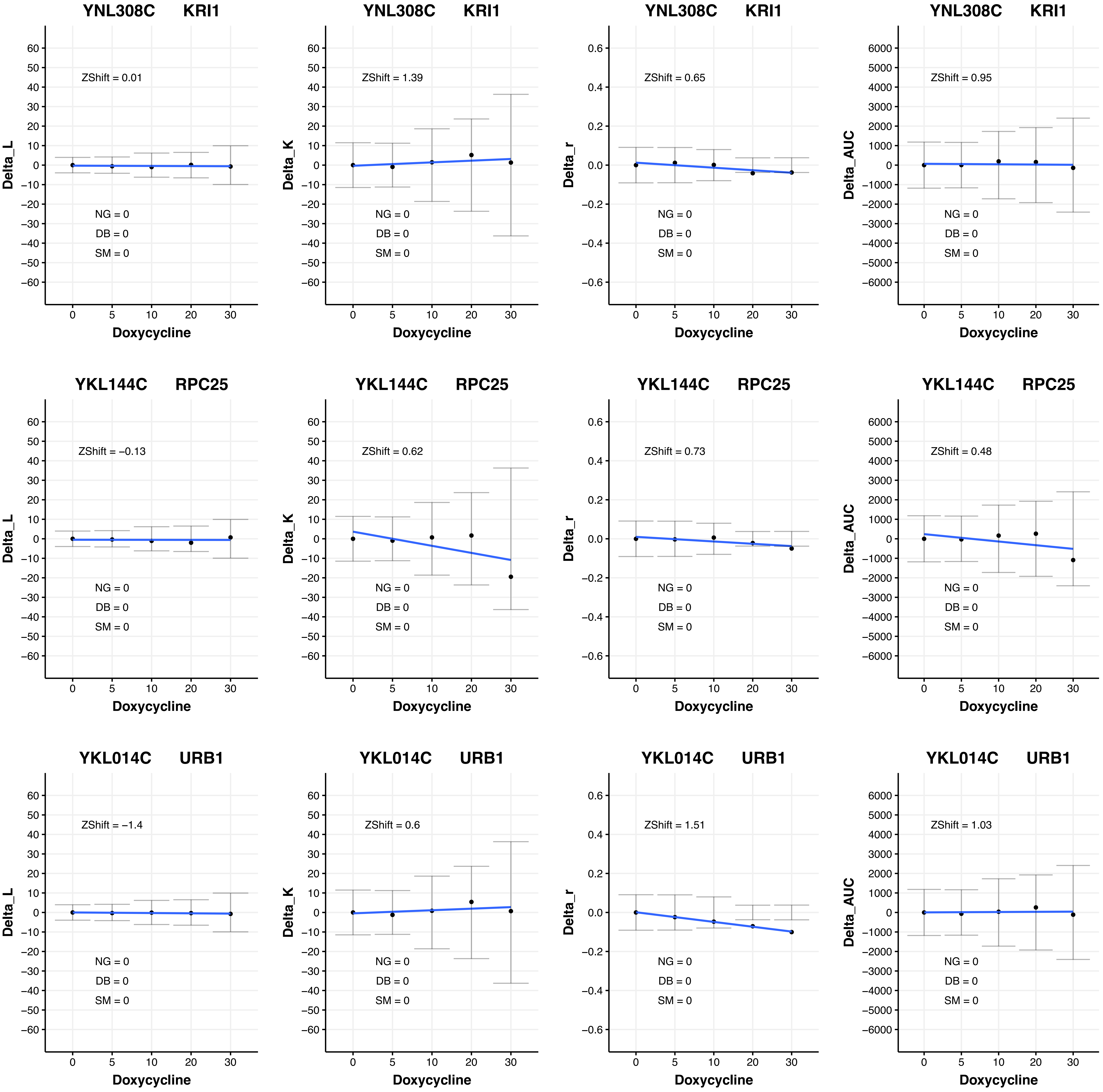
YNL308C KRI1
ZShift = 0.01
Delta_L
NG = 0
DB = 0
SM = 0
Doxycycline
YNL308C KRI1
ZShift = 1.39
Delta_K
NG = 0
DB = 0
SM = 0
Doxycycline
YNL308C KRI1
ZShift = 0.65
Delta_r
NG = 0
DB = 0
SM = 0
Doxycycline
YNL308C KRI1
ZShift = 0.95
Delta_AUC
NG = 0
DB = 0
SM = 0
Doxycycline
YKL144C RPC25
ZShift = −0.13
Delta_L
NG = 0
DB = 0
SM = 0
Doxycycline
YKL144C RPC25
ZShift = 0.62
Delta_K
NG = 0
DB = 0
SM = 0
Doxycycline
YKL144C RPC25
ZShift = 0.73
Delta_r
NG = 0
DB = 0
SM = 0
Doxycycline
YKL144C RPC25
ZShift = 0.48
Delta_AUC
NG = 0
DB = 0
SM = 0
Doxycycline
YKL014C URB1
ZShift = −1.4
Delta_L
NG = 0
DB = 0
SM = 0
Doxycycline
YKL014C URB1
ZShift = 0.6
Delta_K
NG = 0
DB = 0
SM = 0
Doxycycline
YKL014C URB1
ZShift = 1.51
Delta_r
NG = 0
DB = 0
SM = 0
Doxycycline
YKL014C URB1
ZShift = 1.03
Delta_AUC
NG = 0
DB = 0
SM = 0
Doxycycline

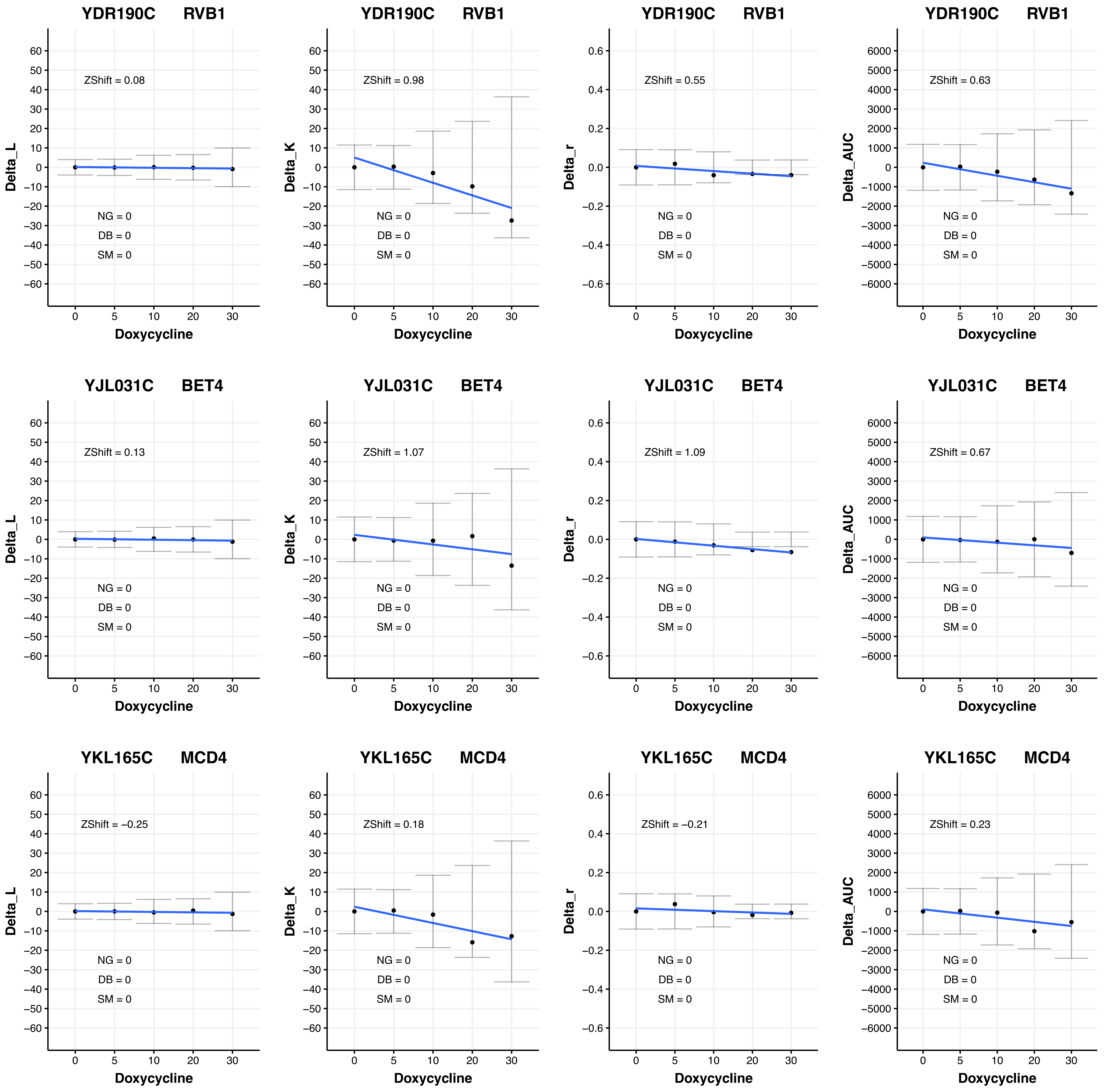

YDR190C RVB1
ZShift = 0.08
NG = 0
DB = 0
SM = 0
Delta_L
Doxycycline
YDR190C RVB1
ZShift = 0.98
NG = 0
DB = 0
SM = 0
Delta_K
Doxycycline
YDR190C RVB1
ZShift = 0.55
NG = 0
DB = 0
SM = 0
Delta_r
Doxycycline
YDR190C RVB1
ZShift = 0.63
NG = 0
DB = 0
SM = 0
Delta_AUC
Doxycycline
YJL031C BET4
ZShift = 0.13
NG = 0
DB = 0
SM = 0
Delta_L
Doxycycline
YJL031C BET4
ZShift = 1.07
NG = 0
DB = 0
SM = 0
Delta_K
Doxycycline
YJL031C BET4
ZShift = 1.09
NG = 0
DB = 0
SM = 0
Delta_r
Doxycycline
YJL031C BET4
ZShift = 0.67
NG = 0
DB = 0
SM = 0
Delta_AUC
Doxycycline
YKL165C MCD4
ZShift = −0.25
NG = 0
DB = 0
SM = 0
Delta_L
Doxycycline
YKL165C MCD4
ZShift = 0.18
NG = 0
DB = 0
SM = 0
Delta_K
Doxycycline
YKL165C MCD4
ZShift = −0.21
NG = 0
DB = 0
SM = 0
Delta_r
Doxycycline
YKL165C MCD4
ZShift = 0.23
NG = 0
DB = 0
SM = 0
Delta_AUC
Doxycycline

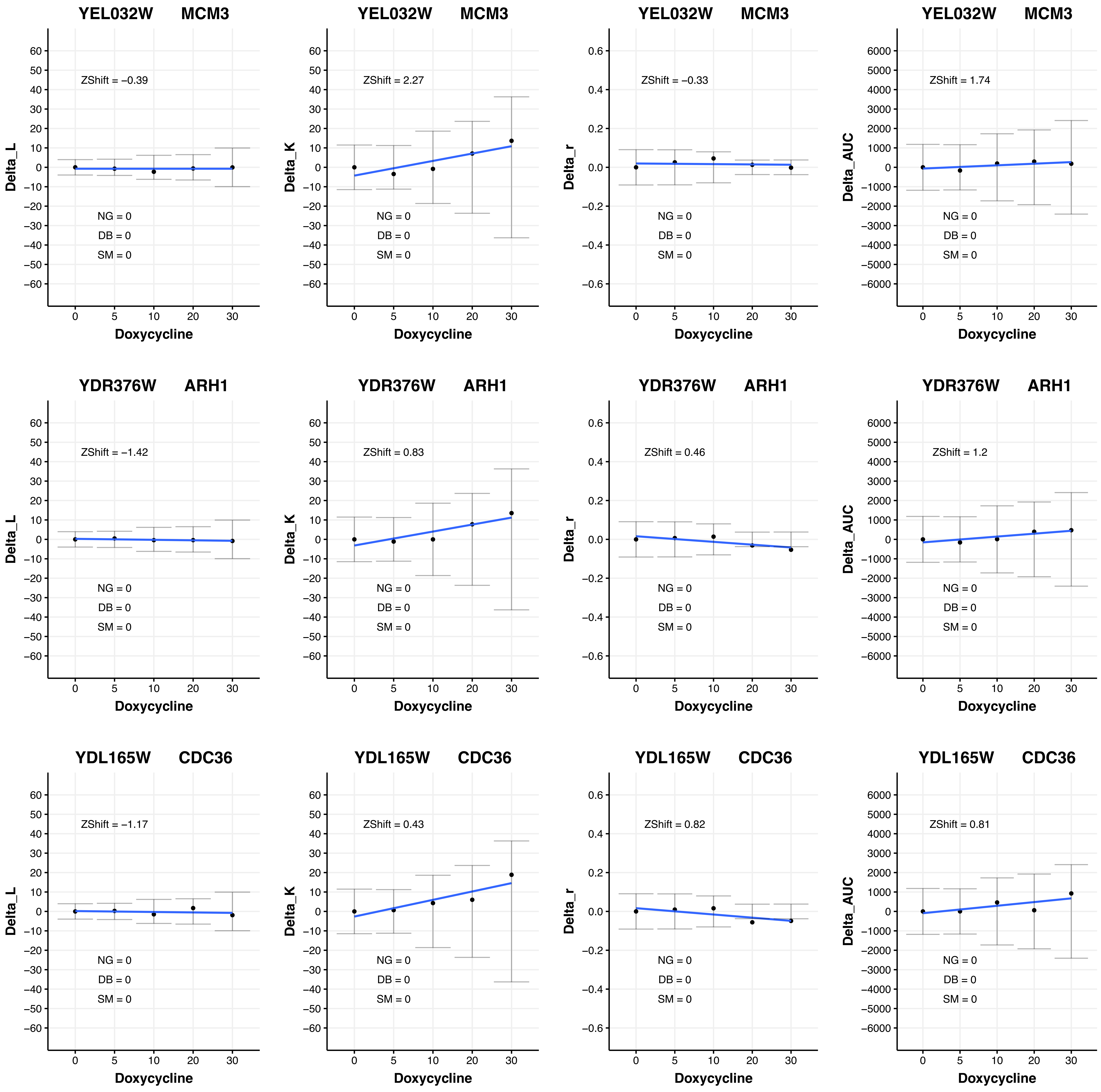
YEL032W MCM3
ZShift = −0.39
NG = 0
DB = 0
SM = 0
Delta_L
Doxycycline
YEL032W MCM3
ZShift = 2.27
NG = 0
DB = 0
SM = 0
Delta_K
Doxycycline
YEL032W MCM3
ZShift = −0.33
NG = 0
DB = 0
SM = 0
Delta_r
Doxycycline
YEL032W MCM3
ZShift = 1.74
NG = 0
DB = 0
SM = 0
Delta_AUC
Doxycycline
YDR376W ARH1
ZShift = −1.42
NG = 0
DB = 0
SM = 0
Delta_L
Doxycycline
YDR376W ARH1
ZShift = 0.83
NG = 0
DB = 0
SM = 0
Delta_K
Doxycycline
YDR376W ARH1
ZShift = 0.46
NG = 0
DB = 0
SM = 0
Delta_r
Doxycycline
YDR376W ARH1
ZShift = 1.2
NG = 0
DB = 0
SM = 0
Delta_AUC
Doxycycline
YDL165W CDC36
ZShift = −1.17
NG = 0
DB = 0
SM = 0
Delta_L
Doxycycline
YDL165W CDC36
ZShift = 0.43
NG = 0
DB = 0
SM = 0
Delta_K
Doxycycline
YDL165W CDC36
ZShift = 0.82
NG = 0
DB = 0
SM = 0
Delta_r
Doxycycline
YDL165W CDC36
ZShift = 0.81
NG = 0
DB = 0
SM = 0
Delta_AUC
Doxycycline

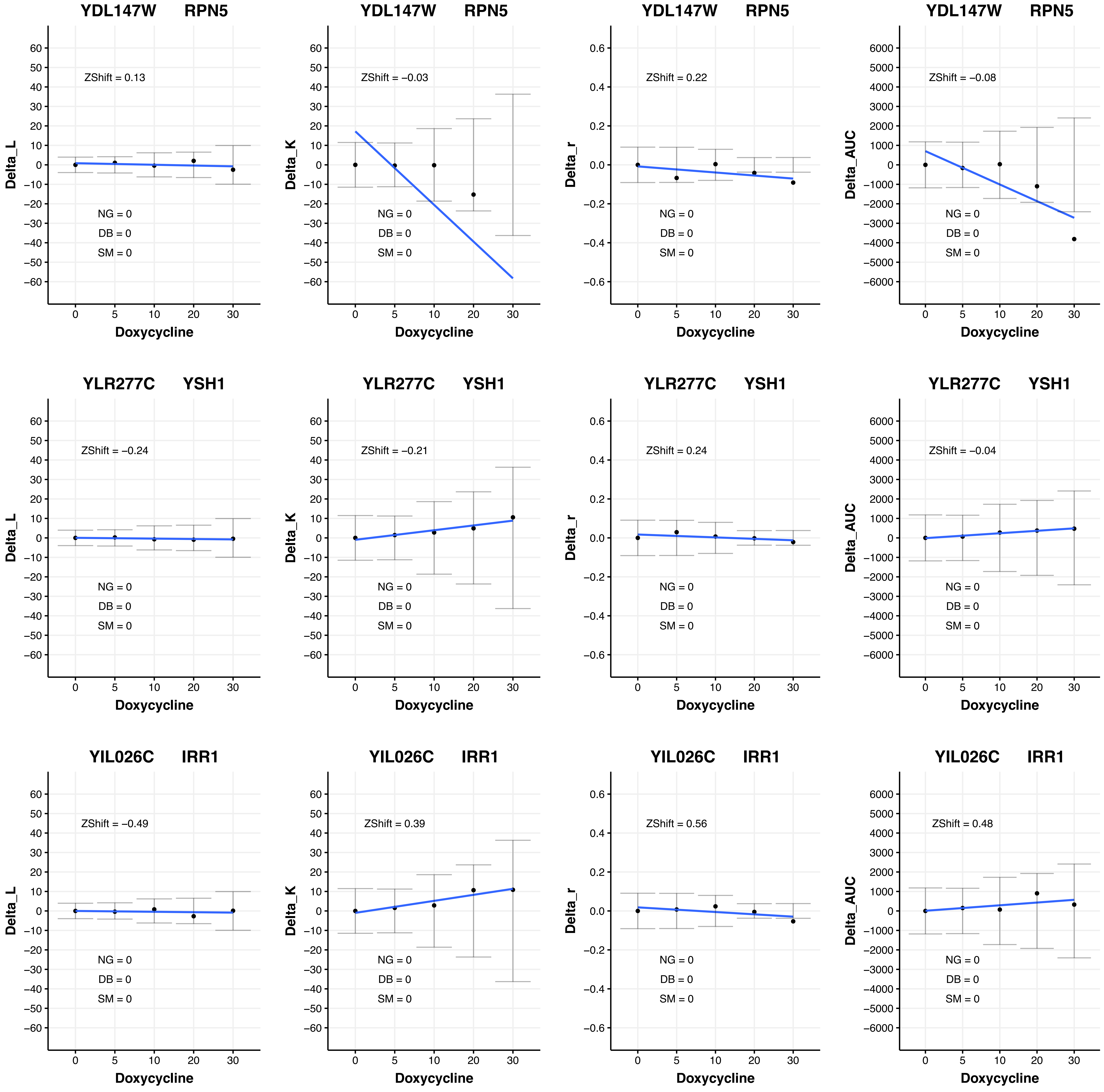
YDL147W RPN5
ZShift = 0.13
NG = 0
DB = 0
SM = 0
Delta_L
Doxycycline
YDL147W RPN5
ZShift = −0.03
NG = 0
DB = 0
SM = 0
Delta_K
Doxycycline
YDL147W RPN5
ZShift = 0.22
NG = 0
DB = 0
SM = 0
Delta_r
Doxycycline
YDL147W RPN5
ZShift = −0.08
NG = 0
DB = 0
SM = 0
Delta_AUC
Doxycycline
YLR277C YSH1
ZShift = −0.24
NG = 0
DB = 0
SM = 0
Delta_L
Doxycycline
YLR277C YSH1
ZShift = −0.21
NG = 0
DB = 0
SM = 0
Delta_K
Doxycycline
YLR277C YSH1
ZShift = 0.24
NG = 0
DB = 0
SM = 0
Delta_r
Doxycycline
YLR277C YSH1
ZShift = −0.04
NG = 0
DB = 0
SM = 0
Delta_AUC
Doxycycline
YIL026C IRR1
ZShift = −0.49
NG = 0
DB = 0
SM = 0
Delta_L
Doxycycline
YIL026C IRR1
ZShift = 0.39
NG = 0
DB = 0
SM = 0
Delta_K
Doxycycline
YIL026C IRR1
ZShift = 0.56
NG = 0
DB = 0
SM = 0
Delta_r
Doxycycline
YIL026C IRR1
ZShift = 0.48
NG = 0
DB = 0
SM = 0
Delta_AUC
Doxycycline

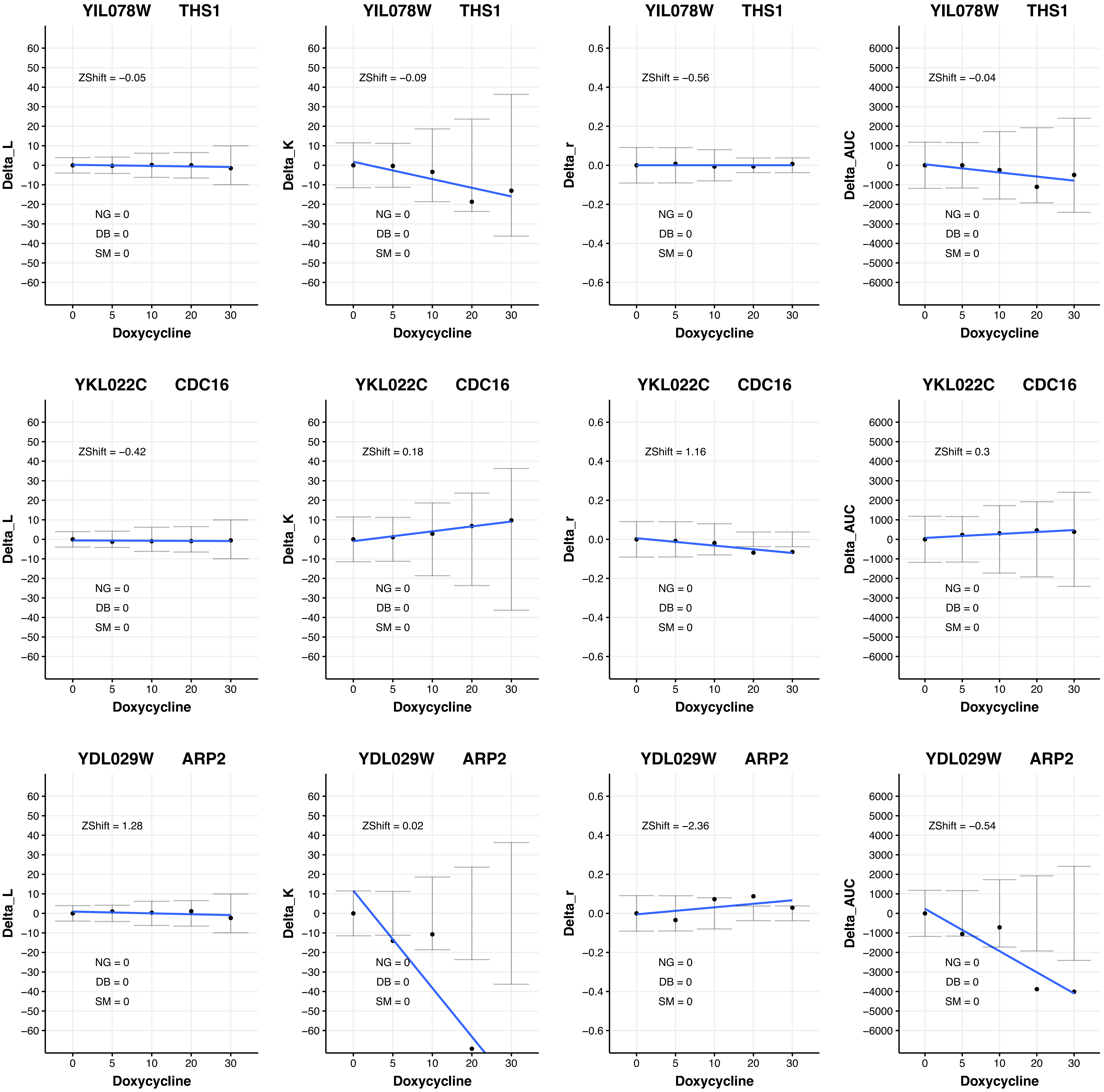

YIL078W THS1
Delta_L
ZShift = −0.05
NG = 0
DB = 0
SM = 0
Doxycycline
YIL078W THS1
Delta_K
ZShift = −0.09
NG = 0
DB = 0
SM = 0
Doxycycline
YIL078W THS1
Delta_r
ZShift = −0.56
NG = 0
DB = 0
SM = 0
Doxycycline
YIL078W THS1
Delta_AUC
ZShift = −0.04
NG = 0
DB = 0
SM = 0
Doxycycline
YKL022C CDC16
Delta_L
ZShift = −0.42
NG = 0
DB = 0
SM = 0
Doxycycline
YKL022C CDC16
Delta_K
ZShift = 0.18
NG = 0
DB = 0
SM = 0
Doxycycline
YKL022C CDC16
Delta_r
ZShift = 1.16
NG = 0
DB = 0
SM = 0
Doxycycline
YKL022C CDC16
Delta_AUC
ZShift = 0.3
NG = 0
DB = 0
SM = 0
Doxycycline
YDL029W ARP2
Delta_L
ZShift = 1.28
NG = 0
DB = 0
SM = 0
Doxycycline
YDL029W ARP2
Delta_K
ZShift = 0.02
NG = 0
DB = 0
SM = 0
Doxycycline
YDL029W ARP2
Delta_r
ZShift = −2.36
NG = 0
DB = 0
SM = 0
Doxycycline
YDL029W ARP2
Delta_AUC
ZShift = −0.54
NG = 0
DB = 0
SM = 0
Doxycycline

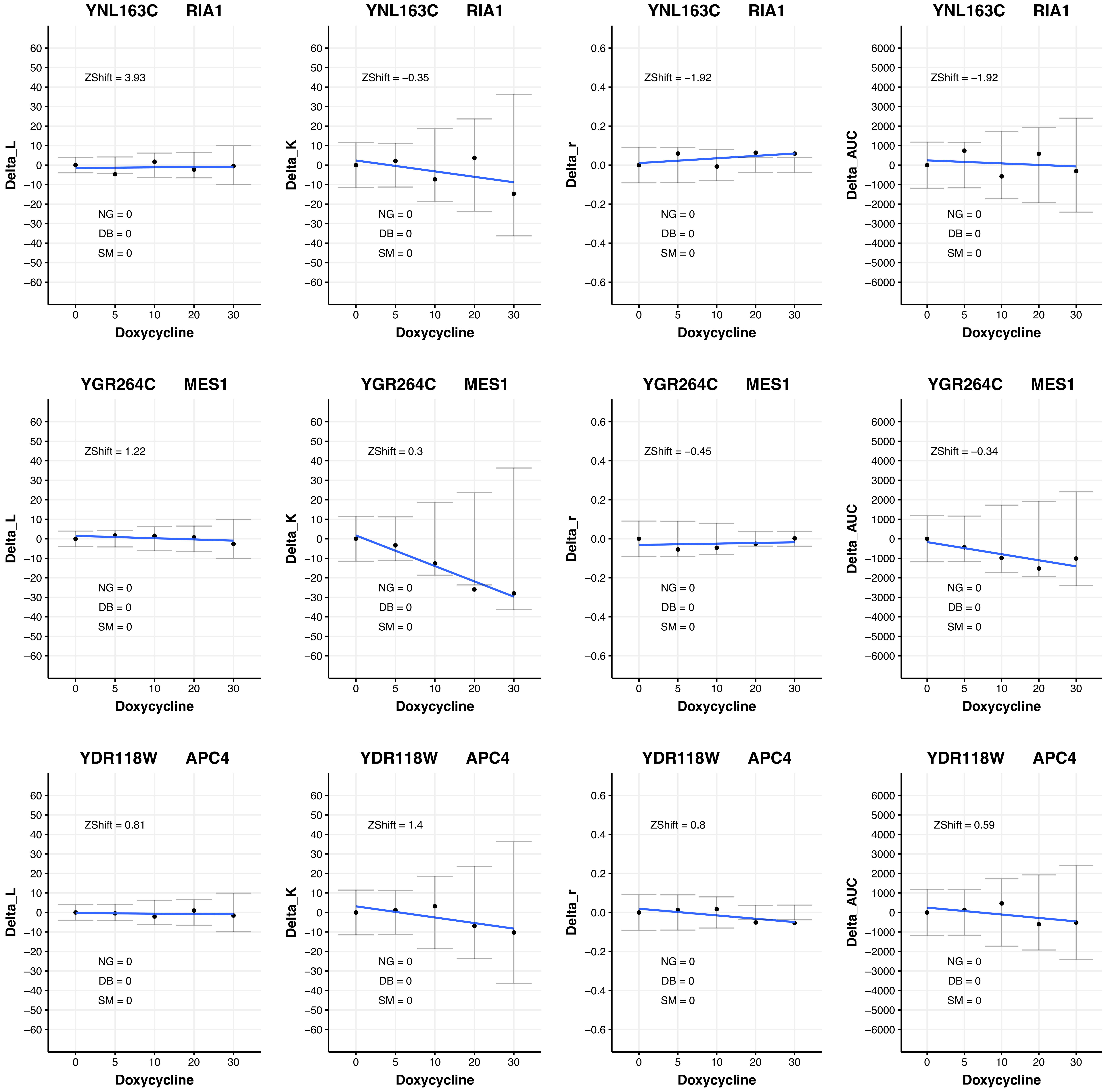

YNL163C RIA1
ZShift = 3.93
NG = 0
DB = 0
SM = 0
Delta_L
Doxycycline
YNL163C RIA1
ZShift = −0.35
NG = 0
DB = 0
SM = 0
Delta_K
Doxycycline
YNL163C RIA1
ZShift = −1.92
NG = 0
DB = 0
SM = 0
Delta_r
Doxycycline
YNL163C RIA1
ZShift = −1.92
NG = 0
DB = 0
SM = 0
Delta_AUC
Doxycycline
YGR264C MES1
ZShift = 1.22
NG = 0
DB = 0
SM = 0
Delta_L
Doxycycline
YGR264C MES1
ZShift = 0.3
NG = 0
DB = 0
SM = 0
Delta_K
Doxycycline
YGR264C MES1
ZShift = −0.45
NG = 0
DB = 0
SM = 0
Delta_r
Doxycycline
YGR264C MES1
ZShift = −0.34
NG = 0
DB = 0
SM = 0
Delta_AUC
Doxycycline
YDR118W APC4
ZShift = 0.81
NG = 0
DB = 0
SM = 0
Delta_L
Doxycycline
YDR118W APC4
ZShift = 1.4
NG = 0
DB = 0
SM = 0
Delta_K
Doxycycline
YDR118W APC4
ZShift = 0.8
NG = 0
DB = 0
SM = 0
Delta_r
Doxycycline
YDR118W APC4
ZShift = 0.59
NG = 0
DB = 0
SM = 0
Delta_AUC
Doxycycline

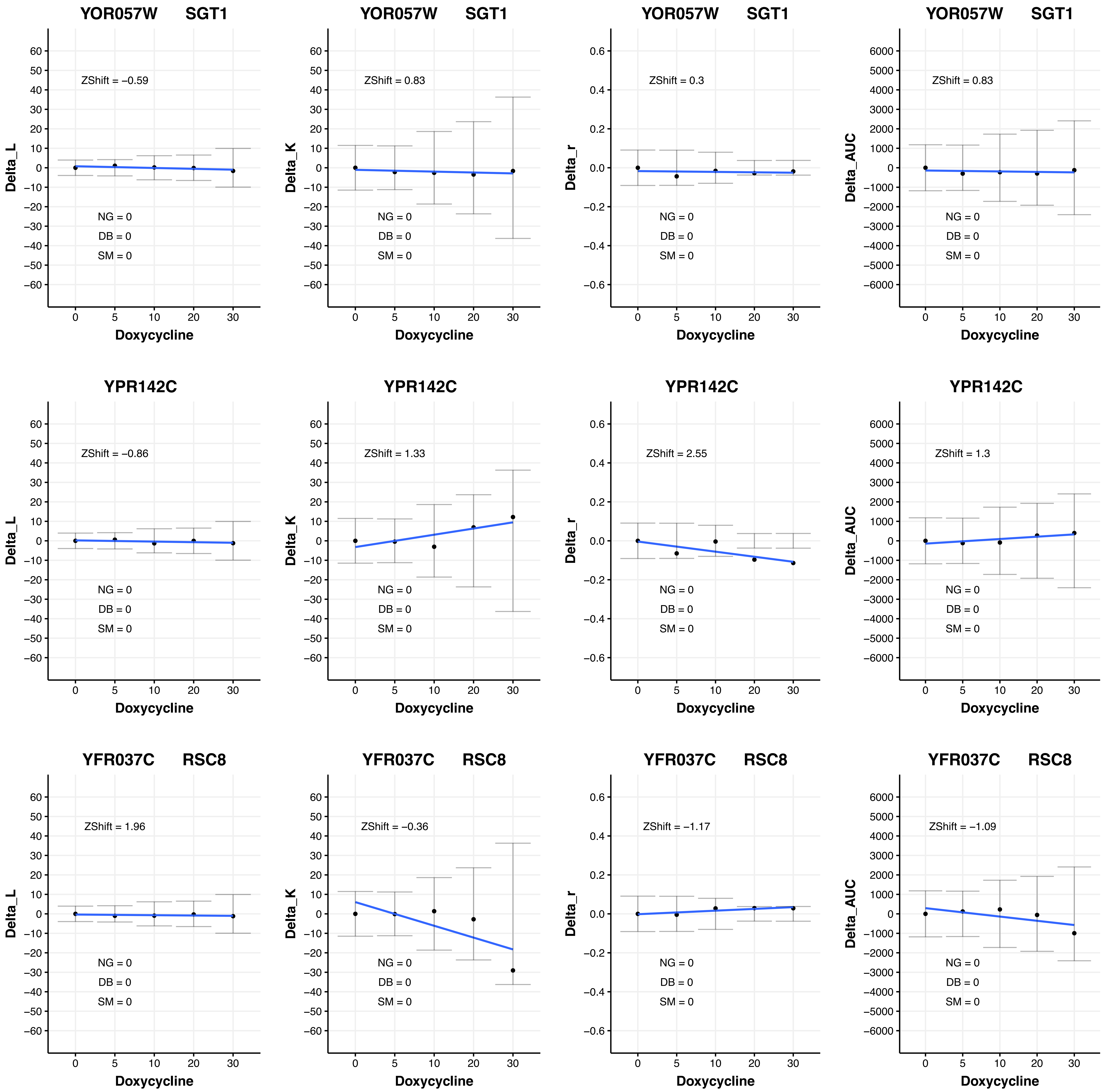

YOR057W SGT1
ZShift = −0.59
NG = 0
DB = 0
SM = 0
Delta_L
Doxycycline
YOR057W SGT1
ZShift = 0.83
NG = 0
DB = 0
SM = 0
Delta_K
Doxycycline
YOR057W SGT1
ZShift = 0.3
NG = 0
DB = 0
SM = 0
Delta_r
Doxycycline
YOR057W SGT1
ZShift = 0.83
NG = 0
DB = 0
SM = 0
Delta_AUC
Doxycycline
YPR142C
ZShift = −0.86
NG = 0
DB = 0
SM = 0
Delta_L
Doxycycline
YPR142C
ZShift = 1.33
NG = 0
DB = 0
SM = 0
Delta_K
Doxycycline
YPR142C
ZShift = 2.55
NG = 0
DB = 0
SM = 0
Delta_r
Doxycycline
YPR142C
ZShift = 1.3
NG = 0
DB = 0
SM = 0
Delta_AUC
Doxycycline
YFR037C RSC8
ZShift = 1.96
NG = 0
DB = 0
SM = 0
Delta_L
Doxycycline
YFR037C RSC8
ZShift = −0.36
NG = 0
DB = 0
SM = 0
Delta_K
Doxycycline
YFR037C RSC8
ZShift = −1.17
NG = 0
DB = 0
SM = 0
Delta_r
Doxycycline
YFR037C RSC8
ZShift = −1.09
NG = 0
DB = 0
SM = 0
Delta_AUC
Doxycycline

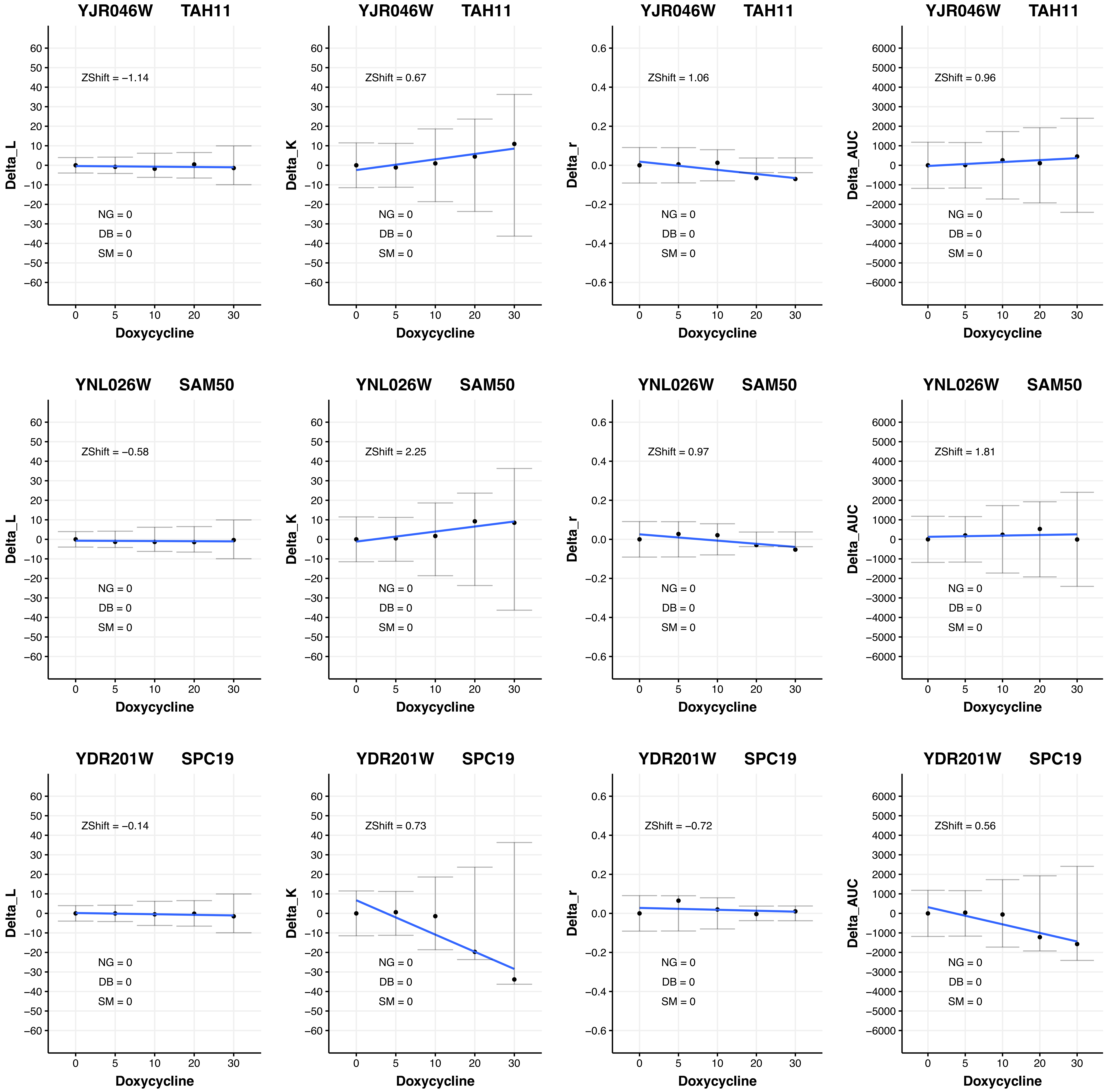
YJR046W TAH11
ZShift = −1.14
Delta_L
NG = 0
DB = 0
SM = 0
Doxycycline
YJR046W TAH11
ZShift = 0.67
Delta_K
NG = 0
DB = 0
SM = 0
Doxycycline
YJR046W TAH11
ZShift = 1.06
Delta_r
NG = 0
DB = 0
SM = 0
Doxycycline
YJR046W TAH11
ZShift = 0.96
Delta_AUC
NG = 0
DB = 0
SM = 0
Doxycycline
YNL026W SAM50
ZShift = −0.58
Delta_L
NG = 0
DB = 0
SM = 0
Doxycycline
YNL026W SAM50
ZShift = 2.25
Delta_K
NG = 0
DB = 0
SM = 0
Doxycycline
YNL026W SAM50
ZShift = 0.97
Delta_r
NG = 0
DB = 0
SM = 0
Doxycycline
YNL026W SAM50
ZShift = 1.81
Delta_AUC
NG = 0
DB = 0
SM = 0
Doxycycline
YDR201W SPC19
ZShift = −0.14
Delta_L
NG = 0
DB = 0
SM = 0
Doxycycline
YDR201W SPC19
ZShift = 0.73
Delta_K
NG = 0
DB = 0
SM = 0
Doxycycline
YDR201W SPC19
ZShift = −0.72
Delta_r
NG = 0
DB = 0
SM = 0
Doxycycline
YDR201W SPC19
ZShift = 0.56
Delta_AUC
NG = 0
DB = 0
SM = 0
Doxycycline

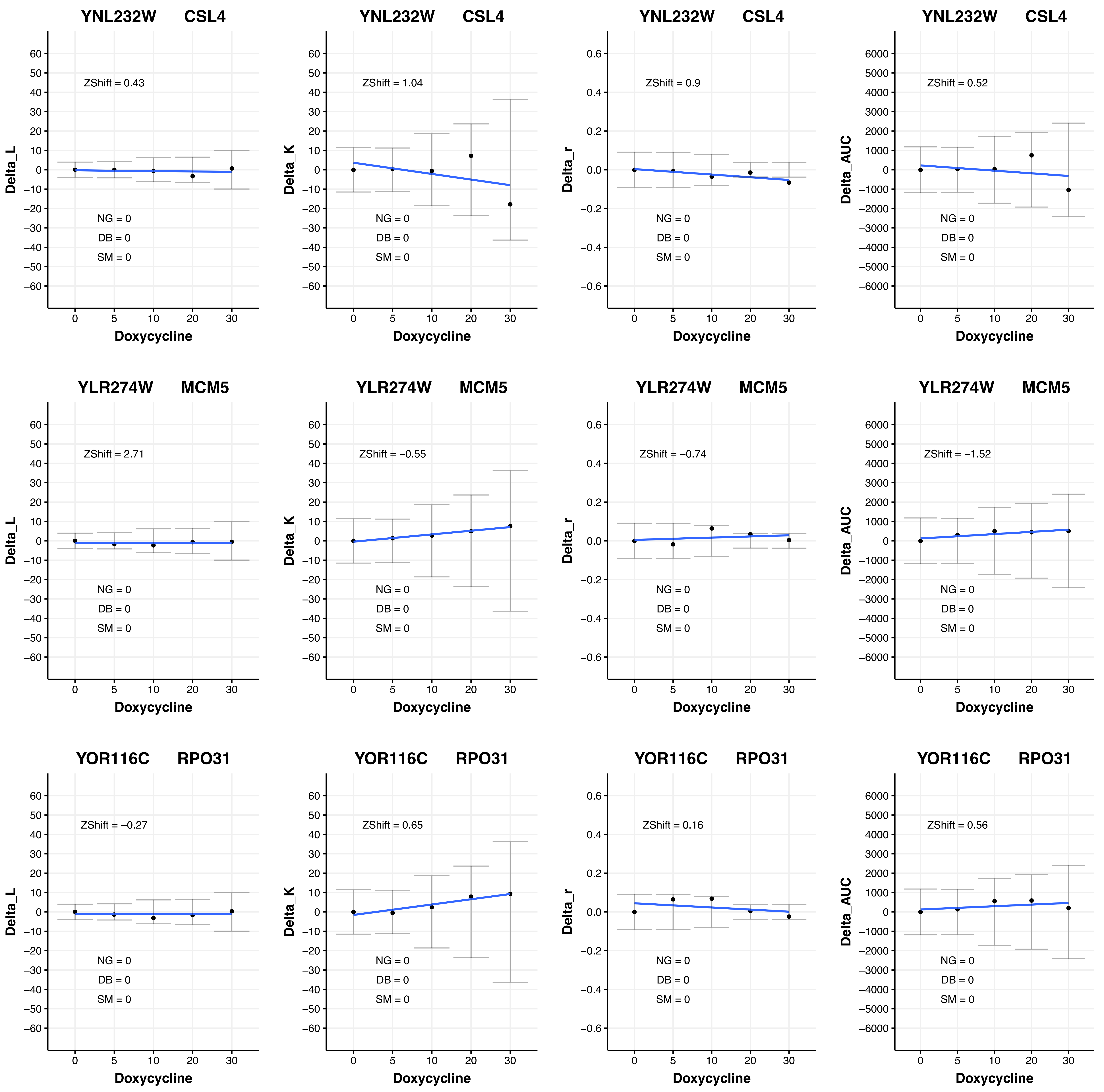
YNL232W CSL4
ZShift = 0.43
ZShift = 1.04
ZShift = 0.9
ZShift = 0.52
YLR274W MCM5
ZShift = 2.71
ZShift = −0.55
ZShift = −0.74
ZShift = −1.52
YOR116C RPO31
ZShift = −0.27
ZShift = 0.65
ZShift = 0.16
ZShift = 0.56
NG = 0
DB = 0
SM = 0
Delta_L
Delta_K
Delta_r
Delta_AUC
Doxycycline

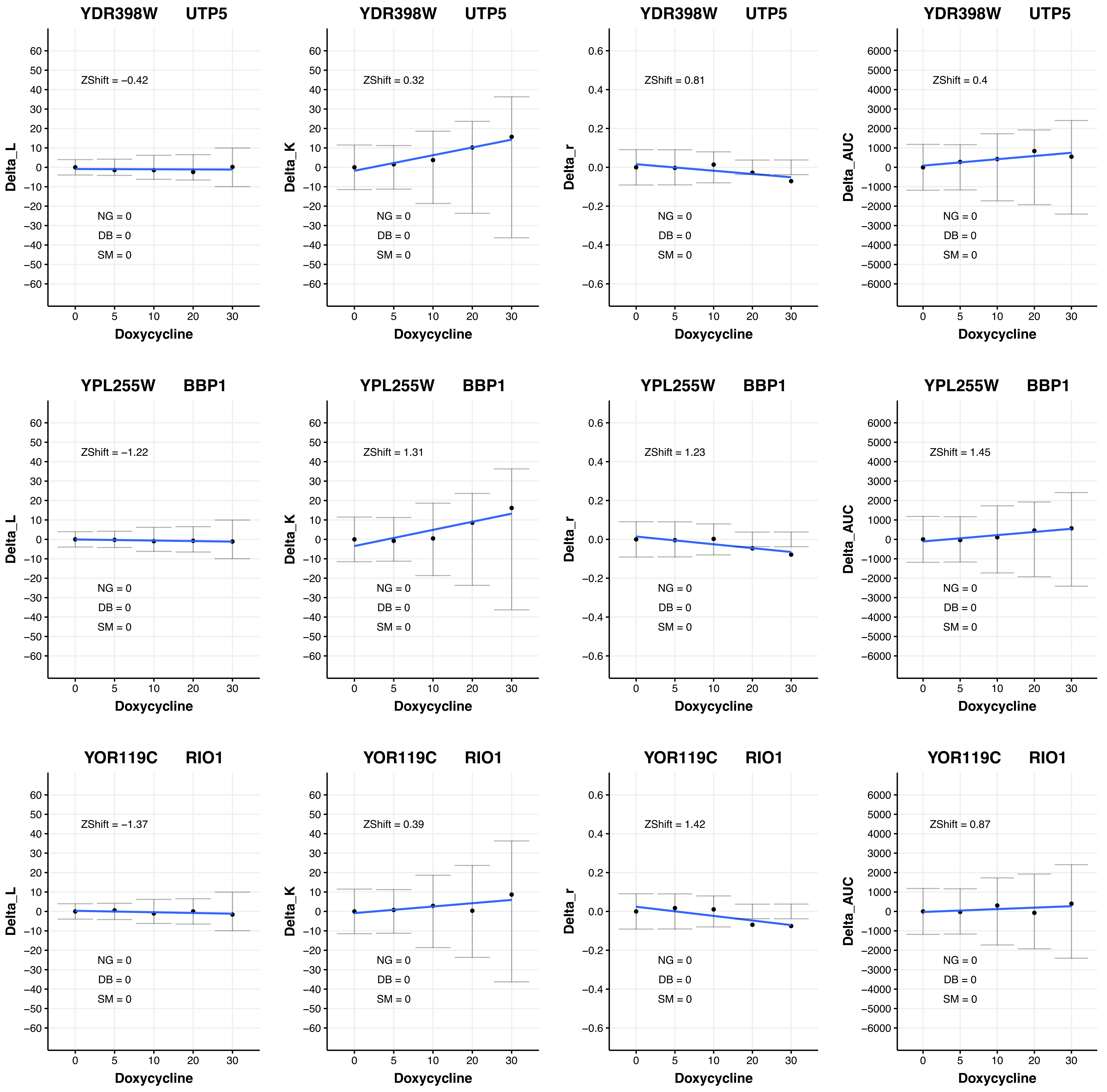
YDR398W UTP5
ZShift = −0.42
NG = 0
DB = 0
SM = 0
Delta_L
Doxycycline
YDR398W UTP5
ZShift = 0.32
NG = 0
DB = 0
SM = 0
Delta_K
Doxycycline
YDR398W UTP5
ZShift = 0.81
NG = 0
DB = 0
SM = 0
Delta_r
Doxycycline
YDR398W UTP5
ZShift = 0.4
NG = 0
DB = 0
SM = 0
Delta_AUC
Doxycycline
YPL255W BBP1
ZShift = −1.22
NG = 0
DB = 0
SM = 0
Delta_L
Doxycycline
YPL255W BBP1
ZShift = 1.31
NG = 0
DB = 0
SM = 0
Delta_K
Doxycycline
YPL255W BBP1
ZShift = 1.23
NG = 0
DB = 0
SM = 0
Delta_r
Doxycycline
YPL255W BBP1
ZShift = 1.45
NG = 0
DB = 0
SM = 0
Delta_AUC
Doxycycline
YOR119C RIO1
ZShift = −1.37
NG = 0
DB = 0
SM = 0
Delta_L
Doxycycline
YOR119C RIO1
ZShift = 0.39
NG = 0
DB = 0
SM = 0
Delta_K
Doxycycline
YOR119C RIO1
ZShift = 1.42
NG = 0
DB = 0
SM = 0
Delta_r
Doxycycline
YOR119C RIO1
ZShift = 0.87
NG = 0
DB = 0
SM = 0
Delta_AUC
Doxycycline

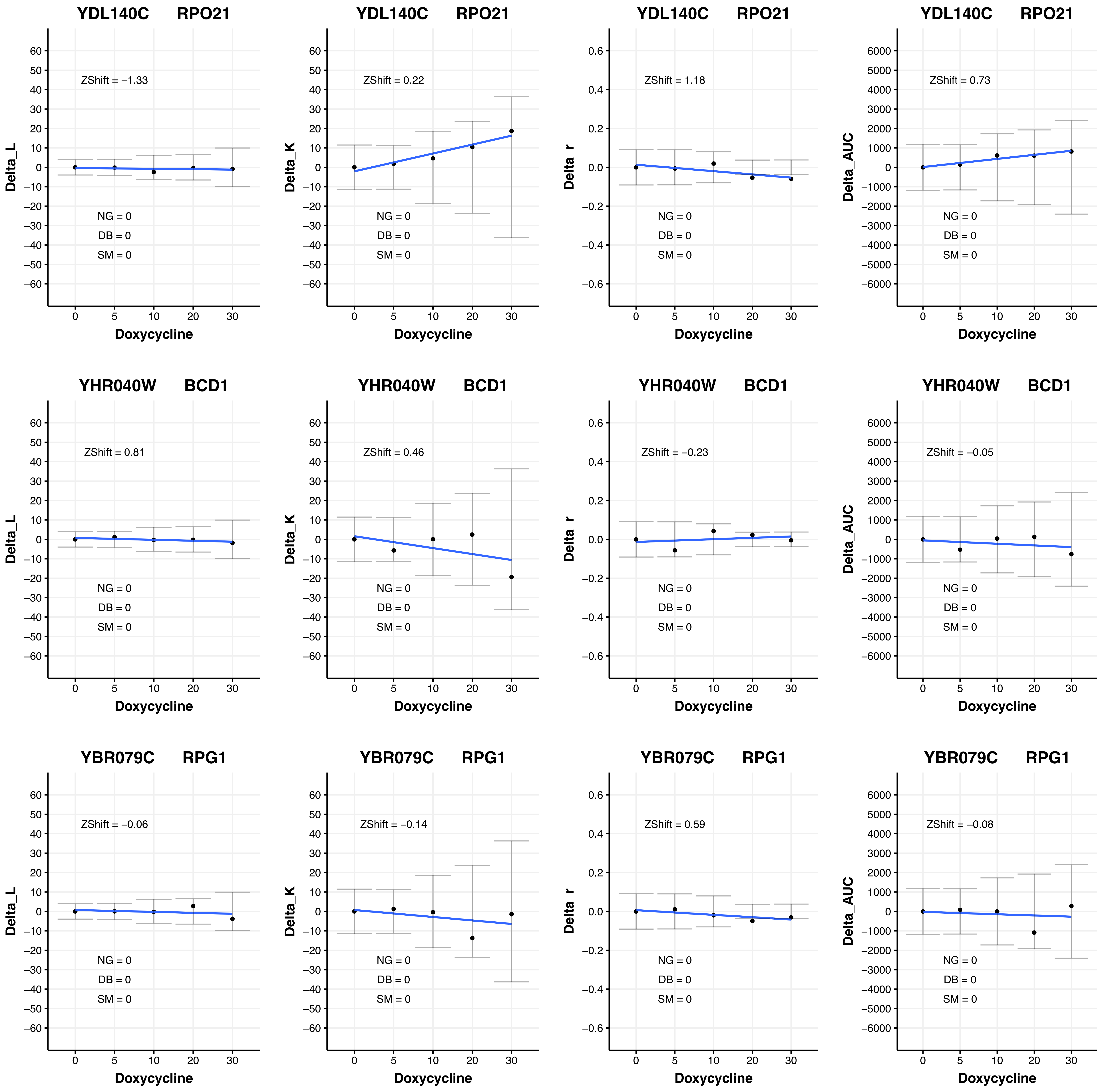
YDL140C RPO21
ZShift = −1.33
NG = 0
DB = 0
SM = 0
Delta_L
Doxycycline
YDL140C RPO21
ZShift = 0.22
NG = 0
DB = 0
SM = 0
Delta_K
Doxycycline
YDL140C RPO21
ZShift = 1.18
NG = 0
DB = 0
SM = 0
Delta_r
Doxycycline
YDL140C RPO21
ZShift = 0.73
NG = 0
DB = 0
SM = 0
Delta_AUC
Doxycycline
YHR040W BCD1
ZShift = 0.81
NG = 0
DB = 0
SM = 0
Delta_L
Doxycycline
YHR040W BCD1
ZShift = 0.46
NG = 0
DB = 0
SM = 0
Delta_K
Doxycycline
YHR040W BCD1
ZShift = −0.23
NG = 0
DB = 0
SM = 0
Delta_r
Doxycycline
YHR040W BCD1
ZShift = −0.05
NG = 0
DB = 0
SM = 0
Delta_AUC
Doxycycline
YBR079C RPG1
ZShift = −0.06
NG = 0
DB = 0
SM = 0
Delta_L
Doxycycline
YBR079C RPG1
ZShift = −0.14
NG = 0
DB = 0
SM = 0
Delta_K
Doxycycline
YBR079C RPG1
ZShift = 0.59
NG = 0
DB = 0
SM = 0
Delta_r
Doxycycline
YBR079C RPG1
ZShift = −0.08
NG = 0
DB = 0
SM = 0
Delta_AUC
Doxycycline

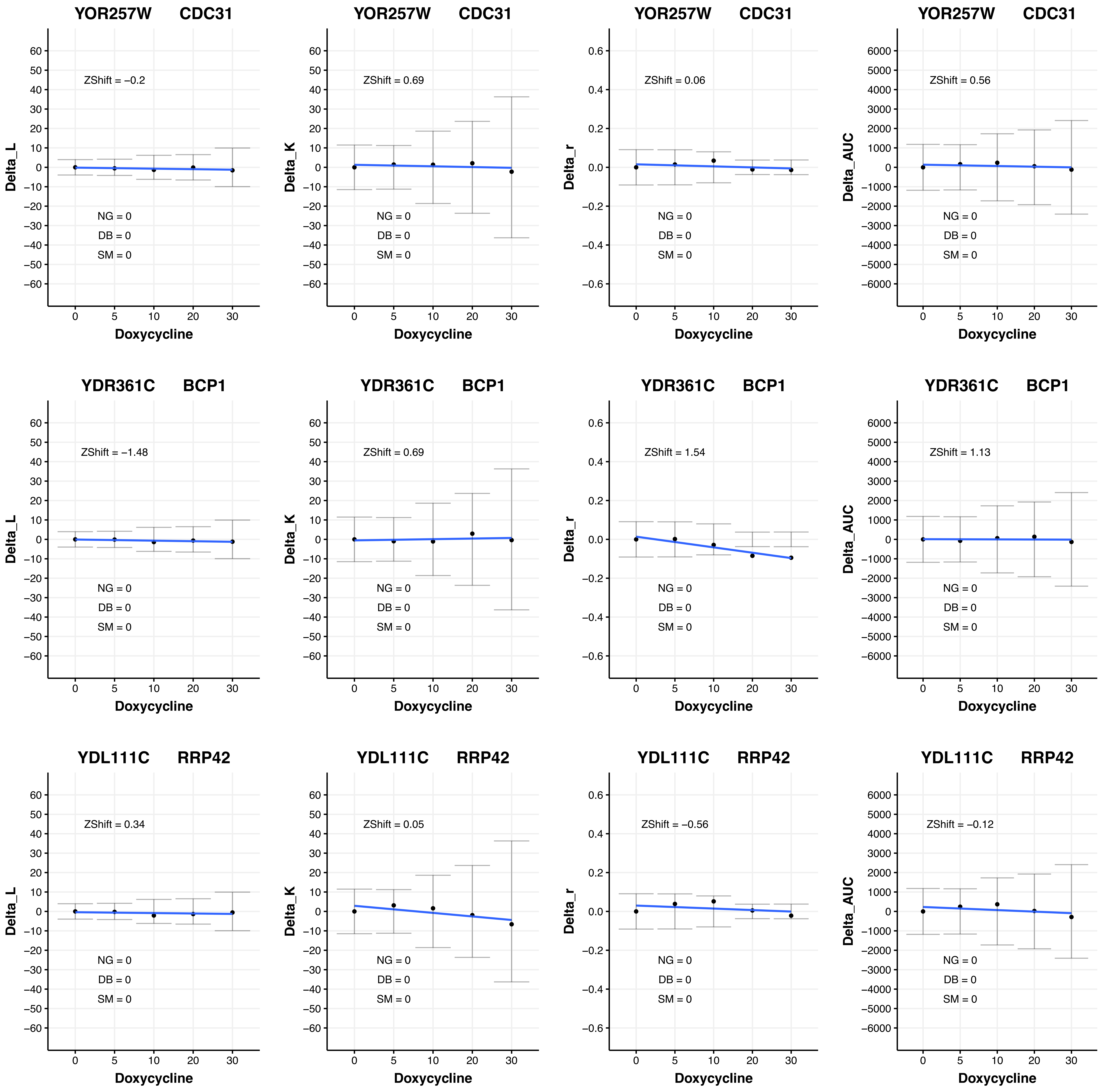
YOR257W CDC31
ZShift = −0.2
NG = 0
DB = 0
SM = 0
Delta_L
Doxycycline
YOR257W CDC31
ZShift = 0.69
NG = 0
DB = 0
SM = 0
Delta_K
Doxycycline
YOR257W CDC31
ZShift = 0.06
NG = 0
DB = 0
SM = 0
Delta_r
Doxycycline
YOR257W CDC31
ZShift = 0.56
NG = 0
DB = 0
SM = 0
Delta_AUC
Doxycycline
YDR361C BCP1
ZShift = −1.48
NG = 0
DB = 0
SM = 0
Delta_L
Doxycycline
YDR361C BCP1
ZShift = 0.69
NG = 0
DB = 0
SM = 0
Delta_K
Doxycycline
YDR361C BCP1
ZShift = 1.54
NG = 0
DB = 0
SM = 0
Delta_r
Doxycycline
YDR361C BCP1
ZShift = 1.13
NG = 0
DB = 0
SM = 0
Delta_AUC
Doxycycline
YDL111C RRP42
ZShift = 0.34
NG = 0
DB = 0
SM = 0
Delta_L
Doxycycline
YDL111C RRP42
ZShift = 0.05
NG = 0
DB = 0
SM = 0
Delta_K
Doxycycline
YDL111C RRP42
ZShift = −0.56
NG = 0
DB = 0
SM = 0
Delta_r
Doxycycline
YDL111C RRP42
ZShift = −0.12
NG = 0
DB = 0
SM = 0
Delta_AUC
Doxycycline

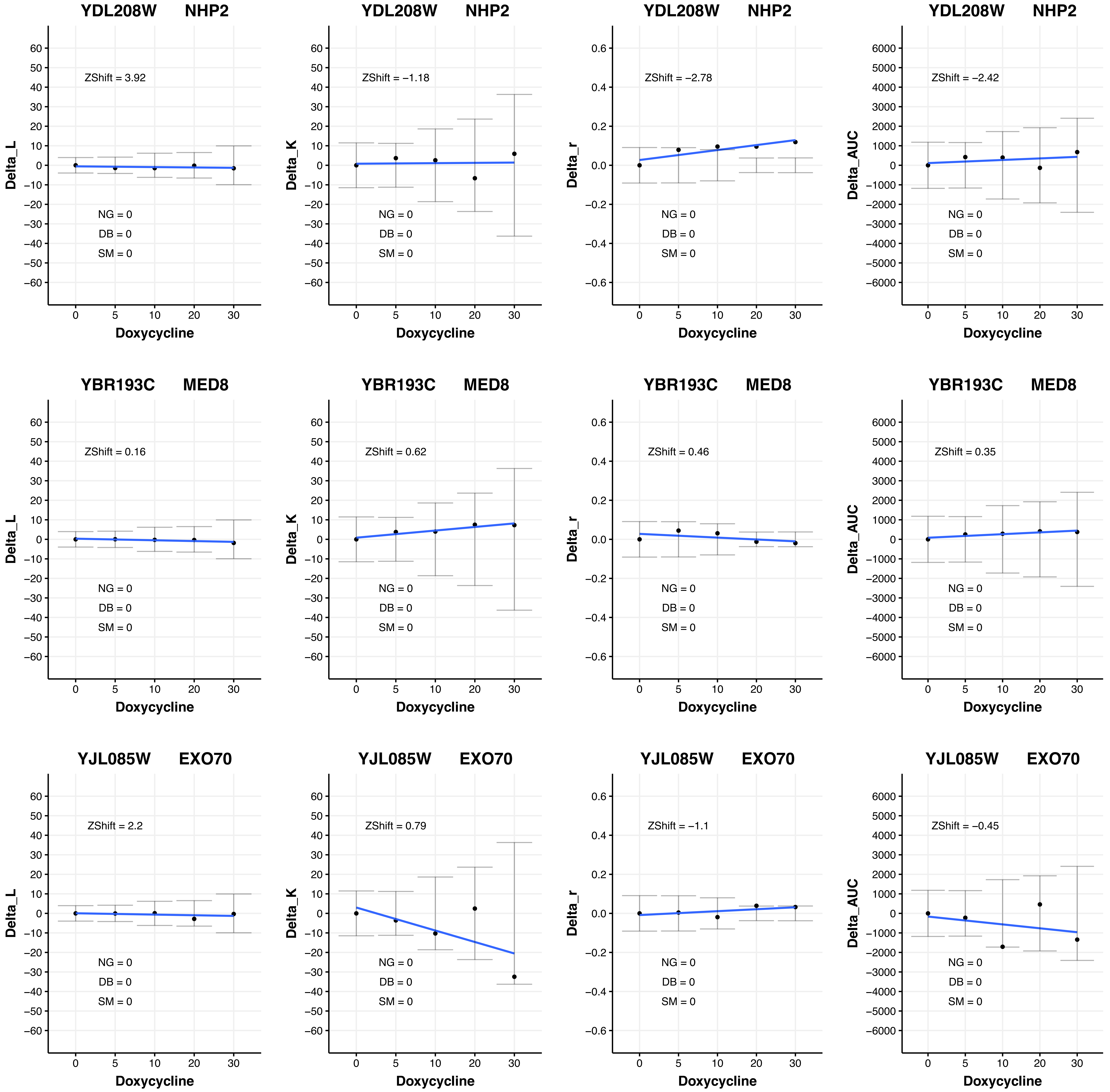
YDL208W NHP2
ZShift = 3.92
Delta_L
Doxycycline
NG = 0
DB = 0
SM = 0
YDL208W NHP2
ZShift = −1.18
Delta_K
Doxycycline
NG = 0
DB = 0
SM = 0
YDL208W NHP2
ZShift = −2.78
Delta_r
Doxycycline
NG = 0
DB = 0
SM = 0
YDL208W NHP2
ZShift = −2.42
Delta_AUC
Doxycycline
NG = 0
DB = 0
SM = 0
YBR193C MED8
ZShift = 0.16
Delta_L
Doxycycline
NG = 0
DB = 0
SM = 0
YBR193C MED8
ZShift = 0.62
Delta_K
Doxycycline
NG = 0
DB = 0
SM = 0
YBR193C MED8
ZShift = 0.46
Delta_r
Doxycycline
NG = 0
DB = 0
SM = 0
YBR193C MED8
ZShift = 0.35
Delta_AUC
Doxycycline
NG = 0
DB = 0
SM = 0
YJL085W EXO70
ZShift = 2.2
Delta_L
Doxycycline
NG = 0
DB = 0
SM = 0
YJL085W EXO70
ZShift = 0.79
Delta_K
Doxycycline
NG = 0
DB = 0
SM = 0
YJL085W EXO70
ZShift = −1.1
Delta_r
Doxycycline
NG = 0
DB = 0
SM = 0
YJL085W EXO70
ZShift = −0.45
Delta_AUC
Doxycycline
NG = 0
DB = 0
SM = 0

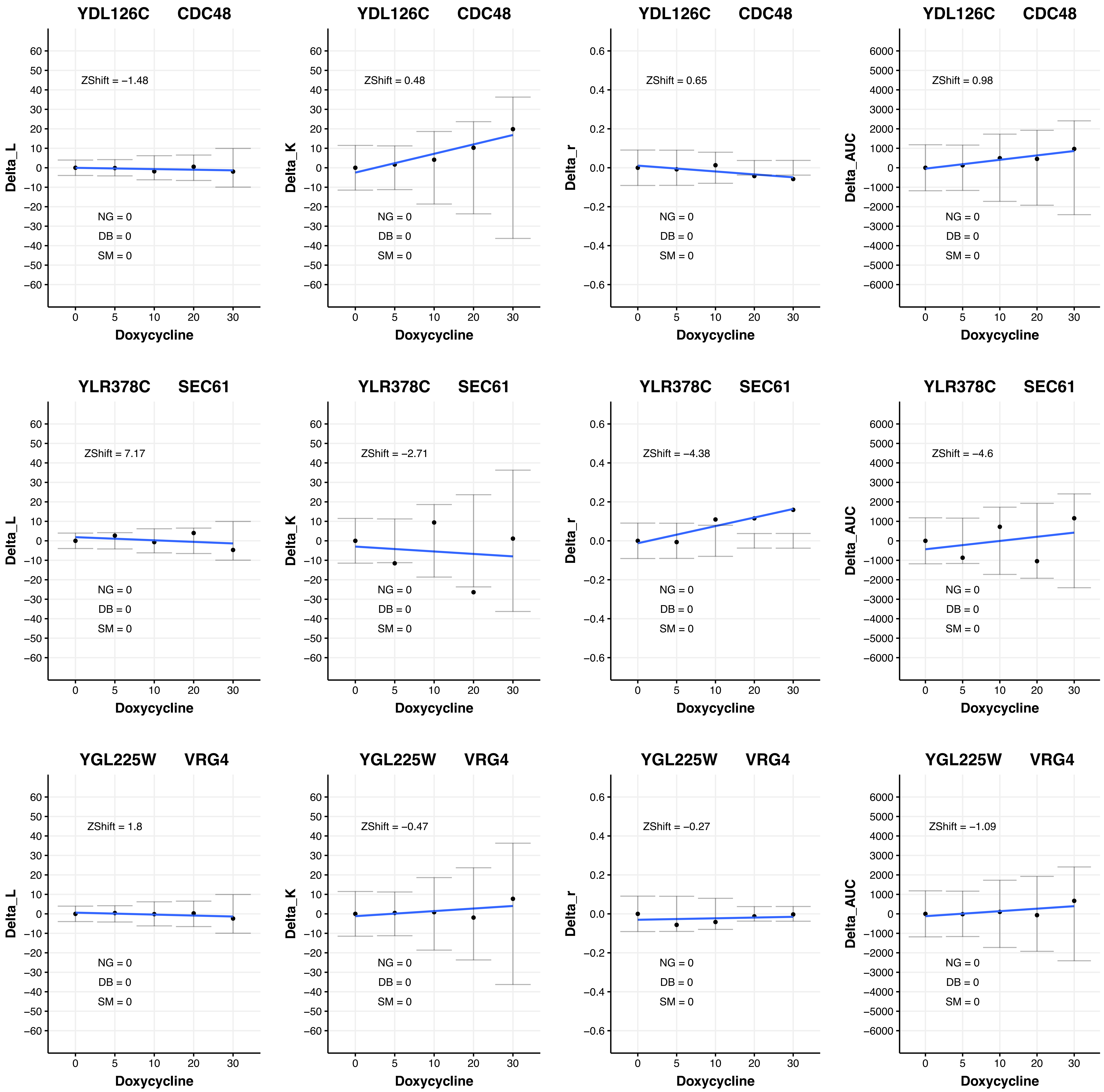
YDL126C CDC48
ZShift = −1.48
ZShift = 0.48
ZShift = 0.65
ZShift = 0.98
YLR378C SEC61
ZShift = 7.17
ZShift = −2.71
ZShift = −4.38
ZShift = −4.6
YGL225W VRG4
ZShift = 1.8
ZShift = −0.47
ZShift = −0.27
ZShift = −1.09
NG = 0
DB = 0
SM = 0
Delta_L
Delta_K
Delta_r
Delta_AUC
Doxycycline

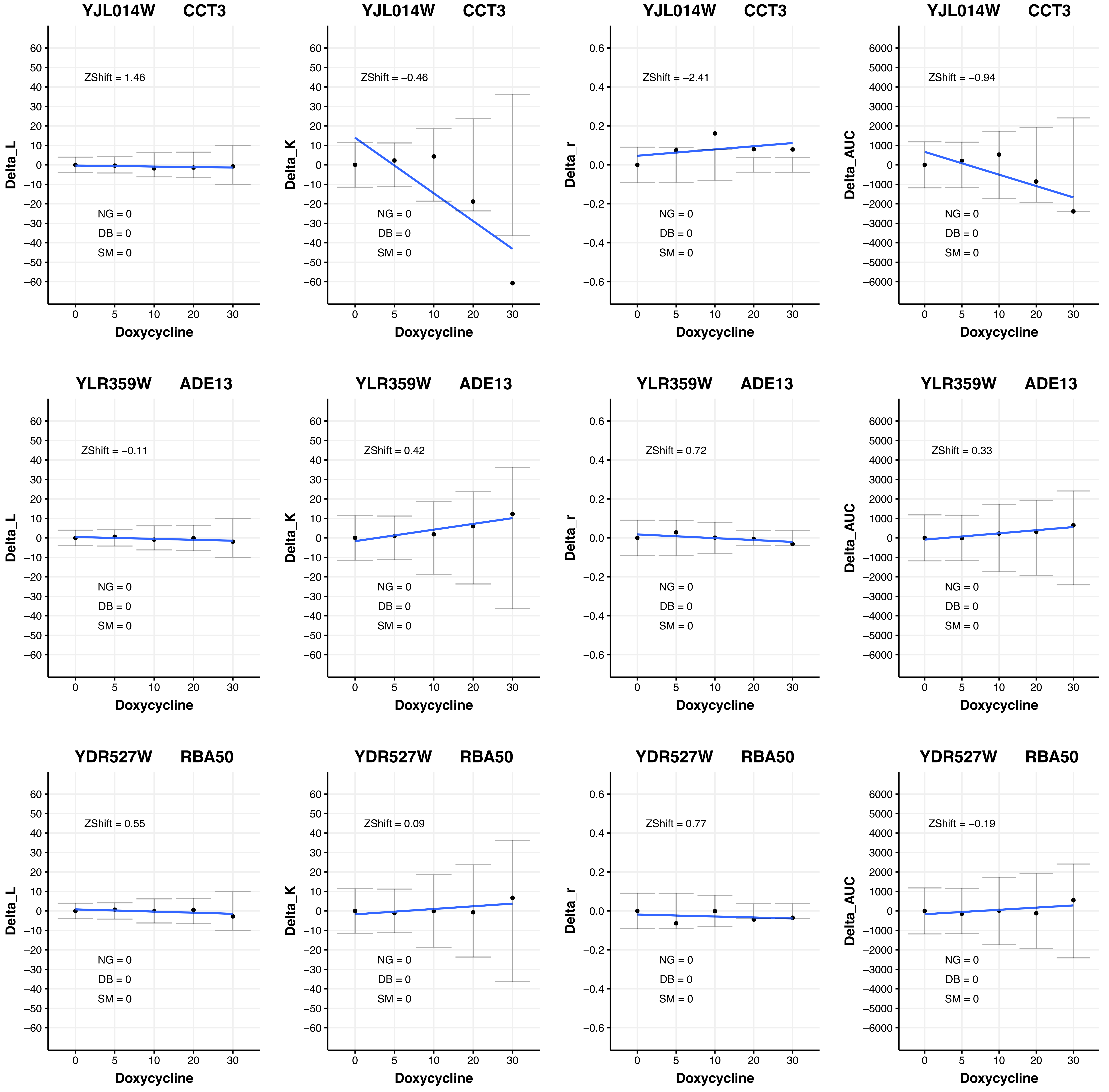
YJL014W CCT3
ZShift = 1.46
NG = 0
DB = 0
SM = 0
Delta_L
Doxycycline
YJL014W CCT3
ZShift = −0.46
NG = 0
DB = 0
SM = 0
Delta_K
Doxycycline
YJL014W CCT3
ZShift = −2.41
NG = 0
DB = 0
SM = 0
Delta_r
Doxycycline
YJL014W CCT3
ZShift = −0.94
NG = 0
DB = 0
SM = 0
Delta_AUC
Doxycycline
YLR359W ADE13
ZShift = −0.11
NG = 0
DB = 0
SM = 0
Delta_L
Doxycycline
YLR359W ADE13
ZShift = 0.42
NG = 0
DB = 0
SM = 0
Delta_K
Doxycycline
YLR359W ADE13
ZShift = 0.72
NG = 0
DB = 0
SM = 0
Delta_r
Doxycycline
YLR359W ADE13
ZShift = 0.33
NG = 0
DB = 0
SM = 0
Delta_AUC
Doxycycline
YDR527W RBA50
ZShift = 0.55
NG = 0
DB = 0
SM = 0
Delta_L
Doxycycline
YDR527W RBA50
ZShift = 0.09
NG = 0
DB = 0
SM = 0
Delta_K
Doxycycline
YDR527W RBA50
ZShift = 0.77
NG = 0
DB = 0
SM = 0
Delta_r
Doxycycline
YDR527W RBA50
ZShift = −0.19
NG = 0
DB = 0
SM = 0
Delta_AUC
Doxycycline

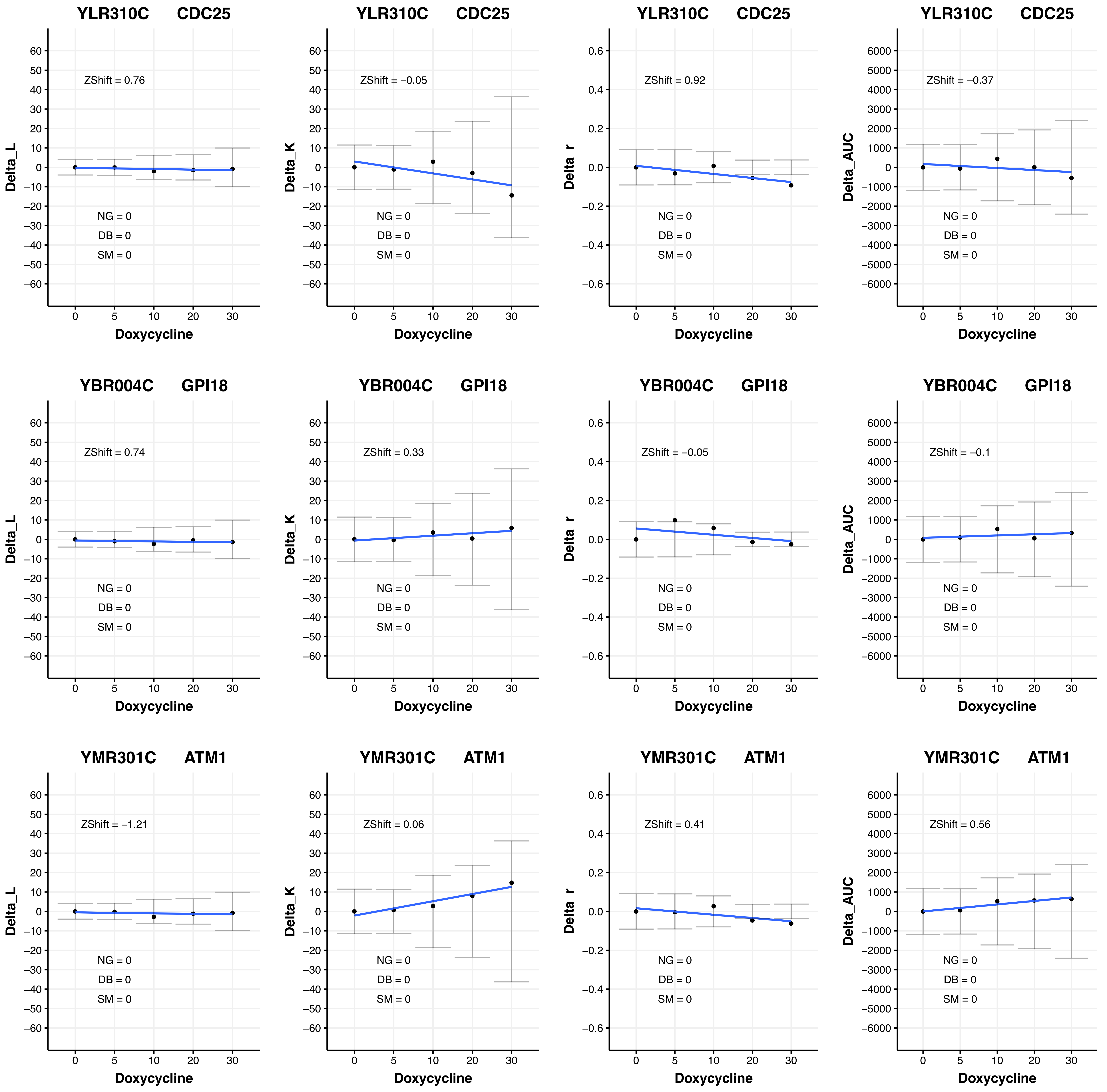

YLR310C CDC25
ZShift = 0.76
Delta_L
NG = 0
DB = 0
SM = 0
Doxycycline
YLR310C CDC25
ZShift = −0.05
Delta_K
NG = 0
DB = 0
SM = 0
Doxycycline
YLR310C CDC25
ZShift = 0.92
Delta_r
NG = 0
DB = 0
SM = 0
Doxycycline
YLR310C CDC25
ZShift = −0.37
Delta_AUC
NG = 0
DB = 0
SM = 0
Doxycycline
YBR004C GPI18
ZShift = 0.74
Delta_L
NG = 0
DB = 0
SM = 0
Doxycycline
YBR004C GPI18
ZShift = 0.33
Delta_K
NG = 0
DB = 0
SM = 0
Doxycycline
YBR004C GPI18
ZShift = −0.05
Delta_r
NG = 0
DB = 0
SM = 0
Doxycycline
YBR004C GPI18
ZShift = −0.1
Delta_AUC
NG = 0
DB = 0
SM = 0
Doxycycline
YMR301C ATM1
ZShift = −1.21
Delta_L
NG = 0
DB = 0
SM = 0
Doxycycline
YMR301C ATM1
ZShift = 0.06
Delta_K
NG = 0
DB = 0
SM = 0
Doxycycline
YMR301C ATM1
ZShift = 0.41
Delta_r
NG = 0
DB = 0
SM = 0
Doxycycline
YMR301C ATM1
ZShift = 0.56
Delta_AUC
NG = 0
DB = 0
SM = 0
Doxycycline

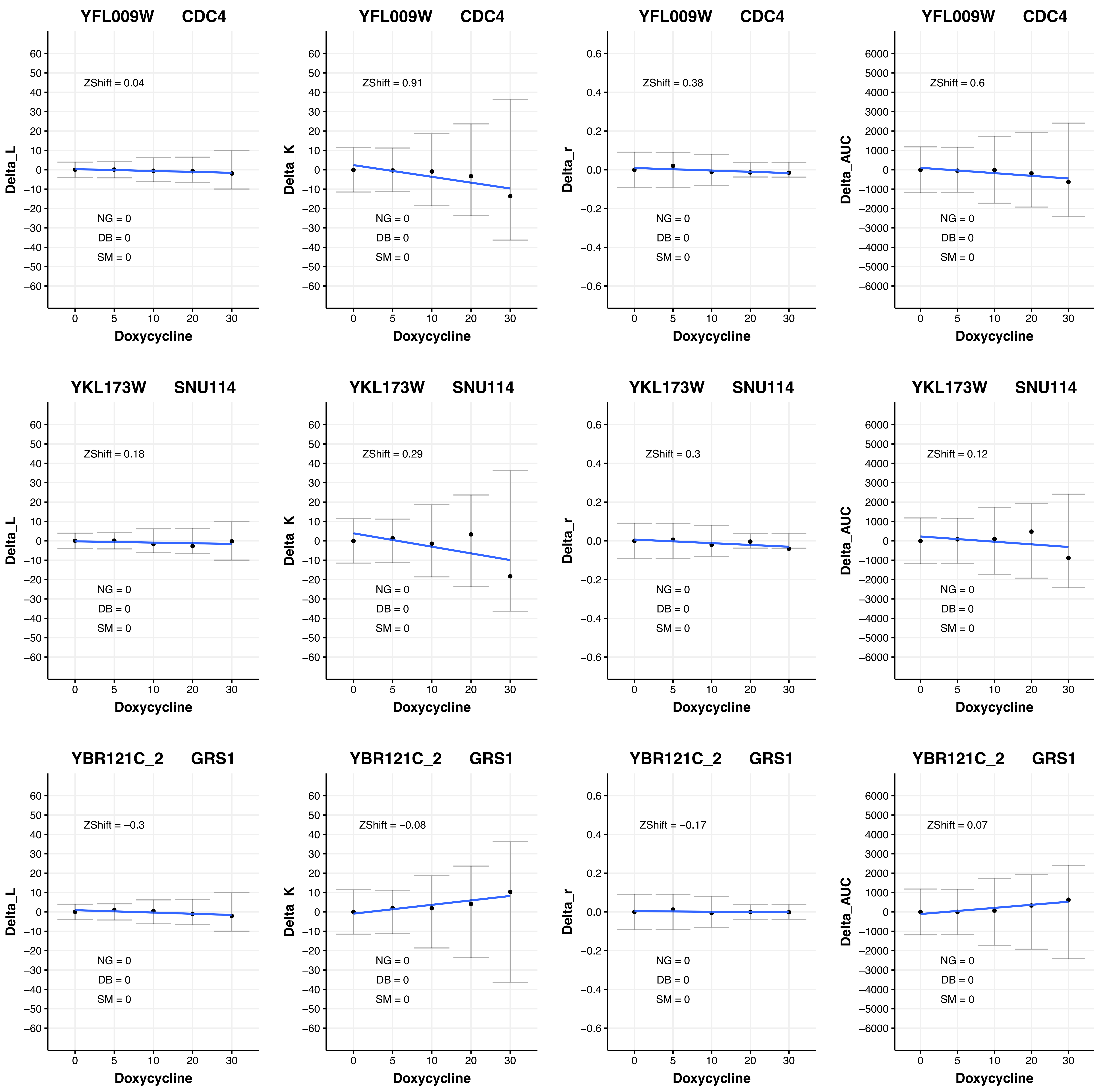
YFL009W CDC4
ZShift = 0.04
ZShift = 0.91
ZShift = 0.38
ZShift = 0.6
YKL173W SNU114
ZShift = 0.18
ZShift = 0.29
ZShift = 0.3
ZShift = 0.12
YBR121C_2 GRS1
ZShift = −0.3
ZShift = −0.08
ZShift = −0.17
ZShift = 0.07
NG = 0
DB = 0
SM = 0
Delta_L
Delta_K
Delta_r
Delta_AUC
Doxycycline
0 5 10 20 30

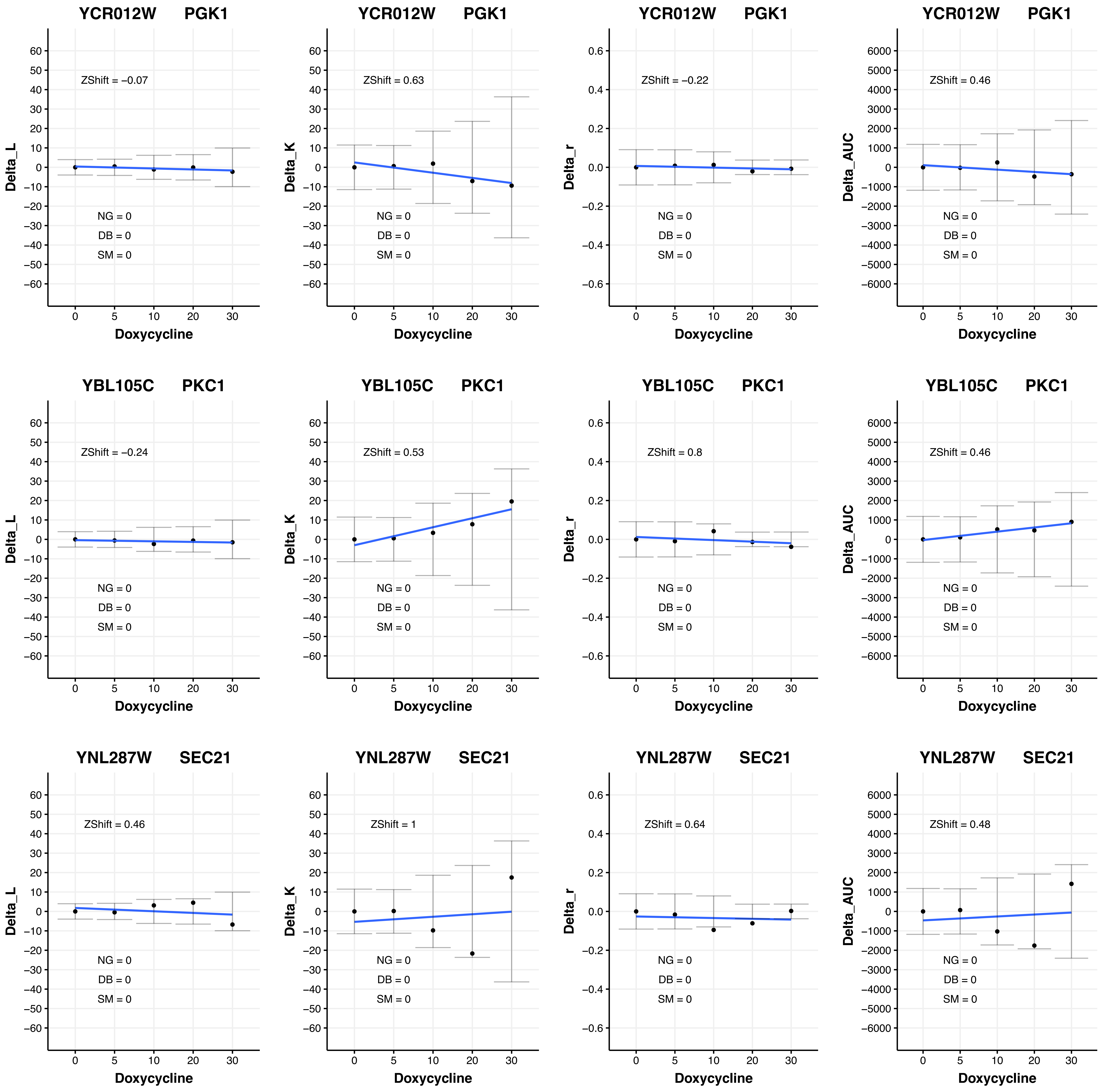
YCR012W PGK1
Delta_L
ZShift = −0.07
NG = 0
DB = 0
SM = 0
Doxycycline
YCR012W PGK1
Delta_K
ZShift = 0.63
NG = 0
DB = 0
SM = 0
Doxycycline
YCR012W PGK1
Delta_r
ZShift = −0.22
NG = 0
DB = 0
SM = 0
Doxycycline
YCR012W PGK1
Delta_AUC
ZShift = 0.46
NG = 0
DB = 0
SM = 0
Doxycycline
YBL105C PKC1
Delta_L
ZShift = −0.24
NG = 0
DB = 0
SM = 0
Doxycycline
YBL105C PKC1
Delta_K
ZShift = 0.53
NG = 0
DB = 0
SM = 0
Doxycycline
YBL105C PKC1
Delta_r
ZShift = 0.8
NG = 0
DB = 0
SM = 0
Doxycycline
YBL105C PKC1
Delta_AUC
ZShift = 0.46
NG = 0
DB = 0
SM = 0
Doxycycline
YNL287W SEC21
Delta_L
ZShift = 0.46
NG = 0
DB = 0
SM = 0
Doxycycline
YNL287W SEC21
Delta_K
ZShift = 1
NG = 0
DB = 0
SM = 0
Doxycycline
YNL287W SEC21
Delta_r
ZShift = 0.64
NG = 0
DB = 0
SM = 0
Doxycycline
YNL287W SEC21
Delta_AUC
ZShift = 0.48
NG = 0
DB = 0
SM = 0
Doxycycline

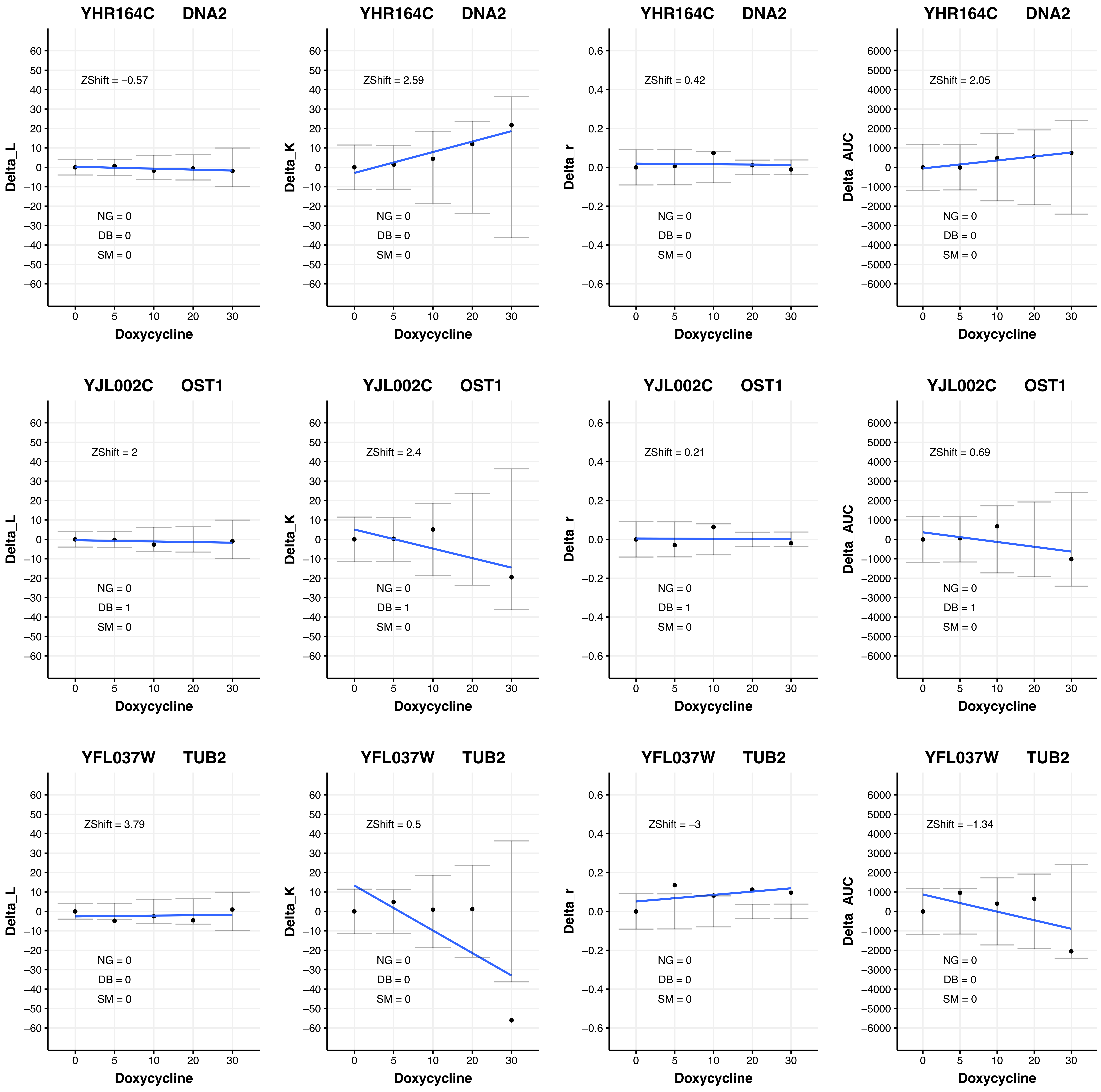

YHR164C DNA2
ZShift = −0.57
ZShift = 2.59
ZShift = 0.42
ZShift = 2.05
NG = 0
DB = 0
SM = 0
YJL002C OST1
ZShift = 2
ZShift = 2.4
ZShift = 0.21
ZShift = 0.69
NG = 0
DB = 1
SM = 0
YFL037W TUB2
ZShift = 3.79
ZShift = 0.5
ZShift = −3
ZShift = −1.34
NG = 0
DB = 0
SM = 0
Delta_L
Delta_K
Delta_r
Delta_AUC
Doxycycline
0
5
10
20
30

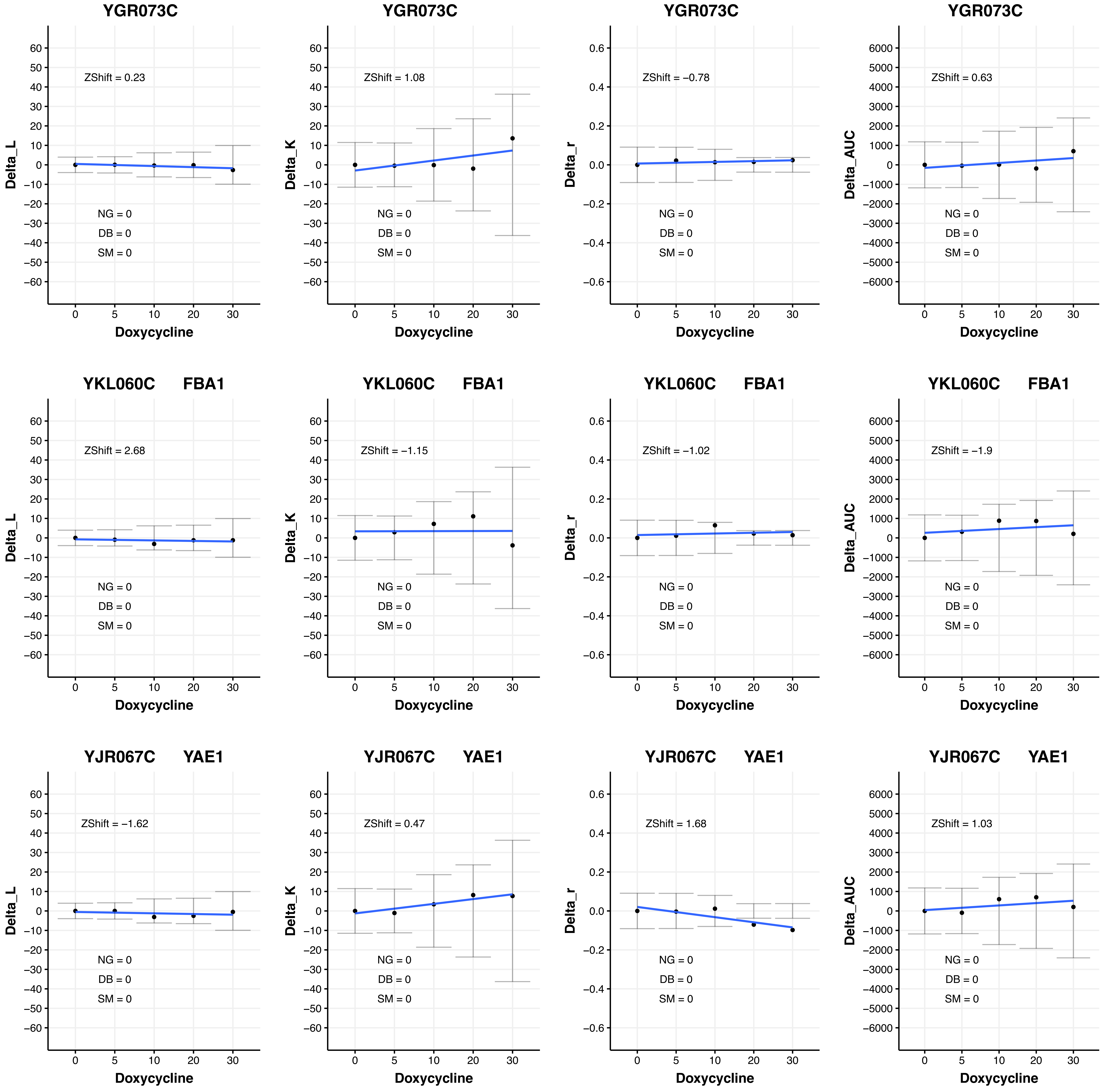
YGR073C
ZShift = 0.23
NG = 0
DB = 0
SM = 0
Delta_L
Doxycycline
YGR073C
ZShift = 1.08
NG = 0
DB = 0
SM = 0
Delta_K
Doxycycline
YGR073C
ZShift = −0.78
NG = 0
DB = 0
SM = 0
Delta_r
Doxycycline
YGR073C
ZShift = 0.63
NG = 0
DB = 0
SM = 0
Delta_AUC
Doxycycline
YKL060C FBA1
ZShift = 2.68
NG = 0
DB = 0
SM = 0
Delta_L
Doxycycline
YKL060C FBA1
ZShift = −1.15
NG = 0
DB = 0
SM = 0
Delta_K
Doxycycline
YKL060C FBA1
ZShift = −1.02
NG = 0
DB = 0
SM = 0
Delta_r
Doxycycline
YKL060C FBA1
ZShift = −1.9
NG = 0
DB = 0
SM = 0
Delta_AUC
Doxycycline
YJR067C YAE1
ZShift = −1.62
NG = 0
DB = 0
SM = 0
Delta_L
Doxycycline
YJR067C YAE1
ZShift = 0.47
NG = 0
DB = 0
SM = 0
Delta_K
Doxycycline
YJR067C YAE1
ZShift = 1.68
NG = 0
DB = 0
SM = 0
Delta_r
Doxycycline
YJR067C YAE1
ZShift = 1.03
NG = 0
DB = 0
SM = 0
Delta_AUC
Doxycycline

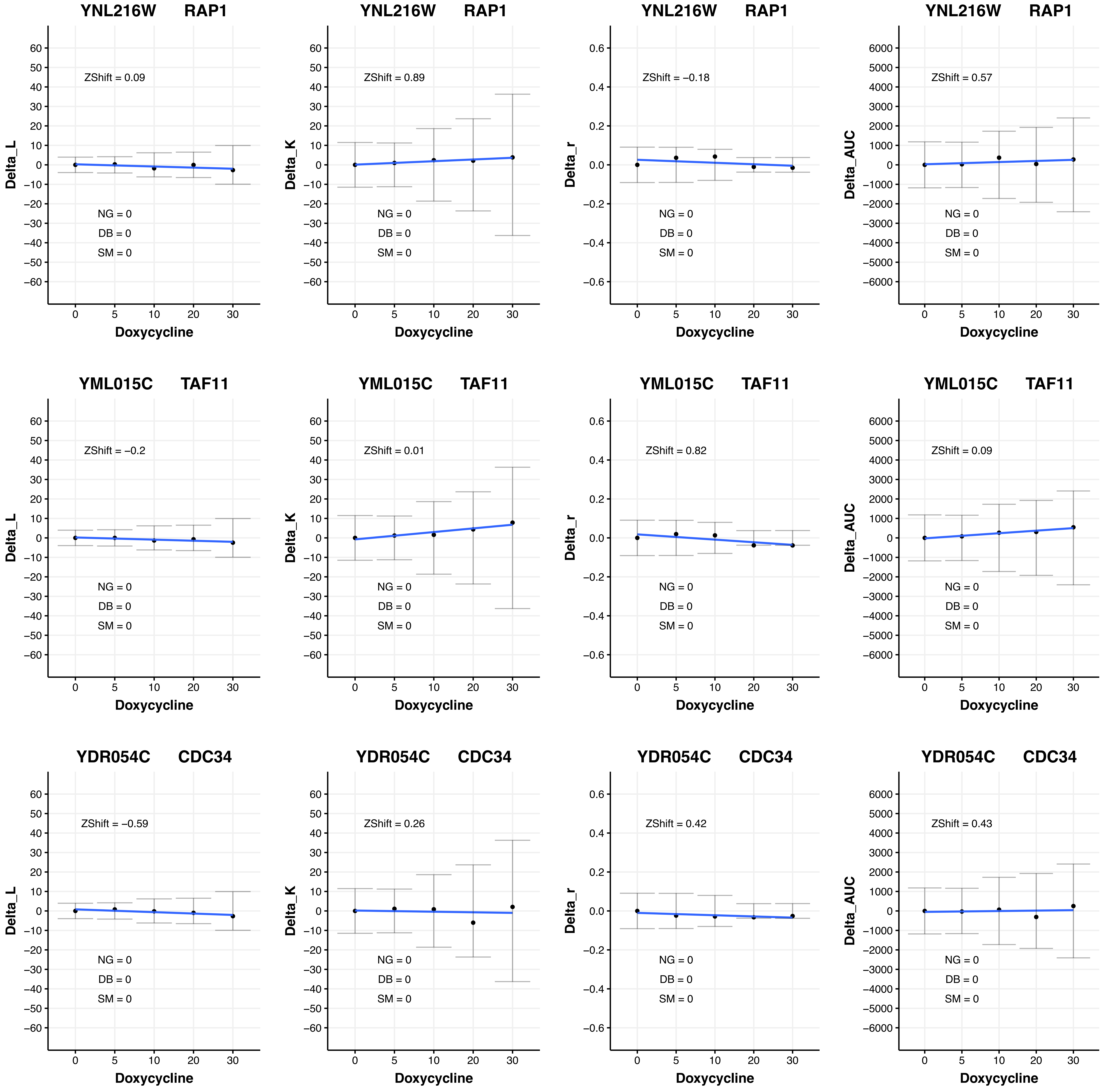
YNL216W RAP1
ZShift = 0.09
NG = 0
DB = 0
SM = 0
Delta_L
Doxycycline
YNL216W RAP1
ZShift = 0.89
NG = 0
DB = 0
SM = 0
Delta_K
Doxycycline
YNL216W RAP1
ZShift = −0.18
NG = 0
DB = 0
SM = 0
Delta_r
Doxycycline
YNL216W RAP1
ZShift = 0.57
NG = 0
DB = 0
SM = 0
Delta_AUC
Doxycycline
YML015C TAF11
ZShift = −0.2
NG = 0
DB = 0
SM = 0
Delta_L
Doxycycline
YML015C TAF11
ZShift = 0.01
NG = 0
DB = 0
SM = 0
Delta_K
Doxycycline
YML015C TAF11
ZShift = 0.82
NG = 0
DB = 0
SM = 0
Delta_r
Doxycycline
YML015C TAF11
ZShift = 0.09
NG = 0
DB = 0
SM = 0
Delta_AUC
Doxycycline
YDR054C CDC34
ZShift = −0.59
NG = 0
DB = 0
SM = 0
Delta_L
Doxycycline
YDR054C CDC34
ZShift = 0.26
NG = 0
DB = 0
SM = 0
Delta_K
Doxycycline
YDR054C CDC34
ZShift = 0.42
NG = 0
DB = 0
SM = 0
Delta_r
Doxycycline
YDR054C CDC34
ZShift = 0.43
NG = 0
DB = 0
SM = 0
Delta_AUC
Doxycycline

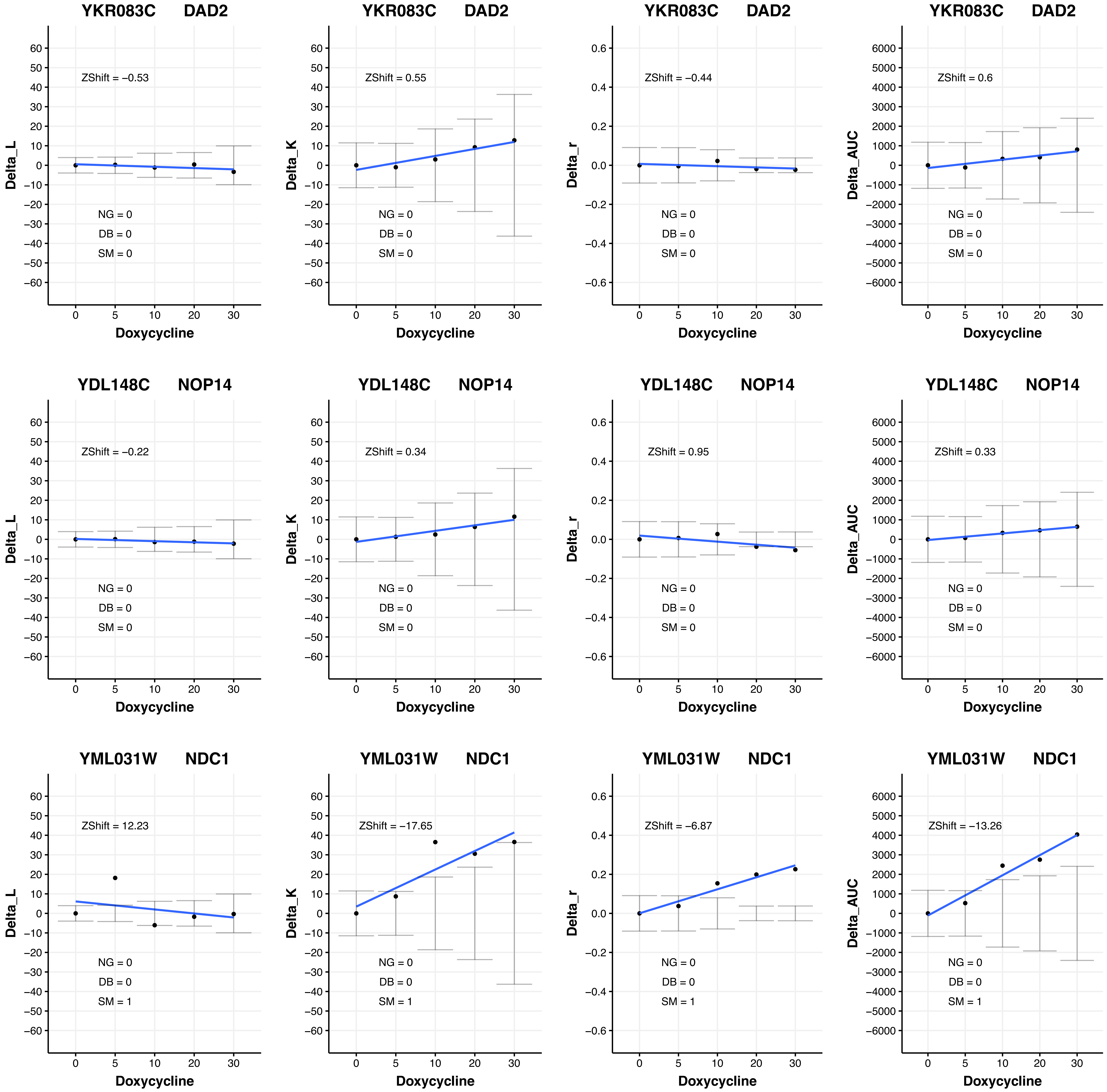

YKR083C DAD2
ZShift = −0.53
NG = 0
DB = 0
SM = 0
Delta_L
Doxycycline
YKR083C DAD2
ZShift = 0.55
NG = 0
DB = 0
SM = 0
Delta_K
Doxycycline
YKR083C DAD2
ZShift = −0.44
NG = 0
DB = 0
SM = 0
Delta_r
Doxycycline
YKR083C DAD2
ZShift = 0.6
NG = 0
DB = 0
SM = 0
Delta_AUC
Doxycycline
YDL148C NOP14
ZShift = −0.22
NG = 0
DB = 0
SM = 0
Delta_L
Doxycycline
YDL148C NOP14
ZShift = 0.34
NG = 0
DB = 0
SM = 0
Delta_K
Doxycycline
YDL148C NOP14
ZShift = 0.95
NG = 0
DB = 0
SM = 0
Delta_r
Doxycycline
YDL148C NOP14
ZShift = 0.33
NG = 0
DB = 0
SM = 0
Delta_AUC
Doxycycline
YML031W NDC1
ZShift = 12.23
NG = 0
DB = 0
SM = 1
Delta_L
Doxycycline
YML031W NDC1
ZShift = −17.65
NG = 0
DB = 0
SM = 1
Delta_K
Doxycycline
YML031W NDC1
ZShift = −6.87
NG = 0
DB = 0
SM = 1
Delta_r
Doxycycline
YML031W NDC1
ZShift = −13.26
NG = 0
DB = 0
SM = 1
Delta_AUC
Doxycycline

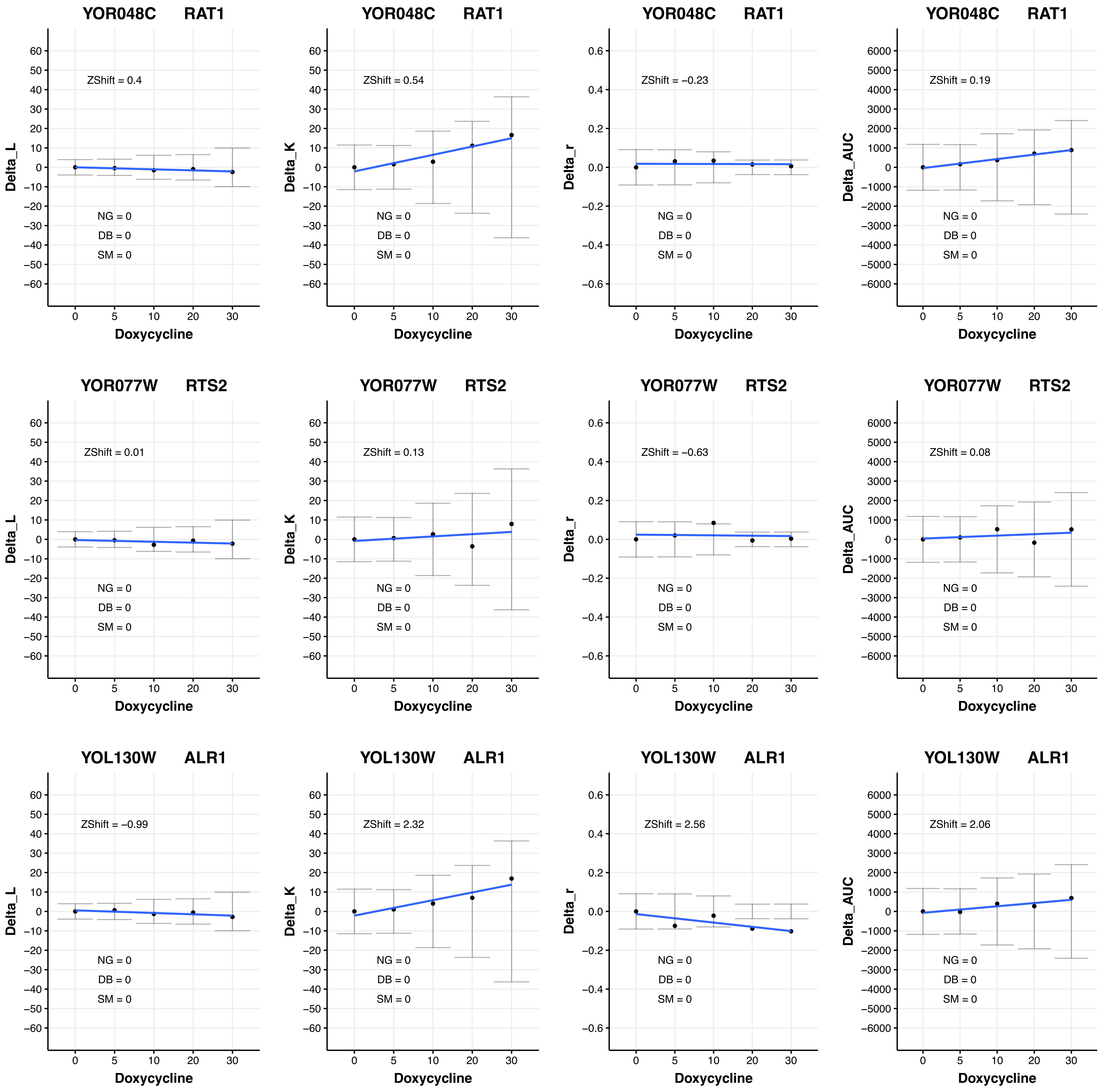
YOR048C RAT1
ZShift = 0.4
NG = 0
DB = 0
SM = 0
Delta_L
Doxycycline
YOR048C RAT1
ZShift = 0.54
NG = 0
DB = 0
SM = 0
Delta_K
Doxycycline
YOR048C RAT1
ZShift = −0.23
NG = 0
DB = 0
SM = 0
Delta_r
Doxycycline
YOR048C RAT1
ZShift = 0.19
NG = 0
DB = 0
SM = 0
Delta_AUC
Doxycycline
YOR077W RTS2
ZShift = 0.01
NG = 0
DB = 0
SM = 0
Delta_L
Doxycycline
YOR077W RTS2
ZShift = 0.13
NG = 0
DB = 0
SM = 0
Delta_K
Doxycycline
YOR077W RTS2
ZShift = −0.63
NG = 0
DB = 0
SM = 0
Delta_r
Doxycycline
YOR077W RTS2
ZShift = 0.08
NG = 0
DB = 0
SM = 0
Delta_AUC
Doxycycline
YOL130W ALR1
ZShift = −0.99
NG = 0
DB = 0
SM = 0
Delta_L
Doxycycline
YOL130W ALR1
ZShift = 2.32
NG = 0
DB = 0
SM = 0
Delta_K
Doxycycline
YOL130W ALR1
ZShift = 2.56
NG = 0
DB = 0
SM = 0
Delta_r
Doxycycline
YOL130W ALR1
ZShift = 2.06
NG = 0
DB = 0
SM = 0
Delta_AUC
Doxycycline

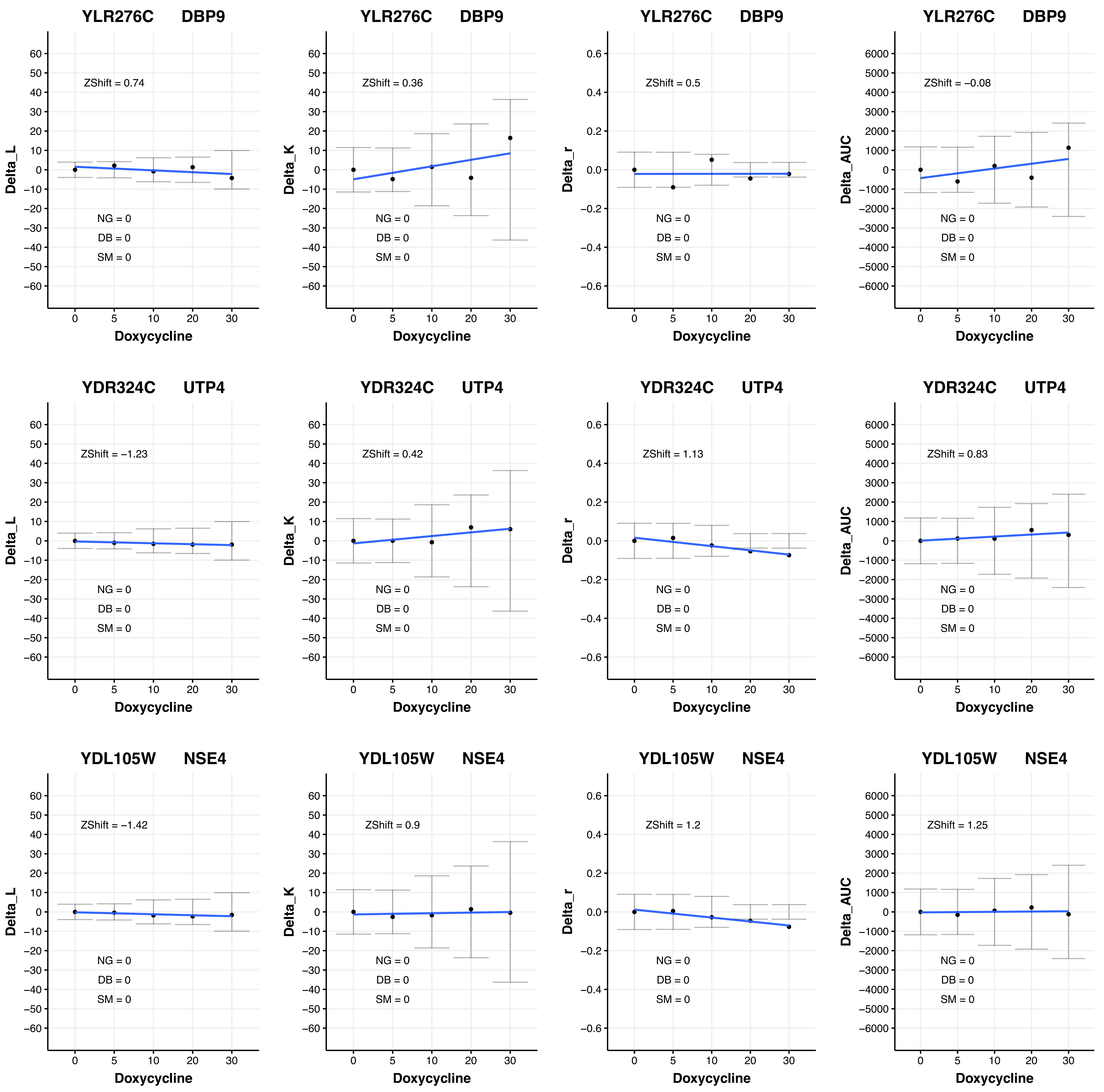
YLR276C DBP9
ZShift = 0.74
NG = 0
DB = 0
SM = 0
Delta_L
Doxycycline
YLR276C DBP9
ZShift = 0.36
NG = 0
DB = 0
SM = 0
Delta_K
Doxycycline
YLR276C DBP9
ZShift = 0.5
NG = 0
DB = 0
SM = 0
Delta_r
Doxycycline
YLR276C DBP9
ZShift = −0.08
NG = 0
DB = 0
SM = 0
Delta_AUC
Doxycycline
YDR324C UTP4
ZShift = −1.23
NG = 0
DB = 0
SM = 0
Delta_L
Doxycycline
YDR324C UTP4
ZShift = 0.42
NG = 0
DB = 0
SM = 0
Delta_K
Doxycycline
YDR324C UTP4
ZShift = 1.13
NG = 0
DB = 0
SM = 0
Delta_r
Doxycycline
YDR324C UTP4
ZShift = 0.83
NG = 0
DB = 0
SM = 0
Delta_AUC
Doxycycline
YDL105W NSE4
ZShift = −1.42
NG = 0
DB = 0
SM = 0
Delta_L
Doxycycline
YDL105W NSE4
ZShift = 0.9
NG = 0
DB = 0
SM = 0
Delta_K
Doxycycline
YDL105W NSE4
ZShift = 1.2
NG = 0
DB = 0
SM = 0
Delta_r
Doxycycline
YDL105W NSE4
ZShift = 1.25
NG = 0
DB = 0
SM = 0
Delta_AUC
Doxycycline

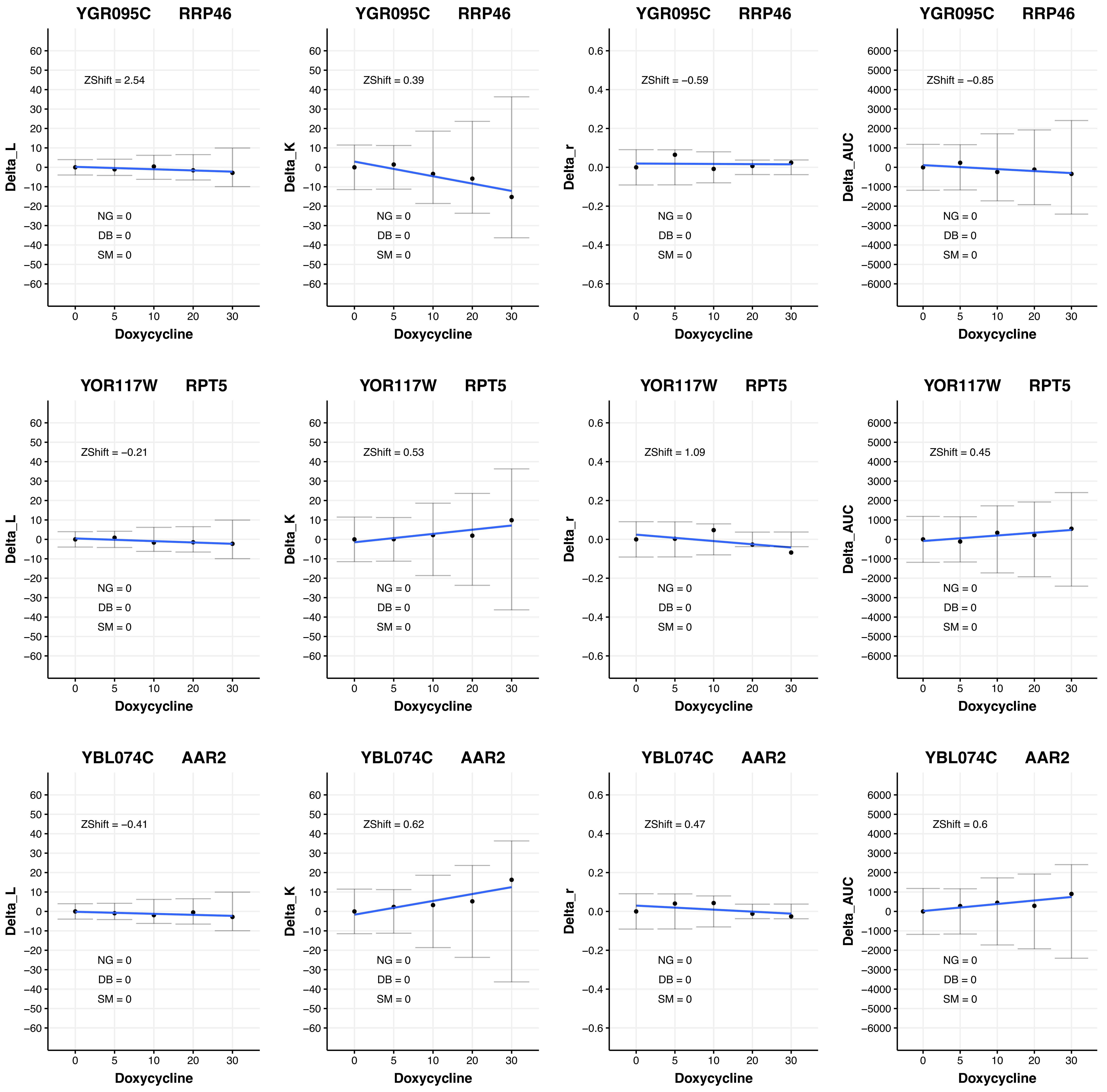
YGR095C RRP46
ZShift = 2.54
Delta_L
NG = 0
DB = 0
SM = 0
Doxycycline
YGR095C RRP46
ZShift = 0.39
Delta_K
NG = 0
DB = 0
SM = 0
Doxycycline
YGR095C RRP46
ZShift = −0.59
Delta_r
NG = 0
DB = 0
SM = 0
Doxycycline
YGR095C RRP46
ZShift = −0.85
Delta_AUC
NG = 0
DB = 0
SM = 0
Doxycycline
YOR117W RPT5
ZShift = −0.21
Delta_L
NG = 0
DB = 0
SM = 0
Doxycycline
YOR117W RPT5
ZShift = 0.53
Delta_K
NG = 0
DB = 0
SM = 0
Doxycycline
YOR117W RPT5
ZShift = 1.09
Delta_r
NG = 0
DB = 0
SM = 0
Doxycycline
YOR117W RPT5
ZShift = 0.45
Delta_AUC
NG = 0
DB = 0
SM = 0
Doxycycline
YBL074C AAR2
ZShift = −0.41
Delta_L
NG = 0
DB = 0
SM = 0
Doxycycline
YBL074C AAR2
ZShift = 0.62
Delta_K
NG = 0
DB = 0
SM = 0
Doxycycline
YBL074C AAR2
ZShift = 0.47
Delta_r
NG = 0
DB = 0
SM = 0
Doxycycline
YBL074C AAR2
ZShift = 0.6
Delta_AUC
NG = 0
DB = 0
SM = 0
Doxycycline

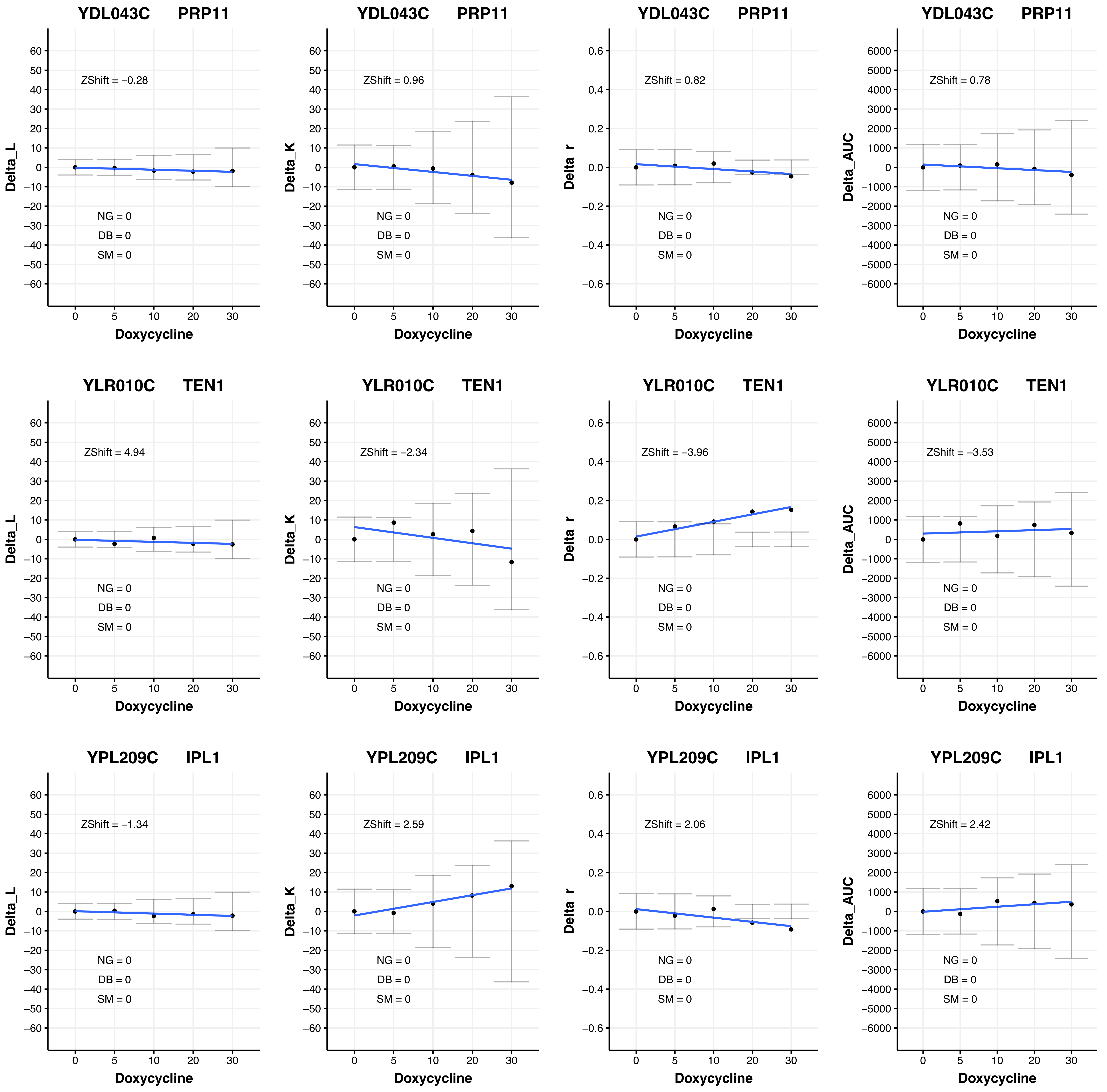
YDL043C PRP11
ZShift = −0.28
NG = 0
DB = 0
SM = 0
Delta_L
Doxycycline
YDL043C PRP11
ZShift = 0.96
NG = 0
DB = 0
SM = 0
Delta_K
Doxycycline
YDL043C PRP11
ZShift = 0.82
NG = 0
DB = 0
SM = 0
Delta_r
Doxycycline
YDL043C PRP11
ZShift = 0.78
NG = 0
DB = 0
SM = 0
Delta_AUC
Doxycycline
YLR010C TEN1
ZShift = 4.94
NG = 0
DB = 0
SM = 0
Delta_L
Doxycycline
YLR010C TEN1
ZShift = −2.34
NG = 0
DB = 0
SM = 0
Delta_K
Doxycycline
YLR010C TEN1
ZShift = −3.96
NG = 0
DB = 0
SM = 0
Delta_r
Doxycycline
YLR010C TEN1
ZShift = −3.53
NG = 0
DB = 0
SM = 0
Delta_AUC
Doxycycline
YPL209C IPL1
ZShift = −1.34
NG = 0
DB = 0
SM = 0
Delta_L
Doxycycline
YPL209C IPL1
ZShift = 2.59
NG = 0
DB = 0
SM = 0
Delta_K
Doxycycline
YPL209C IPL1
ZShift = 2.06
NG = 0
DB = 0
SM = 0
Delta_r
Doxycycline
YPL209C IPL1
ZShift = 2.42
NG = 0
DB = 0
SM = 0
Delta_AUC
Doxycycline

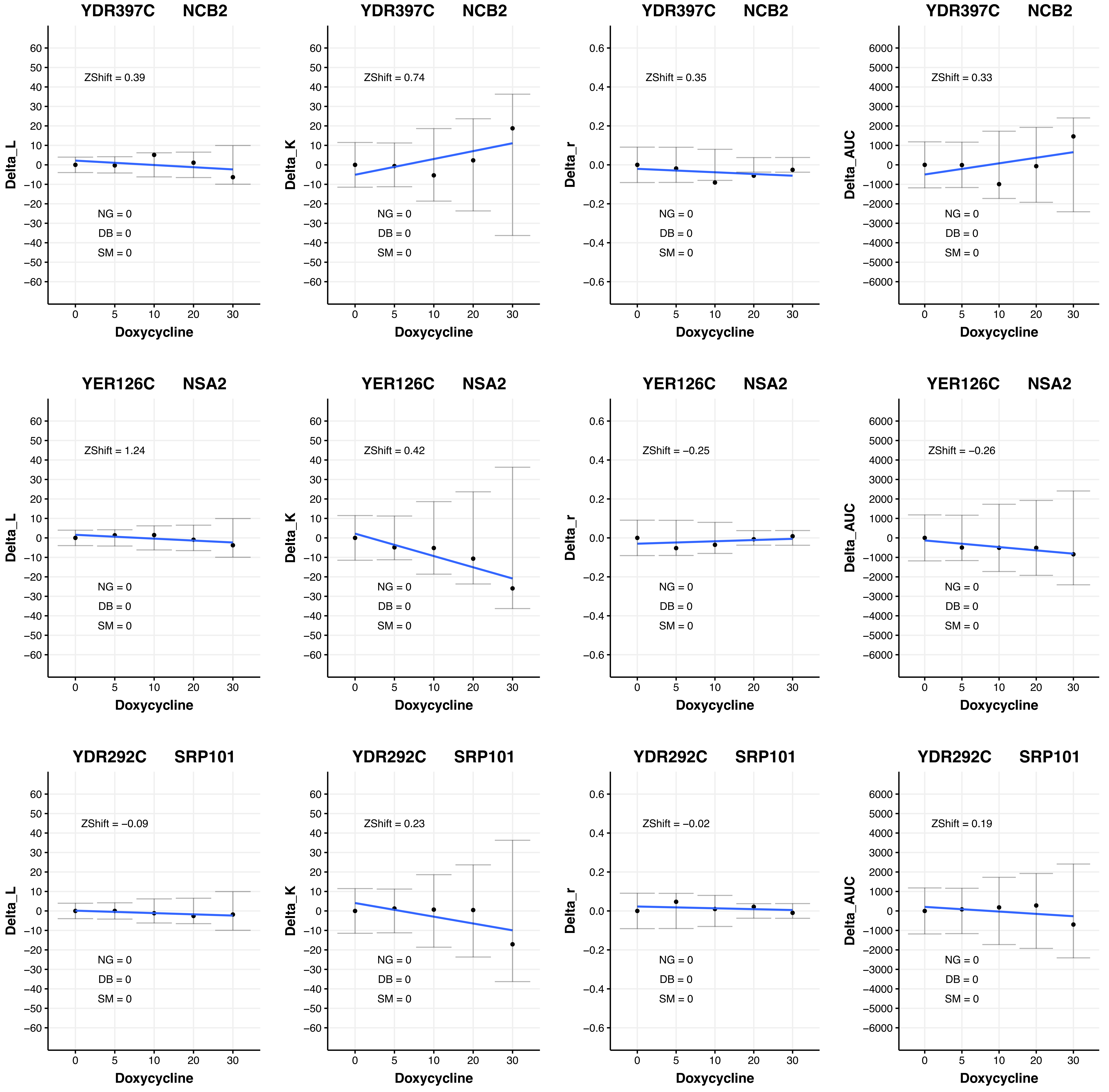
YDR397C NCB2
ZShift = 0.39
NG = 0
DB = 0
SM = 0
Delta_L
Doxycycline
YDR397C NCB2
ZShift = 0.74
NG = 0
DB = 0
SM = 0
Delta_K
Doxycycline
YDR397C NCB2
ZShift = 0.35
NG = 0
DB = 0
SM = 0
Delta_r
Doxycycline
YDR397C NCB2
ZShift = 0.33
NG = 0
DB = 0
SM = 0
Delta_AUC
Doxycycline
YER126C NSA2
ZShift = 1.24
NG = 0
DB = 0
SM = 0
Delta_L
Doxycycline
YER126C NSA2
ZShift = 0.42
NG = 0
DB = 0
SM = 0
Delta_K
Doxycycline
YER126C NSA2
ZShift = −0.25
NG = 0
DB = 0
SM = 0
Delta_r
Doxycycline
YER126C NSA2
ZShift = −0.26
NG = 0
DB = 0
SM = 0
Delta_AUC
Doxycycline
YDR292C SRP101
ZShift = −0.09
NG = 0
DB = 0
SM = 0
Delta_L
Doxycycline
YDR292C SRP101
ZShift = 0.23
NG = 0
DB = 0
SM = 0
Delta_K
Doxycycline
YDR292C SRP101
ZShift = −0.02
NG = 0
DB = 0
SM = 0
Delta_r
Doxycycline
YDR292C SRP101
ZShift = 0.19
NG = 0
DB = 0
SM = 0
Delta_AUC
Doxycycline

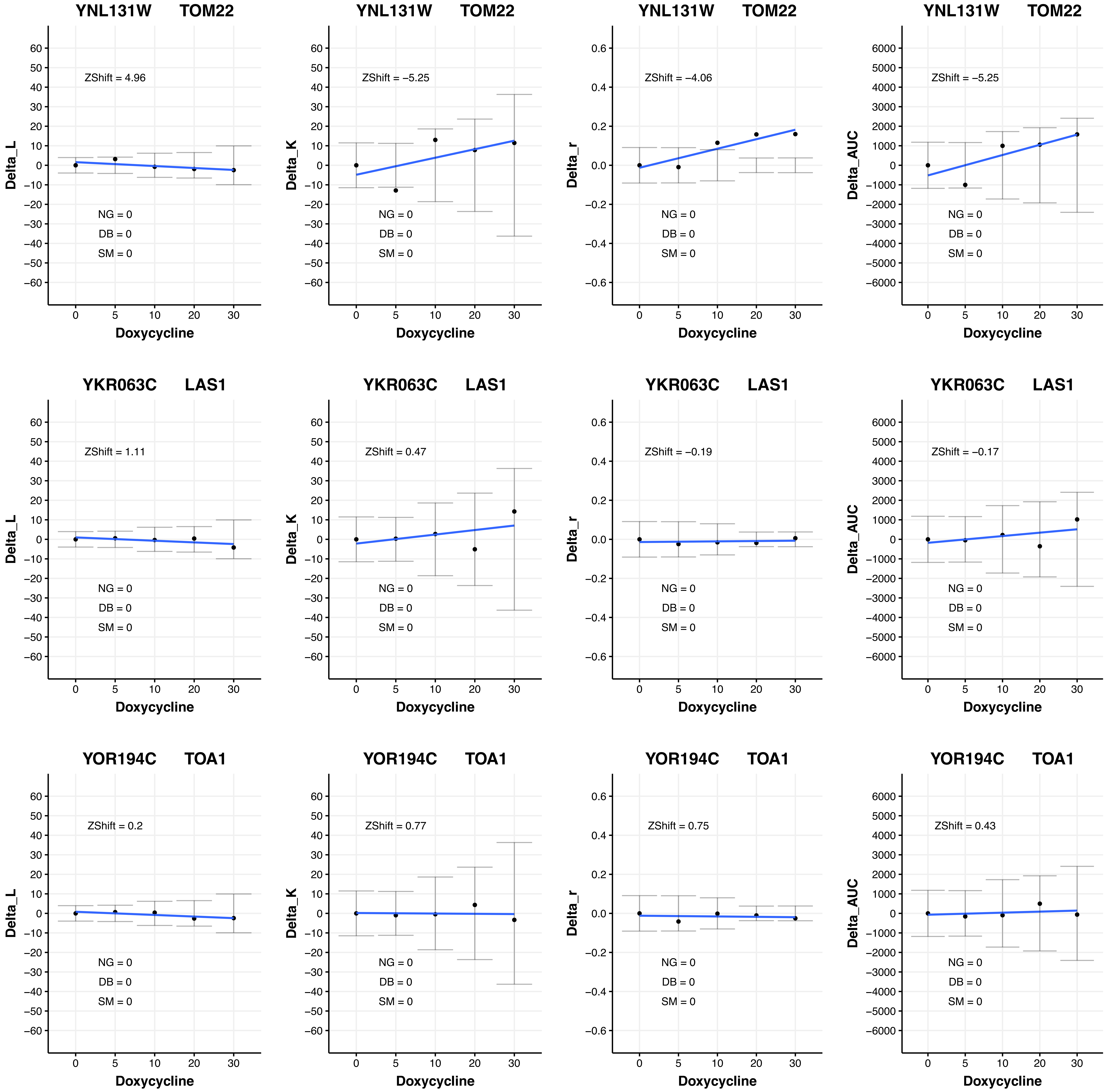
YNL131W TOM22
ZShift = 4.96
NG = 0
DB = 0
SM = 0
Delta_L
Doxycycline
YNL131W TOM22
ZShift = −5.25
NG = 0
DB = 0
SM = 0
Delta_K
Doxycycline
YNL131W TOM22
ZShift = −4.06
NG = 0
DB = 0
SM = 0
Delta_r
Doxycycline
YNL131W TOM22
ZShift = −5.25
NG = 0
DB = 0
SM = 0
Delta_AUC
Doxycycline
YKR063C LAS1
ZShift = 1.11
NG = 0
DB = 0
SM = 0
Delta_L
Doxycycline
YKR063C LAS1
ZShift = 0.47
NG = 0
DB = 0
SM = 0
Delta_K
Doxycycline
YKR063C LAS1
ZShift = −0.19
NG = 0
DB = 0
SM = 0
Delta_r
Doxycycline
YKR063C LAS1
ZShift = −0.17
NG = 0
DB = 0
SM = 0
Delta_AUC
Doxycycline
YOR194C TOA1
ZShift = 0.2
NG = 0
DB = 0
SM = 0
Delta_L
Doxycycline
YOR194C TOA1
ZShift = 0.77
NG = 0
DB = 0
SM = 0
Delta_K
Doxycycline
YOR194C TOA1
ZShift = 0.75
NG = 0
DB = 0
SM = 0
Delta_r
Doxycycline
YOR194C TOA1
ZShift = 0.43
NG = 0
DB = 0
SM = 0
Delta_AUC
Doxycycline

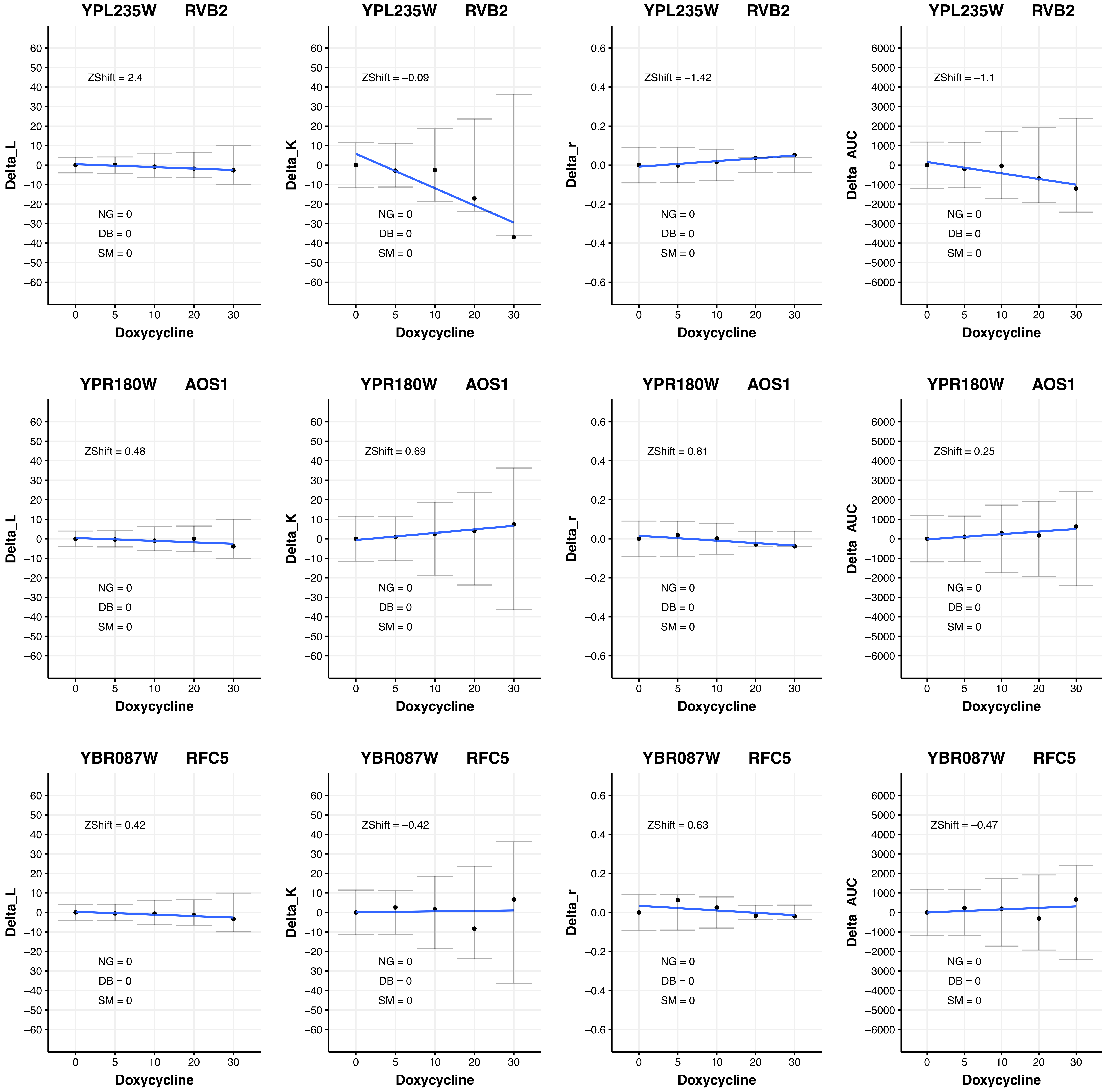
YPL235W RVB2
ZShift = 2.4
NG = 0
DB = 0
SM = 0
Delta_L
Doxycycline
YPL235W RVB2
ZShift = −0.09
NG = 0
DB = 0
SM = 0
Delta_K
Doxycycline
YPL235W RVB2
ZShift = −1.42
NG = 0
DB = 0
SM = 0
Delta_r
Doxycycline
YPL235W RVB2
ZShift = −1.1
NG = 0
DB = 0
SM = 0
Delta_AUC
Doxycycline
YPR180W AOS1
ZShift = 0.48
NG = 0
DB = 0
SM = 0
Delta_L
Doxycycline
YPR180W AOS1
ZShift = 0.69
NG = 0
DB = 0
SM = 0
Delta_K
Doxycycline
YPR180W AOS1
ZShift = 0.81
NG = 0
DB = 0
SM = 0
Delta_r
Doxycycline
YPR180W AOS1
ZShift = 0.25
NG = 0
DB = 0
SM = 0
Delta_AUC
Doxycycline
YBR087W RFC5
ZShift = 0.42
NG = 0
DB = 0
SM = 0
Delta_L
Doxycycline
YBR087W RFC5
ZShift = −0.42
NG = 0
DB = 0
SM = 0
Delta_K
Doxycycline
YBR087W RFC5
ZShift = 0.63
NG = 0
DB = 0
SM = 0
Delta_r
Doxycycline
YBR087W RFC5
ZShift = −0.47
NG = 0
DB = 0
SM = 0
Delta_AUC
Doxycycline

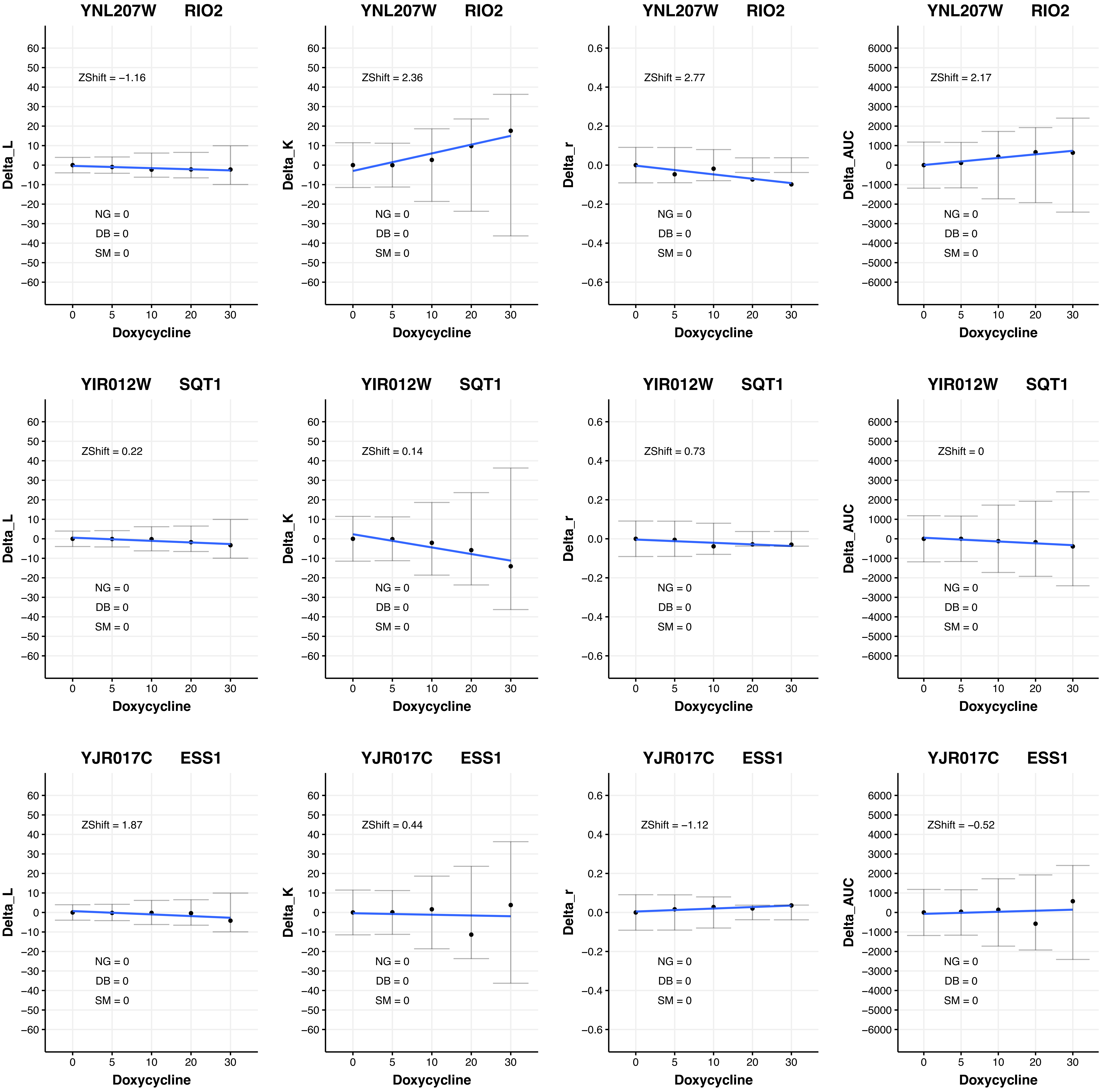
YNL207W RIO2
ZShift = −1.16
Delta_L
NG = 0
DB = 0
SM = 0
Doxycycline
YNL207W RIO2
ZShift = 2.36
Delta_K
NG = 0
DB = 0
SM = 0
Doxycycline
YNL207W RIO2
ZShift = 2.77
Delta_r
NG = 0
DB = 0
SM = 0
Doxycycline
YNL207W RIO2
ZShift = 2.17
Delta_AUC
NG = 0
DB = 0
SM = 0
Doxycycline
YIR012W SQT1
ZShift = 0.22
Delta_L
NG = 0
DB = 0
SM = 0
Doxycycline
YIR012W SQT1
ZShift = 0.14
Delta_K
NG = 0
DB = 0
SM = 0
Doxycycline
YIR012W SQT1
ZShift = 0.73
Delta_r
NG = 0
DB = 0
SM = 0
Doxycycline
YIR012W SQT1
ZShift = 0
Delta_AUC
NG = 0
DB = 0
SM = 0
Doxycycline
YJR017C ESS1
ZShift = 1.87
Delta_L
NG = 0
DB = 0
SM = 0
Doxycycline
YJR017C ESS1
ZShift = 0.44
Delta_K
NG = 0
DB = 0
SM = 0
Doxycycline
YJR017C ESS1
ZShift = −1.12
Delta_r
NG = 0
DB = 0
SM = 0
Doxycycline
YJR017C ESS1
ZShift = −0.52
Delta_AUC
NG = 0
DB = 0
SM = 0
Doxycycline

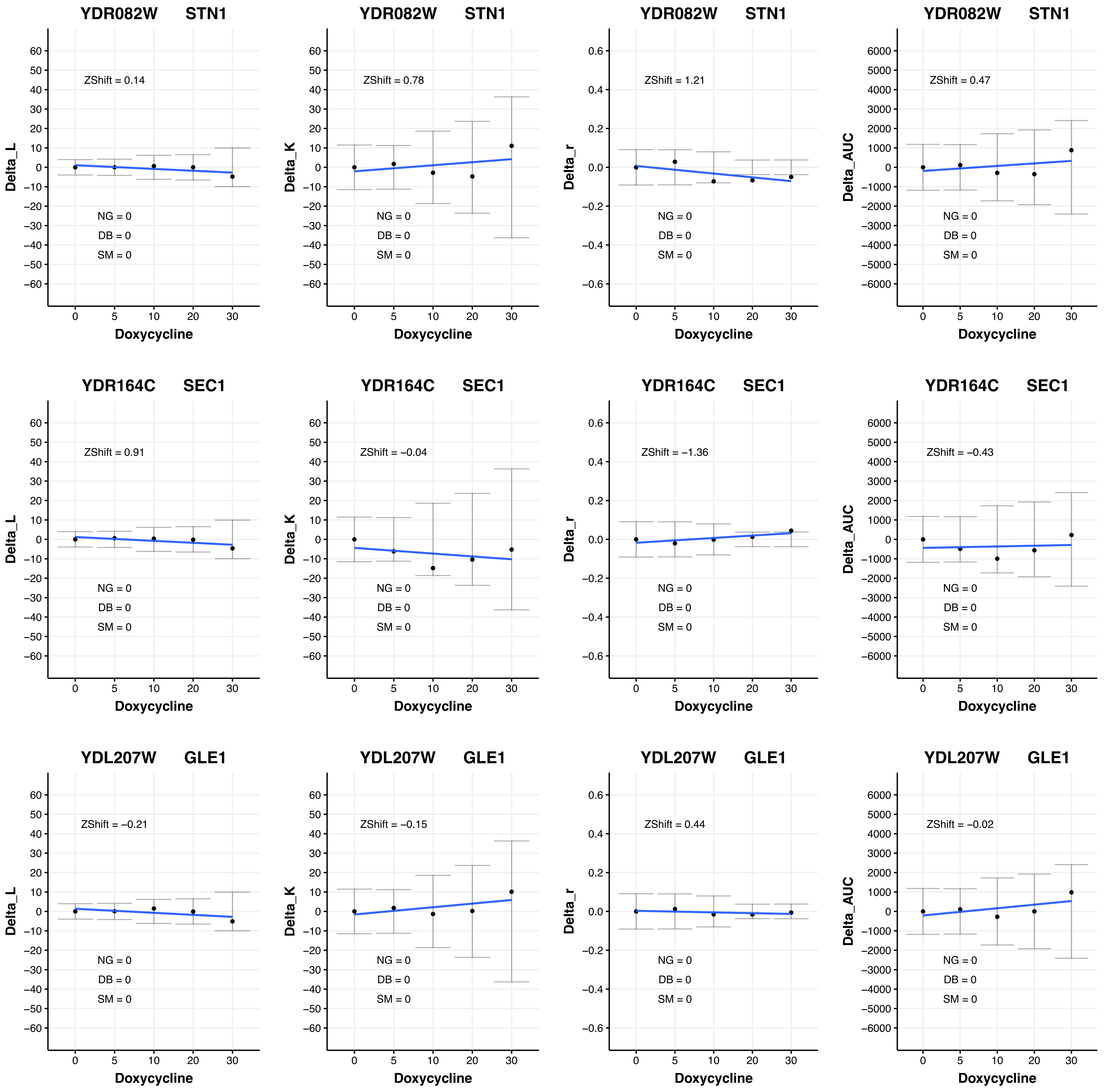

YDR082W STN1
ZShift = 0.14
NG = 0
DB = 0
SM = 0
Delta_L
Doxycycline
YDR082W STN1
ZShift = 0.78
NG = 0
DB = 0
SM = 0
Delta_K
Doxycycline
YDR082W STN1
ZShift = 1.21
NG = 0
DB = 0
SM = 0
Delta_r
Doxycycline
YDR082W STN1
ZShift = 0.47
NG = 0
DB = 0
SM = 0
Delta_AUC
Doxycycline
YDR164C SEC1
ZShift = 0.91
NG = 0
DB = 0
SM = 0
Delta_L
Doxycycline
YDR164C SEC1
ZShift = −0.04
NG = 0
DB = 0
SM = 0
Delta_K
Doxycycline
YDR164C SEC1
ZShift = −1.36
NG = 0
DB = 0
SM = 0
Delta_r
Doxycycline
YDR164C SEC1
ZShift = −0.43
NG = 0
DB = 0
SM = 0
Delta_AUC
Doxycycline
YDL207W GLE1
ZShift = −0.21
NG = 0
DB = 0
SM = 0
Delta_L
Doxycycline
YDL207W GLE1
ZShift = −0.15
NG = 0
DB = 0
SM = 0
Delta_K
Doxycycline
YDL207W GLE1
ZShift = 0.44
NG = 0
DB = 0
SM = 0
Delta_r
Doxycycline
YDL207W GLE1
ZShift = −0.02
NG = 0
DB = 0
SM = 0
Delta_AUC
Doxycycline

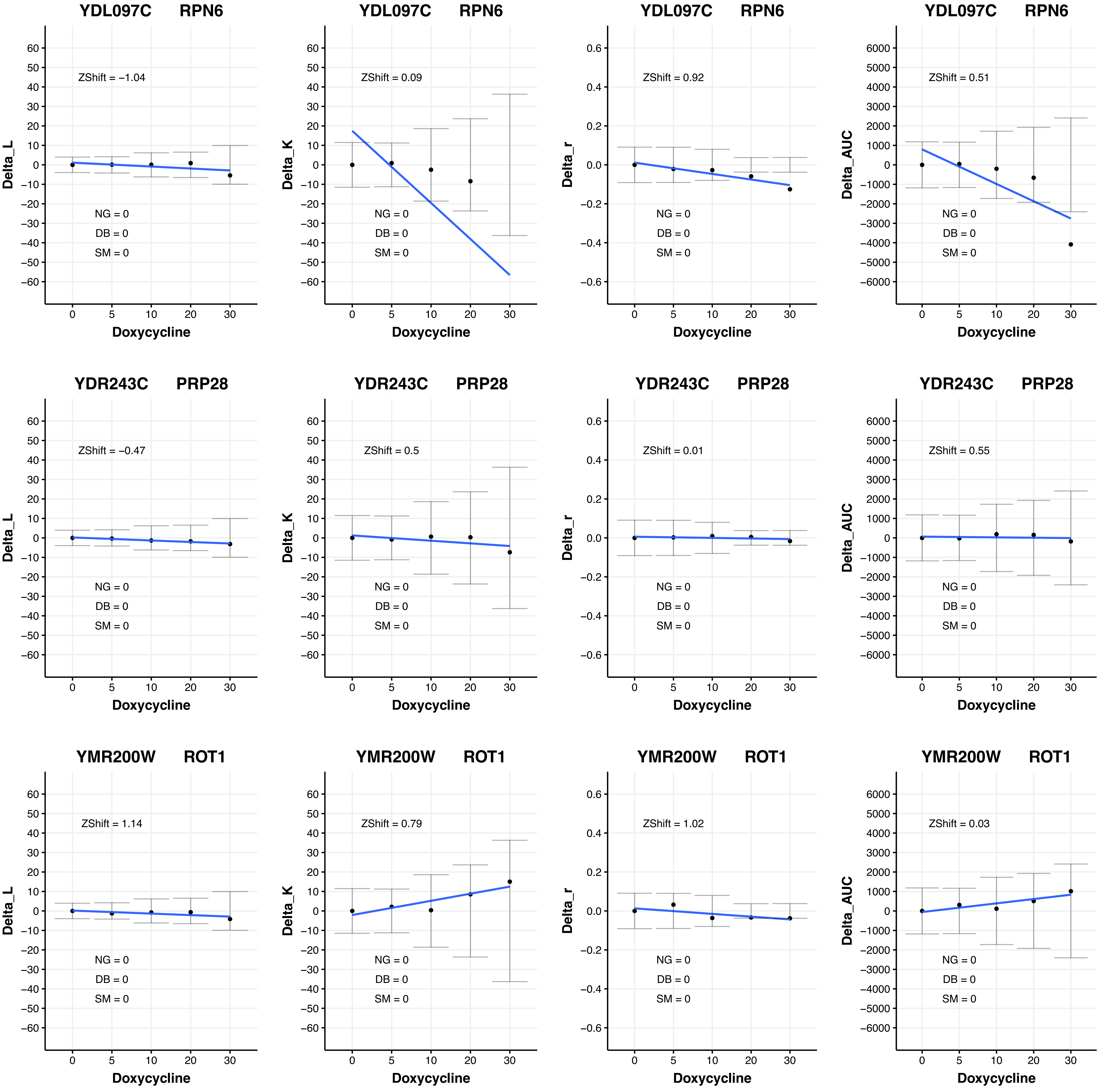

YDL097C RPN6
ZShift = −1.04
NG = 0
DB = 0
SM = 0
Delta_L
Doxycycline
YDL097C RPN6
ZShift = 0.09
NG = 0
DB = 0
SM = 0
Delta_K
Doxycycline
YDL097C RPN6
ZShift = 0.92
NG = 0
DB = 0
SM = 0
Delta_r
Doxycycline
YDL097C RPN6
ZShift = 0.51
NG = 0
DB = 0
SM = 0
Delta_AUC
Doxycycline
YDR243C PRP28
ZShift = −0.47
NG = 0
DB = 0
SM = 0
Delta_L
Doxycycline
YDR243C PRP28
ZShift = 0.5
NG = 0
DB = 0
SM = 0
Delta_K
Doxycycline
YDR243C PRP28
ZShift = 0.01
NG = 0
DB = 0
SM = 0
Delta_r
Doxycycline
YDR243C PRP28
ZShift = 0.55
NG = 0
DB = 0
SM = 0
Delta_AUC
Doxycycline
YMR200W ROT1
ZShift = 1.14
NG = 0
DB = 0
SM = 0
Delta_L
Doxycycline
YMR200W ROT1
ZShift = 0.79
NG = 0
DB = 0
SM = 0
Delta_K
Doxycycline
YMR200W ROT1
ZShift = 1.02
NG = 0
DB = 0
SM = 0
Delta_r
Doxycycline
YMR200W ROT1
ZShift = 0.03
NG = 0
DB = 0
SM = 0
Delta_AUC
Doxycycline

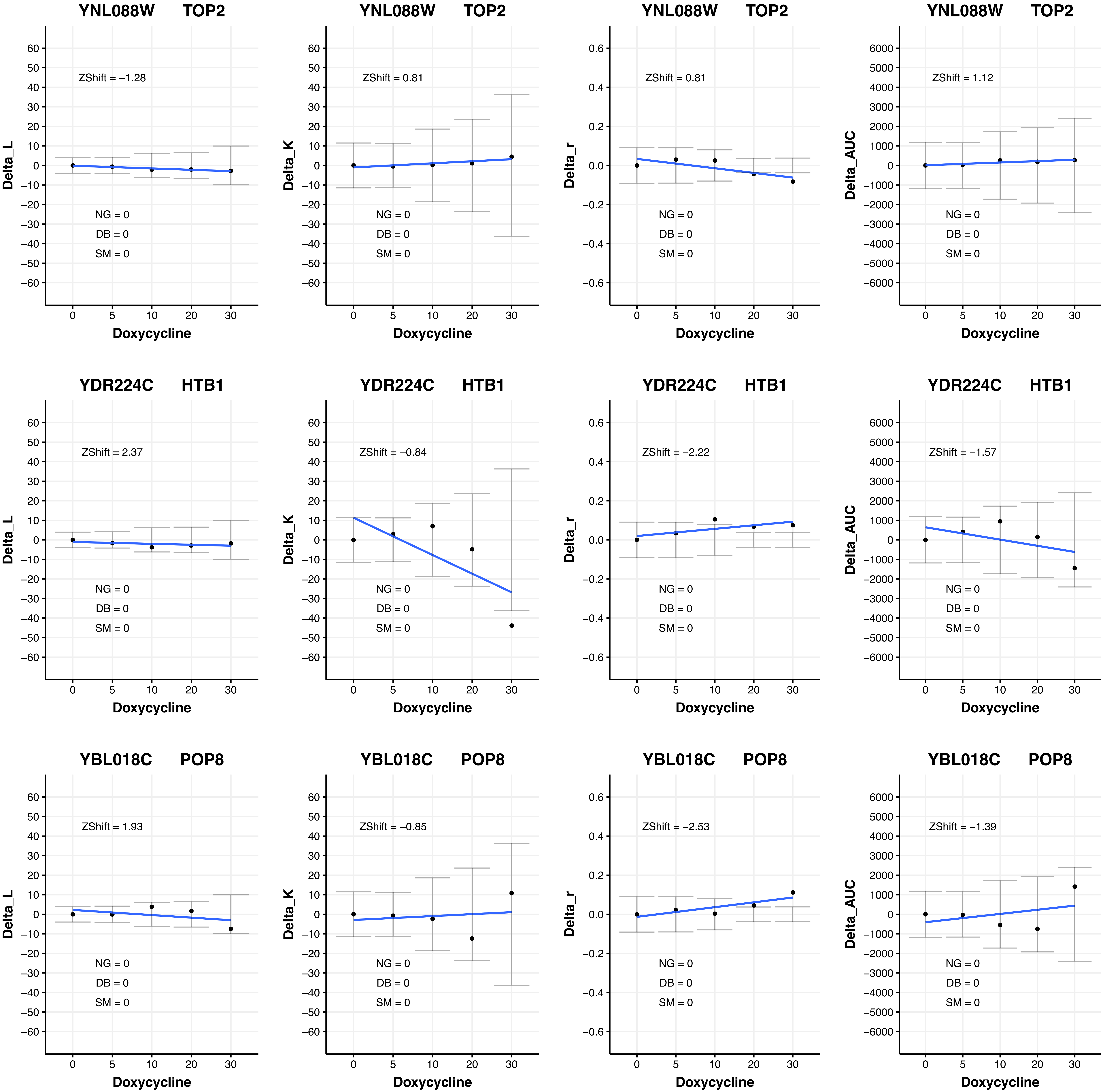
YNL088W TOP2
ZShift = −1.28
NG = 0
DB = 0
SM = 0
Delta_L
Doxycycline
YNL088W TOP2
ZShift = 0.81
NG = 0
DB = 0
SM = 0
Delta_K
Doxycycline
YNL088W TOP2
ZShift = 0.81
NG = 0
DB = 0
SM = 0
Delta_r
Doxycycline
YNL088W TOP2
ZShift = 1.12
NG = 0
DB = 0
SM = 0
Delta_AUC
Doxycycline
YDR224C HTB1
ZShift = 2.37
NG = 0
DB = 0
SM = 0
Delta_L
Doxycycline
YDR224C HTB1
ZShift = −0.84
NG = 0
DB = 0
SM = 0
Delta_K
Doxycycline
YDR224C HTB1
ZShift = −2.22
NG = 0
DB = 0
SM = 0
Delta_r
Doxycycline
YDR224C HTB1
ZShift = −1.57
NG = 0
DB = 0
SM = 0
Delta_AUC
Doxycycline
YBL018C POP8
ZShift = 1.93
NG = 0
DB = 0
SM = 0
Delta_L
Doxycycline
YBL018C POP8
ZShift = −0.85
NG = 0
DB = 0
SM = 0
Delta_K
Doxycycline
YBL018C POP8
ZShift = −2.53
NG = 0
DB = 0
SM = 0
Delta_r
Doxycycline
YBL018C POP8
ZShift = −1.39
NG = 0
DB = 0
SM = 0
Delta_AUC
Doxycycline

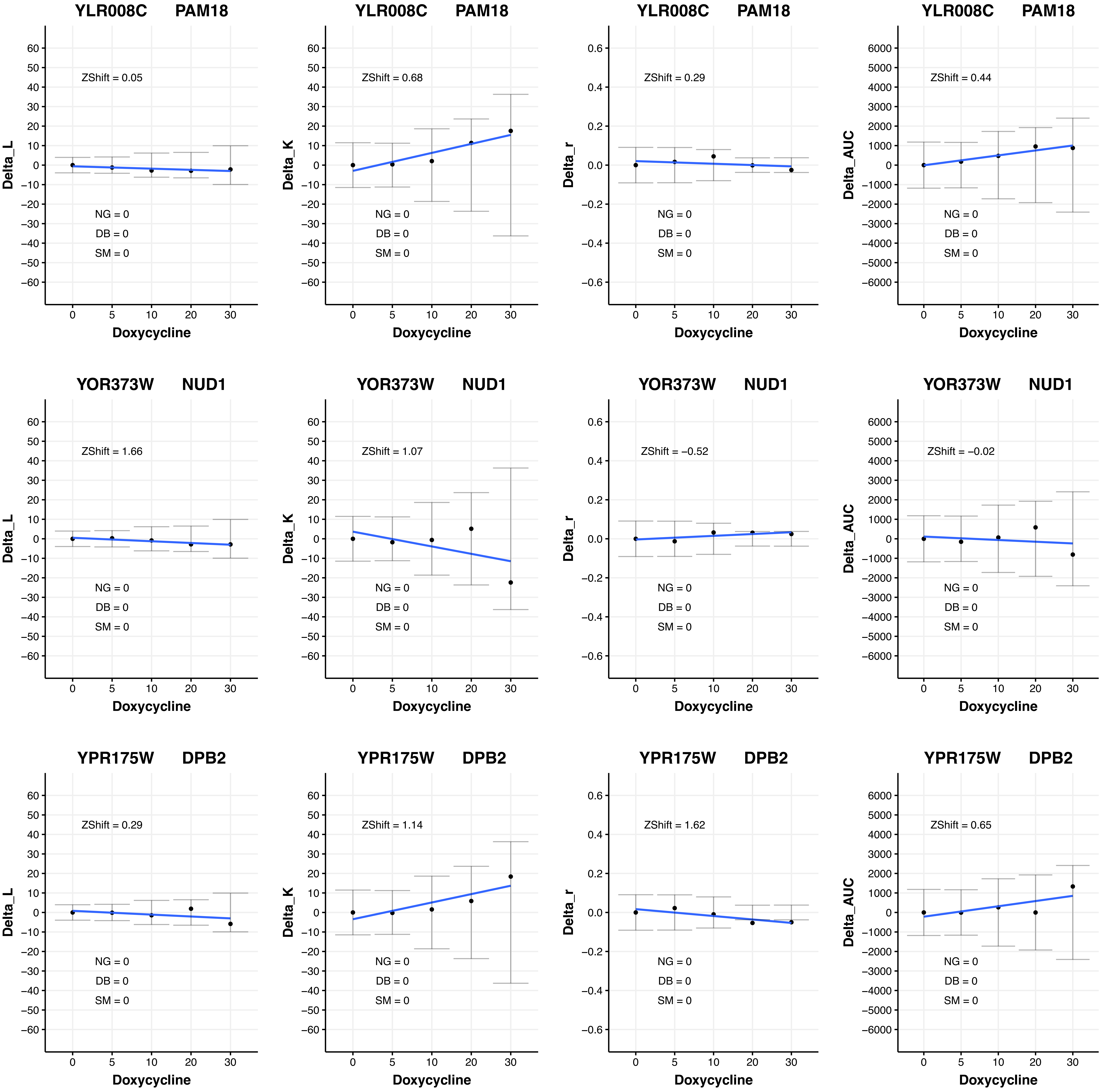
YLR008C PAM18
Delta_L
ZShift = 0.05
NG = 0
DB = 0
SM = 0
Doxycycline
YLR008C PAM18
Delta_K
ZShift = 0.68
NG = 0
DB = 0
SM = 0
Doxycycline
YLR008C PAM18
Delta_r
ZShift = 0.29
NG = 0
DB = 0
SM = 0
Doxycycline
YLR008C PAM18
Delta_AUC
ZShift = 0.44
NG = 0
DB = 0
SM = 0
Doxycycline
YOR373W NUD1
Delta_L
ZShift = 1.66
NG = 0
DB = 0
SM = 0
Doxycycline
YOR373W NUD1
Delta_K
ZShift = 1.07
NG = 0
DB = 0
SM = 0
Doxycycline
YOR373W NUD1
Delta_r
ZShift = −0.52
NG = 0
DB = 0
SM = 0
Doxycycline
YOR373W NUD1
Delta_AUC
ZShift = −0.02
NG = 0
DB = 0
SM = 0
Doxycycline
YPR175W DPB2
Delta_L
ZShift = 0.29
NG = 0
DB = 0
SM = 0
Doxycycline
YPR175W DPB2
Delta_K
ZShift = 1.14
NG = 0
DB = 0
SM = 0
Doxycycline
YPR175W DPB2
Delta_r
ZShift = 1.62
NG = 0
DB = 0
SM = 0
Doxycycline
YPR175W DPB2
Delta_AUC
ZShift = 0.65
NG = 0
DB = 0
SM = 0
Doxycycline

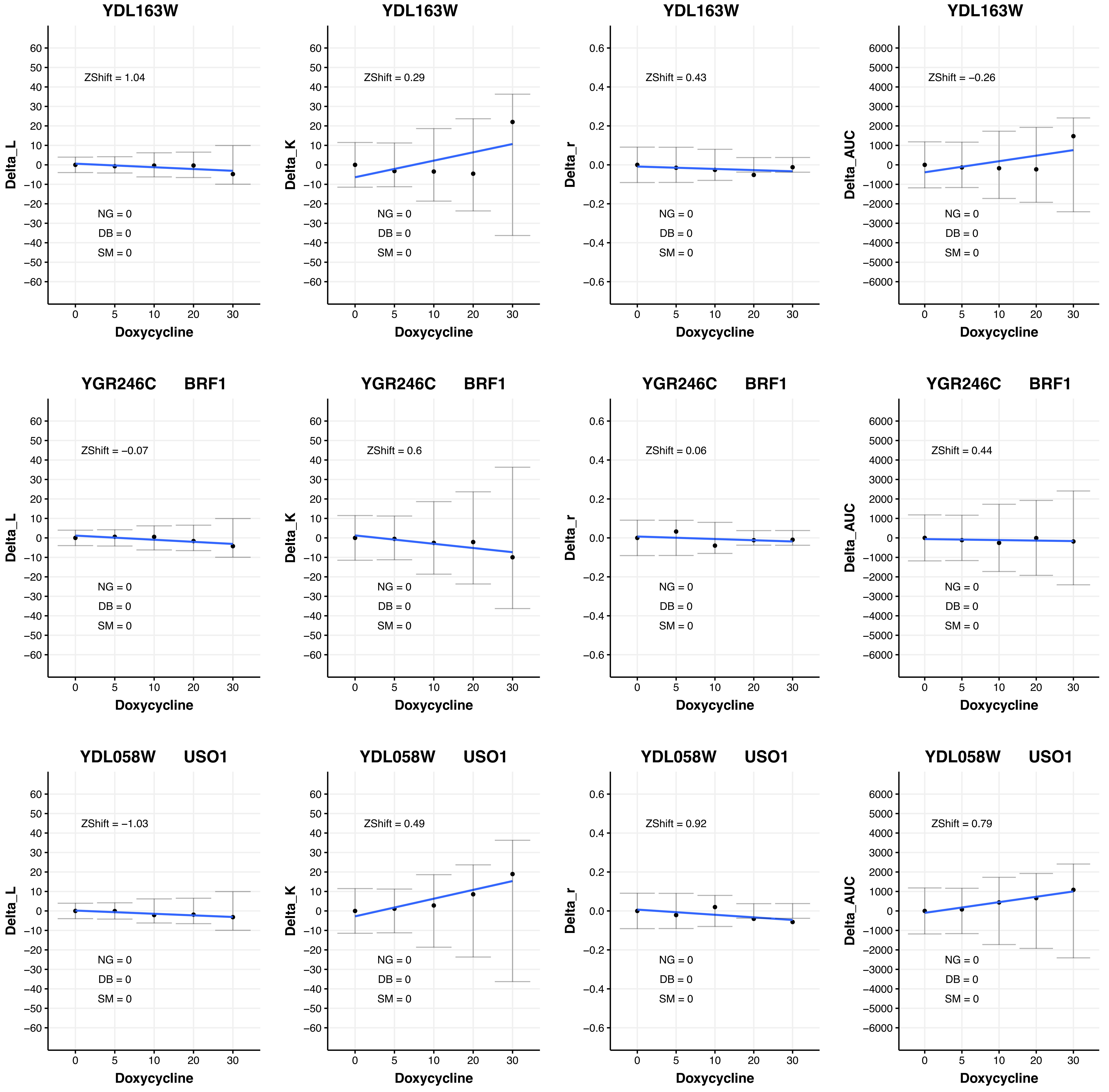
YDL163W
ZShift = 1.04
NG = 0
DB = 0
SM = 0
Delta_L
Doxycycline
YDL163W
ZShift = 0.29
NG = 0
DB = 0
SM = 0
Delta_K
Doxycycline
YDL163W
ZShift = 0.43
NG = 0
DB = 0
SM = 0
Delta_r
Doxycycline
YDL163W
ZShift = −0.26
NG = 0
DB = 0
SM = 0
Delta_AUC
Doxycycline
YGR246C BRF1
ZShift = −0.07
NG = 0
DB = 0
SM = 0
Delta_L
Doxycycline
YGR246C BRF1
ZShift = 0.6
NG = 0
DB = 0
SM = 0
Delta_K
Doxycycline
YGR246C BRF1
ZShift = 0.06
NG = 0
DB = 0
SM = 0
Delta_r
Doxycycline
YGR246C BRF1
ZShift = 0.44
NG = 0
DB = 0
SM = 0
Delta_AUC
Doxycycline
YDL058W USO1
ZShift = −1.03
NG = 0
DB = 0
SM = 0
Delta_L
Doxycycline
YDL058W USO1
ZShift = 0.49
NG = 0
DB = 0
SM = 0
Delta_K
Doxycycline
YDL058W USO1
ZShift = 0.92
NG = 0
DB = 0
SM = 0
Delta_r
Doxycycline
YDL058W USO1
ZShift = 0.79
NG = 0
DB = 0
SM = 0
Delta_AUC
Doxycycline

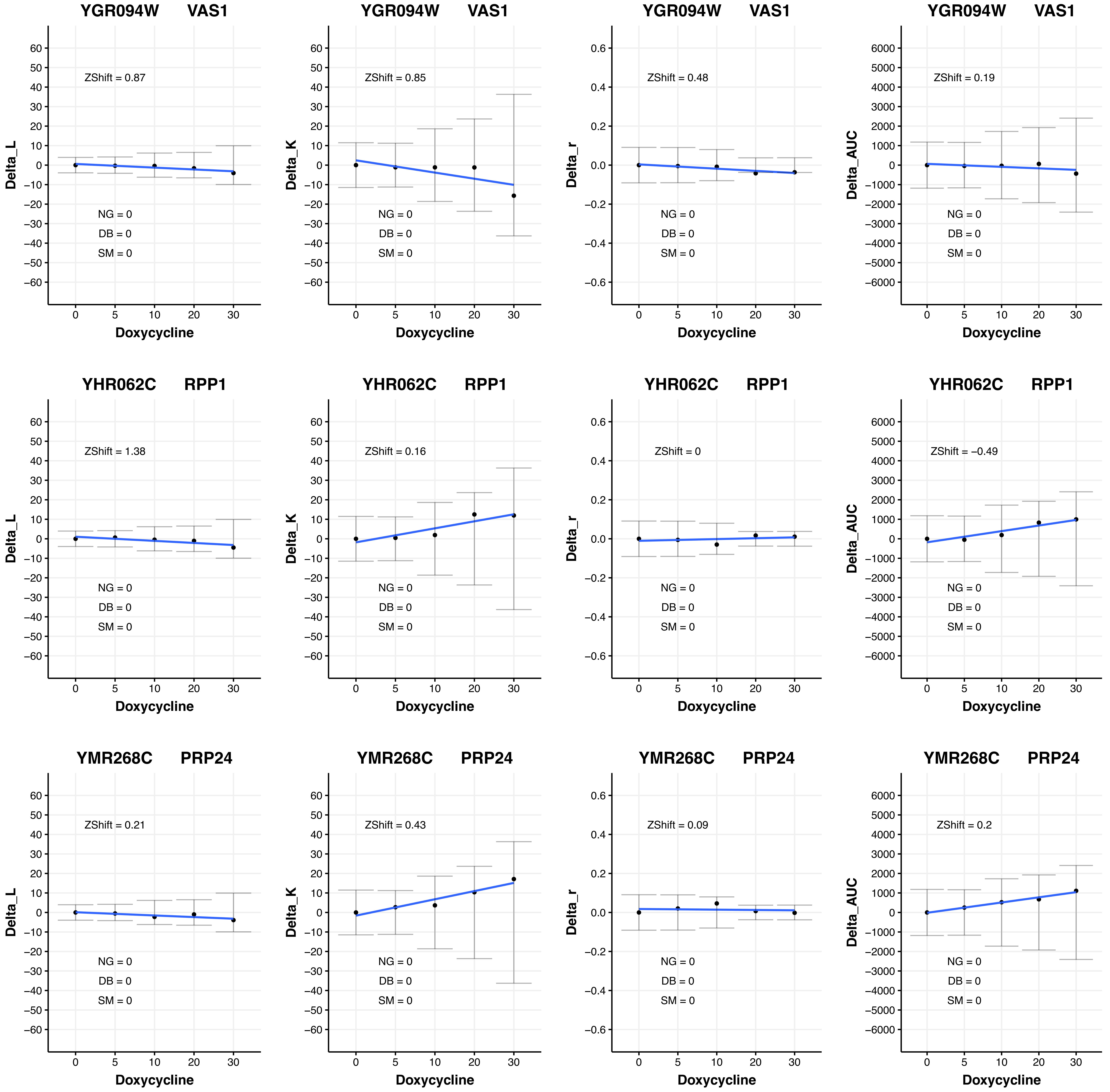
YGR094W VAS1
ZShift = 0.87
NG = 0
DB = 0
SM = 0
Delta_L
Doxycycline
YGR094W VAS1
ZShift = 0.85
NG = 0
DB = 0
SM = 0
Delta_K
Doxycycline
YGR094W VAS1
ZShift = 0.48
NG = 0
DB = 0
SM = 0
Delta_r
Doxycycline
YGR094W VAS1
ZShift = 0.19
NG = 0
DB = 0
SM = 0
Delta_AUC
Doxycycline
YHR062C RPP1
ZShift = 1.38
NG = 0
DB = 0
SM = 0
Delta_L
Doxycycline
YHR062C RPP1
ZShift = 0.16
NG = 0
DB = 0
SM = 0
Delta_K
Doxycycline
YHR062C RPP1
ZShift = 0
NG = 0
DB = 0
SM = 0
Delta_r
Doxycycline
YHR062C RPP1
ZShift = −0.49
NG = 0
DB = 0
SM = 0
Delta_AUC
Doxycycline
YMR268C PRP24
ZShift = 0.21
NG = 0
DB = 0
SM = 0
Delta_L
Doxycycline
YMR268C PRP24
ZShift = 0.43
NG = 0
DB = 0
SM = 0
Delta_K
Doxycycline
YMR268C PRP24
ZShift = 0.09
NG = 0
DB = 0
SM = 0
Delta_r
Doxycycline
YMR268C PRP24
ZShift = 0.2
NG = 0
DB = 0
SM = 0
Delta_AUC
Doxycycline

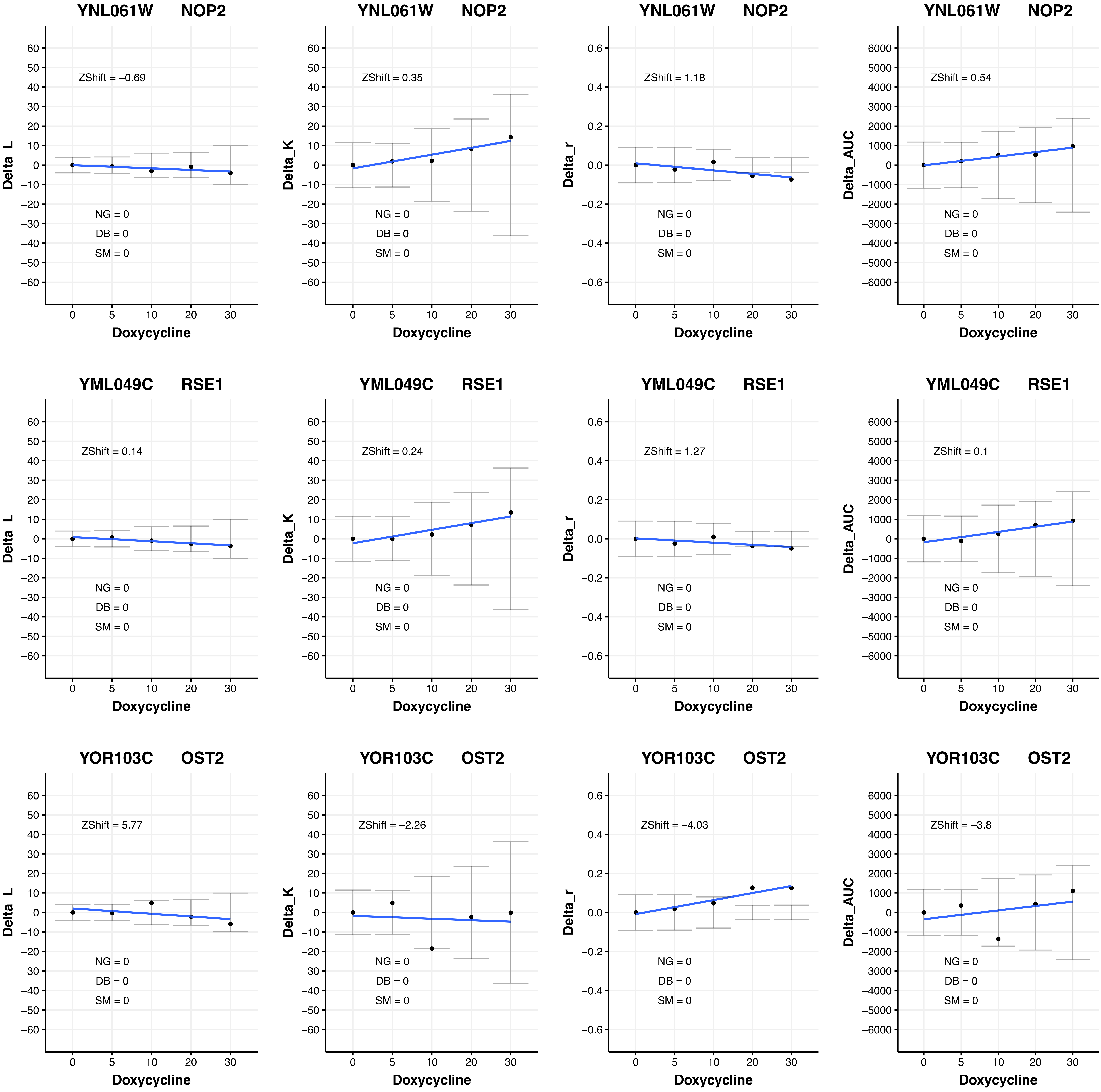
YNL061W NOP2
ZShift = −0.69
Delta_L
NG = 0
DB = 0
SM = 0
Doxycycline
YNL061W NOP2
ZShift = 0.35
Delta_K
NG = 0
DB = 0
SM = 0
Doxycycline
YNL061W NOP2
ZShift = 1.18
Delta_r
NG = 0
DB = 0
SM = 0
Doxycycline
YNL061W NOP2
ZShift = 0.54
Delta_AUC
NG = 0
DB = 0
SM = 0
Doxycycline
YML049C RSE1
ZShift = 0.14
Delta_L
NG = 0
DB = 0
SM = 0
Doxycycline
YML049C RSE1
ZShift = 0.24
Delta_K
NG = 0
DB = 0
SM = 0
Doxycycline
YML049C RSE1
ZShift = 1.27
Delta_r
NG = 0
DB = 0
SM = 0
Doxycycline
YML049C RSE1
ZShift = 0.1
Delta_AUC
NG = 0
DB = 0
SM = 0
Doxycycline
YOR103C OST2
ZShift = 5.77
Delta_L
NG = 0
DB = 0
SM = 0
Doxycycline
YOR103C OST2
ZShift = −2.26
Delta_K
NG = 0
DB = 0
SM = 0
Doxycycline
YOR103C OST2
ZShift = −4.03
Delta_r
NG = 0
DB = 0
SM = 0
Doxycycline
YOR103C OST2
ZShift = −3.8
Delta_AUC
NG = 0
DB = 0
SM = 0
Doxycycline

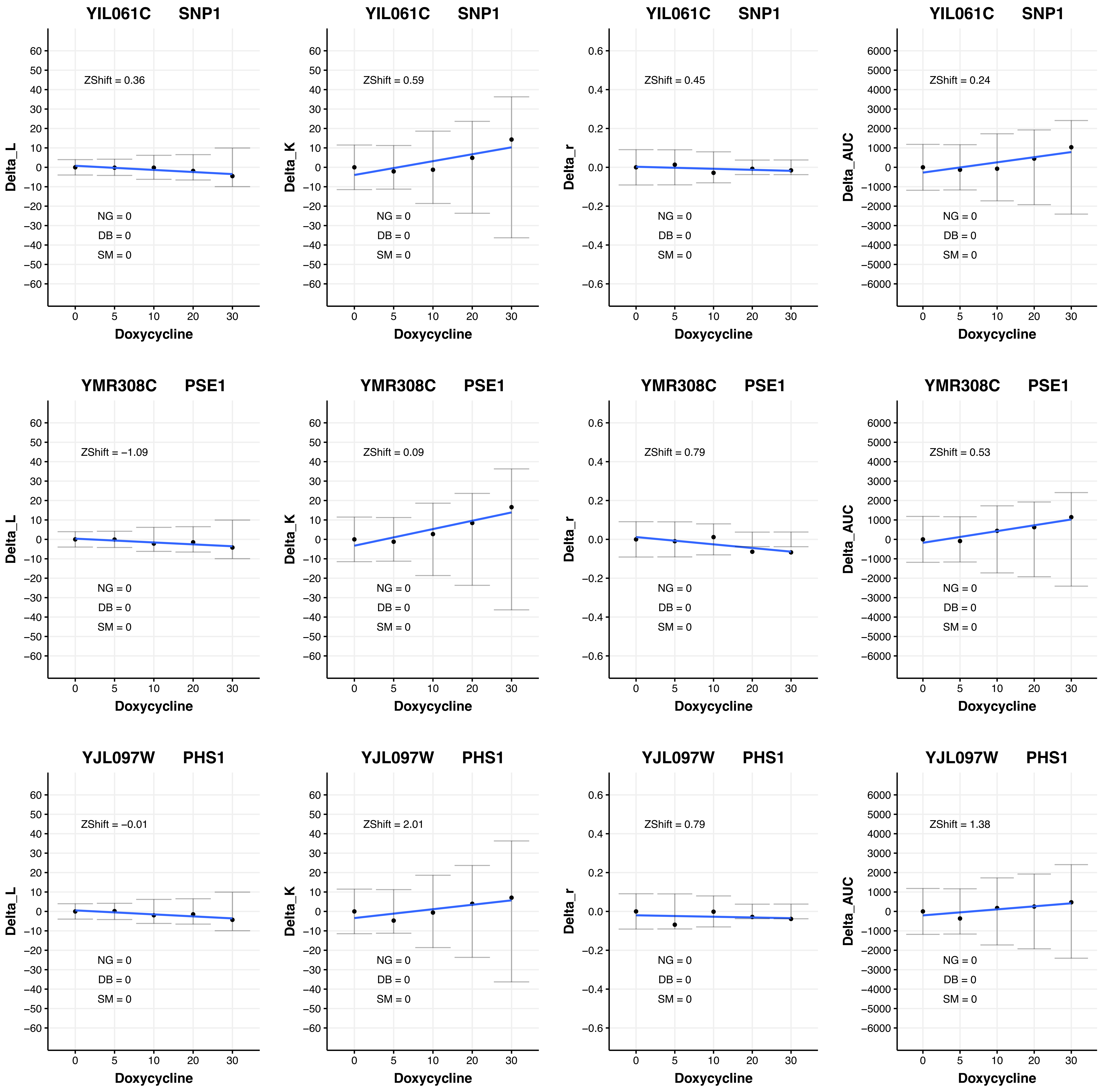
YIL061C SNP1
ZShift = 0.36
NG = 0
DB = 0
SM = 0
Delta_L
Doxycycline
YIL061C SNP1
ZShift = 0.59
NG = 0
DB = 0
SM = 0
Delta_K
Doxycycline
YIL061C SNP1
ZShift = 0.45
NG = 0
DB = 0
SM = 0
Delta_r
Doxycycline
YIL061C SNP1
ZShift = 0.24
NG = 0
DB = 0
SM = 0
Delta_AUC
Doxycycline
YMR308C PSE1
ZShift = −1.09
NG = 0
DB = 0
SM = 0
Delta_L
Doxycycline
YMR308C PSE1
ZShift = 0.09
NG = 0
DB = 0
SM = 0
Delta_K
Doxycycline
YMR308C PSE1
ZShift = 0.79
NG = 0
DB = 0
SM = 0
Delta_r
Doxycycline
YMR308C PSE1
ZShift = 0.53
NG = 0
DB = 0
SM = 0
Delta_AUC
Doxycycline
YJL097W PHS1
ZShift = −0.01
NG = 0
DB = 0
SM = 0
Delta_L
Doxycycline
YJL097W PHS1
ZShift = 2.01
NG = 0
DB = 0
SM = 0
Delta_K
Doxycycline
YJL097W PHS1
ZShift = 0.79
NG = 0
DB = 0
SM = 0
Delta_r
Doxycycline
YJL097W PHS1
ZShift = 1.38
NG = 0
DB = 0
SM = 0
Delta_AUC
Doxycycline

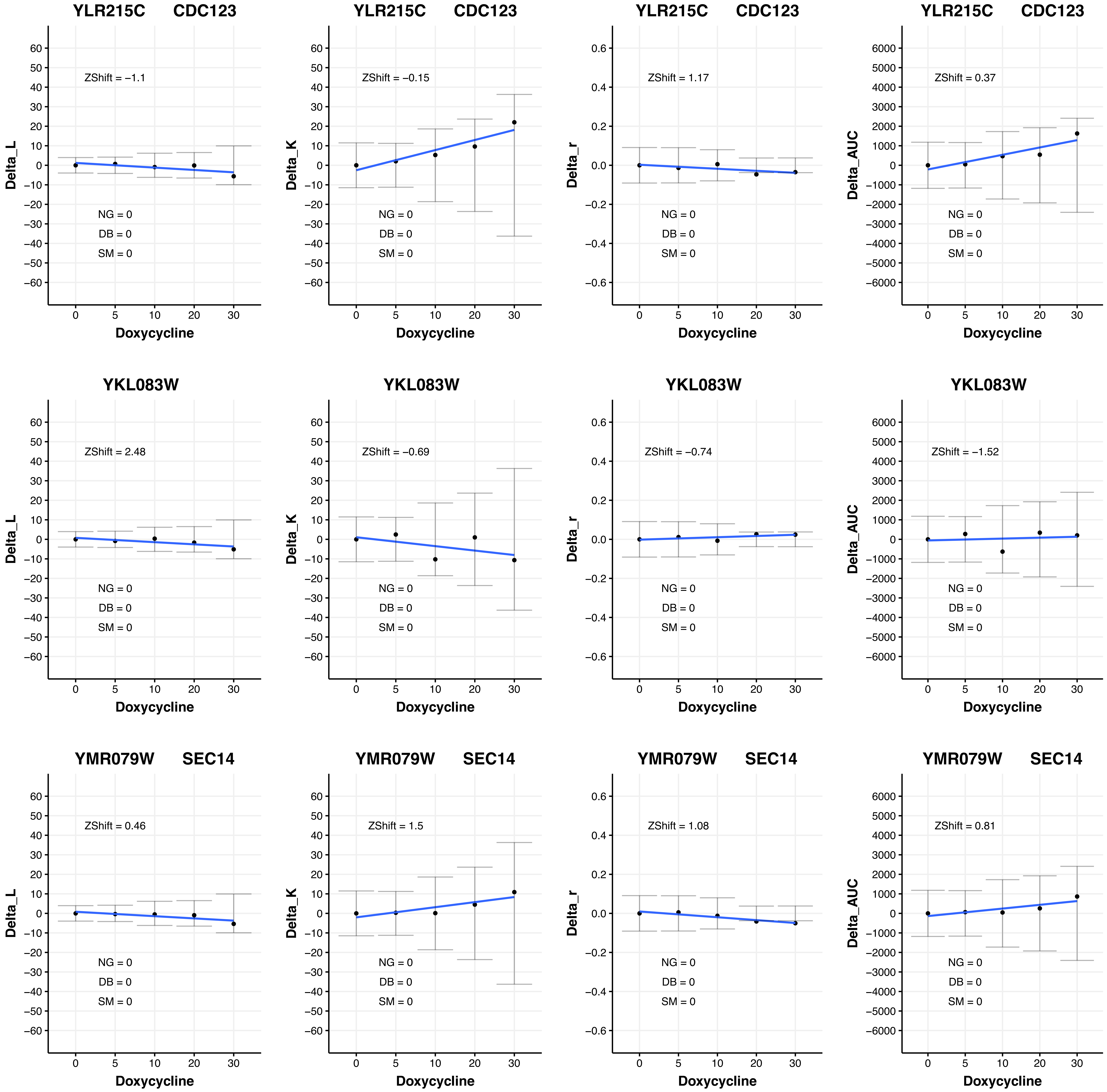

YLR215C CDC123
ZShift = −1.1
NG = 0
DB = 0
SM = 0
Delta_L
Doxycycline
YLR215C CDC123
ZShift = −0.15
NG = 0
DB = 0
SM = 0
Delta_K
Doxycycline
YLR215C CDC123
ZShift = 1.17
NG = 0
DB = 0
SM = 0
Delta_r
Doxycycline
YLR215C CDC123
ZShift = 0.37
NG = 0
DB = 0
SM = 0
Delta_AUC
Doxycycline
YKL083W
ZShift = 2.48
NG = 0
DB = 0
SM = 0
Delta_L
Doxycycline
YKL083W
ZShift = −0.69
NG = 0
DB = 0
SM = 0
Delta_K
Doxycycline
YKL083W
ZShift = −0.74
NG = 0
DB = 0
SM = 0
Delta_r
Doxycycline
YKL083W
ZShift = −1.52
NG = 0
DB = 0
SM = 0
Delta_AUC
Doxycycline
YMR079W SEC14
ZShift = 0.46
NG = 0
DB = 0
SM = 0
Delta_L
Doxycycline
YMR079W SEC14
ZShift = 1.5
NG = 0
DB = 0
SM = 0
Delta_K
Doxycycline
YMR079W SEC14
ZShift = 1.08
NG = 0
DB = 0
SM = 0
Delta_r
Doxycycline
YMR079W SEC14
ZShift = 0.81
NG = 0
DB = 0
SM = 0
Delta_AUC
Doxycycline

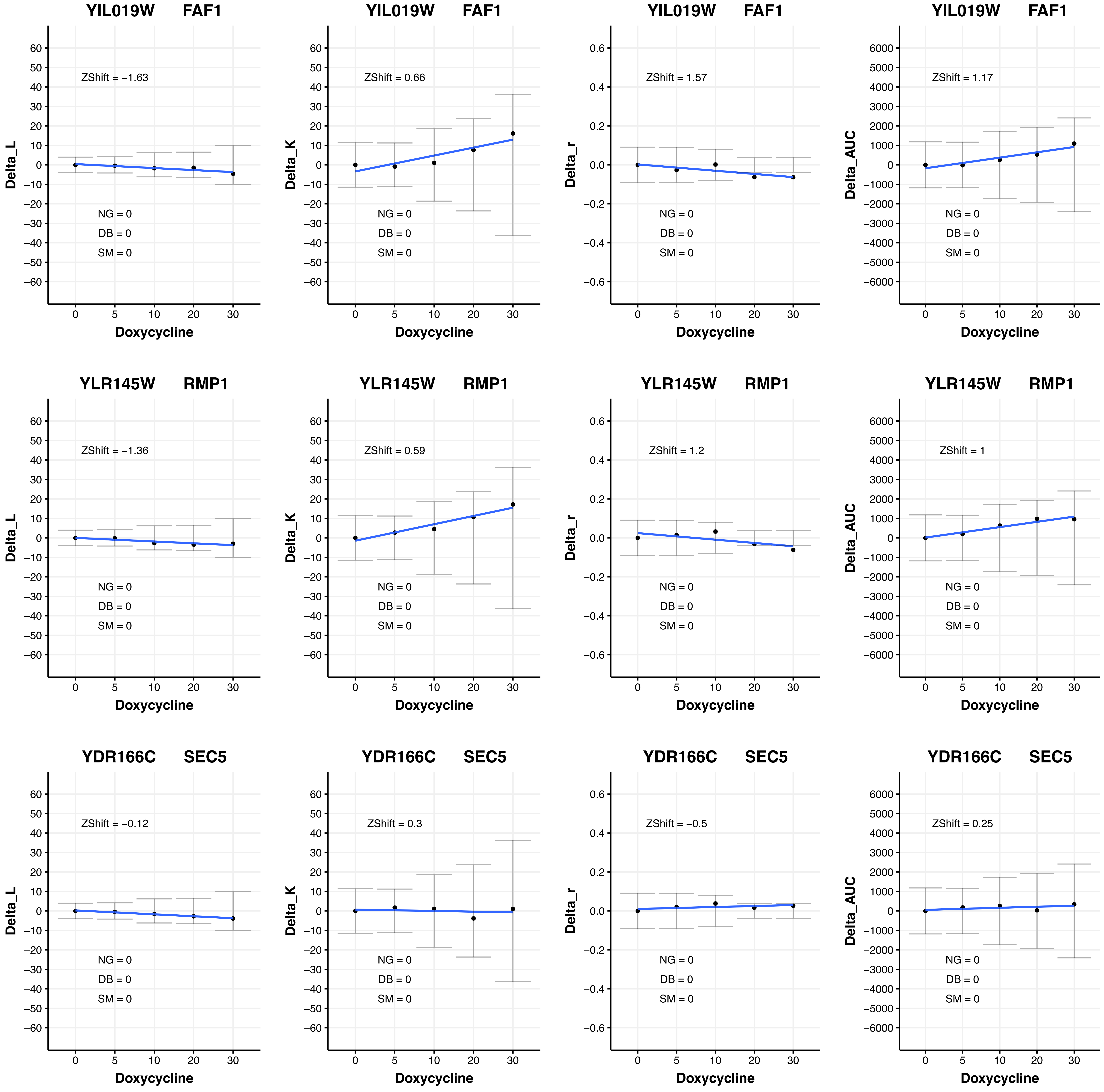
YIL019W FAF1
ZShift = −1.63
ZShift = 0.66
ZShift = 1.57
ZShift = 1.17
YLR145W RMP1
ZShift = −1.36
ZShift = 0.59
ZShift = 1.2
ZShift = 1
YDR166C SEC5
ZShift = −0.12
ZShift = 0.3
ZShift = −0.5
ZShift = 0.25
NG = 0
DB = 0
SM = 0
Delta_L
Delta_K
Delta_r
Delta_AUC
Doxycycline
0
5
10
20
30

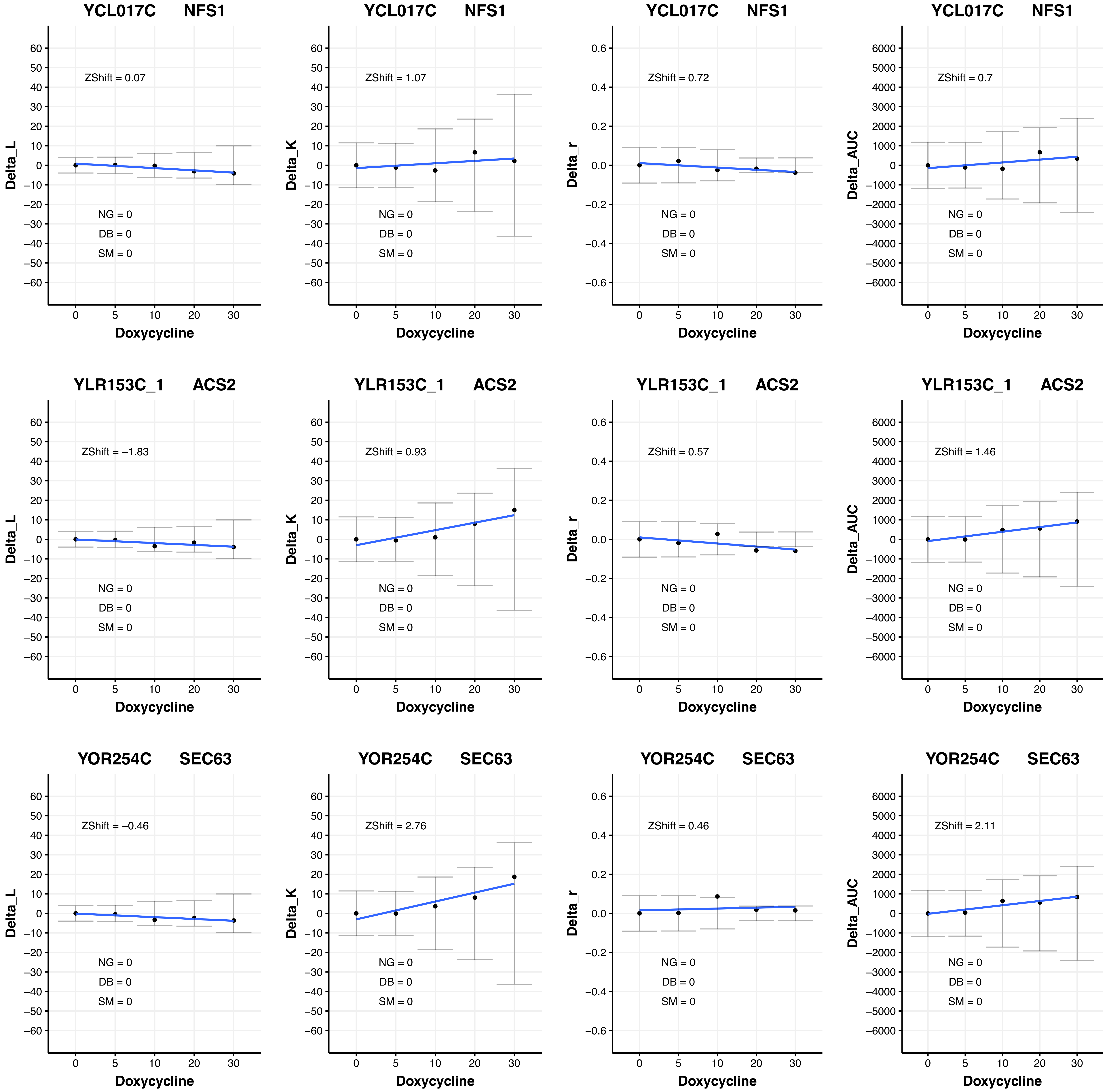

YCL017C NFS1
ZShift = 0.07
NG = 0
DB = 0
SM = 0
Delta_L
Doxycycline
YCL017C NFS1
ZShift = 1.07
NG = 0
DB = 0
SM = 0
Delta_K
Doxycycline
YCL017C NFS1
ZShift = 0.72
NG = 0
DB = 0
SM = 0
Delta_r
Doxycycline
YCL017C NFS1
ZShift = 0.7
NG = 0
DB = 0
SM = 0
Delta_AUC
Doxycycline
YLR153C_1 ACS2
ZShift = −1.83
NG = 0
DB = 0
SM = 0
Delta_L
Doxycycline
YLR153C_1 ACS2
ZShift = 0.93
NG = 0
DB = 0
SM = 0
Delta_K
Doxycycline
YLR153C_1 ACS2
ZShift = 0.57
NG = 0
DB = 0
SM = 0
Delta_r
Doxycycline
YLR153C_1 ACS2
ZShift = 1.46
NG = 0
DB = 0
SM = 0
Delta_AUC
Doxycycline
YOR254C SEC63
ZShift = −0.46
NG = 0
DB = 0
SM = 0
Delta_L
Doxycycline
YOR254C SEC63
ZShift = 2.76
NG = 0
DB = 0
SM = 0
Delta_K
Doxycycline
YOR254C SEC63
ZShift = 0.46
NG = 0
DB = 0
SM = 0
Delta_r
Doxycycline
YOR254C SEC63
ZShift = 2.11
NG = 0
DB = 0
SM = 0
Delta_AUC
Doxycycline

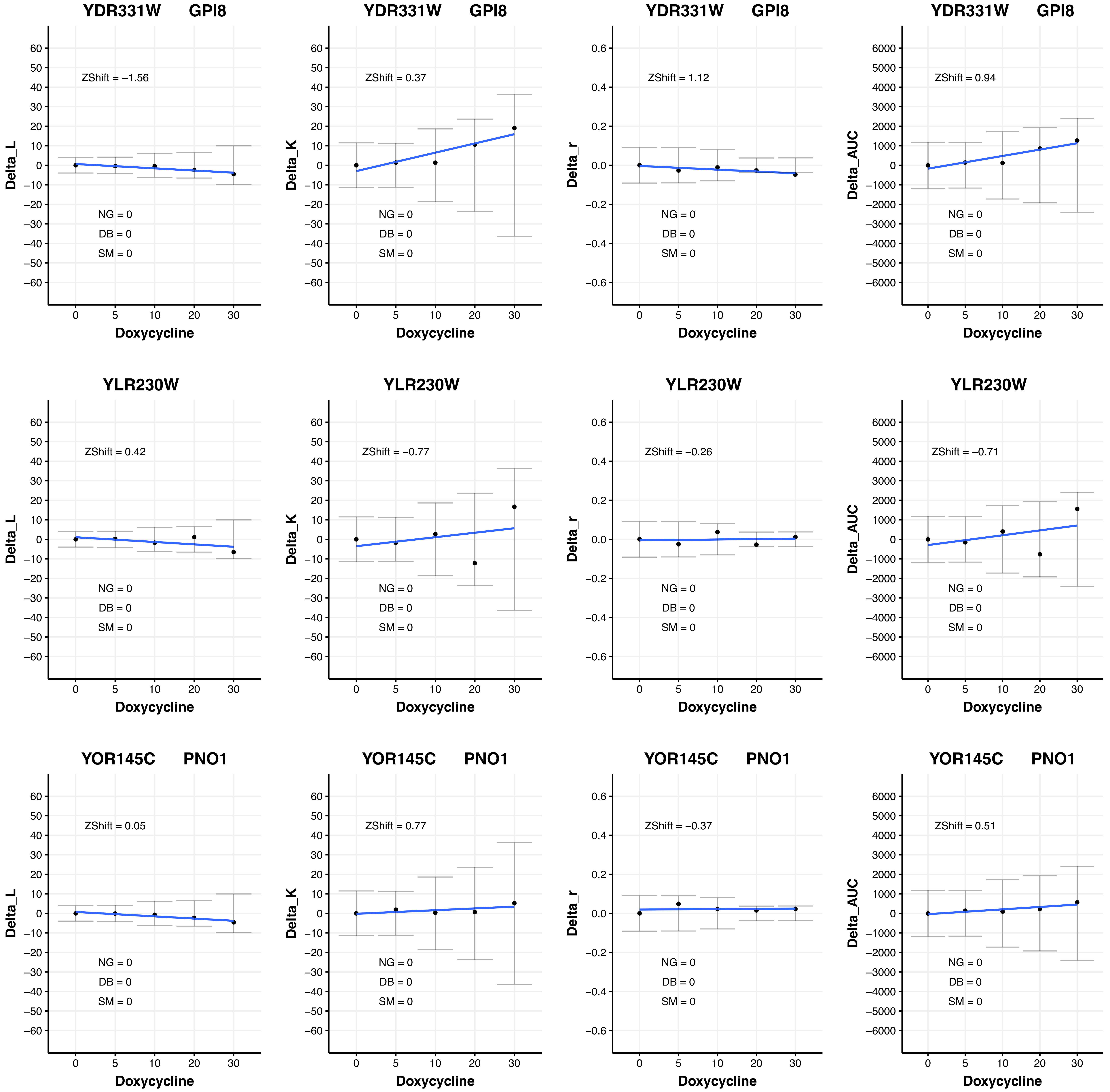
YDR331W GPI8
ZShift = −1.56
ZShift = 0.37
ZShift = 1.12
ZShift = 0.94
YLR230W
ZShift = 0.42
ZShift = −0.77
ZShift = −0.26
ZShift = −0.71
YOR145C PNO1
ZShift = 0.05
ZShift = 0.77
ZShift = −0.37
ZShift = 0.51
NG = 0
DB = 0
SM = 0
Delta_L
Delta_K
Delta_r
Delta_AUC
Doxycycline
0 5 10 20 30

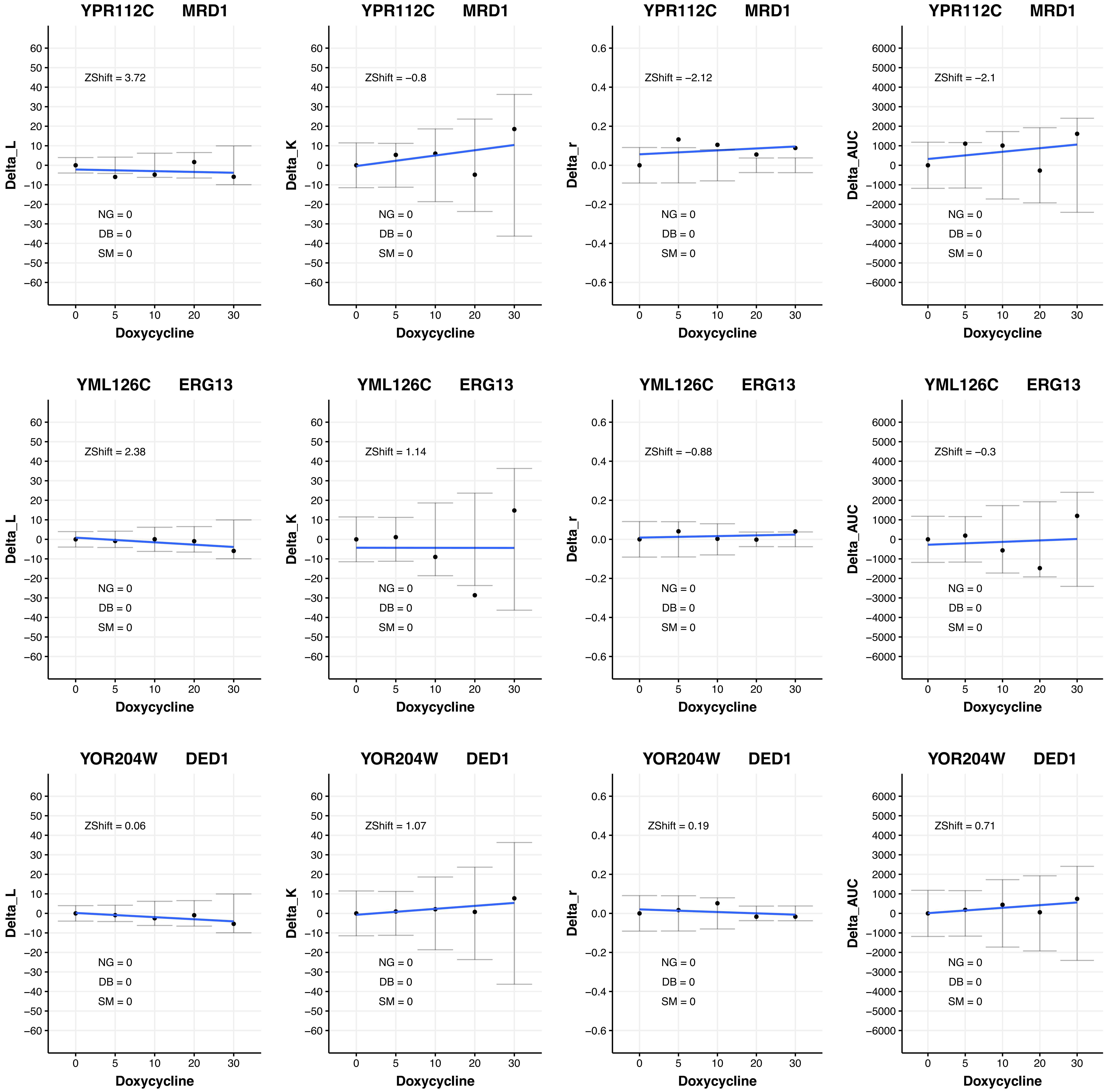
YPR112C MRD1
ZShift = 3.72
ZShift = −0.8
ZShift = −2.12
ZShift = −2.1
YML126C ERG13
ZShift = 2.38
ZShift = 1.14
ZShift = −0.88
ZShift = −0.3
YOR204W DED1
ZShift = 0.06
ZShift = 1.07
ZShift = 0.19
ZShift = 0.71
NG = 0
DB = 0
SM = 0
Delta_L
Delta_K
Delta_r
Delta_AUC
Doxycycline
0
5
10
20
30

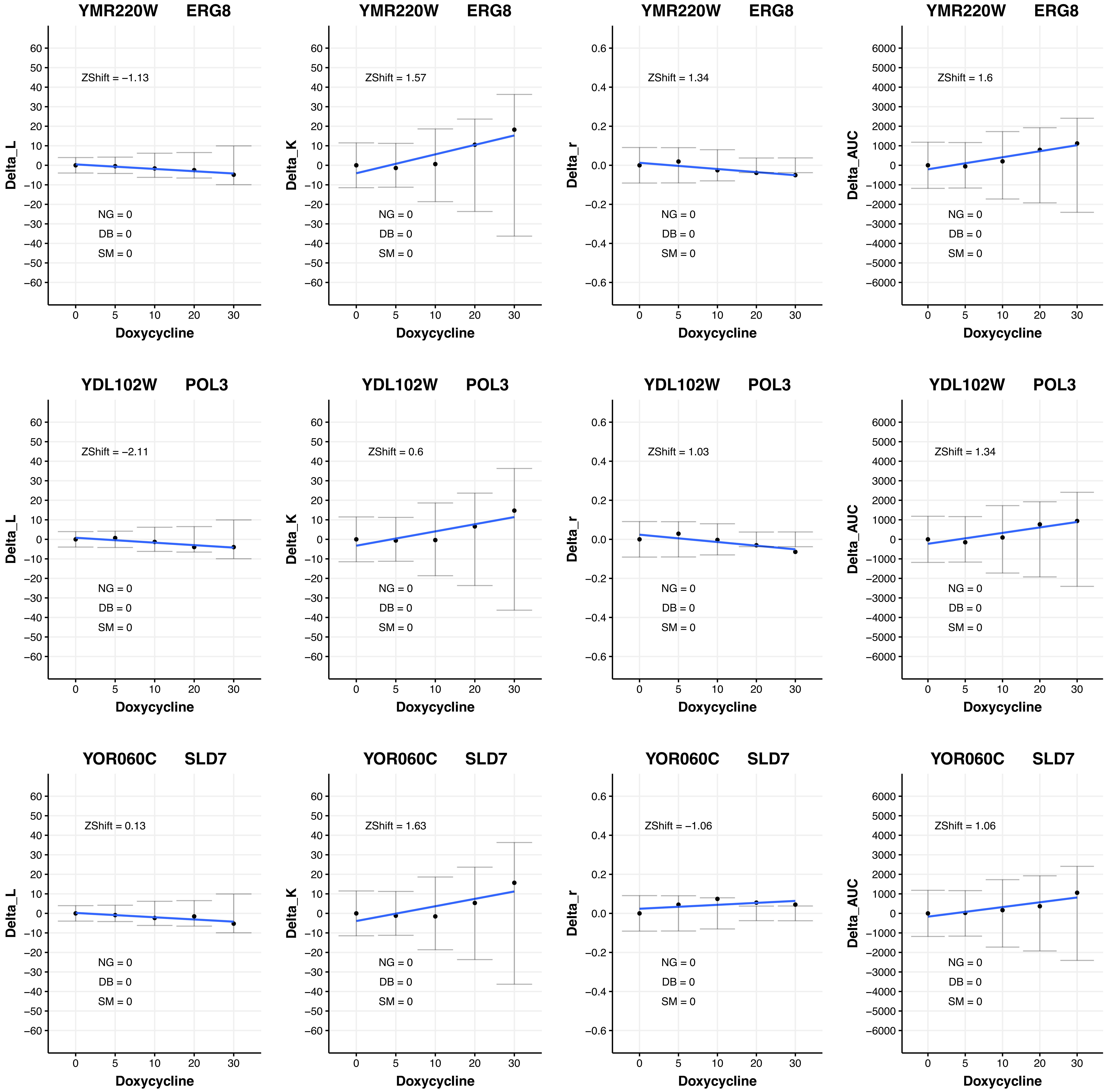
YMR220W ERG8
ZShift = −1.13
Delta_L
NG = 0
DB = 0
SM = 0
Doxycycline
YMR220W ERG8
ZShift = 1.57
Delta_K
NG = 0
DB = 0
SM = 0
Doxycycline
YMR220W ERG8
ZShift = 1.34
Delta_r
NG = 0
DB = 0
SM = 0
Doxycycline
YMR220W ERG8
ZShift = 1.6
Delta_AUC
NG = 0
DB = 0
SM = 0
Doxycycline
YDL102W POL3
ZShift = −2.11
Delta_L
NG = 0
DB = 0
SM = 0
Doxycycline
YDL102W POL3
ZShift = 0.6
Delta_K
NG = 0
DB = 0
SM = 0
Doxycycline
YDL102W POL3
ZShift = 1.03
Delta_r
NG = 0
DB = 0
SM = 0
Doxycycline
YDL102W POL3
ZShift = 1.34
Delta_AUC
NG = 0
DB = 0
SM = 0
Doxycycline
YOR060C SLD7
ZShift = 0.13
Delta_L
NG = 0
DB = 0
SM = 0
Doxycycline
YOR060C SLD7
ZShift = 1.63
Delta_K
NG = 0
DB = 0
SM = 0
Doxycycline
YOR060C SLD7
ZShift = −1.06
Delta_r
NG = 0
DB = 0
SM = 0
Doxycycline
YOR060C SLD7
ZShift = 1.06
Delta_AUC
NG = 0
DB = 0
SM = 0
Doxycycline

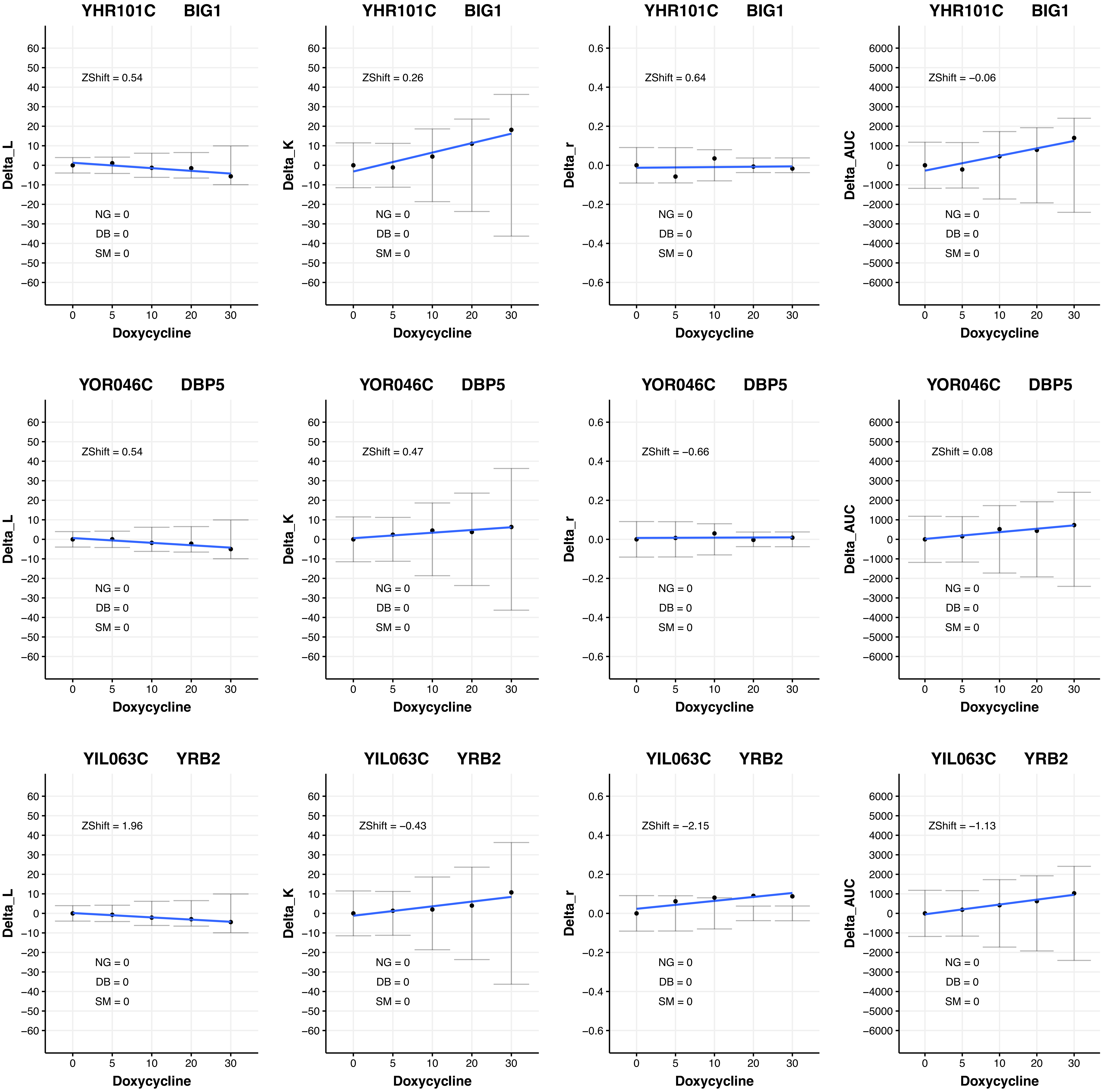
YHR101C BIG1
ZShift = 0.54
Delta_L
ZShift = 0.26
Delta_K
ZShift = 0.64
Delta_r
ZShift = −0.06
Delta_AUC
YOR046C DBP5
ZShift = 0.54
ZShift = 0.47
ZShift = −0.66
ZShift = 0.08
YIL063C YRB2
ZShift = 1.96
ZShift = −0.43
ZShift = −2.15
ZShift = −1.13
NG = 0
DB = 0
SM = 0
Doxycycline
0
5
10
20
30

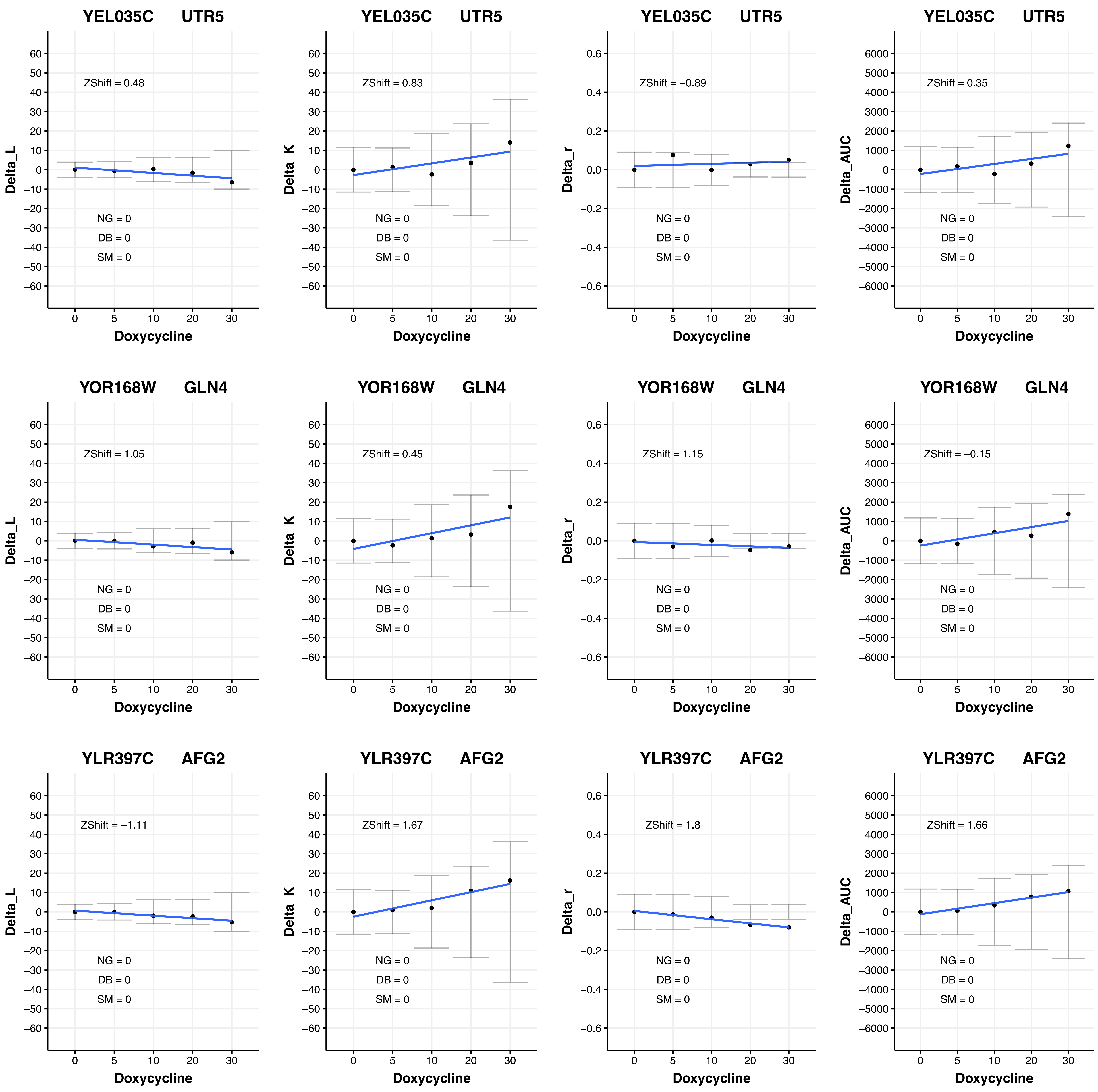

YEL035C UTR5
ZShift = 0.48
Delta_L
NG = 0
DB = 0
SM = 0
Doxycycline
YEL035C UTR5
ZShift = 0.83
Delta_K
NG = 0
DB = 0
SM = 0
Doxycycline
YEL035C UTR5
ZShift = −0.89
Delta_r
NG = 0
DB = 0
SM = 0
Doxycycline
YEL035C UTR5
ZShift = 0.35
Delta_AUC
NG = 0
DB = 0
SM = 0
Doxycycline
YOR168W GLN4
ZShift = 1.05
Delta_L
NG = 0
DB = 0
SM = 0
Doxycycline
YOR168W GLN4
ZShift = 0.45
Delta_K
NG = 0
DB = 0
SM = 0
Doxycycline
YOR168W GLN4
ZShift = 1.15
Delta_r
NG = 0
DB = 0
SM = 0
Doxycycline
YOR168W GLN4
ZShift = −0.15
Delta_AUC
NG = 0
DB = 0
SM = 0
Doxycycline
YLR397C AFG2
ZShift = −1.11
Delta_L
NG = 0
DB = 0
SM = 0
Doxycycline
YLR397C AFG2
ZShift = 1.67
Delta_K
NG = 0
DB = 0
SM = 0
Doxycycline
YLR397C AFG2
ZShift = 1.8
Delta_r
NG = 0
DB = 0
SM = 0
Doxycycline
YLR397C AFG2
ZShift = 1.66
Delta_AUC
NG = 0
DB = 0
SM = 0
Doxycycline

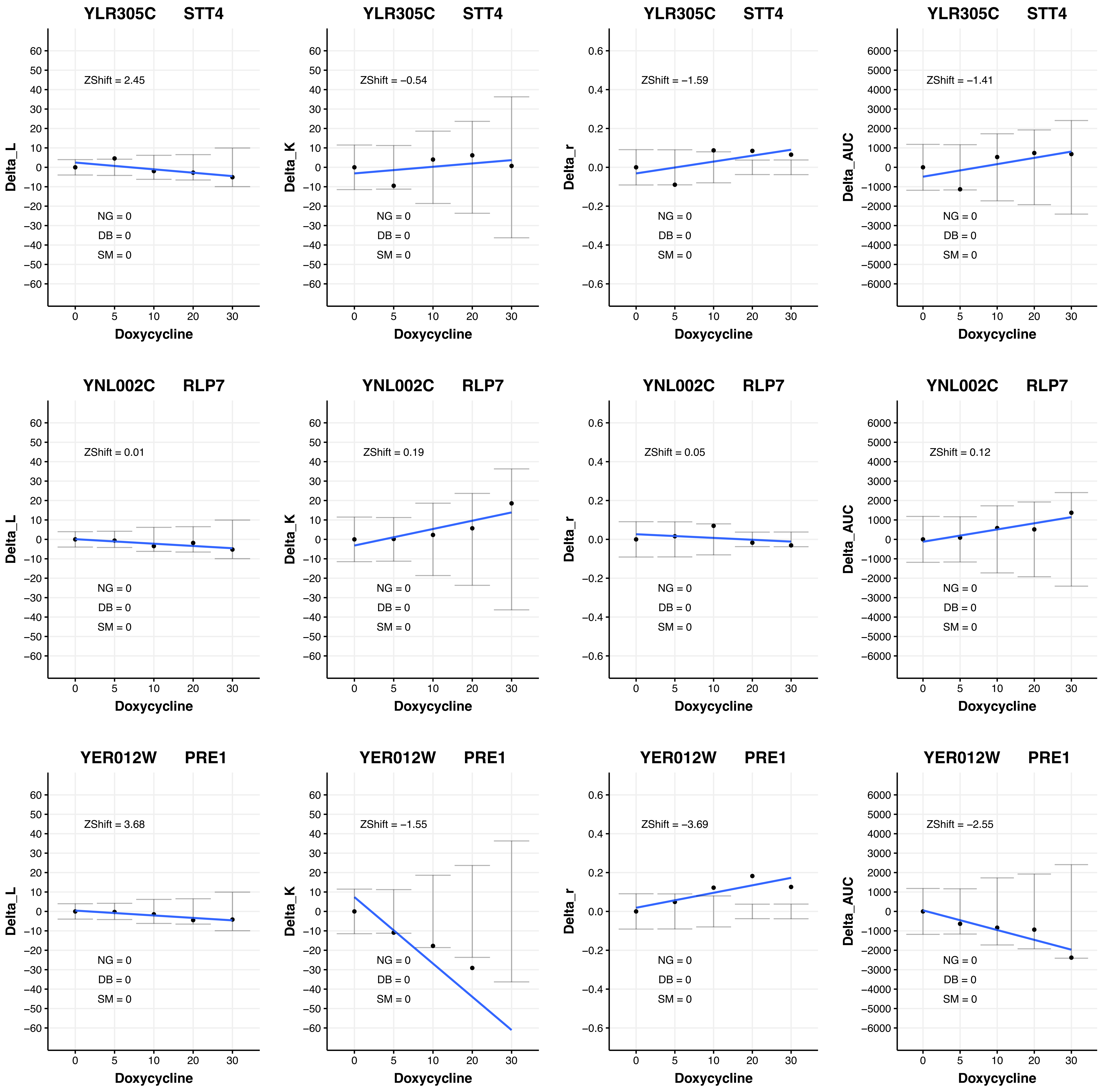
YLR305C STT4
ZShift = 2.45
NG = 0
DB = 0
SM = 0
Delta_L
Doxycycline
YLR305C STT4
ZShift = −0.54
NG = 0
DB = 0
SM = 0
Delta_K
Doxycycline
YLR305C STT4
ZShift = −1.59
NG = 0
DB = 0
SM = 0
Delta_r
Doxycycline
YLR305C STT4
ZShift = −1.41
NG = 0
DB = 0
SM = 0
Delta_AUC
Doxycycline
YNL002C RLP7
ZShift = 0.01
NG = 0
DB = 0
SM = 0
Delta_L
Doxycycline
YNL002C RLP7
ZShift = 0.19
NG = 0
DB = 0
SM = 0
Delta_K
Doxycycline
YNL002C RLP7
ZShift = 0.05
NG = 0
DB = 0
SM = 0
Delta_r
Doxycycline
YNL002C RLP7
ZShift = 0.12
NG = 0
DB = 0
SM = 0
Delta_AUC
Doxycycline
YER012W PRE1
ZShift = 3.68
NG = 0
DB = 0
SM = 0
Delta_L
Doxycycline
YER012W PRE1
ZShift = −1.55
NG = 0
DB = 0
SM = 0
Delta_K
Doxycycline
YER012W PRE1
ZShift = −3.69
NG = 0
DB = 0
SM = 0
Delta_r
Doxycycline
YER012W PRE1
ZShift = −2.55
NG = 0
DB = 0
SM = 0
Delta_AUC
Doxycycline

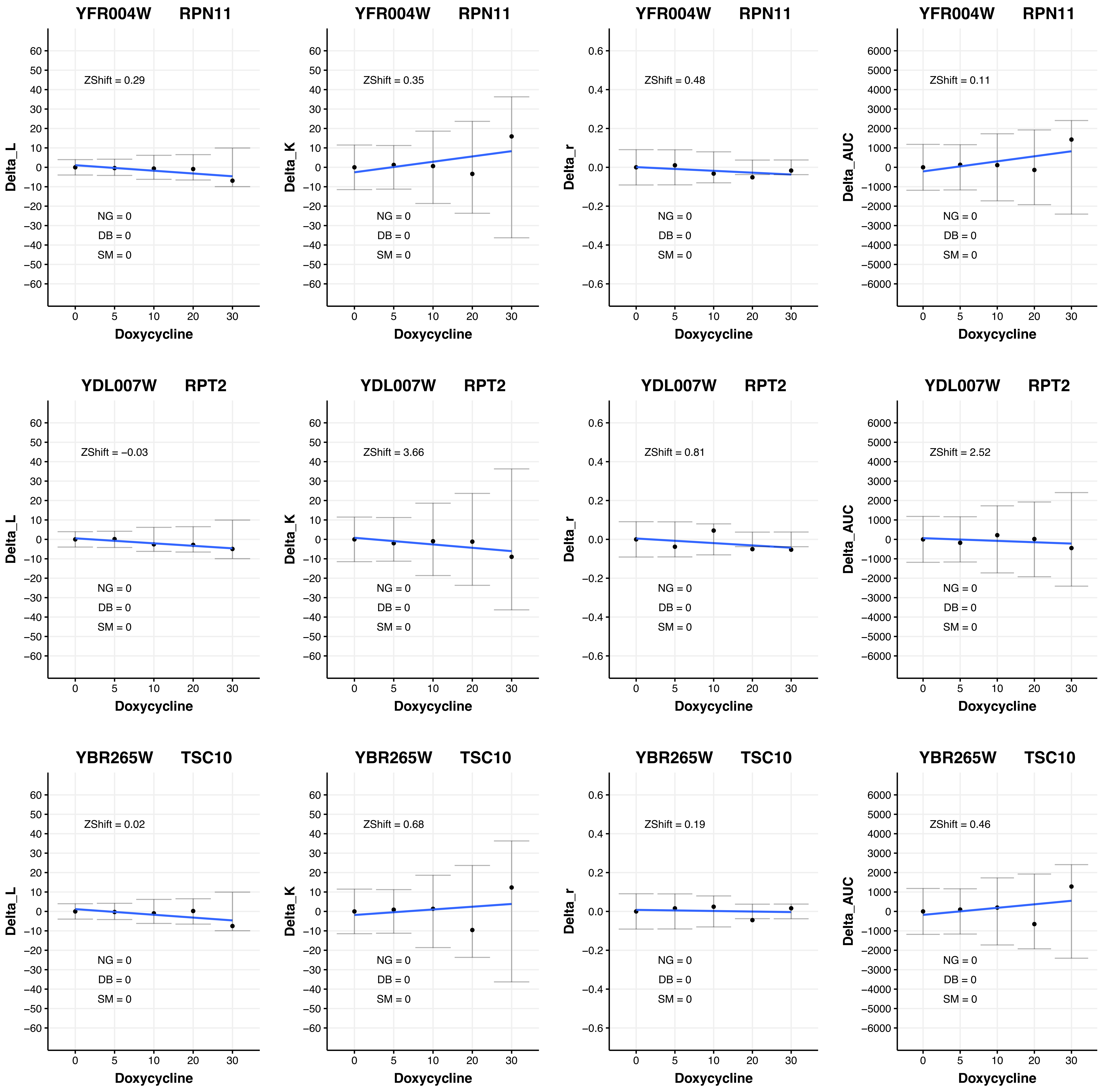
YFR004W RPN11
ZShift = 0.29
NG = 0
DB = 0
SM = 0
Delta_L
Doxycycline
YFR004W RPN11
ZShift = 0.35
NG = 0
DB = 0
SM = 0
Delta_K
Doxycycline
YFR004W RPN11
ZShift = 0.48
NG = 0
DB = 0
SM = 0
Delta_r
Doxycycline
YFR004W RPN11
ZShift = 0.11
NG = 0
DB = 0
SM = 0
Delta_AUC
Doxycycline
YDL007W RPT2
ZShift = −0.03
NG = 0
DB = 0
SM = 0
Delta_L
Doxycycline
YDL007W RPT2
ZShift = 3.66
NG = 0
DB = 0
SM = 0
Delta_K
Doxycycline
YDL007W RPT2
ZShift = 0.81
NG = 0
DB = 0
SM = 0
Delta_r
Doxycycline
YDL007W RPT2
ZShift = 2.52
NG = 0
DB = 0
SM = 0
Delta_AUC
Doxycycline
YBR265W TSC10
ZShift = 0.02
NG = 0
DB = 0
SM = 0
Delta_L
Doxycycline
YBR265W TSC10
ZShift = 0.68
NG = 0
DB = 0
SM = 0
Delta_K
Doxycycline
YBR265W TSC10
ZShift = 0.19
NG = 0
DB = 0
SM = 0
Delta_r
Doxycycline
YBR265W TSC10
ZShift = 0.46
NG = 0
DB = 0
SM = 0
Delta_AUC
Doxycycline

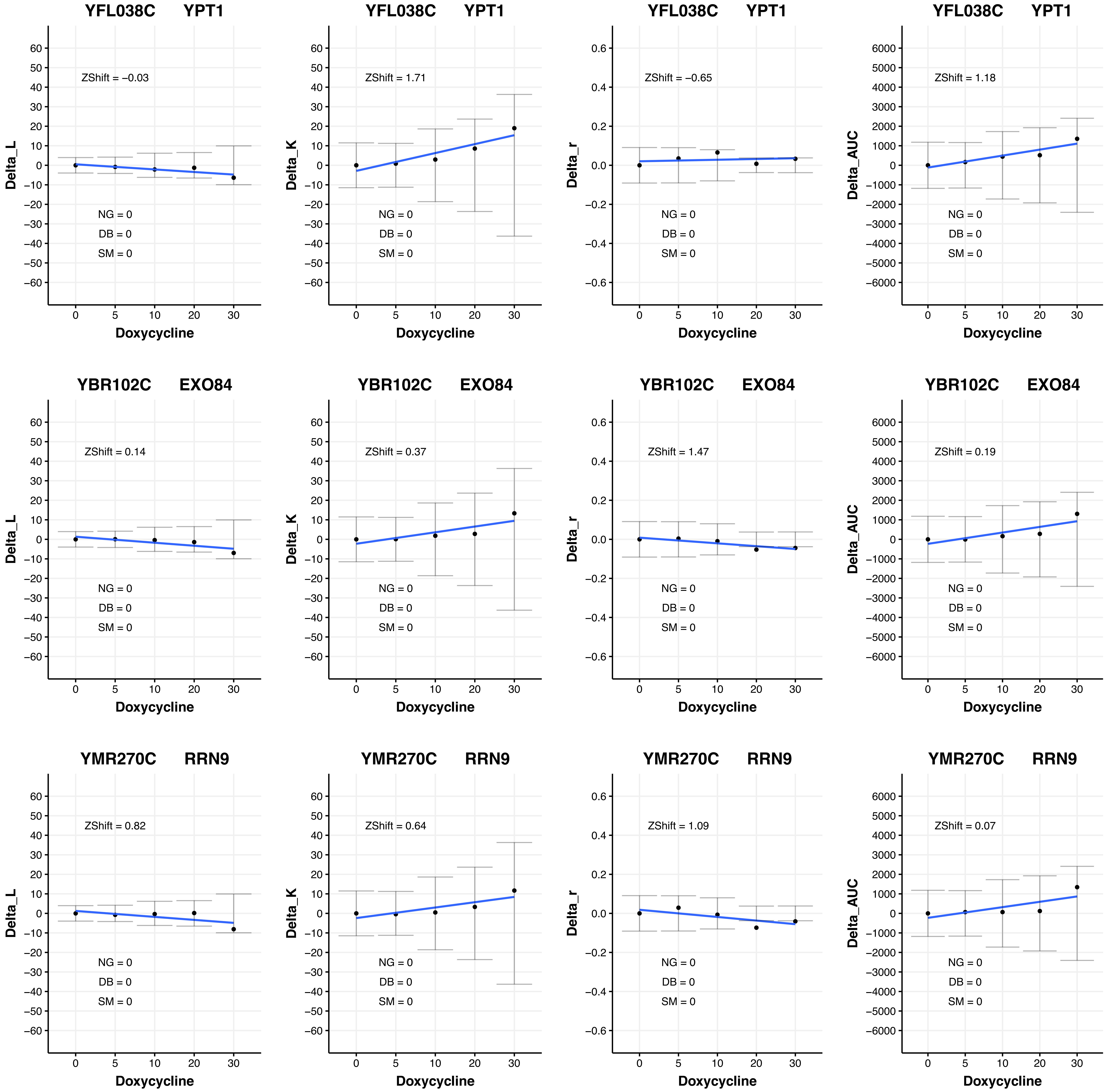
YFL038C YPT1
ZShift = −0.03
NG = 0
DB = 0
SM = 0
Delta_L
YFL038C YPT1
ZShift = 1.71
NG = 0
DB = 0
SM = 0
Delta_K
YFL038C YPT1
ZShift = −0.65
NG = 0
DB = 0
SM = 0
Delta_r
YFL038C YPT1
ZShift = 1.18
NG = 0
DB = 0
SM = 0
Delta_AUC
YBR102C EXO84
ZShift = 0.14
NG = 0
DB = 0
SM = 0
Delta_L
YBR102C EXO84
ZShift = 0.37
NG = 0
DB = 0
SM = 0
Delta_K
YBR102C EXO84
ZShift = 1.47
NG = 0
DB = 0
SM = 0
Delta_r
YBR102C EXO84
ZShift = 0.19
NG = 0
DB = 0
SM = 0
Delta_AUC
YMR270C RRN9
ZShift = 0.82
NG = 0
DB = 0
SM = 0
Delta_L
YMR270C RRN9
ZShift = 0.64
NG = 0
DB = 0
SM = 0
Delta_K
YMR270C RRN9
ZShift = 1.09
NG = 0
DB = 0
SM = 0
Delta_r
YMR270C RRN9
ZShift = 0.07
NG = 0
DB = 0
SM = 0
Delta_AUC
Doxycycline
0 5 10 20 30

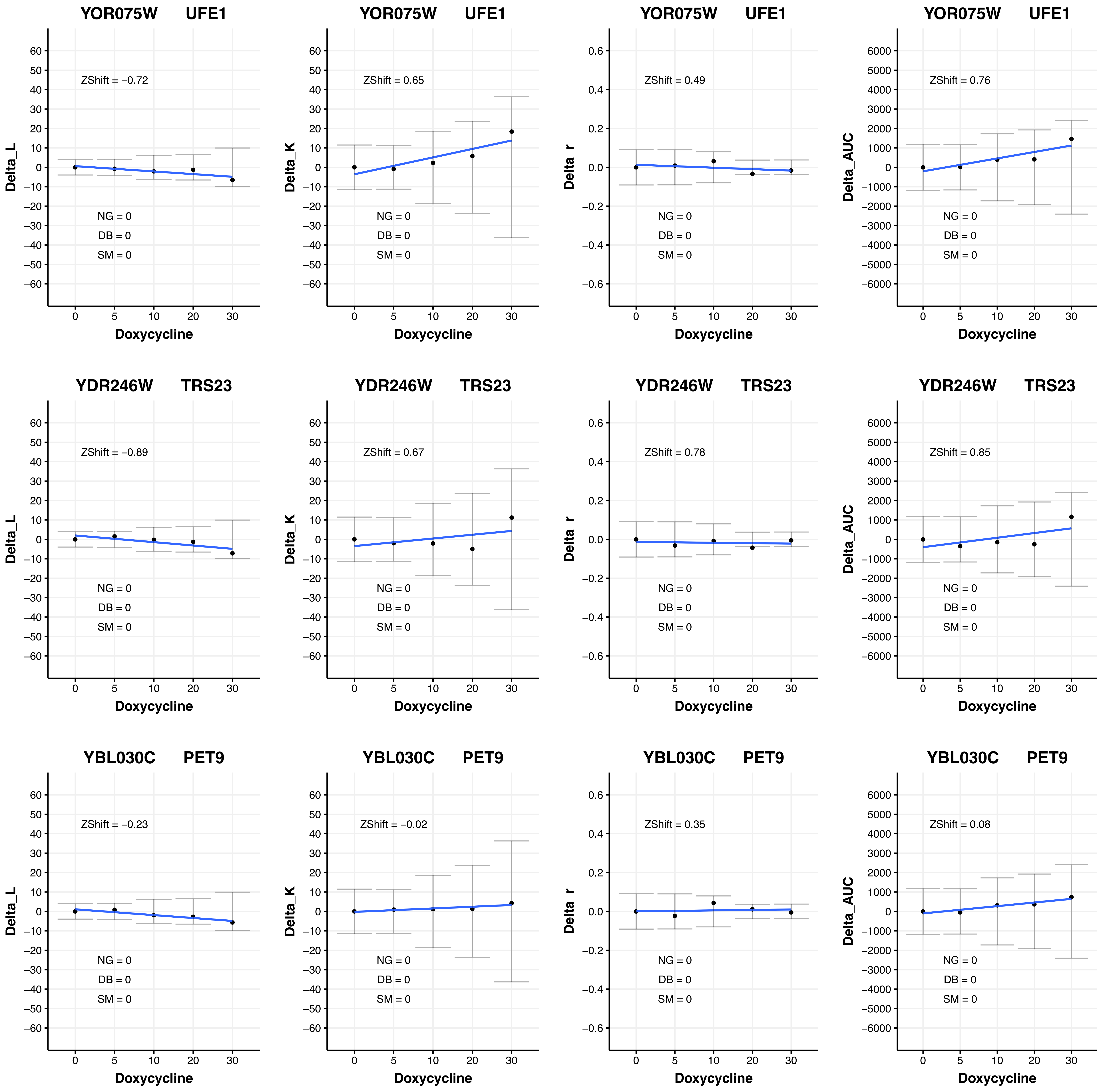
YOR075W UFE1
ZShift = −0.72
NG = 0
DB = 0
SM = 0
Delta_L
Doxycycline
YOR075W UFE1
ZShift = 0.65
NG = 0
DB = 0
SM = 0
Delta_K
Doxycycline
YOR075W UFE1
ZShift = 0.49
NG = 0
DB = 0
SM = 0
Delta_r
Doxycycline
YOR075W UFE1
ZShift = 0.76
NG = 0
DB = 0
SM = 0
Delta_AUC
Doxycycline
YDR246W TRS23
ZShift = −0.89
NG = 0
DB = 0
SM = 0
Delta_L
Doxycycline
YDR246W TRS23
ZShift = 0.67
NG = 0
DB = 0
SM = 0
Delta_K
Doxycycline
YDR246W TRS23
ZShift = 0.78
NG = 0
DB = 0
SM = 0
Delta_r
Doxycycline
YDR246W TRS23
ZShift = 0.85
NG = 0
DB = 0
SM = 0
Delta_AUC
Doxycycline
YBL030C PET9
ZShift = −0.23
NG = 0
DB = 0
SM = 0
Delta_L
Doxycycline
YBL030C PET9
ZShift = −0.02
NG = 0
DB = 0
SM = 0
Delta_K
Doxycycline
YBL030C PET9
ZShift = 0.35
NG = 0
DB = 0
SM = 0
Delta_r
Doxycycline
YBL030C PET9
ZShift = 0.08
NG = 0
DB = 0
SM = 0
Delta_AUC
Doxycycline

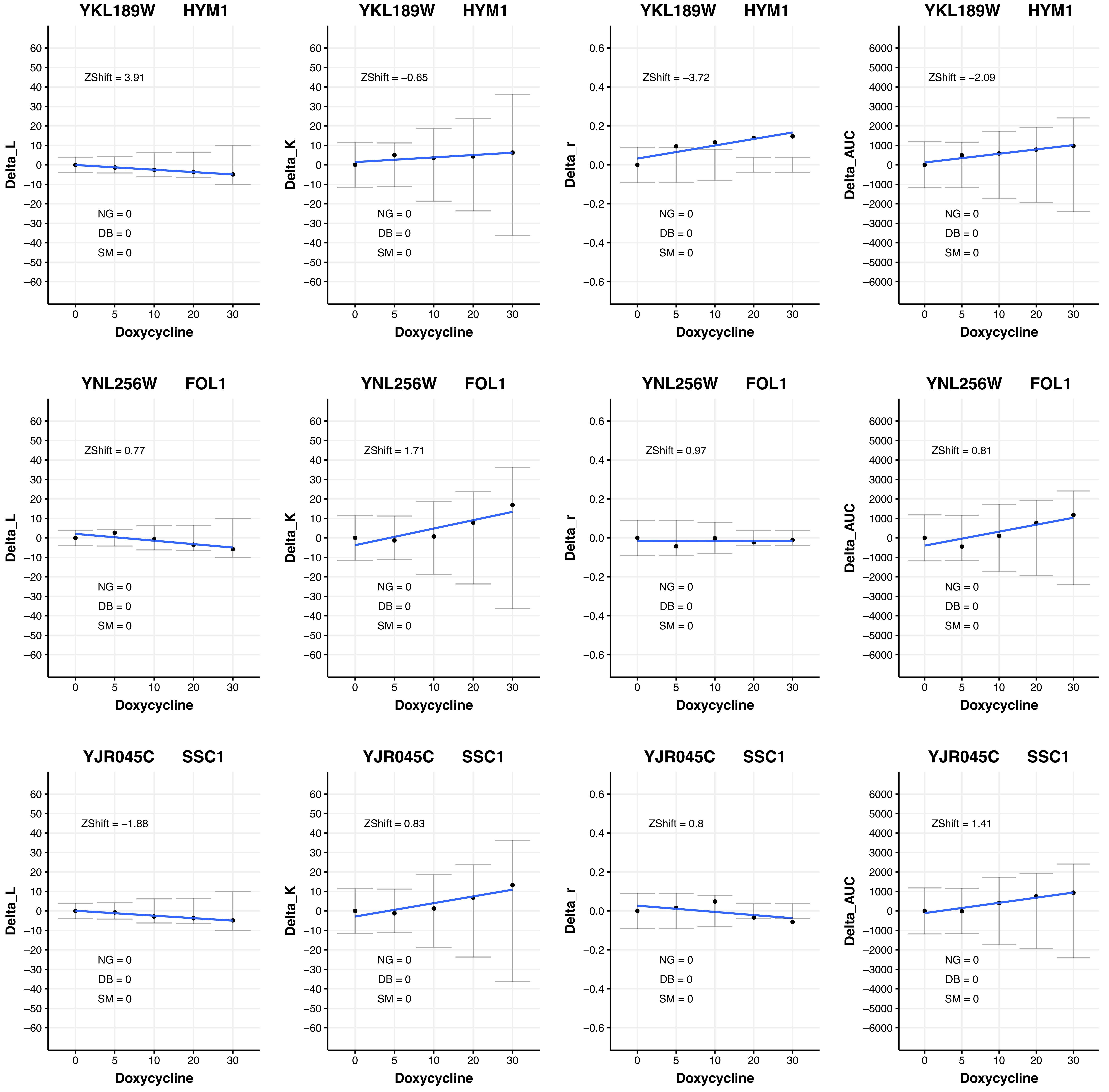
YKL189W HYM1
ZShift = 3.91
NG = 0
DB = 0
SM = 0
Delta_L
Doxycycline
YKL189W HYM1
ZShift = −0.65
NG = 0
DB = 0
SM = 0
Delta_K
Doxycycline
YKL189W HYM1
ZShift = −3.72
NG = 0
DB = 0
SM = 0
Delta_r
Doxycycline
YKL189W HYM1
ZShift = −2.09
NG = 0
DB = 0
SM = 0
Delta_AUC
Doxycycline
YNL256W FOL1
ZShift = 0.77
NG = 0
DB = 0
SM = 0
Delta_L
Doxycycline
YNL256W FOL1
ZShift = 1.71
NG = 0
DB = 0
SM = 0
Delta_K
Doxycycline
YNL256W FOL1
ZShift = 0.97
NG = 0
DB = 0
SM = 0
Delta_r
Doxycycline
YNL256W FOL1
ZShift = 0.81
NG = 0
DB = 0
SM = 0
Delta_AUC
Doxycycline
YJR045C SSC1
ZShift = −1.88
NG = 0
DB = 0
SM = 0
Delta_L
Doxycycline
YJR045C SSC1
ZShift = 0.83
NG = 0
DB = 0
SM = 0
Delta_K
Doxycycline
YJR045C SSC1
ZShift = 0.8
NG = 0
DB = 0
SM = 0
Delta_r
Doxycycline
YJR045C SSC1
ZShift = 1.41
NG = 0
DB = 0
SM = 0
Delta_AUC
Doxycycline

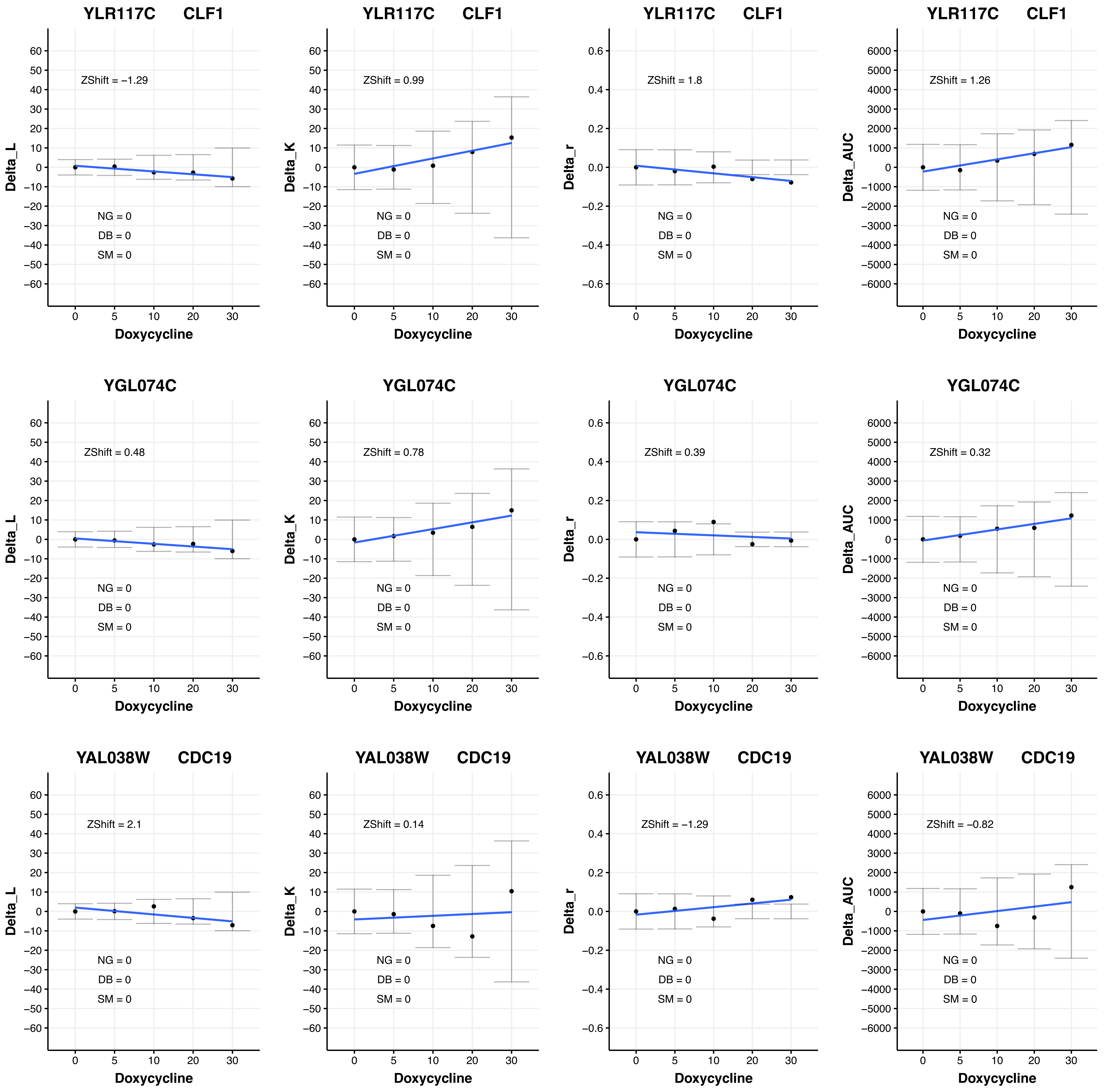
YLR117C CLF1
ZShift = −1.29
ZShift = 0.99
ZShift = 1.8
ZShift = 1.26
YGL074C
ZShift = 0.48
ZShift = 0.78
ZShift = 0.39
ZShift = 0.32
YAL038W CDC19
ZShift = 2.1
ZShift = 0.14
ZShift = −1.29
ZShift = −0.82
NG = 0
DB = 0
SM = 0
Delta_L
Delta_K
Delta_r
Delta_AUC
Doxycycline

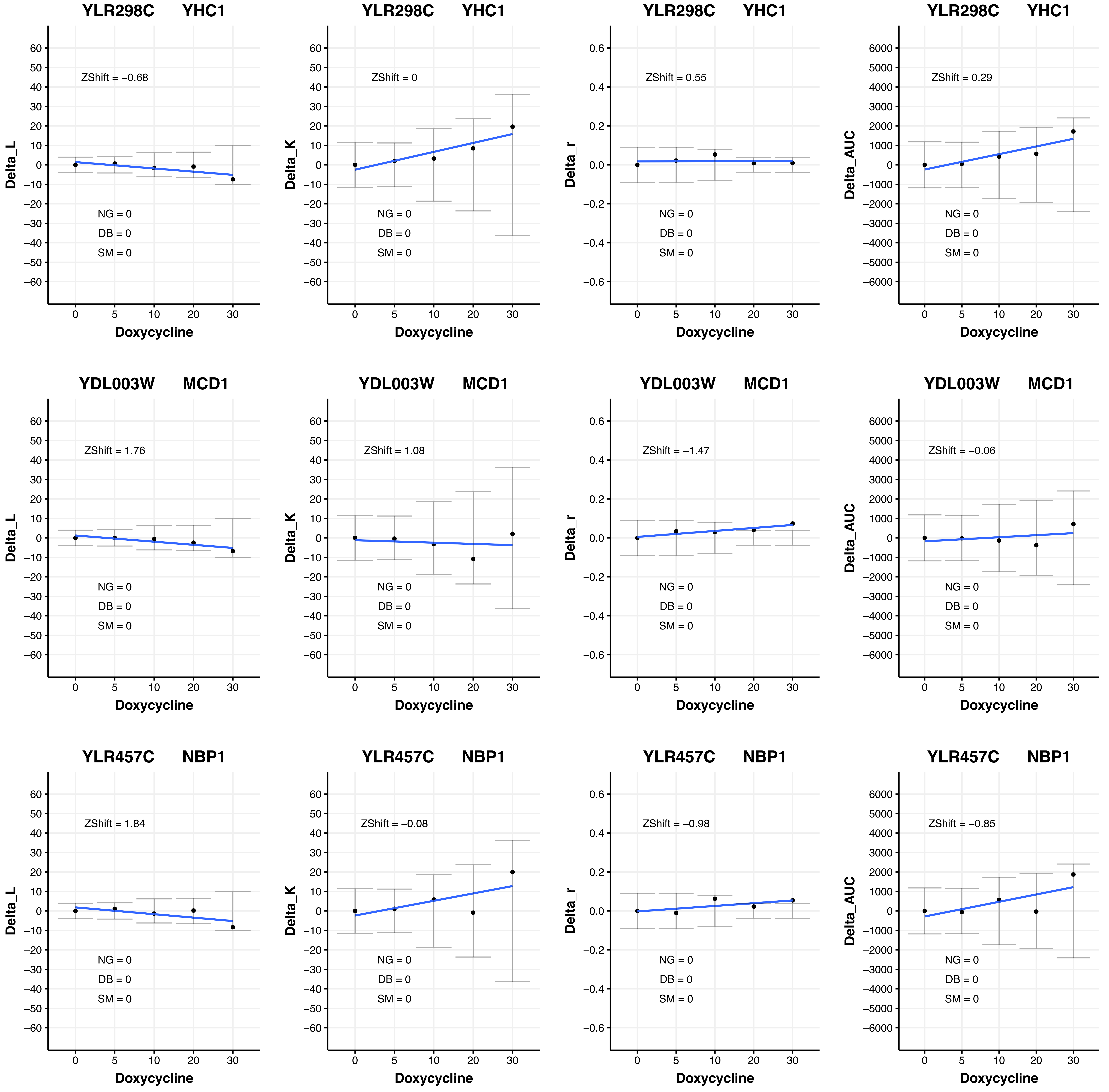

YLR298C YHC1
Delta_L
ZShift = −0.68
NG = 0
DB = 0
SM = 0
Doxycycline
YLR298C YHC1
Delta_K
ZShift = 0
NG = 0
DB = 0
SM = 0
Doxycycline
YLR298C YHC1
Delta_r
ZShift = 0.55
NG = 0
DB = 0
SM = 0
Doxycycline
YLR298C YHC1
Delta_AUC
ZShift = 0.29
NG = 0
DB = 0
SM = 0
Doxycycline
YDL003W MCD1
Delta_L
ZShift = 1.76
NG = 0
DB = 0
SM = 0
Doxycycline
YDL003W MCD1
Delta_K
ZShift = 1.08
NG = 0
DB = 0
SM = 0
Doxycycline
YDL003W MCD1
Delta_r
ZShift = −1.47
NG = 0
DB = 0
SM = 0
Doxycycline
YDL003W MCD1
Delta_AUC
ZShift = −0.06
NG = 0
DB = 0
SM = 0
Doxycycline
YLR457C NBP1
Delta_L
ZShift = 1.84
NG = 0
DB = 0
SM = 0
Doxycycline
YLR457C NBP1
Delta_K
ZShift = −0.08
NG = 0
DB = 0
SM = 0
Doxycycline
YLR457C NBP1
Delta_r
ZShift = −0.98
NG = 0
DB = 0
SM = 0
Doxycycline
YLR457C NBP1
Delta_AUC
ZShift = −0.85
NG = 0
DB = 0
SM = 0
Doxycycline

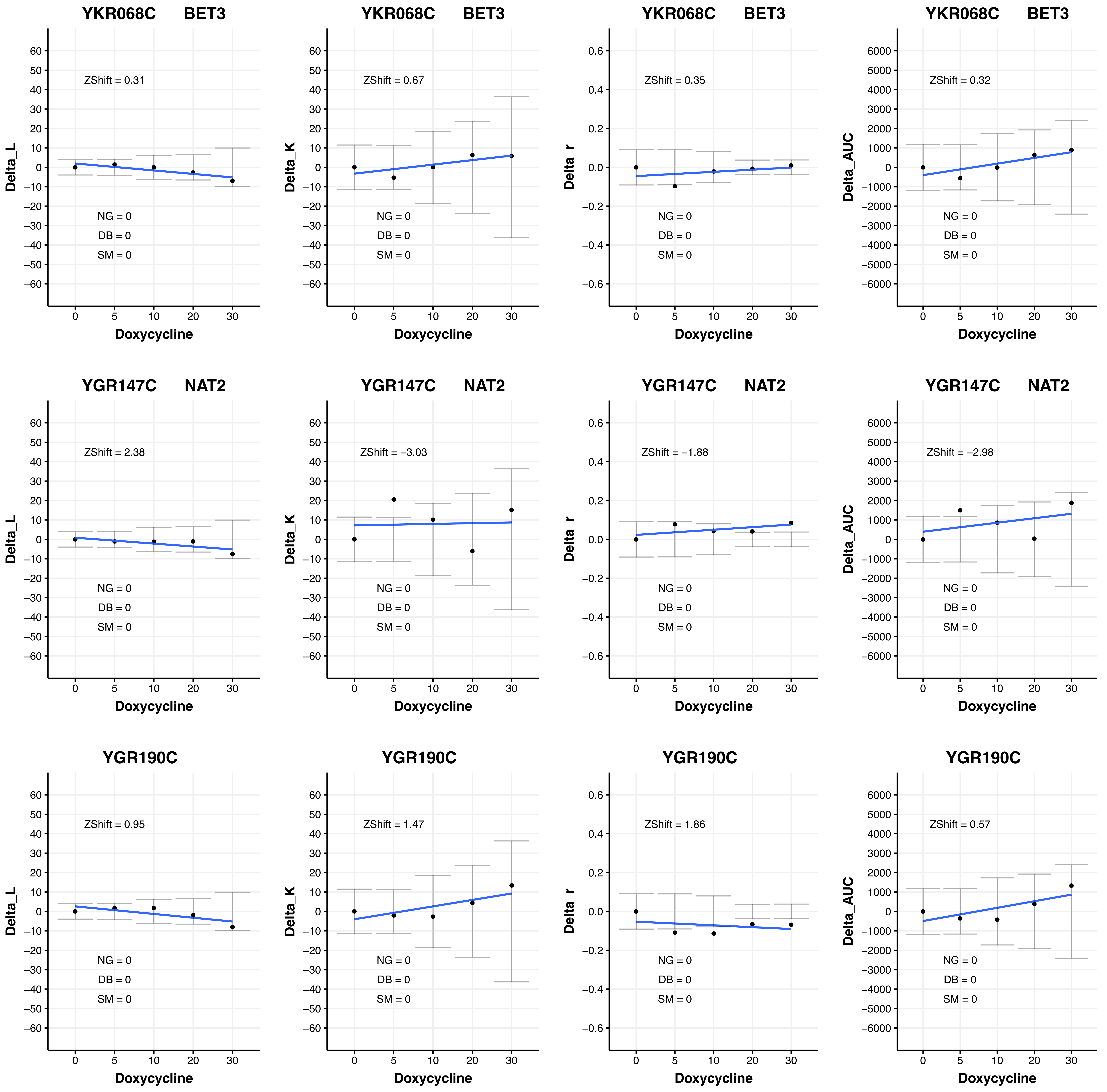
YKR068C BET3
ZShift = 0.31
NG = 0
DB = 0
SM = 0
Delta_L
Doxycycline
YKR068C BET3
ZShift = 0.67
NG = 0
DB = 0
SM = 0
Delta_K
Doxycycline
YKR068C BET3
ZShift = 0.35
NG = 0
DB = 0
SM = 0
Delta_r
Doxycycline
YKR068C BET3
ZShift = 0.32
NG = 0
DB = 0
SM = 0
Delta_AUC
Doxycycline
YGR147C NAT2
ZShift = 2.38
NG = 0
DB = 0
SM = 0
Delta_L
Doxycycline
YGR147C NAT2
ZShift = −3.03
NG = 0
DB = 0
SM = 0
Delta_K
Doxycycline
YGR147C NAT2
ZShift = −1.88
NG = 0
DB = 0
SM = 0
Delta_r
Doxycycline
YGR147C NAT2
ZShift = −2.98
NG = 0
DB = 0
SM = 0
Delta_AUC
Doxycycline
YGR190C
ZShift = 0.95
NG = 0
DB = 0
SM = 0
Delta_L
Doxycycline
YGR190C
ZShift = 1.47
NG = 0
DB = 0
SM = 0
Delta_K
Doxycycline
YGR190C
ZShift = 1.86
NG = 0
DB = 0
SM = 0
Delta_r
Doxycycline
YGR190C
ZShift = 0.57
NG = 0
DB = 0
SM = 0
Delta_AUC
Doxycycline

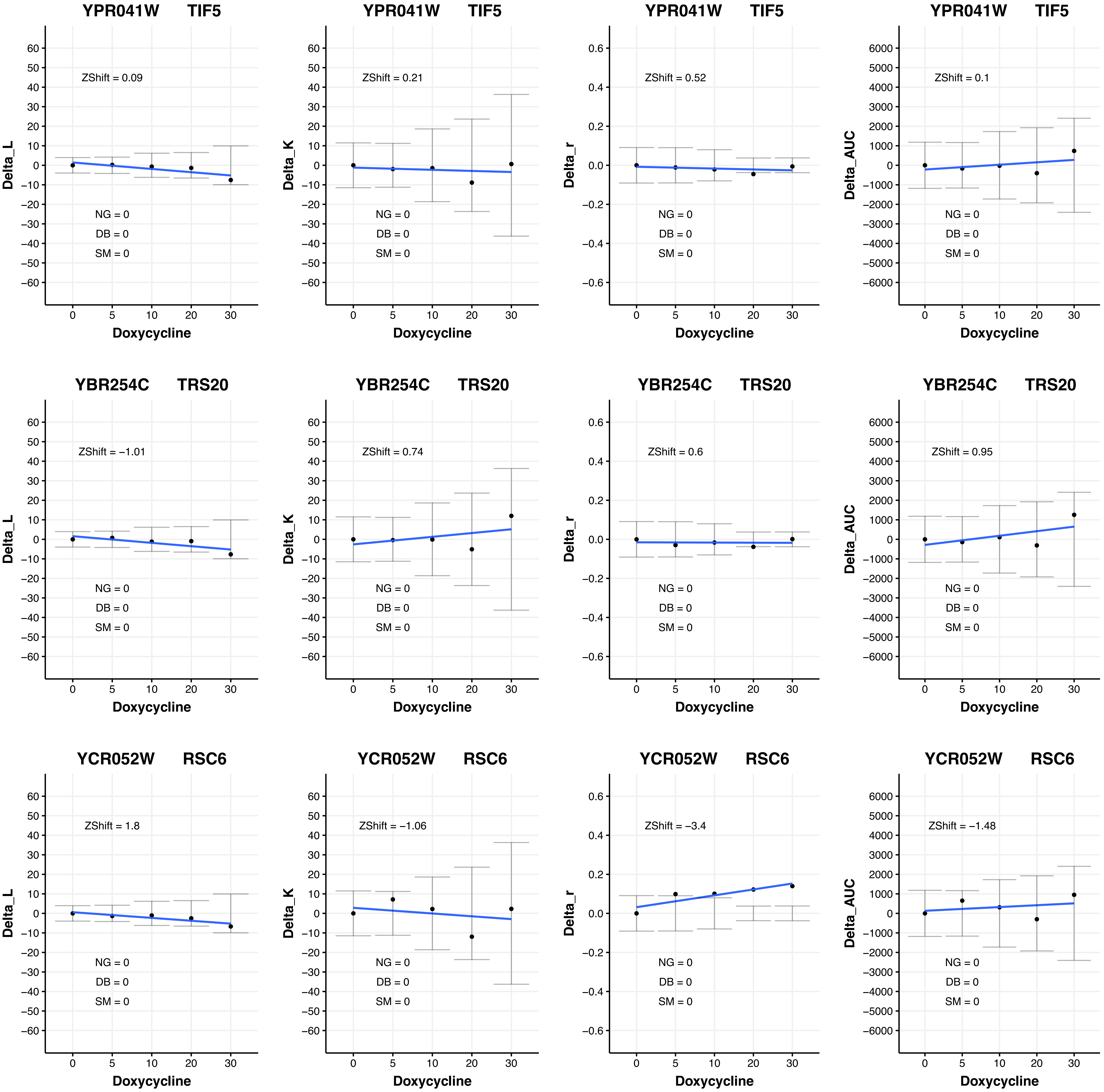

YPR041W TIF5
ZShift = 0.09
NG = 0
DB = 0
SM = 0
Delta_L
Doxycycline
YPR041W TIF5
ZShift = 0.21
NG = 0
DB = 0
SM = 0
Delta_K
Doxycycline
YPR041W TIF5
ZShift = 0.52
NG = 0
DB = 0
SM = 0
Delta_r
Doxycycline
YPR041W TIF5
ZShift = 0.1
NG = 0
DB = 0
SM = 0
Delta_AUC
Doxycycline
YBR254C TRS20
ZShift = −1.01
NG = 0
DB = 0
SM = 0
Delta_L
Doxycycline
YBR254C TRS20
ZShift = 0.74
NG = 0
DB = 0
SM = 0
Delta_K
Doxycycline
YBR254C TRS20
ZShift = 0.6
NG = 0
DB = 0
SM = 0
Delta_r
Doxycycline
YBR254C TRS20
ZShift = 0.95
NG = 0
DB = 0
SM = 0
Delta_AUC
Doxycycline
YCR052W RSC6
ZShift = 1.8
NG = 0
DB = 0
SM = 0
Delta_L
Doxycycline
YCR052W RSC6
ZShift = −1.06
NG = 0
DB = 0
SM = 0
Delta_K
Doxycycline
YCR052W RSC6
ZShift = −3.4
NG = 0
DB = 0
SM = 0
Delta_r
Doxycycline
YCR052W RSC6
ZShift = −1.48
NG = 0
DB = 0
SM = 0
Delta_AUC
Doxycycline

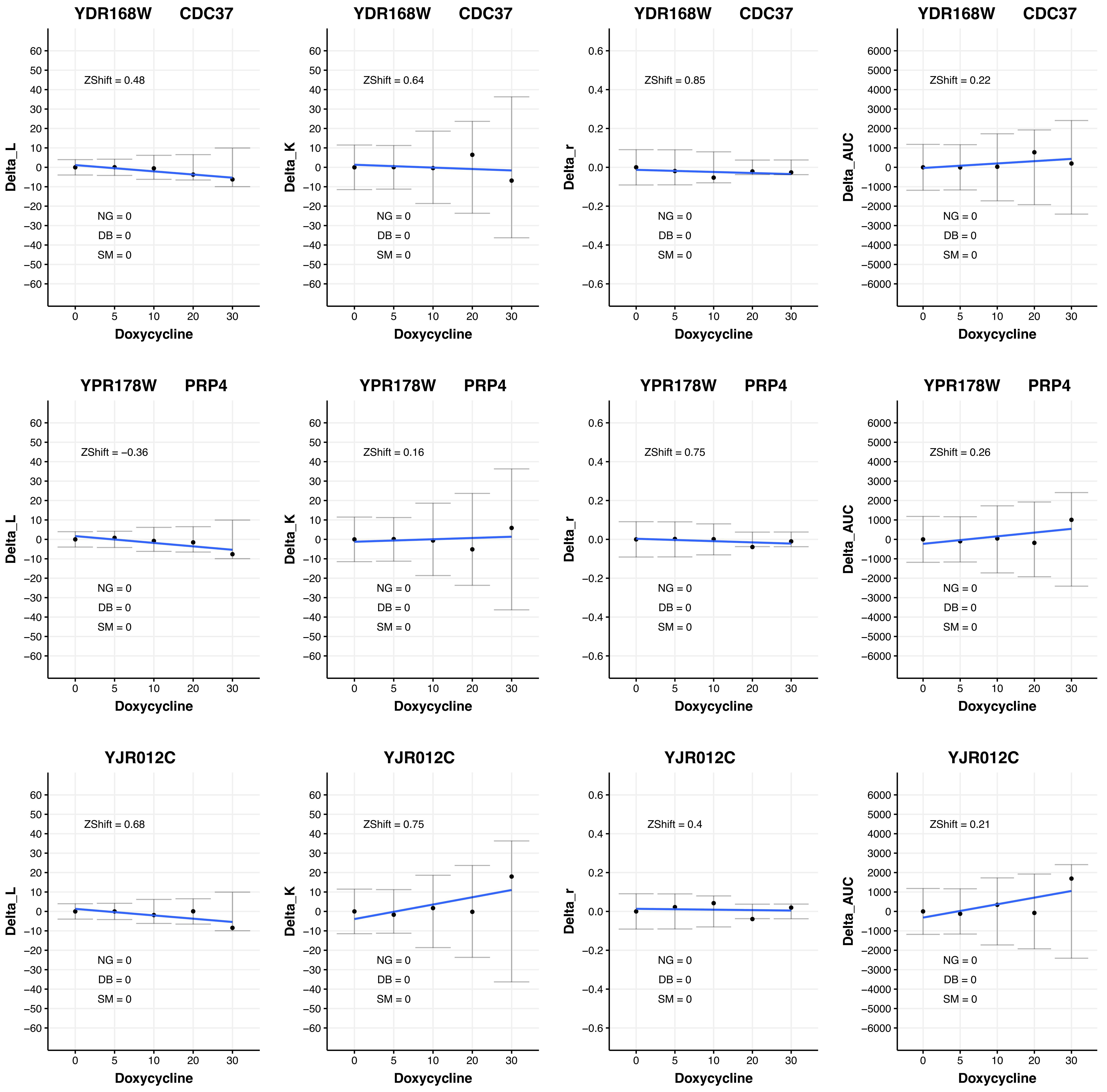

YDR168W CDC37
ZShift = 0.48
NG = 0
DB = 0
SM = 0
Delta_L
Doxycycline
YDR168W CDC37
ZShift = 0.64
NG = 0
DB = 0
SM = 0
Delta_K
Doxycycline
YDR168W CDC37
ZShift = 0.85
NG = 0
DB = 0
SM = 0
Delta_r
Doxycycline
YDR168W CDC37
ZShift = 0.22
NG = 0
DB = 0
SM = 0
Delta_AUC
Doxycycline
YPR178W PRP4
ZShift = −0.36
NG = 0
DB = 0
SM = 0
Delta_L
Doxycycline
YPR178W PRP4
ZShift = 0.16
NG = 0
DB = 0
SM = 0
Delta_K
Doxycycline
YPR178W PRP4
ZShift = 0.75
NG = 0
DB = 0
SM = 0
Delta_r
Doxycycline
YPR178W PRP4
ZShift = 0.26
NG = 0
DB = 0
SM = 0
Delta_AUC
Doxycycline
YJR012C
ZShift = 0.68
NG = 0
DB = 0
SM = 0
Delta_L
Doxycycline
YJR012C
ZShift = 0.75
NG = 0
DB = 0
SM = 0
Delta_K
Doxycycline
YJR012C
ZShift = 0.4
NG = 0
DB = 0
SM = 0
Delta_r
Doxycycline
YJR012C
ZShift = 0.21
NG = 0
DB = 0
SM = 0
Delta_AUC
Doxycycline

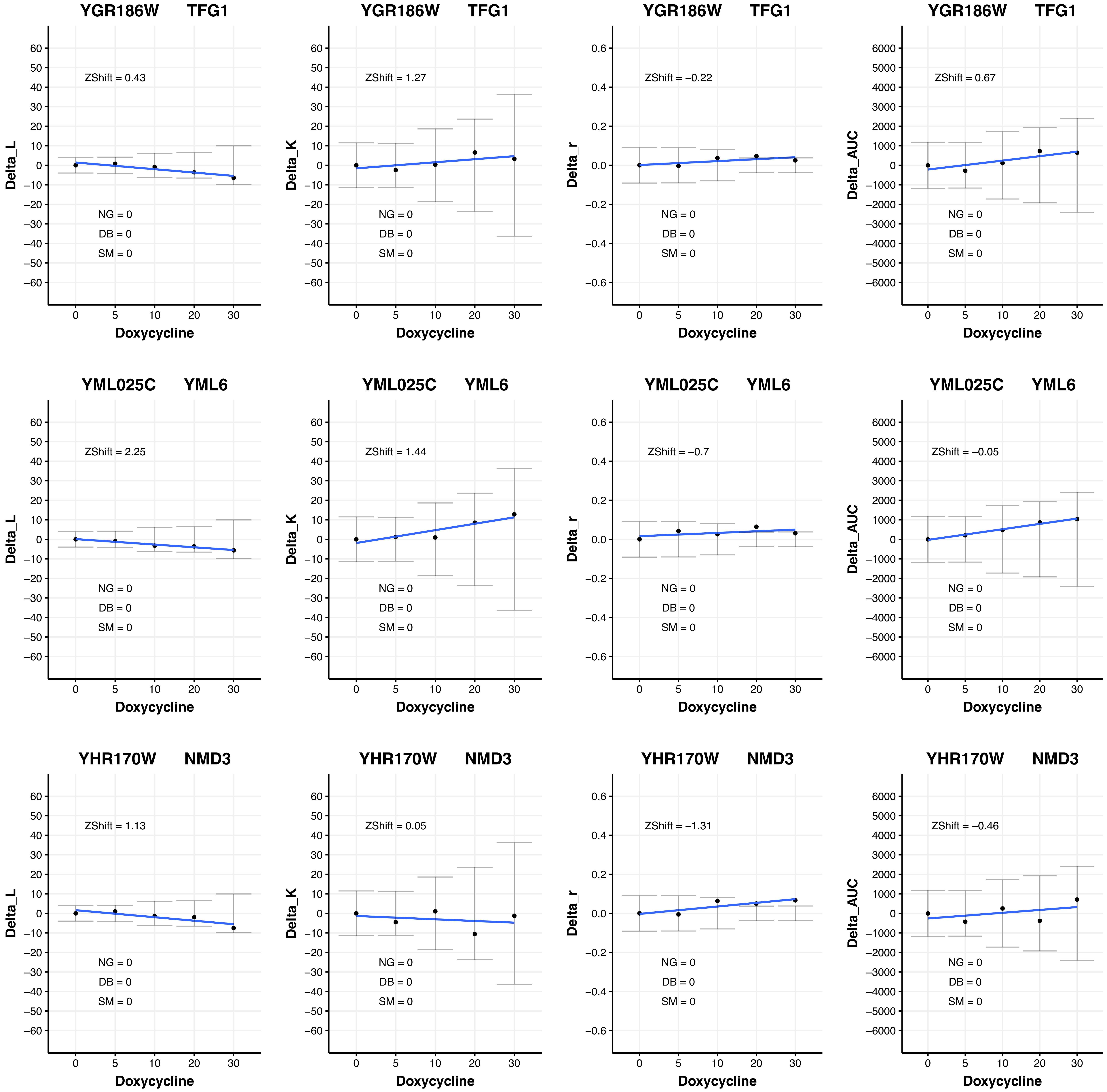
YGR186W TFG1
ZShift = 0.43
NG = 0
DB = 0
SM = 0
Delta_L
Doxycycline
YGR186W TFG1
ZShift = 1.27
NG = 0
DB = 0
SM = 0
Delta_K
Doxycycline
YGR186W TFG1
ZShift = −0.22
NG = 0
DB = 0
SM = 0
Delta_r
Doxycycline
YGR186W TFG1
ZShift = 0.67
NG = 0
DB = 0
SM = 0
Delta_AUC
Doxycycline
YML025C YML6
ZShift = 2.25
NG = 0
DB = 0
SM = 0
Delta_L
Doxycycline
YML025C YML6
ZShift = 1.44
NG = 0
DB = 0
SM = 0
Delta_K
Doxycycline
YML025C YML6
ZShift = −0.7
NG = 0
DB = 0
SM = 0
Delta_r
Doxycycline
YML025C YML6
ZShift = −0.05
NG = 0
DB = 0
SM = 0
Delta_AUC
Doxycycline
YHR170W NMD3
ZShift = 1.13
NG = 0
DB = 0
SM = 0
Delta_L
Doxycycline
YHR170W NMD3
ZShift = 0.05
NG = 0
DB = 0
SM = 0
Delta_K
Doxycycline
YHR170W NMD3
ZShift = −1.31
NG = 0
DB = 0
SM = 0
Delta_r
Doxycycline
YHR170W NMD3
ZShift = −0.46
NG = 0
DB = 0
SM = 0
Delta_AUC
Doxycycline

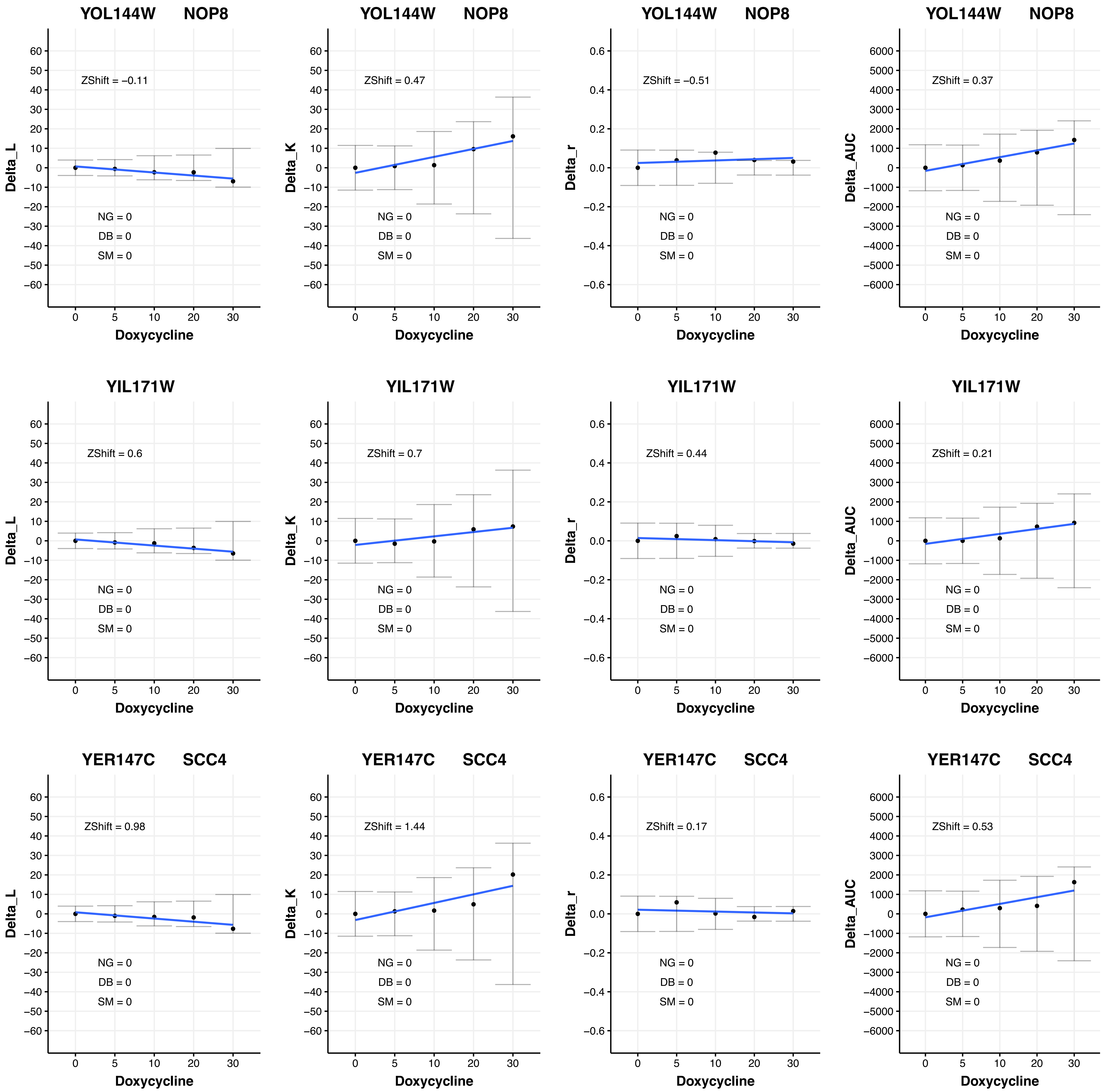
YOL144W NOP8
ZShift = −0.11
NG = 0
DB = 0
SM = 0
Delta_L
Doxycycline
YOL144W NOP8
ZShift = 0.47
NG = 0
DB = 0
SM = 0
Delta_K
Doxycycline
YOL144W NOP8
ZShift = −0.51
NG = 0
DB = 0
SM = 0
Delta_r
Doxycycline
YOL144W NOP8
ZShift = 0.37
NG = 0
DB = 0
SM = 0
Delta_AUC
Doxycycline
YIL171W
ZShift = 0.6
NG = 0
DB = 0
SM = 0
Delta_L
Doxycycline
YIL171W
ZShift = 0.7
NG = 0
DB = 0
SM = 0
Delta_K
Doxycycline
YIL171W
ZShift = 0.44
NG = 0
DB = 0
SM = 0
Delta_r
Doxycycline
YIL171W
ZShift = 0.21
NG = 0
DB = 0
SM = 0
Delta_AUC
Doxycycline
YER147C SCC4
ZShift = 0.98
NG = 0
DB = 0
SM = 0
Delta_L
Doxycycline
YER147C SCC4
ZShift = 1.44
NG = 0
DB = 0
SM = 0
Delta_K
Doxycycline
YER147C SCC4
ZShift = 0.17
NG = 0
DB = 0
SM = 0
Delta_r
Doxycycline
YER147C SCC4
ZShift = 0.53
NG = 0
DB = 0
SM = 0
Delta_AUC
Doxycycline

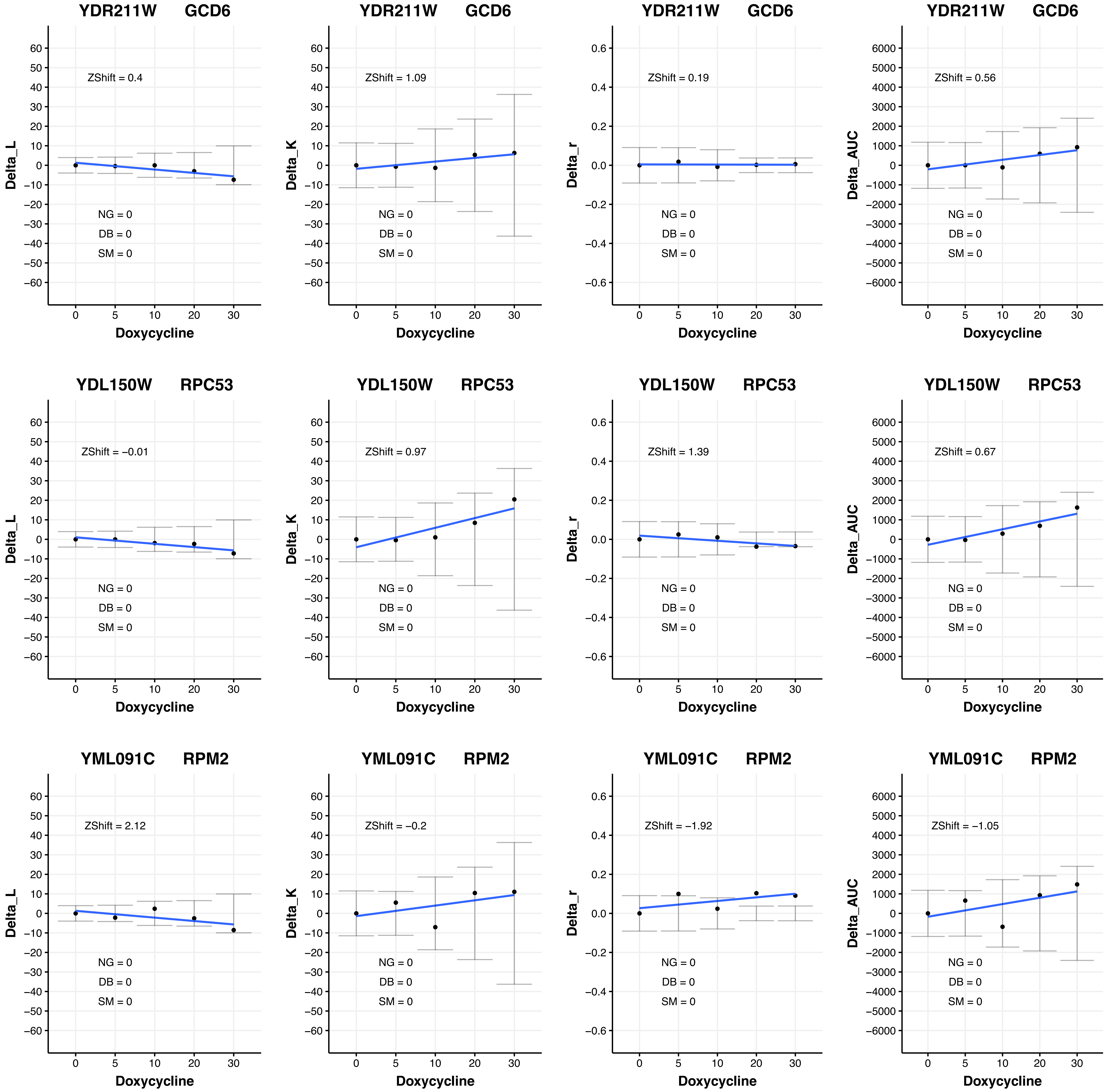

YDR211W GCD6
ZShift = 0.4
NG = 0
DB = 0
SM = 0
Delta_L
Doxycycline
YDR211W GCD6
ZShift = 1.09
NG = 0
DB = 0
SM = 0
Delta_K
Doxycycline
YDR211W GCD6
ZShift = 0.19
NG = 0
DB = 0
SM = 0
Delta_r
Doxycycline
YDR211W GCD6
ZShift = 0.56
NG = 0
DB = 0
SM = 0
Delta_AUC
Doxycycline
YDL150W RPC53
ZShift = −0.01
NG = 0
DB = 0
SM = 0
Delta_L
Doxycycline
YDL150W RPC53
ZShift = 0.97
NG = 0
DB = 0
SM = 0
Delta_K
Doxycycline
YDL150W RPC53
ZShift = 1.39
NG = 0
DB = 0
SM = 0
Delta_r
Doxycycline
YDL150W RPC53
ZShift = 0.67
NG = 0
DB = 0
SM = 0
Delta_AUC
Doxycycline
YML091C RPM2
ZShift = 2.12
NG = 0
DB = 0
SM = 0
Delta_L
Doxycycline
YML091C RPM2
ZShift = −0.2
NG = 0
DB = 0
SM = 0
Delta_K
Doxycycline
YML091C RPM2
ZShift = −1.92
NG = 0
DB = 0
SM = 0
Delta_r
Doxycycline
YML091C RPM2
ZShift = −1.05
NG = 0
DB = 0
SM = 0
Delta_AUC
Doxycycline

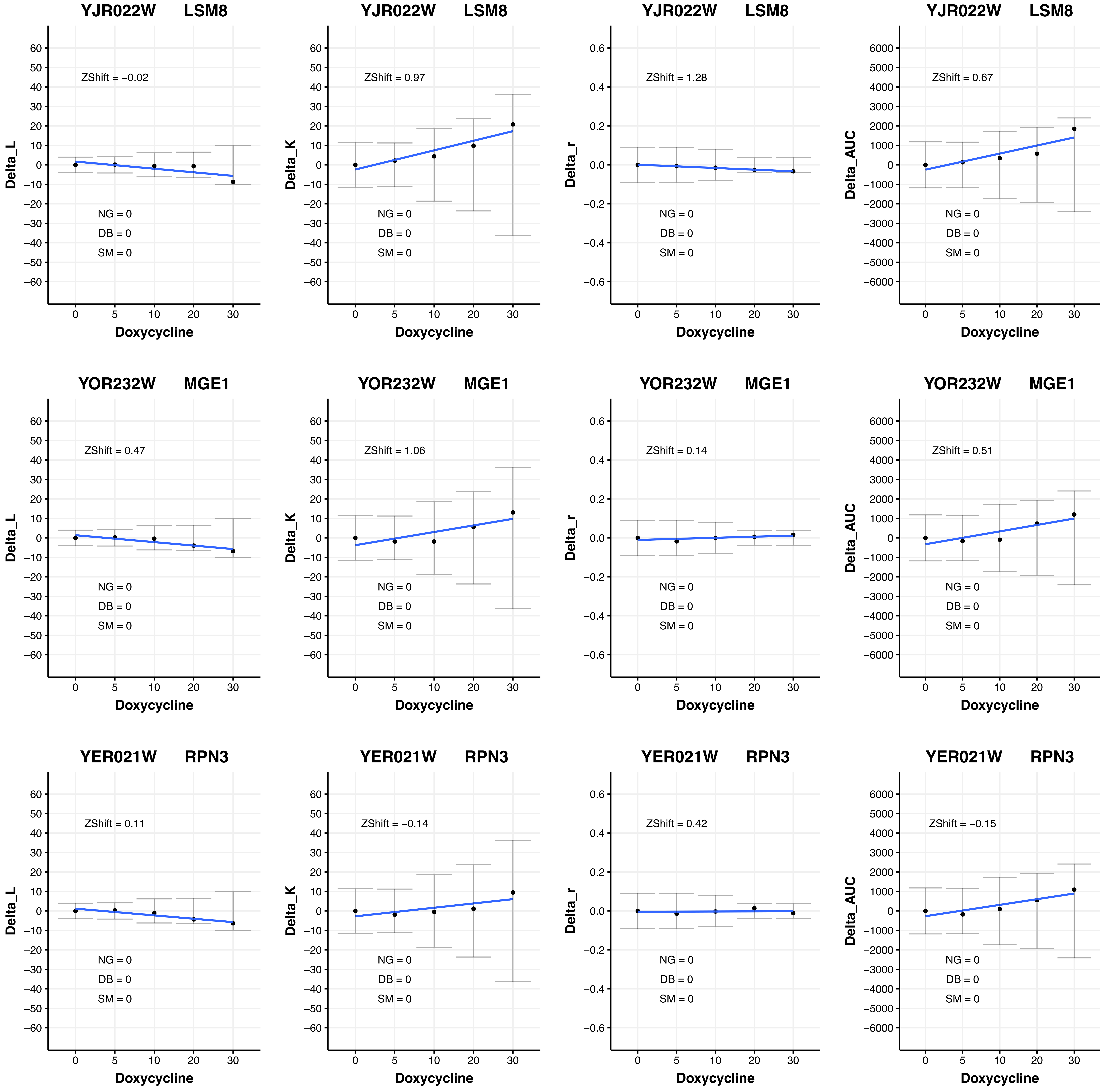
YJR022W LSM8
ZShift = –0.02
NG = 0
DB = 0
SM = 0
Delta_L
Doxycycline
YJR022W LSM8
ZShift = 0.97
NG = 0
DB = 0
SM = 0
Delta_K
Doxycycline
YJR022W LSM8
ZShift = 1.28
NG = 0
DB = 0
SM = 0
Delta_r
Doxycycline
YJR022W LSM8
ZShift = 0.67
NG = 0
DB = 0
SM = 0
Delta_AUC
Doxycycline
YOR232W MGE1
ZShift = 0.47
NG = 0
DB = 0
SM = 0
Delta_L
Doxycycline
YOR232W MGE1
ZShift = 1.06
NG = 0
DB = 0
SM = 0
Delta_K
Doxycycline
YOR232W MGE1
ZShift = 0.14
NG = 0
DB = 0
SM = 0
Delta_r
Doxycycline
YOR232W MGE1
ZShift = 0.51
NG = 0
DB = 0
SM = 0
Delta_AUC
Doxycycline
YER021W RPN3
ZShift = 0.11
NG = 0
DB = 0
SM = 0
Delta_L
Doxycycline
YER021W RPN3
ZShift = –0.14
NG = 0
DB = 0
SM = 0
Delta_K
Doxycycline
YER021W RPN3
ZShift = 0.42
NG = 0
DB = 0
SM = 0
Delta_r
Doxycycline
YER021W RPN3
ZShift = –0.15
NG = 0
DB = 0
SM = 0
Delta_AUC
Doxycycline

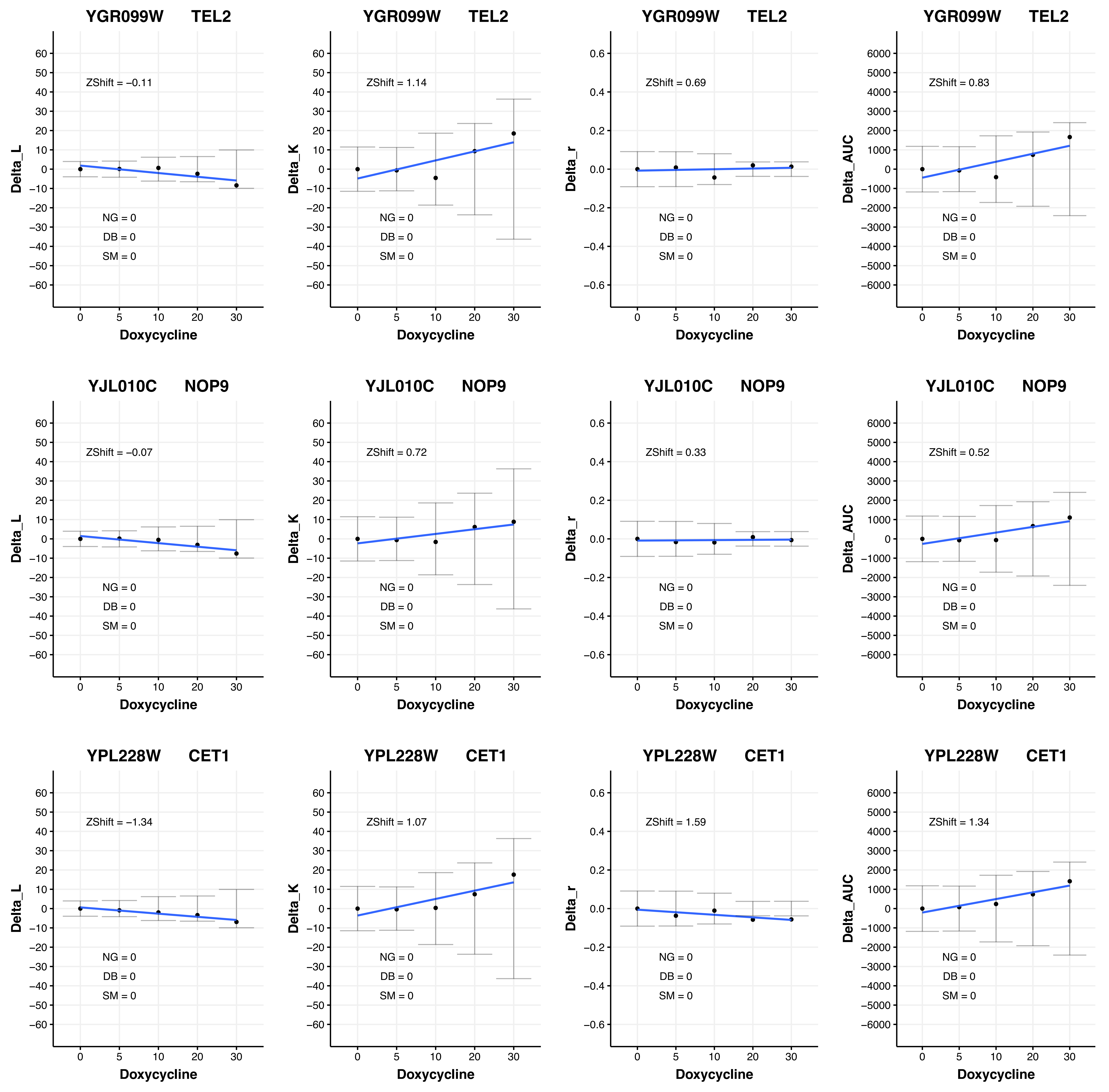

YGR099W TEL2
ZShift = −0.11
NG = 0
DB = 0
SM = 0
Delta_L
Doxycycline
YGR099W TEL2
ZShift = 1.14
NG = 0
DB = 0
SM = 0
Delta_K
Doxycycline
YGR099W TEL2
ZShift = 0.69
NG = 0
DB = 0
SM = 0
Delta_r
Doxycycline
YGR099W TEL2
ZShift = 0.83
NG = 0
DB = 0
SM = 0
Delta_AUC
Doxycycline
YJL010C NOP9
ZShift = −0.07
NG = 0
DB = 0
SM = 0
Delta_L
Doxycycline
YJL010C NOP9
ZShift = 0.72
NG = 0
DB = 0
SM = 0
Delta_K
Doxycycline
YJL010C NOP9
ZShift = 0.33
NG = 0
DB = 0
SM = 0
Delta_r
Doxycycline
YJL010C NOP9
ZShift = 0.52
NG = 0
DB = 0
SM = 0
Delta_AUC
Doxycycline
YPL228W CET1
ZShift = −1.34
NG = 0
DB = 0
SM = 0
Delta_L
Doxycycline
YPL228W CET1
ZShift = 1.07
NG = 0
DB = 0
SM = 0
Delta_K
Doxycycline
YPL228W CET1
ZShift = 1.59
NG = 0
DB = 0
SM = 0
Delta_r
Doxycycline
YPL228W CET1
ZShift = 1.34
NG = 0
DB = 0
SM = 0
Delta_AUC
Doxycycline

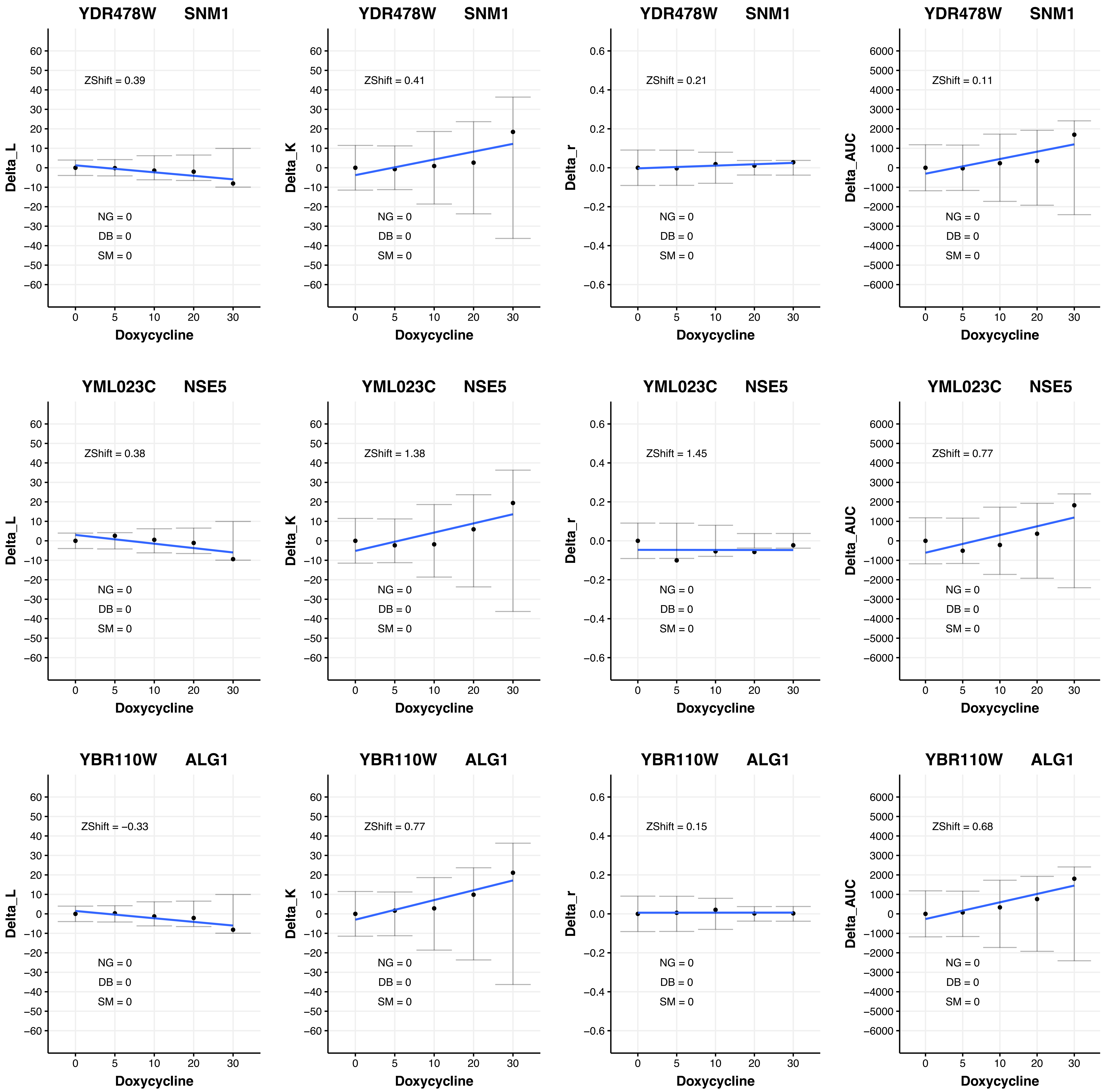

YDR478W SNM1
ZShift = 0.39
ZShift = 0.41
ZShift = 0.21
ZShift = 0.11
YML023C NSE5
ZShift = 0.38
ZShift = 1.38
ZShift = 1.45
ZShift = 0.77
YBR110W ALG1
ZShift = −0.33
ZShift = 0.77
ZShift = 0.15
ZShift = 0.68
NG = 0
DB = 0
SM = 0
Delta_L
Delta_K
Delta_r
Delta_AUC
Doxycycline

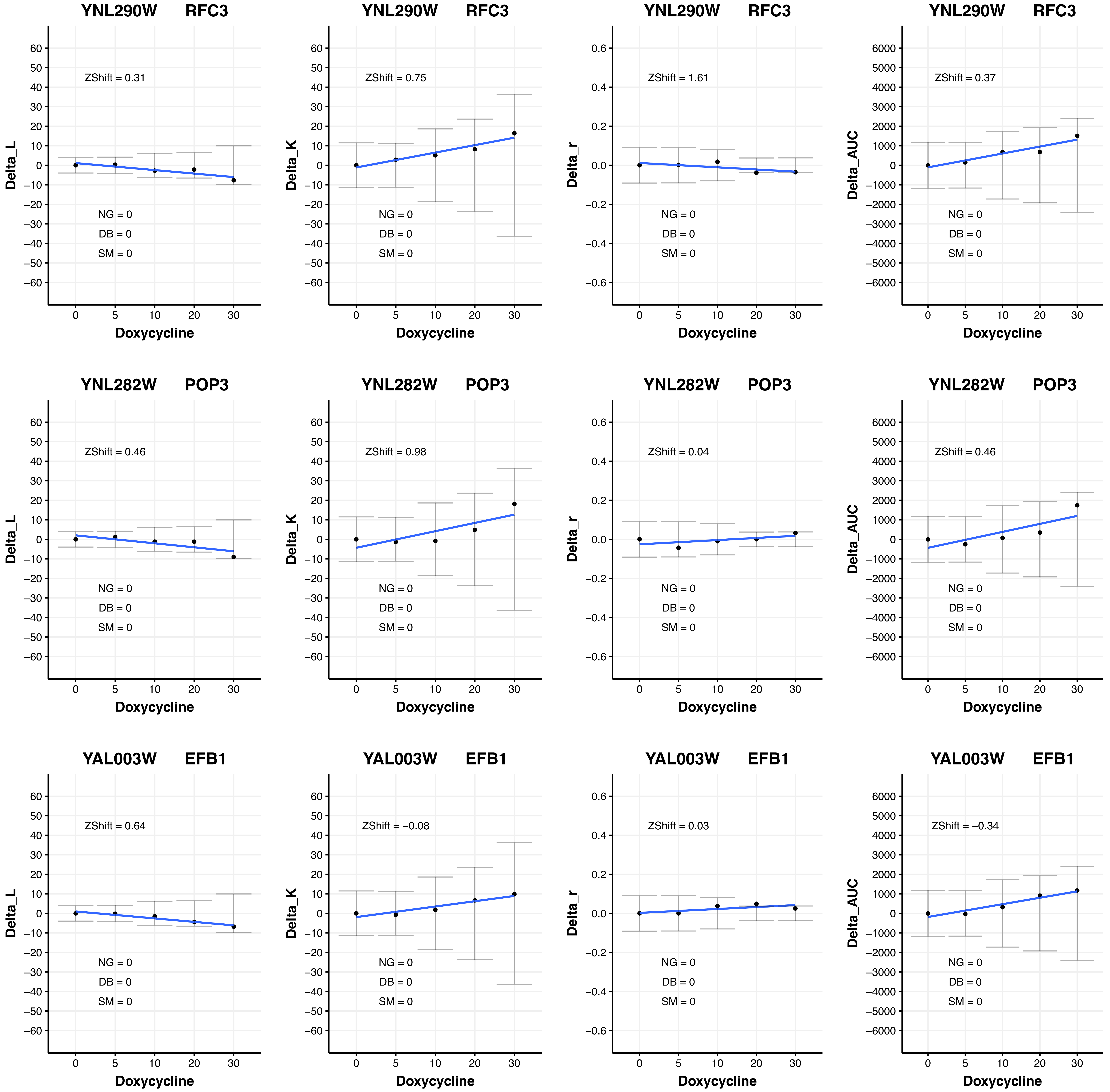

YNL290W RFC3
ZShift = 0.31
NG = 0
DB = 0
SM = 0
Delta_L
Doxycycline
YNL290W RFC3
ZShift = 0.75
NG = 0
DB = 0
SM = 0
Delta_K
Doxycycline
YNL290W RFC3
ZShift = 1.61
NG = 0
DB = 0
SM = 0
Delta_r
Doxycycline
YNL290W RFC3
ZShift = 0.37
NG = 0
DB = 0
SM = 0
Delta_AUC
Doxycycline
YNL282W POP3
ZShift = 0.46
NG = 0
DB = 0
SM = 0
Delta_L
Doxycycline
YNL282W POP3
ZShift = 0.98
NG = 0
DB = 0
SM = 0
Delta_K
Doxycycline
YNL282W POP3
ZShift = 0.04
NG = 0
DB = 0
SM = 0
Delta_r
Doxycycline
YNL282W POP3
ZShift = 0.46
NG = 0
DB = 0
SM = 0
Delta_AUC
Doxycycline
YAL003W EFB1
ZShift = 0.64
NG = 0
DB = 0
SM = 0
Delta_L
Doxycycline
YAL003W EFB1
ZShift = −0.08
NG = 0
DB = 0
SM = 0
Delta_K
Doxycycline
YAL003W EFB1
ZShift = 0.03
NG = 0
DB = 0
SM = 0
Delta_r
Doxycycline
YAL003W EFB1
ZShift = −0.34
NG = 0
DB = 0
SM = 0
Delta_AUC
Doxycycline

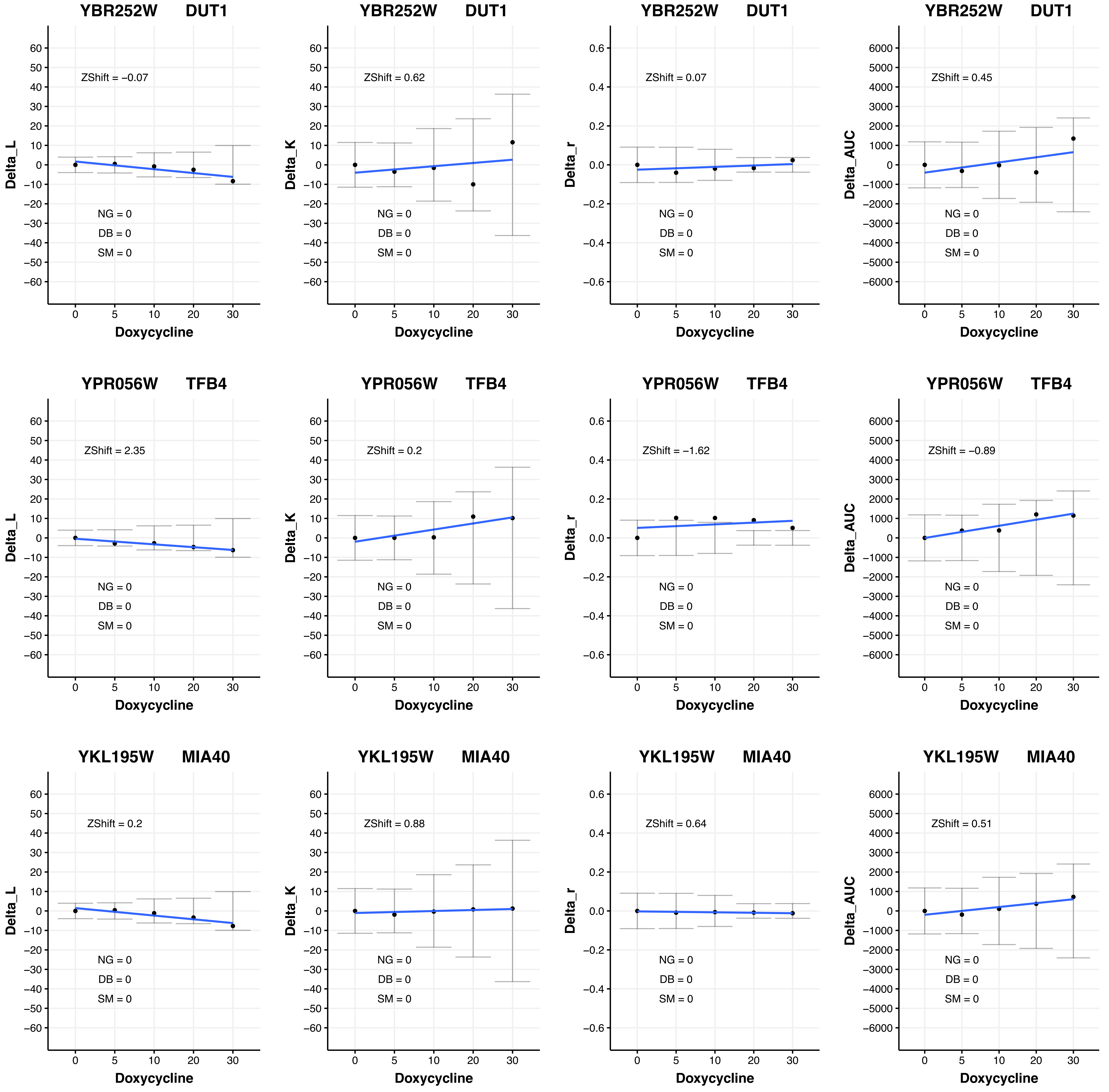

YBR252W DUT1
ZShift = −0.07
NG = 0
DB = 0
SM = 0
Delta_L
Doxycycline
YBR252W DUT1
ZShift = 0.62
NG = 0
DB = 0
SM = 0
Delta_K
Doxycycline
YBR252W DUT1
ZShift = 0.07
NG = 0
DB = 0
SM = 0
Delta_r
Doxycycline
YBR252W DUT1
ZShift = 0.45
NG = 0
DB = 0
SM = 0
Delta_AUC
Doxycycline
YPR056W TFB4
ZShift = 2.35
NG = 0
DB = 0
SM = 0
Delta_L
Doxycycline
YPR056W TFB4
ZShift = 0.2
NG = 0
DB = 0
SM = 0
Delta_K
Doxycycline
YPR056W TFB4
ZShift = −1.62
NG = 0
DB = 0
SM = 0
Delta_r
Doxycycline
YPR056W TFB4
ZShift = −0.89
NG = 0
DB = 0
SM = 0
Delta_AUC
Doxycycline
YKL195W MIA40
ZShift = 0.2
NG = 0
DB = 0
SM = 0
Delta_L
Doxycycline
YKL195W MIA40
ZShift = 0.88
NG = 0
DB = 0
SM = 0
Delta_K
Doxycycline
YKL195W MIA40
ZShift = 0.64
NG = 0
DB = 0
SM = 0
Delta_r
Doxycycline
YKL195W MIA40
ZShift = 0.51
NG = 0
DB = 0
SM = 0
Delta_AUC
Doxycycline

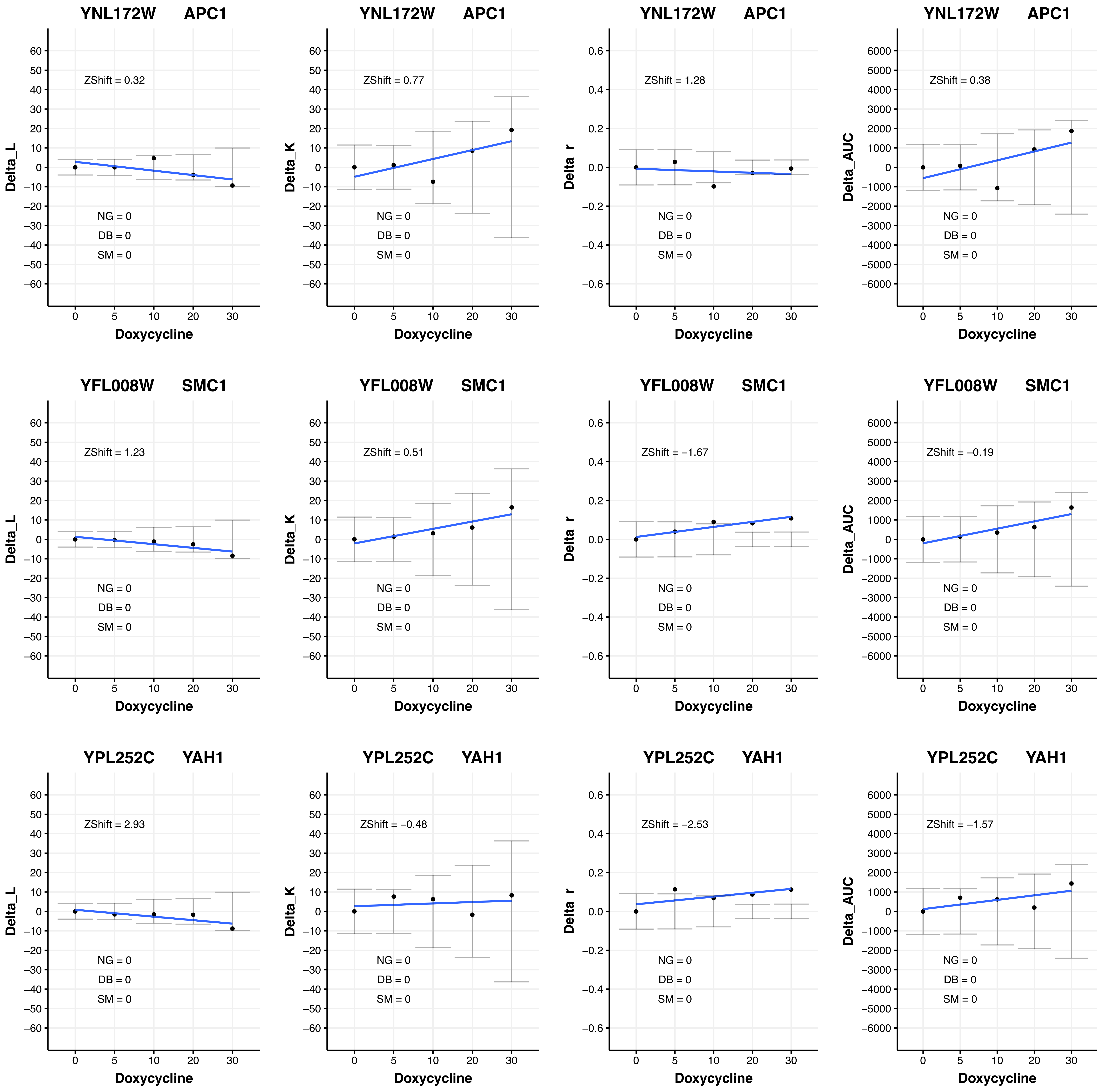
YNL172W APC1
ZShift = 0.32
NG = 0
DB = 0
SM = 0
Delta_L
Doxycycline
YNL172W APC1
ZShift = 0.77
NG = 0
DB = 0
SM = 0
Delta_K
Doxycycline
YNL172W APC1
ZShift = 1.28
NG = 0
DB = 0
SM = 0
Delta_r
Doxycycline
YNL172W APC1
ZShift = 0.38
NG = 0
DB = 0
SM = 0
Delta_AUC
Doxycycline
YFL008W SMC1
ZShift = 1.23
NG = 0
DB = 0
SM = 0
Delta_L
Doxycycline
YFL008W SMC1
ZShift = 0.51
NG = 0
DB = 0
SM = 0
Delta_K
Doxycycline
YFL008W SMC1
ZShift = −1.67
NG = 0
DB = 0
SM = 0
Delta_r
Doxycycline
YFL008W SMC1
ZShift = −0.19
NG = 0
DB = 0
SM = 0
Delta_AUC
Doxycycline
YPL252C YAH1
ZShift = 2.93
NG = 0
DB = 0
SM = 0
Delta_L
Doxycycline
YPL252C YAH1
ZShift = −0.48
NG = 0
DB = 0
SM = 0
Delta_K
Doxycycline
YPL252C YAH1
ZShift = −2.53
NG = 0
DB = 0
SM = 0
Delta_r
Doxycycline
YPL252C YAH1
ZShift = −1.57
NG = 0
DB = 0
SM = 0
Delta_AUC
Doxycycline

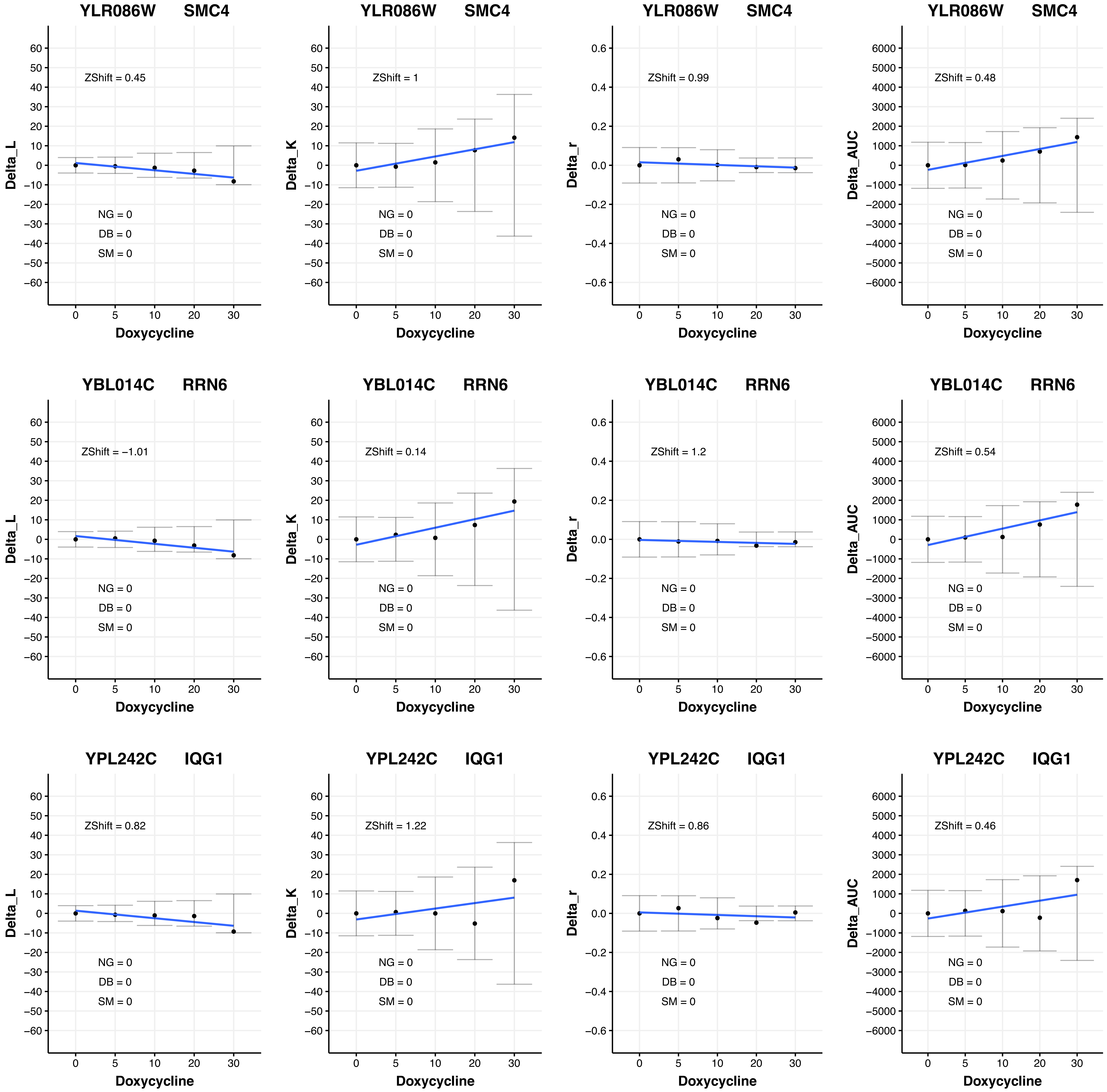
YLR086W SMC4
ZShift = 0.45
NG = 0
DB = 0
SM = 0
Delta_L
Doxycycline
YLR086W SMC4
ZShift = 1
NG = 0
DB = 0
SM = 0
Delta_K
Doxycycline
YLR086W SMC4
ZShift = 0.99
NG = 0
DB = 0
SM = 0
Delta_r
Doxycycline
YLR086W SMC4
ZShift = 0.48
NG = 0
DB = 0
SM = 0
Delta_AUC
Doxycycline
YBL014C RRN6
ZShift = −1.01
NG = 0
DB = 0
SM = 0
Delta_L
Doxycycline
YBL014C RRN6
ZShift = 0.14
NG = 0
DB = 0
SM = 0
Delta_K
Doxycycline
YBL014C RRN6
ZShift = 1.2
NG = 0
DB = 0
SM = 0
Delta_r
Doxycycline
YBL014C RRN6
ZShift = 0.54
NG = 0
DB = 0
SM = 0
Delta_AUC
Doxycycline
YPL242C IQG1
ZShift = 0.82
NG = 0
DB = 0
SM = 0
Delta_L
Doxycycline
YPL242C IQG1
ZShift = 1.22
NG = 0
DB = 0
SM = 0
Delta_K
Doxycycline
YPL242C IQG1
ZShift = 0.86
NG = 0
DB = 0
SM = 0
Delta_r
Doxycycline
YPL242C IQG1
ZShift = 0.46
NG = 0
DB = 0
SM = 0
Delta_AUC
Doxycycline

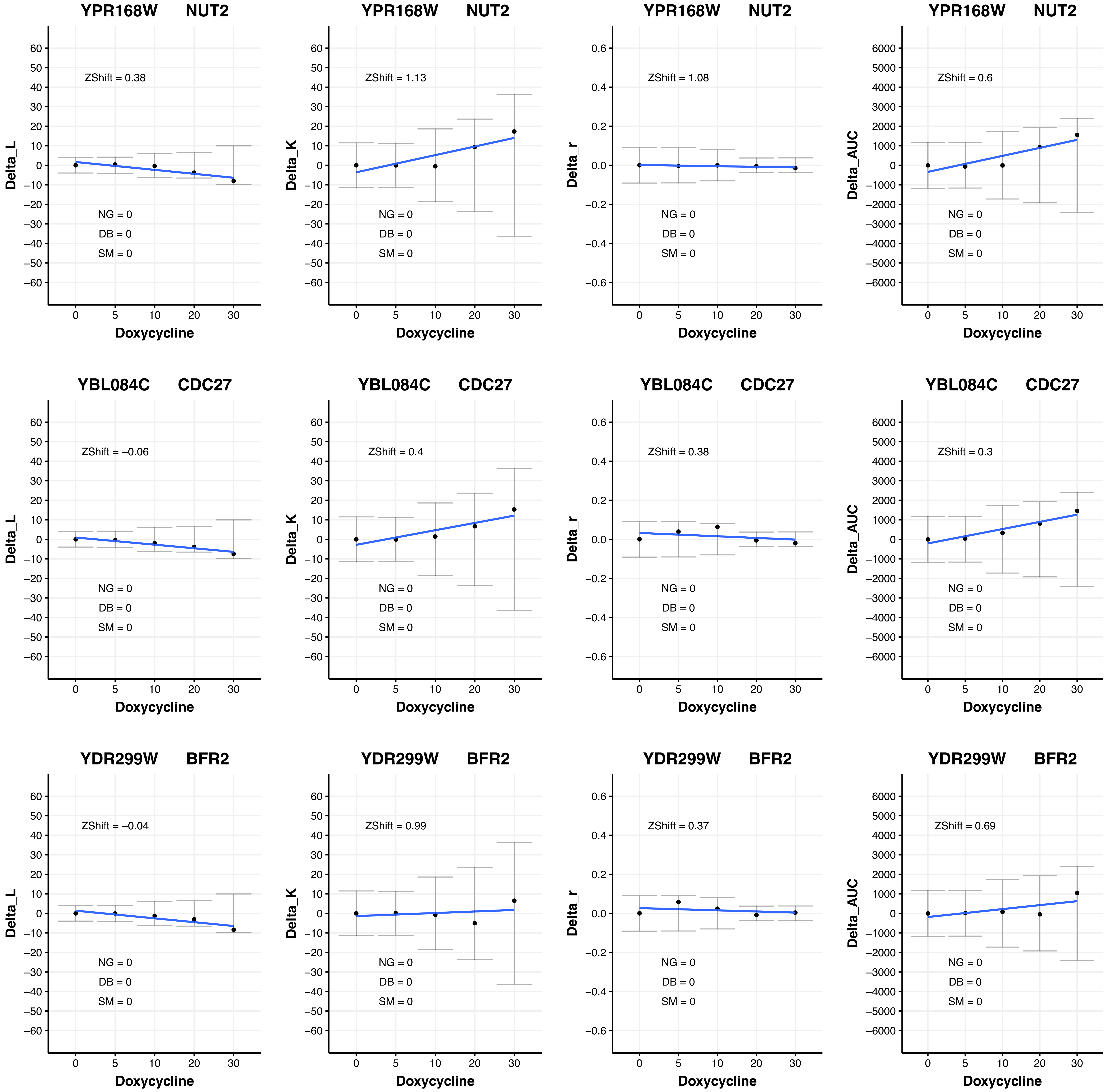
YPR168W NUT2
ZShift = 0.38
ZShift = 1.13
ZShift = 1.08
ZShift = 0.6
YBL084C CDC27
ZShift = −0.06
ZShift = 0.4
ZShift = 0.38
ZShift = 0.3
YDR299W BFR2
ZShift = −0.04
ZShift = 0.99
ZShift = 0.37
ZShift = 0.69
NG = 0
DB = 0
SM = 0
Delta_L
Delta_K
Delta_r
Delta_AUC
Doxycycline

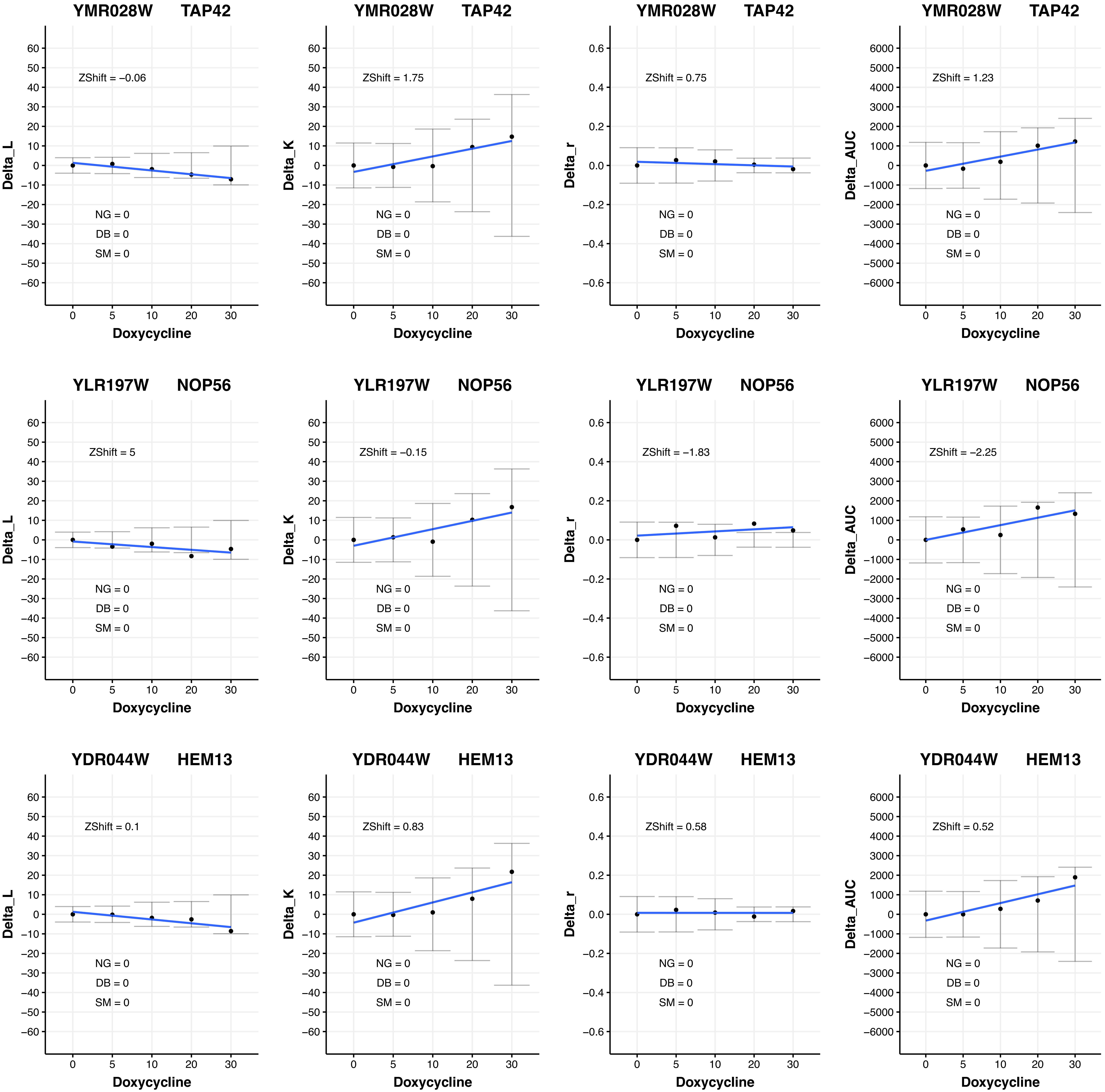
YMR028W TAP42
ZShift = −0.06
NG = 0
DB = 0
SM = 0
Delta_L
Doxycycline
YMR028W TAP42
ZShift = 1.75
NG = 0
DB = 0
SM = 0
Delta_K
Doxycycline
YMR028W TAP42
ZShift = 0.75
NG = 0
DB = 0
SM = 0
Delta_r
Doxycycline
YMR028W TAP42
ZShift = 1.23
NG = 0
DB = 0
SM = 0
Delta_AUC
Doxycycline
YLR197W NOP56
ZShift = 5
NG = 0
DB = 0
SM = 0
Delta_L
Doxycycline
YLR197W NOP56
ZShift = −0.15
NG = 0
DB = 0
SM = 0
Delta_K
Doxycycline
YLR197W NOP56
ZShift = −1.83
NG = 0
DB = 0
SM = 0
Delta_r
Doxycycline
YLR197W NOP56
ZShift = −2.25
NG = 0
DB = 0
SM = 0
Delta_AUC
Doxycycline
YDR044W HEM13
ZShift = 0.1
NG = 0
DB = 0
SM = 0
Delta_L
Doxycycline
YDR044W HEM13
ZShift = 0.83
NG = 0
DB = 0
SM = 0
Delta_K
Doxycycline
YDR044W HEM13
ZShift = 0.58
NG = 0
DB = 0
SM = 0
Delta_r
Doxycycline
YDR044W HEM13
ZShift = 0.52
NG = 0
DB = 0
SM = 0
Delta_AUC
Doxycycline

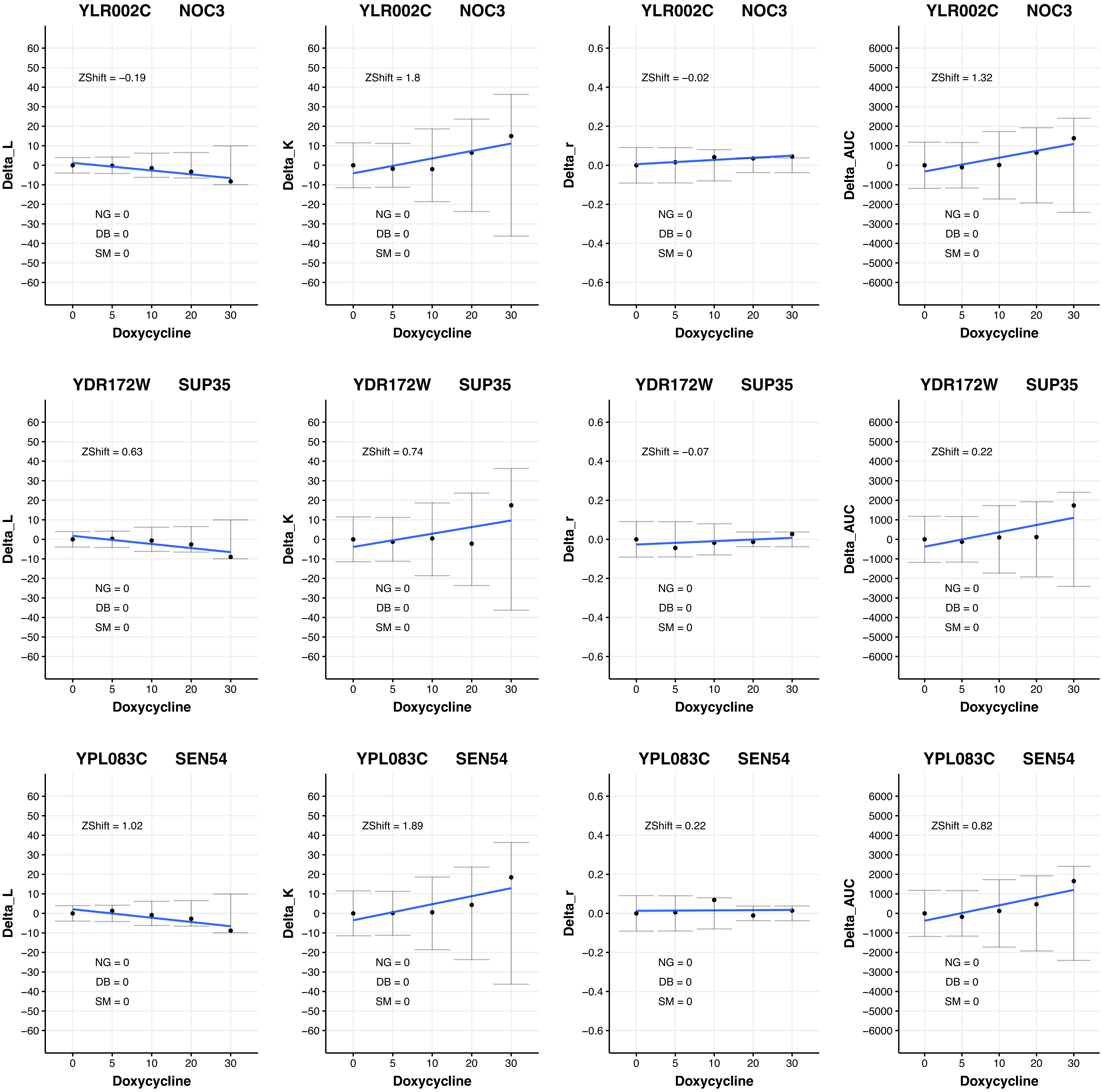
YLR002C NOC3
ZShift = −0.19
NG = 0
DB = 0
SM = 0
Delta_L
Doxycycline
YLR002C NOC3
ZShift = 1.8
NG = 0
DB = 0
SM = 0
Delta_K
Doxycycline
YLR002C NOC3
ZShift = −0.02
NG = 0
DB = 0
SM = 0
Delta_r
Doxycycline
YLR002C NOC3
ZShift = 1.32
NG = 0
DB = 0
SM = 0
Delta_AUC
Doxycycline
YDR172W SUP35
ZShift = 0.63
NG = 0
DB = 0
SM = 0
Delta_L
Doxycycline
YDR172W SUP35
ZShift = 0.74
NG = 0
DB = 0
SM = 0
Delta_K
Doxycycline
YDR172W SUP35
ZShift = −0.07
NG = 0
DB = 0
SM = 0
Delta_r
Doxycycline
YDR172W SUP35
ZShift = 0.22
NG = 0
DB = 0
SM = 0
Delta_AUC
Doxycycline
YPL083C SEN54
ZShift = 1.02
NG = 0
DB = 0
SM = 0
Delta_L
Doxycycline
YPL083C SEN54
ZShift = 1.89
NG = 0
DB = 0
SM = 0
Delta_K
Doxycycline
YPL083C SEN54
ZShift = 0.22
NG = 0
DB = 0
SM = 0
Delta_r
Doxycycline
YPL083C SEN54
ZShift = 0.82
NG = 0
DB = 0
SM = 0
Delta_AUC
Doxycycline

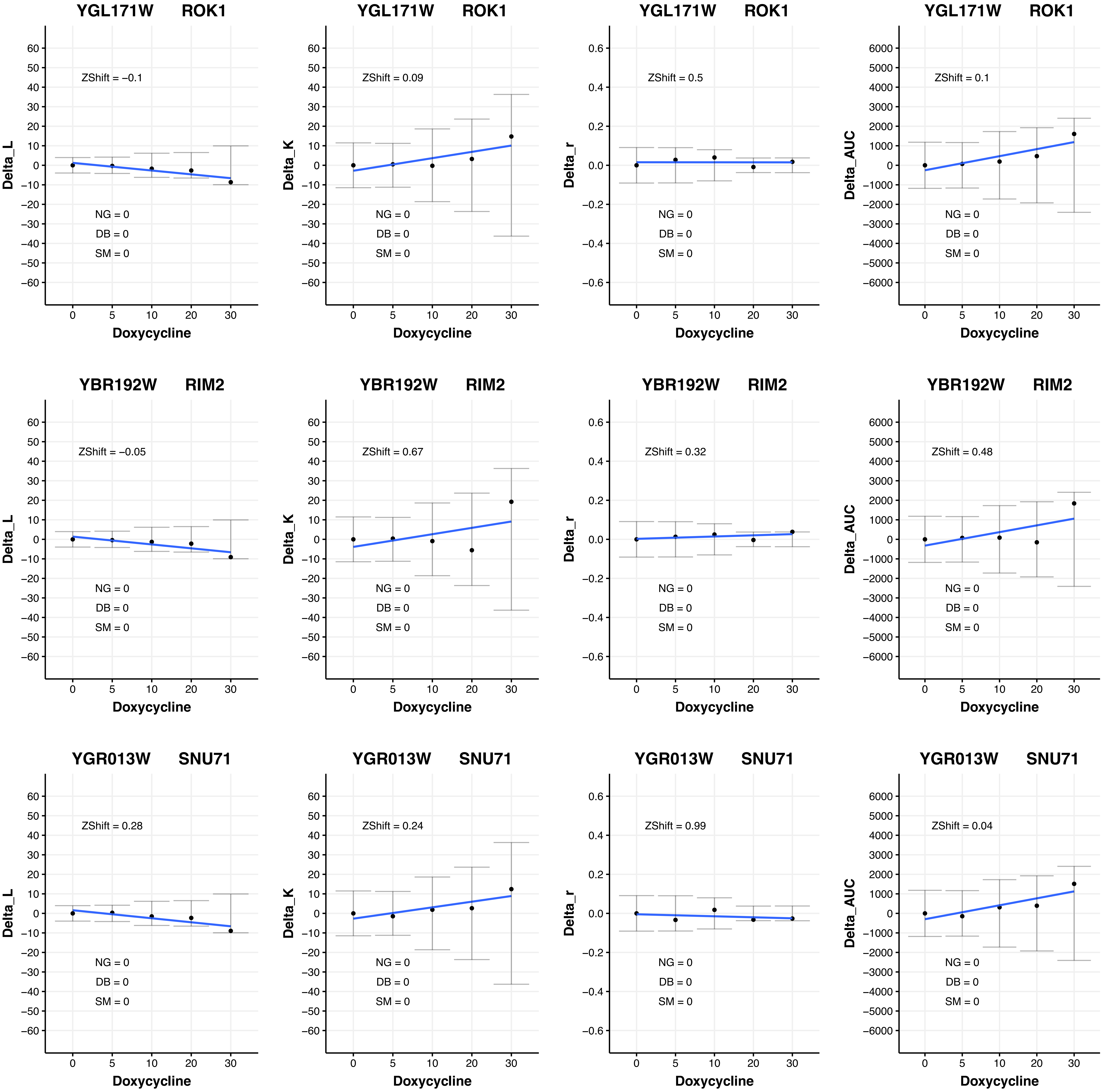
YGL171W ROK1
Delta_L
ZShift = −0.1
NG = 0
DB = 0
SM = 0
Doxycycline
YGL171W ROK1
Delta_K
ZShift = 0.09
NG = 0
DB = 0
SM = 0
Doxycycline
YGL171W ROK1
Delta_r
ZShift = 0.5
NG = 0
DB = 0
SM = 0
Doxycycline
YGL171W ROK1
Delta_AUC
ZShift = 0.1
NG = 0
DB = 0
SM = 0
Doxycycline
YBR192W RIM2
Delta_L
ZShift = −0.05
NG = 0
DB = 0
SM = 0
Doxycycline
YBR192W RIM2
Delta_K
ZShift = 0.67
NG = 0
DB = 0
SM = 0
Doxycycline
YBR192W RIM2
Delta_r
ZShift = 0.32
NG = 0
DB = 0
SM = 0
Doxycycline
YBR192W RIM2
Delta_AUC
ZShift = 0.48
NG = 0
DB = 0
SM = 0
Doxycycline
YGR013W SNU71
Delta_L
ZShift = 0.28
NG = 0
DB = 0
SM = 0
Doxycycline
YGR013W SNU71
Delta_K
ZShift = 0.24
NG = 0
DB = 0
SM = 0
Doxycycline
YGR013W SNU71
Delta_r
ZShift = 0.99
NG = 0
DB = 0
SM = 0
Doxycycline
YGR013W SNU71
Delta_AUC
ZShift = 0.04
NG = 0
DB = 0
SM = 0
Doxycycline

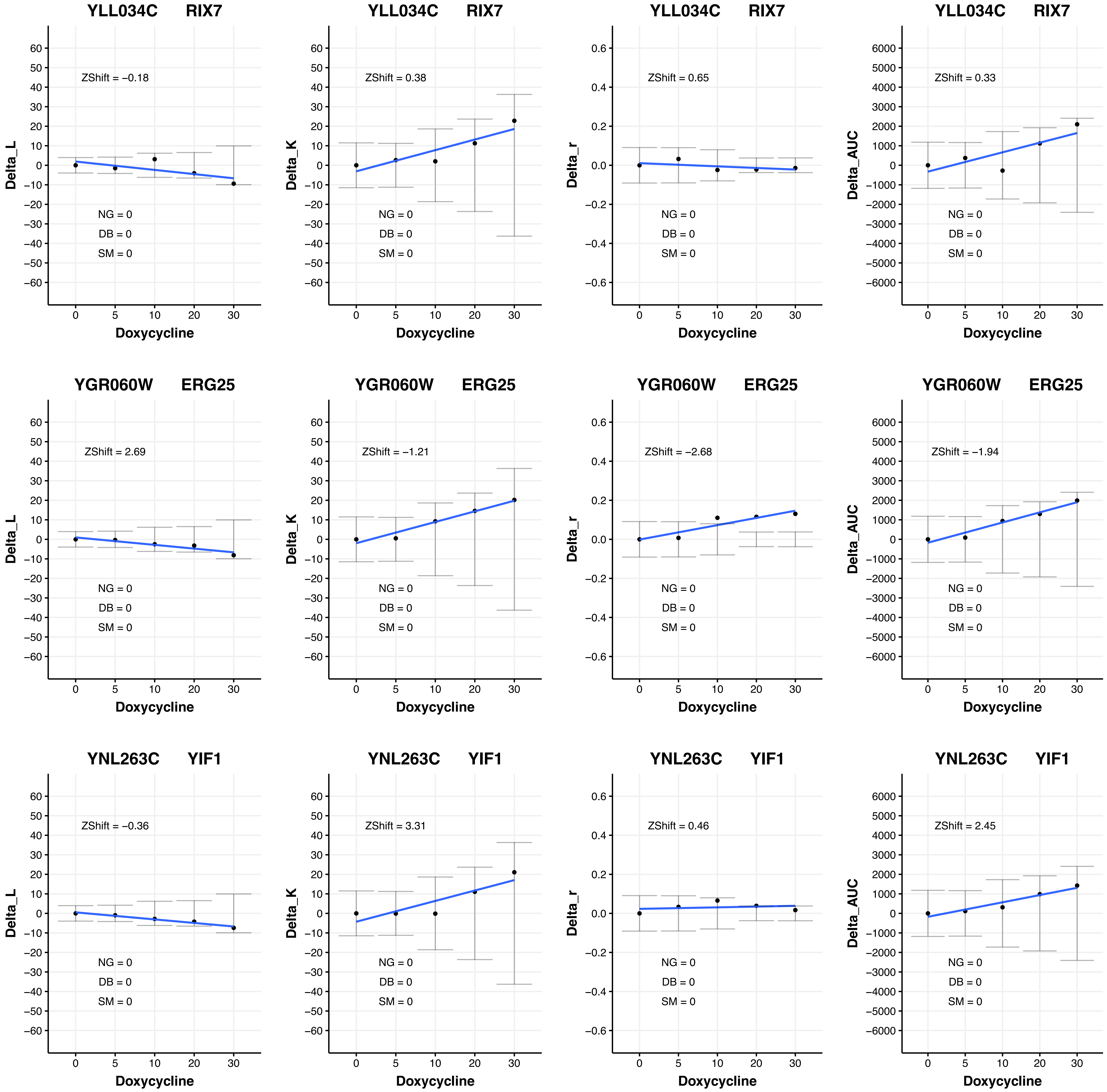

YLL034C RIX7
ZShift = −0.18
NG = 0
DB = 0
SM = 0
Delta_L
Doxycycline
YLL034C RIX7
ZShift = 0.38
NG = 0
DB = 0
SM = 0
Delta_K
Doxycycline
YLL034C RIX7
ZShift = 0.65
NG = 0
DB = 0
SM = 0
Delta_r
Doxycycline
YLL034C RIX7
ZShift = 0.33
NG = 0
DB = 0
SM = 0
Delta_AUC
Doxycycline
YGR060W ERG25
ZShift = 2.69
NG = 0
DB = 0
SM = 0
Delta_L
Doxycycline
YGR060W ERG25
ZShift = −1.21
NG = 0
DB = 0
SM = 0
Delta_K
Doxycycline
YGR060W ERG25
ZShift = −2.68
NG = 0
DB = 0
SM = 0
Delta_r
Doxycycline
YGR060W ERG25
ZShift = −1.94
NG = 0
DB = 0
SM = 0
Delta_AUC
Doxycycline
YNL263C YIF1
ZShift = −0.36
NG = 0
DB = 0
SM = 0
Delta_L
Doxycycline
YNL263C YIF1
ZShift = 3.31
NG = 0
DB = 0
SM = 0
Delta_K
Doxycycline
YNL263C YIF1
ZShift = 0.46
NG = 0
DB = 0
SM = 0
Delta_r
Doxycycline
YNL263C YIF1
ZShift = 2.45
NG = 0
DB = 0
SM = 0
Delta_AUC
Doxycycline

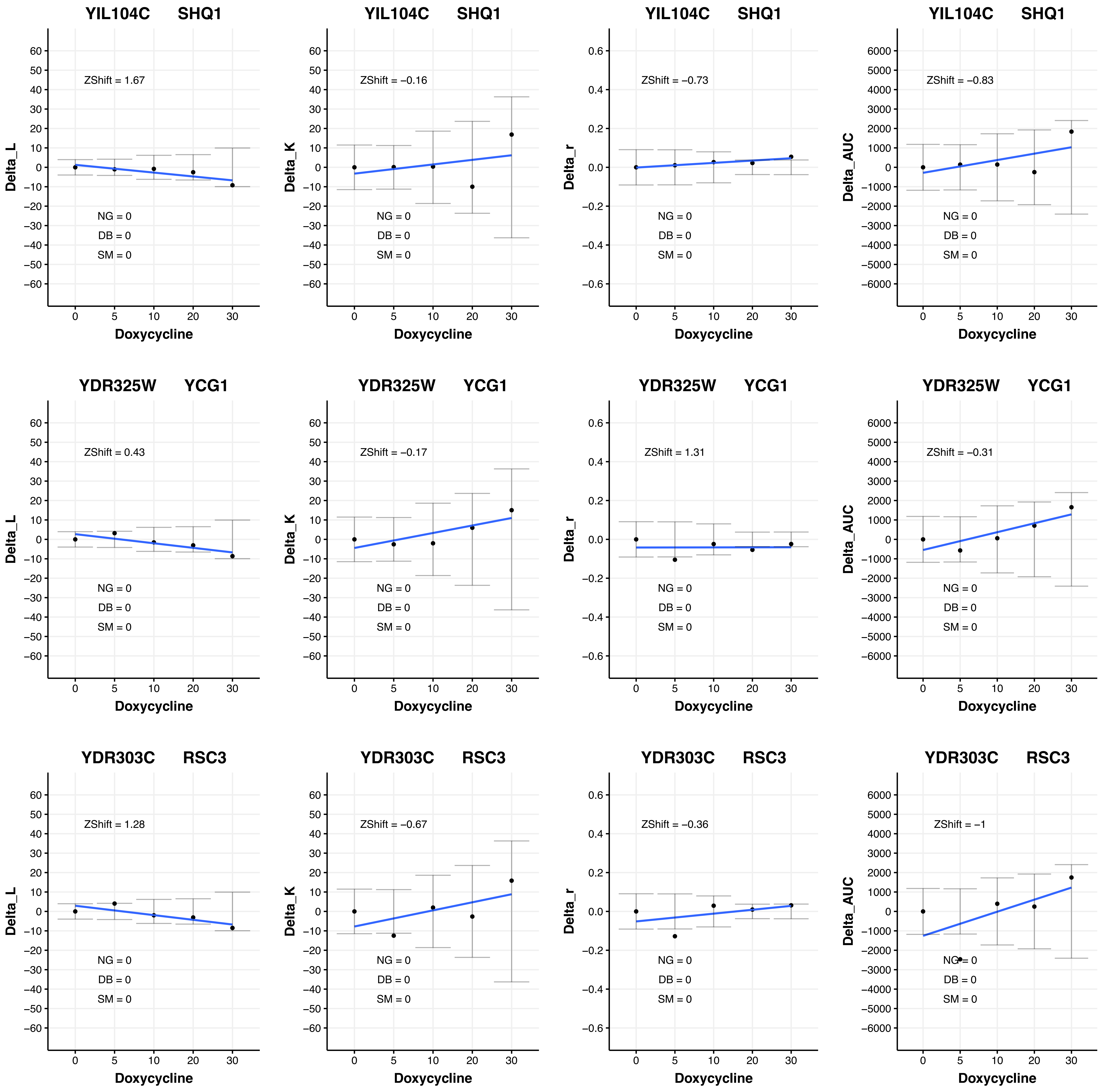
YIL104C SHQ1
ZShift = 1.67
NG = 0
DB = 0
SM = 0
Delta_L
Doxycycline
YIL104C SHQ1
ZShift = −0.16
NG = 0
DB = 0
SM = 0
Delta_K
Doxycycline
YIL104C SHQ1
ZShift = −0.73
NG = 0
DB = 0
SM = 0
Delta_r
Doxycycline
YIL104C SHQ1
ZShift = −0.83
NG = 0
DB = 0
SM = 0
Delta_AUC
Doxycycline
YDR325W YCG1
ZShift = 0.43
NG = 0
DB = 0
SM = 0
Delta_L
Doxycycline
YDR325W YCG1
ZShift = −0.17
NG = 0
DB = 0
SM = 0
Delta_K
Doxycycline
YDR325W YCG1
ZShift = 1.31
NG = 0
DB = 0
SM = 0
Delta_r
Doxycycline
YDR325W YCG1
ZShift = −0.31
NG = 0
DB = 0
SM = 0
Delta_AUC
Doxycycline
YDR303C RSC3
ZShift = 1.28
NG = 0
DB = 0
SM = 0
Delta_L
Doxycycline
YDR303C RSC3
ZShift = −0.67
NG = 0
DB = 0
SM = 0
Delta_K
Doxycycline
YDR303C RSC3
ZShift = −0.36
NG = 0
DB = 0
SM = 0
Delta_r
Doxycycline
YDR303C RSC3
ZShift = −1
NG = 0
DB = 0
SM = 0
Delta_AUC
Doxycycline

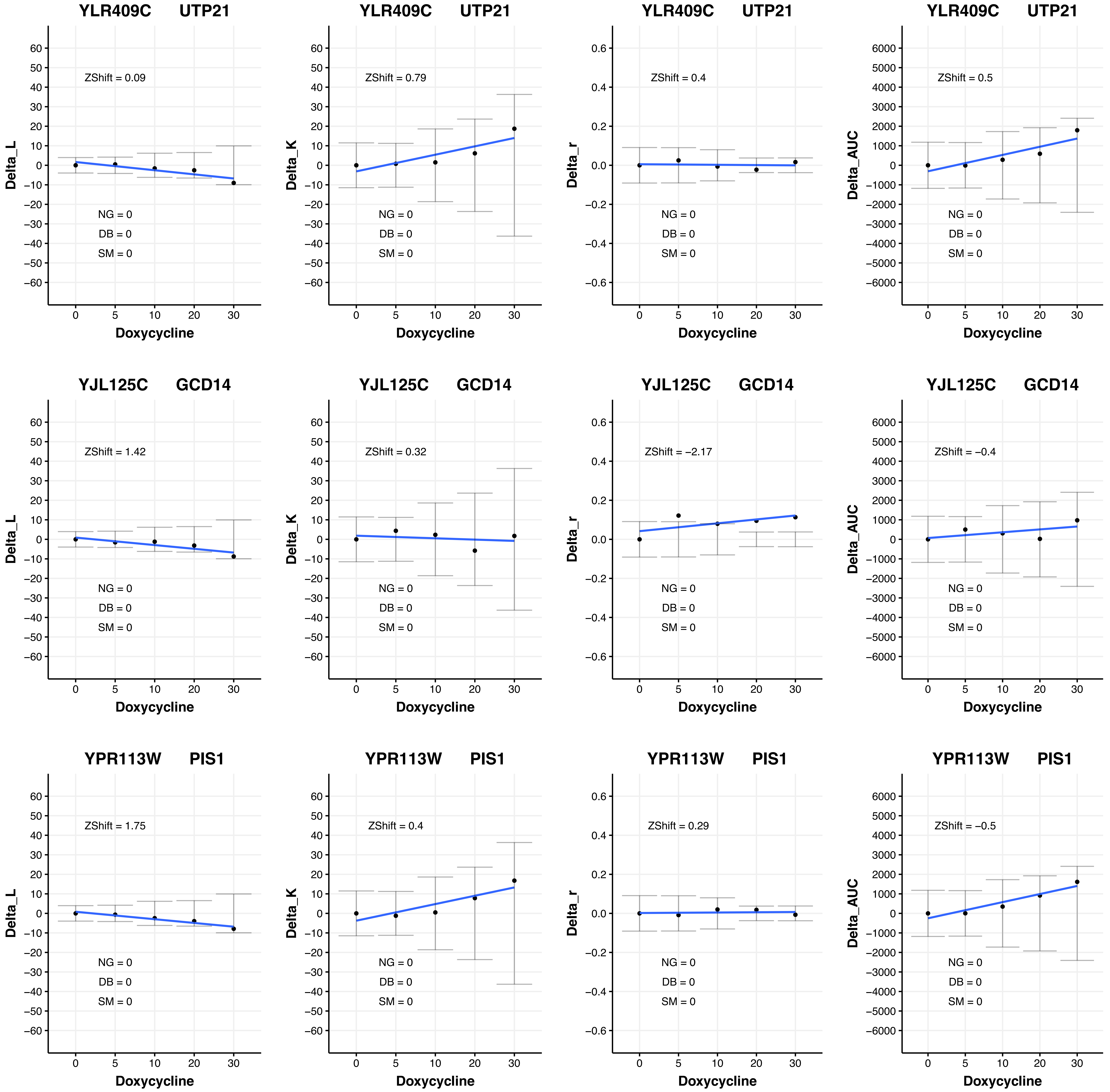

YLR409C UTP21
Delta_L
ZShift = 0.09
NG = 0
DB = 0
SM = 0
Doxycycline
YLR409C UTP21
Delta_K
ZShift = 0.79
NG = 0
DB = 0
SM = 0
Doxycycline
YLR409C UTP21
Delta_r
ZShift = 0.4
NG = 0
DB = 0
SM = 0
Doxycycline
YLR409C UTP21
Delta_AUC
ZShift = 0.5
NG = 0
DB = 0
SM = 0
Doxycycline
YJL125C GCD14
Delta_L
ZShift = 1.42
NG = 0
DB = 0
SM = 0
Doxycycline
YJL125C GCD14
Delta_K
ZShift = 0.32
NG = 0
DB = 0
SM = 0
Doxycycline
YJL125C GCD14
Delta_r
ZShift = −2.17
NG = 0
DB = 0
SM = 0
Doxycycline
YJL125C GCD14
Delta_AUC
ZShift = −0.4
NG = 0
DB = 0
SM = 0
Doxycycline
YPR113W PIS1
Delta_L
ZShift = 1.75
NG = 0
DB = 0
SM = 0
Doxycycline
YPR113W PIS1
Delta_K
ZShift = 0.4
NG = 0
DB = 0
SM = 0
Doxycycline
YPR113W PIS1
Delta_r
ZShift = 0.29
NG = 0
DB = 0
SM = 0
Doxycycline
YPR113W PIS1
Delta_AUC
ZShift = −0.5
NG = 0
DB = 0
SM = 0
Doxycycline

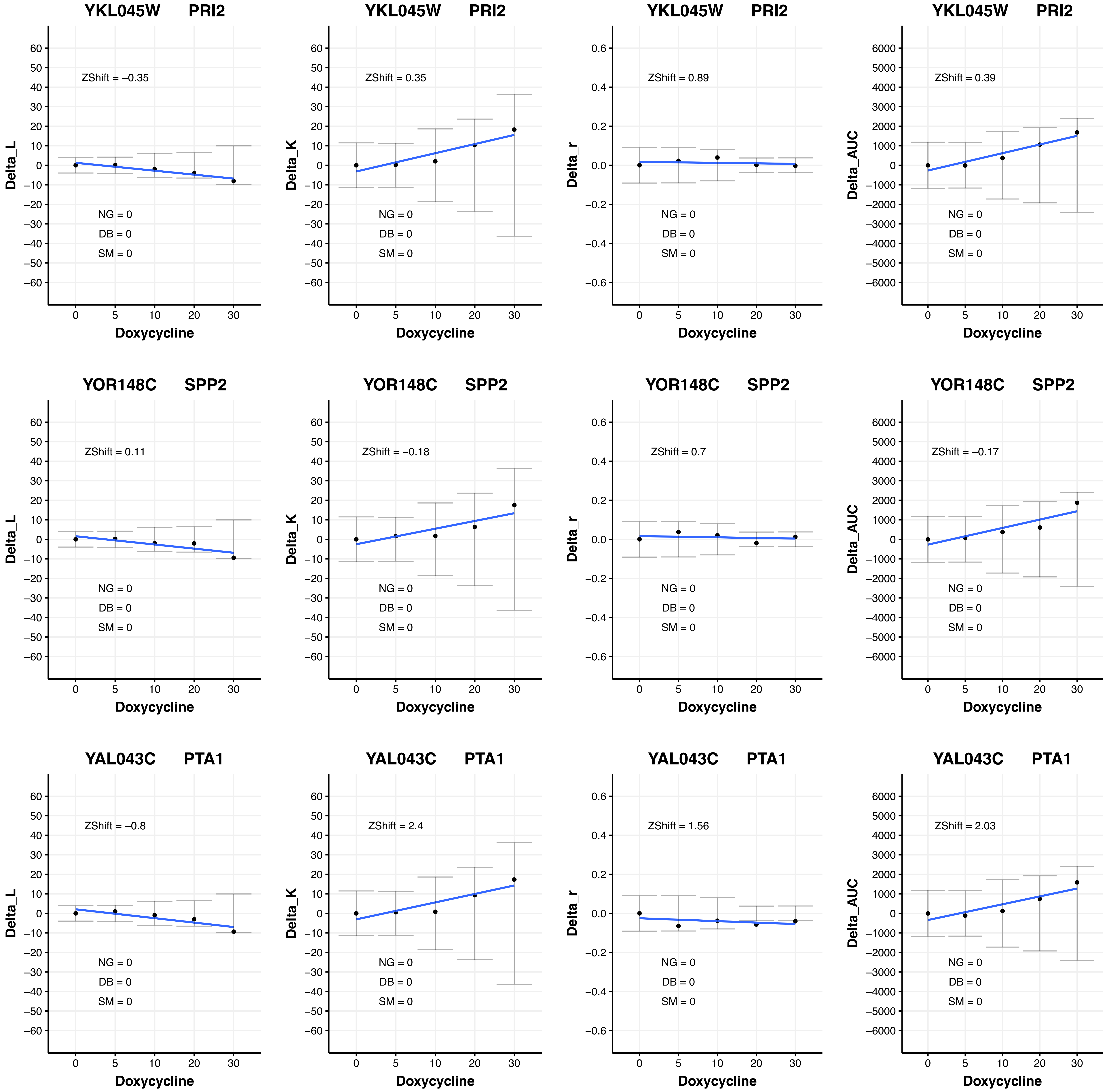
YKL045W PRI2
ZShift = −0.35
NG = 0
DB = 0
SM = 0
Delta_L
Doxycycline
YKL045W PRI2
ZShift = 0.35
NG = 0
DB = 0
SM = 0
Delta_K
Doxycycline
YKL045W PRI2
ZShift = 0.89
NG = 0
DB = 0
SM = 0
Delta_r
Doxycycline
YKL045W PRI2
ZShift = 0.39
NG = 0
DB = 0
SM = 0
Delta_AUC
Doxycycline
YOR148C SPP2
ZShift = 0.11
NG = 0
DB = 0
SM = 0
Delta_L
Doxycycline
YOR148C SPP2
ZShift = −0.18
NG = 0
DB = 0
SM = 0
Delta_K
Doxycycline
YOR148C SPP2
ZShift = 0.7
NG = 0
DB = 0
SM = 0
Delta_r
Doxycycline
YOR148C SPP2
ZShift = −0.17
NG = 0
DB = 0
SM = 0
Delta_AUC
Doxycycline
YAL043C PTA1
ZShift = −0.8
NG = 0
DB = 0
SM = 0
Delta_L
Doxycycline
YAL043C PTA1
ZShift = 2.4
NG = 0
DB = 0
SM = 0
Delta_K
Doxycycline
YAL043C PTA1
ZShift = 1.56
NG = 0
DB = 0
SM = 0
Delta_r
Doxycycline
YAL043C PTA1
ZShift = 2.03
NG = 0
DB = 0
SM = 0
Delta_AUC
Doxycycline

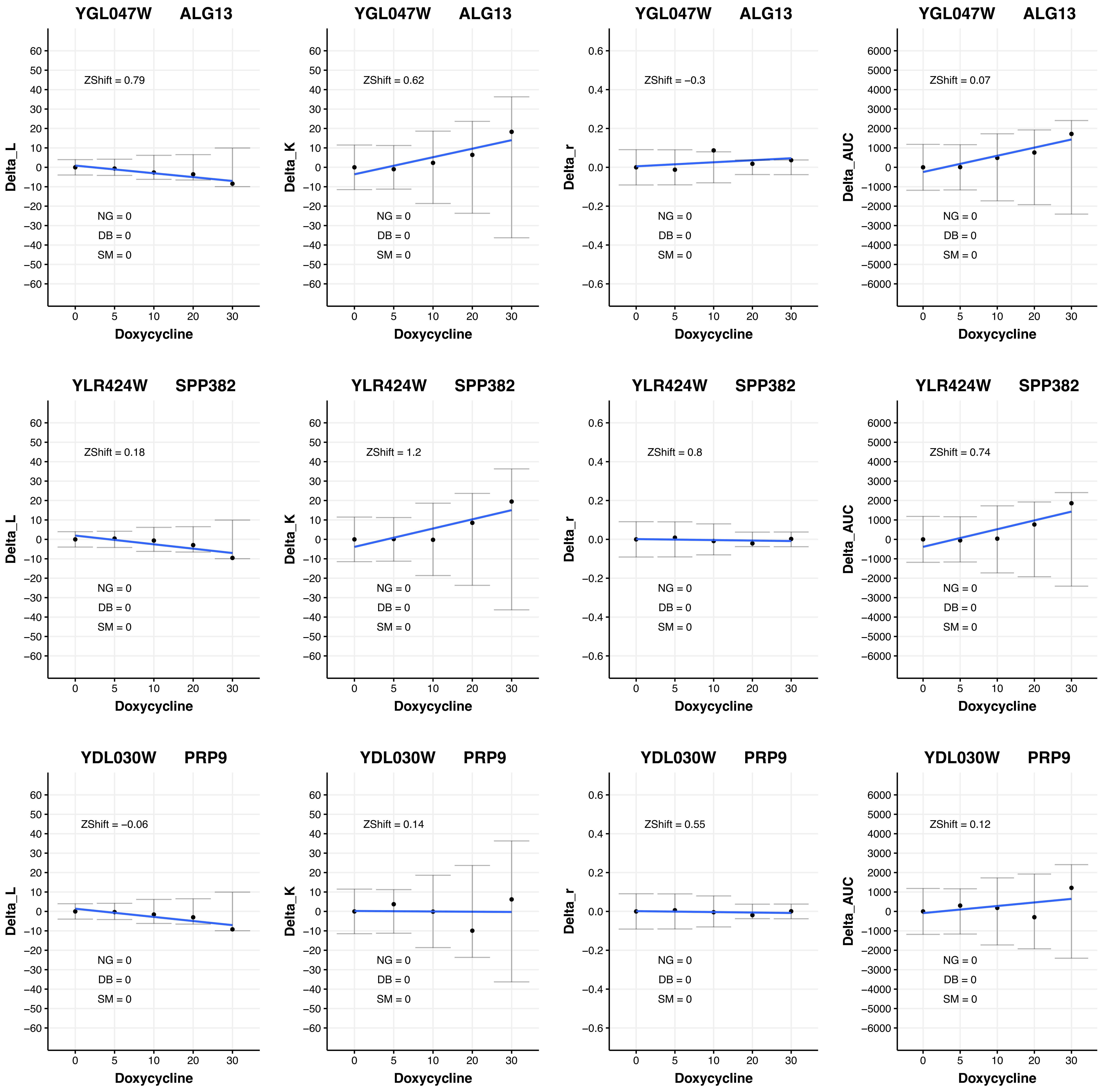
YGL047W ALG13
ZShift = 0.79
Delta_L
NG = 0
DB = 0
SM = 0
Doxycycline
YGL047W ALG13
ZShift = 0.62
Delta_K
NG = 0
DB = 0
SM = 0
Doxycycline
YGL047W ALG13
ZShift = −0.3
Delta_r
NG = 0
DB = 0
SM = 0
Doxycycline
YGL047W ALG13
ZShift = 0.07
Delta_AUC
NG = 0
DB = 0
SM = 0
Doxycycline
YLR424W SPP382
ZShift = 0.18
Delta_L
NG = 0
DB = 0
SM = 0
Doxycycline
YLR424W SPP382
ZShift = 1.2
Delta_K
NG = 0
DB = 0
SM = 0
Doxycycline
YLR424W SPP382
ZShift = 0.8
Delta_r
NG = 0
DB = 0
SM = 0
Doxycycline
YLR424W SPP382
ZShift = 0.74
Delta_AUC
NG = 0
DB = 0
SM = 0
Doxycycline
YDL030W PRP9
ZShift = −0.06
Delta_L
NG = 0
DB = 0
SM = 0
Doxycycline
YDL030W PRP9
ZShift = 0.14
Delta_K
NG = 0
DB = 0
SM = 0
Doxycycline
YDL030W PRP9
ZShift = 0.55
Delta_r
NG = 0
DB = 0
SM = 0
Doxycycline
YDL030W PRP9
ZShift = 0.12
Delta_AUC
NG = 0
DB = 0
SM = 0
Doxycycline

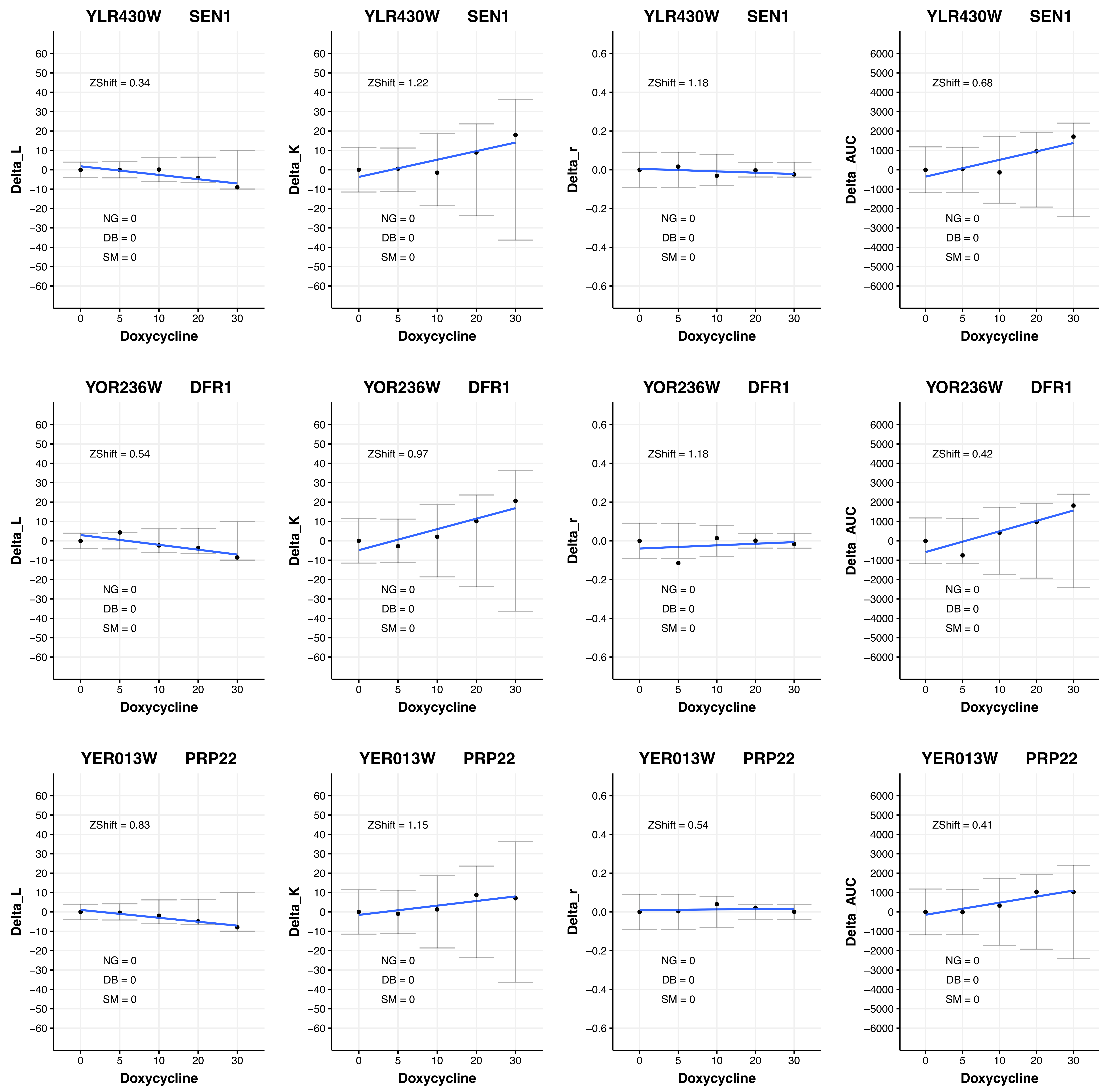
YLR430W SEN1
ZShift = 0.34
NG = 0
DB = 0
SM = 0
Delta_L
Doxycycline
YLR430W SEN1
ZShift = 1.22
NG = 0
DB = 0
SM = 0
Delta_K
Doxycycline
YLR430W SEN1
ZShift = 1.18
NG = 0
DB = 0
SM = 0
Delta_r
Doxycycline
YLR430W SEN1
ZShift = 0.68
NG = 0
DB = 0
SM = 0
Delta_AUC
Doxycycline
YOR236W DFR1
ZShift = 0.54
NG = 0
DB = 0
SM = 0
Delta_L
Doxycycline
YOR236W DFR1
ZShift = 0.97
NG = 0
DB = 0
SM = 0
Delta_K
Doxycycline
YOR236W DFR1
ZShift = 1.18
NG = 0
DB = 0
SM = 0
Delta_r
Doxycycline
YOR236W DFR1
ZShift = 0.42
NG = 0
DB = 0
SM = 0
Delta_AUC
Doxycycline
YER013W PRP22
ZShift = 0.83
NG = 0
DB = 0
SM = 0
Delta_L
Doxycycline
YER013W PRP22
ZShift = 1.15
NG = 0
DB = 0
SM = 0
Delta_K
Doxycycline
YER013W PRP22
ZShift = 0.54
NG = 0
DB = 0
SM = 0
Delta_r
Doxycycline
YER013W PRP22
ZShift = 0.41
NG = 0
DB = 0
SM = 0
Delta_AUC
Doxycycline

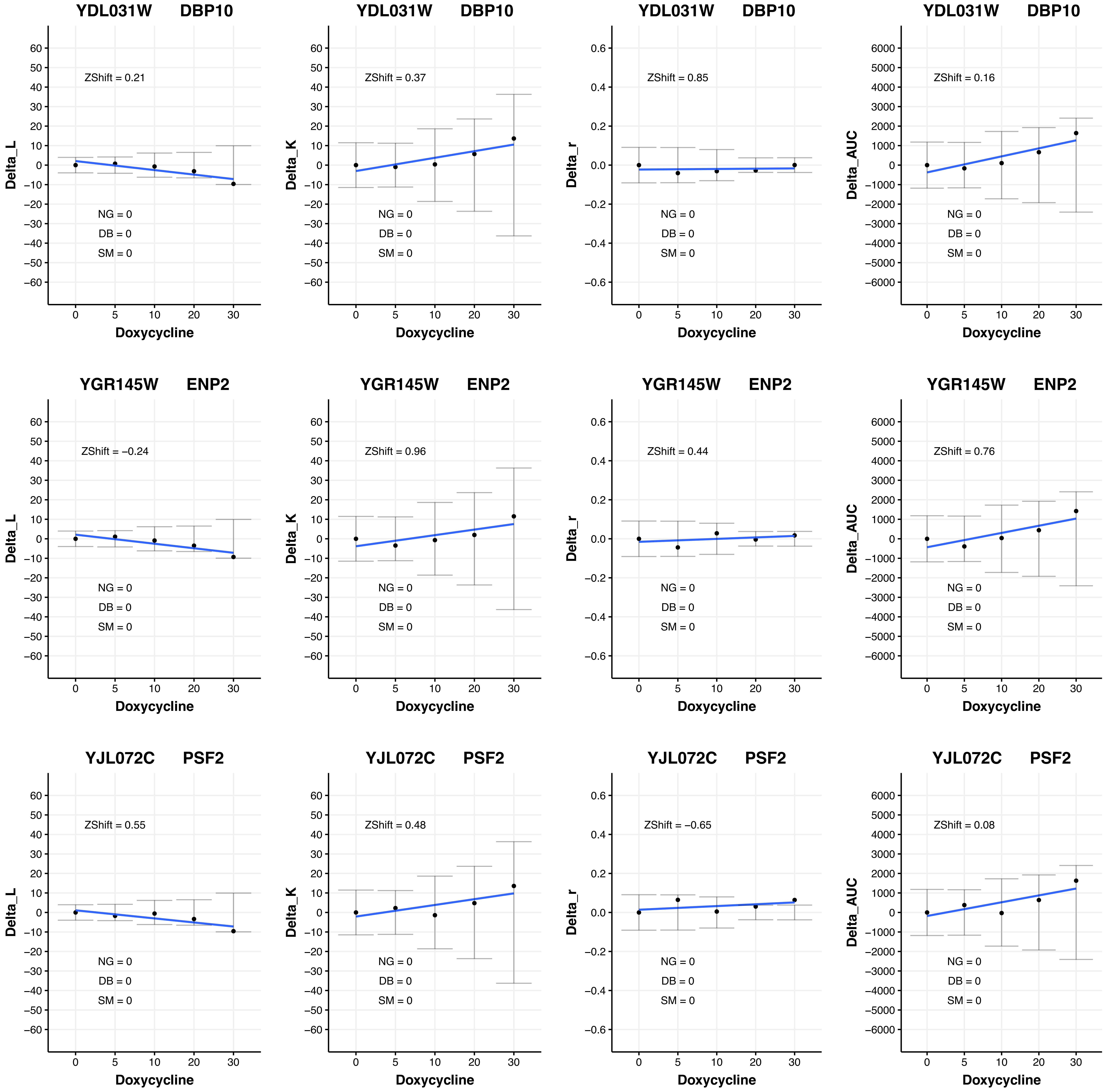
YDL031W DBP10
ZShift = 0.21
NG = 0
DB = 0
SM = 0
Delta_L
Doxycycline
YDL031W DBP10
ZShift = 0.37
NG = 0
DB = 0
SM = 0
Delta_K
Doxycycline
YDL031W DBP10
ZShift = 0.85
NG = 0
DB = 0
SM = 0
Delta_r
Doxycycline
YDL031W DBP10
ZShift = 0.16
NG = 0
DB = 0
SM = 0
Delta_AUC
Doxycycline
YGR145W ENP2
ZShift = −0.24
NG = 0
DB = 0
SM = 0
Delta_L
Doxycycline
YGR145W ENP2
ZShift = 0.96
NG = 0
DB = 0
SM = 0
Delta_K
Doxycycline
YGR145W ENP2
ZShift = 0.44
NG = 0
DB = 0
SM = 0
Delta_r
Doxycycline
YGR145W ENP2
ZShift = 0.76
NG = 0
DB = 0
SM = 0
Delta_AUC
Doxycycline
YJL072C PSF2
ZShift = 0.55
NG = 0
DB = 0
SM = 0
Delta_L
Doxycycline
YJL072C PSF2
ZShift = 0.48
NG = 0
DB = 0
SM = 0
Delta_K
Doxycycline
YJL072C PSF2
ZShift = −0.65
NG = 0
DB = 0
SM = 0
Delta_r
Doxycycline
YJL072C PSF2
ZShift = 0.08
NG = 0
DB = 0
SM = 0
Delta_AUC
Doxycycline

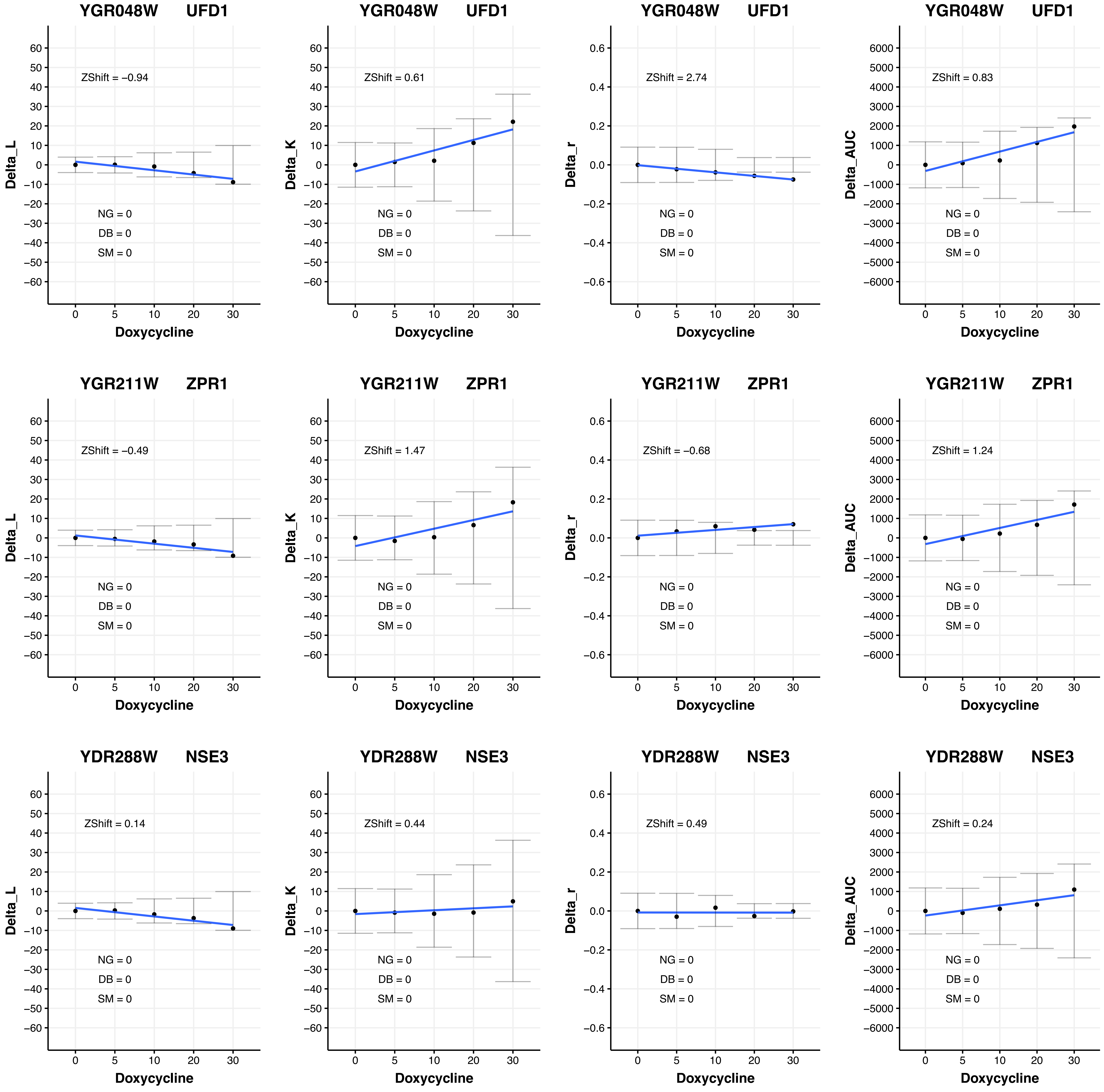
YGR048W UFD1
ZShift = −0.94
NG = 0
DB = 0
SM = 0
Delta_L
Doxycycline
YGR048W UFD1
ZShift = 0.61
NG = 0
DB = 0
SM = 0
Delta_K
Doxycycline
YGR048W UFD1
ZShift = 2.74
NG = 0
DB = 0
SM = 0
Delta_r
Doxycycline
YGR048W UFD1
ZShift = 0.83
NG = 0
DB = 0
SM = 0
Delta_AUC
Doxycycline
YGR211W ZPR1
ZShift = −0.49
NG = 0
DB = 0
SM = 0
Delta_L
Doxycycline
YGR211W ZPR1
ZShift = 1.47
NG = 0
DB = 0
SM = 0
Delta_K
Doxycycline
YGR211W ZPR1
ZShift = −0.68
NG = 0
DB = 0
SM = 0
Delta_r
Doxycycline
YGR211W ZPR1
ZShift = 1.24
NG = 0
DB = 0
SM = 0
Delta_AUC
Doxycycline
YDR288W NSE3
ZShift = 0.14
NG = 0
DB = 0
SM = 0
Delta_L
Doxycycline
YDR288W NSE3
ZShift = 0.44
NG = 0
DB = 0
SM = 0
Delta_K
Doxycycline
YDR288W NSE3
ZShift = 0.49
NG = 0
DB = 0
SM = 0
Delta_r
Doxycycline
YDR288W NSE3
ZShift = 0.24
NG = 0
DB = 0
SM = 0
Delta_AUC
Doxycycline

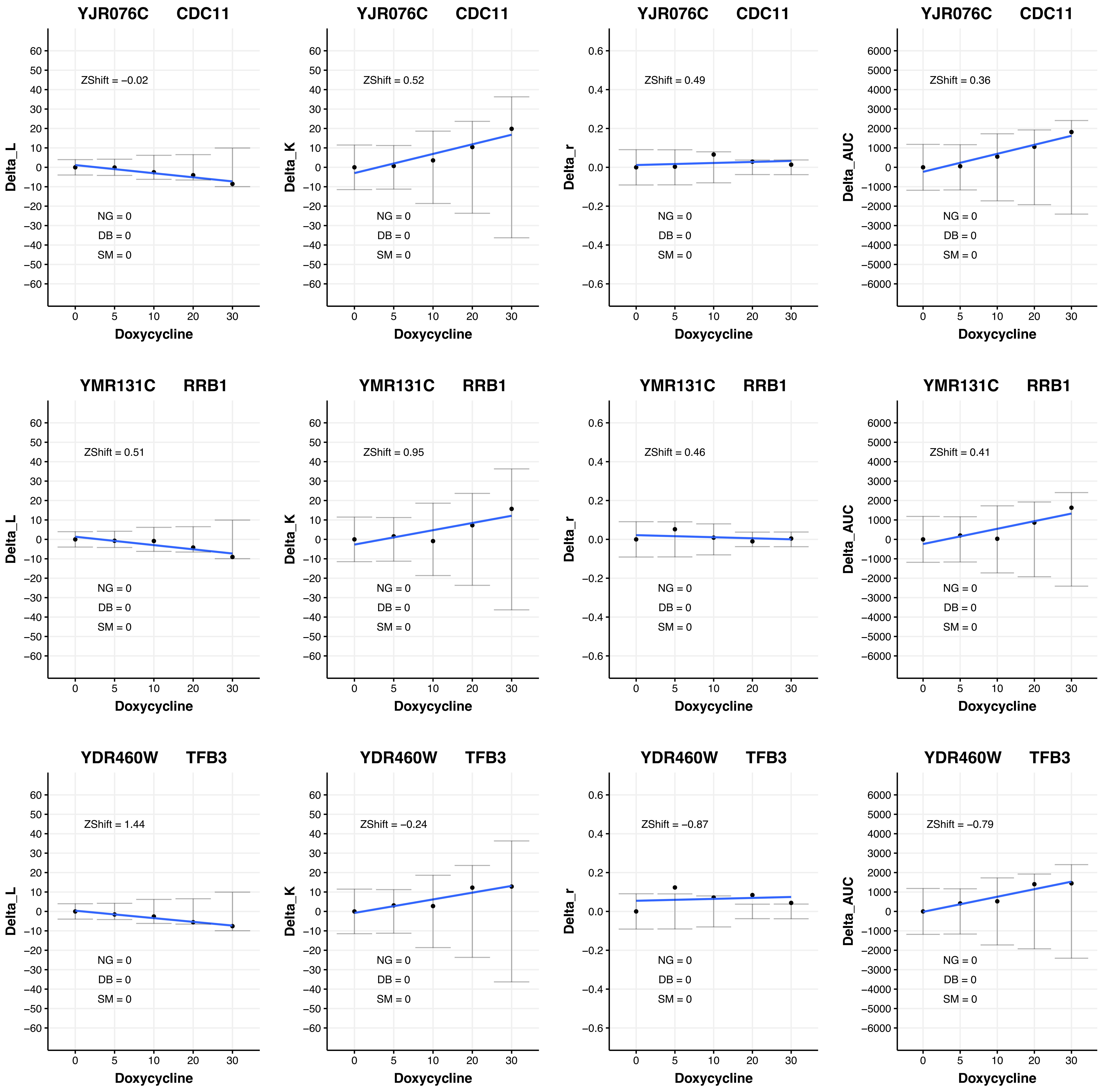
YJR076C CDC11
ZShift = −0.02
NG = 0
DB = 0
SM = 0
Delta_L
Doxycycline
YJR076C CDC11
ZShift = 0.52
NG = 0
DB = 0
SM = 0
Delta_K
Doxycycline
YJR076C CDC11
ZShift = 0.49
NG = 0
DB = 0
SM = 0
Delta_r
Doxycycline
YJR076C CDC11
ZShift = 0.36
NG = 0
DB = 0
SM = 0
Delta_AUC
Doxycycline
YMR131C RRB1
ZShift = 0.51
NG = 0
DB = 0
SM = 0
Delta_L
Doxycycline
YMR131C RRB1
ZShift = 0.95
NG = 0
DB = 0
SM = 0
Delta_K
Doxycycline
YMR131C RRB1
ZShift = 0.46
NG = 0
DB = 0
SM = 0
Delta_r
Doxycycline
YMR131C RRB1
ZShift = 0.41
NG = 0
DB = 0
SM = 0
Delta_AUC
Doxycycline
YDR460W TFB3
ZShift = 1.44
NG = 0
DB = 0
SM = 0
Delta_L
Doxycycline
YDR460W TFB3
ZShift = −0.24
NG = 0
DB = 0
SM = 0
Delta_K
Doxycycline
YDR460W TFB3
ZShift = −0.87
NG = 0
DB = 0
SM = 0
Delta_r
Doxycycline
YDR460W TFB3
ZShift = −0.79
NG = 0
DB = 0
SM = 0
Delta_AUC
Doxycycline

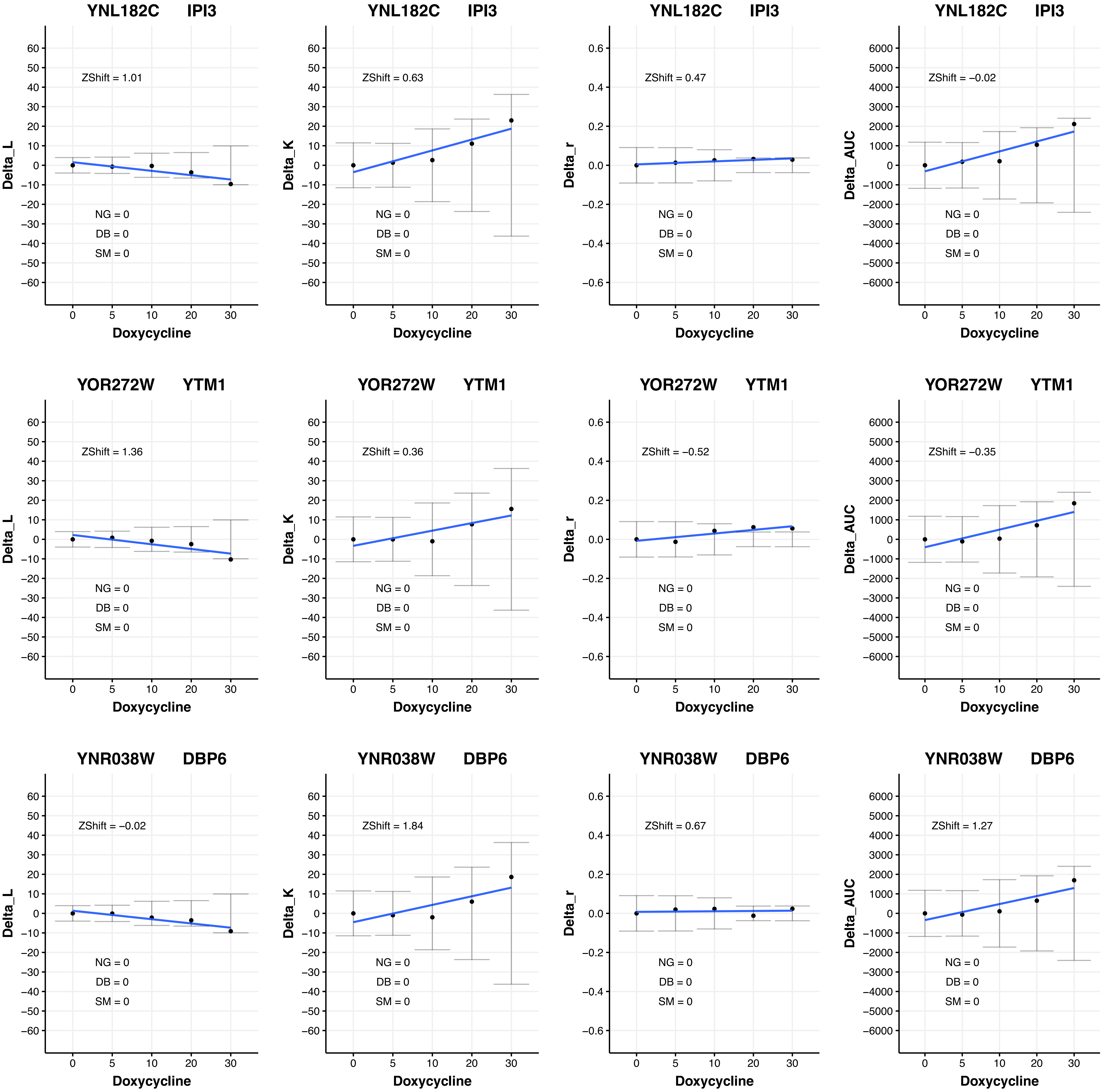
YNL182C IPI3
ZShift = 1.01
Delta_L
NG = 0
DB = 0
SM = 0
Doxycycline
YNL182C IPI3
ZShift = 0.63
Delta_K
NG = 0
DB = 0
SM = 0
Doxycycline
YNL182C IPI3
ZShift = 0.47
Delta_r
NG = 0
DB = 0
SM = 0
Doxycycline
YNL182C IPI3
ZShift = −0.02
Delta_AUC
NG = 0
DB = 0
SM = 0
Doxycycline
YOR272W YTM1
ZShift = 1.36
Delta_L
NG = 0
DB = 0
SM = 0
Doxycycline
YOR272W YTM1
ZShift = 0.36
Delta_K
NG = 0
DB = 0
SM = 0
Doxycycline
YOR272W YTM1
ZShift = −0.52
Delta_r
NG = 0
DB = 0
SM = 0
Doxycycline
YOR272W YTM1
ZShift = −0.35
Delta_AUC
NG = 0
DB = 0
SM = 0
Doxycycline
YNR038W DBP6
ZShift = −0.02
Delta_L
NG = 0
DB = 0
SM = 0
Doxycycline
YNR038W DBP6
ZShift = 1.84
Delta_K
NG = 0
DB = 0
SM = 0
Doxycycline
YNR038W DBP6
ZShift = 0.67
Delta_r
NG = 0
DB = 0
SM = 0
Doxycycline
YNR038W DBP6
ZShift = 1.27
Delta_AUC
NG = 0
DB = 0
SM = 0
Doxycycline

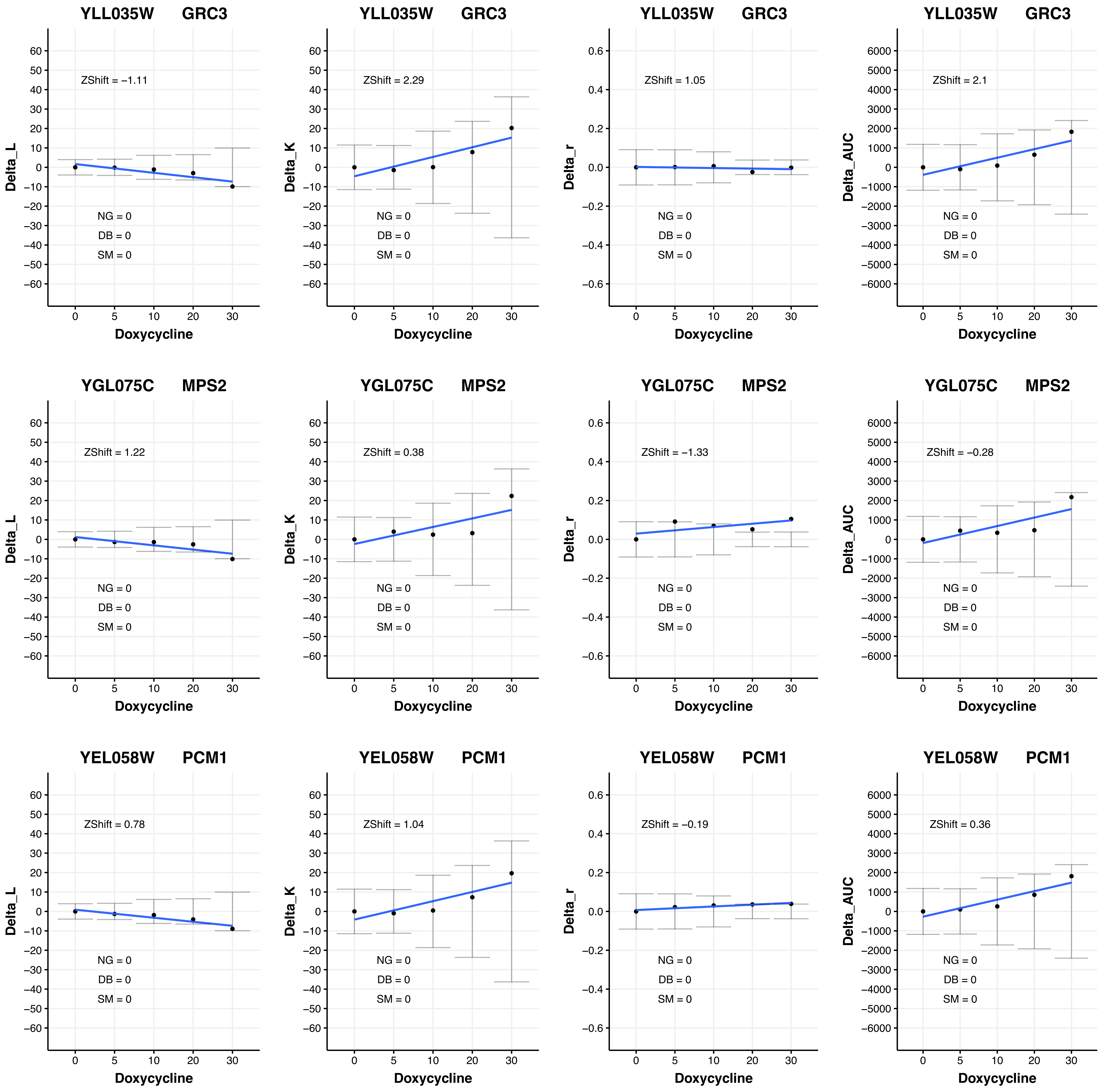

YLL035W GRC3
ZShift = −1.11
NG = 0
DB = 0
SM = 0
Delta_L
Doxycycline
YLL035W GRC3
ZShift = 2.29
NG = 0
DB = 0
SM = 0
Delta_K
Doxycycline
YLL035W GRC3
ZShift = 1.05
NG = 0
DB = 0
SM = 0
Delta_r
Doxycycline
YLL035W GRC3
ZShift = 2.1
NG = 0
DB = 0
SM = 0
Delta_AUC
Doxycycline
YGL075C MPS2
ZShift = 1.22
NG = 0
DB = 0
SM = 0
Delta_L
Doxycycline
YGL075C MPS2
ZShift = 0.38
NG = 0
DB = 0
SM = 0
Delta_K
Doxycycline
YGL075C MPS2
ZShift = −1.33
NG = 0
DB = 0
SM = 0
Delta_r
Doxycycline
YGL075C MPS2
ZShift = −0.28
NG = 0
DB = 0
SM = 0
Delta_AUC
Doxycycline
YEL058W PCM1
ZShift = 0.78
NG = 0
DB = 0
SM = 0
Delta_L
Doxycycline
YEL058W PCM1
ZShift = 1.04
NG = 0
DB = 0
SM = 0
Delta_K
Doxycycline
YEL058W PCM1
ZShift = −0.19
NG = 0
DB = 0
SM = 0
Delta_r
Doxycycline
YEL058W PCM1
ZShift = 0.36
NG = 0
DB = 0
SM = 0
Delta_AUC
Doxycycline

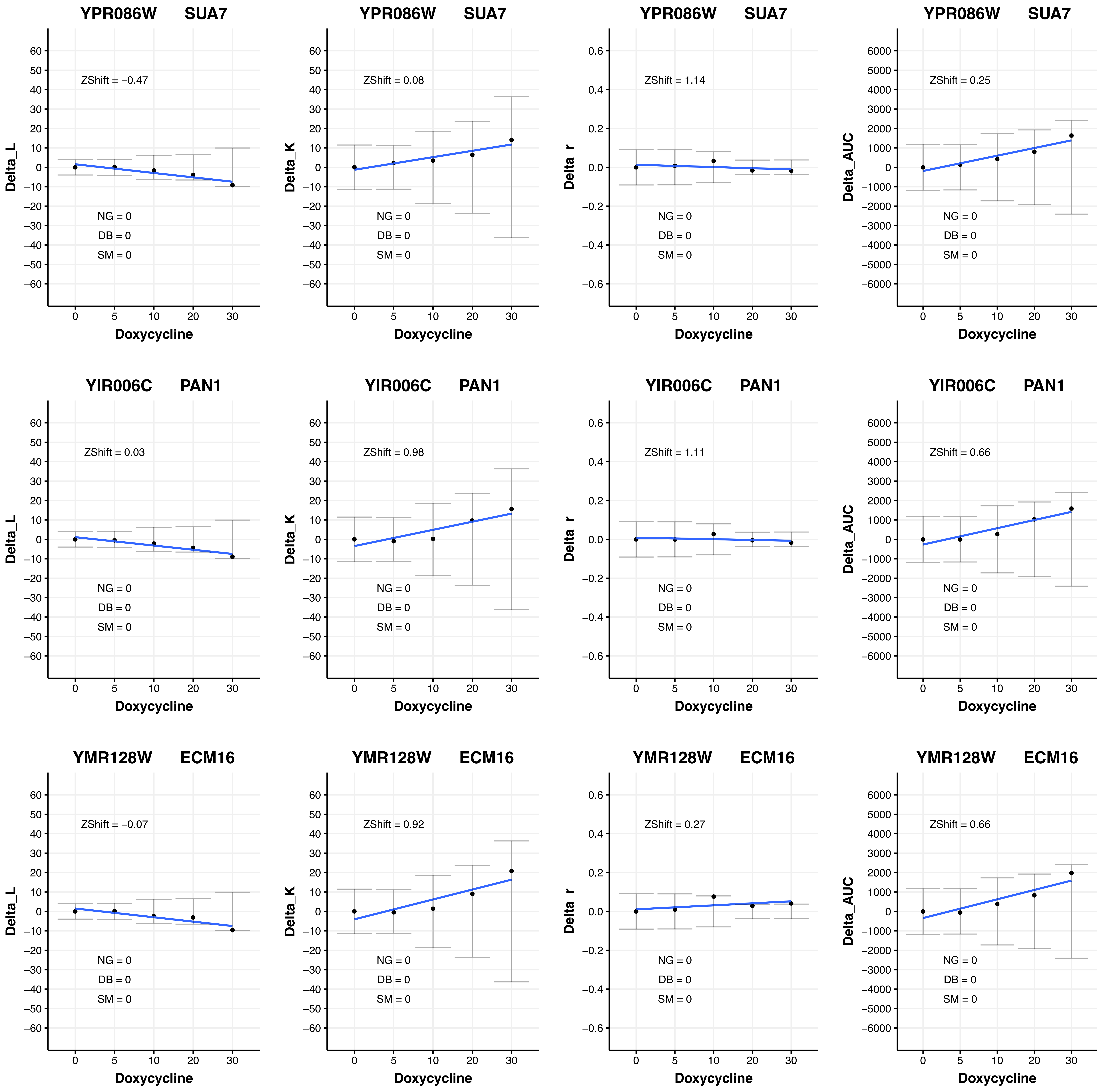

YPR086W SUA7
ZShift = −0.47
Delta_L
NG = 0
DB = 0
SM = 0
Doxycycline
YPR086W SUA7
ZShift = 0.08
Delta_K
NG = 0
DB = 0
SM = 0
Doxycycline
YPR086W SUA7
ZShift = 1.14
Delta_r
NG = 0
DB = 0
SM = 0
Doxycycline
YPR086W SUA7
ZShift = 0.25
Delta_AUC
NG = 0
DB = 0
SM = 0
Doxycycline
YIR006C PAN1
ZShift = 0.03
Delta_L
NG = 0
DB = 0
SM = 0
Doxycycline
YIR006C PAN1
ZShift = 0.98
Delta_K
NG = 0
DB = 0
SM = 0
Doxycycline
YIR006C PAN1
ZShift = 1.11
Delta_r
NG = 0
DB = 0
SM = 0
Doxycycline
YIR006C PAN1
ZShift = 0.66
Delta_AUC
NG = 0
DB = 0
SM = 0
Doxycycline
YMR128W ECM16
ZShift = −0.07
Delta_L
NG = 0
DB = 0
SM = 0
Doxycycline
YMR128W ECM16
ZShift = 0.92
Delta_K
NG = 0
DB = 0
SM = 0
Doxycycline
YMR128W ECM16
ZShift = 0.27
Delta_r
NG = 0
DB = 0
SM = 0
Doxycycline
YMR128W ECM16
ZShift = 0.66
Delta_AUC
NG = 0
DB = 0
SM = 0
Doxycycline

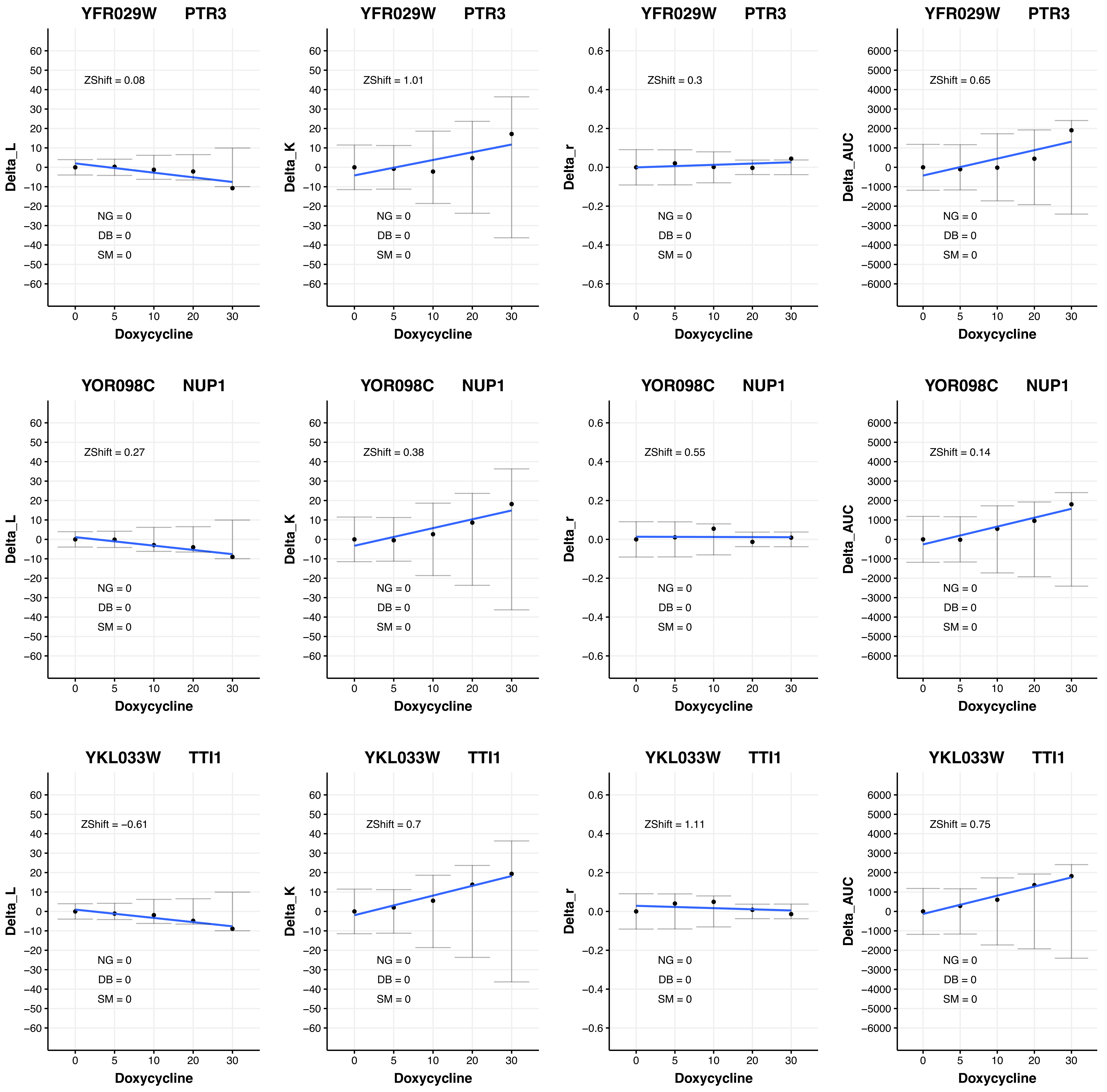

YFR029W PTR3
ZShift = 0.08
NG = 0
DB = 0
SM = 0
Delta_L
Doxycycline
YFR029W PTR3
ZShift = 1.01
NG = 0
DB = 0
SM = 0
Delta_K
Doxycycline
YFR029W PTR3
ZShift = 0.3
NG = 0
DB = 0
SM = 0
Delta_r
Doxycycline
YFR029W PTR3
ZShift = 0.65
NG = 0
DB = 0
SM = 0
Delta_AUC
Doxycycline
YOR098C NUP1
ZShift = 0.27
NG = 0
DB = 0
SM = 0
Delta_L
Doxycycline
YOR098C NUP1
ZShift = 0.38
NG = 0
DB = 0
SM = 0
Delta_K
Doxycycline
YOR098C NUP1
ZShift = 0.55
NG = 0
DB = 0
SM = 0
Delta_r
Doxycycline
YOR098C NUP1
ZShift = 0.14
NG = 0
DB = 0
SM = 0
Delta_AUC
Doxycycline
YKL033W TTI1
ZShift = −0.61
NG = 0
DB = 0
SM = 0
Delta_L
Doxycycline
YKL033W TTI1
ZShift = 0.7
NG = 0
DB = 0
SM = 0
Delta_K
Doxycycline
YKL033W TTI1
ZShift = 1.11
NG = 0
DB = 0
SM = 0
Delta_r
Doxycycline
YKL033W TTI1
ZShift = 0.75
NG = 0
DB = 0
SM = 0
Delta_AUC
Doxycycline

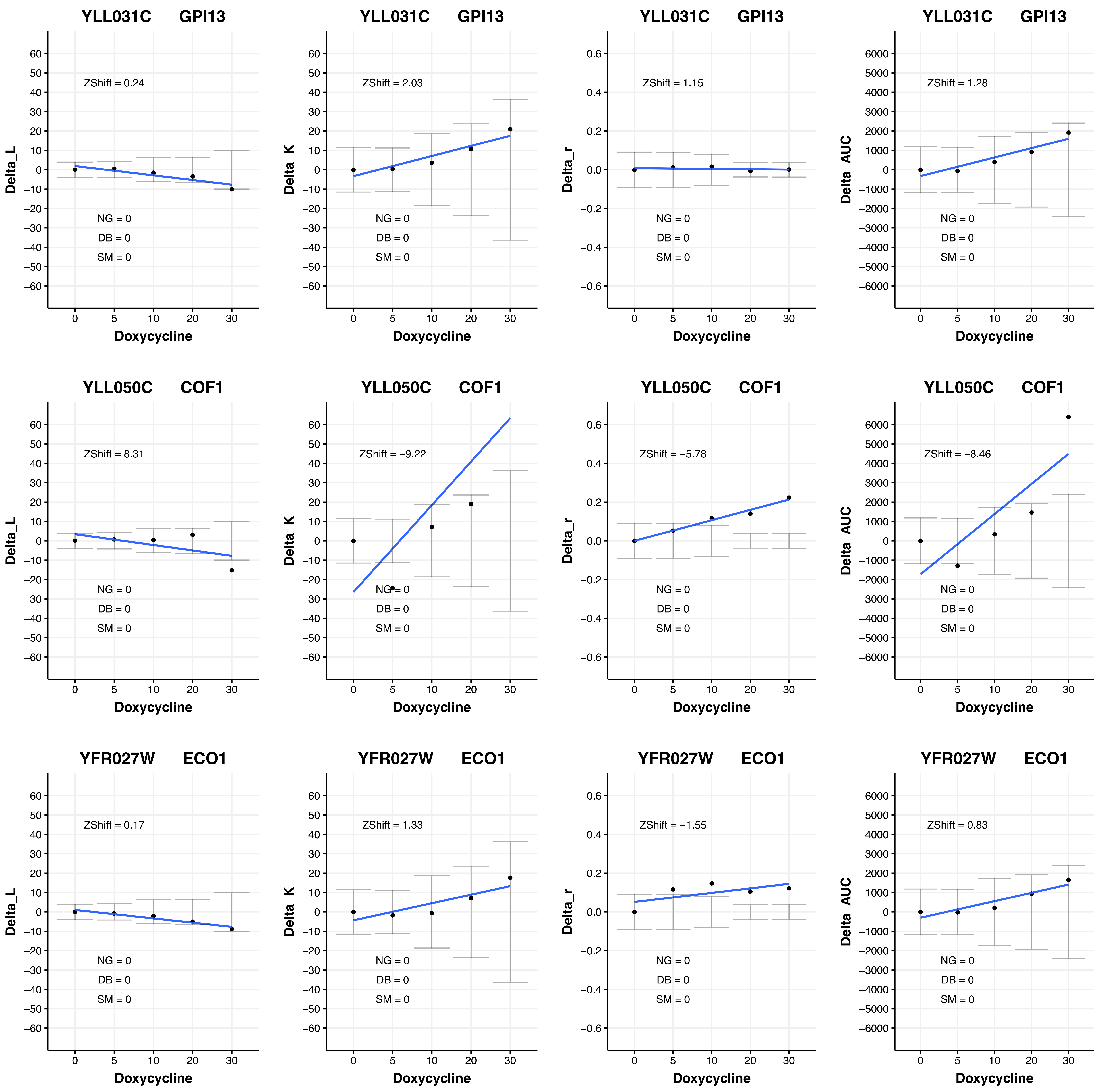
YLL031C GPI13
ZShift = 0.24
NG = 0
DB = 0
SM = 0
Delta_L
Doxycycline
YLL031C GPI13
ZShift = 2.03
NG = 0
DB = 0
SM = 0
Delta_K
Doxycycline
YLL031C GPI13
ZShift = 1.15
NG = 0
DB = 0
SM = 0
Delta_r
Doxycycline
YLL031C GPI13
ZShift = 1.28
NG = 0
DB = 0
SM = 0
Delta_AUC
Doxycycline
YLL050C COF1
ZShift = 8.31
NG = 0
DB = 0
SM = 0
Delta_L
Doxycycline
YLL050C COF1
ZShift = −9.22
NG = 0
DB = 0
SM = 0
Delta_K
Doxycycline
YLL050C COF1
ZShift = −5.78
NG = 0
DB = 0
SM = 0
Delta_r
Doxycycline
YLL050C COF1
ZShift = −8.46
NG = 0
DB = 0
SM = 0
Delta_AUC
Doxycycline
YFR027W ECO1
ZShift = 0.17
NG = 0
DB = 0
SM = 0
Delta_L
Doxycycline
YFR027W ECO1
ZShift = 1.33
NG = 0
DB = 0
SM = 0
Delta_K
Doxycycline
YFR027W ECO1
ZShift = −1.55
NG = 0
DB = 0
SM = 0
Delta_r
Doxycycline
YFR027W ECO1
ZShift = 0.83
NG = 0
DB = 0
SM = 0
Delta_AUC
Doxycycline

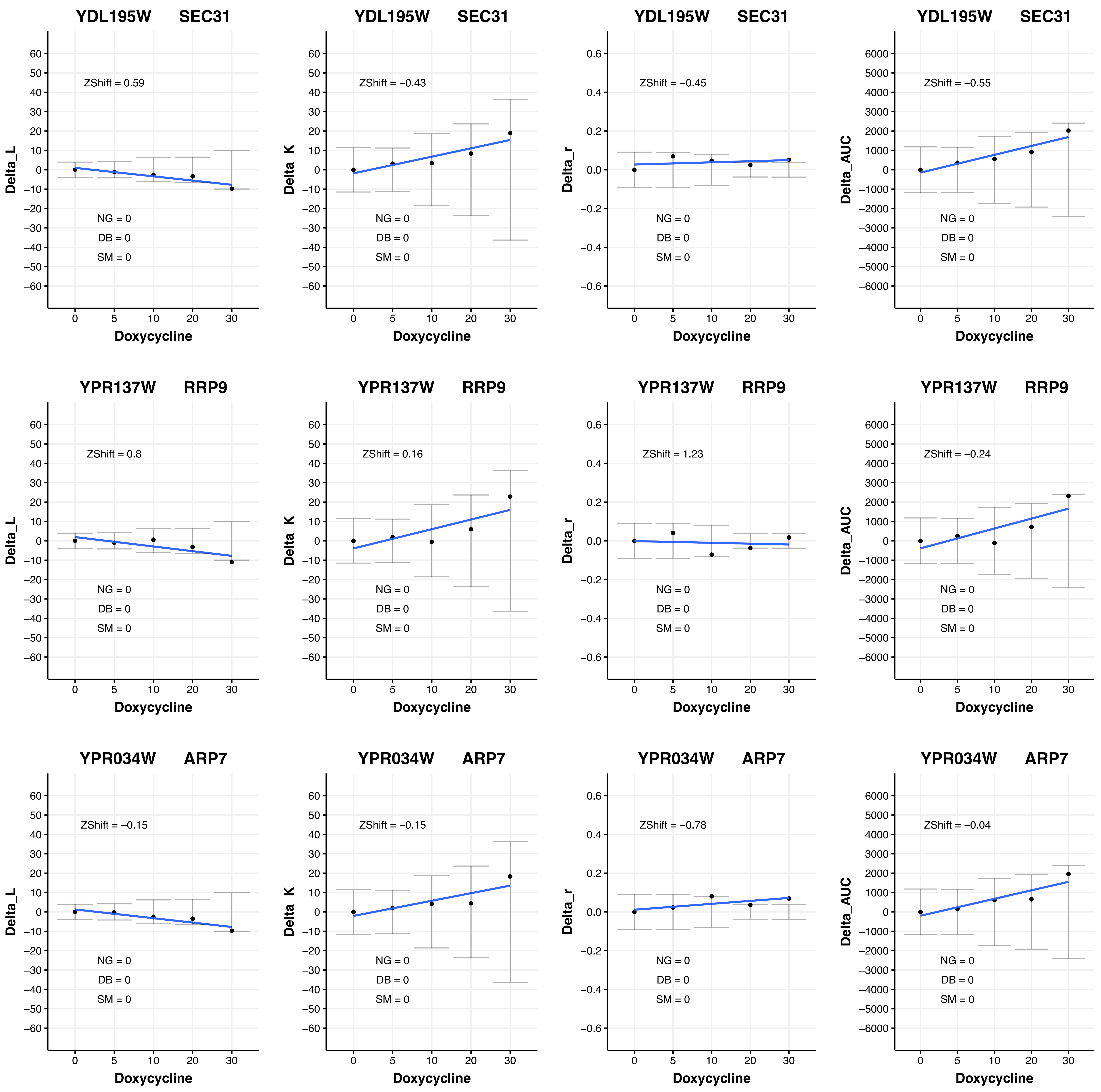
YDL195W SEC31
ZShift = 0.59
NG = 0
DB = 0
SM = 0
Delta_L
Doxycycline
YDL195W SEC31
ZShift = –0.43
NG = 0
DB = 0
SM = 0
Delta_K
Doxycycline
YDL195W SEC31
ZShift = –0.45
NG = 0
DB = 0
SM = 0
Delta_r
Doxycycline
YDL195W SEC31
ZShift = –0.55
NG = 0
DB = 0
SM = 0
Delta_AUC
Doxycycline
YPR137W RRP9
ZShift = 0.8
NG = 0
DB = 0
SM = 0
Delta_L
Doxycycline
YPR137W RRP9
ZShift = 0.16
NG = 0
DB = 0
SM = 0
Delta_K
Doxycycline
YPR137W RRP9
ZShift = 1.23
NG = 0
DB = 0
SM = 0
Delta_r
Doxycycline
YPR137W RRP9
ZShift = –0.24
NG = 0
DB = 0
SM = 0
Delta_AUC
Doxycycline
YPR034W ARP7
ZShift = –0.15
NG = 0
DB = 0
SM = 0
Delta_L
Doxycycline
YPR034W ARP7
ZShift = –0.15
NG = 0
DB = 0
SM = 0
Delta_K
Doxycycline
YPR034W ARP7
ZShift = –0.78
NG = 0
DB = 0
SM = 0
Delta_r
Doxycycline
YPR034W ARP7
ZShift = –0.04
NG = 0
DB = 0
SM = 0
Delta_AUC
Doxycycline

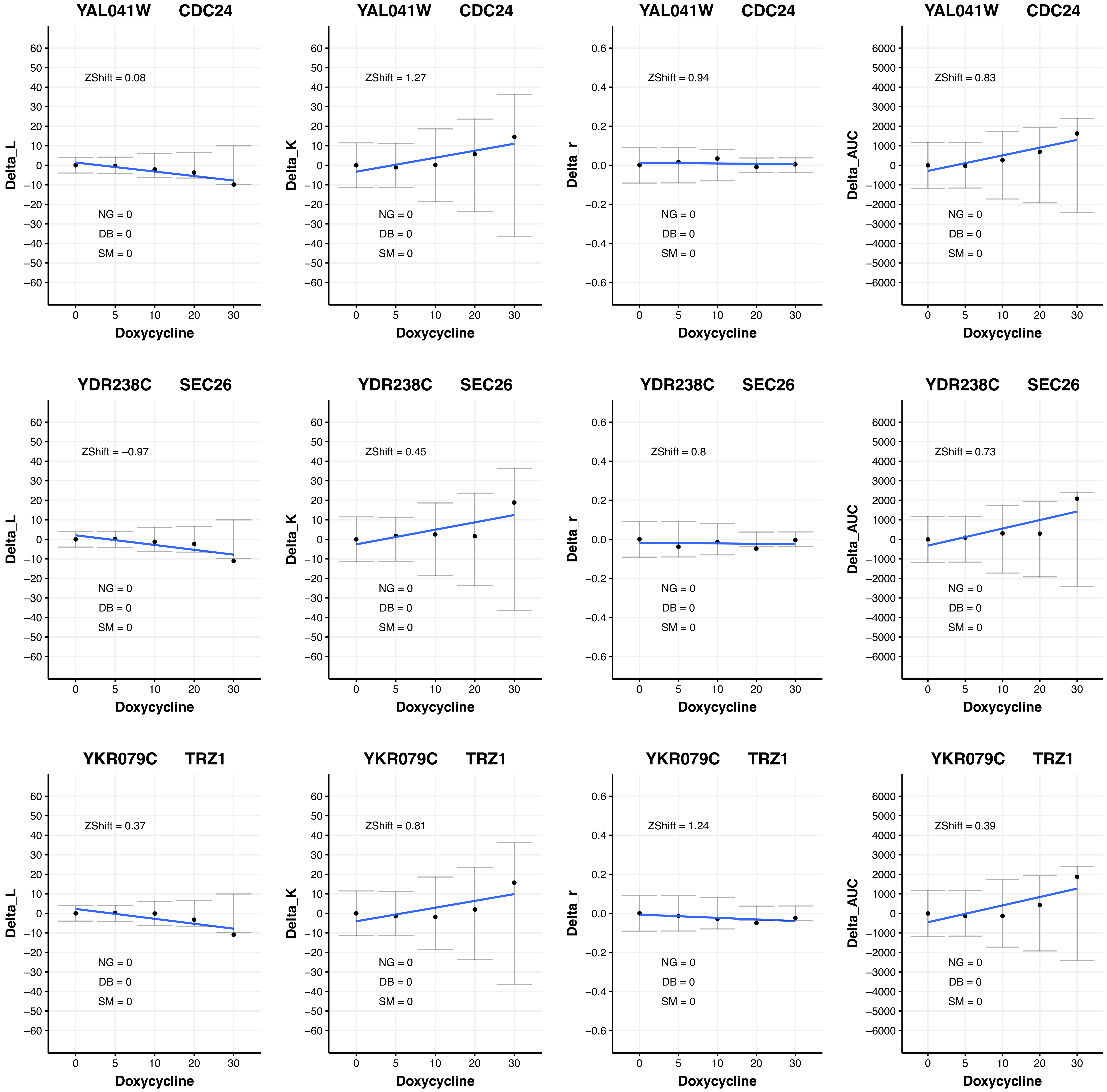
YAL041W CDC24
ZShift = 0.08
ZShift = 1.27
ZShift = 0.94
ZShift = 0.83
YDR238C SEC26
ZShift = −0.97
ZShift = 0.45
ZShift = 0.8
ZShift = 0.73
YKR079C TRZ1
ZShift = 0.37
ZShift = 0.81
ZShift = 1.24
ZShift = 0.39
NG = 0
DB = 0
SM = 0
Delta_L
Delta_K
Delta_r
Delta_AUC
Doxycycline

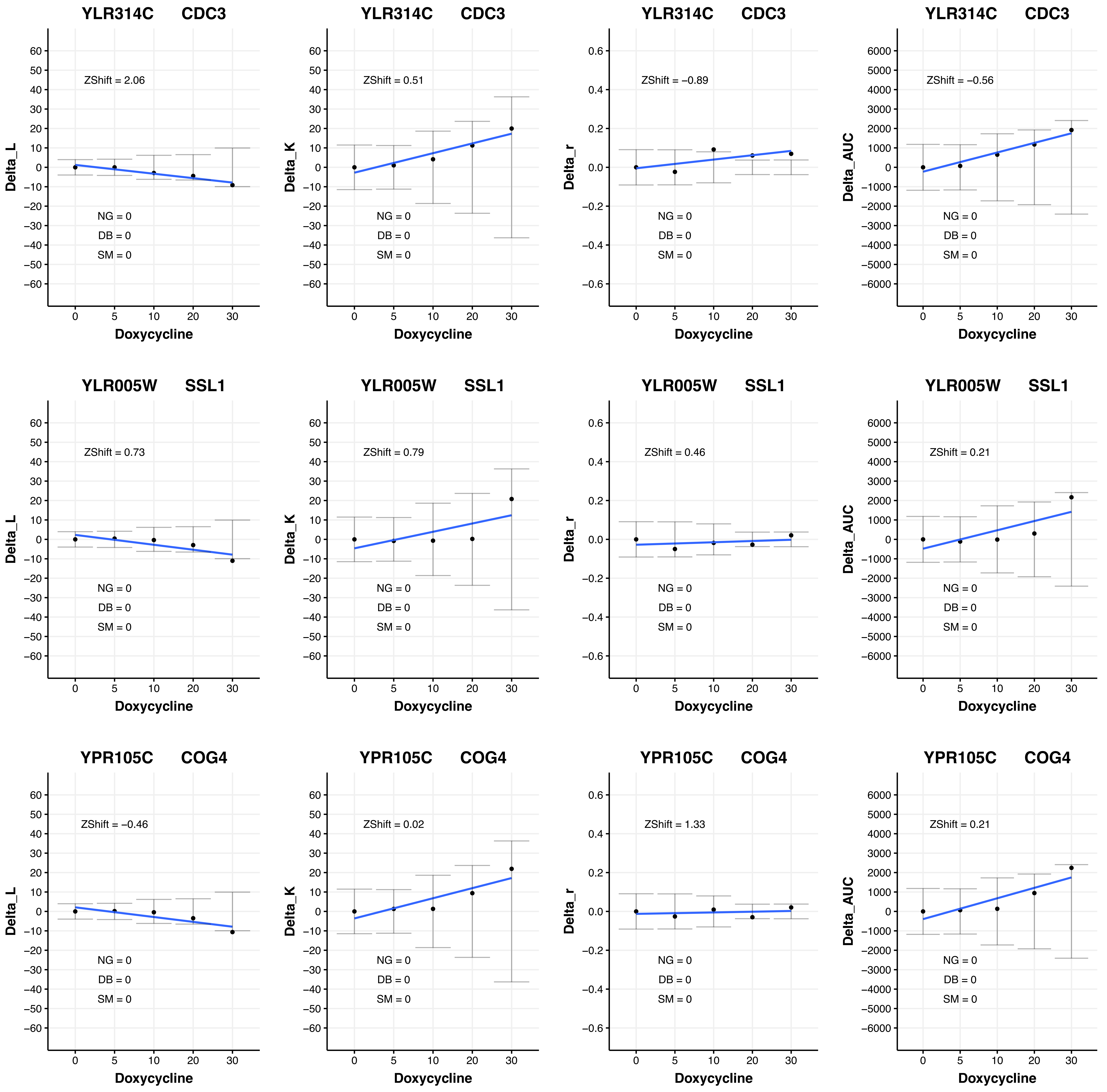
YLR314C CDC3
ZShift = 2.06
NG = 0
DB = 0
SM = 0
Delta_L
Doxycycline
YLR314C CDC3
ZShift = 0.51
NG = 0
DB = 0
SM = 0
Delta_K
Doxycycline
YLR314C CDC3
ZShift = −0.89
NG = 0
DB = 0
SM = 0
Delta_r
Doxycycline
YLR314C CDC3
ZShift = −0.56
NG = 0
DB = 0
SM = 0
Delta_AUC
Doxycycline
YLR005W SSL1
ZShift = 0.73
NG = 0
DB = 0
SM = 0
Delta_L
Doxycycline
YLR005W SSL1
ZShift = 0.79
NG = 0
DB = 0
SM = 0
Delta_K
Doxycycline
YLR005W SSL1
ZShift = 0.46
NG = 0
DB = 0
SM = 0
Delta_r
Doxycycline
YLR005W SSL1
ZShift = 0.21
NG = 0
DB = 0
SM = 0
Delta_AUC
Doxycycline
YPR105C COG4
ZShift = −0.46
NG = 0
DB = 0
SM = 0
Delta_L
Doxycycline
YPR105C COG4
ZShift = 0.02
NG = 0
DB = 0
SM = 0
Delta_K
Doxycycline
YPR105C COG4
ZShift = 1.33
NG = 0
DB = 0
SM = 0
Delta_r
Doxycycline
YPR105C COG4
ZShift = 0.21
NG = 0
DB = 0
SM = 0
Delta_AUC
Doxycycline

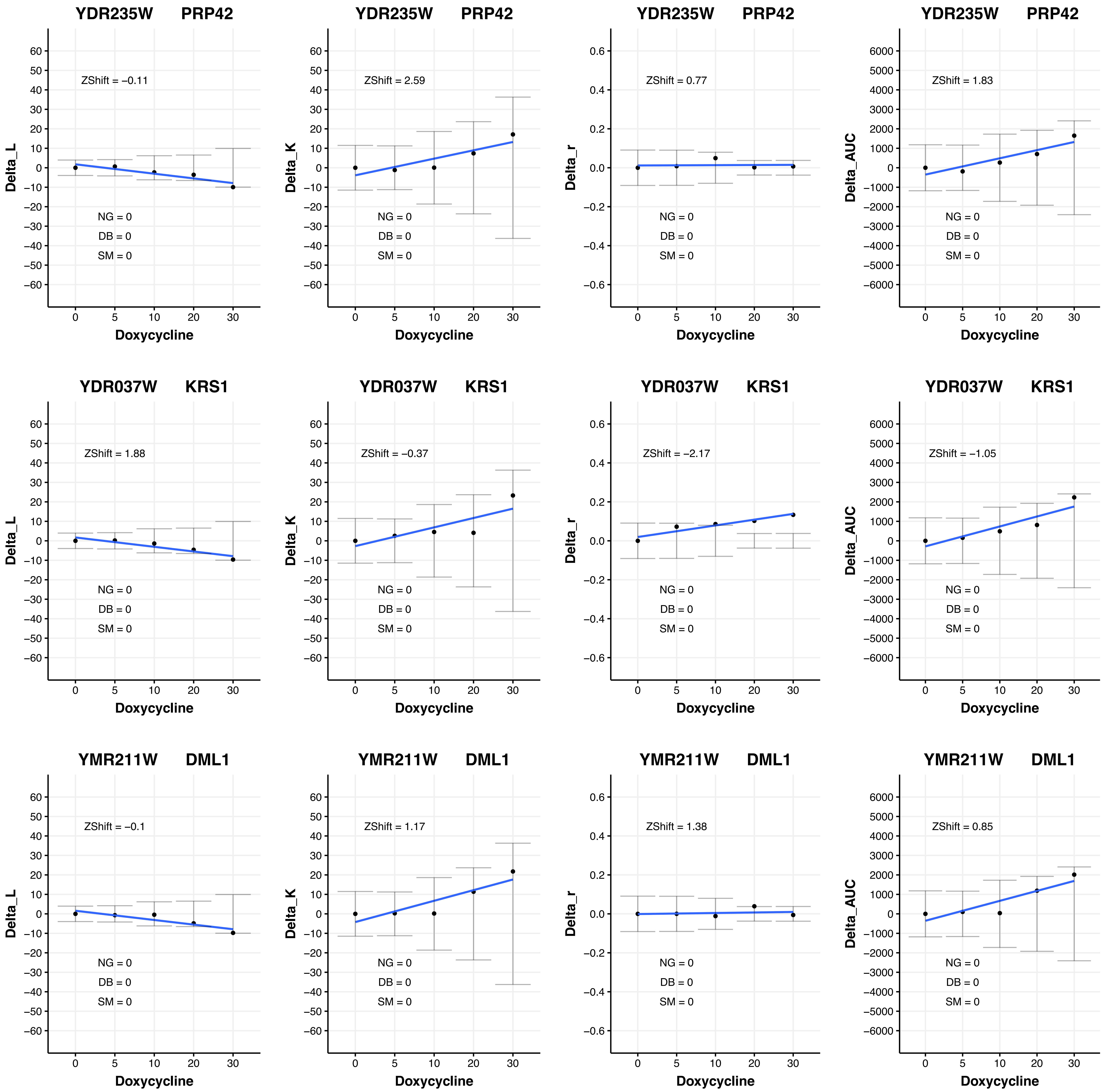
YDR235W PRP42
ZShift = −0.11
NG = 0
DB = 0
SM = 0
Delta_L
Doxycycline
YDR235W PRP42
ZShift = 2.59
NG = 0
DB = 0
SM = 0
Delta_K
Doxycycline
YDR235W PRP42
ZShift = 0.77
NG = 0
DB = 0
SM = 0
Delta_r
Doxycycline
YDR235W PRP42
ZShift = 1.83
NG = 0
DB = 0
SM = 0
Delta_AUC
Doxycycline
YDR037W KRS1
ZShift = 1.88
NG = 0
DB = 0
SM = 0
Delta_L
Doxycycline
YDR037W KRS1
ZShift = −0.37
NG = 0
DB = 0
SM = 0
Delta_K
Doxycycline
YDR037W KRS1
ZShift = −2.17
NG = 0
DB = 0
SM = 0
Delta_r
Doxycycline
YDR037W KRS1
ZShift = −1.05
NG = 0
DB = 0
SM = 0
Delta_AUC
Doxycycline
YMR211W DML1
ZShift = −0.1
NG = 0
DB = 0
SM = 0
Delta_L
Doxycycline
YMR211W DML1
ZShift = 1.17
NG = 0
DB = 0
SM = 0
Delta_K
Doxycycline
YMR211W DML1
ZShift = 1.38
NG = 0
DB = 0
SM = 0
Delta_r
Doxycycline
YMR211W DML1
ZShift = 0.85
NG = 0
DB = 0
SM = 0
Delta_AUC
Doxycycline

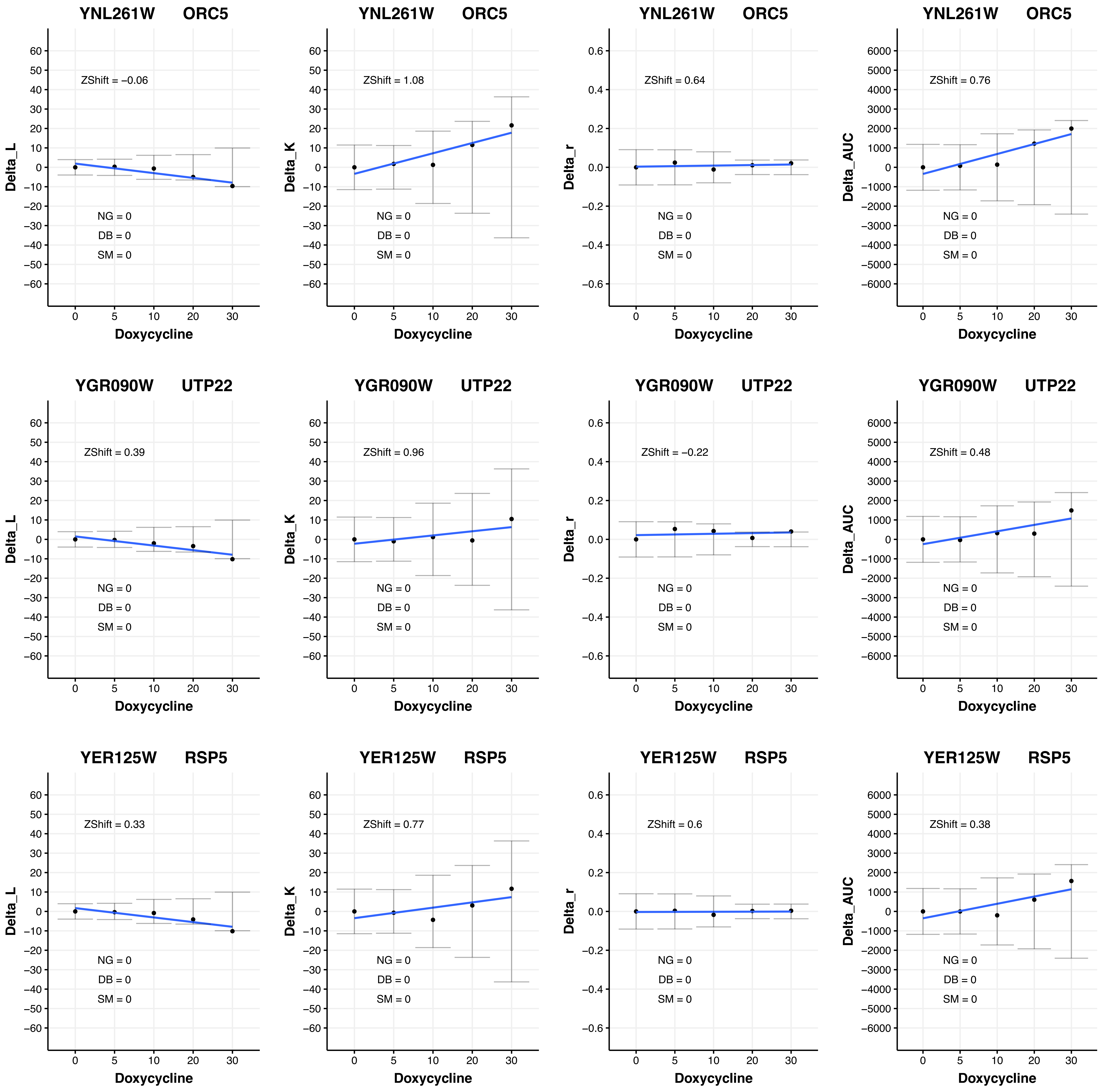
YNL261W ORC5
ZShift = −0.06
NG = 0
DB = 0
SM = 0
Delta_L
Doxycycline
YNL261W ORC5
ZShift = 1.08
NG = 0
DB = 0
SM = 0
Delta_K
Doxycycline
YNL261W ORC5
ZShift = 0.64
NG = 0
DB = 0
SM = 0
Delta_r
Doxycycline
YNL261W ORC5
ZShift = 0.76
NG = 0
DB = 0
SM = 0
Delta_AUC
Doxycycline
YGR090W UTP22
ZShift = 0.39
NG = 0
DB = 0
SM = 0
Delta_L
Doxycycline
YGR090W UTP22
ZShift = 0.96
NG = 0
DB = 0
SM = 0
Delta_K
Doxycycline
YGR090W UTP22
ZShift = −0.22
NG = 0
DB = 0
SM = 0
Delta_r
Doxycycline
YGR090W UTP22
ZShift = 0.48
NG = 0
DB = 0
SM = 0
Delta_AUC
Doxycycline
YER125W RSP5
ZShift = 0.33
NG = 0
DB = 0
SM = 0
Delta_L
Doxycycline
YER125W RSP5
ZShift = 0.77
NG = 0
DB = 0
SM = 0
Delta_K
Doxycycline
YER125W RSP5
ZShift = 0.6
NG = 0
DB = 0
SM = 0
Delta_r
Doxycycline
YER125W RSP5
ZShift = 0.38
NG = 0
DB = 0
SM = 0
Delta_AUC
Doxycycline

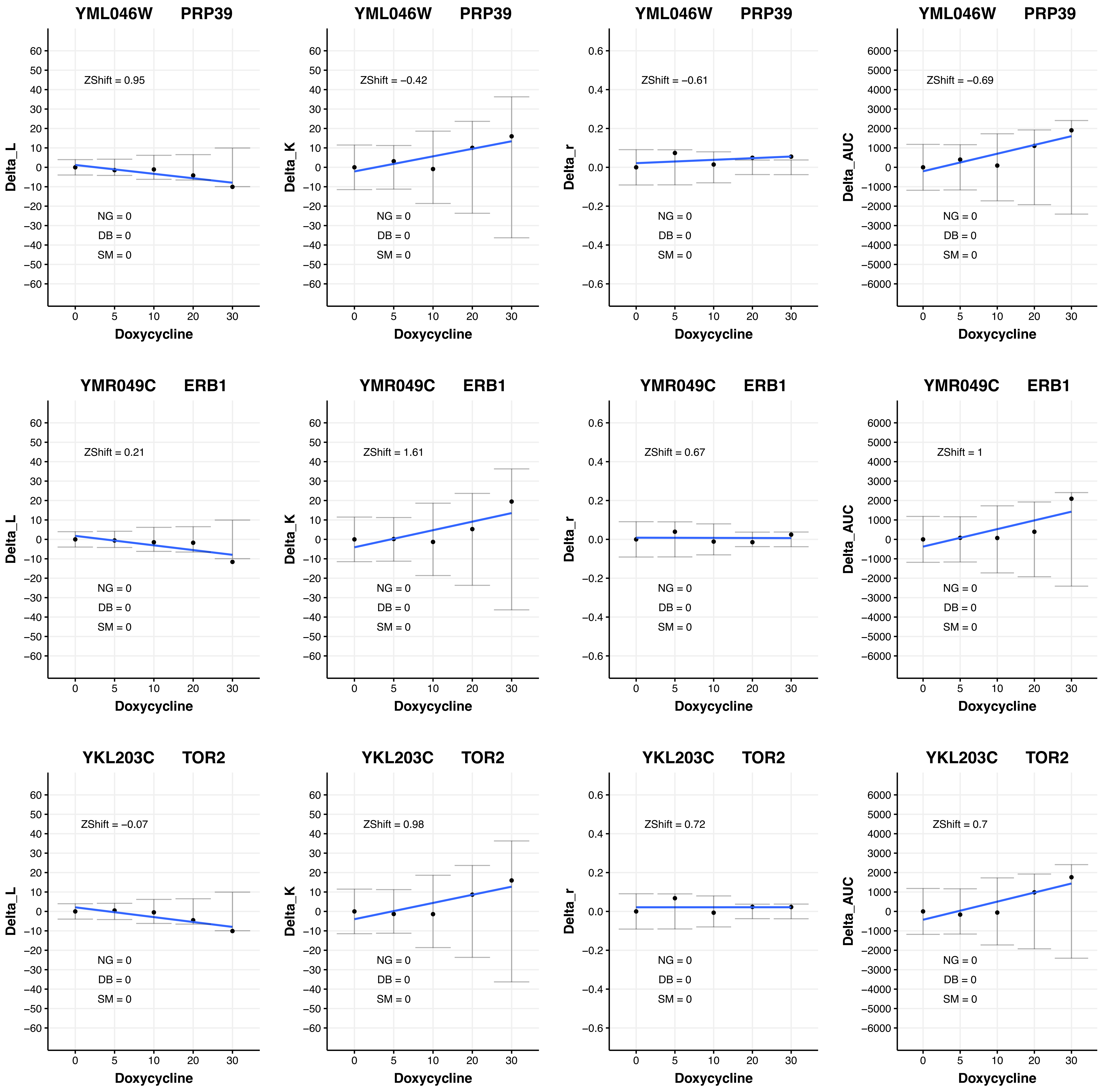
YML046W PRP39
ZShift = 0.95
NG = 0
DB = 0
SM = 0
Delta_L
Doxycycline
YML046W PRP39
ZShift = −0.42
NG = 0
DB = 0
SM = 0
Delta_K
Doxycycline
YML046W PRP39
ZShift = −0.61
NG = 0
DB = 0
SM = 0
Delta_r
Doxycycline
YML046W PRP39
ZShift = −0.69
NG = 0
DB = 0
SM = 0
Delta_AUC
Doxycycline
YMR049C ERB1
ZShift = 0.21
NG = 0
DB = 0
SM = 0
Delta_L
Doxycycline
YMR049C ERB1
ZShift = 1.61
NG = 0
DB = 0
SM = 0
Delta_K
Doxycycline
YMR049C ERB1
ZShift = 0.67
NG = 0
DB = 0
SM = 0
Delta_r
Doxycycline
YMR049C ERB1
ZShift = 1
NG = 0
DB = 0
SM = 0
Delta_AUC
Doxycycline
YKL203C TOR2
ZShift = −0.07
NG = 0
DB = 0
SM = 0
Delta_L
Doxycycline
YKL203C TOR2
ZShift = 0.98
NG = 0
DB = 0
SM = 0
Delta_K
Doxycycline
YKL203C TOR2
ZShift = 0.72
NG = 0
DB = 0
SM = 0
Delta_r
Doxycycline
YKL203C TOR2
ZShift = 0.7
NG = 0
DB = 0
SM = 0
Delta_AUC
Doxycycline

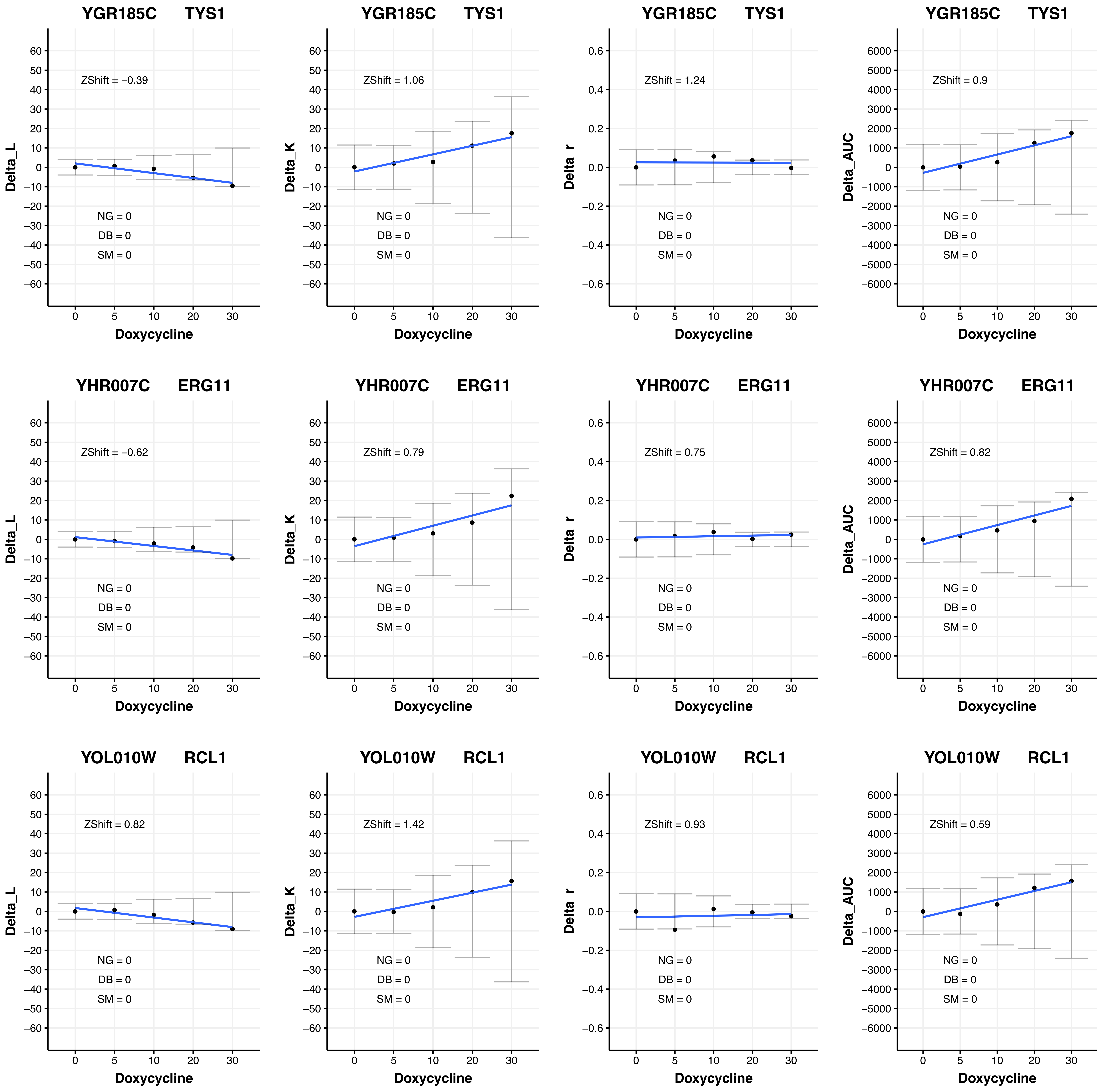
YGR185C TYS1
ZShift = −0.39
NG = 0
DB = 0
SM = 0
Delta_L
Doxycycline
YGR185C TYS1
ZShift = 1.06
NG = 0
DB = 0
SM = 0
Delta_K
Doxycycline
YGR185C TYS1
ZShift = 1.24
NG = 0
DB = 0
SM = 0
Delta_r
Doxycycline
YGR185C TYS1
ZShift = 0.9
NG = 0
DB = 0
SM = 0
Delta_AUC
Doxycycline
YHR007C ERG11
ZShift = −0.62
NG = 0
DB = 0
SM = 0
Delta_L
Doxycycline
YHR007C ERG11
ZShift = 0.79
NG = 0
DB = 0
SM = 0
Delta_K
Doxycycline
YHR007C ERG11
ZShift = 0.75
NG = 0
DB = 0
SM = 0
Delta_r
Doxycycline
YHR007C ERG11
ZShift = 0.82
NG = 0
DB = 0
SM = 0
Delta_AUC
Doxycycline
YOL010W RCL1
ZShift = 0.82
NG = 0
DB = 0
SM = 0
Delta_L
Doxycycline
YOL010W RCL1
ZShift = 1.42
NG = 0
DB = 0
SM = 0
Delta_K
Doxycycline
YOL010W RCL1
ZShift = 0.93
NG = 0
DB = 0
SM = 0
Delta_r
Doxycycline
YOL010W RCL1
ZShift = 0.59
NG = 0
DB = 0
SM = 0
Delta_AUC
Doxycycline

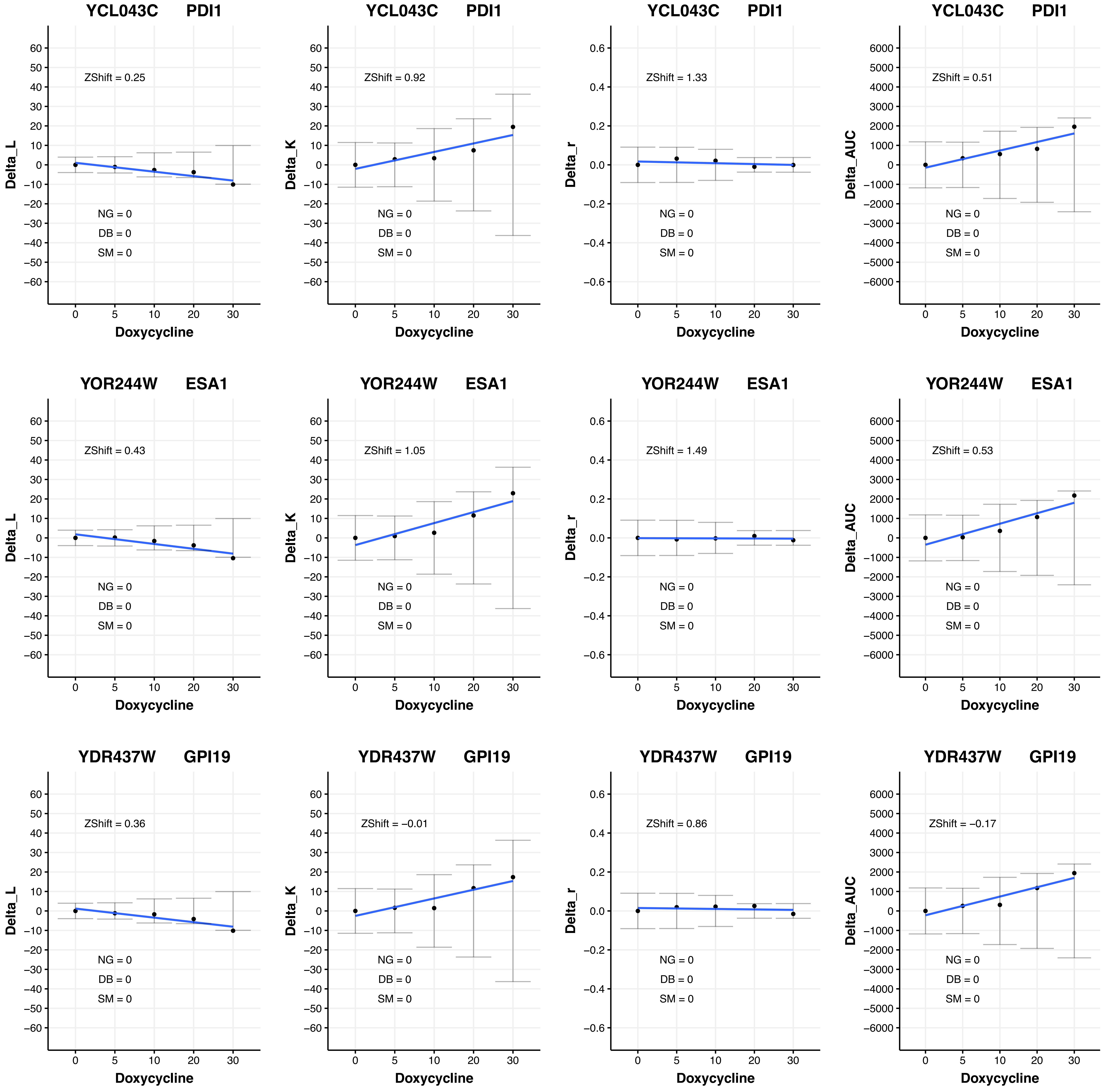
YCL043C PDI1
ZShift = 0.25
NG = 0
DB = 0
SM = 0
Delta_L
Doxycycline
YCL043C PDI1
ZShift = 0.92
NG = 0
DB = 0
SM = 0
Delta_K
Doxycycline
YCL043C PDI1
ZShift = 1.33
NG = 0
DB = 0
SM = 0
Delta_r
Doxycycline
YCL043C PDI1
ZShift = 0.51
NG = 0
DB = 0
SM = 0
Delta_AUC
Doxycycline
YOR244W ESA1
ZShift = 0.43
NG = 0
DB = 0
SM = 0
Delta_L
Doxycycline
YOR244W ESA1
ZShift = 1.05
NG = 0
DB = 0
SM = 0
Delta_K
Doxycycline
YOR244W ESA1
ZShift = 1.49
NG = 0
DB = 0
SM = 0
Delta_r
Doxycycline
YOR244W ESA1
ZShift = 0.53
NG = 0
DB = 0
SM = 0
Delta_AUC
Doxycycline
YDR437W GPI19
ZShift = 0.36
NG = 0
DB = 0
SM = 0
Delta_L
Doxycycline
YDR437W GPI19
ZShift = −0.01
NG = 0
DB = 0
SM = 0
Delta_K
Doxycycline
YDR437W GPI19
ZShift = 0.86
NG = 0
DB = 0
SM = 0
Delta_r
Doxycycline
YDR437W GPI19
ZShift = −0.17
NG = 0
DB = 0
SM = 0
Delta_AUC
Doxycycline

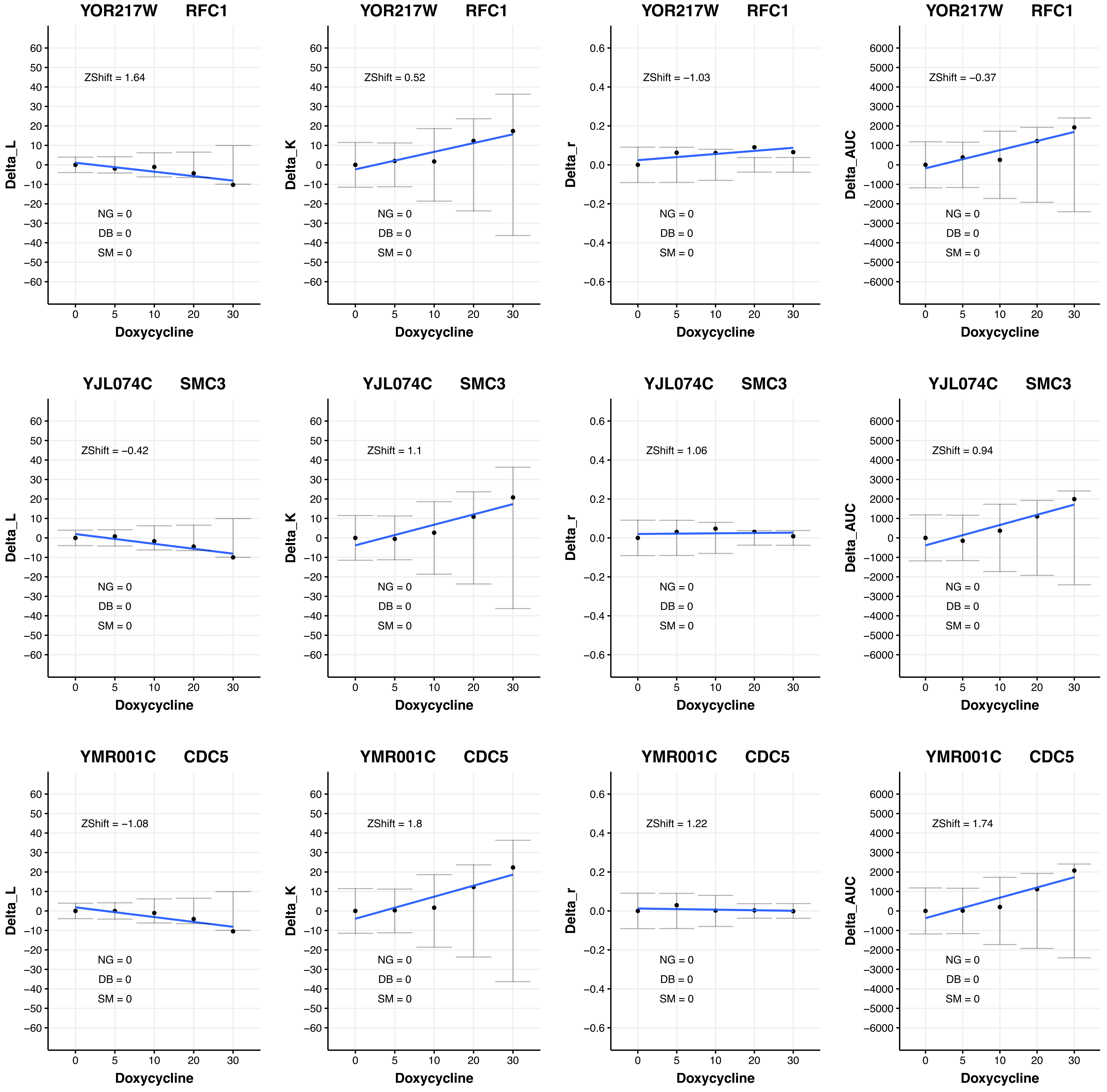

YOR217W RFC1
ZShift = 1.64
NG = 0
DB = 0
SM = 0
Delta_L
Doxycycline
YOR217W RFC1
ZShift = 0.52
NG = 0
DB = 0
SM = 0
Delta_K
Doxycycline
YOR217W RFC1
ZShift = −1.03
NG = 0
DB = 0
SM = 0
Delta_r
Doxycycline
YOR217W RFC1
ZShift = −0.37
NG = 0
DB = 0
SM = 0
Delta_AUC
Doxycycline
YJL074C SMC3
ZShift = −0.42
NG = 0
DB = 0
SM = 0
Delta_L
Doxycycline
YJL074C SMC3
ZShift = 1.1
NG = 0
DB = 0
SM = 0
Delta_K
Doxycycline
YJL074C SMC3
ZShift = 1.06
NG = 0
DB = 0
SM = 0
Delta_r
Doxycycline
YJL074C SMC3
ZShift = 0.94
NG = 0
DB = 0
SM = 0
Delta_AUC
Doxycycline
YMR001C CDC5
ZShift = −1.08
NG = 0
DB = 0
SM = 0
Delta_L
Doxycycline
YMR001C CDC5
ZShift = 1.8
NG = 0
DB = 0
SM = 0
Delta_K
Doxycycline
YMR001C CDC5
ZShift = 1.22
NG = 0
DB = 0
SM = 0
Delta_r
Doxycycline
YMR001C CDC5
ZShift = 1.74
NG = 0
DB = 0
SM = 0
Delta_AUC
Doxycycline

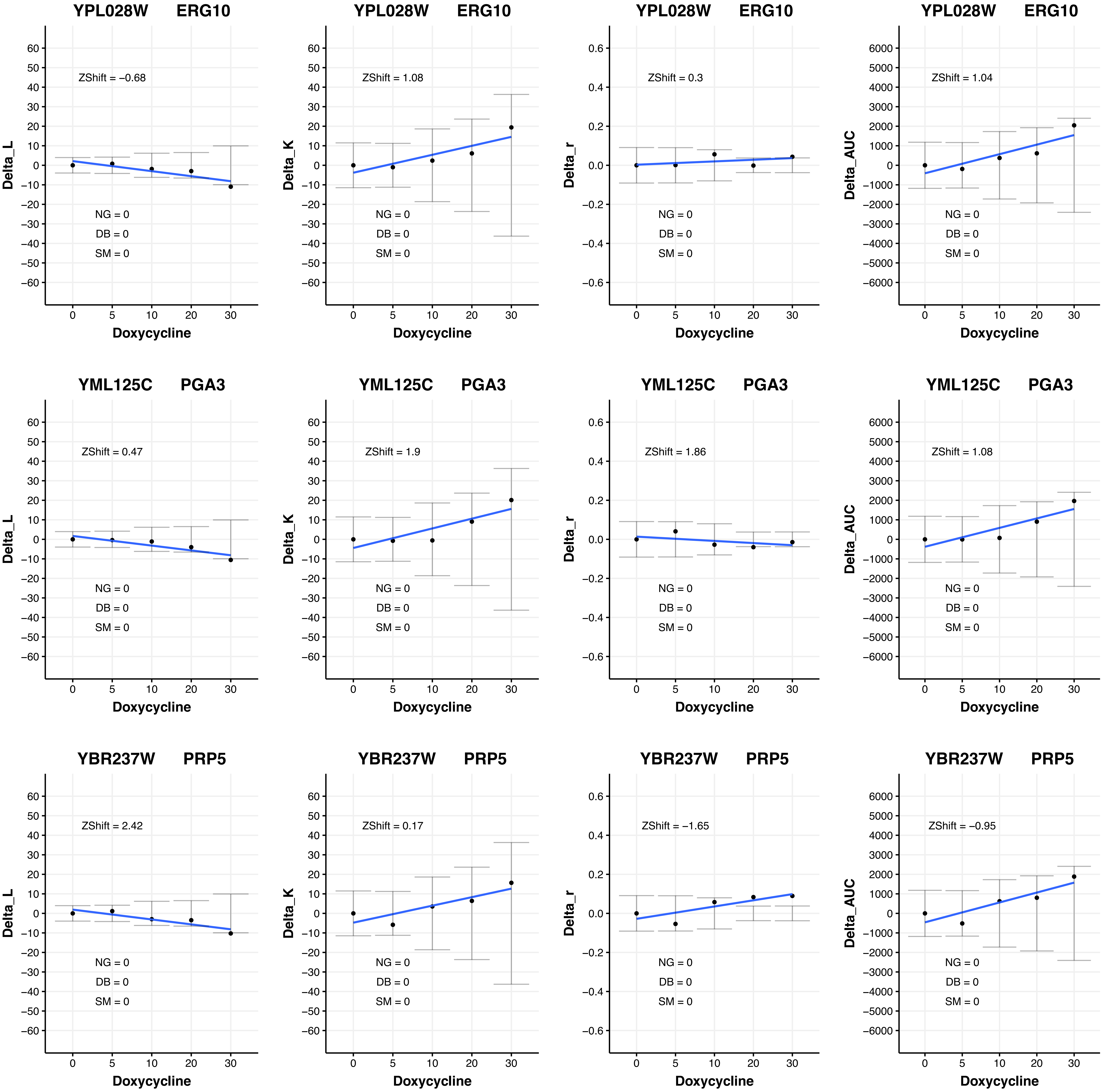
YPL028W ERG10
ZShift = −0.68
NG = 0
DB = 0
SM = 0
Delta_L
Doxycycline
YPL028W ERG10
ZShift = 1.08
NG = 0
DB = 0
SM = 0
Delta_K
Doxycycline
YPL028W ERG10
ZShift = 0.3
NG = 0
DB = 0
SM = 0
Delta_r
Doxycycline
YPL028W ERG10
ZShift = 1.04
NG = 0
DB = 0
SM = 0
Delta_AUC
Doxycycline
YML125C PGA3
ZShift = 0.47
NG = 0
DB = 0
SM = 0
Delta_L
Doxycycline
YML125C PGA3
ZShift = 1.9
NG = 0
DB = 0
SM = 0
Delta_K
Doxycycline
YML125C PGA3
ZShift = 1.86
NG = 0
DB = 0
SM = 0
Delta_r
Doxycycline
YML125C PGA3
ZShift = 1.08
NG = 0
DB = 0
SM = 0
Delta_AUC
Doxycycline
YBR237W PRP5
ZShift = 2.42
NG = 0
DB = 0
SM = 0
Delta_L
Doxycycline
YBR237W PRP5
ZShift = 0.17
NG = 0
DB = 0
SM = 0
Delta_K
Doxycycline
YBR237W PRP5
ZShift = −1.65
NG = 0
DB = 0
SM = 0
Delta_r
Doxycycline
YBR237W PRP5
ZShift = −0.95
NG = 0
DB = 0
SM = 0
Delta_AUC
Doxycycline

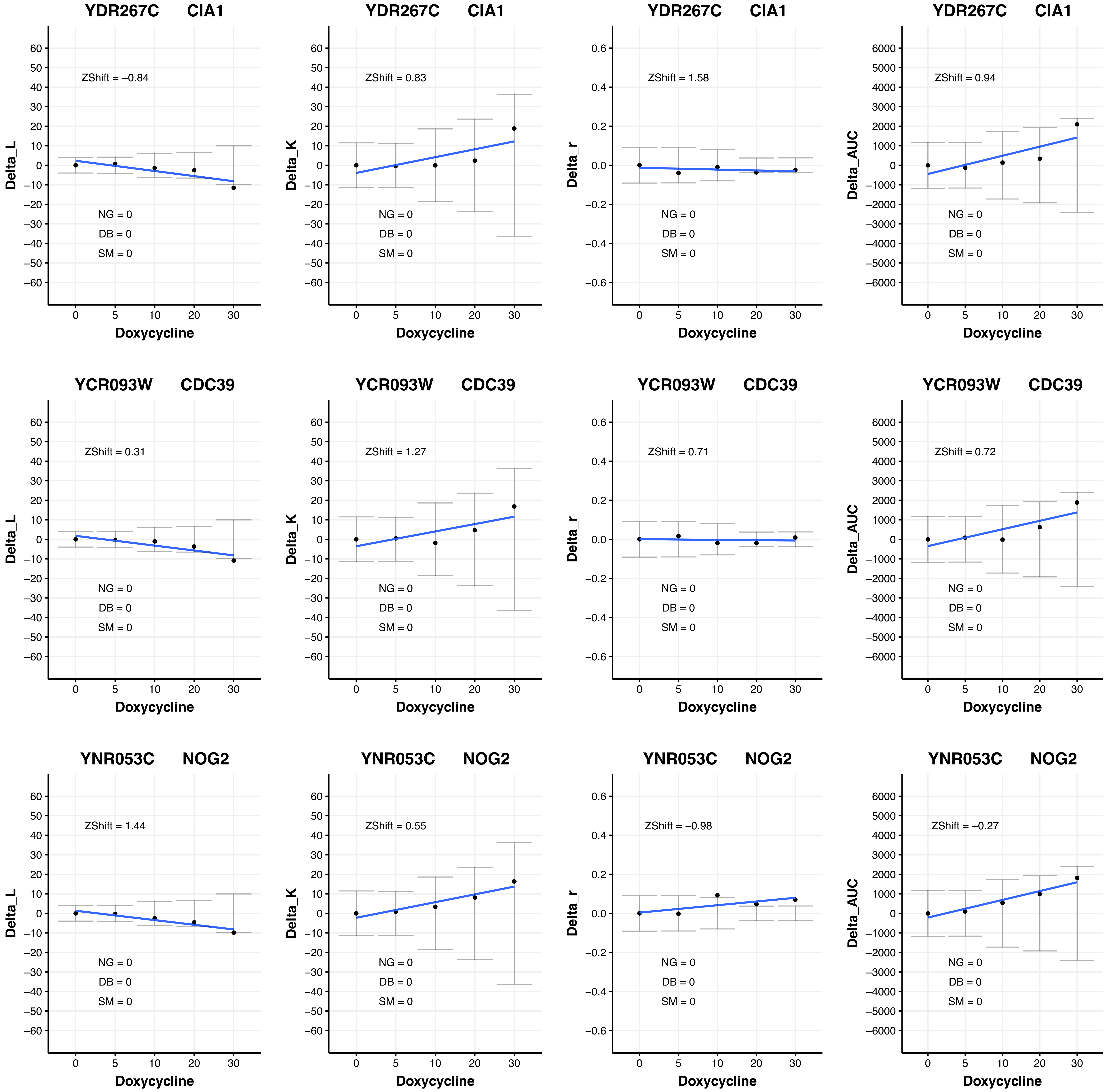
YDR267C CIA1
ZShift = −0.84
NG = 0
DB = 0
SM = 0
Delta_L
Doxycycline
YDR267C CIA1
ZShift = 0.83
NG = 0
DB = 0
SM = 0
Delta_K
Doxycycline
YDR267C CIA1
ZShift = 1.58
NG = 0
DB = 0
SM = 0
Delta_r
Doxycycline
YDR267C CIA1
ZShift = 0.94
NG = 0
DB = 0
SM = 0
Delta_AUC
Doxycycline
YCR093W CDC39
ZShift = 0.31
NG = 0
DB = 0
SM = 0
Delta_L
Doxycycline
YCR093W CDC39
ZShift = 1.27
NG = 0
DB = 0
SM = 0
Delta_K
Doxycycline
YCR093W CDC39
ZShift = 0.71
NG = 0
DB = 0
SM = 0
Delta_r
Doxycycline
YCR093W CDC39
ZShift = 0.72
NG = 0
DB = 0
SM = 0
Delta_AUC
Doxycycline
YNR053C NOG2
ZShift = 1.44
NG = 0
DB = 0
SM = 0
Delta_L
Doxycycline
YNR053C NOG2
ZShift = 0.55
NG = 0
DB = 0
SM = 0
Delta_K
Doxycycline
YNR053C NOG2
ZShift = −0.98
NG = 0
DB = 0
SM = 0
Delta_r
Doxycycline
YNR053C NOG2
ZShift = −0.27
NG = 0
DB = 0
SM = 0
Delta_AUC
Doxycycline

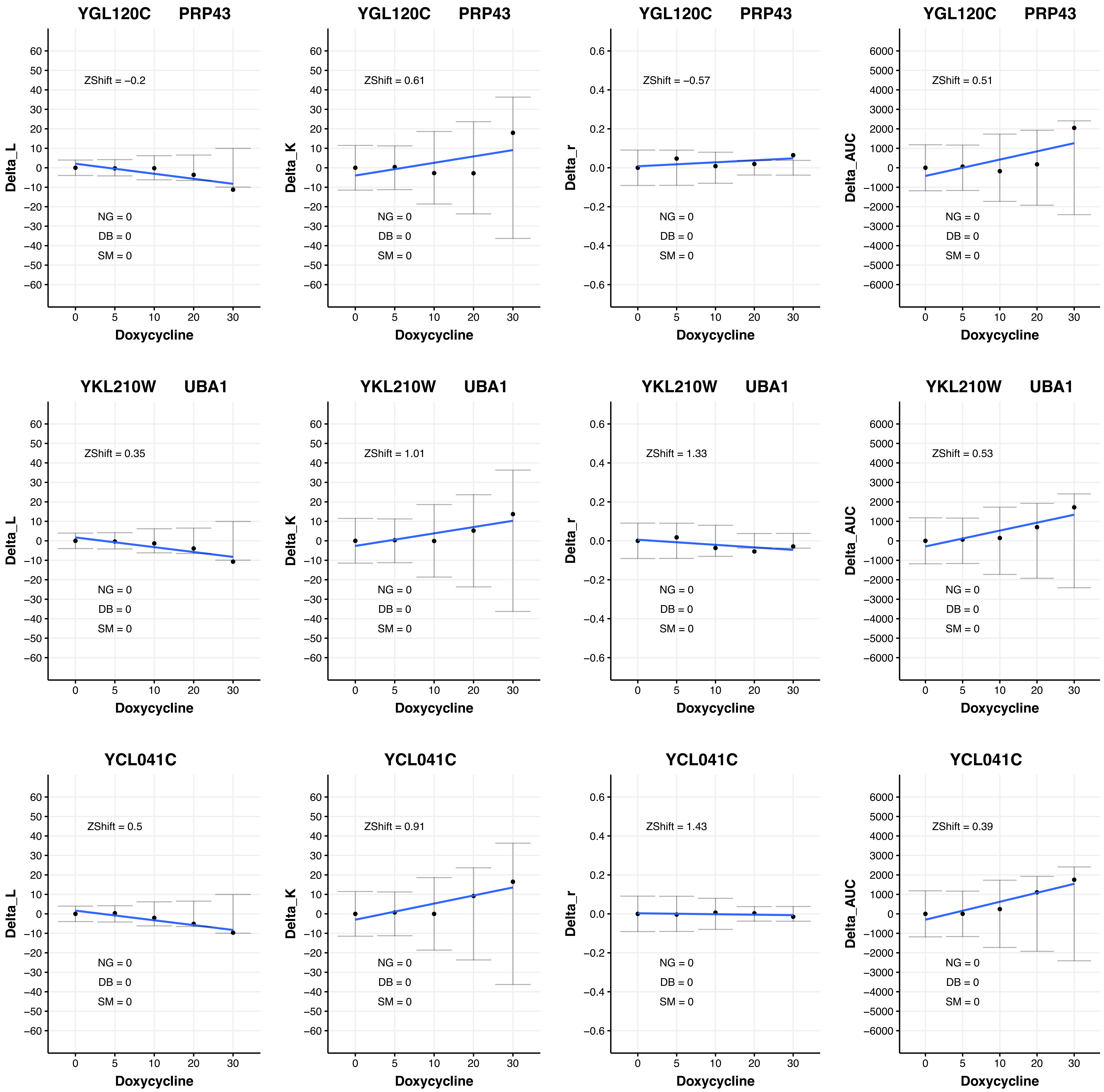

YGL120C PRP43
ZShift = −0.2
NG = 0
DB = 0
SM = 0
Delta_L
Doxycycline
YGL120C PRP43
ZShift = 0.61
NG = 0
DB = 0
SM = 0
Delta_K
Doxycycline
YGL120C PRP43
ZShift = −0.57
NG = 0
DB = 0
SM = 0
Delta_r
Doxycycline
YGL120C PRP43
ZShift = 0.51
NG = 0
DB = 0
SM = 0
Delta_AUC
Doxycycline
YKL210W UBA1
ZShift = 0.35
NG = 0
DB = 0
SM = 0
Delta_L
Doxycycline
YKL210W UBA1
ZShift = 1.01
NG = 0
DB = 0
SM = 0
Delta_K
Doxycycline
YKL210W UBA1
ZShift = 1.33
NG = 0
DB = 0
SM = 0
Delta_r
Doxycycline
YKL210W UBA1
ZShift = 0.53
NG = 0
DB = 0
SM = 0
Delta_AUC
Doxycycline
YCL041C
ZShift = 0.5
NG = 0
DB = 0
SM = 0
Delta_L
Doxycycline
YCL041C
ZShift = 0.91
NG = 0
DB = 0
SM = 0
Delta_K
Doxycycline
YCL041C
ZShift = 1.43
NG = 0
DB = 0
SM = 0
Delta_r
Doxycycline
YCL041C
ZShift = 0.39
NG = 0
DB = 0
SM = 0
Delta_AUC
Doxycycline

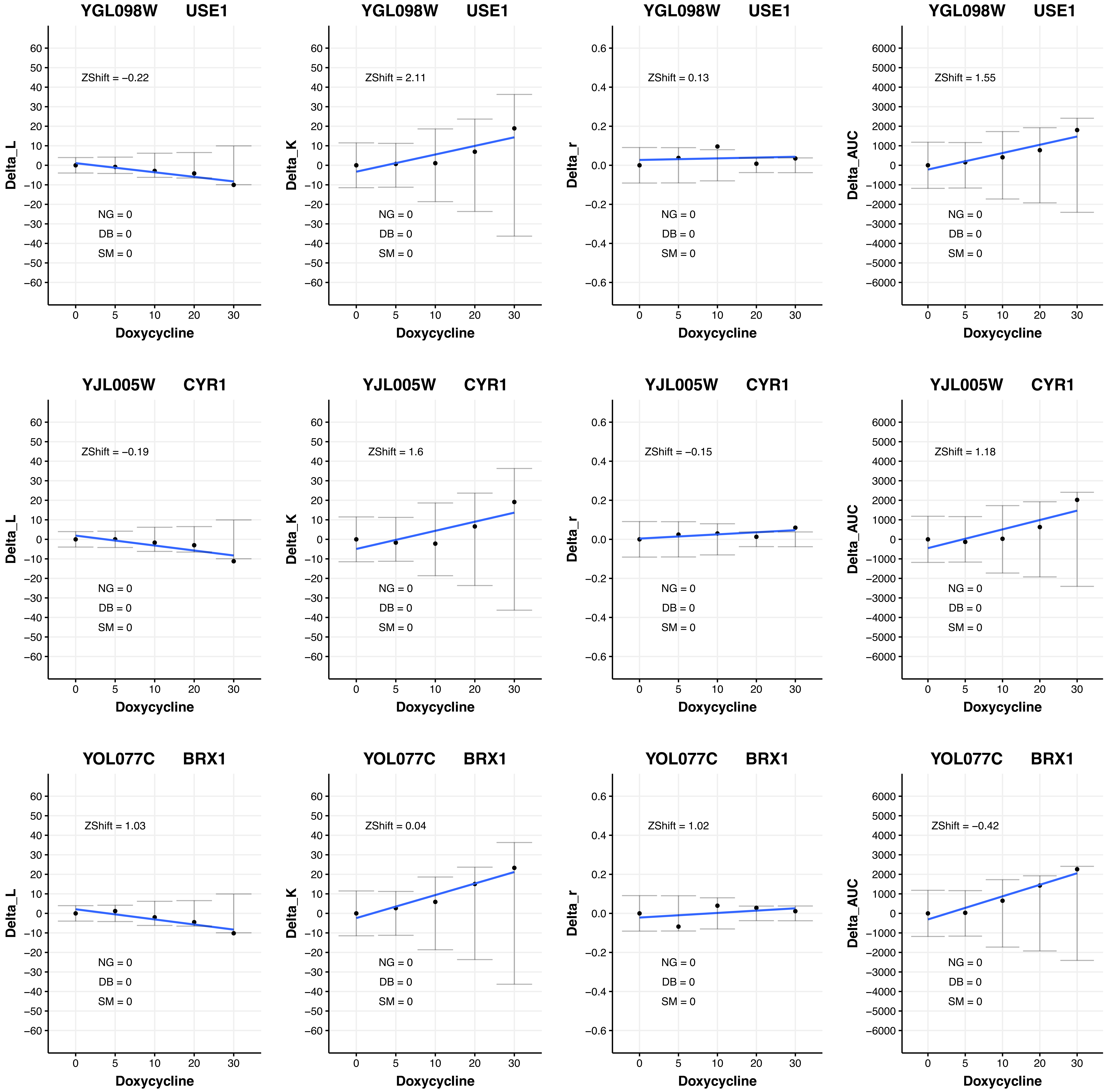

YGL098W USE1
ZShift = −0.22
NG = 0
DB = 0
SM = 0
Delta_L
Doxycycline
YGL098W USE1
ZShift = 2.11
NG = 0
DB = 0
SM = 0
Delta_K
Doxycycline
YGL098W USE1
ZShift = 0.13
NG = 0
DB = 0
SM = 0
Delta_r
Doxycycline
YGL098W USE1
ZShift = 1.55
NG = 0
DB = 0
SM = 0
Delta_AUC
Doxycycline
YJL005W CYR1
ZShift = −0.19
NG = 0
DB = 0
SM = 0
Delta_L
Doxycycline
YJL005W CYR1
ZShift = 1.6
NG = 0
DB = 0
SM = 0
Delta_K
Doxycycline
YJL005W CYR1
ZShift = −0.15
NG = 0
DB = 0
SM = 0
Delta_r
Doxycycline
YJL005W CYR1
ZShift = 1.18
NG = 0
DB = 0
SM = 0
Delta_AUC
Doxycycline
YOL077C BRX1
ZShift = 1.03
NG = 0
DB = 0
SM = 0
Delta_L
Doxycycline
YOL077C BRX1
ZShift = 0.04
NG = 0
DB = 0
SM = 0
Delta_K
Doxycycline
YOL077C BRX1
ZShift = 1.02
NG = 0
DB = 0
SM = 0
Delta_r
Doxycycline
YOL077C BRX1
ZShift = −0.42
NG = 0
DB = 0
SM = 0
Delta_AUC
Doxycycline

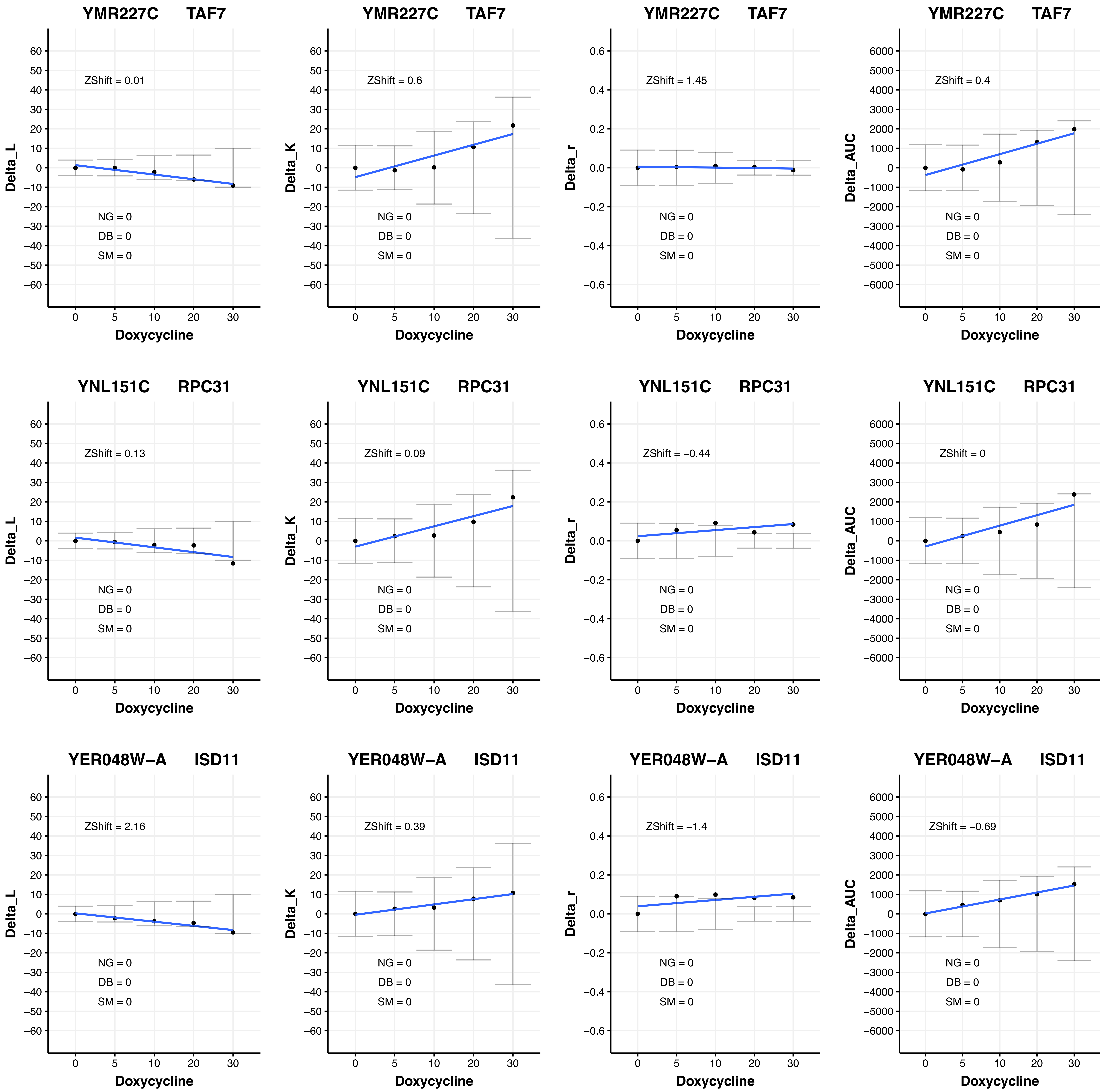
YMR227C TAF7
Delta_L
ZShift = 0.01
NG = 0
DB = 0
SM = 0
Doxycycline
YMR227C TAF7
Delta_K
ZShift = 0.6
NG = 0
DB = 0
SM = 0
Doxycycline
YMR227C TAF7
Delta_r
ZShift = 1.45
NG = 0
DB = 0
SM = 0
Doxycycline
YMR227C TAF7
Delta_AUC
ZShift = 0.4
NG = 0
DB = 0
SM = 0
Doxycycline
YNL151C RPC31
Delta_L
ZShift = 0.13
NG = 0
DB = 0
SM = 0
Doxycycline
YNL151C RPC31
Delta_K
ZShift = 0.09
NG = 0
DB = 0
SM = 0
Doxycycline
YNL151C RPC31
Delta_r
ZShift = −0.44
NG = 0
DB = 0
SM = 0
Doxycycline
YNL151C RPC31
Delta_AUC
ZShift = 0
NG = 0
DB = 0
SM = 0
Doxycycline
YER048W−A ISD11
Delta_L
ZShift = 2.16
NG = 0
DB = 0
SM = 0
Doxycycline
YER048W−A ISD11
Delta_K
ZShift = 0.39
NG = 0
DB = 0
SM = 0
Doxycycline
YER048W−A ISD11
Delta_r
ZShift = −1.4
NG = 0
DB = 0
SM = 0
Doxycycline
YER048W−A ISD11
Delta_AUC
ZShift = −0.69
NG = 0
DB = 0
SM = 0
Doxycycline

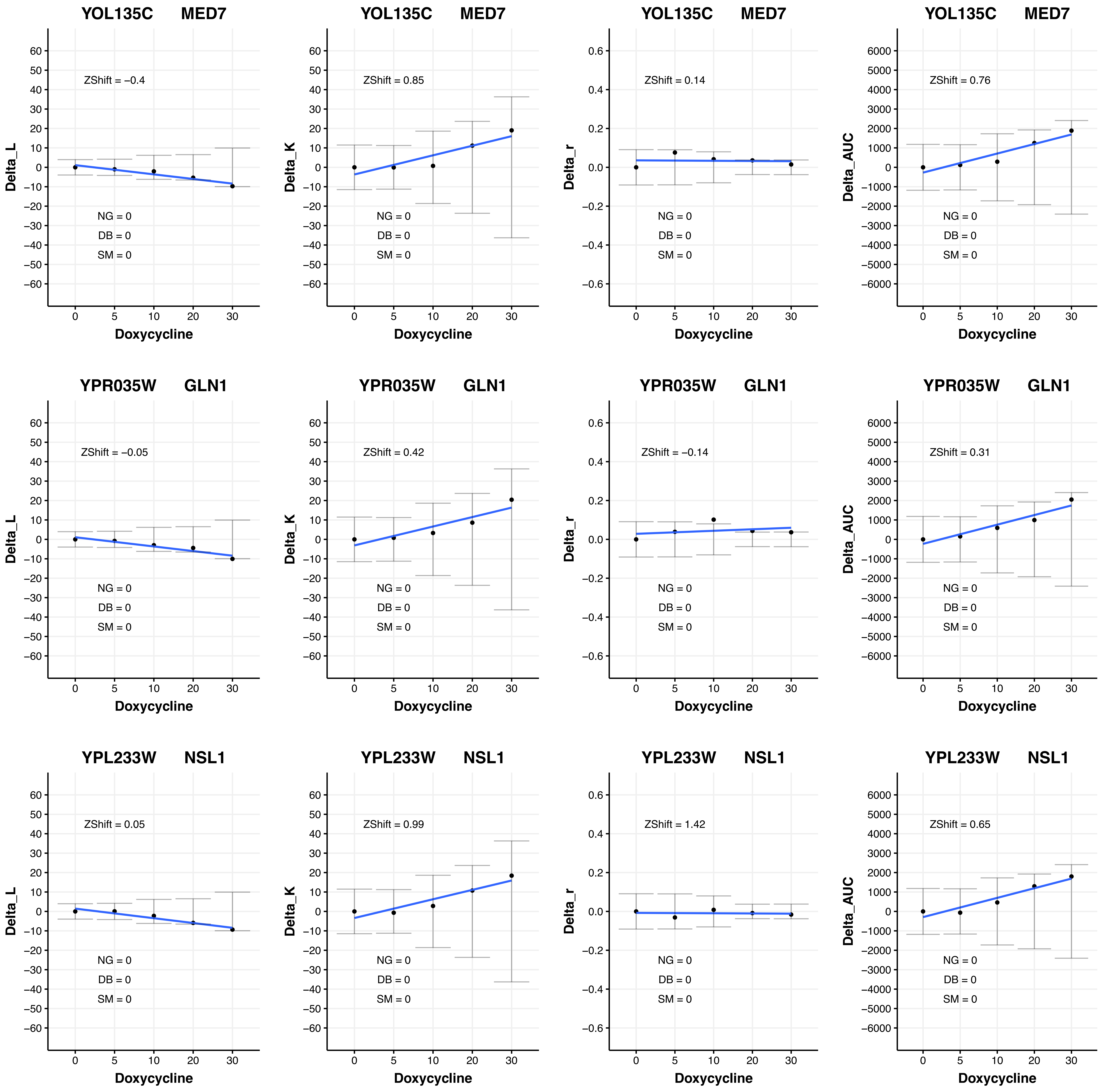

YOL135C MED7
Delta_L
ZShift = −0.4
NG = 0
DB = 0
SM = 0
Doxycycline
YOL135C MED7
Delta_K
ZShift = 0.85
NG = 0
DB = 0
SM = 0
Doxycycline
YOL135C MED7
Delta_r
ZShift = 0.14
NG = 0
DB = 0
SM = 0
Doxycycline
YOL135C MED7
Delta_AUC
ZShift = 0.76
NG = 0
DB = 0
SM = 0
Doxycycline
YPR035W GLN1
Delta_L
ZShift = −0.05
NG = 0
DB = 0
SM = 0
Doxycycline
YPR035W GLN1
Delta_K
ZShift = 0.42
NG = 0
DB = 0
SM = 0
Doxycycline
YPR035W GLN1
Delta_r
ZShift = −0.14
NG = 0
DB = 0
SM = 0
Doxycycline
YPR035W GLN1
Delta_AUC
ZShift = 0.31
NG = 0
DB = 0
SM = 0
Doxycycline
YPL233W NSL1
Delta_L
ZShift = 0.05
NG = 0
DB = 0
SM = 0
Doxycycline
YPL233W NSL1
Delta_K
ZShift = 0.99
NG = 0
DB = 0
SM = 0
Doxycycline
YPL233W NSL1
Delta_r
ZShift = 1.42
NG = 0
DB = 0
SM = 0
Doxycycline
YPL233W NSL1
Delta_AUC
ZShift = 0.65
NG = 0
DB = 0
SM = 0
Doxycycline

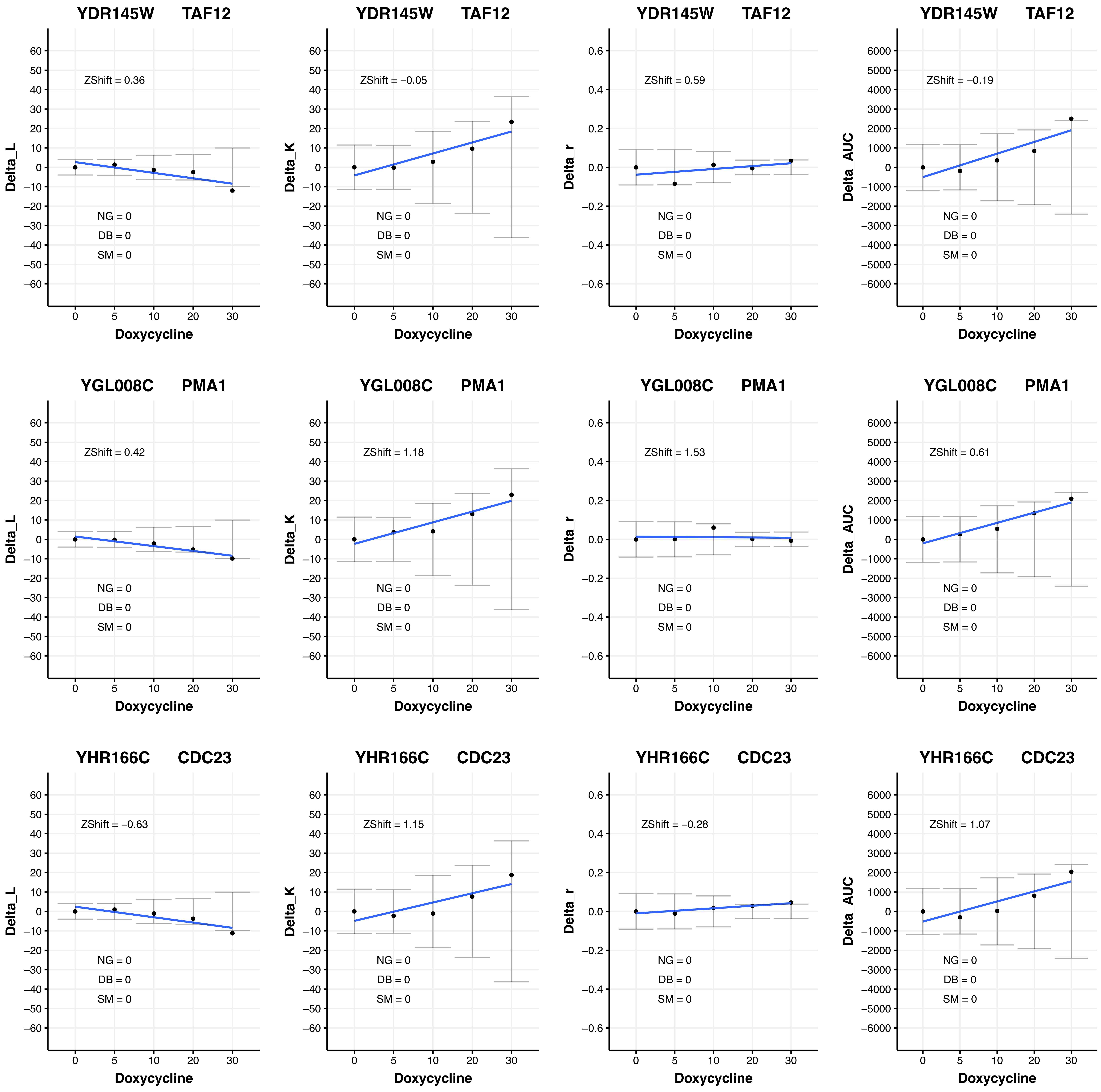
YDR145W TAF12
ZShift = 0.36
Delta_L
NG = 0
DB = 0
SM = 0
Doxycycline
YDR145W TAF12
ZShift = –0.05
Delta_K
NG = 0
DB = 0
SM = 0
Doxycycline
YDR145W TAF12
ZShift = 0.59
Delta_r
NG = 0
DB = 0
SM = 0
Doxycycline
YDR145W TAF12
ZShift = –0.19
Delta_AUC
NG = 0
DB = 0
SM = 0
Doxycycline
YGL008C PMA1
ZShift = 0.42
Delta_L
NG = 0
DB = 0
SM = 0
Doxycycline
YGL008C PMA1
ZShift = 1.18
Delta_K
NG = 0
DB = 0
SM = 0
Doxycycline
YGL008C PMA1
ZShift = 1.53
Delta_r
NG = 0
DB = 0
SM = 0
Doxycycline
YGL008C PMA1
ZShift = 0.61
Delta_AUC
NG = 0
DB = 0
SM = 0
Doxycycline
YHR166C CDC23
ZShift = –0.63
Delta_L
NG = 0
DB = 0
SM = 0
Doxycycline
YHR166C CDC23
ZShift = 1.15
Delta_K
NG = 0
DB = 0
SM = 0
Doxycycline
YHR166C CDC23
ZShift = –0.28
Delta_r
NG = 0
DB = 0
SM = 0
Doxycycline
YHR166C CDC23
ZShift = 1.07
Delta_AUC
NG = 0
DB = 0
SM = 0
Doxycycline

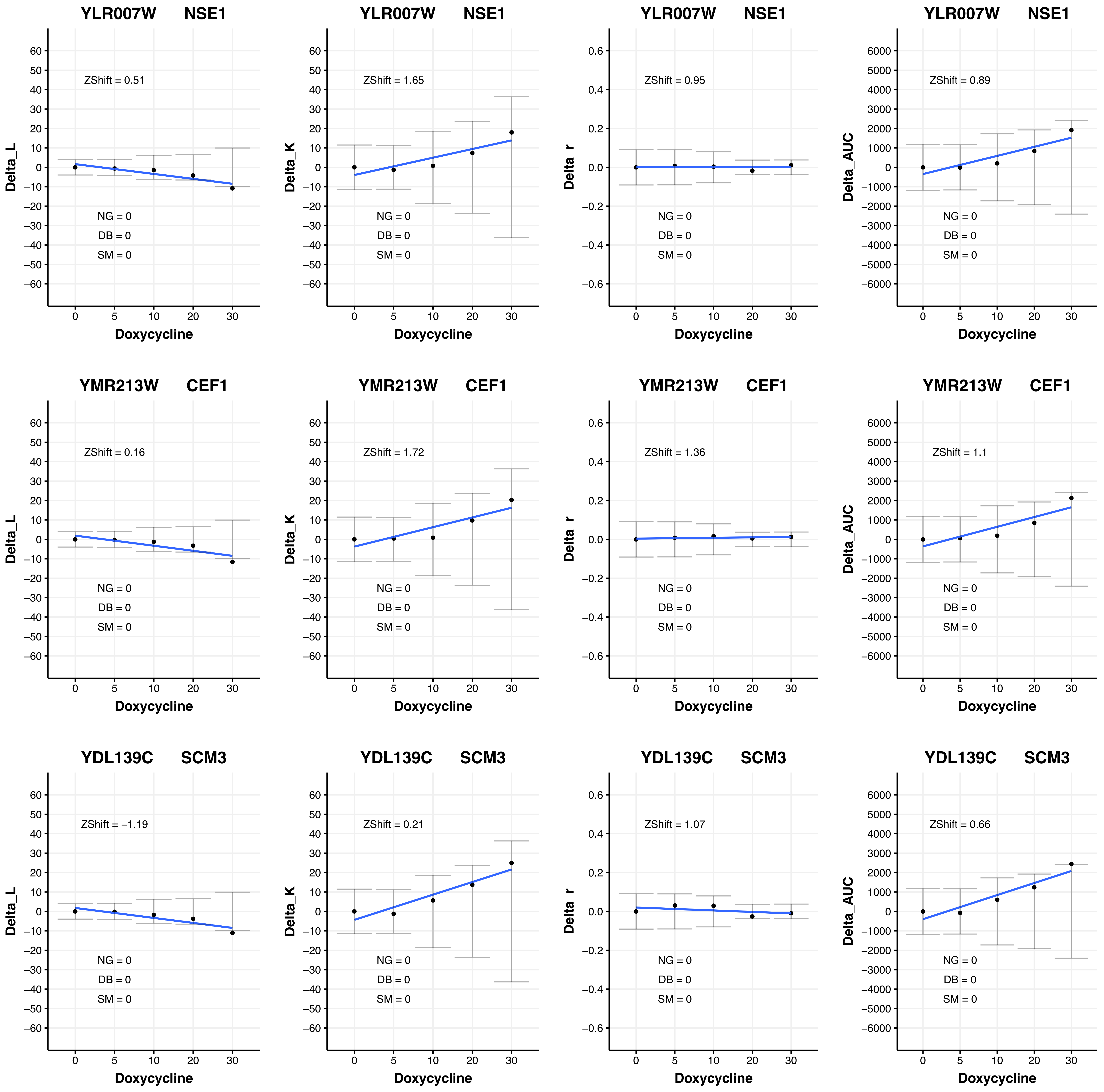
YLR007W NSE1
ZShift = 0.51
NG = 0
DB = 0
SM = 0
Delta_L
Doxycycline
YLR007W NSE1
ZShift = 1.65
NG = 0
DB = 0
SM = 0
Delta_K
Doxycycline
YLR007W NSE1
ZShift = 0.95
NG = 0
DB = 0
SM = 0
Delta_r
Doxycycline
YLR007W NSE1
ZShift = 0.89
NG = 0
DB = 0
SM = 0
Delta_AUC
Doxycycline
YMR213W CEF1
ZShift = 0.16
NG = 0
DB = 0
SM = 0
Delta_L
Doxycycline
YMR213W CEF1
ZShift = 1.72
NG = 0
DB = 0
SM = 0
Delta_K
Doxycycline
YMR213W CEF1
ZShift = 1.36
NG = 0
DB = 0
SM = 0
Delta_r
Doxycycline
YMR213W CEF1
ZShift = 1.1
NG = 0
DB = 0
SM = 0
Delta_AUC
Doxycycline
YDL139C SCM3
ZShift = −1.19
NG = 0
DB = 0
SM = 0
Delta_L
Doxycycline
YDL139C SCM3
ZShift = 0.21
NG = 0
DB = 0
SM = 0
Delta_K
Doxycycline
YDL139C SCM3
ZShift = 1.07
NG = 0
DB = 0
SM = 0
Delta_r
Doxycycline
YDL139C SCM3
ZShift = 0.66
NG = 0
DB = 0
SM = 0
Delta_AUC
Doxycycline
0 5 10 20 30

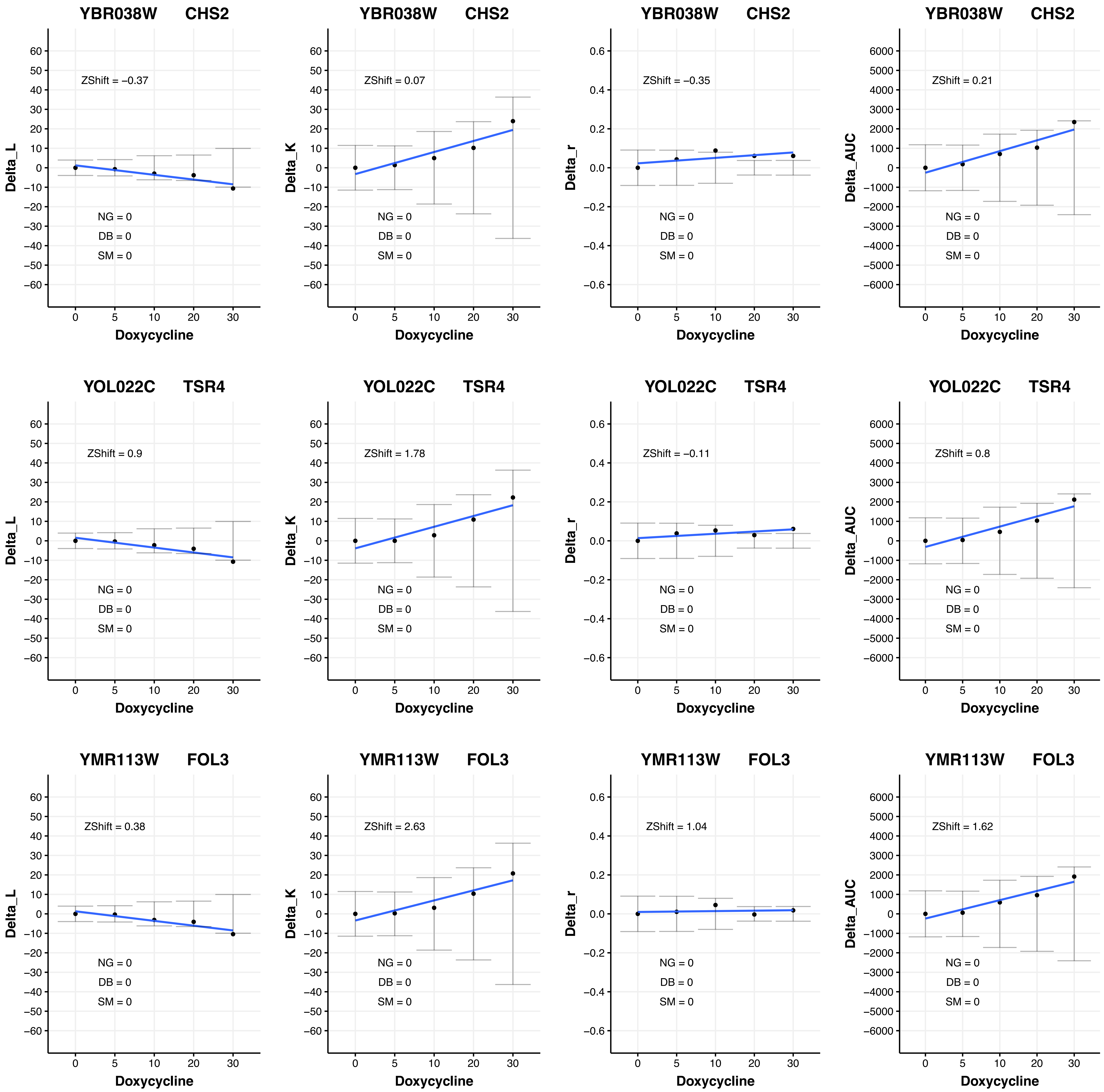
YBR038W CHS2
ZShift = −0.37
Delta_L
NG = 0
DB = 0
SM = 0
Doxycycline
YBR038W CHS2
ZShift = 0.07
Delta_K
NG = 0
DB = 0
SM = 0
Doxycycline
YBR038W CHS2
ZShift = −0.35
Delta_r
NG = 0
DB = 0
SM = 0
Doxycycline
YBR038W CHS2
ZShift = 0.21
Delta_AUC
NG = 0
DB = 0
SM = 0
Doxycycline
YOL022C TSR4
ZShift = 0.9
Delta_L
NG = 0
DB = 0
SM = 0
Doxycycline
YOL022C TSR4
ZShift = 1.78
Delta_K
NG = 0
DB = 0
SM = 0
Doxycycline
YOL022C TSR4
ZShift = −0.11
Delta_r
NG = 0
DB = 0
SM = 0
Doxycycline
YOL022C TSR4
ZShift = 0.8
Delta_AUC
NG = 0
DB = 0
SM = 0
Doxycycline
YMR113W FOL3
ZShift = 0.38
Delta_L
NG = 0
DB = 0
SM = 0
Doxycycline
YMR113W FOL3
ZShift = 2.63
Delta_K
NG = 0
DB = 0
SM = 0
Doxycycline
YMR113W FOL3
ZShift = 1.04
Delta_r
NG = 0
DB = 0
SM = 0
Doxycycline
YMR113W FOL3
ZShift = 1.62
Delta_AUC
NG = 0
DB = 0
SM = 0
Doxycycline

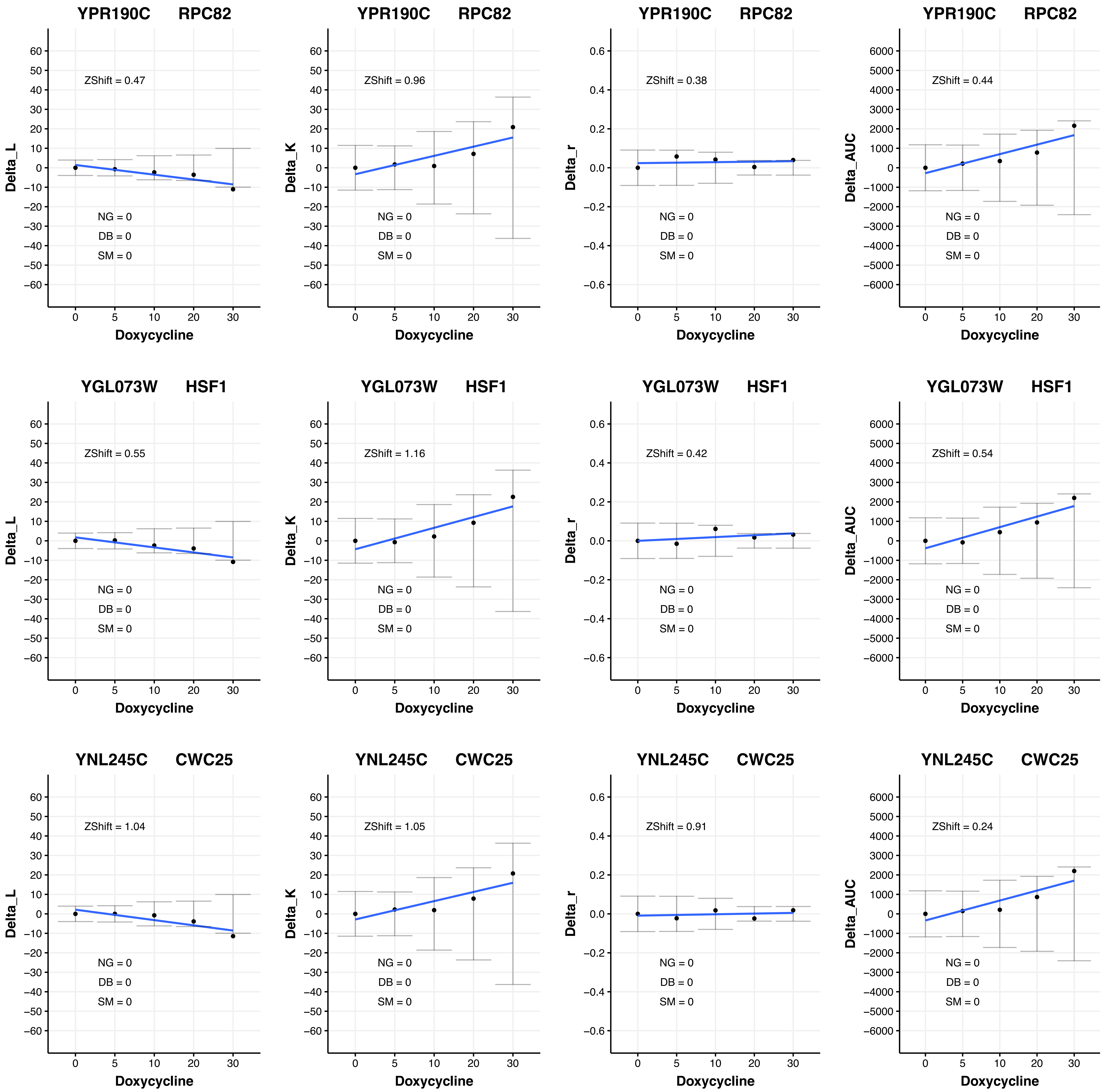
YPR190C RPC82
Delta_L
ZShift = 0.47
NG = 0
DB = 0
SM = 0
Doxycycline
YPR190C RPC82
Delta_K
ZShift = 0.96
NG = 0
DB = 0
SM = 0
Doxycycline
YPR190C RPC82
Delta_r
ZShift = 0.38
NG = 0
DB = 0
SM = 0
Doxycycline
YPR190C RPC82
Delta_AUC
ZShift = 0.44
NG = 0
DB = 0
SM = 0
Doxycycline
YGL073W HSF1
Delta_L
ZShift = 0.55
NG = 0
DB = 0
SM = 0
Doxycycline
YGL073W HSF1
Delta_K
ZShift = 1.16
NG = 0
DB = 0
SM = 0
Doxycycline
YGL073W HSF1
Delta_r
ZShift = 0.42
NG = 0
DB = 0
SM = 0
Doxycycline
YGL073W HSF1
Delta_AUC
ZShift = 0.54
NG = 0
DB = 0
SM = 0
Doxycycline
YNL245C CWC25
Delta_L
ZShift = 1.04
NG = 0
DB = 0
SM = 0
Doxycycline
YNL245C CWC25
Delta_K
ZShift = 1.05
NG = 0
DB = 0
SM = 0
Doxycycline
YNL245C CWC25
Delta_r
ZShift = 0.91
NG = 0
DB = 0
SM = 0
Doxycycline
YNL245C CWC25
Delta_AUC
ZShift = 0.24
NG = 0
DB = 0
SM = 0
Doxycycline

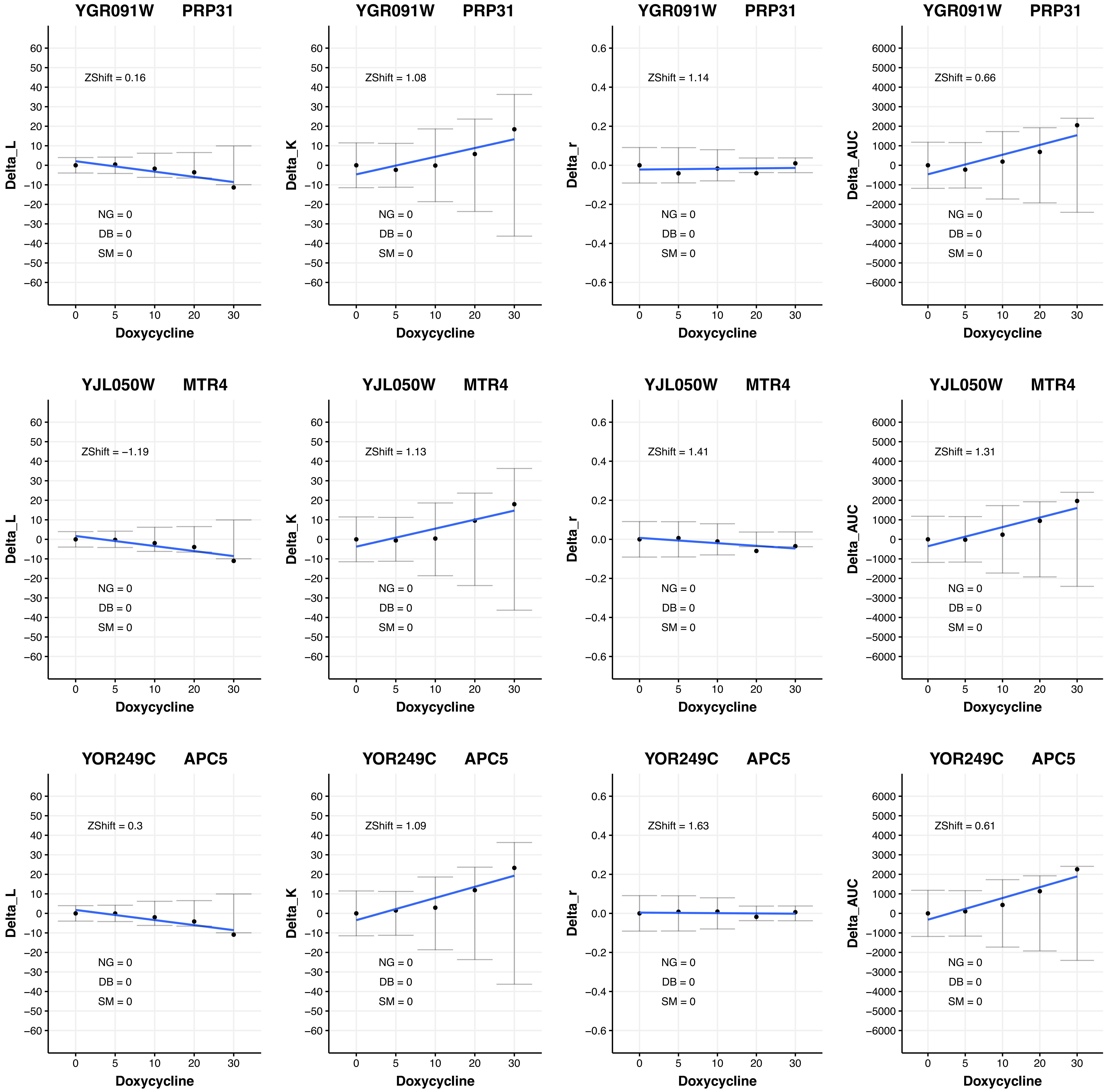

YGR091W PRP31
ZShift = 0.16
NG = 0
DB = 0
SM = 0
Delta_L
Doxycycline
YGR091W PRP31
ZShift = 1.08
NG = 0
DB = 0
SM = 0
Delta_K
Doxycycline
YGR091W PRP31
ZShift = 1.14
NG = 0
DB = 0
SM = 0
Delta_r
Doxycycline
YGR091W PRP31
ZShift = 0.66
NG = 0
DB = 0
SM = 0
Delta_AUC
Doxycycline
YJL050W MTR4
ZShift = −1.19
NG = 0
DB = 0
SM = 0
Delta_L
Doxycycline
YJL050W MTR4
ZShift = 1.13
NG = 0
DB = 0
SM = 0
Delta_K
Doxycycline
YJL050W MTR4
ZShift = 1.41
NG = 0
DB = 0
SM = 0
Delta_r
Doxycycline
YJL050W MTR4
ZShift = 1.31
NG = 0
DB = 0
SM = 0
Delta_AUC
Doxycycline
YOR249C APC5
ZShift = 0.3
NG = 0
DB = 0
SM = 0
Delta_L
Doxycycline
YOR249C APC5
ZShift = 1.09
NG = 0
DB = 0
SM = 0
Delta_K
Doxycycline
YOR249C APC5
ZShift = 1.63
NG = 0
DB = 0
SM = 0
Delta_r
Doxycycline
YOR249C APC5
ZShift = 0.61
NG = 0
DB = 0
SM = 0
Delta_AUC
Doxycycline

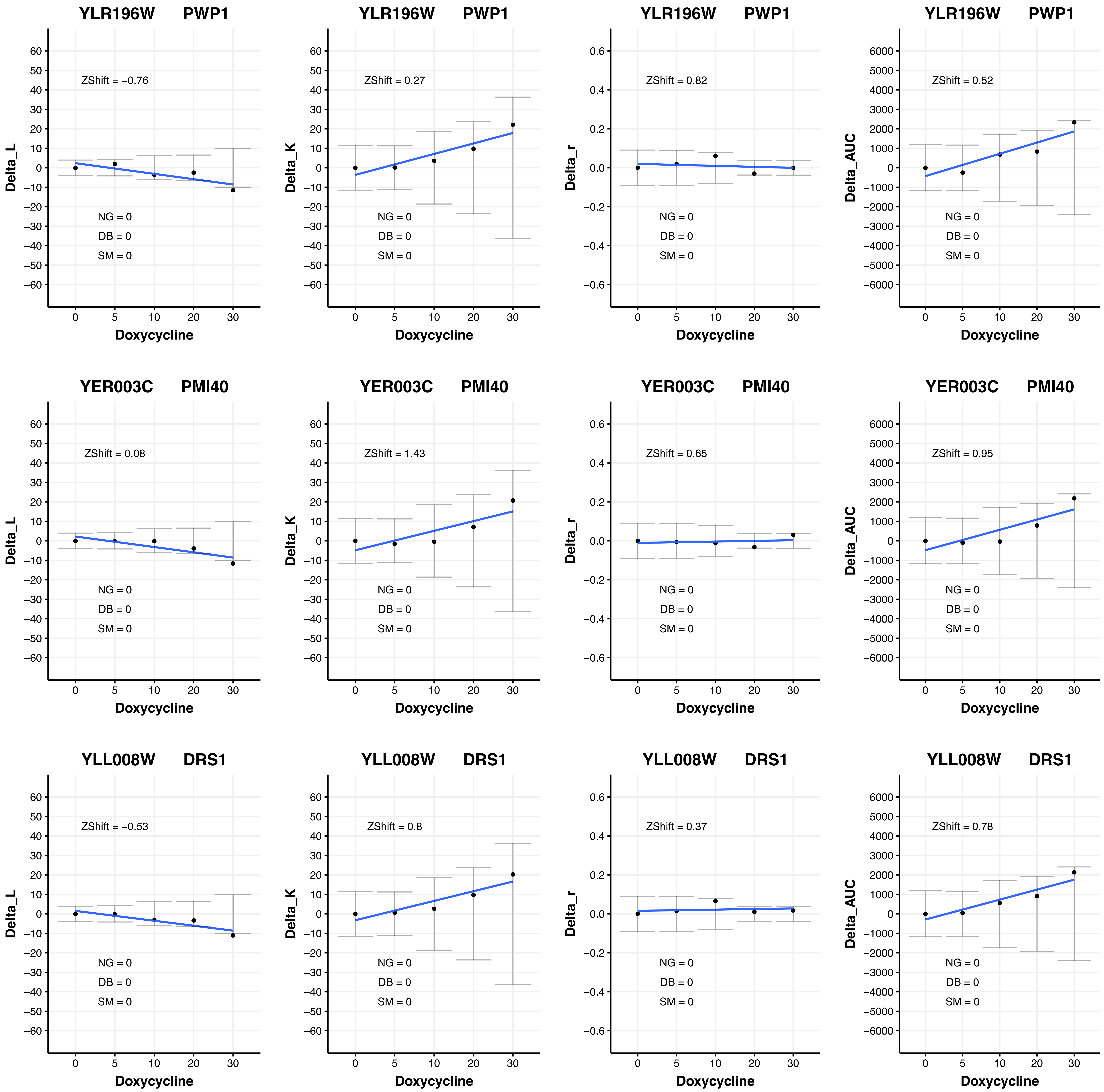
YLR196W PWP1
ZShift = −0.76
NG = 0
DB = 0
SM = 0
Delta_L
Doxycycline
YLR196W PWP1
ZShift = 0.27
NG = 0
DB = 0
SM = 0
Delta_K
Doxycycline
YLR196W PWP1
ZShift = 0.82
NG = 0
DB = 0
SM = 0
Delta_r
Doxycycline
YLR196W PWP1
ZShift = 0.52
NG = 0
DB = 0
SM = 0
Delta_AUC
Doxycycline
YER003C PMI40
ZShift = 0.08
NG = 0
DB = 0
SM = 0
Delta_L
Doxycycline
YER003C PMI40
ZShift = 1.43
NG = 0
DB = 0
SM = 0
Delta_K
Doxycycline
YER003C PMI40
ZShift = 0.65
NG = 0
DB = 0
SM = 0
Delta_r
Doxycycline
YER003C PMI40
ZShift = 0.95
NG = 0
DB = 0
SM = 0
Delta_AUC
Doxycycline
YLL008W DRS1
ZShift = −0.53
NG = 0
DB = 0
SM = 0
Delta_L
Doxycycline
YLL008W DRS1
ZShift = 0.8
NG = 0
DB = 0
SM = 0
Delta_K
Doxycycline
YLL008W DRS1
ZShift = 0.37
NG = 0
DB = 0
SM = 0
Delta_r
Doxycycline
YLL008W DRS1
ZShift = 0.78
NG = 0
DB = 0
SM = 0
Delta_AUC
Doxycycline

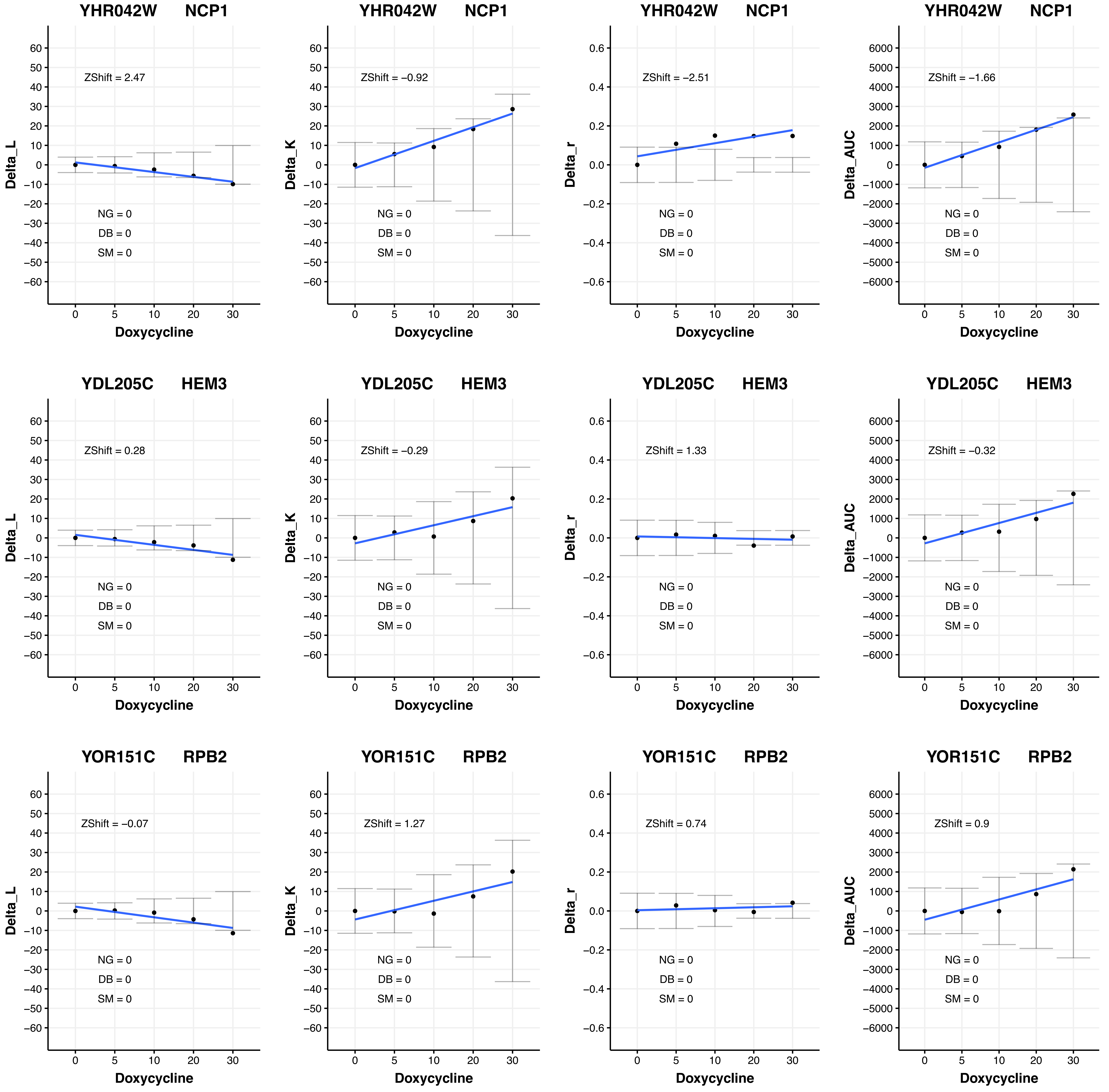
YHR042W NCP1
ZShift = 2.47
Delta_L
NG = 0
DB = 0
SM = 0
Doxycycline
YHR042W NCP1
ZShift = −0.92
Delta_K
NG = 0
DB = 0
SM = 0
Doxycycline
YHR042W NCP1
ZShift = −2.51
Delta_r
NG = 0
DB = 0
SM = 0
Doxycycline
YHR042W NCP1
ZShift = −1.66
Delta_AUC
NG = 0
DB = 0
SM = 0
Doxycycline
YDL205C HEM3
ZShift = 0.28
Delta_L
NG = 0
DB = 0
SM = 0
Doxycycline
YDL205C HEM3
ZShift = −0.29
Delta_K
NG = 0
DB = 0
SM = 0
Doxycycline
YDL205C HEM3
ZShift = 1.33
Delta_r
NG = 0
DB = 0
SM = 0
Doxycycline
YDL205C HEM3
ZShift = −0.32
Delta_AUC
NG = 0
DB = 0
SM = 0
Doxycycline
YOR151C RPB2
ZShift = −0.07
Delta_L
NG = 0
DB = 0
SM = 0
Doxycycline
YOR151C RPB2
ZShift = 1.27
Delta_K
NG = 0
DB = 0
SM = 0
Doxycycline
YOR151C RPB2
ZShift = 0.74
Delta_r
NG = 0
DB = 0
SM = 0
Doxycycline
YOR151C RPB2
ZShift = 0.9
Delta_AUC
NG = 0
DB = 0
SM = 0
Doxycycline

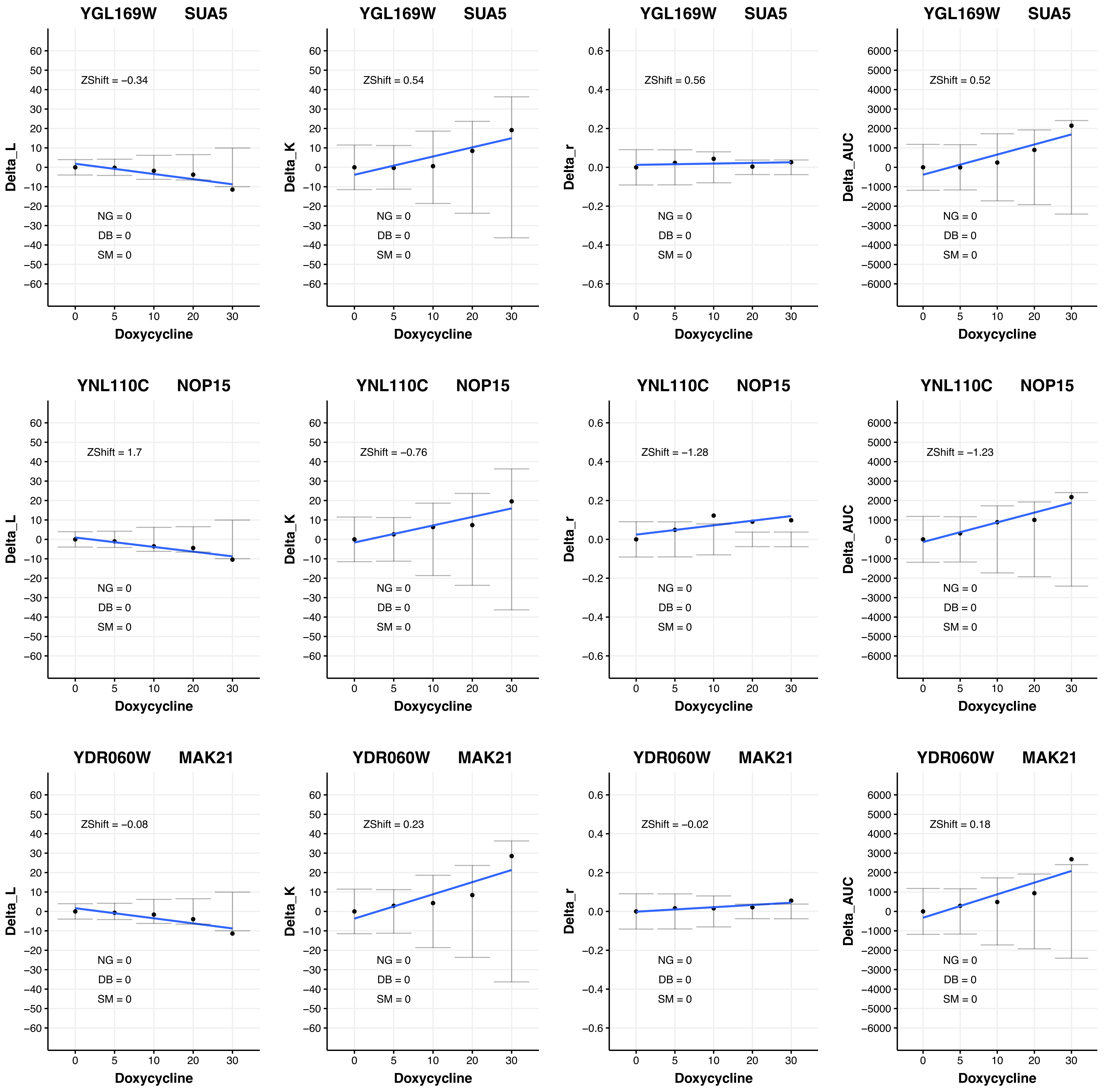

YGL169W SUA5
ZShift = −0.34
NG = 0
DB = 0
SM = 0
Delta_L
Doxycycline
YGL169W SUA5
ZShift = 0.54
NG = 0
DB = 0
SM = 0
Delta_K
Doxycycline
YGL169W SUA5
ZShift = 0.56
NG = 0
DB = 0
SM = 0
Delta_r
Doxycycline
YGL169W SUA5
ZShift = 0.52
NG = 0
DB = 0
SM = 0
Delta_AUC
Doxycycline
YNL110C NOP15
ZShift = 1.7
NG = 0
DB = 0
SM = 0
Delta_L
Doxycycline
YNL110C NOP15
ZShift = −0.76
NG = 0
DB = 0
SM = 0
Delta_K
Doxycycline
YNL110C NOP15
ZShift = −1.28
NG = 0
DB = 0
SM = 0
Delta_r
Doxycycline
YNL110C NOP15
ZShift = −1.23
NG = 0
DB = 0
SM = 0
Delta_AUC
Doxycycline
YDR060W MAK21
ZShift = −0.08
NG = 0
DB = 0
SM = 0
Delta_L
Doxycycline
YDR060W MAK21
ZShift = 0.23
NG = 0
DB = 0
SM = 0
Delta_K
Doxycycline
YDR060W MAK21
ZShift = −0.02
NG = 0
DB = 0
SM = 0
Delta_r
Doxycycline
YDR060W MAK21
ZShift = 0.18
NG = 0
DB = 0
SM = 0
Delta_AUC
Doxycycline

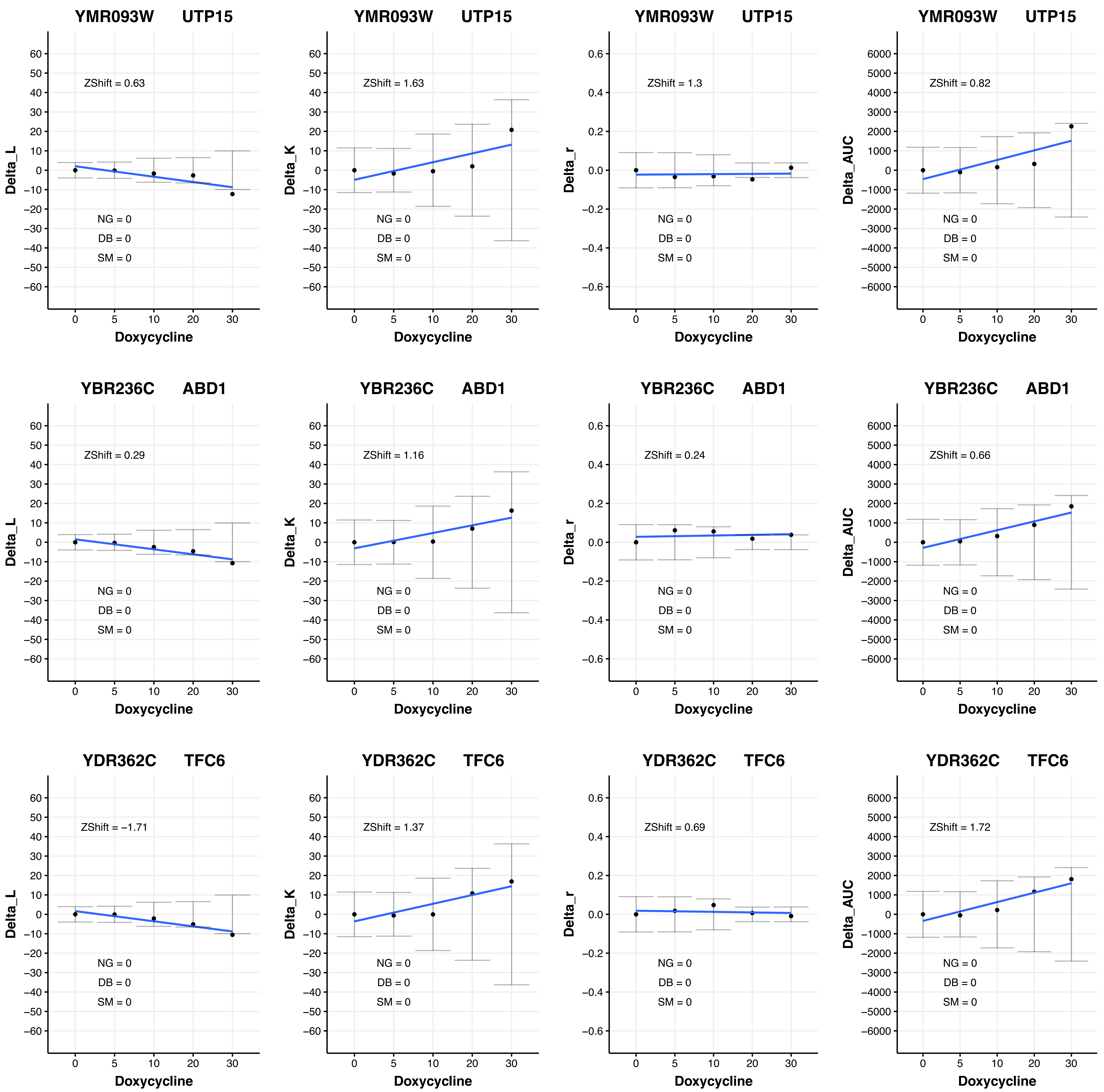
YMR093W UTP15
ZShift = 0.63
NG = 0
DB = 0
SM = 0
Delta_L
Doxycycline
YMR093W UTP15
ZShift = 1.63
NG = 0
DB = 0
SM = 0
Delta_K
Doxycycline
YMR093W UTP15
ZShift = 1.3
NG = 0
DB = 0
SM = 0
Delta_r
Doxycycline
YMR093W UTP15
ZShift = 0.82
NG = 0
DB = 0
SM = 0
Delta_AUC
Doxycycline
YBR236C ABD1
ZShift = 0.29
NG = 0
DB = 0
SM = 0
Delta_L
Doxycycline
YBR236C ABD1
ZShift = 1.16
NG = 0
DB = 0
SM = 0
Delta_K
Doxycycline
YBR236C ABD1
ZShift = 0.24
NG = 0
DB = 0
SM = 0
Delta_r
Doxycycline
YBR236C ABD1
ZShift = 0.66
NG = 0
DB = 0
SM = 0
Delta_AUC
Doxycycline
YDR362C TFC6
ZShift = −1.71
NG = 0
DB = 0
SM = 0
Delta_L
Doxycycline
YDR362C TFC6
ZShift = 1.37
NG = 0
DB = 0
SM = 0
Delta_K
Doxycycline
YDR362C TFC6
ZShift = 0.69
NG = 0
DB = 0
SM = 0
Delta_r
Doxycycline
YDR362C TFC6
ZShift = 1.72
NG = 0
DB = 0
SM = 0
Delta_AUC
Doxycycline

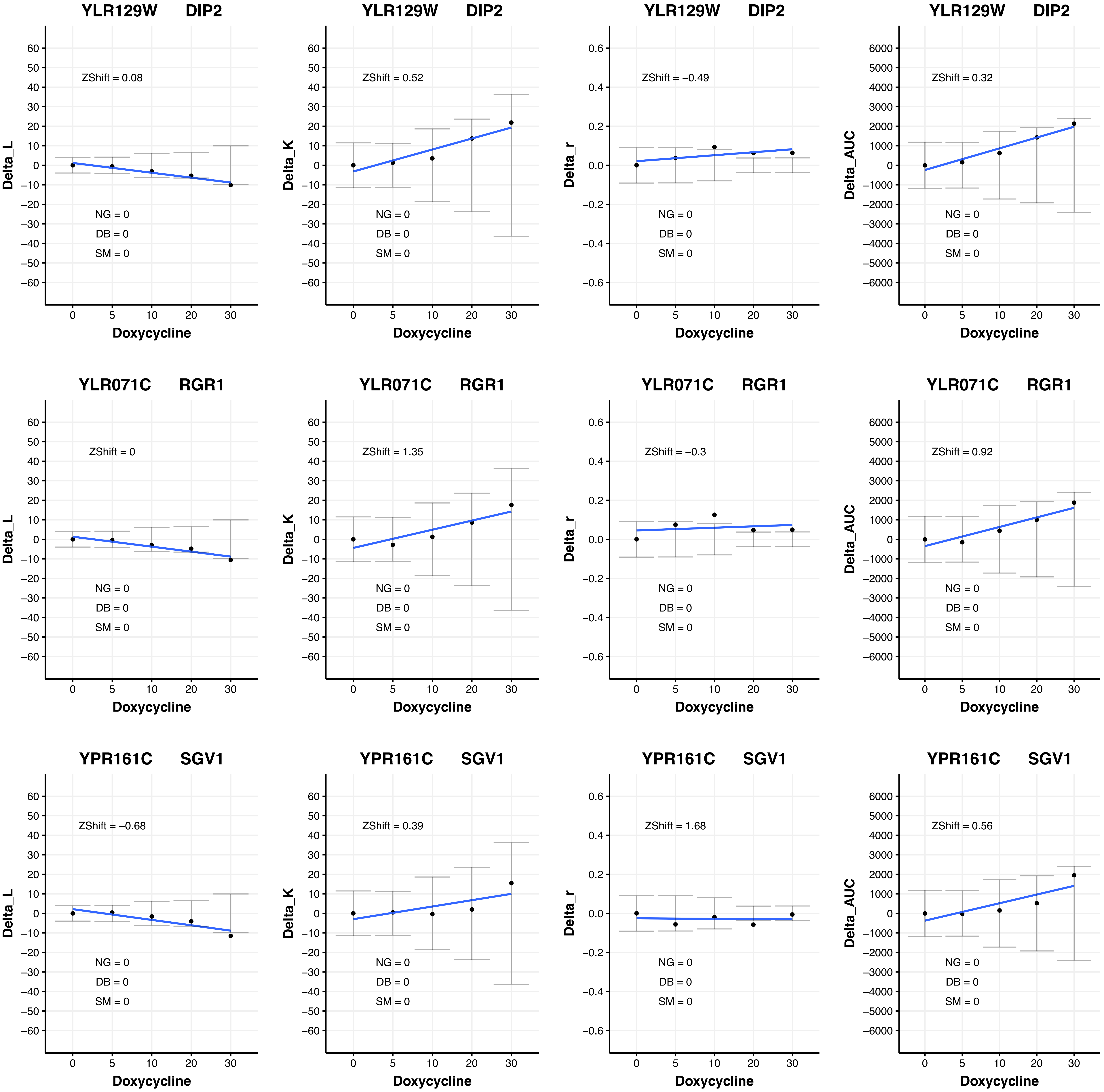
YLR129W DIP2
ZShift = 0.08
NG = 0
DB = 0
SM = 0
Delta_L
Doxycycline
YLR129W DIP2
ZShift = 0.52
NG = 0
DB = 0
SM = 0
Delta_K
Doxycycline
YLR129W DIP2
ZShift = −0.49
NG = 0
DB = 0
SM = 0
Delta_r
Doxycycline
YLR129W DIP2
ZShift = 0.32
NG = 0
DB = 0
SM = 0
Delta_AUC
Doxycycline
YLR071C RGR1
ZShift = 0
NG = 0
DB = 0
SM = 0
Delta_L
Doxycycline
YLR071C RGR1
ZShift = 1.35
NG = 0
DB = 0
SM = 0
Delta_K
Doxycycline
YLR071C RGR1
ZShift = −0.3
NG = 0
DB = 0
SM = 0
Delta_r
Doxycycline
YLR071C RGR1
ZShift = 0.92
NG = 0
DB = 0
SM = 0
Delta_AUC
Doxycycline
YPR161C SGV1
ZShift = −0.68
NG = 0
DB = 0
SM = 0
Delta_L
Doxycycline
YPR161C SGV1
ZShift = 0.39
NG = 0
DB = 0
SM = 0
Delta_K
Doxycycline
YPR161C SGV1
ZShift = 1.68
NG = 0
DB = 0
SM = 0
Delta_r
Doxycycline
YPR161C SGV1
ZShift = 0.56
NG = 0
DB = 0
SM = 0
Delta_AUC
Doxycycline

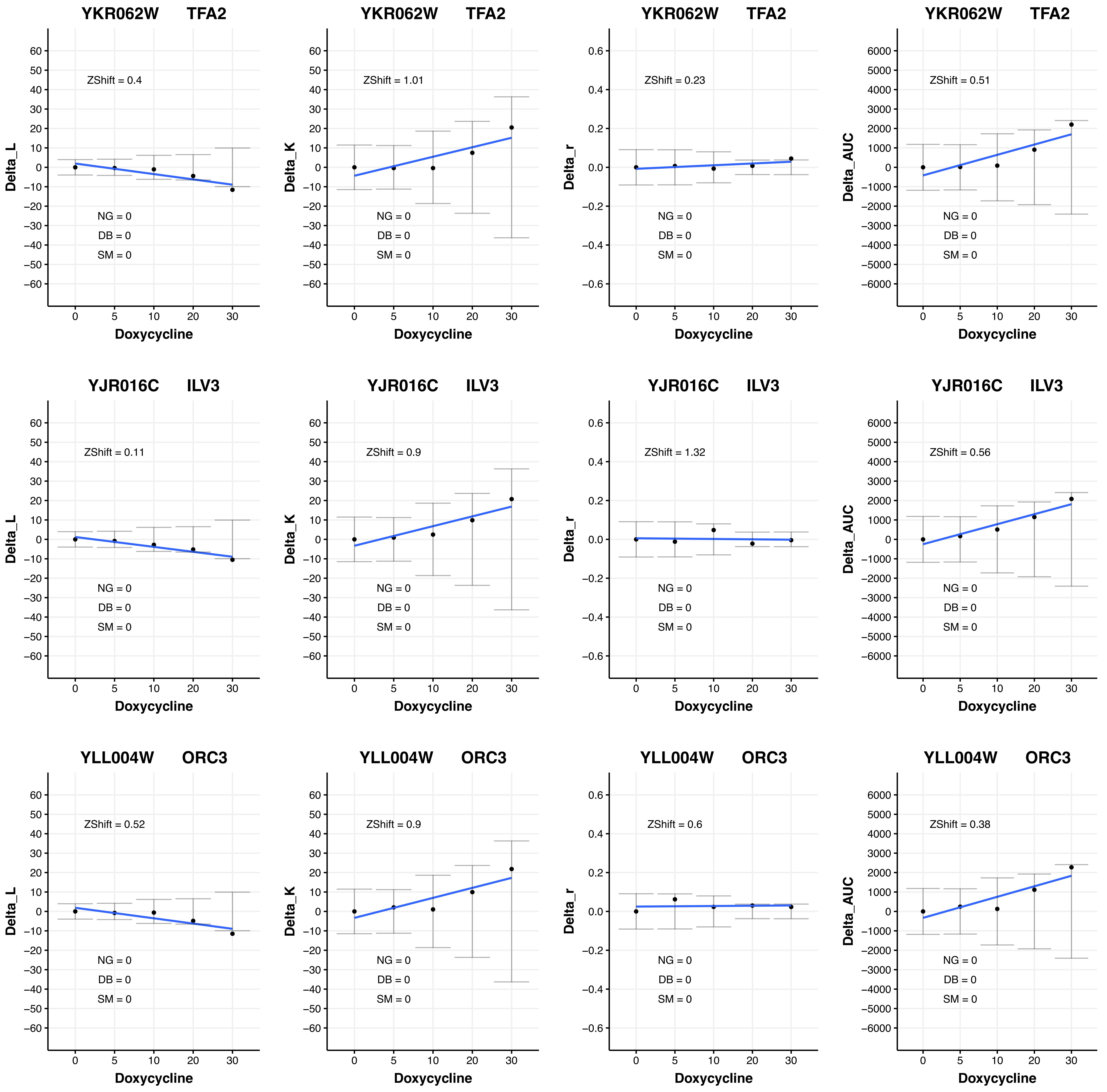

YKR062W TFA2
ZShift = 0.4
NG = 0
DB = 0
SM = 0
Delta_L
Doxycycline
YKR062W TFA2
ZShift = 1.01
NG = 0
DB = 0
SM = 0
Delta_K
Doxycycline
YKR062W TFA2
ZShift = 0.23
NG = 0
DB = 0
SM = 0
Delta_r
Doxycycline
YKR062W TFA2
ZShift = 0.51
NG = 0
DB = 0
SM = 0
Delta_AUC
Doxycycline
YJR016C ILV3
ZShift = 0.11
NG = 0
DB = 0
SM = 0
Delta_L
Doxycycline
YJR016C ILV3
ZShift = 0.9
NG = 0
DB = 0
SM = 0
Delta_K
Doxycycline
YJR016C ILV3
ZShift = 1.32
NG = 0
DB = 0
SM = 0
Delta_r
Doxycycline
YJR016C ILV3
ZShift = 0.56
NG = 0
DB = 0
SM = 0
Delta_AUC
Doxycycline
YLL004W ORC3
ZShift = 0.52
NG = 0
DB = 0
SM = 0
Delta_L
Doxycycline
YLL004W ORC3
ZShift = 0.9
NG = 0
DB = 0
SM = 0
Delta_K
Doxycycline
YLL004W ORC3
ZShift = 0.6
NG = 0
DB = 0
SM = 0
Delta_r
Doxycycline
YLL004W ORC3
ZShift = 0.38
NG = 0
DB = 0
SM = 0
Delta_AUC
Doxycycline

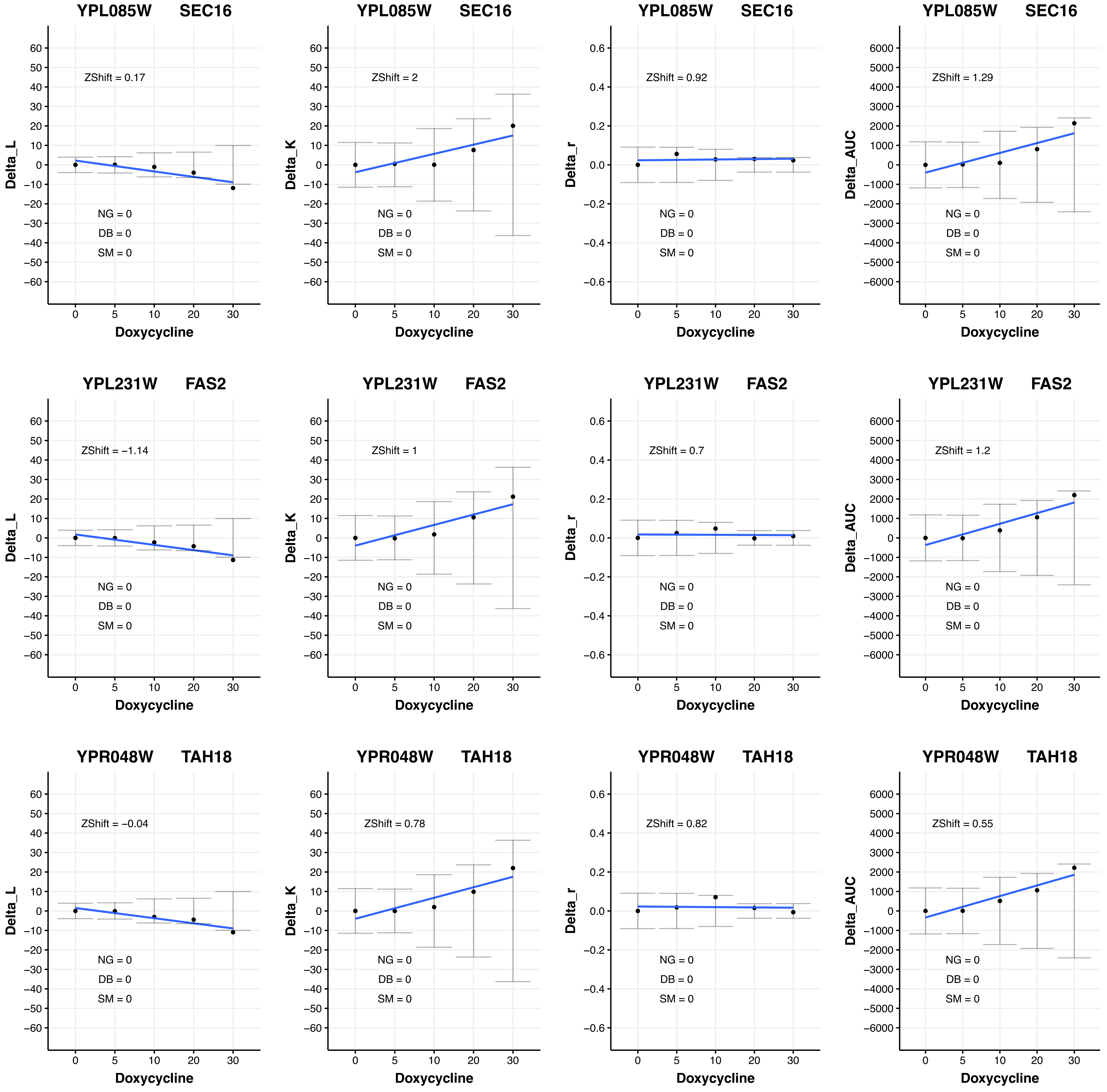
YPL085W SEC16
ZShift = 0.17
NG = 0
DB = 0
SM = 0
Delta_L
Doxycycline
YPL085W SEC16
ZShift = 2
NG = 0
DB = 0
SM = 0
Delta_K
Doxycycline
YPL085W SEC16
ZShift = 0.92
NG = 0
DB = 0
SM = 0
Delta_r
Doxycycline
YPL085W SEC16
ZShift = 1.29
NG = 0
DB = 0
SM = 0
Delta_AUC
Doxycycline
YPL231W FAS2
ZShift = −1.14
NG = 0
DB = 0
SM = 0
Delta_L
Doxycycline
YPL231W FAS2
ZShift = 1
NG = 0
DB = 0
SM = 0
Delta_K
Doxycycline
YPL231W FAS2
ZShift = 0.7
NG = 0
DB = 0
SM = 0
Delta_r
Doxycycline
YPL231W FAS2
ZShift = 1.2
NG = 0
DB = 0
SM = 0
Delta_AUC
Doxycycline
YPR048W TAH18
ZShift = −0.04
NG = 0
DB = 0
SM = 0
Delta_L
Doxycycline
YPR048W TAH18
ZShift = 0.78
NG = 0
DB = 0
SM = 0
Delta_K
Doxycycline
YPR048W TAH18
ZShift = 0.82
NG = 0
DB = 0
SM = 0
Delta_r
Doxycycline
YPR048W TAH18
ZShift = 0.55
NG = 0
DB = 0
SM = 0
Delta_AUC
Doxycycline

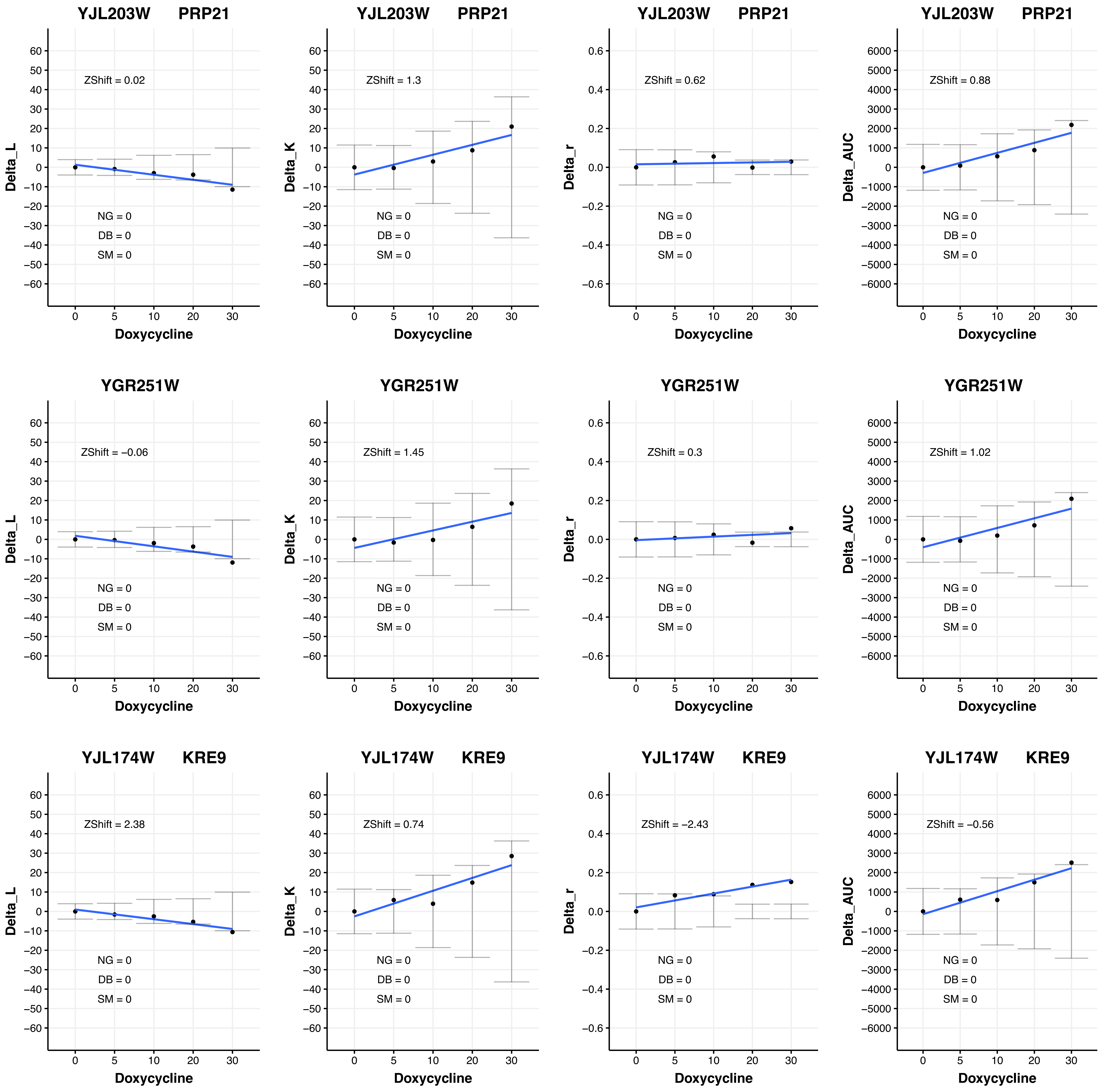
YJL203W PRP21
ZShift = 0.02
Delta_L
ZShift = 1.3
Delta_K
ZShift = 0.62
Delta_r
ZShift = 0.88
Delta_AUC
YGR251W
ZShift = −0.06
ZShift = 1.45
ZShift = 0.3
ZShift = 1.02
YJL174W KRE9
ZShift = 2.38
ZShift = 0.74
ZShift = −2.43
ZShift = −0.56
NG = 0
DB = 0
SM = 0
Doxycycline

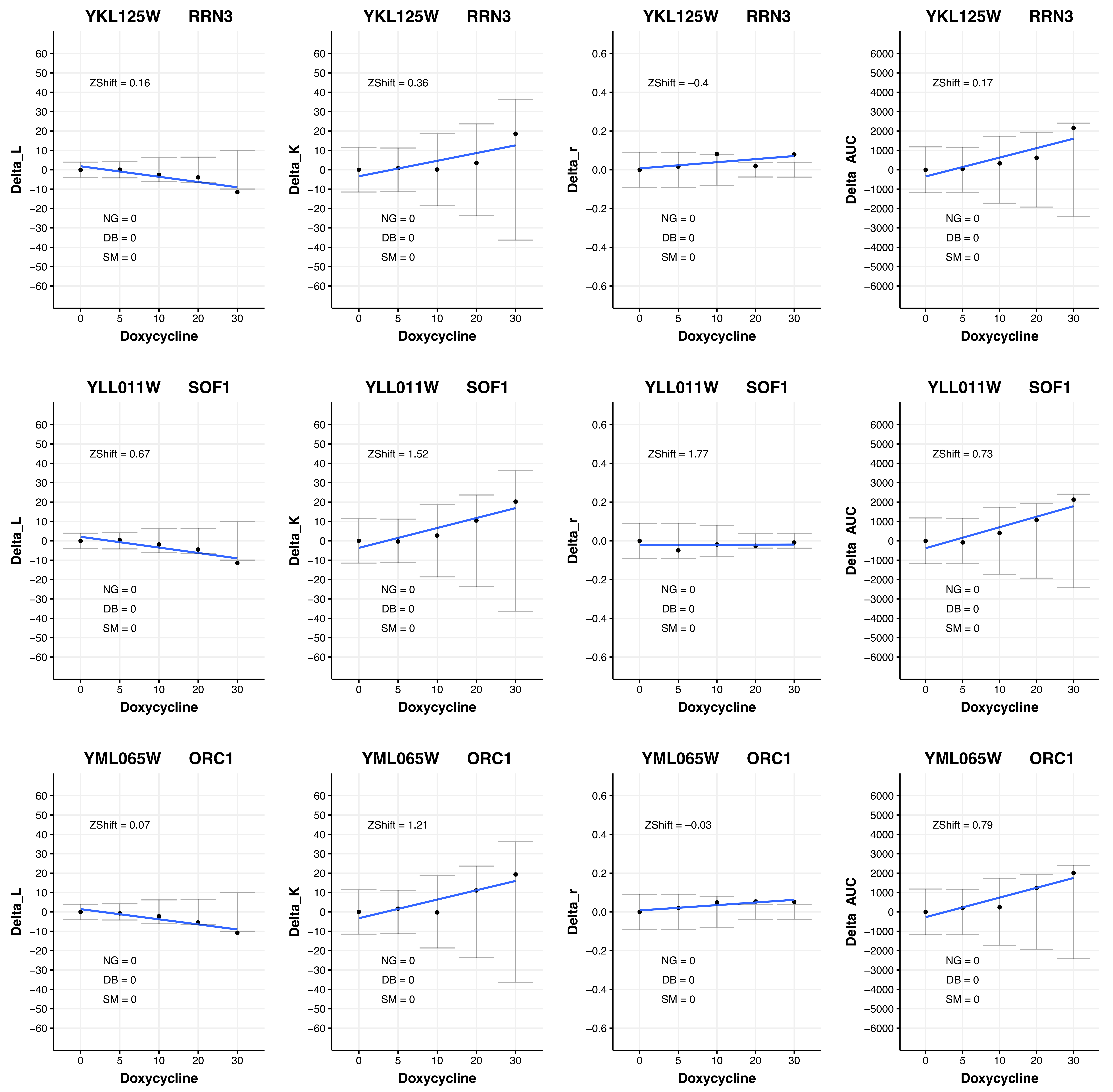
YKL125W RRN3
ZShift = 0.16
NG = 0
DB = 0
SM = 0
Delta_L
Doxycycline
YKL125W RRN3
ZShift = 0.36
NG = 0
DB = 0
SM = 0
Delta_K
Doxycycline
YKL125W RRN3
ZShift = −0.4
NG = 0
DB = 0
SM = 0
Delta_r
Doxycycline
YKL125W RRN3
ZShift = 0.17
NG = 0
DB = 0
SM = 0
Delta_AUC
Doxycycline
YLL011W SOF1
ZShift = 0.67
NG = 0
DB = 0
SM = 0
Delta_L
Doxycycline
YLL011W SOF1
ZShift = 1.52
NG = 0
DB = 0
SM = 0
Delta_K
Doxycycline
YLL011W SOF1
ZShift = 1.77
NG = 0
DB = 0
SM = 0
Delta_r
Doxycycline
YLL011W SOF1
ZShift = 0.73
NG = 0
DB = 0
SM = 0
Delta_AUC
Doxycycline
YML065W ORC1
ZShift = 0.07
NG = 0
DB = 0
SM = 0
Delta_L
Doxycycline
YML065W ORC1
ZShift = 1.21
NG = 0
DB = 0
SM = 0
Delta_K
Doxycycline
YML065W ORC1
ZShift = −0.03
NG = 0
DB = 0
SM = 0
Delta_r
Doxycycline
YML065W ORC1
ZShift = 0.79
NG = 0
DB = 0
SM = 0
Delta_AUC
Doxycycline

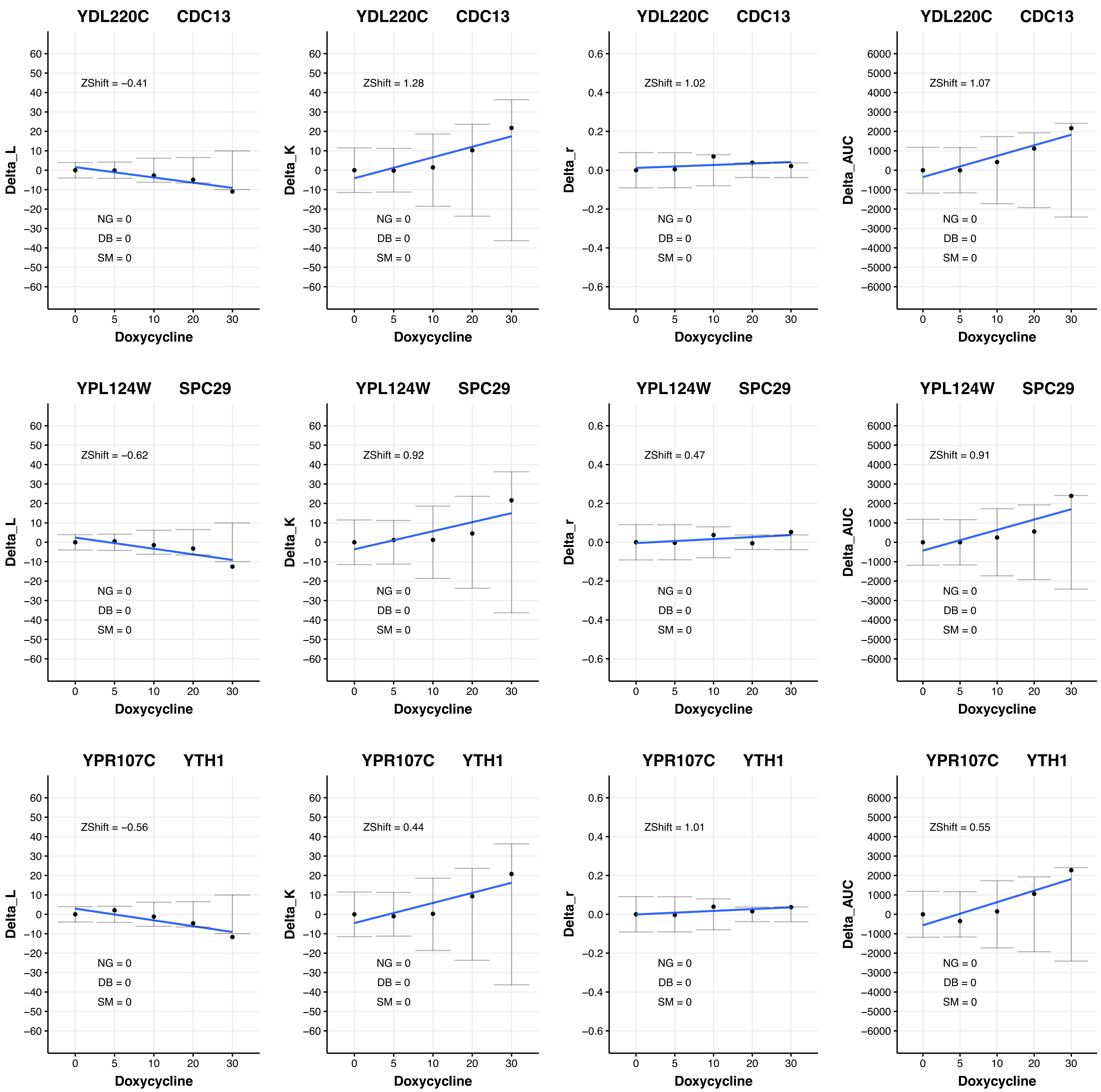
YDL220C CDC13
ZShift = −0.41
NG = 0
DB = 0
SM = 0
Delta_L
Doxycycline
YDL220C CDC13
ZShift = 1.28
NG = 0
DB = 0
SM = 0
Delta_K
Doxycycline
YDL220C CDC13
ZShift = 1.02
NG = 0
DB = 0
SM = 0
Delta_r
Doxycycline
YDL220C CDC13
ZShift = 1.07
NG = 0
DB = 0
SM = 0
Delta_AUC
Doxycycline
YPL124W SPC29
ZShift = −0.62
NG = 0
DB = 0
SM = 0
Delta_L
Doxycycline
YPL124W SPC29
ZShift = 0.92
NG = 0
DB = 0
SM = 0
Delta_K
Doxycycline
YPL124W SPC29
ZShift = 0.47
NG = 0
DB = 0
SM = 0
Delta_r
Doxycycline
YPL124W SPC29
ZShift = 0.91
NG = 0
DB = 0
SM = 0
Delta_AUC
Doxycycline
YPR107C YTH1
ZShift = −0.56
NG = 0
DB = 0
SM = 0
Delta_L
Doxycycline
YPR107C YTH1
ZShift = 0.44
NG = 0
DB = 0
SM = 0
Delta_K
Doxycycline
YPR107C YTH1
ZShift = 1.01
NG = 0
DB = 0
SM = 0
Delta_r
Doxycycline
YPR107C YTH1
ZShift = 0.55
NG = 0
DB = 0
SM = 0
Delta_AUC
Doxycycline

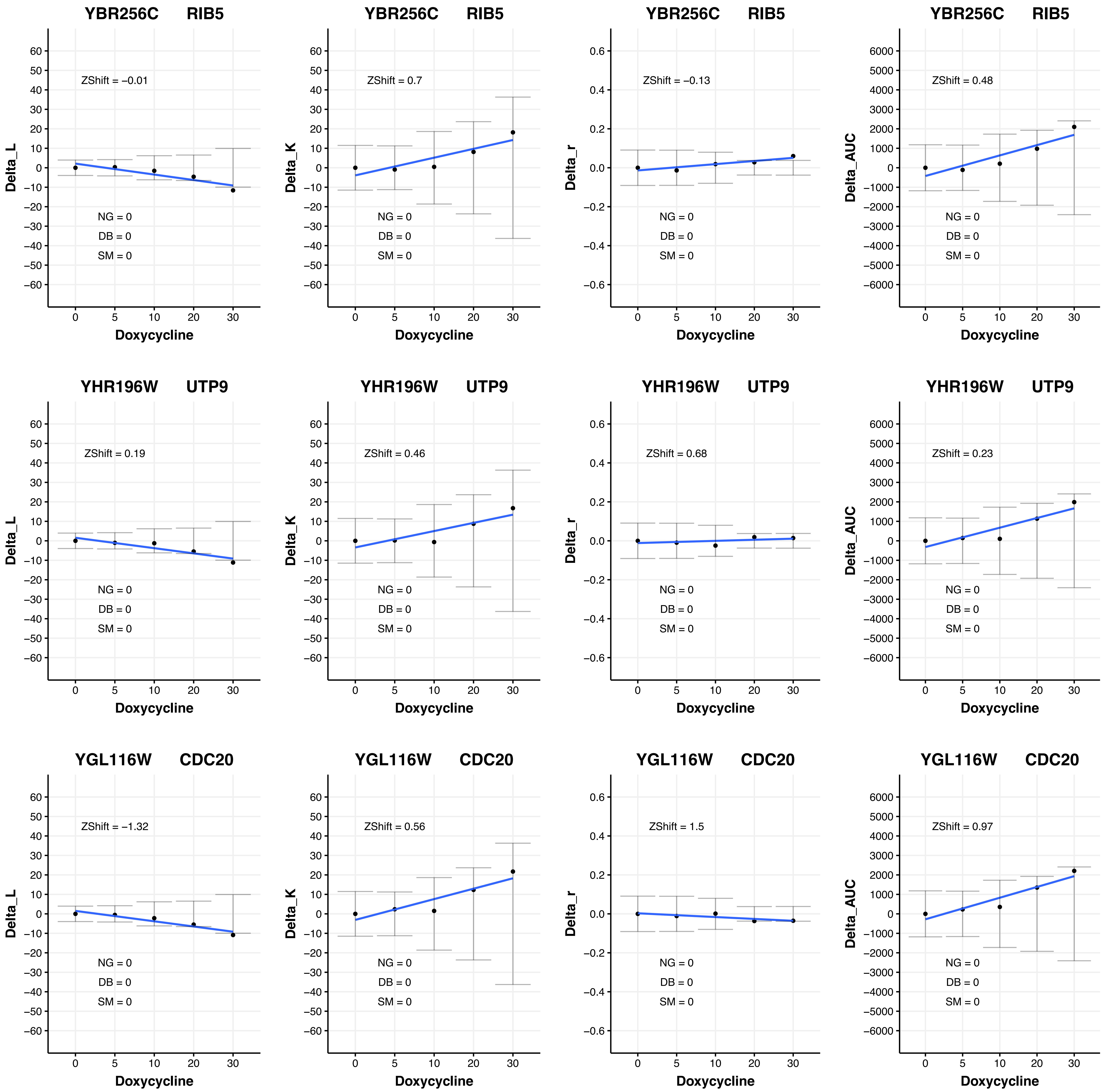

YBR256C RIB5
ZShift = −0.01
NG = 0
DB = 0
SM = 0
Delta_L
Doxycycline
YBR256C RIB5
ZShift = 0.7
NG = 0
DB = 0
SM = 0
Delta_K
Doxycycline
YBR256C RIB5
ZShift = −0.13
NG = 0
DB = 0
SM = 0
Delta_r
Doxycycline
YBR256C RIB5
ZShift = 0.48
NG = 0
DB = 0
SM = 0
Delta_AUC
Doxycycline
YHR196W UTP9
ZShift = 0.19
NG = 0
DB = 0
SM = 0
Delta_L
Doxycycline
YHR196W UTP9
ZShift = 0.46
NG = 0
DB = 0
SM = 0
Delta_K
Doxycycline
YHR196W UTP9
ZShift = 0.68
NG = 0
DB = 0
SM = 0
Delta_r
Doxycycline
YHR196W UTP9
ZShift = 0.23
NG = 0
DB = 0
SM = 0
Delta_AUC
Doxycycline
YGL116W CDC20
ZShift = −1.32
NG = 0
DB = 0
SM = 0
Delta_L
Doxycycline
YGL116W CDC20
ZShift = 0.56
NG = 0
DB = 0
SM = 0
Delta_K
Doxycycline
YGL116W CDC20
ZShift = 1.5
NG = 0
DB = 0
SM = 0
Delta_r
Doxycycline
YGL116W CDC20
ZShift = 0.97
NG = 0
DB = 0
SM = 0
Delta_AUC
Doxycycline

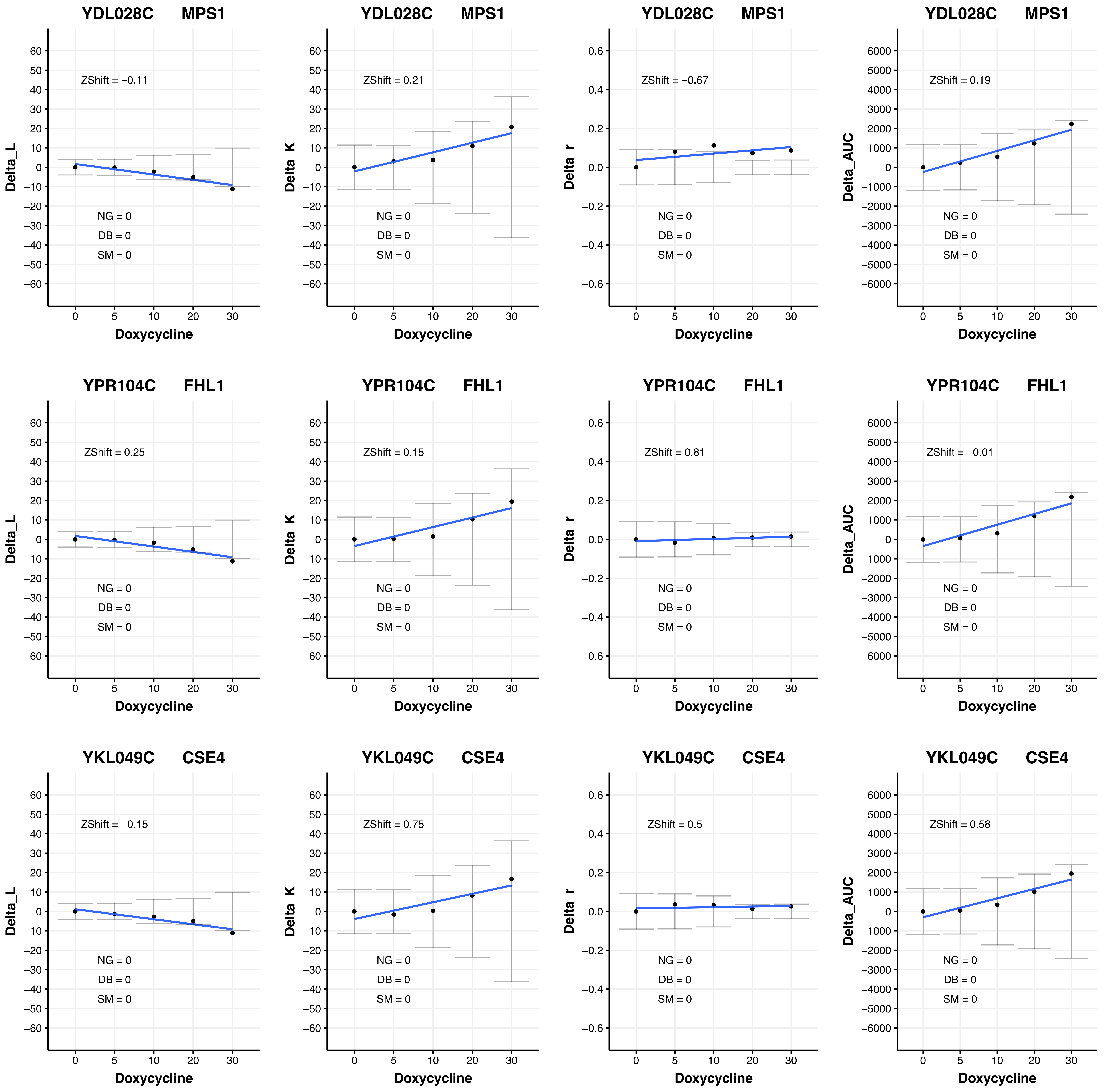
YDL028C MPS1
ZShift = −0.11
NG = 0
DB = 0
SM = 0
Delta_L
Doxycycline
YDL028C MPS1
ZShift = 0.21
NG = 0
DB = 0
SM = 0
Delta_K
Doxycycline
YDL028C MPS1
ZShift = −0.67
NG = 0
DB = 0
SM = 0
Delta_r
Doxycycline
YDL028C MPS1
ZShift = 0.19
NG = 0
DB = 0
SM = 0
Delta_AUC
Doxycycline
YPR104C FHL1
ZShift = 0.25
NG = 0
DB = 0
SM = 0
Delta_L
Doxycycline
YPR104C FHL1
ZShift = 0.15
NG = 0
DB = 0
SM = 0
Delta_K
Doxycycline
YPR104C FHL1
ZShift = 0.81
NG = 0
DB = 0
SM = 0
Delta_r
Doxycycline
YPR104C FHL1
ZShift = −0.01
NG = 0
DB = 0
SM = 0
Delta_AUC
Doxycycline
YKL049C CSE4
ZShift = −0.15
NG = 0
DB = 0
SM = 0
Delta_L
Doxycycline
YKL049C CSE4
ZShift = 0.75
NG = 0
DB = 0
SM = 0
Delta_K
Doxycycline
YKL049C CSE4
ZShift = 0.5
NG = 0
DB = 0
SM = 0
Delta_r
Doxycycline
YKL049C CSE4
ZShift = 0.58
NG = 0
DB = 0
SM = 0
Delta_AUC
Doxycycline

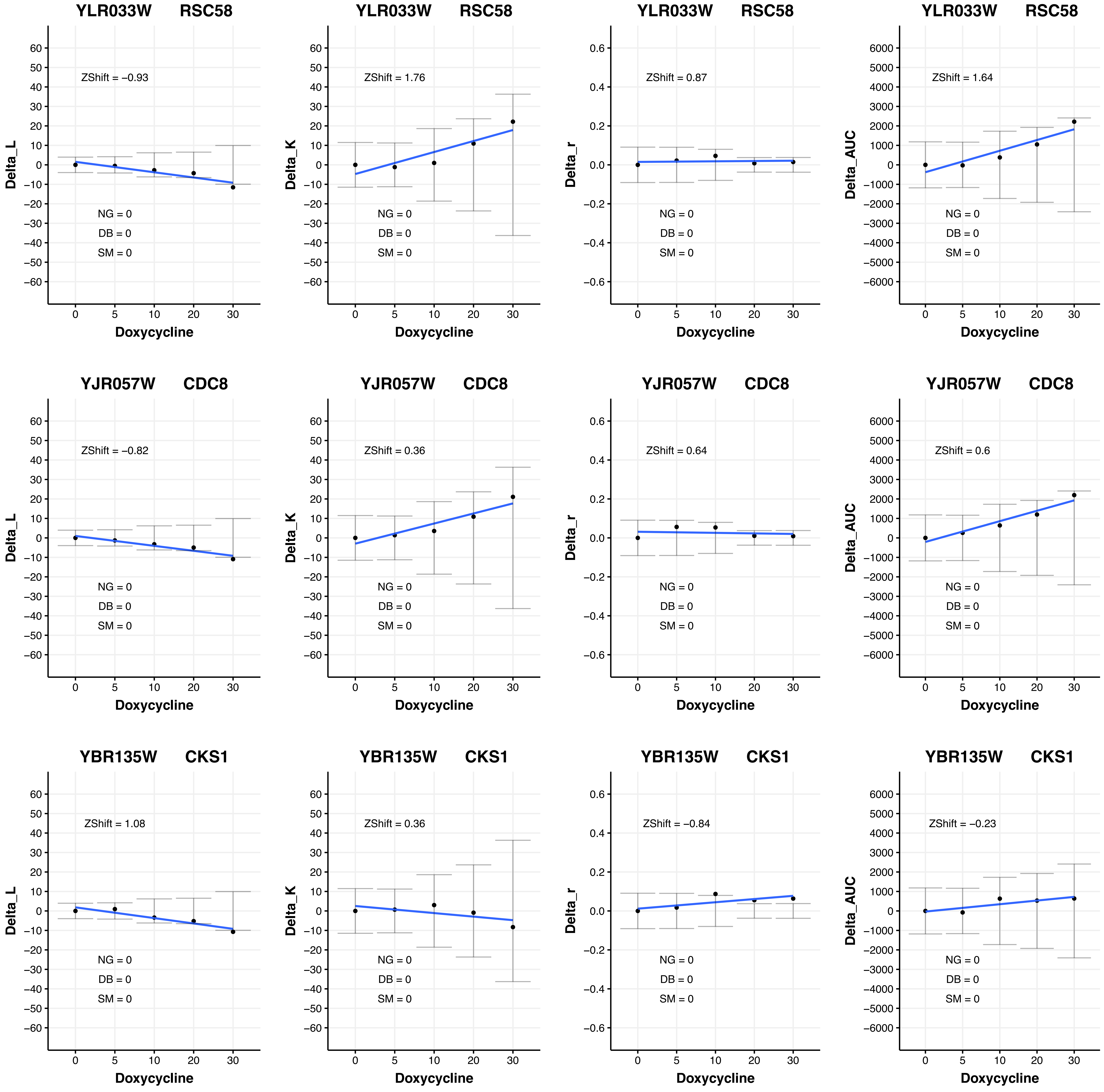
YLR033W RSC58
ZShift = −0.93
NG = 0
DB = 0
SM = 0
Delta_L
Doxycycline
YLR033W RSC58
ZShift = 1.76
NG = 0
DB = 0
SM = 0
Delta_K
Doxycycline
YLR033W RSC58
ZShift = 0.87
NG = 0
DB = 0
SM = 0
Delta_r
Doxycycline
YLR033W RSC58
ZShift = 1.64
NG = 0
DB = 0
SM = 0
Delta_AUC
Doxycycline
YJR057W CDC8
ZShift = −0.82
NG = 0
DB = 0
SM = 0
Delta_L
Doxycycline
YJR057W CDC8
ZShift = 0.36
NG = 0
DB = 0
SM = 0
Delta_K
Doxycycline
YJR057W CDC8
ZShift = 0.64
NG = 0
DB = 0
SM = 0
Delta_r
Doxycycline
YJR057W CDC8
ZShift = 0.6
NG = 0
DB = 0
SM = 0
Delta_AUC
Doxycycline
YBR135W CKS1
ZShift = 1.08
NG = 0
DB = 0
SM = 0
Delta_L
Doxycycline
YBR135W CKS1
ZShift = 0.36
NG = 0
DB = 0
SM = 0
Delta_K
Doxycycline
YBR135W CKS1
ZShift = −0.84
NG = 0
DB = 0
SM = 0
Delta_r
Doxycycline
YBR135W CKS1
ZShift = −0.23
NG = 0
DB = 0
SM = 0
Delta_AUC
Doxycycline

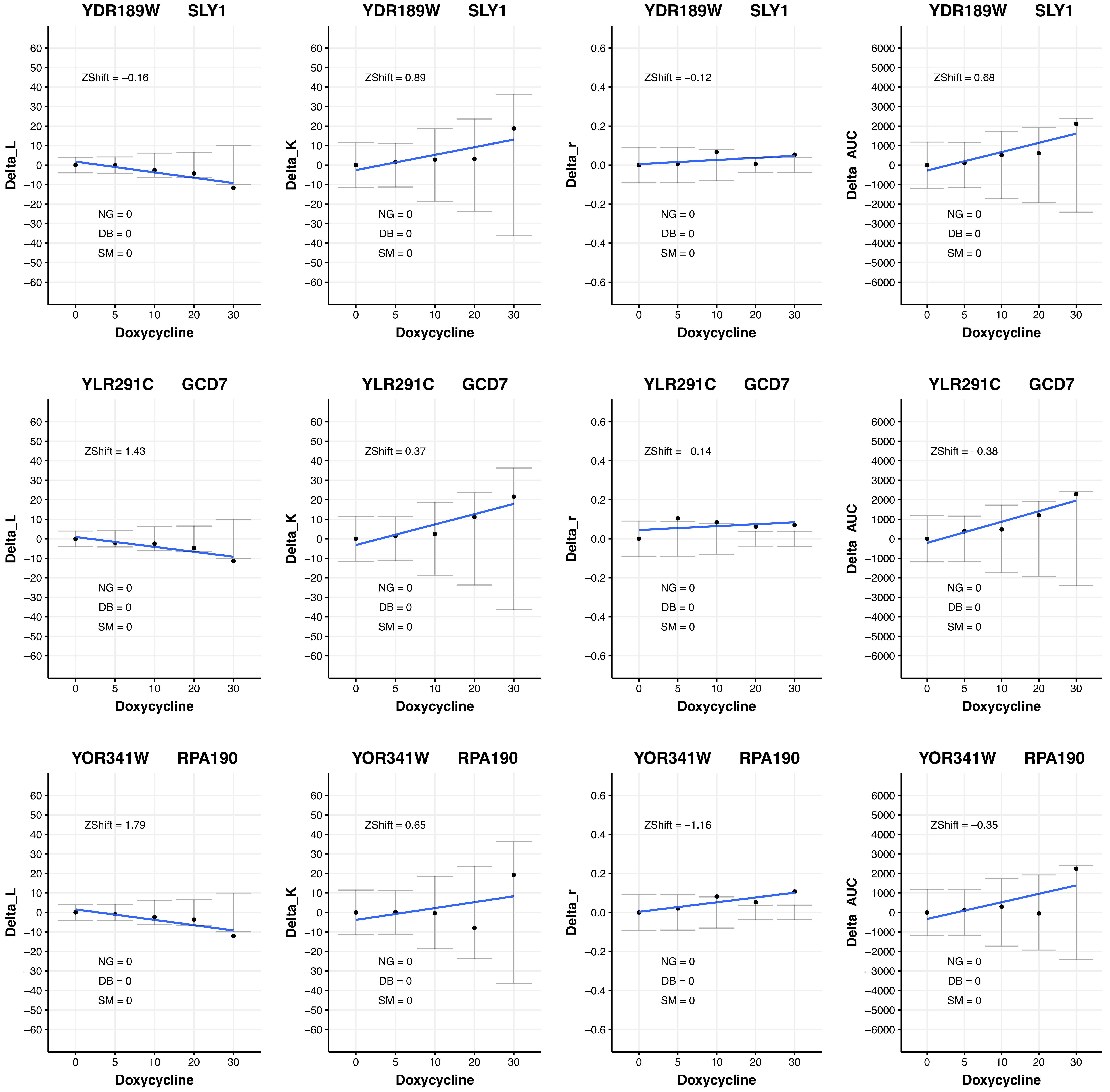

YDR189W SLY1
ZShift = −0.16
NG = 0
DB = 0
SM = 0
Delta_L
Doxycycline
YDR189W SLY1
ZShift = 0.89
NG = 0
DB = 0
SM = 0
Delta_K
Doxycycline
YDR189W SLY1
ZShift = −0.12
NG = 0
DB = 0
SM = 0
Delta_r
Doxycycline
YDR189W SLY1
ZShift = 0.68
NG = 0
DB = 0
SM = 0
Delta_AUC
Doxycycline
YLR291C GCD7
ZShift = 1.43
NG = 0
DB = 0
SM = 0
Delta_L
Doxycycline
YLR291C GCD7
ZShift = 0.37
NG = 0
DB = 0
SM = 0
Delta_K
Doxycycline
YLR291C GCD7
ZShift = −0.14
NG = 0
DB = 0
SM = 0
Delta_r
Doxycycline
YLR291C GCD7
ZShift = −0.38
NG = 0
DB = 0
SM = 0
Delta_AUC
Doxycycline
YOR341W RPA190
ZShift = 1.79
NG = 0
DB = 0
SM = 0
Delta_L
Doxycycline
YOR341W RPA190
ZShift = 0.65
NG = 0
DB = 0
SM = 0
Delta_K
Doxycycline
YOR341W RPA190
ZShift = −1.16
NG = 0
DB = 0
SM = 0
Delta_r
Doxycycline
YOR341W RPA190
ZShift = −0.35
NG = 0
DB = 0
SM = 0
Delta_AUC
Doxycycline

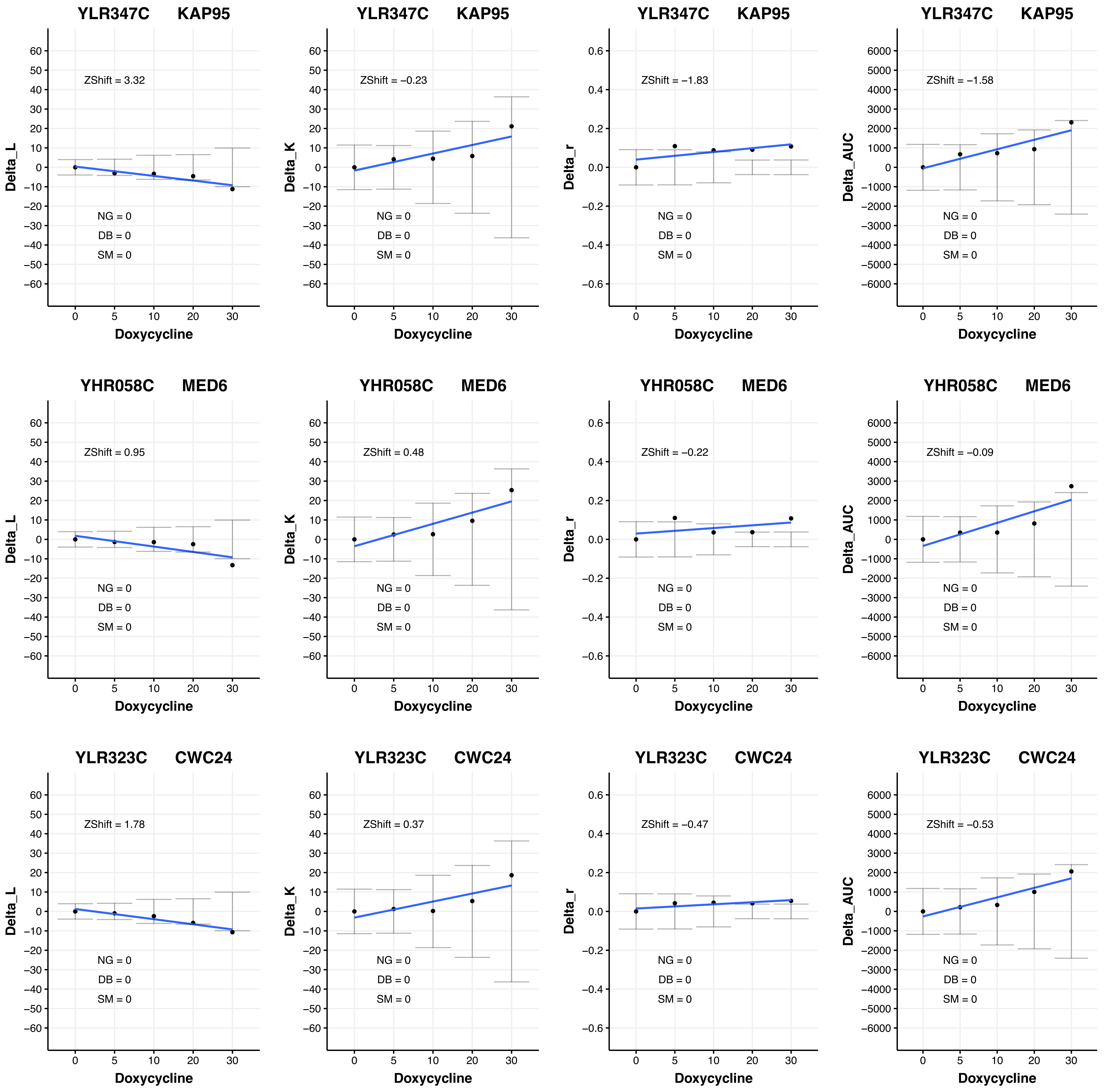
YLR347C KAP95
ZShift = 3.32
NG = 0
DB = 0
SM = 0
Delta_L
Doxycycline
YLR347C KAP95
ZShift = −0.23
NG = 0
DB = 0
SM = 0
Delta_K
Doxycycline
YLR347C KAP95
ZShift = −1.83
NG = 0
DB = 0
SM = 0
Delta_r
Doxycycline
YLR347C KAP95
ZShift = −1.58
NG = 0
DB = 0
SM = 0
Delta_AUC
Doxycycline
YHR058C MED6
ZShift = 0.95
NG = 0
DB = 0
SM = 0
Delta_L
Doxycycline
YHR058C MED6
ZShift = 0.48
NG = 0
DB = 0
SM = 0
Delta_K
Doxycycline
YHR058C MED6
ZShift = −0.22
NG = 0
DB = 0
SM = 0
Delta_r
Doxycycline
YHR058C MED6
ZShift = −0.09
NG = 0
DB = 0
SM = 0
Delta_AUC
Doxycycline
YLR323C CWC24
ZShift = 1.78
NG = 0
DB = 0
SM = 0
Delta_L
Doxycycline
YLR323C CWC24
ZShift = 0.37
NG = 0
DB = 0
SM = 0
Delta_K
Doxycycline
YLR323C CWC24
ZShift = −0.47
NG = 0
DB = 0
SM = 0
Delta_r
Doxycycline
YLR323C CWC24
ZShift = −0.53
NG = 0
DB = 0
SM = 0
Delta_AUC
Doxycycline

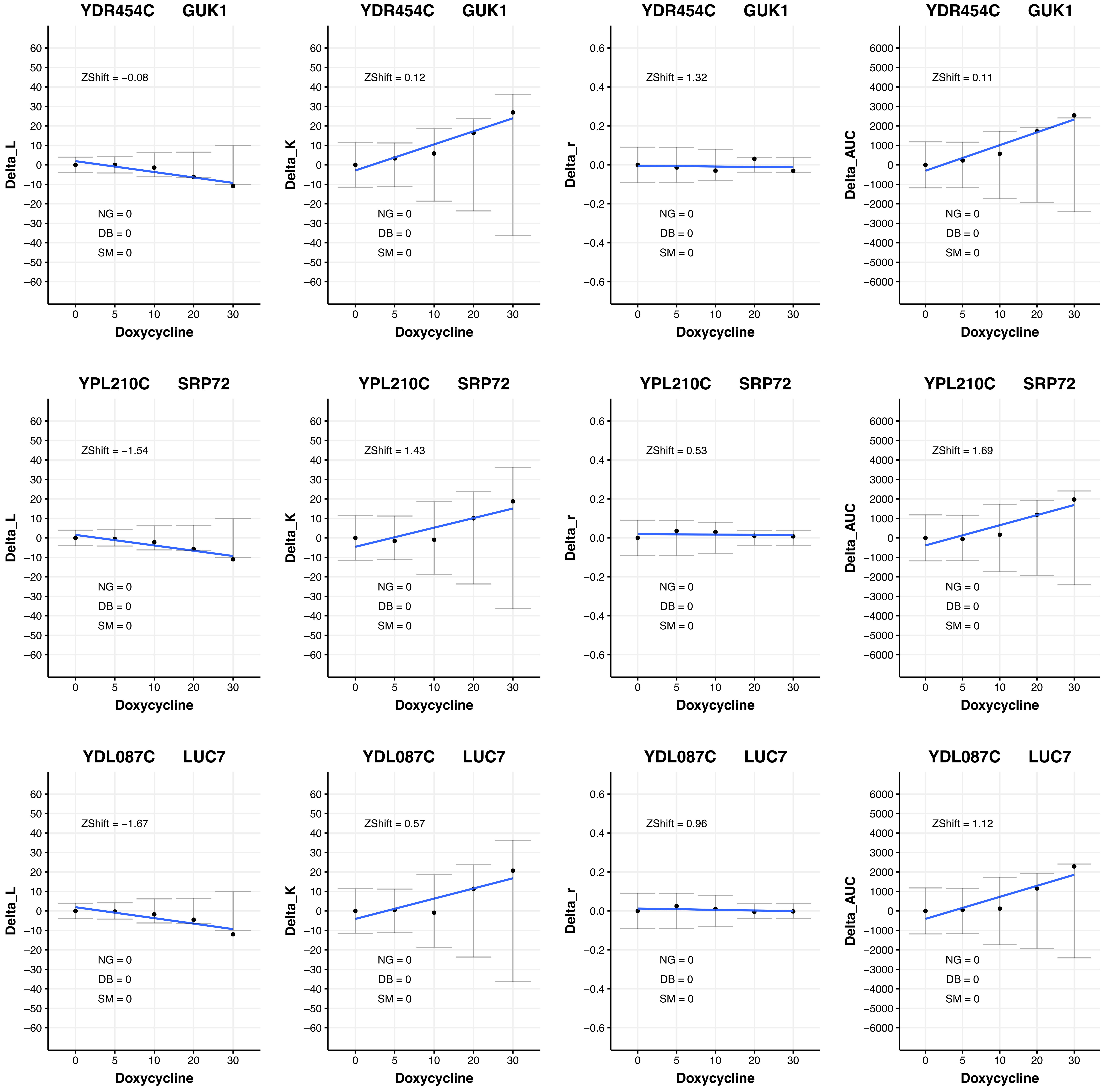

YDR454C GUK1
ZShift = −0.08
NG = 0
DB = 0
SM = 0
Delta_L
Doxycycline
YDR454C GUK1
ZShift = 0.12
NG = 0
DB = 0
SM = 0
Delta_K
Doxycycline
YDR454C GUK1
ZShift = 1.32
NG = 0
DB = 0
SM = 0
Delta_r
Doxycycline
YDR454C GUK1
ZShift = 0.11
NG = 0
DB = 0
SM = 0
Delta_AUC
Doxycycline
YPL210C SRP72
ZShift = −1.54
NG = 0
DB = 0
SM = 0
Delta_L
Doxycycline
YPL210C SRP72
ZShift = 1.43
NG = 0
DB = 0
SM = 0
Delta_K
Doxycycline
YPL210C SRP72
ZShift = 0.53
NG = 0
DB = 0
SM = 0
Delta_r
Doxycycline
YPL210C SRP72
ZShift = 1.69
NG = 0
DB = 0
SM = 0
Delta_AUC
Doxycycline
YDL087C LUC7
ZShift = −1.67
NG = 0
DB = 0
SM = 0
Delta_L
Doxycycline
YDL087C LUC7
ZShift = 0.57
NG = 0
DB = 0
SM = 0
Delta_K
Doxycycline
YDL087C LUC7
ZShift = 0.96
NG = 0
DB = 0
SM = 0
Delta_r
Doxycycline
YDL087C LUC7
ZShift = 1.12
NG = 0
DB = 0
SM = 0
Delta_AUC
Doxycycline

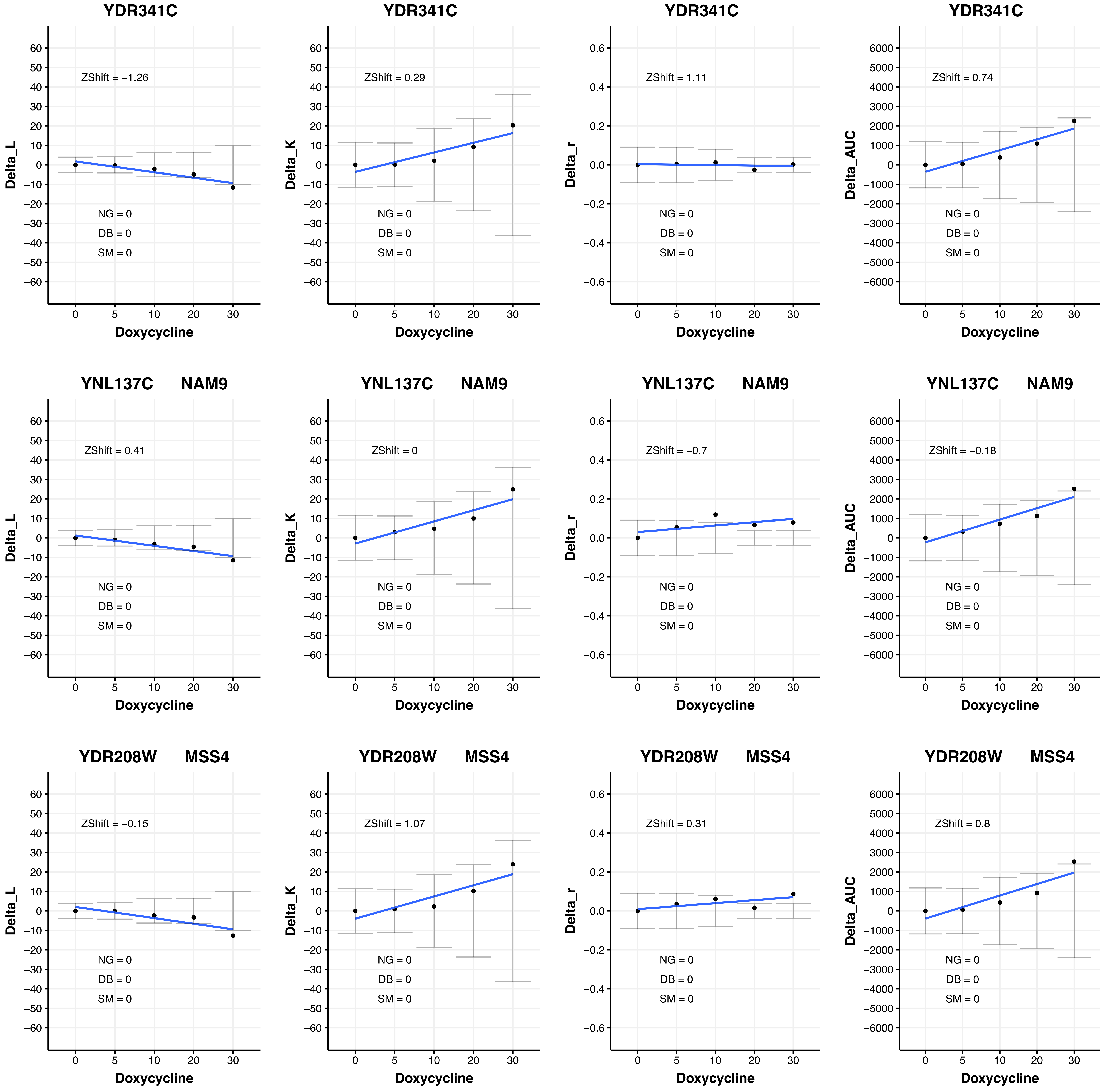

YDR341C
Delta_L
ZShift = −1.26
NG = 0
DB = 0
SM = 0
Doxycycline
YDR341C
Delta_K
ZShift = 0.29
NG = 0
DB = 0
SM = 0
Doxycycline
YDR341C
Delta_r
ZShift = 1.11
NG = 0
DB = 0
SM = 0
Doxycycline
YDR341C
Delta_AUC
ZShift = 0.74
NG = 0
DB = 0
SM = 0
Doxycycline
YNL137C NAM9
Delta_L
ZShift = 0.41
NG = 0
DB = 0
SM = 0
Doxycycline
YNL137C NAM9
Delta_K
ZShift = 0
NG = 0
DB = 0
SM = 0
Doxycycline
YNL137C NAM9
Delta_r
ZShift = −0.7
NG = 0
DB = 0
SM = 0
Doxycycline
YNL137C NAM9
Delta_AUC
ZShift = −0.18
NG = 0
DB = 0
SM = 0
Doxycycline
YDR208W MSS4
Delta_L
ZShift = −0.15
NG = 0
DB = 0
SM = 0
Doxycycline
YDR208W MSS4
Delta_K
ZShift = 1.07
NG = 0
DB = 0
SM = 0
Doxycycline
YDR208W MSS4
Delta_r
ZShift = 0.31
NG = 0
DB = 0
SM = 0
Doxycycline
YDR208W MSS4
Delta_AUC
ZShift = 0.8
NG = 0
DB = 0
SM = 0
Doxycycline

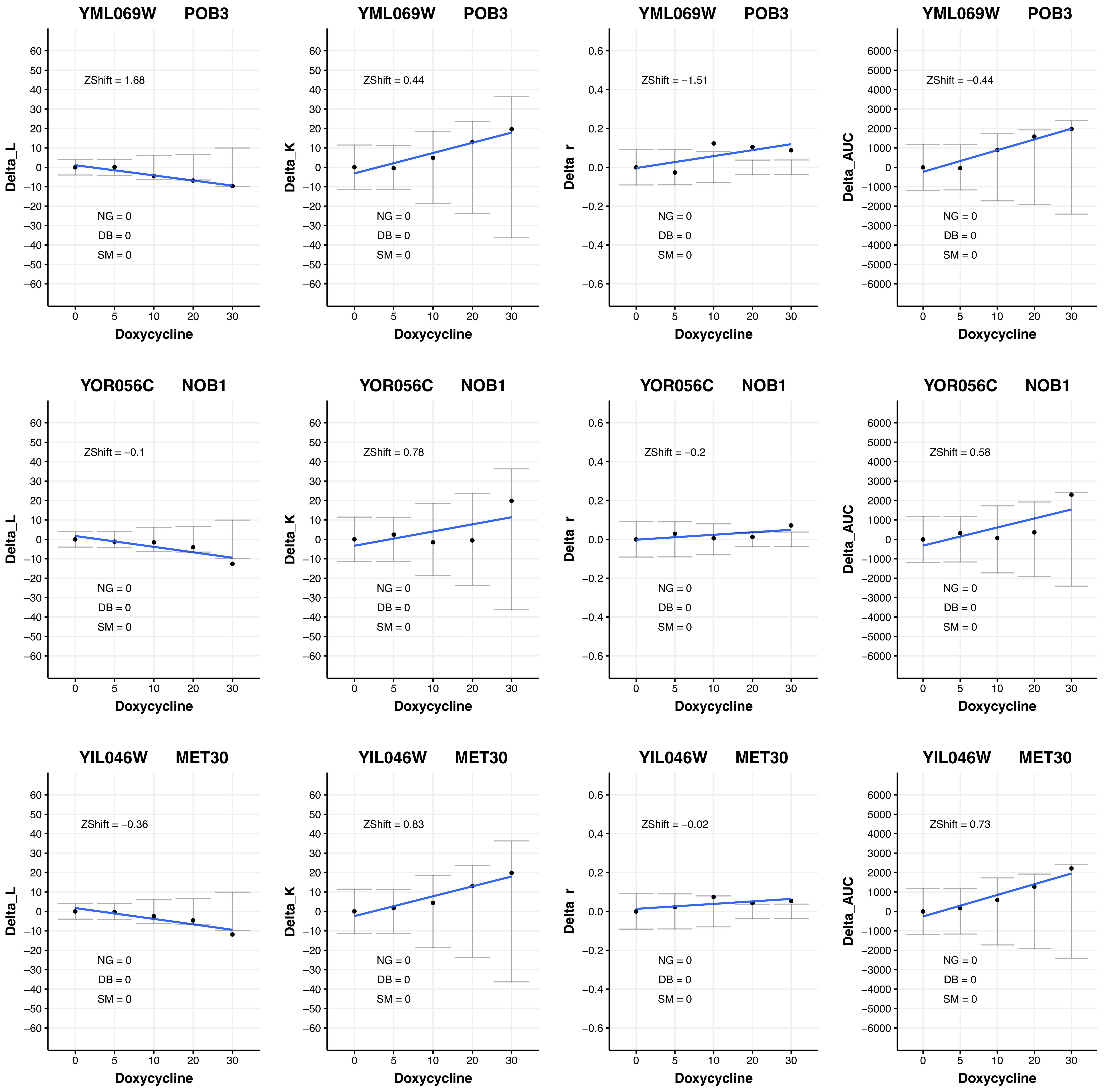
YML069W POB3
ZShift = 1.68
NG = 0
DB = 0
SM = 0
Delta_L
Doxycycline
YML069W POB3
ZShift = 0.44
NG = 0
DB = 0
SM = 0
Delta_K
Doxycycline
YML069W POB3
ZShift = −1.51
NG = 0
DB = 0
SM = 0
Delta_r
Doxycycline
YML069W POB3
ZShift = −0.44
NG = 0
DB = 0
SM = 0
Delta_AUC
Doxycycline
YOR056C NOB1
ZShift = −0.1
NG = 0
DB = 0
SM = 0
Delta_L
Doxycycline
YOR056C NOB1
ZShift = 0.78
NG = 0
DB = 0
SM = 0
Delta_K
Doxycycline
YOR056C NOB1
ZShift = −0.2
NG = 0
DB = 0
SM = 0
Delta_r
Doxycycline
YOR056C NOB1
ZShift = 0.58
NG = 0
DB = 0
SM = 0
Delta_AUC
Doxycycline
YIL046W MET30
ZShift = −0.36
NG = 0
DB = 0
SM = 0
Delta_L
Doxycycline
YIL046W MET30
ZShift = 0.83
NG = 0
DB = 0
SM = 0
Delta_K
Doxycycline
YIL046W MET30
ZShift = −0.02
NG = 0
DB = 0
SM = 0
Delta_r
Doxycycline
YIL046W MET30
ZShift = 0.73
NG = 0
DB = 0
SM = 0
Delta_AUC
Doxycycline

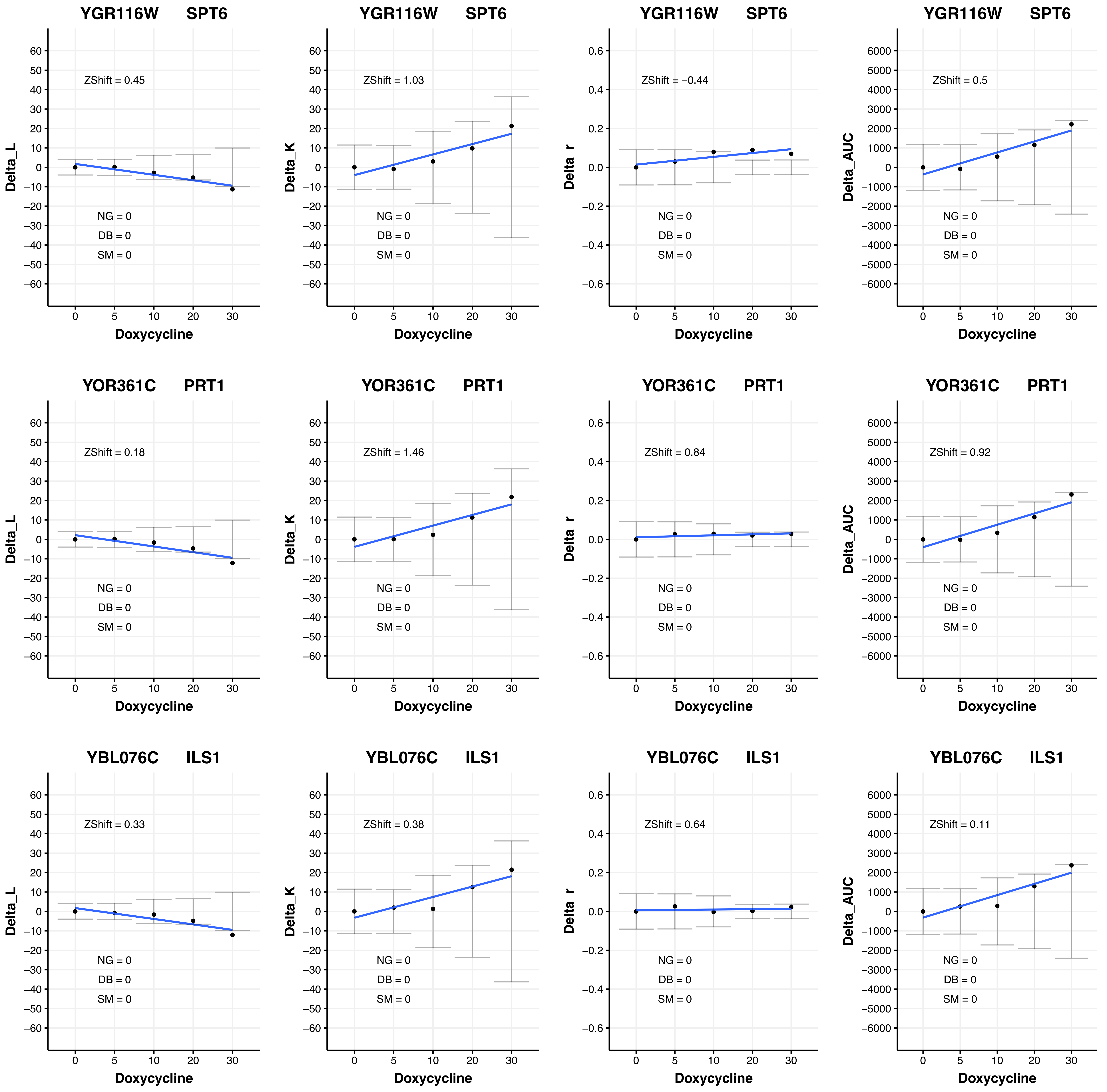

YGR116W SPT6
ZShift = 0.45
NG = 0
DB = 0
SM = 0
Delta_L
Doxycycline
YGR116W SPT6
ZShift = 1.03
NG = 0
DB = 0
SM = 0
Delta_K
Doxycycline
YGR116W SPT6
ZShift = −0.44
NG = 0
DB = 0
SM = 0
Delta_r
Doxycycline
YGR116W SPT6
ZShift = 0.5
NG = 0
DB = 0
SM = 0
Delta_AUC
Doxycycline
YOR361C PRT1
ZShift = 0.18
NG = 0
DB = 0
SM = 0
Delta_L
Doxycycline
YOR361C PRT1
ZShift = 1.46
NG = 0
DB = 0
SM = 0
Delta_K
Doxycycline
YOR361C PRT1
ZShift = 0.84
NG = 0
DB = 0
SM = 0
Delta_r
Doxycycline
YOR361C PRT1
ZShift = 0.92
NG = 0
DB = 0
SM = 0
Delta_AUC
Doxycycline
YBL076C ILS1
ZShift = 0.33
NG = 0
DB = 0
SM = 0
Delta_L
Doxycycline
YBL076C ILS1
ZShift = 0.38
NG = 0
DB = 0
SM = 0
Delta_K
Doxycycline
YBL076C ILS1
ZShift = 0.64
NG = 0
DB = 0
SM = 0
Delta_r
Doxycycline
YBL076C ILS1
ZShift = 0.11
NG = 0
DB = 0
SM = 0
Delta_AUC
Doxycycline

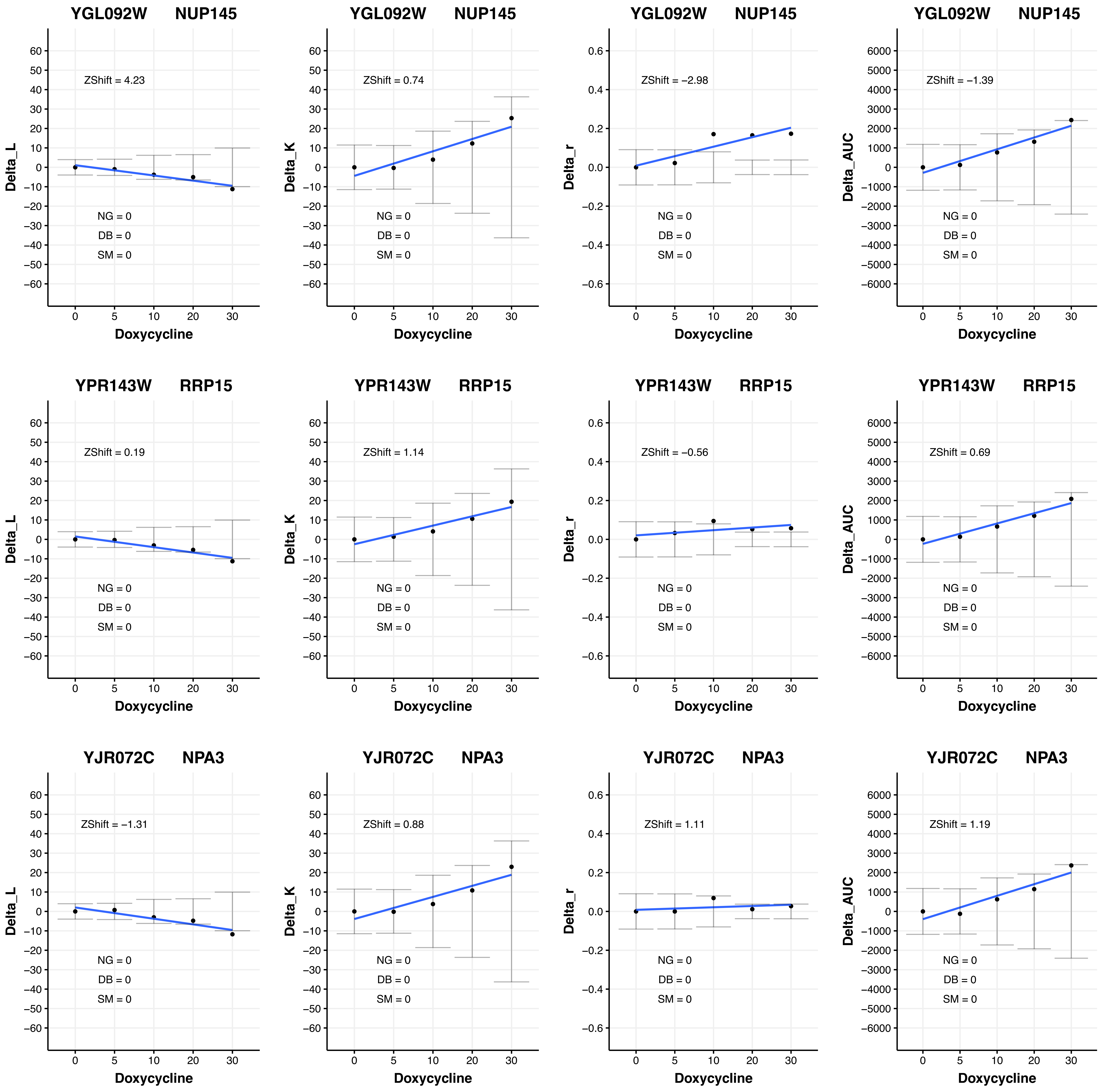
YGL092W NUP145
Delta_L
ZShift = 4.23
NG = 0
DB = 0
SM = 0
Doxycycline
YGL092W NUP145
Delta_K
ZShift = 0.74
NG = 0
DB = 0
SM = 0
Doxycycline
YGL092W NUP145
Delta_r
ZShift = −2.98
NG = 0
DB = 0
SM = 0
Doxycycline
YGL092W NUP145
Delta_AUC
ZShift = −1.39
NG = 0
DB = 0
SM = 0
Doxycycline
YPR143W RRP15
Delta_L
ZShift = 0.19
NG = 0
DB = 0
SM = 0
Doxycycline
YPR143W RRP15
Delta_K
ZShift = 1.14
NG = 0
DB = 0
SM = 0
Doxycycline
YPR143W RRP15
Delta_r
ZShift = −0.56
NG = 0
DB = 0
SM = 0
Doxycycline
YPR143W RRP15
Delta_AUC
ZShift = 0.69
NG = 0
DB = 0
SM = 0
Doxycycline
YJR072C NPA3
Delta_L
ZShift = −1.31
NG = 0
DB = 0
SM = 0
Doxycycline
YJR072C NPA3
Delta_K
ZShift = 0.88
NG = 0
DB = 0
SM = 0
Doxycycline
YJR072C NPA3
Delta_r
ZShift = 1.11
NG = 0
DB = 0
SM = 0
Doxycycline
YJR072C NPA3
Delta_AUC
ZShift = 1.19
NG = 0
DB = 0
SM = 0
Doxycycline

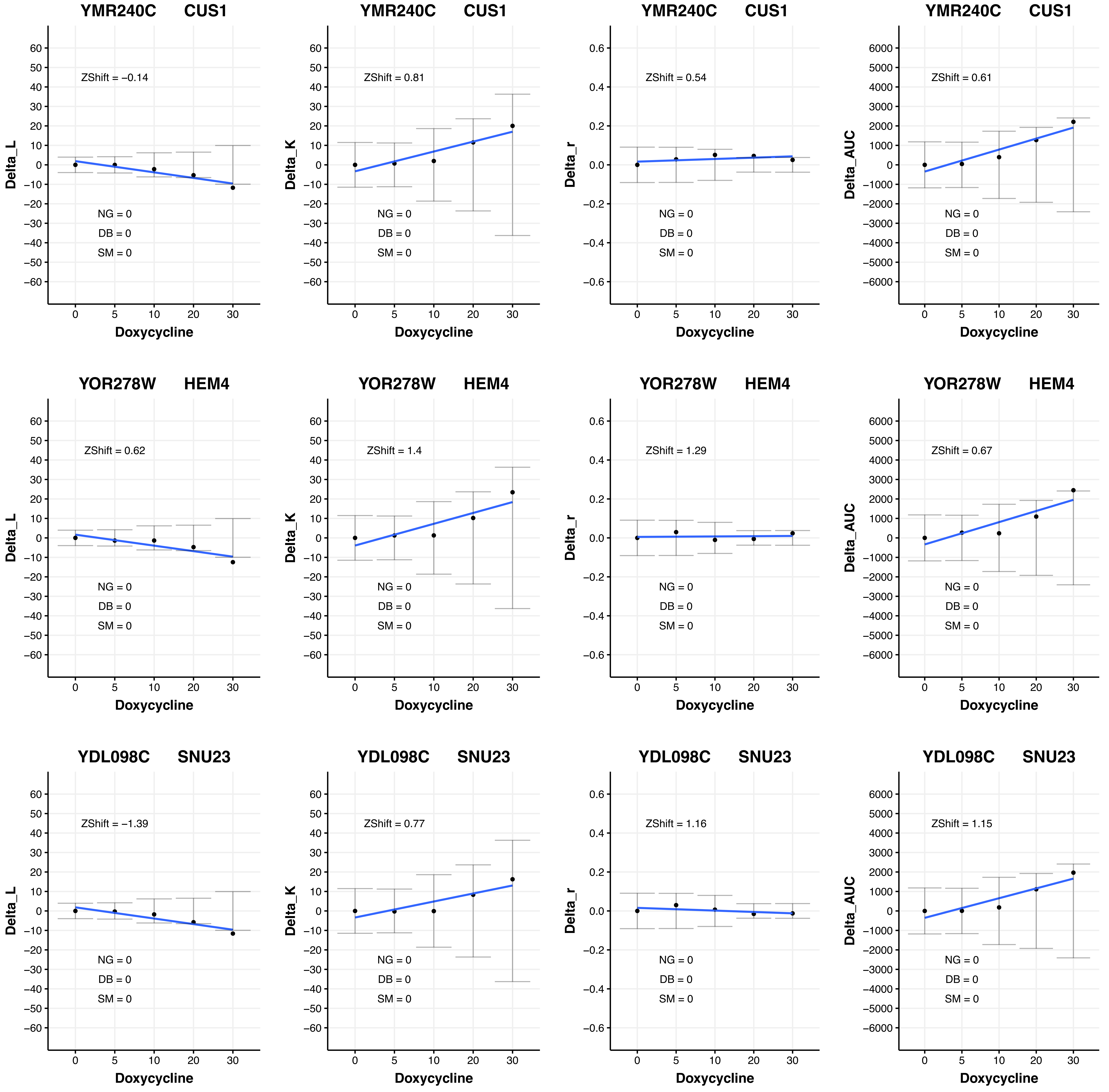
YMR240C CUS1
ZShift = –0.14
NG = 0
DB = 0
SM = 0
Delta_L
Doxycycline
YMR240C CUS1
ZShift = 0.81
NG = 0
DB = 0
SM = 0
Delta_K
Doxycycline
YMR240C CUS1
ZShift = 0.54
NG = 0
DB = 0
SM = 0
Delta_r
Doxycycline
YMR240C CUS1
ZShift = 0.61
NG = 0
DB = 0
SM = 0
Delta_AUC
Doxycycline
YOR278W HEM4
ZShift = 0.62
NG = 0
DB = 0
SM = 0
Delta_L
Doxycycline
YOR278W HEM4
ZShift = 1.4
NG = 0
DB = 0
SM = 0
Delta_K
Doxycycline
YOR278W HEM4
ZShift = 1.29
NG = 0
DB = 0
SM = 0
Delta_r
Doxycycline
YOR278W HEM4
ZShift = 0.67
NG = 0
DB = 0
SM = 0
Delta_AUC
Doxycycline
YDL098C SNU23
ZShift = –1.39
NG = 0
DB = 0
SM = 0
Delta_L
Doxycycline
YDL098C SNU23
ZShift = 0.77
NG = 0
DB = 0
SM = 0
Delta_K
Doxycycline
YDL098C SNU23
ZShift = 1.16
NG = 0
DB = 0
SM = 0
Delta_r
Doxycycline
YDL098C SNU23
ZShift = 1.15
NG = 0
DB = 0
SM = 0
Delta_AUC
Doxycycline

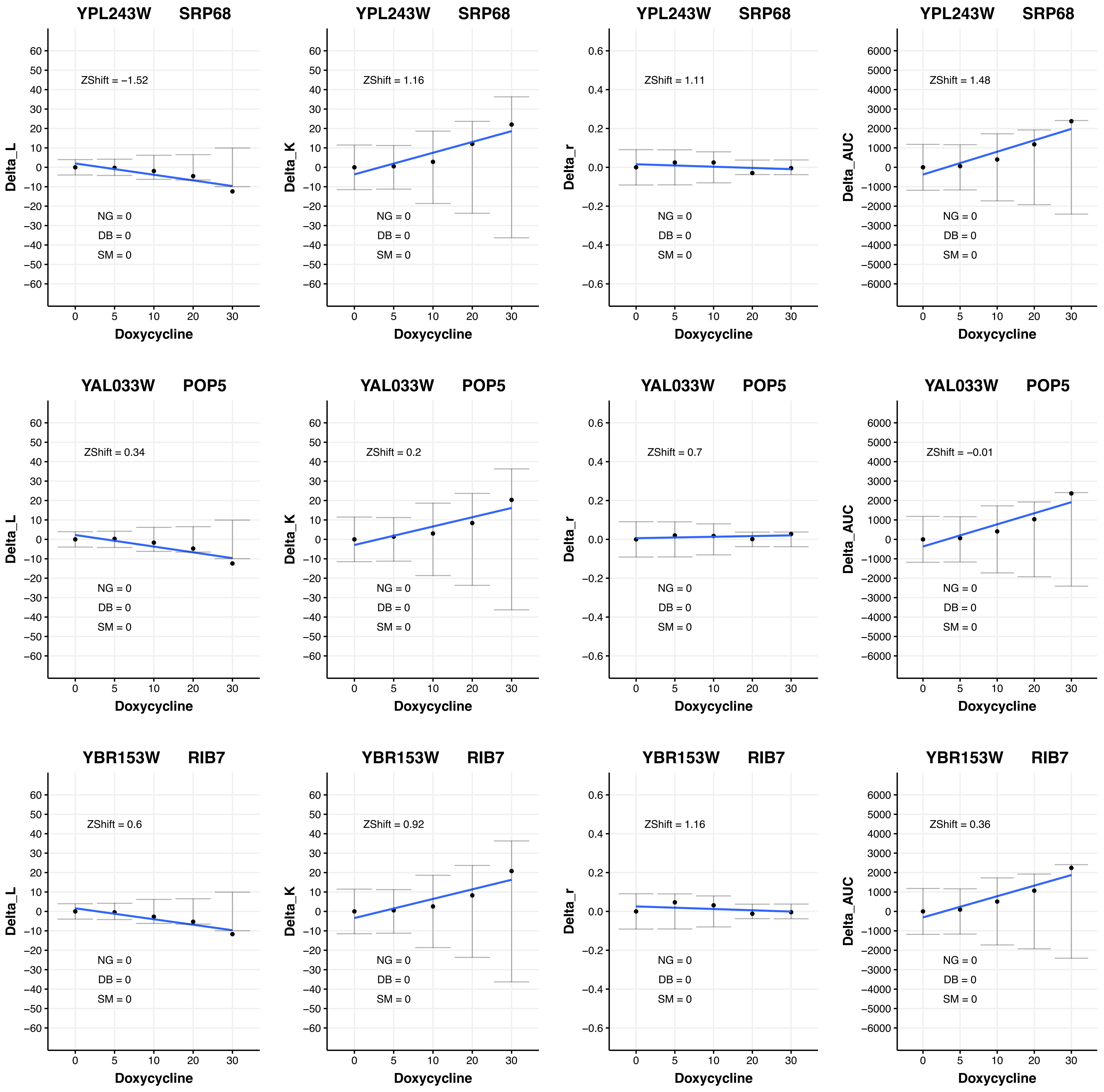
YPL243W SRP68
ZShift = −1.52
NG = 0
DB = 0
SM = 0
Delta_L
Doxycycline
YPL243W SRP68
ZShift = 1.16
NG = 0
DB = 0
SM = 0
Delta_K
Doxycycline
YPL243W SRP68
ZShift = 1.11
NG = 0
DB = 0
SM = 0
Delta_r
Doxycycline
YPL243W SRP68
ZShift = 1.48
NG = 0
DB = 0
SM = 0
Delta_AUC
Doxycycline
YAL033W POP5
ZShift = 0.34
NG = 0
DB = 0
SM = 0
Delta_L
Doxycycline
YAL033W POP5
ZShift = 0.2
NG = 0
DB = 0
SM = 0
Delta_K
Doxycycline
YAL033W POP5
ZShift = 0.7
NG = 0
DB = 0
SM = 0
Delta_r
Doxycycline
YAL033W POP5
ZShift = −0.01
NG = 0
DB = 0
SM = 0
Delta_AUC
Doxycycline
YBR153W RIB7
ZShift = 0.6
NG = 0
DB = 0
SM = 0
Delta_L
Doxycycline
YBR153W RIB7
ZShift = 0.92
NG = 0
DB = 0
SM = 0
Delta_K
Doxycycline
YBR153W RIB7
ZShift = 1.16
NG = 0
DB = 0
SM = 0
Delta_r
Doxycycline
YBR153W RIB7
ZShift = 0.36
NG = 0
DB = 0
SM = 0
Delta_AUC
Doxycycline

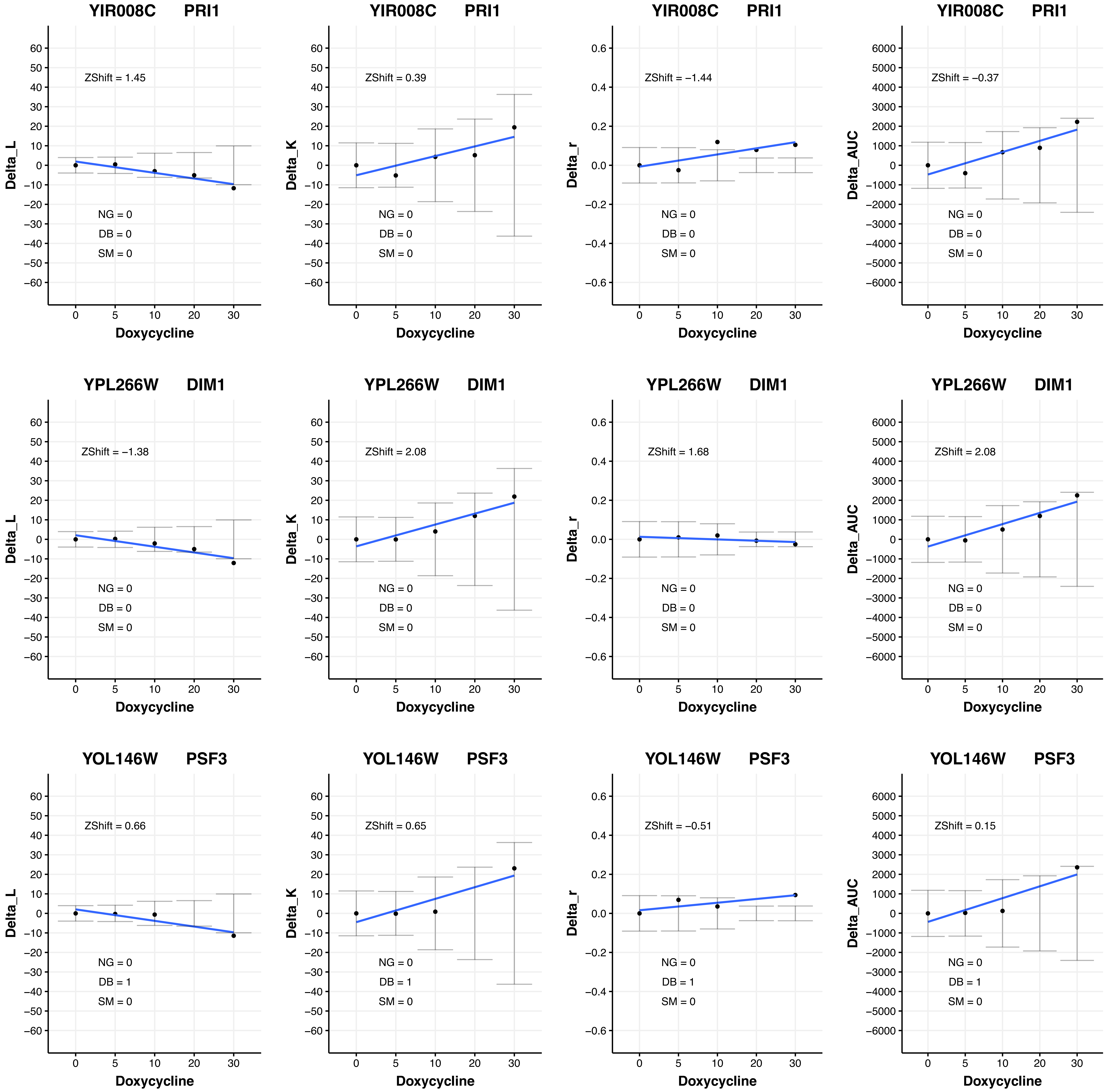

YIR008C PRI1
ZShift = 1.45
NG = 0
DB = 0
SM = 0
Delta_L
Doxycycline
YIR008C PRI1
ZShift = 0.39
NG = 0
DB = 0
SM = 0
Delta_K
Doxycycline
YIR008C PRI1
ZShift = −1.44
NG = 0
DB = 0
SM = 0
Delta_r
Doxycycline
YIR008C PRI1
ZShift = −0.37
NG = 0
DB = 0
SM = 0
Delta_AUC
Doxycycline
YPL266W DIM1
ZShift = −1.38
NG = 0
DB = 0
SM = 0
Delta_L
Doxycycline
YPL266W DIM1
ZShift = 2.08
NG = 0
DB = 0
SM = 0
Delta_K
Doxycycline
YPL266W DIM1
ZShift = 1.68
NG = 0
DB = 0
SM = 0
Delta_r
Doxycycline
YPL266W DIM1
ZShift = 2.08
NG = 0
DB = 0
SM = 0
Delta_AUC
Doxycycline
YOL146W PSF3
ZShift = 0.66
NG = 0
DB = 1
SM = 0
Delta_L
Doxycycline
YOL146W PSF3
ZShift = 0.65
NG = 0
DB = 1
SM = 0
Delta_K
Doxycycline
YOL146W PSF3
ZShift = −0.51
NG = 0
DB = 1
SM = 0
Delta_r
Doxycycline
YOL146W PSF3
ZShift = 0.15
NG = 0
DB = 1
SM = 0
Delta_AUC
Doxycycline

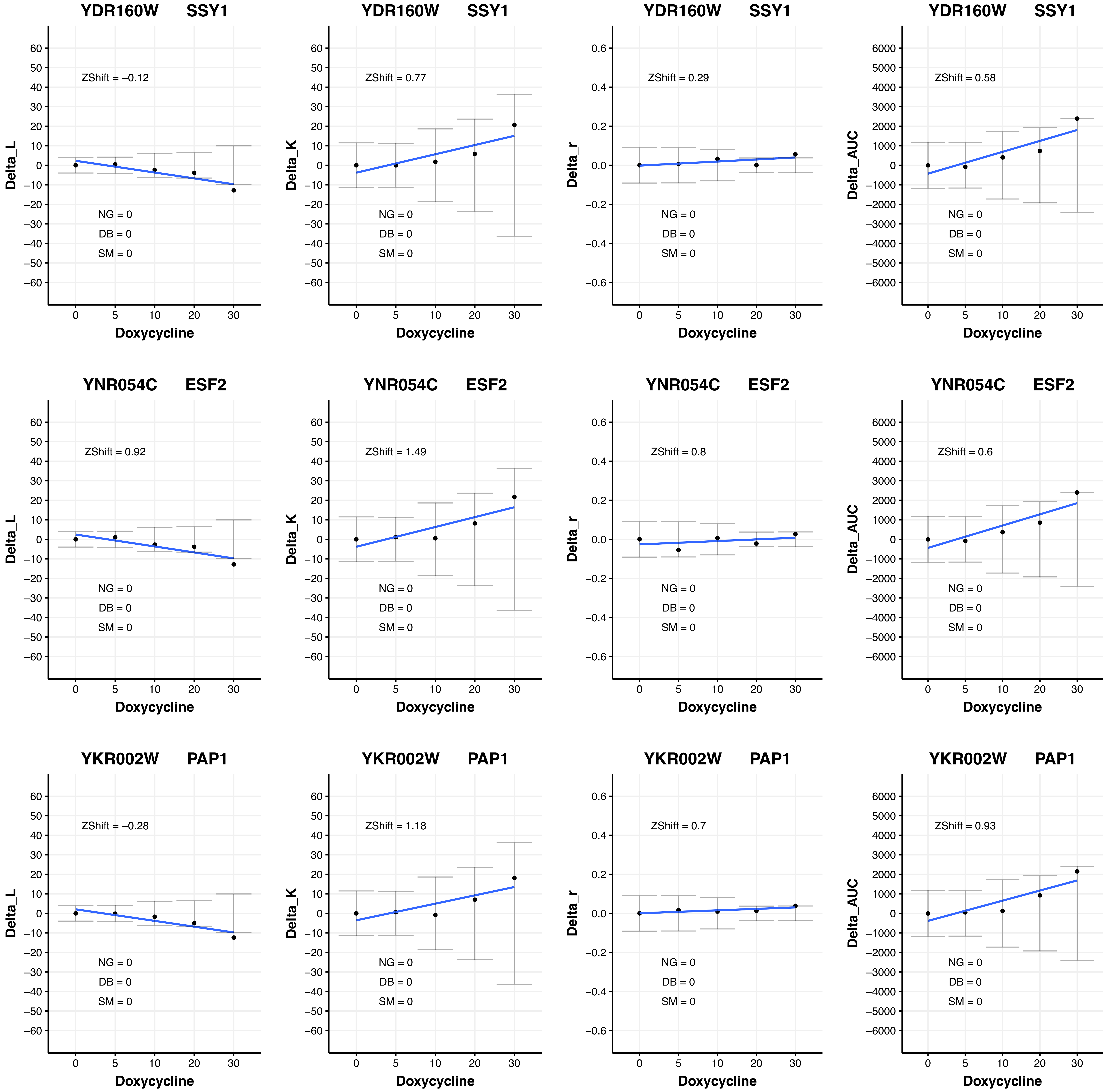

YDR160W SSY1
Delta_L
ZShift = −0.12
NG = 0
DB = 0
SM = 0
Doxycycline
YDR160W SSY1
Delta_K
ZShift = 0.77
NG = 0
DB = 0
SM = 0
Doxycycline
YDR160W SSY1
Delta_r
ZShift = 0.29
NG = 0
DB = 0
SM = 0
Doxycycline
YDR160W SSY1
Delta_AUC
ZShift = 0.58
NG = 0
DB = 0
SM = 0
Doxycycline
YNR054C ESF2
Delta_L
ZShift = 0.92
NG = 0
DB = 0
SM = 0
Doxycycline
YNR054C ESF2
Delta_K
ZShift = 1.49
NG = 0
DB = 0
SM = 0
Doxycycline
YNR054C ESF2
Delta_r
ZShift = 0.8
NG = 0
DB = 0
SM = 0
Doxycycline
YNR054C ESF2
Delta_AUC
ZShift = 0.6
NG = 0
DB = 0
SM = 0
Doxycycline
YKR002W PAP1
Delta_L
ZShift = −0.28
NG = 0
DB = 0
SM = 0
Doxycycline
YKR002W PAP1
Delta_K
ZShift = 1.18
NG = 0
DB = 0
SM = 0
Doxycycline
YKR002W PAP1
Delta_r
ZShift = 0.7
NG = 0
DB = 0
SM = 0
Doxycycline
YKR002W PAP1
Delta_AUC
ZShift = 0.93
NG = 0
DB = 0
SM = 0
Doxycycline

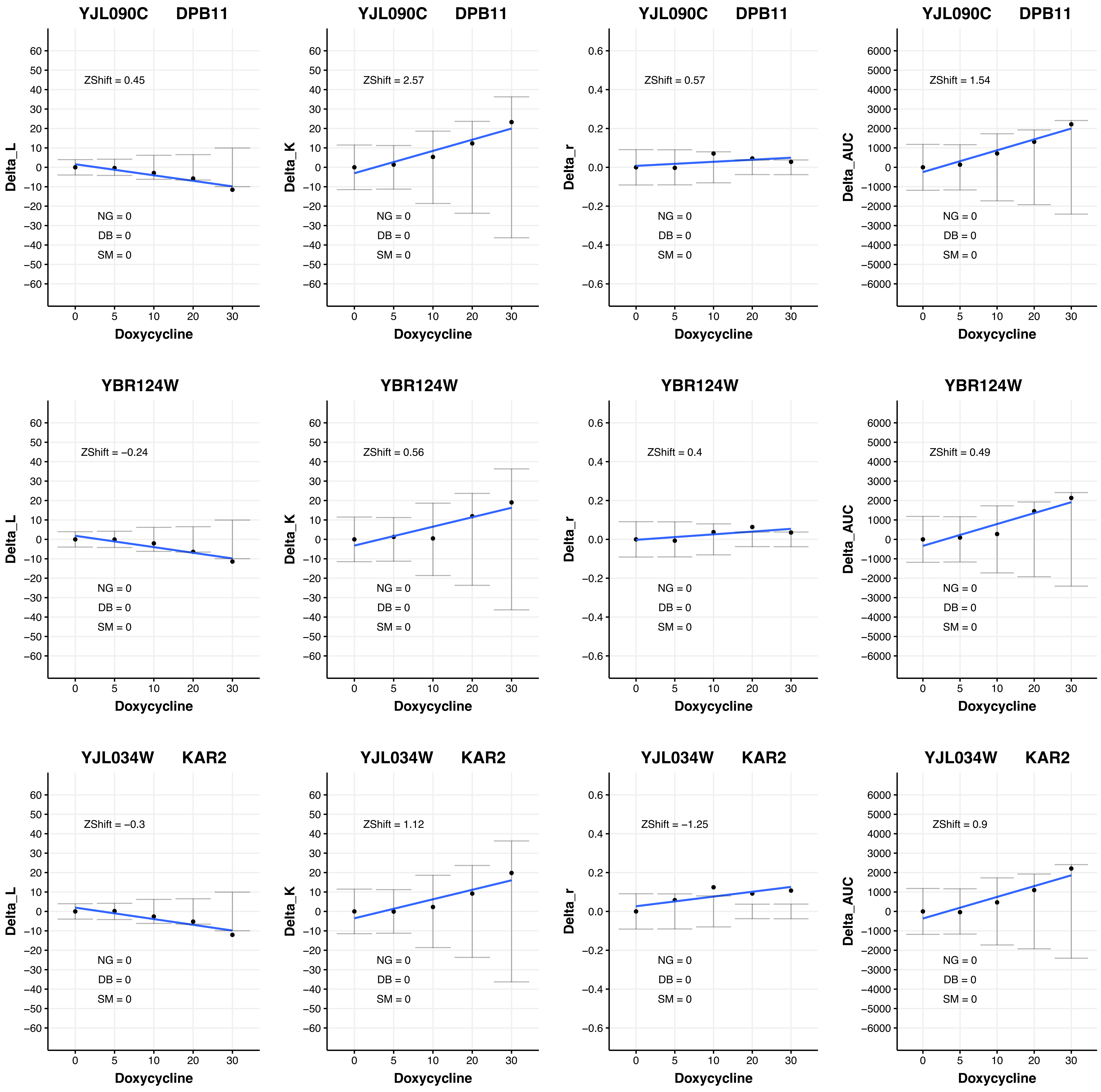
YJL090C DPB11
ZShift = 0.45
NG = 0
DB = 0
SM = 0
Delta_L
Doxycycline
YJL090C DPB11
ZShift = 2.57
NG = 0
DB = 0
SM = 0
Delta_K
Doxycycline
YJL090C DPB11
ZShift = 0.57
NG = 0
DB = 0
SM = 0
Delta_r
Doxycycline
YJL090C DPB11
ZShift = 1.54
NG = 0
DB = 0
SM = 0
Delta_AUC
Doxycycline
YBR124W
ZShift = −0.24
NG = 0
DB = 0
SM = 0
Delta_L
Doxycycline
YBR124W
ZShift = 0.56
NG = 0
DB = 0
SM = 0
Delta_K
Doxycycline
YBR124W
ZShift = 0.4
NG = 0
DB = 0
SM = 0
Delta_r
Doxycycline
YBR124W
ZShift = 0.49
NG = 0
DB = 0
SM = 0
Delta_AUC
Doxycycline
YJL034W KAR2
ZShift = −0.3
NG = 0
DB = 0
SM = 0
Delta_L
Doxycycline
YJL034W KAR2
ZShift = 1.12
NG = 0
DB = 0
SM = 0
Delta_K
Doxycycline
YJL034W KAR2
ZShift = −1.25
NG = 0
DB = 0
SM = 0
Delta_r
Doxycycline
YJL034W KAR2
ZShift = 0.9
NG = 0
DB = 0
SM = 0
Delta_AUC
Doxycycline

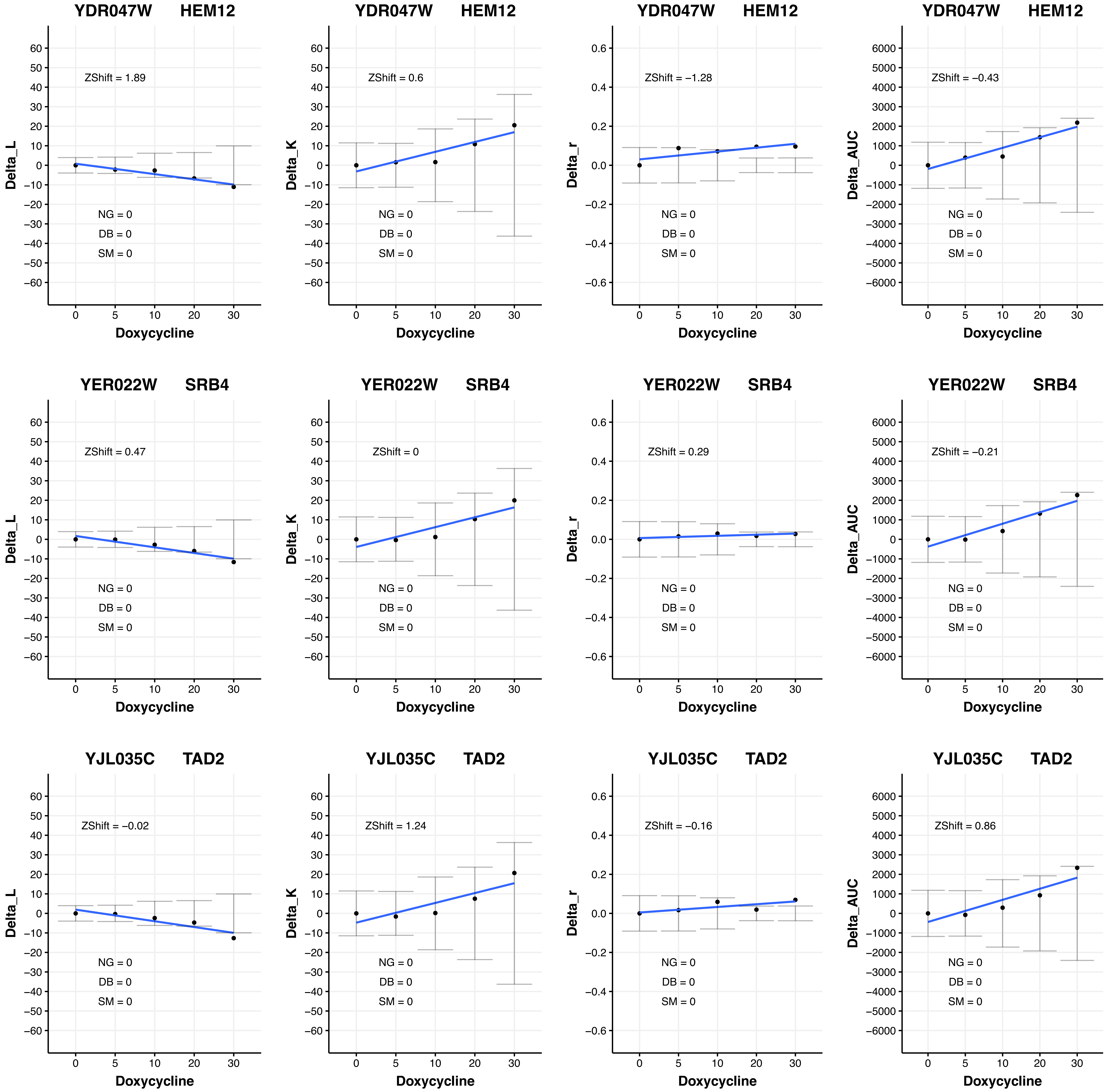

YDR047W HEM12
ZShift = 1.89
NG = 0
DB = 0
SM = 0
Delta_L
Doxycycline
YDR047W HEM12
ZShift = 0.6
NG = 0
DB = 0
SM = 0
Delta_K
Doxycycline
YDR047W HEM12
ZShift = −1.28
NG = 0
DB = 0
SM = 0
Delta_r
Doxycycline
YDR047W HEM12
ZShift = −0.43
NG = 0
DB = 0
SM = 0
Delta_AUC
Doxycycline
YER022W SRB4
ZShift = 0.47
NG = 0
DB = 0
SM = 0
Delta_L
Doxycycline
YER022W SRB4
ZShift = 0
NG = 0
DB = 0
SM = 0
Delta_K
Doxycycline
YER022W SRB4
ZShift = 0.29
NG = 0
DB = 0
SM = 0
Delta_r
Doxycycline
YER022W SRB4
ZShift = −0.21
NG = 0
DB = 0
SM = 0
Delta_AUC
Doxycycline
YJL035C TAD2
ZShift = −0.02
NG = 0
DB = 0
SM = 0
Delta_L
Doxycycline
YJL035C TAD2
ZShift = 1.24
NG = 0
DB = 0
SM = 0
Delta_K
Doxycycline
YJL035C TAD2
ZShift = −0.16
NG = 0
DB = 0
SM = 0
Delta_r
Doxycycline
YJL035C TAD2
ZShift = 0.86
NG = 0
DB = 0
SM = 0
Delta_AUC
Doxycycline

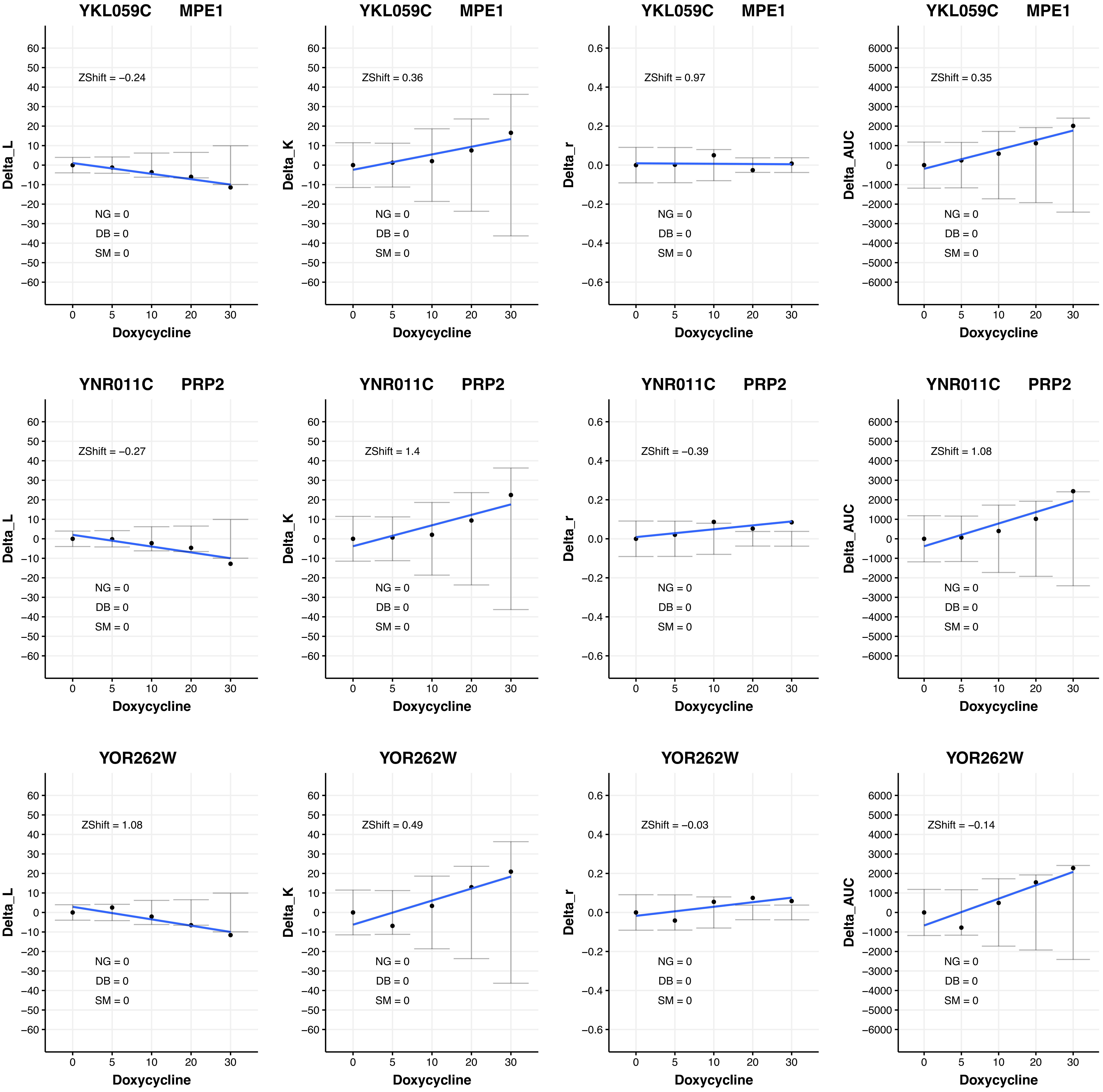

YKL059C MPE1
ZShift = −0.24
Delta_L
NG = 0
DB = 0
SM = 0
Doxycycline
YKL059C MPE1
ZShift = 0.36
Delta_K
NG = 0
DB = 0
SM = 0
Doxycycline
YKL059C MPE1
ZShift = 0.97
Delta_r
NG = 0
DB = 0
SM = 0
Doxycycline
YKL059C MPE1
ZShift = 0.35
Delta_AUC
NG = 0
DB = 0
SM = 0
Doxycycline
YNR011C PRP2
ZShift = −0.27
Delta_L
NG = 0
DB = 0
SM = 0
Doxycycline
YNR011C PRP2
ZShift = 1.4
Delta_K
NG = 0
DB = 0
SM = 0
Doxycycline
YNR011C PRP2
ZShift = −0.39
Delta_r
NG = 0
DB = 0
SM = 0
Doxycycline
YNR011C PRP2
ZShift = 1.08
Delta_AUC
NG = 0
DB = 0
SM = 0
Doxycycline
YOR262W
ZShift = 1.08
Delta_L
NG = 0
DB = 0
SM = 0
Doxycycline
YOR262W
ZShift = 0.49
Delta_K
NG = 0
DB = 0
SM = 0
Doxycycline
YOR262W
ZShift = −0.03
Delta_r
NG = 0
DB = 0
SM = 0
Doxycycline
YOR262W
ZShift = −0.14
Delta_AUC
NG = 0
DB = 0
SM = 0
Doxycycline

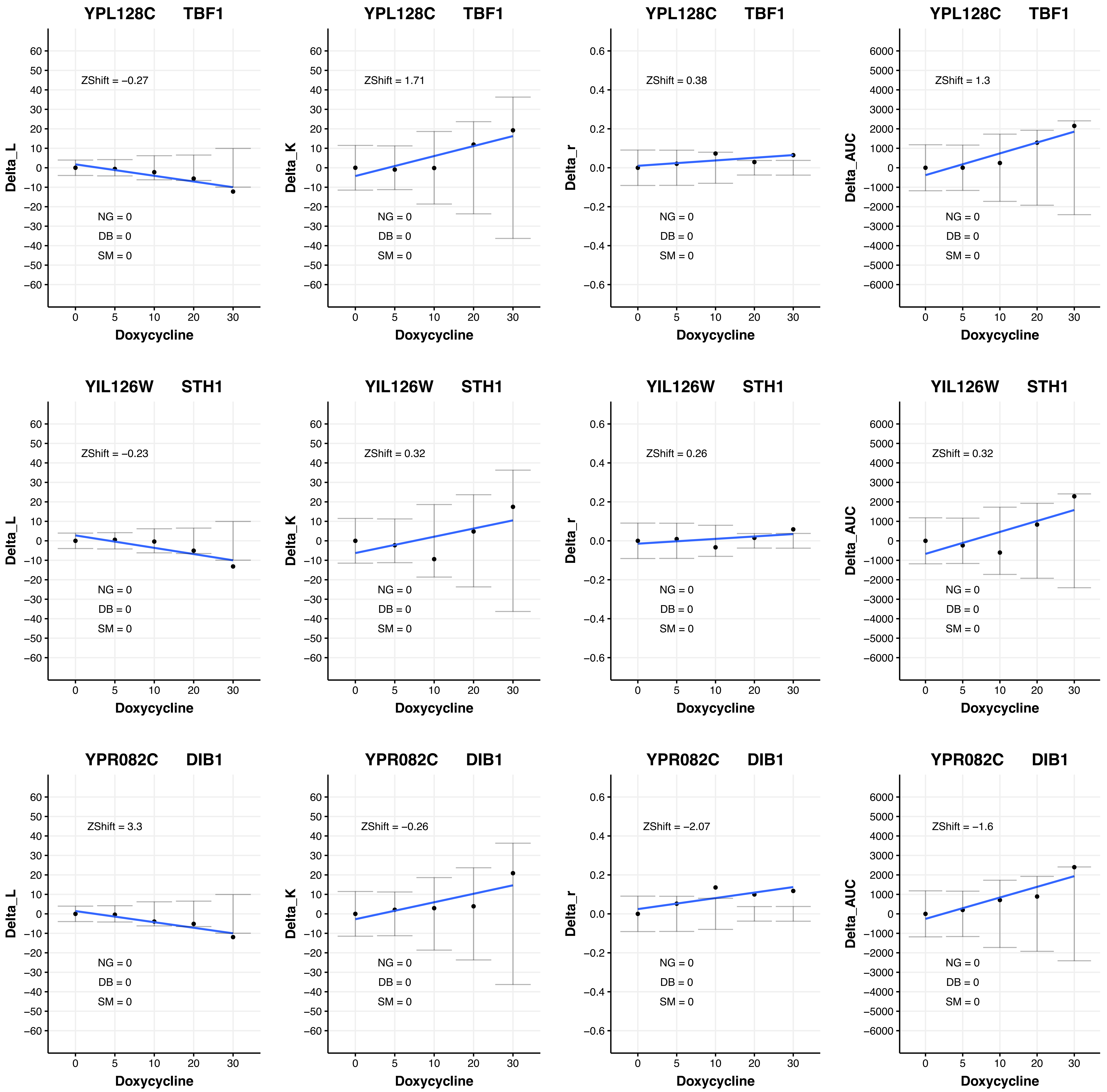
YPL128C TBF1
ZShift = −0.27
NG = 0
DB = 0
SM = 0
Delta_L
Doxycycline
YPL128C TBF1
ZShift = 1.71
NG = 0
DB = 0
SM = 0
Delta_K
Doxycycline
YPL128C TBF1
ZShift = 0.38
NG = 0
DB = 0
SM = 0
Delta_r
Doxycycline
YPL128C TBF1
ZShift = 1.3
NG = 0
DB = 0
SM = 0
Delta_AUC
Doxycycline
YIL126W STH1
ZShift = −0.23
NG = 0
DB = 0
SM = 0
Delta_L
Doxycycline
YIL126W STH1
ZShift = 0.32
NG = 0
DB = 0
SM = 0
Delta_K
Doxycycline
YIL126W STH1
ZShift = 0.26
NG = 0
DB = 0
SM = 0
Delta_r
Doxycycline
YIL126W STH1
ZShift = 0.32
NG = 0
DB = 0
SM = 0
Delta_AUC
Doxycycline
YPR082C DIB1
ZShift = 3.3
NG = 0
DB = 0
SM = 0
Delta_L
Doxycycline
YPR082C DIB1
ZShift = −0.26
NG = 0
DB = 0
SM = 0
Delta_K
Doxycycline
YPR082C DIB1
ZShift = −2.07
NG = 0
DB = 0
SM = 0
Delta_r
Doxycycline
YPR082C DIB1
ZShift = −1.6
NG = 0
DB = 0
SM = 0
Delta_AUC
Doxycycline

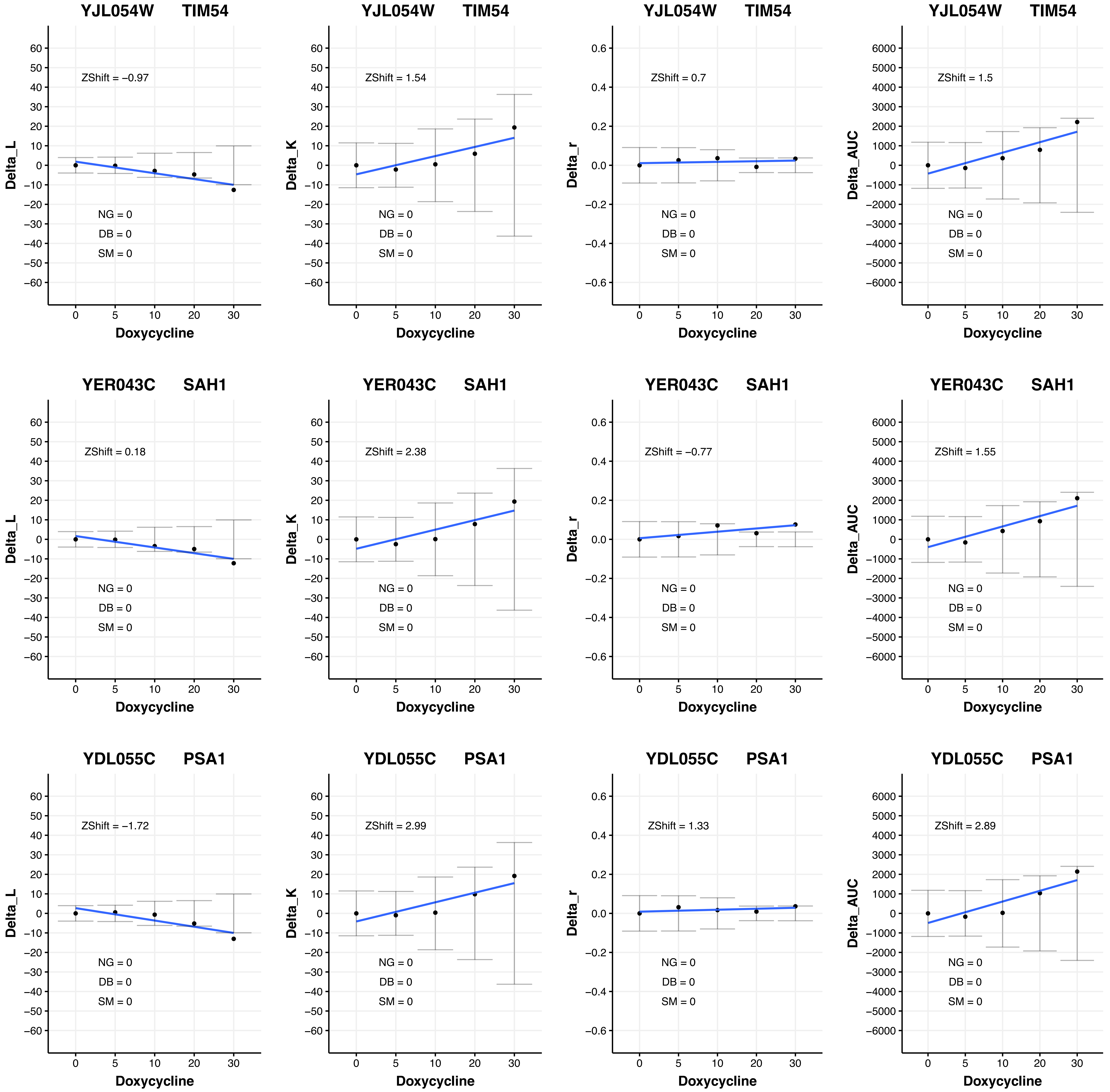
YJL054W TIM54
ZShift = −0.97
NG = 0
DB = 0
SM = 0
Delta_L
Doxycycline
YJL054W TIM54
ZShift = 1.54
NG = 0
DB = 0
SM = 0
Delta_K
Doxycycline
YJL054W TIM54
ZShift = 0.7
NG = 0
DB = 0
SM = 0
Delta_r
Doxycycline
YJL054W TIM54
ZShift = 1.5
NG = 0
DB = 0
SM = 0
Delta_AUC
Doxycycline
YER043C SAH1
ZShift = 0.18
NG = 0
DB = 0
SM = 0
Delta_L
Doxycycline
YER043C SAH1
ZShift = 2.38
NG = 0
DB = 0
SM = 0
Delta_K
Doxycycline
YER043C SAH1
ZShift = −0.77
NG = 0
DB = 0
SM = 0
Delta_r
Doxycycline
YER043C SAH1
ZShift = 1.55
NG = 0
DB = 0
SM = 0
Delta_AUC
Doxycycline
YDL055C PSA1
ZShift = −1.72
NG = 0
DB = 0
SM = 0
Delta_L
Doxycycline
YDL055C PSA1
ZShift = 2.99
NG = 0
DB = 0
SM = 0
Delta_K
Doxycycline
YDL055C PSA1
ZShift = 1.33
NG = 0
DB = 0
SM = 0
Delta_r
Doxycycline
YDL055C PSA1
ZShift = 2.89
NG = 0
DB = 0
SM = 0
Delta_AUC
Doxycycline

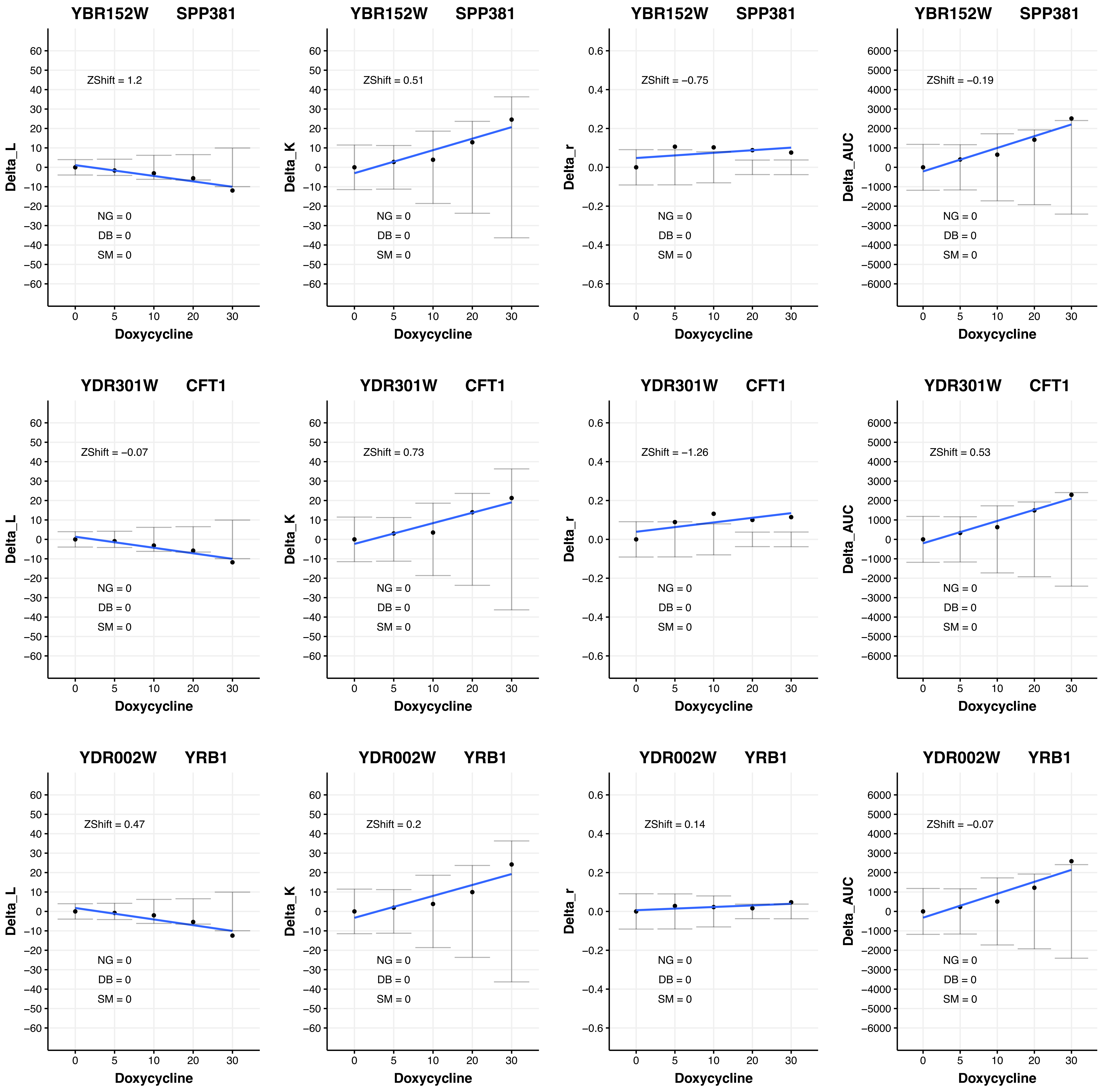

YBR152W SPP381
ZShift = 1.2
Delta_L
ZShift = 0.51
Delta_K
ZShift = −0.75
Delta_r
ZShift = −0.19
Delta_AUC
YDR301W CFT1
ZShift = −0.07
Delta_L
ZShift = 0.73
Delta_K
ZShift = −1.26
Delta_r
ZShift = 0.53
Delta_AUC
YDR002W YRB1
ZShift = 0.47
Delta_L
ZShift = 0.2
Delta_K
ZShift = 0.14
Delta_r
ZShift = −0.07
Delta_AUC
NG = 0
DB = 0
SM = 0
Doxycycline
0 5 10 20 30

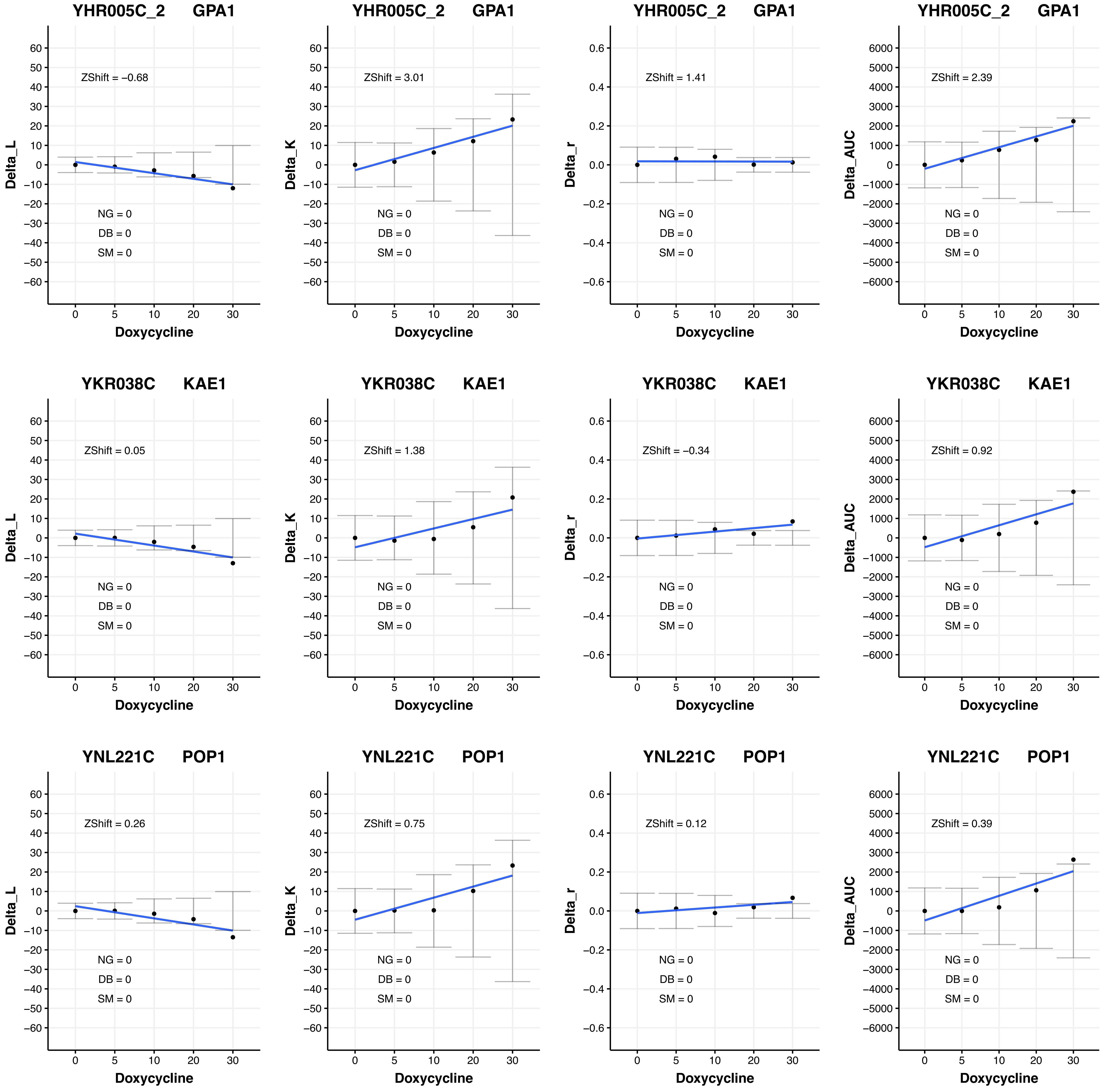

YHR005C_2 GPA1
ZShift = −0.68
NG = 0
DB = 0
SM = 0
Delta_L
Doxycycline
YHR005C_2 GPA1
ZShift = 3.01
NG = 0
DB = 0
SM = 0
Delta_K
Doxycycline
YHR005C_2 GPA1
ZShift = 1.41
NG = 0
DB = 0
SM = 0
Delta_r
Doxycycline
YHR005C_2 GPA1
ZShift = 2.39
NG = 0
DB = 0
SM = 0
Delta_AUC
Doxycycline
YKR038C KAE1
ZShift = 0.05
NG = 0
DB = 0
SM = 0
Delta_L
Doxycycline
YKR038C KAE1
ZShift = 1.38
NG = 0
DB = 0
SM = 0
Delta_K
Doxycycline
YKR038C KAE1
ZShift = −0.34
NG = 0
DB = 0
SM = 0
Delta_r
Doxycycline
YKR038C KAE1
ZShift = 0.92
NG = 0
DB = 0
SM = 0
Delta_AUC
Doxycycline
YNL221C POP1
ZShift = 0.26
NG = 0
DB = 0
SM = 0
Delta_L
Doxycycline
YNL221C POP1
ZShift = 0.75
NG = 0
DB = 0
SM = 0
Delta_K
Doxycycline
YNL221C POP1
ZShift = 0.12
NG = 0
DB = 0
SM = 0
Delta_r
Doxycycline
YNL221C POP1
ZShift = 0.39
NG = 0
DB = 0
SM = 0
Delta_AUC
Doxycycline

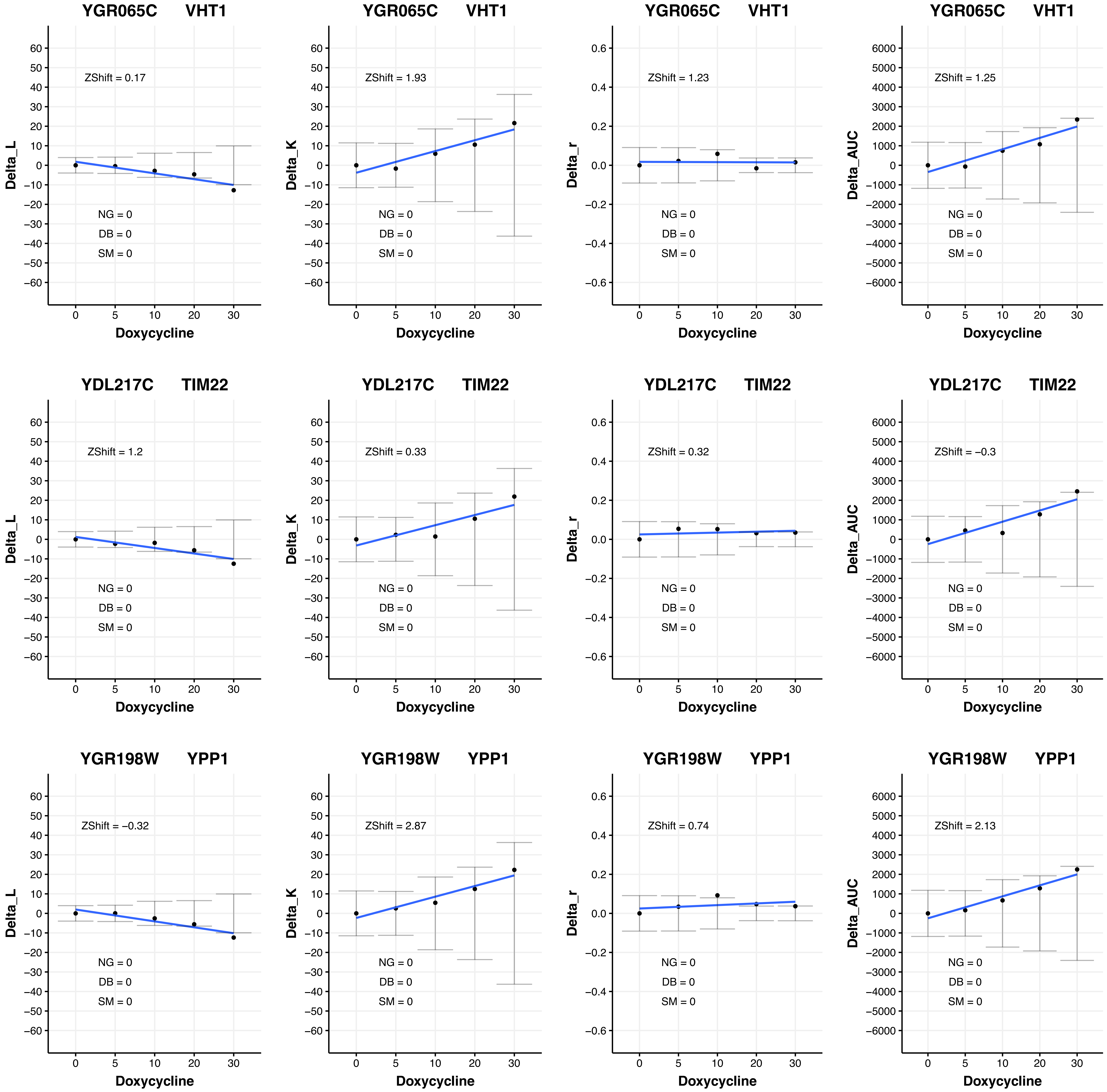

YGR065C VHT1
ZShift = 0.17
NG = 0
DB = 0
SM = 0
Delta_L
Doxycycline
YGR065C VHT1
ZShift = 1.93
NG = 0
DB = 0
SM = 0
Delta_K
Doxycycline
YGR065C VHT1
ZShift = 1.23
NG = 0
DB = 0
SM = 0
Delta_r
Doxycycline
YGR065C VHT1
ZShift = 1.25
NG = 0
DB = 0
SM = 0
Delta_AUC
Doxycycline
YDL217C TIM22
ZShift = 1.2
NG = 0
DB = 0
SM = 0
Delta_L
Doxycycline
YDL217C TIM22
ZShift = 0.33
NG = 0
DB = 0
SM = 0
Delta_K
Doxycycline
YDL217C TIM22
ZShift = 0.32
NG = 0
DB = 0
SM = 0
Delta_r
Doxycycline
YDL217C TIM22
ZShift = −0.3
NG = 0
DB = 0
SM = 0
Delta_AUC
Doxycycline
YGR198W YPP1
ZShift = −0.32
NG = 0
DB = 0
SM = 0
Delta_L
Doxycycline
YGR198W YPP1
ZShift = 2.87
NG = 0
DB = 0
SM = 0
Delta_K
Doxycycline
YGR198W YPP1
ZShift = 0.74
NG = 0
DB = 0
SM = 0
Delta_r
Doxycycline
YGR198W YPP1
ZShift = 2.13
NG = 0
DB = 0
SM = 0
Delta_AUC
Doxycycline

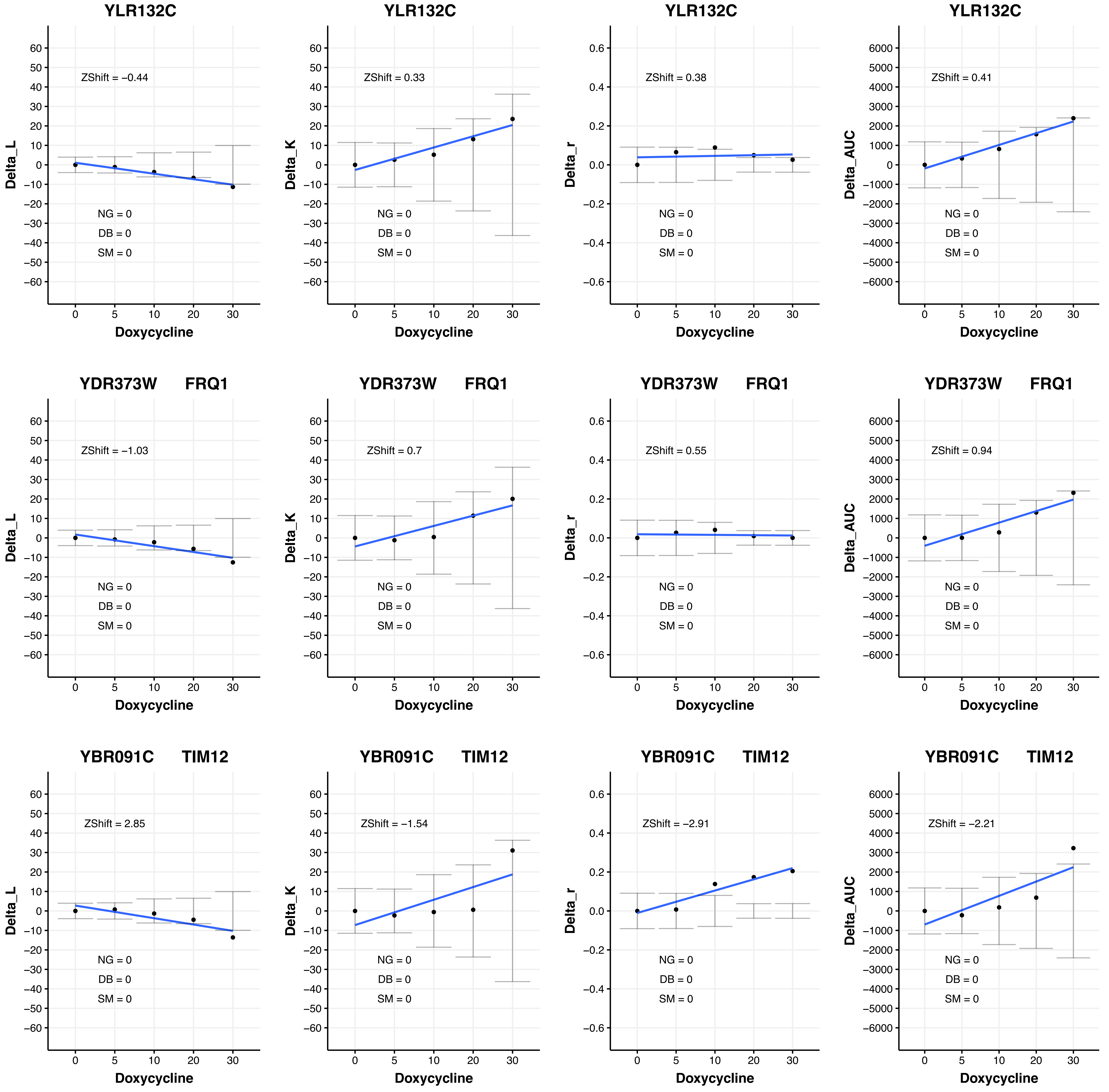
YLR132C
ZShift = −0.44
NG = 0
DB = 0
SM = 0
Delta_L
Doxycycline
YLR132C
ZShift = 0.33
NG = 0
DB = 0
SM = 0
Delta_K
Doxycycline
YLR132C
ZShift = 0.38
NG = 0
DB = 0
SM = 0
Delta_r
Doxycycline
YLR132C
ZShift = 0.41
NG = 0
DB = 0
SM = 0
Delta_AUC
Doxycycline
YDR373W FRQ1
ZShift = −1.03
NG = 0
DB = 0
SM = 0
Delta_L
Doxycycline
YDR373W FRQ1
ZShift = 0.7
NG = 0
DB = 0
SM = 0
Delta_K
Doxycycline
YDR373W FRQ1
ZShift = 0.55
NG = 0
DB = 0
SM = 0
Delta_r
Doxycycline
YDR373W FRQ1
ZShift = 0.94
NG = 0
DB = 0
SM = 0
Delta_AUC
Doxycycline
YBR091C TIM12
ZShift = 2.85
NG = 0
DB = 0
SM = 0
Delta_L
Doxycycline
YBR091C TIM12
ZShift = −1.54
NG = 0
DB = 0
SM = 0
Delta_K
Doxycycline
YBR091C TIM12
ZShift = −2.91
NG = 0
DB = 0
SM = 0
Delta_r
Doxycycline
YBR091C TIM12
ZShift = −2.21
NG = 0
DB = 0
SM = 0
Delta_AUC
Doxycycline

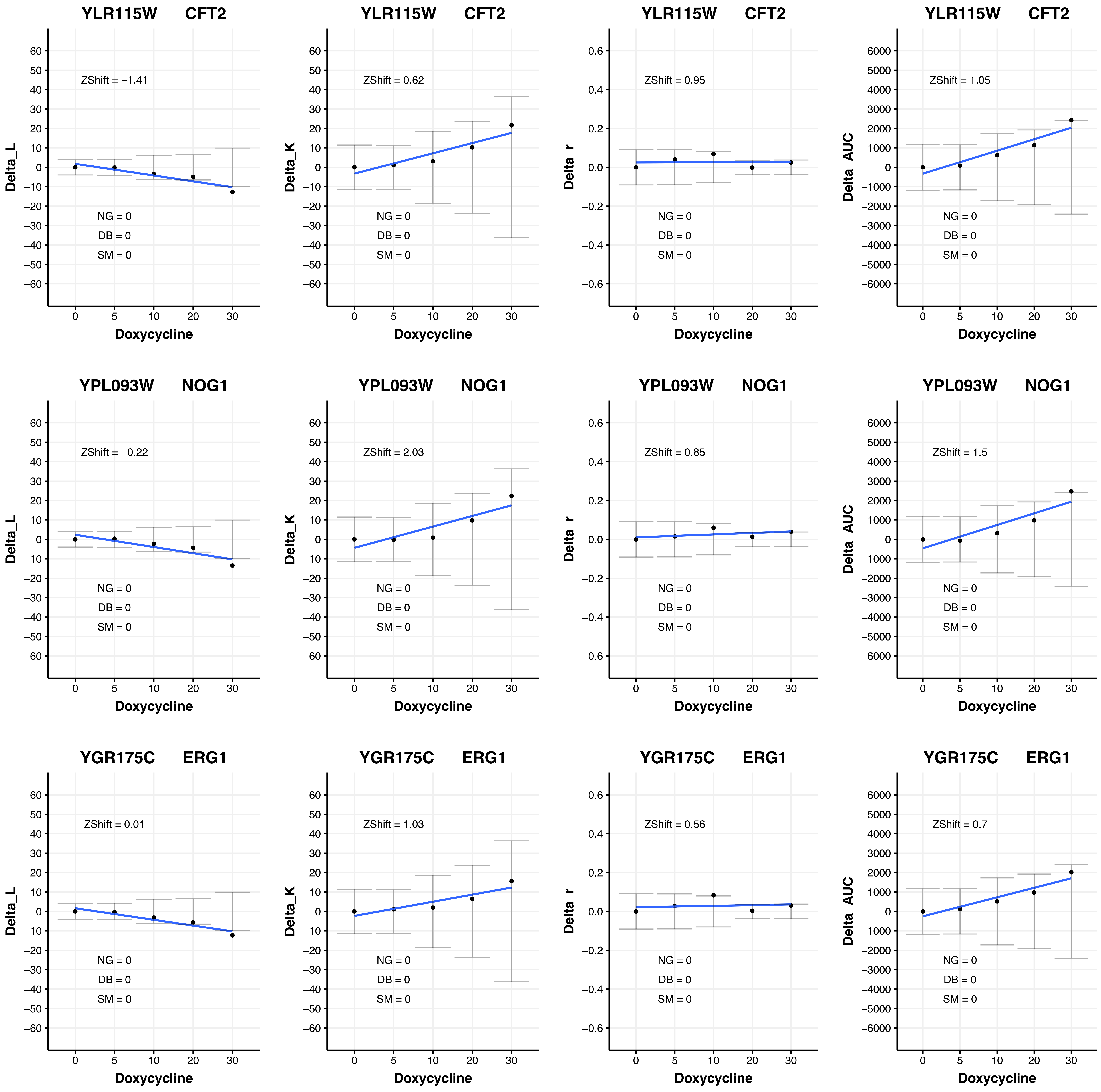
YLR115W CFT2
ZShift = −1.41
NG = 0
DB = 0
SM = 0
Delta_L
Doxycycline
YLR115W CFT2
ZShift = 0.62
NG = 0
DB = 0
SM = 0
Delta_K
Doxycycline
YLR115W CFT2
ZShift = 0.95
NG = 0
DB = 0
SM = 0
Delta_r
Doxycycline
YLR115W CFT2
ZShift = 1.05
NG = 0
DB = 0
SM = 0
Delta_AUC
Doxycycline
YPL093W NOG1
ZShift = −0.22
NG = 0
DB = 0
SM = 0
Delta_L
Doxycycline
YPL093W NOG1
ZShift = 2.03
NG = 0
DB = 0
SM = 0
Delta_K
Doxycycline
YPL093W NOG1
ZShift = 0.85
NG = 0
DB = 0
SM = 0
Delta_r
Doxycycline
YPL093W NOG1
ZShift = 1.5
NG = 0
DB = 0
SM = 0
Delta_AUC
Doxycycline
YGR175C ERG1
ZShift = 0.01
NG = 0
DB = 0
SM = 0
Delta_L
Doxycycline
YGR175C ERG1
ZShift = 1.03
NG = 0
DB = 0
SM = 0
Delta_K
Doxycycline
YGR175C ERG1
ZShift = 0.56
NG = 0
DB = 0
SM = 0
Delta_r
Doxycycline
YGR175C ERG1
ZShift = 0.7
NG = 0
DB = 0
SM = 0
Delta_AUC
Doxycycline

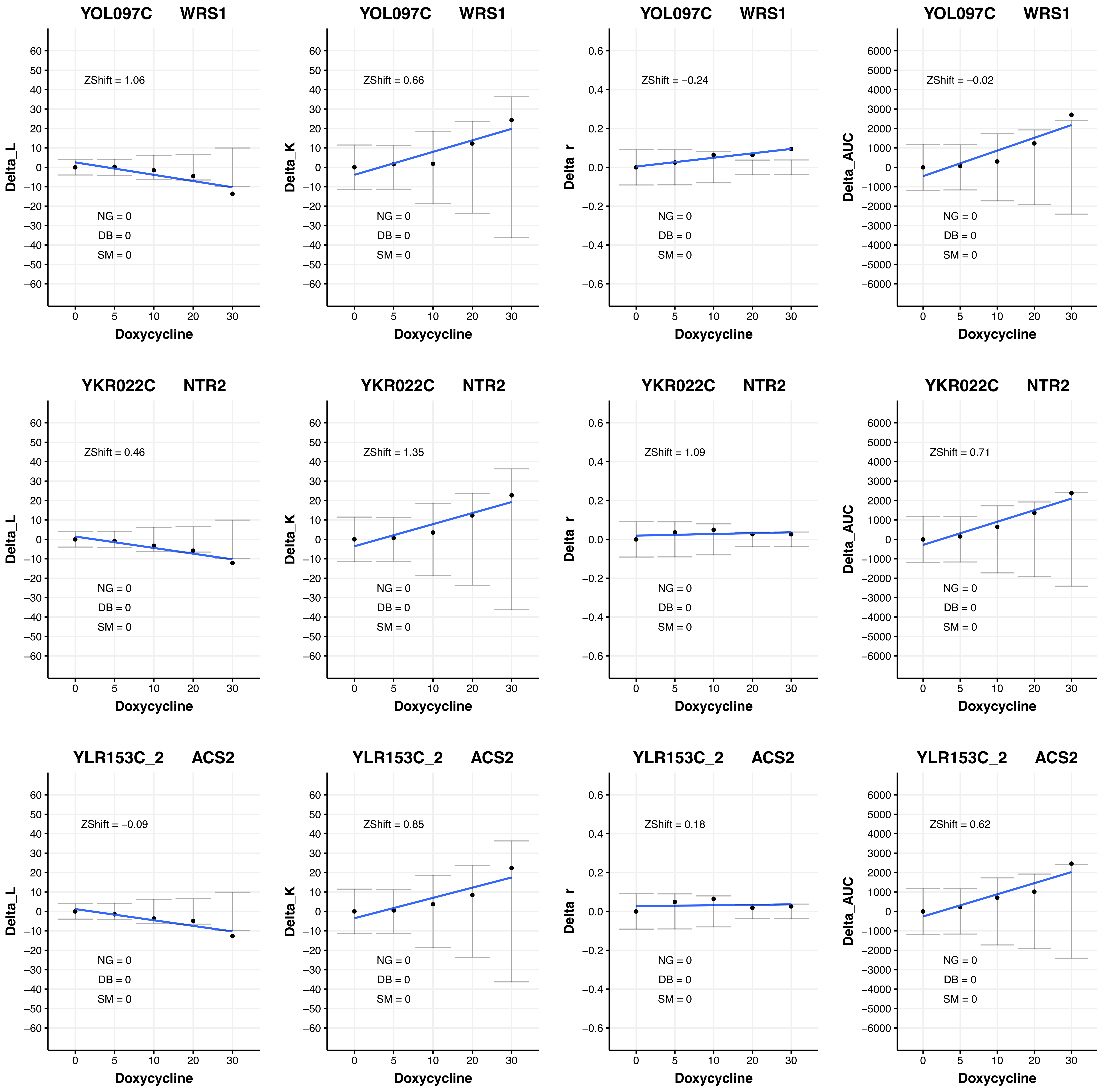

YOL097C WRS1
ZShift = 1.06
NG = 0
DB = 0
SM = 0
Delta_L
Doxycycline
YOL097C WRS1
ZShift = 0.66
NG = 0
DB = 0
SM = 0
Delta_K
Doxycycline
YOL097C WRS1
ZShift = −0.24
NG = 0
DB = 0
SM = 0
Delta_r
Doxycycline
YOL097C WRS1
ZShift = −0.02
NG = 0
DB = 0
SM = 0
Delta_AUC
Doxycycline
YKR022C NTR2
ZShift = 0.46
NG = 0
DB = 0
SM = 0
Delta_L
Doxycycline
YKR022C NTR2
ZShift = 1.35
NG = 0
DB = 0
SM = 0
Delta_K
Doxycycline
YKR022C NTR2
ZShift = 1.09
NG = 0
DB = 0
SM = 0
Delta_r
Doxycycline
YKR022C NTR2
ZShift = 0.71
NG = 0
DB = 0
SM = 0
Delta_AUC
Doxycycline
YLR153C_2 ACS2
ZShift = −0.09
NG = 0
DB = 0
SM = 0
Delta_L
Doxycycline
YLR153C_2 ACS2
ZShift = 0.85
NG = 0
DB = 0
SM = 0
Delta_K
Doxycycline
YLR153C_2 ACS2
ZShift = 0.18
NG = 0
DB = 0
SM = 0
Delta_r
Doxycycline
YLR153C_2 ACS2
ZShift = 0.62
NG = 0
DB = 0
SM = 0
Delta_AUC
Doxycycline

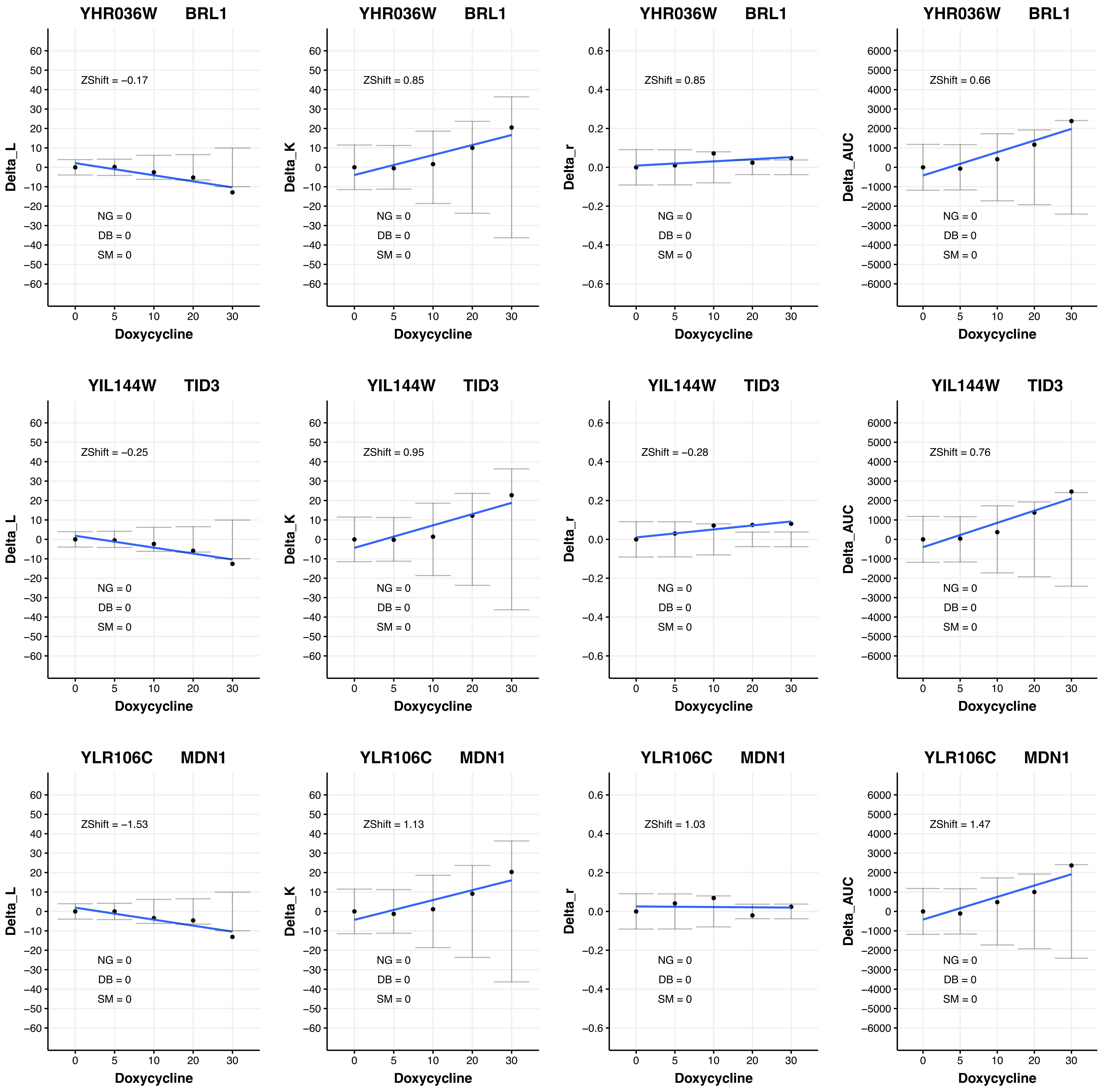
YHR036W BRL1
Delta_L
ZShift = −0.17
NG = 0
DB = 0
SM = 0
Doxycycline
YHR036W BRL1
Delta_K
ZShift = 0.85
NG = 0
DB = 0
SM = 0
Doxycycline
YHR036W BRL1
Delta_r
ZShift = 0.85
NG = 0
DB = 0
SM = 0
Doxycycline
YHR036W BRL1
Delta_AUC
ZShift = 0.66
NG = 0
DB = 0
SM = 0
Doxycycline
YIL144W TID3
Delta_L
ZShift = −0.25
NG = 0
DB = 0
SM = 0
Doxycycline
YIL144W TID3
Delta_K
ZShift = 0.95
NG = 0
DB = 0
SM = 0
Doxycycline
YIL144W TID3
Delta_r
ZShift = −0.28
NG = 0
DB = 0
SM = 0
Doxycycline
YIL144W TID3
Delta_AUC
ZShift = 0.76
NG = 0
DB = 0
SM = 0
Doxycycline
YLR106C MDN1
Delta_L
ZShift = −1.53
NG = 0
DB = 0
SM = 0
Doxycycline
YLR106C MDN1
Delta_K
ZShift = 1.13
NG = 0
DB = 0
SM = 0
Doxycycline
YLR106C MDN1
Delta_r
ZShift = 1.03
NG = 0
DB = 0
SM = 0
Doxycycline
YLR106C MDN1
Delta_AUC
ZShift = 1.47
NG = 0
DB = 0
SM = 0
Doxycycline

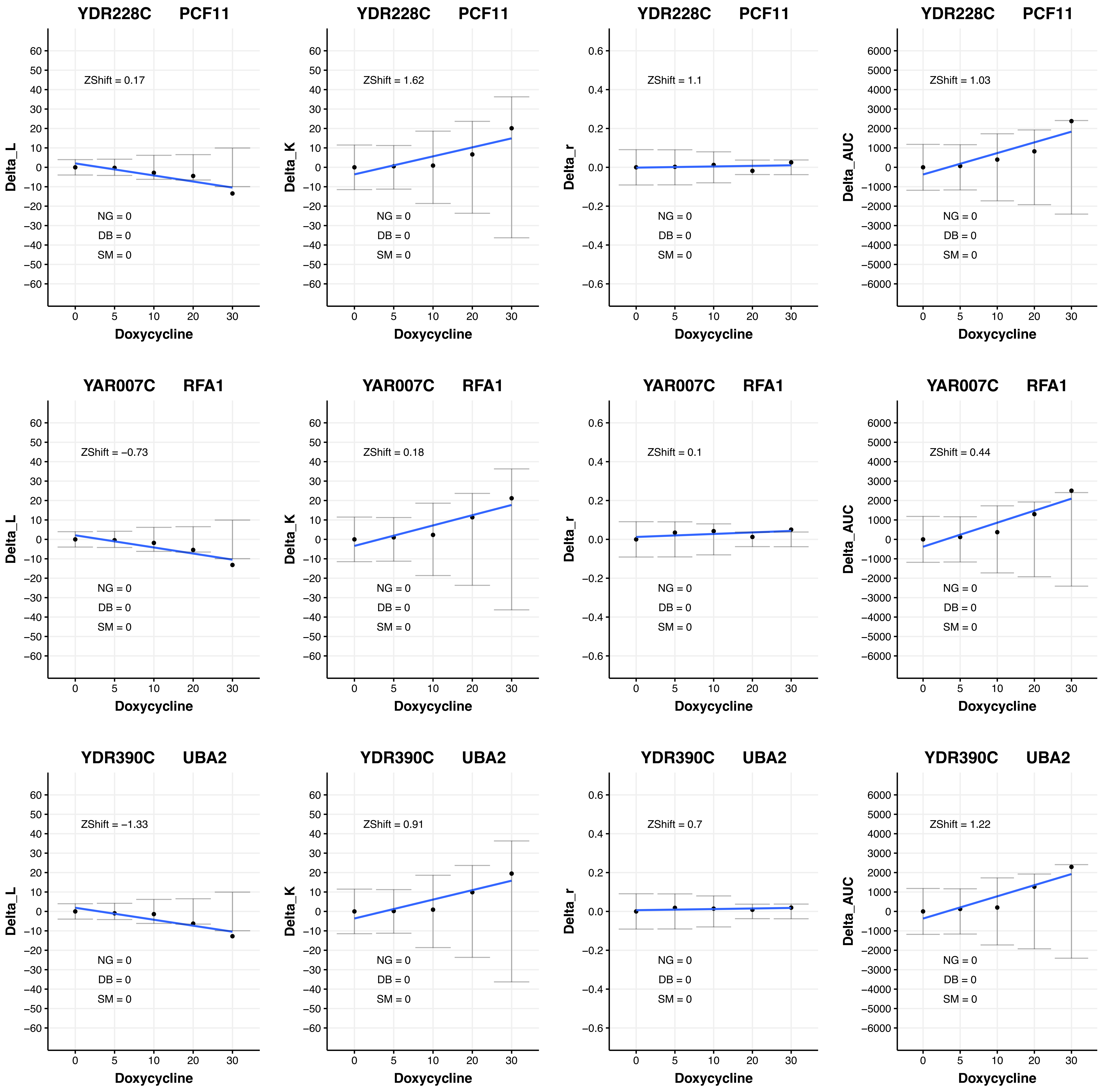

YDR228C PCF11
ZShift = 0.17
Delta_L
NG = 0
DB = 0
SM = 0
Doxycycline
YDR228C PCF11
ZShift = 1.62
Delta_K
NG = 0
DB = 0
SM = 0
Doxycycline
YDR228C PCF11
ZShift = 1.1
Delta_r
NG = 0
DB = 0
SM = 0
Doxycycline
YDR228C PCF11
ZShift = 1.03
Delta_AUC
NG = 0
DB = 0
SM = 0
Doxycycline
YAR007C RFA1
ZShift = −0.73
Delta_L
NG = 0
DB = 0
SM = 0
Doxycycline
YAR007C RFA1
ZShift = 0.18
Delta_K
NG = 0
DB = 0
SM = 0
Doxycycline
YAR007C RFA1
ZShift = 0.1
Delta_r
NG = 0
DB = 0
SM = 0
Doxycycline
YAR007C RFA1
ZShift = 0.44
Delta_AUC
NG = 0
DB = 0
SM = 0
Doxycycline
YDR390C UBA2
ZShift = −1.33
Delta_L
NG = 0
DB = 0
SM = 0
Doxycycline
YDR390C UBA2
ZShift = 0.91
Delta_K
NG = 0
DB = 0
SM = 0
Doxycycline
YDR390C UBA2
ZShift = 0.7
Delta_r
NG = 0
DB = 0
SM = 0
Doxycycline
YDR390C UBA2
ZShift = 1.22
Delta_AUC
NG = 0
DB = 0
SM = 0
Doxycycline

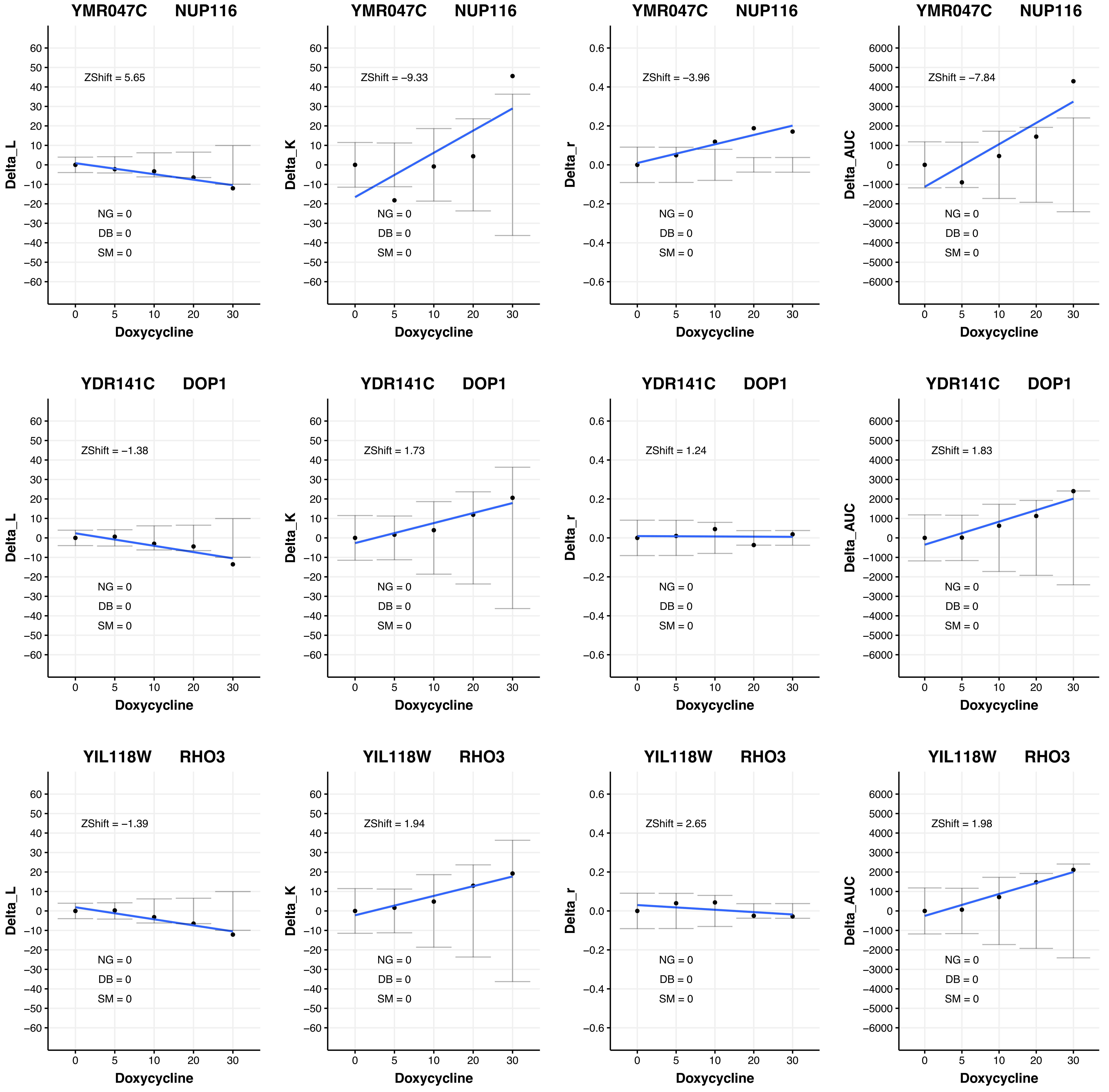

YMR047C NUP116
ZShift = 5.65
NG = 0
DB = 0
SM = 0
Delta_L
Doxycycline
YMR047C NUP116
ZShift = −9.33
NG = 0
DB = 0
SM = 0
Delta_K
Doxycycline
YMR047C NUP116
ZShift = −3.96
NG = 0
DB = 0
SM = 0
Delta_r
Doxycycline
YMR047C NUP116
ZShift = −7.84
NG = 0
DB = 0
SM = 0
Delta_AUC
Doxycycline
YDR141C DOP1
ZShift = −1.38
NG = 0
DB = 0
SM = 0
Delta_L
Doxycycline
YDR141C DOP1
ZShift = 1.73
NG = 0
DB = 0
SM = 0
Delta_K
Doxycycline
YDR141C DOP1
ZShift = 1.24
NG = 0
DB = 0
SM = 0
Delta_r
Doxycycline
YDR141C DOP1
ZShift = 1.83
NG = 0
DB = 0
SM = 0
Delta_AUC
Doxycycline
YIL118W RHO3
ZShift = −1.39
NG = 0
DB = 0
SM = 0
Delta_L
Doxycycline
YIL118W RHO3
ZShift = 1.94
NG = 0
DB = 0
SM = 0
Delta_K
Doxycycline
YIL118W RHO3
ZShift = 2.65
NG = 0
DB = 0
SM = 0
Delta_r
Doxycycline
YIL118W RHO3
ZShift = 1.98
NG = 0
DB = 0
SM = 0
Delta_AUC
Doxycycline

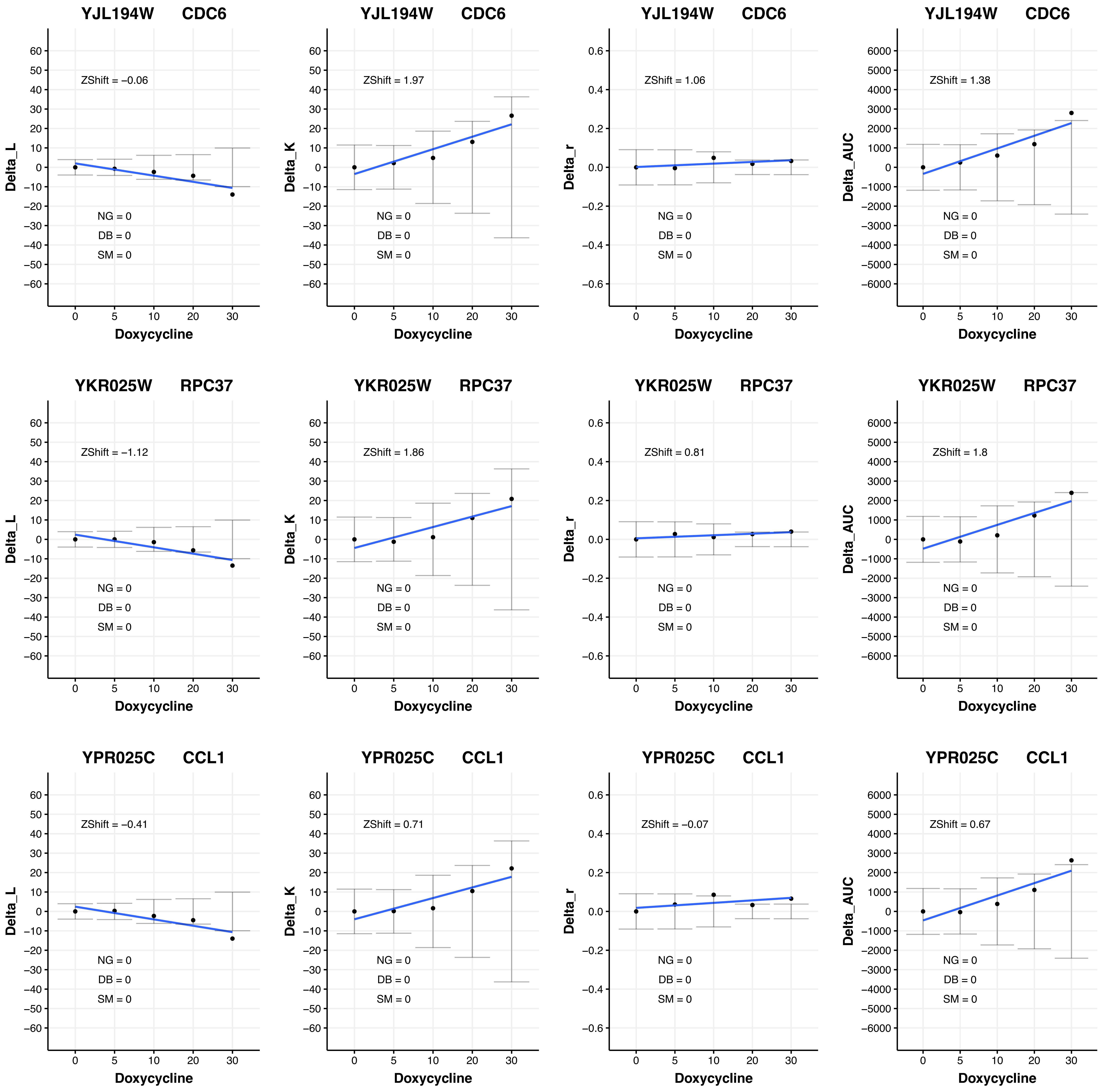

YJL194W CDC6
ZShift = −0.06
Delta_L
ZShift = 1.97
Delta_K
ZShift = 1.06
Delta_r
ZShift = 1.38
Delta_AUC
YKR025W RPC37
ZShift = −1.12
ZShift = 1.86
ZShift = 0.81
ZShift = 1.8
YPR025C CCL1
ZShift = −0.41
ZShift = 0.71
ZShift = −0.07
ZShift = 0.67
NG = 0
DB = 0
SM = 0
Doxycycline
0
5
10
20
30

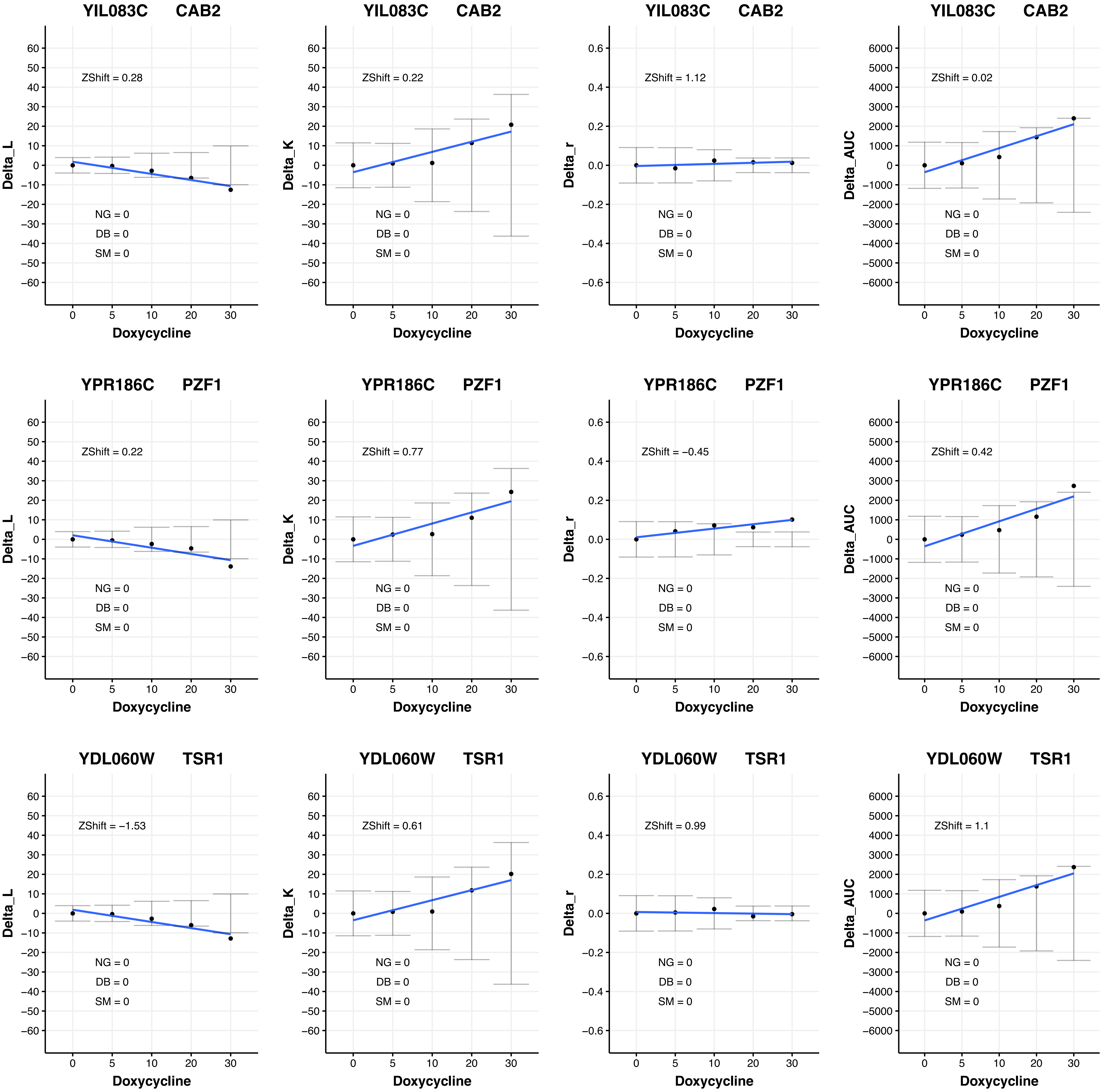
YIL083C CAB2
ZShift = 0.28
NG = 0
DB = 0
SM = 0
Delta_L
Doxycycline
YIL083C CAB2
ZShift = 0.22
NG = 0
DB = 0
SM = 0
Delta_K
Doxycycline
YIL083C CAB2
ZShift = 1.12
NG = 0
DB = 0
SM = 0
Delta_r
Doxycycline
YIL083C CAB2
ZShift = 0.02
NG = 0
DB = 0
SM = 0
Delta_AUC
Doxycycline
YPR186C PZF1
ZShift = 0.22
NG = 0
DB = 0
SM = 0
Delta_L
Doxycycline
YPR186C PZF1
ZShift = 0.77
NG = 0
DB = 0
SM = 0
Delta_K
Doxycycline
YPR186C PZF1
ZShift = −0.45
NG = 0
DB = 0
SM = 0
Delta_r
Doxycycline
YPR186C PZF1
ZShift = 0.42
NG = 0
DB = 0
SM = 0
Delta_AUC
Doxycycline
YDL060W TSR1
ZShift = −1.53
NG = 0
DB = 0
SM = 0
Delta_L
Doxycycline
YDL060W TSR1
ZShift = 0.61
NG = 0
DB = 0
SM = 0
Delta_K
Doxycycline
YDL060W TSR1
ZShift = 0.99
NG = 0
DB = 0
SM = 0
Delta_r
Doxycycline
YDL060W TSR1
ZShift = 1.1
NG = 0
DB = 0
SM = 0
Delta_AUC
Doxycycline

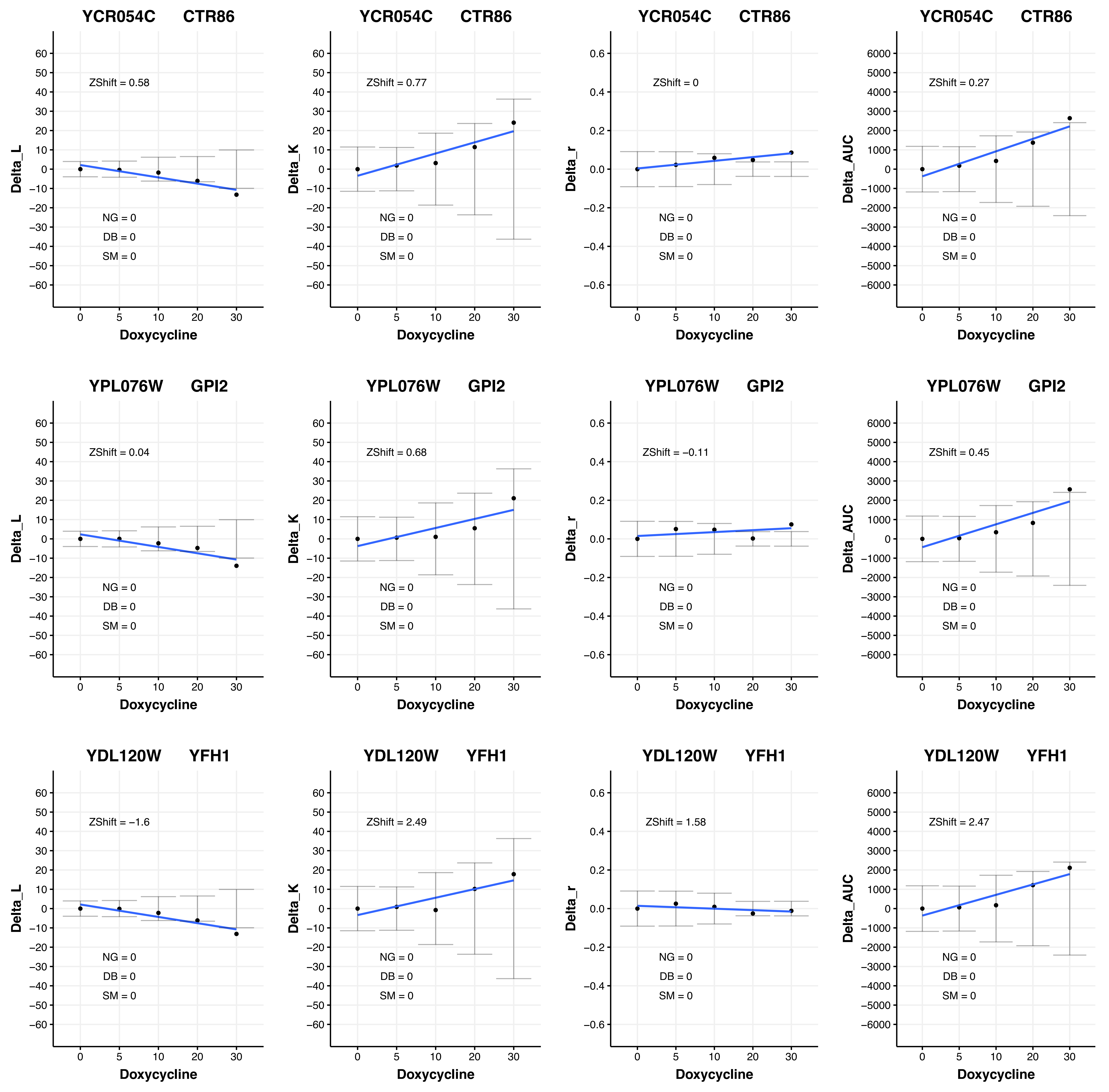
YCR054C CTR86
ZShift = 0.58
NG = 0
DB = 0
SM = 0
Delta_L
Doxycycline
YCR054C CTR86
ZShift = 0.77
NG = 0
DB = 0
SM = 0
Delta_K
Doxycycline
YCR054C CTR86
ZShift = 0
NG = 0
DB = 0
SM = 0
Delta_r
Doxycycline
YCR054C CTR86
ZShift = 0.27
NG = 0
DB = 0
SM = 0
Delta_AUC
Doxycycline
YPL076W GPI2
ZShift = 0.04
NG = 0
DB = 0
SM = 0
Delta_L
Doxycycline
YPL076W GPI2
ZShift = 0.68
NG = 0
DB = 0
SM = 0
Delta_K
Doxycycline
YPL076W GPI2
ZShift = −0.11
NG = 0
DB = 0
SM = 0
Delta_r
Doxycycline
YPL076W GPI2
ZShift = 0.45
NG = 0
DB = 0
SM = 0
Delta_AUC
Doxycycline
YDL120W YFH1
ZShift = −1.6
NG = 0
DB = 0
SM = 0
Delta_L
Doxycycline
YDL120W YFH1
ZShift = 2.49
NG = 0
DB = 0
SM = 0
Delta_K
Doxycycline
YDL120W YFH1
ZShift = 1.58
NG = 0
DB = 0
SM = 0
Delta_r
Doxycycline
YDL120W YFH1
ZShift = 2.47
NG = 0
DB = 0
SM = 0
Delta_AUC
Doxycycline

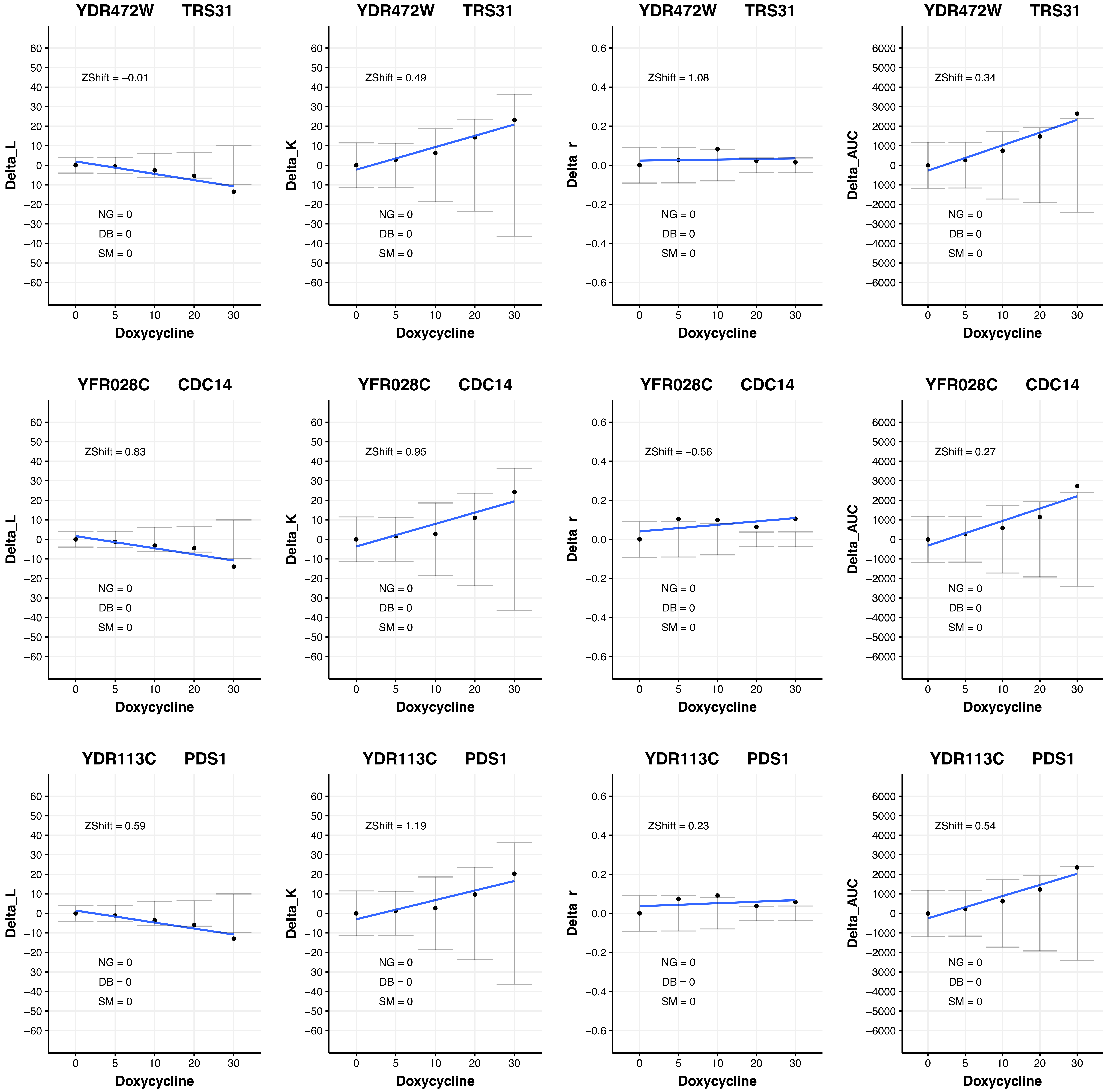

YDR472W TRS31
ZShift = −0.01
NG = 0
DB = 0
SM = 0
Delta_L
Doxycycline
YDR472W TRS31
ZShift = 0.49
NG = 0
DB = 0
SM = 0
Delta_K
Doxycycline
YDR472W TRS31
ZShift = 1.08
NG = 0
DB = 0
SM = 0
Delta_r
Doxycycline
YDR472W TRS31
ZShift = 0.34
NG = 0
DB = 0
SM = 0
Delta_AUC
Doxycycline
YFR028C CDC14
ZShift = 0.83
NG = 0
DB = 0
SM = 0
Delta_L
Doxycycline
YFR028C CDC14
ZShift = 0.95
NG = 0
DB = 0
SM = 0
Delta_K
Doxycycline
YFR028C CDC14
ZShift = −0.56
NG = 0
DB = 0
SM = 0
Delta_r
Doxycycline
YFR028C CDC14
ZShift = 0.27
NG = 0
DB = 0
SM = 0
Delta_AUC
Doxycycline
YDR113C PDS1
ZShift = 0.59
NG = 0
DB = 0
SM = 0
Delta_L
Doxycycline
YDR113C PDS1
ZShift = 1.19
NG = 0
DB = 0
SM = 0
Delta_K
Doxycycline
YDR113C PDS1
ZShift = 0.23
NG = 0
DB = 0
SM = 0
Delta_r
Doxycycline
YDR113C PDS1
ZShift = 0.54
NG = 0
DB = 0
SM = 0
Delta_AUC
Doxycycline

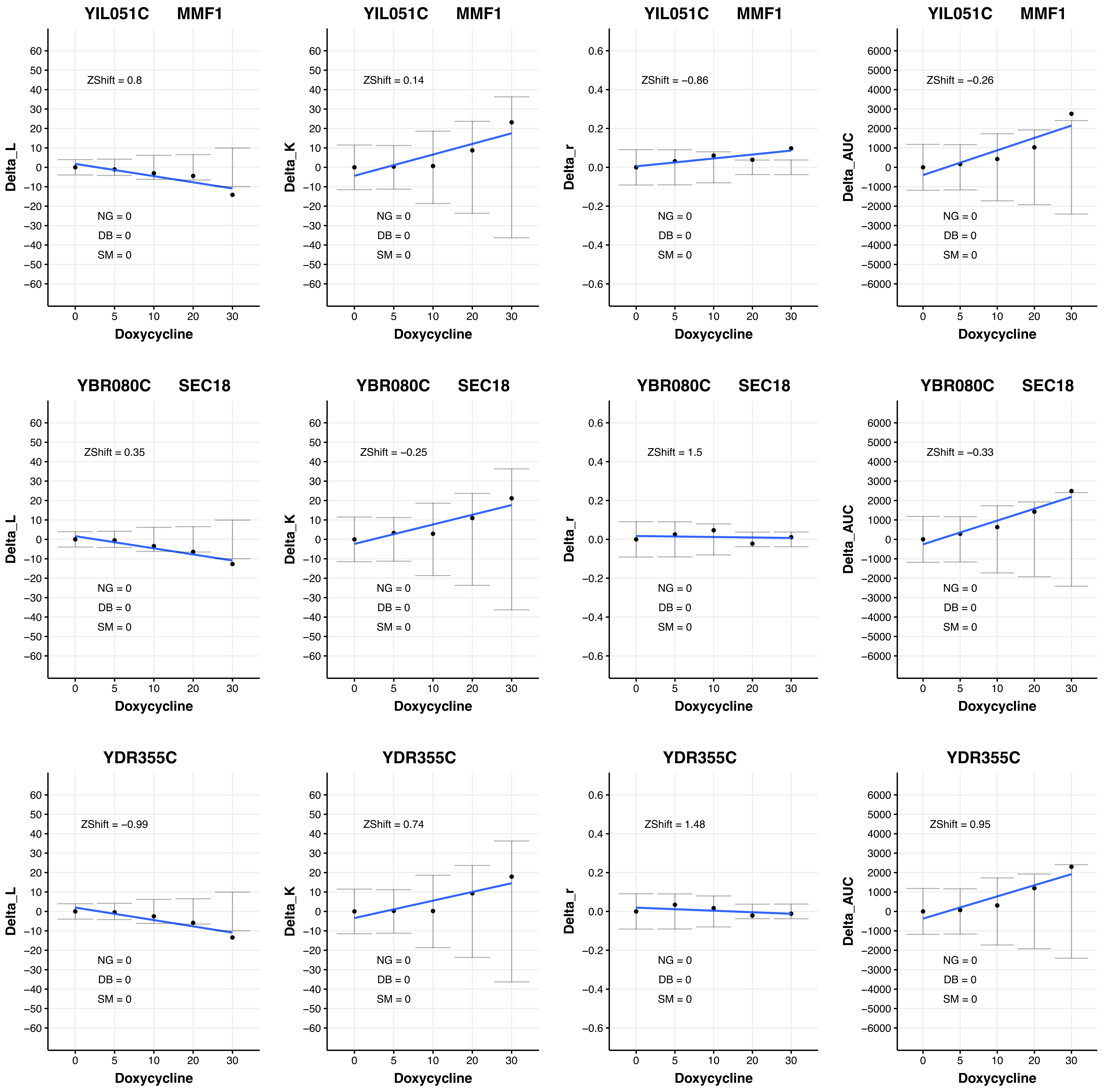
YIL051C MMF1
ZShift = 0.8
NG = 0
DB = 0
SM = 0
Delta_L
Doxycycline
YIL051C MMF1
ZShift = 0.14
NG = 0
DB = 0
SM = 0
Delta_K
Doxycycline
YIL051C MMF1
ZShift = −0.86
NG = 0
DB = 0
SM = 0
Delta_r
Doxycycline
YIL051C MMF1
ZShift = −0.26
NG = 0
DB = 0
SM = 0
Delta_AUC
Doxycycline
YBR080C SEC18
ZShift = 0.35
NG = 0
DB = 0
SM = 0
Delta_L
Doxycycline
YBR080C SEC18
ZShift = −0.25
NG = 0
DB = 0
SM = 0
Delta_K
Doxycycline
YBR080C SEC18
ZShift = 1.5
NG = 0
DB = 0
SM = 0
Delta_r
Doxycycline
YBR080C SEC18
ZShift = −0.33
NG = 0
DB = 0
SM = 0
Delta_AUC
Doxycycline
YDR355C
ZShift = −0.99
NG = 0
DB = 0
SM = 0
Delta_L
Doxycycline
YDR355C
ZShift = 0.74
NG = 0
DB = 0
SM = 0
Delta_K
Doxycycline
YDR355C
ZShift = 1.48
NG = 0
DB = 0
SM = 0
Delta_r
Doxycycline
YDR355C
ZShift = 0.95
NG = 0
DB = 0
SM = 0
Delta_AUC
Doxycycline

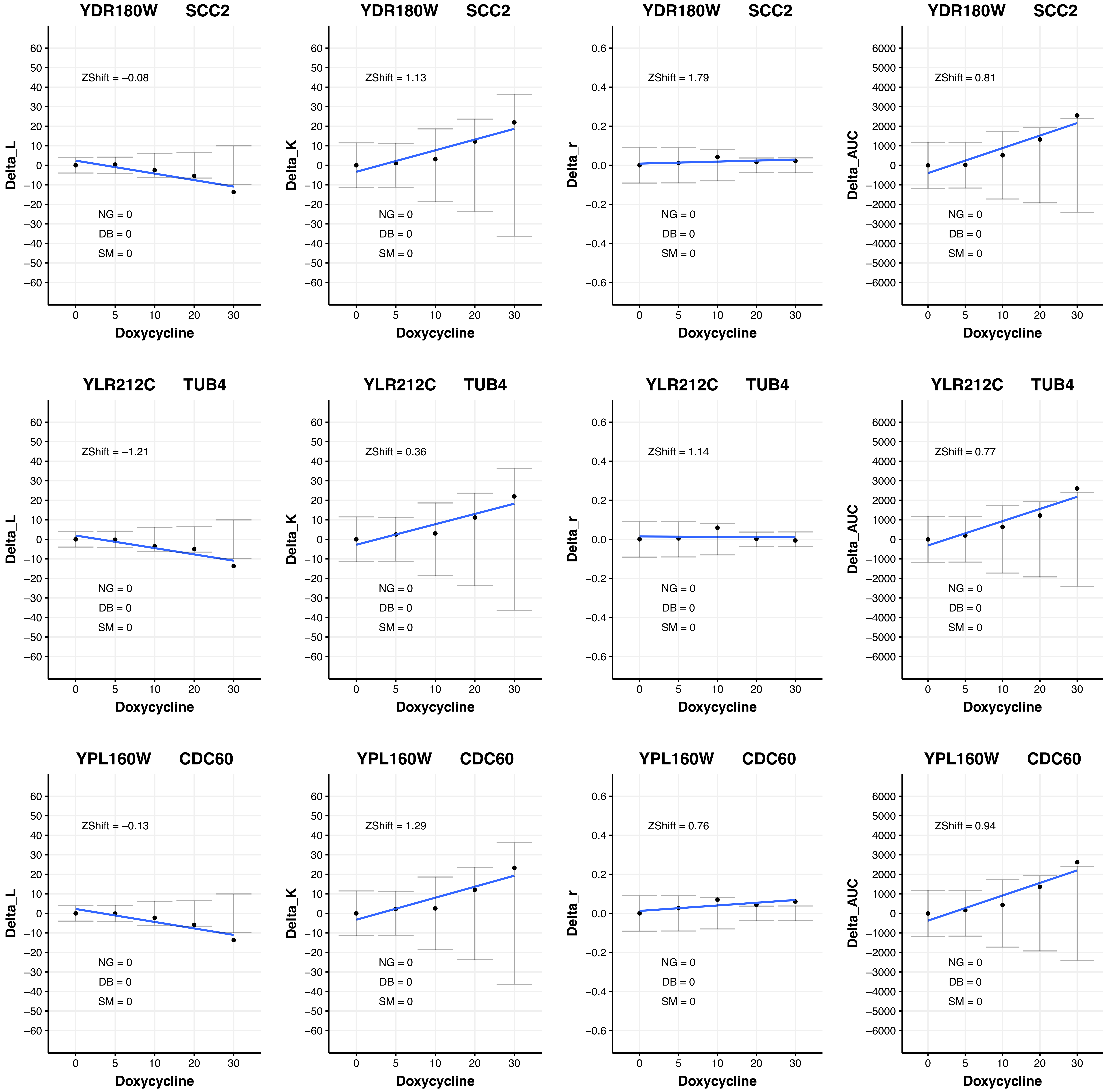
YDR180W SCC2
ZShift = −0.08
ZShift = 1.13
ZShift = 1.79
ZShift = 0.81
YLR212C TUB4
ZShift = −1.21
ZShift = 0.36
ZShift = 1.14
ZShift = 0.77
YPL160W CDC60
ZShift = −0.13
ZShift = 1.29
ZShift = 0.76
ZShift = 0.94
NG = 0
DB = 0
SM = 0
Delta_L
Delta_K
Delta_r
Delta_AUC
Doxycycline

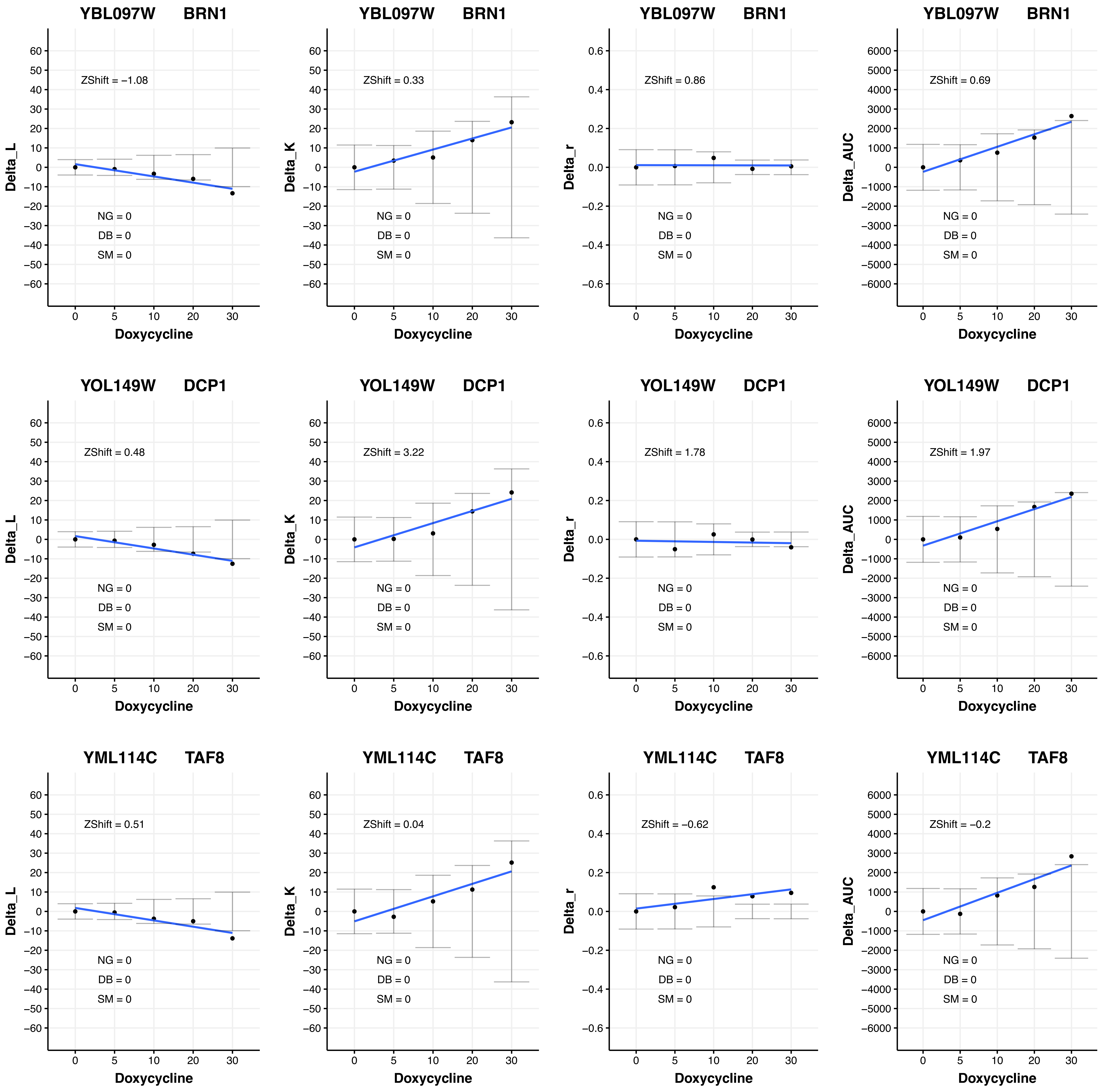

YBL097W BRN1
ZShift = −1.08
NG = 0
DB = 0
SM = 0
Delta_L
Doxycycline
YBL097W BRN1
ZShift = 0.33
NG = 0
DB = 0
SM = 0
Delta_K
Doxycycline
YBL097W BRN1
ZShift = 0.86
NG = 0
DB = 0
SM = 0
Delta_r
Doxycycline
YBL097W BRN1
ZShift = 0.69
NG = 0
DB = 0
SM = 0
Delta_AUC
Doxycycline
YOL149W DCP1
ZShift = 0.48
NG = 0
DB = 0
SM = 0
Delta_L
Doxycycline
YOL149W DCP1
ZShift = 3.22
NG = 0
DB = 0
SM = 0
Delta_K
Doxycycline
YOL149W DCP1
ZShift = 1.78
NG = 0
DB = 0
SM = 0
Delta_r
Doxycycline
YOL149W DCP1
ZShift = 1.97
NG = 0
DB = 0
SM = 0
Delta_AUC
Doxycycline
YML114C TAF8
ZShift = 0.51
NG = 0
DB = 0
SM = 0
Delta_L
Doxycycline
YML114C TAF8
ZShift = 0.04
NG = 0
DB = 0
SM = 0
Delta_K
Doxycycline
YML114C TAF8
ZShift = −0.62
NG = 0
DB = 0
SM = 0
Delta_r
Doxycycline
YML114C TAF8
ZShift = −0.2
NG = 0
DB = 0
SM = 0
Delta_AUC
Doxycycline

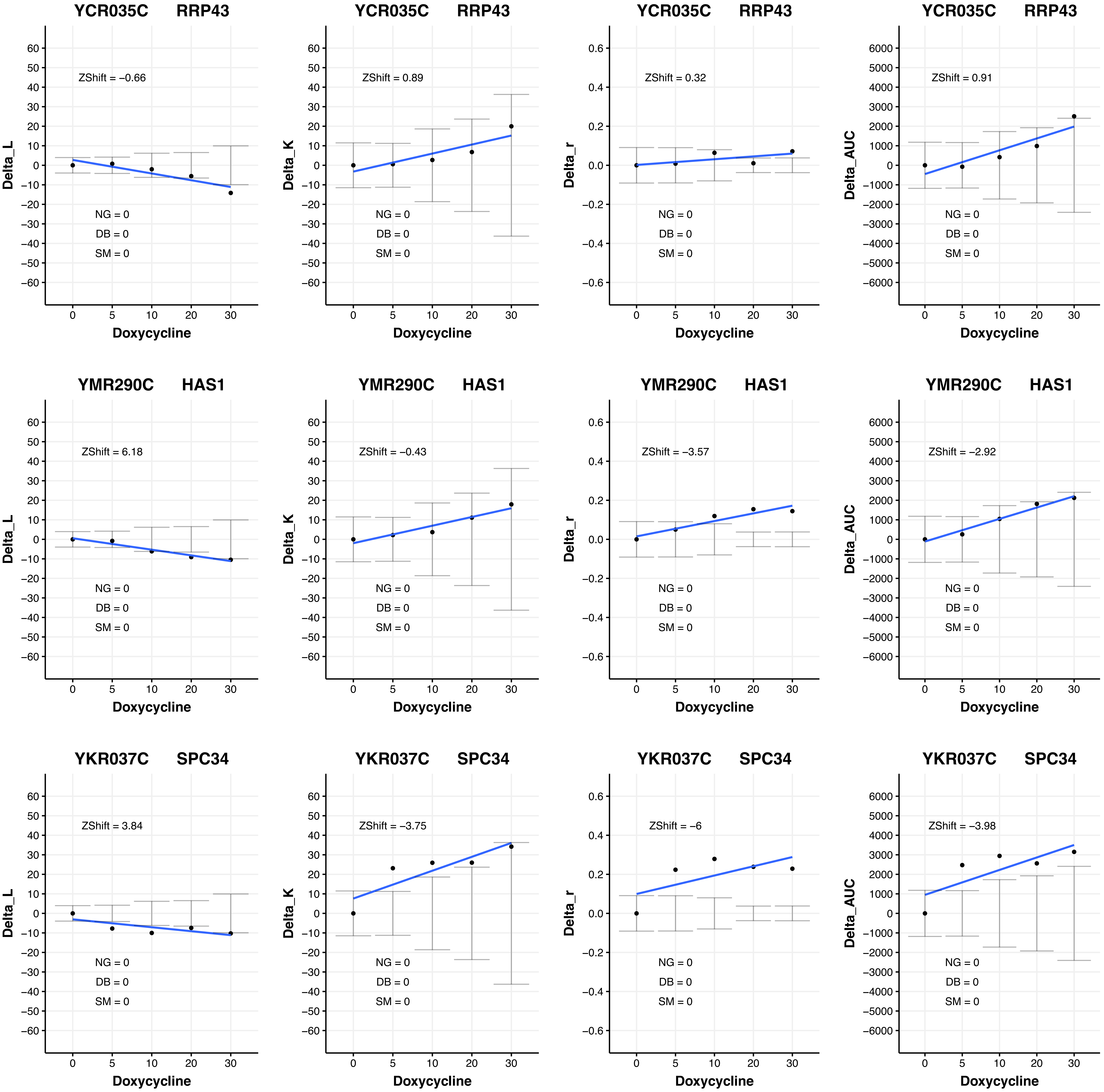
YCR035C RRP43
ZShift = −0.66
NG = 0
DB = 0
SM = 0
Delta_L
Doxycycline
YCR035C RRP43
ZShift = 0.89
NG = 0
DB = 0
SM = 0
Delta_K
Doxycycline
YCR035C RRP43
ZShift = 0.32
NG = 0
DB = 0
SM = 0
Delta_r
Doxycycline
YCR035C RRP43
ZShift = 0.91
NG = 0
DB = 0
SM = 0
Delta_AUC
Doxycycline
YMR290C HAS1
ZShift = 6.18
NG = 0
DB = 0
SM = 0
Delta_L
Doxycycline
YMR290C HAS1
ZShift = −0.43
NG = 0
DB = 0
SM = 0
Delta_K
Doxycycline
YMR290C HAS1
ZShift = −3.57
NG = 0
DB = 0
SM = 0
Delta_r
Doxycycline
YMR290C HAS1
ZShift = −2.92
NG = 0
DB = 0
SM = 0
Delta_AUC
Doxycycline
YKR037C SPC34
ZShift = 3.84
NG = 0
DB = 0
SM = 0
Delta_L
Doxycycline
YKR037C SPC34
ZShift = −3.75
NG = 0
DB = 0
SM = 0
Delta_K
Doxycycline
YKR037C SPC34
ZShift = −6
NG = 0
DB = 0
SM = 0
Delta_r
Doxycycline
YKR037C SPC34
ZShift = −3.98
NG = 0
DB = 0
SM = 0
Delta_AUC
Doxycycline

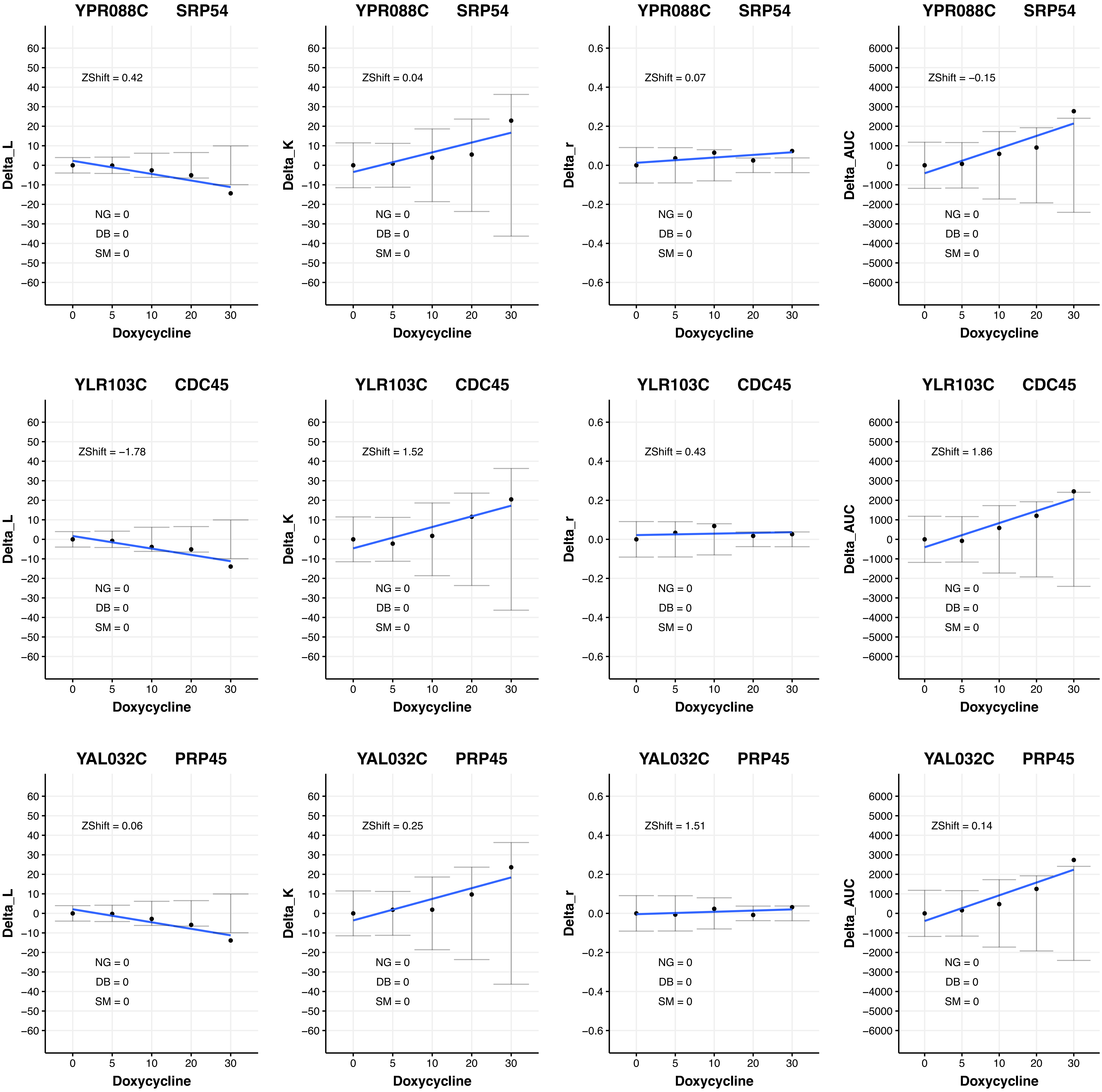
YPR088C SRP54
ZShift = 0.42
NG = 0
DB = 0
SM = 0
Delta_L
Doxycycline
YPR088C SRP54
ZShift = 0.04
NG = 0
DB = 0
SM = 0
Delta_K
Doxycycline
YPR088C SRP54
ZShift = 0.07
NG = 0
DB = 0
SM = 0
Delta_r
Doxycycline
YPR088C SRP54
ZShift = −0.15
NG = 0
DB = 0
SM = 0
Delta_AUC
Doxycycline
YLR103C CDC45
ZShift = −1.78
NG = 0
DB = 0
SM = 0
Delta_L
Doxycycline
YLR103C CDC45
ZShift = 1.52
NG = 0
DB = 0
SM = 0
Delta_K
Doxycycline
YLR103C CDC45
ZShift = 0.43
NG = 0
DB = 0
SM = 0
Delta_r
Doxycycline
YLR103C CDC45
ZShift = 1.86
NG = 0
DB = 0
SM = 0
Delta_AUC
Doxycycline
YAL032C PRP45
ZShift = 0.06
NG = 0
DB = 0
SM = 0
Delta_L
Doxycycline
YAL032C PRP45
ZShift = 0.25
NG = 0
DB = 0
SM = 0
Delta_K
Doxycycline
YAL032C PRP45
ZShift = 1.51
NG = 0
DB = 0
SM = 0
Delta_r
Doxycycline
YAL032C PRP45
ZShift = 0.14
NG = 0
DB = 0
SM = 0
Delta_AUC
Doxycycline

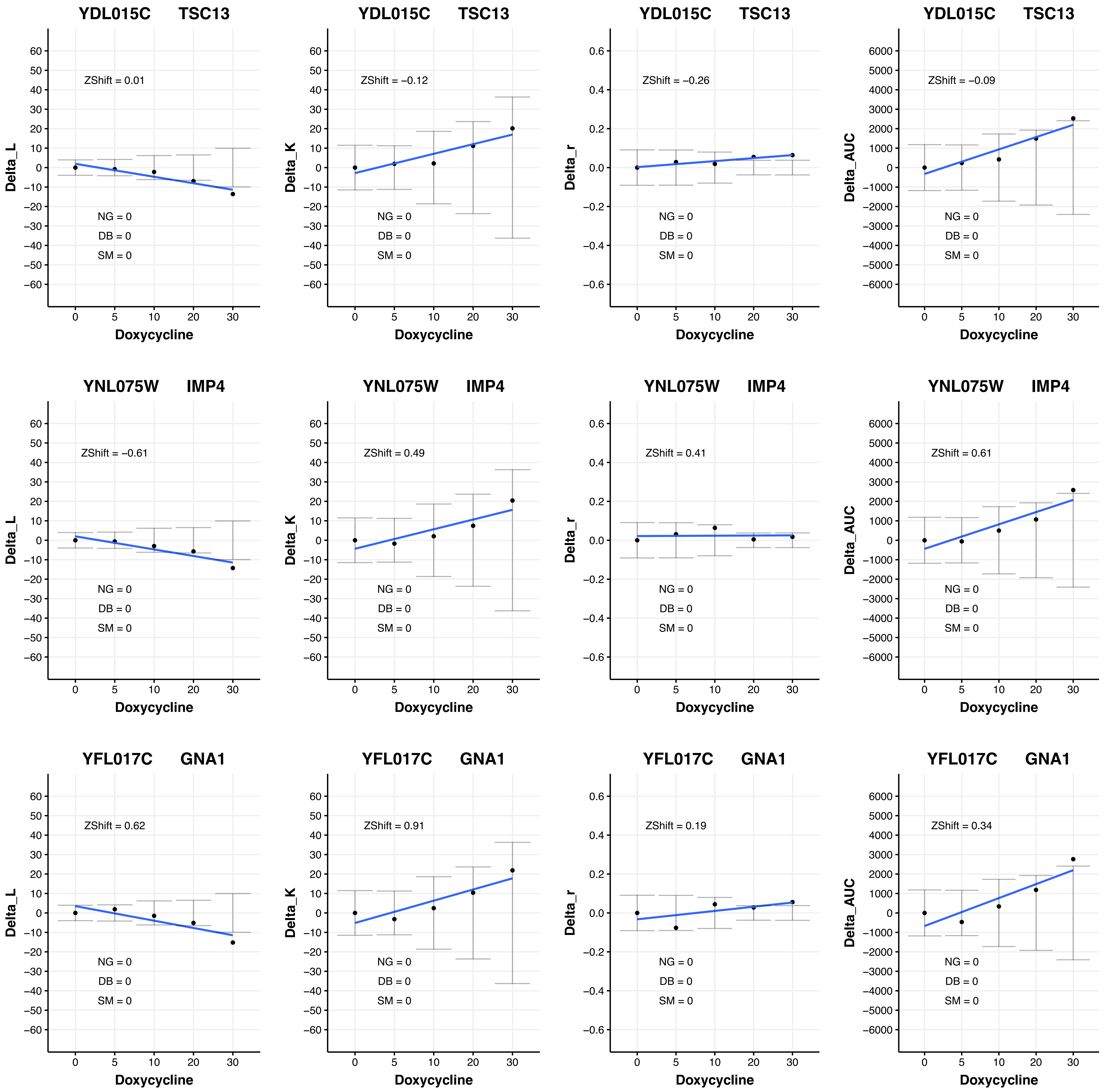

YDL015C TSC13
ZShift = 0.01
Delta_L
NG = 0
DB = 0
SM = 0
Doxycycline
YDL015C TSC13
ZShift = –0.12
Delta_K
NG = 0
DB = 0
SM = 0
Doxycycline
YDL015C TSC13
ZShift = –0.26
Delta_r
NG = 0
DB = 0
SM = 0
Doxycycline
YDL015C TSC13
ZShift = –0.09
Delta_AUC
NG = 0
DB = 0
SM = 0
Doxycycline
YNL075W IMP4
ZShift = –0.61
Delta_L
NG = 0
DB = 0
SM = 0
Doxycycline
YNL075W IMP4
ZShift = 0.49
Delta_K
NG = 0
DB = 0
SM = 0
Doxycycline
YNL075W IMP4
ZShift = 0.41
Delta_r
NG = 0
DB = 0
SM = 0
Doxycycline
YNL075W IMP4
ZShift = 0.61
Delta_AUC
NG = 0
DB = 0
SM = 0
Doxycycline
YFL017C GNA1
ZShift = 0.62
Delta_L
NG = 0
DB = 0
SM = 0
Doxycycline
YFL017C GNA1
ZShift = 0.91
Delta_K
NG = 0
DB = 0
SM = 0
Doxycycline
YFL017C GNA1
ZShift = 0.19
Delta_r
NG = 0
DB = 0
SM = 0
Doxycycline
YFL017C GNA1
ZShift = 0.34
Delta_AUC
NG = 0
DB = 0
SM = 0
Doxycycline

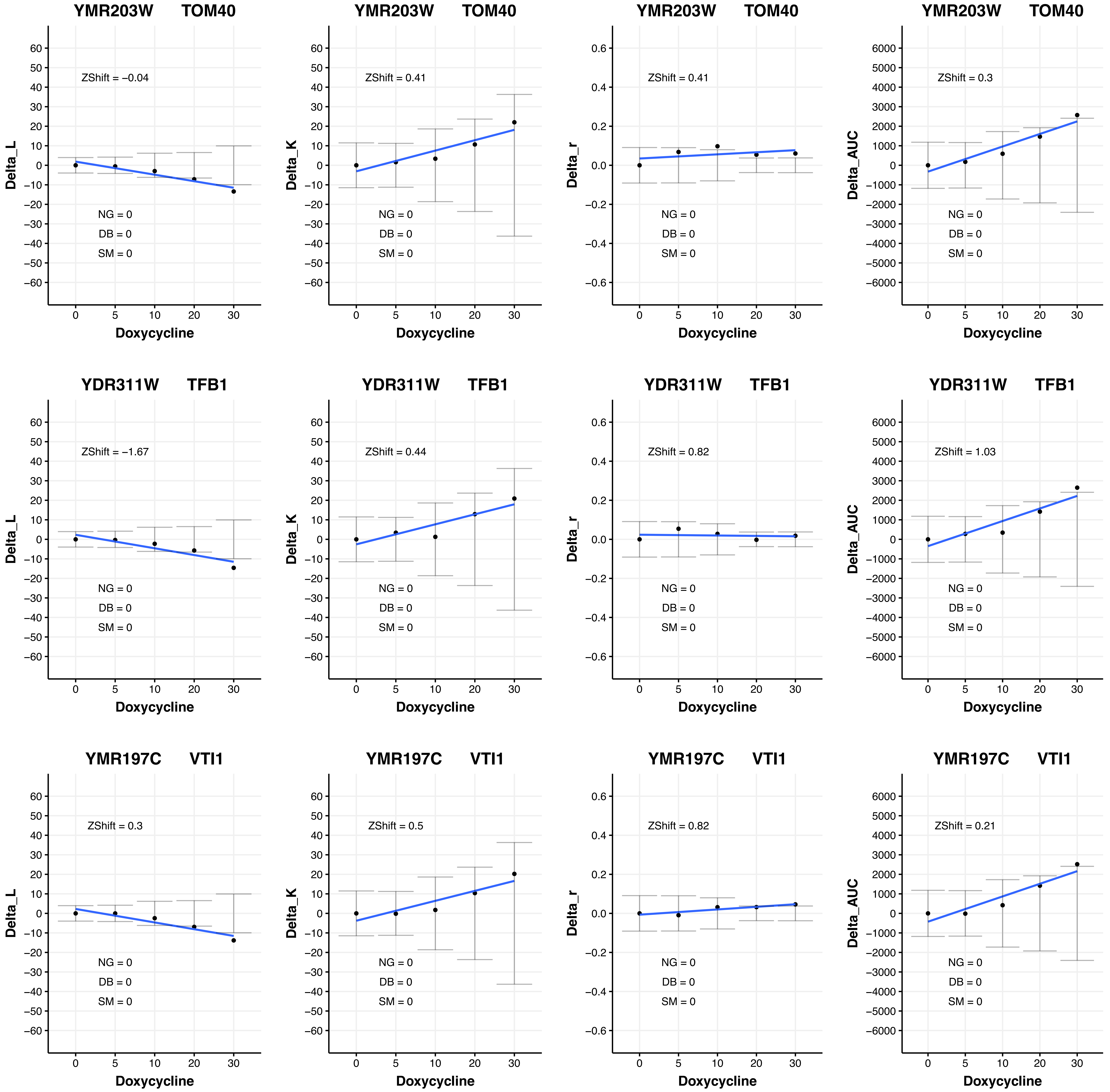
YMR203W TOM40
ZShift = −0.04
NG = 0
DB = 0
SM = 0
Delta_L
Doxycycline
YMR203W TOM40
ZShift = 0.41
NG = 0
DB = 0
SM = 0
Delta_K
Doxycycline
YMR203W TOM40
ZShift = 0.41
NG = 0
DB = 0
SM = 0
Delta_r
Doxycycline
YMR203W TOM40
ZShift = 0.3
NG = 0
DB = 0
SM = 0
Delta_AUC
Doxycycline
YDR311W TFB1
ZShift = −1.67
NG = 0
DB = 0
SM = 0
Delta_L
Doxycycline
YDR311W TFB1
ZShift = 0.44
NG = 0
DB = 0
SM = 0
Delta_K
Doxycycline
YDR311W TFB1
ZShift = 0.82
NG = 0
DB = 0
SM = 0
Delta_r
Doxycycline
YDR311W TFB1
ZShift = 1.03
NG = 0
DB = 0
SM = 0
Delta_AUC
Doxycycline
YMR197C VTI1
ZShift = 0.3
NG = 0
DB = 0
SM = 0
Delta_L
Doxycycline
YMR197C VTI1
ZShift = 0.5
NG = 0
DB = 0
SM = 0
Delta_K
Doxycycline
YMR197C VTI1
ZShift = 0.82
NG = 0
DB = 0
SM = 0
Delta_r
Doxycycline
YMR197C VTI1
ZShift = 0.21
NG = 0
DB = 0
SM = 0
Delta_AUC
Doxycycline

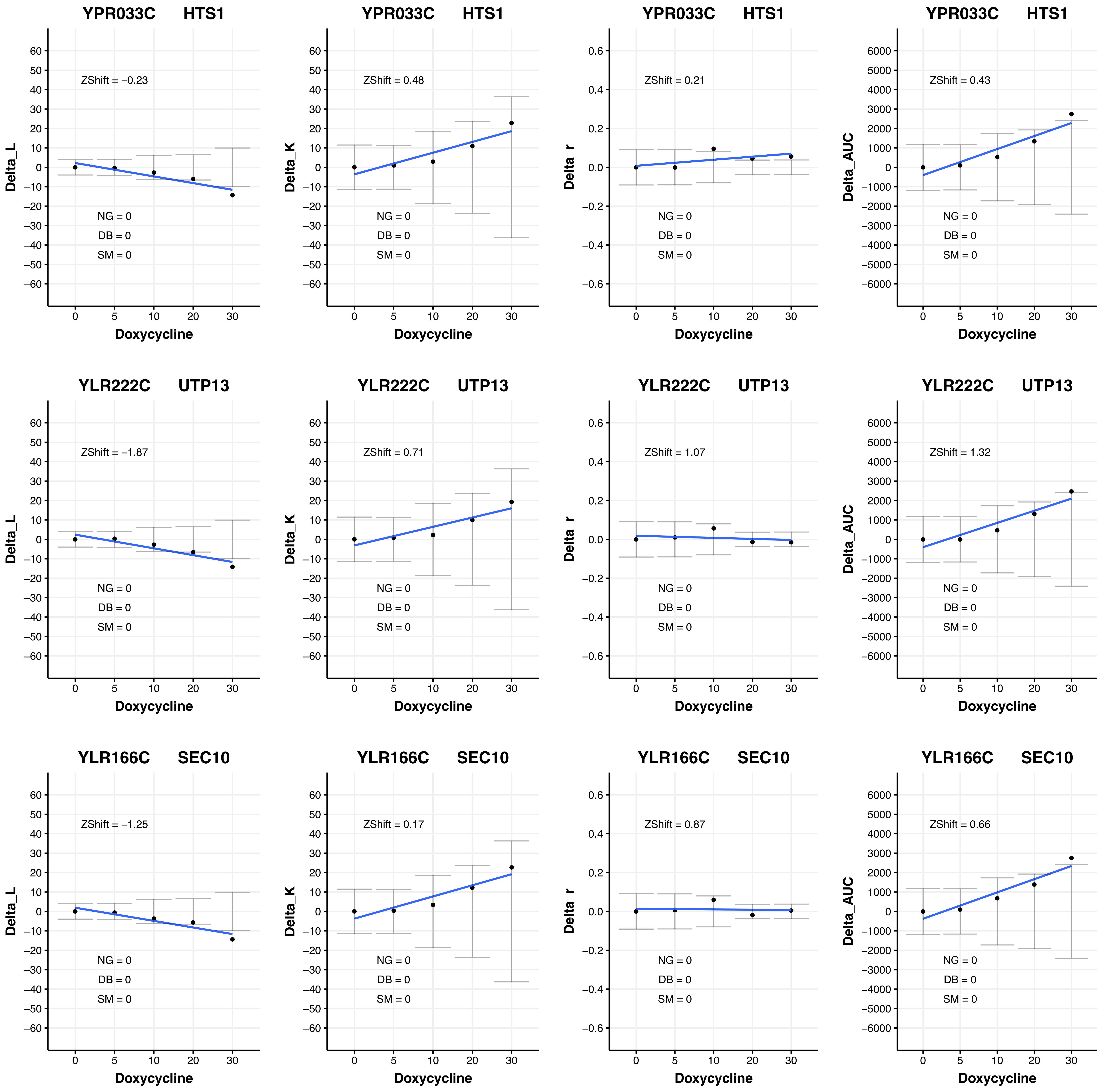

YPR033C HTS1
ZShift = −0.23
NG = 0
DB = 0
SM = 0
Delta_L
Doxycycline
YPR033C HTS1
ZShift = 0.48
NG = 0
DB = 0
SM = 0
Delta_K
Doxycycline
YPR033C HTS1
ZShift = 0.21
NG = 0
DB = 0
SM = 0
Delta_r
Doxycycline
YPR033C HTS1
ZShift = 0.43
NG = 0
DB = 0
SM = 0
Delta_AUC
Doxycycline
YLR222C UTP13
ZShift = −1.87
NG = 0
DB = 0
SM = 0
Delta_L
Doxycycline
YLR222C UTP13
ZShift = 0.71
NG = 0
DB = 0
SM = 0
Delta_K
Doxycycline
YLR222C UTP13
ZShift = 1.07
NG = 0
DB = 0
SM = 0
Delta_r
Doxycycline
YLR222C UTP13
ZShift = 1.32
NG = 0
DB = 0
SM = 0
Delta_AUC
Doxycycline
YLR166C SEC10
ZShift = −1.25
NG = 0
DB = 0
SM = 0
Delta_L
Doxycycline
YLR166C SEC10
ZShift = 0.17
NG = 0
DB = 0
SM = 0
Delta_K
Doxycycline
YLR166C SEC10
ZShift = 0.87
NG = 0
DB = 0
SM = 0
Delta_r
Doxycycline
YLR166C SEC10
ZShift = 0.66
NG = 0
DB = 0
SM = 0
Delta_AUC
Doxycycline

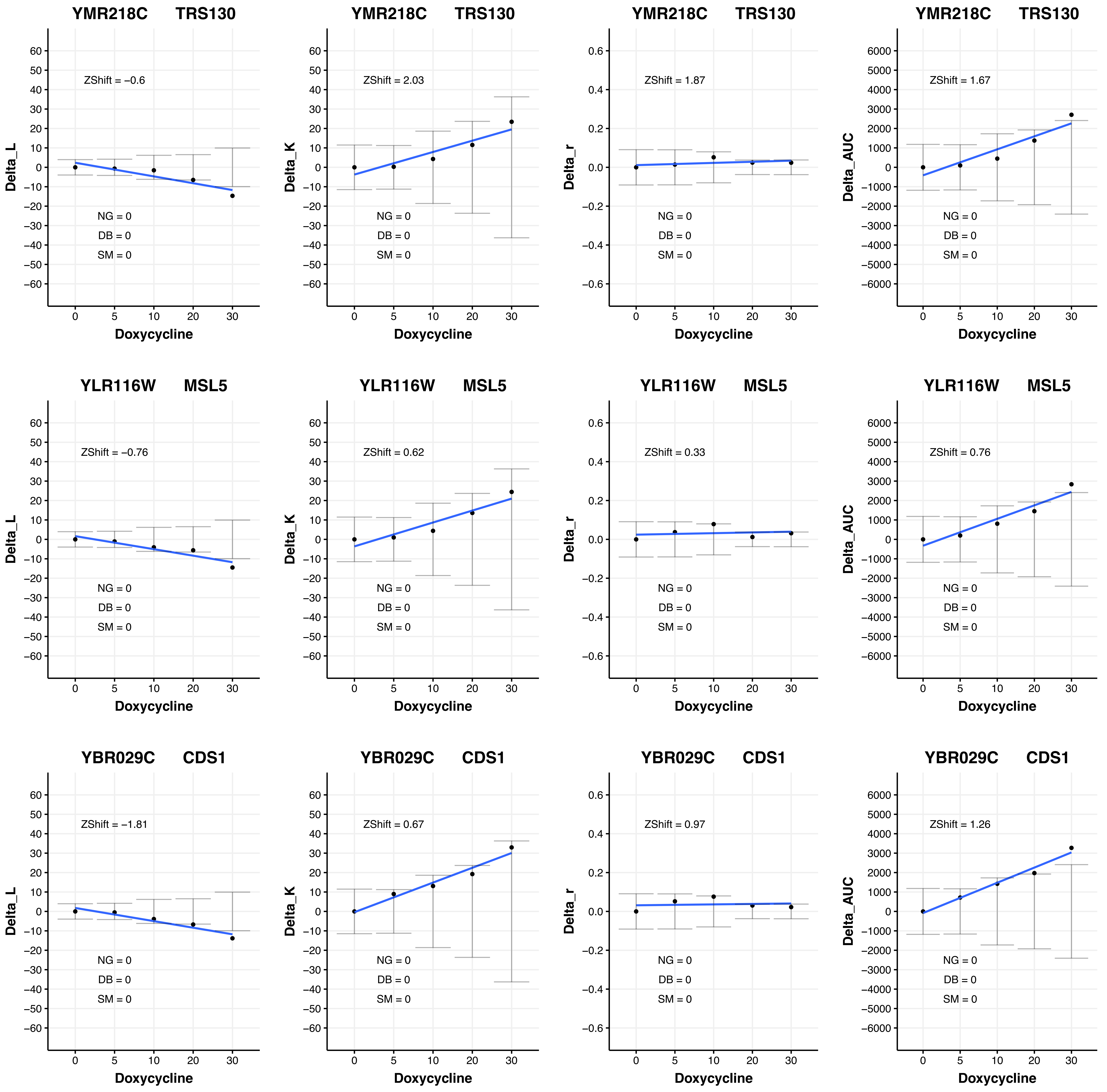

YMR218C TRS130
ZShift = −0.6
NG = 0
DB = 0
SM = 0
Delta_L
Doxycycline
YMR218C TRS130
ZShift = 2.03
NG = 0
DB = 0
SM = 0
Delta_K
Doxycycline
YMR218C TRS130
ZShift = 1.87
NG = 0
DB = 0
SM = 0
Delta_r
Doxycycline
YMR218C TRS130
ZShift = 1.67
NG = 0
DB = 0
SM = 0
Delta_AUC
Doxycycline
YLR116W MSL5
ZShift = −0.76
NG = 0
DB = 0
SM = 0
Delta_L
Doxycycline
YLR116W MSL5
ZShift = 0.62
NG = 0
DB = 0
SM = 0
Delta_K
Doxycycline
YLR116W MSL5
ZShift = 0.33
NG = 0
DB = 0
SM = 0
Delta_r
Doxycycline
YLR116W MSL5
ZShift = 0.76
NG = 0
DB = 0
SM = 0
Delta_AUC
Doxycycline
YBR029C CDS1
ZShift = −1.81
NG = 0
DB = 0
SM = 0
Delta_L
Doxycycline
YBR029C CDS1
ZShift = 0.67
NG = 0
DB = 0
SM = 0
Delta_K
Doxycycline
YBR029C CDS1
ZShift = 0.97
NG = 0
DB = 0
SM = 0
Delta_r
Doxycycline
YBR029C CDS1
ZShift = 1.26
NG = 0
DB = 0
SM = 0
Delta_AUC
Doxycycline

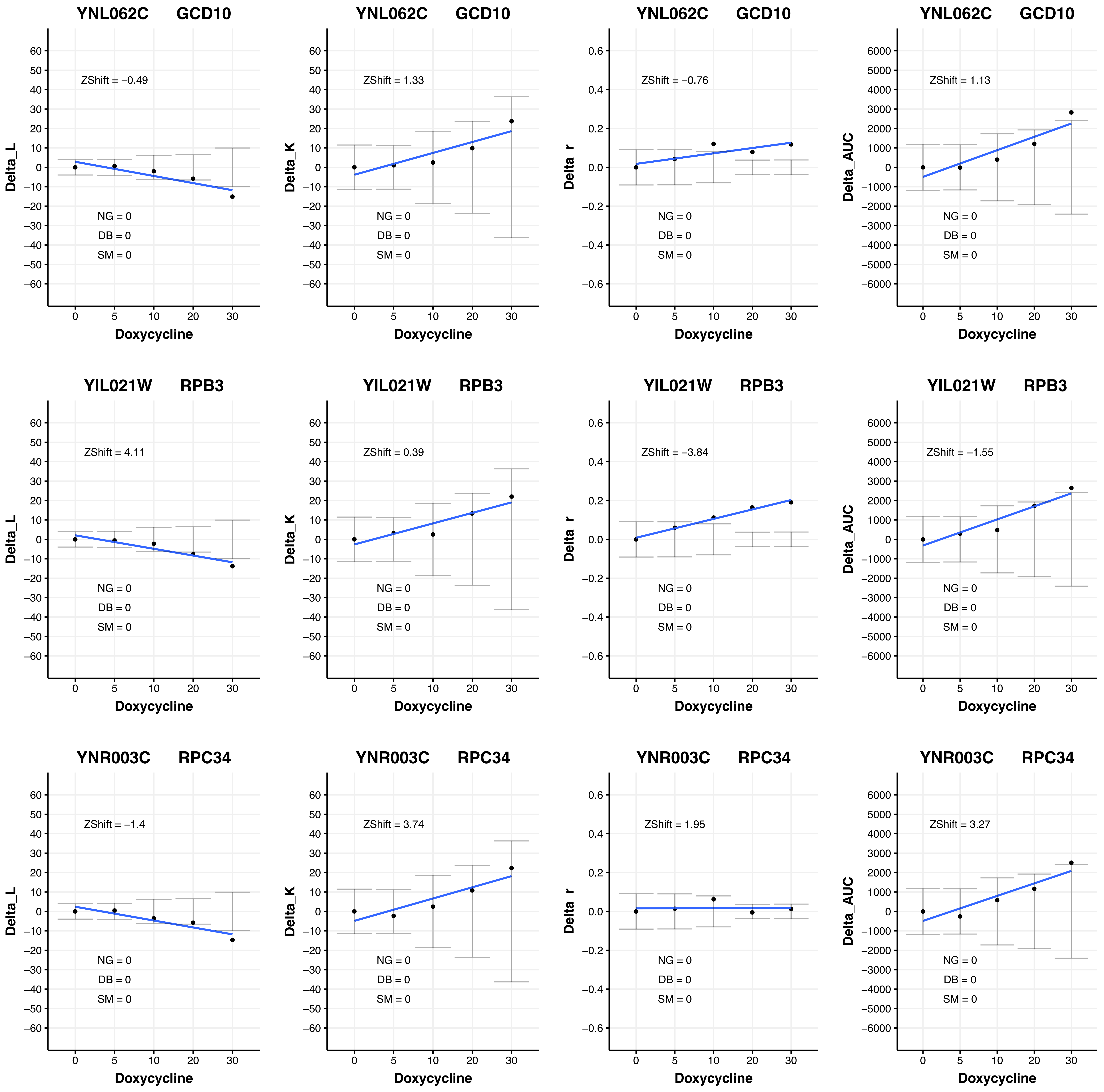
YNL062C GCD10
ZShift = −0.49
NG = 0
DB = 0
SM = 0
Delta_L
Doxycycline
YNL062C GCD10
ZShift = 1.33
NG = 0
DB = 0
SM = 0
Delta_K
Doxycycline
YNL062C GCD10
ZShift = −0.76
NG = 0
DB = 0
SM = 0
Delta_r
Doxycycline
YNL062C GCD10
ZShift = 1.13
NG = 0
DB = 0
SM = 0
Delta_AUC
Doxycycline
YIL021W RPB3
ZShift = 4.11
NG = 0
DB = 0
SM = 0
Delta_L
Doxycycline
YIL021W RPB3
ZShift = 0.39
NG = 0
DB = 0
SM = 0
Delta_K
Doxycycline
YIL021W RPB3
ZShift = −3.84
NG = 0
DB = 0
SM = 0
Delta_r
Doxycycline
YIL021W RPB3
ZShift = −1.55
NG = 0
DB = 0
SM = 0
Delta_AUC
Doxycycline
YNR003C RPC34
ZShift = −1.4
NG = 0
DB = 0
SM = 0
Delta_L
Doxycycline
YNR003C RPC34
ZShift = 3.74
NG = 0
DB = 0
SM = 0
Delta_K
Doxycycline
YNR003C RPC34
ZShift = 1.95
NG = 0
DB = 0
SM = 0
Delta_r
Doxycycline
YNR003C RPC34
ZShift = 3.27
NG = 0
DB = 0
SM = 0
Delta_AUC
Doxycycline

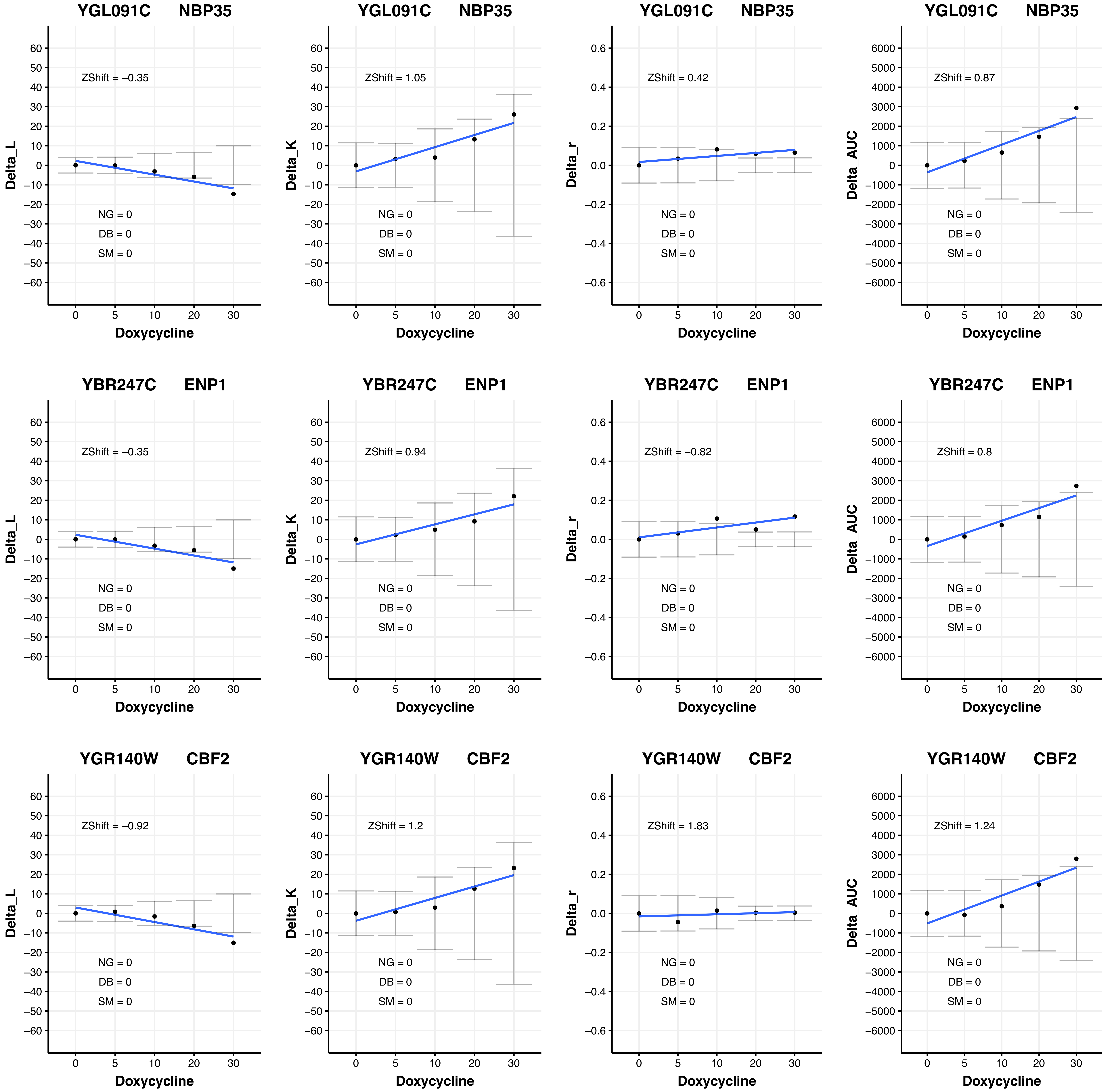
YGL091C NBP35
ZShift = −0.35
NG = 0
DB = 0
SM = 0
Delta_L
YGL091C NBP35
ZShift = 1.05
NG = 0
DB = 0
SM = 0
Delta_K
YGL091C NBP35
ZShift = 0.42
NG = 0
DB = 0
SM = 0
Delta_r
YGL091C NBP35
ZShift = 0.87
NG = 0
DB = 0
SM = 0
Delta_AUC
YBR247C ENP1
ZShift = −0.35
NG = 0
DB = 0
SM = 0
Delta_L
YBR247C ENP1
ZShift = 0.94
NG = 0
DB = 0
SM = 0
Delta_K
YBR247C ENP1
ZShift = −0.82
NG = 0
DB = 0
SM = 0
Delta_r
YBR247C ENP1
ZShift = 0.8
NG = 0
DB = 0
SM = 0
Delta_AUC
YGR140W CBF2
ZShift = −0.92
NG = 0
DB = 0
SM = 0
Delta_L
YGR140W CBF2
ZShift = 1.2
NG = 0
DB = 0
SM = 0
Delta_K
YGR140W CBF2
ZShift = 1.83
NG = 0
DB = 0
SM = 0
Delta_r
YGR140W CBF2
ZShift = 1.24
NG = 0
DB = 0
SM = 0
Delta_AUC
Doxycycline

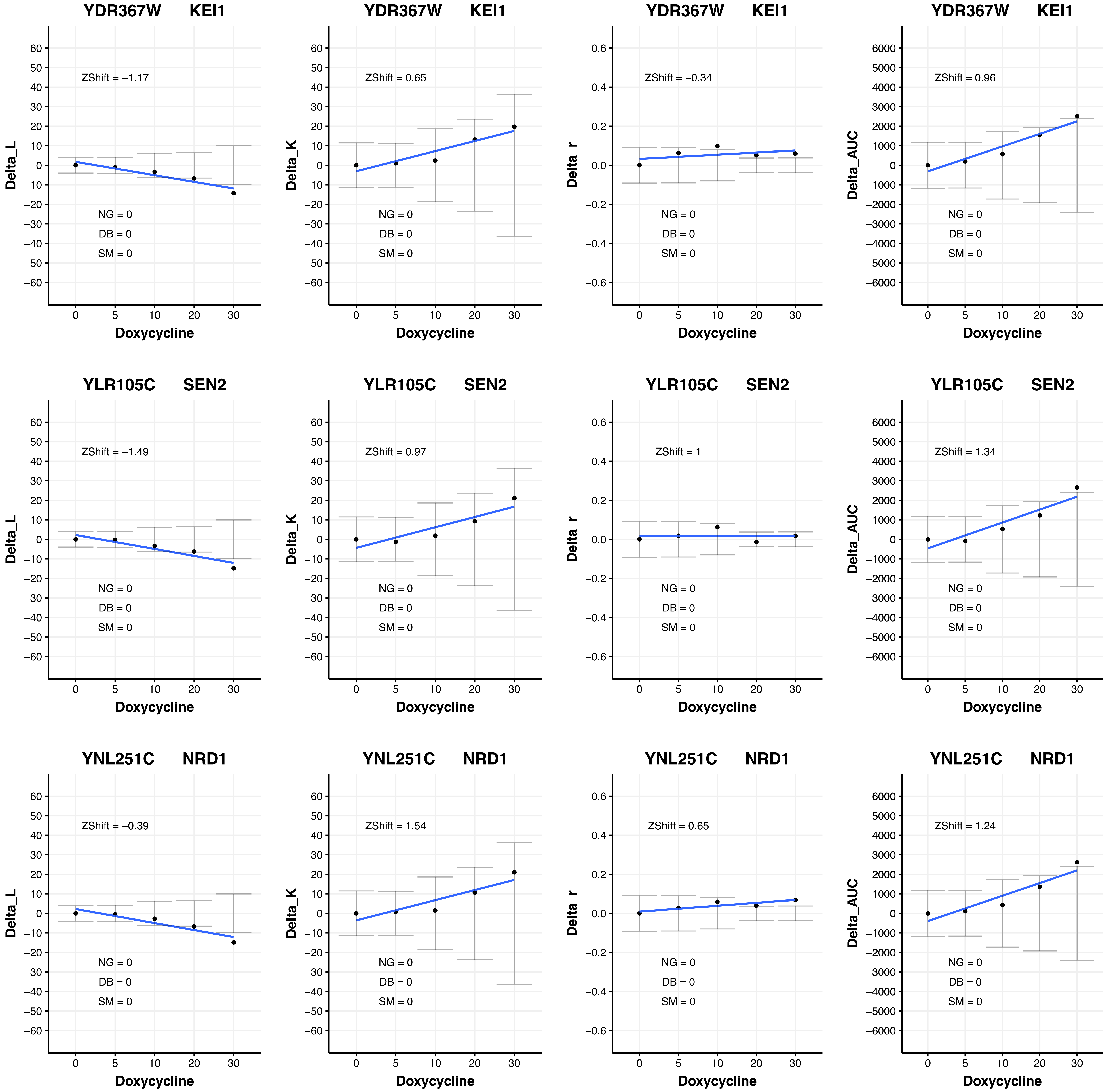
YDR367W KEI1
ZShift = −1.17
ZShift = 0.65
ZShift = −0.34
ZShift = 0.96
YLR105C SEN2
ZShift = −1.49
ZShift = 0.97
ZShift = 1
ZShift = 1.34
YNL251C NRD1
ZShift = −0.39
ZShift = 1.54
ZShift = 0.65
ZShift = 1.24
NG = 0
DB = 0
SM = 0
Delta_L
Delta_K
Delta_r
Delta_AUC
Doxycycline
0
5
10
20
30

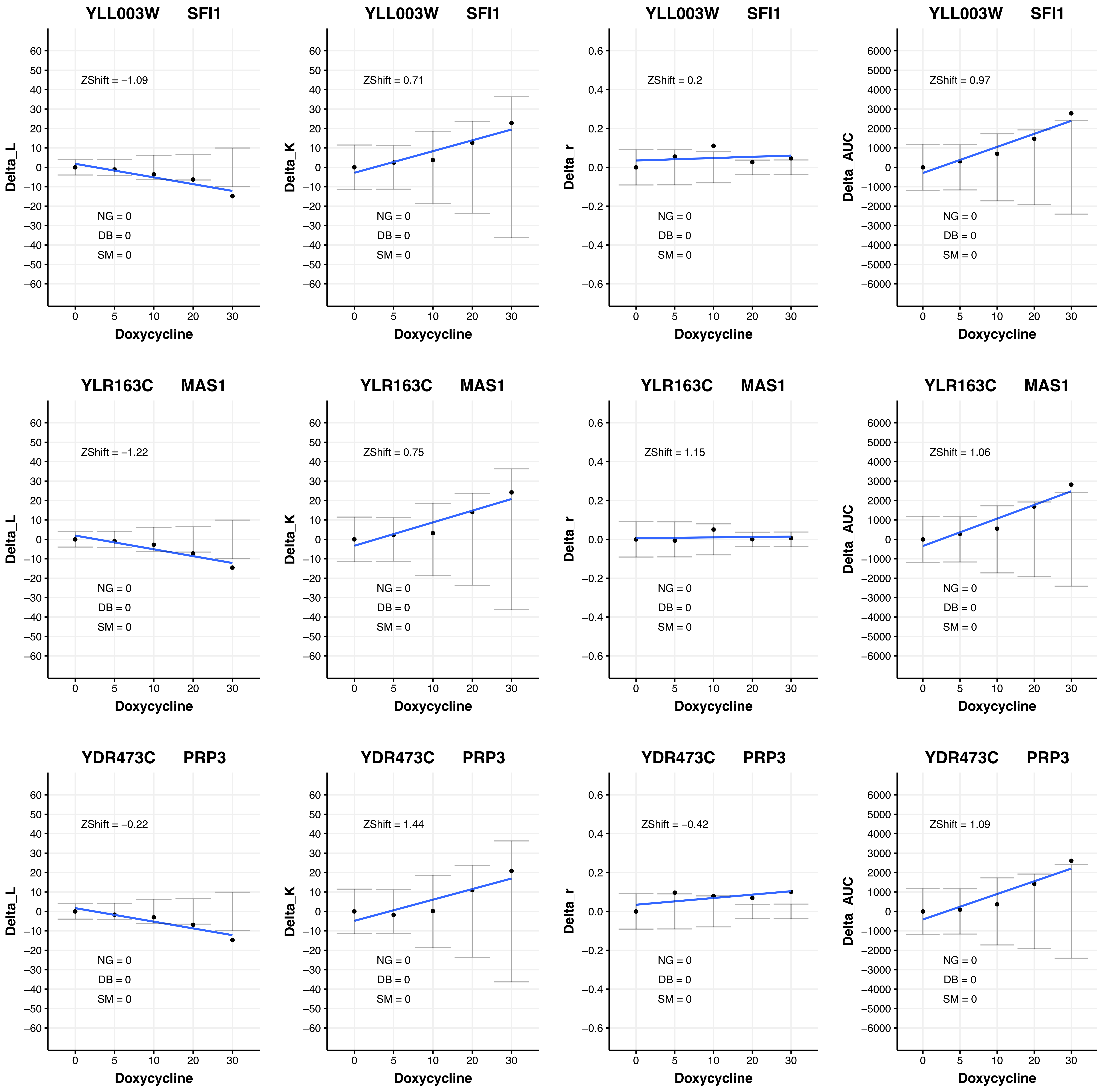
YLL003W SFI1
ZShift = −1.09
NG = 0
DB = 0
SM = 0
Delta_L
Doxycycline
YLL003W SFI1
ZShift = 0.71
NG = 0
DB = 0
SM = 0
Delta_K
Doxycycline
YLL003W SFI1
ZShift = 0.2
NG = 0
DB = 0
SM = 0
Delta_r
Doxycycline
YLL003W SFI1
ZShift = 0.97
NG = 0
DB = 0
SM = 0
Delta_AUC
Doxycycline
YLR163C MAS1
ZShift = −1.22
NG = 0
DB = 0
SM = 0
Delta_L
Doxycycline
YLR163C MAS1
ZShift = 0.75
NG = 0
DB = 0
SM = 0
Delta_K
Doxycycline
YLR163C MAS1
ZShift = 1.15
NG = 0
DB = 0
SM = 0
Delta_r
Doxycycline
YLR163C MAS1
ZShift = 1.06
NG = 0
DB = 0
SM = 0
Delta_AUC
Doxycycline
YDR473C PRP3
ZShift = −0.22
NG = 0
DB = 0
SM = 0
Delta_L
Doxycycline
YDR473C PRP3
ZShift = 1.44
NG = 0
DB = 0
SM = 0
Delta_K
Doxycycline
YDR473C PRP3
ZShift = −0.42
NG = 0
DB = 0
SM = 0
Delta_r
Doxycycline
YDR473C PRP3
ZShift = 1.09
NG = 0
DB = 0
SM = 0
Delta_AUC
Doxycycline

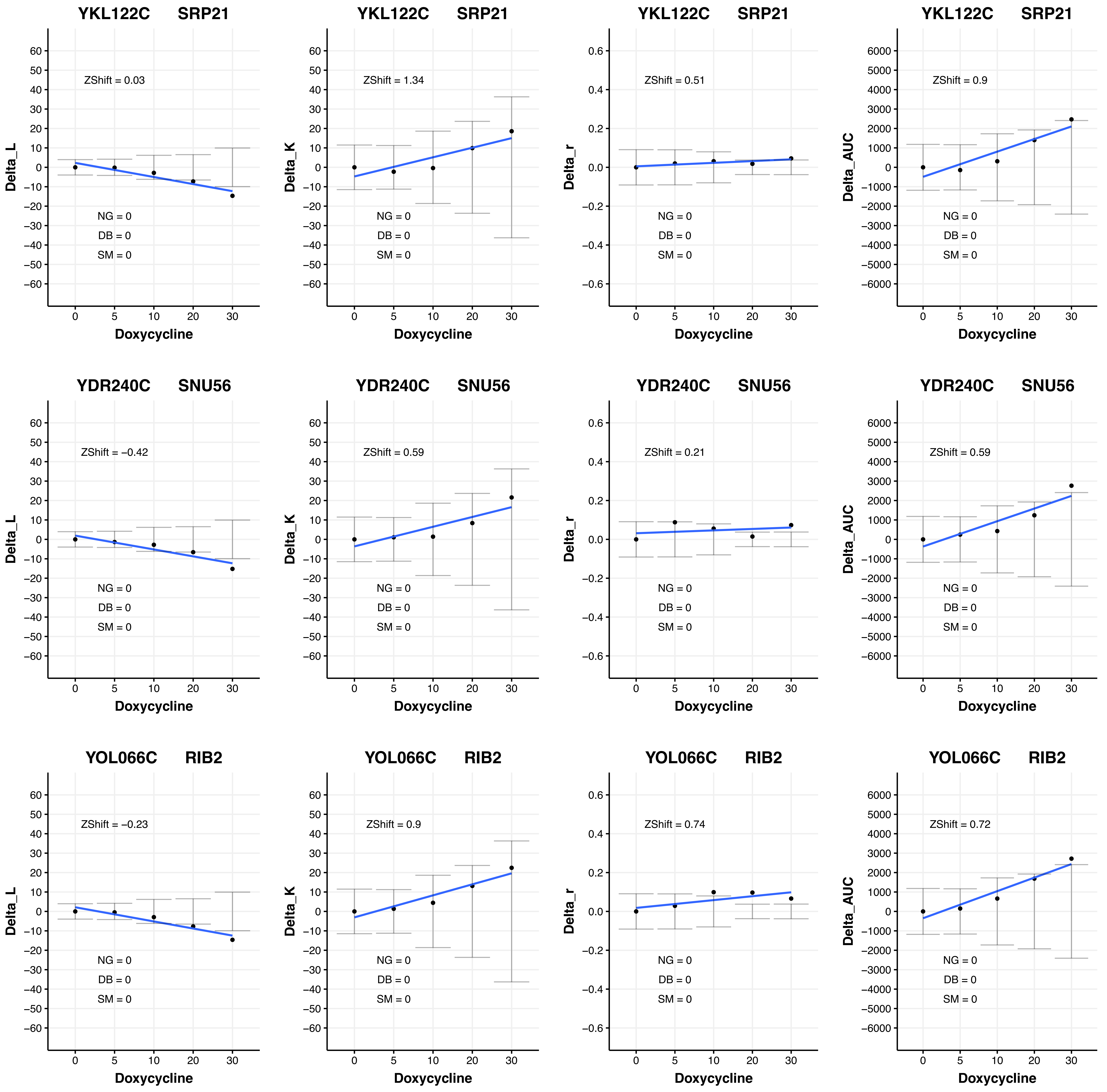

YKL122C SRP21
ZShift = 0.03
Delta_L
NG = 0
DB = 0
SM = 0
Doxycycline
YKL122C SRP21
ZShift = 1.34
Delta_K
NG = 0
DB = 0
SM = 0
Doxycycline
YKL122C SRP21
ZShift = 0.51
Delta_r
NG = 0
DB = 0
SM = 0
Doxycycline
YKL122C SRP21
ZShift = 0.9
Delta_AUC
NG = 0
DB = 0
SM = 0
Doxycycline
YDR240C SNU56
ZShift = −0.42
Delta_L
NG = 0
DB = 0
SM = 0
Doxycycline
YDR240C SNU56
ZShift = 0.59
Delta_K
NG = 0
DB = 0
SM = 0
Doxycycline
YDR240C SNU56
ZShift = 0.21
Delta_r
NG = 0
DB = 0
SM = 0
Doxycycline
YDR240C SNU56
ZShift = 0.59
Delta_AUC
NG = 0
DB = 0
SM = 0
Doxycycline
YOL066C RIB2
ZShift = −0.23
Delta_L
NG = 0
DB = 0
SM = 0
Doxycycline
YOL066C RIB2
ZShift = 0.9
Delta_K
NG = 0
DB = 0
SM = 0
Doxycycline
YOL066C RIB2
ZShift = 0.74
Delta_r
NG = 0
DB = 0
SM = 0
Doxycycline
YOL066C RIB2
ZShift = 0.72
Delta_AUC
NG = 0
DB = 0
SM = 0
Doxycycline

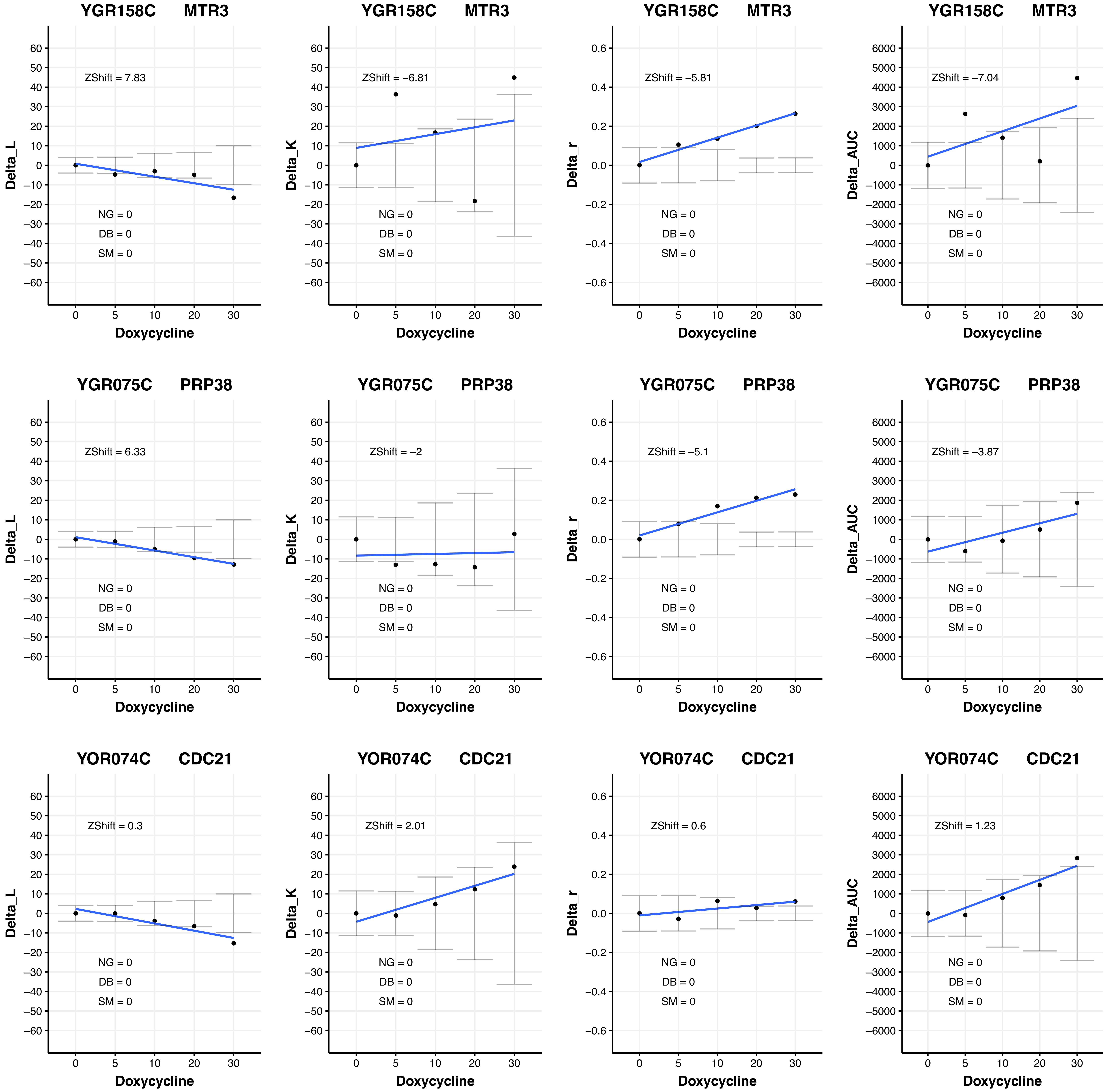
YGR158C MTR3
ZShift = 7.83
NG = 0
DB = 0
SM = 0
Delta_L
Doxycycline
YGR158C MTR3
ZShift = −6.81
NG = 0
DB = 0
SM = 0
Delta_K
Doxycycline
YGR158C MTR3
ZShift = −5.81
NG = 0
DB = 0
SM = 0
Delta_r
Doxycycline
YGR158C MTR3
ZShift = −7.04
NG = 0
DB = 0
SM = 0
Delta_AUC
Doxycycline
YGR075C PRP38
ZShift = 6.33
NG = 0
DB = 0
SM = 0
Delta_L
Doxycycline
YGR075C PRP38
ZShift = −2
NG = 0
DB = 0
SM = 0
Delta_K
Doxycycline
YGR075C PRP38
ZShift = −5.1
NG = 0
DB = 0
SM = 0
Delta_r
Doxycycline
YGR075C PRP38
ZShift = −3.87
NG = 0
DB = 0
SM = 0
Delta_AUC
Doxycycline
YOR074C CDC21
ZShift = 0.3
NG = 0
DB = 0
SM = 0
Delta_L
Doxycycline
YOR074C CDC21
ZShift = 2.01
NG = 0
DB = 0
SM = 0
Delta_K
Doxycycline
YOR074C CDC21
ZShift = 0.6
NG = 0
DB = 0
SM = 0
Delta_r
Doxycycline
YOR074C CDC21
ZShift = 1.23
NG = 0
DB = 0
SM = 0
Delta_AUC
Doxycycline

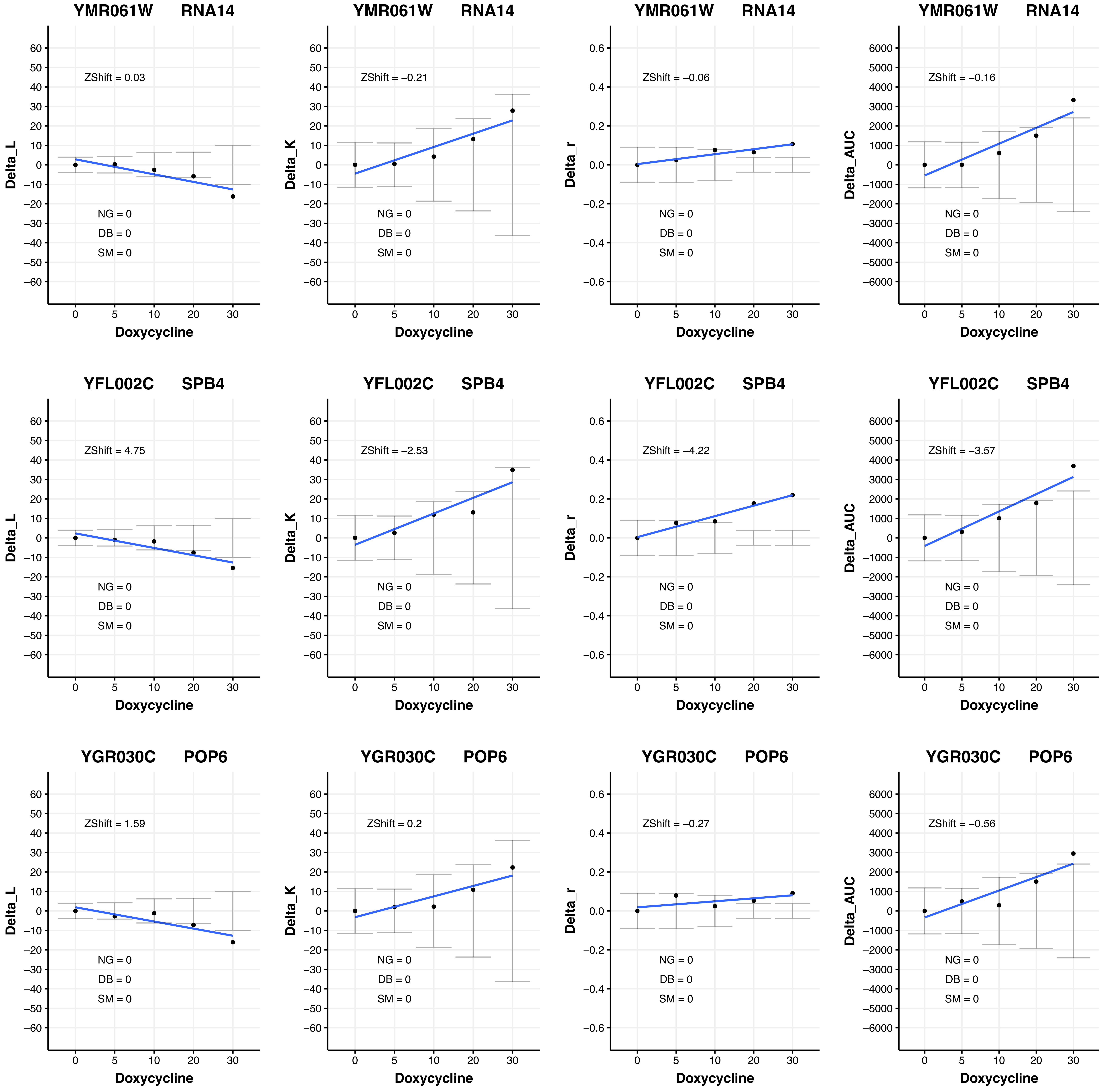

YMR061W RNA14
ZShift = 0.03
Delta_L
NG = 0
DB = 0
SM = 0
Doxycycline
YMR061W RNA14
ZShift = −0.21
Delta_K
NG = 0
DB = 0
SM = 0
Doxycycline
YMR061W RNA14
ZShift = −0.06
Delta_r
NG = 0
DB = 0
SM = 0
Doxycycline
YMR061W RNA14
ZShift = −0.16
Delta_AUC
NG = 0
DB = 0
SM = 0
Doxycycline
YFL002C SPB4
ZShift = 4.75
Delta_L
NG = 0
DB = 0
SM = 0
Doxycycline
YFL002C SPB4
ZShift = −2.53
Delta_K
NG = 0
DB = 0
SM = 0
Doxycycline
YFL002C SPB4
ZShift = −4.22
Delta_r
NG = 0
DB = 0
SM = 0
Doxycycline
YFL002C SPB4
ZShift = −3.57
Delta_AUC
NG = 0
DB = 0
SM = 0
Doxycycline
YGR030C POP6
ZShift = 1.59
Delta_L
NG = 0
DB = 0
SM = 0
Doxycycline
YGR030C POP6
ZShift = 0.2
Delta_K
NG = 0
DB = 0
SM = 0
Doxycycline
YGR030C POP6
ZShift = −0.27
Delta_r
NG = 0
DB = 0
SM = 0
Doxycycline
YGR030C POP6
ZShift = −0.56
Delta_AUC
NG = 0
DB = 0
SM = 0
Doxycycline

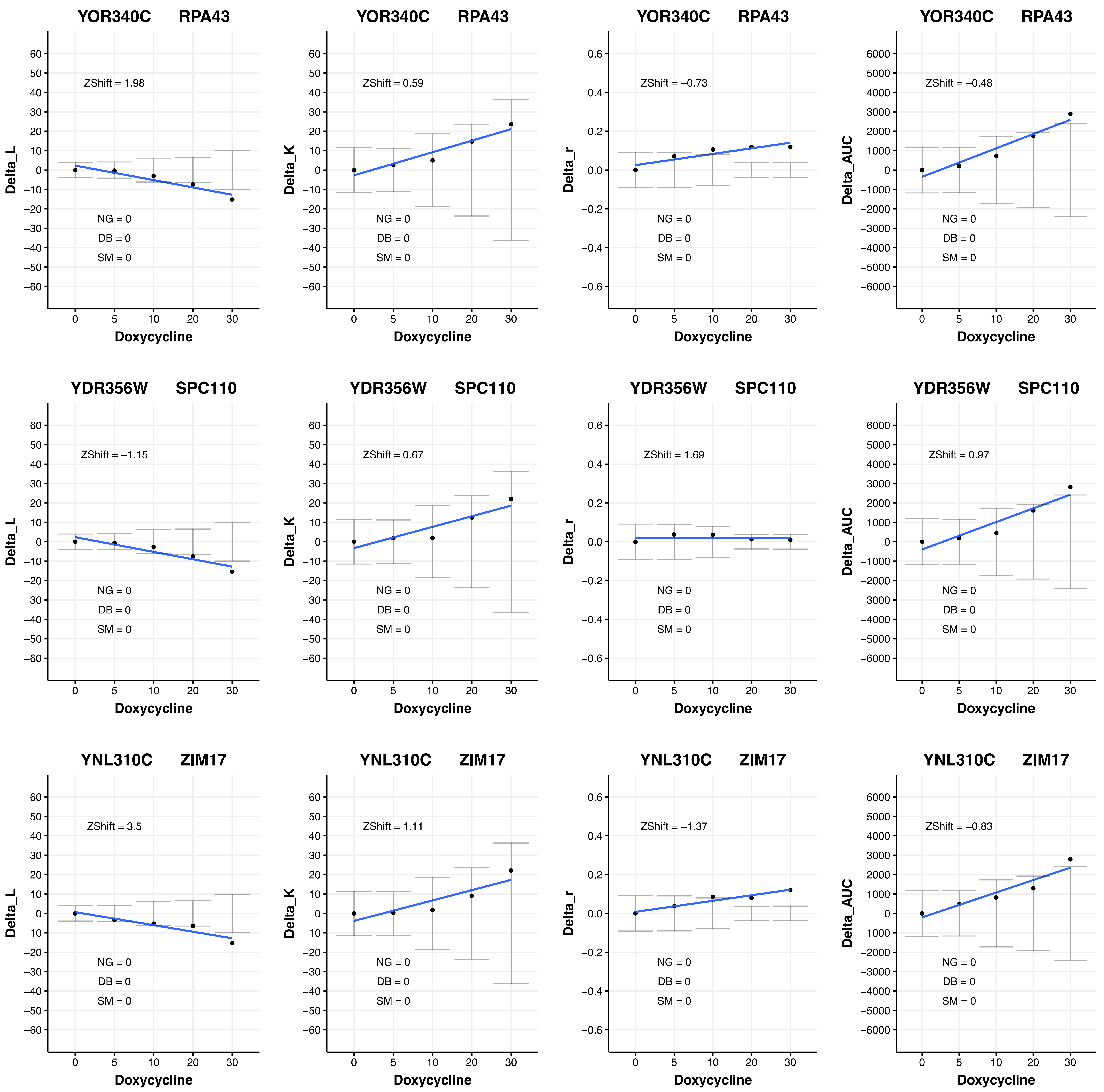
YOR340C RPA43
ZShift = 1.98
NG = 0
DB = 0
SM = 0
Delta_L
Doxycycline
YOR340C RPA43
ZShift = 0.59
NG = 0
DB = 0
SM = 0
Delta_K
Doxycycline
YOR340C RPA43
ZShift = −0.73
NG = 0
DB = 0
SM = 0
Delta_r
Doxycycline
YOR340C RPA43
ZShift = −0.48
NG = 0
DB = 0
SM = 0
Delta_AUC
Doxycycline
YDR356W SPC110
ZShift = −1.15
NG = 0
DB = 0
SM = 0
Delta_L
Doxycycline
YDR356W SPC110
ZShift = 0.67
NG = 0
DB = 0
SM = 0
Delta_K
Doxycycline
YDR356W SPC110
ZShift = 1.69
NG = 0
DB = 0
SM = 0
Delta_r
Doxycycline
YDR356W SPC110
ZShift = 0.97
NG = 0
DB = 0
SM = 0
Delta_AUC
Doxycycline
YNL310C ZIM17
ZShift = 3.5
NG = 0
DB = 0
SM = 0
Delta_L
Doxycycline
YNL310C ZIM17
ZShift = 1.11
NG = 0
DB = 0
SM = 0
Delta_K
Doxycycline
YNL310C ZIM17
ZShift = −1.37
NG = 0
DB = 0
SM = 0
Delta_r
Doxycycline
YNL310C ZIM17
ZShift = −0.83
NG = 0
DB = 0
SM = 0
Delta_AUC
Doxycycline

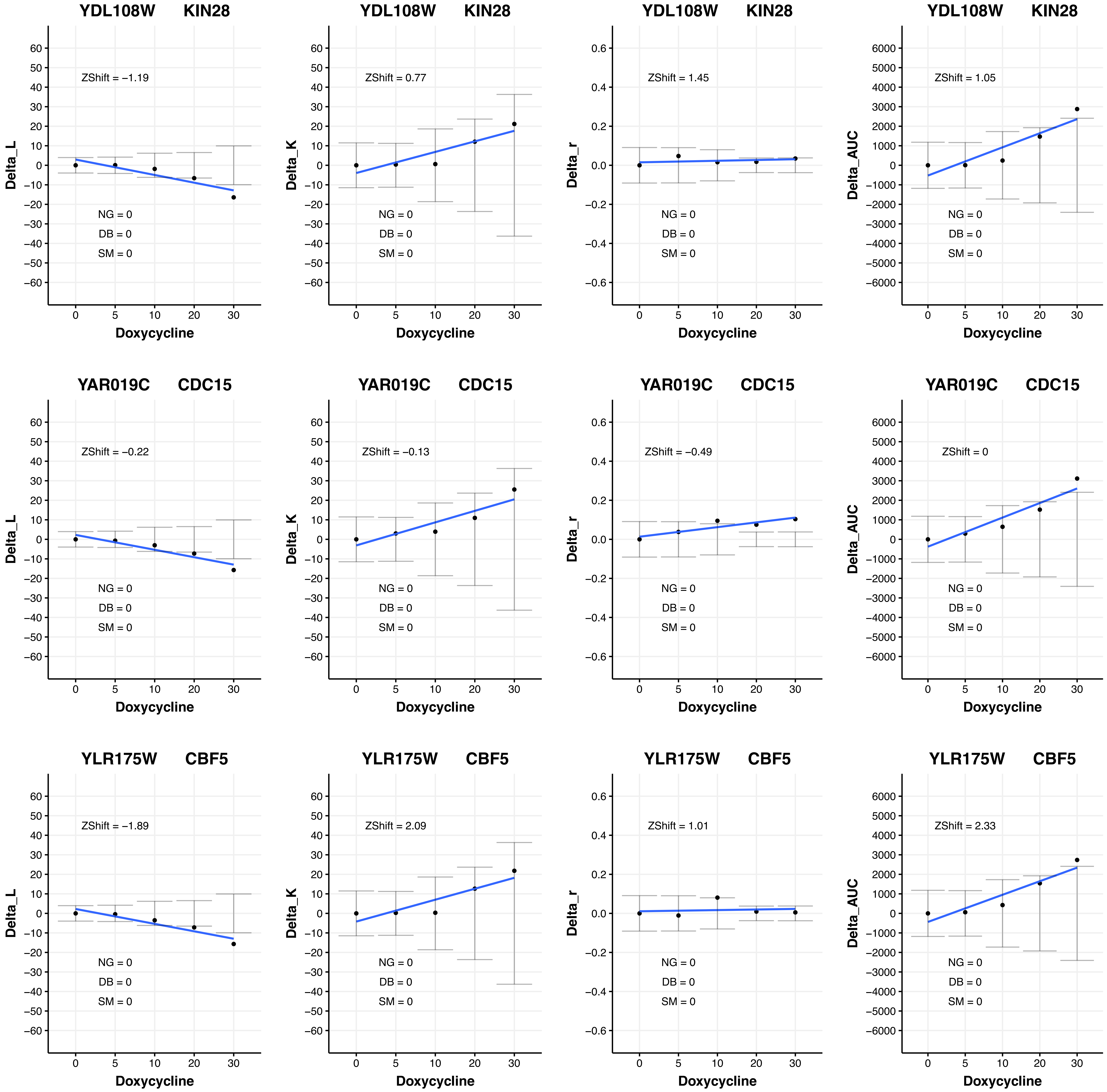

YDL108W KIN28
ZShift = −1.19
NG = 0
DB = 0
SM = 0
Delta_L
Doxycycline
YDL108W KIN28
ZShift = 0.77
NG = 0
DB = 0
SM = 0
Delta_K
Doxycycline
YDL108W KIN28
ZShift = 1.45
NG = 0
DB = 0
SM = 0
Delta_r
Doxycycline
YDL108W KIN28
ZShift = 1.05
NG = 0
DB = 0
SM = 0
Delta_AUC
Doxycycline
YAR019C CDC15
ZShift = −0.22
NG = 0
DB = 0
SM = 0
Delta_L
Doxycycline
YAR019C CDC15
ZShift = −0.13
NG = 0
DB = 0
SM = 0
Delta_K
Doxycycline
YAR019C CDC15
ZShift = −0.49
NG = 0
DB = 0
SM = 0
Delta_r
Doxycycline
YAR019C CDC15
ZShift = 0
NG = 0
DB = 0
SM = 0
Delta_AUC
Doxycycline
YLR175W CBF5
ZShift = −1.89
NG = 0
DB = 0
SM = 0
Delta_L
Doxycycline
YLR175W CBF5
ZShift = 2.09
NG = 0
DB = 0
SM = 0
Delta_K
Doxycycline
YLR175W CBF5
ZShift = 1.01
NG = 0
DB = 0
SM = 0
Delta_r
Doxycycline
YLR175W CBF5
ZShift = 2.33
NG = 0
DB = 0
SM = 0
Delta_AUC
Doxycycline

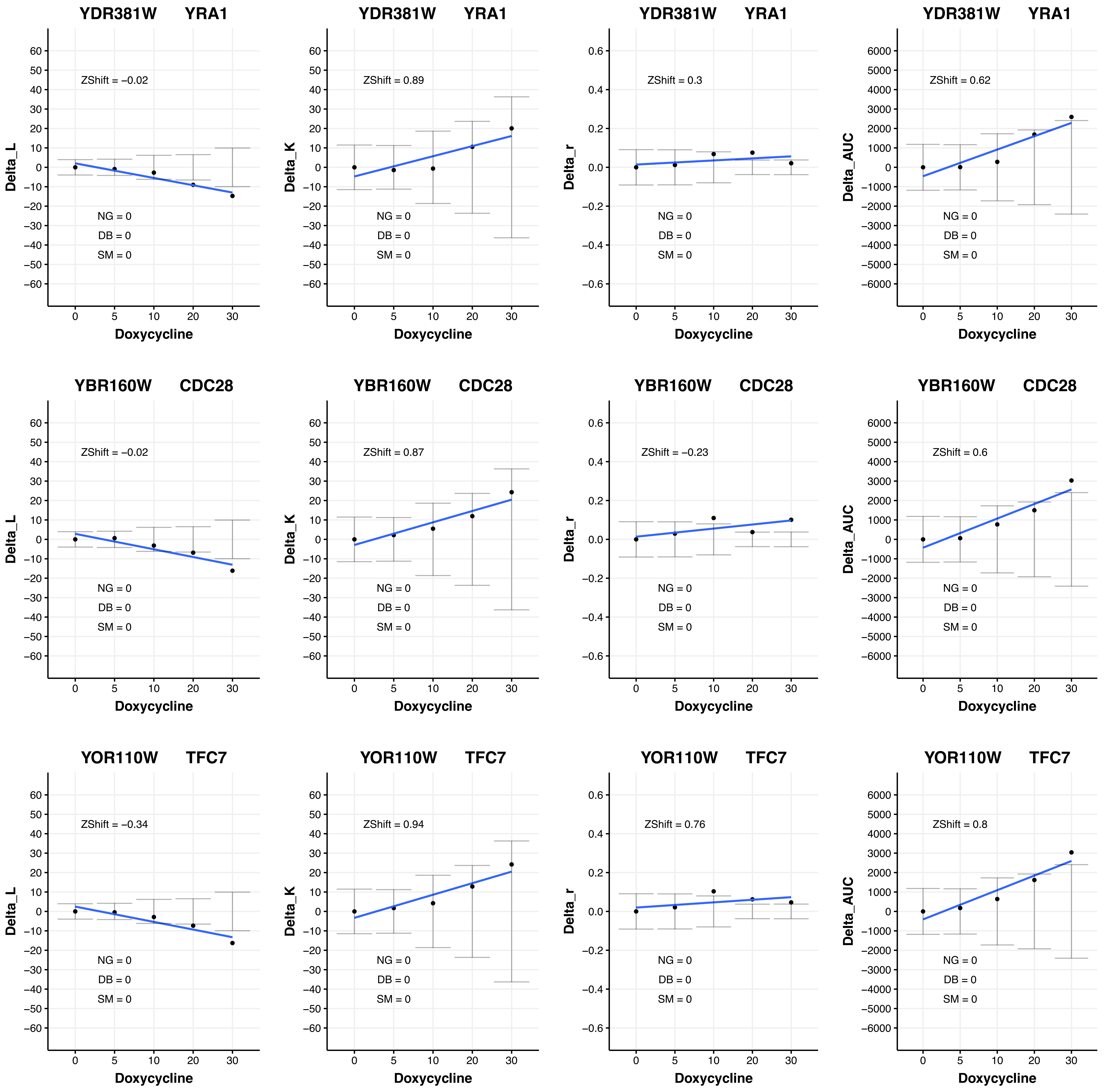
YDR381W YRA1
ZShift = −0.02
NG = 0
DB = 0
SM = 0
Delta_L
Doxycycline
YDR381W YRA1
ZShift = 0.89
NG = 0
DB = 0
SM = 0
Delta_K
Doxycycline
YDR381W YRA1
ZShift = 0.3
NG = 0
DB = 0
SM = 0
Delta_r
Doxycycline
YDR381W YRA1
ZShift = 0.62
NG = 0
DB = 0
SM = 0
Delta_AUC
Doxycycline
YBR160W CDC28
ZShift = −0.02
NG = 0
DB = 0
SM = 0
Delta_L
Doxycycline
YBR160W CDC28
ZShift = 0.87
NG = 0
DB = 0
SM = 0
Delta_K
Doxycycline
YBR160W CDC28
ZShift = −0.23
NG = 0
DB = 0
SM = 0
Delta_r
Doxycycline
YBR160W CDC28
ZShift = 0.6
NG = 0
DB = 0
SM = 0
Delta_AUC
Doxycycline
YOR110W TFC7
ZShift = −0.34
NG = 0
DB = 0
SM = 0
Delta_L
Doxycycline
YOR110W TFC7
ZShift = 0.94
NG = 0
DB = 0
SM = 0
Delta_K
Doxycycline
YOR110W TFC7
ZShift = 0.76
NG = 0
DB = 0
SM = 0
Delta_r
Doxycycline
YOR110W TFC7
ZShift = 0.8
NG = 0
DB = 0
SM = 0
Delta_AUC
Doxycycline

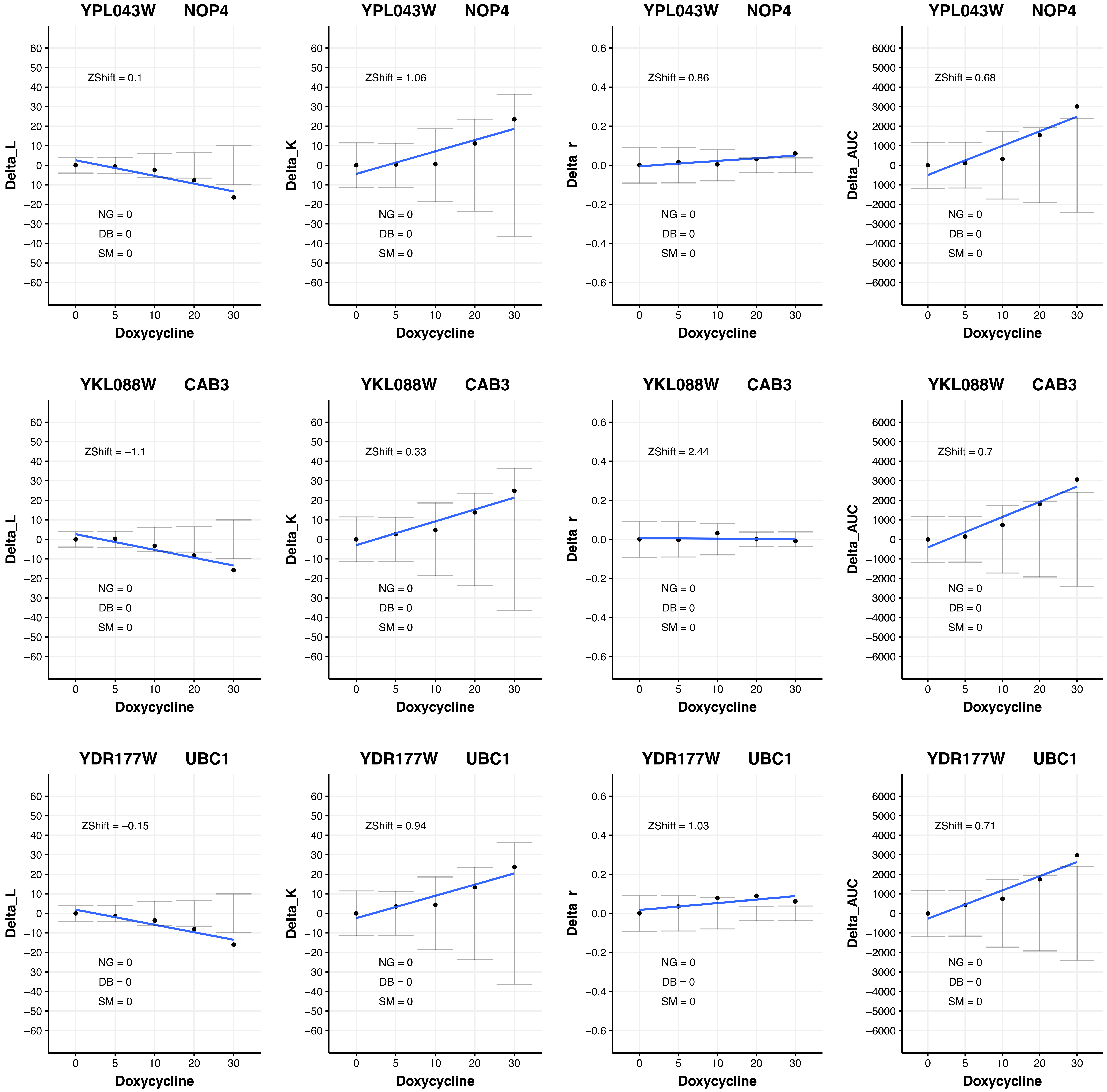
YPL043W NOP4
ZShift = 0.1
NG = 0
DB = 0
SM = 0
Delta_L
Doxycycline
YPL043W NOP4
ZShift = 1.06
NG = 0
DB = 0
SM = 0
Delta_K
Doxycycline
YPL043W NOP4
ZShift = 0.86
NG = 0
DB = 0
SM = 0
Delta_r
Doxycycline
YPL043W NOP4
ZShift = 0.68
NG = 0
DB = 0
SM = 0
Delta_AUC
Doxycycline
YKL088W CAB3
ZShift = −1.1
NG = 0
DB = 0
SM = 0
Delta_L
Doxycycline
YKL088W CAB3
ZShift = 0.33
NG = 0
DB = 0
SM = 0
Delta_K
Doxycycline
YKL088W CAB3
ZShift = 2.44
NG = 0
DB = 0
SM = 0
Delta_r
Doxycycline
YKL088W CAB3
ZShift = 0.7
NG = 0
DB = 0
SM = 0
Delta_AUC
Doxycycline
YDR177W UBC1
ZShift = −0.15
NG = 0
DB = 0
SM = 0
Delta_L
Doxycycline
YDR177W UBC1
ZShift = 0.94
NG = 0
DB = 0
SM = 0
Delta_K
Doxycycline
YDR177W UBC1
ZShift = 1.03
NG = 0
DB = 0
SM = 0
Delta_r
Doxycycline
YDR177W UBC1
ZShift = 0.71
NG = 0
DB = 0
SM = 0
Delta_AUC
Doxycycline

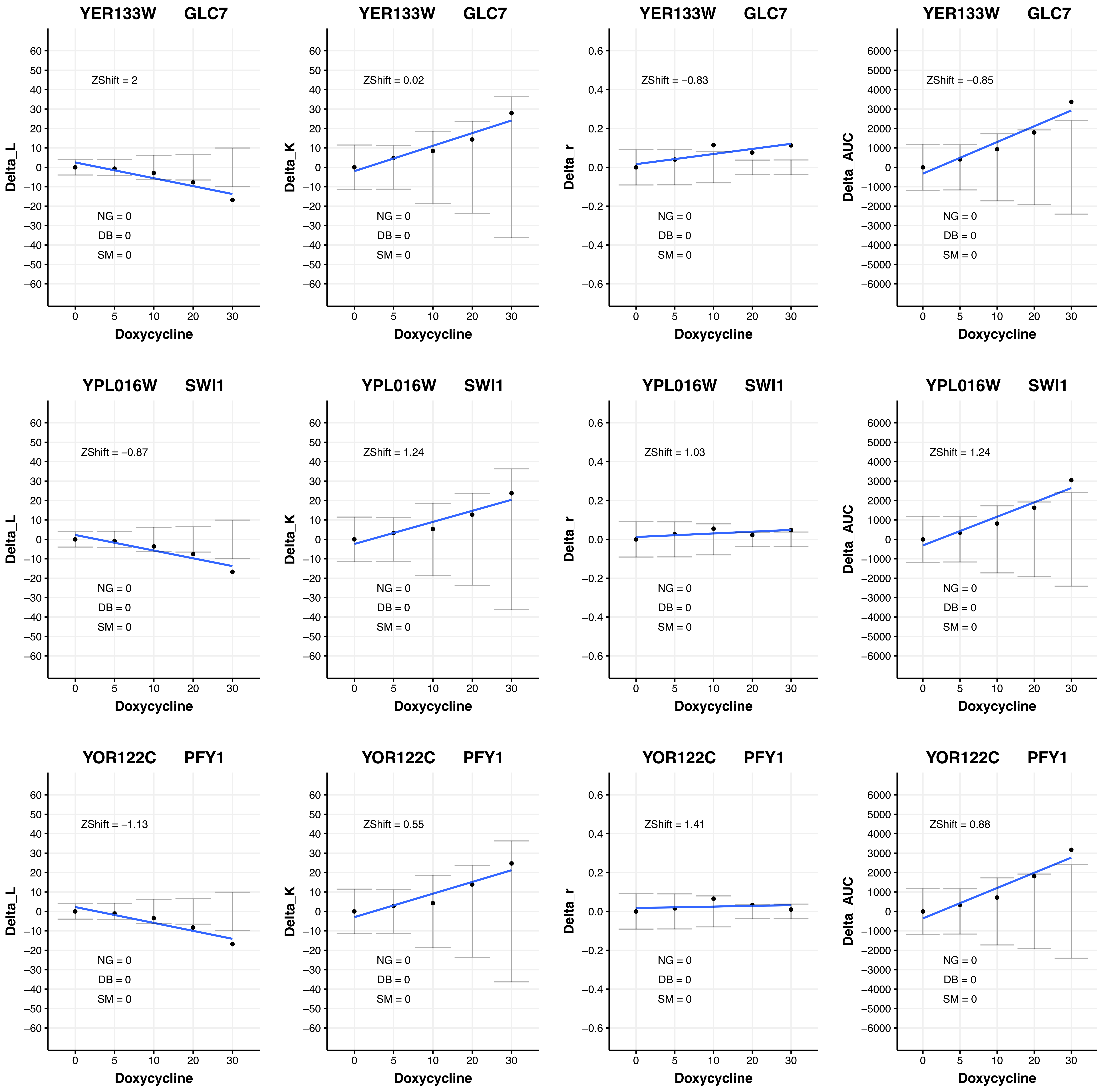
YER133W GLC7
ZShift = 2
NG = 0
DB = 0
SM = 0
Delta_L
Doxycycline
YER133W GLC7
ZShift = 0.02
NG = 0
DB = 0
SM = 0
Delta_K
Doxycycline
YER133W GLC7
ZShift = −0.83
NG = 0
DB = 0
SM = 0
Delta_r
Doxycycline
YER133W GLC7
ZShift = −0.85
NG = 0
DB = 0
SM = 0
Delta_AUC
Doxycycline
YPL016W SWI1
ZShift = −0.87
NG = 0
DB = 0
SM = 0
Delta_L
Doxycycline
YPL016W SWI1
ZShift = 1.24
NG = 0
DB = 0
SM = 0
Delta_K
Doxycycline
YPL016W SWI1
ZShift = 1.03
NG = 0
DB = 0
SM = 0
Delta_r
Doxycycline
YPL016W SWI1
ZShift = 1.24
NG = 0
DB = 0
SM = 0
Delta_AUC
Doxycycline
YOR122C PFY1
ZShift = −1.13
NG = 0
DB = 0
SM = 0
Delta_L
Doxycycline
YOR122C PFY1
ZShift = 0.55
NG = 0
DB = 0
SM = 0
Delta_K
Doxycycline
YOR122C PFY1
ZShift = 1.41
NG = 0
DB = 0
SM = 0
Delta_r
Doxycycline
YOR122C PFY1
ZShift = 0.88
NG = 0
DB = 0
SM = 0
Delta_AUC
Doxycycline

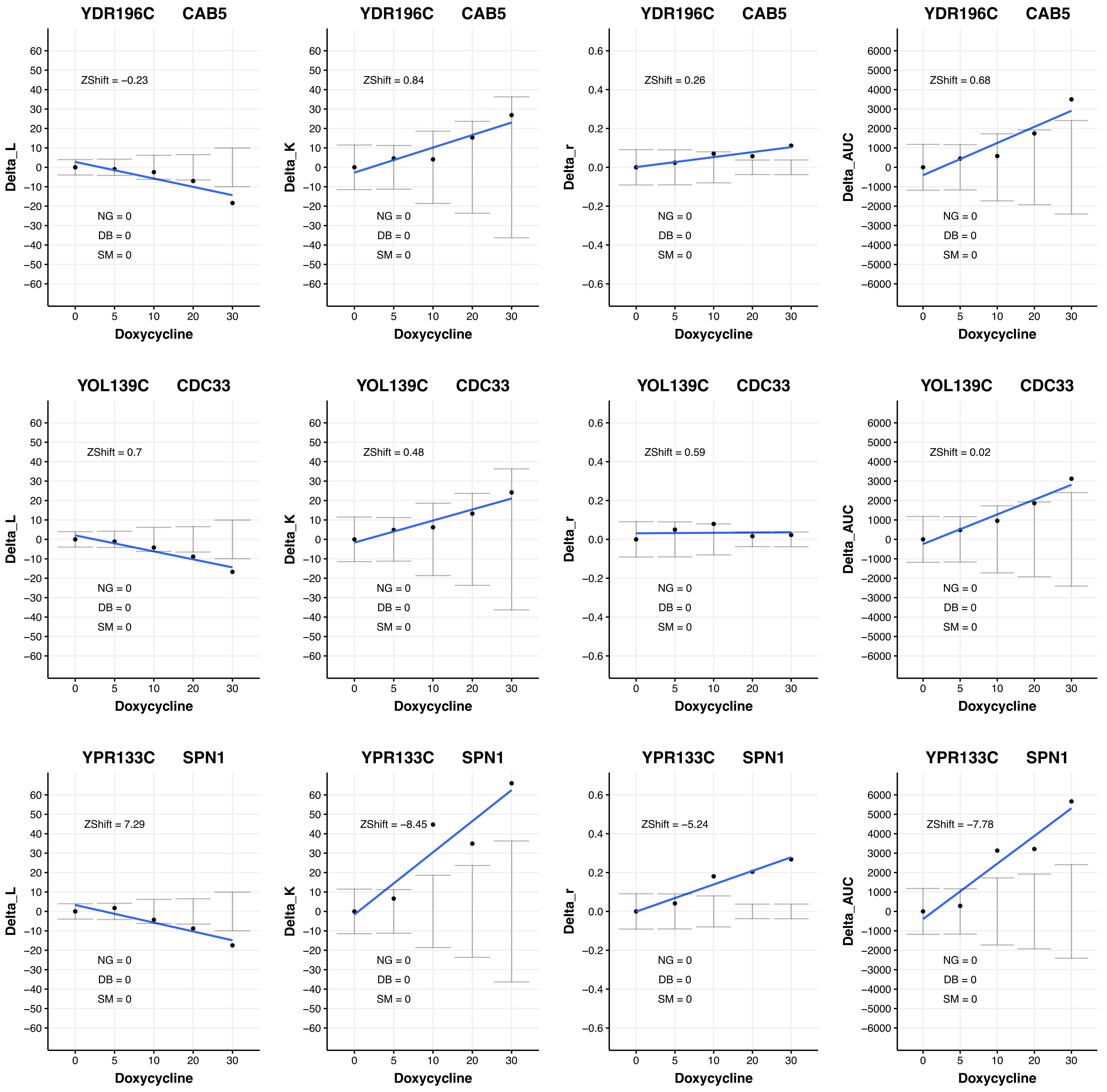
YDR196C CAB5
ZShift = −0.23
NG = 0
DB = 0
SM = 0
Delta_L
Doxycycline
YDR196C CAB5
ZShift = 0.84
NG = 0
DB = 0
SM = 0
Delta_K
Doxycycline
YDR196C CAB5
ZShift = 0.26
NG = 0
DB = 0
SM = 0
Delta_r
Doxycycline
YDR196C CAB5
ZShift = 0.68
NG = 0
DB = 0
SM = 0
Delta_AUC
Doxycycline
YOL139C CDC33
ZShift = 0.7
NG = 0
DB = 0
SM = 0
Delta_L
Doxycycline
YOL139C CDC33
ZShift = 0.48
NG = 0
DB = 0
SM = 0
Delta_K
Doxycycline
YOL139C CDC33
ZShift = 0.59
NG = 0
DB = 0
SM = 0
Delta_r
Doxycycline
YOL139C CDC33
ZShift = 0.02
NG = 0
DB = 0
SM = 0
Delta_AUC
Doxycycline
YPR133C SPN1
ZShift = 7.29
NG = 0
DB = 0
SM = 0
Delta_L
Doxycycline
YPR133C SPN1
ZShift = −8.45
NG = 0
DB = 0
SM = 0
Delta_K
Doxycycline
YPR133C SPN1
ZShift = −5.24
NG = 0
DB = 0
SM = 0
Delta_r
Doxycycline
YPR133C SPN1
ZShift = −7.78
NG = 0
DB = 0
SM = 0
Delta_AUC
Doxycycline

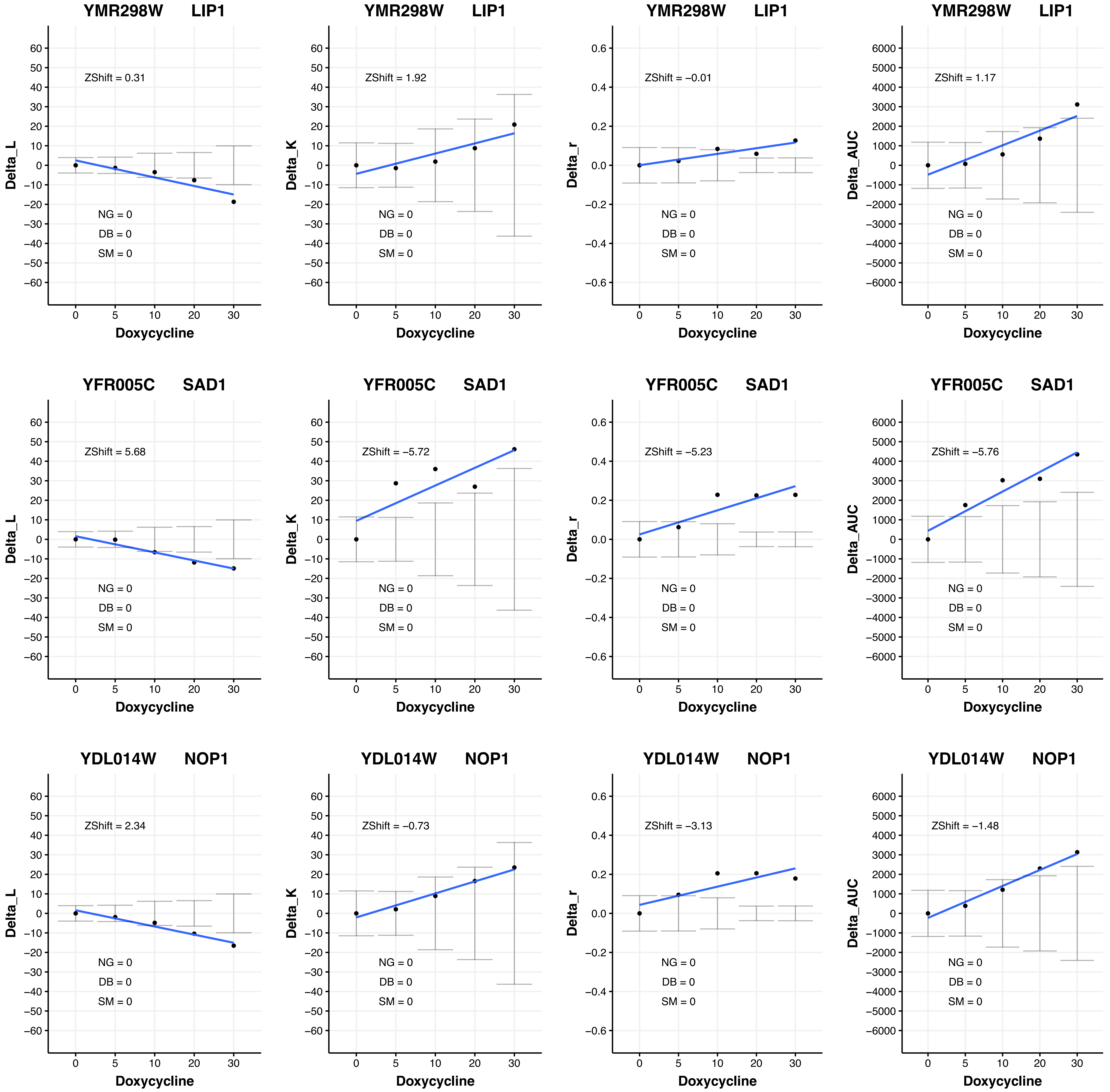

YMR298W LIP1
ZShift = 0.31
NG = 0
DB = 0
SM = 0
Delta_L
Doxycycline
YMR298W LIP1
ZShift = 1.92
NG = 0
DB = 0
SM = 0
Delta_K
Doxycycline
YMR298W LIP1
ZShift = −0.01
NG = 0
DB = 0
SM = 0
Delta_r
Doxycycline
YMR298W LIP1
ZShift = 1.17
NG = 0
DB = 0
SM = 0
Delta_AUC
Doxycycline
YFR005C SAD1
ZShift = 5.68
NG = 0
DB = 0
SM = 0
Delta_L
Doxycycline
YFR005C SAD1
ZShift = −5.72
NG = 0
DB = 0
SM = 0
Delta_K
Doxycycline
YFR005C SAD1
ZShift = −5.23
NG = 0
DB = 0
SM = 0
Delta_r
Doxycycline
YFR005C SAD1
ZShift = −5.76
NG = 0
DB = 0
SM = 0
Delta_AUC
Doxycycline
YDL014W NOP1
ZShift = 2.34
NG = 0
DB = 0
SM = 0
Delta_L
Doxycycline
YDL014W NOP1
ZShift = −0.73
NG = 0
DB = 0
SM = 0
Delta_K
Doxycycline
YDL014W NOP1
ZShift = −3.13
NG = 0
DB = 0
SM = 0
Delta_r
Doxycycline
YDL014W NOP1
ZShift = −1.48
NG = 0
DB = 0
SM = 0
Delta_AUC
Doxycycline

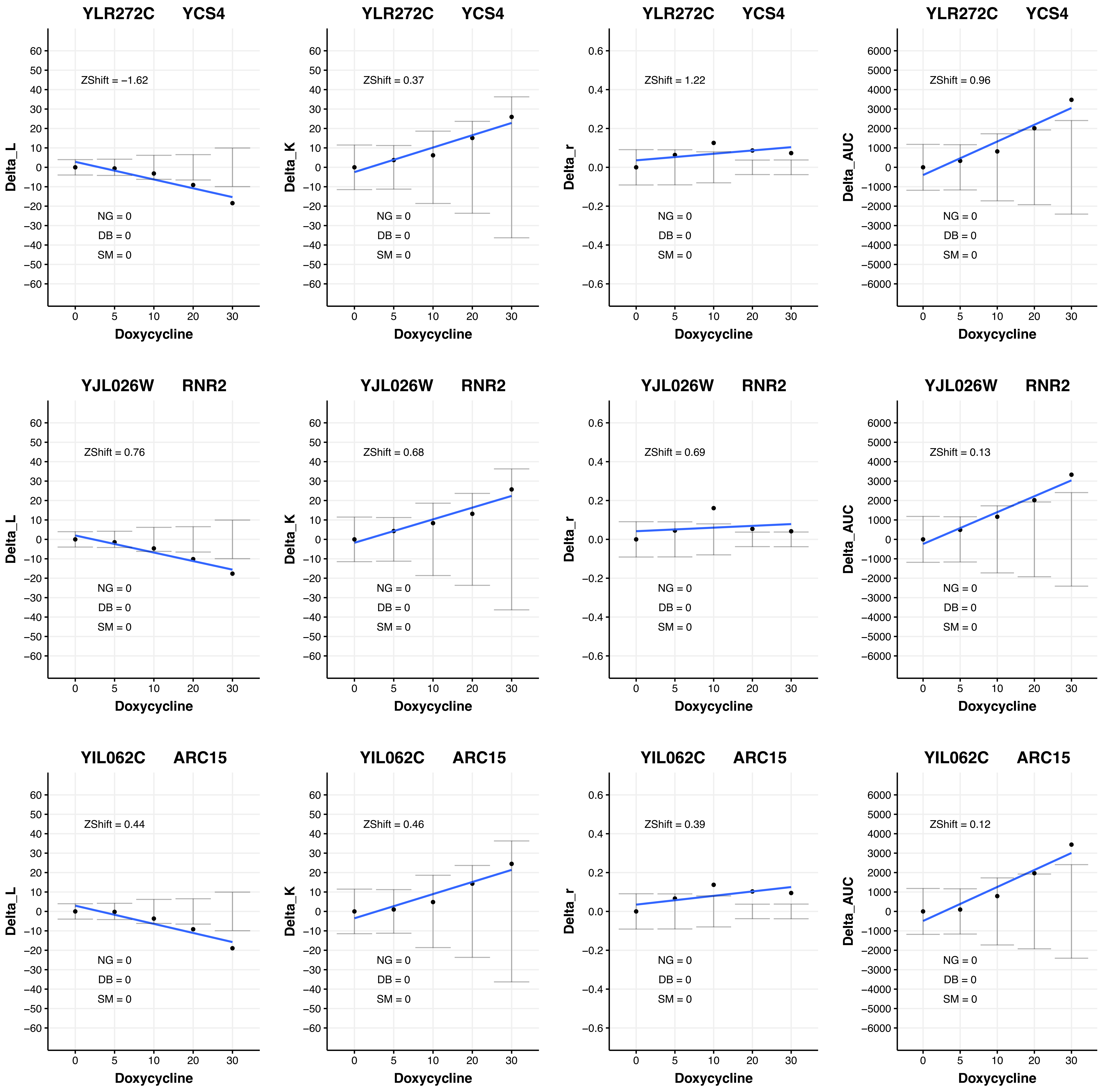

YLR272C YCS4
ZShift = −1.62
NG = 0
DB = 0
SM = 0
Delta_L
Doxycycline
YLR272C YCS4
ZShift = 0.37
NG = 0
DB = 0
SM = 0
Delta_K
Doxycycline
YLR272C YCS4
ZShift = 1.22
NG = 0
DB = 0
SM = 0
Delta_r
Doxycycline
YLR272C YCS4
ZShift = 0.96
NG = 0
DB = 0
SM = 0
Delta_AUC
Doxycycline
YJL026W RNR2
ZShift = 0.76
NG = 0
DB = 0
SM = 0
Delta_L
Doxycycline
YJL026W RNR2
ZShift = 0.68
NG = 0
DB = 0
SM = 0
Delta_K
Doxycycline
YJL026W RNR2
ZShift = 0.69
NG = 0
DB = 0
SM = 0
Delta_r
Doxycycline
YJL026W RNR2
ZShift = 0.13
NG = 0
DB = 0
SM = 0
Delta_AUC
Doxycycline
YIL062C ARC15
ZShift = 0.44
NG = 0
DB = 0
SM = 0
Delta_L
Doxycycline
YIL062C ARC15
ZShift = 0.46
NG = 0
DB = 0
SM = 0
Delta_K
Doxycycline
YIL062C ARC15
ZShift = 0.39
NG = 0
DB = 0
SM = 0
Delta_r
Doxycycline
YIL062C ARC15
ZShift = 0.12
NG = 0
DB = 0
SM = 0
Delta_AUC
Doxycycline

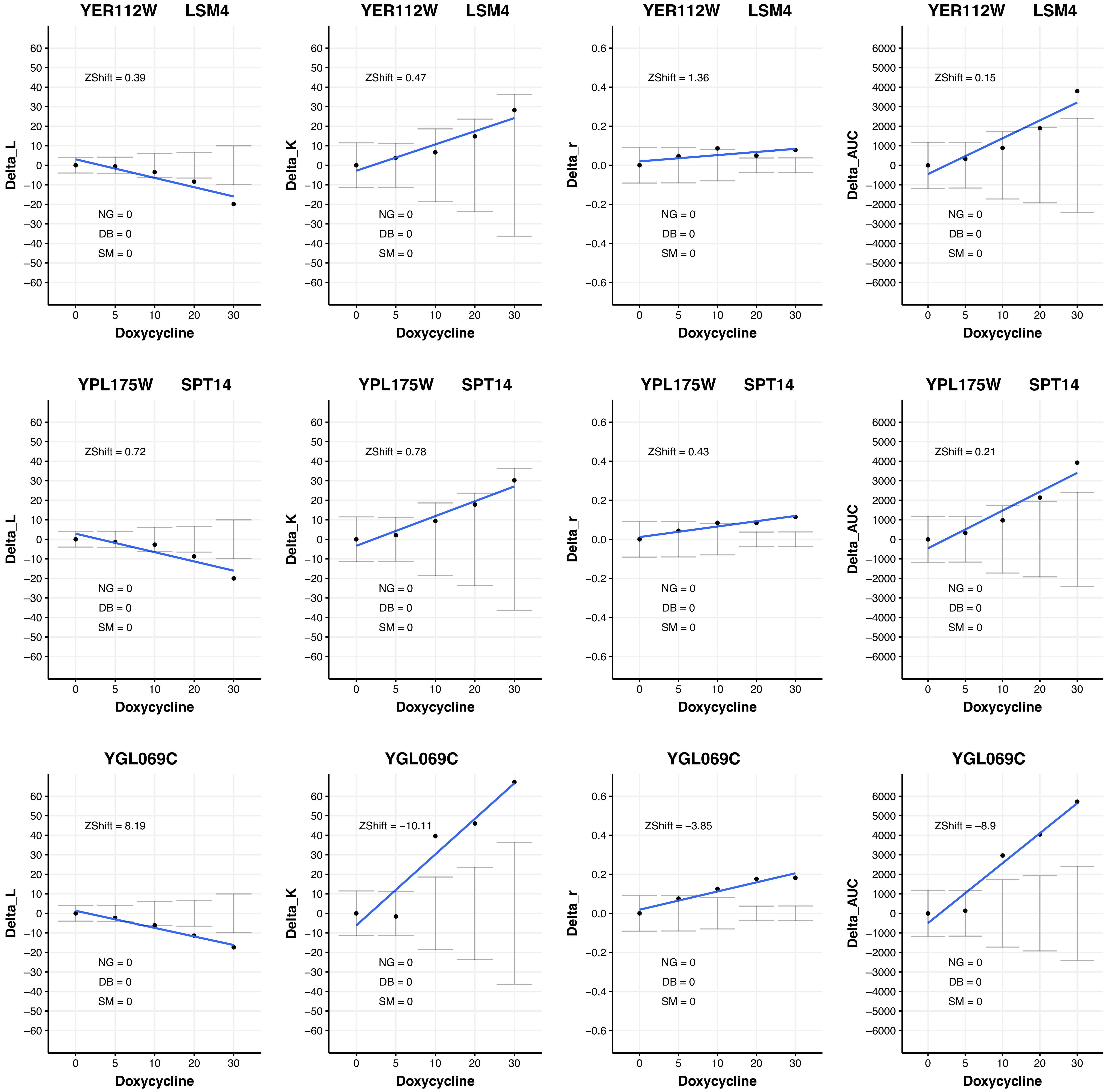
YER112W LSM4
ZShift = 0.39
NG = 0
DB = 0
SM = 0
Delta_L
Doxycycline
YER112W LSM4
ZShift = 0.47
NG = 0
DB = 0
SM = 0
Delta_K
Doxycycline
YER112W LSM4
ZShift = 1.36
NG = 0
DB = 0
SM = 0
Delta_r
Doxycycline
YER112W LSM4
ZShift = 0.15
NG = 0
DB = 0
SM = 0
Delta_AUC
Doxycycline
YPL175W SPT14
ZShift = 0.72
NG = 0
DB = 0
SM = 0
Delta_L
Doxycycline
YPL175W SPT14
ZShift = 0.78
NG = 0
DB = 0
SM = 0
Delta_K
Doxycycline
YPL175W SPT14
ZShift = 0.43
NG = 0
DB = 0
SM = 0
Delta_r
Doxycycline
YPL175W SPT14
ZShift = 0.21
NG = 0
DB = 0
SM = 0
Delta_AUC
Doxycycline
YGL069C
ZShift = 8.19
NG = 0
DB = 0
SM = 0
Delta_L
Doxycycline
YGL069C
ZShift = −10.11
NG = 0
DB = 0
SM = 0
Delta_K
Doxycycline
YGL069C
ZShift = −3.85
NG = 0
DB = 0
SM = 0
Delta_r
Doxycycline
YGL069C
ZShift = −8.9
NG = 0
DB = 0
SM = 0
Delta_AUC
Doxycycline

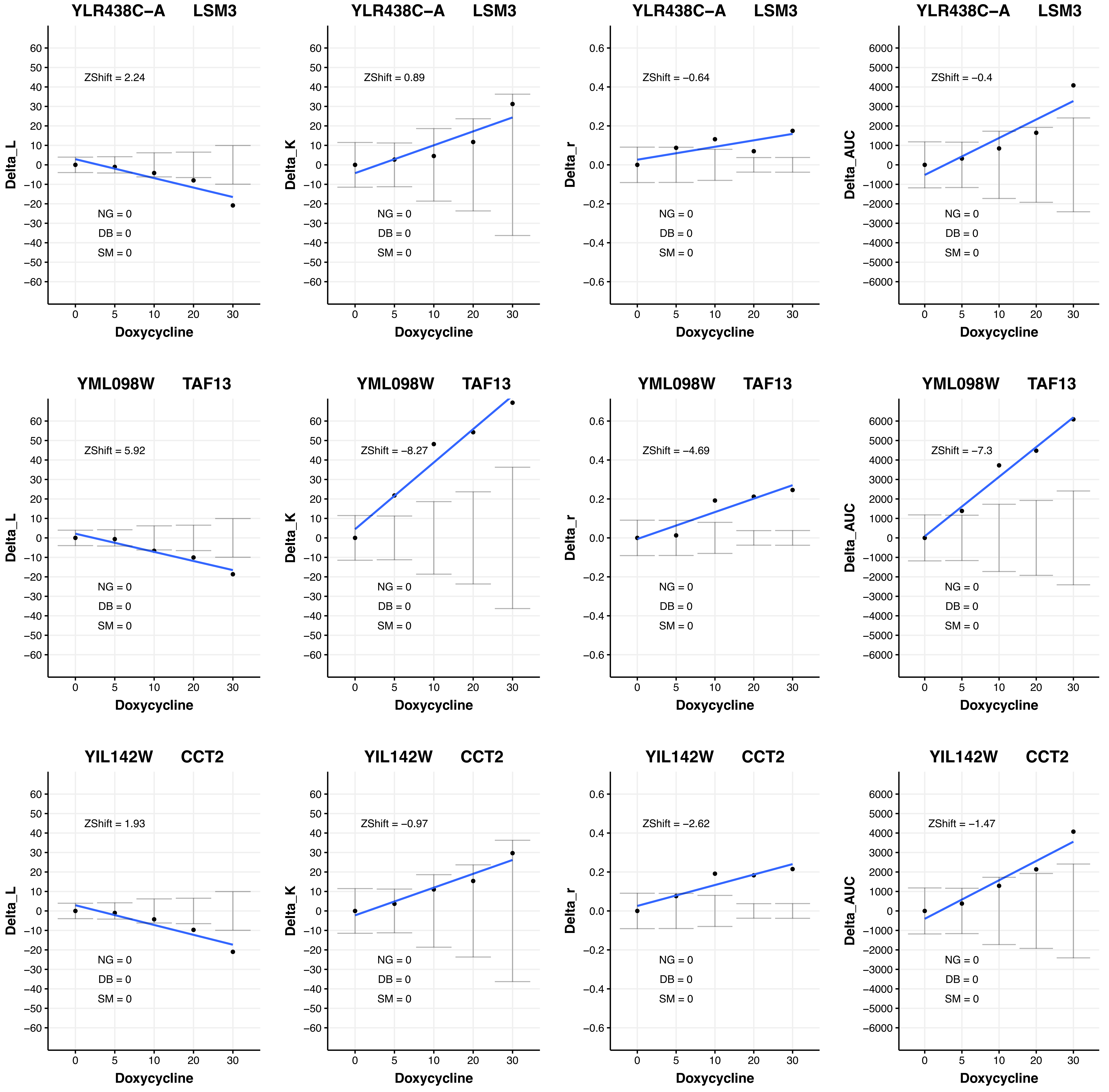
YLR438C−A LSM3
ZShift = 2.24
NG = 0
DB = 0
SM = 0
Delta_L
Doxycycline
YLR438C−A LSM3
ZShift = 0.89
NG = 0
DB = 0
SM = 0
Delta_K
Doxycycline
YLR438C−A LSM3
ZShift = −0.64
NG = 0
DB = 0
SM = 0
Delta_r
Doxycycline
YLR438C−A LSM3
ZShift = −0.4
NG = 0
DB = 0
SM = 0
Delta_AUC
Doxycycline
YML098W TAF13
ZShift = 5.92
NG = 0
DB = 0
SM = 0
Delta_L
Doxycycline
YML098W TAF13
ZShift = −8.27
NG = 0
DB = 0
SM = 0
Delta_K
Doxycycline
YML098W TAF13
ZShift = −4.69
NG = 0
DB = 0
SM = 0
Delta_r
Doxycycline
YML098W TAF13
ZShift = −7.3
NG = 0
DB = 0
SM = 0
Delta_AUC
Doxycycline
YIL142W CCT2
ZShift = 1.93
NG = 0
DB = 0
SM = 0
Delta_L
Doxycycline
YIL142W CCT2
ZShift = −0.97
NG = 0
DB = 0
SM = 0
Delta_K
Doxycycline
YIL142W CCT2
ZShift = −2.62
NG = 0
DB = 0
SM = 0
Delta_r
Doxycycline
YIL142W CCT2
ZShift = −1.47
NG = 0
DB = 0
SM = 0
Delta_AUC
Doxycycline

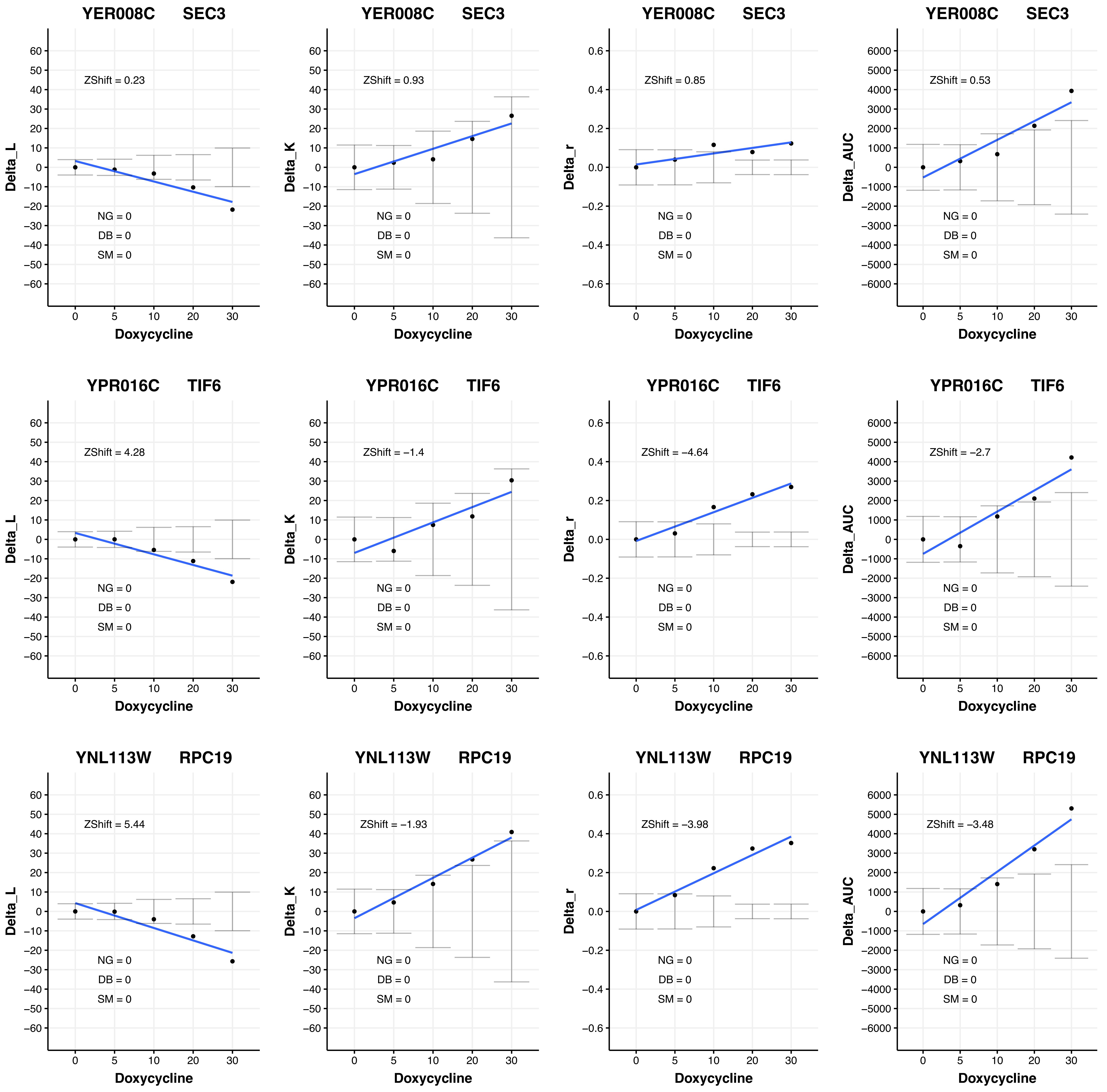
YER008C SEC3
Delta_L
ZShift = 0.23
NG = 0
DB = 0
SM = 0
Doxycycline
YER008C SEC3
Delta_K
ZShift = 0.93
NG = 0
DB = 0
SM = 0
Doxycycline
YER008C SEC3
Delta_r
ZShift = 0.85
NG = 0
DB = 0
SM = 0
Doxycycline
YER008C SEC3
Delta_AUC
ZShift = 0.53
NG = 0
DB = 0
SM = 0
Doxycycline
YPR016C TIF6
Delta_L
ZShift = 4.28
NG = 0
DB = 0
SM = 0
Doxycycline
YPR016C TIF6
Delta_K
ZShift = −1.4
NG = 0
DB = 0
SM = 0
Doxycycline
YPR016C TIF6
Delta_r
ZShift = −4.64
NG = 0
DB = 0
SM = 0
Doxycycline
YPR016C TIF6
Delta_AUC
ZShift = −2.7
NG = 0
DB = 0
SM = 0
Doxycycline
YNL113W RPC19
Delta_L
ZShift = 5.44
NG = 0
DB = 0
SM = 0
Doxycycline
YNL113W RPC19
Delta_K
ZShift = −1.93
NG = 0
DB = 0
SM = 0
Doxycycline
YNL113W RPC19
Delta_r
ZShift = −3.98
NG = 0
DB = 0
SM = 0
Doxycycline
YNL113W RPC19
Delta_AUC
ZShift = −3.48
NG = 0
DB = 0
SM = 0
Doxycycline
