## Additional File 4 for "A humanized yeast phenomic model of deoxycytidine kinase to predict genetic buffering of nucleoside analog cytotoxicity": A - InteractionPlots_Cytarabine.pdf

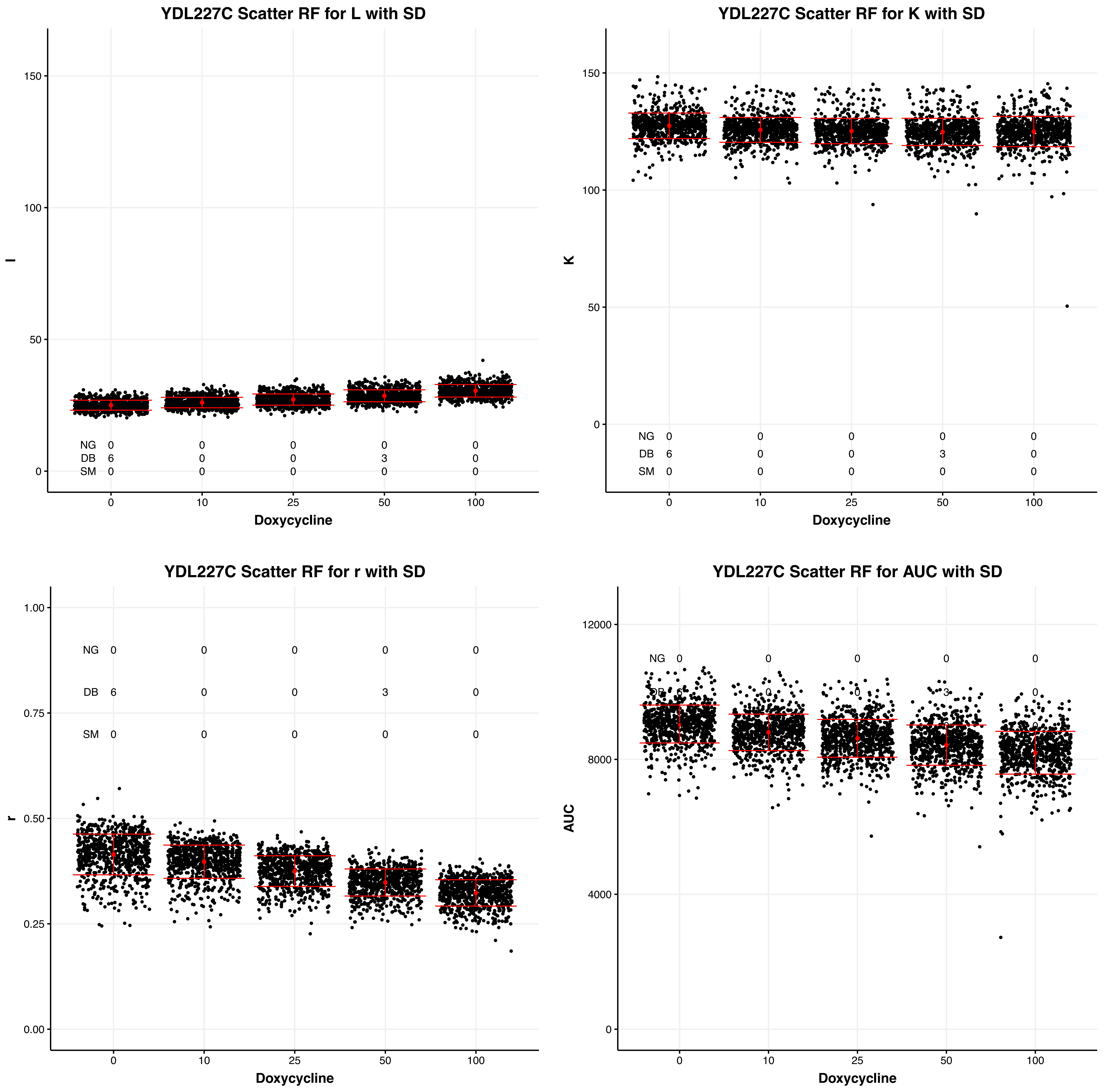

YDL227C Scatter RF for L with SD
l
0
50
100
150
NG 0 0 0 0 0
DB 6 0 0 3 0
SM 0 0 0 0 0
0 10 25 50 100
Doxycycline
YDL227C Scatter RF for K with SD
K
0
50
100
150
NG 0 0 0 0 0
DB 6 0 0 3 0
SM 0 0 0 0 0
0 10 25 50 100
Doxycycline
YDL227C Scatter RF for r with SD
r
0.00
0.25
0.50
0.75
1.00
NG 0 0 0 0 0
DB 6 0 0 3 0
SM 0 0 0 0 0
0 10 25 50 100
Doxycycline
YDL227C Scatter RF for AUC with SD
AUC
0
4000
8000
12000
NG 0 0 0 0 0
0 10 25 50 100
Doxycycline

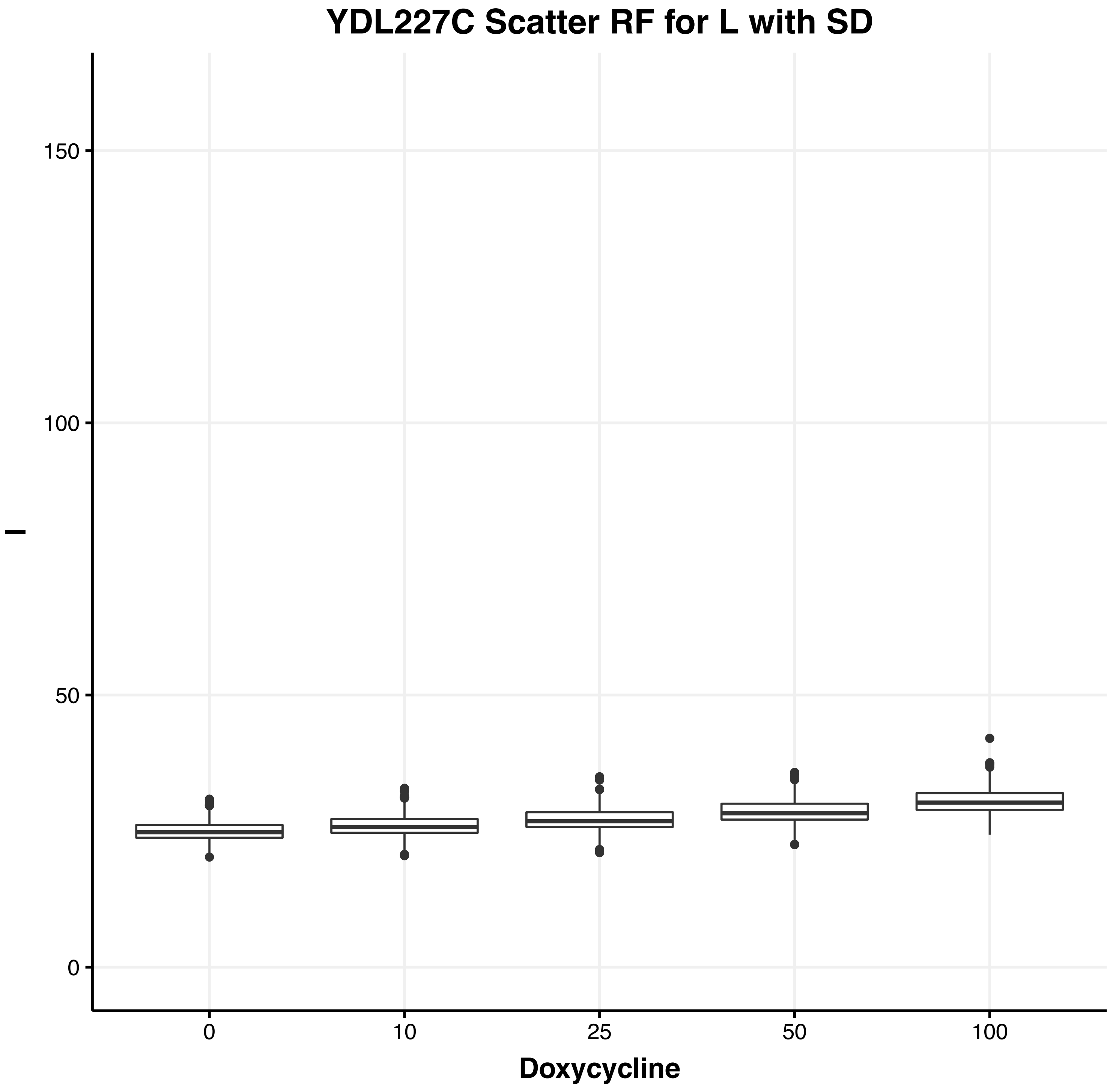
YDL227C Scatter RF for L with SD
l
Doxycycline
0
50
100
150
0
10
25
50
100

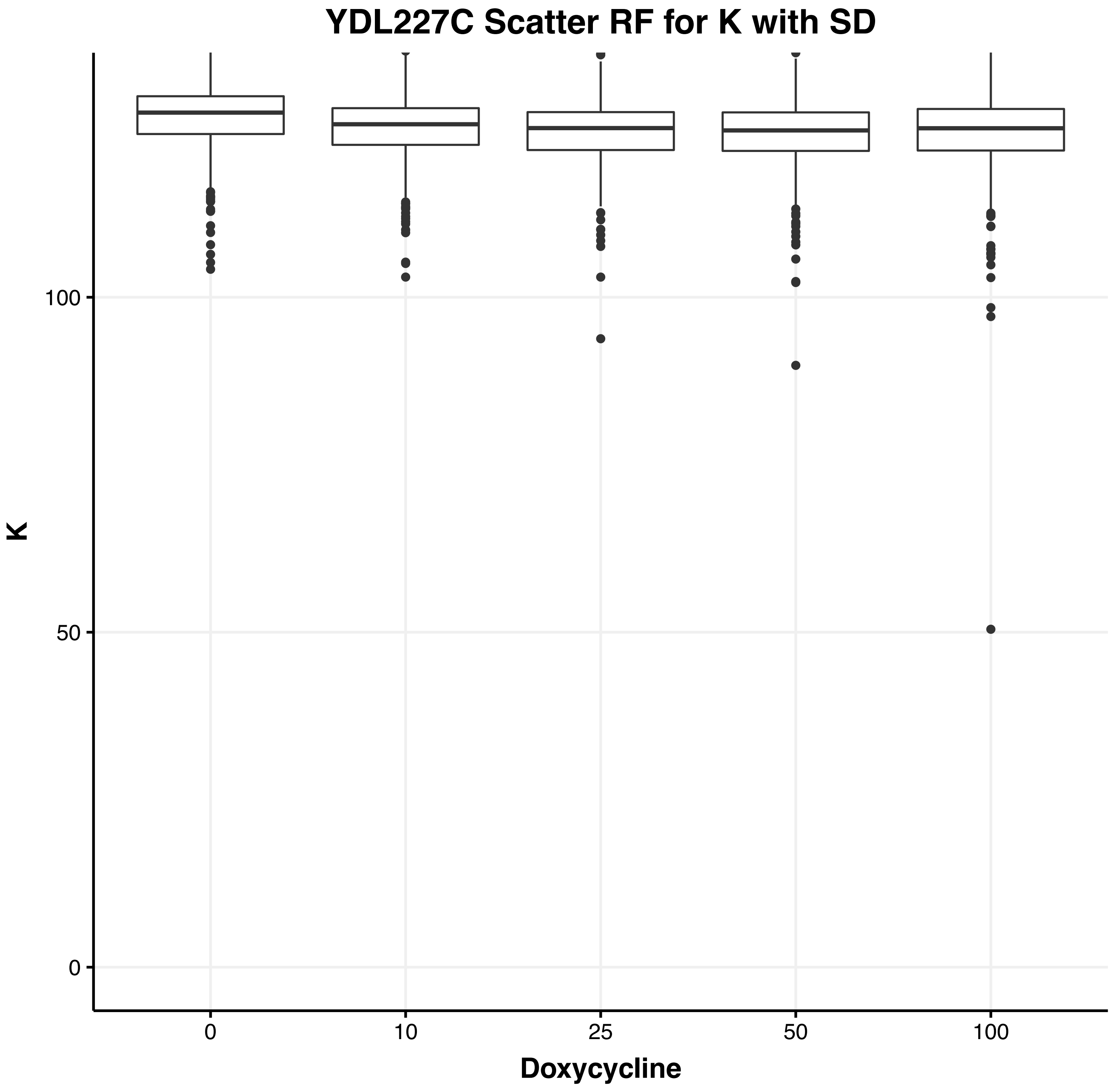
YDL227C Scatter RF for K with SD
K
Doxycycline
0
50
100
0
10
25
50
100

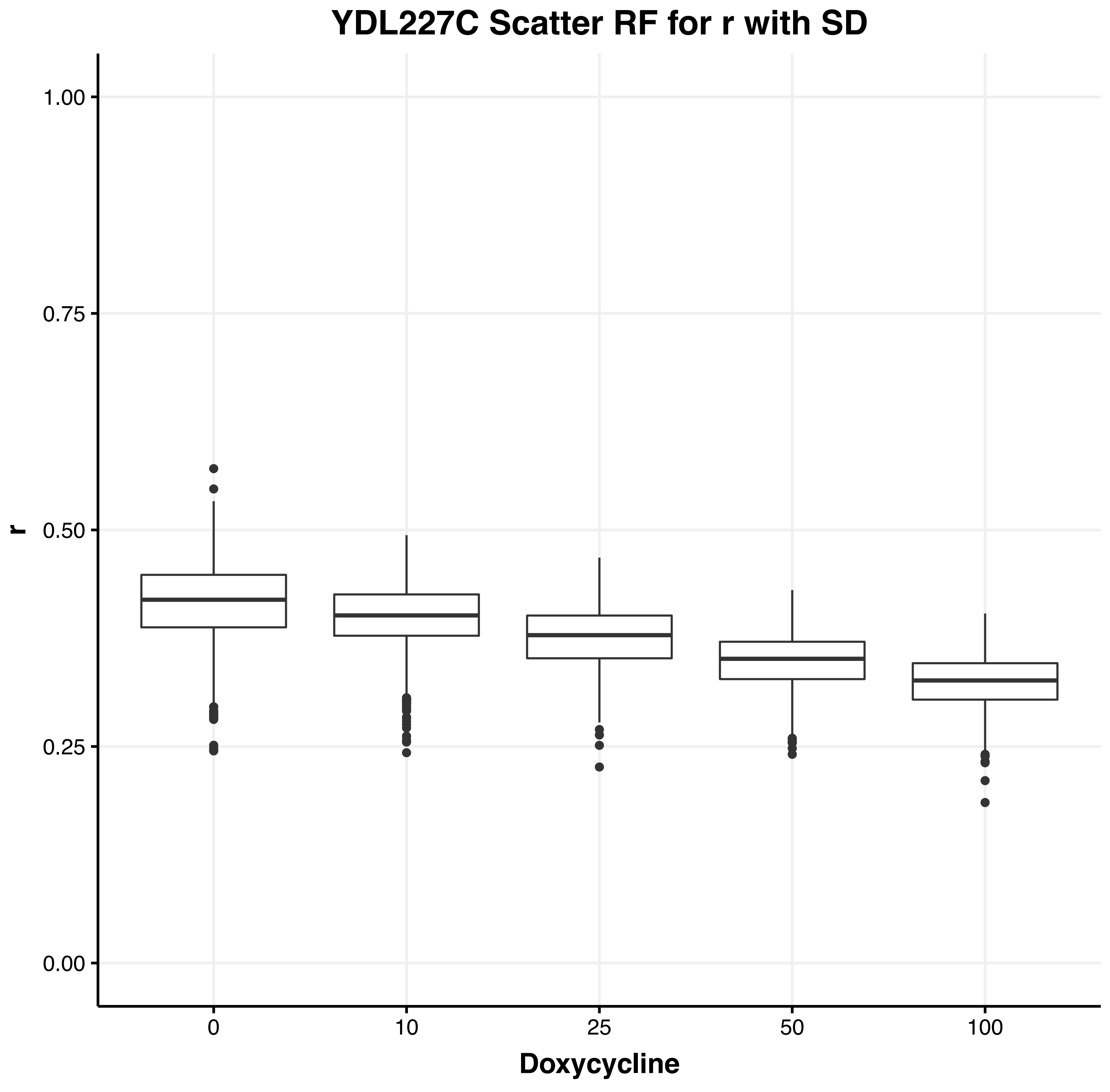
YDL227C Scatter RF for r with SD
r
Doxycycline
0.00
0.25
0.50
0.75
1.00
0
10
25
50
100

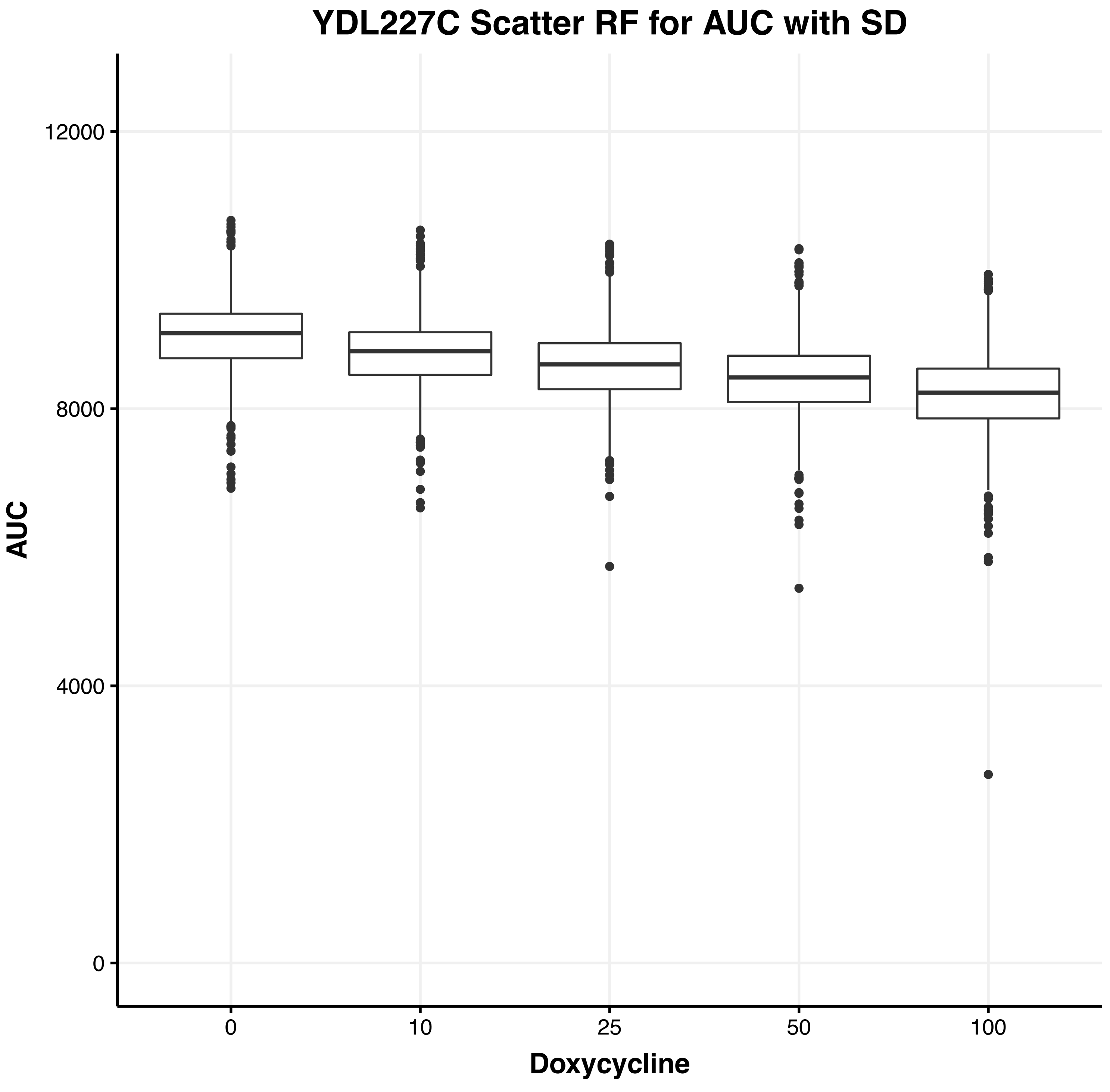
YDL227C Scatter RF for AUC with SD
AUC
Doxycycline
0
4000
8000
12000
0
10
25
50
100

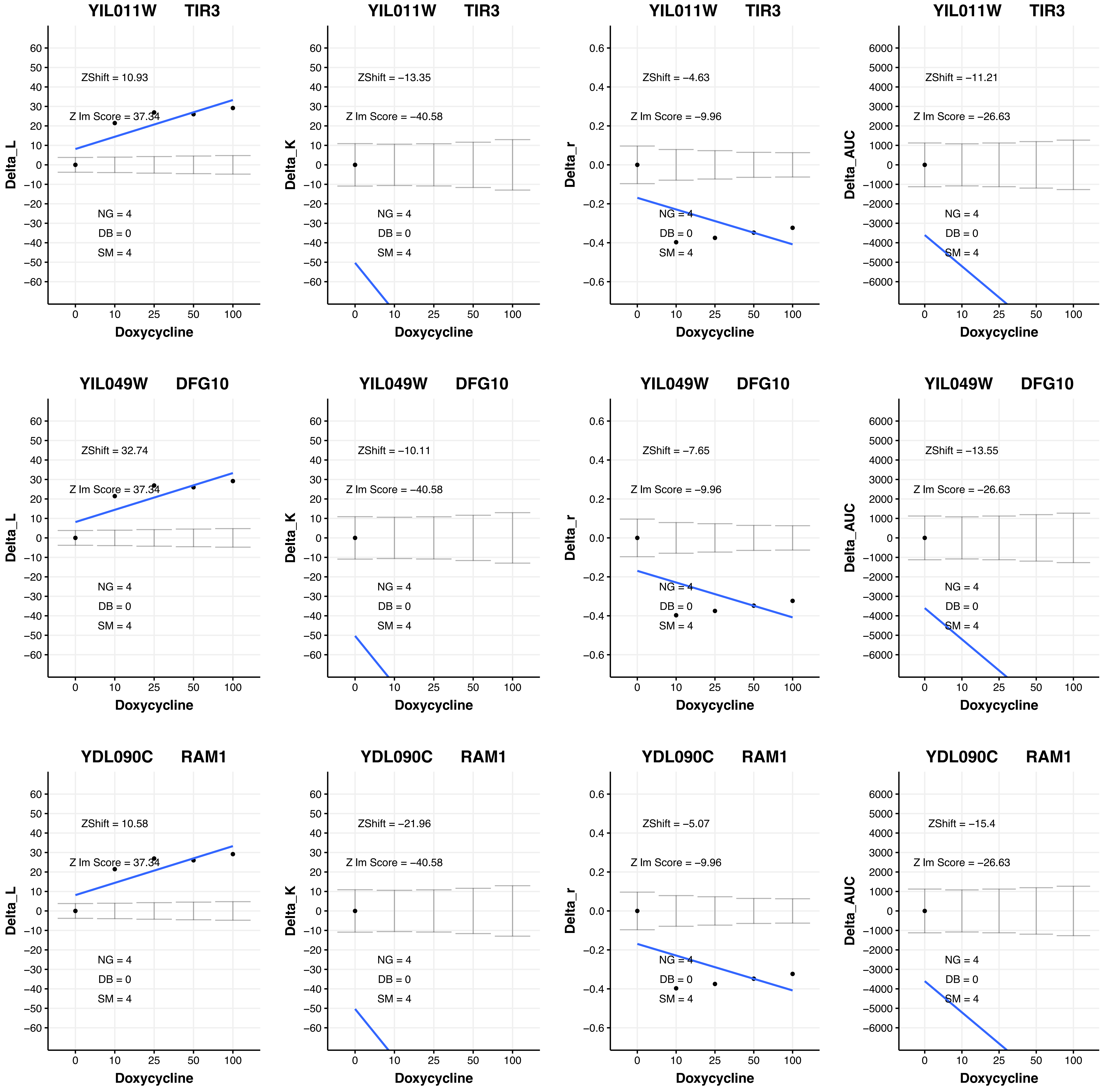
YIL011W TIR3
ZShift = 10.93
Z lm Score = 37.34
NG = 4
DB = 0
SM = 4
Delta_L
Doxycycline
YIL011W TIR3
ZShift = −13.35
Z lm Score = −40.58
NG = 4
DB = 0
SM = 4
Delta_K
Doxycycline
YIL011W TIR3
ZShift = −4.63
Z lm Score = −9.96
NG = 4
DB = 0
SM = 4
Delta_r
Doxycycline
YIL011W TIR3
ZShift = −11.21
Z lm Score = −26.63
NG = 4
DB = 0
SM = 4
Delta_AUC
Doxycycline
YIL049W DFG10
ZShift = 32.74
Z lm Score = 37.34
NG = 4
DB = 0
SM = 4
Delta_L
Doxycycline
YIL049W DFG10
ZShift = −10.11
Z lm Score = −40.58
NG = 4
DB = 0
SM = 4
Delta_K
Doxycycline
YIL049W DFG10
ZShift = −7.65
Z lm Score = −9.96
NG = 4
DB = 0
SM = 4
Delta_r
Doxycycline
YIL049W DFG10
ZShift = −13.55
Z lm Score = −26.63
NG = 4
DB = 0
SM = 4
Delta_AUC
Doxycycline
YDL090C RAM1
ZShift = 10.58
Z lm Score = 37.34
NG = 4
DB = 0
SM = 4
Delta_L
Doxycycline
YDL090C RAM1
ZShift = −21.96
Z lm Score = −40.58
NG = 4
DB = 0
SM = 4
Delta_K
Doxycycline
YDL090C RAM1
ZShift = −5.07
Z lm Score = −9.96
NG = 4
DB = 0
SM = 4
Delta_r
Doxycycline
YDL090C RAM1
ZShift = −15.4
Z lm Score = −26.63
NG = 4
DB = 0
SM = 4
Delta_AUC
Doxycycline

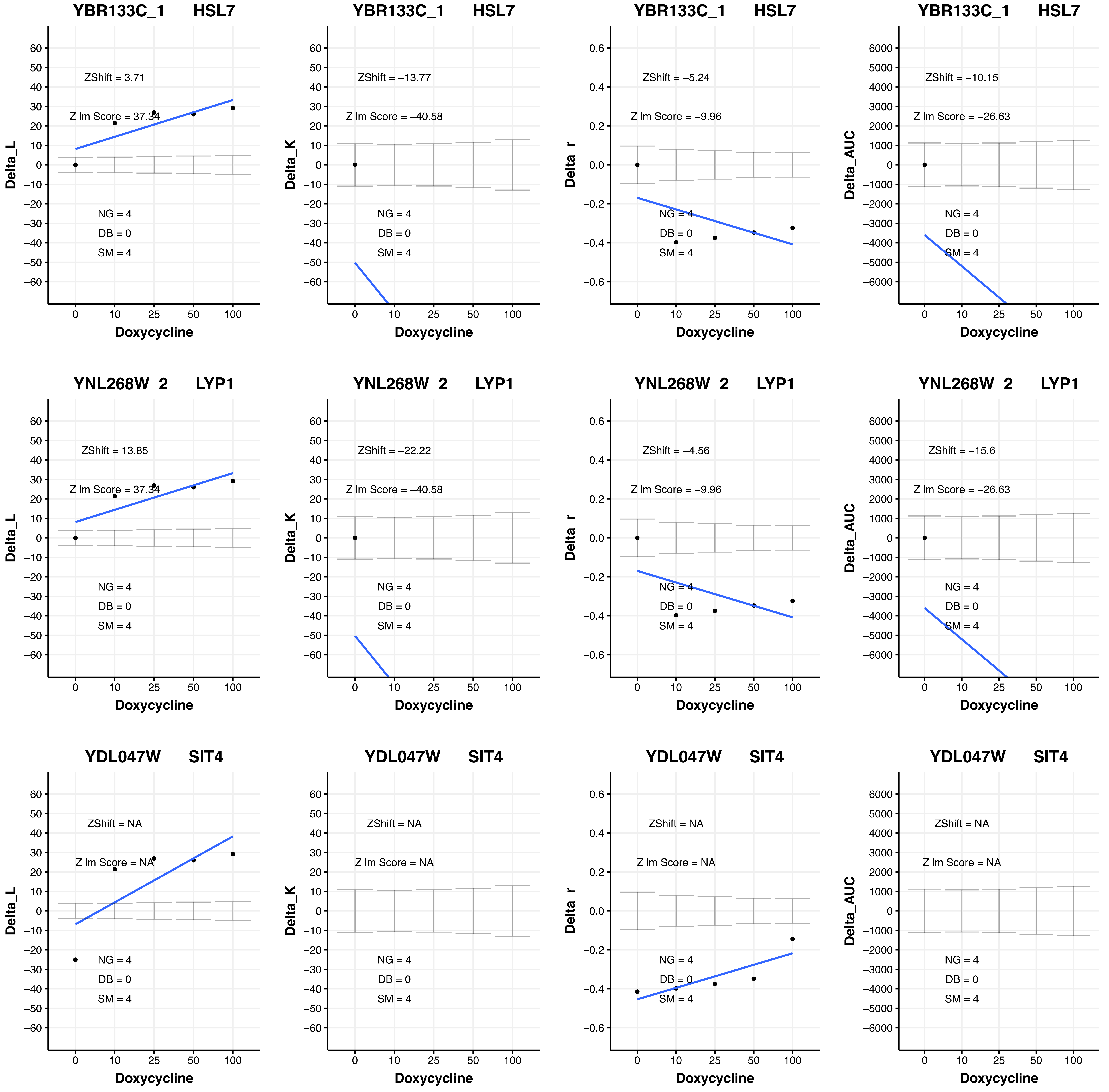
YBR133C_1 HSL7
ZShift = 3.71
Z lm Score = 37.34
NG = 4
DB = 0
SM = 4
Delta_L
Doxycycline
YBR133C_1 HSL7
ZShift = −13.77
Z lm Score = −40.58
NG = 4
DB = 0
SM = 4
Delta_K
Doxycycline
YBR133C_1 HSL7
ZShift = −5.24
Z lm Score = −9.96
NG = 4
DB = 0
SM = 4
Delta_r
Doxycycline
YBR133C_1 HSL7
ZShift = −10.15
Z lm Score = −26.63
NG = 4
DB = 0
SM = 4
Delta_AUC
Doxycycline
YNL268W_2 LYP1
ZShift = 13.85
Z lm Score = 37.34
NG = 4
DB = 0
SM = 4
Delta_L
Doxycycline
YNL268W_2 LYP1
ZShift = −22.22
Z lm Score = −40.58
NG = 4
DB = 0
SM = 4
Delta_K
Doxycycline
YNL268W_2 LYP1
ZShift = −4.56
Z lm Score = −9.96
NG = 4
DB = 0
SM = 4
Delta_r
Doxycycline
YNL268W_2 LYP1
ZShift = −15.6
Z lm Score = −26.63
NG = 4
DB = 0
SM = 4
Delta_AUC
Doxycycline
YDL047W SIT4
ZShift = NA
Z lm Score = NA
NG = 4
DB = 0
SM = 4
Delta_L
Doxycycline
YDL047W SIT4
ZShift = NA
Z lm Score = NA
NG = 4
DB = 0
SM = 4
Delta_K
Doxycycline
YDL047W SIT4
ZShift = NA
Z lm Score = NA
NG = 4
DB = 0
SM = 4
Delta_r
Doxycycline
YDL047W SIT4
ZShift = NA
Z lm Score = NA
NG = 4
DB = 0
SM = 4
Delta_AUC
Doxycycline

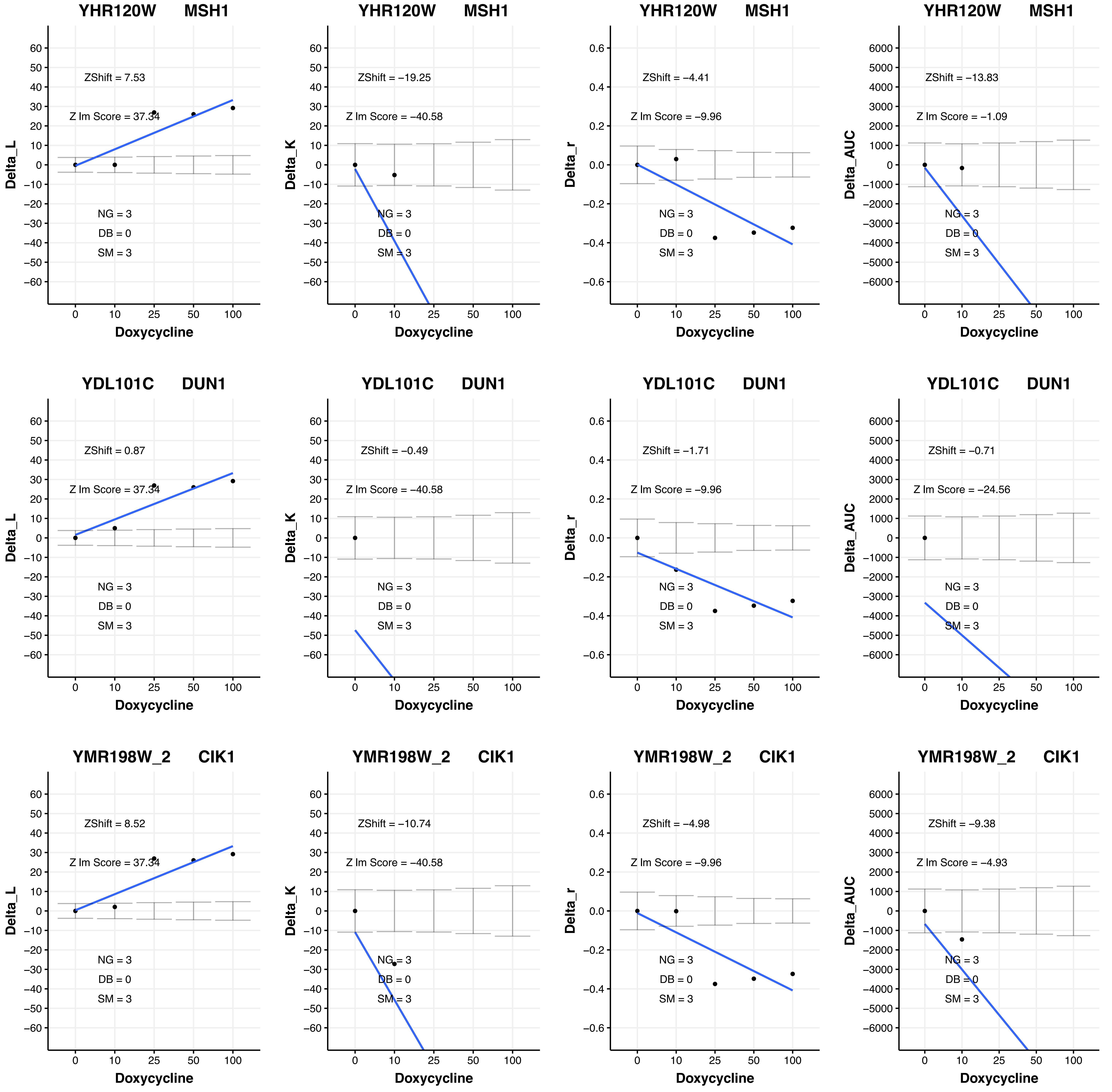
YHR120W MSH1
ZShift = 7.53
Z lm Score = 37.34
NG = 3
DB = 0
SM = 3
Delta_L
Doxycycline
YHR120W MSH1
ZShift = −19.25
Z lm Score = −40.58
NG = 3
DB = 0
SM = 3
Delta_K
Doxycycline
YHR120W MSH1
ZShift = −4.41
Z lm Score = −9.96
NG = 3
DB = 0
SM = 3
Delta_r
Doxycycline
YHR120W MSH1
ZShift = −13.83
Z lm Score = −1.09
NG = 3
DB = 0
SM = 3
Delta_AUC
Doxycycline
YDL101C DUN1
ZShift = 0.87
Z lm Score = 37.34
NG = 3
DB = 0
SM = 3
Delta_L
Doxycycline
YDL101C DUN1
ZShift = −0.49
Z lm Score = −40.58
NG = 3
DB = 0
SM = 3
Delta_K
Doxycycline
YDL101C DUN1
ZShift = −1.71
Z lm Score = −9.96
NG = 3
DB = 0
SM = 3
Delta_r
Doxycycline
YDL101C DUN1
ZShift = −0.71
Z lm Score = −24.56
NG = 3
DB = 0
SM = 3
Delta_AUC
Doxycycline
YMR198W_2 CIK1
ZShift = 8.52
Z lm Score = 37.34
NG = 3
DB = 0
SM = 3
Delta_L
Doxycycline
YMR198W_2 CIK1
ZShift = −10.74
Z lm Score = −40.58
NG = 3
DB = 0
SM = 3
Delta_K
Doxycycline
YMR198W_2 CIK1
ZShift = −4.98
Z lm Score = −9.96
NG = 3
DB = 0
SM = 3
Delta_r
Doxycycline
YMR198W_2 CIK1
ZShift = −9.38
Z lm Score = −4.93
NG = 3
DB = 0
SM = 3
Delta_AUC
Doxycycline

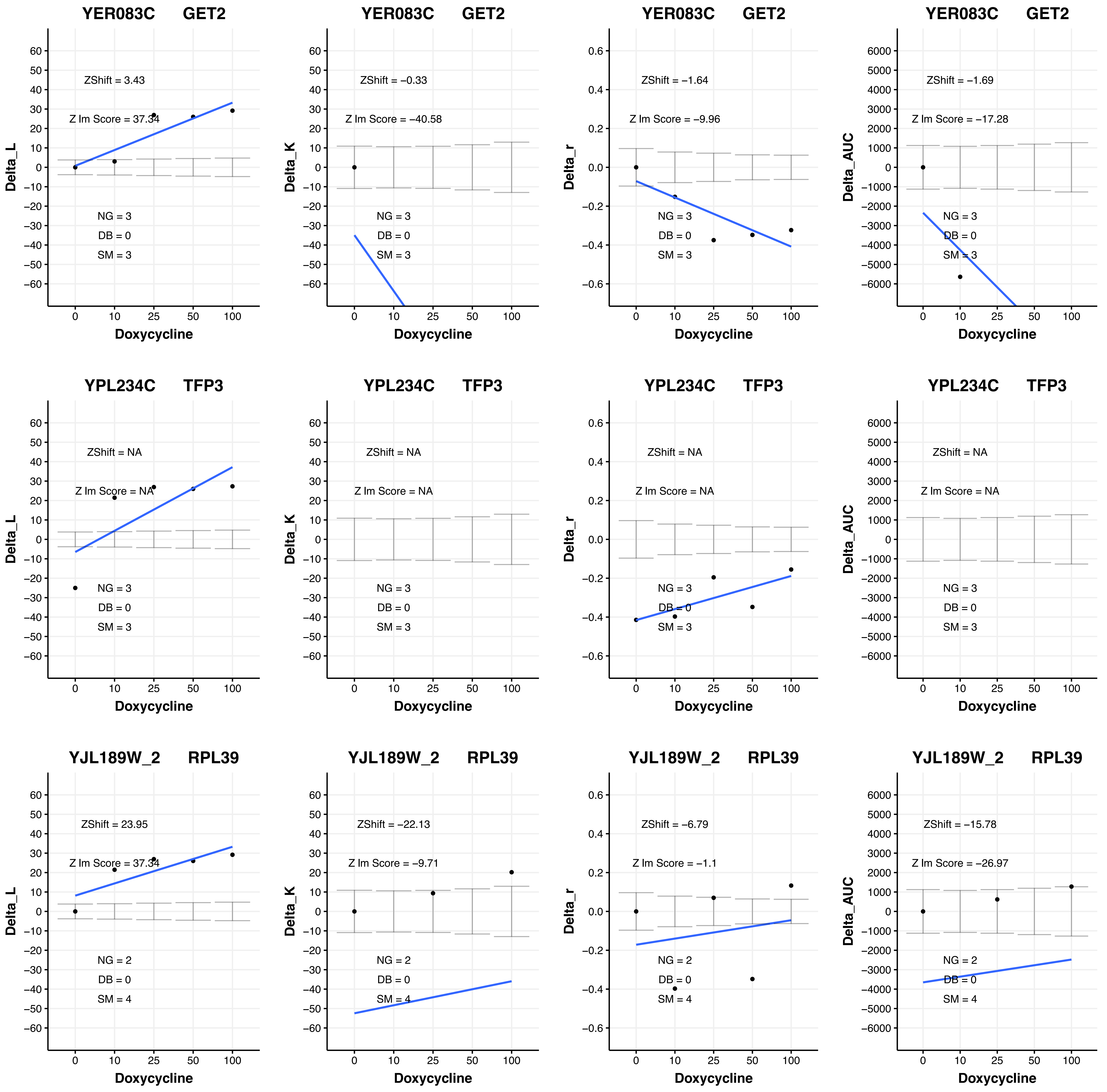
YER083C GET2
ZShift = 3.43
Z lm Score = 37.34
NG = 3
DB = 0
SM = 3
Delta_L
Doxycycline
YER083C GET2
ZShift = −0.33
Z lm Score = −40.58
NG = 3
DB = 0
SM = 3
Delta_K
Doxycycline
YER083C GET2
ZShift = −1.64
Z lm Score = −9.96
NG = 3
DB = 0
SM = 3
Delta_r
Doxycycline
YER083C GET2
ZShift = −1.69
Z lm Score = −17.28
NG = 3
DB = 0
SM = 3
Delta_AUC
Doxycycline
YPL234C TFP3
ZShift = NA
Z lm Score = NA
NG = 3
DB = 0
SM = 3
Delta_L
Doxycycline
YPL234C TFP3
ZShift = NA
Z lm Score = NA
NG = 3
DB = 0
SM = 3
Delta_K
Doxycycline
YPL234C TFP3
ZShift = NA
Z lm Score = NA
NG = 3
DB = 0
SM = 3
Delta_r
Doxycycline
YPL234C TFP3
ZShift = NA
Z lm Score = NA
NG = 3
DB = 0
SM = 3
Delta_AUC
Doxycycline
YJL189W_2 RPL39
ZShift = 23.95
Z lm Score = 37.34
NG = 2
DB = 0
SM = 4
Delta_L
Doxycycline
YJL189W_2 RPL39
ZShift = −22.13
Z lm Score = −9.71
NG = 2
DB = 0
SM = 4
Delta_K
Doxycycline
YJL189W_2 RPL39
ZShift = −6.79
Z lm Score = −1.1
NG = 2
DB = 0
SM = 4
Delta_r
Doxycycline
YJL189W_2 RPL39
ZShift = −15.78
Z lm Score = −26.97
NG = 2
DB = 0
SM = 4
Delta_AUC
Doxycycline

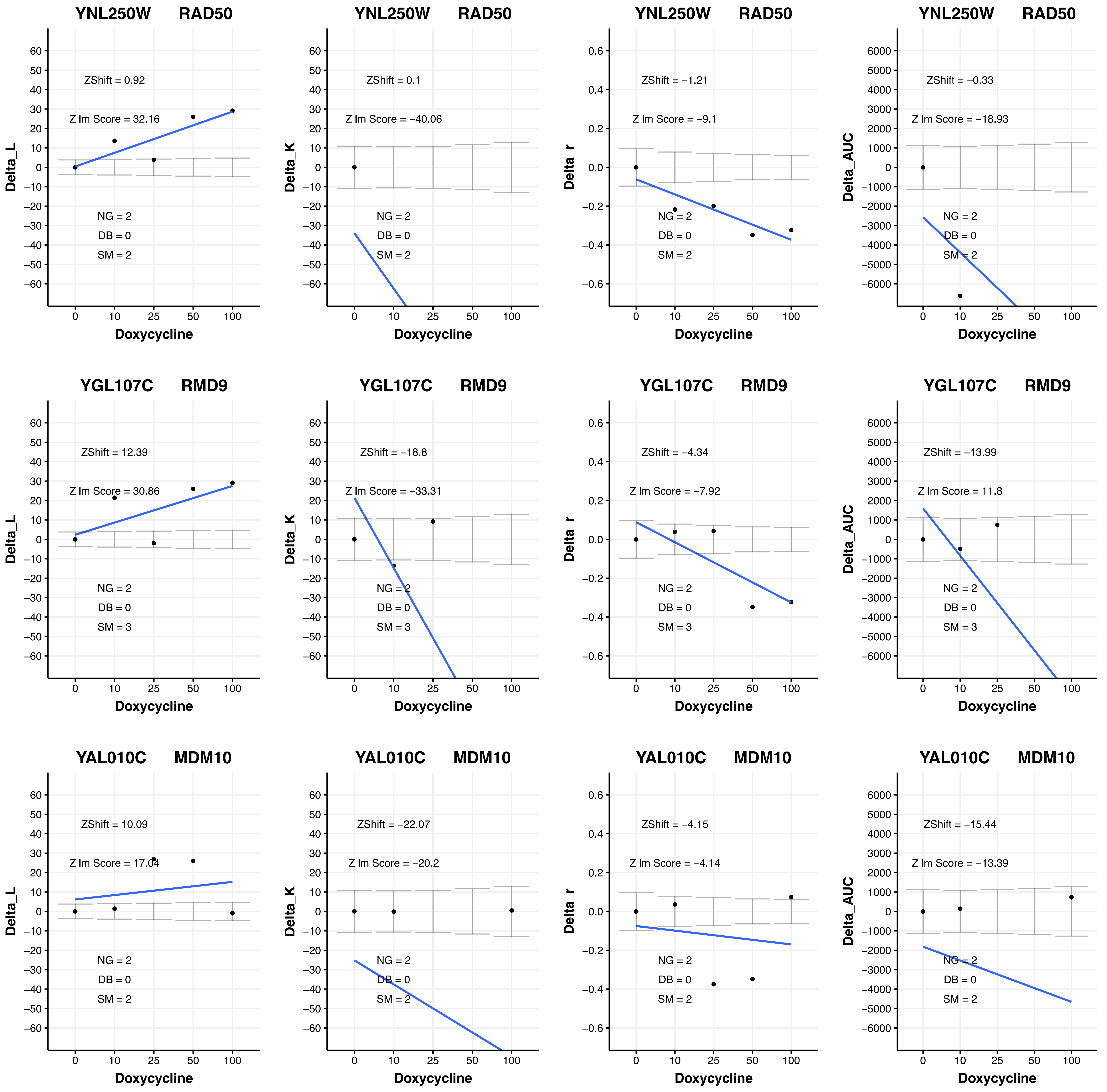
YNL250W RAD50
ZShift = 0.92
Z lm Score = 32.16
NG = 2
DB = 0
SM = 2
Delta_L
Doxycycline
YNL250W RAD50
ZShift = 0.1
Z lm Score = −40.06
NG = 2
DB = 0
SM = 2
Delta_K
Doxycycline
YNL250W RAD50
ZShift = −1.21
Z lm Score = −9.1
NG = 2
DB = 0
SM = 2
Delta_r
Doxycycline
YNL250W RAD50
ZShift = −0.33
Z lm Score = −18.93
NG = 2
DB = 0
SM = 2
Delta_AUC
Doxycycline
YGL107C RMD9
ZShift = 12.39
Z lm Score = 30.86
NG = 2
DB = 0
SM = 3
Delta_L
Doxycycline
YGL107C RMD9
ZShift = −18.8
Z lm Score = −33.31
NG = 2
DB = 0
SM = 3
Delta_K
Doxycycline
YGL107C RMD9
ZShift = −4.34
Z lm Score = −7.92
NG = 2
DB = 0
SM = 3
Delta_r
Doxycycline
YGL107C RMD9
ZShift = −13.99
Z lm Score = 11.8
NG = 2
DB = 0
SM = 3
Delta_AUC
Doxycycline
YAL010C MDM10
ZShift = 10.09
Z lm Score = 17.04
NG = 2
DB = 0
SM = 2
Delta_L
Doxycycline
YAL010C MDM10
ZShift = −22.07
Z lm Score = −20.2
NG = 2
DB = 0
SM = 2
Delta_K
Doxycycline
YAL010C MDM10
ZShift = −4.15
Z lm Score = −4.14
NG = 2
DB = 0
SM = 2
Delta_r
Doxycycline
YAL010C MDM10
ZShift = −15.44
Z lm Score = −13.39
NG = 2
DB = 0
SM = 2
Delta_AUC
Doxycycline

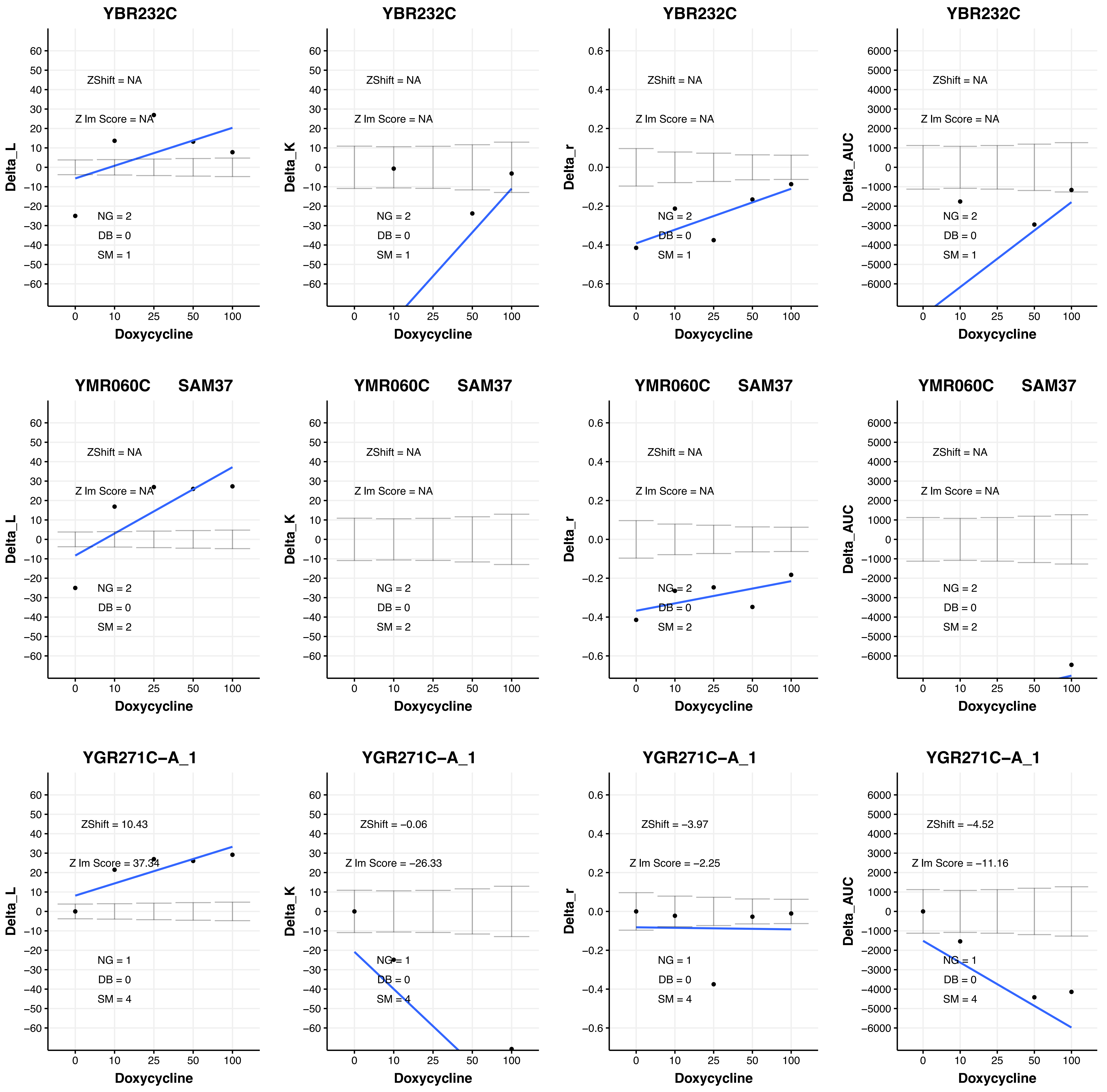

YBR232C
ZShift = NA
Z lm Score = NA
NG = 2
DB = 0
SM = 1
Delta_L
Delta_K
Delta_r
Delta_AUC
Doxycycline
YMR060C SAM37
ZShift = NA
Z lm Score = NA
NG = 2
DB = 0
SM = 2
YGR271C−A_1
ZShift = 10.43
Z lm Score = 37.34
ZShift = −0.06
Z lm Score = −26.33
ZShift = −3.97
Z lm Score = −2.25
ZShift = −4.52
Z lm Score = −11.16
NG = 1
DB = 0
SM = 4

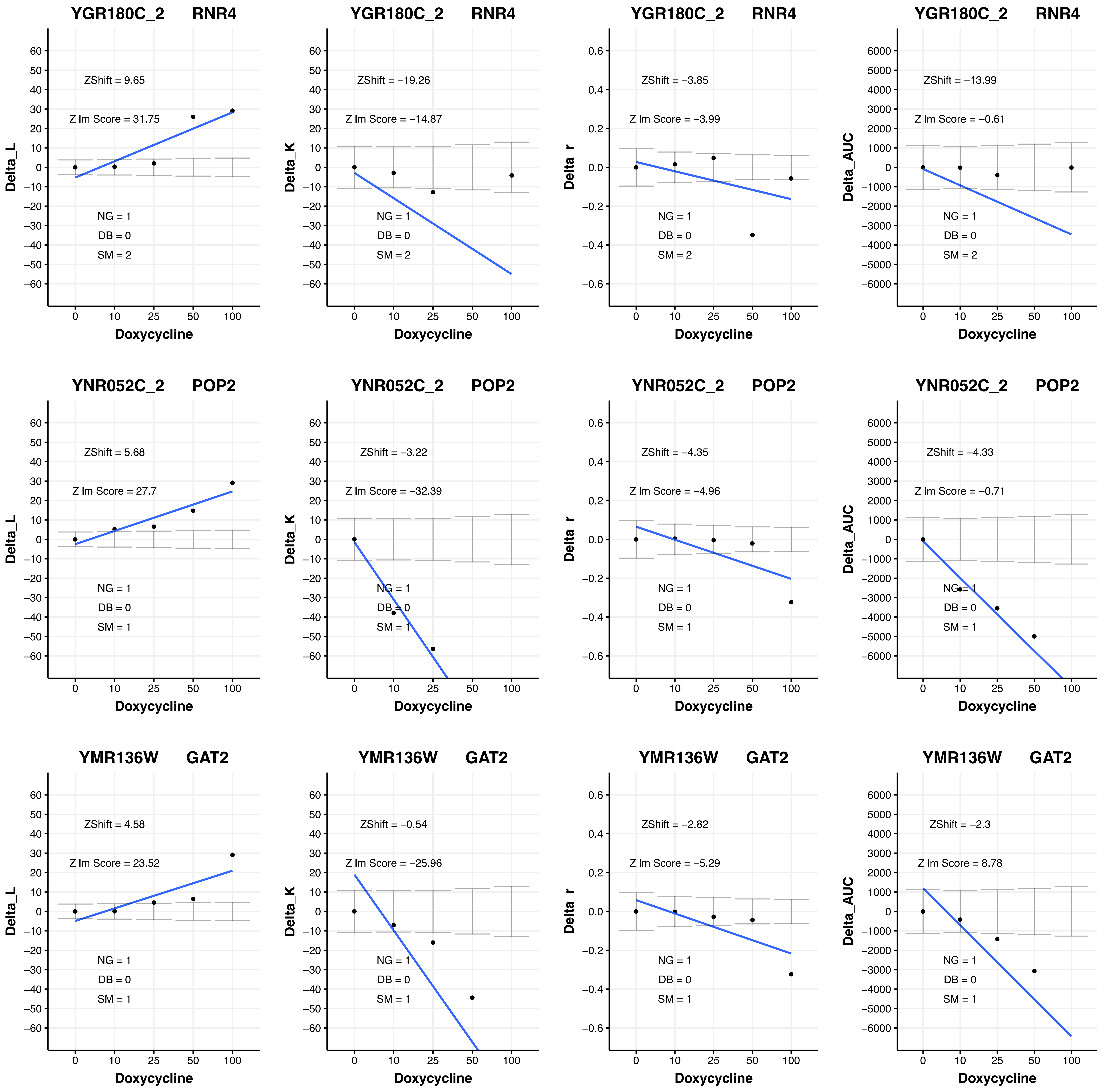

YGR180C_2 RNR4
ZShift = 9.65
Z lm Score = 31.75
NG = 1
DB = 0
SM = 2
Delta_L
Doxycycline
YGR180C_2 RNR4
ZShift = −19.26
Z lm Score = −14.87
NG = 1
DB = 0
SM = 2
Delta_K
Doxycycline
YGR180C_2 RNR4
ZShift = −3.85
Z lm Score = −3.99
NG = 1
DB = 0
SM = 2
Delta_r
Doxycycline
YGR180C_2 RNR4
ZShift = −13.99
Z lm Score = −0.61
NG = 1
DB = 0
SM = 2
Delta_AUC
Doxycycline
YNR052C_2 POP2
ZShift = 5.68
Z lm Score = 27.7
NG = 1
DB = 0
SM = 1
Delta_L
Doxycycline
YNR052C_2 POP2
ZShift = −3.22
Z lm Score = −32.39
NG = 1
DB = 0
SM = 1
Delta_K
Doxycycline
YNR052C_2 POP2
ZShift = −4.35
Z lm Score = −4.96
NG = 1
DB = 0
SM = 1
Delta_r
Doxycycline
YNR052C_2 POP2
ZShift = −4.33
Z lm Score = −0.71
NG = 1
DB = 0
SM = 1
Delta_AUC
Doxycycline
YMR136W GAT2
ZShift = 4.58
Z lm Score = 23.52
NG = 1
DB = 0
SM = 1
Delta_L
Doxycycline
YMR136W GAT2
ZShift = −0.54
Z lm Score = −25.96
NG = 1
DB = 0
SM = 1
Delta_K
Doxycycline
YMR136W GAT2
ZShift = −2.82
Z lm Score = −5.29
NG = 1
DB = 0
SM = 1
Delta_r
Doxycycline
YMR136W GAT2
ZShift = −2.3
Z lm Score = 8.78
NG = 1
DB = 0
SM = 1
Delta_AUC
Doxycycline

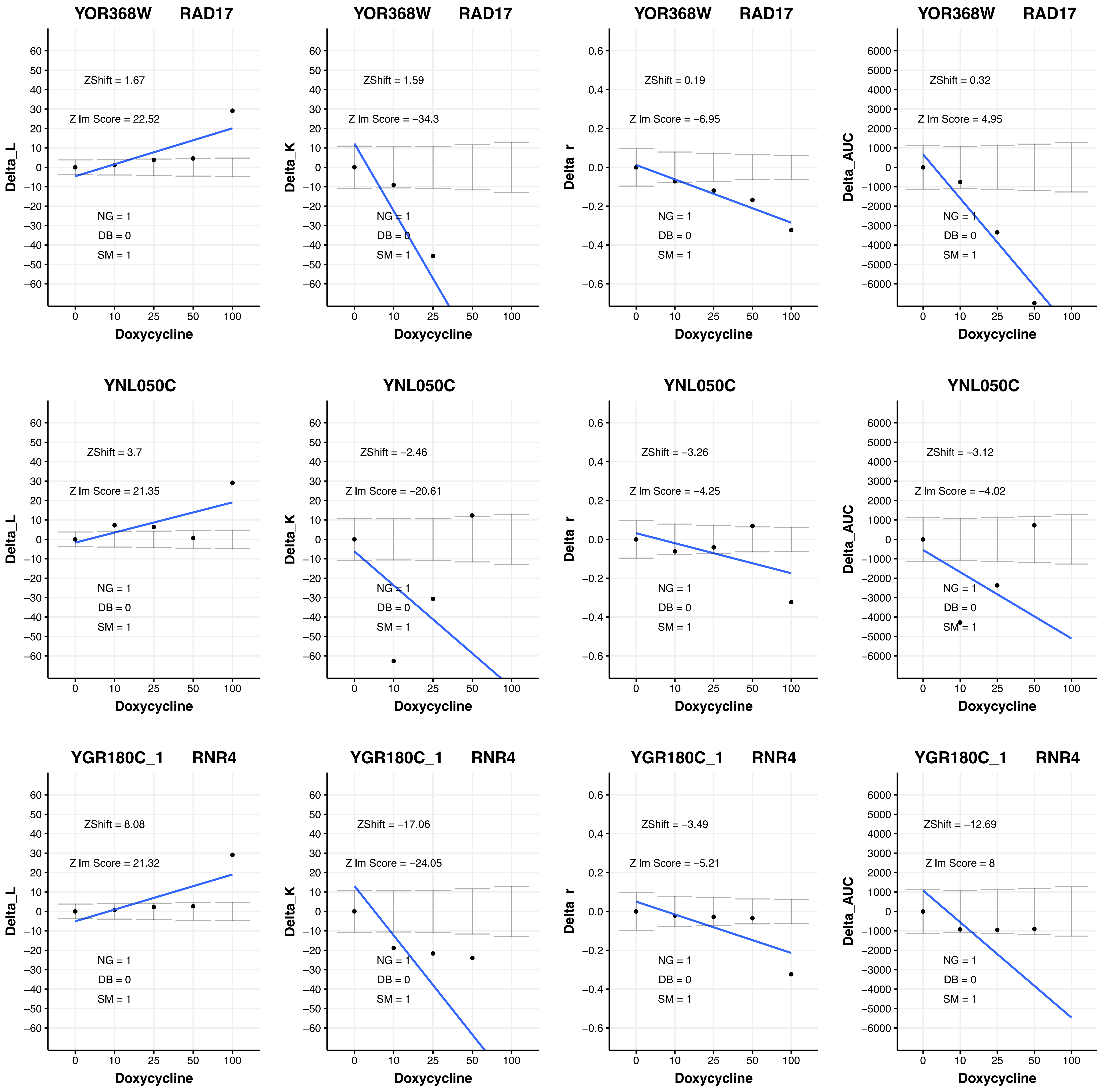
YOR368W RAD17
ZShift = 1.67
Z lm Score = 22.52
NG = 1
DB = 0
SM = 1
Delta_L
Doxycycline
YOR368W RAD17
ZShift = 1.59
Z lm Score = −34.3
NG = 1
DB = 0
SM = 1
Delta_K
Doxycycline
YOR368W RAD17
ZShift = 0.19
Z lm Score = −6.95
NG = 1
DB = 0
SM = 1
Delta_r
Doxycycline
YOR368W RAD17
ZShift = 0.32
Z lm Score = 4.95
NG = 1
DB = 0
SM = 1
Delta_AUC
Doxycycline
YNL050C
ZShift = 3.7
Z lm Score = 21.35
NG = 1
DB = 0
SM = 1
Delta_L
Doxycycline
YNL050C
ZShift = −2.46
Z lm Score = −20.61
NG = 1
DB = 0
SM = 1
Delta_K
Doxycycline
YNL050C
ZShift = −3.26
Z lm Score = −4.25
NG = 1
DB = 0
SM = 1
Delta_r
Doxycycline
YNL050C
ZShift = −3.12
Z lm Score = −4.02
NG = 1
DB = 0
SM = 1
Delta_AUC
Doxycycline
YGR180C_1 RNR4
ZShift = 8.08
Z lm Score = 21.32
NG = 1
DB = 0
SM = 1
Delta_L
Doxycycline
YGR180C_1 RNR4
ZShift = −17.06
Z lm Score = −24.05
NG = 1
DB = 0
SM = 1
Delta_K
Doxycycline
YGR180C_1 RNR4
ZShift = −3.49
Z lm Score = −5.21
NG = 1
DB = 0
SM = 1
Delta_r
Doxycycline
YGR180C_1 RNR4
ZShift = −12.69
Z lm Score = 8
NG = 1
DB = 0
SM = 1
Delta_AUC
Doxycycline

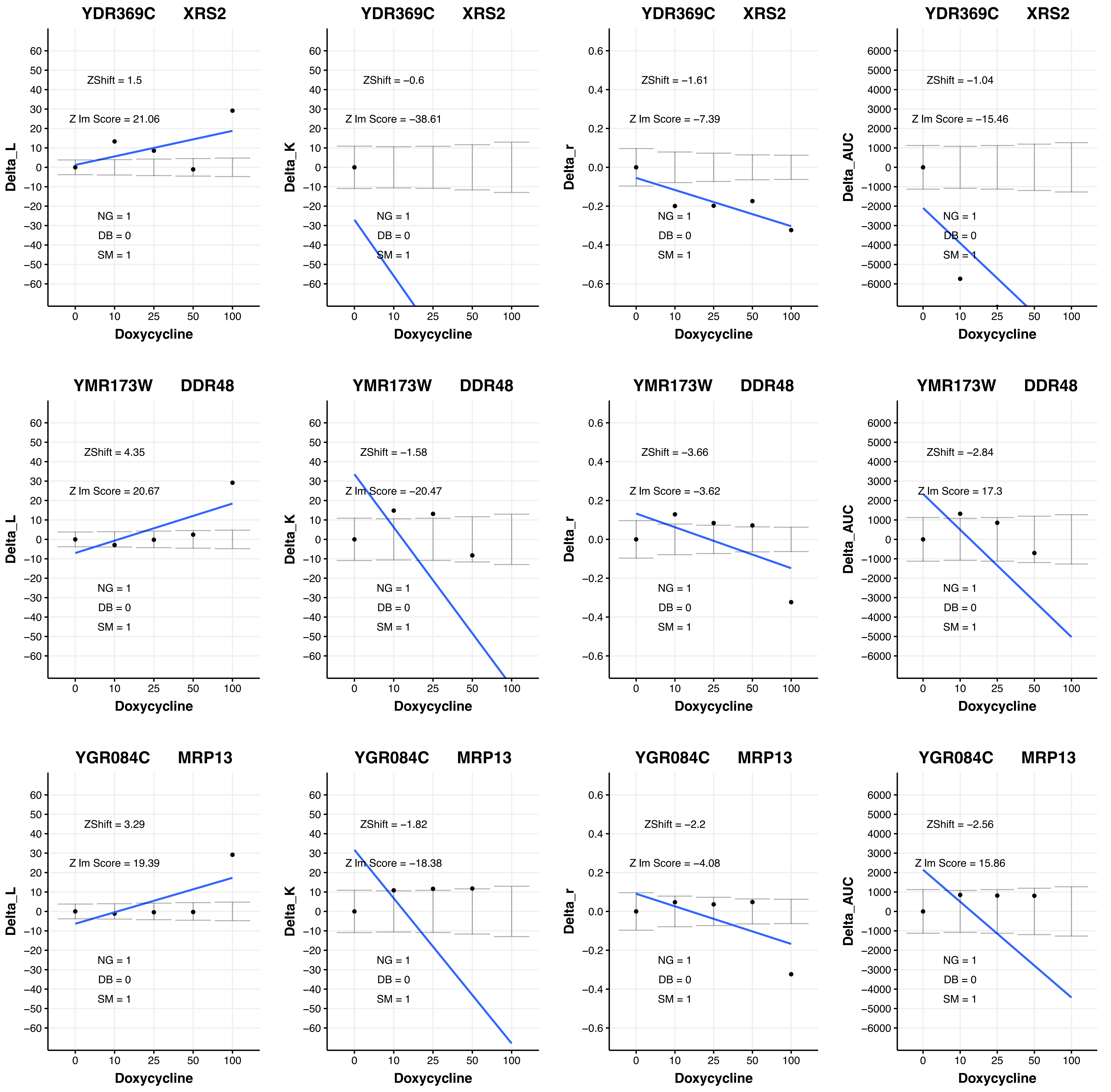
YDR369C XRS2
ZShift = 1.5
Z lm Score = 21.06
NG = 1
DB = 0
SM = 1
Delta_L
Doxycycline
YDR369C XRS2
ZShift = −0.6
Z lm Score = −38.61
NG = 1
DB = 0
SM = 1
Delta_K
Doxycycline
YDR369C XRS2
ZShift = −1.61
Z lm Score = −7.39
NG = 1
DB = 0
SM = 1
Delta_r
Doxycycline
YDR369C XRS2
ZShift = −1.04
Z lm Score = −15.46
NG = 1
DB = 0
SM = 1
Delta_AUC
Doxycycline
YMR173W DDR48
ZShift = 4.35
Z lm Score = 20.67
NG = 1
DB = 0
SM = 1
Delta_L
Doxycycline
YMR173W DDR48
ZShift = −1.58
Z lm Score = −20.47
NG = 1
DB = 0
SM = 1
Delta_K
Doxycycline
YMR173W DDR48
ZShift = −3.66
Z lm Score = −3.62
NG = 1
DB = 0
SM = 1
Delta_r
Doxycycline
YMR173W DDR48
ZShift = −2.84
Z lm Score = 17.3
NG = 1
DB = 0
SM = 1
Delta_AUC
Doxycycline
YGR084C MRP13
ZShift = 3.29
Z lm Score = 19.39
NG = 1
DB = 0
SM = 1
Delta_L
Doxycycline
YGR084C MRP13
ZShift = −1.82
Z lm Score = −18.38
NG = 1
DB = 0
SM = 1
Delta_K
Doxycycline
YGR084C MRP13
ZShift = −2.2
Z lm Score = −4.08
NG = 1
DB = 0
SM = 1
Delta_r
Doxycycline
YGR084C MRP13
ZShift = −2.56
Z lm Score = 15.86
NG = 1
DB = 0
SM = 1
Delta_AUC
Doxycycline

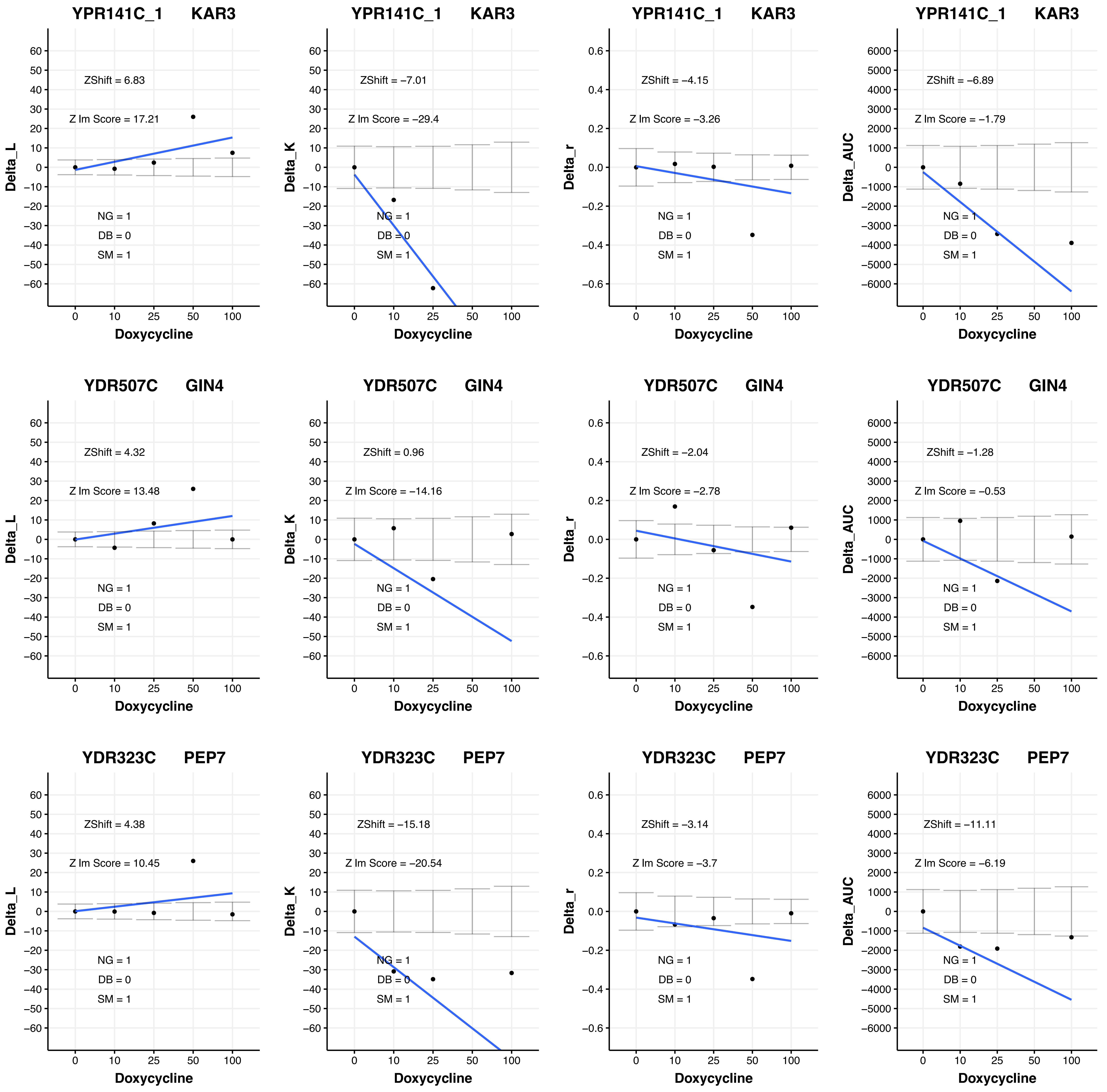

YPR141C_1 KAR3
ZShift = 6.83
Z lm Score = 17.21
NG = 1
DB = 0
SM = 1
Delta_L
Doxycycline
YPR141C_1 KAR3
ZShift = −7.01
Z lm Score = −29.4
NG = 1
DB = 0
SM = 1
Delta_K
Doxycycline
YPR141C_1 KAR3
ZShift = −4.15
Z lm Score = −3.26
NG = 1
DB = 0
SM = 1
Delta_r
Doxycycline
YPR141C_1 KAR3
ZShift = −6.89
Z lm Score = −1.79
NG = 1
DB = 0
SM = 1
Delta_AUC
Doxycycline
YDR507C GIN4
ZShift = 4.32
Z lm Score = 13.48
NG = 1
DB = 0
SM = 1
Delta_L
Doxycycline
YDR507C GIN4
ZShift = 0.96
Z lm Score = −14.16
NG = 1
DB = 0
SM = 1
Delta_K
Doxycycline
YDR507C GIN4
ZShift = −2.04
Z lm Score = −2.78
NG = 1
DB = 0
SM = 1
Delta_r
Doxycycline
YDR507C GIN4
ZShift = −1.28
Z lm Score = −0.53
NG = 1
DB = 0
SM = 1
Delta_AUC
Doxycycline
YDR323C PEP7
ZShift = 4.38
Z lm Score = 10.45
NG = 1
DB = 0
SM = 1
Delta_L
Doxycycline
YDR323C PEP7
ZShift = −15.18
Z lm Score = −20.54
NG = 1
DB = 0
SM = 1
Delta_K
Doxycycline
YDR323C PEP7
ZShift = −3.14
Z lm Score = −3.7
NG = 1
DB = 0
SM = 1
Delta_r
Doxycycline
YDR323C PEP7
ZShift = −11.11
Z lm Score = −6.19
NG = 1
DB = 0
SM = 1
Delta_AUC
Doxycycline

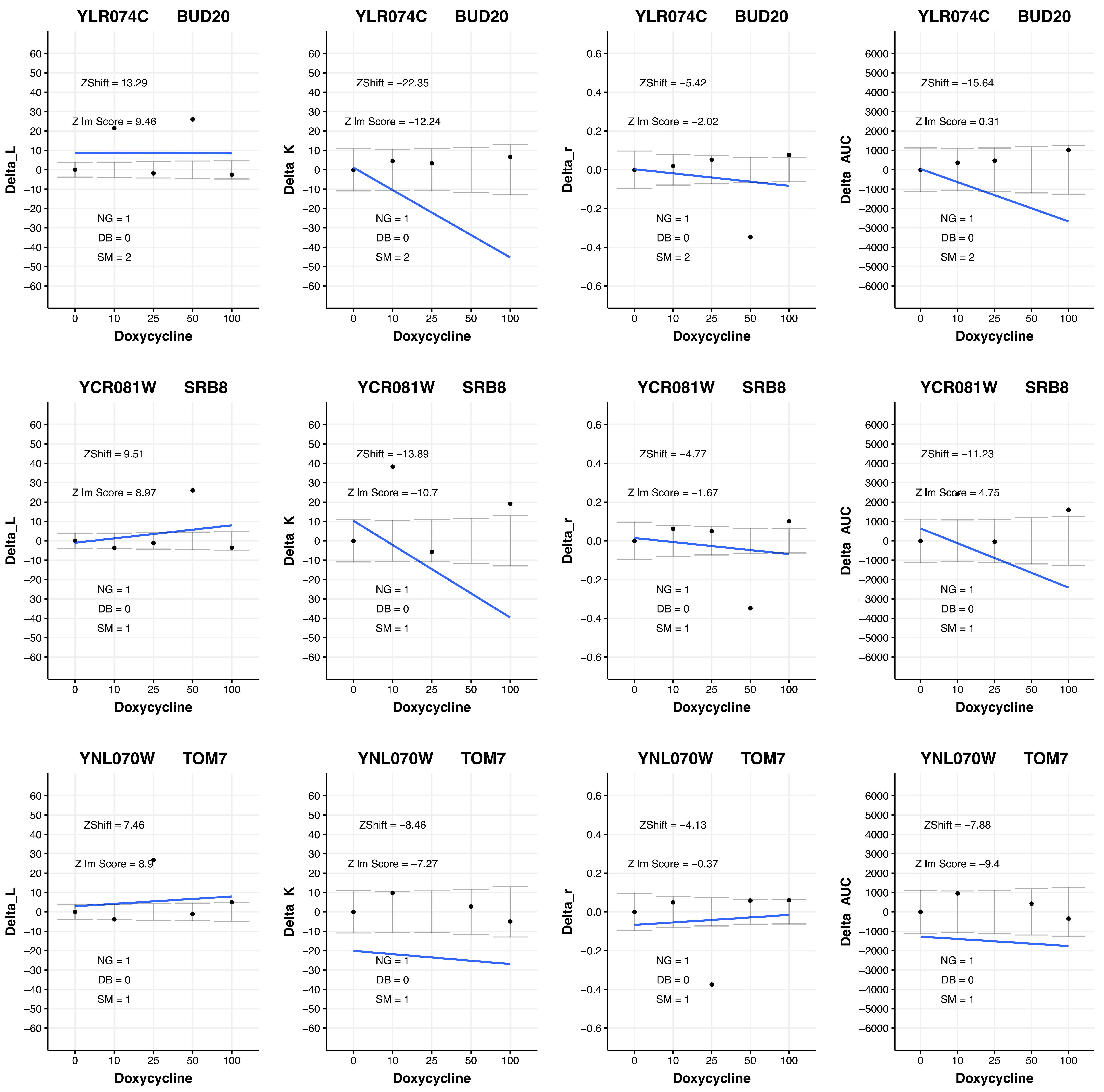

YLR074C BUD20
ZShift = 13.29
Z lm Score = 9.46
NG = 1
DB = 0
SM = 2
Delta_L
Doxycycline
YLR074C BUD20
ZShift = −22.35
Z lm Score = −12.24
NG = 1
DB = 0
SM = 2
Delta_K
Doxycycline
YLR074C BUD20
ZShift = −5.42
Z lm Score = −2.02
NG = 1
DB = 0
SM = 2
Delta_r
Doxycycline
YLR074C BUD20
ZShift = −15.64
Z lm Score = 0.31
NG = 1
DB = 0
SM = 2
Delta_AUC
Doxycycline
YCR081W SRB8
ZShift = 9.51
Z lm Score = 8.97
NG = 1
DB = 0
SM = 1
Delta_L
Doxycycline
YCR081W SRB8
ZShift = −13.89
Z lm Score = −10.7
NG = 1
DB = 0
SM = 1
Delta_K
Doxycycline
YCR081W SRB8
ZShift = −4.77
Z lm Score = −1.67
NG = 1
DB = 0
SM = 1
Delta_r
Doxycycline
YCR081W SRB8
ZShift = −11.23
Z lm Score = 4.75
NG = 1
DB = 0
SM = 1
Delta_AUC
Doxycycline
YNL070W TOM7
ZShift = 7.46
Z lm Score = 8.9
NG = 1
DB = 0
SM = 1
Delta_L
Doxycycline
YNL070W TOM7
ZShift = −8.46
Z lm Score = −7.27
NG = 1
DB = 0
SM = 1
Delta_K
Doxycycline
YNL070W TOM7
ZShift = −4.13
Z lm Score = −0.37
NG = 1
DB = 0
SM = 1
Delta_r
Doxycycline
YNL070W TOM7
ZShift = −7.88
Z lm Score = −9.4
NG = 1
DB = 0
SM = 1
Delta_AUC
Doxycycline

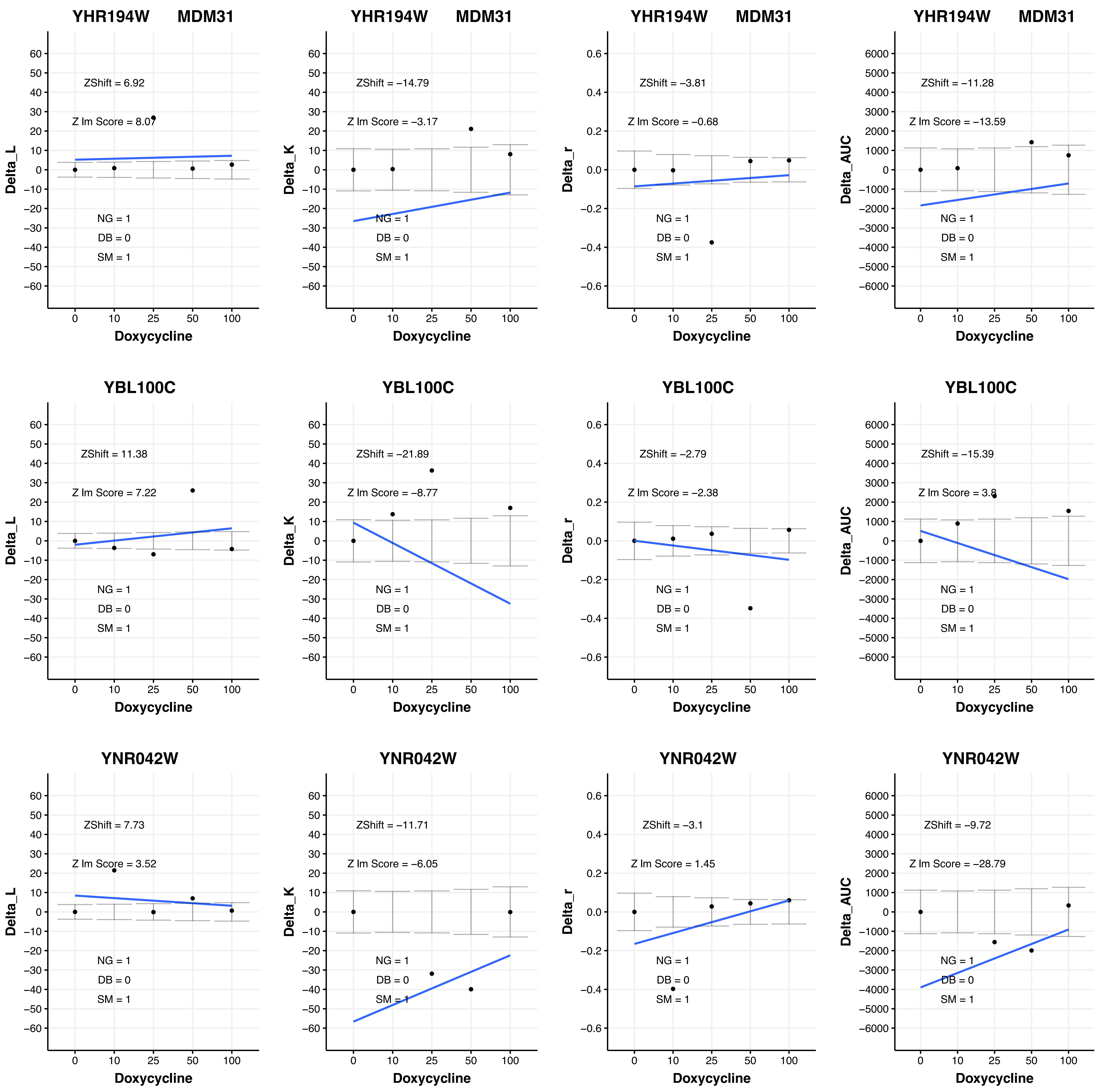
YHR194W MDM31
ZShift = 6.92
Z lm Score = 8.07
NG = 1
DB = 0
SM = 1
Delta_L
Doxycycline
YHR194W MDM31
ZShift = −14.79
Z lm Score = −3.17
NG = 1
DB = 0
SM = 1
Delta_K
Doxycycline
YHR194W MDM31
ZShift = −3.81
Z lm Score = −0.68
NG = 1
DB = 0
SM = 1
Delta_r
Doxycycline
YHR194W MDM31
ZShift = −11.28
Z lm Score = −13.59
NG = 1
DB = 0
SM = 1
Delta_AUC
Doxycycline
YBL100C
ZShift = 11.38
Z lm Score = 7.22
NG = 1
DB = 0
SM = 1
Delta_L
Doxycycline
YBL100C
ZShift = −21.89
Z lm Score = −8.77
NG = 1
DB = 0
SM = 1
Delta_K
Doxycycline
YBL100C
ZShift = −2.79
Z lm Score = −2.38
NG = 1
DB = 0
SM = 1
Delta_r
Doxycycline
YBL100C
ZShift = −15.39
Z lm Score = 3.8
NG = 1
DB = 0
SM = 1
Delta_AUC
Doxycycline
YNR042W
ZShift = 7.73
Z lm Score = 3.52
NG = 1
DB = 0
SM = 1
Delta_L
Doxycycline
YNR042W
ZShift = −11.71
Z lm Score = −6.05
NG = 1
DB = 0
SM = 1
Delta_K
Doxycycline
YNR042W
ZShift = −3.1
Z lm Score = 1.45
NG = 1
DB = 0
SM = 1
Delta_r
Doxycycline
YNR042W
ZShift = −9.72
Z lm Score = −28.79
NG = 1
DB = 0
SM = 1
Delta_AUC
Doxycycline

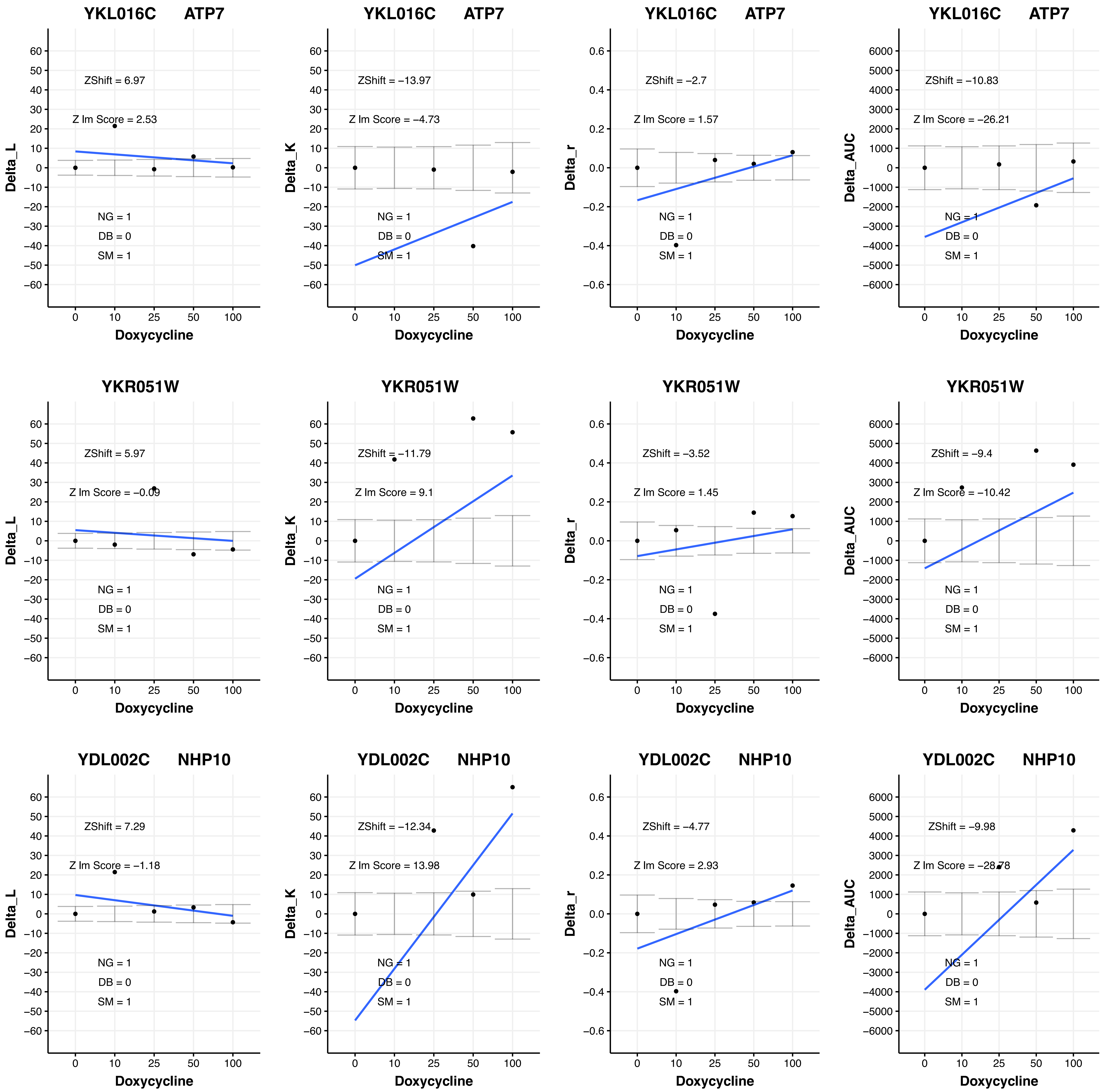

YKL016C ATP7
ZShift = 6.97
Z lm Score = 2.53
NG = 1
DB = 0
SM = 1
Delta_L
Doxycycline
YKL016C ATP7
ZShift = −13.97
Z lm Score = −4.73
NG = 1
DB = 0
SM = 1
Delta_K
Doxycycline
YKL016C ATP7
ZShift = −2.7
Z lm Score = 1.57
NG = 1
DB = 0
SM = 1
Delta_r
Doxycycline
YKL016C ATP7
ZShift = −10.83
Z lm Score = −26.21
NG = 1
DB = 0
SM = 1
Delta_AUC
Doxycycline
YKR051W
ZShift = 5.97
Z lm Score = −0.09
NG = 1
DB = 0
SM = 1
Delta_L
Doxycycline
YKR051W
ZShift = −11.79
Z lm Score = 9.1
NG = 1
DB = 0
SM = 1
Delta_K
Doxycycline
YKR051W
ZShift = −3.52
Z lm Score = 1.45
NG = 1
DB = 0
SM = 1
Delta_r
Doxycycline
YKR051W
ZShift = −9.4
Z lm Score = −10.42
NG = 1
DB = 0
SM = 1
Delta_AUC
Doxycycline
YDL002C NHP10
ZShift = 7.29
Z lm Score = −1.18
NG = 1
DB = 0
SM = 1
Delta_L
Doxycycline
YDL002C NHP10
ZShift = −12.34
Z lm Score = 13.98
NG = 1
DB = 0
SM = 1
Delta_K
Doxycycline
YDL002C NHP10
ZShift = −4.77
Z lm Score = 2.93
NG = 1
DB = 0
SM = 1
Delta_r
Doxycycline
YDL002C NHP10
ZShift = −9.98
Z lm Score = −28.78
NG = 1
DB = 0
SM = 1
Delta_AUC
Doxycycline

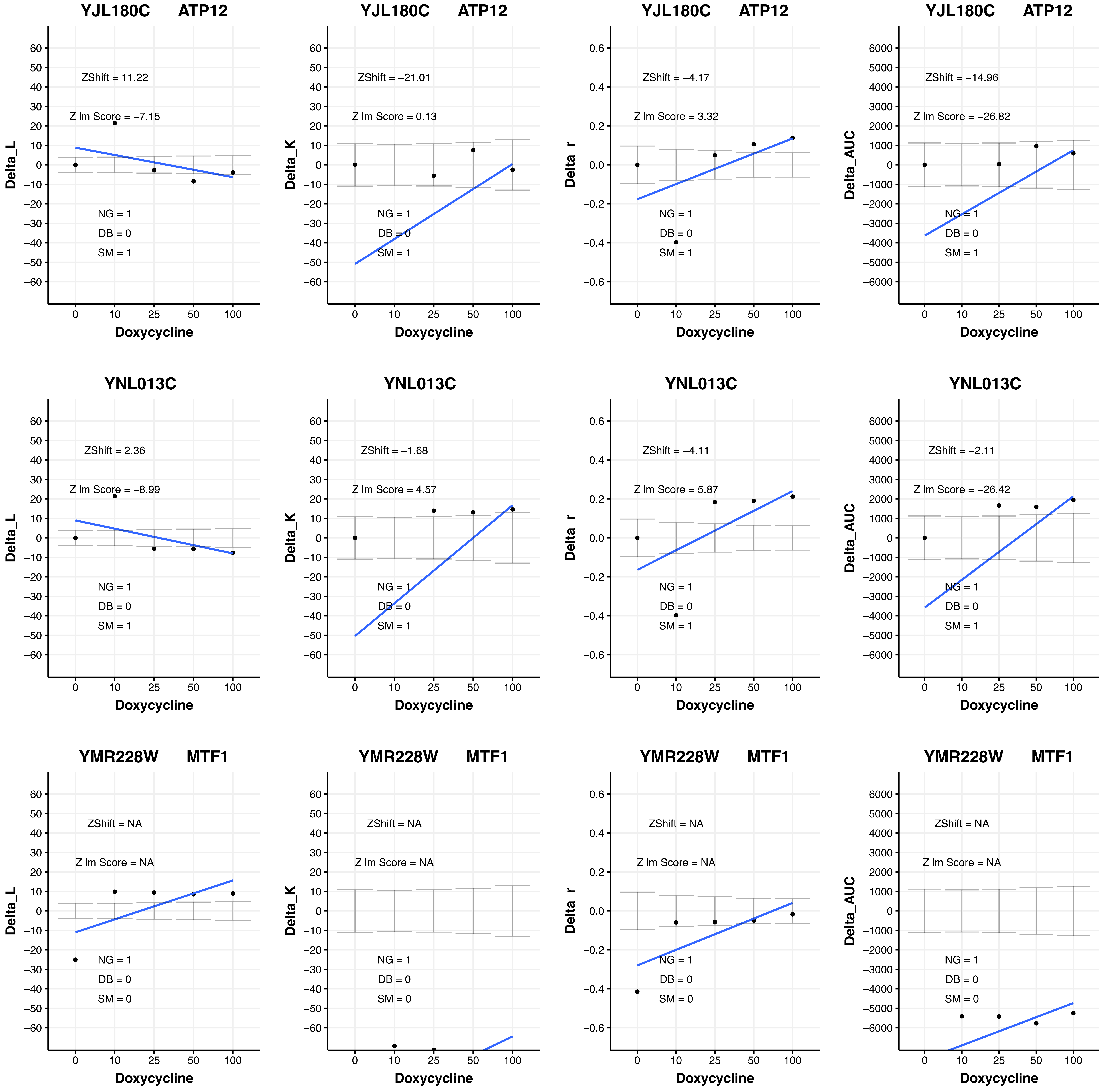

YJL180C ATP12
ZShift = 11.22
Z lm Score = −7.15
NG = 1
DB = 0
SM = 1
Delta_L
Doxycycline
YJL180C ATP12
ZShift = −21.01
Z lm Score = 0.13
NG = 1
DB = 0
SM = 1
Delta_K
Doxycycline
YJL180C ATP12
ZShift = −4.17
Z lm Score = 3.32
NG = 1
DB = 0
SM = 1
Delta_r
Doxycycline
YJL180C ATP12
ZShift = −14.96
Z lm Score = −26.82
NG = 1
DB = 0
SM = 1
Delta_AUC
Doxycycline
YNL013C
ZShift = 2.36
Z lm Score = −8.99
NG = 1
DB = 0
SM = 1
Delta_L
Doxycycline
YNL013C
ZShift = −1.68
Z lm Score = 4.57
NG = 1
DB = 0
SM = 1
Delta_K
Doxycycline
YNL013C
ZShift = −4.11
Z lm Score = 5.87
NG = 1
DB = 0
SM = 1
Delta_r
Doxycycline
YNL013C
ZShift = −2.11
Z lm Score = −26.42
NG = 1
DB = 0
SM = 1
Delta_AUC
Doxycycline
YMR228W MTF1
ZShift = NA
Z lm Score = NA
NG = 1
DB = 0
SM = 0
Delta_L
Doxycycline
YMR228W MTF1
ZShift = NA
Z lm Score = NA
NG = 1
DB = 0
SM = 0
Delta_K
Doxycycline
YMR228W MTF1
ZShift = NA
Z lm Score = NA
NG = 1
DB = 0
SM = 0
Delta_r
Doxycycline
YMR228W MTF1
ZShift = NA
Z lm Score = NA
NG = 1
DB = 0
SM = 0
Delta_AUC
Doxycycline

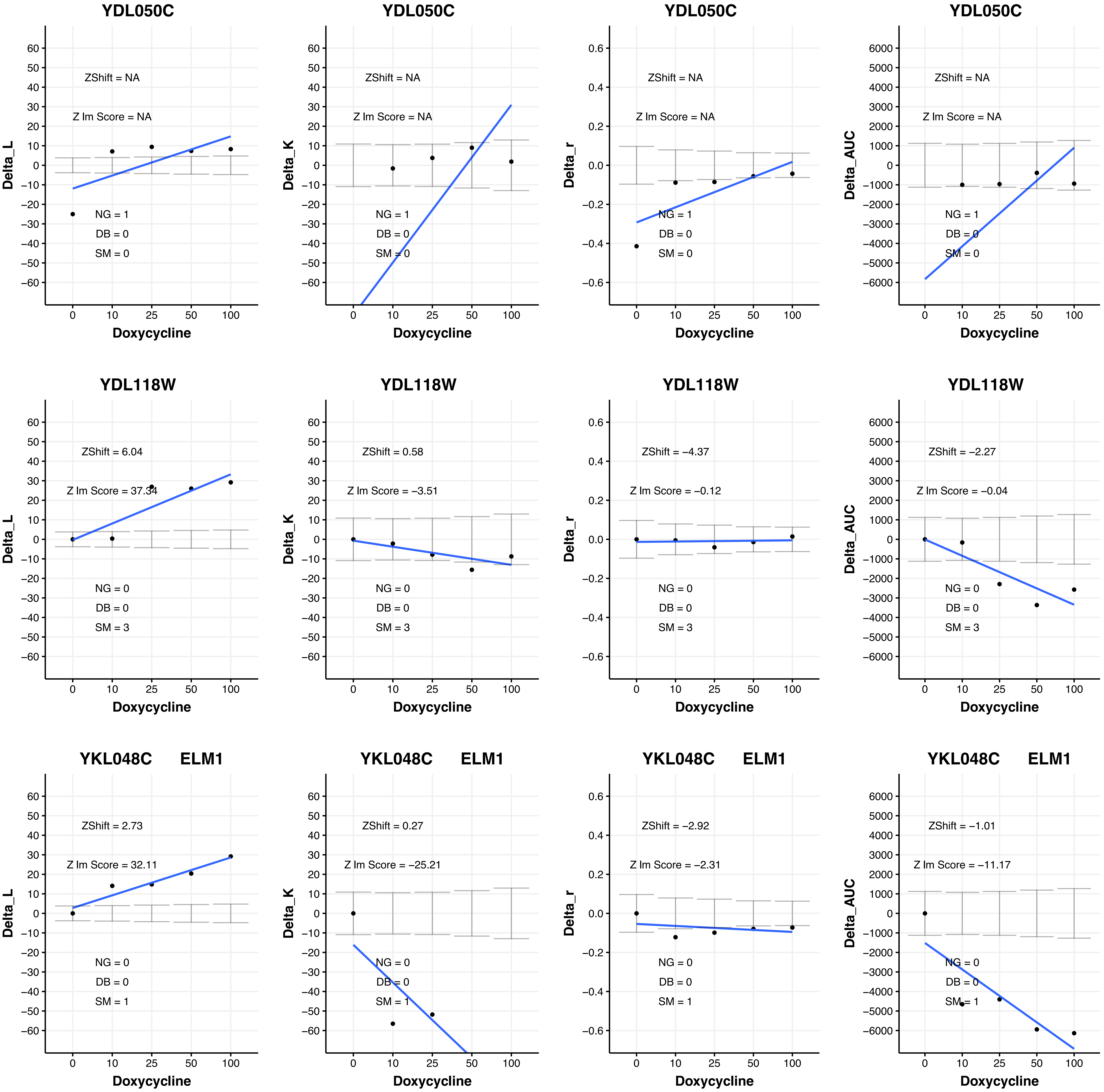

YDL050C
ZShift = NA
Z lm Score = NA
NG = 1
DB = 0
SM = 0
Delta_L
Delta_K
Delta_r
Delta_AUC
Doxycycline
YDL118W
ZShift = 6.04
Z lm Score = 37.34
NG = 0
DB = 0
SM = 3
ZShift = 0.58
Z lm Score = −3.51
ZShift = −4.37
Z lm Score = −0.12
ZShift = −2.27
Z lm Score = −0.04
YKL048C ELM1
ZShift = 2.73
Z lm Score = 32.11
NG = 0
DB = 0
SM = 1
ZShift = 0.27
Z lm Score = −25.21
ZShift = −2.92
Z lm Score = −2.31
ZShift = −1.01
Z lm Score = −11.17

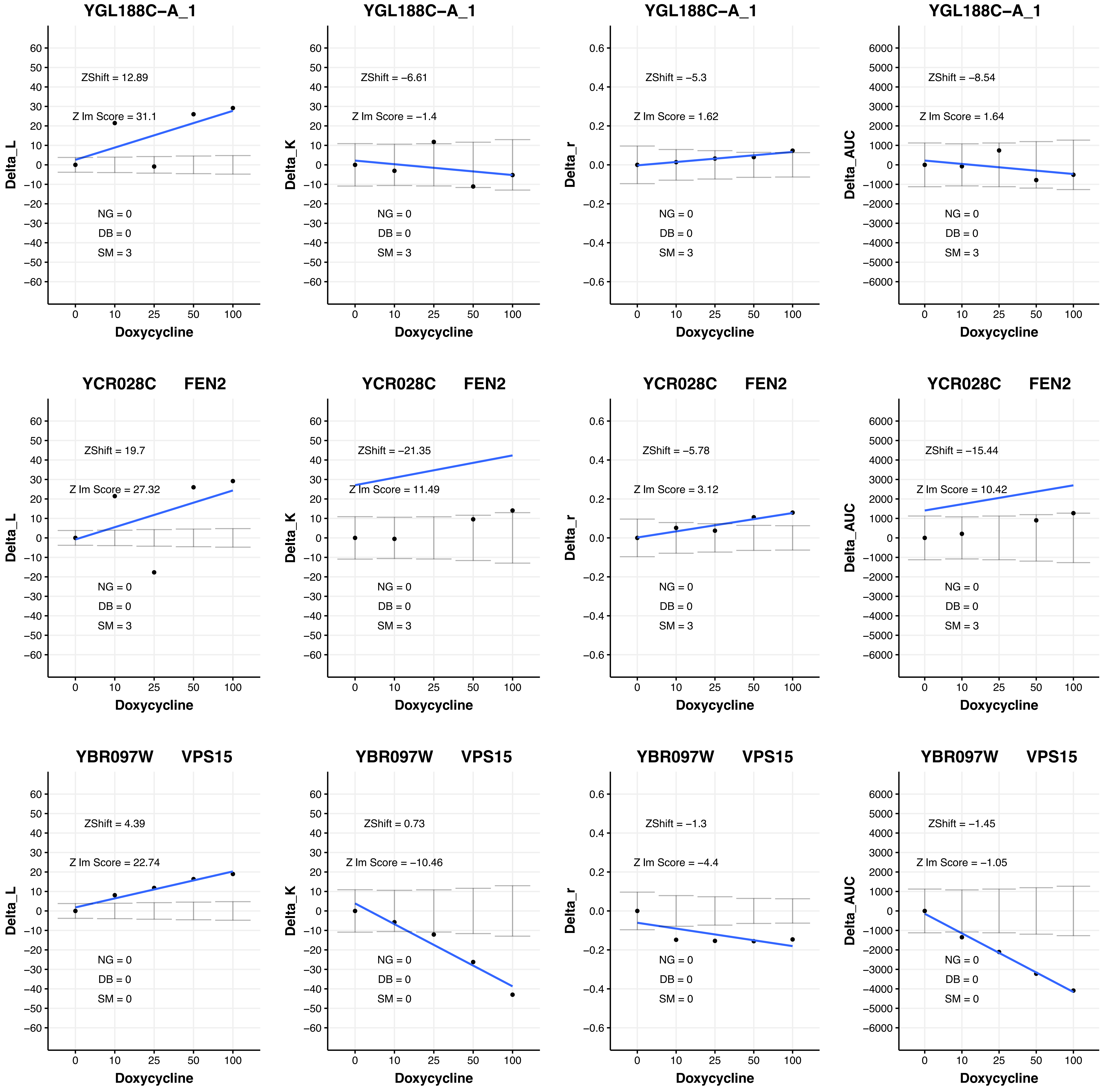
YGL188C−A_1
ZShift = 12.89
Z lm Score = 31.1
NG = 0
DB = 0
SM = 3
Delta_L
Doxycycline
YGL188C−A_1
ZShift = −6.61
Z lm Score = −1.4
NG = 0
DB = 0
SM = 3
Delta_K
Doxycycline
YGL188C−A_1
ZShift = −5.3
Z lm Score = 1.62
NG = 0
DB = 0
SM = 3
Delta_r
Doxycycline
YGL188C−A_1
ZShift = −8.54
Z lm Score = 1.64
NG = 0
DB = 0
SM = 3
Delta_AUC
Doxycycline
YCR028C FEN2
ZShift = 19.7
Z lm Score = 27.32
NG = 0
DB = 0
SM = 3
Delta_L
Doxycycline
YCR028C FEN2
ZShift = −21.35
Z lm Score = 11.49
NG = 0
DB = 0
SM = 3
Delta_K
Doxycycline
YCR028C FEN2
ZShift = −5.78
Z lm Score = 3.12
NG = 0
DB = 0
SM = 3
Delta_r
Doxycycline
YCR028C FEN2
ZShift = −15.44
Z lm Score = 10.42
NG = 0
DB = 0
SM = 3
Delta_AUC
Doxycycline
YBR097W VPS15
ZShift = 4.39
Z lm Score = 22.74
NG = 0
DB = 0
SM = 0
Delta_L
Doxycycline
YBR097W VPS15
ZShift = 0.73
Z lm Score = −10.46
NG = 0
DB = 0
SM = 0
Delta_K
Doxycycline
YBR097W VPS15
ZShift = −1.3
Z lm Score = −4.4
NG = 0
DB = 0
SM = 0
Delta_r
Doxycycline
YBR097W VPS15
ZShift = −1.45
Z lm Score = −1.05
NG = 0
DB = 0
SM = 0
Delta_AUC
Doxycycline

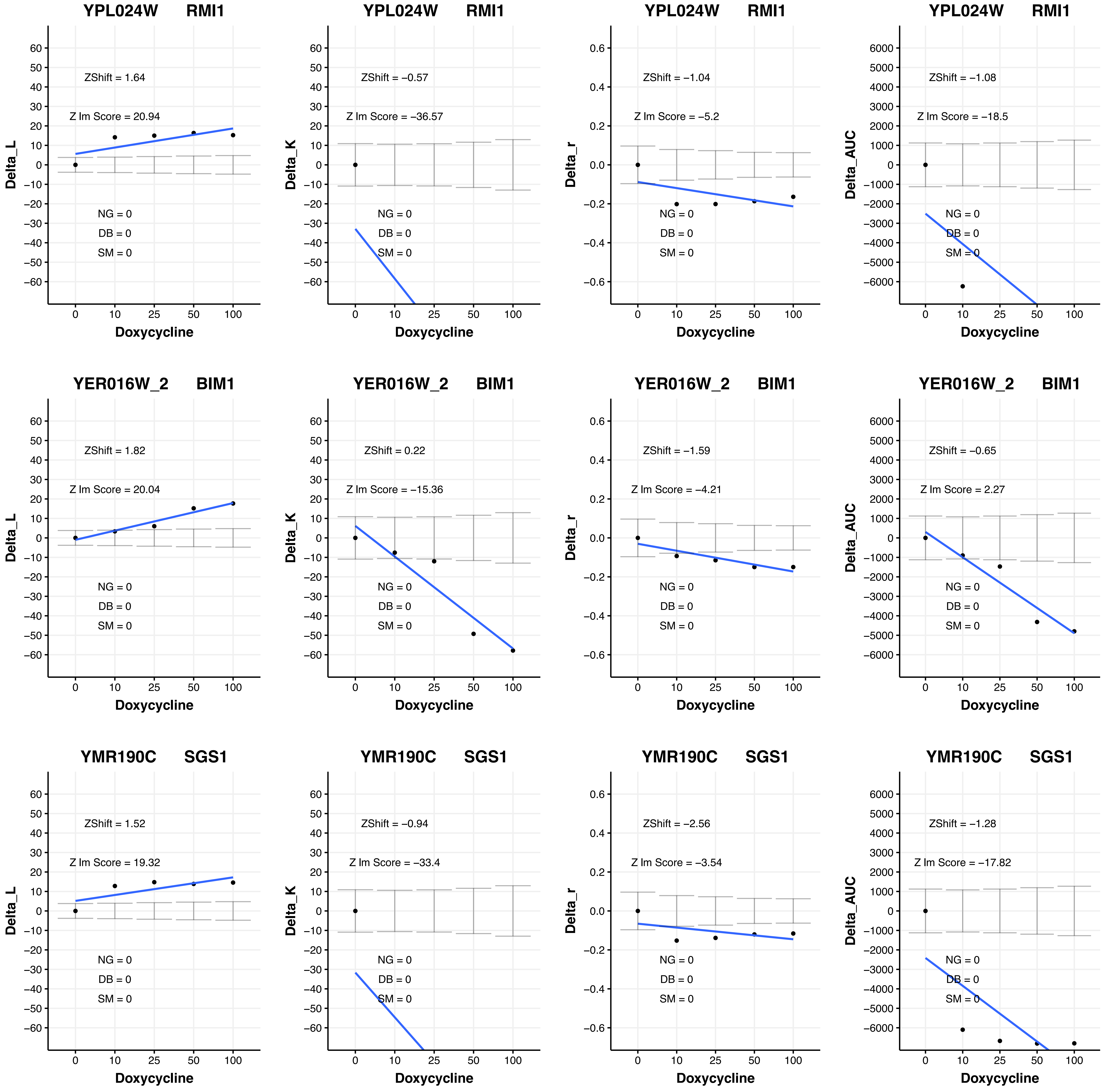
YPL024W RMI1
ZShift = 1.64
Z lm Score = 20.94
NG = 0
DB = 0
SM = 0
Delta_L
Doxycycline
YPL024W RMI1
ZShift = −0.57
Z lm Score = −36.57
NG = 0
DB = 0
SM = 0
Delta_K
Doxycycline
YPL024W RMI1
ZShift = −1.04
Z lm Score = −5.2
NG = 0
DB = 0
SM = 0
Delta_r
Doxycycline
YPL024W RMI1
ZShift = −1.08
Z lm Score = −18.5
NG = 0
DB = 0
SM = 0
Delta_AUC
Doxycycline
YER016W_2 BIM1
ZShift = 1.82
Z lm Score = 20.04
NG = 0
DB = 0
SM = 0
Delta_L
Doxycycline
YER016W_2 BIM1
ZShift = 0.22
Z lm Score = −15.36
NG = 0
DB = 0
SM = 0
Delta_K
Doxycycline
YER016W_2 BIM1
ZShift = −1.59
Z lm Score = −4.21
NG = 0
DB = 0
SM = 0
Delta_r
Doxycycline
YER016W_2 BIM1
ZShift = −0.65
Z lm Score = 2.27
NG = 0
DB = 0
SM = 0
Delta_AUC
Doxycycline
YMR190C SGS1
ZShift = 1.52
Z lm Score = 19.32
NG = 0
DB = 0
SM = 0
Delta_L
Doxycycline
YMR190C SGS1
ZShift = −0.94
Z lm Score = −33.4
NG = 0
DB = 0
SM = 0
Delta_K
Doxycycline
YMR190C SGS1
ZShift = −2.56
Z lm Score = −3.54
NG = 0
DB = 0
SM = 0
Delta_r
Doxycycline
YMR190C SGS1
ZShift = −1.28
Z lm Score = −17.82
NG = 0
DB = 0
SM = 0
Delta_AUC
Doxycycline

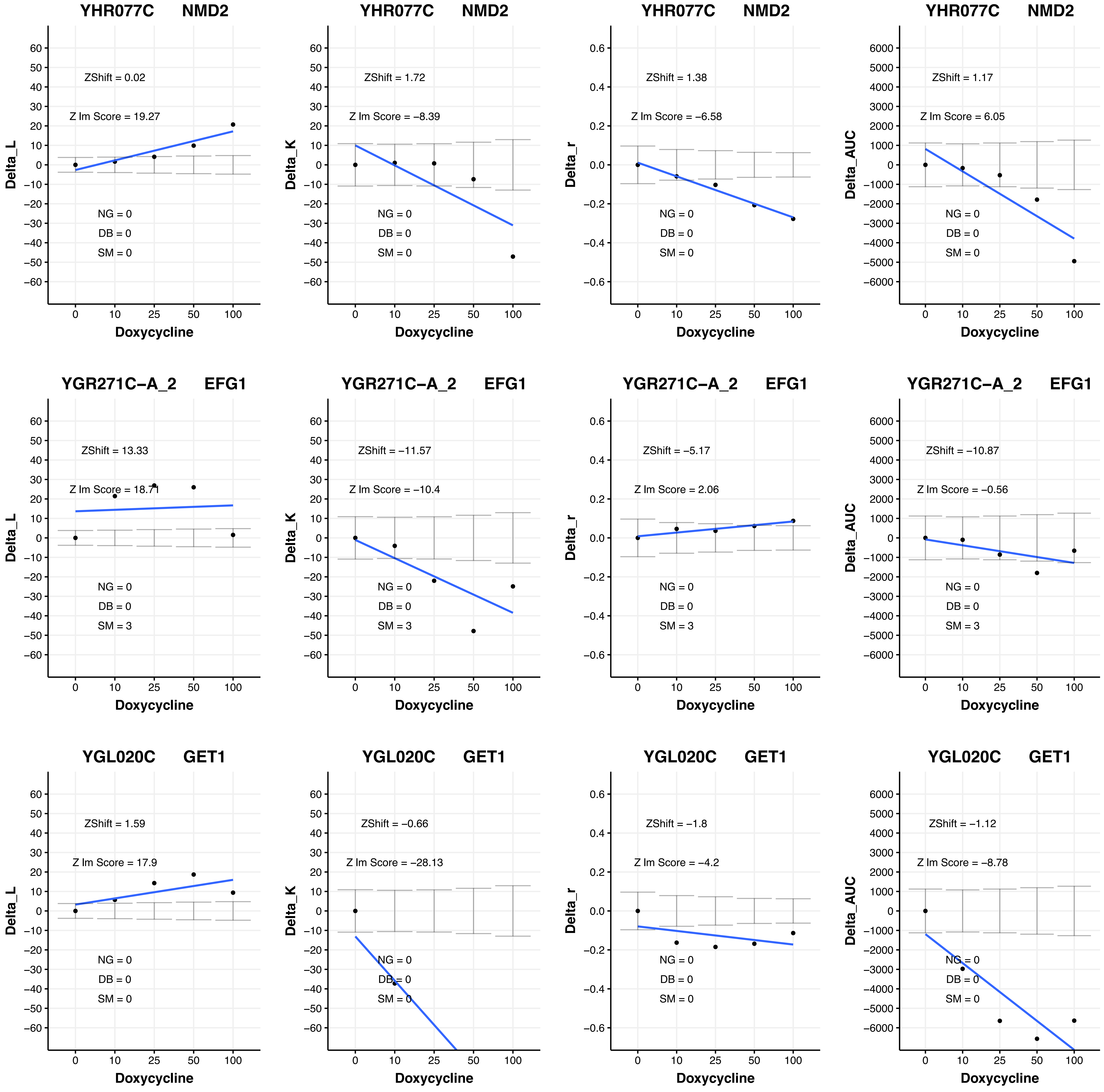

YHR077C NMD2
ZShift = 0.02
Z lm Score = 19.27
Delta_L
NG = 0
DB = 0
SM = 0
Doxycycline
YHR077C NMD2
ZShift = 1.72
Z lm Score = −8.39
Delta_K
NG = 0
DB = 0
SM = 0
Doxycycline
YHR077C NMD2
ZShift = 1.38
Z lm Score = −6.58
Delta_r
NG = 0
DB = 0
SM = 0
Doxycycline
YHR077C NMD2
ZShift = 1.17
Z lm Score = 6.05
Delta_AUC
NG = 0
DB = 0
SM = 0
Doxycycline
YGR271C−A_2 EFG1
ZShift = 13.33
Z lm Score = 18.71
Delta_L
NG = 0
DB = 0
SM = 3
Doxycycline
YGR271C−A_2 EFG1
ZShift = −11.57
Z lm Score = −10.4
Delta_K
NG = 0
DB = 0
SM = 3
Doxycycline
YGR271C−A_2 EFG1
ZShift = −5.17
Z lm Score = 2.06
Delta_r
NG = 0
DB = 0
SM = 3
Doxycycline
YGR271C−A_2 EFG1
ZShift = −10.87
Z lm Score = −0.56
Delta_AUC
NG = 0
DB = 0
SM = 3
Doxycycline
YGL020C GET1
ZShift = 1.59
Z lm Score = 17.9
Delta_L
NG = 0
DB = 0
SM = 0
Doxycycline
YGL020C GET1
ZShift = −0.66
Z lm Score = −28.13
Delta_K
NG = 0
DB = 0
SM = 0
Doxycycline
YGL020C GET1
ZShift = −1.8
Z lm Score = −4.2
Delta_r
NG = 0
DB = 0
SM = 0
Doxycycline
YGL020C GET1
ZShift = −1.12
Z lm Score = −8.78
Delta_AUC
NG = 0
DB = 0
SM = 0
Doxycycline
0
10
25
50
100

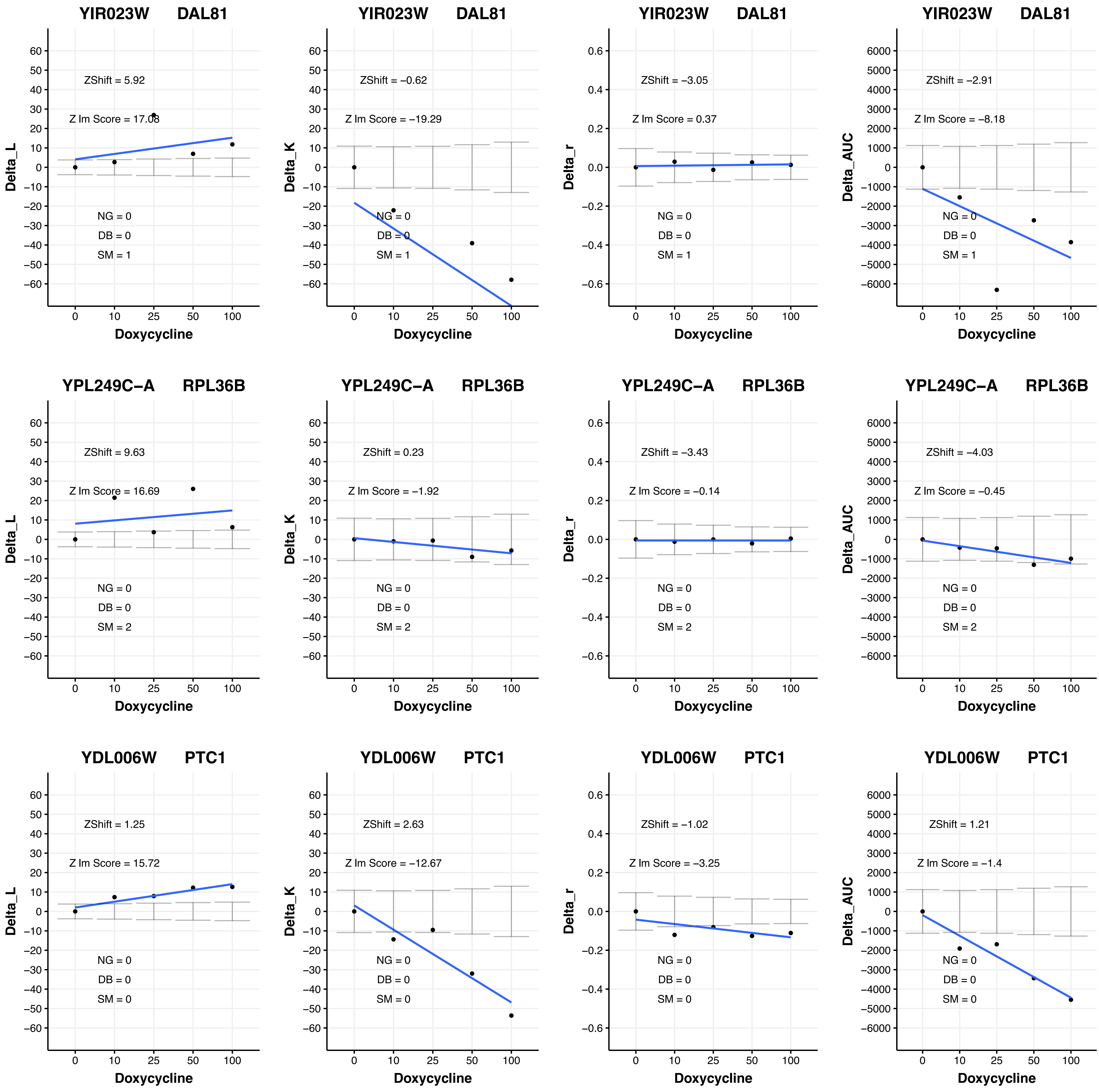
YIR023W DAL81
ZShift = 5.92
Z lm Score = 17.08
NG = 0
DB = 0
SM = 1
Delta_L
Doxycycline
YIR023W DAL81
ZShift = −0.62
Z lm Score = −19.29
NG = 0
DB = 0
SM = 1
Delta_K
Doxycycline
YIR023W DAL81
ZShift = −3.05
Z lm Score = 0.37
NG = 0
DB = 0
SM = 1
Delta_r
Doxycycline
YIR023W DAL81
ZShift = −2.91
Z lm Score = −8.18
NG = 0
DB = 0
SM = 1
Delta_AUC
Doxycycline
YPL249C−A RPL36B
ZShift = 9.63
Z lm Score = 16.69
NG = 0
DB = 0
SM = 2
Delta_L
Doxycycline
YPL249C−A RPL36B
ZShift = 0.23
Z lm Score = −1.92
NG = 0
DB = 0
SM = 2
Delta_K
Doxycycline
YPL249C−A RPL36B
ZShift = −3.43
Z lm Score = −0.14
NG = 0
DB = 0
SM = 2
Delta_r
Doxycycline
YPL249C−A RPL36B
ZShift = −4.03
Z lm Score = −0.45
NG = 0
DB = 0
SM = 2
Delta_AUC
Doxycycline
YDL006W PTC1
ZShift = 1.25
Z lm Score = 15.72
NG = 0
DB = 0
SM = 0
Delta_L
Doxycycline
YDL006W PTC1
ZShift = 2.63
Z lm Score = −12.67
NG = 0
DB = 0
SM = 0
Delta_K
Doxycycline
YDL006W PTC1
ZShift = −1.02
Z lm Score = −3.25
NG = 0
DB = 0
SM = 0
Delta_r
Doxycycline
YDL006W PTC1
ZShift = 1.21
Z lm Score = −1.4
NG = 0
DB = 0
SM = 0
Delta_AUC
Doxycycline

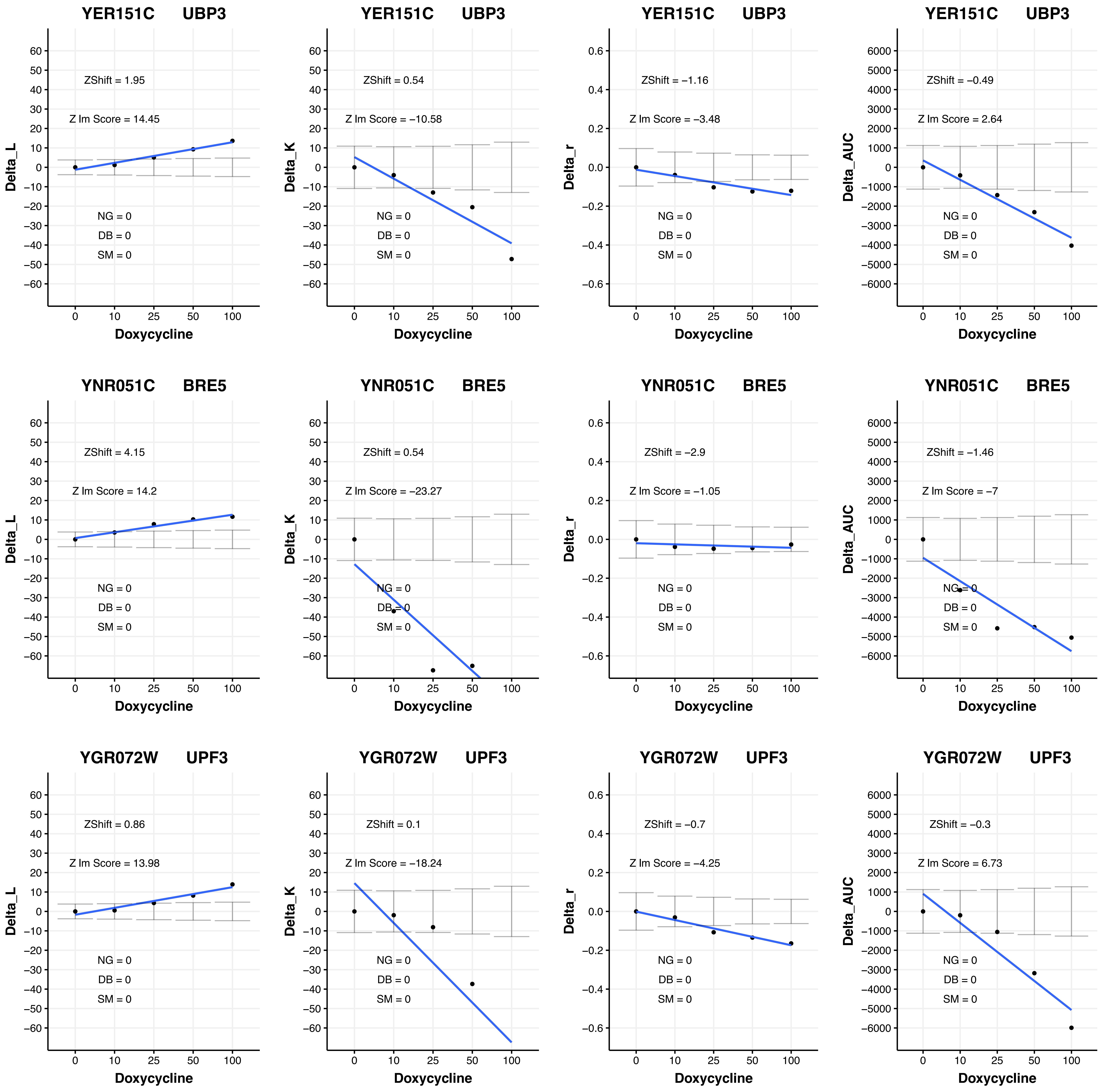
YER151C UBP3
ZShift = 1.95
Z lm Score = 14.45
NG = 0
DB = 0
SM = 0
Delta_L
Doxycycline
YER151C UBP3
ZShift = 0.54
Z lm Score = −10.58
NG = 0
DB = 0
SM = 0
Delta_K
Doxycycline
YER151C UBP3
ZShift = −1.16
Z lm Score = −3.48
NG = 0
DB = 0
SM = 0
Delta_r
Doxycycline
YER151C UBP3
ZShift = −0.49
Z lm Score = 2.64
NG = 0
DB = 0
SM = 0
Delta_AUC
Doxycycline
YNR051C BRE5
ZShift = 4.15
Z lm Score = 14.2
NG = 0
DB = 0
SM = 0
Delta_L
Doxycycline
YNR051C BRE5
ZShift = 0.54
Z lm Score = −23.27
NG = 0
DB = 0
SM = 0
Delta_K
Doxycycline
YNR051C BRE5
ZShift = −2.9
Z lm Score = −1.05
NG = 0
DB = 0
SM = 0
Delta_r
Doxycycline
YNR051C BRE5
ZShift = −1.46
Z lm Score = −7
NG = 0
DB = 0
SM = 0
Delta_AUC
Doxycycline
YGR072W UPF3
ZShift = 0.86
Z lm Score = 13.98
NG = 0
DB = 0
SM = 0
Delta_L
Doxycycline
YGR072W UPF3
ZShift = 0.1
Z lm Score = −18.24
NG = 0
DB = 0
SM = 0
Delta_K
Doxycycline
YGR072W UPF3
ZShift = −0.7
Z lm Score = −4.25
NG = 0
DB = 0
SM = 0
Delta_r
Doxycycline
YGR072W UPF3
ZShift = −0.3
Z lm Score = 6.73
NG = 0
DB = 0
SM = 0
Delta_AUC
Doxycycline

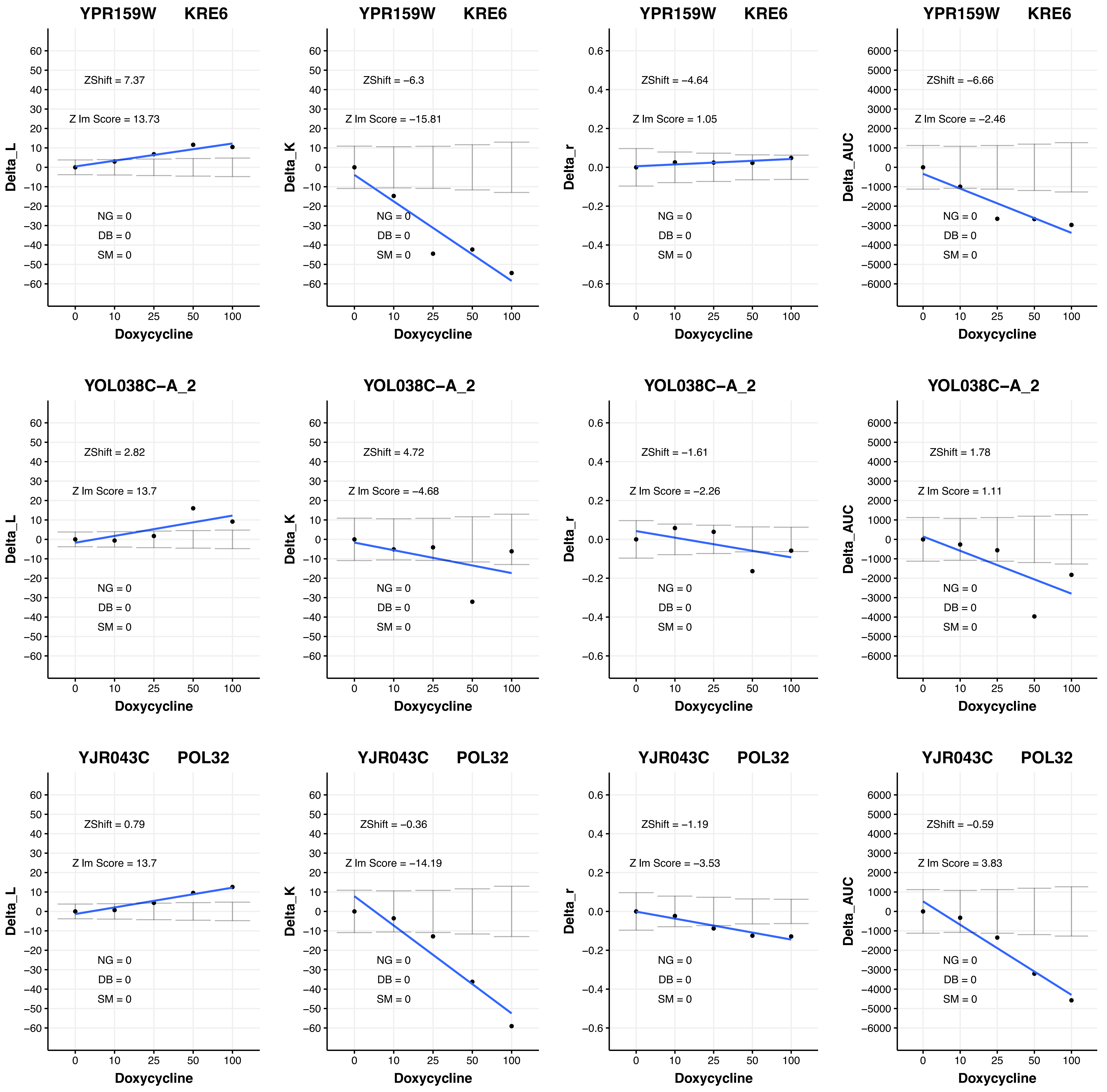

YPR159W KRE6
ZShift = 7.37
Z lm Score = 13.73
NG = 0
DB = 0
SM = 0
Delta_L
Doxycycline
YPR159W KRE6
ZShift = −6.3
Z lm Score = −15.81
NG = 0
DB = 0
SM = 0
Delta_K
Doxycycline
YPR159W KRE6
ZShift = −4.64
Z lm Score = 1.05
NG = 0
DB = 0
SM = 0
Delta_r
Doxycycline
YPR159W KRE6
ZShift = −6.66
Z lm Score = −2.46
NG = 0
DB = 0
SM = 0
Delta_AUC
Doxycycline
YOL038C−A_2
ZShift = 2.82
Z lm Score = 13.7
NG = 0
DB = 0
SM = 0
Delta_L
Doxycycline
YOL038C−A_2
ZShift = 4.72
Z lm Score = −4.68
NG = 0
DB = 0
SM = 0
Delta_K
Doxycycline
YOL038C−A_2
ZShift = −1.61
Z lm Score = −2.26
NG = 0
DB = 0
SM = 0
Delta_r
Doxycycline
YOL038C−A_2
ZShift = 1.78
Z lm Score = 1.11
NG = 0
DB = 0
SM = 0
Delta_AUC
Doxycycline
YJR043C POL32
ZShift = 0.79
Z lm Score = 13.7
NG = 0
DB = 0
SM = 0
Delta_L
Doxycycline
YJR043C POL32
ZShift = −0.36
Z lm Score = −14.19
NG = 0
DB = 0
SM = 0
Delta_K
Doxycycline
YJR043C POL32
ZShift = −1.19
Z lm Score = −3.53
NG = 0
DB = 0
SM = 0
Delta_r
Doxycycline
YJR043C POL32
ZShift = −0.59
Z lm Score = 3.83
NG = 0
DB = 0
SM = 0
Delta_AUC
Doxycycline

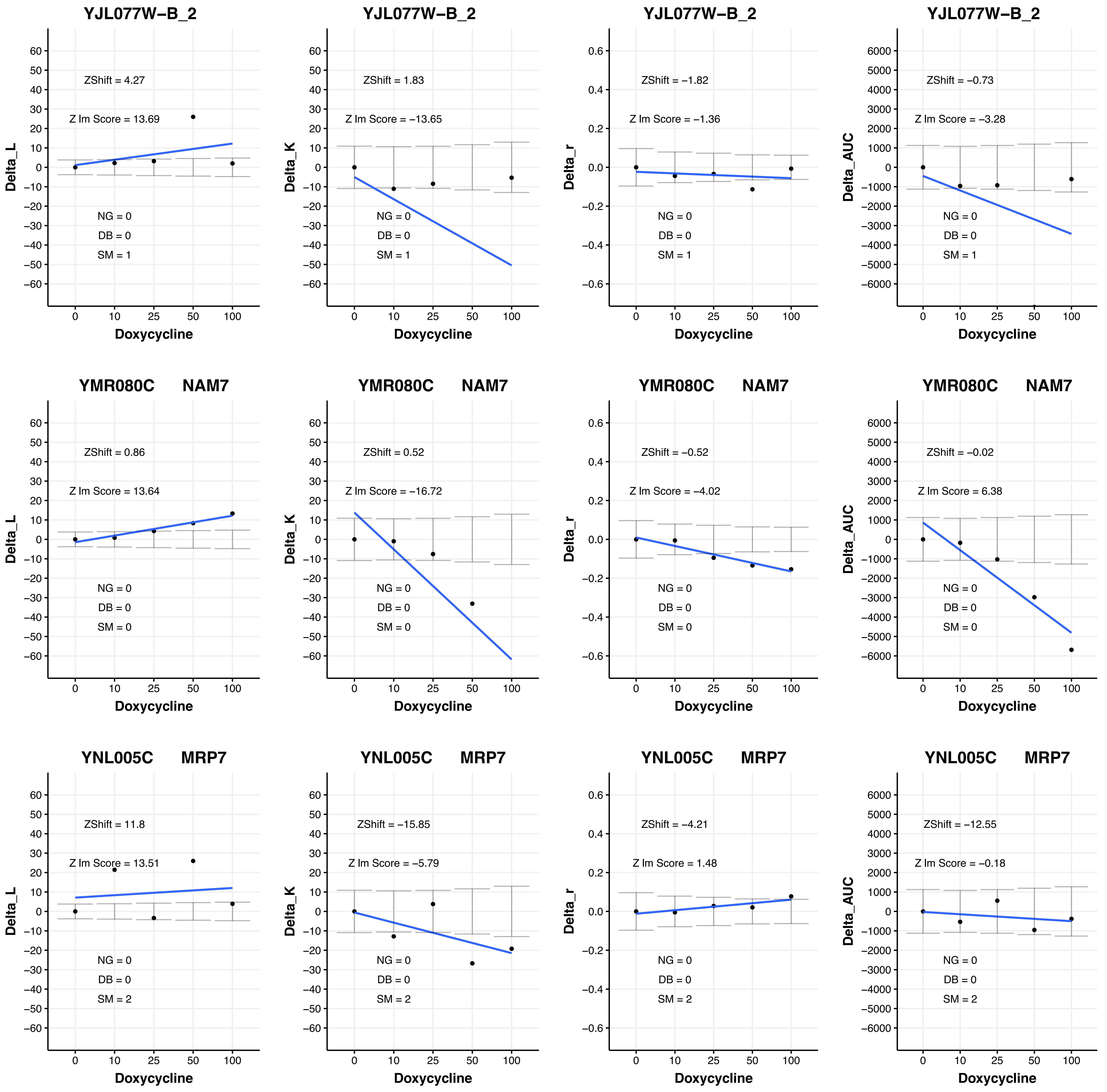

YJL077W−B_2
ZShift = 4.27
Z lm Score = 13.69
NG = 0
DB = 0
SM = 1
Delta_L
Doxycycline
YJL077W−B_2
ZShift = 1.83
Z lm Score = −13.65
NG = 0
DB = 0
SM = 1
Delta_K
Doxycycline
YJL077W−B_2
ZShift = −1.82
Z lm Score = −1.36
NG = 0
DB = 0
SM = 1
Delta_r
Doxycycline
YJL077W−B_2
ZShift = −0.73
Z lm Score = −3.28
NG = 0
DB = 0
SM = 1
Delta_AUC
Doxycycline
YMR080C NAM7
ZShift = 0.86
Z lm Score = 13.64
NG = 0
DB = 0
SM = 0
Delta_L
Doxycycline
YMR080C NAM7
ZShift = 0.52
Z lm Score = −16.72
NG = 0
DB = 0
SM = 0
Delta_K
Doxycycline
YMR080C NAM7
ZShift = −0.52
Z lm Score = −4.02
NG = 0
DB = 0
SM = 0
Delta_r
Doxycycline
YMR080C NAM7
ZShift = −0.02
Z lm Score = 6.38
NG = 0
DB = 0
SM = 0
Delta_AUC
Doxycycline
YNL005C MRP7
ZShift = 11.8
Z lm Score = 13.51
NG = 0
DB = 0
SM = 2
Delta_L
Doxycycline
YNL005C MRP7
ZShift = −15.85
Z lm Score = −5.79
NG = 0
DB = 0
SM = 2
Delta_K
Doxycycline
YNL005C MRP7
ZShift = −4.21
Z lm Score = 1.48
NG = 0
DB = 0
SM = 2
Delta_r
Doxycycline
YNL005C MRP7
ZShift = −12.55
Z lm Score = −0.18
NG = 0
DB = 0
SM = 2
Delta_AUC
Doxycycline

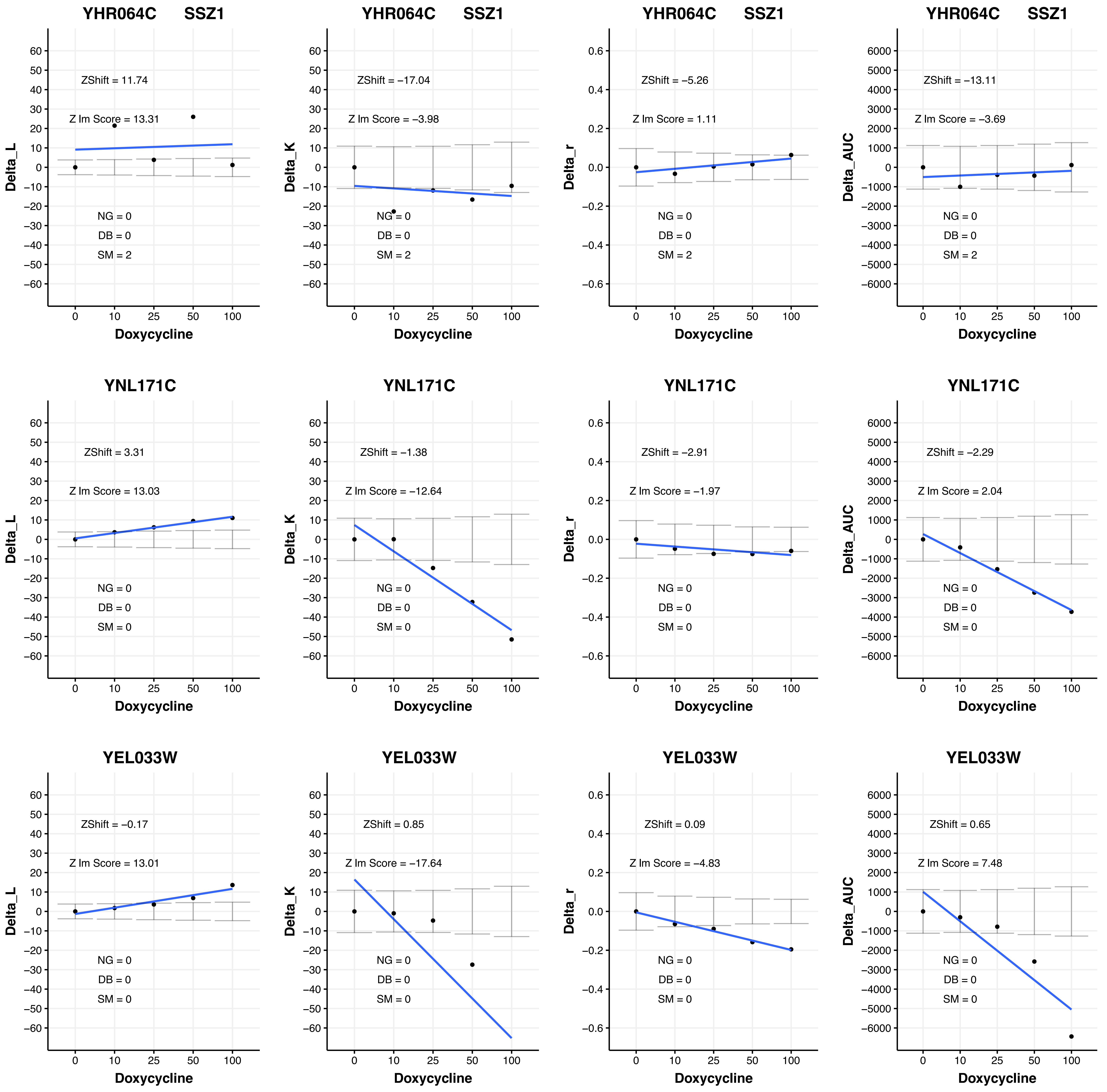

YHR064C SSZ1
ZShift = 11.74
Z lm Score = 13.31
NG = 0
DB = 0
SM = 2
Delta_L
Doxycycline
YHR064C SSZ1
ZShift = −17.04
Z lm Score = −3.98
NG = 0
DB = 0
SM = 2
Delta_K
Doxycycline
YHR064C SSZ1
ZShift = −5.26
Z lm Score = 1.11
NG = 0
DB = 0
SM = 2
Delta_r
Doxycycline
YHR064C SSZ1
ZShift = −13.11
Z lm Score = −3.69
NG = 0
DB = 0
SM = 2
Delta_AUC
Doxycycline
YNL171C
ZShift = 3.31
Z lm Score = 13.03
NG = 0
DB = 0
SM = 0
Delta_L
Doxycycline
YNL171C
ZShift = −1.38
Z lm Score = −12.64
NG = 0
DB = 0
SM = 0
Delta_K
Doxycycline
YNL171C
ZShift = −2.91
Z lm Score = −1.97
NG = 0
DB = 0
SM = 0
Delta_r
Doxycycline
YNL171C
ZShift = −2.29
Z lm Score = 2.04
NG = 0
DB = 0
SM = 0
Delta_AUC
Doxycycline
YEL033W
ZShift = −0.17
Z lm Score = 13.01
NG = 0
DB = 0
SM = 0
Delta_L
Doxycycline
YEL033W
ZShift = 0.85
Z lm Score = −17.64
NG = 0
DB = 0
SM = 0
Delta_K
Doxycycline
YEL033W
ZShift = 0.09
Z lm Score = −4.83
NG = 0
DB = 0
SM = 0
Delta_r
Doxycycline
YEL033W
ZShift = 0.65
Z lm Score = 7.48
NG = 0
DB = 0
SM = 0
Delta_AUC
Doxycycline

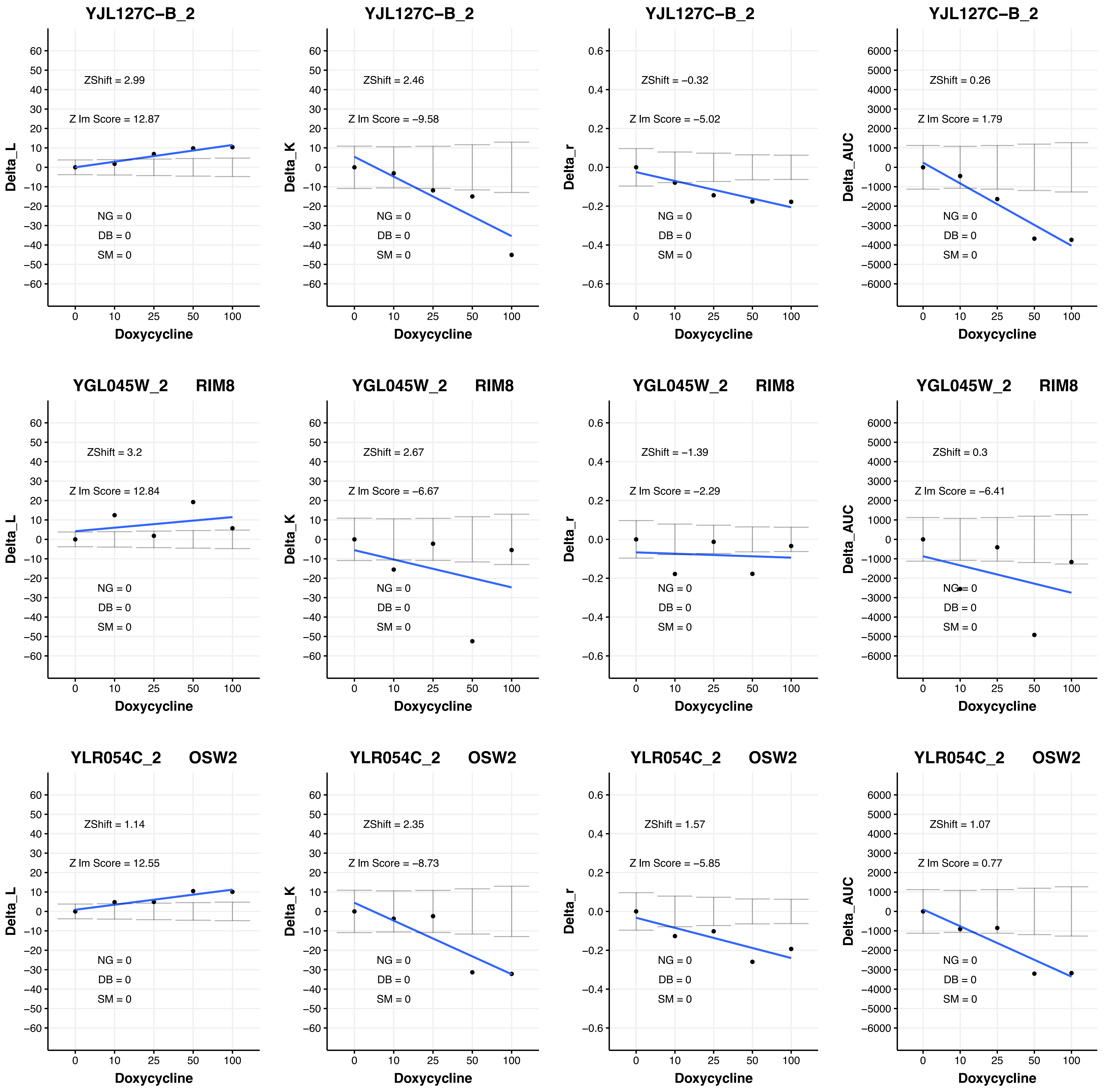

YJL127C–B_2
ZShift = 2.99
Z lm Score = 12.87
NG = 0
DB = 0
SM = 0
Delta_L
Doxycycline
YJL127C–B_2
ZShift = 2.46
Z lm Score = −9.58
NG = 0
DB = 0
SM = 0
Delta_K
Doxycycline
YJL127C–B_2
ZShift = −0.32
Z lm Score = −5.02
NG = 0
DB = 0
SM = 0
Delta_r
Doxycycline
YJL127C–B_2
ZShift = 0.26
Z lm Score = 1.79
NG = 0
DB = 0
SM = 0
Delta_AUC
Doxycycline
YGL045W_2 RIM8
ZShift = 3.2
Z lm Score = 12.84
NG = 0
DB = 0
SM = 0
Delta_L
Doxycycline
YGL045W_2 RIM8
ZShift = 2.67
Z lm Score = −6.67
NG = 0
DB = 0
SM = 0
Delta_K
Doxycycline
YGL045W_2 RIM8
ZShift = −1.39
Z lm Score = −2.29
NG = 0
DB = 0
SM = 0
Delta_r
Doxycycline
YGL045W_2 RIM8
ZShift = 0.3
Z lm Score = −6.41
NG = 0
DB = 0
SM = 0
Delta_AUC
Doxycycline
YLR054C_2 OSW2
ZShift = 1.14
Z lm Score = 12.55
NG = 0
DB = 0
SM = 0
Delta_L
Doxycycline
YLR054C_2 OSW2
ZShift = 2.35
Z lm Score = −8.73
NG = 0
DB = 0
SM = 0
Delta_K
Doxycycline
YLR054C_2 OSW2
ZShift = 1.57
Z lm Score = −5.85
NG = 0
DB = 0
SM = 0
Delta_r
Doxycycline
YLR054C_2 OSW2
ZShift = 1.07
Z lm Score = 0.77
NG = 0
DB = 0
SM = 0
Delta_AUC
Doxycycline

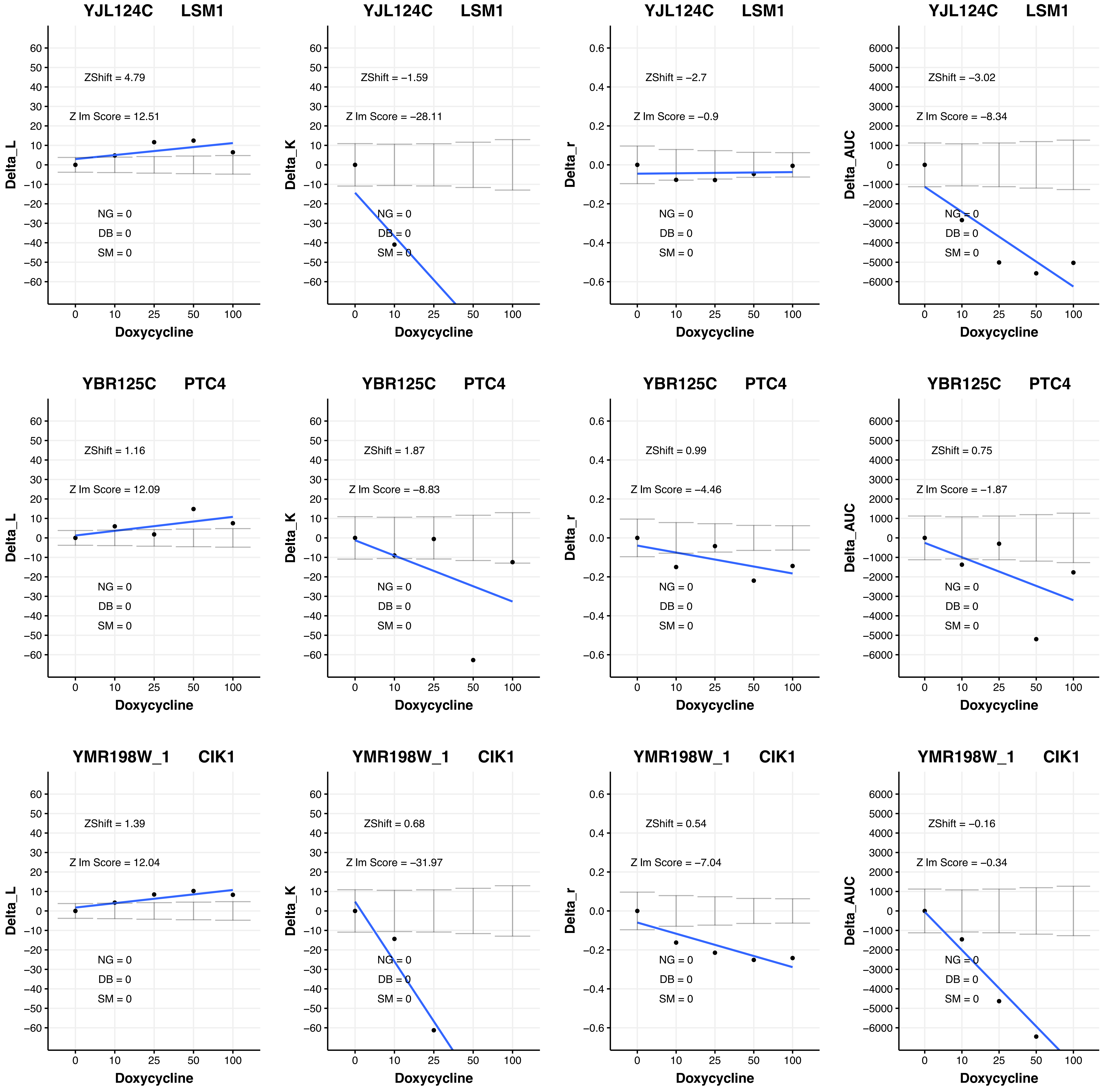
YJL124C LSM1
ZShift = 4.79
Z lm Score = 12.51
NG = 0
DB = 0
SM = 0
Delta_L
Doxycycline
YJL124C LSM1
ZShift = −1.59
Z lm Score = −28.11
NG = 0
DB = 0
SM = 0
Delta_K
Doxycycline
YJL124C LSM1
ZShift = −2.7
Z lm Score = −0.9
NG = 0
DB = 0
SM = 0
Delta_r
Doxycycline
YJL124C LSM1
ZShift = −3.02
Z lm Score = −8.34
NG = 0
DB = 0
SM = 0
Delta_AUC
Doxycycline
YBR125C PTC4
ZShift = 1.16
Z lm Score = 12.09
NG = 0
DB = 0
SM = 0
Delta_L
Doxycycline
YBR125C PTC4
ZShift = 1.87
Z lm Score = −8.83
NG = 0
DB = 0
SM = 0
Delta_K
Doxycycline
YBR125C PTC4
ZShift = 0.99
Z lm Score = −4.46
NG = 0
DB = 0
SM = 0
Delta_r
Doxycycline
YBR125C PTC4
ZShift = 0.75
Z lm Score = −1.87
NG = 0
DB = 0
SM = 0
Delta_AUC
Doxycycline
YMR198W_1 CIK1
ZShift = 1.39
Z lm Score = 12.04
NG = 0
DB = 0
SM = 0
Delta_L
Doxycycline
YMR198W_1 CIK1
ZShift = 0.68
Z lm Score = −31.97
NG = 0
DB = 0
SM = 0
Delta_K
Doxycycline
YMR198W_1 CIK1
ZShift = 0.54
Z lm Score = −7.04
NG = 0
DB = 0
SM = 0
Delta_r
Doxycycline
YMR198W_1 CIK1
ZShift = −0.16
Z lm Score = −0.34
NG = 0
DB = 0
SM = 0
Delta_AUC
Doxycycline

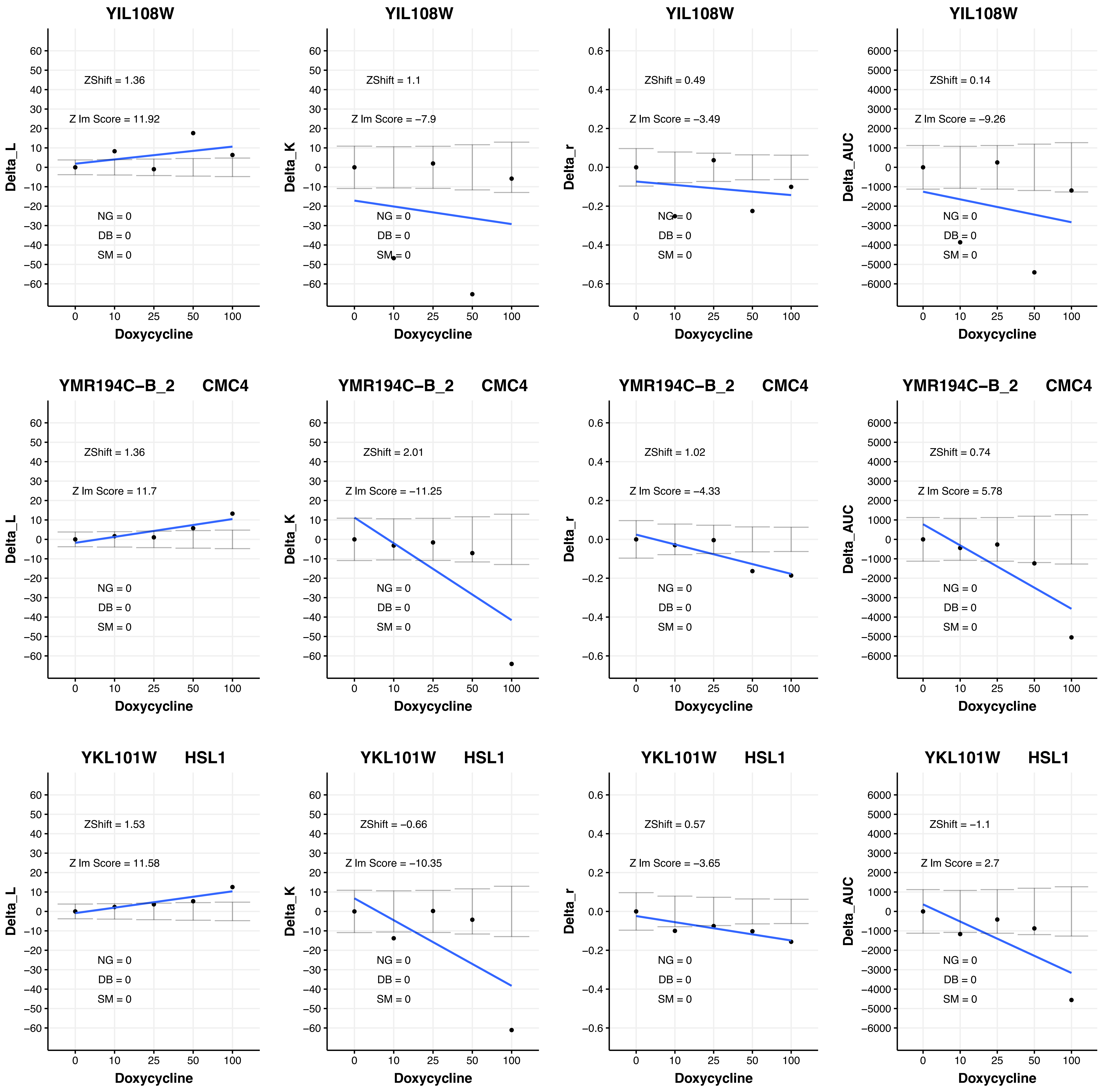
YIL108W
ZShift = 1.36
Z lm Score = 11.92
NG = 0
DB = 0
SM = 0
Delta_L
Doxycycline
YIL108W
ZShift = 1.1
Z lm Score = −7.9
NG = 0
DB = 0
SM = 0
Delta_K
Doxycycline
YIL108W
ZShift = 0.49
Z lm Score = −3.49
NG = 0
DB = 0
SM = 0
Delta_r
Doxycycline
YIL108W
ZShift = 0.14
Z lm Score = −9.26
NG = 0
DB = 0
SM = 0
Delta_AUC
Doxycycline
YMR194C−B_2 CMC4
ZShift = 1.36
Z lm Score = 11.7
NG = 0
DB = 0
SM = 0
Delta_L
Doxycycline
YMR194C−B_2 CMC4
ZShift = 2.01
Z lm Score = −11.25
NG = 0
DB = 0
SM = 0
Delta_K
Doxycycline
YMR194C−B_2 CMC4
ZShift = 1.02
Z lm Score = −4.33
NG = 0
DB = 0
SM = 0
Delta_r
Doxycycline
YMR194C−B_2 CMC4
ZShift = 0.74
Z lm Score = 5.78
NG = 0
DB = 0
SM = 0
Delta_AUC
Doxycycline
YKL101W HSL1
ZShift = 1.53
Z lm Score = 11.58
NG = 0
DB = 0
SM = 0
Delta_L
Doxycycline
YKL101W HSL1
ZShift = −0.66
Z lm Score = −10.35
NG = 0
DB = 0
SM = 0
Delta_K
Doxycycline
YKL101W HSL1
ZShift = 0.57
Z lm Score = −3.65
NG = 0
DB = 0
SM = 0
Delta_r
Doxycycline
YKL101W HSL1
ZShift = −1.1
Z lm Score = 2.7
NG = 0
DB = 0
SM = 0
Delta_AUC
Doxycycline

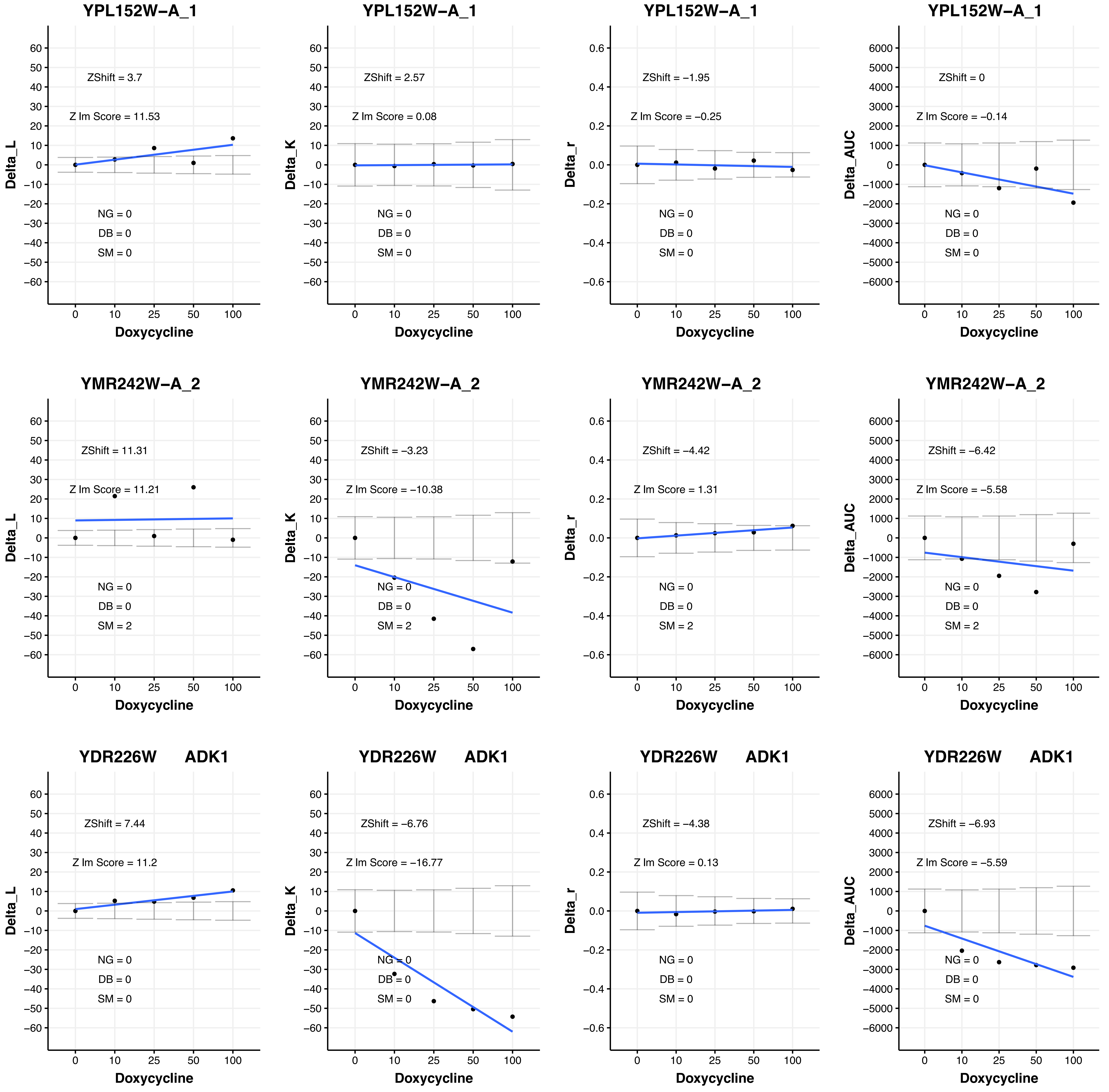

YPL152W−A_1
ZShift = 3.7
Z lm Score = 11.53
NG = 0
DB = 0
SM = 0
Delta_L
Doxycycline
YPL152W−A_1
ZShift = 2.57
Z lm Score = 0.08
NG = 0
DB = 0
SM = 0
Delta_K
Doxycycline
YPL152W−A_1
ZShift = −1.95
Z lm Score = −0.25
NG = 0
DB = 0
SM = 0
Delta_r
Doxycycline
YPL152W−A_1
ZShift = 0
Z lm Score = −0.14
NG = 0
DB = 0
SM = 0
Delta_AUC
Doxycycline
YMR242W−A_2
ZShift = 11.31
Z lm Score = 11.21
NG = 0
DB = 0
SM = 2
Delta_L
Doxycycline
YMR242W−A_2
ZShift = −3.23
Z lm Score = −10.38
NG = 0
DB = 0
SM = 2
Delta_K
Doxycycline
YMR242W−A_2
ZShift = −4.42
Z lm Score = 1.31
NG = 0
DB = 0
SM = 2
Delta_r
Doxycycline
YMR242W−A_2
ZShift = −6.42
Z lm Score = −5.58
NG = 0
DB = 0
SM = 2
Delta_AUC
Doxycycline
YDR226W ADK1
ZShift = 7.44
Z lm Score = 11.2
NG = 0
DB = 0
SM = 0
Delta_L
Doxycycline
YDR226W ADK1
ZShift = −6.76
Z lm Score = −16.77
NG = 0
DB = 0
SM = 0
Delta_K
Doxycycline
YDR226W ADK1
ZShift = −4.38
Z lm Score = 0.13
NG = 0
DB = 0
SM = 0
Delta_r
Doxycycline
YDR226W ADK1
ZShift = −6.93
Z lm Score = −5.59
NG = 0
DB = 0
SM = 0
Delta_AUC
Doxycycline

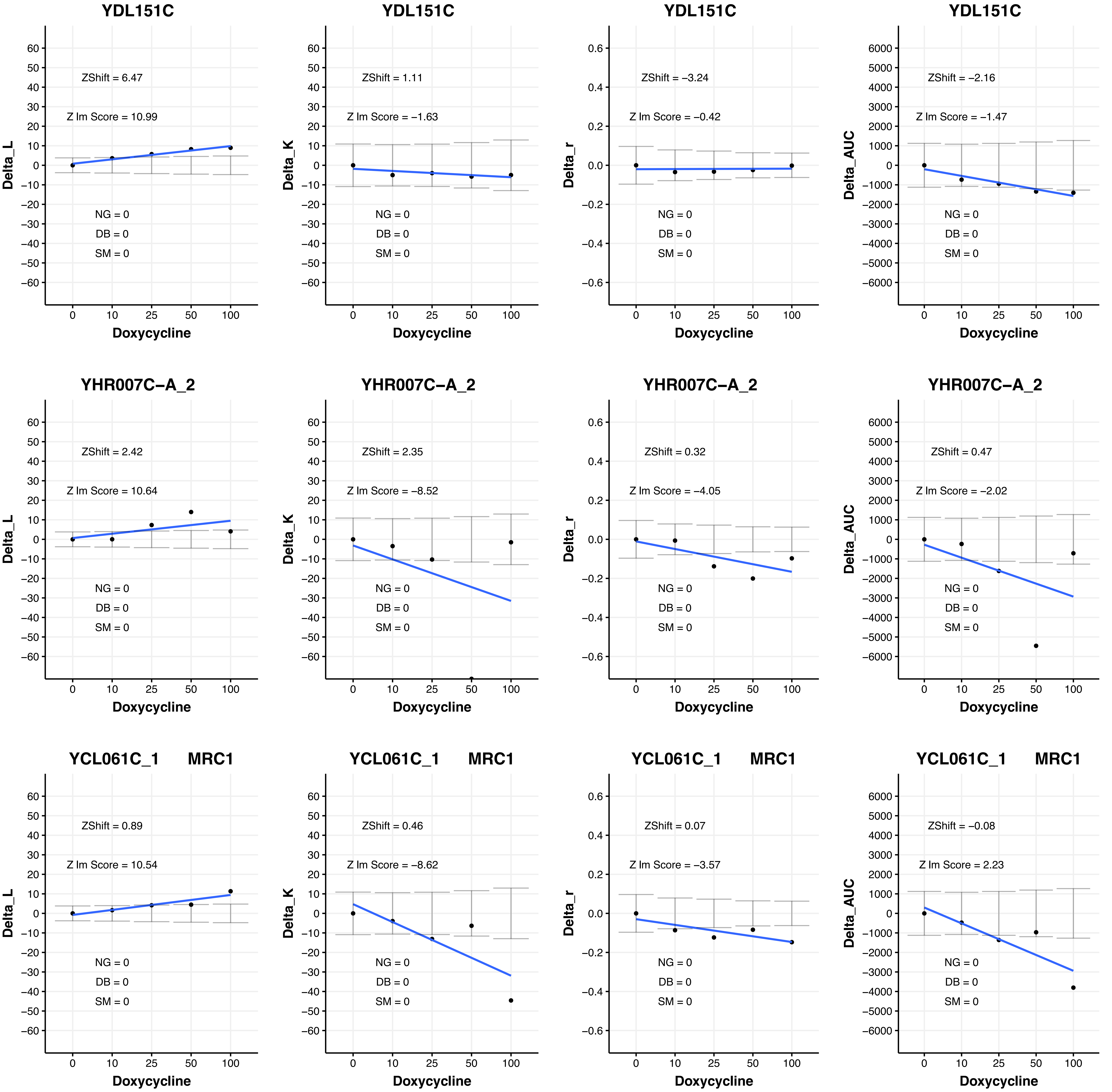
YDL151C
Delta_L
ZShift = 6.47
Z lm Score = 10.99
NG = 0
DB = 0
SM = 0
Doxycycline
YDL151C
Delta_K
ZShift = 1.11
Z lm Score = −1.63
NG = 0
DB = 0
SM = 0
Doxycycline
YDL151C
Delta_r
ZShift = −3.24
Z lm Score = −0.42
NG = 0
DB = 0
SM = 0
Doxycycline
YDL151C
Delta_AUC
ZShift = −2.16
Z lm Score = −1.47
NG = 0
DB = 0
SM = 0
Doxycycline
YHR007C−A_2
Delta_L
ZShift = 2.42
Z lm Score = 10.64
NG = 0
DB = 0
SM = 0
Doxycycline
YHR007C−A_2
Delta_K
ZShift = 2.35
Z lm Score = −8.52
NG = 0
DB = 0
SM = 0
Doxycycline
YHR007C−A_2
Delta_r
ZShift = 0.32
Z lm Score = −4.05
NG = 0
DB = 0
SM = 0
Doxycycline
YHR007C−A_2
Delta_AUC
ZShift = 0.47
Z lm Score = −2.02
NG = 0
DB = 0
SM = 0
Doxycycline
YCL061C_1 MRC1
Delta_L
ZShift = 0.89
Z lm Score = 10.54
NG = 0
DB = 0
SM = 0
Doxycycline
YCL061C_1 MRC1
Delta_K
ZShift = 0.46
Z lm Score = −8.62
NG = 0
DB = 0
SM = 0
Doxycycline
YCL061C_1 MRC1
Delta_r
ZShift = 0.07
Z lm Score = −3.57
NG = 0
DB = 0
SM = 0
Doxycycline
YCL061C_1 MRC1
Delta_AUC
ZShift = −0.08
Z lm Score = 2.23
NG = 0
DB = 0
SM = 0
Doxycycline

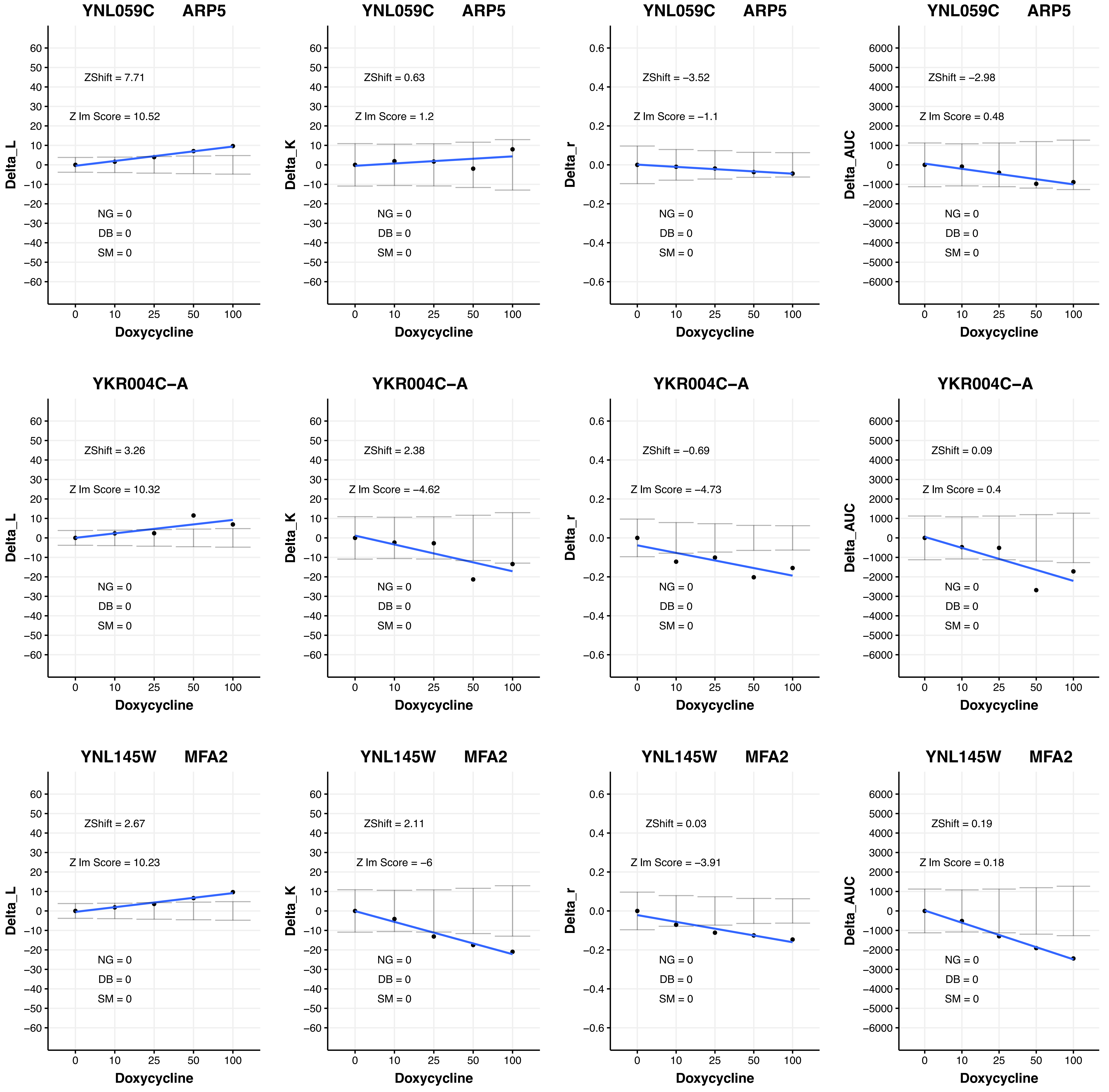

YNL059C ARP5
ZShift = 7.71
Z lm Score = 10.52
NG = 0
DB = 0
SM = 0
Delta_L
Doxycycline
YNL059C ARP5
ZShift = 0.63
Z lm Score = 1.2
NG = 0
DB = 0
SM = 0
Delta_K
Doxycycline
YNL059C ARP5
ZShift = −3.52
Z lm Score = −1.1
NG = 0
DB = 0
SM = 0
Delta_r
Doxycycline
YNL059C ARP5
ZShift = −2.98
Z lm Score = 0.48
NG = 0
DB = 0
SM = 0
Delta_AUC
Doxycycline
YKR004C−A
ZShift = 3.26
Z lm Score = 10.32
NG = 0
DB = 0
SM = 0
Delta_L
Doxycycline
YKR004C−A
ZShift = 2.38
Z lm Score = −4.62
NG = 0
DB = 0
SM = 0
Delta_K
Doxycycline
YKR004C−A
ZShift = −0.69
Z lm Score = −4.73
NG = 0
DB = 0
SM = 0
Delta_r
Doxycycline
YKR004C−A
ZShift = 0.09
Z lm Score = 0.4
NG = 0
DB = 0
SM = 0
Delta_AUC
Doxycycline
YNL145W MFA2
ZShift = 2.67
Z lm Score = 10.23
NG = 0
DB = 0
SM = 0
Delta_L
Doxycycline
YNL145W MFA2
ZShift = 2.11
Z lm Score = −6
NG = 0
DB = 0
SM = 0
Delta_K
Doxycycline
YNL145W MFA2
ZShift = 0.03
Z lm Score = −3.91
NG = 0
DB = 0
SM = 0
Delta_r
Doxycycline
YNL145W MFA2
ZShift = 0.19
Z lm Score = 0.18
NG = 0
DB = 0
SM = 0
Delta_AUC
Doxycycline

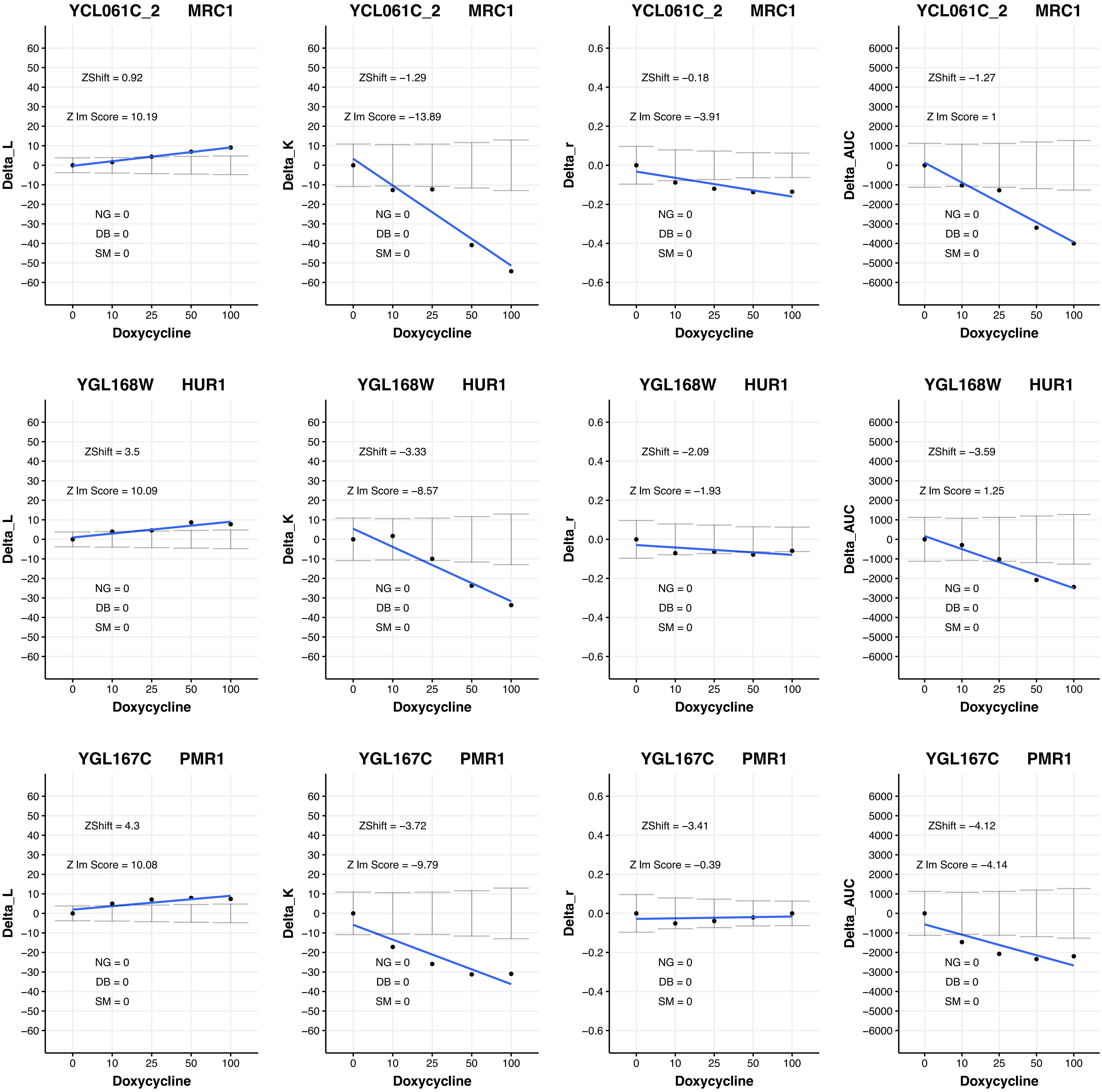
YCL061C_2 MRC1
ZShift = 0.92
Z lm Score = 10.19
NG = 0
DB = 0
SM = 0
Delta_L
Doxycycline
YCL061C_2 MRC1
ZShift = −1.29
Z lm Score = −13.89
NG = 0
DB = 0
SM = 0
Delta_K
Doxycycline
YCL061C_2 MRC1
ZShift = −0.18
Z lm Score = −3.91
NG = 0
DB = 0
SM = 0
Delta_r
Doxycycline
YCL061C_2 MRC1
ZShift = −1.27
Z lm Score = 1
NG = 0
DB = 0
SM = 0
Delta_AUC
Doxycycline
YGL168W HUR1
ZShift = 3.5
Z lm Score = 10.09
NG = 0
DB = 0
SM = 0
Delta_L
Doxycycline
YGL168W HUR1
ZShift = −3.33
Z lm Score = −8.57
NG = 0
DB = 0
SM = 0
Delta_K
Doxycycline
YGL168W HUR1
ZShift = −2.09
Z lm Score = −1.93
NG = 0
DB = 0
SM = 0
Delta_r
Doxycycline
YGL168W HUR1
ZShift = −3.59
Z lm Score = 1.25
NG = 0
DB = 0
SM = 0
Delta_AUC
Doxycycline
YGL167C PMR1
ZShift = 4.3
Z lm Score = 10.08
NG = 0
DB = 0
SM = 0
Delta_L
Doxycycline
YGL167C PMR1
ZShift = −3.72
Z lm Score = −9.79
NG = 0
DB = 0
SM = 0
Delta_K
Doxycycline
YGL167C PMR1
ZShift = −3.41
Z lm Score = −0.39
NG = 0
DB = 0
SM = 0
Delta_r
Doxycycline
YGL167C PMR1
ZShift = −4.12
Z lm Score = −4.14
NG = 0
DB = 0
SM = 0
Delta_AUC
Doxycycline

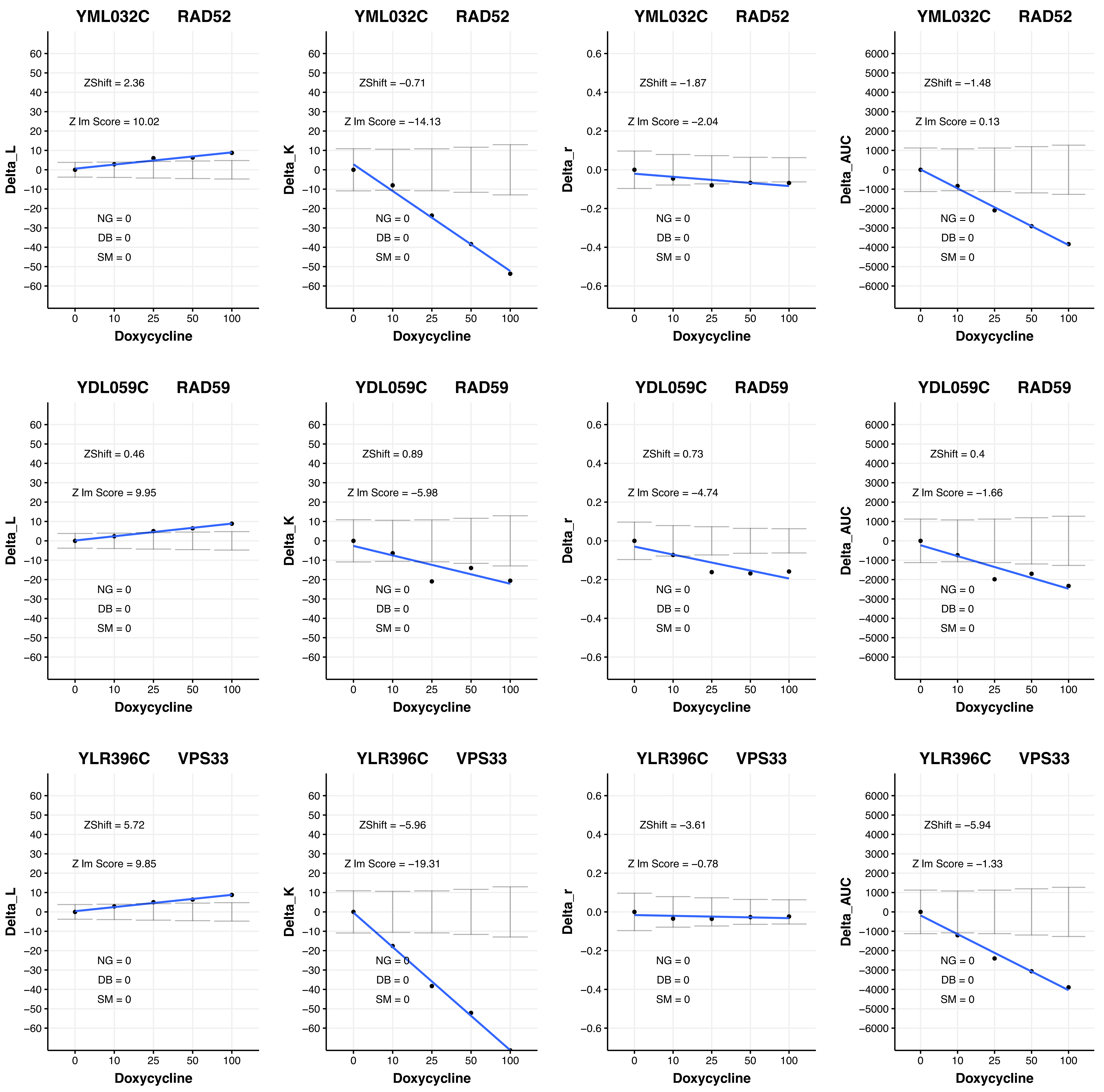

YML032C RAD52
Delta_L
ZShift = 2.36
Z lm Score = 10.02
NG = 0
DB = 0
SM = 0
Doxycycline
YML032C RAD52
Delta_K
ZShift = −0.71
Z lm Score = −14.13
NG = 0
DB = 0
SM = 0
Doxycycline
YML032C RAD52
Delta_r
ZShift = −1.87
Z lm Score = −2.04
NG = 0
DB = 0
SM = 0
Doxycycline
YML032C RAD52
Delta_AUC
ZShift = −1.48
Z lm Score = 0.13
NG = 0
DB = 0
SM = 0
Doxycycline
YDL059C RAD59
Delta_L
ZShift = 0.46
Z lm Score = 9.95
NG = 0
DB = 0
SM = 0
Doxycycline
YDL059C RAD59
Delta_K
ZShift = 0.89
Z lm Score = −5.98
NG = 0
DB = 0
SM = 0
Doxycycline
YDL059C RAD59
Delta_r
ZShift = 0.73
Z lm Score = −4.74
NG = 0
DB = 0
SM = 0
Doxycycline
YDL059C RAD59
Delta_AUC
ZShift = 0.4
Z lm Score = −1.66
NG = 0
DB = 0
SM = 0
Doxycycline
YLR396C VPS33
Delta_L
ZShift = 5.72
Z lm Score = 9.85
NG = 0
DB = 0
SM = 0
Doxycycline
YLR396C VPS33
Delta_K
ZShift = −5.96
Z lm Score = −19.31
NG = 0
DB = 0
SM = 0
Doxycycline
YLR396C VPS33
Delta_r
ZShift = −3.61
Z lm Score = −0.78
NG = 0
DB = 0
SM = 0
Doxycycline
YLR396C VPS33
Delta_AUC
ZShift = −5.94
Z lm Score = −1.33
NG = 0
DB = 0
SM = 0
Doxycycline

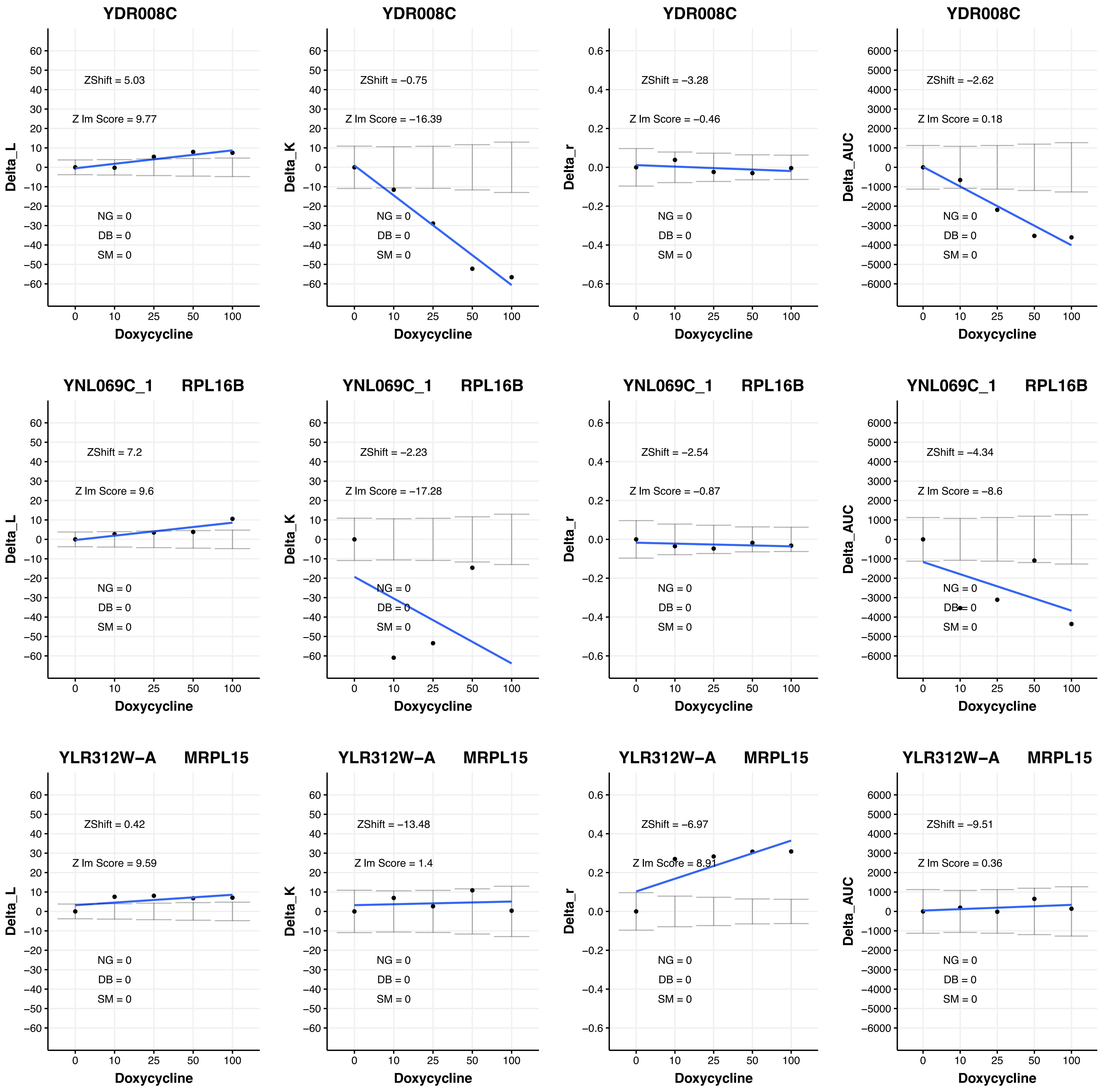
YDR008C
ZShift = 5.03
Z lm Score = 9.77
NG = 0
DB = 0
SM = 0
Delta_L
Doxycycline
YDR008C
ZShift = −0.75
Z lm Score = −16.39
NG = 0
DB = 0
SM = 0
Delta_K
Doxycycline
YDR008C
ZShift = −3.28
Z lm Score = −0.46
NG = 0
DB = 0
SM = 0
Delta_r
Doxycycline
YDR008C
ZShift = −2.62
Z lm Score = 0.18
NG = 0
DB = 0
SM = 0
Delta_AUC
Doxycycline
YNL069C_1 RPL16B
ZShift = 7.2
Z lm Score = 9.6
NG = 0
DB = 0
SM = 0
Delta_L
Doxycycline
YNL069C_1 RPL16B
ZShift = −2.23
Z lm Score = −17.28
NG = 0
DB = 0
SM = 0
Delta_K
Doxycycline
YNL069C_1 RPL16B
ZShift = −2.54
Z lm Score = −0.87
NG = 0
DB = 0
SM = 0
Delta_r
Doxycycline
YNL069C_1 RPL16B
ZShift = −4.34
Z lm Score = −8.6
NG = 0
DB = 0
SM = 0
Delta_AUC
Doxycycline
YLR312W−A MRPL15
ZShift = 0.42
Z lm Score = 9.59
NG = 0
DB = 0
SM = 0
Delta_L
Doxycycline
YLR312W−A MRPL15
ZShift = −13.48
Z lm Score = 1.4
NG = 0
DB = 0
SM = 0
Delta_K
Doxycycline
YLR312W−A MRPL15
ZShift = −6.97
Z lm Score = 8.91
NG = 0
DB = 0
SM = 0
Delta_r
Doxycycline
YLR312W−A MRPL15
ZShift = −9.51
Z lm Score = 0.36
NG = 0
DB = 0
SM = 0
Delta_AUC
Doxycycline

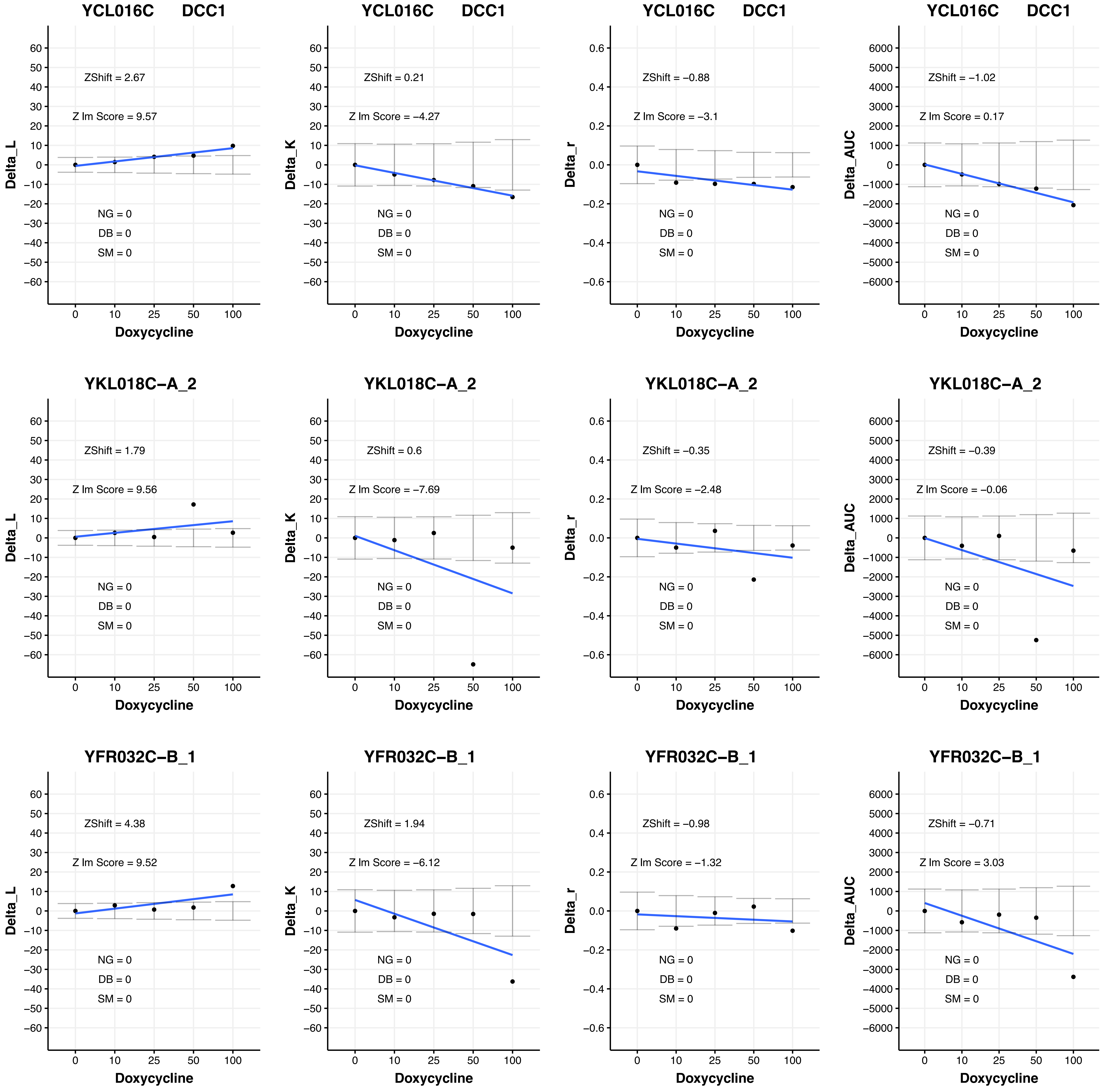
YCL016C DCC1
ZShift = 2.67
Z lm Score = 9.57
NG = 0
DB = 0
SM = 0
Delta_L
Doxycycline
YCL016C DCC1
ZShift = 0.21
Z lm Score = −4.27
NG = 0
DB = 0
SM = 0
Delta_K
Doxycycline
YCL016C DCC1
ZShift = −0.88
Z lm Score = −3.1
NG = 0
DB = 0
SM = 0
Delta_r
Doxycycline
YCL016C DCC1
ZShift = −1.02
Z lm Score = 0.17
NG = 0
DB = 0
SM = 0
Delta_AUC
Doxycycline
YKL018C−A_2
ZShift = 1.79
Z lm Score = 9.56
NG = 0
DB = 0
SM = 0
Delta_L
Doxycycline
YKL018C−A_2
ZShift = 0.6
Z lm Score = −7.69
NG = 0
DB = 0
SM = 0
Delta_K
Doxycycline
YKL018C−A_2
ZShift = −0.35
Z lm Score = −2.48
NG = 0
DB = 0
SM = 0
Delta_r
Doxycycline
YKL018C−A_2
ZShift = −0.39
Z lm Score = −0.06
NG = 0
DB = 0
SM = 0
Delta_AUC
Doxycycline
YFR032C−B_1
ZShift = 4.38
Z lm Score = 9.52
NG = 0
DB = 0
SM = 0
Delta_L
Doxycycline
YFR032C−B_1
ZShift = 1.94
Z lm Score = −6.12
NG = 0
DB = 0
SM = 0
Delta_K
Doxycycline
YFR032C−B_1
ZShift = −0.98
Z lm Score = −1.32
NG = 0
DB = 0
SM = 0
Delta_r
Doxycycline
YFR032C−B_1
ZShift = −0.71
Z lm Score = 3.03
NG = 0
DB = 0
SM = 0
Delta_AUC
Doxycycline

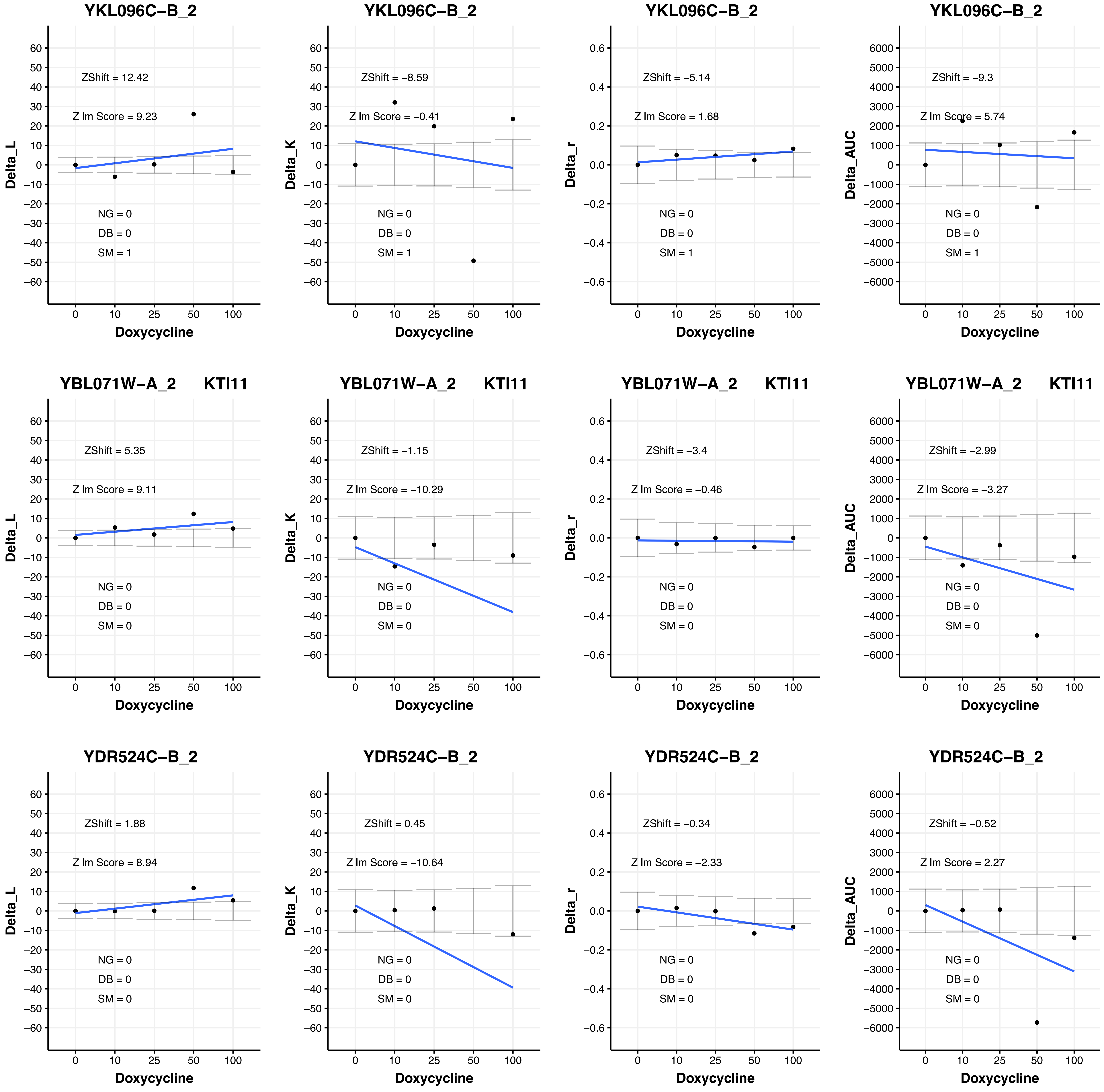
YKL096C−B_2
ZShift = 12.42
Z lm Score = 9.23
NG = 0
DB = 0
SM = 1
Delta_L
Doxycycline
YKL096C−B_2
ZShift = −8.59
Z lm Score = −0.41
NG = 0
DB = 0
SM = 1
Delta_K
Doxycycline
YKL096C−B_2
ZShift = −5.14
Z lm Score = 1.68
NG = 0
DB = 0
SM = 1
Delta_r
Doxycycline
YKL096C−B_2
ZShift = −9.3
Z lm Score = 5.74
NG = 0
DB = 0
SM = 1
Delta_AUC
Doxycycline
YBL071W−A_2 KTI11
ZShift = 5.35
Z lm Score = 9.11
NG = 0
DB = 0
SM = 0
Delta_L
Doxycycline
YBL071W−A_2 KTI11
ZShift = −1.15
Z lm Score = −10.29
NG = 0
DB = 0
SM = 0
Delta_K
Doxycycline
YBL071W−A_2 KTI11
ZShift = −3.4
Z lm Score = −0.46
NG = 0
DB = 0
SM = 0
Delta_r
Doxycycline
YBL071W−A_2 KTI11
ZShift = −2.99
Z lm Score = −3.27
NG = 0
DB = 0
SM = 0
Delta_AUC
Doxycycline
YDR524C−B_2
ZShift = 1.88
Z lm Score = 8.94
NG = 0
DB = 0
SM = 0
Delta_L
Doxycycline
YDR524C−B_2
ZShift = 0.45
Z lm Score = −10.64
NG = 0
DB = 0
SM = 0
Delta_K
Doxycycline
YDR524C−B_2
ZShift = −0.34
Z lm Score = −2.33
NG = 0
DB = 0
SM = 0
Delta_r
Doxycycline
YDR524C−B_2
ZShift = −0.52
Z lm Score = 2.27
NG = 0
DB = 0
SM = 0
Delta_AUC
Doxycycline

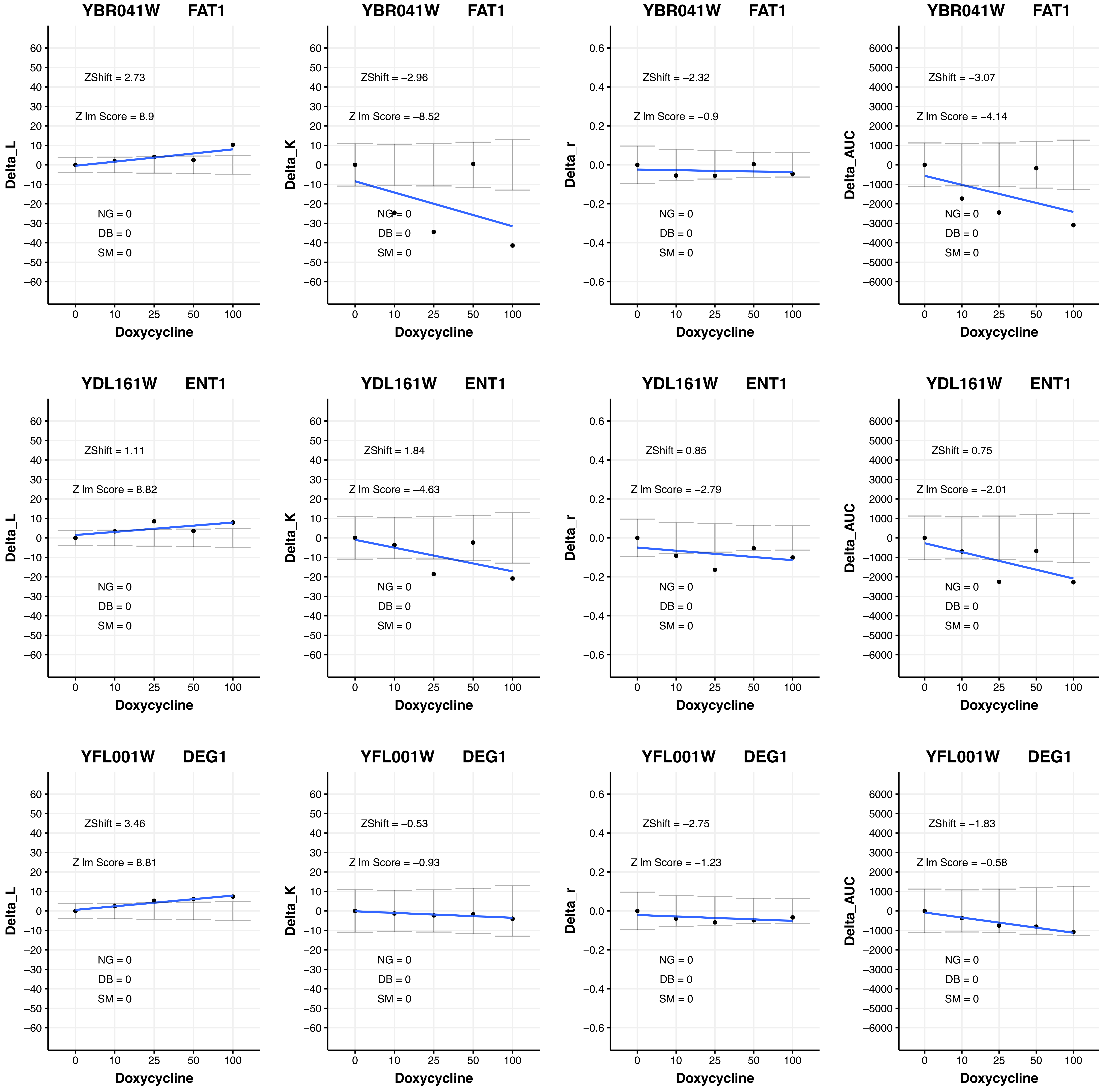
YBR041W FAT1
ZShift = 2.73
Z lm Score = 8.9
NG = 0
DB = 0
SM = 0
Delta_L
Doxycycline
YBR041W FAT1
ZShift = −2.96
Z lm Score = −8.52
NG = 0
DB = 0
SM = 0
Delta_K
Doxycycline
YBR041W FAT1
ZShift = −2.32
Z lm Score = −0.9
NG = 0
DB = 0
SM = 0
Delta_r
Doxycycline
YBR041W FAT1
ZShift = −3.07
Z lm Score = −4.14
NG = 0
DB = 0
SM = 0
Delta_AUC
Doxycycline
YDL161W ENT1
ZShift = 1.11
Z lm Score = 8.82
NG = 0
DB = 0
SM = 0
Delta_L
Doxycycline
YDL161W ENT1
ZShift = 1.84
Z lm Score = −4.63
NG = 0
DB = 0
SM = 0
Delta_K
Doxycycline
YDL161W ENT1
ZShift = 0.85
Z lm Score = −2.79
NG = 0
DB = 0
SM = 0
Delta_r
Doxycycline
YDL161W ENT1
ZShift = 0.75
Z lm Score = −2.01
NG = 0
DB = 0
SM = 0
Delta_AUC
Doxycycline
YFL001W DEG1
ZShift = 3.46
Z lm Score = 8.81
NG = 0
DB = 0
SM = 0
Delta_L
Doxycycline
YFL001W DEG1
ZShift = −0.53
Z lm Score = −0.93
NG = 0
DB = 0
SM = 0
Delta_K
Doxycycline
YFL001W DEG1
ZShift = −2.75
Z lm Score = −1.23
NG = 0
DB = 0
SM = 0
Delta_r
Doxycycline
YFL001W DEG1
ZShift = −1.83
Z lm Score = −0.58
NG = 0
DB = 0
SM = 0
Delta_AUC
Doxycycline

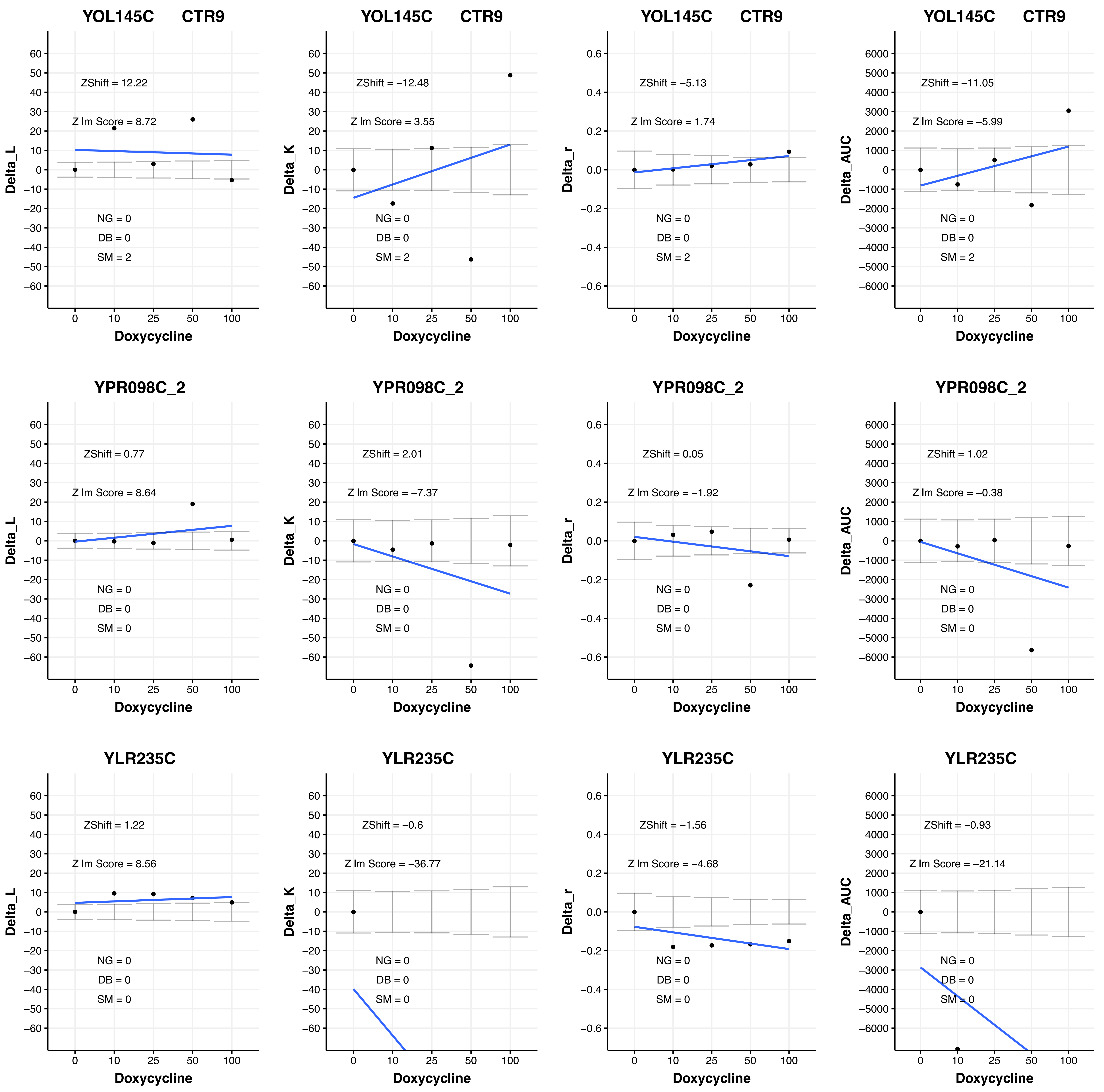
YOL145C CTR9
ZShift = 12.22
Z lm Score = 8.72
NG = 0
DB = 0
SM = 2
Delta_L
Doxycycline
YOL145C CTR9
ZShift = −12.48
Z lm Score = 3.55
NG = 0
DB = 0
SM = 2
Delta_K
Doxycycline
YOL145C CTR9
ZShift = −5.13
Z lm Score = 1.74
NG = 0
DB = 0
SM = 2
Delta_r
Doxycycline
YOL145C CTR9
ZShift = −11.05
Z lm Score = −5.99
NG = 0
DB = 0
SM = 2
Delta_AUC
Doxycycline
YPR098C_2
ZShift = 0.77
Z lm Score = 8.64
NG = 0
DB = 0
SM = 0
Delta_L
Doxycycline
YPR098C_2
ZShift = 2.01
Z lm Score = −7.37
NG = 0
DB = 0
SM = 0
Delta_K
Doxycycline
YPR098C_2
ZShift = 0.05
Z lm Score = −1.92
NG = 0
DB = 0
SM = 0
Delta_r
Doxycycline
YPR098C_2
ZShift = 1.02
Z lm Score = −0.38
NG = 0
DB = 0
SM = 0
Delta_AUC
Doxycycline
YLR235C
ZShift = 1.22
Z lm Score = 8.56
NG = 0
DB = 0
SM = 0
Delta_L
Doxycycline
YLR235C
ZShift = −0.6
Z lm Score = −36.77
NG = 0
DB = 0
SM = 0
Delta_K
Doxycycline
YLR235C
ZShift = −1.56
Z lm Score = −4.68
NG = 0
DB = 0
SM = 0
Delta_r
Doxycycline
YLR235C
ZShift = −0.93
Z lm Score = −21.14
NG = 0
DB = 0
SM = 0
Delta_AUC
Doxycycline

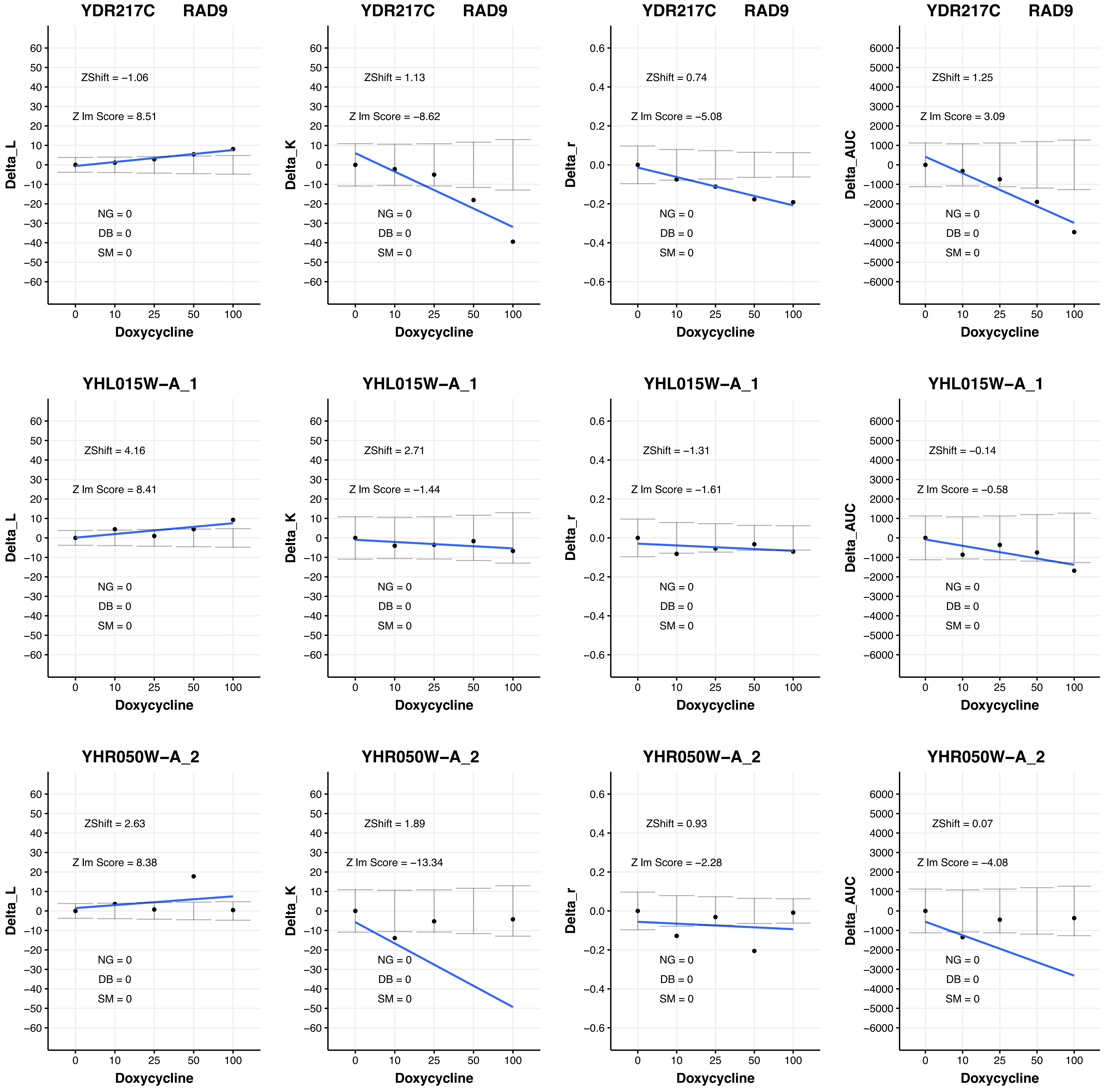
YDR217C RAD9
ZShift = −1.06
Z lm Score = 8.51
NG = 0
DB = 0
SM = 0
Delta_L
Doxycycline
YDR217C RAD9
ZShift = 1.13
Z lm Score = −8.62
NG = 0
DB = 0
SM = 0
Delta_K
Doxycycline
YDR217C RAD9
ZShift = 0.74
Z lm Score = −5.08
NG = 0
DB = 0
SM = 0
Delta_r
Doxycycline
YDR217C RAD9
ZShift = 1.25
Z lm Score = 3.09
NG = 0
DB = 0
SM = 0
Delta_AUC
Doxycycline
YHL015W−A_1
ZShift = 4.16
Z lm Score = 8.41
NG = 0
DB = 0
SM = 0
Delta_L
Doxycycline
YHL015W−A_1
ZShift = 2.71
Z lm Score = −1.44
NG = 0
DB = 0
SM = 0
Delta_K
Doxycycline
YHL015W−A_1
ZShift = −1.31
Z lm Score = −1.61
NG = 0
DB = 0
SM = 0
Delta_r
Doxycycline
YHL015W−A_1
ZShift = −0.14
Z lm Score = −0.58
NG = 0
DB = 0
SM = 0
Delta_AUC
Doxycycline
YHR050W−A_2
ZShift = 2.63
Z lm Score = 8.38
NG = 0
DB = 0
SM = 0
Delta_L
Doxycycline
YHR050W−A_2
ZShift = 1.89
Z lm Score = −13.34
NG = 0
DB = 0
SM = 0
Delta_K
Doxycycline
YHR050W−A_2
ZShift = 0.93
Z lm Score = −2.28
NG = 0
DB = 0
SM = 0
Delta_r
Doxycycline
YHR050W−A_2
ZShift = 0.07
Z lm Score = −4.08
NG = 0
DB = 0
SM = 0
Delta_AUC
Doxycycline

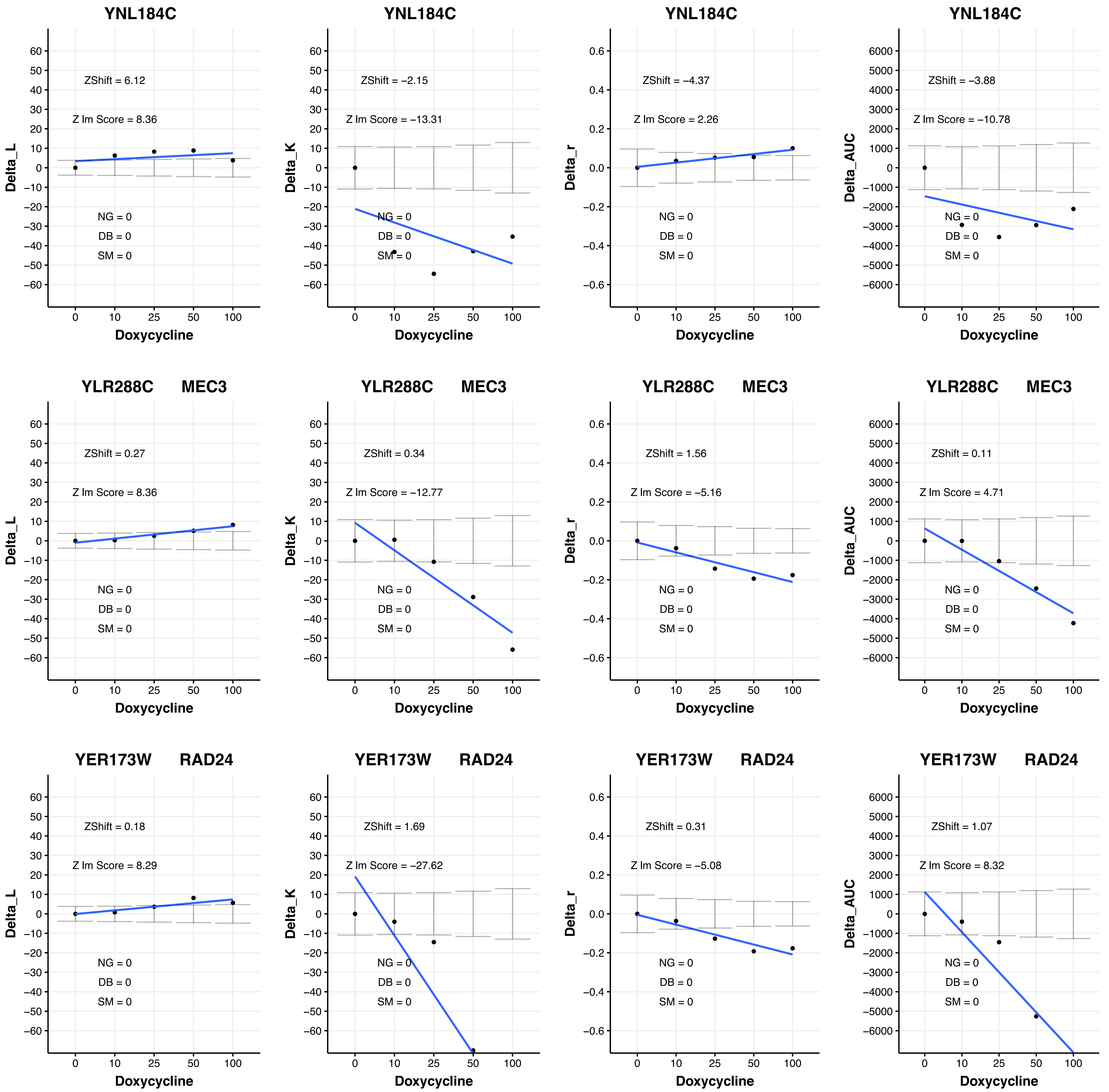
YNL184C
ZShift = 6.12
Z lm Score = 8.36
NG = 0
DB = 0
SM = 0
Delta_L
Doxycycline
YNL184C
ZShift = −2.15
Z lm Score = −13.31
NG = 0
DB = 0
SM = 0
Delta_K
Doxycycline
YNL184C
ZShift = −4.37
Z lm Score = 2.26
NG = 0
DB = 0
SM = 0
Delta_r
Doxycycline
YNL184C
ZShift = −3.88
Z lm Score = −10.78
NG = 0
DB = 0
SM = 0
Delta_AUC
Doxycycline
YLR288C MEC3
ZShift = 0.27
Z lm Score = 8.36
NG = 0
DB = 0
SM = 0
Delta_L
Doxycycline
YLR288C MEC3
ZShift = 0.34
Z lm Score = −12.77
NG = 0
DB = 0
SM = 0
Delta_K
Doxycycline
YLR288C MEC3
ZShift = 1.56
Z lm Score = −5.16
NG = 0
DB = 0
SM = 0
Delta_r
Doxycycline
YLR288C MEC3
ZShift = 0.11
Z lm Score = 4.71
NG = 0
DB = 0
SM = 0
Delta_AUC
Doxycycline
YER173W RAD24
ZShift = 0.18
Z lm Score = 8.29
NG = 0
DB = 0
SM = 0
Delta_L
Doxycycline
YER173W RAD24
ZShift = 1.69
Z lm Score = −27.62
NG = 0
DB = 0
SM = 0
Delta_K
Doxycycline
YER173W RAD24
ZShift = 0.31
Z lm Score = −5.08
NG = 0
DB = 0
SM = 0
Delta_r
Doxycycline
YER173W RAD24
ZShift = 1.07
Z lm Score = 8.32
NG = 0
DB = 0
SM = 0
Delta_AUC
Doxycycline

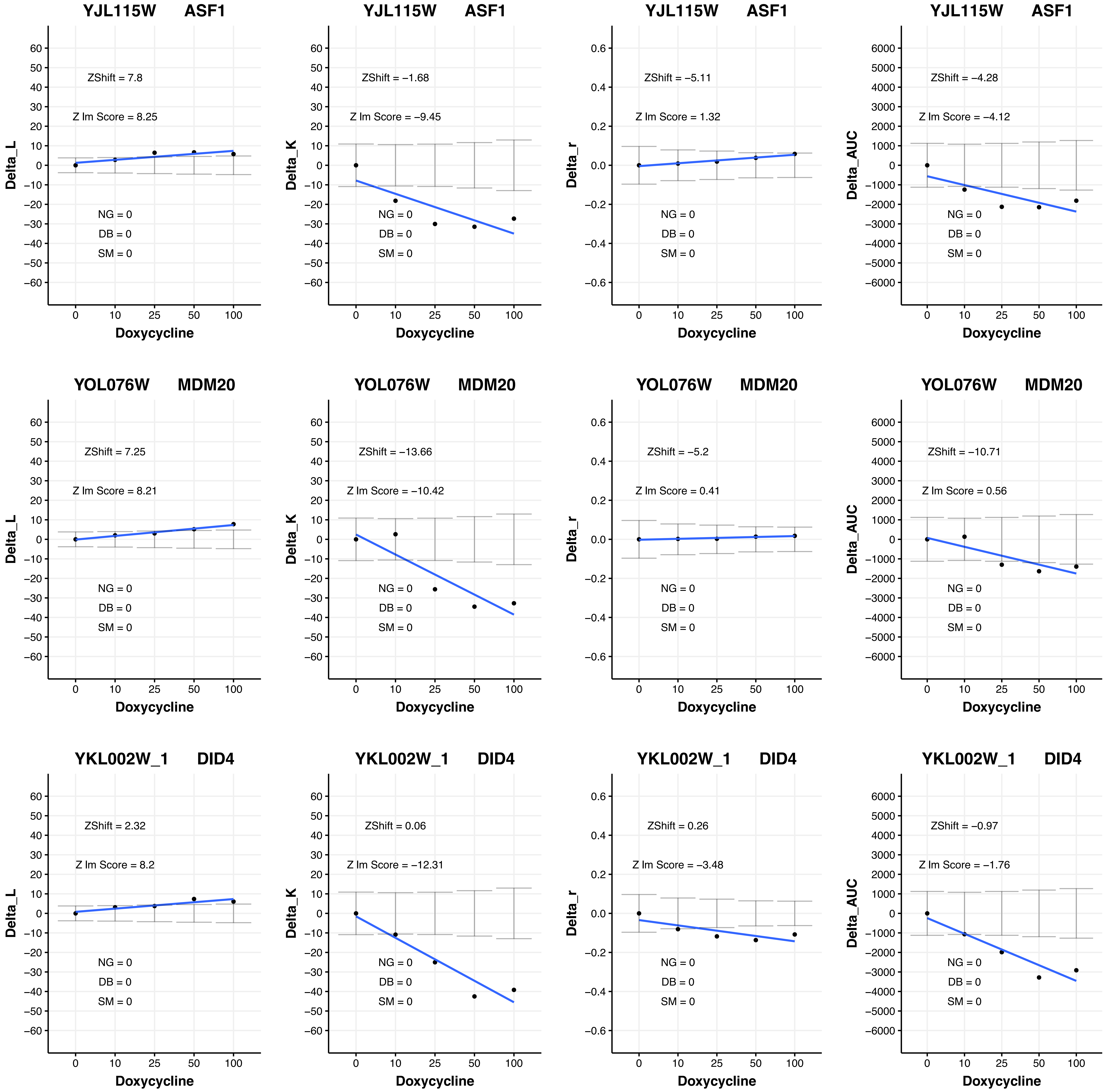

YJL115W ASF1
ZShift = 7.8
Z lm Score = 8.25
NG = 0
DB = 0
SM = 0
Delta_L
Doxycycline
YJL115W ASF1
ZShift = −1.68
Z lm Score = −9.45
NG = 0
DB = 0
SM = 0
Delta_K
Doxycycline
YJL115W ASF1
ZShift = −5.11
Z lm Score = 1.32
NG = 0
DB = 0
SM = 0
Delta_r
Doxycycline
YJL115W ASF1
ZShift = −4.28
Z lm Score = −4.12
NG = 0
DB = 0
SM = 0
Delta_AUC
Doxycycline
YOL076W MDM20
ZShift = 7.25
Z lm Score = 8.21
NG = 0
DB = 0
SM = 0
Delta_L
Doxycycline
YOL076W MDM20
ZShift = −13.66
Z lm Score = −10.42
NG = 0
DB = 0
SM = 0
Delta_K
Doxycycline
YOL076W MDM20
ZShift = −5.2
Z lm Score = 0.41
NG = 0
DB = 0
SM = 0
Delta_r
Doxycycline
YOL076W MDM20
ZShift = −10.71
Z lm Score = 0.56
NG = 0
DB = 0
SM = 0
Delta_AUC
Doxycycline
YKL002W_1 DID4
ZShift = 2.32
Z lm Score = 8.2
NG = 0
DB = 0
SM = 0
Delta_L
Doxycycline
YKL002W_1 DID4
ZShift = 0.06
Z lm Score = −12.31
NG = 0
DB = 0
SM = 0
Delta_K
Doxycycline
YKL002W_1 DID4
ZShift = 0.26
Z lm Score = −3.48
NG = 0
DB = 0
SM = 0
Delta_r
Doxycycline
YKL002W_1 DID4
ZShift = −0.97
Z lm Score = −1.76
NG = 0
DB = 0
SM = 0
Delta_AUC
Doxycycline

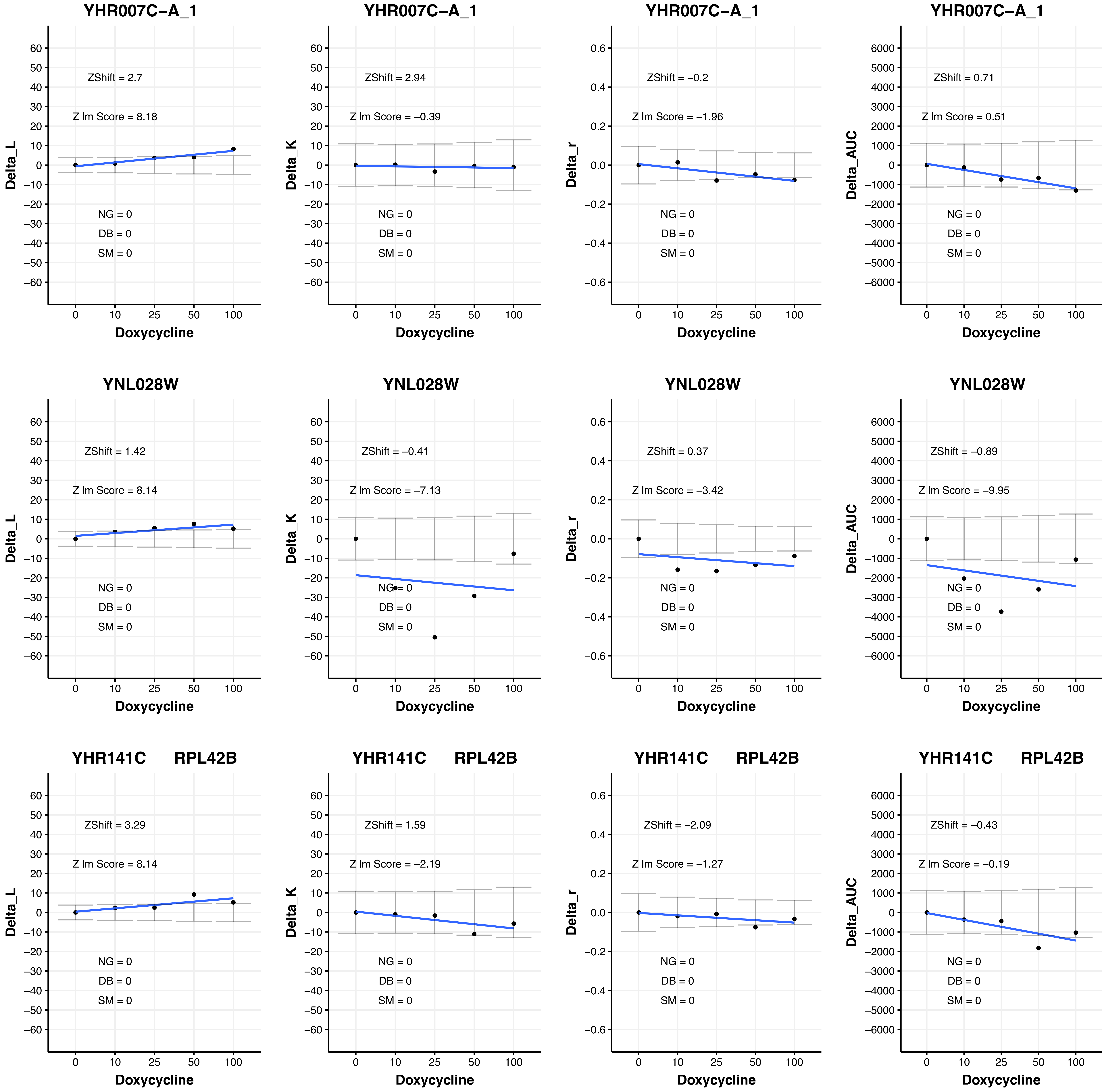

YHR007C−A_1
ZShift = 2.7
Z lm Score = 8.18
NG = 0
DB = 0
SM = 0
Delta_L
Doxycycline
YHR007C−A_1
ZShift = 2.94
Z lm Score = −0.39
NG = 0
DB = 0
SM = 0
Delta_K
Doxycycline
YHR007C−A_1
ZShift = −0.2
Z lm Score = −1.96
NG = 0
DB = 0
SM = 0
Delta_r
Doxycycline
YHR007C−A_1
ZShift = 0.71
Z lm Score = 0.51
NG = 0
DB = 0
SM = 0
Delta_AUC
Doxycycline
YNL028W
ZShift = 1.42
Z lm Score = 8.14
NG = 0
DB = 0
SM = 0
Delta_L
Doxycycline
YNL028W
ZShift = −0.41
Z lm Score = −7.13
NG = 0
DB = 0
SM = 0
Delta_K
Doxycycline
YNL028W
ZShift = 0.37
Z lm Score = −3.42
NG = 0
DB = 0
SM = 0
Delta_r
Doxycycline
YNL028W
ZShift = −0.89
Z lm Score = −9.95
NG = 0
DB = 0
SM = 0
Delta_AUC
Doxycycline
YHR141C RPL42B
ZShift = 3.29
Z lm Score = 8.14
NG = 0
DB = 0
SM = 0
Delta_L
Doxycycline
YHR141C RPL42B
ZShift = 1.59
Z lm Score = −2.19
NG = 0
DB = 0
SM = 0
Delta_K
Doxycycline
YHR141C RPL42B
ZShift = −2.09
Z lm Score = −1.27
NG = 0
DB = 0
SM = 0
Delta_r
Doxycycline
YHR141C RPL42B
ZShift = −0.43
Z lm Score = −0.19
NG = 0
DB = 0
SM = 0
Delta_AUC
Doxycycline

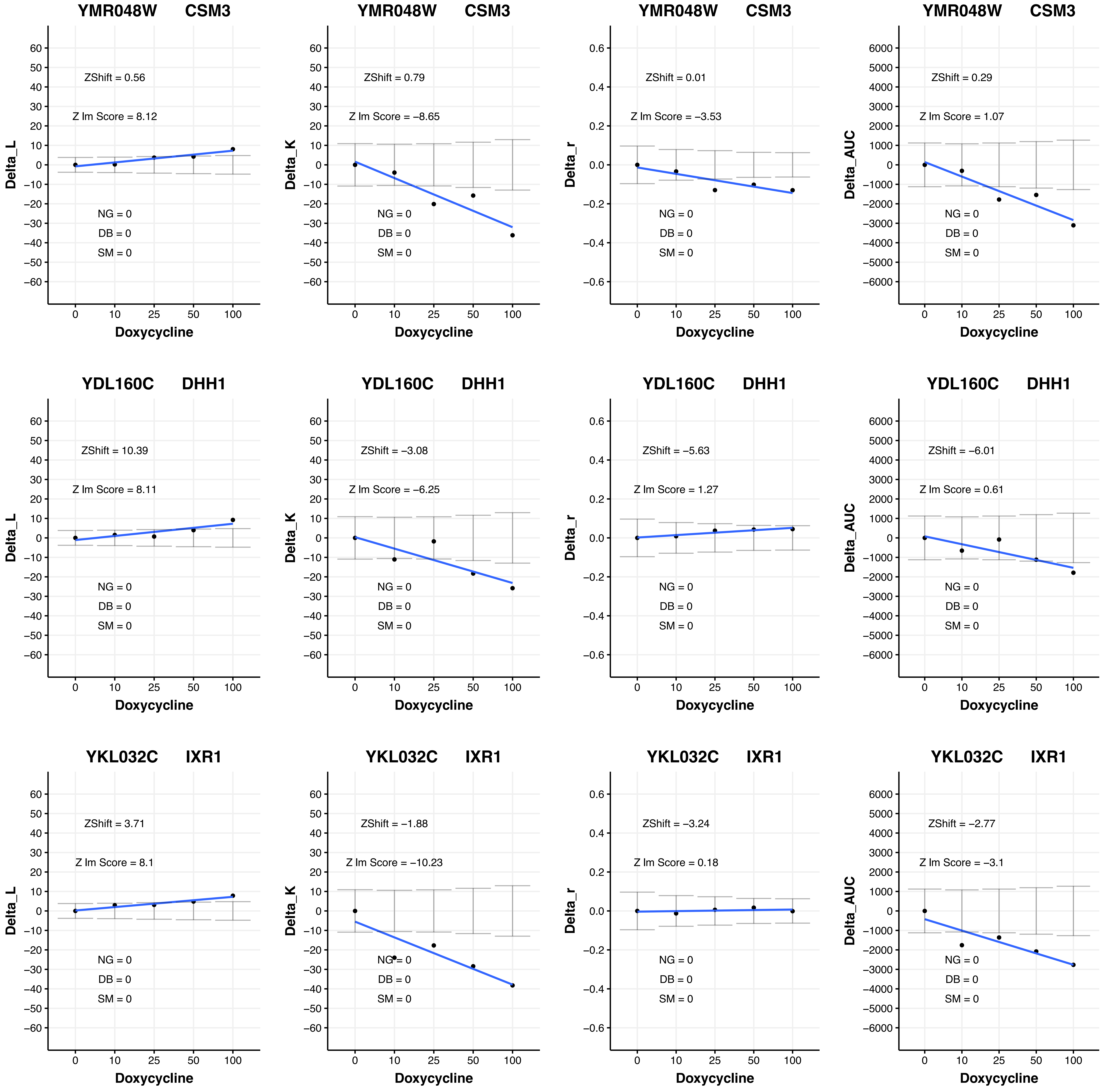

YMR048W CSM3
ZShift = 0.56
Z lm Score = 8.12
NG = 0
DB = 0
SM = 0
Delta_L
Doxycycline
YMR048W CSM3
ZShift = 0.79
Z lm Score = −8.65
NG = 0
DB = 0
SM = 0
Delta_K
Doxycycline
YMR048W CSM3
ZShift = 0.01
Z lm Score = −3.53
NG = 0
DB = 0
SM = 0
Delta_r
Doxycycline
YMR048W CSM3
ZShift = 0.29
Z lm Score = 1.07
NG = 0
DB = 0
SM = 0
Delta_AUC
Doxycycline
YDL160C DHH1
ZShift = 10.39
Z lm Score = 8.11
NG = 0
DB = 0
SM = 0
Delta_L
Doxycycline
YDL160C DHH1
ZShift = −3.08
Z lm Score = −6.25
NG = 0
DB = 0
SM = 0
Delta_K
Doxycycline
YDL160C DHH1
ZShift = −5.63
Z lm Score = 1.27
NG = 0
DB = 0
SM = 0
Delta_r
Doxycycline
YDL160C DHH1
ZShift = −6.01
Z lm Score = 0.61
NG = 0
DB = 0
SM = 0
Delta_AUC
Doxycycline
YKL032C IXR1
ZShift = 3.71
Z lm Score = 8.1
NG = 0
DB = 0
SM = 0
Delta_L
Doxycycline
YKL032C IXR1
ZShift = −1.88
Z lm Score = −10.23
NG = 0
DB = 0
SM = 0
Delta_K
Doxycycline
YKL032C IXR1
ZShift = −3.24
Z lm Score = 0.18
NG = 0
DB = 0
SM = 0
Delta_r
Doxycycline
YKL032C IXR1
ZShift = −2.77
Z lm Score = −3.1
NG = 0
DB = 0
SM = 0
Delta_AUC
Doxycycline

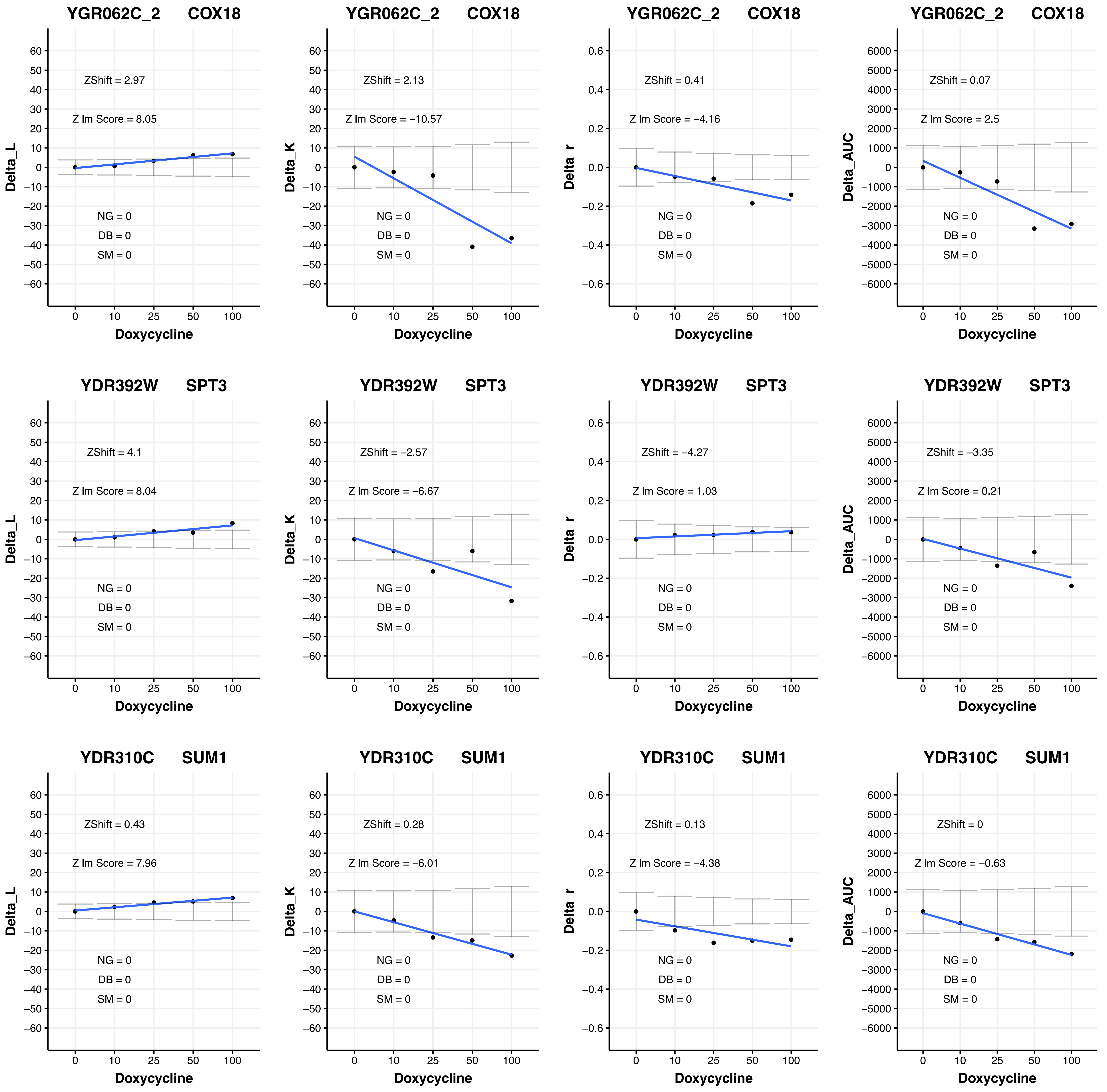

YGR062C_2 COX18
ZShift = 2.97
Z lm Score = 8.05
NG = 0
DB = 0
SM = 0
Delta_L
Doxycycline
YGR062C_2 COX18
ZShift = 2.13
Z lm Score = −10.57
NG = 0
DB = 0
SM = 0
Delta_K
Doxycycline
YGR062C_2 COX18
ZShift = 0.41
Z lm Score = −4.16
NG = 0
DB = 0
SM = 0
Delta_r
Doxycycline
YGR062C_2 COX18
ZShift = 0.07
Z lm Score = 2.5
NG = 0
DB = 0
SM = 0
Delta_AUC
Doxycycline
YDR392W SPT3
ZShift = 4.1
Z lm Score = 8.04
NG = 0
DB = 0
SM = 0
Delta_L
Doxycycline
YDR392W SPT3
ZShift = −2.57
Z lm Score = −6.67
NG = 0
DB = 0
SM = 0
Delta_K
Doxycycline
YDR392W SPT3
ZShift = −4.27
Z lm Score = 1.03
NG = 0
DB = 0
SM = 0
Delta_r
Doxycycline
YDR392W SPT3
ZShift = −3.35
Z lm Score = 0.21
NG = 0
DB = 0
SM = 0
Delta_AUC
Doxycycline
YDR310C SUM1
ZShift = 0.43
Z lm Score = 7.96
NG = 0
DB = 0
SM = 0
Delta_L
Doxycycline
YDR310C SUM1
ZShift = 0.28
Z lm Score = −6.01
NG = 0
DB = 0
SM = 0
Delta_K
Doxycycline
YDR310C SUM1
ZShift = 0.13
Z lm Score = −4.38
NG = 0
DB = 0
SM = 0
Delta_r
Doxycycline
YDR310C SUM1
ZShift = 0
Z lm Score = −0.63
NG = 0
DB = 0
SM = 0
Delta_AUC
Doxycycline

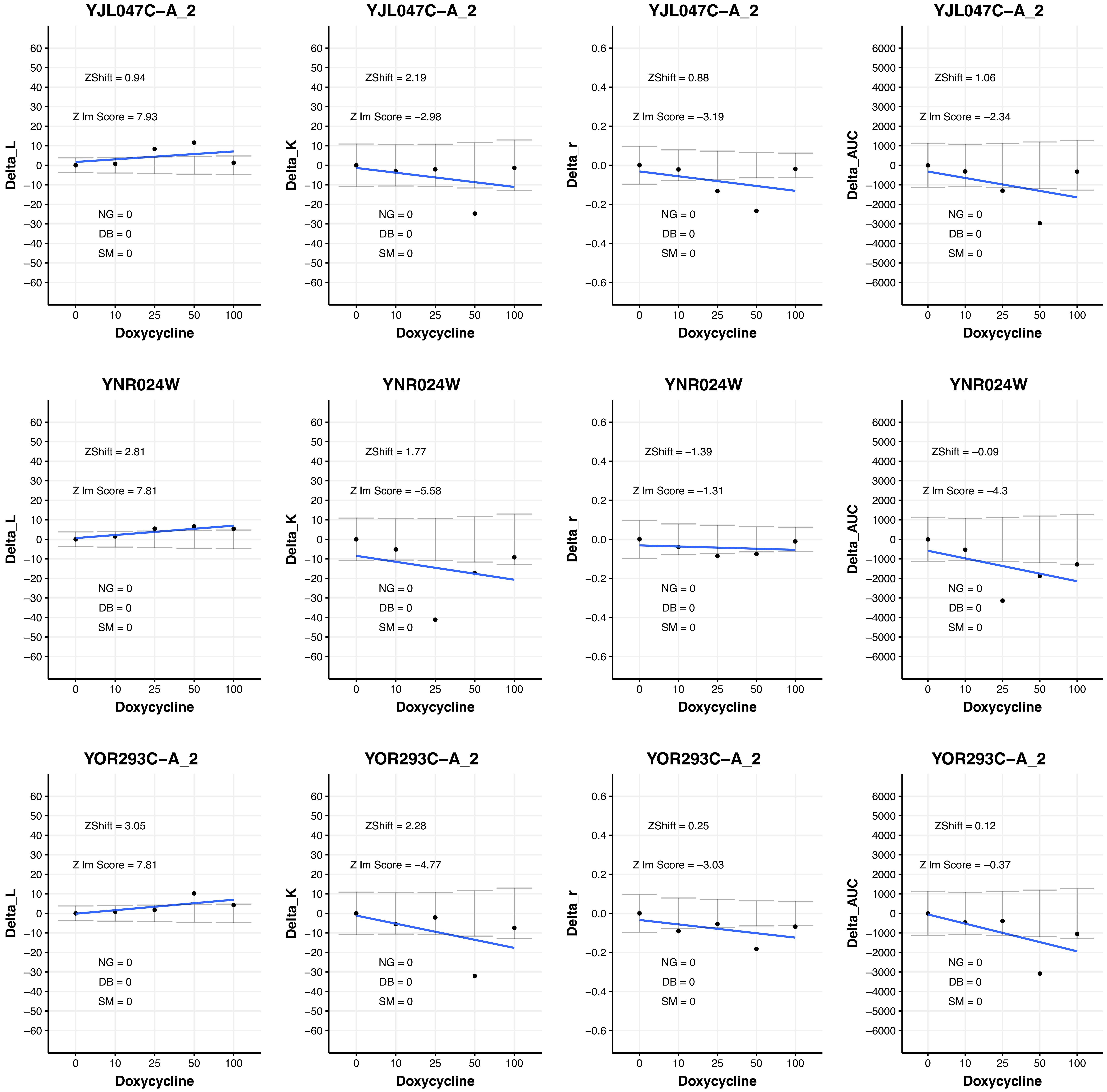

YJL047C−A_2
ZShift = 0.94
Z lm Score = 7.93
NG = 0
DB = 0
SM = 0
Delta_L
Doxycycline
YJL047C−A_2
ZShift = 2.19
Z lm Score = −2.98
NG = 0
DB = 0
SM = 0
Delta_K
Doxycycline
YJL047C−A_2
ZShift = 0.88
Z lm Score = −3.19
NG = 0
DB = 0
SM = 0
Delta_r
Doxycycline
YJL047C−A_2
ZShift = 1.06
Z lm Score = −2.34
NG = 0
DB = 0
SM = 0
Delta_AUC
Doxycycline
YNR024W
ZShift = 2.81
Z lm Score = 7.81
NG = 0
DB = 0
SM = 0
Delta_L
Doxycycline
YNR024W
ZShift = 1.77
Z lm Score = −5.58
NG = 0
DB = 0
SM = 0
Delta_K
Doxycycline
YNR024W
ZShift = −1.39
Z lm Score = −1.31
NG = 0
DB = 0
SM = 0
Delta_r
Doxycycline
YNR024W
ZShift = −0.09
Z lm Score = −4.3
NG = 0
DB = 0
SM = 0
Delta_AUC
Doxycycline
YOR293C−A_2
ZShift = 3.05
Z lm Score = 7.81
NG = 0
DB = 0
SM = 0
Delta_L
Doxycycline
YOR293C−A_2
ZShift = 2.28
Z lm Score = −4.77
NG = 0
DB = 0
SM = 0
Delta_K
Doxycycline
YOR293C−A_2
ZShift = 0.25
Z lm Score = −3.03
NG = 0
DB = 0
SM = 0
Delta_r
Doxycycline
YOR293C−A_2
ZShift = 0.12
Z lm Score = −0.37
NG = 0
DB = 0
SM = 0
Delta_AUC
Doxycycline

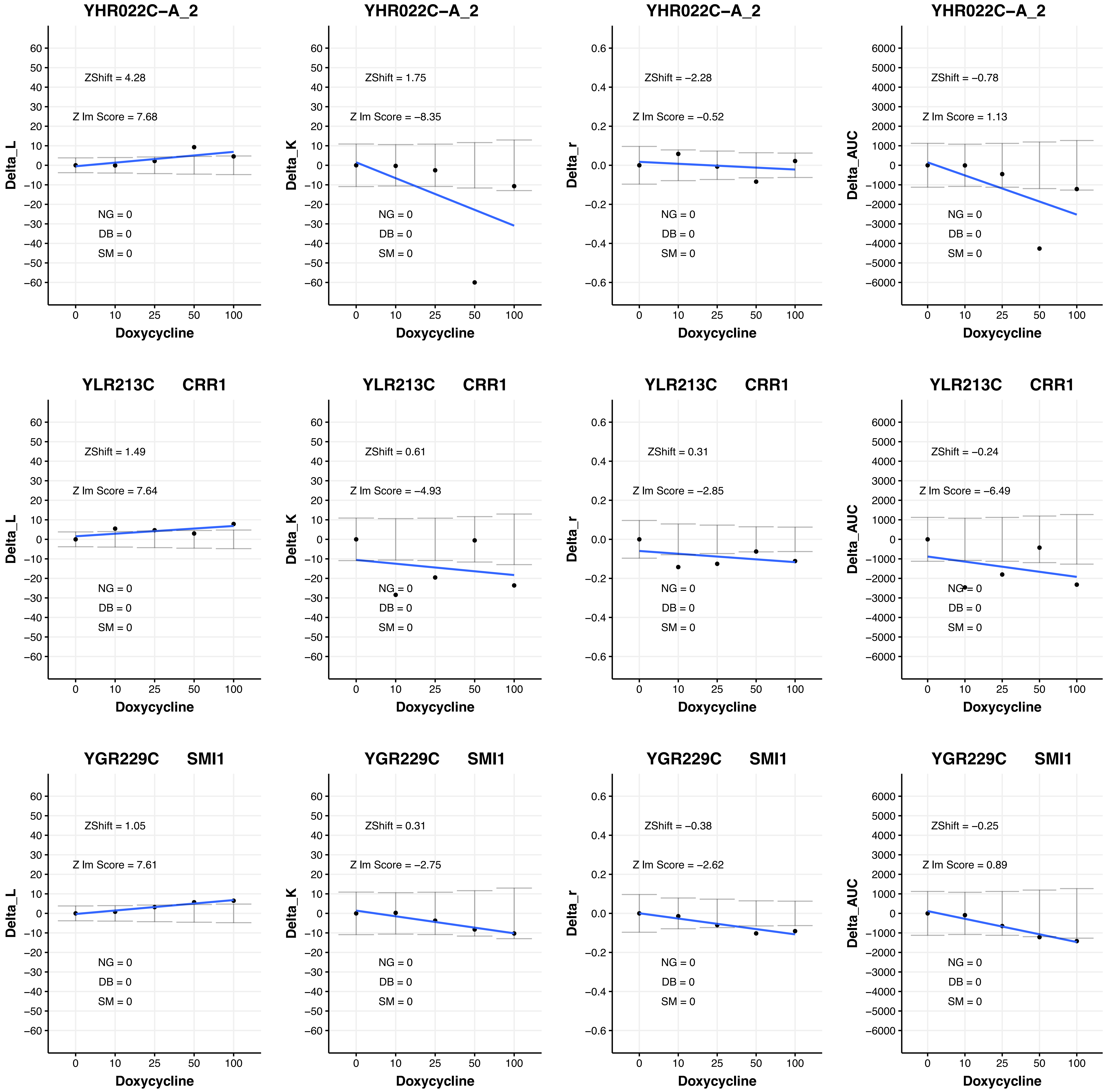
YHR022C−A_2
ZShift = 4.28
Z lm Score = 7.68
NG = 0
DB = 0
SM = 0
Delta_L
Doxycycline
YHR022C−A_2
ZShift = 1.75
Z lm Score = −8.35
NG = 0
DB = 0
SM = 0
Delta_K
Doxycycline
YHR022C−A_2
ZShift = −2.28
Z lm Score = −0.52
NG = 0
DB = 0
SM = 0
Delta_r
Doxycycline
YHR022C−A_2
ZShift = −0.78
Z lm Score = 1.13
NG = 0
DB = 0
SM = 0
Delta_AUC
Doxycycline
YLR213C CRR1
ZShift = 1.49
Z lm Score = 7.64
NG = 0
DB = 0
SM = 0
Delta_L
Doxycycline
YLR213C CRR1
ZShift = 0.61
Z lm Score = −4.93
NG = 0
DB = 0
SM = 0
Delta_K
Doxycycline
YLR213C CRR1
ZShift = 0.31
Z lm Score = −2.85
NG = 0
DB = 0
SM = 0
Delta_r
Doxycycline
YLR213C CRR1
ZShift = −0.24
Z lm Score = −6.49
NG = 0
DB = 0
SM = 0
Delta_AUC
Doxycycline
YGR229C SMI1
ZShift = 1.05
Z lm Score = 7.61
NG = 0
DB = 0
SM = 0
Delta_L
Doxycycline
YGR229C SMI1
ZShift = 0.31
Z lm Score = −2.75
NG = 0
DB = 0
SM = 0
Delta_K
Doxycycline
YGR229C SMI1
ZShift = −0.38
Z lm Score = −2.62
NG = 0
DB = 0
SM = 0
Delta_r
Doxycycline
YGR229C SMI1
ZShift = −0.25
Z lm Score = 0.89
NG = 0
DB = 0
SM = 0
Delta_AUC
Doxycycline

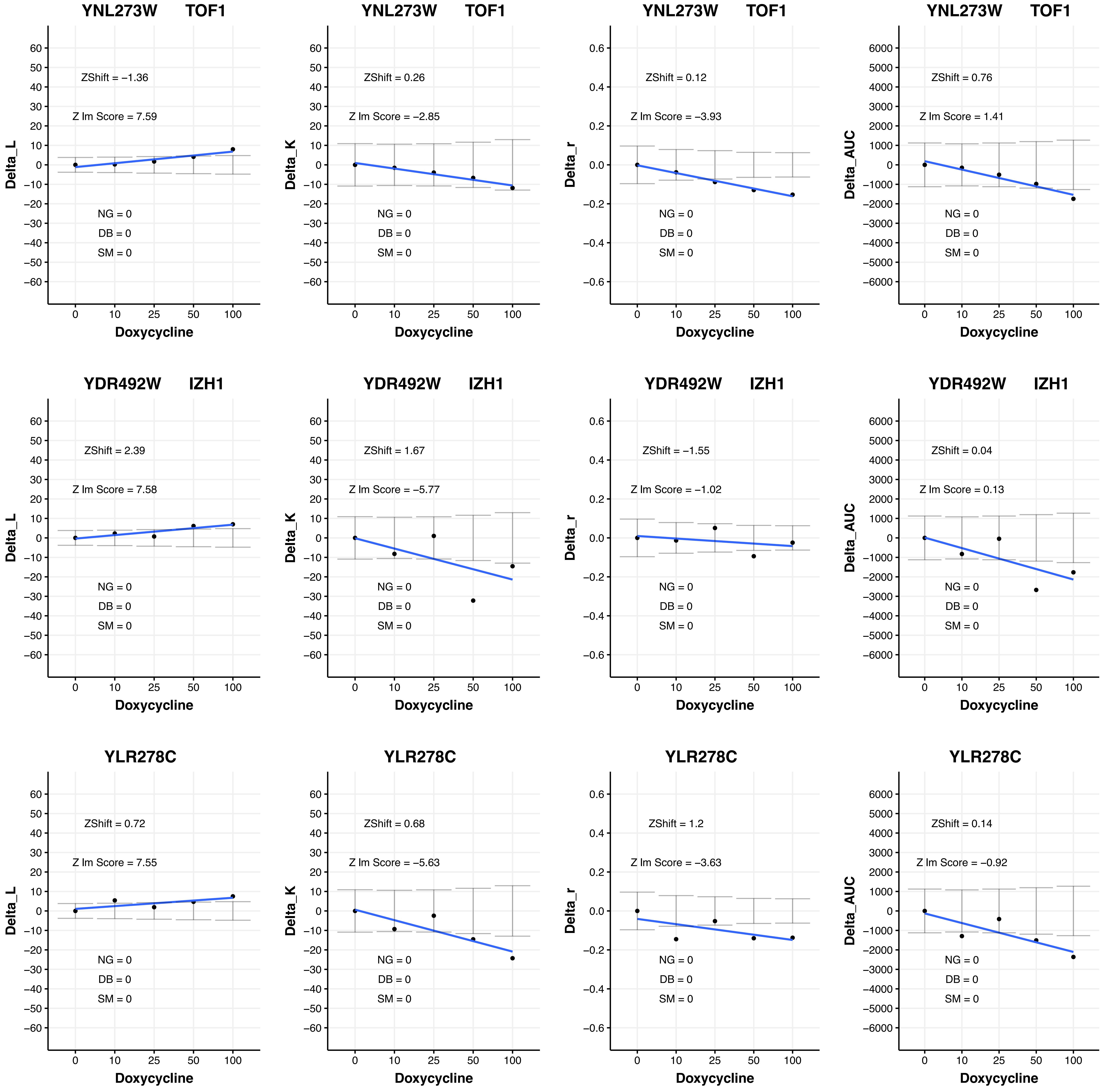

YNL273W TOF1
ZShift = −1.36
Z lm Score = 7.59
NG = 0
DB = 0
SM = 0
Delta_L
Doxycycline
YNL273W TOF1
ZShift = 0.26
Z lm Score = −2.85
NG = 0
DB = 0
SM = 0
Delta_K
Doxycycline
YNL273W TOF1
ZShift = 0.12
Z lm Score = −3.93
NG = 0
DB = 0
SM = 0
Delta_r
Doxycycline
YNL273W TOF1
ZShift = 0.76
Z lm Score = 1.41
NG = 0
DB = 0
SM = 0
Delta_AUC
Doxycycline
YDR492W IZH1
ZShift = 2.39
Z lm Score = 7.58
NG = 0
DB = 0
SM = 0
Delta_L
Doxycycline
YDR492W IZH1
ZShift = 1.67
Z lm Score = −5.77
NG = 0
DB = 0
SM = 0
Delta_K
Doxycycline
YDR492W IZH1
ZShift = −1.55
Z lm Score = −1.02
NG = 0
DB = 0
SM = 0
Delta_r
Doxycycline
YDR492W IZH1
ZShift = 0.04
Z lm Score = 0.13
NG = 0
DB = 0
SM = 0
Delta_AUC
Doxycycline
YLR278C
ZShift = 0.72
Z lm Score = 7.55
NG = 0
DB = 0
SM = 0
Delta_L
Doxycycline
YLR278C
ZShift = 0.68
Z lm Score = −5.63
NG = 0
DB = 0
SM = 0
Delta_K
Doxycycline
YLR278C
ZShift = 1.2
Z lm Score = −3.63
NG = 0
DB = 0
SM = 0
Delta_r
Doxycycline
YLR278C
ZShift = 0.14
Z lm Score = −0.92
NG = 0
DB = 0
SM = 0
Delta_AUC
Doxycycline

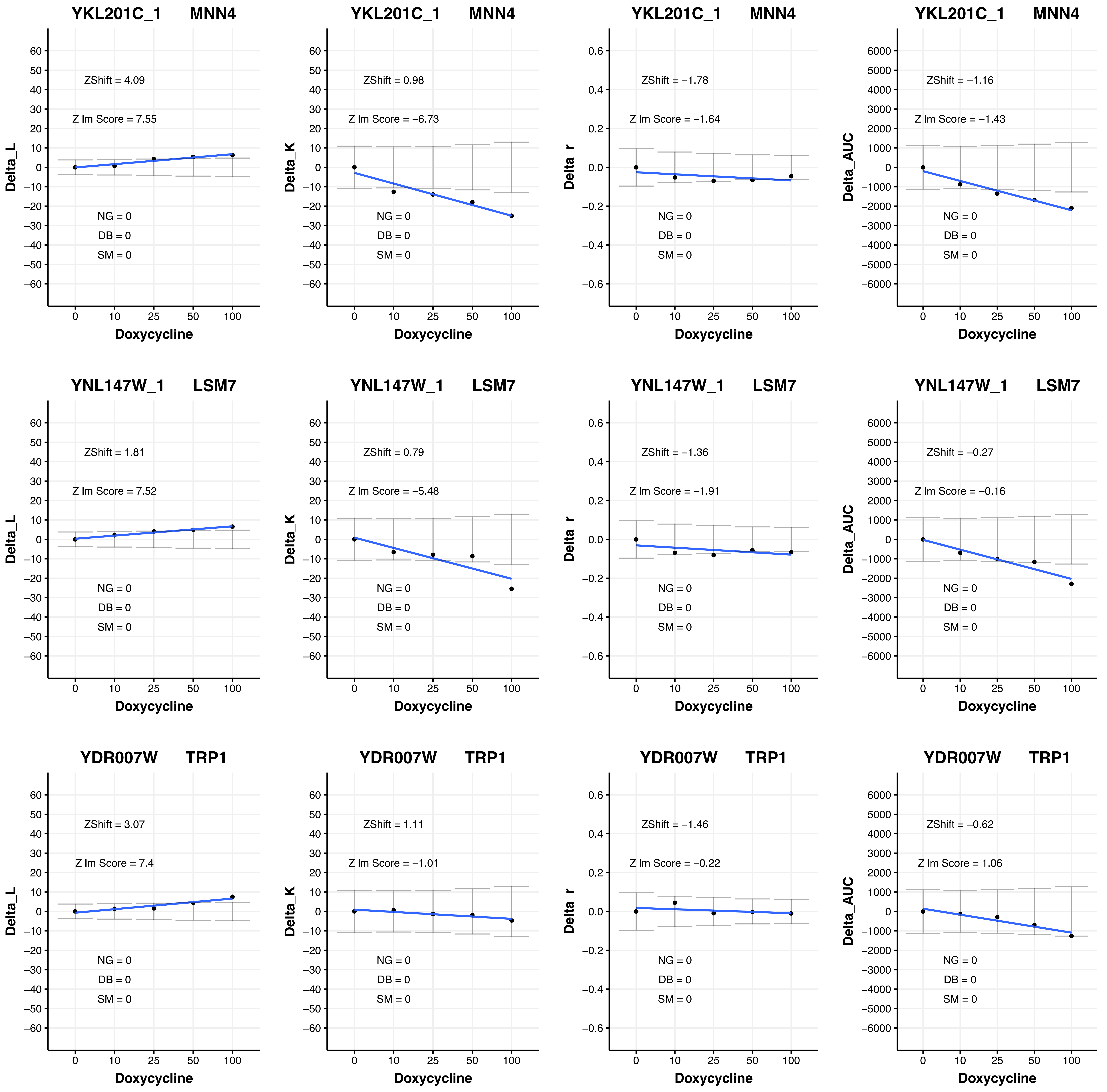
YKL201C_1 MNN4
ZShift = 4.09
Z lm Score = 7.55
NG = 0
DB = 0
SM = 0
Delta_L
Doxycycline
YKL201C_1 MNN4
ZShift = 0.98
Z lm Score = −6.73
NG = 0
DB = 0
SM = 0
Delta_K
Doxycycline
YKL201C_1 MNN4
ZShift = −1.78
Z lm Score = −1.64
NG = 0
DB = 0
SM = 0
Delta_r
Doxycycline
YKL201C_1 MNN4
ZShift = −1.16
Z lm Score = −1.43
NG = 0
DB = 0
SM = 0
Delta_AUC
Doxycycline
YNL147W_1 LSM7
ZShift = 1.81
Z lm Score = 7.52
NG = 0
DB = 0
SM = 0
Delta_L
Doxycycline
YNL147W_1 LSM7
ZShift = 0.79
Z lm Score = −5.48
NG = 0
DB = 0
SM = 0
Delta_K
Doxycycline
YNL147W_1 LSM7
ZShift = −1.36
Z lm Score = −1.91
NG = 0
DB = 0
SM = 0
Delta_r
Doxycycline
YNL147W_1 LSM7
ZShift = −0.27
Z lm Score = −0.16
NG = 0
DB = 0
SM = 0
Delta_AUC
Doxycycline
YDR007W TRP1
ZShift = 3.07
Z lm Score = 7.4
NG = 0
DB = 0
SM = 0
Delta_L
Doxycycline
YDR007W TRP1
ZShift = 1.11
Z lm Score = −1.01
NG = 0
DB = 0
SM = 0
Delta_K
Doxycycline
YDR007W TRP1
ZShift = −1.46
Z lm Score = −0.22
NG = 0
DB = 0
SM = 0
Delta_r
Doxycycline
YDR007W TRP1
ZShift = −0.62
Z lm Score = 1.06
NG = 0
DB = 0
SM = 0
Delta_AUC
Doxycycline

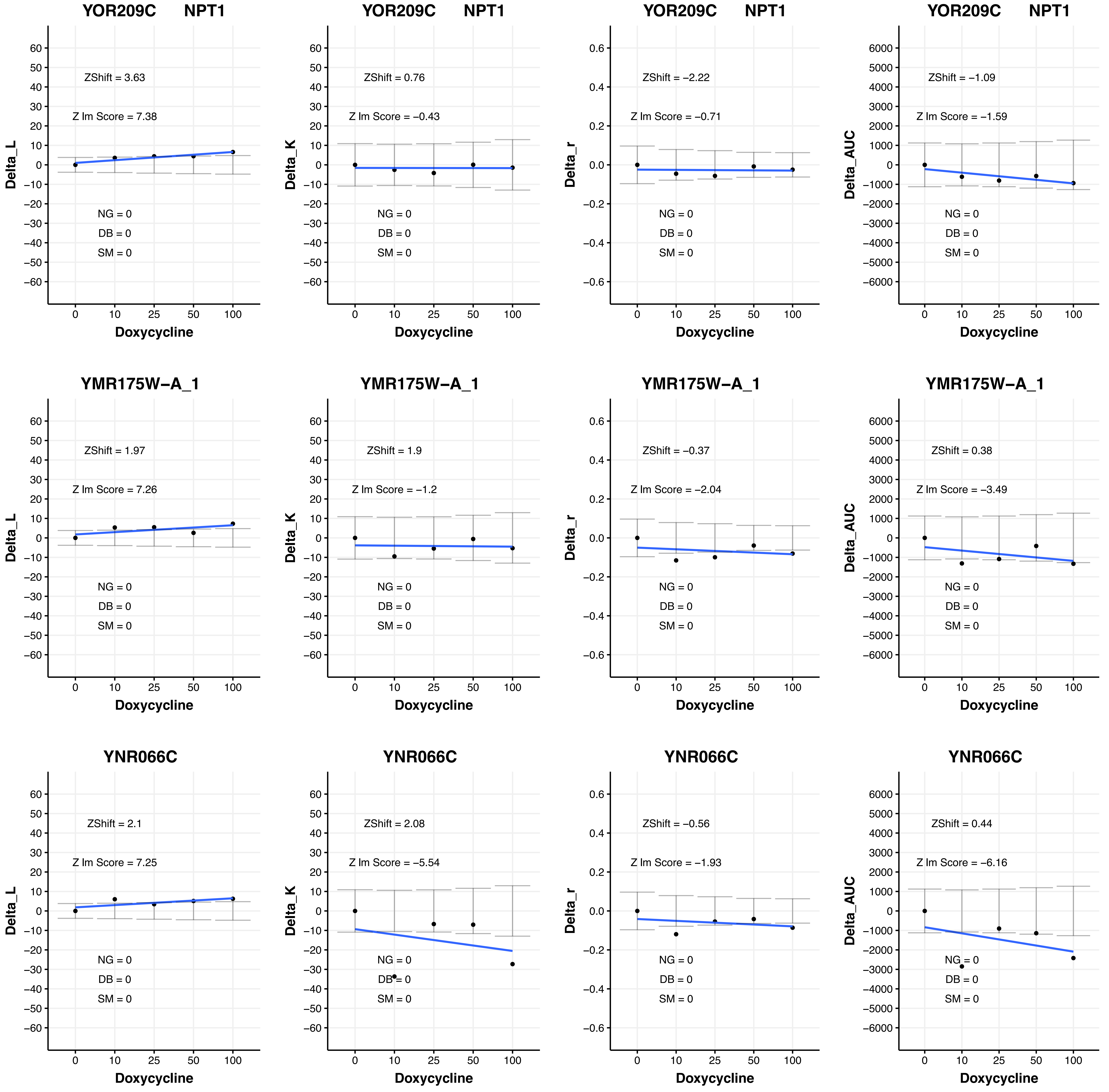

YOR209C NPT1
ZShift = 3.63
Z lm Score = 7.38
NG = 0
DB = 0
SM = 0
Delta_L
Doxycycline
YOR209C NPT1
ZShift = 0.76
Z lm Score = −0.43
NG = 0
DB = 0
SM = 0
Delta_K
Doxycycline
YOR209C NPT1
ZShift = −2.22
Z lm Score = −0.71
NG = 0
DB = 0
SM = 0
Delta_r
Doxycycline
YOR209C NPT1
ZShift = −1.09
Z lm Score = −1.59
NG = 0
DB = 0
SM = 0
Delta_AUC
Doxycycline
YMR175W−A_1
ZShift = 1.97
Z lm Score = 7.26
NG = 0
DB = 0
SM = 0
Delta_L
Doxycycline
YMR175W−A_1
ZShift = 1.9
Z lm Score = −1.2
NG = 0
DB = 0
SM = 0
Delta_K
Doxycycline
YMR175W−A_1
ZShift = −0.37
Z lm Score = −2.04
NG = 0
DB = 0
SM = 0
Delta_r
Doxycycline
YMR175W−A_1
ZShift = 0.38
Z lm Score = −3.49
NG = 0
DB = 0
SM = 0
Delta_AUC
Doxycycline
YNR066C
ZShift = 2.1
Z lm Score = 7.25
NG = 0
DB = 0
SM = 0
Delta_L
Doxycycline
YNR066C
ZShift = 2.08
Z lm Score = −5.54
NG = 0
DB = 0
SM = 0
Delta_K
Doxycycline
YNR066C
ZShift = −0.56
Z lm Score = −1.93
NG = 0
DB = 0
SM = 0
Delta_r
Doxycycline
YNR066C
ZShift = 0.44
Z lm Score = −6.16
NG = 0
DB = 0
SM = 0
Delta_AUC
Doxycycline

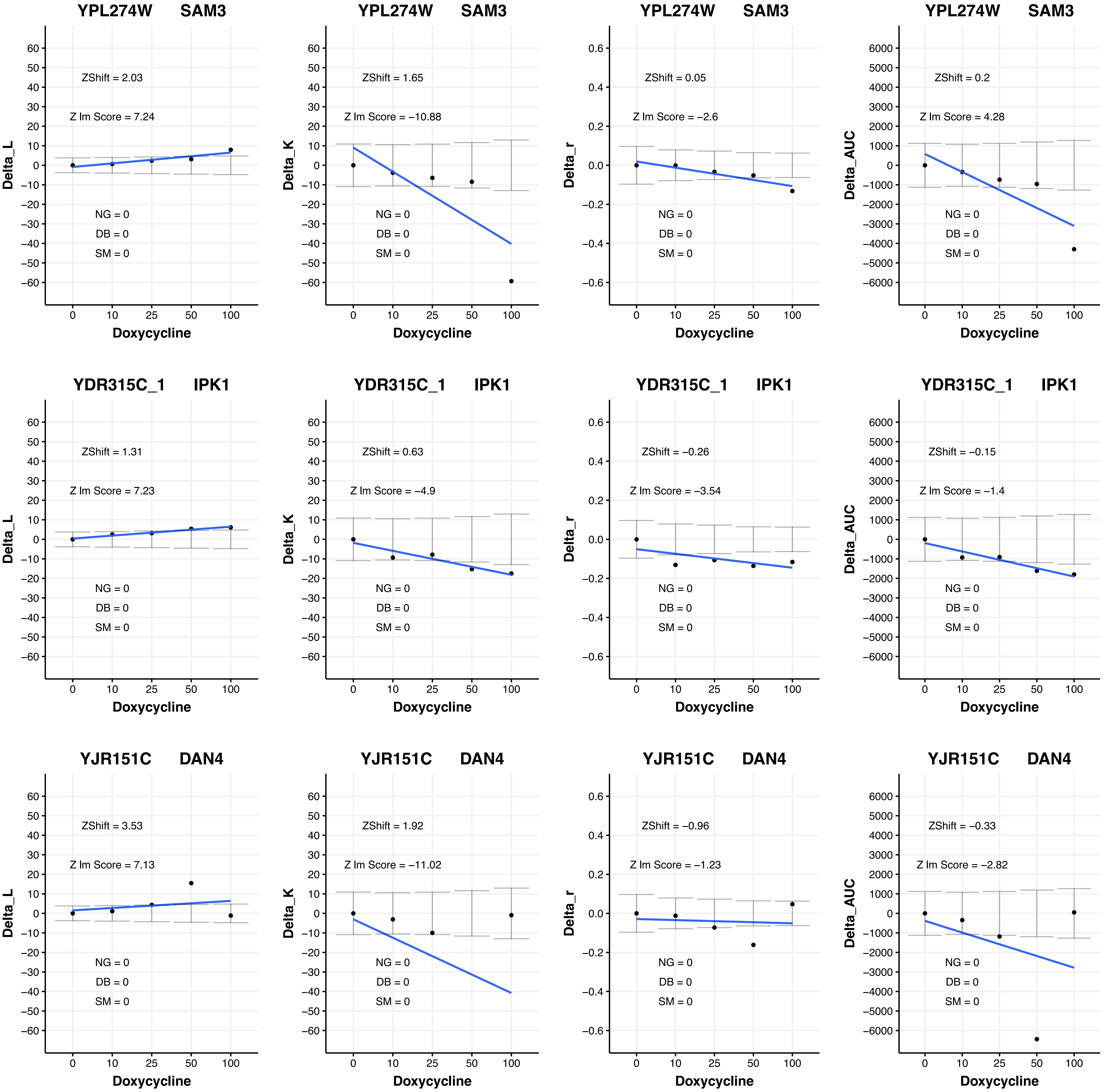

YPL274W SAM3
ZShift = 2.03
Z lm Score = 7.24
NG = 0
DB = 0
SM = 0
Delta_L
Doxycycline
YPL274W SAM3
ZShift = 1.65
Z lm Score = −10.88
NG = 0
DB = 0
SM = 0
Delta_K
Doxycycline
YPL274W SAM3
ZShift = 0.05
Z lm Score = −2.6
NG = 0
DB = 0
SM = 0
Delta_r
Doxycycline
YPL274W SAM3
ZShift = 0.2
Z lm Score = 4.28
NG = 0
DB = 0
SM = 0
Delta_AUC
Doxycycline
YDR315C_1 IPK1
ZShift = 1.31
Z lm Score = 7.23
NG = 0
DB = 0
SM = 0
Delta_L
Doxycycline
YDR315C_1 IPK1
ZShift = 0.63
Z lm Score = −4.9
NG = 0
DB = 0
SM = 0
Delta_K
Doxycycline
YDR315C_1 IPK1
ZShift = −0.26
Z lm Score = −3.54
NG = 0
DB = 0
SM = 0
Delta_r
Doxycycline
YDR315C_1 IPK1
ZShift = −0.15
Z lm Score = −1.4
NG = 0
DB = 0
SM = 0
Delta_AUC
Doxycycline
YJR151C DAN4
ZShift = 3.53
Z lm Score = 7.13
NG = 0
DB = 0
SM = 0
Delta_L
Doxycycline
YJR151C DAN4
ZShift = 1.92
Z lm Score = −11.02
NG = 0
DB = 0
SM = 0
Delta_K
Doxycycline
YJR151C DAN4
ZShift = −0.96
Z lm Score = −1.23
NG = 0
DB = 0
SM = 0
Delta_r
Doxycycline
YJR151C DAN4
ZShift = −0.33
Z lm Score = −2.82
NG = 0
DB = 0
SM = 0
Delta_AUC
Doxycycline

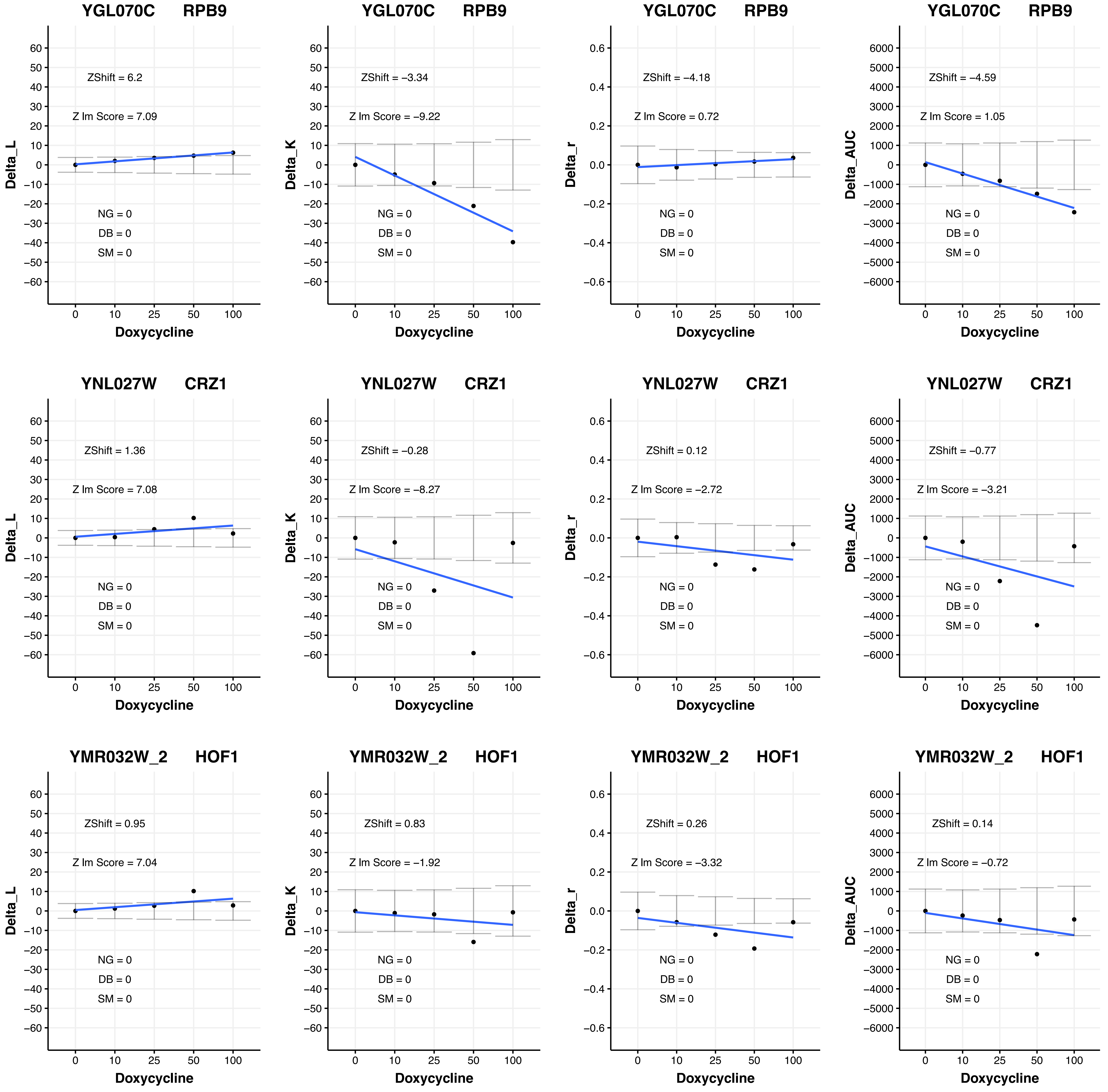

YGL070C RPB9
ZShift = 6.2
Z lm Score = 7.09
NG = 0
DB = 0
SM = 0
Delta_L
Doxycycline
YGL070C RPB9
ZShift = −3.34
Z lm Score = −9.22
NG = 0
DB = 0
SM = 0
Delta_K
Doxycycline
YGL070C RPB9
ZShift = −4.18
Z lm Score = 0.72
NG = 0
DB = 0
SM = 0
Delta_r
Doxycycline
YGL070C RPB9
ZShift = −4.59
Z lm Score = 1.05
NG = 0
DB = 0
SM = 0
Delta_AUC
Doxycycline
YNL027W CRZ1
ZShift = 1.36
Z lm Score = 7.08
NG = 0
DB = 0
SM = 0
Delta_L
Doxycycline
YNL027W CRZ1
ZShift = −0.28
Z lm Score = −8.27
NG = 0
DB = 0
SM = 0
Delta_K
Doxycycline
YNL027W CRZ1
ZShift = 0.12
Z lm Score = −2.72
NG = 0
DB = 0
SM = 0
Delta_r
Doxycycline
YNL027W CRZ1
ZShift = −0.77
Z lm Score = −3.21
NG = 0
DB = 0
SM = 0
Delta_AUC
Doxycycline
YMR032W_2 HOF1
ZShift = 0.95
Z lm Score = 7.04
NG = 0
DB = 0
SM = 0
Delta_L
Doxycycline
YMR032W_2 HOF1
ZShift = 0.83
Z lm Score = −1.92
NG = 0
DB = 0
SM = 0
Delta_K
Doxycycline
YMR032W_2 HOF1
ZShift = 0.26
Z lm Score = −3.32
NG = 0
DB = 0
SM = 0
Delta_r
Doxycycline
YMR032W_2 HOF1
ZShift = 0.14
Z lm Score = −0.72
NG = 0
DB = 0
SM = 0
Delta_AUC
Doxycycline

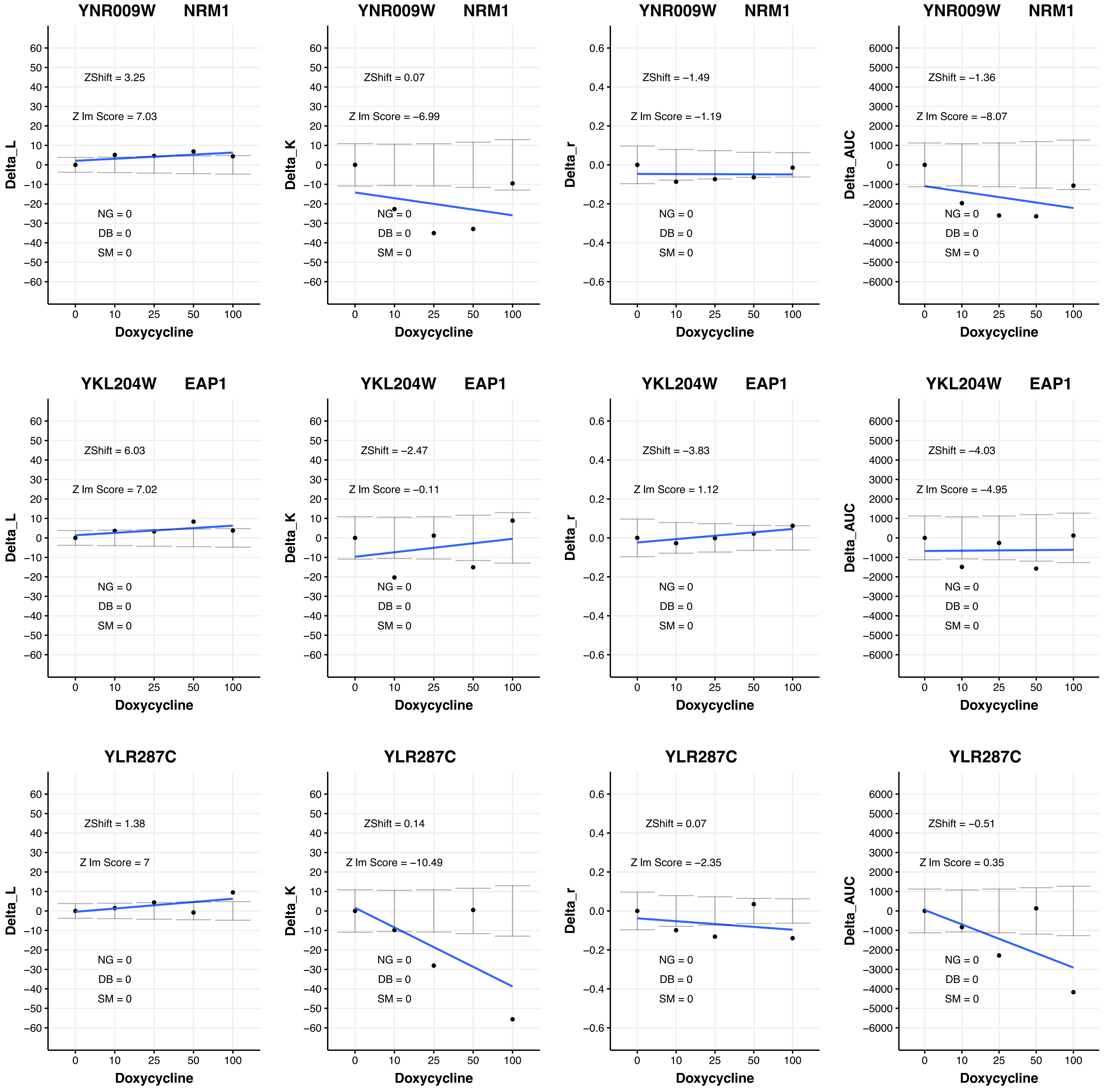
YNR009W NRM1
ZShift = 3.25
Z lm Score = 7.03
NG = 0
DB = 0
SM = 0
Delta_L
Doxycycline
YNR009W NRM1
ZShift = 0.07
Z lm Score = −6.99
NG = 0
DB = 0
SM = 0
Delta_K
Doxycycline
YNR009W NRM1
ZShift = −1.49
Z lm Score = −1.19
NG = 0
DB = 0
SM = 0
Delta_r
Doxycycline
YNR009W NRM1
ZShift = −1.36
Z lm Score = −8.07
NG = 0
DB = 0
SM = 0
Delta_AUC
Doxycycline
YKL204W EAP1
ZShift = 6.03
Z lm Score = 7.02
NG = 0
DB = 0
SM = 0
Delta_L
Doxycycline
YKL204W EAP1
ZShift = −2.47
Z lm Score = −0.11
NG = 0
DB = 0
SM = 0
Delta_K
Doxycycline
YKL204W EAP1
ZShift = −3.83
Z lm Score = 1.12
NG = 0
DB = 0
SM = 0
Delta_r
Doxycycline
YKL204W EAP1
ZShift = −4.03
Z lm Score = −4.95
NG = 0
DB = 0
SM = 0
Delta_AUC
Doxycycline
YLR287C
ZShift = 1.38
Z lm Score = 7
NG = 0
DB = 0
SM = 0
Delta_L
Doxycycline
YLR287C
ZShift = 0.14
Z lm Score = −10.49
NG = 0
DB = 0
SM = 0
Delta_K
Doxycycline
YLR287C
ZShift = 0.07
Z lm Score = −2.35
NG = 0
DB = 0
SM = 0
Delta_r
Doxycycline
YLR287C
ZShift = −0.51
Z lm Score = 0.35
NG = 0
DB = 0
SM = 0
Delta_AUC
Doxycycline

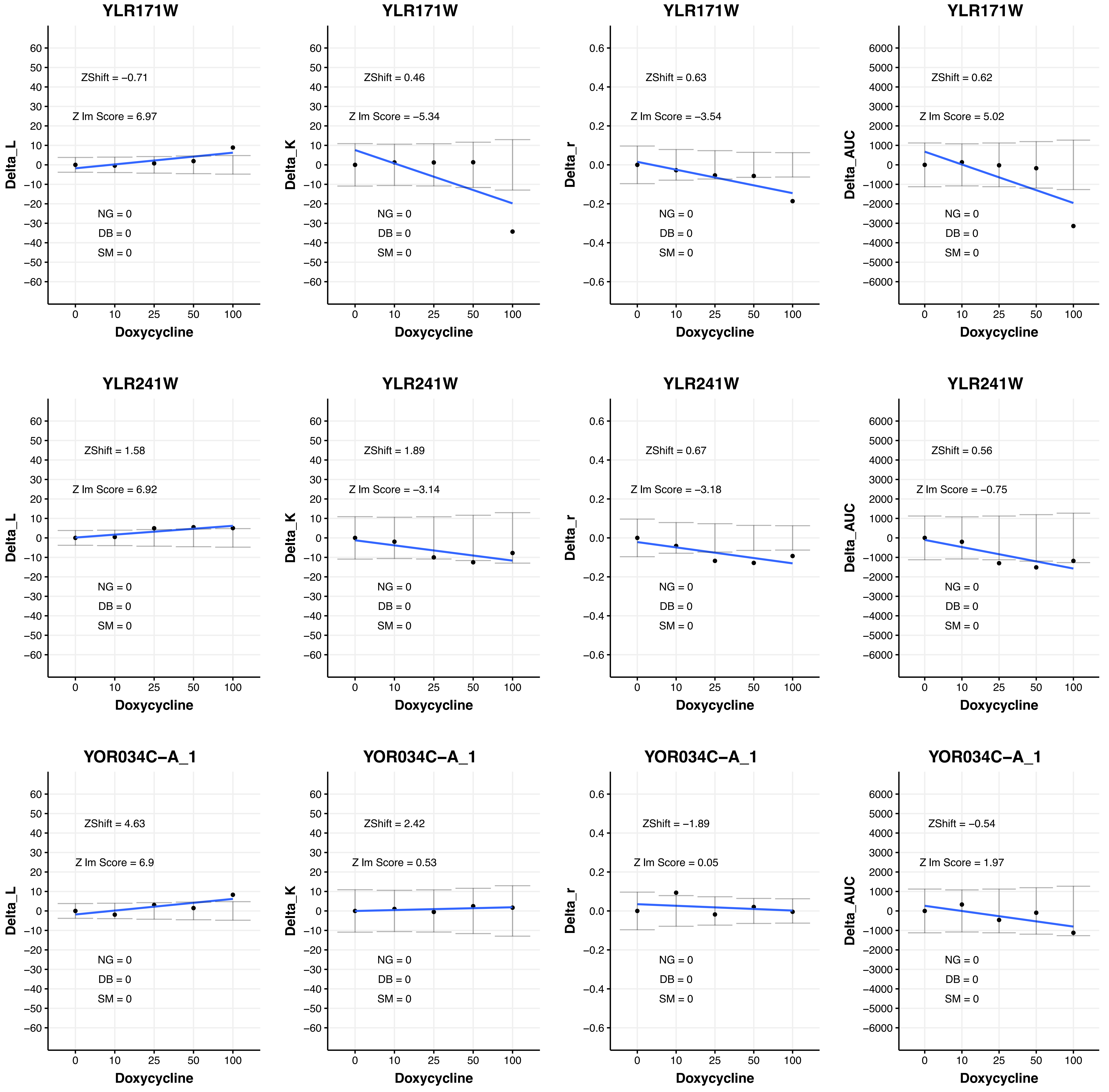

YLR171W
Delta_L
ZShift = −0.71
Z lm Score = 6.97
NG = 0
DB = 0
SM = 0
YLR171W
Delta_K
ZShift = 0.46
Z lm Score = −5.34
NG = 0
DB = 0
SM = 0
YLR171W
Delta_r
ZShift = 0.63
Z lm Score = −3.54
NG = 0
DB = 0
SM = 0
YLR171W
Delta_AUC
ZShift = 0.62
Z lm Score = 5.02
NG = 0
DB = 0
SM = 0
YLR241W
Delta_L
ZShift = 1.58
Z lm Score = 6.92
NG = 0
DB = 0
SM = 0
YLR241W
Delta_K
ZShift = 1.89
Z lm Score = −3.14
NG = 0
DB = 0
SM = 0
YLR241W
Delta_r
ZShift = 0.67
Z lm Score = −3.18
NG = 0
DB = 0
SM = 0
YLR241W
Delta_AUC
ZShift = 0.56
Z lm Score = −0.75
NG = 0
DB = 0
SM = 0
YOR034C−A_1
Delta_L
ZShift = 4.63
Z lm Score = 6.9
NG = 0
DB = 0
SM = 0
YOR034C−A_1
Delta_K
ZShift = 2.42
Z lm Score = 0.53
NG = 0
DB = 0
SM = 0
YOR034C−A_1
Delta_r
ZShift = −1.89
Z lm Score = 0.05
NG = 0
DB = 0
SM = 0
YOR034C−A_1
Delta_AUC
ZShift = −0.54
Z lm Score = 1.97
NG = 0
DB = 0
SM = 0
Doxycycline
0
10
25
50
100

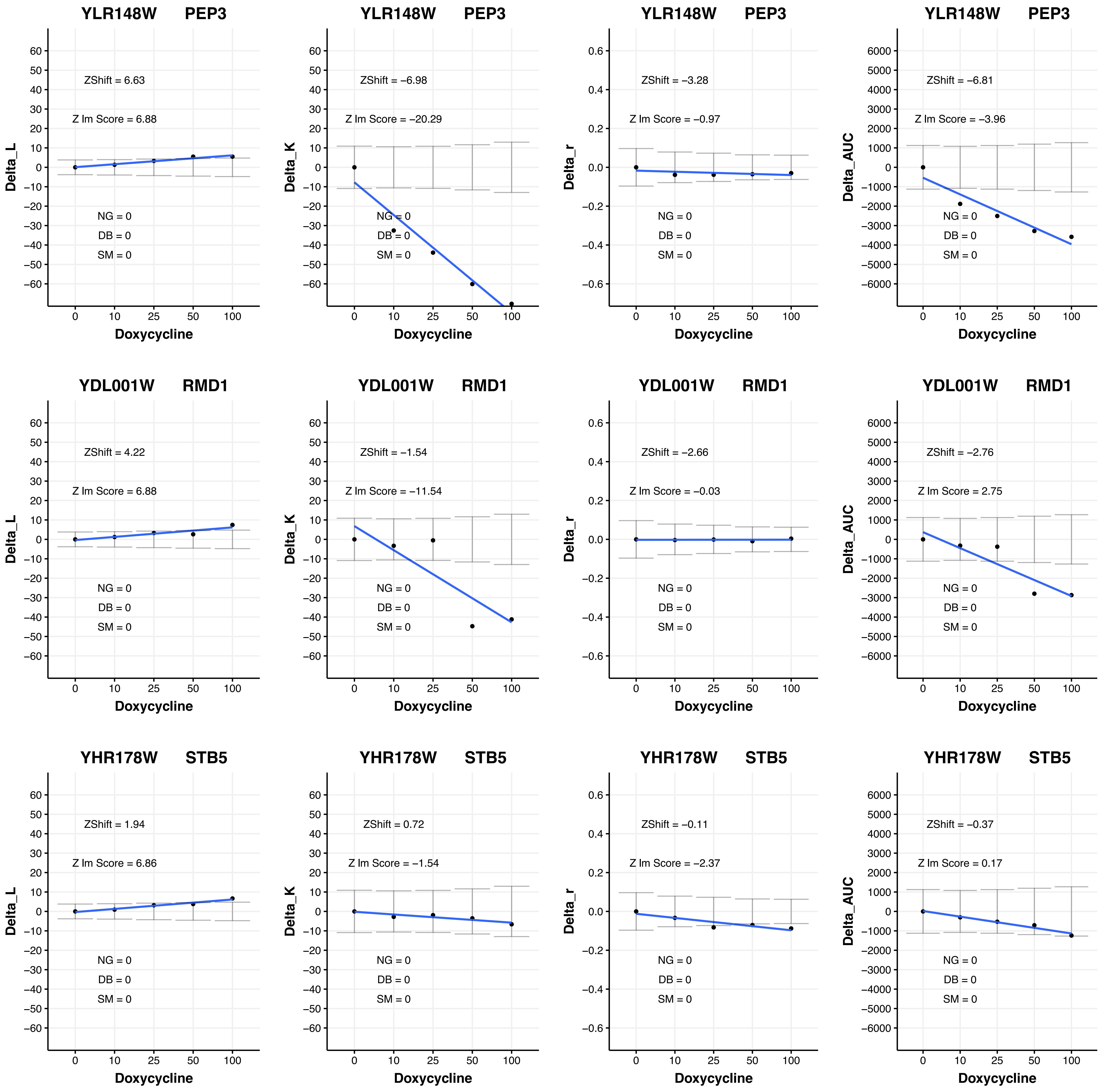
YLR148W PEP3
ZShift = 6.63
Z lm Score = 6.88
NG = 0
DB = 0
SM = 0
Delta_L
Doxycycline
YLR148W PEP3
ZShift = −6.98
Z lm Score = −20.29
NG = 0
DB = 0
SM = 0
Delta_K
Doxycycline
YLR148W PEP3
ZShift = −3.28
Z lm Score = −0.97
NG = 0
DB = 0
SM = 0
Delta_r
Doxycycline
YLR148W PEP3
ZShift = −6.81
Z lm Score = −3.96
NG = 0
DB = 0
SM = 0
Delta_AUC
Doxycycline
YDL001W RMD1
ZShift = 4.22
Z lm Score = 6.88
NG = 0
DB = 0
SM = 0
Delta_L
Doxycycline
YDL001W RMD1
ZShift = −1.54
Z lm Score = −11.54
NG = 0
DB = 0
SM = 0
Delta_K
Doxycycline
YDL001W RMD1
ZShift = −2.66
Z lm Score = −0.03
NG = 0
DB = 0
SM = 0
Delta_r
Doxycycline
YDL001W RMD1
ZShift = −2.76
Z lm Score = 2.75
NG = 0
DB = 0
SM = 0
Delta_AUC
Doxycycline
YHR178W STB5
ZShift = 1.94
Z lm Score = 6.86
NG = 0
DB = 0
SM = 0
Delta_L
Doxycycline
YHR178W STB5
ZShift = 0.72
Z lm Score = −1.54
NG = 0
DB = 0
SM = 0
Delta_K
Doxycycline
YHR178W STB5
ZShift = −0.11
Z lm Score = −2.37
NG = 0
DB = 0
SM = 0
Delta_r
Doxycycline
YHR178W STB5
ZShift = −0.37
Z lm Score = 0.17
NG = 0
DB = 0
SM = 0
Delta_AUC
Doxycycline

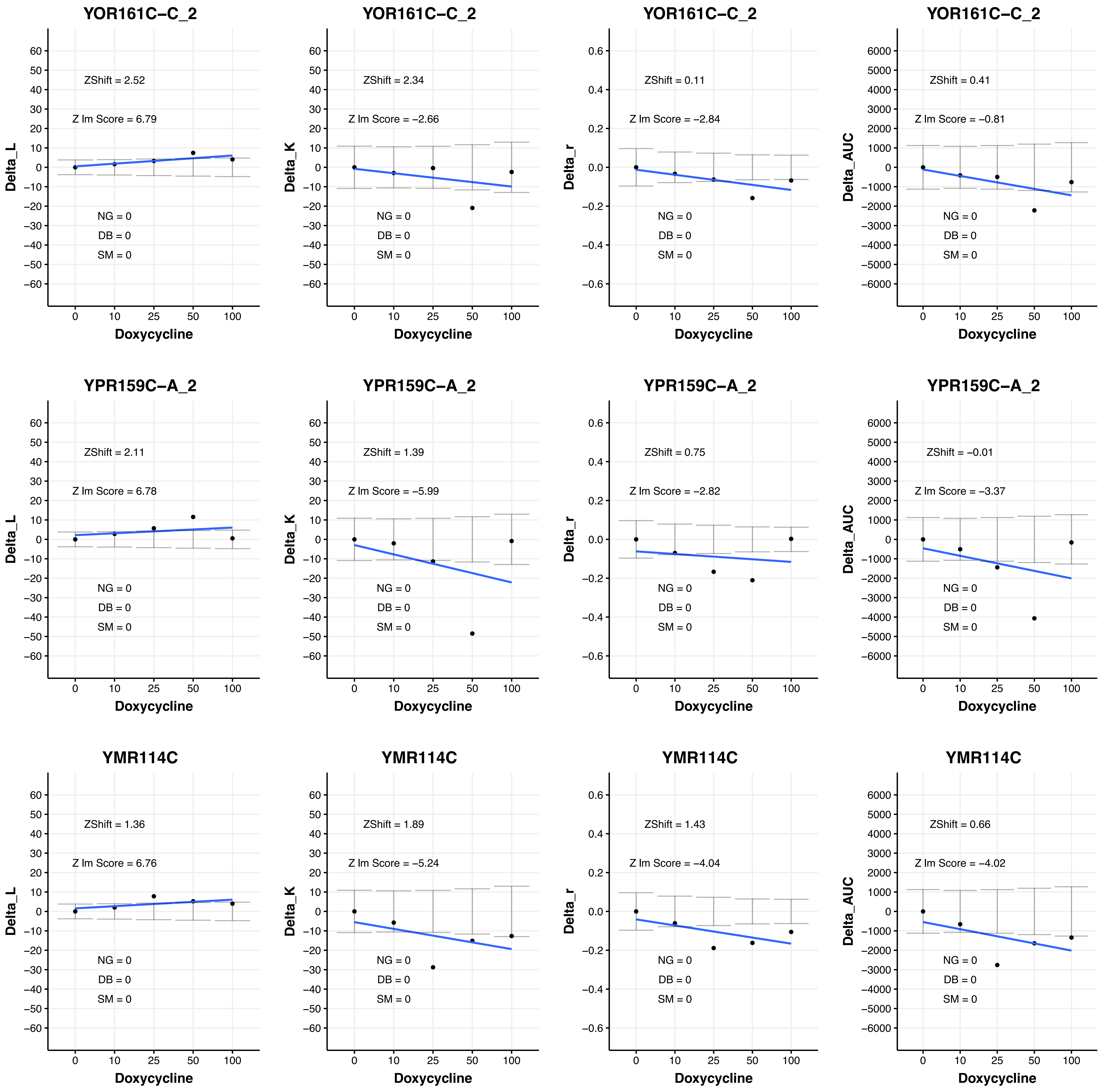

YOR161C–C_2
ZShift = 2.52
Z lm Score = 6.79
NG = 0
DB = 0
SM = 0
Delta_L
Doxycycline
YOR161C–C_2
ZShift = 2.34
Z lm Score = –2.66
NG = 0
DB = 0
SM = 0
Delta_K
Doxycycline
YOR161C–C_2
ZShift = 0.11
Z lm Score = –2.84
NG = 0
DB = 0
SM = 0
Delta_r
Doxycycline
YOR161C–C_2
ZShift = 0.41
Z lm Score = –0.81
NG = 0
DB = 0
SM = 0
Delta_AUC
Doxycycline
YPR159C–A_2
ZShift = 2.11
Z lm Score = 6.78
NG = 0
DB = 0
SM = 0
Delta_L
Doxycycline
YPR159C–A_2
ZShift = 1.39
Z lm Score = –5.99
NG = 0
DB = 0
SM = 0
Delta_K
Doxycycline
YPR159C–A_2
ZShift = 0.75
Z lm Score = –2.82
NG = 0
DB = 0
SM = 0
Delta_r
Doxycycline
YPR159C–A_2
ZShift = –0.01
Z lm Score = –3.37
NG = 0
DB = 0
SM = 0
Delta_AUC
Doxycycline
YMR114C
ZShift = 1.36
Z lm Score = 6.76
NG = 0
DB = 0
SM = 0
Delta_L
Doxycycline
YMR114C
ZShift = 1.89
Z lm Score = –5.24
NG = 0
DB = 0
SM = 0
Delta_K
Doxycycline
YMR114C
ZShift = 1.43
Z lm Score = –4.04
NG = 0
DB = 0
SM = 0
Delta_r
Doxycycline
YMR114C
ZShift = 0.66
Z lm Score = –4.02
NG = 0
DB = 0
SM = 0
Delta_AUC
Doxycycline

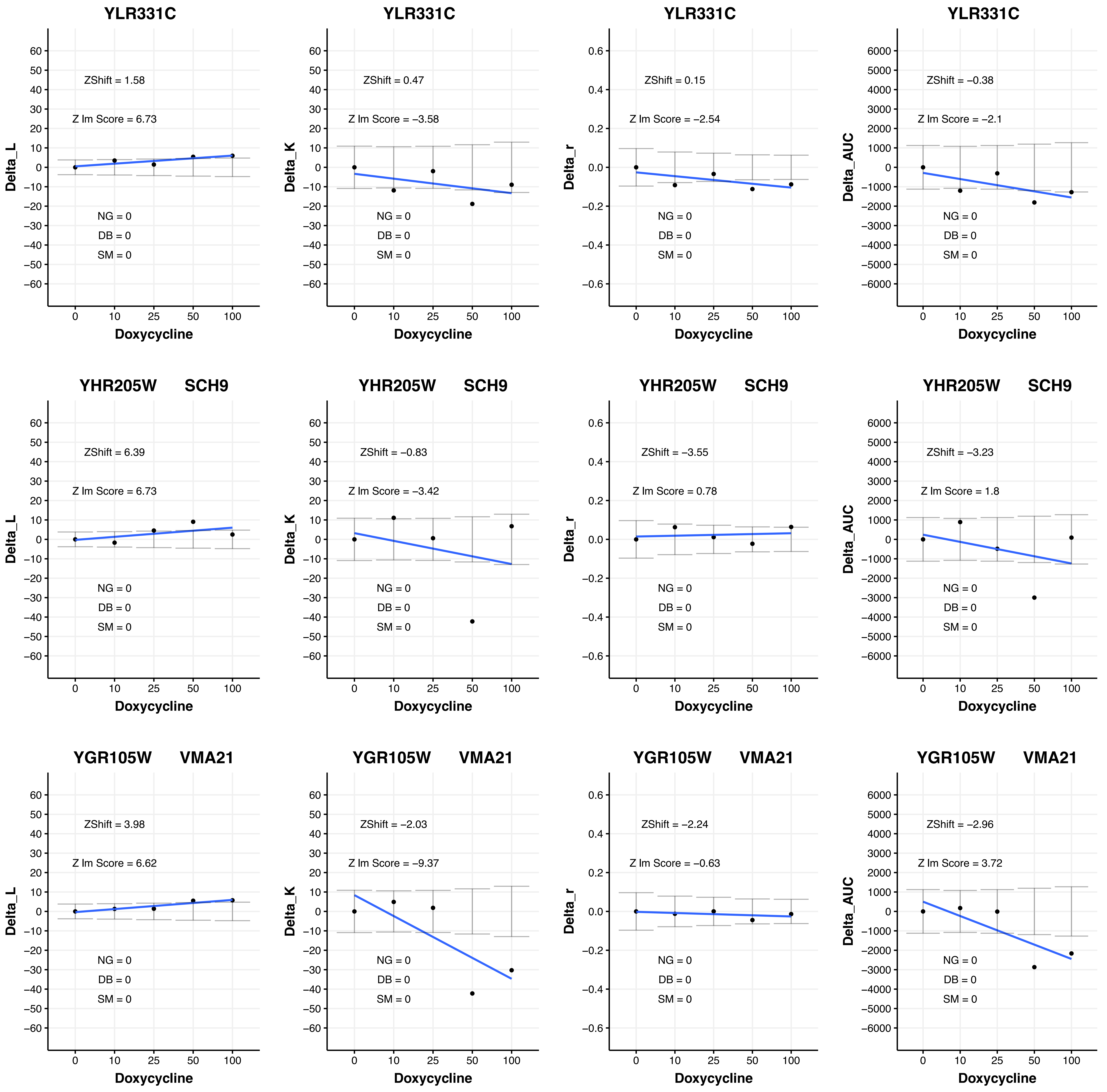
YLR331C
ZShift = 1.58
Z lm Score = 6.73
NG = 0
DB = 0
SM = 0
Delta_L
Doxycycline
YLR331C
ZShift = 0.47
Z lm Score = −3.58
NG = 0
DB = 0
SM = 0
Delta_K
Doxycycline
YLR331C
ZShift = 0.15
Z lm Score = −2.54
NG = 0
DB = 0
SM = 0
Delta_r
Doxycycline
YLR331C
ZShift = −0.38
Z lm Score = −2.1
NG = 0
DB = 0
SM = 0
Delta_AUC
Doxycycline
YHR205W SCH9
ZShift = 6.39
Z lm Score = 6.73
NG = 0
DB = 0
SM = 0
Delta_L
Doxycycline
YHR205W SCH9
ZShift = −0.83
Z lm Score = −3.42
NG = 0
DB = 0
SM = 0
Delta_K
Doxycycline
YHR205W SCH9
ZShift = −3.55
Z lm Score = 0.78
NG = 0
DB = 0
SM = 0
Delta_r
Doxycycline
YHR205W SCH9
ZShift = −3.23
Z lm Score = 1.8
NG = 0
DB = 0
SM = 0
Delta_AUC
Doxycycline
YGR105W VMA21
ZShift = 3.98
Z lm Score = 6.62
NG = 0
DB = 0
SM = 0
Delta_L
Doxycycline
YGR105W VMA21
ZShift = −2.03
Z lm Score = −9.37
NG = 0
DB = 0
SM = 0
Delta_K
Doxycycline
YGR105W VMA21
ZShift = −2.24
Z lm Score = −0.63
NG = 0
DB = 0
SM = 0
Delta_r
Doxycycline
YGR105W VMA21
ZShift = −2.96
Z lm Score = 3.72
NG = 0
DB = 0
SM = 0
Delta_AUC
Doxycycline

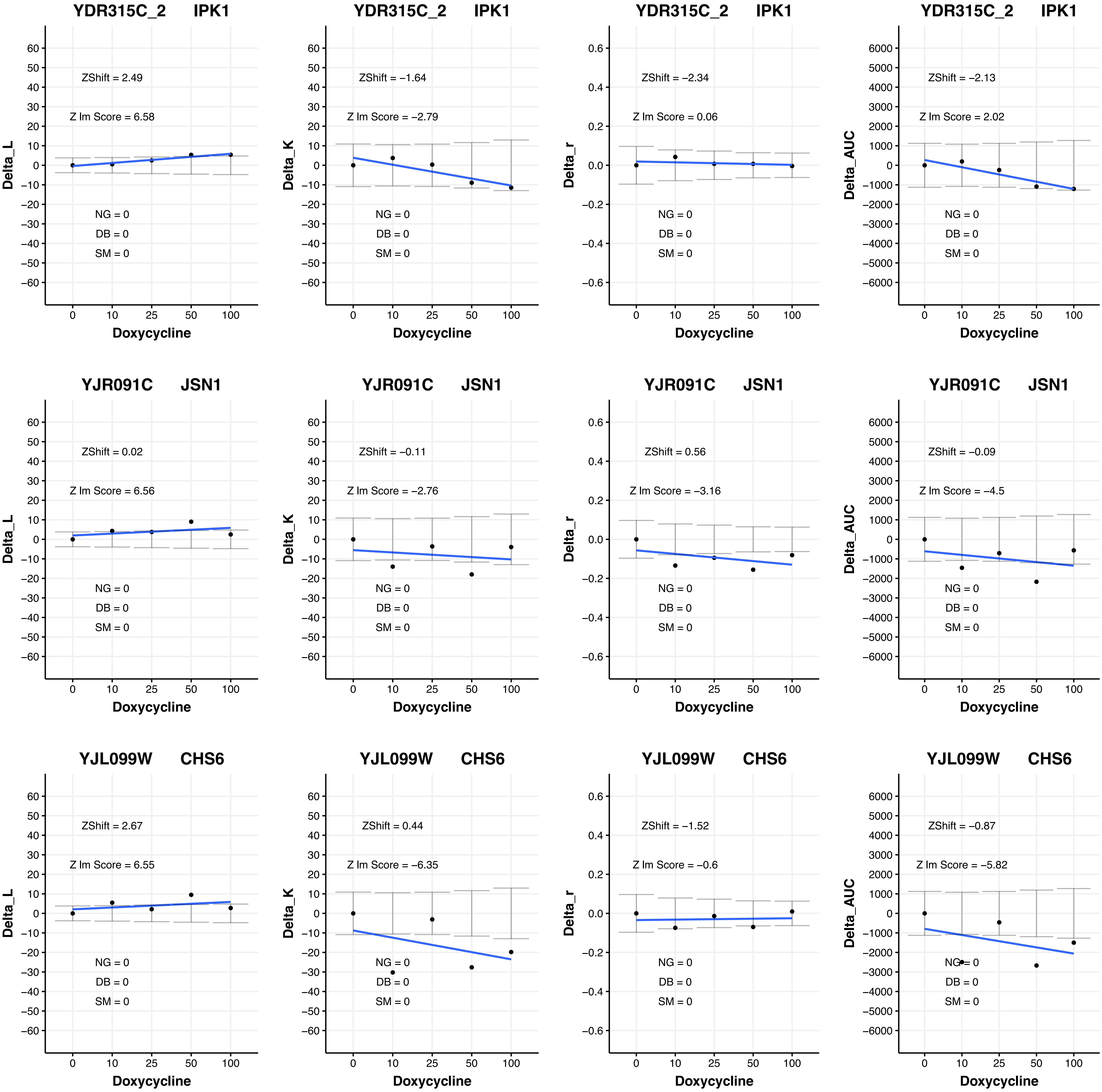
YDR315C_2 IPK1
ZShift = 2.49
Z lm Score = 6.58
NG = 0
DB = 0
SM = 0
Delta_L
Doxycycline
YDR315C_2 IPK1
ZShift = −1.64
Z lm Score = −2.79
NG = 0
DB = 0
SM = 0
Delta_K
Doxycycline
YDR315C_2 IPK1
ZShift = −2.34
Z lm Score = 0.06
NG = 0
DB = 0
SM = 0
Delta_r
Doxycycline
YDR315C_2 IPK1
ZShift = −2.13
Z lm Score = 2.02
NG = 0
DB = 0
SM = 0
Delta_AUC
Doxycycline
YJR091C JSN1
ZShift = 0.02
Z lm Score = 6.56
NG = 0
DB = 0
SM = 0
Delta_L
Doxycycline
YJR091C JSN1
ZShift = −0.11
Z lm Score = −2.76
NG = 0
DB = 0
SM = 0
Delta_K
Doxycycline
YJR091C JSN1
ZShift = 0.56
Z lm Score = −3.16
NG = 0
DB = 0
SM = 0
Delta_r
Doxycycline
YJR091C JSN1
ZShift = −0.09
Z lm Score = −4.5
NG = 0
DB = 0
SM = 0
Delta_AUC
Doxycycline
YJL099W CHS6
ZShift = 2.67
Z lm Score = 6.55
NG = 0
DB = 0
SM = 0
Delta_L
Doxycycline
YJL099W CHS6
ZShift = 0.44
Z lm Score = −6.35
NG = 0
DB = 0
SM = 0
Delta_K
Doxycycline
YJL099W CHS6
ZShift = −1.52
Z lm Score = −0.6
NG = 0
DB = 0
SM = 0
Delta_r
Doxycycline
YJL099W CHS6
ZShift = −0.87
Z lm Score = −5.82
NG = 0
DB = 0
SM = 0
Delta_AUC
Doxycycline

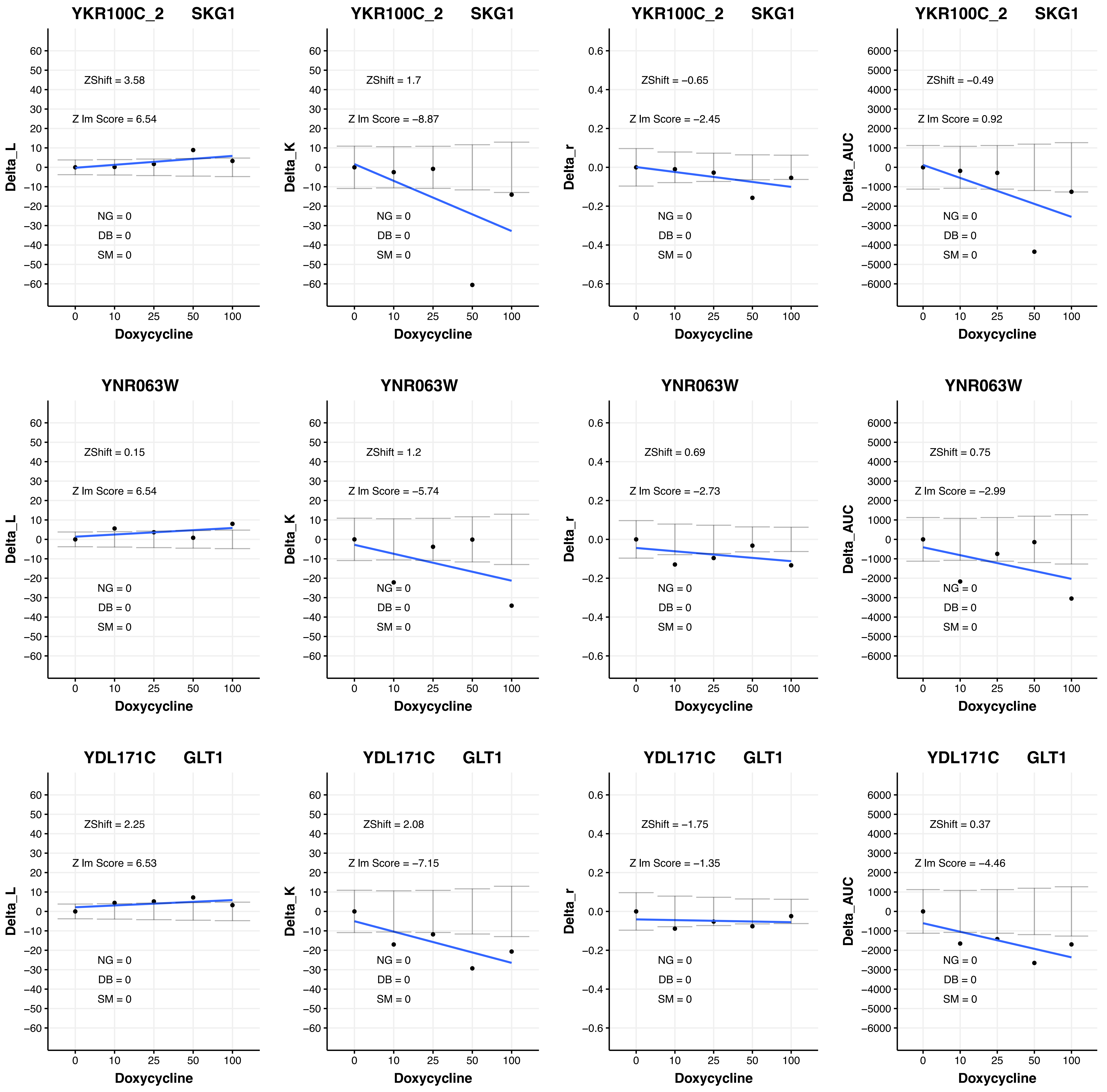
YKR100C_2 SKG1
ZShift = 3.58
Z lm Score = 6.54
NG = 0
DB = 0
SM = 0
Delta_L
Doxycycline
0 10 25 50 100
YKR100C_2 SKG1
ZShift = 1.7
Z lm Score = −8.87
NG = 0
DB = 0
SM = 0
Delta_K
Doxycycline
YKR100C_2 SKG1
ZShift = −0.65
Z lm Score = −2.45
NG = 0
DB = 0
SM = 0
Delta_r
Doxycycline
YKR100C_2 SKG1
ZShift = −0.49
Z lm Score = 0.92
NG = 0
DB = 0
SM = 0
Delta_AUC
Doxycycline
YNR063W
ZShift = 0.15
Z lm Score = 6.54
NG = 0
DB = 0
SM = 0
Delta_L
Doxycycline
YNR063W
ZShift = 1.2
Z lm Score = −5.74
NG = 0
DB = 0
SM = 0
Delta_K
Doxycycline
YNR063W
ZShift = 0.69
Z lm Score = −2.73
NG = 0
DB = 0
SM = 0
Delta_r
Doxycycline
YNR063W
ZShift = 0.75
Z lm Score = −2.99
NG = 0
DB = 0
SM = 0
Delta_AUC
Doxycycline
YDL171C GLT1
ZShift = 2.25
Z lm Score = 6.53
NG = 0
DB = 0
SM = 0
Delta_L
Doxycycline
YDL171C GLT1
ZShift = 2.08
Z lm Score = −7.15
NG = 0
DB = 0
SM = 0
Delta_K
Doxycycline
YDL171C GLT1
ZShift = −1.75
Z lm Score = −1.35
NG = 0
DB = 0
SM = 0
Delta_r
Doxycycline
YDL171C GLT1
ZShift = 0.37
Z lm Score = −4.46
NG = 0
DB = 0
SM = 0
Delta_AUC
Doxycycline

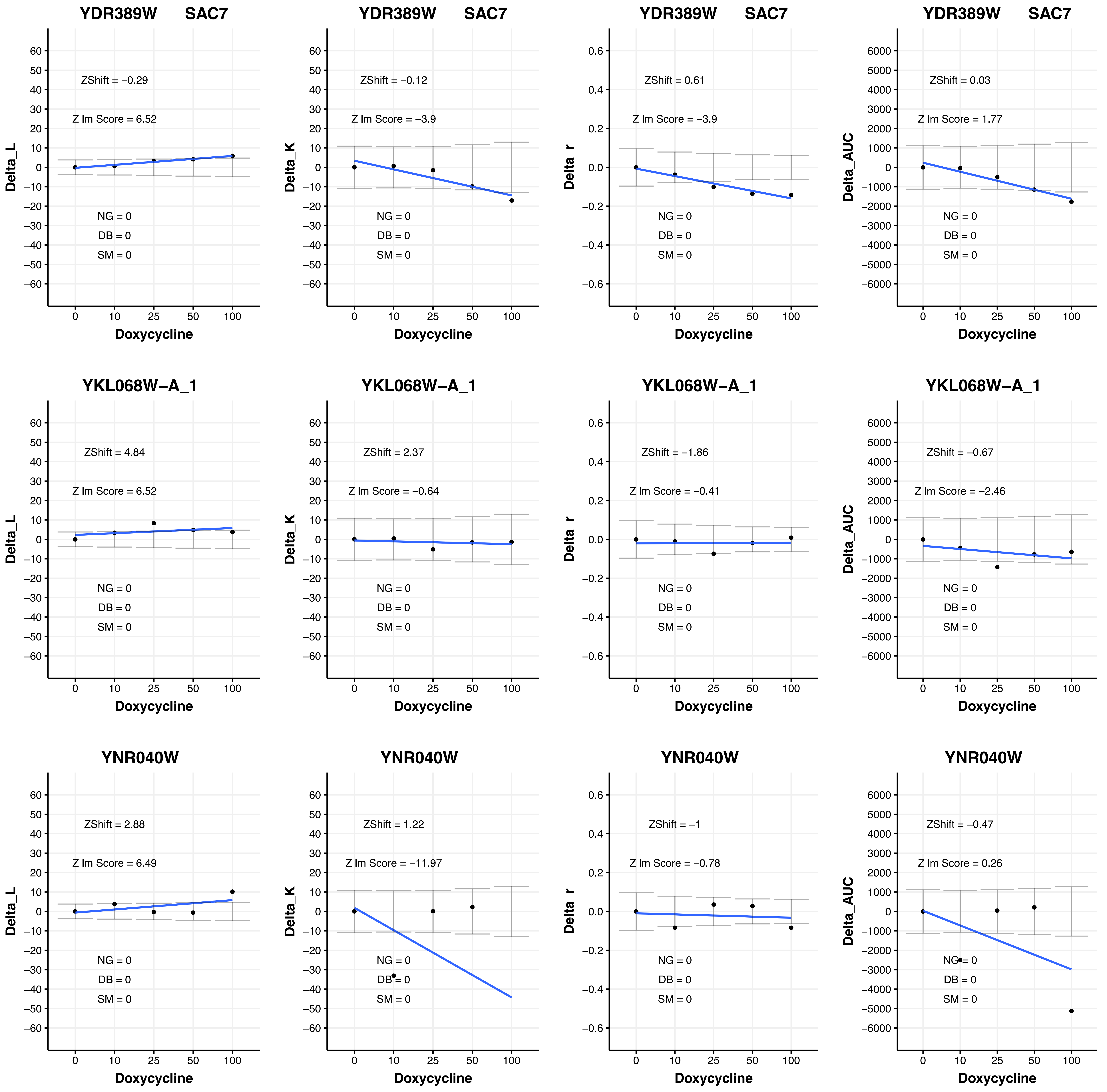
YDR389W SAC7
ZShift = −0.29
Z lm Score = 6.52
NG = 0
DB = 0
SM = 0
Delta_L
Doxycycline
YDR389W SAC7
ZShift = −0.12
Z lm Score = −3.9
NG = 0
DB = 0
SM = 0
Delta_K
Doxycycline
YDR389W SAC7
ZShift = 0.61
Z lm Score = −3.9
NG = 0
DB = 0
SM = 0
Delta_r
Doxycycline
YDR389W SAC7
ZShift = 0.03
Z lm Score = 1.77
NG = 0
DB = 0
SM = 0
Delta_AUC
Doxycycline
YKL068W−A_1
ZShift = 4.84
Z lm Score = 6.52
NG = 0
DB = 0
SM = 0
Delta_L
Doxycycline
YKL068W−A_1
ZShift = 2.37
Z lm Score = −0.64
NG = 0
DB = 0
SM = 0
Delta_K
Doxycycline
YKL068W−A_1
ZShift = −1.86
Z lm Score = −0.41
NG = 0
DB = 0
SM = 0
Delta_r
Doxycycline
YKL068W−A_1
ZShift = −0.67
Z lm Score = −2.46
NG = 0
DB = 0
SM = 0
Delta_AUC
Doxycycline
YNR040W
ZShift = 2.88
Z lm Score = 6.49
NG = 0
DB = 0
SM = 0
Delta_L
Doxycycline
YNR040W
ZShift = 1.22
Z lm Score = −11.97
NG = 0
DB = 0
SM = 0
Delta_K
Doxycycline
YNR040W
ZShift = −1
Z lm Score = −0.78
NG = 0
DB = 0
SM = 0
Delta_r
Doxycycline
YNR040W
ZShift = −0.47
Z lm Score = 0.26
NG = 0
DB = 0
SM = 0
Delta_AUC
Doxycycline

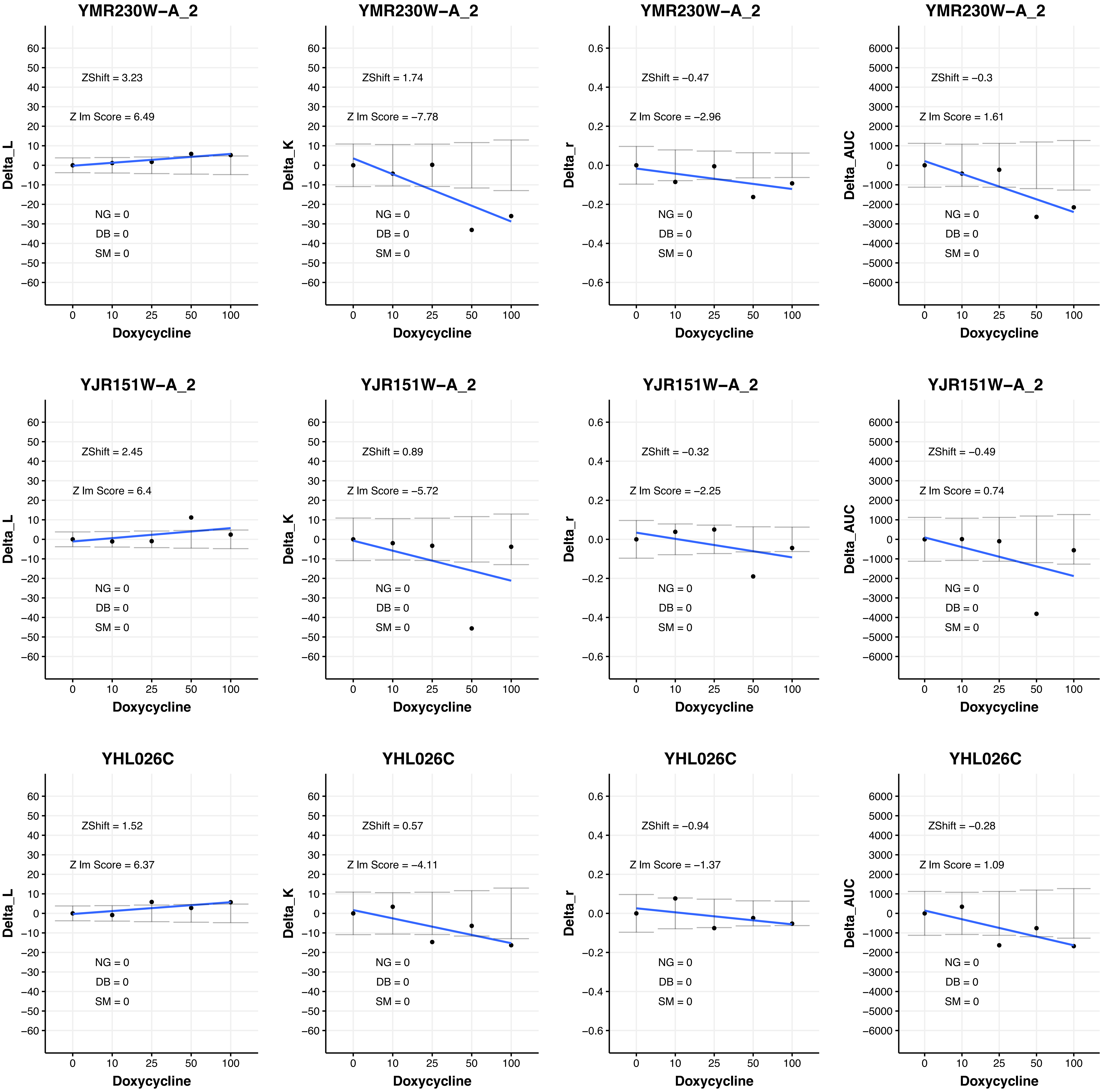
YMR230W−A_2
ZShift = 3.23
Z lm Score = 6.49
NG = 0
DB = 0
SM = 0
Delta_L
Doxycycline
YMR230W−A_2
ZShift = 1.74
Z lm Score = −7.78
NG = 0
DB = 0
SM = 0
Delta_K
Doxycycline
YMR230W−A_2
ZShift = −0.47
Z lm Score = −2.96
NG = 0
DB = 0
SM = 0
Delta_r
Doxycycline
YMR230W−A_2
ZShift = −0.3
Z lm Score = 1.61
NG = 0
DB = 0
SM = 0
Delta_AUC
Doxycycline
YJR151W−A_2
ZShift = 2.45
Z lm Score = 6.4
NG = 0
DB = 0
SM = 0
Delta_L
Doxycycline
YJR151W−A_2
ZShift = 0.89
Z lm Score = −5.72
NG = 0
DB = 0
SM = 0
Delta_K
Doxycycline
YJR151W−A_2
ZShift = −0.32
Z lm Score = −2.25
NG = 0
DB = 0
SM = 0
Delta_r
Doxycycline
YJR151W−A_2
ZShift = −0.49
Z lm Score = 0.74
NG = 0
DB = 0
SM = 0
Delta_AUC
Doxycycline
YHL026C
ZShift = 1.52
Z lm Score = 6.37
NG = 0
DB = 0
SM = 0
Delta_L
Doxycycline
YHL026C
ZShift = 0.57
Z lm Score = −4.11
NG = 0
DB = 0
SM = 0
Delta_K
Doxycycline
YHL026C
ZShift = −0.94
Z lm Score = −1.37
NG = 0
DB = 0
SM = 0
Delta_r
Doxycycline
YHL026C
ZShift = −0.28
Z lm Score = 1.09
NG = 0
DB = 0
SM = 0
Delta_AUC
Doxycycline

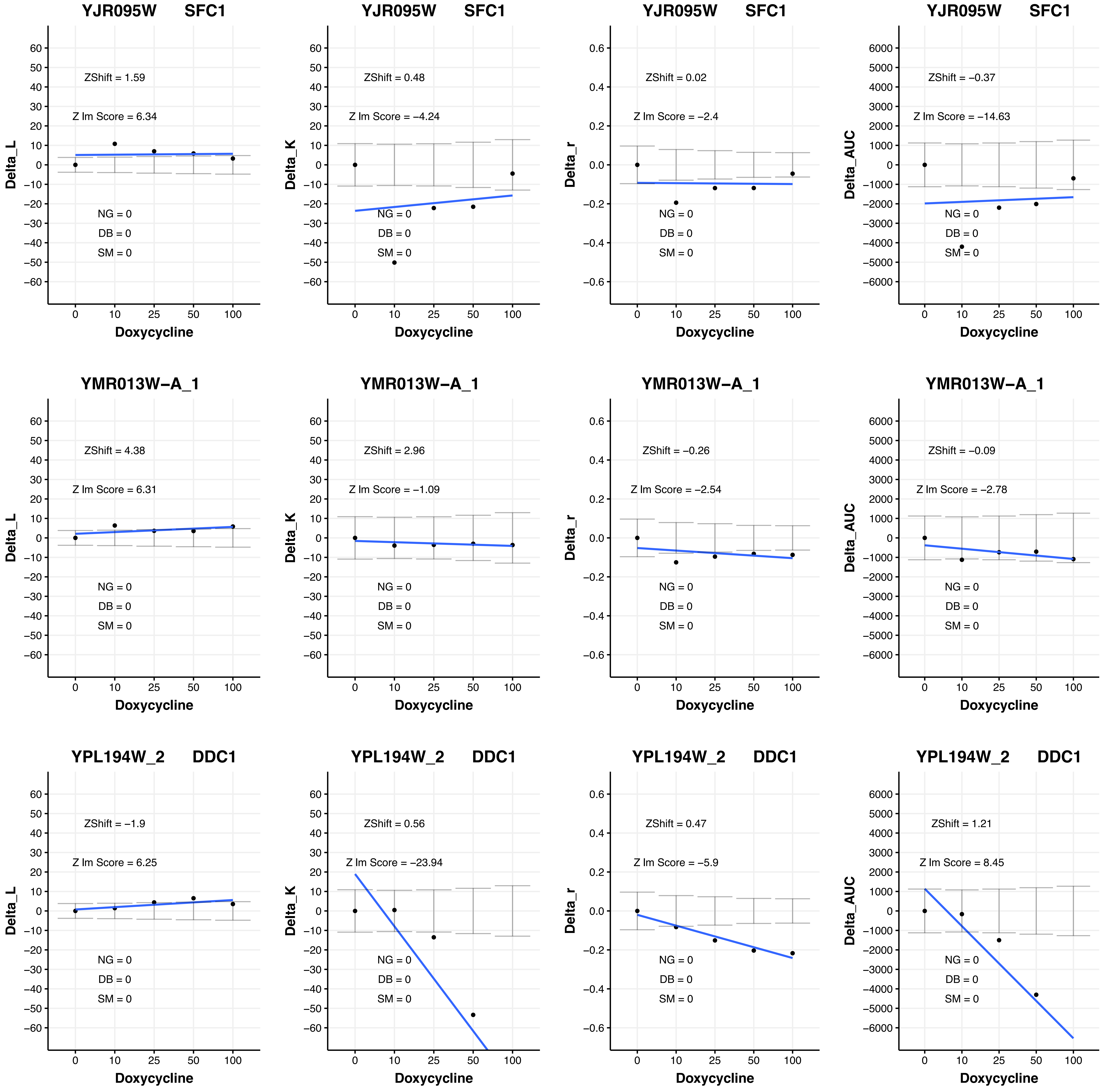

YJR095W SFC1
ZShift = 1.59
Z lm Score = 6.34
NG = 0
DB = 0
SM = 0
Delta_L
Doxycycline
YJR095W SFC1
ZShift = 0.48
Z lm Score = −4.24
NG = 0
DB = 0
SM = 0
Delta_K
Doxycycline
YJR095W SFC1
ZShift = 0.02
Z lm Score = −2.4
NG = 0
DB = 0
SM = 0
Delta_r
Doxycycline
YJR095W SFC1
ZShift = −0.37
Z lm Score = −14.63
NG = 0
DB = 0
SM = 0
Delta_AUC
Doxycycline
YMR013W−A_1
ZShift = 4.38
Z lm Score = 6.31
NG = 0
DB = 0
SM = 0
Delta_L
Doxycycline
YMR013W−A_1
ZShift = 2.96
Z lm Score = −1.09
NG = 0
DB = 0
SM = 0
Delta_K
Doxycycline
YMR013W−A_1
ZShift = −0.26
Z lm Score = −2.54
NG = 0
DB = 0
SM = 0
Delta_r
Doxycycline
YMR013W−A_1
ZShift = −0.09
Z lm Score = −2.78
NG = 0
DB = 0
SM = 0
Delta_AUC
Doxycycline
YPL194W_2 DDC1
ZShift = −1.9
Z lm Score = 6.25
NG = 0
DB = 0
SM = 0
Delta_L
Doxycycline
YPL194W_2 DDC1
ZShift = 0.56
Z lm Score = −23.94
NG = 0
DB = 0
SM = 0
Delta_K
Doxycycline
YPL194W_2 DDC1
ZShift = 0.47
Z lm Score = −5.9
NG = 0
DB = 0
SM = 0
Delta_r
Doxycycline
YPL194W_2 DDC1
ZShift = 1.21
Z lm Score = 8.45
NG = 0
DB = 0
SM = 0
Delta_AUC
Doxycycline

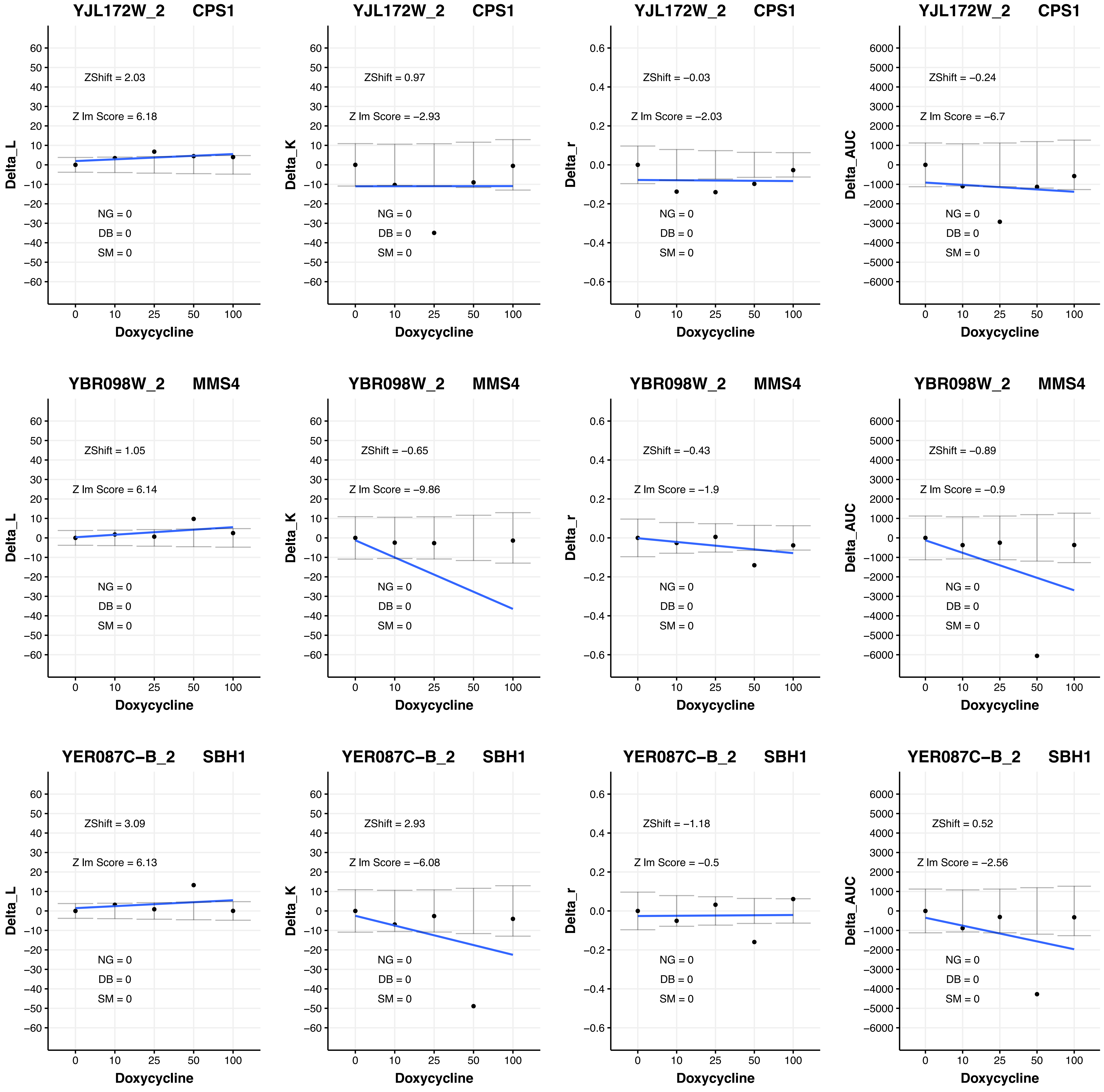

YJL172W_2 CPS1
ZShift = 2.03
Z lm Score = 6.18
NG = 0
DB = 0
SM = 0
Delta_L
Doxycycline
YJL172W_2 CPS1
ZShift = 0.97
Z lm Score = −2.93
NG = 0
DB = 0
SM = 0
Delta_K
Doxycycline
YJL172W_2 CPS1
ZShift = −0.03
Z lm Score = −2.03
NG = 0
DB = 0
SM = 0
Delta_r
Doxycycline
YJL172W_2 CPS1
ZShift = −0.24
Z lm Score = −6.7
NG = 0
DB = 0
SM = 0
Delta_AUC
Doxycycline
YBR098W_2 MMS4
ZShift = 1.05
Z lm Score = 6.14
NG = 0
DB = 0
SM = 0
Delta_L
Doxycycline
YBR098W_2 MMS4
ZShift = −0.65
Z lm Score = −9.86
NG = 0
DB = 0
SM = 0
Delta_K
Doxycycline
YBR098W_2 MMS4
ZShift = −0.43
Z lm Score = −1.9
NG = 0
DB = 0
SM = 0
Delta_r
Doxycycline
YBR098W_2 MMS4
ZShift = −0.89
Z lm Score = −0.9
NG = 0
DB = 0
SM = 0
Delta_AUC
Doxycycline
YER087C−B_2 SBH1
ZShift = 3.09
Z lm Score = 6.13
NG = 0
DB = 0
SM = 0
Delta_L
Doxycycline
YER087C−B_2 SBH1
ZShift = 2.93
Z lm Score = −6.08
NG = 0
DB = 0
SM = 0
Delta_K
Doxycycline
YER087C−B_2 SBH1
ZShift = −1.18
Z lm Score = −0.5
NG = 0
DB = 0
SM = 0
Delta_r
Doxycycline
YER087C−B_2 SBH1
ZShift = 0.52
Z lm Score = −2.56
NG = 0
DB = 0
SM = 0
Delta_AUC
Doxycycline
0 10 25 50 100

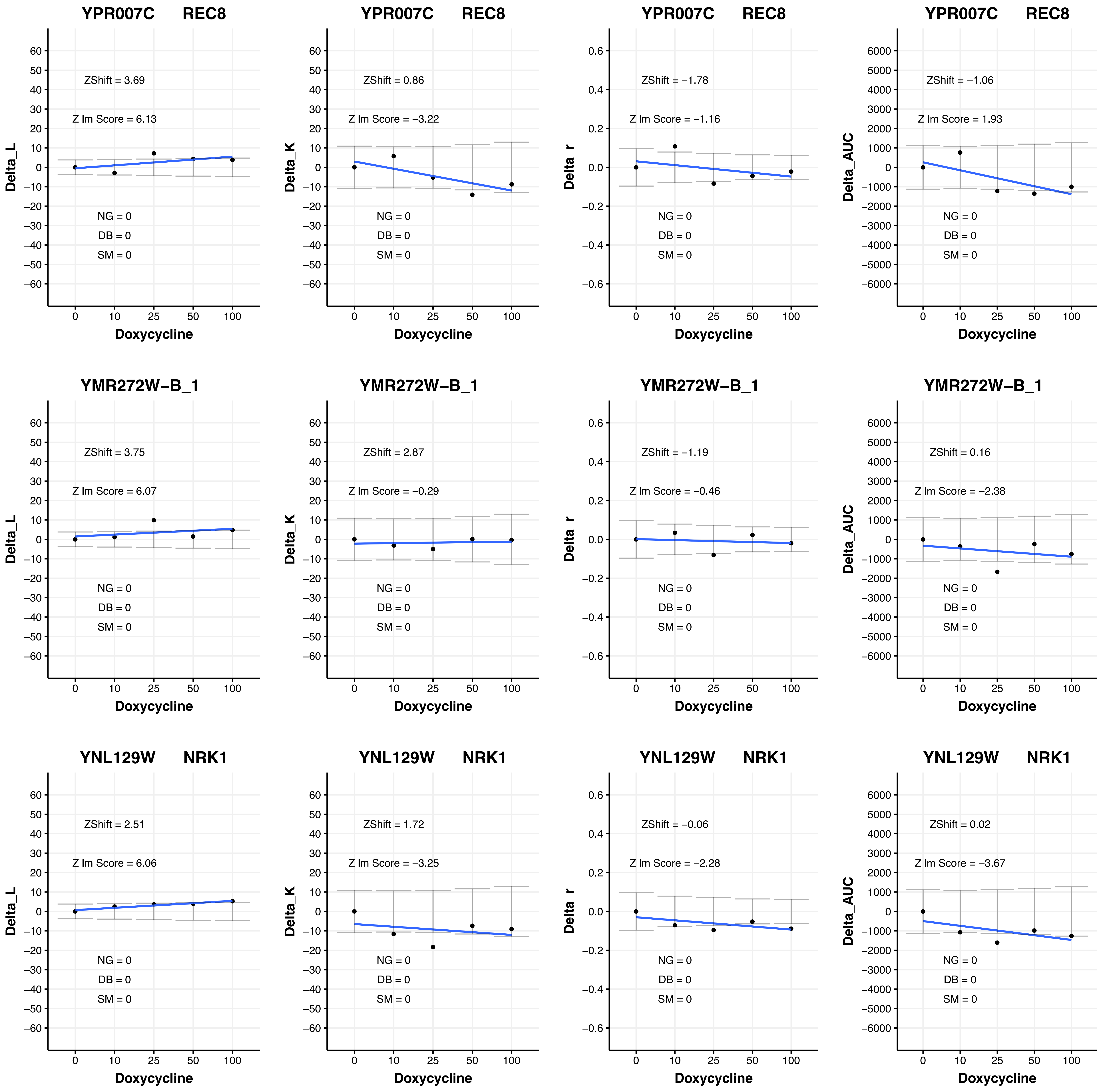

YPR007C REC8
ZShift = 3.69
Z lm Score = 6.13
NG = 0
DB = 0
SM = 0
Delta_L
Doxycycline
YPR007C REC8
ZShift = 0.86
Z lm Score = −3.22
NG = 0
DB = 0
SM = 0
Delta_K
Doxycycline
YPR007C REC8
ZShift = −1.78
Z lm Score = −1.16
NG = 0
DB = 0
SM = 0
Delta_r
Doxycycline
YPR007C REC8
ZShift = −1.06
Z lm Score = 1.93
NG = 0
DB = 0
SM = 0
Delta_AUC
Doxycycline
YMR272W−B_1
ZShift = 3.75
Z lm Score = 6.07
NG = 0
DB = 0
SM = 0
Delta_L
Doxycycline
YMR272W−B_1
ZShift = 2.87
Z lm Score = −0.29
NG = 0
DB = 0
SM = 0
Delta_K
Doxycycline
YMR272W−B_1
ZShift = −1.19
Z lm Score = −0.46
NG = 0
DB = 0
SM = 0
Delta_r
Doxycycline
YMR272W−B_1
ZShift = 0.16
Z lm Score = −2.38
NG = 0
DB = 0
SM = 0
Delta_AUC
Doxycycline
YNL129W NRK1
ZShift = 2.51
Z lm Score = 6.06
NG = 0
DB = 0
SM = 0
Delta_L
Doxycycline
YNL129W NRK1
ZShift = 1.72
Z lm Score = −3.25
NG = 0
DB = 0
SM = 0
Delta_K
Doxycycline
YNL129W NRK1
ZShift = −0.06
Z lm Score = −2.28
NG = 0
DB = 0
SM = 0
Delta_r
Doxycycline
YNL129W NRK1
ZShift = 0.02
Z lm Score = −3.67
NG = 0
DB = 0
SM = 0
Delta_AUC
Doxycycline

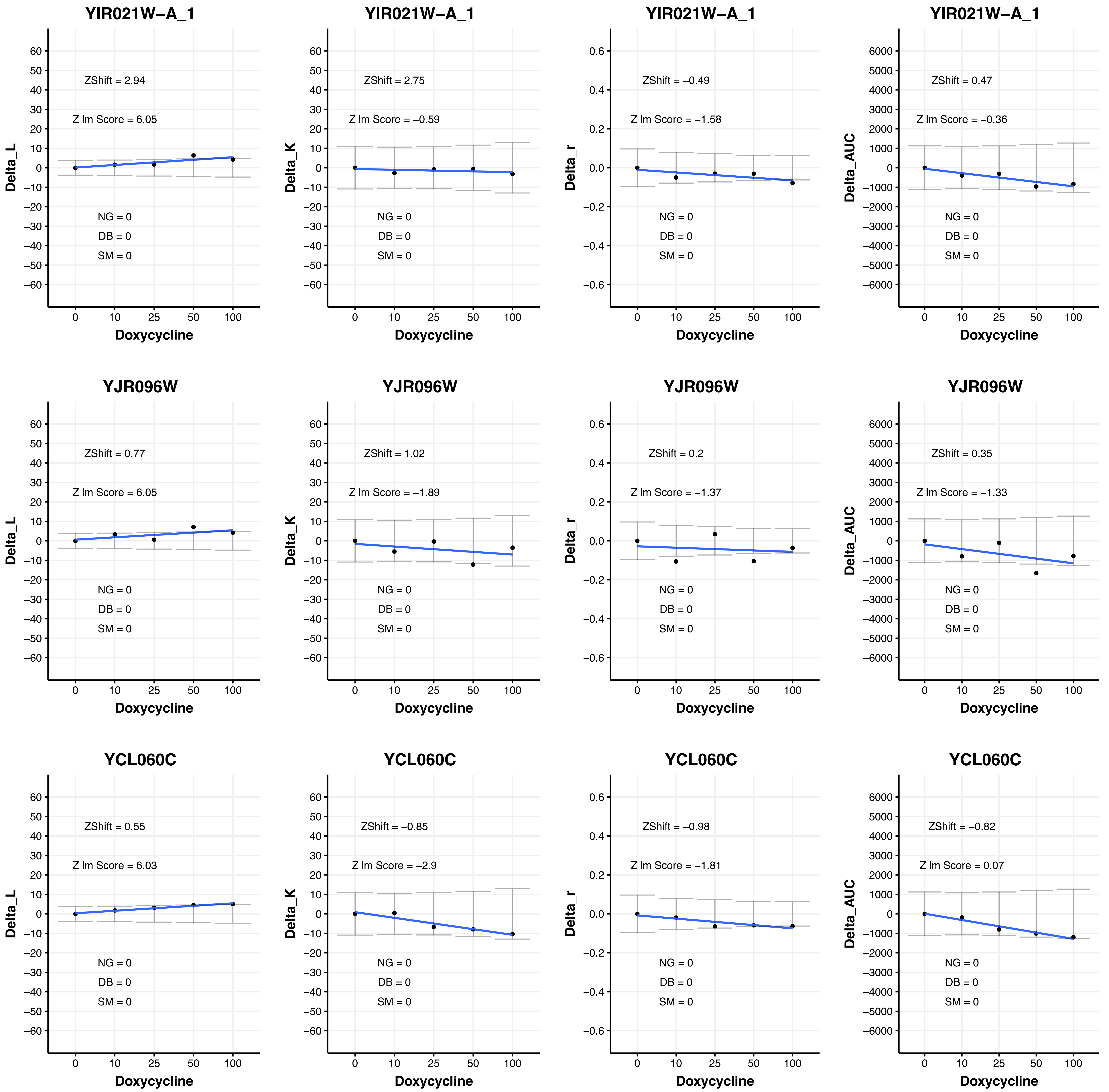
YIR021W−A_1
ZShift = 2.94
Z lm Score = 6.05
NG = 0
DB = 0
SM = 0
Delta_L
Doxycycline
YIR021W−A_1
ZShift = 2.75
Z lm Score = −0.59
NG = 0
DB = 0
SM = 0
Delta_K
Doxycycline
YIR021W−A_1
ZShift = −0.49
Z lm Score = −1.58
NG = 0
DB = 0
SM = 0
Delta_r
Doxycycline
YIR021W−A_1
ZShift = 0.47
Z lm Score = −0.36
NG = 0
DB = 0
SM = 0
Delta_AUC
Doxycycline
YJR096W
ZShift = 0.77
Z lm Score = 6.05
NG = 0
DB = 0
SM = 0
Delta_L
Doxycycline
YJR096W
ZShift = 1.02
Z lm Score = −1.89
NG = 0
DB = 0
SM = 0
Delta_K
Doxycycline
YJR096W
ZShift = 0.2
Z lm Score = −1.37
NG = 0
DB = 0
SM = 0
Delta_r
Doxycycline
YJR096W
ZShift = 0.35
Z lm Score = −1.33
NG = 0
DB = 0
SM = 0
Delta_AUC
Doxycycline
YCL060C
ZShift = 0.55
Z lm Score = 6.03
NG = 0
DB = 0
SM = 0
Delta_L
Doxycycline
YCL060C
ZShift = −0.85
Z lm Score = −2.9
NG = 0
DB = 0
SM = 0
Delta_K
Doxycycline
YCL060C
ZShift = −0.98
Z lm Score = −1.81
NG = 0
DB = 0
SM = 0
Delta_r
Doxycycline
YCL060C
ZShift = −0.82
Z lm Score = 0.07
NG = 0
DB = 0
SM = 0
Delta_AUC
Doxycycline

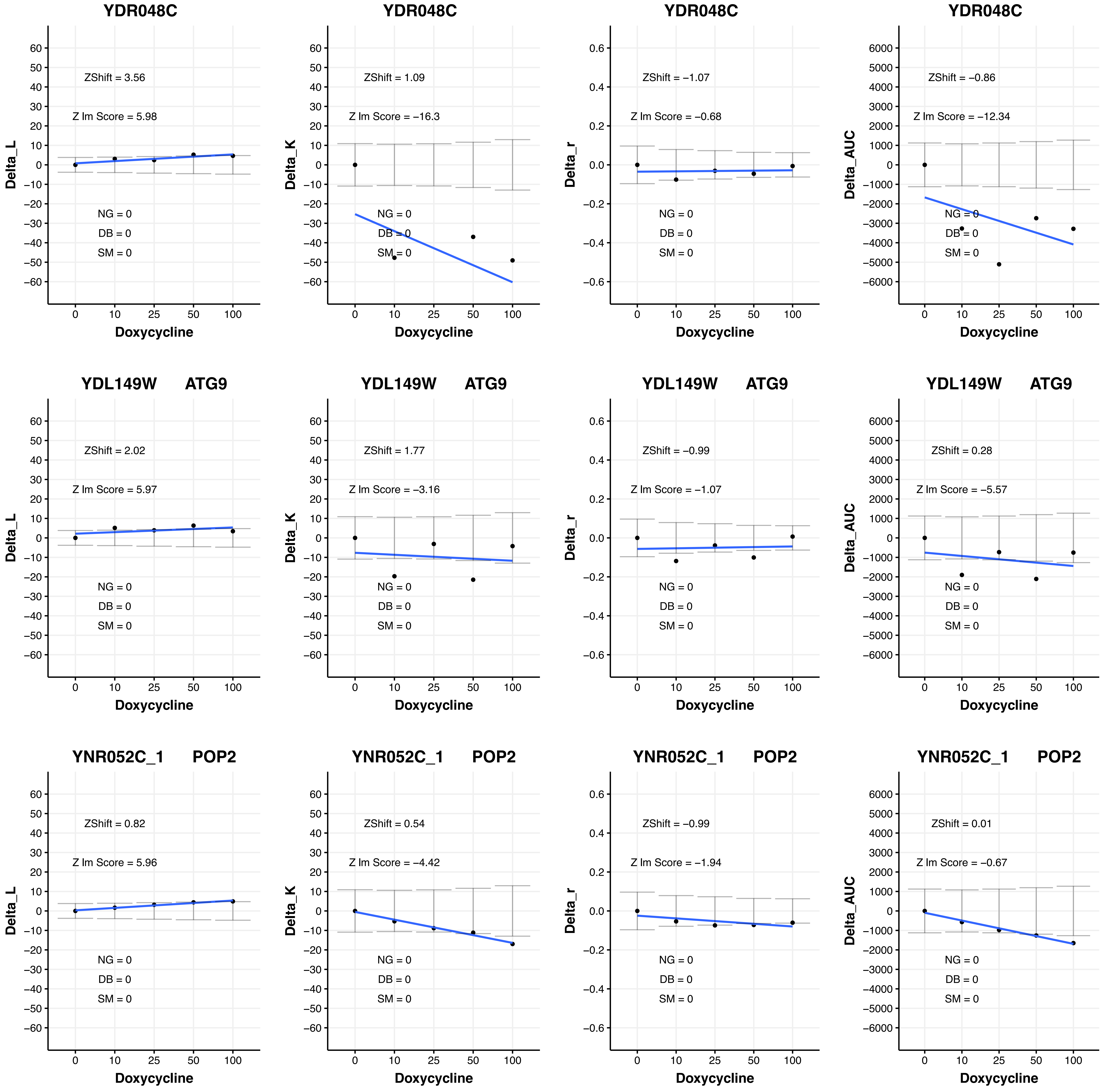
YDR048C
Delta_L
ZShift = 3.56
Z lm Score = 5.98
NG = 0
DB = 0
SM = 0
Doxycycline
YDR048C
Delta_K
ZShift = 1.09
Z lm Score = −16.3
NG = 0
DB = 0
SM = 0
Doxycycline
YDR048C
Delta_r
ZShift = −1.07
Z lm Score = −0.68
NG = 0
DB = 0
SM = 0
Doxycycline
YDR048C
Delta_AUC
ZShift = −0.86
Z lm Score = −12.34
NG = 0
DB = 0
SM = 0
Doxycycline
YDL149W ATG9
Delta_L
ZShift = 2.02
Z lm Score = 5.97
NG = 0
DB = 0
SM = 0
Doxycycline
YDL149W ATG9
Delta_K
ZShift = 1.77
Z lm Score = −3.16
NG = 0
DB = 0
SM = 0
Doxycycline
YDL149W ATG9
Delta_r
ZShift = −0.99
Z lm Score = −1.07
NG = 0
DB = 0
SM = 0
Doxycycline
YDL149W ATG9
Delta_AUC
ZShift = 0.28
Z lm Score = −5.57
NG = 0
DB = 0
SM = 0
Doxycycline
YNR052C_1 POP2
Delta_L
ZShift = 0.82
Z lm Score = 5.96
NG = 0
DB = 0
SM = 0
Doxycycline
YNR052C_1 POP2
Delta_K
ZShift = 0.54
Z lm Score = −4.42
NG = 0
DB = 0
SM = 0
Doxycycline
YNR052C_1 POP2
Delta_r
ZShift = −0.99
Z lm Score = −1.94
NG = 0
DB = 0
SM = 0
Doxycycline
YNR052C_1 POP2
Delta_AUC
ZShift = 0.01
Z lm Score = −0.67
NG = 0
DB = 0
SM = 0
Doxycycline

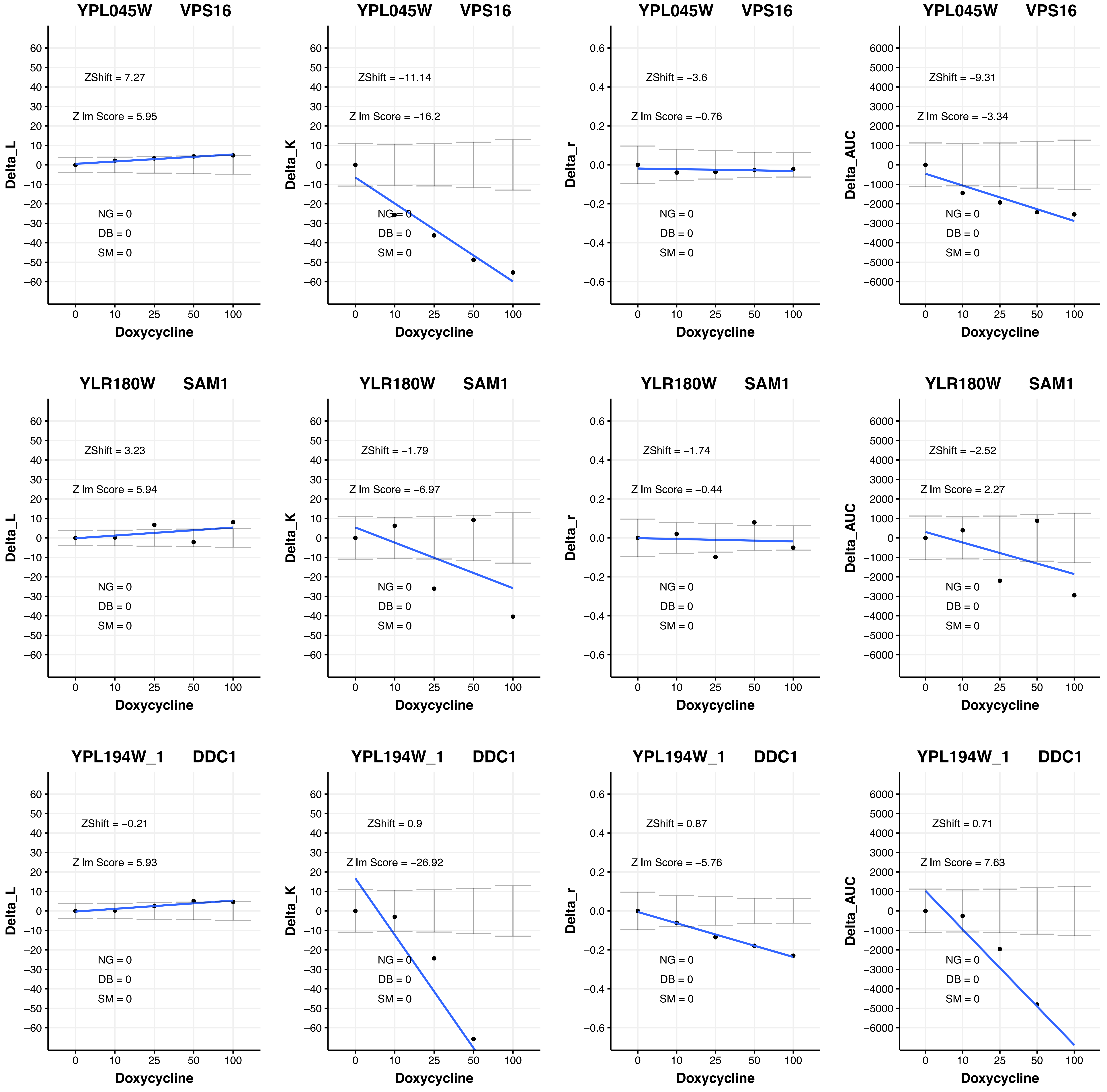
YPL045W VPS16
ZShift = 7.27
Z lm Score = 5.95
NG = 0
DB = 0
SM = 0
Delta_L
Doxycycline
YPL045W VPS16
ZShift = −11.14
Z lm Score = −16.2
NG = 0
DB = 0
SM = 0
Delta_K
Doxycycline
YPL045W VPS16
ZShift = −3.6
Z lm Score = −0.76
NG = 0
DB = 0
SM = 0
Delta_r
Doxycycline
YPL045W VPS16
ZShift = −9.31
Z lm Score = −3.34
NG = 0
DB = 0
SM = 0
Delta_AUC
Doxycycline
YLR180W SAM1
ZShift = 3.23
Z lm Score = 5.94
NG = 0
DB = 0
SM = 0
Delta_L
Doxycycline
YLR180W SAM1
ZShift = −1.79
Z lm Score = −6.97
NG = 0
DB = 0
SM = 0
Delta_K
Doxycycline
YLR180W SAM1
ZShift = −1.74
Z lm Score = −0.44
NG = 0
DB = 0
SM = 0
Delta_r
Doxycycline
YLR180W SAM1
ZShift = −2.52
Z lm Score = 2.27
NG = 0
DB = 0
SM = 0
Delta_AUC
Doxycycline
YPL194W_1 DDC1
ZShift = −0.21
Z lm Score = 5.93
NG = 0
DB = 0
SM = 0
Delta_L
Doxycycline
YPL194W_1 DDC1
ZShift = 0.9
Z lm Score = −26.92
NG = 0
DB = 0
SM = 0
Delta_K
Doxycycline
YPL194W_1 DDC1
ZShift = 0.87
Z lm Score = −5.76
NG = 0
DB = 0
SM = 0
Delta_r
Doxycycline
YPL194W_1 DDC1
ZShift = 0.71
Z lm Score = 7.63
NG = 0
DB = 0
SM = 0
Delta_AUC
Doxycycline

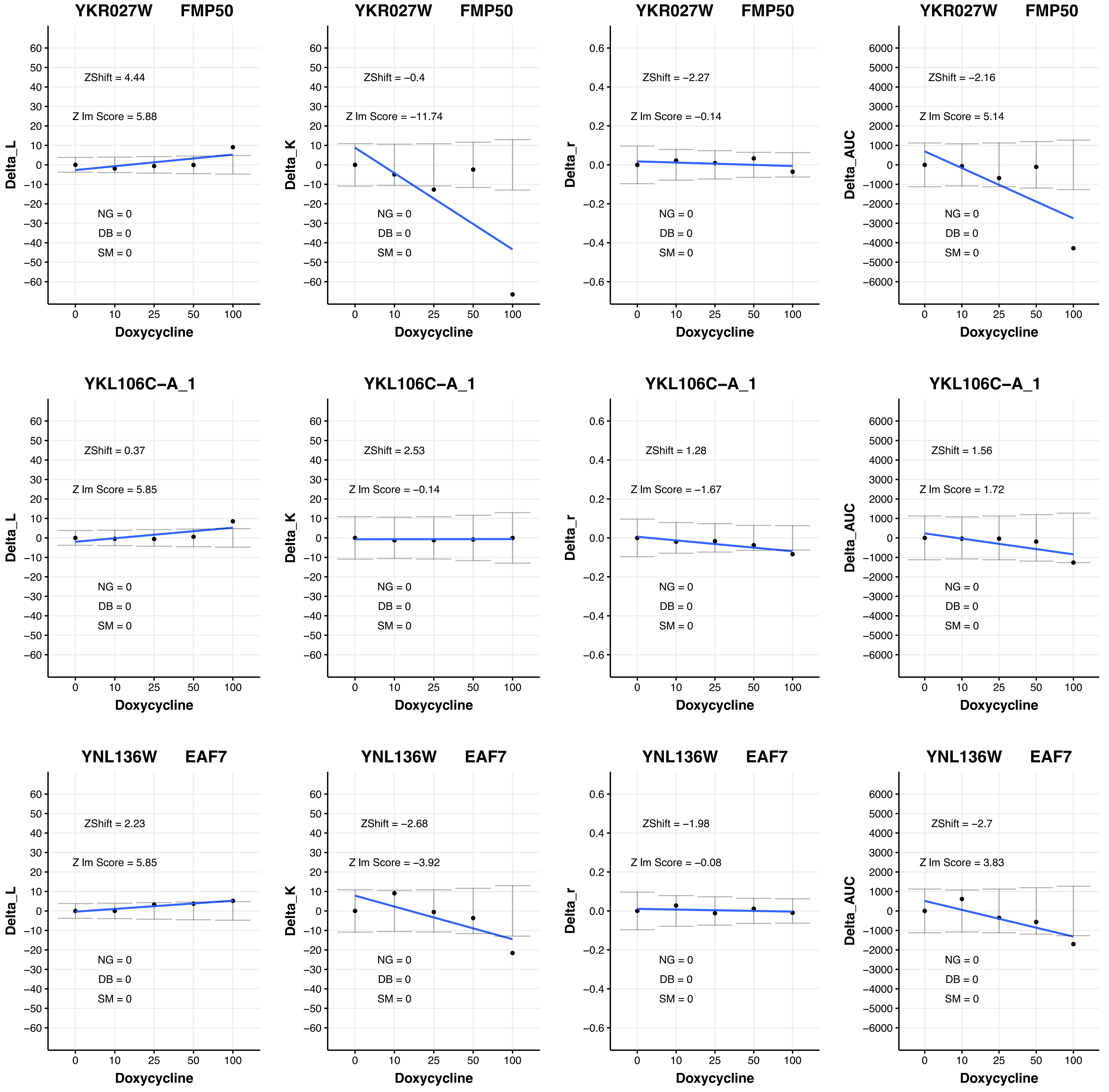

YKR027W FMP50
ZShift = 4.44
Z lm Score = 5.88
NG = 0
DB = 0
SM = 0
Delta_L
Doxycycline
YKR027W FMP50
ZShift = −0.4
Z lm Score = −11.74
NG = 0
DB = 0
SM = 0
Delta_K
Doxycycline
YKR027W FMP50
ZShift = −2.27
Z lm Score = −0.14
NG = 0
DB = 0
SM = 0
Delta_r
Doxycycline
YKR027W FMP50
ZShift = −2.16
Z lm Score = 5.14
NG = 0
DB = 0
SM = 0
Delta_AUC
Doxycycline
YKL106C−A_1
ZShift = 0.37
Z lm Score = 5.85
NG = 0
DB = 0
SM = 0
Delta_L
Doxycycline
YKL106C−A_1
ZShift = 2.53
Z lm Score = −0.14
NG = 0
DB = 0
SM = 0
Delta_K
Doxycycline
YKL106C−A_1
ZShift = 1.28
Z lm Score = −1.67
NG = 0
DB = 0
SM = 0
Delta_r
Doxycycline
YKL106C−A_1
ZShift = 1.56
Z lm Score = 1.72
NG = 0
DB = 0
SM = 0
Delta_AUC
Doxycycline
YNL136W EAF7
ZShift = 2.23
Z lm Score = 5.85
NG = 0
DB = 0
SM = 0
Delta_L
Doxycycline
YNL136W EAF7
ZShift = −2.68
Z lm Score = −3.92
NG = 0
DB = 0
SM = 0
Delta_K
Doxycycline
YNL136W EAF7
ZShift = −1.98
Z lm Score = −0.08
NG = 0
DB = 0
SM = 0
Delta_r
Doxycycline
YNL136W EAF7
ZShift = −2.7
Z lm Score = 3.83
NG = 0
DB = 0
SM = 0
Delta_AUC
Doxycycline

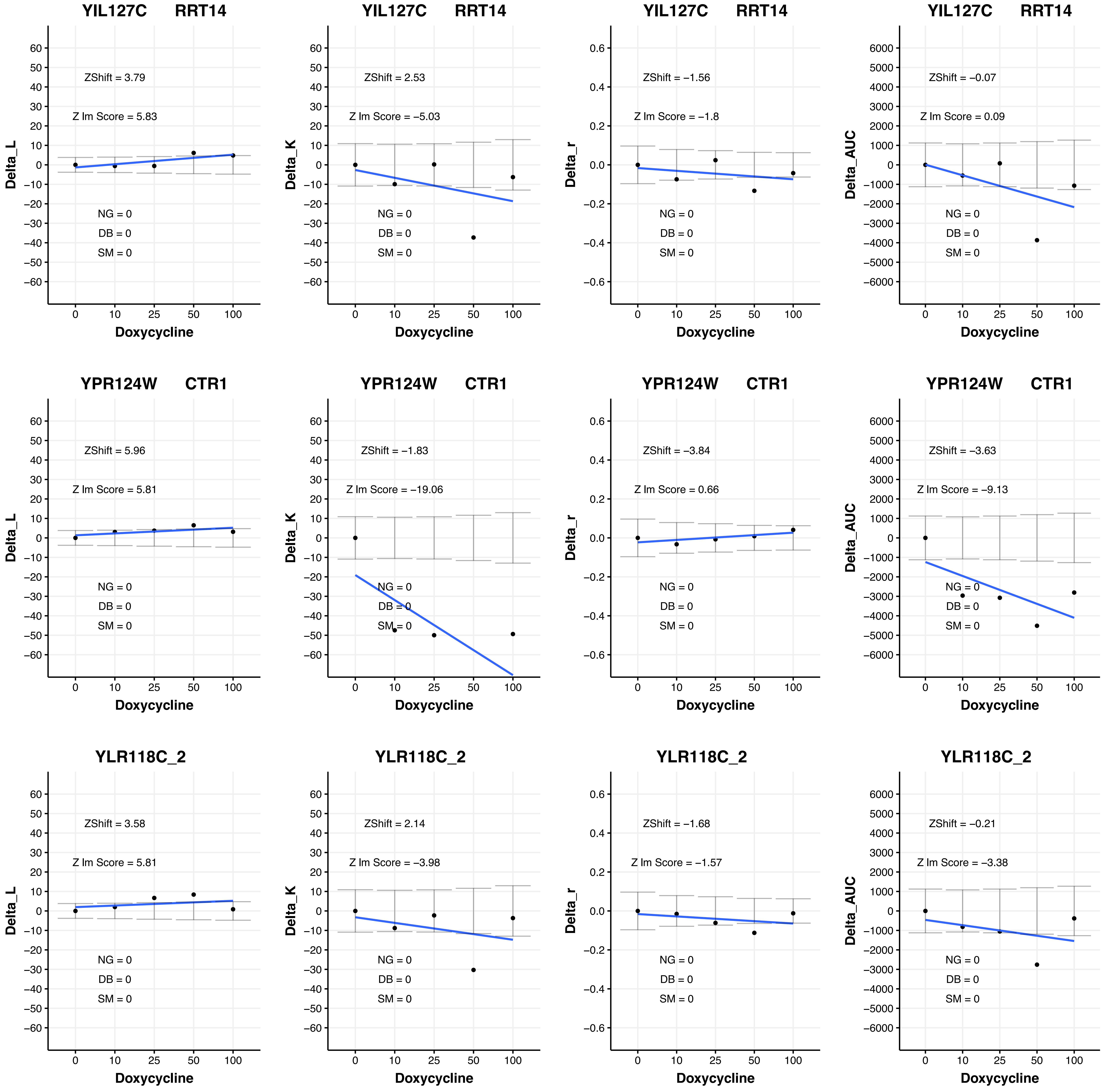

YIL127C RRT14
ZShift = 3.79
Z lm Score = 5.83
NG = 0
DB = 0
SM = 0
Delta_L
Doxycycline
YIL127C RRT14
ZShift = 2.53
Z lm Score = −5.03
Delta_K
YIL127C RRT14
ZShift = −1.56
Z lm Score = −1.8
Delta_r
YIL127C RRT14
ZShift = −0.07
Z lm Score = 0.09
Delta_AUC
YPR124W CTR1
ZShift = 5.96
Z lm Score = 5.81
YPR124W CTR1
ZShift = −1.83
Z lm Score = −19.06
YPR124W CTR1
ZShift = −3.84
Z lm Score = 0.66
YPR124W CTR1
ZShift = −3.63
Z lm Score = −9.13
YLR118C_2
ZShift = 3.58
Z lm Score = 5.81
YLR118C_2
ZShift = 2.14
Z lm Score = −3.98
YLR118C_2
ZShift = −1.68
Z lm Score = −1.57
YLR118C_2
ZShift = −0.21
Z lm Score = −3.38

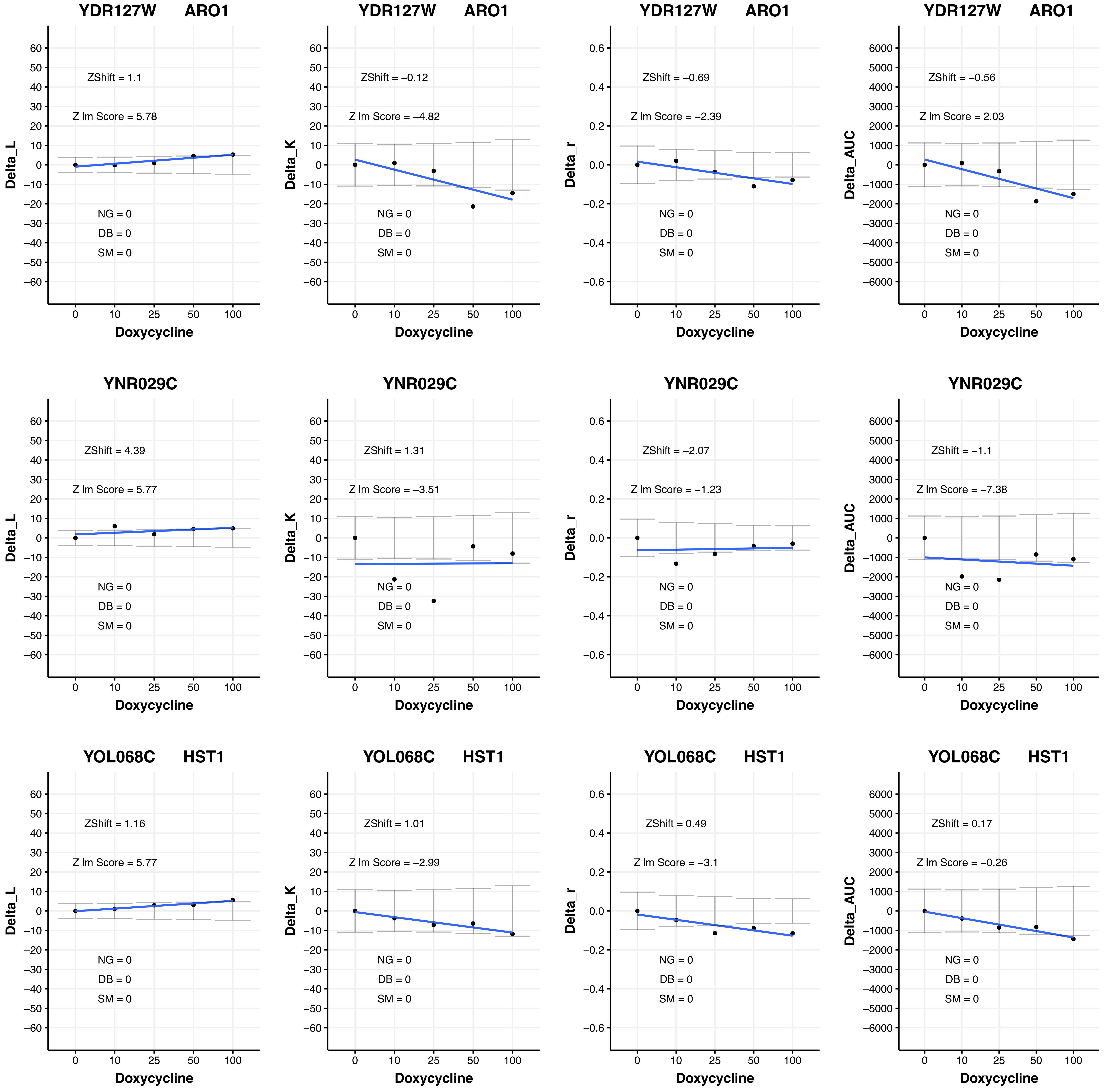

YDR127W ARO1
ZShift = 1.1
Z lm Score = 5.78
NG = 0
DB = 0
SM = 0
Delta_L
Doxycycline
YDR127W ARO1
ZShift = −0.12
Z lm Score = −4.82
NG = 0
DB = 0
SM = 0
Delta_K
Doxycycline
YDR127W ARO1
ZShift = −0.69
Z lm Score = −2.39
NG = 0
DB = 0
SM = 0
Delta_r
Doxycycline
YDR127W ARO1
ZShift = −0.56
Z lm Score = 2.03
NG = 0
DB = 0
SM = 0
Delta_AUC
Doxycycline
YNR029C
ZShift = 4.39
Z lm Score = 5.77
NG = 0
DB = 0
SM = 0
Delta_L
Doxycycline
YNR029C
ZShift = 1.31
Z lm Score = −3.51
NG = 0
DB = 0
SM = 0
Delta_K
Doxycycline
YNR029C
ZShift = −2.07
Z lm Score = −1.23
NG = 0
DB = 0
SM = 0
Delta_r
Doxycycline
YNR029C
ZShift = −1.1
Z lm Score = −7.38
NG = 0
DB = 0
SM = 0
Delta_AUC
Doxycycline
YOL068C HST1
ZShift = 1.16
Z lm Score = 5.77
NG = 0
DB = 0
SM = 0
Delta_L
Doxycycline
YOL068C HST1
ZShift = 1.01
Z lm Score = −2.99
NG = 0
DB = 0
SM = 0
Delta_K
Doxycycline
YOL068C HST1
ZShift = 0.49
Z lm Score = −3.1
NG = 0
DB = 0
SM = 0
Delta_r
Doxycycline
YOL068C HST1
ZShift = 0.17
Z lm Score = −0.26
NG = 0
DB = 0
SM = 0
Delta_AUC
Doxycycline

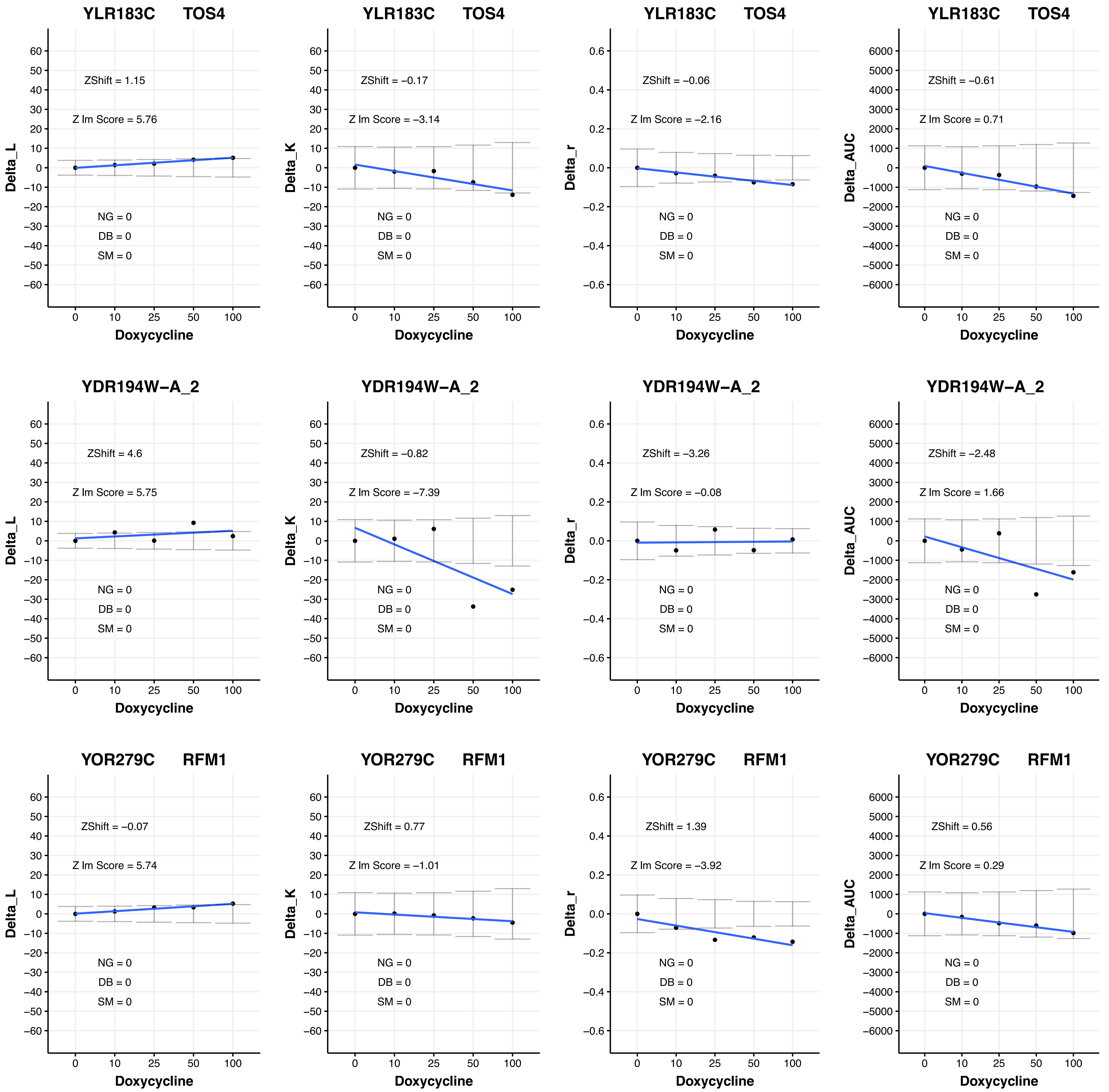
YLR183C TOS4
ZShift = 1.15
Z lm Score = 5.76
NG = 0
DB = 0
SM = 0
Delta_L
Doxycycline
YLR183C TOS4
ZShift = −0.17
Z lm Score = −3.14
NG = 0
DB = 0
SM = 0
Delta_K
Doxycycline
YLR183C TOS4
ZShift = −0.06
Z lm Score = −2.16
NG = 0
DB = 0
SM = 0
Delta_r
Doxycycline
YLR183C TOS4
ZShift = −0.61
Z lm Score = 0.71
NG = 0
DB = 0
SM = 0
Delta_AUC
Doxycycline
YDR194W−A_2
ZShift = 4.6
Z lm Score = 5.75
NG = 0
DB = 0
SM = 0
Delta_L
Doxycycline
YDR194W−A_2
ZShift = −0.82
Z lm Score = −7.39
NG = 0
DB = 0
SM = 0
Delta_K
Doxycycline
YDR194W−A_2
ZShift = −3.26
Z lm Score = −0.08
NG = 0
DB = 0
SM = 0
Delta_r
Doxycycline
YDR194W−A_2
ZShift = −2.48
Z lm Score = 1.66
NG = 0
DB = 0
SM = 0
Delta_AUC
Doxycycline
YOR279C RFM1
ZShift = −0.07
Z lm Score = 5.74
NG = 0
DB = 0
SM = 0
Delta_L
Doxycycline
YOR279C RFM1
ZShift = 0.77
Z lm Score = −1.01
NG = 0
DB = 0
SM = 0
Delta_K
Doxycycline
YOR279C RFM1
ZShift = 1.39
Z lm Score = −3.92
NG = 0
DB = 0
SM = 0
Delta_r
Doxycycline
YOR279C RFM1
ZShift = 0.56
Z lm Score = 0.29
NG = 0
DB = 0
SM = 0
Delta_AUC
Doxycycline

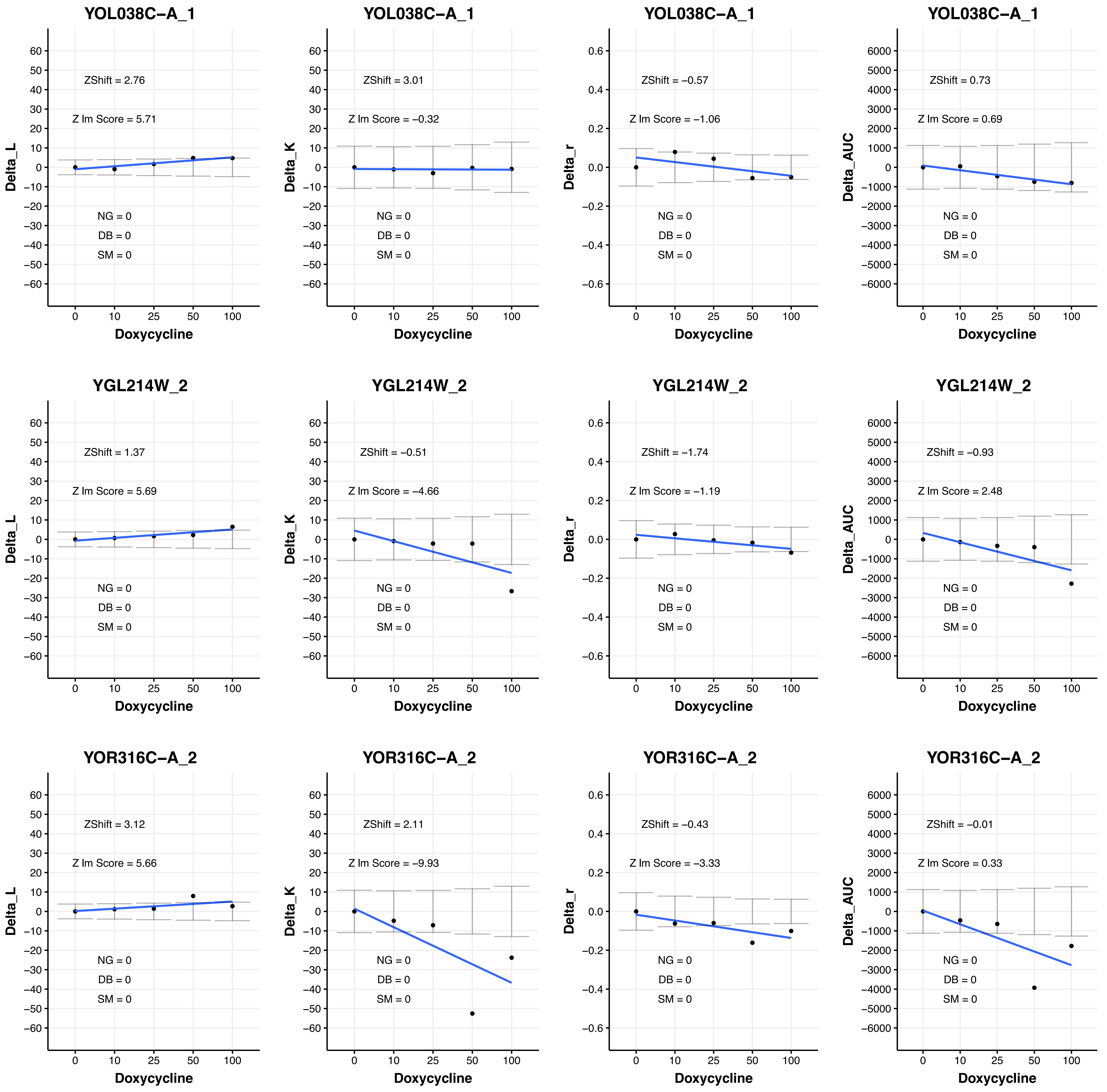
YOL038C−A_1
ZShift = 2.76
Z lm Score = 5.71
NG = 0
DB = 0
SM = 0
Delta_L
Doxycycline
YOL038C−A_1
ZShift = 3.01
Z lm Score = −0.32
NG = 0
DB = 0
SM = 0
Delta_K
Doxycycline
YOL038C−A_1
ZShift = −0.57
Z lm Score = −1.06
NG = 0
DB = 0
SM = 0
Delta_r
Doxycycline
YOL038C−A_1
ZShift = 0.73
Z lm Score = 0.69
NG = 0
DB = 0
SM = 0
Delta_AUC
Doxycycline
YGL214W_2
ZShift = 1.37
Z lm Score = 5.69
NG = 0
DB = 0
SM = 0
Delta_L
Doxycycline
YGL214W_2
ZShift = −0.51
Z lm Score = −4.66
NG = 0
DB = 0
SM = 0
Delta_K
Doxycycline
YGL214W_2
ZShift = −1.74
Z lm Score = −1.19
NG = 0
DB = 0
SM = 0
Delta_r
Doxycycline
YGL214W_2
ZShift = −0.93
Z lm Score = 2.48
NG = 0
DB = 0
SM = 0
Delta_AUC
Doxycycline
YOR316C−A_2
ZShift = 3.12
Z lm Score = 5.66
NG = 0
DB = 0
SM = 0
Delta_L
Doxycycline
YOR316C−A_2
ZShift = 2.11
Z lm Score = −9.93
NG = 0
DB = 0
SM = 0
Delta_K
Doxycycline
YOR316C−A_2
ZShift = −0.43
Z lm Score = −3.33
NG = 0
DB = 0
SM = 0
Delta_r
Doxycycline
YOR316C−A_2
ZShift = −0.01
Z lm Score = 0.33
NG = 0
DB = 0
SM = 0
Delta_AUC
Doxycycline

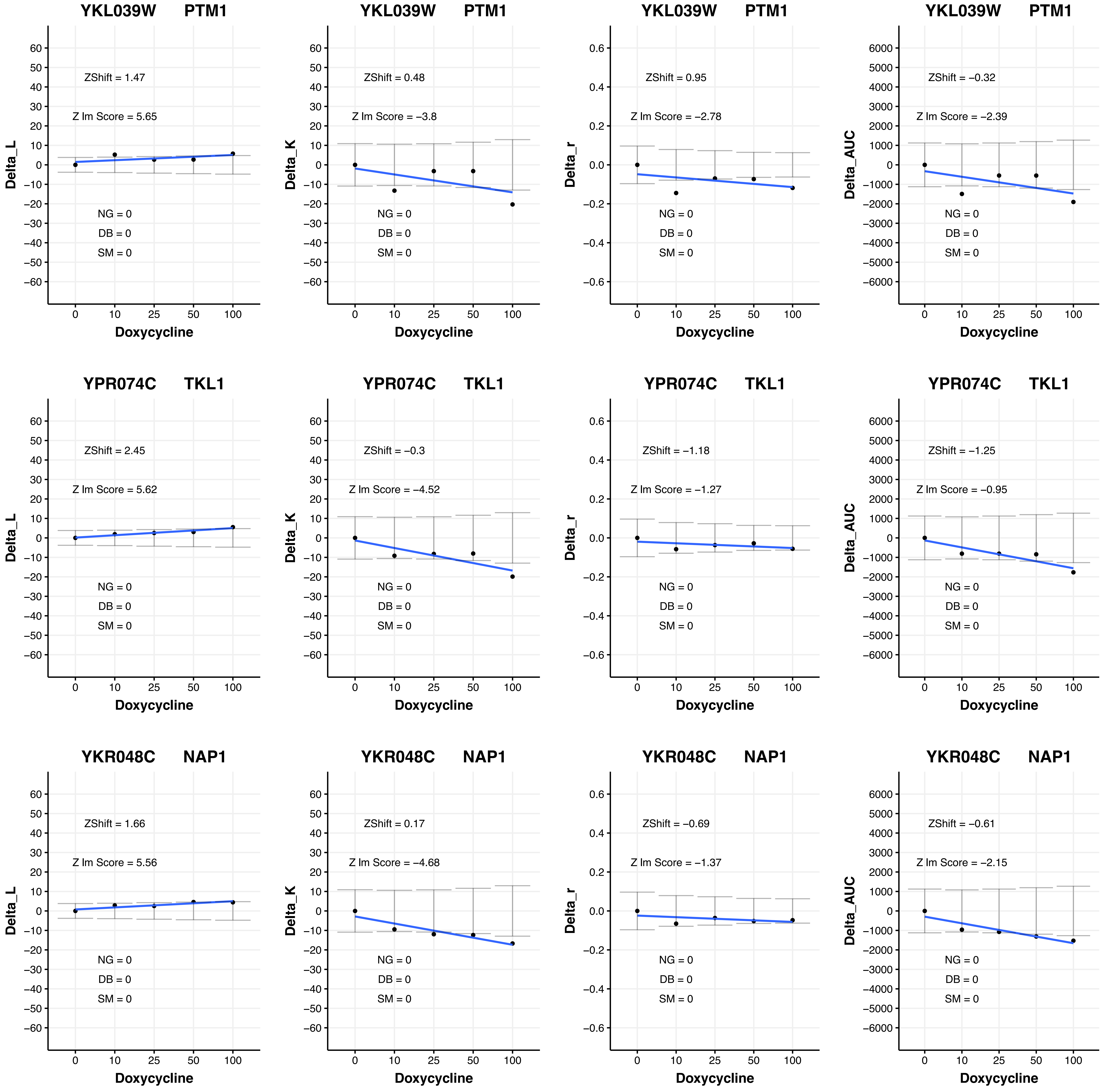

YKL039W PTM1
ZShift = 1.47
Z lm Score = 5.65
NG = 0
DB = 0
SM = 0
Delta_L
Doxycycline
YKL039W PTM1
ZShift = 0.48
Z lm Score = −3.8
NG = 0
DB = 0
SM = 0
Delta_K
Doxycycline
YKL039W PTM1
ZShift = 0.95
Z lm Score = −2.78
NG = 0
DB = 0
SM = 0
Delta_r
Doxycycline
YKL039W PTM1
ZShift = −0.32
Z lm Score = −2.39
NG = 0
DB = 0
SM = 0
Delta_AUC
Doxycycline
YPR074C TKL1
ZShift = 2.45
Z lm Score = 5.62
NG = 0
DB = 0
SM = 0
Delta_L
Doxycycline
YPR074C TKL1
ZShift = −0.3
Z lm Score = −4.52
NG = 0
DB = 0
SM = 0
Delta_K
Doxycycline
YPR074C TKL1
ZShift = −1.18
Z lm Score = −1.27
NG = 0
DB = 0
SM = 0
Delta_r
Doxycycline
YPR074C TKL1
ZShift = −1.25
Z lm Score = −0.95
NG = 0
DB = 0
SM = 0
Delta_AUC
Doxycycline
YKR048C NAP1
ZShift = 1.66
Z lm Score = 5.56
NG = 0
DB = 0
SM = 0
Delta_L
Doxycycline
YKR048C NAP1
ZShift = 0.17
Z lm Score = −4.68
NG = 0
DB = 0
SM = 0
Delta_K
Doxycycline
YKR048C NAP1
ZShift = −0.69
Z lm Score = −1.37
NG = 0
DB = 0
SM = 0
Delta_r
Doxycycline
YKR048C NAP1
ZShift = −0.61
Z lm Score = −2.15
NG = 0
DB = 0
SM = 0
Delta_AUC
Doxycycline

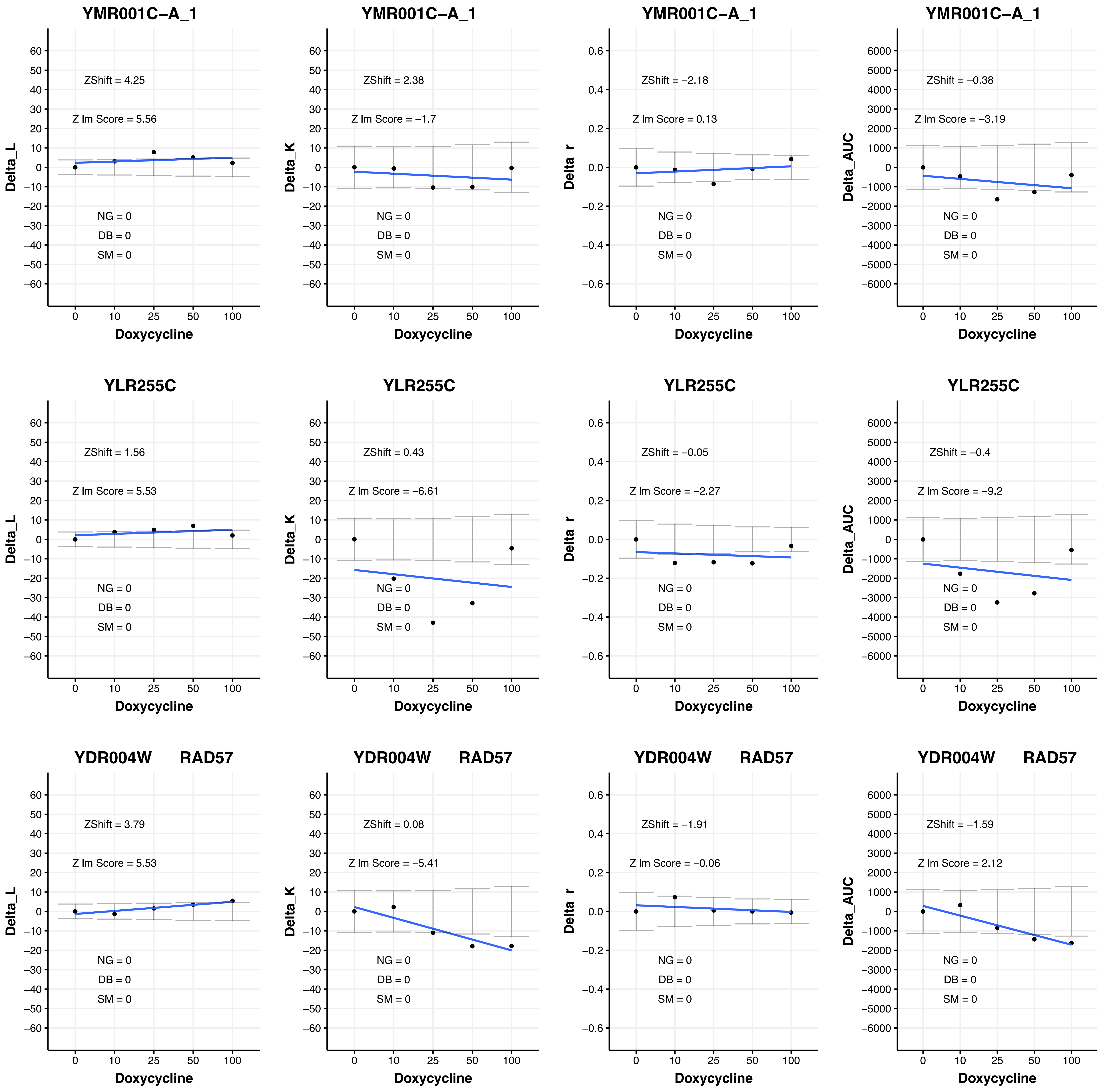
YMR001C−A_1
ZShift = 4.25
Z lm Score = 5.56
NG = 0
DB = 0
SM = 0
Delta_L
Doxycycline
YMR001C−A_1
ZShift = 2.38
Z lm Score = −1.7
NG = 0
DB = 0
SM = 0
Delta_K
Doxycycline
YMR001C−A_1
ZShift = −2.18
Z lm Score = 0.13
NG = 0
DB = 0
SM = 0
Delta_r
Doxycycline
YMR001C−A_1
ZShift = −0.38
Z lm Score = −3.19
NG = 0
DB = 0
SM = 0
Delta_AUC
Doxycycline
YLR255C
ZShift = 1.56
Z lm Score = 5.53
NG = 0
DB = 0
SM = 0
Delta_L
Doxycycline
YLR255C
ZShift = 0.43
Z lm Score = −6.61
NG = 0
DB = 0
SM = 0
Delta_K
Doxycycline
YLR255C
ZShift = −0.05
Z lm Score = −2.27
NG = 0
DB = 0
SM = 0
Delta_r
Doxycycline
YLR255C
ZShift = −0.4
Z lm Score = −9.2
NG = 0
DB = 0
SM = 0
Delta_AUC
Doxycycline
YDR004W RAD57
ZShift = 3.79
Z lm Score = 5.53
NG = 0
DB = 0
SM = 0
Delta_L
Doxycycline
YDR004W RAD57
ZShift = 0.08
Z lm Score = −5.41
NG = 0
DB = 0
SM = 0
Delta_K
Doxycycline
YDR004W RAD57
ZShift = −1.91
Z lm Score = −0.06
NG = 0
DB = 0
SM = 0
Delta_r
Doxycycline
YDR004W RAD57
ZShift = −1.59
Z lm Score = 2.12
NG = 0
DB = 0
SM = 0
Delta_AUC
Doxycycline

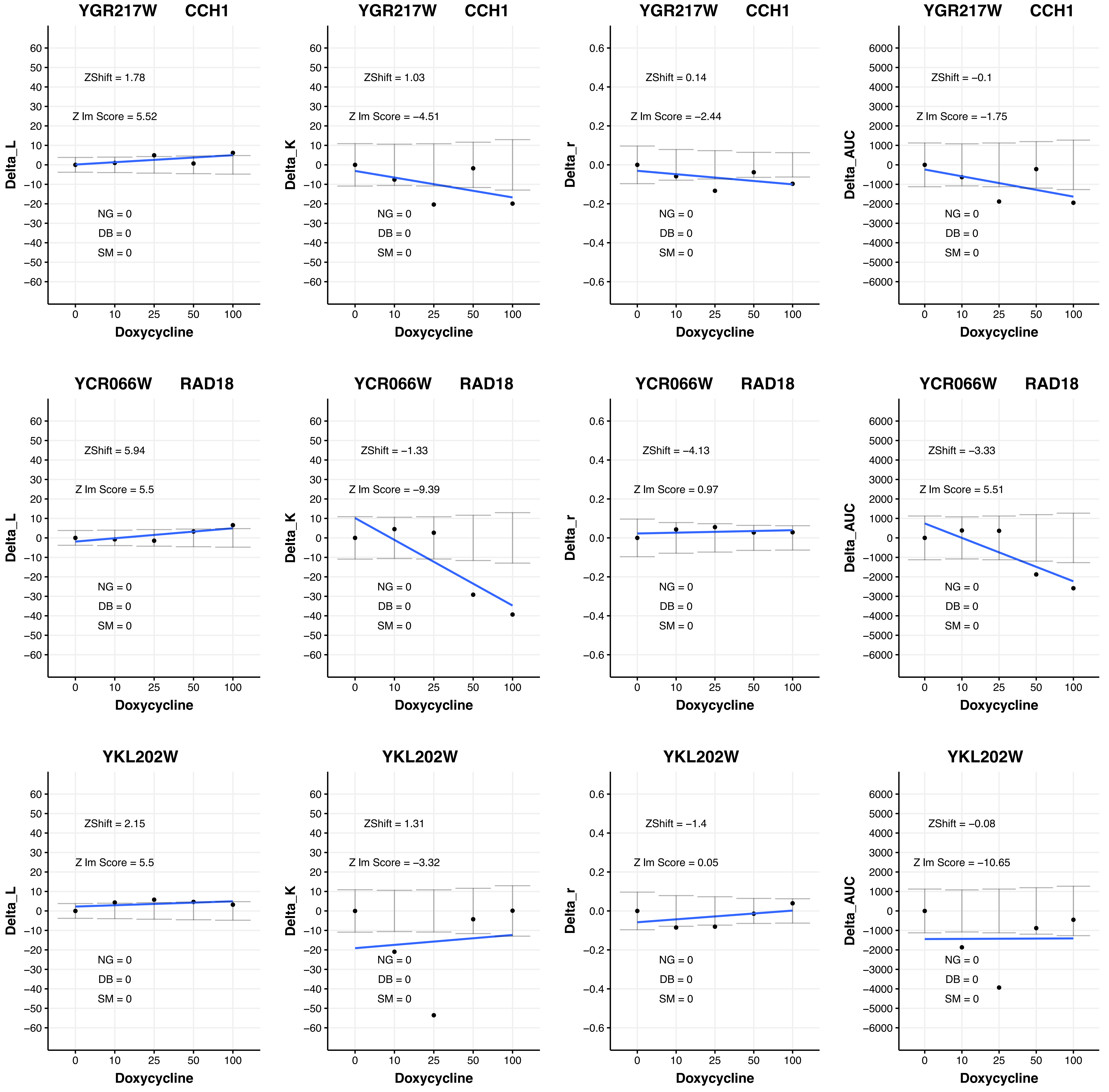

YGR217W CCH1
ZShift = 1.78
Z lm Score = 5.52
NG = 0
DB = 0
SM = 0
Delta_L
Doxycycline
YGR217W CCH1
ZShift = 1.03
Z lm Score = −4.51
NG = 0
DB = 0
SM = 0
Delta_K
Doxycycline
YGR217W CCH1
ZShift = 0.14
Z lm Score = −2.44
NG = 0
DB = 0
SM = 0
Delta_r
Doxycycline
YGR217W CCH1
ZShift = −0.1
Z lm Score = −1.75
NG = 0
DB = 0
SM = 0
Delta_AUC
Doxycycline
YCR066W RAD18
ZShift = 5.94
Z lm Score = 5.5
NG = 0
DB = 0
SM = 0
Delta_L
Doxycycline
YCR066W RAD18
ZShift = −1.33
Z lm Score = −9.39
NG = 0
DB = 0
SM = 0
Delta_K
Doxycycline
YCR066W RAD18
ZShift = −4.13
Z lm Score = 0.97
NG = 0
DB = 0
SM = 0
Delta_r
Doxycycline
YCR066W RAD18
ZShift = −3.33
Z lm Score = 5.51
NG = 0
DB = 0
SM = 0
Delta_AUC
Doxycycline
YKL202W
ZShift = 2.15
Z lm Score = 5.5
NG = 0
DB = 0
SM = 0
Delta_L
Doxycycline
YKL202W
ZShift = 1.31
Z lm Score = −3.32
NG = 0
DB = 0
SM = 0
Delta_K
Doxycycline
YKL202W
ZShift = −1.4
Z lm Score = 0.05
NG = 0
DB = 0
SM = 0
Delta_r
Doxycycline
YKL202W
ZShift = −0.08
Z lm Score = −10.65
NG = 0
DB = 0
SM = 0
Delta_AUC
Doxycycline

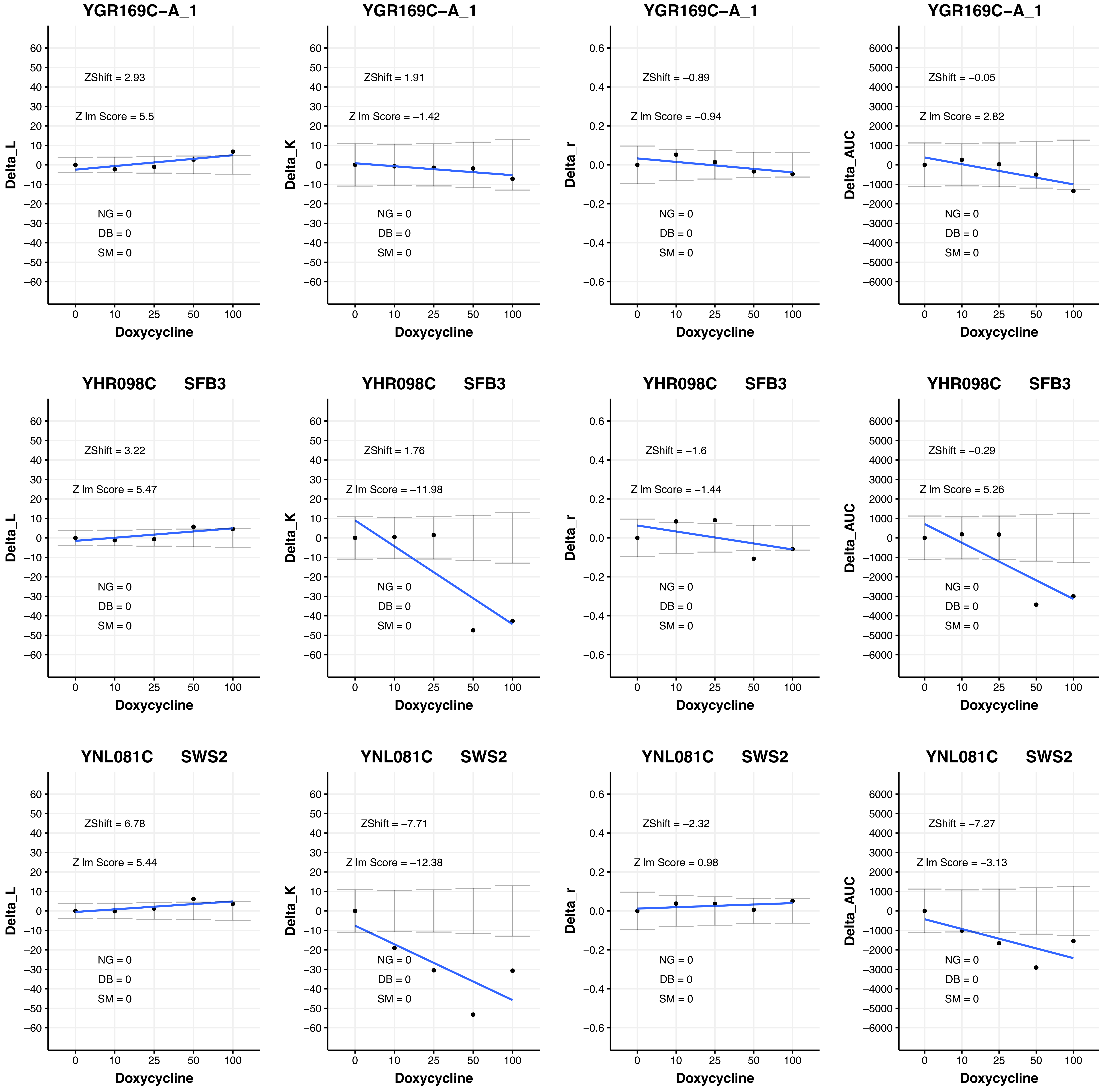

YGR169C−A_1
ZShift = 2.93
Z lm Score = 5.5
NG = 0
DB = 0
SM = 0
Delta_L
Doxycycline
YGR169C−A_1
ZShift = 1.91
Z lm Score = −1.42
NG = 0
DB = 0
SM = 0
Delta_K
Doxycycline
YGR169C−A_1
ZShift = −0.89
Z lm Score = −0.94
NG = 0
DB = 0
SM = 0
Delta_r
Doxycycline
YGR169C−A_1
ZShift = −0.05
Z lm Score = 2.82
NG = 0
DB = 0
SM = 0
Delta_AUC
Doxycycline
YHR098C SFB3
ZShift = 3.22
Z lm Score = 5.47
NG = 0
DB = 0
SM = 0
Delta_L
Doxycycline
YHR098C SFB3
ZShift = 1.76
Z lm Score = −11.98
NG = 0
DB = 0
SM = 0
Delta_K
Doxycycline
YHR098C SFB3
ZShift = −1.6
Z lm Score = −1.44
NG = 0
DB = 0
SM = 0
Delta_r
Doxycycline
YHR098C SFB3
ZShift = −0.29
Z lm Score = 5.26
NG = 0
DB = 0
SM = 0
Delta_AUC
Doxycycline
YNL081C SWS2
ZShift = 6.78
Z lm Score = 5.44
NG = 0
DB = 0
SM = 0
Delta_L
Doxycycline
YNL081C SWS2
ZShift = −7.71
Z lm Score = −12.38
NG = 0
DB = 0
SM = 0
Delta_K
Doxycycline
YNL081C SWS2
ZShift = −2.32
Z lm Score = 0.98
NG = 0
DB = 0
SM = 0
Delta_r
Doxycycline
YNL081C SWS2
ZShift = −7.27
Z lm Score = −3.13
NG = 0
DB = 0
SM = 0
Delta_AUC
Doxycycline

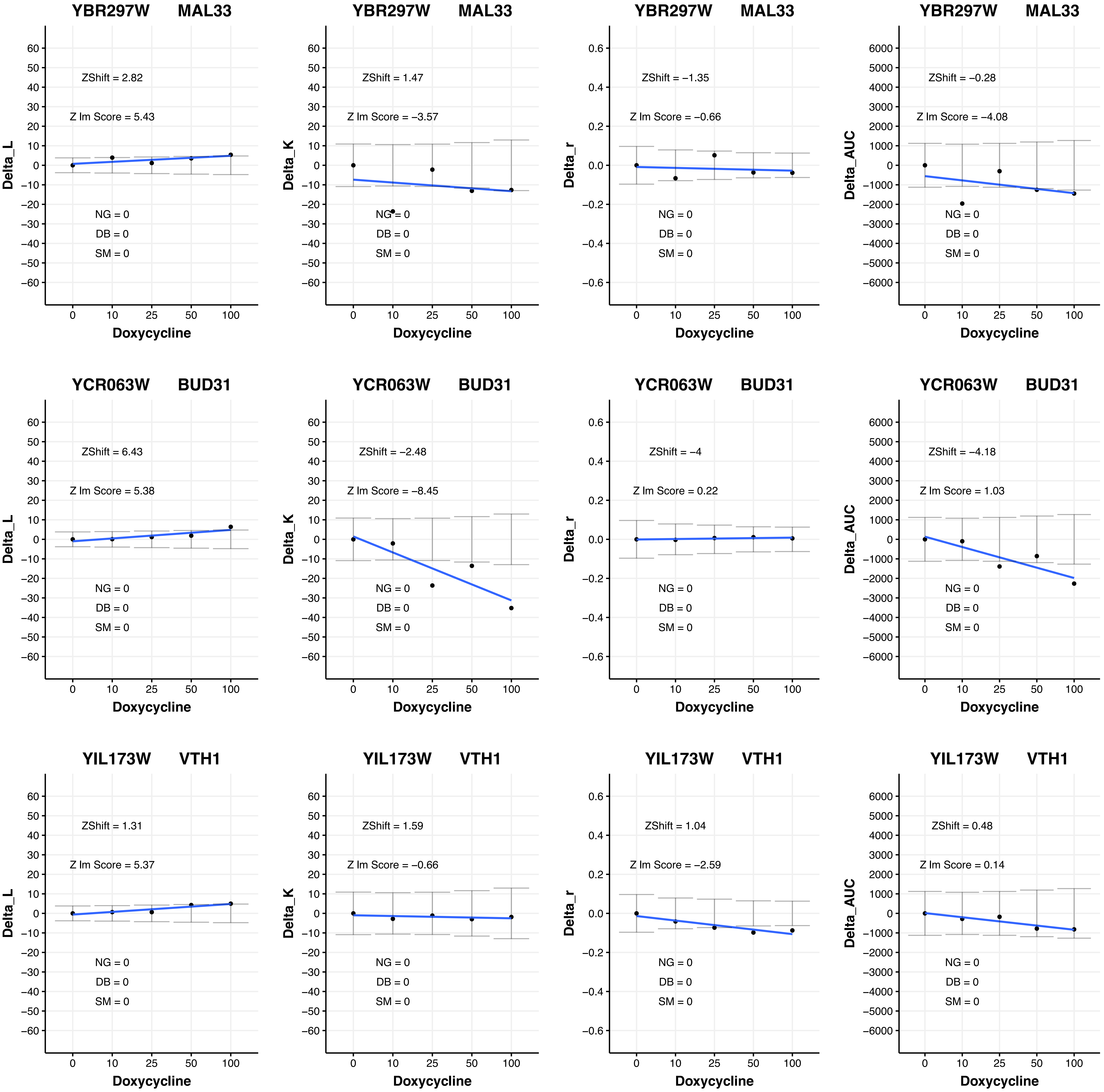
YBR297W MAL33
ZShift = 2.82
Z lm Score = 5.43
NG = 0
DB = 0
SM = 0
Delta_L
Doxycycline
YBR297W MAL33
ZShift = 1.47
Z lm Score = −3.57
NG = 0
DB = 0
SM = 0
Delta_K
Doxycycline
YBR297W MAL33
ZShift = −1.35
Z lm Score = −0.66
NG = 0
DB = 0
SM = 0
Delta_r
Doxycycline
YBR297W MAL33
ZShift = −0.28
Z lm Score = −4.08
NG = 0
DB = 0
SM = 0
Delta_AUC
Doxycycline
YCR063W BUD31
ZShift = 6.43
Z lm Score = 5.38
NG = 0
DB = 0
SM = 0
Delta_L
Doxycycline
YCR063W BUD31
ZShift = −2.48
Z lm Score = −8.45
NG = 0
DB = 0
SM = 0
Delta_K
Doxycycline
YCR063W BUD31
ZShift = −4
Z lm Score = 0.22
NG = 0
DB = 0
SM = 0
Delta_r
Doxycycline
YCR063W BUD31
ZShift = −4.18
Z lm Score = 1.03
NG = 0
DB = 0
SM = 0
Delta_AUC
Doxycycline
YIL173W VTH1
ZShift = 1.31
Z lm Score = 5.37
NG = 0
DB = 0
SM = 0
Delta_L
Doxycycline
YIL173W VTH1
ZShift = 1.59
Z lm Score = −0.66
NG = 0
DB = 0
SM = 0
Delta_K
Doxycycline
YIL173W VTH1
ZShift = 1.04
Z lm Score = −2.59
NG = 0
DB = 0
SM = 0
Delta_r
Doxycycline
YIL173W VTH1
ZShift = 0.48
Z lm Score = 0.14
NG = 0
DB = 0
SM = 0
Delta_AUC
Doxycycline

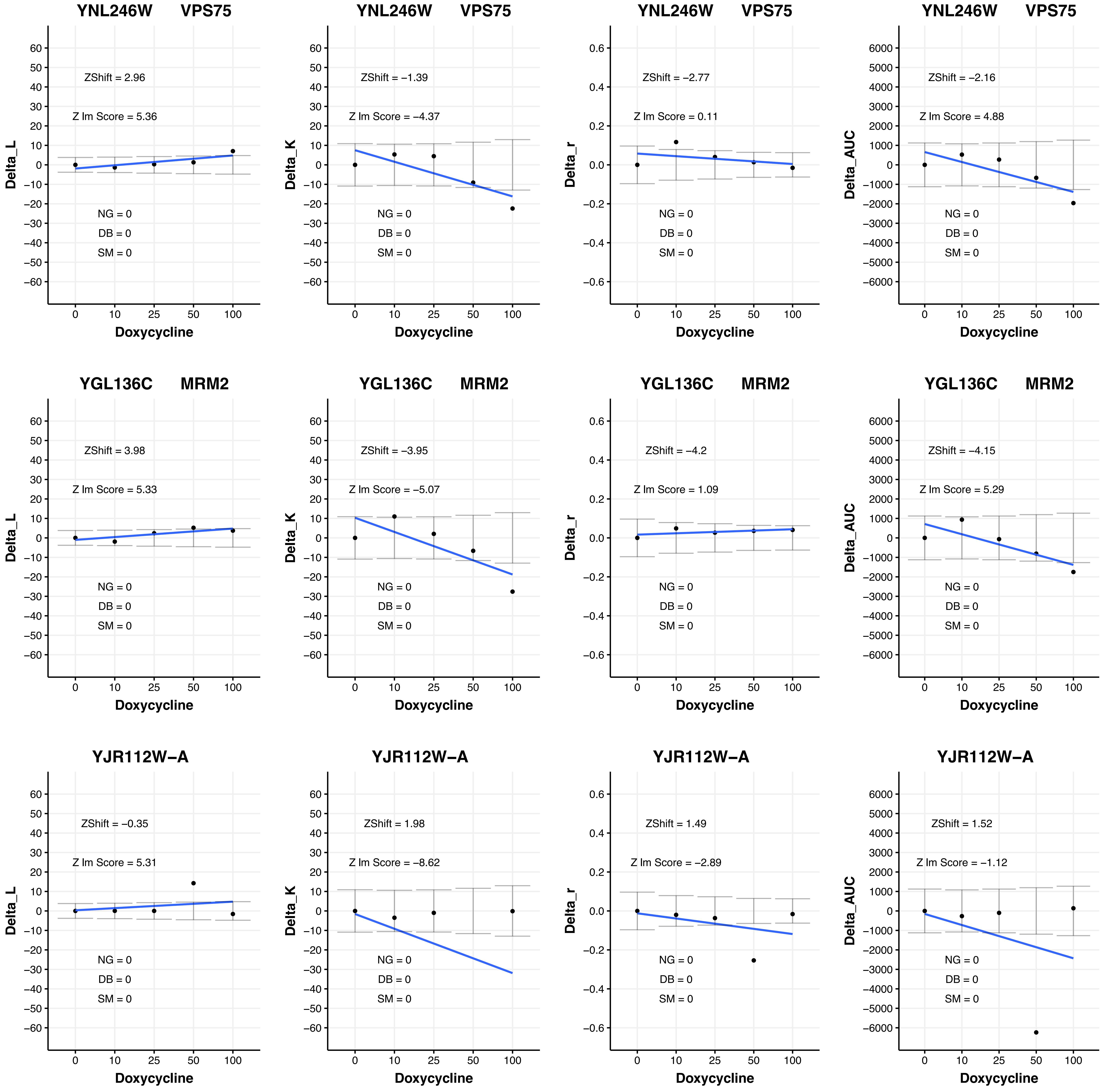

YNL246W VPS75
ZShift = 2.96
Z lm Score = 5.36
NG = 0
DB = 0
SM = 0
Delta_L
Doxycycline
YNL246W VPS75
ZShift = −1.39
Z lm Score = −4.37
NG = 0
DB = 0
SM = 0
Delta_K
Doxycycline
YNL246W VPS75
ZShift = −2.77
Z lm Score = 0.11
NG = 0
DB = 0
SM = 0
Delta_r
Doxycycline
YNL246W VPS75
ZShift = −2.16
Z lm Score = 4.88
NG = 0
DB = 0
SM = 0
Delta_AUC
Doxycycline
YGL136C MRM2
ZShift = 3.98
Z lm Score = 5.33
NG = 0
DB = 0
SM = 0
Delta_L
Doxycycline
YGL136C MRM2
ZShift = −3.95
Z lm Score = −5.07
NG = 0
DB = 0
SM = 0
Delta_K
Doxycycline
YGL136C MRM2
ZShift = −4.2
Z lm Score = 1.09
NG = 0
DB = 0
SM = 0
Delta_r
Doxycycline
YGL136C MRM2
ZShift = −4.15
Z lm Score = 5.29
NG = 0
DB = 0
SM = 0
Delta_AUC
Doxycycline
YJR112W−A
ZShift = −0.35
Z lm Score = 5.31
NG = 0
DB = 0
SM = 0
Delta_L
Doxycycline
YJR112W−A
ZShift = 1.98
Z lm Score = −8.62
NG = 0
DB = 0
SM = 0
Delta_K
Doxycycline
YJR112W−A
ZShift = 1.49
Z lm Score = −2.89
NG = 0
DB = 0
SM = 0
Delta_r
Doxycycline
YJR112W−A
ZShift = 1.52
Z lm Score = −1.12
NG = 0
DB = 0
SM = 0
Delta_AUC
Doxycycline

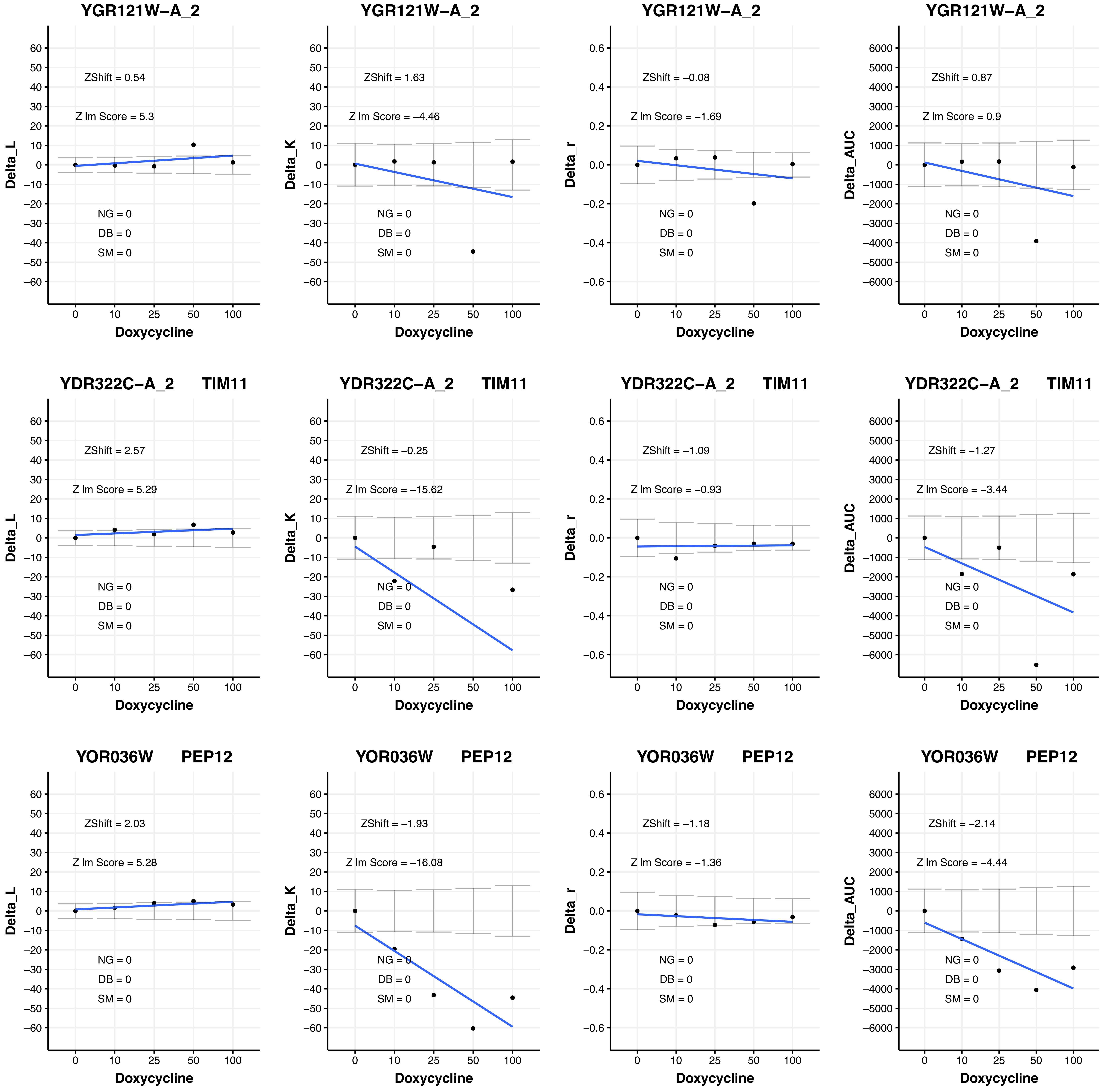

YGR121W−A_2
ZShift = 0.54
Z lm Score = 5.3
NG = 0
DB = 0
SM = 0
Delta_L
Doxycycline
YGR121W−A_2
ZShift = 1.63
Z lm Score = −4.46
NG = 0
DB = 0
SM = 0
Delta_K
Doxycycline
YGR121W−A_2
ZShift = −0.08
Z lm Score = −1.69
NG = 0
DB = 0
SM = 0
Delta_r
Doxycycline
YGR121W−A_2
ZShift = 0.87
Z lm Score = 0.9
NG = 0
DB = 0
SM = 0
Delta_AUC
Doxycycline
YDR322C−A_2 TIM11
ZShift = 2.57
Z lm Score = 5.29
NG = 0
DB = 0
SM = 0
Delta_L
Doxycycline
YDR322C−A_2 TIM11
ZShift = −0.25
Z lm Score = −15.62
NG = 0
DB = 0
SM = 0
Delta_K
Doxycycline
YDR322C−A_2 TIM11
ZShift = −1.09
Z lm Score = −0.93
NG = 0
DB = 0
SM = 0
Delta_r
Doxycycline
YDR322C−A_2 TIM11
ZShift = −1.27
Z lm Score = −3.44
NG = 0
DB = 0
SM = 0
Delta_AUC
Doxycycline
YOR036W PEP12
ZShift = 2.03
Z lm Score = 5.28
NG = 0
DB = 0
SM = 0
Delta_L
Doxycycline
YOR036W PEP12
ZShift = −1.93
Z lm Score = −16.08
NG = 0
DB = 0
SM = 0
Delta_K
Doxycycline
YOR036W PEP12
ZShift = −1.18
Z lm Score = −1.36
NG = 0
DB = 0
SM = 0
Delta_r
Doxycycline
YOR036W PEP12
ZShift = −2.14
Z lm Score = −4.44
NG = 0
DB = 0
SM = 0
Delta_AUC
Doxycycline

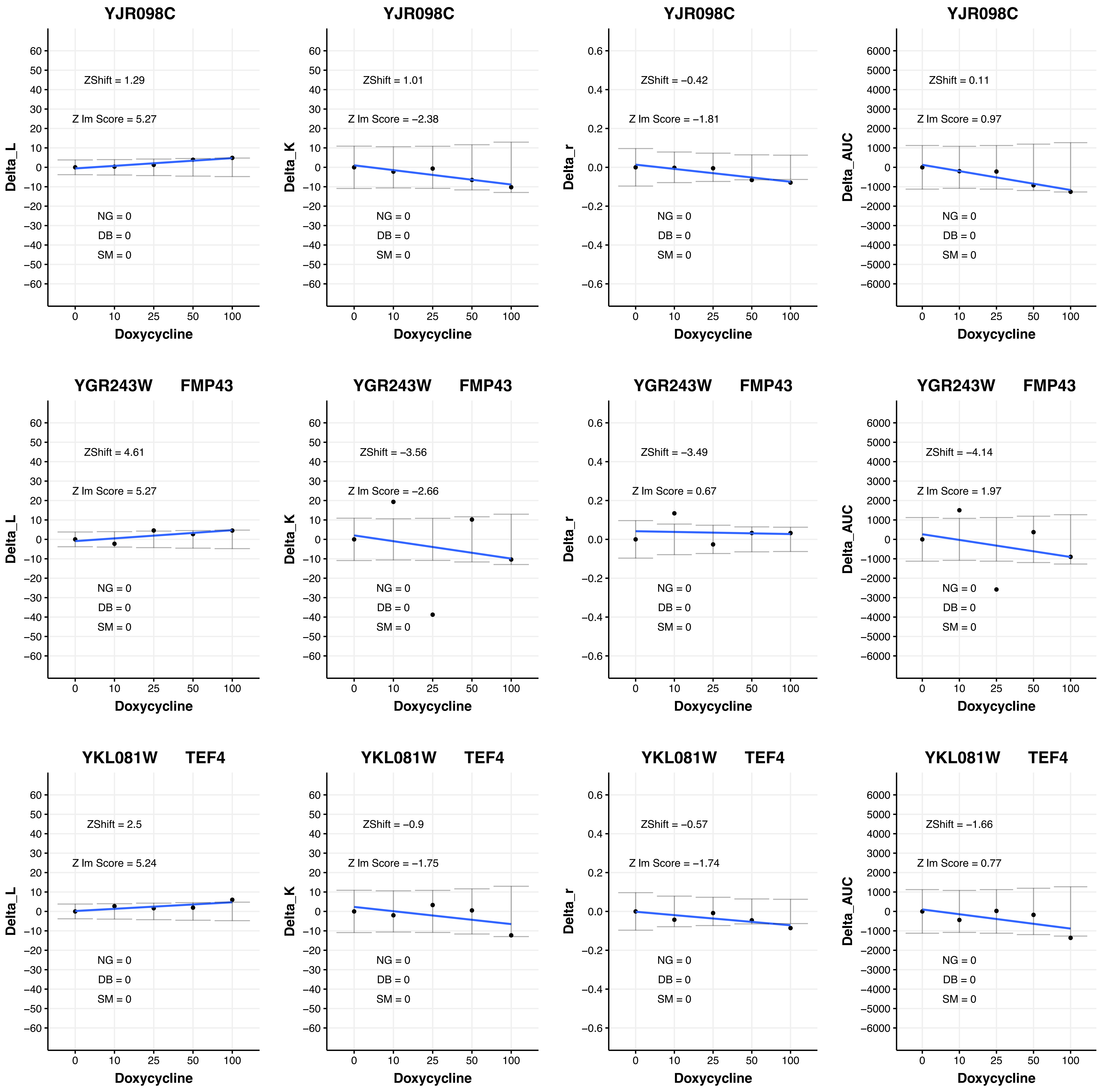
YJR098C
ZShift = 1.29
Z lm Score = 5.27
NG = 0
DB = 0
SM = 0
Delta_L
Doxycycline
0 10 25 50 100
60 50 40 30 20 10 0 −10 −20 −30 −40 −50 −60
YJR098C
ZShift = 1.01
Z lm Score = −2.38
NG = 0
DB = 0
SM = 0
Delta_K
Doxycycline
YJR098C
ZShift = −0.42
Z lm Score = −1.81
NG = 0
DB = 0
SM = 0
Delta_r
Doxycycline
0.6 0.4 0.2 0.0 −0.2 −0.4 −0.6
YJR098C
ZShift = 0.11
Z lm Score = 0.97
NG = 0
DB = 0
SM = 0
Delta_AUC
Doxycycline
6000 5000 4000 3000 2000 1000 0 −1000 −2000 −3000 −4000 −5000 −6000
YGR243W FMP43
ZShift = 4.61
Z lm Score = 5.27
NG = 0
DB = 0
SM = 0
Delta_L
Doxycycline
YGR243W FMP43
ZShift = −3.56
Z lm Score = −2.66
NG = 0
DB = 0
SM = 0
Delta_K
Doxycycline
YGR243W FMP43
ZShift = −3.49
Z lm Score = 0.67
NG = 0
DB = 0
SM = 0
Delta_r
Doxycycline
YGR243W FMP43
ZShift = −4.14
Z lm Score = 1.97
NG = 0
DB = 0
SM = 0
Delta_AUC
Doxycycline
YKL081W TEF4
ZShift = 2.5
Z lm Score = 5.24
NG = 0
DB = 0
SM = 0
Delta_L
Doxycycline
YKL081W TEF4
ZShift = −0.9
Z lm Score = −1.75
NG = 0
DB = 0
SM = 0
Delta_K
Doxycycline
YKL081W TEF4
ZShift = −0.57
Z lm Score = −1.74
NG = 0
DB = 0
SM = 0
Delta_r
Doxycycline
YKL081W TEF4
ZShift = −1.66
Z lm Score = 0.77
NG = 0
DB = 0
SM = 0
Delta_AUC
Doxycycline

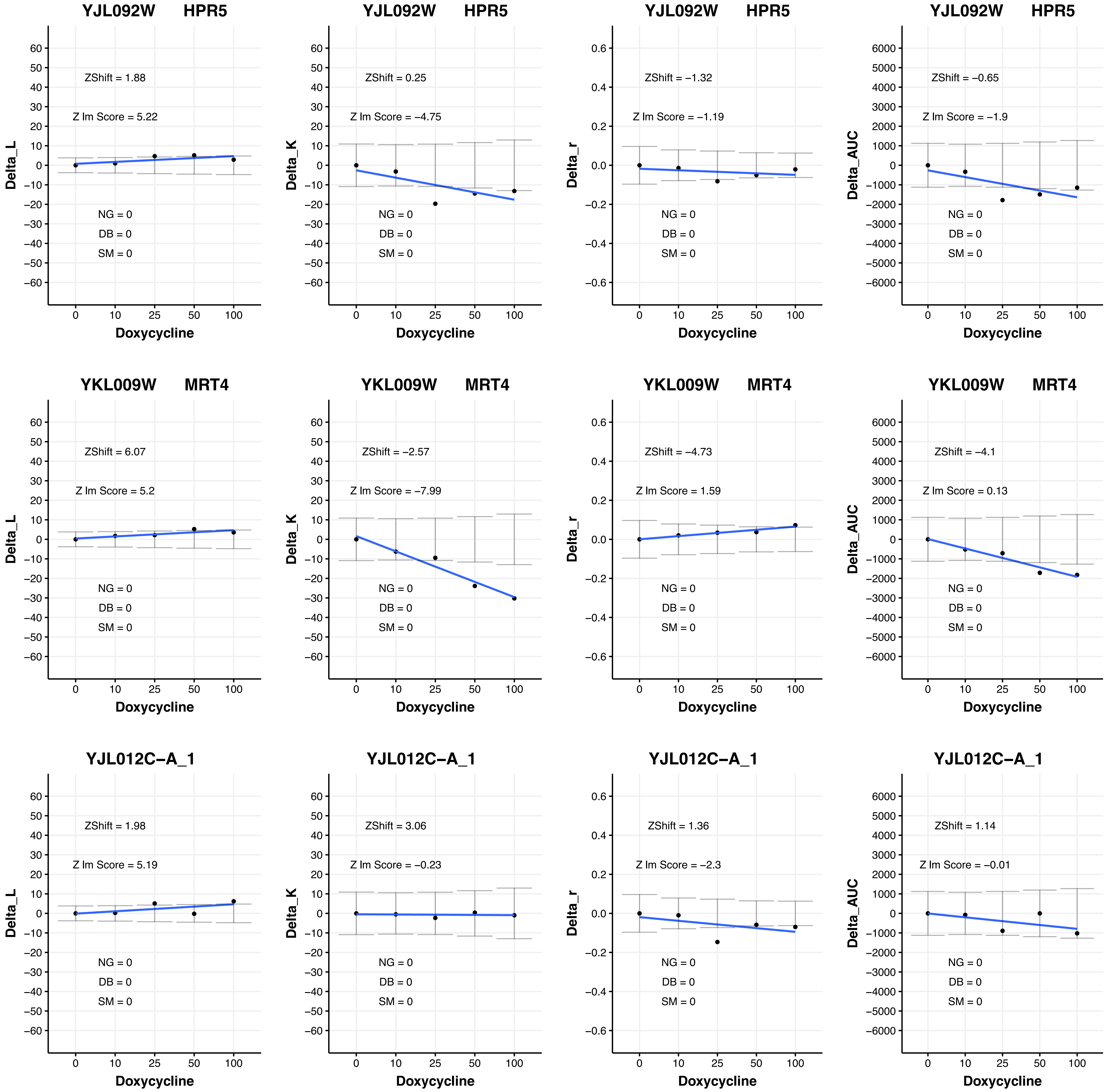
YJL092W HPR5
ZShift = 1.88
Z lm Score = 5.22
NG = 0
DB = 0
SM = 0
Delta_L
Doxycycline
YJL092W HPR5
ZShift = 0.25
Z lm Score = −4.75
NG = 0
DB = 0
SM = 0
Delta_K
Doxycycline
YJL092W HPR5
ZShift = −1.32
Z lm Score = −1.19
NG = 0
DB = 0
SM = 0
Delta_r
Doxycycline
YJL092W HPR5
ZShift = −0.65
Z lm Score = −1.9
NG = 0
DB = 0
SM = 0
Delta_AUC
Doxycycline
YKL009W MRT4
ZShift = 6.07
Z lm Score = 5.2
NG = 0
DB = 0
SM = 0
Delta_L
Doxycycline
YKL009W MRT4
ZShift = −2.57
Z lm Score = −7.99
NG = 0
DB = 0
SM = 0
Delta_K
Doxycycline
YKL009W MRT4
ZShift = −4.73
Z lm Score = 1.59
NG = 0
DB = 0
SM = 0
Delta_r
Doxycycline
YKL009W MRT4
ZShift = −4.1
Z lm Score = 0.13
NG = 0
DB = 0
SM = 0
Delta_AUC
Doxycycline
YJL012C−A_1
ZShift = 1.98
Z lm Score = 5.19
NG = 0
DB = 0
SM = 0
Delta_L
Doxycycline
YJL012C−A_1
ZShift = 3.06
Z lm Score = −0.23
NG = 0
DB = 0
SM = 0
Delta_K
Doxycycline
YJL012C−A_1
ZShift = 1.36
Z lm Score = −2.3
NG = 0
DB = 0
SM = 0
Delta_r
Doxycycline
YJL012C−A_1
ZShift = 1.14
Z lm Score = −0.01
NG = 0
DB = 0
SM = 0
Delta_AUC
Doxycycline

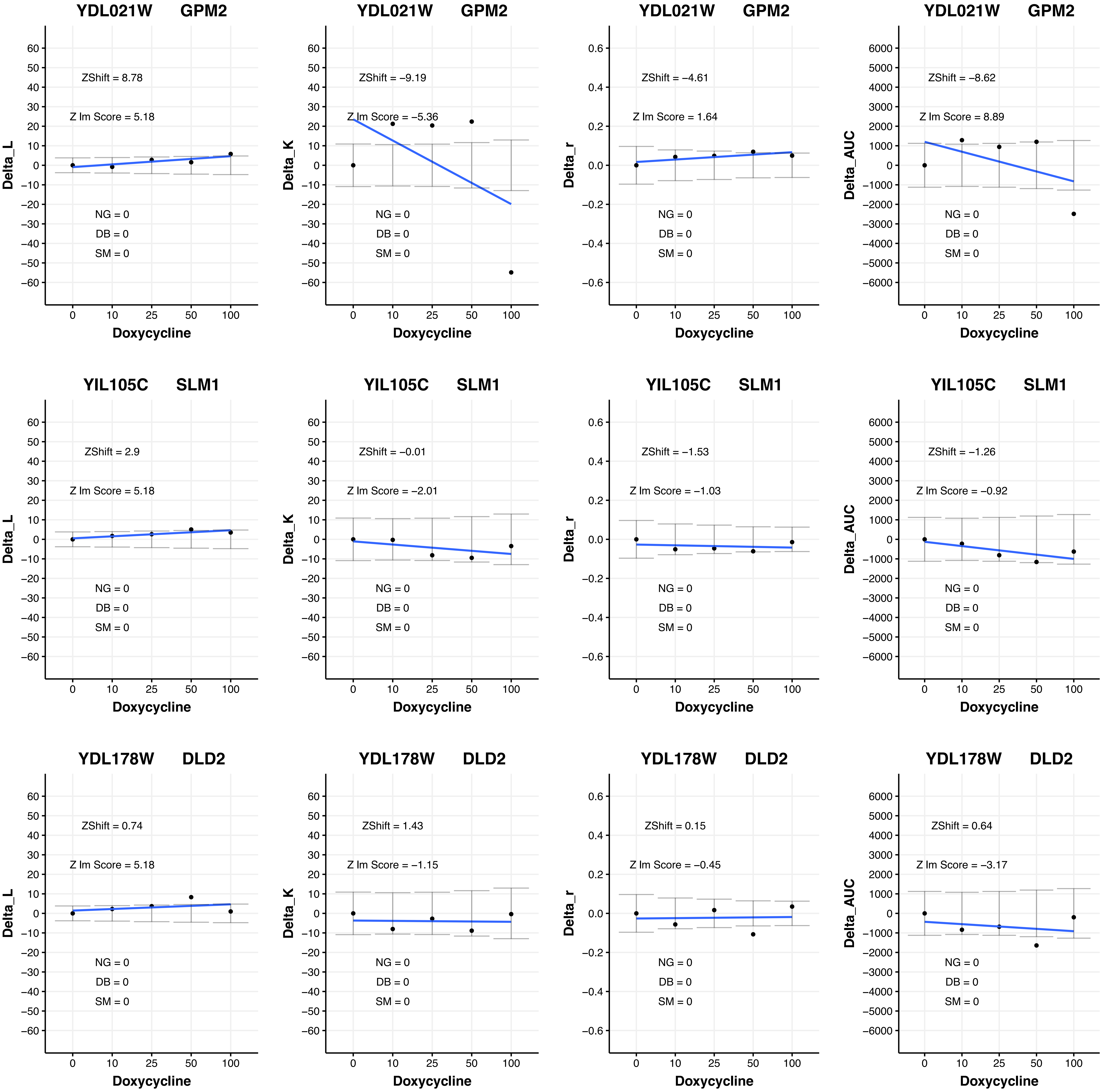

YDL021W GPM2
ZShift = 8.78
Z lm Score = 5.18
NG = 0
DB = 0
SM = 0
Delta_L
Doxycycline
YDL021W GPM2
ZShift = −9.19
Z lm Score = −5.36
NG = 0
DB = 0
SM = 0
Delta_K
Doxycycline
YDL021W GPM2
ZShift = −4.61
Z lm Score = 1.64
NG = 0
DB = 0
SM = 0
Delta_r
Doxycycline
YDL021W GPM2
ZShift = −8.62
Z lm Score = 8.89
NG = 0
DB = 0
SM = 0
Delta_AUC
Doxycycline
YIL105C SLM1
ZShift = 2.9
Z lm Score = 5.18
NG = 0
DB = 0
SM = 0
Delta_L
Doxycycline
YIL105C SLM1
ZShift = −0.01
Z lm Score = −2.01
NG = 0
DB = 0
SM = 0
Delta_K
Doxycycline
YIL105C SLM1
ZShift = −1.53
Z lm Score = −1.03
NG = 0
DB = 0
SM = 0
Delta_r
Doxycycline
YIL105C SLM1
ZShift = −1.26
Z lm Score = −0.92
NG = 0
DB = 0
SM = 0
Delta_AUC
Doxycycline
YDL178W DLD2
ZShift = 0.74
Z lm Score = 5.18
NG = 0
DB = 0
SM = 0
Delta_L
Doxycycline
YDL178W DLD2
ZShift = 1.43
Z lm Score = −1.15
NG = 0
DB = 0
SM = 0
Delta_K
Doxycycline
YDL178W DLD2
ZShift = 0.15
Z lm Score = −0.45
NG = 0
DB = 0
SM = 0
Delta_r
Doxycycline
YDL178W DLD2
ZShift = 0.64
Z lm Score = −3.17
NG = 0
DB = 0
SM = 0
Delta_AUC
Doxycycline

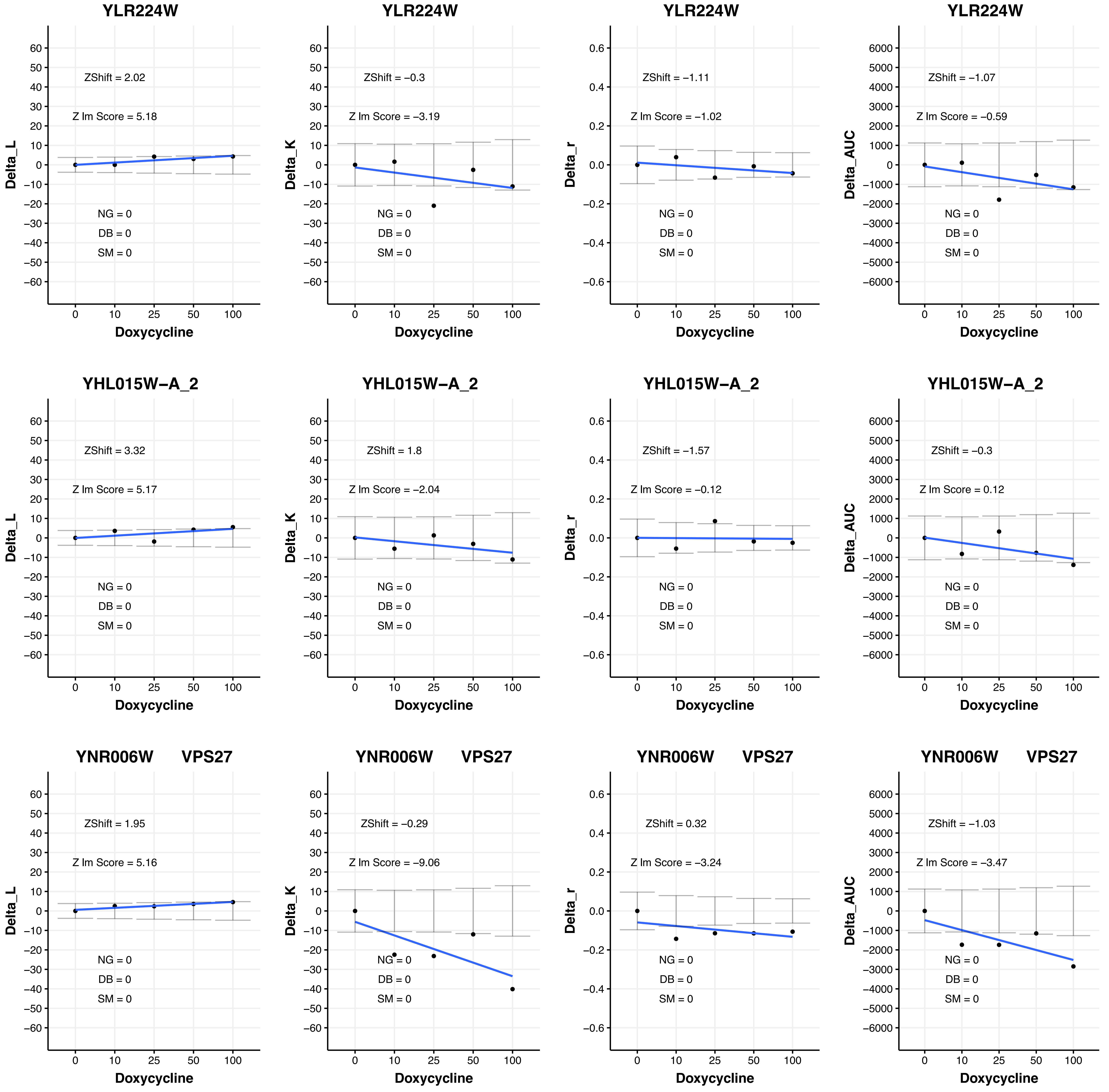
YLR224W
ZShift = 2.02
Z lm Score = 5.18
NG = 0
DB = 0
SM = 0
Delta_L
Doxycycline
YLR224W
ZShift = −0.3
Z lm Score = −3.19
NG = 0
DB = 0
SM = 0
Delta_K
Doxycycline
YLR224W
ZShift = −1.11
Z lm Score = −1.02
NG = 0
DB = 0
SM = 0
Delta_r
Doxycycline
YLR224W
ZShift = −1.07
Z lm Score = −0.59
NG = 0
DB = 0
SM = 0
Delta_AUC
Doxycycline
YHL015W−A_2
ZShift = 3.32
Z lm Score = 5.17
NG = 0
DB = 0
SM = 0
Delta_L
Doxycycline
YHL015W−A_2
ZShift = 1.8
Z lm Score = −2.04
NG = 0
DB = 0
SM = 0
Delta_K
Doxycycline
YHL015W−A_2
ZShift = −1.57
Z lm Score = −0.12
NG = 0
DB = 0
SM = 0
Delta_r
Doxycycline
YHL015W−A_2
ZShift = −0.3
Z lm Score = 0.12
NG = 0
DB = 0
SM = 0
Delta_AUC
Doxycycline
YNR006W VPS27
ZShift = 1.95
Z lm Score = 5.16
NG = 0
DB = 0
SM = 0
Delta_L
Doxycycline
YNR006W VPS27
ZShift = −0.29
Z lm Score = −9.06
NG = 0
DB = 0
SM = 0
Delta_K
Doxycycline
YNR006W VPS27
ZShift = 0.32
Z lm Score = −3.24
NG = 0
DB = 0
SM = 0
Delta_r
Doxycycline
YNR006W VPS27
ZShift = −1.03
Z lm Score = −3.47
NG = 0
DB = 0
SM = 0
Delta_AUC
Doxycycline

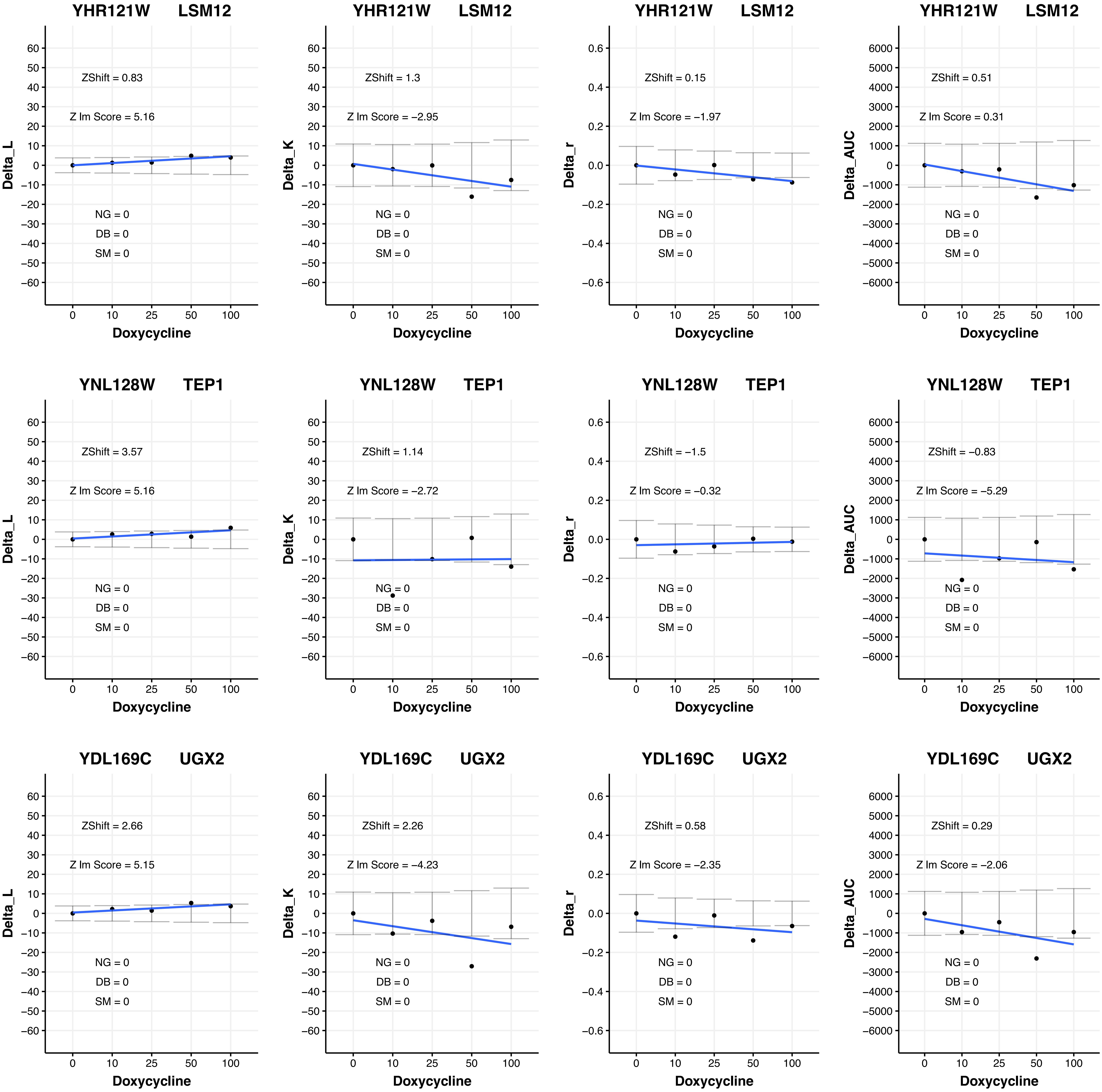
YHR121W LSM12
ZShift = 0.83
Z lm Score = 5.16
NG = 0
DB = 0
SM = 0
Delta_L
Doxycycline
YHR121W LSM12
ZShift = 1.3
Z lm Score = −2.95
NG = 0
DB = 0
SM = 0
Delta_K
Doxycycline
YHR121W LSM12
ZShift = 0.15
Z lm Score = −1.97
NG = 0
DB = 0
SM = 0
Delta_r
Doxycycline
YHR121W LSM12
ZShift = 0.51
Z lm Score = 0.31
NG = 0
DB = 0
SM = 0
Delta_AUC
Doxycycline
YNL128W TEP1
ZShift = 3.57
Z lm Score = 5.16
NG = 0
DB = 0
SM = 0
Delta_L
Doxycycline
YNL128W TEP1
ZShift = 1.14
Z lm Score = −2.72
NG = 0
DB = 0
SM = 0
Delta_K
Doxycycline
YNL128W TEP1
ZShift = −1.5
Z lm Score = −0.32
NG = 0
DB = 0
SM = 0
Delta_r
Doxycycline
YNL128W TEP1
ZShift = −0.83
Z lm Score = −5.29
NG = 0
DB = 0
SM = 0
Delta_AUC
Doxycycline
YDL169C UGX2
ZShift = 2.66
Z lm Score = 5.15
NG = 0
DB = 0
SM = 0
Delta_L
Doxycycline
YDL169C UGX2
ZShift = 2.26
Z lm Score = −4.23
NG = 0
DB = 0
SM = 0
Delta_K
Doxycycline
YDL169C UGX2
ZShift = 0.58
Z lm Score = −2.35
NG = 0
DB = 0
SM = 0
Delta_r
Doxycycline
YDL169C UGX2
ZShift = 0.29
Z lm Score = −2.06
NG = 0
DB = 0
SM = 0
Delta_AUC
Doxycycline

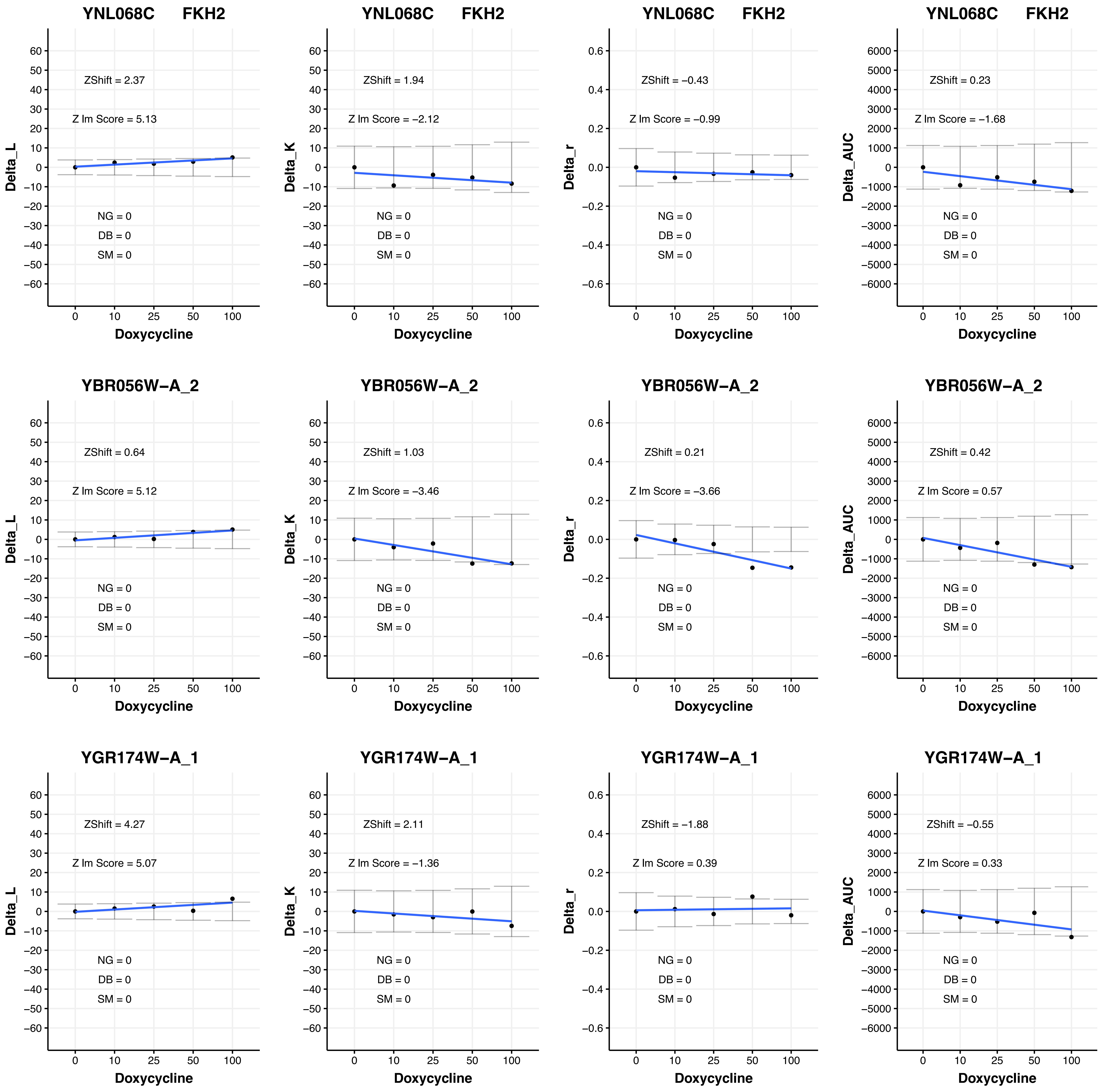

YNL068C FKH2
ZShift = 2.37
Z lm Score = 5.13
NG = 0
DB = 0
SM = 0
Delta_L
Doxycycline
YNL068C FKH2
ZShift = 1.94
Z lm Score = −2.12
NG = 0
DB = 0
SM = 0
Delta_K
Doxycycline
YNL068C FKH2
ZShift = −0.43
Z lm Score = −0.99
NG = 0
DB = 0
SM = 0
Delta_r
Doxycycline
YNL068C FKH2
ZShift = 0.23
Z lm Score = −1.68
NG = 0
DB = 0
SM = 0
Delta_AUC
Doxycycline
YBR056W−A_2
ZShift = 0.64
Z lm Score = 5.12
NG = 0
DB = 0
SM = 0
Delta_L
Doxycycline
YBR056W−A_2
ZShift = 1.03
Z lm Score = −3.46
NG = 0
DB = 0
SM = 0
Delta_K
Doxycycline
YBR056W−A_2
ZShift = 0.21
Z lm Score = −3.66
NG = 0
DB = 0
SM = 0
Delta_r
Doxycycline
YBR056W−A_2
ZShift = 0.42
Z lm Score = 0.57
NG = 0
DB = 0
SM = 0
Delta_AUC
Doxycycline
YGR174W−A_1
ZShift = 4.27
Z lm Score = 5.07
NG = 0
DB = 0
SM = 0
Delta_L
Doxycycline
YGR174W−A_1
ZShift = 2.11
Z lm Score = −1.36
NG = 0
DB = 0
SM = 0
Delta_K
Doxycycline
YGR174W−A_1
ZShift = −1.88
Z lm Score = 0.39
NG = 0
DB = 0
SM = 0
Delta_r
Doxycycline
YGR174W−A_1
ZShift = −0.55
Z lm Score = 0.33
NG = 0
DB = 0
SM = 0
Delta_AUC
Doxycycline

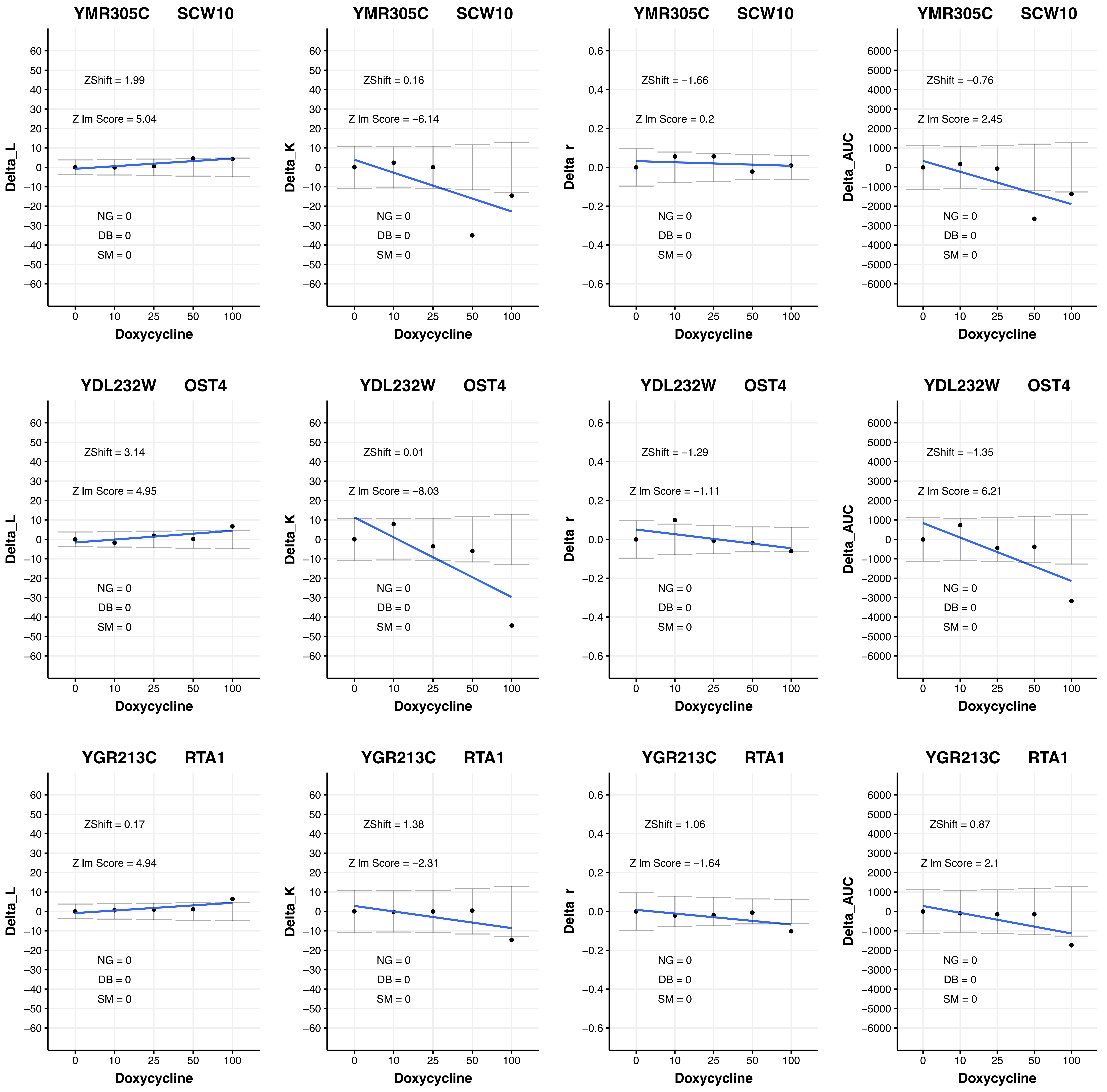

YMR305C SCW10
ZShift = 1.99
Z lm Score = 5.04
NG = 0
DB = 0
SM = 0
Delta_L
Doxycycline
YMR305C SCW10
ZShift = 0.16
Z lm Score = −6.14
NG = 0
DB = 0
SM = 0
Delta_K
Doxycycline
YMR305C SCW10
ZShift = −1.66
Z lm Score = 0.2
NG = 0
DB = 0
SM = 0
Delta_r
Doxycycline
YMR305C SCW10
ZShift = −0.76
Z lm Score = 2.45
NG = 0
DB = 0
SM = 0
Delta_AUC
Doxycycline
YDL232W OST4
ZShift = 3.14
Z lm Score = 4.95
NG = 0
DB = 0
SM = 0
Delta_L
Doxycycline
YDL232W OST4
ZShift = 0.01
Z lm Score = −8.03
NG = 0
DB = 0
SM = 0
Delta_K
Doxycycline
YDL232W OST4
ZShift = −1.29
Z lm Score = −1.11
NG = 0
DB = 0
SM = 0
Delta_r
Doxycycline
YDL232W OST4
ZShift = −1.35
Z lm Score = 6.21
NG = 0
DB = 0
SM = 0
Delta_AUC
Doxycycline
YGR213C RTA1
ZShift = 0.17
Z lm Score = 4.94
NG = 0
DB = 0
SM = 0
Delta_L
Doxycycline
YGR213C RTA1
ZShift = 1.38
Z lm Score = −2.31
NG = 0
DB = 0
SM = 0
Delta_K
Doxycycline
YGR213C RTA1
ZShift = 1.06
Z lm Score = −1.64
NG = 0
DB = 0
SM = 0
Delta_r
Doxycycline
YGR213C RTA1
ZShift = 0.87
Z lm Score = 2.1
NG = 0
DB = 0
SM = 0
Delta_AUC
Doxycycline

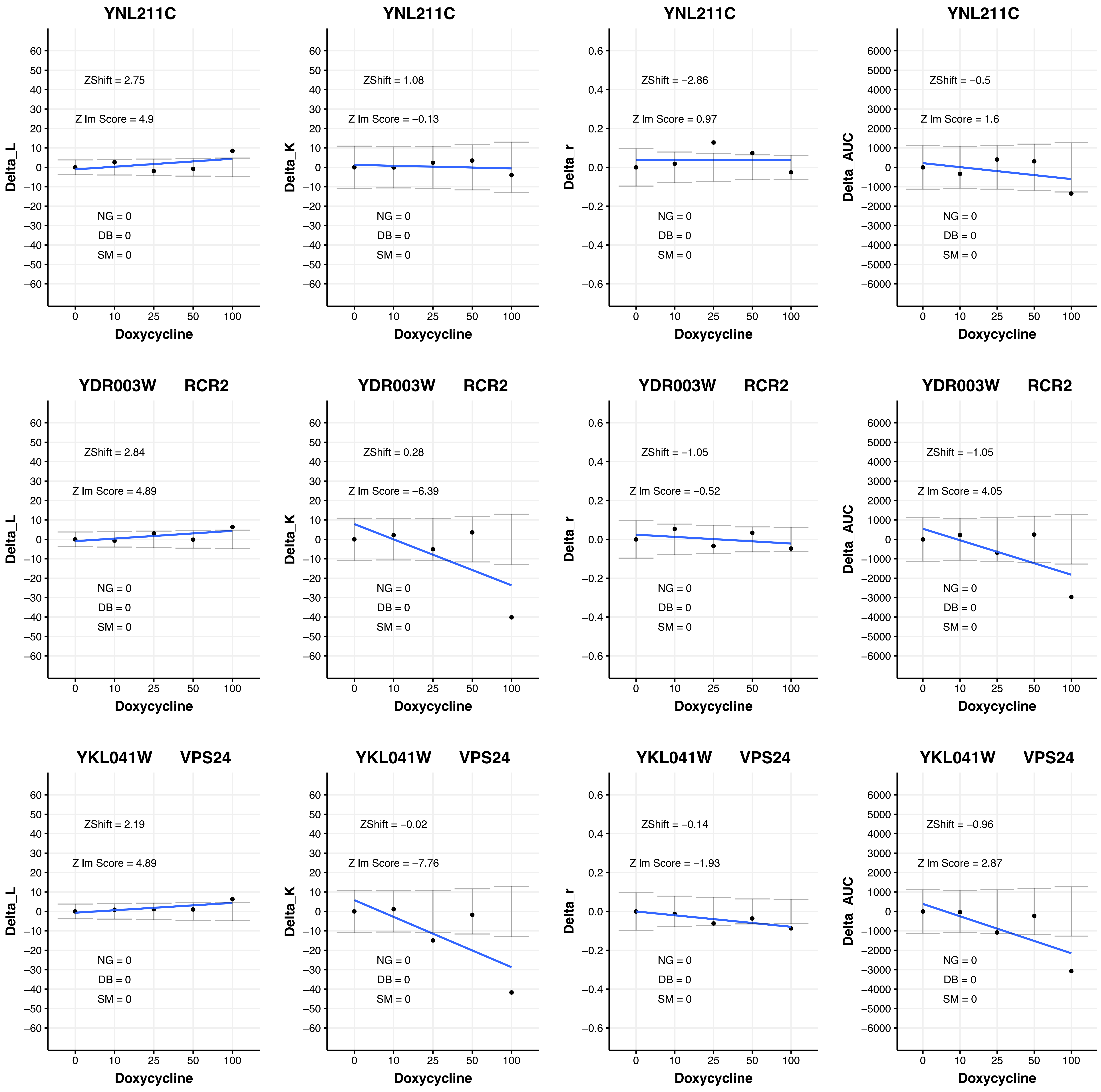
YNL211C
ZShift = 2.75
Z lm Score = 4.9
NG = 0
DB = 0
SM = 0
Delta_L
Doxycycline
YNL211C
ZShift = 1.08
Z lm Score = −0.13
NG = 0
DB = 0
SM = 0
Delta_K
Doxycycline
YNL211C
ZShift = −2.86
Z lm Score = 0.97
NG = 0
DB = 0
SM = 0
Delta_r
Doxycycline
YNL211C
ZShift = −0.5
Z lm Score = 1.6
NG = 0
DB = 0
SM = 0
Delta_AUC
Doxycycline
YDR003W RCR2
ZShift = 2.84
Z lm Score = 4.89
NG = 0
DB = 0
SM = 0
Delta_L
Doxycycline
YDR003W RCR2
ZShift = 0.28
Z lm Score = −6.39
NG = 0
DB = 0
SM = 0
Delta_K
Doxycycline
YDR003W RCR2
ZShift = −1.05
Z lm Score = −0.52
NG = 0
DB = 0
SM = 0
Delta_r
Doxycycline
YDR003W RCR2
ZShift = −1.05
Z lm Score = 4.05
NG = 0
DB = 0
SM = 0
Delta_AUC
Doxycycline
YKL041W VPS24
ZShift = 2.19
Z lm Score = 4.89
NG = 0
DB = 0
SM = 0
Delta_L
Doxycycline
YKL041W VPS24
ZShift = −0.02
Z lm Score = −7.76
NG = 0
DB = 0
SM = 0
Delta_K
Doxycycline
YKL041W VPS24
ZShift = −0.14
Z lm Score = −1.93
NG = 0
DB = 0
SM = 0
Delta_r
Doxycycline
YKL041W VPS24
ZShift = −0.96
Z lm Score = 2.87
NG = 0
DB = 0
SM = 0
Delta_AUC
Doxycycline

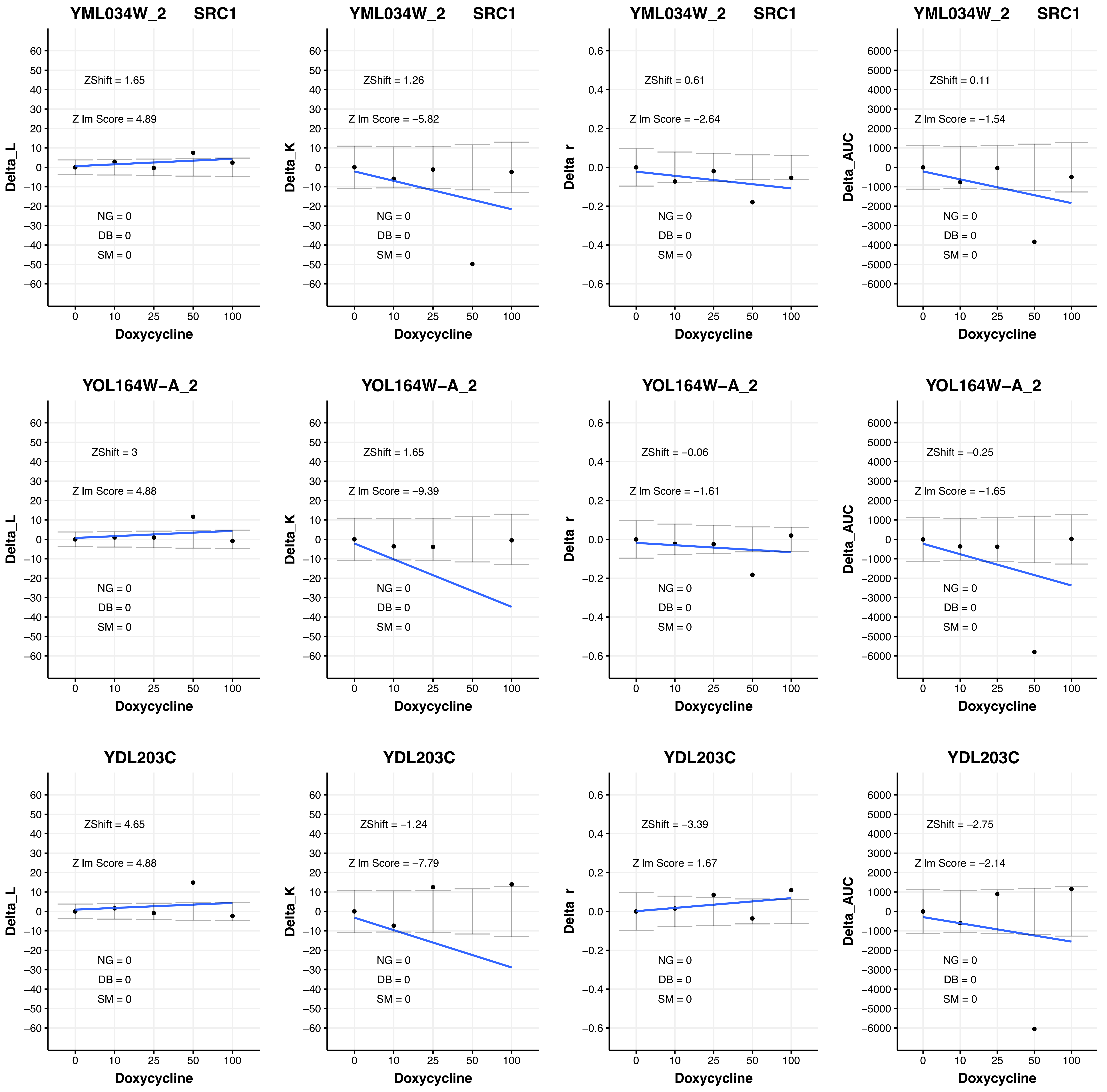

YML034W_2 SRC1
ZShift = 1.65
Z lm Score = 4.89
NG = 0
DB = 0
SM = 0
Delta_L
Doxycycline
YML034W_2 SRC1
ZShift = 1.26
Z lm Score = −5.82
NG = 0
DB = 0
SM = 0
Delta_K
Doxycycline
YML034W_2 SRC1
ZShift = 0.61
Z lm Score = −2.64
NG = 0
DB = 0
SM = 0
Delta_r
Doxycycline
YML034W_2 SRC1
ZShift = 0.11
Z lm Score = −1.54
NG = 0
DB = 0
SM = 0
Delta_AUC
Doxycycline
YOL164W−A_2
ZShift = 3
Z lm Score = 4.88
NG = 0
DB = 0
SM = 0
Delta_L
Doxycycline
YOL164W−A_2
ZShift = 1.65
Z lm Score = −9.39
NG = 0
DB = 0
SM = 0
Delta_K
Doxycycline
YOL164W−A_2
ZShift = −0.06
Z lm Score = −1.61
NG = 0
DB = 0
SM = 0
Delta_r
Doxycycline
YOL164W−A_2
ZShift = −0.25
Z lm Score = −1.65
NG = 0
DB = 0
SM = 0
Delta_AUC
Doxycycline
YDL203C
ZShift = 4.65
Z lm Score = 4.88
NG = 0
DB = 0
SM = 0
Delta_L
Doxycycline
YDL203C
ZShift = −1.24
Z lm Score = −7.79
NG = 0
DB = 0
SM = 0
Delta_K
Doxycycline
YDL203C
ZShift = −3.39
Z lm Score = 1.67
NG = 0
DB = 0
SM = 0
Delta_r
Doxycycline
YDL203C
ZShift = −2.75
Z lm Score = −2.14
NG = 0
DB = 0
SM = 0
Delta_AUC
Doxycycline

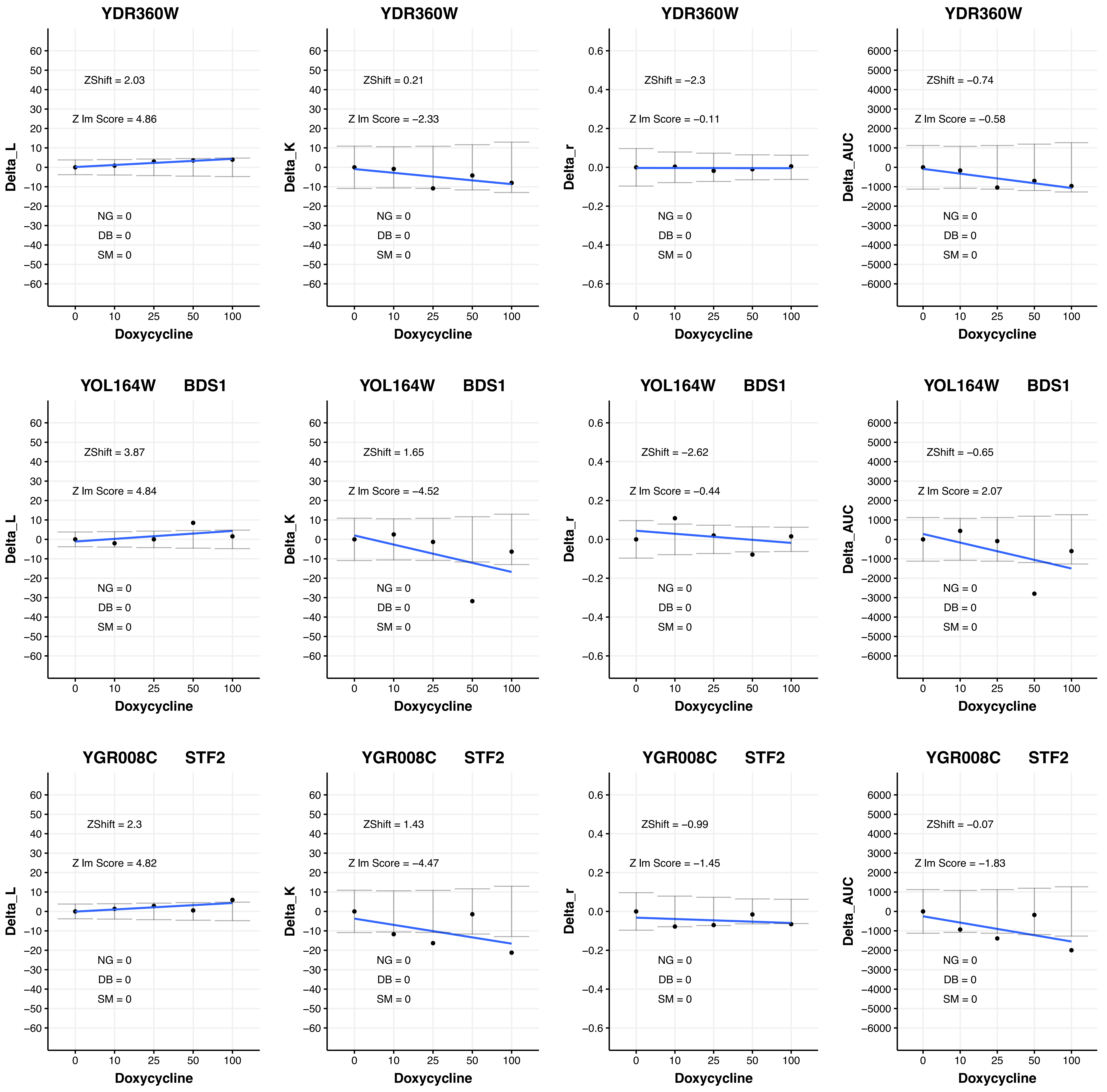

YDR360W
ZShift = 2.03
Z lm Score = 4.86
NG = 0
DB = 0
SM = 0
Delta_L
Doxycycline
YDR360W
ZShift = 0.21
Z lm Score = −2.33
NG = 0
DB = 0
SM = 0
Delta_K
Doxycycline
YDR360W
ZShift = −2.3
Z lm Score = −0.11
NG = 0
DB = 0
SM = 0
Delta_r
Doxycycline
YDR360W
ZShift = −0.74
Z lm Score = −0.58
NG = 0
DB = 0
SM = 0
Delta_AUC
Doxycycline
YOL164W BDS1
ZShift = 3.87
Z lm Score = 4.84
NG = 0
DB = 0
SM = 0
Delta_L
Doxycycline
YOL164W BDS1
ZShift = 1.65
Z lm Score = −4.52
NG = 0
DB = 0
SM = 0
Delta_K
Doxycycline
YOL164W BDS1
ZShift = −2.62
Z lm Score = −0.44
NG = 0
DB = 0
SM = 0
Delta_r
Doxycycline
YOL164W BDS1
ZShift = −0.65
Z lm Score = 2.07
NG = 0
DB = 0
SM = 0
Delta_AUC
Doxycycline
YGR008C STF2
ZShift = 2.3
Z lm Score = 4.82
NG = 0
DB = 0
SM = 0
Delta_L
Doxycycline
YGR008C STF2
ZShift = 1.43
Z lm Score = −4.47
NG = 0
DB = 0
SM = 0
Delta_K
Doxycycline
YGR008C STF2
ZShift = −0.99
Z lm Score = −1.45
NG = 0
DB = 0
SM = 0
Delta_r
Doxycycline
YGR008C STF2
ZShift = −0.07
Z lm Score = −1.83
NG = 0
DB = 0
SM = 0
Delta_AUC
Doxycycline

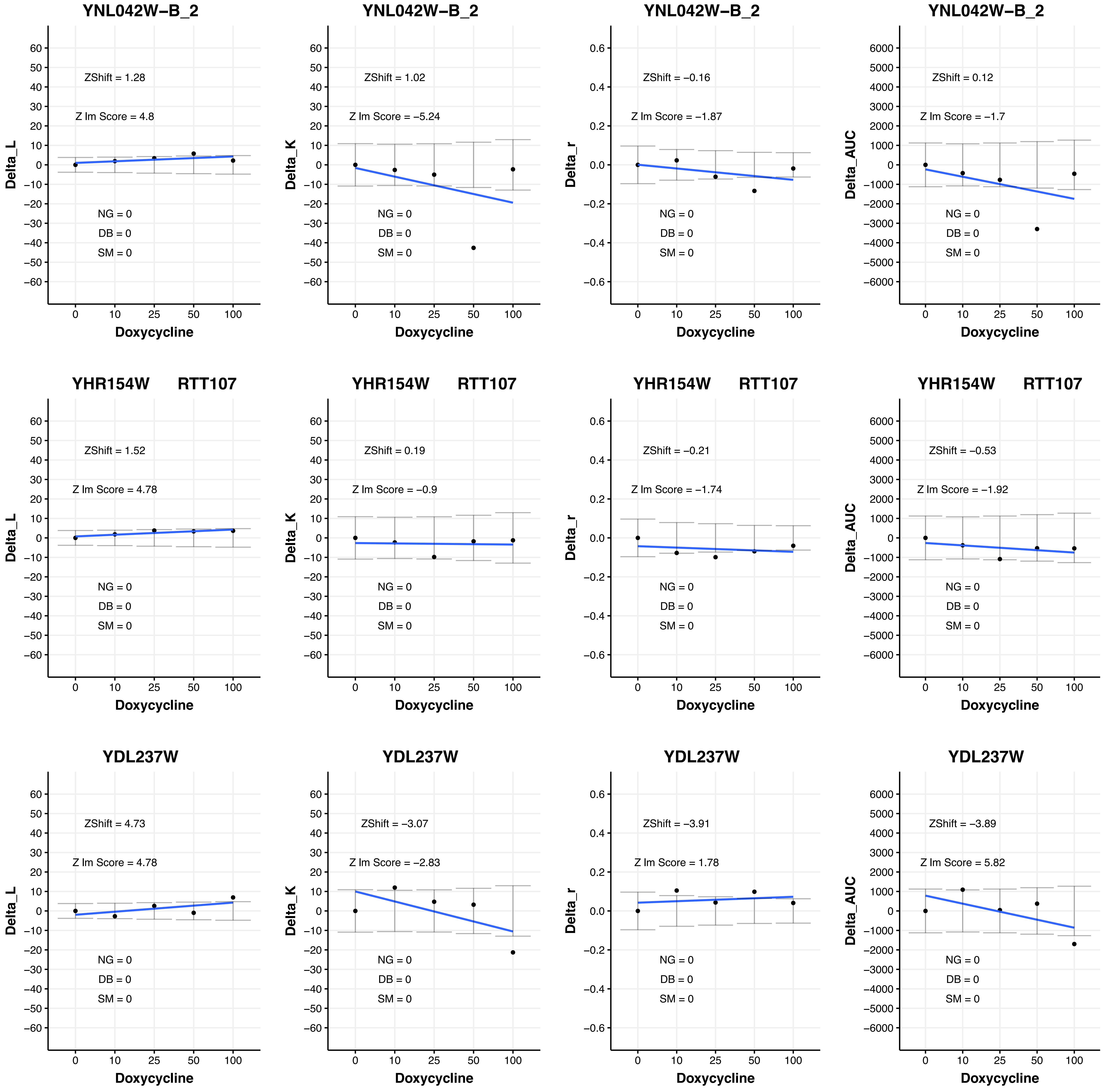

YNL042W−B_2
ZShift = 1.28
Z lm Score = 4.8
NG = 0
DB = 0
SM = 0
Delta_L
Doxycycline
YNL042W−B_2
ZShift = 1.02
Z lm Score = −5.24
NG = 0
DB = 0
SM = 0
Delta_K
Doxycycline
YNL042W−B_2
ZShift = −0.16
Z lm Score = −1.87
NG = 0
DB = 0
SM = 0
Delta_r
Doxycycline
YNL042W−B_2
ZShift = 0.12
Z lm Score = −1.7
NG = 0
DB = 0
SM = 0
Delta_AUC
Doxycycline
YHR154W RTT107
ZShift = 1.52
Z lm Score = 4.78
NG = 0
DB = 0
SM = 0
Delta_L
Doxycycline
YHR154W RTT107
ZShift = 0.19
Z lm Score = −0.9
NG = 0
DB = 0
SM = 0
Delta_K
Doxycycline
YHR154W RTT107
ZShift = −0.21
Z lm Score = −1.74
NG = 0
DB = 0
SM = 0
Delta_r
Doxycycline
YHR154W RTT107
ZShift = −0.53
Z lm Score = −1.92
NG = 0
DB = 0
SM = 0
Delta_AUC
Doxycycline
YDL237W
ZShift = 4.73
Z lm Score = 4.78
NG = 0
DB = 0
SM = 0
Delta_L
Doxycycline
YDL237W
ZShift = −3.07
Z lm Score = −2.83
NG = 0
DB = 0
SM = 0
Delta_K
Doxycycline
YDL237W
ZShift = −3.91
Z lm Score = 1.78
NG = 0
DB = 0
SM = 0
Delta_r
Doxycycline
YDL237W
ZShift = −3.89
Z lm Score = 5.82
NG = 0
DB = 0
SM = 0
Delta_AUC
Doxycycline

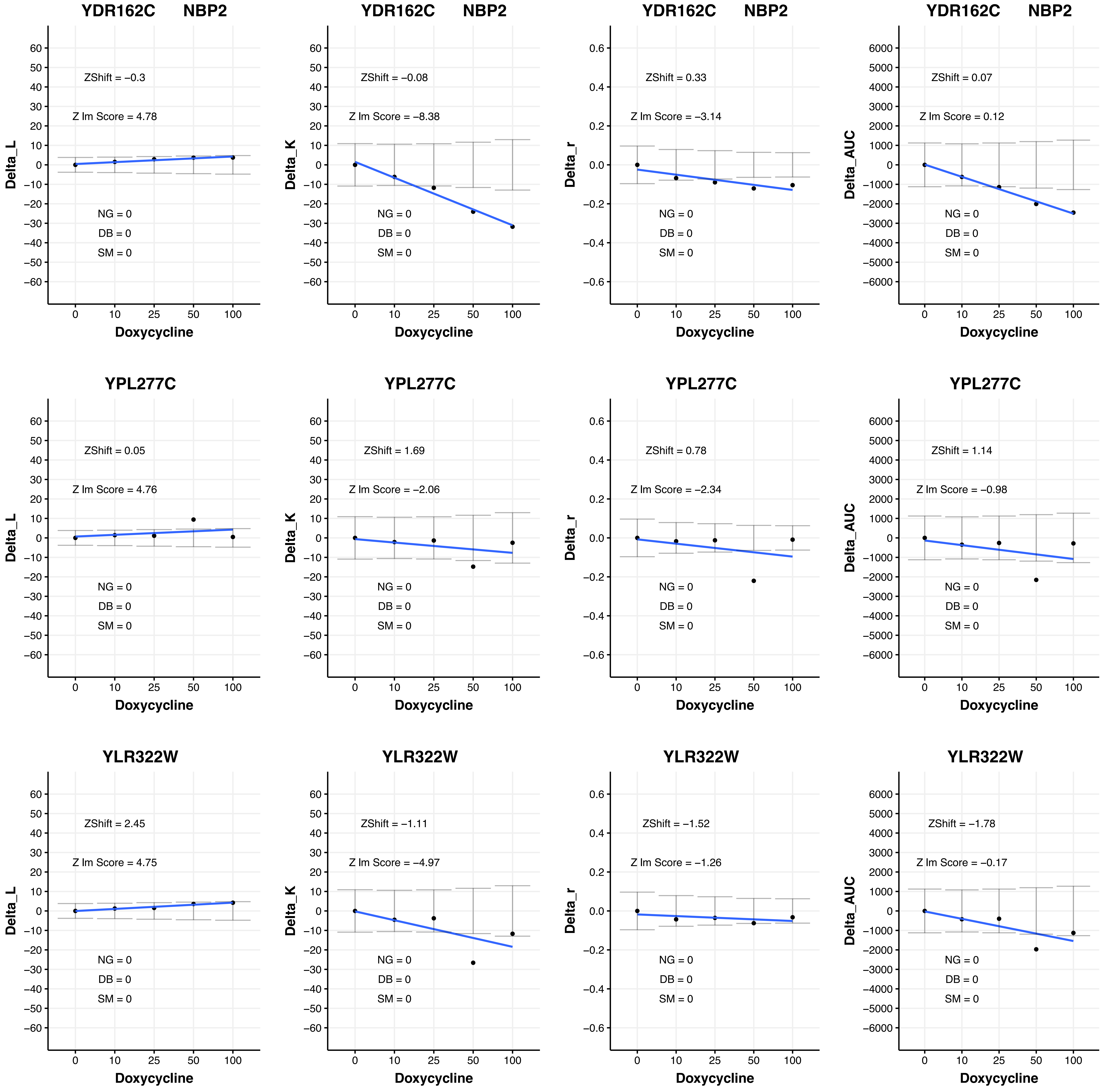

YDR162C NBP2
ZShift = −0.3
Z lm Score = 4.78
NG = 0
DB = 0
SM = 0
Delta_L
Doxycycline
YDR162C NBP2
ZShift = −0.08
Z lm Score = −8.38
NG = 0
DB = 0
SM = 0
Delta_K
Doxycycline
YDR162C NBP2
ZShift = 0.33
Z lm Score = −3.14
NG = 0
DB = 0
SM = 0
Delta_r
Doxycycline
YDR162C NBP2
ZShift = 0.07
Z lm Score = 0.12
NG = 0
DB = 0
SM = 0
Delta_AUC
Doxycycline
YPL277C
ZShift = 0.05
Z lm Score = 4.76
NG = 0
DB = 0
SM = 0
Delta_L
Doxycycline
YPL277C
ZShift = 1.69
Z lm Score = −2.06
NG = 0
DB = 0
SM = 0
Delta_K
Doxycycline
YPL277C
ZShift = 0.78
Z lm Score = −2.34
NG = 0
DB = 0
SM = 0
Delta_r
Doxycycline
YPL277C
ZShift = 1.14
Z lm Score = −0.98
NG = 0
DB = 0
SM = 0
Delta_AUC
Doxycycline
YLR322W
ZShift = 2.45
Z lm Score = 4.75
NG = 0
DB = 0
SM = 0
Delta_L
Doxycycline
YLR322W
ZShift = −1.11
Z lm Score = −4.97
NG = 0
DB = 0
SM = 0
Delta_K
Doxycycline
YLR322W
ZShift = −1.52
Z lm Score = −1.26
NG = 0
DB = 0
SM = 0
Delta_r
Doxycycline
YLR322W
ZShift = −1.78
Z lm Score = −0.17
NG = 0
DB = 0
SM = 0
Delta_AUC
Doxycycline

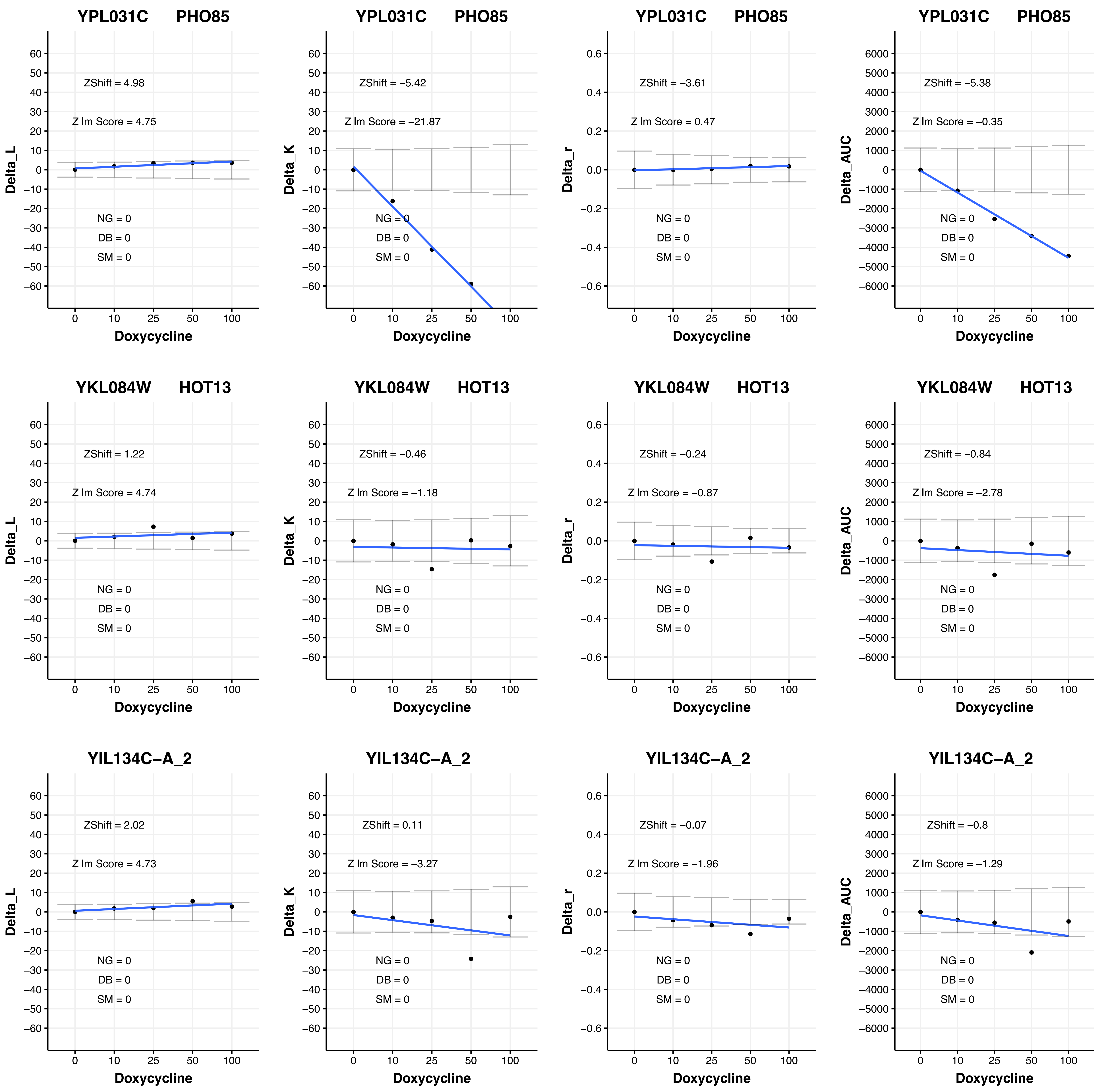

YPL031C PHO85
ZShift = 4.98
Z lm Score = 4.75
NG = 0
DB = 0
SM = 0
Delta_L
Doxycycline
YPL031C PHO85
ZShift = −5.42
Z lm Score = −21.87
NG = 0
DB = 0
SM = 0
Delta_K
Doxycycline
YPL031C PHO85
ZShift = −3.61
Z lm Score = 0.47
NG = 0
DB = 0
SM = 0
Delta_r
Doxycycline
YPL031C PHO85
ZShift = −5.38
Z lm Score = −0.35
NG = 0
DB = 0
SM = 0
Delta_AUC
Doxycycline
YKL084W HOT13
ZShift = 1.22
Z lm Score = 4.74
NG = 0
DB = 0
SM = 0
Delta_L
Doxycycline
YKL084W HOT13
ZShift = −0.46
Z lm Score = −1.18
NG = 0
DB = 0
SM = 0
Delta_K
Doxycycline
YKL084W HOT13
ZShift = −0.24
Z lm Score = −0.87
NG = 0
DB = 0
SM = 0
Delta_r
Doxycycline
YKL084W HOT13
ZShift = −0.84
Z lm Score = −2.78
NG = 0
DB = 0
SM = 0
Delta_AUC
Doxycycline
YIL134C−A_2
ZShift = 2.02
Z lm Score = 4.73
NG = 0
DB = 0
SM = 0
Delta_L
Doxycycline
YIL134C−A_2
ZShift = 0.11
Z lm Score = −3.27
NG = 0
DB = 0
SM = 0
Delta_K
Doxycycline
YIL134C−A_2
ZShift = −0.07
Z lm Score = −1.96
NG = 0
DB = 0
SM = 0
Delta_r
Doxycycline
YIL134C−A_2
ZShift = −0.8
Z lm Score = −1.29
NG = 0
DB = 0
SM = 0
Delta_AUC
Doxycycline

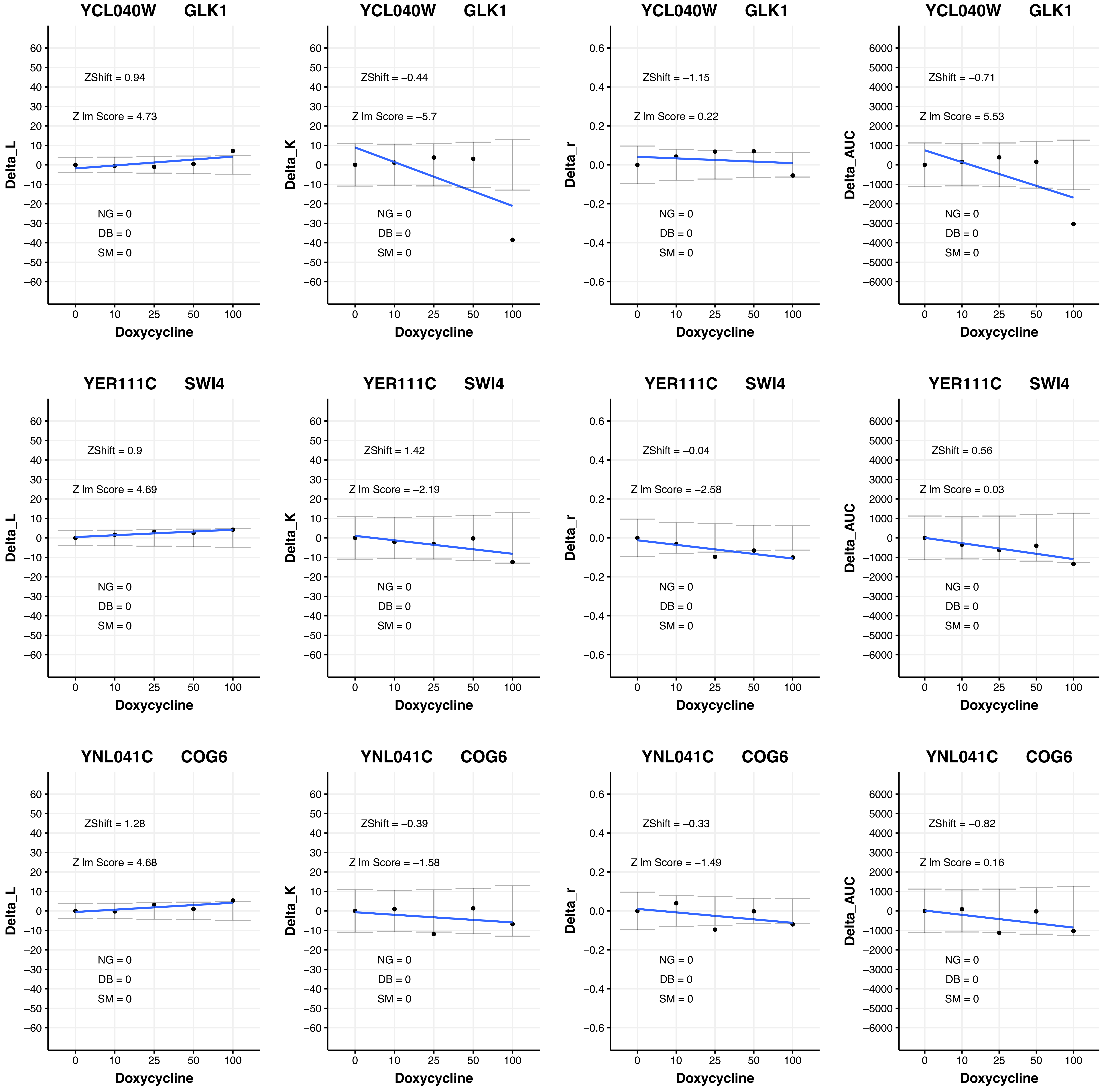
YCL040W GLK1
ZShift = 0.94
Z lm Score = 4.73
NG = 0
DB = 0
SM = 0
Delta_L
Doxycycline
YCL040W GLK1
ZShift = −0.44
Z lm Score = −5.7
NG = 0
DB = 0
SM = 0
Delta_K
Doxycycline
YCL040W GLK1
ZShift = −1.15
Z lm Score = 0.22
NG = 0
DB = 0
SM = 0
Delta_r
Doxycycline
YCL040W GLK1
ZShift = −0.71
Z lm Score = 5.53
NG = 0
DB = 0
SM = 0
Delta_AUC
Doxycycline
YER111C SWI4
ZShift = 0.9
Z lm Score = 4.69
NG = 0
DB = 0
SM = 0
Delta_L
Doxycycline
YER111C SWI4
ZShift = 1.42
Z lm Score = −2.19
NG = 0
DB = 0
SM = 0
Delta_K
Doxycycline
YER111C SWI4
ZShift = −0.04
Z lm Score = −2.58
NG = 0
DB = 0
SM = 0
Delta_r
Doxycycline
YER111C SWI4
ZShift = 0.56
Z lm Score = 0.03
NG = 0
DB = 0
SM = 0
Delta_AUC
Doxycycline
YNL041C COG6
ZShift = 1.28
Z lm Score = 4.68
NG = 0
DB = 0
SM = 0
Delta_L
Doxycycline
YNL041C COG6
ZShift = −0.39
Z lm Score = −1.58
NG = 0
DB = 0
SM = 0
Delta_K
Doxycycline
YNL041C COG6
ZShift = −0.33
Z lm Score = −1.49
NG = 0
DB = 0
SM = 0
Delta_r
Doxycycline
YNL041C COG6
ZShift = −0.82
Z lm Score = 0.16
NG = 0
DB = 0
SM = 0
Delta_AUC
Doxycycline

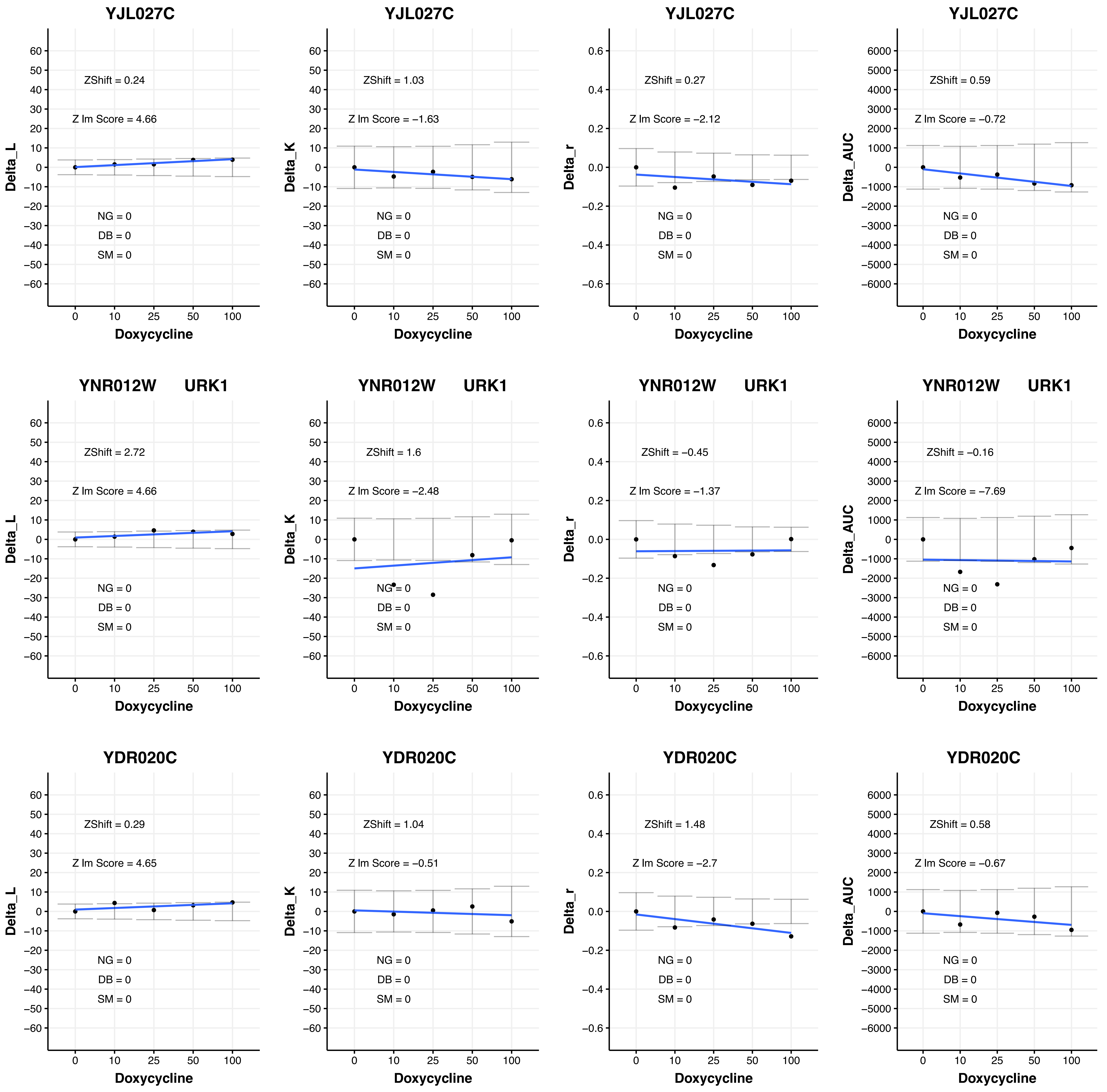

YJL027C
ZShift = 0.24
Z lm Score = 4.66
NG = 0
DB = 0
SM = 0
Delta_L
Doxycycline
YJL027C
ZShift = 1.03
Z lm Score = −1.63
NG = 0
DB = 0
SM = 0
Delta_K
Doxycycline
YJL027C
ZShift = 0.27
Z lm Score = −2.12
NG = 0
DB = 0
SM = 0
Delta_r
Doxycycline
YJL027C
ZShift = 0.59
Z lm Score = −0.72
NG = 0
DB = 0
SM = 0
Delta_AUC
Doxycycline
YNR012W URK1
ZShift = 2.72
Z lm Score = 4.66
NG = 0
DB = 0
SM = 0
Delta_L
Doxycycline
YNR012W URK1
ZShift = 1.6
Z lm Score = −2.48
NG = 0
DB = 0
SM = 0
Delta_K
Doxycycline
YNR012W URK1
ZShift = −0.45
Z lm Score = −1.37
NG = 0
DB = 0
SM = 0
Delta_r
Doxycycline
YNR012W URK1
ZShift = −0.16
Z lm Score = −7.69
NG = 0
DB = 0
SM = 0
Delta_AUC
Doxycycline
YDR020C
ZShift = 0.29
Z lm Score = 4.65
NG = 0
DB = 0
SM = 0
Delta_L
Doxycycline
YDR020C
ZShift = 1.04
Z lm Score = −0.51
NG = 0
DB = 0
SM = 0
Delta_K
Doxycycline
YDR020C
ZShift = 1.48
Z lm Score = −2.7
NG = 0
DB = 0
SM = 0
Delta_r
Doxycycline
YDR020C
ZShift = 0.58
Z lm Score = −0.67
NG = 0
DB = 0
SM = 0
Delta_AUC
Doxycycline

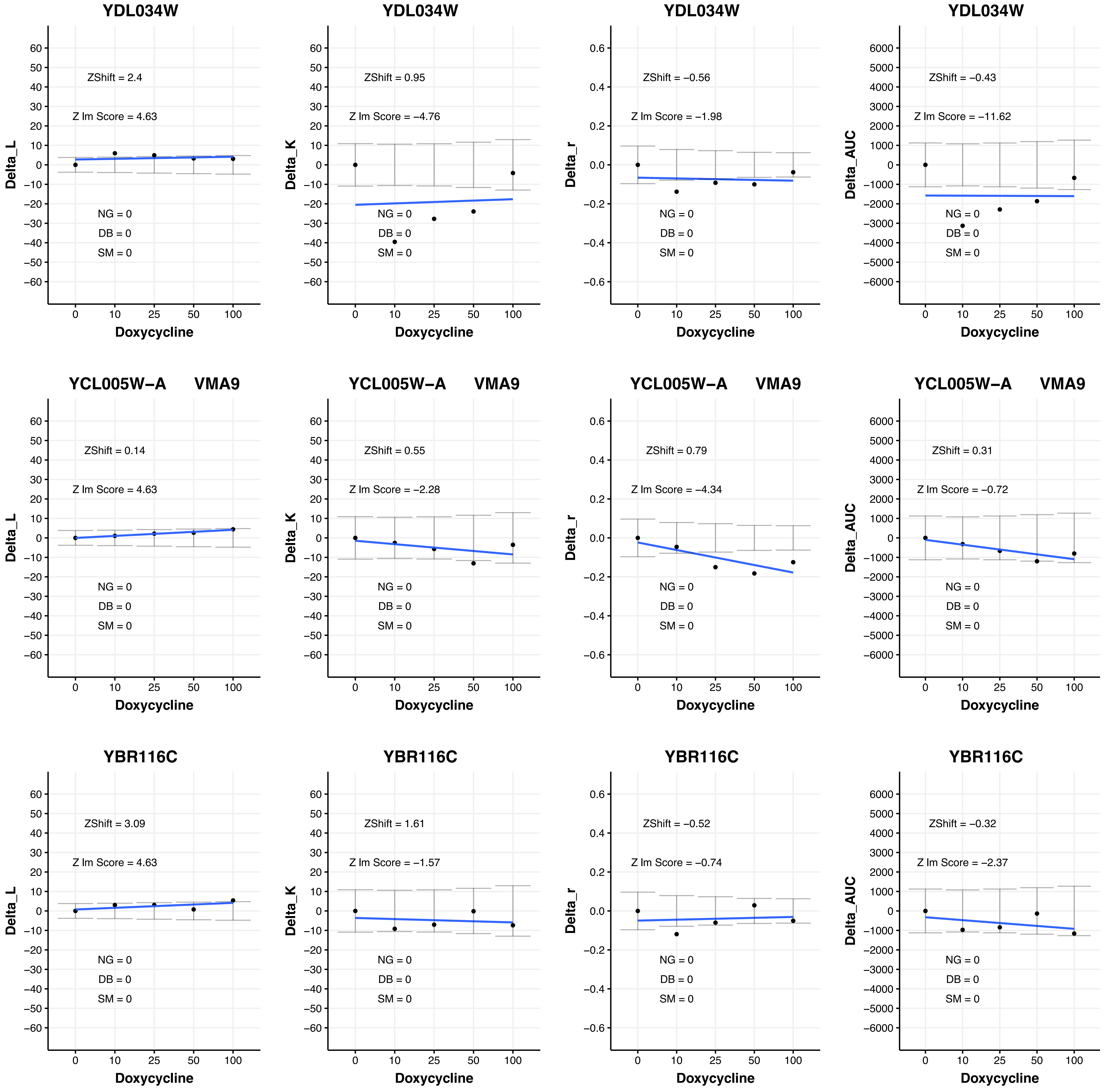
YDL034W
ZShift = 2.4
Z lm Score = 4.63
NG = 0
DB = 0
SM = 0
Delta_L
Doxycycline
YDL034W
ZShift = 0.95
Z lm Score = −4.76
NG = 0
DB = 0
SM = 0
Delta_K
Doxycycline
YDL034W
ZShift = −0.56
Z lm Score = −1.98
NG = 0
DB = 0
SM = 0
Delta_r
Doxycycline
YDL034W
ZShift = −0.43
Z lm Score = −11.62
NG = 0
DB = 0
SM = 0
Delta_AUC
Doxycycline
YCL005W−A VMA9
ZShift = 0.14
Z lm Score = 4.63
NG = 0
DB = 0
SM = 0
Delta_L
Doxycycline
YCL005W−A VMA9
ZShift = 0.55
Z lm Score = −2.28
NG = 0
DB = 0
SM = 0
Delta_K
Doxycycline
YCL005W−A VMA9
ZShift = 0.79
Z lm Score = −4.34
NG = 0
DB = 0
SM = 0
Delta_r
Doxycycline
YCL005W−A VMA9
ZShift = 0.31
Z lm Score = −0.72
NG = 0
DB = 0
SM = 0
Delta_AUC
Doxycycline
YBR116C
ZShift = 3.09
Z lm Score = 4.63
NG = 0
DB = 0
SM = 0
Delta_L
Doxycycline
YBR116C
ZShift = 1.61
Z lm Score = −1.57
NG = 0
DB = 0
SM = 0
Delta_K
Doxycycline
YBR116C
ZShift = −0.52
Z lm Score = −0.74
NG = 0
DB = 0
SM = 0
Delta_r
Doxycycline
YBR116C
ZShift = −0.32
Z lm Score = −2.37
NG = 0
DB = 0
SM = 0
Delta_AUC
Doxycycline
0
10
25
50
100

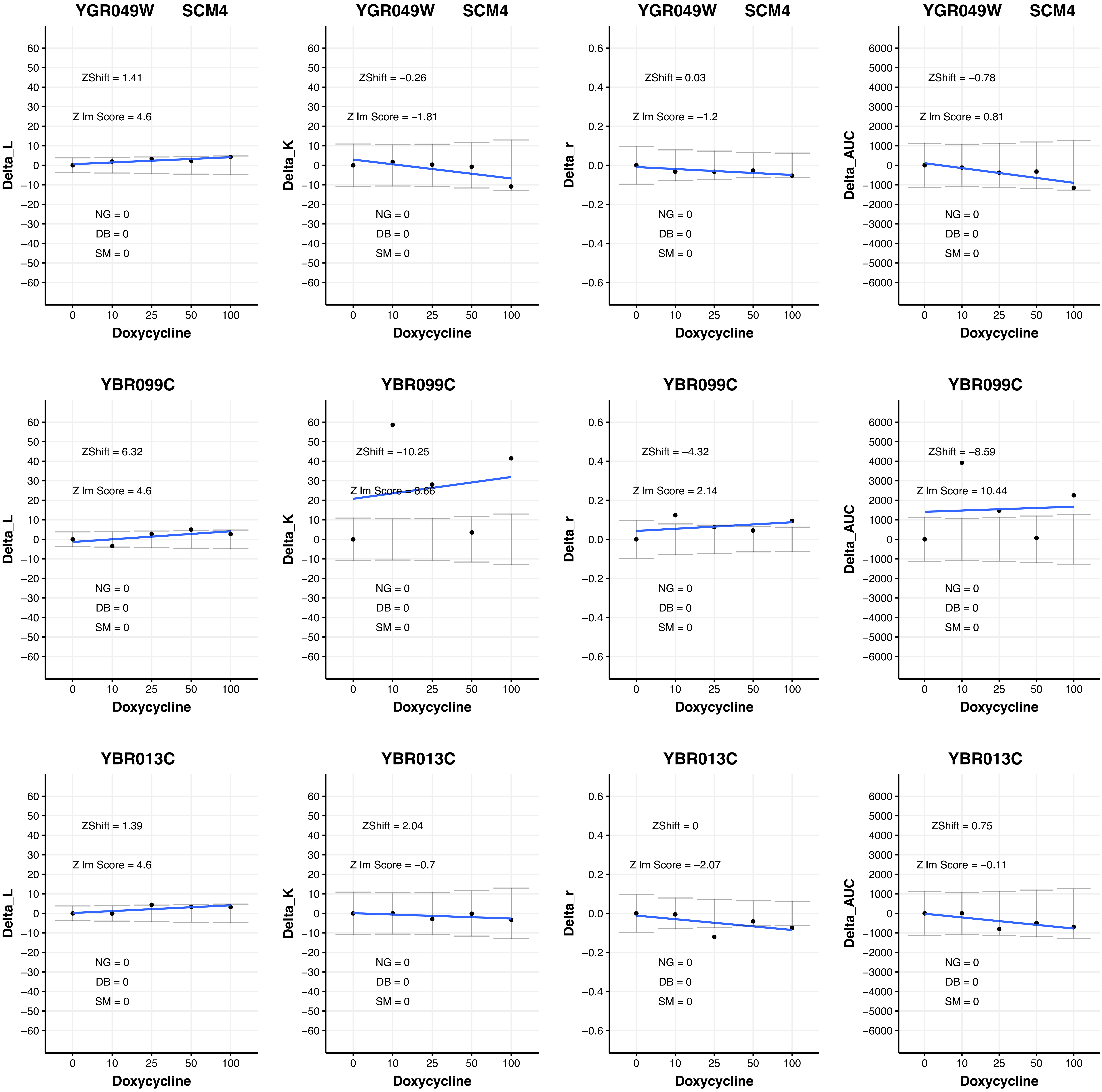

YGR049W SCM4
ZShift = 1.41
Z lm Score = 4.6
NG = 0
DB = 0
SM = 0
Delta_L
Doxycycline
YGR049W SCM4
ZShift = −0.26
Z lm Score = −1.81
NG = 0
DB = 0
SM = 0
Delta_K
Doxycycline
YGR049W SCM4
ZShift = 0.03
Z lm Score = −1.2
NG = 0
DB = 0
SM = 0
Delta_r
Doxycycline
YGR049W SCM4
ZShift = −0.78
Z lm Score = 0.81
NG = 0
DB = 0
SM = 0
Delta_AUC
Doxycycline
YBR099C
ZShift = 6.32
Z lm Score = 4.6
NG = 0
DB = 0
SM = 0
Delta_L
Doxycycline
YBR099C
ZShift = −10.25
Z lm Score = 8.66
NG = 0
DB = 0
SM = 0
Delta_K
Doxycycline
YBR099C
ZShift = −4.32
Z lm Score = 2.14
NG = 0
DB = 0
SM = 0
Delta_r
Doxycycline
YBR099C
ZShift = −8.59
Z lm Score = 10.44
NG = 0
DB = 0
SM = 0
Delta_AUC
Doxycycline
YBR013C
ZShift = 1.39
Z lm Score = 4.6
NG = 0
DB = 0
SM = 0
Delta_L
Doxycycline
YBR013C
ZShift = 2.04
Z lm Score = −0.7
NG = 0
DB = 0
SM = 0
Delta_K
Doxycycline
YBR013C
ZShift = 0
Z lm Score = −2.07
NG = 0
DB = 0
SM = 0
Delta_r
Doxycycline
YBR013C
ZShift = 0.75
Z lm Score = −0.11
NG = 0
DB = 0
SM = 0
Delta_AUC
Doxycycline

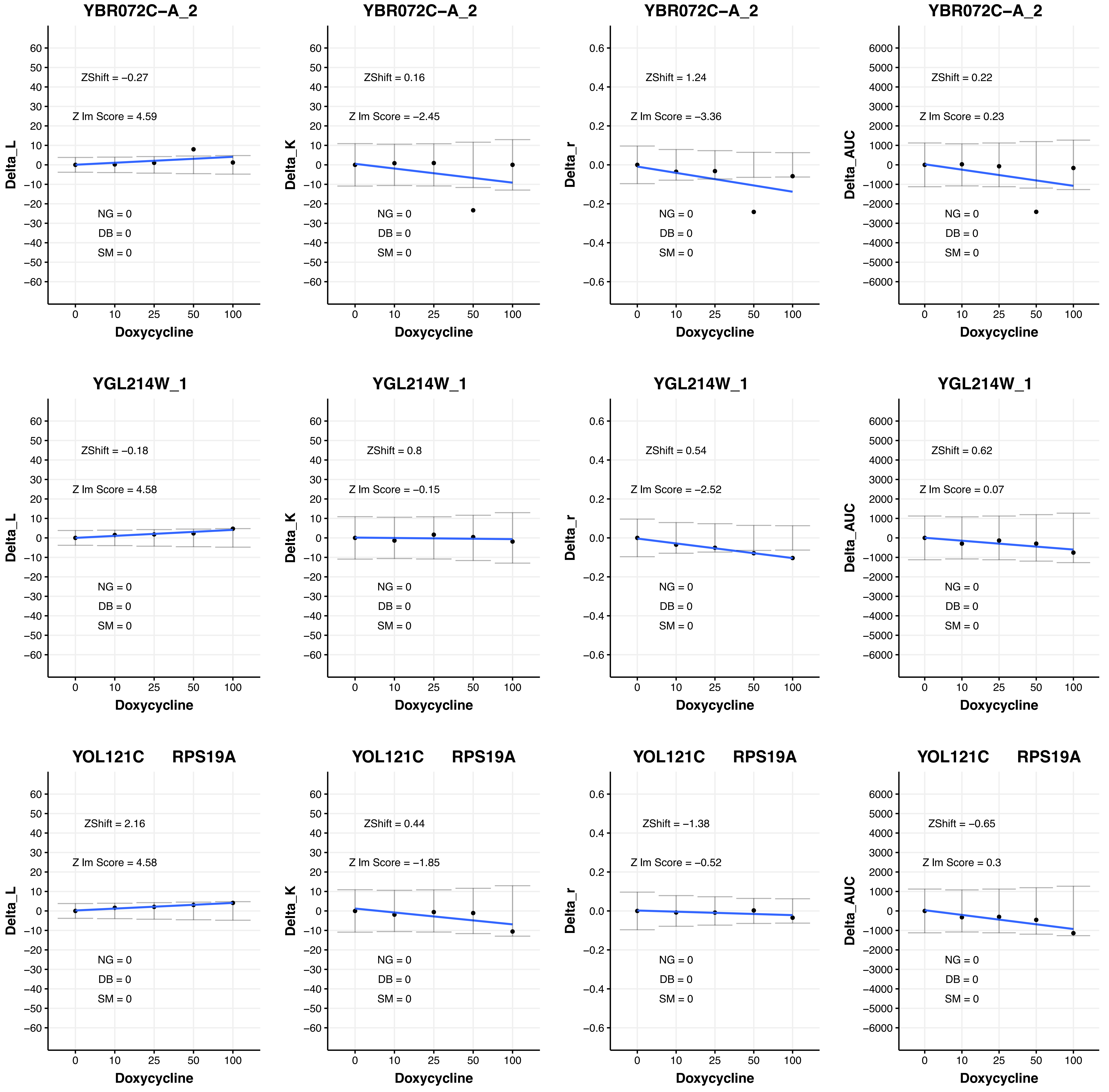

YBR072C−A_2
ZShift = −0.27
Z lm Score = 4.59
NG = 0
DB = 0
SM = 0
Delta_L
Doxycycline
YBR072C−A_2
ZShift = 0.16
Z lm Score = −2.45
NG = 0
DB = 0
SM = 0
Delta_K
Doxycycline
YBR072C−A_2
ZShift = 1.24
Z lm Score = −3.36
NG = 0
DB = 0
SM = 0
Delta_r
Doxycycline
YBR072C−A_2
ZShift = 0.22
Z lm Score = 0.23
NG = 0
DB = 0
SM = 0
Delta_AUC
Doxycycline
YGL214W_1
ZShift = −0.18
Z lm Score = 4.58
NG = 0
DB = 0
SM = 0
Delta_L
Doxycycline
YGL214W_1
ZShift = 0.8
Z lm Score = −0.15
NG = 0
DB = 0
SM = 0
Delta_K
Doxycycline
YGL214W_1
ZShift = 0.54
Z lm Score = −2.52
NG = 0
DB = 0
SM = 0
Delta_r
Doxycycline
YGL214W_1
ZShift = 0.62
Z lm Score = 0.07
NG = 0
DB = 0
SM = 0
Delta_AUC
Doxycycline
YOL121C RPS19A
ZShift = 2.16
Z lm Score = 4.58
NG = 0
DB = 0
SM = 0
Delta_L
Doxycycline
YOL121C RPS19A
ZShift = 0.44
Z lm Score = −1.85
NG = 0
DB = 0
SM = 0
Delta_K
Doxycycline
YOL121C RPS19A
ZShift = −1.38
Z lm Score = −0.52
NG = 0
DB = 0
SM = 0
Delta_r
Doxycycline
YOL121C RPS19A
ZShift = −0.65
Z lm Score = 0.3
NG = 0
DB = 0
SM = 0
Delta_AUC
Doxycycline

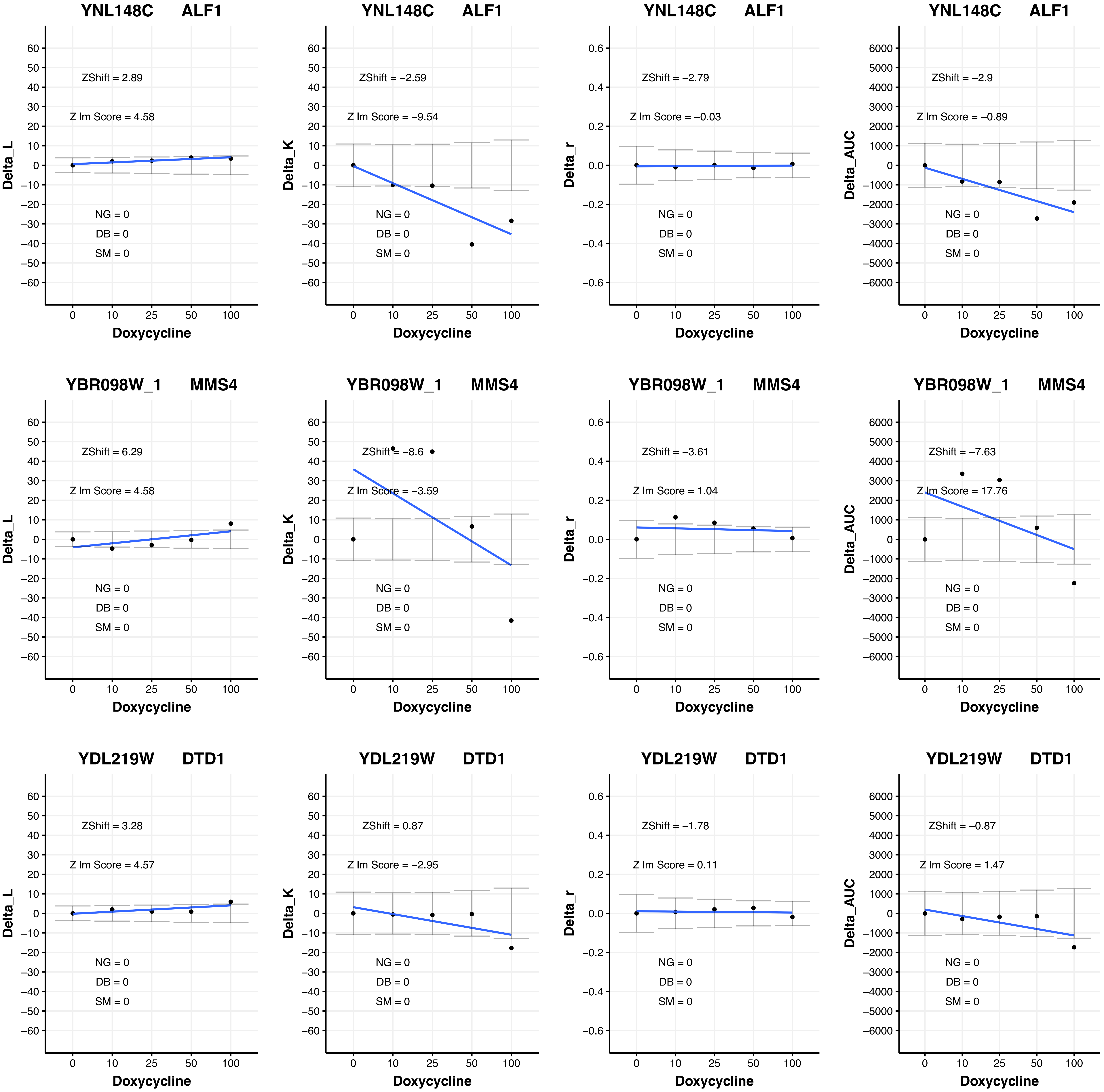
YNL148C ALF1
ZShift = 2.89
Z lm Score = 4.58
NG = 0
DB = 0
SM = 0
Delta_L
Doxycycline
YNL148C ALF1
ZShift = −2.59
Z lm Score = −9.54
NG = 0
DB = 0
SM = 0
Delta_K
Doxycycline
YNL148C ALF1
ZShift = −2.79
Z lm Score = −0.03
NG = 0
DB = 0
SM = 0
Delta_r
Doxycycline
YNL148C ALF1
ZShift = −2.9
Z lm Score = −0.89
NG = 0
DB = 0
SM = 0
Delta_AUC
Doxycycline
YBR098W_1 MMS4
ZShift = 6.29
Z lm Score = 4.58
NG = 0
DB = 0
SM = 0
Delta_L
Doxycycline
YBR098W_1 MMS4
ZShift = −8.6
Z lm Score = −3.59
NG = 0
DB = 0
SM = 0
Delta_K
Doxycycline
YBR098W_1 MMS4
ZShift = −3.61
Z lm Score = 1.04
NG = 0
DB = 0
SM = 0
Delta_r
Doxycycline
YBR098W_1 MMS4
ZShift = −7.63
Z lm Score = 17.76
NG = 0
DB = 0
SM = 0
Delta_AUC
Doxycycline
YDL219W DTD1
ZShift = 3.28
Z lm Score = 4.57
NG = 0
DB = 0
SM = 0
Delta_L
Doxycycline
YDL219W DTD1
ZShift = 0.87
Z lm Score = −2.95
NG = 0
DB = 0
SM = 0
Delta_K
Doxycycline
YDL219W DTD1
ZShift = −1.78
Z lm Score = 0.11
NG = 0
DB = 0
SM = 0
Delta_r
Doxycycline
YDL219W DTD1
ZShift = −0.87
Z lm Score = 1.47
NG = 0
DB = 0
SM = 0
Delta_AUC
Doxycycline

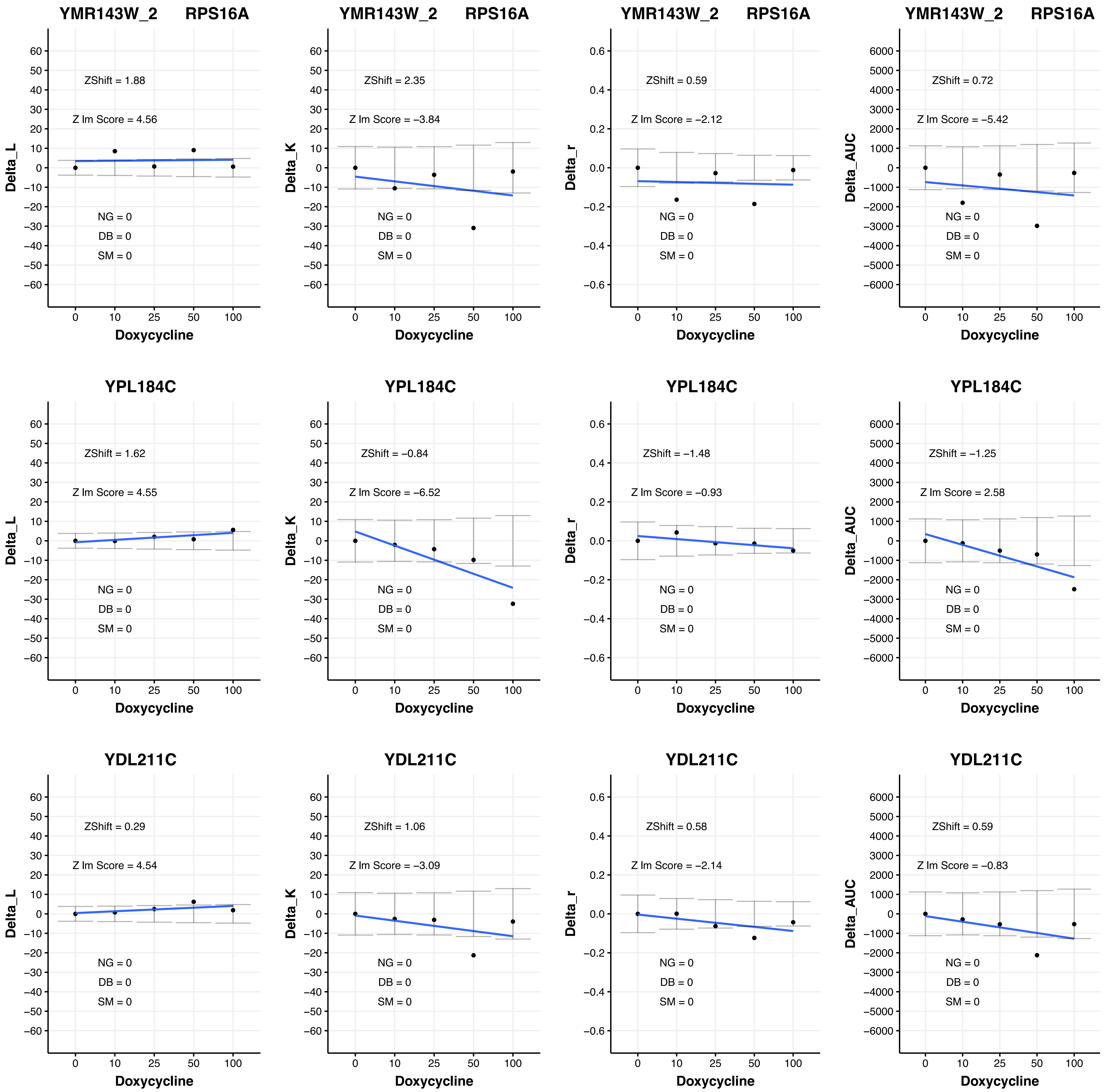

YMR143W_2 RPS16A
ZShift = 1.88
Z lm Score = 4.56
NG = 0
DB = 0
SM = 0
Delta_L
Doxycycline
YMR143W_2 RPS16A
ZShift = 2.35
Z lm Score = −3.84
NG = 0
DB = 0
SM = 0
Delta_K
Doxycycline
YMR143W_2 RPS16A
ZShift = 0.59
Z lm Score = −2.12
NG = 0
DB = 0
SM = 0
Delta_r
Doxycycline
YMR143W_2 RPS16A
ZShift = 0.72
Z lm Score = −5.42
NG = 0
DB = 0
SM = 0
Delta_AUC
Doxycycline
YPL184C
ZShift = 1.62
Z lm Score = 4.55
NG = 0
DB = 0
SM = 0
Delta_L
Doxycycline
YPL184C
ZShift = −0.84
Z lm Score = −6.52
NG = 0
DB = 0
SM = 0
Delta_K
Doxycycline
YPL184C
ZShift = −1.48
Z lm Score = −0.93
NG = 0
DB = 0
SM = 0
Delta_r
Doxycycline
YPL184C
ZShift = −1.25
Z lm Score = 2.58
NG = 0
DB = 0
SM = 0
Delta_AUC
Doxycycline
YDL211C
ZShift = 0.29
Z lm Score = 4.54
NG = 0
DB = 0
SM = 0
Delta_L
Doxycycline
YDL211C
ZShift = 1.06
Z lm Score = −3.09
NG = 0
DB = 0
SM = 0
Delta_K
Doxycycline
YDL211C
ZShift = 0.58
Z lm Score = −2.14
NG = 0
DB = 0
SM = 0
Delta_r
Doxycycline
YDL211C
ZShift = 0.59
Z lm Score = −0.83
NG = 0
DB = 0
SM = 0
Delta_AUC
Doxycycline

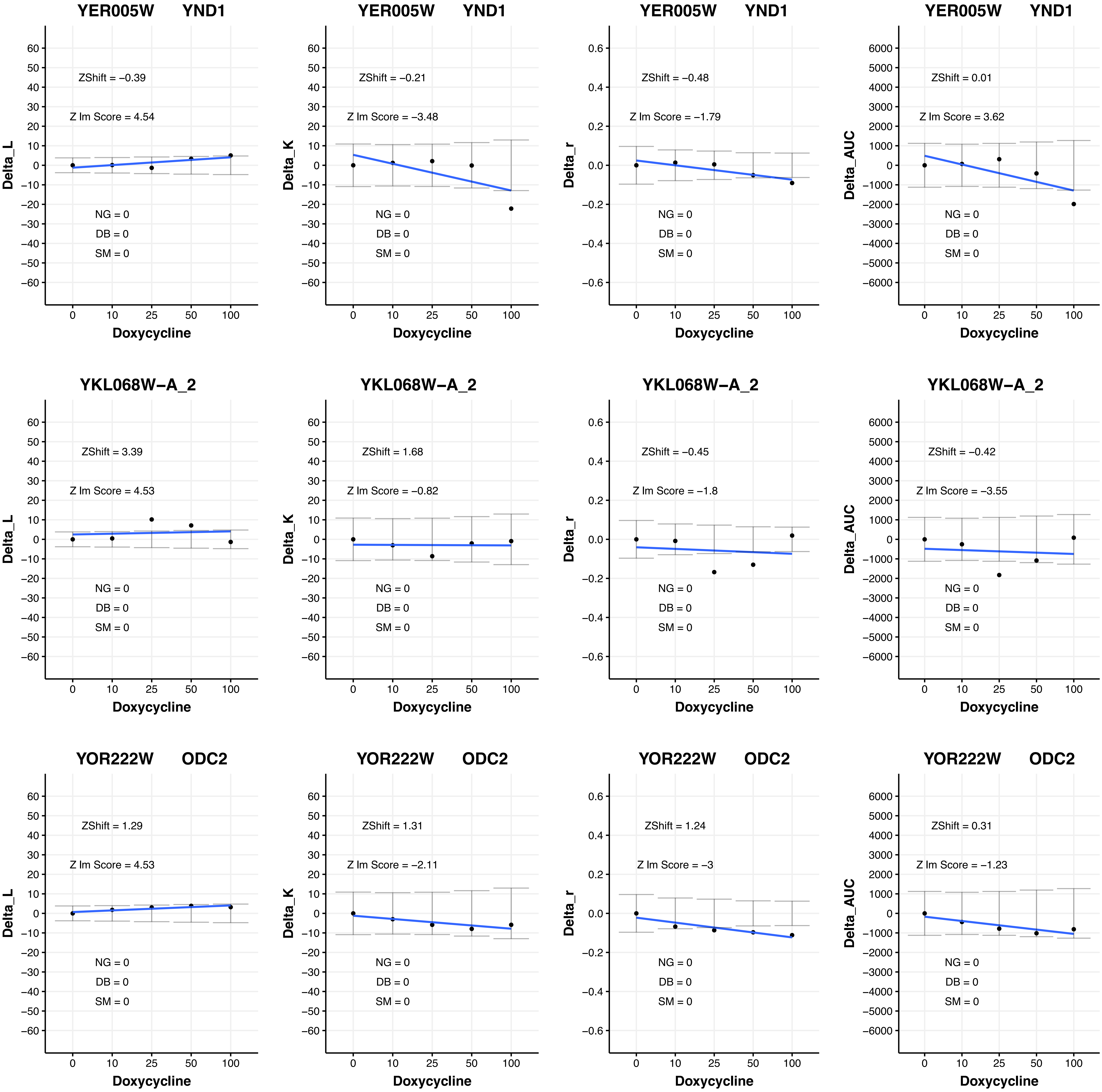

YER005W YND1
ZShift = −0.39
Z lm Score = 4.54
NG = 0
DB = 0
SM = 0
Delta_L
Doxycycline
YER005W YND1
ZShift = −0.21
Z lm Score = −3.48
NG = 0
DB = 0
SM = 0
Delta_K
Doxycycline
YER005W YND1
ZShift = −0.48
Z lm Score = −1.79
NG = 0
DB = 0
SM = 0
Delta_r
Doxycycline
YER005W YND1
ZShift = 0.01
Z lm Score = 3.62
NG = 0
DB = 0
SM = 0
Delta_AUC
Doxycycline
YKL068W−A_2
ZShift = 3.39
Z lm Score = 4.53
NG = 0
DB = 0
SM = 0
Delta_L
Doxycycline
YKL068W−A_2
ZShift = 1.68
Z lm Score = −0.82
NG = 0
DB = 0
SM = 0
Delta_K
Doxycycline
YKL068W−A_2
ZShift = −0.45
Z lm Score = −1.8
NG = 0
DB = 0
SM = 0
Delta_r
Doxycycline
YKL068W−A_2
ZShift = −0.42
Z lm Score = −3.55
NG = 0
DB = 0
SM = 0
Delta_AUC
Doxycycline
YOR222W ODC2
ZShift = 1.29
Z lm Score = 4.53
NG = 0
DB = 0
SM = 0
Delta_L
Doxycycline
YOR222W ODC2
ZShift = 1.31
Z lm Score = −2.11
NG = 0
DB = 0
SM = 0
Delta_K
Doxycycline
YOR222W ODC2
ZShift = 1.24
Z lm Score = −3
NG = 0
DB = 0
SM = 0
Delta_r
Doxycycline
YOR222W ODC2
ZShift = 0.31
Z lm Score = −1.23
NG = 0
DB = 0
SM = 0
Delta_AUC
Doxycycline

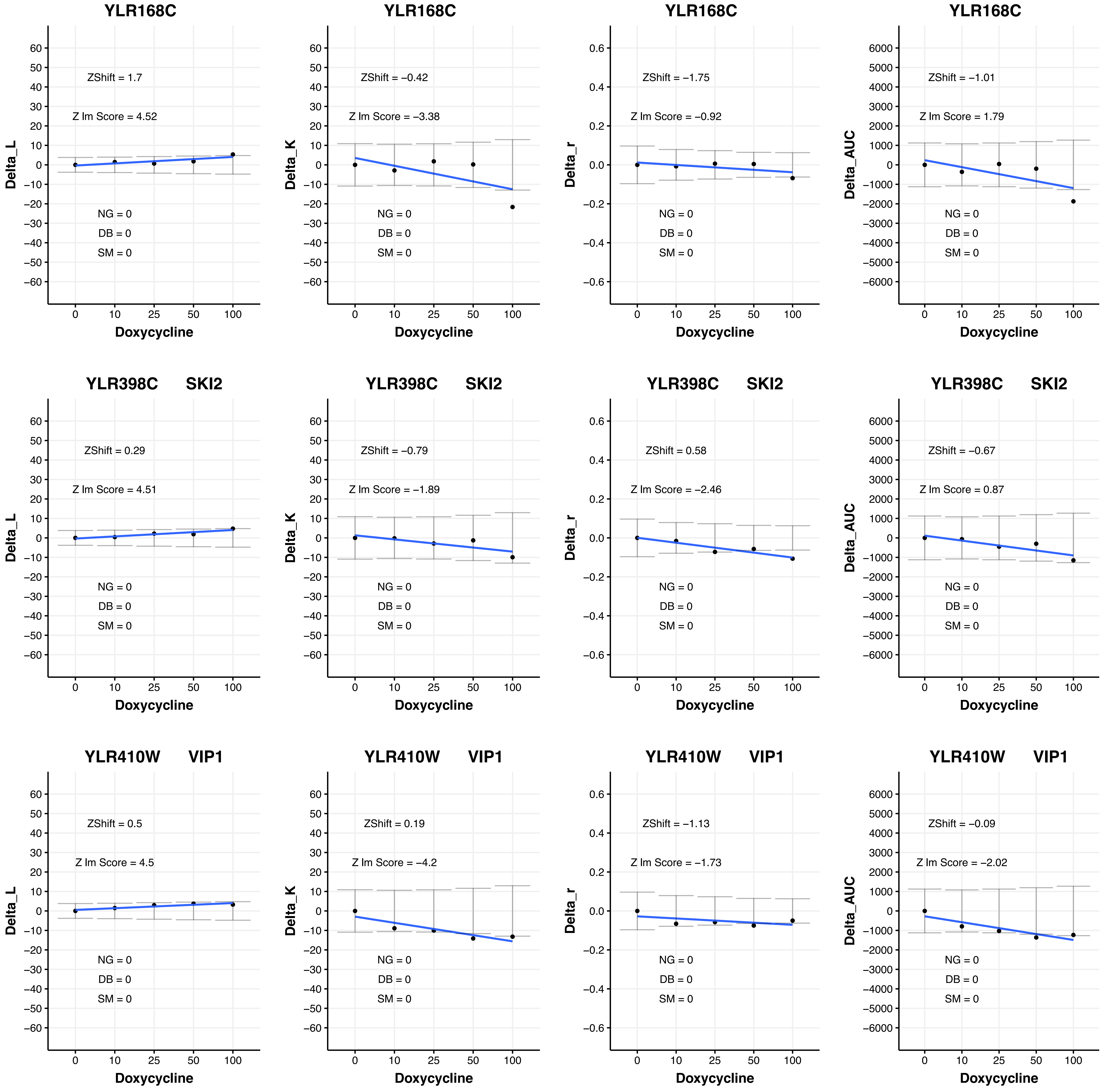

YLR168C
ZShift = 1.7
Z lm Score = 4.52
NG = 0
DB = 0
SM = 0
Delta_L
Doxycycline
YLR168C
ZShift = −0.42
Z lm Score = −3.38
NG = 0
DB = 0
SM = 0
Delta_K
Doxycycline
YLR168C
ZShift = −1.75
Z lm Score = −0.92
NG = 0
DB = 0
SM = 0
Delta_r
Doxycycline
YLR168C
ZShift = −1.01
Z lm Score = 1.79
NG = 0
DB = 0
SM = 0
Delta_AUC
Doxycycline
YLR398C SKI2
ZShift = 0.29
Z lm Score = 4.51
NG = 0
DB = 0
SM = 0
Delta_L
Doxycycline
YLR398C SKI2
ZShift = −0.79
Z lm Score = −1.89
NG = 0
DB = 0
SM = 0
Delta_K
Doxycycline
YLR398C SKI2
ZShift = 0.58
Z lm Score = −2.46
NG = 0
DB = 0
SM = 0
Delta_r
Doxycycline
YLR398C SKI2
ZShift = −0.67
Z lm Score = 0.87
NG = 0
DB = 0
SM = 0
Delta_AUC
Doxycycline
YLR410W VIP1
ZShift = 0.5
Z lm Score = 4.5
NG = 0
DB = 0
SM = 0
Delta_L
Doxycycline
YLR410W VIP1
ZShift = 0.19
Z lm Score = −4.2
NG = 0
DB = 0
SM = 0
Delta_K
Doxycycline
YLR410W VIP1
ZShift = −1.13
Z lm Score = −1.73
NG = 0
DB = 0
SM = 0
Delta_r
Doxycycline
YLR410W VIP1
ZShift = −0.09
Z lm Score = −2.02
NG = 0
DB = 0
SM = 0
Delta_AUC
Doxycycline

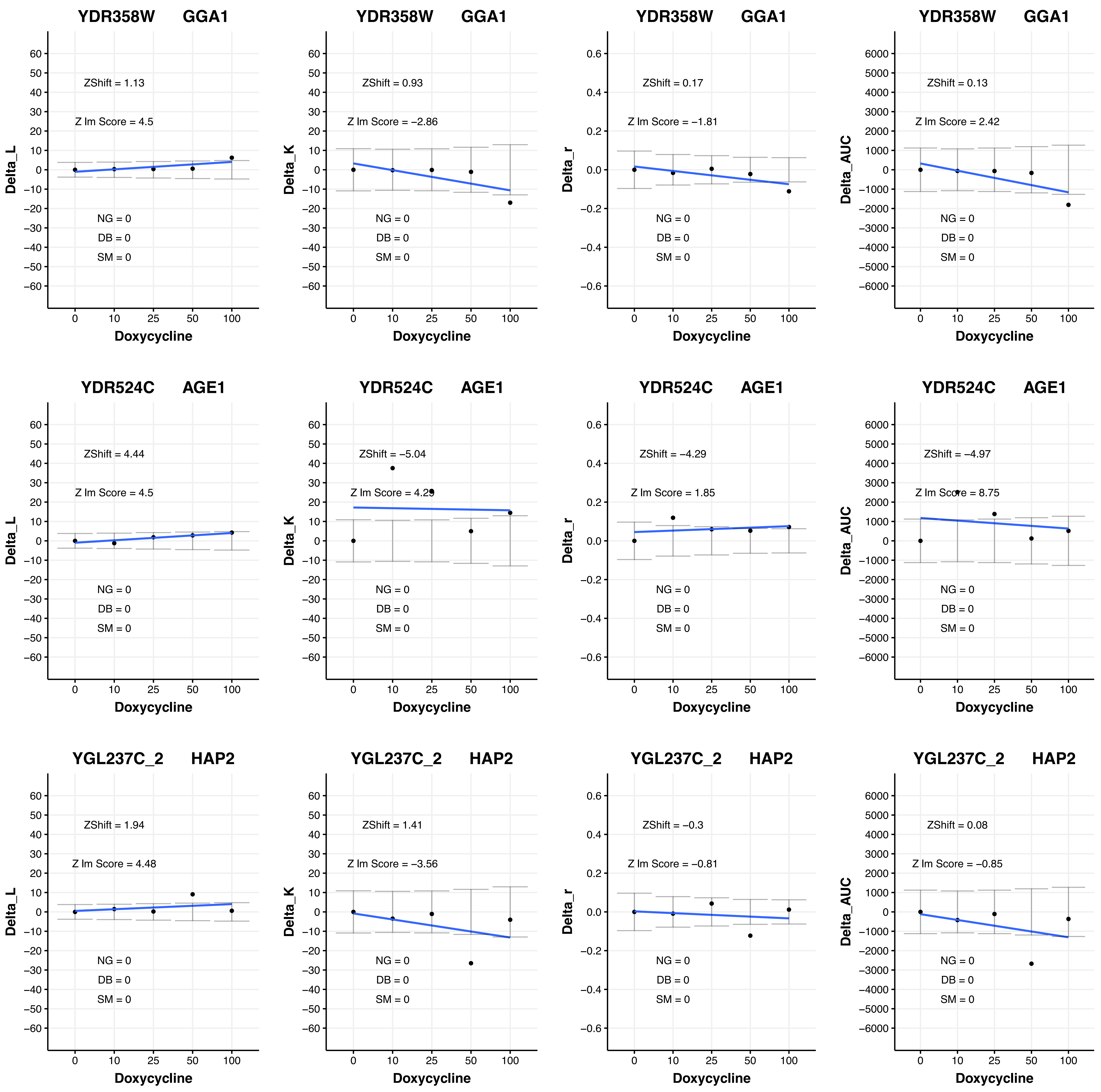

YDR358W GGA1
ZShift = 1.13
Z lm Score = 4.5
NG = 0
DB = 0
SM = 0
Delta_L
Doxycycline
YDR358W GGA1
ZShift = 0.93
Z lm Score = −2.86
NG = 0
DB = 0
SM = 0
Delta_K
Doxycycline
YDR358W GGA1
ZShift = 0.17
Z lm Score = −1.81
NG = 0
DB = 0
SM = 0
Delta_r
Doxycycline
YDR358W GGA1
ZShift = 0.13
Z lm Score = 2.42
NG = 0
DB = 0
SM = 0
Delta_AUC
Doxycycline
YDR524C AGE1
ZShift = 4.44
Z lm Score = 4.5
NG = 0
DB = 0
SM = 0
Delta_L
Doxycycline
YDR524C AGE1
ZShift = −5.04
Z lm Score = 4.29
NG = 0
DB = 0
SM = 0
Delta_K
Doxycycline
YDR524C AGE1
ZShift = −4.29
Z lm Score = 1.85
NG = 0
DB = 0
SM = 0
Delta_r
Doxycycline
YDR524C AGE1
ZShift = −4.97
Z lm Score = 8.75
NG = 0
DB = 0
SM = 0
Delta_AUC
Doxycycline
YGL237C_2 HAP2
ZShift = 1.94
Z lm Score = 4.48
NG = 0
DB = 0
SM = 0
Delta_L
Doxycycline
YGL237C_2 HAP2
ZShift = 1.41
Z lm Score = −3.56
NG = 0
DB = 0
SM = 0
Delta_K
Doxycycline
YGL237C_2 HAP2
ZShift = −0.3
Z lm Score = −0.81
NG = 0
DB = 0
SM = 0
Delta_r
Doxycycline
YGL237C_2 HAP2
ZShift = 0.08
Z lm Score = −0.85
NG = 0
DB = 0
SM = 0
Delta_AUC
Doxycycline

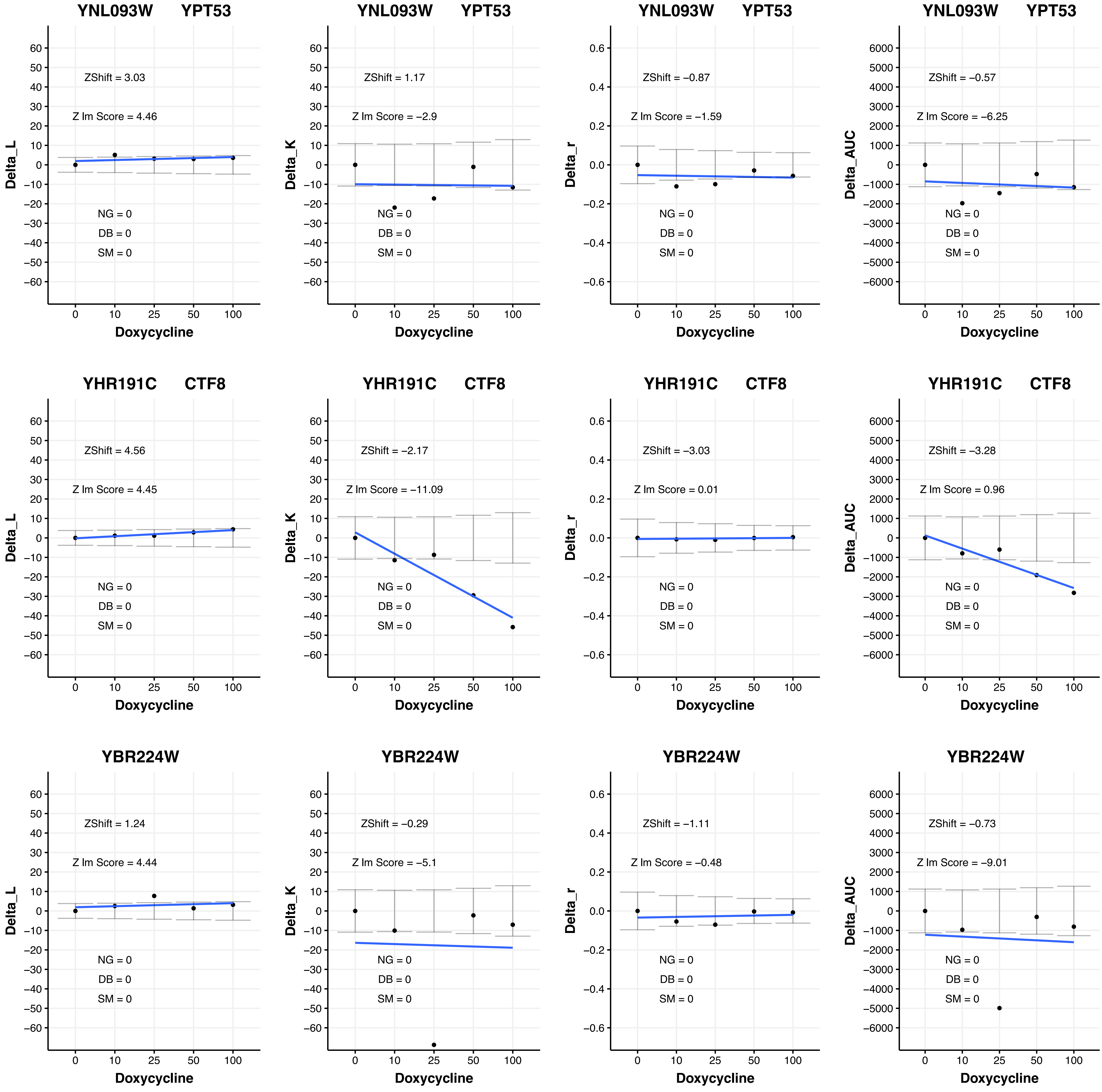

YNL093W YPT53
ZShift = 3.03
Z lm Score = 4.46
NG = 0
DB = 0
SM = 0
Delta_L
Doxycycline
0 10 25 50 100
YNL093W YPT53
ZShift = 1.17
Z lm Score = −2.9
NG = 0
DB = 0
SM = 0
Delta_K
Doxycycline
YNL093W YPT53
ZShift = −0.87
Z lm Score = −1.59
NG = 0
DB = 0
SM = 0
Delta_r
Doxycycline
YNL093W YPT53
ZShift = −0.57
Z lm Score = −6.25
NG = 0
DB = 0
SM = 0
Delta_AUC
Doxycycline
YHR191C CTF8
ZShift = 4.56
Z lm Score = 4.45
NG = 0
DB = 0
SM = 0
Delta_L
Doxycycline
YHR191C CTF8
ZShift = −2.17
Z lm Score = −11.09
NG = 0
DB = 0
SM = 0
Delta_K
Doxycycline
YHR191C CTF8
ZShift = −3.03
Z lm Score = 0.01
NG = 0
DB = 0
SM = 0
Delta_r
Doxycycline
YHR191C CTF8
ZShift = −3.28
Z lm Score = 0.96
NG = 0
DB = 0
SM = 0
Delta_AUC
Doxycycline
YBR224W
ZShift = 1.24
Z lm Score = 4.44
NG = 0
DB = 0
SM = 0
Delta_L
Doxycycline
YBR224W
ZShift = −0.29
Z lm Score = −5.1
NG = 0
DB = 0
SM = 0
Delta_K
Doxycycline
YBR224W
ZShift = −1.11
Z lm Score = −0.48
NG = 0
DB = 0
SM = 0
Delta_r
Doxycycline
YBR224W
ZShift = −0.73
Z lm Score = −9.01
NG = 0
DB = 0
SM = 0
Delta_AUC
Doxycycline

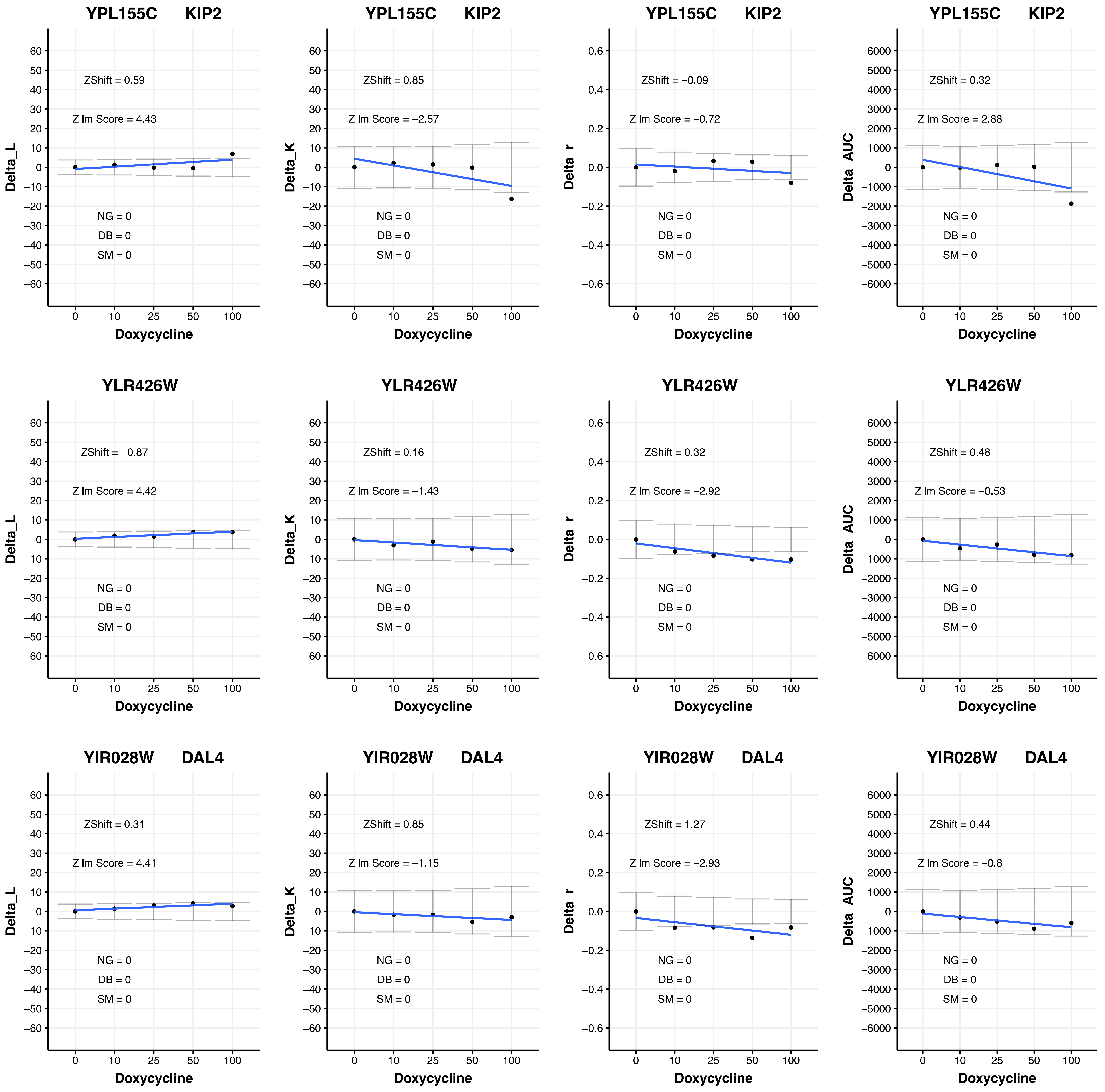

YPL155C KIP2
ZShift = 0.59
Z lm Score = 4.43
NG = 0
DB = 0
SM = 0
Delta_L
Doxycycline
YPL155C KIP2
ZShift = 0.85
Z lm Score = −2.57
NG = 0
DB = 0
SM = 0
Delta_K
Doxycycline
YPL155C KIP2
ZShift = −0.09
Z lm Score = −0.72
NG = 0
DB = 0
SM = 0
Delta_r
Doxycycline
YPL155C KIP2
ZShift = 0.32
Z lm Score = 2.88
NG = 0
DB = 0
SM = 0
Delta_AUC
Doxycycline
YLR426W
ZShift = −0.87
Z lm Score = 4.42
NG = 0
DB = 0
SM = 0
Delta_L
Doxycycline
YLR426W
ZShift = 0.16
Z lm Score = −1.43
NG = 0
DB = 0
SM = 0
Delta_K
Doxycycline
YLR426W
ZShift = 0.32
Z lm Score = −2.92
NG = 0
DB = 0
SM = 0
Delta_r
Doxycycline
YLR426W
ZShift = 0.48
Z lm Score = −0.53
NG = 0
DB = 0
SM = 0
Delta_AUC
Doxycycline
YIR028W DAL4
ZShift = 0.31
Z lm Score = 4.41
NG = 0
DB = 0
SM = 0
Delta_L
Doxycycline
YIR028W DAL4
ZShift = 0.85
Z lm Score = −1.15
NG = 0
DB = 0
SM = 0
Delta_K
Doxycycline
YIR028W DAL4
ZShift = 1.27
Z lm Score = −2.93
NG = 0
DB = 0
SM = 0
Delta_r
Doxycycline
YIR028W DAL4
ZShift = 0.44
Z lm Score = −0.8
NG = 0
DB = 0
SM = 0
Delta_AUC
Doxycycline

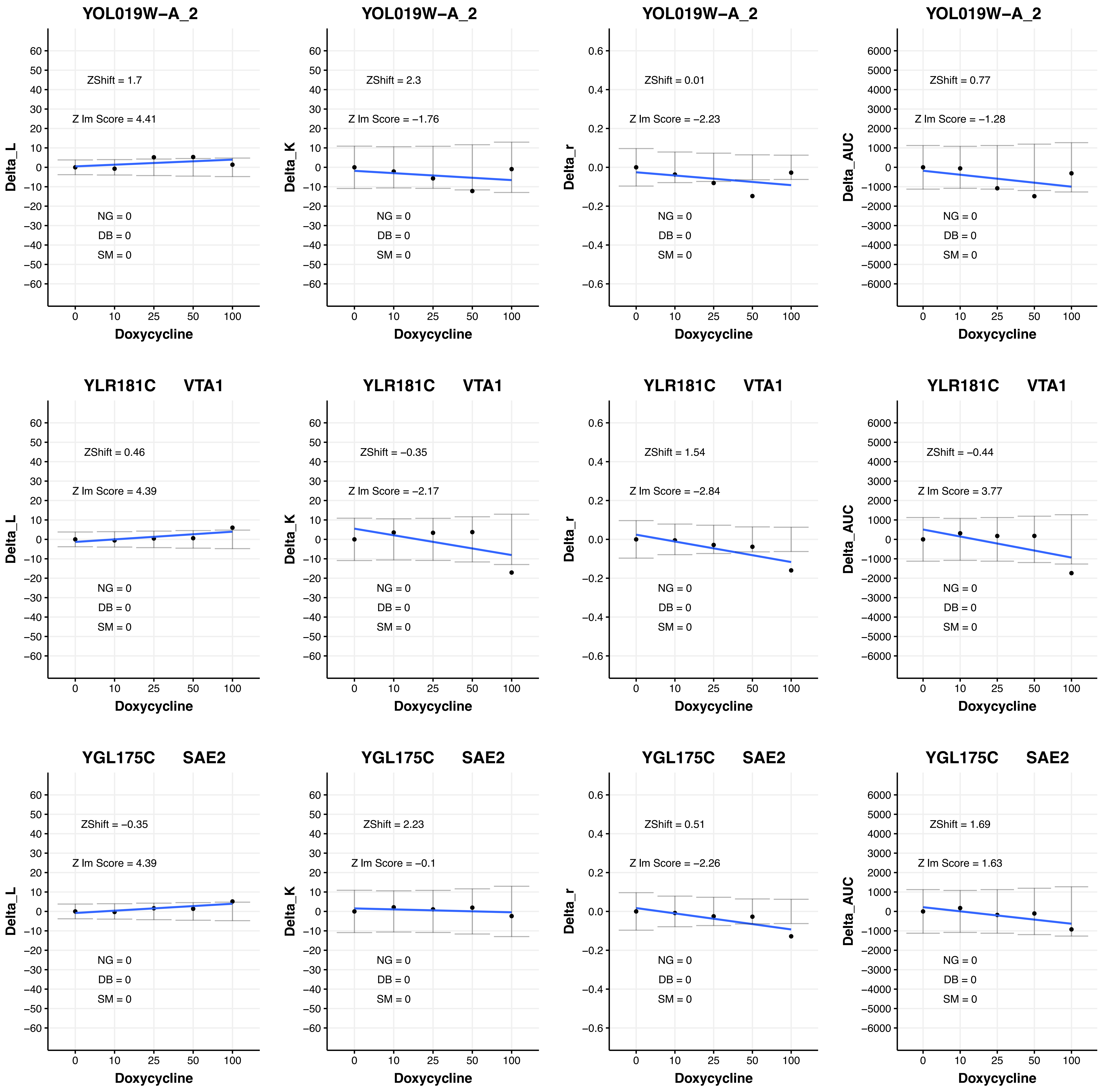

YOL019W−A_2
ZShift = 1.7
Z lm Score = 4.41
NG = 0
DB = 0
SM = 0
Delta_L
Doxycycline
YOL019W−A_2
ZShift = 2.3
Z lm Score = −1.76
NG = 0
DB = 0
SM = 0
Delta_K
Doxycycline
YOL019W−A_2
ZShift = 0.01
Z lm Score = −2.23
NG = 0
DB = 0
SM = 0
Delta_r
Doxycycline
YOL019W−A_2
ZShift = 0.77
Z lm Score = −1.28
NG = 0
DB = 0
SM = 0
Delta_AUC
Doxycycline
YLR181C VTA1
ZShift = 0.46
Z lm Score = 4.39
NG = 0
DB = 0
SM = 0
Delta_L
Doxycycline
YLR181C VTA1
ZShift = −0.35
Z lm Score = −2.17
NG = 0
DB = 0
SM = 0
Delta_K
Doxycycline
YLR181C VTA1
ZShift = 1.54
Z lm Score = −2.84
NG = 0
DB = 0
SM = 0
Delta_r
Doxycycline
YLR181C VTA1
ZShift = −0.44
Z lm Score = 3.77
NG = 0
DB = 0
SM = 0
Delta_AUC
Doxycycline
YGL175C SAE2
ZShift = −0.35
Z lm Score = 4.39
NG = 0
DB = 0
SM = 0
Delta_L
Doxycycline
YGL175C SAE2
ZShift = 2.23
Z lm Score = −0.1
NG = 0
DB = 0
SM = 0
Delta_K
Doxycycline
YGL175C SAE2
ZShift = 0.51
Z lm Score = −2.26
NG = 0
DB = 0
SM = 0
Delta_r
Doxycycline
YGL175C SAE2
ZShift = 1.69
Z lm Score = 1.63
NG = 0
DB = 0
SM = 0
Delta_AUC
Doxycycline

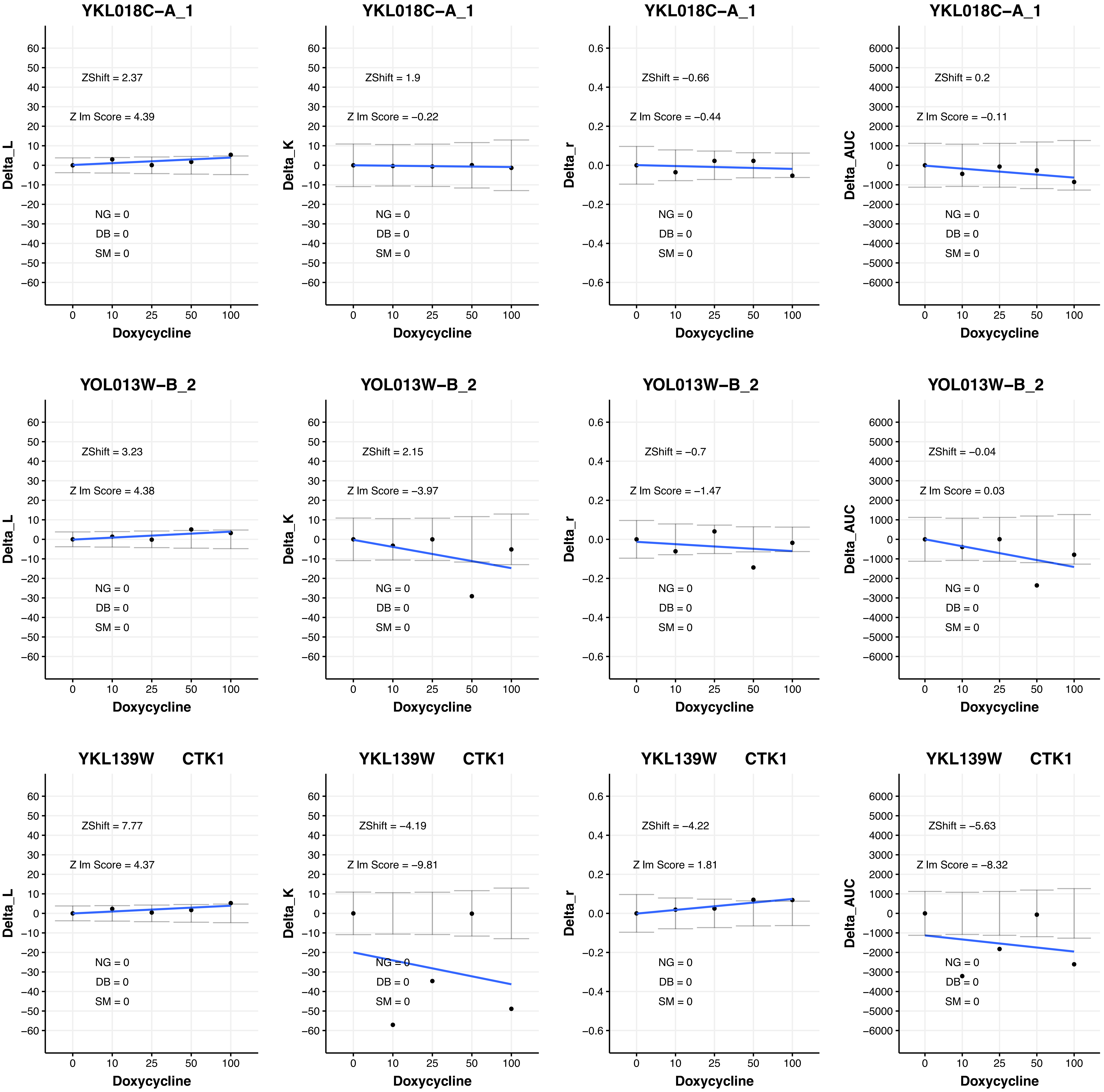
YKL018C−A_1
ZShift = 2.37
Z lm Score = 4.39
NG = 0
DB = 0
SM = 0
Delta_L
Doxycycline
YKL018C−A_1
ZShift = 1.9
Z lm Score = −0.22
NG = 0
DB = 0
SM = 0
Delta_K
Doxycycline
YKL018C−A_1
ZShift = −0.66
Z lm Score = −0.44
NG = 0
DB = 0
SM = 0
Delta_r
Doxycycline
YKL018C−A_1
ZShift = 0.2
Z lm Score = −0.11
NG = 0
DB = 0
SM = 0
Delta_AUC
Doxycycline
YOL013W−B_2
ZShift = 3.23
Z lm Score = 4.38
NG = 0
DB = 0
SM = 0
Delta_L
Doxycycline
YOL013W−B_2
ZShift = 2.15
Z lm Score = −3.97
NG = 0
DB = 0
SM = 0
Delta_K
Doxycycline
YOL013W−B_2
ZShift = −0.7
Z lm Score = −1.47
NG = 0
DB = 0
SM = 0
Delta_r
Doxycycline
YOL013W−B_2
ZShift = −0.04
Z lm Score = 0.03
NG = 0
DB = 0
SM = 0
Delta_AUC
Doxycycline
YKL139W CTK1
ZShift = 7.77
Z lm Score = 4.37
NG = 0
DB = 0
SM = 0
Delta_L
Doxycycline
YKL139W CTK1
ZShift = −4.19
Z lm Score = −9.81
NG = 0
DB = 0
SM = 0
Delta_K
Doxycycline
YKL139W CTK1
ZShift = −4.22
Z lm Score = 1.81
NG = 0
DB = 0
SM = 0
Delta_r
Doxycycline
YKL139W CTK1
ZShift = −5.63
Z lm Score = −8.32
NG = 0
DB = 0
SM = 0
Delta_AUC
Doxycycline

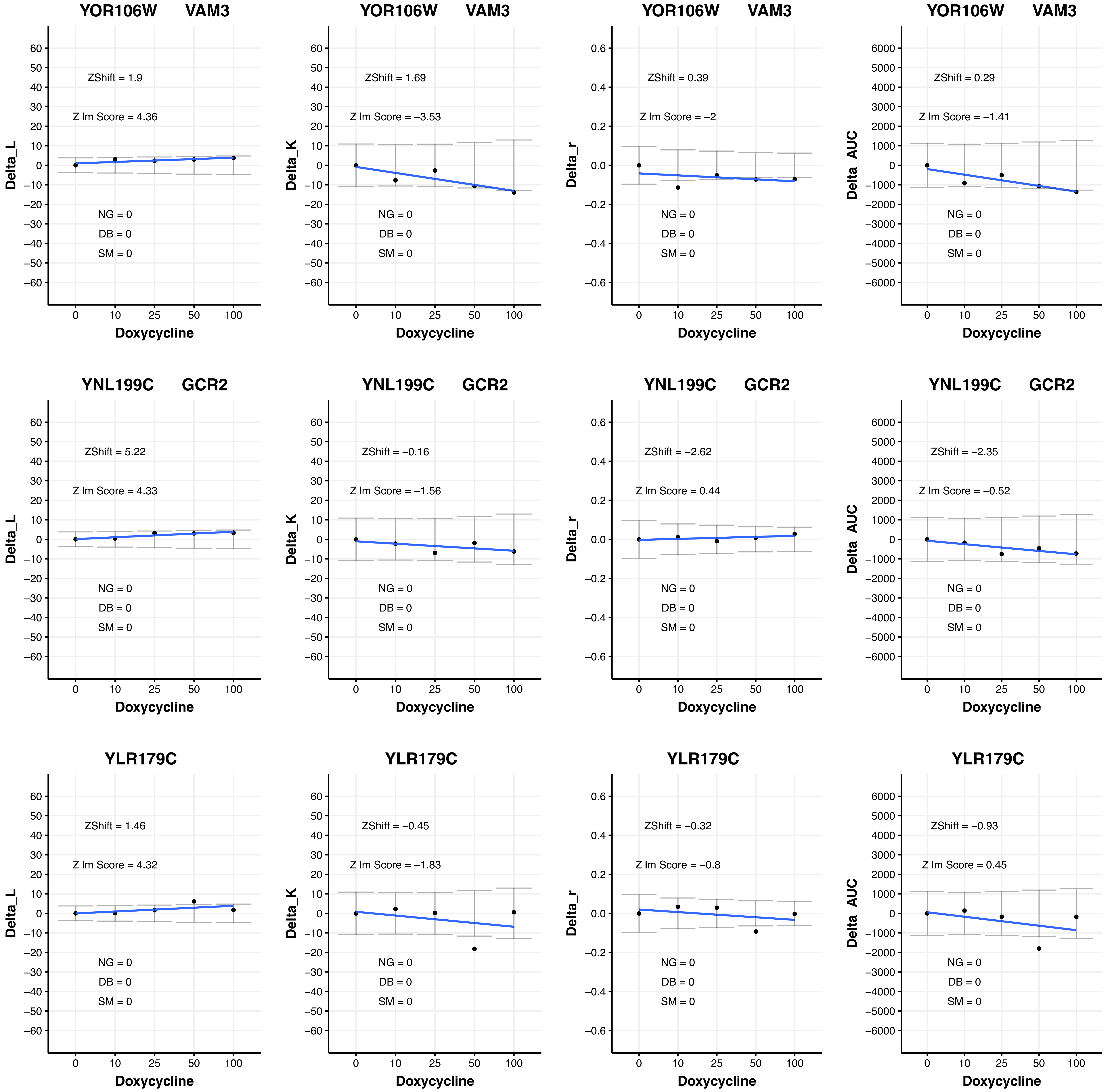
YOR106W VAM3
ZShift = 1.9
Z lm Score = 4.36
NG = 0
DB = 0
SM = 0
Delta_L
Doxycycline
YOR106W VAM3
ZShift = 1.69
Z lm Score = −3.53
NG = 0
DB = 0
SM = 0
Delta_K
Doxycycline
YOR106W VAM3
ZShift = 0.39
Z lm Score = −2
NG = 0
DB = 0
SM = 0
Delta_r
Doxycycline
YOR106W VAM3
ZShift = 0.29
Z lm Score = −1.41
NG = 0
DB = 0
SM = 0
Delta_AUC
Doxycycline
YNL199C GCR2
ZShift = 5.22
Z lm Score = 4.33
NG = 0
DB = 0
SM = 0
Delta_L
Doxycycline
YNL199C GCR2
ZShift = −0.16
Z lm Score = −1.56
NG = 0
DB = 0
SM = 0
Delta_K
Doxycycline
YNL199C GCR2
ZShift = −2.62
Z lm Score = 0.44
NG = 0
DB = 0
SM = 0
Delta_r
Doxycycline
YNL199C GCR2
ZShift = −2.35
Z lm Score = −0.52
NG = 0
DB = 0
SM = 0
Delta_AUC
Doxycycline
YLR179C
ZShift = 1.46
Z lm Score = 4.32
NG = 0
DB = 0
SM = 0
Delta_L
Doxycycline
YLR179C
ZShift = −0.45
Z lm Score = −1.83
NG = 0
DB = 0
SM = 0
Delta_K
Doxycycline
YLR179C
ZShift = −0.32
Z lm Score = −0.8
NG = 0
DB = 0
SM = 0
Delta_r
Doxycycline
YLR179C
ZShift = −0.93
Z lm Score = 0.45
NG = 0
DB = 0
SM = 0
Delta_AUC
Doxycycline

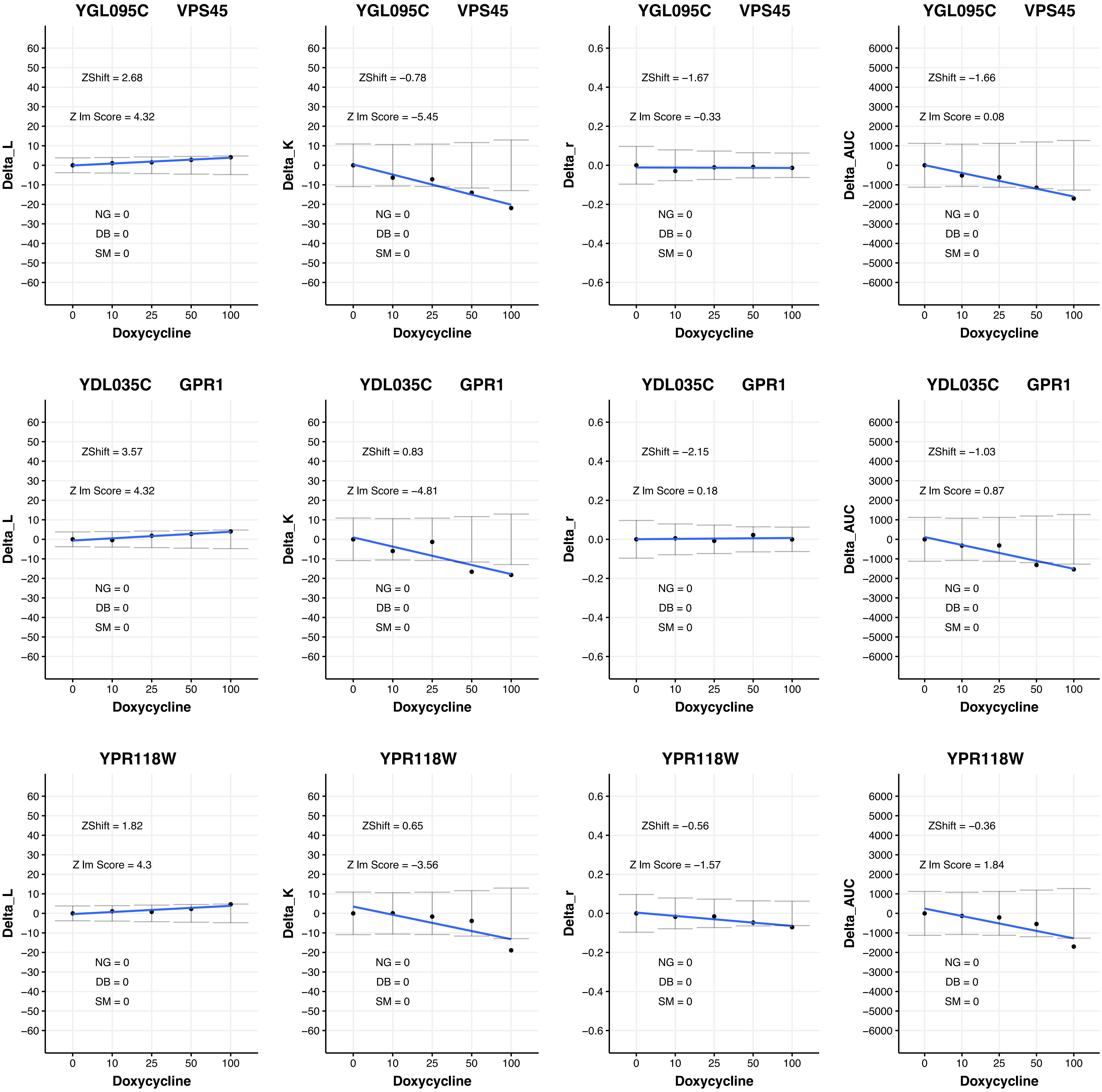
YGL095C VPS45
ZShift = 2.68
Z lm Score = 4.32
NG = 0
DB = 0
SM = 0
Delta_L
Doxycycline
YGL095C VPS45
ZShift = −0.78
Z lm Score = −5.45
NG = 0
DB = 0
SM = 0
Delta_K
Doxycycline
YGL095C VPS45
ZShift = −1.67
Z lm Score = −0.33
NG = 0
DB = 0
SM = 0
Delta_r
Doxycycline
YGL095C VPS45
ZShift = −1.66
Z lm Score = 0.08
NG = 0
DB = 0
SM = 0
Delta_AUC
Doxycycline
YDL035C GPR1
ZShift = 3.57
Z lm Score = 4.32
NG = 0
DB = 0
SM = 0
Delta_L
Doxycycline
YDL035C GPR1
ZShift = 0.83
Z lm Score = −4.81
NG = 0
DB = 0
SM = 0
Delta_K
Doxycycline
YDL035C GPR1
ZShift = −2.15
Z lm Score = 0.18
NG = 0
DB = 0
SM = 0
Delta_r
Doxycycline
YDL035C GPR1
ZShift = −1.03
Z lm Score = 0.87
NG = 0
DB = 0
SM = 0
Delta_AUC
Doxycycline
YPR118W
ZShift = 1.82
Z lm Score = 4.3
NG = 0
DB = 0
SM = 0
Delta_L
Doxycycline
YPR118W
ZShift = 0.65
Z lm Score = −3.56
NG = 0
DB = 0
SM = 0
Delta_K
Doxycycline
YPR118W
ZShift = −0.56
Z lm Score = −1.57
NG = 0
DB = 0
SM = 0
Delta_r
Doxycycline
YPR118W
ZShift = −0.36
Z lm Score = 1.84
NG = 0
DB = 0
SM = 0
Delta_AUC
Doxycycline
0
10
25
50
100

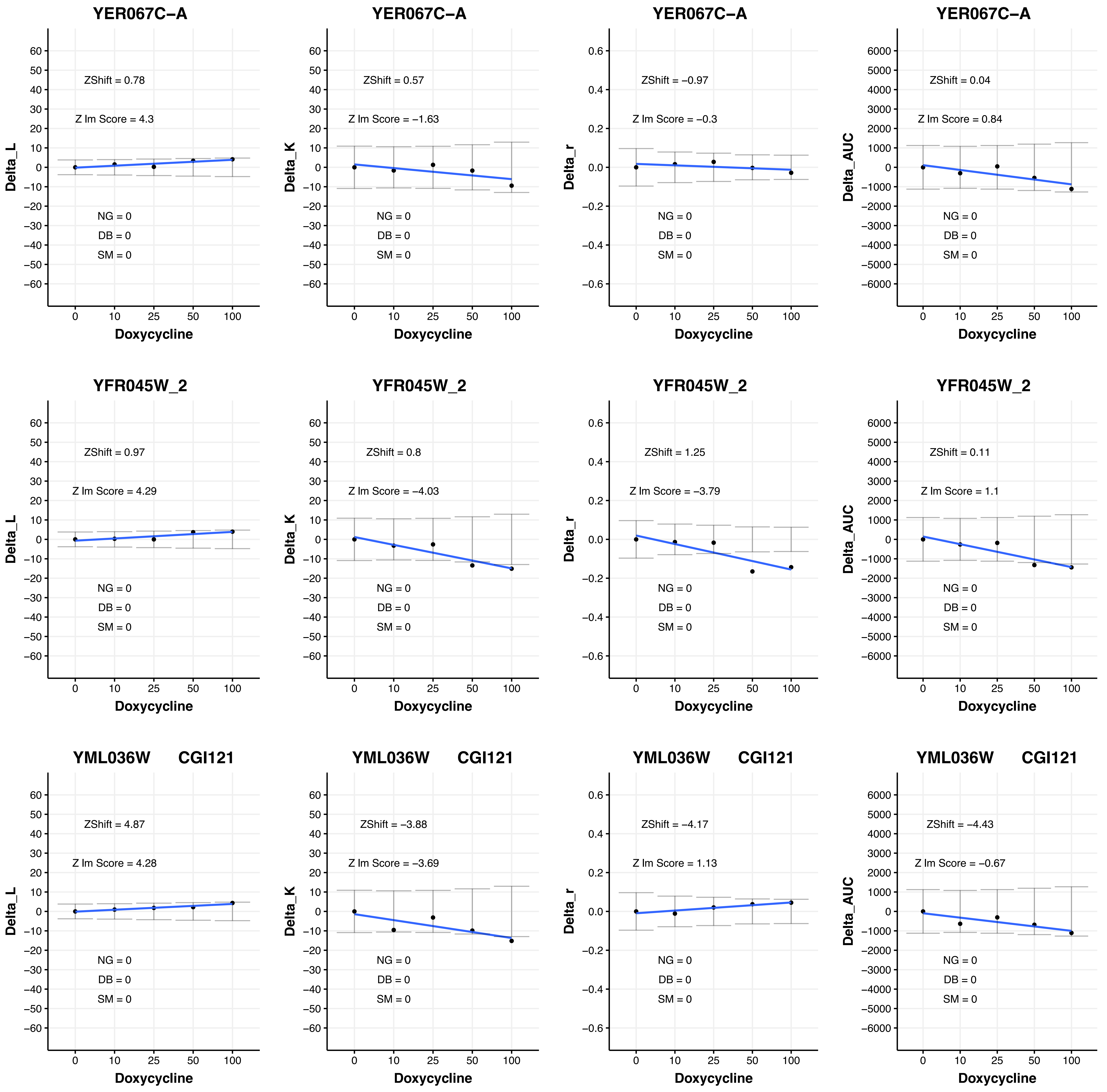

YER067C−A
ZShift = 0.78
Z lm Score = 4.3
NG = 0
DB = 0
SM = 0
Delta_L
Doxycycline
YER067C−A
ZShift = 0.57
Z lm Score = −1.63
NG = 0
DB = 0
SM = 0
Delta_K
Doxycycline
YER067C−A
ZShift = −0.97
Z lm Score = −0.3
NG = 0
DB = 0
SM = 0
Delta_r
Doxycycline
YER067C−A
ZShift = 0.04
Z lm Score = 0.84
NG = 0
DB = 0
SM = 0
Delta_AUC
Doxycycline
YFR045W_2
ZShift = 0.97
Z lm Score = 4.29
NG = 0
DB = 0
SM = 0
Delta_L
Doxycycline
YFR045W_2
ZShift = 0.8
Z lm Score = −4.03
NG = 0
DB = 0
SM = 0
Delta_K
Doxycycline
YFR045W_2
ZShift = 1.25
Z lm Score = −3.79
NG = 0
DB = 0
SM = 0
Delta_r
Doxycycline
YFR045W_2
ZShift = 0.11
Z lm Score = 1.1
NG = 0
DB = 0
SM = 0
Delta_AUC
Doxycycline
YML036W CGI121
ZShift = 4.87
Z lm Score = 4.28
NG = 0
DB = 0
SM = 0
Delta_L
Doxycycline
YML036W CGI121
ZShift = −3.88
Z lm Score = −3.69
NG = 0
DB = 0
SM = 0
Delta_K
Doxycycline
YML036W CGI121
ZShift = −4.17
Z lm Score = 1.13
NG = 0
DB = 0
SM = 0
Delta_r
Doxycycline
YML036W CGI121
ZShift = −4.43
Z lm Score = −0.67
NG = 0
DB = 0
SM = 0
Delta_AUC
Doxycycline

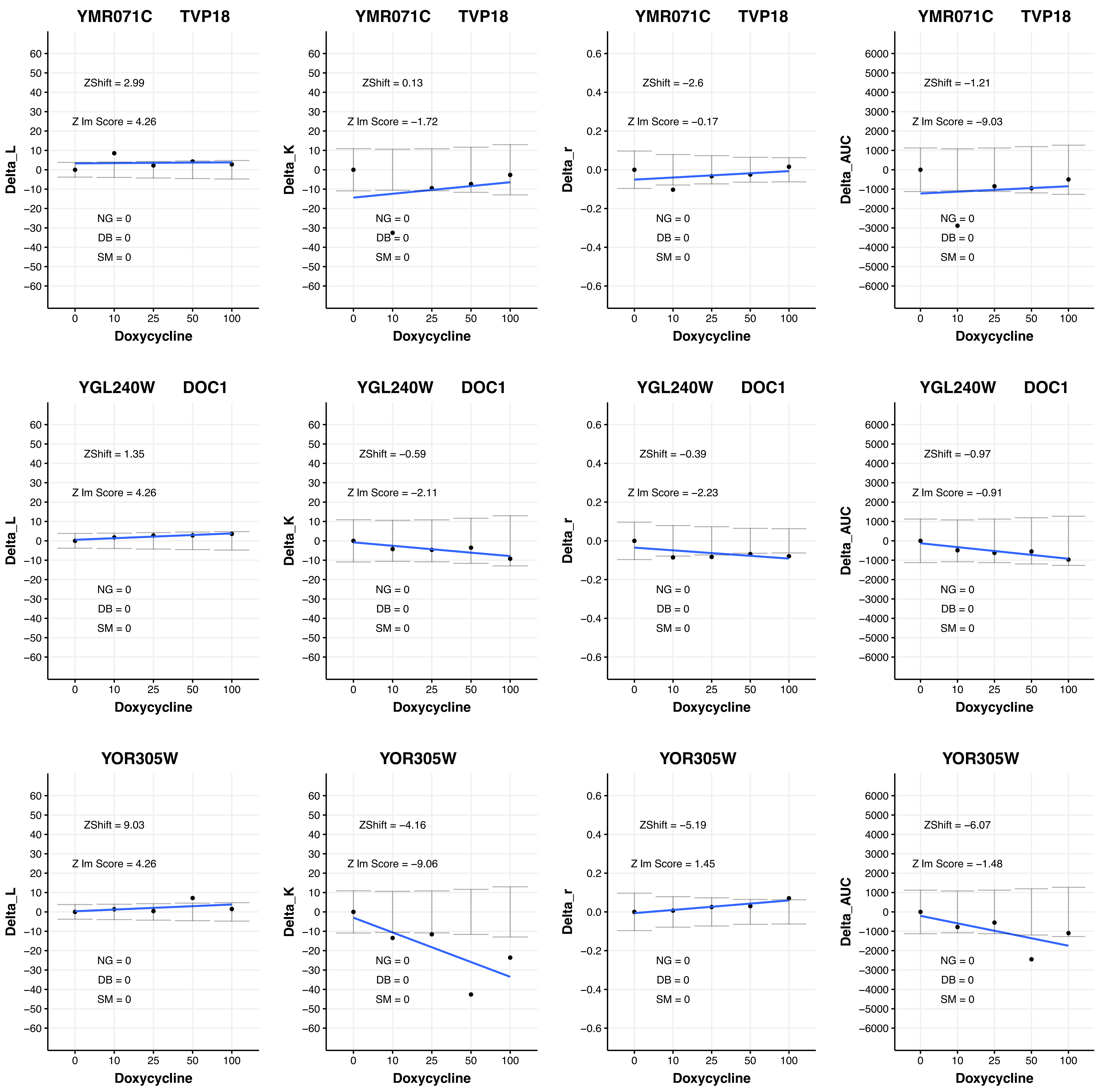
YMR071C TVP18
ZShift = 2.99
Z lm Score = 4.26
NG = 0
DB = 0
SM = 0
Delta_L
Doxycycline
YMR071C TVP18
ZShift = 0.13
Z lm Score = −1.72
NG = 0
DB = 0
SM = 0
Delta_K
Doxycycline
YMR071C TVP18
ZShift = −2.6
Z lm Score = −0.17
NG = 0
DB = 0
SM = 0
Delta_r
Doxycycline
YMR071C TVP18
ZShift = −1.21
Z lm Score = −9.03
NG = 0
DB = 0
SM = 0
Delta_AUC
Doxycycline
YGL240W DOC1
ZShift = 1.35
Z lm Score = 4.26
NG = 0
DB = 0
SM = 0
Delta_L
Doxycycline
YGL240W DOC1
ZShift = −0.59
Z lm Score = −2.11
NG = 0
DB = 0
SM = 0
Delta_K
Doxycycline
YGL240W DOC1
ZShift = −0.39
Z lm Score = −2.23
NG = 0
DB = 0
SM = 0
Delta_r
Doxycycline
YGL240W DOC1
ZShift = −0.97
Z lm Score = −0.91
NG = 0
DB = 0
SM = 0
Delta_AUC
Doxycycline
YOR305W
ZShift = 9.03
Z lm Score = 4.26
NG = 0
DB = 0
SM = 0
Delta_L
Doxycycline
YOR305W
ZShift = −4.16
Z lm Score = −9.06
NG = 0
DB = 0
SM = 0
Delta_K
Doxycycline
YOR305W
ZShift = −5.19
Z lm Score = 1.45
NG = 0
DB = 0
SM = 0
Delta_r
Doxycycline
YOR305W
ZShift = −6.07
Z lm Score = −1.48
NG = 0
DB = 0
SM = 0
Delta_AUC
Doxycycline

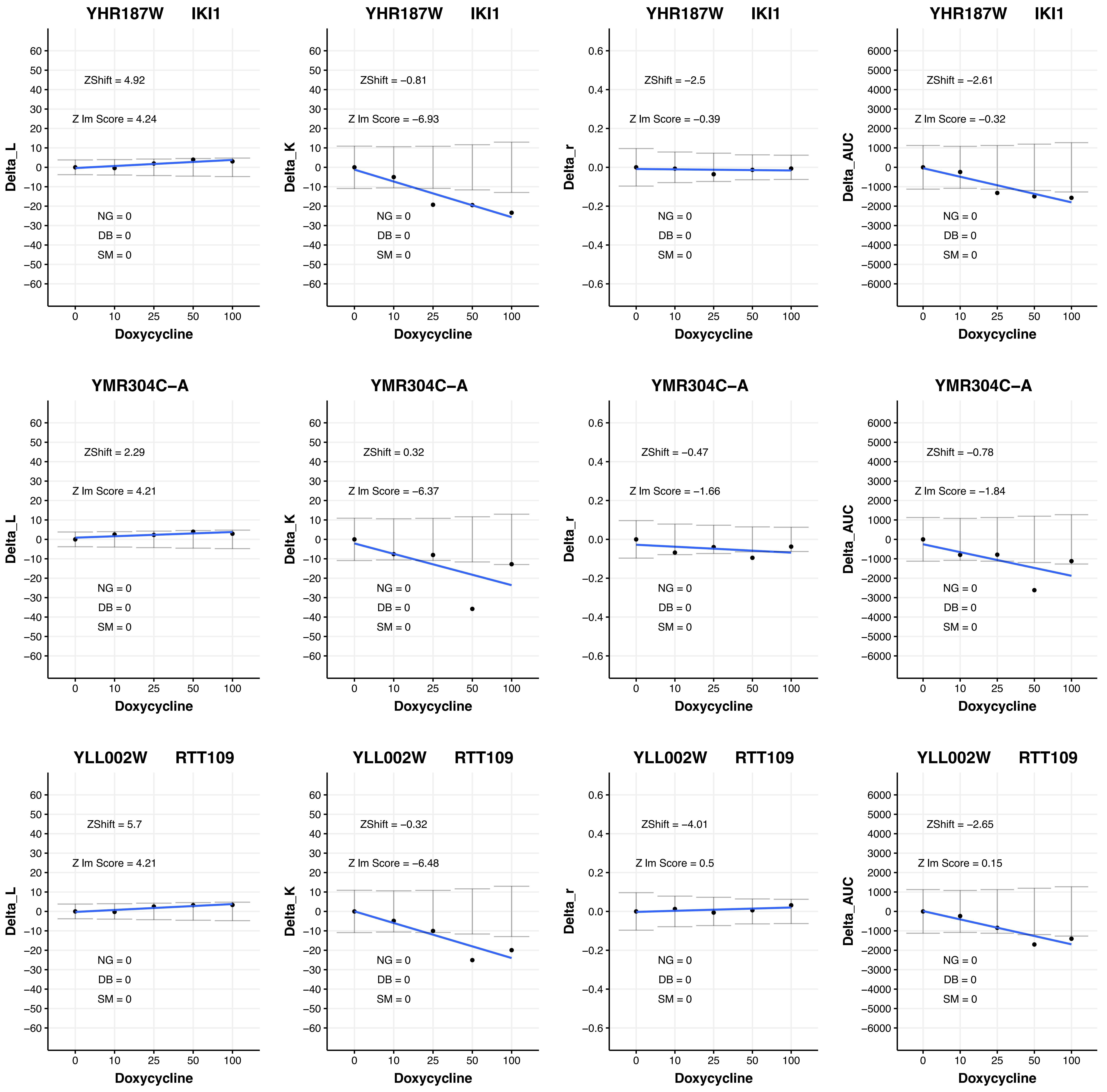
YHR187W IKI1
ZShift = 4.92
Z lm Score = 4.24
NG = 0
DB = 0
SM = 0
Delta_L
Doxycycline
YHR187W IKI1
ZShift = −0.81
Z lm Score = −6.93
NG = 0
DB = 0
SM = 0
Delta_K
Doxycycline
YHR187W IKI1
ZShift = −2.5
Z lm Score = −0.39
NG = 0
DB = 0
SM = 0
Delta_r
Doxycycline
YHR187W IKI1
ZShift = −2.61
Z lm Score = −0.32
NG = 0
DB = 0
SM = 0
Delta_AUC
Doxycycline
YMR304C−A
ZShift = 2.29
Z lm Score = 4.21
NG = 0
DB = 0
SM = 0
Delta_L
Doxycycline
YMR304C−A
ZShift = 0.32
Z lm Score = −6.37
NG = 0
DB = 0
SM = 0
Delta_K
Doxycycline
YMR304C−A
ZShift = −0.47
Z lm Score = −1.66
NG = 0
DB = 0
SM = 0
Delta_r
Doxycycline
YMR304C−A
ZShift = −0.78
Z lm Score = −1.84
NG = 0
DB = 0
SM = 0
Delta_AUC
Doxycycline
YLL002W RTT109
ZShift = 5.7
Z lm Score = 4.21
NG = 0
DB = 0
SM = 0
Delta_L
Doxycycline
YLL002W RTT109
ZShift = −0.32
Z lm Score = −6.48
NG = 0
DB = 0
SM = 0
Delta_K
Doxycycline
YLL002W RTT109
ZShift = −4.01
Z lm Score = 0.5
NG = 0
DB = 0
SM = 0
Delta_r
Doxycycline
YLL002W RTT109
ZShift = −2.65
Z lm Score = 0.15
NG = 0
DB = 0
SM = 0
Delta_AUC
Doxycycline

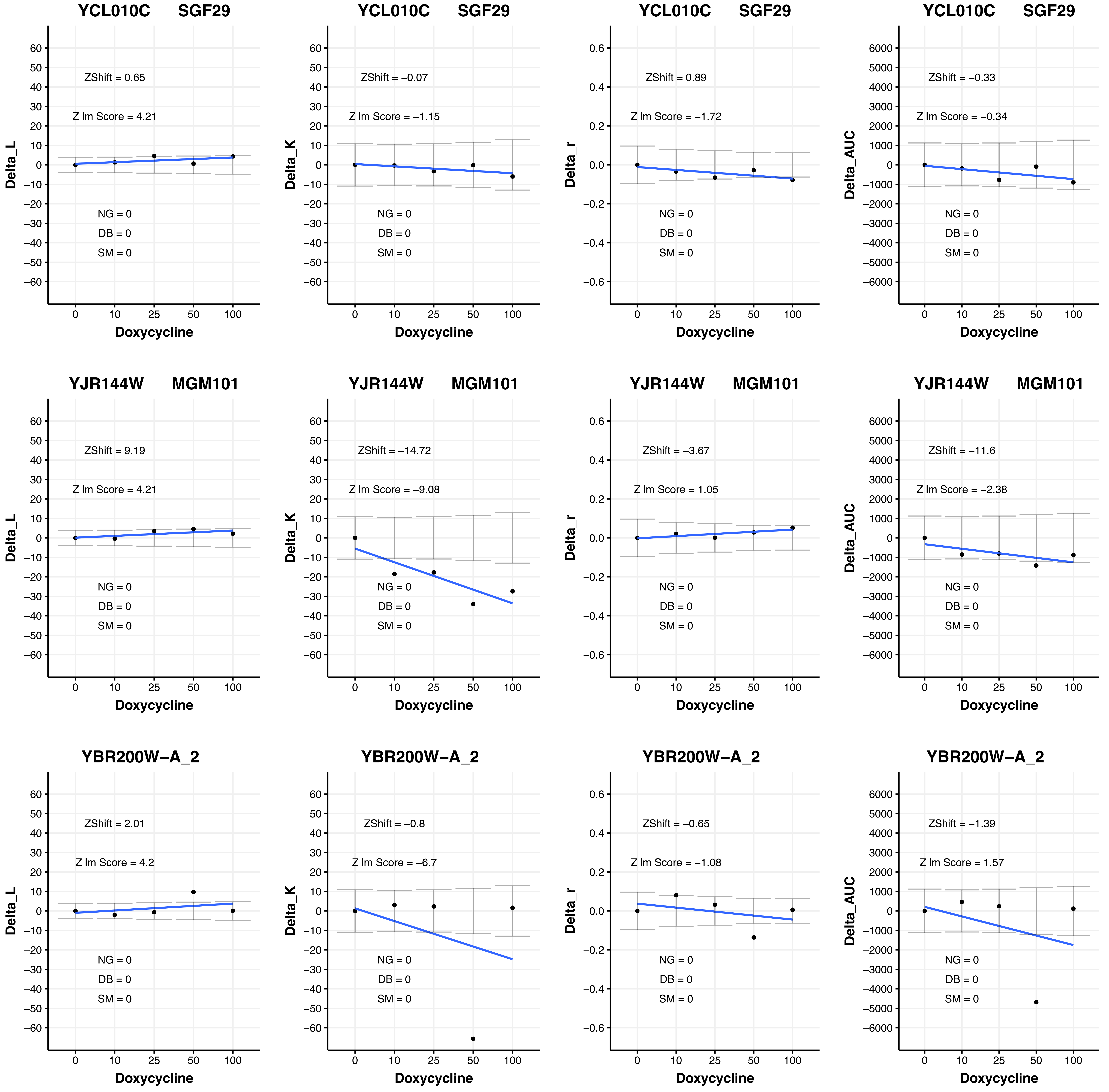
YCL010C SGF29
ZShift = 0.65
Z lm Score = 4.21
NG = 0
DB = 0
SM = 0
Delta_L
Doxycycline
YCL010C SGF29
ZShift = −0.07
Z lm Score = −1.15
NG = 0
DB = 0
SM = 0
Delta_K
Doxycycline
YCL010C SGF29
ZShift = 0.89
Z lm Score = −1.72
NG = 0
DB = 0
SM = 0
Delta_r
Doxycycline
YCL010C SGF29
ZShift = −0.33
Z lm Score = −0.34
NG = 0
DB = 0
SM = 0
Delta_AUC
Doxycycline
YJR144W MGM101
ZShift = 9.19
Z lm Score = 4.21
NG = 0
DB = 0
SM = 0
Delta_L
Doxycycline
YJR144W MGM101
ZShift = −14.72
Z lm Score = −9.08
NG = 0
DB = 0
SM = 0
Delta_K
Doxycycline
YJR144W MGM101
ZShift = −3.67
Z lm Score = 1.05
NG = 0
DB = 0
SM = 0
Delta_r
Doxycycline
YJR144W MGM101
ZShift = −11.6
Z lm Score = −2.38
NG = 0
DB = 0
SM = 0
Delta_AUC
Doxycycline
YBR200W−A_2
ZShift = 2.01
Z lm Score = 4.2
NG = 0
DB = 0
SM = 0
Delta_L
Doxycycline
YBR200W−A_2
ZShift = −0.8
Z lm Score = −6.7
NG = 0
DB = 0
SM = 0
Delta_K
Doxycycline
YBR200W−A_2
ZShift = −0.65
Z lm Score = −1.08
NG = 0
DB = 0
SM = 0
Delta_r
Doxycycline
YBR200W−A_2
ZShift = −1.39
Z lm Score = 1.57
NG = 0
DB = 0
SM = 0
Delta_AUC
Doxycycline

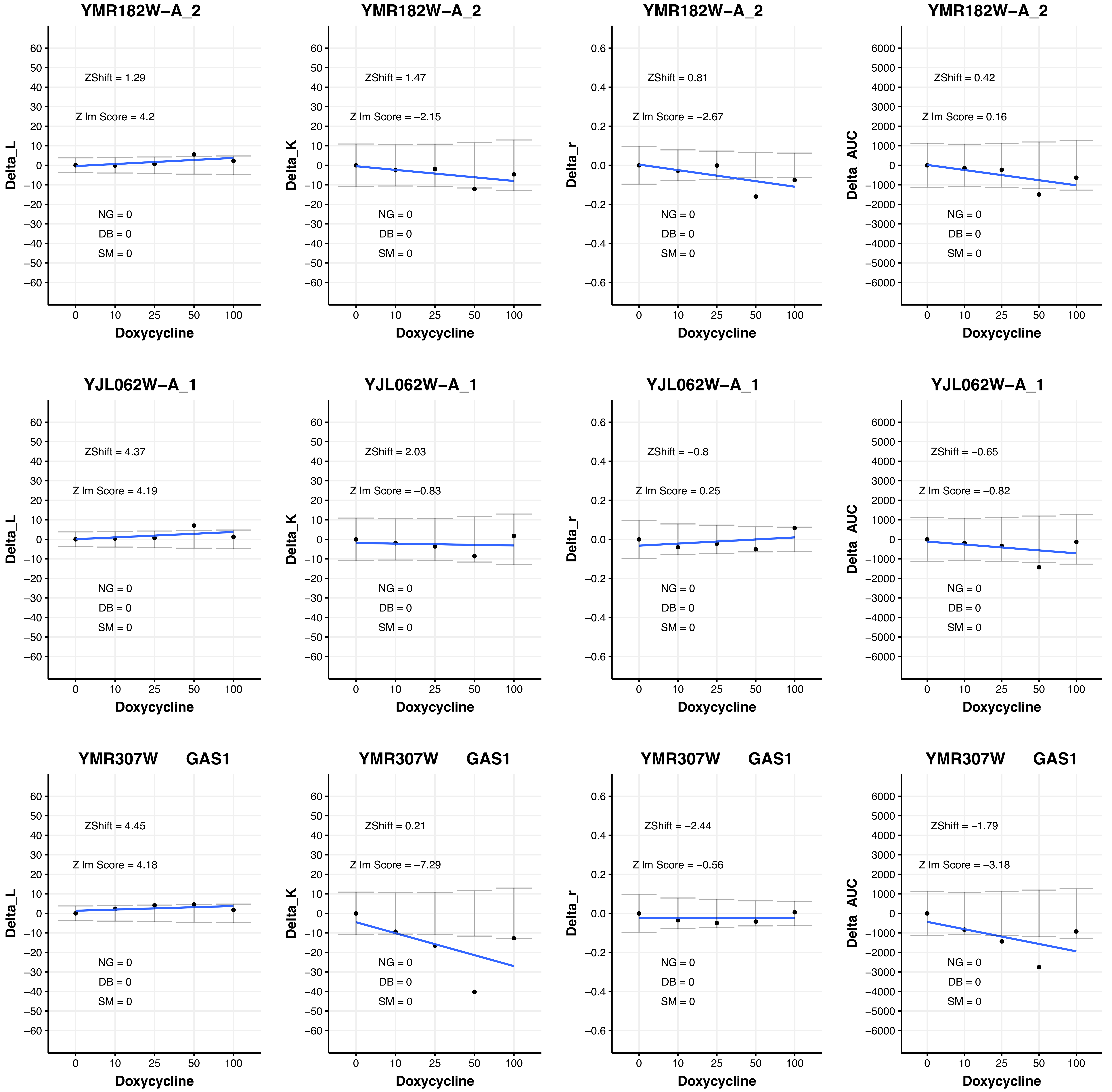
YMR182W−A_2
ZShift = 1.29
Z lm Score = 4.2
NG = 0
DB = 0
SM = 0
Delta_L
Doxycycline
YMR182W−A_2
ZShift = 1.47
Z lm Score = −2.15
NG = 0
DB = 0
SM = 0
Delta_K
Doxycycline
YMR182W−A_2
ZShift = 0.81
Z lm Score = −2.67
NG = 0
DB = 0
SM = 0
Delta_r
Doxycycline
YMR182W−A_2
ZShift = 0.42
Z lm Score = 0.16
NG = 0
DB = 0
SM = 0
Delta_AUC
Doxycycline
YJL062W−A_1
ZShift = 4.37
Z lm Score = 4.19
NG = 0
DB = 0
SM = 0
Delta_L
Doxycycline
YJL062W−A_1
ZShift = 2.03
Z lm Score = −0.83
NG = 0
DB = 0
SM = 0
Delta_K
Doxycycline
YJL062W−A_1
ZShift = −0.8
Z lm Score = 0.25
NG = 0
DB = 0
SM = 0
Delta_r
Doxycycline
YJL062W−A_1
ZShift = −0.65
Z lm Score = −0.82
NG = 0
DB = 0
SM = 0
Delta_AUC
Doxycycline
YMR307W GAS1
ZShift = 4.45
Z lm Score = 4.18
NG = 0
DB = 0
SM = 0
Delta_L
Doxycycline
YMR307W GAS1
ZShift = 0.21
Z lm Score = −7.29
NG = 0
DB = 0
SM = 0
Delta_K
Doxycycline
YMR307W GAS1
ZShift = −2.44
Z lm Score = −0.56
NG = 0
DB = 0
SM = 0
Delta_r
Doxycycline
YMR307W GAS1
ZShift = −1.79
Z lm Score = −3.18
NG = 0
DB = 0
SM = 0
Delta_AUC
Doxycycline

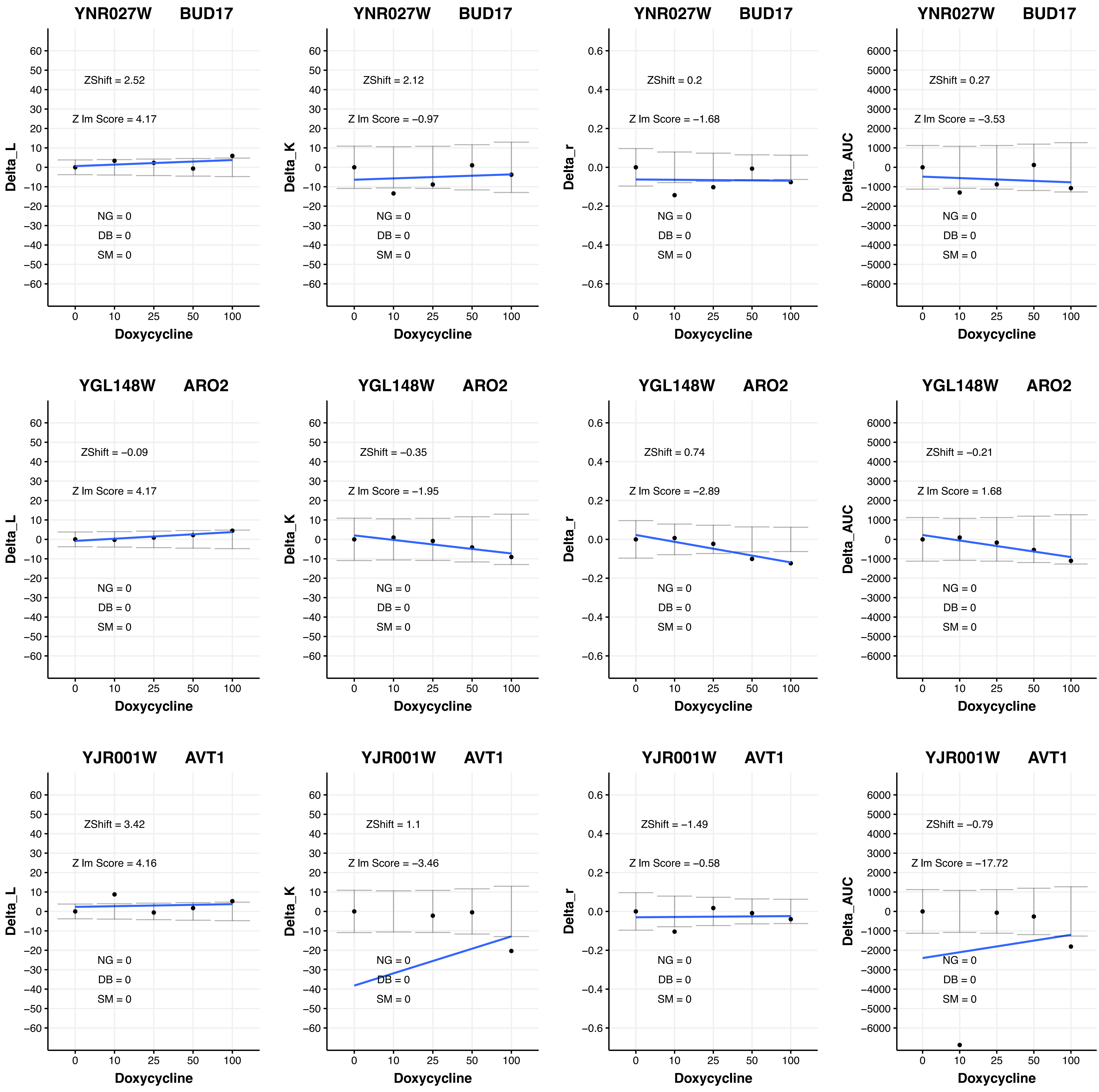
YNR027W BUD17
ZShift = 2.52
Z lm Score = 4.17
NG = 0
DB = 0
SM = 0
Delta_L
Doxycycline
YNR027W BUD17
ZShift = 2.12
Z lm Score = −0.97
NG = 0
DB = 0
SM = 0
Delta_K
Doxycycline
YNR027W BUD17
ZShift = 0.2
Z lm Score = −1.68
NG = 0
DB = 0
SM = 0
Delta_r
Doxycycline
YNR027W BUD17
ZShift = 0.27
Z lm Score = −3.53
NG = 0
DB = 0
SM = 0
Delta_AUC
Doxycycline
YGL148W ARO2
ZShift = −0.09
Z lm Score = 4.17
NG = 0
DB = 0
SM = 0
Delta_L
Doxycycline
YGL148W ARO2
ZShift = −0.35
Z lm Score = −1.95
NG = 0
DB = 0
SM = 0
Delta_K
Doxycycline
YGL148W ARO2
ZShift = 0.74
Z lm Score = −2.89
NG = 0
DB = 0
SM = 0
Delta_r
Doxycycline
YGL148W ARO2
ZShift = −0.21
Z lm Score = 1.68
NG = 0
DB = 0
SM = 0
Delta_AUC
Doxycycline
YJR001W AVT1
ZShift = 3.42
Z lm Score = 4.16
NG = 0
DB = 0
SM = 0
Delta_L
Doxycycline
YJR001W AVT1
ZShift = 1.1
Z lm Score = −3.46
NG = 0
DB = 0
SM = 0
Delta_K
Doxycycline
YJR001W AVT1
ZShift = −1.49
Z lm Score = −0.58
NG = 0
DB = 0
SM = 0
Delta_r
Doxycycline
YJR001W AVT1
ZShift = −0.79
Z lm Score = −17.72
NG = 0
DB = 0
SM = 0
Delta_AUC
Doxycycline

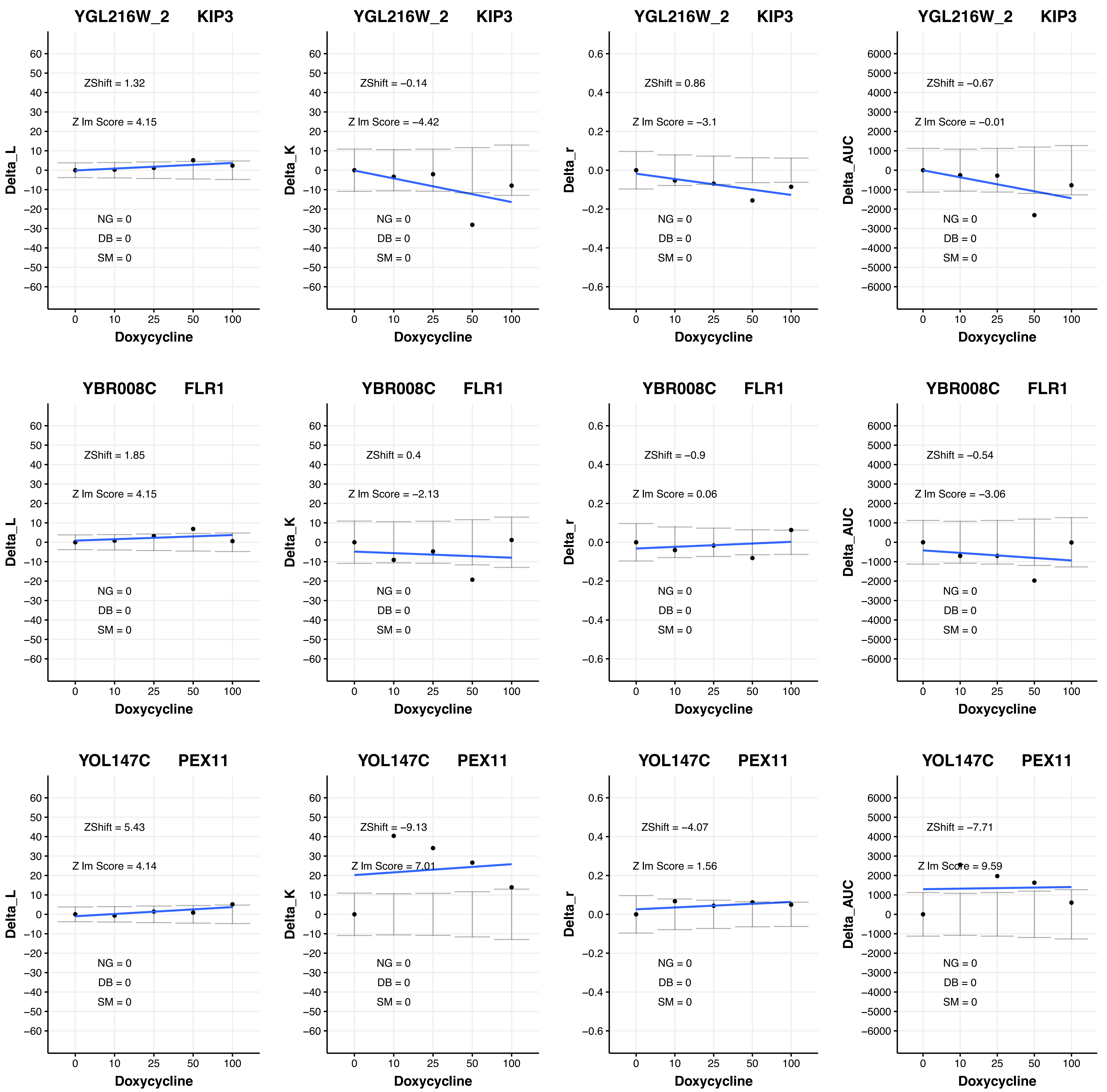

YGL216W_2 KIP3
ZShift = 1.32
Z lm Score = 4.15
Delta_L
NG = 0
DB = 0
SM = 0
Doxycycline
YGL216W_2 KIP3
ZShift = −0.14
Z lm Score = −4.42
Delta_K
YGL216W_2 KIP3
ZShift = 0.86
Z lm Score = −3.1
Delta_r
YGL216W_2 KIP3
ZShift = −0.67
Z lm Score = −0.01
Delta_AUC
YBR008C FLR1
ZShift = 1.85
Z lm Score = 4.15
YBR008C FLR1
ZShift = 0.4
Z lm Score = −2.13
YBR008C FLR1
ZShift = −0.9
Z lm Score = 0.06
YBR008C FLR1
ZShift = −0.54
Z lm Score = −3.06
YOL147C PEX11
ZShift = 5.43
Z lm Score = 4.14
YOL147C PEX11
ZShift = −9.13
Z lm Score = 7.01
YOL147C PEX11
ZShift = −4.07
Z lm Score = 1.56
YOL147C PEX11
ZShift = −7.71
Z lm Score = 9.59

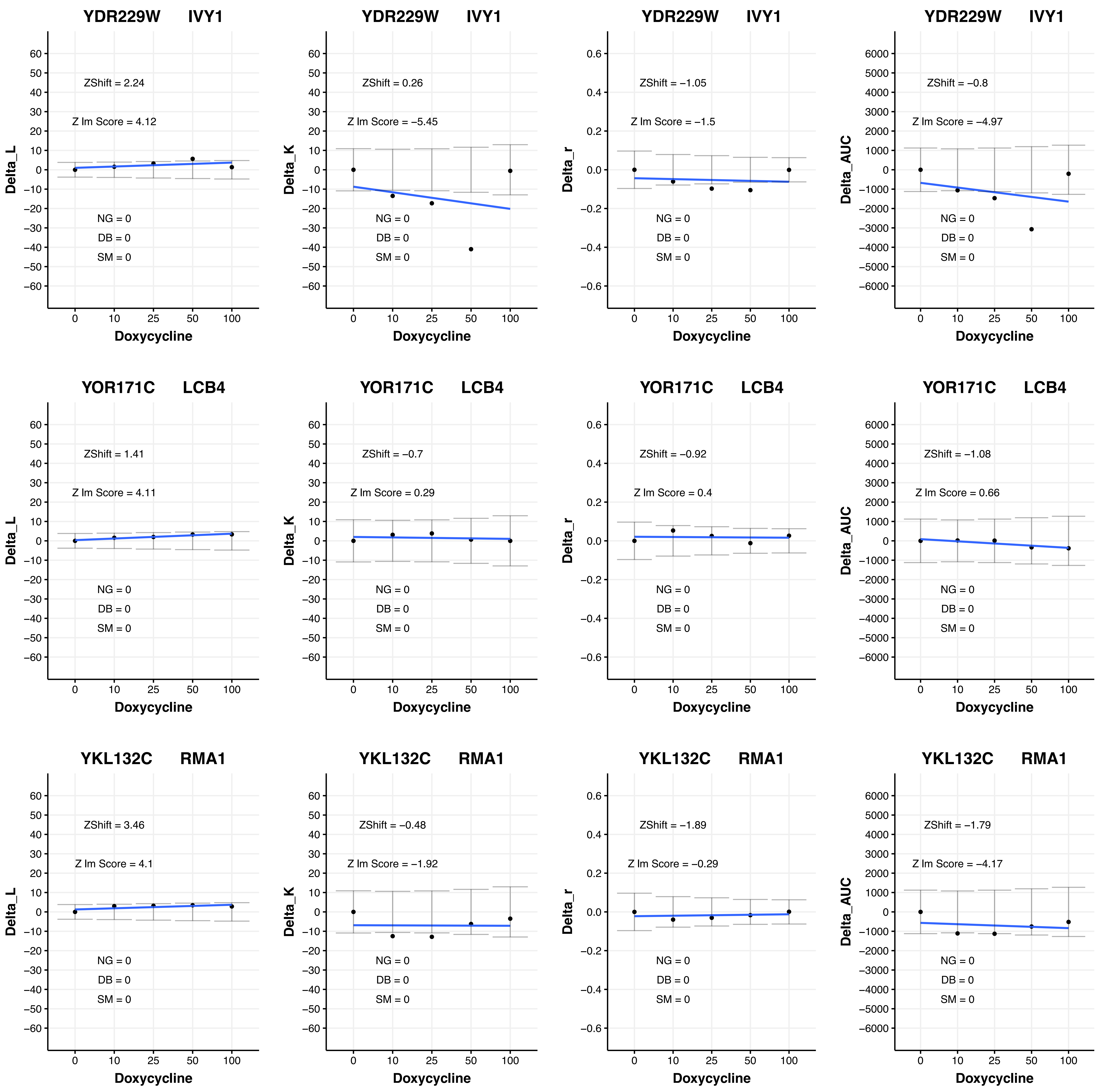
YDR229W IVY1
Delta_L
ZShift = 2.24
Z lm Score = 4.12
NG = 0
DB = 0
SM = 0
Doxycycline
YDR229W IVY1
Delta_K
ZShift = 0.26
Z lm Score = −5.45
NG = 0
DB = 0
SM = 0
Doxycycline
YDR229W IVY1
Delta_r
ZShift = −1.05
Z lm Score = −1.5
NG = 0
DB = 0
SM = 0
Doxycycline
YDR229W IVY1
Delta_AUC
ZShift = −0.8
Z lm Score = −4.97
NG = 0
DB = 0
SM = 0
Doxycycline
YOR171C LCB4
Delta_L
ZShift = 1.41
Z lm Score = 4.11
NG = 0
DB = 0
SM = 0
Doxycycline
YOR171C LCB4
Delta_K
ZShift = −0.7
Z lm Score = 0.29
NG = 0
DB = 0
SM = 0
Doxycycline
YOR171C LCB4
Delta_r
ZShift = −0.92
Z lm Score = 0.4
NG = 0
DB = 0
SM = 0
Doxycycline
YOR171C LCB4
Delta_AUC
ZShift = −1.08
Z lm Score = 0.66
NG = 0
DB = 0
SM = 0
Doxycycline
YKL132C RMA1
Delta_L
ZShift = 3.46
Z lm Score = 4.1
NG = 0
DB = 0
SM = 0
Doxycycline
YKL132C RMA1
Delta_K
ZShift = −0.48
Z lm Score = −1.92
NG = 0
DB = 0
SM = 0
Doxycycline
YKL132C RMA1
Delta_r
ZShift = −1.89
Z lm Score = −0.29
NG = 0
DB = 0
SM = 0
Doxycycline
YKL132C RMA1
Delta_AUC
ZShift = −1.79
Z lm Score = −4.17
NG = 0
DB = 0
SM = 0
Doxycycline

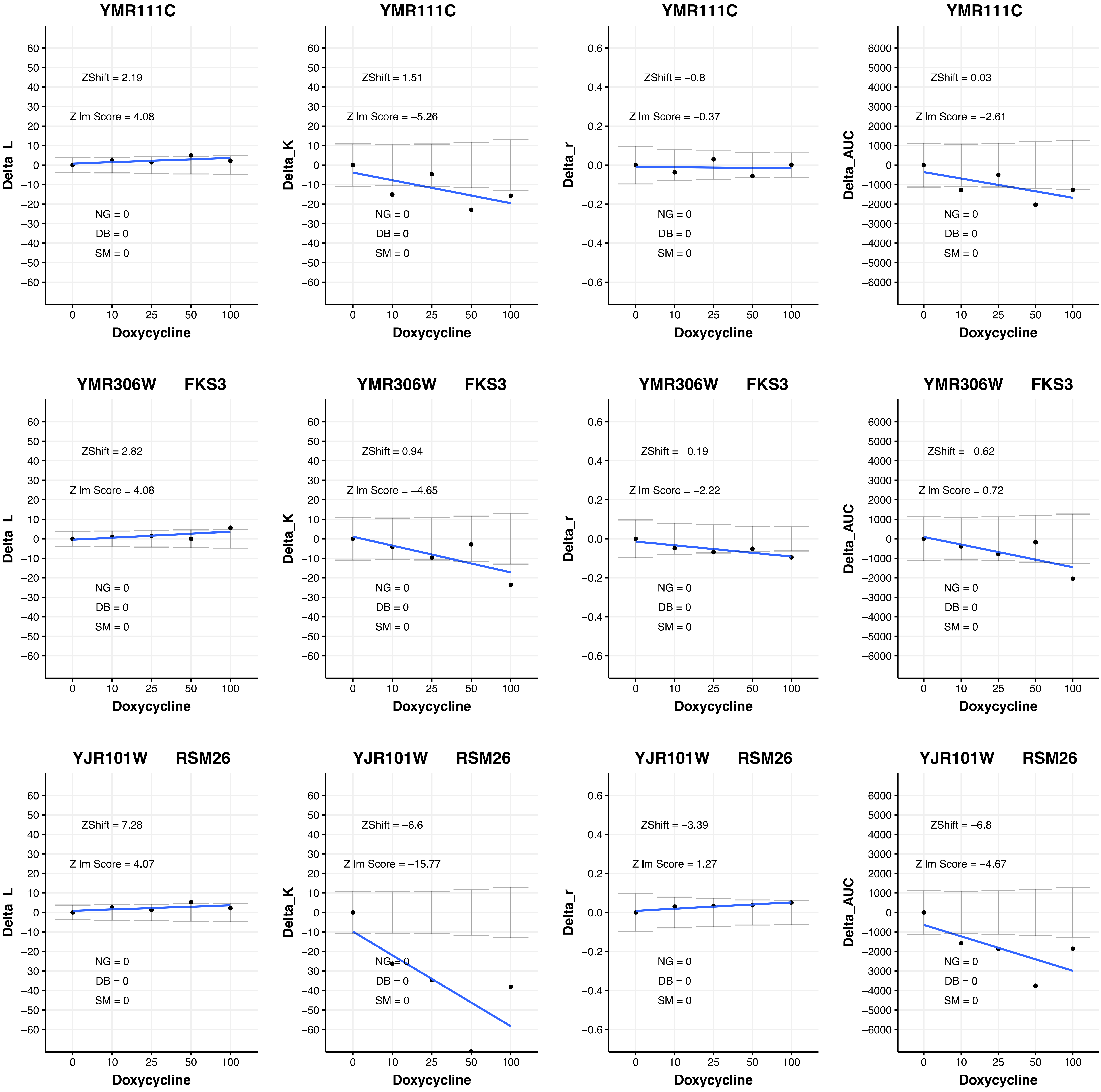

YMR111C
ZShift = 2.19
Z lm Score = 4.08
NG = 0
DB = 0
SM = 0
Delta_L
Doxycycline
YMR111C
ZShift = 1.51
Z lm Score = −5.26
NG = 0
DB = 0
SM = 0
Delta_K
Doxycycline
YMR111C
ZShift = −0.8
Z lm Score = −0.37
NG = 0
DB = 0
SM = 0
Delta_r
Doxycycline
YMR111C
ZShift = 0.03
Z lm Score = −2.61
NG = 0
DB = 0
SM = 0
Delta_AUC
Doxycycline
YMR306W FKS3
ZShift = 2.82
Z lm Score = 4.08
NG = 0
DB = 0
SM = 0
Delta_L
Doxycycline
YMR306W FKS3
ZShift = 0.94
Z lm Score = −4.65
NG = 0
DB = 0
SM = 0
Delta_K
Doxycycline
YMR306W FKS3
ZShift = −0.19
Z lm Score = −2.22
NG = 0
DB = 0
SM = 0
Delta_r
Doxycycline
YMR306W FKS3
ZShift = −0.62
Z lm Score = 0.72
NG = 0
DB = 0
SM = 0
Delta_AUC
Doxycycline
YJR101W RSM26
ZShift = 7.28
Z lm Score = 4.07
NG = 0
DB = 0
SM = 0
Delta_L
Doxycycline
YJR101W RSM26
ZShift = −6.6
Z lm Score = −15.77
NG = 0
DB = 0
SM = 0
Delta_K
Doxycycline
YJR101W RSM26
ZShift = −3.39
Z lm Score = 1.27
NG = 0
DB = 0
SM = 0
Delta_r
Doxycycline
YJR101W RSM26
ZShift = −6.8
Z lm Score = −4.67
NG = 0
DB = 0
SM = 0
Delta_AUC
Doxycycline

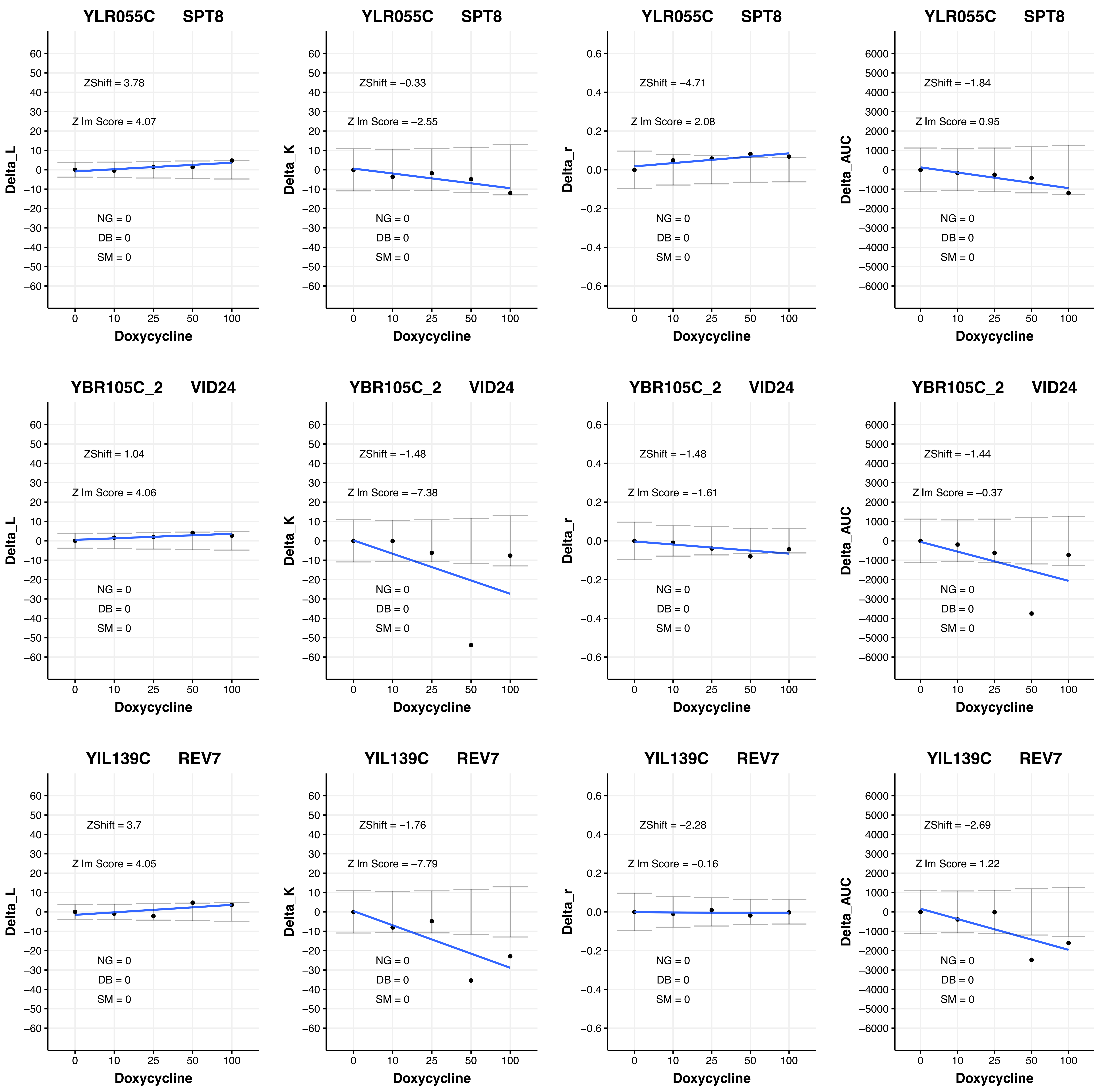
YLR055C SPT8
ZShift = 3.78
Z lm Score = 4.07
NG = 0
DB = 0
SM = 0
Delta_L
Doxycycline
YLR055C SPT8
ZShift = −0.33
Z lm Score = −2.55
NG = 0
DB = 0
SM = 0
Delta_K
Doxycycline
YLR055C SPT8
ZShift = −4.71
Z lm Score = 2.08
NG = 0
DB = 0
SM = 0
Delta_r
Doxycycline
YLR055C SPT8
ZShift = −1.84
Z lm Score = 0.95
NG = 0
DB = 0
SM = 0
Delta_AUC
Doxycycline
YBR105C_2 VID24
ZShift = 1.04
Z lm Score = 4.06
NG = 0
DB = 0
SM = 0
Delta_L
Doxycycline
YBR105C_2 VID24
ZShift = −1.48
Z lm Score = −7.38
NG = 0
DB = 0
SM = 0
Delta_K
Doxycycline
YBR105C_2 VID24
ZShift = −1.48
Z lm Score = −1.61
NG = 0
DB = 0
SM = 0
Delta_r
Doxycycline
YBR105C_2 VID24
ZShift = −1.44
Z lm Score = −0.37
NG = 0
DB = 0
SM = 0
Delta_AUC
Doxycycline
YIL139C REV7
ZShift = 3.7
Z lm Score = 4.05
NG = 0
DB = 0
SM = 0
Delta_L
Doxycycline
YIL139C REV7
ZShift = −1.76
Z lm Score = −7.79
NG = 0
DB = 0
SM = 0
Delta_K
Doxycycline
YIL139C REV7
ZShift = −2.28
Z lm Score = −0.16
NG = 0
DB = 0
SM = 0
Delta_r
Doxycycline
YIL139C REV7
ZShift = −2.69
Z lm Score = 1.22
NG = 0
DB = 0
SM = 0
Delta_AUC
Doxycycline

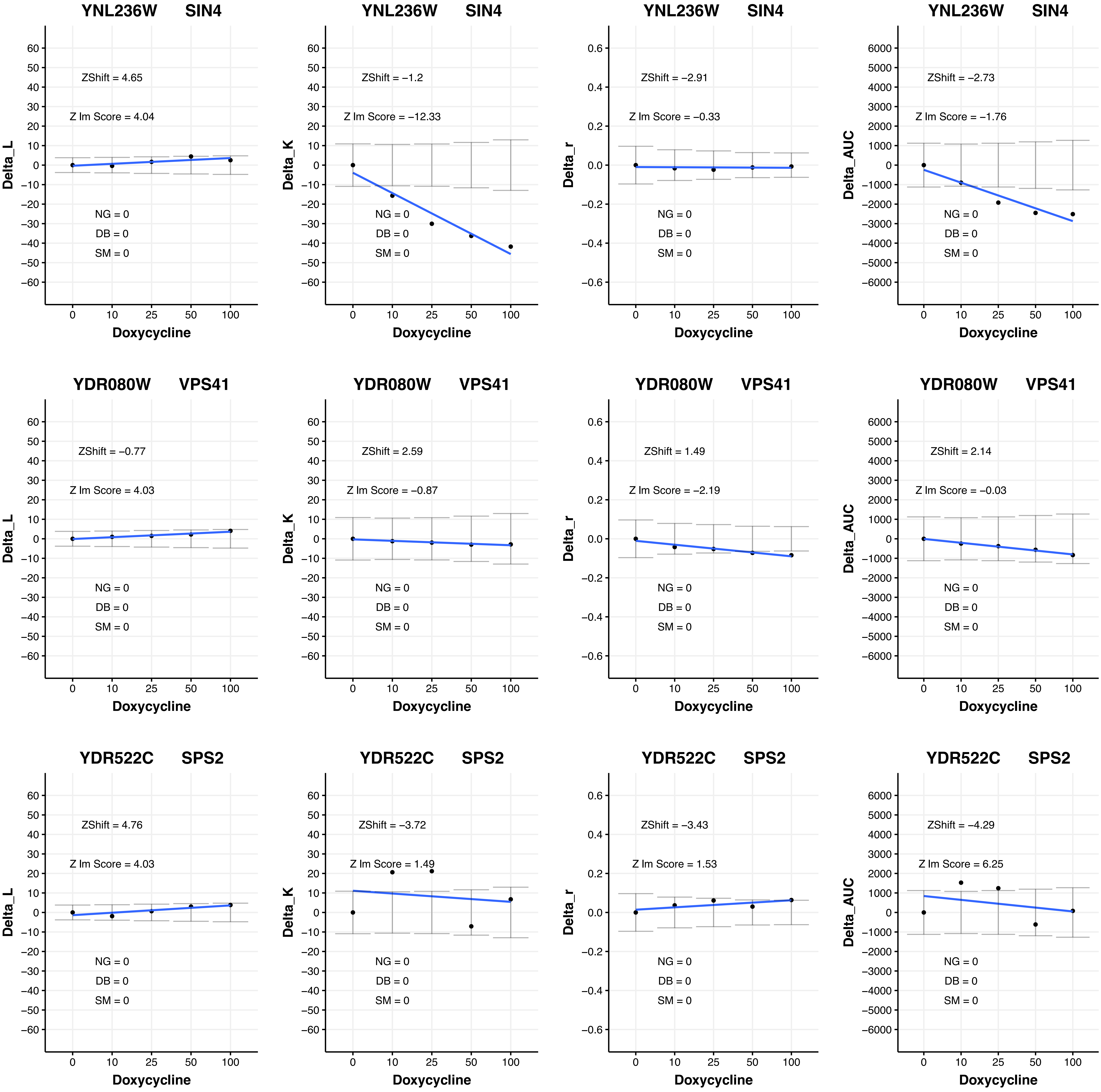
YNL236W SIN4
ZShift = 4.65
Z lm Score = 4.04
NG = 0
DB = 0
SM = 0
Delta_L
Doxycycline
YNL236W SIN4
ZShift = −1.2
Z lm Score = −12.33
NG = 0
DB = 0
SM = 0
Delta_K
Doxycycline
YNL236W SIN4
ZShift = −2.91
Z lm Score = −0.33
NG = 0
DB = 0
SM = 0
Delta_r
Doxycycline
YNL236W SIN4
ZShift = −2.73
Z lm Score = −1.76
NG = 0
DB = 0
SM = 0
Delta_AUC
Doxycycline
YDR080W VPS41
ZShift = −0.77
Z lm Score = 4.03
NG = 0
DB = 0
SM = 0
Delta_L
Doxycycline
YDR080W VPS41
ZShift = 2.59
Z lm Score = −0.87
NG = 0
DB = 0
SM = 0
Delta_K
Doxycycline
YDR080W VPS41
ZShift = 1.49
Z lm Score = −2.19
NG = 0
DB = 0
SM = 0
Delta_r
Doxycycline
YDR080W VPS41
ZShift = 2.14
Z lm Score = −0.03
NG = 0
DB = 0
SM = 0
Delta_AUC
Doxycycline
YDR522C SPS2
ZShift = 4.76
Z lm Score = 4.03
NG = 0
DB = 0
SM = 0
Delta_L
Doxycycline
YDR522C SPS2
ZShift = −3.72
Z lm Score = 1.49
NG = 0
DB = 0
SM = 0
Delta_K
Doxycycline
YDR522C SPS2
ZShift = −3.43
Z lm Score = 1.53
NG = 0
DB = 0
SM = 0
Delta_r
Doxycycline
YDR522C SPS2
ZShift = −4.29
Z lm Score = 6.25
NG = 0
DB = 0
SM = 0
Delta_AUC
Doxycycline

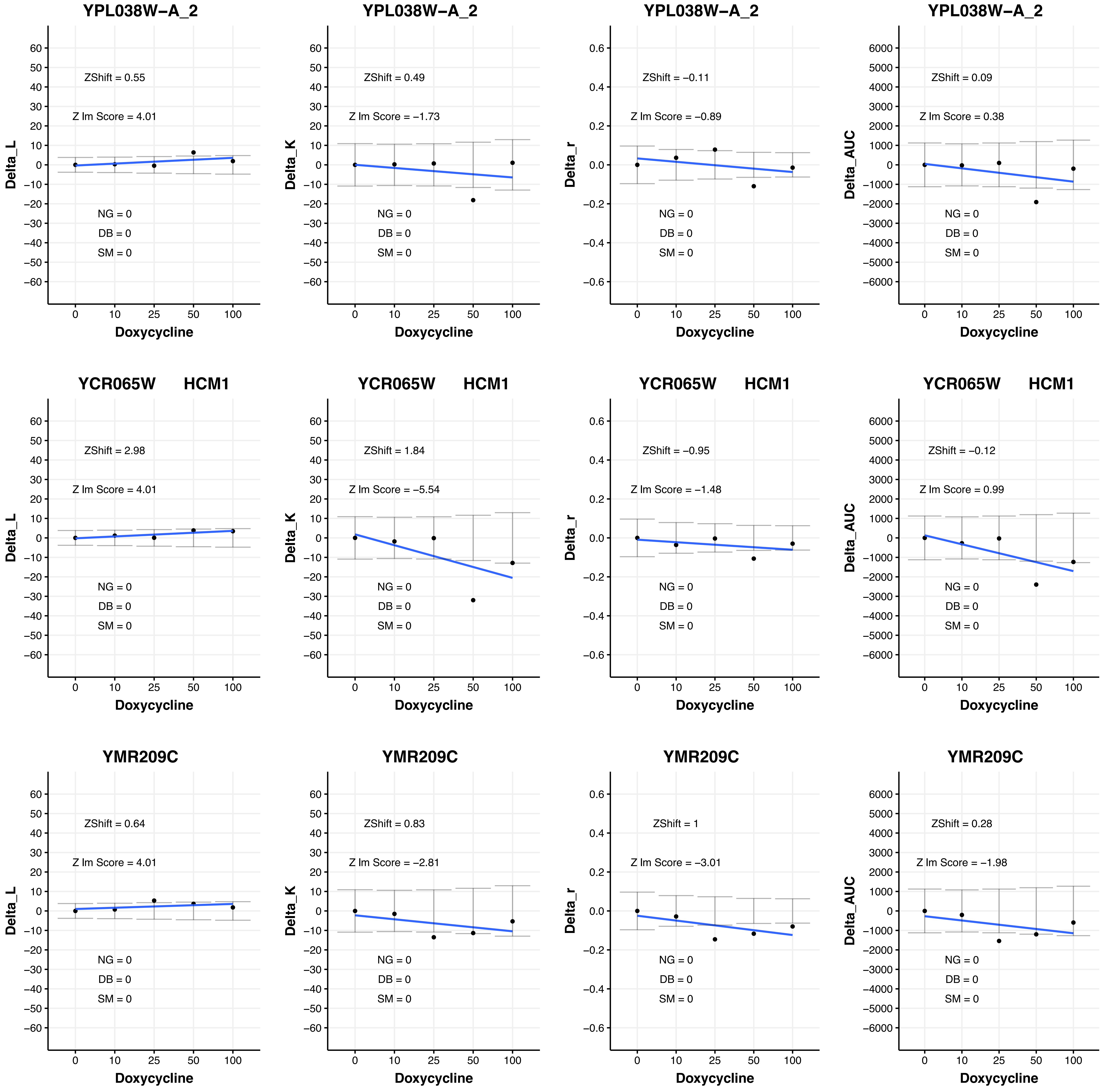

YPL038W−A_2
ZShift = 0.55
Z lm Score = 4.01
NG = 0
DB = 0
SM = 0
Delta_L
Doxycycline
YPL038W−A_2
ZShift = 0.49
Z lm Score = −1.73
NG = 0
DB = 0
SM = 0
Delta_K
Doxycycline
YPL038W−A_2
ZShift = −0.11
Z lm Score = −0.89
NG = 0
DB = 0
SM = 0
Delta_r
Doxycycline
YPL038W−A_2
ZShift = 0.09
Z lm Score = 0.38
NG = 0
DB = 0
SM = 0
Delta_AUC
Doxycycline
YCR065W HCM1
ZShift = 2.98
Z lm Score = 4.01
NG = 0
DB = 0
SM = 0
Delta_L
Doxycycline
YCR065W HCM1
ZShift = 1.84
Z lm Score = −5.54
NG = 0
DB = 0
SM = 0
Delta_K
Doxycycline
YCR065W HCM1
ZShift = −0.95
Z lm Score = −1.48
NG = 0
DB = 0
SM = 0
Delta_r
Doxycycline
YCR065W HCM1
ZShift = −0.12
Z lm Score = 0.99
NG = 0
DB = 0
SM = 0
Delta_AUC
Doxycycline
YMR209C
ZShift = 0.64
Z lm Score = 4.01
NG = 0
DB = 0
SM = 0
Delta_L
Doxycycline
YMR209C
ZShift = 0.83
Z lm Score = −2.81
NG = 0
DB = 0
SM = 0
Delta_K
Doxycycline
YMR209C
ZShift = 1
Z lm Score = −3.01
NG = 0
DB = 0
SM = 0
Delta_r
Doxycycline
YMR209C
ZShift = 0.28
Z lm Score = −1.98
NG = 0
DB = 0
SM = 0
Delta_AUC
Doxycycline

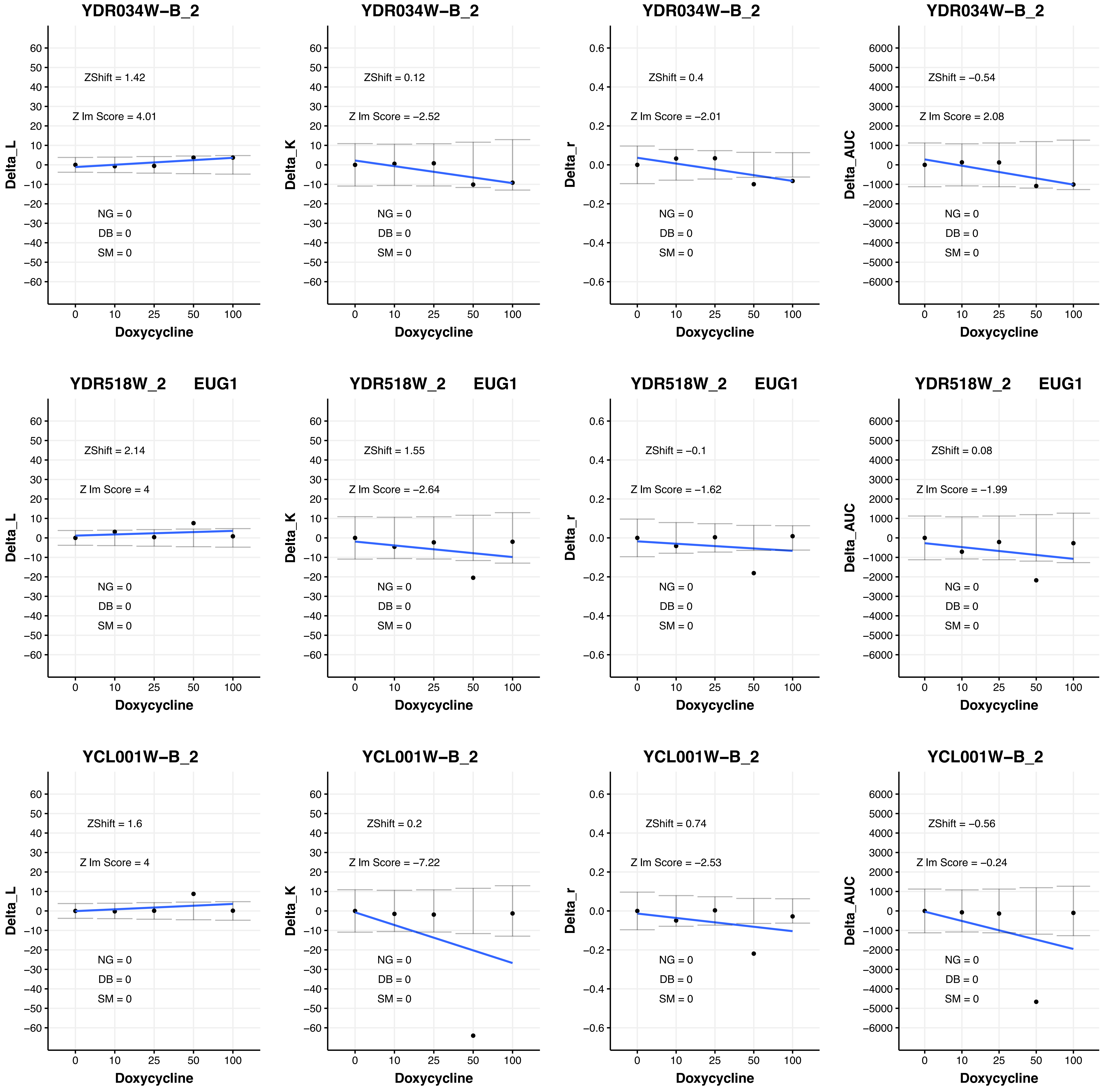

YDR034W−B_2
ZShift = 1.42
Z lm Score = 4.01
NG = 0
DB = 0
SM = 0
Delta_L
Doxycycline
YDR034W−B_2
ZShift = 0.12
Z lm Score = −2.52
NG = 0
DB = 0
SM = 0
Delta_K
Doxycycline
YDR034W−B_2
ZShift = 0.4
Z lm Score = −2.01
NG = 0
DB = 0
SM = 0
Delta_r
Doxycycline
YDR034W−B_2
ZShift = −0.54
Z lm Score = 2.08
NG = 0
DB = 0
SM = 0
Delta_AUC
Doxycycline
YDR518W_2 EUG1
ZShift = 2.14
Z lm Score = 4
NG = 0
DB = 0
SM = 0
Delta_L
Doxycycline
YDR518W_2 EUG1
ZShift = 1.55
Z lm Score = −2.64
NG = 0
DB = 0
SM = 0
Delta_K
Doxycycline
YDR518W_2 EUG1
ZShift = −0.1
Z lm Score = −1.62
NG = 0
DB = 0
SM = 0
Delta_r
Doxycycline
YDR518W_2 EUG1
ZShift = 0.08
Z lm Score = −1.99
NG = 0
DB = 0
SM = 0
Delta_AUC
Doxycycline
YCL001W−B_2
ZShift = 1.6
Z lm Score = 4
NG = 0
DB = 0
SM = 0
Delta_L
Doxycycline
YCL001W−B_2
ZShift = 0.2
Z lm Score = −7.22
NG = 0
DB = 0
SM = 0
Delta_K
Doxycycline
YCL001W−B_2
ZShift = 0.74
Z lm Score = −2.53
NG = 0
DB = 0
SM = 0
Delta_r
Doxycycline
YCL001W−B_2
ZShift = −0.56
Z lm Score = −0.24
NG = 0
DB = 0
SM = 0
Delta_AUC
Doxycycline

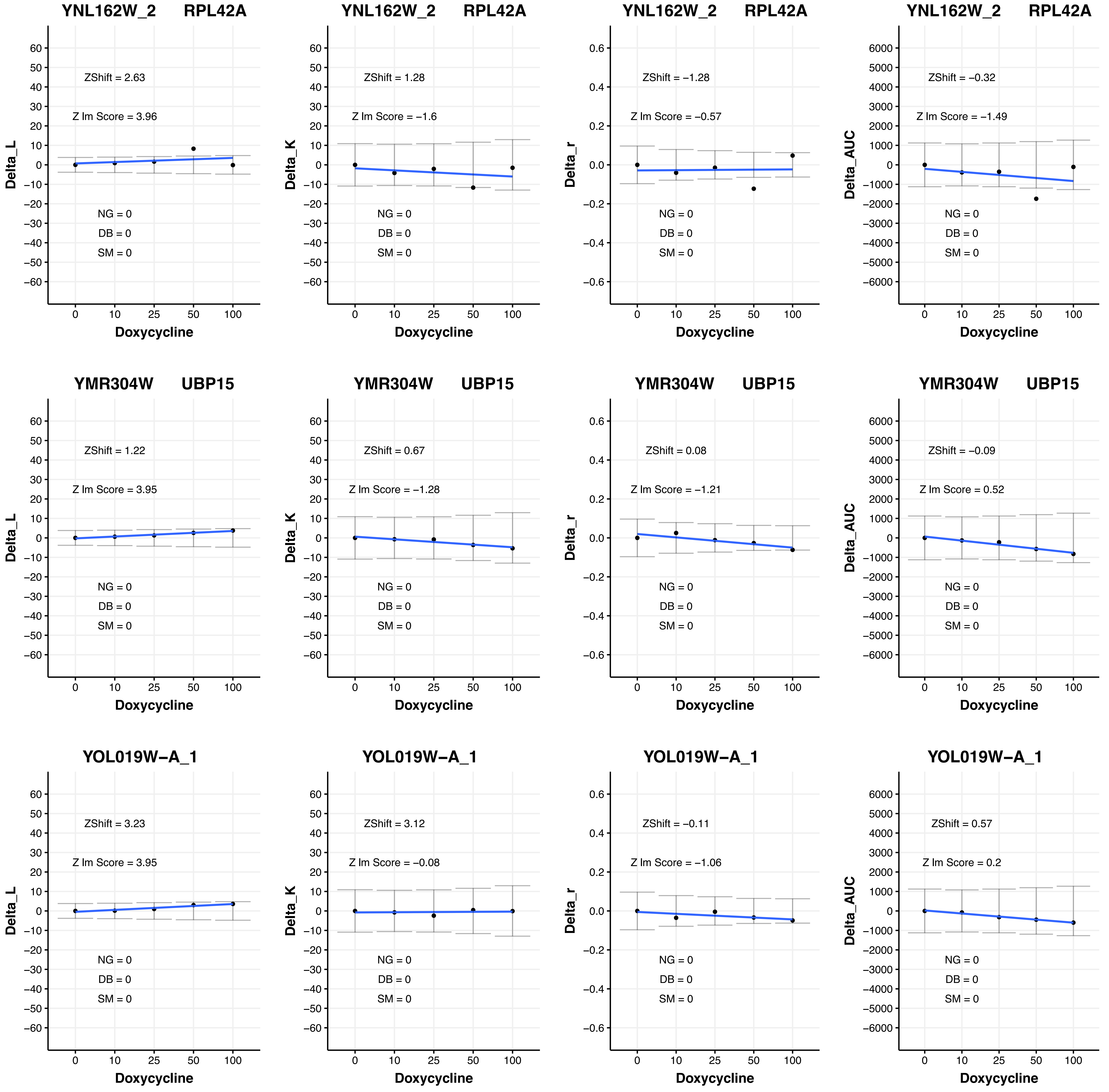

YNL162W_2 RPL42A
ZShift = 2.63
Z lm Score = 3.96
NG = 0
DB = 0
SM = 0
Delta_L
Doxycycline
YNL162W_2 RPL42A
ZShift = 1.28
Z lm Score = −1.6
NG = 0
DB = 0
SM = 0
Delta_K
Doxycycline
YNL162W_2 RPL42A
ZShift = −1.28
Z lm Score = −0.57
NG = 0
DB = 0
SM = 0
Delta_r
Doxycycline
YNL162W_2 RPL42A
ZShift = −0.32
Z lm Score = −1.49
NG = 0
DB = 0
SM = 0
Delta_AUC
Doxycycline
YMR304W UBP15
ZShift = 1.22
Z lm Score = 3.95
NG = 0
DB = 0
SM = 0
Delta_L
Doxycycline
YMR304W UBP15
ZShift = 0.67
Z lm Score = −1.28
NG = 0
DB = 0
SM = 0
Delta_K
Doxycycline
YMR304W UBP15
ZShift = 0.08
Z lm Score = −1.21
NG = 0
DB = 0
SM = 0
Delta_r
Doxycycline
YMR304W UBP15
ZShift = −0.09
Z lm Score = 0.52
NG = 0
DB = 0
SM = 0
Delta_AUC
Doxycycline
YOL019W−A_1
ZShift = 3.23
Z lm Score = 3.95
NG = 0
DB = 0
SM = 0
Delta_L
Doxycycline
YOL019W−A_1
ZShift = 3.12
Z lm Score = −0.08
NG = 0
DB = 0
SM = 0
Delta_K
Doxycycline
YOL019W−A_1
ZShift = −0.11
Z lm Score = −1.06
NG = 0
DB = 0
SM = 0
Delta_r
Doxycycline
YOL019W−A_1
ZShift = 0.57
Z lm Score = 0.2
NG = 0
DB = 0
SM = 0
Delta_AUC
Doxycycline

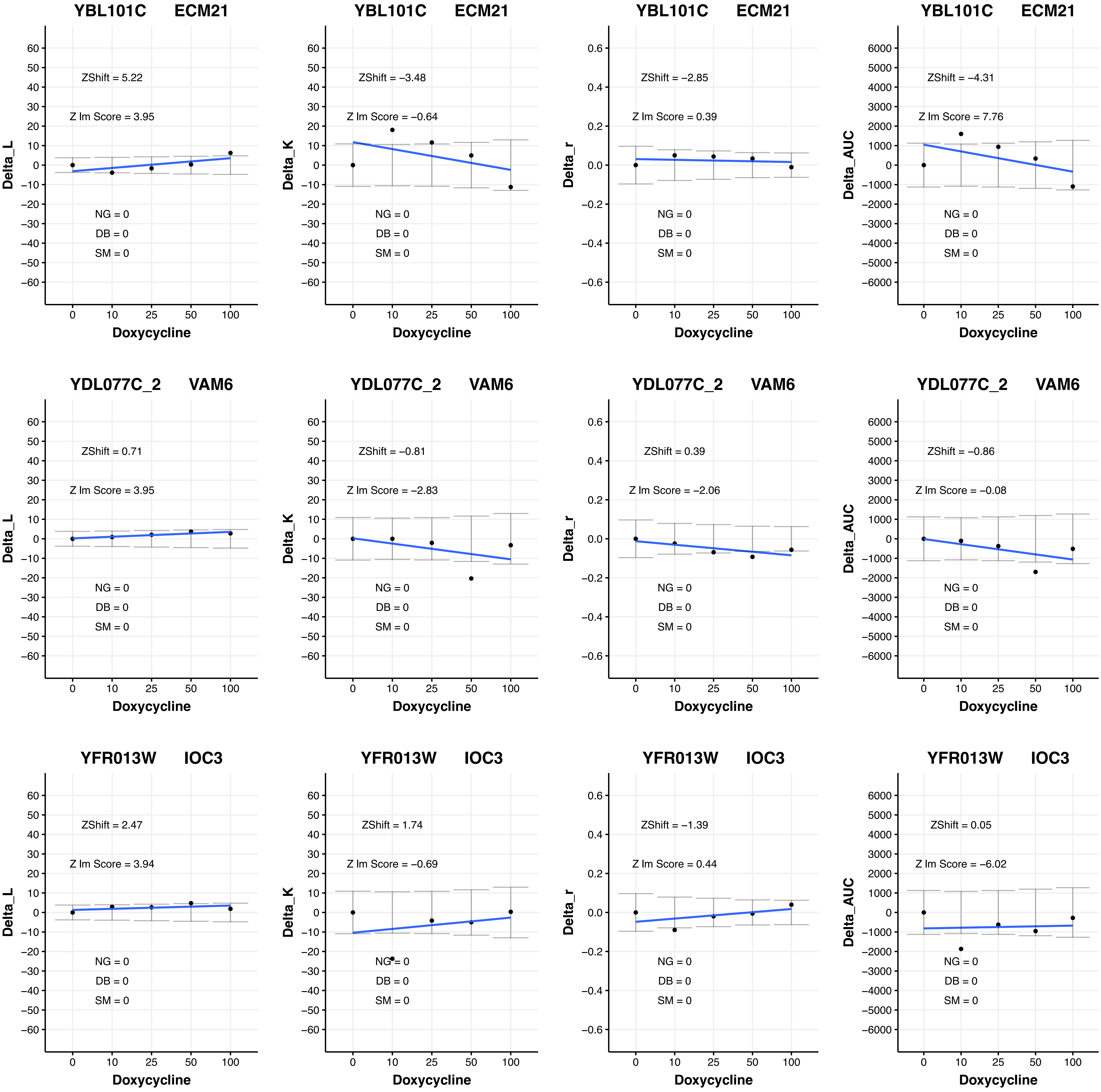
YBL101C ECM21
ZShift = 5.22
Z lm Score = 3.95
NG = 0
DB = 0
SM = 0
Delta_L
Doxycycline
YBL101C ECM21
ZShift = −3.48
Z lm Score = −0.64
NG = 0
DB = 0
SM = 0
Delta_K
Doxycycline
YBL101C ECM21
ZShift = −2.85
Z lm Score = 0.39
NG = 0
DB = 0
SM = 0
Delta_r
Doxycycline
YBL101C ECM21
ZShift = −4.31
Z lm Score = 7.76
NG = 0
DB = 0
SM = 0
Delta_AUC
Doxycycline
YDL077C_2 VAM6
ZShift = 0.71
Z lm Score = 3.95
NG = 0
DB = 0
SM = 0
Delta_L
Doxycycline
YDL077C_2 VAM6
ZShift = −0.81
Z lm Score = −2.83
NG = 0
DB = 0
SM = 0
Delta_K
Doxycycline
YDL077C_2 VAM6
ZShift = 0.39
Z lm Score = −2.06
NG = 0
DB = 0
SM = 0
Delta_r
Doxycycline
YDL077C_2 VAM6
ZShift = −0.86
Z lm Score = −0.08
NG = 0
DB = 0
SM = 0
Delta_AUC
Doxycycline
YFR013W IOC3
ZShift = 2.47
Z lm Score = 3.94
NG = 0
DB = 0
SM = 0
Delta_L
Doxycycline
YFR013W IOC3
ZShift = 1.74
Z lm Score = −0.69
NG = 0
DB = 0
SM = 0
Delta_K
Doxycycline
YFR013W IOC3
ZShift = −1.39
Z lm Score = 0.44
NG = 0
DB = 0
SM = 0
Delta_r
Doxycycline
YFR013W IOC3
ZShift = 0.05
Z lm Score = −6.02
NG = 0
DB = 0
SM = 0
Delta_AUC
Doxycycline

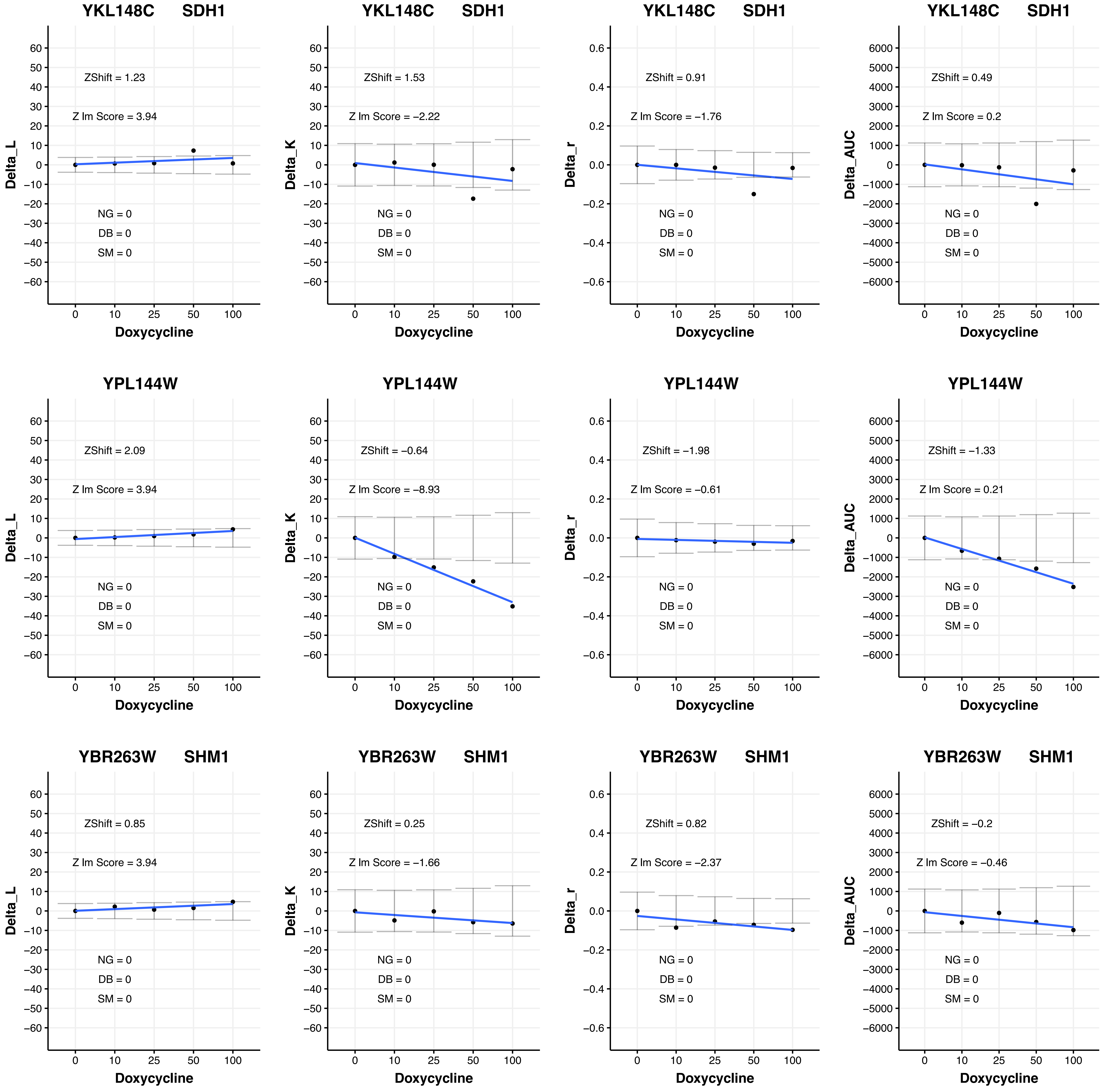
YKL148C SDH1
ZShift = 1.23
Z lm Score = 3.94
NG = 0
DB = 0
SM = 0
Delta_L
Doxycycline
YKL148C SDH1
ZShift = 1.53
Z lm Score = −2.22
NG = 0
DB = 0
SM = 0
Delta_K
Doxycycline
YKL148C SDH1
ZShift = 0.91
Z lm Score = −1.76
NG = 0
DB = 0
SM = 0
Delta_r
Doxycycline
YKL148C SDH1
ZShift = 0.49
Z lm Score = 0.2
NG = 0
DB = 0
SM = 0
Delta_AUC
Doxycycline
YPL144W
ZShift = 2.09
Z lm Score = 3.94
NG = 0
DB = 0
SM = 0
Delta_L
Doxycycline
YPL144W
ZShift = −0.64
Z lm Score = −8.93
NG = 0
DB = 0
SM = 0
Delta_K
Doxycycline
YPL144W
ZShift = −1.98
Z lm Score = −0.61
NG = 0
DB = 0
SM = 0
Delta_r
Doxycycline
YPL144W
ZShift = −1.33
Z lm Score = 0.21
NG = 0
DB = 0
SM = 0
Delta_AUC
Doxycycline
YBR263W SHM1
ZShift = 0.85
Z lm Score = 3.94
NG = 0
DB = 0
SM = 0
Delta_L
Doxycycline
YBR263W SHM1
ZShift = 0.25
Z lm Score = −1.66
NG = 0
DB = 0
SM = 0
Delta_K
Doxycycline
YBR263W SHM1
ZShift = 0.82
Z lm Score = −2.37
NG = 0
DB = 0
SM = 0
Delta_r
Doxycycline
YBR263W SHM1
ZShift = −0.2
Z lm Score = −0.46
NG = 0
DB = 0
SM = 0
Delta_AUC
Doxycycline

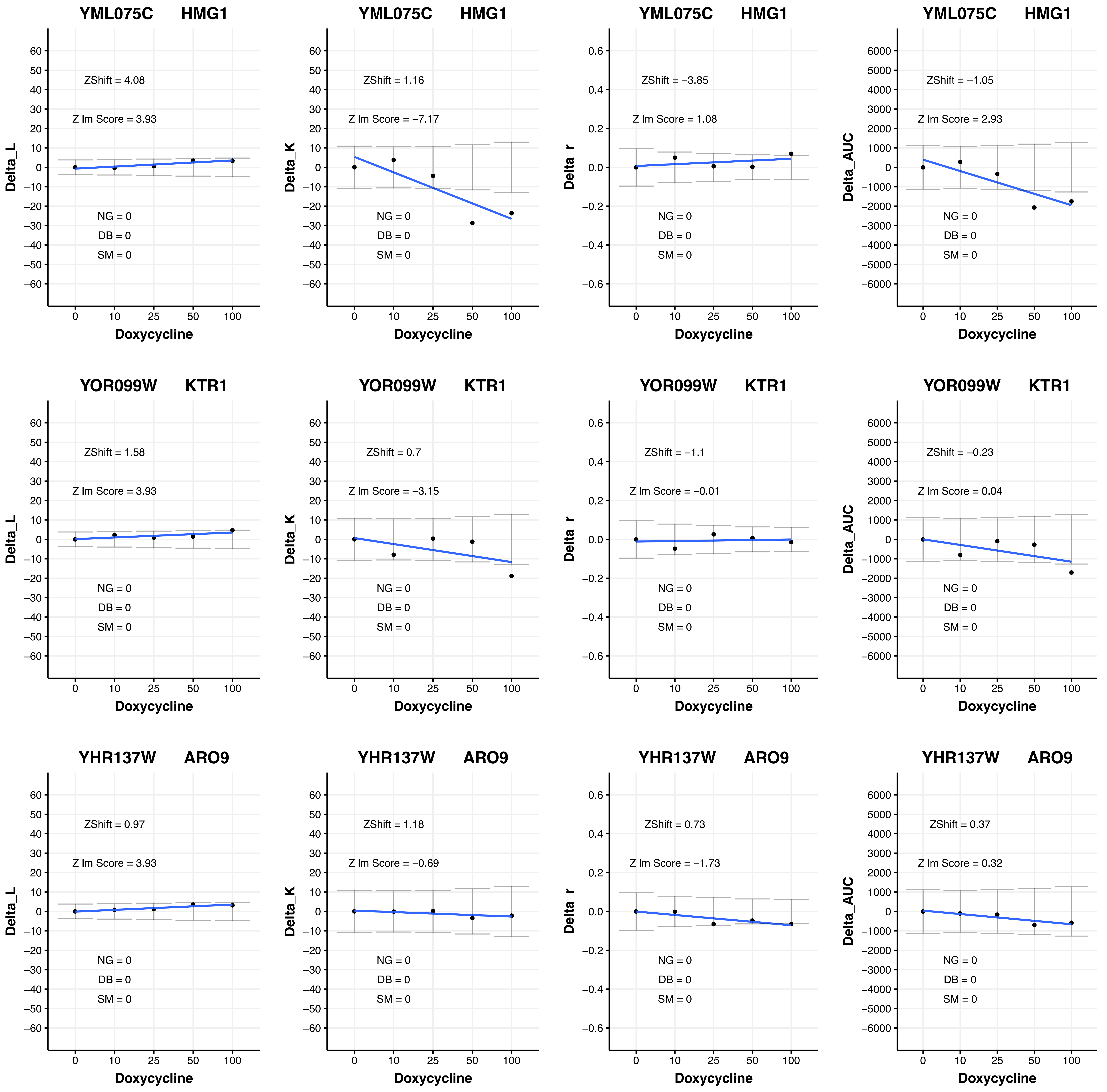

YML075C HMG1
ZShift = 4.08
Z lm Score = 3.93
NG = 0
DB = 0
SM = 0
Delta_L
Doxycycline
YML075C HMG1
ZShift = 1.16
Z lm Score = −7.17
NG = 0
DB = 0
SM = 0
Delta_K
Doxycycline
YML075C HMG1
ZShift = −3.85
Z lm Score = 1.08
NG = 0
DB = 0
SM = 0
Delta_r
Doxycycline
YML075C HMG1
ZShift = −1.05
Z lm Score = 2.93
NG = 0
DB = 0
SM = 0
Delta_AUC
Doxycycline
YOR099W KTR1
ZShift = 1.58
Z lm Score = 3.93
NG = 0
DB = 0
SM = 0
Delta_L
Doxycycline
YOR099W KTR1
ZShift = 0.7
Z lm Score = −3.15
NG = 0
DB = 0
SM = 0
Delta_K
Doxycycline
YOR099W KTR1
ZShift = −1.1
Z lm Score = −0.01
NG = 0
DB = 0
SM = 0
Delta_r
Doxycycline
YOR099W KTR1
ZShift = −0.23
Z lm Score = 0.04
NG = 0
DB = 0
SM = 0
Delta_AUC
Doxycycline
YHR137W ARO9
ZShift = 0.97
Z lm Score = 3.93
NG = 0
DB = 0
SM = 0
Delta_L
Doxycycline
YHR137W ARO9
ZShift = 1.18
Z lm Score = −0.69
NG = 0
DB = 0
SM = 0
Delta_K
Doxycycline
YHR137W ARO9
ZShift = 0.73
Z lm Score = −1.73
NG = 0
DB = 0
SM = 0
Delta_r
Doxycycline
YHR137W ARO9
ZShift = 0.37
Z lm Score = 0.32
NG = 0
DB = 0
SM = 0
Delta_AUC
Doxycycline

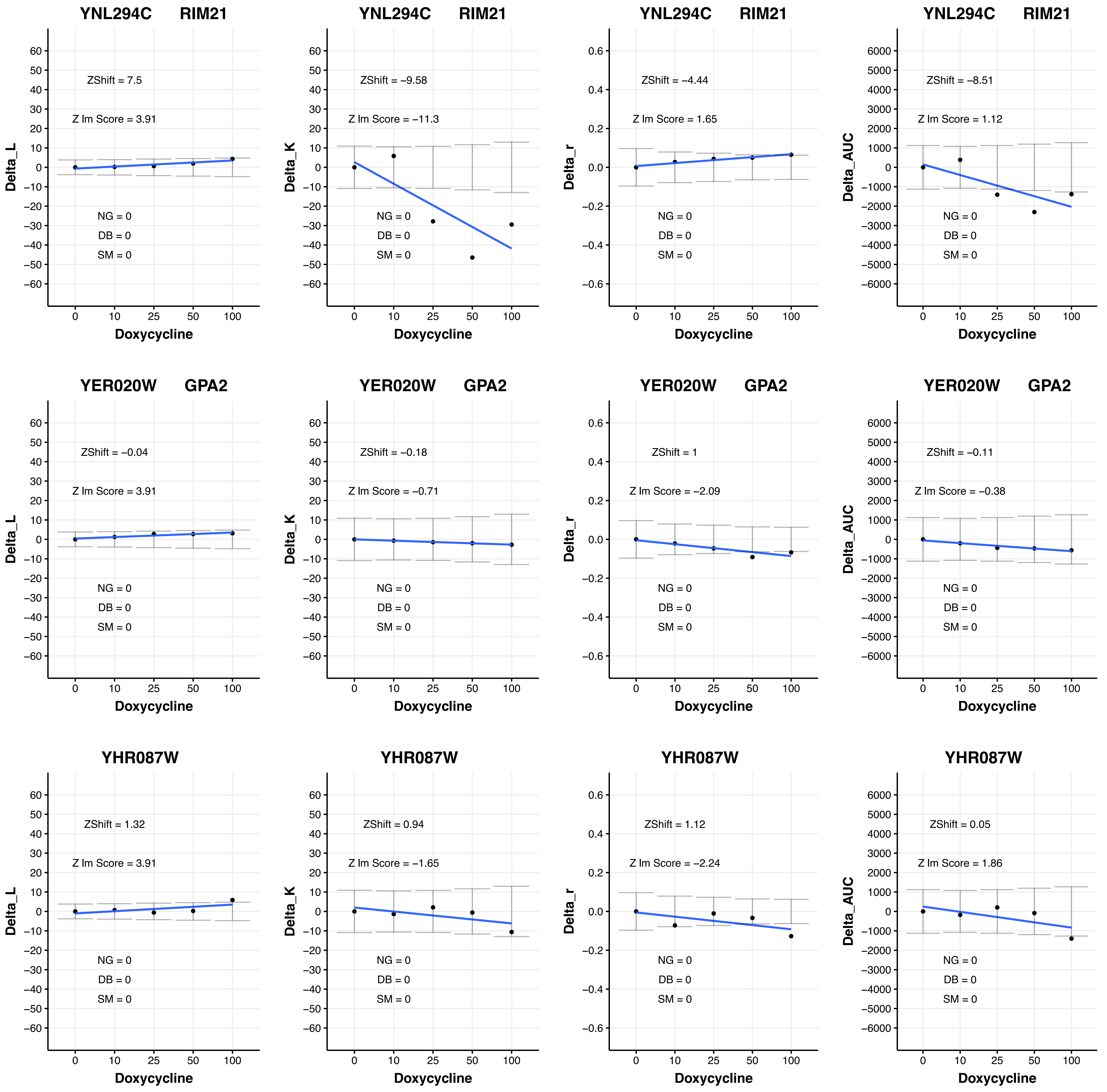
YNL294C RIM21
ZShift = 7.5
Z lm Score = 3.91
NG = 0
DB = 0
SM = 0
Delta_L
Doxycycline
YNL294C RIM21
ZShift = −9.58
Z lm Score = −11.3
NG = 0
DB = 0
SM = 0
Delta_K
Doxycycline
YNL294C RIM21
ZShift = −4.44
Z lm Score = 1.65
NG = 0
DB = 0
SM = 0
Delta_r
Doxycycline
YNL294C RIM21
ZShift = −8.51
Z lm Score = 1.12
NG = 0
DB = 0
SM = 0
Delta_AUC
Doxycycline
YER020W GPA2
ZShift = −0.04
Z lm Score = 3.91
NG = 0
DB = 0
SM = 0
Delta_L
Doxycycline
YER020W GPA2
ZShift = −0.18
Z lm Score = −0.71
NG = 0
DB = 0
SM = 0
Delta_K
Doxycycline
YER020W GPA2
ZShift = 1
Z lm Score = −2.09
NG = 0
DB = 0
SM = 0
Delta_r
Doxycycline
YER020W GPA2
ZShift = −0.11
Z lm Score = −0.38
NG = 0
DB = 0
SM = 0
Delta_AUC
Doxycycline
YHR087W
ZShift = 1.32
Z lm Score = 3.91
NG = 0
DB = 0
SM = 0
Delta_L
Doxycycline
YHR087W
ZShift = 0.94
Z lm Score = −1.65
NG = 0
DB = 0
SM = 0
Delta_K
Doxycycline
YHR087W
ZShift = 1.12
Z lm Score = −2.24
NG = 0
DB = 0
SM = 0
Delta_r
Doxycycline
YHR087W
ZShift = 0.05
Z lm Score = 1.86
NG = 0
DB = 0
SM = 0
Delta_AUC
Doxycycline

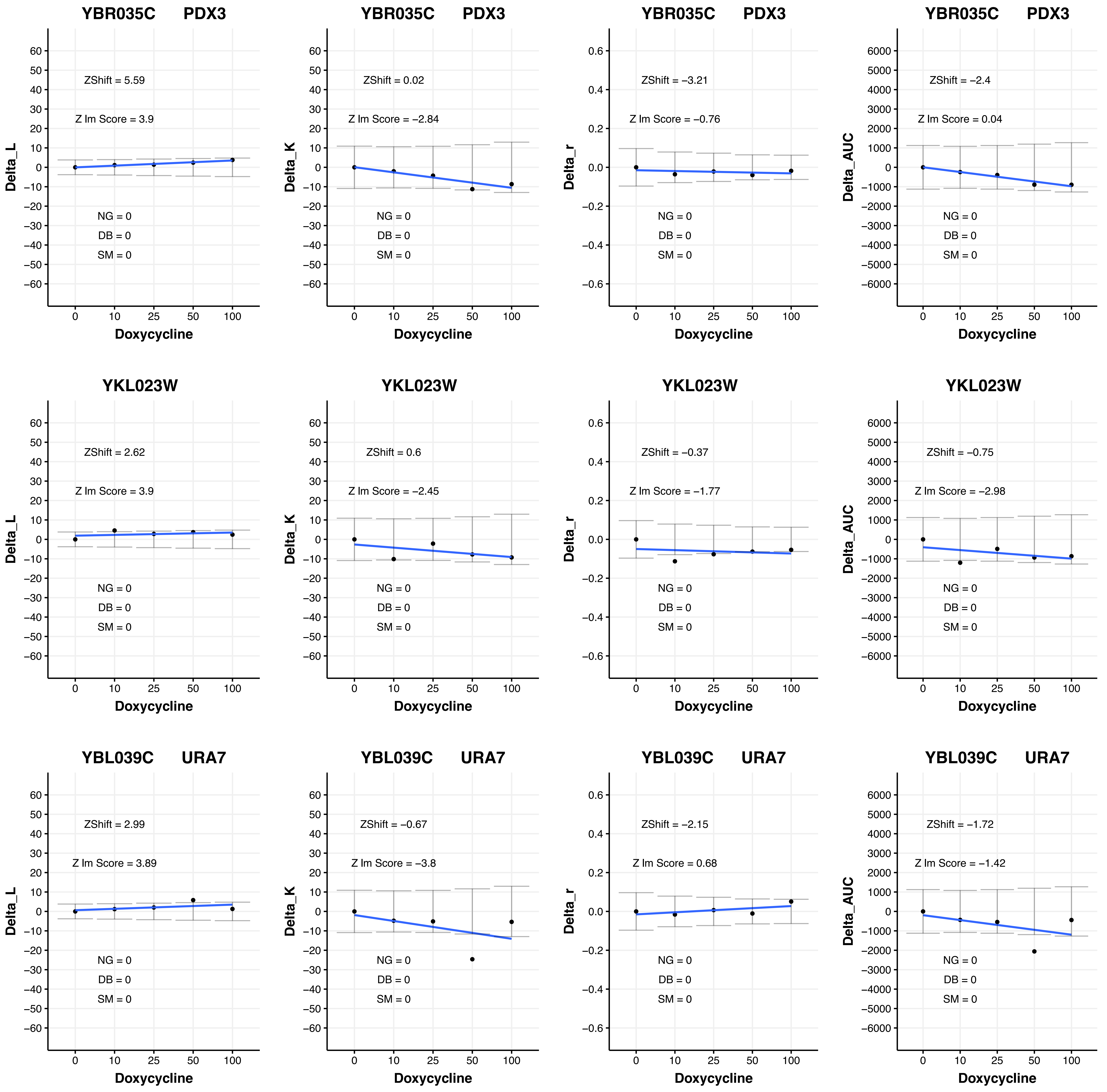

YBR035C PDX3
ZShift = 5.59
Z lm Score = 3.9
NG = 0
DB = 0
SM = 0
Delta_L
Doxycycline
YBR035C PDX3
ZShift = 0.02
Z lm Score = −2.84
NG = 0
DB = 0
SM = 0
Delta_K
Doxycycline
YBR035C PDX3
ZShift = −3.21
Z lm Score = −0.76
NG = 0
DB = 0
SM = 0
Delta_r
Doxycycline
YBR035C PDX3
ZShift = −2.4
Z lm Score = 0.04
NG = 0
DB = 0
SM = 0
Delta_AUC
Doxycycline
YKL023W
ZShift = 2.62
Z lm Score = 3.9
NG = 0
DB = 0
SM = 0
Delta_L
Doxycycline
YKL023W
ZShift = 0.6
Z lm Score = −2.45
NG = 0
DB = 0
SM = 0
Delta_K
Doxycycline
YKL023W
ZShift = −0.37
Z lm Score = −1.77
NG = 0
DB = 0
SM = 0
Delta_r
Doxycycline
YKL023W
ZShift = −0.75
Z lm Score = −2.98
NG = 0
DB = 0
SM = 0
Delta_AUC
Doxycycline
YBL039C URA7
ZShift = 2.99
Z lm Score = 3.89
NG = 0
DB = 0
SM = 0
Delta_L
Doxycycline
YBL039C URA7
ZShift = −0.67
Z lm Score = −3.8
NG = 0
DB = 0
SM = 0
Delta_K
Doxycycline
YBL039C URA7
ZShift = −2.15
Z lm Score = 0.68
NG = 0
DB = 0
SM = 0
Delta_r
Doxycycline
YBL039C URA7
ZShift = −1.72
Z lm Score = −1.42
NG = 0
DB = 0
SM = 0
Delta_AUC
Doxycycline

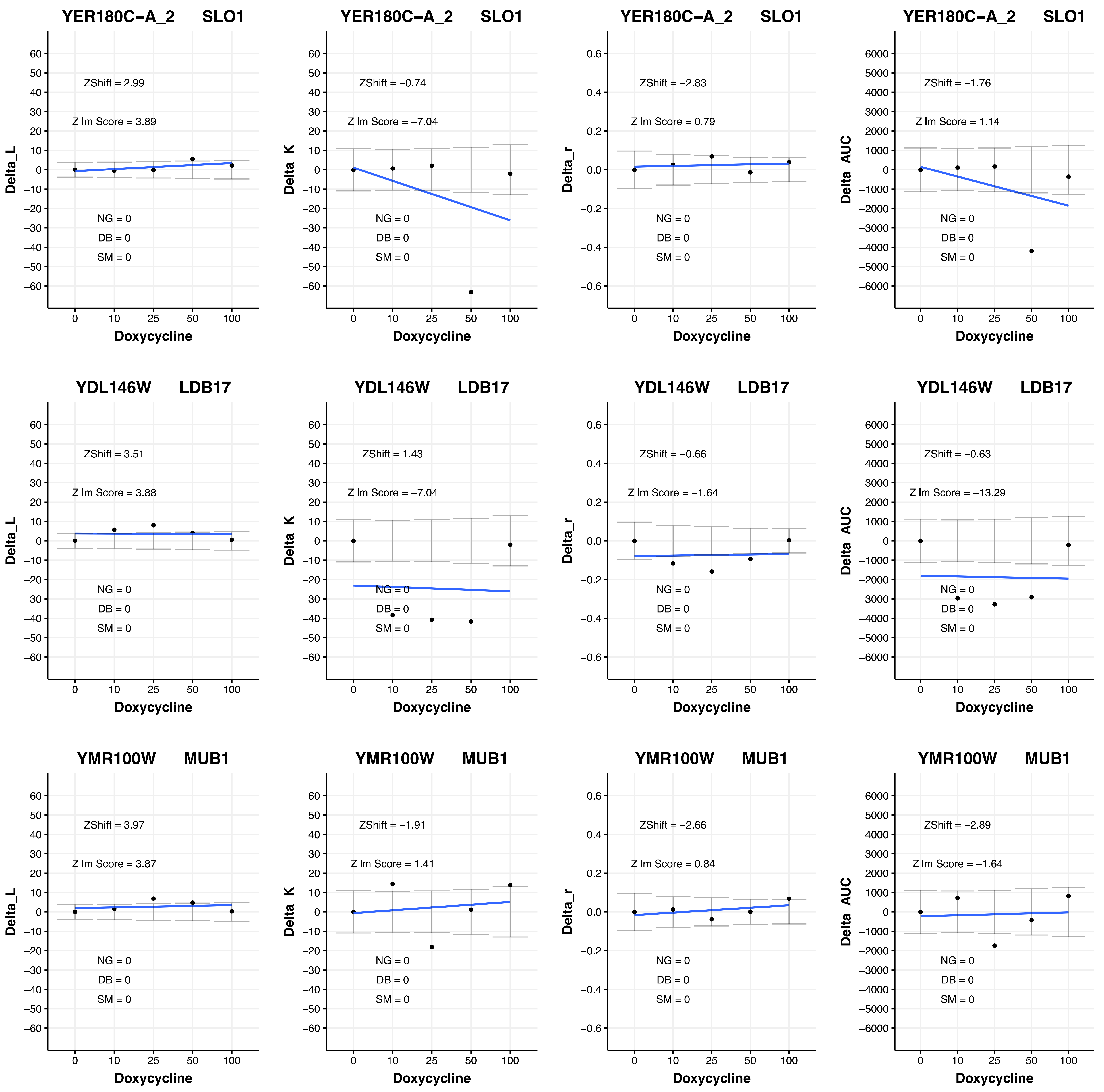

YER180C−A_2 SLO1
ZShift = 2.99
Z lm Score = 3.89
NG = 0
DB = 0
SM = 0
Delta_L
Doxycycline
YER180C−A_2 SLO1
ZShift = −0.74
Z lm Score = −7.04
NG = 0
DB = 0
SM = 0
Delta_K
Doxycycline
YER180C−A_2 SLO1
ZShift = −2.83
Z lm Score = 0.79
NG = 0
DB = 0
SM = 0
Delta_r
Doxycycline
YER180C−A_2 SLO1
ZShift = −1.76
Z lm Score = 1.14
NG = 0
DB = 0
SM = 0
Delta_AUC
Doxycycline
YDL146W LDB17
ZShift = 3.51
Z lm Score = 3.88
NG = 0
DB = 0
SM = 0
Delta_L
Doxycycline
YDL146W LDB17
ZShift = 1.43
Z lm Score = −7.04
NG = 0
DB = 0
SM = 0
Delta_K
Doxycycline
YDL146W LDB17
ZShift = −0.66
Z lm Score = −1.64
NG = 0
DB = 0
SM = 0
Delta_r
Doxycycline
YDL146W LDB17
ZShift = −0.63
Z lm Score = −13.29
NG = 0
DB = 0
SM = 0
Delta_AUC
Doxycycline
YMR100W MUB1
ZShift = 3.97
Z lm Score = 3.87
NG = 0
DB = 0
SM = 0
Delta_L
Doxycycline
YMR100W MUB1
ZShift = −1.91
Z lm Score = 1.41
NG = 0
DB = 0
SM = 0
Delta_K
Doxycycline
YMR100W MUB1
ZShift = −2.66
Z lm Score = 0.84
NG = 0
DB = 0
SM = 0
Delta_r
Doxycycline
YMR100W MUB1
ZShift = −2.89
Z lm Score = −1.64
NG = 0
DB = 0
SM = 0
Delta_AUC
Doxycycline

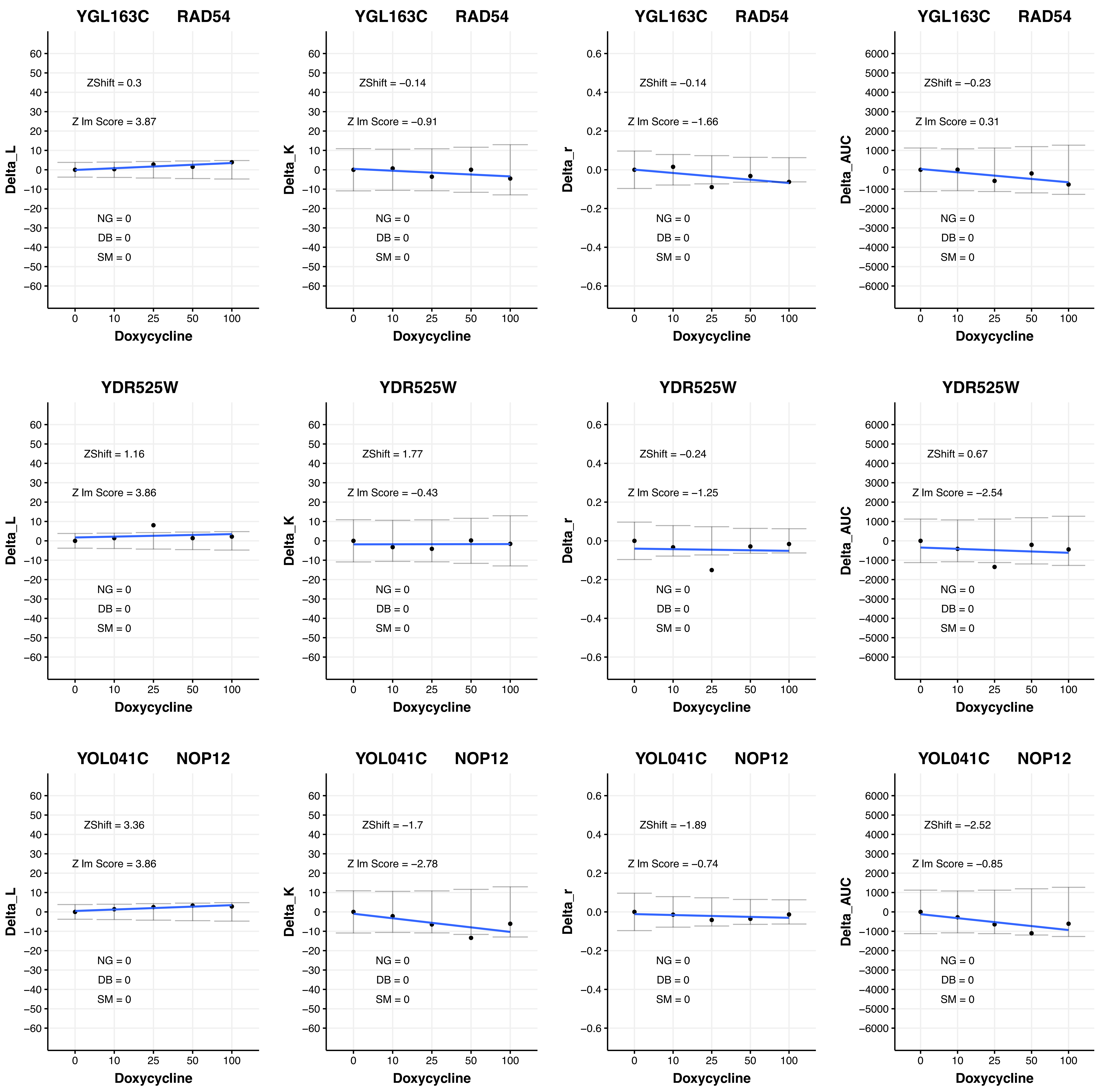

YGL163C RAD54
ZShift = 0.3
Z lm Score = 3.87
Delta_L
YGL163C RAD54
ZShift = −0.14
Z lm Score = −0.91
Delta_K
YGL163C RAD54
ZShift = −0.14
Z lm Score = −1.66
Delta_r
YGL163C RAD54
ZShift = −0.23
Z lm Score = 0.31
Delta_AUC
YDR525W
ZShift = 1.16
Z lm Score = 3.86
Delta_L
YDR525W
ZShift = 1.77
Z lm Score = −0.43
Delta_K
YDR525W
ZShift = −0.24
Z lm Score = −1.25
Delta_r
YDR525W
ZShift = 0.67
Z lm Score = −2.54
Delta_AUC
YOL041C NOP12
ZShift = 3.36
Z lm Score = 3.86
Delta_L
YOL041C NOP12
ZShift = −1.7
Z lm Score = −2.78
Delta_K
YOL041C NOP12
ZShift = −1.89
Z lm Score = −0.74
Delta_r
YOL041C NOP12
ZShift = −2.52
Z lm Score = −0.85
Delta_AUC
NG = 0
DB = 0
SM = 0
Doxycycline
0 10 25 50 100

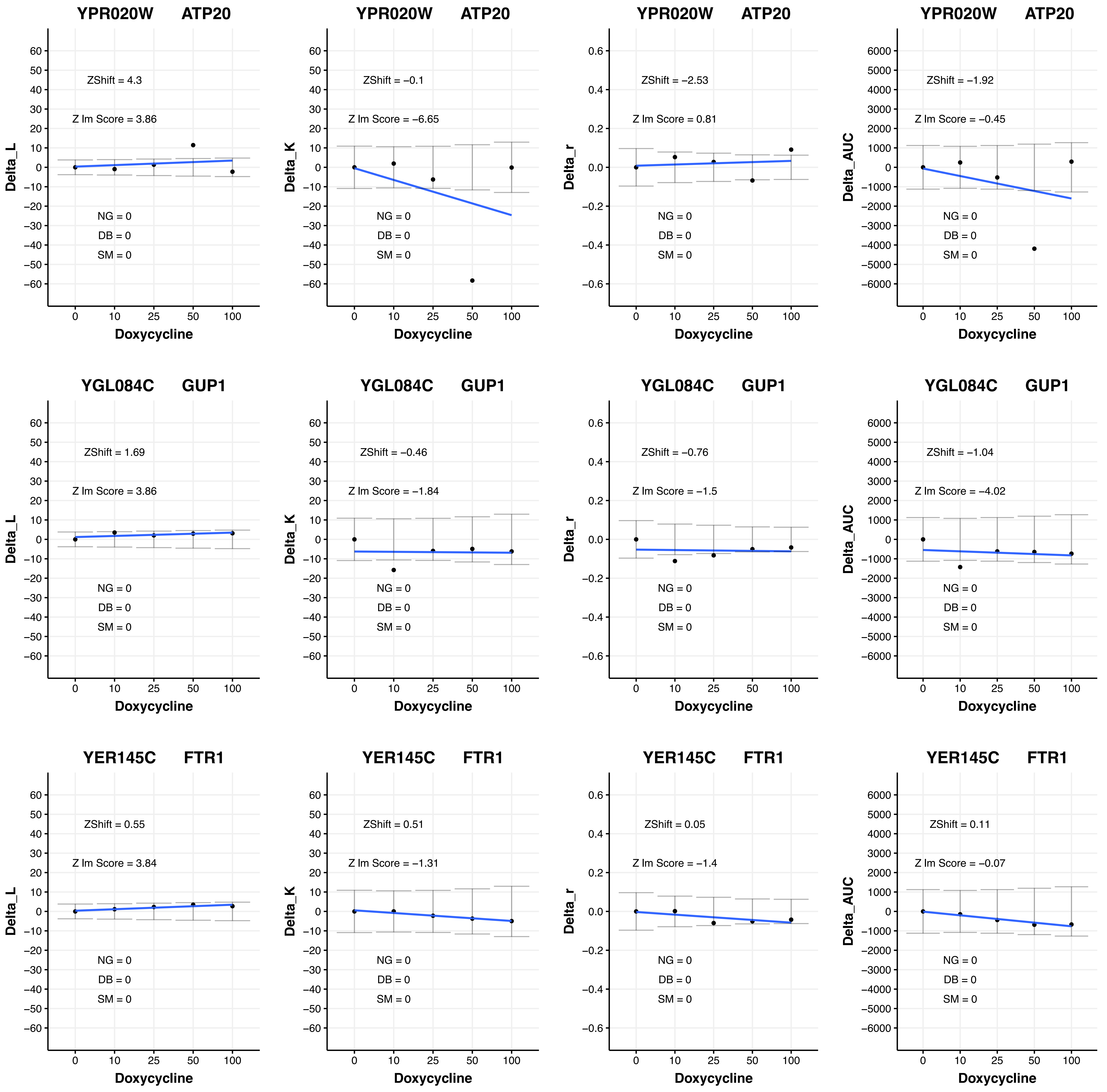
YPR020W ATP20
ZShift = 4.3
Z lm Score = 3.86
NG = 0
DB = 0
SM = 0
Delta_L
Doxycycline
YPR020W ATP20
ZShift = −0.1
Z lm Score = −6.65
NG = 0
DB = 0
SM = 0
Delta_K
Doxycycline
YPR020W ATP20
ZShift = −2.53
Z lm Score = 0.81
NG = 0
DB = 0
SM = 0
Delta_r
Doxycycline
YPR020W ATP20
ZShift = −1.92
Z lm Score = −0.45
NG = 0
DB = 0
SM = 0
Delta_AUC
Doxycycline
YGL084C GUP1
ZShift = 1.69
Z lm Score = 3.86
NG = 0
DB = 0
SM = 0
Delta_L
Doxycycline
YGL084C GUP1
ZShift = −0.46
Z lm Score = −1.84
NG = 0
DB = 0
SM = 0
Delta_K
Doxycycline
YGL084C GUP1
ZShift = −0.76
Z lm Score = −1.5
NG = 0
DB = 0
SM = 0
Delta_r
Doxycycline
YGL084C GUP1
ZShift = −1.04
Z lm Score = −4.02
NG = 0
DB = 0
SM = 0
Delta_AUC
Doxycycline
YER145C FTR1
ZShift = 0.55
Z lm Score = 3.84
NG = 0
DB = 0
SM = 0
Delta_L
Doxycycline
YER145C FTR1
ZShift = 0.51
Z lm Score = −1.31
NG = 0
DB = 0
SM = 0
Delta_K
Doxycycline
YER145C FTR1
ZShift = 0.05
Z lm Score = −1.4
NG = 0
DB = 0
SM = 0
Delta_r
Doxycycline
YER145C FTR1
ZShift = 0.11
Z lm Score = −0.07
NG = 0
DB = 0
SM = 0
Delta_AUC
Doxycycline

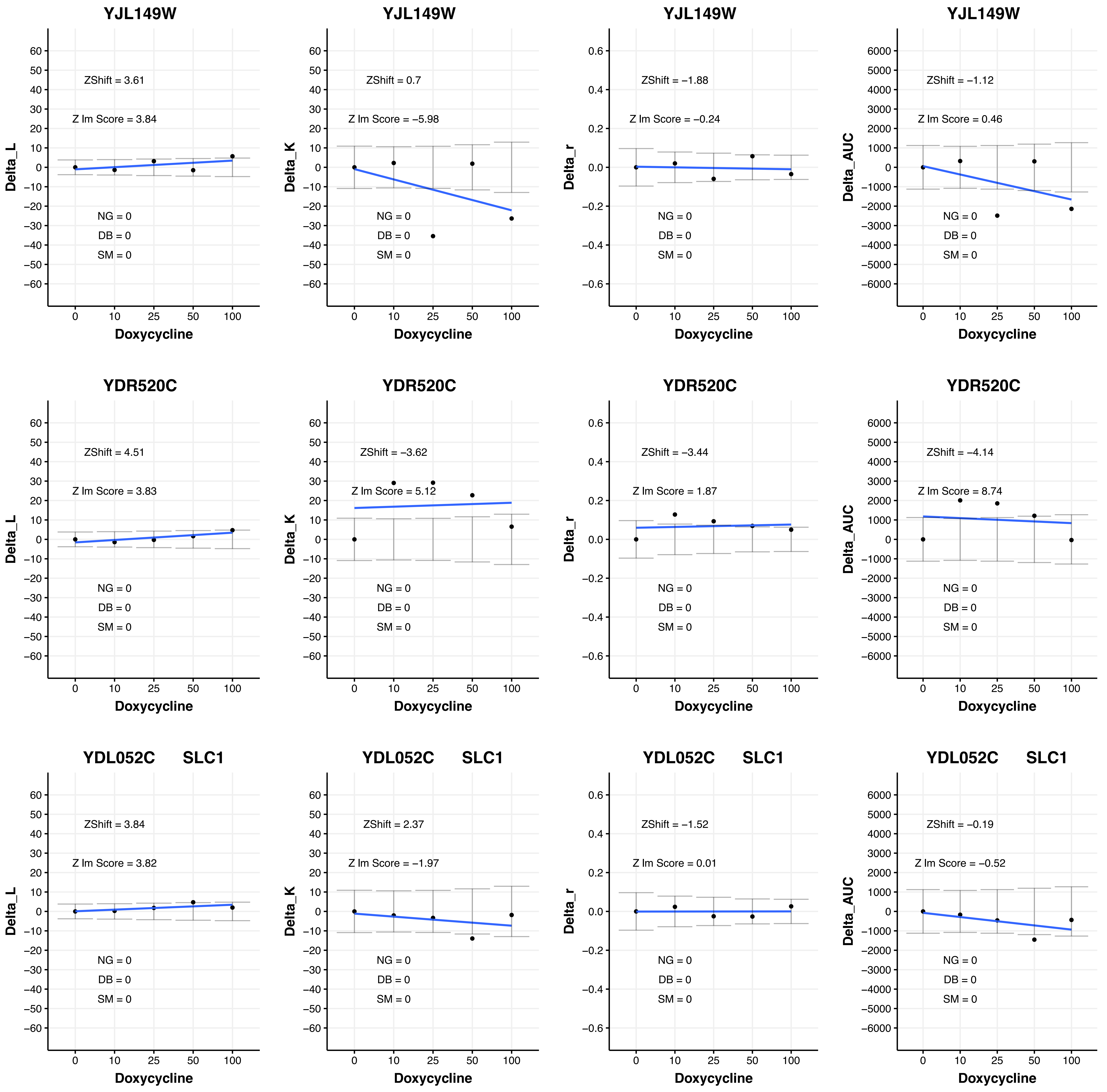

YJL149W
ZShift = 3.61
Z lm Score = 3.84
NG = 0
DB = 0
SM = 0
Delta_L
Doxycycline
YJL149W
ZShift = 0.7
Z lm Score = −5.98
NG = 0
DB = 0
SM = 0
Delta_K
Doxycycline
YJL149W
ZShift = −1.88
Z lm Score = −0.24
NG = 0
DB = 0
SM = 0
Delta_r
Doxycycline
YJL149W
ZShift = −1.12
Z lm Score = 0.46
NG = 0
DB = 0
SM = 0
Delta_AUC
Doxycycline
YDR520C
ZShift = 4.51
Z lm Score = 3.83
NG = 0
DB = 0
SM = 0
Delta_L
Doxycycline
YDR520C
ZShift = −3.62
Z lm Score = 5.12
NG = 0
DB = 0
SM = 0
Delta_K
Doxycycline
YDR520C
ZShift = −3.44
Z lm Score = 1.87
NG = 0
DB = 0
SM = 0
Delta_r
Doxycycline
YDR520C
ZShift = −4.14
Z lm Score = 8.74
NG = 0
DB = 0
SM = 0
Delta_AUC
Doxycycline
YDL052C SLC1
ZShift = 3.84
Z lm Score = 3.82
NG = 0
DB = 0
SM = 0
Delta_L
Doxycycline
YDL052C SLC1
ZShift = 2.37
Z lm Score = −1.97
NG = 0
DB = 0
SM = 0
Delta_K
Doxycycline
YDL052C SLC1
ZShift = −1.52
Z lm Score = 0.01
NG = 0
DB = 0
SM = 0
Delta_r
Doxycycline
YDL052C SLC1
ZShift = −0.19
Z lm Score = −0.52
NG = 0
DB = 0
SM = 0
Delta_AUC
Doxycycline

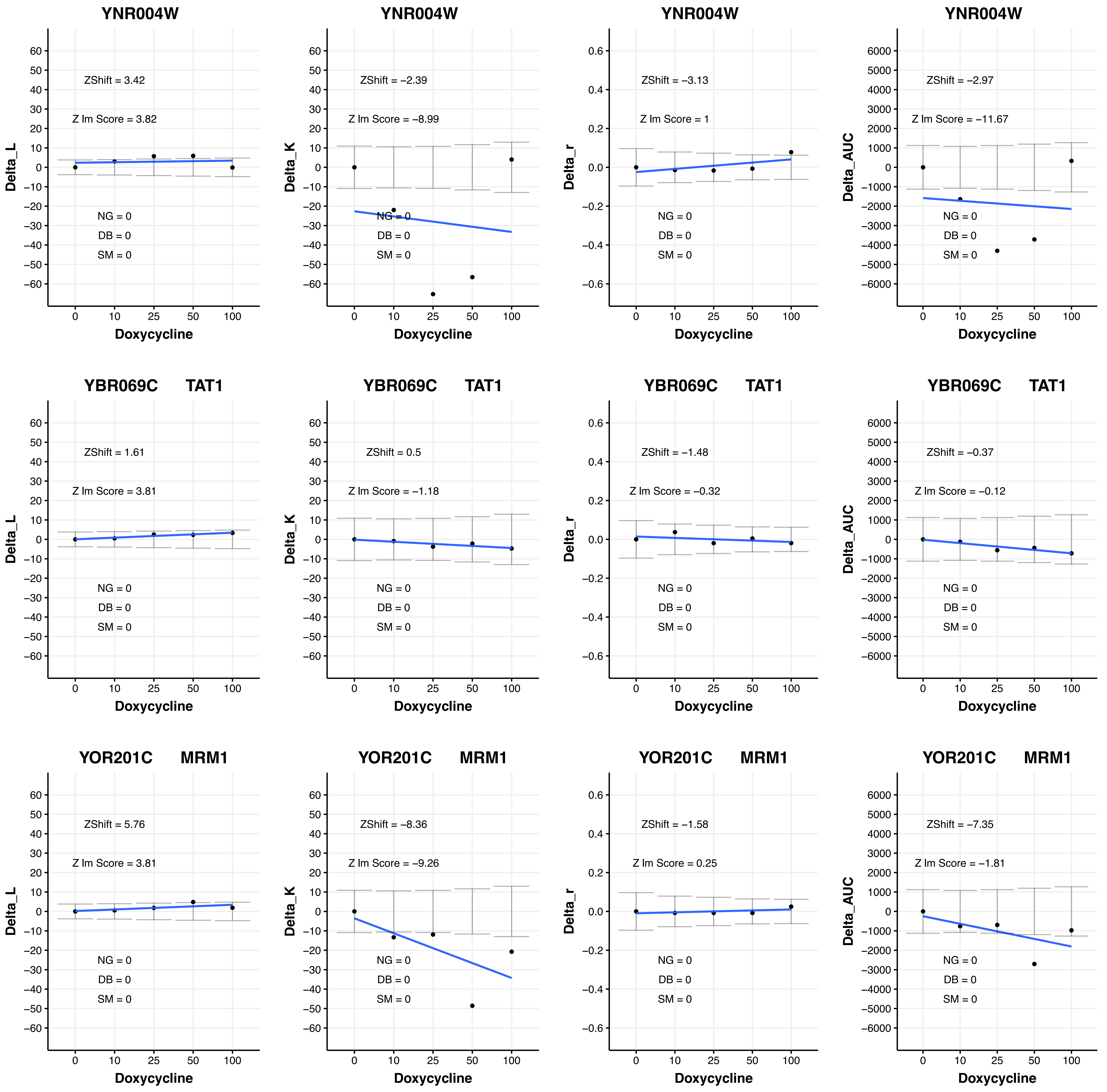
YNR004W
ZShift = 3.42
Z lm Score = 3.82
NG = 0
DB = 0
SM = 0
Delta_L
Doxycycline
YNR004W
ZShift = −2.39
Z lm Score = −8.99
NG = 0
DB = 0
SM = 0
Delta_K
Doxycycline
YNR004W
ZShift = −3.13
Z lm Score = 1
NG = 0
DB = 0
SM = 0
Delta_r
Doxycycline
YNR004W
ZShift = −2.97
Z lm Score = −11.67
NG = 0
DB = 0
SM = 0
Delta_AUC
Doxycycline
YBR069C TAT1
ZShift = 1.61
Z lm Score = 3.81
NG = 0
DB = 0
SM = 0
Delta_L
Doxycycline
YBR069C TAT1
ZShift = 0.5
Z lm Score = −1.18
NG = 0
DB = 0
SM = 0
Delta_K
Doxycycline
YBR069C TAT1
ZShift = −1.48
Z lm Score = −0.32
NG = 0
DB = 0
SM = 0
Delta_r
Doxycycline
YBR069C TAT1
ZShift = −0.37
Z lm Score = −0.12
NG = 0
DB = 0
SM = 0
Delta_AUC
Doxycycline
YOR201C MRM1
ZShift = 5.76
Z lm Score = 3.81
NG = 0
DB = 0
SM = 0
Delta_L
Doxycycline
YOR201C MRM1
ZShift = −8.36
Z lm Score = −9.26
NG = 0
DB = 0
SM = 0
Delta_K
Doxycycline
YOR201C MRM1
ZShift = −1.58
Z lm Score = 0.25
NG = 0
DB = 0
SM = 0
Delta_r
Doxycycline
YOR201C MRM1
ZShift = −7.35
Z lm Score = −1.81
NG = 0
DB = 0
SM = 0
Delta_AUC
Doxycycline

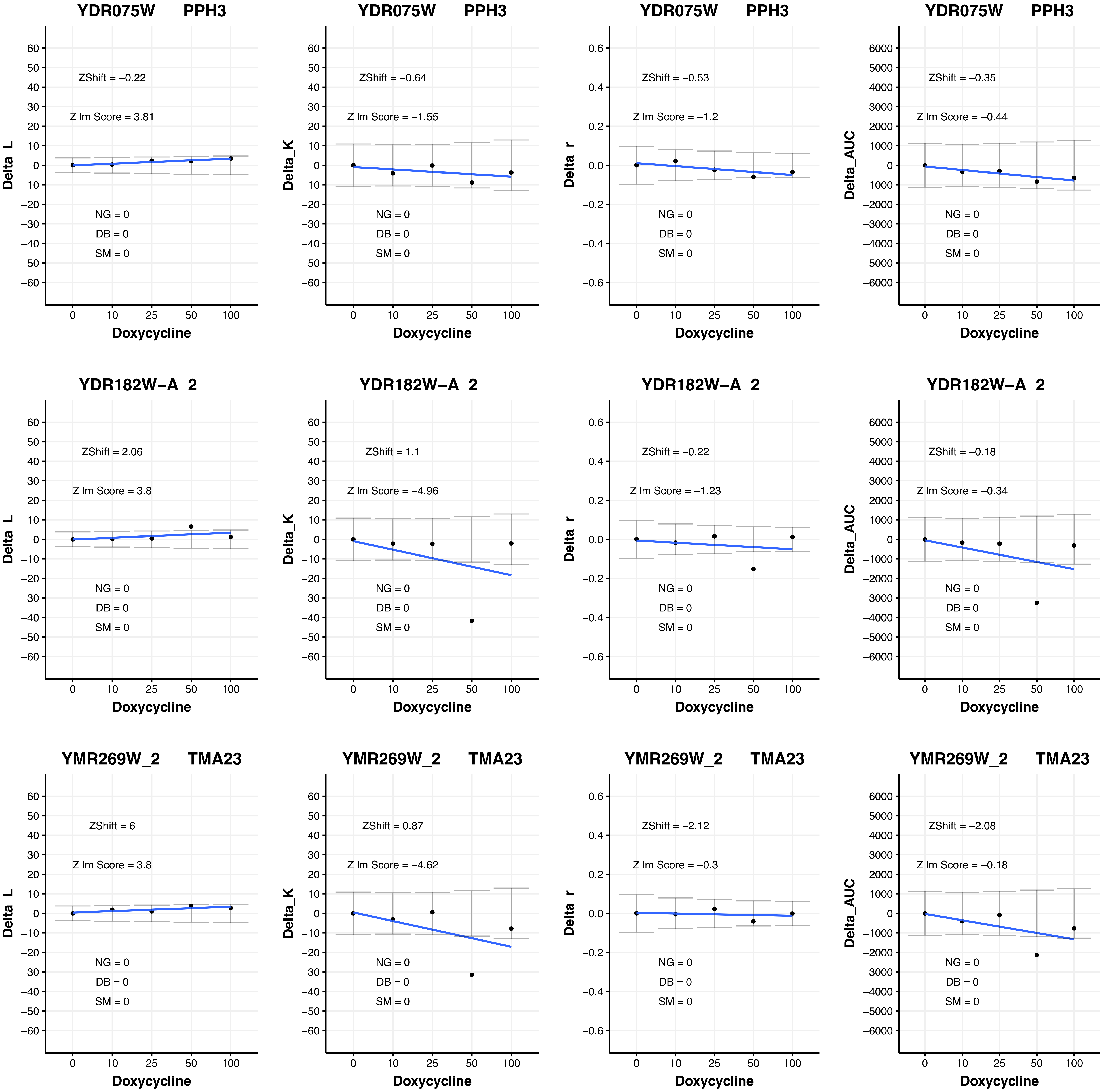

YDR075W PPH3
ZShift = −0.22
Z lm Score = 3.81
NG = 0
DB = 0
SM = 0
Delta_L
Doxycycline
YDR075W PPH3
ZShift = −0.64
Z lm Score = −1.55
NG = 0
DB = 0
SM = 0
Delta_K
Doxycycline
YDR075W PPH3
ZShift = −0.53
Z lm Score = −1.2
NG = 0
DB = 0
SM = 0
Delta_r
Doxycycline
YDR075W PPH3
ZShift = −0.35
Z lm Score = −0.44
NG = 0
DB = 0
SM = 0
Delta_AUC
Doxycycline
YDR182W−A_2
ZShift = 2.06
Z lm Score = 3.8
NG = 0
DB = 0
SM = 0
Delta_L
Doxycycline
YDR182W−A_2
ZShift = 1.1
Z lm Score = −4.96
NG = 0
DB = 0
SM = 0
Delta_K
Doxycycline
YDR182W−A_2
ZShift = −0.22
Z lm Score = −1.23
NG = 0
DB = 0
SM = 0
Delta_r
Doxycycline
YDR182W−A_2
ZShift = −0.18
Z lm Score = −0.34
NG = 0
DB = 0
SM = 0
Delta_AUC
Doxycycline
YMR269W_2 TMA23
ZShift = 6
Z lm Score = 3.8
NG = 0
DB = 0
SM = 0
Delta_L
Doxycycline
YMR269W_2 TMA23
ZShift = 0.87
Z lm Score = −4.62
NG = 0
DB = 0
SM = 0
Delta_K
Doxycycline
YMR269W_2 TMA23
ZShift = −2.12
Z lm Score = −0.3
NG = 0
DB = 0
SM = 0
Delta_r
Doxycycline
YMR269W_2 TMA23
ZShift = −2.08
Z lm Score = −0.18
NG = 0
DB = 0
SM = 0
Delta_AUC
Doxycycline

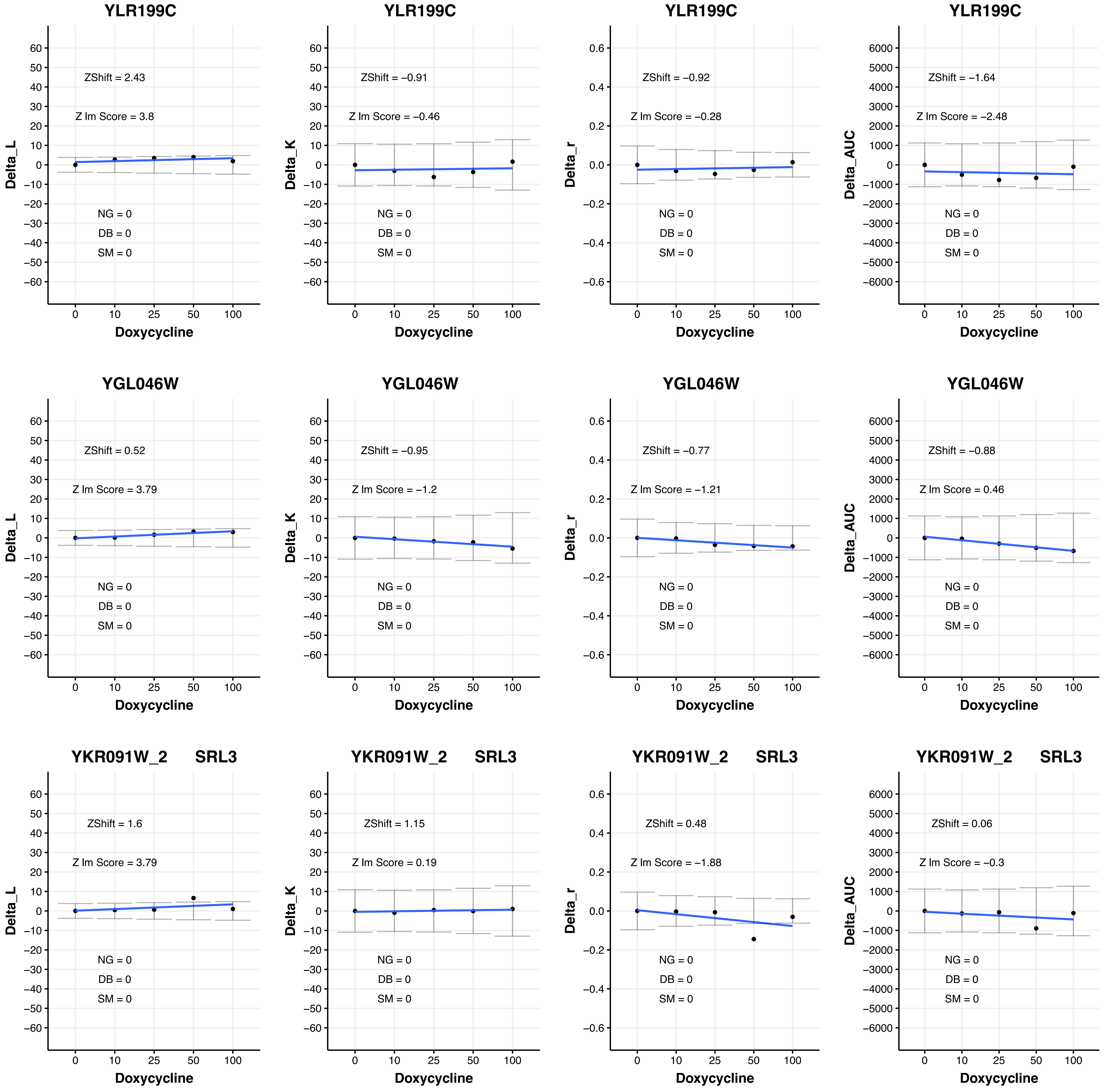
YLR199C
ZShift = 2.43
Z lm Score = 3.8
NG = 0
DB = 0
SM = 0
Delta_L
Doxycycline
YLR199C
ZShift = −0.91
Z lm Score = −0.46
NG = 0
DB = 0
SM = 0
Delta_K
Doxycycline
YLR199C
ZShift = −0.92
Z lm Score = −0.28
NG = 0
DB = 0
SM = 0
Delta_r
Doxycycline
YLR199C
ZShift = −1.64
Z lm Score = −2.48
NG = 0
DB = 0
SM = 0
Delta_AUC
Doxycycline
YGL046W
ZShift = 0.52
Z lm Score = 3.79
NG = 0
DB = 0
SM = 0
Delta_L
Doxycycline
YGL046W
ZShift = −0.95
Z lm Score = −1.2
NG = 0
DB = 0
SM = 0
Delta_K
Doxycycline
YGL046W
ZShift = −0.77
Z lm Score = −1.21
NG = 0
DB = 0
SM = 0
Delta_r
Doxycycline
YGL046W
ZShift = −0.88
Z lm Score = 0.46
NG = 0
DB = 0
SM = 0
Delta_AUC
Doxycycline
YKR091W_2 SRL3
ZShift = 1.6
Z lm Score = 3.79
NG = 0
DB = 0
SM = 0
Delta_L
Doxycycline
YKR091W_2 SRL3
ZShift = 1.15
Z lm Score = 0.19
NG = 0
DB = 0
SM = 0
Delta_K
Doxycycline
YKR091W_2 SRL3
ZShift = 0.48
Z lm Score = −1.88
NG = 0
DB = 0
SM = 0
Delta_r
Doxycycline
YKR091W_2 SRL3
ZShift = 0.06
Z lm Score = −0.3
NG = 0
DB = 0
SM = 0
Delta_AUC
Doxycycline

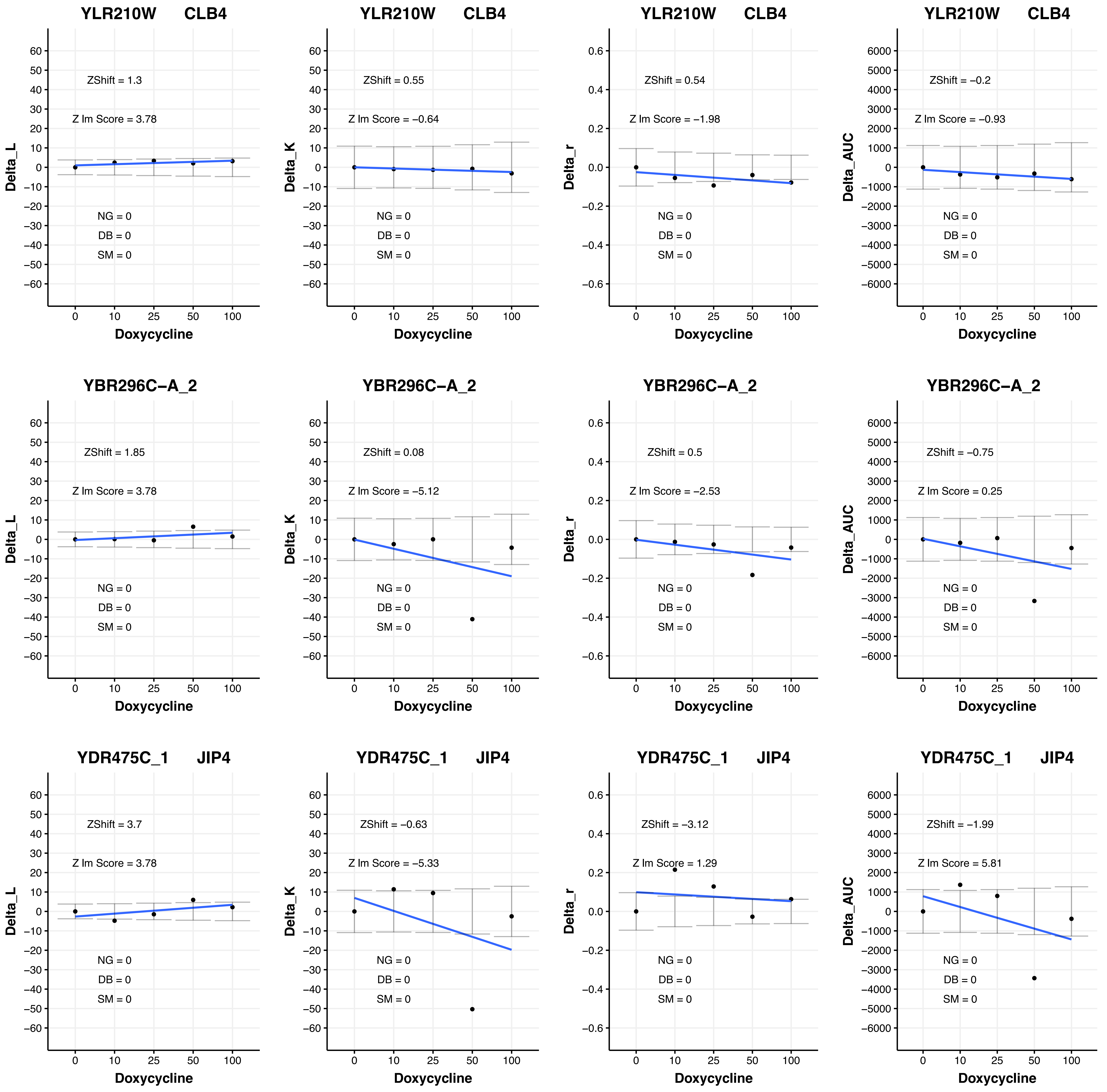

YLR210W CLB4
ZShift = 1.3
Z lm Score = 3.78
NG = 0
DB = 0
SM = 0
Delta_L
Doxycycline
YLR210W CLB4
ZShift = 0.55
Z lm Score = −0.64
NG = 0
DB = 0
SM = 0
Delta_K
Doxycycline
YLR210W CLB4
ZShift = 0.54
Z lm Score = −1.98
NG = 0
DB = 0
SM = 0
Delta_r
Doxycycline
YLR210W CLB4
ZShift = −0.2
Z lm Score = −0.93
NG = 0
DB = 0
SM = 0
Delta_AUC
Doxycycline
YBR296C−A_2
ZShift = 1.85
Z lm Score = 3.78
NG = 0
DB = 0
SM = 0
Delta_L
Doxycycline
YBR296C−A_2
ZShift = 0.08
Z lm Score = −5.12
NG = 0
DB = 0
SM = 0
Delta_K
Doxycycline
YBR296C−A_2
ZShift = 0.5
Z lm Score = −2.53
NG = 0
DB = 0
SM = 0
Delta_r
Doxycycline
YBR296C−A_2
ZShift = −0.75
Z lm Score = 0.25
NG = 0
DB = 0
SM = 0
Delta_AUC
Doxycycline
YDR475C_1 JIP4
ZShift = 3.7
Z lm Score = 3.78
NG = 0
DB = 0
SM = 0
Delta_L
Doxycycline
YDR475C_1 JIP4
ZShift = −0.63
Z lm Score = −5.33
NG = 0
DB = 0
SM = 0
Delta_K
Doxycycline
YDR475C_1 JIP4
ZShift = −3.12
Z lm Score = 1.29
NG = 0
DB = 0
SM = 0
Delta_r
Doxycycline
YDR475C_1 JIP4
ZShift = −1.99
Z lm Score = 5.81
NG = 0
DB = 0
SM = 0
Delta_AUC
Doxycycline

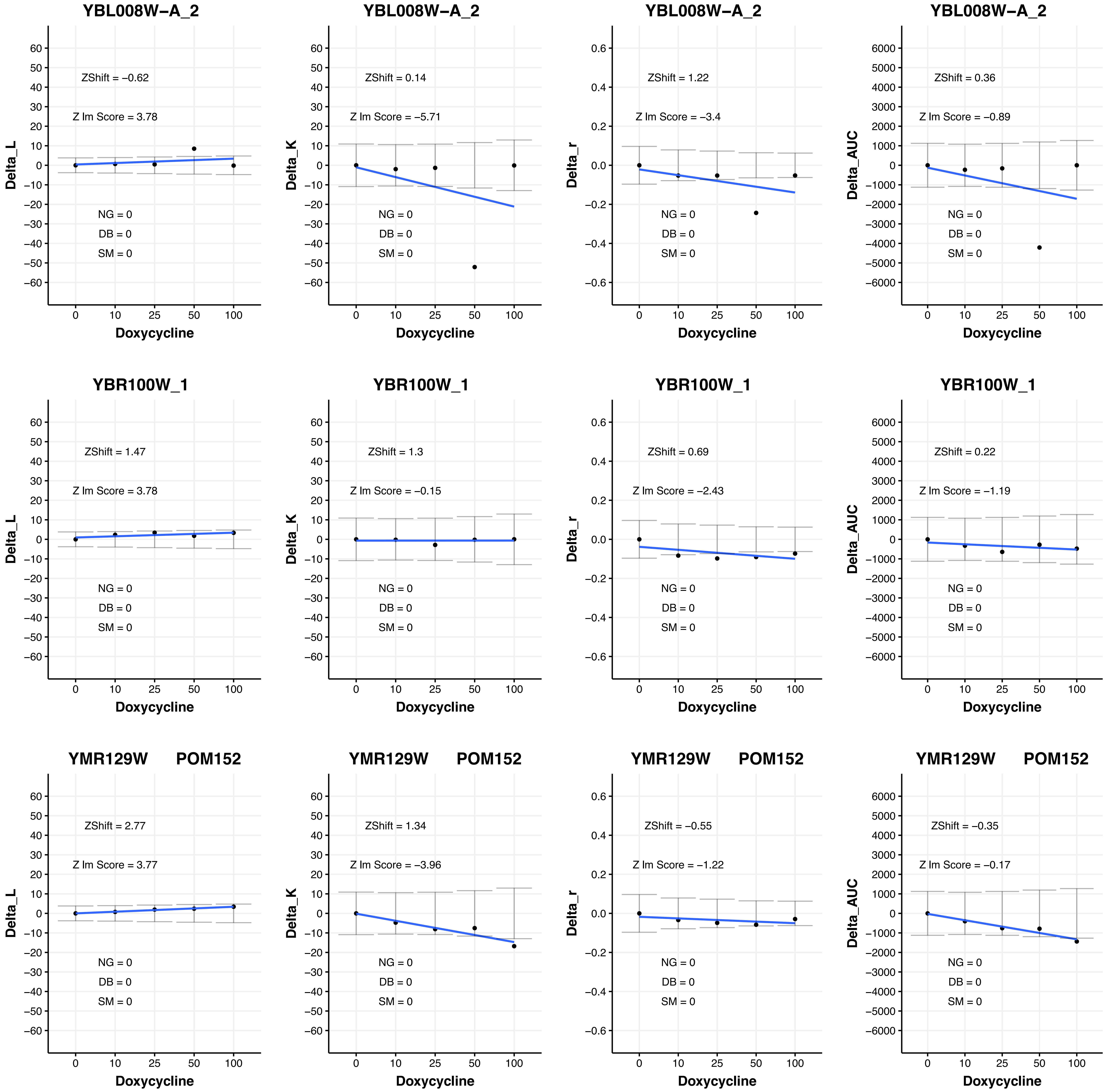

YBL008W−A_2
ZShift = −0.62
Z lm Score = 3.78
NG = 0
DB = 0
SM = 0
Delta_L
Doxycycline
YBL008W−A_2
ZShift = 0.14
Z lm Score = −5.71
NG = 0
DB = 0
SM = 0
Delta_K
Doxycycline
YBL008W−A_2
ZShift = 1.22
Z lm Score = −3.4
NG = 0
DB = 0
SM = 0
Delta_r
Doxycycline
YBL008W−A_2
ZShift = 0.36
Z lm Score = −0.89
NG = 0
DB = 0
SM = 0
Delta_AUC
Doxycycline
YBR100W_1
ZShift = 1.47
Z lm Score = 3.78
NG = 0
DB = 0
SM = 0
Delta_L
Doxycycline
YBR100W_1
ZShift = 1.3
Z lm Score = −0.15
NG = 0
DB = 0
SM = 0
Delta_K
Doxycycline
YBR100W_1
ZShift = 0.69
Z lm Score = −2.43
NG = 0
DB = 0
SM = 0
Delta_r
Doxycycline
YBR100W_1
ZShift = 0.22
Z lm Score = −1.19
NG = 0
DB = 0
SM = 0
Delta_AUC
Doxycycline
YMR129W POM152
ZShift = 2.77
Z lm Score = 3.77
NG = 0
DB = 0
SM = 0
Delta_L
Doxycycline
YMR129W POM152
ZShift = 1.34
Z lm Score = −3.96
NG = 0
DB = 0
SM = 0
Delta_K
Doxycycline
YMR129W POM152
ZShift = −0.55
Z lm Score = −1.22
NG = 0
DB = 0
SM = 0
Delta_r
Doxycycline
YMR129W POM152
ZShift = −0.35
Z lm Score = −0.17
NG = 0
DB = 0
SM = 0
Delta_AUC
Doxycycline

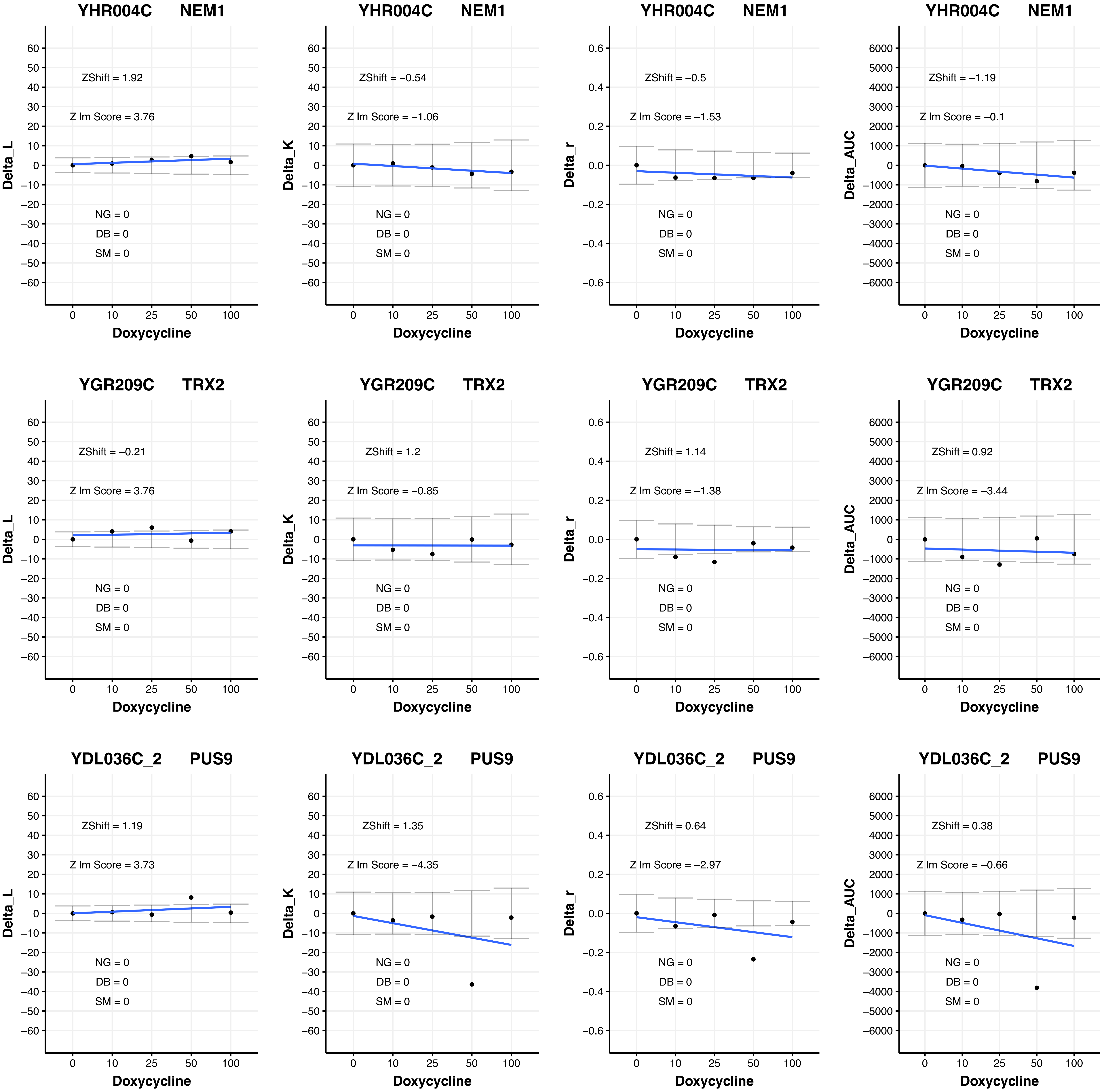

YHR004C NEM1
ZShift = 1.92
Z lm Score = 3.76
NG = 0
DB = 0
SM = 0
Delta_L
Doxycycline
YHR004C NEM1
ZShift = −0.54
Z lm Score = −1.06
NG = 0
DB = 0
SM = 0
Delta_K
Doxycycline
YHR004C NEM1
ZShift = −0.5
Z lm Score = −1.53
NG = 0
DB = 0
SM = 0
Delta_r
Doxycycline
YHR004C NEM1
ZShift = −1.19
Z lm Score = −0.1
NG = 0
DB = 0
SM = 0
Delta_AUC
Doxycycline
YGR209C TRX2
ZShift = −0.21
Z lm Score = 3.76
NG = 0
DB = 0
SM = 0
Delta_L
Doxycycline
YGR209C TRX2
ZShift = 1.2
Z lm Score = −0.85
NG = 0
DB = 0
SM = 0
Delta_K
Doxycycline
YGR209C TRX2
ZShift = 1.14
Z lm Score = −1.38
NG = 0
DB = 0
SM = 0
Delta_r
Doxycycline
YGR209C TRX2
ZShift = 0.92
Z lm Score = −3.44
NG = 0
DB = 0
SM = 0
Delta_AUC
Doxycycline
YDL036C_2 PUS9
ZShift = 1.19
Z lm Score = 3.73
NG = 0
DB = 0
SM = 0
Delta_L
Doxycycline
YDL036C_2 PUS9
ZShift = 1.35
Z lm Score = −4.35
NG = 0
DB = 0
SM = 0
Delta_K
Doxycycline
YDL036C_2 PUS9
ZShift = 0.64
Z lm Score = −2.97
NG = 0
DB = 0
SM = 0
Delta_r
Doxycycline
YDL036C_2 PUS9
ZShift = 0.38
Z lm Score = −0.66
NG = 0
DB = 0
SM = 0
Delta_AUC
Doxycycline

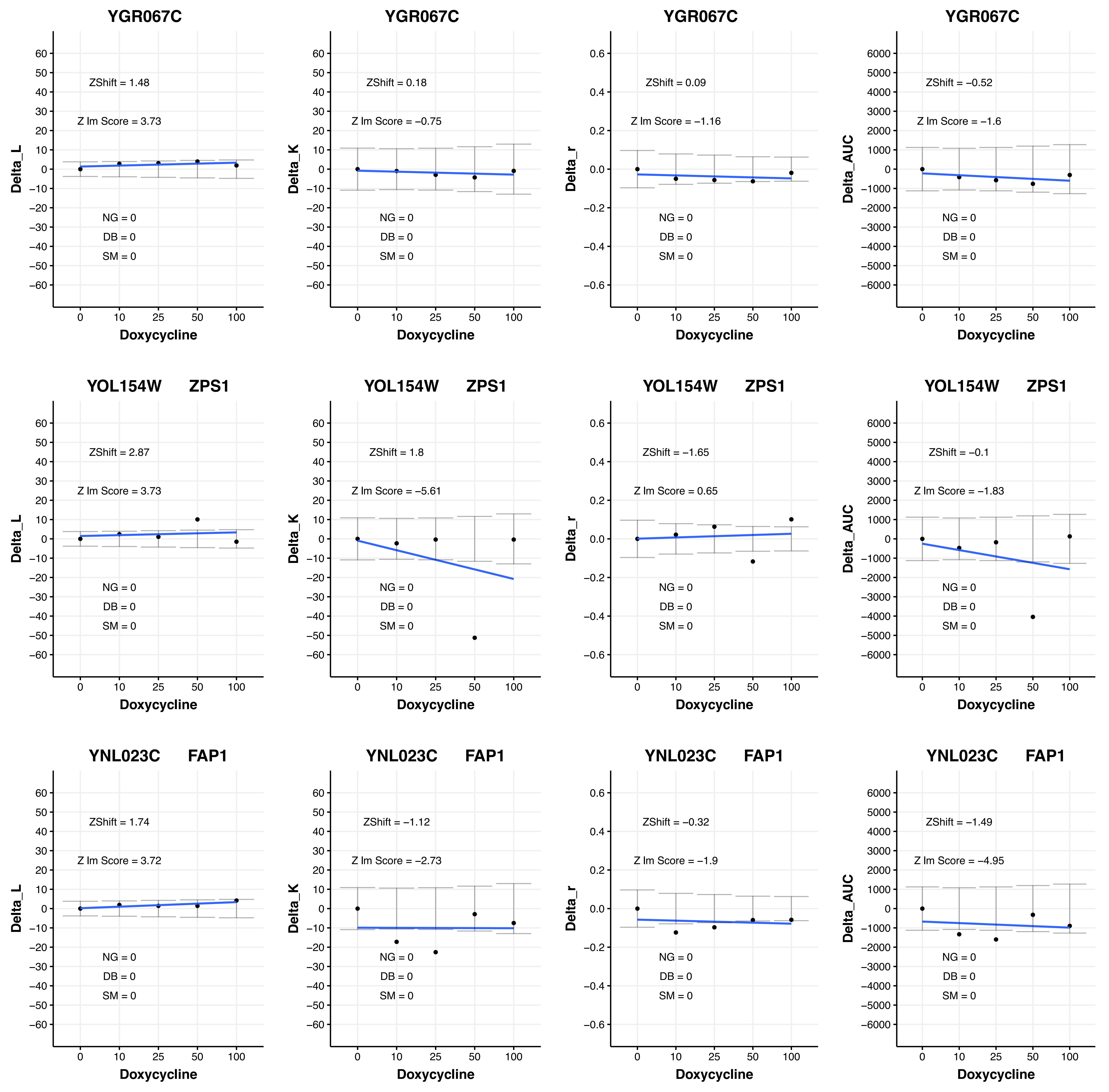
YGR067C
ZShift = 1.48
Z lm Score = 3.73
NG = 0
DB = 0
SM = 0
Delta_L
Doxycycline
YGR067C
ZShift = 0.18
Z lm Score = −0.75
NG = 0
DB = 0
SM = 0
Delta_K
Doxycycline
YGR067C
ZShift = 0.09
Z lm Score = −1.16
NG = 0
DB = 0
SM = 0
Delta_r
Doxycycline
YGR067C
ZShift = −0.52
Z lm Score = −1.6
NG = 0
DB = 0
SM = 0
Delta_AUC
Doxycycline
YOL154W ZPS1
ZShift = 2.87
Z lm Score = 3.73
NG = 0
DB = 0
SM = 0
Delta_L
Doxycycline
YOL154W ZPS1
ZShift = 1.8
Z lm Score = −5.61
NG = 0
DB = 0
SM = 0
Delta_K
Doxycycline
YOL154W ZPS1
ZShift = −1.65
Z lm Score = 0.65
NG = 0
DB = 0
SM = 0
Delta_r
Doxycycline
YOL154W ZPS1
ZShift = −0.1
Z lm Score = −1.83
NG = 0
DB = 0
SM = 0
Delta_AUC
Doxycycline
YNL023C FAP1
ZShift = 1.74
Z lm Score = 3.72
NG = 0
DB = 0
SM = 0
Delta_L
Doxycycline
YNL023C FAP1
ZShift = −1.12
Z lm Score = −2.73
NG = 0
DB = 0
SM = 0
Delta_K
Doxycycline
YNL023C FAP1
ZShift = −0.32
Z lm Score = −1.9
NG = 0
DB = 0
SM = 0
Delta_r
Doxycycline
YNL023C FAP1
ZShift = −1.49
Z lm Score = −4.95
NG = 0
DB = 0
SM = 0
Delta_AUC
Doxycycline

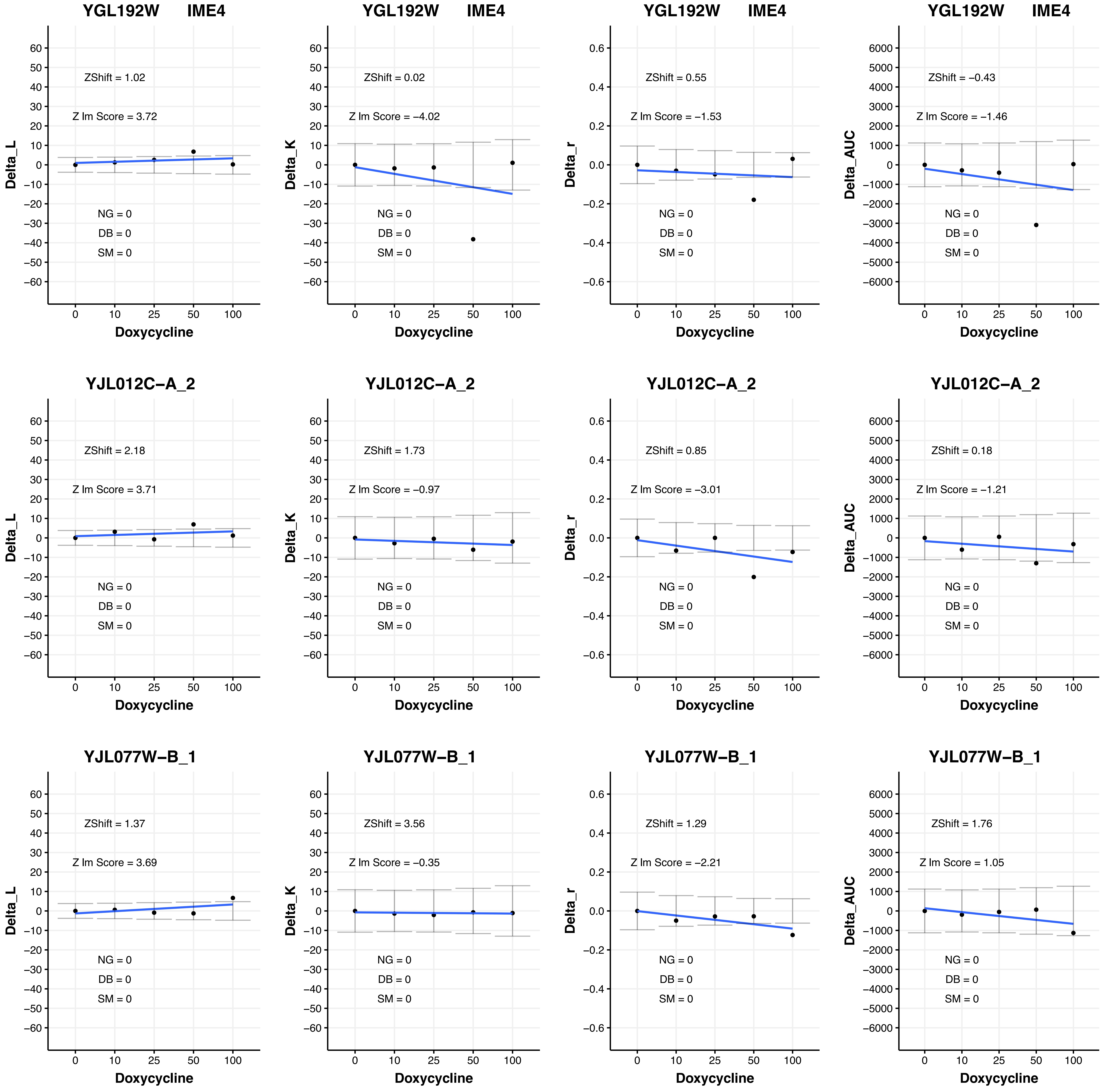
YGL192W IME4
ZShift = 1.02
Z lm Score = 3.72
NG = 0
DB = 0
SM = 0
Delta_L
Doxycycline
YGL192W IME4
ZShift = 0.02
Z lm Score = −4.02
NG = 0
DB = 0
SM = 0
Delta_K
Doxycycline
YGL192W IME4
ZShift = 0.55
Z lm Score = −1.53
NG = 0
DB = 0
SM = 0
Delta_r
Doxycycline
YGL192W IME4
ZShift = −0.43
Z lm Score = −1.46
NG = 0
DB = 0
SM = 0
Delta_AUC
Doxycycline
YJL012C−A_2
ZShift = 2.18
Z lm Score = 3.71
NG = 0
DB = 0
SM = 0
Delta_L
Doxycycline
YJL012C−A_2
ZShift = 1.73
Z lm Score = −0.97
NG = 0
DB = 0
SM = 0
Delta_K
Doxycycline
YJL012C−A_2
ZShift = 0.85
Z lm Score = −3.01
NG = 0
DB = 0
SM = 0
Delta_r
Doxycycline
YJL012C−A_2
ZShift = 0.18
Z lm Score = −1.21
NG = 0
DB = 0
SM = 0
Delta_AUC
Doxycycline
YJL077W−B_1
ZShift = 1.37
Z lm Score = 3.69
NG = 0
DB = 0
SM = 0
Delta_L
Doxycycline
YJL077W−B_1
ZShift = 3.56
Z lm Score = −0.35
NG = 0
DB = 0
SM = 0
Delta_K
Doxycycline
YJL077W−B_1
ZShift = 1.29
Z lm Score = −2.21
NG = 0
DB = 0
SM = 0
Delta_r
Doxycycline
YJL077W−B_1
ZShift = 1.76
Z lm Score = 1.05
NG = 0
DB = 0
SM = 0
Delta_AUC
Doxycycline

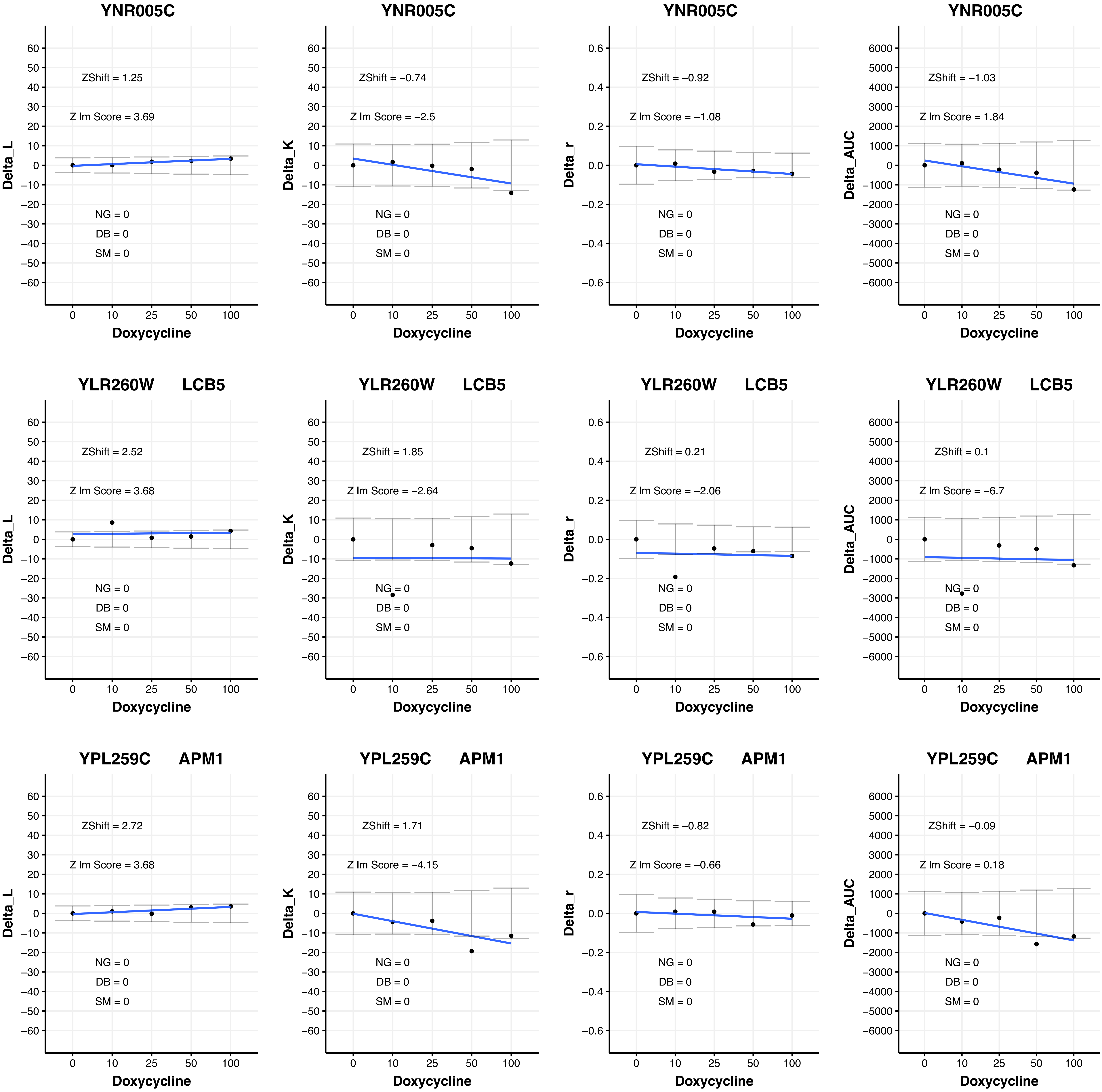

YNR005C
Delta_L
ZShift = 1.25
Z lm Score = 3.69
NG = 0
DB = 0
SM = 0
Doxycycline
YNR005C
Delta_K
ZShift = −0.74
Z lm Score = −2.5
NG = 0
DB = 0
SM = 0
Doxycycline
YNR005C
Delta_r
ZShift = −0.92
Z lm Score = −1.08
NG = 0
DB = 0
SM = 0
Doxycycline
YNR005C
Delta_AUC
ZShift = −1.03
Z lm Score = 1.84
NG = 0
DB = 0
SM = 0
Doxycycline
YLR260W LCB5
Delta_L
ZShift = 2.52
Z lm Score = 3.68
NG = 0
DB = 0
SM = 0
Doxycycline
YLR260W LCB5
Delta_K
ZShift = 1.85
Z lm Score = −2.64
NG = 0
DB = 0
SM = 0
Doxycycline
YLR260W LCB5
Delta_r
ZShift = 0.21
Z lm Score = −2.06
NG = 0
DB = 0
SM = 0
Doxycycline
YLR260W LCB5
Delta_AUC
ZShift = 0.1
Z lm Score = −6.7
NG = 0
DB = 0
SM = 0
Doxycycline
YPL259C APM1
Delta_L
ZShift = 2.72
Z lm Score = 3.68
NG = 0
DB = 0
SM = 0
Doxycycline
YPL259C APM1
Delta_K
ZShift = 1.71
Z lm Score = −4.15
NG = 0
DB = 0
SM = 0
Doxycycline
YPL259C APM1
Delta_r
ZShift = −0.82
Z lm Score = −0.66
NG = 0
DB = 0
SM = 0
Doxycycline
YPL259C APM1
Delta_AUC
ZShift = −0.09
Z lm Score = 0.18
NG = 0
DB = 0
SM = 0
Doxycycline

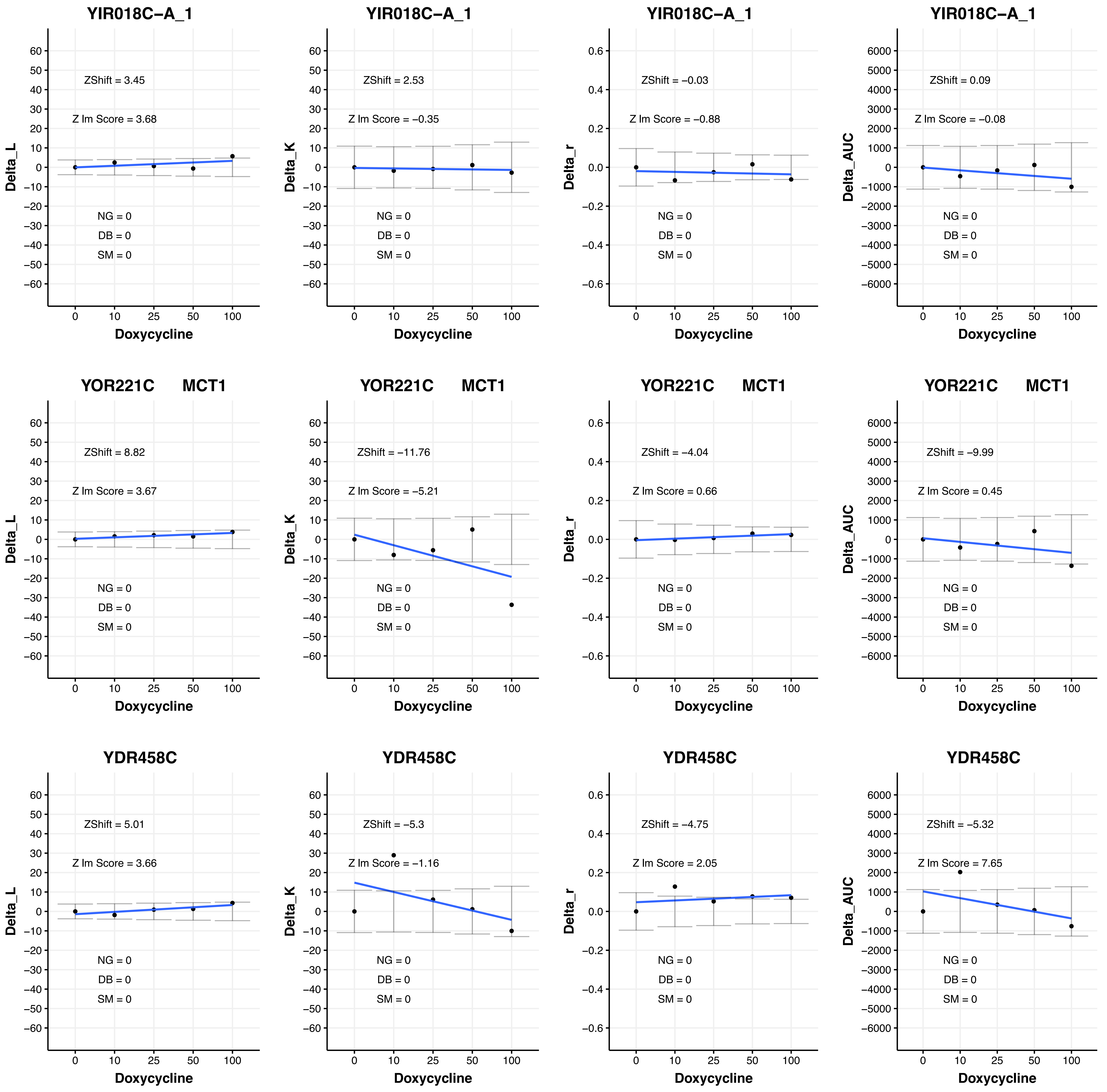

YIR018C−A_1
ZShift = 3.45
Z lm Score = 3.68
NG = 0
DB = 0
SM = 0
Delta_L
Doxycycline
YIR018C−A_1
ZShift = 2.53
Z lm Score = −0.35
NG = 0
DB = 0
SM = 0
Delta_K
Doxycycline
YIR018C−A_1
ZShift = −0.03
Z lm Score = −0.88
NG = 0
DB = 0
SM = 0
Delta_r
Doxycycline
YIR018C−A_1
ZShift = 0.09
Z lm Score = −0.08
NG = 0
DB = 0
SM = 0
Delta_AUC
Doxycycline
YOR221C MCT1
ZShift = 8.82
Z lm Score = 3.67
NG = 0
DB = 0
SM = 0
Delta_L
Doxycycline
YOR221C MCT1
ZShift = −11.76
Z lm Score = −5.21
NG = 0
DB = 0
SM = 0
Delta_K
Doxycycline
YOR221C MCT1
ZShift = −4.04
Z lm Score = 0.66
NG = 0
DB = 0
SM = 0
Delta_r
Doxycycline
YOR221C MCT1
ZShift = −9.99
Z lm Score = 0.45
NG = 0
DB = 0
SM = 0
Delta_AUC
Doxycycline
YDR458C
ZShift = 5.01
Z lm Score = 3.66
NG = 0
DB = 0
SM = 0
Delta_L
Doxycycline
YDR458C
ZShift = −5.3
Z lm Score = −1.16
NG = 0
DB = 0
SM = 0
Delta_K
Doxycycline
YDR458C
ZShift = −4.75
Z lm Score = 2.05
NG = 0
DB = 0
SM = 0
Delta_r
Doxycycline
YDR458C
ZShift = −5.32
Z lm Score = 7.65
NG = 0
DB = 0
SM = 0
Delta_AUC
Doxycycline

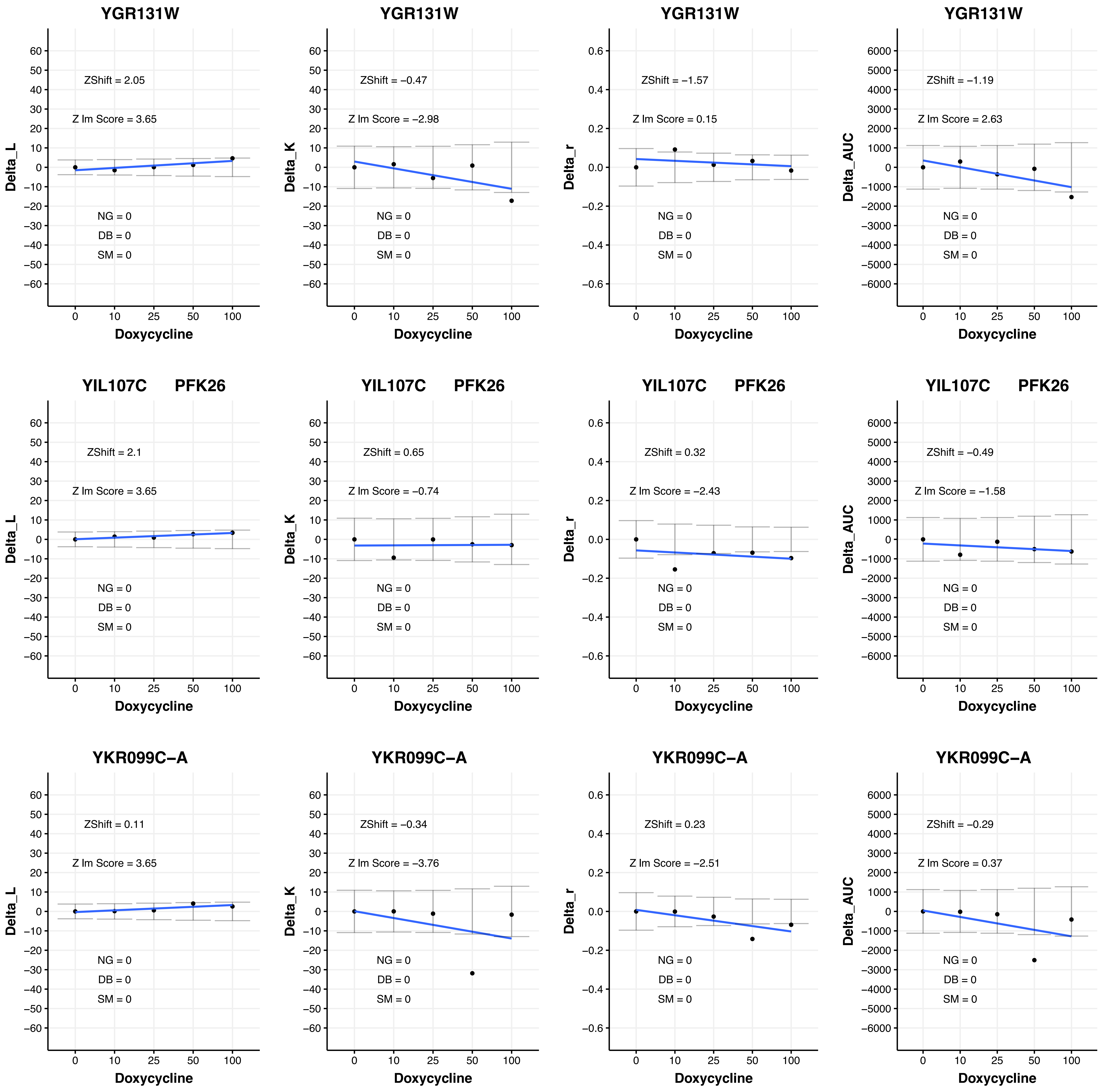

YGR131W
ZShift = 2.05
Z lm Score = 3.65
NG = 0
DB = 0
SM = 0
Delta_L
Doxycycline
YGR131W
ZShift = −0.47
Z lm Score = −2.98
NG = 0
DB = 0
SM = 0
Delta_K
Doxycycline
YGR131W
ZShift = −1.57
Z lm Score = 0.15
NG = 0
DB = 0
SM = 0
Delta_r
Doxycycline
YGR131W
ZShift = −1.19
Z lm Score = 2.63
NG = 0
DB = 0
SM = 0
Delta_AUC
Doxycycline
YIL107C PFK26
ZShift = 2.1
Z lm Score = 3.65
NG = 0
DB = 0
SM = 0
Delta_L
Doxycycline
YIL107C PFK26
ZShift = 0.65
Z lm Score = −0.74
NG = 0
DB = 0
SM = 0
Delta_K
Doxycycline
YIL107C PFK26
ZShift = 0.32
Z lm Score = −2.43
NG = 0
DB = 0
SM = 0
Delta_r
Doxycycline
YIL107C PFK26
ZShift = −0.49
Z lm Score = −1.58
NG = 0
DB = 0
SM = 0
Delta_AUC
Doxycycline
YKR099C−A
ZShift = 0.11
Z lm Score = 3.65
NG = 0
DB = 0
SM = 0
Delta_L
Doxycycline
YKR099C−A
ZShift = −0.34
Z lm Score = −3.76
NG = 0
DB = 0
SM = 0
Delta_K
Doxycycline
YKR099C−A
ZShift = 0.23
Z lm Score = −2.51
NG = 0
DB = 0
SM = 0
Delta_r
Doxycycline
YKR099C−A
ZShift = −0.29
Z lm Score = 0.37
NG = 0
DB = 0
SM = 0
Delta_AUC
Doxycycline

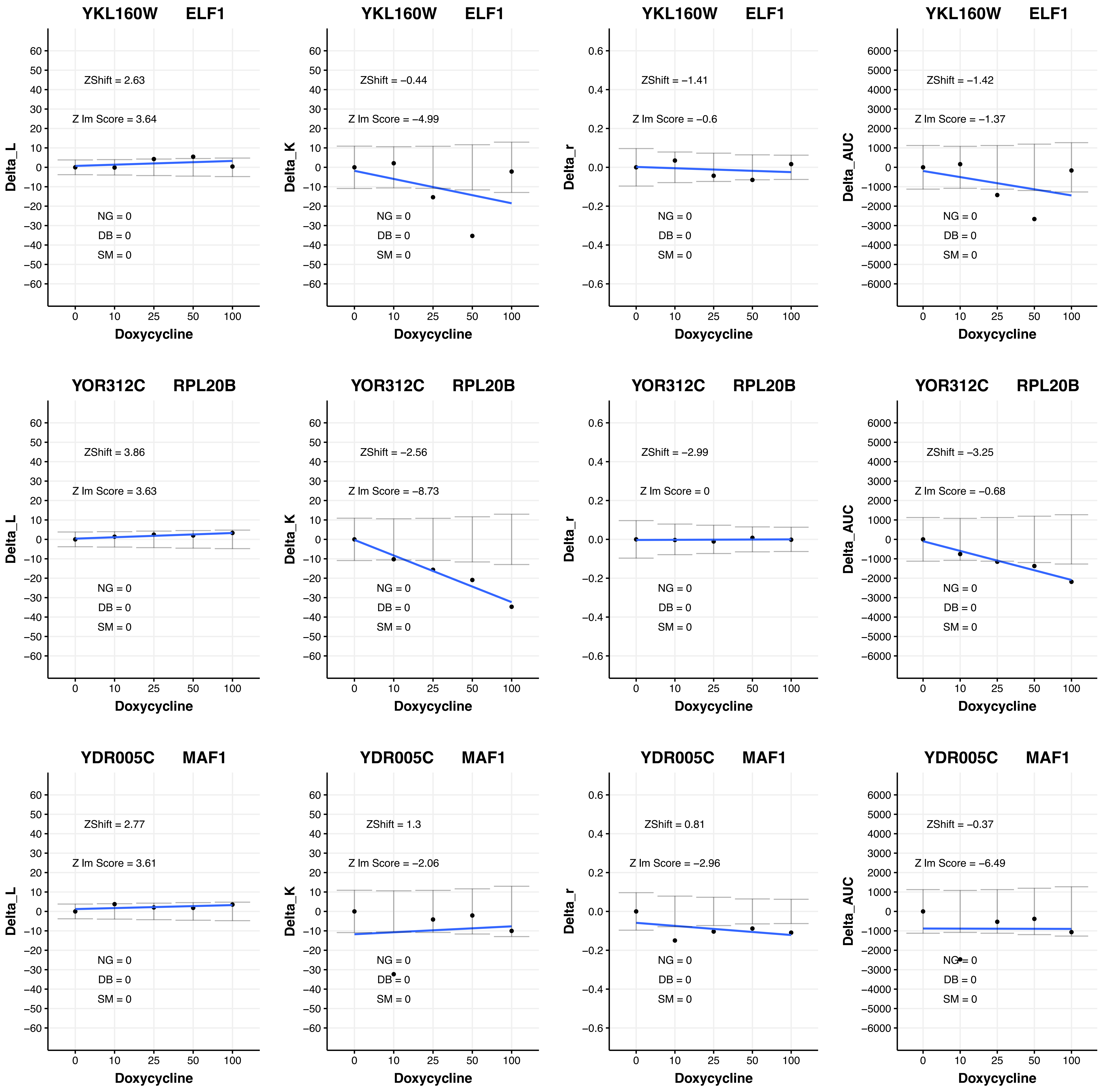

YKL160W ELF1
ZShift = 2.63
Z lm Score = 3.64
NG = 0
DB = 0
SM = 0
Delta_L
Doxycycline
YKL160W ELF1
ZShift = −0.44
Z lm Score = −4.99
NG = 0
DB = 0
SM = 0
Delta_K
Doxycycline
YKL160W ELF1
ZShift = −1.41
Z lm Score = −0.6
NG = 0
DB = 0
SM = 0
Delta_r
Doxycycline
YKL160W ELF1
ZShift = −1.42
Z lm Score = −1.37
NG = 0
DB = 0
SM = 0
Delta_AUC
Doxycycline
YOR312C RPL20B
ZShift = 3.86
Z lm Score = 3.63
NG = 0
DB = 0
SM = 0
Delta_L
Doxycycline
YOR312C RPL20B
ZShift = −2.56
Z lm Score = −8.73
NG = 0
DB = 0
SM = 0
Delta_K
Doxycycline
YOR312C RPL20B
ZShift = −2.99
Z lm Score = 0
NG = 0
DB = 0
SM = 0
Delta_r
Doxycycline
YOR312C RPL20B
ZShift = −3.25
Z lm Score = −0.68
NG = 0
DB = 0
SM = 0
Delta_AUC
Doxycycline
YDR005C MAF1
ZShift = 2.77
Z lm Score = 3.61
NG = 0
DB = 0
SM = 0
Delta_L
Doxycycline
YDR005C MAF1
ZShift = 1.3
Z lm Score = −2.06
NG = 0
DB = 0
SM = 0
Delta_K
Doxycycline
YDR005C MAF1
ZShift = 0.81
Z lm Score = −2.96
NG = 0
DB = 0
SM = 0
Delta_r
Doxycycline
YDR005C MAF1
ZShift = −0.37
Z lm Score = −6.49
NG = 0
DB = 0
SM = 0
Delta_AUC
Doxycycline

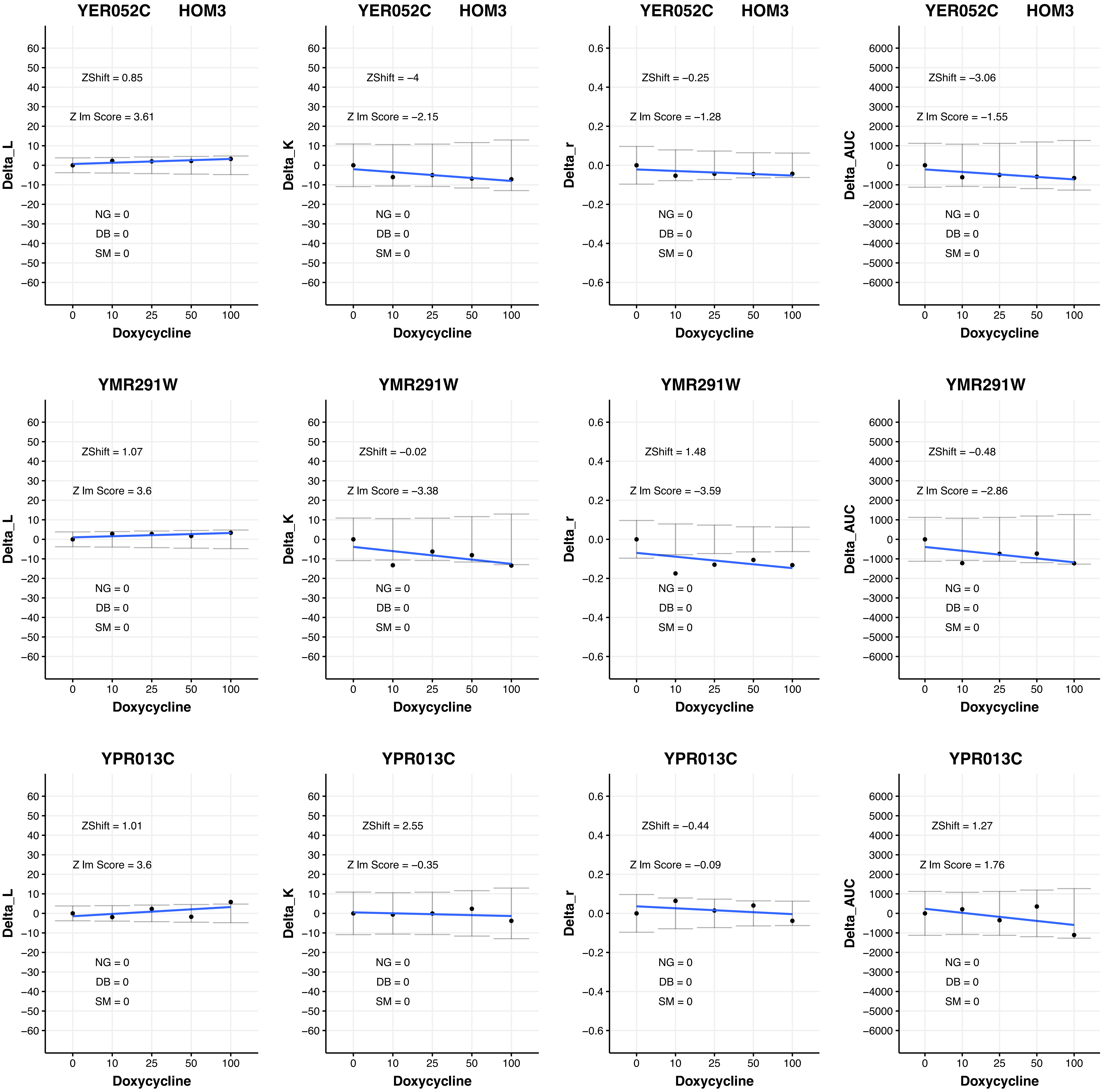

YER052C HOM3
ZShift = 0.85
Z lm Score = 3.61
NG = 0
DB = 0
SM = 0
Delta_L
Doxycycline
YER052C HOM3
ZShift = −4
Z lm Score = −2.15
NG = 0
DB = 0
SM = 0
Delta_K
Doxycycline
YER052C HOM3
ZShift = −0.25
Z lm Score = −1.28
NG = 0
DB = 0
SM = 0
Delta_r
Doxycycline
YER052C HOM3
ZShift = −3.06
Z lm Score = −1.55
NG = 0
DB = 0
SM = 0
Delta_AUC
Doxycycline
YMR291W
ZShift = 1.07
Z lm Score = 3.6
NG = 0
DB = 0
SM = 0
Delta_L
Doxycycline
YMR291W
ZShift = −0.02
Z lm Score = −3.38
NG = 0
DB = 0
SM = 0
Delta_K
Doxycycline
YMR291W
ZShift = 1.48
Z lm Score = −3.59
NG = 0
DB = 0
SM = 0
Delta_r
Doxycycline
YMR291W
ZShift = −0.48
Z lm Score = −2.86
NG = 0
DB = 0
SM = 0
Delta_AUC
Doxycycline
YPR013C
ZShift = 1.01
Z lm Score = 3.6
NG = 0
DB = 0
SM = 0
Delta_L
Doxycycline
YPR013C
ZShift = 2.55
Z lm Score = −0.35
NG = 0
DB = 0
SM = 0
Delta_K
Doxycycline
YPR013C
ZShift = −0.44
Z lm Score = −0.09
NG = 0
DB = 0
SM = 0
Delta_r
Doxycycline
YPR013C
ZShift = 1.27
Z lm Score = 1.76
NG = 0
DB = 0
SM = 0
Delta_AUC
Doxycycline

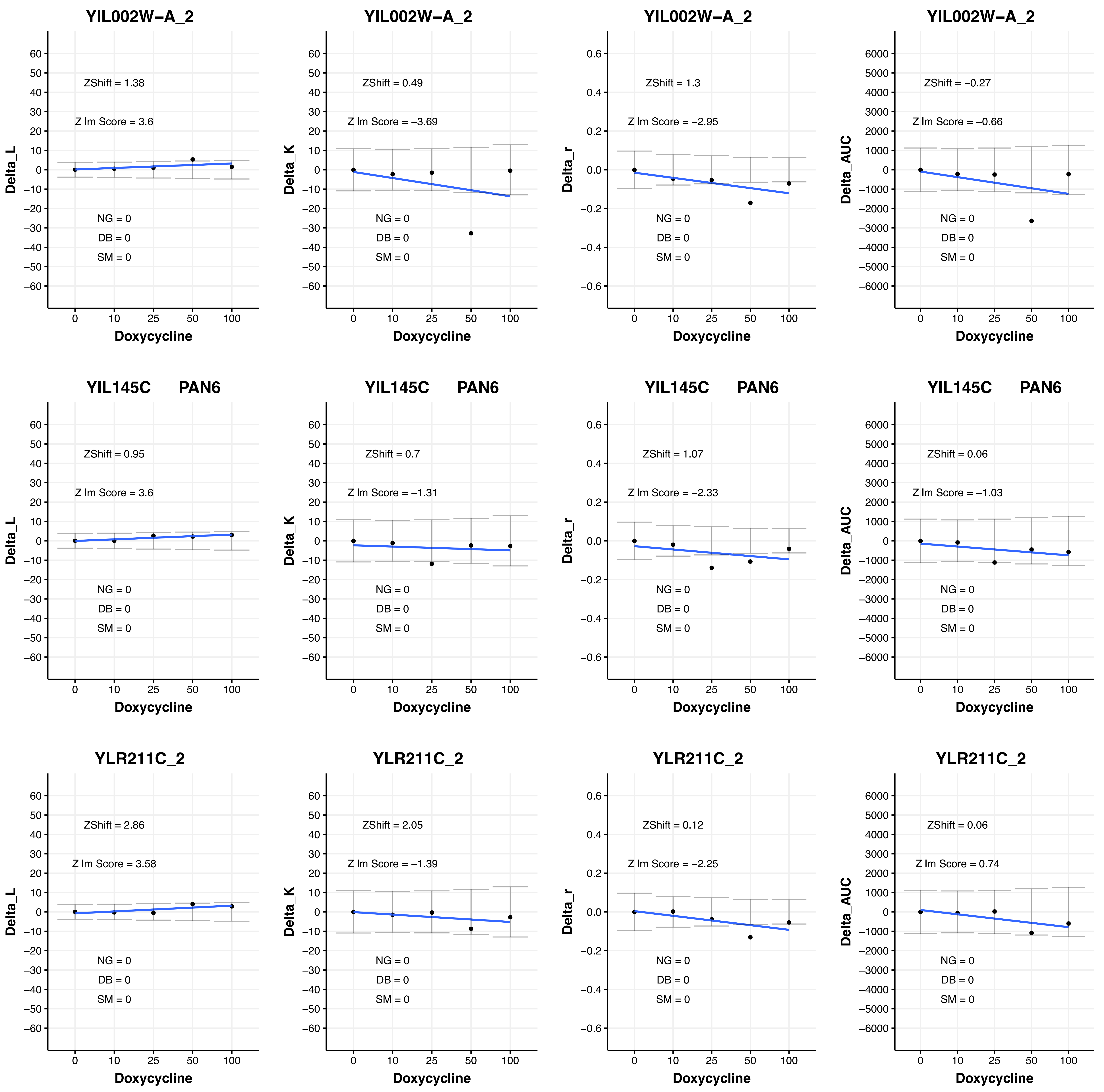

YIL002W−A_2
ZShift = 1.38
Z lm Score = 3.6
NG = 0
DB = 0
SM = 0
Delta_L
Doxycycline
YIL002W−A_2
ZShift = 0.49
Z lm Score = −3.69
NG = 0
DB = 0
SM = 0
Delta_K
Doxycycline
YIL002W−A_2
ZShift = 1.3
Z lm Score = −2.95
NG = 0
DB = 0
SM = 0
Delta_r
Doxycycline
YIL002W−A_2
ZShift = −0.27
Z lm Score = −0.66
NG = 0
DB = 0
SM = 0
Delta_AUC
Doxycycline
YIL145C PAN6
ZShift = 0.95
Z lm Score = 3.6
NG = 0
DB = 0
SM = 0
Delta_L
Doxycycline
YIL145C PAN6
ZShift = 0.7
Z lm Score = −1.31
NG = 0
DB = 0
SM = 0
Delta_K
Doxycycline
YIL145C PAN6
ZShift = 1.07
Z lm Score = −2.33
NG = 0
DB = 0
SM = 0
Delta_r
Doxycycline
YIL145C PAN6
ZShift = 0.06
Z lm Score = −1.03
NG = 0
DB = 0
SM = 0
Delta_AUC
Doxycycline
YLR211C_2
ZShift = 2.86
Z lm Score = 3.58
NG = 0
DB = 0
SM = 0
Delta_L
Doxycycline
YLR211C_2
ZShift = 2.05
Z lm Score = −1.39
NG = 0
DB = 0
SM = 0
Delta_K
Doxycycline
YLR211C_2
ZShift = 0.12
Z lm Score = −2.25
NG = 0
DB = 0
SM = 0
Delta_r
Doxycycline
YLR211C_2
ZShift = 0.06
Z lm Score = 0.74
NG = 0
DB = 0
SM = 0
Delta_AUC
Doxycycline

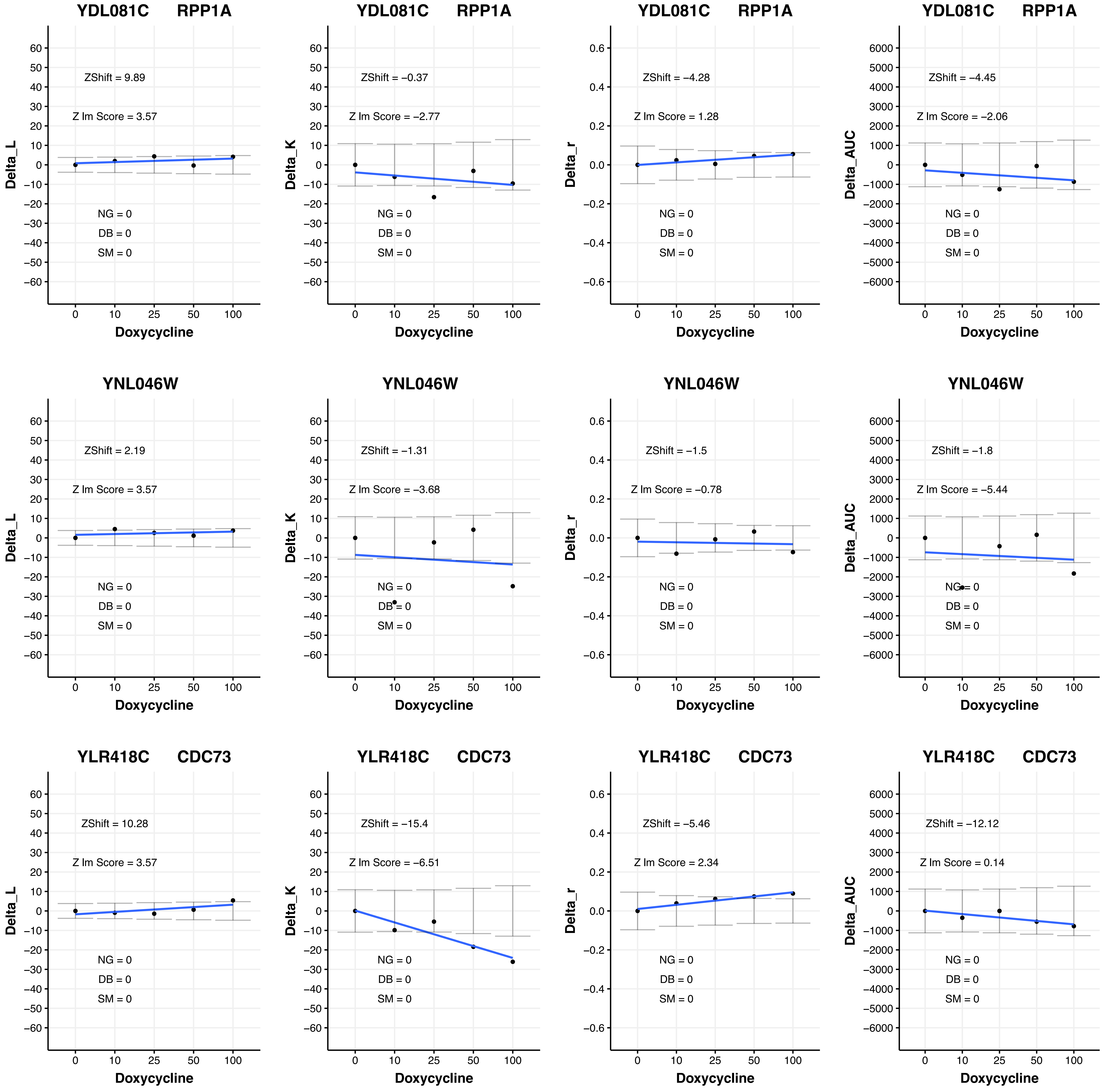

YDL081C RPP1A
ZShift = 9.89
Z lm Score = 3.57
NG = 0
DB = 0
SM = 0
Delta_L
Doxycycline
YDL081C RPP1A
ZShift = −0.37
Z lm Score = −2.77
NG = 0
DB = 0
SM = 0
Delta_K
Doxycycline
YDL081C RPP1A
ZShift = −4.28
Z lm Score = 1.28
NG = 0
DB = 0
SM = 0
Delta_r
Doxycycline
YDL081C RPP1A
ZShift = −4.45
Z lm Score = −2.06
NG = 0
DB = 0
SM = 0
Delta_AUC
Doxycycline
YNL046W
ZShift = 2.19
Z lm Score = 3.57
NG = 0
DB = 0
SM = 0
Delta_L
Doxycycline
YNL046W
ZShift = −1.31
Z lm Score = −3.68
NG = 0
DB = 0
SM = 0
Delta_K
Doxycycline
YNL046W
ZShift = −1.5
Z lm Score = −0.78
NG = 0
DB = 0
SM = 0
Delta_r
Doxycycline
YNL046W
ZShift = −1.8
Z lm Score = −5.44
NG = 0
DB = 0
SM = 0
Delta_AUC
Doxycycline
YLR418C CDC73
ZShift = 10.28
Z lm Score = 3.57
NG = 0
DB = 0
SM = 0
Delta_L
Doxycycline
YLR418C CDC73
ZShift = −15.4
Z lm Score = −6.51
NG = 0
DB = 0
SM = 0
Delta_K
Doxycycline
YLR418C CDC73
ZShift = −5.46
Z lm Score = 2.34
NG = 0
DB = 0
SM = 0
Delta_r
Doxycycline
YLR418C CDC73
ZShift = −12.12
Z lm Score = 0.14
NG = 0
DB = 0
SM = 0
Delta_AUC
Doxycycline

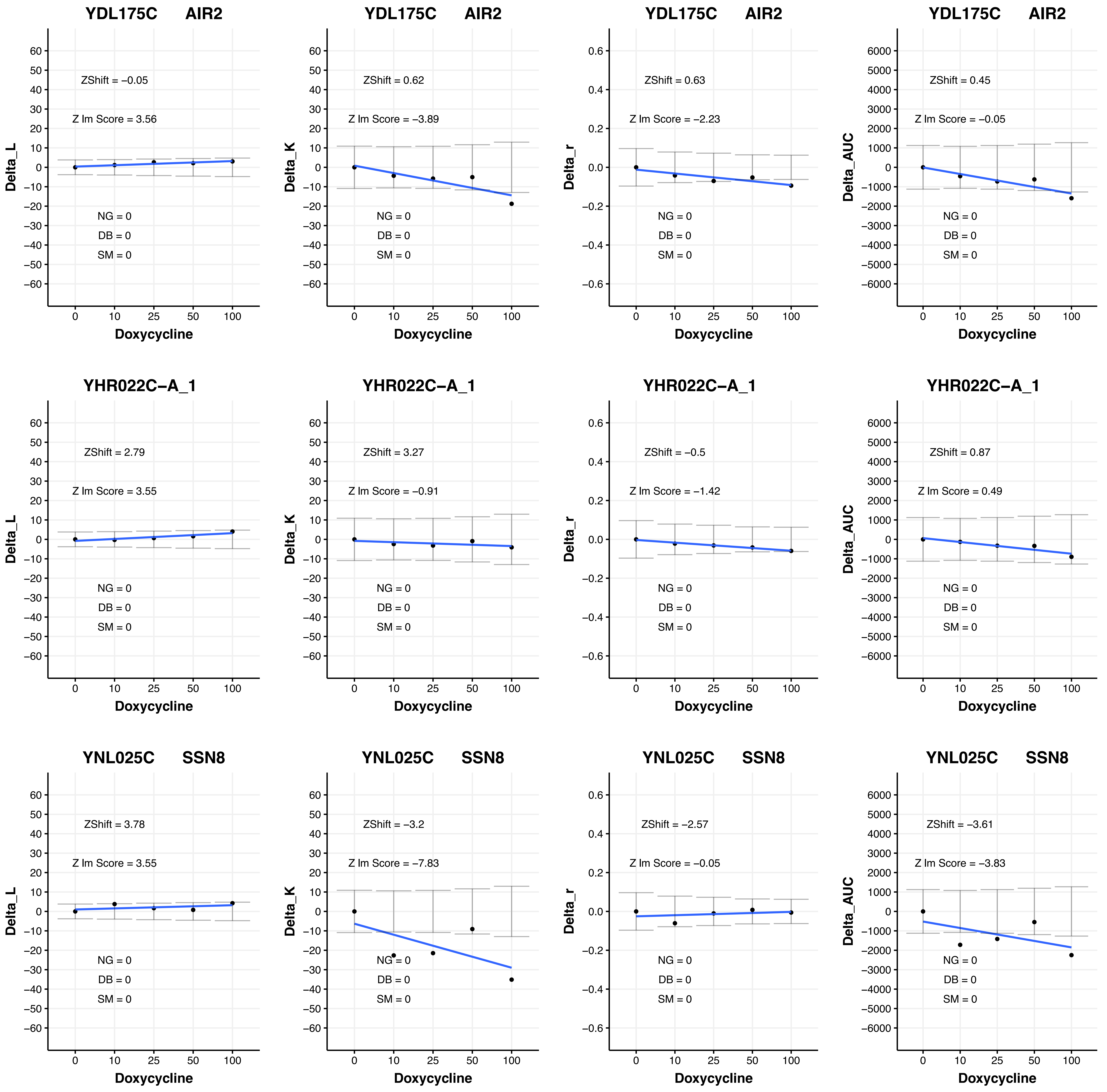
YDL175C AIR2
ZShift = −0.05
Z lm Score = 3.56
NG = 0
DB = 0
SM = 0
Delta_L
Doxycycline
YDL175C AIR2
ZShift = 0.62
Z lm Score = −3.89
NG = 0
DB = 0
SM = 0
Delta_K
Doxycycline
YDL175C AIR2
ZShift = 0.63
Z lm Score = −2.23
NG = 0
DB = 0
SM = 0
Delta_r
Doxycycline
YDL175C AIR2
ZShift = 0.45
Z lm Score = −0.05
NG = 0
DB = 0
SM = 0
Delta_AUC
Doxycycline
YHR022C−A_1
ZShift = 2.79
Z lm Score = 3.55
NG = 0
DB = 0
SM = 0
Delta_L
Doxycycline
YHR022C−A_1
ZShift = 3.27
Z lm Score = −0.91
NG = 0
DB = 0
SM = 0
Delta_K
Doxycycline
YHR022C−A_1
ZShift = −0.5
Z lm Score = −1.42
NG = 0
DB = 0
SM = 0
Delta_r
Doxycycline
YHR022C−A_1
ZShift = 0.87
Z lm Score = 0.49
NG = 0
DB = 0
SM = 0
Delta_AUC
Doxycycline
YNL025C SSN8
ZShift = 3.78
Z lm Score = 3.55
NG = 0
DB = 0
SM = 0
Delta_L
Doxycycline
YNL025C SSN8
ZShift = −3.2
Z lm Score = −7.83
NG = 0
DB = 0
SM = 0
Delta_K
Doxycycline
YNL025C SSN8
ZShift = −2.57
Z lm Score = −0.05
NG = 0
DB = 0
SM = 0
Delta_r
Doxycycline
YNL025C SSN8
ZShift = −3.61
Z lm Score = −3.83
NG = 0
DB = 0
SM = 0
Delta_AUC
Doxycycline

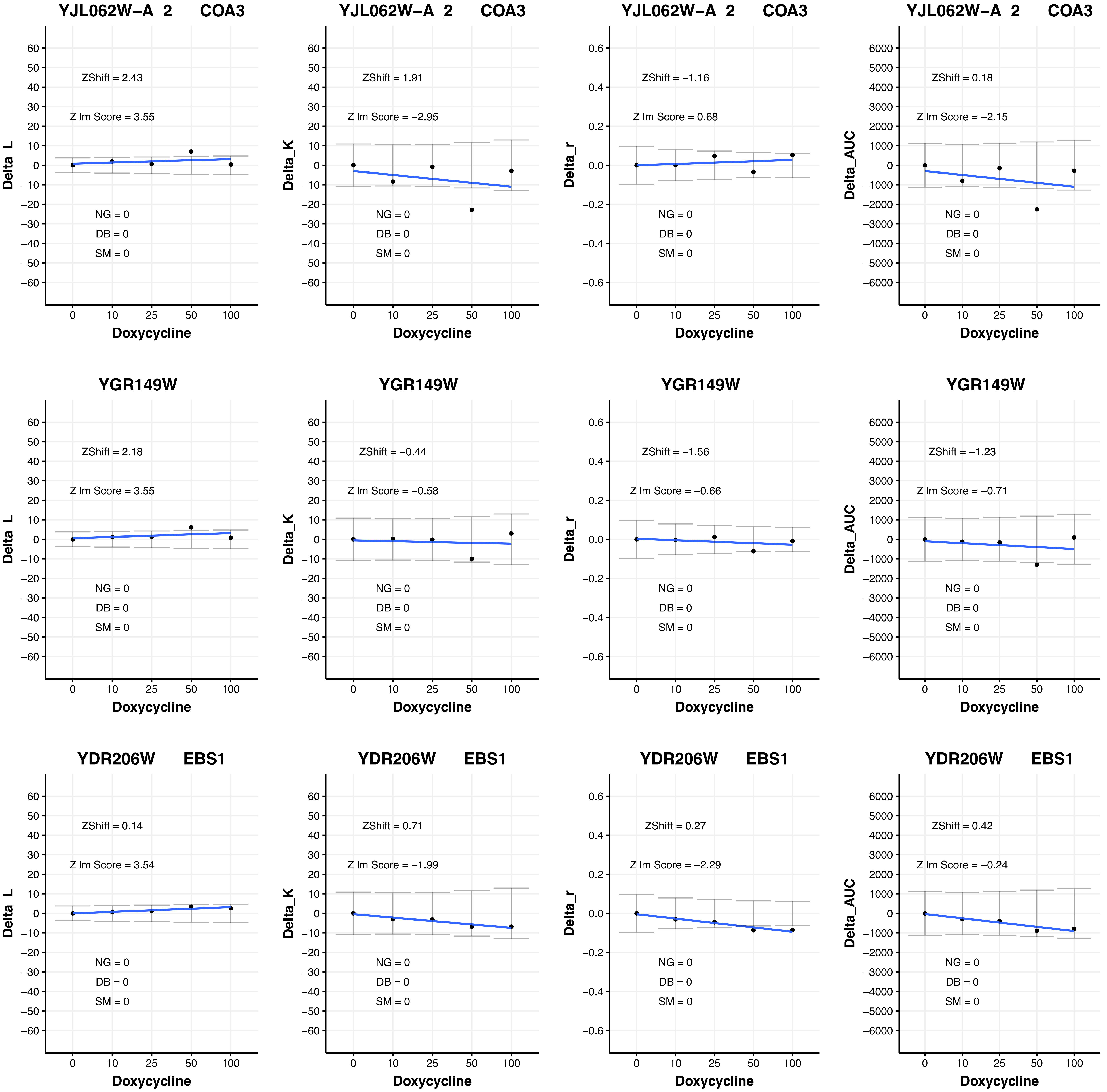

YJL062W-A_2 COA3
ZShift = 2.43
Z lm Score = 3.55
NG = 0
DB = 0
SM = 0
Delta_L
Doxycycline
YJL062W-A_2 COA3
ZShift = 1.91
Z lm Score = −2.95
NG = 0
DB = 0
SM = 0
Delta_K
Doxycycline
YJL062W-A_2 COA3
ZShift = −1.16
Z lm Score = 0.68
NG = 0
DB = 0
SM = 0
Delta_r
Doxycycline
YJL062W-A_2 COA3
ZShift = 0.18
Z lm Score = −2.15
NG = 0
DB = 0
SM = 0
Delta_AUC
Doxycycline
YGR149W
ZShift = 2.18
Z lm Score = 3.55
NG = 0
DB = 0
SM = 0
Delta_L
Doxycycline
YGR149W
ZShift = −0.44
Z lm Score = −0.58
NG = 0
DB = 0
SM = 0
Delta_K
Doxycycline
YGR149W
ZShift = −1.56
Z lm Score = −0.66
NG = 0
DB = 0
SM = 0
Delta_r
Doxycycline
YGR149W
ZShift = −1.23
Z lm Score = −0.71
NG = 0
DB = 0
SM = 0
Delta_AUC
Doxycycline
YDR206W EBS1
ZShift = 0.14
Z lm Score = 3.54
NG = 0
DB = 0
SM = 0
Delta_L
Doxycycline
YDR206W EBS1
ZShift = 0.71
Z lm Score = −1.99
NG = 0
DB = 0
SM = 0
Delta_K
Doxycycline
YDR206W EBS1
ZShift = 0.27
Z lm Score = −2.29
NG = 0
DB = 0
SM = 0
Delta_r
Doxycycline
YDR206W EBS1
ZShift = 0.42
Z lm Score = −0.24
NG = 0
DB = 0
SM = 0
Delta_AUC
Doxycycline

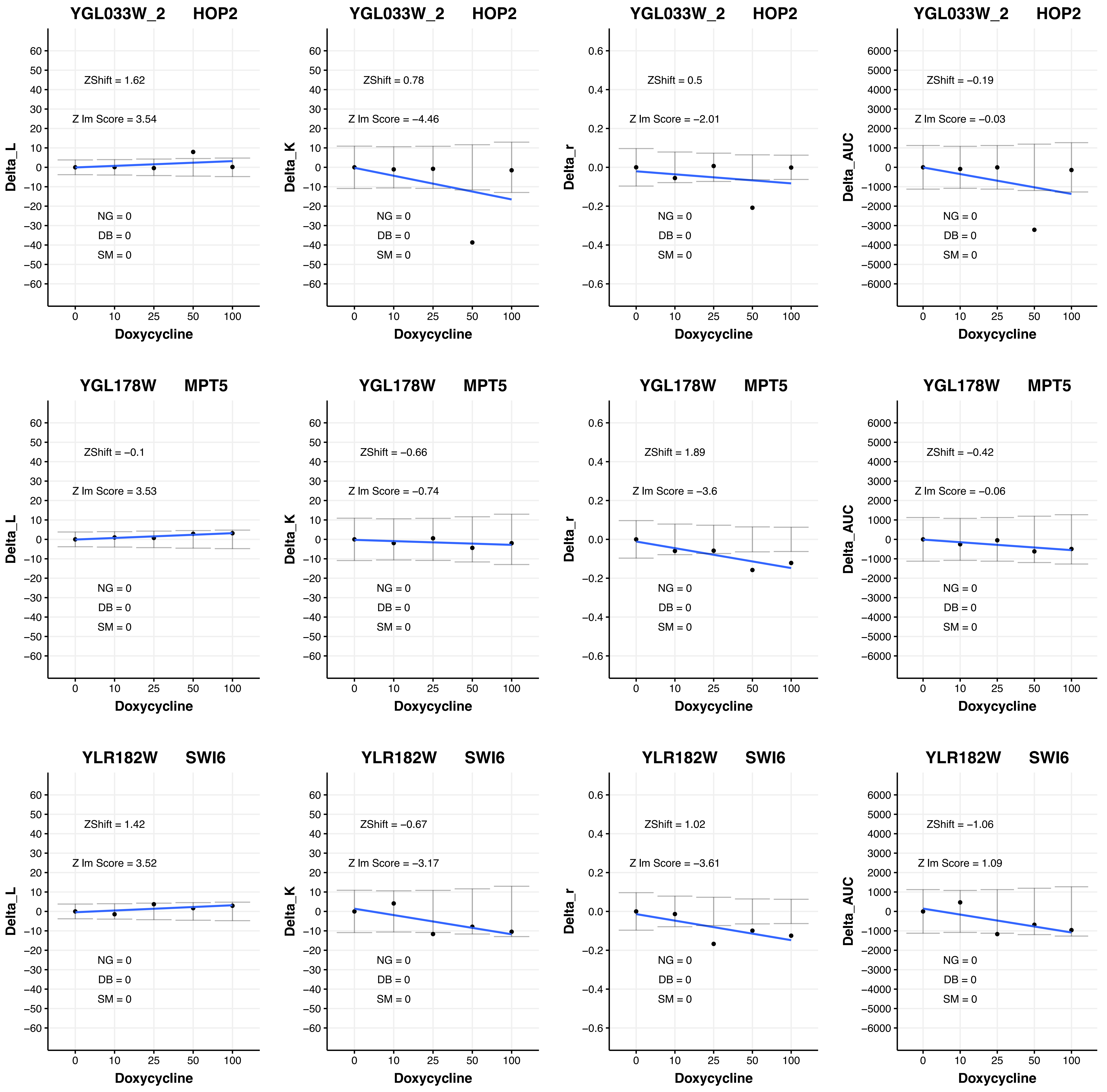
YGL033W_2 HOP2
ZShift = 1.62
Z lm Score = 3.54
NG = 0
DB = 0
SM = 0
Delta_L
Doxycycline
YGL033W_2 HOP2
ZShift = 0.78
Z lm Score = −4.46
NG = 0
DB = 0
SM = 0
Delta_K
Doxycycline
YGL033W_2 HOP2
ZShift = 0.5
Z lm Score = −2.01
NG = 0
DB = 0
SM = 0
Delta_r
Doxycycline
YGL033W_2 HOP2
ZShift = −0.19
Z lm Score = −0.03
NG = 0
DB = 0
SM = 0
Delta_AUC
Doxycycline
YGL178W MPT5
ZShift = −0.1
Z lm Score = 3.53
NG = 0
DB = 0
SM = 0
Delta_L
Doxycycline
YGL178W MPT5
ZShift = −0.66
Z lm Score = −0.74
NG = 0
DB = 0
SM = 0
Delta_K
Doxycycline
YGL178W MPT5
ZShift = 1.89
Z lm Score = −3.6
NG = 0
DB = 0
SM = 0
Delta_r
Doxycycline
YGL178W MPT5
ZShift = −0.42
Z lm Score = −0.06
NG = 0
DB = 0
SM = 0
Delta_AUC
Doxycycline
YLR182W SWI6
ZShift = 1.42
Z lm Score = 3.52
NG = 0
DB = 0
SM = 0
Delta_L
Doxycycline
YLR182W SWI6
ZShift = −0.67
Z lm Score = −3.17
NG = 0
DB = 0
SM = 0
Delta_K
Doxycycline
YLR182W SWI6
ZShift = 1.02
Z lm Score = −3.61
NG = 0
DB = 0
SM = 0
Delta_r
Doxycycline
YLR182W SWI6
ZShift = −1.06
Z lm Score = 1.09
NG = 0
DB = 0
SM = 0
Delta_AUC
Doxycycline

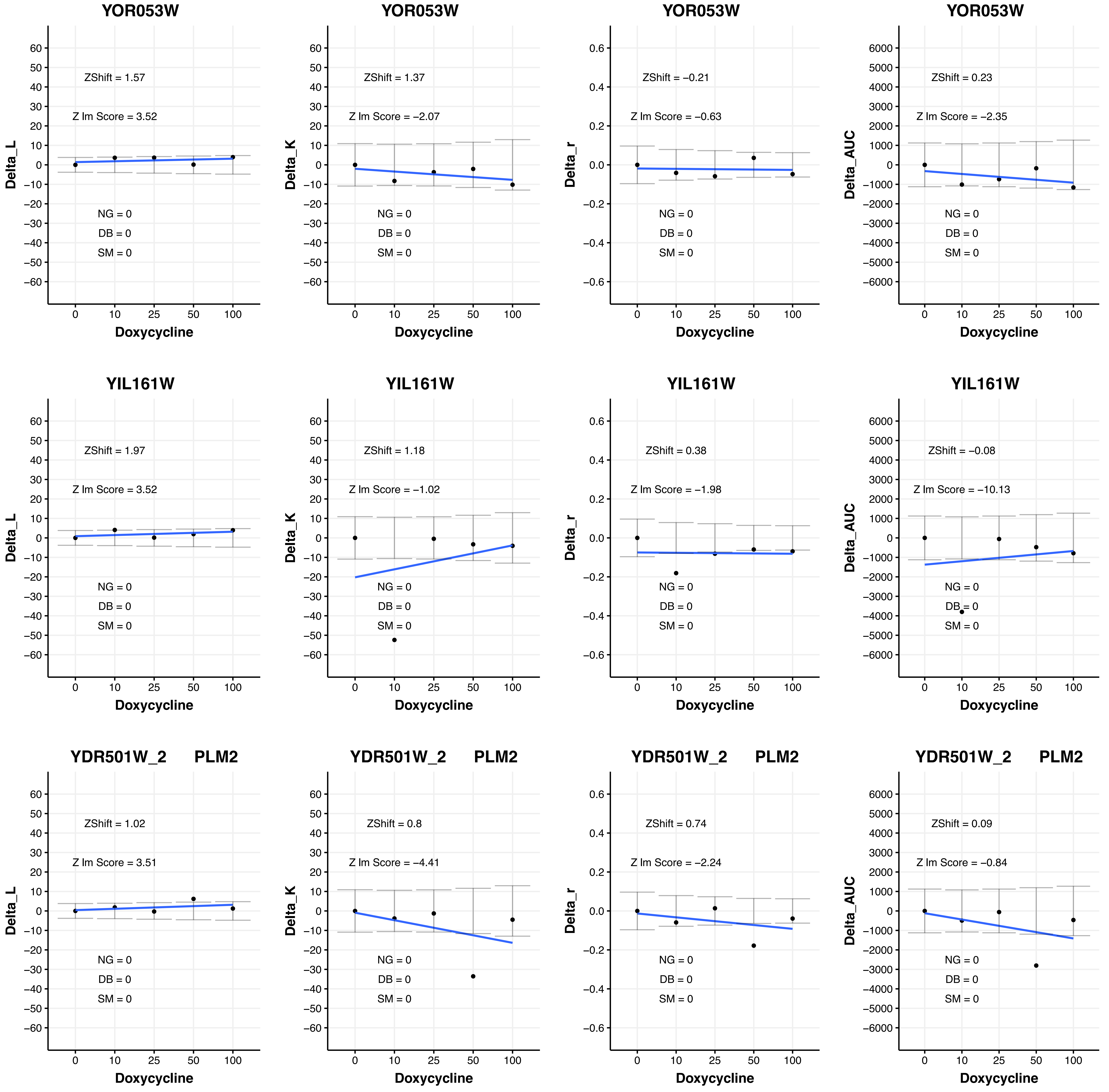
YOR053W
ZShift = 1.57
Z lm Score = 3.52
NG = 0
DB = 0
SM = 0
Delta_L
Doxycycline
YOR053W
ZShift = 1.37
Z lm Score = −2.07
NG = 0
DB = 0
SM = 0
Delta_K
Doxycycline
YOR053W
ZShift = −0.21
Z lm Score = −0.63
NG = 0
DB = 0
SM = 0
Delta_r
Doxycycline
YOR053W
ZShift = 0.23
Z lm Score = −2.35
NG = 0
DB = 0
SM = 0
Delta_AUC
Doxycycline
YIL161W
ZShift = 1.97
Z lm Score = 3.52
NG = 0
DB = 0
SM = 0
Delta_L
Doxycycline
YIL161W
ZShift = 1.18
Z lm Score = −1.02
NG = 0
DB = 0
SM = 0
Delta_K
Doxycycline
YIL161W
ZShift = 0.38
Z lm Score = −1.98
NG = 0
DB = 0
SM = 0
Delta_r
Doxycycline
YIL161W
ZShift = −0.08
Z lm Score = −10.13
NG = 0
DB = 0
SM = 0
Delta_AUC
Doxycycline
YDR501W_2 PLM2
ZShift = 1.02
Z lm Score = 3.51
NG = 0
DB = 0
SM = 0
Delta_L
Doxycycline
YDR501W_2 PLM2
ZShift = 0.8
Z lm Score = −4.41
NG = 0
DB = 0
SM = 0
Delta_K
Doxycycline
YDR501W_2 PLM2
ZShift = 0.74
Z lm Score = −2.24
NG = 0
DB = 0
SM = 0
Delta_r
Doxycycline
YDR501W_2 PLM2
ZShift = 0.09
Z lm Score = −0.84
NG = 0
DB = 0
SM = 0
Delta_AUC
Doxycycline

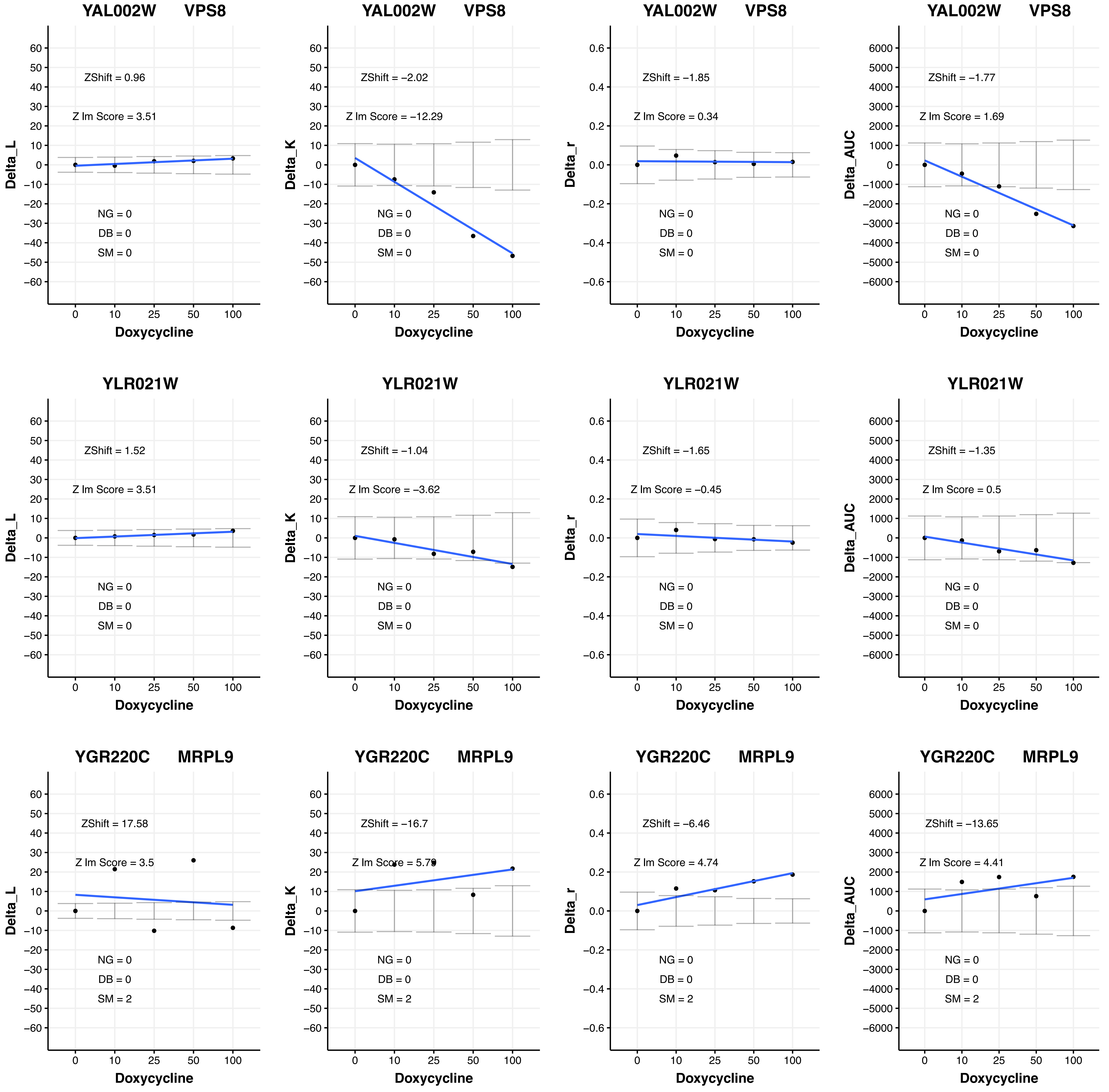
YAL002W VPS8
ZShift = 0.96
Z lm Score = 3.51
NG = 0
DB = 0
SM = 0
Delta_L
Doxycycline
YAL002W VPS8
ZShift = −2.02
Z lm Score = −12.29
NG = 0
DB = 0
SM = 0
Delta_K
Doxycycline
YAL002W VPS8
ZShift = −1.85
Z lm Score = 0.34
NG = 0
DB = 0
SM = 0
Delta_r
Doxycycline
YAL002W VPS8
ZShift = −1.77
Z lm Score = 1.69
NG = 0
DB = 0
SM = 0
Delta_AUC
Doxycycline
YLR021W
ZShift = 1.52
Z lm Score = 3.51
NG = 0
DB = 0
SM = 0
Delta_L
Doxycycline
YLR021W
ZShift = −1.04
Z lm Score = −3.62
NG = 0
DB = 0
SM = 0
Delta_K
Doxycycline
YLR021W
ZShift = −1.65
Z lm Score = −0.45
NG = 0
DB = 0
SM = 0
Delta_r
Doxycycline
YLR021W
ZShift = −1.35
Z lm Score = 0.5
NG = 0
DB = 0
SM = 0
Delta_AUC
Doxycycline
YGR220C MRPL9
ZShift = 17.58
Z lm Score = 3.5
NG = 0
DB = 0
SM = 2
Delta_L
Doxycycline
YGR220C MRPL9
ZShift = −16.7
Z lm Score = 5.79
NG = 0
DB = 0
SM = 2
Delta_K
Doxycycline
YGR220C MRPL9
ZShift = −6.46
Z lm Score = 4.74
NG = 0
DB = 0
SM = 2
Delta_r
Doxycycline
YGR220C MRPL9
ZShift = −13.65
Z lm Score = 4.41
NG = 0
DB = 0
SM = 2
Delta_AUC
Doxycycline

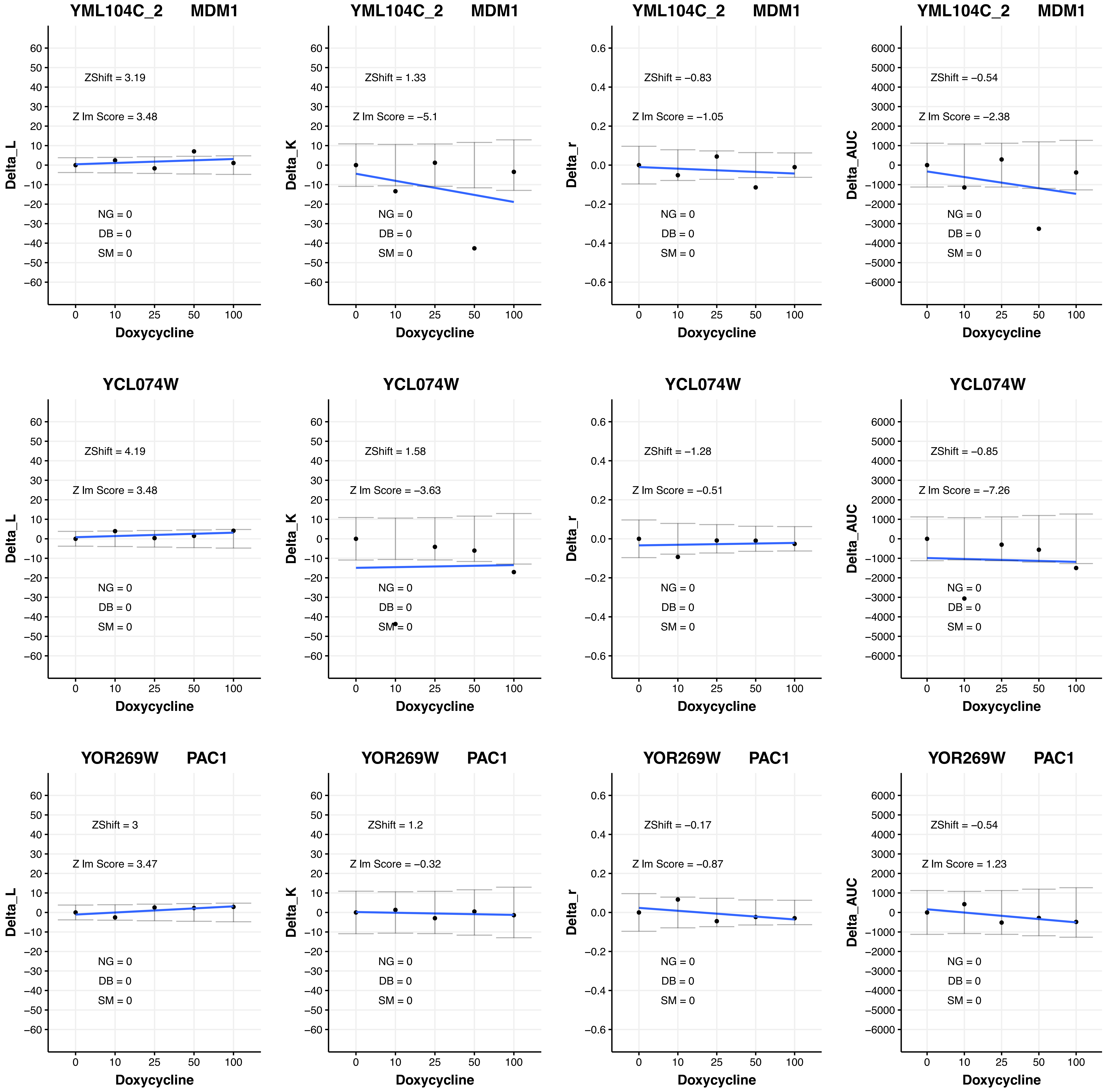
YML104C_2 MDM1
ZShift = 3.19
Z lm Score = 3.48
NG = 0
DB = 0
SM = 0
Delta_L
Doxycycline
YML104C_2 MDM1
ZShift = 1.33
Z lm Score = −5.1
NG = 0
DB = 0
SM = 0
Delta_K
Doxycycline
YML104C_2 MDM1
ZShift = −0.83
Z lm Score = −1.05
NG = 0
DB = 0
SM = 0
Delta_r
Doxycycline
YML104C_2 MDM1
ZShift = −0.54
Z lm Score = −2.38
NG = 0
DB = 0
SM = 0
Delta_AUC
Doxycycline
YCL074W
ZShift = 4.19
Z lm Score = 3.48
NG = 0
DB = 0
SM = 0
Delta_L
Doxycycline
YCL074W
ZShift = 1.58
Z lm Score = −3.63
NG = 0
DB = 0
SM = 0
Delta_K
Doxycycline
YCL074W
ZShift = −1.28
Z lm Score = −0.51
NG = 0
DB = 0
SM = 0
Delta_r
Doxycycline
YCL074W
ZShift = −0.85
Z lm Score = −7.26
NG = 0
DB = 0
SM = 0
Delta_AUC
Doxycycline
YOR269W PAC1
ZShift = 3
Z lm Score = 3.47
NG = 0
DB = 0
SM = 0
Delta_L
Doxycycline
YOR269W PAC1
ZShift = 1.2
Z lm Score = −0.32
NG = 0
DB = 0
SM = 0
Delta_K
Doxycycline
YOR269W PAC1
ZShift = −0.17
Z lm Score = −0.87
NG = 0
DB = 0
SM = 0
Delta_r
Doxycycline
YOR269W PAC1
ZShift = −0.54
Z lm Score = 1.23
NG = 0
DB = 0
SM = 0
Delta_AUC
Doxycycline

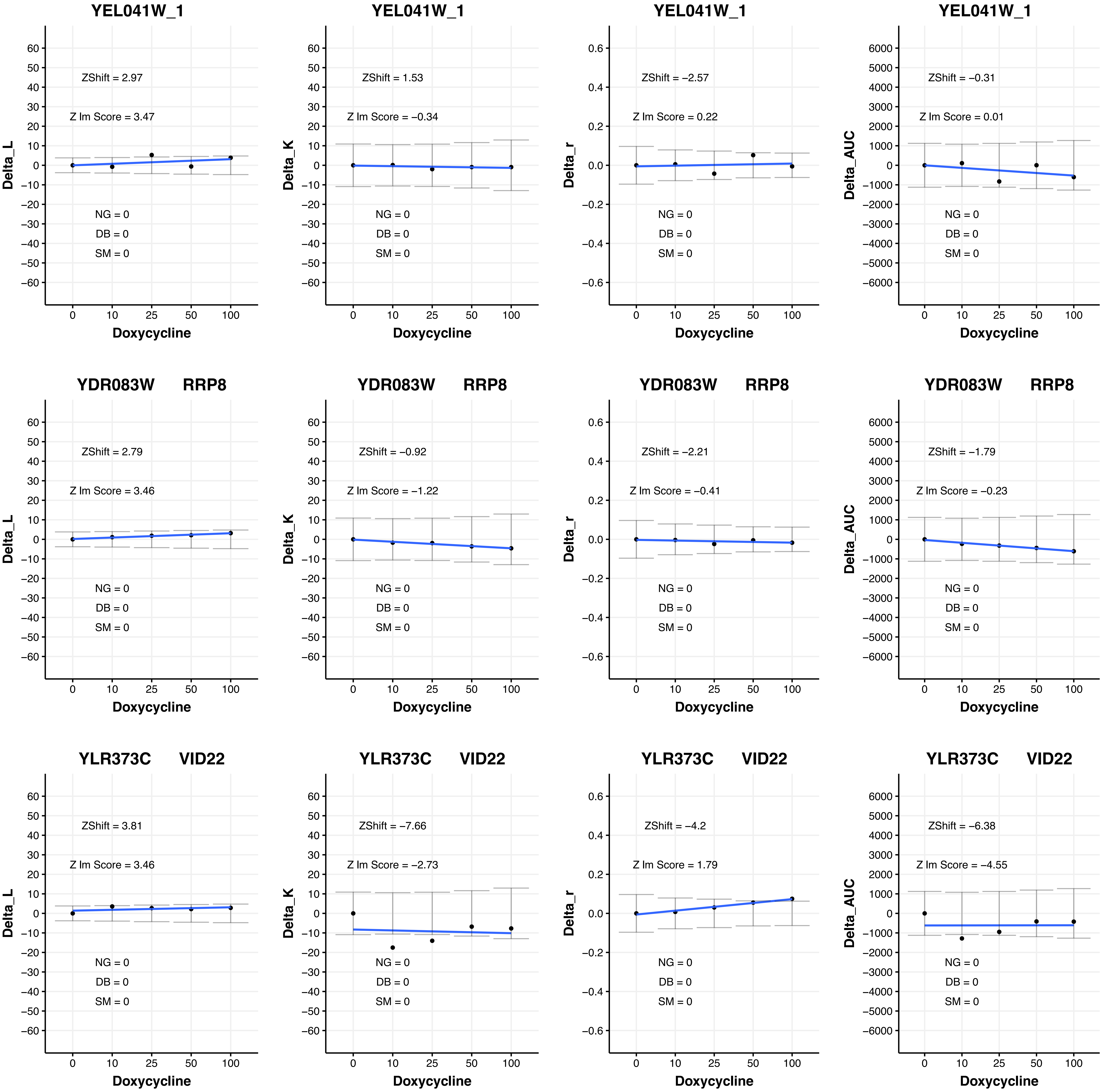

YEL041W_1
ZShift = 2.97
Z lm Score = 3.47
Delta_L
NG = 0
DB = 0
SM = 0
Doxycycline
YEL041W_1
ZShift = 1.53
Z lm Score = −0.34
Delta_K
NG = 0
DB = 0
SM = 0
Doxycycline
YEL041W_1
ZShift = −2.57
Z lm Score = 0.22
Delta_r
NG = 0
DB = 0
SM = 0
Doxycycline
YEL041W_1
ZShift = −0.31
Z lm Score = 0.01
Delta_AUC
NG = 0
DB = 0
SM = 0
Doxycycline
YDR083W RRP8
ZShift = 2.79
Z lm Score = 3.46
Delta_L
NG = 0
DB = 0
SM = 0
Doxycycline
YDR083W RRP8
ZShift = −0.92
Z lm Score = −1.22
Delta_K
NG = 0
DB = 0
SM = 0
Doxycycline
YDR083W RRP8
ZShift = −2.21
Z lm Score = −0.41
Delta_r
NG = 0
DB = 0
SM = 0
Doxycycline
YDR083W RRP8
ZShift = −1.79
Z lm Score = −0.23
Delta_AUC
NG = 0
DB = 0
SM = 0
Doxycycline
YLR373C VID22
ZShift = 3.81
Z lm Score = 3.46
Delta_L
NG = 0
DB = 0
SM = 0
Doxycycline
YLR373C VID22
ZShift = −7.66
Z lm Score = −2.73
Delta_K
NG = 0
DB = 0
SM = 0
Doxycycline
YLR373C VID22
ZShift = −4.2
Z lm Score = 1.79
Delta_r
NG = 0
DB = 0
SM = 0
Doxycycline
YLR373C VID22
ZShift = −6.38
Z lm Score = −4.55
Delta_AUC
NG = 0
DB = 0
SM = 0
Doxycycline

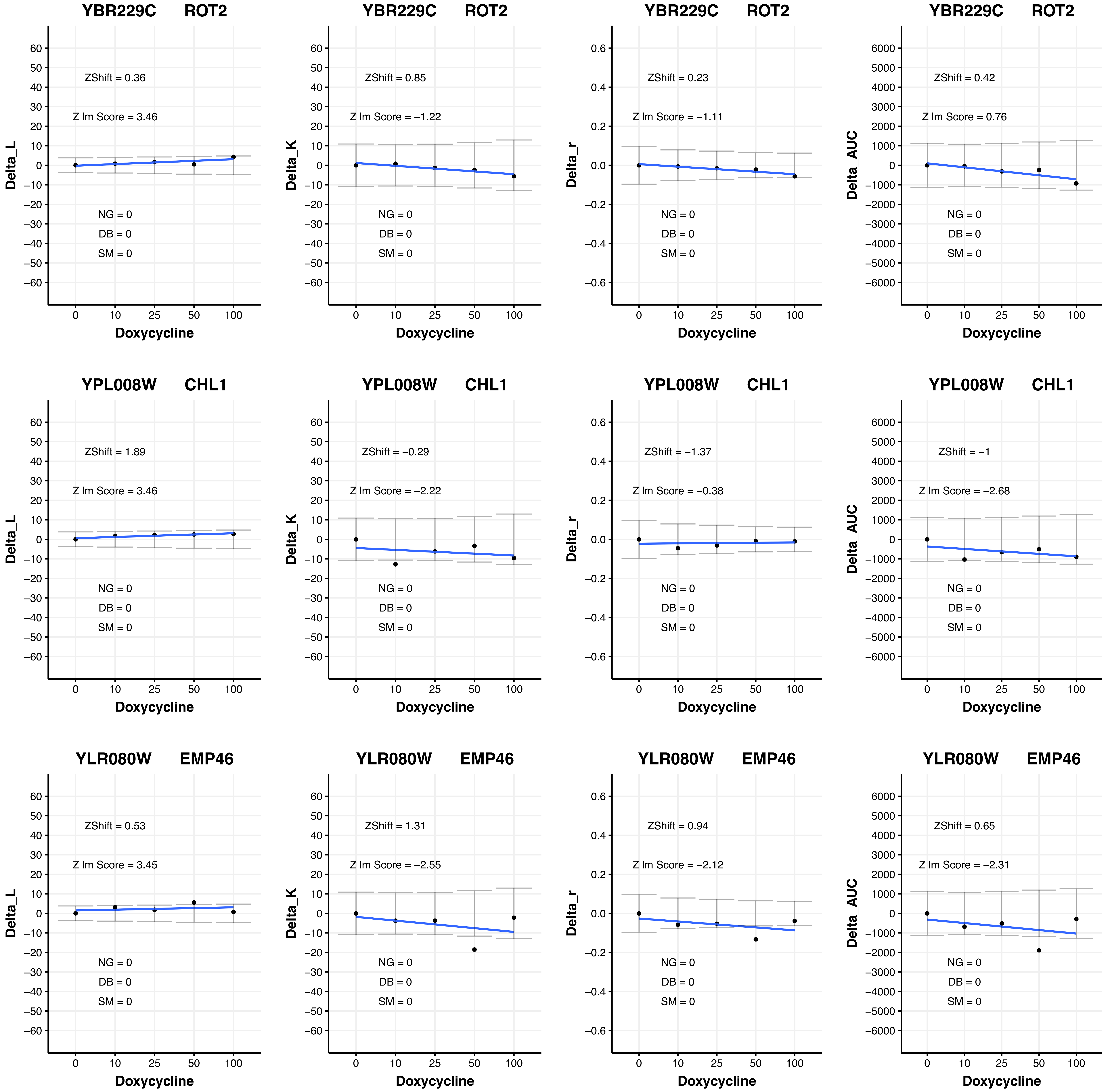

YBR229C ROT2
ZShift = 0.36
Z lm Score = 3.46
ZShift = 0.85
Z lm Score = −1.22
ZShift = 0.23
Z lm Score = −1.11
ZShift = 0.42
Z lm Score = 0.76
YPL008W CHL1
ZShift = 1.89
Z lm Score = 3.46
ZShift = −0.29
Z lm Score = −2.22
ZShift = −1.37
Z lm Score = −0.38
ZShift = −1
Z lm Score = −2.68
YLR080W EMP46
ZShift = 0.53
Z lm Score = 3.45
ZShift = 1.31
Z lm Score = −2.55
ZShift = 0.94
Z lm Score = −2.12
ZShift = 0.65
Z lm Score = −2.31
NG = 0
DB = 0
SM = 0
Delta_L
Delta_K
Delta_r
Delta_AUC
Doxycycline
0 10 25 50 100

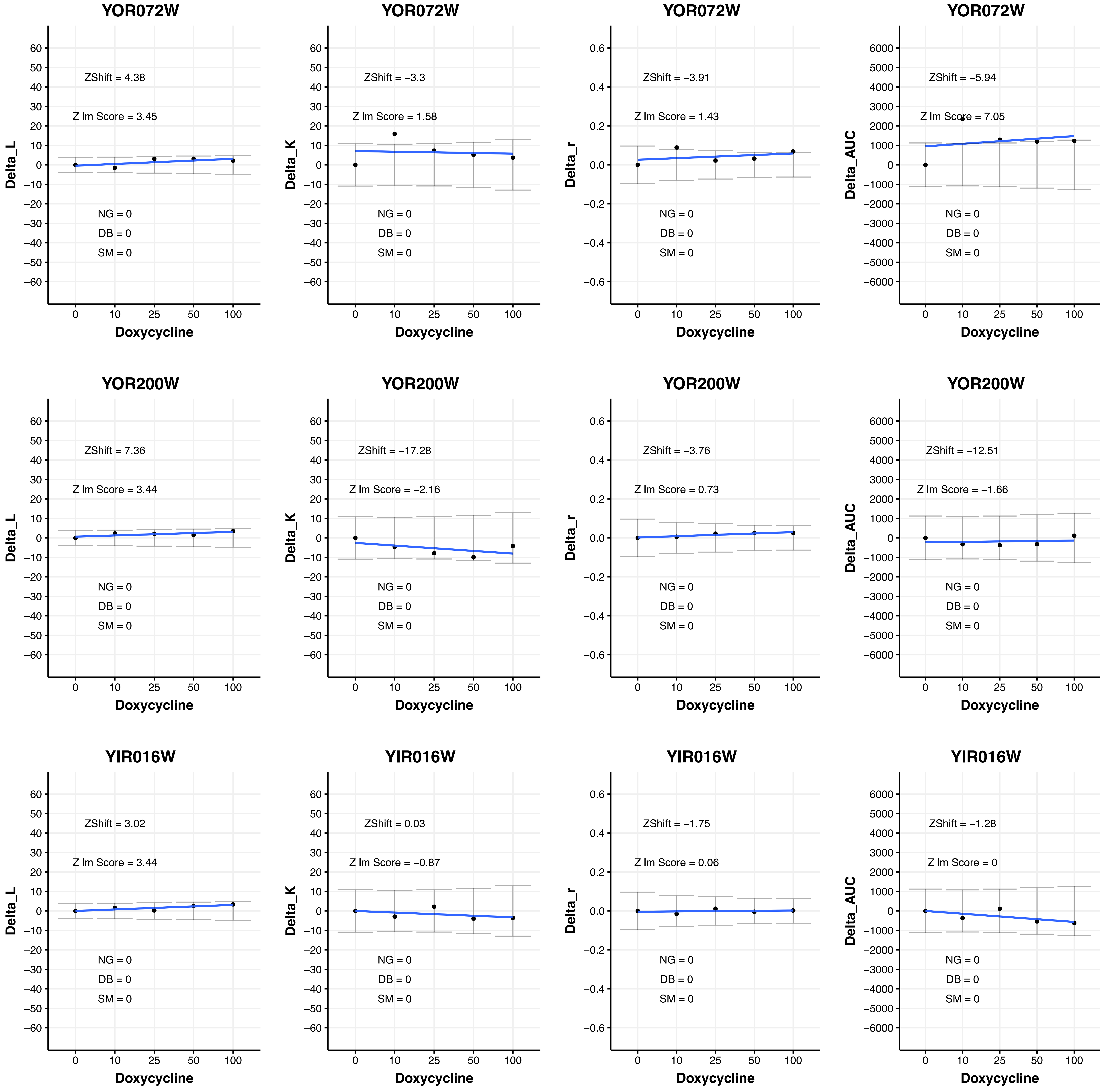

YOR072W
ZShift = 4.38
Z lm Score = 3.45
NG = 0
DB = 0
SM = 0
Delta_L
Doxycycline
YOR072W
ZShift = −3.3
Z lm Score = 1.58
NG = 0
DB = 0
SM = 0
Delta_K
Doxycycline
YOR072W
ZShift = −3.91
Z lm Score = 1.43
NG = 0
DB = 0
SM = 0
Delta_r
Doxycycline
YOR072W
ZShift = −5.94
Z lm Score = 7.05
NG = 0
DB = 0
SM = 0
Delta_AUC
Doxycycline
YOR200W
ZShift = 7.36
Z lm Score = 3.44
NG = 0
DB = 0
SM = 0
Delta_L
Doxycycline
YOR200W
ZShift = −17.28
Z lm Score = −2.16
NG = 0
DB = 0
SM = 0
Delta_K
Doxycycline
YOR200W
ZShift = −3.76
Z lm Score = 0.73
NG = 0
DB = 0
SM = 0
Delta_r
Doxycycline
YOR200W
ZShift = −12.51
Z lm Score = −1.66
NG = 0
DB = 0
SM = 0
Delta_AUC
Doxycycline
YIR016W
ZShift = 3.02
Z lm Score = 3.44
NG = 0
DB = 0
SM = 0
Delta_L
Doxycycline
YIR016W
ZShift = 0.03
Z lm Score = −0.87
NG = 0
DB = 0
SM = 0
Delta_K
Doxycycline
YIR016W
ZShift = −1.75
Z lm Score = 0.06
NG = 0
DB = 0
SM = 0
Delta_r
Doxycycline
YIR016W
ZShift = −1.28
Z lm Score = 0
NG = 0
DB = 0
SM = 0
Delta_AUC
Doxycycline

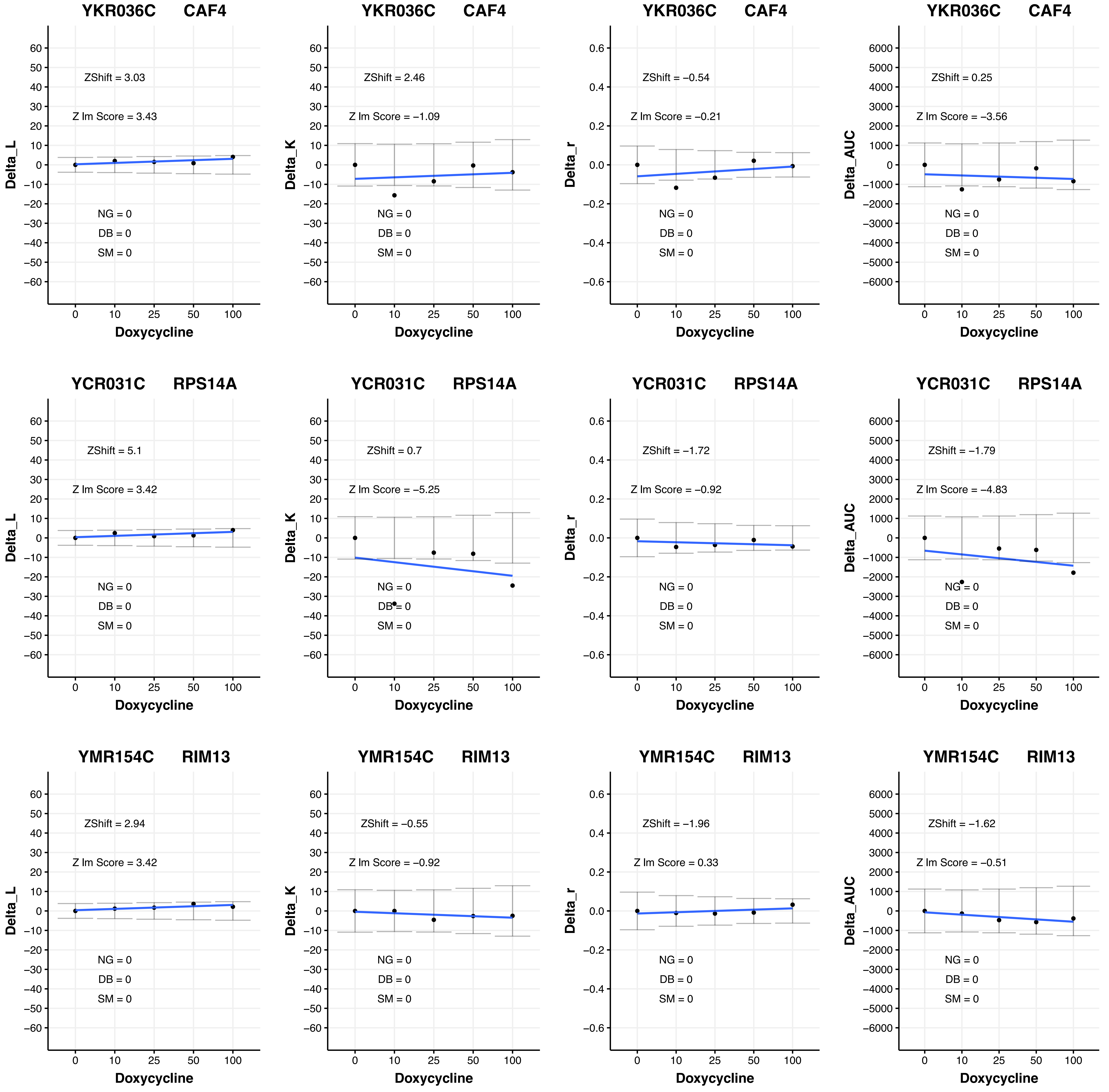
YKR036C CAF4
ZShift = 3.03
Z lm Score = 3.43
NG = 0
DB = 0
SM = 0
Delta_L
Doxycycline
YKR036C CAF4
ZShift = 2.46
Z lm Score = −1.09
NG = 0
DB = 0
SM = 0
Delta_K
Doxycycline
YKR036C CAF4
ZShift = −0.54
Z lm Score = −0.21
NG = 0
DB = 0
SM = 0
Delta_r
Doxycycline
YKR036C CAF4
ZShift = 0.25
Z lm Score = −3.56
NG = 0
DB = 0
SM = 0
Delta_AUC
Doxycycline
YCR031C RPS14A
ZShift = 5.1
Z lm Score = 3.42
NG = 0
DB = 0
SM = 0
Delta_L
Doxycycline
YCR031C RPS14A
ZShift = 0.7
Z lm Score = −5.25
NG = 0
DB = 0
SM = 0
Delta_K
Doxycycline
YCR031C RPS14A
ZShift = −1.72
Z lm Score = −0.92
NG = 0
DB = 0
SM = 0
Delta_r
Doxycycline
YCR031C RPS14A
ZShift = −1.79
Z lm Score = −4.83
NG = 0
DB = 0
SM = 0
Delta_AUC
Doxycycline
YMR154C RIM13
ZShift = 2.94
Z lm Score = 3.42
NG = 0
DB = 0
SM = 0
Delta_L
Doxycycline
YMR154C RIM13
ZShift = −0.55
Z lm Score = −0.92
NG = 0
DB = 0
SM = 0
Delta_K
Doxycycline
YMR154C RIM13
ZShift = −1.96
Z lm Score = 0.33
NG = 0
DB = 0
SM = 0
Delta_r
Doxycycline
YMR154C RIM13
ZShift = −1.62
Z lm Score = −0.51
NG = 0
DB = 0
SM = 0
Delta_AUC
Doxycycline

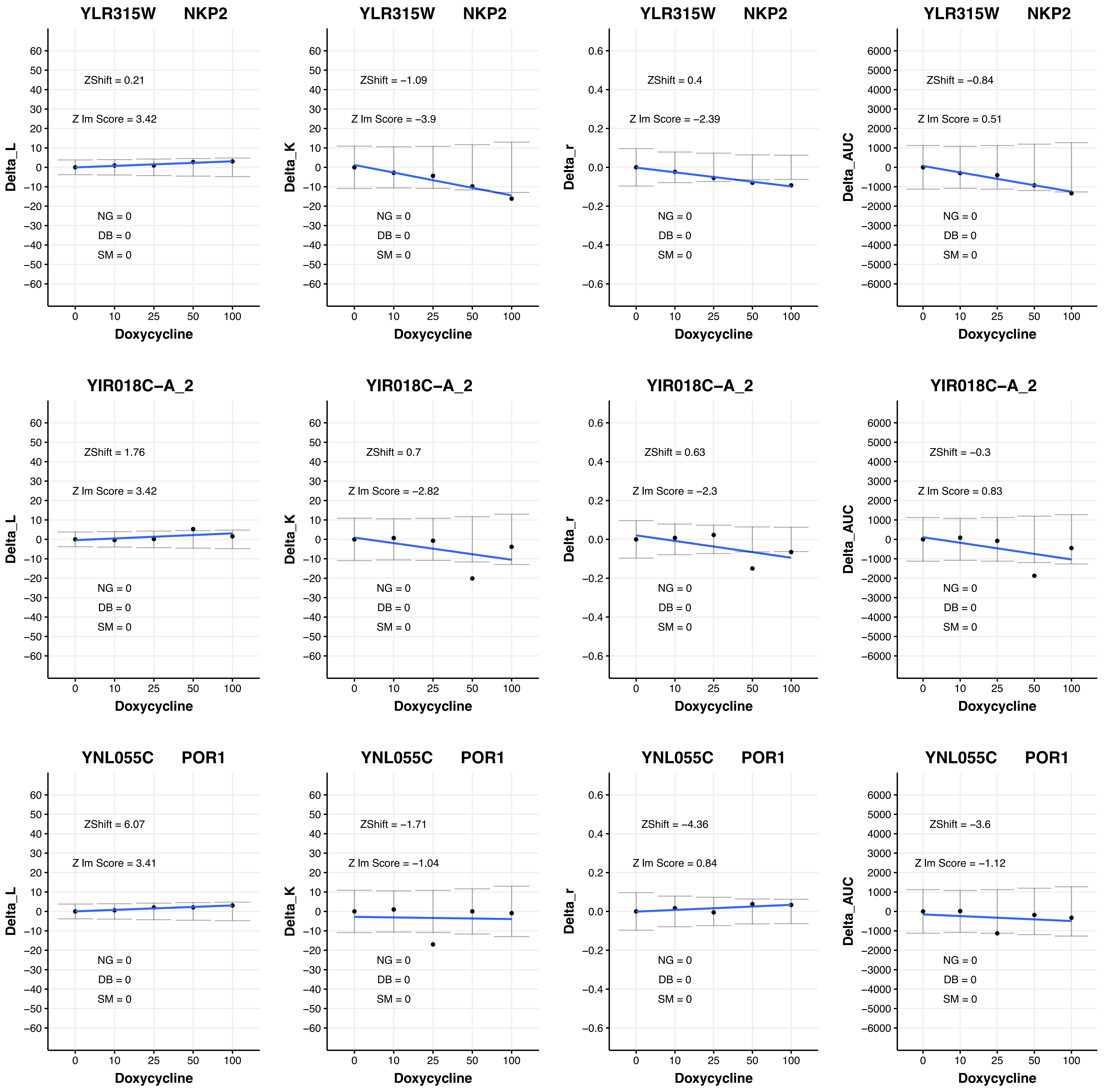

YLR315W NKP2
ZShift = 0.21
Z lm Score = 3.42
NG = 0
DB = 0
SM = 0
Delta_L
Doxycycline
YLR315W NKP2
ZShift = −1.09
Z lm Score = −3.9
NG = 0
DB = 0
SM = 0
Delta_K
Doxycycline
YLR315W NKP2
ZShift = 0.4
Z lm Score = −2.39
NG = 0
DB = 0
SM = 0
Delta_r
Doxycycline
YLR315W NKP2
ZShift = −0.84
Z lm Score = 0.51
NG = 0
DB = 0
SM = 0
Delta_AUC
Doxycycline
YIR018C−A_2
ZShift = 1.76
Z lm Score = 3.42
NG = 0
DB = 0
SM = 0
Delta_L
Doxycycline
YIR018C−A_2
ZShift = 0.7
Z lm Score = −2.82
NG = 0
DB = 0
SM = 0
Delta_K
Doxycycline
YIR018C−A_2
ZShift = 0.63
Z lm Score = −2.3
NG = 0
DB = 0
SM = 0
Delta_r
Doxycycline
YIR018C−A_2
ZShift = −0.3
Z lm Score = 0.83
NG = 0
DB = 0
SM = 0
Delta_AUC
Doxycycline
YNL055C POR1
ZShift = 6.07
Z lm Score = 3.41
NG = 0
DB = 0
SM = 0
Delta_L
Doxycycline
YNL055C POR1
ZShift = −1.71
Z lm Score = −1.04
NG = 0
DB = 0
SM = 0
Delta_K
Doxycycline
YNL055C POR1
ZShift = −4.36
Z lm Score = 0.84
NG = 0
DB = 0
SM = 0
Delta_r
Doxycycline
YNL055C POR1
ZShift = −3.6
Z lm Score = −1.12
NG = 0
DB = 0
SM = 0
Delta_AUC
Doxycycline

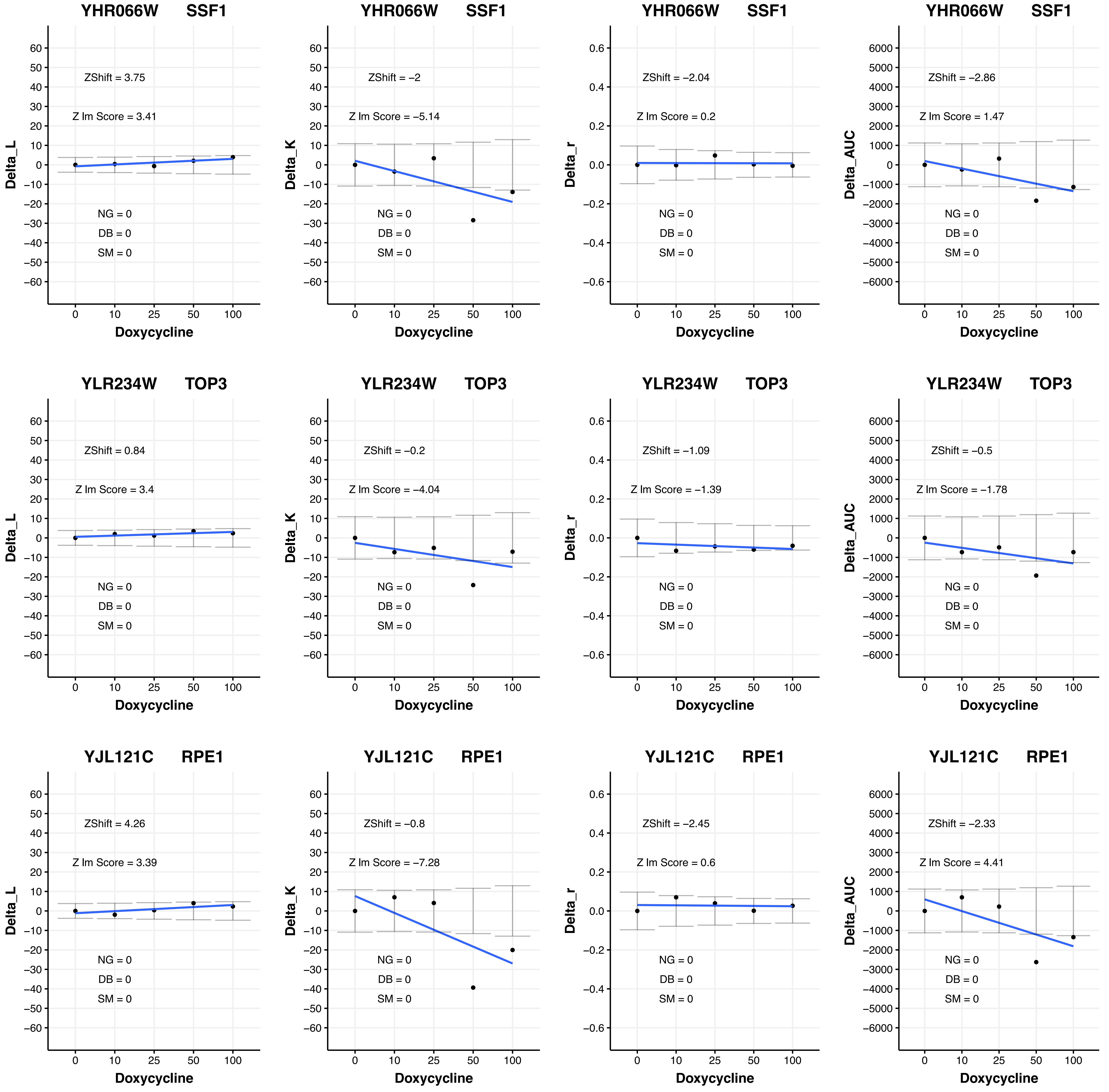
YHR066W SSF1
ZShift = 3.75
Z lm Score = 3.41
NG = 0
DB = 0
SM = 0
Delta_L
Doxycycline
YHR066W SSF1
ZShift = −2
Z lm Score = −5.14
NG = 0
DB = 0
SM = 0
Delta_K
Doxycycline
YHR066W SSF1
ZShift = −2.04
Z lm Score = 0.2
NG = 0
DB = 0
SM = 0
Delta_r
Doxycycline
YHR066W SSF1
ZShift = −2.86
Z lm Score = 1.47
NG = 0
DB = 0
SM = 0
Delta_AUC
Doxycycline
YLR234W TOP3
ZShift = 0.84
Z lm Score = 3.4
NG = 0
DB = 0
SM = 0
Delta_L
Doxycycline
YLR234W TOP3
ZShift = −0.2
Z lm Score = −4.04
NG = 0
DB = 0
SM = 0
Delta_K
Doxycycline
YLR234W TOP3
ZShift = −1.09
Z lm Score = −1.39
NG = 0
DB = 0
SM = 0
Delta_r
Doxycycline
YLR234W TOP3
ZShift = −0.5
Z lm Score = −1.78
NG = 0
DB = 0
SM = 0
Delta_AUC
Doxycycline
YJL121C RPE1
ZShift = 4.26
Z lm Score = 3.39
NG = 0
DB = 0
SM = 0
Delta_L
Doxycycline
YJL121C RPE1
ZShift = −0.8
Z lm Score = −7.28
NG = 0
DB = 0
SM = 0
Delta_K
Doxycycline
YJL121C RPE1
ZShift = −2.45
Z lm Score = 0.6
NG = 0
DB = 0
SM = 0
Delta_r
Doxycycline
YJL121C RPE1
ZShift = −2.33
Z lm Score = 4.41
NG = 0
DB = 0
SM = 0
Delta_AUC
Doxycycline

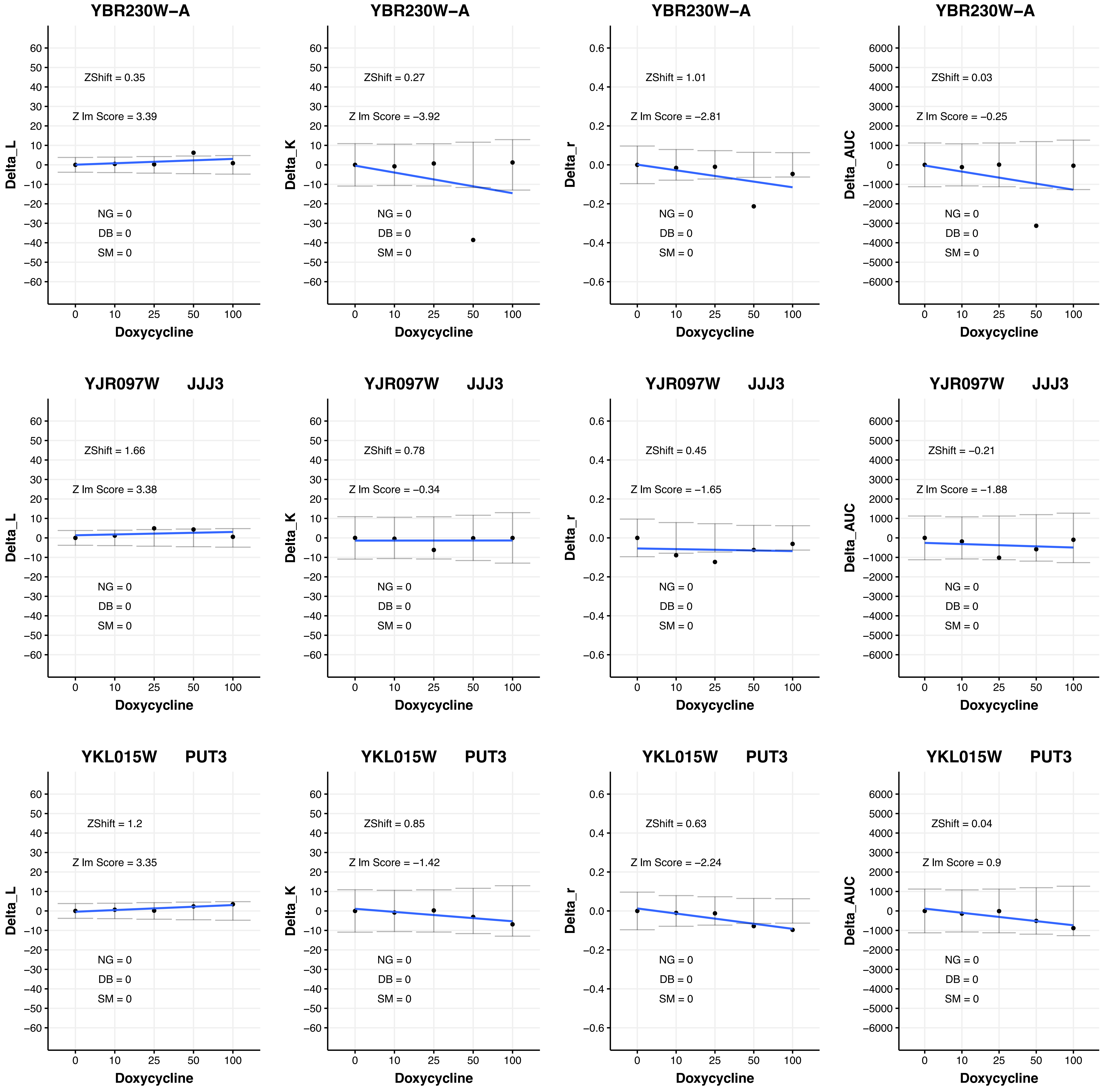
YBR230W−A
ZShift = 0.35
Z lm Score = 3.39
NG = 0
DB = 0
SM = 0
Delta_L
Doxycycline
YBR230W−A
ZShift = 0.27
Z lm Score = −3.92
NG = 0
DB = 0
SM = 0
Delta_K
Doxycycline
YBR230W−A
ZShift = 1.01
Z lm Score = −2.81
NG = 0
DB = 0
SM = 0
Delta_r
Doxycycline
YBR230W−A
ZShift = 0.03
Z lm Score = −0.25
NG = 0
DB = 0
SM = 0
Delta_AUC
Doxycycline
YJR097W JJJ3
ZShift = 1.66
Z lm Score = 3.38
NG = 0
DB = 0
SM = 0
Delta_L
Doxycycline
YJR097W JJJ3
ZShift = 0.78
Z lm Score = −0.34
NG = 0
DB = 0
SM = 0
Delta_K
Doxycycline
YJR097W JJJ3
ZShift = 0.45
Z lm Score = −1.65
NG = 0
DB = 0
SM = 0
Delta_r
Doxycycline
YJR097W JJJ3
ZShift = −0.21
Z lm Score = −1.88
NG = 0
DB = 0
SM = 0
Delta_AUC
Doxycycline
YKL015W PUT3
ZShift = 1.2
Z lm Score = 3.35
NG = 0
DB = 0
SM = 0
Delta_L
Doxycycline
YKL015W PUT3
ZShift = 0.85
Z lm Score = −1.42
NG = 0
DB = 0
SM = 0
Delta_K
Doxycycline
YKL015W PUT3
ZShift = 0.63
Z lm Score = −2.24
NG = 0
DB = 0
SM = 0
Delta_r
Doxycycline
YKL015W PUT3
ZShift = 0.04
Z lm Score = 0.9
NG = 0
DB = 0
SM = 0
Delta_AUC
Doxycycline

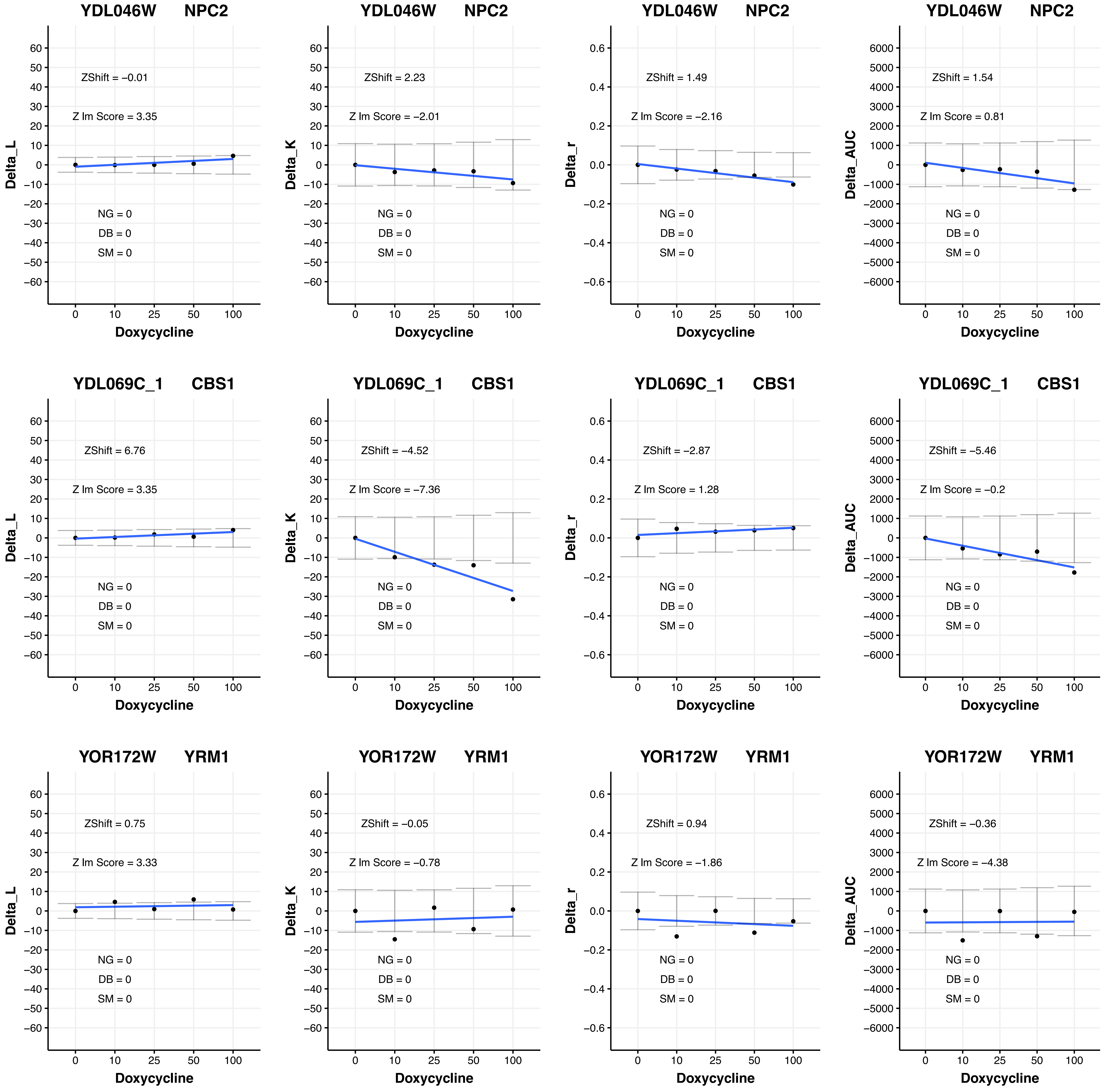

YDL046W NPC2
Delta_L
ZShift = −0.01
Z lm Score = 3.35
NG = 0
DB = 0
SM = 0
Doxycycline
YDL046W NPC2
Delta_K
ZShift = 2.23
Z lm Score = −2.01
NG = 0
DB = 0
SM = 0
Doxycycline
YDL046W NPC2
Delta_r
ZShift = 1.49
Z lm Score = −2.16
NG = 0
DB = 0
SM = 0
Doxycycline
YDL046W NPC2
Delta_AUC
ZShift = 1.54
Z lm Score = 0.81
NG = 0
DB = 0
SM = 0
Doxycycline
YDL069C_1 CBS1
Delta_L
ZShift = 6.76
Z lm Score = 3.35
NG = 0
DB = 0
SM = 0
Doxycycline
YDL069C_1 CBS1
Delta_K
ZShift = −4.52
Z lm Score = −7.36
NG = 0
DB = 0
SM = 0
Doxycycline
YDL069C_1 CBS1
Delta_r
ZShift = −2.87
Z lm Score = 1.28
NG = 0
DB = 0
SM = 0
Doxycycline
YDL069C_1 CBS1
Delta_AUC
ZShift = −5.46
Z lm Score = −0.2
NG = 0
DB = 0
SM = 0
Doxycycline
YOR172W YRM1
Delta_L
ZShift = 0.75
Z lm Score = 3.33
NG = 0
DB = 0
SM = 0
Doxycycline
YOR172W YRM1
Delta_K
ZShift = −0.05
Z lm Score = −0.78
NG = 0
DB = 0
SM = 0
Doxycycline
YOR172W YRM1
Delta_r
ZShift = 0.94
Z lm Score = −1.86
NG = 0
DB = 0
SM = 0
Doxycycline
YOR172W YRM1
Delta_AUC
ZShift = −0.36
Z lm Score = −4.38
NG = 0
DB = 0
SM = 0
Doxycycline

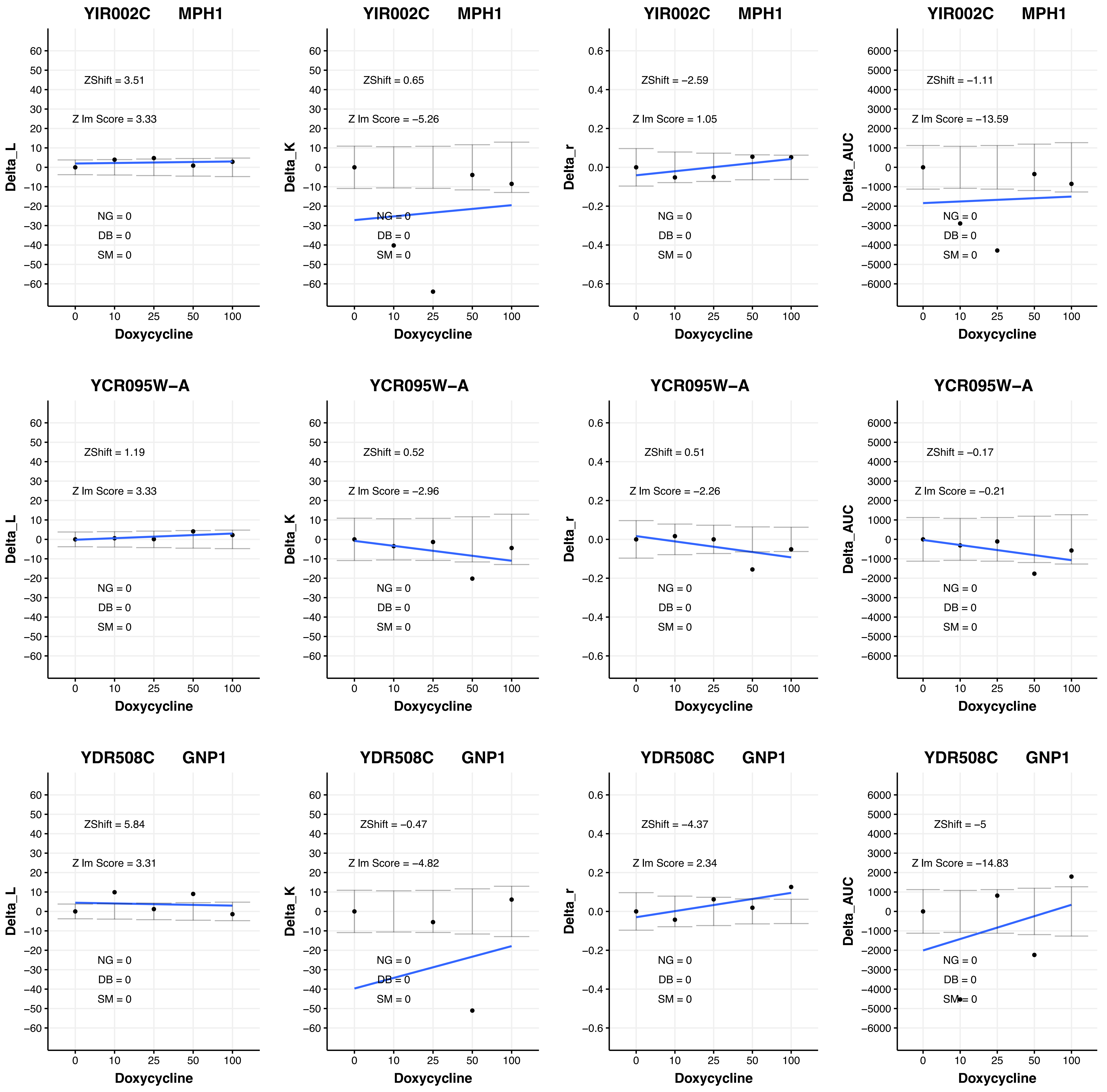
YIR002C MPH1
ZShift = 3.51
Z lm Score = 3.33
NG = 0
DB = 0
SM = 0
Delta_L
Doxycycline
YIR002C MPH1
ZShift = 0.65
Z lm Score = −5.26
NG = 0
DB = 0
SM = 0
Delta_K
Doxycycline
YIR002C MPH1
ZShift = −2.59
Z lm Score = 1.05
NG = 0
DB = 0
SM = 0
Delta_r
Doxycycline
YIR002C MPH1
ZShift = −1.11
Z lm Score = −13.59
NG = 0
DB = 0
SM = 0
Delta_AUC
Doxycycline
YCR095W−A
ZShift = 1.19
Z lm Score = 3.33
NG = 0
DB = 0
SM = 0
Delta_L
Doxycycline
YCR095W−A
ZShift = 0.52
Z lm Score = −2.96
NG = 0
DB = 0
SM = 0
Delta_K
Doxycycline
YCR095W−A
ZShift = 0.51
Z lm Score = −2.26
NG = 0
DB = 0
SM = 0
Delta_r
Doxycycline
YCR095W−A
ZShift = −0.17
Z lm Score = −0.21
NG = 0
DB = 0
SM = 0
Delta_AUC
Doxycycline
YDR508C GNP1
ZShift = 5.84
Z lm Score = 3.31
NG = 0
DB = 0
SM = 0
Delta_L
Doxycycline
YDR508C GNP1
ZShift = −0.47
Z lm Score = −4.82
NG = 0
DB = 0
SM = 0
Delta_K
Doxycycline
YDR508C GNP1
ZShift = −4.37
Z lm Score = 2.34
NG = 0
DB = 0
SM = 0
Delta_r
Doxycycline
YDR508C GNP1
ZShift = −5
Z lm Score = −14.83
NG = 0
DB = 0
SM = 0
Delta_AUC
Doxycycline

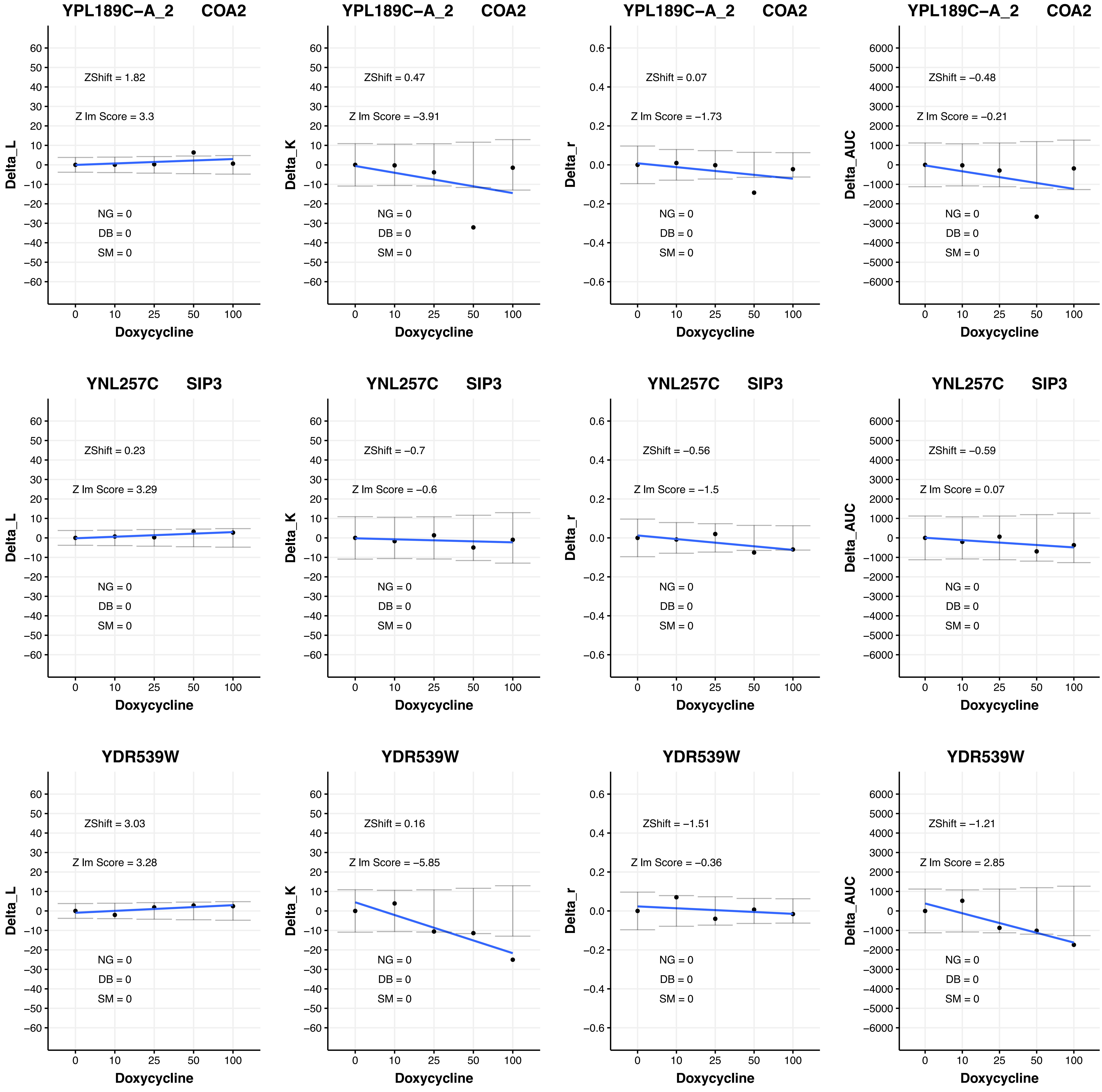

YPL189C−A_2 COA2
ZShift = 1.82
Z lm Score = 3.3
NG = 0
DB = 0
SM = 0
Delta_L
Doxycycline
YPL189C−A_2 COA2
ZShift = 0.47
Z lm Score = −3.91
NG = 0
DB = 0
SM = 0
Delta_K
Doxycycline
YPL189C−A_2 COA2
ZShift = 0.07
Z lm Score = −1.73
NG = 0
DB = 0
SM = 0
Delta_r
Doxycycline
YPL189C−A_2 COA2
ZShift = −0.48
Z lm Score = −0.21
NG = 0
DB = 0
SM = 0
Delta_AUC
Doxycycline
YNL257C SIP3
ZShift = 0.23
Z lm Score = 3.29
NG = 0
DB = 0
SM = 0
Delta_L
Doxycycline
YNL257C SIP3
ZShift = −0.7
Z lm Score = −0.6
NG = 0
DB = 0
SM = 0
Delta_K
Doxycycline
YNL257C SIP3
ZShift = −0.56
Z lm Score = −1.5
NG = 0
DB = 0
SM = 0
Delta_r
Doxycycline
YNL257C SIP3
ZShift = −0.59
Z lm Score = 0.07
NG = 0
DB = 0
SM = 0
Delta_AUC
Doxycycline
YDR539W
ZShift = 3.03
Z lm Score = 3.28
NG = 0
DB = 0
SM = 0
Delta_L
Doxycycline
YDR539W
ZShift = 0.16
Z lm Score = −5.85
NG = 0
DB = 0
SM = 0
Delta_K
Doxycycline
YDR539W
ZShift = −1.51
Z lm Score = −0.36
NG = 0
DB = 0
SM = 0
Delta_r
Doxycycline
YDR539W
ZShift = −1.21
Z lm Score = 2.85
NG = 0
DB = 0
SM = 0
Delta_AUC
Doxycycline

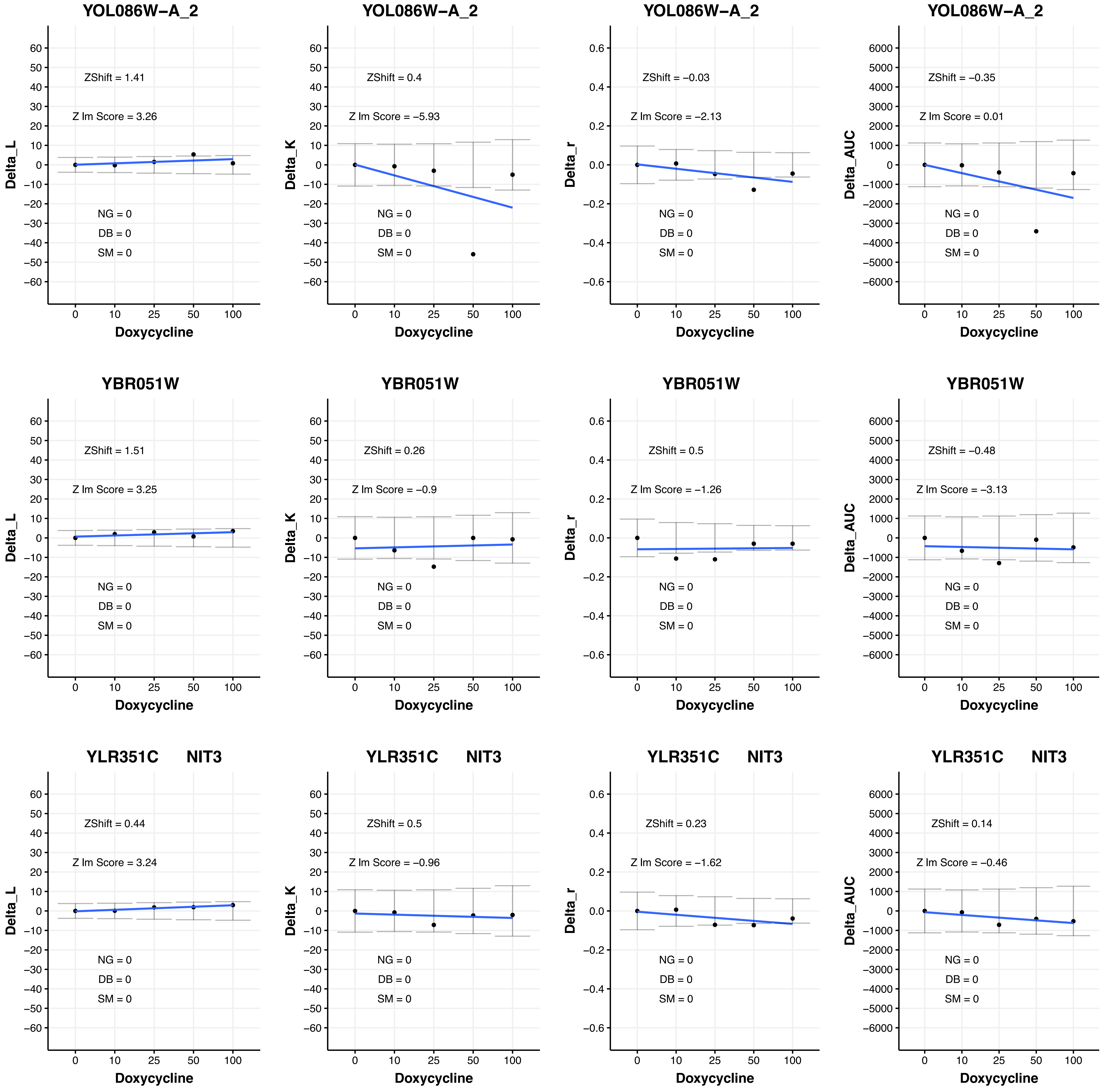

YOL086W−A_2
ZShift = 1.41
Z lm Score = 3.26
NG = 0
DB = 0
SM = 0
Delta_L
Doxycycline
YOL086W−A_2
ZShift = 0.4
Z lm Score = −5.93
NG = 0
DB = 0
SM = 0
Delta_K
Doxycycline
YOL086W−A_2
ZShift = −0.03
Z lm Score = −2.13
NG = 0
DB = 0
SM = 0
Delta_r
Doxycycline
YOL086W−A_2
ZShift = −0.35
Z lm Score = 0.01
NG = 0
DB = 0
SM = 0
Delta_AUC
Doxycycline
YBR051W
ZShift = 1.51
Z lm Score = 3.25
NG = 0
DB = 0
SM = 0
Delta_L
Doxycycline
YBR051W
ZShift = 0.26
Z lm Score = −0.9
NG = 0
DB = 0
SM = 0
Delta_K
Doxycycline
YBR051W
ZShift = 0.5
Z lm Score = −1.26
NG = 0
DB = 0
SM = 0
Delta_r
Doxycycline
YBR051W
ZShift = −0.48
Z lm Score = −3.13
NG = 0
DB = 0
SM = 0
Delta_AUC
Doxycycline
YLR351C NIT3
ZShift = 0.44
Z lm Score = 3.24
NG = 0
DB = 0
SM = 0
Delta_L
Doxycycline
YLR351C NIT3
ZShift = 0.5
Z lm Score = −0.96
NG = 0
DB = 0
SM = 0
Delta_K
Doxycycline
YLR351C NIT3
ZShift = 0.23
Z lm Score = −1.62
NG = 0
DB = 0
SM = 0
Delta_r
Doxycycline
YLR351C NIT3
ZShift = 0.14
Z lm Score = −0.46
NG = 0
DB = 0
SM = 0
Delta_AUC
Doxycycline

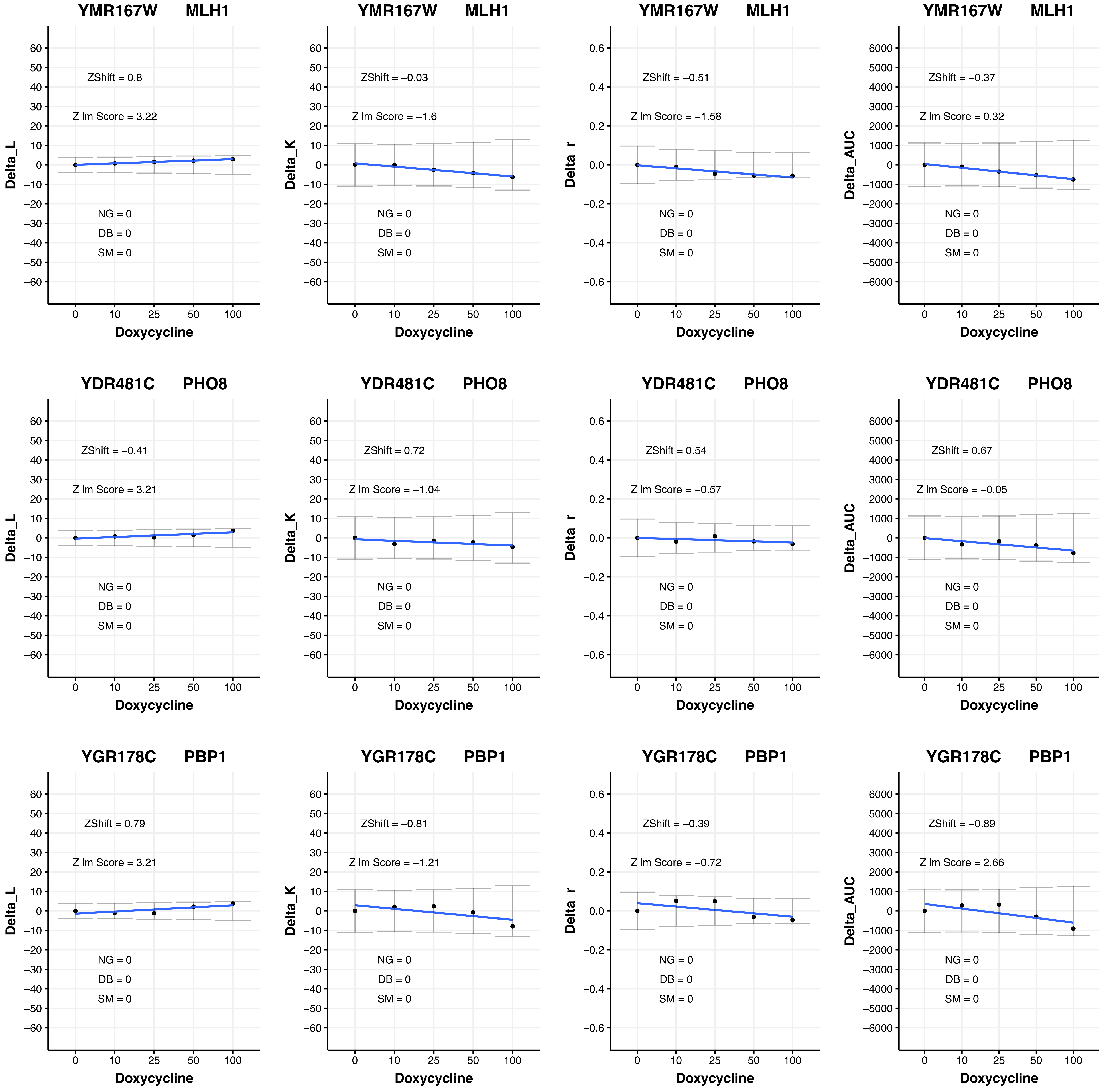
YMR167W MLH1
ZShift = 0.8
Z lm Score = 3.22
NG = 0
DB = 0
SM = 0
Delta_L
Doxycycline
YMR167W MLH1
ZShift = −0.03
Z lm Score = −1.6
NG = 0
DB = 0
SM = 0
Delta_K
Doxycycline
YMR167W MLH1
ZShift = −0.51
Z lm Score = −1.58
NG = 0
DB = 0
SM = 0
Delta_r
Doxycycline
YMR167W MLH1
ZShift = −0.37
Z lm Score = 0.32
NG = 0
DB = 0
SM = 0
Delta_AUC
Doxycycline
YDR481C PHO8
ZShift = −0.41
Z lm Score = 3.21
NG = 0
DB = 0
SM = 0
Delta_L
Doxycycline
YDR481C PHO8
ZShift = 0.72
Z lm Score = −1.04
NG = 0
DB = 0
SM = 0
Delta_K
Doxycycline
YDR481C PHO8
ZShift = 0.54
Z lm Score = −0.57
NG = 0
DB = 0
SM = 0
Delta_r
Doxycycline
YDR481C PHO8
ZShift = 0.67
Z lm Score = −0.05
NG = 0
DB = 0
SM = 0
Delta_AUC
Doxycycline
YGR178C PBP1
ZShift = 0.79
Z lm Score = 3.21
NG = 0
DB = 0
SM = 0
Delta_L
Doxycycline
YGR178C PBP1
ZShift = −0.81
Z lm Score = −1.21
NG = 0
DB = 0
SM = 0
Delta_K
Doxycycline
YGR178C PBP1
ZShift = −0.39
Z lm Score = −0.72
NG = 0
DB = 0
SM = 0
Delta_r
Doxycycline
YGR178C PBP1
ZShift = −0.89
Z lm Score = 2.66
NG = 0
DB = 0
SM = 0
Delta_AUC
Doxycycline

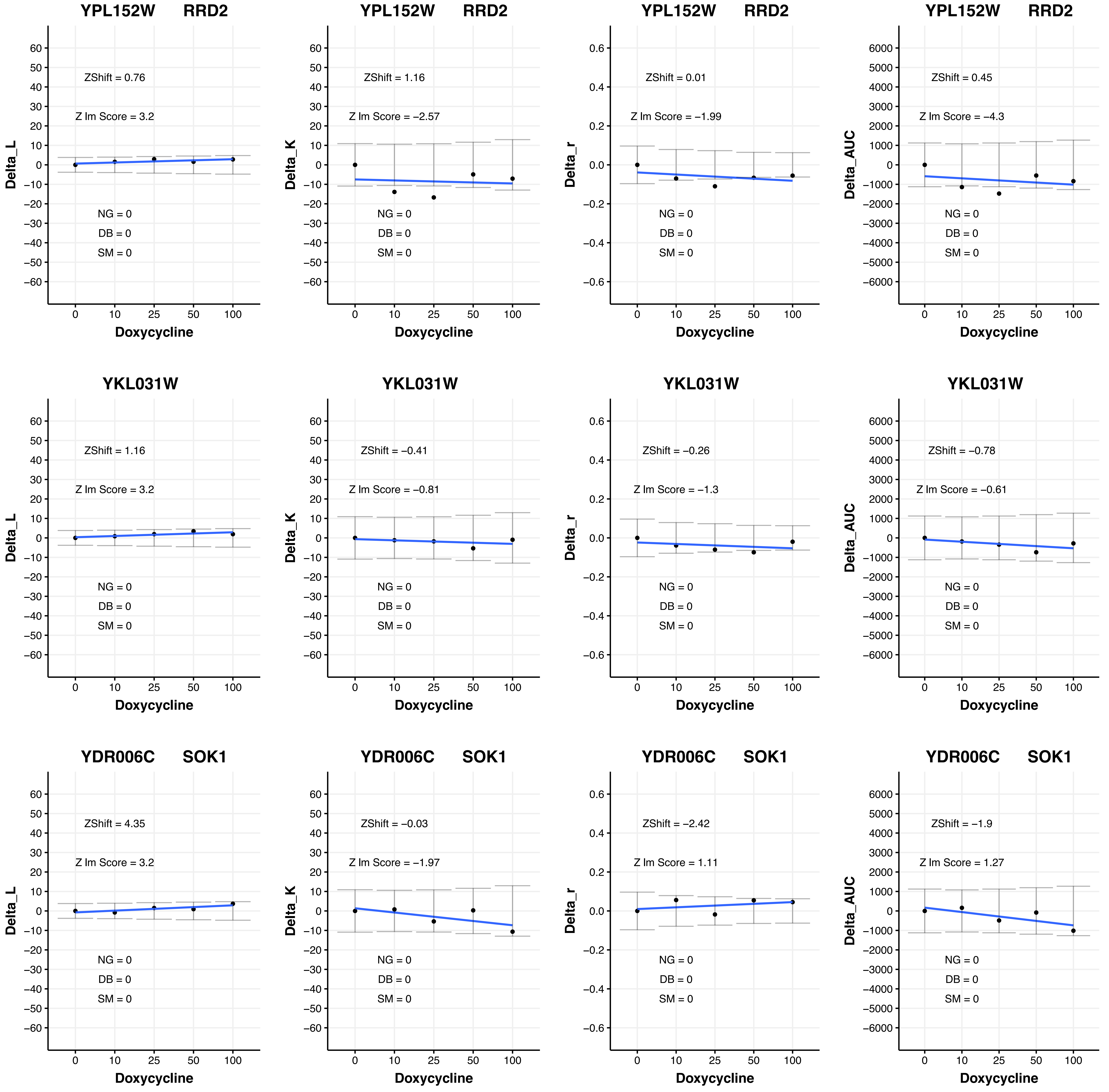
YPL152W RRD2
ZShift = 0.76
Z lm Score = 3.2
NG = 0
DB = 0
SM = 0
Delta_L
Doxycycline
YPL152W RRD2
ZShift = 1.16
Z lm Score = −2.57
NG = 0
DB = 0
SM = 0
Delta_K
Doxycycline
YPL152W RRD2
ZShift = 0.01
Z lm Score = −1.99
NG = 0
DB = 0
SM = 0
Delta_r
Doxycycline
YPL152W RRD2
ZShift = 0.45
Z lm Score = −4.3
NG = 0
DB = 0
SM = 0
Delta_AUC
Doxycycline
YKL031W
ZShift = 1.16
Z lm Score = 3.2
NG = 0
DB = 0
SM = 0
Delta_L
Doxycycline
YKL031W
ZShift = −0.41
Z lm Score = −0.81
NG = 0
DB = 0
SM = 0
Delta_K
Doxycycline
YKL031W
ZShift = −0.26
Z lm Score = −1.3
NG = 0
DB = 0
SM = 0
Delta_r
Doxycycline
YKL031W
ZShift = −0.78
Z lm Score = −0.61
NG = 0
DB = 0
SM = 0
Delta_AUC
Doxycycline
YDR006C SOK1
ZShift = 4.35
Z lm Score = 3.2
NG = 0
DB = 0
SM = 0
Delta_L
Doxycycline
YDR006C SOK1
ZShift = −0.03
Z lm Score = −1.97
NG = 0
DB = 0
SM = 0
Delta_K
Doxycycline
YDR006C SOK1
ZShift = −2.42
Z lm Score = 1.11
NG = 0
DB = 0
SM = 0
Delta_r
Doxycycline
YDR006C SOK1
ZShift = −1.9
Z lm Score = 1.27
NG = 0
DB = 0
SM = 0
Delta_AUC
Doxycycline

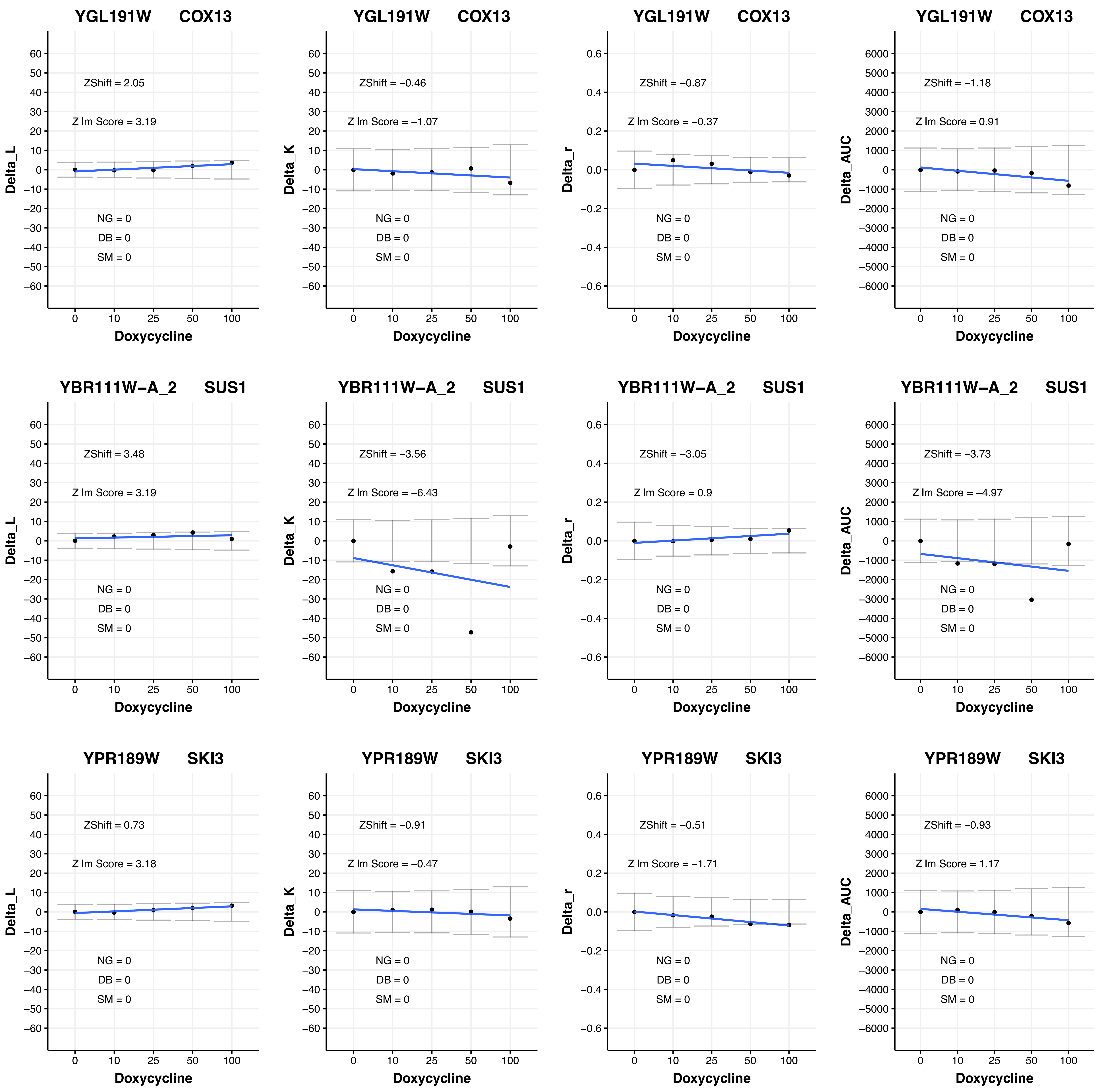
YGL191W COX13
ZShift = 2.05
Z lm Score = 3.19
NG = 0
DB = 0
SM = 0
Delta_L
Doxycycline
YGL191W COX13
ZShift = −0.46
Z lm Score = −1.07
NG = 0
DB = 0
SM = 0
Delta_K
Doxycycline
YGL191W COX13
ZShift = −0.87
Z lm Score = −0.37
NG = 0
DB = 0
SM = 0
Delta_r
Doxycycline
YGL191W COX13
ZShift = −1.18
Z lm Score = 0.91
NG = 0
DB = 0
SM = 0
Delta_AUC
Doxycycline
YBR111W−A_2 SUS1
ZShift = 3.48
Z lm Score = 3.19
NG = 0
DB = 0
SM = 0
Delta_L
Doxycycline
YBR111W−A_2 SUS1
ZShift = −3.56
Z lm Score = −6.43
NG = 0
DB = 0
SM = 0
Delta_K
Doxycycline
YBR111W−A_2 SUS1
ZShift = −3.05
Z lm Score = 0.9
NG = 0
DB = 0
SM = 0
Delta_r
Doxycycline
YBR111W−A_2 SUS1
ZShift = −3.73
Z lm Score = −4.97
NG = 0
DB = 0
SM = 0
Delta_AUC
Doxycycline
YPR189W SKI3
ZShift = 0.73
Z lm Score = 3.18
NG = 0
DB = 0
SM = 0
Delta_L
Doxycycline
YPR189W SKI3
ZShift = −0.91
Z lm Score = −0.47
NG = 0
DB = 0
SM = 0
Delta_K
Doxycycline
YPR189W SKI3
ZShift = −0.51
Z lm Score = −1.71
NG = 0
DB = 0
SM = 0
Delta_r
Doxycycline
YPR189W SKI3
ZShift = −0.93
Z lm Score = 1.17
NG = 0
DB = 0
SM = 0
Delta_AUC
Doxycycline

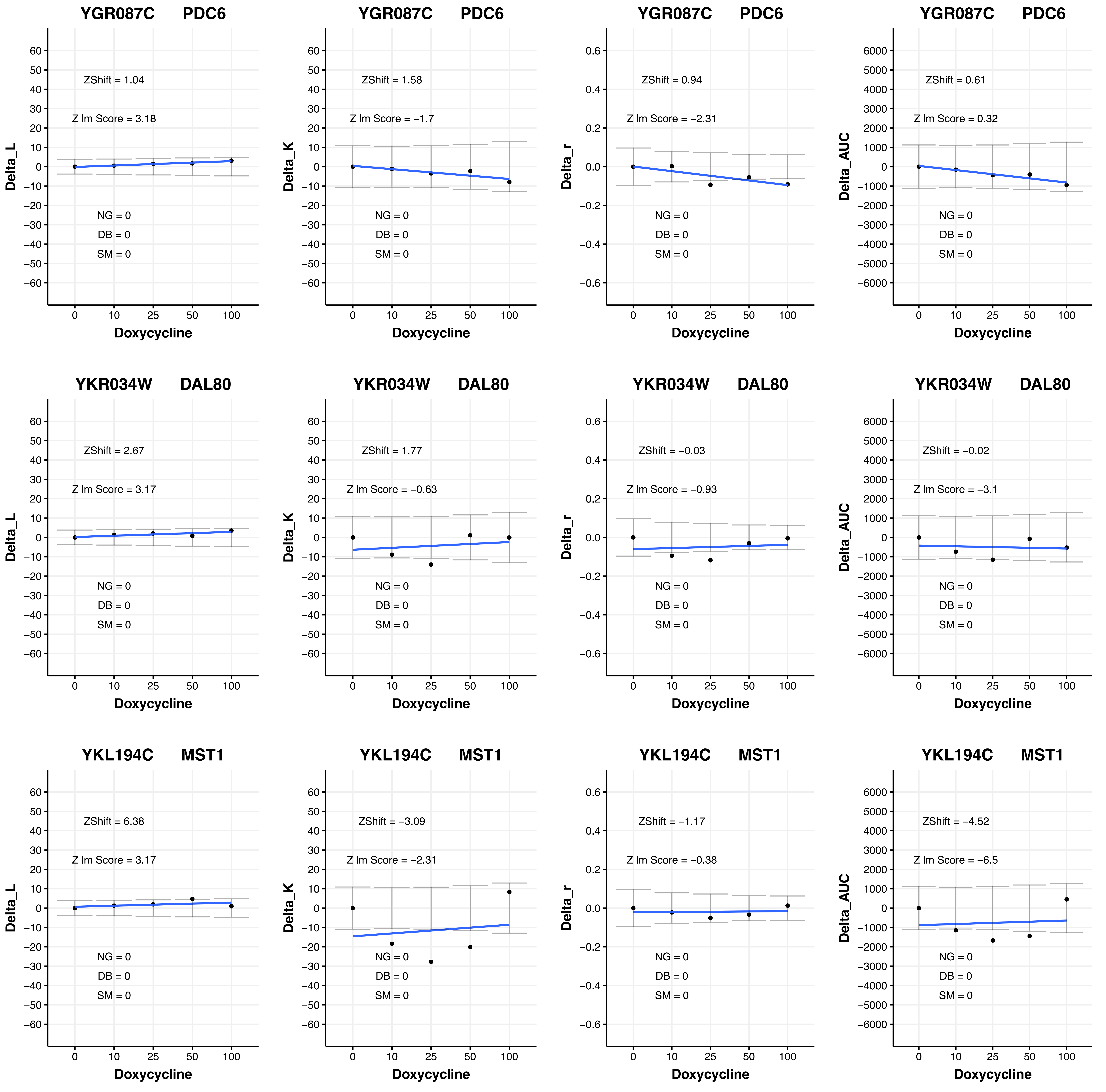
YGR087C PDC6
ZShift = 1.04
Z lm Score = 3.18
NG = 0
DB = 0
SM = 0
Delta_L
Doxycycline
YGR087C PDC6
ZShift = 1.58
Z lm Score = −1.7
NG = 0
DB = 0
SM = 0
Delta_K
Doxycycline
YGR087C PDC6
ZShift = 0.94
Z lm Score = −2.31
NG = 0
DB = 0
SM = 0
Delta_r
Doxycycline
YGR087C PDC6
ZShift = 0.61
Z lm Score = 0.32
NG = 0
DB = 0
SM = 0
Delta_AUC
Doxycycline
YKR034W DAL80
ZShift = 2.67
Z lm Score = 3.17
NG = 0
DB = 0
SM = 0
Delta_L
Doxycycline
YKR034W DAL80
ZShift = 1.77
Z lm Score = −0.63
NG = 0
DB = 0
SM = 0
Delta_K
Doxycycline
YKR034W DAL80
ZShift = −0.03
Z lm Score = −0.93
NG = 0
DB = 0
SM = 0
Delta_r
Doxycycline
YKR034W DAL80
ZShift = −0.02
Z lm Score = −3.1
NG = 0
DB = 0
SM = 0
Delta_AUC
Doxycycline
YKL194C MST1
ZShift = 6.38
Z lm Score = 3.17
NG = 0
DB = 0
SM = 0
Delta_L
Doxycycline
YKL194C MST1
ZShift = −3.09
Z lm Score = −2.31
NG = 0
DB = 0
SM = 0
Delta_K
Doxycycline
YKL194C MST1
ZShift = −1.17
Z lm Score = −0.38
NG = 0
DB = 0
SM = 0
Delta_r
Doxycycline
YKL194C MST1
ZShift = −4.52
Z lm Score = −6.5
NG = 0
DB = 0
SM = 0
Delta_AUC
Doxycycline

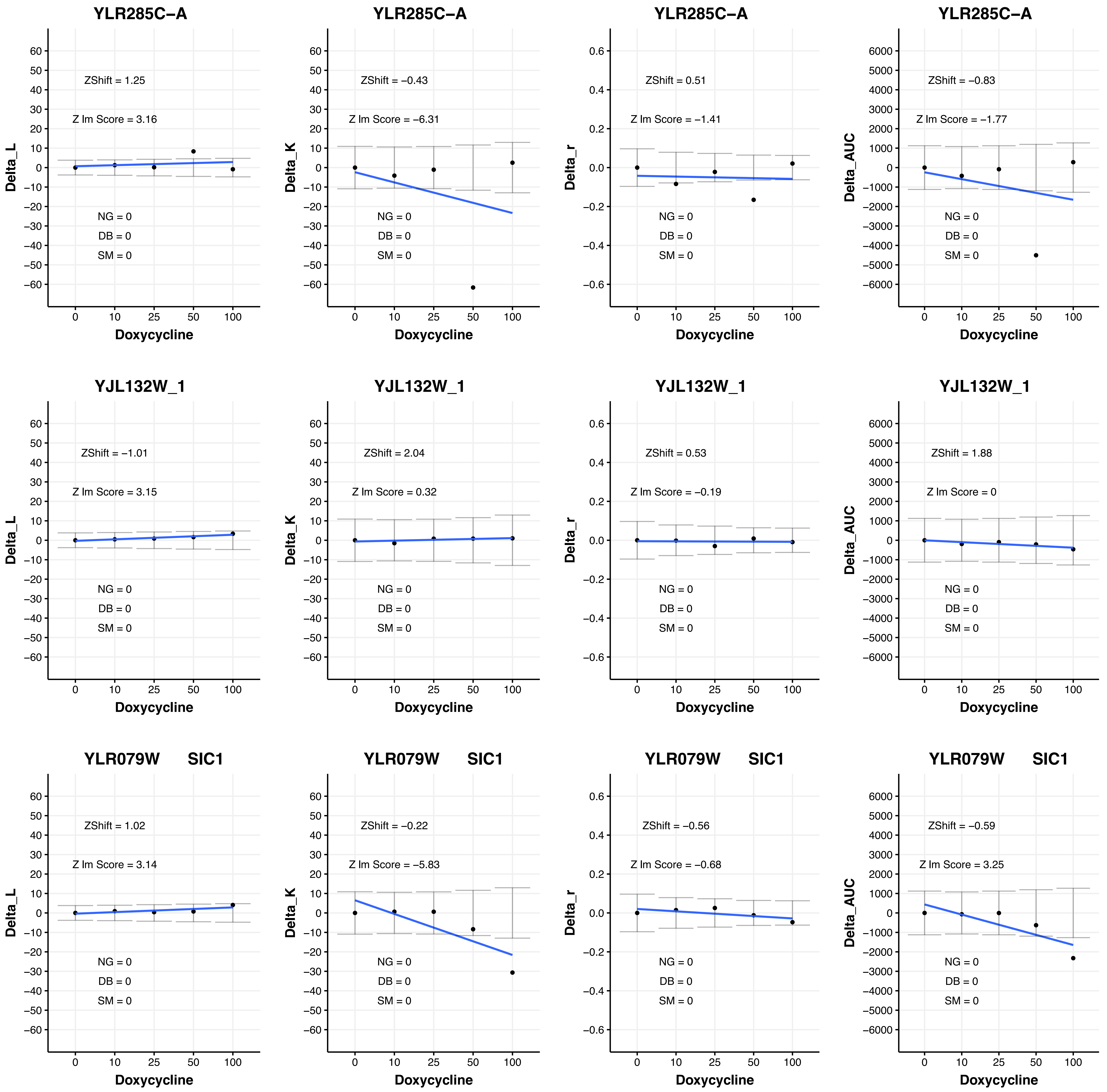

YLR285C−A
ZShift = 1.25
Z lm Score = 3.16
NG = 0
DB = 0
SM = 0
Delta_L
Doxycycline
YLR285C−A
ZShift = −0.43
Z lm Score = −6.31
NG = 0
DB = 0
SM = 0
Delta_K
Doxycycline
YLR285C−A
ZShift = 0.51
Z lm Score = −1.41
NG = 0
DB = 0
SM = 0
Delta_r
Doxycycline
YLR285C−A
ZShift = −0.83
Z lm Score = −1.77
NG = 0
DB = 0
SM = 0
Delta_AUC
Doxycycline
YJL132W_1
ZShift = −1.01
Z lm Score = 3.15
NG = 0
DB = 0
SM = 0
Delta_L
Doxycycline
YJL132W_1
ZShift = 2.04
Z lm Score = 0.32
NG = 0
DB = 0
SM = 0
Delta_K
Doxycycline
YJL132W_1
ZShift = 0.53
Z lm Score = −0.19
NG = 0
DB = 0
SM = 0
Delta_r
Doxycycline
YJL132W_1
ZShift = 1.88
Z lm Score = 0
NG = 0
DB = 0
SM = 0
Delta_AUC
Doxycycline
YLR079W SIC1
ZShift = 1.02
Z lm Score = 3.14
NG = 0
DB = 0
SM = 0
Delta_L
Doxycycline
YLR079W SIC1
ZShift = −0.22
Z lm Score = −5.83
NG = 0
DB = 0
SM = 0
Delta_K
Doxycycline
YLR079W SIC1
ZShift = −0.56
Z lm Score = −0.68
NG = 0
DB = 0
SM = 0
Delta_r
Doxycycline
YLR079W SIC1
ZShift = −0.59
Z lm Score = 3.25
NG = 0
DB = 0
SM = 0
Delta_AUC
Doxycycline

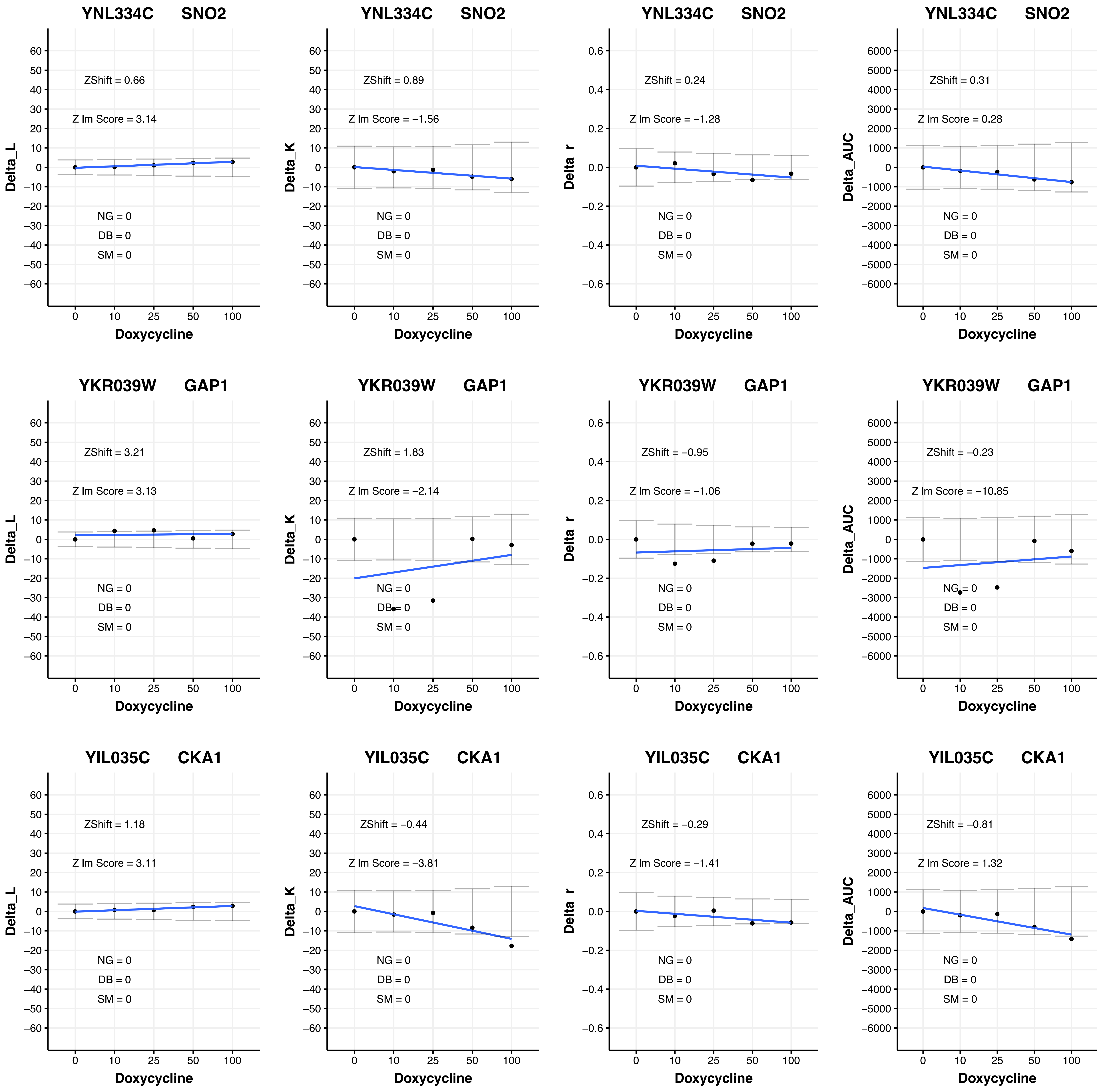

YNL334C SNO2
ZShift = 0.66
Z lm Score = 3.14
NG = 0
DB = 0
SM = 0
Delta_L
Doxycycline
YNL334C SNO2
ZShift = 0.89
Z lm Score = −1.56
NG = 0
DB = 0
SM = 0
Delta_K
Doxycycline
YNL334C SNO2
ZShift = 0.24
Z lm Score = −1.28
NG = 0
DB = 0
SM = 0
Delta_r
Doxycycline
YNL334C SNO2
ZShift = 0.31
Z lm Score = 0.28
NG = 0
DB = 0
SM = 0
Delta_AUC
Doxycycline
YKR039W GAP1
ZShift = 3.21
Z lm Score = 3.13
NG = 0
DB = 0
SM = 0
Delta_L
Doxycycline
YKR039W GAP1
ZShift = 1.83
Z lm Score = −2.14
NG = 0
DB = 0
SM = 0
Delta_K
Doxycycline
YKR039W GAP1
ZShift = −0.95
Z lm Score = −1.06
NG = 0
DB = 0
SM = 0
Delta_r
Doxycycline
YKR039W GAP1
ZShift = −0.23
Z lm Score = −10.85
NG = 0
DB = 0
SM = 0
Delta_AUC
Doxycycline
YIL035C CKA1
ZShift = 1.18
Z lm Score = 3.11
NG = 0
DB = 0
SM = 0
Delta_L
Doxycycline
YIL035C CKA1
ZShift = −0.44
Z lm Score = −3.81
NG = 0
DB = 0
SM = 0
Delta_K
Doxycycline
YIL035C CKA1
ZShift = −0.29
Z lm Score = −1.41
NG = 0
DB = 0
SM = 0
Delta_r
Doxycycline
YIL035C CKA1
ZShift = −0.81
Z lm Score = 1.32
NG = 0
DB = 0
SM = 0
Delta_AUC
Doxycycline

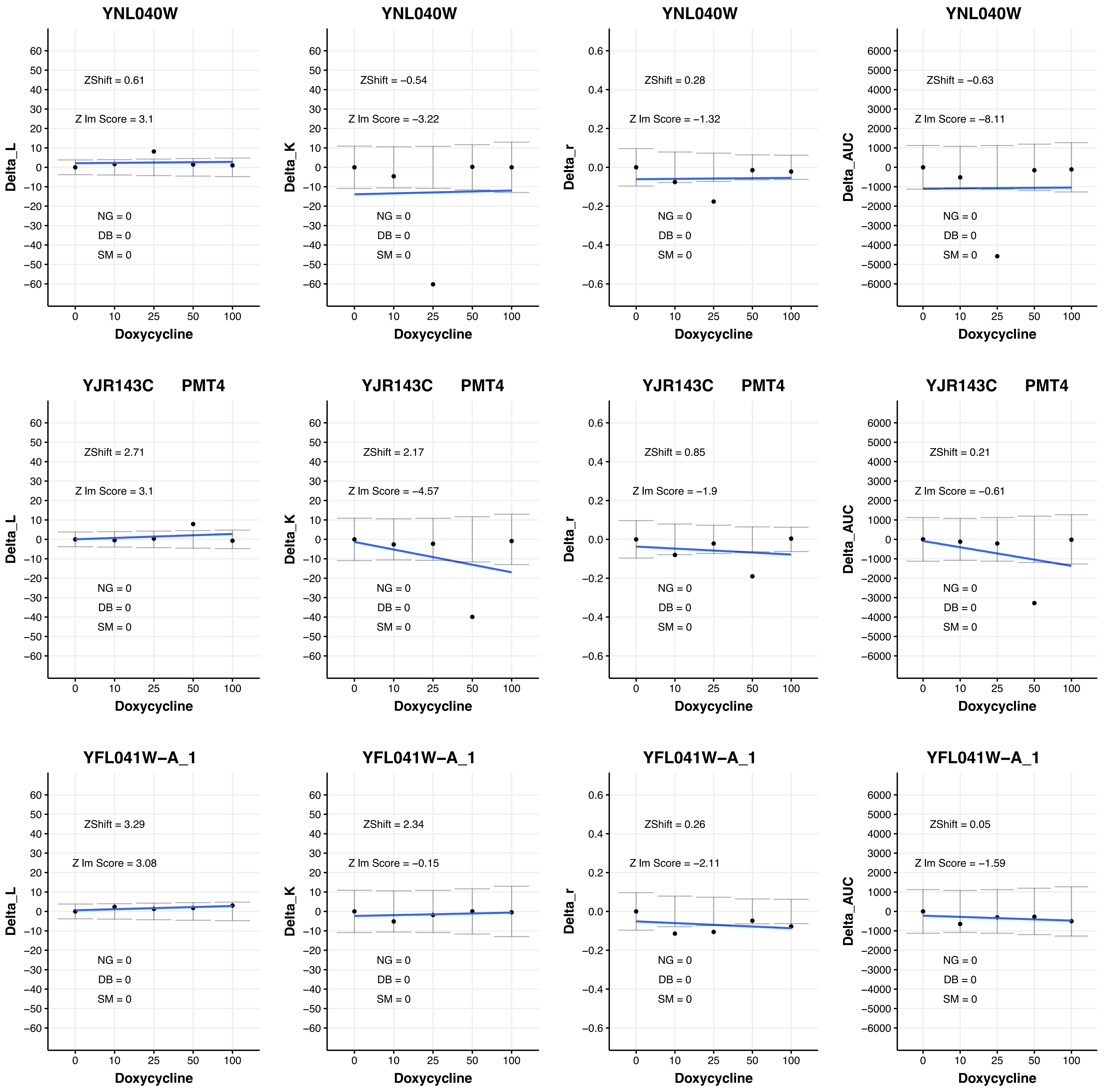

YNL040W
ZShift = 0.61
Z lm Score = 3.1
NG = 0
DB = 0
SM = 0
Delta_L
Doxycycline
YNL040W
ZShift = −0.54
Z lm Score = −3.22
NG = 0
DB = 0
SM = 0
Delta_K
Doxycycline
YNL040W
ZShift = 0.28
Z lm Score = −1.32
NG = 0
DB = 0
SM = 0
Delta_r
Doxycycline
YNL040W
ZShift = −0.63
Z lm Score = −8.11
NG = 0
DB = 0
SM = 0
Delta_AUC
Doxycycline
YJR143C PMT4
ZShift = 2.71
Z lm Score = 3.1
NG = 0
DB = 0
SM = 0
Delta_L
Doxycycline
YJR143C PMT4
ZShift = 2.17
Z lm Score = −4.57
NG = 0
DB = 0
SM = 0
Delta_K
Doxycycline
YJR143C PMT4
ZShift = 0.85
Z lm Score = −1.9
NG = 0
DB = 0
SM = 0
Delta_r
Doxycycline
YJR143C PMT4
ZShift = 0.21
Z lm Score = −0.61
NG = 0
DB = 0
SM = 0
Delta_AUC
Doxycycline
YFL041W−A_1
ZShift = 3.29
Z lm Score = 3.08
NG = 0
DB = 0
SM = 0
Delta_L
Doxycycline
YFL041W−A_1
ZShift = 2.34
Z lm Score = −0.15
NG = 0
DB = 0
SM = 0
Delta_K
Doxycycline
YFL041W−A_1
ZShift = 0.26
Z lm Score = −2.11
NG = 0
DB = 0
SM = 0
Delta_r
Doxycycline
YFL041W−A_1
ZShift = 0.05
Z lm Score = −1.59
NG = 0
DB = 0
SM = 0
Delta_AUC
Doxycycline

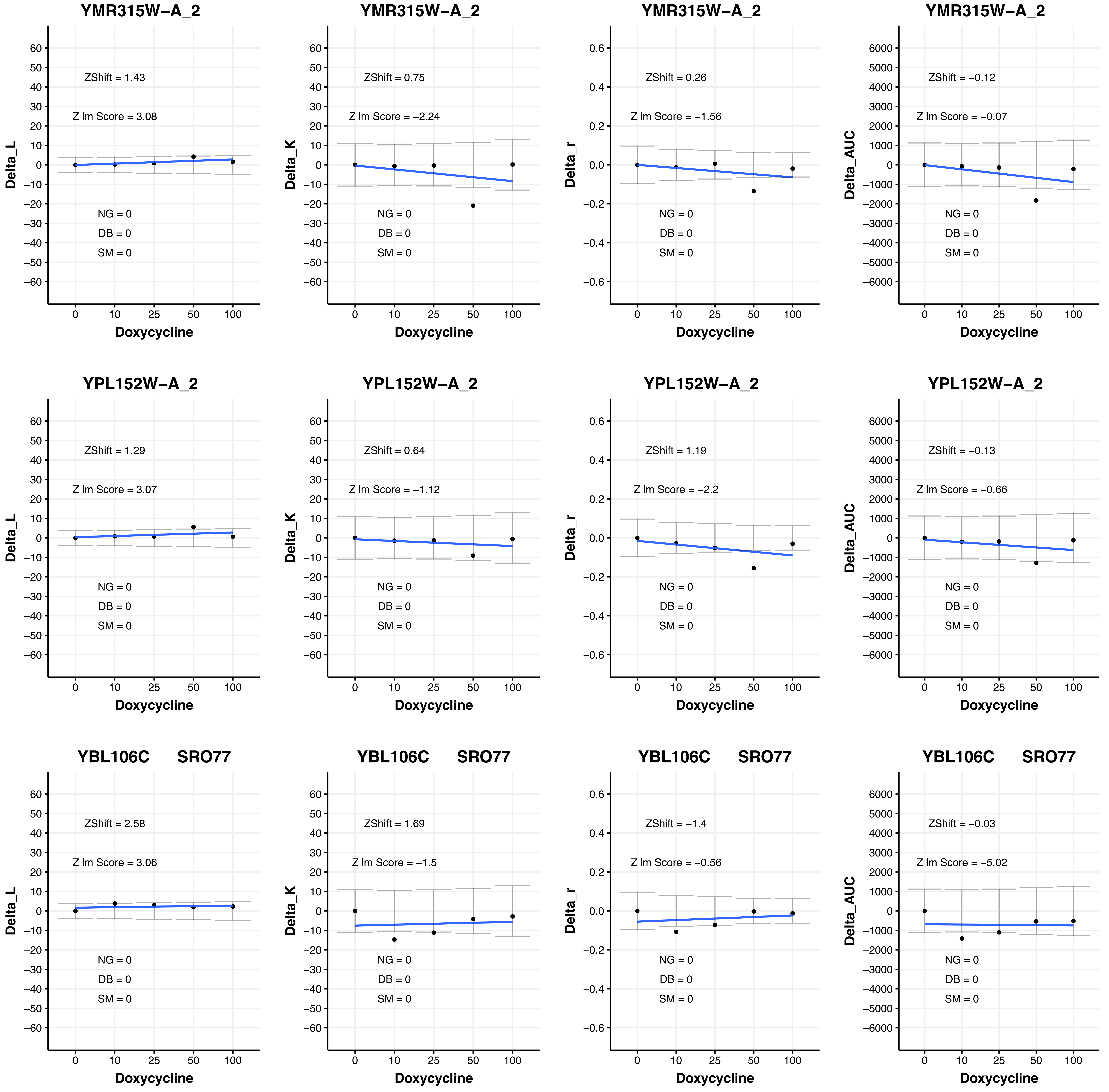

YMR315W−A_2
ZShift = 1.43
Z lm Score = 3.08
NG = 0
DB = 0
SM = 0
Delta_L
Doxycycline
YMR315W−A_2
ZShift = 0.75
Z lm Score = −2.24
NG = 0
DB = 0
SM = 0
Delta_K
Doxycycline
YMR315W−A_2
ZShift = 0.26
Z lm Score = −1.56
NG = 0
DB = 0
SM = 0
Delta_r
Doxycycline
YMR315W−A_2
ZShift = −0.12
Z lm Score = −0.07
NG = 0
DB = 0
SM = 0
Delta_AUC
Doxycycline
YPL152W−A_2
ZShift = 1.29
Z lm Score = 3.07
NG = 0
DB = 0
SM = 0
Delta_L
Doxycycline
YPL152W−A_2
ZShift = 0.64
Z lm Score = −1.12
NG = 0
DB = 0
SM = 0
Delta_K
Doxycycline
YPL152W−A_2
ZShift = 1.19
Z lm Score = −2.2
NG = 0
DB = 0
SM = 0
Delta_r
Doxycycline
YPL152W−A_2
ZShift = −0.13
Z lm Score = −0.66
NG = 0
DB = 0
SM = 0
Delta_AUC
Doxycycline
YBL106C SRO77
ZShift = 2.58
Z lm Score = 3.06
NG = 0
DB = 0
SM = 0
Delta_L
Doxycycline
YBL106C SRO77
ZShift = 1.69
Z lm Score = −1.5
NG = 0
DB = 0
SM = 0
Delta_K
Doxycycline
YBL106C SRO77
ZShift = −1.4
Z lm Score = −0.56
NG = 0
DB = 0
SM = 0
Delta_r
Doxycycline
YBL106C SRO77
ZShift = −0.03
Z lm Score = −5.02
NG = 0
DB = 0
SM = 0
Delta_AUC
Doxycycline

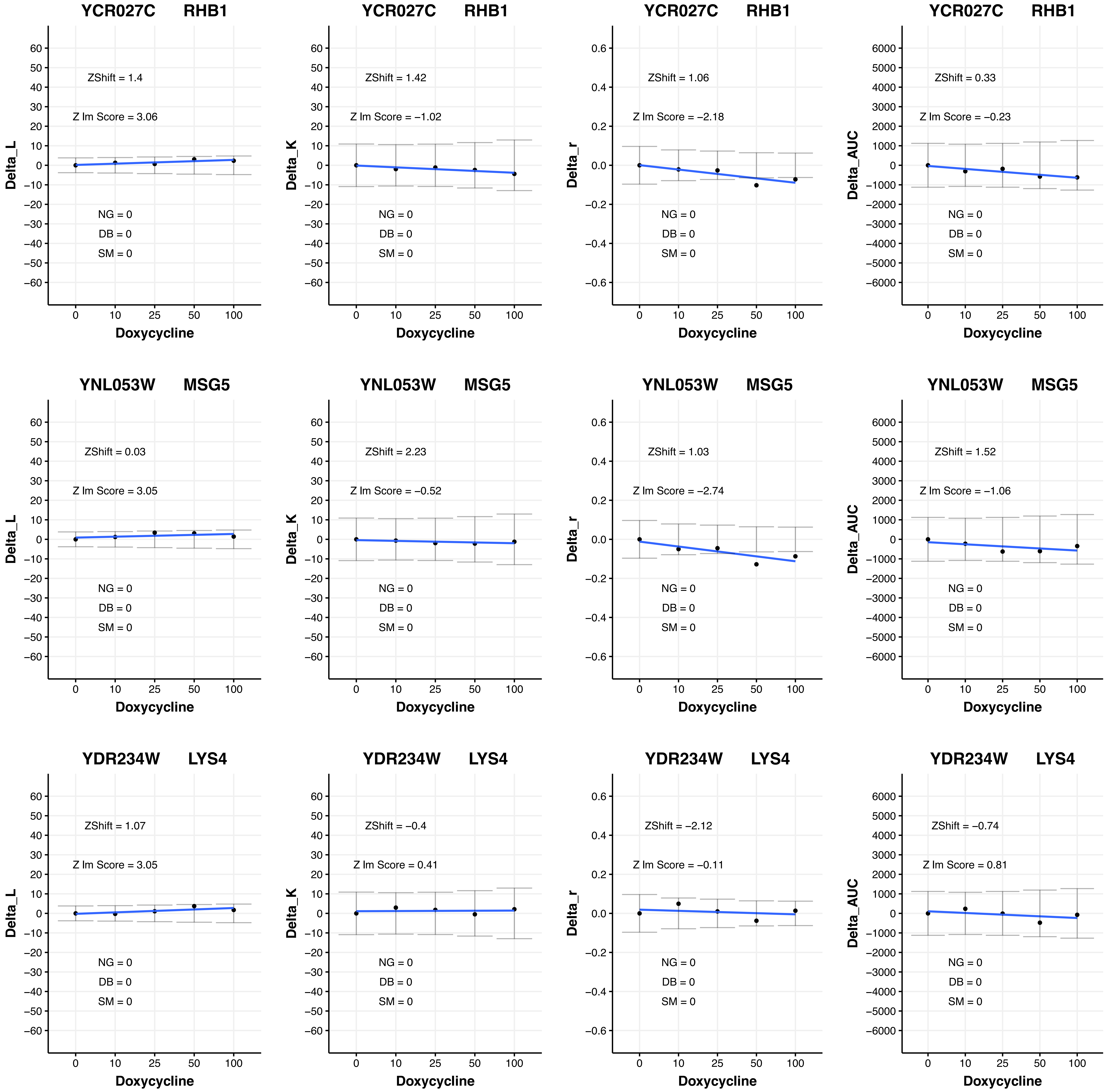
YCR027C RHB1
Delta_L
ZShift = 1.4
Z lm Score = 3.06
NG = 0
DB = 0
SM = 0
Doxycycline
YCR027C RHB1
Delta_K
ZShift = 1.42
Z lm Score = −1.02
NG = 0
DB = 0
SM = 0
Doxycycline
YCR027C RHB1
Delta_r
ZShift = 1.06
Z lm Score = −2.18
NG = 0
DB = 0
SM = 0
Doxycycline
YCR027C RHB1
Delta_AUC
ZShift = 0.33
Z lm Score = −0.23
NG = 0
DB = 0
SM = 0
Doxycycline
YNL053W MSG5
Delta_L
ZShift = 0.03
Z lm Score = 3.05
NG = 0
DB = 0
SM = 0
Doxycycline
YNL053W MSG5
Delta_K
ZShift = 2.23
Z lm Score = −0.52
NG = 0
DB = 0
SM = 0
Doxycycline
YNL053W MSG5
Delta_r
ZShift = 1.03
Z lm Score = −2.74
NG = 0
DB = 0
SM = 0
Doxycycline
YNL053W MSG5
Delta_AUC
ZShift = 1.52
Z lm Score = −1.06
NG = 0
DB = 0
SM = 0
Doxycycline
YDR234W LYS4
Delta_L
ZShift = 1.07
Z lm Score = 3.05
NG = 0
DB = 0
SM = 0
Doxycycline
YDR234W LYS4
Delta_K
ZShift = −0.4
Z lm Score = 0.41
NG = 0
DB = 0
SM = 0
Doxycycline
YDR234W LYS4
Delta_r
ZShift = −2.12
Z lm Score = −0.11
NG = 0
DB = 0
SM = 0
Doxycycline
YDR234W LYS4
Delta_AUC
ZShift = −0.74
Z lm Score = 0.81
NG = 0
DB = 0
SM = 0
Doxycycline

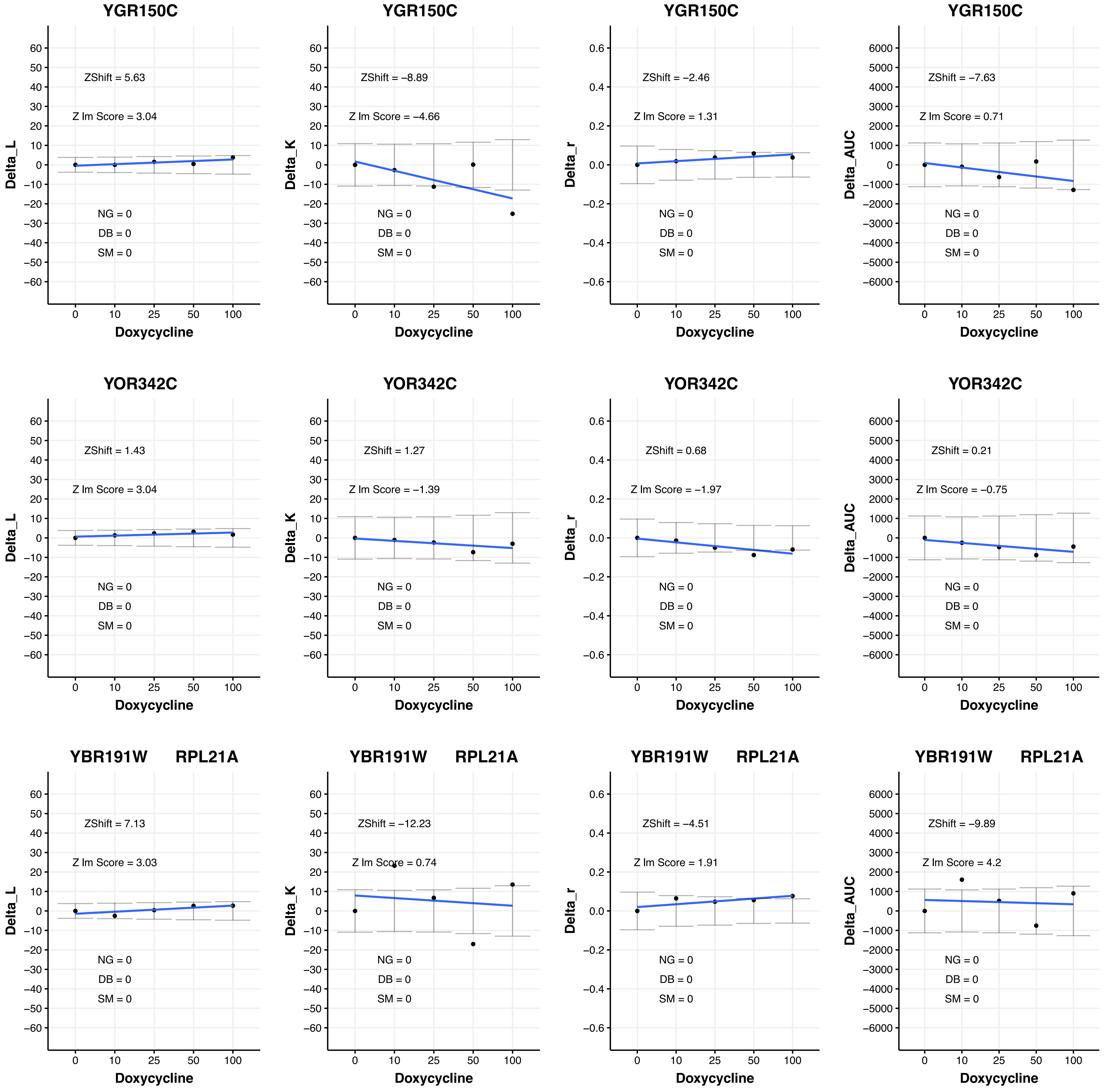

YGR150C
ZShift = 5.63
Z lm Score = 3.04
NG = 0
DB = 0
SM = 0
Delta_L
Doxycycline
YGR150C
ZShift = −8.89
Z lm Score = −4.66
NG = 0
DB = 0
SM = 0
Delta_K
Doxycycline
YGR150C
ZShift = −2.46
Z lm Score = 1.31
NG = 0
DB = 0
SM = 0
Delta_r
Doxycycline
YGR150C
ZShift = −7.63
Z lm Score = 0.71
NG = 0
DB = 0
SM = 0
Delta_AUC
Doxycycline
YOR342C
ZShift = 1.43
Z lm Score = 3.04
NG = 0
DB = 0
SM = 0
Delta_L
Doxycycline
YOR342C
ZShift = 1.27
Z lm Score = −1.39
NG = 0
DB = 0
SM = 0
Delta_K
Doxycycline
YOR342C
ZShift = 0.68
Z lm Score = −1.97
NG = 0
DB = 0
SM = 0
Delta_r
Doxycycline
YOR342C
ZShift = 0.21
Z lm Score = −0.75
NG = 0
DB = 0
SM = 0
Delta_AUC
Doxycycline
YBR191W RPL21A
ZShift = 7.13
Z lm Score = 3.03
NG = 0
DB = 0
SM = 0
Delta_L
Doxycycline
YBR191W RPL21A
ZShift = −12.23
Z lm Score = 0.74
NG = 0
DB = 0
SM = 0
Delta_K
Doxycycline
YBR191W RPL21A
ZShift = −4.51
Z lm Score = 1.91
NG = 0
DB = 0
SM = 0
Delta_r
Doxycycline
YBR191W RPL21A
ZShift = −9.89
Z lm Score = 4.2
NG = 0
DB = 0
SM = 0
Delta_AUC
Doxycycline

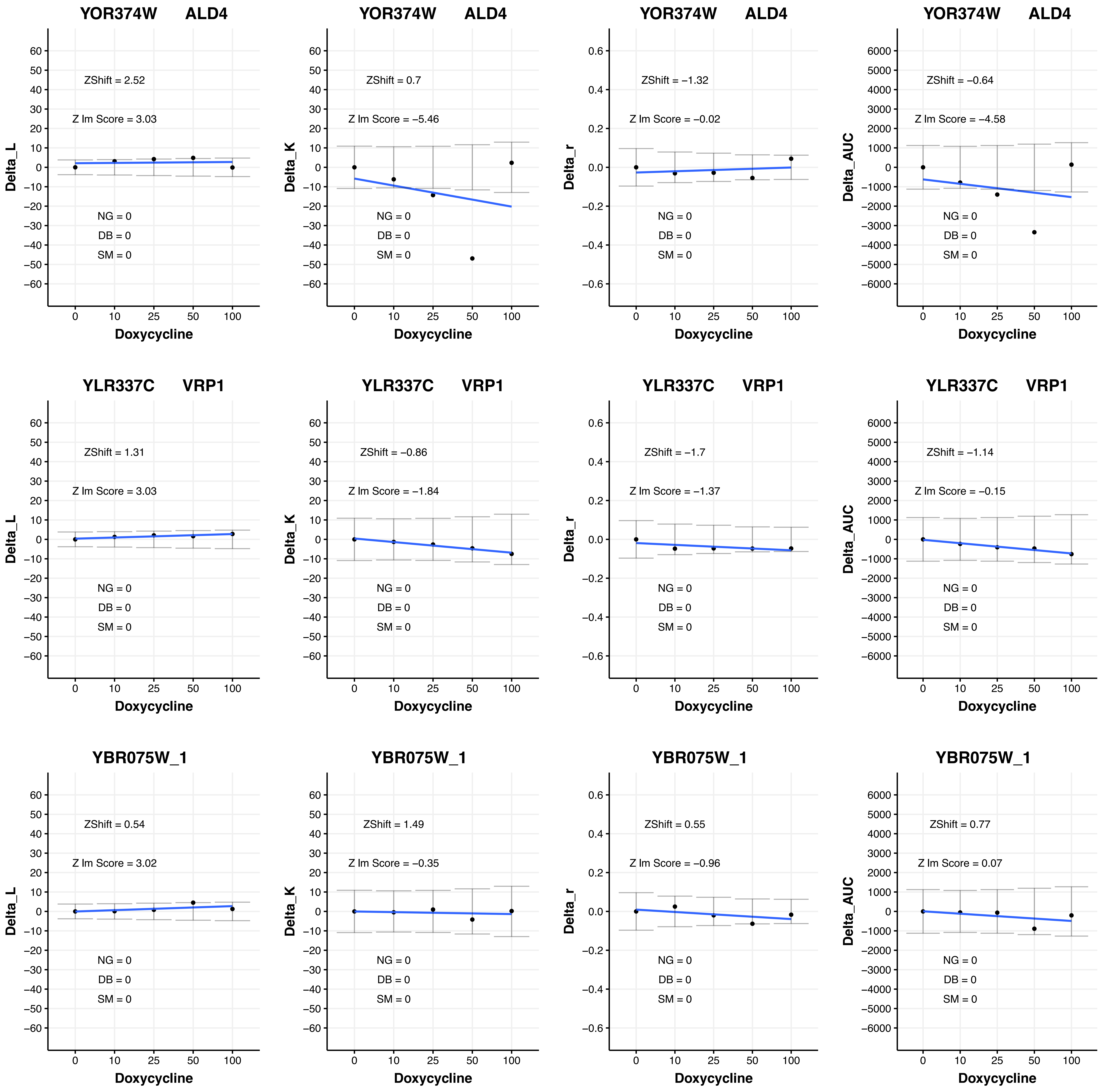

YOR374W ALD4
ZShift = 2.52
Z lm Score = 3.03
NG = 0
DB = 0
SM = 0
Delta_L
Doxycycline
YOR374W ALD4
ZShift = 0.7
Z lm Score = −5.46
NG = 0
DB = 0
SM = 0
Delta_K
Doxycycline
YOR374W ALD4
ZShift = −1.32
Z lm Score = −0.02
NG = 0
DB = 0
SM = 0
Delta_r
Doxycycline
YOR374W ALD4
ZShift = −0.64
Z lm Score = −4.58
NG = 0
DB = 0
SM = 0
Delta_AUC
Doxycycline
YLR337C VRP1
ZShift = 1.31
Z lm Score = 3.03
NG = 0
DB = 0
SM = 0
Delta_L
Doxycycline
YLR337C VRP1
ZShift = −0.86
Z lm Score = −1.84
NG = 0
DB = 0
SM = 0
Delta_K
Doxycycline
YLR337C VRP1
ZShift = −1.7
Z lm Score = −1.37
NG = 0
DB = 0
SM = 0
Delta_r
Doxycycline
YLR337C VRP1
ZShift = −1.14
Z lm Score = −0.15
NG = 0
DB = 0
SM = 0
Delta_AUC
Doxycycline
YBR075W_1
ZShift = 0.54
Z lm Score = 3.02
NG = 0
DB = 0
SM = 0
Delta_L
Doxycycline
YBR075W_1
ZShift = 1.49
Z lm Score = −0.35
NG = 0
DB = 0
SM = 0
Delta_K
Doxycycline
YBR075W_1
ZShift = 0.55
Z lm Score = −0.96
NG = 0
DB = 0
SM = 0
Delta_r
Doxycycline
YBR075W_1
ZShift = 0.77
Z lm Score = 0.07
NG = 0
DB = 0
SM = 0
Delta_AUC
Doxycycline

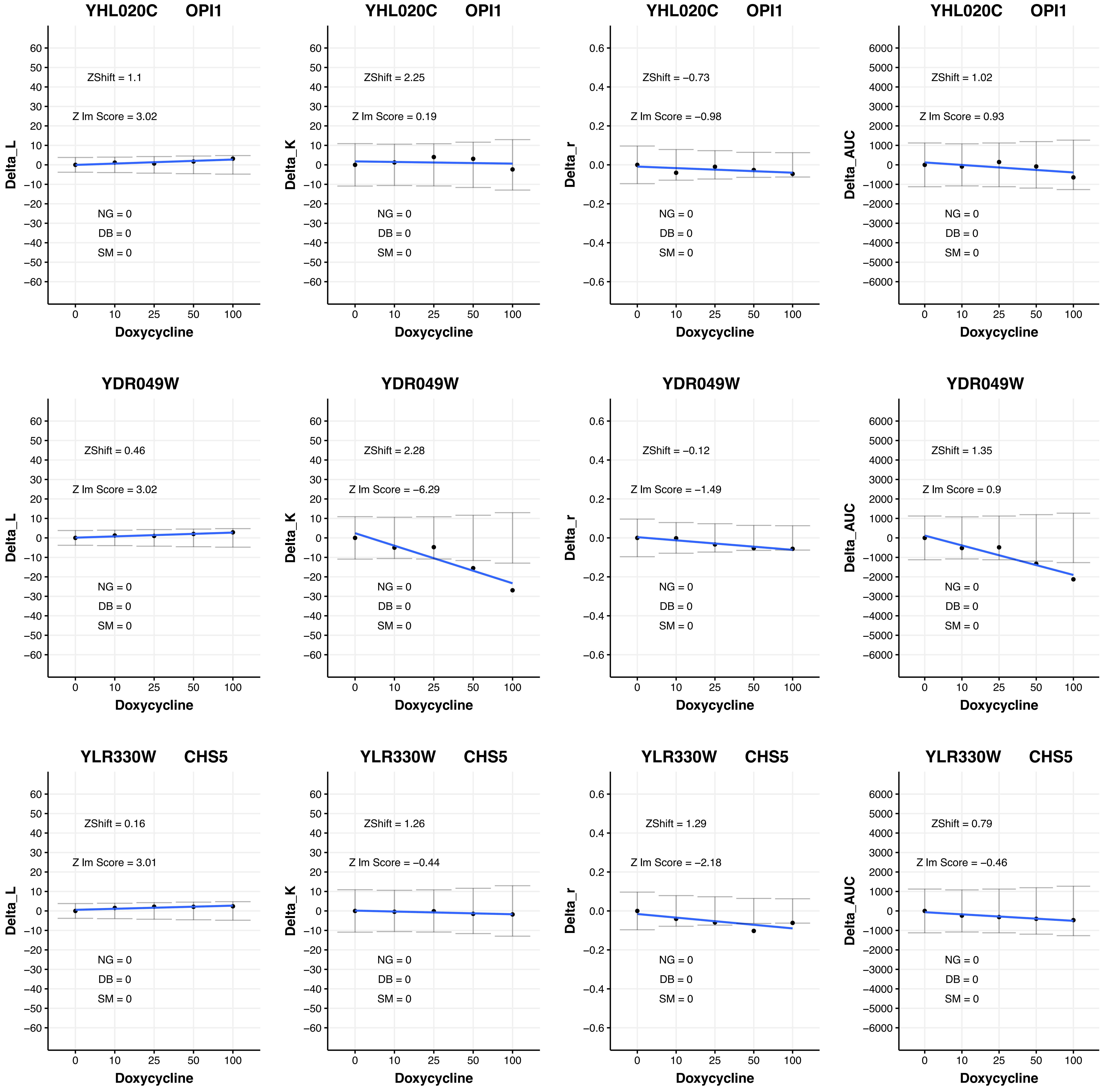

YHL020C OPI1
ZShift = 1.1
Z lm Score = 3.02
NG = 0
DB = 0
SM = 0
Delta_L
Doxycycline
YHL020C OPI1
ZShift = 2.25
Z lm Score = 0.19
NG = 0
DB = 0
SM = 0
Delta_K
Doxycycline
YHL020C OPI1
ZShift = −0.73
Z lm Score = −0.98
NG = 0
DB = 0
SM = 0
Delta_r
Doxycycline
YHL020C OPI1
ZShift = 1.02
Z lm Score = 0.93
NG = 0
DB = 0
SM = 0
Delta_AUC
Doxycycline
YDR049W
ZShift = 0.46
Z lm Score = 3.02
NG = 0
DB = 0
SM = 0
Delta_L
Doxycycline
YDR049W
ZShift = 2.28
Z lm Score = −6.29
NG = 0
DB = 0
SM = 0
Delta_K
Doxycycline
YDR049W
ZShift = −0.12
Z lm Score = −1.49
NG = 0
DB = 0
SM = 0
Delta_r
Doxycycline
YDR049W
ZShift = 1.35
Z lm Score = 0.9
NG = 0
DB = 0
SM = 0
Delta_AUC
Doxycycline
YLR330W CHS5
ZShift = 0.16
Z lm Score = 3.01
NG = 0
DB = 0
SM = 0
Delta_L
Doxycycline
YLR330W CHS5
ZShift = 1.26
Z lm Score = −0.44
NG = 0
DB = 0
SM = 0
Delta_K
Doxycycline
YLR330W CHS5
ZShift = 1.29
Z lm Score = −2.18
NG = 0
DB = 0
SM = 0
Delta_r
Doxycycline
YLR330W CHS5
ZShift = 0.79
Z lm Score = −0.46
NG = 0
DB = 0
SM = 0
Delta_AUC
Doxycycline

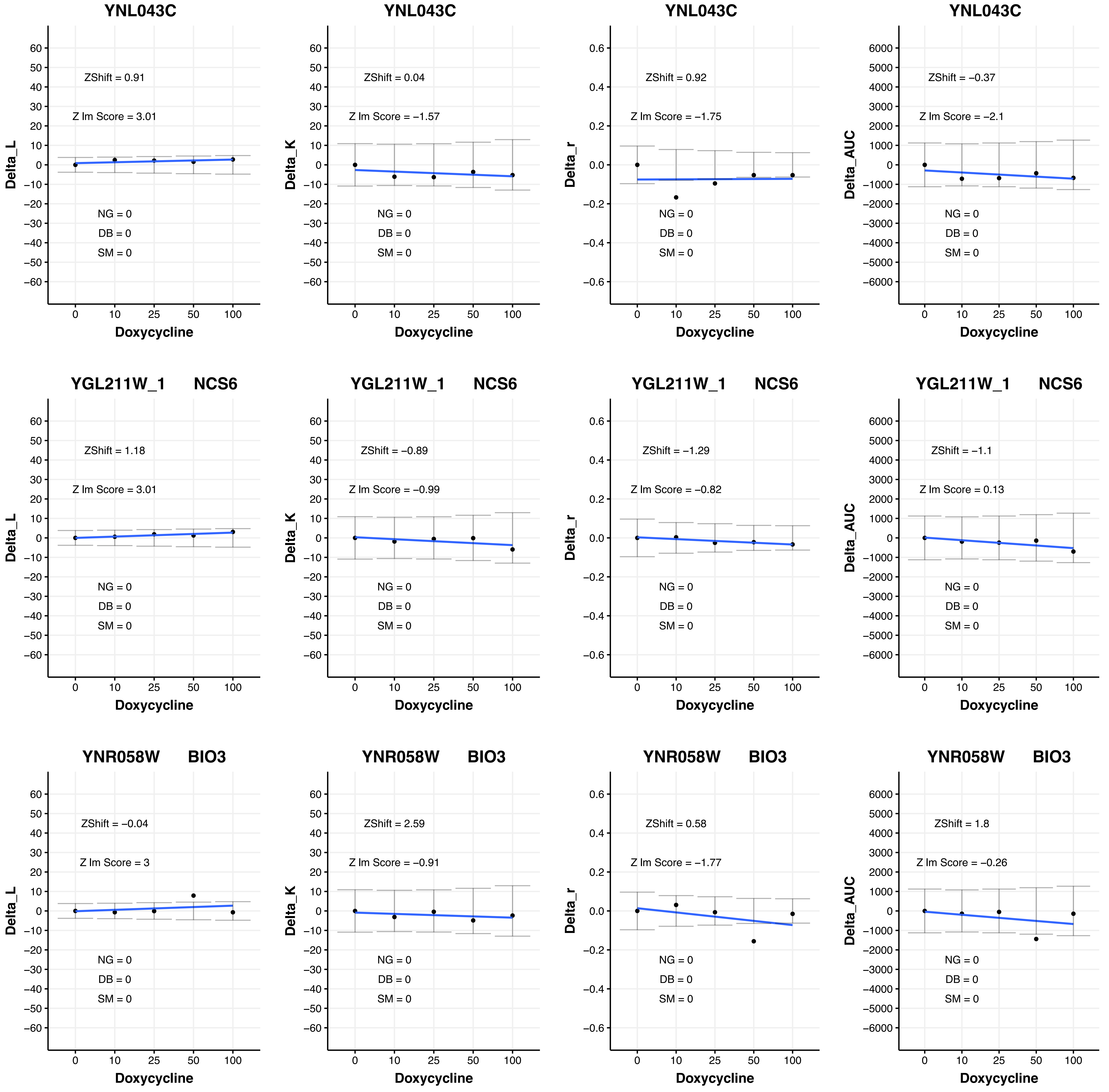

YNL043C
ZShift = 0.91
Z lm Score = 3.01
NG = 0
DB = 0
SM = 0
Delta_L
Doxycycline
YNL043C
ZShift = 0.04
Z lm Score = −1.57
NG = 0
DB = 0
SM = 0
Delta_K
Doxycycline
YNL043C
ZShift = 0.92
Z lm Score = −1.75
NG = 0
DB = 0
SM = 0
Delta_r
Doxycycline
YNL043C
ZShift = −0.37
Z lm Score = −2.1
NG = 0
DB = 0
SM = 0
Delta_AUC
Doxycycline
YGL211W_1 NCS6
ZShift = 1.18
Z lm Score = 3.01
NG = 0
DB = 0
SM = 0
Delta_L
Doxycycline
YGL211W_1 NCS6
ZShift = −0.89
Z lm Score = −0.99
NG = 0
DB = 0
SM = 0
Delta_K
Doxycycline
YGL211W_1 NCS6
ZShift = −1.29
Z lm Score = −0.82
NG = 0
DB = 0
SM = 0
Delta_r
Doxycycline
YGL211W_1 NCS6
ZShift = −1.1
Z lm Score = 0.13
NG = 0
DB = 0
SM = 0
Delta_AUC
Doxycycline
YNR058W BIO3
ZShift = −0.04
Z lm Score = 3
NG = 0
DB = 0
SM = 0
Delta_L
Doxycycline
YNR058W BIO3
ZShift = 2.59
Z lm Score = −0.91
NG = 0
DB = 0
SM = 0
Delta_K
Doxycycline
YNR058W BIO3
ZShift = 0.58
Z lm Score = −1.77
NG = 0
DB = 0
SM = 0
Delta_r
Doxycycline
YNR058W BIO3
ZShift = 1.8
Z lm Score = −0.26
NG = 0
DB = 0
SM = 0
Delta_AUC
Doxycycline

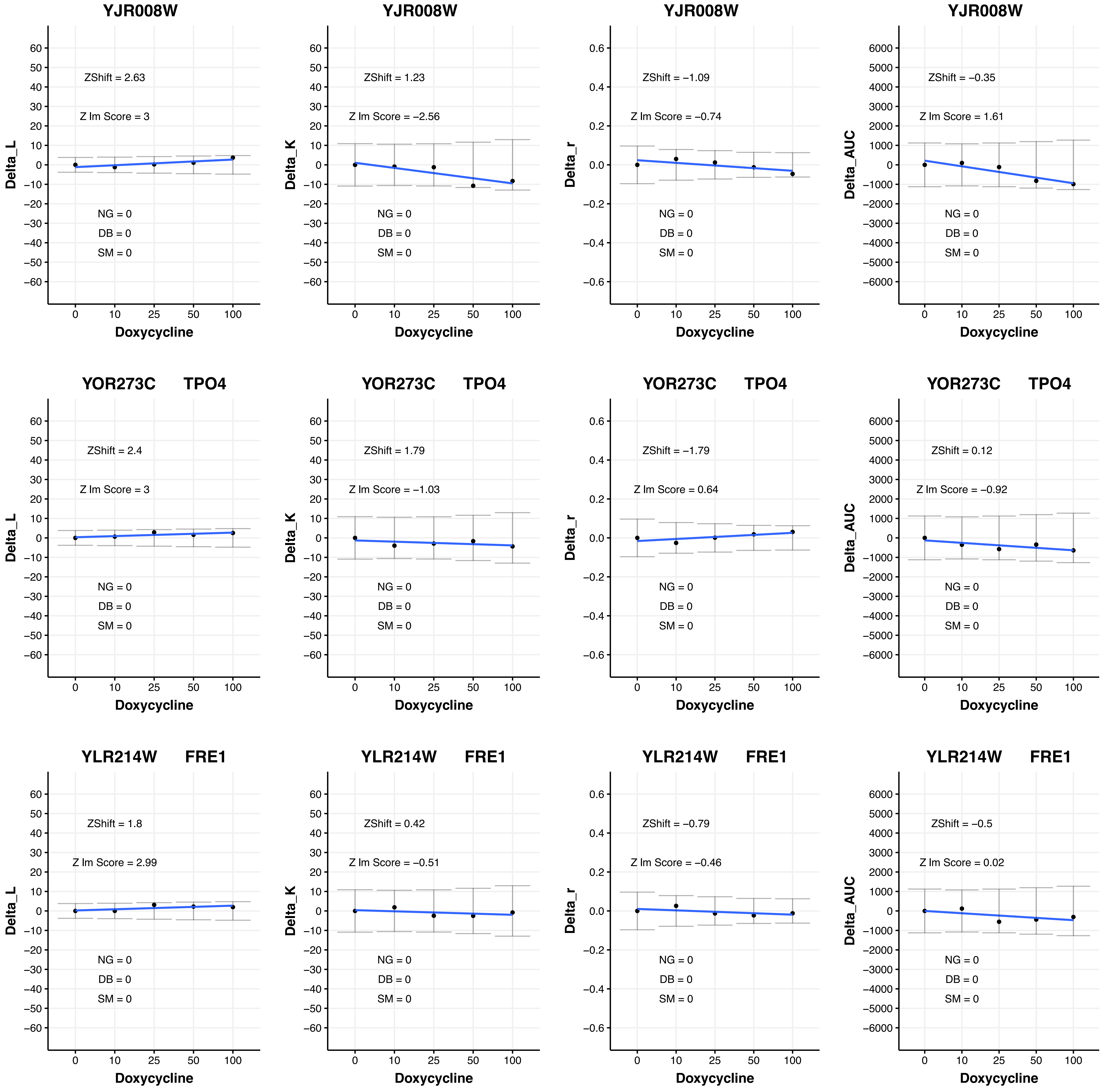
YJR008W
ZShift = 2.63
Z lm Score = 3
NG = 0
DB = 0
SM = 0
Delta_L
Doxycycline
YJR008W
ZShift = 1.23
Z lm Score = −2.56
NG = 0
DB = 0
SM = 0
Delta_K
Doxycycline
YJR008W
ZShift = −1.09
Z lm Score = −0.74
NG = 0
DB = 0
SM = 0
Delta_r
Doxycycline
YJR008W
ZShift = −0.35
Z lm Score = 1.61
NG = 0
DB = 0
SM = 0
Delta_AUC
Doxycycline
YOR273C TPO4
ZShift = 2.4
Z lm Score = 3
NG = 0
DB = 0
SM = 0
Delta_L
Doxycycline
YOR273C TPO4
ZShift = 1.79
Z lm Score = −1.03
NG = 0
DB = 0
SM = 0
Delta_K
Doxycycline
YOR273C TPO4
ZShift = −1.79
Z lm Score = 0.64
NG = 0
DB = 0
SM = 0
Delta_r
Doxycycline
YOR273C TPO4
ZShift = 0.12
Z lm Score = −0.92
NG = 0
DB = 0
SM = 0
Delta_AUC
Doxycycline
YLR214W FRE1
ZShift = 1.8
Z lm Score = 2.99
NG = 0
DB = 0
SM = 0
Delta_L
Doxycycline
YLR214W FRE1
ZShift = 0.42
Z lm Score = −0.51
NG = 0
DB = 0
SM = 0
Delta_K
Doxycycline
YLR214W FRE1
ZShift = −0.79
Z lm Score = −0.46
NG = 0
DB = 0
SM = 0
Delta_r
Doxycycline
YLR214W FRE1
ZShift = −0.5
Z lm Score = 0.02
NG = 0
DB = 0
SM = 0
Delta_AUC
Doxycycline

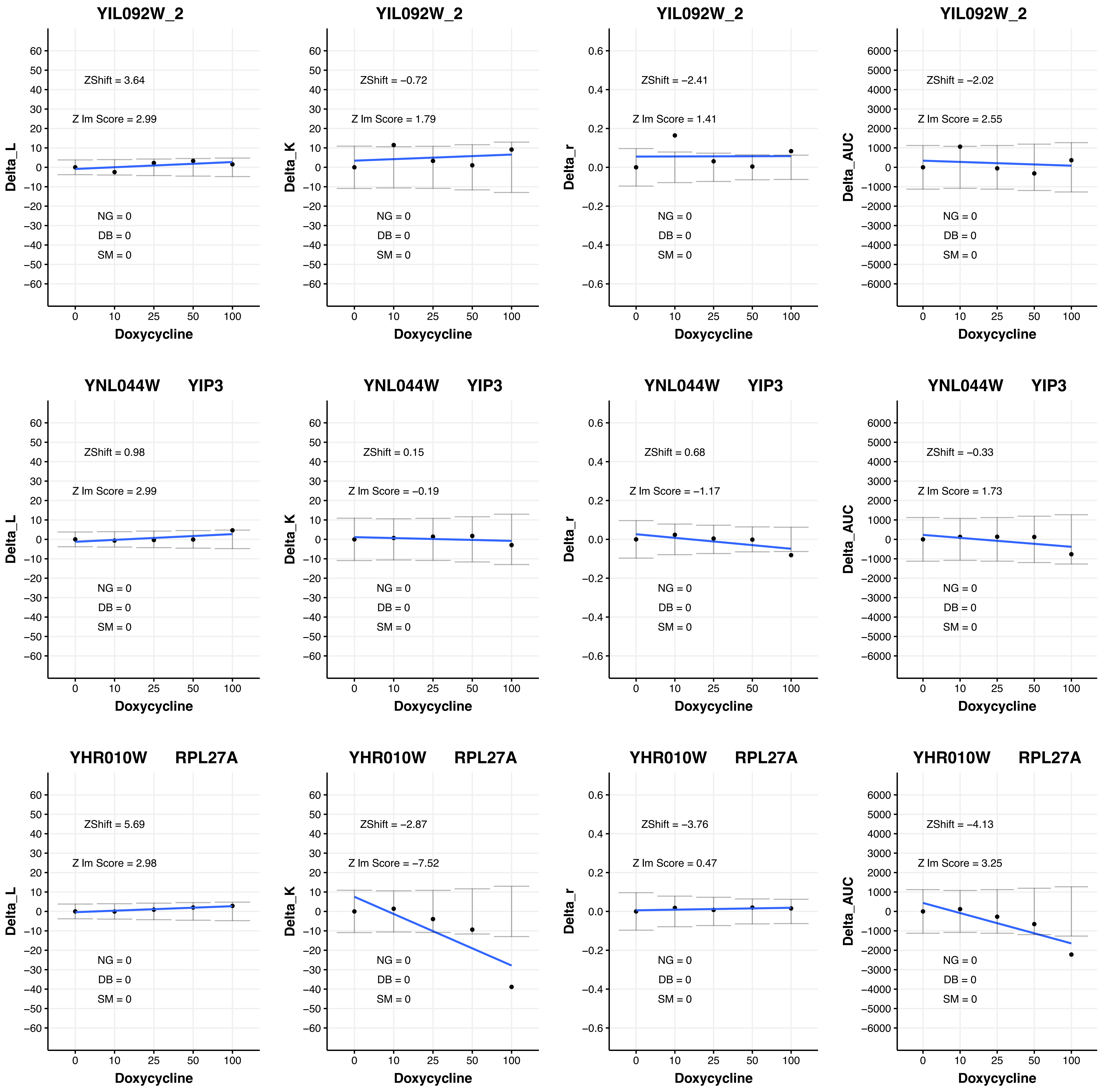
YIL092W_2
ZShift = 3.64
Z lm Score = 2.99
Delta_L
NG = 0
DB = 0
SM = 0
Doxycycline
YIL092W_2
ZShift = −0.72
Z lm Score = 1.79
Delta_K
NG = 0
DB = 0
SM = 0
Doxycycline
YIL092W_2
ZShift = −2.41
Z lm Score = 1.41
Delta_r
NG = 0
DB = 0
SM = 0
Doxycycline
YIL092W_2
ZShift = −2.02
Z lm Score = 2.55
Delta_AUC
NG = 0
DB = 0
SM = 0
Doxycycline
YNL044W YIP3
ZShift = 0.98
Z lm Score = 2.99
Delta_L
NG = 0
DB = 0
SM = 0
Doxycycline
YNL044W YIP3
ZShift = 0.15
Z lm Score = −0.19
Delta_K
NG = 0
DB = 0
SM = 0
Doxycycline
YNL044W YIP3
ZShift = 0.68
Z lm Score = −1.17
Delta_r
NG = 0
DB = 0
SM = 0
Doxycycline
YNL044W YIP3
ZShift = −0.33
Z lm Score = 1.73
Delta_AUC
NG = 0
DB = 0
SM = 0
Doxycycline
YHR010W RPL27A
ZShift = 5.69
Z lm Score = 2.98
Delta_L
NG = 0
DB = 0
SM = 0
Doxycycline
YHR010W RPL27A
ZShift = −2.87
Z lm Score = −7.52
Delta_K
NG = 0
DB = 0
SM = 0
Doxycycline
YHR010W RPL27A
ZShift = −3.76
Z lm Score = 0.47
Delta_r
NG = 0
DB = 0
SM = 0
Doxycycline
YHR010W RPL27A
ZShift = −4.13
Z lm Score = 3.25
Delta_AUC
NG = 0
DB = 0
SM = 0
Doxycycline

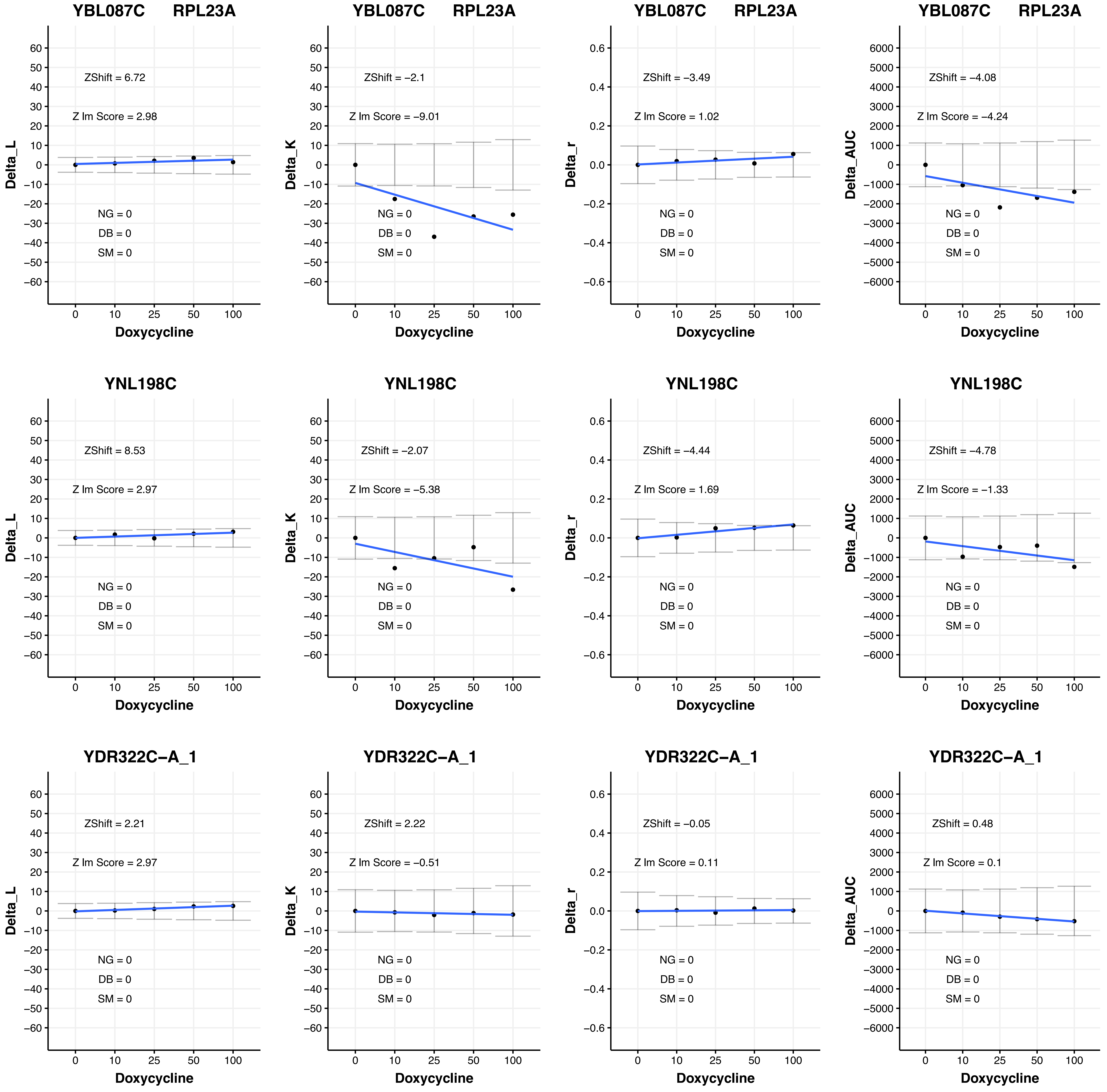

YBL087C RPL23A
ZShift = 6.72
Z lm Score = 2.98
NG = 0
DB = 0
SM = 0
Delta_L
Doxycycline
YBL087C RPL23A
ZShift = −2.1
Z lm Score = −9.01
NG = 0
DB = 0
SM = 0
Delta_K
Doxycycline
YBL087C RPL23A
ZShift = −3.49
Z lm Score = 1.02
NG = 0
DB = 0
SM = 0
Delta_r
Doxycycline
YBL087C RPL23A
ZShift = −4.08
Z lm Score = −4.24
NG = 0
DB = 0
SM = 0
Delta_AUC
Doxycycline
YNL198C
ZShift = 8.53
Z lm Score = 2.97
NG = 0
DB = 0
SM = 0
Delta_L
Doxycycline
YNL198C
ZShift = −2.07
Z lm Score = −5.38
NG = 0
DB = 0
SM = 0
Delta_K
Doxycycline
YNL198C
ZShift = −4.44
Z lm Score = 1.69
NG = 0
DB = 0
SM = 0
Delta_r
Doxycycline
YNL198C
ZShift = −4.78
Z lm Score = −1.33
NG = 0
DB = 0
SM = 0
Delta_AUC
Doxycycline
YDR322C−A_1
ZShift = 2.21
Z lm Score = 2.97
NG = 0
DB = 0
SM = 0
Delta_L
Doxycycline
YDR322C−A_1
ZShift = 2.22
Z lm Score = −0.51
NG = 0
DB = 0
SM = 0
Delta_K
Doxycycline
YDR322C−A_1
ZShift = −0.05
Z lm Score = 0.11
NG = 0
DB = 0
SM = 0
Delta_r
Doxycycline
YDR322C−A_1
ZShift = 0.48
Z lm Score = 0.1
NG = 0
DB = 0
SM = 0
Delta_AUC
Doxycycline

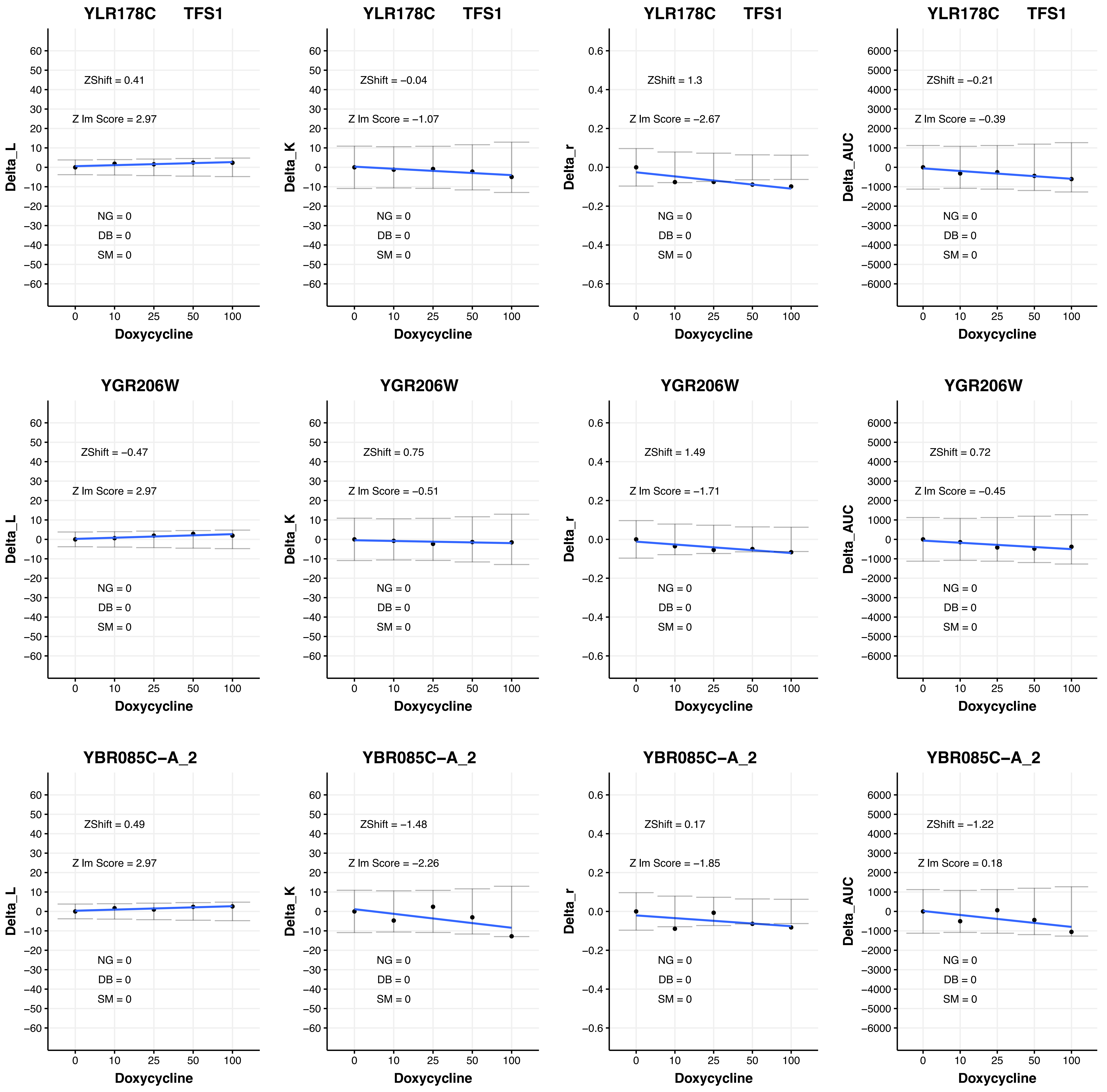

YLR178C TFS1
ZShift = 0.41
Z lm Score = 2.97
NG = 0
DB = 0
SM = 0
Delta_L
Doxycycline
YLR178C TFS1
ZShift = −0.04
Z lm Score = −1.07
NG = 0
DB = 0
SM = 0
Delta_K
Doxycycline
YLR178C TFS1
ZShift = 1.3
Z lm Score = −2.67
NG = 0
DB = 0
SM = 0
Delta_r
Doxycycline
YLR178C TFS1
ZShift = −0.21
Z lm Score = −0.39
NG = 0
DB = 0
SM = 0
Delta_AUC
Doxycycline
YGR206W
ZShift = −0.47
Z lm Score = 2.97
NG = 0
DB = 0
SM = 0
Delta_L
Doxycycline
YGR206W
ZShift = 0.75
Z lm Score = −0.51
NG = 0
DB = 0
SM = 0
Delta_K
Doxycycline
YGR206W
ZShift = 1.49
Z lm Score = −1.71
NG = 0
DB = 0
SM = 0
Delta_r
Doxycycline
YGR206W
ZShift = 0.72
Z lm Score = −0.45
NG = 0
DB = 0
SM = 0
Delta_AUC
Doxycycline
YBR085C−A_2
ZShift = 0.49
Z lm Score = 2.97
NG = 0
DB = 0
SM = 0
Delta_L
Doxycycline
YBR085C−A_2
ZShift = −1.48
Z lm Score = −2.26
NG = 0
DB = 0
SM = 0
Delta_K
Doxycycline
YBR085C−A_2
ZShift = 0.17
Z lm Score = −1.85
NG = 0
DB = 0
SM = 0
Delta_r
Doxycycline
YBR085C−A_2
ZShift = −1.22
Z lm Score = 0.18
NG = 0
DB = 0
SM = 0
Delta_AUC
Doxycycline
0
10
25
50
100

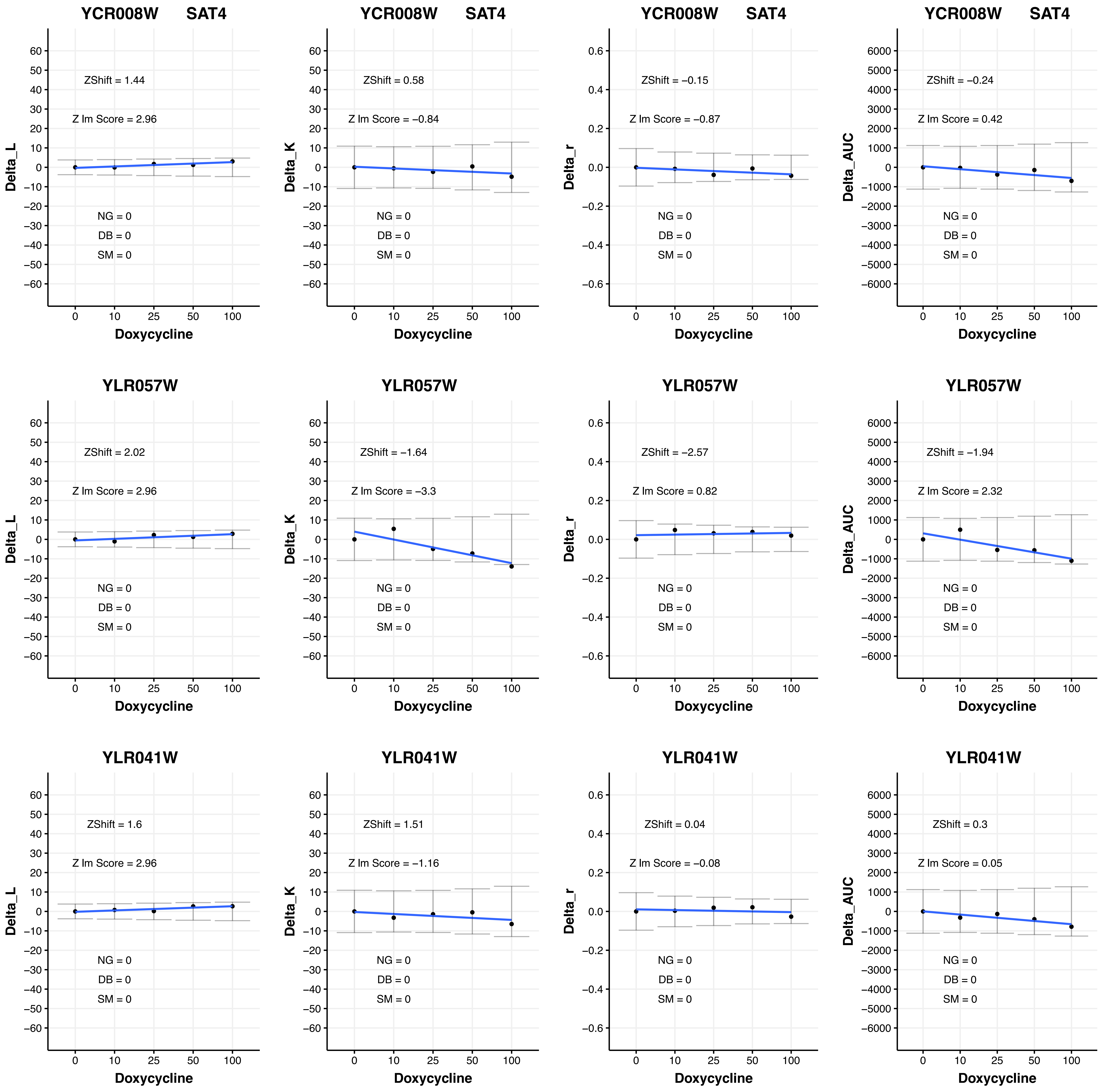
YCR008W SAT4
ZShift = 1.44
Z lm Score = 2.96
NG = 0
DB = 0
SM = 0
Delta_L
Doxycycline
YCR008W SAT4
ZShift = 0.58
Z lm Score = −0.84
NG = 0
DB = 0
SM = 0
Delta_K
Doxycycline
YCR008W SAT4
ZShift = −0.15
Z lm Score = −0.87
NG = 0
DB = 0
SM = 0
Delta_r
Doxycycline
YCR008W SAT4
ZShift = −0.24
Z lm Score = 0.42
NG = 0
DB = 0
SM = 0
Delta_AUC
Doxycycline
YLR057W
ZShift = 2.02
Z lm Score = 2.96
NG = 0
DB = 0
SM = 0
Delta_L
Doxycycline
YLR057W
ZShift = −1.64
Z lm Score = −3.3
NG = 0
DB = 0
SM = 0
Delta_K
Doxycycline
YLR057W
ZShift = −2.57
Z lm Score = 0.82
NG = 0
DB = 0
SM = 0
Delta_r
Doxycycline
YLR057W
ZShift = −1.94
Z lm Score = 2.32
NG = 0
DB = 0
SM = 0
Delta_AUC
Doxycycline
YLR041W
ZShift = 1.6
Z lm Score = 2.96
NG = 0
DB = 0
SM = 0
Delta_L
Doxycycline
YLR041W
ZShift = 1.51
Z lm Score = −1.16
NG = 0
DB = 0
SM = 0
Delta_K
Doxycycline
YLR041W
ZShift = 0.04
Z lm Score = −0.08
NG = 0
DB = 0
SM = 0
Delta_r
Doxycycline
YLR041W
ZShift = 0.3
Z lm Score = 0.05
NG = 0
DB = 0
SM = 0
Delta_AUC
Doxycycline

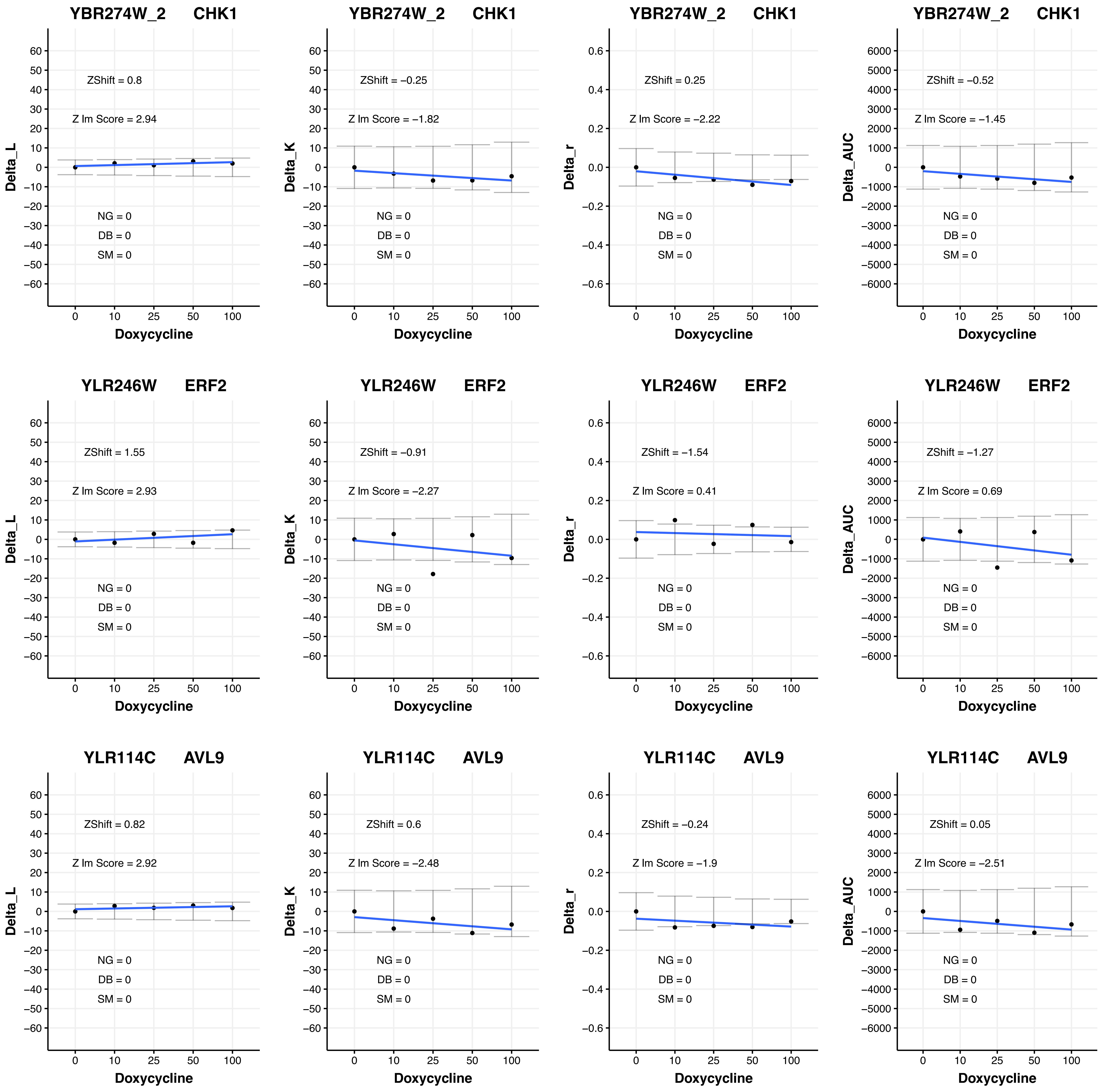

YBR274W_2 CHK1
ZShift = 0.8
Z lm Score = 2.94
NG = 0
DB = 0
SM = 0
Delta_L
Doxycycline
YBR274W_2 CHK1
ZShift = −0.25
Z lm Score = −1.82
NG = 0
DB = 0
SM = 0
Delta_K
Doxycycline
YBR274W_2 CHK1
ZShift = 0.25
Z lm Score = −2.22
NG = 0
DB = 0
SM = 0
Delta_r
Doxycycline
YBR274W_2 CHK1
ZShift = −0.52
Z lm Score = −1.45
NG = 0
DB = 0
SM = 0
Delta_AUC
Doxycycline
YLR246W ERF2
ZShift = 1.55
Z lm Score = 2.93
NG = 0
DB = 0
SM = 0
Delta_L
Doxycycline
YLR246W ERF2
ZShift = −0.91
Z lm Score = −2.27
NG = 0
DB = 0
SM = 0
Delta_K
Doxycycline
YLR246W ERF2
ZShift = −1.54
Z lm Score = 0.41
NG = 0
DB = 0
SM = 0
Delta_r
Doxycycline
YLR246W ERF2
ZShift = −1.27
Z lm Score = 0.69
NG = 0
DB = 0
SM = 0
Delta_AUC
Doxycycline
YLR114C AVL9
ZShift = 0.82
Z lm Score = 2.92
NG = 0
DB = 0
SM = 0
Delta_L
Doxycycline
YLR114C AVL9
ZShift = 0.6
Z lm Score = −2.48
NG = 0
DB = 0
SM = 0
Delta_K
Doxycycline
YLR114C AVL9
ZShift = −0.24
Z lm Score = −1.9
NG = 0
DB = 0
SM = 0
Delta_r
Doxycycline
YLR114C AVL9
ZShift = 0.05
Z lm Score = −2.51
NG = 0
DB = 0
SM = 0
Delta_AUC
Doxycycline

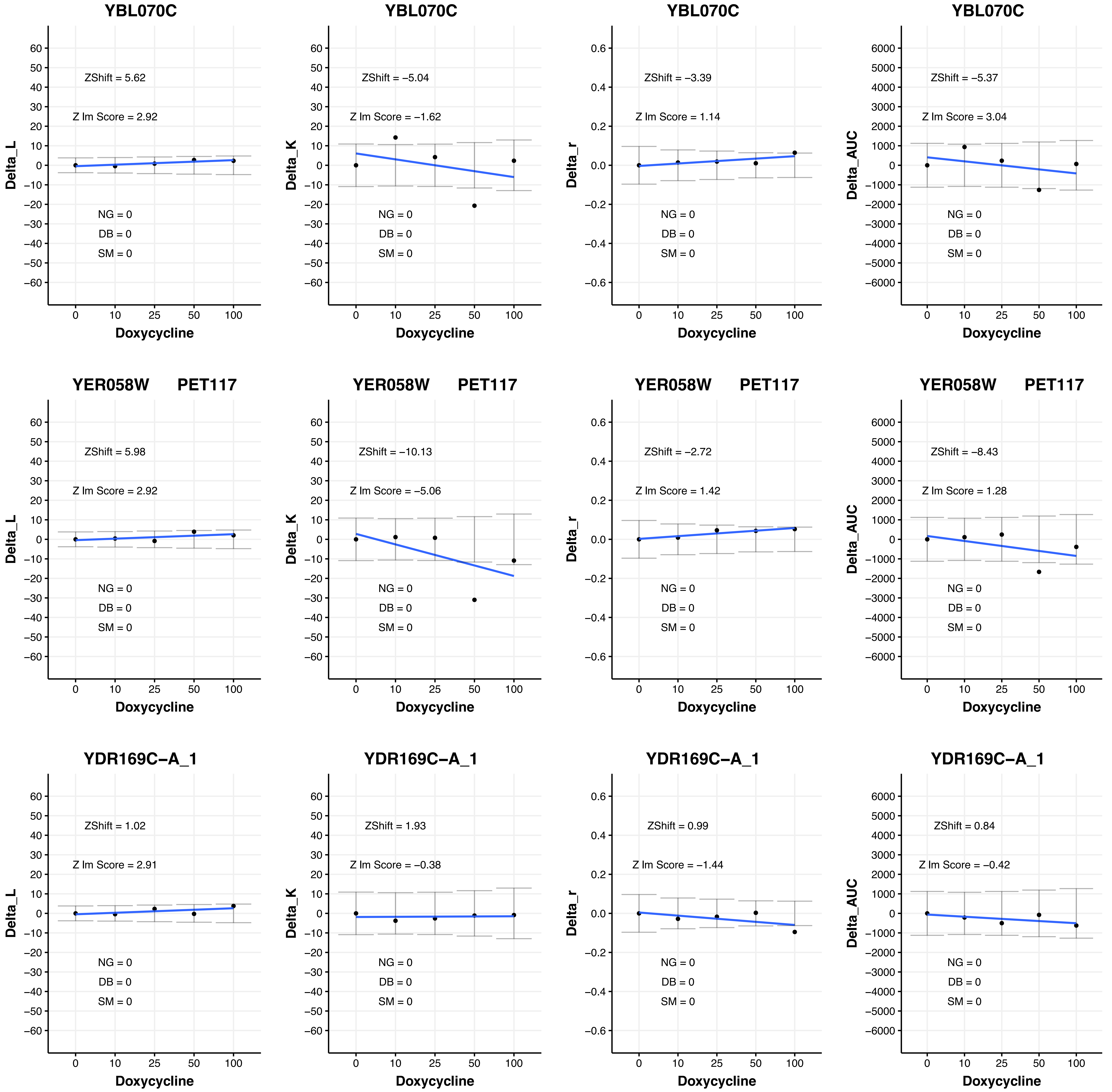
YBL070C
ZShift = 5.62
Z lm Score = 2.92
NG = 0
DB = 0
SM = 0
Delta_L
Doxycycline
YBL070C
ZShift = −5.04
Z lm Score = −1.62
NG = 0
DB = 0
SM = 0
Delta_K
Doxycycline
YBL070C
ZShift = −3.39
Z lm Score = 1.14
NG = 0
DB = 0
SM = 0
Delta_r
Doxycycline
YBL070C
ZShift = −5.37
Z lm Score = 3.04
NG = 0
DB = 0
SM = 0
Delta_AUC
Doxycycline
YER058W PET117
ZShift = 5.98
Z lm Score = 2.92
NG = 0
DB = 0
SM = 0
Delta_L
Doxycycline
YER058W PET117
ZShift = −10.13
Z lm Score = −5.06
NG = 0
DB = 0
SM = 0
Delta_K
Doxycycline
YER058W PET117
ZShift = −2.72
Z lm Score = 1.42
NG = 0
DB = 0
SM = 0
Delta_r
Doxycycline
YER058W PET117
ZShift = −8.43
Z lm Score = 1.28
NG = 0
DB = 0
SM = 0
Delta_AUC
Doxycycline
YDR169C−A_1
ZShift = 1.02
Z lm Score = 2.91
NG = 0
DB = 0
SM = 0
Delta_L
Doxycycline
YDR169C−A_1
ZShift = 1.93
Z lm Score = −0.38
NG = 0
DB = 0
SM = 0
Delta_K
Doxycycline
YDR169C−A_1
ZShift = 0.99
Z lm Score = −1.44
NG = 0
DB = 0
SM = 0
Delta_r
Doxycycline
YDR169C−A_1
ZShift = 0.84
Z lm Score = −0.42
NG = 0
DB = 0
SM = 0
Delta_AUC
Doxycycline

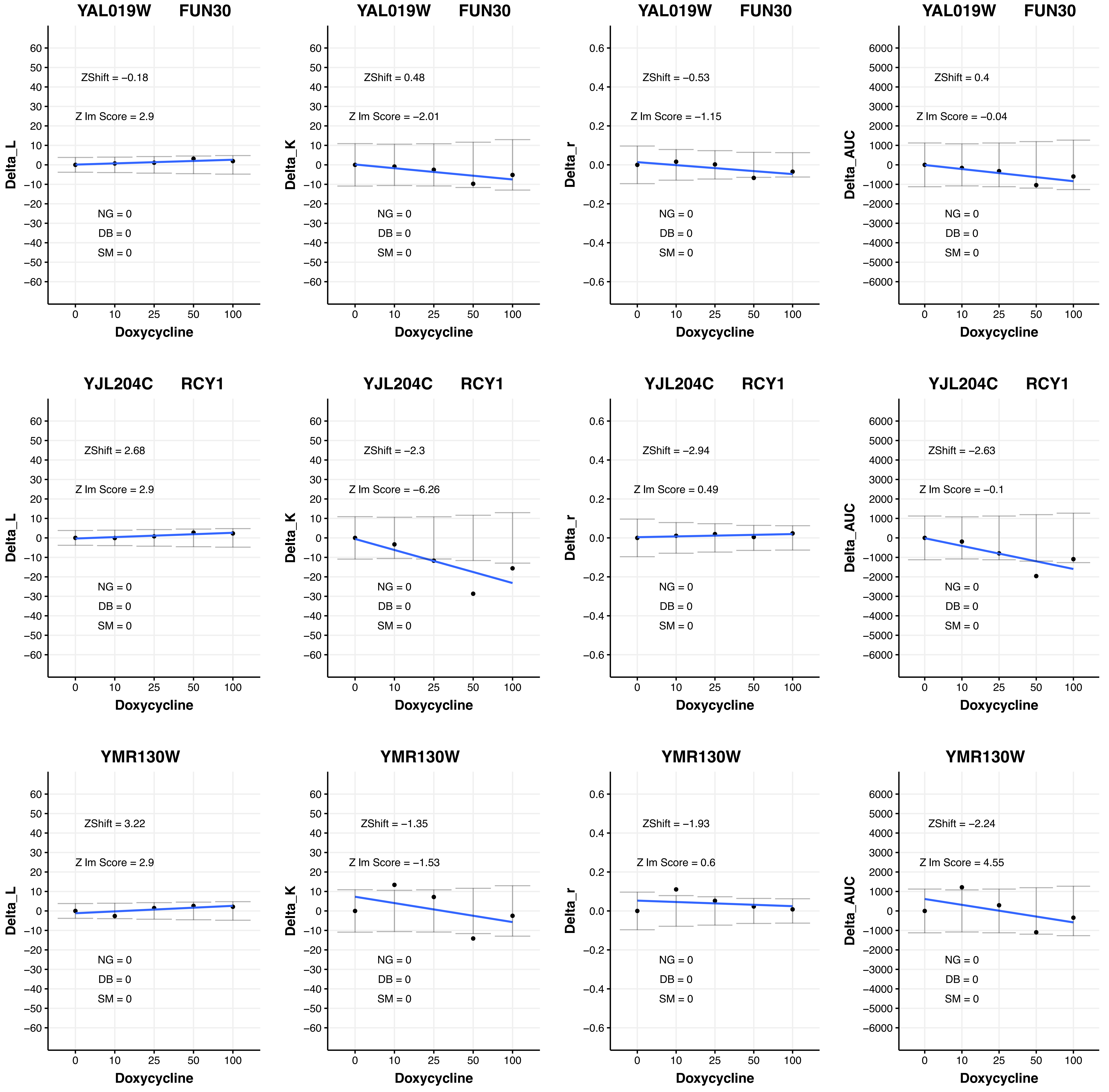
YAL019W FUN30
ZShift = −0.18
Z lm Score = 2.9
NG = 0
DB = 0
SM = 0
Delta_L
Doxycycline
YAL019W FUN30
ZShift = 0.48
Z lm Score = −2.01
NG = 0
DB = 0
SM = 0
Delta_K
Doxycycline
YAL019W FUN30
ZShift = −0.53
Z lm Score = −1.15
NG = 0
DB = 0
SM = 0
Delta_r
Doxycycline
YAL019W FUN30
ZShift = 0.4
Z lm Score = −0.04
NG = 0
DB = 0
SM = 0
Delta_AUC
Doxycycline
YJL204C RCY1
ZShift = 2.68
Z lm Score = 2.9
NG = 0
DB = 0
SM = 0
Delta_L
Doxycycline
YJL204C RCY1
ZShift = −2.3
Z lm Score = −6.26
NG = 0
DB = 0
SM = 0
Delta_K
Doxycycline
YJL204C RCY1
ZShift = −2.94
Z lm Score = 0.49
NG = 0
DB = 0
SM = 0
Delta_r
Doxycycline
YJL204C RCY1
ZShift = −2.63
Z lm Score = −0.1
NG = 0
DB = 0
SM = 0
Delta_AUC
Doxycycline
YMR130W
ZShift = 3.22
Z lm Score = 2.9
NG = 0
DB = 0
SM = 0
Delta_L
Doxycycline
YMR130W
ZShift = −1.35
Z lm Score = −1.53
NG = 0
DB = 0
SM = 0
Delta_K
Doxycycline
YMR130W
ZShift = −1.93
Z lm Score = 0.6
NG = 0
DB = 0
SM = 0
Delta_r
Doxycycline
YMR130W
ZShift = −2.24
Z lm Score = 4.55
NG = 0
DB = 0
SM = 0
Delta_AUC
Doxycycline

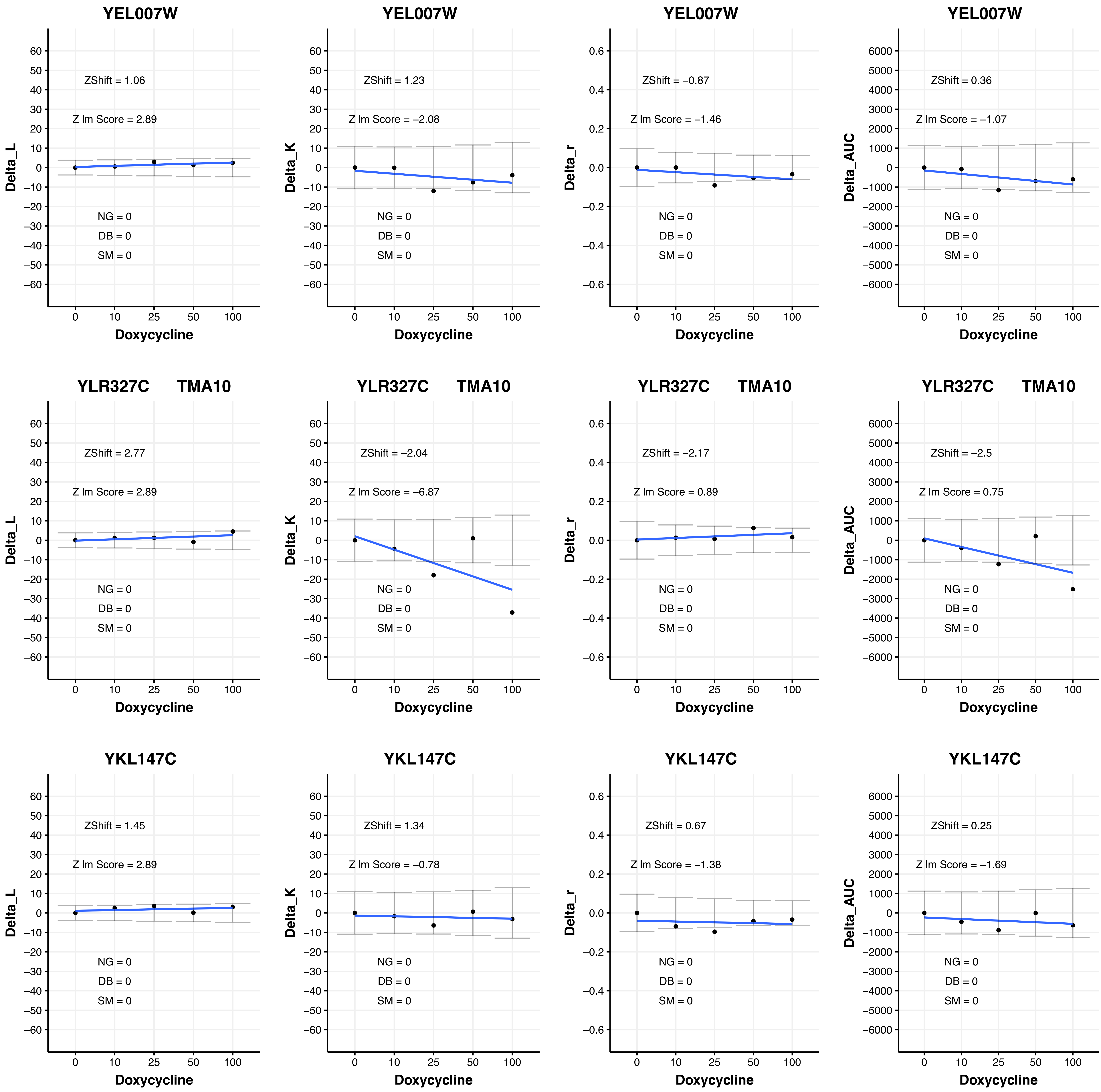

YEL007W
ZShift = 1.06
Z lm Score = 2.89
NG = 0
DB = 0
SM = 0
Delta_L
Doxycycline
YEL007W
ZShift = 1.23
Z lm Score = −2.08
NG = 0
DB = 0
SM = 0
Delta_K
Doxycycline
YEL007W
ZShift = −0.87
Z lm Score = −1.46
NG = 0
DB = 0
SM = 0
Delta_r
Doxycycline
YEL007W
ZShift = 0.36
Z lm Score = −1.07
NG = 0
DB = 0
SM = 0
Delta_AUC
Doxycycline
YLR327C TMA10
ZShift = 2.77
Z lm Score = 2.89
NG = 0
DB = 0
SM = 0
Delta_L
Doxycycline
YLR327C TMA10
ZShift = −2.04
Z lm Score = −6.87
NG = 0
DB = 0
SM = 0
Delta_K
Doxycycline
YLR327C TMA10
ZShift = −2.17
Z lm Score = 0.89
NG = 0
DB = 0
SM = 0
Delta_r
Doxycycline
YLR327C TMA10
ZShift = −2.5
Z lm Score = 0.75
NG = 0
DB = 0
SM = 0
Delta_AUC
Doxycycline
YKL147C
ZShift = 1.45
Z lm Score = 2.89
NG = 0
DB = 0
SM = 0
Delta_L
Doxycycline
YKL147C
ZShift = 1.34
Z lm Score = −0.78
NG = 0
DB = 0
SM = 0
Delta_K
Doxycycline
YKL147C
ZShift = 0.67
Z lm Score = −1.38
NG = 0
DB = 0
SM = 0
Delta_r
Doxycycline
YKL147C
ZShift = 0.25
Z lm Score = −1.69
NG = 0
DB = 0
SM = 0
Delta_AUC
Doxycycline

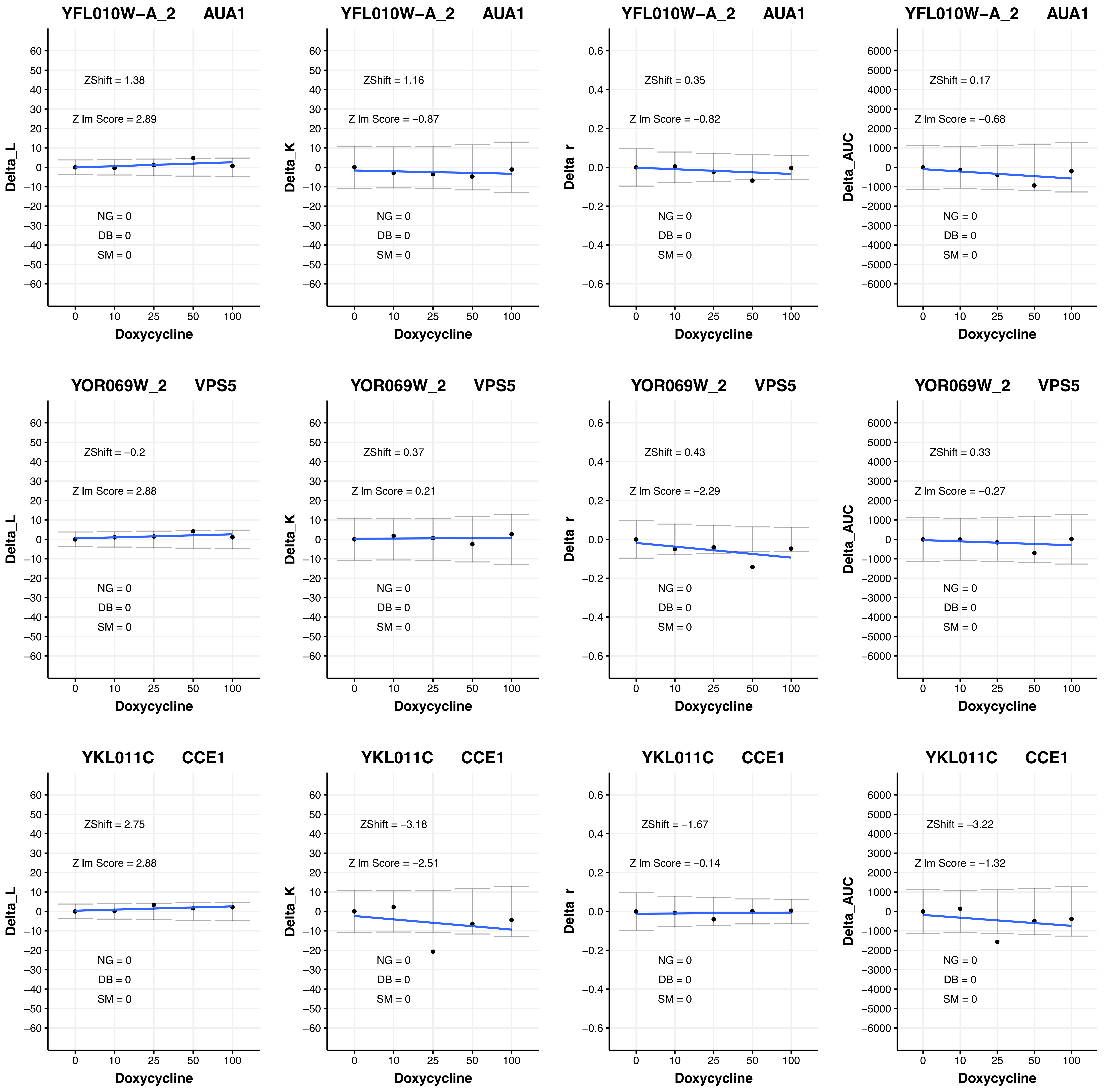

YFL010W−A_2 AUA1
ZShift = 1.38
Z lm Score = 2.89
NG = 0
DB = 0
SM = 0
Delta_L
YFL010W−A_2 AUA1
ZShift = 1.16
Z lm Score = −0.87
NG = 0
DB = 0
SM = 0
Delta_K
YFL010W−A_2 AUA1
ZShift = 0.35
Z lm Score = −0.82
NG = 0
DB = 0
SM = 0
Delta_r
YFL010W−A_2 AUA1
ZShift = 0.17
Z lm Score = −0.68
NG = 0
DB = 0
SM = 0
Delta_AUC
YOR069W_2 VPS5
ZShift = −0.2
Z lm Score = 2.88
NG = 0
DB = 0
SM = 0
Delta_L
YOR069W_2 VPS5
ZShift = 0.37
Z lm Score = 0.21
NG = 0
DB = 0
SM = 0
Delta_K
YOR069W_2 VPS5
ZShift = 0.43
Z lm Score = −2.29
NG = 0
DB = 0
SM = 0
Delta_r
YOR069W_2 VPS5
ZShift = 0.33
Z lm Score = −0.27
NG = 0
DB = 0
SM = 0
Delta_AUC
YKL011C CCE1
ZShift = 2.75
Z lm Score = 2.88
NG = 0
DB = 0
SM = 0
Delta_L
YKL011C CCE1
ZShift = −3.18
Z lm Score = −2.51
NG = 0
DB = 0
SM = 0
Delta_K
YKL011C CCE1
ZShift = −1.67
Z lm Score = −0.14
NG = 0
DB = 0
SM = 0
Delta_r
YKL011C CCE1
ZShift = −3.22
Z lm Score = −1.32
NG = 0
DB = 0
SM = 0
Delta_AUC
Doxycycline
0
10
25
50
100

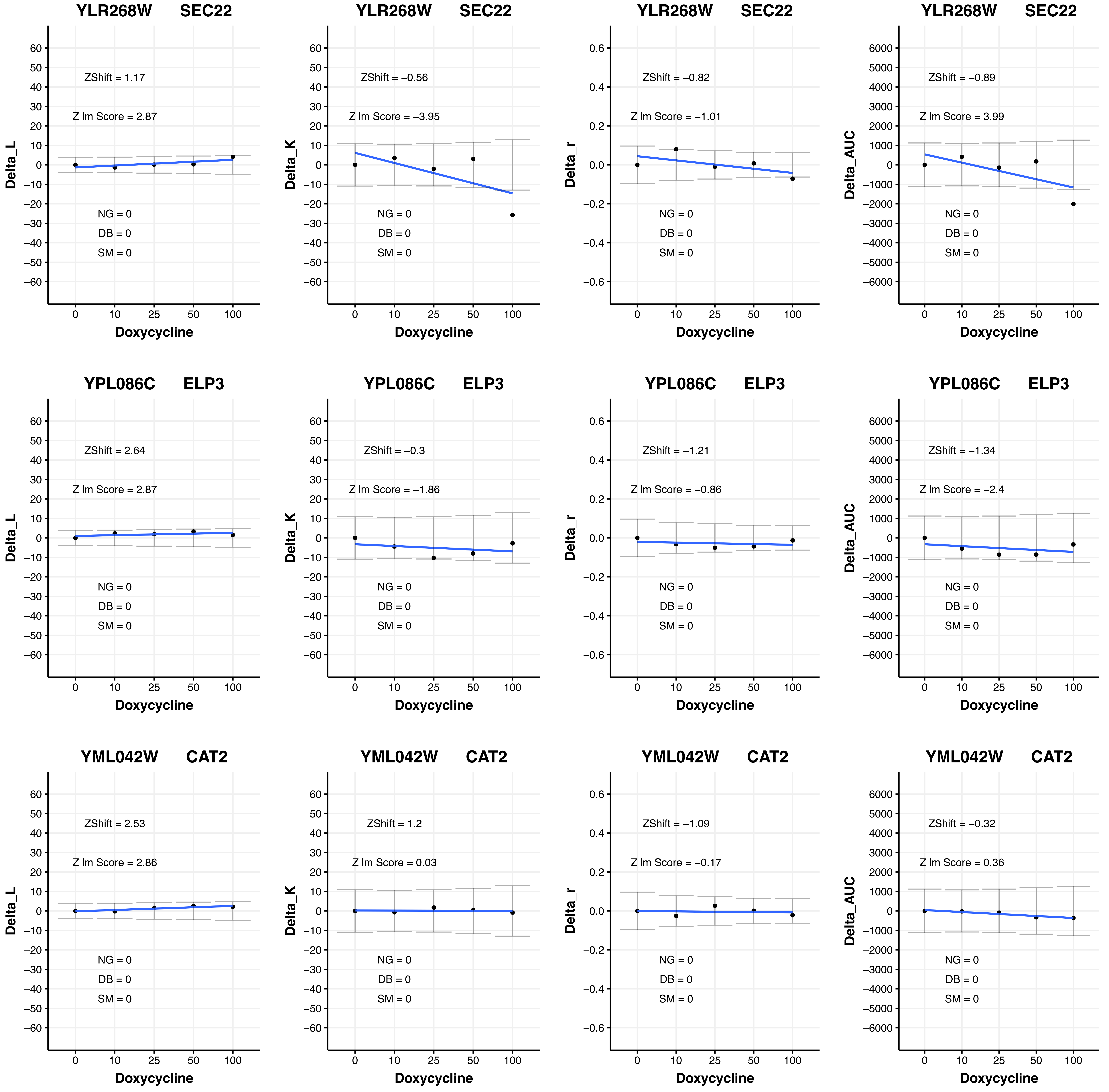

YLR268W SEC22
ZShift = 1.17
Z lm Score = 2.87
NG = 0
DB = 0
SM = 0
Delta_L
Doxycycline
YLR268W SEC22
ZShift = −0.56
Z lm Score = −3.95
NG = 0
DB = 0
SM = 0
Delta_K
Doxycycline
YLR268W SEC22
ZShift = −0.82
Z lm Score = −1.01
NG = 0
DB = 0
SM = 0
Delta_r
Doxycycline
YLR268W SEC22
ZShift = −0.89
Z lm Score = 3.99
NG = 0
DB = 0
SM = 0
Delta_AUC
Doxycycline
YPL086C ELP3
ZShift = 2.64
Z lm Score = 2.87
NG = 0
DB = 0
SM = 0
Delta_L
Doxycycline
YPL086C ELP3
ZShift = −0.3
Z lm Score = −1.86
NG = 0
DB = 0
SM = 0
Delta_K
Doxycycline
YPL086C ELP3
ZShift = −1.21
Z lm Score = −0.86
NG = 0
DB = 0
SM = 0
Delta_r
Doxycycline
YPL086C ELP3
ZShift = −1.34
Z lm Score = −2.4
NG = 0
DB = 0
SM = 0
Delta_AUC
Doxycycline
YML042W CAT2
ZShift = 2.53
Z lm Score = 2.86
NG = 0
DB = 0
SM = 0
Delta_L
Doxycycline
YML042W CAT2
ZShift = 1.2
Z lm Score = 0.03
NG = 0
DB = 0
SM = 0
Delta_K
Doxycycline
YML042W CAT2
ZShift = −1.09
Z lm Score = −0.17
NG = 0
DB = 0
SM = 0
Delta_r
Doxycycline
YML042W CAT2
ZShift = −0.32
Z lm Score = 0.36
NG = 0
DB = 0
SM = 0
Delta_AUC
Doxycycline

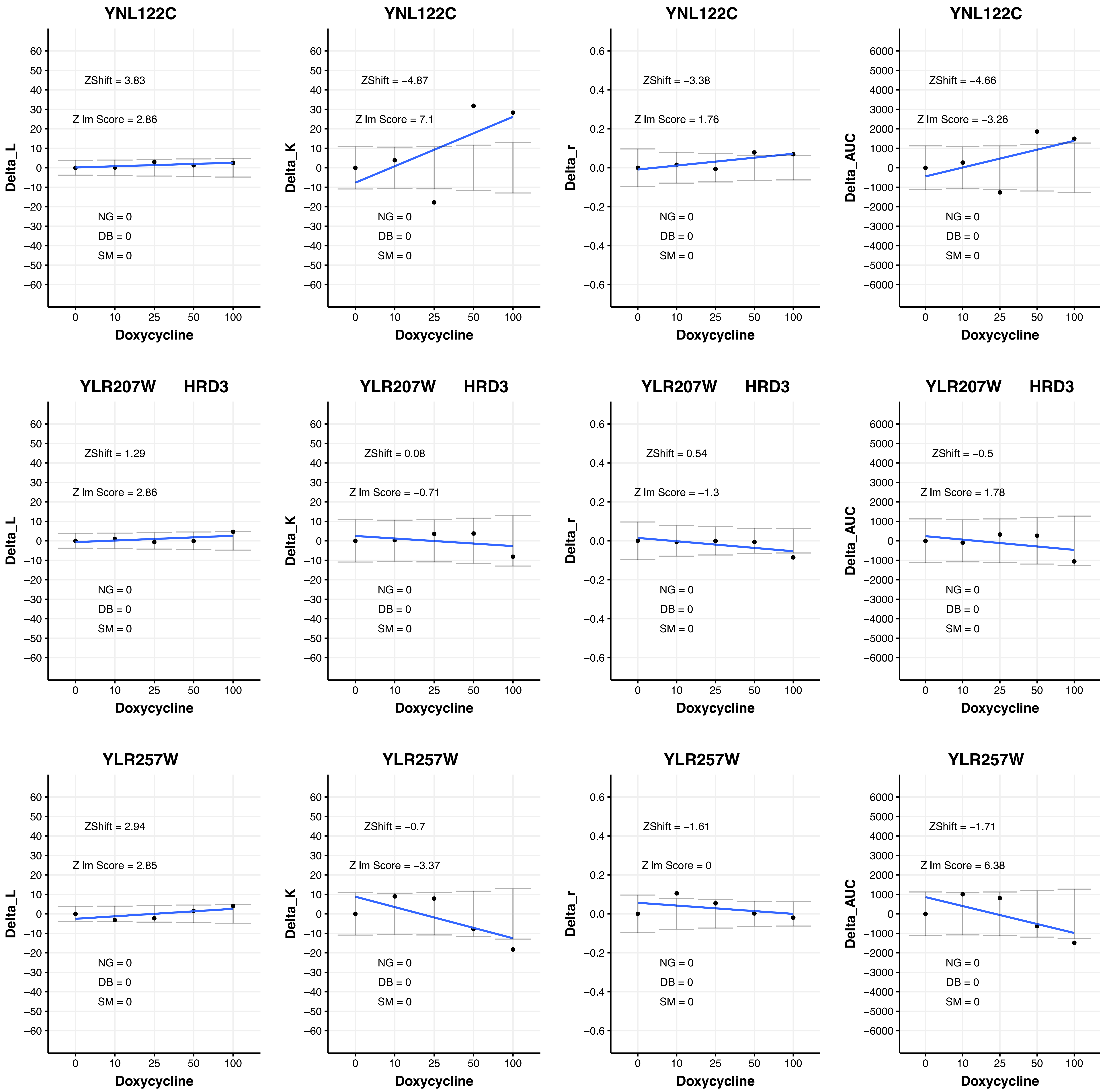

YNL122C
ZShift = 3.83
Z lm Score = 2.86
NG = 0
DB = 0
SM = 0
Delta_L
Doxycycline
YNL122C
ZShift = −4.87
Z lm Score = 7.1
NG = 0
DB = 0
SM = 0
Delta_K
Doxycycline
YNL122C
ZShift = −3.38
Z lm Score = 1.76
NG = 0
DB = 0
SM = 0
Delta_r
Doxycycline
YNL122C
ZShift = −4.66
Z lm Score = −3.26
NG = 0
DB = 0
SM = 0
Delta_AUC
Doxycycline
YLR207W HRD3
ZShift = 1.29
Z lm Score = 2.86
NG = 0
DB = 0
SM = 0
Delta_L
Doxycycline
YLR207W HRD3
ZShift = 0.08
Z lm Score = −0.71
NG = 0
DB = 0
SM = 0
Delta_K
Doxycycline
YLR207W HRD3
ZShift = 0.54
Z lm Score = −1.3
NG = 0
DB = 0
SM = 0
Delta_r
Doxycycline
YLR207W HRD3
ZShift = −0.5
Z lm Score = 1.78
NG = 0
DB = 0
SM = 0
Delta_AUC
Doxycycline
YLR257W
ZShift = 2.94
Z lm Score = 2.85
NG = 0
DB = 0
SM = 0
Delta_L
Doxycycline
YLR257W
ZShift = −0.7
Z lm Score = −3.37
NG = 0
DB = 0
SM = 0
Delta_K
Doxycycline
YLR257W
ZShift = −1.61
Z lm Score = 0
NG = 0
DB = 0
SM = 0
Delta_r
Doxycycline
YLR257W
ZShift = −1.71
Z lm Score = 6.38
NG = 0
DB = 0
SM = 0
Delta_AUC
Doxycycline

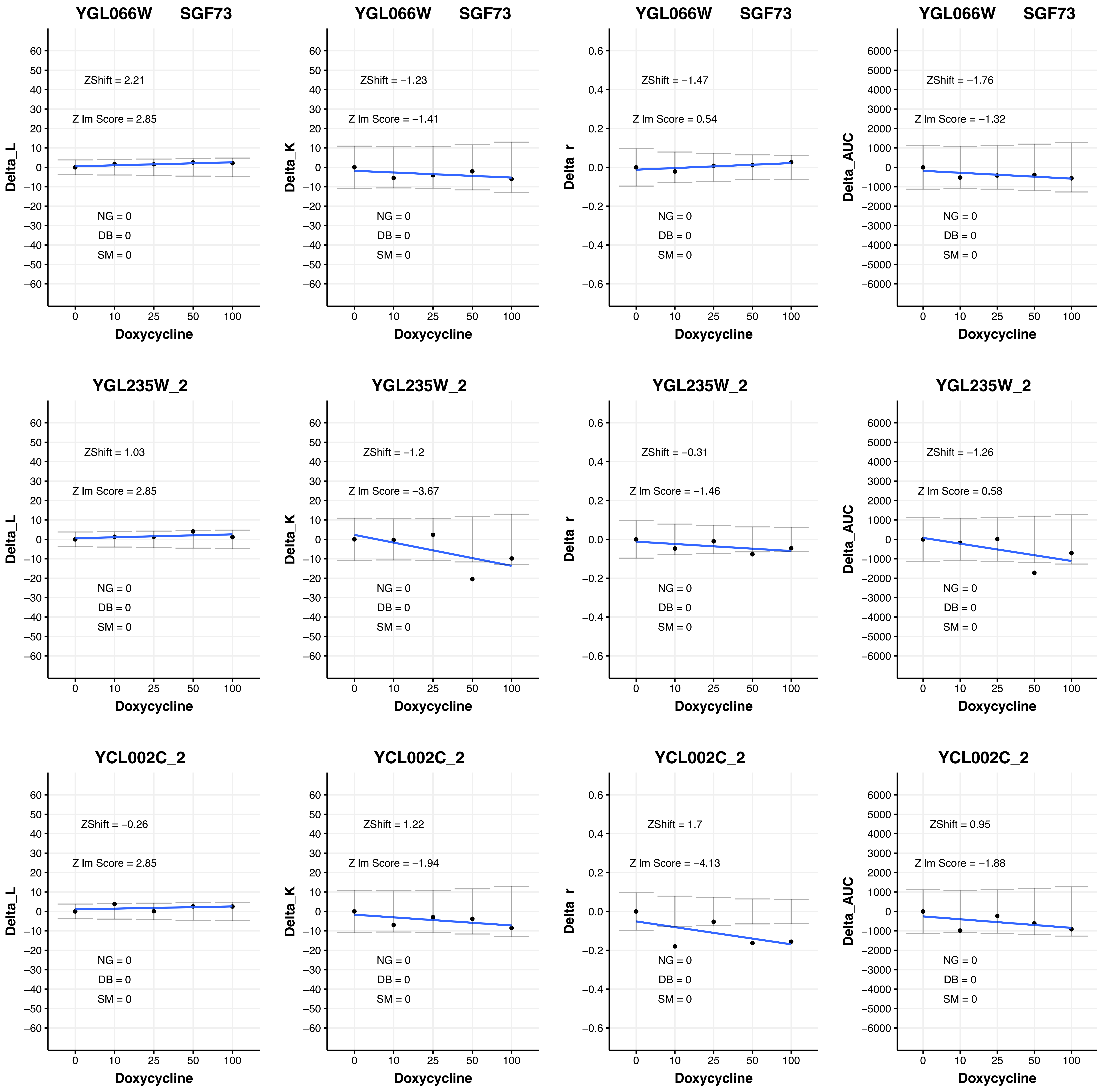

YGL066W SGF73
ZShift = 2.21
Z lm Score = 2.85
NG = 0
DB = 0
SM = 0
Delta_L
Doxycycline
YGL066W SGF73
ZShift = −1.23
Z lm Score = −1.41
NG = 0
DB = 0
SM = 0
Delta_K
Doxycycline
YGL066W SGF73
ZShift = −1.47
Z lm Score = 0.54
NG = 0
DB = 0
SM = 0
Delta_r
Doxycycline
YGL066W SGF73
ZShift = −1.76
Z lm Score = −1.32
NG = 0
DB = 0
SM = 0
Delta_AUC
Doxycycline
YGL235W_2
ZShift = 1.03
Z lm Score = 2.85
NG = 0
DB = 0
SM = 0
Delta_L
Doxycycline
YGL235W_2
ZShift = −1.2
Z lm Score = −3.67
NG = 0
DB = 0
SM = 0
Delta_K
Doxycycline
YGL235W_2
ZShift = −0.31
Z lm Score = −1.46
NG = 0
DB = 0
SM = 0
Delta_r
Doxycycline
YGL235W_2
ZShift = −1.26
Z lm Score = 0.58
NG = 0
DB = 0
SM = 0
Delta_AUC
Doxycycline
YCL002C_2
ZShift = −0.26
Z lm Score = 2.85
NG = 0
DB = 0
SM = 0
Delta_L
Doxycycline
YCL002C_2
ZShift = 1.22
Z lm Score = −1.94
NG = 0
DB = 0
SM = 0
Delta_K
Doxycycline
YCL002C_2
ZShift = 1.7
Z lm Score = −4.13
NG = 0
DB = 0
SM = 0
Delta_r
Doxycycline
YCL002C_2
ZShift = 0.95
Z lm Score = −1.88
NG = 0
DB = 0
SM = 0
Delta_AUC
Doxycycline

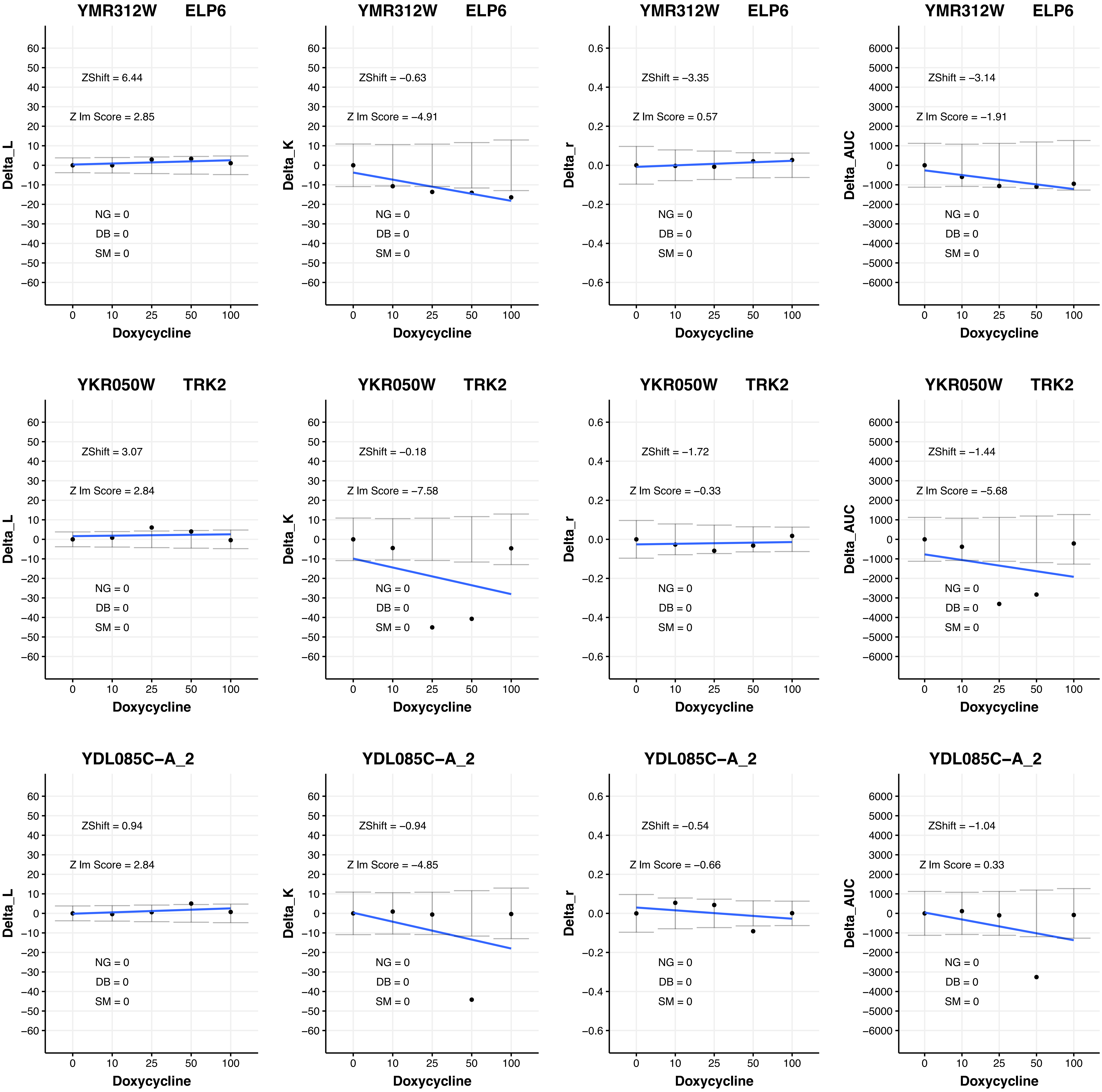
YMR312W ELP6
ZShift = 6.44
Z lm Score = 2.85
NG = 0
DB = 0
SM = 0
Delta_L
Doxycycline
YMR312W ELP6
ZShift = −0.63
Z lm Score = −4.91
NG = 0
DB = 0
SM = 0
Delta_K
Doxycycline
YMR312W ELP6
ZShift = −3.35
Z lm Score = 0.57
NG = 0
DB = 0
SM = 0
Delta_r
Doxycycline
YMR312W ELP6
ZShift = −3.14
Z lm Score = −1.91
NG = 0
DB = 0
SM = 0
Delta_AUC
Doxycycline
YKR050W TRK2
ZShift = 3.07
Z lm Score = 2.84
NG = 0
DB = 0
SM = 0
Delta_L
Doxycycline
YKR050W TRK2
ZShift = −0.18
Z lm Score = −7.58
NG = 0
DB = 0
SM = 0
Delta_K
Doxycycline
YKR050W TRK2
ZShift = −1.72
Z lm Score = −0.33
NG = 0
DB = 0
SM = 0
Delta_r
Doxycycline
YKR050W TRK2
ZShift = −1.44
Z lm Score = −5.68
NG = 0
DB = 0
SM = 0
Delta_AUC
Doxycycline
YDL085C−A_2
ZShift = 0.94
Z lm Score = 2.84
NG = 0
DB = 0
SM = 0
Delta_L
Doxycycline
YDL085C−A_2
ZShift = −0.94
Z lm Score = −4.85
NG = 0
DB = 0
SM = 0
Delta_K
Doxycycline
YDL085C−A_2
ZShift = −0.54
Z lm Score = −0.66
NG = 0
DB = 0
SM = 0
Delta_r
Doxycycline
YDL085C−A_2
ZShift = −1.04
Z lm Score = 0.33
NG = 0
DB = 0
SM = 0
Delta_AUC
Doxycycline

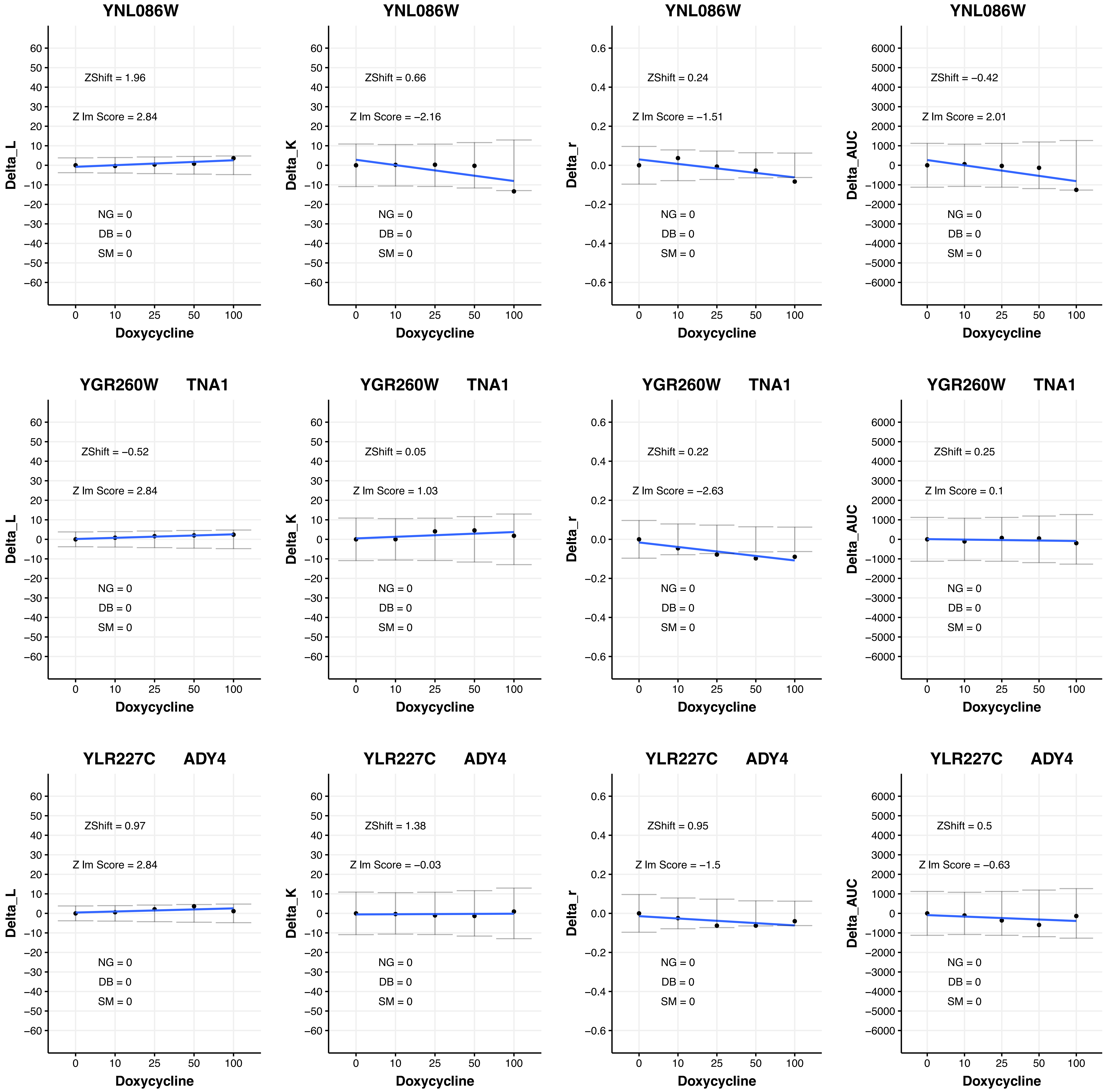
YNL086W
ZShift = 1.96
Z lm Score = 2.84
NG = 0
DB = 0
SM = 0
Delta_L
Doxycycline
YNL086W
ZShift = 0.66
Z lm Score = −2.16
NG = 0
DB = 0
SM = 0
Delta_K
Doxycycline
YNL086W
ZShift = 0.24
Z lm Score = −1.51
NG = 0
DB = 0
SM = 0
Delta_r
Doxycycline
YNL086W
ZShift = −0.42
Z lm Score = 2.01
NG = 0
DB = 0
SM = 0
Delta_AUC
Doxycycline
YGR260W TNA1
ZShift = −0.52
Z lm Score = 2.84
NG = 0
DB = 0
SM = 0
Delta_L
Doxycycline
YGR260W TNA1
ZShift = 0.05
Z lm Score = 1.03
NG = 0
DB = 0
SM = 0
Delta_K
Doxycycline
YGR260W TNA1
ZShift = 0.22
Z lm Score = −2.63
NG = 0
DB = 0
SM = 0
Delta_r
Doxycycline
YGR260W TNA1
ZShift = 0.25
Z lm Score = 0.1
NG = 0
DB = 0
SM = 0
Delta_AUC
Doxycycline
YLR227C ADY4
ZShift = 0.97
Z lm Score = 2.84
NG = 0
DB = 0
SM = 0
Delta_L
Doxycycline
YLR227C ADY4
ZShift = 1.38
Z lm Score = −0.03
NG = 0
DB = 0
SM = 0
Delta_K
Doxycycline
YLR227C ADY4
ZShift = 0.95
Z lm Score = −1.5
NG = 0
DB = 0
SM = 0
Delta_r
Doxycycline
YLR227C ADY4
ZShift = 0.5
Z lm Score = −0.63
NG = 0
DB = 0
SM = 0
Delta_AUC
Doxycycline

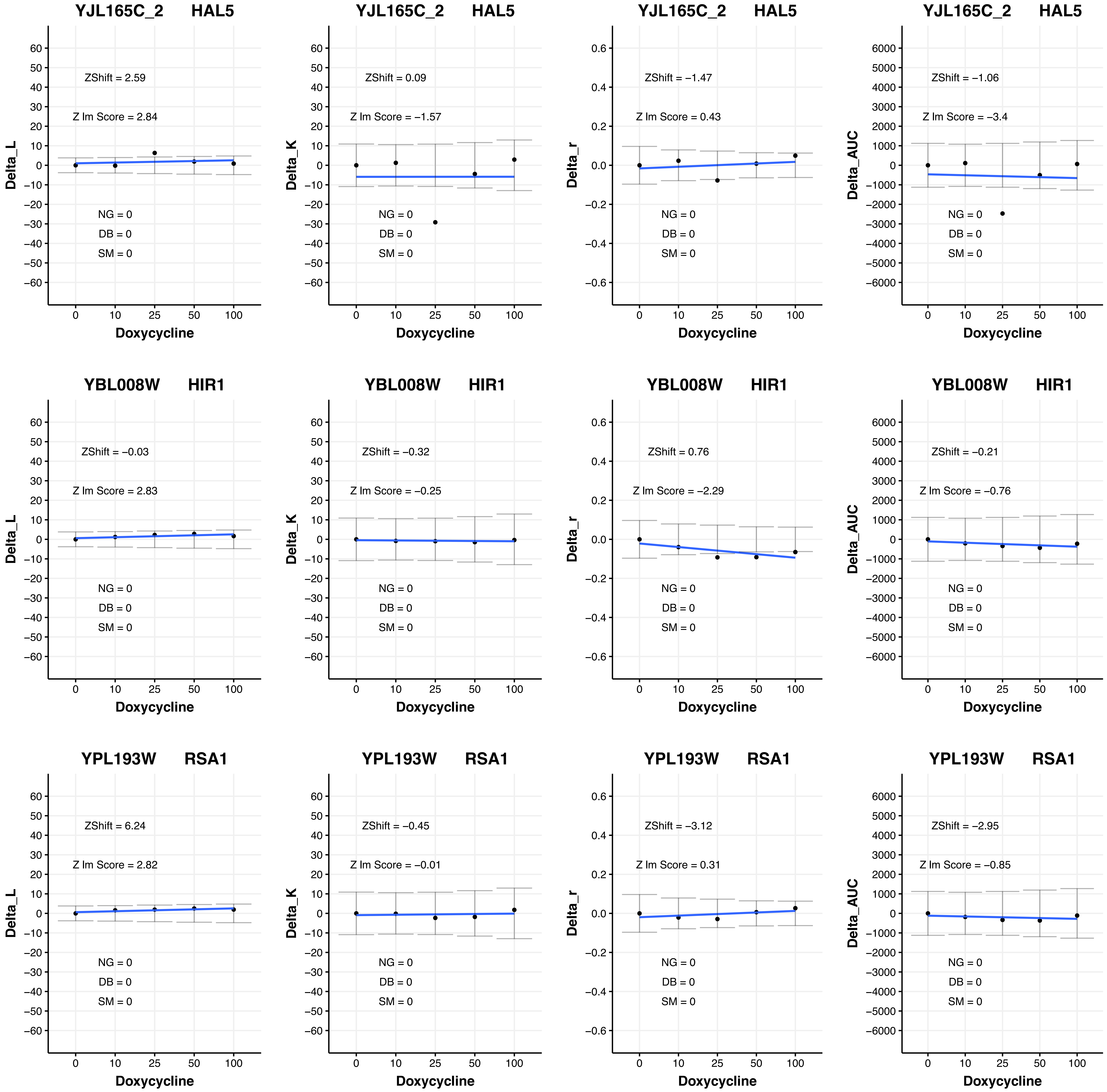
YJL165C_2 HAL5
ZShift = 2.59
Z lm Score = 2.84
NG = 0
DB = 0
SM = 0
Delta_L
Doxycycline
YJL165C_2 HAL5
ZShift = 0.09
Z lm Score = −1.57
NG = 0
DB = 0
SM = 0
Delta_K
Doxycycline
YJL165C_2 HAL5
ZShift = −1.47
Z lm Score = 0.43
NG = 0
DB = 0
SM = 0
Delta_r
Doxycycline
YJL165C_2 HAL5
ZShift = −1.06
Z lm Score = −3.4
NG = 0
DB = 0
SM = 0
Delta_AUC
Doxycycline
YBL008W HIR1
ZShift = −0.03
Z lm Score = 2.83
NG = 0
DB = 0
SM = 0
Delta_L
Doxycycline
YBL008W HIR1
ZShift = −0.32
Z lm Score = −0.25
NG = 0
DB = 0
SM = 0
Delta_K
Doxycycline
YBL008W HIR1
ZShift = 0.76
Z lm Score = −2.29
NG = 0
DB = 0
SM = 0
Delta_r
Doxycycline
YBL008W HIR1
ZShift = −0.21
Z lm Score = −0.76
NG = 0
DB = 0
SM = 0
Delta_AUC
Doxycycline
YPL193W RSA1
ZShift = 6.24
Z lm Score = 2.82
NG = 0
DB = 0
SM = 0
Delta_L
Doxycycline
YPL193W RSA1
ZShift = −0.45
Z lm Score = −0.01
NG = 0
DB = 0
SM = 0
Delta_K
Doxycycline
YPL193W RSA1
ZShift = −3.12
Z lm Score = 0.31
NG = 0
DB = 0
SM = 0
Delta_r
Doxycycline
YPL193W RSA1
ZShift = −2.95
Z lm Score = −0.85
NG = 0
DB = 0
SM = 0
Delta_AUC
Doxycycline

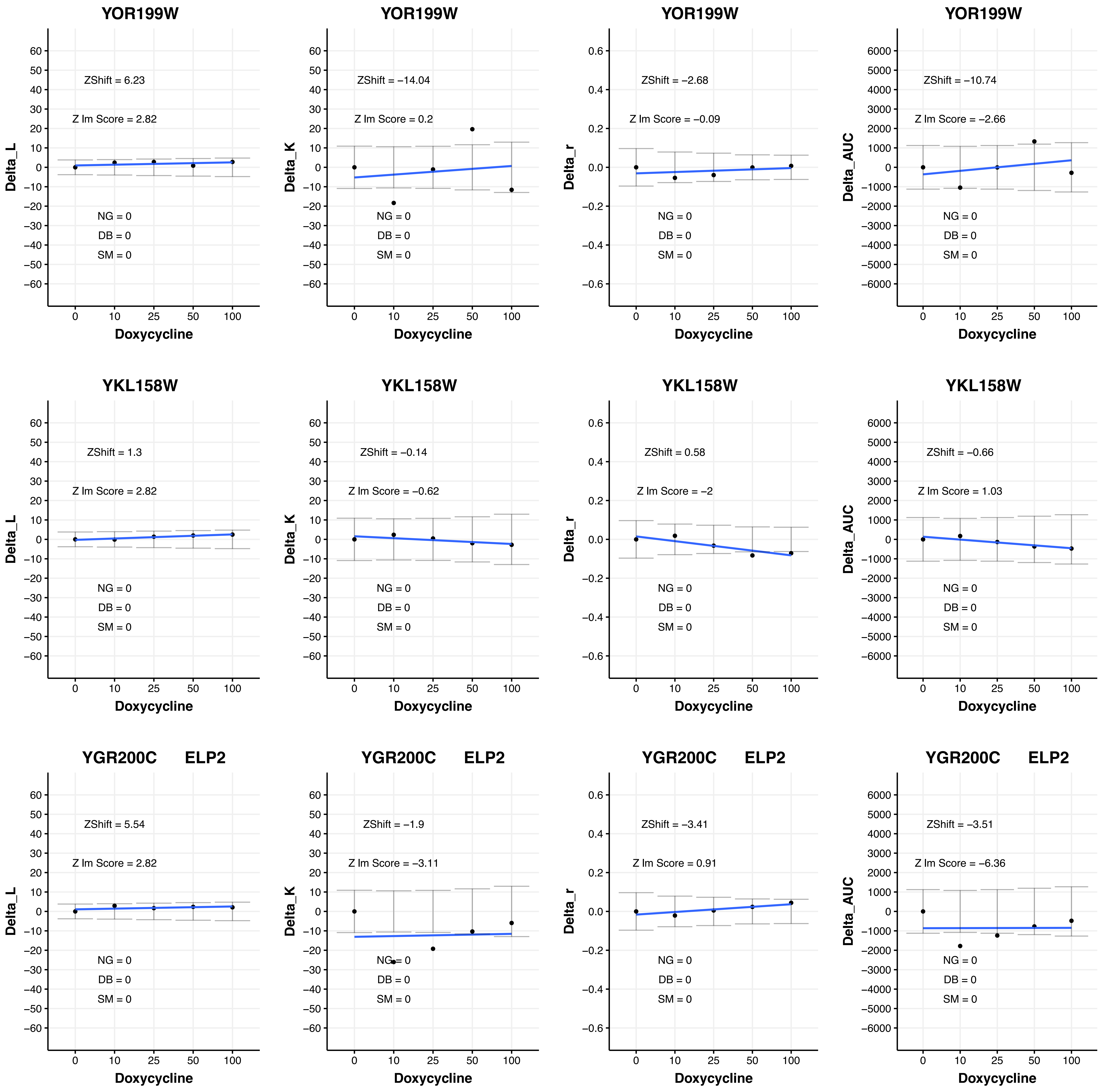

YOR199W
ZShift = 6.23
Z lm Score = 2.82
NG = 0
DB = 0
SM = 0
Delta_L
Doxycycline
YOR199W
ZShift = −14.04
Z lm Score = 0.2
NG = 0
DB = 0
SM = 0
Delta_K
Doxycycline
YOR199W
ZShift = −2.68
Z lm Score = −0.09
NG = 0
DB = 0
SM = 0
Delta_r
Doxycycline
YOR199W
ZShift = −10.74
Z lm Score = −2.66
NG = 0
DB = 0
SM = 0
Delta_AUC
Doxycycline
YKL158W
ZShift = 1.3
Z lm Score = 2.82
NG = 0
DB = 0
SM = 0
Delta_L
Doxycycline
YKL158W
ZShift = −0.14
Z lm Score = −0.62
NG = 0
DB = 0
SM = 0
Delta_K
Doxycycline
YKL158W
ZShift = 0.58
Z lm Score = −2
NG = 0
DB = 0
SM = 0
Delta_r
Doxycycline
YKL158W
ZShift = −0.66
Z lm Score = 1.03
NG = 0
DB = 0
SM = 0
Delta_AUC
Doxycycline
YGR200C ELP2
ZShift = 5.54
Z lm Score = 2.82
NG = 0
DB = 0
SM = 0
Delta_L
Doxycycline
YGR200C ELP2
ZShift = −1.9
Z lm Score = −3.11
NG = 0
DB = 0
SM = 0
Delta_K
Doxycycline
YGR200C ELP2
ZShift = −3.41
Z lm Score = 0.91
NG = 0
DB = 0
SM = 0
Delta_r
Doxycycline
YGR200C ELP2
ZShift = −3.51
Z lm Score = −6.36
NG = 0
DB = 0
SM = 0
Delta_AUC
Doxycycline

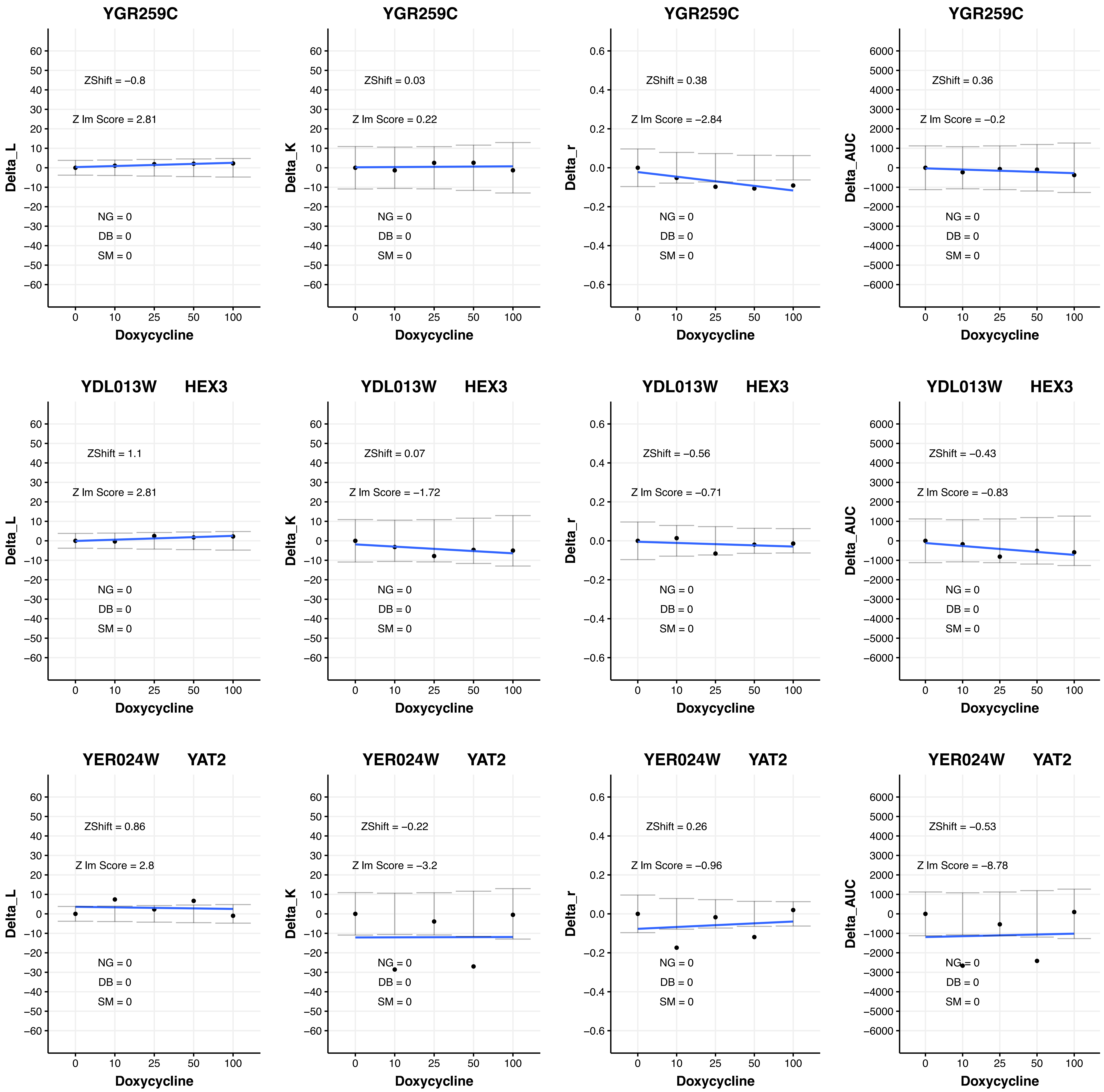

YGR259C
ZShift = −0.8
Z lm Score = 2.81
NG = 0
DB = 0
SM = 0
Delta_L
Doxycycline
YGR259C
ZShift = 0.03
Z lm Score = 0.22
NG = 0
DB = 0
SM = 0
Delta_K
Doxycycline
YGR259C
ZShift = 0.38
Z lm Score = −2.84
NG = 0
DB = 0
SM = 0
Delta_r
Doxycycline
YGR259C
ZShift = 0.36
Z lm Score = −0.2
NG = 0
DB = 0
SM = 0
Delta_AUC
Doxycycline
YDL013W HEX3
ZShift = 1.1
Z lm Score = 2.81
NG = 0
DB = 0
SM = 0
Delta_L
Doxycycline
YDL013W HEX3
ZShift = 0.07
Z lm Score = −1.72
NG = 0
DB = 0
SM = 0
Delta_K
Doxycycline
YDL013W HEX3
ZShift = −0.56
Z lm Score = −0.71
NG = 0
DB = 0
SM = 0
Delta_r
Doxycycline
YDL013W HEX3
ZShift = −0.43
Z lm Score = −0.83
NG = 0
DB = 0
SM = 0
Delta_AUC
Doxycycline
YER024W YAT2
ZShift = 0.86
Z lm Score = 2.8
NG = 0
DB = 0
SM = 0
Delta_L
Doxycycline
YER024W YAT2
ZShift = −0.22
Z lm Score = −3.2
NG = 0
DB = 0
SM = 0
Delta_K
Doxycycline
YER024W YAT2
ZShift = 0.26
Z lm Score = −0.96
NG = 0
DB = 0
SM = 0
Delta_r
Doxycycline
YER024W YAT2
ZShift = −0.53
Z lm Score = −8.78
NG = 0
DB = 0
SM = 0
Delta_AUC
Doxycycline

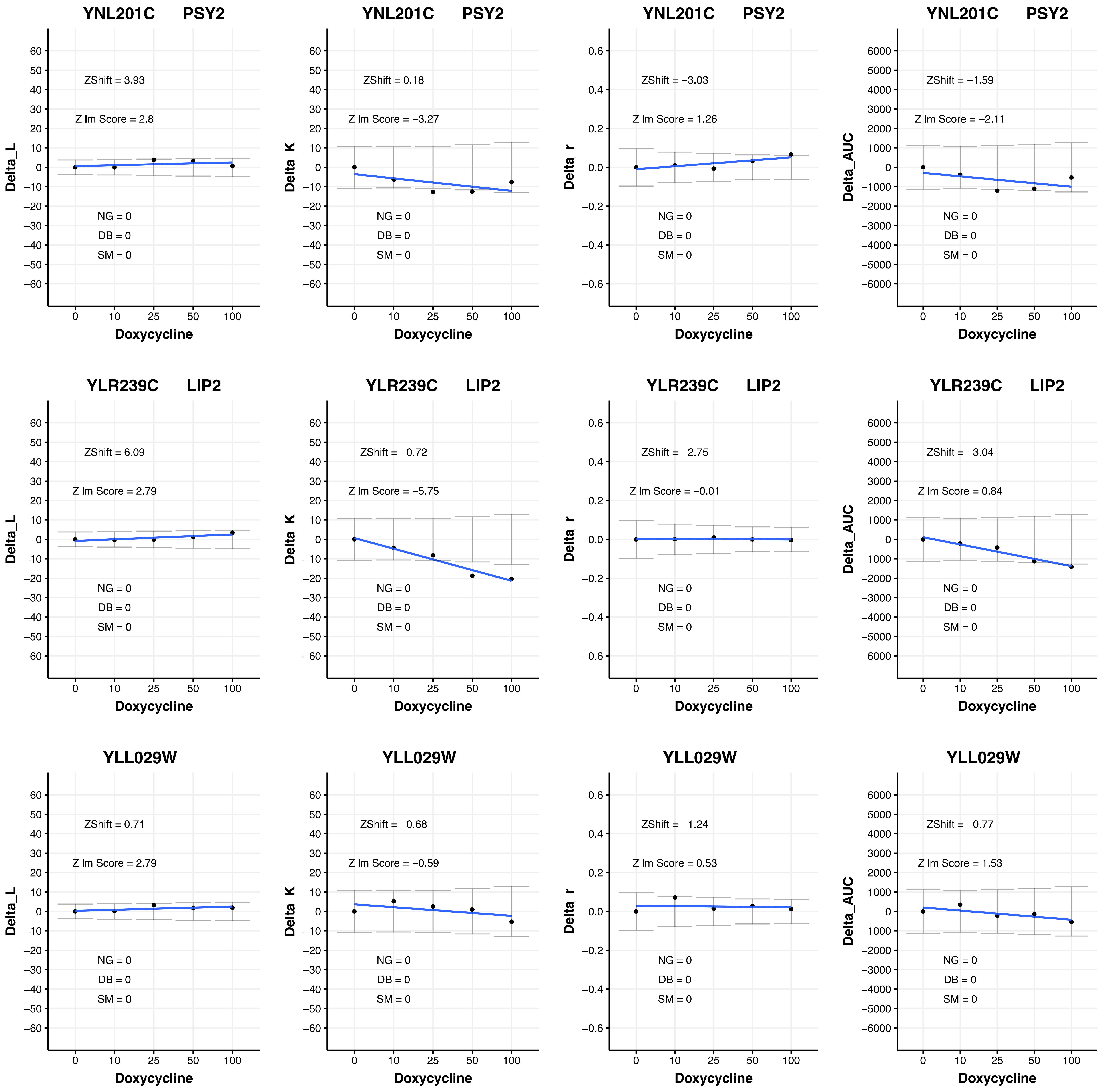

YNL201C PSY2
ZShift = 3.93
Z lm Score = 2.8
NG = 0
DB = 0
SM = 0
Delta_L
Doxycycline
YNL201C PSY2
ZShift = 0.18
Z lm Score = −3.27
NG = 0
DB = 0
SM = 0
Delta_K
Doxycycline
YNL201C PSY2
ZShift = −3.03
Z lm Score = 1.26
NG = 0
DB = 0
SM = 0
Delta_r
Doxycycline
YNL201C PSY2
ZShift = −1.59
Z lm Score = −2.11
NG = 0
DB = 0
SM = 0
Delta_AUC
Doxycycline
YLR239C LIP2
ZShift = 6.09
Z lm Score = 2.79
NG = 0
DB = 0
SM = 0
Delta_L
Doxycycline
YLR239C LIP2
ZShift = −0.72
Z lm Score = −5.75
NG = 0
DB = 0
SM = 0
Delta_K
Doxycycline
YLR239C LIP2
ZShift = −2.75
Z lm Score = −0.01
NG = 0
DB = 0
SM = 0
Delta_r
Doxycycline
YLR239C LIP2
ZShift = −3.04
Z lm Score = 0.84
NG = 0
DB = 0
SM = 0
Delta_AUC
Doxycycline
YLL029W
ZShift = 0.71
Z lm Score = 2.79
NG = 0
DB = 0
SM = 0
Delta_L
Doxycycline
YLL029W
ZShift = −0.68
Z lm Score = −0.59
NG = 0
DB = 0
SM = 0
Delta_K
Doxycycline
YLL029W
ZShift = −1.24
Z lm Score = 0.53
NG = 0
DB = 0
SM = 0
Delta_r
Doxycycline
YLL029W
ZShift = −0.77
Z lm Score = 1.53
NG = 0
DB = 0
SM = 0
Delta_AUC
Doxycycline

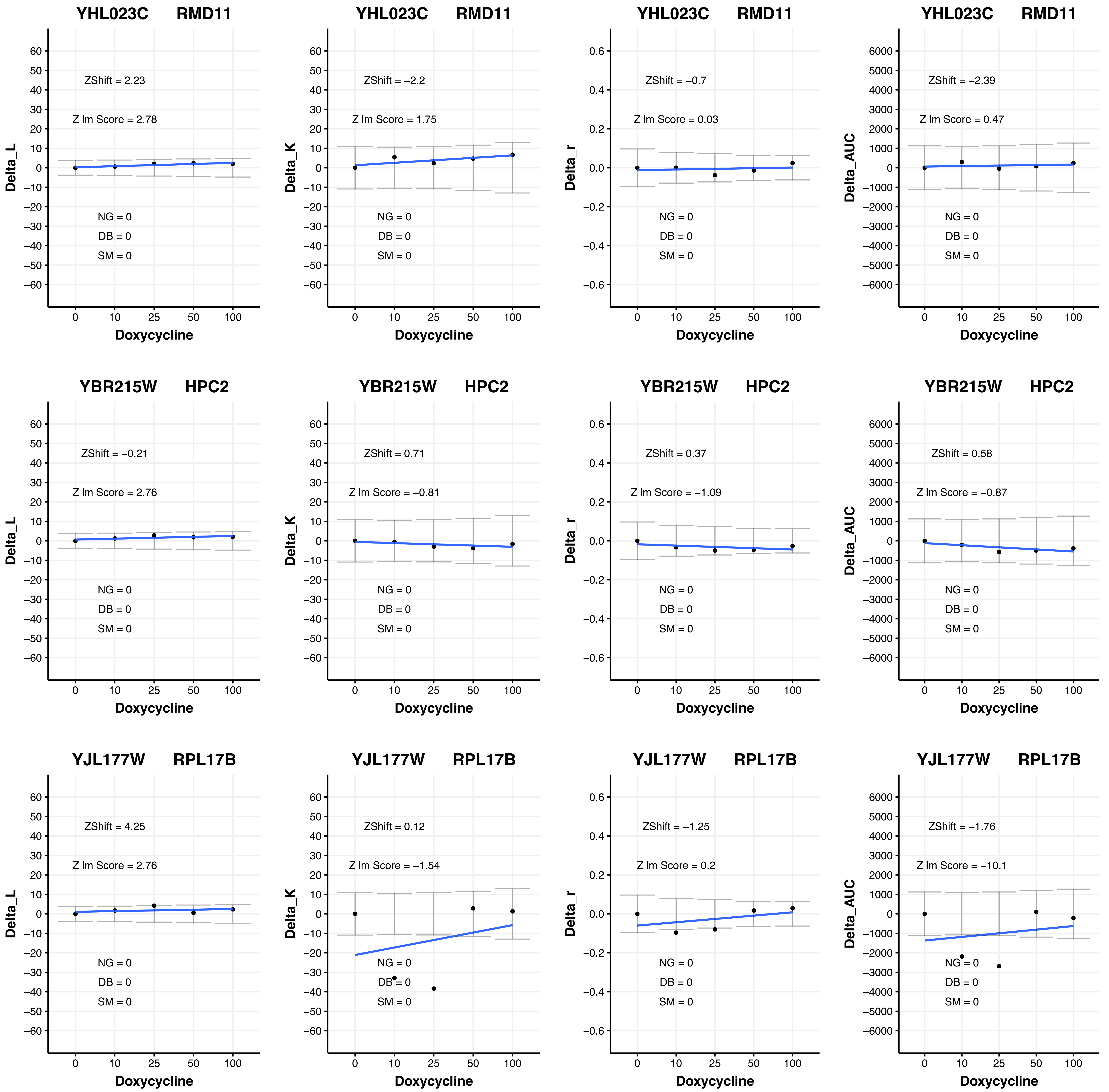
YHL023C RMD11
ZShift = 2.23
Z lm Score = 2.78
NG = 0
DB = 0
SM = 0
Delta_L
Doxycycline
YHL023C RMD11
ZShift = −2.2
Z lm Score = 1.75
NG = 0
DB = 0
SM = 0
Delta_K
Doxycycline
YHL023C RMD11
ZShift = −0.7
Z lm Score = 0.03
NG = 0
DB = 0
SM = 0
Delta_r
Doxycycline
YHL023C RMD11
ZShift = −2.39
Z lm Score = 0.47
NG = 0
DB = 0
SM = 0
Delta_AUC
Doxycycline
YBR215W HPC2
ZShift = −0.21
Z lm Score = 2.76
NG = 0
DB = 0
SM = 0
Delta_L
Doxycycline
YBR215W HPC2
ZShift = 0.71
Z lm Score = −0.81
NG = 0
DB = 0
SM = 0
Delta_K
Doxycycline
YBR215W HPC2
ZShift = 0.37
Z lm Score = −1.09
NG = 0
DB = 0
SM = 0
Delta_r
Doxycycline
YBR215W HPC2
ZShift = 0.58
Z lm Score = −0.87
NG = 0
DB = 0
SM = 0
Delta_AUC
Doxycycline
YJL177W RPL17B
ZShift = 4.25
Z lm Score = 2.76
NG = 0
DB = 0
SM = 0
Delta_L
Doxycycline
YJL177W RPL17B
ZShift = 0.12
Z lm Score = −1.54
NG = 0
DB = 0
SM = 0
Delta_K
Doxycycline
YJL177W RPL17B
ZShift = −1.25
Z lm Score = 0.2
NG = 0
DB = 0
SM = 0
Delta_r
Doxycycline
YJL177W RPL17B
ZShift = −1.76
Z lm Score = −10.1
NG = 0
DB = 0
SM = 0
Delta_AUC
Doxycycline

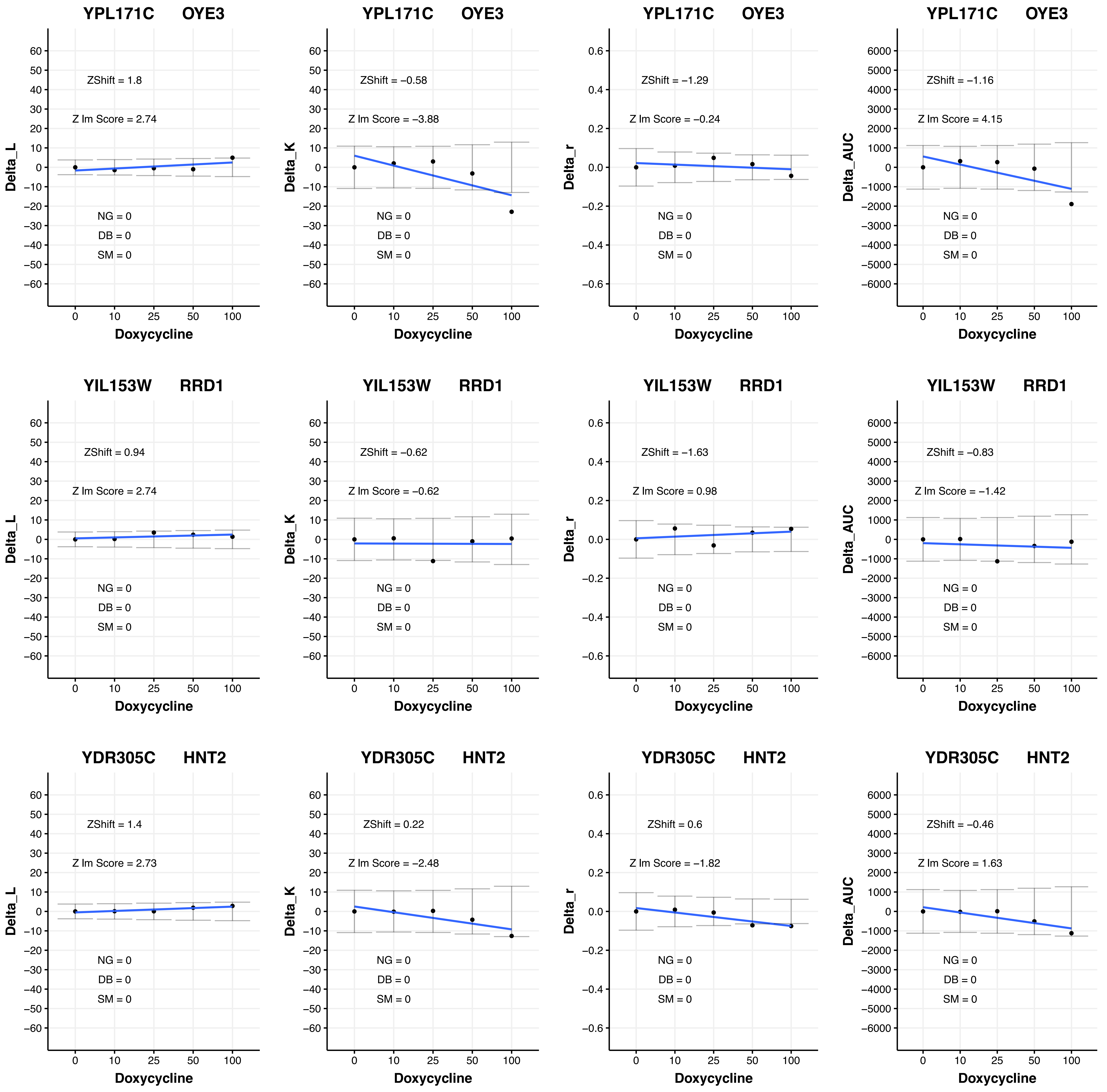
YPL171C OYE3
ZShift = 1.8
Z lm Score = 2.74
NG = 0
DB = 0
SM = 0
Delta_L
Doxycycline
YPL171C OYE3
ZShift = −0.58
Z lm Score = −3.88
NG = 0
DB = 0
SM = 0
Delta_K
Doxycycline
YPL171C OYE3
ZShift = −1.29
Z lm Score = −0.24
NG = 0
DB = 0
SM = 0
Delta_r
Doxycycline
YPL171C OYE3
ZShift = −1.16
Z lm Score = 4.15
NG = 0
DB = 0
SM = 0
Delta_AUC
Doxycycline
YIL153W RRD1
ZShift = 0.94
Z lm Score = 2.74
NG = 0
DB = 0
SM = 0
Delta_L
Doxycycline
YIL153W RRD1
ZShift = −0.62
Z lm Score = −0.62
NG = 0
DB = 0
SM = 0
Delta_K
Doxycycline
YIL153W RRD1
ZShift = −1.63
Z lm Score = 0.98
NG = 0
DB = 0
SM = 0
Delta_r
Doxycycline
YIL153W RRD1
ZShift = −0.83
Z lm Score = −1.42
NG = 0
DB = 0
SM = 0
Delta_AUC
Doxycycline
YDR305C HNT2
ZShift = 1.4
Z lm Score = 2.73
NG = 0
DB = 0
SM = 0
Delta_L
Doxycycline
YDR305C HNT2
ZShift = 0.22
Z lm Score = −2.48
NG = 0
DB = 0
SM = 0
Delta_K
Doxycycline
YDR305C HNT2
ZShift = 0.6
Z lm Score = −1.82
NG = 0
DB = 0
SM = 0
Delta_r
Doxycycline
YDR305C HNT2
ZShift = −0.46
Z lm Score = 1.63
NG = 0
DB = 0
SM = 0
Delta_AUC
Doxycycline

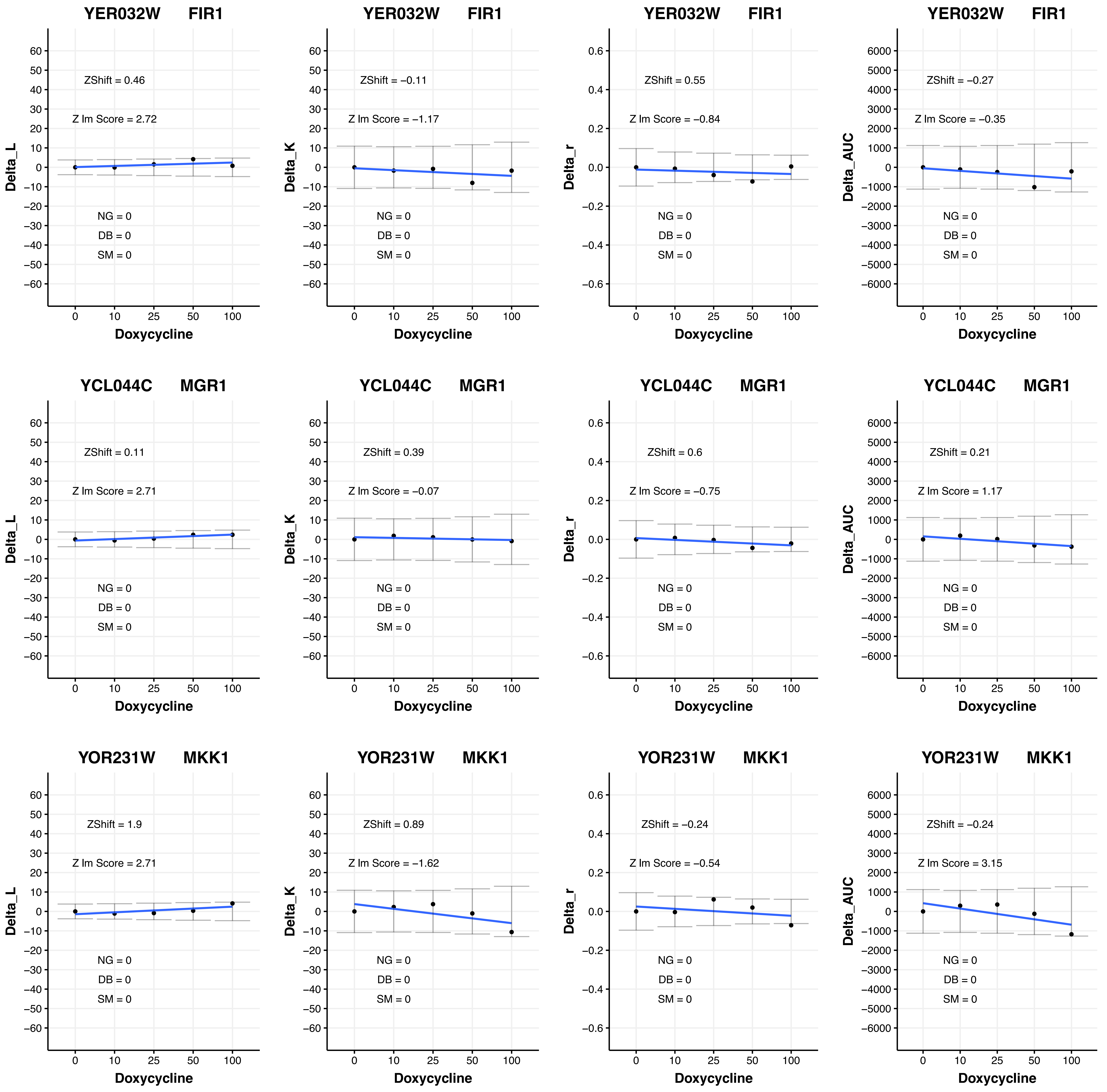

YER032W FIR1
ZShift = 0.46
Z lm Score = 2.72
NG = 0
DB = 0
SM = 0
Delta_L
Doxycycline
YER032W FIR1
ZShift = −0.11
Z lm Score = −1.17
NG = 0
DB = 0
SM = 0
Delta_K
Doxycycline
YER032W FIR1
ZShift = 0.55
Z lm Score = −0.84
NG = 0
DB = 0
SM = 0
Delta_r
Doxycycline
YER032W FIR1
ZShift = −0.27
Z lm Score = −0.35
NG = 0
DB = 0
SM = 0
Delta_AUC
Doxycycline
YCL044C MGR1
ZShift = 0.11
Z lm Score = 2.71
NG = 0
DB = 0
SM = 0
Delta_L
Doxycycline
YCL044C MGR1
ZShift = 0.39
Z lm Score = −0.07
NG = 0
DB = 0
SM = 0
Delta_K
Doxycycline
YCL044C MGR1
ZShift = 0.6
Z lm Score = −0.75
NG = 0
DB = 0
SM = 0
Delta_r
Doxycycline
YCL044C MGR1
ZShift = 0.21
Z lm Score = 1.17
NG = 0
DB = 0
SM = 0
Delta_AUC
Doxycycline
YOR231W MKK1
ZShift = 1.9
Z lm Score = 2.71
NG = 0
DB = 0
SM = 0
Delta_L
Doxycycline
YOR231W MKK1
ZShift = 0.89
Z lm Score = −1.62
NG = 0
DB = 0
SM = 0
Delta_K
Doxycycline
YOR231W MKK1
ZShift = −0.24
Z lm Score = −0.54
NG = 0
DB = 0
SM = 0
Delta_r
Doxycycline
YOR231W MKK1
ZShift = −0.24
Z lm Score = 3.15
NG = 0
DB = 0
SM = 0
Delta_AUC
Doxycycline

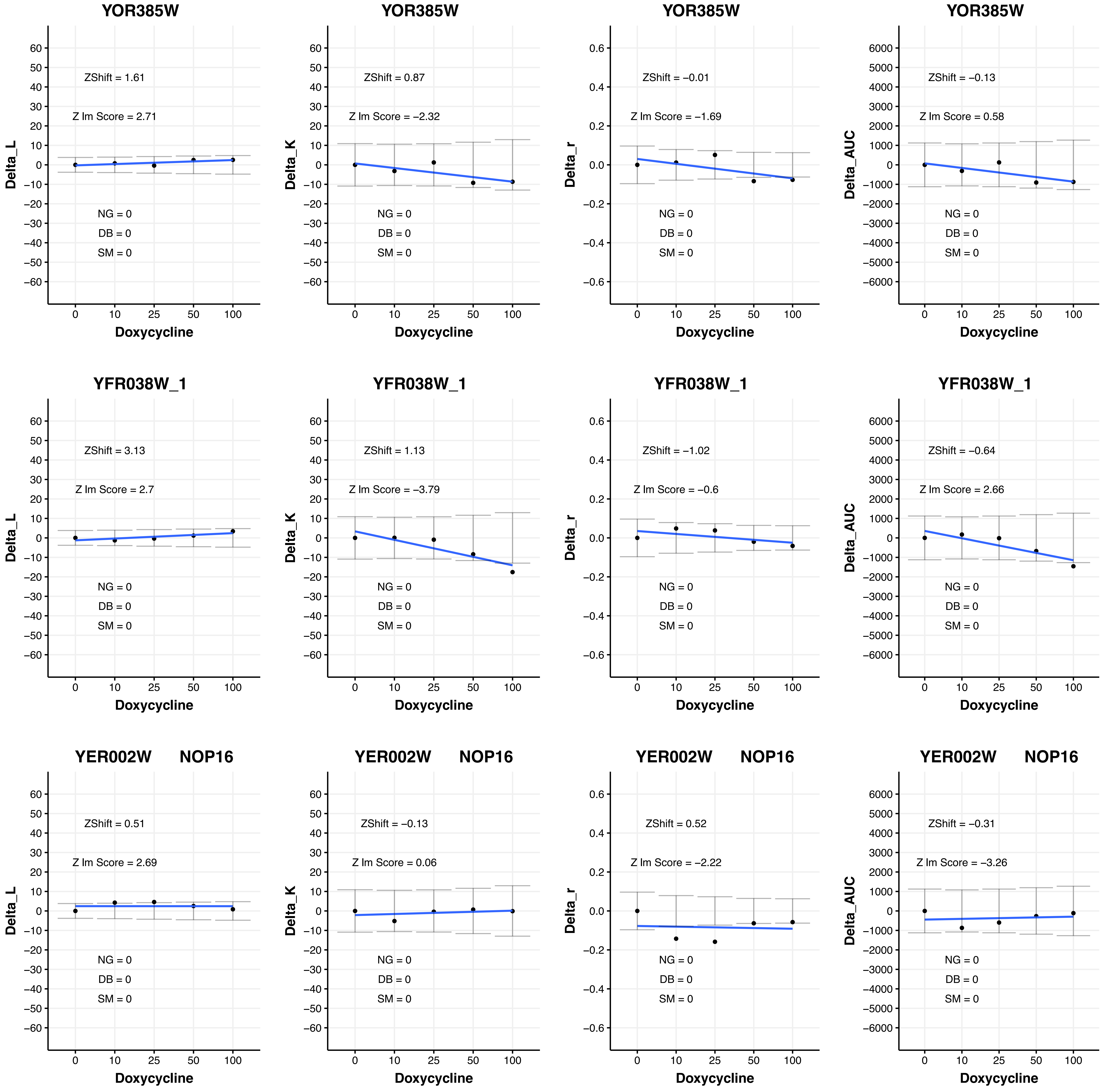
YOR385W
ZShift = 1.61
Z lm Score = 2.71
NG = 0
DB = 0
SM = 0
Delta_L
Doxycycline
YOR385W
ZShift = 0.87
Z lm Score = −2.32
NG = 0
DB = 0
SM = 0
Delta_K
Doxycycline
YOR385W
ZShift = −0.01
Z lm Score = −1.69
NG = 0
DB = 0
SM = 0
Delta_r
Doxycycline
YOR385W
ZShift = −0.13
Z lm Score = 0.58
NG = 0
DB = 0
SM = 0
Delta_AUC
Doxycycline
YFR038W_1
ZShift = 3.13
Z lm Score = 2.7
NG = 0
DB = 0
SM = 0
Delta_L
Doxycycline
YFR038W_1
ZShift = 1.13
Z lm Score = −3.79
NG = 0
DB = 0
SM = 0
Delta_K
Doxycycline
YFR038W_1
ZShift = −1.02
Z lm Score = −0.6
NG = 0
DB = 0
SM = 0
Delta_r
Doxycycline
YFR038W_1
ZShift = −0.64
Z lm Score = 2.66
NG = 0
DB = 0
SM = 0
Delta_AUC
Doxycycline
YER002W NOP16
ZShift = 0.51
Z lm Score = 2.69
NG = 0
DB = 0
SM = 0
Delta_L
Doxycycline
YER002W NOP16
ZShift = −0.13
Z lm Score = 0.06
NG = 0
DB = 0
SM = 0
Delta_K
Doxycycline
YER002W NOP16
ZShift = 0.52
Z lm Score = −2.22
NG = 0
DB = 0
SM = 0
Delta_r
Doxycycline
YER002W NOP16
ZShift = −0.31
Z lm Score = −3.26
NG = 0
DB = 0
SM = 0
Delta_AUC
Doxycycline

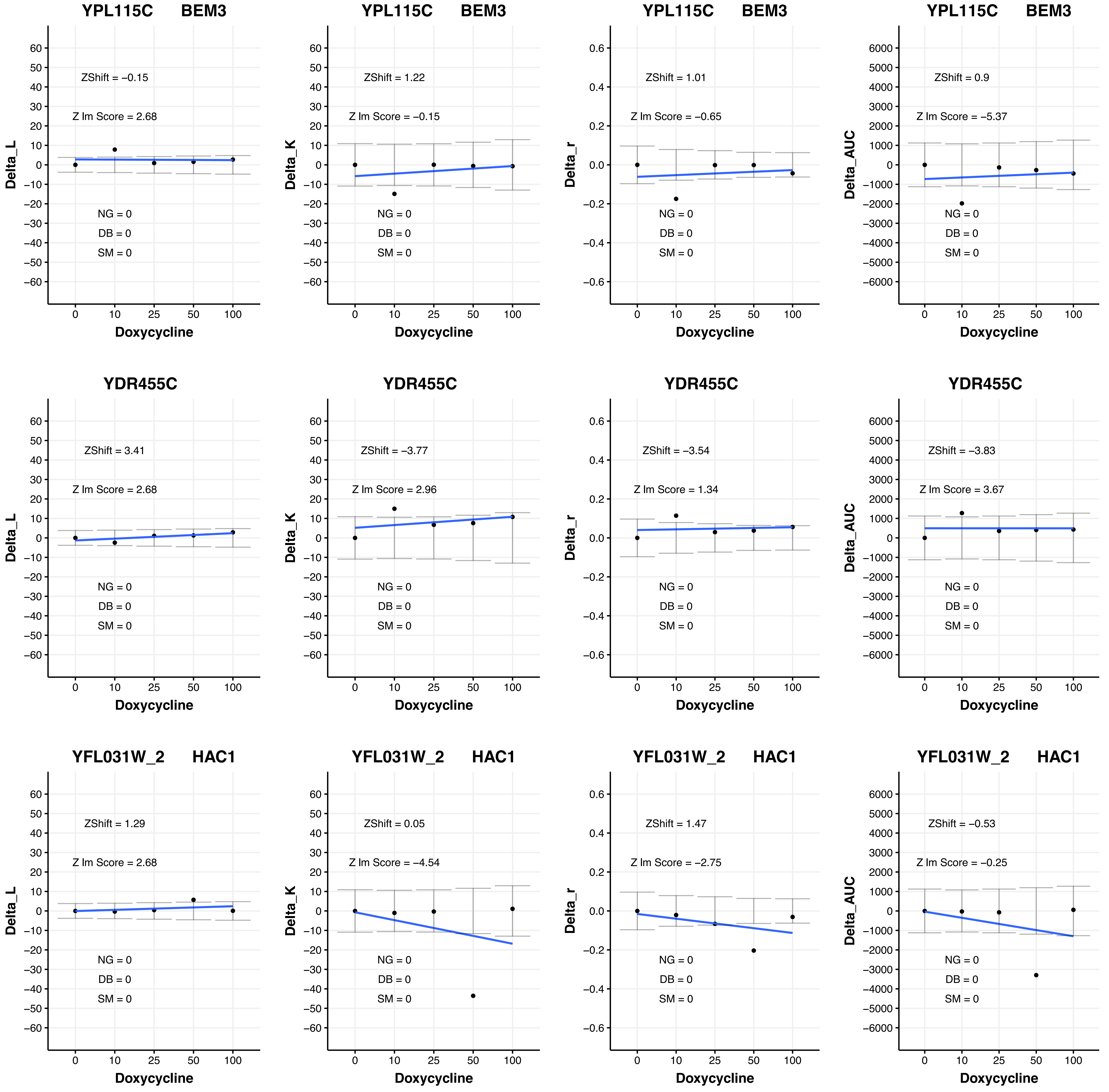

YPL115C BEM3
ZShift = −0.15
Z lm Score = 2.68
NG = 0
DB = 0
SM = 0
Delta_L
Doxycycline
YPL115C BEM3
ZShift = 1.22
Z lm Score = −0.15
NG = 0
DB = 0
SM = 0
Delta_K
Doxycycline
YPL115C BEM3
ZShift = 1.01
Z lm Score = −0.65
NG = 0
DB = 0
SM = 0
Delta_r
Doxycycline
YPL115C BEM3
ZShift = 0.9
Z lm Score = −5.37
NG = 0
DB = 0
SM = 0
Delta_AUC
Doxycycline
YDR455C
ZShift = 3.41
Z lm Score = 2.68
NG = 0
DB = 0
SM = 0
Delta_L
Doxycycline
YDR455C
ZShift = −3.77
Z lm Score = 2.96
NG = 0
DB = 0
SM = 0
Delta_K
Doxycycline
YDR455C
ZShift = −3.54
Z lm Score = 1.34
NG = 0
DB = 0
SM = 0
Delta_r
Doxycycline
YDR455C
ZShift = −3.83
Z lm Score = 3.67
NG = 0
DB = 0
SM = 0
Delta_AUC
Doxycycline
YFL031W_2 HAC1
ZShift = 1.29
Z lm Score = 2.68
NG = 0
DB = 0
SM = 0
Delta_L
Doxycycline
YFL031W_2 HAC1
ZShift = 0.05
Z lm Score = −4.54
NG = 0
DB = 0
SM = 0
Delta_K
Doxycycline
YFL031W_2 HAC1
ZShift = 1.47
Z lm Score = −2.75
NG = 0
DB = 0
SM = 0
Delta_r
Doxycycline
YFL031W_2 HAC1
ZShift = −0.53
Z lm Score = −0.25
NG = 0
DB = 0
SM = 0
Delta_AUC
Doxycycline

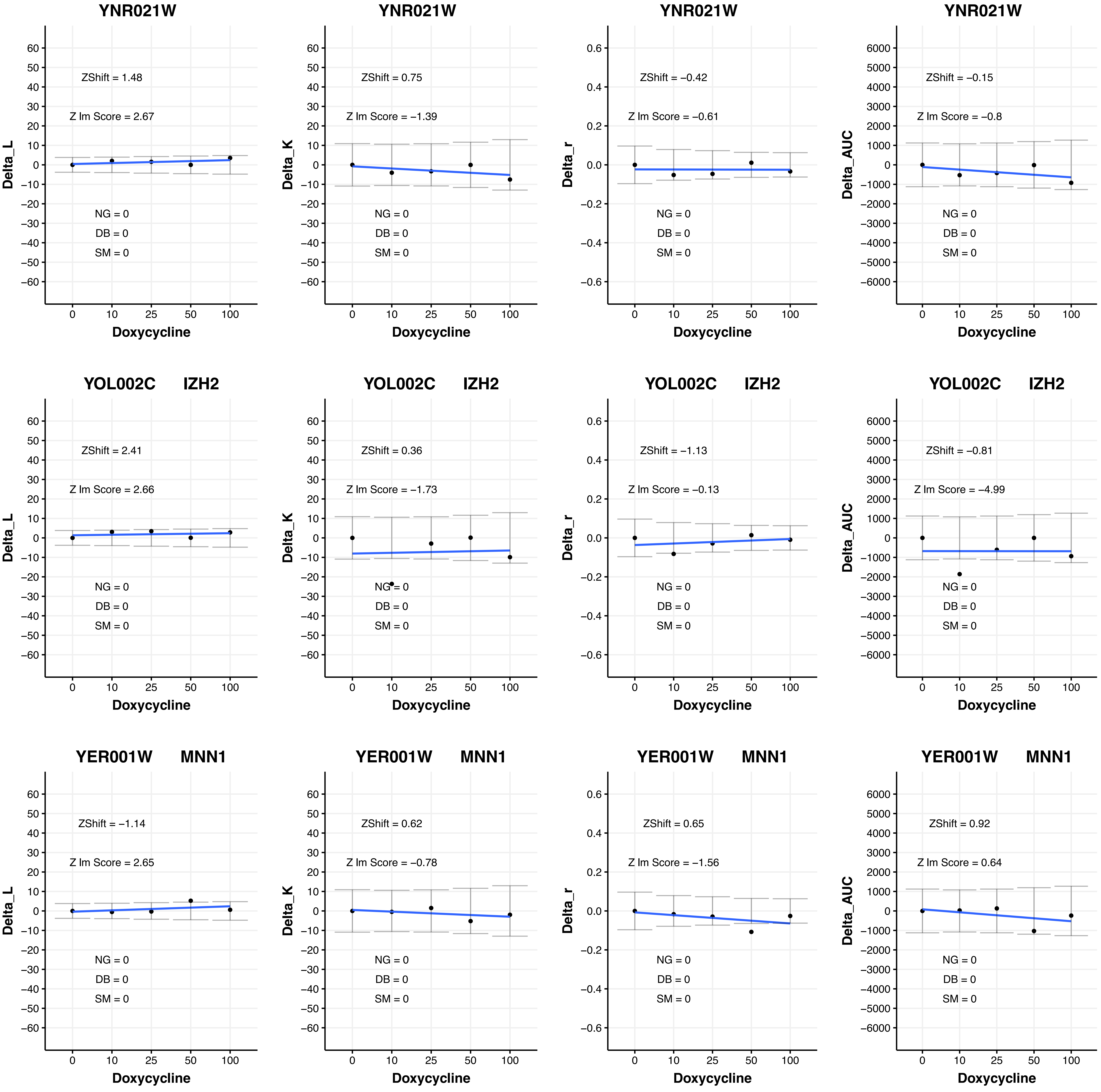
YNR021W
ZShift = 1.48
Z lm Score = 2.67
NG = 0
DB = 0
SM = 0
Delta_L
Doxycycline
YNR021W
ZShift = 0.75
Z lm Score = −1.39
NG = 0
DB = 0
SM = 0
Delta_K
Doxycycline
YNR021W
ZShift = −0.42
Z lm Score = −0.61
NG = 0
DB = 0
SM = 0
Delta_r
Doxycycline
YNR021W
ZShift = −0.15
Z lm Score = −0.8
NG = 0
DB = 0
SM = 0
Delta_AUC
Doxycycline
YOL002C IZH2
ZShift = 2.41
Z lm Score = 2.66
NG = 0
DB = 0
SM = 0
Delta_L
Doxycycline
YOL002C IZH2
ZShift = 0.36
Z lm Score = −1.73
NG = 0
DB = 0
SM = 0
Delta_K
Doxycycline
YOL002C IZH2
ZShift = −1.13
Z lm Score = −0.13
NG = 0
DB = 0
SM = 0
Delta_r
Doxycycline
YOL002C IZH2
ZShift = −0.81
Z lm Score = −4.99
NG = 0
DB = 0
SM = 0
Delta_AUC
Doxycycline
YER001W MNN1
ZShift = −1.14
Z lm Score = 2.65
NG = 0
DB = 0
SM = 0
Delta_L
Doxycycline
YER001W MNN1
ZShift = 0.62
Z lm Score = −0.78
NG = 0
DB = 0
SM = 0
Delta_K
Doxycycline
YER001W MNN1
ZShift = 0.65
Z lm Score = −1.56
NG = 0
DB = 0
SM = 0
Delta_r
Doxycycline
YER001W MNN1
ZShift = 0.92
Z lm Score = 0.64
NG = 0
DB = 0
SM = 0
Delta_AUC
Doxycycline

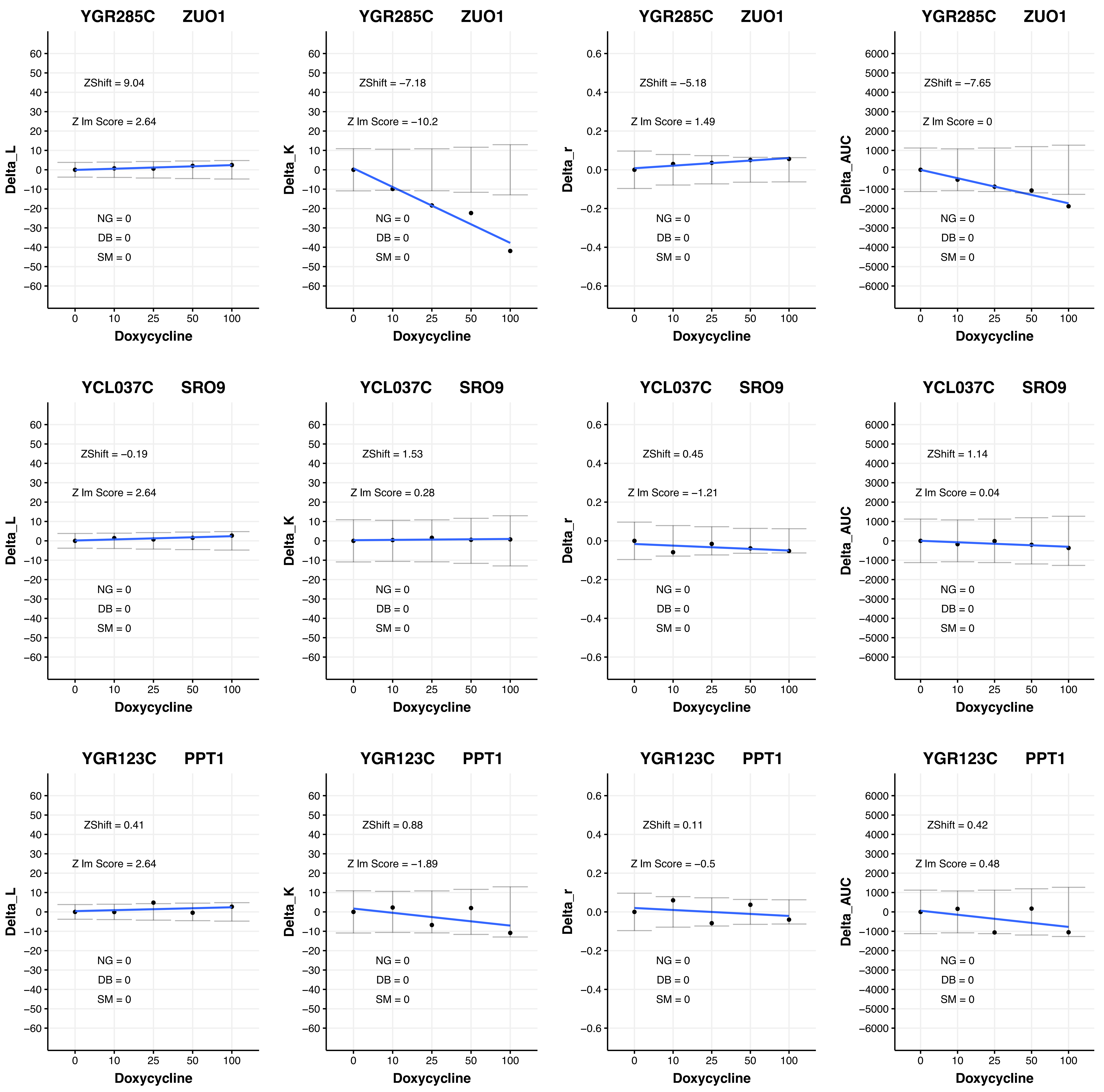
YGR285C ZUO1
ZShift = 9.04
Z lm Score = 2.64
NG = 0
DB = 0
SM = 0
Delta_L
Doxycycline
YGR285C ZUO1
ZShift = −7.18
Z lm Score = −10.2
NG = 0
DB = 0
SM = 0
Delta_K
Doxycycline
YGR285C ZUO1
ZShift = −5.18
Z lm Score = 1.49
NG = 0
DB = 0
SM = 0
Delta_r
Doxycycline
YGR285C ZUO1
ZShift = −7.65
Z lm Score = 0
NG = 0
DB = 0
SM = 0
Delta_AUC
Doxycycline
YCL037C SRO9
ZShift = −0.19
Z lm Score = 2.64
NG = 0
DB = 0
SM = 0
Delta_L
Doxycycline
YCL037C SRO9
ZShift = 1.53
Z lm Score = 0.28
NG = 0
DB = 0
SM = 0
Delta_K
Doxycycline
YCL037C SRO9
ZShift = 0.45
Z lm Score = −1.21
NG = 0
DB = 0
SM = 0
Delta_r
Doxycycline
YCL037C SRO9
ZShift = 1.14
Z lm Score = 0.04
NG = 0
DB = 0
SM = 0
Delta_AUC
Doxycycline
YGR123C PPT1
ZShift = 0.41
Z lm Score = 2.64
NG = 0
DB = 0
SM = 0
Delta_L
Doxycycline
YGR123C PPT1
ZShift = 0.88
Z lm Score = −1.89
NG = 0
DB = 0
SM = 0
Delta_K
Doxycycline
YGR123C PPT1
ZShift = 0.11
Z lm Score = −0.5
NG = 0
DB = 0
SM = 0
Delta_r
Doxycycline
YGR123C PPT1
ZShift = 0.42
Z lm Score = 0.48
NG = 0
DB = 0
SM = 0
Delta_AUC
Doxycycline

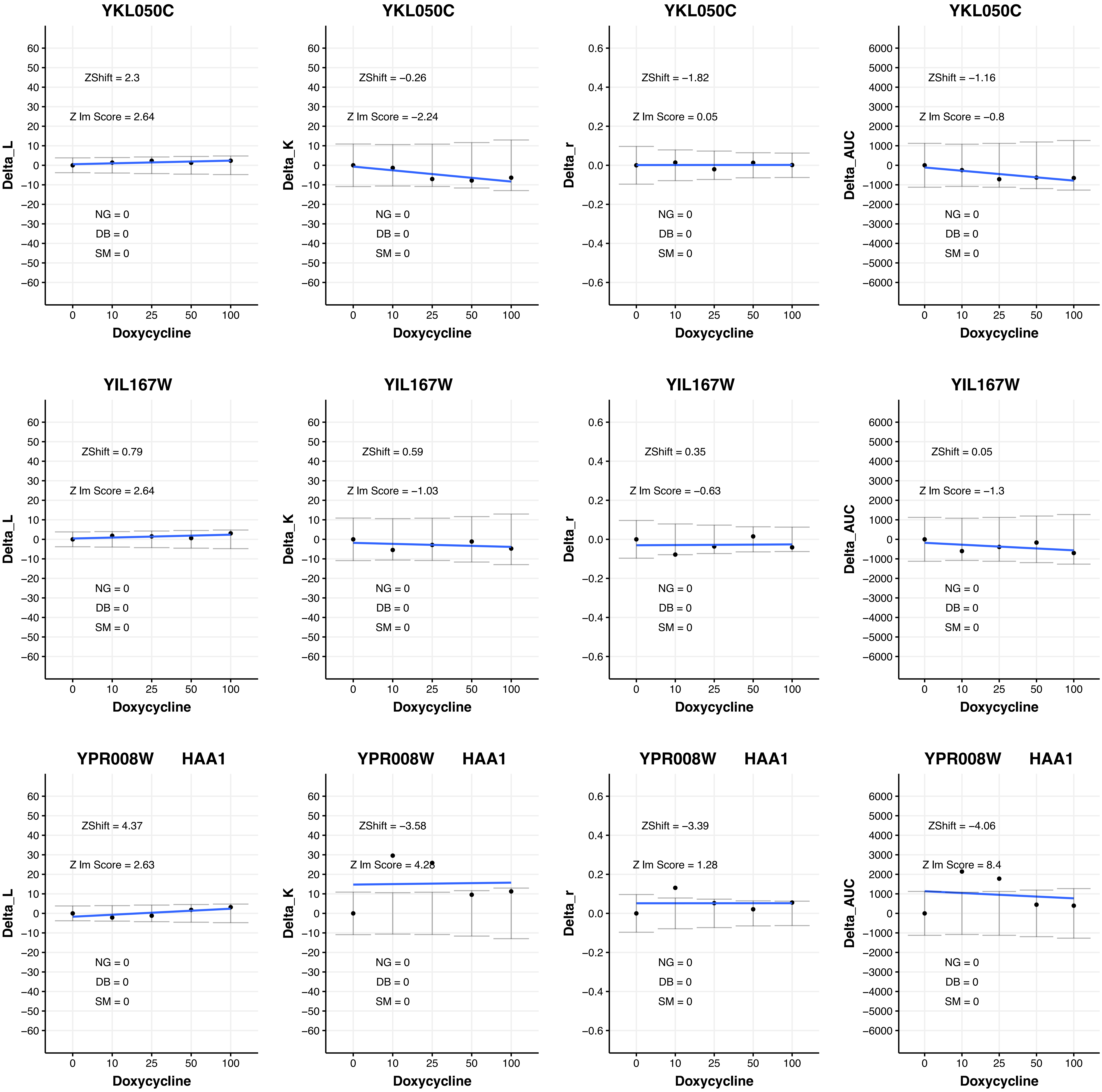

YKL050C
ZShift = 2.3
Z lm Score = 2.64
NG = 0
DB = 0
SM = 0
Delta_L
Doxycycline
YKL050C
ZShift = −0.26
Z lm Score = −2.24
NG = 0
DB = 0
SM = 0
Delta_K
Doxycycline
YKL050C
ZShift = −1.82
Z lm Score = 0.05
NG = 0
DB = 0
SM = 0
Delta_r
Doxycycline
YKL050C
ZShift = −1.16
Z lm Score = −0.8
NG = 0
DB = 0
SM = 0
Delta_AUC
Doxycycline
YIL167W
ZShift = 0.79
Z lm Score = 2.64
NG = 0
DB = 0
SM = 0
Delta_L
Doxycycline
YIL167W
ZShift = 0.59
Z lm Score = −1.03
NG = 0
DB = 0
SM = 0
Delta_K
Doxycycline
YIL167W
ZShift = 0.35
Z lm Score = −0.63
NG = 0
DB = 0
SM = 0
Delta_r
Doxycycline
YIL167W
ZShift = 0.05
Z lm Score = −1.3
NG = 0
DB = 0
SM = 0
Delta_AUC
Doxycycline
YPR008W HAA1
ZShift = 4.37
Z lm Score = 2.63
NG = 0
DB = 0
SM = 0
Delta_L
Doxycycline
YPR008W HAA1
ZShift = −3.58
Z lm Score = 4.28
NG = 0
DB = 0
SM = 0
Delta_K
Doxycycline
YPR008W HAA1
ZShift = −3.39
Z lm Score = 1.28
NG = 0
DB = 0
SM = 0
Delta_r
Doxycycline
YPR008W HAA1
ZShift = −4.06
Z lm Score = 8.4
NG = 0
DB = 0
SM = 0
Delta_AUC
Doxycycline

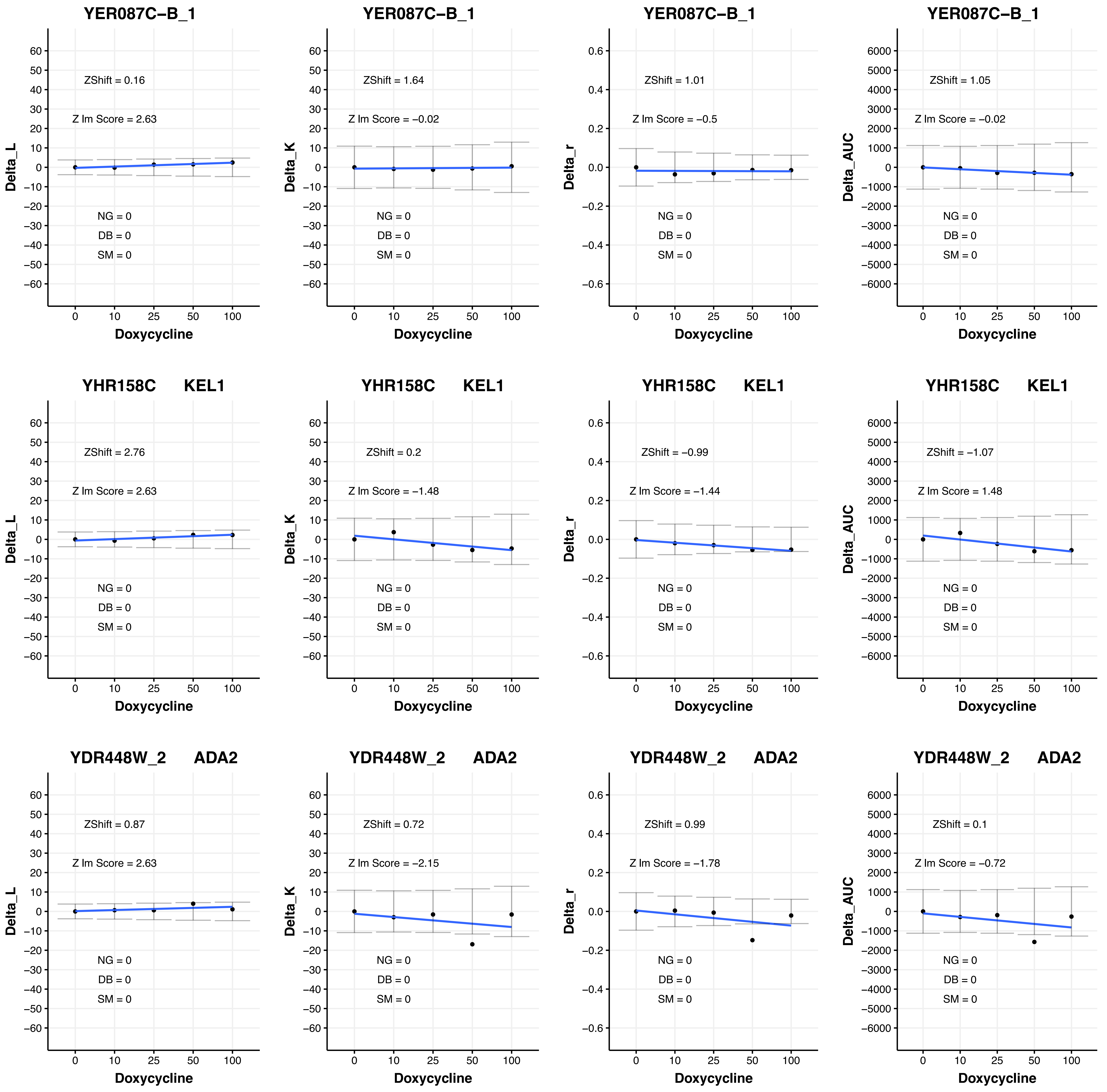

YER087C–B_1
ZShift = 0.16
Z lm Score = 2.63
NG = 0
DB = 0
SM = 0
Delta_L
Doxycycline
YER087C–B_1
ZShift = 1.64
Z lm Score = −0.02
NG = 0
DB = 0
SM = 0
Delta_K
Doxycycline
YER087C–B_1
ZShift = 1.01
Z lm Score = −0.5
NG = 0
DB = 0
SM = 0
Delta_r
Doxycycline
YER087C–B_1
ZShift = 1.05
Z lm Score = −0.02
NG = 0
DB = 0
SM = 0
Delta_AUC
Doxycycline
YHR158C KEL1
ZShift = 2.76
Z lm Score = 2.63
NG = 0
DB = 0
SM = 0
Delta_L
Doxycycline
YHR158C KEL1
ZShift = 0.2
Z lm Score = −1.48
NG = 0
DB = 0
SM = 0
Delta_K
Doxycycline
YHR158C KEL1
ZShift = −0.99
Z lm Score = −1.44
NG = 0
DB = 0
SM = 0
Delta_r
Doxycycline
YHR158C KEL1
ZShift = −1.07
Z lm Score = 1.48
NG = 0
DB = 0
SM = 0
Delta_AUC
Doxycycline
YDR448W_2 ADA2
ZShift = 0.87
Z lm Score = 2.63
NG = 0
DB = 0
SM = 0
Delta_L
Doxycycline
YDR448W_2 ADA2
ZShift = 0.72
Z lm Score = −2.15
NG = 0
DB = 0
SM = 0
Delta_K
Doxycycline
YDR448W_2 ADA2
ZShift = 0.99
Z lm Score = −1.78
NG = 0
DB = 0
SM = 0
Delta_r
Doxycycline
YDR448W_2 ADA2
ZShift = 0.1
Z lm Score = −0.72
NG = 0
DB = 0
SM = 0
Delta_AUC
Doxycycline

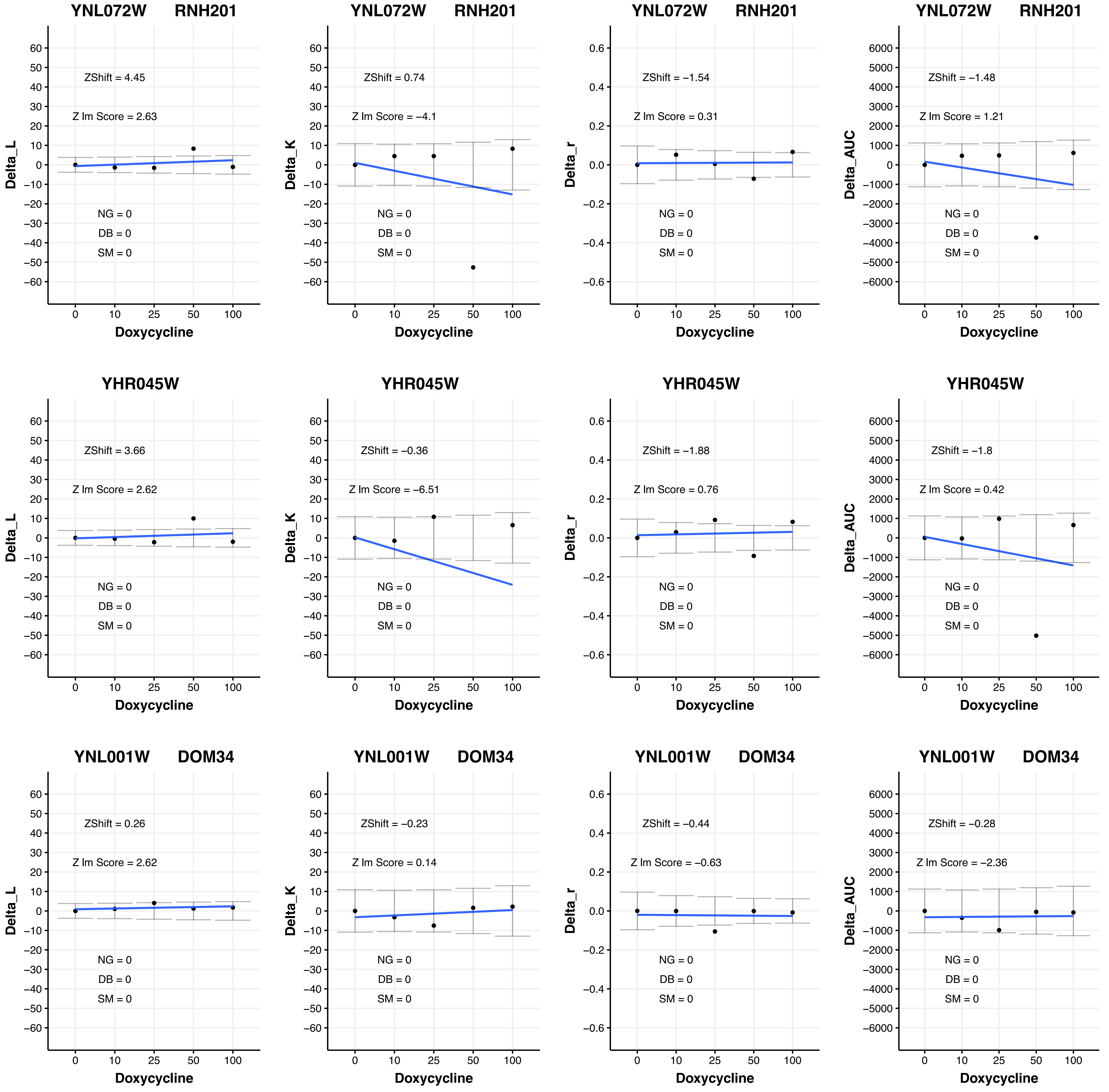

YNL072W RNH201
ZShift = 4.45
Z lm Score = 2.63
NG = 0
DB = 0
SM = 0
Delta_L
Doxycycline
YNL072W RNH201
ZShift = 0.74
Z lm Score = −4.1
Delta_K
YNL072W RNH201
ZShift = −1.54
Z lm Score = 0.31
Delta_r
YNL072W RNH201
ZShift = −1.48
Z lm Score = 1.21
Delta_AUC
YHR045W
ZShift = 3.66
Z lm Score = 2.62
ZShift = −0.36
Z lm Score = −6.51
ZShift = −1.88
Z lm Score = 0.76
ZShift = −1.8
Z lm Score = 0.42
YNL001W DOM34
ZShift = 0.26
Z lm Score = 2.62
ZShift = −0.23
Z lm Score = 0.14
ZShift = −0.44
Z lm Score = −0.63
ZShift = −0.28
Z lm Score = −2.36

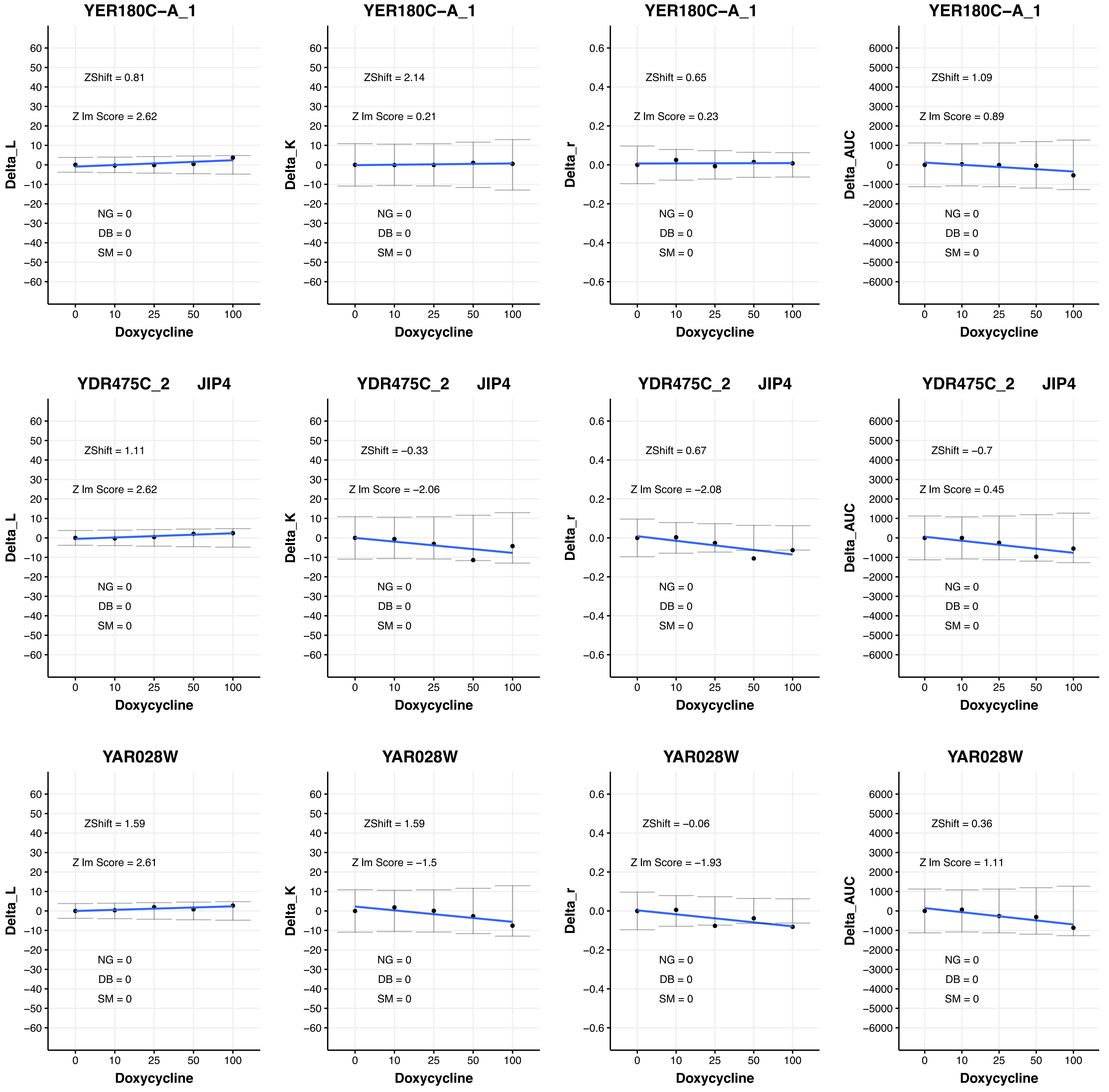

YER180C−A_1
ZShift = 0.81
Z lm Score = 2.62
NG = 0
DB = 0
SM = 0
Delta_L
Doxycycline
YER180C−A_1
ZShift = 2.14
Z lm Score = 0.21
NG = 0
DB = 0
SM = 0
Delta_K
Doxycycline
YER180C−A_1
ZShift = 0.65
Z lm Score = 0.23
NG = 0
DB = 0
SM = 0
Delta_r
Doxycycline
YER180C−A_1
ZShift = 1.09
Z lm Score = 0.89
NG = 0
DB = 0
SM = 0
Delta_AUC
Doxycycline
YDR475C_2 JIP4
ZShift = 1.11
Z lm Score = 2.62
NG = 0
DB = 0
SM = 0
Delta_L
Doxycycline
YDR475C_2 JIP4
ZShift = −0.33
Z lm Score = −2.06
NG = 0
DB = 0
SM = 0
Delta_K
Doxycycline
YDR475C_2 JIP4
ZShift = 0.67
Z lm Score = −2.08
NG = 0
DB = 0
SM = 0
Delta_r
Doxycycline
YDR475C_2 JIP4
ZShift = −0.7
Z lm Score = 0.45
NG = 0
DB = 0
SM = 0
Delta_AUC
Doxycycline
YAR028W
ZShift = 1.59
Z lm Score = 2.61
NG = 0
DB = 0
SM = 0
Delta_L
Doxycycline
YAR028W
ZShift = 1.59
Z lm Score = −1.5
NG = 0
DB = 0
SM = 0
Delta_K
Doxycycline
YAR028W
ZShift = −0.06
Z lm Score = −1.93
NG = 0
DB = 0
SM = 0
Delta_r
Doxycycline
YAR028W
ZShift = 0.36
Z lm Score = 1.11
NG = 0
DB = 0
SM = 0
Delta_AUC
Doxycycline

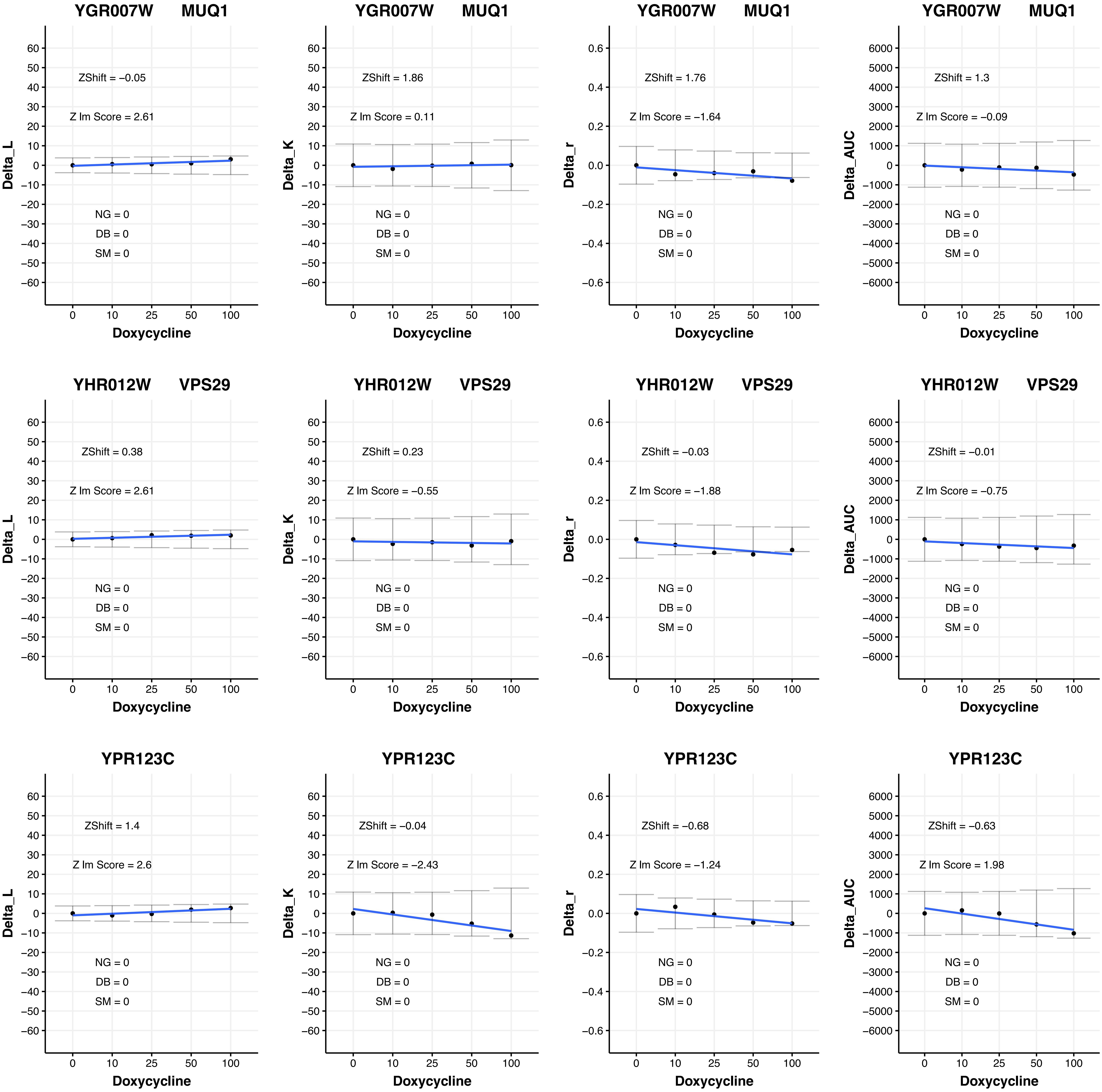

YGR007W MUQ1
ZShift = −0.05
Z lm Score = 2.61
Delta_L
NG = 0
DB = 0
SM = 0
Doxycycline
YGR007W MUQ1
ZShift = 1.86
Z lm Score = 0.11
Delta_K
NG = 0
DB = 0
SM = 0
Doxycycline
YGR007W MUQ1
ZShift = 1.76
Z lm Score = −1.64
Delta_r
NG = 0
DB = 0
SM = 0
Doxycycline
YGR007W MUQ1
ZShift = 1.3
Z lm Score = −0.09
Delta_AUC
NG = 0
DB = 0
SM = 0
Doxycycline
YHR012W VPS29
ZShift = 0.38
Z lm Score = 2.61
Delta_L
NG = 0
DB = 0
SM = 0
Doxycycline
YHR012W VPS29
ZShift = 0.23
Z lm Score = −0.55
Delta_K
NG = 0
DB = 0
SM = 0
Doxycycline
YHR012W VPS29
ZShift = −0.03
Z lm Score = −1.88
Delta_r
NG = 0
DB = 0
SM = 0
Doxycycline
YHR012W VPS29
ZShift = −0.01
Z lm Score = −0.75
Delta_AUC
NG = 0
DB = 0
SM = 0
Doxycycline
YPR123C
ZShift = 1.4
Z lm Score = 2.6
Delta_L
NG = 0
DB = 0
SM = 0
Doxycycline
YPR123C
ZShift = −0.04
Z lm Score = −2.43
Delta_K
NG = 0
DB = 0
SM = 0
Doxycycline
YPR123C
ZShift = −0.68
Z lm Score = −1.24
Delta_r
NG = 0
DB = 0
SM = 0
Doxycycline
YPR123C
ZShift = −0.63
Z lm Score = 1.98
Delta_AUC
NG = 0
DB = 0
SM = 0
Doxycycline

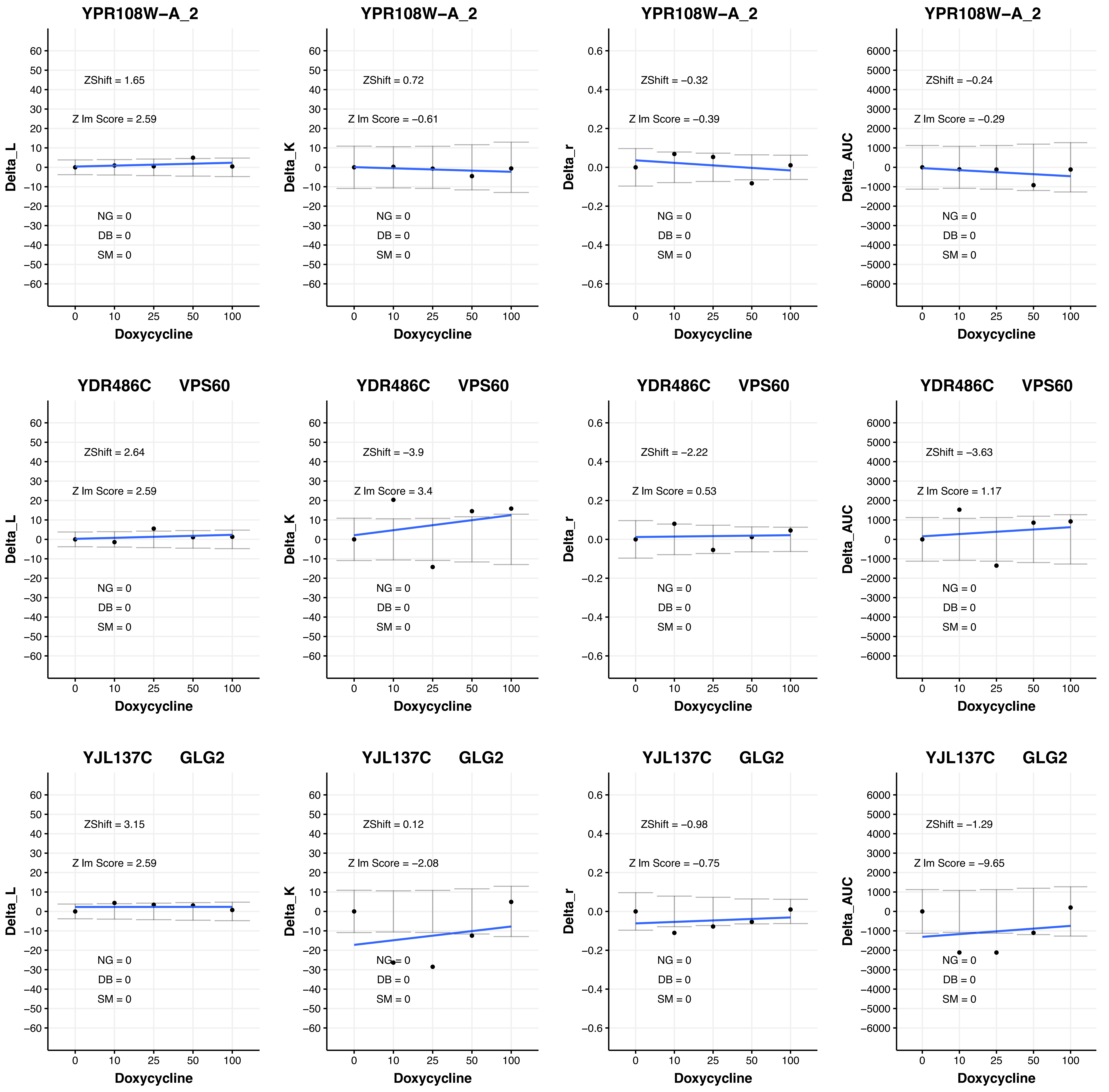
YPR108W−A_2
ZShift = 1.65
Z lm Score = 2.59
NG = 0
DB = 0
SM = 0
Delta_L
Doxycycline
YPR108W−A_2
ZShift = 0.72
Z lm Score = −0.61
NG = 0
DB = 0
SM = 0
Delta_K
Doxycycline
YPR108W−A_2
ZShift = −0.32
Z lm Score = −0.39
NG = 0
DB = 0
SM = 0
Delta_r
Doxycycline
YPR108W−A_2
ZShift = −0.24
Z lm Score = −0.29
NG = 0
DB = 0
SM = 0
Delta_AUC
Doxycycline
YDR486C VPS60
ZShift = 2.64
Z lm Score = 2.59
NG = 0
DB = 0
SM = 0
Delta_L
Doxycycline
YDR486C VPS60
ZShift = −3.9
Z lm Score = 3.4
NG = 0
DB = 0
SM = 0
Delta_K
Doxycycline
YDR486C VPS60
ZShift = −2.22
Z lm Score = 0.53
NG = 0
DB = 0
SM = 0
Delta_r
Doxycycline
YDR486C VPS60
ZShift = −3.63
Z lm Score = 1.17
NG = 0
DB = 0
SM = 0
Delta_AUC
Doxycycline
YJL137C GLG2
ZShift = 3.15
Z lm Score = 2.59
NG = 0
DB = 0
SM = 0
Delta_L
Doxycycline
YJL137C GLG2
ZShift = 0.12
Z lm Score = −2.08
NG = 0
DB = 0
SM = 0
Delta_K
Doxycycline
YJL137C GLG2
ZShift = −0.98
Z lm Score = −0.75
NG = 0
DB = 0
SM = 0
Delta_r
Doxycycline
YJL137C GLG2
ZShift = −1.29
Z lm Score = −9.65
NG = 0
DB = 0
SM = 0
Delta_AUC
Doxycycline

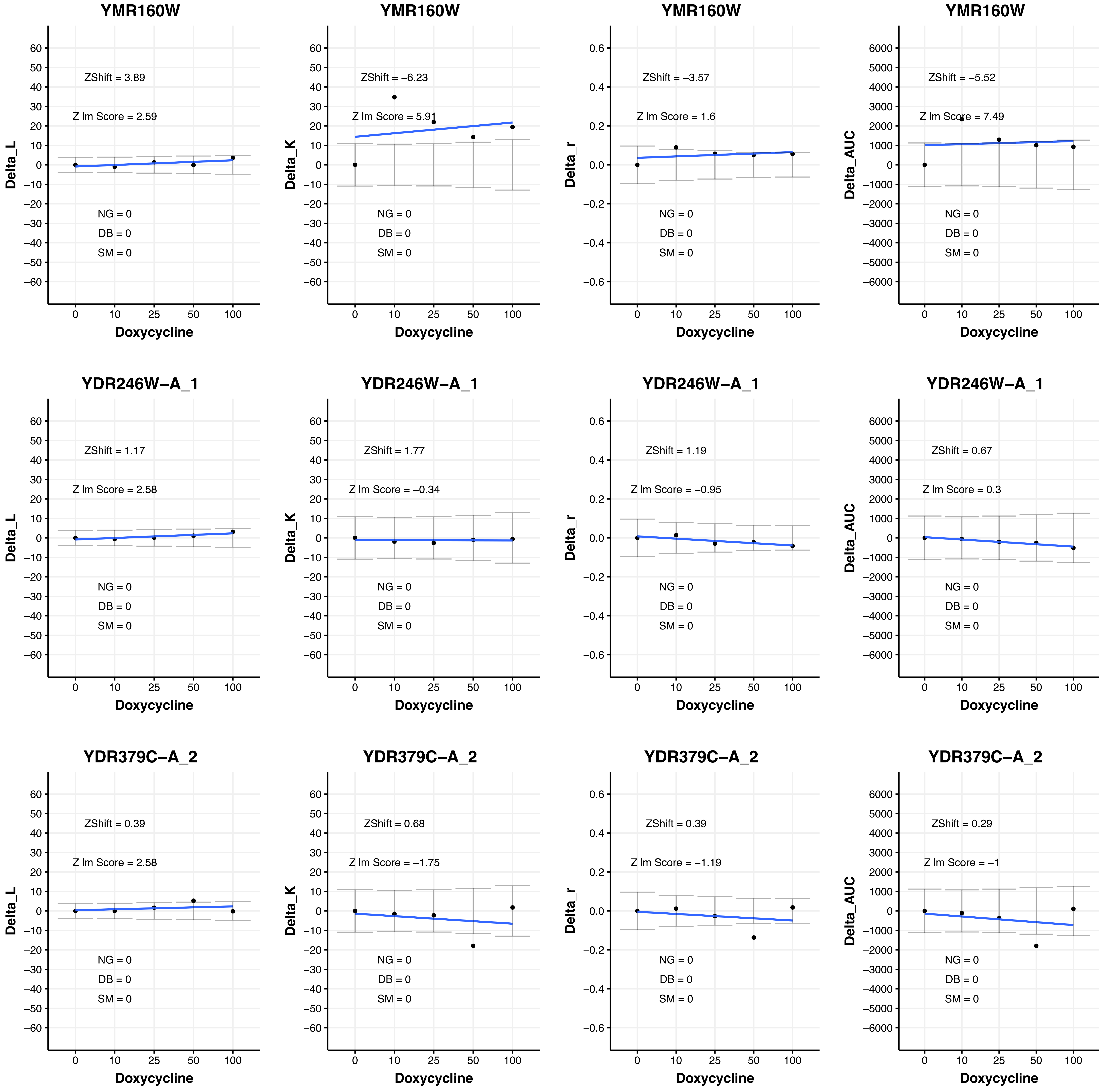
YMR160W
ZShift = 3.89
Z lm Score = 2.59
NG = 0
DB = 0
SM = 0
Delta_L
Doxycycline
YMR160W
ZShift = −6.23
Z lm Score = 5.91
NG = 0
DB = 0
SM = 0
Delta_K
Doxycycline
YMR160W
ZShift = −3.57
Z lm Score = 1.6
NG = 0
DB = 0
SM = 0
Delta_r
Doxycycline
YMR160W
ZShift = −5.52
Z lm Score = 7.49
NG = 0
DB = 0
SM = 0
Delta_AUC
Doxycycline
YDR246W−A_1
ZShift = 1.17
Z lm Score = 2.58
NG = 0
DB = 0
SM = 0
Delta_L
Doxycycline
YDR246W−A_1
ZShift = 1.77
Z lm Score = −0.34
NG = 0
DB = 0
SM = 0
Delta_K
Doxycycline
YDR246W−A_1
ZShift = 1.19
Z lm Score = −0.95
NG = 0
DB = 0
SM = 0
Delta_r
Doxycycline
YDR246W−A_1
ZShift = 0.67
Z lm Score = 0.3
NG = 0
DB = 0
SM = 0
Delta_AUC
Doxycycline
YDR379C−A_2
ZShift = 0.39
Z lm Score = 2.58
NG = 0
DB = 0
SM = 0
Delta_L
Doxycycline
YDR379C−A_2
ZShift = 0.68
Z lm Score = −1.75
NG = 0
DB = 0
SM = 0
Delta_K
Doxycycline
YDR379C−A_2
ZShift = 0.39
Z lm Score = −1.19
NG = 0
DB = 0
SM = 0
Delta_r
Doxycycline
YDR379C−A_2
ZShift = 0.29
Z lm Score = −1
NG = 0
DB = 0
SM = 0
Delta_AUC
Doxycycline

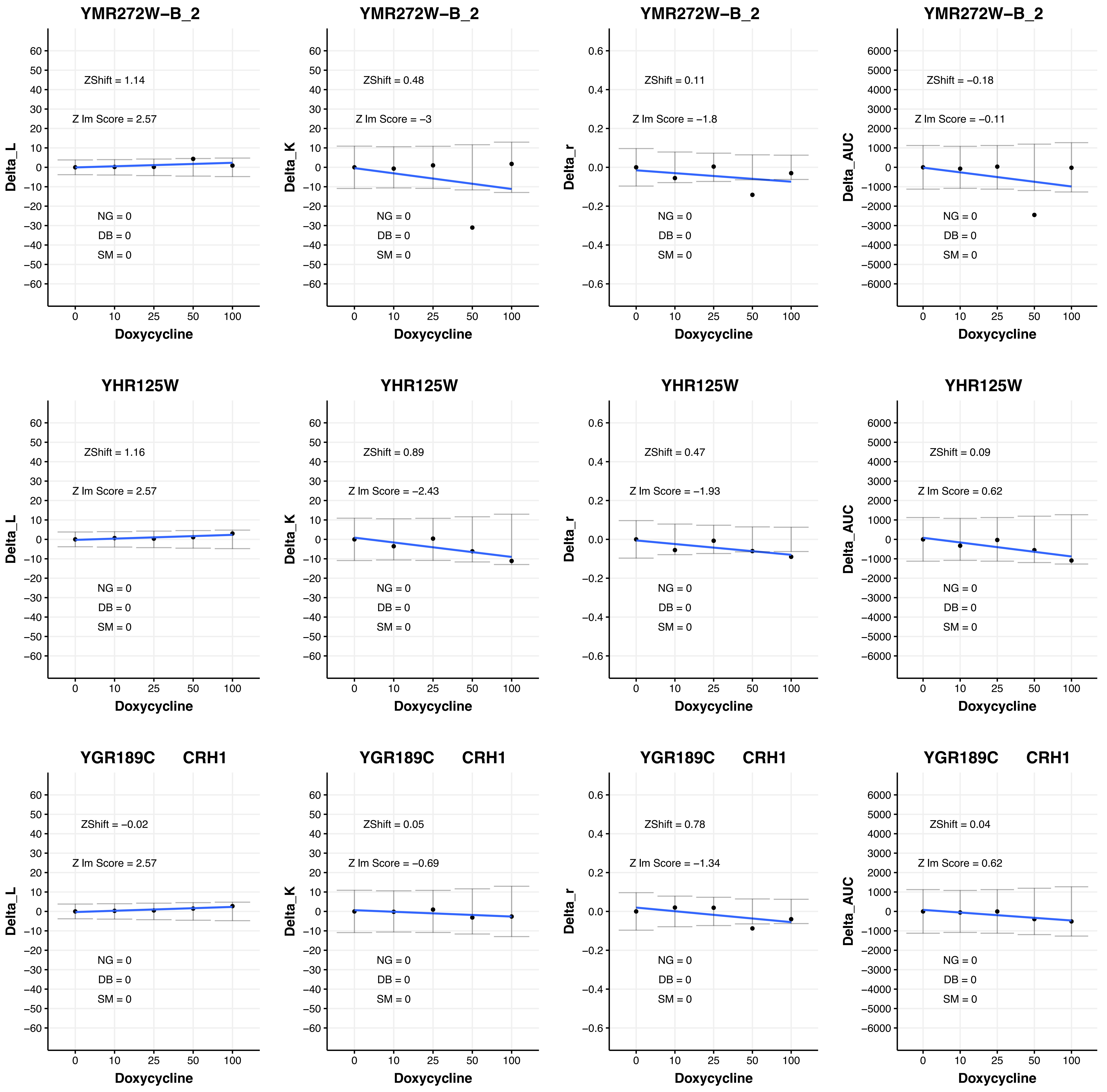
YMR272W−B_2
ZShift = 1.14
Z lm Score = 2.57
NG = 0
DB = 0
SM = 0
Delta_L
Doxycycline
YMR272W−B_2
ZShift = 0.48
Z lm Score = −3
NG = 0
DB = 0
SM = 0
Delta_K
Doxycycline
YMR272W−B_2
ZShift = 0.11
Z lm Score = −1.8
NG = 0
DB = 0
SM = 0
Delta_r
Doxycycline
YMR272W−B_2
ZShift = −0.18
Z lm Score = −0.11
NG = 0
DB = 0
SM = 0
Delta_AUC
Doxycycline
YHR125W
ZShift = 1.16
Z lm Score = 2.57
NG = 0
DB = 0
SM = 0
Delta_L
Doxycycline
YHR125W
ZShift = 0.89
Z lm Score = −2.43
NG = 0
DB = 0
SM = 0
Delta_K
Doxycycline
YHR125W
ZShift = 0.47
Z lm Score = −1.93
NG = 0
DB = 0
SM = 0
Delta_r
Doxycycline
YHR125W
ZShift = 0.09
Z lm Score = 0.62
NG = 0
DB = 0
SM = 0
Delta_AUC
Doxycycline
YGR189C CRH1
ZShift = −0.02
Z lm Score = 2.57
NG = 0
DB = 0
SM = 0
Delta_L
Doxycycline
YGR189C CRH1
ZShift = 0.05
Z lm Score = −0.69
NG = 0
DB = 0
SM = 0
Delta_K
Doxycycline
YGR189C CRH1
ZShift = 0.78
Z lm Score = −1.34
NG = 0
DB = 0
SM = 0
Delta_r
Doxycycline
YGR189C CRH1
ZShift = 0.04
Z lm Score = 0.62
NG = 0
DB = 0
SM = 0
Delta_AUC
Doxycycline

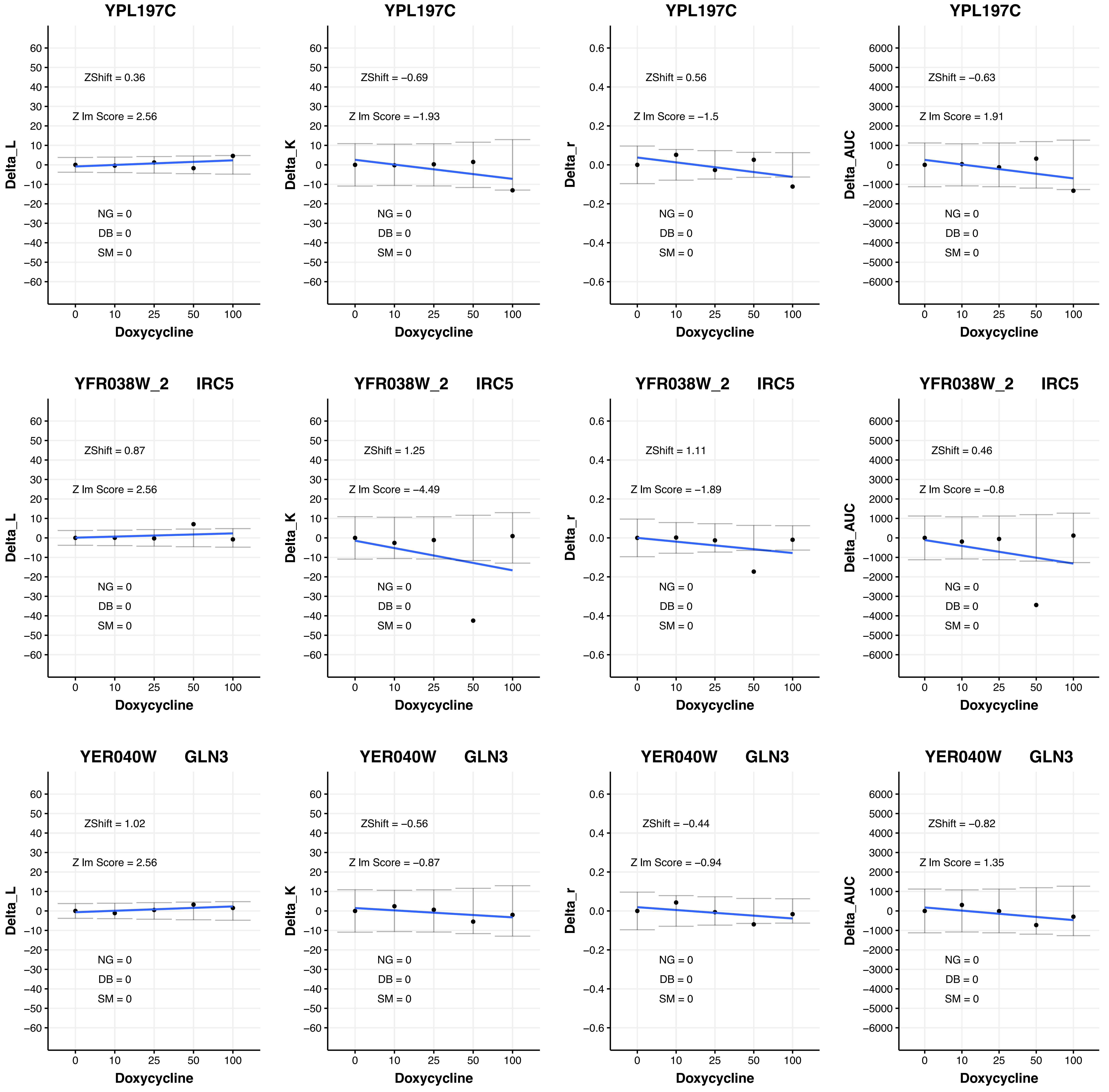

YPL197C
ZShift = 0.36
Z lm Score = 2.56
NG = 0
DB = 0
SM = 0
Delta_L
Doxycycline
YPL197C
ZShift = −0.69
Z lm Score = −1.93
NG = 0
DB = 0
SM = 0
Delta_K
Doxycycline
YPL197C
ZShift = 0.56
Z lm Score = −1.5
NG = 0
DB = 0
SM = 0
Delta_r
Doxycycline
YPL197C
ZShift = −0.63
Z lm Score = 1.91
NG = 0
DB = 0
SM = 0
Delta_AUC
Doxycycline
YFR038W_2 IRC5
ZShift = 0.87
Z lm Score = 2.56
NG = 0
DB = 0
SM = 0
Delta_L
Doxycycline
YFR038W_2 IRC5
ZShift = 1.25
Z lm Score = −4.49
NG = 0
DB = 0
SM = 0
Delta_K
Doxycycline
YFR038W_2 IRC5
ZShift = 1.11
Z lm Score = −1.89
NG = 0
DB = 0
SM = 0
Delta_r
Doxycycline
YFR038W_2 IRC5
ZShift = 0.46
Z lm Score = −0.8
NG = 0
DB = 0
SM = 0
Delta_AUC
Doxycycline
YER040W GLN3
ZShift = 1.02
Z lm Score = 2.56
NG = 0
DB = 0
SM = 0
Delta_L
Doxycycline
YER040W GLN3
ZShift = −0.56
Z lm Score = −0.87
NG = 0
DB = 0
SM = 0
Delta_K
Doxycycline
YER040W GLN3
ZShift = −0.44
Z lm Score = −0.94
NG = 0
DB = 0
SM = 0
Delta_r
Doxycycline
YER040W GLN3
ZShift = −0.82
Z lm Score = 1.35
NG = 0
DB = 0
SM = 0
Delta_AUC
Doxycycline

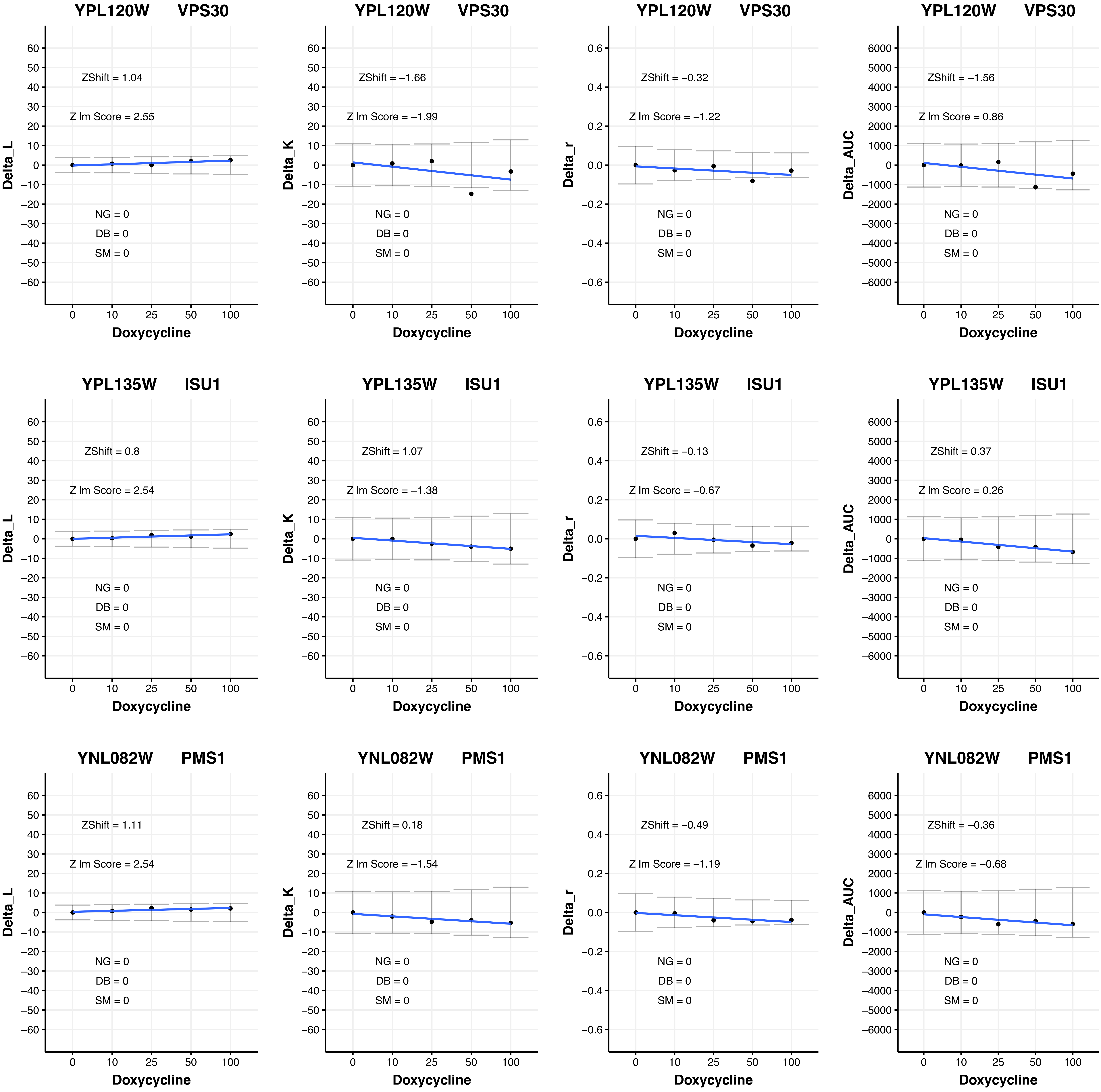

YPL120W VPS30
ZShift = 1.04
Z lm Score = 2.55
NG = 0
DB = 0
SM = 0
Delta_L
Doxycycline
YPL120W VPS30
ZShift = −1.66
Z lm Score = −1.99
NG = 0
DB = 0
SM = 0
Delta_K
Doxycycline
YPL120W VPS30
ZShift = −0.32
Z lm Score = −1.22
NG = 0
DB = 0
SM = 0
Delta_r
Doxycycline
YPL120W VPS30
ZShift = −1.56
Z lm Score = 0.86
NG = 0
DB = 0
SM = 0
Delta_AUC
Doxycycline
YPL135W ISU1
ZShift = 0.8
Z lm Score = 2.54
NG = 0
DB = 0
SM = 0
Delta_L
Doxycycline
YPL135W ISU1
ZShift = 1.07
Z lm Score = −1.38
NG = 0
DB = 0
SM = 0
Delta_K
Doxycycline
YPL135W ISU1
ZShift = −0.13
Z lm Score = −0.67
NG = 0
DB = 0
SM = 0
Delta_r
Doxycycline
YPL135W ISU1
ZShift = 0.37
Z lm Score = 0.26
NG = 0
DB = 0
SM = 0
Delta_AUC
Doxycycline
YNL082W PMS1
ZShift = 1.11
Z lm Score = 2.54
NG = 0
DB = 0
SM = 0
Delta_L
Doxycycline
YNL082W PMS1
ZShift = 0.18
Z lm Score = −1.54
NG = 0
DB = 0
SM = 0
Delta_K
Doxycycline
YNL082W PMS1
ZShift = −0.49
Z lm Score = −1.19
NG = 0
DB = 0
SM = 0
Delta_r
Doxycycline
YNL082W PMS1
ZShift = −0.36
Z lm Score = −0.68
NG = 0
DB = 0
SM = 0
Delta_AUC
Doxycycline

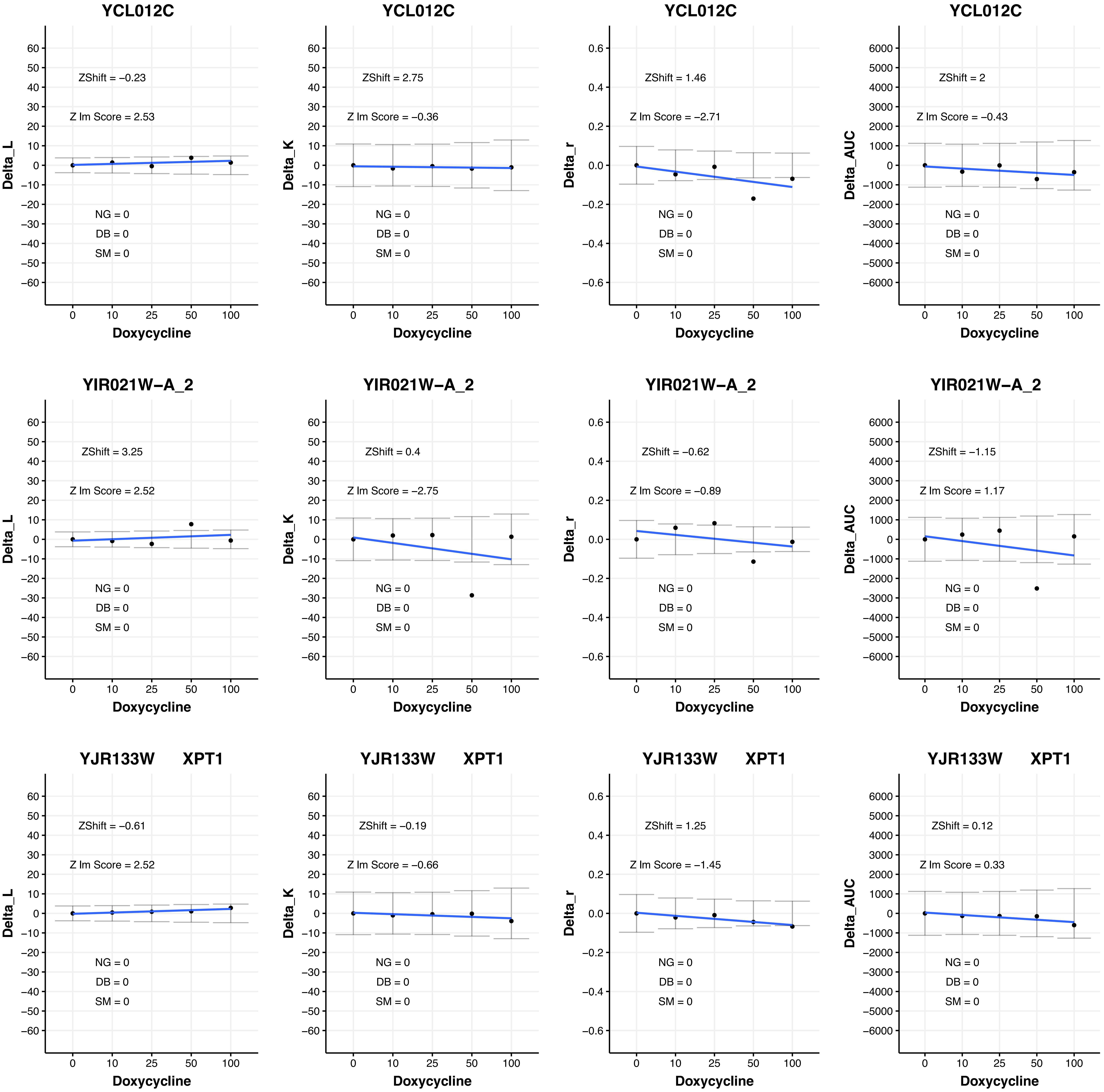
YCL012C
ZShift = −0.23
Z lm Score = 2.53
NG = 0
DB = 0
SM = 0
Delta_L
Doxycycline
YCL012C
ZShift = 2.75
Z lm Score = −0.36
NG = 0
DB = 0
SM = 0
Delta_K
Doxycycline
YCL012C
ZShift = 1.46
Z lm Score = −2.71
NG = 0
DB = 0
SM = 0
Delta_r
Doxycycline
YCL012C
ZShift = 2
Z lm Score = −0.43
NG = 0
DB = 0
SM = 0
Delta_AUC
Doxycycline
YIR021W−A_2
ZShift = 3.25
Z lm Score = 2.52
NG = 0
DB = 0
SM = 0
Delta_L
Doxycycline
YIR021W−A_2
ZShift = 0.4
Z lm Score = −2.75
NG = 0
DB = 0
SM = 0
Delta_K
Doxycycline
YIR021W−A_2
ZShift = −0.62
Z lm Score = −0.89
NG = 0
DB = 0
SM = 0
Delta_r
Doxycycline
YIR021W−A_2
ZShift = −1.15
Z lm Score = 1.17
NG = 0
DB = 0
SM = 0
Delta_AUC
Doxycycline
YJR133W XPT1
ZShift = −0.61
Z lm Score = 2.52
NG = 0
DB = 0
SM = 0
Delta_L
Doxycycline
YJR133W XPT1
ZShift = −0.19
Z lm Score = −0.66
NG = 0
DB = 0
SM = 0
Delta_K
Doxycycline
YJR133W XPT1
ZShift = 1.25
Z lm Score = −1.45
NG = 0
DB = 0
SM = 0
Delta_r
Doxycycline
YJR133W XPT1
ZShift = 0.12
Z lm Score = 0.33
NG = 0
DB = 0
SM = 0
Delta_AUC
Doxycycline

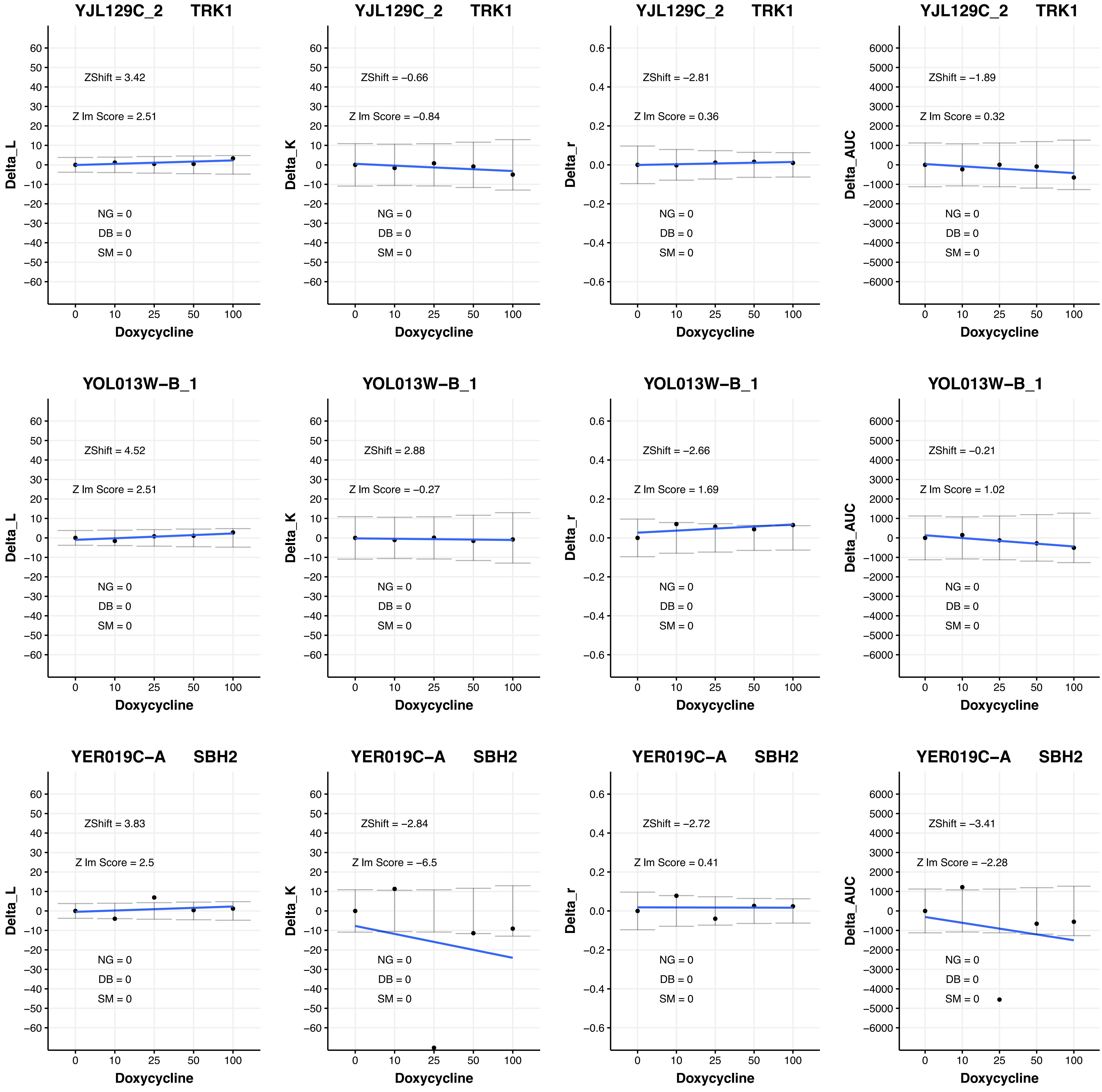
YJL129C_2 TRK1
ZShift = 3.42
Z lm Score = 2.51
NG = 0
DB = 0
SM = 0
Delta_L
Doxycycline
YJL129C_2 TRK1
ZShift = −0.66
Z lm Score = −0.84
NG = 0
DB = 0
SM = 0
Delta_K
Doxycycline
YJL129C_2 TRK1
ZShift = −2.81
Z lm Score = 0.36
NG = 0
DB = 0
SM = 0
Delta_r
Doxycycline
YJL129C_2 TRK1
ZShift = −1.89
Z lm Score = 0.32
NG = 0
DB = 0
SM = 0
Delta_AUC
Doxycycline
YOL013W−B_1
ZShift = 4.52
Z lm Score = 2.51
NG = 0
DB = 0
SM = 0
Delta_L
Doxycycline
YOL013W−B_1
ZShift = 2.88
Z lm Score = −0.27
NG = 0
DB = 0
SM = 0
Delta_K
Doxycycline
YOL013W−B_1
ZShift = −2.66
Z lm Score = 1.69
NG = 0
DB = 0
SM = 0
Delta_r
Doxycycline
YOL013W−B_1
ZShift = −0.21
Z lm Score = 1.02
NG = 0
DB = 0
SM = 0
Delta_AUC
Doxycycline
YER019C−A SBH2
ZShift = 3.83
Z lm Score = 2.5
NG = 0
DB = 0
SM = 0
Delta_L
Doxycycline
YER019C−A SBH2
ZShift = −2.84
Z lm Score = −6.5
NG = 0
DB = 0
SM = 0
Delta_K
Doxycycline
YER019C−A SBH2
ZShift = −2.72
Z lm Score = 0.41
NG = 0
DB = 0
SM = 0
Delta_r
Doxycycline
YER019C−A SBH2
ZShift = −3.41
Z lm Score = −2.28
NG = 0
DB = 0
SM = 0
Delta_AUC
Doxycycline

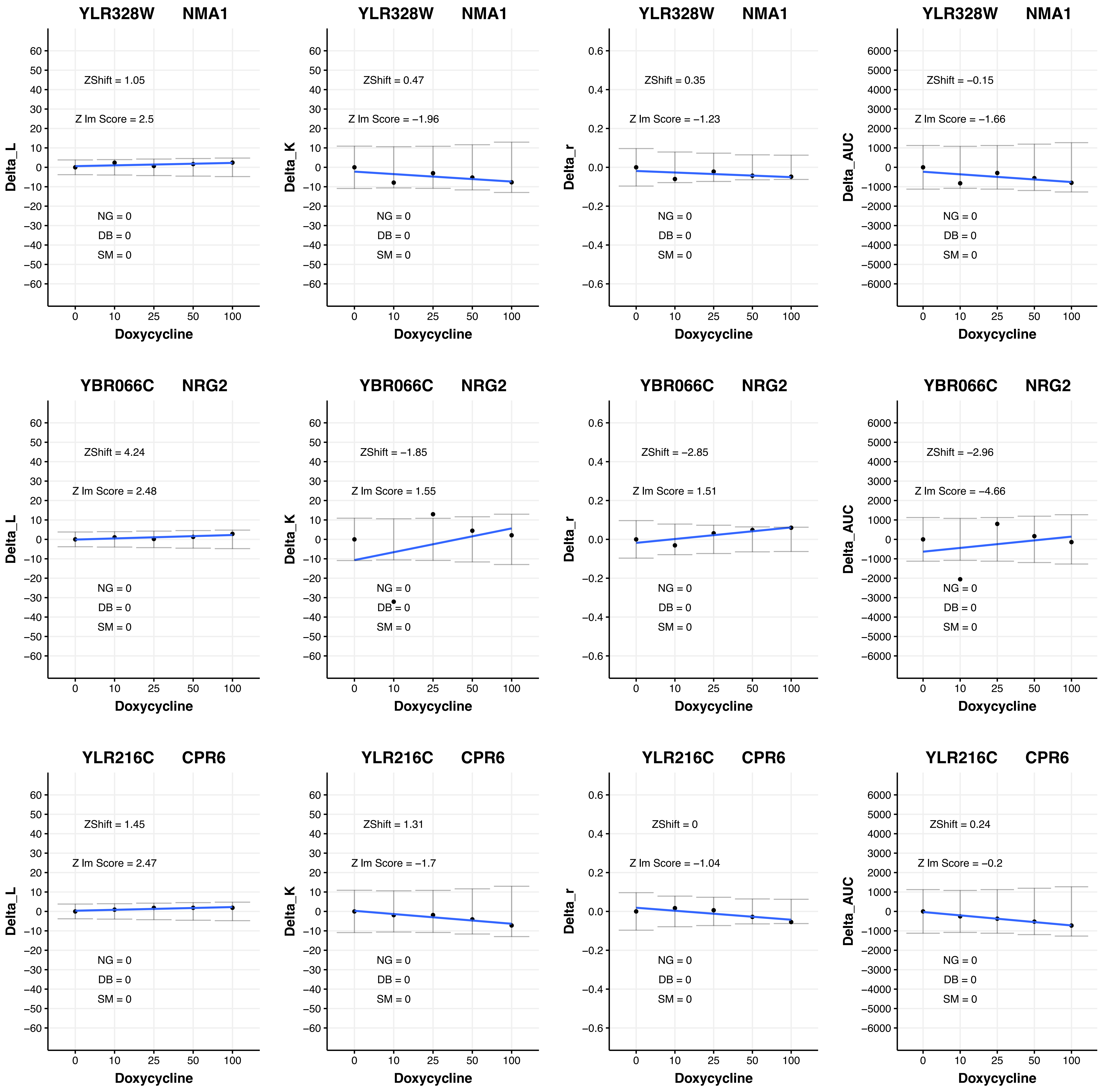

YLR328W NMA1
ZShift = 1.05
Z lm Score = 2.5
NG = 0
DB = 0
SM = 0
Delta_L
Doxycycline
YLR328W NMA1
ZShift = 0.47
Z lm Score = −1.96
NG = 0
DB = 0
SM = 0
Delta_K
Doxycycline
YLR328W NMA1
ZShift = 0.35
Z lm Score = −1.23
NG = 0
DB = 0
SM = 0
Delta_r
Doxycycline
YLR328W NMA1
ZShift = −0.15
Z lm Score = −1.66
NG = 0
DB = 0
SM = 0
Delta_AUC
Doxycycline
YBR066C NRG2
ZShift = 4.24
Z lm Score = 2.48
NG = 0
DB = 0
SM = 0
Delta_L
Doxycycline
YBR066C NRG2
ZShift = −1.85
Z lm Score = 1.55
NG = 0
DB = 0
SM = 0
Delta_K
Doxycycline
YBR066C NRG2
ZShift = −2.85
Z lm Score = 1.51
NG = 0
DB = 0
SM = 0
Delta_r
Doxycycline
YBR066C NRG2
ZShift = −2.96
Z lm Score = −4.66
NG = 0
DB = 0
SM = 0
Delta_AUC
Doxycycline
YLR216C CPR6
ZShift = 1.45
Z lm Score = 2.47
NG = 0
DB = 0
SM = 0
Delta_L
Doxycycline
YLR216C CPR6
ZShift = 1.31
Z lm Score = −1.7
NG = 0
DB = 0
SM = 0
Delta_K
Doxycycline
YLR216C CPR6
ZShift = 0
Z lm Score = −1.04
NG = 0
DB = 0
SM = 0
Delta_r
Doxycycline
YLR216C CPR6
ZShift = 0.24
Z lm Score = −0.2
NG = 0
DB = 0
SM = 0
Delta_AUC
Doxycycline

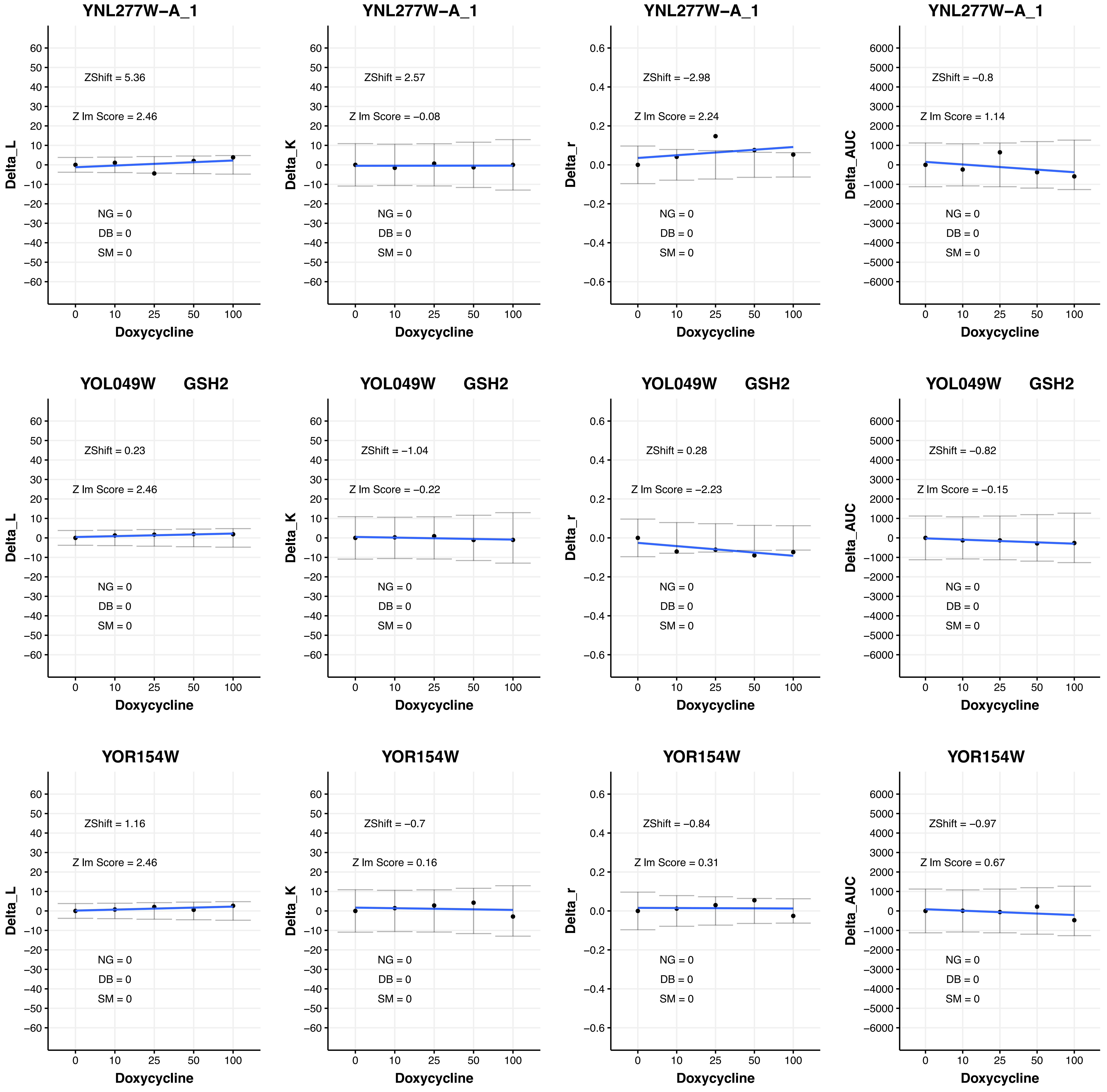
YNL277W−A_1
ZShift = 5.36
Z lm Score = 2.46
NG = 0
DB = 0
SM = 0
Delta_L
Doxycycline
YNL277W−A_1
ZShift = 2.57
Z lm Score = −0.08
NG = 0
DB = 0
SM = 0
Delta_K
Doxycycline
YNL277W−A_1
ZShift = −2.98
Z lm Score = 2.24
NG = 0
DB = 0
SM = 0
Delta_r
Doxycycline
YNL277W−A_1
ZShift = −0.8
Z lm Score = 1.14
NG = 0
DB = 0
SM = 0
Delta_AUC
Doxycycline
YOL049W GSH2
ZShift = 0.23
Z lm Score = 2.46
NG = 0
DB = 0
SM = 0
Delta_L
Doxycycline
YOL049W GSH2
ZShift = −1.04
Z lm Score = −0.22
NG = 0
DB = 0
SM = 0
Delta_K
Doxycycline
YOL049W GSH2
ZShift = 0.28
Z lm Score = −2.23
NG = 0
DB = 0
SM = 0
Delta_r
Doxycycline
YOL049W GSH2
ZShift = −0.82
Z lm Score = −0.15
NG = 0
DB = 0
SM = 0
Delta_AUC
Doxycycline
YOR154W
ZShift = 1.16
Z lm Score = 2.46
NG = 0
DB = 0
SM = 0
Delta_L
Doxycycline
YOR154W
ZShift = −0.7
Z lm Score = 0.16
NG = 0
DB = 0
SM = 0
Delta_K
Doxycycline
YOR154W
ZShift = −0.84
Z lm Score = 0.31
NG = 0
DB = 0
SM = 0
Delta_r
Doxycycline
YOR154W
ZShift = −0.97
Z lm Score = 0.67
NG = 0
DB = 0
SM = 0
Delta_AUC
Doxycycline

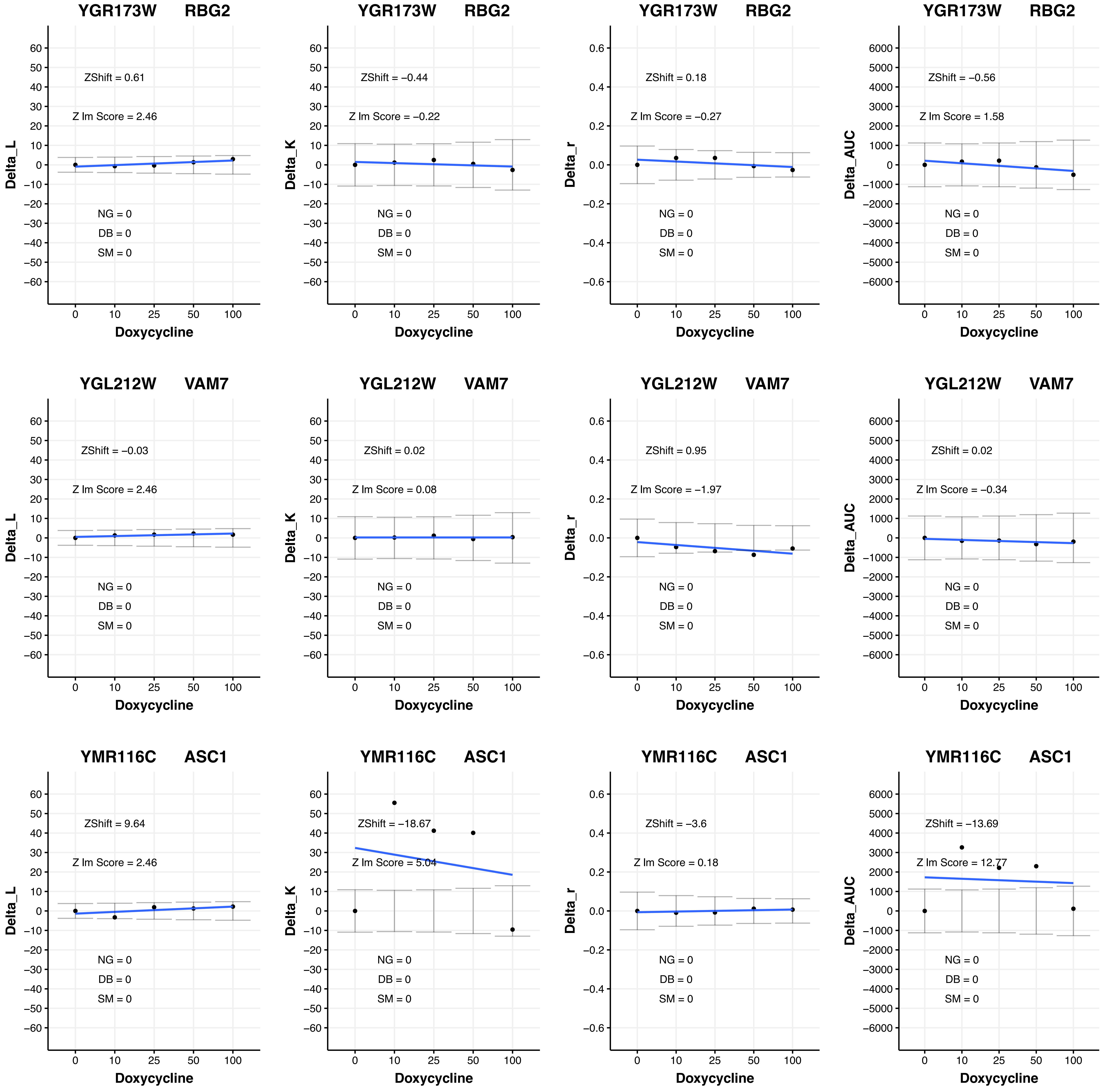

YGR173W RBG2
ZShift = 0.61
Z lm Score = 2.46
NG = 0
DB = 0
SM = 0
Delta_L
Doxycycline
YGR173W RBG2
ZShift = −0.44
Z lm Score = −0.22
NG = 0
DB = 0
SM = 0
Delta_K
Doxycycline
YGR173W RBG2
ZShift = 0.18
Z lm Score = −0.27
NG = 0
DB = 0
SM = 0
Delta_r
Doxycycline
YGR173W RBG2
ZShift = −0.56
Z lm Score = 1.58
NG = 0
DB = 0
SM = 0
Delta_AUC
Doxycycline
YGL212W VAM7
ZShift = −0.03
Z lm Score = 2.46
NG = 0
DB = 0
SM = 0
Delta_L
Doxycycline
YGL212W VAM7
ZShift = 0.02
Z lm Score = 0.08
NG = 0
DB = 0
SM = 0
Delta_K
Doxycycline
YGL212W VAM7
ZShift = 0.95
Z lm Score = −1.97
NG = 0
DB = 0
SM = 0
Delta_r
Doxycycline
YGL212W VAM7
ZShift = 0.02
Z lm Score = −0.34
NG = 0
DB = 0
SM = 0
Delta_AUC
Doxycycline
YMR116C ASC1
ZShift = 9.64
Z lm Score = 2.46
NG = 0
DB = 0
SM = 0
Delta_L
Doxycycline
YMR116C ASC1
ZShift = −18.67
Z lm Score = 5.04
NG = 0
DB = 0
SM = 0
Delta_K
Doxycycline
YMR116C ASC1
ZShift = −3.6
Z lm Score = 0.18
NG = 0
DB = 0
SM = 0
Delta_r
Doxycycline
YMR116C ASC1
ZShift = −13.69
Z lm Score = 12.77
NG = 0
DB = 0
SM = 0
Delta_AUC
Doxycycline

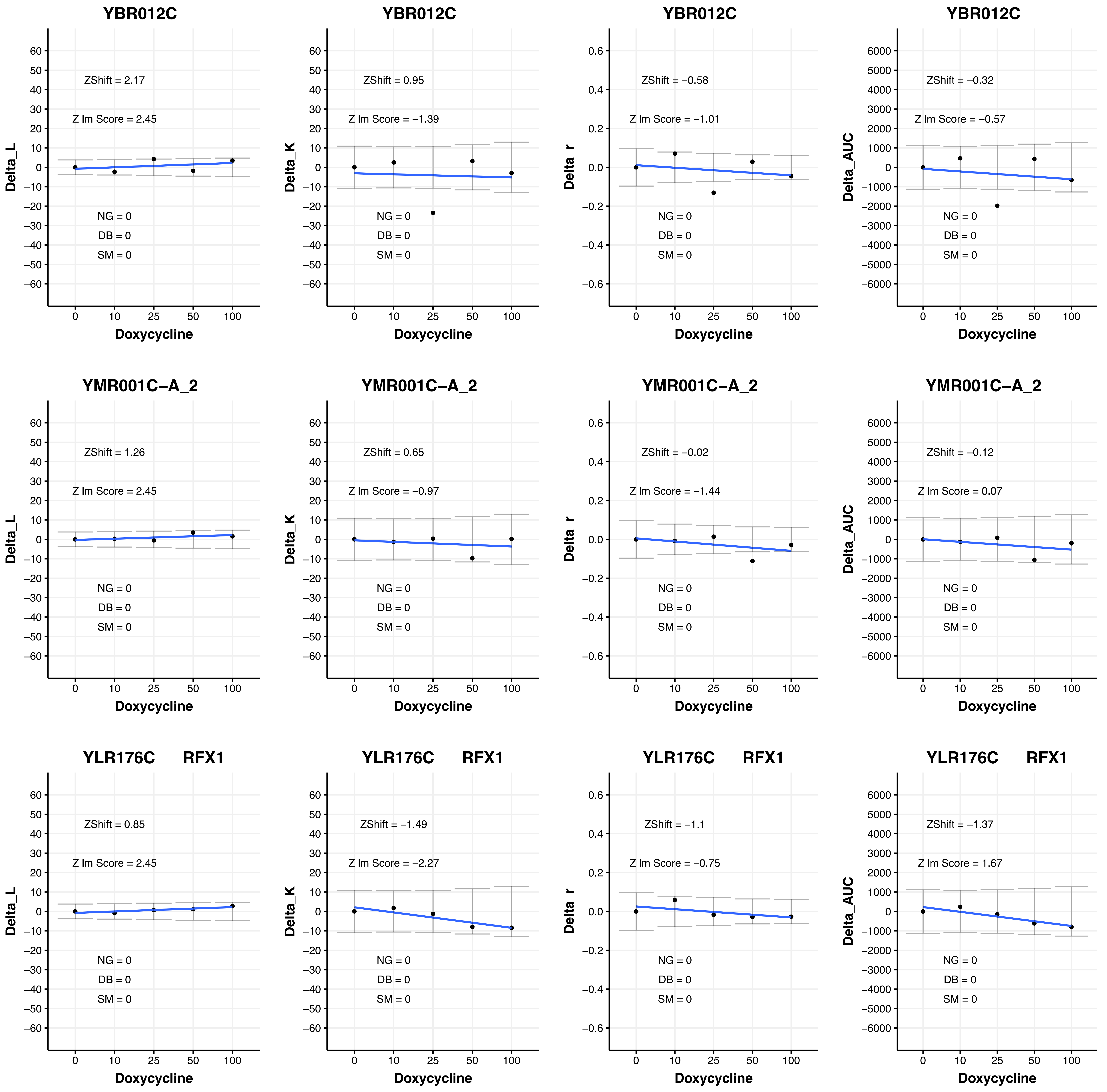
YBR012C
ZShift = 2.17
Z lm Score = 2.45
NG = 0
DB = 0
SM = 0
Delta_L
Doxycycline
YBR012C
ZShift = 0.95
Z lm Score = −1.39
NG = 0
DB = 0
SM = 0
Delta_K
Doxycycline
YBR012C
ZShift = −0.58
Z lm Score = −1.01
NG = 0
DB = 0
SM = 0
Delta_r
Doxycycline
YBR012C
ZShift = −0.32
Z lm Score = −0.57
NG = 0
DB = 0
SM = 0
Delta_AUC
Doxycycline
YMR001C−A_2
ZShift = 1.26
Z lm Score = 2.45
NG = 0
DB = 0
SM = 0
Delta_L
Doxycycline
YMR001C−A_2
ZShift = 0.65
Z lm Score = −0.97
NG = 0
DB = 0
SM = 0
Delta_K
Doxycycline
YMR001C−A_2
ZShift = −0.02
Z lm Score = −1.44
NG = 0
DB = 0
SM = 0
Delta_r
Doxycycline
YMR001C−A_2
ZShift = −0.12
Z lm Score = 0.07
NG = 0
DB = 0
SM = 0
Delta_AUC
Doxycycline
YLR176C RFX1
ZShift = 0.85
Z lm Score = 2.45
NG = 0
DB = 0
SM = 0
Delta_L
Doxycycline
YLR176C RFX1
ZShift = −1.49
Z lm Score = −2.27
NG = 0
DB = 0
SM = 0
Delta_K
Doxycycline
YLR176C RFX1
ZShift = −1.1
Z lm Score = −0.75
NG = 0
DB = 0
SM = 0
Delta_r
Doxycycline
YLR176C RFX1
ZShift = −1.37
Z lm Score = 1.67
NG = 0
DB = 0
SM = 0
Delta_AUC
Doxycycline

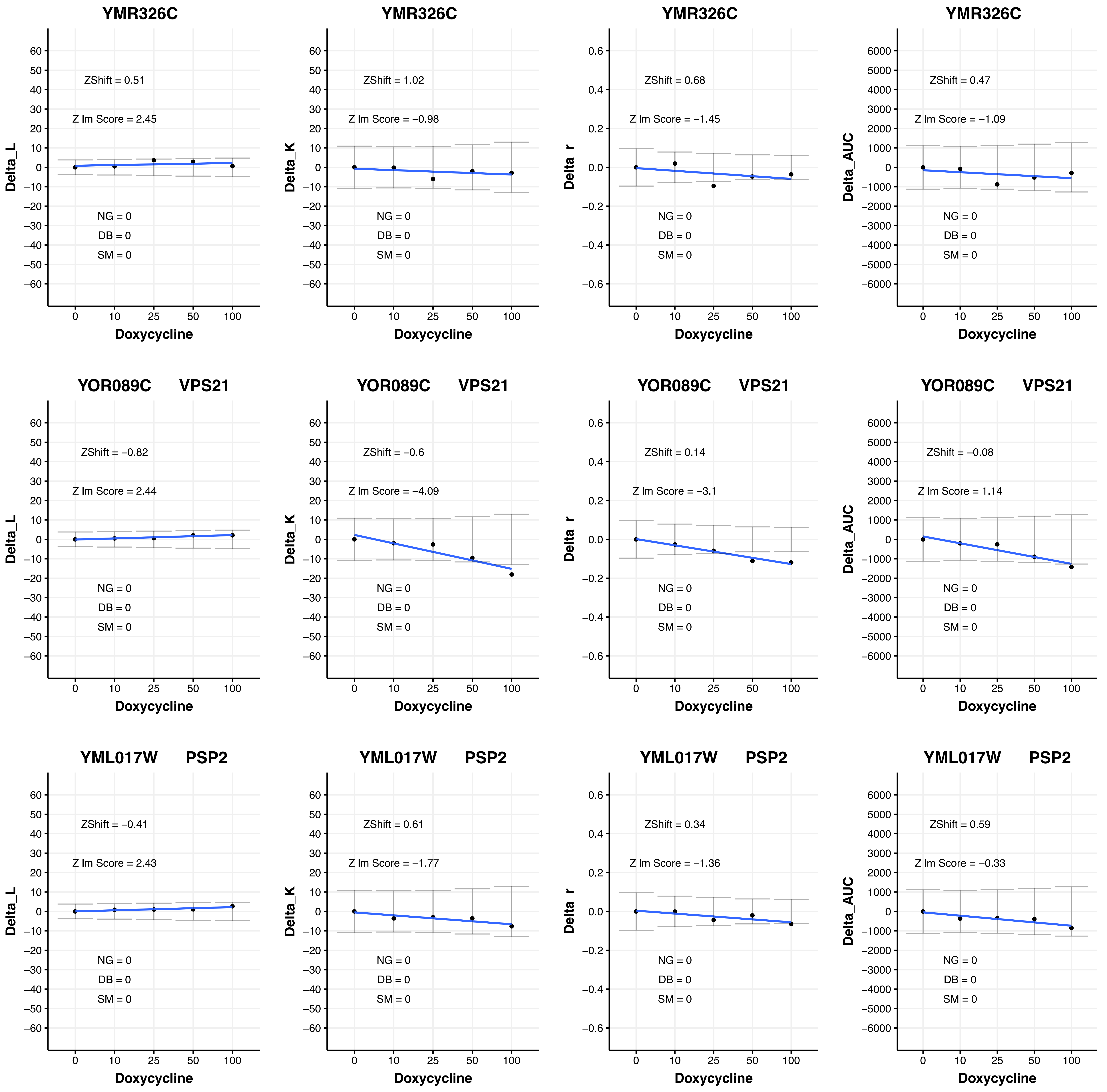
YMR326C
ZShift = 0.51
Z lm Score = 2.45
NG = 0
DB = 0
SM = 0
Delta_L
Doxycycline
YMR326C
ZShift = 1.02
Z lm Score = −0.98
NG = 0
DB = 0
SM = 0
Delta_K
Doxycycline
YMR326C
ZShift = 0.68
Z lm Score = −1.45
NG = 0
DB = 0
SM = 0
Delta_r
Doxycycline
YMR326C
ZShift = 0.47
Z lm Score = −1.09
NG = 0
DB = 0
SM = 0
Delta_AUC
Doxycycline
YOR089C VPS21
ZShift = −0.82
Z lm Score = 2.44
NG = 0
DB = 0
SM = 0
Delta_L
Doxycycline
YOR089C VPS21
ZShift = −0.6
Z lm Score = −4.09
NG = 0
DB = 0
SM = 0
Delta_K
Doxycycline
YOR089C VPS21
ZShift = 0.14
Z lm Score = −3.1
NG = 0
DB = 0
SM = 0
Delta_r
Doxycycline
YOR089C VPS21
ZShift = −0.08
Z lm Score = 1.14
NG = 0
DB = 0
SM = 0
Delta_AUC
Doxycycline
YML017W PSP2
ZShift = −0.41
Z lm Score = 2.43
NG = 0
DB = 0
SM = 0
Delta_L
Doxycycline
YML017W PSP2
ZShift = 0.61
Z lm Score = −1.77
NG = 0
DB = 0
SM = 0
Delta_K
Doxycycline
YML017W PSP2
ZShift = 0.34
Z lm Score = −1.36
NG = 0
DB = 0
SM = 0
Delta_r
Doxycycline
YML017W PSP2
ZShift = 0.59
Z lm Score = −0.33
NG = 0
DB = 0
SM = 0
Delta_AUC
Doxycycline

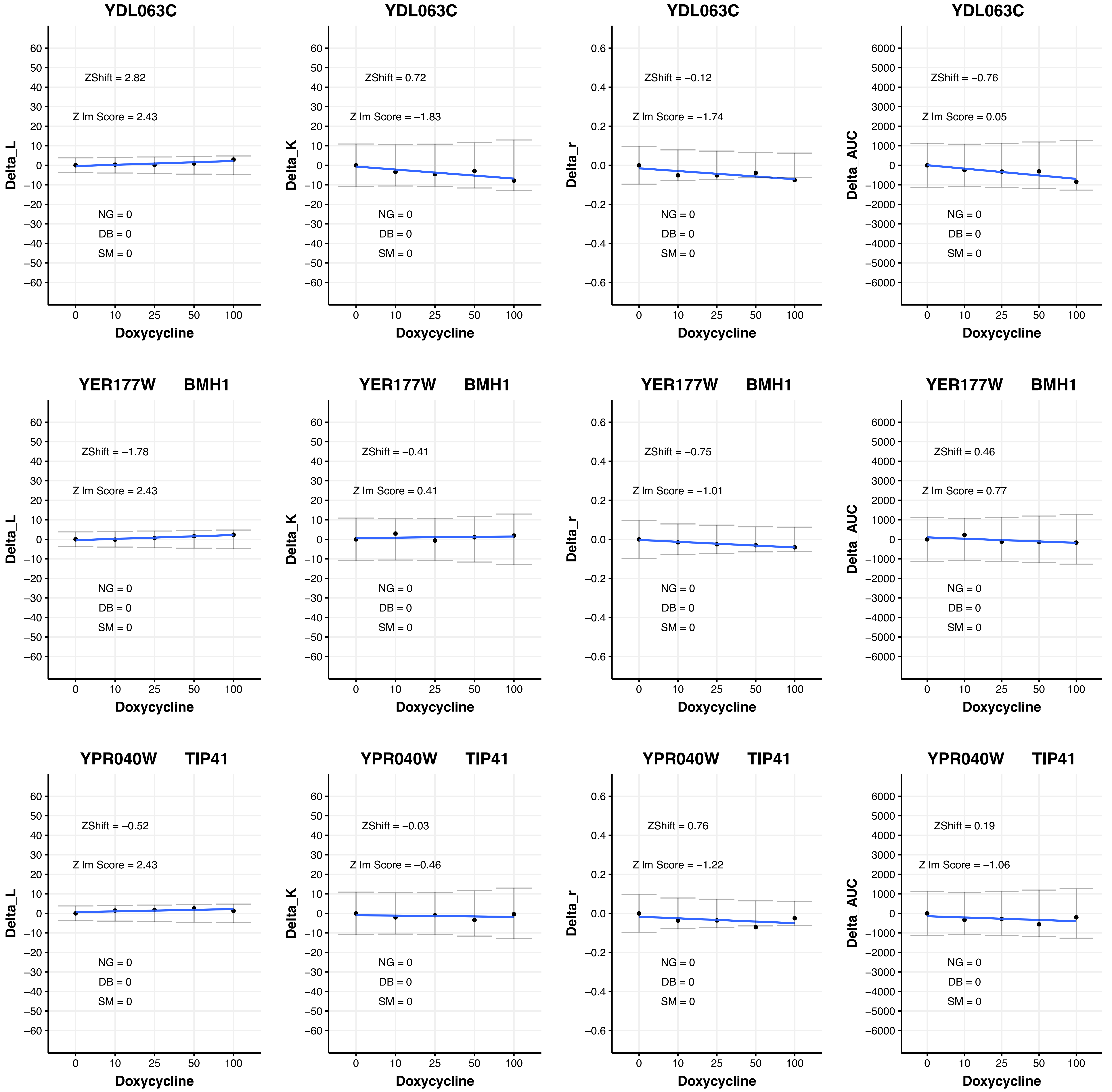
YDL063C
ZShift = 2.82
Z lm Score = 2.43
NG = 0
DB = 0
SM = 0
Delta_L
Doxycycline
YDL063C
ZShift = 0.72
Z lm Score = −1.83
NG = 0
DB = 0
SM = 0
Delta_K
Doxycycline
YDL063C
ZShift = −0.12
Z lm Score = −1.74
NG = 0
DB = 0
SM = 0
Delta_r
Doxycycline
YDL063C
ZShift = −0.76
Z lm Score = 0.05
NG = 0
DB = 0
SM = 0
Delta_AUC
Doxycycline
YER177W BMH1
ZShift = −1.78
Z lm Score = 2.43
NG = 0
DB = 0
SM = 0
Delta_L
Doxycycline
YER177W BMH1
ZShift = −0.41
Z lm Score = 0.41
NG = 0
DB = 0
SM = 0
Delta_K
Doxycycline
YER177W BMH1
ZShift = −0.75
Z lm Score = −1.01
NG = 0
DB = 0
SM = 0
Delta_r
Doxycycline
YER177W BMH1
ZShift = 0.46
Z lm Score = 0.77
NG = 0
DB = 0
SM = 0
Delta_AUC
Doxycycline
YPR040W TIP41
ZShift = −0.52
Z lm Score = 2.43
NG = 0
DB = 0
SM = 0
Delta_L
Doxycycline
YPR040W TIP41
ZShift = −0.03
Z lm Score = −0.46
NG = 0
DB = 0
SM = 0
Delta_K
Doxycycline
YPR040W TIP41
ZShift = 0.76
Z lm Score = −1.22
NG = 0
DB = 0
SM = 0
Delta_r
Doxycycline
YPR040W TIP41
ZShift = 0.19
Z lm Score = −1.06
NG = 0
DB = 0
SM = 0
Delta_AUC
Doxycycline

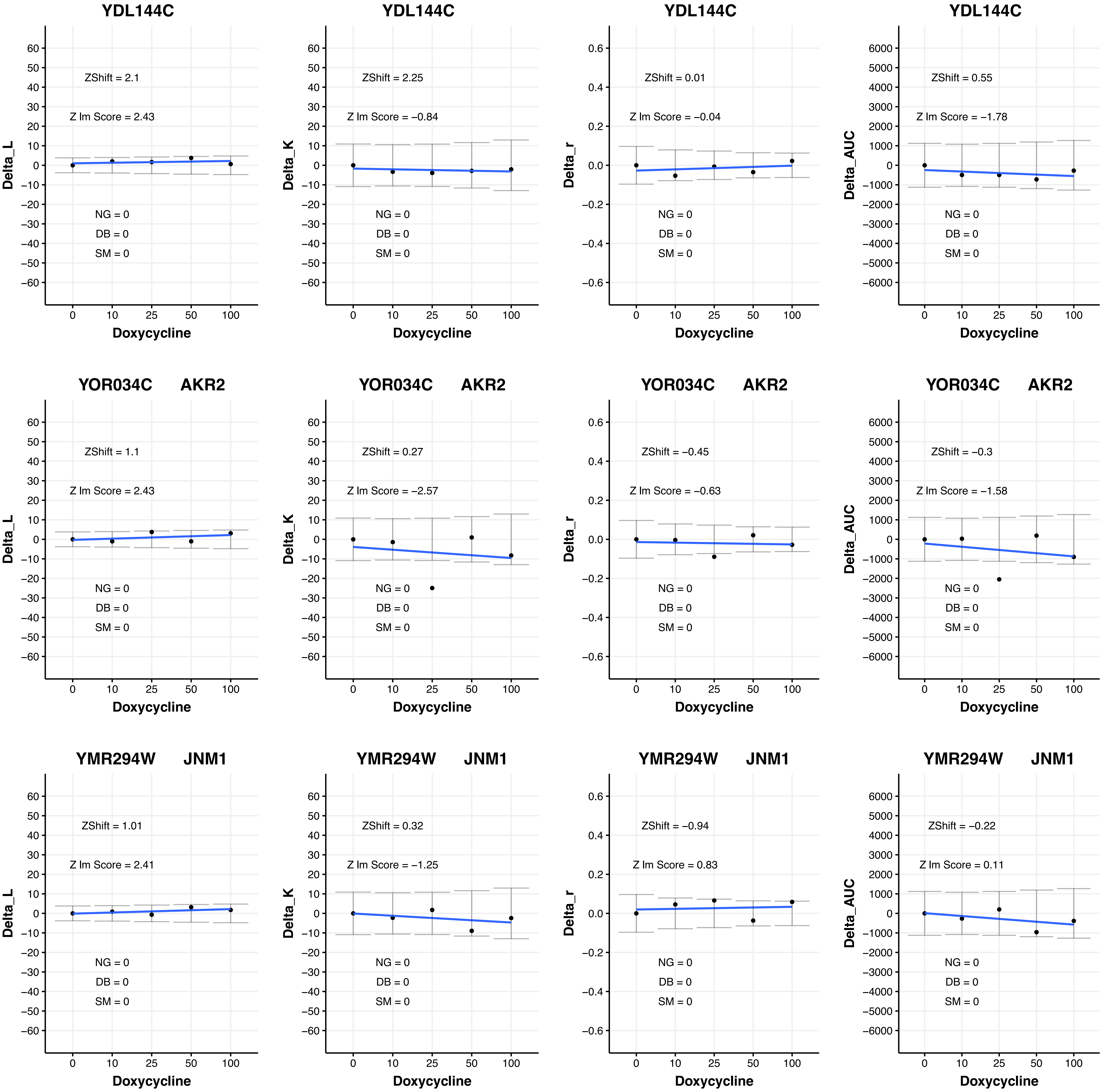
YDL144C
Delta_L
ZShift = 2.1
Z lm Score = 2.43
NG = 0
DB = 0
SM = 0
Doxycycline
YDL144C
Delta_K
ZShift = 2.25
Z lm Score = −0.84
NG = 0
DB = 0
SM = 0
Doxycycline
YDL144C
Delta_r
ZShift = 0.01
Z lm Score = −0.04
NG = 0
DB = 0
SM = 0
Doxycycline
YDL144C
Delta_AUC
ZShift = 0.55
Z lm Score = −1.78
NG = 0
DB = 0
SM = 0
Doxycycline
YOR034C AKR2
Delta_L
ZShift = 1.1
Z lm Score = 2.43
NG = 0
DB = 0
SM = 0
Doxycycline
YOR034C AKR2
Delta_K
ZShift = 0.27
Z lm Score = −2.57
NG = 0
DB = 0
SM = 0
Doxycycline
YOR034C AKR2
Delta_r
ZShift = −0.45
Z lm Score = −0.63
NG = 0
DB = 0
SM = 0
Doxycycline
YOR034C AKR2
Delta_AUC
ZShift = −0.3
Z lm Score = −1.58
NG = 0
DB = 0
SM = 0
Doxycycline
YMR294W JNM1
Delta_L
ZShift = 1.01
Z lm Score = 2.41
NG = 0
DB = 0
SM = 0
Doxycycline
YMR294W JNM1
Delta_K
ZShift = 0.32
Z lm Score = −1.25
NG = 0
DB = 0
SM = 0
Doxycycline
YMR294W JNM1
Delta_r
ZShift = −0.94
Z lm Score = 0.83
NG = 0
DB = 0
SM = 0
Doxycycline
YMR294W JNM1
Delta_AUC
ZShift = −0.22
Z lm Score = 0.11
NG = 0
DB = 0
SM = 0
Doxycycline

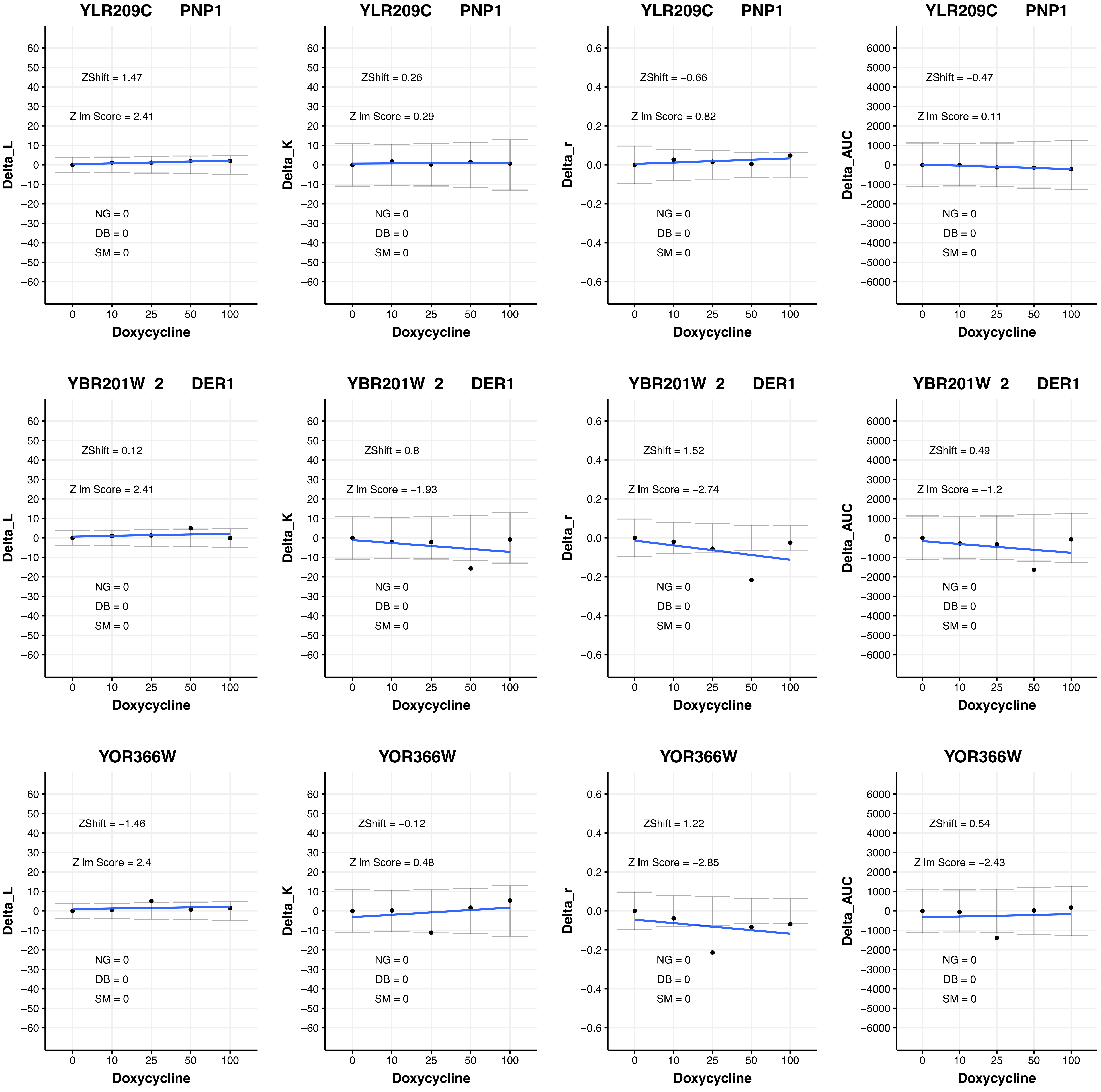
YLR209C PNP1
ZShift = 1.47
Z lm Score = 2.41
NG = 0
DB = 0
SM = 0
Delta_L
Doxycycline
YLR209C PNP1
ZShift = 0.26
Z lm Score = 0.29
NG = 0
DB = 0
SM = 0
Delta_K
Doxycycline
YLR209C PNP1
ZShift = −0.66
Z lm Score = 0.82
NG = 0
DB = 0
SM = 0
Delta_r
Doxycycline
YLR209C PNP1
ZShift = −0.47
Z lm Score = 0.11
NG = 0
DB = 0
SM = 0
Delta_AUC
Doxycycline
YBR201W_2 DER1
ZShift = 0.12
Z lm Score = 2.41
NG = 0
DB = 0
SM = 0
Delta_L
Doxycycline
YBR201W_2 DER1
ZShift = 0.8
Z lm Score = −1.93
NG = 0
DB = 0
SM = 0
Delta_K
Doxycycline
YBR201W_2 DER1
ZShift = 1.52
Z lm Score = −2.74
NG = 0
DB = 0
SM = 0
Delta_r
Doxycycline
YBR201W_2 DER1
ZShift = 0.49
Z lm Score = −1.2
NG = 0
DB = 0
SM = 0
Delta_AUC
Doxycycline
YOR366W
ZShift = −1.46
Z lm Score = 2.4
NG = 0
DB = 0
SM = 0
Delta_L
Doxycycline
YOR366W
ZShift = −0.12
Z lm Score = 0.48
NG = 0
DB = 0
SM = 0
Delta_K
Doxycycline
YOR366W
ZShift = 1.22
Z lm Score = −2.85
NG = 0
DB = 0
SM = 0
Delta_r
Doxycycline
YOR366W
ZShift = 0.54
Z lm Score = −2.43
NG = 0
DB = 0
SM = 0
Delta_AUC
Doxycycline

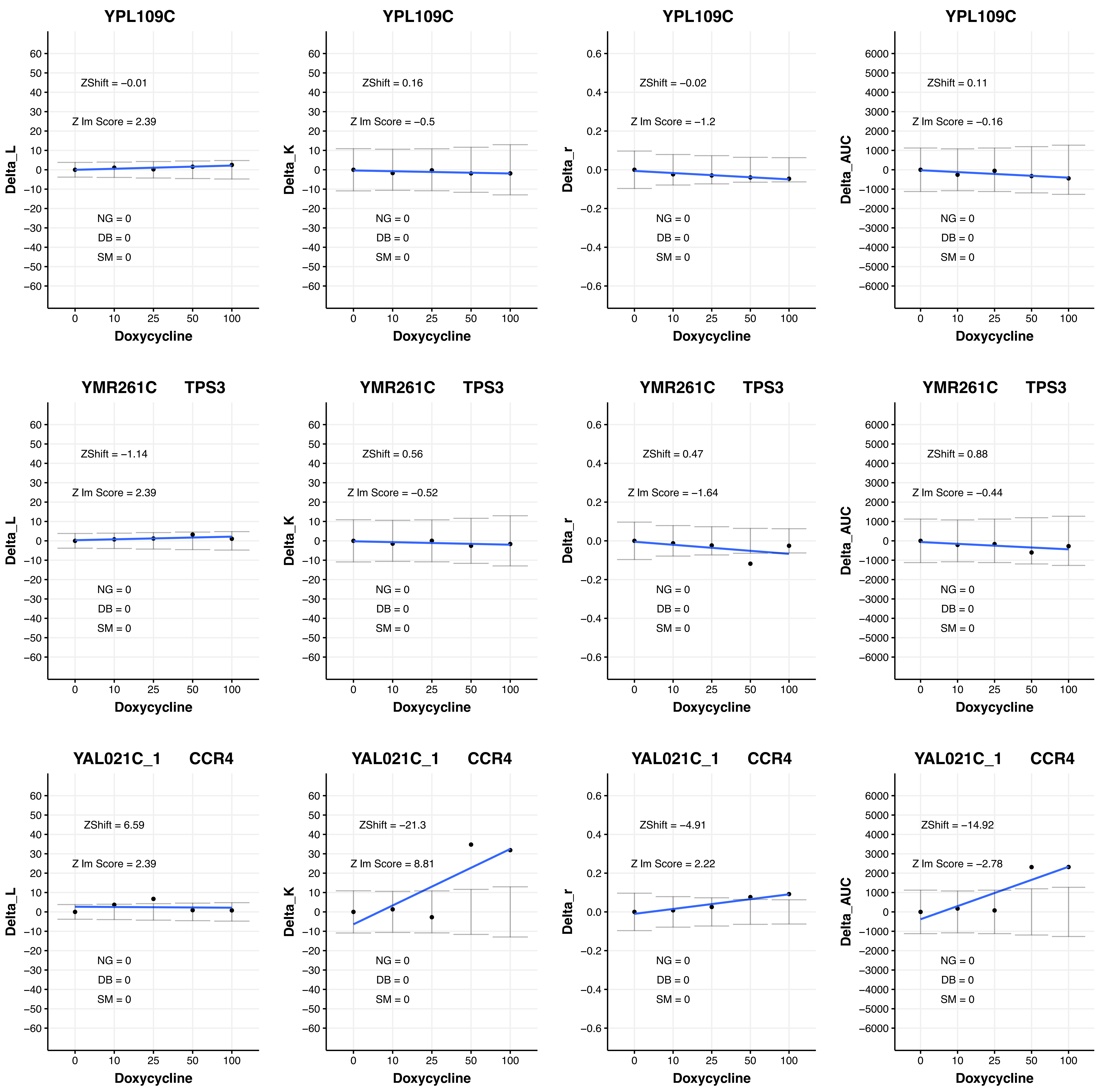

YPL109C
ZShift = −0.01
Z lm Score = 2.39
NG = 0
DB = 0
SM = 0
Delta_L
Doxycycline
YPL109C
ZShift = 0.16
Z lm Score = −0.5
NG = 0
DB = 0
SM = 0
Delta_K
Doxycycline
YPL109C
ZShift = −0.02
Z lm Score = −1.2
NG = 0
DB = 0
SM = 0
Delta_r
Doxycycline
YPL109C
ZShift = 0.11
Z lm Score = −0.16
NG = 0
DB = 0
SM = 0
Delta_AUC
Doxycycline
YMR261C TPS3
ZShift = −1.14
Z lm Score = 2.39
NG = 0
DB = 0
SM = 0
Delta_L
Doxycycline
YMR261C TPS3
ZShift = 0.56
Z lm Score = −0.52
NG = 0
DB = 0
SM = 0
Delta_K
Doxycycline
YMR261C TPS3
ZShift = 0.47
Z lm Score = −1.64
NG = 0
DB = 0
SM = 0
Delta_r
Doxycycline
YMR261C TPS3
ZShift = 0.88
Z lm Score = −0.44
NG = 0
DB = 0
SM = 0
Delta_AUC
Doxycycline
YAL021C_1 CCR4
ZShift = 6.59
Z lm Score = 2.39
NG = 0
DB = 0
SM = 0
Delta_L
Doxycycline
YAL021C_1 CCR4
ZShift = −21.3
Z lm Score = 8.81
NG = 0
DB = 0
SM = 0
Delta_K
Doxycycline
YAL021C_1 CCR4
ZShift = −4.91
Z lm Score = 2.22
NG = 0
DB = 0
SM = 0
Delta_r
Doxycycline
YAL021C_1 CCR4
ZShift = −14.92
Z lm Score = −2.78
NG = 0
DB = 0
SM = 0
Delta_AUC
Doxycycline

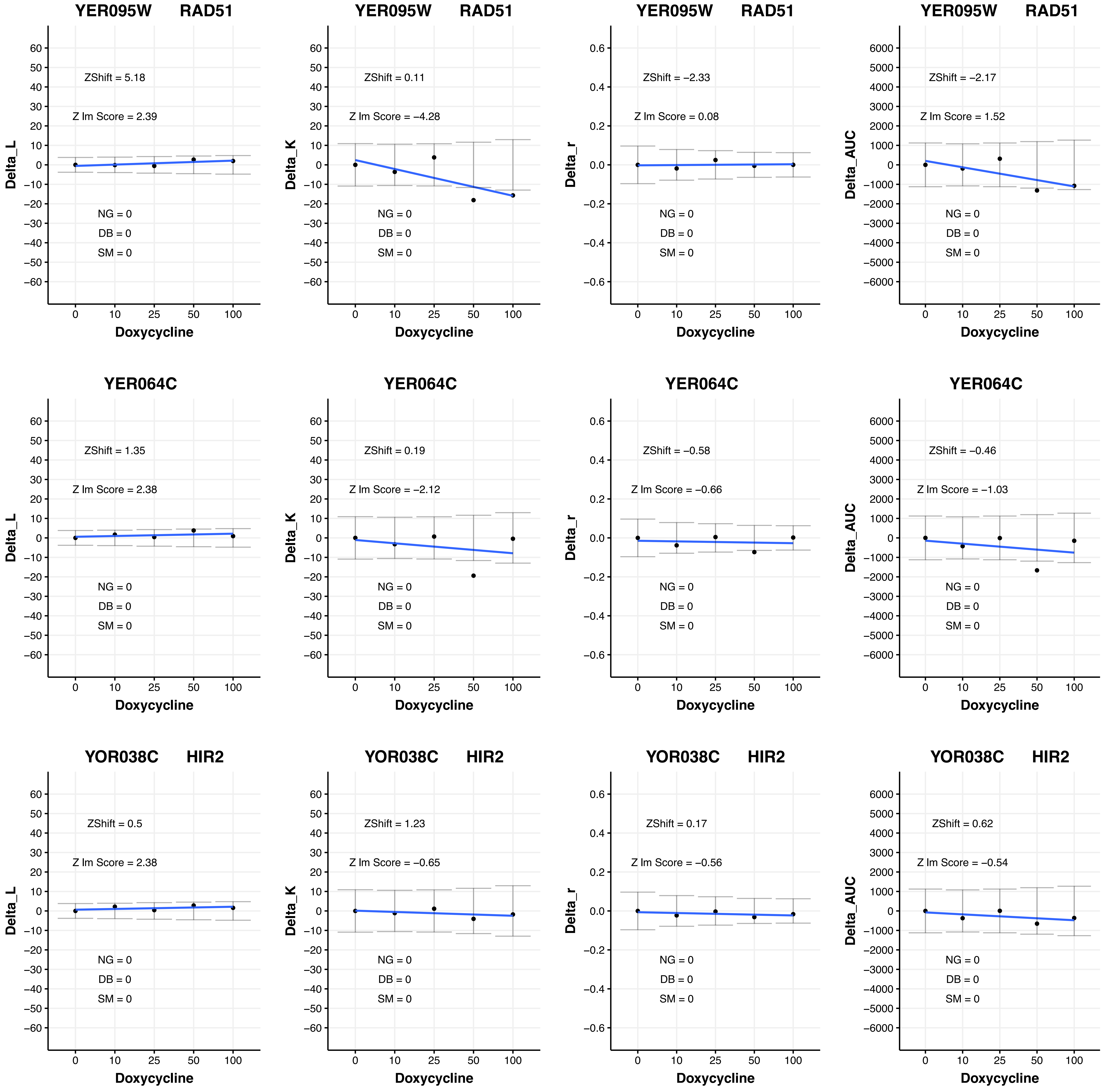
YER095W RAD51
ZShift = 5.18
Z lm Score = 2.39
NG = 0
DB = 0
SM = 0
Delta_L
Doxycycline
YER095W RAD51
ZShift = 0.11
Z lm Score = −4.28
NG = 0
DB = 0
SM = 0
Delta_K
Doxycycline
YER095W RAD51
ZShift = −2.33
Z lm Score = 0.08
NG = 0
DB = 0
SM = 0
Delta_r
Doxycycline
YER095W RAD51
ZShift = −2.17
Z lm Score = 1.52
NG = 0
DB = 0
SM = 0
Delta_AUC
Doxycycline
YER064C
ZShift = 1.35
Z lm Score = 2.38
NG = 0
DB = 0
SM = 0
Delta_L
Doxycycline
YER064C
ZShift = 0.19
Z lm Score = −2.12
NG = 0
DB = 0
SM = 0
Delta_K
Doxycycline
YER064C
ZShift = −0.58
Z lm Score = −0.66
NG = 0
DB = 0
SM = 0
Delta_r
Doxycycline
YER064C
ZShift = −0.46
Z lm Score = −1.03
NG = 0
DB = 0
SM = 0
Delta_AUC
Doxycycline
YOR038C HIR2
ZShift = 0.5
Z lm Score = 2.38
NG = 0
DB = 0
SM = 0
Delta_L
Doxycycline
YOR038C HIR2
ZShift = 1.23
Z lm Score = −0.65
NG = 0
DB = 0
SM = 0
Delta_K
Doxycycline
YOR038C HIR2
ZShift = 0.17
Z lm Score = −0.56
NG = 0
DB = 0
SM = 0
Delta_r
Doxycycline
YOR038C HIR2
ZShift = 0.62
Z lm Score = −0.54
NG = 0
DB = 0
SM = 0
Delta_AUC
Doxycycline

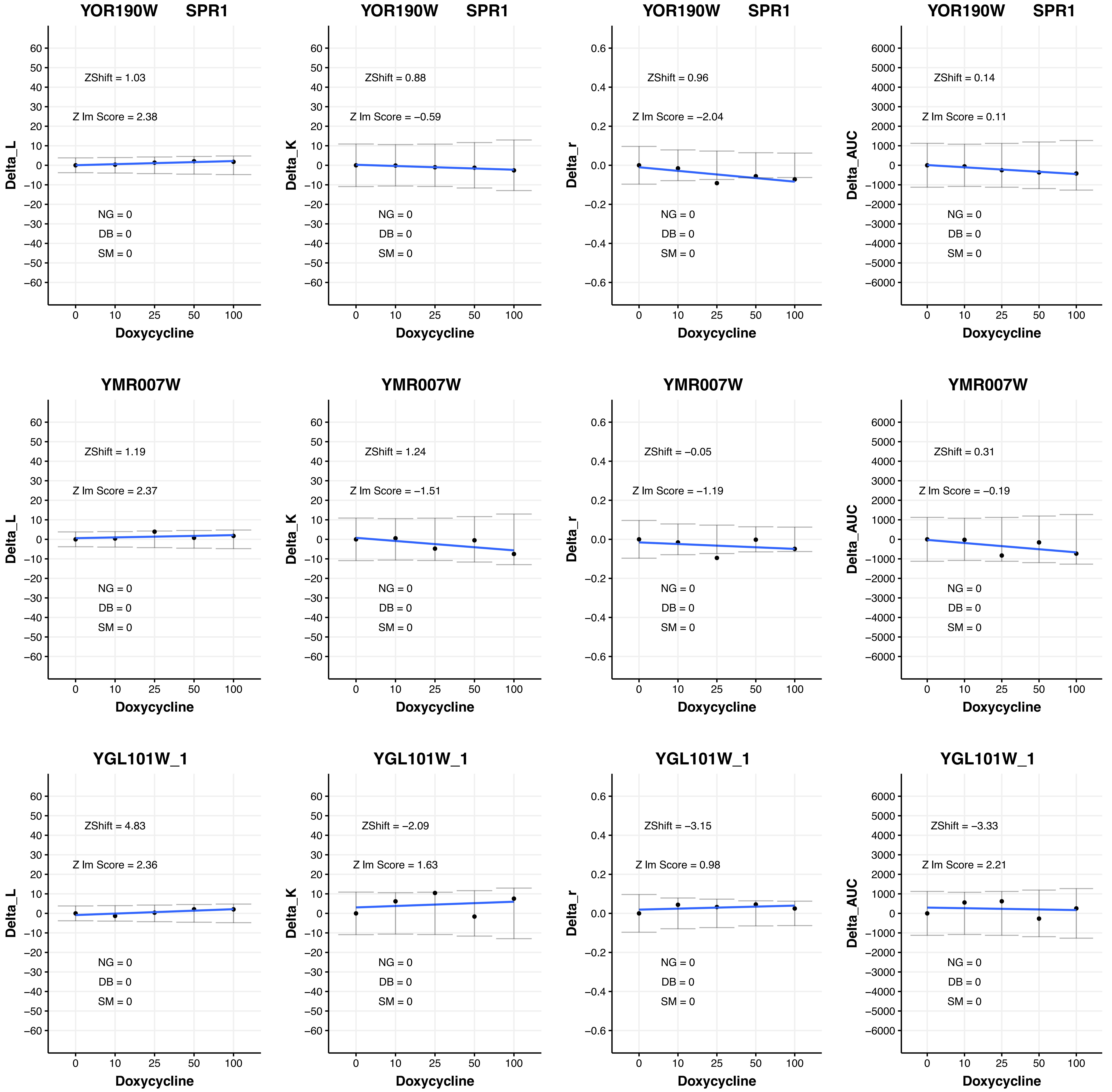
YOR190W SPR1
ZShift = 1.03
Z lm Score = 2.38
NG = 0
DB = 0
SM = 0
Delta_L
Doxycycline
YOR190W SPR1
ZShift = 0.88
Z lm Score = −0.59
NG = 0
DB = 0
SM = 0
Delta_K
Doxycycline
YOR190W SPR1
ZShift = 0.96
Z lm Score = −2.04
NG = 0
DB = 0
SM = 0
Delta_r
Doxycycline
YOR190W SPR1
ZShift = 0.14
Z lm Score = 0.11
NG = 0
DB = 0
SM = 0
Delta_AUC
Doxycycline
YMR007W
ZShift = 1.19
Z lm Score = 2.37
NG = 0
DB = 0
SM = 0
Delta_L
Doxycycline
YMR007W
ZShift = 1.24
Z lm Score = −1.51
NG = 0
DB = 0
SM = 0
Delta_K
Doxycycline
YMR007W
ZShift = −0.05
Z lm Score = −1.19
NG = 0
DB = 0
SM = 0
Delta_r
Doxycycline
YMR007W
ZShift = 0.31
Z lm Score = −0.19
NG = 0
DB = 0
SM = 0
Delta_AUC
Doxycycline
YGL101W_1
ZShift = 4.83
Z lm Score = 2.36
NG = 0
DB = 0
SM = 0
Delta_L
Doxycycline
YGL101W_1
ZShift = −2.09
Z lm Score = 1.63
NG = 0
DB = 0
SM = 0
Delta_K
Doxycycline
YGL101W_1
ZShift = −3.15
Z lm Score = 0.98
NG = 0
DB = 0
SM = 0
Delta_r
Doxycycline
YGL101W_1
ZShift = −3.33
Z lm Score = 2.21
NG = 0
DB = 0
SM = 0
Delta_AUC
Doxycycline

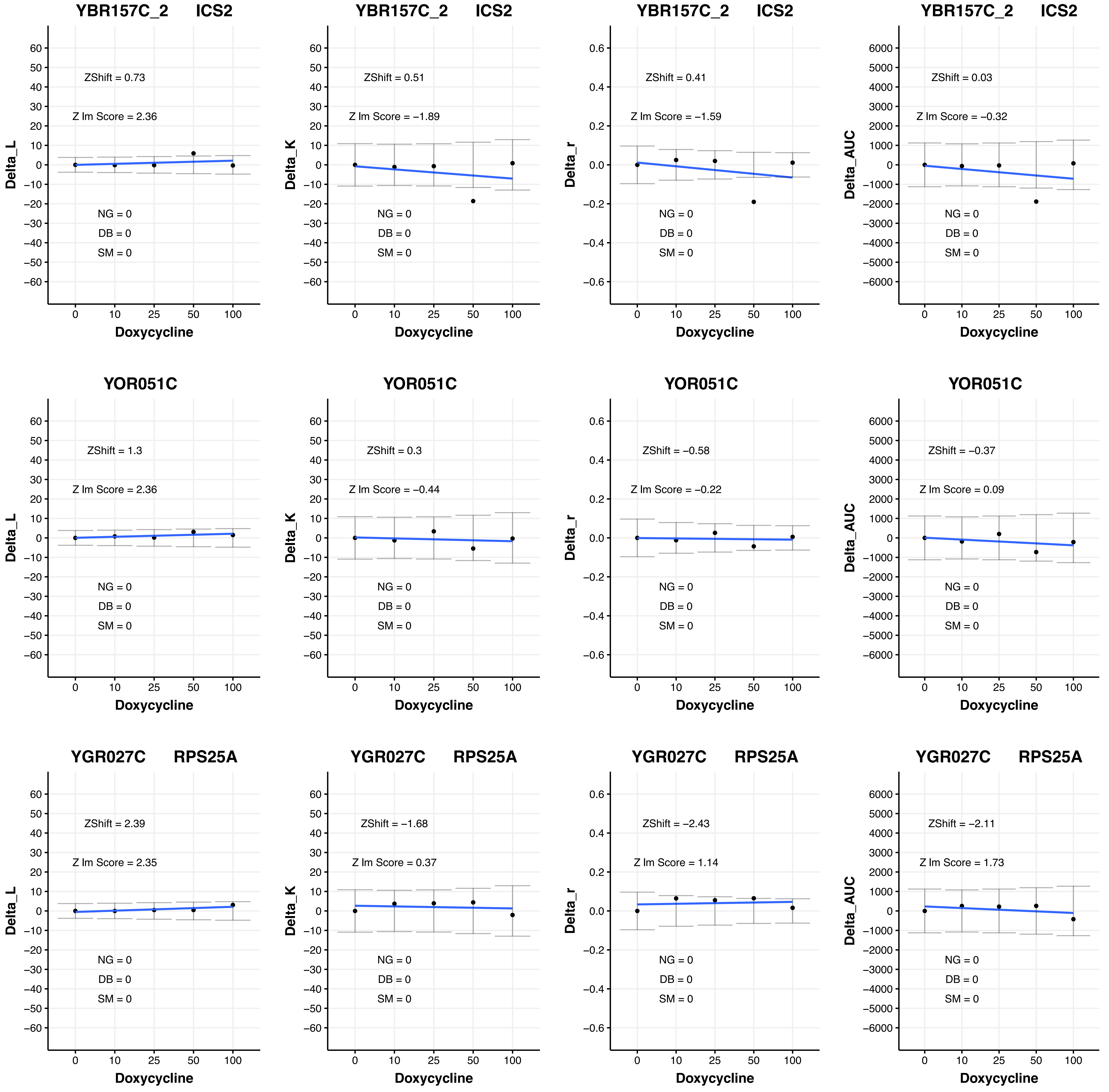

YBR157C_2 ICS2
ZShift = 0.73
Z lm Score = 2.36
NG = 0
DB = 0
SM = 0
Delta_L
Doxycycline
YBR157C_2 ICS2
ZShift = 0.51
Z lm Score = −1.89
NG = 0
DB = 0
SM = 0
Delta_K
Doxycycline
YBR157C_2 ICS2
ZShift = 0.41
Z lm Score = −1.59
NG = 0
DB = 0
SM = 0
Delta_r
Doxycycline
YBR157C_2 ICS2
ZShift = 0.03
Z lm Score = −0.32
NG = 0
DB = 0
SM = 0
Delta_AUC
Doxycycline
YOR051C
ZShift = 1.3
Z lm Score = 2.36
NG = 0
DB = 0
SM = 0
Delta_L
Doxycycline
YOR051C
ZShift = 0.3
Z lm Score = −0.44
NG = 0
DB = 0
SM = 0
Delta_K
Doxycycline
YOR051C
ZShift = −0.58
Z lm Score = −0.22
NG = 0
DB = 0
SM = 0
Delta_r
Doxycycline
YOR051C
ZShift = −0.37
Z lm Score = 0.09
NG = 0
DB = 0
SM = 0
Delta_AUC
Doxycycline
YGR027C RPS25A
ZShift = 2.39
Z lm Score = 2.35
NG = 0
DB = 0
SM = 0
Delta_L
Doxycycline
YGR027C RPS25A
ZShift = −1.68
Z lm Score = 0.37
NG = 0
DB = 0
SM = 0
Delta_K
Doxycycline
YGR027C RPS25A
ZShift = −2.43
Z lm Score = 1.14
NG = 0
DB = 0
SM = 0
Delta_r
Doxycycline
YGR027C RPS25A
ZShift = −2.11
Z lm Score = 1.73
NG = 0
DB = 0
SM = 0
Delta_AUC
Doxycycline

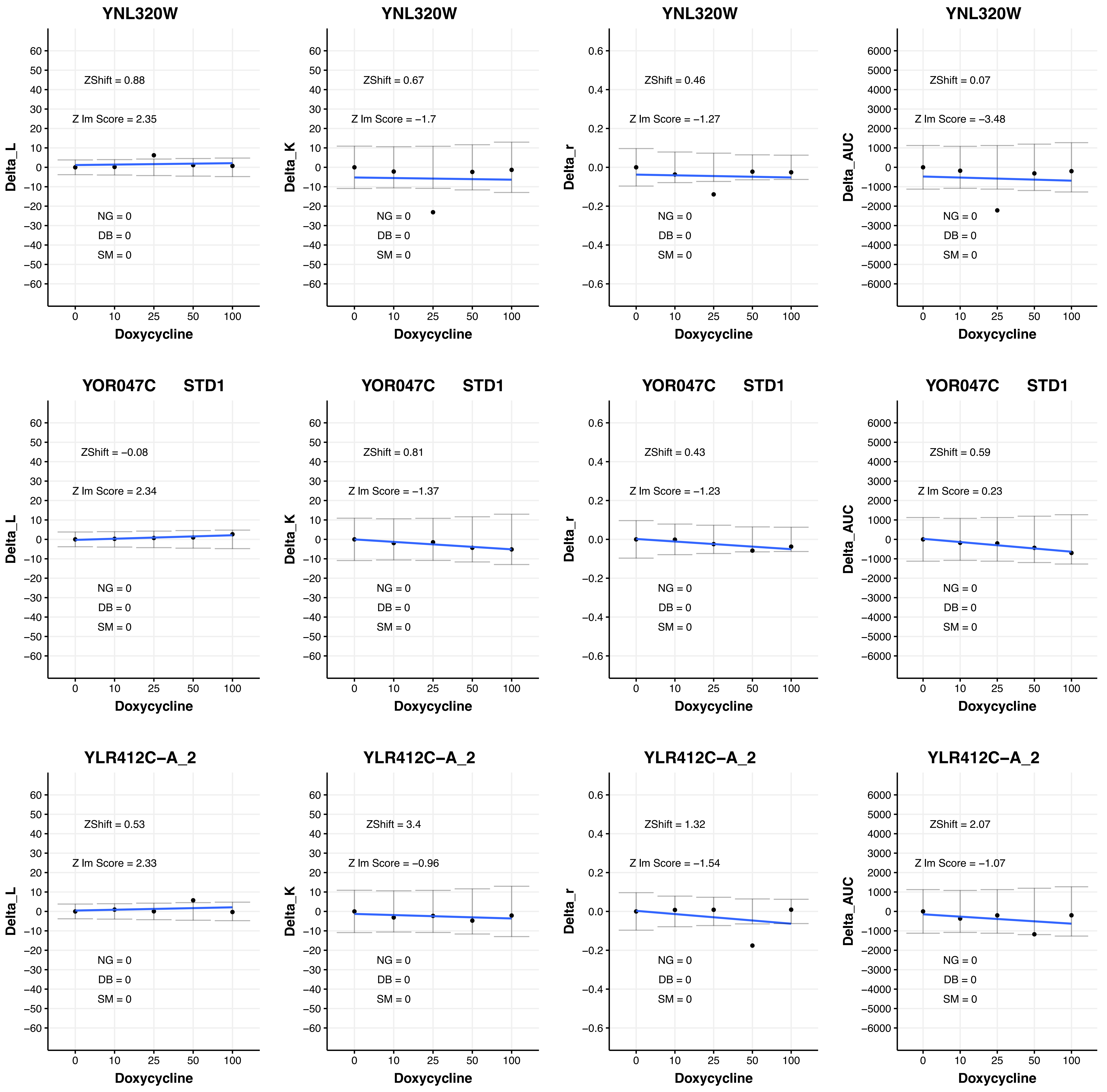
YNL320W
ZShift = 0.88
Z lm Score = 2.35
NG = 0
DB = 0
SM = 0
Delta_L
Doxycycline
YNL320W
ZShift = 0.67
Z lm Score = −1.7
NG = 0
DB = 0
SM = 0
Delta_K
Doxycycline
YNL320W
ZShift = 0.46
Z lm Score = −1.27
NG = 0
DB = 0
SM = 0
Delta_r
Doxycycline
YNL320W
ZShift = 0.07
Z lm Score = −3.48
NG = 0
DB = 0
SM = 0
Delta_AUC
Doxycycline
YOR047C STD1
ZShift = −0.08
Z lm Score = 2.34
NG = 0
DB = 0
SM = 0
Delta_L
Doxycycline
YOR047C STD1
ZShift = 0.81
Z lm Score = −1.37
NG = 0
DB = 0
SM = 0
Delta_K
Doxycycline
YOR047C STD1
ZShift = 0.43
Z lm Score = −1.23
NG = 0
DB = 0
SM = 0
Delta_r
Doxycycline
YOR047C STD1
ZShift = 0.59
Z lm Score = 0.23
NG = 0
DB = 0
SM = 0
Delta_AUC
Doxycycline
YLR412C−A_2
ZShift = 0.53
Z lm Score = 2.33
NG = 0
DB = 0
SM = 0
Delta_L
Doxycycline
YLR412C−A_2
ZShift = 3.4
Z lm Score = −0.96
NG = 0
DB = 0
SM = 0
Delta_K
Doxycycline
YLR412C−A_2
ZShift = 1.32
Z lm Score = −1.54
NG = 0
DB = 0
SM = 0
Delta_r
Doxycycline
YLR412C−A_2
ZShift = 2.07
Z lm Score = −1.07
NG = 0
DB = 0
SM = 0
Delta_AUC
Doxycycline

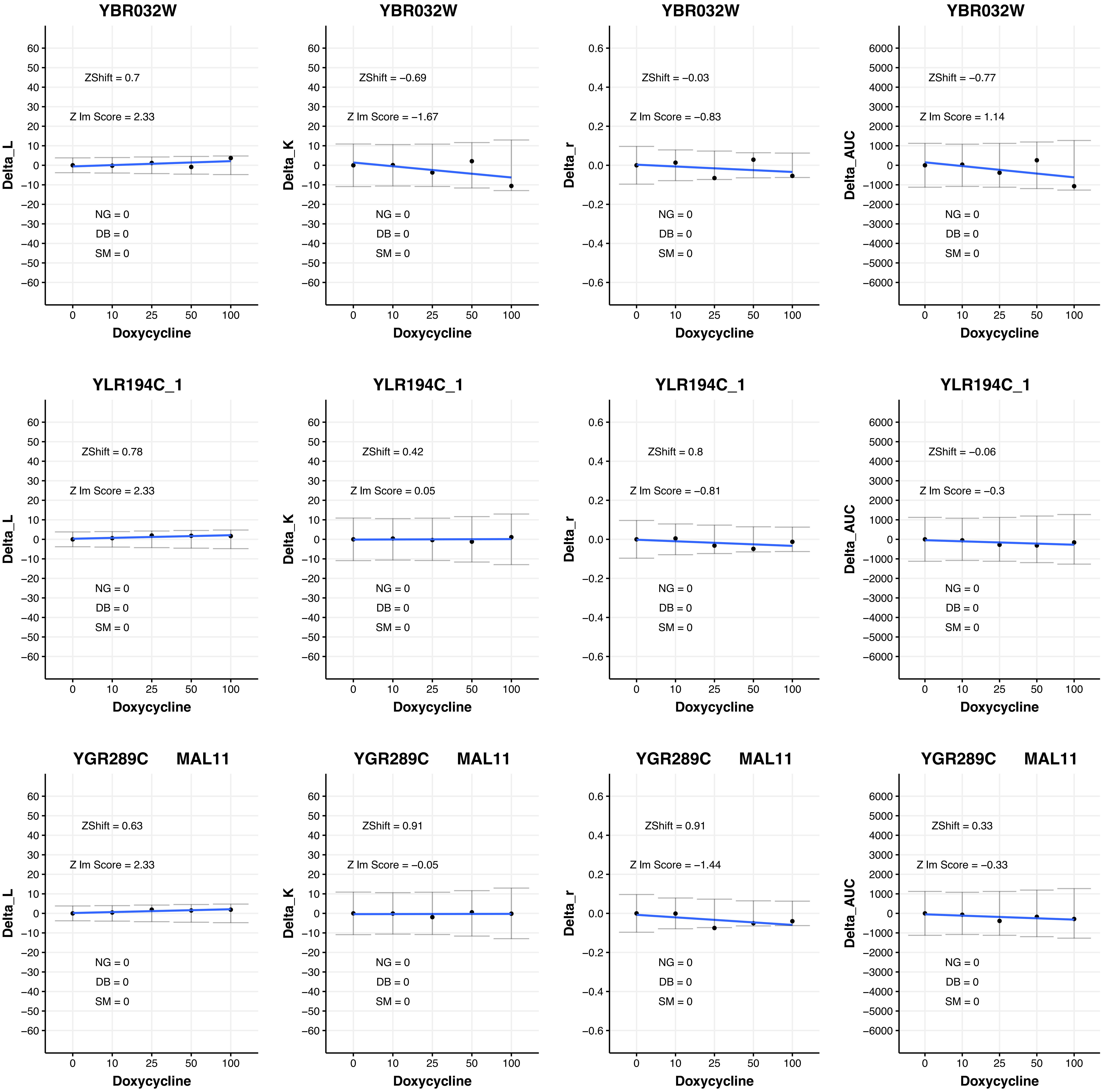

YBR032W
ZShift = 0.7
Z lm Score = 2.33
NG = 0
DB = 0
SM = 0
Delta_L
Doxycycline
YBR032W
ZShift = −0.69
Z lm Score = −1.67
NG = 0
DB = 0
SM = 0
Delta_K
Doxycycline
YBR032W
ZShift = −0.03
Z lm Score = −0.83
NG = 0
DB = 0
SM = 0
Delta_r
Doxycycline
YBR032W
ZShift = −0.77
Z lm Score = 1.14
NG = 0
DB = 0
SM = 0
Delta_AUC
Doxycycline
YLR194C_1
ZShift = 0.78
Z lm Score = 2.33
NG = 0
DB = 0
SM = 0
Delta_L
Doxycycline
YLR194C_1
ZShift = 0.42
Z lm Score = 0.05
NG = 0
DB = 0
SM = 0
Delta_K
Doxycycline
YLR194C_1
ZShift = 0.8
Z lm Score = −0.81
NG = 0
DB = 0
SM = 0
Delta_r
Doxycycline
YLR194C_1
ZShift = −0.06
Z lm Score = −0.3
NG = 0
DB = 0
SM = 0
Delta_AUC
Doxycycline
YGR289C MAL11
ZShift = 0.63
Z lm Score = 2.33
NG = 0
DB = 0
SM = 0
Delta_L
Doxycycline
YGR289C MAL11
ZShift = 0.91
Z lm Score = −0.05
NG = 0
DB = 0
SM = 0
Delta_K
Doxycycline
YGR289C MAL11
ZShift = 0.91
Z lm Score = −1.44
NG = 0
DB = 0
SM = 0
Delta_r
Doxycycline
YGR289C MAL11
ZShift = 0.33
Z lm Score = −0.33
NG = 0
DB = 0
SM = 0
Delta_AUC
Doxycycline

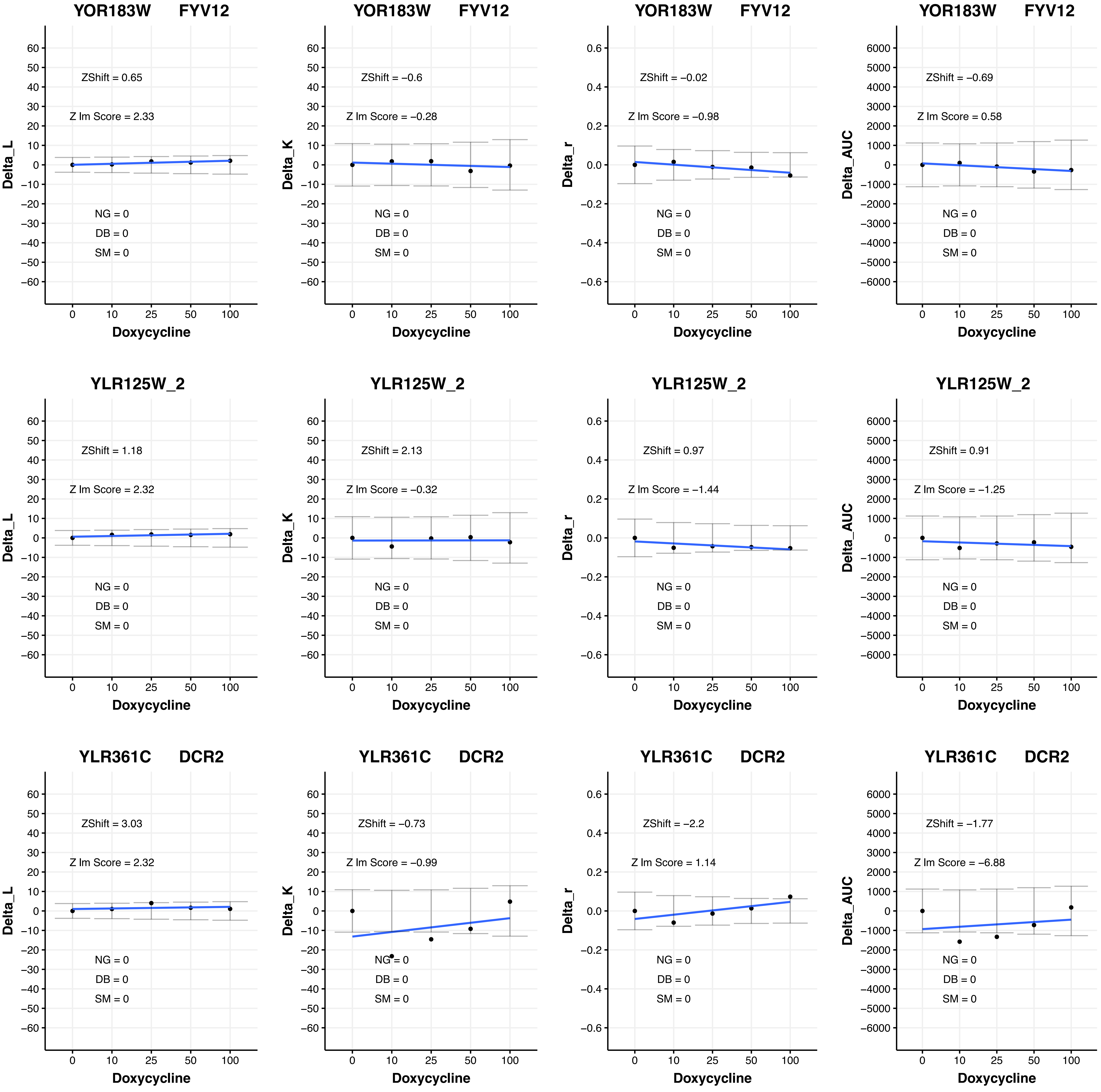

YOR183W FYV12
ZShift = 0.65
Z lm Score = 2.33
NG = 0
DB = 0
SM = 0
Delta_L
Doxycycline
YOR183W FYV12
ZShift = −0.6
Z lm Score = −0.28
NG = 0
DB = 0
SM = 0
Delta_K
Doxycycline
YOR183W FYV12
ZShift = −0.02
Z lm Score = −0.98
NG = 0
DB = 0
SM = 0
Delta_r
Doxycycline
YOR183W FYV12
ZShift = −0.69
Z lm Score = 0.58
NG = 0
DB = 0
SM = 0
Delta_AUC
Doxycycline
YLR125W_2
ZShift = 1.18
Z lm Score = 2.32
NG = 0
DB = 0
SM = 0
Delta_L
Doxycycline
YLR125W_2
ZShift = 2.13
Z lm Score = −0.32
NG = 0
DB = 0
SM = 0
Delta_K
Doxycycline
YLR125W_2
ZShift = 0.97
Z lm Score = −1.44
NG = 0
DB = 0
SM = 0
Delta_r
Doxycycline
YLR125W_2
ZShift = 0.91
Z lm Score = −1.25
NG = 0
DB = 0
SM = 0
Delta_AUC
Doxycycline
YLR361C DCR2
ZShift = 3.03
Z lm Score = 2.32
NG = 0
DB = 0
SM = 0
Delta_L
Doxycycline
YLR361C DCR2
ZShift = −0.73
Z lm Score = −0.99
NG = 0
DB = 0
SM = 0
Delta_K
Doxycycline
YLR361C DCR2
ZShift = −2.2
Z lm Score = 1.14
NG = 0
DB = 0
SM = 0
Delta_r
Doxycycline
YLR361C DCR2
ZShift = −1.77
Z lm Score = −6.88
NG = 0
DB = 0
SM = 0
Delta_AUC
Doxycycline

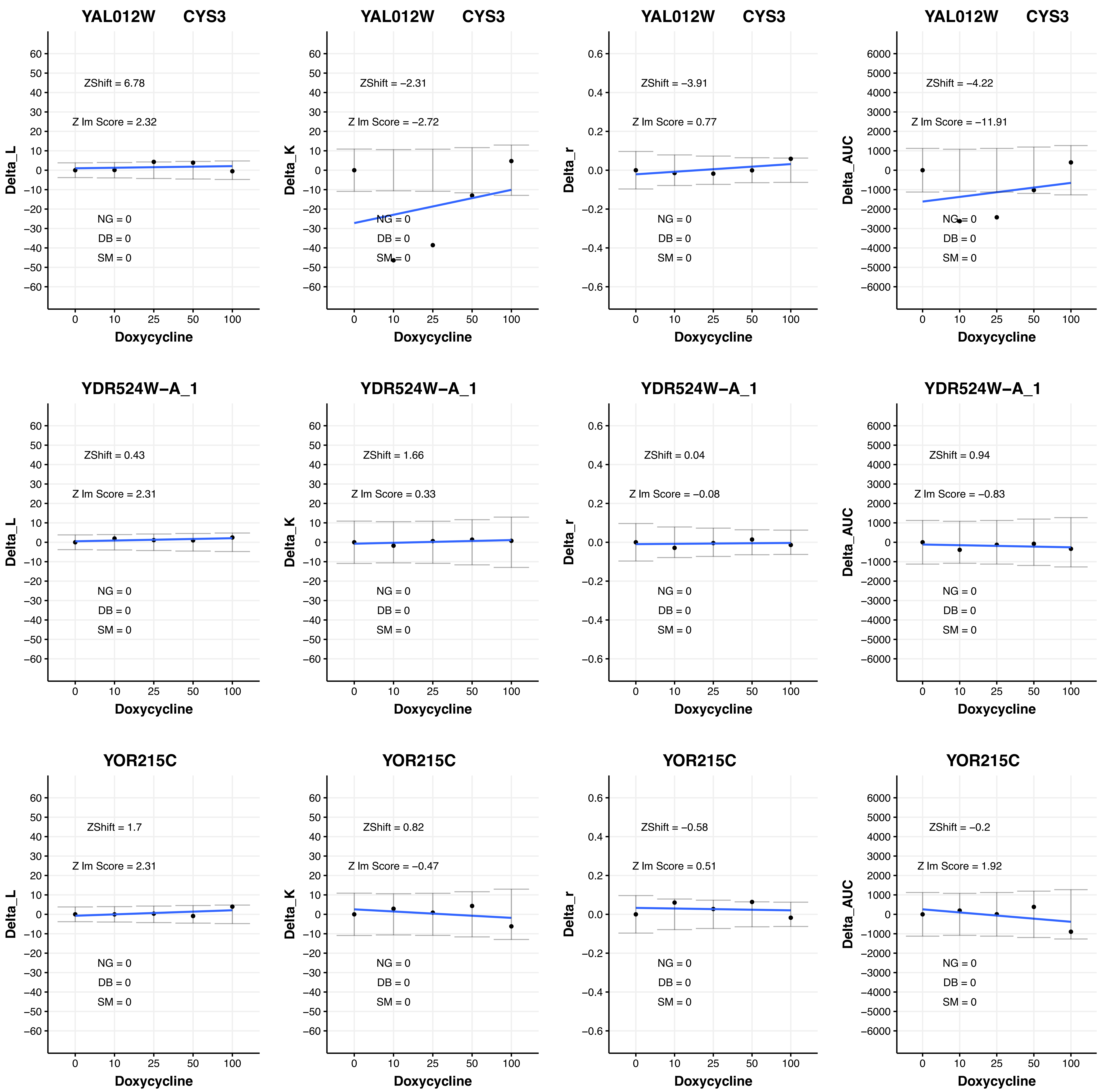

YAL012W CYS3
ZShift = 6.78
Z lm Score = 2.32
NG = 0
DB = 0
SM = 0
Delta_L
Doxycycline
YAL012W CYS3
ZShift = −2.31
Z lm Score = −2.72
NG = 0
DB = 0
SM = 0
Delta_K
Doxycycline
YAL012W CYS3
ZShift = −3.91
Z lm Score = 0.77
NG = 0
DB = 0
SM = 0
Delta_r
Doxycycline
YAL012W CYS3
ZShift = −4.22
Z lm Score = −11.91
NG = 0
DB = 0
SM = 0
Delta_AUC
Doxycycline
YDR524W−A_1
ZShift = 0.43
Z lm Score = 2.31
NG = 0
DB = 0
SM = 0
Delta_L
Doxycycline
YDR524W−A_1
ZShift = 1.66
Z lm Score = 0.33
NG = 0
DB = 0
SM = 0
Delta_K
Doxycycline
YDR524W−A_1
ZShift = 0.04
Z lm Score = −0.08
NG = 0
DB = 0
SM = 0
Delta_r
Doxycycline
YDR524W−A_1
ZShift = 0.94
Z lm Score = −0.83
NG = 0
DB = 0
SM = 0
Delta_AUC
Doxycycline
YOR215C
ZShift = 1.7
Z lm Score = 2.31
NG = 0
DB = 0
SM = 0
Delta_L
Doxycycline
YOR215C
ZShift = 0.82
Z lm Score = −0.47
NG = 0
DB = 0
SM = 0
Delta_K
Doxycycline
YOR215C
ZShift = −0.58
Z lm Score = 0.51
NG = 0
DB = 0
SM = 0
Delta_r
Doxycycline
YOR215C
ZShift = −0.2
Z lm Score = 1.92
NG = 0
DB = 0
SM = 0
Delta_AUC
Doxycycline
0 10 25 50 100

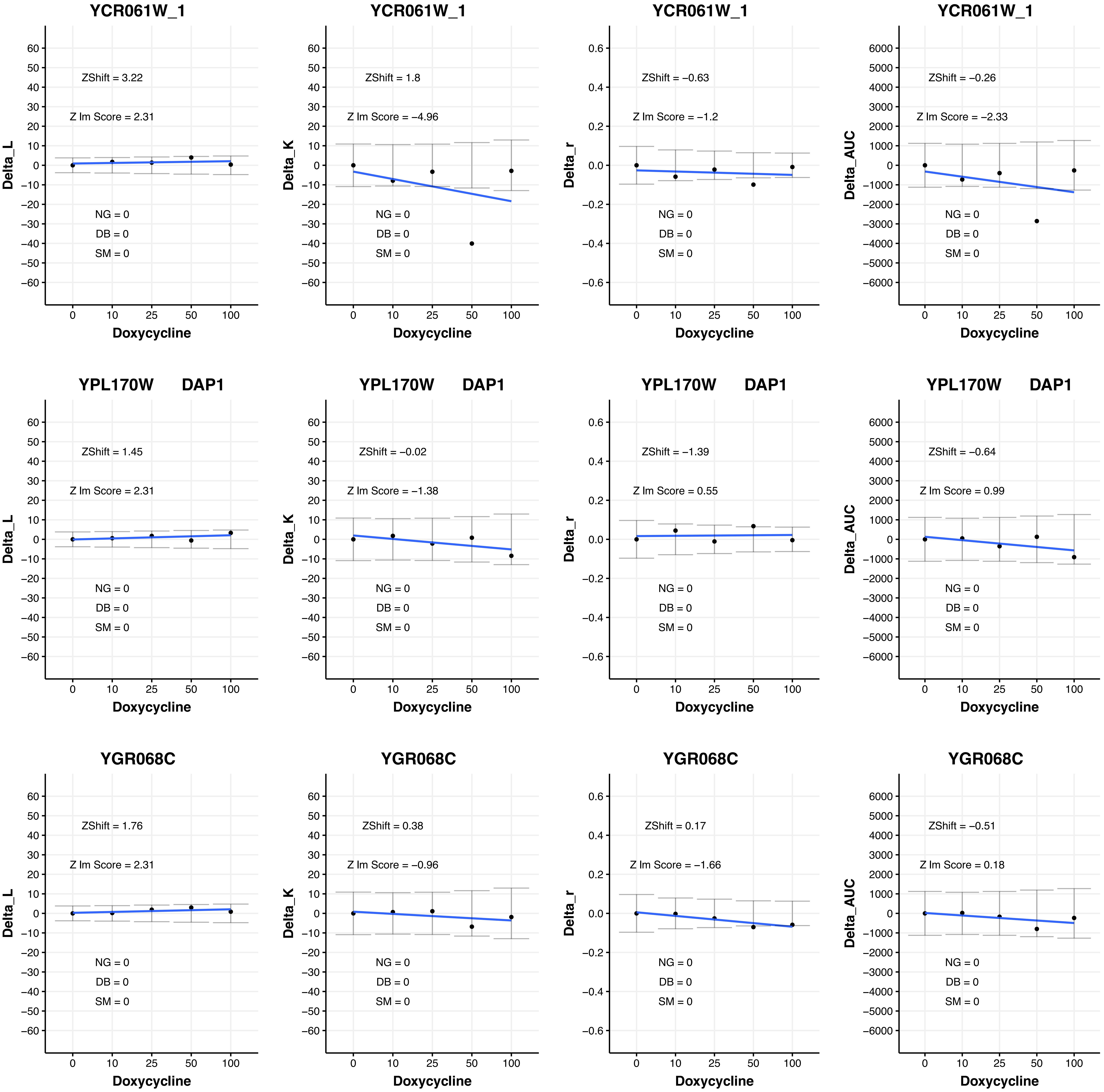

YCR061W_1
Delta_L
ZShift = 3.22
Z lm Score = 2.31
NG = 0
DB = 0
SM = 0
Doxycycline
YCR061W_1
Delta_K
ZShift = 1.8
Z lm Score = −4.96
NG = 0
DB = 0
SM = 0
Doxycycline
YCR061W_1
Delta_r
ZShift = −0.63
Z lm Score = −1.2
NG = 0
DB = 0
SM = 0
Doxycycline
YCR061W_1
Delta_AUC
ZShift = −0.26
Z lm Score = −2.33
NG = 0
DB = 0
SM = 0
Doxycycline
YPL170W DAP1
Delta_L
ZShift = 1.45
Z lm Score = 2.31
NG = 0
DB = 0
SM = 0
Doxycycline
YPL170W DAP1
Delta_K
ZShift = −0.02
Z lm Score = −1.38
NG = 0
DB = 0
SM = 0
Doxycycline
YPL170W DAP1
Delta_r
ZShift = −1.39
Z lm Score = 0.55
NG = 0
DB = 0
SM = 0
Doxycycline
YPL170W DAP1
Delta_AUC
ZShift = −0.64
Z lm Score = 0.99
NG = 0
DB = 0
SM = 0
Doxycycline
YGR068C
Delta_L
ZShift = 1.76
Z lm Score = 2.31
NG = 0
DB = 0
SM = 0
Doxycycline
YGR068C
Delta_K
ZShift = 0.38
Z lm Score = −0.96
NG = 0
DB = 0
SM = 0
Doxycycline
YGR068C
Delta_r
ZShift = 0.17
Z lm Score = −1.66
NG = 0
DB = 0
SM = 0
Doxycycline
YGR068C
Delta_AUC
ZShift = −0.51
Z lm Score = 0.18
NG = 0
DB = 0
SM = 0
Doxycycline

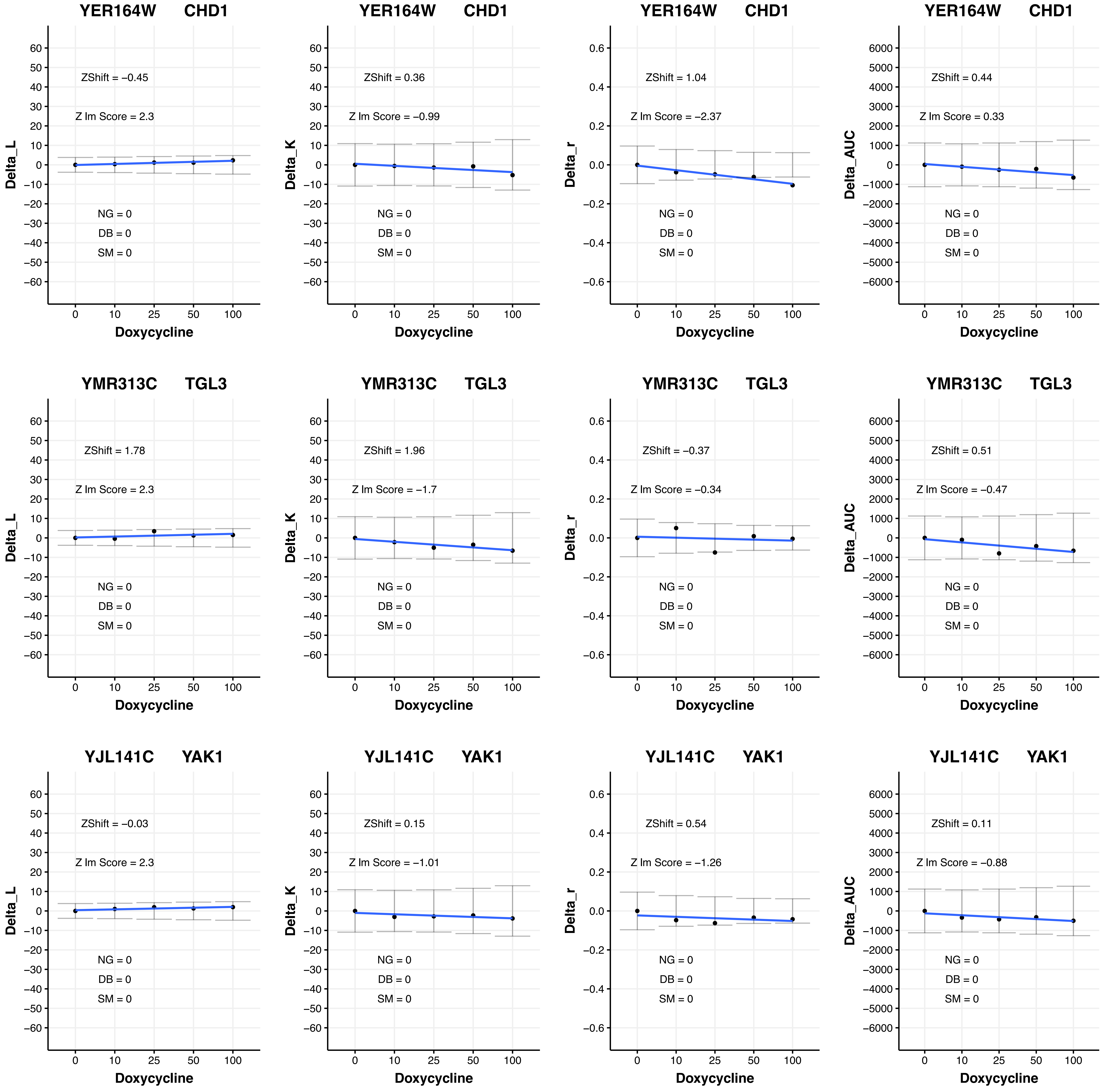

YER164W CHD1
ZShift = −0.45
Z lm Score = 2.3
NG = 0
DB = 0
SM = 0
Delta_L
Doxycycline
YER164W CHD1
ZShift = 0.36
Z lm Score = −0.99
NG = 0
DB = 0
SM = 0
Delta_K
Doxycycline
YER164W CHD1
ZShift = 1.04
Z lm Score = −2.37
NG = 0
DB = 0
SM = 0
Delta_r
Doxycycline
YER164W CHD1
ZShift = 0.44
Z lm Score = 0.33
NG = 0
DB = 0
SM = 0
Delta_AUC
Doxycycline
YMR313C TGL3
ZShift = 1.78
Z lm Score = 2.3
NG = 0
DB = 0
SM = 0
Delta_L
Doxycycline
YMR313C TGL3
ZShift = 1.96
Z lm Score = −1.7
NG = 0
DB = 0
SM = 0
Delta_K
Doxycycline
YMR313C TGL3
ZShift = −0.37
Z lm Score = −0.34
NG = 0
DB = 0
SM = 0
Delta_r
Doxycycline
YMR313C TGL3
ZShift = 0.51
Z lm Score = −0.47
NG = 0
DB = 0
SM = 0
Delta_AUC
Doxycycline
YJL141C YAK1
ZShift = −0.03
Z lm Score = 2.3
NG = 0
DB = 0
SM = 0
Delta_L
Doxycycline
YJL141C YAK1
ZShift = 0.15
Z lm Score = −1.01
NG = 0
DB = 0
SM = 0
Delta_K
Doxycycline
YJL141C YAK1
ZShift = 0.54
Z lm Score = −1.26
NG = 0
DB = 0
SM = 0
Delta_r
Doxycycline
YJL141C YAK1
ZShift = 0.11
Z lm Score = −0.88
NG = 0
DB = 0
SM = 0
Delta_AUC
Doxycycline

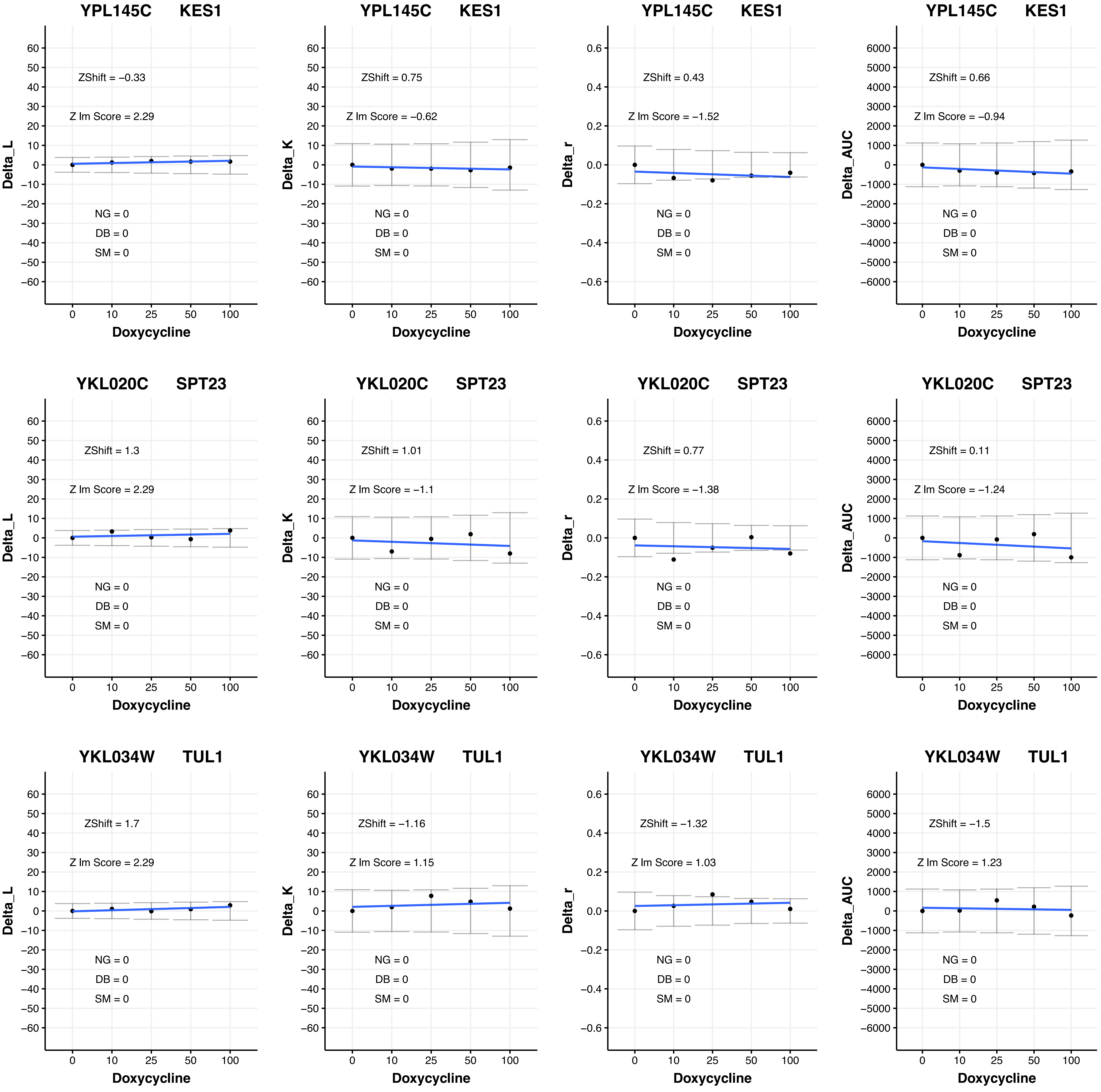
YPL145C KES1
ZShift = −0.33
Z lm Score = 2.29
NG = 0
DB = 0
SM = 0
Delta_L
Doxycycline
YPL145C KES1
ZShift = 0.75
Z lm Score = −0.62
NG = 0
DB = 0
SM = 0
Delta_K
Doxycycline
YPL145C KES1
ZShift = 0.43
Z lm Score = −1.52
NG = 0
DB = 0
SM = 0
Delta_r
Doxycycline
YPL145C KES1
ZShift = 0.66
Z lm Score = −0.94
NG = 0
DB = 0
SM = 0
Delta_AUC
Doxycycline
YKL020C SPT23
ZShift = 1.3
Z lm Score = 2.29
NG = 0
DB = 0
SM = 0
Delta_L
Doxycycline
YKL020C SPT23
ZShift = 1.01
Z lm Score = −1.1
NG = 0
DB = 0
SM = 0
Delta_K
Doxycycline
YKL020C SPT23
ZShift = 0.77
Z lm Score = −1.38
NG = 0
DB = 0
SM = 0
Delta_r
Doxycycline
YKL020C SPT23
ZShift = 0.11
Z lm Score = −1.24
NG = 0
DB = 0
SM = 0
Delta_AUC
Doxycycline
YKL034W TUL1
ZShift = 1.7
Z lm Score = 2.29
NG = 0
DB = 0
SM = 0
Delta_L
Doxycycline
YKL034W TUL1
ZShift = −1.16
Z lm Score = 1.15
NG = 0
DB = 0
SM = 0
Delta_K
Doxycycline
YKL034W TUL1
ZShift = −1.32
Z lm Score = 1.03
NG = 0
DB = 0
SM = 0
Delta_r
Doxycycline
YKL034W TUL1
ZShift = −1.5
Z lm Score = 1.23
NG = 0
DB = 0
SM = 0
Delta_AUC
Doxycycline

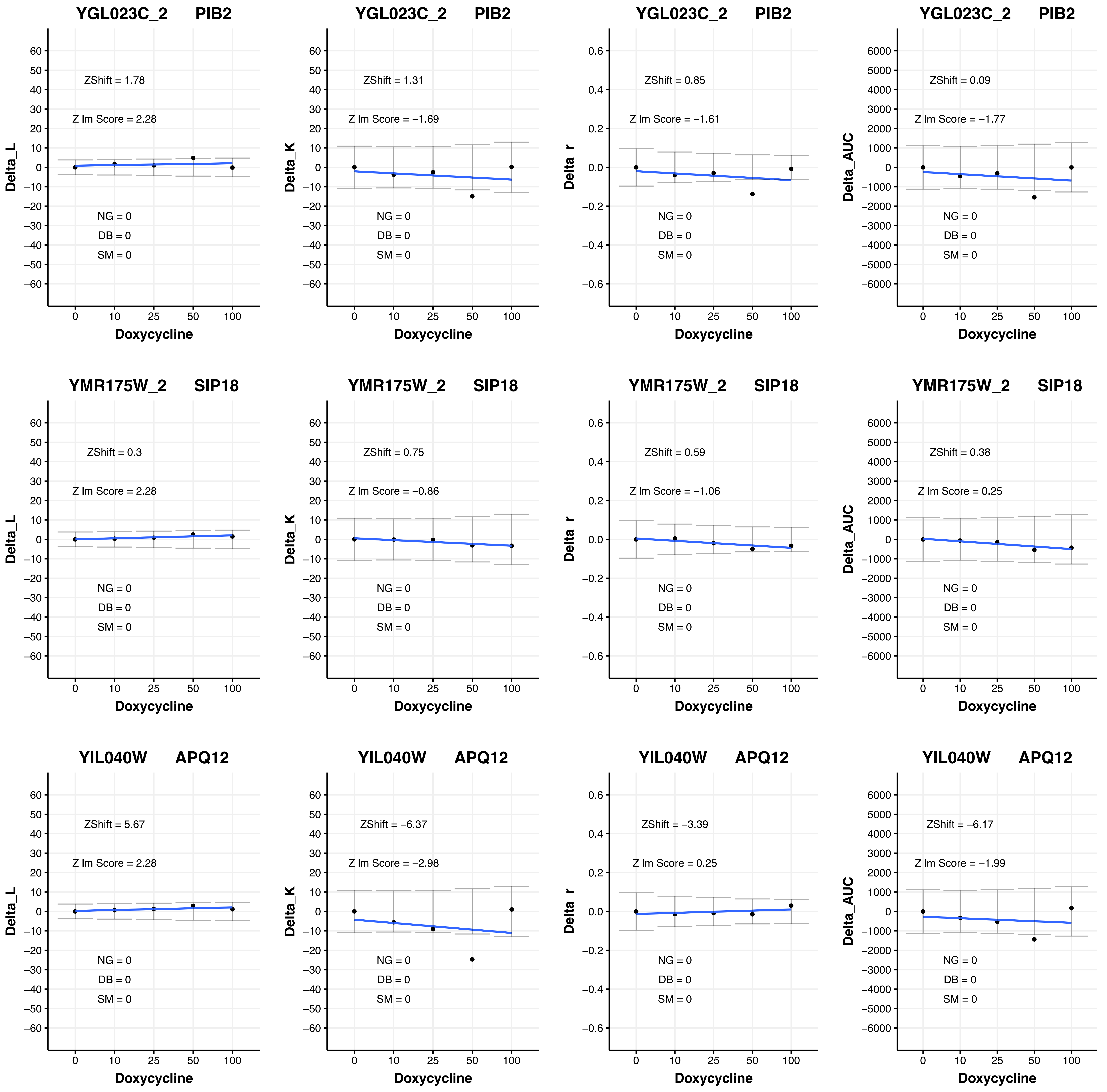

YGL023C_2 PIB2
ZShift = 1.78
Z lm Score = 2.28
NG = 0
DB = 0
SM = 0
Delta_L
Doxycycline
YGL023C_2 PIB2
ZShift = 1.31
Z lm Score = −1.69
NG = 0
DB = 0
SM = 0
Delta_K
Doxycycline
YGL023C_2 PIB2
ZShift = 0.85
Z lm Score = −1.61
NG = 0
DB = 0
SM = 0
Delta_r
Doxycycline
YGL023C_2 PIB2
ZShift = 0.09
Z lm Score = −1.77
NG = 0
DB = 0
SM = 0
Delta_AUC
Doxycycline
YMR175W_2 SIP18
ZShift = 0.3
Z lm Score = 2.28
NG = 0
DB = 0
SM = 0
Delta_L
Doxycycline
YMR175W_2 SIP18
ZShift = 0.75
Z lm Score = −0.86
NG = 0
DB = 0
SM = 0
Delta_K
Doxycycline
YMR175W_2 SIP18
ZShift = 0.59
Z lm Score = −1.06
NG = 0
DB = 0
SM = 0
Delta_r
Doxycycline
YMR175W_2 SIP18
ZShift = 0.38
Z lm Score = 0.25
NG = 0
DB = 0
SM = 0
Delta_AUC
Doxycycline
YIL040W APQ12
ZShift = 5.67
Z lm Score = 2.28
NG = 0
DB = 0
SM = 0
Delta_L
Doxycycline
YIL040W APQ12
ZShift = −6.37
Z lm Score = −2.98
NG = 0
DB = 0
SM = 0
Delta_K
Doxycycline
YIL040W APQ12
ZShift = −3.39
Z lm Score = 0.25
NG = 0
DB = 0
SM = 0
Delta_r
Doxycycline
YIL040W APQ12
ZShift = −6.17
Z lm Score = −1.99
NG = 0
DB = 0
SM = 0
Delta_AUC
Doxycycline

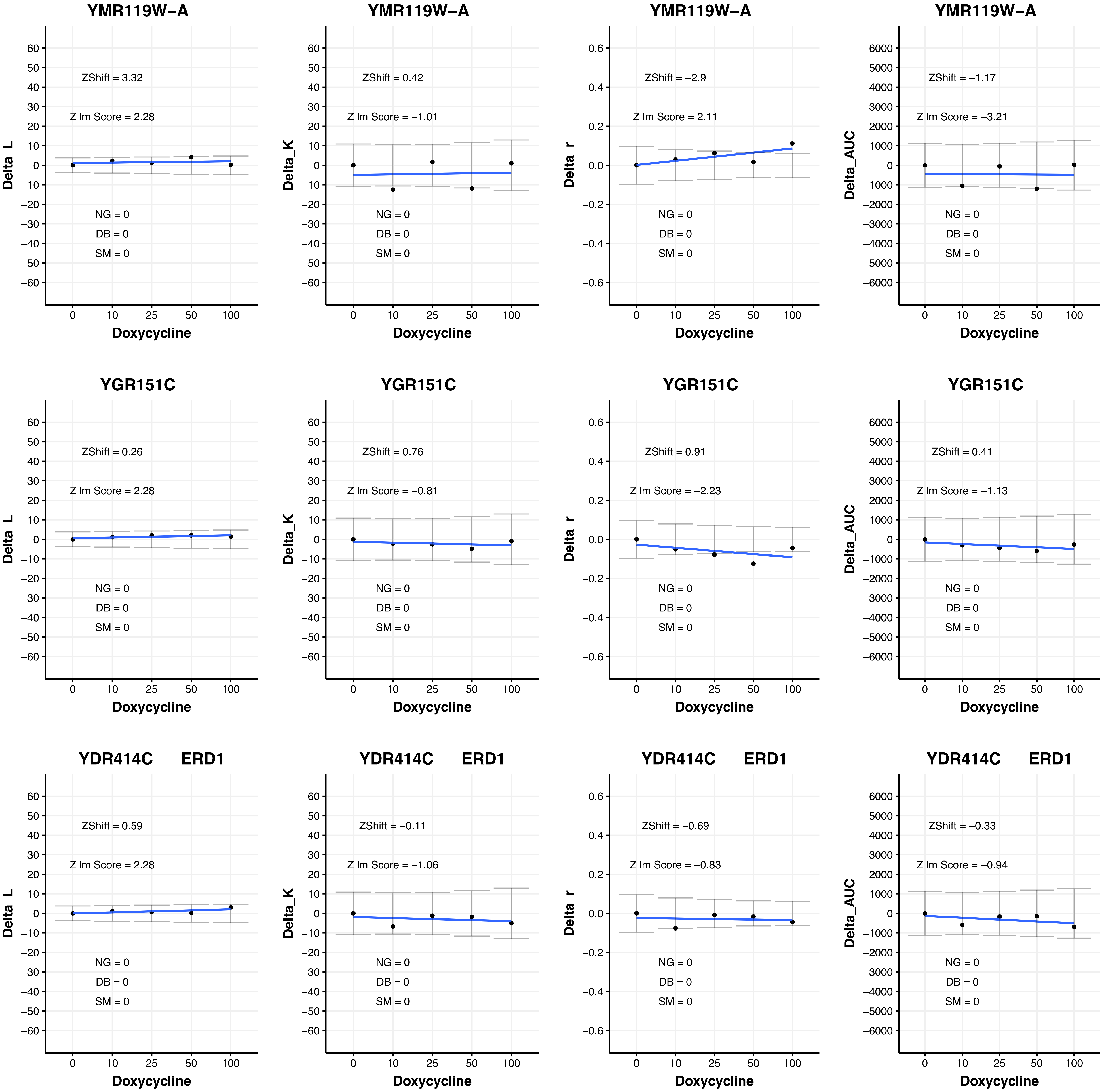
YMR119W−A
ZShift = 3.32
Z lm Score = 2.28
NG = 0
DB = 0
SM = 0
Delta_L
Doxycycline
YMR119W−A
ZShift = 0.42
Z lm Score = −1.01
NG = 0
DB = 0
SM = 0
Delta_K
Doxycycline
YMR119W−A
ZShift = −2.9
Z lm Score = 2.11
NG = 0
DB = 0
SM = 0
Delta_r
Doxycycline
YMR119W−A
ZShift = −1.17
Z lm Score = −3.21
NG = 0
DB = 0
SM = 0
Delta_AUC
Doxycycline
YGR151C
ZShift = 0.26
Z lm Score = 2.28
NG = 0
DB = 0
SM = 0
Delta_L
Doxycycline
YGR151C
ZShift = 0.76
Z lm Score = −0.81
NG = 0
DB = 0
SM = 0
Delta_K
Doxycycline
YGR151C
ZShift = 0.91
Z lm Score = −2.23
NG = 0
DB = 0
SM = 0
Delta_r
Doxycycline
YGR151C
ZShift = 0.41
Z lm Score = −1.13
NG = 0
DB = 0
SM = 0
Delta_AUC
Doxycycline
YDR414C ERD1
ZShift = 0.59
Z lm Score = 2.28
NG = 0
DB = 0
SM = 0
Delta_L
Doxycycline
YDR414C ERD1
ZShift = −0.11
Z lm Score = −1.06
NG = 0
DB = 0
SM = 0
Delta_K
Doxycycline
YDR414C ERD1
ZShift = −0.69
Z lm Score = −0.83
NG = 0
DB = 0
SM = 0
Delta_r
Doxycycline
YDR414C ERD1
ZShift = −0.33
Z lm Score = −0.94
NG = 0
DB = 0
SM = 0
Delta_AUC
Doxycycline

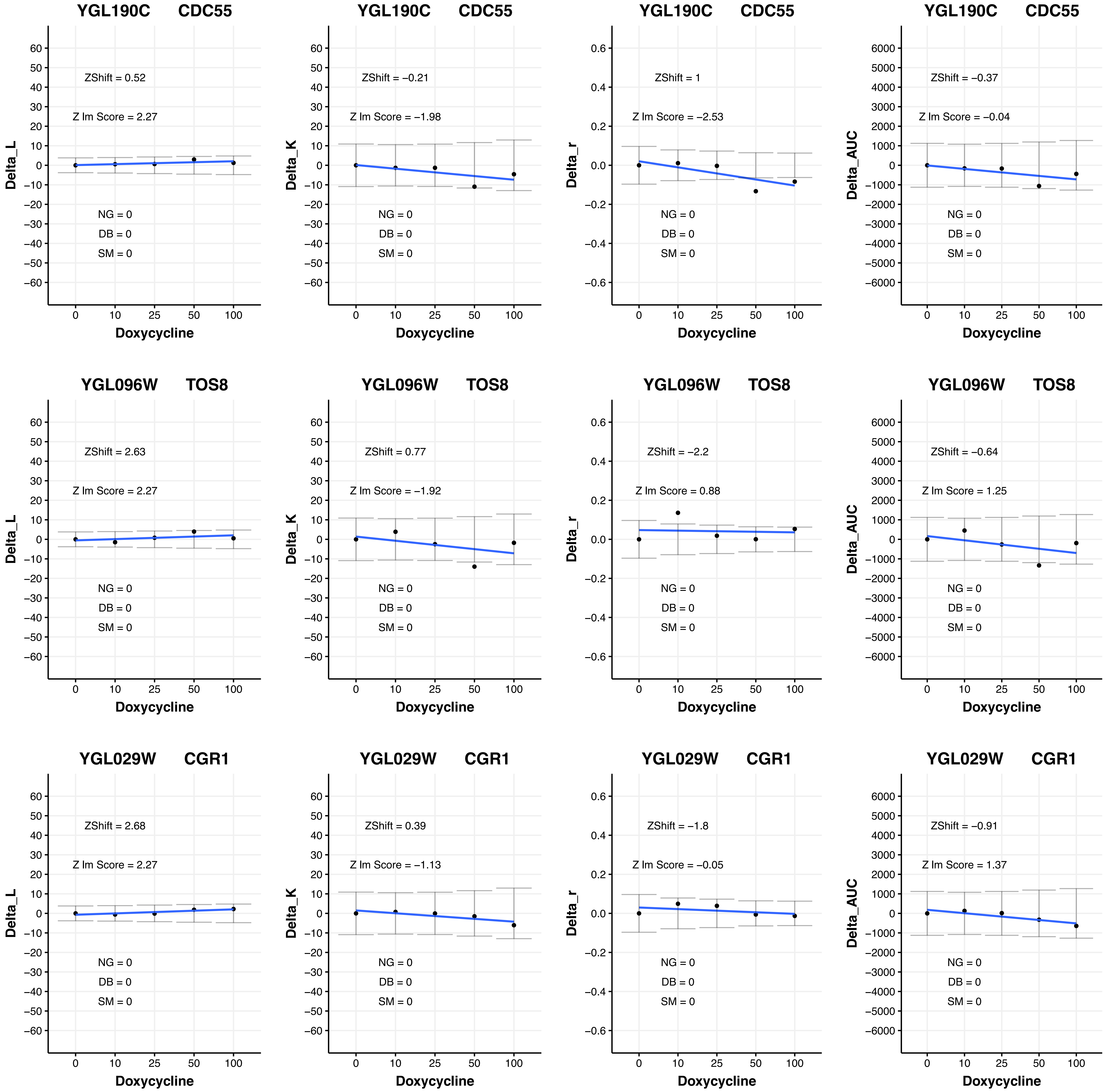
YGL190C CDC55
ZShift = 0.52
Z lm Score = 2.27
NG = 0
DB = 0
SM = 0
Delta_L
Doxycycline
YGL190C CDC55
ZShift = −0.21
Z lm Score = −1.98
NG = 0
DB = 0
SM = 0
Delta_K
Doxycycline
YGL190C CDC55
ZShift = 1
Z lm Score = −2.53
NG = 0
DB = 0
SM = 0
Delta_r
Doxycycline
YGL190C CDC55
ZShift = −0.37
Z lm Score = −0.04
NG = 0
DB = 0
SM = 0
Delta_AUC
Doxycycline
YGL096W TOS8
ZShift = 2.63
Z lm Score = 2.27
NG = 0
DB = 0
SM = 0
Delta_L
Doxycycline
YGL096W TOS8
ZShift = 0.77
Z lm Score = −1.92
NG = 0
DB = 0
SM = 0
Delta_K
Doxycycline
YGL096W TOS8
ZShift = −2.2
Z lm Score = 0.88
NG = 0
DB = 0
SM = 0
Delta_r
Doxycycline
YGL096W TOS8
ZShift = −0.64
Z lm Score = 1.25
NG = 0
DB = 0
SM = 0
Delta_AUC
Doxycycline
YGL029W CGR1
ZShift = 2.68
Z lm Score = 2.27
NG = 0
DB = 0
SM = 0
Delta_L
Doxycycline
YGL029W CGR1
ZShift = 0.39
Z lm Score = −1.13
NG = 0
DB = 0
SM = 0
Delta_K
Doxycycline
YGL029W CGR1
ZShift = −1.8
Z lm Score = −0.05
NG = 0
DB = 0
SM = 0
Delta_r
Doxycycline
YGL029W CGR1
ZShift = −0.91
Z lm Score = 1.37
NG = 0
DB = 0
SM = 0
Delta_AUC
Doxycycline

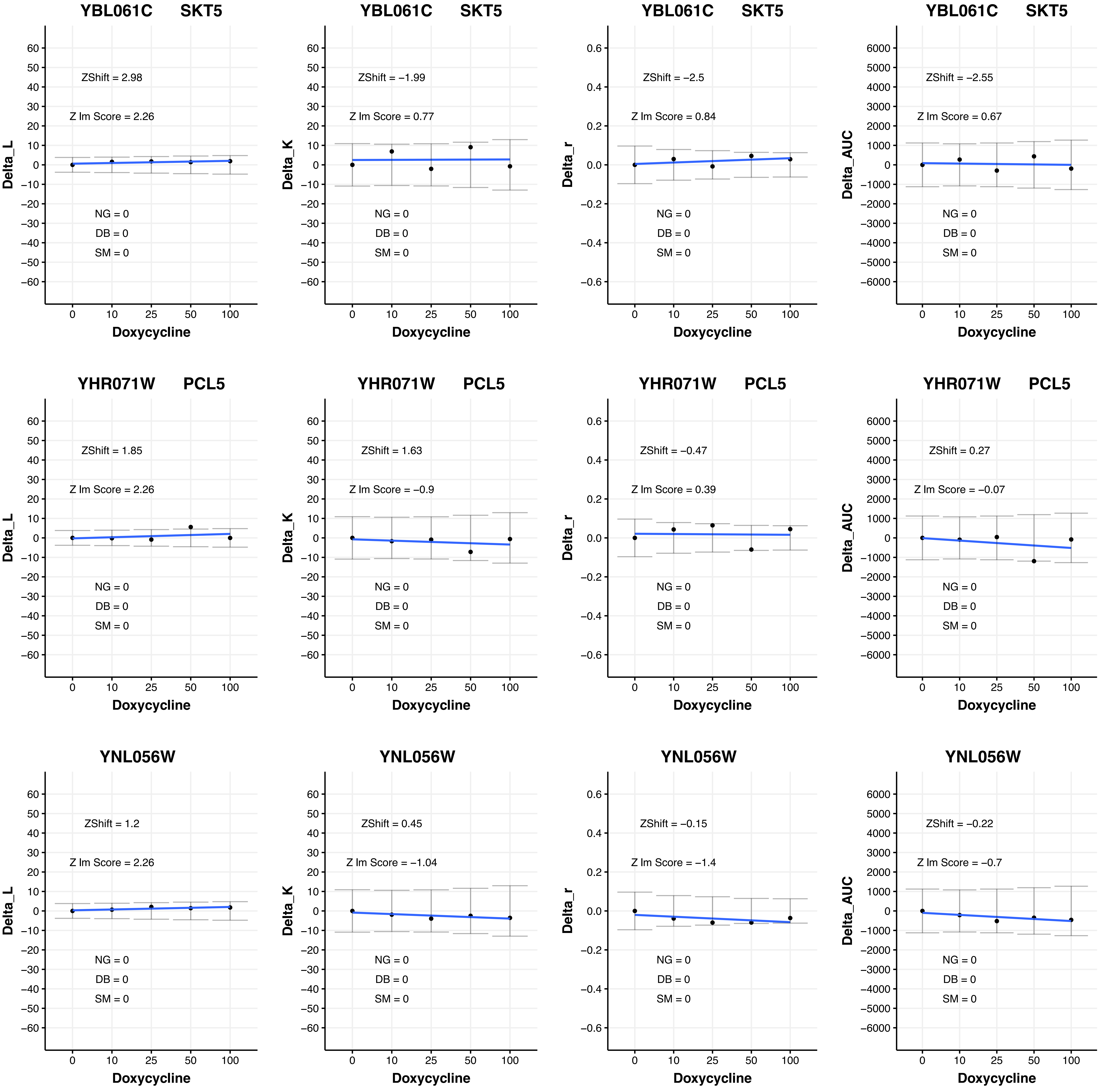

YBL061C SKT5
ZShift = 2.98
Z lm Score = 2.26
NG = 0
DB = 0
SM = 0
Delta_L
Doxycycline
YBL061C SKT5
ZShift = −1.99
Z lm Score = 0.77
NG = 0
DB = 0
SM = 0
Delta_K
Doxycycline
YBL061C SKT5
ZShift = −2.5
Z lm Score = 0.84
NG = 0
DB = 0
SM = 0
Delta_r
Doxycycline
YBL061C SKT5
ZShift = −2.55
Z lm Score = 0.67
NG = 0
DB = 0
SM = 0
Delta_AUC
Doxycycline
YHR071W PCL5
ZShift = 1.85
Z lm Score = 2.26
NG = 0
DB = 0
SM = 0
Delta_L
Doxycycline
YHR071W PCL5
ZShift = 1.63
Z lm Score = −0.9
NG = 0
DB = 0
SM = 0
Delta_K
Doxycycline
YHR071W PCL5
ZShift = −0.47
Z lm Score = 0.39
NG = 0
DB = 0
SM = 0
Delta_r
Doxycycline
YHR071W PCL5
ZShift = 0.27
Z lm Score = −0.07
NG = 0
DB = 0
SM = 0
Delta_AUC
Doxycycline
YNL056W
ZShift = 1.2
Z lm Score = 2.26
NG = 0
DB = 0
SM = 0
Delta_L
Doxycycline
YNL056W
ZShift = 0.45
Z lm Score = −1.04
NG = 0
DB = 0
SM = 0
Delta_K
Doxycycline
YNL056W
ZShift = −0.15
Z lm Score = −1.4
NG = 0
DB = 0
SM = 0
Delta_r
Doxycycline
YNL056W
ZShift = −0.22
Z lm Score = −0.7
NG = 0
DB = 0
SM = 0
Delta_AUC
Doxycycline

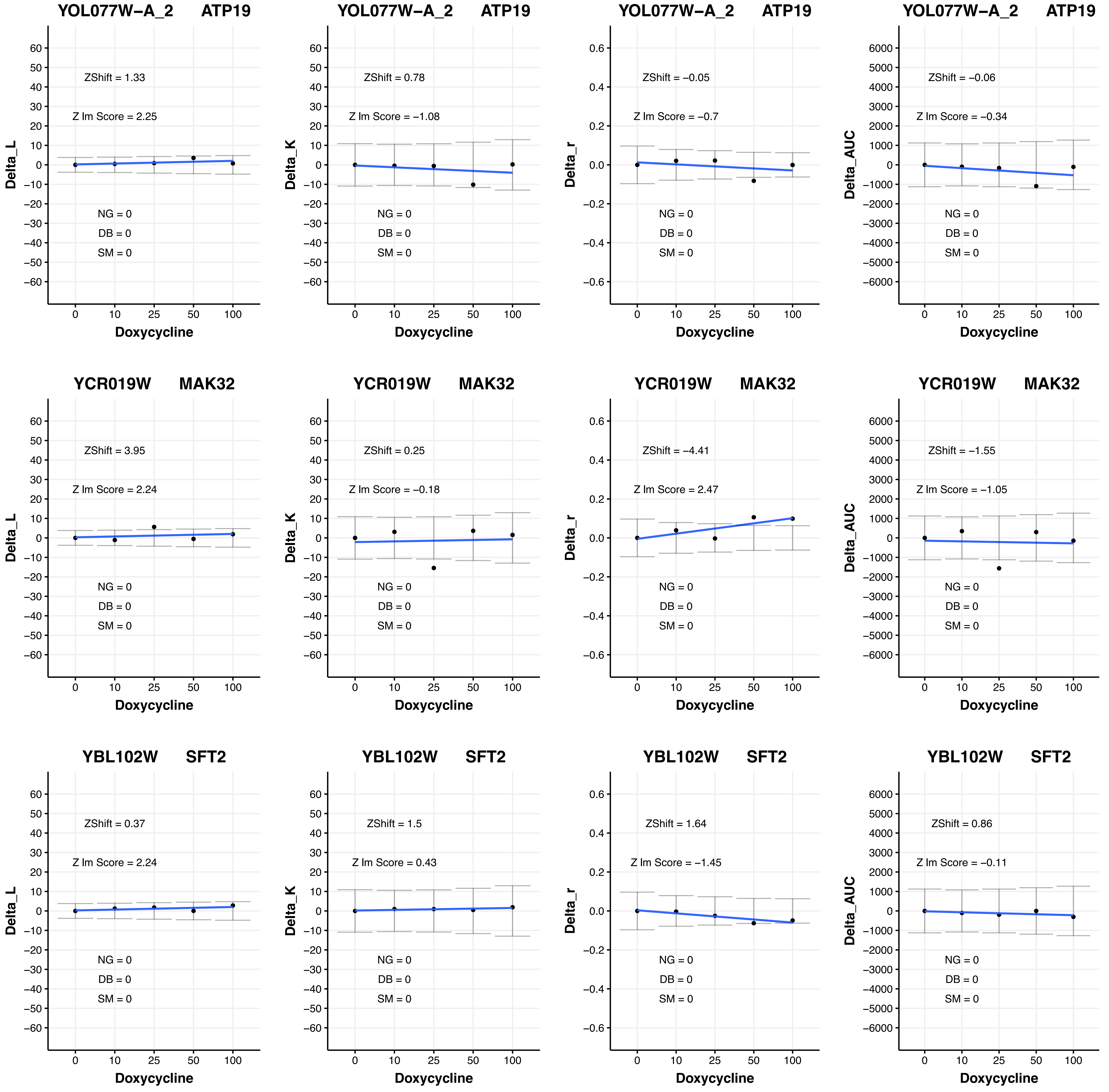

YOL077W−A_2 ATP19
ZShift = 1.33
Z lm Score = 2.25
NG = 0
DB = 0
SM = 0
Delta_L
Doxycycline
YOL077W−A_2 ATP19
ZShift = 0.78
Z lm Score = −1.08
NG = 0
DB = 0
SM = 0
Delta_K
Doxycycline
YOL077W−A_2 ATP19
ZShift = −0.05
Z lm Score = −0.7
NG = 0
DB = 0
SM = 0
Delta_r
Doxycycline
YOL077W−A_2 ATP19
ZShift = −0.06
Z lm Score = −0.34
NG = 0
DB = 0
SM = 0
Delta_AUC
Doxycycline
YCR019W MAK32
ZShift = 3.95
Z lm Score = 2.24
NG = 0
DB = 0
SM = 0
Delta_L
Doxycycline
YCR019W MAK32
ZShift = 0.25
Z lm Score = −0.18
NG = 0
DB = 0
SM = 0
Delta_K
Doxycycline
YCR019W MAK32
ZShift = −4.41
Z lm Score = 2.47
NG = 0
DB = 0
SM = 0
Delta_r
Doxycycline
YCR019W MAK32
ZShift = −1.55
Z lm Score = −1.05
NG = 0
DB = 0
SM = 0
Delta_AUC
Doxycycline
YBL102W SFT2
ZShift = 0.37
Z lm Score = 2.24
NG = 0
DB = 0
SM = 0
Delta_L
Doxycycline
YBL102W SFT2
ZShift = 1.5
Z lm Score = 0.43
NG = 0
DB = 0
SM = 0
Delta_K
Doxycycline
YBL102W SFT2
ZShift = 1.64
Z lm Score = −1.45
NG = 0
DB = 0
SM = 0
Delta_r
Doxycycline
YBL102W SFT2
ZShift = 0.86
Z lm Score = −0.11
NG = 0
DB = 0
SM = 0
Delta_AUC
Doxycycline

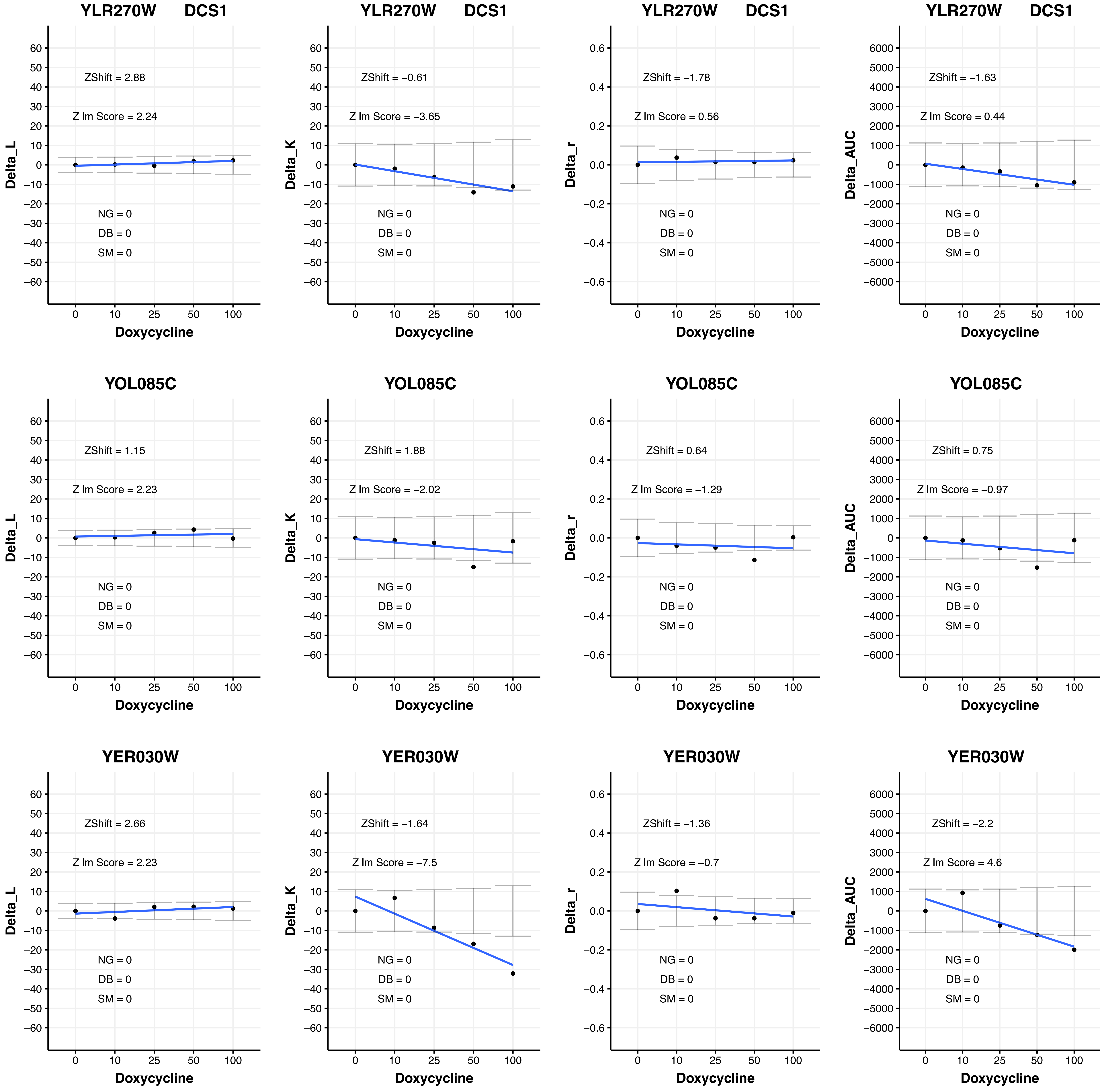

YLR270W DCS1
ZShift = 2.88
Z lm Score = 2.24
NG = 0
DB = 0
SM = 0
Delta_L
Doxycycline
YLR270W DCS1
ZShift = −0.61
Z lm Score = −3.65
NG = 0
DB = 0
SM = 0
Delta_K
Doxycycline
YLR270W DCS1
ZShift = −1.78
Z lm Score = 0.56
NG = 0
DB = 0
SM = 0
Delta_r
Doxycycline
YLR270W DCS1
ZShift = −1.63
Z lm Score = 0.44
NG = 0
DB = 0
SM = 0
Delta_AUC
Doxycycline
YOL085C
ZShift = 1.15
Z lm Score = 2.23
NG = 0
DB = 0
SM = 0
Delta_L
Doxycycline
YOL085C
ZShift = 1.88
Z lm Score = −2.02
NG = 0
DB = 0
SM = 0
Delta_K
Doxycycline
YOL085C
ZShift = 0.64
Z lm Score = −1.29
NG = 0
DB = 0
SM = 0
Delta_r
Doxycycline
YOL085C
ZShift = 0.75
Z lm Score = −0.97
NG = 0
DB = 0
SM = 0
Delta_AUC
Doxycycline
YER030W
ZShift = 2.66
Z lm Score = 2.23
NG = 0
DB = 0
SM = 0
Delta_L
Doxycycline
YER030W
ZShift = −1.64
Z lm Score = −7.5
NG = 0
DB = 0
SM = 0
Delta_K
Doxycycline
YER030W
ZShift = −1.36
Z lm Score = −0.7
NG = 0
DB = 0
SM = 0
Delta_r
Doxycycline
YER030W
ZShift = −2.2
Z lm Score = 4.6
NG = 0
DB = 0
SM = 0
Delta_AUC
Doxycycline

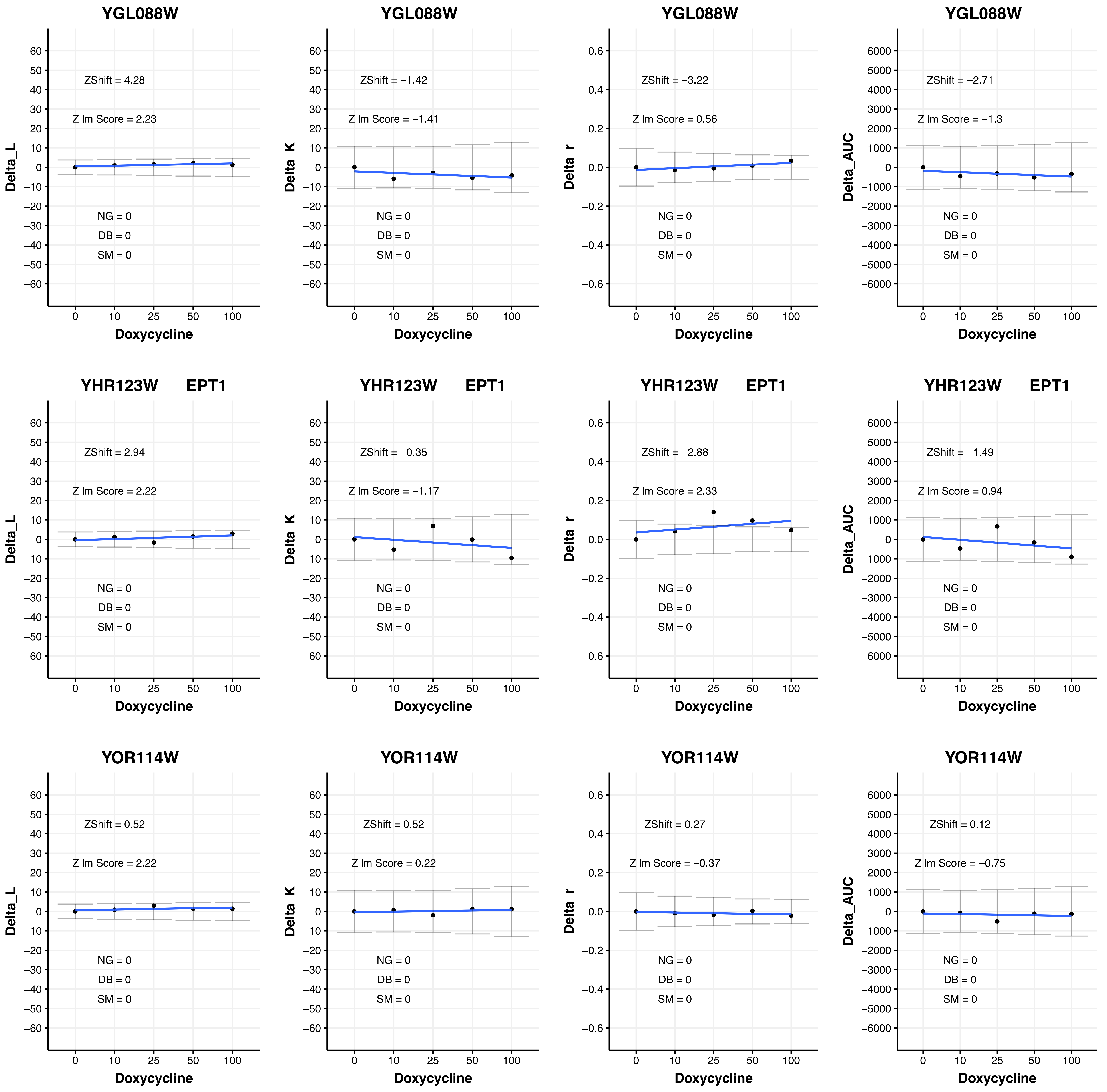
YGL088W
ZShift = 4.28
Z lm Score = 2.23
NG = 0
DB = 0
SM = 0
Delta_L
Doxycycline
YGL088W
ZShift = −1.42
Z lm Score = −1.41
NG = 0
DB = 0
SM = 0
Delta_K
Doxycycline
YGL088W
ZShift = −3.22
Z lm Score = 0.56
NG = 0
DB = 0
SM = 0
Delta_r
Doxycycline
YGL088W
ZShift = −2.71
Z lm Score = −1.3
NG = 0
DB = 0
SM = 0
Delta_AUC
Doxycycline
YHR123W EPT1
ZShift = 2.94
Z lm Score = 2.22
NG = 0
DB = 0
SM = 0
Delta_L
Doxycycline
YHR123W EPT1
ZShift = −0.35
Z lm Score = −1.17
NG = 0
DB = 0
SM = 0
Delta_K
Doxycycline
YHR123W EPT1
ZShift = −2.88
Z lm Score = 2.33
NG = 0
DB = 0
SM = 0
Delta_r
Doxycycline
YHR123W EPT1
ZShift = −1.49
Z lm Score = 0.94
NG = 0
DB = 0
SM = 0
Delta_AUC
Doxycycline
YOR114W
ZShift = 0.52
Z lm Score = 2.22
NG = 0
DB = 0
SM = 0
Delta_L
Doxycycline
YOR114W
ZShift = 0.52
Z lm Score = 0.22
NG = 0
DB = 0
SM = 0
Delta_K
Doxycycline
YOR114W
ZShift = 0.27
Z lm Score = −0.37
NG = 0
DB = 0
SM = 0
Delta_r
Doxycycline
YOR114W
ZShift = 0.12
Z lm Score = −0.75
NG = 0
DB = 0
SM = 0
Delta_AUC
Doxycycline

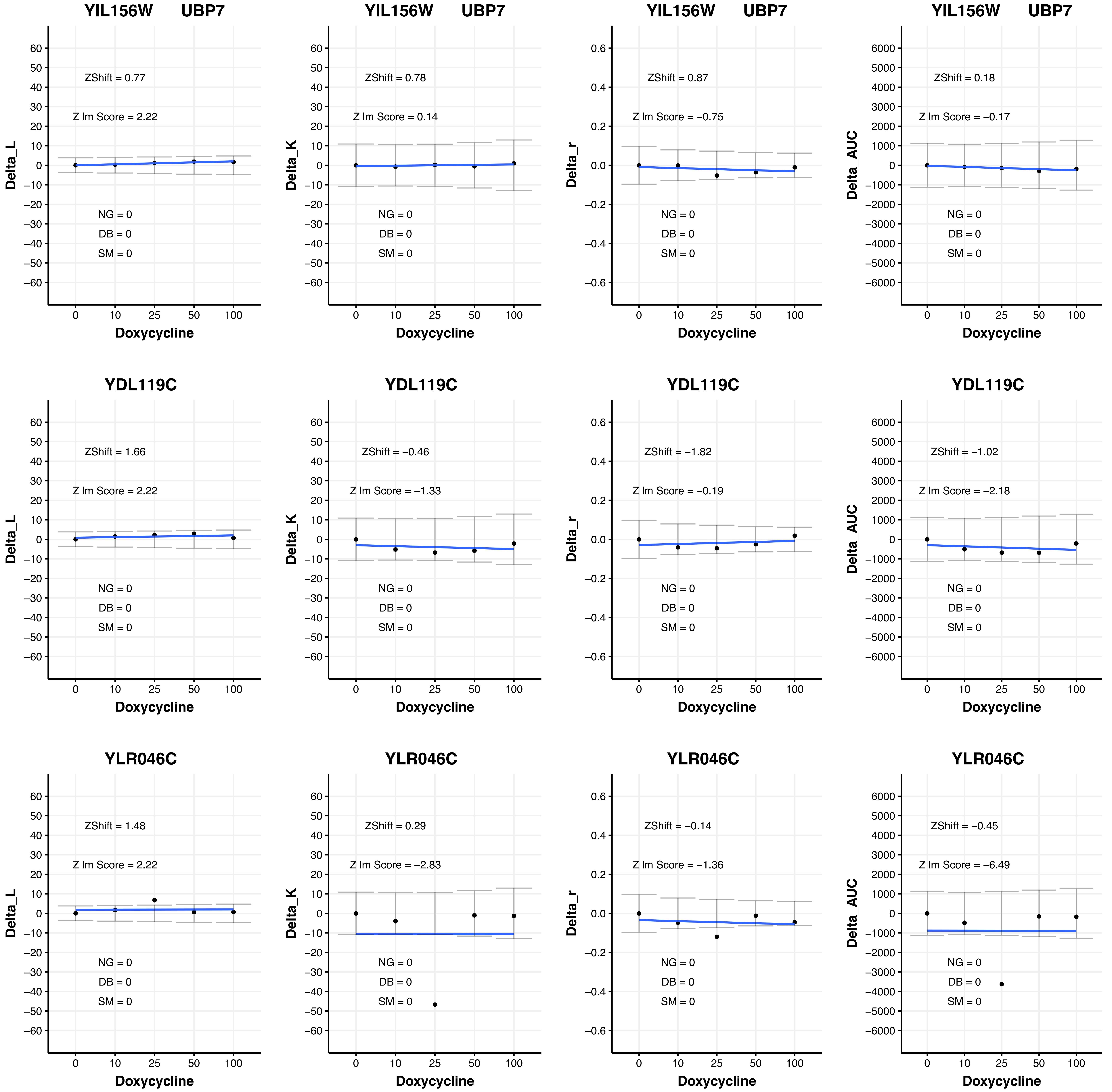

YIL156W UBP7
ZShift = 0.77
Z lm Score = 2.22
NG = 0
DB = 0
SM = 0
Delta_L
Doxycycline
YIL156W UBP7
ZShift = 0.78
Z lm Score = 0.14
NG = 0
DB = 0
SM = 0
Delta_K
Doxycycline
YIL156W UBP7
ZShift = 0.87
Z lm Score = −0.75
NG = 0
DB = 0
SM = 0
Delta_r
Doxycycline
YIL156W UBP7
ZShift = 0.18
Z lm Score = −0.17
NG = 0
DB = 0
SM = 0
Delta_AUC
Doxycycline
YDL119C
ZShift = 1.66
Z lm Score = 2.22
NG = 0
DB = 0
SM = 0
Delta_L
Doxycycline
YDL119C
ZShift = −0.46
Z lm Score = −1.33
NG = 0
DB = 0
SM = 0
Delta_K
Doxycycline
YDL119C
ZShift = −1.82
Z lm Score = −0.19
NG = 0
DB = 0
SM = 0
Delta_r
Doxycycline
YDL119C
ZShift = −1.02
Z lm Score = −2.18
NG = 0
DB = 0
SM = 0
Delta_AUC
Doxycycline
YLR046C
ZShift = 1.48
Z lm Score = 2.22
NG = 0
DB = 0
SM = 0
Delta_L
Doxycycline
YLR046C
ZShift = 0.29
Z lm Score = −2.83
NG = 0
DB = 0
SM = 0
Delta_K
Doxycycline
YLR046C
ZShift = −0.14
Z lm Score = −1.36
NG = 0
DB = 0
SM = 0
Delta_r
Doxycycline
YLR046C
ZShift = −0.45
Z lm Score = −6.49
NG = 0
DB = 0
SM = 0
Delta_AUC
Doxycycline

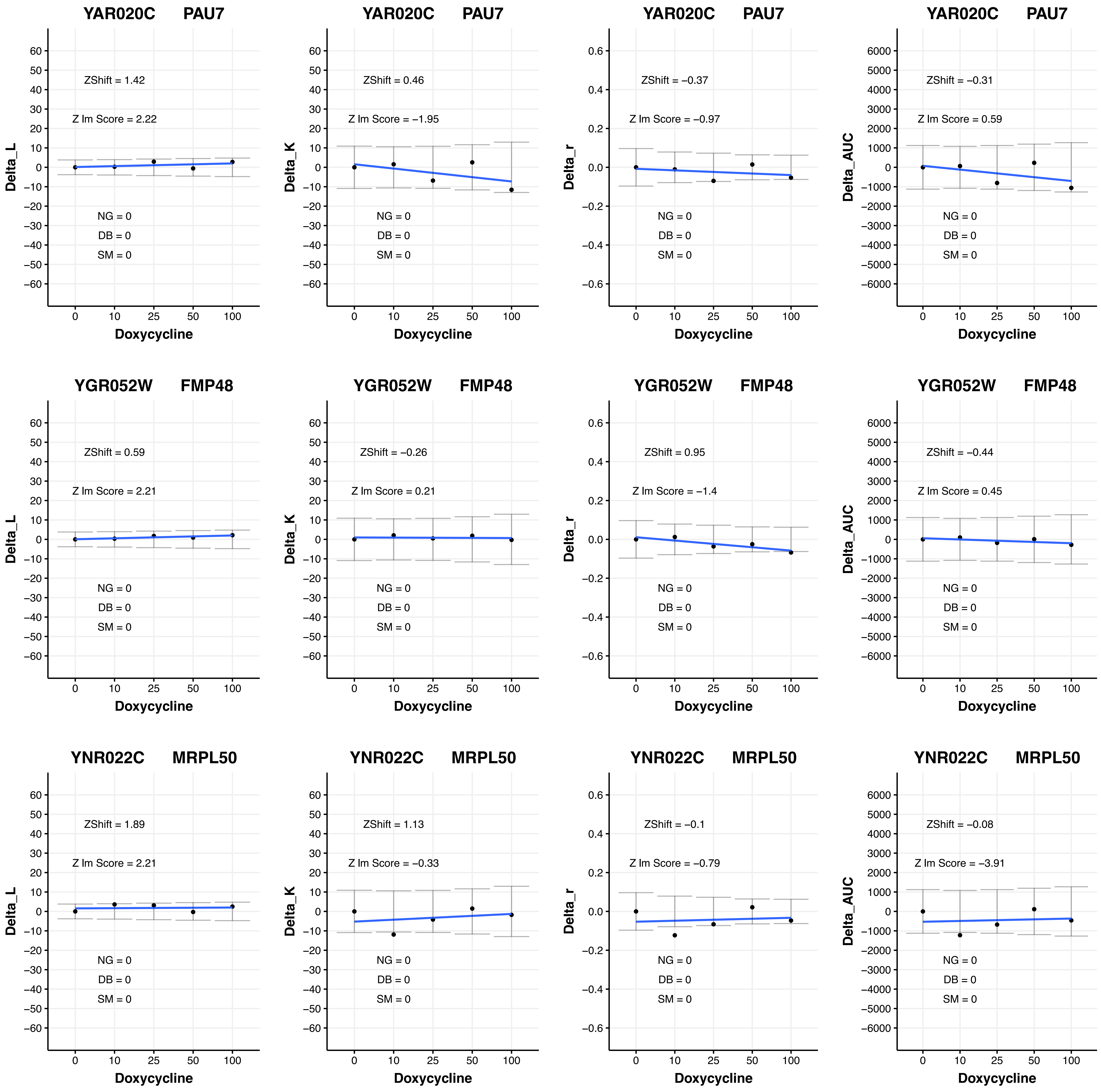

YAR020C PAU7
ZShift = 1.42
Z lm Score = 2.22
NG = 0
DB = 0
SM = 0
Delta_L
Doxycycline
YAR020C PAU7
ZShift = 0.46
Z lm Score = −1.95
NG = 0
DB = 0
SM = 0
Delta_K
Doxycycline
YAR020C PAU7
ZShift = −0.37
Z lm Score = −0.97
NG = 0
DB = 0
SM = 0
Delta_r
Doxycycline
YAR020C PAU7
ZShift = −0.31
Z lm Score = 0.59
NG = 0
DB = 0
SM = 0
Delta_AUC
Doxycycline
YGR052W FMP48
ZShift = 0.59
Z lm Score = 2.21
NG = 0
DB = 0
SM = 0
Delta_L
Doxycycline
YGR052W FMP48
ZShift = −0.26
Z lm Score = 0.21
NG = 0
DB = 0
SM = 0
Delta_K
Doxycycline
YGR052W FMP48
ZShift = 0.95
Z lm Score = −1.4
NG = 0
DB = 0
SM = 0
Delta_r
Doxycycline
YGR052W FMP48
ZShift = −0.44
Z lm Score = 0.45
NG = 0
DB = 0
SM = 0
Delta_AUC
Doxycycline
YNR022C MRPL50
ZShift = 1.89
Z lm Score = 2.21
NG = 0
DB = 0
SM = 0
Delta_L
Doxycycline
YNR022C MRPL50
ZShift = 1.13
Z lm Score = −0.33
NG = 0
DB = 0
SM = 0
Delta_K
Doxycycline
YNR022C MRPL50
ZShift = −0.1
Z lm Score = −0.79
NG = 0
DB = 0
SM = 0
Delta_r
Doxycycline
YNR022C MRPL50
ZShift = −0.08
Z lm Score = −3.91
NG = 0
DB = 0
SM = 0
Delta_AUC
Doxycycline

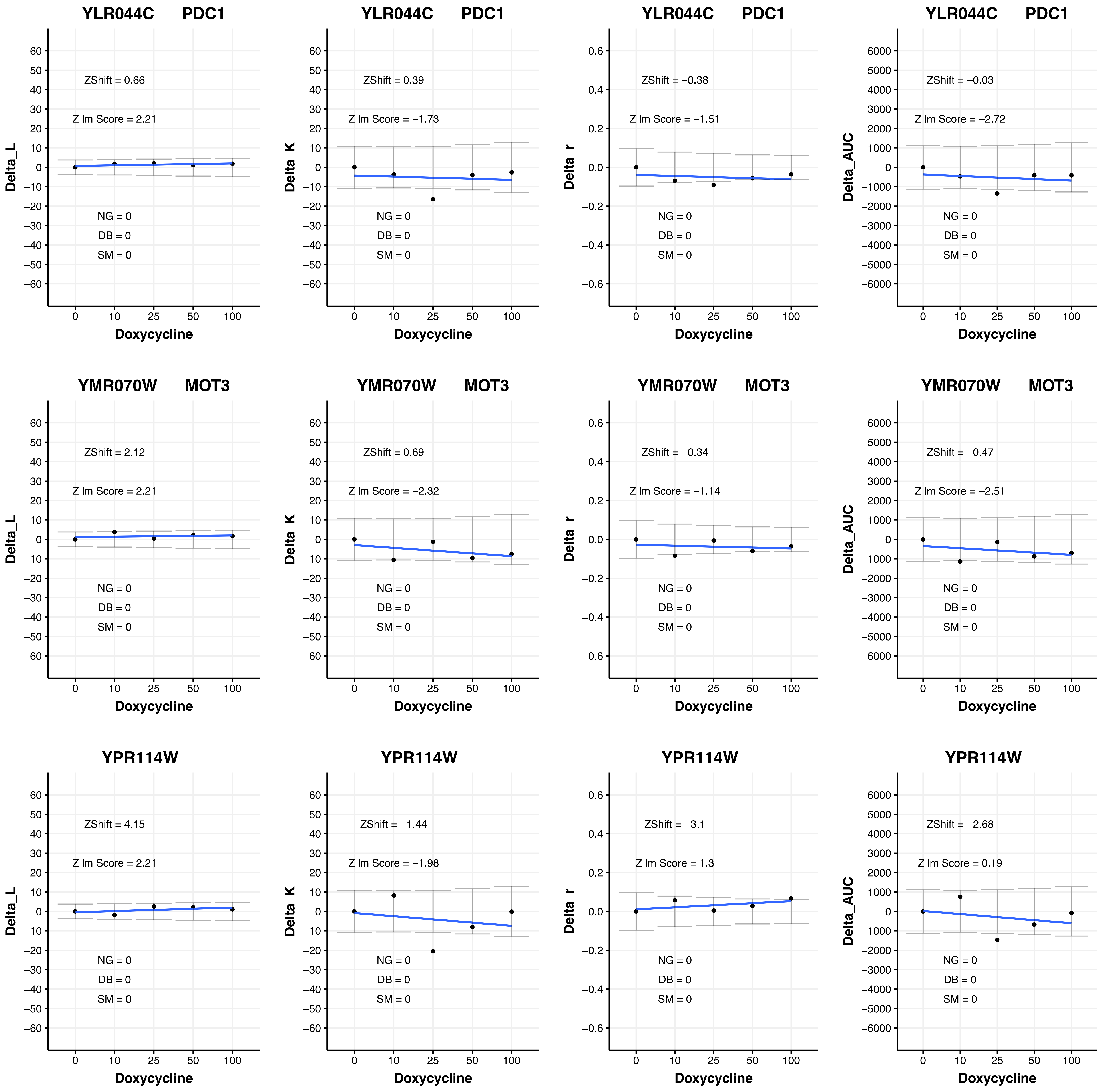
YLR044C PDC1
ZShift = 0.66
Z lm Score = 2.21
NG = 0
DB = 0
SM = 0
Delta_L
Doxycycline
YLR044C PDC1
ZShift = 0.39
Z lm Score = −1.73
NG = 0
DB = 0
SM = 0
Delta_K
Doxycycline
YLR044C PDC1
ZShift = −0.38
Z lm Score = −1.51
NG = 0
DB = 0
SM = 0
Delta_r
Doxycycline
YLR044C PDC1
ZShift = −0.03
Z lm Score = −2.72
NG = 0
DB = 0
SM = 0
Delta_AUC
Doxycycline
YMR070W MOT3
ZShift = 2.12
Z lm Score = 2.21
NG = 0
DB = 0
SM = 0
Delta_L
Doxycycline
YMR070W MOT3
ZShift = 0.69
Z lm Score = −2.32
NG = 0
DB = 0
SM = 0
Delta_K
Doxycycline
YMR070W MOT3
ZShift = −0.34
Z lm Score = −1.14
NG = 0
DB = 0
SM = 0
Delta_r
Doxycycline
YMR070W MOT3
ZShift = −0.47
Z lm Score = −2.51
NG = 0
DB = 0
SM = 0
Delta_AUC
Doxycycline
YPR114W
ZShift = 4.15
Z lm Score = 2.21
NG = 0
DB = 0
SM = 0
Delta_L
Doxycycline
YPR114W
ZShift = −1.44
Z lm Score = −1.98
NG = 0
DB = 0
SM = 0
Delta_K
Doxycycline
YPR114W
ZShift = −3.1
Z lm Score = 1.3
NG = 0
DB = 0
SM = 0
Delta_r
Doxycycline
YPR114W
ZShift = −2.68
Z lm Score = 0.19
NG = 0
DB = 0
SM = 0
Delta_AUC
Doxycycline

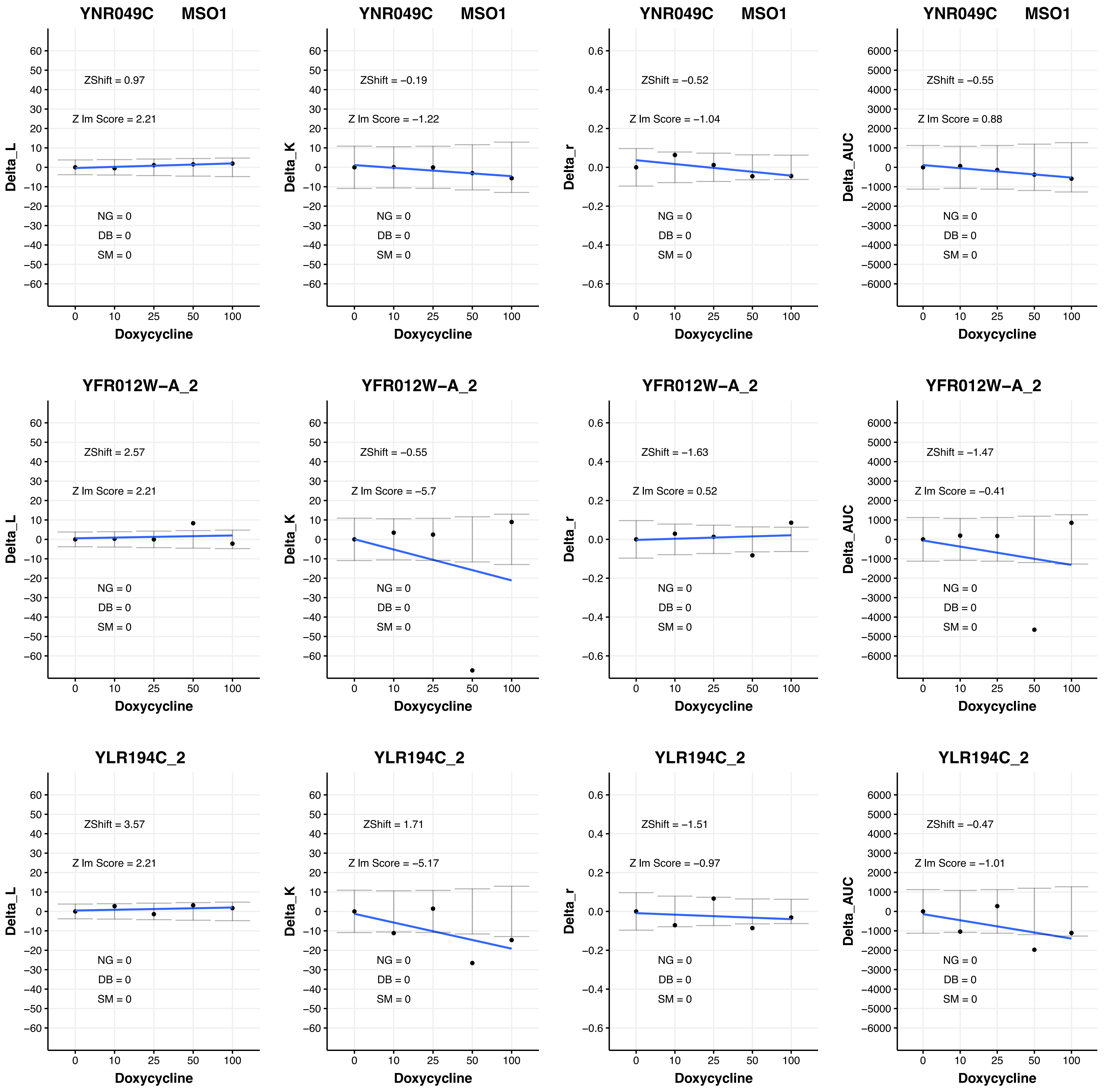

YNR049C MSO1
ZShift = 0.97
Z lm Score = 2.21
NG = 0
DB = 0
SM = 0
Delta_L
Doxycycline
YNR049C MSO1
ZShift = −0.19
Z lm Score = −1.22
NG = 0
DB = 0
SM = 0
Delta_K
Doxycycline
YNR049C MSO1
ZShift = −0.52
Z lm Score = −1.04
NG = 0
DB = 0
SM = 0
Delta_r
Doxycycline
YNR049C MSO1
ZShift = −0.55
Z lm Score = 0.88
NG = 0
DB = 0
SM = 0
Delta_AUC
Doxycycline
YFR012W−A_2
ZShift = 2.57
Z lm Score = 2.21
NG = 0
DB = 0
SM = 0
Delta_L
Doxycycline
YFR012W−A_2
ZShift = −0.55
Z lm Score = −5.7
NG = 0
DB = 0
SM = 0
Delta_K
Doxycycline
YFR012W−A_2
ZShift = −1.63
Z lm Score = 0.52
NG = 0
DB = 0
SM = 0
Delta_r
Doxycycline
YFR012W−A_2
ZShift = −1.47
Z lm Score = −0.41
NG = 0
DB = 0
SM = 0
Delta_AUC
Doxycycline
YLR194C_2
ZShift = 3.57
Z lm Score = 2.21
NG = 0
DB = 0
SM = 0
Delta_L
Doxycycline
YLR194C_2
ZShift = 1.71
Z lm Score = −5.17
NG = 0
DB = 0
SM = 0
Delta_K
Doxycycline
YLR194C_2
ZShift = −1.51
Z lm Score = −0.97
NG = 0
DB = 0
SM = 0
Delta_r
Doxycycline
YLR194C_2
ZShift = −0.47
Z lm Score = −1.01
NG = 0
DB = 0
SM = 0
Delta_AUC
Doxycycline

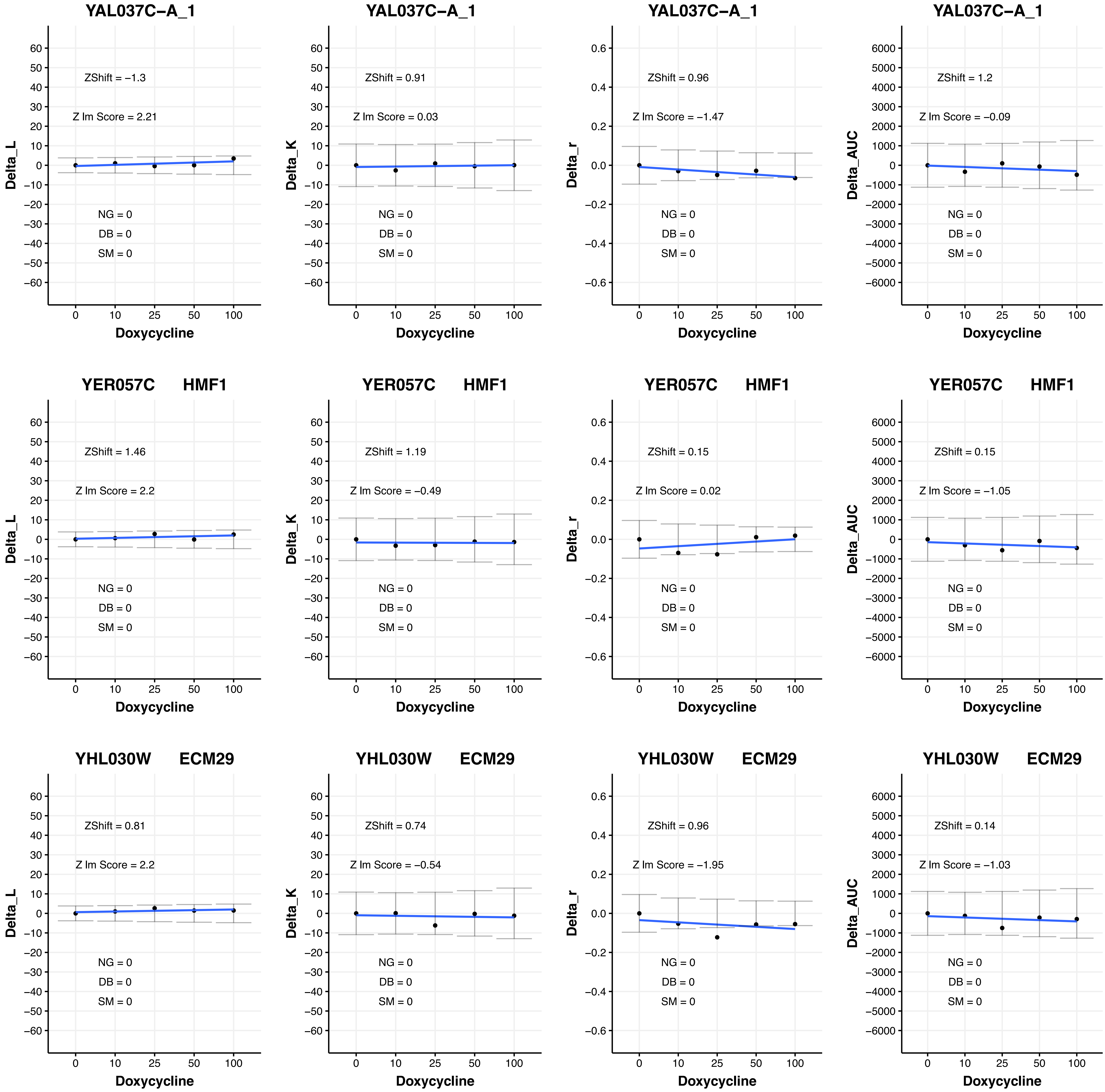

YAL037C−A_1
ZShift = −1.3
Z lm Score = 2.21
NG = 0
DB = 0
SM = 0
Delta_L
Doxycycline
YAL037C−A_1
ZShift = 0.91
Z lm Score = 0.03
NG = 0
DB = 0
SM = 0
Delta_K
Doxycycline
YAL037C−A_1
ZShift = 0.96
Z lm Score = −1.47
NG = 0
DB = 0
SM = 0
Delta_r
Doxycycline
YAL037C−A_1
ZShift = 1.2
Z lm Score = −0.09
NG = 0
DB = 0
SM = 0
Delta_AUC
Doxycycline
YER057C HMF1
ZShift = 1.46
Z lm Score = 2.2
NG = 0
DB = 0
SM = 0
Delta_L
Doxycycline
YER057C HMF1
ZShift = 1.19
Z lm Score = −0.49
NG = 0
DB = 0
SM = 0
Delta_K
Doxycycline
YER057C HMF1
ZShift = 0.15
Z lm Score = 0.02
NG = 0
DB = 0
SM = 0
Delta_r
Doxycycline
YER057C HMF1
ZShift = 0.15
Z lm Score = −1.05
NG = 0
DB = 0
SM = 0
Delta_AUC
Doxycycline
YHL030W ECM29
ZShift = 0.81
Z lm Score = 2.2
NG = 0
DB = 0
SM = 0
Delta_L
Doxycycline
YHL030W ECM29
ZShift = 0.74
Z lm Score = −0.54
NG = 0
DB = 0
SM = 0
Delta_K
Doxycycline
YHL030W ECM29
ZShift = 0.96
Z lm Score = −1.95
NG = 0
DB = 0
SM = 0
Delta_r
Doxycycline
YHL030W ECM29
ZShift = 0.14
Z lm Score = −1.03
NG = 0
DB = 0
SM = 0
Delta_AUC
Doxycycline

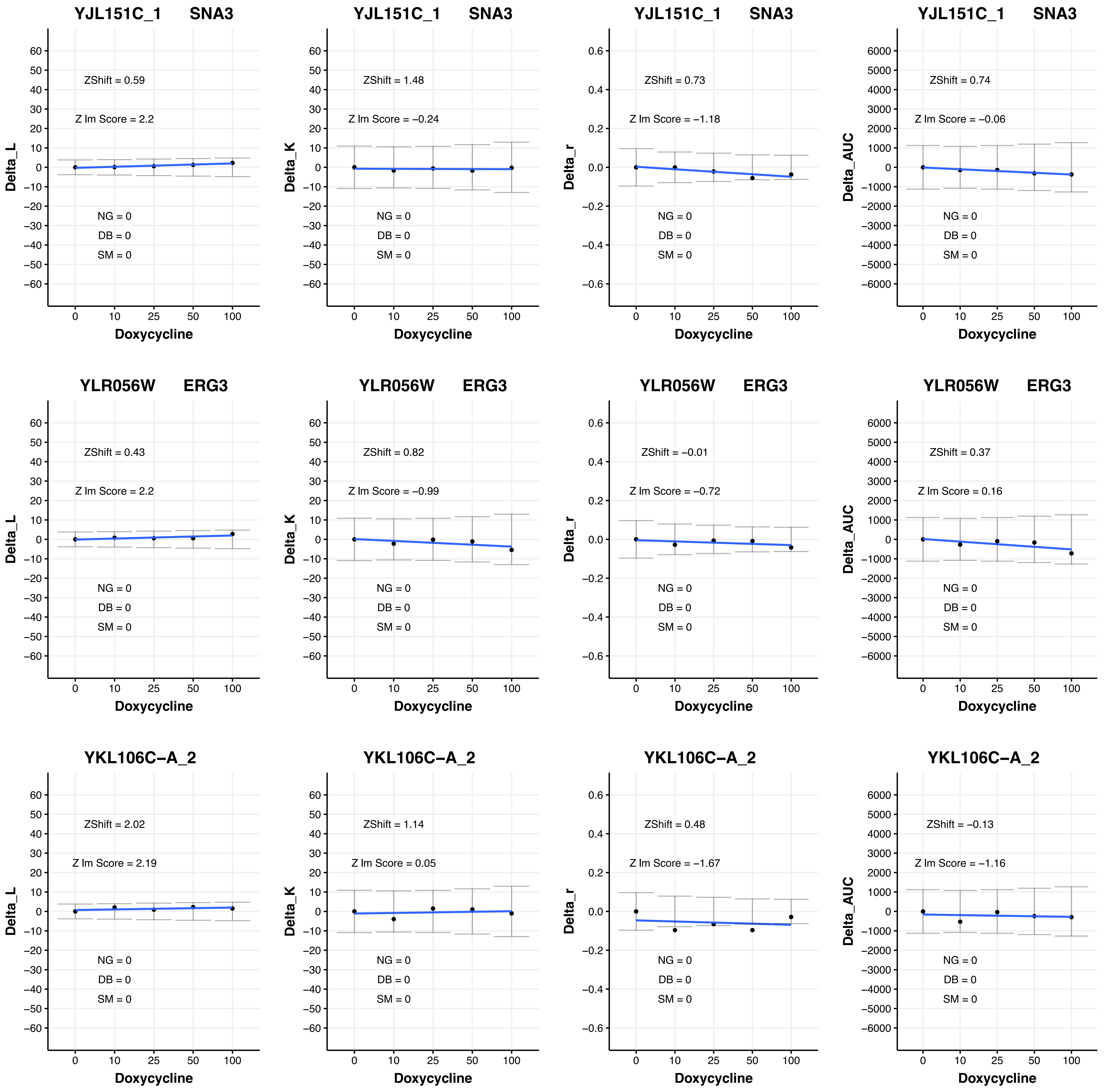

YJL151C_1 SNA3
ZShift = 0.59
Z lm Score = 2.2
Delta_L
NG = 0
DB = 0
SM = 0
Doxycycline
YJL151C_1 SNA3
ZShift = 1.48
Z lm Score = −0.24
Delta_K
NG = 0
DB = 0
SM = 0
Doxycycline
YJL151C_1 SNA3
ZShift = 0.73
Z lm Score = −1.18
Delta_r
NG = 0
DB = 0
SM = 0
Doxycycline
YJL151C_1 SNA3
ZShift = 0.74
Z lm Score = −0.06
Delta_AUC
NG = 0
DB = 0
SM = 0
Doxycycline
YLR056W ERG3
ZShift = 0.43
Z lm Score = 2.2
Delta_L
NG = 0
DB = 0
SM = 0
Doxycycline
YLR056W ERG3
ZShift = 0.82
Z lm Score = −0.99
Delta_K
NG = 0
DB = 0
SM = 0
Doxycycline
YLR056W ERG3
ZShift = −0.01
Z lm Score = −0.72
Delta_r
NG = 0
DB = 0
SM = 0
Doxycycline
YLR056W ERG3
ZShift = 0.37
Z lm Score = 0.16
Delta_AUC
NG = 0
DB = 0
SM = 0
Doxycycline
YKL106C−A_2
ZShift = 2.02
Z lm Score = 2.19
Delta_L
NG = 0
DB = 0
SM = 0
Doxycycline
YKL106C−A_2
ZShift = 1.14
Z lm Score = 0.05
Delta_K
NG = 0
DB = 0
SM = 0
Doxycycline
YKL106C−A_2
ZShift = 0.48
Z lm Score = −1.67
Delta_r
NG = 0
DB = 0
SM = 0
Doxycycline
YKL106C−A_2
ZShift = −0.13
Z lm Score = −1.16
Delta_AUC
NG = 0
DB = 0
SM = 0
Doxycycline

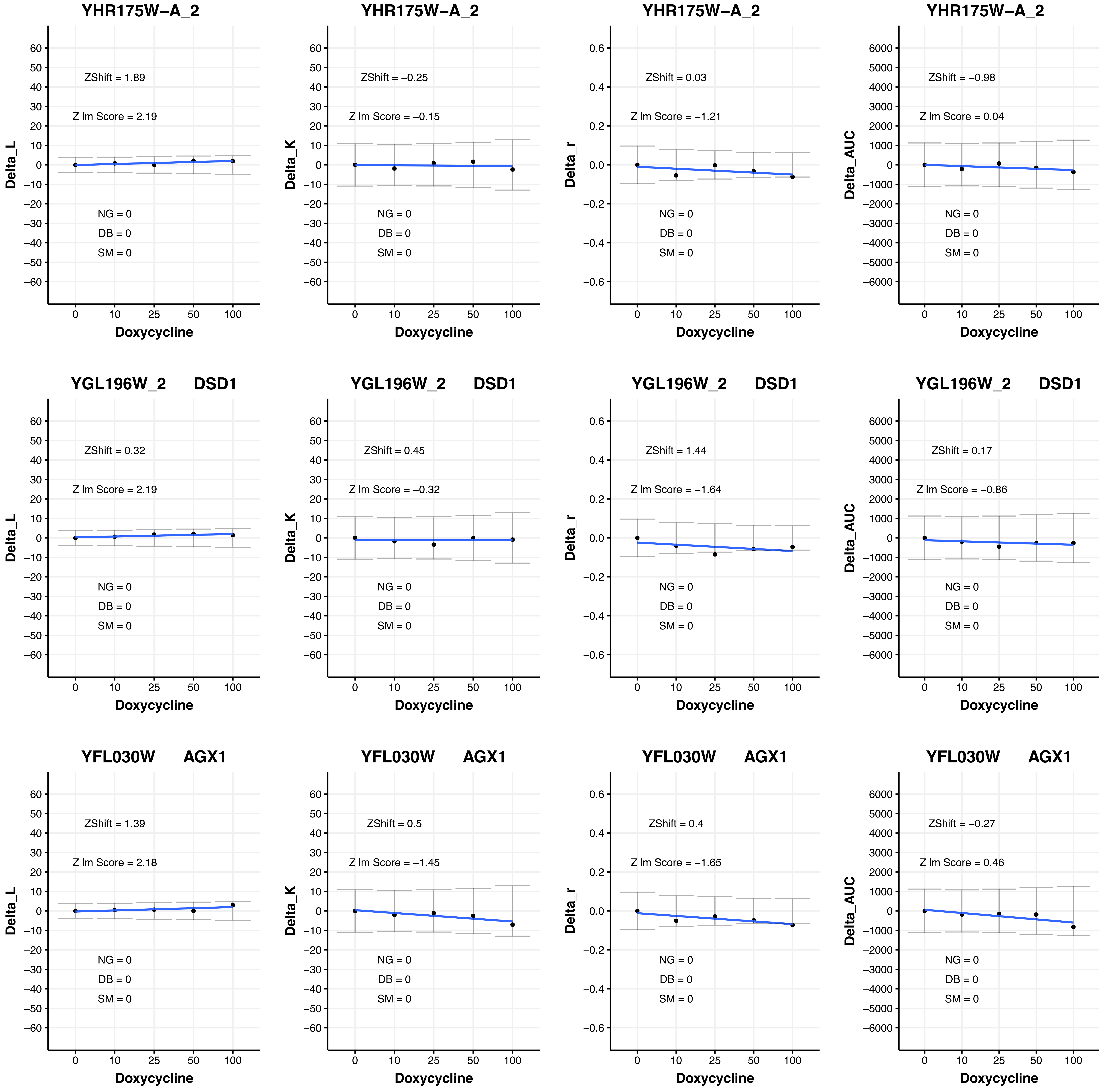
YHR175W−A_2
ZShift = 1.89
Z lm Score = 2.19
NG = 0
DB = 0
SM = 0
Delta_L
Doxycycline
YHR175W−A_2
ZShift = −0.25
Z lm Score = −0.15
NG = 0
DB = 0
SM = 0
Delta_K
Doxycycline
YHR175W−A_2
ZShift = 0.03
Z lm Score = −1.21
NG = 0
DB = 0
SM = 0
Delta_r
Doxycycline
YHR175W−A_2
ZShift = −0.98
Z lm Score = 0.04
NG = 0
DB = 0
SM = 0
Delta_AUC
Doxycycline
YGL196W_2 DSD1
ZShift = 0.32
Z lm Score = 2.19
NG = 0
DB = 0
SM = 0
Delta_L
Doxycycline
YGL196W_2 DSD1
ZShift = 0.45
Z lm Score = −0.32
NG = 0
DB = 0
SM = 0
Delta_K
Doxycycline
YGL196W_2 DSD1
ZShift = 1.44
Z lm Score = −1.64
NG = 0
DB = 0
SM = 0
Delta_r
Doxycycline
YGL196W_2 DSD1
ZShift = 0.17
Z lm Score = −0.86
NG = 0
DB = 0
SM = 0
Delta_AUC
Doxycycline
YFL030W AGX1
ZShift = 1.39
Z lm Score = 2.18
NG = 0
DB = 0
SM = 0
Delta_L
Doxycycline
YFL030W AGX1
ZShift = 0.5
Z lm Score = −1.45
NG = 0
DB = 0
SM = 0
Delta_K
Doxycycline
YFL030W AGX1
ZShift = 0.4
Z lm Score = −1.65
NG = 0
DB = 0
SM = 0
Delta_r
Doxycycline
YFL030W AGX1
ZShift = −0.27
Z lm Score = 0.46
NG = 0
DB = 0
SM = 0
Delta_AUC
Doxycycline

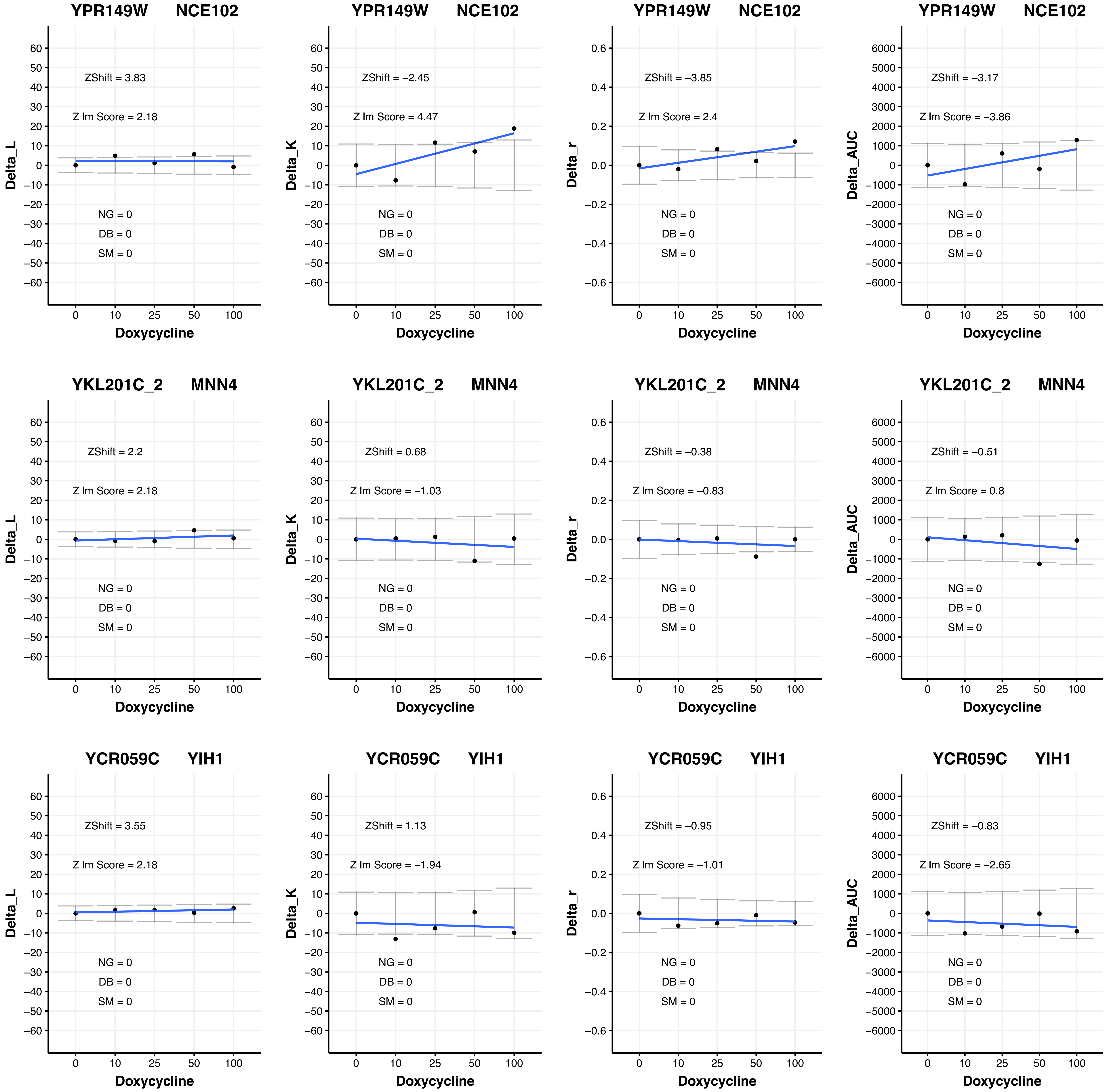

YPR149W NCE102
ZShift = 3.83
Z lm Score = 2.18
NG = 0
DB = 0
SM = 0
Delta_L
Doxycycline
YPR149W NCE102
ZShift = −2.45
Z lm Score = 4.47
NG = 0
DB = 0
SM = 0
Delta_K
Doxycycline
YPR149W NCE102
ZShift = −3.85
Z lm Score = 2.4
NG = 0
DB = 0
SM = 0
Delta_r
Doxycycline
YPR149W NCE102
ZShift = −3.17
Z lm Score = −3.86
NG = 0
DB = 0
SM = 0
Delta_AUC
Doxycycline
YKL201C_2 MNN4
ZShift = 2.2
Z lm Score = 2.18
NG = 0
DB = 0
SM = 0
Delta_L
Doxycycline
YKL201C_2 MNN4
ZShift = 0.68
Z lm Score = −1.03
NG = 0
DB = 0
SM = 0
Delta_K
Doxycycline
YKL201C_2 MNN4
ZShift = −0.38
Z lm Score = −0.83
NG = 0
DB = 0
SM = 0
Delta_r
Doxycycline
YKL201C_2 MNN4
ZShift = −0.51
Z lm Score = 0.8
NG = 0
DB = 0
SM = 0
Delta_AUC
Doxycycline
YCR059C YIH1
ZShift = 3.55
Z lm Score = 2.18
NG = 0
DB = 0
SM = 0
Delta_L
Doxycycline
YCR059C YIH1
ZShift = 1.13
Z lm Score = −1.94
NG = 0
DB = 0
SM = 0
Delta_K
Doxycycline
YCR059C YIH1
ZShift = −0.95
Z lm Score = −1.01
NG = 0
DB = 0
SM = 0
Delta_r
Doxycycline
YCR059C YIH1
ZShift = −0.83
Z lm Score = −2.65
NG = 0
DB = 0
SM = 0
Delta_AUC
Doxycycline

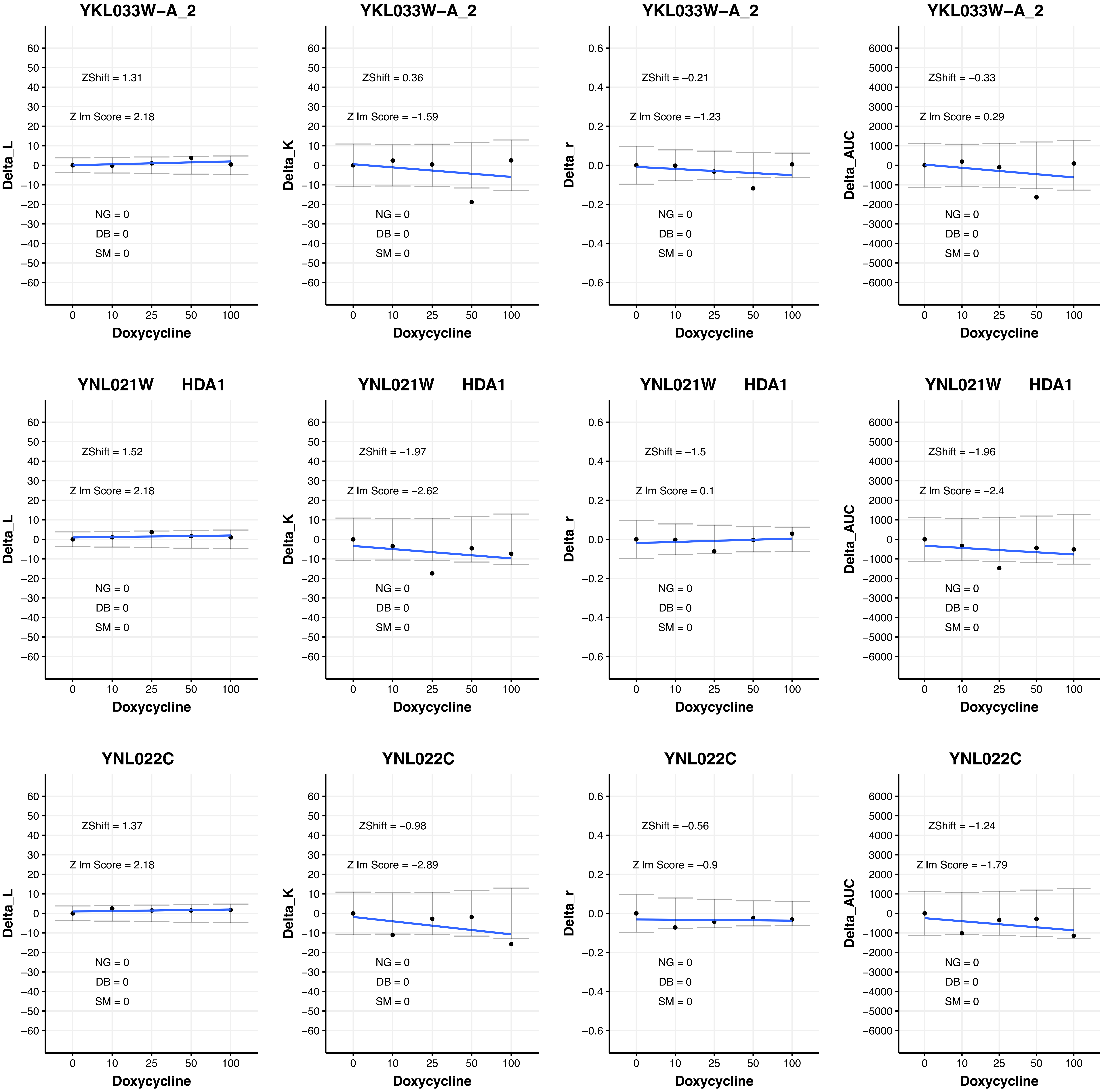

YKL033W−A_2
ZShift = 1.31
Z lm Score = 2.18
NG = 0
DB = 0
SM = 0
Delta_L
Doxycycline
YKL033W−A_2
ZShift = 0.36
Z lm Score = −1.59
NG = 0
DB = 0
SM = 0
Delta_K
Doxycycline
YKL033W−A_2
ZShift = −0.21
Z lm Score = −1.23
NG = 0
DB = 0
SM = 0
Delta_r
Doxycycline
YKL033W−A_2
ZShift = −0.33
Z lm Score = 0.29
NG = 0
DB = 0
SM = 0
Delta_AUC
Doxycycline
YNL021W HDA1
ZShift = 1.52
Z lm Score = 2.18
NG = 0
DB = 0
SM = 0
Delta_L
Doxycycline
YNL021W HDA1
ZShift = −1.97
Z lm Score = −2.62
NG = 0
DB = 0
SM = 0
Delta_K
Doxycycline
YNL021W HDA1
ZShift = −1.5
Z lm Score = 0.1
NG = 0
DB = 0
SM = 0
Delta_r
Doxycycline
YNL021W HDA1
ZShift = −1.96
Z lm Score = −2.4
NG = 0
DB = 0
SM = 0
Delta_AUC
Doxycycline
YNL022C
ZShift = 1.37
Z lm Score = 2.18
NG = 0
DB = 0
SM = 0
Delta_L
Doxycycline
YNL022C
ZShift = −0.98
Z lm Score = −2.89
NG = 0
DB = 0
SM = 0
Delta_K
Doxycycline
YNL022C
ZShift = −0.56
Z lm Score = −0.9
NG = 0
DB = 0
SM = 0
Delta_r
Doxycycline
YNL022C
ZShift = −1.24
Z lm Score = −1.79
NG = 0
DB = 0
SM = 0
Delta_AUC
Doxycycline

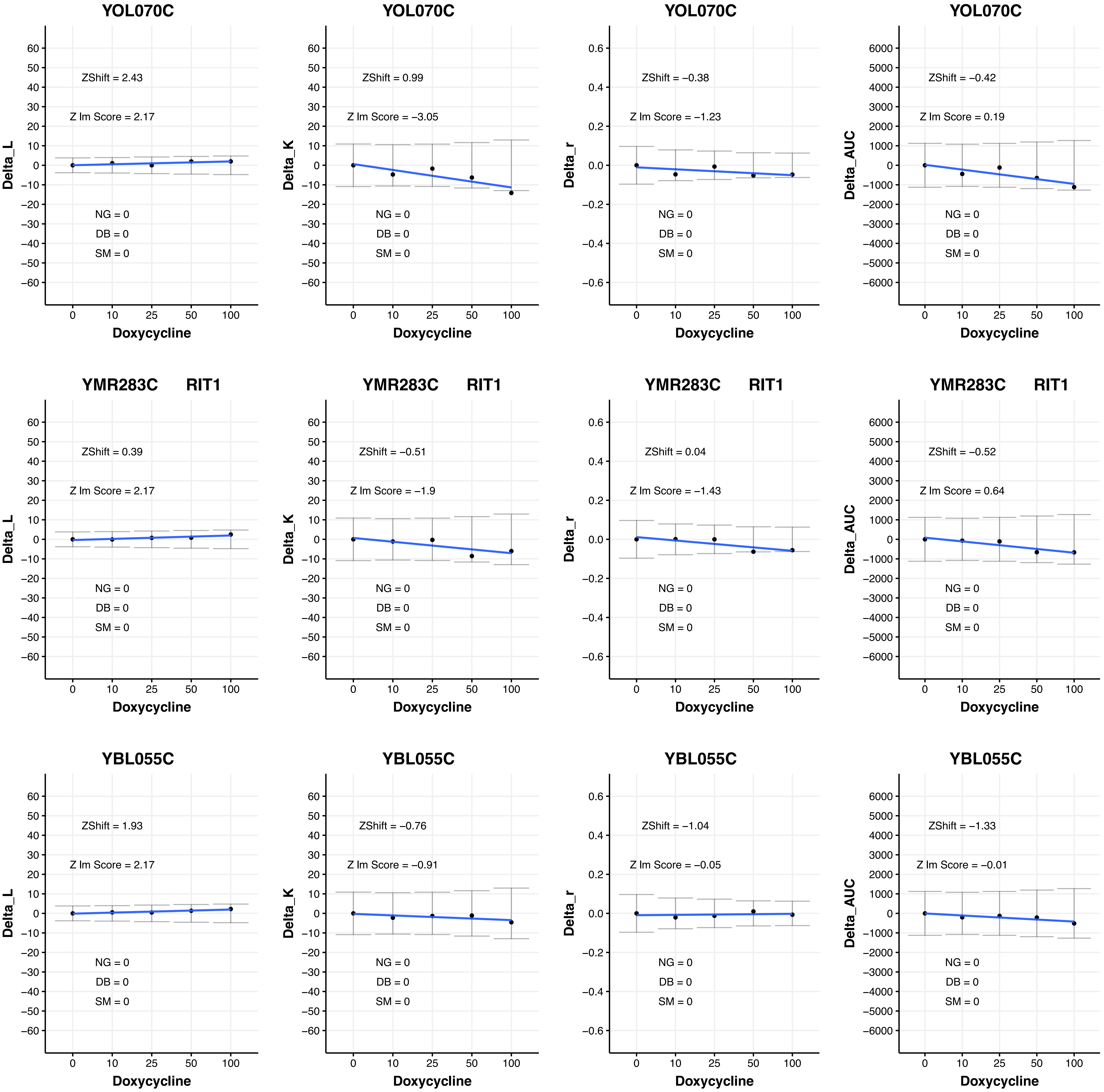
YOL070C
ZShift = 2.43
Z lm Score = 2.17
NG = 0
DB = 0
SM = 0
Delta_L
Doxycycline
YOL070C
ZShift = 0.99
Z lm Score = −3.05
NG = 0
DB = 0
SM = 0
Delta_K
Doxycycline
YOL070C
ZShift = −0.38
Z lm Score = −1.23
NG = 0
DB = 0
SM = 0
Delta_r
Doxycycline
YOL070C
ZShift = −0.42
Z lm Score = 0.19
NG = 0
DB = 0
SM = 0
Delta_AUC
Doxycycline
YMR283C RIT1
ZShift = 0.39
Z lm Score = 2.17
NG = 0
DB = 0
SM = 0
Delta_L
Doxycycline
YMR283C RIT1
ZShift = −0.51
Z lm Score = −1.9
NG = 0
DB = 0
SM = 0
Delta_K
Doxycycline
YMR283C RIT1
ZShift = 0.04
Z lm Score = −1.43
NG = 0
DB = 0
SM = 0
Delta_r
Doxycycline
YMR283C RIT1
ZShift = −0.52
Z lm Score = 0.64
NG = 0
DB = 0
SM = 0
Delta_AUC
Doxycycline
YBL055C
ZShift = 1.93
Z lm Score = 2.17
NG = 0
DB = 0
SM = 0
Delta_L
Doxycycline
YBL055C
ZShift = −0.76
Z lm Score = −0.91
NG = 0
DB = 0
SM = 0
Delta_K
Doxycycline
YBL055C
ZShift = −1.04
Z lm Score = −0.05
NG = 0
DB = 0
SM = 0
Delta_r
Doxycycline
YBL055C
ZShift = −1.33
Z lm Score = −0.01
NG = 0
DB = 0
SM = 0
Delta_AUC
Doxycycline

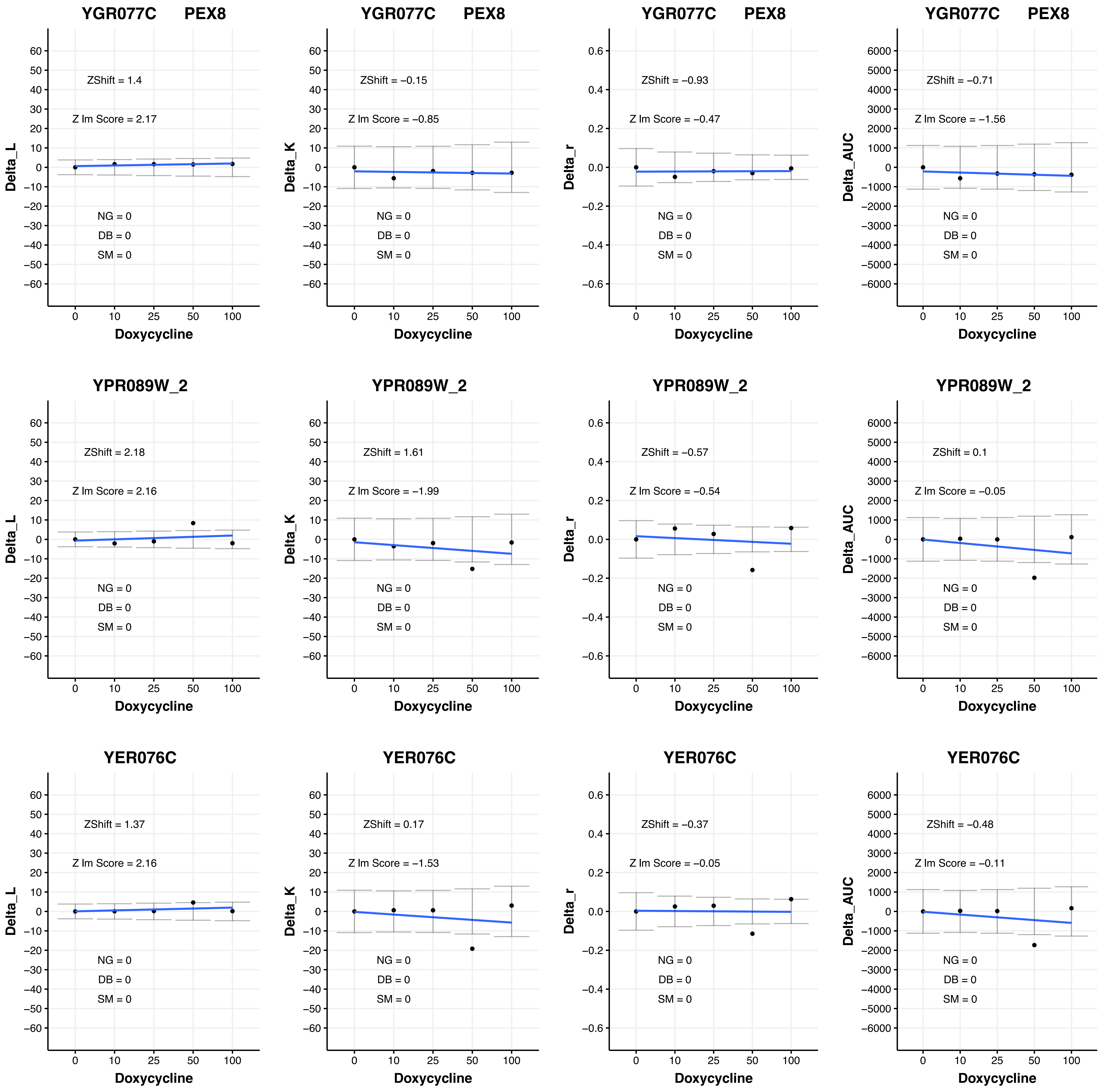
YGR077C PEX8
ZShift = 1.4
Z lm Score = 2.17
NG = 0
DB = 0
SM = 0
Delta_L
Doxycycline
YGR077C PEX8
ZShift = −0.15
Z lm Score = −0.85
NG = 0
DB = 0
SM = 0
Delta_K
Doxycycline
YGR077C PEX8
ZShift = −0.93
Z lm Score = −0.47
NG = 0
DB = 0
SM = 0
Delta_r
Doxycycline
YGR077C PEX8
ZShift = −0.71
Z lm Score = −1.56
NG = 0
DB = 0
SM = 0
Delta_AUC
Doxycycline
YPR089W_2
ZShift = 2.18
Z lm Score = 2.16
NG = 0
DB = 0
SM = 0
Delta_L
Doxycycline
YPR089W_2
ZShift = 1.61
Z lm Score = −1.99
NG = 0
DB = 0
SM = 0
Delta_K
Doxycycline
YPR089W_2
ZShift = −0.57
Z lm Score = −0.54
NG = 0
DB = 0
SM = 0
Delta_r
Doxycycline
YPR089W_2
ZShift = 0.1
Z lm Score = −0.05
NG = 0
DB = 0
SM = 0
Delta_AUC
Doxycycline
YER076C
ZShift = 1.37
Z lm Score = 2.16
NG = 0
DB = 0
SM = 0
Delta_L
Doxycycline
YER076C
ZShift = 0.17
Z lm Score = −1.53
NG = 0
DB = 0
SM = 0
Delta_K
Doxycycline
YER076C
ZShift = −0.37
Z lm Score = −0.05
NG = 0
DB = 0
SM = 0
Delta_r
Doxycycline
YER076C
ZShift = −0.48
Z lm Score = −0.11
NG = 0
DB = 0
SM = 0
Delta_AUC
Doxycycline
0
10
25
50
100

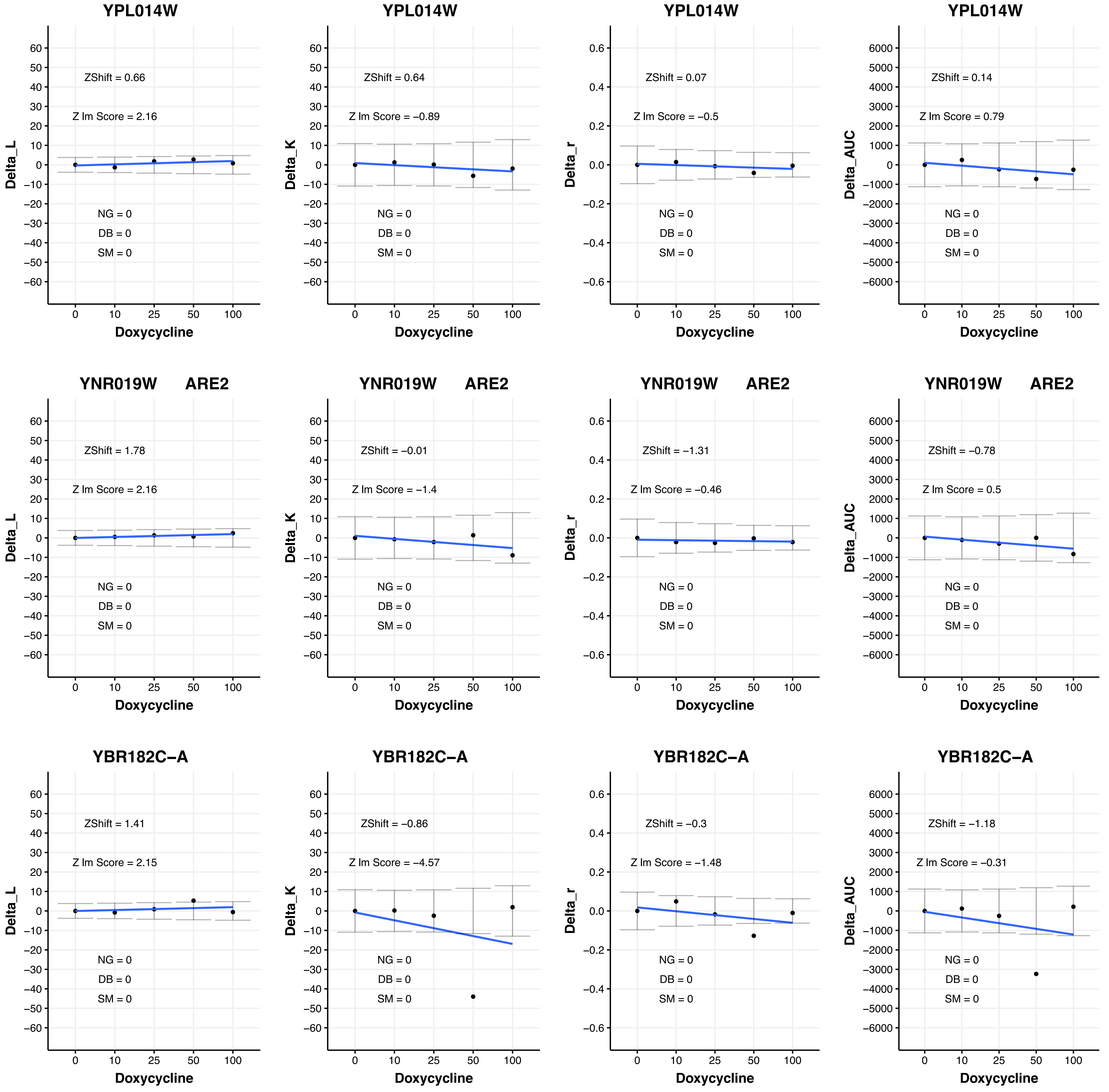
YPL014W
ZShift = 0.66
Z lm Score = 2.16
NG = 0
DB = 0
SM = 0
Delta_L
Doxycycline
YPL014W
ZShift = 0.64
Z lm Score = −0.89
NG = 0
DB = 0
SM = 0
Delta_K
Doxycycline
YPL014W
ZShift = 0.07
Z lm Score = −0.5
NG = 0
DB = 0
SM = 0
Delta_r
Doxycycline
YPL014W
ZShift = 0.14
Z lm Score = 0.79
NG = 0
DB = 0
SM = 0
Delta_AUC
Doxycycline
YNR019W ARE2
ZShift = 1.78
Z lm Score = 2.16
NG = 0
DB = 0
SM = 0
Delta_L
Doxycycline
YNR019W ARE2
ZShift = −0.01
Z lm Score = −1.4
NG = 0
DB = 0
SM = 0
Delta_K
Doxycycline
YNR019W ARE2
ZShift = −1.31
Z lm Score = −0.46
NG = 0
DB = 0
SM = 0
Delta_r
Doxycycline
YNR019W ARE2
ZShift = −0.78
Z lm Score = 0.5
NG = 0
DB = 0
SM = 0
Delta_AUC
Doxycycline
YBR182C−A
ZShift = 1.41
Z lm Score = 2.15
NG = 0
DB = 0
SM = 0
Delta_L
Doxycycline
YBR182C−A
ZShift = −0.86
Z lm Score = −4.57
NG = 0
DB = 0
SM = 0
Delta_K
Doxycycline
YBR182C−A
ZShift = −0.3
Z lm Score = −1.48
NG = 0
DB = 0
SM = 0
Delta_r
Doxycycline
YBR182C−A
ZShift = −1.18
Z lm Score = −0.31
NG = 0
DB = 0
SM = 0
Delta_AUC
Doxycycline

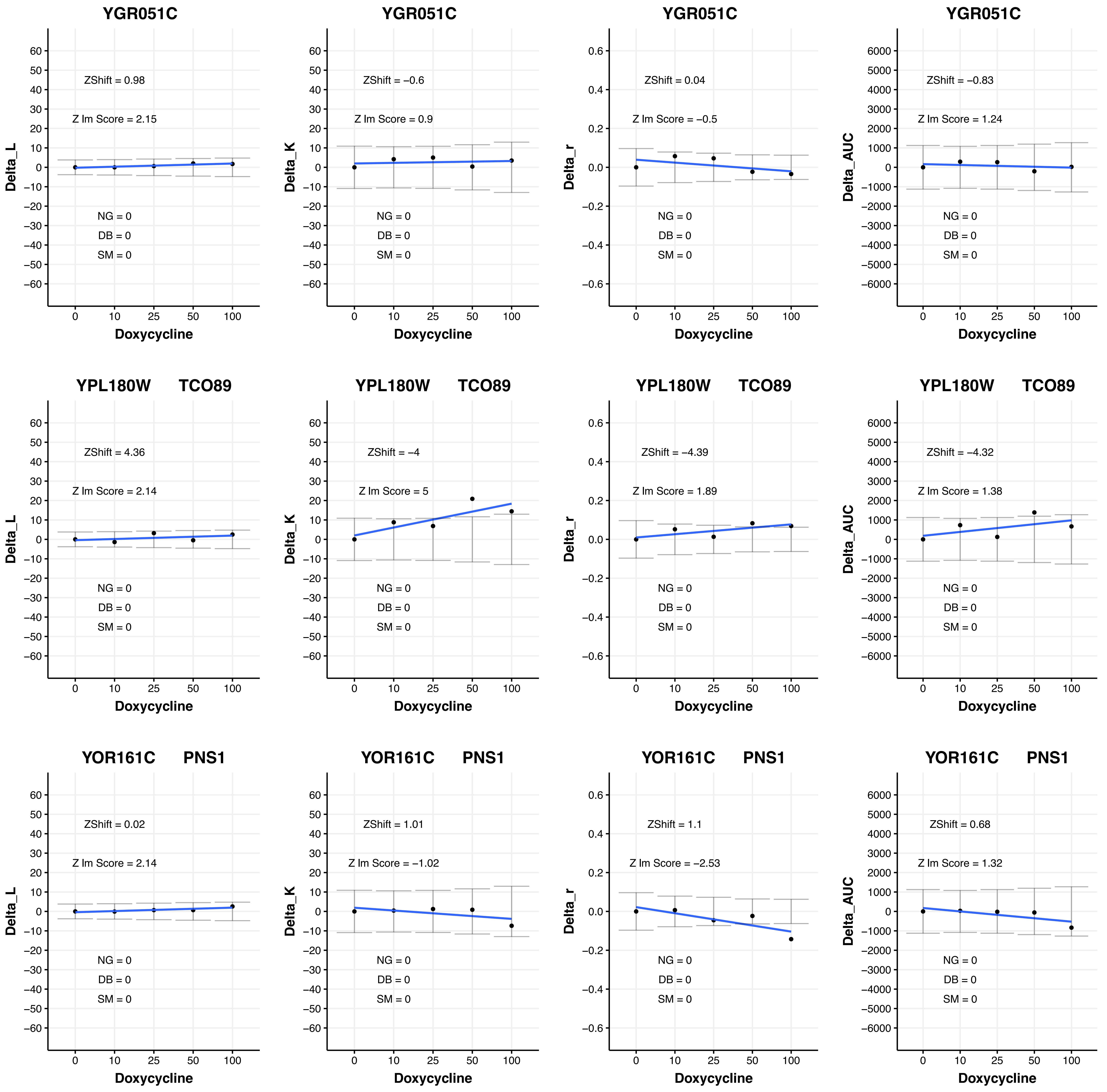
YGR051C
ZShift = 0.98
Z lm Score = 2.15
NG = 0
DB = 0
SM = 0
Delta_L
Doxycycline
YGR051C
ZShift = −0.6
Z lm Score = 0.9
NG = 0
DB = 0
SM = 0
Delta_K
Doxycycline
YGR051C
ZShift = 0.04
Z lm Score = −0.5
NG = 0
DB = 0
SM = 0
Delta_r
Doxycycline
YGR051C
ZShift = −0.83
Z lm Score = 1.24
NG = 0
DB = 0
SM = 0
Delta_AUC
Doxycycline
YPL180W TCO89
ZShift = 4.36
Z lm Score = 2.14
NG = 0
DB = 0
SM = 0
Delta_L
Doxycycline
YPL180W TCO89
ZShift = −4
Z lm Score = 5
NG = 0
DB = 0
SM = 0
Delta_K
Doxycycline
YPL180W TCO89
ZShift = −4.39
Z lm Score = 1.89
NG = 0
DB = 0
SM = 0
Delta_r
Doxycycline
YPL180W TCO89
ZShift = −4.32
Z lm Score = 1.38
NG = 0
DB = 0
SM = 0
Delta_AUC
Doxycycline
YOR161C PNS1
ZShift = 0.02
Z lm Score = 2.14
NG = 0
DB = 0
SM = 0
Delta_L
Doxycycline
YOR161C PNS1
ZShift = 1.01
Z lm Score = −1.02
NG = 0
DB = 0
SM = 0
Delta_K
Doxycycline
YOR161C PNS1
ZShift = 1.1
Z lm Score = −2.53
NG = 0
DB = 0
SM = 0
Delta_r
Doxycycline
YOR161C PNS1
ZShift = 0.68
Z lm Score = 1.32
NG = 0
DB = 0
SM = 0
Delta_AUC
Doxycycline

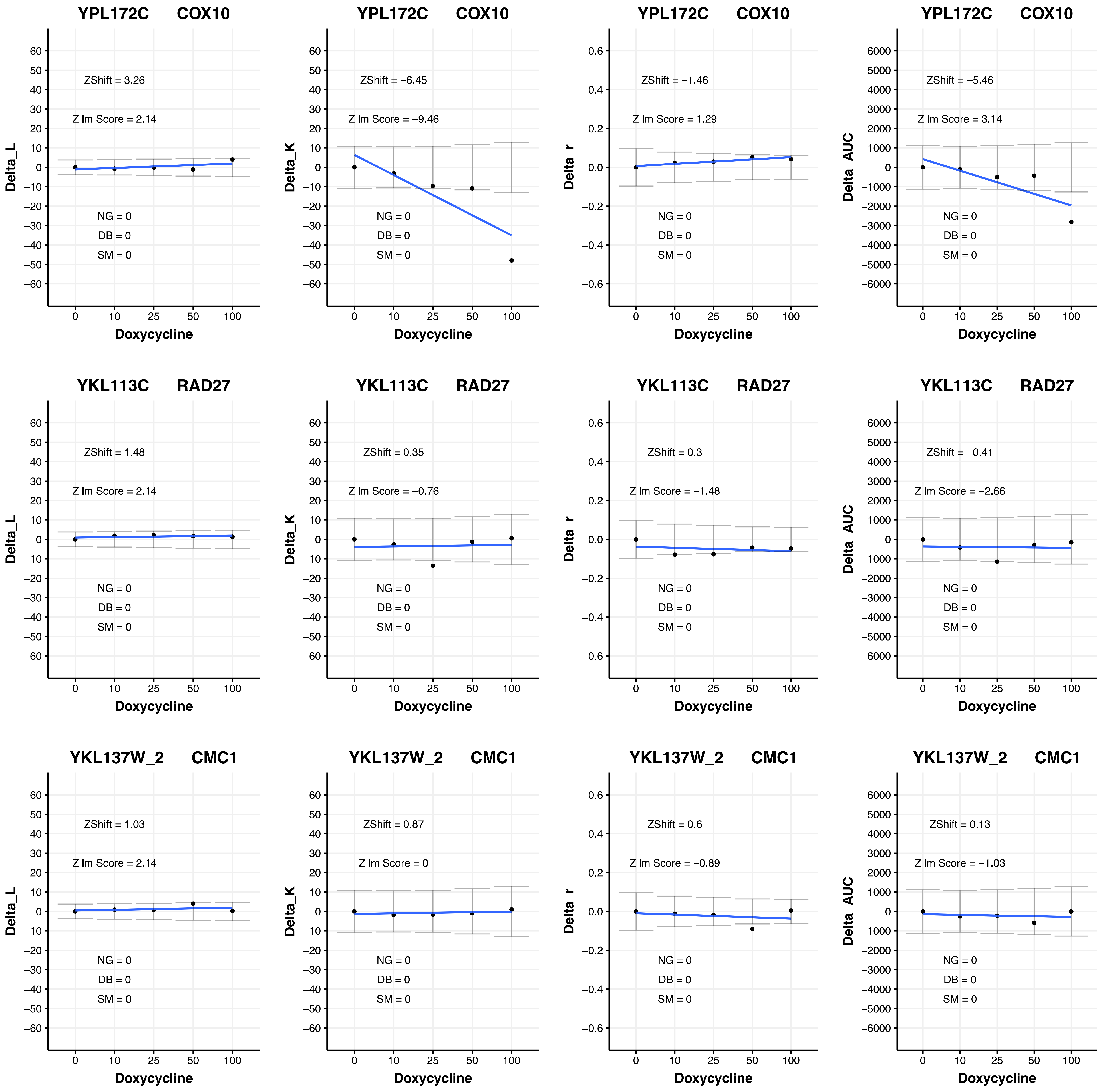
YPL172C COX10
ZShift = 3.26
Z lm Score = 2.14
NG = 0
DB = 0
SM = 0
Delta_L
Doxycycline
YPL172C COX10
ZShift = −6.45
Z lm Score = −9.46
NG = 0
DB = 0
SM = 0
Delta_K
Doxycycline
YPL172C COX10
ZShift = −1.46
Z lm Score = 1.29
NG = 0
DB = 0
SM = 0
Delta_r
Doxycycline
YPL172C COX10
ZShift = −5.46
Z lm Score = 3.14
NG = 0
DB = 0
SM = 0
Delta_AUC
Doxycycline
YKL113C RAD27
ZShift = 1.48
Z lm Score = 2.14
NG = 0
DB = 0
SM = 0
Delta_L
Doxycycline
YKL113C RAD27
ZShift = 0.35
Z lm Score = −0.76
NG = 0
DB = 0
SM = 0
Delta_K
Doxycycline
YKL113C RAD27
ZShift = 0.3
Z lm Score = −1.48
NG = 0
DB = 0
SM = 0
Delta_r
Doxycycline
YKL113C RAD27
ZShift = −0.41
Z lm Score = −2.66
NG = 0
DB = 0
SM = 0
Delta_AUC
Doxycycline
YKL137W_2 CMC1
ZShift = 1.03
Z lm Score = 2.14
NG = 0
DB = 0
SM = 0
Delta_L
Doxycycline
YKL137W_2 CMC1
ZShift = 0.87
Z lm Score = 0
NG = 0
DB = 0
SM = 0
Delta_K
Doxycycline
YKL137W_2 CMC1
ZShift = 0.6
Z lm Score = −0.89
NG = 0
DB = 0
SM = 0
Delta_r
Doxycycline
YKL137W_2 CMC1
ZShift = 0.13
Z lm Score = −1.03
NG = 0
DB = 0
SM = 0
Delta_AUC
Doxycycline

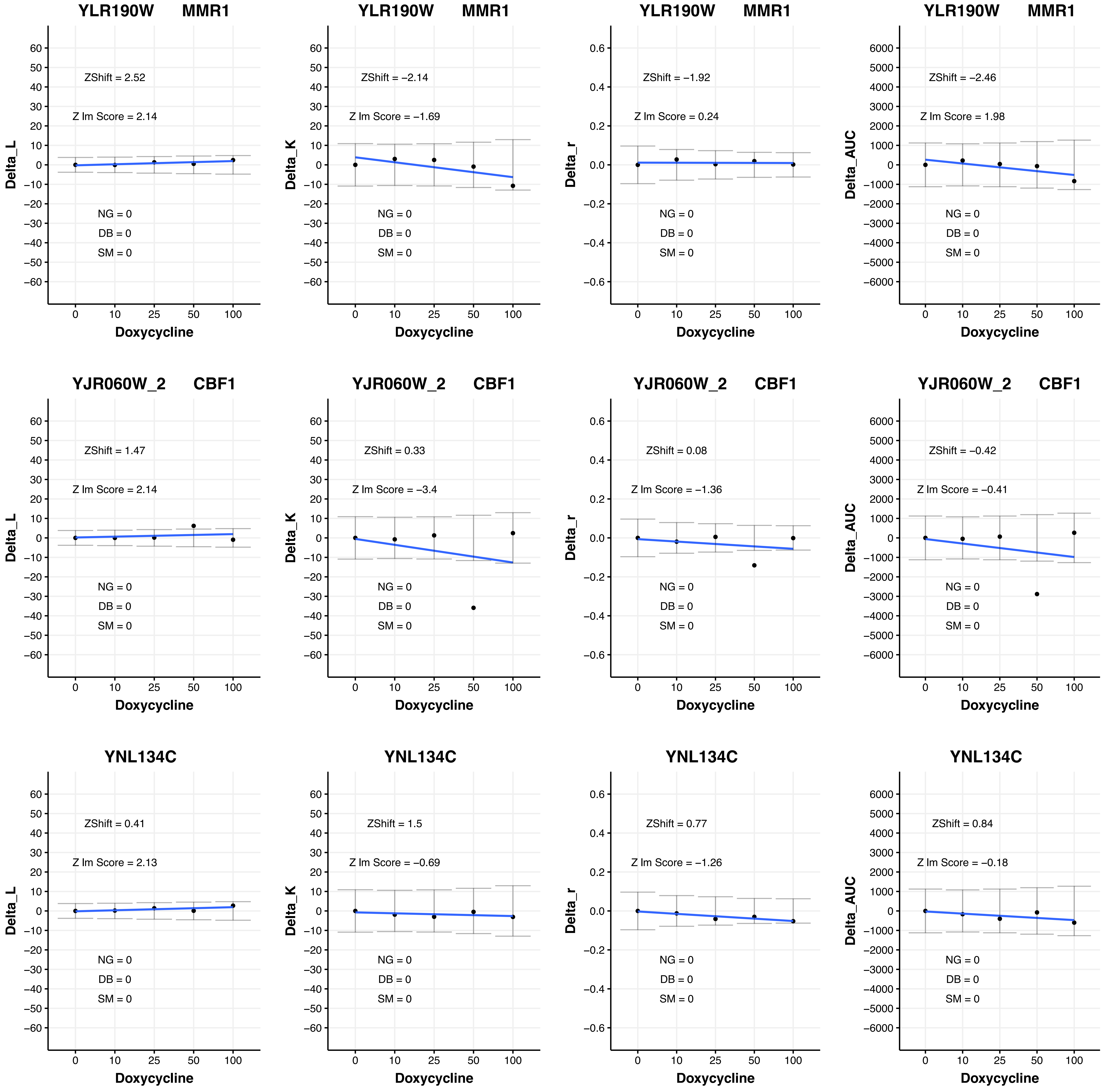
YLR190W MMR1
ZShift = 2.52
Z lm Score = 2.14
NG = 0
DB = 0
SM = 0
Delta_L
Doxycycline
YLR190W MMR1
ZShift = −2.14
Z lm Score = −1.69
NG = 0
DB = 0
SM = 0
Delta_K
Doxycycline
YLR190W MMR1
ZShift = −1.92
Z lm Score = 0.24
NG = 0
DB = 0
SM = 0
Delta_r
Doxycycline
YLR190W MMR1
ZShift = −2.46
Z lm Score = 1.98
NG = 0
DB = 0
SM = 0
Delta_AUC
Doxycycline
YJR060W_2 CBF1
ZShift = 1.47
Z lm Score = 2.14
NG = 0
DB = 0
SM = 0
Delta_L
Doxycycline
YJR060W_2 CBF1
ZShift = 0.33
Z lm Score = −3.4
NG = 0
DB = 0
SM = 0
Delta_K
Doxycycline
YJR060W_2 CBF1
ZShift = 0.08
Z lm Score = −1.36
NG = 0
DB = 0
SM = 0
Delta_r
Doxycycline
YJR060W_2 CBF1
ZShift = −0.42
Z lm Score = −0.41
NG = 0
DB = 0
SM = 0
Delta_AUC
Doxycycline
YNL134C
ZShift = 0.41
Z lm Score = 2.13
NG = 0
DB = 0
SM = 0
Delta_L
Doxycycline
YNL134C
ZShift = 1.5
Z lm Score = −0.69
NG = 0
DB = 0
SM = 0
Delta_K
Doxycycline
YNL134C
ZShift = 0.77
Z lm Score = −1.26
NG = 0
DB = 0
SM = 0
Delta_r
Doxycycline
YNL134C
ZShift = 0.84
Z lm Score = −0.18
NG = 0
DB = 0
SM = 0
Delta_AUC
Doxycycline

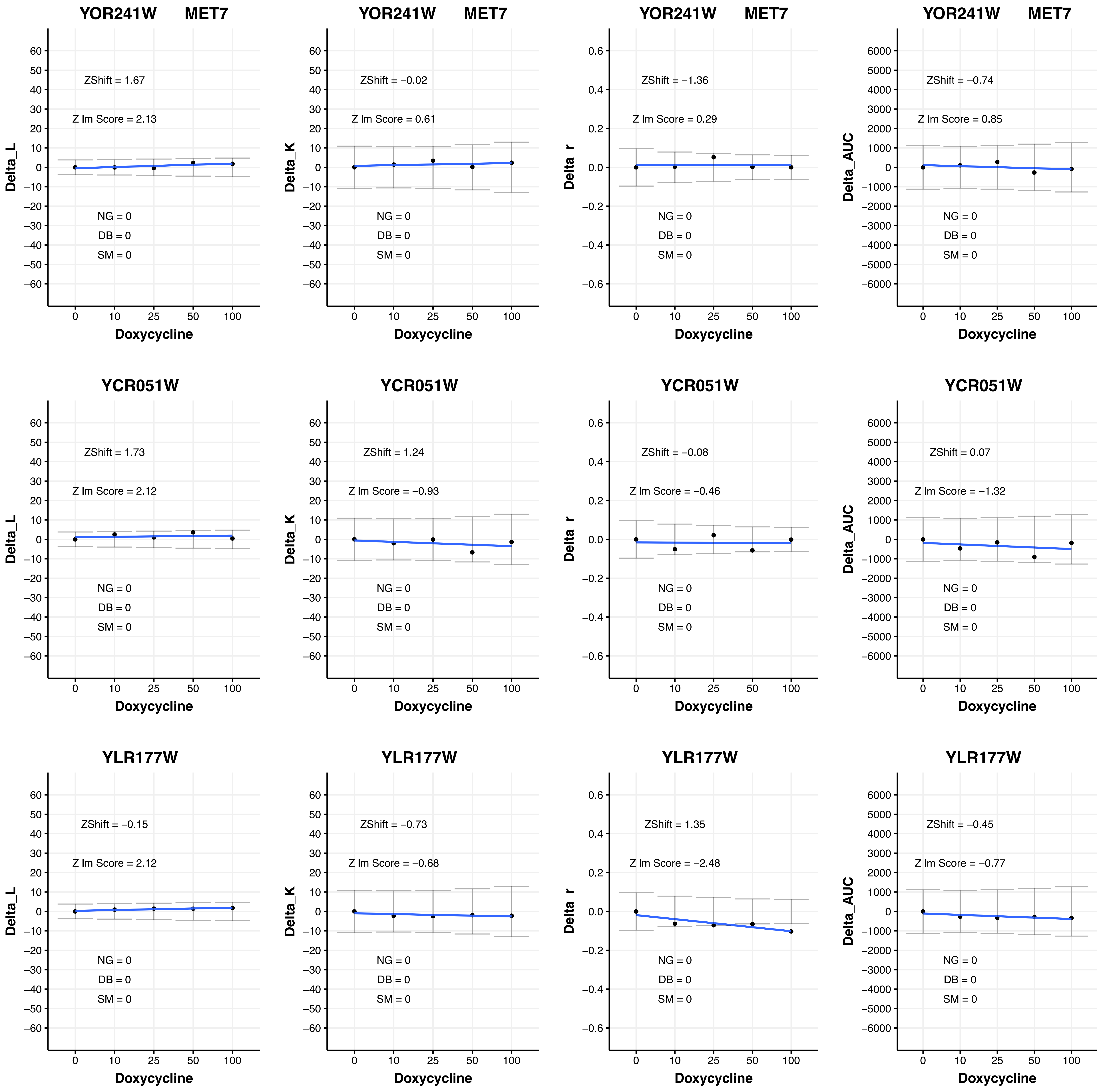
YOR241W MET7
Delta_L
ZShift = 1.67
Z lm Score = 2.13
NG = 0
DB = 0
SM = 0
Doxycycline
YOR241W MET7
Delta_K
ZShift = −0.02
Z lm Score = 0.61
NG = 0
DB = 0
SM = 0
Doxycycline
YOR241W MET7
Delta_r
ZShift = −1.36
Z lm Score = 0.29
NG = 0
DB = 0
SM = 0
Doxycycline
YOR241W MET7
Delta_AUC
ZShift = −0.74
Z lm Score = 0.85
NG = 0
DB = 0
SM = 0
Doxycycline
YCR051W
Delta_L
ZShift = 1.73
Z lm Score = 2.12
NG = 0
DB = 0
SM = 0
Doxycycline
YCR051W
Delta_K
ZShift = 1.24
Z lm Score = −0.93
NG = 0
DB = 0
SM = 0
Doxycycline
YCR051W
Delta_r
ZShift = −0.08
Z lm Score = −0.46
NG = 0
DB = 0
SM = 0
Doxycycline
YCR051W
Delta_AUC
ZShift = 0.07
Z lm Score = −1.32
NG = 0
DB = 0
SM = 0
Doxycycline
YLR177W
Delta_L
ZShift = −0.15
Z lm Score = 2.12
NG = 0
DB = 0
SM = 0
Doxycycline
YLR177W
Delta_K
ZShift = −0.73
Z lm Score = −0.68
NG = 0
DB = 0
SM = 0
Doxycycline
YLR177W
Delta_r
ZShift = 1.35
Z lm Score = −2.48
NG = 0
DB = 0
SM = 0
Doxycycline
YLR177W
Delta_AUC
ZShift = −0.45
Z lm Score = −0.77
NG = 0
DB = 0
SM = 0
Doxycycline

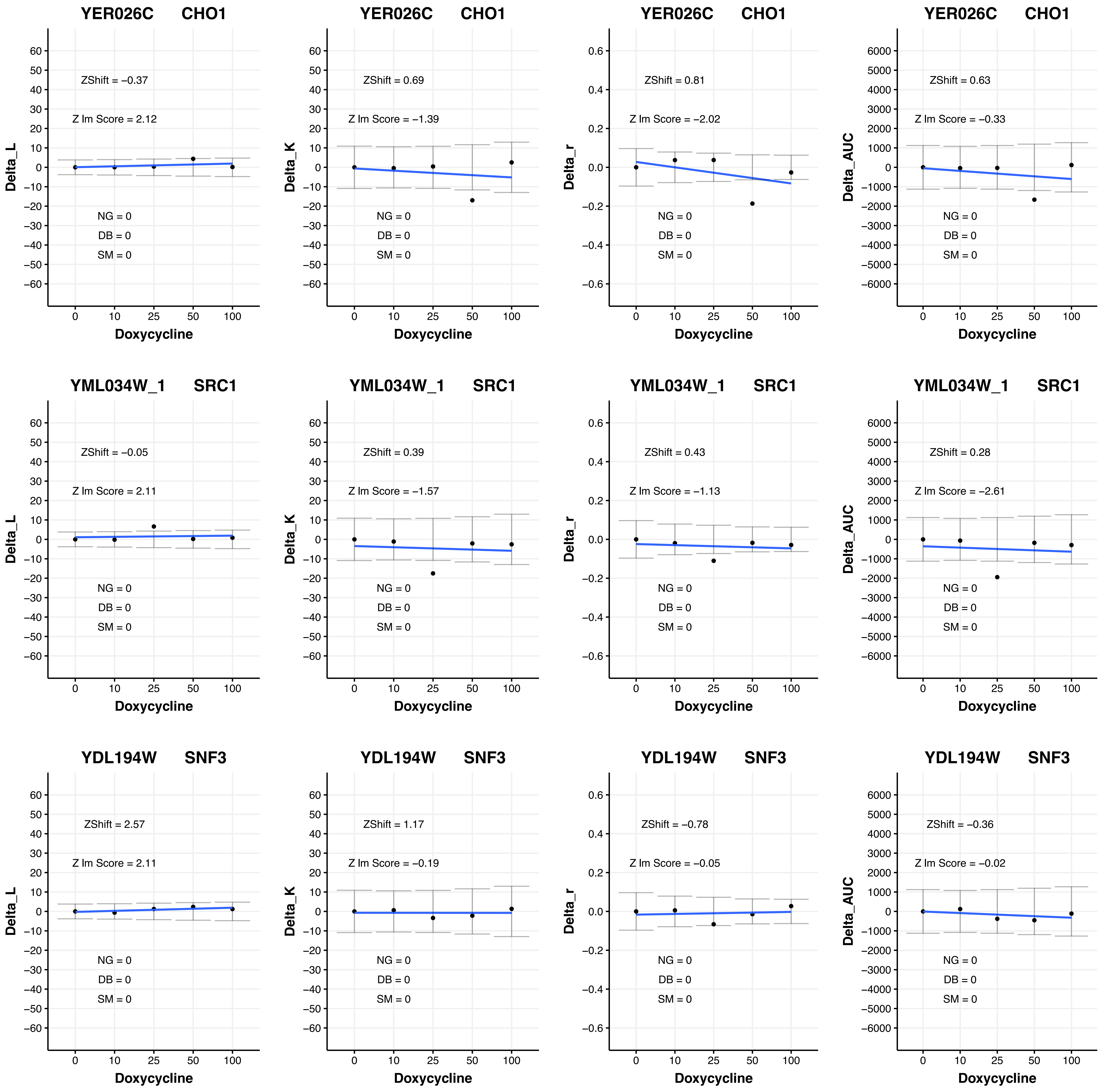
YER026C CHO1
ZShift = −0.37
Z lm Score = 2.12
NG = 0
DB = 0
SM = 0
Delta_L
Doxycycline
YER026C CHO1
ZShift = 0.69
Z lm Score = −1.39
NG = 0
DB = 0
SM = 0
Delta_K
Doxycycline
YER026C CHO1
ZShift = 0.81
Z lm Score = −2.02
NG = 0
DB = 0
SM = 0
Delta_r
Doxycycline
YER026C CHO1
ZShift = 0.63
Z lm Score = −0.33
NG = 0
DB = 0
SM = 0
Delta_AUC
Doxycycline
YML034W_1 SRC1
ZShift = −0.05
Z lm Score = 2.11
NG = 0
DB = 0
SM = 0
Delta_L
Doxycycline
YML034W_1 SRC1
ZShift = 0.39
Z lm Score = −1.57
NG = 0
DB = 0
SM = 0
Delta_K
Doxycycline
YML034W_1 SRC1
ZShift = 0.43
Z lm Score = −1.13
NG = 0
DB = 0
SM = 0
Delta_r
Doxycycline
YML034W_1 SRC1
ZShift = 0.28
Z lm Score = −2.61
NG = 0
DB = 0
SM = 0
Delta_AUC
Doxycycline
YDL194W SNF3
ZShift = 2.57
Z lm Score = 2.11
NG = 0
DB = 0
SM = 0
Delta_L
Doxycycline
YDL194W SNF3
ZShift = 1.17
Z lm Score = −0.19
NG = 0
DB = 0
SM = 0
Delta_K
Doxycycline
YDL194W SNF3
ZShift = −0.78
Z lm Score = −0.05
NG = 0
DB = 0
SM = 0
Delta_r
Doxycycline
YDL194W SNF3
ZShift = −0.36
Z lm Score = −0.02
NG = 0
DB = 0
SM = 0
Delta_AUC
Doxycycline
0 10 25 50 100

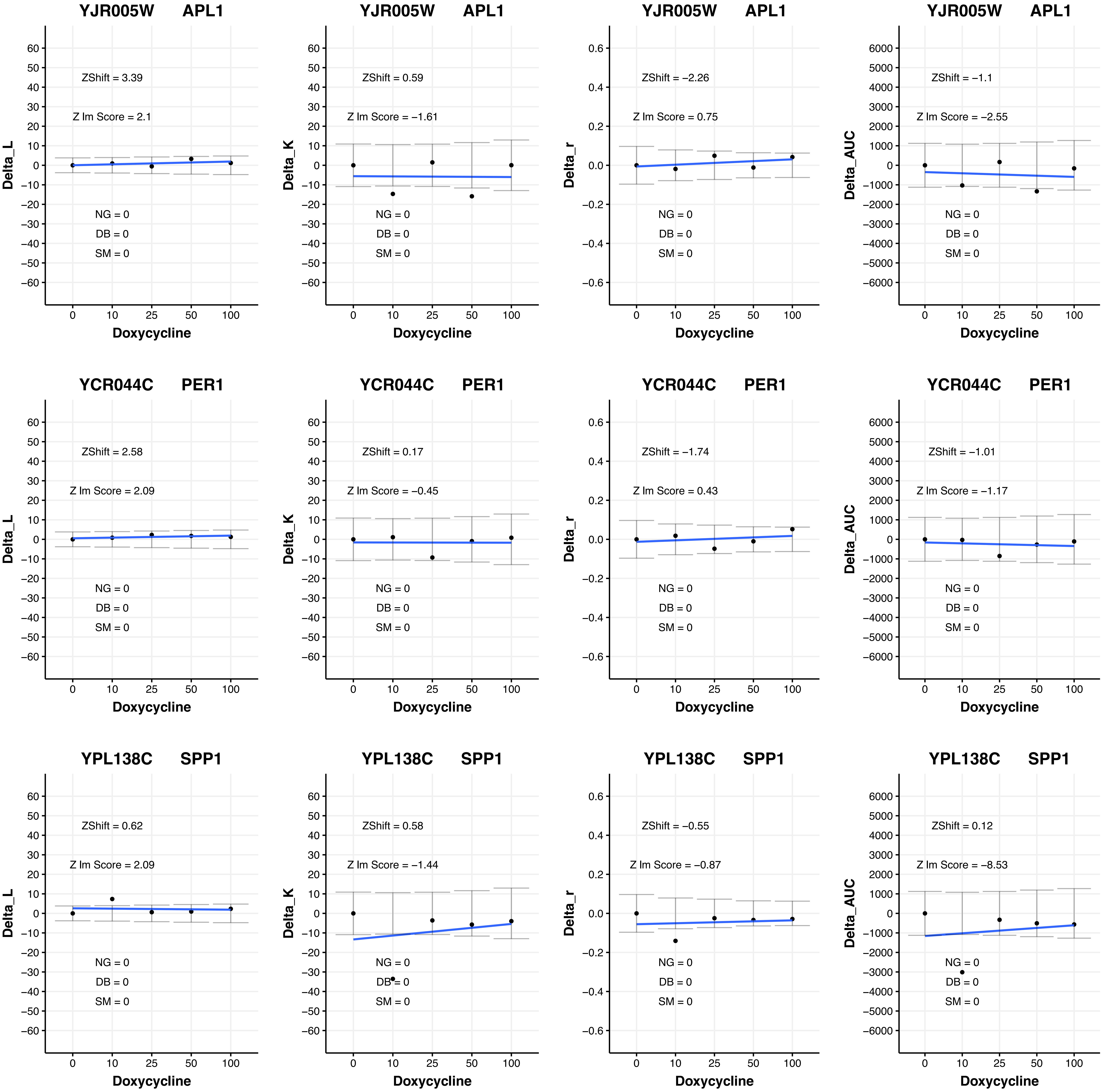

YJR005W APL1
ZShift = 3.39
Z lm Score = 2.1
NG = 0
DB = 0
SM = 0
Delta_L
Doxycycline
YJR005W APL1
ZShift = 0.59
Z lm Score = −1.61
NG = 0
DB = 0
SM = 0
Delta_K
Doxycycline
YJR005W APL1
ZShift = −2.26
Z lm Score = 0.75
NG = 0
DB = 0
SM = 0
Delta_r
Doxycycline
YJR005W APL1
ZShift = −1.1
Z lm Score = −2.55
NG = 0
DB = 0
SM = 0
Delta_AUC
Doxycycline
YCR044C PER1
ZShift = 2.58
Z lm Score = 2.09
NG = 0
DB = 0
SM = 0
Delta_L
Doxycycline
YCR044C PER1
ZShift = 0.17
Z lm Score = −0.45
NG = 0
DB = 0
SM = 0
Delta_K
Doxycycline
YCR044C PER1
ZShift = −1.74
Z lm Score = 0.43
NG = 0
DB = 0
SM = 0
Delta_r
Doxycycline
YCR044C PER1
ZShift = −1.01
Z lm Score = −1.17
NG = 0
DB = 0
SM = 0
Delta_AUC
Doxycycline
YPL138C SPP1
ZShift = 0.62
Z lm Score = 2.09
NG = 0
DB = 0
SM = 0
Delta_L
Doxycycline
YPL138C SPP1
ZShift = 0.58
Z lm Score = −1.44
NG = 0
DB = 0
SM = 0
Delta_K
Doxycycline
YPL138C SPP1
ZShift = −0.55
Z lm Score = −0.87
NG = 0
DB = 0
SM = 0
Delta_r
Doxycycline
YPL138C SPP1
ZShift = 0.12
Z lm Score = −8.53
NG = 0
DB = 0
SM = 0
Delta_AUC
Doxycycline

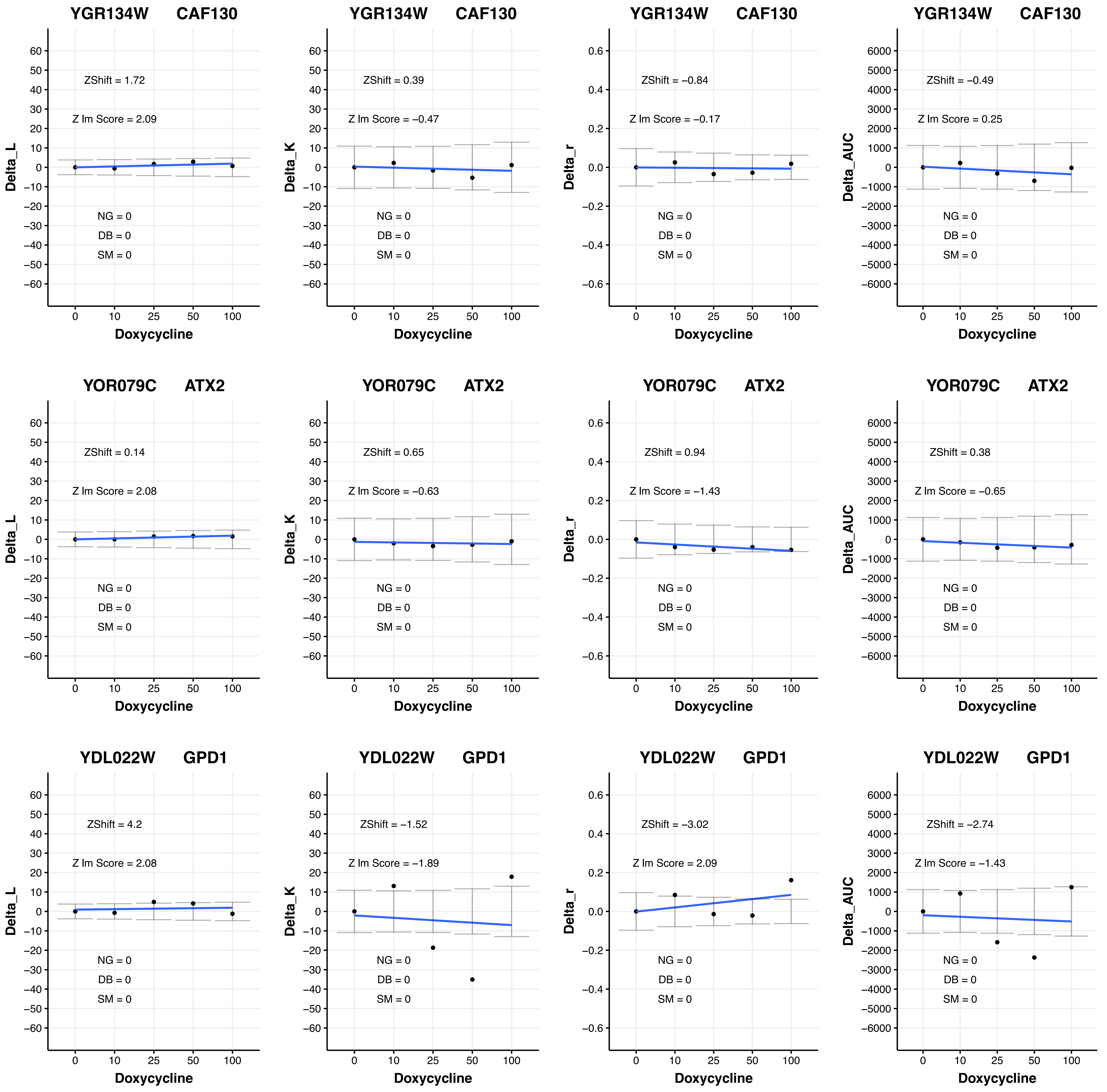
YGR134W CAF130
Delta_L
ZShift = 1.72
Z lm Score = 2.09
NG = 0
DB = 0
SM = 0
Doxycycline
YGR134W CAF130
Delta_K
ZShift = 0.39
Z lm Score = −0.47
NG = 0
DB = 0
SM = 0
Doxycycline
YGR134W CAF130
Delta_r
ZShift = −0.84
Z lm Score = −0.17
NG = 0
DB = 0
SM = 0
Doxycycline
YGR134W CAF130
Delta_AUC
ZShift = −0.49
Z lm Score = 0.25
NG = 0
DB = 0
SM = 0
Doxycycline
YOR079C ATX2
Delta_L
ZShift = 0.14
Z lm Score = 2.08
NG = 0
DB = 0
SM = 0
Doxycycline
YOR079C ATX2
Delta_K
ZShift = 0.65
Z lm Score = −0.63
NG = 0
DB = 0
SM = 0
Doxycycline
YOR079C ATX2
Delta_r
ZShift = 0.94
Z lm Score = −1.43
NG = 0
DB = 0
SM = 0
Doxycycline
YOR079C ATX2
Delta_AUC
ZShift = 0.38
Z lm Score = −0.65
NG = 0
DB = 0
SM = 0
Doxycycline
YDL022W GPD1
Delta_L
ZShift = 4.2
Z lm Score = 2.08
NG = 0
DB = 0
SM = 0
Doxycycline
YDL022W GPD1
Delta_K
ZShift = −1.52
Z lm Score = −1.89
NG = 0
DB = 0
SM = 0
Doxycycline
YDL022W GPD1
Delta_r
ZShift = −3.02
Z lm Score = 2.09
NG = 0
DB = 0
SM = 0
Doxycycline
YDL022W GPD1
Delta_AUC
ZShift = −2.74
Z lm Score = −1.43
NG = 0
DB = 0
SM = 0
Doxycycline

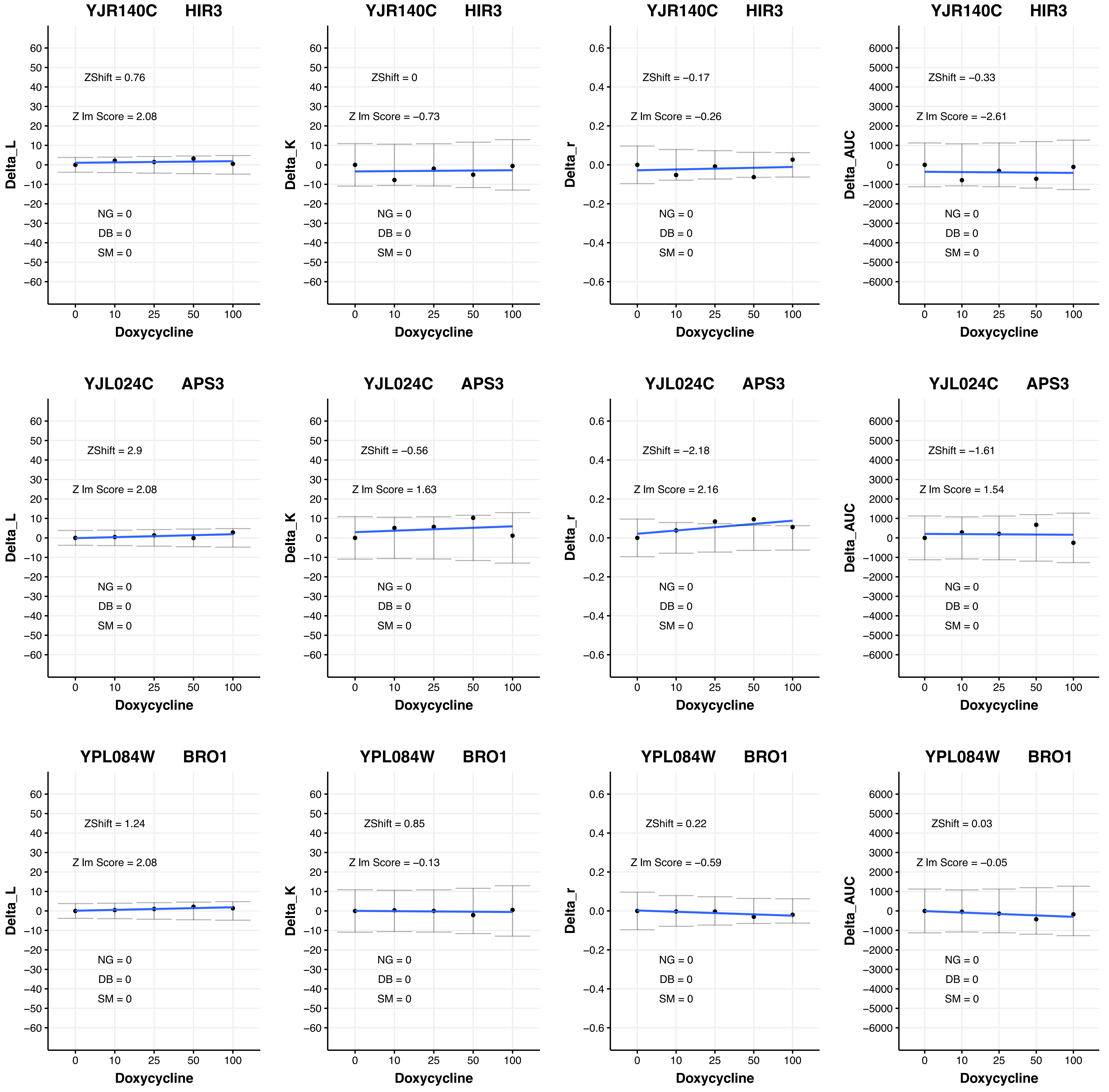
YJR140C HIR3
ZShift = 0.76
Z lm Score = 2.08
NG = 0
DB = 0
SM = 0
Delta_L
Doxycycline
YJR140C HIR3
ZShift = 0
Z lm Score = −0.73
NG = 0
DB = 0
SM = 0
Delta_K
Doxycycline
YJR140C HIR3
ZShift = −0.17
Z lm Score = −0.26
NG = 0
DB = 0
SM = 0
Delta_r
Doxycycline
YJR140C HIR3
ZShift = −0.33
Z lm Score = −2.61
NG = 0
DB = 0
SM = 0
Delta_AUC
Doxycycline
YJL024C APS3
ZShift = 2.9
Z lm Score = 2.08
NG = 0
DB = 0
SM = 0
Delta_L
Doxycycline
YJL024C APS3
ZShift = −0.56
Z lm Score = 1.63
NG = 0
DB = 0
SM = 0
Delta_K
Doxycycline
YJL024C APS3
ZShift = −2.18
Z lm Score = 2.16
NG = 0
DB = 0
SM = 0
Delta_r
Doxycycline
YJL024C APS3
ZShift = −1.61
Z lm Score = 1.54
NG = 0
DB = 0
SM = 0
Delta_AUC
Doxycycline
YPL084W BRO1
ZShift = 1.24
Z lm Score = 2.08
NG = 0
DB = 0
SM = 0
Delta_L
Doxycycline
YPL084W BRO1
ZShift = 0.85
Z lm Score = −0.13
NG = 0
DB = 0
SM = 0
Delta_K
Doxycycline
YPL084W BRO1
ZShift = 0.22
Z lm Score = −0.59
NG = 0
DB = 0
SM = 0
Delta_r
Doxycycline
YPL084W BRO1
ZShift = 0.03
Z lm Score = −0.05
NG = 0
DB = 0
SM = 0
Delta_AUC
Doxycycline

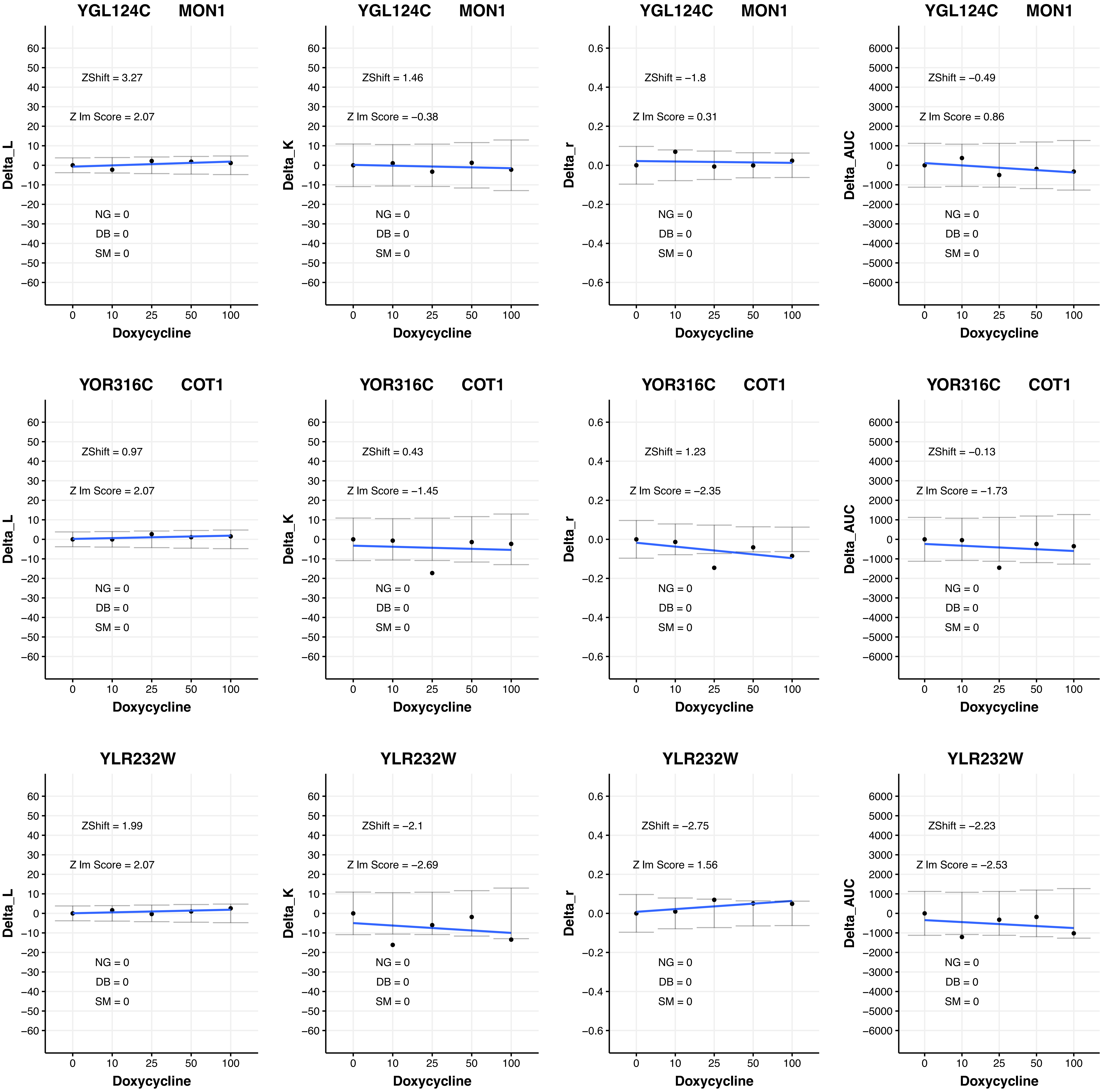

YGL124C MON1
ZShift = 3.27
Z lm Score = 2.07
NG = 0
DB = 0
SM = 0
Delta_L
Doxycycline
YGL124C MON1
ZShift = 1.46
Z lm Score = −0.38
NG = 0
DB = 0
SM = 0
Delta_K
Doxycycline
YGL124C MON1
ZShift = −1.8
Z lm Score = 0.31
NG = 0
DB = 0
SM = 0
Delta_r
Doxycycline
YGL124C MON1
ZShift = −0.49
Z lm Score = 0.86
NG = 0
DB = 0
SM = 0
Delta_AUC
Doxycycline
YOR316C COT1
ZShift = 0.97
Z lm Score = 2.07
NG = 0
DB = 0
SM = 0
Delta_L
Doxycycline
YOR316C COT1
ZShift = 0.43
Z lm Score = −1.45
NG = 0
DB = 0
SM = 0
Delta_K
Doxycycline
YOR316C COT1
ZShift = 1.23
Z lm Score = −2.35
NG = 0
DB = 0
SM = 0
Delta_r
Doxycycline
YOR316C COT1
ZShift = −0.13
Z lm Score = −1.73
NG = 0
DB = 0
SM = 0
Delta_AUC
Doxycycline
YLR232W
ZShift = 1.99
Z lm Score = 2.07
NG = 0
DB = 0
SM = 0
Delta_L
Doxycycline
YLR232W
ZShift = −2.1
Z lm Score = −2.69
NG = 0
DB = 0
SM = 0
Delta_K
Doxycycline
YLR232W
ZShift = −2.75
Z lm Score = 1.56
NG = 0
DB = 0
SM = 0
Delta_r
Doxycycline
YLR232W
ZShift = −2.23
Z lm Score = −2.53
NG = 0
DB = 0
SM = 0
Delta_AUC
Doxycycline

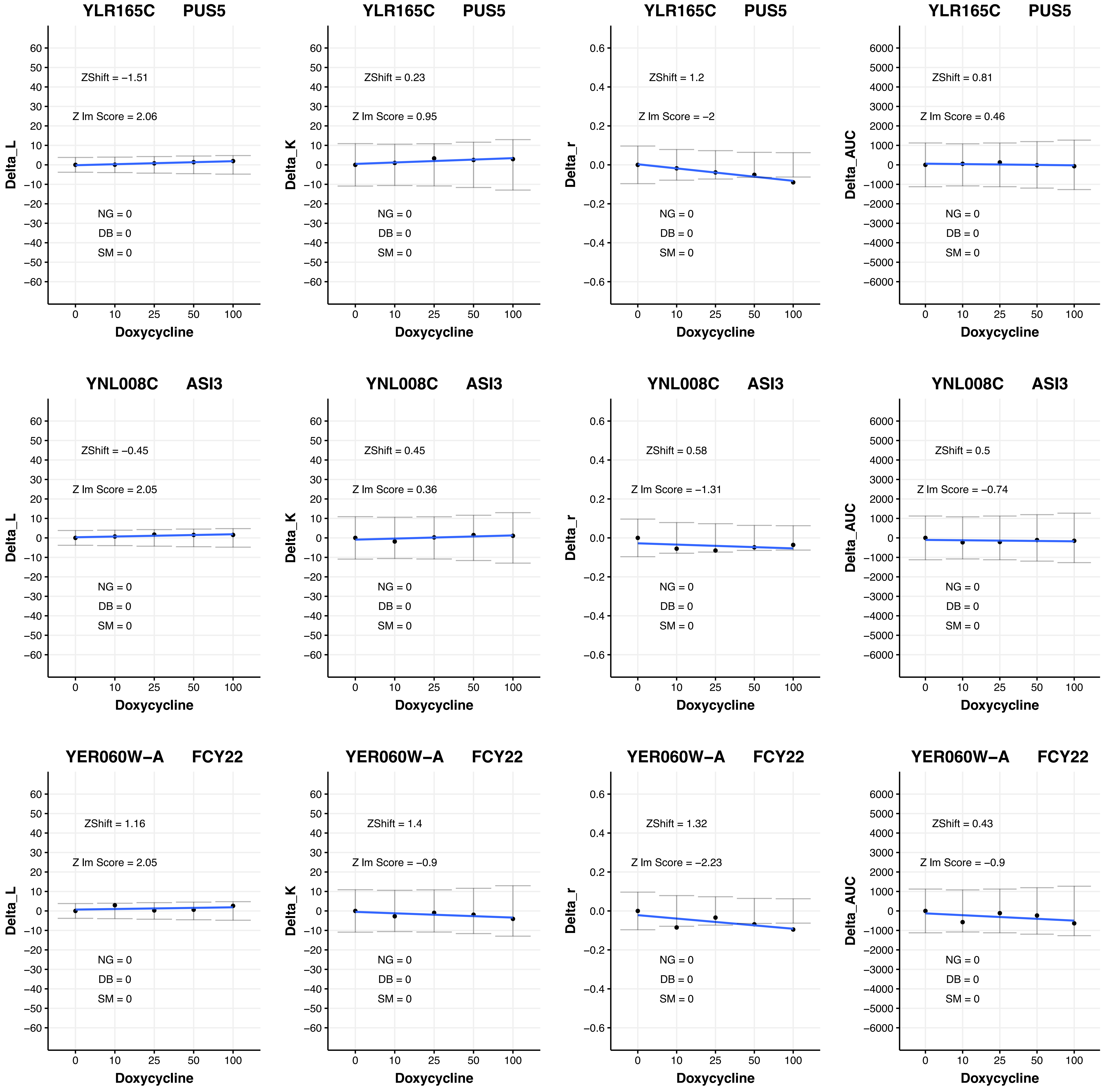

YLR165C PUS5
ZShift = −1.51
Z lm Score = 2.06
Delta_L
NG = 0
DB = 0
SM = 0
Doxycycline
YLR165C PUS5
ZShift = 0.23
Z lm Score = 0.95
Delta_K
NG = 0
DB = 0
SM = 0
Doxycycline
YLR165C PUS5
ZShift = 1.2
Z lm Score = −2
Delta_r
NG = 0
DB = 0
SM = 0
Doxycycline
YLR165C PUS5
ZShift = 0.81
Z lm Score = 0.46
Delta_AUC
NG = 0
DB = 0
SM = 0
Doxycycline
YNL008C ASI3
ZShift = −0.45
Z lm Score = 2.05
Delta_L
NG = 0
DB = 0
SM = 0
Doxycycline
YNL008C ASI3
ZShift = 0.45
Z lm Score = 0.36
Delta_K
NG = 0
DB = 0
SM = 0
Doxycycline
YNL008C ASI3
ZShift = 0.58
Z lm Score = −1.31
Delta_r
NG = 0
DB = 0
SM = 0
Doxycycline
YNL008C ASI3
ZShift = 0.5
Z lm Score = −0.74
Delta_AUC
NG = 0
DB = 0
SM = 0
Doxycycline
YER060W−A FCY22
ZShift = 1.16
Z lm Score = 2.05
Delta_L
NG = 0
DB = 0
SM = 0
Doxycycline
YER060W−A FCY22
ZShift = 1.4
Z lm Score = −0.9
Delta_K
NG = 0
DB = 0
SM = 0
Doxycycline
YER060W−A FCY22
ZShift = 1.32
Z lm Score = −2.23
Delta_r
NG = 0
DB = 0
SM = 0
Doxycycline
YER060W−A FCY22
ZShift = 0.43
Z lm Score = −0.9
Delta_AUC
NG = 0
DB = 0
SM = 0
Doxycycline

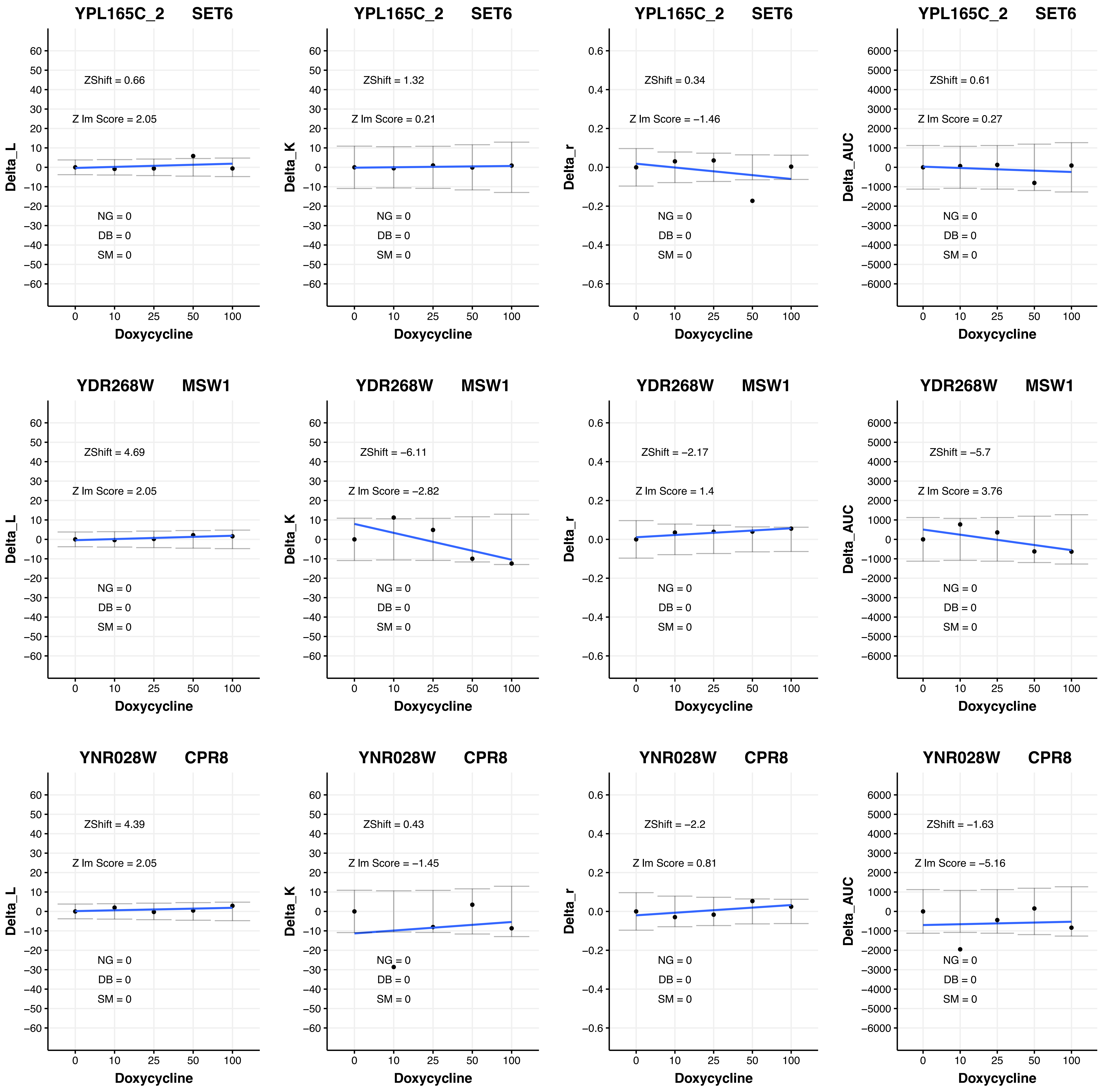
YPL165C_2 SET6
ZShift = 0.66
Z lm Score = 2.05
NG = 0
DB = 0
SM = 0
Delta_L
Doxycycline
YPL165C_2 SET6
ZShift = 1.32
Z lm Score = 0.21
NG = 0
DB = 0
SM = 0
Delta_K
Doxycycline
YPL165C_2 SET6
ZShift = 0.34
Z lm Score = −1.46
NG = 0
DB = 0
SM = 0
Delta_r
Doxycycline
YPL165C_2 SET6
ZShift = 0.61
Z lm Score = 0.27
NG = 0
DB = 0
SM = 0
Delta_AUC
Doxycycline
YDR268W MSW1
ZShift = 4.69
Z lm Score = 2.05
NG = 0
DB = 0
SM = 0
Delta_L
Doxycycline
YDR268W MSW1
ZShift = −6.11
Z lm Score = −2.82
NG = 0
DB = 0
SM = 0
Delta_K
Doxycycline
YDR268W MSW1
ZShift = −2.17
Z lm Score = 1.4
NG = 0
DB = 0
SM = 0
Delta_r
Doxycycline
YDR268W MSW1
ZShift = −5.7
Z lm Score = 3.76
NG = 0
DB = 0
SM = 0
Delta_AUC
Doxycycline
YNR028W CPR8
ZShift = 4.39
Z lm Score = 2.05
NG = 0
DB = 0
SM = 0
Delta_L
Doxycycline
YNR028W CPR8
ZShift = 0.43
Z lm Score = −1.45
NG = 0
DB = 0
SM = 0
Delta_K
Doxycycline
YNR028W CPR8
ZShift = −2.2
Z lm Score = 0.81
NG = 0
DB = 0
SM = 0
Delta_r
Doxycycline
YNR028W CPR8
ZShift = −1.63
Z lm Score = −5.16
NG = 0
DB = 0
SM = 0
Delta_AUC
Doxycycline

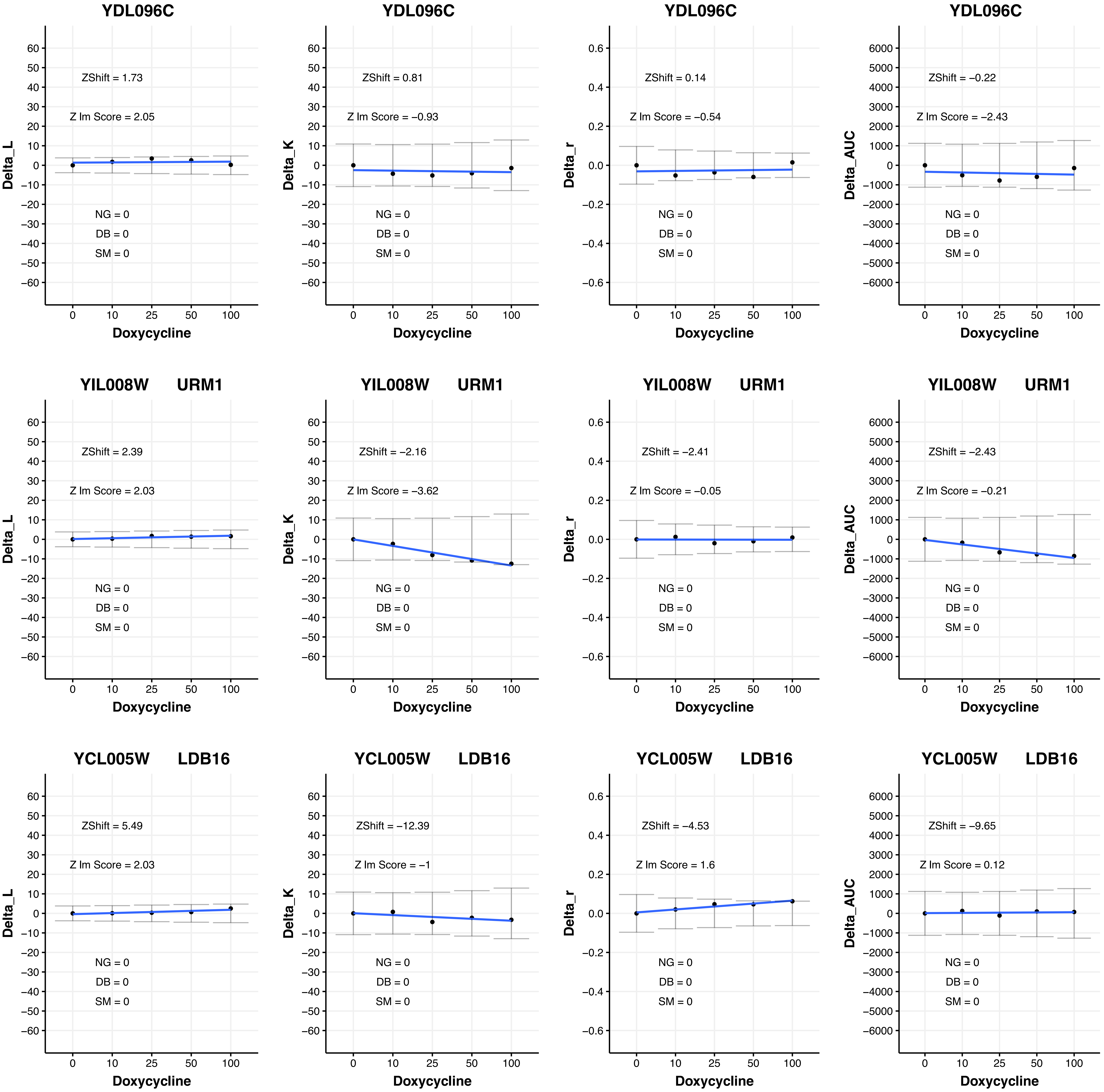

YDL096C
ZShift = 1.73
Z lm Score = 2.05
NG = 0
DB = 0
SM = 0
Delta_L
Doxycycline
YDL096C
ZShift = 0.81
Z lm Score = −0.93
NG = 0
DB = 0
SM = 0
Delta_K
Doxycycline
YDL096C
ZShift = 0.14
Z lm Score = −0.54
NG = 0
DB = 0
SM = 0
Delta_r
Doxycycline
YDL096C
ZShift = −0.22
Z lm Score = −2.43
NG = 0
DB = 0
SM = 0
Delta_AUC
Doxycycline
YIL008W URM1
ZShift = 2.39
Z lm Score = 2.03
NG = 0
DB = 0
SM = 0
Delta_L
Doxycycline
YIL008W URM1
ZShift = −2.16
Z lm Score = −3.62
NG = 0
DB = 0
SM = 0
Delta_K
Doxycycline
YIL008W URM1
ZShift = −2.41
Z lm Score = −0.05
NG = 0
DB = 0
SM = 0
Delta_r
Doxycycline
YIL008W URM1
ZShift = −2.43
Z lm Score = −0.21
NG = 0
DB = 0
SM = 0
Delta_AUC
Doxycycline
YCL005W LDB16
ZShift = 5.49
Z lm Score = 2.03
NG = 0
DB = 0
SM = 0
Delta_L
Doxycycline
YCL005W LDB16
ZShift = −12.39
Z lm Score = −1
NG = 0
DB = 0
SM = 0
Delta_K
Doxycycline
YCL005W LDB16
ZShift = −4.53
Z lm Score = 1.6
NG = 0
DB = 0
SM = 0
Delta_r
Doxycycline
YCL005W LDB16
ZShift = −9.65
Z lm Score = 0.12
NG = 0
DB = 0
SM = 0
Delta_AUC
Doxycycline

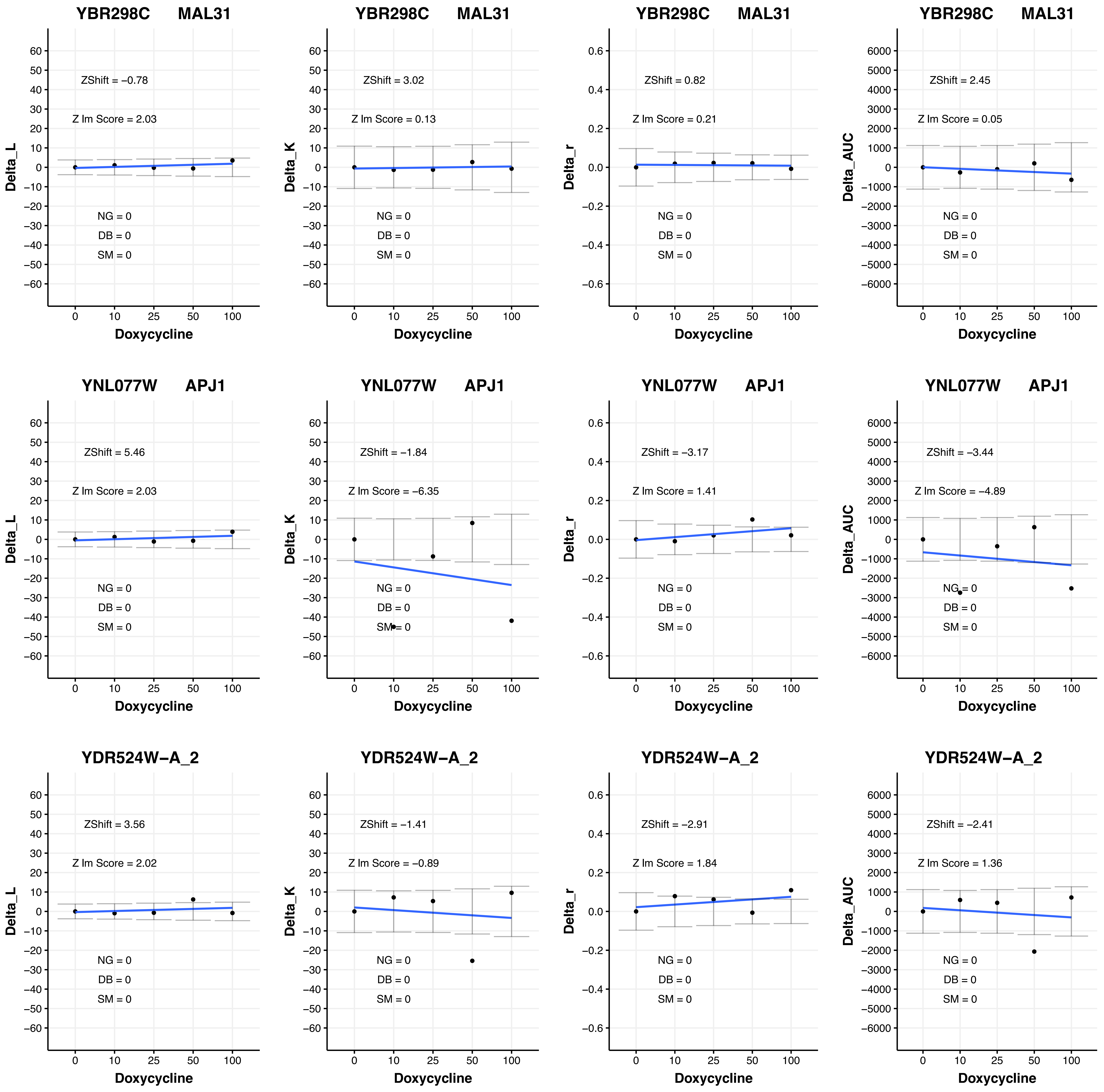
YBR298C MAL31
ZShift = −0.78
Z lm Score = 2.03
NG = 0
DB = 0
SM = 0
Delta_L
Doxycycline
YBR298C MAL31
ZShift = 3.02
Z lm Score = 0.13
NG = 0
DB = 0
SM = 0
Delta_K
Doxycycline
YBR298C MAL31
ZShift = 0.82
Z lm Score = 0.21
NG = 0
DB = 0
SM = 0
Delta_r
Doxycycline
YBR298C MAL31
ZShift = 2.45
Z lm Score = 0.05
NG = 0
DB = 0
SM = 0
Delta_AUC
Doxycycline
YNL077W APJ1
ZShift = 5.46
Z lm Score = 2.03
NG = 0
DB = 0
SM = 0
Delta_L
Doxycycline
YNL077W APJ1
ZShift = −1.84
Z lm Score = −6.35
NG = 0
DB = 0
SM = 0
Delta_K
Doxycycline
YNL077W APJ1
ZShift = −3.17
Z lm Score = 1.41
NG = 0
DB = 0
SM = 0
Delta_r
Doxycycline
YNL077W APJ1
ZShift = −3.44
Z lm Score = −4.89
NG = 0
DB = 0
SM = 0
Delta_AUC
Doxycycline
YDR524W−A_2
ZShift = 3.56
Z lm Score = 2.02
NG = 0
DB = 0
SM = 0
Delta_L
Doxycycline
YDR524W−A_2
ZShift = −1.41
Z lm Score = −0.89
NG = 0
DB = 0
SM = 0
Delta_K
Doxycycline
YDR524W−A_2
ZShift = −2.91
Z lm Score = 1.84
NG = 0
DB = 0
SM = 0
Delta_r
Doxycycline
YDR524W−A_2
ZShift = −2.41
Z lm Score = 1.36
NG = 0
DB = 0
SM = 0
Delta_AUC
Doxycycline

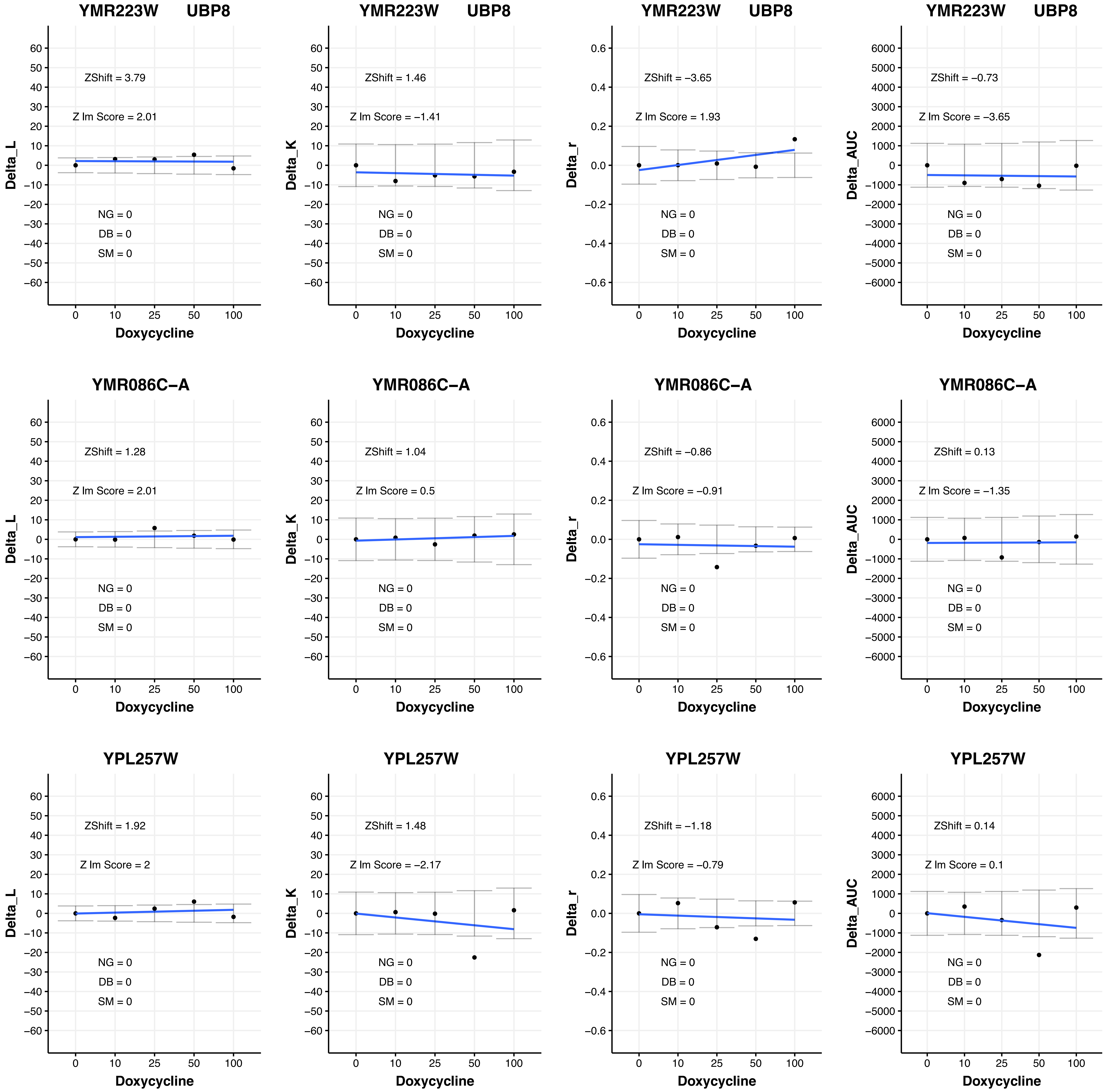

YMR223W UBP8
ZShift = 3.79
Z lm Score = 2.01
NG = 0
DB = 0
SM = 0
Delta_L
Doxycycline
YMR223W UBP8
ZShift = 1.46
Z lm Score = −1.41
NG = 0
DB = 0
SM = 0
Delta_K
Doxycycline
YMR223W UBP8
ZShift = −3.65
Z lm Score = 1.93
NG = 0
DB = 0
SM = 0
Delta_r
Doxycycline
YMR223W UBP8
ZShift = −0.73
Z lm Score = −3.65
NG = 0
DB = 0
SM = 0
Delta_AUC
Doxycycline
YMR086C−A
ZShift = 1.28
Z lm Score = 2.01
NG = 0
DB = 0
SM = 0
Delta_L
Doxycycline
YMR086C−A
ZShift = 1.04
Z lm Score = 0.5
NG = 0
DB = 0
SM = 0
Delta_K
Doxycycline
YMR086C−A
ZShift = −0.86
Z lm Score = −0.91
NG = 0
DB = 0
SM = 0
Delta_r
Doxycycline
YMR086C−A
ZShift = 0.13
Z lm Score = −1.35
NG = 0
DB = 0
SM = 0
Delta_AUC
Doxycycline
YPL257W
ZShift = 1.92
Z lm Score = 2
NG = 0
DB = 0
SM = 0
Delta_L
Doxycycline
YPL257W
ZShift = 1.48
Z lm Score = −2.17
NG = 0
DB = 0
SM = 0
Delta_K
Doxycycline
YPL257W
ZShift = −1.18
Z lm Score = −0.79
NG = 0
DB = 0
SM = 0
Delta_r
Doxycycline
YPL257W
ZShift = 0.14
Z lm Score = 0.1
NG = 0
DB = 0
SM = 0
Delta_AUC
Doxycycline

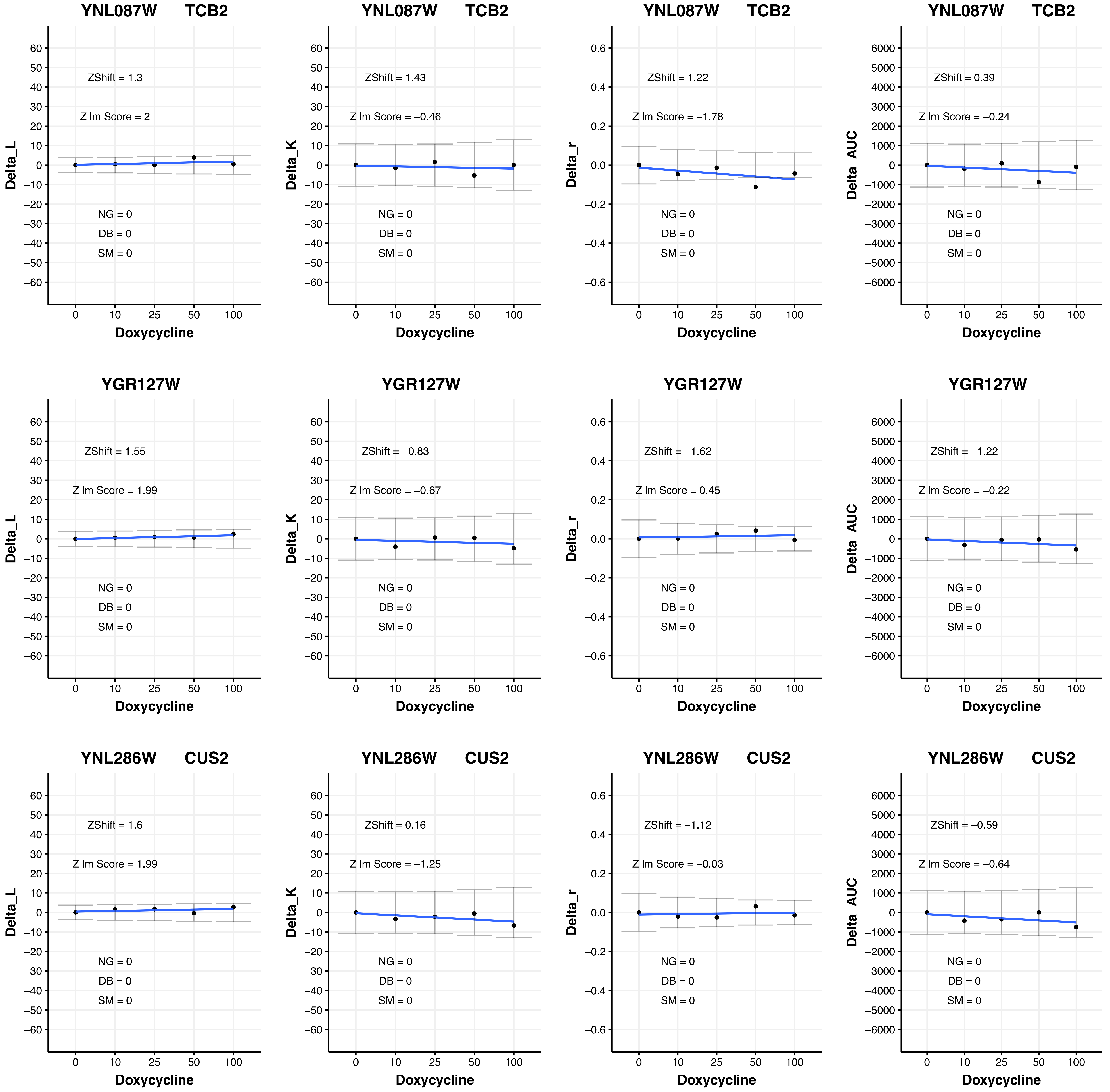

YNL087W TCB2
ZShift = 1.3
Z lm Score = 2
NG = 0
DB = 0
SM = 0
Delta_L
Doxycycline
YNL087W TCB2
ZShift = 1.43
Z lm Score = −0.46
NG = 0
DB = 0
SM = 0
Delta_K
Doxycycline
YNL087W TCB2
ZShift = 1.22
Z lm Score = −1.78
NG = 0
DB = 0
SM = 0
Delta_r
Doxycycline
YNL087W TCB2
ZShift = 0.39
Z lm Score = −0.24
NG = 0
DB = 0
SM = 0
Delta_AUC
Doxycycline
YGR127W
ZShift = 1.55
Z lm Score = 1.99
NG = 0
DB = 0
SM = 0
Delta_L
Doxycycline
YGR127W
ZShift = −0.83
Z lm Score = −0.67
NG = 0
DB = 0
SM = 0
Delta_K
Doxycycline
YGR127W
ZShift = −1.62
Z lm Score = 0.45
NG = 0
DB = 0
SM = 0
Delta_r
Doxycycline
YGR127W
ZShift = −1.22
Z lm Score = −0.22
NG = 0
DB = 0
SM = 0
Delta_AUC
Doxycycline
YNL286W CUS2
ZShift = 1.6
Z lm Score = 1.99
NG = 0
DB = 0
SM = 0
Delta_L
Doxycycline
YNL286W CUS2
ZShift = 0.16
Z lm Score = −1.25
NG = 0
DB = 0
SM = 0
Delta_K
Doxycycline
YNL286W CUS2
ZShift = −1.12
Z lm Score = −0.03
NG = 0
DB = 0
SM = 0
Delta_r
Doxycycline
YNL286W CUS2
ZShift = −0.59
Z lm Score = −0.64
NG = 0
DB = 0
SM = 0
Delta_AUC
Doxycycline

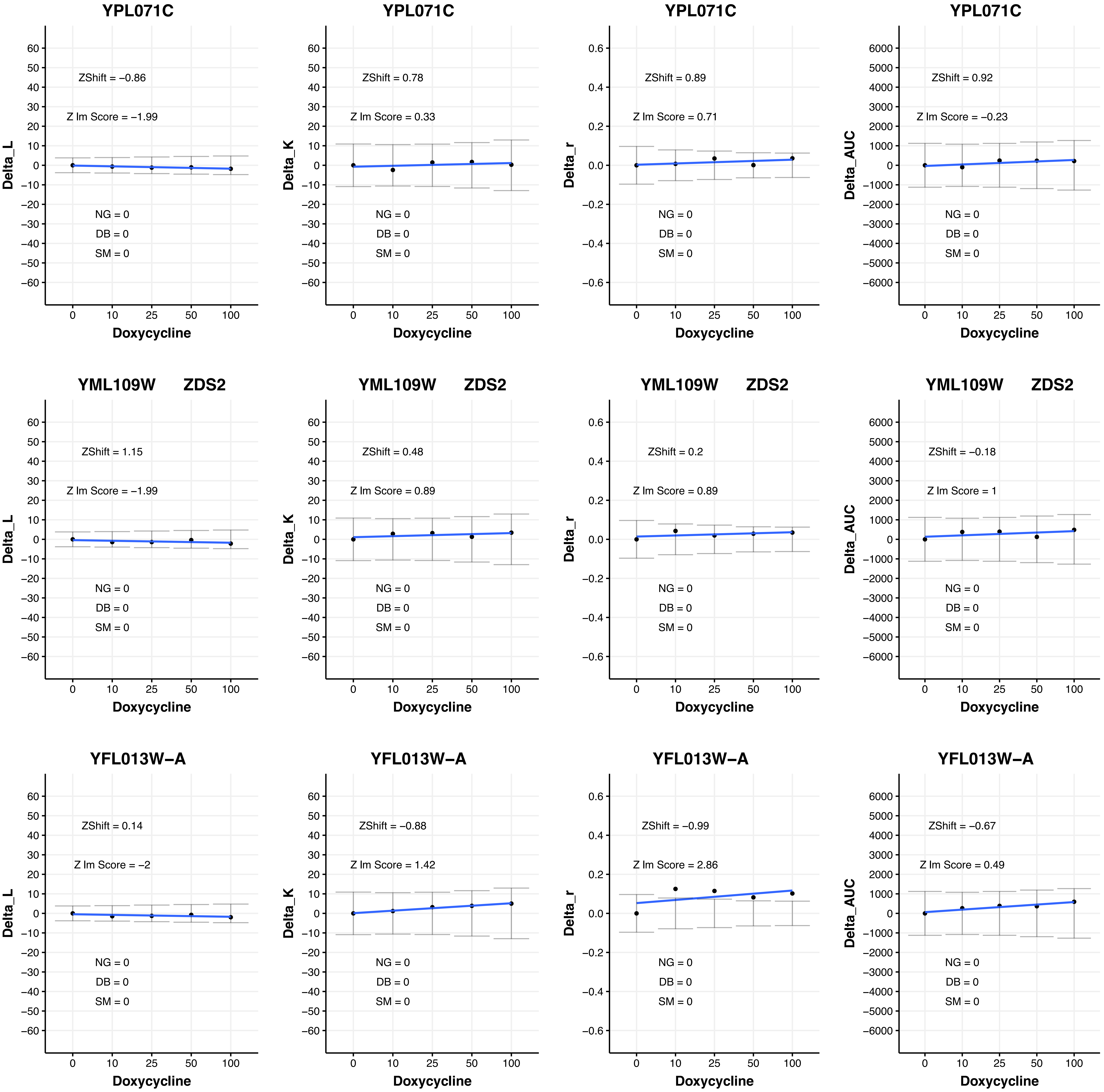

YPL071C
ZShift = −0.86
Z lm Score = −1.99
NG = 0
DB = 0
SM = 0
Delta_L
Doxycycline
YPL071C
ZShift = 0.78
Z lm Score = 0.33
NG = 0
DB = 0
SM = 0
Delta_K
Doxycycline
YPL071C
ZShift = 0.89
Z lm Score = 0.71
NG = 0
DB = 0
SM = 0
Delta_r
Doxycycline
YPL071C
ZShift = 0.92
Z lm Score = −0.23
NG = 0
DB = 0
SM = 0
Delta_AUC
Doxycycline
YML109W ZDS2
ZShift = 1.15
Z lm Score = −1.99
NG = 0
DB = 0
SM = 0
Delta_L
Doxycycline
YML109W ZDS2
ZShift = 0.48
Z lm Score = 0.89
NG = 0
DB = 0
SM = 0
Delta_K
Doxycycline
YML109W ZDS2
ZShift = 0.2
Z lm Score = 0.89
NG = 0
DB = 0
SM = 0
Delta_r
Doxycycline
YML109W ZDS2
ZShift = −0.18
Z lm Score = 1
NG = 0
DB = 0
SM = 0
Delta_AUC
Doxycycline
YFL013W−A
ZShift = 0.14
Z lm Score = −2
NG = 0
DB = 0
SM = 0
Delta_L
Doxycycline
YFL013W−A
ZShift = −0.88
Z lm Score = 1.42
NG = 0
DB = 0
SM = 0
Delta_K
Doxycycline
YFL013W−A
ZShift = −0.99
Z lm Score = 2.86
NG = 0
DB = 0
SM = 0
Delta_r
Doxycycline
YFL013W−A
ZShift = −0.67
Z lm Score = 0.49
NG = 0
DB = 0
SM = 0
Delta_AUC
Doxycycline

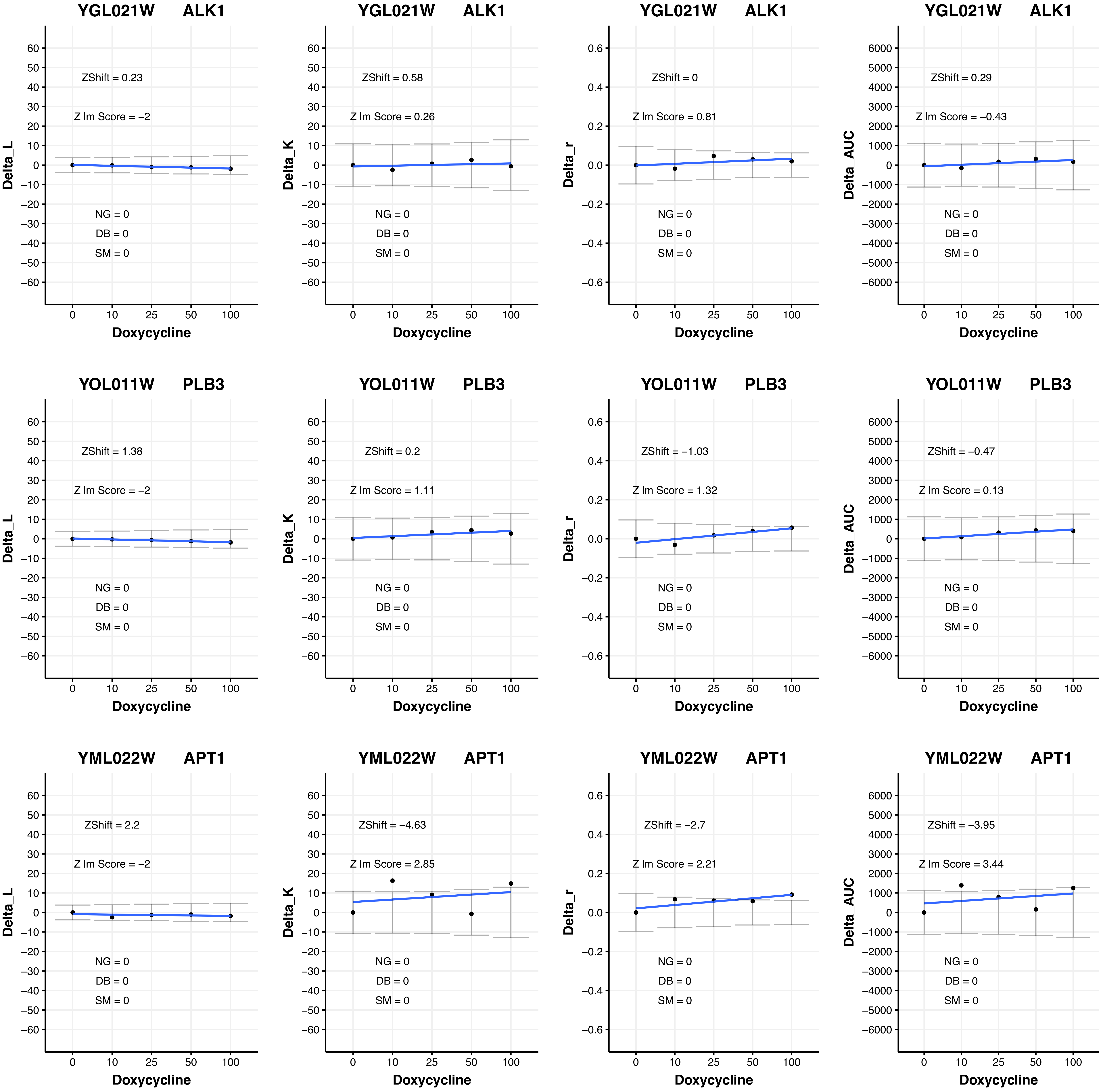

YGL021W ALK1
ZShift = 0.23
Z lm Score = −2
NG = 0
DB = 0
SM = 0
Delta_L
YGL021W ALK1
ZShift = 0.58
Z lm Score = 0.26
NG = 0
DB = 0
SM = 0
Delta_K
YGL021W ALK1
ZShift = 0
Z lm Score = 0.81
NG = 0
DB = 0
SM = 0
Delta_r
YGL021W ALK1
ZShift = 0.29
Z lm Score = −0.43
NG = 0
DB = 0
SM = 0
Delta_AUC
YOL011W PLB3
ZShift = 1.38
Z lm Score = −2
NG = 0
DB = 0
SM = 0
Delta_L
YOL011W PLB3
ZShift = 0.2
Z lm Score = 1.11
NG = 0
DB = 0
SM = 0
Delta_K
YOL011W PLB3
ZShift = −1.03
Z lm Score = 1.32
NG = 0
DB = 0
SM = 0
Delta_r
YOL011W PLB3
ZShift = −0.47
Z lm Score = 0.13
NG = 0
DB = 0
SM = 0
Delta_AUC
YML022W APT1
ZShift = 2.2
Z lm Score = −2
NG = 0
DB = 0
SM = 0
Delta_L
YML022W APT1
ZShift = −4.63
Z lm Score = 2.85
NG = 0
DB = 0
SM = 0
Delta_K
YML022W APT1
ZShift = −2.7
Z lm Score = 2.21
NG = 0
DB = 0
SM = 0
Delta_r
YML022W APT1
ZShift = −3.95
Z lm Score = 3.44
NG = 0
DB = 0
SM = 0
Delta_AUC
Doxycycline
0
10
25
50
100

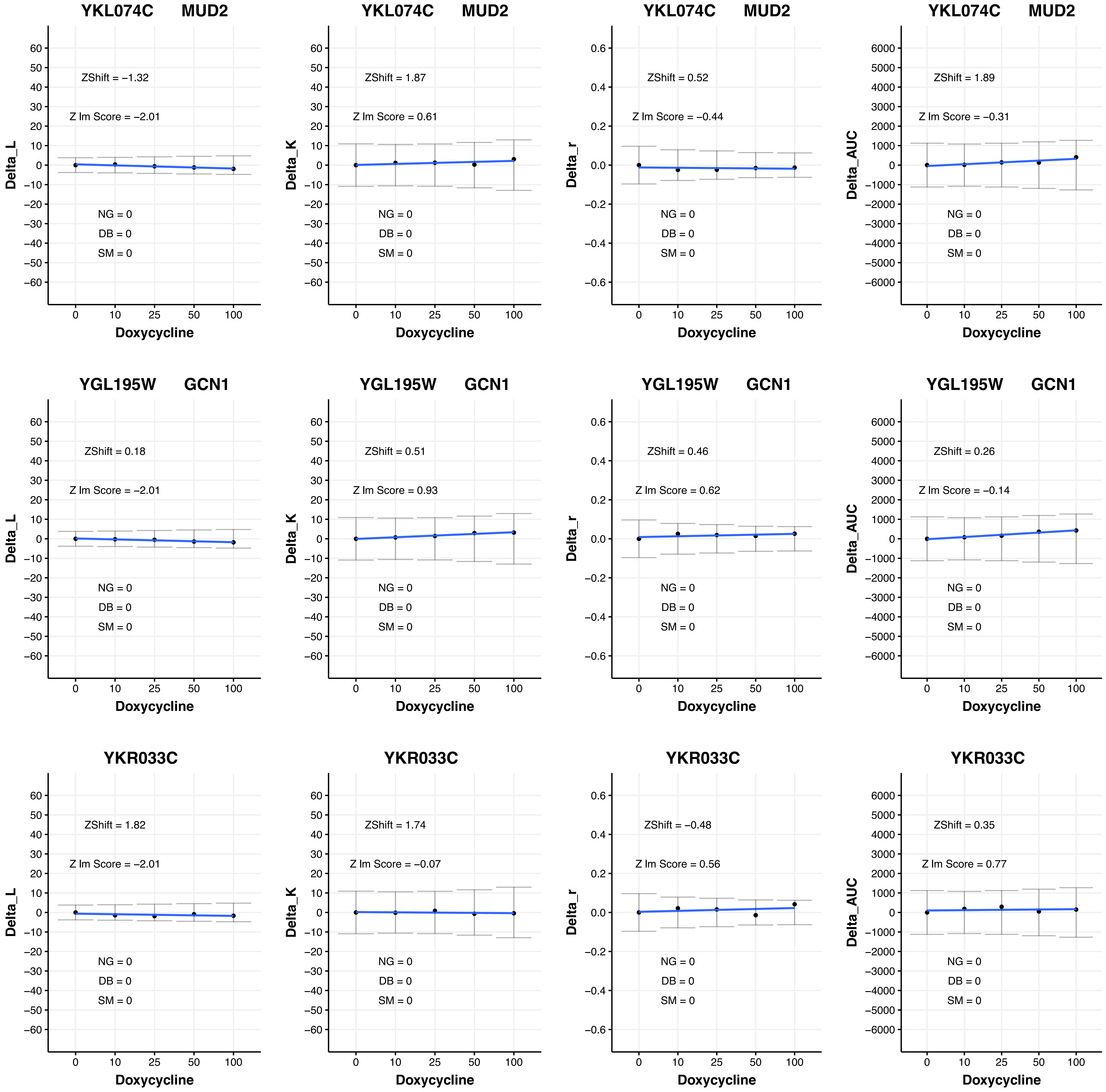
YKL074C MUD2
ZShift = −1.32
Z lm Score = −2.01
NG = 0
DB = 0
SM = 0
Delta_L
Doxycycline
YKL074C MUD2
ZShift = 1.87
Z lm Score = 0.61
NG = 0
DB = 0
SM = 0
Delta_K
Doxycycline
YKL074C MUD2
ZShift = 0.52
Z lm Score = −0.44
NG = 0
DB = 0
SM = 0
Delta_r
Doxycycline
YKL074C MUD2
ZShift = 1.89
Z lm Score = −0.31
NG = 0
DB = 0
SM = 0
Delta_AUC
Doxycycline
YGL195W GCN1
ZShift = 0.18
Z lm Score = −2.01
NG = 0
DB = 0
SM = 0
Delta_L
Doxycycline
YGL195W GCN1
ZShift = 0.51
Z lm Score = 0.93
NG = 0
DB = 0
SM = 0
Delta_K
Doxycycline
YGL195W GCN1
ZShift = 0.46
Z lm Score = 0.62
NG = 0
DB = 0
SM = 0
Delta_r
Doxycycline
YGL195W GCN1
ZShift = 0.26
Z lm Score = −0.14
NG = 0
DB = 0
SM = 0
Delta_AUC
Doxycycline
YKR033C
ZShift = 1.82
Z lm Score = −2.01
NG = 0
DB = 0
SM = 0
Delta_L
Doxycycline
YKR033C
ZShift = 1.74
Z lm Score = −0.07
NG = 0
DB = 0
SM = 0
Delta_K
Doxycycline
YKR033C
ZShift = −0.48
Z lm Score = 0.56
NG = 0
DB = 0
SM = 0
Delta_r
Doxycycline
YKR033C
ZShift = 0.35
Z lm Score = 0.77
NG = 0
DB = 0
SM = 0
Delta_AUC
Doxycycline

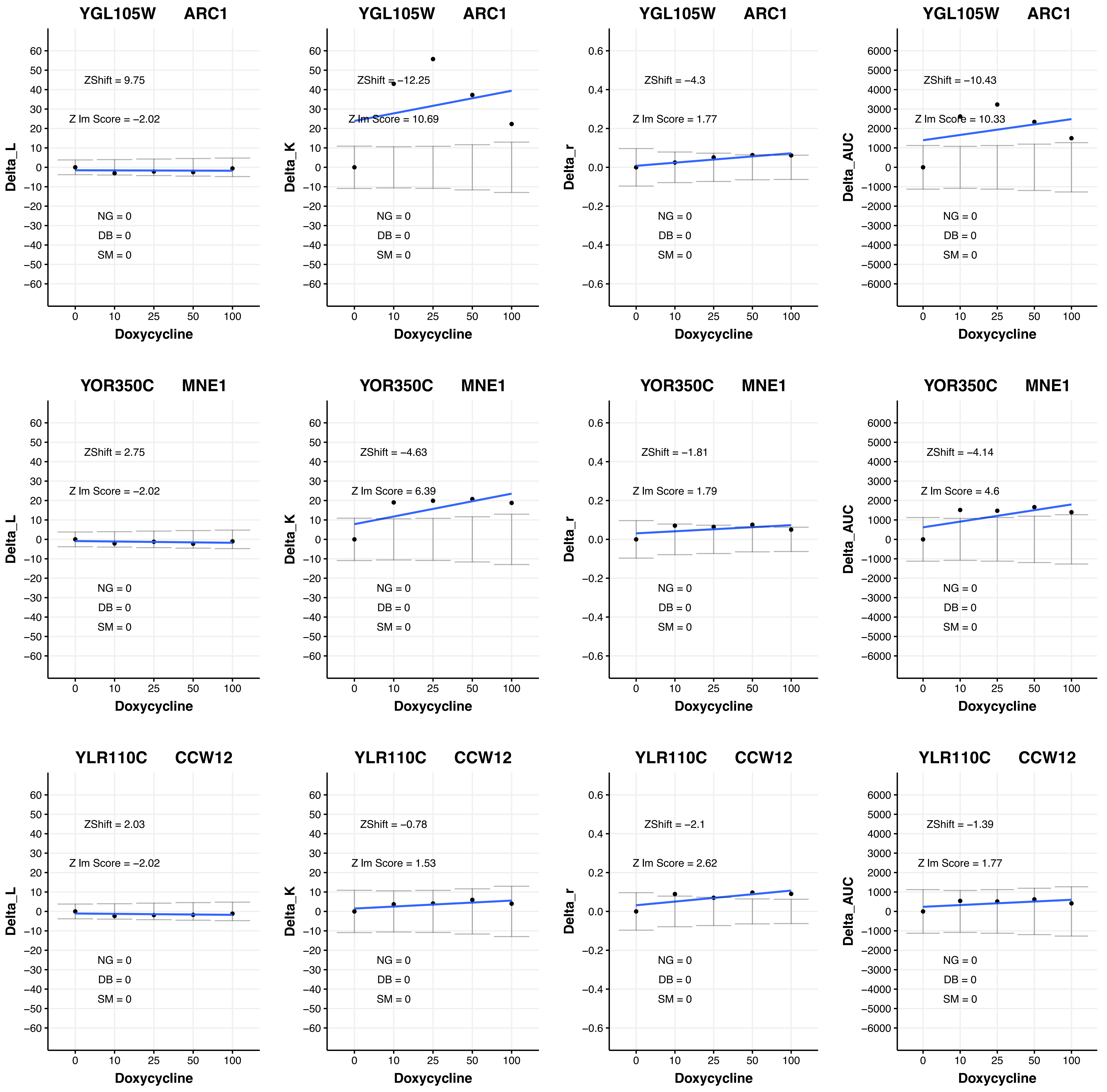
YGL105W ARC1
ZShift = 9.75
Z lm Score = −2.02
NG = 0
DB = 0
SM = 0
Delta_L
Doxycycline
YGL105W ARC1
ZShift = −12.25
Z lm Score = 10.69
NG = 0
DB = 0
SM = 0
Delta_K
Doxycycline
YGL105W ARC1
ZShift = −4.3
Z lm Score = 1.77
NG = 0
DB = 0
SM = 0
Delta_r
Doxycycline
YGL105W ARC1
ZShift = −10.43
Z lm Score = 10.33
NG = 0
DB = 0
SM = 0
Delta_AUC
Doxycycline
YOR350C MNE1
ZShift = 2.75
Z lm Score = −2.02
NG = 0
DB = 0
SM = 0
Delta_L
Doxycycline
YOR350C MNE1
ZShift = −4.63
Z lm Score = 6.39
NG = 0
DB = 0
SM = 0
Delta_K
Doxycycline
YOR350C MNE1
ZShift = −1.81
Z lm Score = 1.79
NG = 0
DB = 0
SM = 0
Delta_r
Doxycycline
YOR350C MNE1
ZShift = −4.14
Z lm Score = 4.6
NG = 0
DB = 0
SM = 0
Delta_AUC
Doxycycline
YLR110C CCW12
ZShift = 2.03
Z lm Score = −2.02
NG = 0
DB = 0
SM = 0
Delta_L
Doxycycline
YLR110C CCW12
ZShift = −0.78
Z lm Score = 1.53
NG = 0
DB = 0
SM = 0
Delta_K
Doxycycline
YLR110C CCW12
ZShift = −2.1
Z lm Score = 2.62
NG = 0
DB = 0
SM = 0
Delta_r
Doxycycline
YLR110C CCW12
ZShift = −1.39
Z lm Score = 1.77
NG = 0
DB = 0
SM = 0
Delta_AUC
Doxycycline

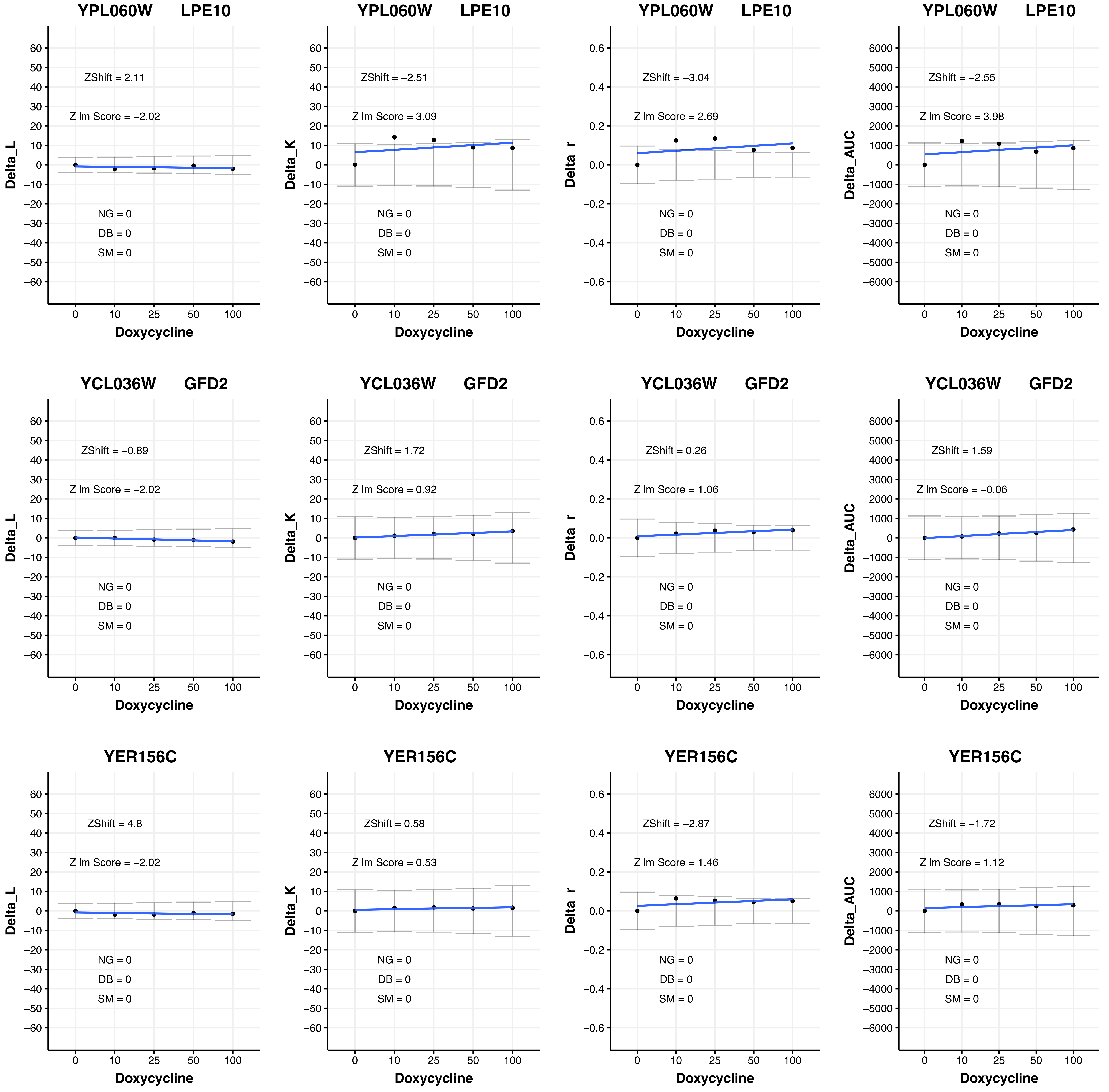

YPL060W LPE10
ZShift = 2.11
Z lm Score = −2.02
NG = 0
DB = 0
SM = 0
Delta_L
Doxycycline
YPL060W LPE10
ZShift = −2.51
Z lm Score = 3.09
NG = 0
DB = 0
SM = 0
Delta_K
Doxycycline
YPL060W LPE10
ZShift = −3.04
Z lm Score = 2.69
NG = 0
DB = 0
SM = 0
Delta_r
Doxycycline
YPL060W LPE10
ZShift = −2.55
Z lm Score = 3.98
NG = 0
DB = 0
SM = 0
Delta_AUC
Doxycycline
YCL036W GFD2
ZShift = −0.89
Z lm Score = −2.02
NG = 0
DB = 0
SM = 0
Delta_L
Doxycycline
YCL036W GFD2
ZShift = 1.72
Z lm Score = 0.92
NG = 0
DB = 0
SM = 0
Delta_K
Doxycycline
YCL036W GFD2
ZShift = 0.26
Z lm Score = 1.06
NG = 0
DB = 0
SM = 0
Delta_r
Doxycycline
YCL036W GFD2
ZShift = 1.59
Z lm Score = −0.06
NG = 0
DB = 0
SM = 0
Delta_AUC
Doxycycline
YER156C
ZShift = 4.8
Z lm Score = −2.02
NG = 0
DB = 0
SM = 0
Delta_L
Doxycycline
YER156C
ZShift = 0.58
Z lm Score = 0.53
NG = 0
DB = 0
SM = 0
Delta_K
Doxycycline
YER156C
ZShift = −2.87
Z lm Score = 1.46
NG = 0
DB = 0
SM = 0
Delta_r
Doxycycline
YER156C
ZShift = −1.72
Z lm Score = 1.12
NG = 0
DB = 0
SM = 0
Delta_AUC
Doxycycline

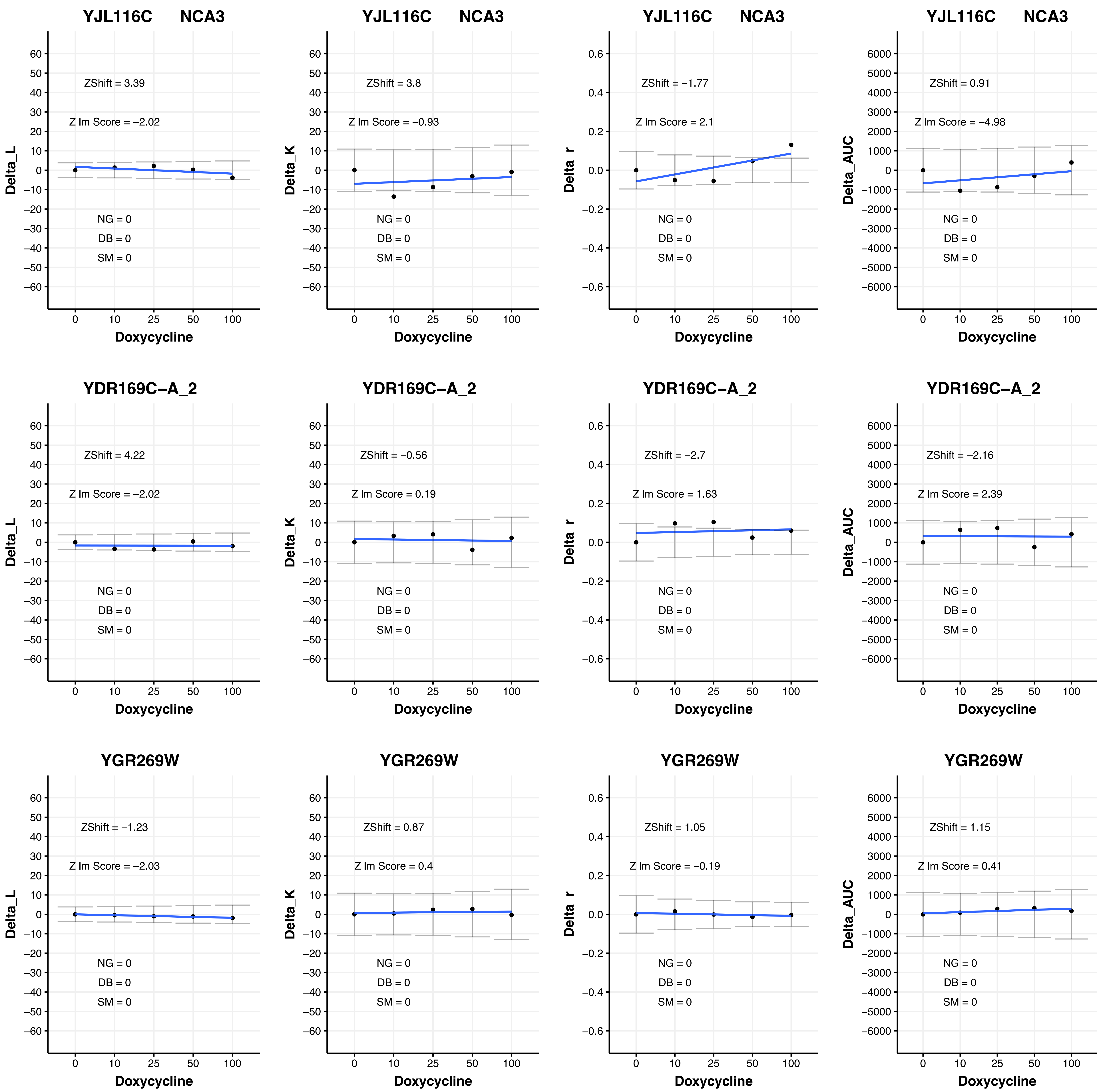

YJL116C NCA3
ZShift = 3.39
Z lm Score = −2.02
NG = 0
DB = 0
SM = 0
Delta_L
Doxycycline
YJL116C NCA3
ZShift = 3.8
Z lm Score = −0.93
NG = 0
DB = 0
SM = 0
Delta_K
Doxycycline
YJL116C NCA3
ZShift = −1.77
Z lm Score = 2.1
NG = 0
DB = 0
SM = 0
Delta_r
Doxycycline
YJL116C NCA3
ZShift = 0.91
Z lm Score = −4.98
NG = 0
DB = 0
SM = 0
Delta_AUC
Doxycycline
YDR169C−A_2
ZShift = 4.22
Z lm Score = −2.02
NG = 0
DB = 0
SM = 0
Delta_L
Doxycycline
YDR169C−A_2
ZShift = −0.56
Z lm Score = 0.19
NG = 0
DB = 0
SM = 0
Delta_K
Doxycycline
YDR169C−A_2
ZShift = −2.7
Z lm Score = 1.63
NG = 0
DB = 0
SM = 0
Delta_r
Doxycycline
YDR169C−A_2
ZShift = −2.16
Z lm Score = 2.39
NG = 0
DB = 0
SM = 0
Delta_AUC
Doxycycline
YGR269W
ZShift = −1.23
Z lm Score = −2.03
NG = 0
DB = 0
SM = 0
Delta_L
Doxycycline
YGR269W
ZShift = 0.87
Z lm Score = 0.4
NG = 0
DB = 0
SM = 0
Delta_K
Doxycycline
YGR269W
ZShift = 1.05
Z lm Score = −0.19
NG = 0
DB = 0
SM = 0
Delta_r
Doxycycline
YGR269W
ZShift = 1.15
Z lm Score = 0.41
NG = 0
DB = 0
SM = 0
Delta_AUC
Doxycycline

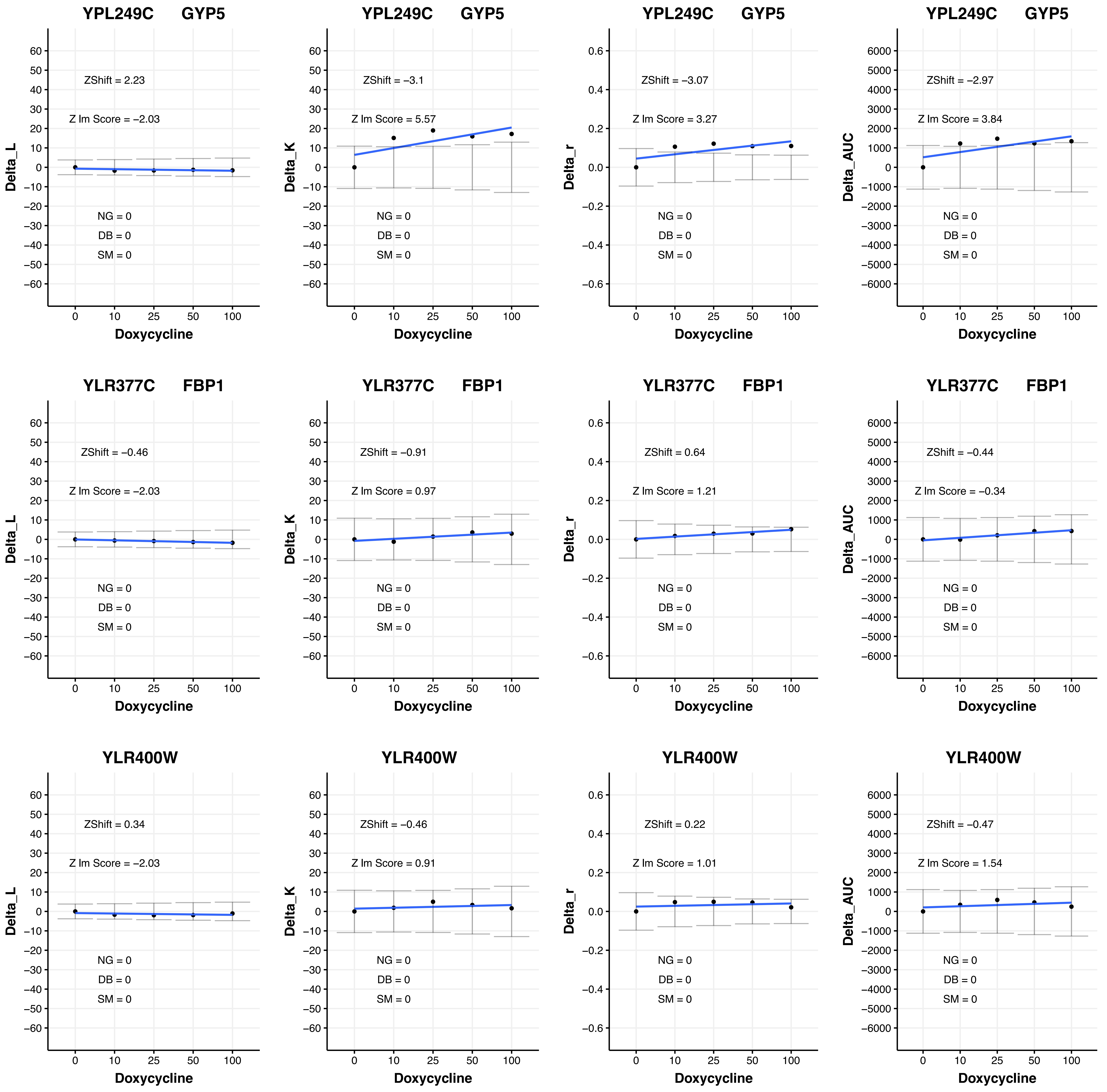
YPL249C GYP5
ZShift = 2.23
Z lm Score = −2.03
NG = 0
DB = 0
SM = 0
Delta_L
Doxycycline
YPL249C GYP5
ZShift = −3.1
Z lm Score = 5.57
NG = 0
DB = 0
SM = 0
Delta_K
Doxycycline
YPL249C GYP5
ZShift = −3.07
Z lm Score = 3.27
NG = 0
DB = 0
SM = 0
Delta_r
Doxycycline
YPL249C GYP5
ZShift = −2.97
Z lm Score = 3.84
NG = 0
DB = 0
SM = 0
Delta_AUC
Doxycycline
YLR377C FBP1
ZShift = −0.46
Z lm Score = −2.03
NG = 0
DB = 0
SM = 0
Delta_L
Doxycycline
YLR377C FBP1
ZShift = −0.91
Z lm Score = 0.97
NG = 0
DB = 0
SM = 0
Delta_K
Doxycycline
YLR377C FBP1
ZShift = 0.64
Z lm Score = 1.21
NG = 0
DB = 0
SM = 0
Delta_r
Doxycycline
YLR377C FBP1
ZShift = −0.44
Z lm Score = −0.34
NG = 0
DB = 0
SM = 0
Delta_AUC
Doxycycline
YLR400W
ZShift = 0.34
Z lm Score = −2.03
NG = 0
DB = 0
SM = 0
Delta_L
Doxycycline
YLR400W
ZShift = −0.46
Z lm Score = 0.91
NG = 0
DB = 0
SM = 0
Delta_K
Doxycycline
YLR400W
ZShift = 0.22
Z lm Score = 1.01
NG = 0
DB = 0
SM = 0
Delta_r
Doxycycline
YLR400W
ZShift = −0.47
Z lm Score = 1.54
NG = 0
DB = 0
SM = 0
Delta_AUC
Doxycycline

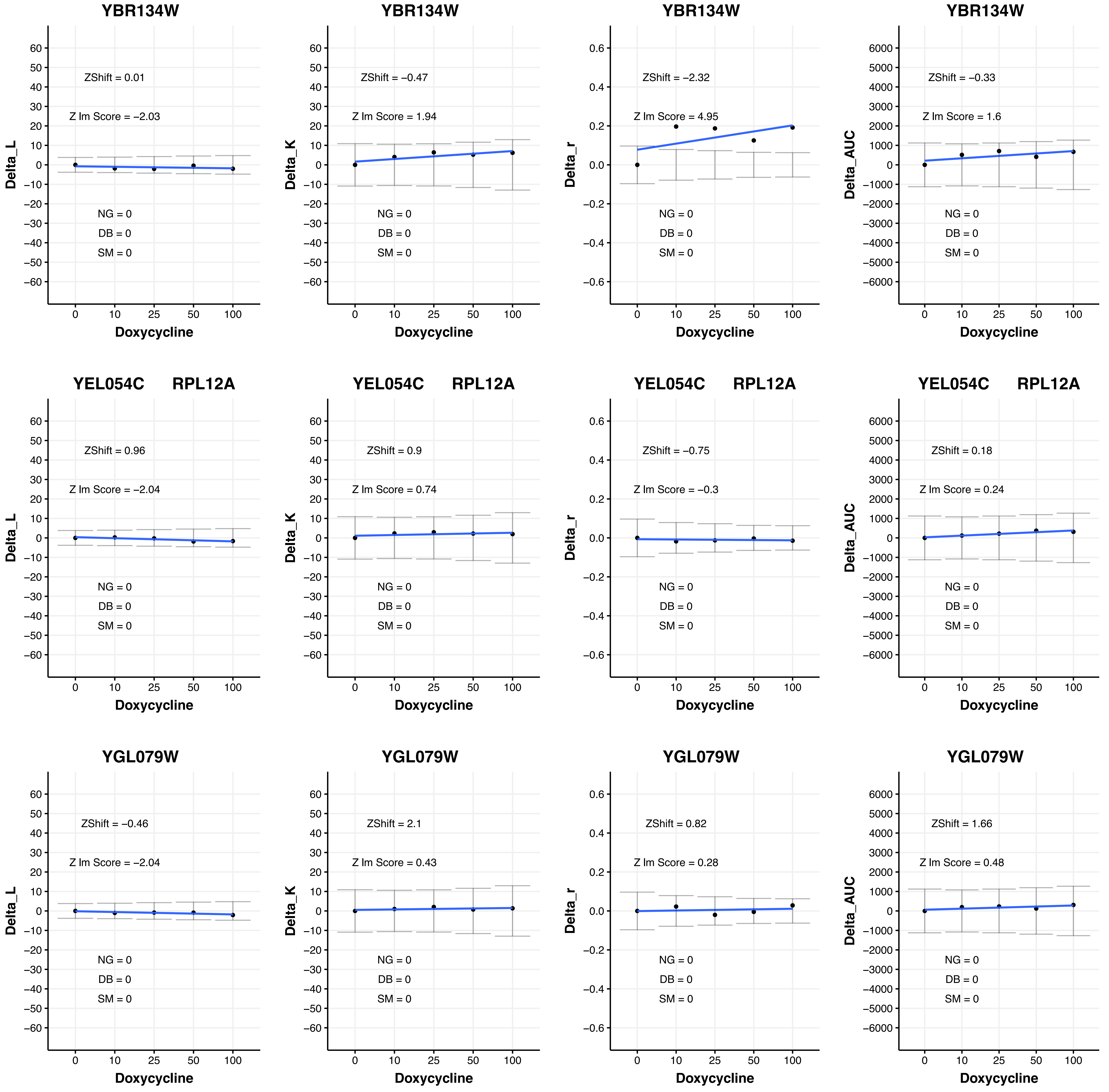

YBR134W
ZShift = 0.01
Z lm Score = −2.03
NG = 0
DB = 0
SM = 0
Delta_L
Doxycycline
YBR134W
ZShift = −0.47
Z lm Score = 1.94
NG = 0
DB = 0
SM = 0
Delta_K
Doxycycline
YBR134W
ZShift = −2.32
Z lm Score = 4.95
NG = 0
DB = 0
SM = 0
Delta_r
Doxycycline
YBR134W
ZShift = −0.33
Z lm Score = 1.6
NG = 0
DB = 0
SM = 0
Delta_AUC
Doxycycline
YEL054C RPL12A
ZShift = 0.96
Z lm Score = −2.04
NG = 0
DB = 0
SM = 0
Delta_L
Doxycycline
YEL054C RPL12A
ZShift = 0.9
Z lm Score = 0.74
NG = 0
DB = 0
SM = 0
Delta_K
Doxycycline
YEL054C RPL12A
ZShift = −0.75
Z lm Score = −0.3
NG = 0
DB = 0
SM = 0
Delta_r
Doxycycline
YEL054C RPL12A
ZShift = 0.18
Z lm Score = 0.24
NG = 0
DB = 0
SM = 0
Delta_AUC
Doxycycline
YGL079W
ZShift = −0.46
Z lm Score = −2.04
NG = 0
DB = 0
SM = 0
Delta_L
Doxycycline
YGL079W
ZShift = 2.1
Z lm Score = 0.43
NG = 0
DB = 0
SM = 0
Delta_K
Doxycycline
YGL079W
ZShift = 0.82
Z lm Score = 0.28
NG = 0
DB = 0
SM = 0
Delta_r
Doxycycline
YGL079W
ZShift = 1.66
Z lm Score = 0.48
NG = 0
DB = 0
SM = 0
Delta_AUC
Doxycycline

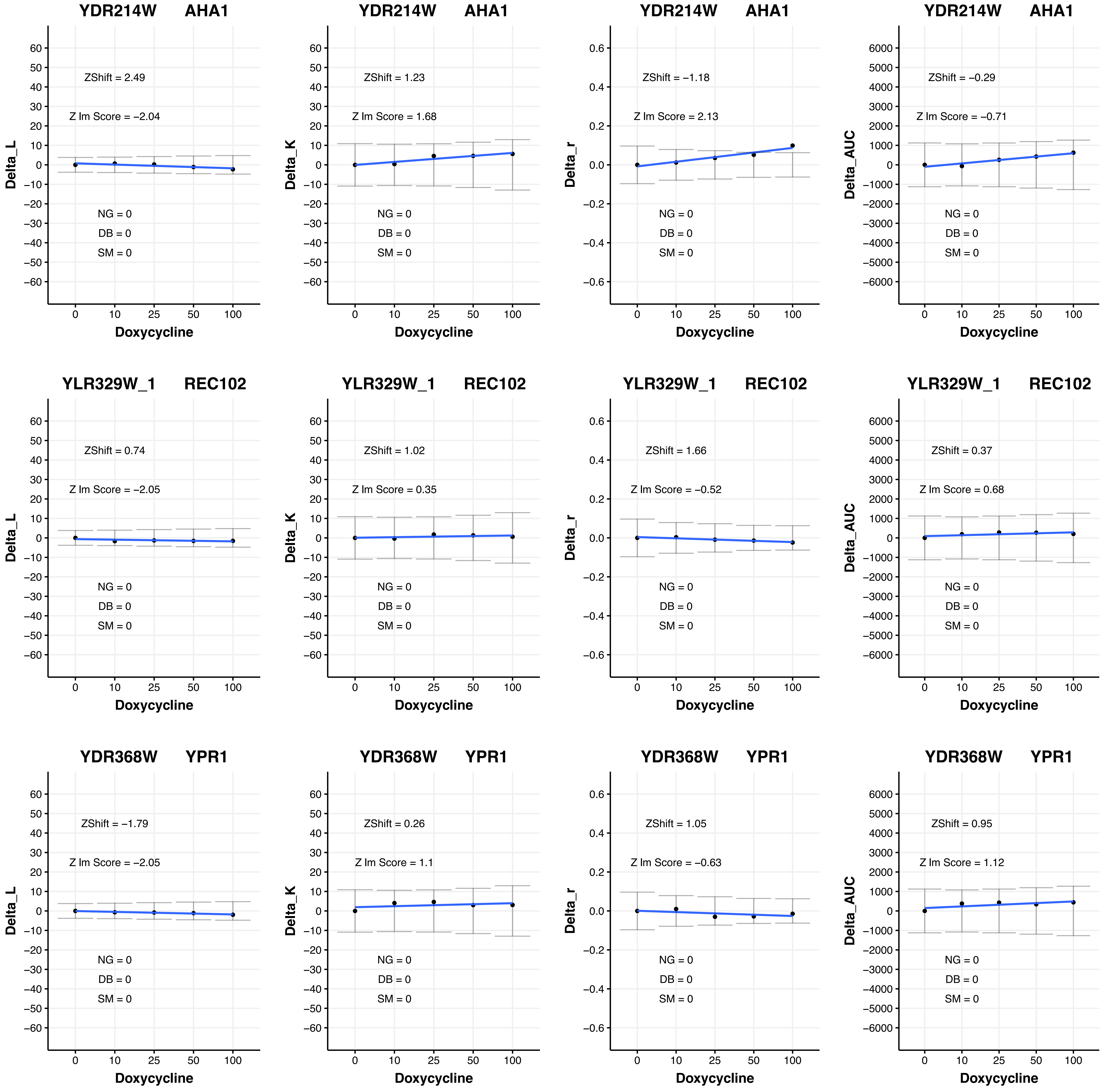

YDR214W AHA1
Delta_L
ZShift = 2.49
Z lm Score = −2.04
NG = 0
DB = 0
SM = 0
Doxycycline
YDR214W AHA1
Delta_K
ZShift = 1.23
Z lm Score = 1.68
NG = 0
DB = 0
SM = 0
Doxycycline
YDR214W AHA1
Delta_r
ZShift = −1.18
Z lm Score = 2.13
NG = 0
DB = 0
SM = 0
Doxycycline
YDR214W AHA1
Delta_AUC
ZShift = −0.29
Z lm Score = −0.71
NG = 0
DB = 0
SM = 0
Doxycycline
YLR329W_1 REC102
Delta_L
ZShift = 0.74
Z lm Score = −2.05
NG = 0
DB = 0
SM = 0
Doxycycline
YLR329W_1 REC102
Delta_K
ZShift = 1.02
Z lm Score = 0.35
NG = 0
DB = 0
SM = 0
Doxycycline
YLR329W_1 REC102
Delta_r
ZShift = 1.66
Z lm Score = −0.52
NG = 0
DB = 0
SM = 0
Doxycycline
YLR329W_1 REC102
Delta_AUC
ZShift = 0.37
Z lm Score = 0.68
NG = 0
DB = 0
SM = 0
Doxycycline
YDR368W YPR1
Delta_L
ZShift = −1.79
Z lm Score = −2.05
NG = 0
DB = 0
SM = 0
Doxycycline
YDR368W YPR1
Delta_K
ZShift = 0.26
Z lm Score = 1.1
NG = 0
DB = 0
SM = 0
Doxycycline
YDR368W YPR1
Delta_r
ZShift = 1.05
Z lm Score = −0.63
NG = 0
DB = 0
SM = 0
Doxycycline
YDR368W YPR1
Delta_AUC
ZShift = 0.95
Z lm Score = 1.12
NG = 0
DB = 0
SM = 0
Doxycycline

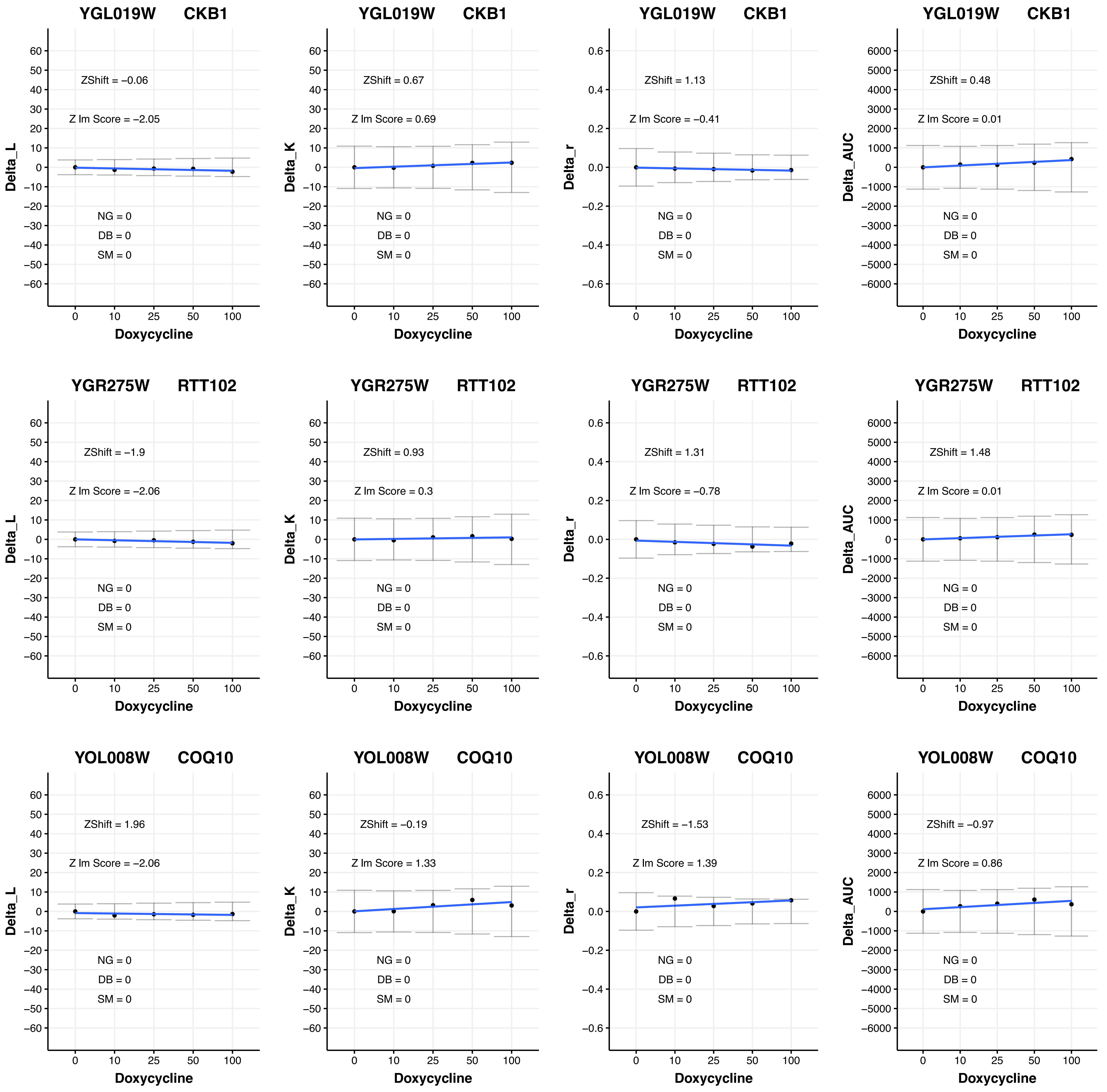
YGL019W CKB1
ZShift = −0.06
Z lm Score = −2.05
NG = 0
DB = 0
SM = 0
Delta_L
Doxycycline
YGL019W CKB1
ZShift = 0.67
Z lm Score = 0.69
NG = 0
DB = 0
SM = 0
Delta_K
Doxycycline
YGL019W CKB1
ZShift = 1.13
Z lm Score = −0.41
NG = 0
DB = 0
SM = 0
Delta_r
Doxycycline
YGL019W CKB1
ZShift = 0.48
Z lm Score = 0.01
NG = 0
DB = 0
SM = 0
Delta_AUC
Doxycycline
YGR275W RTT102
ZShift = −1.9
Z lm Score = −2.06
NG = 0
DB = 0
SM = 0
Delta_L
Doxycycline
YGR275W RTT102
ZShift = 0.93
Z lm Score = 0.3
NG = 0
DB = 0
SM = 0
Delta_K
Doxycycline
YGR275W RTT102
ZShift = 1.31
Z lm Score = −0.78
NG = 0
DB = 0
SM = 0
Delta_r
Doxycycline
YGR275W RTT102
ZShift = 1.48
Z lm Score = 0.01
NG = 0
DB = 0
SM = 0
Delta_AUC
Doxycycline
YOL008W COQ10
ZShift = 1.96
Z lm Score = −2.06
NG = 0
DB = 0
SM = 0
Delta_L
Doxycycline
YOL008W COQ10
ZShift = −0.19
Z lm Score = 1.33
NG = 0
DB = 0
SM = 0
Delta_K
Doxycycline
YOL008W COQ10
ZShift = −1.53
Z lm Score = 1.39
NG = 0
DB = 0
SM = 0
Delta_r
Doxycycline
YOL008W COQ10
ZShift = −0.97
Z lm Score = 0.86
NG = 0
DB = 0
SM = 0
Delta_AUC
Doxycycline

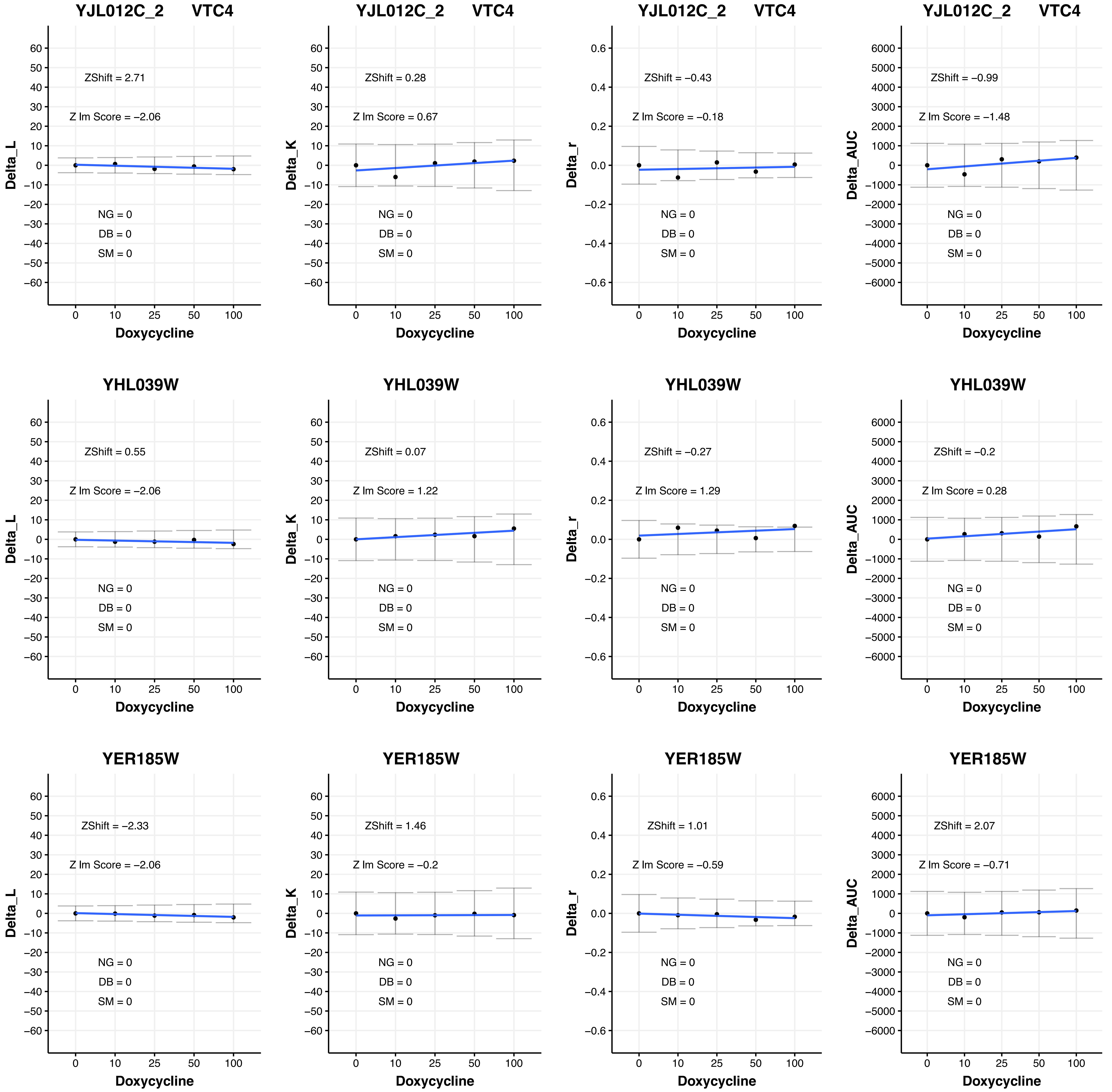

YJL012C_2 VTC4
ZShift = 2.71
Z lm Score = −2.06
NG = 0
DB = 0
SM = 0
Delta_L
Doxycycline
YJL012C_2 VTC4
ZShift = 0.28
Z lm Score = 0.67
NG = 0
DB = 0
SM = 0
Delta_K
Doxycycline
YJL012C_2 VTC4
ZShift = −0.43
Z lm Score = −0.18
NG = 0
DB = 0
SM = 0
Delta_r
Doxycycline
YJL012C_2 VTC4
ZShift = −0.99
Z lm Score = −1.48
NG = 0
DB = 0
SM = 0
Delta_AUC
Doxycycline
YHL039W
ZShift = 0.55
Z lm Score = −2.06
NG = 0
DB = 0
SM = 0
Delta_L
Doxycycline
YHL039W
ZShift = 0.07
Z lm Score = 1.22
NG = 0
DB = 0
SM = 0
Delta_K
Doxycycline
YHL039W
ZShift = −0.27
Z lm Score = 1.29
NG = 0
DB = 0
SM = 0
Delta_r
Doxycycline
YHL039W
ZShift = −0.2
Z lm Score = 0.28
NG = 0
DB = 0
SM = 0
Delta_AUC
Doxycycline
YER185W
ZShift = −2.33
Z lm Score = −2.06
NG = 0
DB = 0
SM = 0
Delta_L
Doxycycline
YER185W
ZShift = 1.46
Z lm Score = −0.2
NG = 0
DB = 0
SM = 0
Delta_K
Doxycycline
YER185W
ZShift = 1.01
Z lm Score = −0.59
NG = 0
DB = 0
SM = 0
Delta_r
Doxycycline
YER185W
ZShift = 2.07
Z lm Score = −0.71
NG = 0
DB = 0
SM = 0
Delta_AUC
Doxycycline

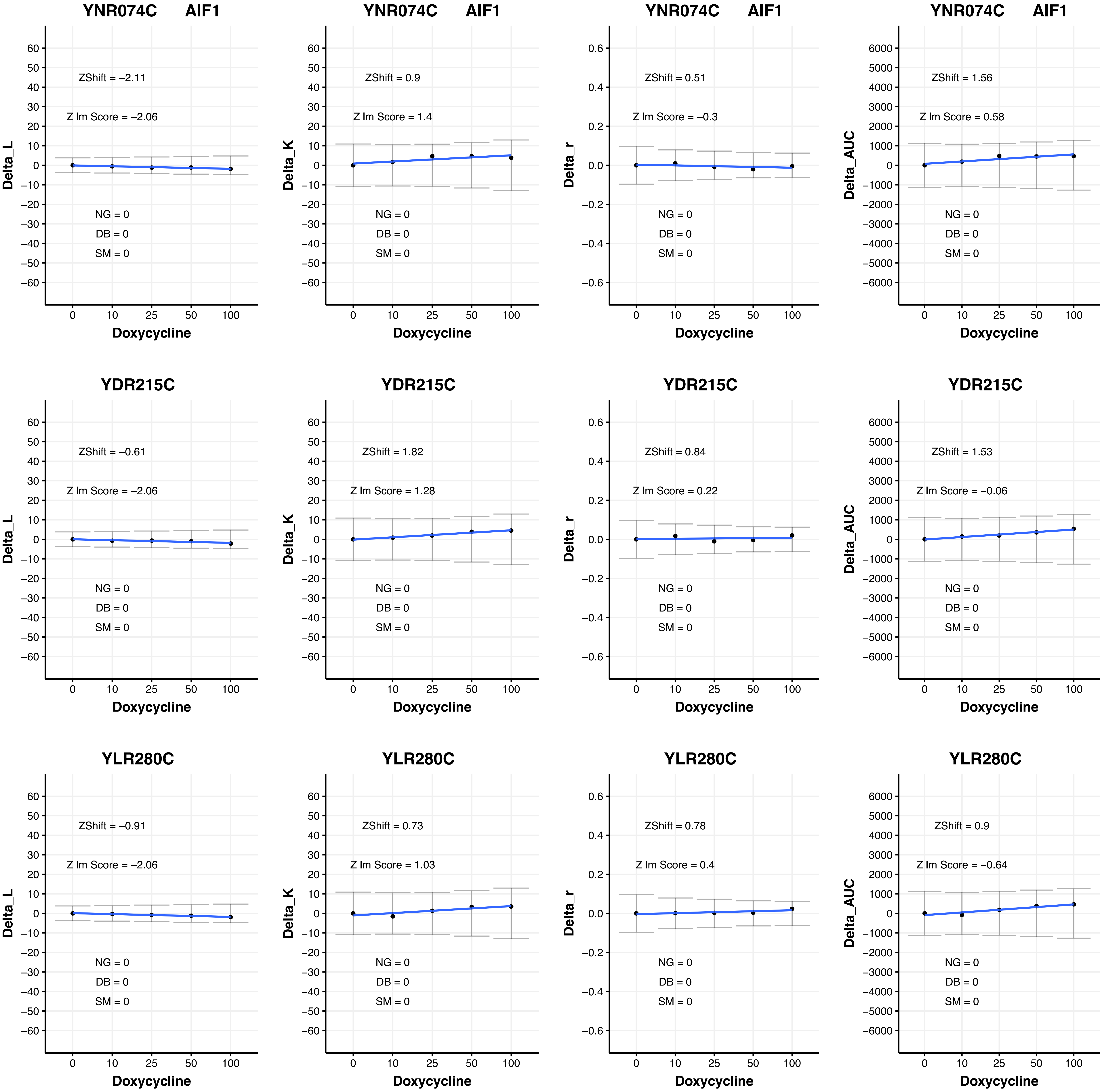
YNR074C AIF1
ZShift = −2.11
Z lm Score = −2.06
NG = 0
DB = 0
SM = 0
Delta_L
Doxycycline
YNR074C AIF1
ZShift = 0.9
Z lm Score = 1.4
NG = 0
DB = 0
SM = 0
Delta_K
Doxycycline
YNR074C AIF1
ZShift = 0.51
Z lm Score = −0.3
NG = 0
DB = 0
SM = 0
Delta_r
Doxycycline
YNR074C AIF1
ZShift = 1.56
Z lm Score = 0.58
NG = 0
DB = 0
SM = 0
Delta_AUC
Doxycycline
YDR215C
ZShift = −0.61
Z lm Score = −2.06
NG = 0
DB = 0
SM = 0
Delta_L
Doxycycline
YDR215C
ZShift = 1.82
Z lm Score = 1.28
NG = 0
DB = 0
SM = 0
Delta_K
Doxycycline
YDR215C
ZShift = 0.84
Z lm Score = 0.22
NG = 0
DB = 0
SM = 0
Delta_r
Doxycycline
YDR215C
ZShift = 1.53
Z lm Score = −0.06
NG = 0
DB = 0
SM = 0
Delta_AUC
Doxycycline
YLR280C
ZShift = −0.91
Z lm Score = −2.06
NG = 0
DB = 0
SM = 0
Delta_L
Doxycycline
YLR280C
ZShift = 0.73
Z lm Score = 1.03
NG = 0
DB = 0
SM = 0
Delta_K
Doxycycline
YLR280C
ZShift = 0.78
Z lm Score = 0.4
NG = 0
DB = 0
SM = 0
Delta_r
Doxycycline
YLR280C
ZShift = 0.9
Z lm Score = −0.64
NG = 0
DB = 0
SM = 0
Delta_AUC
Doxycycline

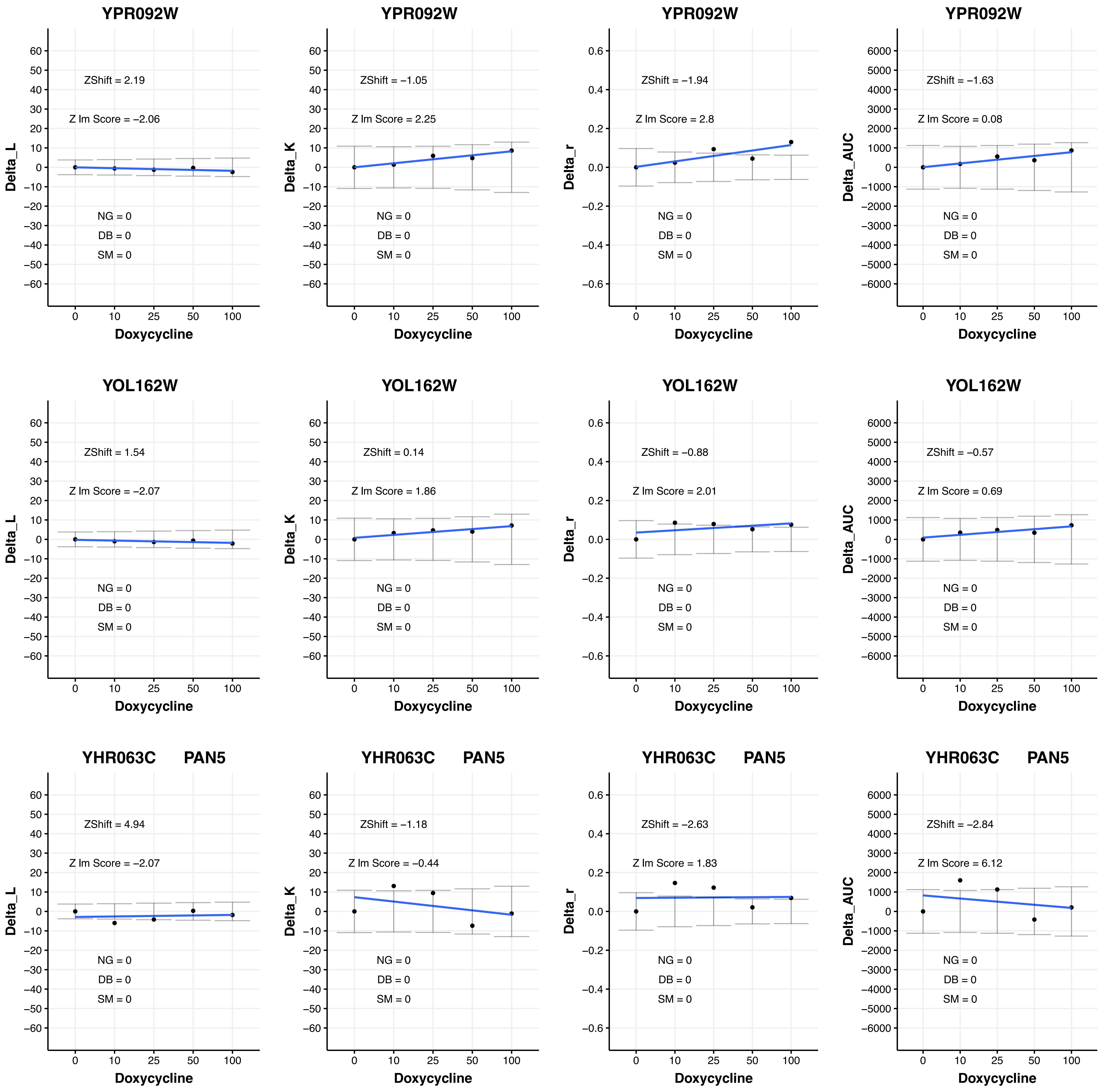
YPR092W
ZShift = 2.19
Z lm Score = −2.06
Delta_L
NG = 0
DB = 0
SM = 0
Doxycycline
YPR092W
ZShift = −1.05
Z lm Score = 2.25
Delta_K
NG = 0
DB = 0
SM = 0
Doxycycline
YPR092W
ZShift = −1.94
Z lm Score = 2.8
Delta_r
NG = 0
DB = 0
SM = 0
Doxycycline
YPR092W
ZShift = −1.63
Z lm Score = 0.08
Delta_AUC
NG = 0
DB = 0
SM = 0
Doxycycline
YOL162W
ZShift = 1.54
Z lm Score = −2.07
Delta_L
NG = 0
DB = 0
SM = 0
Doxycycline
YOL162W
ZShift = 0.14
Z lm Score = 1.86
Delta_K
NG = 0
DB = 0
SM = 0
Doxycycline
YOL162W
ZShift = −0.88
Z lm Score = 2.01
Delta_r
NG = 0
DB = 0
SM = 0
Doxycycline
YOL162W
ZShift = −0.57
Z lm Score = 0.69
Delta_AUC
NG = 0
DB = 0
SM = 0
Doxycycline
YHR063C PAN5
ZShift = 4.94
Z lm Score = −2.07
Delta_L
NG = 0
DB = 0
SM = 0
Doxycycline
YHR063C PAN5
ZShift = −1.18
Z lm Score = −0.44
Delta_K
NG = 0
DB = 0
SM = 0
Doxycycline
YHR063C PAN5
ZShift = −2.63
Z lm Score = 1.83
Delta_r
NG = 0
DB = 0
SM = 0
Doxycycline
YHR063C PAN5
ZShift = −2.84
Z lm Score = 6.12
Delta_AUC
NG = 0
DB = 0
SM = 0
Doxycycline

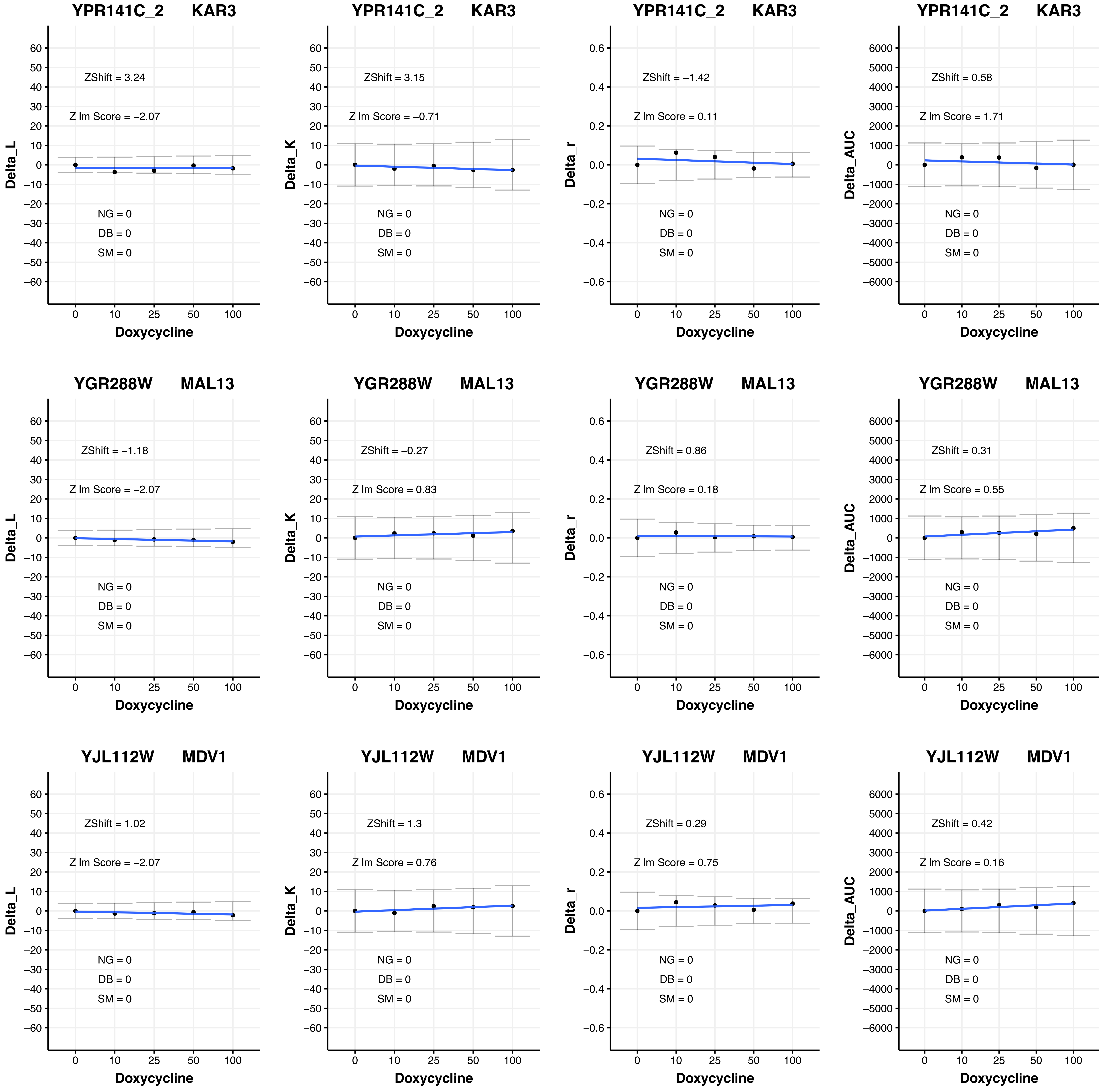
YPR141C_2 KAR3
ZShift = 3.24
Z lm Score = −2.07
NG = 0
DB = 0
SM = 0
Delta_L
Doxycycline
YPR141C_2 KAR3
ZShift = 3.15
Z lm Score = −0.71
NG = 0
DB = 0
SM = 0
Delta_K
Doxycycline
YPR141C_2 KAR3
ZShift = −1.42
Z lm Score = 0.11
NG = 0
DB = 0
SM = 0
Delta_r
Doxycycline
YPR141C_2 KAR3
ZShift = 0.58
Z lm Score = 1.71
NG = 0
DB = 0
SM = 0
Delta_AUC
Doxycycline
YGR288W MAL13
ZShift = −1.18
Z lm Score = −2.07
NG = 0
DB = 0
SM = 0
Delta_L
Doxycycline
YGR288W MAL13
ZShift = −0.27
Z lm Score = 0.83
NG = 0
DB = 0
SM = 0
Delta_K
Doxycycline
YGR288W MAL13
ZShift = 0.86
Z lm Score = 0.18
NG = 0
DB = 0
SM = 0
Delta_r
Doxycycline
YGR288W MAL13
ZShift = 0.31
Z lm Score = 0.55
NG = 0
DB = 0
SM = 0
Delta_AUC
Doxycycline
YJL112W MDV1
ZShift = 1.02
Z lm Score = −2.07
NG = 0
DB = 0
SM = 0
Delta_L
Doxycycline
YJL112W MDV1
ZShift = 1.3
Z lm Score = 0.76
NG = 0
DB = 0
SM = 0
Delta_K
Doxycycline
YJL112W MDV1
ZShift = 0.29
Z lm Score = 0.75
NG = 0
DB = 0
SM = 0
Delta_r
Doxycycline
YJL112W MDV1
ZShift = 0.42
Z lm Score = 0.16
NG = 0
DB = 0
SM = 0
Delta_AUC
Doxycycline

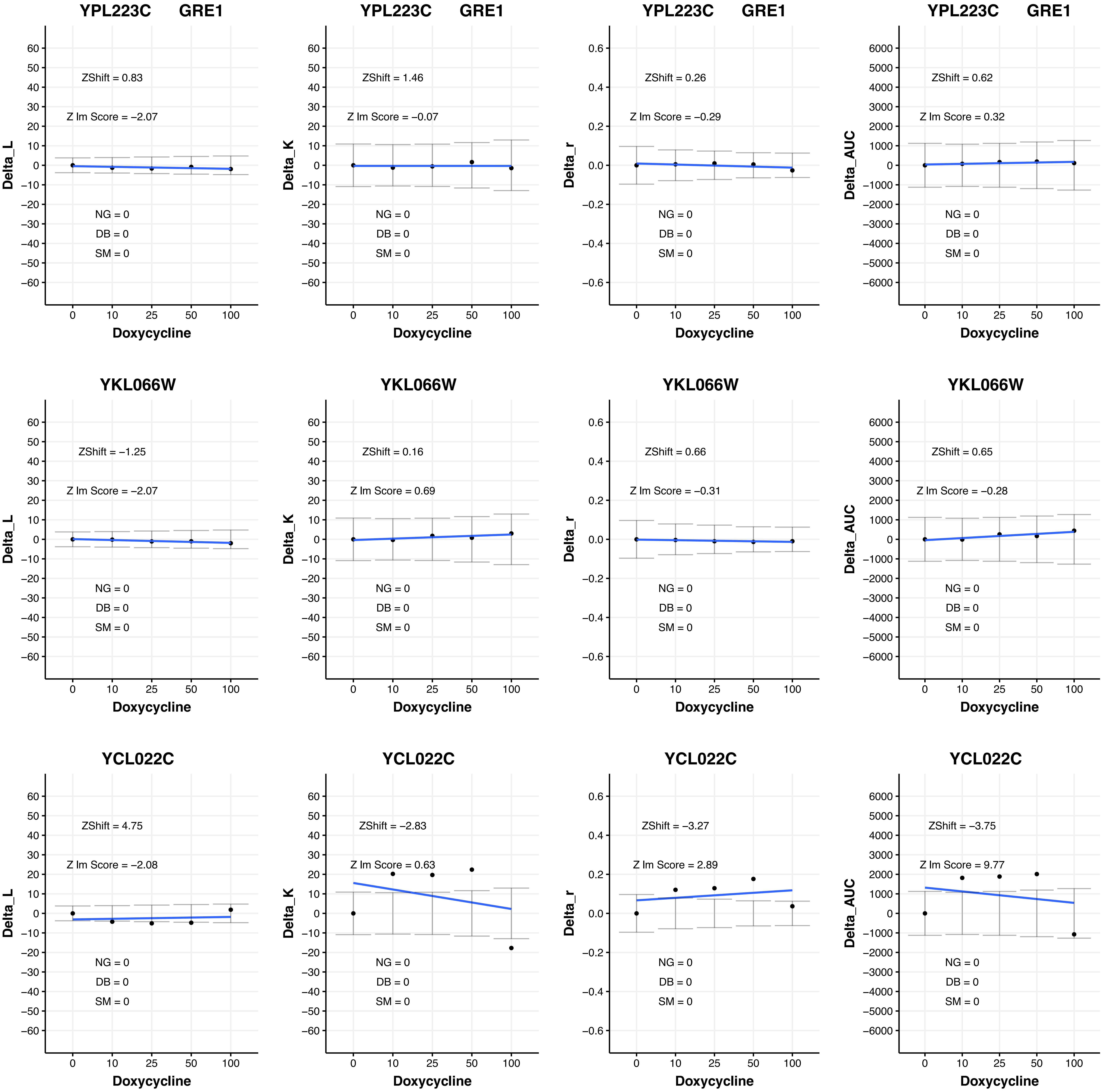

YPL223C GRE1
ZShift = 0.83
Z lm Score = −2.07
NG = 0
DB = 0
SM = 0
Delta_L
Doxycycline
YPL223C GRE1
ZShift = 1.46
Z lm Score = −0.07
NG = 0
DB = 0
SM = 0
Delta_K
Doxycycline
YPL223C GRE1
ZShift = 0.26
Z lm Score = −0.29
NG = 0
DB = 0
SM = 0
Delta_r
Doxycycline
YPL223C GRE1
ZShift = 0.62
Z lm Score = 0.32
NG = 0
DB = 0
SM = 0
Delta_AUC
Doxycycline
YKL066W
ZShift = −1.25
Z lm Score = −2.07
NG = 0
DB = 0
SM = 0
Delta_L
Doxycycline
YKL066W
ZShift = 0.16
Z lm Score = 0.69
NG = 0
DB = 0
SM = 0
Delta_K
Doxycycline
YKL066W
ZShift = 0.66
Z lm Score = −0.31
NG = 0
DB = 0
SM = 0
Delta_r
Doxycycline
YKL066W
ZShift = 0.65
Z lm Score = −0.28
NG = 0
DB = 0
SM = 0
Delta_AUC
Doxycycline
YCL022C
ZShift = 4.75
Z lm Score = −2.08
NG = 0
DB = 0
SM = 0
Delta_L
Doxycycline
YCL022C
ZShift = −2.83
Z lm Score = 0.63
NG = 0
DB = 0
SM = 0
Delta_K
Doxycycline
YCL022C
ZShift = −3.27
Z lm Score = 2.89
NG = 0
DB = 0
SM = 0
Delta_r
Doxycycline
YCL022C
ZShift = −3.75
Z lm Score = 9.77
NG = 0
DB = 0
SM = 0
Delta_AUC
Doxycycline

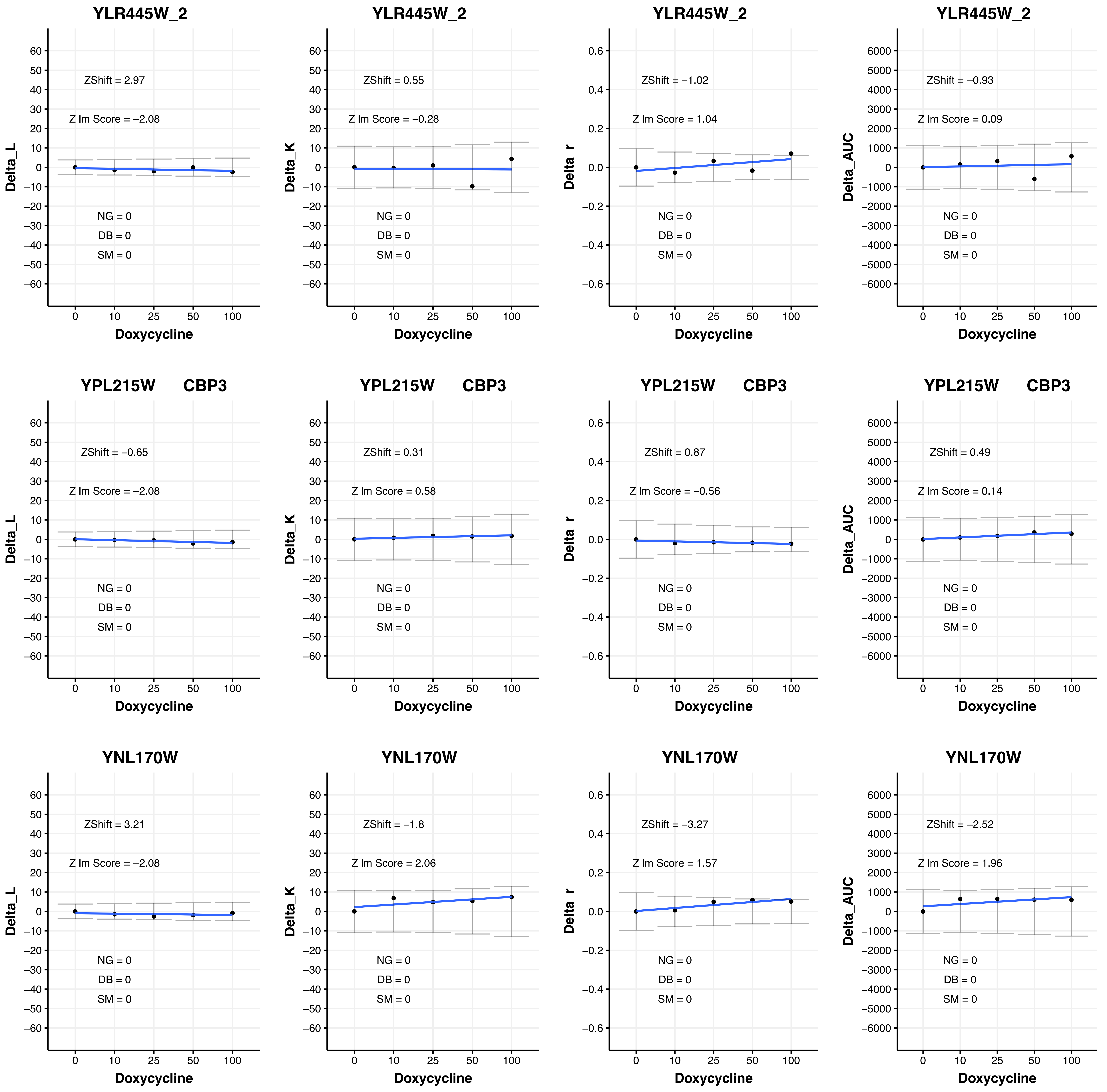

YLR445W_2
ZShift = 2.97
Z lm Score = −2.08
NG = 0
DB = 0
SM = 0
Delta_L
Doxycycline
YLR445W_2
ZShift = 0.55
Z lm Score = −0.28
NG = 0
DB = 0
SM = 0
Delta_K
Doxycycline
YLR445W_2
ZShift = −1.02
Z lm Score = 1.04
NG = 0
DB = 0
SM = 0
Delta_r
Doxycycline
YLR445W_2
ZShift = −0.93
Z lm Score = 0.09
NG = 0
DB = 0
SM = 0
Delta_AUC
Doxycycline
YPL215W CBP3
ZShift = −0.65
Z lm Score = −2.08
NG = 0
DB = 0
SM = 0
Delta_L
Doxycycline
YPL215W CBP3
ZShift = 0.31
Z lm Score = 0.58
NG = 0
DB = 0
SM = 0
Delta_K
Doxycycline
YPL215W CBP3
ZShift = 0.87
Z lm Score = −0.56
NG = 0
DB = 0
SM = 0
Delta_r
Doxycycline
YPL215W CBP3
ZShift = 0.49
Z lm Score = 0.14
NG = 0
DB = 0
SM = 0
Delta_AUC
Doxycycline
YNL170W
ZShift = 3.21
Z lm Score = −2.08
NG = 0
DB = 0
SM = 0
Delta_L
Doxycycline
YNL170W
ZShift = −1.8
Z lm Score = 2.06
NG = 0
DB = 0
SM = 0
Delta_K
Doxycycline
YNL170W
ZShift = −3.27
Z lm Score = 1.57
NG = 0
DB = 0
SM = 0
Delta_r
Doxycycline
YNL170W
ZShift = −2.52
Z lm Score = 1.96
NG = 0
DB = 0
SM = 0
Delta_AUC
Doxycycline

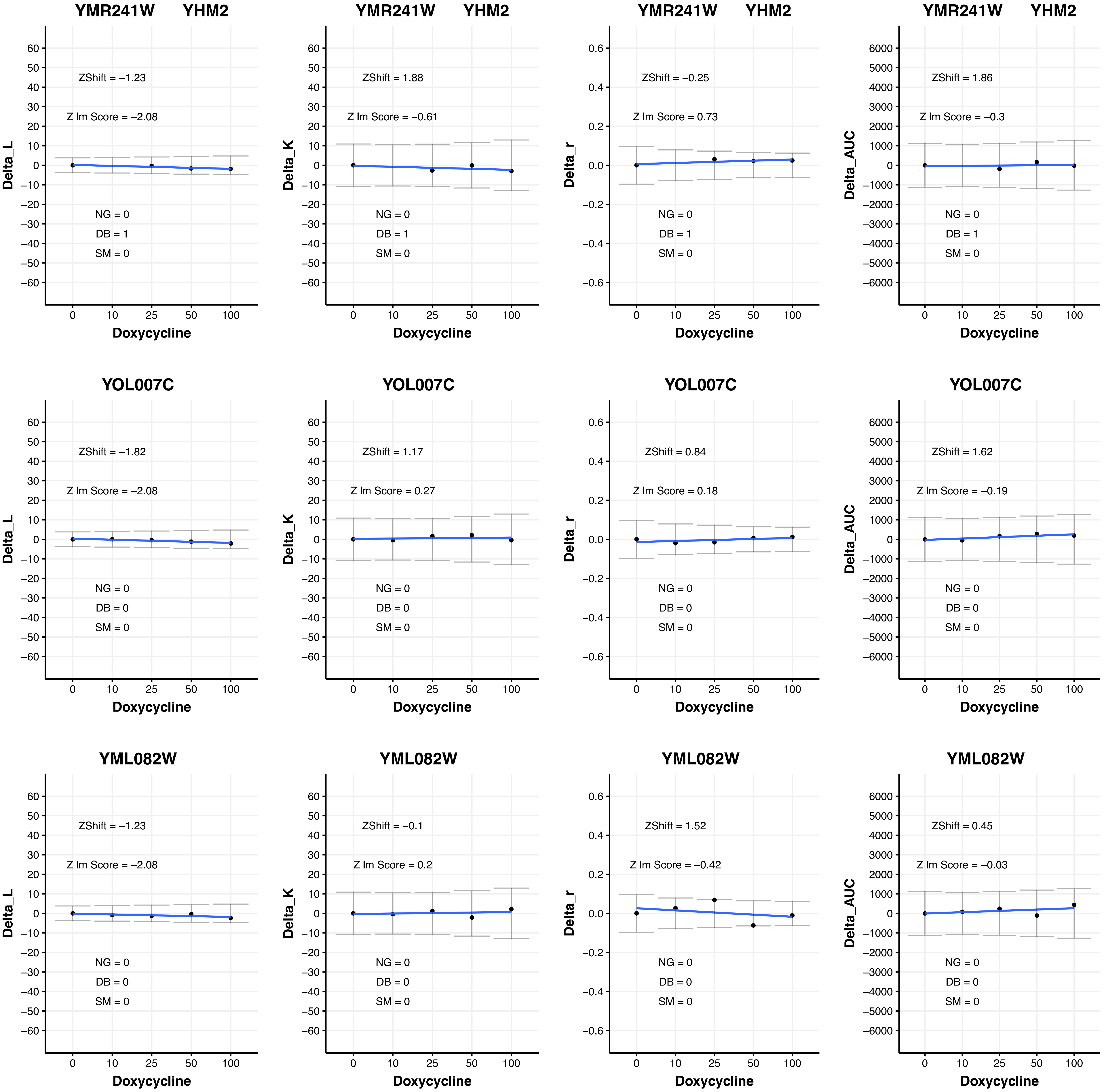

YMR241W YHM2
ZShift = −1.23
Z lm Score = −2.08
NG = 0
DB = 1
SM = 0
Delta_L
Doxycycline
0 10 25 50 100
60 50 40 30 20 10 0 −10 −20 −30 −40 −50 −60
YMR241W YHM2
ZShift = 1.88
Z lm Score = −0.61
NG = 0
DB = 1
SM = 0
Delta_K
Doxycycline
YMR241W YHM2
ZShift = −0.25
Z lm Score = 0.73
NG = 0
DB = 1
SM = 0
Delta_r
Doxycycline
0.6 0.4 0.2 0.0 −0.2 −0.4 −0.6
YMR241W YHM2
ZShift = 1.86
Z lm Score = −0.3
NG = 0
DB = 1
SM = 0
Delta_AUC
Doxycycline
6000 5000 4000 3000 2000 1000 0 −1000 −2000 −3000 −4000 −5000 −6000
YOL007C
ZShift = −1.82
Z lm Score = −2.08
NG = 0
DB = 0
SM = 0
Delta_L
Doxycycline
YOL007C
ZShift = 1.17
Z lm Score = 0.27
NG = 0
DB = 0
SM = 0
Delta_K
Doxycycline
YOL007C
ZShift = 0.84
Z lm Score = 0.18
NG = 0
DB = 0
SM = 0
Delta_r
Doxycycline
YOL007C
ZShift = 1.62
Z lm Score = −0.19
NG = 0
DB = 0
SM = 0
Delta_AUC
Doxycycline
YML082W
ZShift = −1.23
Z lm Score = −2.08
NG = 0
DB = 0
SM = 0
Delta_L
Doxycycline
YML082W
ZShift = −0.1
Z lm Score = 0.2
NG = 0
DB = 0
SM = 0
Delta_K
Doxycycline
YML082W
ZShift = 1.52
Z lm Score = −0.42
NG = 0
DB = 0
SM = 0
Delta_r
Doxycycline
YML082W
ZShift = 0.45
Z lm Score = −0.03
NG = 0
DB = 0
SM = 0
Delta_AUC
Doxycycline

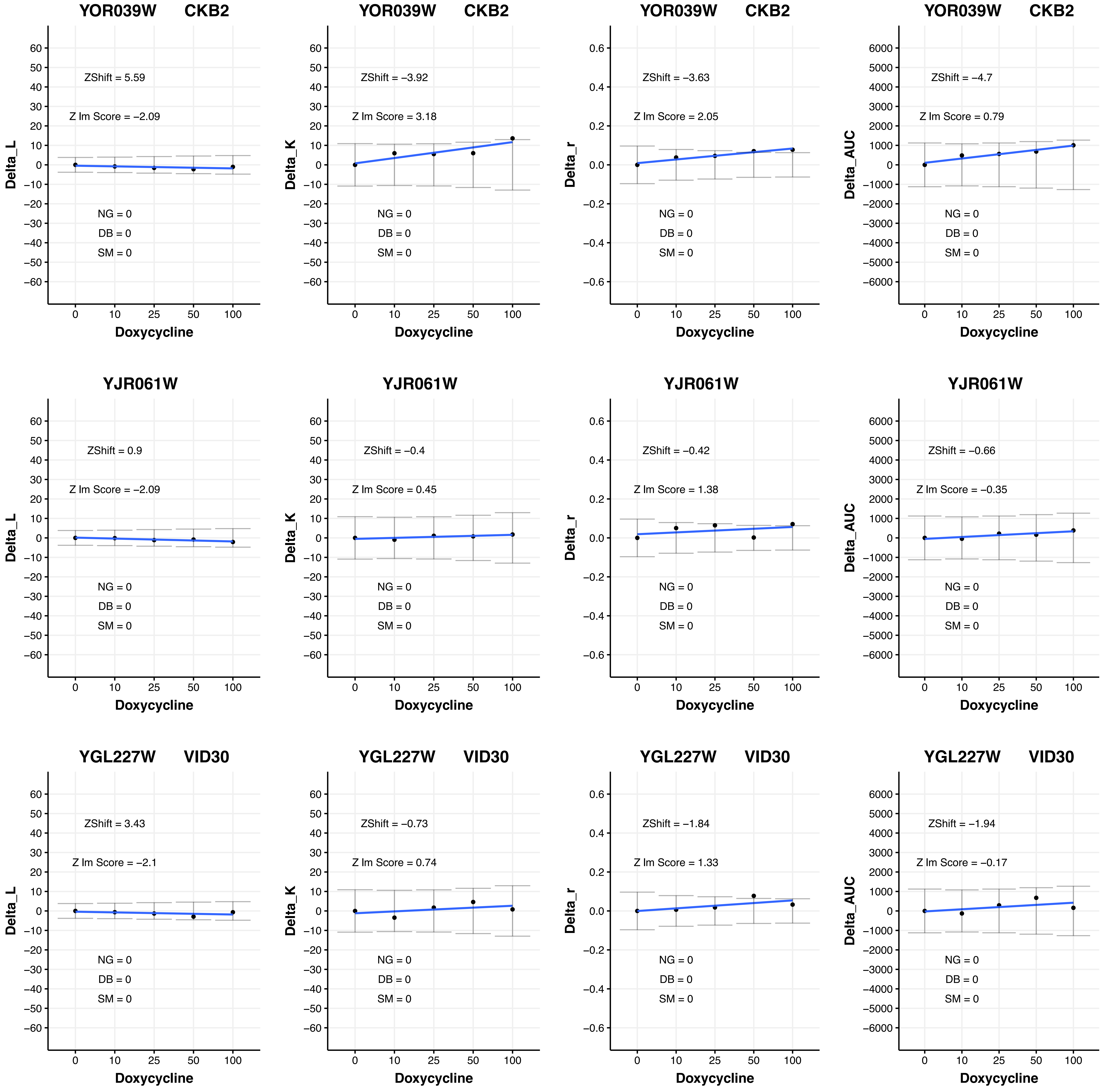

YOR039W CKB2
ZShift = 5.59
Z lm Score = −2.09
NG = 0
DB = 0
SM = 0
Delta_L
Doxycycline
YOR039W CKB2
ZShift = −3.92
Z lm Score = 3.18
NG = 0
DB = 0
SM = 0
Delta_K
Doxycycline
YOR039W CKB2
ZShift = −3.63
Z lm Score = 2.05
NG = 0
DB = 0
SM = 0
Delta_r
Doxycycline
YOR039W CKB2
ZShift = −4.7
Z lm Score = 0.79
NG = 0
DB = 0
SM = 0
Delta_AUC
Doxycycline
YJR061W
ZShift = 0.9
Z lm Score = −2.09
NG = 0
DB = 0
SM = 0
Delta_L
Doxycycline
YJR061W
ZShift = −0.4
Z lm Score = 0.45
NG = 0
DB = 0
SM = 0
Delta_K
Doxycycline
YJR061W
ZShift = −0.42
Z lm Score = 1.38
NG = 0
DB = 0
SM = 0
Delta_r
Doxycycline
YJR061W
ZShift = −0.66
Z lm Score = −0.35
NG = 0
DB = 0
SM = 0
Delta_AUC
Doxycycline
YGL227W VID30
ZShift = 3.43
Z lm Score = −2.1
NG = 0
DB = 0
SM = 0
Delta_L
Doxycycline
YGL227W VID30
ZShift = −0.73
Z lm Score = 0.74
NG = 0
DB = 0
SM = 0
Delta_K
Doxycycline
YGL227W VID30
ZShift = −1.84
Z lm Score = 1.33
NG = 0
DB = 0
SM = 0
Delta_r
Doxycycline
YGL227W VID30
ZShift = −1.94
Z lm Score = −0.17
NG = 0
DB = 0
SM = 0
Delta_AUC
Doxycycline

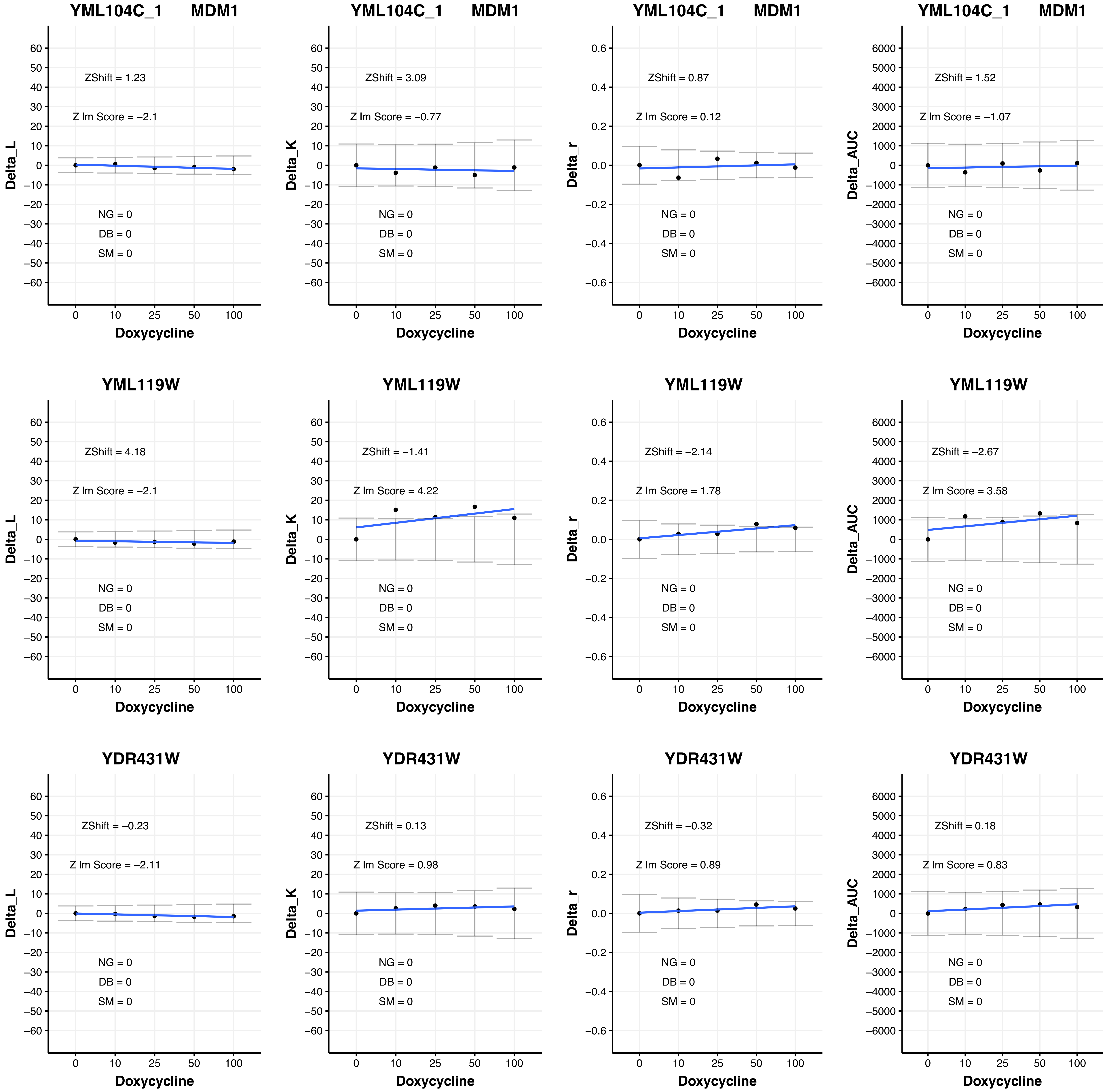

YML104C_1 MDM1
ZShift = 1.23
Z lm Score = −2.1
NG = 0
DB = 0
SM = 0
Delta_L
Doxycycline
YML104C_1 MDM1
ZShift = 3.09
Z lm Score = −0.77
NG = 0
DB = 0
SM = 0
Delta_K
Doxycycline
YML104C_1 MDM1
ZShift = 0.87
Z lm Score = 0.12
NG = 0
DB = 0
SM = 0
Delta_r
Doxycycline
YML104C_1 MDM1
ZShift = 1.52
Z lm Score = −1.07
NG = 0
DB = 0
SM = 0
Delta_AUC
Doxycycline
YML119W
ZShift = 4.18
Z lm Score = −2.1
NG = 0
DB = 0
SM = 0
Delta_L
Doxycycline
YML119W
ZShift = −1.41
Z lm Score = 4.22
NG = 0
DB = 0
SM = 0
Delta_K
Doxycycline
YML119W
ZShift = −2.14
Z lm Score = 1.78
NG = 0
DB = 0
SM = 0
Delta_r
Doxycycline
YML119W
ZShift = −2.67
Z lm Score = 3.58
NG = 0
DB = 0
SM = 0
Delta_AUC
Doxycycline
YDR431W
ZShift = −0.23
Z lm Score = −2.11
NG = 0
DB = 0
SM = 0
Delta_L
Doxycycline
YDR431W
ZShift = 0.13
Z lm Score = 0.98
NG = 0
DB = 0
SM = 0
Delta_K
Doxycycline
YDR431W
ZShift = −0.32
Z lm Score = 0.89
NG = 0
DB = 0
SM = 0
Delta_r
Doxycycline
YDR431W
ZShift = 0.18
Z lm Score = 0.83
NG = 0
DB = 0
SM = 0
Delta_AUC
Doxycycline

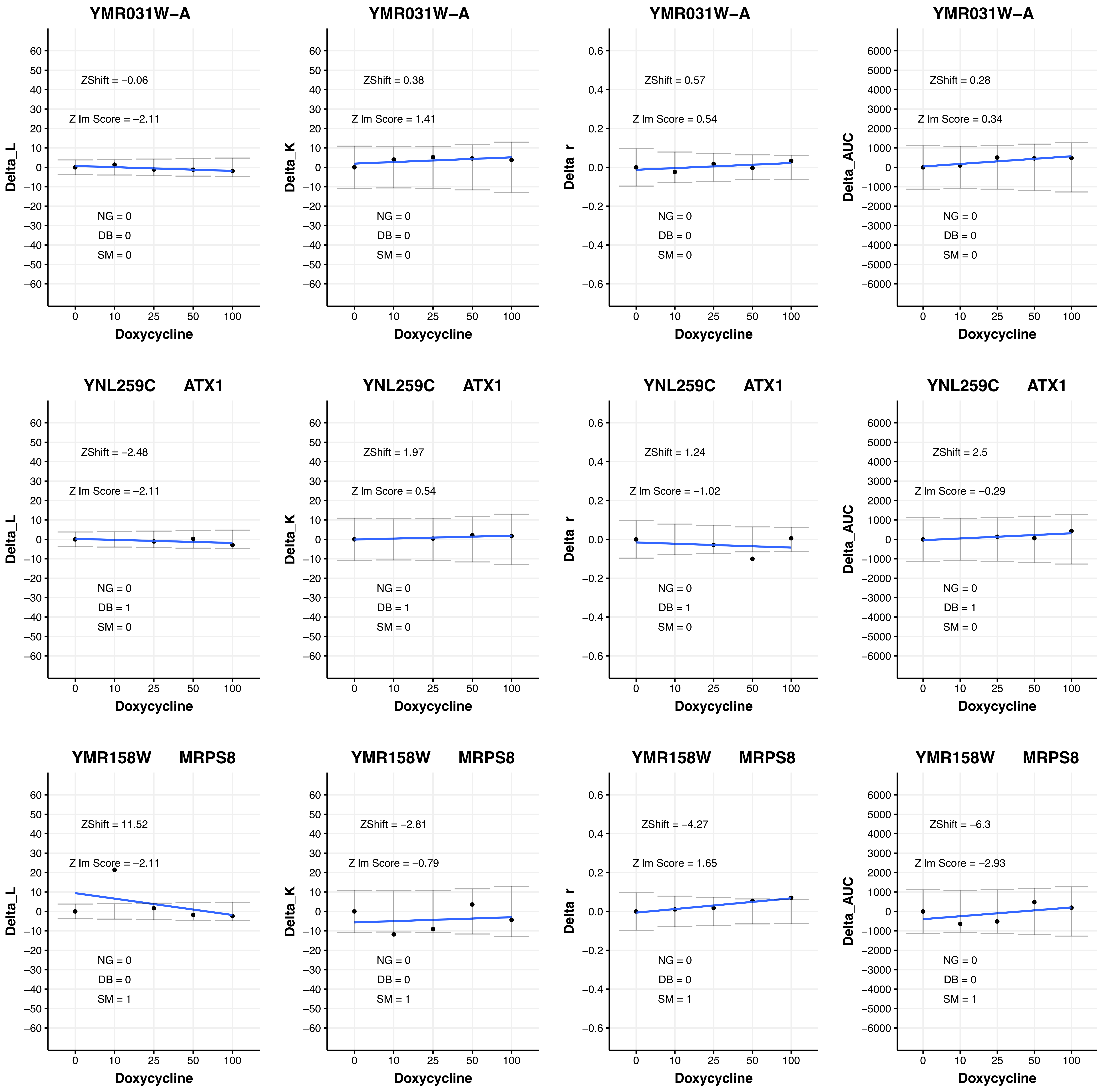

YMR031W−A
ZShift = −0.06
Z lm Score = −2.11
NG = 0
DB = 0
SM = 0
Delta_L
Doxycycline
YMR031W−A
ZShift = 0.38
Z lm Score = 1.41
NG = 0
DB = 0
SM = 0
Delta_K
Doxycycline
YMR031W−A
ZShift = 0.57
Z lm Score = 0.54
NG = 0
DB = 0
SM = 0
Delta_r
Doxycycline
YMR031W−A
ZShift = 0.28
Z lm Score = 0.34
NG = 0
DB = 0
SM = 0
Delta_AUC
Doxycycline
YNL259C ATX1
ZShift = −2.48
Z lm Score = −2.11
NG = 0
DB = 1
SM = 0
Delta_L
Doxycycline
YNL259C ATX1
ZShift = 1.97
Z lm Score = 0.54
NG = 0
DB = 1
SM = 0
Delta_K
Doxycycline
YNL259C ATX1
ZShift = 1.24
Z lm Score = −1.02
NG = 0
DB = 1
SM = 0
Delta_r
Doxycycline
YNL259C ATX1
ZShift = 2.5
Z lm Score = −0.29
NG = 0
DB = 1
SM = 0
Delta_AUC
Doxycycline
YMR158W MRPS8
ZShift = 11.52
Z lm Score = −2.11
NG = 0
DB = 0
SM = 1
Delta_L
Doxycycline
YMR158W MRPS8
ZShift = −2.81
Z lm Score = −0.79
NG = 0
DB = 0
SM = 1
Delta_K
Doxycycline
YMR158W MRPS8
ZShift = −4.27
Z lm Score = 1.65
NG = 0
DB = 0
SM = 1
Delta_r
Doxycycline
YMR158W MRPS8
ZShift = −6.3
Z lm Score = −2.93
NG = 0
DB = 0
SM = 1
Delta_AUC
Doxycycline

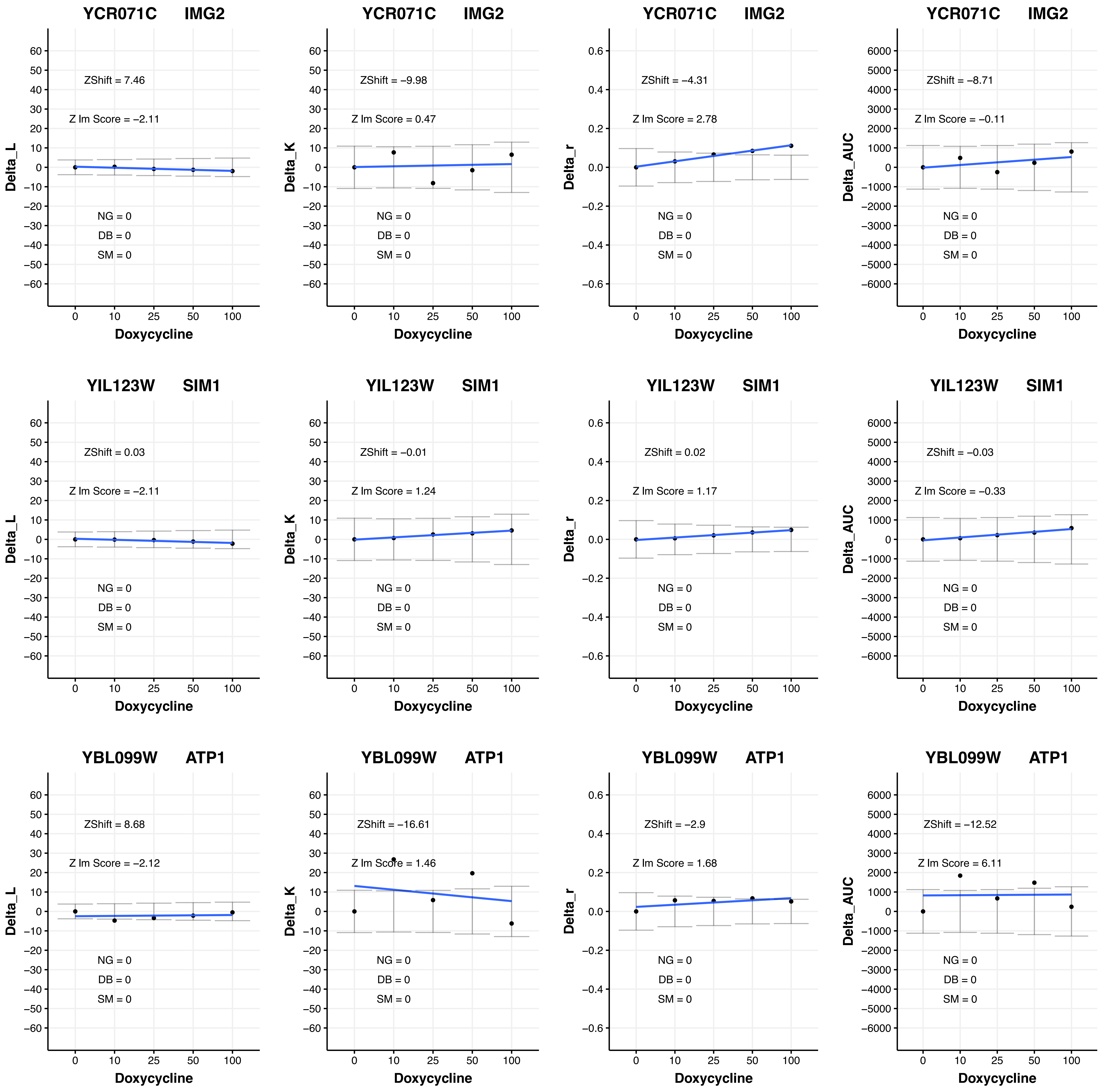

YCR071C IMG2
ZShift = 7.46
Z lm Score = −2.11
NG = 0
DB = 0
SM = 0
Delta_L
Doxycycline
YCR071C IMG2
ZShift = −9.98
Z lm Score = 0.47
NG = 0
DB = 0
SM = 0
Delta_K
Doxycycline
YCR071C IMG2
ZShift = −4.31
Z lm Score = 2.78
NG = 0
DB = 0
SM = 0
Delta_r
Doxycycline
YCR071C IMG2
ZShift = −8.71
Z lm Score = −0.11
NG = 0
DB = 0
SM = 0
Delta_AUC
Doxycycline
YIL123W SIM1
ZShift = 0.03
Z lm Score = −2.11
NG = 0
DB = 0
SM = 0
Delta_L
Doxycycline
YIL123W SIM1
ZShift = −0.01
Z lm Score = 1.24
NG = 0
DB = 0
SM = 0
Delta_K
Doxycycline
YIL123W SIM1
ZShift = 0.02
Z lm Score = 1.17
NG = 0
DB = 0
SM = 0
Delta_r
Doxycycline
YIL123W SIM1
ZShift = −0.03
Z lm Score = −0.33
NG = 0
DB = 0
SM = 0
Delta_AUC
Doxycycline
YBL099W ATP1
ZShift = 8.68
Z lm Score = −2.12
NG = 0
DB = 0
SM = 0
Delta_L
Doxycycline
YBL099W ATP1
ZShift = −16.61
Z lm Score = 1.46
NG = 0
DB = 0
SM = 0
Delta_K
Doxycycline
YBL099W ATP1
ZShift = −2.9
Z lm Score = 1.68
NG = 0
DB = 0
SM = 0
Delta_r
Doxycycline
YBL099W ATP1
ZShift = −12.52
Z lm Score = 6.11
NG = 0
DB = 0
SM = 0
Delta_AUC
Doxycycline

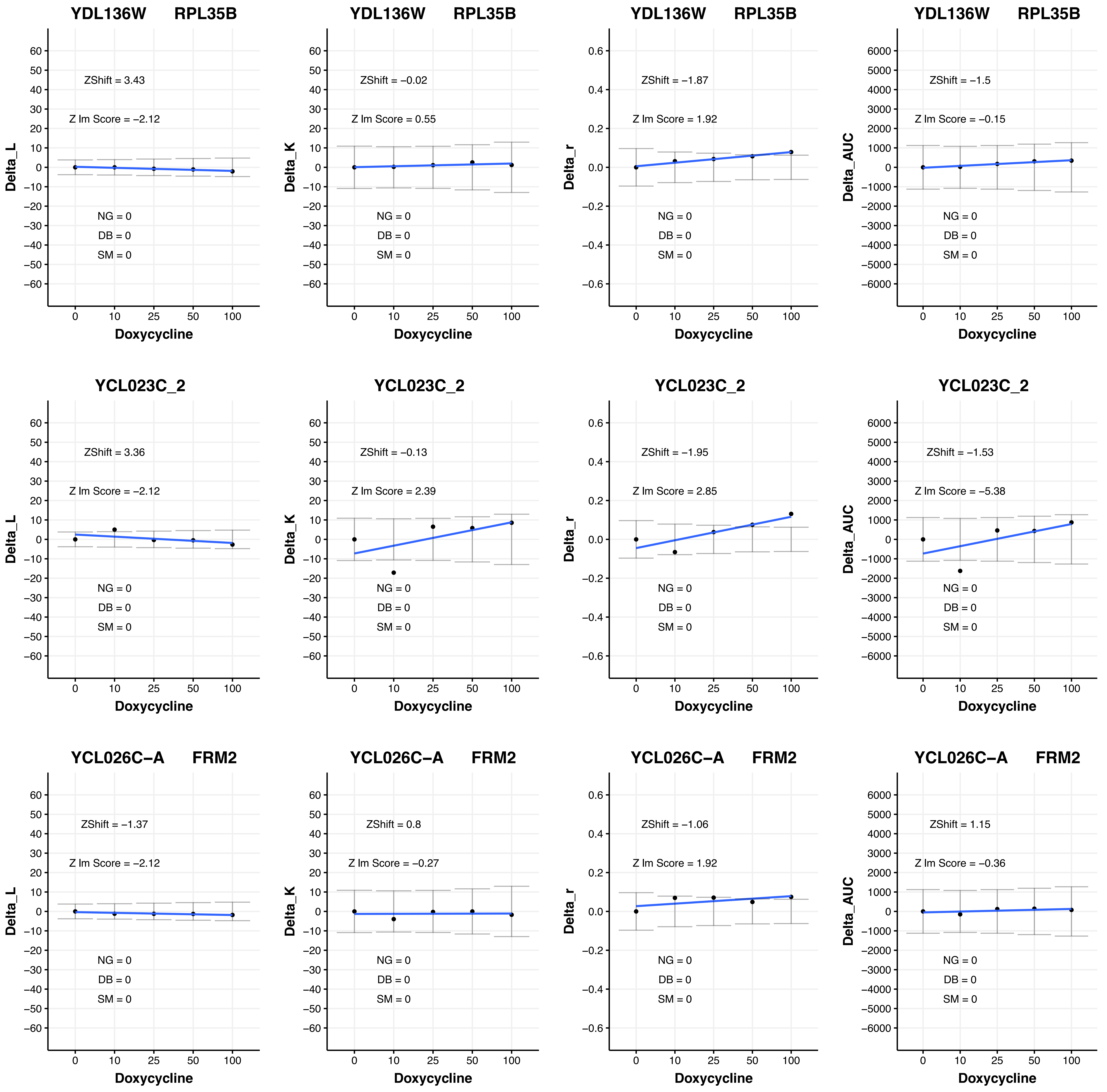
YDL136W RPL35B
ZShift = 3.43
Z lm Score = −2.12
NG = 0
DB = 0
SM = 0
Delta_L
Doxycycline
YDL136W RPL35B
ZShift = −0.02
Z lm Score = 0.55
NG = 0
DB = 0
SM = 0
Delta_K
Doxycycline
YDL136W RPL35B
ZShift = −1.87
Z lm Score = 1.92
NG = 0
DB = 0
SM = 0
Delta_r
Doxycycline
YDL136W RPL35B
ZShift = −1.5
Z lm Score = −0.15
NG = 0
DB = 0
SM = 0
Delta_AUC
Doxycycline
YCL023C_2
ZShift = 3.36
Z lm Score = −2.12
NG = 0
DB = 0
SM = 0
Delta_L
Doxycycline
YCL023C_2
ZShift = −0.13
Z lm Score = 2.39
NG = 0
DB = 0
SM = 0
Delta_K
Doxycycline
YCL023C_2
ZShift = −1.95
Z lm Score = 2.85
NG = 0
DB = 0
SM = 0
Delta_r
Doxycycline
YCL023C_2
ZShift = −1.53
Z lm Score = −5.38
NG = 0
DB = 0
SM = 0
Delta_AUC
Doxycycline
YCL026C−A FRM2
ZShift = −1.37
Z lm Score = −2.12
NG = 0
DB = 0
SM = 0
Delta_L
Doxycycline
YCL026C−A FRM2
ZShift = 0.8
Z lm Score = −0.27
NG = 0
DB = 0
SM = 0
Delta_K
Doxycycline
YCL026C−A FRM2
ZShift = −1.06
Z lm Score = 1.92
NG = 0
DB = 0
SM = 0
Delta_r
Doxycycline
YCL026C−A FRM2
ZShift = 1.15
Z lm Score = −0.36
NG = 0
DB = 0
SM = 0
Delta_AUC
Doxycycline

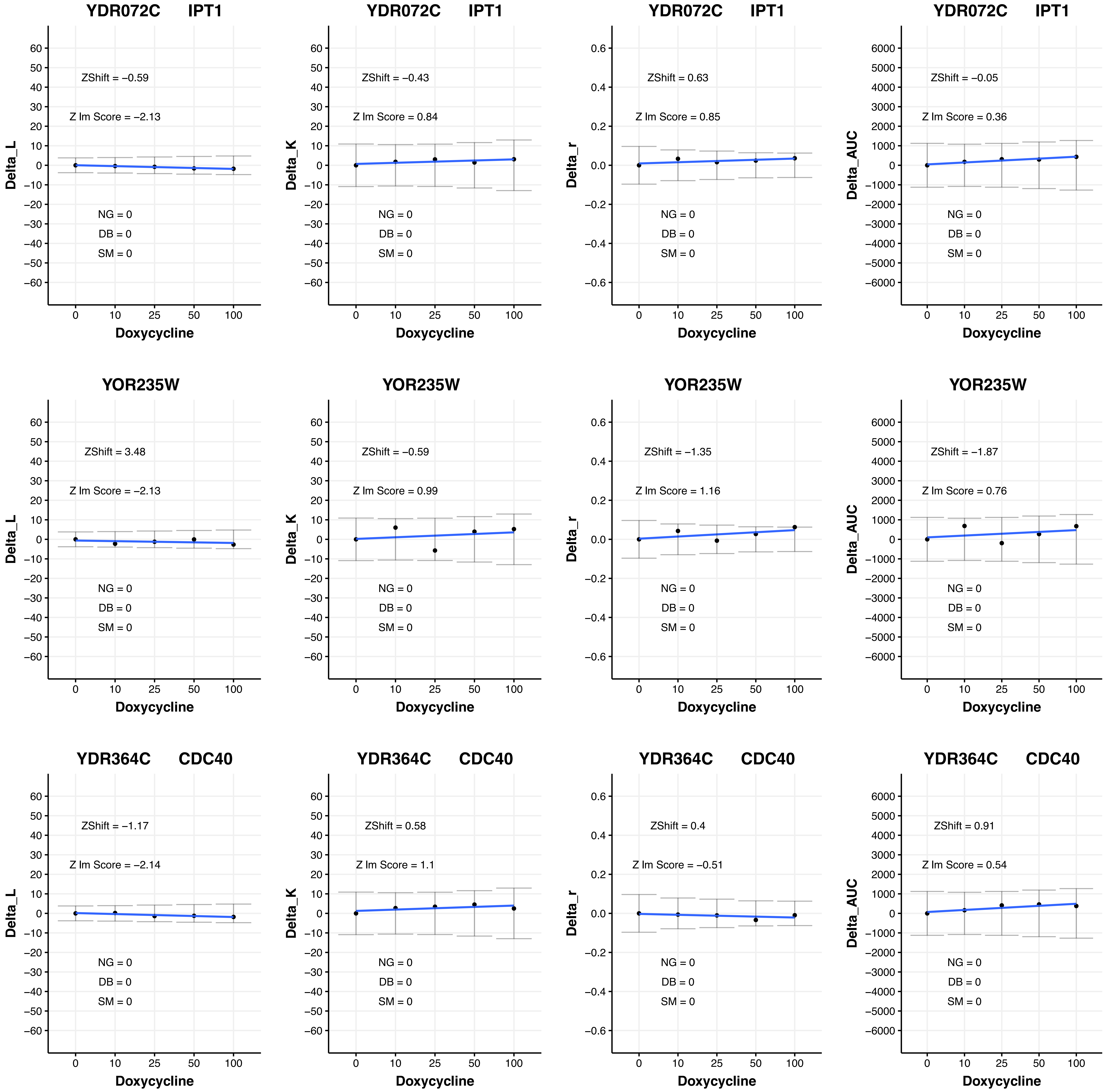
YDR072C IPT1
Delta_L
ZShift = −0.59
Z lm Score = −2.13
NG = 0
DB = 0
SM = 0
Doxycycline
YDR072C IPT1
Delta_K
ZShift = −0.43
Z lm Score = 0.84
NG = 0
DB = 0
SM = 0
Doxycycline
YDR072C IPT1
Delta_r
ZShift = 0.63
Z lm Score = 0.85
NG = 0
DB = 0
SM = 0
Doxycycline
YDR072C IPT1
Delta_AUC
ZShift = −0.05
Z lm Score = 0.36
NG = 0
DB = 0
SM = 0
Doxycycline
YOR235W
Delta_L
ZShift = 3.48
Z lm Score = −2.13
NG = 0
DB = 0
SM = 0
Doxycycline
YOR235W
Delta_K
ZShift = −0.59
Z lm Score = 0.99
NG = 0
DB = 0
SM = 0
Doxycycline
YOR235W
Delta_r
ZShift = −1.35
Z lm Score = 1.16
NG = 0
DB = 0
SM = 0
Doxycycline
YOR235W
Delta_AUC
ZShift = −1.87
Z lm Score = 0.76
NG = 0
DB = 0
SM = 0
Doxycycline
YDR364C CDC40
Delta_L
ZShift = −1.17
Z lm Score = −2.14
NG = 0
DB = 0
SM = 0
Doxycycline
YDR364C CDC40
Delta_K
ZShift = 0.58
Z lm Score = 1.1
NG = 0
DB = 0
SM = 0
Doxycycline
YDR364C CDC40
Delta_r
ZShift = 0.4
Z lm Score = −0.51
NG = 0
DB = 0
SM = 0
Doxycycline
YDR364C CDC40
Delta_AUC
ZShift = 0.91
Z lm Score = 0.54
NG = 0
DB = 0
SM = 0
Doxycycline

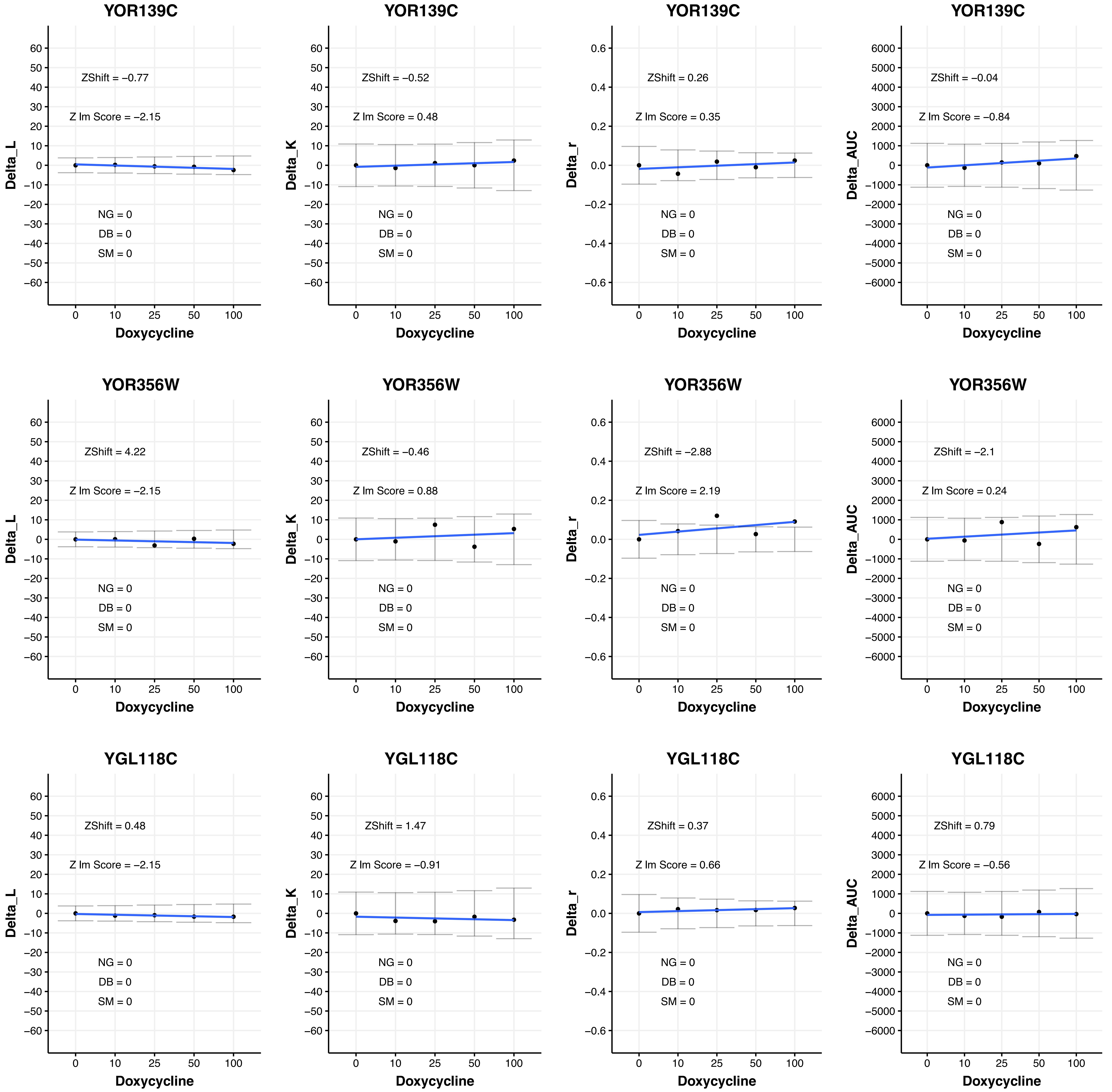

YOR139C
Delta_L
ZShift = −0.77
Z lm Score = −2.15
NG = 0
DB = 0
SM = 0
Doxycycline
YOR139C
Delta_K
ZShift = −0.52
Z lm Score = 0.48
NG = 0
DB = 0
SM = 0
Doxycycline
YOR139C
Delta_r
ZShift = 0.26
Z lm Score = 0.35
NG = 0
DB = 0
SM = 0
Doxycycline
YOR139C
Delta_AUC
ZShift = −0.04
Z lm Score = −0.84
NG = 0
DB = 0
SM = 0
Doxycycline
YOR356W
Delta_L
ZShift = 4.22
Z lm Score = −2.15
NG = 0
DB = 0
SM = 0
Doxycycline
YOR356W
Delta_K
ZShift = −0.46
Z lm Score = 0.88
NG = 0
DB = 0
SM = 0
Doxycycline
YOR356W
Delta_r
ZShift = −2.88
Z lm Score = 2.19
NG = 0
DB = 0
SM = 0
Doxycycline
YOR356W
Delta_AUC
ZShift = −2.1
Z lm Score = 0.24
NG = 0
DB = 0
SM = 0
Doxycycline
YGL118C
Delta_L
ZShift = 0.48
Z lm Score = −2.15
NG = 0
DB = 0
SM = 0
Doxycycline
YGL118C
Delta_K
ZShift = 1.47
Z lm Score = −0.91
NG = 0
DB = 0
SM = 0
Doxycycline
YGL118C
Delta_r
ZShift = 0.37
Z lm Score = 0.66
NG = 0
DB = 0
SM = 0
Doxycycline
YGL118C
Delta_AUC
ZShift = 0.79
Z lm Score = −0.56
NG = 0
DB = 0
SM = 0
Doxycycline

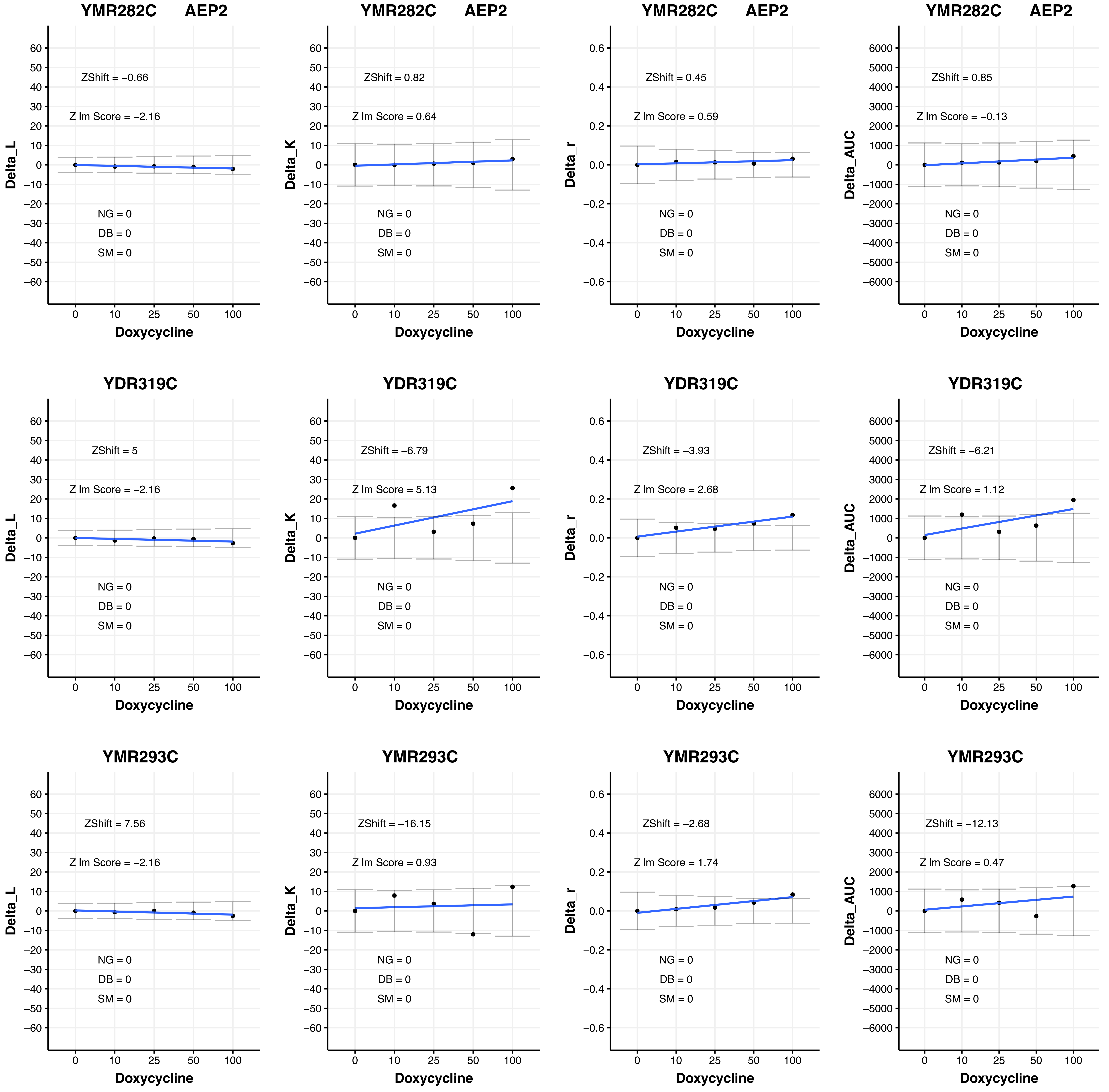

YMR282C AEP2
ZShift = −0.66
Z lm Score = −2.16
NG = 0
DB = 0
SM = 0
Delta_L
Doxycycline
YMR282C AEP2
ZShift = 0.82
Z lm Score = 0.64
NG = 0
DB = 0
SM = 0
Delta_K
Doxycycline
YMR282C AEP2
ZShift = 0.45
Z lm Score = 0.59
NG = 0
DB = 0
SM = 0
Delta_r
Doxycycline
YMR282C AEP2
ZShift = 0.85
Z lm Score = −0.13
NG = 0
DB = 0
SM = 0
Delta_AUC
Doxycycline
YDR319C
ZShift = 5
Z lm Score = −2.16
NG = 0
DB = 0
SM = 0
Delta_L
Doxycycline
YDR319C
ZShift = −6.79
Z lm Score = 5.13
NG = 0
DB = 0
SM = 0
Delta_K
Doxycycline
YDR319C
ZShift = −3.93
Z lm Score = 2.68
NG = 0
DB = 0
SM = 0
Delta_r
Doxycycline
YDR319C
ZShift = −6.21
Z lm Score = 1.12
NG = 0
DB = 0
SM = 0
Delta_AUC
Doxycycline
YMR293C
ZShift = 7.56
Z lm Score = −2.16
NG = 0
DB = 0
SM = 0
Delta_L
Doxycycline
YMR293C
ZShift = −16.15
Z lm Score = 0.93
NG = 0
DB = 0
SM = 0
Delta_K
Doxycycline
YMR293C
ZShift = −2.68
Z lm Score = 1.74
NG = 0
DB = 0
SM = 0
Delta_r
Doxycycline
YMR293C
ZShift = −12.13
Z lm Score = 0.47
NG = 0
DB = 0
SM = 0
Delta_AUC
Doxycycline

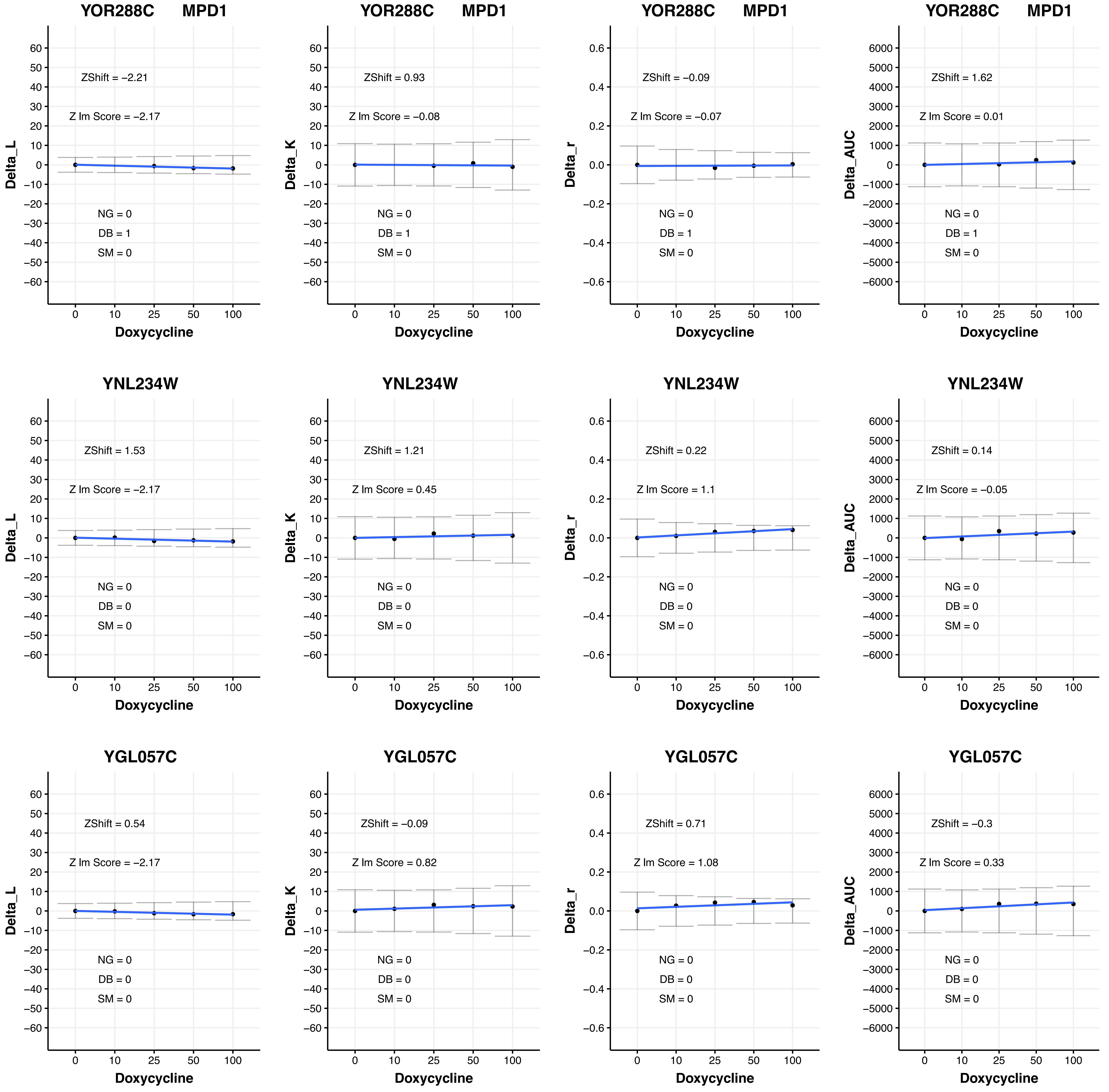
YOR288C MPD1
ZShift = −2.21
Z lm Score = −2.17
NG = 0
DB = 1
SM = 0
Delta_L
Doxycycline
YOR288C MPD1
ZShift = 0.93
Z lm Score = −0.08
NG = 0
DB = 1
SM = 0
Delta_K
Doxycycline
YOR288C MPD1
ZShift = −0.09
Z lm Score = −0.07
NG = 0
DB = 1
SM = 0
Delta_r
Doxycycline
YOR288C MPD1
ZShift = 1.62
Z lm Score = 0.01
NG = 0
DB = 1
SM = 0
Delta_AUC
Doxycycline
YNL234W
ZShift = 1.53
Z lm Score = −2.17
NG = 0
DB = 0
SM = 0
Delta_L
Doxycycline
YNL234W
ZShift = 1.21
Z lm Score = 0.45
NG = 0
DB = 0
SM = 0
Delta_K
Doxycycline
YNL234W
ZShift = 0.22
Z lm Score = 1.1
NG = 0
DB = 0
SM = 0
Delta_r
Doxycycline
YNL234W
ZShift = 0.14
Z lm Score = −0.05
NG = 0
DB = 0
SM = 0
Delta_AUC
Doxycycline
YGL057C
ZShift = 0.54
Z lm Score = −2.17
NG = 0
DB = 0
SM = 0
Delta_L
Doxycycline
YGL057C
ZShift = −0.09
Z lm Score = 0.82
NG = 0
DB = 0
SM = 0
Delta_K
Doxycycline
YGL057C
ZShift = 0.71
Z lm Score = 1.08
NG = 0
DB = 0
SM = 0
Delta_r
Doxycycline
YGL057C
ZShift = −0.3
Z lm Score = 0.33
NG = 0
DB = 0
SM = 0
Delta_AUC
Doxycycline

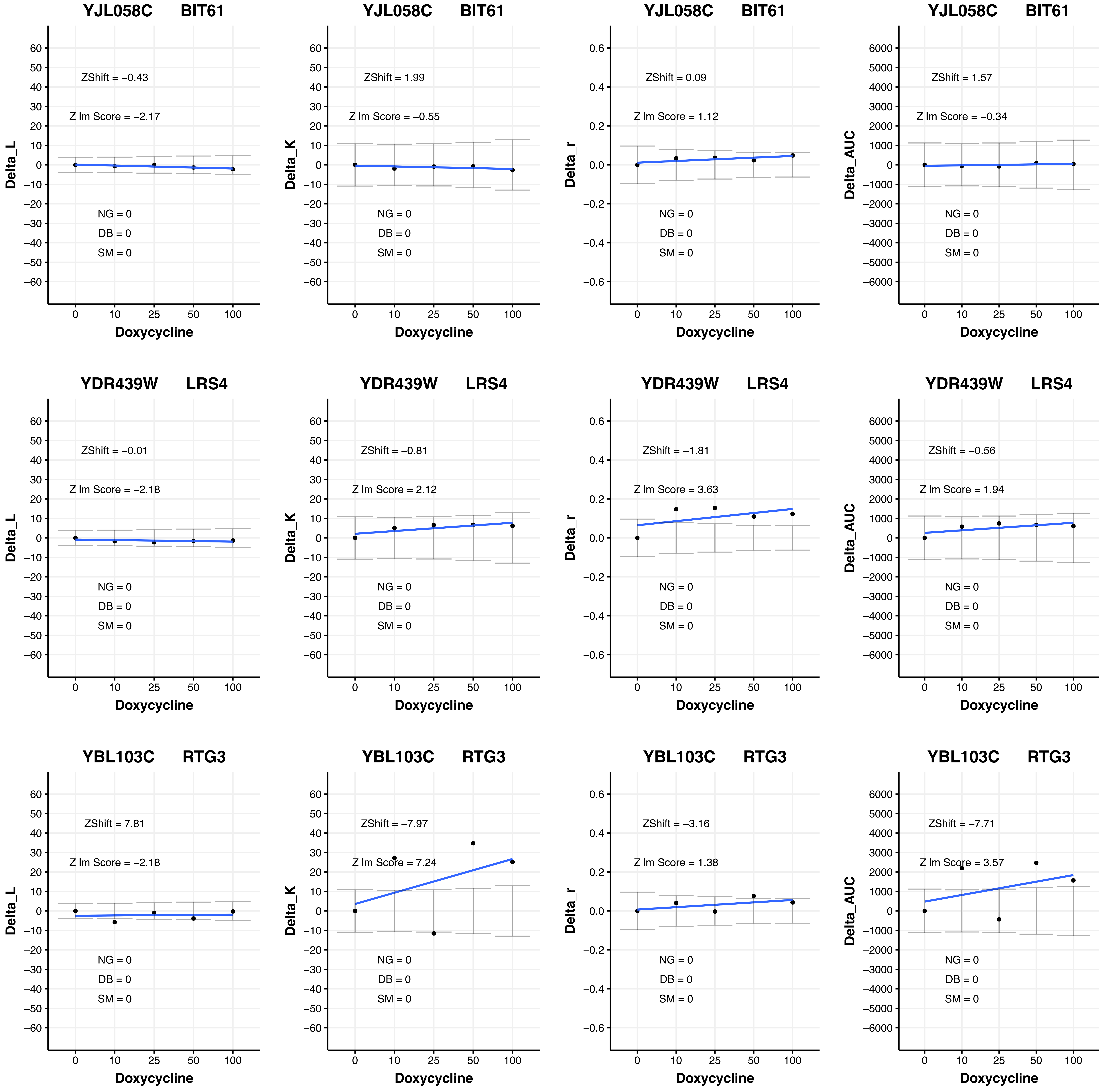
YJL058C BIT61
ZShift = −0.43
Z lm Score = −2.17
NG = 0
DB = 0
SM = 0
Delta_L
Doxycycline
YJL058C BIT61
ZShift = 1.99
Z lm Score = −0.55
NG = 0
DB = 0
SM = 0
Delta_K
Doxycycline
YJL058C BIT61
ZShift = 0.09
Z lm Score = 1.12
NG = 0
DB = 0
SM = 0
Delta_r
Doxycycline
YJL058C BIT61
ZShift = 1.57
Z lm Score = −0.34
NG = 0
DB = 0
SM = 0
Delta_AUC
Doxycycline
YDR439W LRS4
ZShift = −0.01
Z lm Score = −2.18
NG = 0
DB = 0
SM = 0
Delta_L
Doxycycline
YDR439W LRS4
ZShift = −0.81
Z lm Score = 2.12
NG = 0
DB = 0
SM = 0
Delta_K
Doxycycline
YDR439W LRS4
ZShift = −1.81
Z lm Score = 3.63
NG = 0
DB = 0
SM = 0
Delta_r
Doxycycline
YDR439W LRS4
ZShift = −0.56
Z lm Score = 1.94
NG = 0
DB = 0
SM = 0
Delta_AUC
Doxycycline
YBL103C RTG3
ZShift = 7.81
Z lm Score = −2.18
NG = 0
DB = 0
SM = 0
Delta_L
Doxycycline
YBL103C RTG3
ZShift = −7.97
Z lm Score = 7.24
NG = 0
DB = 0
SM = 0
Delta_K
Doxycycline
YBL103C RTG3
ZShift = −3.16
Z lm Score = 1.38
NG = 0
DB = 0
SM = 0
Delta_r
Doxycycline
YBL103C RTG3
ZShift = −7.71
Z lm Score = 3.57
NG = 0
DB = 0
SM = 0
Delta_AUC
Doxycycline

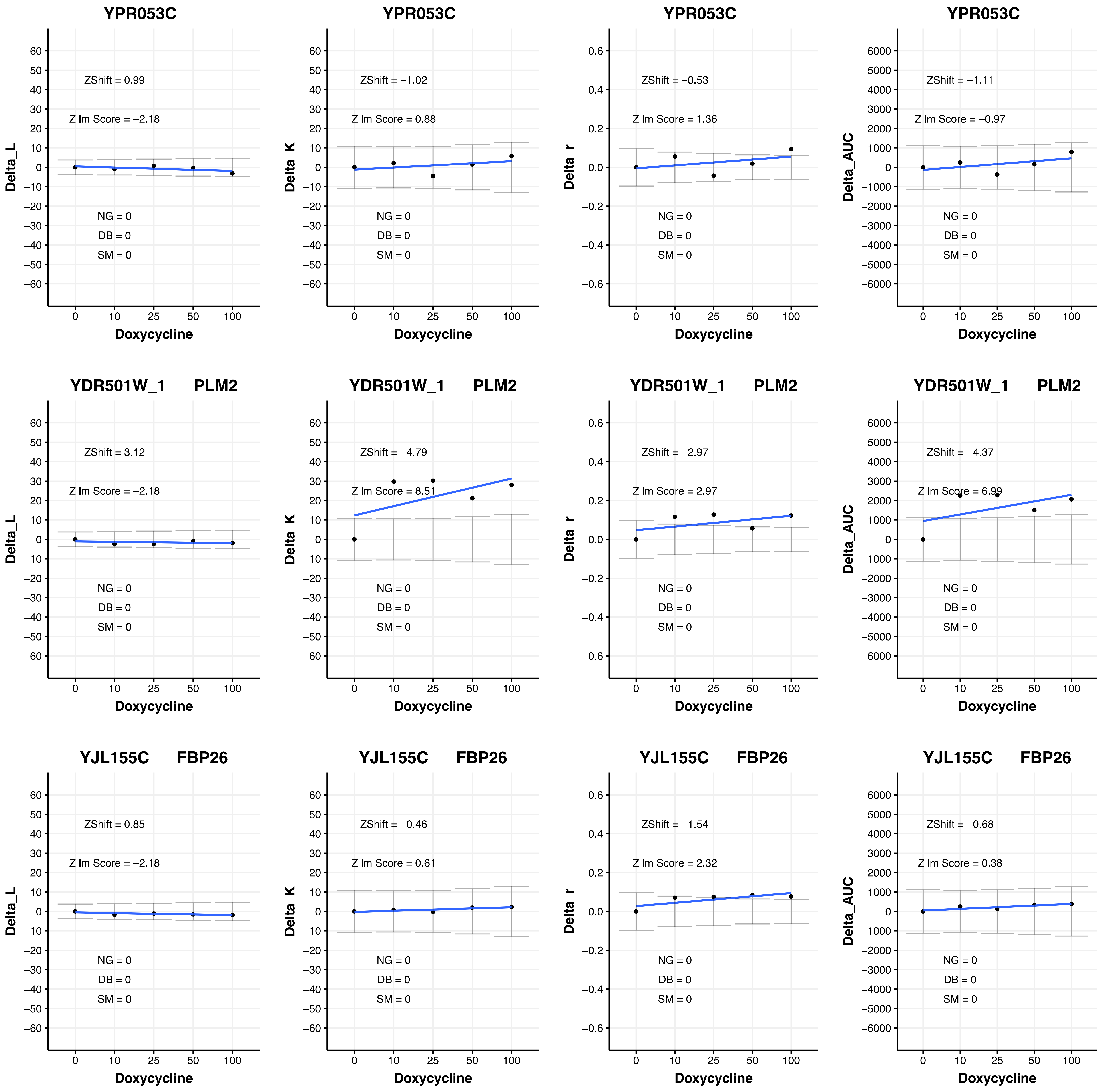

YPR053C
ZShift = 0.99
Z lm Score = −2.18
NG = 0
DB = 0
SM = 0
Delta_L
Doxycycline
YPR053C
ZShift = −1.02
Z lm Score = 0.88
NG = 0
DB = 0
SM = 0
Delta_K
Doxycycline
YPR053C
ZShift = −0.53
Z lm Score = 1.36
NG = 0
DB = 0
SM = 0
Delta_r
Doxycycline
YPR053C
ZShift = −1.11
Z lm Score = −0.97
NG = 0
DB = 0
SM = 0
Delta_AUC
Doxycycline
YDR501W_1 PLM2
ZShift = 3.12
Z lm Score = −2.18
NG = 0
DB = 0
SM = 0
Delta_L
Doxycycline
YDR501W_1 PLM2
ZShift = −4.79
Z lm Score = 8.51
NG = 0
DB = 0
SM = 0
Delta_K
Doxycycline
YDR501W_1 PLM2
ZShift = −2.97
Z lm Score = 2.97
NG = 0
DB = 0
SM = 0
Delta_r
Doxycycline
YDR501W_1 PLM2
ZShift = −4.37
Z lm Score = 6.99
NG = 0
DB = 0
SM = 0
Delta_AUC
Doxycycline
YJL155C FBP26
ZShift = 0.85
Z lm Score = −2.18
NG = 0
DB = 0
SM = 0
Delta_L
Doxycycline
YJL155C FBP26
ZShift = −0.46
Z lm Score = 0.61
NG = 0
DB = 0
SM = 0
Delta_K
Doxycycline
YJL155C FBP26
ZShift = −1.54
Z lm Score = 2.32
NG = 0
DB = 0
SM = 0
Delta_r
Doxycycline
YJL155C FBP26
ZShift = −0.68
Z lm Score = 0.38
NG = 0
DB = 0
SM = 0
Delta_AUC
Doxycycline

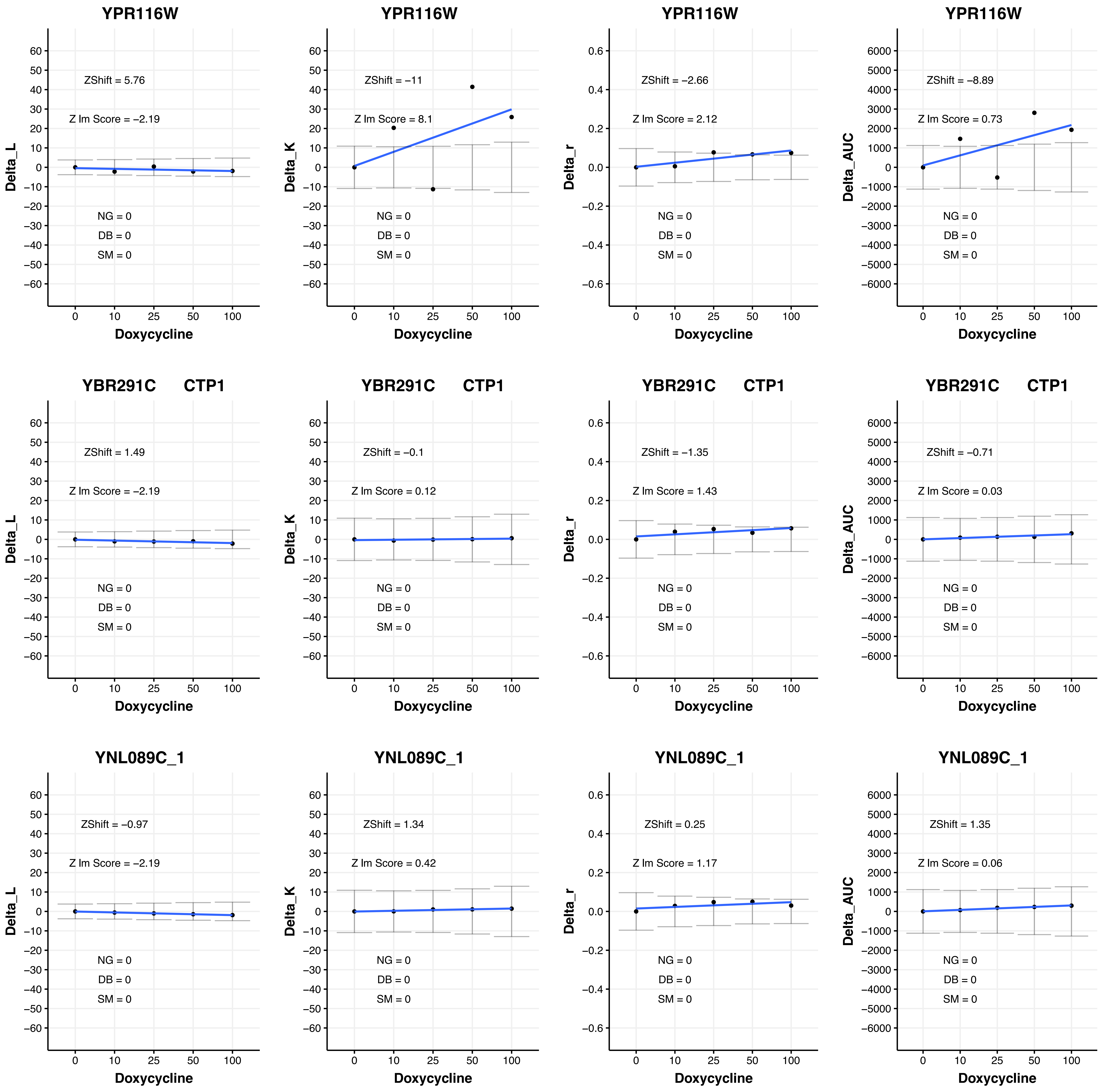

YPR116W
Delta_L
ZShift = 5.76
Z lm Score = −2.19
NG = 0
DB = 0
SM = 0
Doxycycline
YPR116W
Delta_K
ZShift = −11
Z lm Score = 8.1
NG = 0
DB = 0
SM = 0
Doxycycline
YPR116W
Delta_r
ZShift = −2.66
Z lm Score = 2.12
NG = 0
DB = 0
SM = 0
Doxycycline
YPR116W
Delta_AUC
ZShift = −8.89
Z lm Score = 0.73
NG = 0
DB = 0
SM = 0
Doxycycline
YBR291C CTP1
Delta_L
ZShift = 1.49
Z lm Score = −2.19
NG = 0
DB = 0
SM = 0
Doxycycline
YBR291C CTP1
Delta_K
ZShift = −0.1
Z lm Score = 0.12
NG = 0
DB = 0
SM = 0
Doxycycline
YBR291C CTP1
Delta_r
ZShift = −1.35
Z lm Score = 1.43
NG = 0
DB = 0
SM = 0
Doxycycline
YBR291C CTP1
Delta_AUC
ZShift = −0.71
Z lm Score = 0.03
NG = 0
DB = 0
SM = 0
Doxycycline
YNL089C_1
Delta_L
ZShift = −0.97
Z lm Score = −2.19
NG = 0
DB = 0
SM = 0
Doxycycline
YNL089C_1
Delta_K
ZShift = 1.34
Z lm Score = 0.42
NG = 0
DB = 0
SM = 0
Doxycycline
YNL089C_1
Delta_r
ZShift = 0.25
Z lm Score = 1.17
NG = 0
DB = 0
SM = 0
Doxycycline
YNL089C_1
Delta_AUC
ZShift = 1.35
Z lm Score = 0.06
NG = 0
DB = 0
SM = 0
Doxycycline

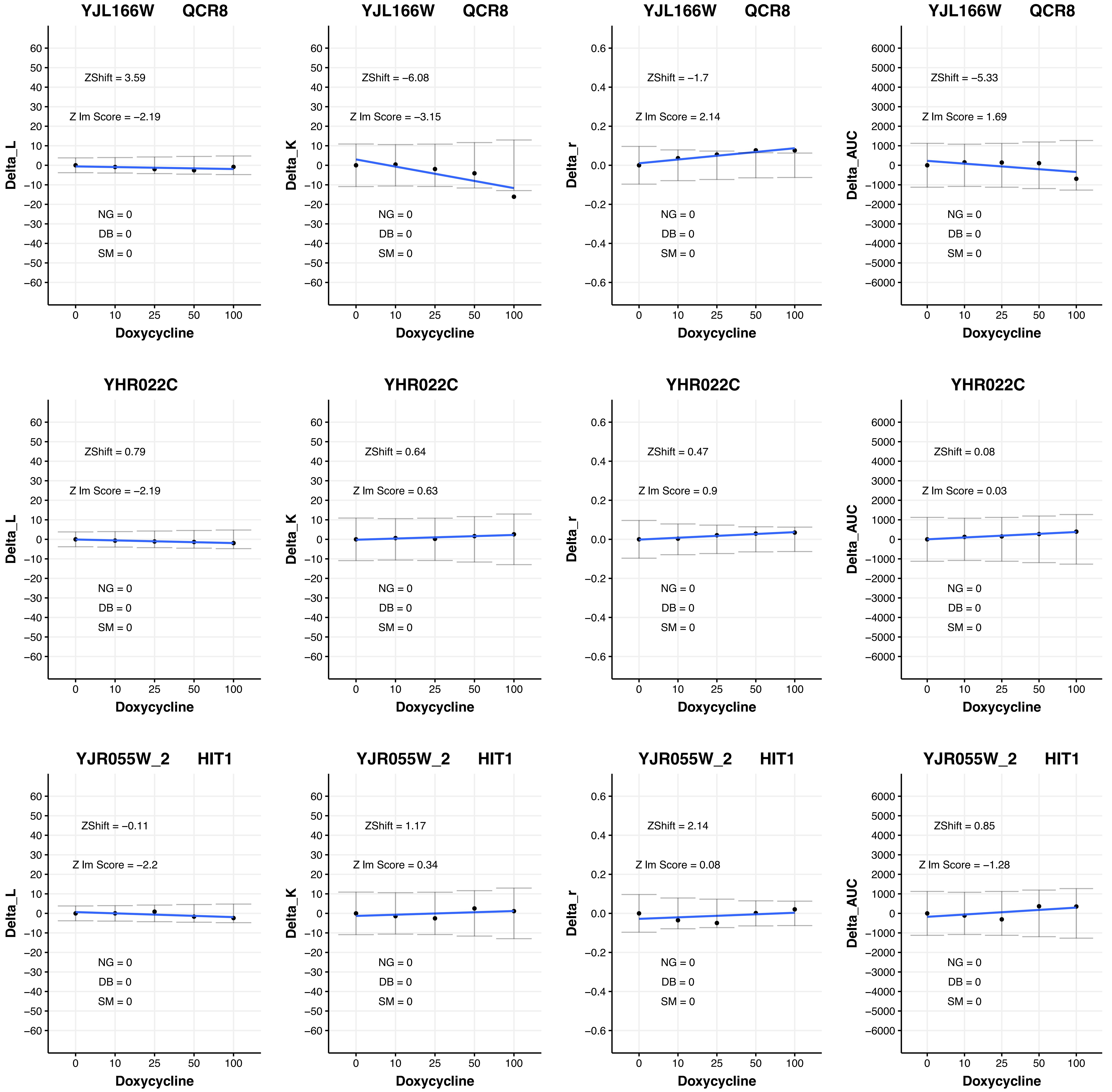

YJL166W QCR8
ZShift = 3.59
Z lm Score = −2.19
NG = 0
DB = 0
SM = 0
Delta_L
Doxycycline
YJL166W QCR8
ZShift = −6.08
Z lm Score = −3.15
NG = 0
DB = 0
SM = 0
Delta_K
Doxycycline
YJL166W QCR8
ZShift = −1.7
Z lm Score = 2.14
NG = 0
DB = 0
SM = 0
Delta_r
Doxycycline
YJL166W QCR8
ZShift = −5.33
Z lm Score = 1.69
NG = 0
DB = 0
SM = 0
Delta_AUC
Doxycycline
YHR022C
ZShift = 0.79
Z lm Score = −2.19
NG = 0
DB = 0
SM = 0
Delta_L
Doxycycline
YHR022C
ZShift = 0.64
Z lm Score = 0.63
NG = 0
DB = 0
SM = 0
Delta_K
Doxycycline
YHR022C
ZShift = 0.47
Z lm Score = 0.9
NG = 0
DB = 0
SM = 0
Delta_r
Doxycycline
YHR022C
ZShift = 0.08
Z lm Score = 0.03
NG = 0
DB = 0
SM = 0
Delta_AUC
Doxycycline
YJR055W_2 HIT1
ZShift = −0.11
Z lm Score = −2.2
NG = 0
DB = 0
SM = 0
Delta_L
Doxycycline
YJR055W_2 HIT1
ZShift = 1.17
Z lm Score = 0.34
NG = 0
DB = 0
SM = 0
Delta_K
Doxycycline
YJR055W_2 HIT1
ZShift = 2.14
Z lm Score = 0.08
NG = 0
DB = 0
SM = 0
Delta_r
Doxycycline
YJR055W_2 HIT1
ZShift = 0.85
Z lm Score = −1.28
NG = 0
DB = 0
SM = 0
Delta_AUC
Doxycycline

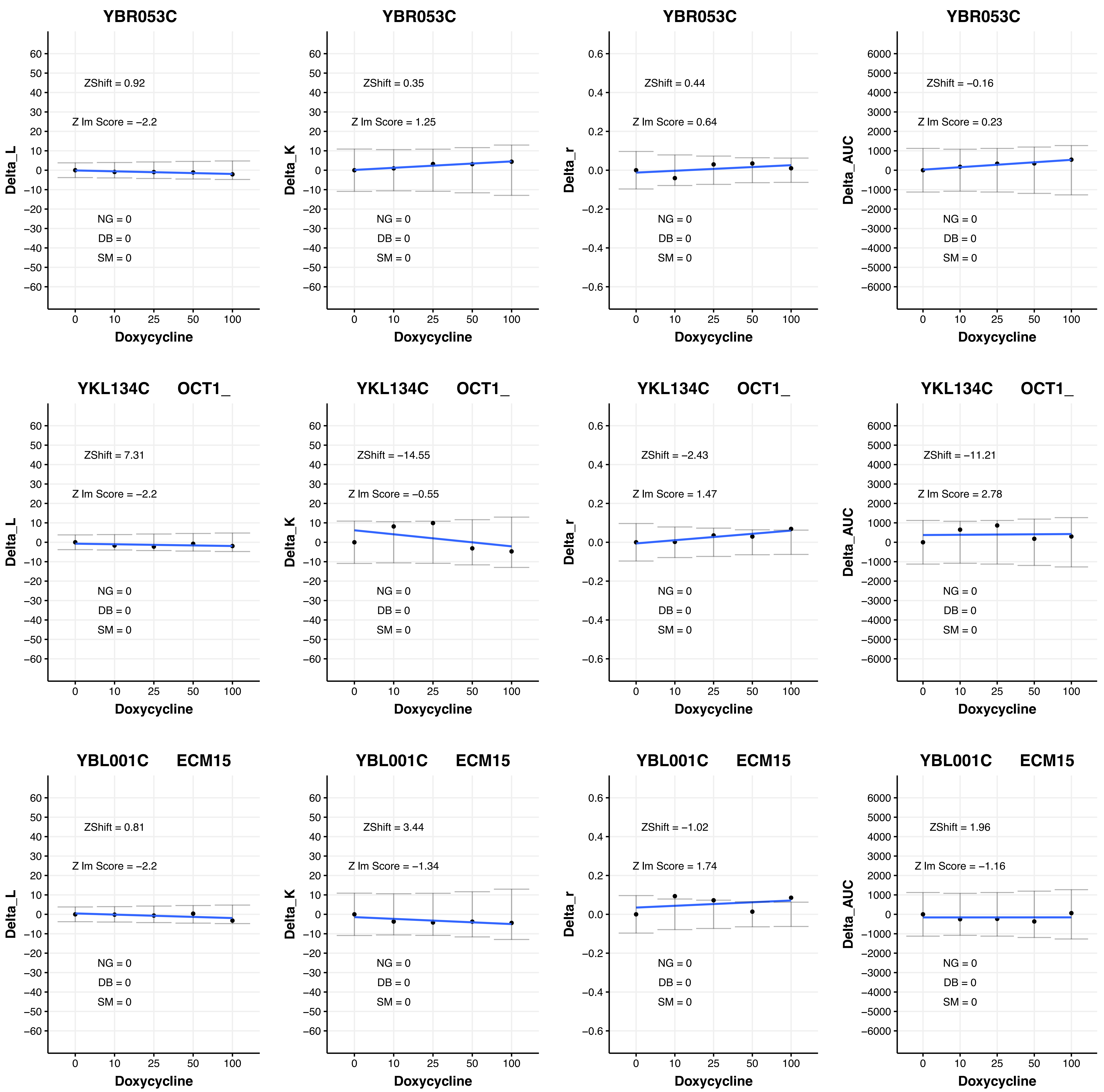
YBR053C
ZShift = 0.92
Z lm Score = −2.2
NG = 0
DB = 0
SM = 0
Delta_L
Doxycycline
YBR053C
ZShift = 0.35
Z lm Score = 1.25
NG = 0
DB = 0
SM = 0
Delta_K
Doxycycline
YBR053C
ZShift = 0.44
Z lm Score = 0.64
NG = 0
DB = 0
SM = 0
Delta_r
Doxycycline
YBR053C
ZShift = −0.16
Z lm Score = 0.23
NG = 0
DB = 0
SM = 0
Delta_AUC
Doxycycline
YKL134C OCT1_
ZShift = 7.31
Z lm Score = −2.2
NG = 0
DB = 0
SM = 0
Delta_L
Doxycycline
YKL134C OCT1_
ZShift = −14.55
Z lm Score = −0.55
NG = 0
DB = 0
SM = 0
Delta_K
Doxycycline
YKL134C OCT1_
ZShift = −2.43
Z lm Score = 1.47
NG = 0
DB = 0
SM = 0
Delta_r
Doxycycline
YKL134C OCT1_
ZShift = −11.21
Z lm Score = 2.78
NG = 0
DB = 0
SM = 0
Delta_AUC
Doxycycline
YBL001C ECM15
ZShift = 0.81
Z lm Score = −2.2
NG = 0
DB = 0
SM = 0
Delta_L
Doxycycline
YBL001C ECM15
ZShift = 3.44
Z lm Score = −1.34
NG = 0
DB = 0
SM = 0
Delta_K
Doxycycline
YBL001C ECM15
ZShift = −1.02
Z lm Score = 1.74
NG = 0
DB = 0
SM = 0
Delta_r
Doxycycline
YBL001C ECM15
ZShift = 1.96
Z lm Score = −1.16
NG = 0
DB = 0
SM = 0
Delta_AUC
Doxycycline

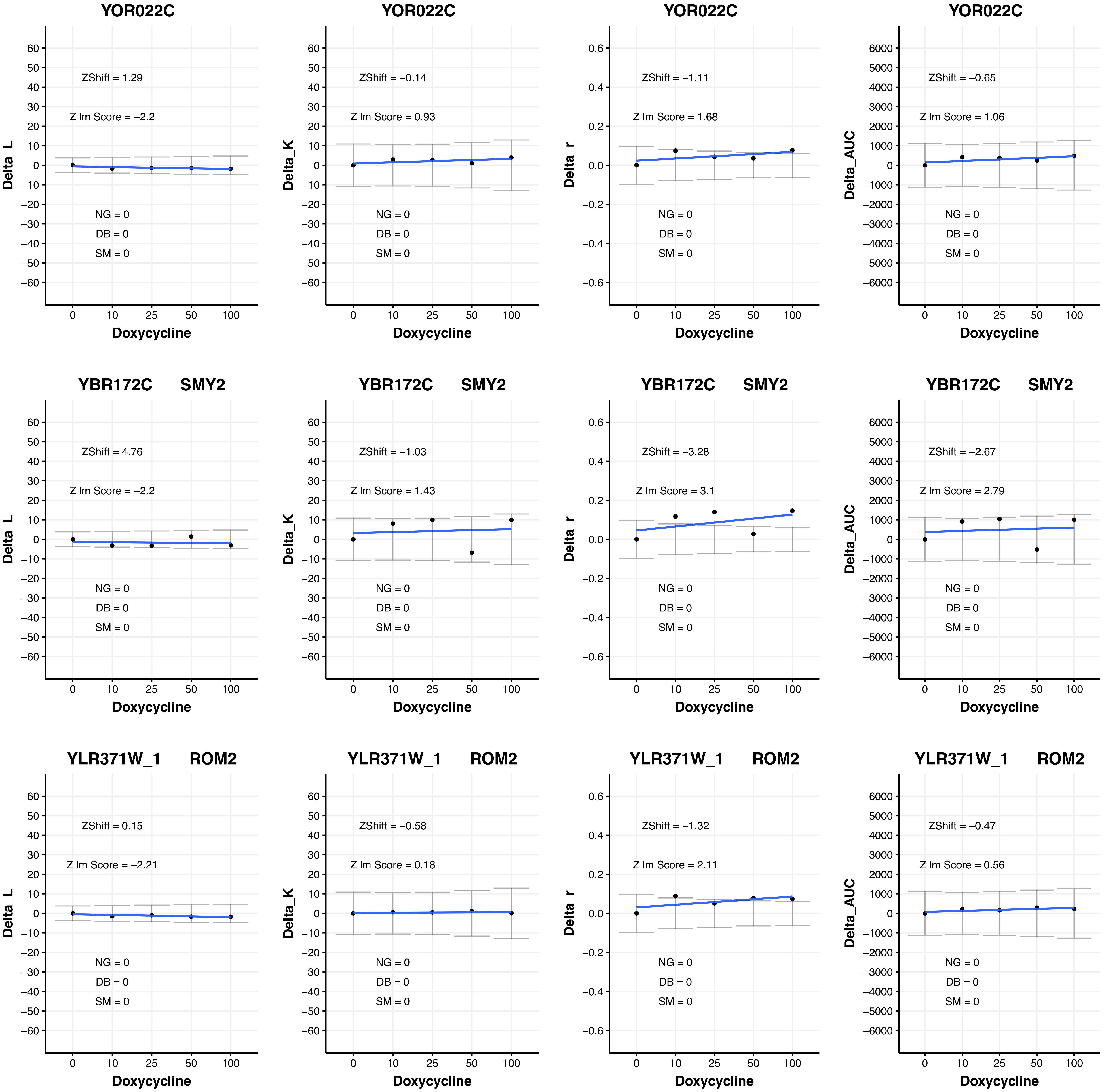

YOR022C
ZShift = 1.29
Z lm Score = −2.2
NG = 0
DB = 0
SM = 0
Delta_L
Doxycycline
YOR022C
ZShift = −0.14
Z lm Score = 0.93
NG = 0
DB = 0
SM = 0
Delta_K
Doxycycline
YOR022C
ZShift = −1.11
Z lm Score = 1.68
NG = 0
DB = 0
SM = 0
Delta_r
Doxycycline
YOR022C
ZShift = −0.65
Z lm Score = 1.06
NG = 0
DB = 0
SM = 0
Delta_AUC
Doxycycline
YBR172C SMY2
ZShift = 4.76
Z lm Score = −2.2
NG = 0
DB = 0
SM = 0
Delta_L
Doxycycline
YBR172C SMY2
ZShift = −1.03
Z lm Score = 1.43
NG = 0
DB = 0
SM = 0
Delta_K
Doxycycline
YBR172C SMY2
ZShift = −3.28
Z lm Score = 3.1
NG = 0
DB = 0
SM = 0
Delta_r
Doxycycline
YBR172C SMY2
ZShift = −2.67
Z lm Score = 2.79
NG = 0
DB = 0
SM = 0
Delta_AUC
Doxycycline
YLR371W_1 ROM2
ZShift = 0.15
Z lm Score = −2.21
NG = 0
DB = 0
SM = 0
Delta_L
Doxycycline
YLR371W_1 ROM2
ZShift = −0.58
Z lm Score = 0.18
NG = 0
DB = 0
SM = 0
Delta_K
Doxycycline
YLR371W_1 ROM2
ZShift = −1.32
Z lm Score = 2.11
NG = 0
DB = 0
SM = 0
Delta_r
Doxycycline
YLR371W_1 ROM2
ZShift = −0.47
Z lm Score = 0.56
NG = 0
DB = 0
SM = 0
Delta_AUC
Doxycycline

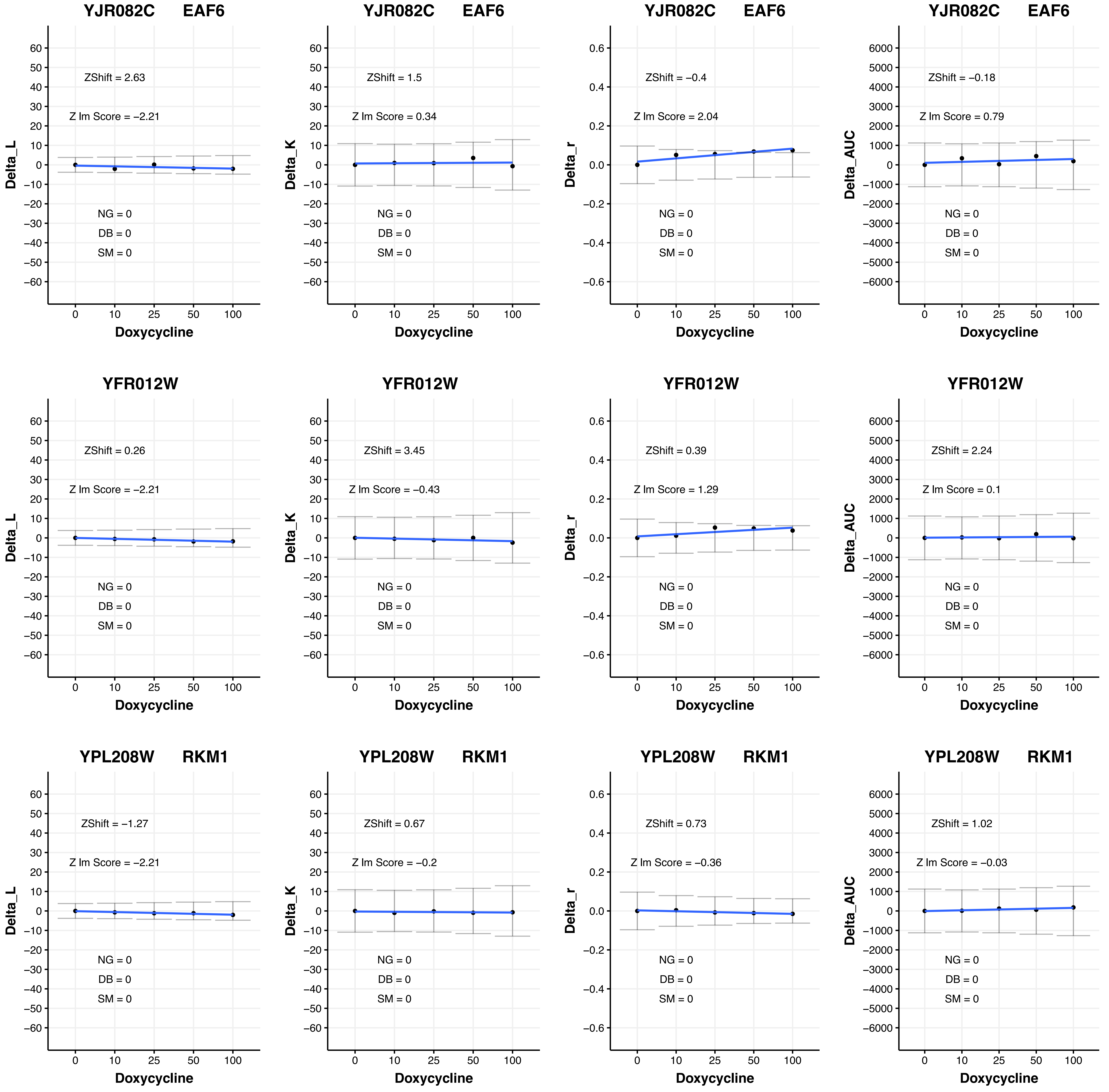

YJR082C EAF6
ZShift = 2.63
Z lm Score = −2.21
NG = 0
DB = 0
SM = 0
Delta_L
Doxycycline
YJR082C EAF6
ZShift = 1.5
Z lm Score = 0.34
NG = 0
DB = 0
SM = 0
Delta_K
Doxycycline
YJR082C EAF6
ZShift = −0.4
Z lm Score = 2.04
NG = 0
DB = 0
SM = 0
Delta_r
Doxycycline
YJR082C EAF6
ZShift = −0.18
Z lm Score = 0.79
NG = 0
DB = 0
SM = 0
Delta_AUC
Doxycycline
YFR012W
ZShift = 0.26
Z lm Score = −2.21
NG = 0
DB = 0
SM = 0
Delta_L
Doxycycline
YFR012W
ZShift = 3.45
Z lm Score = −0.43
NG = 0
DB = 0
SM = 0
Delta_K
Doxycycline
YFR012W
ZShift = 0.39
Z lm Score = 1.29
NG = 0
DB = 0
SM = 0
Delta_r
Doxycycline
YFR012W
ZShift = 2.24
Z lm Score = 0.1
NG = 0
DB = 0
SM = 0
Delta_AUC
Doxycycline
YPL208W RKM1
ZShift = −1.27
Z lm Score = −2.21
NG = 0
DB = 0
SM = 0
Delta_L
Doxycycline
YPL208W RKM1
ZShift = 0.67
Z lm Score = −0.2
NG = 0
DB = 0
SM = 0
Delta_K
Doxycycline
YPL208W RKM1
ZShift = 0.73
Z lm Score = −0.36
NG = 0
DB = 0
SM = 0
Delta_r
Doxycycline
YPL208W RKM1
ZShift = 1.02
Z lm Score = −0.03
NG = 0
DB = 0
SM = 0
Delta_AUC
Doxycycline

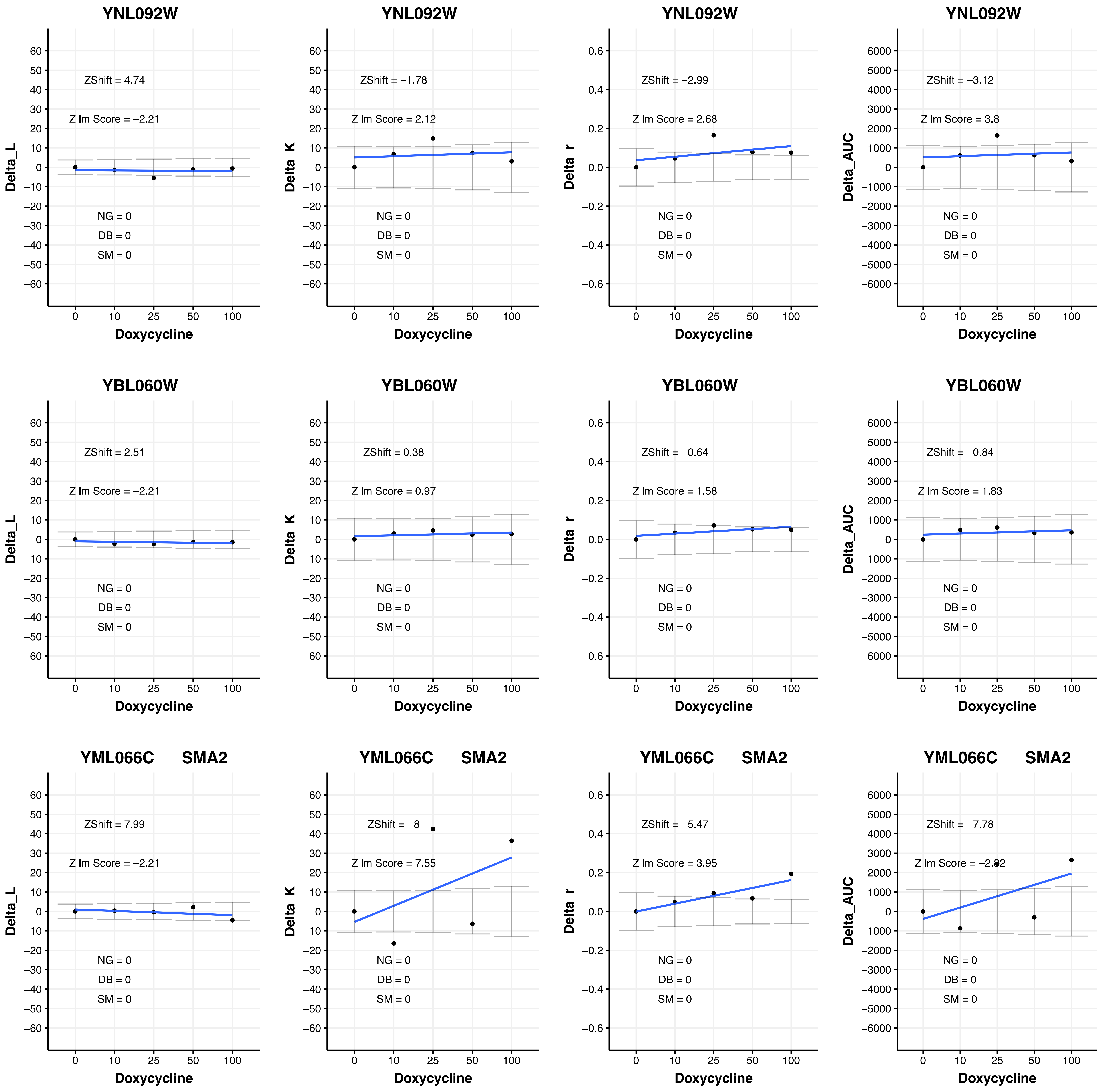
YNL092W
ZShift = 4.74
Z lm Score = −2.21
NG = 0
DB = 0
SM = 0
Delta_L
Doxycycline
YNL092W
ZShift = −1.78
Z lm Score = 2.12
NG = 0
DB = 0
SM = 0
Delta_K
Doxycycline
YNL092W
ZShift = −2.99
Z lm Score = 2.68
NG = 0
DB = 0
SM = 0
Delta_r
Doxycycline
YNL092W
ZShift = −3.12
Z lm Score = 3.8
NG = 0
DB = 0
SM = 0
Delta_AUC
Doxycycline
YBL060W
ZShift = 2.51
Z lm Score = −2.21
NG = 0
DB = 0
SM = 0
Delta_L
Doxycycline
YBL060W
ZShift = 0.38
Z lm Score = 0.97
NG = 0
DB = 0
SM = 0
Delta_K
Doxycycline
YBL060W
ZShift = −0.64
Z lm Score = 1.58
NG = 0
DB = 0
SM = 0
Delta_r
Doxycycline
YBL060W
ZShift = −0.84
Z lm Score = 1.83
NG = 0
DB = 0
SM = 0
Delta_AUC
Doxycycline
YML066C SMA2
ZShift = 7.99
Z lm Score = −2.21
NG = 0
DB = 0
SM = 0
Delta_L
Doxycycline
YML066C SMA2
ZShift = −8
Z lm Score = 7.55
NG = 0
DB = 0
SM = 0
Delta_K
Doxycycline
YML066C SMA2
ZShift = −5.47
Z lm Score = 3.95
NG = 0
DB = 0
SM = 0
Delta_r
Doxycycline
YML066C SMA2
ZShift = −7.78
Z lm Score = −2.82
NG = 0
DB = 0
SM = 0
Delta_AUC
Doxycycline

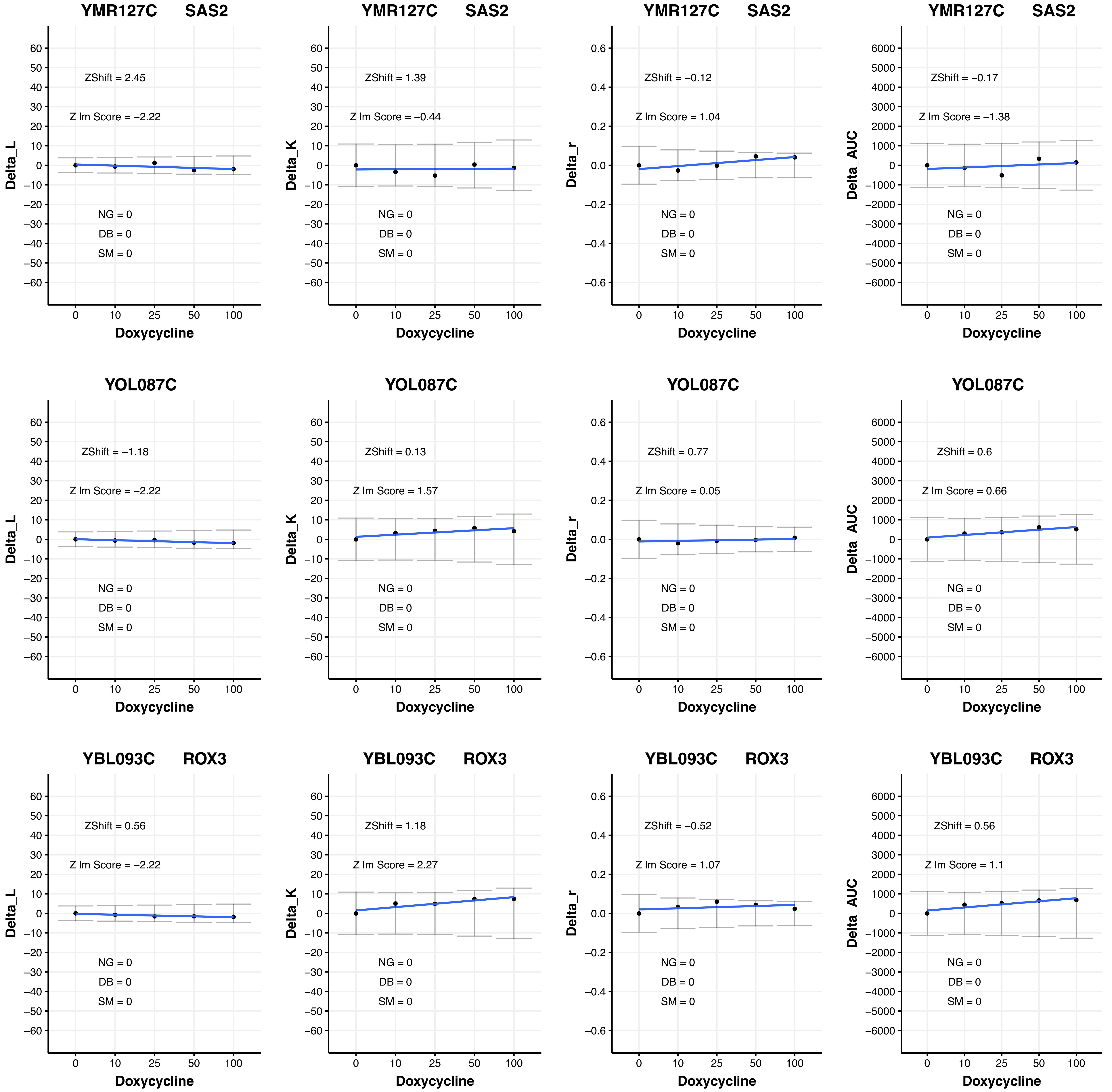

YMR127C SAS2
ZShift = 2.45
Z lm Score = −2.22
NG = 0
DB = 0
SM = 0
Delta_L
Doxycycline
YMR127C SAS2
ZShift = 1.39
Z lm Score = −0.44
NG = 0
DB = 0
SM = 0
Delta_K
Doxycycline
YMR127C SAS2
ZShift = −0.12
Z lm Score = 1.04
NG = 0
DB = 0
SM = 0
Delta_r
Doxycycline
YMR127C SAS2
ZShift = −0.17
Z lm Score = −1.38
NG = 0
DB = 0
SM = 0
Delta_AUC
Doxycycline
YOL087C
ZShift = −1.18
Z lm Score = −2.22
NG = 0
DB = 0
SM = 0
Delta_L
Doxycycline
YOL087C
ZShift = 0.13
Z lm Score = 1.57
NG = 0
DB = 0
SM = 0
Delta_K
Doxycycline
YOL087C
ZShift = 0.77
Z lm Score = 0.05
NG = 0
DB = 0
SM = 0
Delta_r
Doxycycline
YOL087C
ZShift = 0.6
Z lm Score = 0.66
NG = 0
DB = 0
SM = 0
Delta_AUC
Doxycycline
YBL093C ROX3
ZShift = 0.56
Z lm Score = −2.22
NG = 0
DB = 0
SM = 0
Delta_L
Doxycycline
YBL093C ROX3
ZShift = 1.18
Z lm Score = 2.27
NG = 0
DB = 0
SM = 0
Delta_K
Doxycycline
YBL093C ROX3
ZShift = −0.52
Z lm Score = 1.07
NG = 0
DB = 0
SM = 0
Delta_r
Doxycycline
YBL093C ROX3
ZShift = 0.56
Z lm Score = 1.1
NG = 0
DB = 0
SM = 0
Delta_AUC
Doxycycline

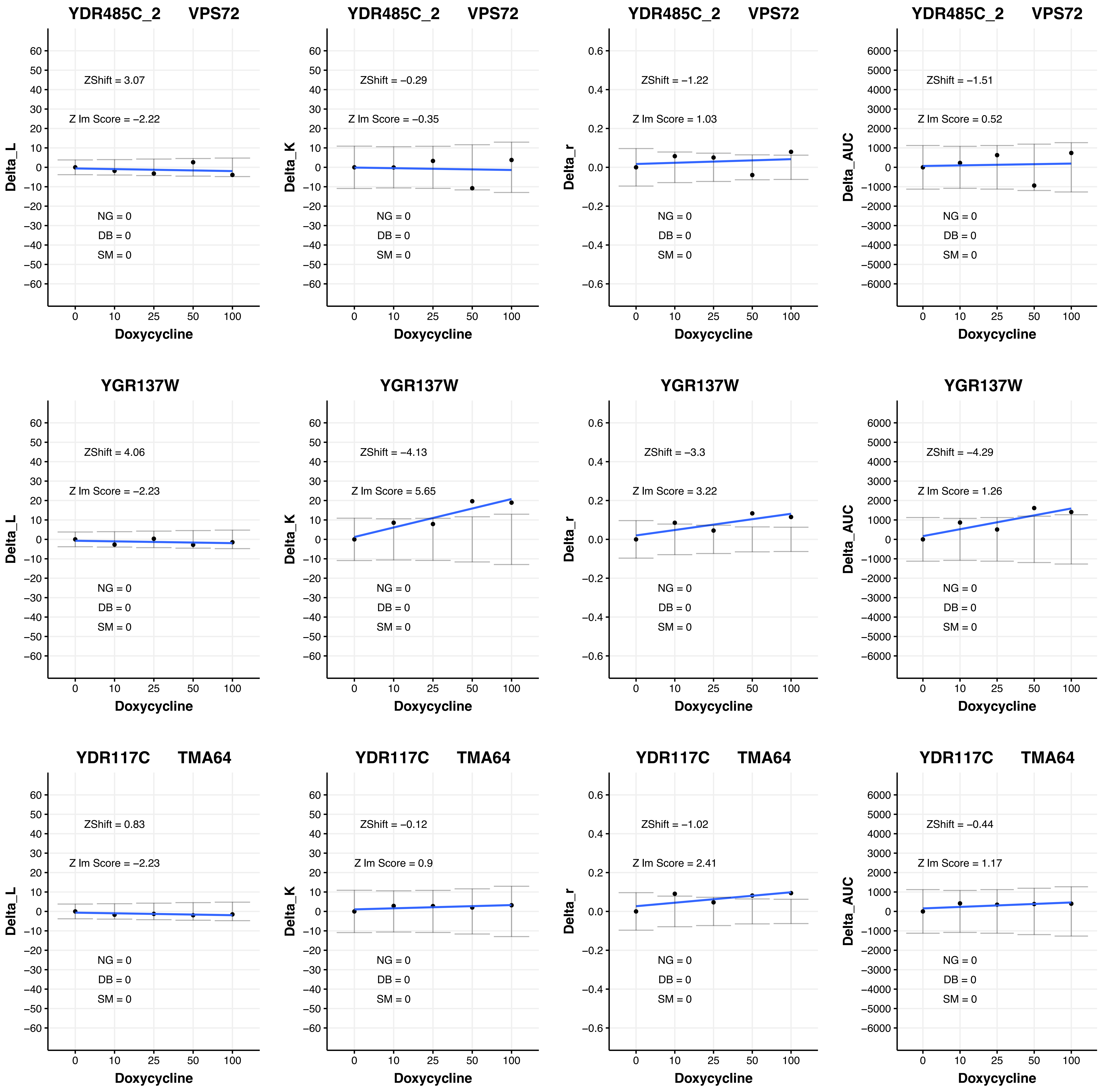

YDR485C_2 VPS72
ZShift = 3.07
Z lm Score = −2.22
NG = 0
DB = 0
SM = 0
Delta_L
Doxycycline
YDR485C_2 VPS72
ZShift = −0.29
Z lm Score = −0.35
NG = 0
DB = 0
SM = 0
Delta_K
Doxycycline
YDR485C_2 VPS72
ZShift = −1.22
Z lm Score = 1.03
NG = 0
DB = 0
SM = 0
Delta_r
Doxycycline
YDR485C_2 VPS72
ZShift = −1.51
Z lm Score = 0.52
NG = 0
DB = 0
SM = 0
Delta_AUC
Doxycycline
YGR137W
ZShift = 4.06
Z lm Score = −2.23
NG = 0
DB = 0
SM = 0
Delta_L
Doxycycline
YGR137W
ZShift = −4.13
Z lm Score = 5.65
NG = 0
DB = 0
SM = 0
Delta_K
Doxycycline
YGR137W
ZShift = −3.3
Z lm Score = 3.22
NG = 0
DB = 0
SM = 0
Delta_r
Doxycycline
YGR137W
ZShift = −4.29
Z lm Score = 1.26
NG = 0
DB = 0
SM = 0
Delta_AUC
Doxycycline
YDR117C TMA64
ZShift = 0.83
Z lm Score = −2.23
NG = 0
DB = 0
SM = 0
Delta_L
Doxycycline
YDR117C TMA64
ZShift = −0.12
Z lm Score = 0.9
NG = 0
DB = 0
SM = 0
Delta_K
Doxycycline
YDR117C TMA64
ZShift = −1.02
Z lm Score = 2.41
NG = 0
DB = 0
SM = 0
Delta_r
Doxycycline
YDR117C TMA64
ZShift = −0.44
Z lm Score = 1.17
NG = 0
DB = 0
SM = 0
Delta_AUC
Doxycycline

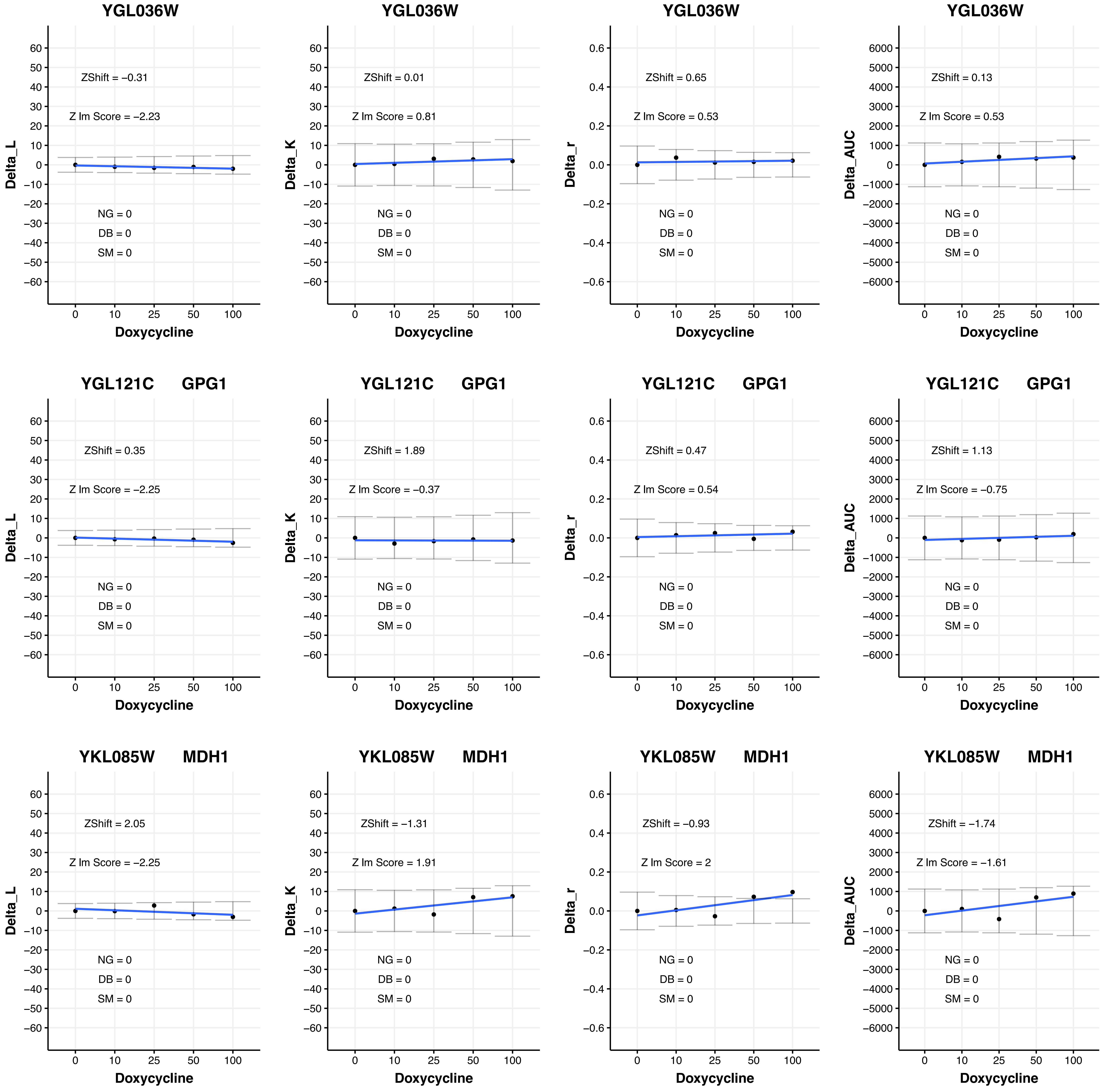

YGL036W
ZShift = −0.31
Z lm Score = −2.23
NG = 0
DB = 0
SM = 0
Delta_L
Doxycycline
YGL036W
ZShift = 0.01
Z lm Score = 0.81
NG = 0
DB = 0
SM = 0
Delta_K
Doxycycline
YGL036W
ZShift = 0.65
Z lm Score = 0.53
NG = 0
DB = 0
SM = 0
Delta_r
Doxycycline
YGL036W
ZShift = 0.13
Z lm Score = 0.53
NG = 0
DB = 0
SM = 0
Delta_AUC
Doxycycline
YGL121C GPG1
ZShift = 0.35
Z lm Score = −2.25
NG = 0
DB = 0
SM = 0
Delta_L
Doxycycline
YGL121C GPG1
ZShift = 1.89
Z lm Score = −0.37
NG = 0
DB = 0
SM = 0
Delta_K
Doxycycline
YGL121C GPG1
ZShift = 0.47
Z lm Score = 0.54
NG = 0
DB = 0
SM = 0
Delta_r
Doxycycline
YGL121C GPG1
ZShift = 1.13
Z lm Score = −0.75
NG = 0
DB = 0
SM = 0
Delta_AUC
Doxycycline
YKL085W MDH1
ZShift = 2.05
Z lm Score = −2.25
NG = 0
DB = 0
SM = 0
Delta_L
Doxycycline
YKL085W MDH1
ZShift = −1.31
Z lm Score = 1.91
NG = 0
DB = 0
SM = 0
Delta_K
Doxycycline
YKL085W MDH1
ZShift = −0.93
Z lm Score = 2
NG = 0
DB = 0
SM = 0
Delta_r
Doxycycline
YKL085W MDH1
ZShift = −1.74
Z lm Score = −1.61
NG = 0
DB = 0
SM = 0
Delta_AUC
Doxycycline

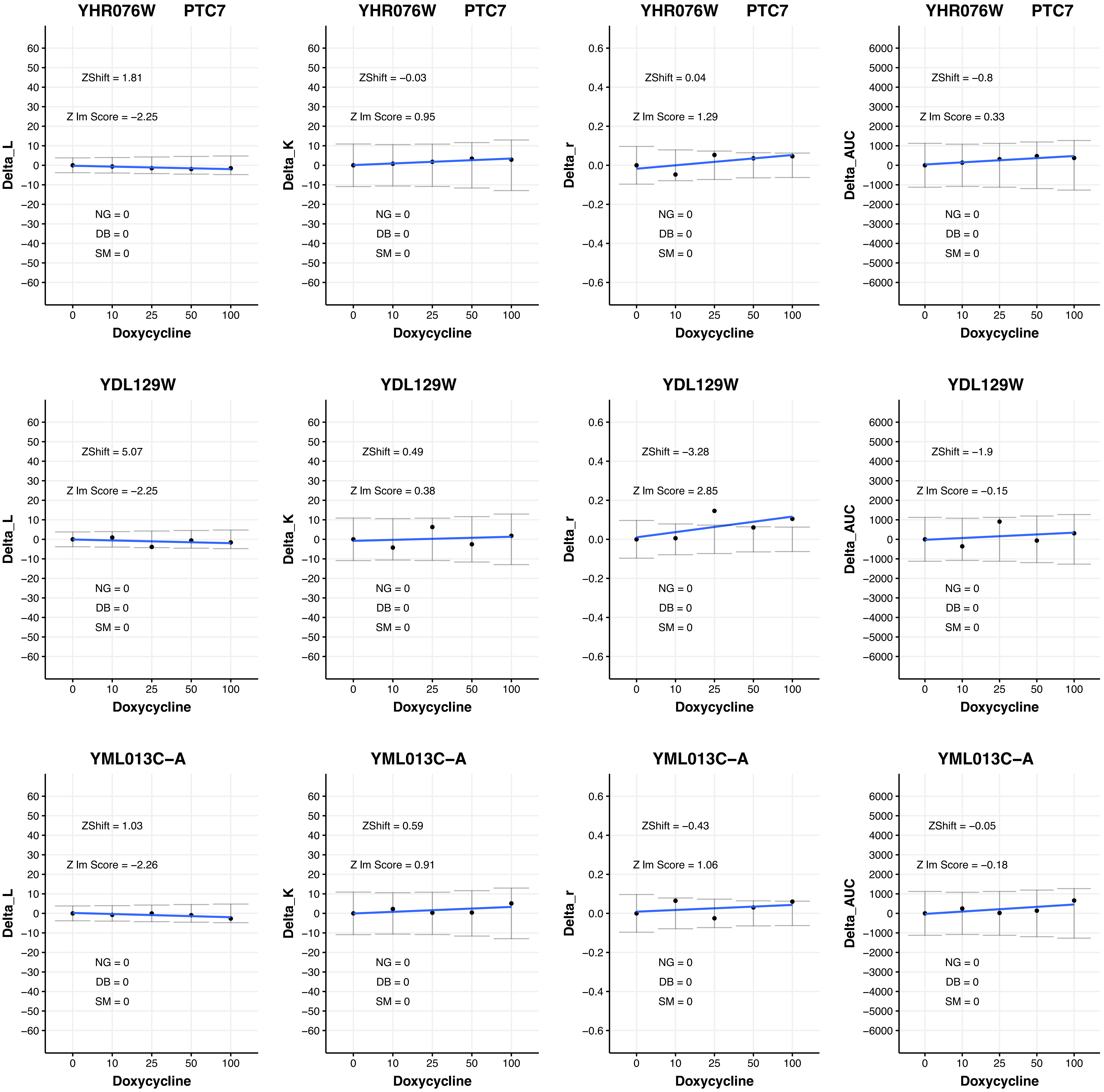

YHR076W PTC7
ZShift = 1.81
Z lm Score = −2.25
Delta_L
NG = 0
DB = 0
SM = 0
Doxycycline
YHR076W PTC7
ZShift = −0.03
Z lm Score = 0.95
Delta_K
NG = 0
DB = 0
SM = 0
Doxycycline
YHR076W PTC7
ZShift = 0.04
Z lm Score = 1.29
Delta_r
NG = 0
DB = 0
SM = 0
Doxycycline
YHR076W PTC7
ZShift = −0.8
Z lm Score = 0.33
Delta_AUC
NG = 0
DB = 0
SM = 0
Doxycycline
YDL129W
ZShift = 5.07
Z lm Score = −2.25
Delta_L
NG = 0
DB = 0
SM = 0
Doxycycline
YDL129W
ZShift = 0.49
Z lm Score = 0.38
Delta_K
NG = 0
DB = 0
SM = 0
Doxycycline
YDL129W
ZShift = −3.28
Z lm Score = 2.85
Delta_r
NG = 0
DB = 0
SM = 0
Doxycycline
YDL129W
ZShift = −1.9
Z lm Score = −0.15
Delta_AUC
NG = 0
DB = 0
SM = 0
Doxycycline
YML013C−A
ZShift = 1.03
Z lm Score = −2.26
Delta_L
NG = 0
DB = 0
SM = 0
Doxycycline
YML013C−A
ZShift = 0.59
Z lm Score = 0.91
Delta_K
NG = 0
DB = 0
SM = 0
Doxycycline
YML013C−A
ZShift = −0.43
Z lm Score = 1.06
Delta_r
NG = 0
DB = 0
SM = 0
Doxycycline
YML013C−A
ZShift = −0.05
Z lm Score = −0.18
Delta_AUC
NG = 0
DB = 0
SM = 0
Doxycycline
0 10 25 50 100

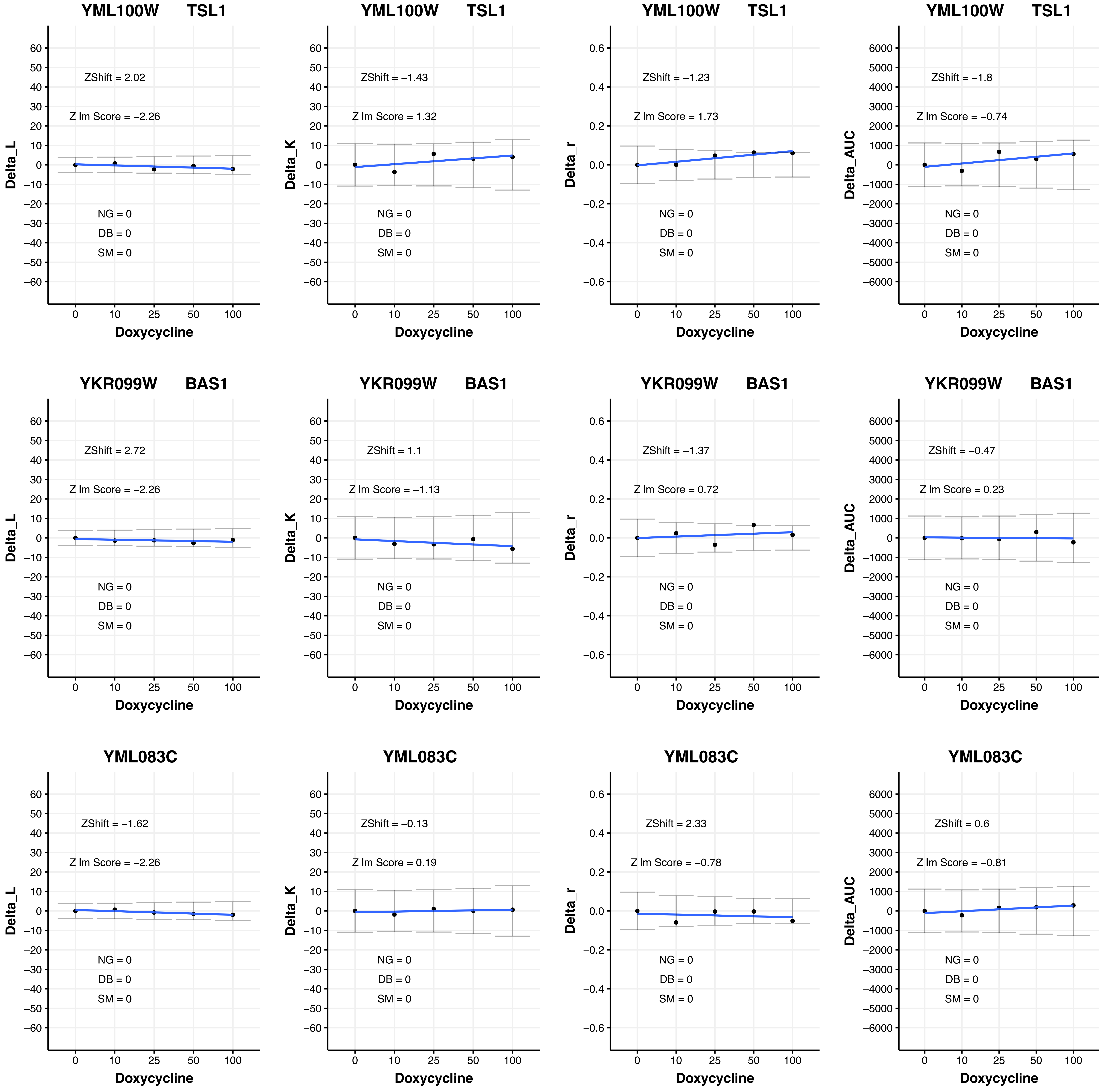

YML100W TSL1
ZShift = 2.02
Z lm Score = −2.26
NG = 0
DB = 0
SM = 0
Delta_L
Doxycycline
YML100W TSL1
ZShift = −1.43
Z lm Score = 1.32
NG = 0
DB = 0
SM = 0
Delta_K
Doxycycline
YML100W TSL1
ZShift = −1.23
Z lm Score = 1.73
NG = 0
DB = 0
SM = 0
Delta_r
Doxycycline
YML100W TSL1
ZShift = −1.8
Z lm Score = −0.74
NG = 0
DB = 0
SM = 0
Delta_AUC
Doxycycline
YKR099W BAS1
ZShift = 2.72
Z lm Score = −2.26
NG = 0
DB = 0
SM = 0
Delta_L
Doxycycline
YKR099W BAS1
ZShift = 1.1
Z lm Score = −1.13
NG = 0
DB = 0
SM = 0
Delta_K
Doxycycline
YKR099W BAS1
ZShift = −1.37
Z lm Score = 0.72
NG = 0
DB = 0
SM = 0
Delta_r
Doxycycline
YKR099W BAS1
ZShift = −0.47
Z lm Score = 0.23
NG = 0
DB = 0
SM = 0
Delta_AUC
Doxycycline
YML083C
ZShift = −1.62
Z lm Score = −2.26
NG = 0
DB = 0
SM = 0
Delta_L
Doxycycline
YML083C
ZShift = −0.13
Z lm Score = 0.19
NG = 0
DB = 0
SM = 0
Delta_K
Doxycycline
YML083C
ZShift = 2.33
Z lm Score = −0.78
NG = 0
DB = 0
SM = 0
Delta_r
Doxycycline
YML083C
ZShift = 0.6
Z lm Score = −0.81
NG = 0
DB = 0
SM = 0
Delta_AUC
Doxycycline

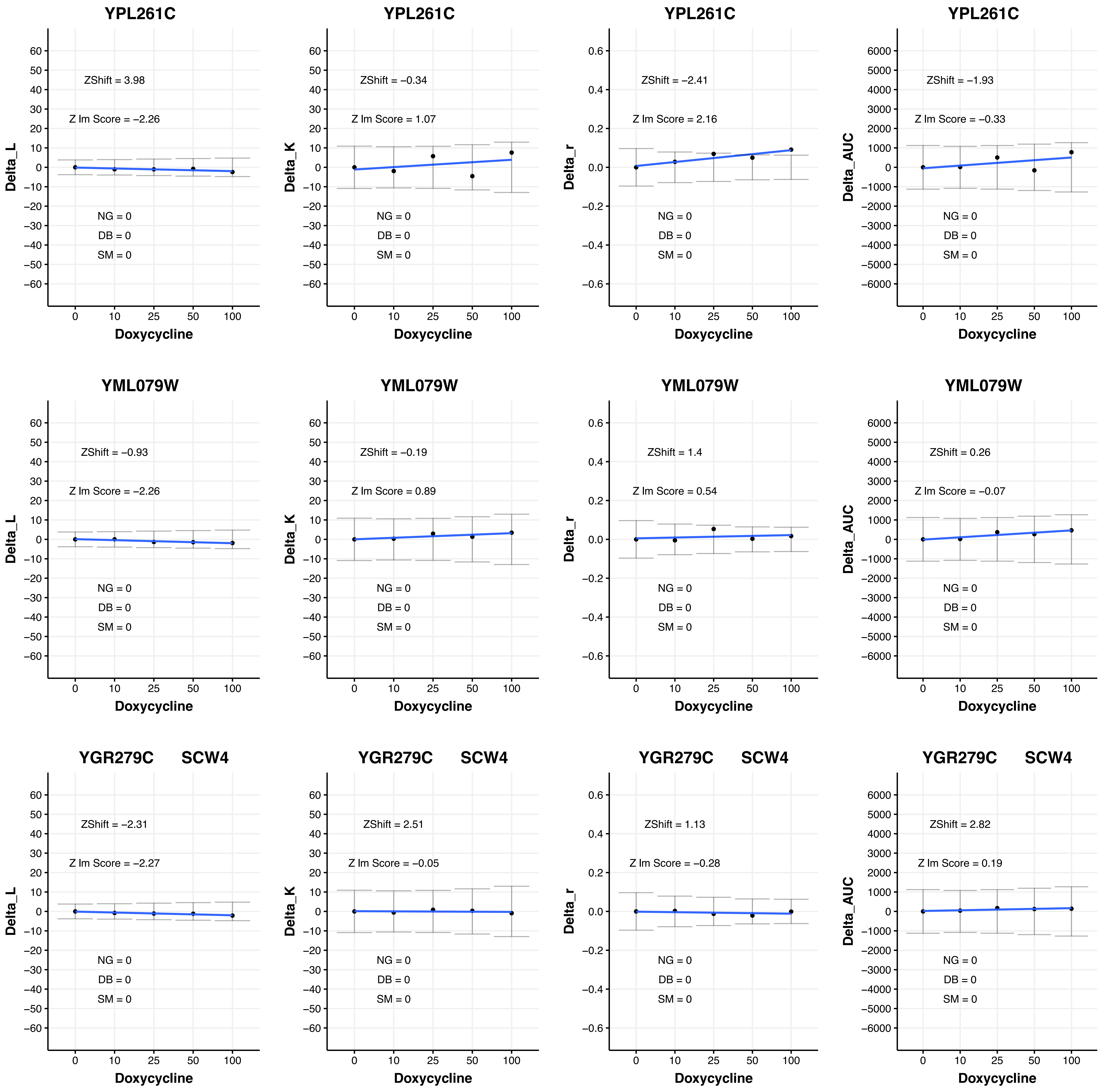
YPL261C
ZShift = 3.98
Z lm Score = −2.26
NG = 0
DB = 0
SM = 0
Delta_L
Doxycycline
YPL261C
ZShift = −0.34
Z lm Score = 1.07
NG = 0
DB = 0
SM = 0
Delta_K
Doxycycline
YPL261C
ZShift = −2.41
Z lm Score = 2.16
NG = 0
DB = 0
SM = 0
Delta_r
Doxycycline
YPL261C
ZShift = −1.93
Z lm Score = −0.33
NG = 0
DB = 0
SM = 0
Delta_AUC
Doxycycline
YML079W
ZShift = −0.93
Z lm Score = −2.26
NG = 0
DB = 0
SM = 0
Delta_L
Doxycycline
YML079W
ZShift = −0.19
Z lm Score = 0.89
NG = 0
DB = 0
SM = 0
Delta_K
Doxycycline
YML079W
ZShift = 1.4
Z lm Score = 0.54
NG = 0
DB = 0
SM = 0
Delta_r
Doxycycline
YML079W
ZShift = 0.26
Z lm Score = −0.07
NG = 0
DB = 0
SM = 0
Delta_AUC
Doxycycline
YGR279C SCW4
ZShift = −2.31
Z lm Score = −2.27
NG = 0
DB = 0
SM = 0
Delta_L
Doxycycline
YGR279C SCW4
ZShift = 2.51
Z lm Score = −0.05
NG = 0
DB = 0
SM = 0
Delta_K
Doxycycline
YGR279C SCW4
ZShift = 1.13
Z lm Score = −0.28
NG = 0
DB = 0
SM = 0
Delta_r
Doxycycline
YGR279C SCW4
ZShift = 2.82
Z lm Score = 0.19
NG = 0
DB = 0
SM = 0
Delta_AUC
Doxycycline

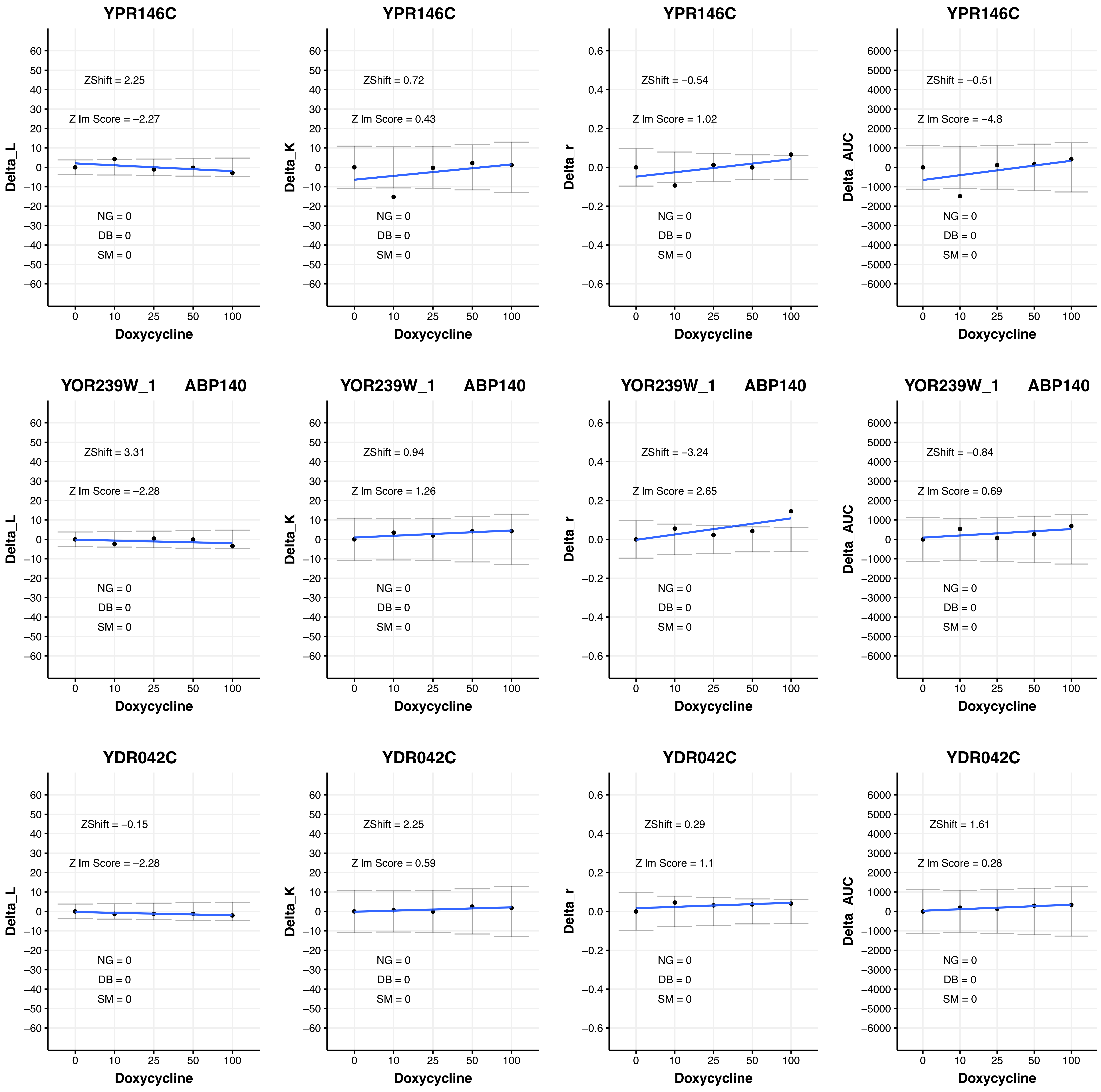

YPR146C
Delta_L
ZShift = 2.25
Z lm Score = −2.27
NG = 0
DB = 0
SM = 0
Doxycycline
YPR146C
Delta_K
ZShift = 0.72
Z lm Score = 0.43
NG = 0
DB = 0
SM = 0
Doxycycline
YPR146C
Delta_r
ZShift = −0.54
Z lm Score = 1.02
NG = 0
DB = 0
SM = 0
Doxycycline
YPR146C
Delta_AUC
ZShift = −0.51
Z lm Score = −4.8
NG = 0
DB = 0
SM = 0
Doxycycline
YOR239W_1 ABP140
Delta_L
ZShift = 3.31
Z lm Score = −2.28
NG = 0
DB = 0
SM = 0
Doxycycline
YOR239W_1 ABP140
Delta_K
ZShift = 0.94
Z lm Score = 1.26
NG = 0
DB = 0
SM = 0
Doxycycline
YOR239W_1 ABP140
Delta_r
ZShift = −3.24
Z lm Score = 2.65
NG = 0
DB = 0
SM = 0
Doxycycline
YOR239W_1 ABP140
Delta_AUC
ZShift = −0.84
Z lm Score = 0.69
NG = 0
DB = 0
SM = 0
Doxycycline
YDR042C
Delta_L
ZShift = −0.15
Z lm Score = −2.28
NG = 0
DB = 0
SM = 0
Doxycycline
YDR042C
Delta_K
ZShift = 2.25
Z lm Score = 0.59
NG = 0
DB = 0
SM = 0
Doxycycline
YDR042C
Delta_r
ZShift = 0.29
Z lm Score = 1.1
NG = 0
DB = 0
SM = 0
Doxycycline
YDR042C
Delta_AUC
ZShift = 1.61
Z lm Score = 0.28
NG = 0
DB = 0
SM = 0
Doxycycline

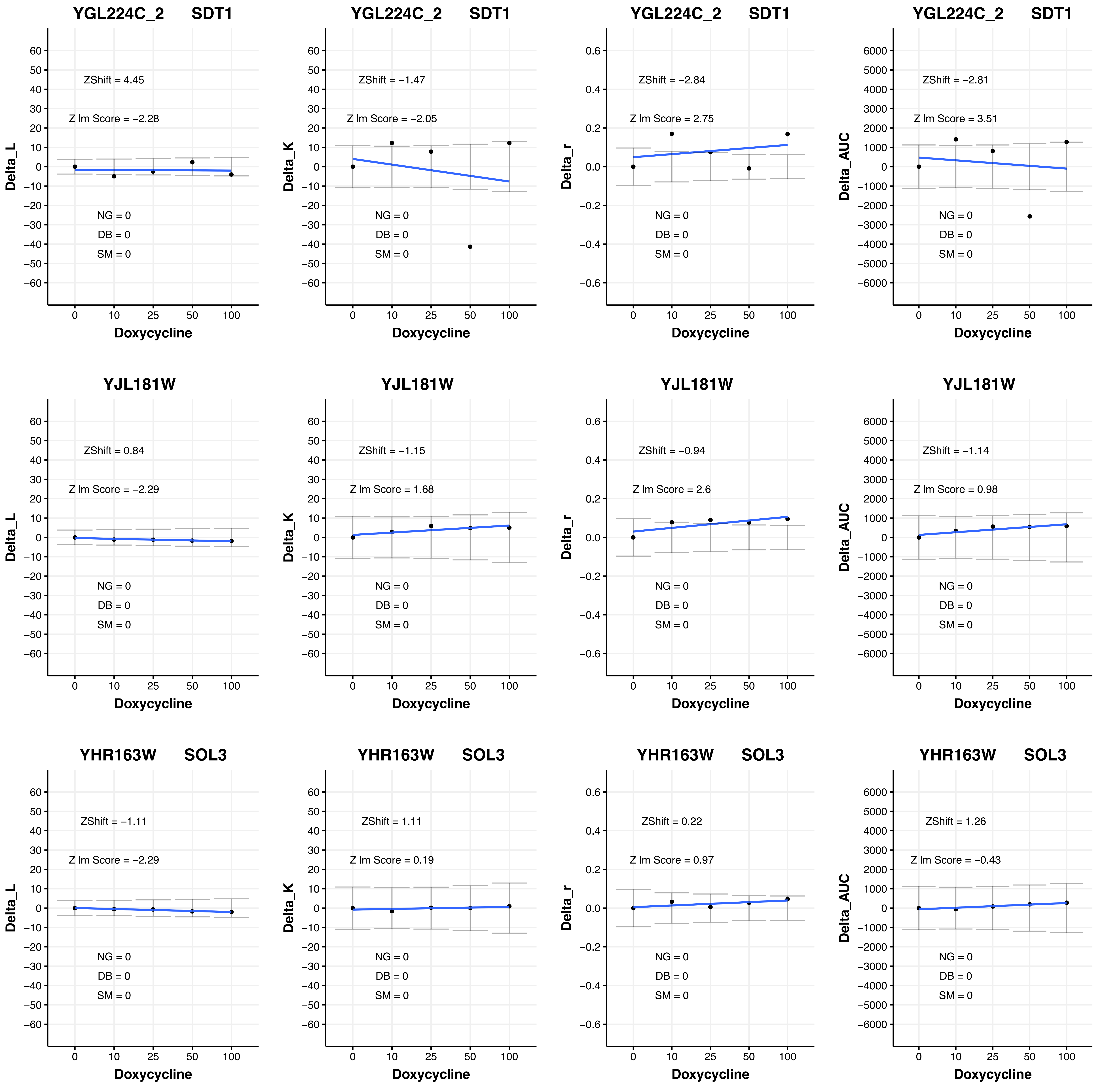

YGL224C_2 SDT1
ZShift = 4.45
Z lm Score = −2.28
NG = 0
DB = 0
SM = 0
Delta_L
Doxycycline
YGL224C_2 SDT1
ZShift = −1.47
Z lm Score = −2.05
NG = 0
DB = 0
SM = 0
Delta_K
Doxycycline
YGL224C_2 SDT1
ZShift = −2.84
Z lm Score = 2.75
NG = 0
DB = 0
SM = 0
Delta_r
Doxycycline
YGL224C_2 SDT1
ZShift = −2.81
Z lm Score = 3.51
NG = 0
DB = 0
SM = 0
Delta_AUC
Doxycycline
YJL181W
ZShift = 0.84
Z lm Score = −2.29
NG = 0
DB = 0
SM = 0
Delta_L
Doxycycline
YJL181W
ZShift = −1.15
Z lm Score = 1.68
NG = 0
DB = 0
SM = 0
Delta_K
Doxycycline
YJL181W
ZShift = −0.94
Z lm Score = 2.6
NG = 0
DB = 0
SM = 0
Delta_r
Doxycycline
YJL181W
ZShift = −1.14
Z lm Score = 0.98
NG = 0
DB = 0
SM = 0
Delta_AUC
Doxycycline
YHR163W SOL3
ZShift = −1.11
Z lm Score = −2.29
NG = 0
DB = 0
SM = 0
Delta_L
Doxycycline
YHR163W SOL3
ZShift = 1.11
Z lm Score = 0.19
NG = 0
DB = 0
SM = 0
Delta_K
Doxycycline
YHR163W SOL3
ZShift = 0.22
Z lm Score = 0.97
NG = 0
DB = 0
SM = 0
Delta_r
Doxycycline
YHR163W SOL3
ZShift = 1.26
Z lm Score = −0.43
NG = 0
DB = 0
SM = 0
Delta_AUC
Doxycycline

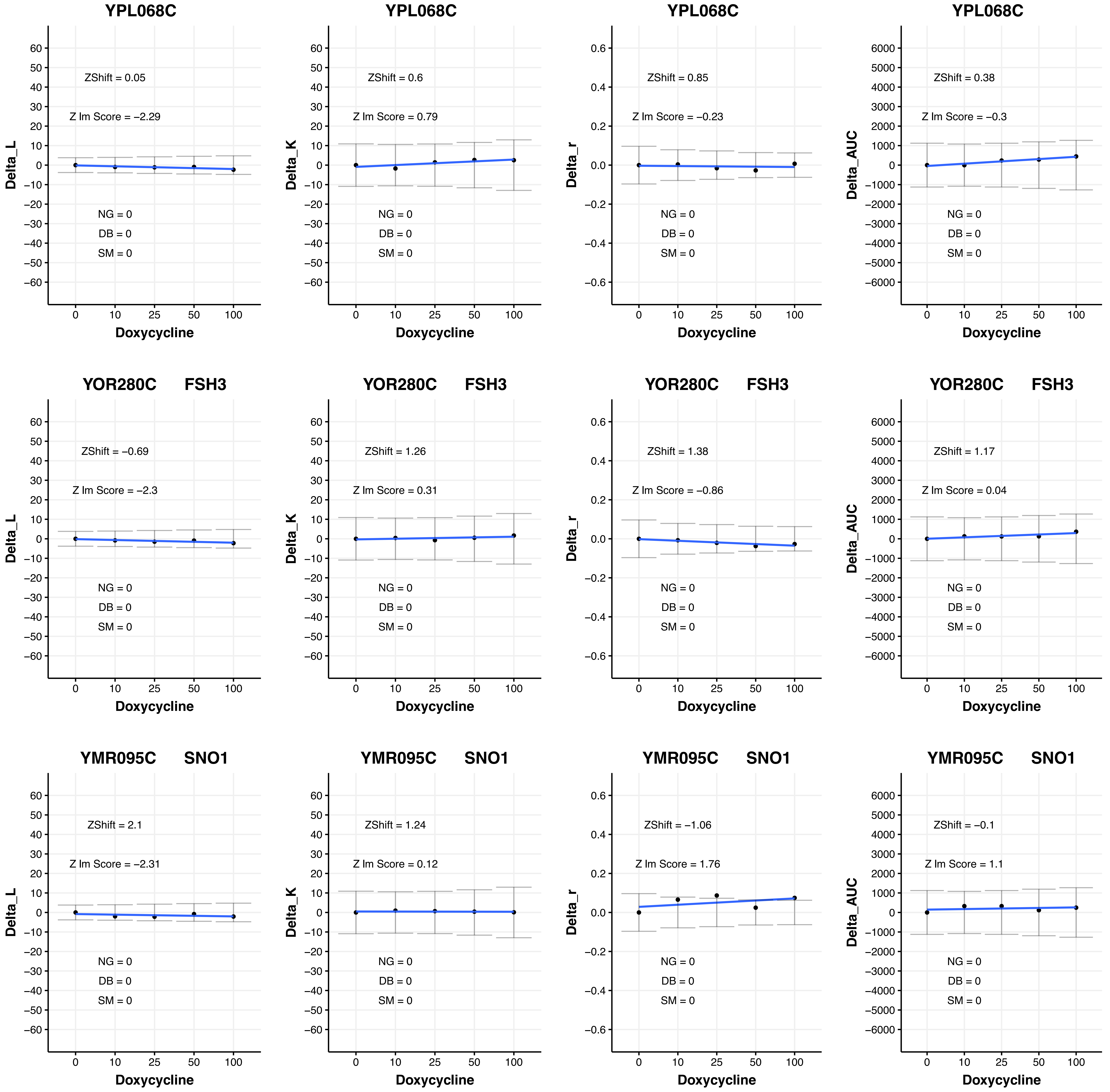
YPL068C
ZShift = 0.05
Z lm Score = −2.29
Delta_L
NG = 0
DB = 0
SM = 0
Doxycycline
YPL068C
ZShift = 0.6
Z lm Score = 0.79
Delta_K
NG = 0
DB = 0
SM = 0
Doxycycline
YPL068C
ZShift = 0.85
Z lm Score = −0.23
Delta_r
NG = 0
DB = 0
SM = 0
Doxycycline
YPL068C
ZShift = 0.38
Z lm Score = −0.3
Delta_AUC
NG = 0
DB = 0
SM = 0
Doxycycline
YOR280C FSH3
ZShift = −0.69
Z lm Score = −2.3
Delta_L
NG = 0
DB = 0
SM = 0
Doxycycline
YOR280C FSH3
ZShift = 1.26
Z lm Score = 0.31
Delta_K
NG = 0
DB = 0
SM = 0
Doxycycline
YOR280C FSH3
ZShift = 1.38
Z lm Score = −0.86
Delta_r
NG = 0
DB = 0
SM = 0
Doxycycline
YOR280C FSH3
ZShift = 1.17
Z lm Score = 0.04
Delta_AUC
NG = 0
DB = 0
SM = 0
Doxycycline
YMR095C SNO1
ZShift = 2.1
Z lm Score = −2.31
Delta_L
NG = 0
DB = 0
SM = 0
Doxycycline
YMR095C SNO1
ZShift = 1.24
Z lm Score = 0.12
Delta_K
NG = 0
DB = 0
SM = 0
Doxycycline
YMR095C SNO1
ZShift = −1.06
Z lm Score = 1.76
Delta_r
NG = 0
DB = 0
SM = 0
Doxycycline
YMR095C SNO1
ZShift = −0.1
Z lm Score = 1.1
Delta_AUC
NG = 0
DB = 0
SM = 0
Doxycycline

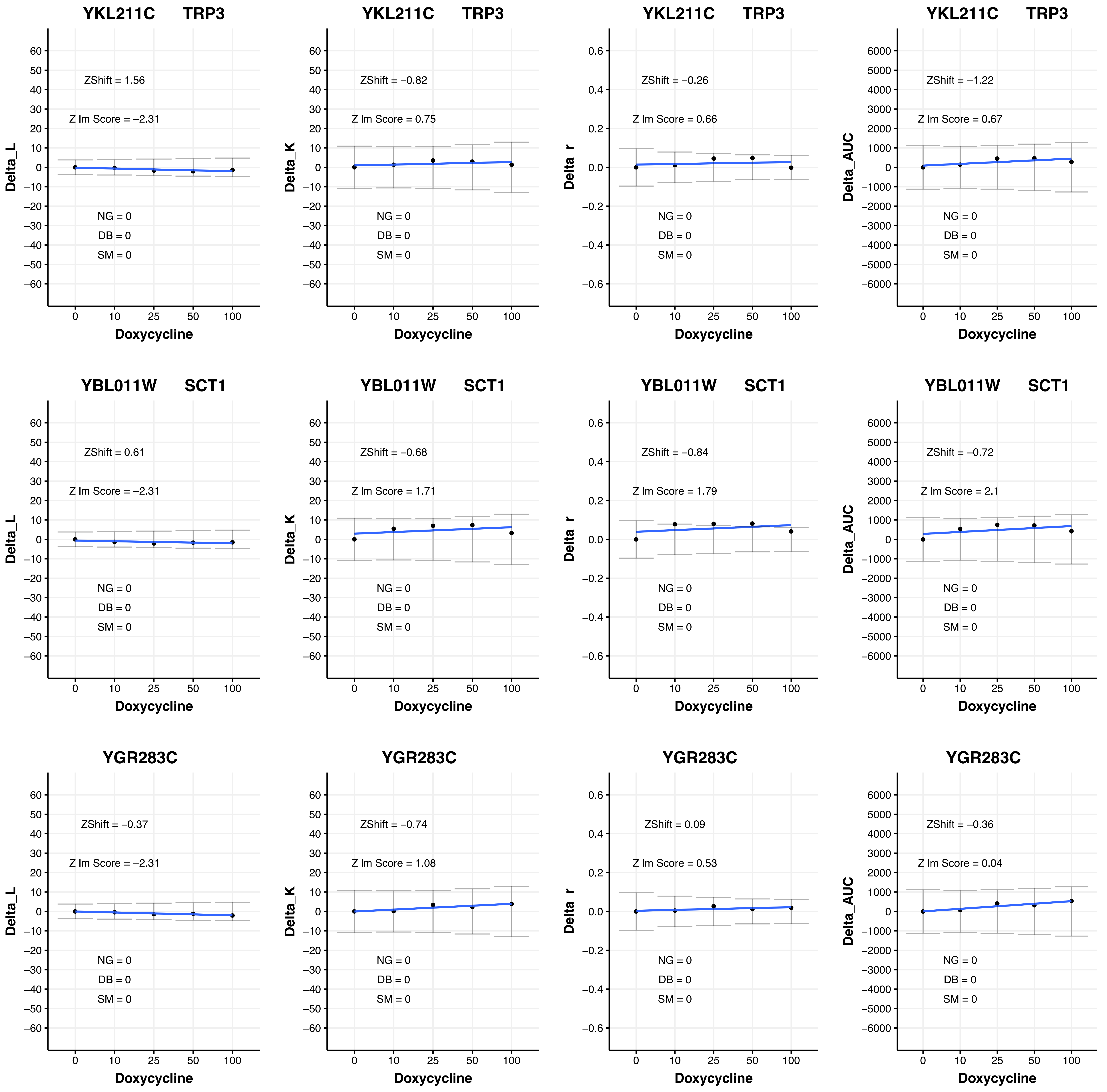
YKL211C TRP3
ZShift = 1.56
Z lm Score = −2.31
NG = 0
DB = 0
SM = 0
Delta_L
Doxycycline
YKL211C TRP3
ZShift = −0.82
Z lm Score = 0.75
NG = 0
DB = 0
SM = 0
Delta_K
Doxycycline
YKL211C TRP3
ZShift = −0.26
Z lm Score = 0.66
NG = 0
DB = 0
SM = 0
Delta_r
Doxycycline
YKL211C TRP3
ZShift = −1.22
Z lm Score = 0.67
NG = 0
DB = 0
SM = 0
Delta_AUC
Doxycycline
YBL011W SCT1
ZShift = 0.61
Z lm Score = −2.31
NG = 0
DB = 0
SM = 0
Delta_L
Doxycycline
YBL011W SCT1
ZShift = −0.68
Z lm Score = 1.71
NG = 0
DB = 0
SM = 0
Delta_K
Doxycycline
YBL011W SCT1
ZShift = −0.84
Z lm Score = 1.79
NG = 0
DB = 0
SM = 0
Delta_r
Doxycycline
YBL011W SCT1
ZShift = −0.72
Z lm Score = 2.1
NG = 0
DB = 0
SM = 0
Delta_AUC
Doxycycline
YGR283C
ZShift = −0.37
Z lm Score = −2.31
NG = 0
DB = 0
SM = 0
Delta_L
Doxycycline
YGR283C
ZShift = −0.74
Z lm Score = 1.08
NG = 0
DB = 0
SM = 0
Delta_K
Doxycycline
YGR283C
ZShift = 0.09
Z lm Score = 0.53
NG = 0
DB = 0
SM = 0
Delta_r
Doxycycline
YGR283C
ZShift = −0.36
Z lm Score = 0.04
NG = 0
DB = 0
SM = 0
Delta_AUC
Doxycycline

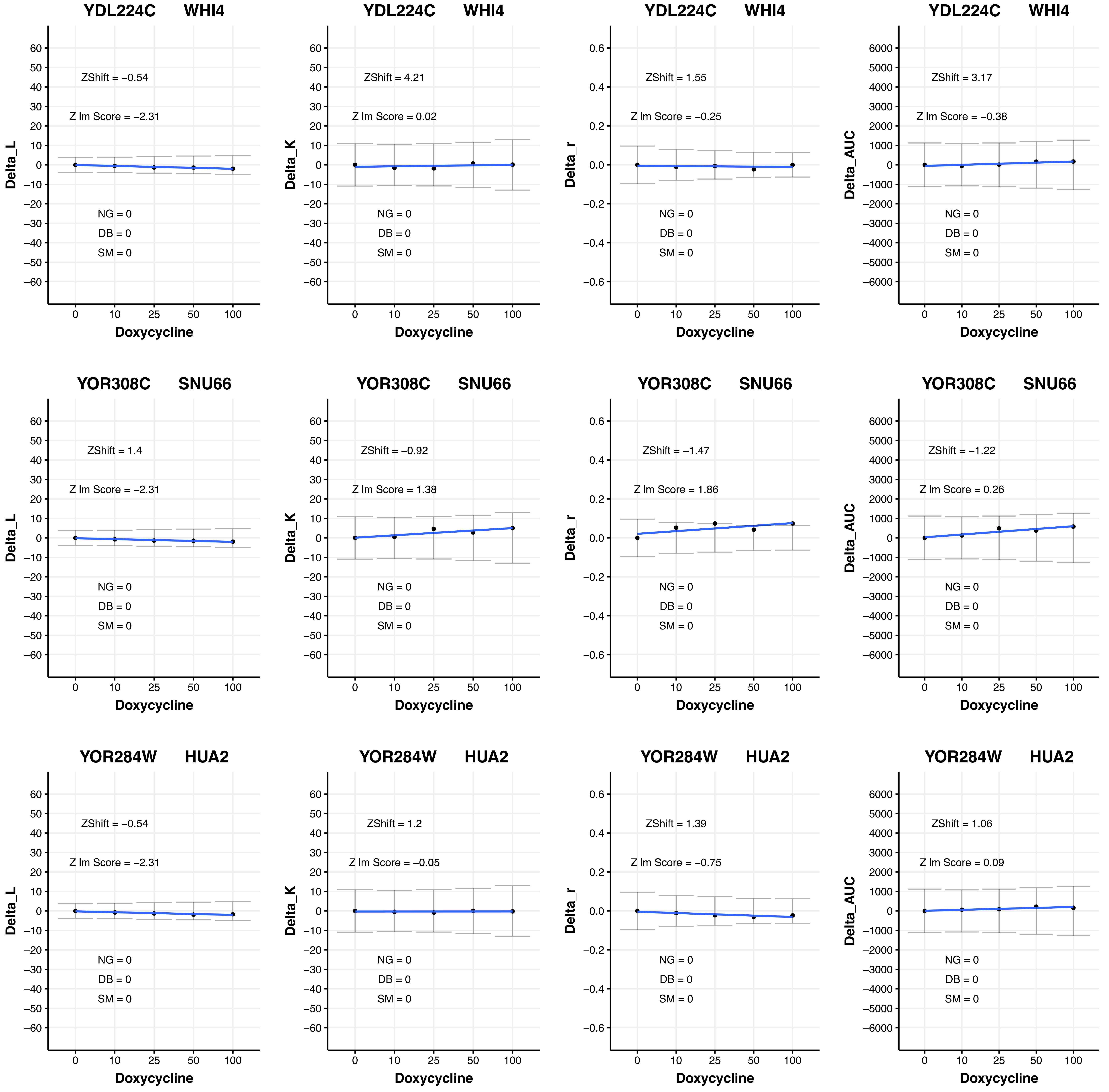
YDL224C WHI4
ZShift = −0.54
Z lm Score = −2.31
NG = 0
DB = 0
SM = 0
Delta_L
Doxycycline
YDL224C WHI4
ZShift = 4.21
Z lm Score = 0.02
NG = 0
DB = 0
SM = 0
Delta_K
Doxycycline
YDL224C WHI4
ZShift = 1.55
Z lm Score = −0.25
NG = 0
DB = 0
SM = 0
Delta_r
Doxycycline
YDL224C WHI4
ZShift = 3.17
Z lm Score = −0.38
NG = 0
DB = 0
SM = 0
Delta_AUC
Doxycycline
YOR308C SNU66
ZShift = 1.4
Z lm Score = −2.31
NG = 0
DB = 0
SM = 0
Delta_L
Doxycycline
YOR308C SNU66
ZShift = −0.92
Z lm Score = 1.38
NG = 0
DB = 0
SM = 0
Delta_K
Doxycycline
YOR308C SNU66
ZShift = −1.47
Z lm Score = 1.86
NG = 0
DB = 0
SM = 0
Delta_r
Doxycycline
YOR308C SNU66
ZShift = −1.22
Z lm Score = 0.26
NG = 0
DB = 0
SM = 0
Delta_AUC
Doxycycline
YOR284W HUA2
ZShift = −0.54
Z lm Score = −2.31
NG = 0
DB = 0
SM = 0
Delta_L
Doxycycline
YOR284W HUA2
ZShift = 1.2
Z lm Score = −0.05
NG = 0
DB = 0
SM = 0
Delta_K
Doxycycline
YOR284W HUA2
ZShift = 1.39
Z lm Score = −0.75
NG = 0
DB = 0
SM = 0
Delta_r
Doxycycline
YOR284W HUA2
ZShift = 1.06
Z lm Score = 0.09
NG = 0
DB = 0
SM = 0
Delta_AUC
Doxycycline

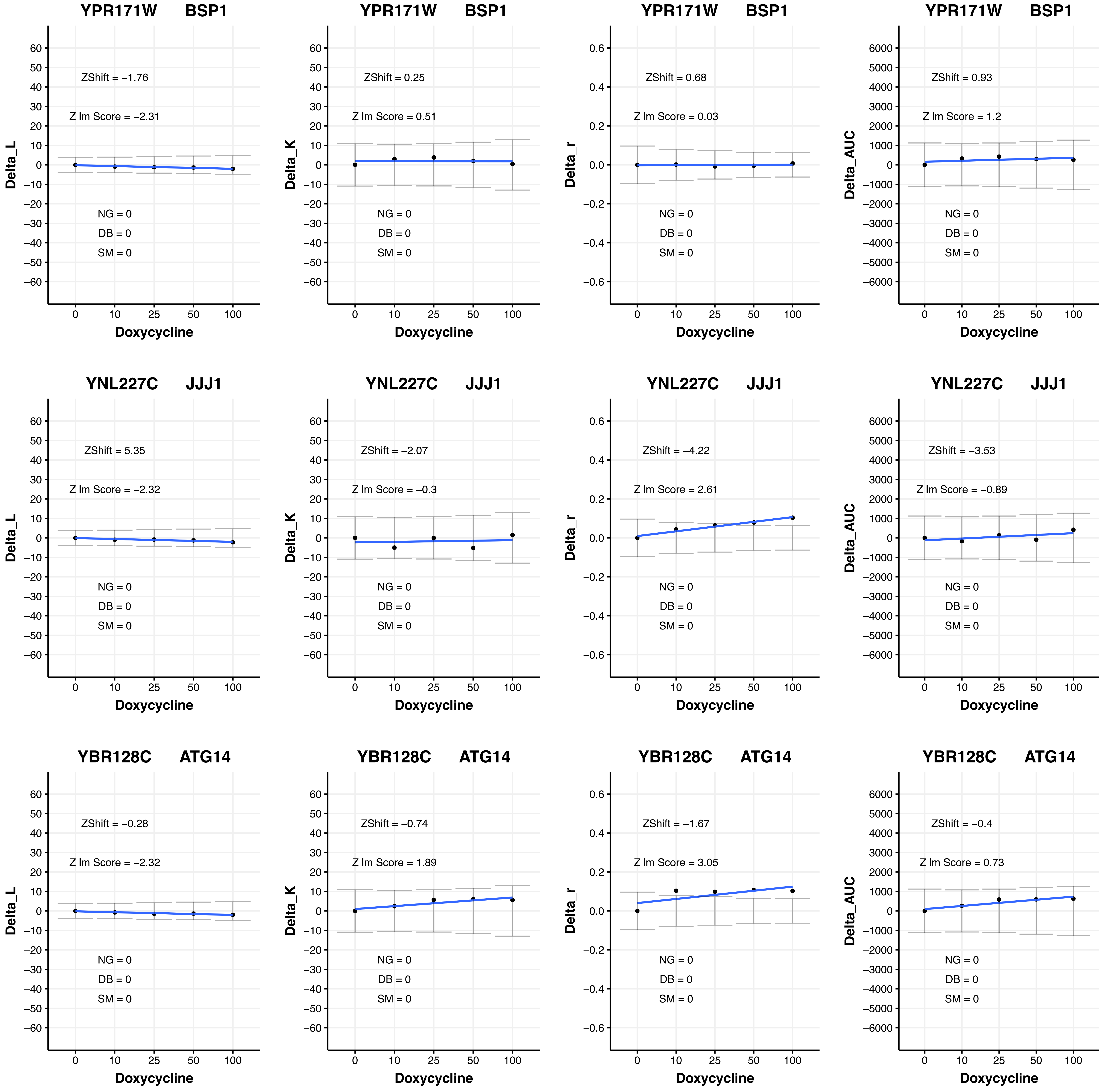
YPR171W BSP1
ZShift = −1.76
Z lm Score = −2.31
NG = 0
DB = 0
SM = 0
Delta_L
Doxycycline
YPR171W BSP1
ZShift = 0.25
Z lm Score = 0.51
NG = 0
DB = 0
SM = 0
Delta_K
Doxycycline
YPR171W BSP1
ZShift = 0.68
Z lm Score = 0.03
NG = 0
DB = 0
SM = 0
Delta_r
Doxycycline
YPR171W BSP1
ZShift = 0.93
Z lm Score = 1.2
NG = 0
DB = 0
SM = 0
Delta_AUC
Doxycycline
YNL227C JJJ1
ZShift = 5.35
Z lm Score = −2.32
NG = 0
DB = 0
SM = 0
Delta_L
Doxycycline
YNL227C JJJ1
ZShift = −2.07
Z lm Score = −0.3
NG = 0
DB = 0
SM = 0
Delta_K
Doxycycline
YNL227C JJJ1
ZShift = −4.22
Z lm Score = 2.61
NG = 0
DB = 0
SM = 0
Delta_r
Doxycycline
YNL227C JJJ1
ZShift = −3.53
Z lm Score = −0.89
NG = 0
DB = 0
SM = 0
Delta_AUC
Doxycycline
YBR128C ATG14
ZShift = −0.28
Z lm Score = −2.32
NG = 0
DB = 0
SM = 0
Delta_L
Doxycycline
YBR128C ATG14
ZShift = −0.74
Z lm Score = 1.89
NG = 0
DB = 0
SM = 0
Delta_K
Doxycycline
YBR128C ATG14
ZShift = −1.67
Z lm Score = 3.05
NG = 0
DB = 0
SM = 0
Delta_r
Doxycycline
YBR128C ATG14
ZShift = −0.4
Z lm Score = 0.73
NG = 0
DB = 0
SM = 0
Delta_AUC
Doxycycline

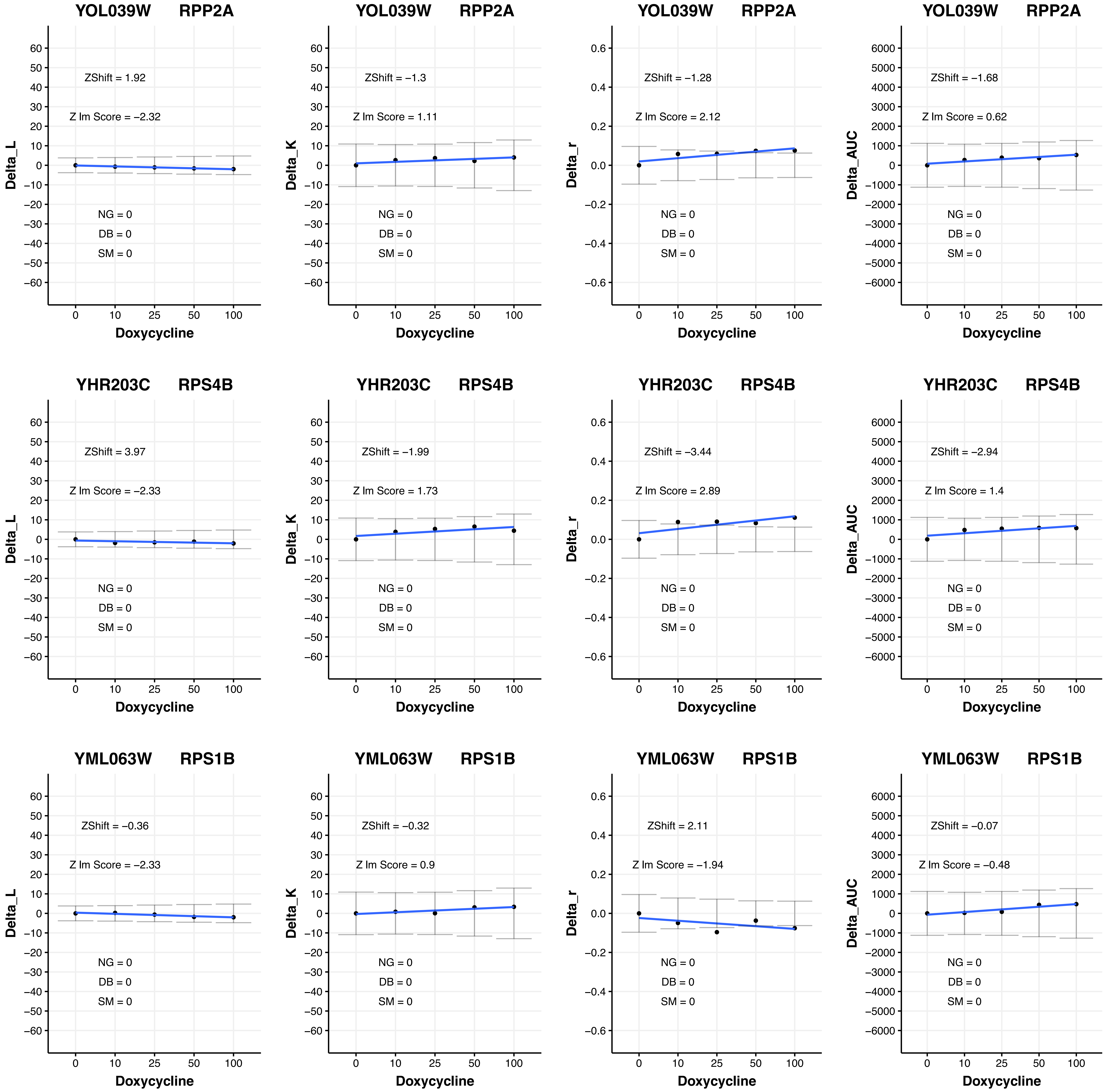
YOL039W RPP2A
ZShift = 1.92
Z lm Score = −2.32
NG = 0
DB = 0
SM = 0
Delta_L
Doxycycline
YOL039W RPP2A
ZShift = −1.3
Z lm Score = 1.11
NG = 0
DB = 0
SM = 0
Delta_K
Doxycycline
YOL039W RPP2A
ZShift = −1.28
Z lm Score = 2.12
NG = 0
DB = 0
SM = 0
Delta_r
Doxycycline
YOL039W RPP2A
ZShift = −1.68
Z lm Score = 0.62
NG = 0
DB = 0
SM = 0
Delta_AUC
Doxycycline
YHR203C RPS4B
ZShift = 3.97
Z lm Score = −2.33
NG = 0
DB = 0
SM = 0
Delta_L
Doxycycline
YHR203C RPS4B
ZShift = −1.99
Z lm Score = 1.73
NG = 0
DB = 0
SM = 0
Delta_K
Doxycycline
YHR203C RPS4B
ZShift = −3.44
Z lm Score = 2.89
NG = 0
DB = 0
SM = 0
Delta_r
Doxycycline
YHR203C RPS4B
ZShift = −2.94
Z lm Score = 1.4
NG = 0
DB = 0
SM = 0
Delta_AUC
Doxycycline
YML063W RPS1B
ZShift = −0.36
Z lm Score = −2.33
NG = 0
DB = 0
SM = 0
Delta_L
Doxycycline
YML063W RPS1B
ZShift = −0.32
Z lm Score = 0.9
NG = 0
DB = 0
SM = 0
Delta_K
Doxycycline
YML063W RPS1B
ZShift = 2.11
Z lm Score = −1.94
NG = 0
DB = 0
SM = 0
Delta_r
Doxycycline
YML063W RPS1B
ZShift = −0.07
Z lm Score = −0.48
NG = 0
DB = 0
SM = 0
Delta_AUC
Doxycycline

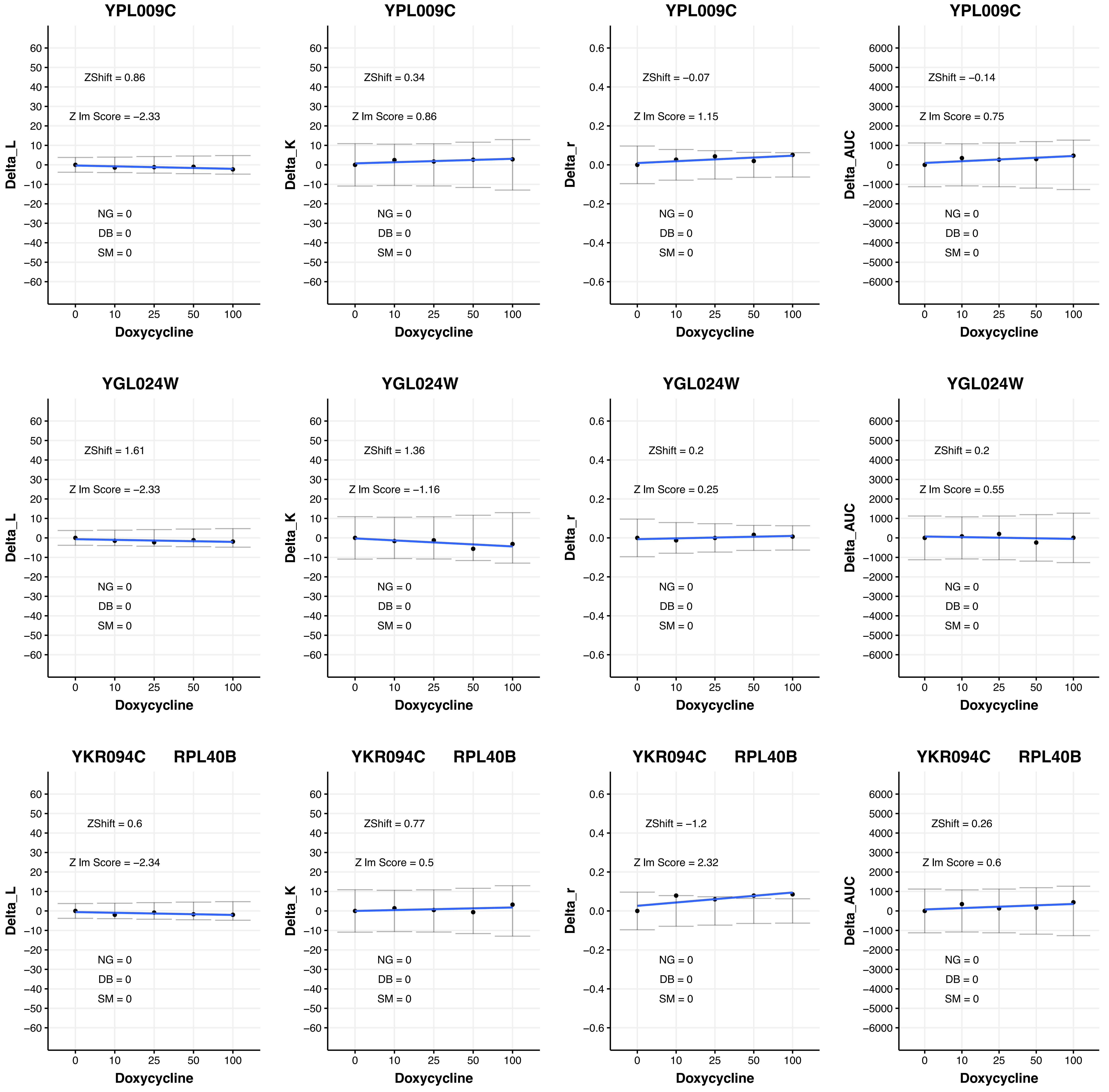
YPL009C
ZShift = 0.86
Z lm Score = −2.33
NG = 0
DB = 0
SM = 0
Delta_L
Doxycycline
YPL009C
ZShift = 0.34
Z lm Score = 0.86
NG = 0
DB = 0
SM = 0
Delta_K
Doxycycline
YPL009C
ZShift = −0.07
Z lm Score = 1.15
NG = 0
DB = 0
SM = 0
Delta_r
Doxycycline
YPL009C
ZShift = −0.14
Z lm Score = 0.75
NG = 0
DB = 0
SM = 0
Delta_AUC
Doxycycline
YGL024W
ZShift = 1.61
Z lm Score = −2.33
NG = 0
DB = 0
SM = 0
Delta_L
Doxycycline
YGL024W
ZShift = 1.36
Z lm Score = −1.16
NG = 0
DB = 0
SM = 0
Delta_K
Doxycycline
YGL024W
ZShift = 0.2
Z lm Score = 0.25
NG = 0
DB = 0
SM = 0
Delta_r
Doxycycline
YGL024W
ZShift = 0.2
Z lm Score = 0.55
NG = 0
DB = 0
SM = 0
Delta_AUC
Doxycycline
YKR094C RPL40B
ZShift = 0.6
Z lm Score = −2.34
NG = 0
DB = 0
SM = 0
Delta_L
Doxycycline
YKR094C RPL40B
ZShift = 0.77
Z lm Score = 0.5
NG = 0
DB = 0
SM = 0
Delta_K
Doxycycline
YKR094C RPL40B
ZShift = −1.2
Z lm Score = 2.32
NG = 0
DB = 0
SM = 0
Delta_r
Doxycycline
YKR094C RPL40B
ZShift = 0.26
Z lm Score = 0.6
NG = 0
DB = 0
SM = 0
Delta_AUC
Doxycycline

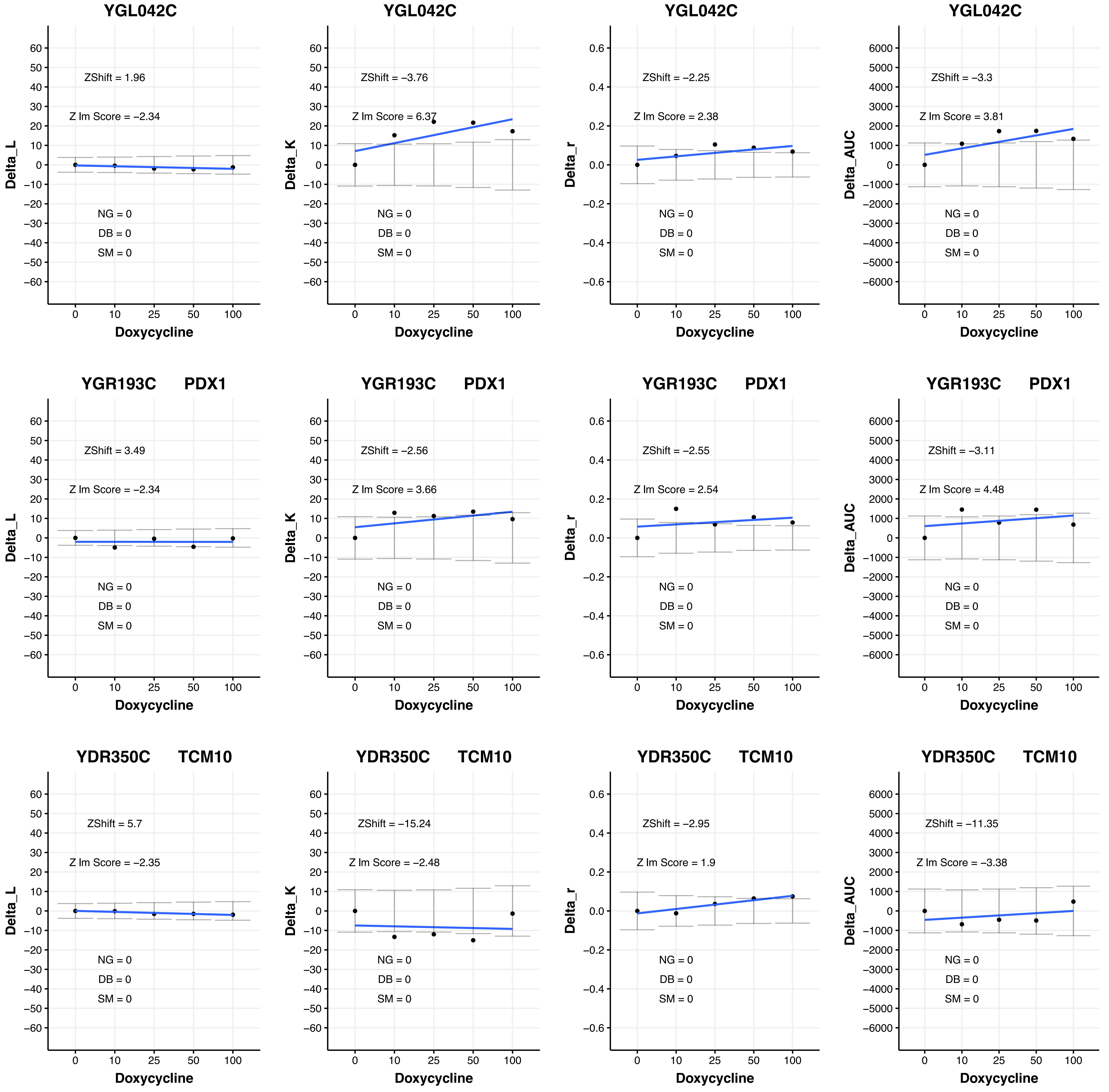
YGL042C
ZShift = 1.96
Z lm Score = −2.34
NG = 0
DB = 0
SM = 0
Delta_L
Doxycycline
YGL042C
ZShift = −3.76
Z lm Score = 6.37
NG = 0
DB = 0
SM = 0
Delta_K
Doxycycline
YGL042C
ZShift = −2.25
Z lm Score = 2.38
NG = 0
DB = 0
SM = 0
Delta_r
Doxycycline
YGL042C
ZShift = −3.3
Z lm Score = 3.81
NG = 0
DB = 0
SM = 0
Delta_AUC
Doxycycline
YGR193C PDX1
ZShift = 3.49
Z lm Score = −2.34
NG = 0
DB = 0
SM = 0
Delta_L
Doxycycline
YGR193C PDX1
ZShift = −2.56
Z lm Score = 3.66
NG = 0
DB = 0
SM = 0
Delta_K
Doxycycline
YGR193C PDX1
ZShift = −2.55
Z lm Score = 2.54
NG = 0
DB = 0
SM = 0
Delta_r
Doxycycline
YGR193C PDX1
ZShift = −3.11
Z lm Score = 4.48
NG = 0
DB = 0
SM = 0
Delta_AUC
Doxycycline
YDR350C TCM10
ZShift = 5.7
Z lm Score = −2.35
NG = 0
DB = 0
SM = 0
Delta_L
Doxycycline
YDR350C TCM10
ZShift = −15.24
Z lm Score = −2.48
NG = 0
DB = 0
SM = 0
Delta_K
Doxycycline
YDR350C TCM10
ZShift = −2.95
Z lm Score = 1.9
NG = 0
DB = 0
SM = 0
Delta_r
Doxycycline
YDR350C TCM10
ZShift = −11.35
Z lm Score = −3.38
NG = 0
DB = 0
SM = 0
Delta_AUC
Doxycycline

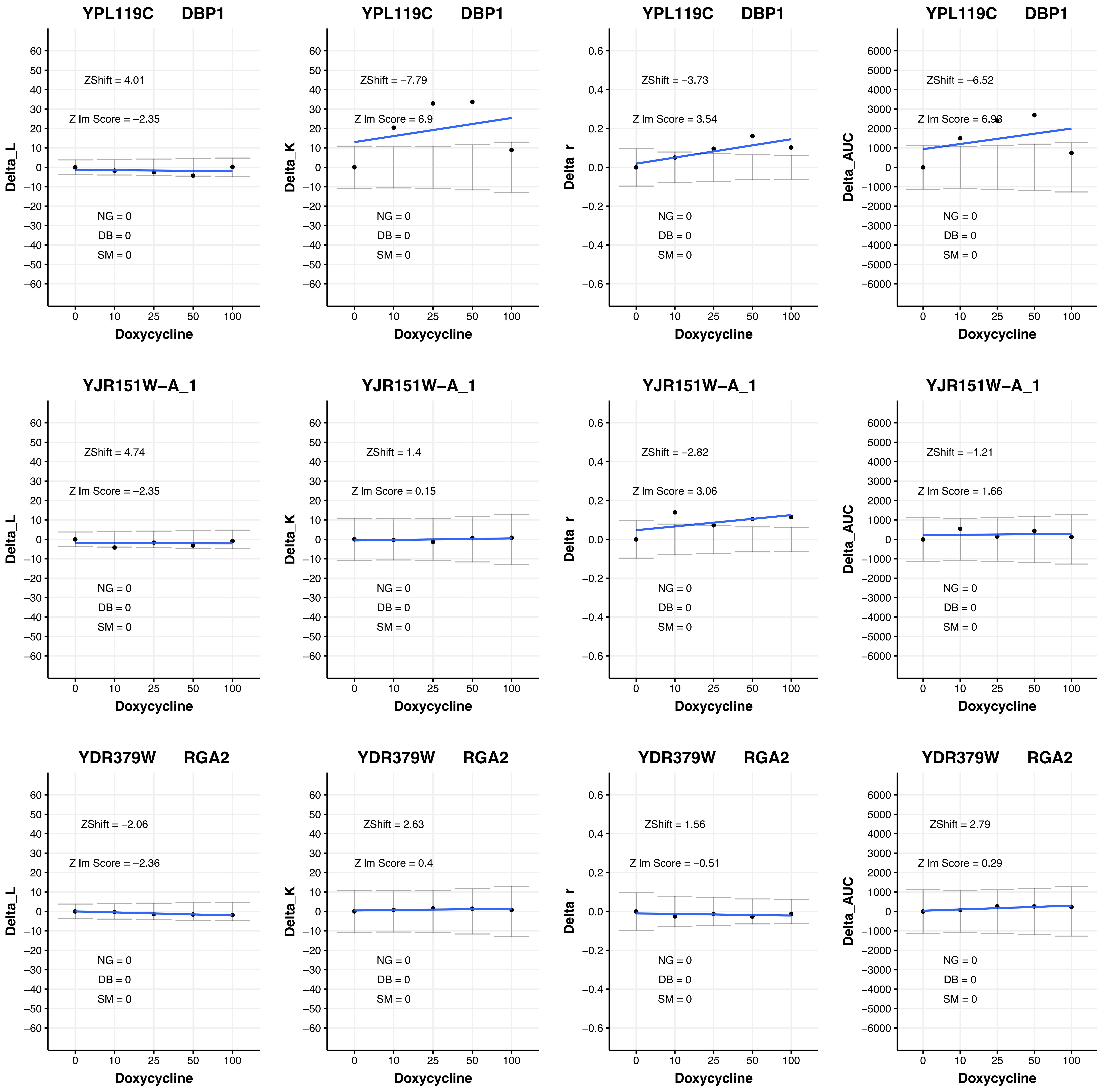

YPL119C DBP1
ZShift = 4.01
Z lm Score = −2.35
NG = 0
DB = 0
SM = 0
Delta_L
Doxycycline
YPL119C DBP1
ZShift = −7.79
Z lm Score = 6.9
NG = 0
DB = 0
SM = 0
Delta_K
Doxycycline
YPL119C DBP1
ZShift = −3.73
Z lm Score = 3.54
NG = 0
DB = 0
SM = 0
Delta_r
Doxycycline
YPL119C DBP1
ZShift = −6.52
Z lm Score = 6.93
NG = 0
DB = 0
SM = 0
Delta_AUC
Doxycycline
YJR151W−A_1
ZShift = 4.74
Z lm Score = −2.35
NG = 0
DB = 0
SM = 0
Delta_L
Doxycycline
YJR151W−A_1
ZShift = 1.4
Z lm Score = 0.15
NG = 0
DB = 0
SM = 0
Delta_K
Doxycycline
YJR151W−A_1
ZShift = −2.82
Z lm Score = 3.06
NG = 0
DB = 0
SM = 0
Delta_r
Doxycycline
YJR151W−A_1
ZShift = −1.21
Z lm Score = 1.66
NG = 0
DB = 0
SM = 0
Delta_AUC
Doxycycline
YDR379W RGA2
ZShift = −2.06
Z lm Score = −2.36
NG = 0
DB = 0
SM = 0
Delta_L
Doxycycline
YDR379W RGA2
ZShift = 2.63
Z lm Score = 0.4
NG = 0
DB = 0
SM = 0
Delta_K
Doxycycline
YDR379W RGA2
ZShift = 1.56
Z lm Score = −0.51
NG = 0
DB = 0
SM = 0
Delta_r
Doxycycline
YDR379W RGA2
ZShift = 2.79
Z lm Score = 0.29
NG = 0
DB = 0
SM = 0
Delta_AUC
Doxycycline

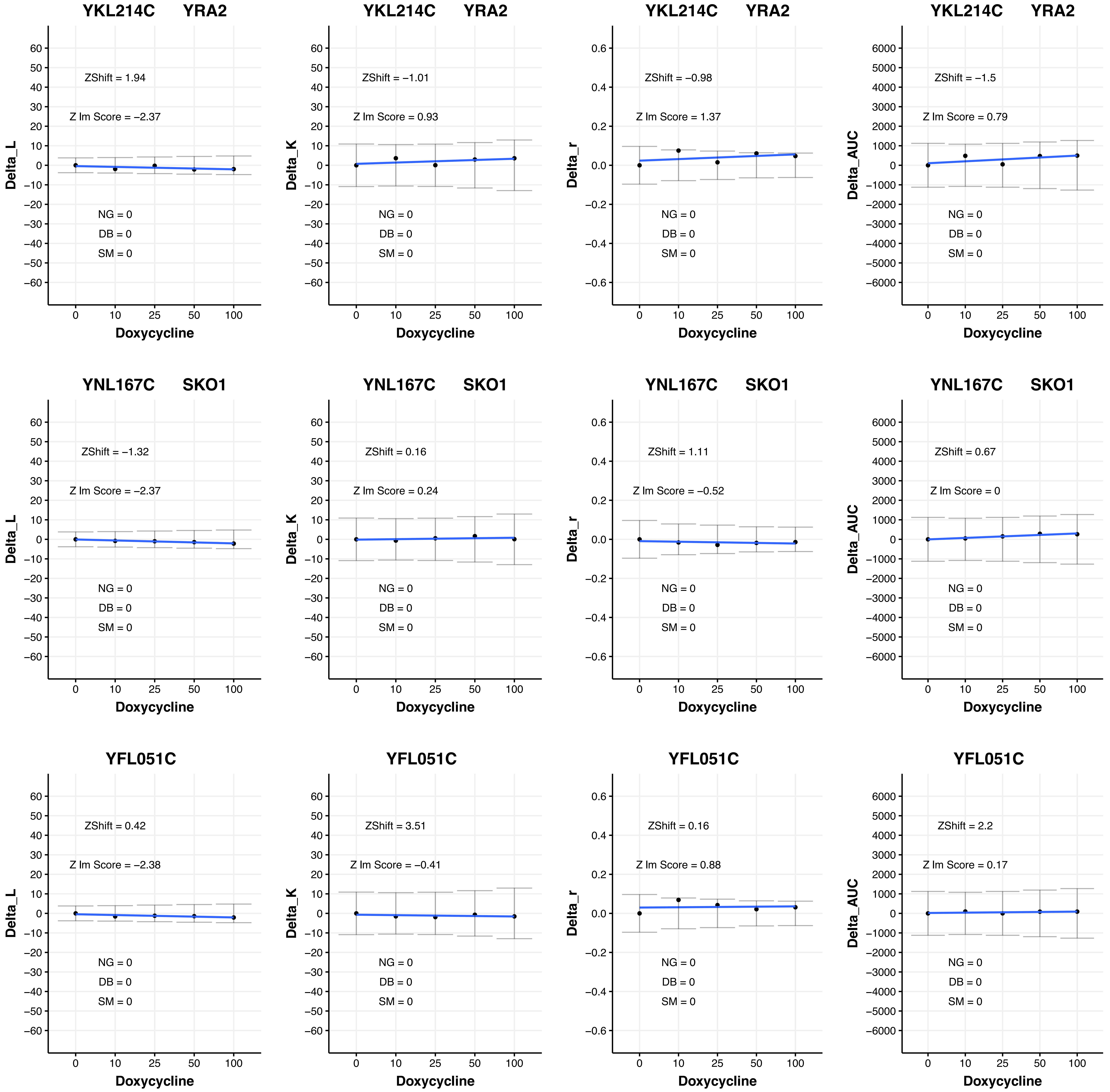

YKL214C YRA2
ZShift = 1.94
Z lm Score = −2.37
NG = 0
DB = 0
SM = 0
Delta_L
Doxycycline
YKL214C YRA2
ZShift = −1.01
Z lm Score = 0.93
NG = 0
DB = 0
SM = 0
Delta_K
Doxycycline
YKL214C YRA2
ZShift = −0.98
Z lm Score = 1.37
NG = 0
DB = 0
SM = 0
Delta_r
Doxycycline
YKL214C YRA2
ZShift = −1.5
Z lm Score = 0.79
NG = 0
DB = 0
SM = 0
Delta_AUC
Doxycycline
YNL167C SKO1
ZShift = −1.32
Z lm Score = −2.37
NG = 0
DB = 0
SM = 0
Delta_L
Doxycycline
YNL167C SKO1
ZShift = 0.16
Z lm Score = 0.24
NG = 0
DB = 0
SM = 0
Delta_K
Doxycycline
YNL167C SKO1
ZShift = 1.11
Z lm Score = −0.52
NG = 0
DB = 0
SM = 0
Delta_r
Doxycycline
YNL167C SKO1
ZShift = 0.67
Z lm Score = 0
NG = 0
DB = 0
SM = 0
Delta_AUC
Doxycycline
YFL051C
ZShift = 0.42
Z lm Score = −2.38
NG = 0
DB = 0
SM = 0
Delta_L
Doxycycline
YFL051C
ZShift = 3.51
Z lm Score = −0.41
NG = 0
DB = 0
SM = 0
Delta_K
Doxycycline
YFL051C
ZShift = 0.16
Z lm Score = 0.88
NG = 0
DB = 0
SM = 0
Delta_r
Doxycycline
YFL051C
ZShift = 2.2
Z lm Score = 0.17
NG = 0
DB = 0
SM = 0
Delta_AUC
Doxycycline

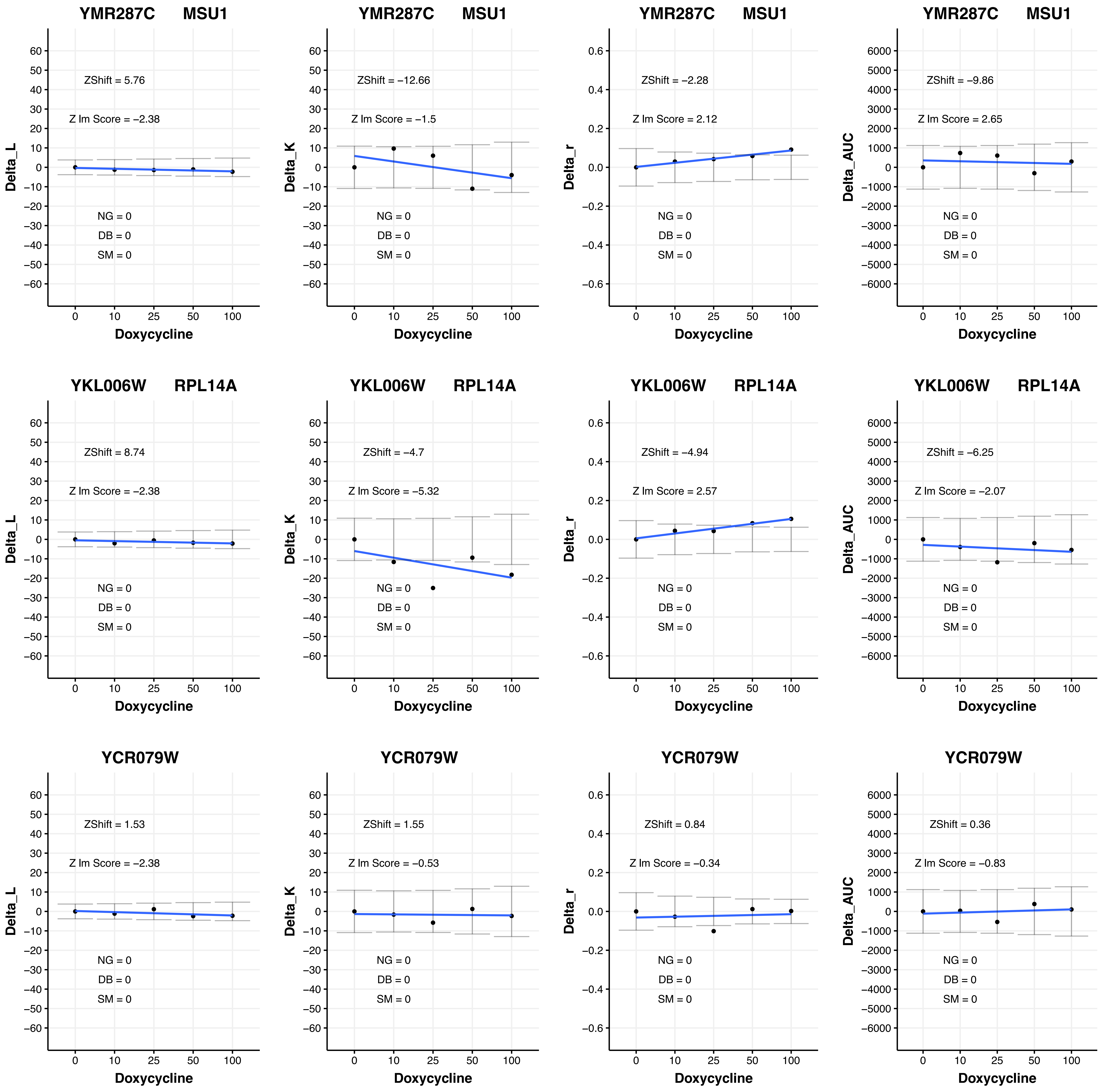
YMR287C MSU1
ZShift = 5.76
Z lm Score = −2.38
NG = 0
DB = 0
SM = 0
Delta_L
Doxycycline
YMR287C MSU1
ZShift = −12.66
Z lm Score = −1.5
NG = 0
DB = 0
SM = 0
Delta_K
Doxycycline
YMR287C MSU1
ZShift = −2.28
Z lm Score = 2.12
NG = 0
DB = 0
SM = 0
Delta_r
Doxycycline
YMR287C MSU1
ZShift = −9.86
Z lm Score = 2.65
NG = 0
DB = 0
SM = 0
Delta_AUC
Doxycycline
YKL006W RPL14A
ZShift = 8.74
Z lm Score = −2.38
NG = 0
DB = 0
SM = 0
Delta_L
Doxycycline
YKL006W RPL14A
ZShift = −4.7
Z lm Score = −5.32
NG = 0
DB = 0
SM = 0
Delta_K
Doxycycline
YKL006W RPL14A
ZShift = −4.94
Z lm Score = 2.57
NG = 0
DB = 0
SM = 0
Delta_r
Doxycycline
YKL006W RPL14A
ZShift = −6.25
Z lm Score = −2.07
NG = 0
DB = 0
SM = 0
Delta_AUC
Doxycycline
YCR079W
ZShift = 1.53
Z lm Score = −2.38
NG = 0
DB = 0
SM = 0
Delta_L
Doxycycline
YCR079W
ZShift = 1.55
Z lm Score = −0.53
NG = 0
DB = 0
SM = 0
Delta_K
Doxycycline
YCR079W
ZShift = 0.84
Z lm Score = −0.34
NG = 0
DB = 0
SM = 0
Delta_r
Doxycycline
YCR079W
ZShift = 0.36
Z lm Score = −0.83
NG = 0
DB = 0
SM = 0
Delta_AUC
Doxycycline

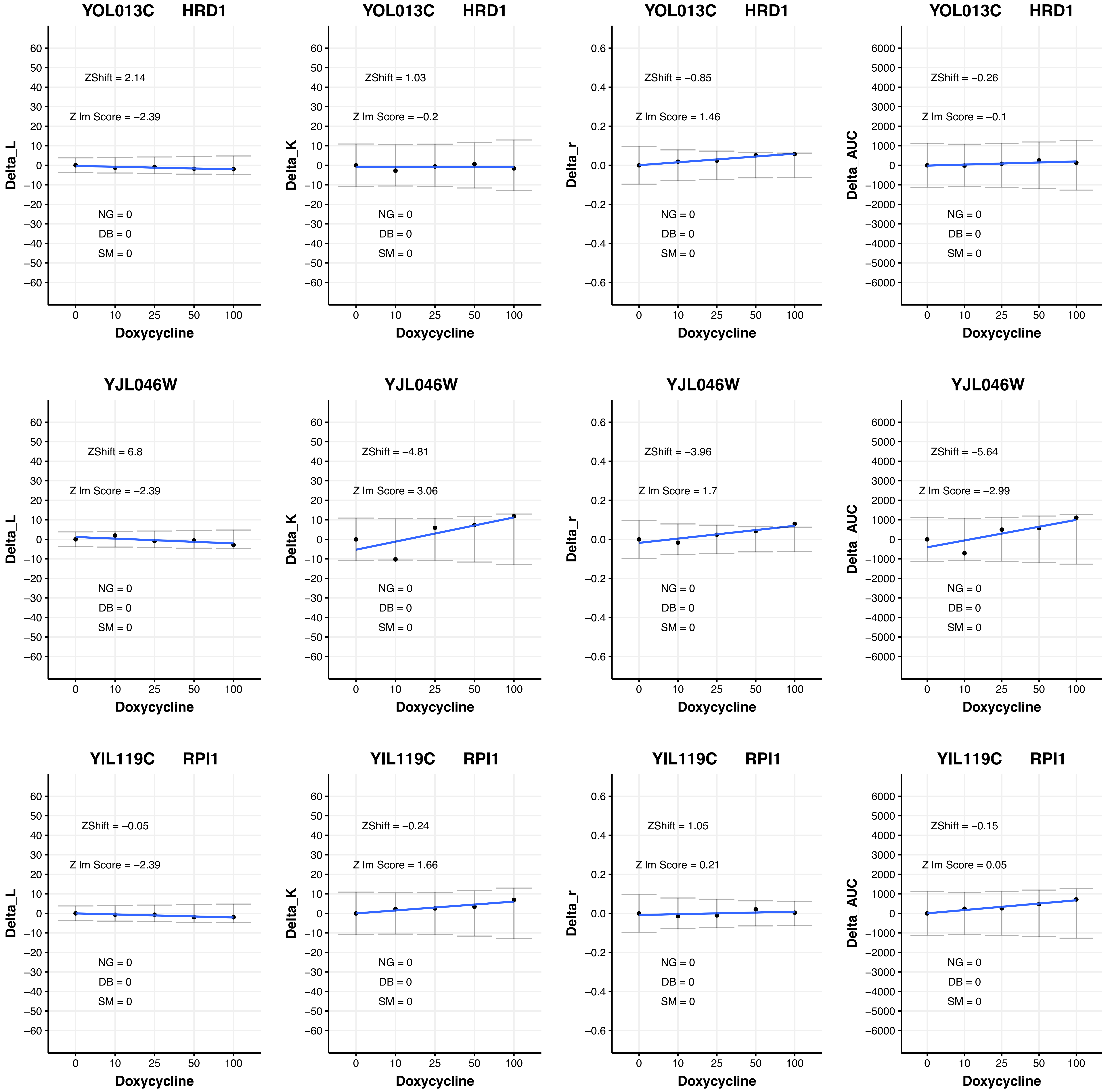

YOL013C HRD1
ZShift = 2.14
Z lm Score = −2.39
NG = 0
DB = 0
SM = 0
Delta_L
Doxycycline
YOL013C HRD1
ZShift = 1.03
Z lm Score = −0.2
NG = 0
DB = 0
SM = 0
Delta_K
Doxycycline
YOL013C HRD1
ZShift = −0.85
Z lm Score = 1.46
NG = 0
DB = 0
SM = 0
Delta_r
Doxycycline
YOL013C HRD1
ZShift = −0.26
Z lm Score = −0.1
NG = 0
DB = 0
SM = 0
Delta_AUC
Doxycycline
YJL046W
ZShift = 6.8
Z lm Score = −2.39
NG = 0
DB = 0
SM = 0
Delta_L
Doxycycline
YJL046W
ZShift = −4.81
Z lm Score = 3.06
NG = 0
DB = 0
SM = 0
Delta_K
Doxycycline
YJL046W
ZShift = −3.96
Z lm Score = 1.7
NG = 0
DB = 0
SM = 0
Delta_r
Doxycycline
YJL046W
ZShift = −5.64
Z lm Score = −2.99
NG = 0
DB = 0
SM = 0
Delta_AUC
Doxycycline
YIL119C RPI1
ZShift = −0.05
Z lm Score = −2.39
NG = 0
DB = 0
SM = 0
Delta_L
Doxycycline
YIL119C RPI1
ZShift = −0.24
Z lm Score = 1.66
NG = 0
DB = 0
SM = 0
Delta_K
Doxycycline
YIL119C RPI1
ZShift = 1.05
Z lm Score = 0.21
NG = 0
DB = 0
SM = 0
Delta_r
Doxycycline
YIL119C RPI1
ZShift = −0.15
Z lm Score = 0.05
NG = 0
DB = 0
SM = 0
Delta_AUC
Doxycycline

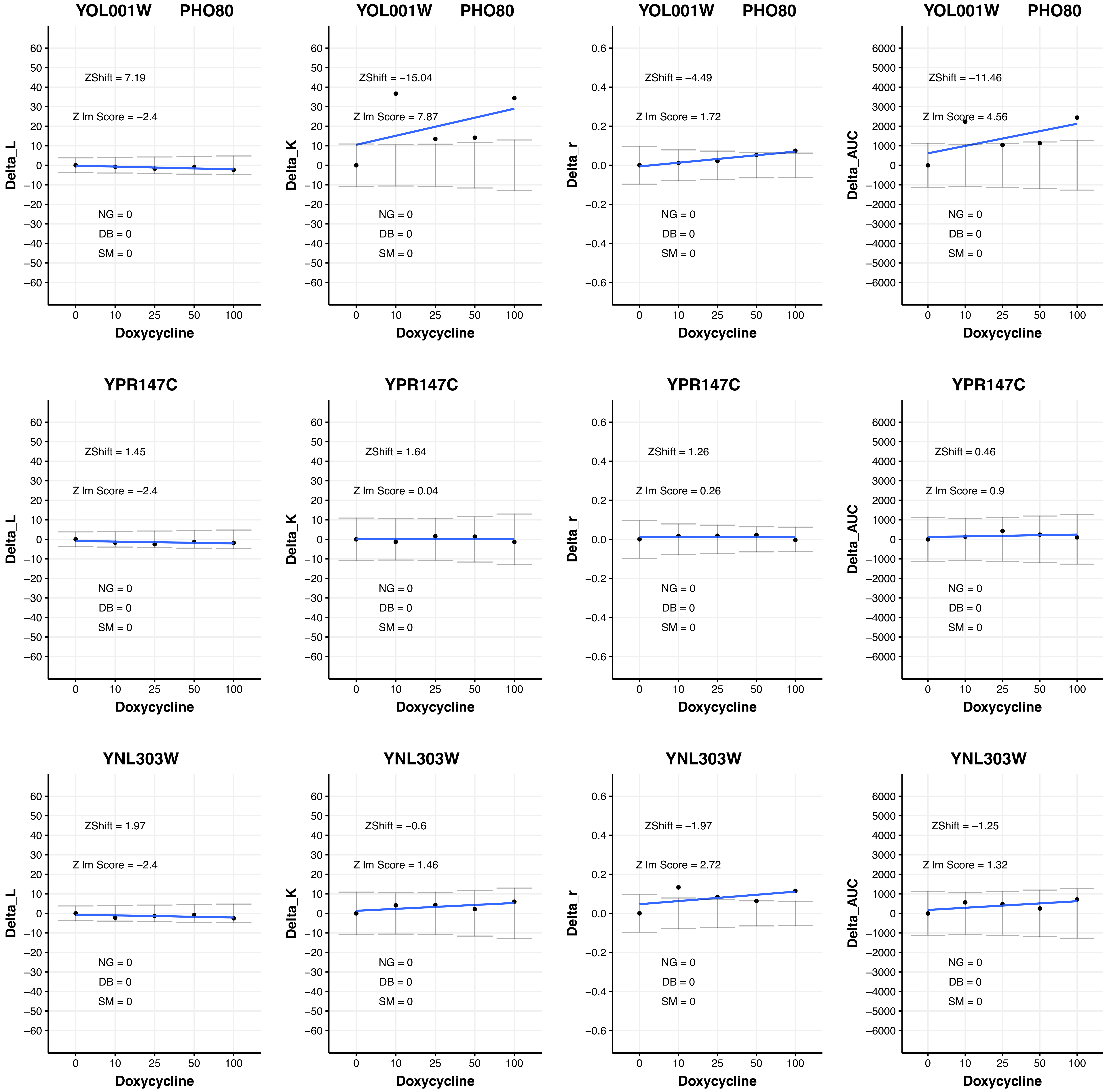
YOL001W PHO80
ZShift = 7.19
Z lm Score = −2.4
NG = 0
DB = 0
SM = 0
Delta_L
Doxycycline
YOL001W PHO80
ZShift = −15.04
Z lm Score = 7.87
NG = 0
DB = 0
SM = 0
Delta_K
Doxycycline
YOL001W PHO80
ZShift = −4.49
Z lm Score = 1.72
NG = 0
DB = 0
SM = 0
Delta_r
Doxycycline
YOL001W PHO80
ZShift = −11.46
Z lm Score = 4.56
NG = 0
DB = 0
SM = 0
Delta_AUC
Doxycycline
YPR147C
ZShift = 1.45
Z lm Score = −2.4
NG = 0
DB = 0
SM = 0
Delta_L
Doxycycline
YPR147C
ZShift = 1.64
Z lm Score = 0.04
NG = 0
DB = 0
SM = 0
Delta_K
Doxycycline
YPR147C
ZShift = 1.26
Z lm Score = 0.26
NG = 0
DB = 0
SM = 0
Delta_r
Doxycycline
YPR147C
ZShift = 0.46
Z lm Score = 0.9
NG = 0
DB = 0
SM = 0
Delta_AUC
Doxycycline
YNL303W
ZShift = 1.97
Z lm Score = −2.4
NG = 0
DB = 0
SM = 0
Delta_L
Doxycycline
YNL303W
ZShift = −0.6
Z lm Score = 1.46
NG = 0
DB = 0
SM = 0
Delta_K
Doxycycline
YNL303W
ZShift = −1.97
Z lm Score = 2.72
NG = 0
DB = 0
SM = 0
Delta_r
Doxycycline
YNL303W
ZShift = −1.25
Z lm Score = 1.32
NG = 0
DB = 0
SM = 0
Delta_AUC
Doxycycline

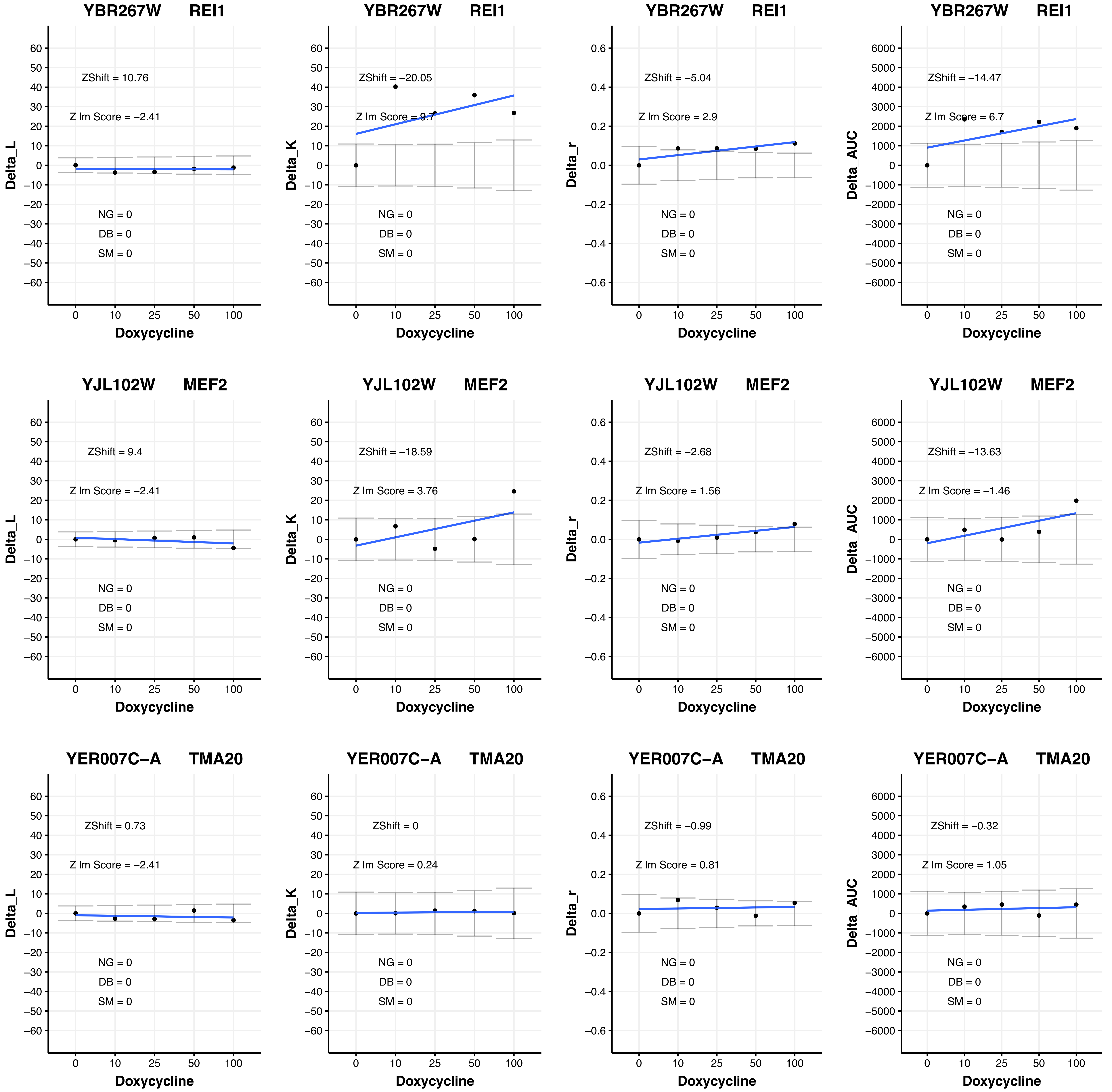

YBR267W REI1
ZShift = 10.76
Z lm Score = −2.41
NG = 0
DB = 0
SM = 0
Delta_L
Doxycycline
YBR267W REI1
ZShift = −20.05
Z lm Score = 9.7
NG = 0
DB = 0
SM = 0
Delta_K
Doxycycline
YBR267W REI1
ZShift = −5.04
Z lm Score = 2.9
NG = 0
DB = 0
SM = 0
Delta_r
Doxycycline
YBR267W REI1
ZShift = −14.47
Z lm Score = 6.7
NG = 0
DB = 0
SM = 0
Delta_AUC
Doxycycline
YJL102W MEF2
ZShift = 9.4
Z lm Score = −2.41
NG = 0
DB = 0
SM = 0
Delta_L
Doxycycline
YJL102W MEF2
ZShift = −18.59
Z lm Score = 3.76
NG = 0
DB = 0
SM = 0
Delta_K
Doxycycline
YJL102W MEF2
ZShift = −2.68
Z lm Score = 1.56
NG = 0
DB = 0
SM = 0
Delta_r
Doxycycline
YJL102W MEF2
ZShift = −13.63
Z lm Score = −1.46
NG = 0
DB = 0
SM = 0
Delta_AUC
Doxycycline
YER007C−A TMA20
ZShift = 0.73
Z lm Score = −2.41
NG = 0
DB = 0
SM = 0
Delta_L
Doxycycline
YER007C−A TMA20
ZShift = 0
Z lm Score = 0.24
NG = 0
DB = 0
SM = 0
Delta_K
Doxycycline
YER007C−A TMA20
ZShift = −0.99
Z lm Score = 0.81
NG = 0
DB = 0
SM = 0
Delta_r
Doxycycline
YER007C−A TMA20
ZShift = −0.32
Z lm Score = 1.05
NG = 0
DB = 0
SM = 0
Delta_AUC
Doxycycline

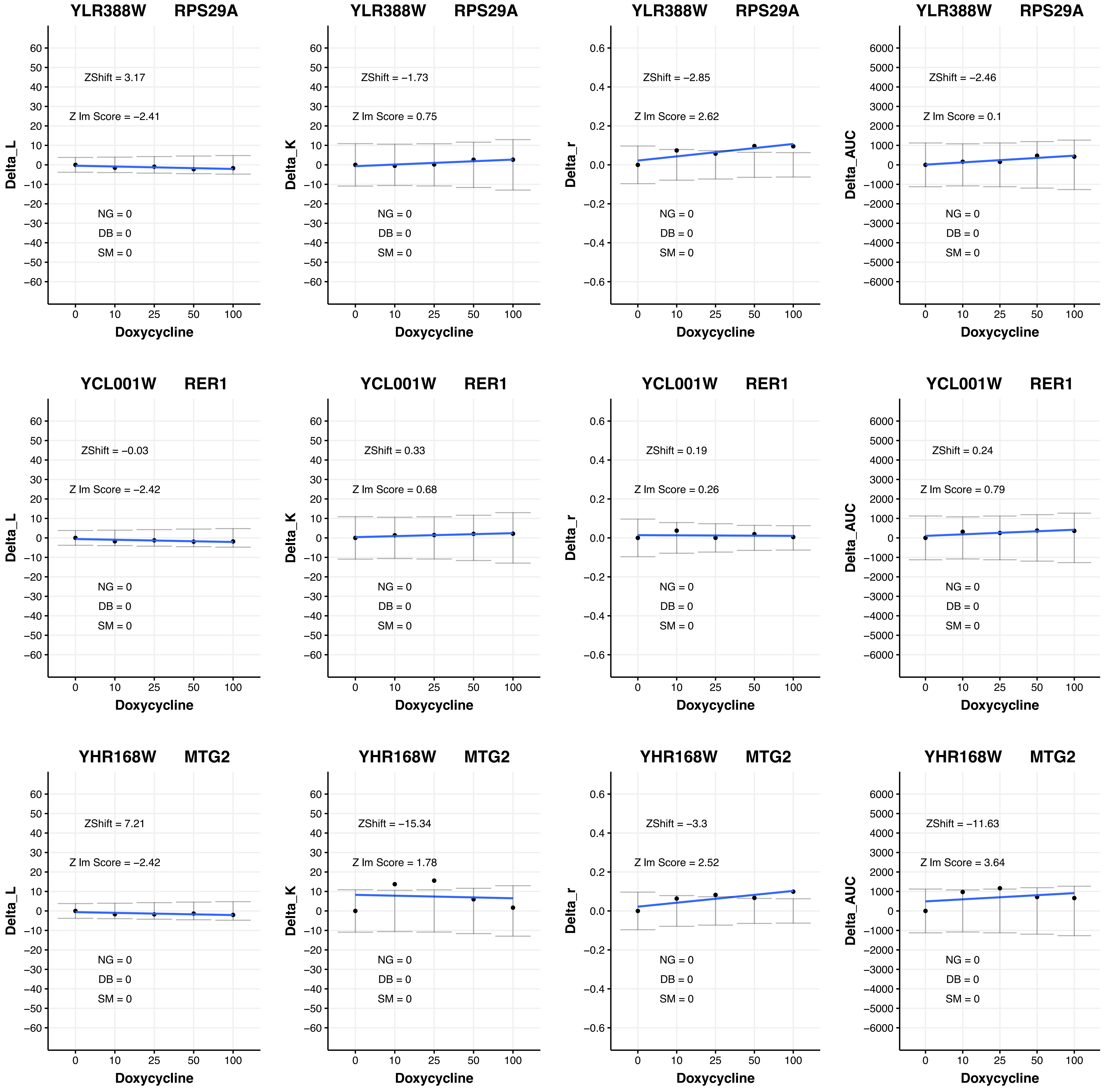

YLR388W RPS29A
ZShift = 3.17
Z lm Score = −2.41
NG = 0
DB = 0
SM = 0
Delta_L
Doxycycline
YLR388W RPS29A
ZShift = −1.73
Z lm Score = 0.75
NG = 0
DB = 0
SM = 0
Delta_K
Doxycycline
YLR388W RPS29A
ZShift = −2.85
Z lm Score = 2.62
NG = 0
DB = 0
SM = 0
Delta_r
Doxycycline
YLR388W RPS29A
ZShift = −2.46
Z lm Score = 0.1
NG = 0
DB = 0
SM = 0
Delta_AUC
Doxycycline
YCL001W RER1
ZShift = −0.03
Z lm Score = −2.42
NG = 0
DB = 0
SM = 0
Delta_L
Doxycycline
YCL001W RER1
ZShift = 0.33
Z lm Score = 0.68
NG = 0
DB = 0
SM = 0
Delta_K
Doxycycline
YCL001W RER1
ZShift = 0.19
Z lm Score = 0.26
NG = 0
DB = 0
SM = 0
Delta_r
Doxycycline
YCL001W RER1
ZShift = 0.24
Z lm Score = 0.79
NG = 0
DB = 0
SM = 0
Delta_AUC
Doxycycline
YHR168W MTG2
ZShift = 7.21
Z lm Score = −2.42
NG = 0
DB = 0
SM = 0
Delta_L
Doxycycline
YHR168W MTG2
ZShift = −15.34
Z lm Score = 1.78
NG = 0
DB = 0
SM = 0
Delta_K
Doxycycline
YHR168W MTG2
ZShift = −3.3
Z lm Score = 2.52
NG = 0
DB = 0
SM = 0
Delta_r
Doxycycline
YHR168W MTG2
ZShift = −11.63
Z lm Score = 3.64
NG = 0
DB = 0
SM = 0
Delta_AUC
Doxycycline

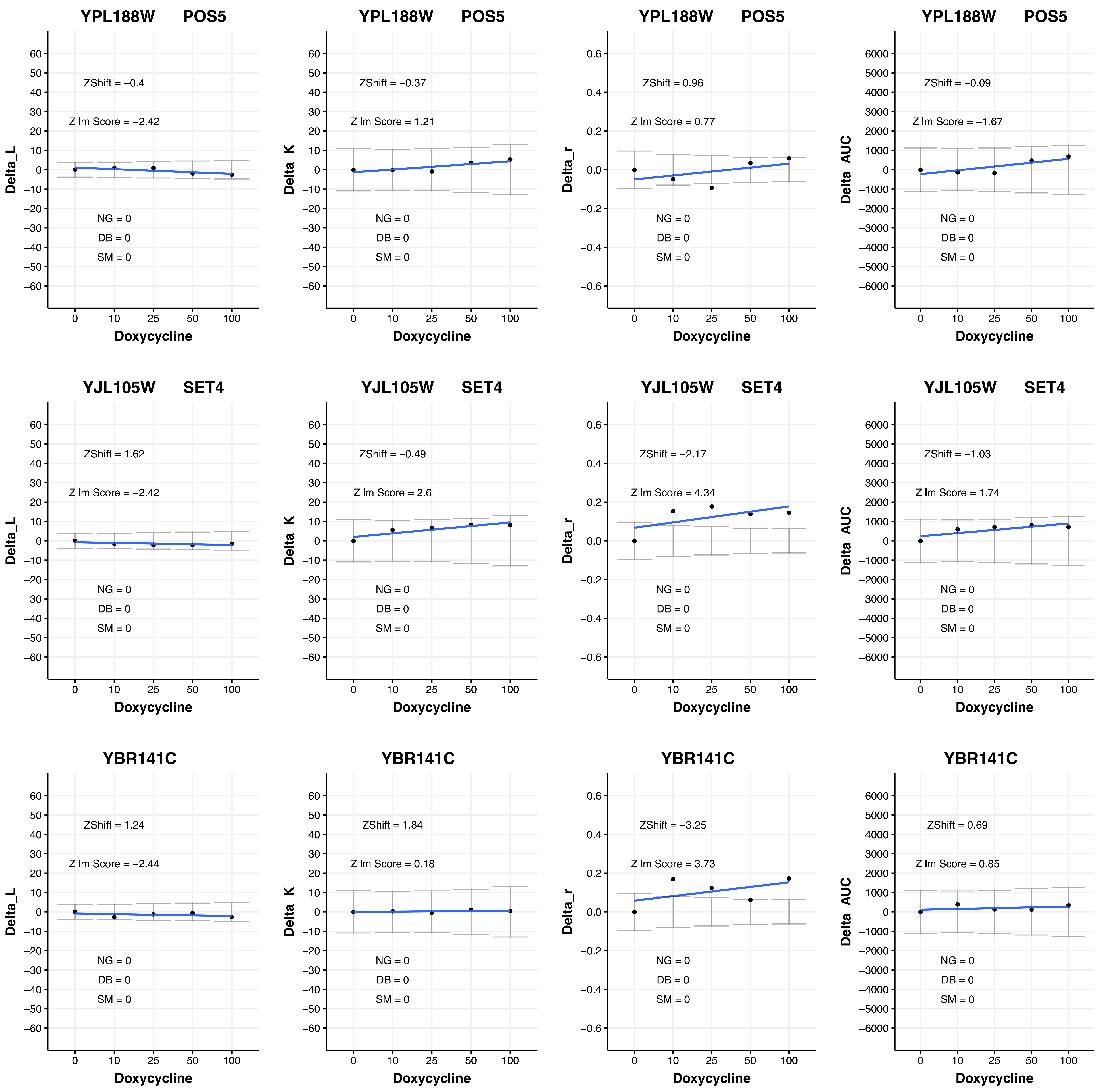
YPL188W POS5
ZShift = −0.4
Z lm Score = −2.42
NG = 0
DB = 0
SM = 0
Delta_L
Doxycycline
YPL188W POS5
ZShift = −0.37
Z lm Score = 1.21
NG = 0
DB = 0
SM = 0
Delta_K
Doxycycline
YPL188W POS5
ZShift = 0.96
Z lm Score = 0.77
NG = 0
DB = 0
SM = 0
Delta_r
Doxycycline
YPL188W POS5
ZShift = −0.09
Z lm Score = −1.67
NG = 0
DB = 0
SM = 0
Delta_AUC
Doxycycline
YJL105W SET4
ZShift = 1.62
Z lm Score = −2.42
NG = 0
DB = 0
SM = 0
Delta_L
Doxycycline
YJL105W SET4
ZShift = −0.49
Z lm Score = 2.6
NG = 0
DB = 0
SM = 0
Delta_K
Doxycycline
YJL105W SET4
ZShift = −2.17
Z lm Score = 4.34
NG = 0
DB = 0
SM = 0
Delta_r
Doxycycline
YJL105W SET4
ZShift = −1.03
Z lm Score = 1.74
NG = 0
DB = 0
SM = 0
Delta_AUC
Doxycycline
YBR141C
ZShift = 1.24
Z lm Score = −2.44
NG = 0
DB = 0
SM = 0
Delta_L
Doxycycline
YBR141C
ZShift = 1.84
Z lm Score = 0.18
NG = 0
DB = 0
SM = 0
Delta_K
Doxycycline
YBR141C
ZShift = −3.25
Z lm Score = 3.73
NG = 0
DB = 0
SM = 0
Delta_r
Doxycycline
YBR141C
ZShift = 0.69
Z lm Score = 0.85
NG = 0
DB = 0
SM = 0
Delta_AUC
Doxycycline

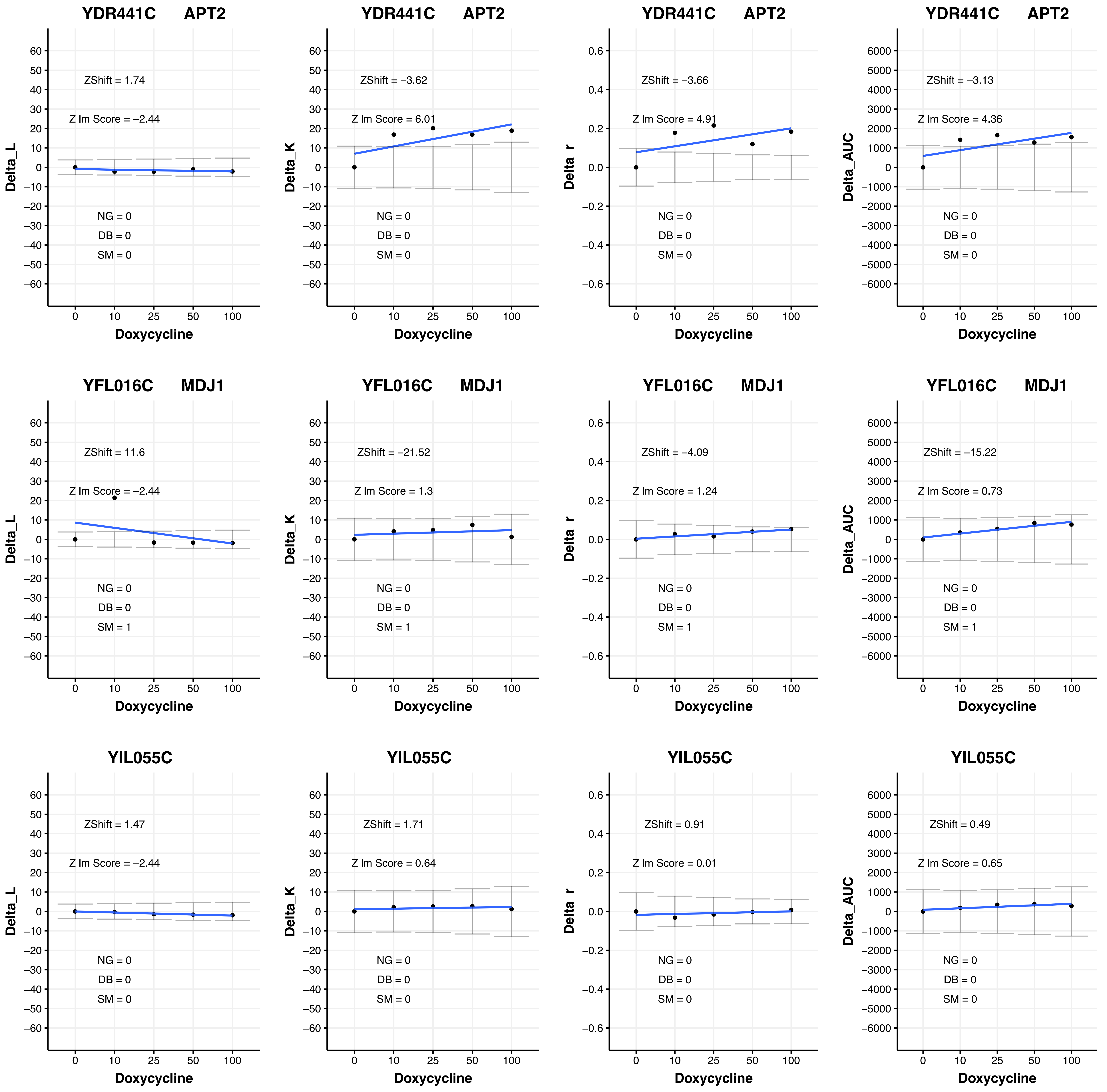

YDR441C APT2
ZShift = 1.74
Z lm Score = −2.44
NG = 0
DB = 0
SM = 0
Delta_L
Doxycycline
YDR441C APT2
ZShift = −3.62
Z lm Score = 6.01
NG = 0
DB = 0
SM = 0
Delta_K
Doxycycline
YDR441C APT2
ZShift = −3.66
Z lm Score = 4.91
NG = 0
DB = 0
SM = 0
Delta_r
Doxycycline
YDR441C APT2
ZShift = −3.13
Z lm Score = 4.36
NG = 0
DB = 0
SM = 0
Delta_AUC
Doxycycline
YFL016C MDJ1
ZShift = 11.6
Z lm Score = −2.44
NG = 0
DB = 0
SM = 1
Delta_L
Doxycycline
YFL016C MDJ1
ZShift = −21.52
Z lm Score = 1.3
NG = 0
DB = 0
SM = 1
Delta_K
Doxycycline
YFL016C MDJ1
ZShift = −4.09
Z lm Score = 1.24
NG = 0
DB = 0
SM = 1
Delta_r
Doxycycline
YFL016C MDJ1
ZShift = −15.22
Z lm Score = 0.73
NG = 0
DB = 0
SM = 1
Delta_AUC
Doxycycline
YIL055C
ZShift = 1.47
Z lm Score = −2.44
NG = 0
DB = 0
SM = 0
Delta_L
Doxycycline
YIL055C
ZShift = 1.71
Z lm Score = 0.64
NG = 0
DB = 0
SM = 0
Delta_K
Doxycycline
YIL055C
ZShift = 0.91
Z lm Score = 0.01
NG = 0
DB = 0
SM = 0
Delta_r
Doxycycline
YIL055C
ZShift = 0.49
Z lm Score = 0.65
NG = 0
DB = 0
SM = 0
Delta_AUC
Doxycycline

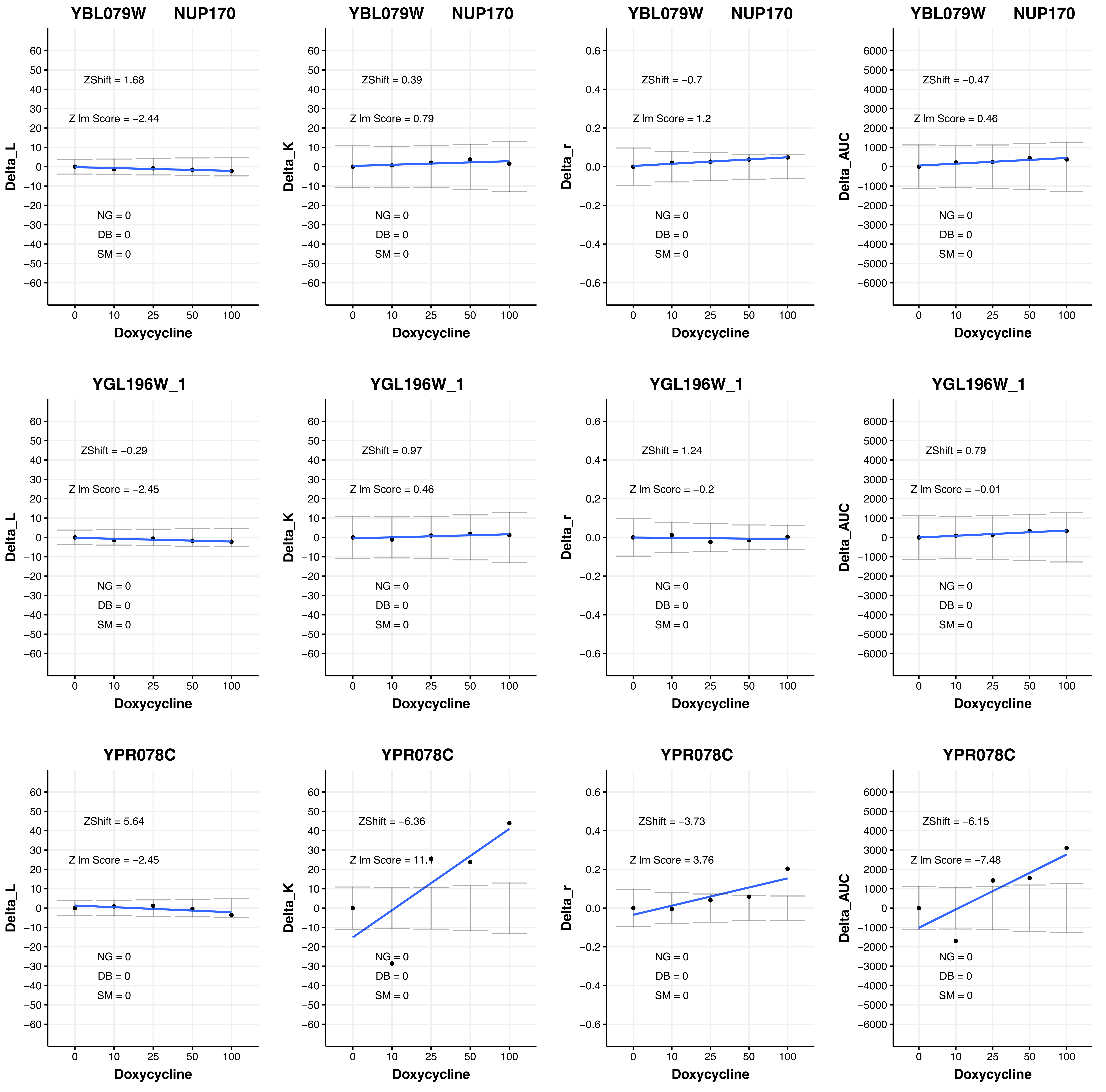

YBL079W NUP170
ZShift = 1.68
Z lm Score = −2.44
NG = 0
DB = 0
SM = 0
Delta_L
Doxycycline
YBL079W NUP170
ZShift = 0.39
Z lm Score = 0.79
NG = 0
DB = 0
SM = 0
Delta_K
Doxycycline
YBL079W NUP170
ZShift = −0.7
Z lm Score = 1.2
NG = 0
DB = 0
SM = 0
Delta_r
Doxycycline
YBL079W NUP170
ZShift = −0.47
Z lm Score = 0.46
NG = 0
DB = 0
SM = 0
Delta_AUC
Doxycycline
YGL196W_1
ZShift = −0.29
Z lm Score = −2.45
NG = 0
DB = 0
SM = 0
Delta_L
Doxycycline
YGL196W_1
ZShift = 0.97
Z lm Score = 0.46
NG = 0
DB = 0
SM = 0
Delta_K
Doxycycline
YGL196W_1
ZShift = 1.24
Z lm Score = −0.2
NG = 0
DB = 0
SM = 0
Delta_r
Doxycycline
YGL196W_1
ZShift = 0.79
Z lm Score = −0.01
NG = 0
DB = 0
SM = 0
Delta_AUC
Doxycycline
YPR078C
ZShift = 5.64
Z lm Score = −2.45
NG = 0
DB = 0
SM = 0
Delta_L
Doxycycline
YPR078C
ZShift = −6.36
Z lm Score = 11.
NG = 0
DB = 0
SM = 0
Delta_K
Doxycycline
YPR078C
ZShift = −3.73
Z lm Score = 3.76
NG = 0
DB = 0
SM = 0
Delta_r
Doxycycline
YPR078C
ZShift = −6.15
Z lm Score = −7.48
NG = 0
DB = 0
SM = 0
Delta_AUC
Doxycycline

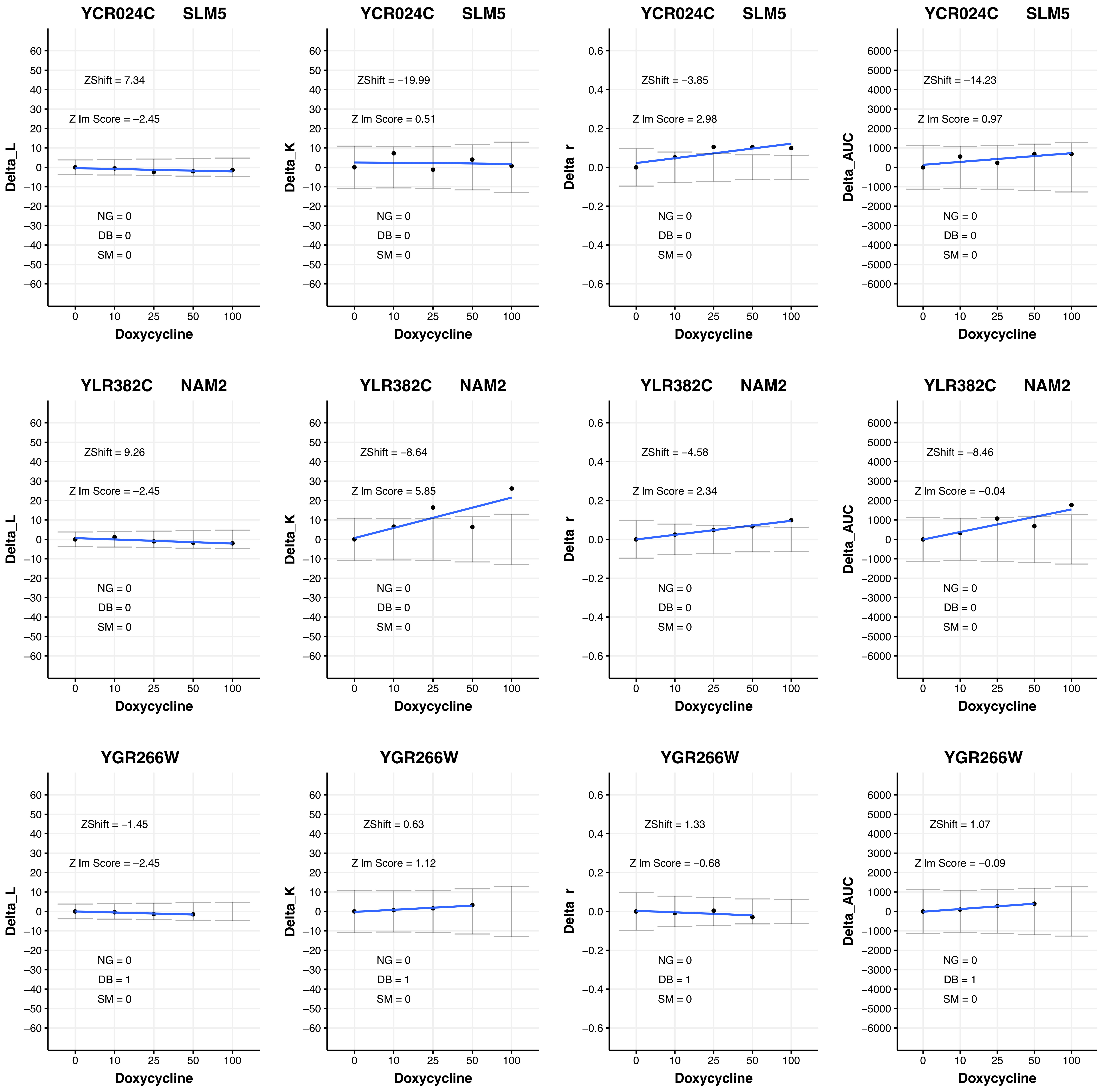

YCR024C SLM5
ZShift = 7.34
Z lm Score = −2.45
NG = 0
DB = 0
SM = 0
Delta_L
Doxycycline
YCR024C SLM5
ZShift = −19.99
Z lm Score = 0.51
NG = 0
DB = 0
SM = 0
Delta_K
Doxycycline
YCR024C SLM5
ZShift = −3.85
Z lm Score = 2.98
NG = 0
DB = 0
SM = 0
Delta_r
Doxycycline
YCR024C SLM5
ZShift = −14.23
Z lm Score = 0.97
NG = 0
DB = 0
SM = 0
Delta_AUC
Doxycycline
YLR382C NAM2
ZShift = 9.26
Z lm Score = −2.45
NG = 0
DB = 0
SM = 0
Delta_L
Doxycycline
YLR382C NAM2
ZShift = −8.64
Z lm Score = 5.85
NG = 0
DB = 0
SM = 0
Delta_K
Doxycycline
YLR382C NAM2
ZShift = −4.58
Z lm Score = 2.34
NG = 0
DB = 0
SM = 0
Delta_r
Doxycycline
YLR382C NAM2
ZShift = −8.46
Z lm Score = −0.04
NG = 0
DB = 0
SM = 0
Delta_AUC
Doxycycline
YGR266W
ZShift = −1.45
Z lm Score = −2.45
NG = 0
DB = 1
SM = 0
Delta_L
Doxycycline
YGR266W
ZShift = 0.63
Z lm Score = 1.12
NG = 0
DB = 1
SM = 0
Delta_K
Doxycycline
YGR266W
ZShift = 1.33
Z lm Score = −0.68
NG = 0
DB = 1
SM = 0
Delta_r
Doxycycline
YGR266W
ZShift = 1.07
Z lm Score = −0.09
NG = 0
DB = 1
SM = 0
Delta_AUC
Doxycycline

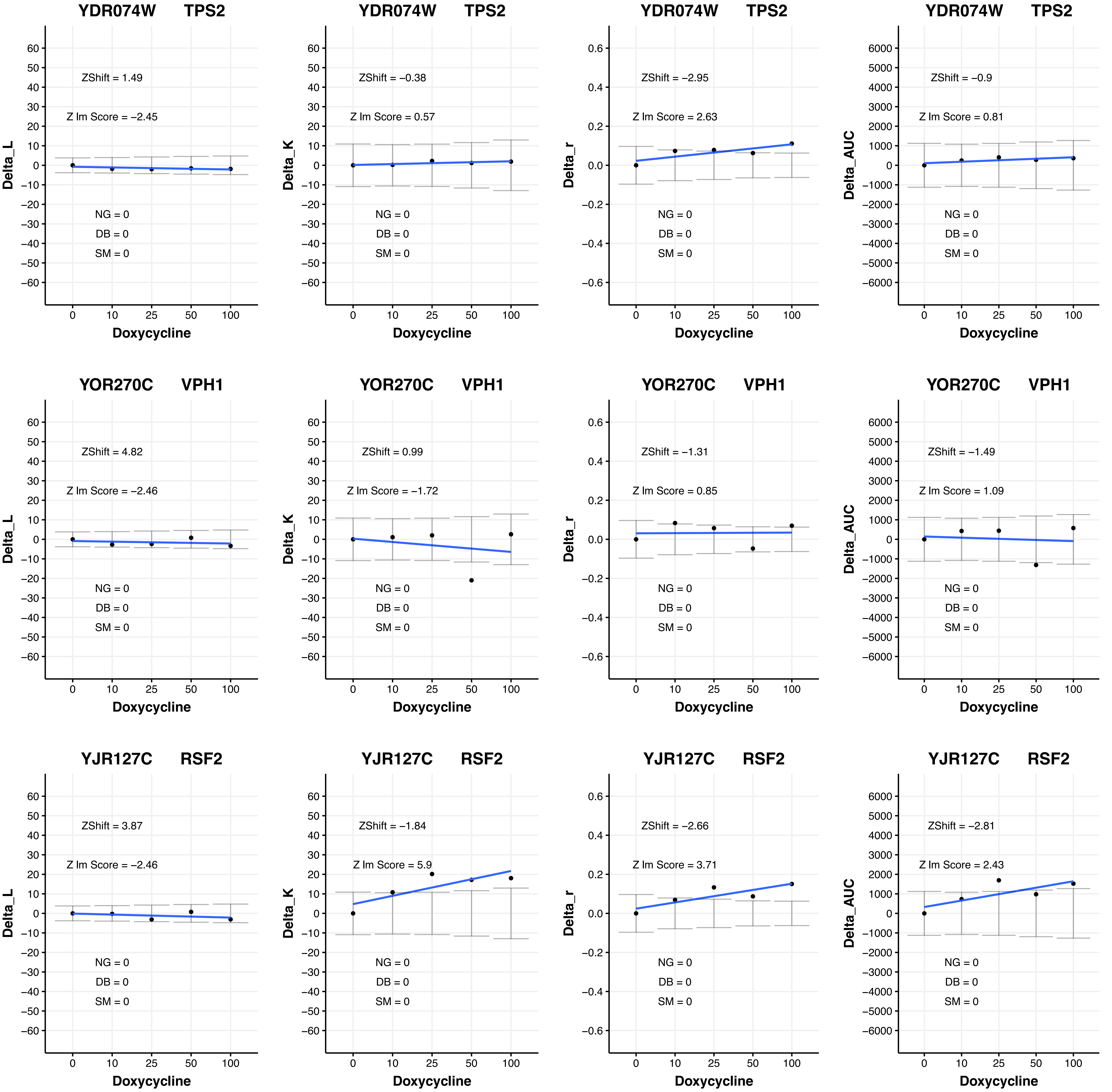

YDR074W TPS2
ZShift = 1.49
Z lm Score = −2.45
NG = 0
DB = 0
SM = 0
Delta_L
Doxycycline
YDR074W TPS2
ZShift = −0.38
Z lm Score = 0.57
NG = 0
DB = 0
SM = 0
Delta_K
Doxycycline
YDR074W TPS2
ZShift = −2.95
Z lm Score = 2.63
NG = 0
DB = 0
SM = 0
Delta_r
Doxycycline
YDR074W TPS2
ZShift = −0.9
Z lm Score = 0.81
NG = 0
DB = 0
SM = 0
Delta_AUC
Doxycycline
YOR270C VPH1
ZShift = 4.82
Z lm Score = −2.46
NG = 0
DB = 0
SM = 0
Delta_L
Doxycycline
YOR270C VPH1
ZShift = 0.99
Z lm Score = −1.72
NG = 0
DB = 0
SM = 0
Delta_K
Doxycycline
YOR270C VPH1
ZShift = −1.31
Z lm Score = 0.85
NG = 0
DB = 0
SM = 0
Delta_r
Doxycycline
YOR270C VPH1
ZShift = −1.49
Z lm Score = 1.09
NG = 0
DB = 0
SM = 0
Delta_AUC
Doxycycline
YJR127C RSF2
ZShift = 3.87
Z lm Score = −2.46
NG = 0
DB = 0
SM = 0
Delta_L
Doxycycline
YJR127C RSF2
ZShift = −1.84
Z lm Score = 5.9
NG = 0
DB = 0
SM = 0
Delta_K
Doxycycline
YJR127C RSF2
ZShift = −2.66
Z lm Score = 3.71
NG = 0
DB = 0
SM = 0
Delta_r
Doxycycline
YJR127C RSF2
ZShift = −2.81
Z lm Score = 2.43
NG = 0
DB = 0
SM = 0
Delta_AUC
Doxycycline

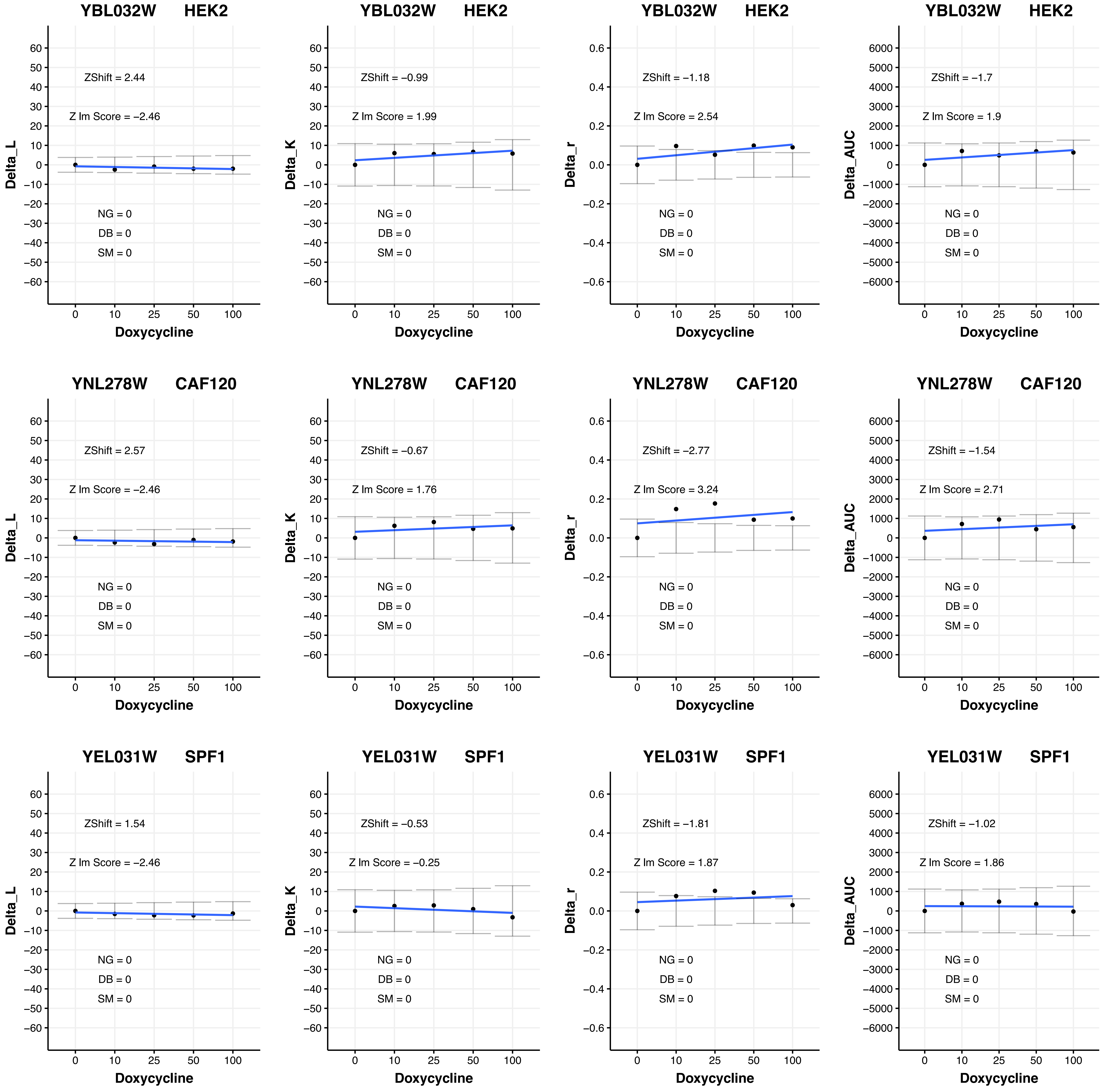
YBL032W HEK2
ZShift = 2.44
Z lm Score = −2.46
NG = 0
DB = 0
SM = 0
Delta_L
Doxycycline
YBL032W HEK2
ZShift = −0.99
Z lm Score = 1.99
NG = 0
DB = 0
SM = 0
Delta_K
Doxycycline
YBL032W HEK2
ZShift = −1.18
Z lm Score = 2.54
NG = 0
DB = 0
SM = 0
Delta_r
Doxycycline
YBL032W HEK2
ZShift = −1.7
Z lm Score = 1.9
NG = 0
DB = 0
SM = 0
Delta_AUC
Doxycycline
YNL278W CAF120
ZShift = 2.57
Z lm Score = −2.46
NG = 0
DB = 0
SM = 0
Delta_L
Doxycycline
YNL278W CAF120
ZShift = −0.67
Z lm Score = 1.76
NG = 0
DB = 0
SM = 0
Delta_K
Doxycycline
YNL278W CAF120
ZShift = −2.77
Z lm Score = 3.24
NG = 0
DB = 0
SM = 0
Delta_r
Doxycycline
YNL278W CAF120
ZShift = −1.54
Z lm Score = 2.71
NG = 0
DB = 0
SM = 0
Delta_AUC
Doxycycline
YEL031W SPF1
ZShift = 1.54
Z lm Score = −2.46
NG = 0
DB = 0
SM = 0
Delta_L
Doxycycline
YEL031W SPF1
ZShift = −0.53
Z lm Score = −0.25
NG = 0
DB = 0
SM = 0
Delta_K
Doxycycline
YEL031W SPF1
ZShift = −1.81
Z lm Score = 1.87
NG = 0
DB = 0
SM = 0
Delta_r
Doxycycline
YEL031W SPF1
ZShift = −1.02
Z lm Score = 1.86
NG = 0
DB = 0
SM = 0
Delta_AUC
Doxycycline

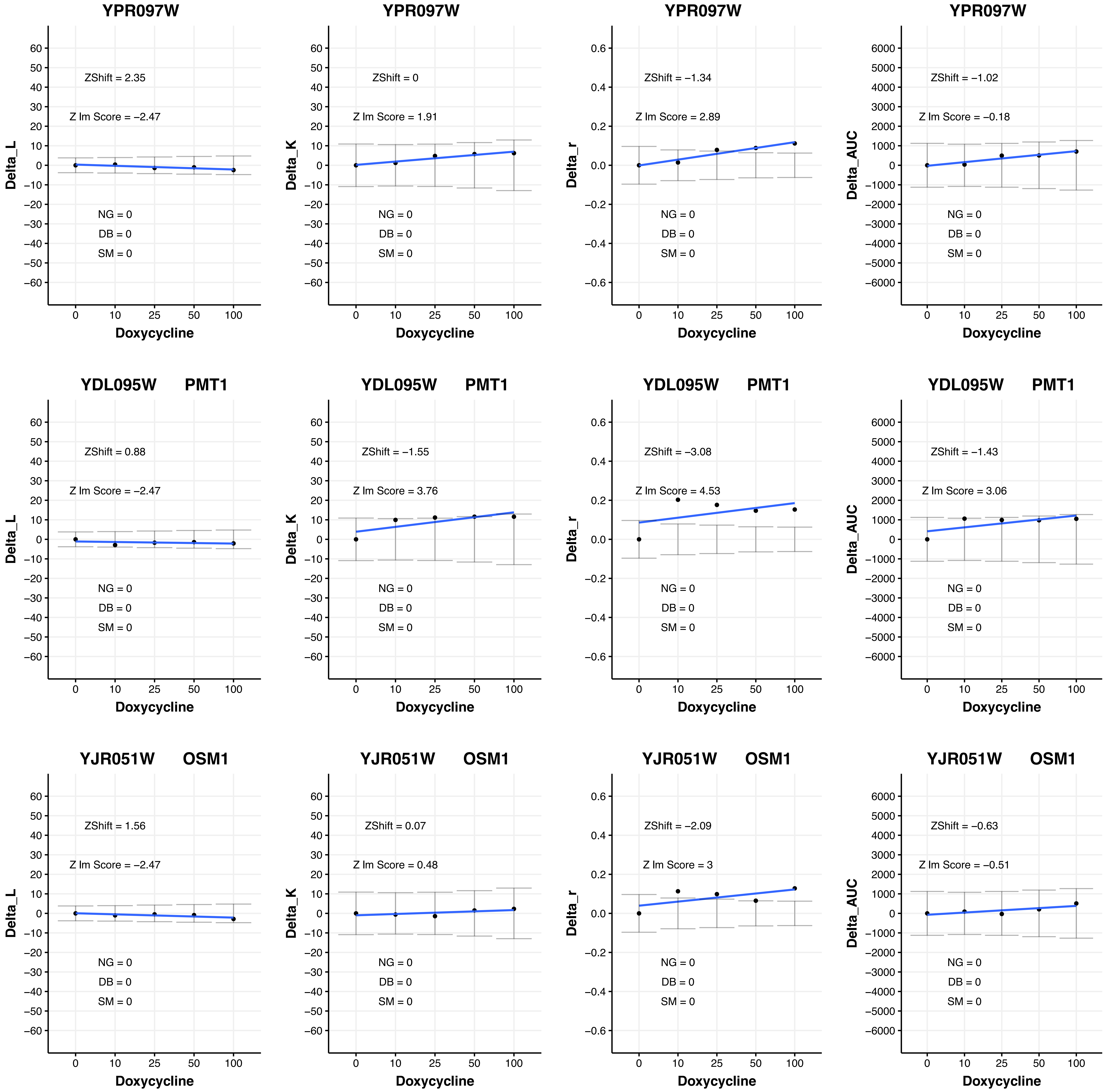

YPR097W
ZShift = 2.35
Z lm Score = −2.47
NG = 0
DB = 0
SM = 0
Delta_L
Doxycycline
YPR097W
ZShift = 0
Z lm Score = 1.91
NG = 0
DB = 0
SM = 0
Delta_K
Doxycycline
YPR097W
ZShift = −1.34
Z lm Score = 2.89
NG = 0
DB = 0
SM = 0
Delta_r
Doxycycline
YPR097W
ZShift = −1.02
Z lm Score = −0.18
NG = 0
DB = 0
SM = 0
Delta_AUC
Doxycycline
YDL095W PMT1
ZShift = 0.88
Z lm Score = −2.47
NG = 0
DB = 0
SM = 0
Delta_L
Doxycycline
YDL095W PMT1
ZShift = −1.55
Z lm Score = 3.76
NG = 0
DB = 0
SM = 0
Delta_K
Doxycycline
YDL095W PMT1
ZShift = −3.08
Z lm Score = 4.53
NG = 0
DB = 0
SM = 0
Delta_r
Doxycycline
YDL095W PMT1
ZShift = −1.43
Z lm Score = 3.06
NG = 0
DB = 0
SM = 0
Delta_AUC
Doxycycline
YJR051W OSM1
ZShift = 1.56
Z lm Score = −2.47
NG = 0
DB = 0
SM = 0
Delta_L
Doxycycline
YJR051W OSM1
ZShift = 0.07
Z lm Score = 0.48
NG = 0
DB = 0
SM = 0
Delta_K
Doxycycline
YJR051W OSM1
ZShift = −2.09
Z lm Score = 3
NG = 0
DB = 0
SM = 0
Delta_r
Doxycycline
YJR051W OSM1
ZShift = −0.63
Z lm Score = −0.51
NG = 0
DB = 0
SM = 0
Delta_AUC
Doxycycline

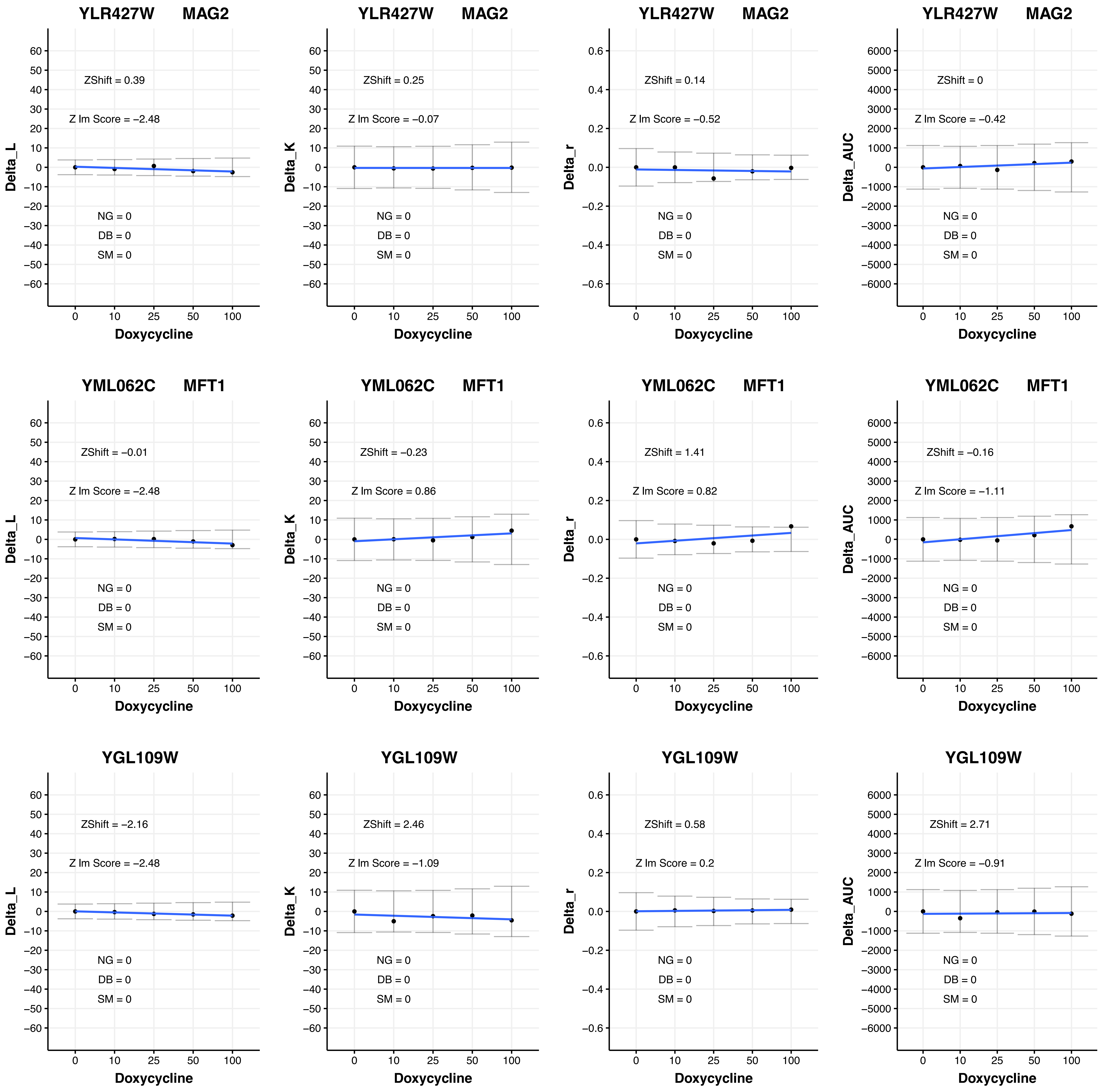
YLR427W MAG2
ZShift = 0.39
Z lm Score = −2.48
NG = 0
DB = 0
SM = 0
Delta_L
Doxycycline
YLR427W MAG2
ZShift = 0.25
Z lm Score = −0.07
NG = 0
DB = 0
SM = 0
Delta_K
Doxycycline
YLR427W MAG2
ZShift = 0.14
Z lm Score = −0.52
NG = 0
DB = 0
SM = 0
Delta_r
Doxycycline
YLR427W MAG2
ZShift = 0
Z lm Score = −0.42
NG = 0
DB = 0
SM = 0
Delta_AUC
Doxycycline
YML062C MFT1
ZShift = −0.01
Z lm Score = −2.48
NG = 0
DB = 0
SM = 0
Delta_L
Doxycycline
YML062C MFT1
ZShift = −0.23
Z lm Score = 0.86
NG = 0
DB = 0
SM = 0
Delta_K
Doxycycline
YML062C MFT1
ZShift = 1.41
Z lm Score = 0.82
NG = 0
DB = 0
SM = 0
Delta_r
Doxycycline
YML062C MFT1
ZShift = −0.16
Z lm Score = −1.11
NG = 0
DB = 0
SM = 0
Delta_AUC
Doxycycline
YGL109W
ZShift = −2.16
Z lm Score = −2.48
NG = 0
DB = 0
SM = 0
Delta_L
Doxycycline
YGL109W
ZShift = 2.46
Z lm Score = −1.09
NG = 0
DB = 0
SM = 0
Delta_K
Doxycycline
YGL109W
ZShift = 0.58
Z lm Score = 0.2
NG = 0
DB = 0
SM = 0
Delta_r
Doxycycline
YGL109W
ZShift = 2.71
Z lm Score = −0.91
NG = 0
DB = 0
SM = 0
Delta_AUC
Doxycycline

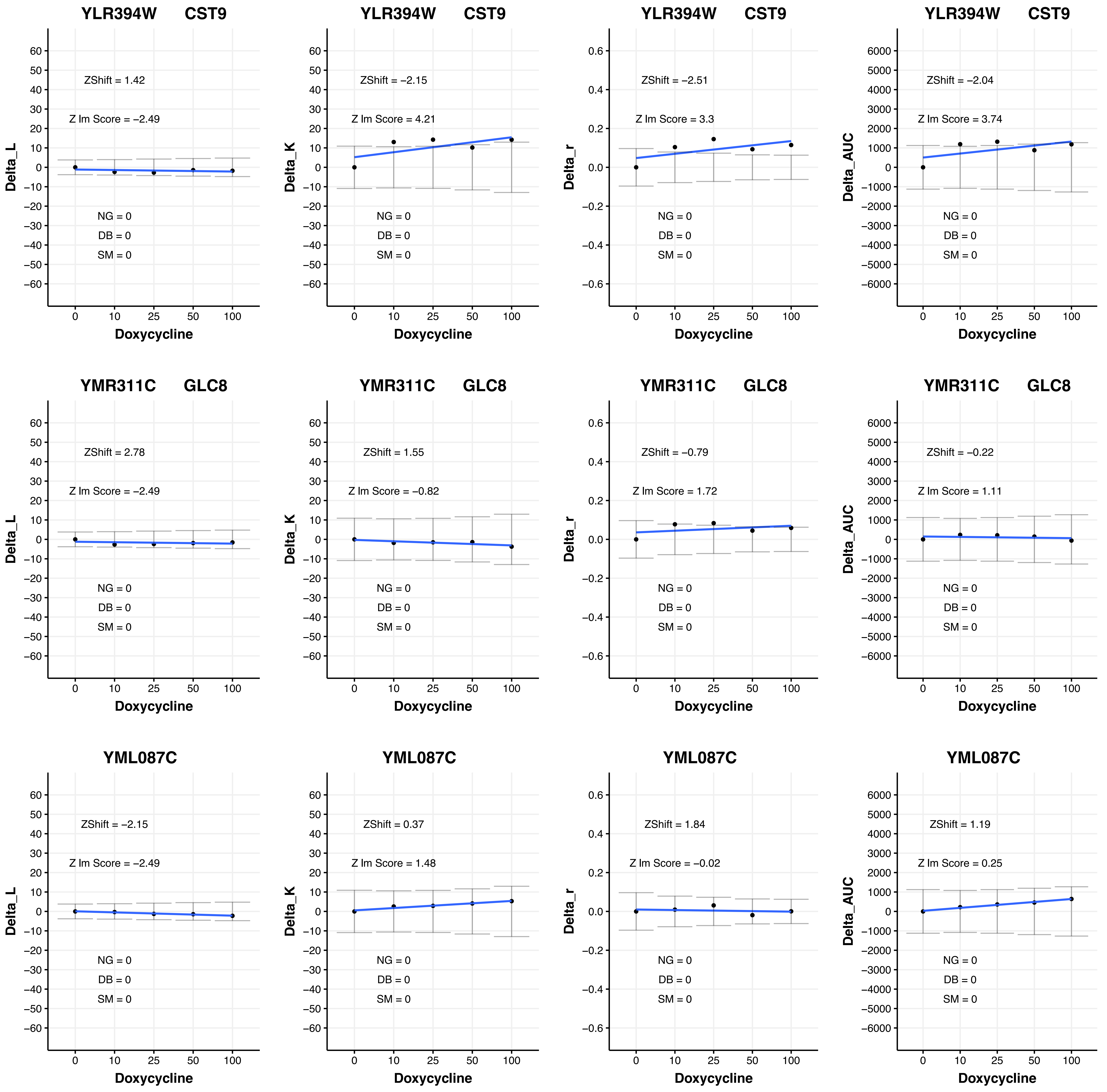

YLR394W CST9
ZShift = 1.42
Z lm Score = −2.49
NG = 0
DB = 0
SM = 0
Delta_L
Doxycycline
YLR394W CST9
ZShift = −2.15
Z lm Score = 4.21
NG = 0
DB = 0
SM = 0
Delta_K
Doxycycline
YLR394W CST9
ZShift = −2.51
Z lm Score = 3.3
NG = 0
DB = 0
SM = 0
Delta_r
Doxycycline
YLR394W CST9
ZShift = −2.04
Z lm Score = 3.74
NG = 0
DB = 0
SM = 0
Delta_AUC
Doxycycline
YMR311C GLC8
ZShift = 2.78
Z lm Score = −2.49
NG = 0
DB = 0
SM = 0
Delta_L
Doxycycline
YMR311C GLC8
ZShift = 1.55
Z lm Score = −0.82
NG = 0
DB = 0
SM = 0
Delta_K
Doxycycline
YMR311C GLC8
ZShift = −0.79
Z lm Score = 1.72
NG = 0
DB = 0
SM = 0
Delta_r
Doxycycline
YMR311C GLC8
ZShift = −0.22
Z lm Score = 1.11
NG = 0
DB = 0
SM = 0
Delta_AUC
Doxycycline
YML087C
ZShift = −2.15
Z lm Score = −2.49
NG = 0
DB = 0
SM = 0
Delta_L
Doxycycline
YML087C
ZShift = 0.37
Z lm Score = 1.48
NG = 0
DB = 0
SM = 0
Delta_K
Doxycycline
YML087C
ZShift = 1.84
Z lm Score = −0.02
NG = 0
DB = 0
SM = 0
Delta_r
Doxycycline
YML087C
ZShift = 1.19
Z lm Score = 0.25
NG = 0
DB = 0
SM = 0
Delta_AUC
Doxycycline

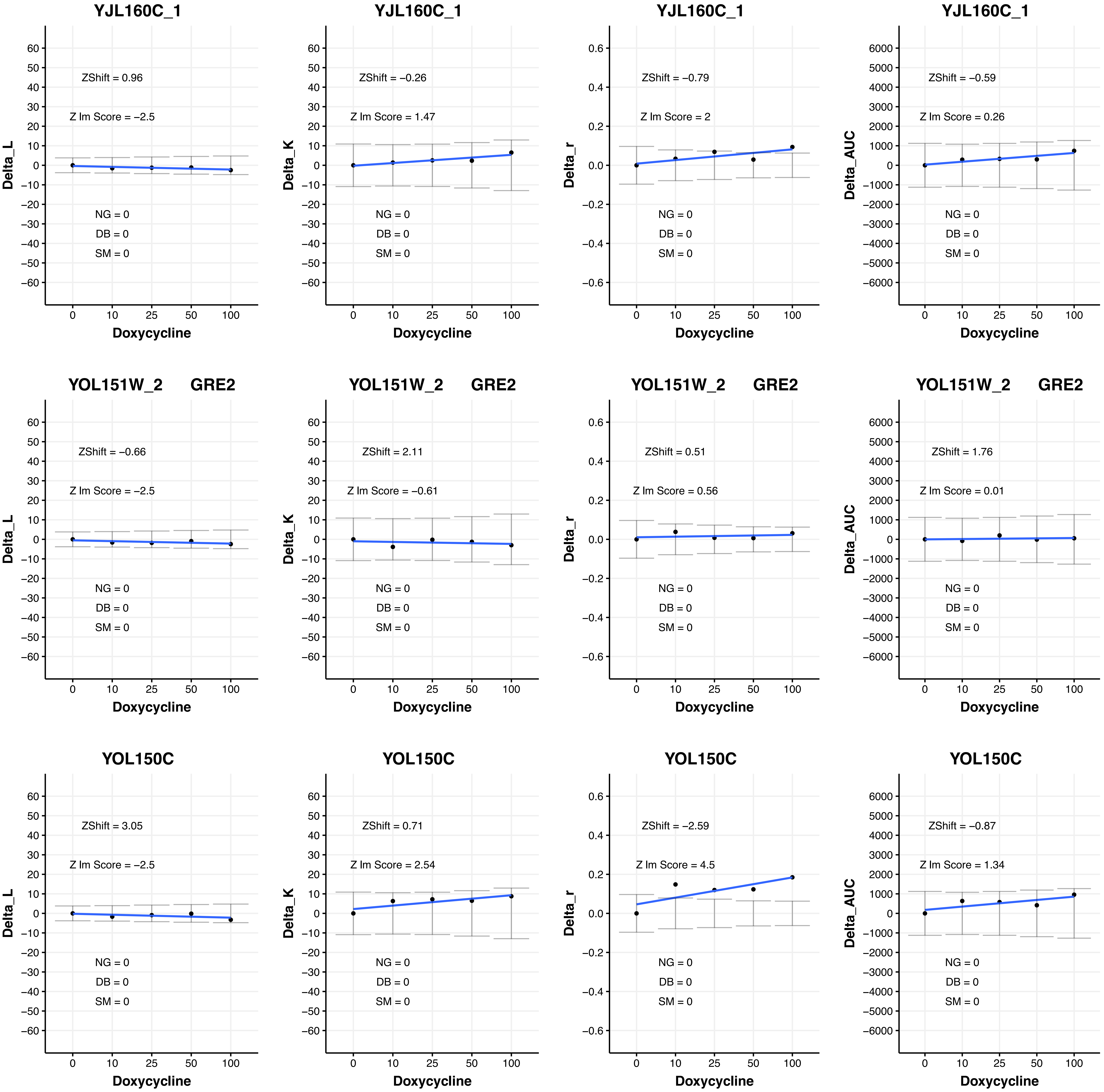
YJL160C_1
ZShift = 0.96
Z lm Score = −2.5
NG = 0
DB = 0
SM = 0
Delta_L
Doxycycline
YJL160C_1
ZShift = −0.26
Z lm Score = 1.47
NG = 0
DB = 0
SM = 0
Delta_K
Doxycycline
YJL160C_1
ZShift = −0.79
Z lm Score = 2
NG = 0
DB = 0
SM = 0
Delta_r
Doxycycline
YJL160C_1
ZShift = −0.59
Z lm Score = 0.26
NG = 0
DB = 0
SM = 0
Delta_AUC
Doxycycline
YOL151W_2 GRE2
ZShift = −0.66
Z lm Score = −2.5
NG = 0
DB = 0
SM = 0
Delta_L
Doxycycline
YOL151W_2 GRE2
ZShift = 2.11
Z lm Score = −0.61
NG = 0
DB = 0
SM = 0
Delta_K
Doxycycline
YOL151W_2 GRE2
ZShift = 0.51
Z lm Score = 0.56
NG = 0
DB = 0
SM = 0
Delta_r
Doxycycline
YOL151W_2 GRE2
ZShift = 1.76
Z lm Score = 0.01
NG = 0
DB = 0
SM = 0
Delta_AUC
Doxycycline
YOL150C
ZShift = 3.05
Z lm Score = −2.5
NG = 0
DB = 0
SM = 0
Delta_L
Doxycycline
YOL150C
ZShift = 0.71
Z lm Score = 2.54
NG = 0
DB = 0
SM = 0
Delta_K
Doxycycline
YOL150C
ZShift = −2.59
Z lm Score = 4.5
NG = 0
DB = 0
SM = 0
Delta_r
Doxycycline
YOL150C
ZShift = −0.87
Z lm Score = 1.34
NG = 0
DB = 0
SM = 0
Delta_AUC
Doxycycline

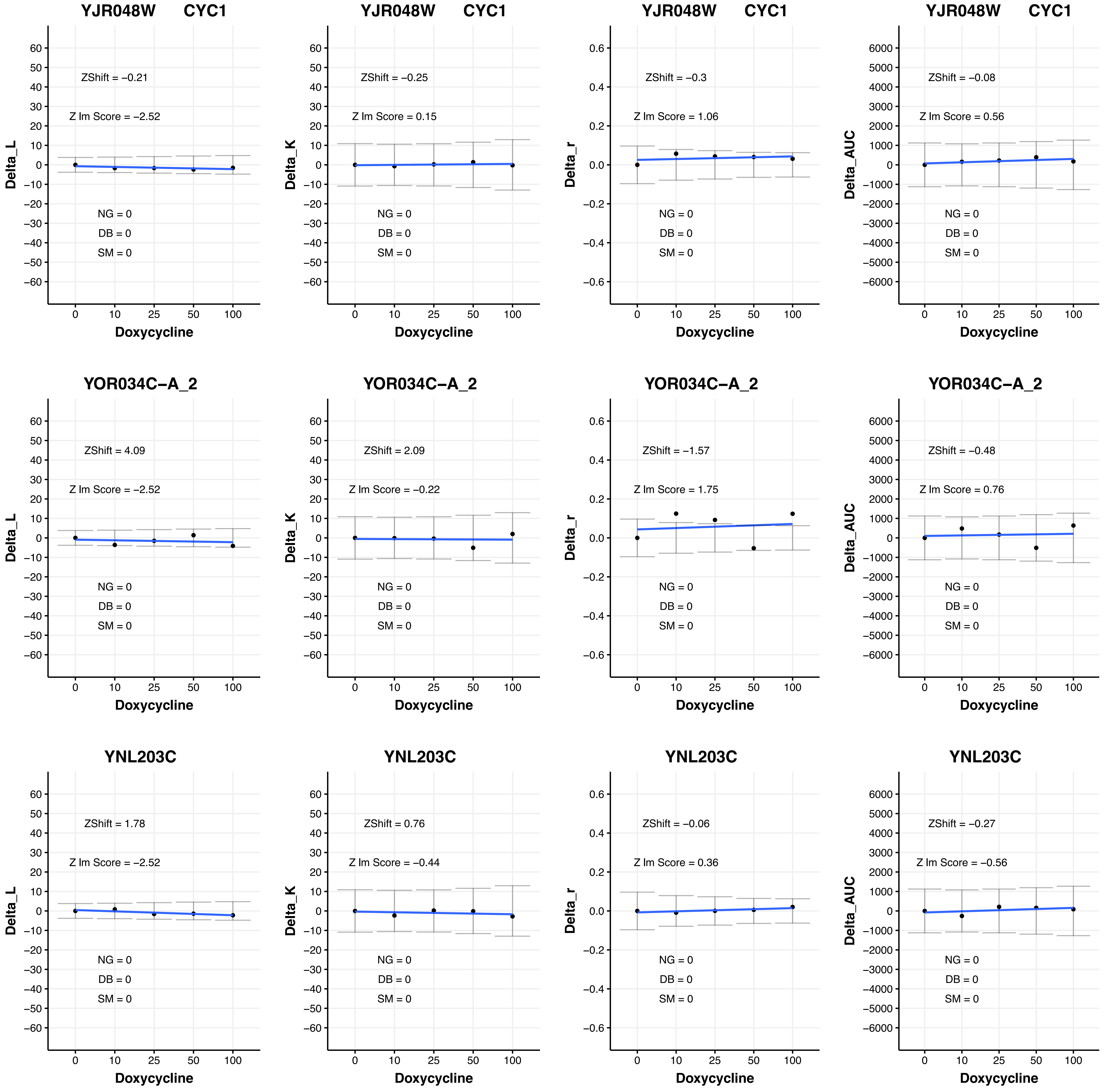
YJR048W CYC1
ZShift = −0.21
Z lm Score = −2.52
NG = 0
DB = 0
SM = 0
Delta_L
Doxycycline
YJR048W CYC1
ZShift = −0.25
Z lm Score = 0.15
NG = 0
DB = 0
SM = 0
Delta_K
Doxycycline
YJR048W CYC1
ZShift = −0.3
Z lm Score = 1.06
NG = 0
DB = 0
SM = 0
Delta_r
Doxycycline
YJR048W CYC1
ZShift = −0.08
Z lm Score = 0.56
NG = 0
DB = 0
SM = 0
Delta_AUC
Doxycycline
YOR034C−A_2
ZShift = 4.09
Z lm Score = −2.52
NG = 0
DB = 0
SM = 0
Delta_L
Doxycycline
YOR034C−A_2
ZShift = 2.09
Z lm Score = −0.22
NG = 0
DB = 0
SM = 0
Delta_K
Doxycycline
YOR034C−A_2
ZShift = −1.57
Z lm Score = 1.75
NG = 0
DB = 0
SM = 0
Delta_r
Doxycycline
YOR034C−A_2
ZShift = −0.48
Z lm Score = 0.76
NG = 0
DB = 0
SM = 0
Delta_AUC
Doxycycline
YNL203C
ZShift = 1.78
Z lm Score = −2.52
NG = 0
DB = 0
SM = 0
Delta_L
Doxycycline
YNL203C
ZShift = 0.76
Z lm Score = −0.44
NG = 0
DB = 0
SM = 0
Delta_K
Doxycycline
YNL203C
ZShift = −0.06
Z lm Score = 0.36
NG = 0
DB = 0
SM = 0
Delta_r
Doxycycline
YNL203C
ZShift = −0.27
Z lm Score = −0.56
NG = 0
DB = 0
SM = 0
Delta_AUC
Doxycycline

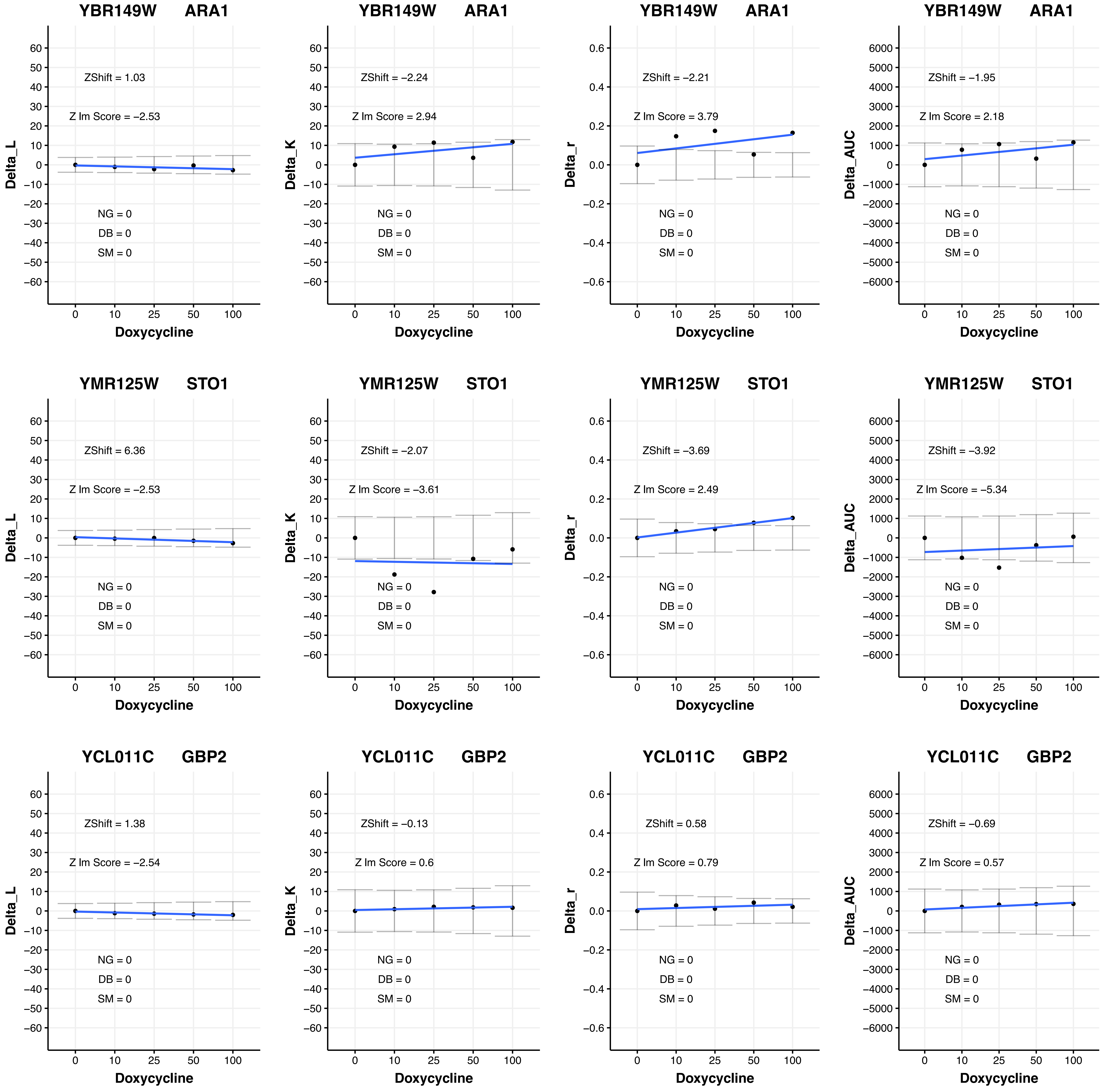

YBR149W ARA1
ZShift = 1.03
Z lm Score = −2.53
ZShift = −2.24
Z lm Score = 2.94
ZShift = −2.21
Z lm Score = 3.79
ZShift = −1.95
Z lm Score = 2.18
YMR125W STO1
ZShift = 6.36
Z lm Score = −2.53
ZShift = −2.07
Z lm Score = −3.61
ZShift = −3.69
Z lm Score = 2.49
ZShift = −3.92
Z lm Score = −5.34
YCL011C GBP2
ZShift = 1.38
Z lm Score = −2.54
ZShift = −0.13
Z lm Score = 0.6
ZShift = 0.58
Z lm Score = 0.79
ZShift = −0.69
Z lm Score = 0.57
NG = 0
DB = 0
SM = 0
Delta_L
Delta_K
Delta_r
Delta_AUC
Doxycycline
0 10 25 50 100

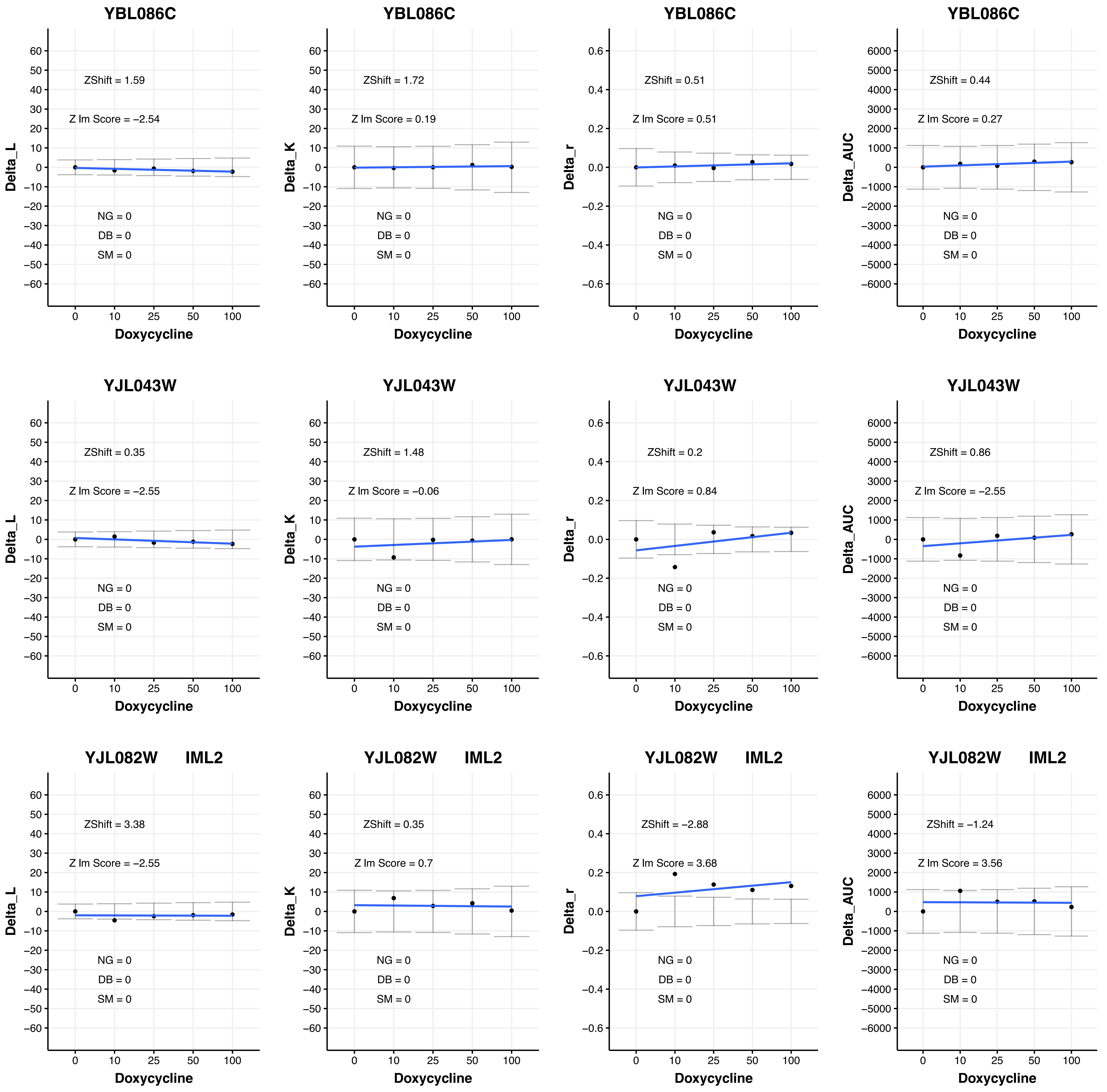

YBL086C
Delta_L
ZShift = 1.59
Z lm Score = −2.54
NG = 0
DB = 0
SM = 0
Doxycycline
YBL086C
Delta_K
ZShift = 1.72
Z lm Score = 0.19
NG = 0
DB = 0
SM = 0
Doxycycline
YBL086C
Delta_r
ZShift = 0.51
Z lm Score = 0.51
NG = 0
DB = 0
SM = 0
Doxycycline
YBL086C
Delta_AUC
ZShift = 0.44
Z lm Score = 0.27
NG = 0
DB = 0
SM = 0
Doxycycline
YJL043W
Delta_L
ZShift = 0.35
Z lm Score = −2.55
NG = 0
DB = 0
SM = 0
Doxycycline
YJL043W
Delta_K
ZShift = 1.48
Z lm Score = −0.06
NG = 0
DB = 0
SM = 0
Doxycycline
YJL043W
Delta_r
ZShift = 0.2
Z lm Score = 0.84
NG = 0
DB = 0
SM = 0
Doxycycline
YJL043W
Delta_AUC
ZShift = 0.86
Z lm Score = −2.55
NG = 0
DB = 0
SM = 0
Doxycycline
YJL082W IML2
Delta_L
ZShift = 3.38
Z lm Score = −2.55
NG = 0
DB = 0
SM = 0
Doxycycline
YJL082W IML2
Delta_K
ZShift = 0.35
Z lm Score = 0.7
NG = 0
DB = 0
SM = 0
Doxycycline
YJL082W IML2
Delta_r
ZShift = −2.88
Z lm Score = 3.68
NG = 0
DB = 0
SM = 0
Doxycycline
YJL082W IML2
Delta_AUC
ZShift = −1.24
Z lm Score = 3.56
NG = 0
DB = 0
SM = 0
Doxycycline

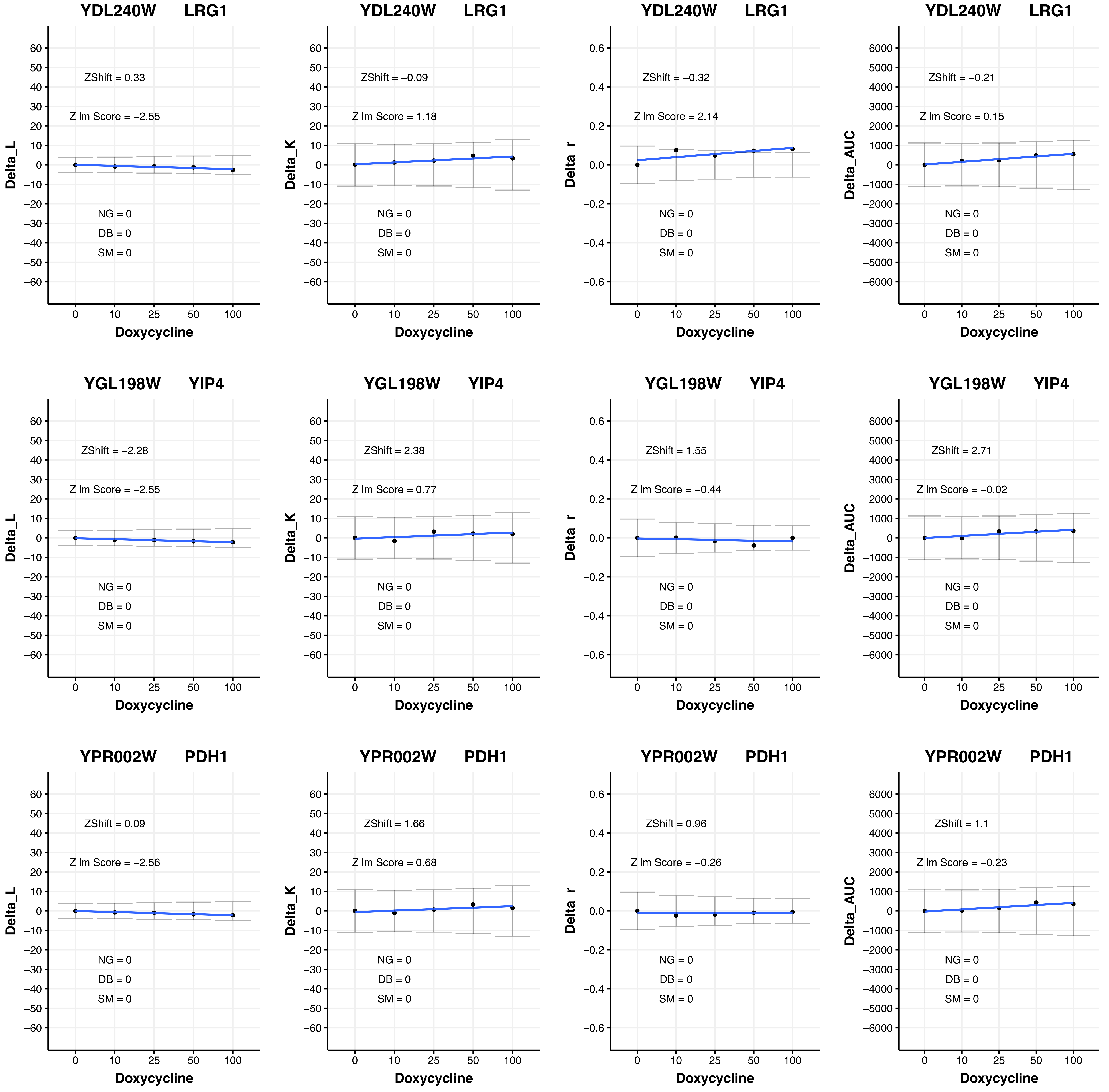

YDL240W LRG1
ZShift = 0.33
Z lm Score = −2.55
NG = 0
DB = 0
SM = 0
Delta_L
Doxycycline
YDL240W LRG1
ZShift = −0.09
Z lm Score = 1.18
NG = 0
DB = 0
SM = 0
Delta_K
Doxycycline
YDL240W LRG1
ZShift = −0.32
Z lm Score = 2.14
NG = 0
DB = 0
SM = 0
Delta_r
Doxycycline
YDL240W LRG1
ZShift = −0.21
Z lm Score = 0.15
NG = 0
DB = 0
SM = 0
Delta_AUC
Doxycycline
YGL198W YIP4
ZShift = −2.28
Z lm Score = −2.55
NG = 0
DB = 0
SM = 0
Delta_L
Doxycycline
YGL198W YIP4
ZShift = 2.38
Z lm Score = 0.77
NG = 0
DB = 0
SM = 0
Delta_K
Doxycycline
YGL198W YIP4
ZShift = 1.55
Z lm Score = −0.44
NG = 0
DB = 0
SM = 0
Delta_r
Doxycycline
YGL198W YIP4
ZShift = 2.71
Z lm Score = −0.02
NG = 0
DB = 0
SM = 0
Delta_AUC
Doxycycline
YPR002W PDH1
ZShift = 0.09
Z lm Score = −2.56
NG = 0
DB = 0
SM = 0
Delta_L
Doxycycline
YPR002W PDH1
ZShift = 1.66
Z lm Score = 0.68
NG = 0
DB = 0
SM = 0
Delta_K
Doxycycline
YPR002W PDH1
ZShift = 0.96
Z lm Score = −0.26
NG = 0
DB = 0
SM = 0
Delta_r
Doxycycline
YPR002W PDH1
ZShift = 1.1
Z lm Score = −0.23
NG = 0
DB = 0
SM = 0
Delta_AUC
Doxycycline

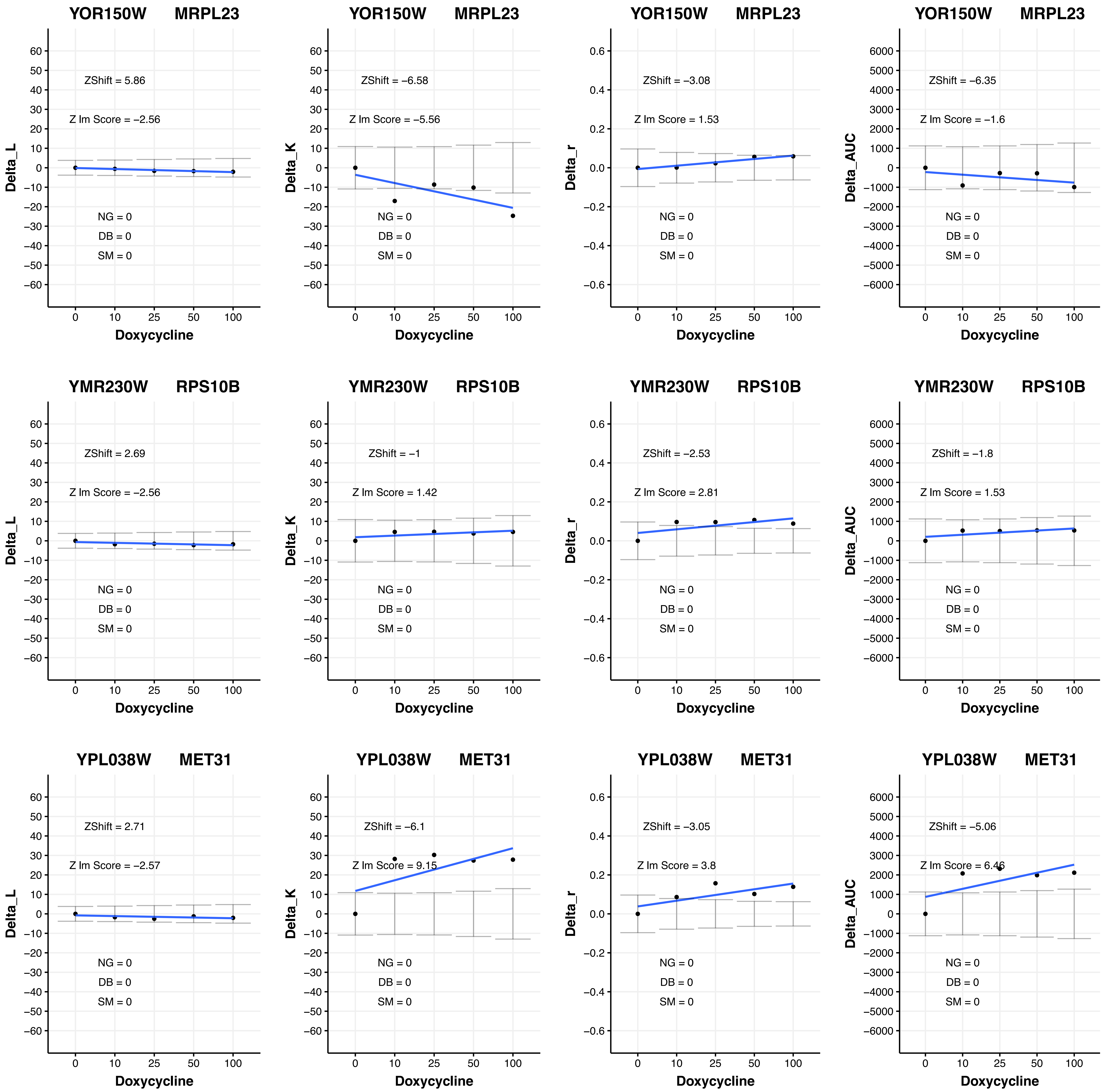
YOR150W MRPL23
ZShift = 5.86
Z lm Score = −2.56
NG = 0
DB = 0
SM = 0
Delta_L
Doxycycline
YOR150W MRPL23
ZShift = −6.58
Z lm Score = −5.56
NG = 0
DB = 0
SM = 0
Delta_K
Doxycycline
YOR150W MRPL23
ZShift = −3.08
Z lm Score = 1.53
NG = 0
DB = 0
SM = 0
Delta_r
Doxycycline
YOR150W MRPL23
ZShift = −6.35
Z lm Score = −1.6
NG = 0
DB = 0
SM = 0
Delta_AUC
Doxycycline
YMR230W RPS10B
ZShift = 2.69
Z lm Score = −2.56
NG = 0
DB = 0
SM = 0
Delta_L
Doxycycline
YMR230W RPS10B
ZShift = −1
Z lm Score = 1.42
NG = 0
DB = 0
SM = 0
Delta_K
Doxycycline
YMR230W RPS10B
ZShift = −2.53
Z lm Score = 2.81
NG = 0
DB = 0
SM = 0
Delta_r
Doxycycline
YMR230W RPS10B
ZShift = −1.8
Z lm Score = 1.53
NG = 0
DB = 0
SM = 0
Delta_AUC
Doxycycline
YPL038W MET31
ZShift = 2.71
Z lm Score = −2.57
NG = 0
DB = 0
SM = 0
Delta_L
Doxycycline
YPL038W MET31
ZShift = −6.1
Z lm Score = 9.15
NG = 0
DB = 0
SM = 0
Delta_K
Doxycycline
YPL038W MET31
ZShift = −3.05
Z lm Score = 3.8
NG = 0
DB = 0
SM = 0
Delta_r
Doxycycline
YPL038W MET31
ZShift = −5.06
Z lm Score = 6.46
NG = 0
DB = 0
SM = 0
Delta_AUC
Doxycycline

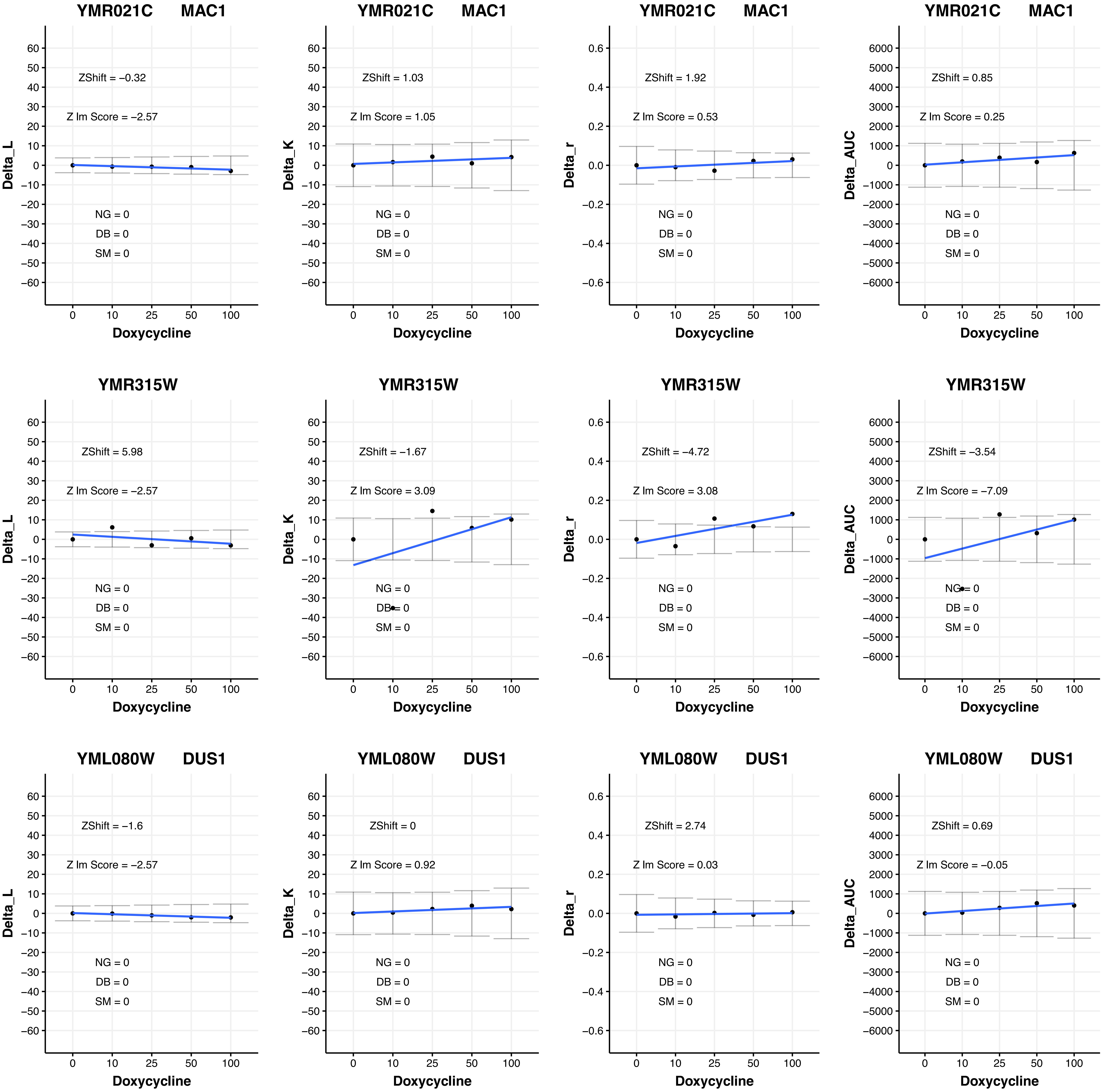
YMR021C MAC1
ZShift = −0.32
Z lm Score = −2.57
NG = 0
DB = 0
SM = 0
Delta_L
Doxycycline
YMR021C MAC1
ZShift = 1.03
Z lm Score = 1.05
NG = 0
DB = 0
SM = 0
Delta_K
Doxycycline
YMR021C MAC1
ZShift = 1.92
Z lm Score = 0.53
NG = 0
DB = 0
SM = 0
Delta_r
Doxycycline
YMR021C MAC1
ZShift = 0.85
Z lm Score = 0.25
NG = 0
DB = 0
SM = 0
Delta_AUC
Doxycycline
YMR315W
ZShift = 5.98
Z lm Score = −2.57
NG = 0
DB = 0
SM = 0
Delta_L
Doxycycline
YMR315W
ZShift = −1.67
Z lm Score = 3.09
NG = 0
DB = 0
SM = 0
Delta_K
Doxycycline
YMR315W
ZShift = −4.72
Z lm Score = 3.08
NG = 0
DB = 0
SM = 0
Delta_r
Doxycycline
YMR315W
ZShift = −3.54
Z lm Score = −7.09
NG = 0
DB = 0
SM = 0
Delta_AUC
Doxycycline
YML080W DUS1
ZShift = −1.6
Z lm Score = −2.57
NG = 0
DB = 0
SM = 0
Delta_L
Doxycycline
YML080W DUS1
ZShift = 0
Z lm Score = 0.92
NG = 0
DB = 0
SM = 0
Delta_K
Doxycycline
YML080W DUS1
ZShift = 2.74
Z lm Score = 0.03
NG = 0
DB = 0
SM = 0
Delta_r
Doxycycline
YML080W DUS1
ZShift = 0.69
Z lm Score = −0.05
NG = 0
DB = 0
SM = 0
Delta_AUC
Doxycycline

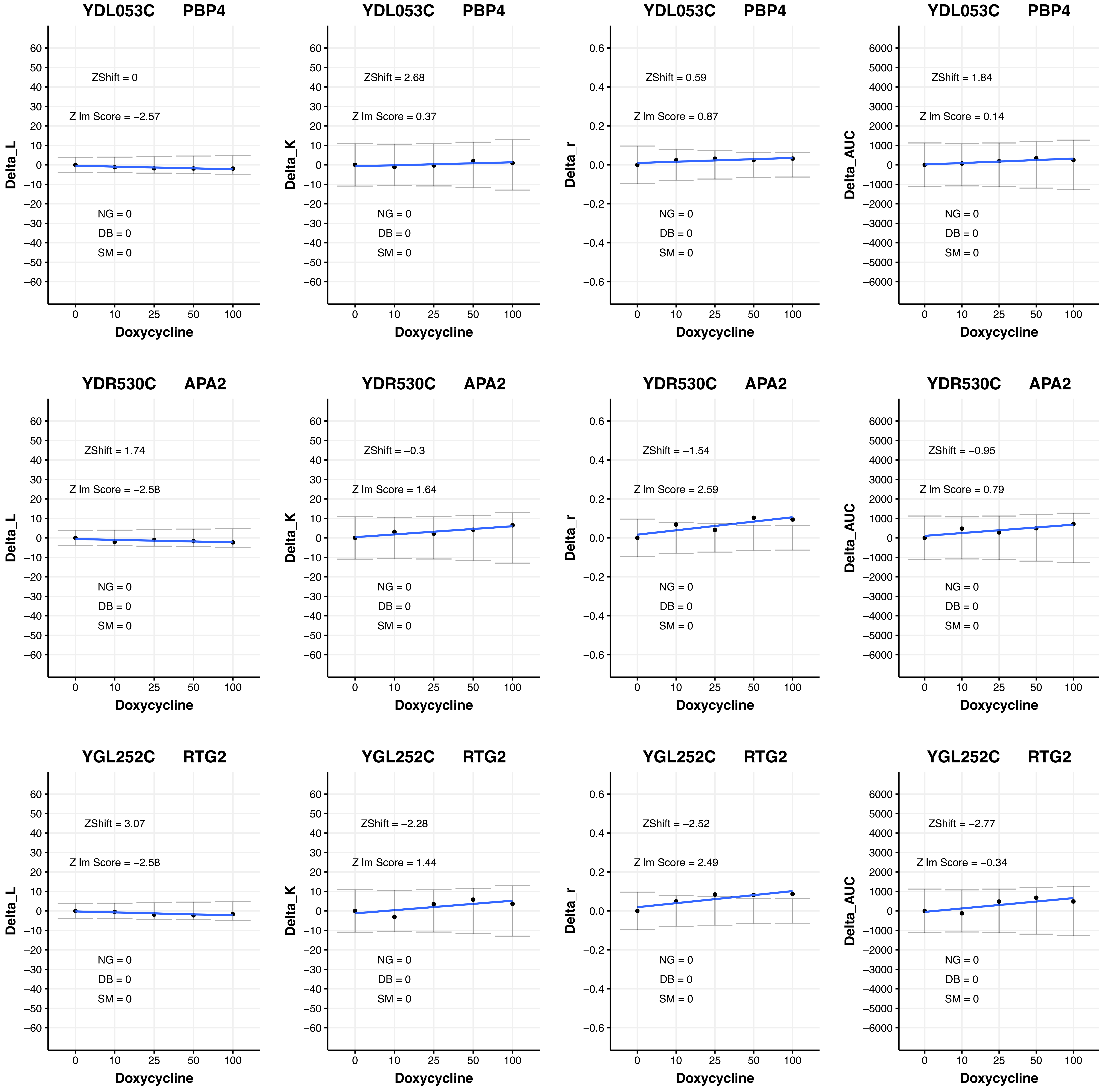

YDL053C PBP4
ZShift = 0
Z lm Score = −2.57
NG = 0
DB = 0
SM = 0
Delta_L
Doxycycline
YDL053C PBP4
ZShift = 2.68
Z lm Score = 0.37
NG = 0
DB = 0
SM = 0
Delta_K
Doxycycline
YDL053C PBP4
ZShift = 0.59
Z lm Score = 0.87
NG = 0
DB = 0
SM = 0
Delta_r
Doxycycline
YDL053C PBP4
ZShift = 1.84
Z lm Score = 0.14
NG = 0
DB = 0
SM = 0
Delta_AUC
Doxycycline
YDR530C APA2
ZShift = 1.74
Z lm Score = −2.58
NG = 0
DB = 0
SM = 0
Delta_L
Doxycycline
YDR530C APA2
ZShift = −0.3
Z lm Score = 1.64
NG = 0
DB = 0
SM = 0
Delta_K
Doxycycline
YDR530C APA2
ZShift = −1.54
Z lm Score = 2.59
NG = 0
DB = 0
SM = 0
Delta_r
Doxycycline
YDR530C APA2
ZShift = −0.95
Z lm Score = 0.79
NG = 0
DB = 0
SM = 0
Delta_AUC
Doxycycline
YGL252C RTG2
ZShift = 3.07
Z lm Score = −2.58
NG = 0
DB = 0
SM = 0
Delta_L
Doxycycline
YGL252C RTG2
ZShift = −2.28
Z lm Score = 1.44
NG = 0
DB = 0
SM = 0
Delta_K
Doxycycline
YGL252C RTG2
ZShift = −2.52
Z lm Score = 2.49
NG = 0
DB = 0
SM = 0
Delta_r
Doxycycline
YGL252C RTG2
ZShift = −2.77
Z lm Score = −0.34
NG = 0
DB = 0
SM = 0
Delta_AUC
Doxycycline

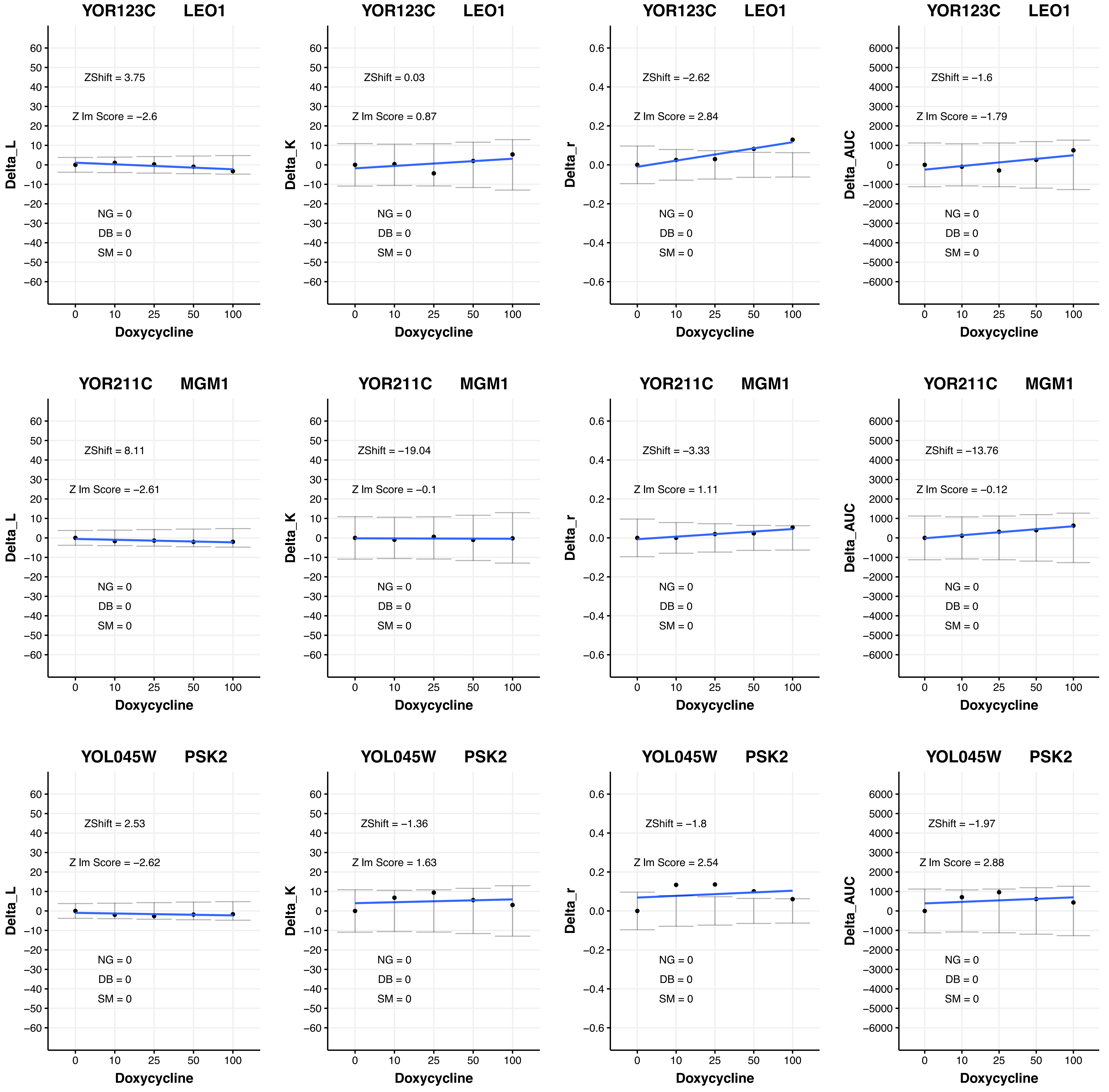

YOR123C LEO1
ZShift = 3.75
Z lm Score = −2.6
Delta_L
NG = 0
DB = 0
SM = 0
Doxycycline
YOR123C LEO1
ZShift = 0.03
Z lm Score = 0.87
Delta_K
NG = 0
DB = 0
SM = 0
Doxycycline
YOR123C LEO1
ZShift = −2.62
Z lm Score = 2.84
Delta_r
NG = 0
DB = 0
SM = 0
Doxycycline
YOR123C LEO1
ZShift = −1.6
Z lm Score = −1.79
Delta_AUC
NG = 0
DB = 0
SM = 0
Doxycycline
YOR211C MGM1
ZShift = 8.11
Z lm Score = −2.61
Delta_L
NG = 0
DB = 0
SM = 0
Doxycycline
YOR211C MGM1
ZShift = −19.04
Z lm Score = −0.1
Delta_K
NG = 0
DB = 0
SM = 0
Doxycycline
YOR211C MGM1
ZShift = −3.33
Z lm Score = 1.11
Delta_r
NG = 0
DB = 0
SM = 0
Doxycycline
YOR211C MGM1
ZShift = −13.76
Z lm Score = −0.12
Delta_AUC
NG = 0
DB = 0
SM = 0
Doxycycline
YOL045W PSK2
ZShift = 2.53
Z lm Score = −2.62
Delta_L
NG = 0
DB = 0
SM = 0
Doxycycline
YOL045W PSK2
ZShift = −1.36
Z lm Score = 1.63
Delta_K
NG = 0
DB = 0
SM = 0
Doxycycline
YOL045W PSK2
ZShift = −1.8
Z lm Score = 2.54
Delta_r
NG = 0
DB = 0
SM = 0
Doxycycline
YOL045W PSK2
ZShift = −1.97
Z lm Score = 2.88
Delta_AUC
NG = 0
DB = 0
SM = 0
Doxycycline

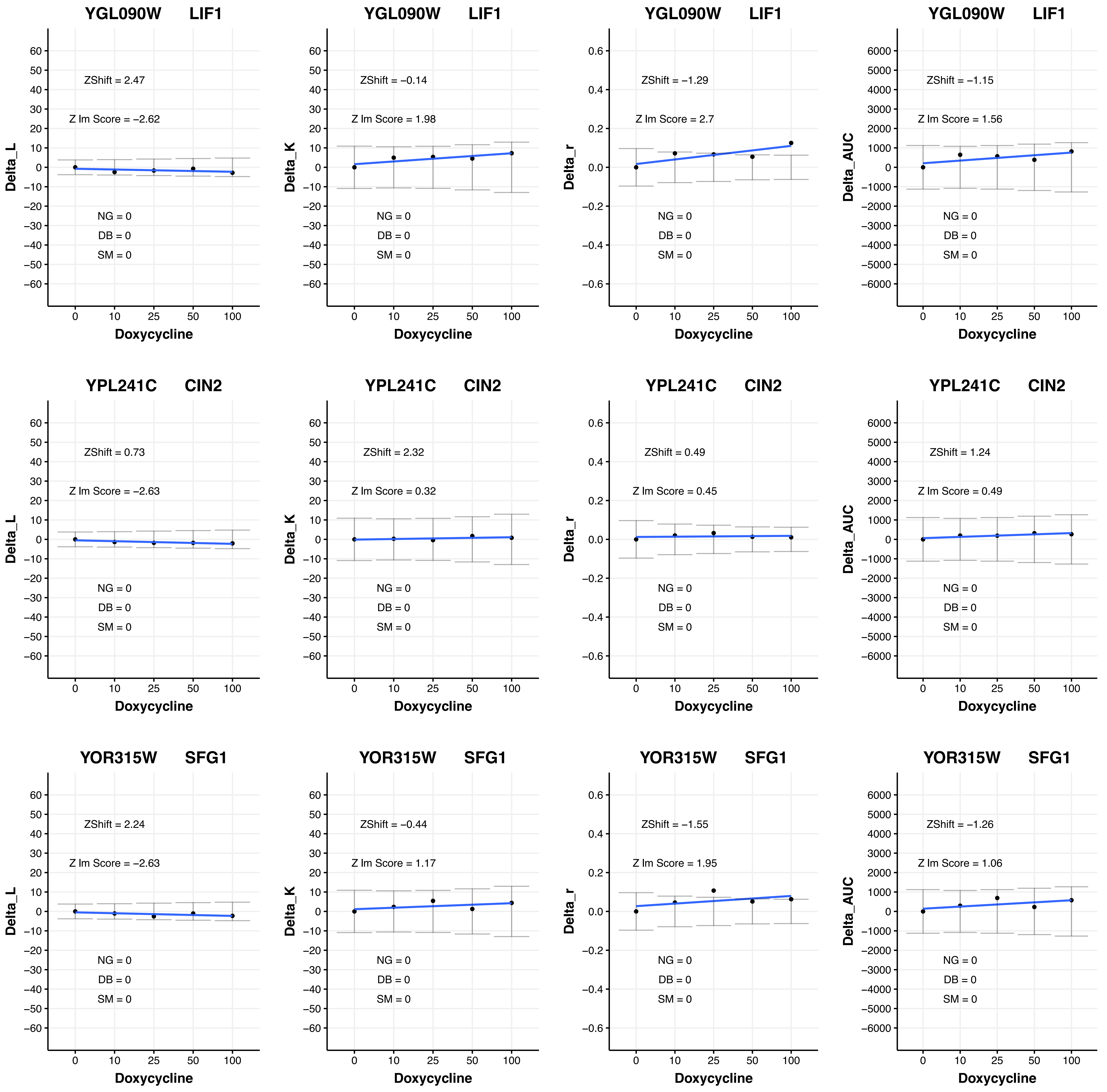

YGL090W LIF1
Delta_L
ZShift = 2.47
Z lm Score = −2.62
NG = 0
DB = 0
SM = 0
Doxycycline
YGL090W LIF1
Delta_K
ZShift = −0.14
Z lm Score = 1.98
NG = 0
DB = 0
SM = 0
Doxycycline
YGL090W LIF1
Delta_r
ZShift = −1.29
Z lm Score = 2.7
NG = 0
DB = 0
SM = 0
Doxycycline
YGL090W LIF1
Delta_AUC
ZShift = −1.15
Z lm Score = 1.56
NG = 0
DB = 0
SM = 0
Doxycycline
YPL241C CIN2
Delta_L
ZShift = 0.73
Z lm Score = −2.63
NG = 0
DB = 0
SM = 0
Doxycycline
YPL241C CIN2
Delta_K
ZShift = 2.32
Z lm Score = 0.32
NG = 0
DB = 0
SM = 0
Doxycycline
YPL241C CIN2
Delta_r
ZShift = 0.49
Z lm Score = 0.45
NG = 0
DB = 0
SM = 0
Doxycycline
YPL241C CIN2
Delta_AUC
ZShift = 1.24
Z lm Score = 0.49
NG = 0
DB = 0
SM = 0
Doxycycline
YOR315W SFG1
Delta_L
ZShift = 2.24
Z lm Score = −2.63
NG = 0
DB = 0
SM = 0
Doxycycline
YOR315W SFG1
Delta_K
ZShift = −0.44
Z lm Score = 1.17
NG = 0
DB = 0
SM = 0
Doxycycline
YOR315W SFG1
Delta_r
ZShift = −1.55
Z lm Score = 1.95
NG = 0
DB = 0
SM = 0
Doxycycline
YOR315W SFG1
Delta_AUC
ZShift = −1.26
Z lm Score = 1.06
NG = 0
DB = 0
SM = 0
Doxycycline

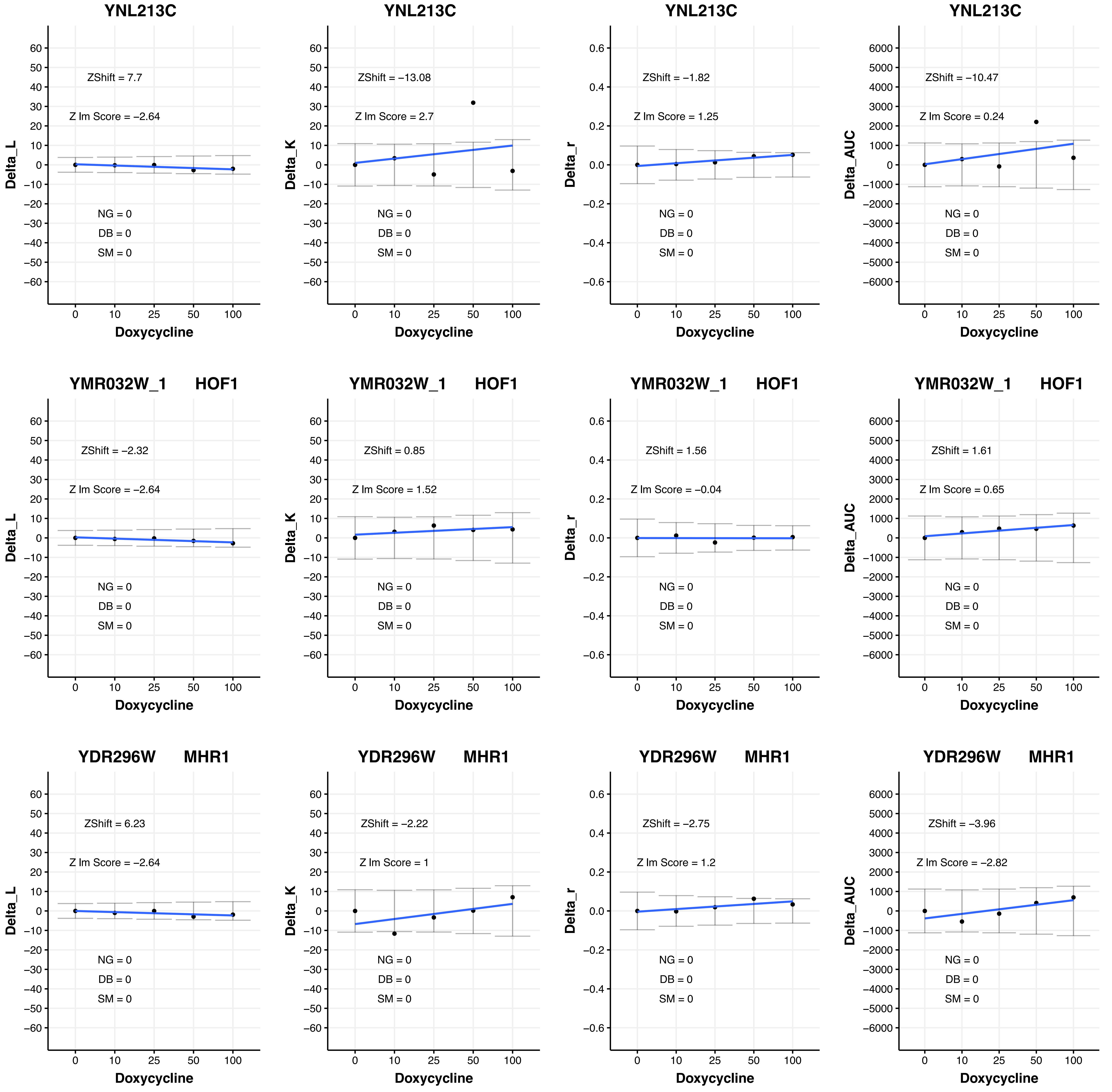

YNL213C
ZShift = −7.7
Z lm Score = −2.64
NG = 0
DB = 0
SM = 0
Delta_L
Doxycycline
YNL213C
ZShift = −13.08
Z lm Score = 2.7
NG = 0
DB = 0
SM = 0
Delta_K
Doxycycline
YNL213C
ZShift = −1.82
Z lm Score = 1.25
NG = 0
DB = 0
SM = 0
Delta_r
Doxycycline
YNL213C
ZShift = −10.47
Z lm Score = 0.24
NG = 0
DB = 0
SM = 0
Delta_AUC
Doxycycline
YMR032W_1 HOF1
ZShift = −2.32
Z lm Score = −2.64
NG = 0
DB = 0
SM = 0
Delta_L
Doxycycline
YMR032W_1 HOF1
ZShift = 0.85
Z lm Score = 1.52
NG = 0
DB = 0
SM = 0
Delta_K
Doxycycline
YMR032W_1 HOF1
ZShift = 1.56
Z lm Score = −0.04
NG = 0
DB = 0
SM = 0
Delta_r
Doxycycline
YMR032W_1 HOF1
ZShift = 1.61
Z lm Score = 0.65
NG = 0
DB = 0
SM = 0
Delta_AUC
Doxycycline
YDR296W MHR1
ZShift = 6.23
Z lm Score = −2.64
NG = 0
DB = 0
SM = 0
Delta_L
Doxycycline
YDR296W MHR1
ZShift = −2.22
Z lm Score = 1
NG = 0
DB = 0
SM = 0
Delta_K
Doxycycline
YDR296W MHR1
ZShift = −2.75
Z lm Score = 1.2
NG = 0
DB = 0
SM = 0
Delta_r
Doxycycline
YDR296W MHR1
ZShift = −3.96
Z lm Score = −2.82
NG = 0
DB = 0
SM = 0
Delta_AUC
Doxycycline

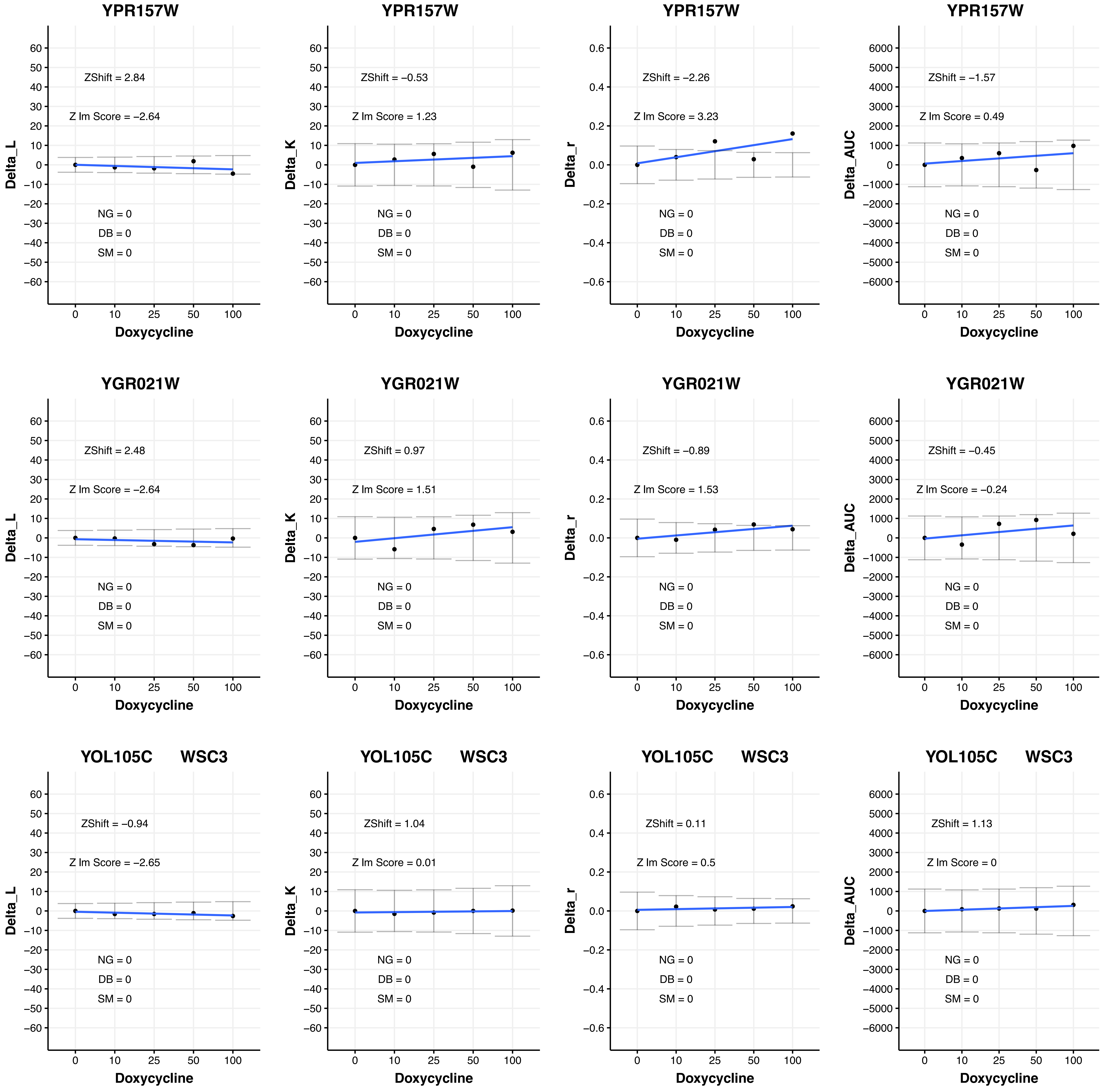
YPR157W
ZShift = 2.84
Z lm Score = −2.64
NG = 0
DB = 0
SM = 0
Delta_L
Doxycycline
YPR157W
ZShift = −0.53
Z lm Score = 1.23
NG = 0
DB = 0
SM = 0
Delta_K
Doxycycline
YPR157W
ZShift = −2.26
Z lm Score = 3.23
NG = 0
DB = 0
SM = 0
Delta_r
Doxycycline
YPR157W
ZShift = −1.57
Z lm Score = 0.49
NG = 0
DB = 0
SM = 0
Delta_AUC
Doxycycline
YGR021W
ZShift = 2.48
Z lm Score = −2.64
NG = 0
DB = 0
SM = 0
Delta_L
Doxycycline
YGR021W
ZShift = 0.97
Z lm Score = 1.51
NG = 0
DB = 0
SM = 0
Delta_K
Doxycycline
YGR021W
ZShift = −0.89
Z lm Score = 1.53
NG = 0
DB = 0
SM = 0
Delta_r
Doxycycline
YGR021W
ZShift = −0.45
Z lm Score = −0.24
NG = 0
DB = 0
SM = 0
Delta_AUC
Doxycycline
YOL105C WSC3
ZShift = −0.94
Z lm Score = −2.65
NG = 0
DB = 0
SM = 0
Delta_L
Doxycycline
YOL105C WSC3
ZShift = 1.04
Z lm Score = 0.01
NG = 0
DB = 0
SM = 0
Delta_K
Doxycycline
YOL105C WSC3
ZShift = 0.11
Z lm Score = 0.5
NG = 0
DB = 0
SM = 0
Delta_r
Doxycycline
YOL105C WSC3
ZShift = 1.13
Z lm Score = 0
NG = 0
DB = 0
SM = 0
Delta_AUC
Doxycycline

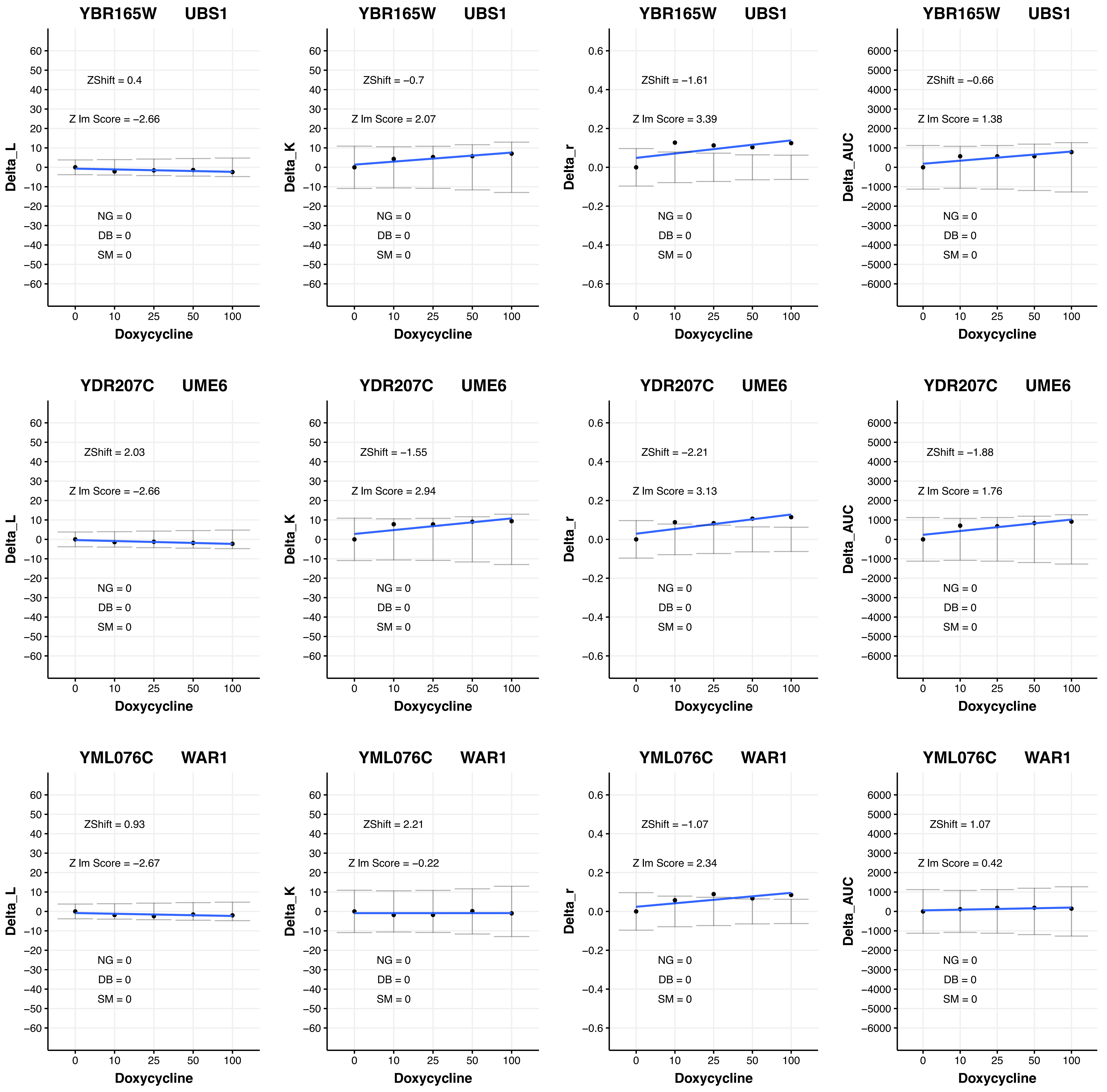
YBR165W UBS1
ZShift = 0.4
Z lm Score = −2.66
NG = 0
DB = 0
SM = 0
Delta_L
Doxycycline
YBR165W UBS1
ZShift = −0.7
Z lm Score = 2.07
NG = 0
DB = 0
SM = 0
Delta_K
Doxycycline
YBR165W UBS1
ZShift = −1.61
Z lm Score = 3.39
NG = 0
DB = 0
SM = 0
Delta_r
Doxycycline
YBR165W UBS1
ZShift = −0.66
Z lm Score = 1.38
NG = 0
DB = 0
SM = 0
Delta_AUC
Doxycycline
YDR207C UME6
ZShift = 2.03
Z lm Score = −2.66
NG = 0
DB = 0
SM = 0
Delta_L
Doxycycline
YDR207C UME6
ZShift = −1.55
Z lm Score = 2.94
NG = 0
DB = 0
SM = 0
Delta_K
Doxycycline
YDR207C UME6
ZShift = −2.21
Z lm Score = 3.13
NG = 0
DB = 0
SM = 0
Delta_r
Doxycycline
YDR207C UME6
ZShift = −1.88
Z lm Score = 1.76
NG = 0
DB = 0
SM = 0
Delta_AUC
Doxycycline
YML076C WAR1
ZShift = 0.93
Z lm Score = −2.67
NG = 0
DB = 0
SM = 0
Delta_L
Doxycycline
YML076C WAR1
ZShift = 2.21
Z lm Score = −0.22
NG = 0
DB = 0
SM = 0
Delta_K
Doxycycline
YML076C WAR1
ZShift = −1.07
Z lm Score = 2.34
NG = 0
DB = 0
SM = 0
Delta_r
Doxycycline
YML076C WAR1
ZShift = 1.07
Z lm Score = 0.42
NG = 0
DB = 0
SM = 0
Delta_AUC
Doxycycline

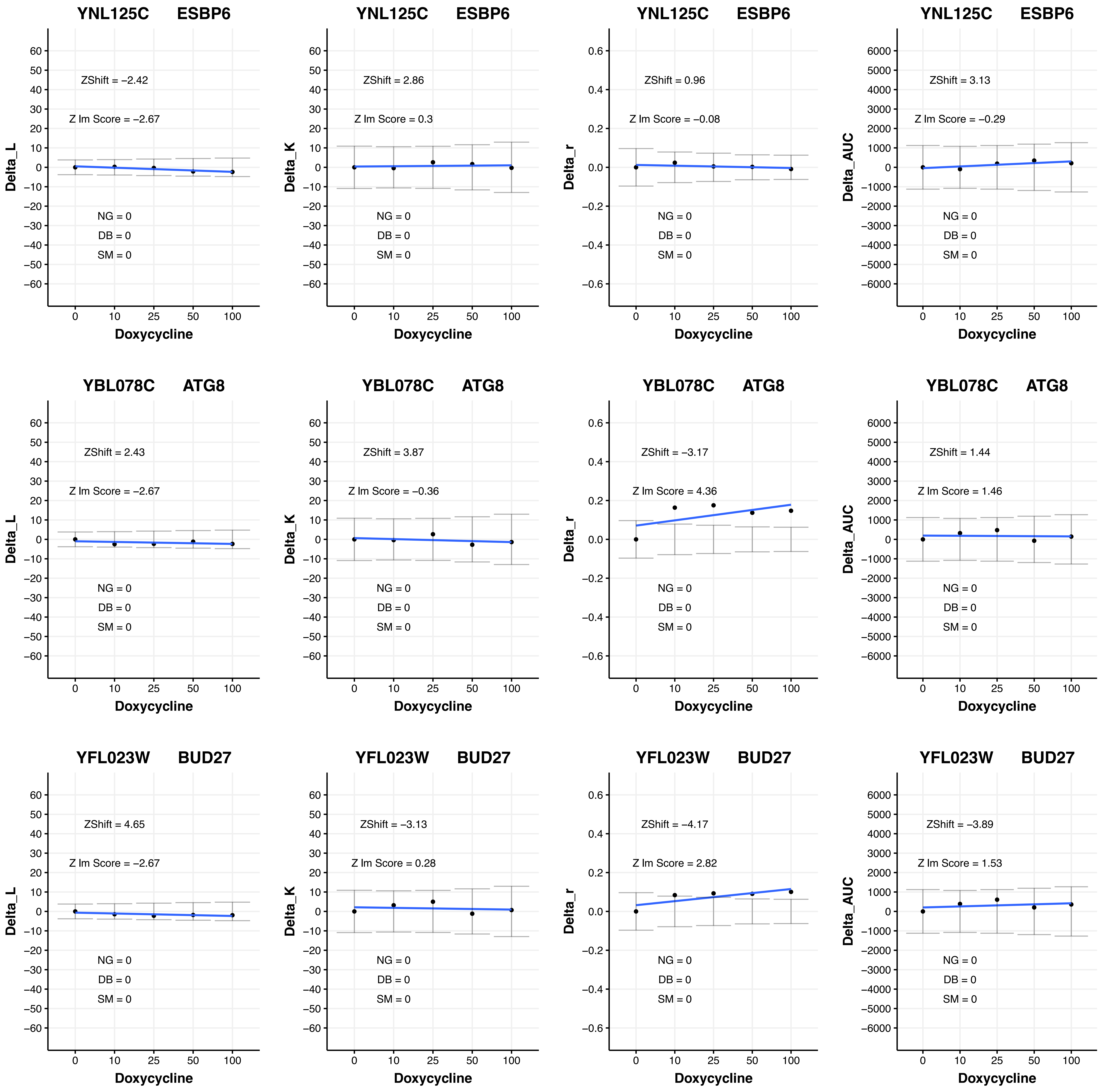

YNL125C ESBP6
ZShift = −2.42
Z Im Score = −2.67
NG = 0
DB = 0
SM = 0
Delta_L
Doxycycline
YNL125C ESBP6
ZShift = 2.86
Z Im Score = 0.3
NG = 0
DB = 0
SM = 0
Delta_K
Doxycycline
YNL125C ESBP6
ZShift = 0.96
Z Im Score = −0.08
NG = 0
DB = 0
SM = 0
Delta_r
Doxycycline
YNL125C ESBP6
ZShift = 3.13
Z Im Score = −0.29
NG = 0
DB = 0
SM = 0
Delta_AUC
Doxycycline
YBL078C ATG8
ZShift = 2.43
Z Im Score = −2.67
NG = 0
DB = 0
SM = 0
Delta_L
Doxycycline
YBL078C ATG8
ZShift = 3.87
Z Im Score = −0.36
NG = 0
DB = 0
SM = 0
Delta_K
Doxycycline
YBL078C ATG8
ZShift = −3.17
Z Im Score = 4.36
NG = 0
DB = 0
SM = 0
Delta_r
Doxycycline
YBL078C ATG8
ZShift = 1.44
Z Im Score = 1.46
NG = 0
DB = 0
SM = 0
Delta_AUC
Doxycycline
YFL023W BUD27
ZShift = 4.65
Z Im Score = −2.67
NG = 0
DB = 0
SM = 0
Delta_L
Doxycycline
YFL023W BUD27
ZShift = −3.13
Z Im Score = 0.28
NG = 0
DB = 0
SM = 0
Delta_K
Doxycycline
YFL023W BUD27
ZShift = −4.17
Z Im Score = 2.82
NG = 0
DB = 0
SM = 0
Delta_r
Doxycycline
YFL023W BUD27
ZShift = −3.89
Z Im Score = 1.53
NG = 0
DB = 0
SM = 0
Delta_AUC
Doxycycline
0
10
25
50
100

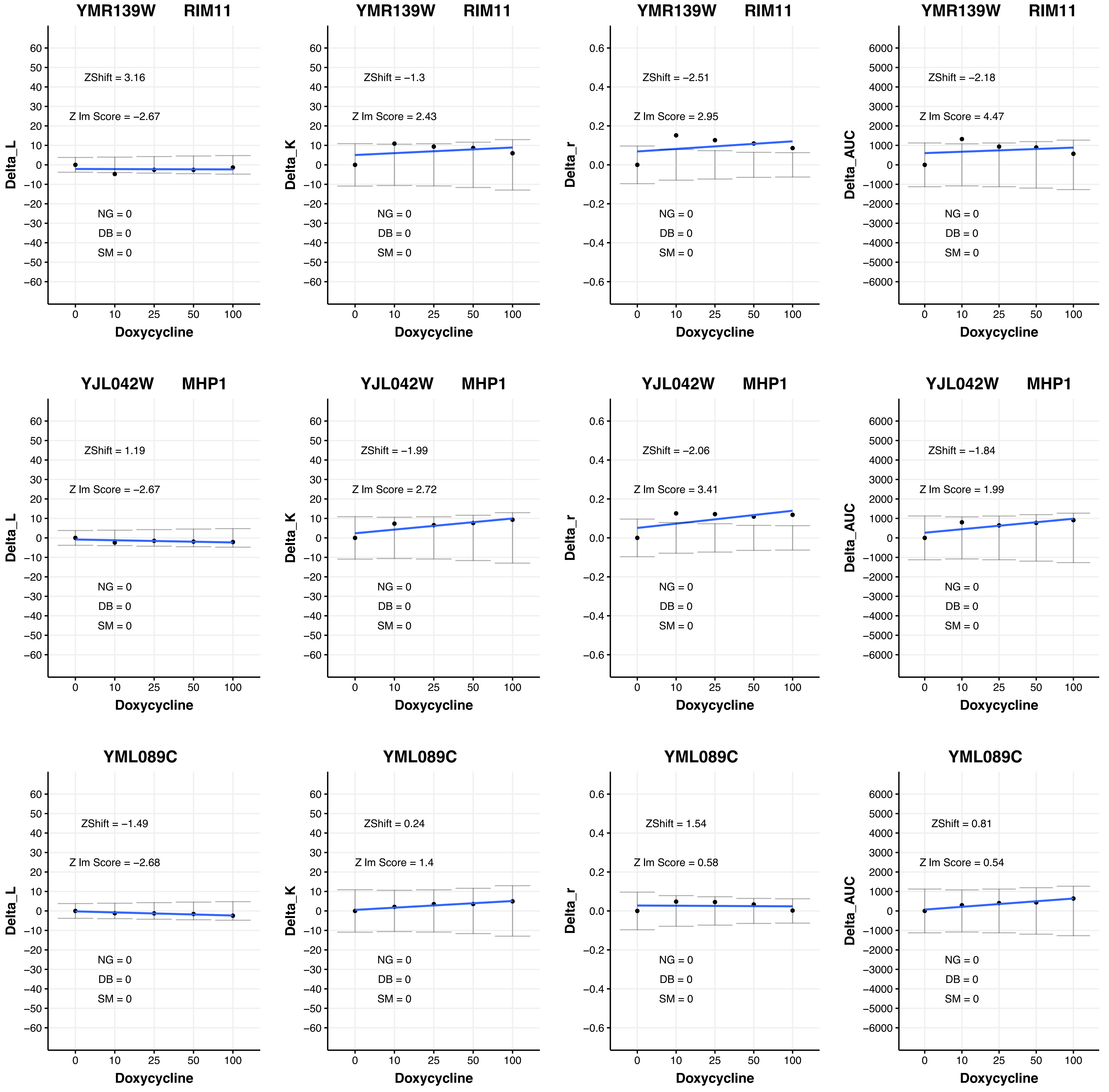
YMR139W RIM11
ZShift = 3.16
Z lm Score = −2.67
NG = 0
DB = 0
SM = 0
Delta_L
Doxycycline
YMR139W RIM11
ZShift = −1.3
Z lm Score = 2.43
NG = 0
DB = 0
SM = 0
Delta_K
Doxycycline
YMR139W RIM11
ZShift = −2.51
Z lm Score = 2.95
NG = 0
DB = 0
SM = 0
Delta_r
Doxycycline
YMR139W RIM11
ZShift = −2.18
Z lm Score = 4.47
NG = 0
DB = 0
SM = 0
Delta_AUC
Doxycycline
YJL042W MHP1
ZShift = 1.19
Z lm Score = −2.67
NG = 0
DB = 0
SM = 0
Delta_L
Doxycycline
YJL042W MHP1
ZShift = −1.99
Z lm Score = 2.72
NG = 0
DB = 0
SM = 0
Delta_K
Doxycycline
YJL042W MHP1
ZShift = −2.06
Z lm Score = 3.41
NG = 0
DB = 0
SM = 0
Delta_r
Doxycycline
YJL042W MHP1
ZShift = −1.84
Z lm Score = 1.99
NG = 0
DB = 0
SM = 0
Delta_AUC
Doxycycline
YML089C
ZShift = −1.49
Z lm Score = −2.68
NG = 0
DB = 0
SM = 0
Delta_L
Doxycycline
YML089C
ZShift = 0.24
Z lm Score = 1.4
NG = 0
DB = 0
SM = 0
Delta_K
Doxycycline
YML089C
ZShift = 1.54
Z lm Score = 0.58
NG = 0
DB = 0
SM = 0
Delta_r
Doxycycline
YML089C
ZShift = 0.81
Z lm Score = 0.54
NG = 0
DB = 0
SM = 0
Delta_AUC
Doxycycline

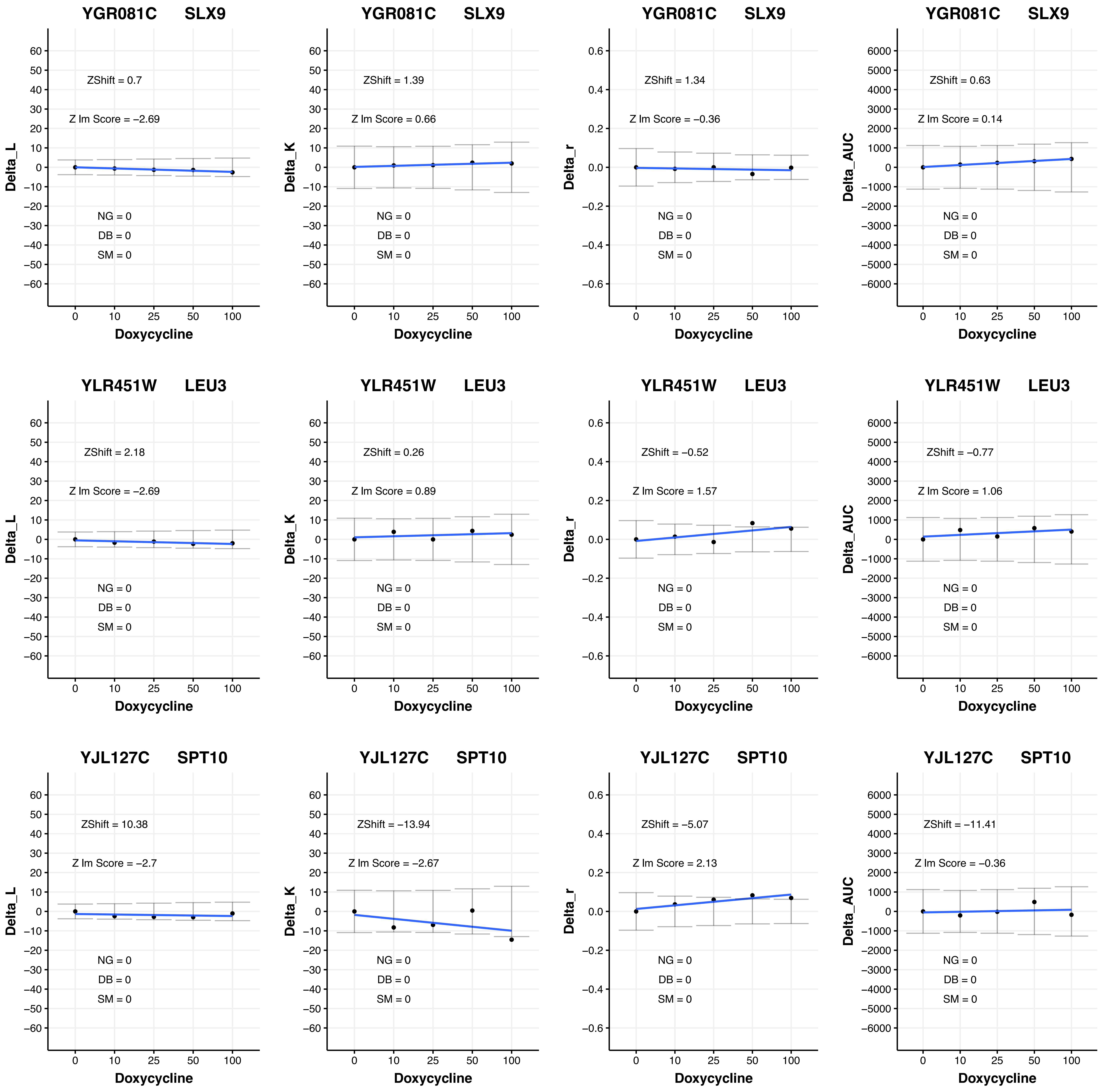
YGR081C SLX9
ZShift = 0.7
Z lm Score = −2.69
NG = 0
DB = 0
SM = 0
Delta_L
Doxycycline
YGR081C SLX9
ZShift = 1.39
Z lm Score = 0.66
NG = 0
DB = 0
SM = 0
Delta_K
Doxycycline
YGR081C SLX9
ZShift = 1.34
Z lm Score = −0.36
NG = 0
DB = 0
SM = 0
Delta_r
Doxycycline
YGR081C SLX9
ZShift = 0.63
Z lm Score = 0.14
NG = 0
DB = 0
SM = 0
Delta_AUC
Doxycycline
YLR451W LEU3
ZShift = 2.18
Z lm Score = −2.69
NG = 0
DB = 0
SM = 0
Delta_L
Doxycycline
YLR451W LEU3
ZShift = 0.26
Z lm Score = 0.89
NG = 0
DB = 0
SM = 0
Delta_K
Doxycycline
YLR451W LEU3
ZShift = −0.52
Z lm Score = 1.57
NG = 0
DB = 0
SM = 0
Delta_r
Doxycycline
YLR451W LEU3
ZShift = −0.77
Z lm Score = 1.06
NG = 0
DB = 0
SM = 0
Delta_AUC
Doxycycline
YJL127C SPT10
ZShift = 10.38
Z lm Score = −2.7
NG = 0
DB = 0
SM = 0
Delta_L
Doxycycline
YJL127C SPT10
ZShift = −13.94
Z lm Score = −2.67
NG = 0
DB = 0
SM = 0
Delta_K
Doxycycline
YJL127C SPT10
ZShift = −5.07
Z lm Score = 2.13
NG = 0
DB = 0
SM = 0
Delta_r
Doxycycline
YJL127C SPT10
ZShift = −11.41
Z lm Score = −0.36
NG = 0
DB = 0
SM = 0
Delta_AUC
Doxycycline

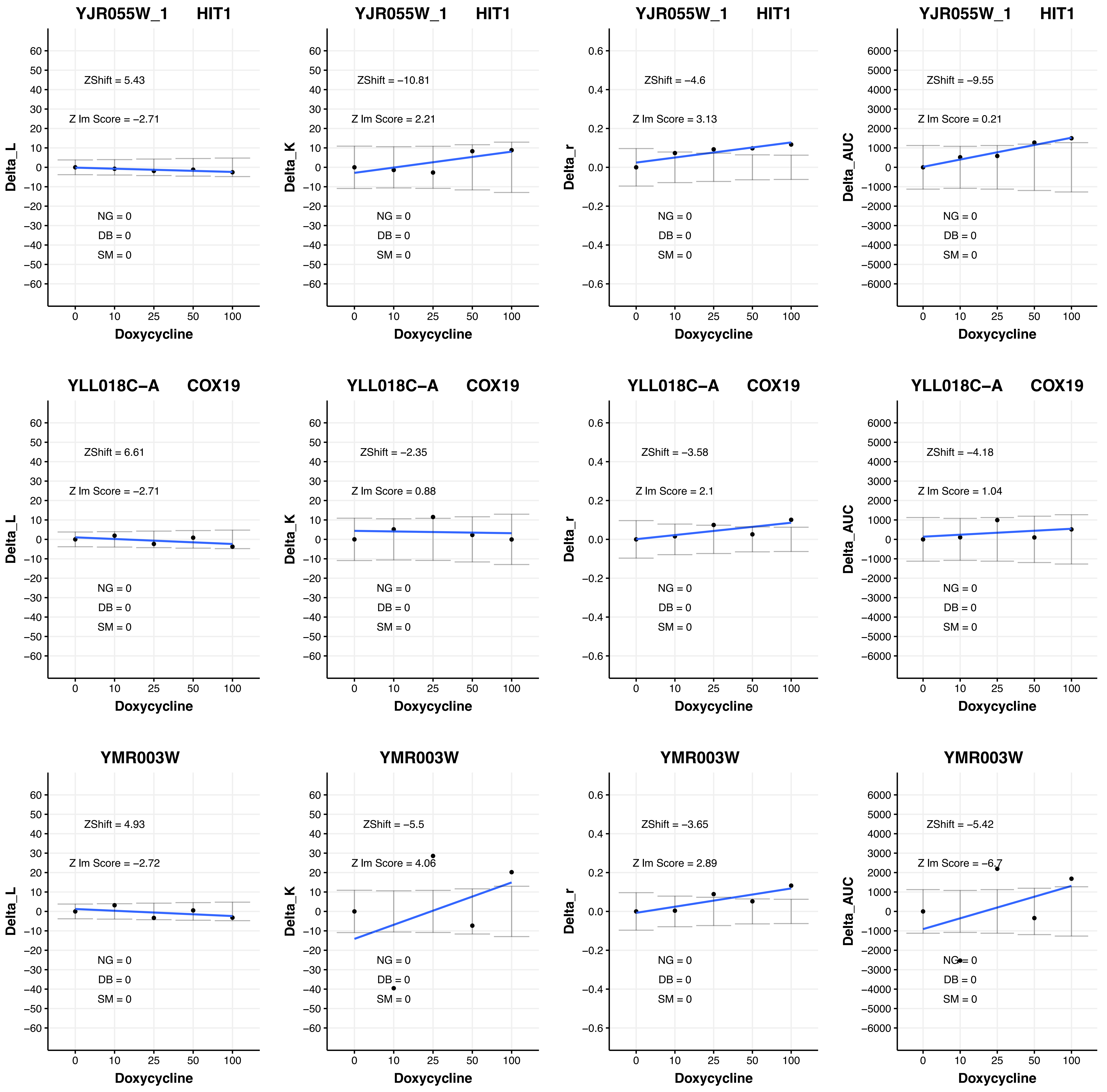
YJR055W_1 HIT1
ZShift = 5.43
Z lm Score = −2.71
NG = 0
DB = 0
SM = 0
Delta_L
Doxycycline
YJR055W_1 HIT1
ZShift = −10.81
Z lm Score = 2.21
NG = 0
DB = 0
SM = 0
Delta_K
Doxycycline
YJR055W_1 HIT1
ZShift = −4.6
Z lm Score = 3.13
NG = 0
DB = 0
SM = 0
Delta_r
Doxycycline
YJR055W_1 HIT1
ZShift = −9.55
Z lm Score = 0.21
NG = 0
DB = 0
SM = 0
Delta_AUC
Doxycycline
YLL018C−A COX19
ZShift = 6.61
Z lm Score = −2.71
NG = 0
DB = 0
SM = 0
Delta_L
Doxycycline
YLL018C−A COX19
ZShift = −2.35
Z lm Score = 0.88
NG = 0
DB = 0
SM = 0
Delta_K
Doxycycline
YLL018C−A COX19
ZShift = −3.58
Z lm Score = 2.1
NG = 0
DB = 0
SM = 0
Delta_r
Doxycycline
YLL018C−A COX19
ZShift = −4.18
Z lm Score = 1.04
NG = 0
DB = 0
SM = 0
Delta_AUC
Doxycycline
YMR003W
ZShift = 4.93
Z lm Score = −2.72
NG = 0
DB = 0
SM = 0
Delta_L
Doxycycline
YMR003W
ZShift = −5.5
Z lm Score = 4.06
NG = 0
DB = 0
SM = 0
Delta_K
Doxycycline
YMR003W
ZShift = −3.65
Z lm Score = 2.89
NG = 0
DB = 0
SM = 0
Delta_r
Doxycycline
YMR003W
ZShift = −5.42
Z lm Score = −6.7
NG = 0
DB = 0
SM = 0
Delta_AUC
Doxycycline

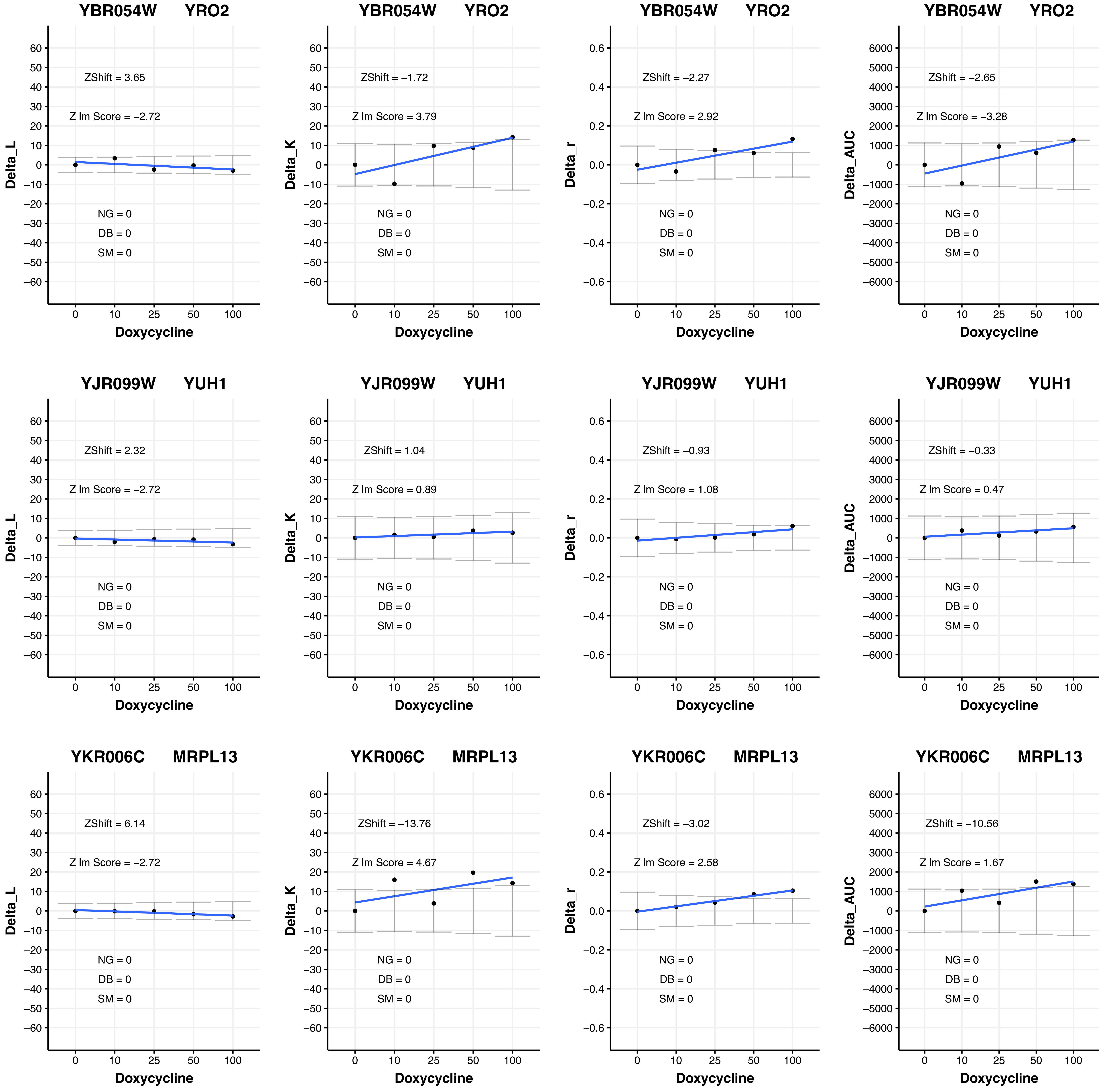
YBR054W YRO2
ZShift = 3.65
Z lm Score = −2.72
NG = 0
DB = 0
SM = 0
Delta_L
Doxycycline
YBR054W YRO2
ZShift = −1.72
Z lm Score = 3.79
NG = 0
DB = 0
SM = 0
Delta_K
Doxycycline
YBR054W YRO2
ZShift = −2.27
Z lm Score = 2.92
NG = 0
DB = 0
SM = 0
Delta_r
Doxycycline
YBR054W YRO2
ZShift = −2.65
Z lm Score = −3.28
NG = 0
DB = 0
SM = 0
Delta_AUC
Doxycycline
YJR099W YUH1
ZShift = 2.32
Z lm Score = −2.72
NG = 0
DB = 0
SM = 0
Delta_L
Doxycycline
YJR099W YUH1
ZShift = 1.04
Z lm Score = 0.89
NG = 0
DB = 0
SM = 0
Delta_K
Doxycycline
YJR099W YUH1
ZShift = −0.93
Z lm Score = 1.08
NG = 0
DB = 0
SM = 0
Delta_r
Doxycycline
YJR099W YUH1
ZShift = −0.33
Z lm Score = 0.47
NG = 0
DB = 0
SM = 0
Delta_AUC
Doxycycline
YKR006C MRPL13
ZShift = 6.14
Z lm Score = −2.72
NG = 0
DB = 0
SM = 0
Delta_L
Doxycycline
YKR006C MRPL13
ZShift = −13.76
Z lm Score = 4.67
NG = 0
DB = 0
SM = 0
Delta_K
Doxycycline
YKR006C MRPL13
ZShift = −3.02
Z lm Score = 2.58
NG = 0
DB = 0
SM = 0
Delta_r
Doxycycline
YKR006C MRPL13
ZShift = −10.56
Z lm Score = 1.67
NG = 0
DB = 0
SM = 0
Delta_AUC
Doxycycline

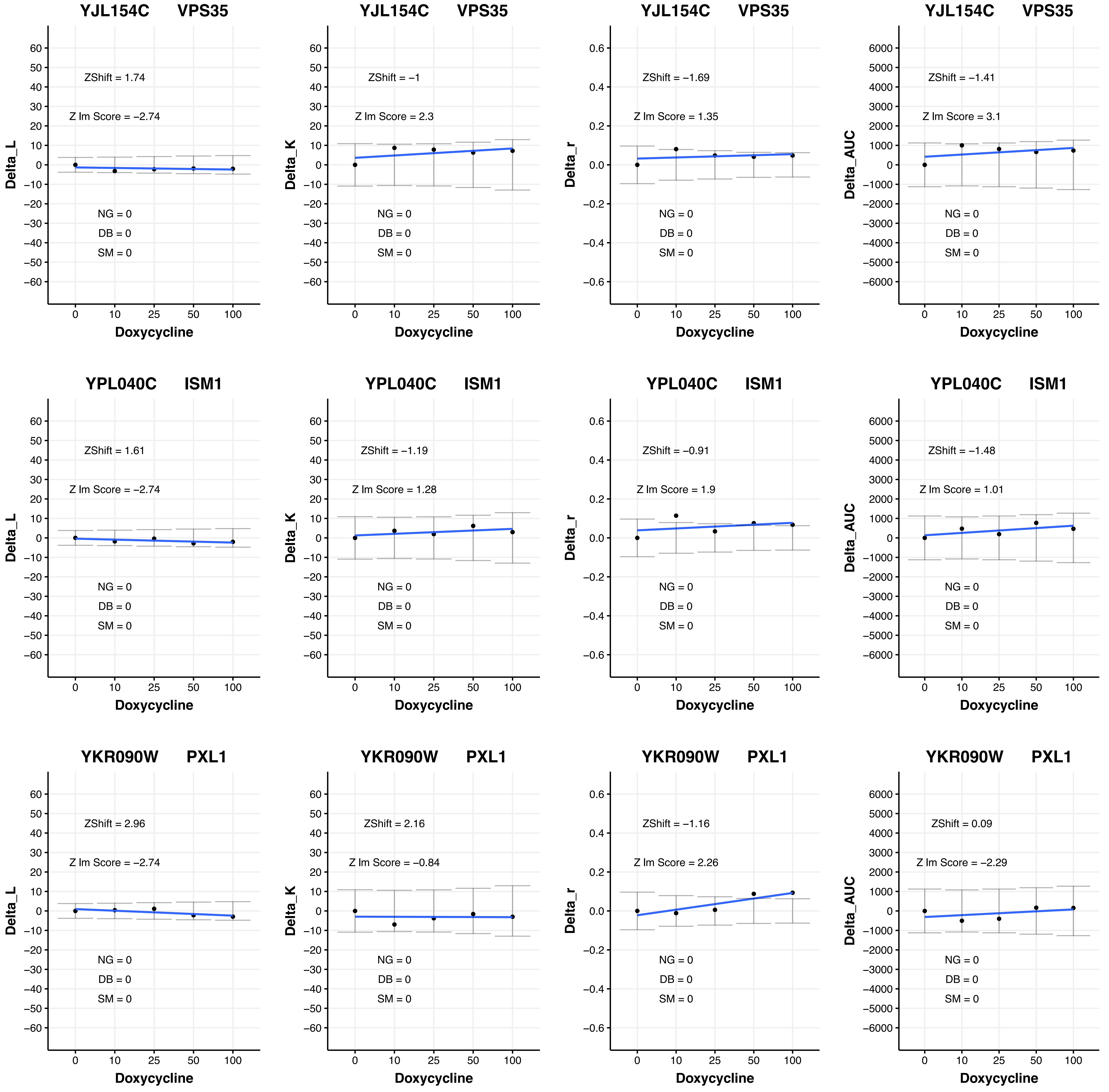

YJL154C VPS35
ZShift = 1.74
Z lm Score = −2.74
NG = 0
DB = 0
SM = 0
Delta_L
Doxycycline
YJL154C VPS35
ZShift = −1
Z lm Score = 2.3
NG = 0
DB = 0
SM = 0
Delta_K
Doxycycline
YJL154C VPS35
ZShift = −1.69
Z lm Score = 1.35
NG = 0
DB = 0
SM = 0
Delta_r
Doxycycline
YJL154C VPS35
ZShift = −1.41
Z lm Score = 3.1
NG = 0
DB = 0
SM = 0
Delta_AUC
Doxycycline
YPL040C ISM1
ZShift = 1.61
Z lm Score = −2.74
NG = 0
DB = 0
SM = 0
Delta_L
Doxycycline
YPL040C ISM1
ZShift = −1.19
Z lm Score = 1.28
NG = 0
DB = 0
SM = 0
Delta_K
Doxycycline
YPL040C ISM1
ZShift = −0.91
Z lm Score = 1.9
NG = 0
DB = 0
SM = 0
Delta_r
Doxycycline
YPL040C ISM1
ZShift = −1.48
Z lm Score = 1.01
NG = 0
DB = 0
SM = 0
Delta_AUC
Doxycycline
YKR090W PXL1
ZShift = 2.96
Z lm Score = −2.74
NG = 0
DB = 0
SM = 0
Delta_L
Doxycycline
YKR090W PXL1
ZShift = 2.16
Z lm Score = −0.84
NG = 0
DB = 0
SM = 0
Delta_K
Doxycycline
YKR090W PXL1
ZShift = −1.16
Z lm Score = 2.26
NG = 0
DB = 0
SM = 0
Delta_r
Doxycycline
YKR090W PXL1
ZShift = 0.09
Z lm Score = −2.29
NG = 0
DB = 0
SM = 0
Delta_AUC
Doxycycline

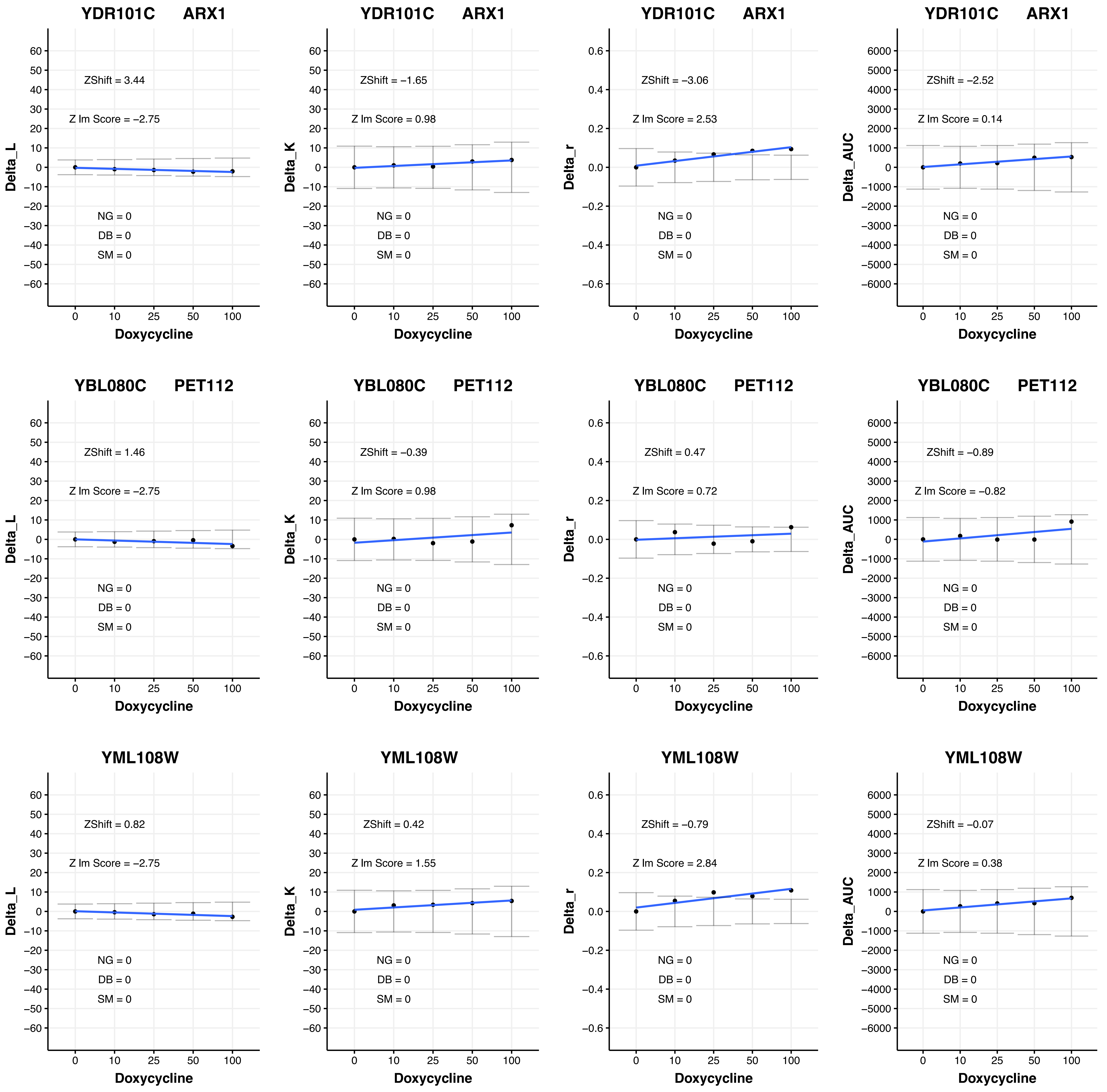

YDR101C ARX1
ZShift = 3.44
Z lm Score = −2.75
NG = 0
DB = 0
SM = 0
Delta_L
Doxycycline
YDR101C ARX1
ZShift = −1.65
Z lm Score = 0.98
NG = 0
DB = 0
SM = 0
Delta_K
Doxycycline
YDR101C ARX1
ZShift = −3.06
Z lm Score = 2.53
NG = 0
DB = 0
SM = 0
Delta_r
Doxycycline
YDR101C ARX1
ZShift = −2.52
Z lm Score = 0.14
NG = 0
DB = 0
SM = 0
Delta_AUC
Doxycycline
YBL080C PET112
ZShift = 1.46
Z lm Score = −2.75
NG = 0
DB = 0
SM = 0
Delta_L
Doxycycline
YBL080C PET112
ZShift = −0.39
Z lm Score = 0.98
NG = 0
DB = 0
SM = 0
Delta_K
Doxycycline
YBL080C PET112
ZShift = 0.47
Z lm Score = 0.72
NG = 0
DB = 0
SM = 0
Delta_r
Doxycycline
YBL080C PET112
ZShift = −0.89
Z lm Score = −0.82
NG = 0
DB = 0
SM = 0
Delta_AUC
Doxycycline
YML108W
ZShift = 0.82
Z lm Score = −2.75
NG = 0
DB = 0
SM = 0
Delta_L
Doxycycline
YML108W
ZShift = 0.42
Z lm Score = 1.55
NG = 0
DB = 0
SM = 0
Delta_K
Doxycycline
YML108W
ZShift = −0.79
Z lm Score = 2.84
NG = 0
DB = 0
SM = 0
Delta_r
Doxycycline
YML108W
ZShift = −0.07
Z lm Score = 0.38
NG = 0
DB = 0
SM = 0
Delta_AUC
Doxycycline

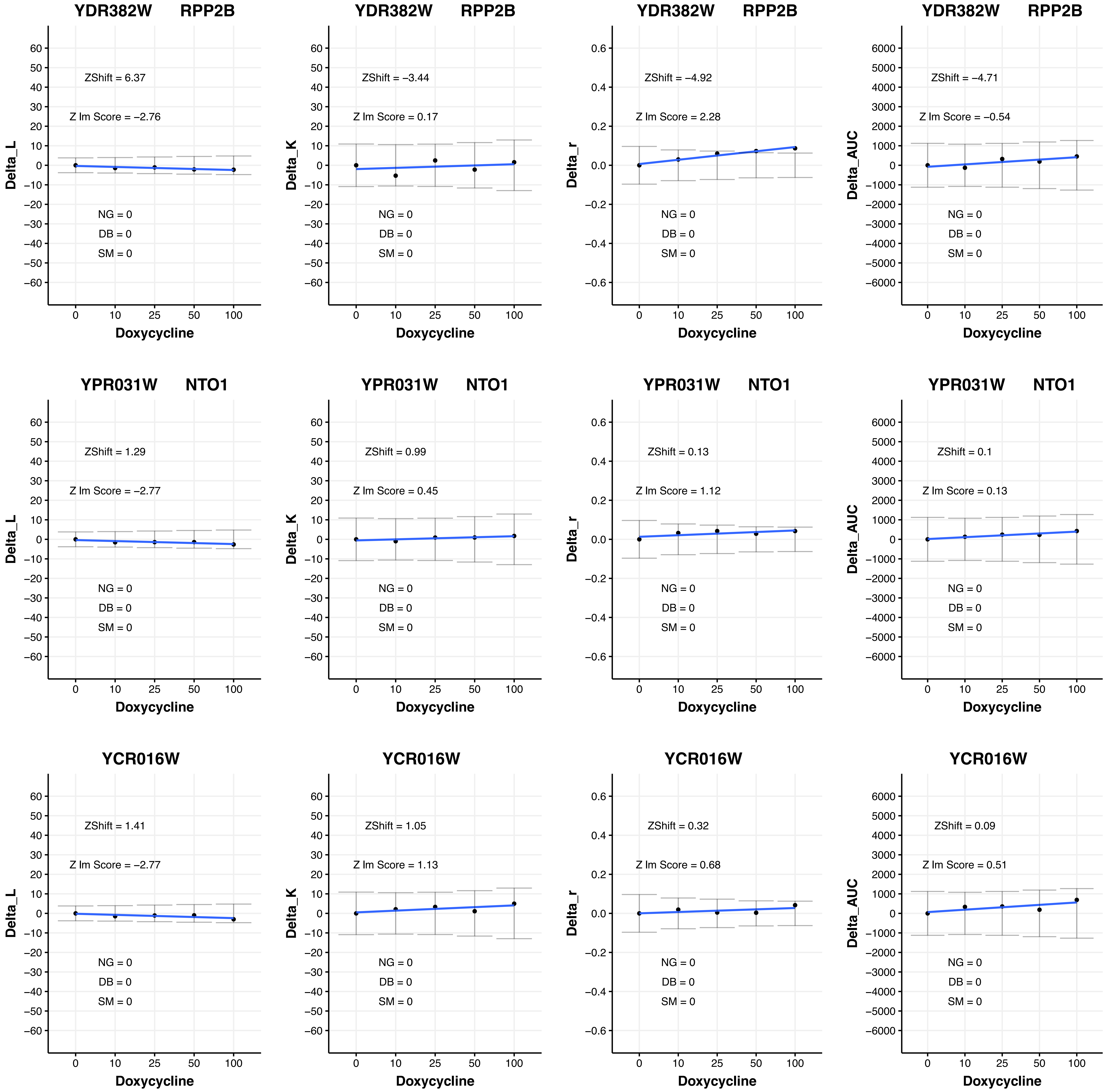

YDR382W RPP2B
ZShift = 6.37
Z lm Score = −2.76
NG = 0
DB = 0
SM = 0
Delta_L
Doxycycline
YDR382W RPP2B
ZShift = −3.44
Z lm Score = 0.17
NG = 0
DB = 0
SM = 0
Delta_K
Doxycycline
YDR382W RPP2B
ZShift = −4.92
Z lm Score = 2.28
NG = 0
DB = 0
SM = 0
Delta_r
Doxycycline
YDR382W RPP2B
ZShift = −4.71
Z lm Score = −0.54
NG = 0
DB = 0
SM = 0
Delta_AUC
Doxycycline
YPR031W NTO1
ZShift = 1.29
Z lm Score = −2.77
NG = 0
DB = 0
SM = 0
Delta_L
Doxycycline
YPR031W NTO1
ZShift = 0.99
Z lm Score = 0.45
NG = 0
DB = 0
SM = 0
Delta_K
Doxycycline
YPR031W NTO1
ZShift = 0.13
Z lm Score = 1.12
NG = 0
DB = 0
SM = 0
Delta_r
Doxycycline
YPR031W NTO1
ZShift = 0.1
Z lm Score = 0.13
NG = 0
DB = 0
SM = 0
Delta_AUC
Doxycycline
YCR016W
ZShift = 1.41
Z lm Score = −2.77
NG = 0
DB = 0
SM = 0
Delta_L
Doxycycline
YCR016W
ZShift = 1.05
Z lm Score = 1.13
NG = 0
DB = 0
SM = 0
Delta_K
Doxycycline
YCR016W
ZShift = 0.32
Z lm Score = 0.68
NG = 0
DB = 0
SM = 0
Delta_r
Doxycycline
YCR016W
ZShift = 0.09
Z lm Score = 0.51
NG = 0
DB = 0
SM = 0
Delta_AUC
Doxycycline

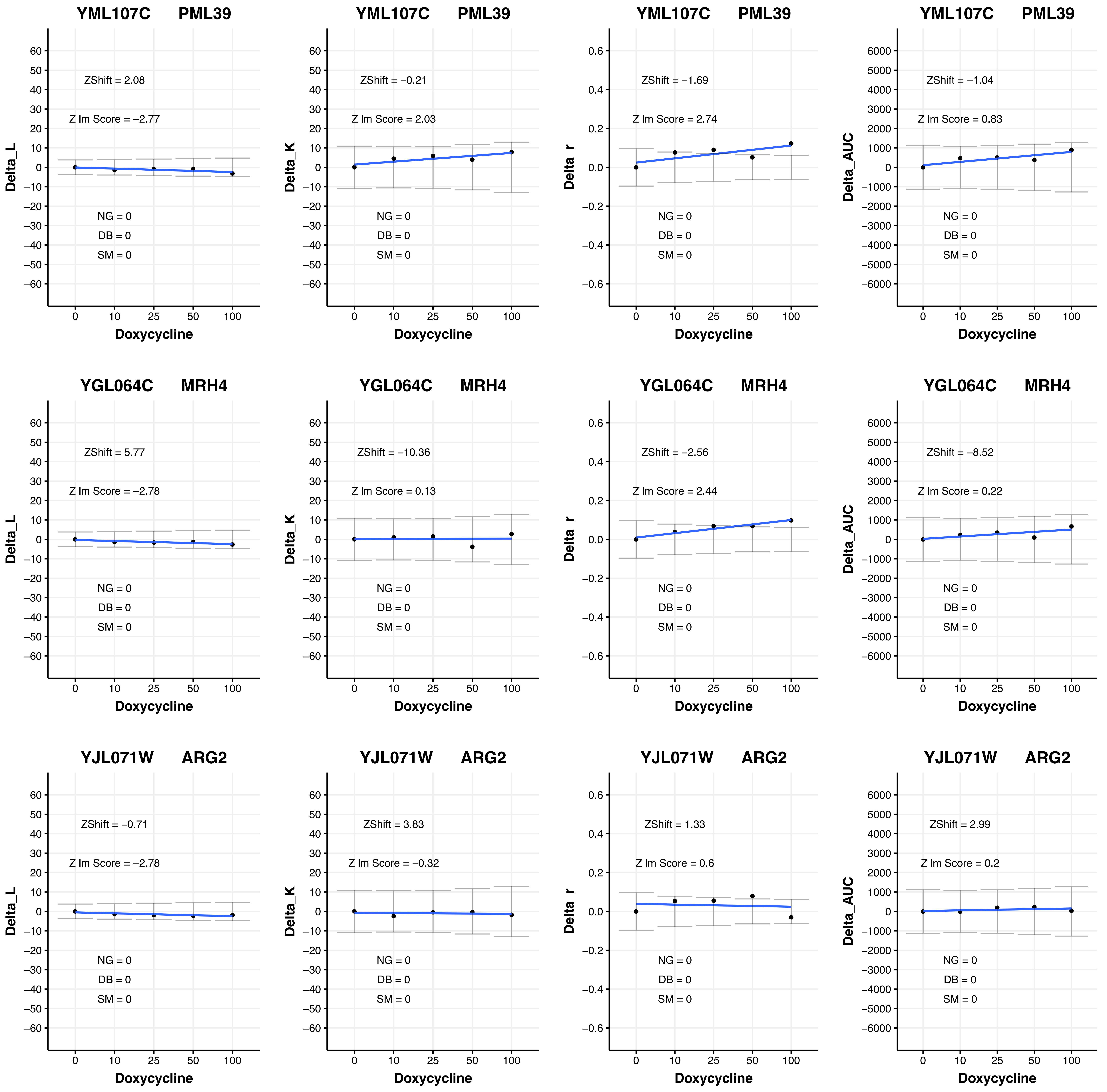

YML107C PML39
ZShift = 2.08
Z lm Score = −2.77
NG = 0
DB = 0
SM = 0
Delta_L
Doxycycline
YML107C PML39
ZShift = −0.21
Z lm Score = 2.03
NG = 0
DB = 0
SM = 0
Delta_K
Doxycycline
YML107C PML39
ZShift = −1.69
Z lm Score = 2.74
NG = 0
DB = 0
SM = 0
Delta_r
Doxycycline
YML107C PML39
ZShift = −1.04
Z lm Score = 0.83
NG = 0
DB = 0
SM = 0
Delta_AUC
Doxycycline
YGL064C MRH4
ZShift = 5.77
Z lm Score = −2.78
NG = 0
DB = 0
SM = 0
Delta_L
Doxycycline
YGL064C MRH4
ZShift = −10.36
Z lm Score = 0.13
NG = 0
DB = 0
SM = 0
Delta_K
Doxycycline
YGL064C MRH4
ZShift = −2.56
Z lm Score = 2.44
NG = 0
DB = 0
SM = 0
Delta_r
Doxycycline
YGL064C MRH4
ZShift = −8.52
Z lm Score = 0.22
NG = 0
DB = 0
SM = 0
Delta_AUC
Doxycycline
YJL071W ARG2
ZShift = −0.71
Z lm Score = −2.78
NG = 0
DB = 0
SM = 0
Delta_L
Doxycycline
YJL071W ARG2
ZShift = 3.83
Z lm Score = −0.32
NG = 0
DB = 0
SM = 0
Delta_K
Doxycycline
YJL071W ARG2
ZShift = 1.33
Z lm Score = 0.6
NG = 0
DB = 0
SM = 0
Delta_r
Doxycycline
YJL071W ARG2
ZShift = 2.99
Z lm Score = 0.2
NG = 0
DB = 0
SM = 0
Delta_AUC
Doxycycline

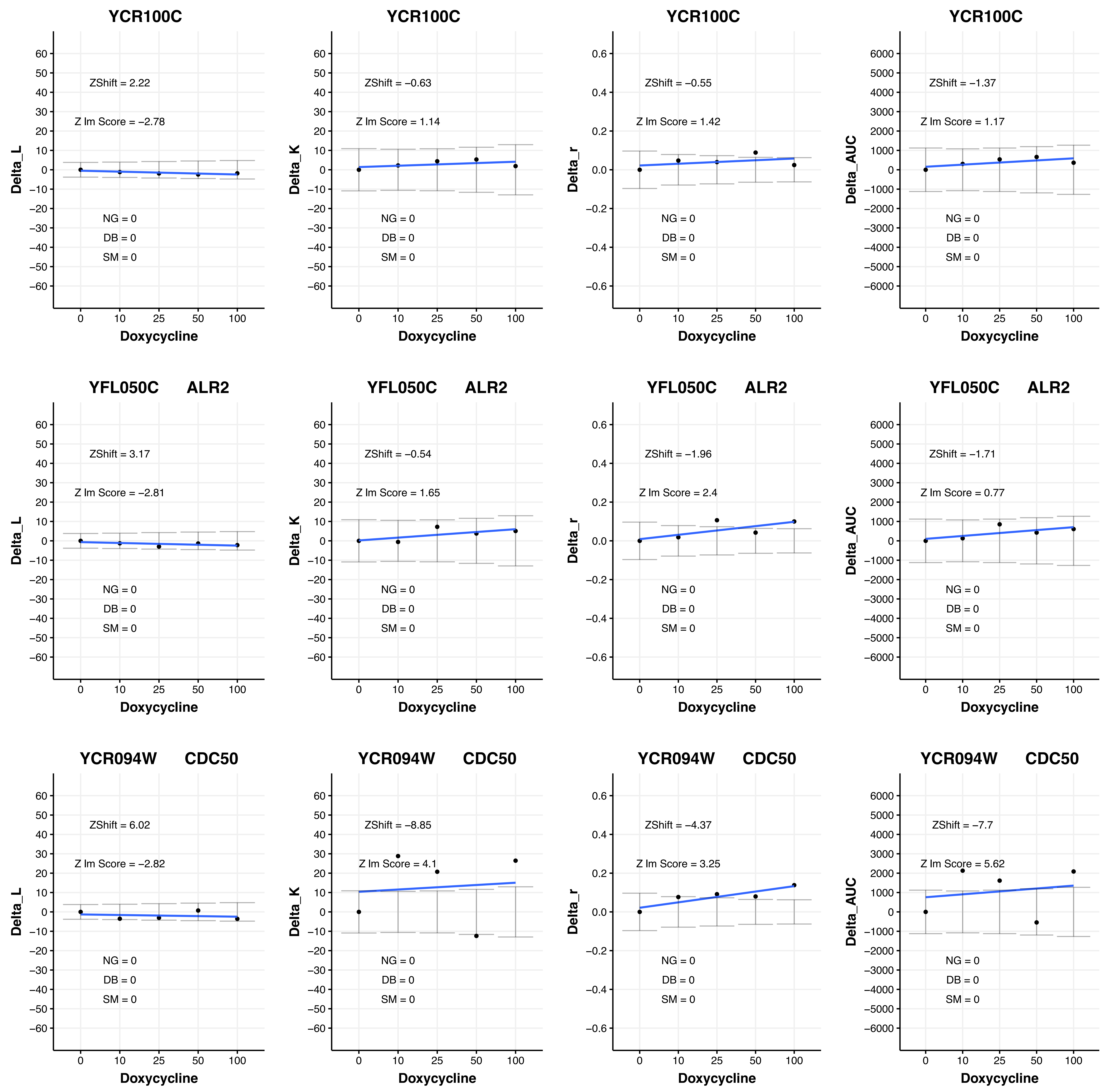

YCR100C
Delta_L
ZShift = 2.22
Z lm Score = −2.78
NG = 0
DB = 0
SM = 0
Doxycycline
YCR100C
Delta_K
ZShift = −0.63
Z lm Score = 1.14
NG = 0
DB = 0
SM = 0
Doxycycline
YCR100C
Delta_r
ZShift = −0.55
Z lm Score = 1.42
NG = 0
DB = 0
SM = 0
Doxycycline
YCR100C
Delta_AUC
ZShift = −1.37
Z lm Score = 1.17
NG = 0
DB = 0
SM = 0
Doxycycline
YFL050C ALR2
Delta_L
ZShift = 3.17
Z lm Score = −2.81
NG = 0
DB = 0
SM = 0
Doxycycline
YFL050C ALR2
Delta_K
ZShift = −0.54
Z lm Score = 1.65
NG = 0
DB = 0
SM = 0
Doxycycline
YFL050C ALR2
Delta_r
ZShift = −1.96
Z lm Score = 2.4
NG = 0
DB = 0
SM = 0
Doxycycline
YFL050C ALR2
Delta_AUC
ZShift = −1.71
Z lm Score = 0.77
NG = 0
DB = 0
SM = 0
Doxycycline
YCR094W CDC50
Delta_L
ZShift = 6.02
Z lm Score = −2.82
NG = 0
DB = 0
SM = 0
Doxycycline
YCR094W CDC50
Delta_K
ZShift = −8.85
Z lm Score = 4.1
NG = 0
DB = 0
SM = 0
Doxycycline
YCR094W CDC50
Delta_r
ZShift = −4.37
Z lm Score = 3.25
NG = 0
DB = 0
SM = 0
Doxycycline
YCR094W CDC50
Delta_AUC
ZShift = −7.7
Z lm Score = 5.62
NG = 0
DB = 0
SM = 0
Doxycycline

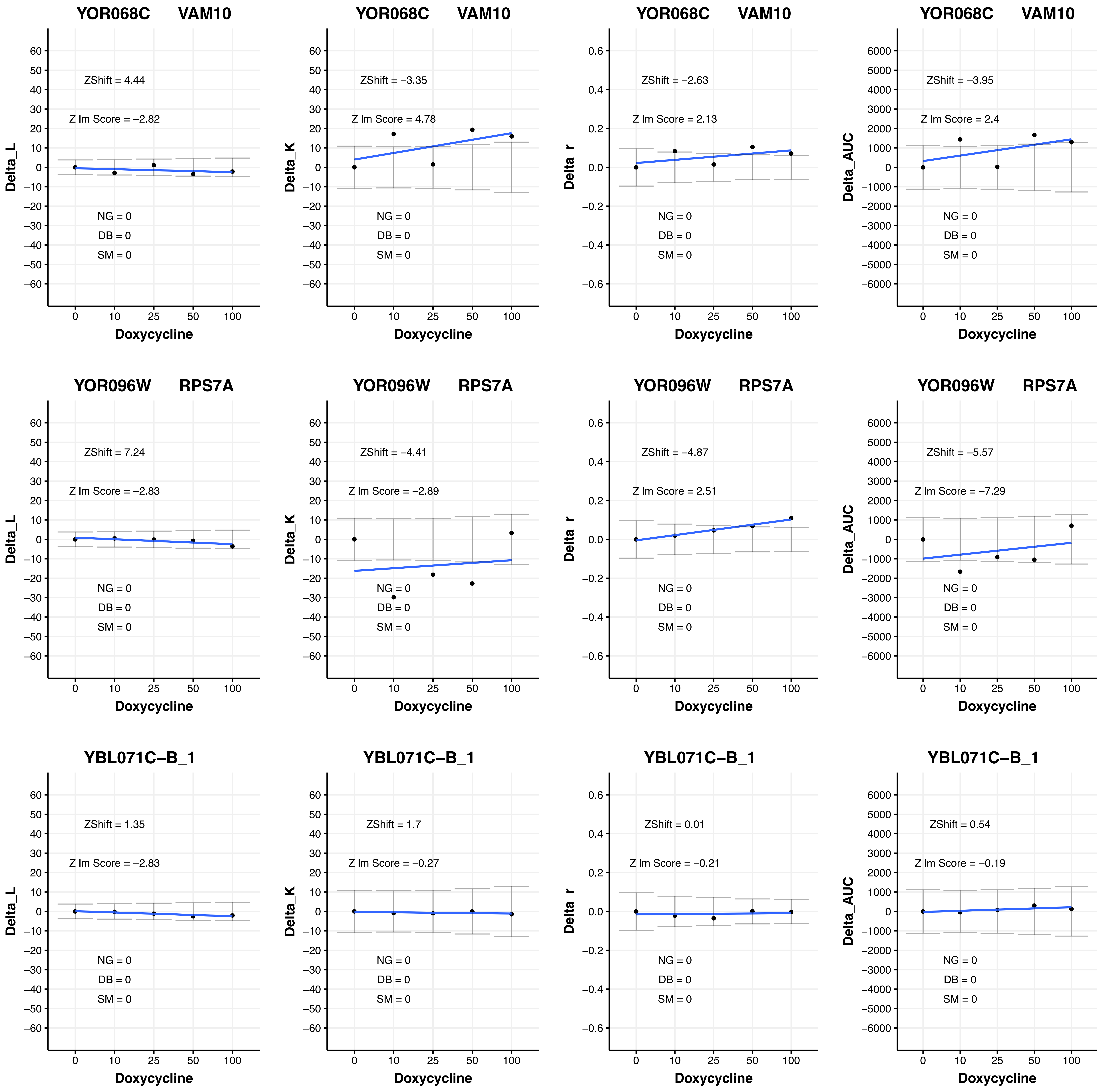
YOR068C VAM10
ZShift = 4.44
Z lm Score = −2.82
NG = 0
DB = 0
SM = 0
Delta_L
Doxycycline
YOR068C VAM10
ZShift = −3.35
Z lm Score = 4.78
NG = 0
DB = 0
SM = 0
Delta_K
Doxycycline
YOR068C VAM10
ZShift = −2.63
Z lm Score = 2.13
NG = 0
DB = 0
SM = 0
Delta_r
Doxycycline
YOR068C VAM10
ZShift = −3.95
Z lm Score = 2.4
NG = 0
DB = 0
SM = 0
Delta_AUC
Doxycycline
YOR096W RPS7A
ZShift = 7.24
Z lm Score = −2.83
NG = 0
DB = 0
SM = 0
Delta_L
Doxycycline
YOR096W RPS7A
ZShift = −4.41
Z lm Score = −2.89
NG = 0
DB = 0
SM = 0
Delta_K
Doxycycline
YOR096W RPS7A
ZShift = −4.87
Z lm Score = 2.51
NG = 0
DB = 0
SM = 0
Delta_r
Doxycycline
YOR096W RPS7A
ZShift = −5.57
Z lm Score = −7.29
NG = 0
DB = 0
SM = 0
Delta_AUC
Doxycycline
YBL071C−B_1
ZShift = 1.35
Z lm Score = −2.83
NG = 0
DB = 0
SM = 0
Delta_L
Doxycycline
YBL071C−B_1
ZShift = 1.7
Z lm Score = −0.27
NG = 0
DB = 0
SM = 0
Delta_K
Doxycycline
YBL071C−B_1
ZShift = 0.01
Z lm Score = −0.21
NG = 0
DB = 0
SM = 0
Delta_r
Doxycycline
YBL071C−B_1
ZShift = 0.54
Z lm Score = −0.19
NG = 0
DB = 0
SM = 0
Delta_AUC
Doxycycline
0 10 25 50 100
60 50 40 30 20 10 0 −10 −20 −30 −40 −50 −60
0.6 0.4 0.2 0.0 −0.2 −0.4 −0.6
6000 5000 4000 3000 2000 1000 0 −1000 −2000 −3000 −4000 −5000 −6000

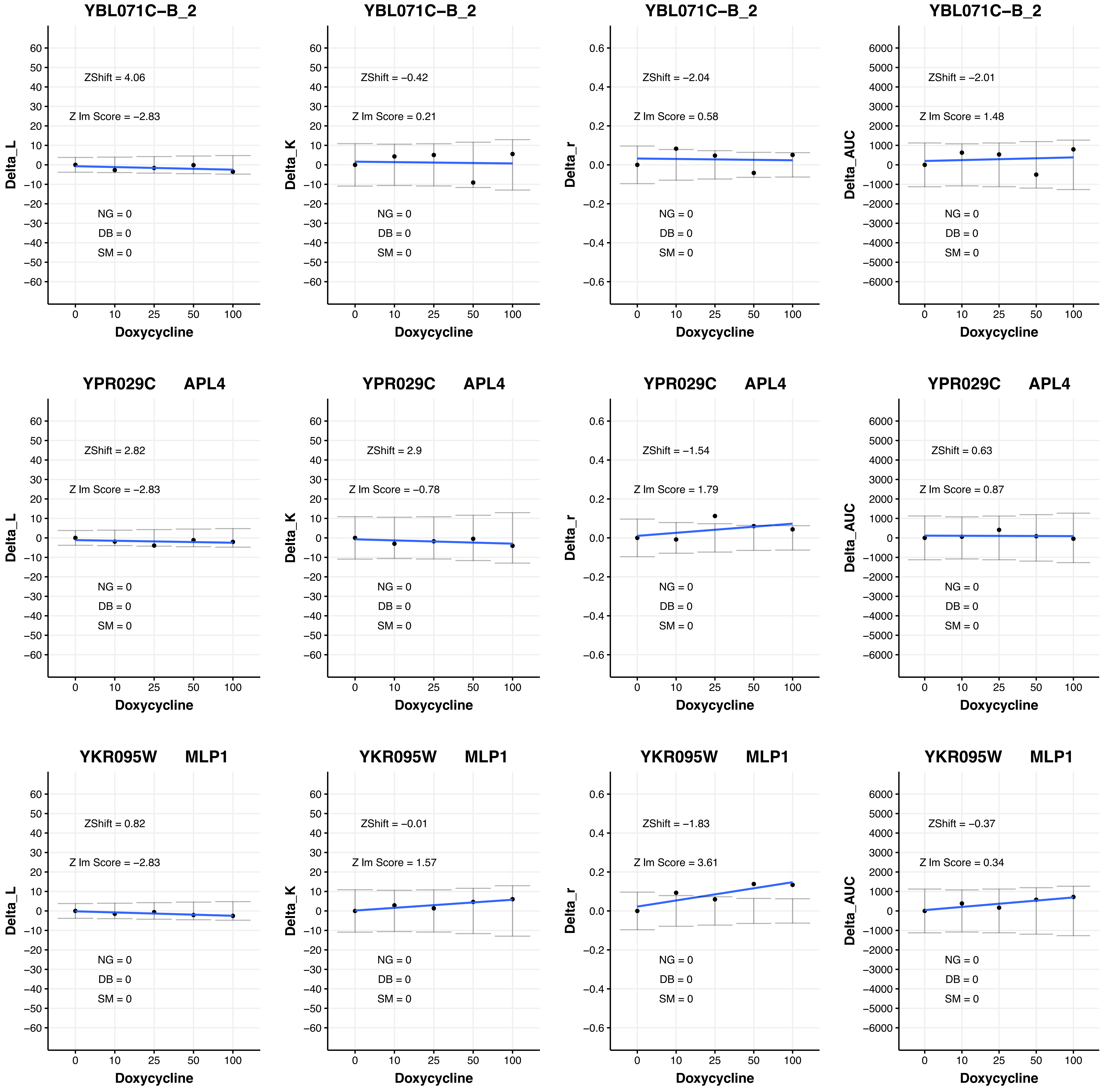
YBL071C–B_2
ZShift = 4.06
Z lm Score = –2.83
NG = 0
DB = 0
SM = 0
Delta_L
Doxycycline
YBL071C–B_2
ZShift = –0.42
Z lm Score = 0.21
NG = 0
DB = 0
SM = 0
Delta_K
Doxycycline
YBL071C–B_2
ZShift = –2.04
Z lm Score = 0.58
NG = 0
DB = 0
SM = 0
Delta_r
Doxycycline
YBL071C–B_2
ZShift = –2.01
Z lm Score = 1.48
NG = 0
DB = 0
SM = 0
Delta_AUC
Doxycycline
YPR029C APL4
ZShift = 2.82
Z lm Score = –2.83
NG = 0
DB = 0
SM = 0
Delta_L
Doxycycline
YPR029C APL4
ZShift = 2.9
Z lm Score = –0.78
NG = 0
DB = 0
SM = 0
Delta_K
Doxycycline
YPR029C APL4
ZShift = –1.54
Z lm Score = 1.79
NG = 0
DB = 0
SM = 0
Delta_r
Doxycycline
YPR029C APL4
ZShift = 0.63
Z lm Score = 0.87
NG = 0
DB = 0
SM = 0
Delta_AUC
Doxycycline
YKR095W MLP1
ZShift = 0.82
Z lm Score = –2.83
NG = 0
DB = 0
SM = 0
Delta_L
Doxycycline
YKR095W MLP1
ZShift = –0.01
Z lm Score = 1.57
NG = 0
DB = 0
SM = 0
Delta_K
Doxycycline
YKR095W MLP1
ZShift = –1.83
Z lm Score = 3.61
NG = 0
DB = 0
SM = 0
Delta_r
Doxycycline
YKR095W MLP1
ZShift = –0.37
Z lm Score = 0.34
NG = 0
DB = 0
SM = 0
Delta_AUC
Doxycycline

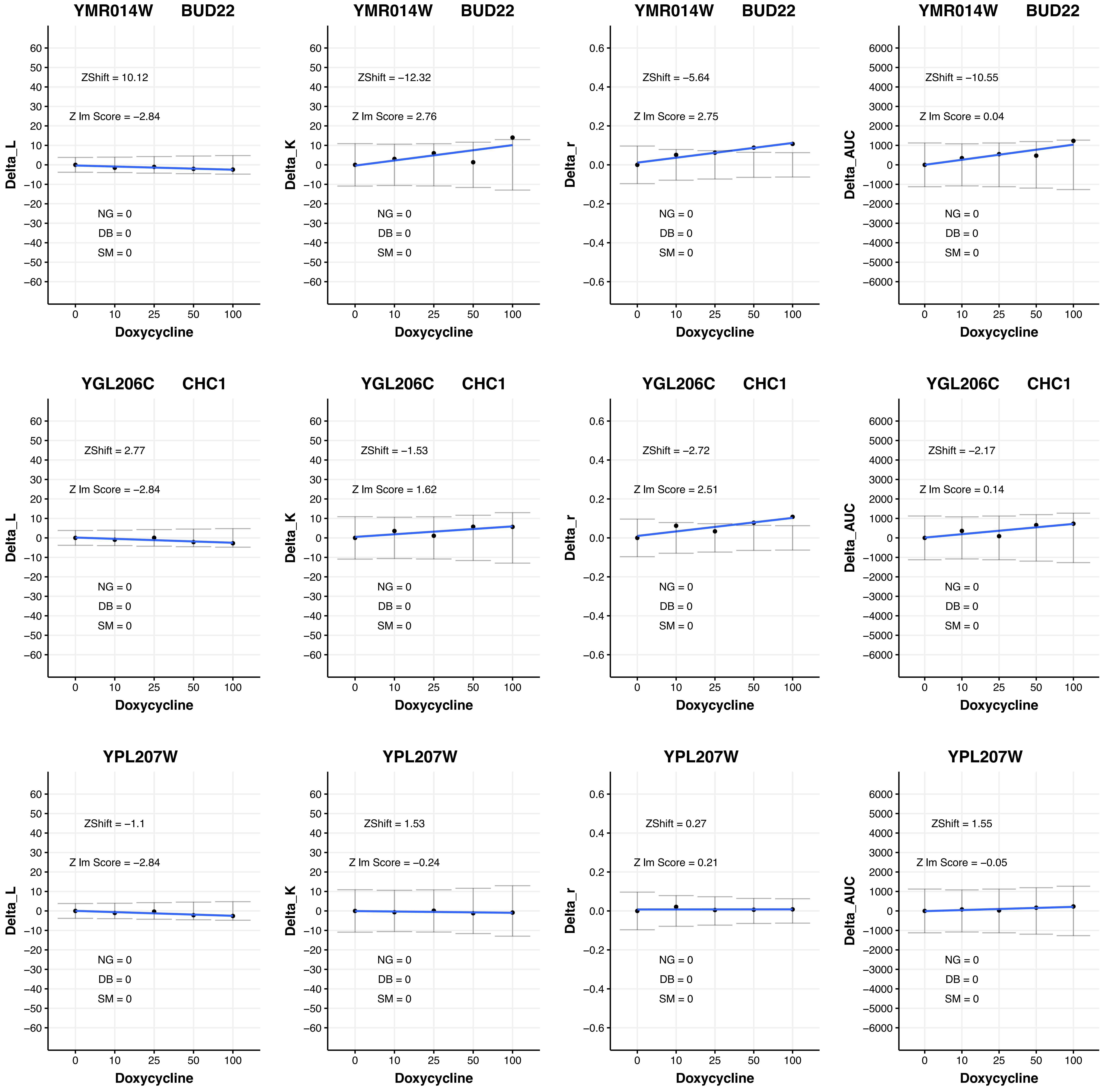
YMR014W BUD22
ZShift = 10.12
Z lm Score = −2.84
NG = 0
DB = 0
SM = 0
Delta_L
Doxycycline
YMR014W BUD22
ZShift = −12.32
Z lm Score = 2.76
NG = 0
DB = 0
SM = 0
Delta_K
Doxycycline
YMR014W BUD22
ZShift = −5.64
Z lm Score = 2.75
NG = 0
DB = 0
SM = 0
Delta_r
Doxycycline
YMR014W BUD22
ZShift = −10.55
Z lm Score = 0.04
NG = 0
DB = 0
SM = 0
Delta_AUC
Doxycycline
YGL206C CHC1
ZShift = 2.77
Z lm Score = −2.84
NG = 0
DB = 0
SM = 0
Delta_L
Doxycycline
YGL206C CHC1
ZShift = −1.53
Z lm Score = 1.62
NG = 0
DB = 0
SM = 0
Delta_K
Doxycycline
YGL206C CHC1
ZShift = −2.72
Z lm Score = 2.51
NG = 0
DB = 0
SM = 0
Delta_r
Doxycycline
YGL206C CHC1
ZShift = −2.17
Z lm Score = 0.14
NG = 0
DB = 0
SM = 0
Delta_AUC
Doxycycline
YPL207W
ZShift = −1.1
Z lm Score = −2.84
NG = 0
DB = 0
SM = 0
Delta_L
Doxycycline
YPL207W
ZShift = 1.53
Z lm Score = −0.24
NG = 0
DB = 0
SM = 0
Delta_K
Doxycycline
YPL207W
ZShift = 0.27
Z lm Score = 0.21
NG = 0
DB = 0
SM = 0
Delta_r
Doxycycline
YPL207W
ZShift = 1.55
Z lm Score = −0.05
NG = 0
DB = 0
SM = 0
Delta_AUC
Doxycycline

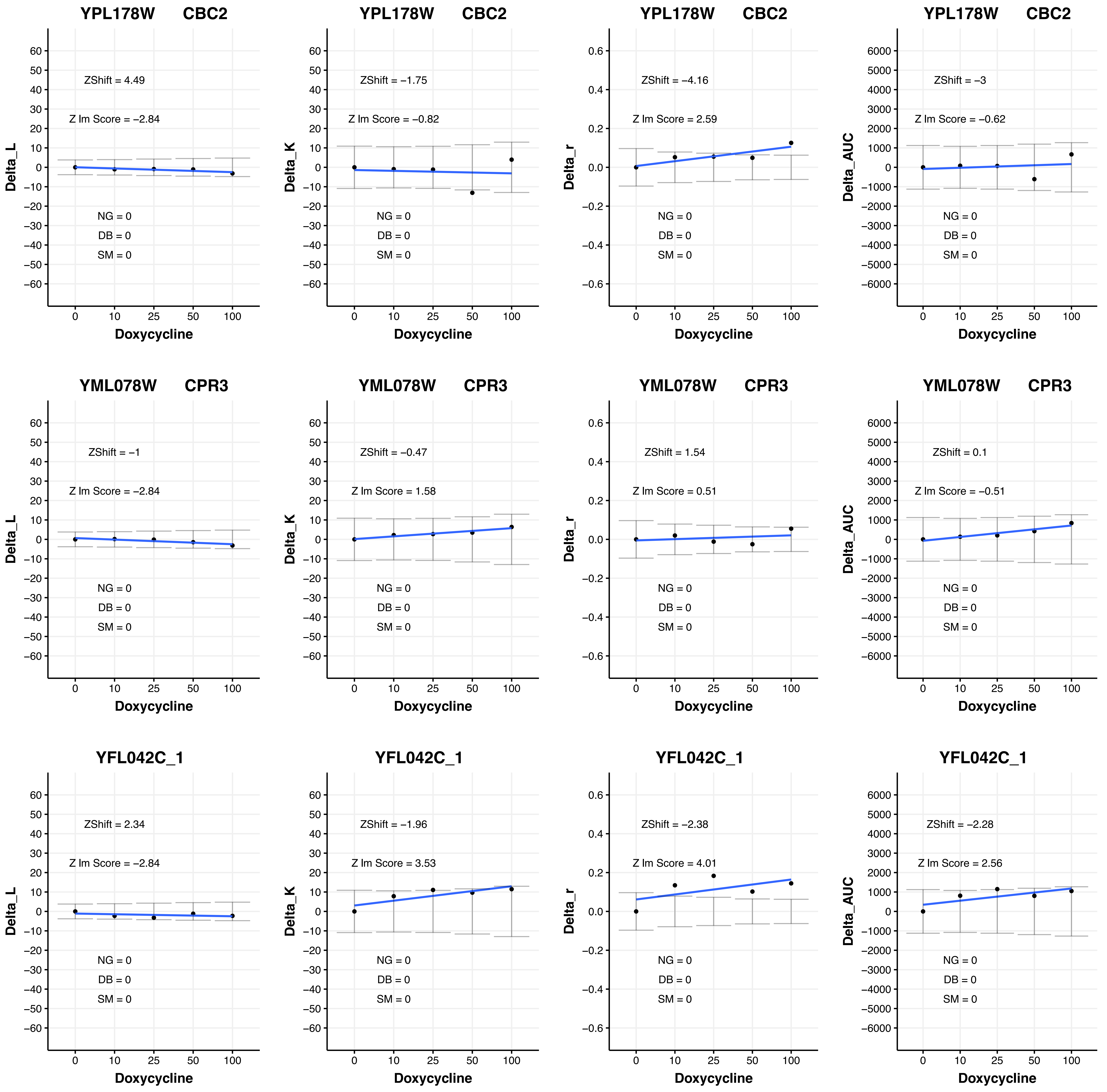
YPL178W CBC2
ZShift = 4.49
Z lm Score = −2.84
NG = 0
DB = 0
SM = 0
Delta_L
Doxycycline
YPL178W CBC2
ZShift = −1.75
Z lm Score = −0.82
NG = 0
DB = 0
SM = 0
Delta_K
Doxycycline
YPL178W CBC2
ZShift = −4.16
Z lm Score = 2.59
NG = 0
DB = 0
SM = 0
Delta_r
Doxycycline
YPL178W CBC2
ZShift = −3
Z lm Score = −0.62
NG = 0
DB = 0
SM = 0
Delta_AUC
Doxycycline
YML078W CPR3
ZShift = −1
Z lm Score = −2.84
NG = 0
DB = 0
SM = 0
Delta_L
Doxycycline
YML078W CPR3
ZShift = −0.47
Z lm Score = 1.58
NG = 0
DB = 0
SM = 0
Delta_K
Doxycycline
YML078W CPR3
ZShift = 1.54
Z lm Score = 0.51
NG = 0
DB = 0
SM = 0
Delta_r
Doxycycline
YML078W CPR3
ZShift = 0.1
Z lm Score = −0.51
NG = 0
DB = 0
SM = 0
Delta_AUC
Doxycycline
YFL042C_1
ZShift = 2.34
Z lm Score = −2.84
NG = 0
DB = 0
SM = 0
Delta_L
Doxycycline
YFL042C_1
ZShift = −1.96
Z lm Score = 3.53
NG = 0
DB = 0
SM = 0
Delta_K
Doxycycline
YFL042C_1
ZShift = −2.38
Z lm Score = 4.01
NG = 0
DB = 0
SM = 0
Delta_r
Doxycycline
YFL042C_1
ZShift = −2.28
Z lm Score = 2.56
NG = 0
DB = 0
SM = 0
Delta_AUC
Doxycycline

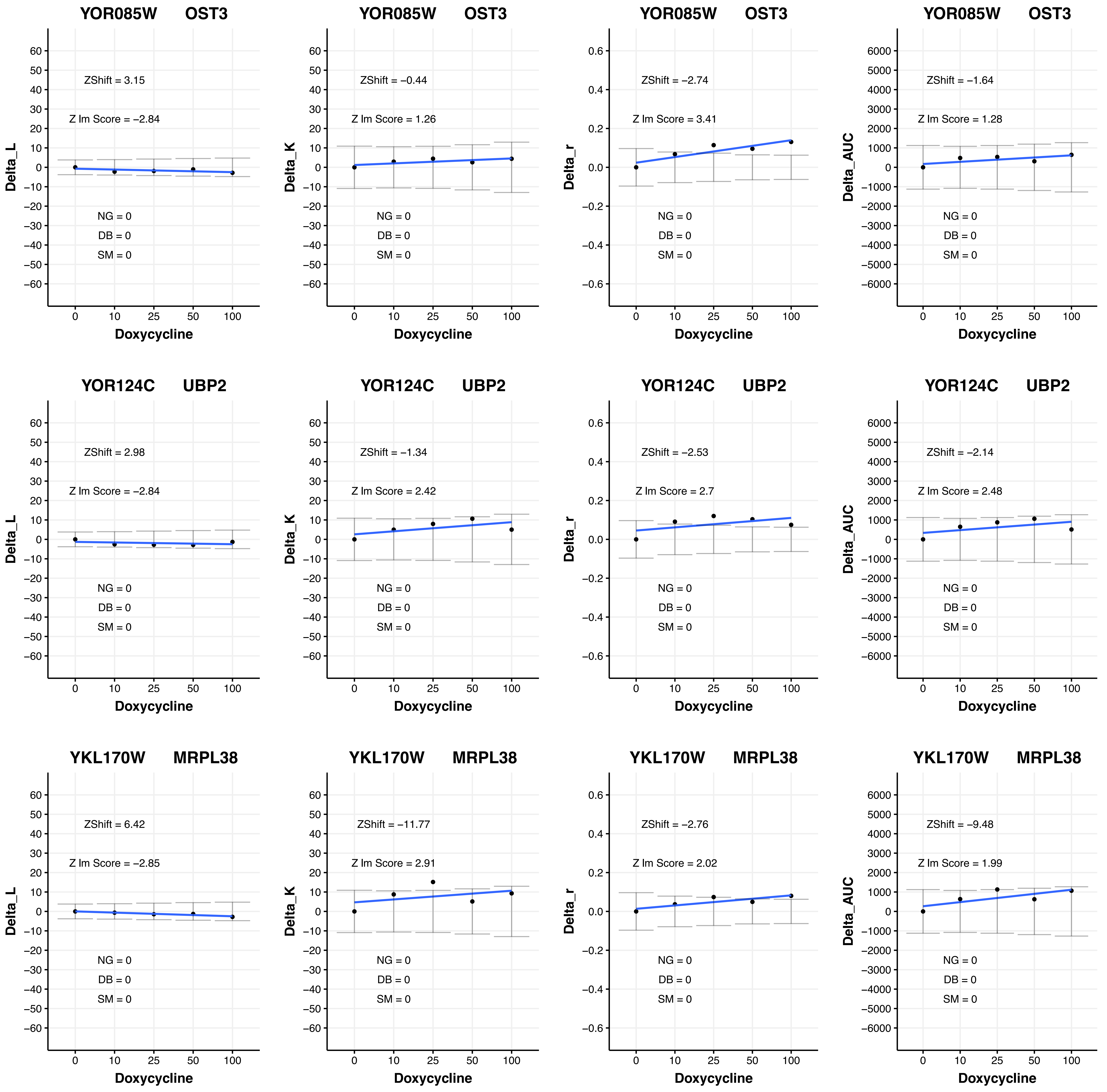

YOR085W OST3
ZShift = 3.15
Z lm Score = −2.84
NG = 0
DB = 0
SM = 0
Delta_L
Doxycycline
YOR085W OST3
ZShift = −0.44
Z lm Score = 1.26
NG = 0
DB = 0
SM = 0
Delta_K
Doxycycline
YOR085W OST3
ZShift = −2.74
Z lm Score = 3.41
NG = 0
DB = 0
SM = 0
Delta_r
Doxycycline
YOR085W OST3
ZShift = −1.64
Z lm Score = 1.28
NG = 0
DB = 0
SM = 0
Delta_AUC
Doxycycline
YOR124C UBP2
ZShift = 2.98
Z lm Score = −2.84
NG = 0
DB = 0
SM = 0
Delta_L
Doxycycline
YOR124C UBP2
ZShift = −1.34
Z lm Score = 2.42
NG = 0
DB = 0
SM = 0
Delta_K
Doxycycline
YOR124C UBP2
ZShift = −2.53
Z lm Score = 2.7
NG = 0
DB = 0
SM = 0
Delta_r
Doxycycline
YOR124C UBP2
ZShift = −2.14
Z lm Score = 2.48
NG = 0
DB = 0
SM = 0
Delta_AUC
Doxycycline
YKL170W MRPL38
ZShift = 6.42
Z lm Score = −2.85
NG = 0
DB = 0
SM = 0
Delta_L
Doxycycline
YKL170W MRPL38
ZShift = −11.77
Z lm Score = 2.91
NG = 0
DB = 0
SM = 0
Delta_K
Doxycycline
YKL170W MRPL38
ZShift = −2.76
Z lm Score = 2.02
NG = 0
DB = 0
SM = 0
Delta_r
Doxycycline
YKL170W MRPL38
ZShift = −9.48
Z lm Score = 1.99
NG = 0
DB = 0
SM = 0
Delta_AUC
Doxycycline

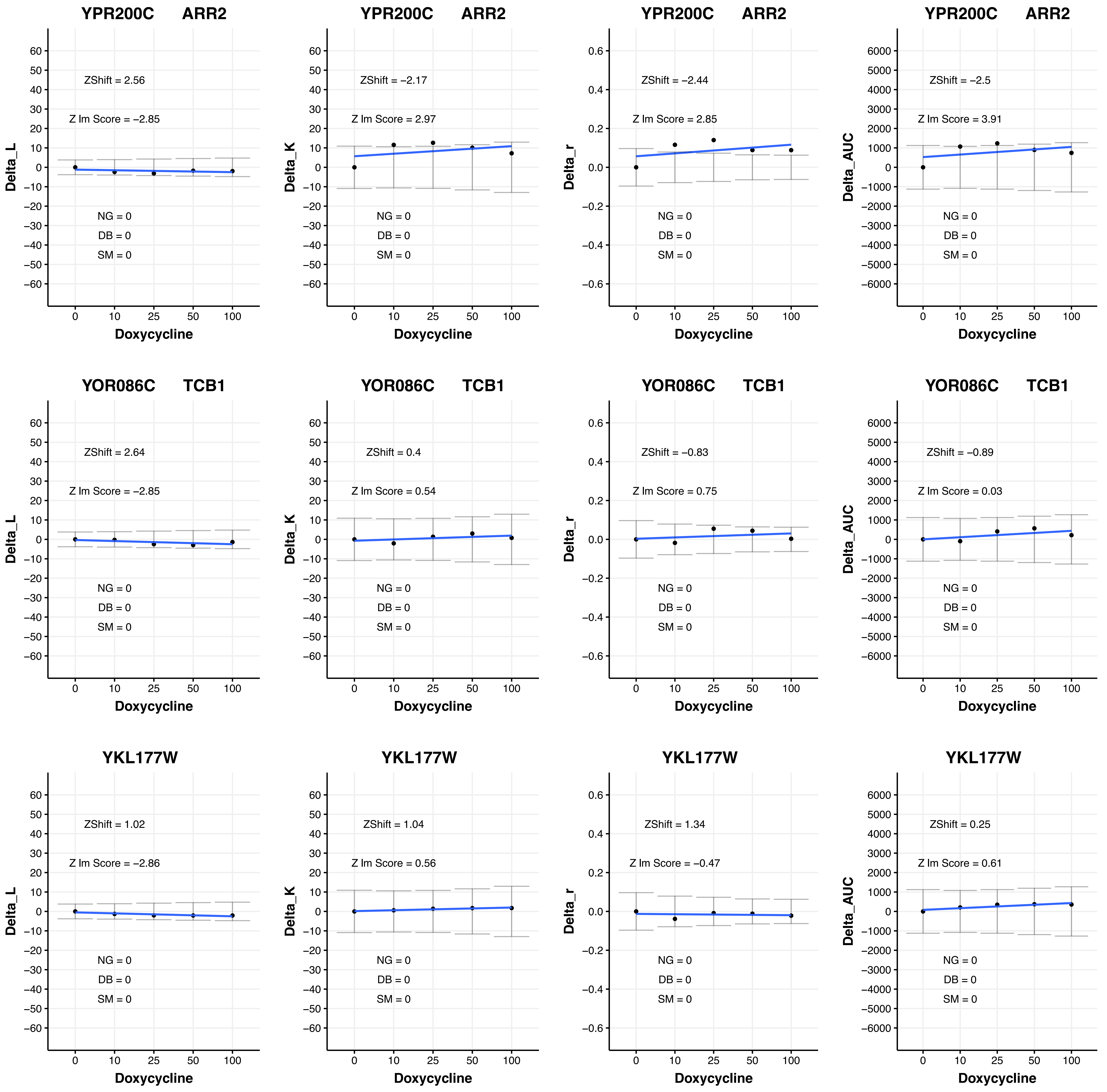

YPR200C ARR2
ZShift = 2.56
Z lm Score = −2.85
NG = 0
DB = 0
SM = 0
Delta_L
Doxycycline
YPR200C ARR2
ZShift = −2.17
Z lm Score = 2.97
Delta_K
YPR200C ARR2
ZShift = −2.44
Z lm Score = 2.85
Delta_r
YPR200C ARR2
ZShift = −2.5
Z lm Score = 3.91
Delta_AUC
YOR086C TCB1
ZShift = 2.64
Z lm Score = −2.85
YOR086C TCB1
ZShift = 0.4
Z lm Score = 0.54
YOR086C TCB1
ZShift = −0.83
Z lm Score = 0.75
YOR086C TCB1
ZShift = −0.89
Z lm Score = 0.03
YKL177W
ZShift = 1.02
Z lm Score = −2.86
YKL177W
ZShift = 1.04
Z lm Score = 0.56
YKL177W
ZShift = 1.34
Z lm Score = −0.47
YKL177W
ZShift = 0.25
Z lm Score = 0.61

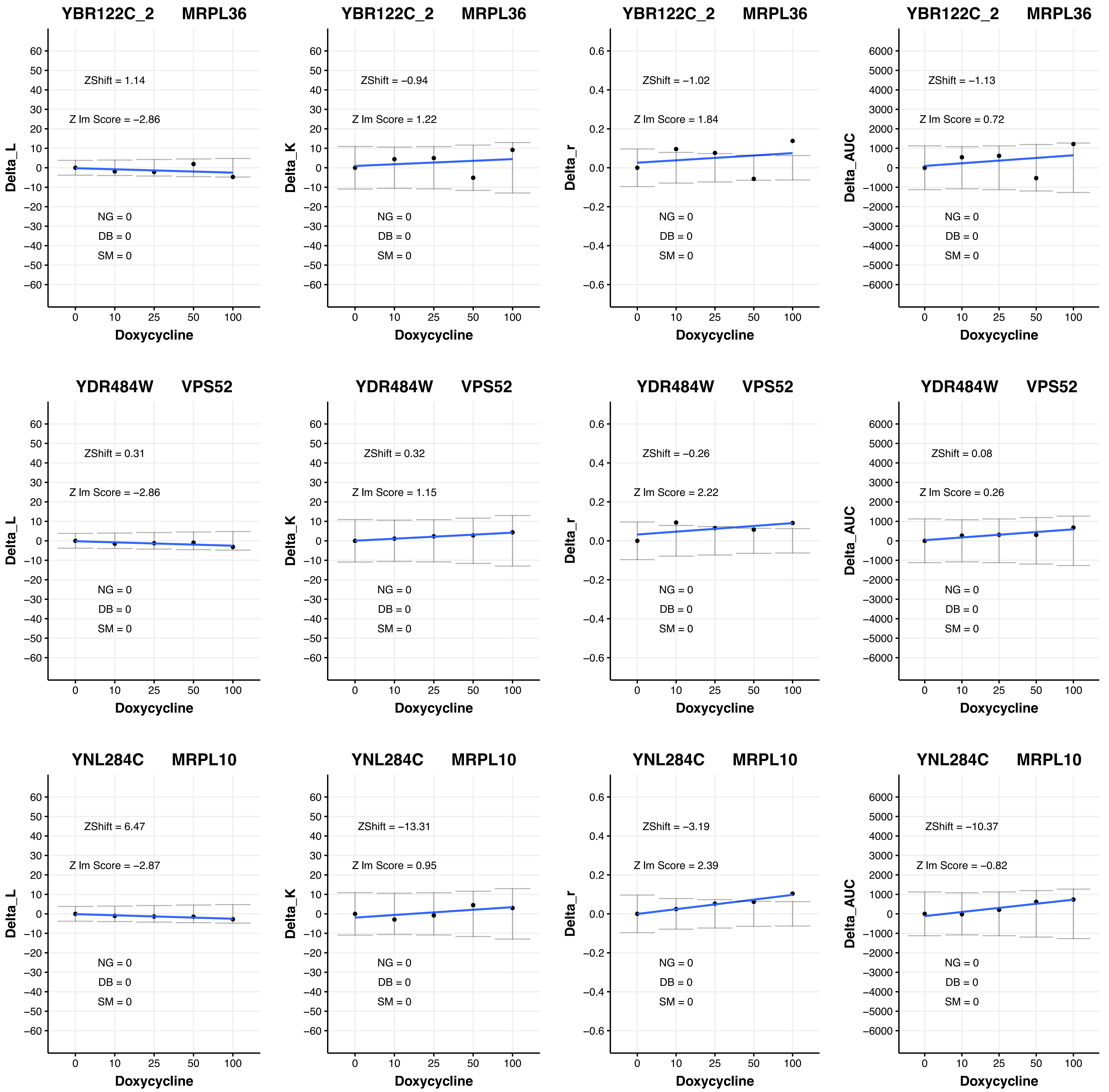
YBR122C_2 MRPL36
ZShift = 1.14
Z lm Score = −2.86
NG = 0
DB = 0
SM = 0
Delta_L
Doxycycline
YBR122C_2 MRPL36
ZShift = −0.94
Z lm Score = 1.22
NG = 0
DB = 0
SM = 0
Delta_K
Doxycycline
YBR122C_2 MRPL36
ZShift = −1.02
Z lm Score = 1.84
NG = 0
DB = 0
SM = 0
Delta_r
Doxycycline
YBR122C_2 MRPL36
ZShift = −1.13
Z lm Score = 0.72
NG = 0
DB = 0
SM = 0
Delta_AUC
Doxycycline
YDR484W VPS52
ZShift = 0.31
Z lm Score = −2.86
NG = 0
DB = 0
SM = 0
Delta_L
Doxycycline
YDR484W VPS52
ZShift = 0.32
Z lm Score = 1.15
NG = 0
DB = 0
SM = 0
Delta_K
Doxycycline
YDR484W VPS52
ZShift = −0.26
Z lm Score = 2.22
NG = 0
DB = 0
SM = 0
Delta_r
Doxycycline
YDR484W VPS52
ZShift = 0.08
Z lm Score = 0.26
NG = 0
DB = 0
SM = 0
Delta_AUC
Doxycycline
YNL284C MRPL10
ZShift = 6.47
Z lm Score = −2.87
NG = 0
DB = 0
SM = 0
Delta_L
Doxycycline
YNL284C MRPL10
ZShift = −13.31
Z lm Score = 0.95
NG = 0
DB = 0
SM = 0
Delta_K
Doxycycline
YNL284C MRPL10
ZShift = −3.19
Z lm Score = 2.39
NG = 0
DB = 0
SM = 0
Delta_r
Doxycycline
YNL284C MRPL10
ZShift = −10.37
Z lm Score = −0.82
NG = 0
DB = 0
SM = 0
Delta_AUC
Doxycycline

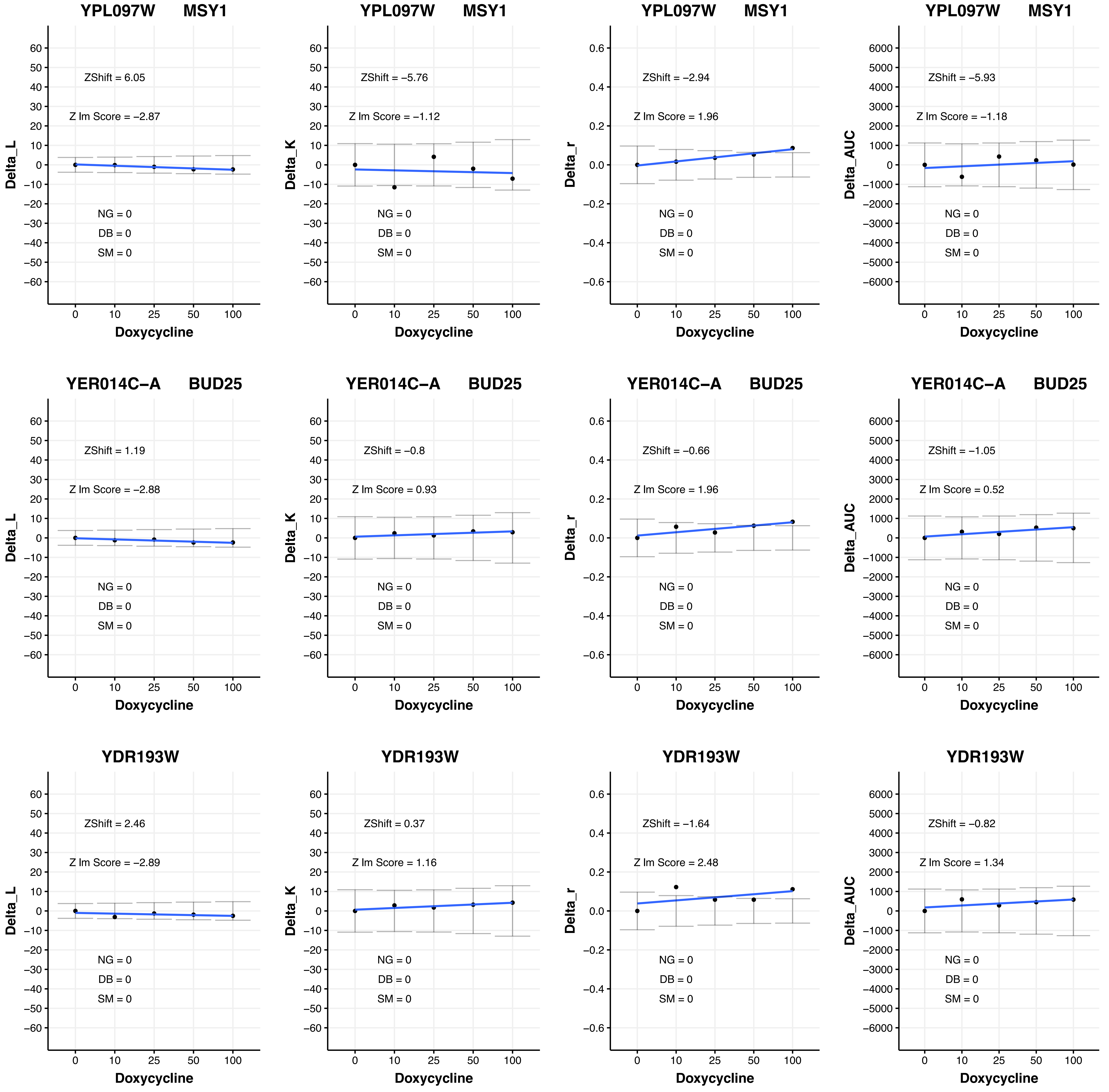
YPL097W MSY1
ZShift = 6.05
Z lm Score = −2.87
NG = 0
DB = 0
SM = 0
Delta_L
YPL097W MSY1
ZShift = −5.76
Z lm Score = −1.12
NG = 0
DB = 0
SM = 0
Delta_K
YPL097W MSY1
ZShift = −2.94
Z lm Score = 1.96
NG = 0
DB = 0
SM = 0
Delta_r
YPL097W MSY1
ZShift = −5.93
Z lm Score = −1.18
NG = 0
DB = 0
SM = 0
Delta_AUC
YER014C−A BUD25
ZShift = 1.19
Z lm Score = −2.88
NG = 0
DB = 0
SM = 0
Delta_L
YER014C−A BUD25
ZShift = −0.8
Z lm Score = 0.93
NG = 0
DB = 0
SM = 0
Delta_K
YER014C−A BUD25
ZShift = −0.66
Z lm Score = 1.96
NG = 0
DB = 0
SM = 0
Delta_r
YER014C−A BUD25
ZShift = −1.05
Z lm Score = 0.52
NG = 0
DB = 0
SM = 0
Delta_AUC
YDR193W
ZShift = 2.46
Z lm Score = −2.89
NG = 0
DB = 0
SM = 0
Delta_L
YDR193W
ZShift = 0.37
Z lm Score = 1.16
NG = 0
DB = 0
SM = 0
Delta_K
YDR193W
ZShift = −1.64
Z lm Score = 2.48
NG = 0
DB = 0
SM = 0
Delta_r
YDR193W
ZShift = −0.82
Z lm Score = 1.34
NG = 0
DB = 0
SM = 0
Delta_AUC
Doxycycline
0 10 25 50 100

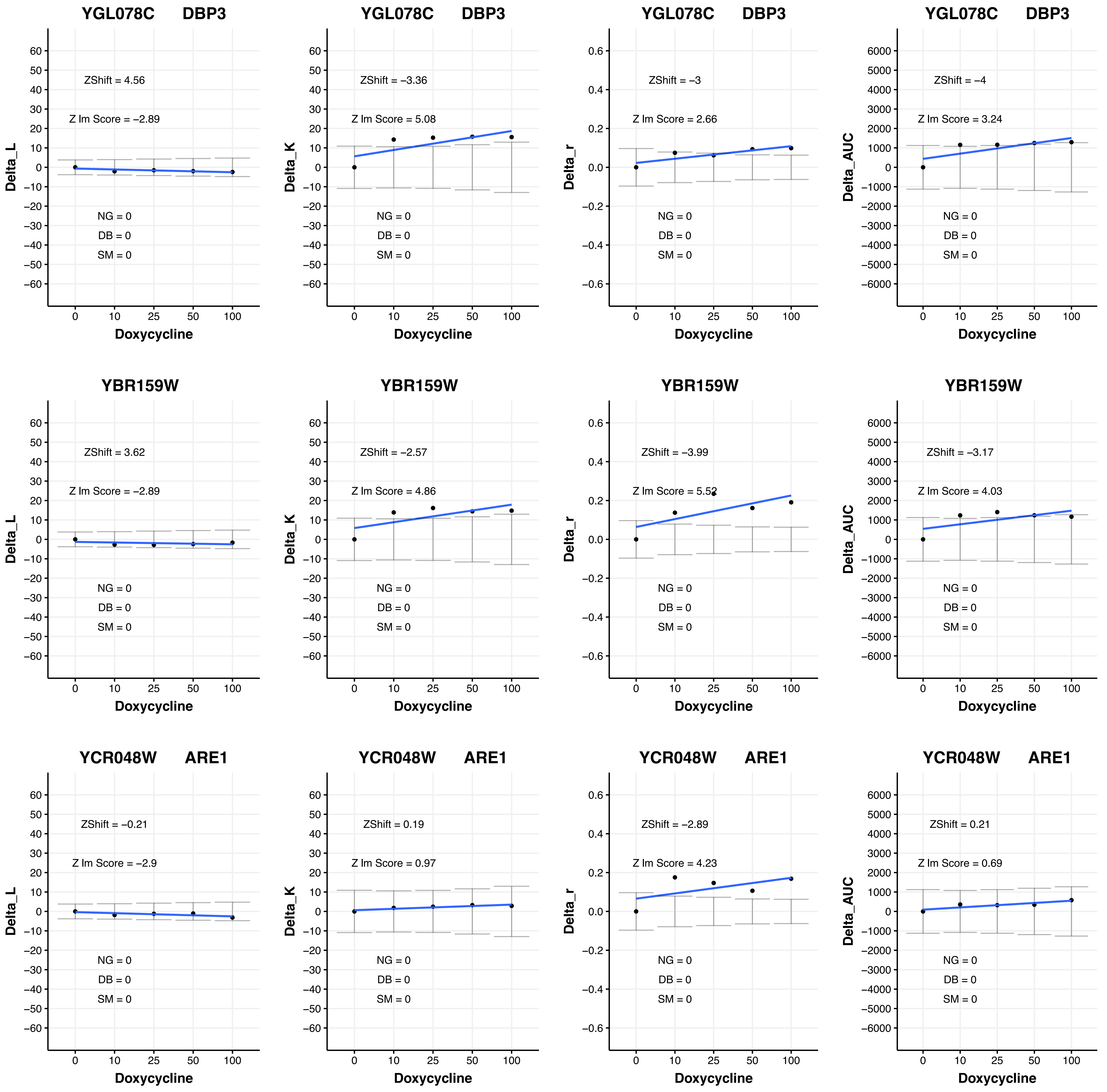
YGL078C DBP3
ZShift = 4.56
Z lm Score = −2.89
NG = 0
DB = 0
SM = 0
Delta_L
Doxycycline
YGL078C DBP3
ZShift = −3.36
Z lm Score = 5.08
NG = 0
DB = 0
SM = 0
Delta_K
Doxycycline
YGL078C DBP3
ZShift = −3
Z lm Score = 2.66
NG = 0
DB = 0
SM = 0
Delta_r
Doxycycline
YGL078C DBP3
ZShift = −4
Z lm Score = 3.24
NG = 0
DB = 0
SM = 0
Delta_AUC
Doxycycline
YBR159W
ZShift = 3.62
Z lm Score = −2.89
NG = 0
DB = 0
SM = 0
Delta_L
Doxycycline
YBR159W
ZShift = −2.57
Z lm Score = 4.86
NG = 0
DB = 0
SM = 0
Delta_K
Doxycycline
YBR159W
ZShift = −3.99
Z lm Score = 5.52
NG = 0
DB = 0
SM = 0
Delta_r
Doxycycline
YBR159W
ZShift = −3.17
Z lm Score = 4.03
NG = 0
DB = 0
SM = 0
Delta_AUC
Doxycycline
YCR048W ARE1
ZShift = −0.21
Z lm Score = −2.9
NG = 0
DB = 0
SM = 0
Delta_L
Doxycycline
YCR048W ARE1
ZShift = 0.19
Z lm Score = 0.97
NG = 0
DB = 0
SM = 0
Delta_K
Doxycycline
YCR048W ARE1
ZShift = −2.89
Z lm Score = 4.23
NG = 0
DB = 0
SM = 0
Delta_r
Doxycycline
YCR048W ARE1
ZShift = 0.21
Z lm Score = 0.69
NG = 0
DB = 0
SM = 0
Delta_AUC
Doxycycline

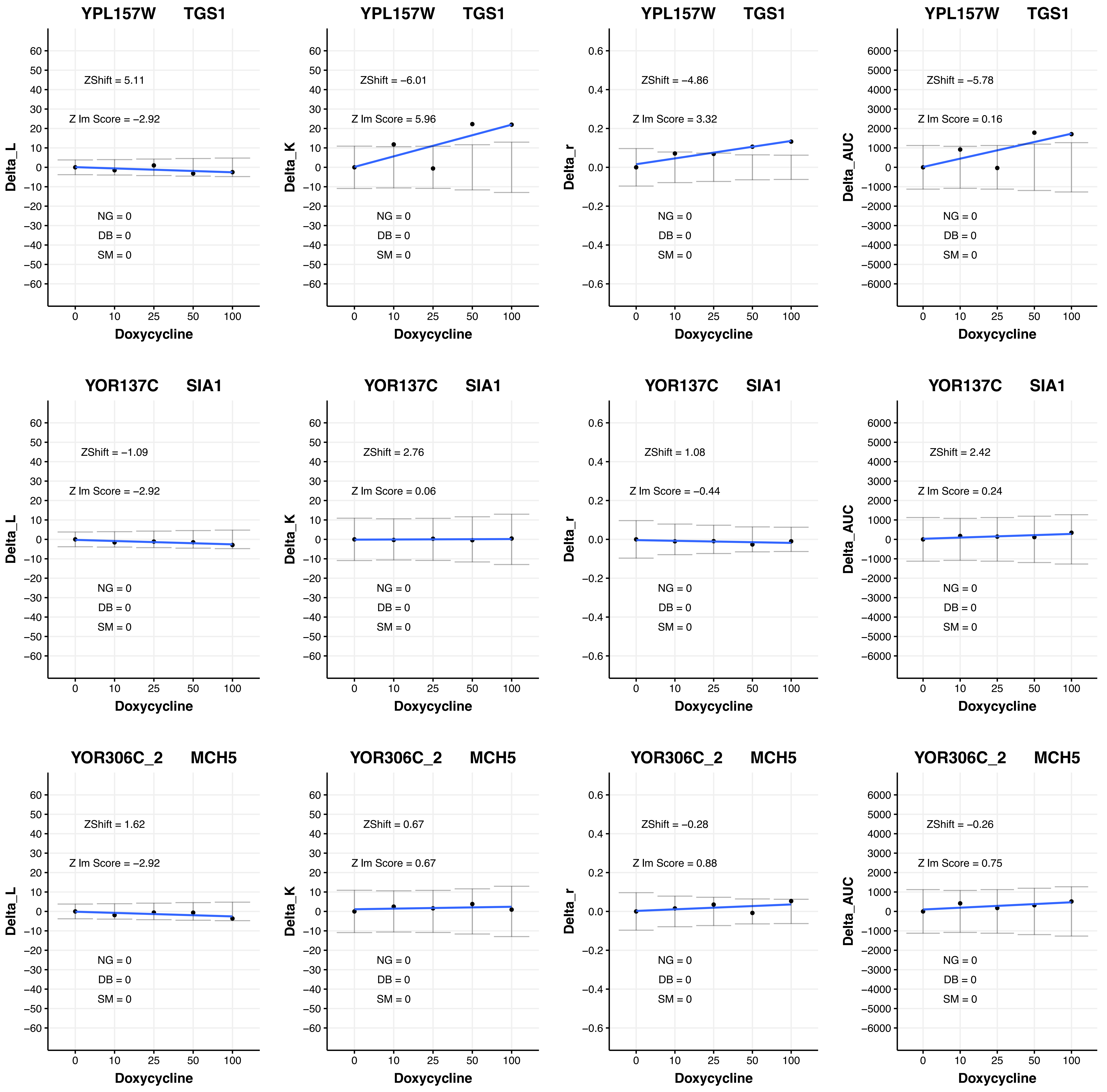
YPL157W TGS1
ZShift = 5.11
Z lm Score = −2.92
NG = 0
DB = 0
SM = 0
Delta_L
Doxycycline
YPL157W TGS1
ZShift = −6.01
Z lm Score = 5.96
NG = 0
DB = 0
SM = 0
Delta_K
Doxycycline
YPL157W TGS1
ZShift = −4.86
Z lm Score = 3.32
NG = 0
DB = 0
SM = 0
Delta_r
Doxycycline
YPL157W TGS1
ZShift = −5.78
Z lm Score = 0.16
NG = 0
DB = 0
SM = 0
Delta_AUC
Doxycycline
YOR137C SIA1
ZShift = −1.09
Z lm Score = −2.92
NG = 0
DB = 0
SM = 0
Delta_L
Doxycycline
YOR137C SIA1
ZShift = 2.76
Z lm Score = 0.06
NG = 0
DB = 0
SM = 0
Delta_K
Doxycycline
YOR137C SIA1
ZShift = 1.08
Z lm Score = −0.44
NG = 0
DB = 0
SM = 0
Delta_r
Doxycycline
YOR137C SIA1
ZShift = 2.42
Z lm Score = 0.24
NG = 0
DB = 0
SM = 0
Delta_AUC
Doxycycline
YOR306C_2 MCH5
ZShift = 1.62
Z lm Score = −2.92
NG = 0
DB = 0
SM = 0
Delta_L
Doxycycline
YOR306C_2 MCH5
ZShift = 0.67
Z lm Score = 0.67
NG = 0
DB = 0
SM = 0
Delta_K
Doxycycline
YOR306C_2 MCH5
ZShift = −0.28
Z lm Score = 0.88
NG = 0
DB = 0
SM = 0
Delta_r
Doxycycline
YOR306C_2 MCH5
ZShift = −0.26
Z lm Score = 0.75
NG = 0
DB = 0
SM = 0
Delta_AUC
Doxycycline

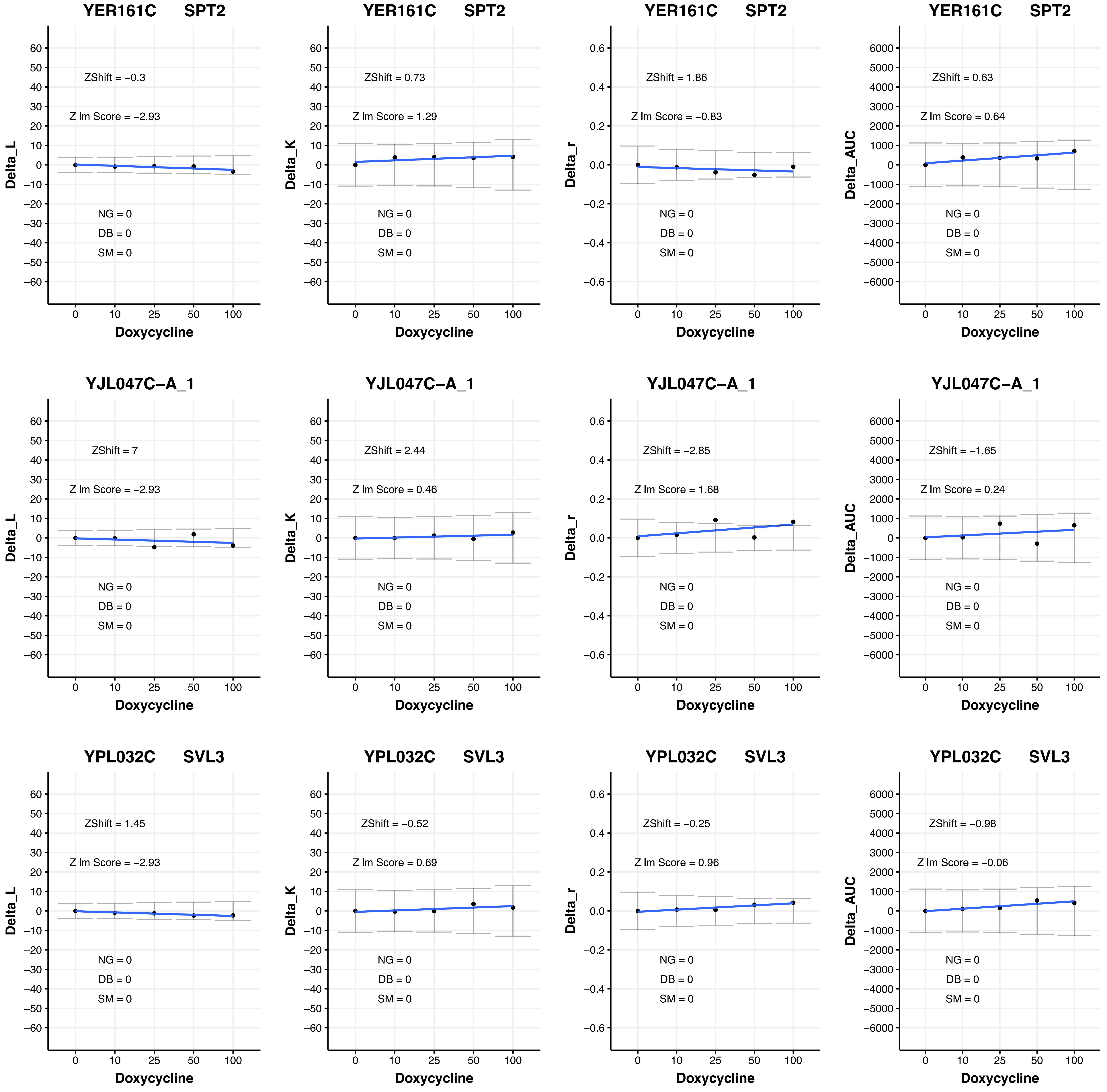

YER161C SPT2
ZShift = −0.3
Z lm Score = −2.93
NG = 0
DB = 0
SM = 0
Delta_L
Doxycycline
YER161C SPT2
ZShift = 0.73
Z lm Score = 1.29
NG = 0
DB = 0
SM = 0
Delta_K
Doxycycline
YER161C SPT2
ZShift = 1.86
Z lm Score = −0.83
NG = 0
DB = 0
SM = 0
Delta_r
Doxycycline
YER161C SPT2
ZShift = 0.63
Z lm Score = 0.64
NG = 0
DB = 0
SM = 0
Delta_AUC
Doxycycline
YJL047C−A_1
ZShift = 7
Z lm Score = −2.93
NG = 0
DB = 0
SM = 0
Delta_L
Doxycycline
YJL047C−A_1
ZShift = 2.44
Z lm Score = 0.46
NG = 0
DB = 0
SM = 0
Delta_K
Doxycycline
YJL047C−A_1
ZShift = −2.85
Z lm Score = 1.68
NG = 0
DB = 0
SM = 0
Delta_r
Doxycycline
YJL047C−A_1
ZShift = −1.65
Z lm Score = 0.24
NG = 0
DB = 0
SM = 0
Delta_AUC
Doxycycline
YPL032C SVL3
ZShift = 1.45
Z lm Score = −2.93
NG = 0
DB = 0
SM = 0
Delta_L
Doxycycline
YPL032C SVL3
ZShift = −0.52
Z lm Score = 0.69
NG = 0
DB = 0
SM = 0
Delta_K
Doxycycline
YPL032C SVL3
ZShift = −0.25
Z lm Score = 0.96
NG = 0
DB = 0
SM = 0
Delta_r
Doxycycline
YPL032C SVL3
ZShift = −0.98
Z lm Score = −0.06
NG = 0
DB = 0
SM = 0
Delta_AUC
Doxycycline

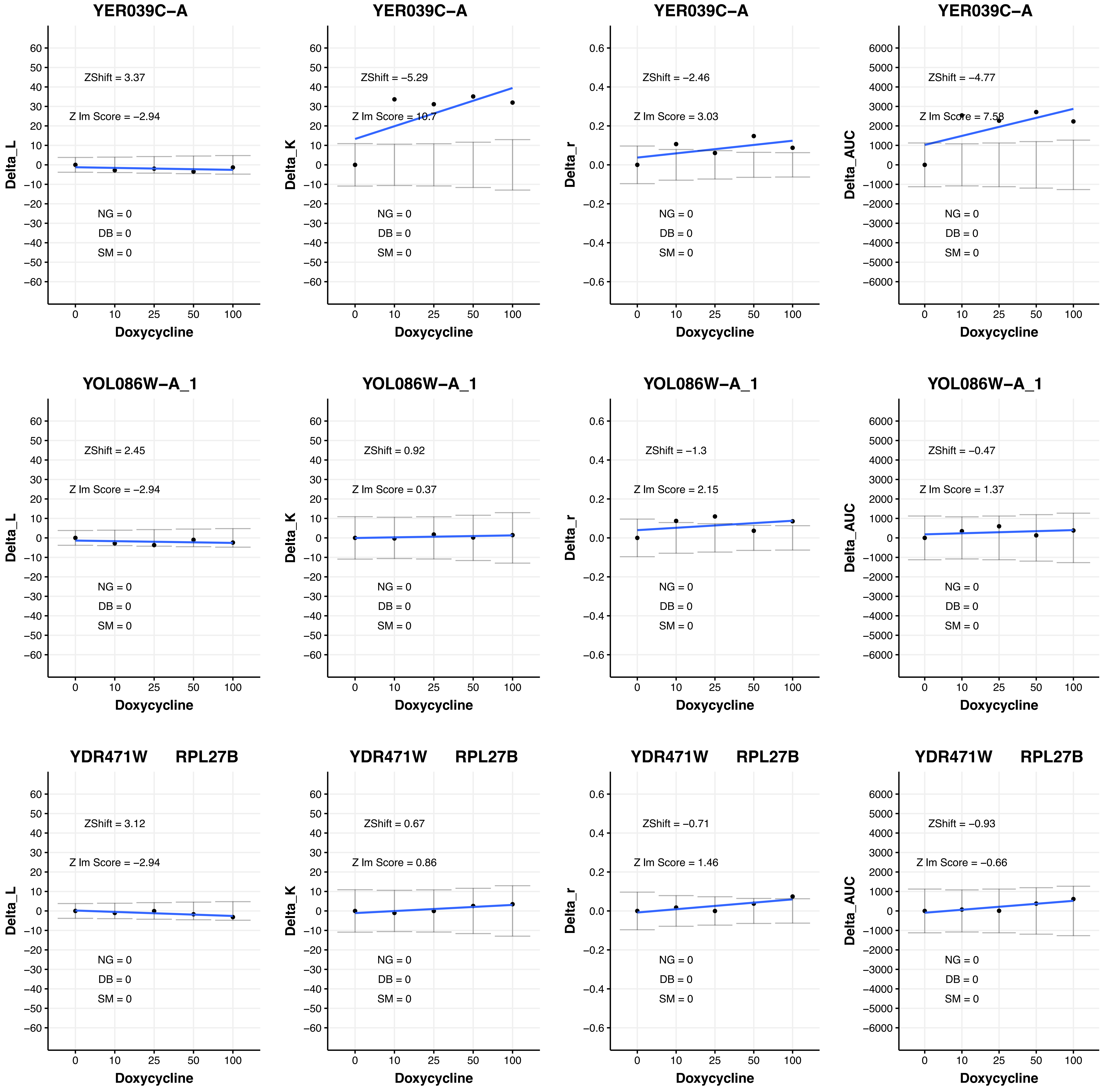

YER039C−A
ZShift = 3.37
Z lm Score = −2.94
NG = 0
DB = 0
SM = 0
Delta_L
Doxycycline
YER039C−A
ZShift = −5.29
Z lm Score = 10.7
NG = 0
DB = 0
SM = 0
Delta_K
Doxycycline
YER039C−A
ZShift = −2.46
Z lm Score = 3.03
NG = 0
DB = 0
SM = 0
Delta_r
Doxycycline
YER039C−A
ZShift = −4.77
Z lm Score = 7.58
NG = 0
DB = 0
SM = 0
Delta_AUC
Doxycycline
YOL086W−A_1
ZShift = 2.45
Z lm Score = −2.94
NG = 0
DB = 0
SM = 0
Delta_L
Doxycycline
YOL086W−A_1
ZShift = 0.92
Z lm Score = 0.37
NG = 0
DB = 0
SM = 0
Delta_K
Doxycycline
YOL086W−A_1
ZShift = −1.3
Z lm Score = 2.15
NG = 0
DB = 0
SM = 0
Delta_r
Doxycycline
YOL086W−A_1
ZShift = −0.47
Z lm Score = 1.37
NG = 0
DB = 0
SM = 0
Delta_AUC
Doxycycline
YDR471W RPL27B
ZShift = 3.12
Z lm Score = −2.94
NG = 0
DB = 0
SM = 0
Delta_L
Doxycycline
YDR471W RPL27B
ZShift = 0.67
Z lm Score = 0.86
NG = 0
DB = 0
SM = 0
Delta_K
Doxycycline
YDR471W RPL27B
ZShift = −0.71
Z lm Score = 1.46
NG = 0
DB = 0
SM = 0
Delta_r
Doxycycline
YDR471W RPL27B
ZShift = −0.93
Z lm Score = −0.66
NG = 0
DB = 0
SM = 0
Delta_AUC
Doxycycline

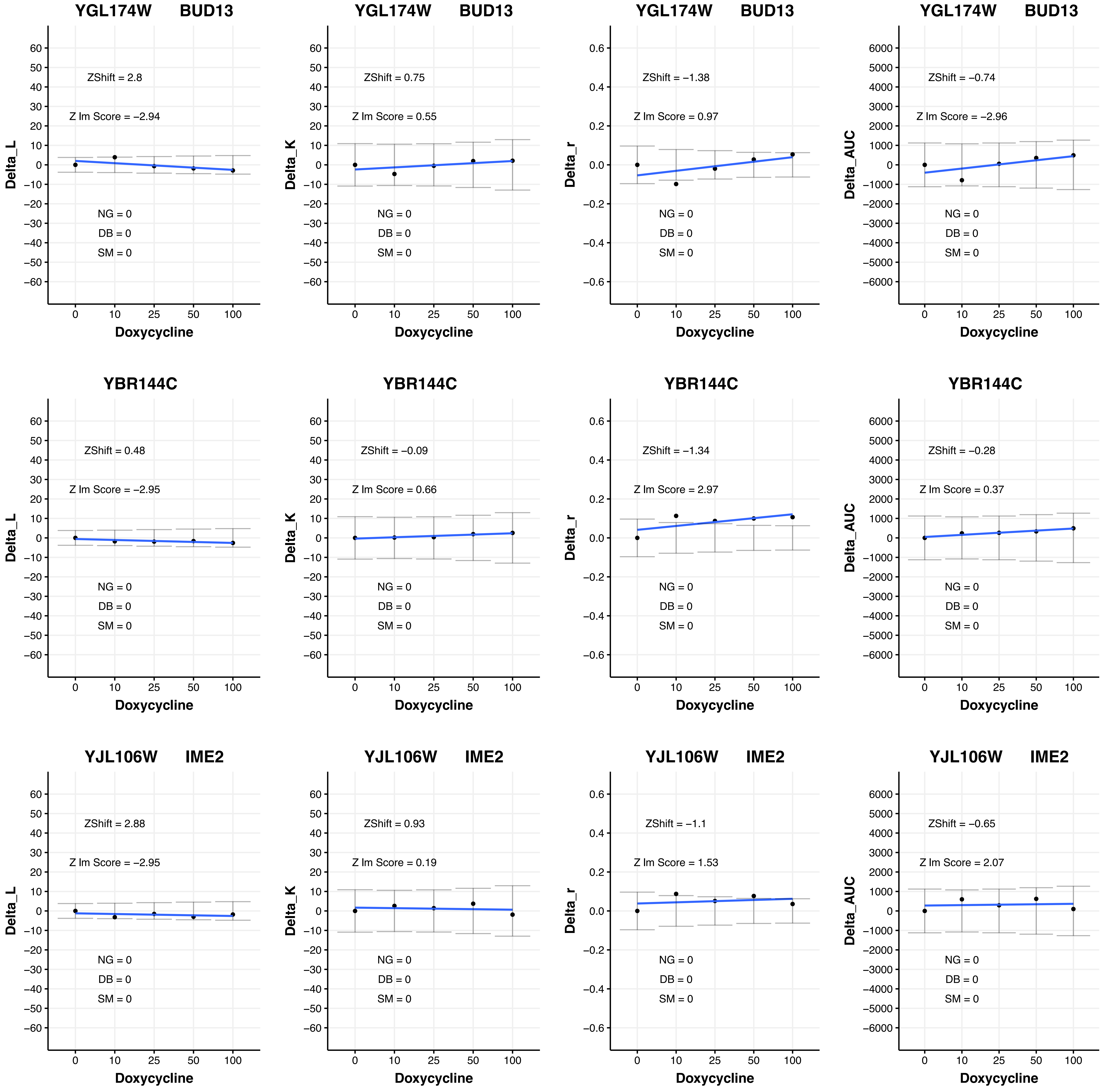
YGL174W BUD13
ZShift = 2.8
Z lm Score = −2.94
NG = 0
DB = 0
SM = 0
Delta_L
Doxycycline
YGL174W BUD13
ZShift = 0.75
Z lm Score = 0.55
NG = 0
DB = 0
SM = 0
Delta_K
Doxycycline
YGL174W BUD13
ZShift = −1.38
Z lm Score = 0.97
NG = 0
DB = 0
SM = 0
Delta_r
Doxycycline
YGL174W BUD13
ZShift = −0.74
Z lm Score = −2.96
NG = 0
DB = 0
SM = 0
Delta_AUC
Doxycycline
YBR144C
ZShift = 0.48
Z lm Score = −2.95
NG = 0
DB = 0
SM = 0
Delta_L
Doxycycline
YBR144C
ZShift = −0.09
Z lm Score = 0.66
NG = 0
DB = 0
SM = 0
Delta_K
Doxycycline
YBR144C
ZShift = −1.34
Z lm Score = 2.97
NG = 0
DB = 0
SM = 0
Delta_r
Doxycycline
YBR144C
ZShift = −0.28
Z lm Score = 0.37
NG = 0
DB = 0
SM = 0
Delta_AUC
Doxycycline
YJL106W IME2
ZShift = 2.88
Z lm Score = −2.95
NG = 0
DB = 0
SM = 0
Delta_L
Doxycycline
YJL106W IME2
ZShift = 0.93
Z lm Score = 0.19
NG = 0
DB = 0
SM = 0
Delta_K
Doxycycline
YJL106W IME2
ZShift = −1.1
Z lm Score = 1.53
NG = 0
DB = 0
SM = 0
Delta_r
Doxycycline
YJL106W IME2
ZShift = −0.65
Z lm Score = 2.07
NG = 0
DB = 0
SM = 0
Delta_AUC
Doxycycline

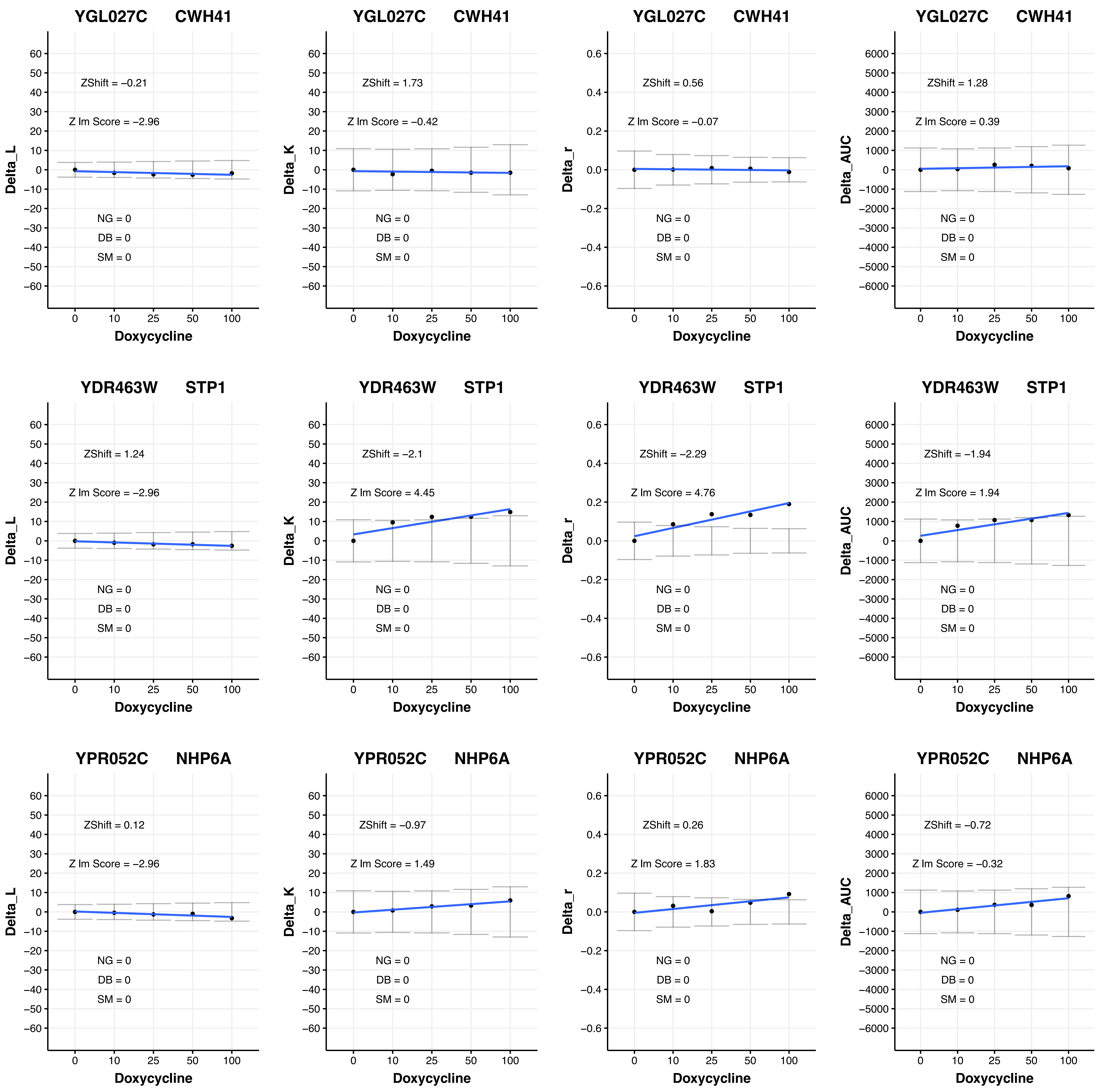
YGL027C CWH41
ZShift = −0.21
Z lm Score = −2.96
NG = 0
DB = 0
SM = 0
Delta_L
Doxycycline
YGL027C CWH41
ZShift = 1.73
Z lm Score = −0.42
NG = 0
DB = 0
SM = 0
Delta_K
Doxycycline
YGL027C CWH41
ZShift = 0.56
Z lm Score = −0.07
NG = 0
DB = 0
SM = 0
Delta_r
Doxycycline
YGL027C CWH41
ZShift = 1.28
Z lm Score = 0.39
NG = 0
DB = 0
SM = 0
Delta_AUC
Doxycycline
YDR463W STP1
ZShift = 1.24
Z lm Score = −2.96
NG = 0
DB = 0
SM = 0
Delta_L
Doxycycline
YDR463W STP1
ZShift = −2.1
Z lm Score = 4.45
NG = 0
DB = 0
SM = 0
Delta_K
Doxycycline
YDR463W STP1
ZShift = −2.29
Z lm Score = 4.76
NG = 0
DB = 0
SM = 0
Delta_r
Doxycycline
YDR463W STP1
ZShift = −1.94
Z lm Score = 1.94
NG = 0
DB = 0
SM = 0
Delta_AUC
Doxycycline
YPR052C NHP6A
ZShift = 0.12
Z lm Score = −2.96
NG = 0
DB = 0
SM = 0
Delta_L
Doxycycline
YPR052C NHP6A
ZShift = −0.97
Z lm Score = 1.49
NG = 0
DB = 0
SM = 0
Delta_K
Doxycycline
YPR052C NHP6A
ZShift = 0.26
Z lm Score = 1.83
NG = 0
DB = 0
SM = 0
Delta_r
Doxycycline
YPR052C NHP6A
ZShift = −0.72
Z lm Score = −0.32
NG = 0
DB = 0
SM = 0
Delta_AUC
Doxycycline

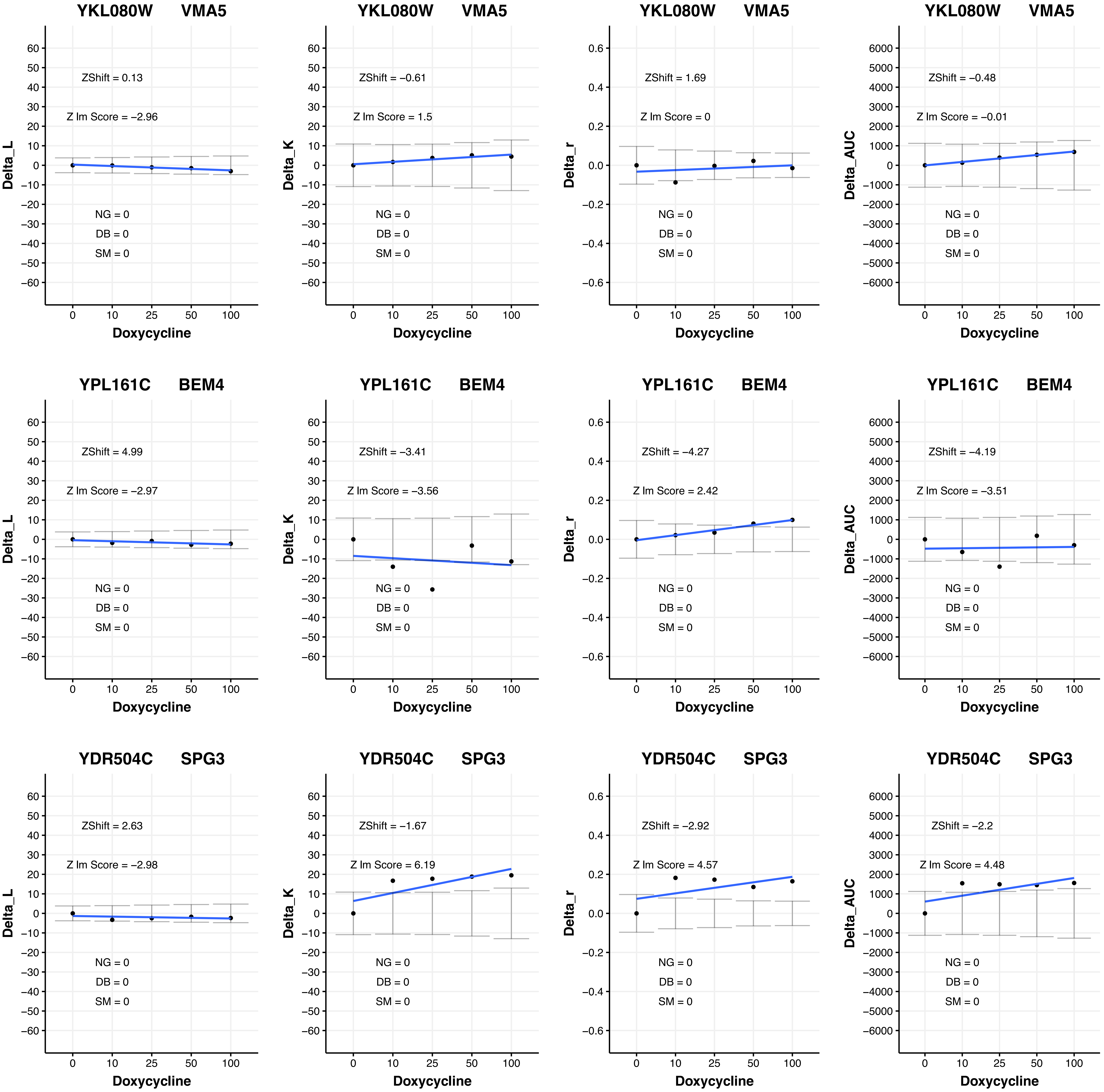
YKL080W VMA5
ZShift = 0.13
Z lm Score = −2.96
NG = 0
DB = 0
SM = 0
Delta_L
Doxycycline
YKL080W VMA5
ZShift = −0.61
Z lm Score = 1.5
NG = 0
DB = 0
SM = 0
Delta_K
Doxycycline
YKL080W VMA5
ZShift = 1.69
Z lm Score = 0
NG = 0
DB = 0
SM = 0
Delta_r
Doxycycline
YKL080W VMA5
ZShift = −0.48
Z lm Score = −0.01
NG = 0
DB = 0
SM = 0
Delta_AUC
Doxycycline
YPL161C BEM4
ZShift = 4.99
Z lm Score = −2.97
NG = 0
DB = 0
SM = 0
Delta_L
Doxycycline
YPL161C BEM4
ZShift = −3.41
Z lm Score = −3.56
NG = 0
DB = 0
SM = 0
Delta_K
Doxycycline
YPL161C BEM4
ZShift = −4.27
Z lm Score = 2.42
NG = 0
DB = 0
SM = 0
Delta_r
Doxycycline
YPL161C BEM4
ZShift = −4.19
Z lm Score = −3.51
NG = 0
DB = 0
SM = 0
Delta_AUC
Doxycycline
YDR504C SPG3
ZShift = 2.63
Z lm Score = −2.98
NG = 0
DB = 0
SM = 0
Delta_L
Doxycycline
YDR504C SPG3
ZShift = −1.67
Z lm Score = 6.19
NG = 0
DB = 0
SM = 0
Delta_K
Doxycycline
YDR504C SPG3
ZShift = −2.92
Z lm Score = 4.57
NG = 0
DB = 0
SM = 0
Delta_r
Doxycycline
YDR504C SPG3
ZShift = −2.2
Z lm Score = 4.48
NG = 0
DB = 0
SM = 0
Delta_AUC
Doxycycline

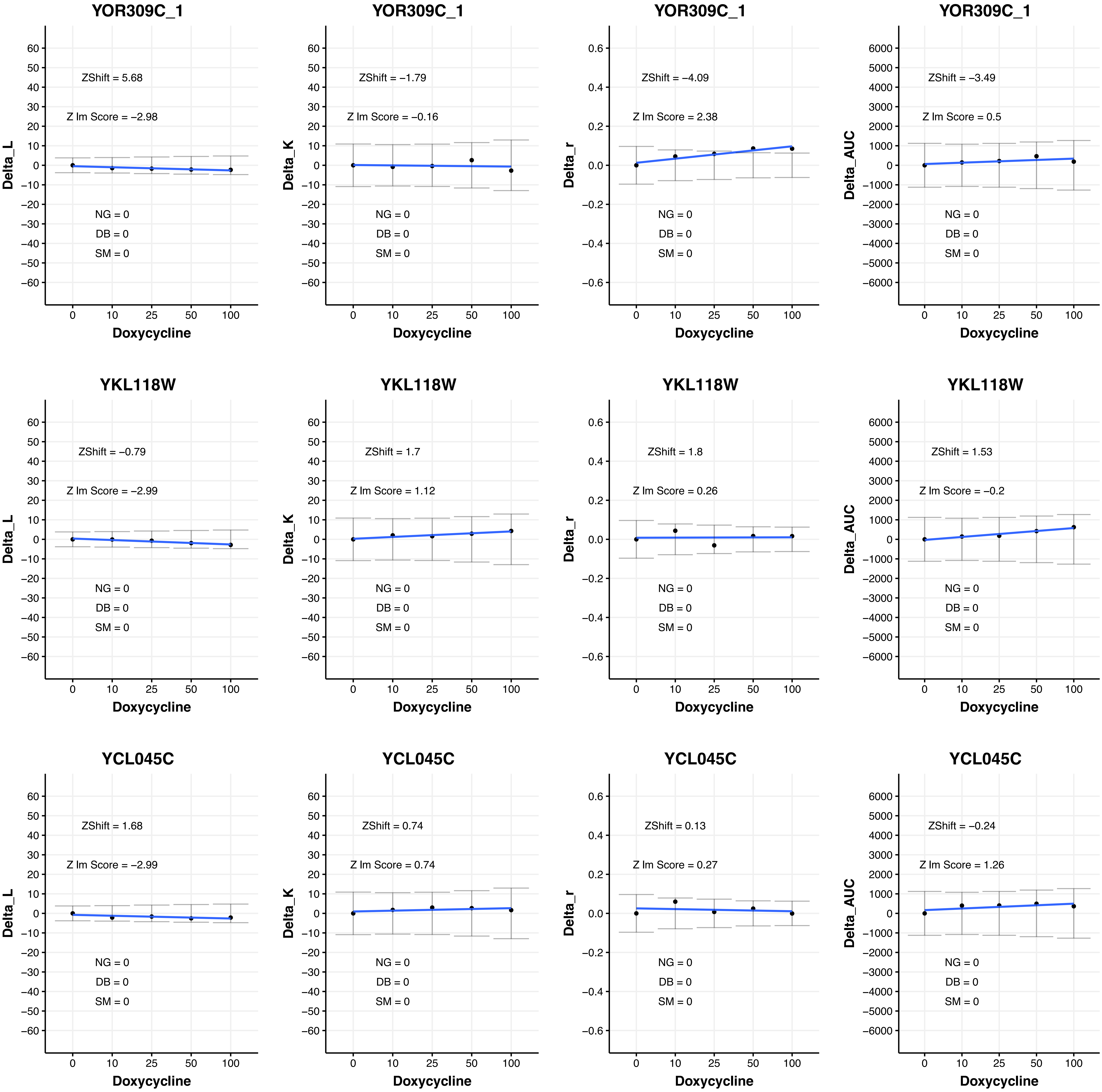
YOR309C_1
ZShift = 5.68
Z lm Score = −2.98
NG = 0
DB = 0
SM = 0
Delta_L
Doxycycline
YOR309C_1
ZShift = −1.79
Z lm Score = −0.16
NG = 0
DB = 0
SM = 0
Delta_K
Doxycycline
YOR309C_1
ZShift = −4.09
Z lm Score = 2.38
NG = 0
DB = 0
SM = 0
Delta_r
Doxycycline
YOR309C_1
ZShift = −3.49
Z lm Score = 0.5
NG = 0
DB = 0
SM = 0
Delta_AUC
Doxycycline
YKL118W
ZShift = −0.79
Z lm Score = −2.99
NG = 0
DB = 0
SM = 0
Delta_L
Doxycycline
YKL118W
ZShift = 1.7
Z lm Score = 1.12
NG = 0
DB = 0
SM = 0
Delta_K
Doxycycline
YKL118W
ZShift = 1.8
Z lm Score = 0.26
NG = 0
DB = 0
SM = 0
Delta_r
Doxycycline
YKL118W
ZShift = 1.53
Z lm Score = −0.2
NG = 0
DB = 0
SM = 0
Delta_AUC
Doxycycline
YCL045C
ZShift = 1.68
Z lm Score = −2.99
NG = 0
DB = 0
SM = 0
Delta_L
Doxycycline
YCL045C
ZShift = 0.74
Z lm Score = 0.74
NG = 0
DB = 0
SM = 0
Delta_K
Doxycycline
YCL045C
ZShift = 0.13
Z lm Score = 0.27
NG = 0
DB = 0
SM = 0
Delta_r
Doxycycline
YCL045C
ZShift = −0.24
Z lm Score = 1.26
NG = 0
DB = 0
SM = 0
Delta_AUC
Doxycycline

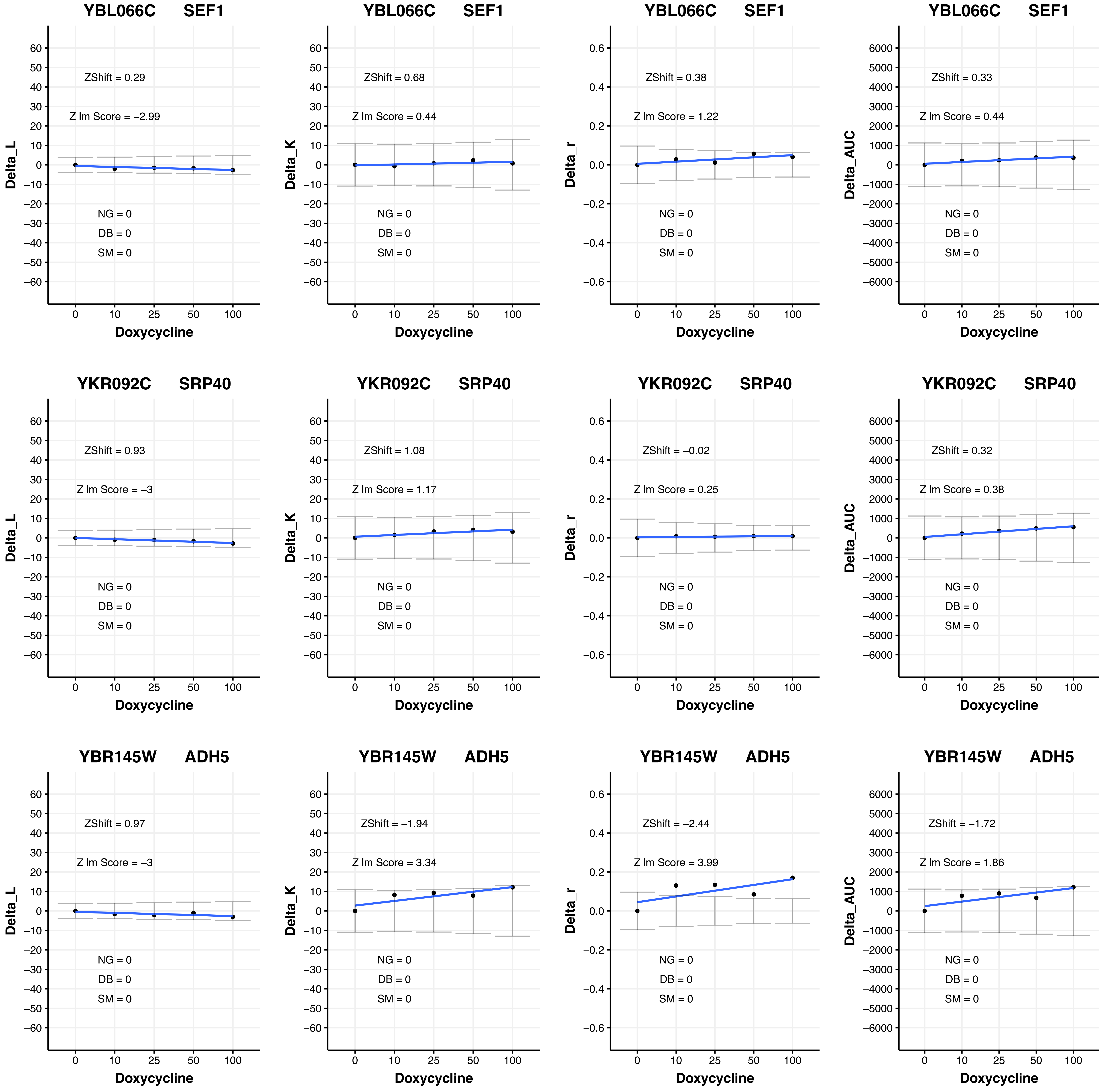
YBL066C SEF1
ZShift = 0.29
Z lm Score = −2.99
NG = 0
DB = 0
SM = 0
Delta_L
Doxycycline
YBL066C SEF1
ZShift = 0.68
Z lm Score = 0.44
NG = 0
DB = 0
SM = 0
Delta_K
Doxycycline
YBL066C SEF1
ZShift = 0.38
Z lm Score = 1.22
NG = 0
DB = 0
SM = 0
Delta_r
Doxycycline
YBL066C SEF1
ZShift = 0.33
Z lm Score = 0.44
NG = 0
DB = 0
SM = 0
Delta_AUC
Doxycycline
YKR092C SRP40
ZShift = 0.93
Z lm Score = −3
NG = 0
DB = 0
SM = 0
Delta_L
Doxycycline
YKR092C SRP40
ZShift = 1.08
Z lm Score = 1.17
NG = 0
DB = 0
SM = 0
Delta_K
Doxycycline
YKR092C SRP40
ZShift = −0.02
Z lm Score = 0.25
NG = 0
DB = 0
SM = 0
Delta_r
Doxycycline
YKR092C SRP40
ZShift = 0.32
Z lm Score = 0.38
NG = 0
DB = 0
SM = 0
Delta_AUC
Doxycycline
YBR145W ADH5
ZShift = 0.97
Z lm Score = −3
NG = 0
DB = 0
SM = 0
Delta_L
Doxycycline
YBR145W ADH5
ZShift = −1.94
Z lm Score = 3.34
NG = 0
DB = 0
SM = 0
Delta_K
Doxycycline
YBR145W ADH5
ZShift = −2.44
Z lm Score = 3.99
NG = 0
DB = 0
SM = 0
Delta_r
Doxycycline
YBR145W ADH5
ZShift = −1.72
Z lm Score = 1.86
NG = 0
DB = 0
SM = 0
Delta_AUC
Doxycycline

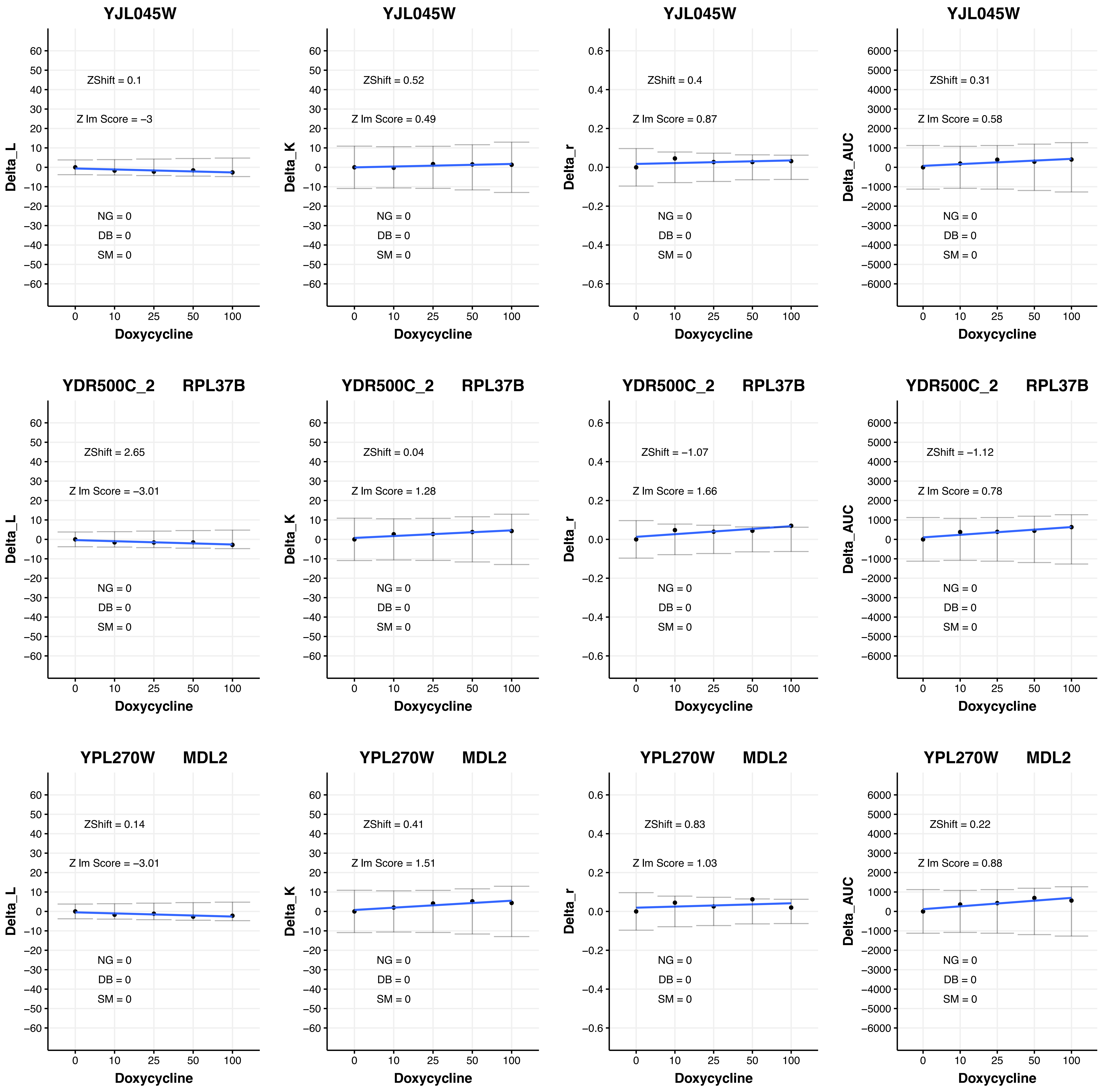
YJL045W
ZShift = 0.1
Z lm Score = −3
NG = 0
DB = 0
SM = 0
Delta_L
Doxycycline
YJL045W
ZShift = 0.52
Z lm Score = 0.49
NG = 0
DB = 0
SM = 0
Delta_K
Doxycycline
YJL045W
ZShift = 0.4
Z lm Score = 0.87
NG = 0
DB = 0
SM = 0
Delta_r
Doxycycline
YJL045W
ZShift = 0.31
Z lm Score = 0.58
NG = 0
DB = 0
SM = 0
Delta_AUC
Doxycycline
YDR500C_2 RPL37B
ZShift = 2.65
Z lm Score = −3.01
NG = 0
DB = 0
SM = 0
Delta_L
Doxycycline
YDR500C_2 RPL37B
ZShift = 0.04
Z lm Score = 1.28
NG = 0
DB = 0
SM = 0
Delta_K
Doxycycline
YDR500C_2 RPL37B
ZShift = −1.07
Z lm Score = 1.66
NG = 0
DB = 0
SM = 0
Delta_r
Doxycycline
YDR500C_2 RPL37B
ZShift = −1.12
Z lm Score = 0.78
NG = 0
DB = 0
SM = 0
Delta_AUC
Doxycycline
YPL270W MDL2
ZShift = 0.14
Z lm Score = −3.01
NG = 0
DB = 0
SM = 0
Delta_L
Doxycycline
YPL270W MDL2
ZShift = 0.41
Z lm Score = 1.51
NG = 0
DB = 0
SM = 0
Delta_K
Doxycycline
YPL270W MDL2
ZShift = 0.83
Z lm Score = 1.03
NG = 0
DB = 0
SM = 0
Delta_r
Doxycycline
YPL270W MDL2
ZShift = 0.22
Z lm Score = 0.88
NG = 0
DB = 0
SM = 0
Delta_AUC
Doxycycline

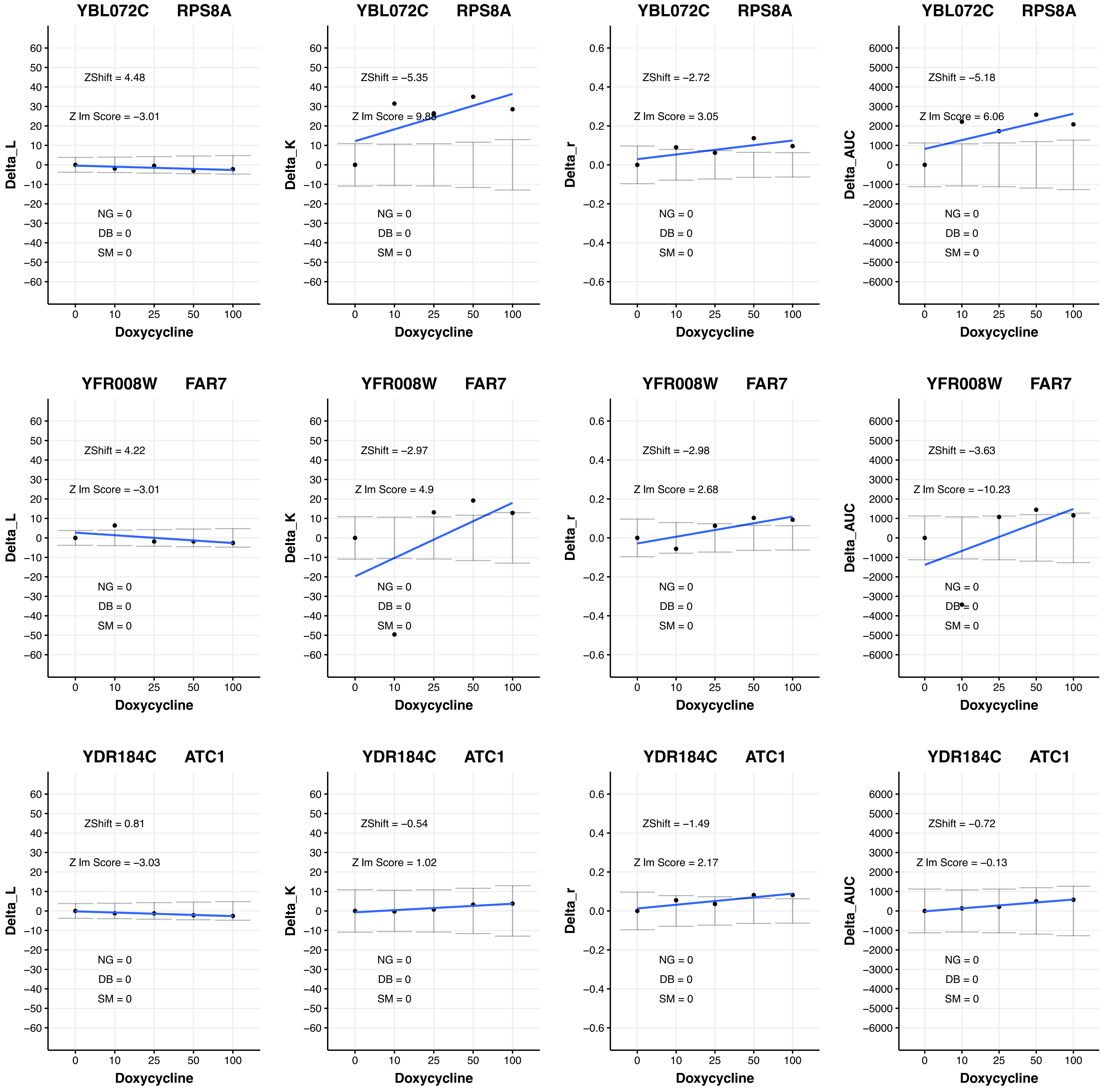

YBL072C RPS8A
ZShift = 4.48
Z lm Score = −3.01
NG = 0
DB = 0
SM = 0
Delta_L
Doxycycline
YBL072C RPS8A
ZShift = −5.35
Z lm Score = 9.88
NG = 0
DB = 0
SM = 0
Delta_K
Doxycycline
YBL072C RPS8A
ZShift = −2.72
Z lm Score = 3.05
NG = 0
DB = 0
SM = 0
Delta_r
Doxycycline
YBL072C RPS8A
ZShift = −5.18
Z lm Score = 6.06
NG = 0
DB = 0
SM = 0
Delta_AUC
Doxycycline
YFR008W FAR7
ZShift = 4.22
Z lm Score = −3.01
NG = 0
DB = 0
SM = 0
Delta_L
Doxycycline
YFR008W FAR7
ZShift = −2.97
Z lm Score = 4.9
NG = 0
DB = 0
SM = 0
Delta_K
Doxycycline
YFR008W FAR7
ZShift = −2.98
Z lm Score = 2.68
NG = 0
DB = 0
SM = 0
Delta_r
Doxycycline
YFR008W FAR7
ZShift = −3.63
Z lm Score = −10.23
NG = 0
DB = 0
SM = 0
Delta_AUC
Doxycycline
YDR184C ATC1
ZShift = 0.81
Z lm Score = −3.03
NG = 0
DB = 0
SM = 0
Delta_L
Doxycycline
YDR184C ATC1
ZShift = −0.54
Z lm Score = 1.02
NG = 0
DB = 0
SM = 0
Delta_K
Doxycycline
YDR184C ATC1
ZShift = −1.49
Z lm Score = 2.17
NG = 0
DB = 0
SM = 0
Delta_r
Doxycycline
YDR184C ATC1
ZShift = −0.72
Z lm Score = −0.13
NG = 0
DB = 0
SM = 0
Delta_AUC
Doxycycline

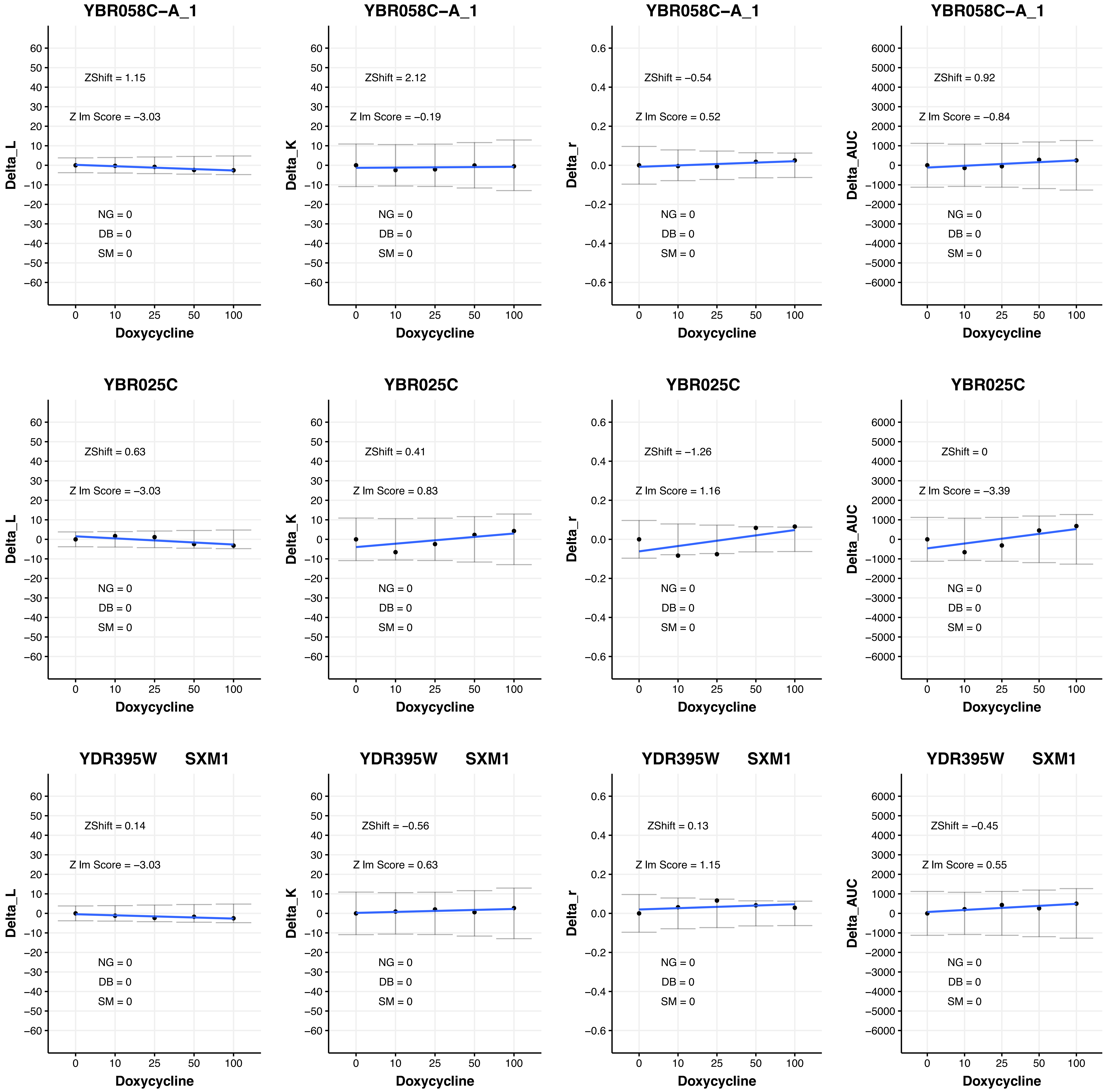
YBR058C−A_1
ZShift = 1.15
Z lm Score = −3.03
NG = 0
DB = 0
SM = 0
Delta_L
Doxycycline
YBR058C−A_1
ZShift = 2.12
Z lm Score = −0.19
NG = 0
DB = 0
SM = 0
Delta_K
Doxycycline
YBR058C−A_1
ZShift = −0.54
Z lm Score = 0.52
NG = 0
DB = 0
SM = 0
Delta_r
Doxycycline
YBR058C−A_1
ZShift = 0.92
Z lm Score = −0.84
NG = 0
DB = 0
SM = 0
Delta_AUC
Doxycycline
YBR025C
ZShift = 0.63
Z lm Score = −3.03
NG = 0
DB = 0
SM = 0
Delta_L
Doxycycline
YBR025C
ZShift = 0.41
Z lm Score = 0.83
NG = 0
DB = 0
SM = 0
Delta_K
Doxycycline
YBR025C
ZShift = −1.26
Z lm Score = 1.16
NG = 0
DB = 0
SM = 0
Delta_r
Doxycycline
YBR025C
ZShift = 0
Z lm Score = −3.39
NG = 0
DB = 0
SM = 0
Delta_AUC
Doxycycline
YDR395W SXM1
ZShift = 0.14
Z lm Score = −3.03
NG = 0
DB = 0
SM = 0
Delta_L
Doxycycline
YDR395W SXM1
ZShift = −0.56
Z lm Score = 0.63
NG = 0
DB = 0
SM = 0
Delta_K
Doxycycline
YDR395W SXM1
ZShift = 0.13
Z lm Score = 1.15
NG = 0
DB = 0
SM = 0
Delta_r
Doxycycline
YDR395W SXM1
ZShift = −0.45
Z lm Score = 0.55
NG = 0
DB = 0
SM = 0
Delta_AUC
Doxycycline

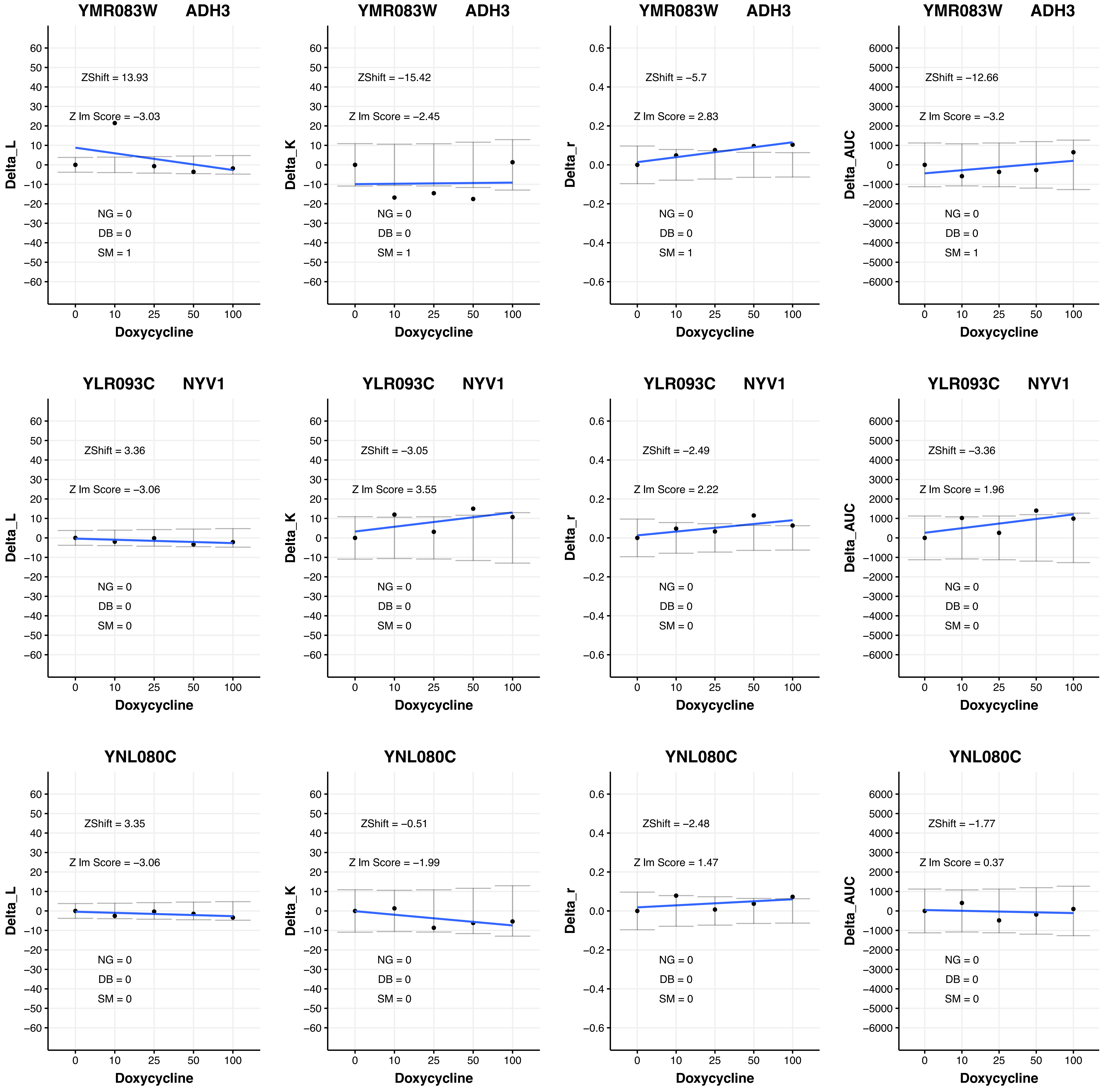
YMR083W ADH3
ZShift = 13.93
Z lm Score = −3.03
NG = 0
DB = 0
SM = 1
Delta_L
Doxycycline
YMR083W ADH3
ZShift = −15.42
Z lm Score = −2.45
NG = 0
DB = 0
SM = 1
Delta_K
Doxycycline
YMR083W ADH3
ZShift = −5.7
Z lm Score = 2.83
NG = 0
DB = 0
SM = 1
Delta_r
Doxycycline
YMR083W ADH3
ZShift = −12.66
Z lm Score = −3.2
NG = 0
DB = 0
SM = 1
Delta_AUC
Doxycycline
YLR093C NYV1
ZShift = 3.36
Z lm Score = −3.06
NG = 0
DB = 0
SM = 0
Delta_L
Doxycycline
YLR093C NYV1
ZShift = −3.05
Z lm Score = 3.55
NG = 0
DB = 0
SM = 0
Delta_K
Doxycycline
YLR093C NYV1
ZShift = −2.49
Z lm Score = 2.22
NG = 0
DB = 0
SM = 0
Delta_r
Doxycycline
YLR093C NYV1
ZShift = −3.36
Z lm Score = 1.96
NG = 0
DB = 0
SM = 0
Delta_AUC
Doxycycline
YNL080C
ZShift = 3.35
Z lm Score = −3.06
NG = 0
DB = 0
SM = 0
Delta_L
Doxycycline
YNL080C
ZShift = −0.51
Z lm Score = −1.99
NG = 0
DB = 0
SM = 0
Delta_K
Doxycycline
YNL080C
ZShift = −2.48
Z lm Score = 1.47
NG = 0
DB = 0
SM = 0
Delta_r
Doxycycline
YNL080C
ZShift = −1.77
Z lm Score = 0.37
NG = 0
DB = 0
SM = 0
Delta_AUC
Doxycycline
0 10 25 50 100

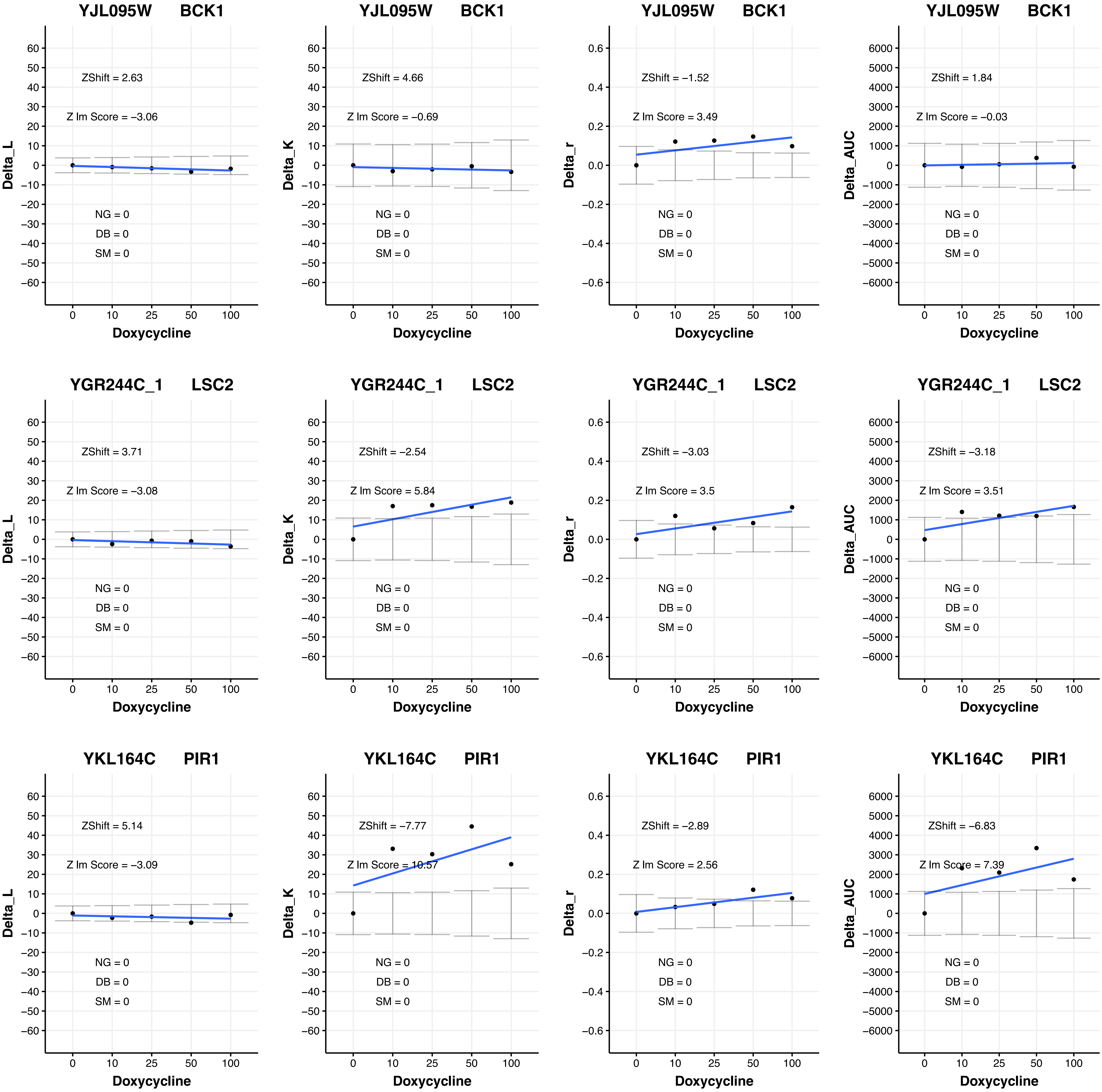
YJL095W BCK1
ZShift = 2.63
Z lm Score = −3.06
NG = 0
DB = 0
SM = 0
Delta_L
Doxycycline
YJL095W BCK1
ZShift = 4.66
Z lm Score = −0.69
NG = 0
DB = 0
SM = 0
Delta_K
Doxycycline
YJL095W BCK1
ZShift = −1.52
Z lm Score = 3.49
NG = 0
DB = 0
SM = 0
Delta_r
Doxycycline
YJL095W BCK1
ZShift = 1.84
Z lm Score = −0.03
NG = 0
DB = 0
SM = 0
Delta_AUC
Doxycycline
YGR244C_1 LSC2
ZShift = 3.71
Z lm Score = −3.08
NG = 0
DB = 0
SM = 0
Delta_L
Doxycycline
YGR244C_1 LSC2
ZShift = −2.54
Z lm Score = 5.84
NG = 0
DB = 0
SM = 0
Delta_K
Doxycycline
YGR244C_1 LSC2
ZShift = −3.03
Z lm Score = 3.5
NG = 0
DB = 0
SM = 0
Delta_r
Doxycycline
YGR244C_1 LSC2
ZShift = −3.18
Z lm Score = 3.51
NG = 0
DB = 0
SM = 0
Delta_AUC
Doxycycline
YKL164C PIR1
ZShift = 5.14
Z lm Score = −3.09
NG = 0
DB = 0
SM = 0
Delta_L
Doxycycline
YKL164C PIR1
ZShift = −7.77
Z lm Score = 10.57
NG = 0
DB = 0
SM = 0
Delta_K
Doxycycline
YKL164C PIR1
ZShift = −2.89
Z lm Score = 2.56
NG = 0
DB = 0
SM = 0
Delta_r
Doxycycline
YKL164C PIR1
ZShift = −6.83
Z lm Score = 7.39
NG = 0
DB = 0
SM = 0
Delta_AUC
Doxycycline

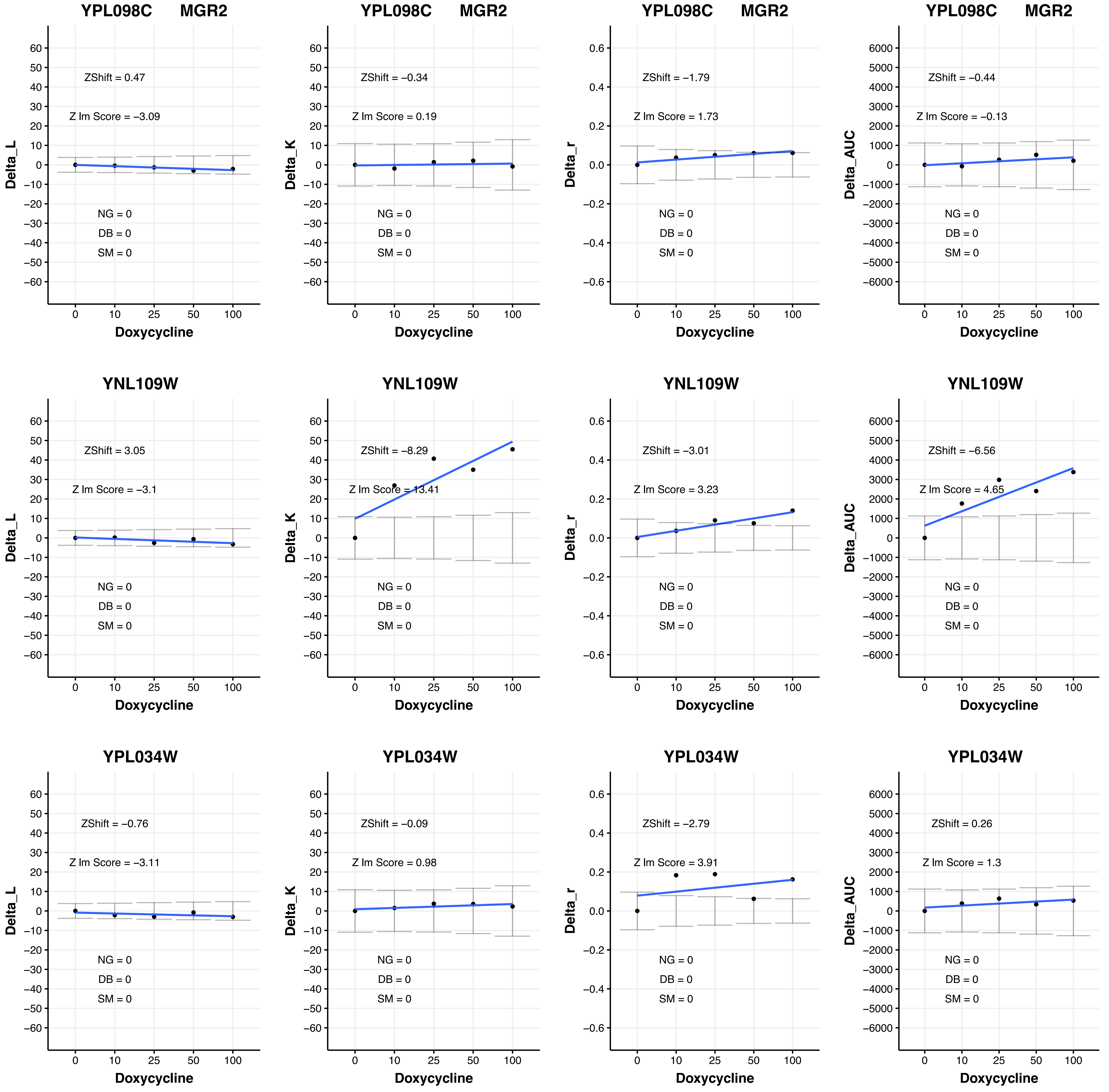
YPL098C MGR2
ZShift = 0.47
Z lm Score = −3.09
NG = 0
DB = 0
SM = 0
Delta_L
Doxycycline
YPL098C MGR2
ZShift = −0.34
Z lm Score = 0.19
NG = 0
DB = 0
SM = 0
Delta_K
Doxycycline
YPL098C MGR2
ZShift = −1.79
Z lm Score = 1.73
NG = 0
DB = 0
SM = 0
Delta_r
Doxycycline
YPL098C MGR2
ZShift = −0.44
Z lm Score = −0.13
NG = 0
DB = 0
SM = 0
Delta_AUC
Doxycycline
YNL109W
ZShift = 3.05
Z lm Score = −3.1
NG = 0
DB = 0
SM = 0
Delta_L
Doxycycline
YNL109W
ZShift = −8.29
Z lm Score = 13.41
NG = 0
DB = 0
SM = 0
Delta_K
Doxycycline
YNL109W
ZShift = −3.01
Z lm Score = 3.23
NG = 0
DB = 0
SM = 0
Delta_r
Doxycycline
YNL109W
ZShift = −6.56
Z lm Score = 4.65
NG = 0
DB = 0
SM = 0
Delta_AUC
Doxycycline
YPL034W
ZShift = −0.76
Z lm Score = −3.11
NG = 0
DB = 0
SM = 0
Delta_L
Doxycycline
YPL034W
ZShift = −0.09
Z lm Score = 0.98
NG = 0
DB = 0
SM = 0
Delta_K
Doxycycline
YPL034W
ZShift = −2.79
Z lm Score = 3.91
NG = 0
DB = 0
SM = 0
Delta_r
Doxycycline
YPL034W
ZShift = 0.26
Z lm Score = 1.3
NG = 0
DB = 0
SM = 0
Delta_AUC
Doxycycline

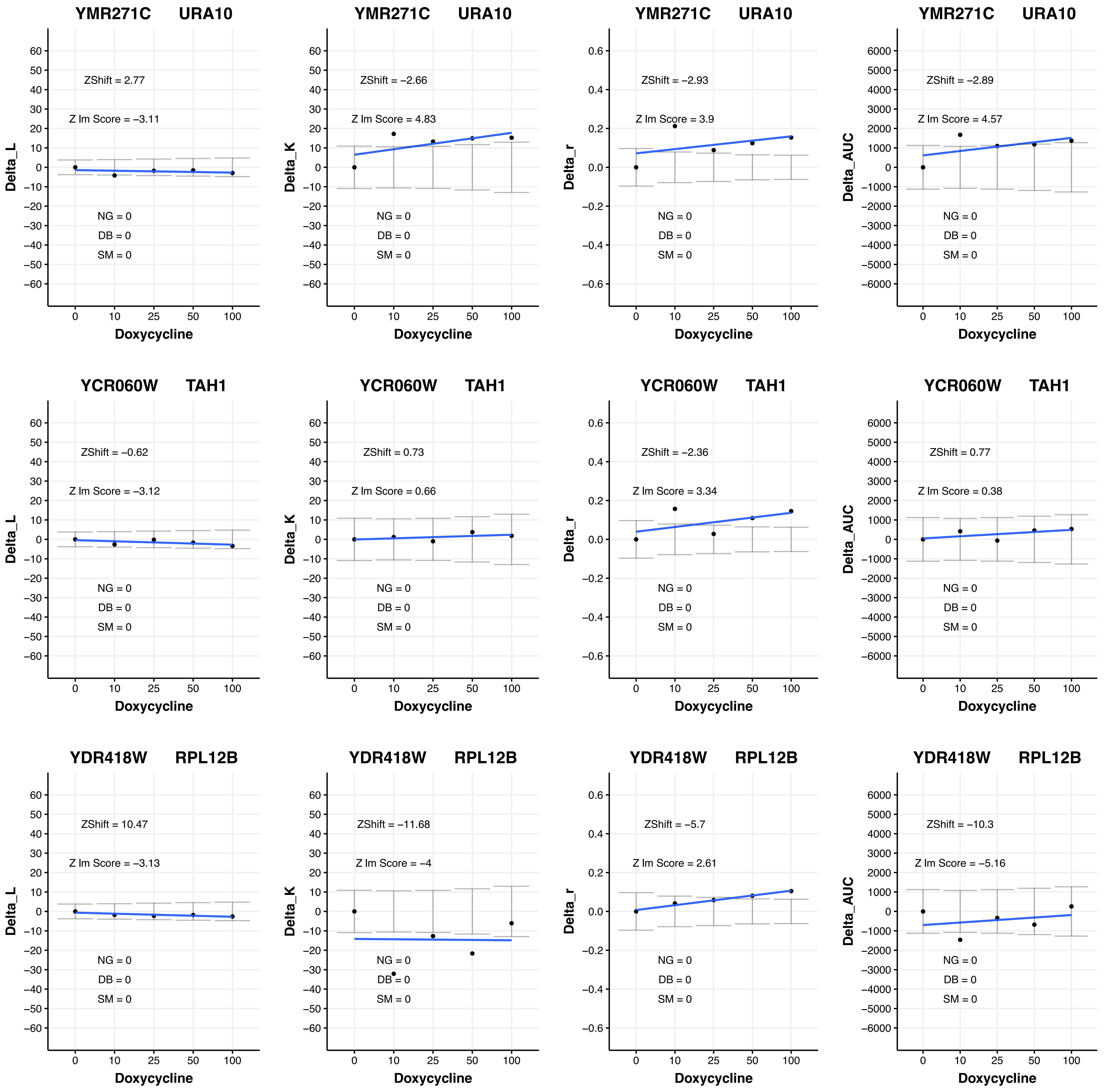

YMR271C URA10
ZShift = 2.77
Z lm Score = −3.11
NG = 0
DB = 0
SM = 0
Delta_L
Doxycycline
YMR271C URA10
ZShift = −2.66
Z lm Score = 4.83
NG = 0
DB = 0
SM = 0
Delta_K
Doxycycline
YMR271C URA10
ZShift = −2.93
Z lm Score = 3.9
NG = 0
DB = 0
SM = 0
Delta_r
Doxycycline
YMR271C URA10
ZShift = −2.89
Z lm Score = 4.57
NG = 0
DB = 0
SM = 0
Delta_AUC
Doxycycline
YCR060W TAH1
ZShift = −0.62
Z lm Score = −3.12
NG = 0
DB = 0
SM = 0
Delta_L
Doxycycline
YCR060W TAH1
ZShift = 0.73
Z lm Score = 0.66
NG = 0
DB = 0
SM = 0
Delta_K
Doxycycline
YCR060W TAH1
ZShift = −2.36
Z lm Score = 3.34
NG = 0
DB = 0
SM = 0
Delta_r
Doxycycline
YCR060W TAH1
ZShift = 0.77
Z lm Score = 0.38
NG = 0
DB = 0
SM = 0
Delta_AUC
Doxycycline
YDR418W RPL12B
ZShift = 10.47
Z lm Score = −3.13
NG = 0
DB = 0
SM = 0
Delta_L
Doxycycline
YDR418W RPL12B
ZShift = −11.68
Z lm Score = −4
NG = 0
DB = 0
SM = 0
Delta_K
Doxycycline
YDR418W RPL12B
ZShift = −5.7
Z lm Score = 2.61
NG = 0
DB = 0
SM = 0
Delta_r
Doxycycline
YDR418W RPL12B
ZShift = −10.3
Z lm Score = −5.16
NG = 0
DB = 0
SM = 0
Delta_AUC
Doxycycline

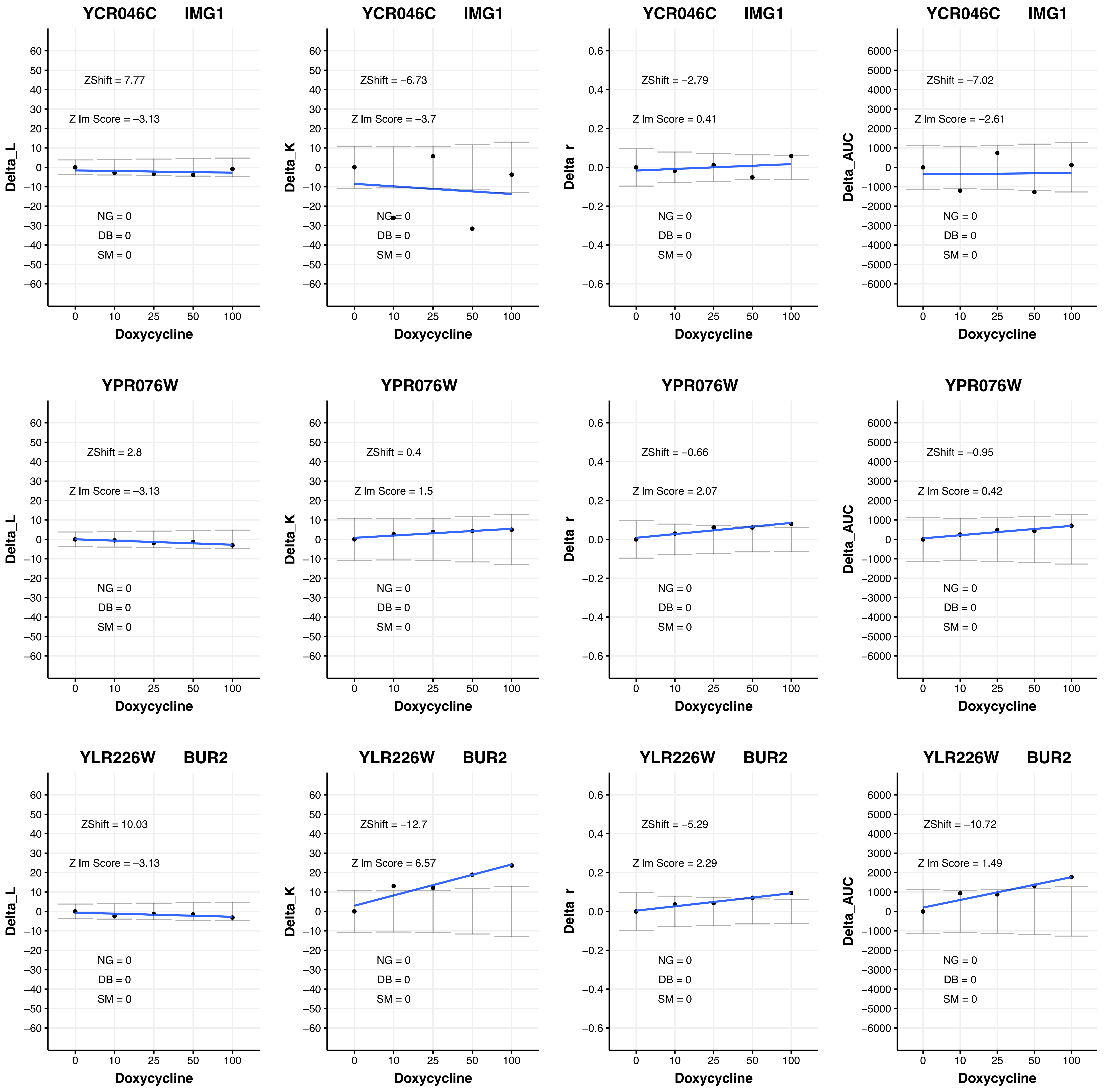
YCR046C IMG1
ZShift = 7.77
Z lm Score = −3.13
NG = 0
DB = 0
SM = 0
Delta_L
Doxycycline
YCR046C IMG1
ZShift = −6.73
Z lm Score = −3.7
NG = 0
DB = 0
SM = 0
Delta_K
Doxycycline
YCR046C IMG1
ZShift = −2.79
Z lm Score = 0.41
NG = 0
DB = 0
SM = 0
Delta_r
Doxycycline
YCR046C IMG1
ZShift = −7.02
Z lm Score = −2.61
NG = 0
DB = 0
SM = 0
Delta_AUC
Doxycycline
YPR076W
ZShift = 2.8
Z lm Score = −3.13
NG = 0
DB = 0
SM = 0
Delta_L
Doxycycline
YPR076W
ZShift = 0.4
Z lm Score = 1.5
NG = 0
DB = 0
SM = 0
Delta_K
Doxycycline
YPR076W
ZShift = −0.66
Z lm Score = 2.07
NG = 0
DB = 0
SM = 0
Delta_r
Doxycycline
YPR076W
ZShift = −0.95
Z lm Score = 0.42
NG = 0
DB = 0
SM = 0
Delta_AUC
Doxycycline
YLR226W BUR2
ZShift = 10.03
Z lm Score = −3.13
NG = 0
DB = 0
SM = 0
Delta_L
Doxycycline
YLR226W BUR2
ZShift = −12.7
Z lm Score = 6.57
NG = 0
DB = 0
SM = 0
Delta_K
Doxycycline
YLR226W BUR2
ZShift = −5.29
Z lm Score = 2.29
NG = 0
DB = 0
SM = 0
Delta_r
Doxycycline
YLR226W BUR2
ZShift = −10.72
Z lm Score = 1.49
NG = 0
DB = 0
SM = 0
Delta_AUC
Doxycycline

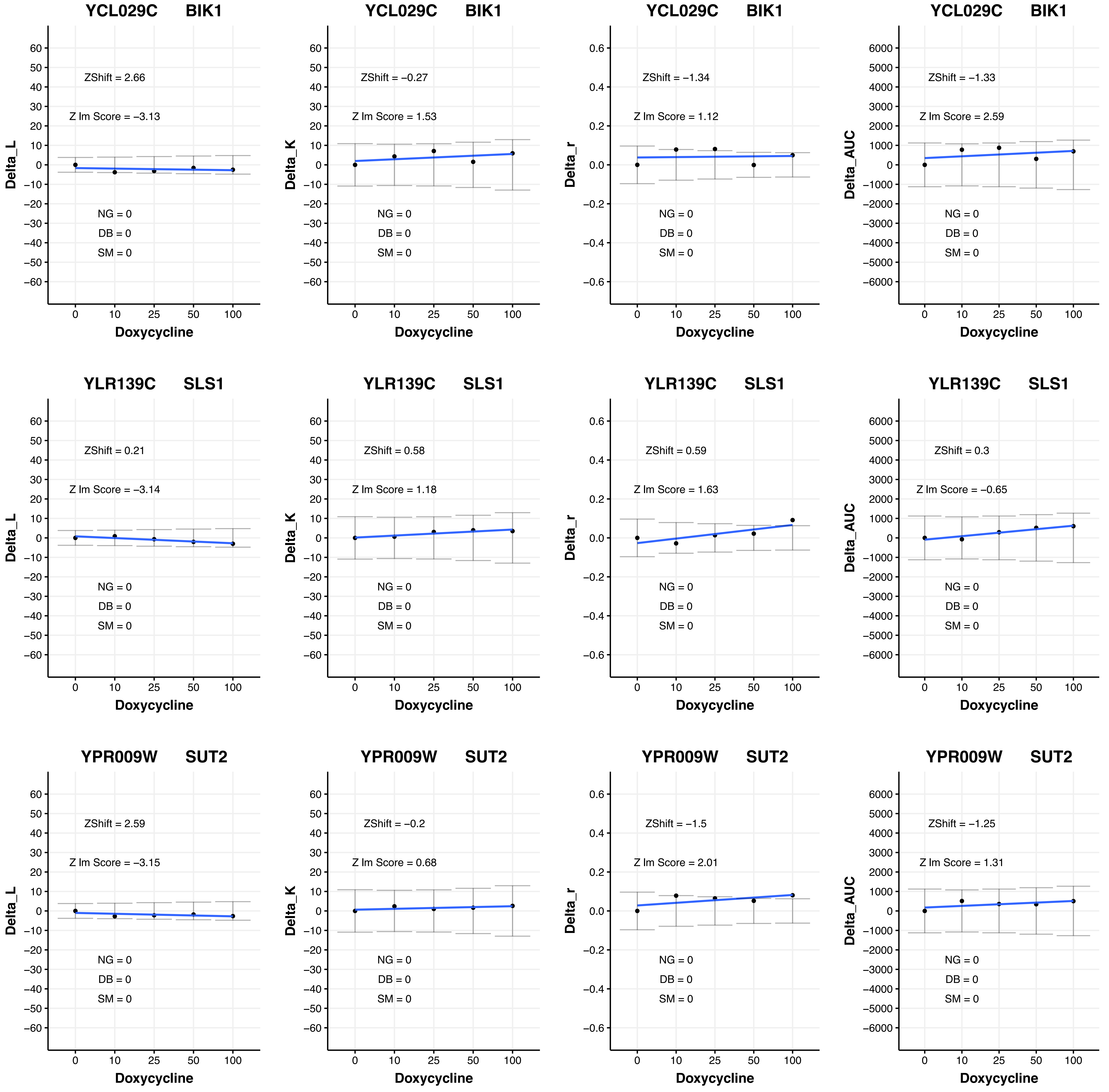

YCL029C BIK1
ZShift = 2.66
Z lm Score = −3.13
NG = 0
DB = 0
SM = 0
Delta_L
Doxycycline
YCL029C BIK1
ZShift = −0.27
Z lm Score = 1.53
NG = 0
DB = 0
SM = 0
Delta_K
Doxycycline
YCL029C BIK1
ZShift = −1.34
Z lm Score = 1.12
NG = 0
DB = 0
SM = 0
Delta_r
Doxycycline
YCL029C BIK1
ZShift = −1.33
Z lm Score = 2.59
NG = 0
DB = 0
SM = 0
Delta_AUC
Doxycycline
YLR139C SLS1
ZShift = 0.21
Z lm Score = −3.14
NG = 0
DB = 0
SM = 0
Delta_L
Doxycycline
YLR139C SLS1
ZShift = 0.58
Z lm Score = 1.18
NG = 0
DB = 0
SM = 0
Delta_K
Doxycycline
YLR139C SLS1
ZShift = 0.59
Z lm Score = 1.63
NG = 0
DB = 0
SM = 0
Delta_r
Doxycycline
YLR139C SLS1
ZShift = 0.3
Z lm Score = −0.65
NG = 0
DB = 0
SM = 0
Delta_AUC
Doxycycline
YPR009W SUT2
ZShift = 2.59
Z lm Score = −3.15
NG = 0
DB = 0
SM = 0
Delta_L
Doxycycline
YPR009W SUT2
ZShift = −0.2
Z lm Score = 0.68
NG = 0
DB = 0
SM = 0
Delta_K
Doxycycline
YPR009W SUT2
ZShift = −1.5
Z lm Score = 2.01
NG = 0
DB = 0
SM = 0
Delta_r
Doxycycline
YPR009W SUT2
ZShift = −1.25
Z lm Score = 1.31
NG = 0
DB = 0
SM = 0
Delta_AUC
Doxycycline

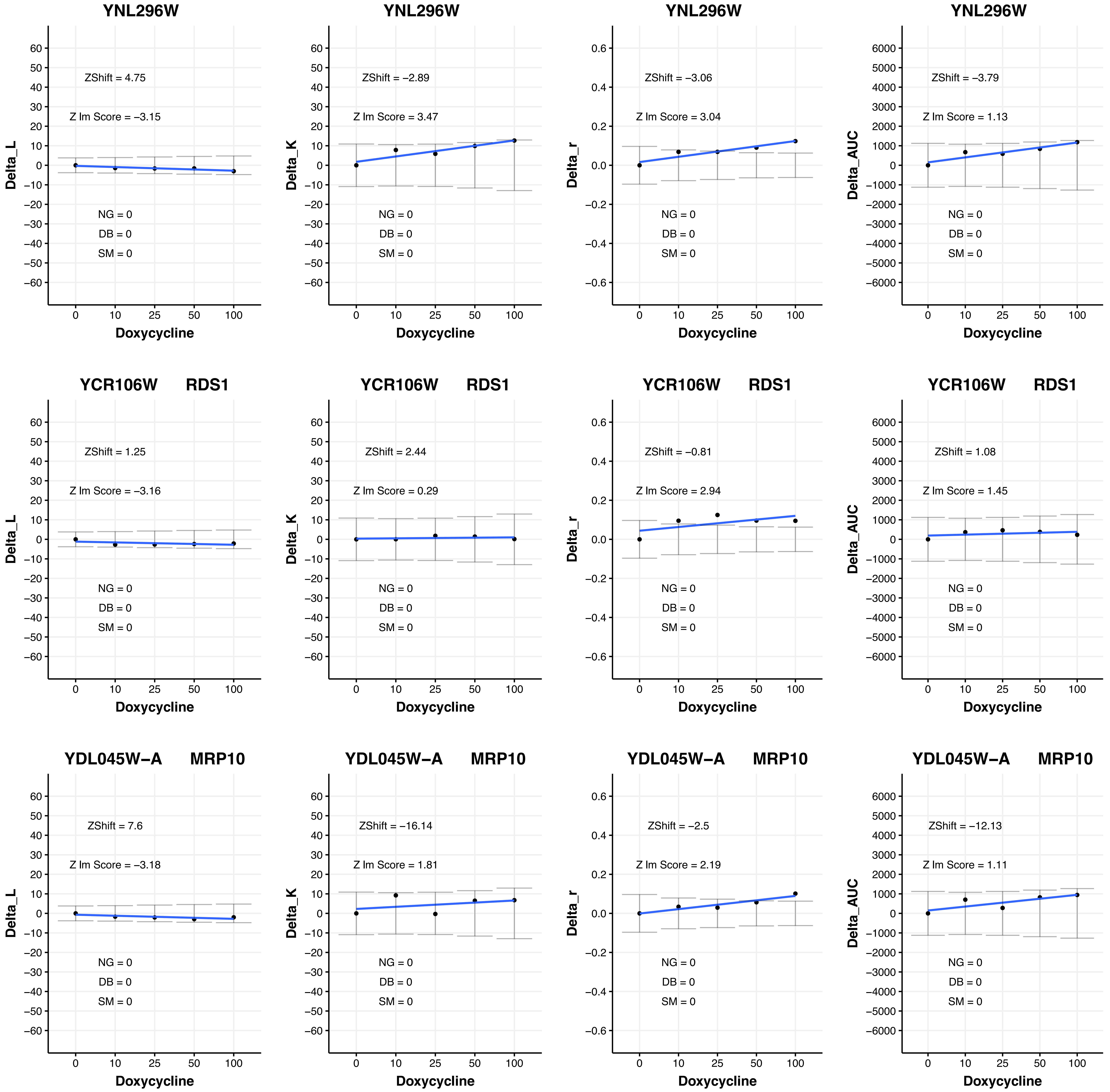
YNL296W
ZShift = 4.75
Z lm Score = −3.15
NG = 0
DB = 0
SM = 0
Delta_L
Doxycycline
YNL296W
ZShift = −2.89
Z lm Score = 3.47
NG = 0
DB = 0
SM = 0
Delta_K
Doxycycline
YNL296W
ZShift = −3.06
Z lm Score = 3.04
NG = 0
DB = 0
SM = 0
Delta_r
Doxycycline
YNL296W
ZShift = −3.79
Z lm Score = 1.13
NG = 0
DB = 0
SM = 0
Delta_AUC
Doxycycline
YCR106W RDS1
ZShift = 1.25
Z lm Score = −3.16
NG = 0
DB = 0
SM = 0
Delta_L
Doxycycline
YCR106W RDS1
ZShift = 2.44
Z lm Score = 0.29
NG = 0
DB = 0
SM = 0
Delta_K
Doxycycline
YCR106W RDS1
ZShift = −0.81
Z lm Score = 2.94
NG = 0
DB = 0
SM = 0
Delta_r
Doxycycline
YCR106W RDS1
ZShift = 1.08
Z lm Score = 1.45
NG = 0
DB = 0
SM = 0
Delta_AUC
Doxycycline
YDL045W−A MRP10
ZShift = 7.6
Z lm Score = −3.18
NG = 0
DB = 0
SM = 0
Delta_L
Doxycycline
YDL045W−A MRP10
ZShift = −16.14
Z lm Score = 1.81
NG = 0
DB = 0
SM = 0
Delta_K
Doxycycline
YDL045W−A MRP10
ZShift = −2.5
Z lm Score = 2.19
NG = 0
DB = 0
SM = 0
Delta_r
Doxycycline
YDL045W−A MRP10
ZShift = −12.13
Z lm Score = 1.11
NG = 0
DB = 0
SM = 0
Delta_AUC
Doxycycline

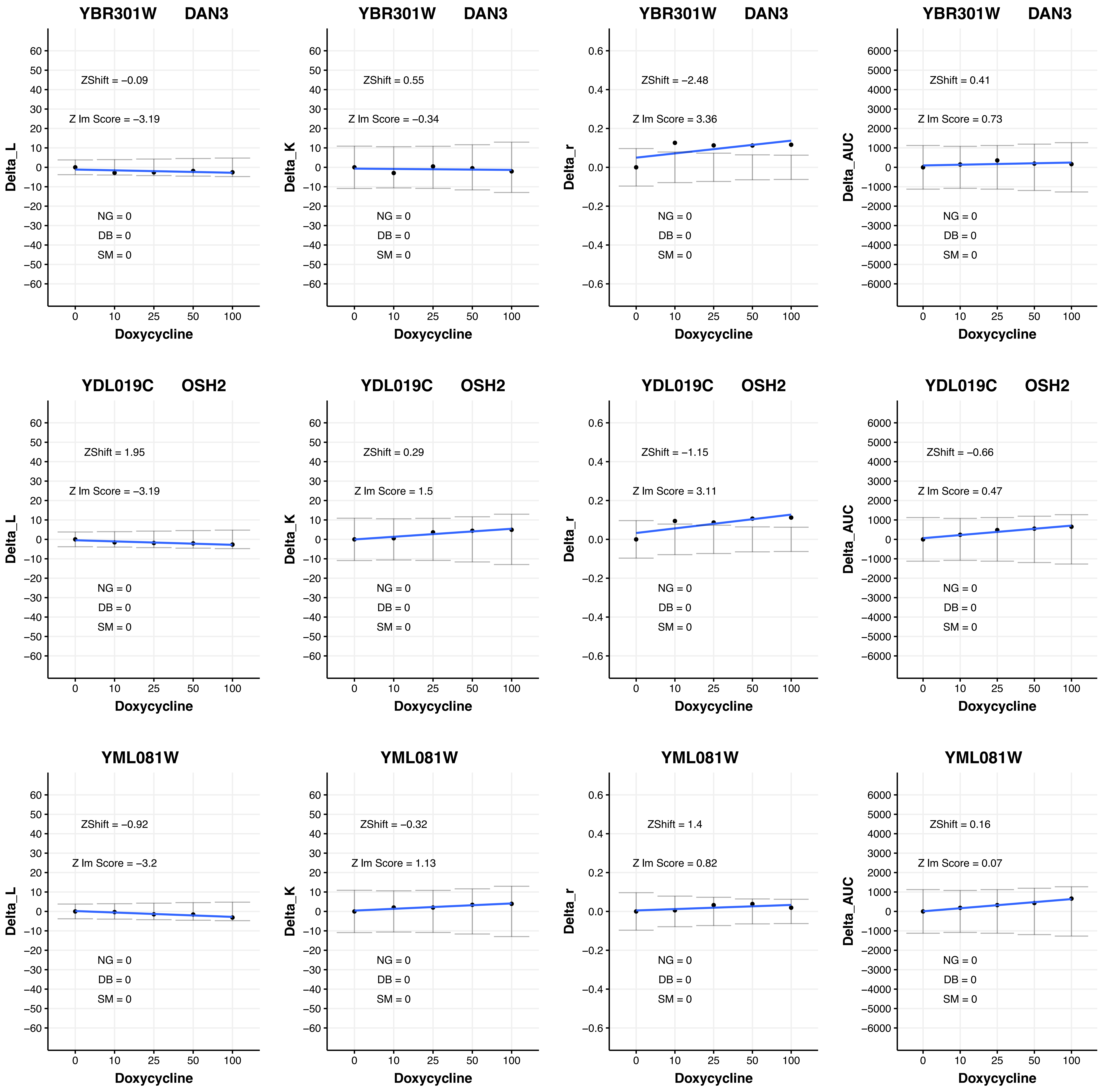
YBR301W DAN3
ZShift = −0.09
Z lm Score = −3.19
NG = 0
DB = 0
SM = 0
Delta_L
Doxycycline
YBR301W DAN3
ZShift = 0.55
Z lm Score = −0.34
NG = 0
DB = 0
SM = 0
Delta_K
Doxycycline
YBR301W DAN3
ZShift = −2.48
Z lm Score = 3.36
NG = 0
DB = 0
SM = 0
Delta_r
Doxycycline
YBR301W DAN3
ZShift = 0.41
Z lm Score = 0.73
NG = 0
DB = 0
SM = 0
Delta_AUC
Doxycycline
YDL019C OSH2
ZShift = 1.95
Z lm Score = −3.19
NG = 0
DB = 0
SM = 0
Delta_L
Doxycycline
YDL019C OSH2
ZShift = 0.29
Z lm Score = 1.5
NG = 0
DB = 0
SM = 0
Delta_K
Doxycycline
YDL019C OSH2
ZShift = −1.15
Z lm Score = 3.11
NG = 0
DB = 0
SM = 0
Delta_r
Doxycycline
YDL019C OSH2
ZShift = −0.66
Z lm Score = 0.47
NG = 0
DB = 0
SM = 0
Delta_AUC
Doxycycline
YML081W
ZShift = −0.92
Z lm Score = −3.2
NG = 0
DB = 0
SM = 0
Delta_L
Doxycycline
YML081W
ZShift = −0.32
Z lm Score = 1.13
NG = 0
DB = 0
SM = 0
Delta_K
Doxycycline
YML081W
ZShift = 1.4
Z lm Score = 0.82
NG = 0
DB = 0
SM = 0
Delta_r
Doxycycline
YML081W
ZShift = 0.16
Z lm Score = 0.07
NG = 0
DB = 0
SM = 0
Delta_AUC
Doxycycline

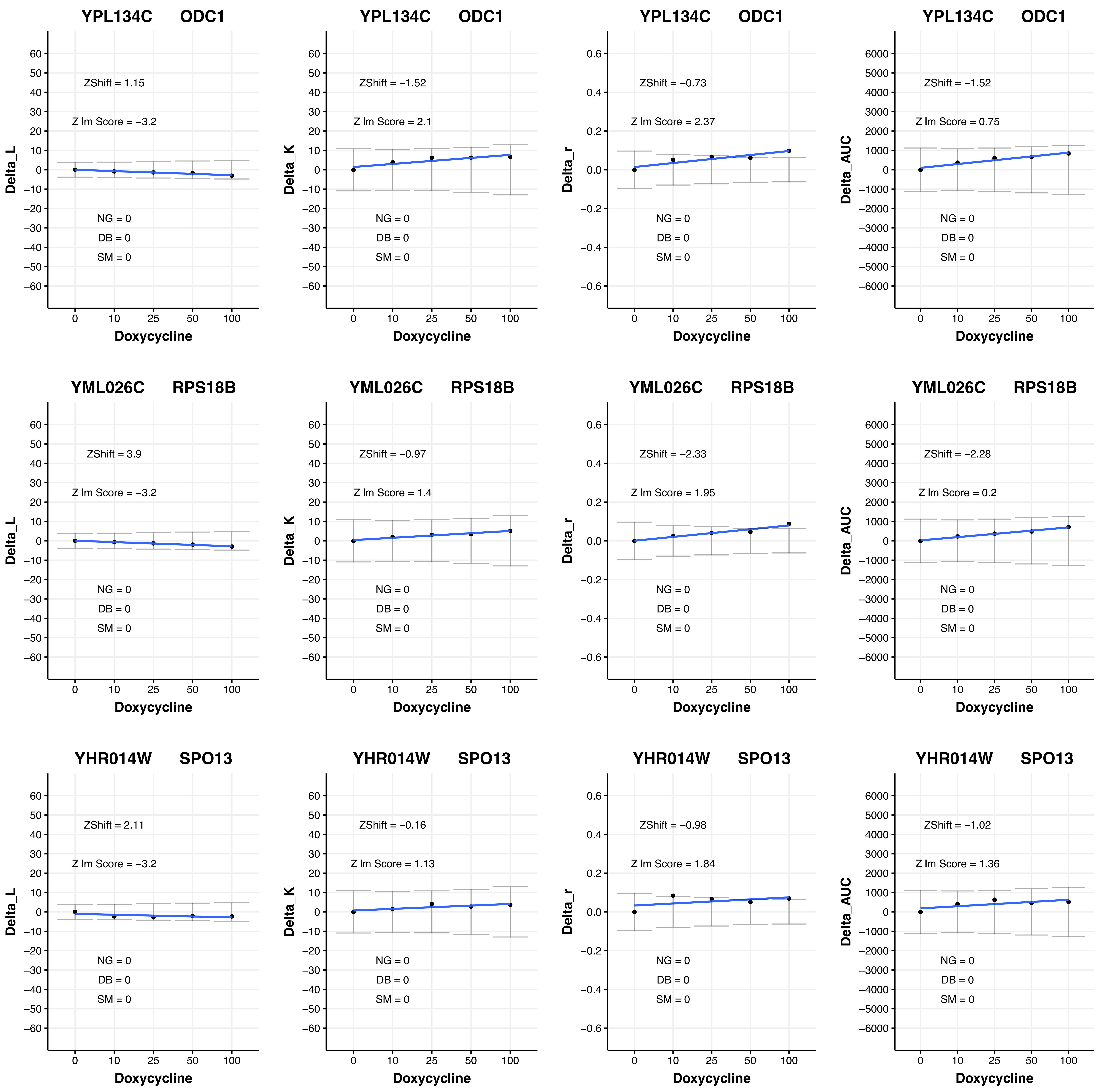
YPL134C ODC1
ZShift = 1.15
Z lm Score = −3.2
NG = 0
DB = 0
SM = 0
Delta_L
Doxycycline
YPL134C ODC1
ZShift = −1.52
Z lm Score = 2.1
NG = 0
DB = 0
SM = 0
Delta_K
Doxycycline
YPL134C ODC1
ZShift = −0.73
Z lm Score = 2.37
NG = 0
DB = 0
SM = 0
Delta_r
Doxycycline
YPL134C ODC1
ZShift = −1.52
Z lm Score = 0.75
NG = 0
DB = 0
SM = 0
Delta_AUC
Doxycycline
YML026C RPS18B
ZShift = 3.9
Z lm Score = −3.2
NG = 0
DB = 0
SM = 0
Delta_L
Doxycycline
YML026C RPS18B
ZShift = −0.97
Z lm Score = 1.4
NG = 0
DB = 0
SM = 0
Delta_K
Doxycycline
YML026C RPS18B
ZShift = −2.33
Z lm Score = 1.95
NG = 0
DB = 0
SM = 0
Delta_r
Doxycycline
YML026C RPS18B
ZShift = −2.28
Z lm Score = 0.2
NG = 0
DB = 0
SM = 0
Delta_AUC
Doxycycline
YHR014W SPO13
ZShift = 2.11
Z lm Score = −3.2
NG = 0
DB = 0
SM = 0
Delta_L
Doxycycline
YHR014W SPO13
ZShift = −0.16
Z lm Score = 1.13
NG = 0
DB = 0
SM = 0
Delta_K
Doxycycline
YHR014W SPO13
ZShift = −0.98
Z lm Score = 1.84
NG = 0
DB = 0
SM = 0
Delta_r
Doxycycline
YHR014W SPO13
ZShift = −1.02
Z lm Score = 1.36
NG = 0
DB = 0
SM = 0
Delta_AUC
Doxycycline

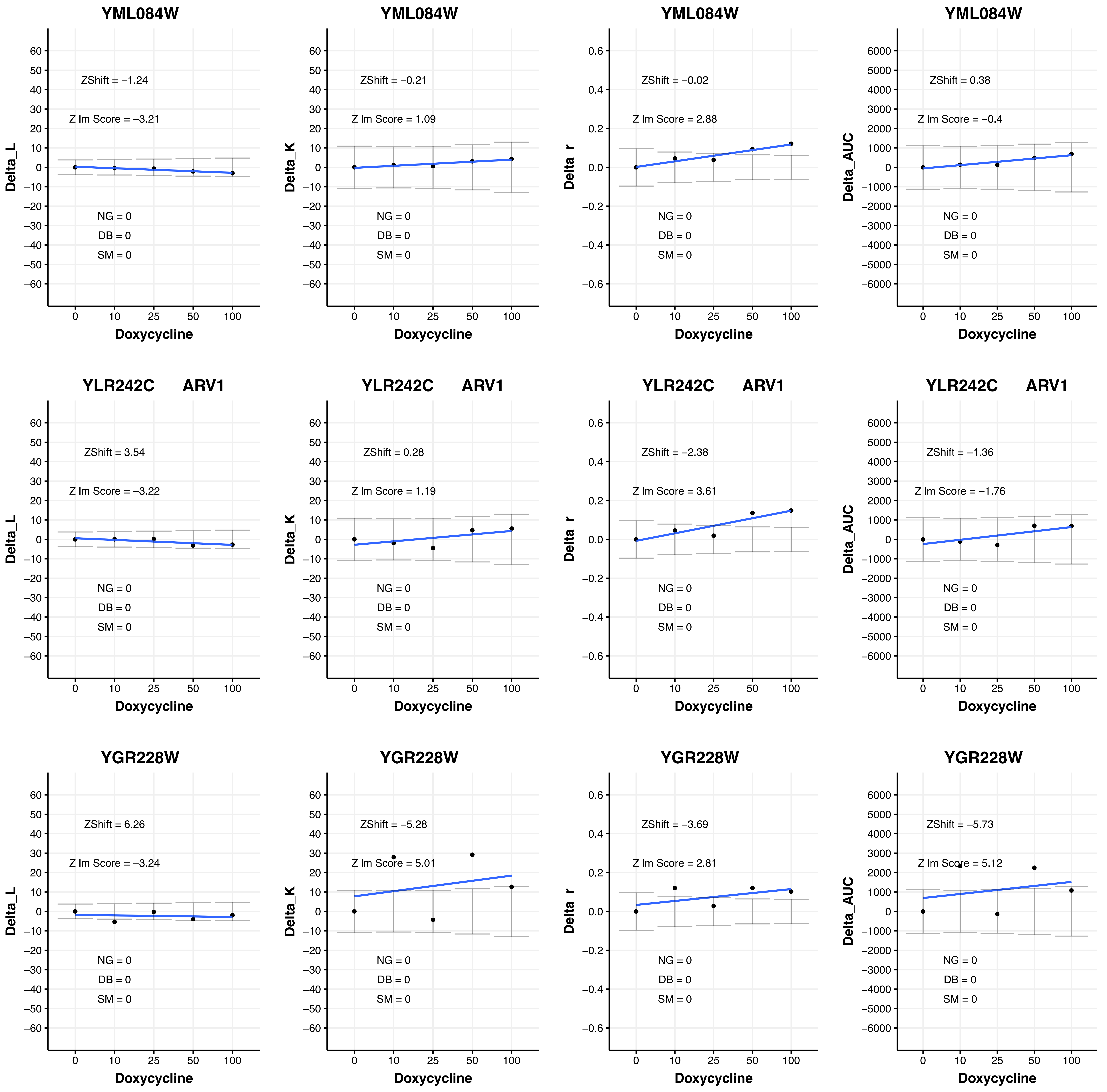

YML084W
ZShift = −1.24
Z lm Score = −3.21
NG = 0
DB = 0
SM = 0
Delta_L
Doxycycline
YML084W
ZShift = −0.21
Z lm Score = 1.09
NG = 0
DB = 0
SM = 0
Delta_K
Doxycycline
YML084W
ZShift = −0.02
Z lm Score = 2.88
NG = 0
DB = 0
SM = 0
Delta_r
Doxycycline
YML084W
ZShift = 0.38
Z lm Score = −0.4
NG = 0
DB = 0
SM = 0
Delta_AUC
Doxycycline
YLR242C ARV1
ZShift = 3.54
Z lm Score = −3.22
NG = 0
DB = 0
SM = 0
Delta_L
Doxycycline
YLR242C ARV1
ZShift = 0.28
Z lm Score = 1.19
NG = 0
DB = 0
SM = 0
Delta_K
Doxycycline
YLR242C ARV1
ZShift = −2.38
Z lm Score = 3.61
NG = 0
DB = 0
SM = 0
Delta_r
Doxycycline
YLR242C ARV1
ZShift = −1.36
Z lm Score = −1.76
NG = 0
DB = 0
SM = 0
Delta_AUC
Doxycycline
YGR228W
ZShift = 6.26
Z lm Score = −3.24
NG = 0
DB = 0
SM = 0
Delta_L
Doxycycline
YGR228W
ZShift = −5.28
Z lm Score = 5.01
NG = 0
DB = 0
SM = 0
Delta_K
Doxycycline
YGR228W
ZShift = −3.69
Z lm Score = 2.81
NG = 0
DB = 0
SM = 0
Delta_r
Doxycycline
YGR228W
ZShift = −5.73
Z lm Score = 5.12
NG = 0
DB = 0
SM = 0
Delta_AUC
Doxycycline

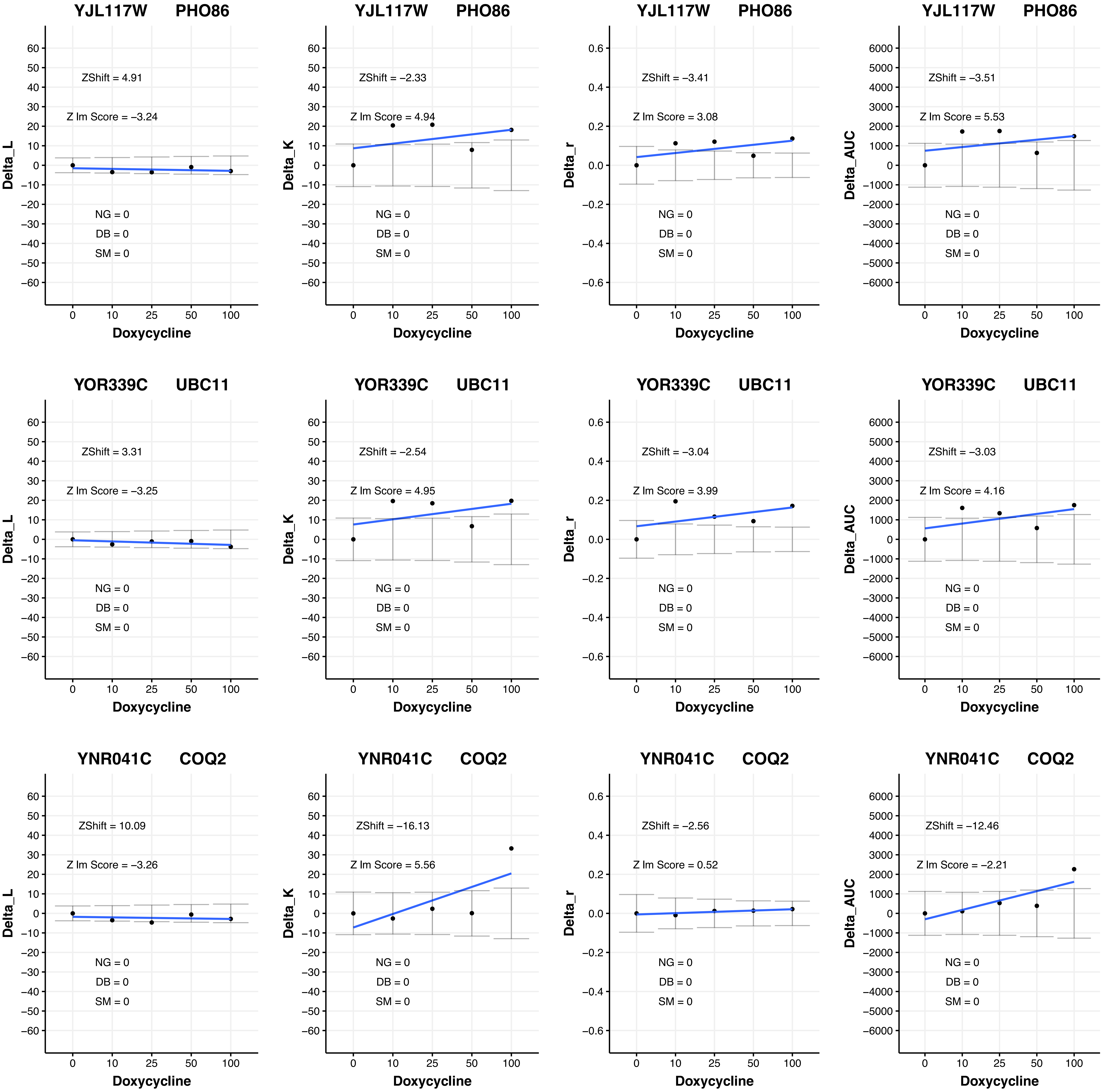

YJL117W PHO86
ZShift = 4.91
Z lm Score = −3.24
NG = 0
DB = 0
SM = 0
Delta_L
YJL117W PHO86
ZShift = −2.33
Z lm Score = 4.94
NG = 0
DB = 0
SM = 0
Delta_K
YJL117W PHO86
ZShift = −3.41
Z lm Score = 3.08
NG = 0
DB = 0
SM = 0
Delta_r
YJL117W PHO86
ZShift = −3.51
Z lm Score = 5.53
NG = 0
DB = 0
SM = 0
Delta_AUC
YOR339C UBC11
ZShift = 3.31
Z lm Score = −3.25
NG = 0
DB = 0
SM = 0
Delta_L
YOR339C UBC11
ZShift = −2.54
Z lm Score = 4.95
NG = 0
DB = 0
SM = 0
Delta_K
YOR339C UBC11
ZShift = −3.04
Z lm Score = 3.99
NG = 0
DB = 0
SM = 0
Delta_r
YOR339C UBC11
ZShift = −3.03
Z lm Score = 4.16
NG = 0
DB = 0
SM = 0
Delta_AUC
YNR041C COQ2
ZShift = 10.09
Z lm Score = −3.26
NG = 0
DB = 0
SM = 0
Delta_L
YNR041C COQ2
ZShift = −16.13
Z lm Score = 5.56
NG = 0
DB = 0
SM = 0
Delta_K
YNR041C COQ2
ZShift = −2.56
Z lm Score = 0.52
NG = 0
DB = 0
SM = 0
Delta_r
YNR041C COQ2
ZShift = −12.46
Z lm Score = −2.21
NG = 0
DB = 0
SM = 0
Delta_AUC
Doxycycline
0 10 25 50 100

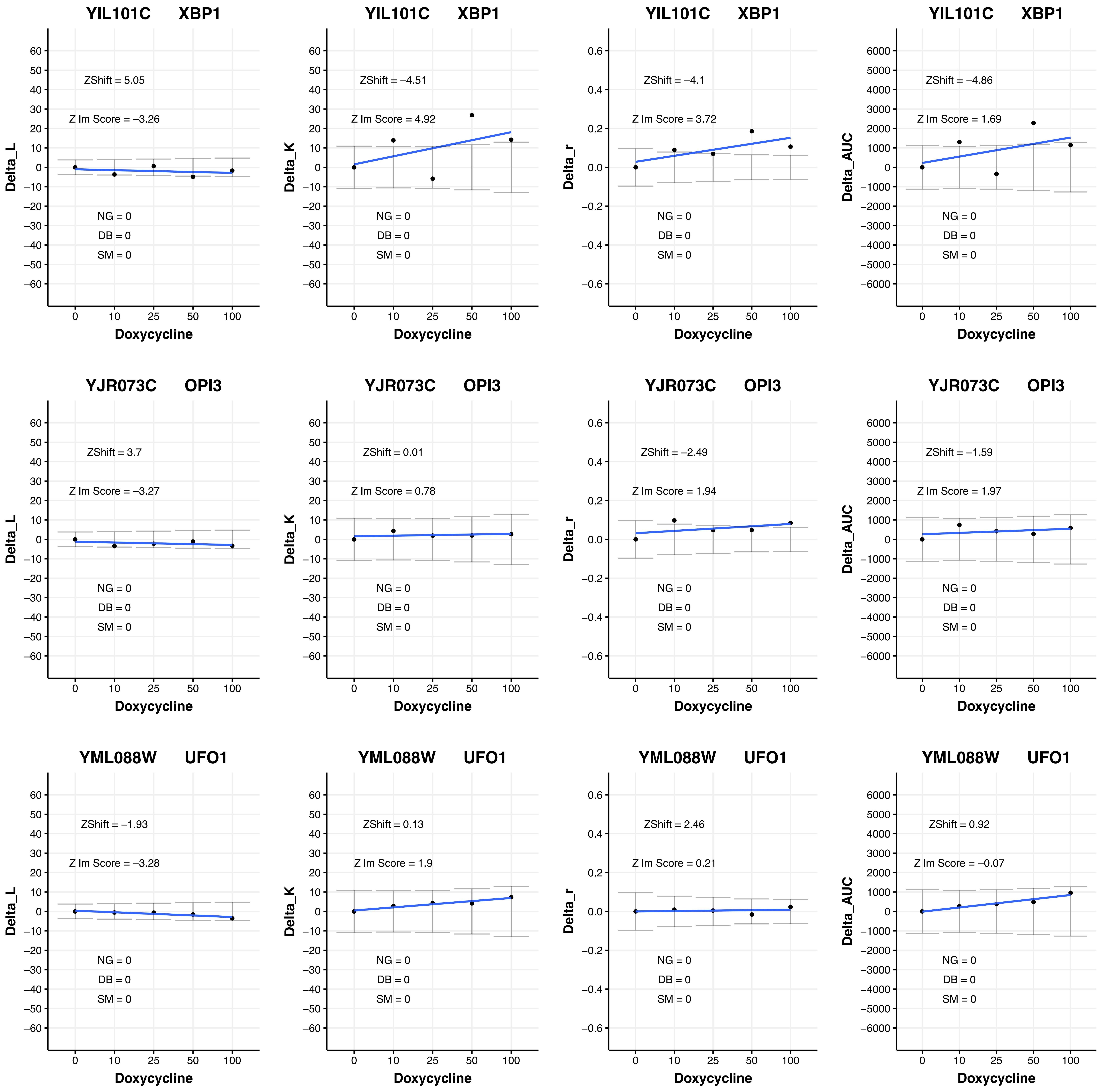

YIL101C XBP1
ZShift = 5.05
Z lm Score = −3.26
NG = 0
DB = 0
SM = 0
Delta_L
Doxycycline
YIL101C XBP1
ZShift = −4.51
Z lm Score = 4.92
NG = 0
DB = 0
SM = 0
Delta_K
Doxycycline
YIL101C XBP1
ZShift = −4.1
Z lm Score = 3.72
NG = 0
DB = 0
SM = 0
Delta_r
Doxycycline
YIL101C XBP1
ZShift = −4.86
Z lm Score = 1.69
NG = 0
DB = 0
SM = 0
Delta_AUC
Doxycycline
YJR073C OPI3
ZShift = 3.7
Z lm Score = −3.27
NG = 0
DB = 0
SM = 0
Delta_L
Doxycycline
YJR073C OPI3
ZShift = 0.01
Z lm Score = 0.78
NG = 0
DB = 0
SM = 0
Delta_K
Doxycycline
YJR073C OPI3
ZShift = −2.49
Z lm Score = 1.94
NG = 0
DB = 0
SM = 0
Delta_r
Doxycycline
YJR073C OPI3
ZShift = −1.59
Z lm Score = 1.97
NG = 0
DB = 0
SM = 0
Delta_AUC
Doxycycline
YML088W UFO1
ZShift = −1.93
Z lm Score = −3.28
NG = 0
DB = 0
SM = 0
Delta_L
Doxycycline
YML088W UFO1
ZShift = 0.13
Z lm Score = 1.9
NG = 0
DB = 0
SM = 0
Delta_K
Doxycycline
YML088W UFO1
ZShift = 2.46
Z lm Score = 0.21
NG = 0
DB = 0
SM = 0
Delta_r
Doxycycline
YML088W UFO1
ZShift = 0.92
Z lm Score = −0.07
NG = 0
DB = 0
SM = 0
Delta_AUC
Doxycycline

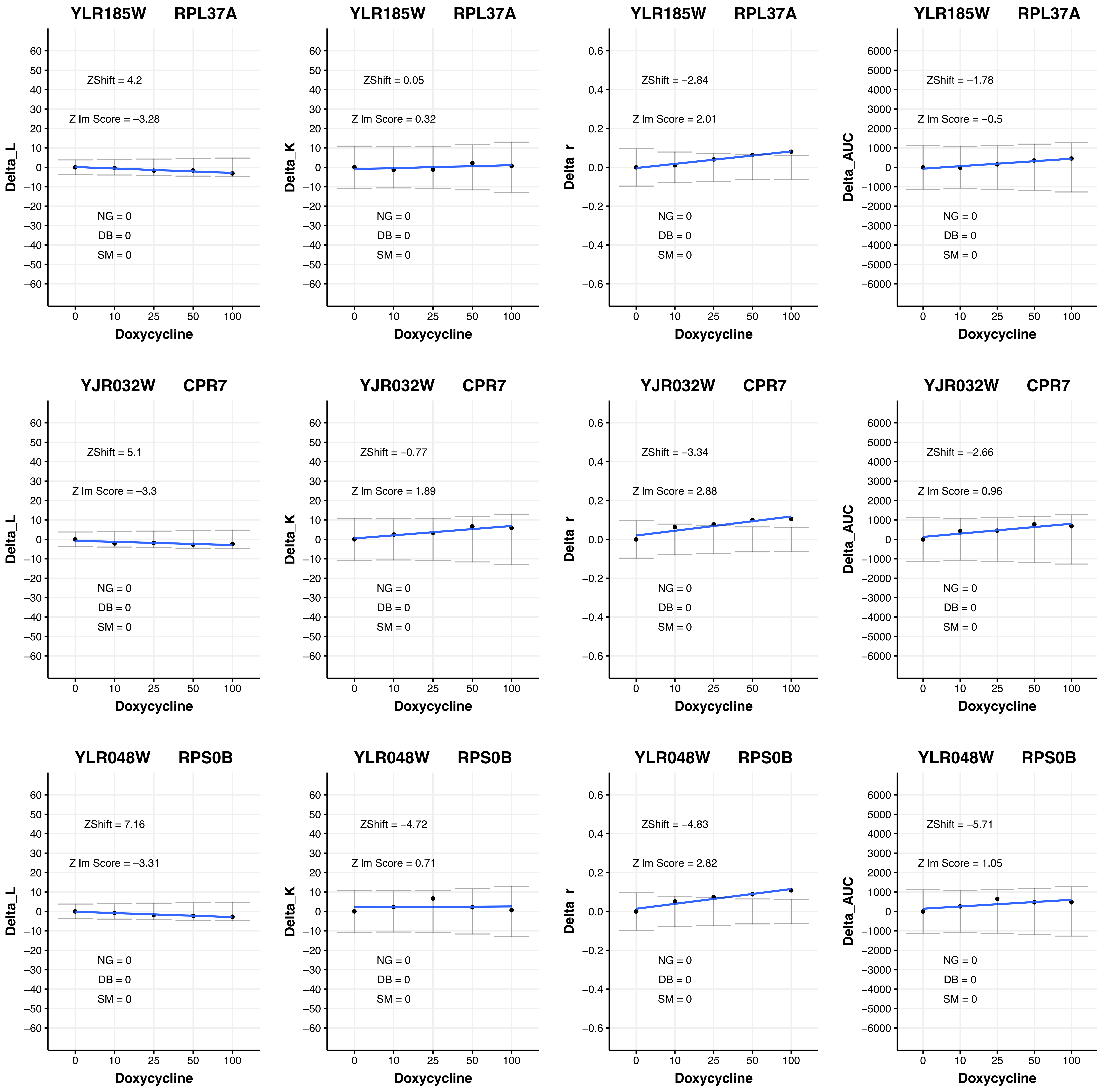

YLR185W RPL37A
ZShift = 4.2
Z lm Score = −3.28
NG = 0
DB = 0
SM = 0
Delta_L
Doxycycline
YLR185W RPL37A
ZShift = 0.05
Z lm Score = 0.32
NG = 0
DB = 0
SM = 0
Delta_K
Doxycycline
YLR185W RPL37A
ZShift = −2.84
Z lm Score = 2.01
NG = 0
DB = 0
SM = 0
Delta_r
Doxycycline
YLR185W RPL37A
ZShift = −1.78
Z lm Score = −0.5
NG = 0
DB = 0
SM = 0
Delta_AUC
Doxycycline
YJR032W CPR7
ZShift = 5.1
Z lm Score = −3.3
NG = 0
DB = 0
SM = 0
Delta_L
Doxycycline
YJR032W CPR7
ZShift = −0.77
Z lm Score = 1.89
NG = 0
DB = 0
SM = 0
Delta_K
Doxycycline
YJR032W CPR7
ZShift = −3.34
Z lm Score = 2.88
NG = 0
DB = 0
SM = 0
Delta_r
Doxycycline
YJR032W CPR7
ZShift = −2.66
Z lm Score = 0.96
NG = 0
DB = 0
SM = 0
Delta_AUC
Doxycycline
YLR048W RPS0B
ZShift = 7.16
Z lm Score = −3.31
NG = 0
DB = 0
SM = 0
Delta_L
Doxycycline
YLR048W RPS0B
ZShift = −4.72
Z lm Score = 0.71
NG = 0
DB = 0
SM = 0
Delta_K
Doxycycline
YLR048W RPS0B
ZShift = −4.83
Z lm Score = 2.82
NG = 0
DB = 0
SM = 0
Delta_r
Doxycycline
YLR048W RPS0B
ZShift = −5.71
Z lm Score = 1.05
NG = 0
DB = 0
SM = 0
Delta_AUC
Doxycycline

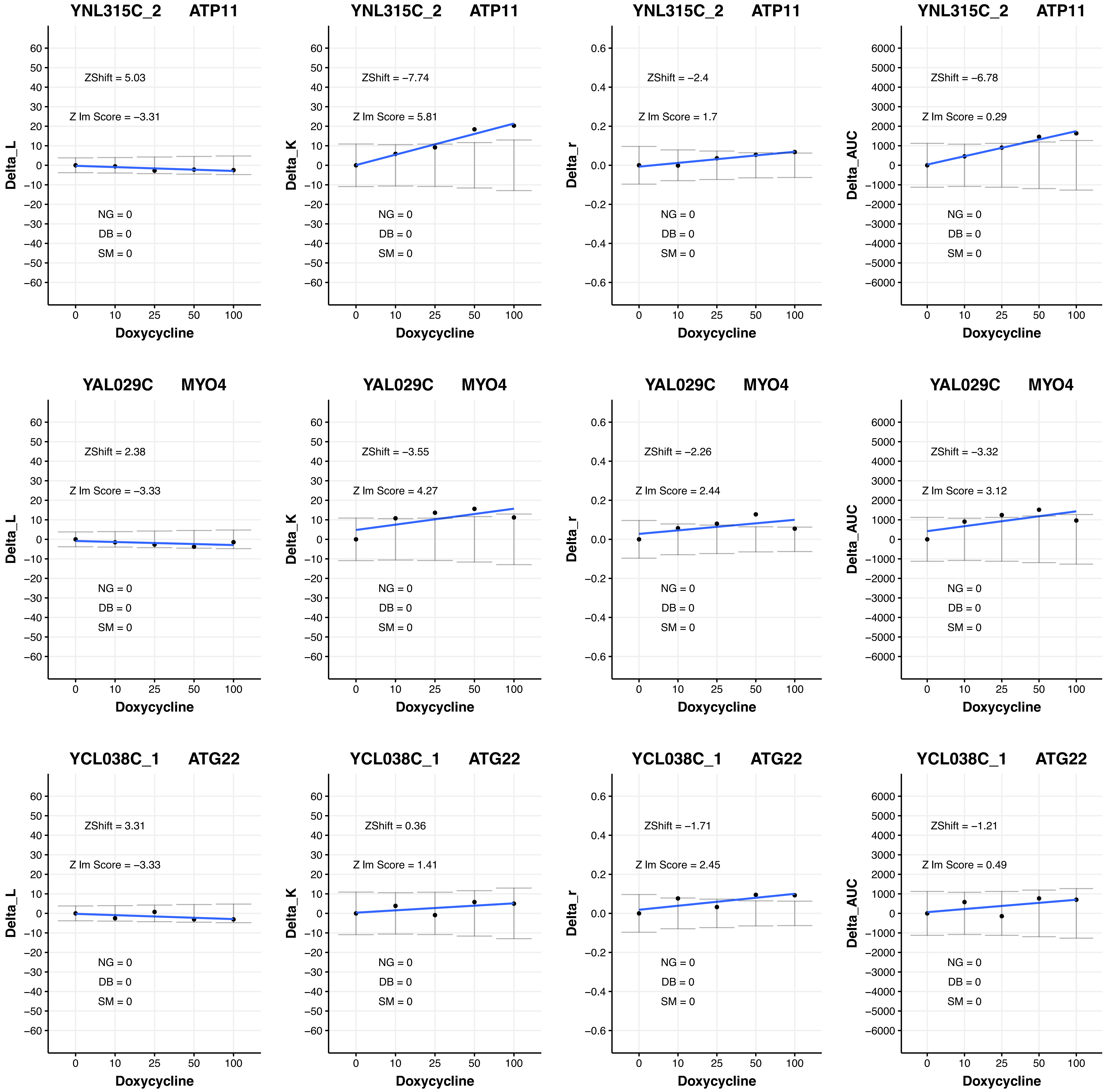
YNL315C_2 ATP11
ZShift = 5.03
Z lm Score = −3.31
NG = 0
DB = 0
SM = 0
Delta_L
Doxycycline
YNL315C_2 ATP11
ZShift = −7.74
Z lm Score = 5.81
NG = 0
DB = 0
SM = 0
Delta_K
Doxycycline
YNL315C_2 ATP11
ZShift = −2.4
Z lm Score = 1.7
NG = 0
DB = 0
SM = 0
Delta_r
Doxycycline
YNL315C_2 ATP11
ZShift = −6.78
Z lm Score = 0.29
NG = 0
DB = 0
SM = 0
Delta_AUC
Doxycycline
YAL029C MYO4
ZShift = 2.38
Z lm Score = −3.33
NG = 0
DB = 0
SM = 0
Delta_L
Doxycycline
YAL029C MYO4
ZShift = −3.55
Z lm Score = 4.27
NG = 0
DB = 0
SM = 0
Delta_K
Doxycycline
YAL029C MYO4
ZShift = −2.26
Z lm Score = 2.44
NG = 0
DB = 0
SM = 0
Delta_r
Doxycycline
YAL029C MYO4
ZShift = −3.32
Z lm Score = 3.12
NG = 0
DB = 0
SM = 0
Delta_AUC
Doxycycline
YCL038C_1 ATG22
ZShift = 3.31
Z lm Score = −3.33
NG = 0
DB = 0
SM = 0
Delta_L
Doxycycline
YCL038C_1 ATG22
ZShift = 0.36
Z lm Score = 1.41
NG = 0
DB = 0
SM = 0
Delta_K
Doxycycline
YCL038C_1 ATG22
ZShift = −1.71
Z lm Score = 2.45
NG = 0
DB = 0
SM = 0
Delta_r
Doxycycline
YCL038C_1 ATG22
ZShift = −1.21
Z lm Score = 0.49
NG = 0
DB = 0
SM = 0
Delta_AUC
Doxycycline

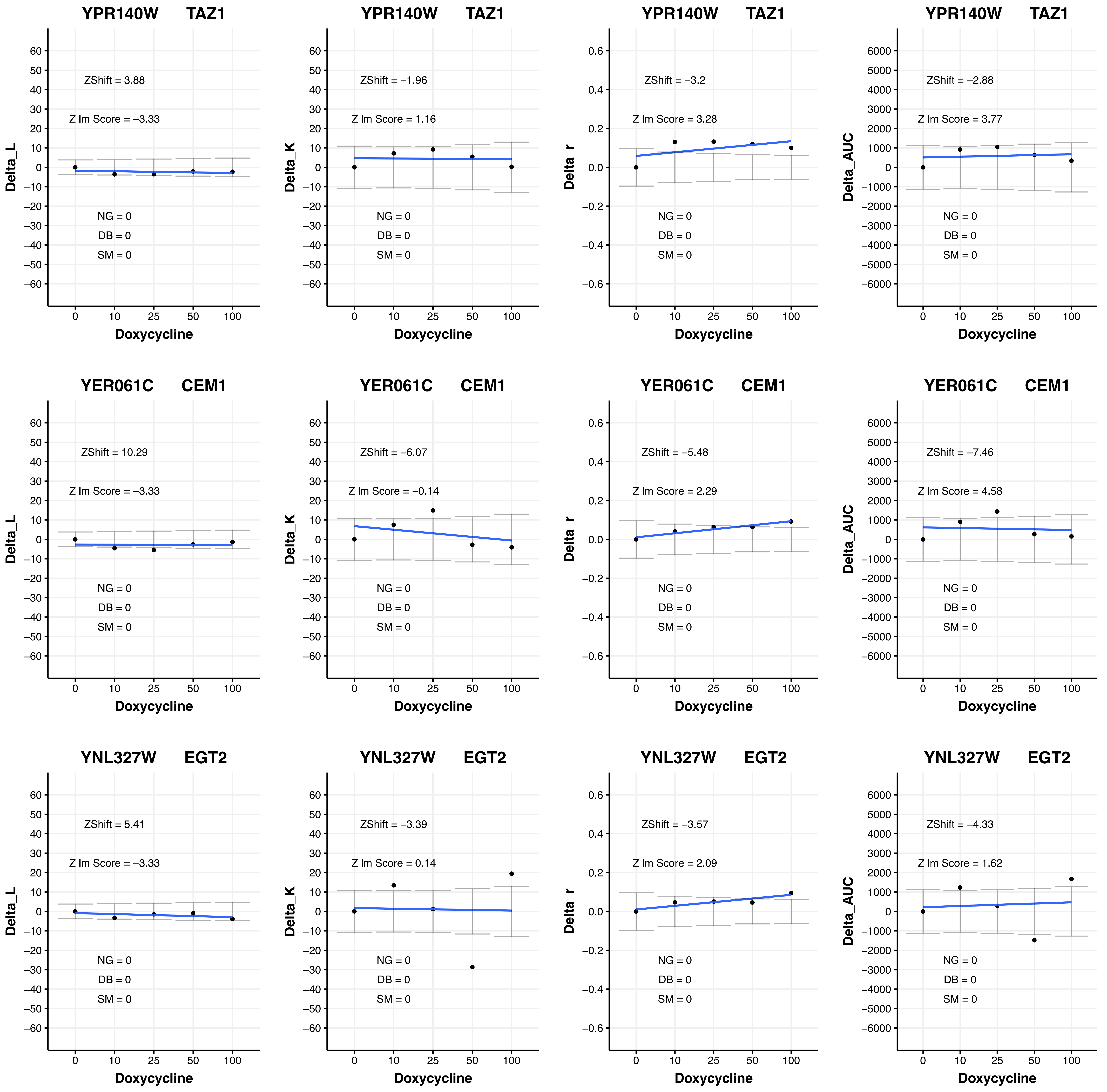
YPR140W TAZ1
ZShift = 3.88
Z lm Score = −3.33
NG = 0
DB = 0
SM = 0
Delta_L
Doxycycline
YPR140W TAZ1
ZShift = −1.96
Z lm Score = 1.16
NG = 0
DB = 0
SM = 0
Delta_K
Doxycycline
YPR140W TAZ1
ZShift = −3.2
Z lm Score = 3.28
NG = 0
DB = 0
SM = 0
Delta_r
Doxycycline
YPR140W TAZ1
ZShift = −2.88
Z lm Score = 3.77
NG = 0
DB = 0
SM = 0
Delta_AUC
Doxycycline
YER061C CEM1
ZShift = 10.29
Z lm Score = −3.33
NG = 0
DB = 0
SM = 0
Delta_L
Doxycycline
YER061C CEM1
ZShift = −6.07
Z lm Score = −0.14
NG = 0
DB = 0
SM = 0
Delta_K
Doxycycline
YER061C CEM1
ZShift = −5.48
Z lm Score = 2.29
NG = 0
DB = 0
SM = 0
Delta_r
Doxycycline
YER061C CEM1
ZShift = −7.46
Z lm Score = 4.58
NG = 0
DB = 0
SM = 0
Delta_AUC
Doxycycline
YNL327W EGT2
ZShift = 5.41
Z lm Score = −3.33
NG = 0
DB = 0
SM = 0
Delta_L
Doxycycline
YNL327W EGT2
ZShift = −3.39
Z lm Score = 0.14
NG = 0
DB = 0
SM = 0
Delta_K
Doxycycline
YNL327W EGT2
ZShift = −3.57
Z lm Score = 2.09
NG = 0
DB = 0
SM = 0
Delta_r
Doxycycline
YNL327W EGT2
ZShift = −4.33
Z lm Score = 1.62
NG = 0
DB = 0
SM = 0
Delta_AUC
Doxycycline

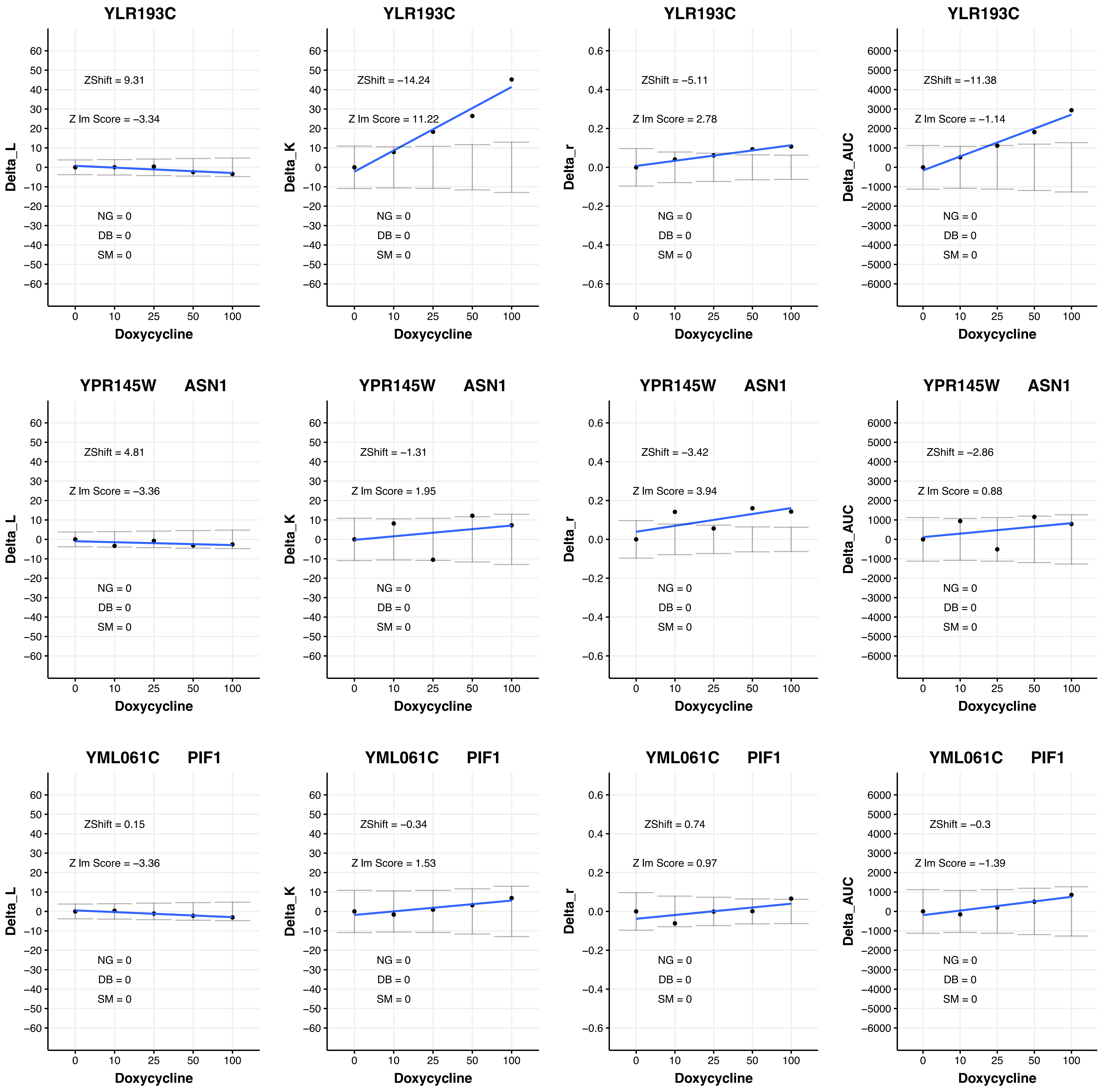

YLR193C
ZShift = 9.31
Z lm Score = −3.34
NG = 0
DB = 0
SM = 0
Delta_L
Doxycycline
YLR193C
ZShift = −14.24
Z lm Score = 11.22
NG = 0
DB = 0
SM = 0
Delta_K
Doxycycline
YLR193C
ZShift = −5.11
Z lm Score = 2.78
NG = 0
DB = 0
SM = 0
Delta_r
Doxycycline
YLR193C
ZShift = −11.38
Z lm Score = −1.14
NG = 0
DB = 0
SM = 0
Delta_AUC
Doxycycline
YPR145W ASN1
ZShift = 4.81
Z lm Score = −3.36
NG = 0
DB = 0
SM = 0
Delta_L
Doxycycline
YPR145W ASN1
ZShift = −1.31
Z lm Score = 1.95
NG = 0
DB = 0
SM = 0
Delta_K
Doxycycline
YPR145W ASN1
ZShift = −3.42
Z lm Score = 3.94
NG = 0
DB = 0
SM = 0
Delta_r
Doxycycline
YPR145W ASN1
ZShift = −2.86
Z lm Score = 0.88
NG = 0
DB = 0
SM = 0
Delta_AUC
Doxycycline
YML061C PIF1
ZShift = 0.15
Z lm Score = −3.36
NG = 0
DB = 0
SM = 0
Delta_L
Doxycycline
YML061C PIF1
ZShift = −0.34
Z lm Score = 1.53
NG = 0
DB = 0
SM = 0
Delta_K
Doxycycline
YML061C PIF1
ZShift = 0.74
Z lm Score = 0.97
NG = 0
DB = 0
SM = 0
Delta_r
Doxycycline
YML061C PIF1
ZShift = −0.3
Z lm Score = −1.39
NG = 0
DB = 0
SM = 0
Delta_AUC
Doxycycline

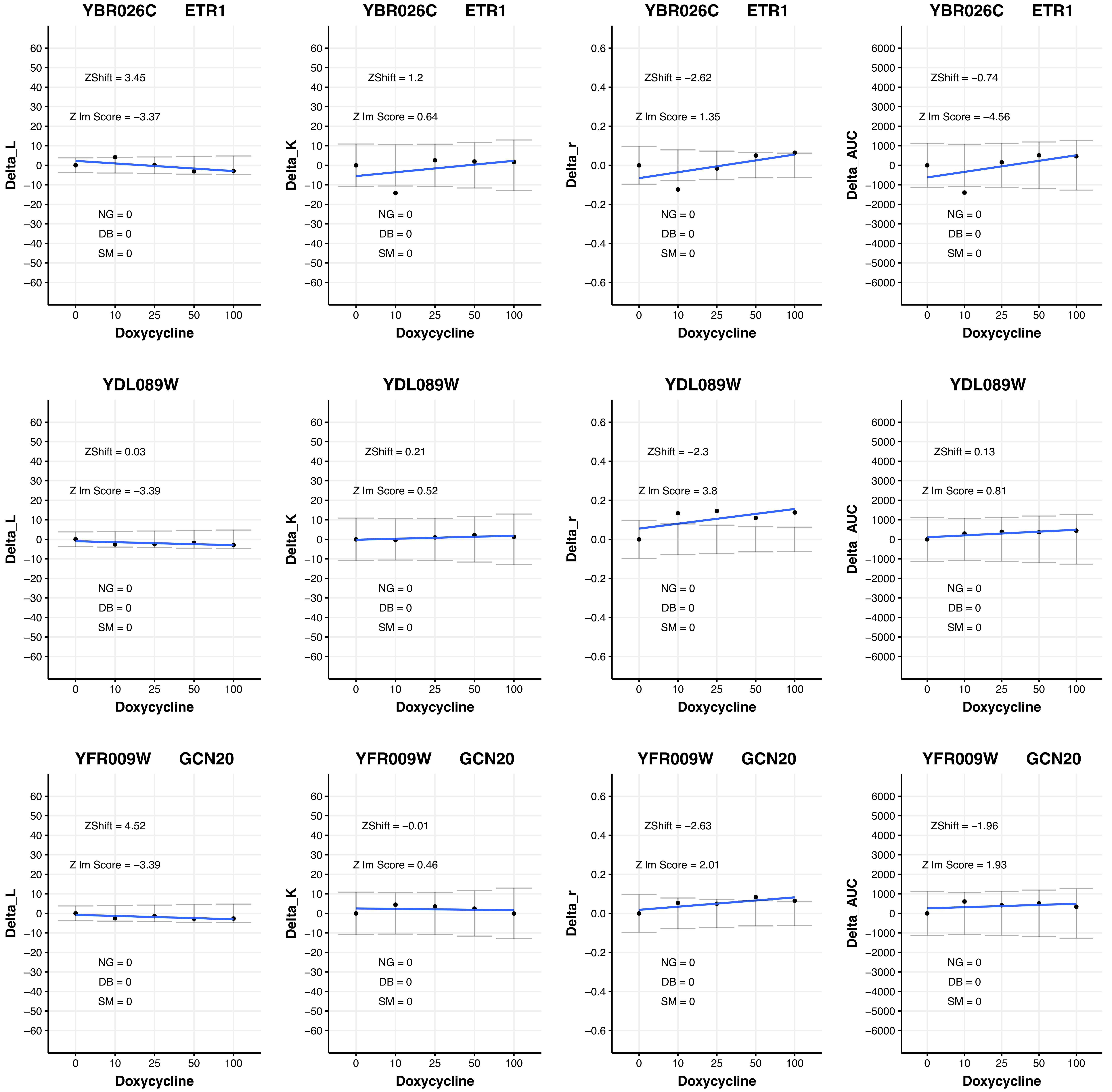

YBR026C ETR1
ZShift = 3.45
Z lm Score = −3.37
NG = 0
DB = 0
SM = 0
Delta_L
Doxycycline
YBR026C ETR1
ZShift = 1.2
Z lm Score = 0.64
NG = 0
DB = 0
SM = 0
Delta_K
Doxycycline
YBR026C ETR1
ZShift = −2.62
Z lm Score = 1.35
NG = 0
DB = 0
SM = 0
Delta_r
Doxycycline
YBR026C ETR1
ZShift = −0.74
Z lm Score = −4.56
NG = 0
DB = 0
SM = 0
Delta_AUC
Doxycycline
YDL089W
ZShift = 0.03
Z lm Score = −3.39
NG = 0
DB = 0
SM = 0
Delta_L
Doxycycline
YDL089W
ZShift = 0.21
Z lm Score = 0.52
NG = 0
DB = 0
SM = 0
Delta_K
Doxycycline
YDL089W
ZShift = −2.3
Z lm Score = 3.8
NG = 0
DB = 0
SM = 0
Delta_r
Doxycycline
YDL089W
ZShift = 0.13
Z lm Score = 0.81
NG = 0
DB = 0
SM = 0
Delta_AUC
Doxycycline
YFR009W GCN20
ZShift = 4.52
Z lm Score = −3.39
NG = 0
DB = 0
SM = 0
Delta_L
Doxycycline
YFR009W GCN20
ZShift = −0.01
Z lm Score = 0.46
NG = 0
DB = 0
SM = 0
Delta_K
Doxycycline
YFR009W GCN20
ZShift = −2.63
Z lm Score = 2.01
NG = 0
DB = 0
SM = 0
Delta_r
Doxycycline
YFR009W GCN20
ZShift = −1.96
Z lm Score = 1.93
NG = 0
DB = 0
SM = 0
Delta_AUC
Doxycycline

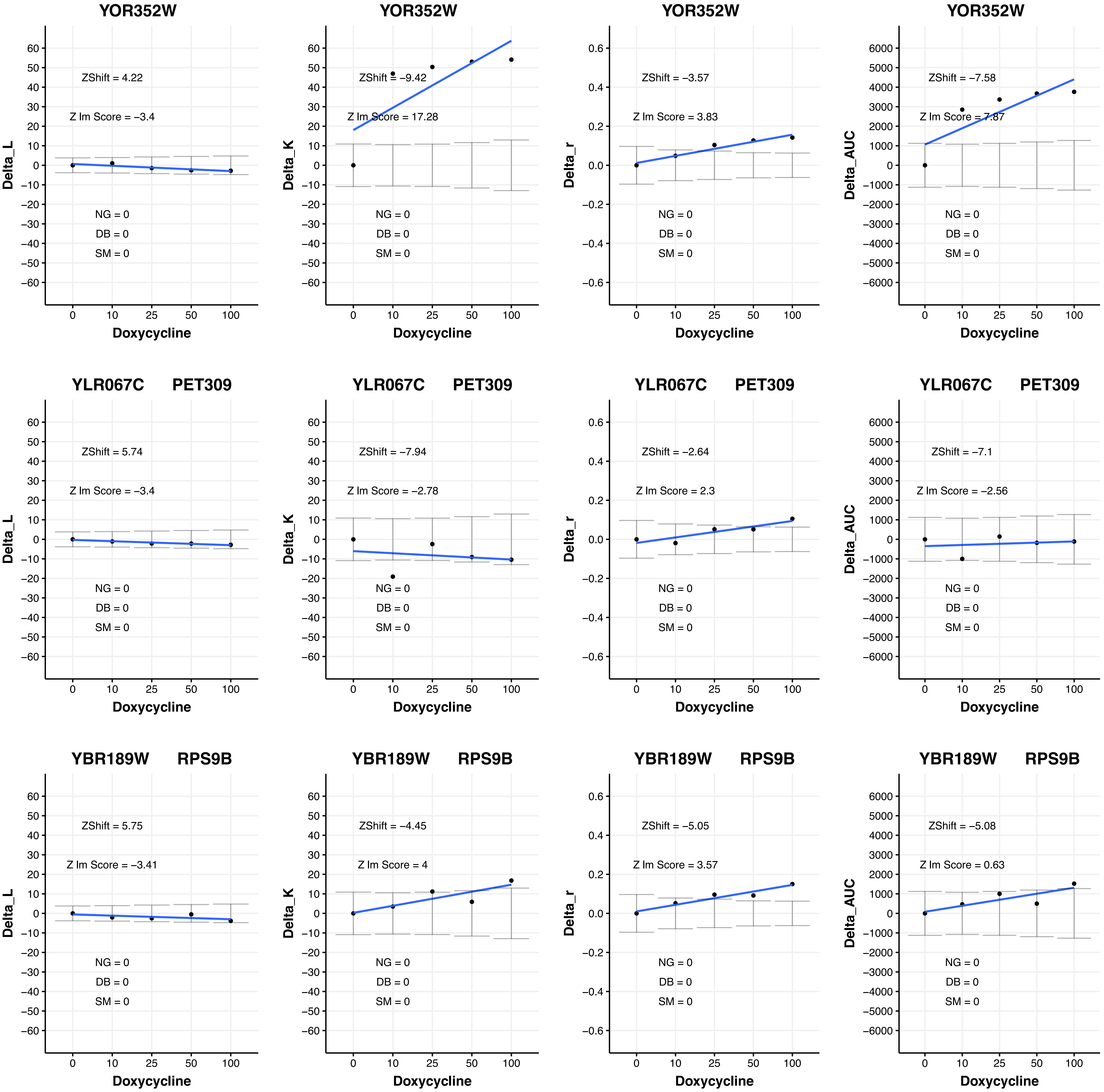

YOR352W
ZShift = 4.22
Z lm Score = −3.4
NG = 0
DB = 0
SM = 0
Delta_L
Doxycycline
YOR352W
ZShift = −9.42
Z lm Score = 17.28
NG = 0
DB = 0
SM = 0
Delta_K
Doxycycline
YOR352W
ZShift = −3.57
Z lm Score = 3.83
NG = 0
DB = 0
SM = 0
Delta_r
Doxycycline
YOR352W
ZShift = −7.58
Z lm Score = 7.87
NG = 0
DB = 0
SM = 0
Delta_AUC
Doxycycline
YLR067C PET309
ZShift = 5.74
Z lm Score = −3.4
NG = 0
DB = 0
SM = 0
Delta_L
Doxycycline
YLR067C PET309
ZShift = −7.94
Z lm Score = −2.78
NG = 0
DB = 0
SM = 0
Delta_K
Doxycycline
YLR067C PET309
ZShift = −2.64
Z lm Score = 2.3
NG = 0
DB = 0
SM = 0
Delta_r
Doxycycline
YLR067C PET309
ZShift = −7.1
Z lm Score = −2.56
NG = 0
DB = 0
SM = 0
Delta_AUC
Doxycycline
YBR189W RPS9B
ZShift = 5.75
Z lm Score = −3.41
NG = 0
DB = 0
SM = 0
Delta_L
Doxycycline
YBR189W RPS9B
ZShift = −4.45
Z lm Score = 4
NG = 0
DB = 0
SM = 0
Delta_K
Doxycycline
YBR189W RPS9B
ZShift = −5.05
Z lm Score = 3.57
NG = 0
DB = 0
SM = 0
Delta_r
Doxycycline
YBR189W RPS9B
ZShift = −5.08
Z lm Score = 0.63
NG = 0
DB = 0
SM = 0
Delta_AUC
Doxycycline

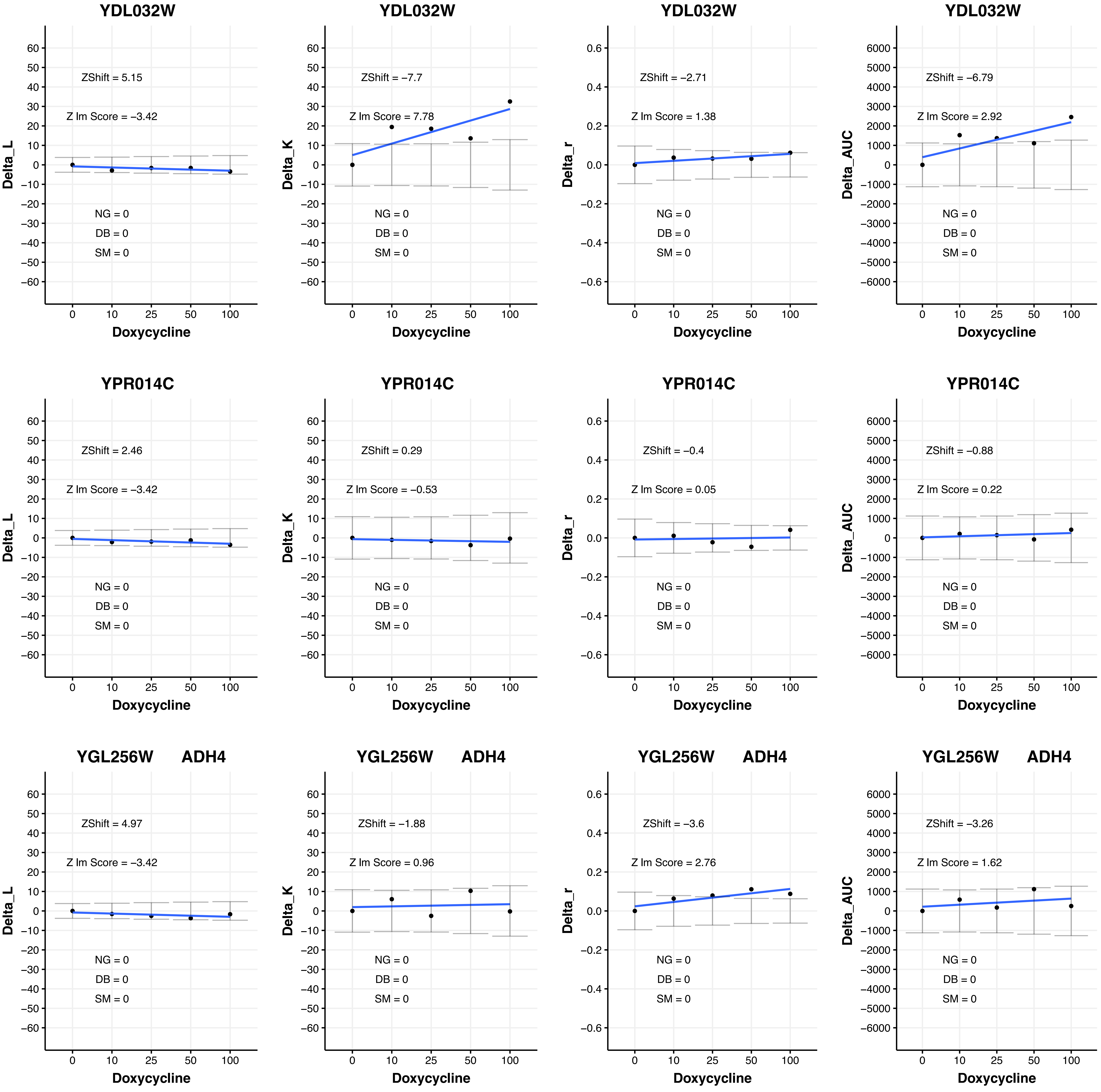

YDL032W
ZShift = 5.15
Z lm Score = −3.42
NG = 0
DB = 0
SM = 0
Delta_L
Doxycycline
YDL032W
ZShift = −7.7
Z lm Score = 7.78
NG = 0
DB = 0
SM = 0
Delta_K
Doxycycline
YDL032W
ZShift = −2.71
Z lm Score = 1.38
NG = 0
DB = 0
SM = 0
Delta_r
Doxycycline
YDL032W
ZShift = −6.79
Z lm Score = 2.92
NG = 0
DB = 0
SM = 0
Delta_AUC
Doxycycline
YPR014C
ZShift = 2.46
Z lm Score = −3.42
NG = 0
DB = 0
SM = 0
Delta_L
Doxycycline
YPR014C
ZShift = 0.29
Z lm Score = −0.53
NG = 0
DB = 0
SM = 0
Delta_K
Doxycycline
YPR014C
ZShift = −0.4
Z lm Score = 0.05
NG = 0
DB = 0
SM = 0
Delta_r
Doxycycline
YPR014C
ZShift = −0.88
Z lm Score = 0.22
NG = 0
DB = 0
SM = 0
Delta_AUC
Doxycycline
YGL256W ADH4
ZShift = 4.97
Z lm Score = −3.42
NG = 0
DB = 0
SM = 0
Delta_L
Doxycycline
YGL256W ADH4
ZShift = −1.88
Z lm Score = 0.96
NG = 0
DB = 0
SM = 0
Delta_K
Doxycycline
YGL256W ADH4
ZShift = −3.6
Z lm Score = 2.76
NG = 0
DB = 0
SM = 0
Delta_r
Doxycycline
YGL256W ADH4
ZShift = −3.26
Z lm Score = 1.62
NG = 0
DB = 0
SM = 0
Delta_AUC
Doxycycline
0 10 25 50 100

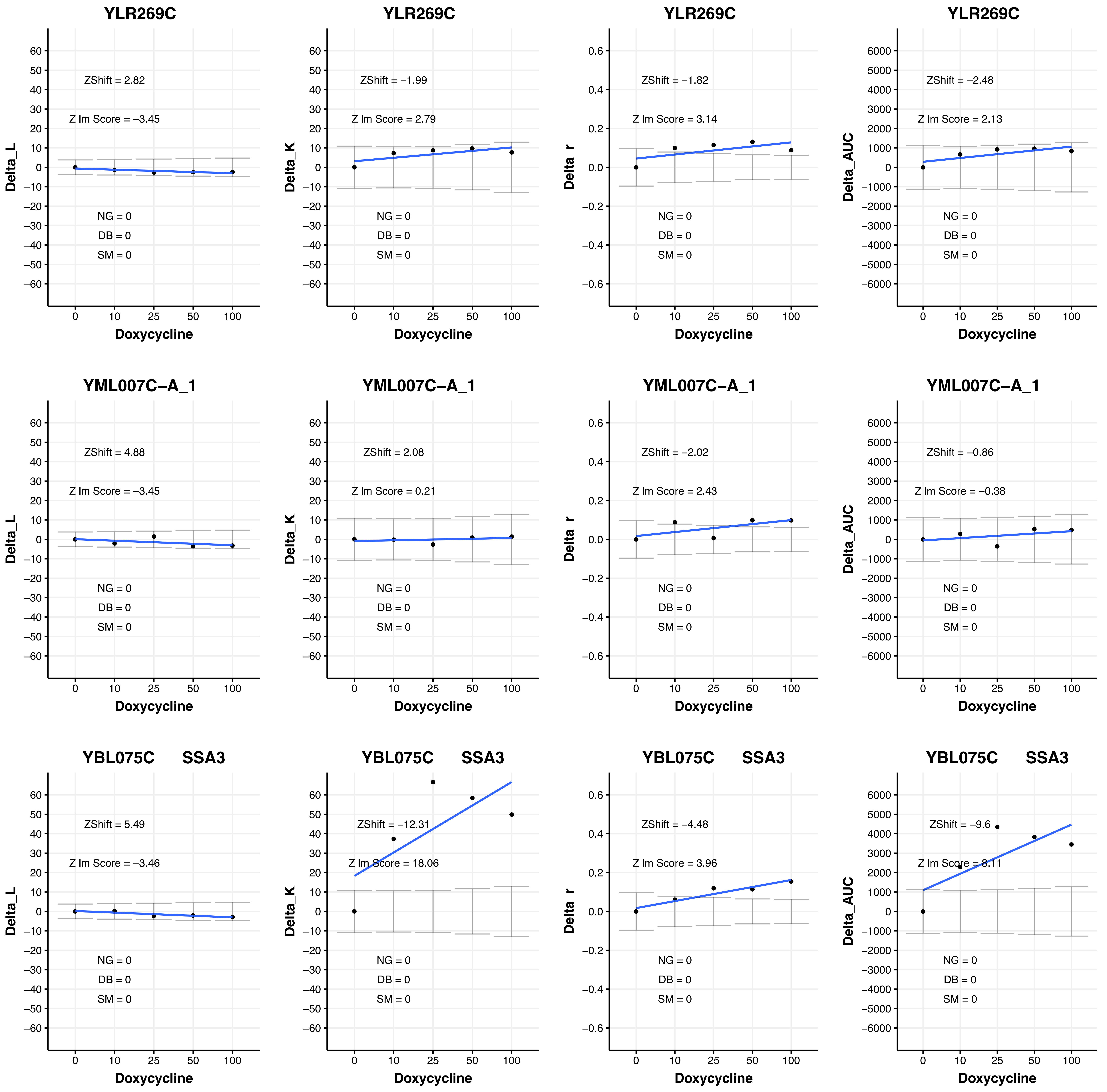

YLR269C
ZShift = 2.82
Z lm Score = −3.45
NG = 0
DB = 0
SM = 0
Delta_L
Doxycycline
YLR269C
ZShift = −1.99
Z lm Score = 2.79
NG = 0
DB = 0
SM = 0
Delta_K
Doxycycline
YLR269C
ZShift = −1.82
Z lm Score = 3.14
NG = 0
DB = 0
SM = 0
Delta_r
Doxycycline
YLR269C
ZShift = −2.48
Z lm Score = 2.13
NG = 0
DB = 0
SM = 0
Delta_AUC
Doxycycline
YML007C−A_1
ZShift = 4.88
Z lm Score = −3.45
NG = 0
DB = 0
SM = 0
Delta_L
Doxycycline
YML007C−A_1
ZShift = 2.08
Z lm Score = 0.21
NG = 0
DB = 0
SM = 0
Delta_K
Doxycycline
YML007C−A_1
ZShift = −2.02
Z lm Score = 2.43
NG = 0
DB = 0
SM = 0
Delta_r
Doxycycline
YML007C−A_1
ZShift = −0.86
Z lm Score = −0.38
NG = 0
DB = 0
SM = 0
Delta_AUC
Doxycycline
YBL075C SSA3
ZShift = 5.49
Z lm Score = −3.46
NG = 0
DB = 0
SM = 0
Delta_L
Doxycycline
YBL075C SSA3
ZShift = −12.31
Z lm Score = 18.06
NG = 0
DB = 0
SM = 0
Delta_K
Doxycycline
YBL075C SSA3
ZShift = −4.48
Z lm Score = 3.96
NG = 0
DB = 0
SM = 0
Delta_r
Doxycycline
YBL075C SSA3
ZShift = −9.6
Z lm Score = 8.11
NG = 0
DB = 0
SM = 0
Delta_AUC
Doxycycline

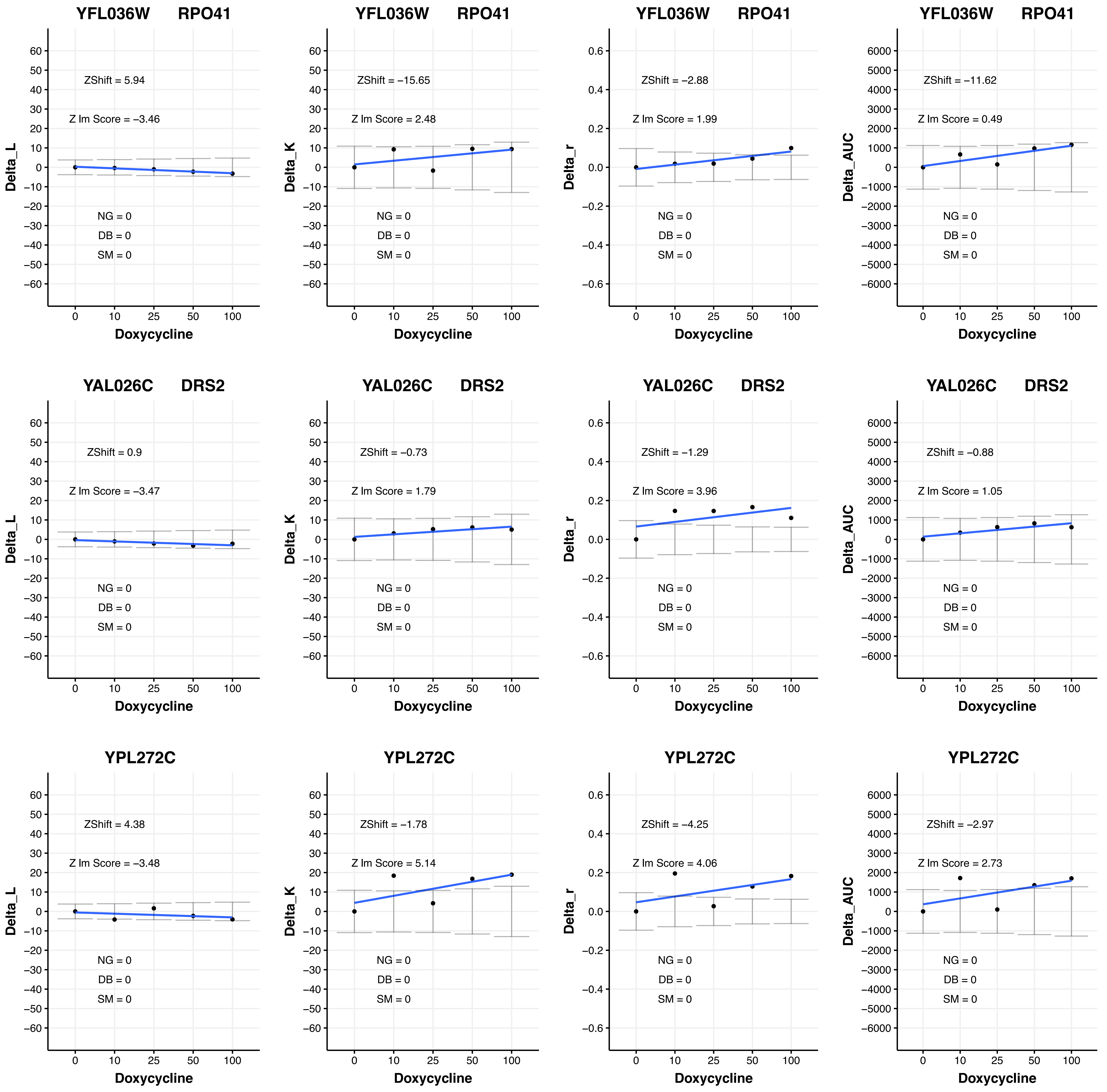
YFL036W RPO41
ZShift = 5.94
Z lm Score = −3.46
NG = 0
DB = 0
SM = 0
Delta_L
Doxycycline
YFL036W RPO41
ZShift = −15.65
Z lm Score = 2.48
NG = 0
DB = 0
SM = 0
Delta_K
Doxycycline
YFL036W RPO41
ZShift = −2.88
Z lm Score = 1.99
NG = 0
DB = 0
SM = 0
Delta_r
Doxycycline
YFL036W RPO41
ZShift = −11.62
Z lm Score = 0.49
NG = 0
DB = 0
SM = 0
Delta_AUC
Doxycycline
YAL026C DRS2
ZShift = 0.9
Z lm Score = −3.47
NG = 0
DB = 0
SM = 0
Delta_L
Doxycycline
YAL026C DRS2
ZShift = −0.73
Z lm Score = 1.79
NG = 0
DB = 0
SM = 0
Delta_K
Doxycycline
YAL026C DRS2
ZShift = −1.29
Z lm Score = 3.96
NG = 0
DB = 0
SM = 0
Delta_r
Doxycycline
YAL026C DRS2
ZShift = −0.88
Z lm Score = 1.05
NG = 0
DB = 0
SM = 0
Delta_AUC
Doxycycline
YPL272C
ZShift = 4.38
Z lm Score = −3.48
NG = 0
DB = 0
SM = 0
Delta_L
Doxycycline
YPL272C
ZShift = −1.78
Z lm Score = 5.14
NG = 0
DB = 0
SM = 0
Delta_K
Doxycycline
YPL272C
ZShift = −4.25
Z lm Score = 4.06
NG = 0
DB = 0
SM = 0
Delta_r
Doxycycline
YPL272C
ZShift = −2.97
Z lm Score = 2.73
NG = 0
DB = 0
SM = 0
Delta_AUC
Doxycycline

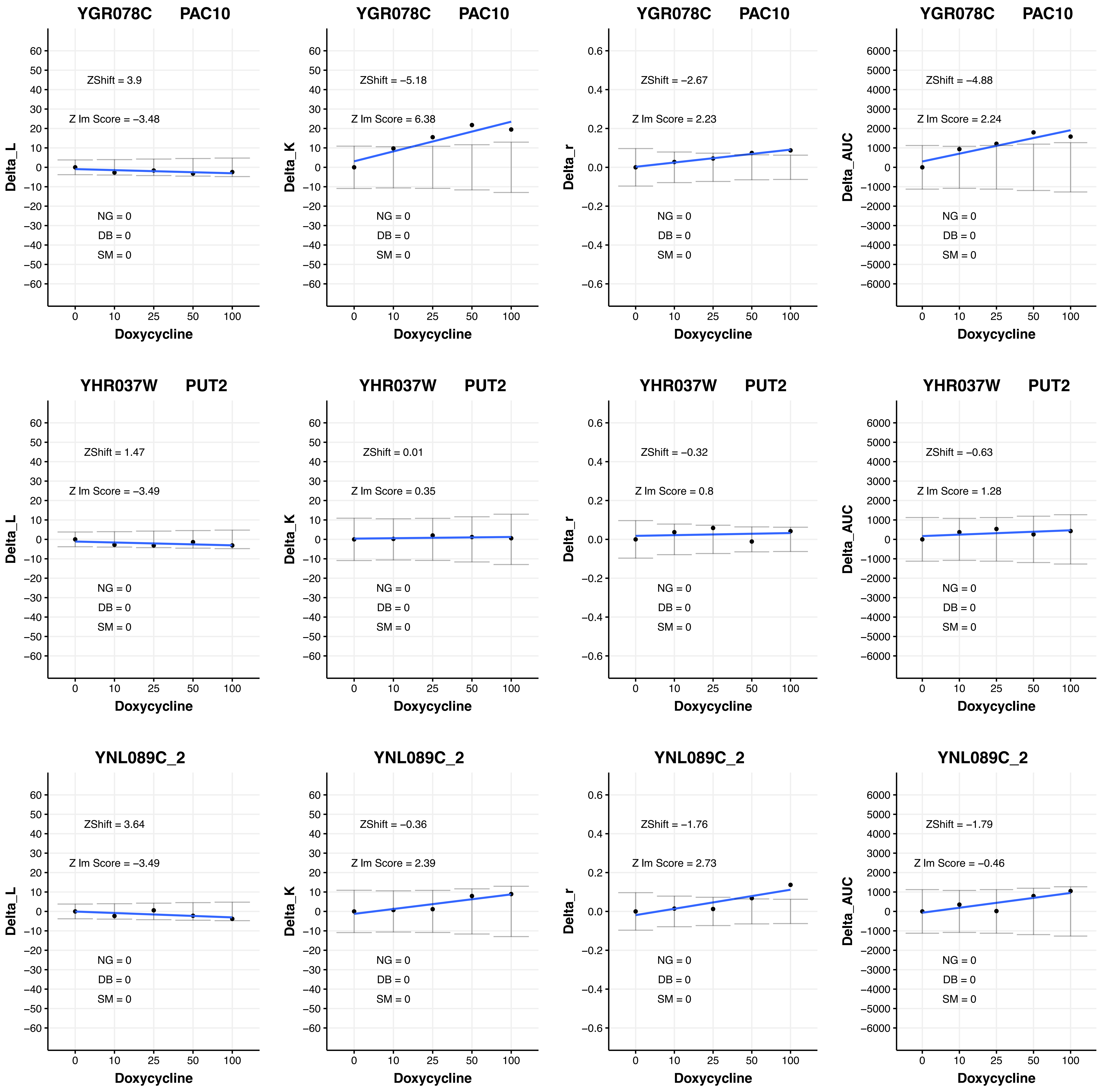

YGR078C PAC10
ZShift = 3.9
Z lm Score = −3.48
NG = 0
DB = 0
SM = 0
Delta_L
Doxycycline
YGR078C PAC10
ZShift = −5.18
Z lm Score = 6.38
NG = 0
DB = 0
SM = 0
Delta_K
Doxycycline
YGR078C PAC10
ZShift = −2.67
Z lm Score = 2.23
NG = 0
DB = 0
SM = 0
Delta_r
Doxycycline
YGR078C PAC10
ZShift = −4.88
Z lm Score = 2.24
NG = 0
DB = 0
SM = 0
Delta_AUC
Doxycycline
YHR037W PUT2
ZShift = 1.47
Z lm Score = −3.49
NG = 0
DB = 0
SM = 0
Delta_L
Doxycycline
YHR037W PUT2
ZShift = 0.01
Z lm Score = 0.35
NG = 0
DB = 0
SM = 0
Delta_K
Doxycycline
YHR037W PUT2
ZShift = −0.32
Z lm Score = 0.8
NG = 0
DB = 0
SM = 0
Delta_r
Doxycycline
YHR037W PUT2
ZShift = −0.63
Z lm Score = 1.28
NG = 0
DB = 0
SM = 0
Delta_AUC
Doxycycline
YNL089C_2
ZShift = 3.64
Z lm Score = −3.49
NG = 0
DB = 0
SM = 0
Delta_L
Doxycycline
YNL089C_2
ZShift = −0.36
Z lm Score = 2.39
NG = 0
DB = 0
SM = 0
Delta_K
Doxycycline
YNL089C_2
ZShift = −1.76
Z lm Score = 2.73
NG = 0
DB = 0
SM = 0
Delta_r
Doxycycline
YNL089C_2
ZShift = −1.79
Z lm Score = −0.46
NG = 0
DB = 0
SM = 0
Delta_AUC
Doxycycline

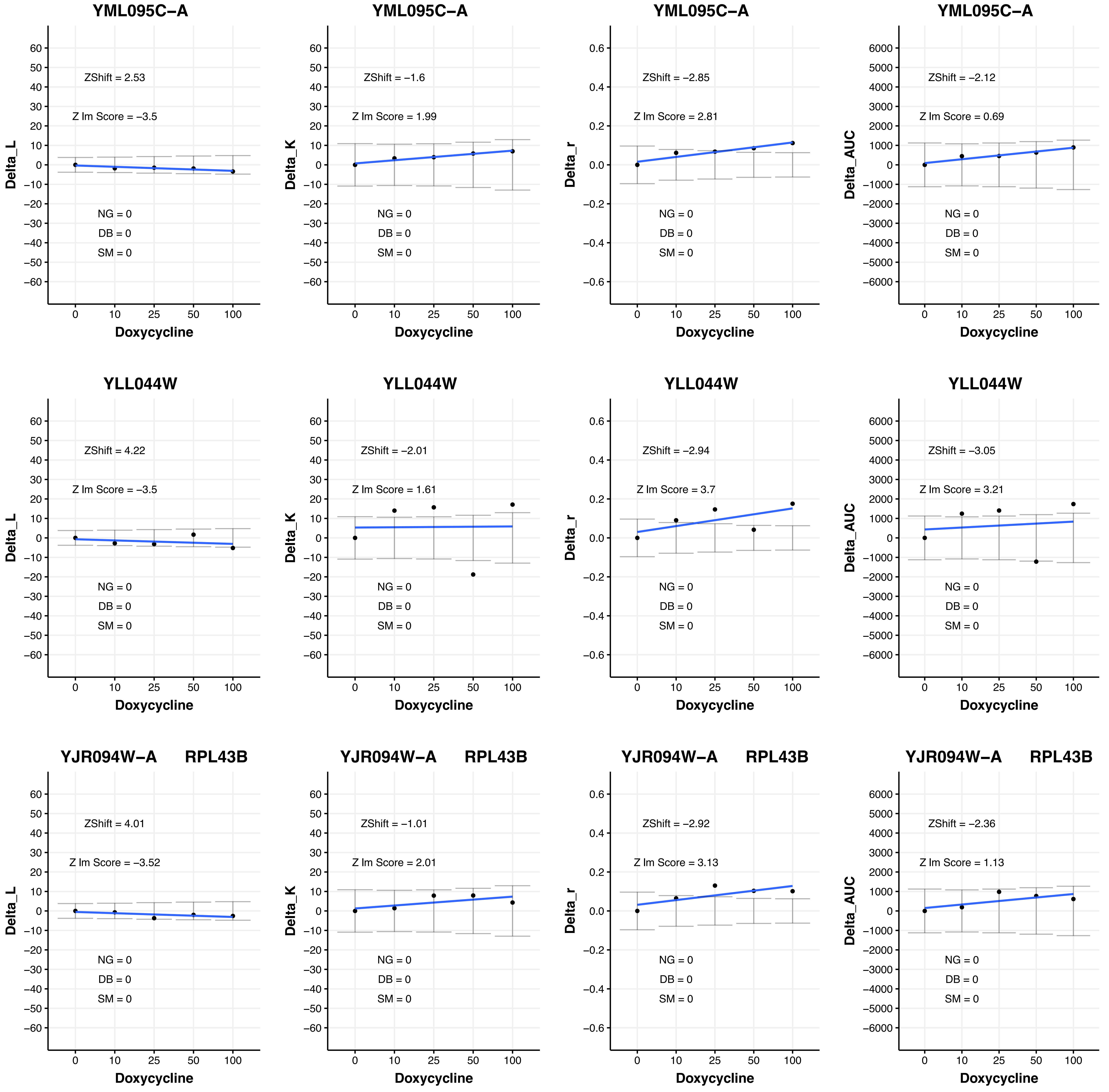

YML095C−A
ZShift = 2.53
Z lm Score = −3.5
NG = 0
DB = 0
SM = 0
Delta_L
Doxycycline
YML095C−A
ZShift = −1.6
Z lm Score = 1.99
NG = 0
DB = 0
SM = 0
Delta_K
Doxycycline
YML095C−A
ZShift = −2.85
Z lm Score = 2.81
NG = 0
DB = 0
SM = 0
Delta_r
Doxycycline
YML095C−A
ZShift = −2.12
Z lm Score = 0.69
NG = 0
DB = 0
SM = 0
Delta_AUC
Doxycycline
YLL044W
ZShift = 4.22
Z lm Score = −3.5
NG = 0
DB = 0
SM = 0
Delta_L
Doxycycline
YLL044W
ZShift = −2.01
Z lm Score = 1.61
NG = 0
DB = 0
SM = 0
Delta_K
Doxycycline
YLL044W
ZShift = −2.94
Z lm Score = 3.7
NG = 0
DB = 0
SM = 0
Delta_r
Doxycycline
YLL044W
ZShift = −3.05
Z lm Score = 3.21
NG = 0
DB = 0
SM = 0
Delta_AUC
Doxycycline
YJR094W−A RPL43B
ZShift = 4.01
Z lm Score = −3.52
NG = 0
DB = 0
SM = 0
Delta_L
Doxycycline
YJR094W−A RPL43B
ZShift = −1.01
Z lm Score = 2.01
NG = 0
DB = 0
SM = 0
Delta_K
Doxycycline
YJR094W−A RPL43B
ZShift = −2.92
Z lm Score = 3.13
NG = 0
DB = 0
SM = 0
Delta_r
Doxycycline
YJR094W−A RPL43B
ZShift = −2.36
Z lm Score = 1.13
NG = 0
DB = 0
SM = 0
Delta_AUC
Doxycycline
0 10 25 50 100

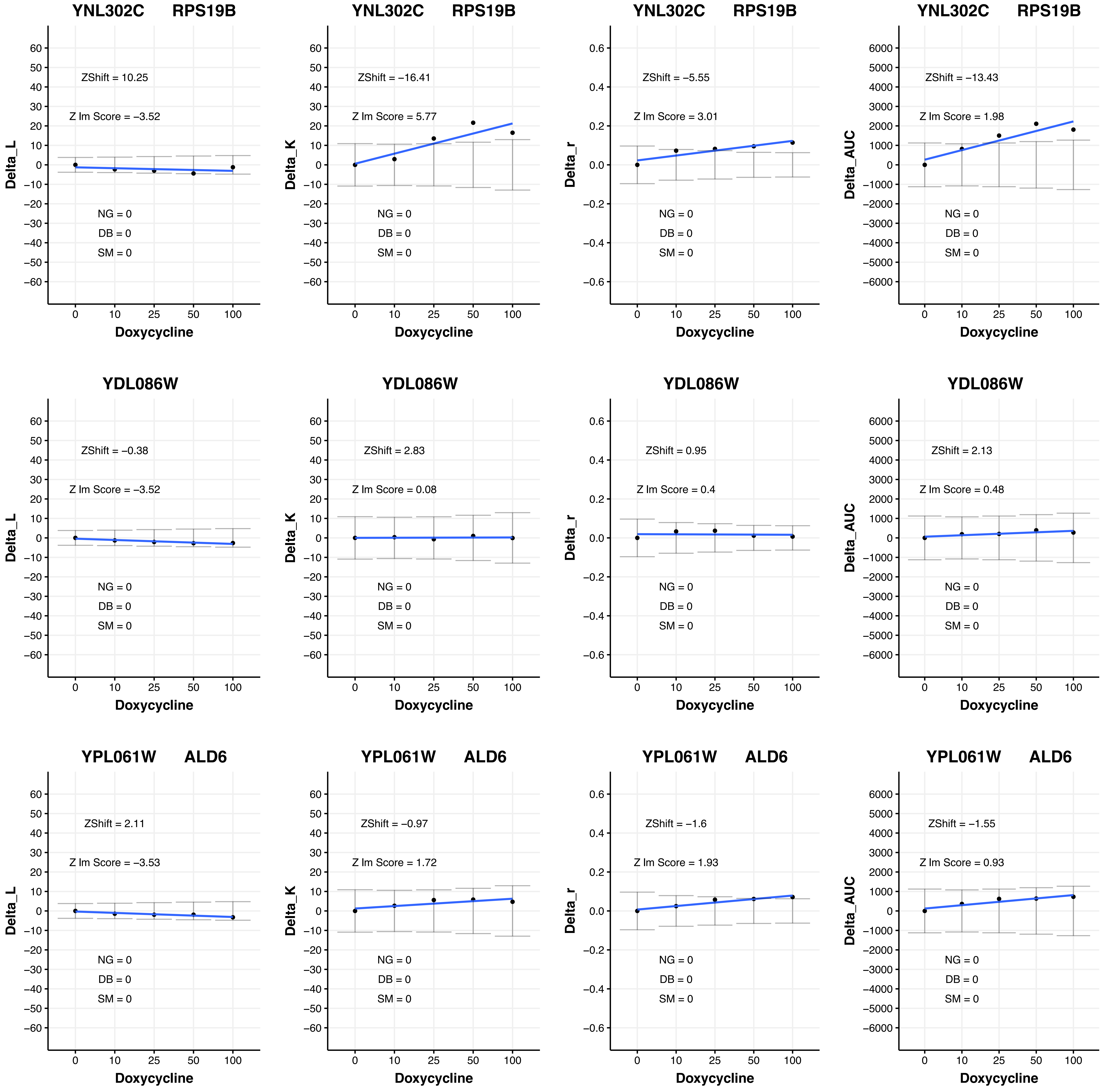

YNL302C RPS19B
ZShift = 10.25
Z lm Score = −3.52
NG = 0
DB = 0
SM = 0
Delta_L
Doxycycline
YNL302C RPS19B
ZShift = −16.41
Z lm Score = 5.77
NG = 0
DB = 0
SM = 0
Delta_K
Doxycycline
YNL302C RPS19B
ZShift = −5.55
Z lm Score = 3.01
NG = 0
DB = 0
SM = 0
Delta_r
Doxycycline
YNL302C RPS19B
ZShift = −13.43
Z lm Score = 1.98
NG = 0
DB = 0
SM = 0
Delta_AUC
Doxycycline
YDL086W
ZShift = −0.38
Z lm Score = −3.52
NG = 0
DB = 0
SM = 0
Delta_L
Doxycycline
YDL086W
ZShift = 2.83
Z lm Score = 0.08
NG = 0
DB = 0
SM = 0
Delta_K
Doxycycline
YDL086W
ZShift = 0.95
Z lm Score = 0.4
NG = 0
DB = 0
SM = 0
Delta_r
Doxycycline
YDL086W
ZShift = 2.13
Z lm Score = 0.48
NG = 0
DB = 0
SM = 0
Delta_AUC
Doxycycline
YPL061W ALD6
ZShift = 2.11
Z lm Score = −3.53
NG = 0
DB = 0
SM = 0
Delta_L
Doxycycline
YPL061W ALD6
ZShift = −0.97
Z lm Score = 1.72
NG = 0
DB = 0
SM = 0
Delta_K
Doxycycline
YPL061W ALD6
ZShift = −1.6
Z lm Score = 1.93
NG = 0
DB = 0
SM = 0
Delta_r
Doxycycline
YPL061W ALD6
ZShift = −1.55
Z lm Score = 0.93
NG = 0
DB = 0
SM = 0
Delta_AUC
Doxycycline

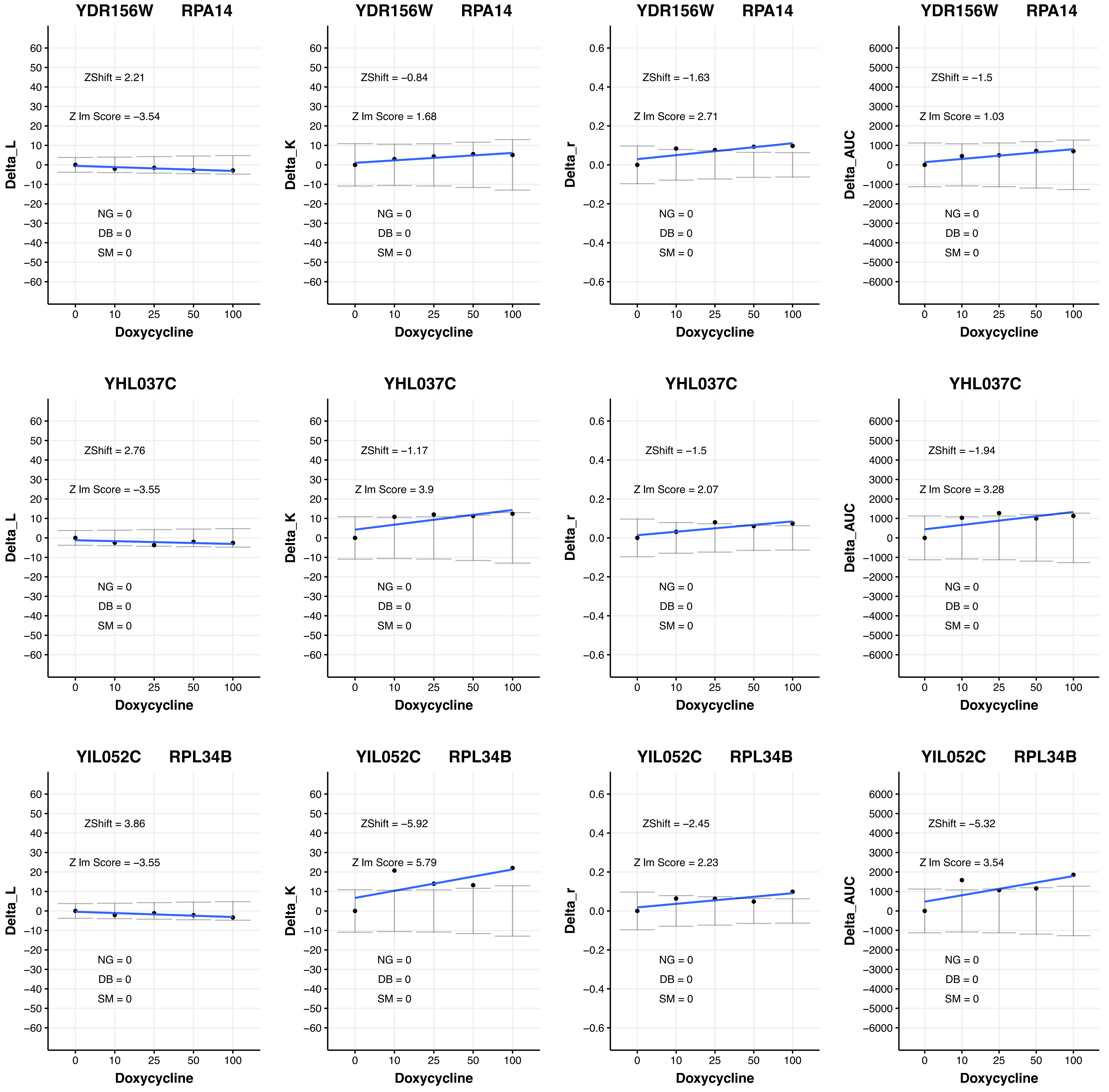

YDR156W RPA14
ZShift = 2.21
Z lm Score = −3.54
Delta_L
NG = 0
DB = 0
SM = 0
Doxycycline
YDR156W RPA14
ZShift = −0.84
Z lm Score = 1.68
Delta_K
NG = 0
DB = 0
SM = 0
Doxycycline
YDR156W RPA14
ZShift = −1.63
Z lm Score = 2.71
Delta_r
NG = 0
DB = 0
SM = 0
Doxycycline
YDR156W RPA14
ZShift = −1.5
Z lm Score = 1.03
Delta_AUC
NG = 0
DB = 0
SM = 0
Doxycycline
YHL037C
ZShift = 2.76
Z lm Score = −3.55
Delta_L
NG = 0
DB = 0
SM = 0
Doxycycline
YHL037C
ZShift = −1.17
Z lm Score = 3.9
Delta_K
NG = 0
DB = 0
SM = 0
Doxycycline
YHL037C
ZShift = −1.5
Z lm Score = 2.07
Delta_r
NG = 0
DB = 0
SM = 0
Doxycycline
YHL037C
ZShift = −1.94
Z lm Score = 3.28
Delta_AUC
NG = 0
DB = 0
SM = 0
Doxycycline
YIL052C RPL34B
ZShift = 3.86
Z lm Score = −3.55
Delta_L
NG = 0
DB = 0
SM = 0
Doxycycline
YIL052C RPL34B
ZShift = −5.92
Z lm Score = 5.79
Delta_K
NG = 0
DB = 0
SM = 0
Doxycycline
YIL052C RPL34B
ZShift = −2.45
Z lm Score = 2.23
Delta_r
NG = 0
DB = 0
SM = 0
Doxycycline
YIL052C RPL34B
ZShift = −5.32
Z lm Score = 3.54
Delta_AUC
NG = 0
DB = 0
SM = 0
Doxycycline

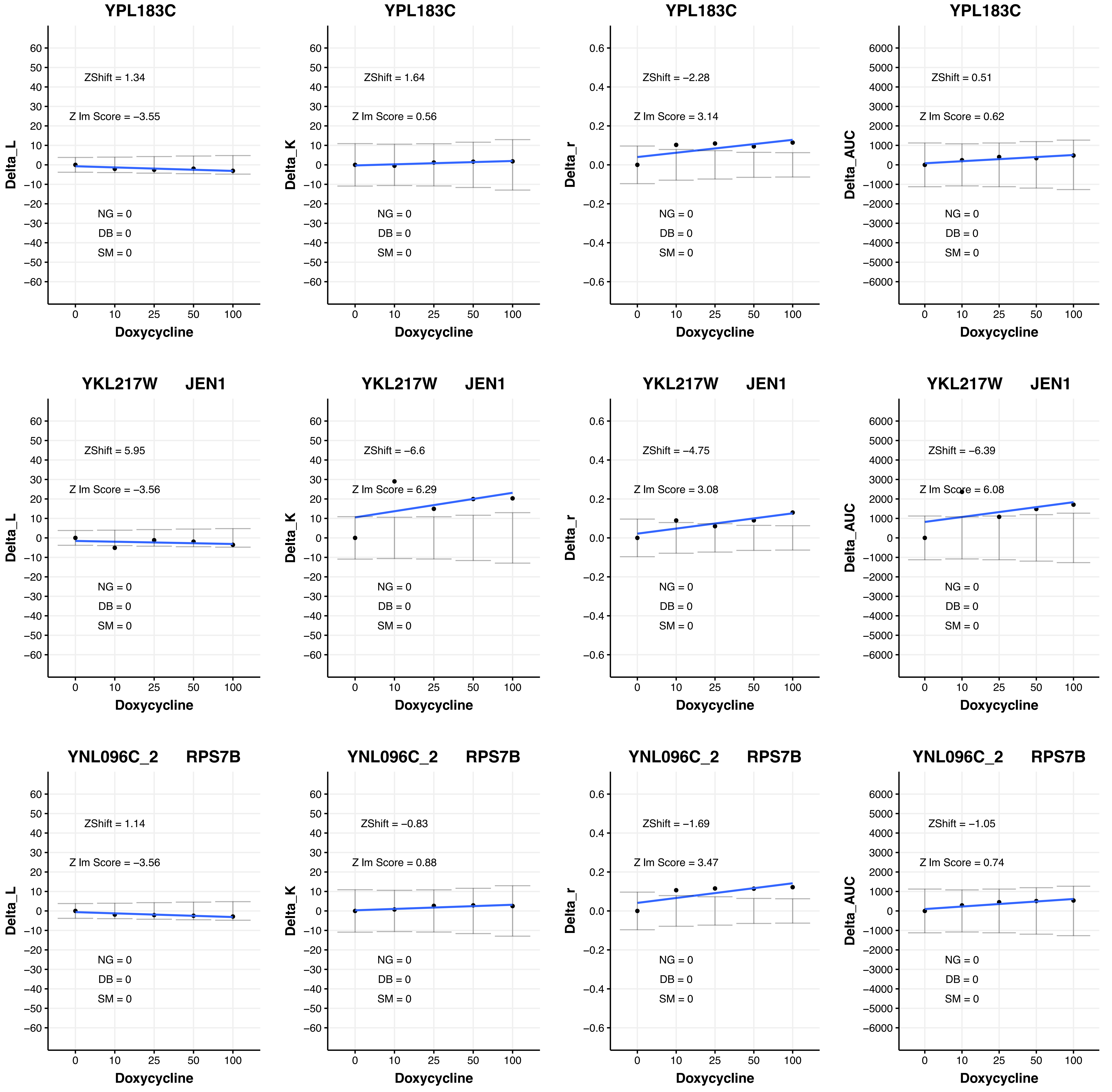
YPL183C
ZShift = 1.34
Z lm Score = −3.55
NG = 0
DB = 0
SM = 0
Delta_L
Doxycycline
YPL183C
ZShift = 1.64
Z lm Score = 0.56
NG = 0
DB = 0
SM = 0
Delta_K
Doxycycline
YPL183C
ZShift = −2.28
Z lm Score = 3.14
NG = 0
DB = 0
SM = 0
Delta_r
Doxycycline
YPL183C
ZShift = 0.51
Z lm Score = 0.62
NG = 0
DB = 0
SM = 0
Delta_AUC
Doxycycline
YKL217W JEN1
ZShift = 5.95
Z lm Score = −3.56
NG = 0
DB = 0
SM = 0
Delta_L
Doxycycline
YKL217W JEN1
ZShift = −6.6
Z lm Score = 6.29
NG = 0
DB = 0
SM = 0
Delta_K
Doxycycline
YKL217W JEN1
ZShift = −4.75
Z lm Score = 3.08
NG = 0
DB = 0
SM = 0
Delta_r
Doxycycline
YKL217W JEN1
ZShift = −6.39
Z lm Score = 6.08
NG = 0
DB = 0
SM = 0
Delta_AUC
Doxycycline
YNL096C_2 RPS7B
ZShift = 1.14
Z lm Score = −3.56
NG = 0
DB = 0
SM = 0
Delta_L
Doxycycline
YNL096C_2 RPS7B
ZShift = −0.83
Z lm Score = 0.88
NG = 0
DB = 0
SM = 0
Delta_K
Doxycycline
YNL096C_2 RPS7B
ZShift = −1.69
Z lm Score = 3.47
NG = 0
DB = 0
SM = 0
Delta_r
Doxycycline
YNL096C_2 RPS7B
ZShift = −1.05
Z lm Score = 0.74
NG = 0
DB = 0
SM = 0
Delta_AUC
Doxycycline

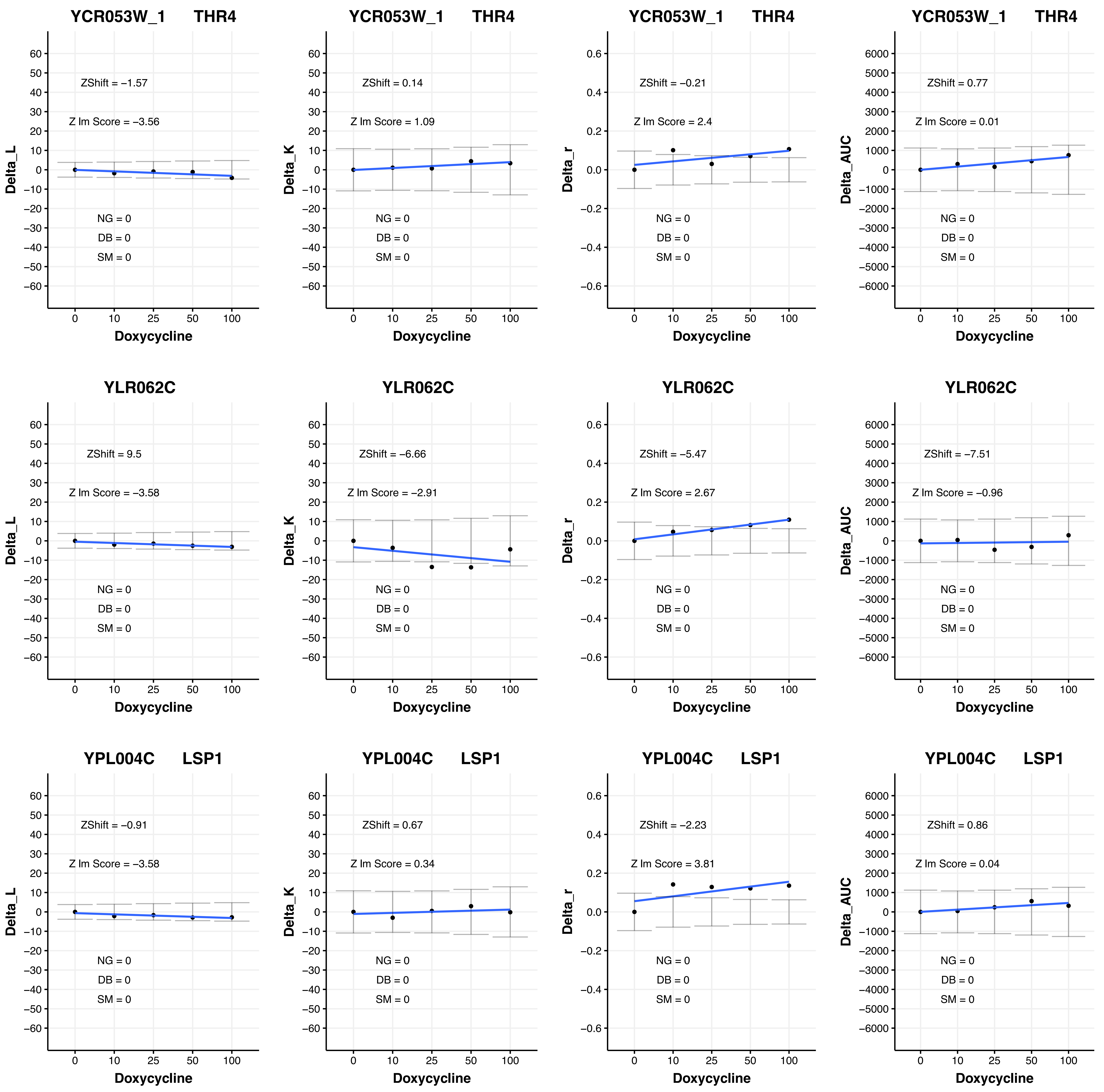
YCR053W_1 THR4
ZShift = −1.57
Z lm Score = −3.56
NG = 0
DB = 0
SM = 0
Delta_L
Doxycycline
YCR053W_1 THR4
ZShift = 0.14
Z lm Score = 1.09
NG = 0
DB = 0
SM = 0
Delta_K
Doxycycline
YCR053W_1 THR4
ZShift = −0.21
Z lm Score = 2.4
NG = 0
DB = 0
SM = 0
Delta_r
Doxycycline
YCR053W_1 THR4
ZShift = 0.77
Z lm Score = 0.01
NG = 0
DB = 0
SM = 0
Delta_AUC
Doxycycline
YLR062C
ZShift = 9.5
Z lm Score = −3.58
NG = 0
DB = 0
SM = 0
Delta_L
Doxycycline
YLR062C
ZShift = −6.66
Z lm Score = −2.91
NG = 0
DB = 0
SM = 0
Delta_K
Doxycycline
YLR062C
ZShift = −5.47
Z lm Score = 2.67
NG = 0
DB = 0
SM = 0
Delta_r
Doxycycline
YLR062C
ZShift = −7.51
Z lm Score = −0.96
NG = 0
DB = 0
SM = 0
Delta_AUC
Doxycycline
YPL004C LSP1
ZShift = −0.91
Z lm Score = −3.58
NG = 0
DB = 0
SM = 0
Delta_L
Doxycycline
YPL004C LSP1
ZShift = 0.67
Z lm Score = 0.34
NG = 0
DB = 0
SM = 0
Delta_K
Doxycycline
YPL004C LSP1
ZShift = −2.23
Z lm Score = 3.81
NG = 0
DB = 0
SM = 0
Delta_r
Doxycycline
YPL004C LSP1
ZShift = 0.86
Z lm Score = 0.04
NG = 0
DB = 0
SM = 0
Delta_AUC
Doxycycline

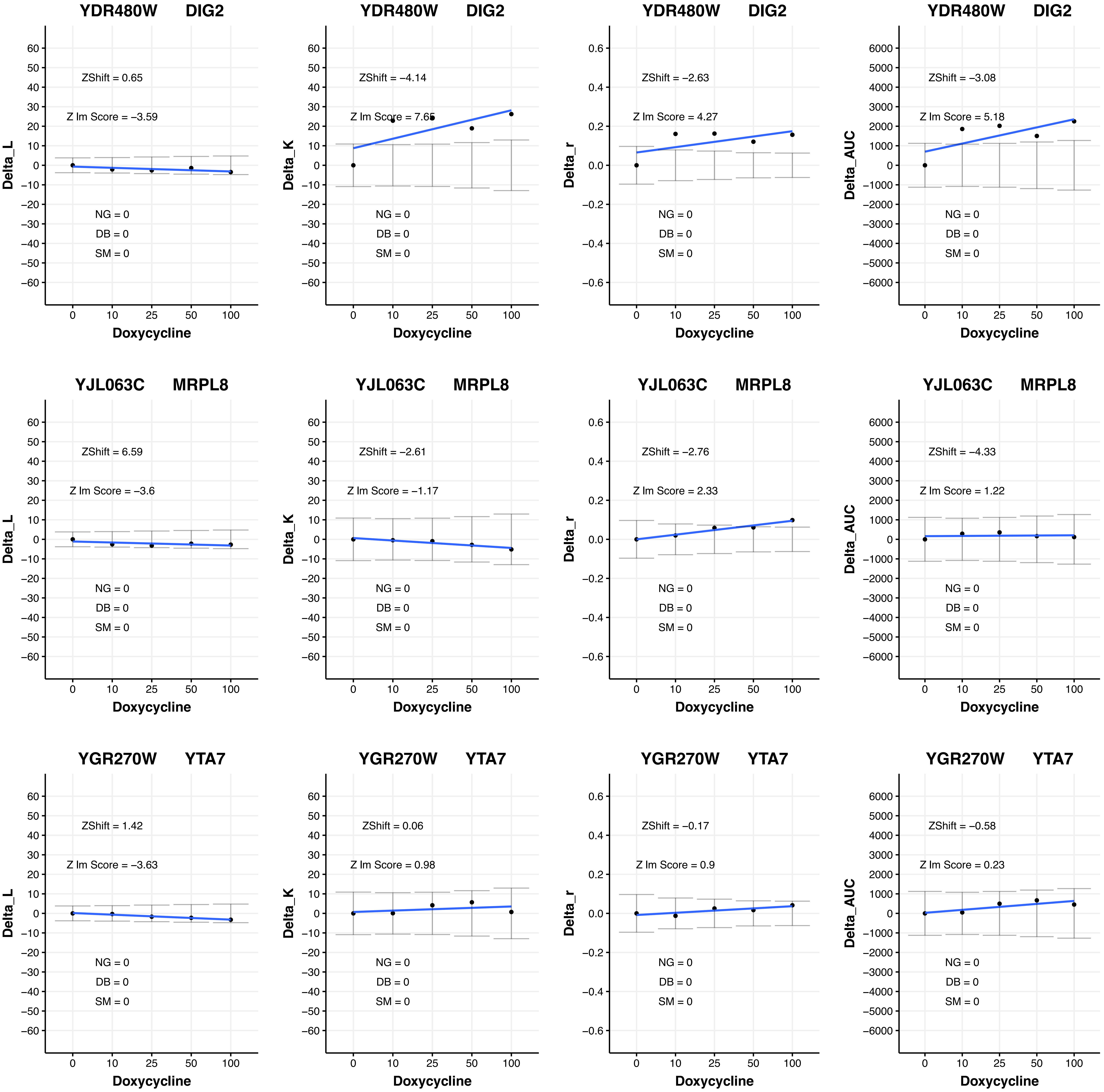

YDR480W DIG2
ZShift = 0.65
Z lm Score = −3.59
NG = 0
DB = 0
SM = 0
Delta_L
Doxycycline
YDR480W DIG2
ZShift = −4.14
Z lm Score = 7.65
NG = 0
DB = 0
SM = 0
Delta_K
Doxycycline
YDR480W DIG2
ZShift = −2.63
Z lm Score = 4.27
NG = 0
DB = 0
SM = 0
Delta_r
Doxycycline
YDR480W DIG2
ZShift = −3.08
Z lm Score = 5.18
NG = 0
DB = 0
SM = 0
Delta_AUC
Doxycycline
YJL063C MRPL8
ZShift = 6.59
Z lm Score = −3.6
NG = 0
DB = 0
SM = 0
Delta_L
Doxycycline
YJL063C MRPL8
ZShift = −2.61
Z lm Score = −1.17
NG = 0
DB = 0
SM = 0
Delta_K
Doxycycline
YJL063C MRPL8
ZShift = −2.76
Z lm Score = 2.33
NG = 0
DB = 0
SM = 0
Delta_r
Doxycycline
YJL063C MRPL8
ZShift = −4.33
Z lm Score = 1.22
NG = 0
DB = 0
SM = 0
Delta_AUC
Doxycycline
YGR270W YTA7
ZShift = 1.42
Z lm Score = −3.63
NG = 0
DB = 0
SM = 0
Delta_L
Doxycycline
YGR270W YTA7
ZShift = 0.06
Z lm Score = 0.98
NG = 0
DB = 0
SM = 0
Delta_K
Doxycycline
YGR270W YTA7
ZShift = −0.17
Z lm Score = 0.9
NG = 0
DB = 0
SM = 0
Delta_r
Doxycycline
YGR270W YTA7
ZShift = −0.58
Z lm Score = 0.23
NG = 0
DB = 0
SM = 0
Delta_AUC
Doxycycline

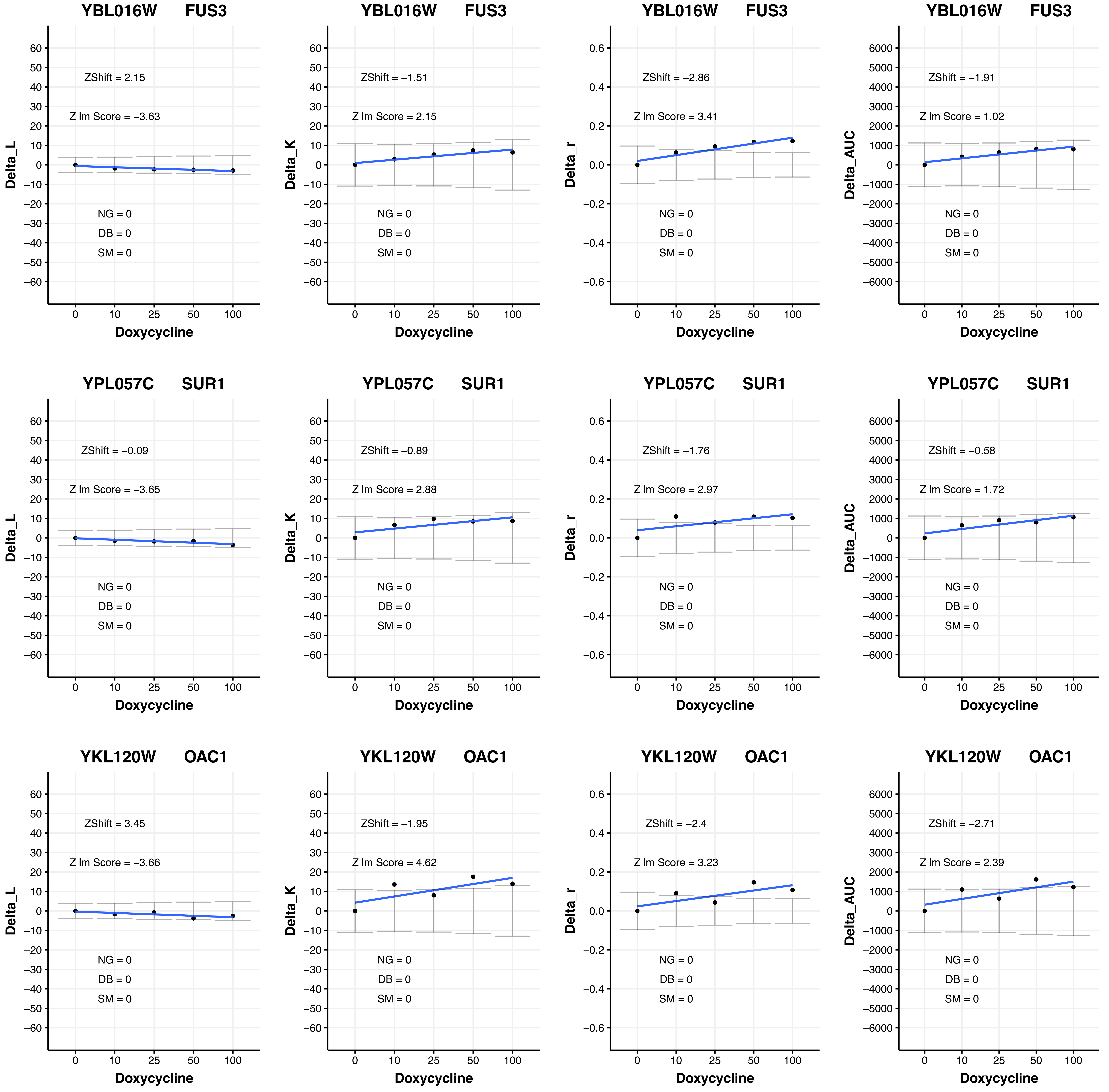

YBL016W FUS3
ZShift = 2.15
Z lm Score = −3.63
NG = 0
DB = 0
SM = 0
Delta_L
Doxycycline
YBL016W FUS3
ZShift = −1.51
Z lm Score = 2.15
NG = 0
DB = 0
SM = 0
Delta_K
Doxycycline
YBL016W FUS3
ZShift = −2.86
Z lm Score = 3.41
NG = 0
DB = 0
SM = 0
Delta_r
Doxycycline
YBL016W FUS3
ZShift = −1.91
Z lm Score = 1.02
NG = 0
DB = 0
SM = 0
Delta_AUC
Doxycycline
YPL057C SUR1
ZShift = −0.09
Z lm Score = −3.65
NG = 0
DB = 0
SM = 0
Delta_L
Doxycycline
YPL057C SUR1
ZShift = −0.89
Z lm Score = 2.88
NG = 0
DB = 0
SM = 0
Delta_K
Doxycycline
YPL057C SUR1
ZShift = −1.76
Z lm Score = 2.97
NG = 0
DB = 0
SM = 0
Delta_r
Doxycycline
YPL057C SUR1
ZShift = −0.58
Z lm Score = 1.72
NG = 0
DB = 0
SM = 0
Delta_AUC
Doxycycline
YKL120W OAC1
ZShift = 3.45
Z lm Score = −3.66
NG = 0
DB = 0
SM = 0
Delta_L
Doxycycline
YKL120W OAC1
ZShift = −1.95
Z lm Score = 4.62
NG = 0
DB = 0
SM = 0
Delta_K
Doxycycline
YKL120W OAC1
ZShift = −2.4
Z lm Score = 3.23
NG = 0
DB = 0
SM = 0
Delta_r
Doxycycline
YKL120W OAC1
ZShift = −2.71
Z lm Score = 2.39
NG = 0
DB = 0
SM = 0
Delta_AUC
Doxycycline

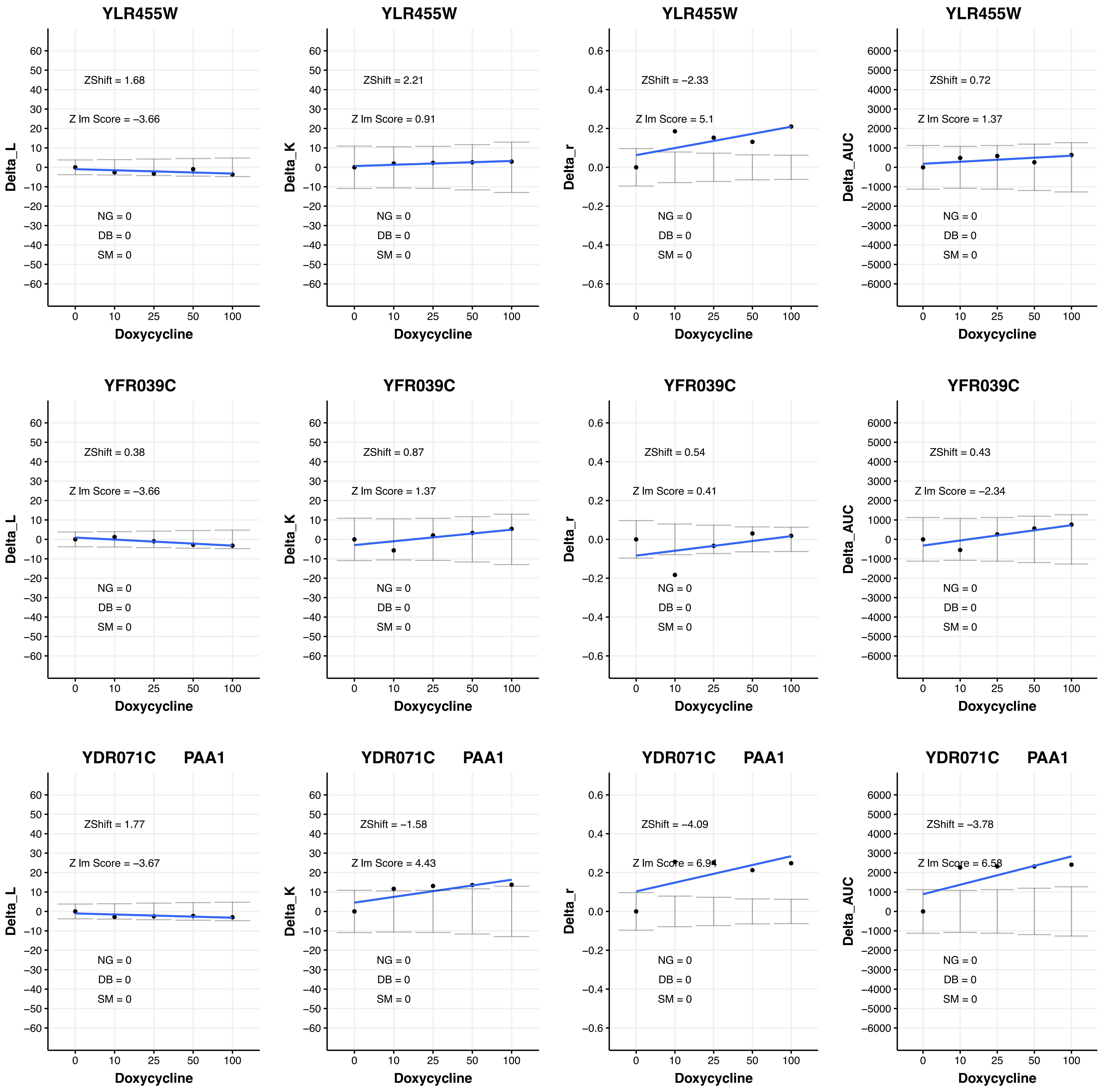

YLR455W
ZShift = 1.68
Z lm Score = −3.66
NG = 0
DB = 0
SM = 0
Delta_L
Doxycycline
YLR455W
ZShift = 2.21
Z lm Score = 0.91
NG = 0
DB = 0
SM = 0
Delta_K
Doxycycline
YLR455W
ZShift = −2.33
Z lm Score = 5.1
NG = 0
DB = 0
SM = 0
Delta_r
Doxycycline
YLR455W
ZShift = 0.72
Z lm Score = 1.37
NG = 0
DB = 0
SM = 0
Delta_AUC
Doxycycline
YFR039C
ZShift = 0.38
Z lm Score = −3.66
NG = 0
DB = 0
SM = 0
Delta_L
Doxycycline
YFR039C
ZShift = 0.87
Z lm Score = 1.37
NG = 0
DB = 0
SM = 0
Delta_K
Doxycycline
YFR039C
ZShift = 0.54
Z lm Score = 0.41
NG = 0
DB = 0
SM = 0
Delta_r
Doxycycline
YFR039C
ZShift = 0.43
Z lm Score = −2.34
NG = 0
DB = 0
SM = 0
Delta_AUC
Doxycycline
YDR071C PAA1
ZShift = 1.77
Z lm Score = −3.67
NG = 0
DB = 0
SM = 0
Delta_L
Doxycycline
YDR071C PAA1
ZShift = −1.58
Z lm Score = 4.43
NG = 0
DB = 0
SM = 0
Delta_K
Doxycycline
YDR071C PAA1
ZShift = −4.09
Z lm Score = 6.94
NG = 0
DB = 0
SM = 0
Delta_r
Doxycycline
YDR071C PAA1
ZShift = −3.78
Z lm Score = 6.58
NG = 0
DB = 0
SM = 0
Delta_AUC
Doxycycline

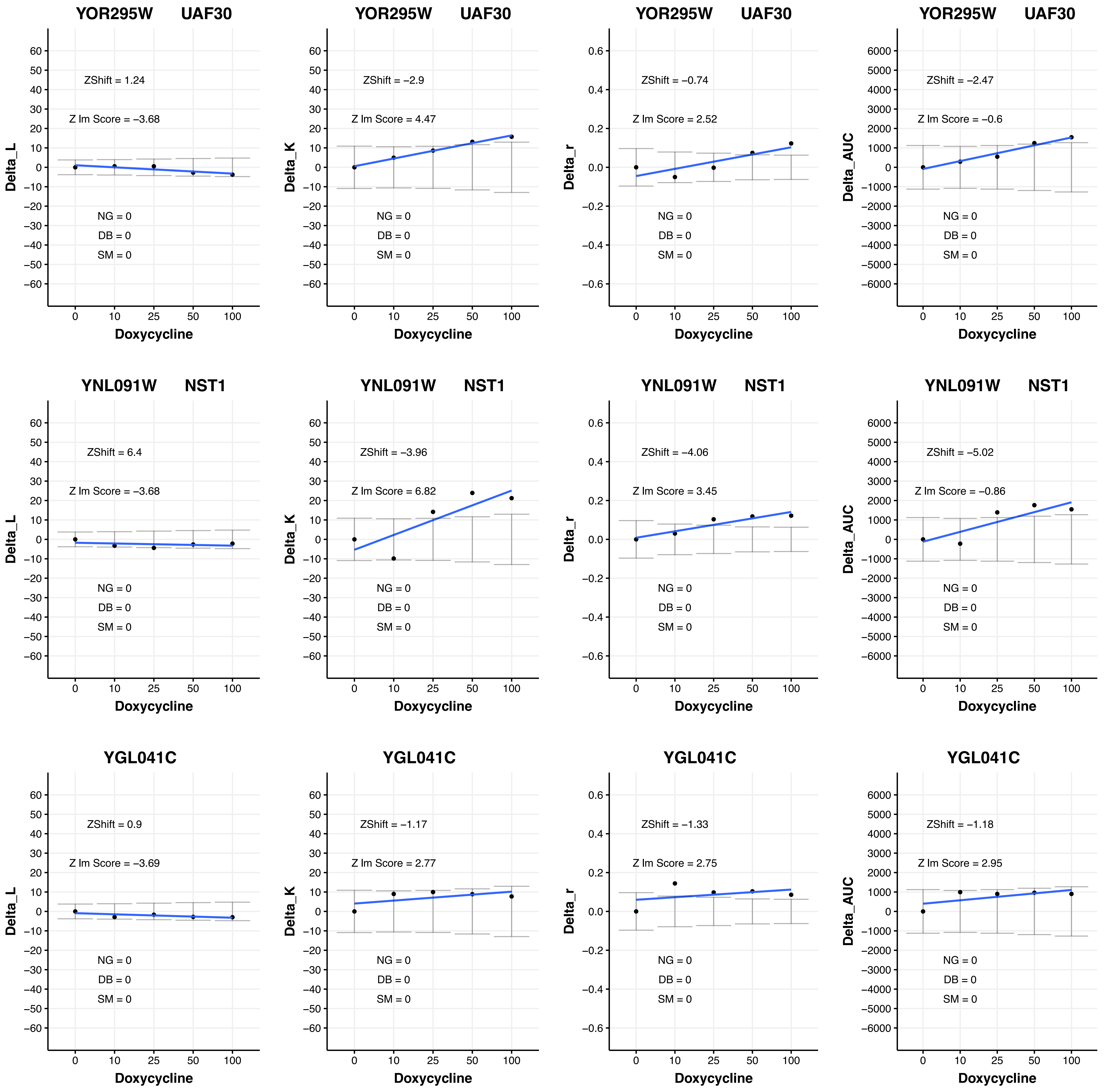
YOR295W UAF30
ZShift = 1.24
Z lm Score = −3.68
NG = 0
DB = 0
SM = 0
Delta_L
Doxycycline
YOR295W UAF30
ZShift = −2.9
Z lm Score = 4.47
NG = 0
DB = 0
SM = 0
Delta_K
Doxycycline
YOR295W UAF30
ZShift = −0.74
Z lm Score = 2.52
NG = 0
DB = 0
SM = 0
Delta_r
Doxycycline
YOR295W UAF30
ZShift = −2.47
Z lm Score = −0.6
NG = 0
DB = 0
SM = 0
Delta_AUC
Doxycycline
YNL091W NST1
ZShift = 6.4
Z lm Score = −3.68
NG = 0
DB = 0
SM = 0
Delta_L
Doxycycline
YNL091W NST1
ZShift = −3.96
Z lm Score = 6.82
NG = 0
DB = 0
SM = 0
Delta_K
Doxycycline
YNL091W NST1
ZShift = −4.06
Z lm Score = 3.45
NG = 0
DB = 0
SM = 0
Delta_r
Doxycycline
YNL091W NST1
ZShift = −5.02
Z lm Score = −0.86
NG = 0
DB = 0
SM = 0
Delta_AUC
Doxycycline
YGL041C
ZShift = 0.9
Z lm Score = −3.69
NG = 0
DB = 0
SM = 0
Delta_L
Doxycycline
YGL041C
ZShift = −1.17
Z lm Score = 2.77
NG = 0
DB = 0
SM = 0
Delta_K
Doxycycline
YGL041C
ZShift = −1.33
Z lm Score = 2.75
NG = 0
DB = 0
SM = 0
Delta_r
Doxycycline
YGL041C
ZShift = −1.18
Z lm Score = 2.95
NG = 0
DB = 0
SM = 0
Delta_AUC
Doxycycline

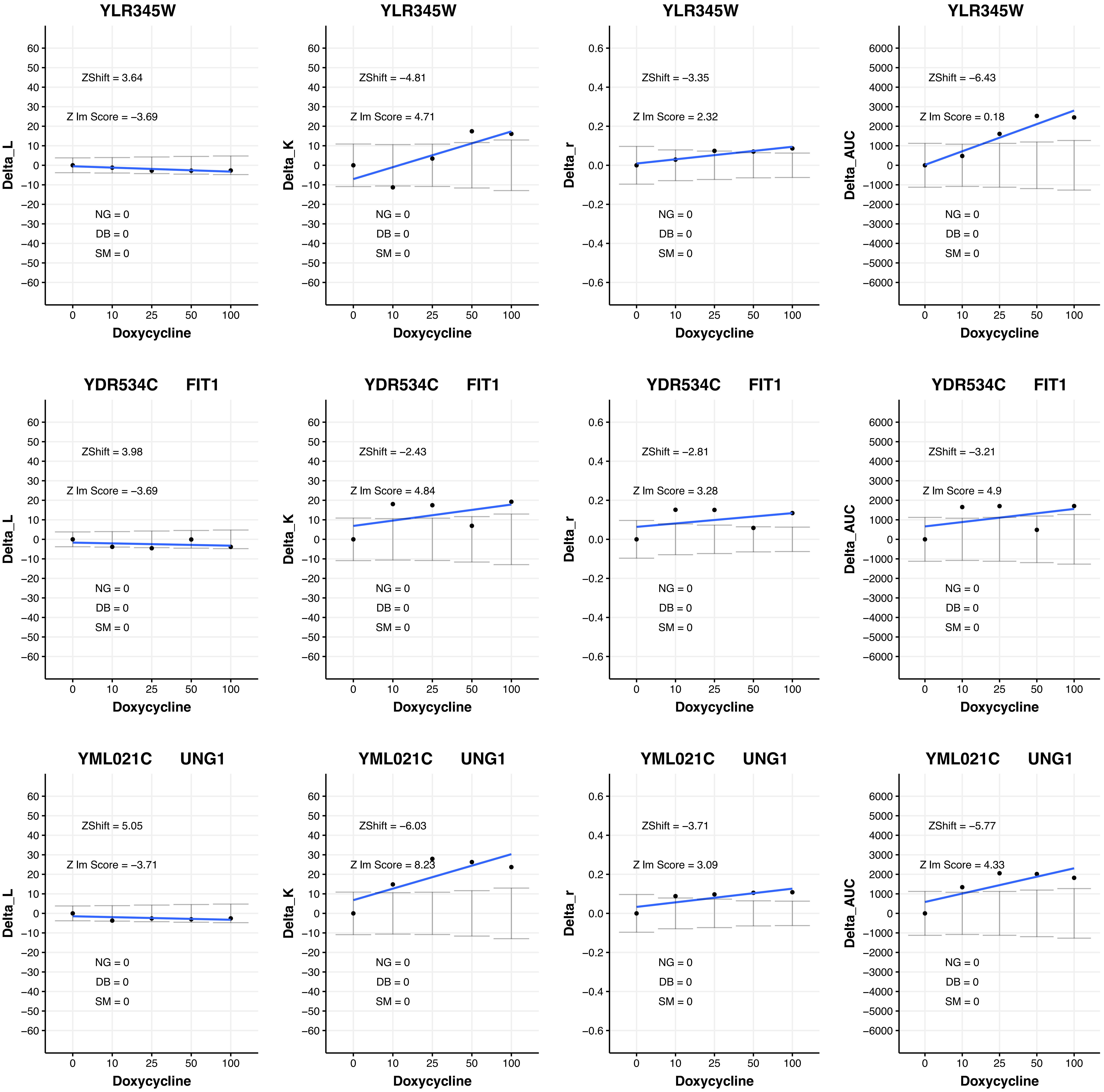

YLR345W
ZShift = 3.64
Z lm Score = −3.69
NG = 0
DB = 0
SM = 0
Delta_L
Doxycycline
YLR345W
ZShift = −4.81
Z lm Score = 4.71
NG = 0
DB = 0
SM = 0
Delta_K
Doxycycline
YLR345W
ZShift = −3.35
Z lm Score = 2.32
NG = 0
DB = 0
SM = 0
Delta_r
Doxycycline
YLR345W
ZShift = −6.43
Z lm Score = 0.18
NG = 0
DB = 0
SM = 0
Delta_AUC
Doxycycline
YDR534C FIT1
ZShift = 3.98
Z lm Score = −3.69
NG = 0
DB = 0
SM = 0
Delta_L
Doxycycline
YDR534C FIT1
ZShift = −2.43
Z lm Score = 4.84
NG = 0
DB = 0
SM = 0
Delta_K
Doxycycline
YDR534C FIT1
ZShift = −2.81
Z lm Score = 3.28
NG = 0
DB = 0
SM = 0
Delta_r
Doxycycline
YDR534C FIT1
ZShift = −3.21
Z lm Score = 4.9
NG = 0
DB = 0
SM = 0
Delta_AUC
Doxycycline
YML021C UNG1
ZShift = 5.05
Z lm Score = −3.71
NG = 0
DB = 0
SM = 0
Delta_L
Doxycycline
YML021C UNG1
ZShift = −6.03
Z lm Score = 8.23
NG = 0
DB = 0
SM = 0
Delta_K
Doxycycline
YML021C UNG1
ZShift = −3.71
Z lm Score = 3.09
NG = 0
DB = 0
SM = 0
Delta_r
Doxycycline
YML021C UNG1
ZShift = −5.77
Z lm Score = 4.33
NG = 0
DB = 0
SM = 0
Delta_AUC
Doxycycline

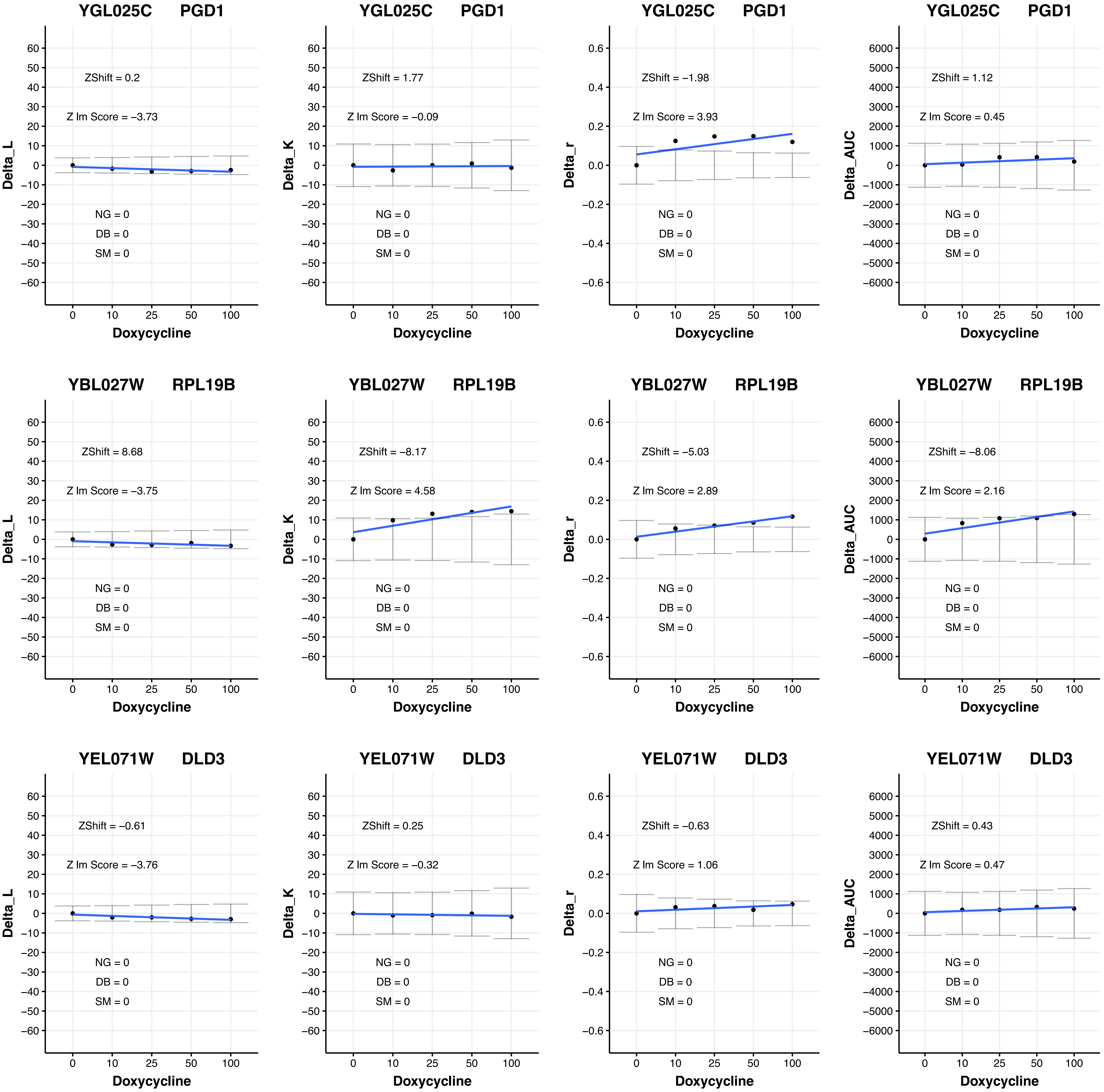
YGL025C PGD1
ZShift = 0.2
Z lm Score = −3.73
NG = 0
DB = 0
SM = 0
Delta_L
Doxycycline
YGL025C PGD1
ZShift = 1.77
Z lm Score = −0.09
NG = 0
DB = 0
SM = 0
Delta_K
Doxycycline
YGL025C PGD1
ZShift = −1.98
Z lm Score = 3.93
NG = 0
DB = 0
SM = 0
Delta_r
Doxycycline
YGL025C PGD1
ZShift = 1.12
Z lm Score = 0.45
NG = 0
DB = 0
SM = 0
Delta_AUC
Doxycycline
YBL027W RPL19B
ZShift = 8.68
Z lm Score = −3.75
NG = 0
DB = 0
SM = 0
Delta_L
Doxycycline
YBL027W RPL19B
ZShift = −8.17
Z lm Score = 4.58
NG = 0
DB = 0
SM = 0
Delta_K
Doxycycline
YBL027W RPL19B
ZShift = −5.03
Z lm Score = 2.89
NG = 0
DB = 0
SM = 0
Delta_r
Doxycycline
YBL027W RPL19B
ZShift = −8.06
Z lm Score = 2.16
NG = 0
DB = 0
SM = 0
Delta_AUC
Doxycycline
YEL071W DLD3
ZShift = −0.61
Z lm Score = −3.76
NG = 0
DB = 0
SM = 0
Delta_L
Doxycycline
YEL071W DLD3
ZShift = 0.25
Z lm Score = −0.32
NG = 0
DB = 0
SM = 0
Delta_K
Doxycycline
YEL071W DLD3
ZShift = −0.63
Z lm Score = 1.06
NG = 0
DB = 0
SM = 0
Delta_r
Doxycycline
YEL071W DLD3
ZShift = 0.43
Z lm Score = 0.47
NG = 0
DB = 0
SM = 0
Delta_AUC
Doxycycline

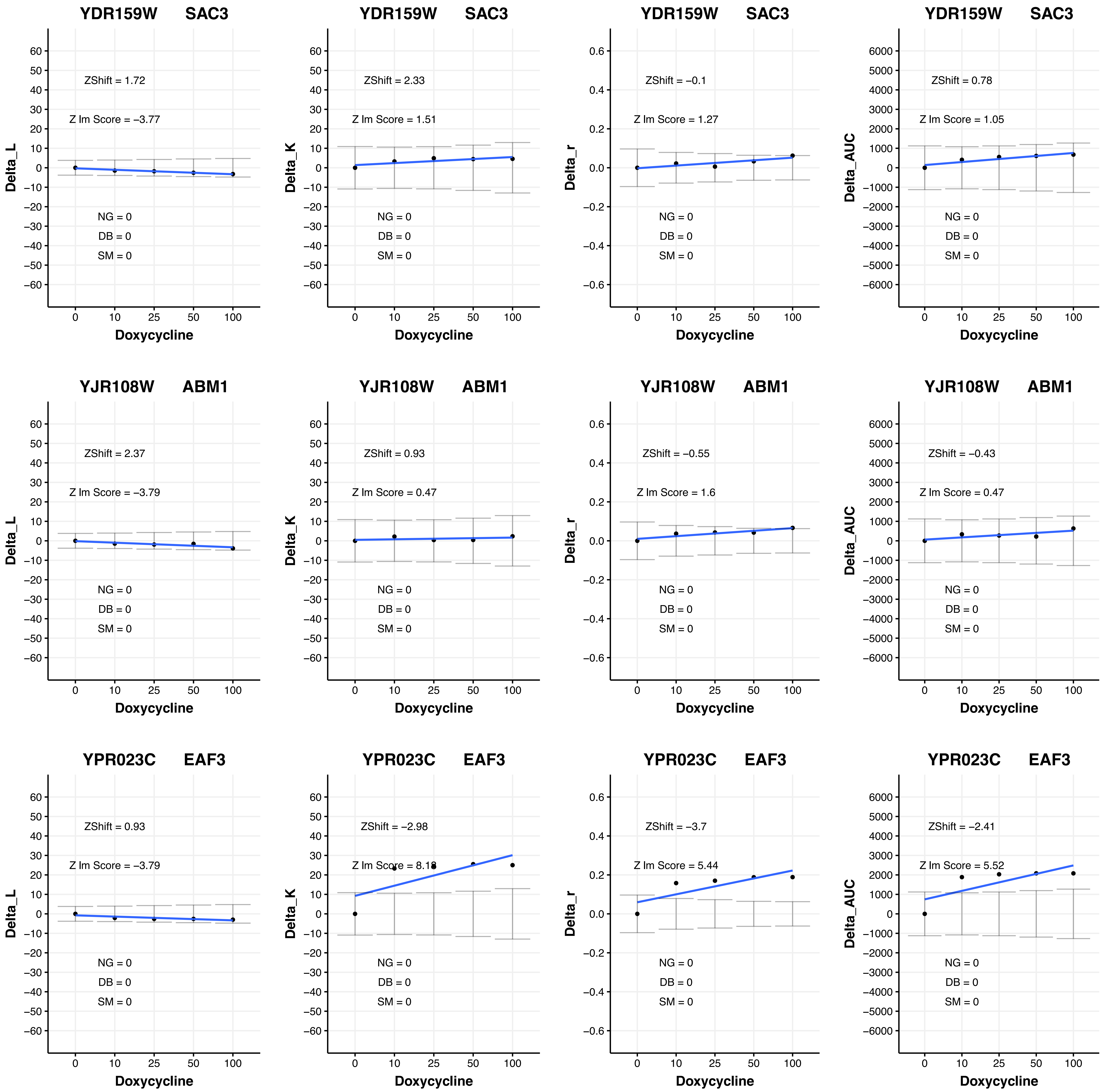
YDR159W SAC3
ZShift = 1.72
Z lm Score = −3.77
NG = 0
DB = 0
SM = 0
Delta_L
Doxycycline
YDR159W SAC3
ZShift = 2.33
Z lm Score = 1.51
NG = 0
DB = 0
SM = 0
Delta_K
Doxycycline
YDR159W SAC3
ZShift = −0.1
Z lm Score = 1.27
NG = 0
DB = 0
SM = 0
Delta_r
Doxycycline
YDR159W SAC3
ZShift = 0.78
Z lm Score = 1.05
NG = 0
DB = 0
SM = 0
Delta_AUC
Doxycycline
YJR108W ABM1
ZShift = 2.37
Z lm Score = −3.79
NG = 0
DB = 0
SM = 0
Delta_L
Doxycycline
YJR108W ABM1
ZShift = 0.93
Z lm Score = 0.47
NG = 0
DB = 0
SM = 0
Delta_K
Doxycycline
YJR108W ABM1
ZShift = −0.55
Z lm Score = 1.6
NG = 0
DB = 0
SM = 0
Delta_r
Doxycycline
YJR108W ABM1
ZShift = −0.43
Z lm Score = 0.47
NG = 0
DB = 0
SM = 0
Delta_AUC
Doxycycline
YPR023C EAF3
ZShift = 0.93
Z lm Score = −3.79
NG = 0
DB = 0
SM = 0
Delta_L
Doxycycline
YPR023C EAF3
ZShift = −2.98
Z lm Score = 8.18
NG = 0
DB = 0
SM = 0
Delta_K
Doxycycline
YPR023C EAF3
ZShift = −3.7
Z lm Score = 5.44
NG = 0
DB = 0
SM = 0
Delta_r
Doxycycline
YPR023C EAF3
ZShift = −2.41
Z lm Score = 5.52
NG = 0
DB = 0
SM = 0
Delta_AUC
Doxycycline

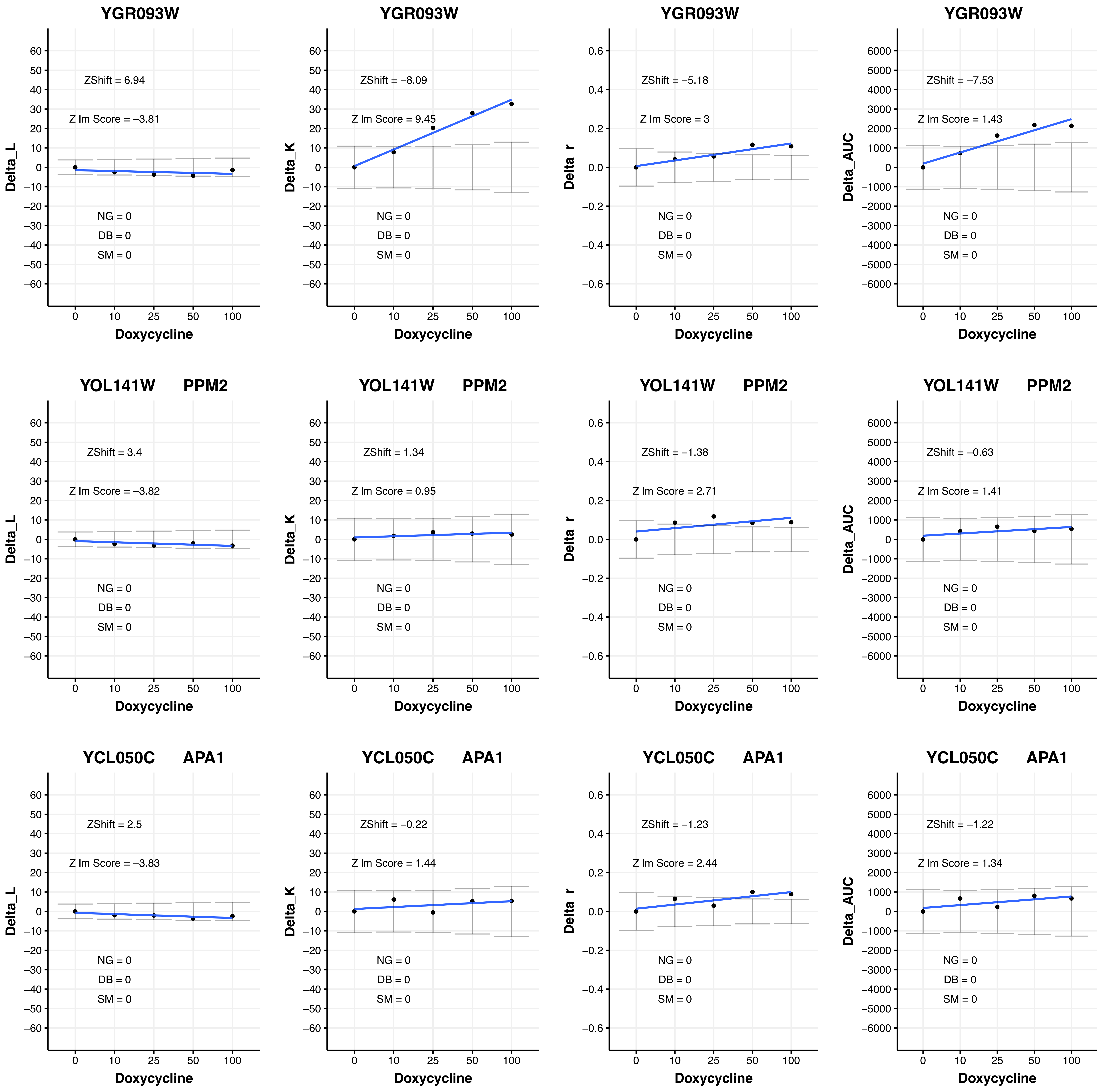

YGR093W
ZShift = 6.94
Z lm Score = −3.81
NG = 0
DB = 0
SM = 0
Delta_L
Doxycycline
YGR093W
ZShift = −8.09
Z lm Score = 9.45
NG = 0
DB = 0
SM = 0
Delta_K
Doxycycline
YGR093W
ZShift = −5.18
Z lm Score = 3
NG = 0
DB = 0
SM = 0
Delta_r
Doxycycline
YGR093W
ZShift = −7.53
Z lm Score = 1.43
NG = 0
DB = 0
SM = 0
Delta_AUC
Doxycycline
YOL141W PPM2
ZShift = 3.4
Z lm Score = −3.82
NG = 0
DB = 0
SM = 0
Delta_L
Doxycycline
YOL141W PPM2
ZShift = 1.34
Z lm Score = 0.95
NG = 0
DB = 0
SM = 0
Delta_K
Doxycycline
YOL141W PPM2
ZShift = −1.38
Z lm Score = 2.71
NG = 0
DB = 0
SM = 0
Delta_r
Doxycycline
YOL141W PPM2
ZShift = −0.63
Z lm Score = 1.41
NG = 0
DB = 0
SM = 0
Delta_AUC
Doxycycline
YCL050C APA1
ZShift = 2.5
Z lm Score = −3.83
NG = 0
DB = 0
SM = 0
Delta_L
Doxycycline
YCL050C APA1
ZShift = −0.22
Z lm Score = 1.44
NG = 0
DB = 0
SM = 0
Delta_K
Doxycycline
YCL050C APA1
ZShift = −1.23
Z lm Score = 2.44
NG = 0
DB = 0
SM = 0
Delta_r
Doxycycline
YCL050C APA1
ZShift = −1.22
Z lm Score = 1.34
NG = 0
DB = 0
SM = 0
Delta_AUC
Doxycycline

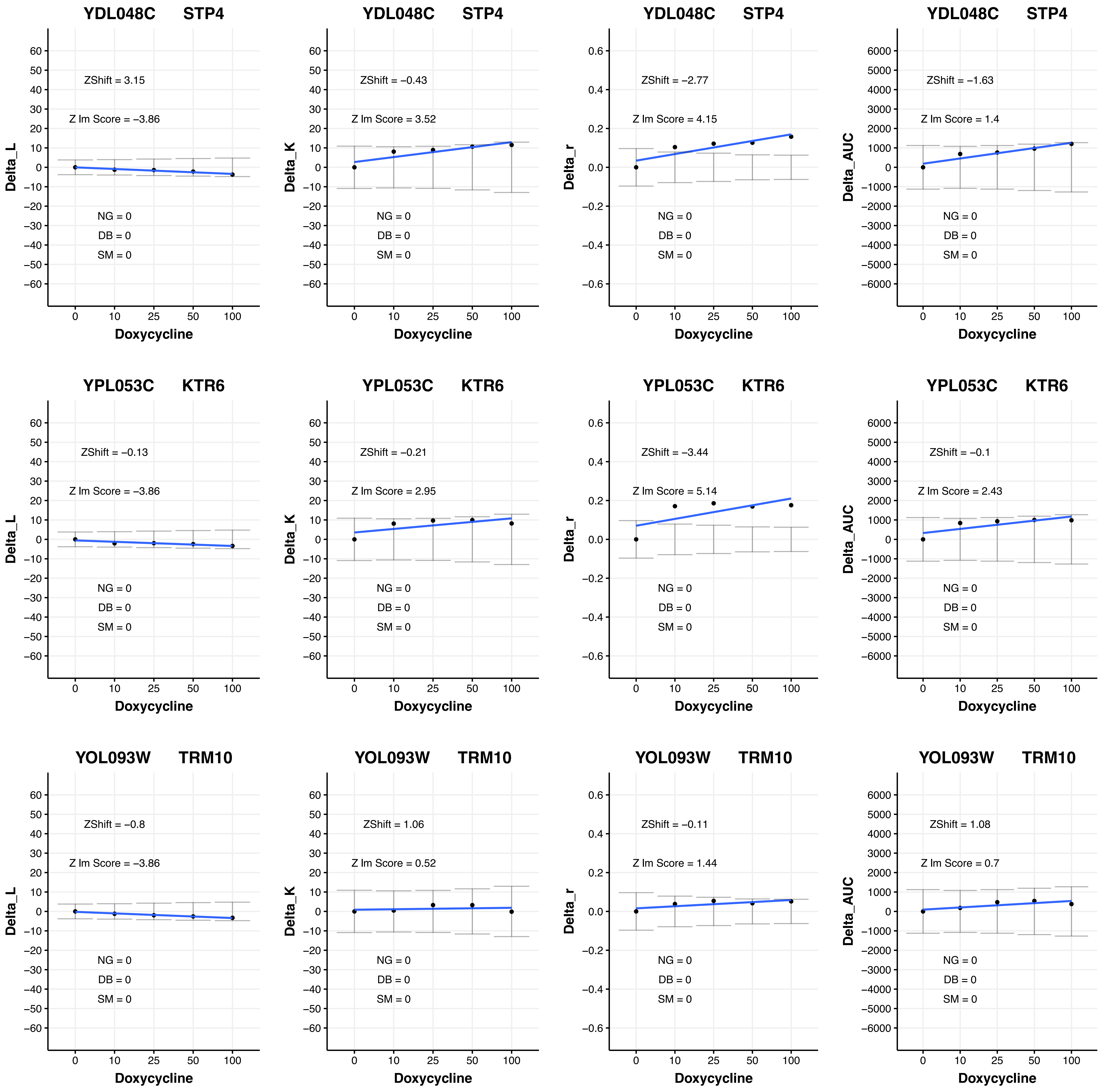

YDL048C STP4
ZShift = 3.15
Z lm Score = −3.86
NG = 0
DB = 0
SM = 0
Delta_L
Doxycycline
YDL048C STP4
ZShift = −0.43
Z lm Score = 3.52
NG = 0
DB = 0
SM = 0
Delta_K
Doxycycline
YDL048C STP4
ZShift = −2.77
Z lm Score = 4.15
NG = 0
DB = 0
SM = 0
Delta_r
Doxycycline
YDL048C STP4
ZShift = −1.63
Z lm Score = 1.4
NG = 0
DB = 0
SM = 0
Delta_AUC
Doxycycline
YPL053C KTR6
ZShift = −0.13
Z lm Score = −3.86
NG = 0
DB = 0
SM = 0
Delta_L
Doxycycline
YPL053C KTR6
ZShift = −0.21
Z lm Score = 2.95
NG = 0
DB = 0
SM = 0
Delta_K
Doxycycline
YPL053C KTR6
ZShift = −3.44
Z lm Score = 5.14
NG = 0
DB = 0
SM = 0
Delta_r
Doxycycline
YPL053C KTR6
ZShift = −0.1
Z lm Score = 2.43
NG = 0
DB = 0
SM = 0
Delta_AUC
Doxycycline
YOL093W TRM10
ZShift = −0.8
Z lm Score = −3.86
NG = 0
DB = 0
SM = 0
Delta_L
Doxycycline
YOL093W TRM10
ZShift = 1.06
Z lm Score = 0.52
NG = 0
DB = 0
SM = 0
Delta_K
Doxycycline
YOL093W TRM10
ZShift = −0.11
Z lm Score = 1.44
NG = 0
DB = 0
SM = 0
Delta_r
Doxycycline
YOL093W TRM10
ZShift = 1.08
Z lm Score = 0.7
NG = 0
DB = 0
SM = 0
Delta_AUC
Doxycycline

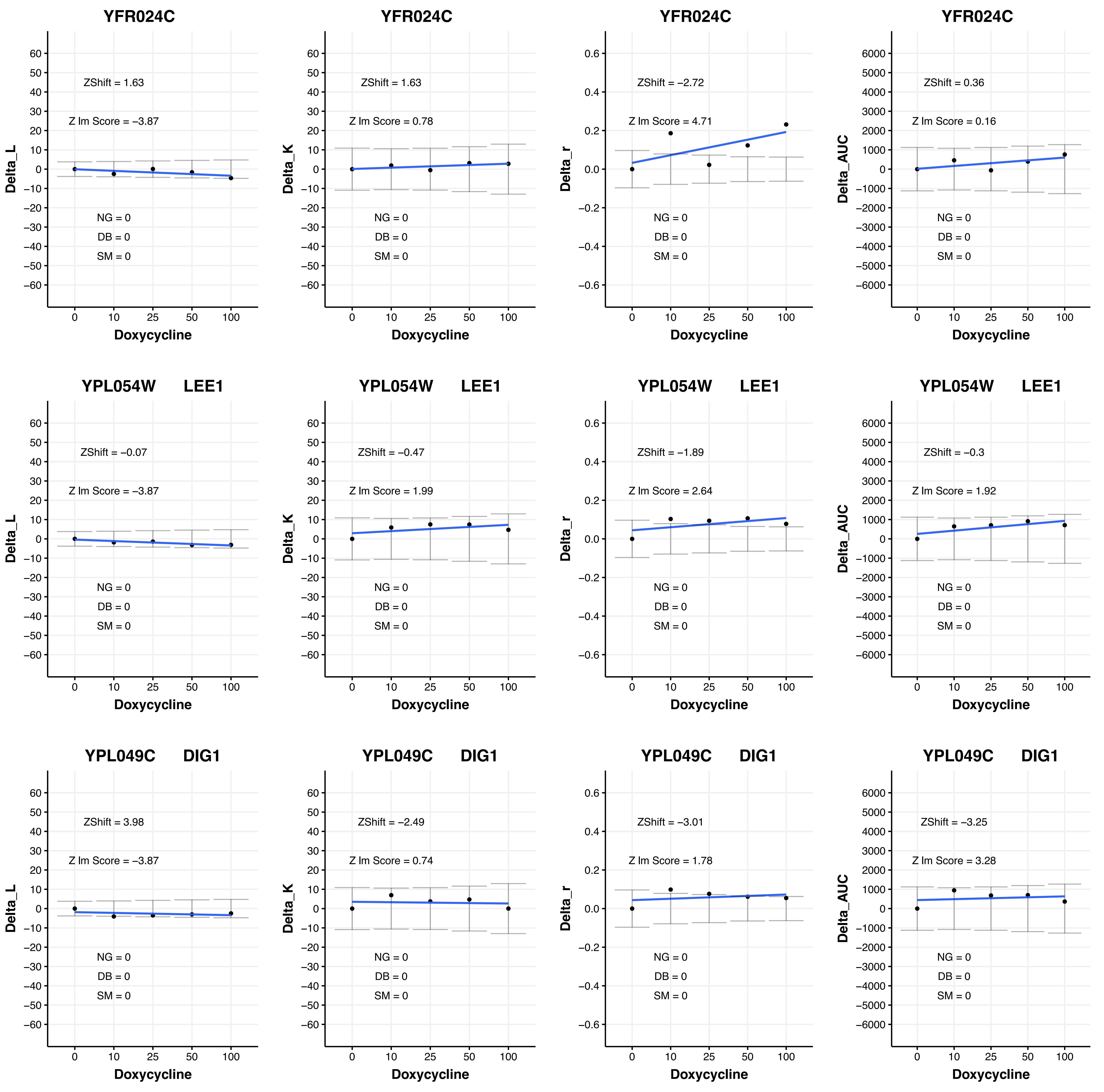

YFR024C
ZShift = 1.63
Z lm Score = −3.87
NG = 0
DB = 0
SM = 0
Delta_L
Doxycycline
YFR024C
ZShift = 1.63
Z lm Score = 0.78
NG = 0
DB = 0
SM = 0
Delta_K
Doxycycline
YFR024C
ZShift = −2.72
Z lm Score = 4.71
NG = 0
DB = 0
SM = 0
Delta_r
Doxycycline
YFR024C
ZShift = 0.36
Z lm Score = 0.16
NG = 0
DB = 0
SM = 0
Delta_AUC
Doxycycline
YPL054W LEE1
ZShift = −0.07
Z lm Score = −3.87
NG = 0
DB = 0
SM = 0
Delta_L
Doxycycline
YPL054W LEE1
ZShift = −0.47
Z lm Score = 1.99
NG = 0
DB = 0
SM = 0
Delta_K
Doxycycline
YPL054W LEE1
ZShift = −1.89
Z lm Score = 2.64
NG = 0
DB = 0
SM = 0
Delta_r
Doxycycline
YPL054W LEE1
ZShift = −0.3
Z lm Score = 1.92
NG = 0
DB = 0
SM = 0
Delta_AUC
Doxycycline
YPL049C DIG1
ZShift = 3.98
Z lm Score = −3.87
NG = 0
DB = 0
SM = 0
Delta_L
Doxycycline
YPL049C DIG1
ZShift = −2.49
Z lm Score = 0.74
NG = 0
DB = 0
SM = 0
Delta_K
Doxycycline
YPL049C DIG1
ZShift = −3.01
Z lm Score = 1.78
NG = 0
DB = 0
SM = 0
Delta_r
Doxycycline
YPL049C DIG1
ZShift = −3.25
Z lm Score = 3.28
NG = 0
DB = 0
SM = 0
Delta_AUC
Doxycycline

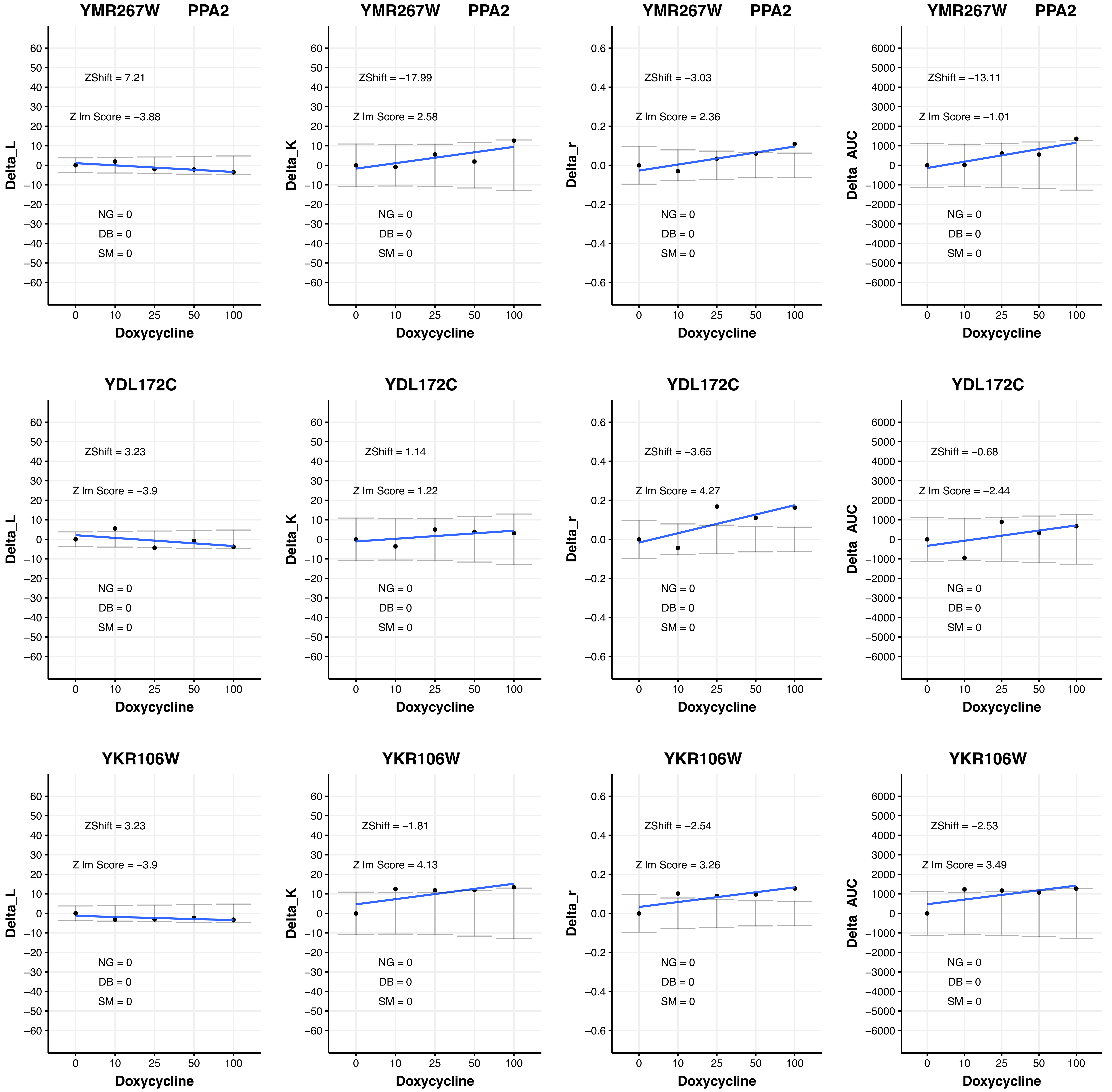

YMR267W PPA2
ZShift = 7.21
Z lm Score = −3.88
NG = 0
DB = 0
SM = 0
Delta_L
Doxycycline
YMR267W PPA2
ZShift = −17.99
Z lm Score = 2.58
NG = 0
DB = 0
SM = 0
Delta_K
Doxycycline
YMR267W PPA2
ZShift = −3.03
Z lm Score = 2.36
NG = 0
DB = 0
SM = 0
Delta_r
Doxycycline
YMR267W PPA2
ZShift = −13.11
Z lm Score = −1.01
NG = 0
DB = 0
SM = 0
Delta_AUC
Doxycycline
YDL172C
ZShift = 3.23
Z lm Score = −3.9
NG = 0
DB = 0
SM = 0
Delta_L
Doxycycline
YDL172C
ZShift = 1.14
Z lm Score = 1.22
NG = 0
DB = 0
SM = 0
Delta_K
Doxycycline
YDL172C
ZShift = −3.65
Z lm Score = 4.27
NG = 0
DB = 0
SM = 0
Delta_r
Doxycycline
YDL172C
ZShift = −0.68
Z lm Score = −2.44
NG = 0
DB = 0
SM = 0
Delta_AUC
Doxycycline
YKR106W
ZShift = 3.23
Z lm Score = −3.9
NG = 0
DB = 0
SM = 0
Delta_L
Doxycycline
YKR106W
ZShift = −1.81
Z lm Score = 4.13
NG = 0
DB = 0
SM = 0
Delta_K
Doxycycline
YKR106W
ZShift = −2.54
Z lm Score = 3.26
NG = 0
DB = 0
SM = 0
Delta_r
Doxycycline
YKR106W
ZShift = −2.53
Z lm Score = 3.49
NG = 0
DB = 0
SM = 0
Delta_AUC
Doxycycline

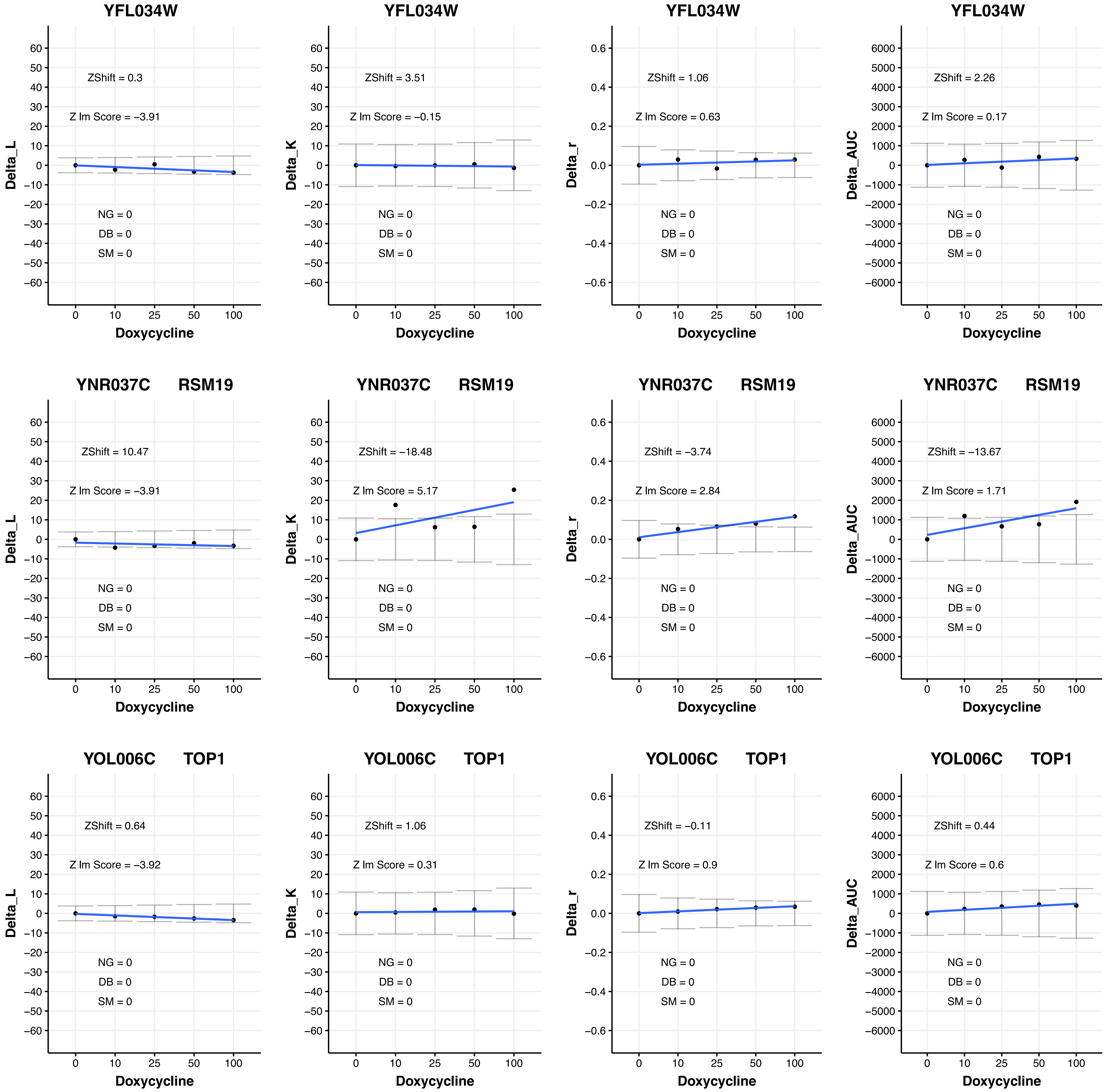

YFL034W
ZShift = 0.3
Z lm Score = −3.91
NG = 0
DB = 0
SM = 0
Delta_L
Doxycycline
YFL034W
ZShift = 3.51
Z lm Score = −0.15
NG = 0
DB = 0
SM = 0
Delta_K
Doxycycline
YFL034W
ZShift = 1.06
Z lm Score = 0.63
NG = 0
DB = 0
SM = 0
Delta_r
Doxycycline
YFL034W
ZShift = 2.26
Z lm Score = 0.17
NG = 0
DB = 0
SM = 0
Delta_AUC
Doxycycline
YNR037C RSM19
ZShift = 10.47
Z lm Score = −3.91
NG = 0
DB = 0
SM = 0
Delta_L
Doxycycline
YNR037C RSM19
ZShift = −18.48
Z lm Score = 5.17
NG = 0
DB = 0
SM = 0
Delta_K
Doxycycline
YNR037C RSM19
ZShift = −3.74
Z lm Score = 2.84
NG = 0
DB = 0
SM = 0
Delta_r
Doxycycline
YNR037C RSM19
ZShift = −13.67
Z lm Score = 1.71
NG = 0
DB = 0
SM = 0
Delta_AUC
Doxycycline
YOL006C TOP1
ZShift = 0.64
Z lm Score = −3.92
NG = 0
DB = 0
SM = 0
Delta_L
Doxycycline
YOL006C TOP1
ZShift = 1.06
Z lm Score = 0.31
NG = 0
DB = 0
SM = 0
Delta_K
Doxycycline
YOL006C TOP1
ZShift = −0.11
Z lm Score = 0.9
NG = 0
DB = 0
SM = 0
Delta_r
Doxycycline
YOL006C TOP1
ZShift = 0.44
Z lm Score = 0.6
NG = 0
DB = 0
SM = 0
Delta_AUC
Doxycycline

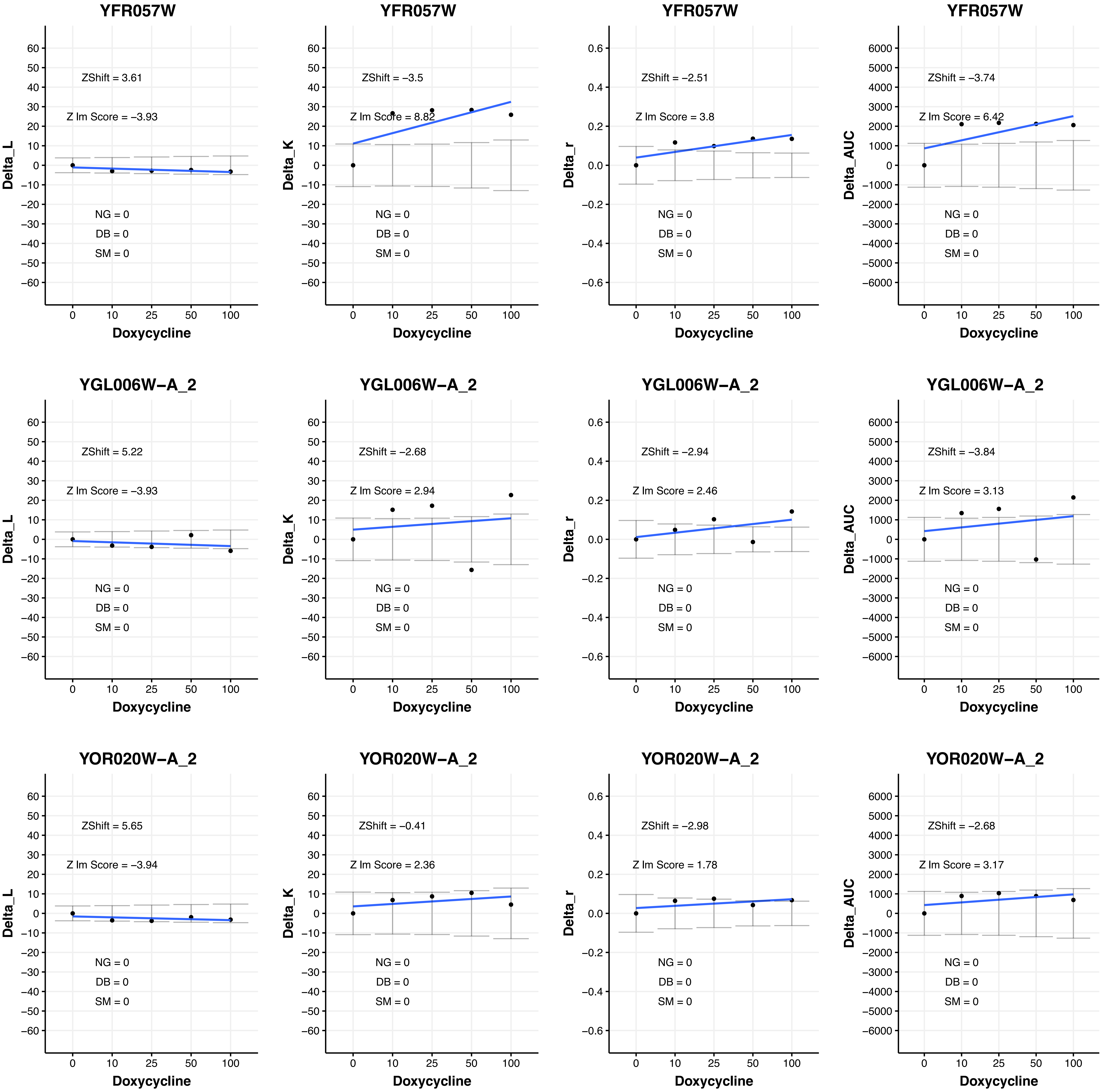
YFR057W
ZShift = 3.61
Z lm Score = −3.93
NG = 0
DB = 0
SM = 0
Delta_L
Doxycycline
YFR057W
ZShift = −3.5
Z lm Score = 8.82
NG = 0
DB = 0
SM = 0
Delta_K
Doxycycline
YFR057W
ZShift = −2.51
Z lm Score = 3.8
NG = 0
DB = 0
SM = 0
Delta_r
Doxycycline
YFR057W
ZShift = −3.74
Z lm Score = 6.42
NG = 0
DB = 0
SM = 0
Delta_AUC
Doxycycline
YGL006W−A_2
ZShift = 5.22
Z lm Score = −3.93
NG = 0
DB = 0
SM = 0
Delta_L
Doxycycline
YGL006W−A_2
ZShift = −2.68
Z lm Score = 2.94
NG = 0
DB = 0
SM = 0
Delta_K
Doxycycline
YGL006W−A_2
ZShift = −2.94
Z lm Score = 2.46
NG = 0
DB = 0
SM = 0
Delta_r
Doxycycline
YGL006W−A_2
ZShift = −3.84
Z lm Score = 3.13
NG = 0
DB = 0
SM = 0
Delta_AUC
Doxycycline
YOR020W−A_2
ZShift = 5.65
Z lm Score = −3.94
NG = 0
DB = 0
SM = 0
Delta_L
Doxycycline
YOR020W−A_2
ZShift = −0.41
Z lm Score = 2.36
NG = 0
DB = 0
SM = 0
Delta_K
Doxycycline
YOR020W−A_2
ZShift = −2.98
Z lm Score = 1.78
NG = 0
DB = 0
SM = 0
Delta_r
Doxycycline
YOR020W−A_2
ZShift = −2.68
Z lm Score = 3.17
NG = 0
DB = 0
SM = 0
Delta_AUC
Doxycycline

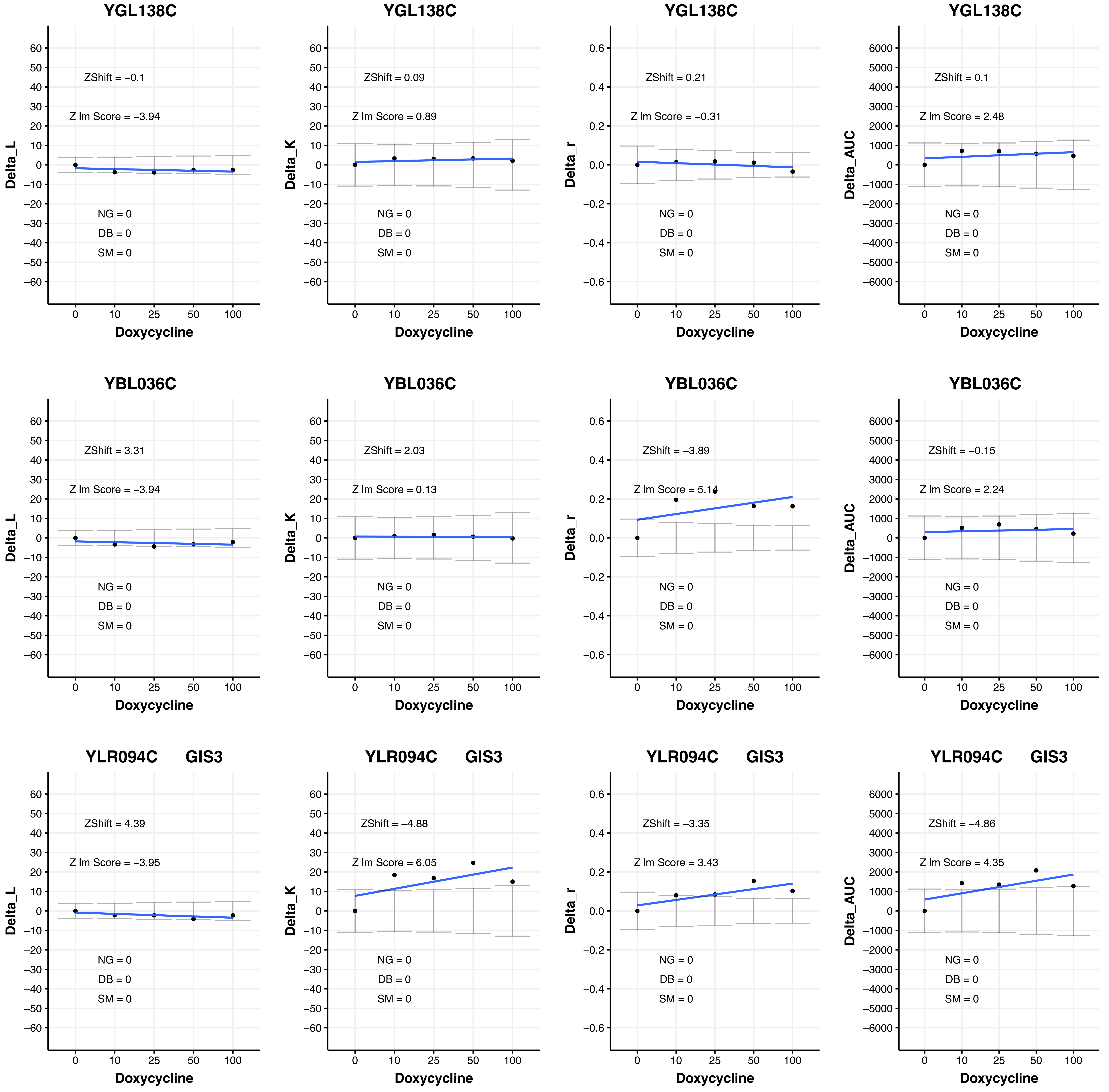
YGL138C
ZShift = –0.1
Z lm Score = –3.94
NG = 0
DB = 0
SM = 0
Delta_L
Doxycycline
YGL138C
ZShift = 0.09
Z lm Score = 0.89
NG = 0
DB = 0
SM = 0
Delta_K
Doxycycline
YGL138C
ZShift = 0.21
Z lm Score = –0.31
NG = 0
DB = 0
SM = 0
Delta_r
Doxycycline
YGL138C
ZShift = 0.1
Z lm Score = 2.48
NG = 0
DB = 0
SM = 0
Delta_AUC
Doxycycline
YBL036C
ZShift = 3.31
Z lm Score = –3.94
NG = 0
DB = 0
SM = 0
Delta_L
Doxycycline
YBL036C
ZShift = 2.03
Z lm Score = 0.13
NG = 0
DB = 0
SM = 0
Delta_K
Doxycycline
YBL036C
ZShift = –3.89
Z lm Score = 5.14
NG = 0
DB = 0
SM = 0
Delta_r
Doxycycline
YBL036C
ZShift = –0.15
Z lm Score = 2.24
NG = 0
DB = 0
SM = 0
Delta_AUC
Doxycycline
YLR094C GIS3
ZShift = 4.39
Z lm Score = –3.95
NG = 0
DB = 0
SM = 0
Delta_L
Doxycycline
YLR094C GIS3
ZShift = –4.88
Z lm Score = 6.05
NG = 0
DB = 0
SM = 0
Delta_K
Doxycycline
YLR094C GIS3
ZShift = –3.35
Z lm Score = 3.43
NG = 0
DB = 0
SM = 0
Delta_r
Doxycycline
YLR094C GIS3
ZShift = –4.86
Z lm Score = 4.35
NG = 0
DB = 0
SM = 0
Delta_AUC
Doxycycline

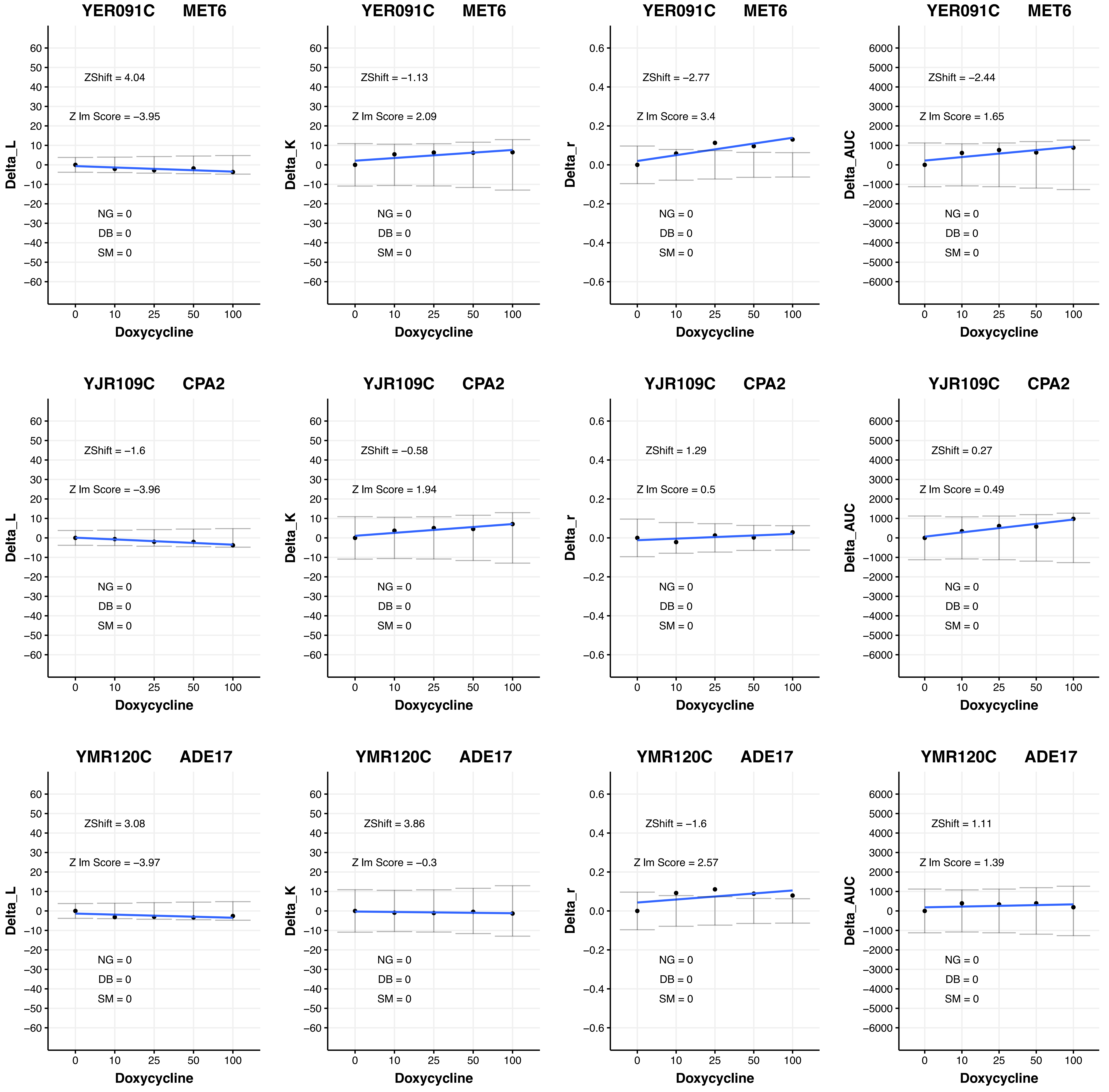
YER091C MET6
ZShift = 4.04
Z lm Score = −3.95
NG = 0
DB = 0
SM = 0
Delta_L
Doxycycline
YER091C MET6
ZShift = −1.13
Z lm Score = 2.09
NG = 0
DB = 0
SM = 0
Delta_K
Doxycycline
YER091C MET6
ZShift = −2.77
Z lm Score = 3.4
NG = 0
DB = 0
SM = 0
Delta_r
Doxycycline
YER091C MET6
ZShift = −2.44
Z lm Score = 1.65
NG = 0
DB = 0
SM = 0
Delta_AUC
Doxycycline
YJR109C CPA2
ZShift = −1.6
Z lm Score = −3.96
NG = 0
DB = 0
SM = 0
Delta_L
Doxycycline
YJR109C CPA2
ZShift = −0.58
Z lm Score = 1.94
NG = 0
DB = 0
SM = 0
Delta_K
Doxycycline
YJR109C CPA2
ZShift = 1.29
Z lm Score = 0.5
NG = 0
DB = 0
SM = 0
Delta_r
Doxycycline
YJR109C CPA2
ZShift = 0.27
Z lm Score = 0.49
NG = 0
DB = 0
SM = 0
Delta_AUC
Doxycycline
YMR120C ADE17
ZShift = 3.08
Z lm Score = −3.97
NG = 0
DB = 0
SM = 0
Delta_L
Doxycycline
YMR120C ADE17
ZShift = 3.86
Z lm Score = −0.3
NG = 0
DB = 0
SM = 0
Delta_K
Doxycycline
YMR120C ADE17
ZShift = −1.6
Z lm Score = 2.57
NG = 0
DB = 0
SM = 0
Delta_r
Doxycycline
YMR120C ADE17
ZShift = 1.11
Z lm Score = 1.39
NG = 0
DB = 0
SM = 0
Delta_AUC
Doxycycline

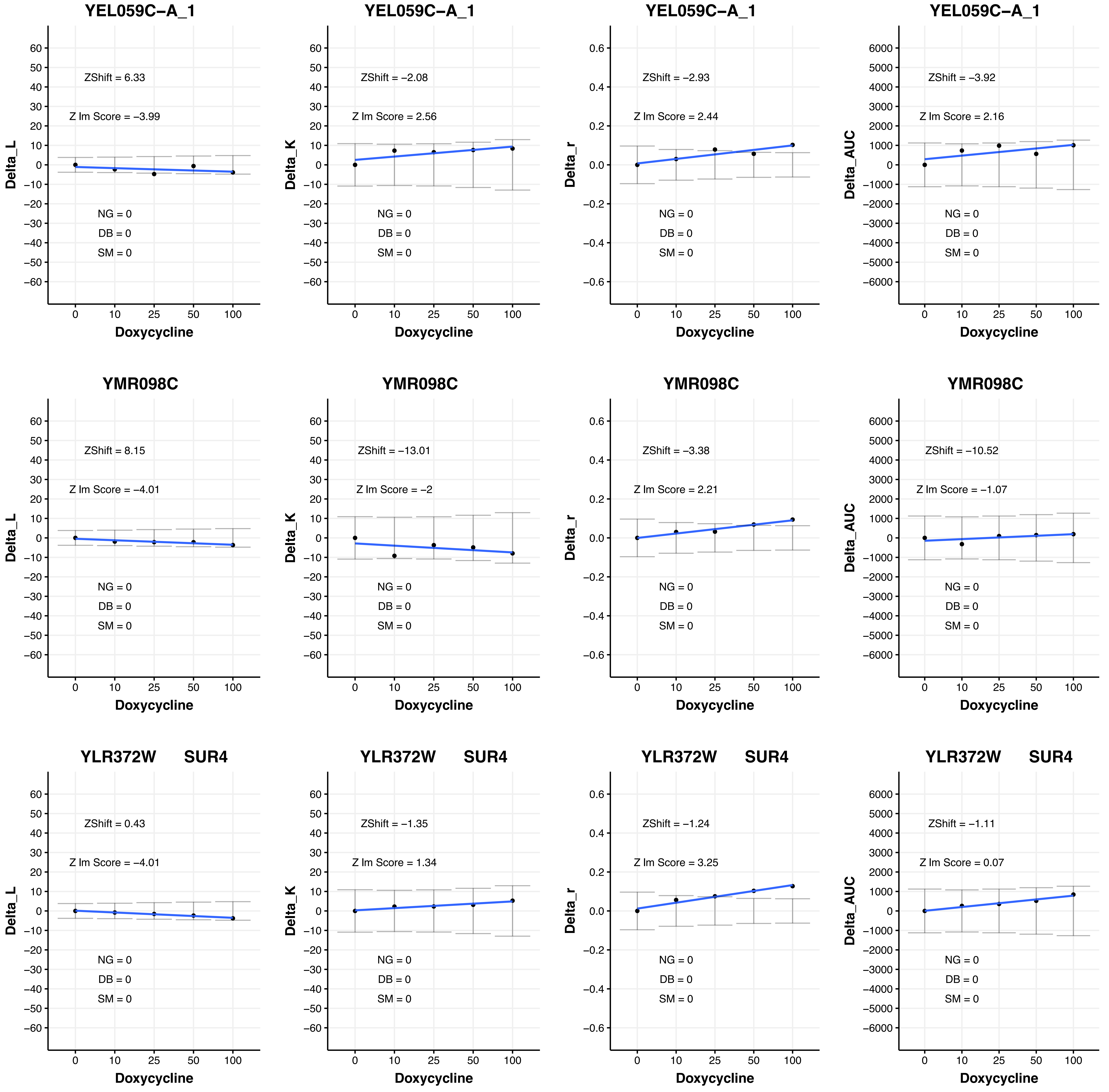

YEL059C−A_1
ZShift = 6.33
Z lm Score = −3.99
Delta_L
NG = 0
DB = 0
SM = 0
Doxycycline
YEL059C−A_1
ZShift = −2.08
Z lm Score = 2.56
Delta_K
NG = 0
DB = 0
SM = 0
Doxycycline
YEL059C−A_1
ZShift = −2.93
Z lm Score = 2.44
Delta_r
NG = 0
DB = 0
SM = 0
Doxycycline
YEL059C−A_1
ZShift = −3.92
Z lm Score = 2.16
Delta_AUC
NG = 0
DB = 0
SM = 0
Doxycycline
YMR098C
ZShift = 8.15
Z lm Score = −4.01
Delta_L
NG = 0
DB = 0
SM = 0
Doxycycline
YMR098C
ZShift = −13.01
Z lm Score = −2
Delta_K
NG = 0
DB = 0
SM = 0
Doxycycline
YMR098C
ZShift = −3.38
Z lm Score = 2.21
Delta_r
NG = 0
DB = 0
SM = 0
Doxycycline
YMR098C
ZShift = −10.52
Z lm Score = −1.07
Delta_AUC
NG = 0
DB = 0
SM = 0
Doxycycline
YLR372W SUR4
ZShift = 0.43
Z lm Score = −4.01
Delta_L
NG = 0
DB = 0
SM = 0
Doxycycline
YLR372W SUR4
ZShift = −1.35
Z lm Score = 1.34
Delta_K
NG = 0
DB = 0
SM = 0
Doxycycline
YLR372W SUR4
ZShift = −1.24
Z lm Score = 3.25
Delta_r
NG = 0
DB = 0
SM = 0
Doxycycline
YLR372W SUR4
ZShift = −1.11
Z lm Score = 0.07
Delta_AUC
NG = 0
DB = 0
SM = 0
Doxycycline

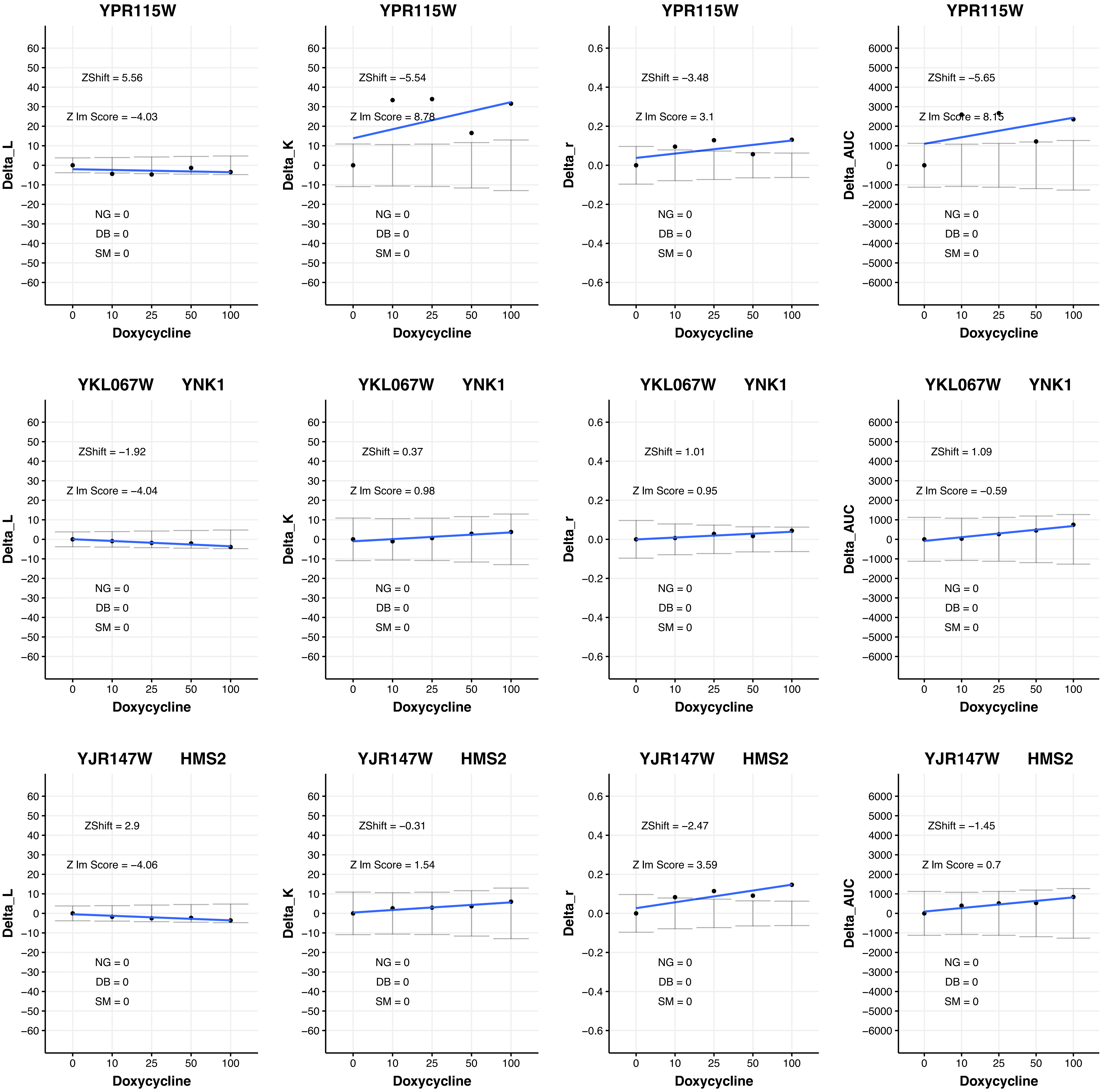
YPR115W
ZShift = 5.56
Z lm Score = −4.03
NG = 0
DB = 0
SM = 0
Delta_L
Doxycycline
YPR115W
ZShift = −5.54
Z lm Score = 8.78
NG = 0
DB = 0
SM = 0
Delta_K
Doxycycline
YPR115W
ZShift = −3.48
Z lm Score = 3.1
NG = 0
DB = 0
SM = 0
Delta_r
Doxycycline
YPR115W
ZShift = −5.65
Z lm Score = 8.15
NG = 0
DB = 0
SM = 0
Delta_AUC
Doxycycline
YKL067W YNK1
ZShift = −1.92
Z lm Score = −4.04
NG = 0
DB = 0
SM = 0
Delta_L
Doxycycline
YKL067W YNK1
ZShift = 0.37
Z lm Score = 0.98
NG = 0
DB = 0
SM = 0
Delta_K
Doxycycline
YKL067W YNK1
ZShift = 1.01
Z lm Score = 0.95
NG = 0
DB = 0
SM = 0
Delta_r
Doxycycline
YKL067W YNK1
ZShift = 1.09
Z lm Score = −0.59
NG = 0
DB = 0
SM = 0
Delta_AUC
Doxycycline
YJR147W HMS2
ZShift = 2.9
Z lm Score = −4.06
NG = 0
DB = 0
SM = 0
Delta_L
Doxycycline
YJR147W HMS2
ZShift = −0.31
Z lm Score = 1.54
NG = 0
DB = 0
SM = 0
Delta_K
Doxycycline
YJR147W HMS2
ZShift = −2.47
Z lm Score = 3.59
NG = 0
DB = 0
SM = 0
Delta_r
Doxycycline
YJR147W HMS2
ZShift = −1.45
Z lm Score = 0.7
NG = 0
DB = 0
SM = 0
Delta_AUC
Doxycycline

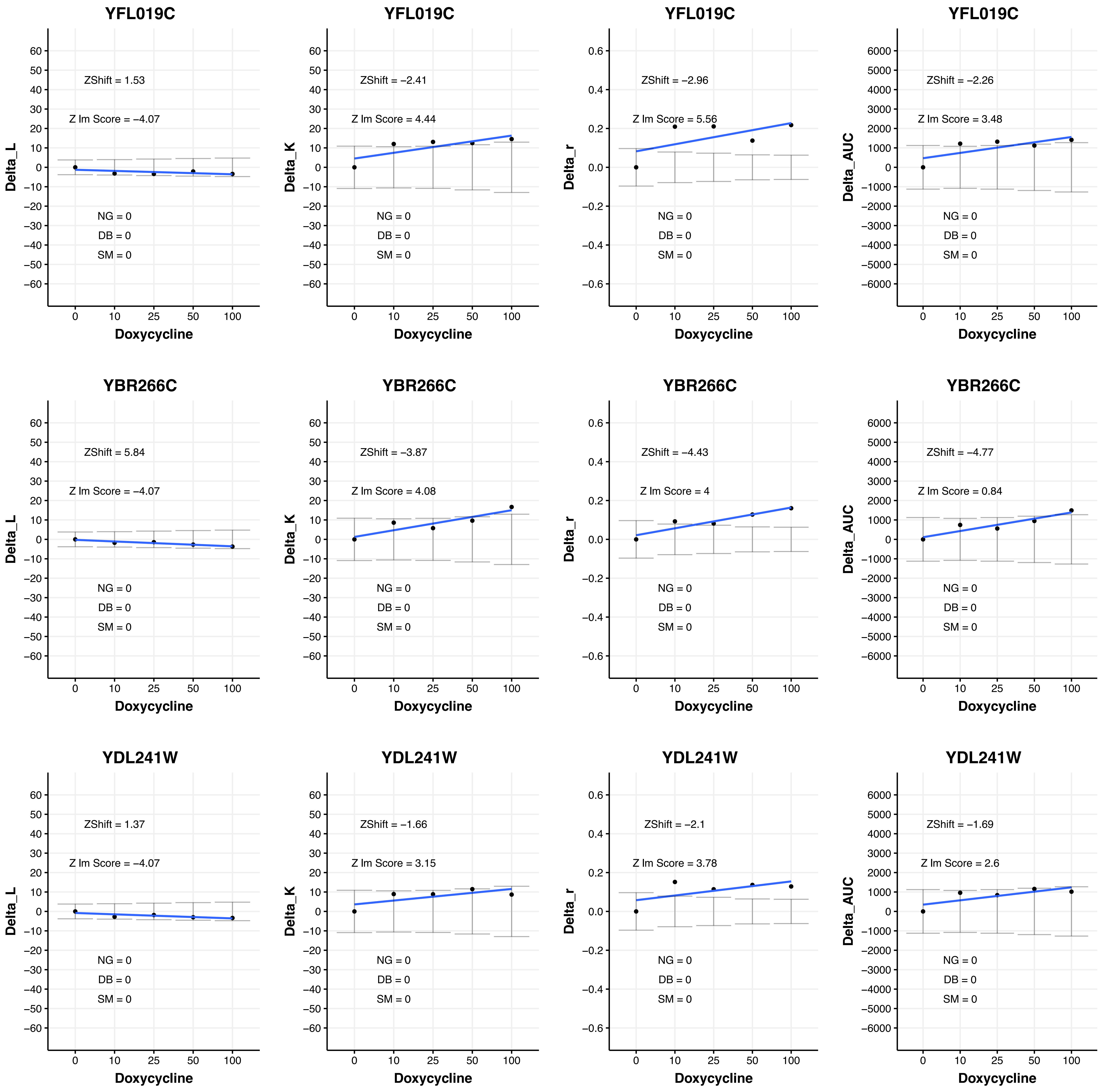

YFL019C
ZShift = 1.53
Z lm Score = −4.07
NG = 0
DB = 0
SM = 0
Delta_L
Doxycycline
YFL019C
ZShift = −2.41
Z lm Score = 4.44
NG = 0
DB = 0
SM = 0
Delta_K
Doxycycline
YFL019C
ZShift = −2.96
Z lm Score = 5.56
NG = 0
DB = 0
SM = 0
Delta_r
Doxycycline
YFL019C
ZShift = −2.26
Z lm Score = 3.48
NG = 0
DB = 0
SM = 0
Delta_AUC
Doxycycline
YBR266C
ZShift = 5.84
Z lm Score = −4.07
NG = 0
DB = 0
SM = 0
Delta_L
Doxycycline
YBR266C
ZShift = −3.87
Z lm Score = 4.08
NG = 0
DB = 0
SM = 0
Delta_K
Doxycycline
YBR266C
ZShift = −4.43
Z lm Score = 4
NG = 0
DB = 0
SM = 0
Delta_r
Doxycycline
YBR266C
ZShift = −4.77
Z lm Score = 0.84
NG = 0
DB = 0
SM = 0
Delta_AUC
Doxycycline
YDL241W
ZShift = 1.37
Z lm Score = −4.07
NG = 0
DB = 0
SM = 0
Delta_L
Doxycycline
YDL241W
ZShift = −1.66
Z lm Score = 3.15
NG = 0
DB = 0
SM = 0
Delta_K
Doxycycline
YDL241W
ZShift = −2.1
Z lm Score = 3.78
NG = 0
DB = 0
SM = 0
Delta_r
Doxycycline
YDL241W
ZShift = −1.69
Z lm Score = 2.6
NG = 0
DB = 0
SM = 0
Delta_AUC
Doxycycline

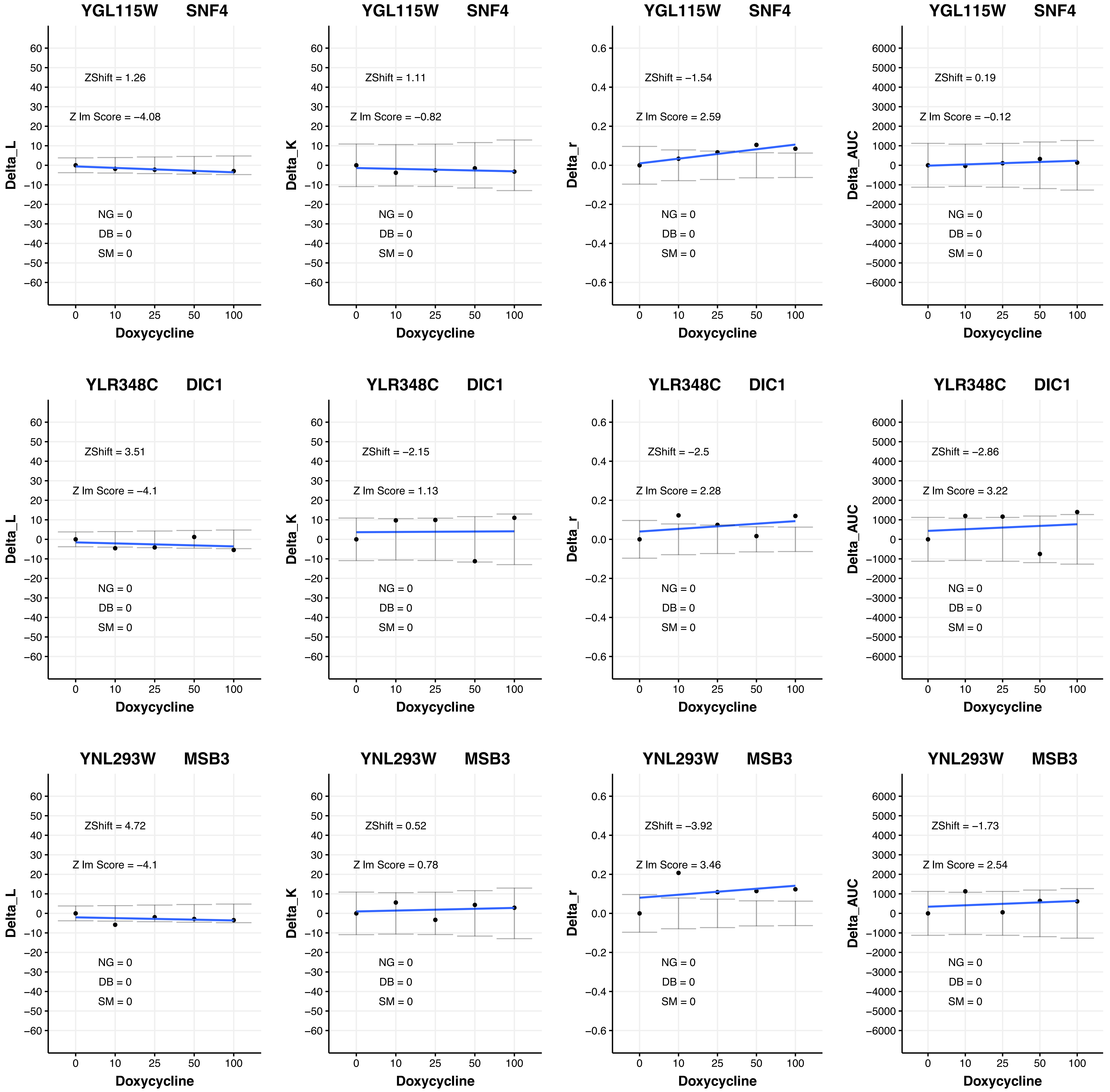

YGL115W SNF4
ZShift = 1.26
Z lm Score = −4.08
NG = 0
DB = 0
SM = 0
Delta_L
Doxycycline
YGL115W SNF4
ZShift = 1.11
Z lm Score = −0.82
NG = 0
DB = 0
SM = 0
Delta_K
Doxycycline
YGL115W SNF4
ZShift = −1.54
Z lm Score = 2.59
NG = 0
DB = 0
SM = 0
Delta_r
Doxycycline
YGL115W SNF4
ZShift = 0.19
Z lm Score = −0.12
NG = 0
DB = 0
SM = 0
Delta_AUC
Doxycycline
YLR348C DIC1
ZShift = 3.51
Z lm Score = −4.1
NG = 0
DB = 0
SM = 0
Delta_L
Doxycycline
YLR348C DIC1
ZShift = −2.15
Z lm Score = 1.13
NG = 0
DB = 0
SM = 0
Delta_K
Doxycycline
YLR348C DIC1
ZShift = −2.5
Z lm Score = 2.28
NG = 0
DB = 0
SM = 0
Delta_r
Doxycycline
YLR348C DIC1
ZShift = −2.86
Z lm Score = 3.22
NG = 0
DB = 0
SM = 0
Delta_AUC
Doxycycline
YNL293W MSB3
ZShift = 4.72
Z lm Score = −4.1
NG = 0
DB = 0
SM = 0
Delta_L
Doxycycline
YNL293W MSB3
ZShift = 0.52
Z lm Score = 0.78
NG = 0
DB = 0
SM = 0
Delta_K
Doxycycline
YNL293W MSB3
ZShift = −3.92
Z lm Score = 3.46
NG = 0
DB = 0
SM = 0
Delta_r
Doxycycline
YNL293W MSB3
ZShift = −1.73
Z lm Score = 2.54
NG = 0
DB = 0
SM = 0
Delta_AUC
Doxycycline

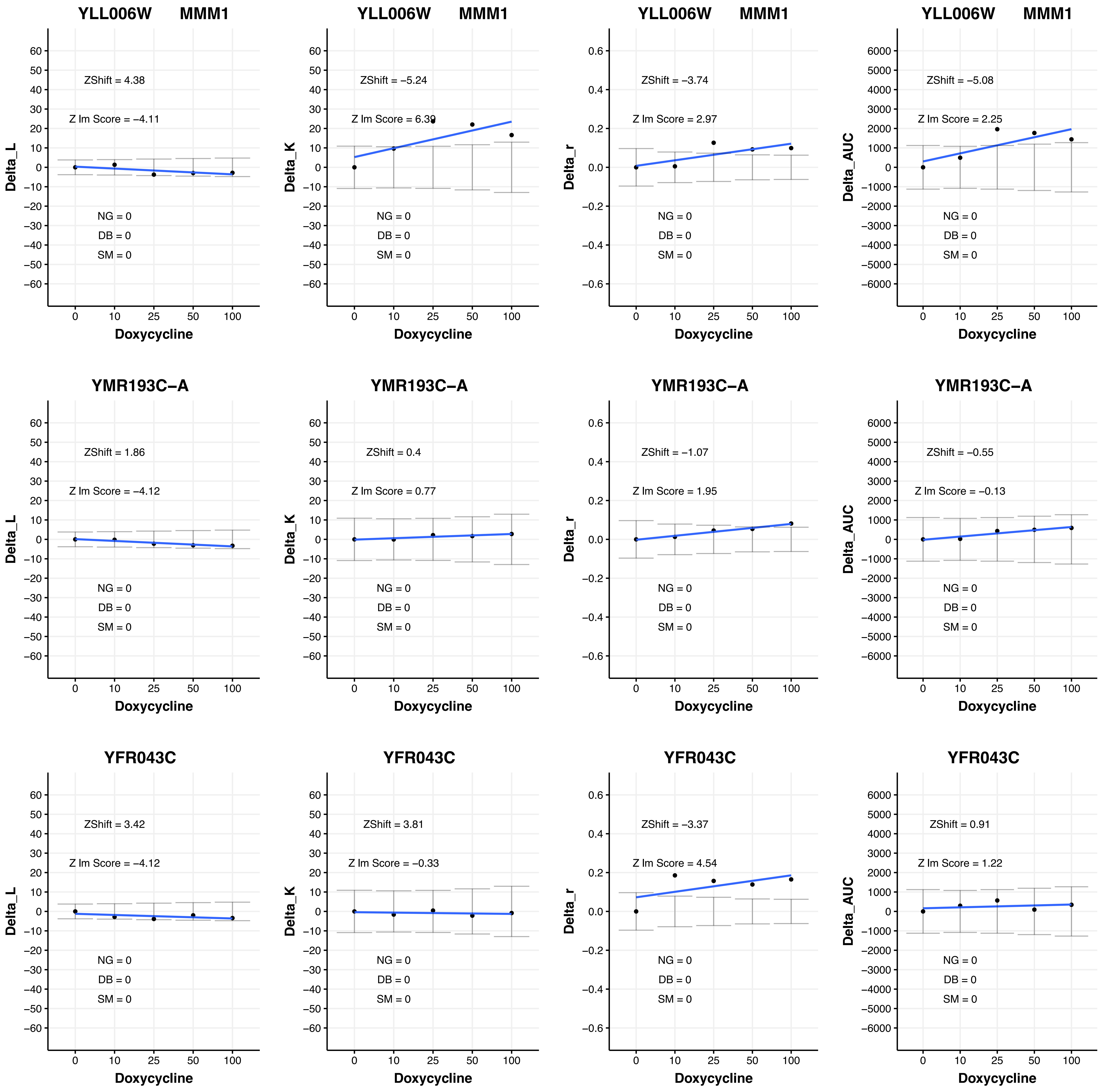

YLL006W MMM1
ZShift = 4.38
Z lm Score = −4.11
NG = 0
DB = 0
SM = 0
Delta_L
Doxycycline
YLL006W MMM1
ZShift = −5.24
Z lm Score = 6.39
NG = 0
DB = 0
SM = 0
Delta_K
Doxycycline
YLL006W MMM1
ZShift = −3.74
Z lm Score = 2.97
NG = 0
DB = 0
SM = 0
Delta_r
Doxycycline
YLL006W MMM1
ZShift = −5.08
Z lm Score = 2.25
NG = 0
DB = 0
SM = 0
Delta_AUC
Doxycycline
YMR193C−A
ZShift = 1.86
Z lm Score = −4.12
NG = 0
DB = 0
SM = 0
Delta_L
Doxycycline
YMR193C−A
ZShift = 0.4
Z lm Score = 0.77
NG = 0
DB = 0
SM = 0
Delta_K
Doxycycline
YMR193C−A
ZShift = −1.07
Z lm Score = 1.95
NG = 0
DB = 0
SM = 0
Delta_r
Doxycycline
YMR193C−A
ZShift = −0.55
Z lm Score = −0.13
NG = 0
DB = 0
SM = 0
Delta_AUC
Doxycycline
YFR043C
ZShift = 3.42
Z lm Score = −4.12
NG = 0
DB = 0
SM = 0
Delta_L
Doxycycline
YFR043C
ZShift = 3.81
Z lm Score = −0.33
NG = 0
DB = 0
SM = 0
Delta_K
Doxycycline
YFR043C
ZShift = −3.37
Z lm Score = 4.54
NG = 0
DB = 0
SM = 0
Delta_r
Doxycycline
YFR043C
ZShift = 0.91
Z lm Score = 1.22
NG = 0
DB = 0
SM = 0
Delta_AUC
Doxycycline

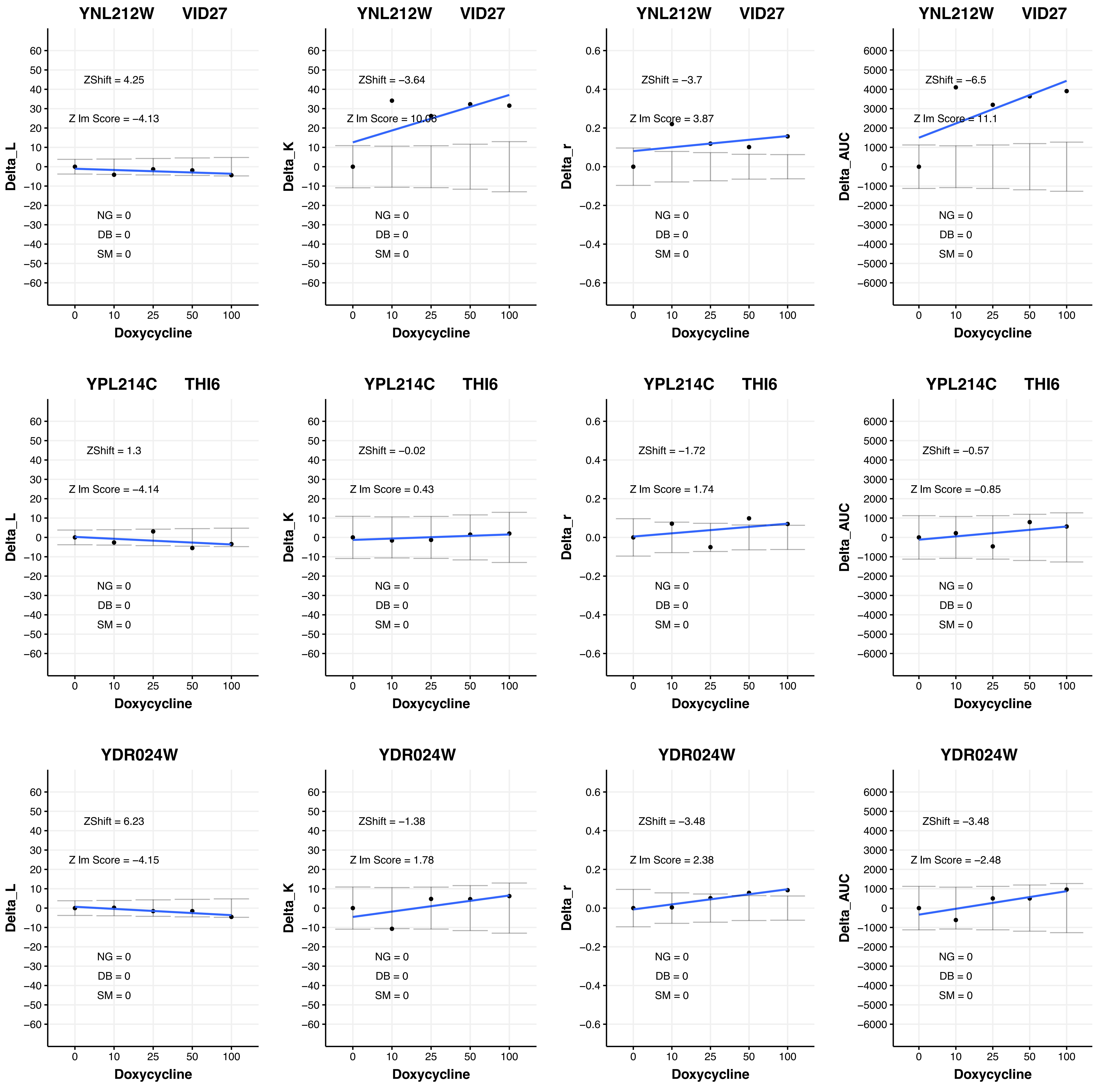

YNL212W VID27
ZShift = 4.25
Z lm Score = −4.13
NG = 0
DB = 0
SM = 0
Delta_L
Doxycycline
YNL212W VID27
ZShift = −3.64
Z lm Score = 10.06
NG = 0
DB = 0
SM = 0
Delta_K
Doxycycline
YNL212W VID27
ZShift = −3.7
Z lm Score = 3.87
NG = 0
DB = 0
SM = 0
Delta_r
Doxycycline
YNL212W VID27
ZShift = −6.5
Z lm Score = 11.1
NG = 0
DB = 0
SM = 0
Delta_AUC
Doxycycline
YPL214C THI6
ZShift = 1.3
Z lm Score = −4.14
NG = 0
DB = 0
SM = 0
Delta_L
Doxycycline
YPL214C THI6
ZShift = −0.02
Z lm Score = 0.43
NG = 0
DB = 0
SM = 0
Delta_K
Doxycycline
YPL214C THI6
ZShift = −1.72
Z lm Score = 1.74
NG = 0
DB = 0
SM = 0
Delta_r
Doxycycline
YPL214C THI6
ZShift = −0.57
Z lm Score = −0.85
NG = 0
DB = 0
SM = 0
Delta_AUC
Doxycycline
YDR024W
ZShift = 6.23
Z lm Score = −4.15
NG = 0
DB = 0
SM = 0
Delta_L
Doxycycline
YDR024W
ZShift = −1.38
Z lm Score = 1.78
NG = 0
DB = 0
SM = 0
Delta_K
Doxycycline
YDR024W
ZShift = −3.48
Z lm Score = 2.38
NG = 0
DB = 0
SM = 0
Delta_r
Doxycycline
YDR024W
ZShift = −3.48
Z lm Score = −2.48
NG = 0
DB = 0
SM = 0
Delta_AUC
Doxycycline

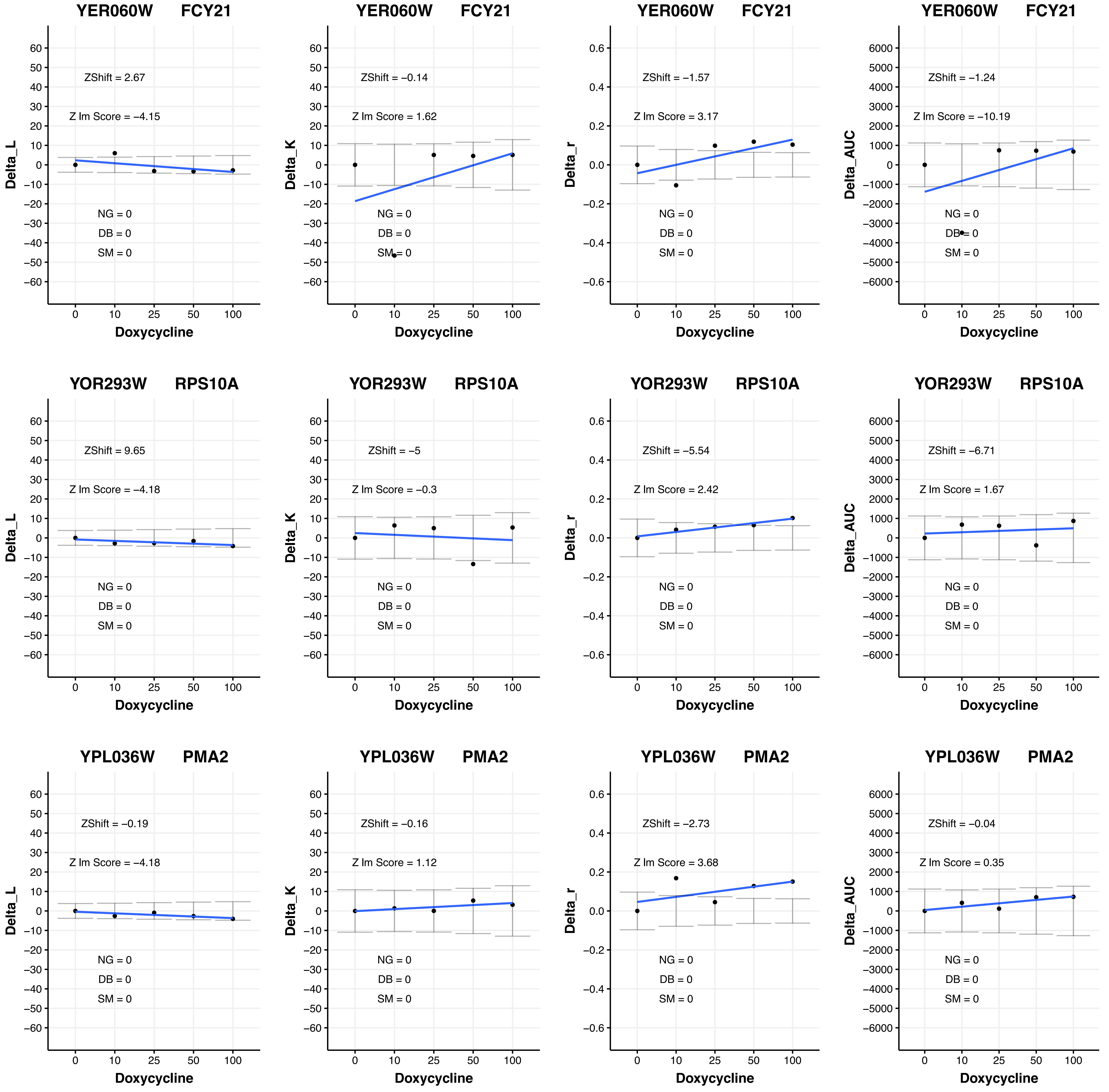
YER060W FCY21
ZShift = 2.67
Z lm Score = −4.15
NG = 0
DB = 0
SM = 0
Delta_L
Doxycycline
YER060W FCY21
ZShift = −0.14
Z lm Score = 1.62
NG = 0
DB = 0
SM = 0
Delta_K
Doxycycline
YER060W FCY21
ZShift = −1.57
Z lm Score = 3.17
NG = 0
DB = 0
SM = 0
Delta_r
Doxycycline
YER060W FCY21
ZShift = −1.24
Z lm Score = −10.19
NG = 0
DB = 0
SM = 0
Delta_AUC
Doxycycline
YOR293W RPS10A
ZShift = 9.65
Z lm Score = −4.18
NG = 0
DB = 0
SM = 0
Delta_L
Doxycycline
YOR293W RPS10A
ZShift = −5
Z lm Score = −0.3
NG = 0
DB = 0
SM = 0
Delta_K
Doxycycline
YOR293W RPS10A
ZShift = −5.54
Z lm Score = 2.42
NG = 0
DB = 0
SM = 0
Delta_r
Doxycycline
YOR293W RPS10A
ZShift = −6.71
Z lm Score = 1.67
NG = 0
DB = 0
SM = 0
Delta_AUC
Doxycycline
YPL036W PMA2
ZShift = −0.19
Z lm Score = −4.18
NG = 0
DB = 0
SM = 0
Delta_L
Doxycycline
YPL036W PMA2
ZShift = −0.16
Z lm Score = 1.12
NG = 0
DB = 0
SM = 0
Delta_K
Doxycycline
YPL036W PMA2
ZShift = −2.73
Z lm Score = 3.68
NG = 0
DB = 0
SM = 0
Delta_r
Doxycycline
YPL036W PMA2
ZShift = −0.04
Z lm Score = 0.35
NG = 0
DB = 0
SM = 0
Delta_AUC
Doxycycline

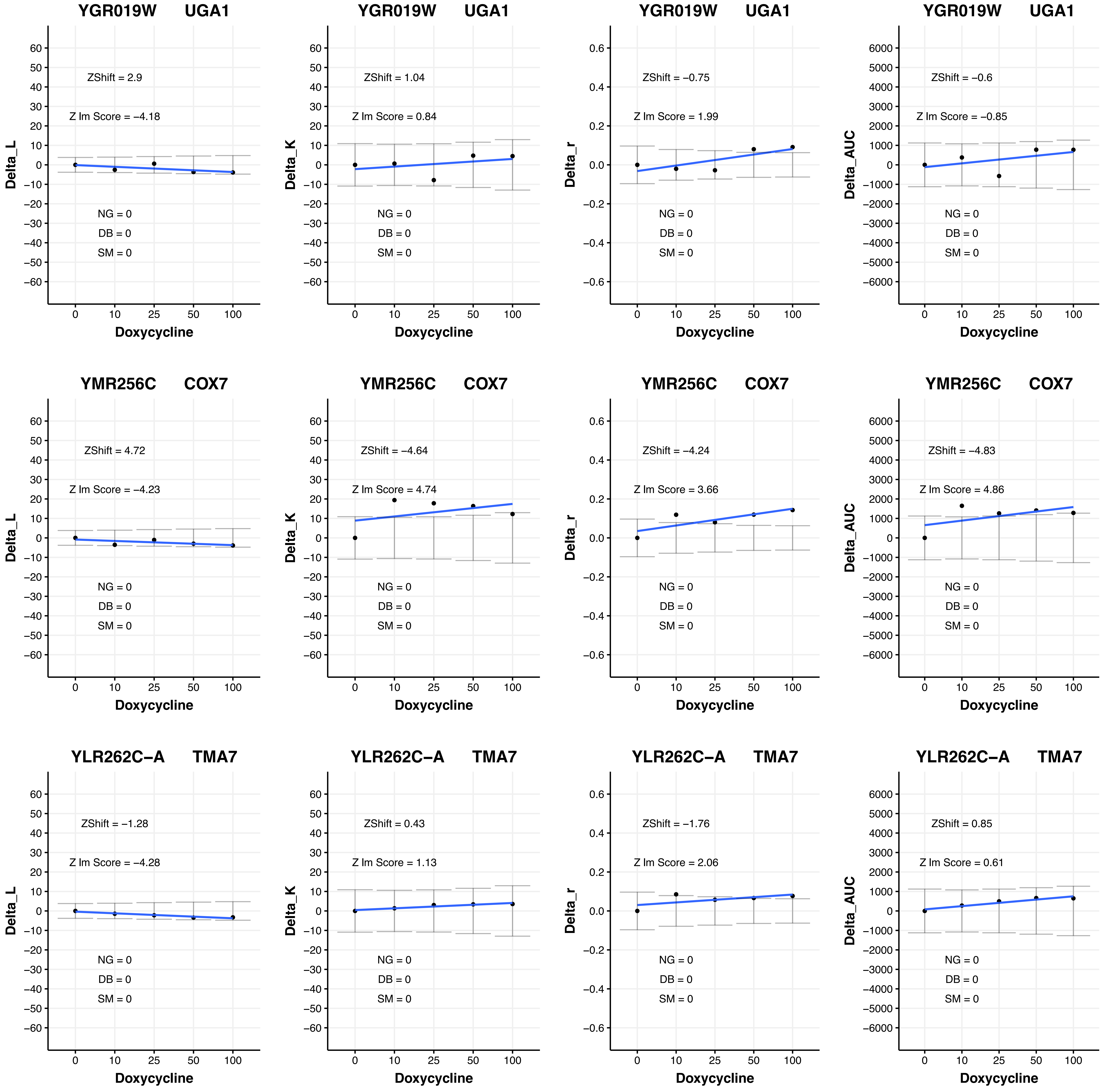

YGR019W UGA1
ZShift = 2.9
Z lm Score = −4.18
NG = 0
DB = 0
SM = 0
Delta_L
Doxycycline
YGR019W UGA1
ZShift = 1.04
Z lm Score = 0.84
NG = 0
DB = 0
SM = 0
Delta_K
Doxycycline
YGR019W UGA1
ZShift = −0.75
Z lm Score = 1.99
NG = 0
DB = 0
SM = 0
Delta_r
Doxycycline
YGR019W UGA1
ZShift = −0.6
Z lm Score = −0.85
NG = 0
DB = 0
SM = 0
Delta_AUC
Doxycycline
YMR256C COX7
ZShift = 4.72
Z lm Score = −4.23
NG = 0
DB = 0
SM = 0
Delta_L
Doxycycline
YMR256C COX7
ZShift = −4.64
Z lm Score = 4.74
NG = 0
DB = 0
SM = 0
Delta_K
Doxycycline
YMR256C COX7
ZShift = −4.24
Z lm Score = 3.66
NG = 0
DB = 0
SM = 0
Delta_r
Doxycycline
YMR256C COX7
ZShift = −4.83
Z lm Score = 4.86
NG = 0
DB = 0
SM = 0
Delta_AUC
Doxycycline
YLR262C−A TMA7
ZShift = −1.28
Z lm Score = −4.28
NG = 0
DB = 0
SM = 0
Delta_L
Doxycycline
YLR262C−A TMA7
ZShift = 0.43
Z lm Score = 1.13
NG = 0
DB = 0
SM = 0
Delta_K
Doxycycline
YLR262C−A TMA7
ZShift = −1.76
Z lm Score = 2.06
NG = 0
DB = 0
SM = 0
Delta_r
Doxycycline
YLR262C−A TMA7
ZShift = 0.85
Z lm Score = 0.61
NG = 0
DB = 0
SM = 0
Delta_AUC
Doxycycline

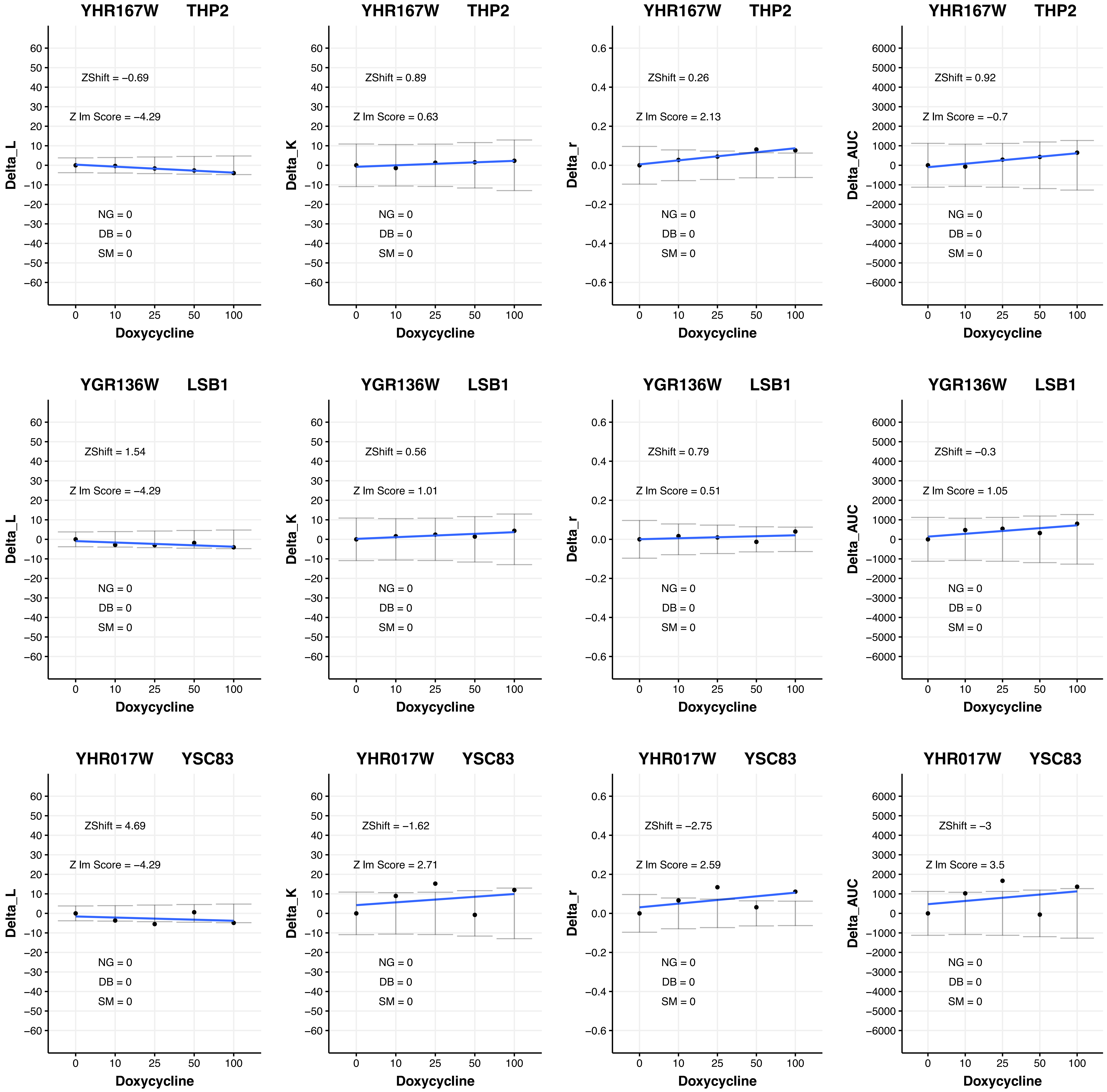

YHR167W THP2
ZShift = −0.69
Z lm Score = −4.29
Delta_L
NG = 0
DB = 0
SM = 0
Doxycycline
YHR167W THP2
ZShift = 0.89
Z lm Score = 0.63
Delta_K
NG = 0
DB = 0
SM = 0
Doxycycline
YHR167W THP2
ZShift = 0.26
Z lm Score = 2.13
Delta_r
NG = 0
DB = 0
SM = 0
Doxycycline
YHR167W THP2
ZShift = 0.92
Z lm Score = −0.7
Delta_AUC
NG = 0
DB = 0
SM = 0
Doxycycline
YGR136W LSB1
ZShift = 1.54
Z lm Score = −4.29
Delta_L
NG = 0
DB = 0
SM = 0
Doxycycline
YGR136W LSB1
ZShift = 0.56
Z lm Score = 1.01
Delta_K
NG = 0
DB = 0
SM = 0
Doxycycline
YGR136W LSB1
ZShift = 0.79
Z lm Score = 0.51
Delta_r
NG = 0
DB = 0
SM = 0
Doxycycline
YGR136W LSB1
ZShift = −0.3
Z lm Score = 1.05
Delta_AUC
NG = 0
DB = 0
SM = 0
Doxycycline
YHR017W YSC83
ZShift = 4.69
Z lm Score = −4.29
Delta_L
NG = 0
DB = 0
SM = 0
Doxycycline
YHR017W YSC83
ZShift = −1.62
Z lm Score = 2.71
Delta_K
NG = 0
DB = 0
SM = 0
Doxycycline
YHR017W YSC83
ZShift = −2.75
Z lm Score = 2.59
Delta_r
NG = 0
DB = 0
SM = 0
Doxycycline
YHR017W YSC83
ZShift = −3
Z lm Score = 3.5
Delta_AUC
NG = 0
DB = 0
SM = 0
Doxycycline

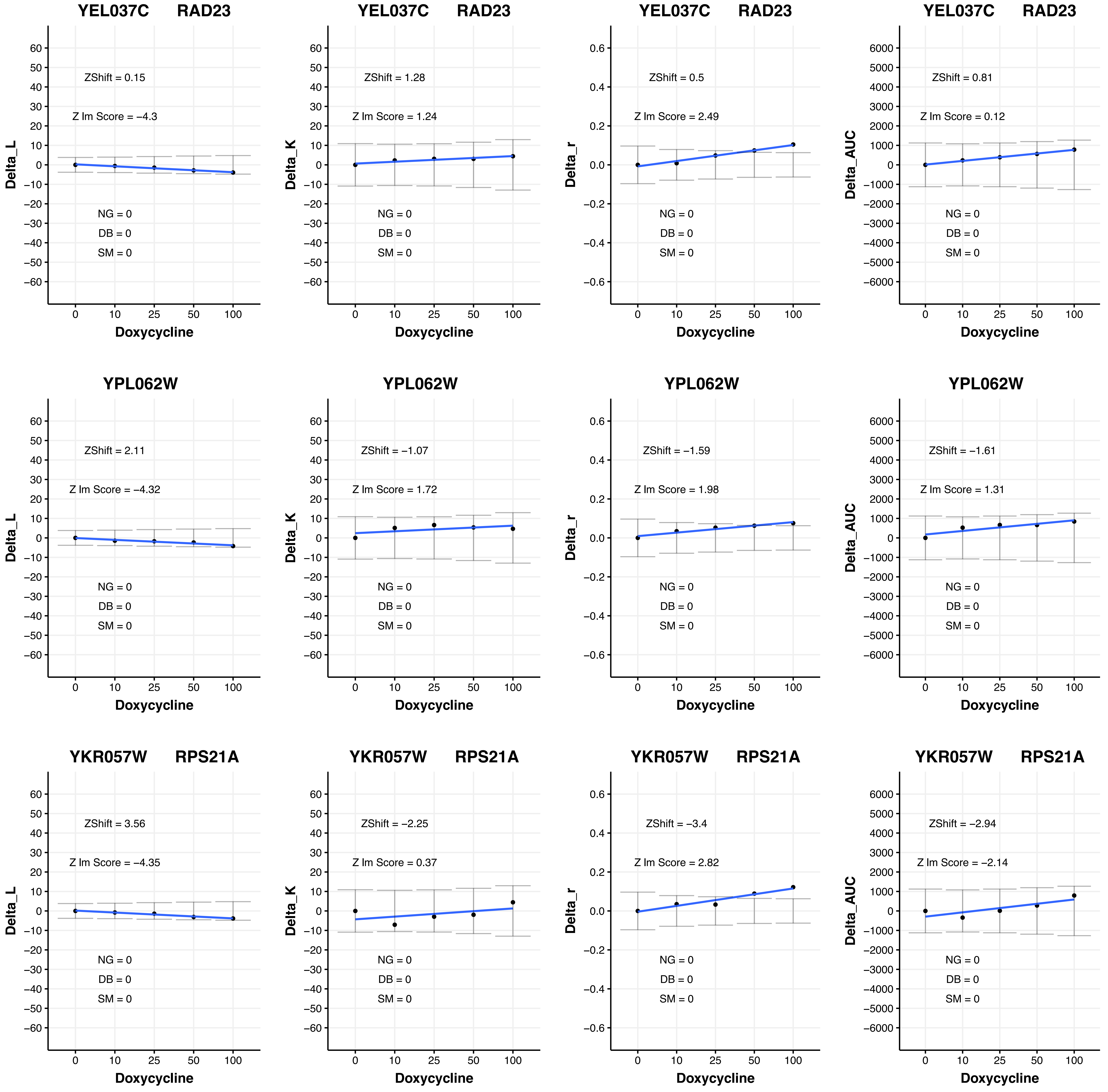

YEL037C RAD23
ZShift = 0.15
Z lm Score = −4.3
NG = 0
DB = 0
SM = 0
Delta_L
Doxycycline
YEL037C RAD23
ZShift = 1.28
Z lm Score = 1.24
NG = 0
DB = 0
SM = 0
Delta_K
Doxycycline
YEL037C RAD23
ZShift = 0.5
Z lm Score = 2.49
NG = 0
DB = 0
SM = 0
Delta_r
Doxycycline
YEL037C RAD23
ZShift = 0.81
Z lm Score = 0.12
NG = 0
DB = 0
SM = 0
Delta_AUC
Doxycycline
YPL062W
ZShift = 2.11
Z lm Score = −4.32
NG = 0
DB = 0
SM = 0
Delta_L
Doxycycline
YPL062W
ZShift = −1.07
Z lm Score = 1.72
NG = 0
DB = 0
SM = 0
Delta_K
Doxycycline
YPL062W
ZShift = −1.59
Z lm Score = 1.98
NG = 0
DB = 0
SM = 0
Delta_r
Doxycycline
YPL062W
ZShift = −1.61
Z lm Score = 1.31
NG = 0
DB = 0
SM = 0
Delta_AUC
Doxycycline
YKR057W RPS21A
ZShift = 3.56
Z lm Score = −4.35
NG = 0
DB = 0
SM = 0
Delta_L
Doxycycline
YKR057W RPS21A
ZShift = −2.25
Z lm Score = 0.37
NG = 0
DB = 0
SM = 0
Delta_K
Doxycycline
YKR057W RPS21A
ZShift = −3.4
Z lm Score = 2.82
NG = 0
DB = 0
SM = 0
Delta_r
Doxycycline
YKR057W RPS21A
ZShift = −2.94
Z lm Score = −2.14
NG = 0
DB = 0
SM = 0
Delta_AUC
Doxycycline

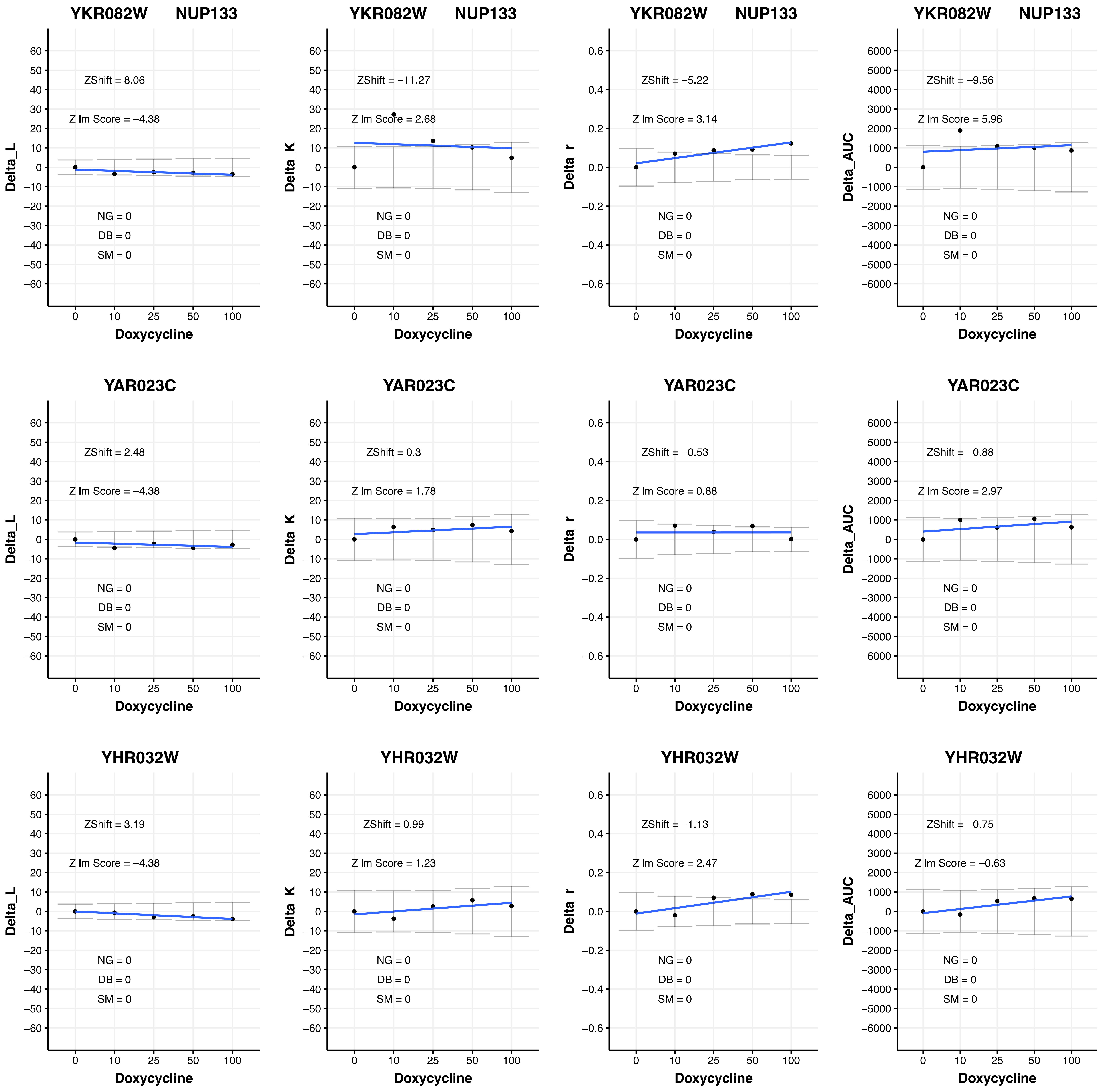
YKR082W NUP133
ZShift = 8.06
Z lm Score = −4.38
NG = 0
DB = 0
SM = 0
Delta_L
Doxycycline
YKR082W NUP133
ZShift = −11.27
Z lm Score = 2.68
NG = 0
DB = 0
SM = 0
Delta_K
Doxycycline
YKR082W NUP133
ZShift = −5.22
Z lm Score = 3.14
NG = 0
DB = 0
SM = 0
Delta_r
Doxycycline
YKR082W NUP133
ZShift = −9.56
Z lm Score = 5.96
NG = 0
DB = 0
SM = 0
Delta_AUC
Doxycycline
YAR023C
ZShift = 2.48
Z lm Score = −4.38
NG = 0
DB = 0
SM = 0
Delta_L
Doxycycline
YAR023C
ZShift = 0.3
Z lm Score = 1.78
NG = 0
DB = 0
SM = 0
Delta_K
Doxycycline
YAR023C
ZShift = −0.53
Z lm Score = 0.88
NG = 0
DB = 0
SM = 0
Delta_r
Doxycycline
YAR023C
ZShift = −0.88
Z lm Score = 2.97
NG = 0
DB = 0
SM = 0
Delta_AUC
Doxycycline
YHR032W
ZShift = 3.19
Z lm Score = −4.38
NG = 0
DB = 0
SM = 0
Delta_L
Doxycycline
YHR032W
ZShift = 0.99
Z lm Score = 1.23
NG = 0
DB = 0
SM = 0
Delta_K
Doxycycline
YHR032W
ZShift = −1.13
Z lm Score = 2.47
NG = 0
DB = 0
SM = 0
Delta_r
Doxycycline
YHR032W
ZShift = −0.75
Z lm Score = −0.63
NG = 0
DB = 0
SM = 0
Delta_AUC
Doxycycline

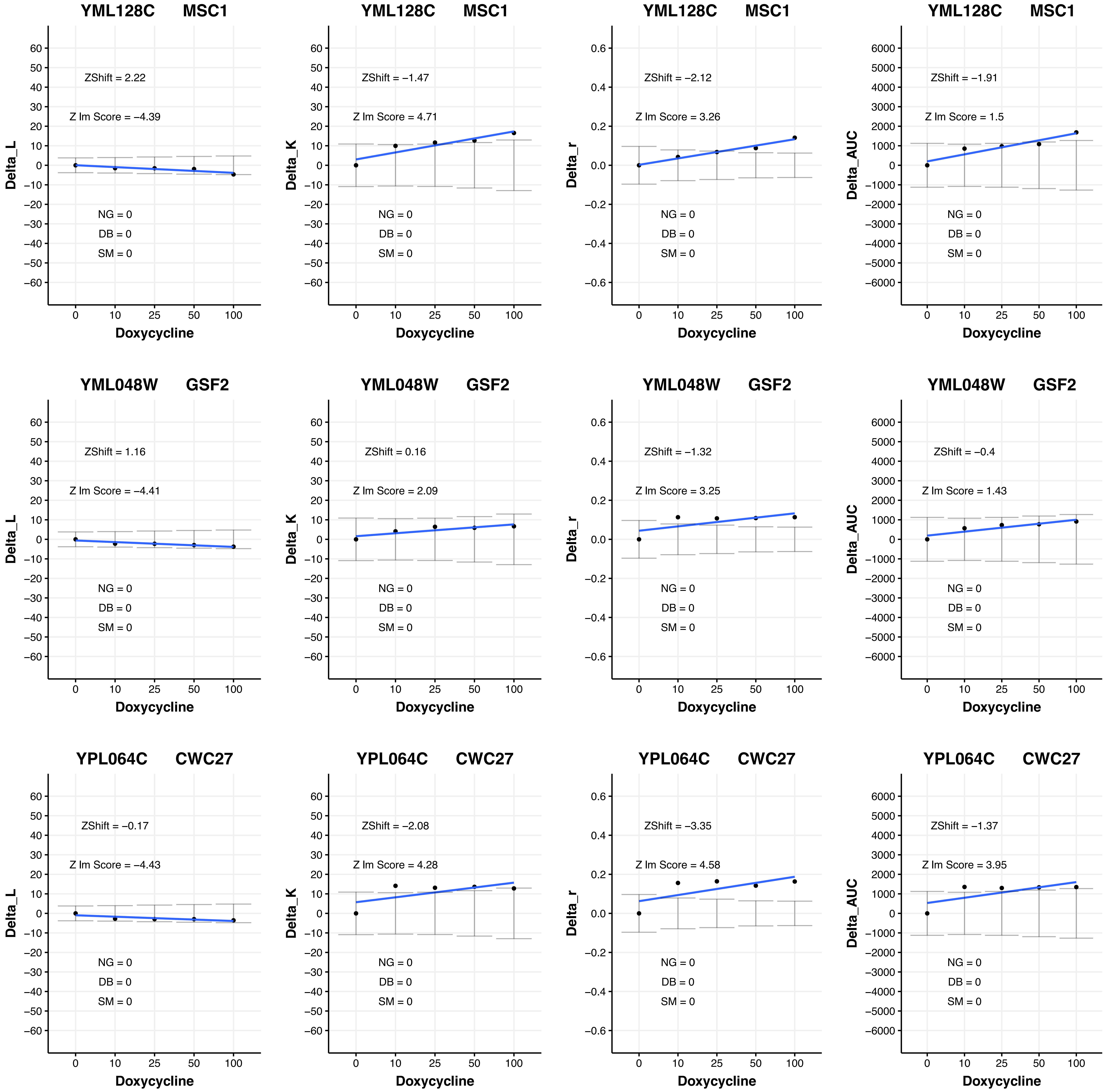

YML128C MSC1
ZShift = 2.22
Z lm Score = −4.39
NG = 0
DB = 0
SM = 0
Delta_L
Doxycycline
YML128C MSC1
ZShift = −1.47
Z lm Score = 4.71
NG = 0
DB = 0
SM = 0
Delta_K
Doxycycline
YML128C MSC1
ZShift = −2.12
Z lm Score = 3.26
NG = 0
DB = 0
SM = 0
Delta_r
Doxycycline
YML128C MSC1
ZShift = −1.91
Z lm Score = 1.5
NG = 0
DB = 0
SM = 0
Delta_AUC
Doxycycline
YML048W GSF2
ZShift = 1.16
Z lm Score = −4.41
NG = 0
DB = 0
SM = 0
Delta_L
Doxycycline
YML048W GSF2
ZShift = 0.16
Z lm Score = 2.09
NG = 0
DB = 0
SM = 0
Delta_K
Doxycycline
YML048W GSF2
ZShift = −1.32
Z lm Score = 3.25
NG = 0
DB = 0
SM = 0
Delta_r
Doxycycline
YML048W GSF2
ZShift = −0.4
Z lm Score = 1.43
NG = 0
DB = 0
SM = 0
Delta_AUC
Doxycycline
YPL064C CWC27
ZShift = −0.17
Z lm Score = −4.43
NG = 0
DB = 0
SM = 0
Delta_L
Doxycycline
YPL064C CWC27
ZShift = −2.08
Z lm Score = 4.28
NG = 0
DB = 0
SM = 0
Delta_K
Doxycycline
YPL064C CWC27
ZShift = −3.35
Z lm Score = 4.58
NG = 0
DB = 0
SM = 0
Delta_r
Doxycycline
YPL064C CWC27
ZShift = −1.37
Z lm Score = 3.95
NG = 0
DB = 0
SM = 0
Delta_AUC
Doxycycline

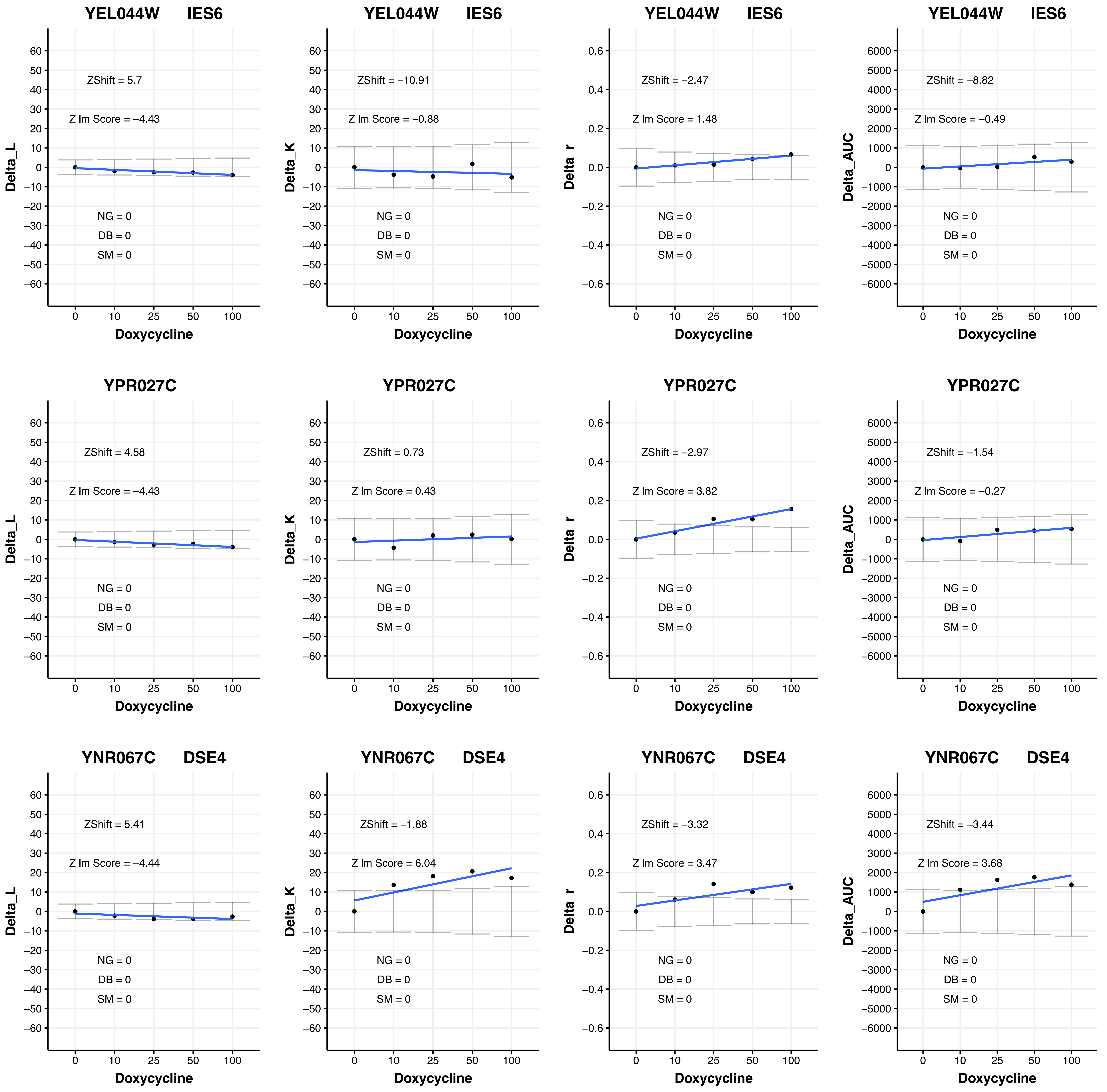

YEL044W IES6
ZShift = 5.7
Z lm Score = −4.43
NG = 0
DB = 0
SM = 0
Delta_L
Doxycycline
YEL044W IES6
ZShift = −10.91
Z lm Score = −0.88
NG = 0
DB = 0
SM = 0
Delta_K
Doxycycline
YEL044W IES6
ZShift = −2.47
Z lm Score = 1.48
NG = 0
DB = 0
SM = 0
Delta_r
Doxycycline
YEL044W IES6
ZShift = −8.82
Z lm Score = −0.49
NG = 0
DB = 0
SM = 0
Delta_AUC
Doxycycline
YPR027C
ZShift = 4.58
Z lm Score = −4.43
NG = 0
DB = 0
SM = 0
Delta_L
Doxycycline
YPR027C
ZShift = 0.73
Z lm Score = 0.43
NG = 0
DB = 0
SM = 0
Delta_K
Doxycycline
YPR027C
ZShift = −2.97
Z lm Score = 3.82
NG = 0
DB = 0
SM = 0
Delta_r
Doxycycline
YPR027C
ZShift = −1.54
Z lm Score = −0.27
NG = 0
DB = 0
SM = 0
Delta_AUC
Doxycycline
YNR067C DSE4
ZShift = 5.41
Z lm Score = −4.44
NG = 0
DB = 0
SM = 0
Delta_L
Doxycycline
YNR067C DSE4
ZShift = −1.88
Z lm Score = 6.04
NG = 0
DB = 0
SM = 0
Delta_K
Doxycycline
YNR067C DSE4
ZShift = −3.32
Z lm Score = 3.47
NG = 0
DB = 0
SM = 0
Delta_r
Doxycycline
YNR067C DSE4
ZShift = −3.44
Z lm Score = 3.68
NG = 0
DB = 0
SM = 0
Delta_AUC
Doxycycline

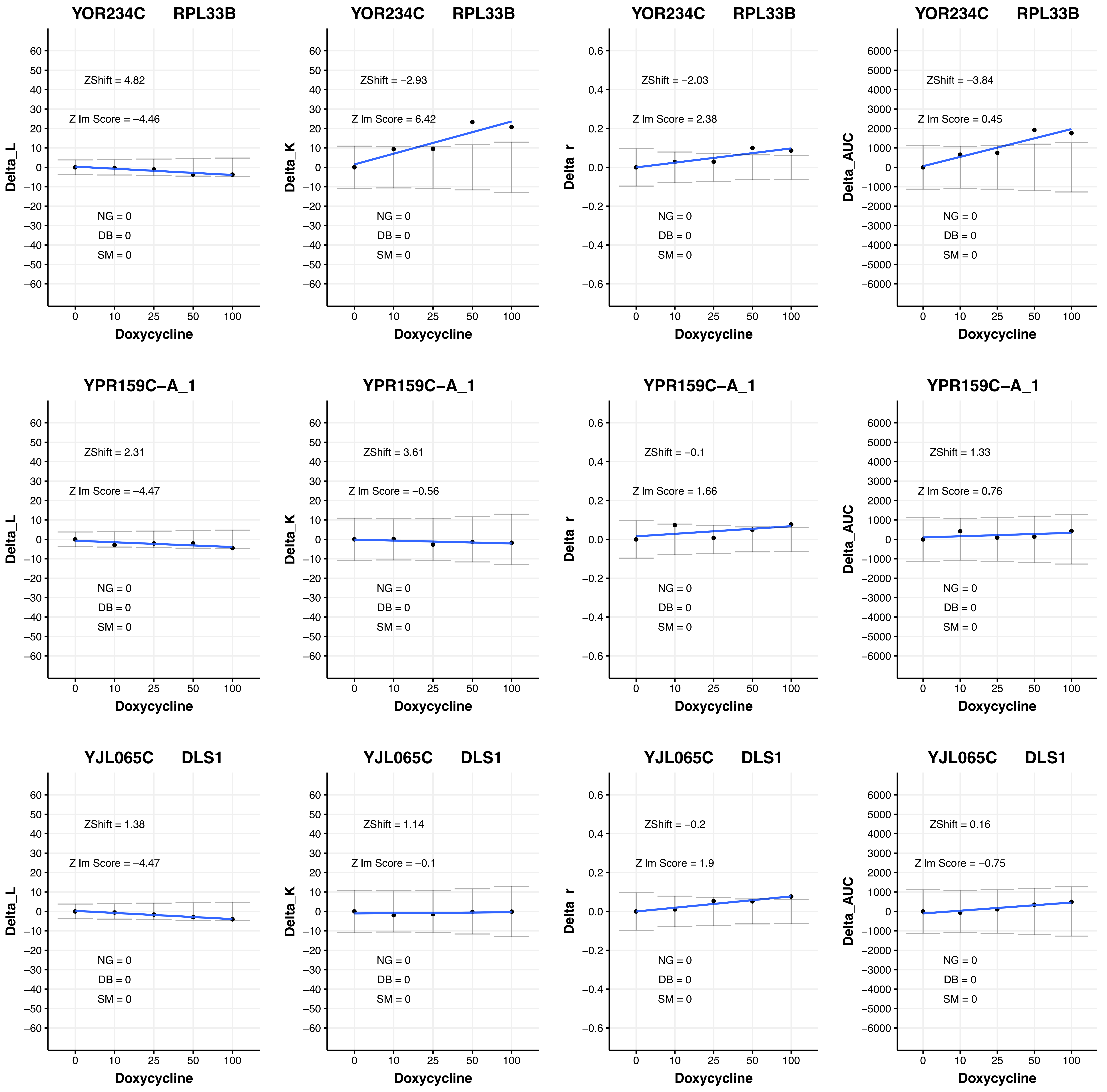
YOR234C RPL33B
ZShift = 4.82
Z lm Score = −4.46
NG = 0
DB = 0
SM = 0
Delta_L
Doxycycline
YOR234C RPL33B
ZShift = −2.93
Z lm Score = 6.42
NG = 0
DB = 0
SM = 0
Delta_K
Doxycycline
YOR234C RPL33B
ZShift = −2.03
Z lm Score = 2.38
NG = 0
DB = 0
SM = 0
Delta_r
Doxycycline
YOR234C RPL33B
ZShift = −3.84
Z lm Score = 0.45
NG = 0
DB = 0
SM = 0
Delta_AUC
Doxycycline
YPR159C−A_1
ZShift = 2.31
Z lm Score = −4.47
NG = 0
DB = 0
SM = 0
Delta_L
Doxycycline
YPR159C−A_1
ZShift = 3.61
Z lm Score = −0.56
NG = 0
DB = 0
SM = 0
Delta_K
Doxycycline
YPR159C−A_1
ZShift = −0.1
Z lm Score = 1.66
NG = 0
DB = 0
SM = 0
Delta_r
Doxycycline
YPR159C−A_1
ZShift = 1.33
Z lm Score = 0.76
NG = 0
DB = 0
SM = 0
Delta_AUC
Doxycycline
YJL065C DLS1
ZShift = 1.38
Z lm Score = −4.47
NG = 0
DB = 0
SM = 0
Delta_L
Doxycycline
YJL065C DLS1
ZShift = 1.14
Z lm Score = −0.1
NG = 0
DB = 0
SM = 0
Delta_K
Doxycycline
YJL065C DLS1
ZShift = −0.2
Z lm Score = 1.9
NG = 0
DB = 0
SM = 0
Delta_r
Doxycycline
YJL065C DLS1
ZShift = 0.16
Z lm Score = −0.75
NG = 0
DB = 0
SM = 0
Delta_AUC
Doxycycline

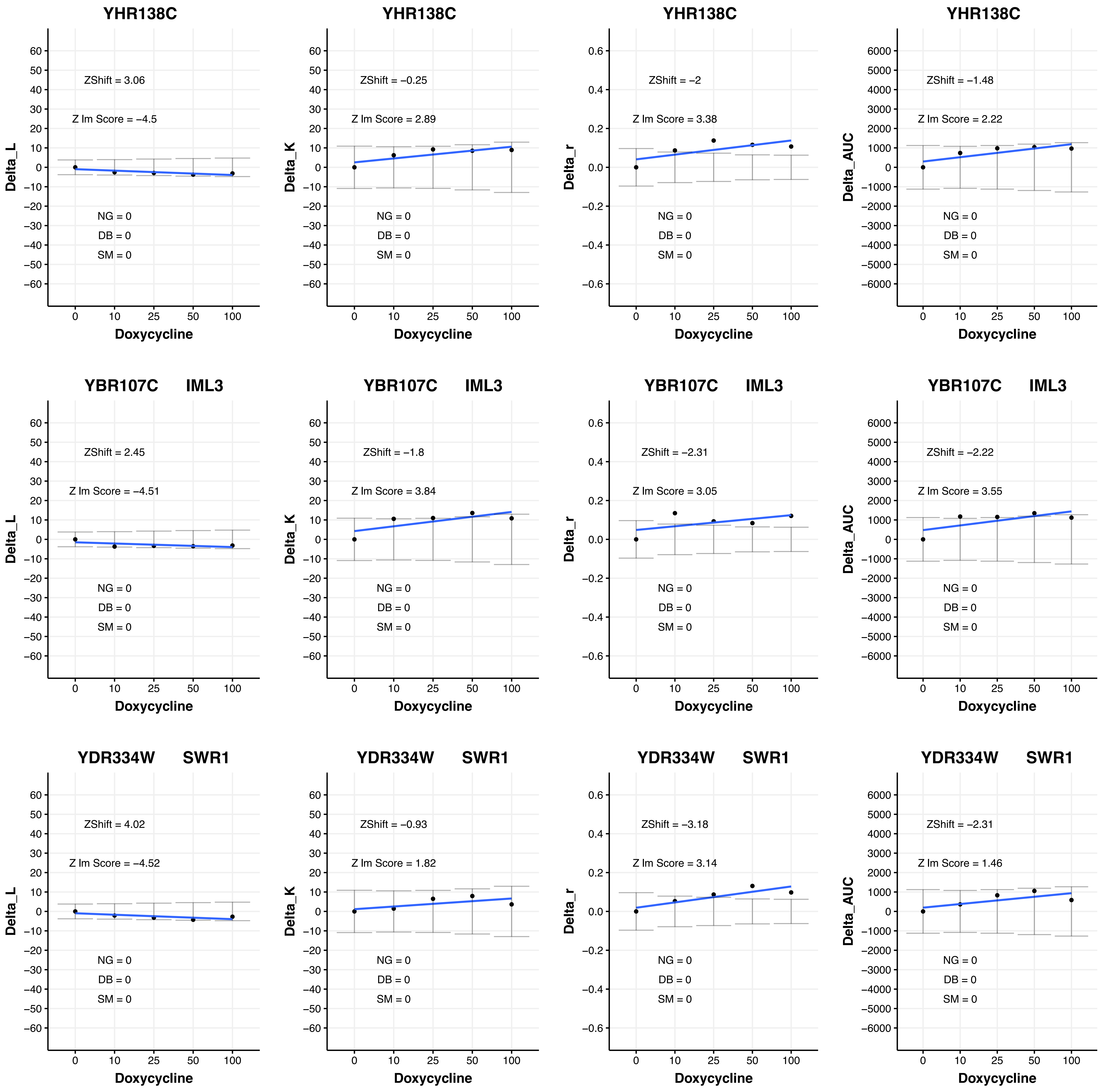

YHR138C
ZShift = 3.06
Z lm Score = −4.5
NG = 0
DB = 0
SM = 0
Delta_L
Doxycycline
YHR138C
ZShift = −0.25
Z lm Score = 2.89
NG = 0
DB = 0
SM = 0
Delta_K
Doxycycline
YHR138C
ZShift = −2
Z lm Score = 3.38
NG = 0
DB = 0
SM = 0
Delta_r
Doxycycline
YHR138C
ZShift = −1.48
Z lm Score = 2.22
NG = 0
DB = 0
SM = 0
Delta_AUC
Doxycycline
YBR107C IML3
ZShift = 2.45
Z lm Score = −4.51
NG = 0
DB = 0
SM = 0
Delta_L
Doxycycline
YBR107C IML3
ZShift = −1.8
Z lm Score = 3.84
NG = 0
DB = 0
SM = 0
Delta_K
Doxycycline
YBR107C IML3
ZShift = −2.31
Z lm Score = 3.05
NG = 0
DB = 0
SM = 0
Delta_r
Doxycycline
YBR107C IML3
ZShift = −2.22
Z lm Score = 3.55
NG = 0
DB = 0
SM = 0
Delta_AUC
Doxycycline
YDR334W SWR1
ZShift = 4.02
Z lm Score = −4.52
NG = 0
DB = 0
SM = 0
Delta_L
Doxycycline
YDR334W SWR1
ZShift = −0.93
Z lm Score = 1.82
NG = 0
DB = 0
SM = 0
Delta_K
Doxycycline
YDR334W SWR1
ZShift = −3.18
Z lm Score = 3.14
NG = 0
DB = 0
SM = 0
Delta_r
Doxycycline
YDR334W SWR1
ZShift = −2.31
Z lm Score = 1.46
NG = 0
DB = 0
SM = 0
Delta_AUC
Doxycycline

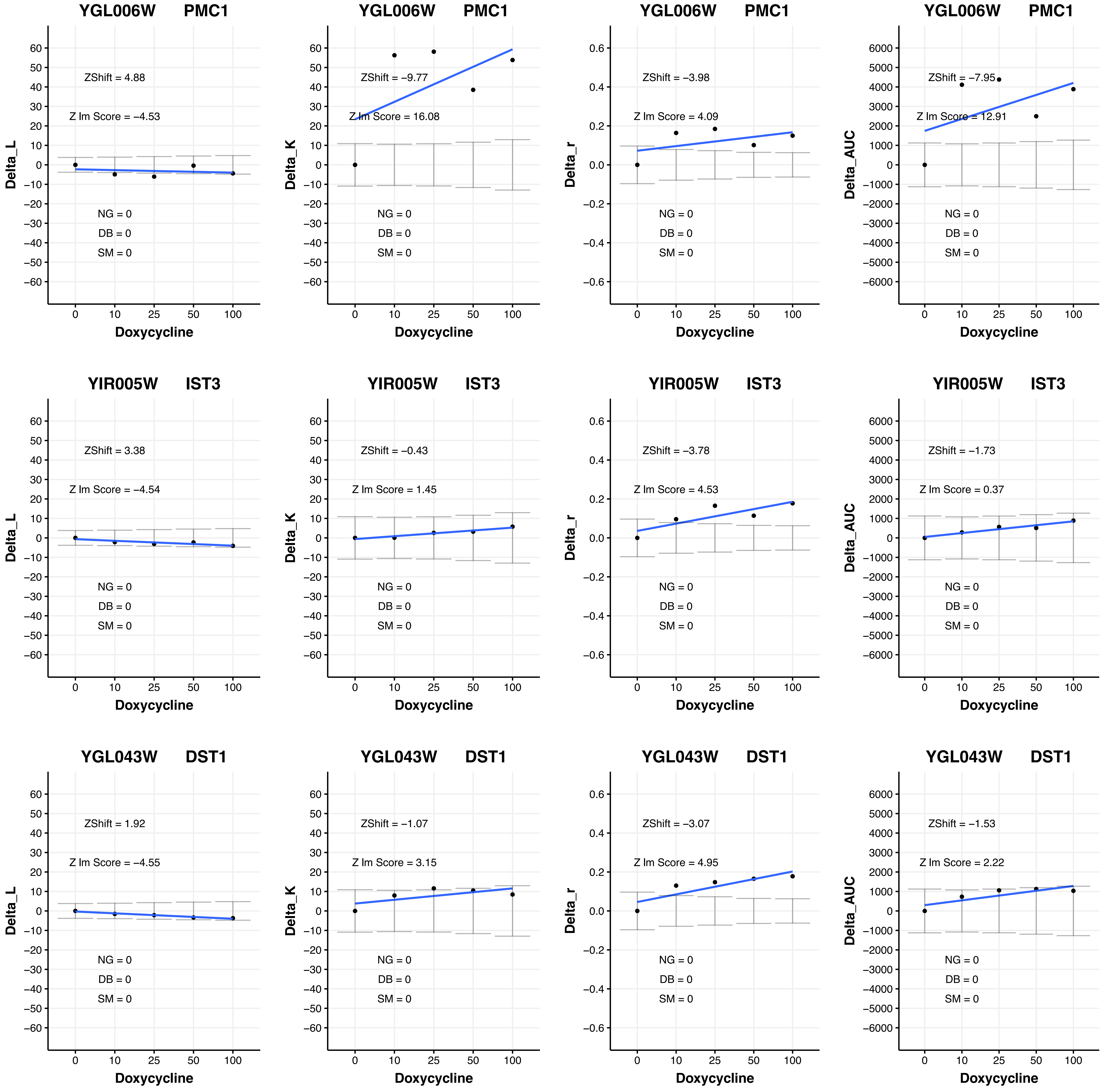

YGL006W PMC1
ZShift = 4.88
Z lm Score = −4.53
ZShift = −9.77
Z lm Score = 16.08
ZShift = −3.98
Z lm Score = 4.09
ZShift = −7.95
Z lm Score = 12.91
YIR005W IST3
ZShift = 3.38
Z lm Score = −4.54
ZShift = −0.43
Z lm Score = 1.45
ZShift = −3.78
Z lm Score = 4.53
ZShift = −1.73
Z lm Score = 0.37
YGL043W DST1
ZShift = 1.92
Z lm Score = −4.55
ZShift = −1.07
Z lm Score = 3.15
ZShift = −3.07
Z lm Score = 4.95
ZShift = −1.53
Z lm Score = 2.22
NG = 0
DB = 0
SM = 0
Delta_L
Delta_K
Delta_r
Delta_AUC
Doxycycline

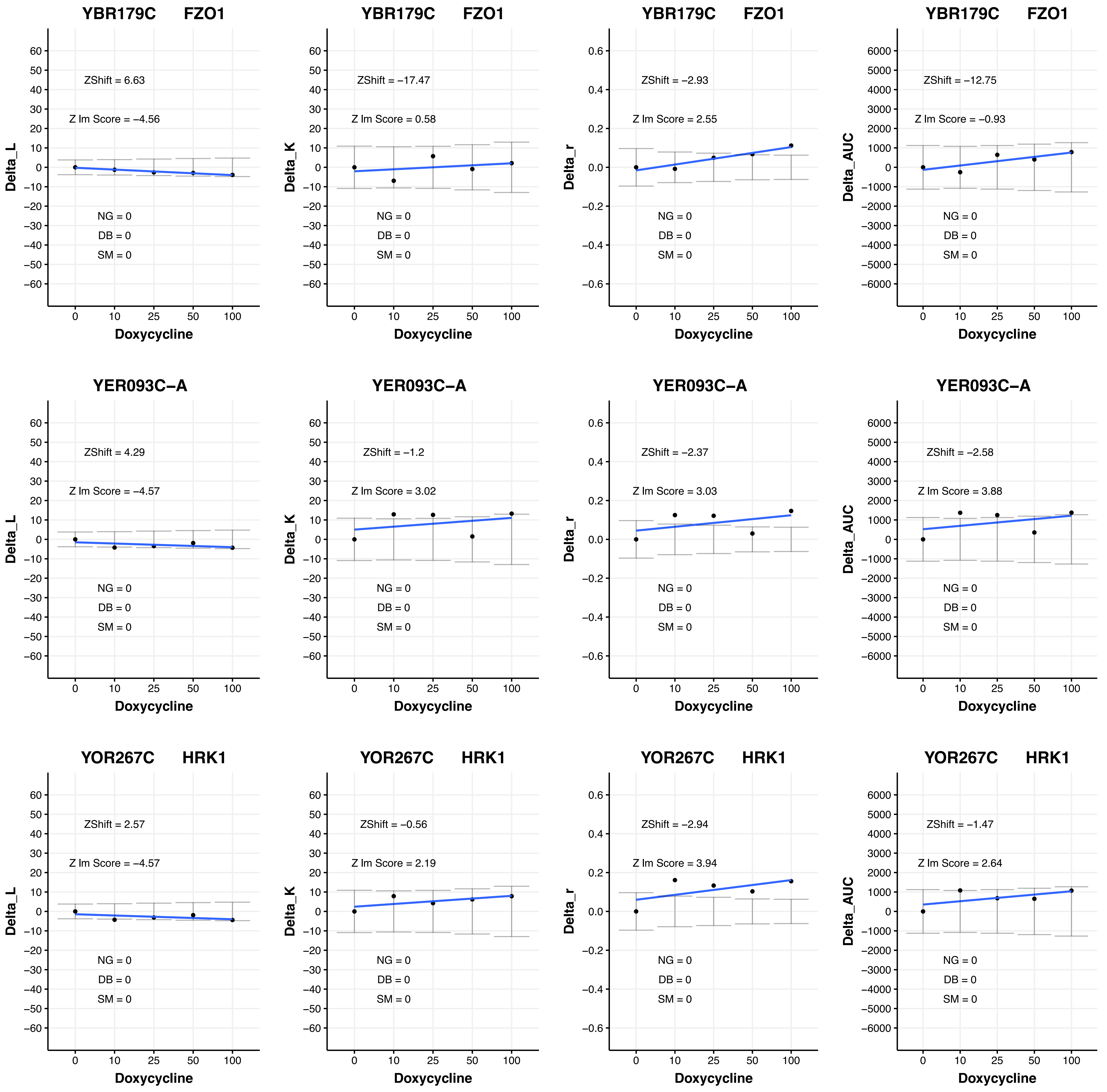

YBR179C FZO1
ZShift = 6.63
Z lm Score = −4.56
NG = 0
DB = 0
SM = 0
Delta_L
Doxycycline
YBR179C FZO1
ZShift = −17.47
Z lm Score = 0.58
NG = 0
DB = 0
SM = 0
Delta_K
Doxycycline
YBR179C FZO1
ZShift = −2.93
Z lm Score = 2.55
NG = 0
DB = 0
SM = 0
Delta_r
Doxycycline
YBR179C FZO1
ZShift = −12.75
Z lm Score = −0.93
NG = 0
DB = 0
SM = 0
Delta_AUC
Doxycycline
YER093C−A
ZShift = 4.29
Z lm Score = −4.57
NG = 0
DB = 0
SM = 0
Delta_L
Doxycycline
YER093C−A
ZShift = −1.2
Z lm Score = 3.02
NG = 0
DB = 0
SM = 0
Delta_K
Doxycycline
YER093C−A
ZShift = −2.37
Z lm Score = 3.03
NG = 0
DB = 0
SM = 0
Delta_r
Doxycycline
YER093C−A
ZShift = −2.58
Z lm Score = 3.88
NG = 0
DB = 0
SM = 0
Delta_AUC
Doxycycline
YOR267C HRK1
ZShift = 2.57
Z lm Score = −4.57
NG = 0
DB = 0
SM = 0
Delta_L
Doxycycline
YOR267C HRK1
ZShift = −0.56
Z lm Score = 2.19
NG = 0
DB = 0
SM = 0
Delta_K
Doxycycline
YOR267C HRK1
ZShift = −2.94
Z lm Score = 3.94
NG = 0
DB = 0
SM = 0
Delta_r
Doxycycline
YOR267C HRK1
ZShift = −1.47
Z lm Score = 2.64
NG = 0
DB = 0
SM = 0
Delta_AUC
Doxycycline

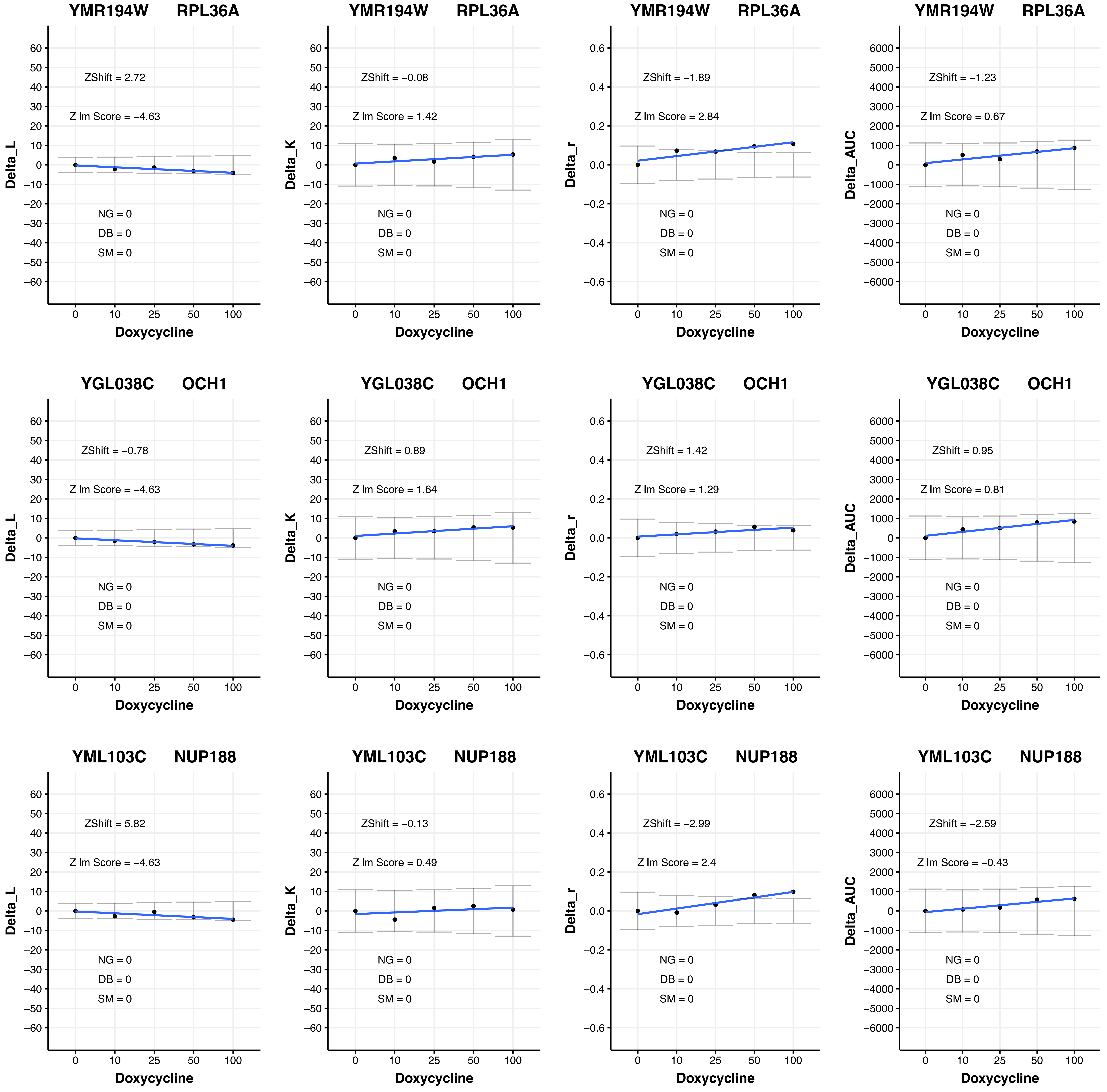
YMR194W RPL36A
ZShift = 2.72
Z lm Score = −4.63
NG = 0
DB = 0
SM = 0
Delta_L
Doxycycline
YMR194W RPL36A
ZShift = −0.08
Z lm Score = 1.42
NG = 0
DB = 0
SM = 0
Delta_K
Doxycycline
YMR194W RPL36A
ZShift = −1.89
Z lm Score = 2.84
NG = 0
DB = 0
SM = 0
Delta_r
Doxycycline
YMR194W RPL36A
ZShift = −1.23
Z lm Score = 0.67
NG = 0
DB = 0
SM = 0
Delta_AUC
Doxycycline
YGL038C OCH1
ZShift = −0.78
Z lm Score = −4.63
NG = 0
DB = 0
SM = 0
Delta_L
Doxycycline
YGL038C OCH1
ZShift = 0.89
Z lm Score = 1.64
NG = 0
DB = 0
SM = 0
Delta_K
Doxycycline
YGL038C OCH1
ZShift = 1.42
Z lm Score = 1.29
NG = 0
DB = 0
SM = 0
Delta_r
Doxycycline
YGL038C OCH1
ZShift = 0.95
Z lm Score = 0.81
NG = 0
DB = 0
SM = 0
Delta_AUC
Doxycycline
YML103C NUP188
ZShift = 5.82
Z lm Score = −4.63
NG = 0
DB = 0
SM = 0
Delta_L
Doxycycline
YML103C NUP188
ZShift = −0.13
Z lm Score = 0.49
NG = 0
DB = 0
SM = 0
Delta_K
Doxycycline
YML103C NUP188
ZShift = −2.99
Z lm Score = 2.4
NG = 0
DB = 0
SM = 0
Delta_r
Doxycycline
YML103C NUP188
ZShift = −2.59
Z lm Score = −0.43
NG = 0
DB = 0
SM = 0
Delta_AUC
Doxycycline

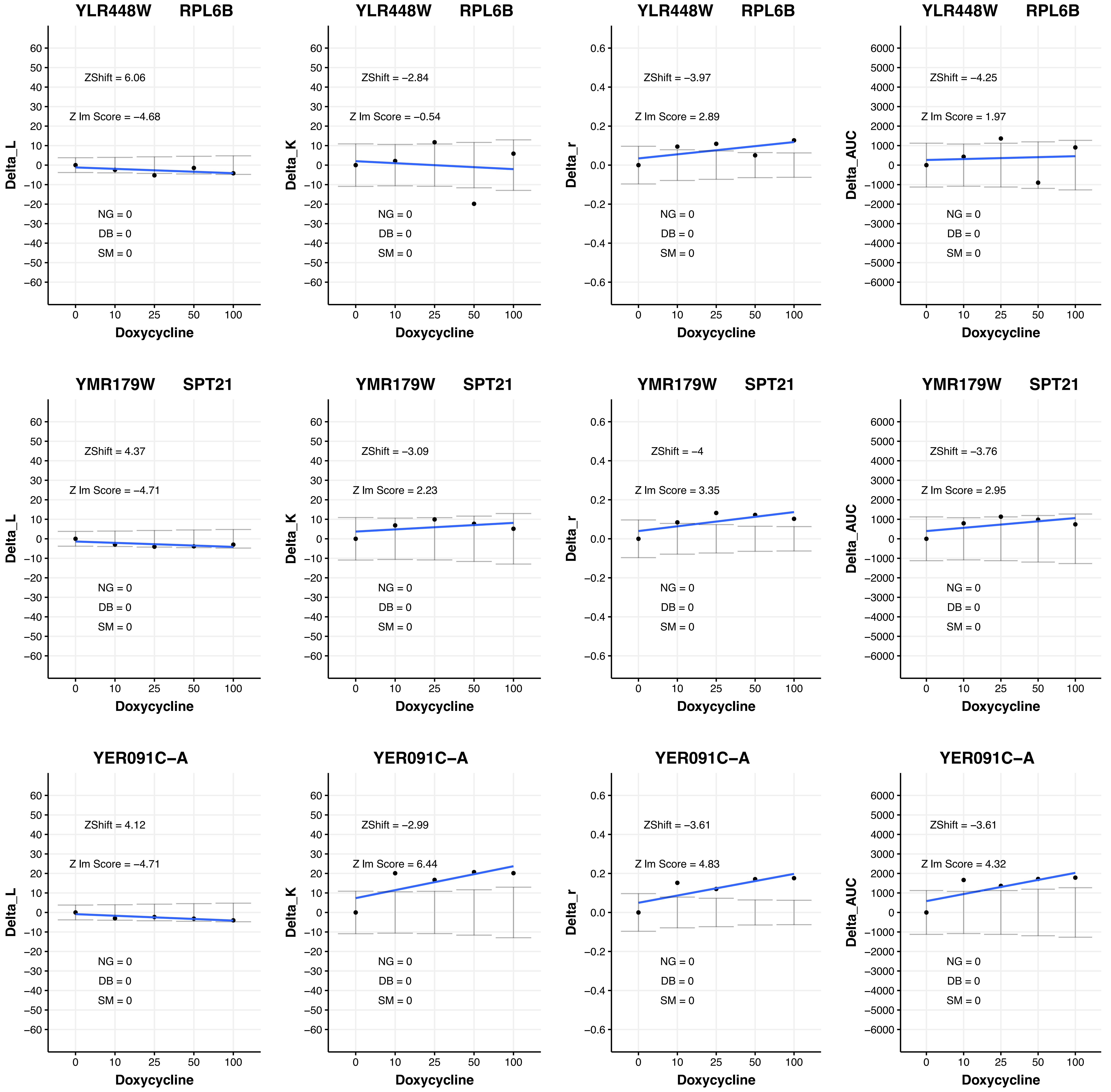

YLR448W RPL6B
ZShift = 6.06
Z lm Score = −4.68
NG = 0
DB = 0
SM = 0
Delta_L
Doxycycline
YLR448W RPL6B
ZShift = −2.84
Z lm Score = −0.54
NG = 0
DB = 0
SM = 0
Delta_K
Doxycycline
YLR448W RPL6B
ZShift = −3.97
Z lm Score = 2.89
NG = 0
DB = 0
SM = 0
Delta_r
Doxycycline
YLR448W RPL6B
ZShift = −4.25
Z lm Score = 1.97
NG = 0
DB = 0
SM = 0
Delta_AUC
Doxycycline
YMR179W SPT21
ZShift = 4.37
Z lm Score = −4.71
NG = 0
DB = 0
SM = 0
Delta_L
Doxycycline
YMR179W SPT21
ZShift = −3.09
Z lm Score = 2.23
NG = 0
DB = 0
SM = 0
Delta_K
Doxycycline
YMR179W SPT21
ZShift = −4
Z lm Score = 3.35
NG = 0
DB = 0
SM = 0
Delta_r
Doxycycline
YMR179W SPT21
ZShift = −3.76
Z lm Score = 2.95
NG = 0
DB = 0
SM = 0
Delta_AUC
Doxycycline
YER091C−A
ZShift = 4.12
Z lm Score = −4.71
NG = 0
DB = 0
SM = 0
Delta_L
Doxycycline
YER091C−A
ZShift = −2.99
Z lm Score = 6.44
NG = 0
DB = 0
SM = 0
Delta_K
Doxycycline
YER091C−A
ZShift = −3.61
Z lm Score = 4.83
NG = 0
DB = 0
SM = 0
Delta_r
Doxycycline
YER091C−A
ZShift = −3.61
Z lm Score = 4.32
NG = 0
DB = 0
SM = 0
Delta_AUC
Doxycycline

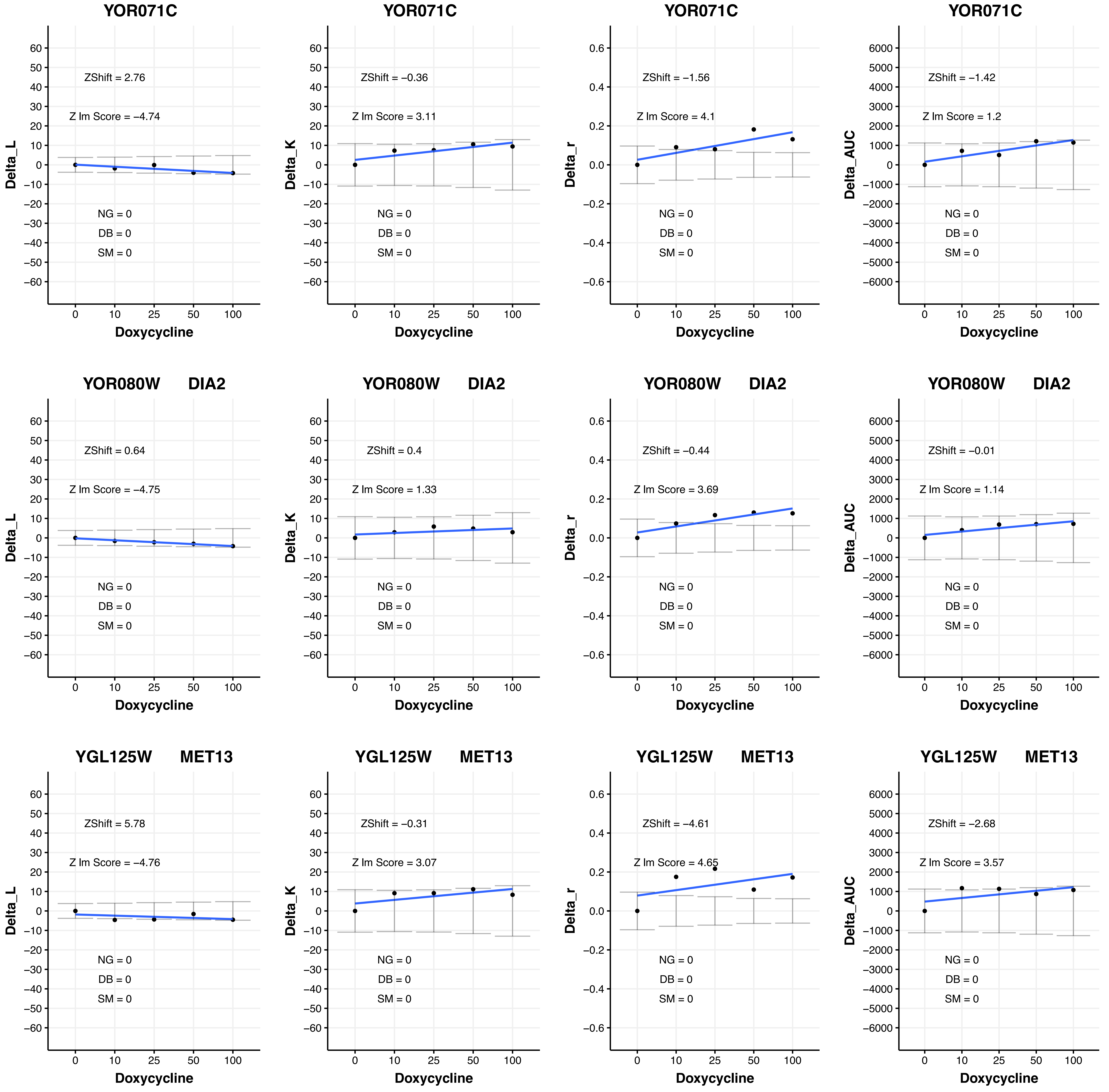

YOR071C
ZShift = 2.76
Z lm Score = −4.74
NG = 0
DB = 0
SM = 0
Delta_L
Doxycycline
YOR071C
ZShift = −0.36
Z lm Score = 3.11
NG = 0
DB = 0
SM = 0
Delta_K
Doxycycline
YOR071C
ZShift = −1.56
Z lm Score = 4.1
NG = 0
DB = 0
SM = 0
Delta_r
Doxycycline
YOR071C
ZShift = −1.42
Z lm Score = 1.2
NG = 0
DB = 0
SM = 0
Delta_AUC
Doxycycline
YOR080W DIA2
ZShift = 0.64
Z lm Score = −4.75
NG = 0
DB = 0
SM = 0
Delta_L
Doxycycline
YOR080W DIA2
ZShift = 0.4
Z lm Score = 1.33
NG = 0
DB = 0
SM = 0
Delta_K
Doxycycline
YOR080W DIA2
ZShift = −0.44
Z lm Score = 3.69
NG = 0
DB = 0
SM = 0
Delta_r
Doxycycline
YOR080W DIA2
ZShift = −0.01
Z lm Score = 1.14
NG = 0
DB = 0
SM = 0
Delta_AUC
Doxycycline
YGL125W MET13
ZShift = 5.78
Z lm Score = −4.76
NG = 0
DB = 0
SM = 0
Delta_L
Doxycycline
YGL125W MET13
ZShift = −0.31
Z lm Score = 3.07
NG = 0
DB = 0
SM = 0
Delta_K
Doxycycline
YGL125W MET13
ZShift = −4.61
Z lm Score = 4.65
NG = 0
DB = 0
SM = 0
Delta_r
Doxycycline
YGL125W MET13
ZShift = −2.68
Z lm Score = 3.57
NG = 0
DB = 0
SM = 0
Delta_AUC
Doxycycline

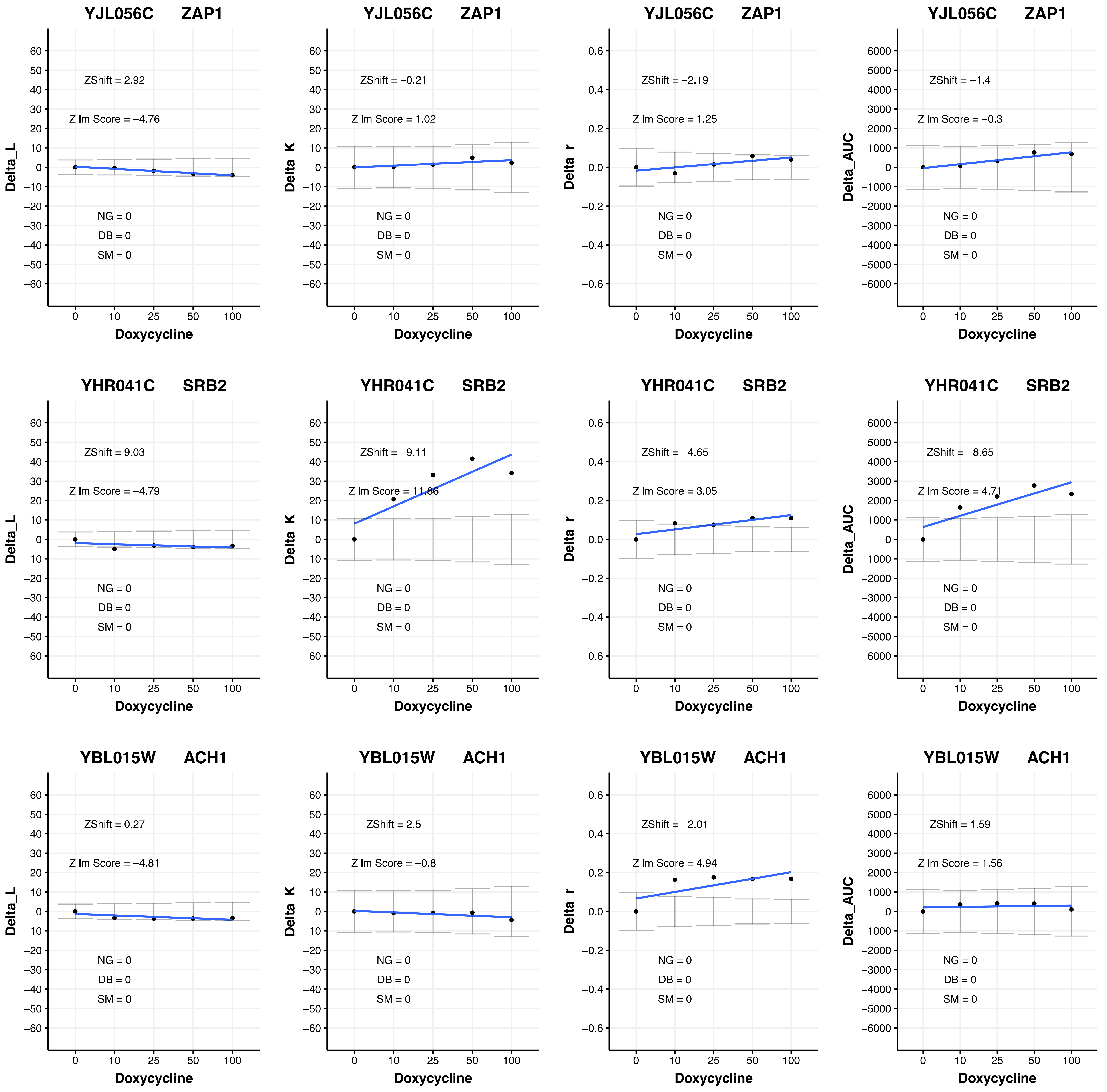
YJL056C ZAP1
ZShift = 2.92
Z lm Score = −4.76
NG = 0
DB = 0
SM = 0
Delta_L
Doxycycline
YJL056C ZAP1
ZShift = −0.21
Z lm Score = 1.02
NG = 0
DB = 0
SM = 0
Delta_K
Doxycycline
YJL056C ZAP1
ZShift = −2.19
Z lm Score = 1.25
NG = 0
DB = 0
SM = 0
Delta_r
Doxycycline
YJL056C ZAP1
ZShift = −1.4
Z lm Score = −0.3
NG = 0
DB = 0
SM = 0
Delta_AUC
Doxycycline
YHR041C SRB2
ZShift = 9.03
Z lm Score = −4.79
NG = 0
DB = 0
SM = 0
Delta_L
Doxycycline
YHR041C SRB2
ZShift = −9.11
Z lm Score = 11.86
NG = 0
DB = 0
SM = 0
Delta_K
Doxycycline
YHR041C SRB2
ZShift = −4.65
Z lm Score = 3.05
NG = 0
DB = 0
SM = 0
Delta_r
Doxycycline
YHR041C SRB2
ZShift = −8.65
Z lm Score = 4.71
NG = 0
DB = 0
SM = 0
Delta_AUC
Doxycycline
YBL015W ACH1
ZShift = 0.27
Z lm Score = −4.81
NG = 0
DB = 0
SM = 0
Delta_L
Doxycycline
YBL015W ACH1
ZShift = 2.5
Z lm Score = −0.8
NG = 0
DB = 0
SM = 0
Delta_K
Doxycycline
YBL015W ACH1
ZShift = −2.01
Z lm Score = 4.94
NG = 0
DB = 0
SM = 0
Delta_r
Doxycycline
YBL015W ACH1
ZShift = 1.59
Z lm Score = 1.56
NG = 0
DB = 0
SM = 0
Delta_AUC
Doxycycline

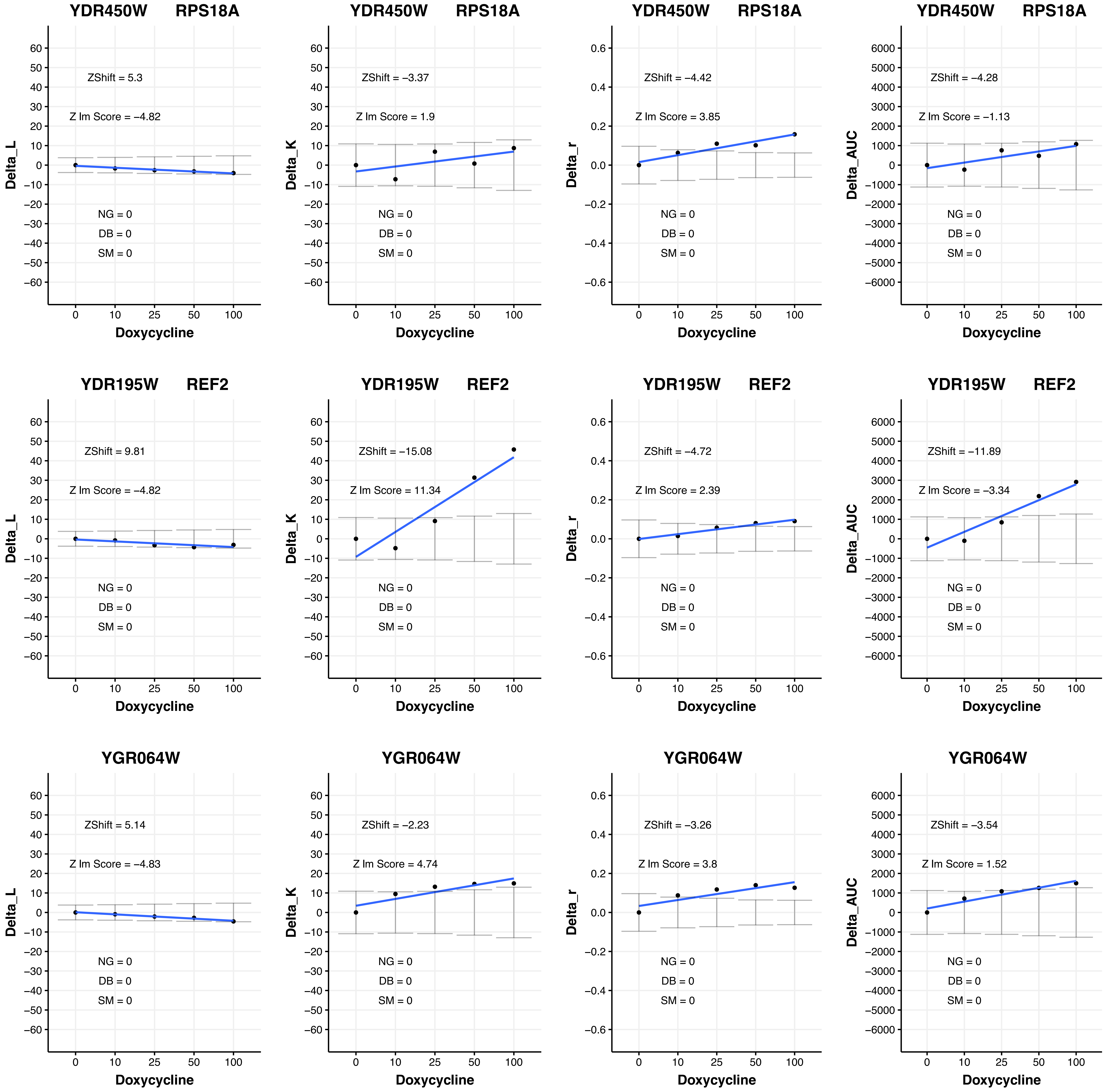

YDR450W RPS18A
ZShift = 5.3
Z lm Score = −4.82
NG = 0
DB = 0
SM = 0
Delta_L
Doxycycline
YDR450W RPS18A
ZShift = −3.37
Z lm Score = 1.9
NG = 0
DB = 0
SM = 0
Delta_K
Doxycycline
YDR450W RPS18A
ZShift = −4.42
Z lm Score = 3.85
NG = 0
DB = 0
SM = 0
Delta_r
Doxycycline
YDR450W RPS18A
ZShift = −4.28
Z lm Score = −1.13
NG = 0
DB = 0
SM = 0
Delta_AUC
Doxycycline
YDR195W REF2
ZShift = 9.81
Z lm Score = −4.82
NG = 0
DB = 0
SM = 0
Delta_L
Doxycycline
YDR195W REF2
ZShift = −15.08
Z lm Score = 11.34
NG = 0
DB = 0
SM = 0
Delta_K
Doxycycline
YDR195W REF2
ZShift = −4.72
Z lm Score = 2.39
NG = 0
DB = 0
SM = 0
Delta_r
Doxycycline
YDR195W REF2
ZShift = −11.89
Z lm Score = −3.34
NG = 0
DB = 0
SM = 0
Delta_AUC
Doxycycline
YGR064W
ZShift = 5.14
Z lm Score = −4.83
NG = 0
DB = 0
SM = 0
Delta_L
Doxycycline
YGR064W
ZShift = −2.23
Z lm Score = 4.74
NG = 0
DB = 0
SM = 0
Delta_K
Doxycycline
YGR064W
ZShift = −3.26
Z lm Score = 3.8
NG = 0
DB = 0
SM = 0
Delta_r
Doxycycline
YGR064W
ZShift = −3.54
Z lm Score = 1.52
NG = 0
DB = 0
SM = 0
Delta_AUC
Doxycycline

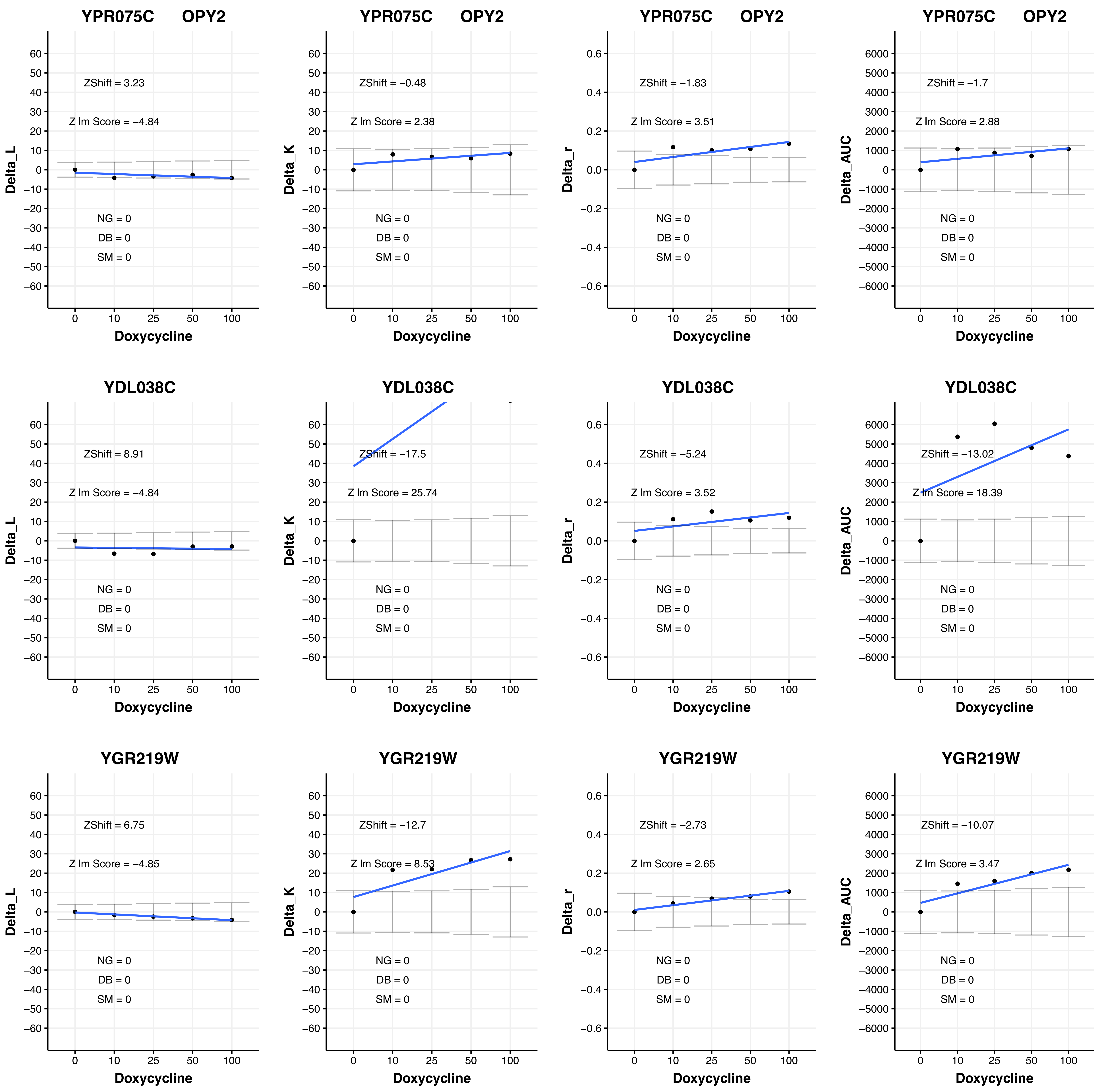

YPR075C OPY2
ZShift = 3.23
Z lm Score = −4.84
NG = 0
DB = 0
SM = 0
Delta_L
Doxycycline
YPR075C OPY2
ZShift = −0.48
Z lm Score = 2.38
NG = 0
DB = 0
SM = 0
Delta_K
Doxycycline
YPR075C OPY2
ZShift = −1.83
Z lm Score = 3.51
NG = 0
DB = 0
SM = 0
Delta_r
Doxycycline
YPR075C OPY2
ZShift = −1.7
Z lm Score = 2.88
NG = 0
DB = 0
SM = 0
Delta_AUC
Doxycycline
YDL038C
ZShift = 8.91
Z lm Score = −4.84
NG = 0
DB = 0
SM = 0
Delta_L
Doxycycline
YDL038C
ZShift = −17.5
Z lm Score = 25.74
NG = 0
DB = 0
SM = 0
Delta_K
Doxycycline
YDL038C
ZShift = −5.24
Z lm Score = 3.52
NG = 0
DB = 0
SM = 0
Delta_r
Doxycycline
YDL038C
ZShift = −13.02
Z lm Score = 18.39
NG = 0
DB = 0
SM = 0
Delta_AUC
Doxycycline
YGR219W
ZShift = 6.75
Z lm Score = −4.85
NG = 0
DB = 0
SM = 0
Delta_L
Doxycycline
YGR219W
ZShift = −12.7
Z lm Score = 8.53
NG = 0
DB = 0
SM = 0
Delta_K
Doxycycline
YGR219W
ZShift = −2.73
Z lm Score = 2.65
NG = 0
DB = 0
SM = 0
Delta_r
Doxycycline
YGR219W
ZShift = −10.07
Z lm Score = 3.47
NG = 0
DB = 0
SM = 0
Delta_AUC
Doxycycline

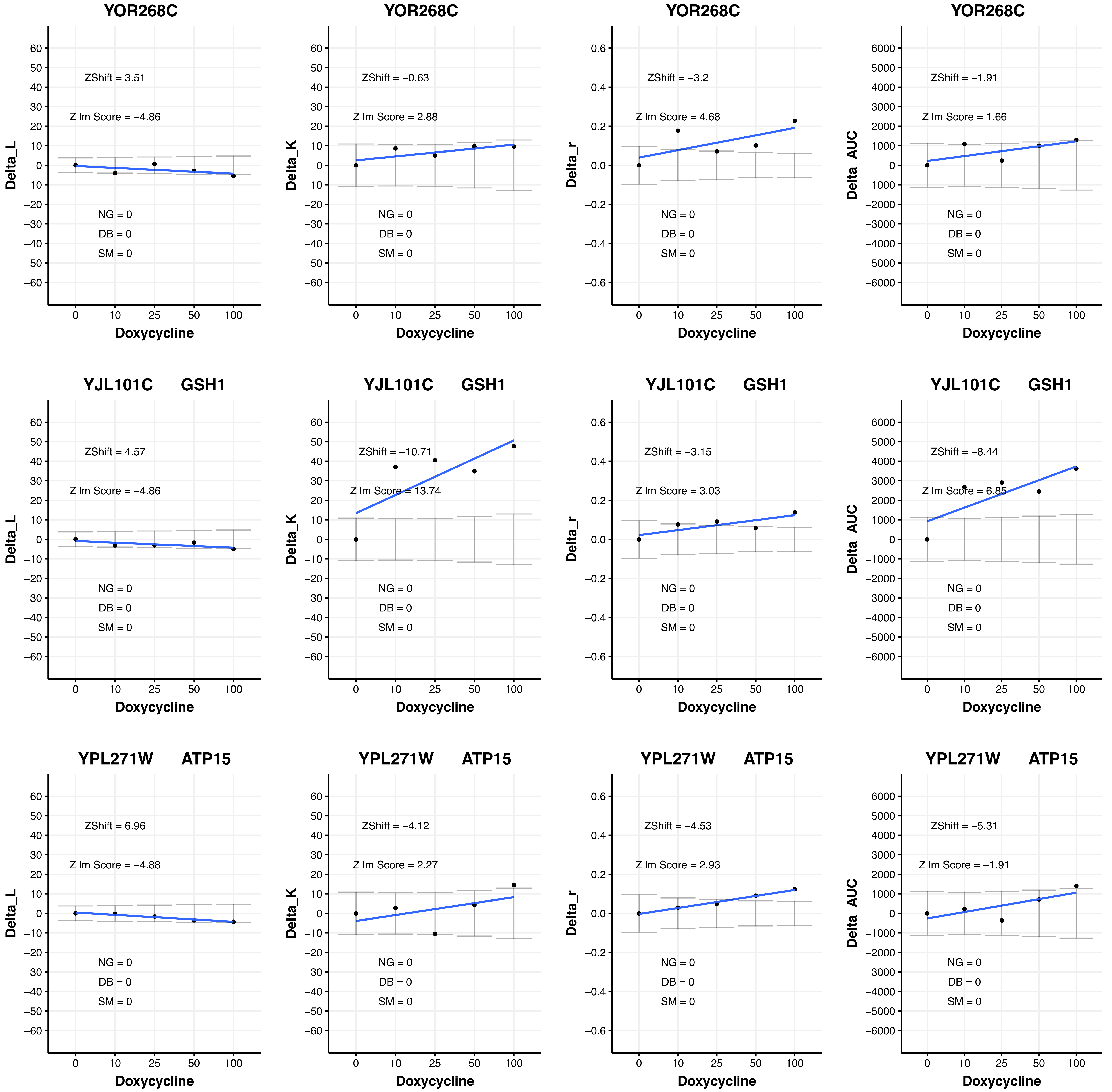

YOR268C
ZShift = 3.51
Z lm Score = −4.86
NG = 0
DB = 0
SM = 0
Delta_L
Doxycycline
YOR268C
ZShift = −0.63
Z lm Score = 2.88
NG = 0
DB = 0
SM = 0
Delta_K
Doxycycline
YOR268C
ZShift = −3.2
Z lm Score = 4.68
NG = 0
DB = 0
SM = 0
Delta_r
Doxycycline
YOR268C
ZShift = −1.91
Z lm Score = 1.66
NG = 0
DB = 0
SM = 0
Delta_AUC
Doxycycline
YJL101C GSH1
ZShift = 4.57
Z lm Score = −4.86
NG = 0
DB = 0
SM = 0
Delta_L
Doxycycline
YJL101C GSH1
ZShift = −10.71
Z lm Score = 13.74
NG = 0
DB = 0
SM = 0
Delta_K
Doxycycline
YJL101C GSH1
ZShift = −3.15
Z lm Score = 3.03
NG = 0
DB = 0
SM = 0
Delta_r
Doxycycline
YJL101C GSH1
ZShift = −8.44
Z lm Score = 6.85
NG = 0
DB = 0
SM = 0
Delta_AUC
Doxycycline
YPL271W ATP15
ZShift = 6.96
Z lm Score = −4.88
NG = 0
DB = 0
SM = 0
Delta_L
Doxycycline
YPL271W ATP15
ZShift = −4.12
Z lm Score = 2.27
NG = 0
DB = 0
SM = 0
Delta_K
Doxycycline
YPL271W ATP15
ZShift = −4.53
Z lm Score = 2.93
NG = 0
DB = 0
SM = 0
Delta_r
Doxycycline
YPL271W ATP15
ZShift = −5.31
Z lm Score = −1.91
NG = 0
DB = 0
SM = 0
Delta_AUC
Doxycycline

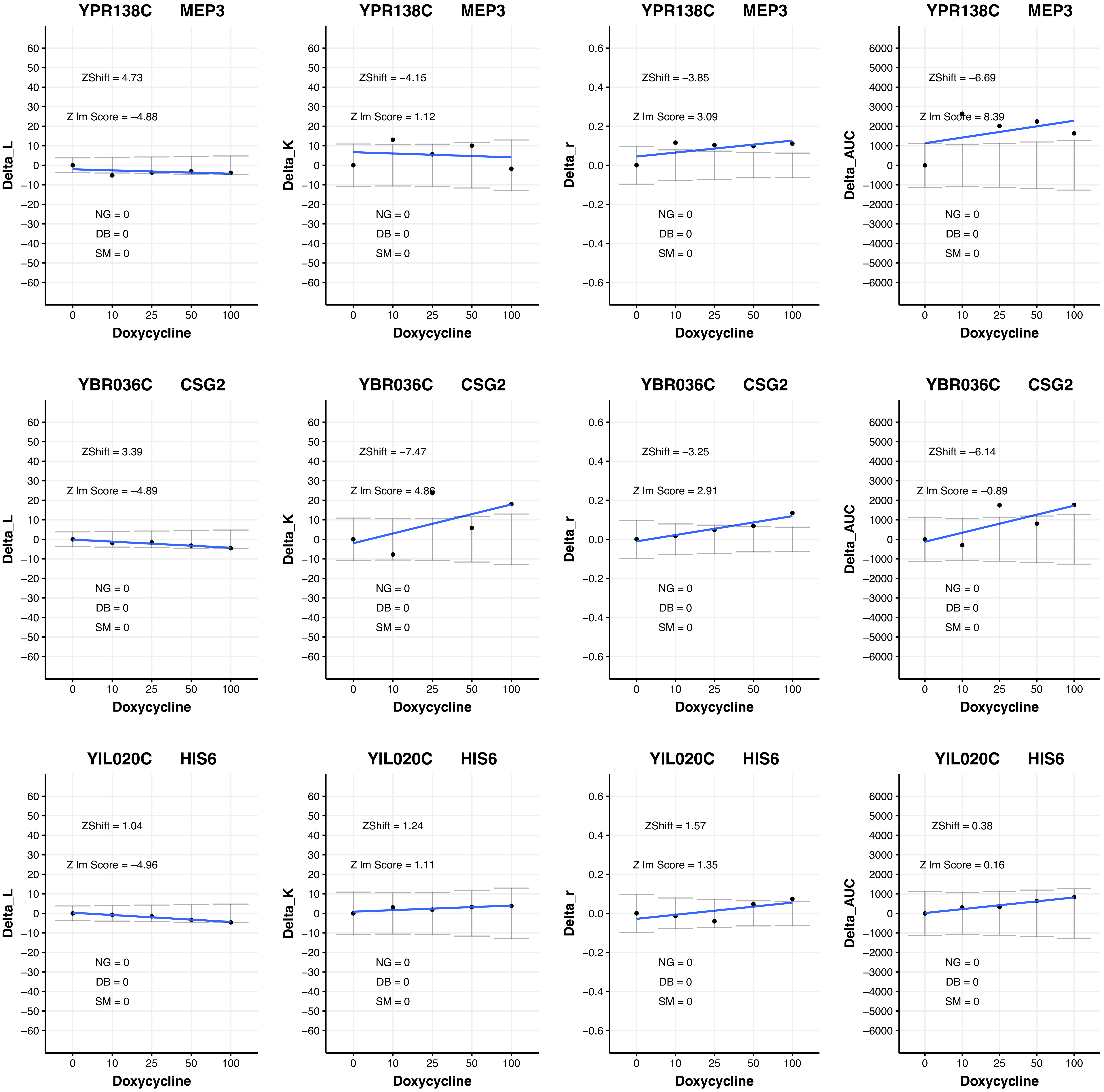
YPR138C MEP3
ZShift = 4.73
Z lm Score = −4.88
NG = 0
DB = 0
SM = 0
Delta_L
Doxycycline
YPR138C MEP3
ZShift = −4.15
Z lm Score = 1.12
NG = 0
DB = 0
SM = 0
Delta_K
Doxycycline
YPR138C MEP3
ZShift = −3.85
Z lm Score = 3.09
NG = 0
DB = 0
SM = 0
Delta_r
Doxycycline
YPR138C MEP3
ZShift = −6.69
Z lm Score = 8.39
NG = 0
DB = 0
SM = 0
Delta_AUC
Doxycycline
YBR036C CSG2
ZShift = 3.39
Z lm Score = −4.89
NG = 0
DB = 0
SM = 0
Delta_L
Doxycycline
YBR036C CSG2
ZShift = −7.47
Z lm Score = 4.86
NG = 0
DB = 0
SM = 0
Delta_K
Doxycycline
YBR036C CSG2
ZShift = −3.25
Z lm Score = 2.91
NG = 0
DB = 0
SM = 0
Delta_r
Doxycycline
YBR036C CSG2
ZShift = −6.14
Z lm Score = −0.89
NG = 0
DB = 0
SM = 0
Delta_AUC
Doxycycline
YIL020C HIS6
ZShift = 1.04
Z lm Score = −4.96
NG = 0
DB = 0
SM = 0
Delta_L
Doxycycline
YIL020C HIS6
ZShift = 1.24
Z lm Score = 1.11
NG = 0
DB = 0
SM = 0
Delta_K
Doxycycline
YIL020C HIS6
ZShift = 1.57
Z lm Score = 1.35
NG = 0
DB = 0
SM = 0
Delta_r
Doxycycline
YIL020C HIS6
ZShift = 0.38
Z lm Score = 0.16
NG = 0
DB = 0
SM = 0
Delta_AUC
Doxycycline

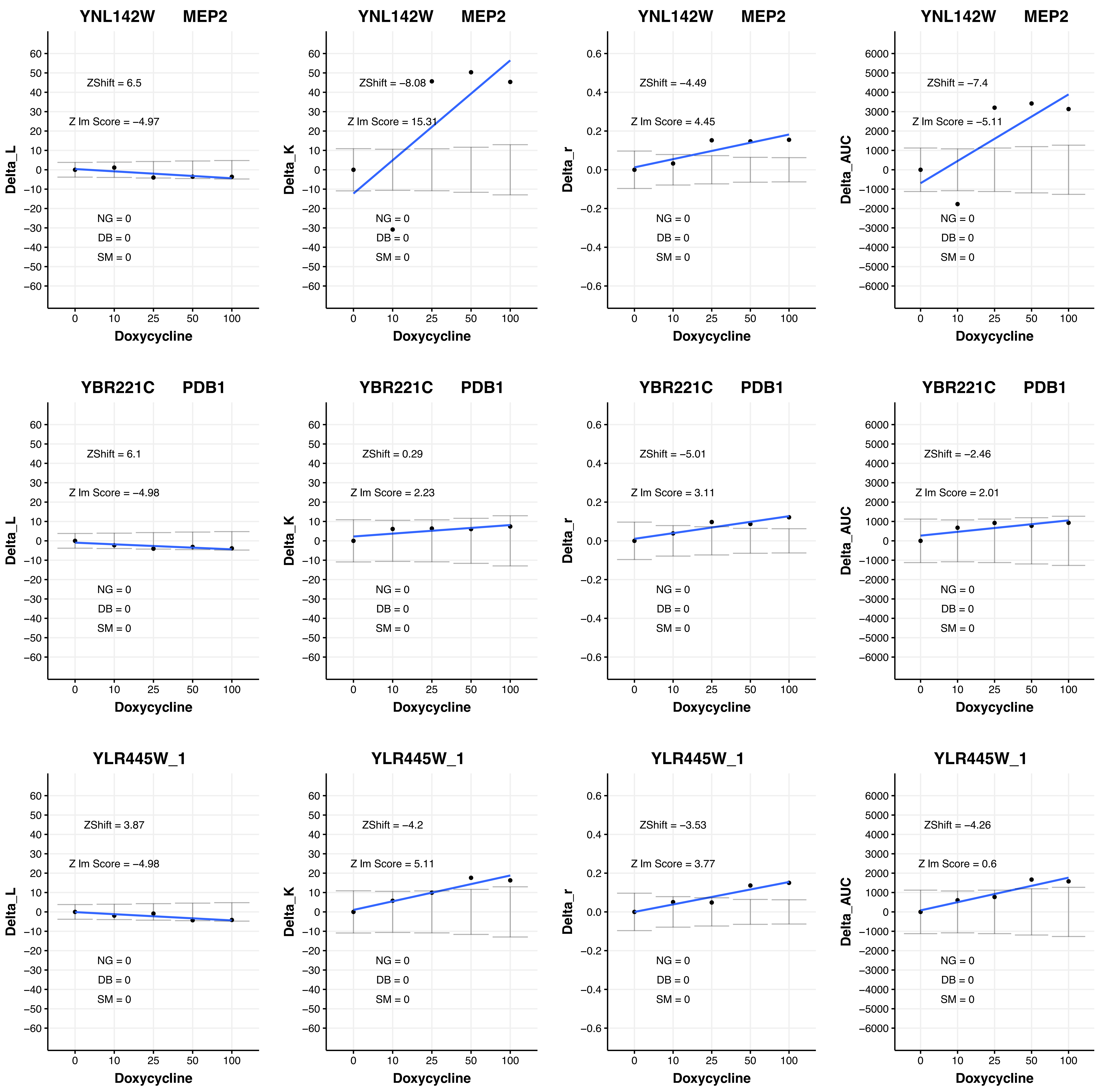
YNL142W MEP2
ZShift = 6.5
Z lm Score = −4.97
NG = 0
DB = 0
SM = 0
Delta_L
Doxycycline
YNL142W MEP2
ZShift = −8.08
Z lm Score = 15.31
NG = 0
DB = 0
SM = 0
Delta_K
Doxycycline
YNL142W MEP2
ZShift = −4.49
Z lm Score = 4.45
NG = 0
DB = 0
SM = 0
Delta_r
Doxycycline
YNL142W MEP2
ZShift = −7.4
Z lm Score = −5.11
NG = 0
DB = 0
SM = 0
Delta_AUC
Doxycycline
YBR221C PDB1
ZShift = 6.1
Z lm Score = −4.98
NG = 0
DB = 0
SM = 0
Delta_L
Doxycycline
YBR221C PDB1
ZShift = 0.29
Z lm Score = 2.23
NG = 0
DB = 0
SM = 0
Delta_K
Doxycycline
YBR221C PDB1
ZShift = −5.01
Z lm Score = 3.11
NG = 0
DB = 0
SM = 0
Delta_r
Doxycycline
YBR221C PDB1
ZShift = −2.46
Z lm Score = 2.01
NG = 0
DB = 0
SM = 0
Delta_AUC
Doxycycline
YLR445W_1
ZShift = 3.87
Z lm Score = −4.98
NG = 0
DB = 0
SM = 0
Delta_L
Doxycycline
YLR445W_1
ZShift = −4.2
Z lm Score = 5.11
NG = 0
DB = 0
SM = 0
Delta_K
Doxycycline
YLR445W_1
ZShift = −3.53
Z lm Score = 3.77
NG = 0
DB = 0
SM = 0
Delta_r
Doxycycline
YLR445W_1
ZShift = −4.26
Z lm Score = 0.6
NG = 0
DB = 0
SM = 0
Delta_AUC
Doxycycline

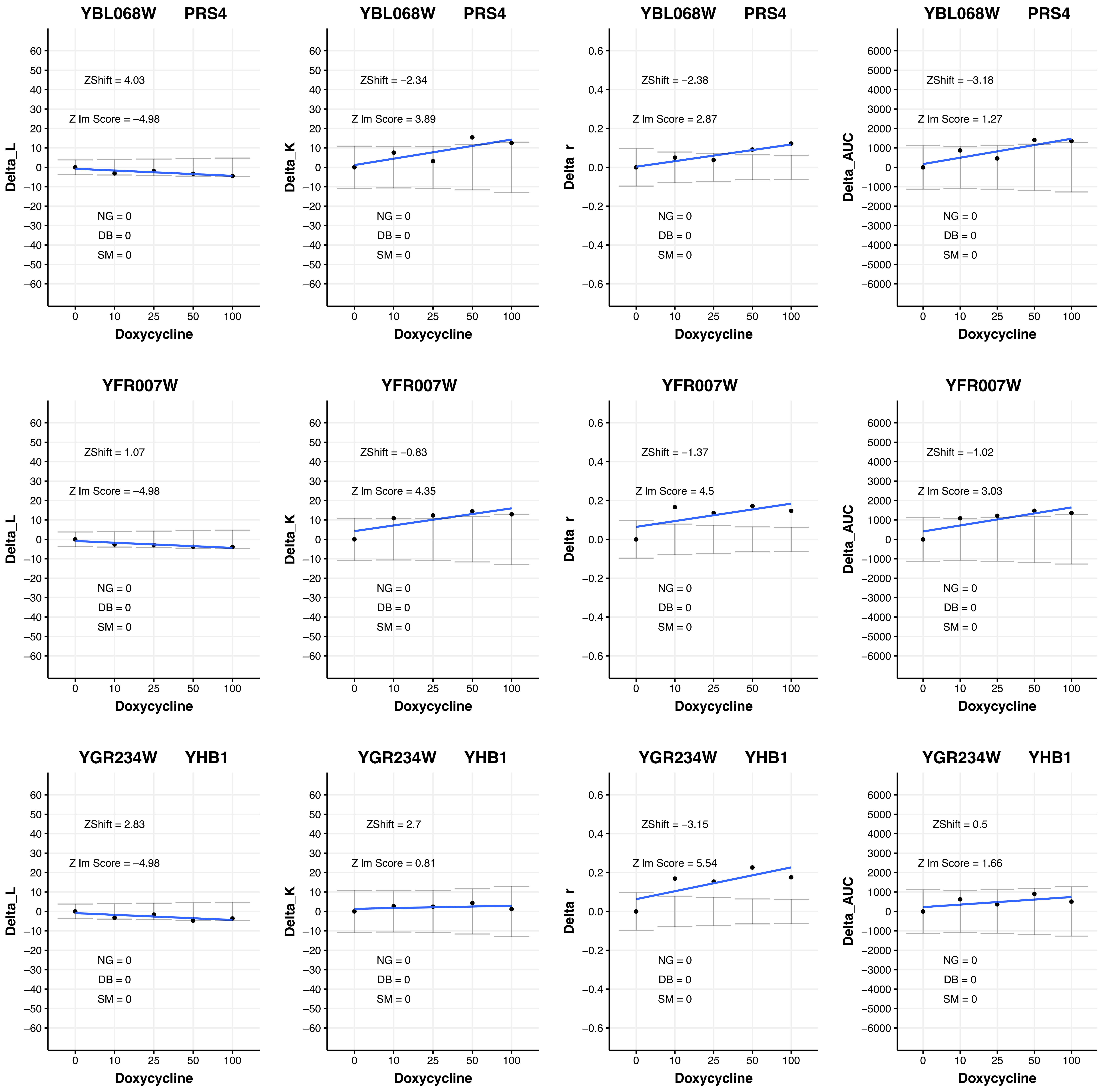
YBL068W PRS4
ZShift = 4.03
Z lm Score = −4.98
NG = 0
DB = 0
SM = 0
Delta_L
Doxycycline
YBL068W PRS4
ZShift = −2.34
Z lm Score = 3.89
NG = 0
DB = 0
SM = 0
Delta_K
Doxycycline
YBL068W PRS4
ZShift = −2.38
Z lm Score = 2.87
NG = 0
DB = 0
SM = 0
Delta_r
Doxycycline
YBL068W PRS4
ZShift = −3.18
Z lm Score = 1.27
NG = 0
DB = 0
SM = 0
Delta_AUC
Doxycycline
YFR007W
ZShift = 1.07
Z lm Score = −4.98
NG = 0
DB = 0
SM = 0
Delta_L
Doxycycline
YFR007W
ZShift = −0.83
Z lm Score = 4.35
NG = 0
DB = 0
SM = 0
Delta_K
Doxycycline
YFR007W
ZShift = −1.37
Z lm Score = 4.5
NG = 0
DB = 0
SM = 0
Delta_r
Doxycycline
YFR007W
ZShift = −1.02
Z lm Score = 3.03
NG = 0
DB = 0
SM = 0
Delta_AUC
Doxycycline
YGR234W YHB1
ZShift = 2.83
Z lm Score = −4.98
NG = 0
DB = 0
SM = 0
Delta_L
Doxycycline
YGR234W YHB1
ZShift = 2.7
Z lm Score = 0.81
NG = 0
DB = 0
SM = 0
Delta_K
Doxycycline
YGR234W YHB1
ZShift = −3.15
Z lm Score = 5.54
NG = 0
DB = 0
SM = 0
Delta_r
Doxycycline
YGR234W YHB1
ZShift = 0.5
Z lm Score = 1.66
NG = 0
DB = 0
SM = 0
Delta_AUC
Doxycycline

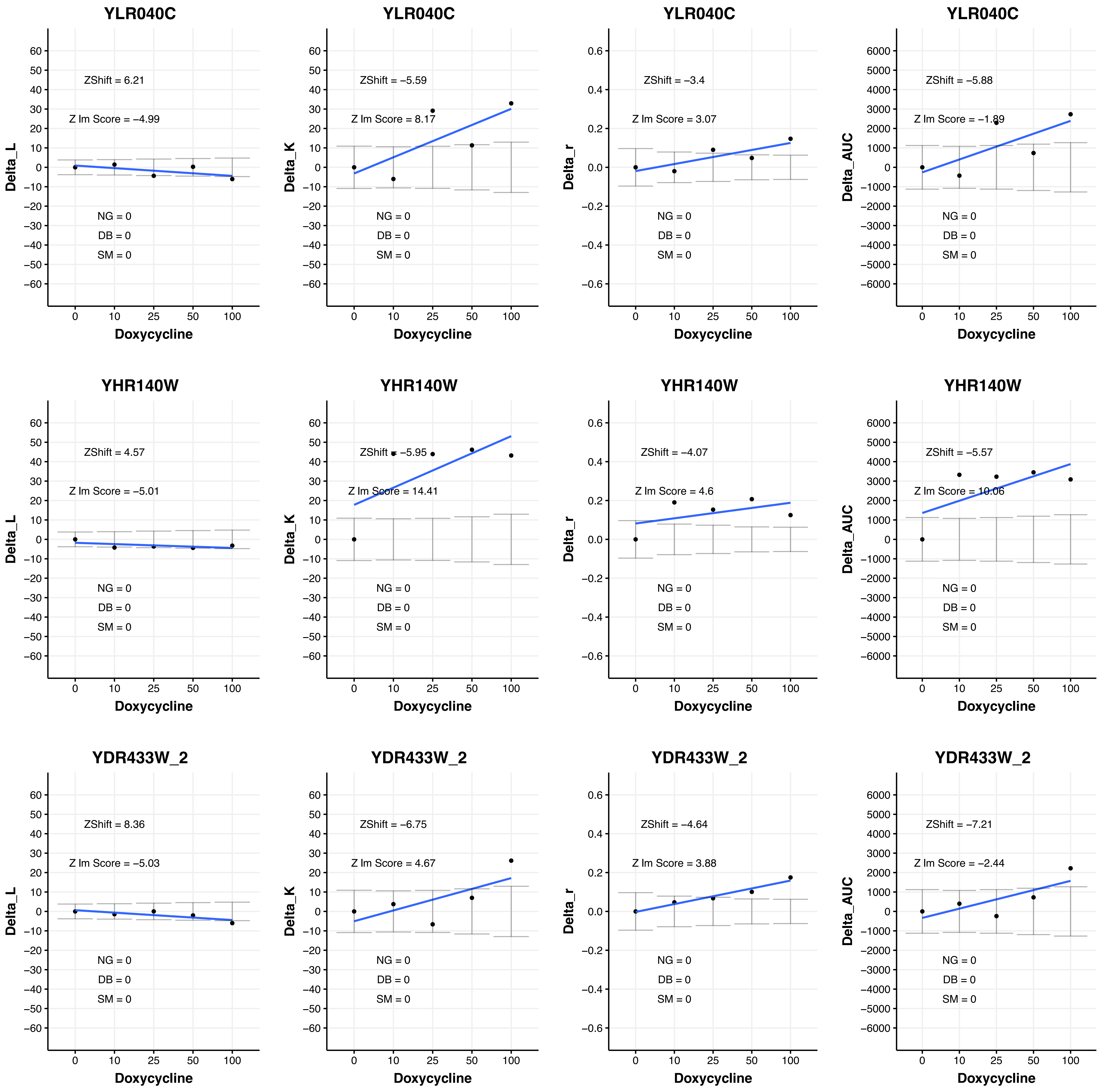

YLR040C
ZShift = 6.21
Z lm Score = −4.99
NG = 0
DB = 0
SM = 0
Delta_L
Doxycycline
YLR040C
ZShift = −5.59
Z lm Score = 8.17
NG = 0
DB = 0
SM = 0
Delta_K
Doxycycline
YLR040C
ZShift = −3.4
Z lm Score = 3.07
NG = 0
DB = 0
SM = 0
Delta_r
Doxycycline
YLR040C
ZShift = −5.88
Z lm Score = −1.89
NG = 0
DB = 0
SM = 0
Delta_AUC
Doxycycline
YHR140W
ZShift = 4.57
Z lm Score = −5.01
NG = 0
DB = 0
SM = 0
Delta_L
Doxycycline
YHR140W
ZShift = −5.95
Z lm Score = 14.41
NG = 0
DB = 0
SM = 0
Delta_K
Doxycycline
YHR140W
ZShift = −4.07
Z lm Score = 4.6
NG = 0
DB = 0
SM = 0
Delta_r
Doxycycline
YHR140W
ZShift = −5.57
Z lm Score = 10.06
NG = 0
DB = 0
SM = 0
Delta_AUC
Doxycycline
YDR433W_2
ZShift = 8.36
Z lm Score = −5.03
NG = 0
DB = 0
SM = 0
Delta_L
Doxycycline
YDR433W_2
ZShift = −6.75
Z lm Score = 4.67
NG = 0
DB = 0
SM = 0
Delta_K
Doxycycline
YDR433W_2
ZShift = −4.64
Z lm Score = 3.88
NG = 0
DB = 0
SM = 0
Delta_r
Doxycycline
YDR433W_2
ZShift = −7.21
Z lm Score = −2.44
NG = 0
DB = 0
SM = 0
Delta_AUC
Doxycycline

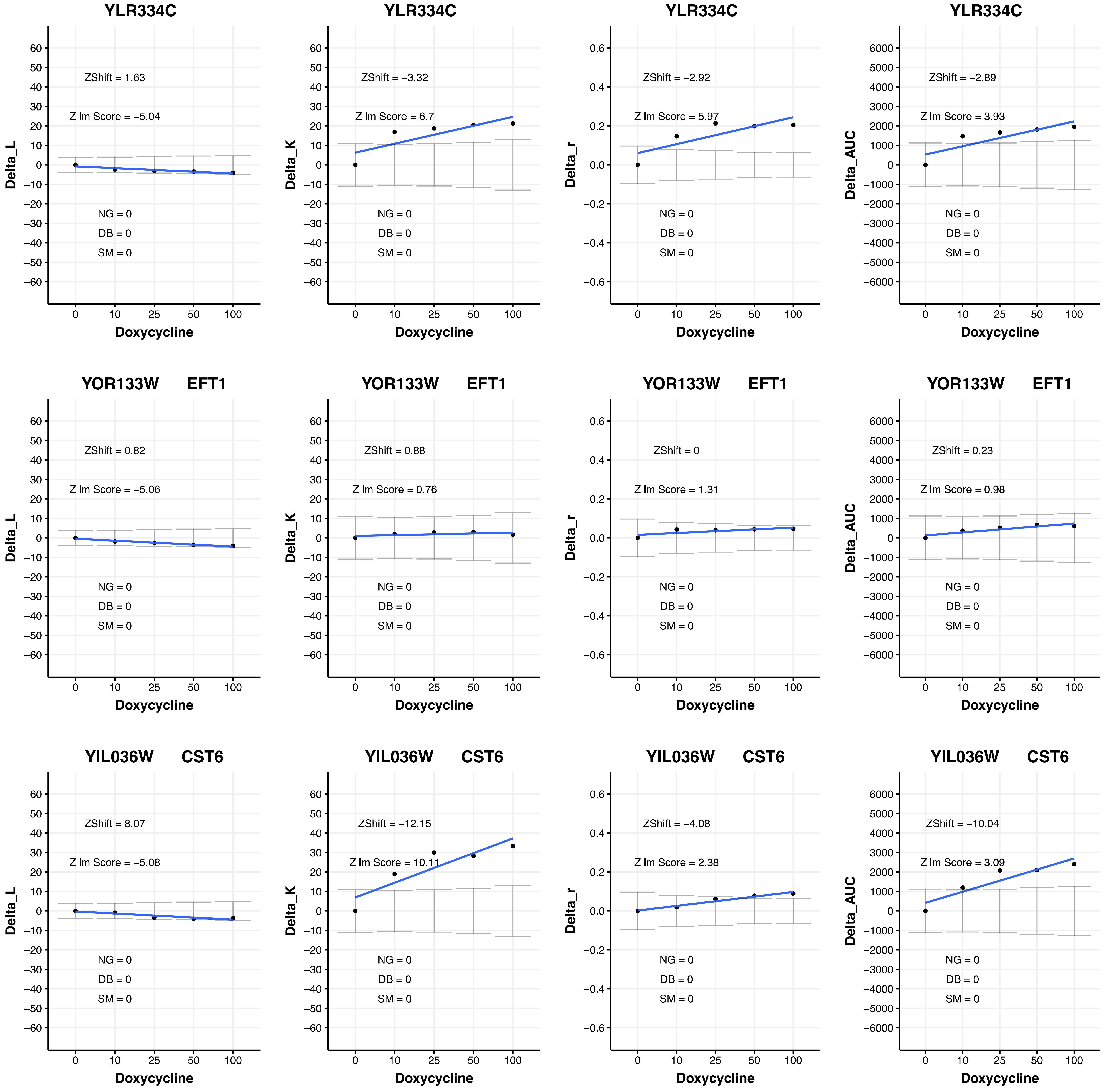
YLR334C
ZShift = 1.63
Z lm Score = −5.04
NG = 0
DB = 0
SM = 0
Delta_L
Doxycycline
YLR334C
ZShift = −3.32
Z lm Score = 6.7
NG = 0
DB = 0
SM = 0
Delta_K
Doxycycline
YLR334C
ZShift = −2.92
Z lm Score = 5.97
NG = 0
DB = 0
SM = 0
Delta_r
Doxycycline
YLR334C
ZShift = −2.89
Z lm Score = 3.93
NG = 0
DB = 0
SM = 0
Delta_AUC
Doxycycline
YOR133W EFT1
ZShift = 0.82
Z lm Score = −5.06
NG = 0
DB = 0
SM = 0
Delta_L
Doxycycline
YOR133W EFT1
ZShift = 0.88
Z lm Score = 0.76
NG = 0
DB = 0
SM = 0
Delta_K
Doxycycline
YOR133W EFT1
ZShift = 0
Z lm Score = 1.31
NG = 0
DB = 0
SM = 0
Delta_r
Doxycycline
YOR133W EFT1
ZShift = 0.23
Z lm Score = 0.98
NG = 0
DB = 0
SM = 0
Delta_AUC
Doxycycline
YIL036W CST6
ZShift = 8.07
Z lm Score = −5.08
NG = 0
DB = 0
SM = 0
Delta_L
Doxycycline
YIL036W CST6
ZShift = −12.15
Z lm Score = 10.11
NG = 0
DB = 0
SM = 0
Delta_K
Doxycycline
YIL036W CST6
ZShift = −4.08
Z lm Score = 2.38
NG = 0
DB = 0
SM = 0
Delta_r
Doxycycline
YIL036W CST6
ZShift = −10.04
Z lm Score = 3.09
NG = 0
DB = 0
SM = 0
Delta_AUC
Doxycycline

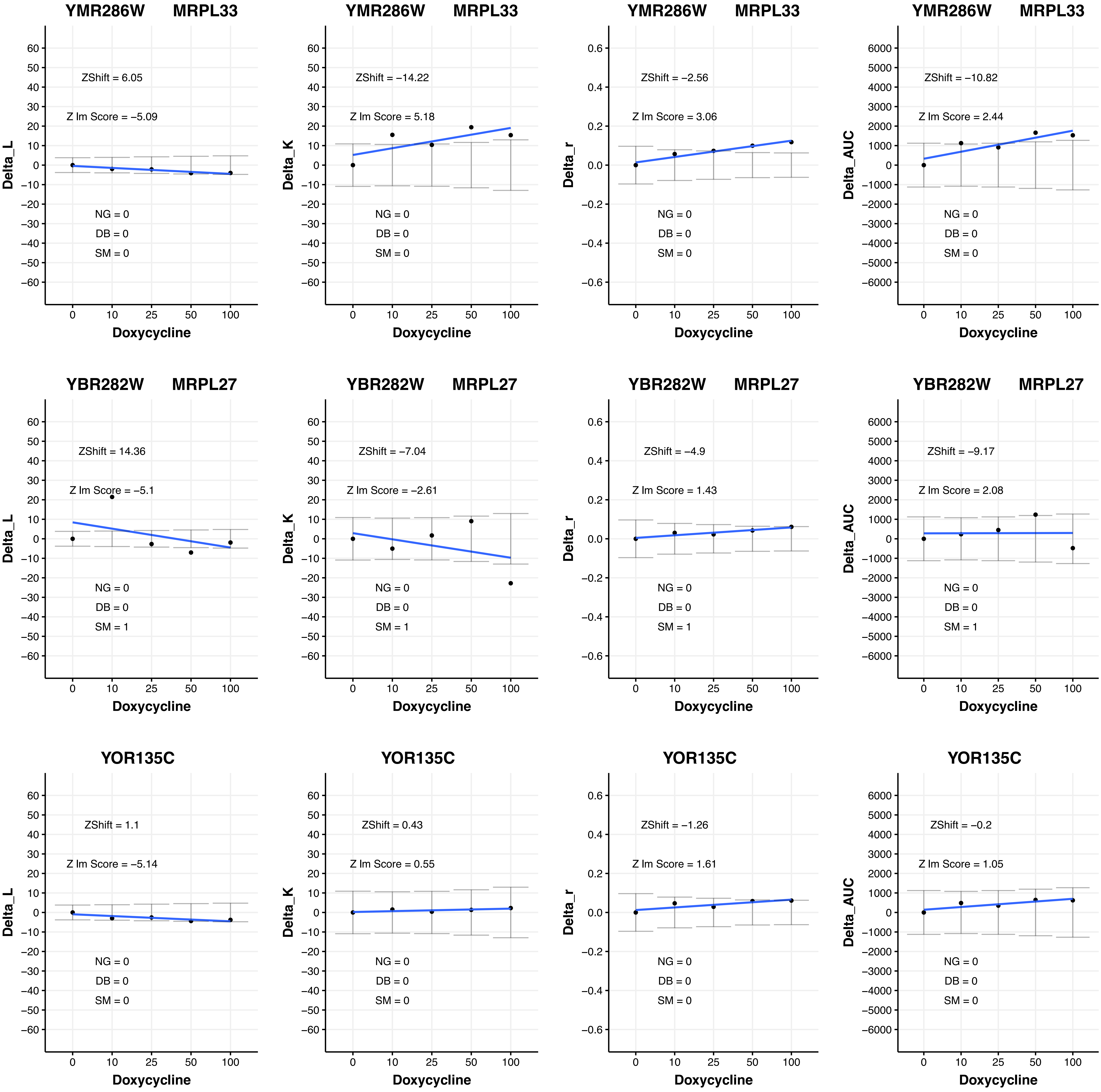

YMR286W MRPL33
ZShift = 6.05
Z lm Score = −5.09
Delta_L
NG = 0
DB = 0
SM = 0
Doxycycline
YMR286W MRPL33
ZShift = −14.22
Z lm Score = 5.18
Delta_K
NG = 0
DB = 0
SM = 0
Doxycycline
YMR286W MRPL33
ZShift = −2.56
Z lm Score = 3.06
Delta_r
NG = 0
DB = 0
SM = 0
Doxycycline
YMR286W MRPL33
ZShift = −10.82
Z lm Score = 2.44
Delta_AUC
NG = 0
DB = 0
SM = 0
Doxycycline
YBR282W MRPL27
ZShift = 14.36
Z lm Score = −5.1
Delta_L
NG = 0
DB = 0
SM = 1
Doxycycline
YBR282W MRPL27
ZShift = −7.04
Z lm Score = −2.61
Delta_K
NG = 0
DB = 0
SM = 1
Doxycycline
YBR282W MRPL27
ZShift = −4.9
Z lm Score = 1.43
Delta_r
NG = 0
DB = 0
SM = 1
Doxycycline
YBR282W MRPL27
ZShift = −9.17
Z lm Score = 2.08
Delta_AUC
NG = 0
DB = 0
SM = 1
Doxycycline
YOR135C
ZShift = 1.1
Z lm Score = −5.14
Delta_L
NG = 0
DB = 0
SM = 0
Doxycycline
YOR135C
ZShift = 0.43
Z lm Score = 0.55
Delta_K
NG = 0
DB = 0
SM = 0
Doxycycline
YOR135C
ZShift = −1.26
Z lm Score = 1.61
Delta_r
NG = 0
DB = 0
SM = 0
Doxycycline
YOR135C
ZShift = −0.2
Z lm Score = 1.05
Delta_AUC
NG = 0
DB = 0
SM = 0
Doxycycline

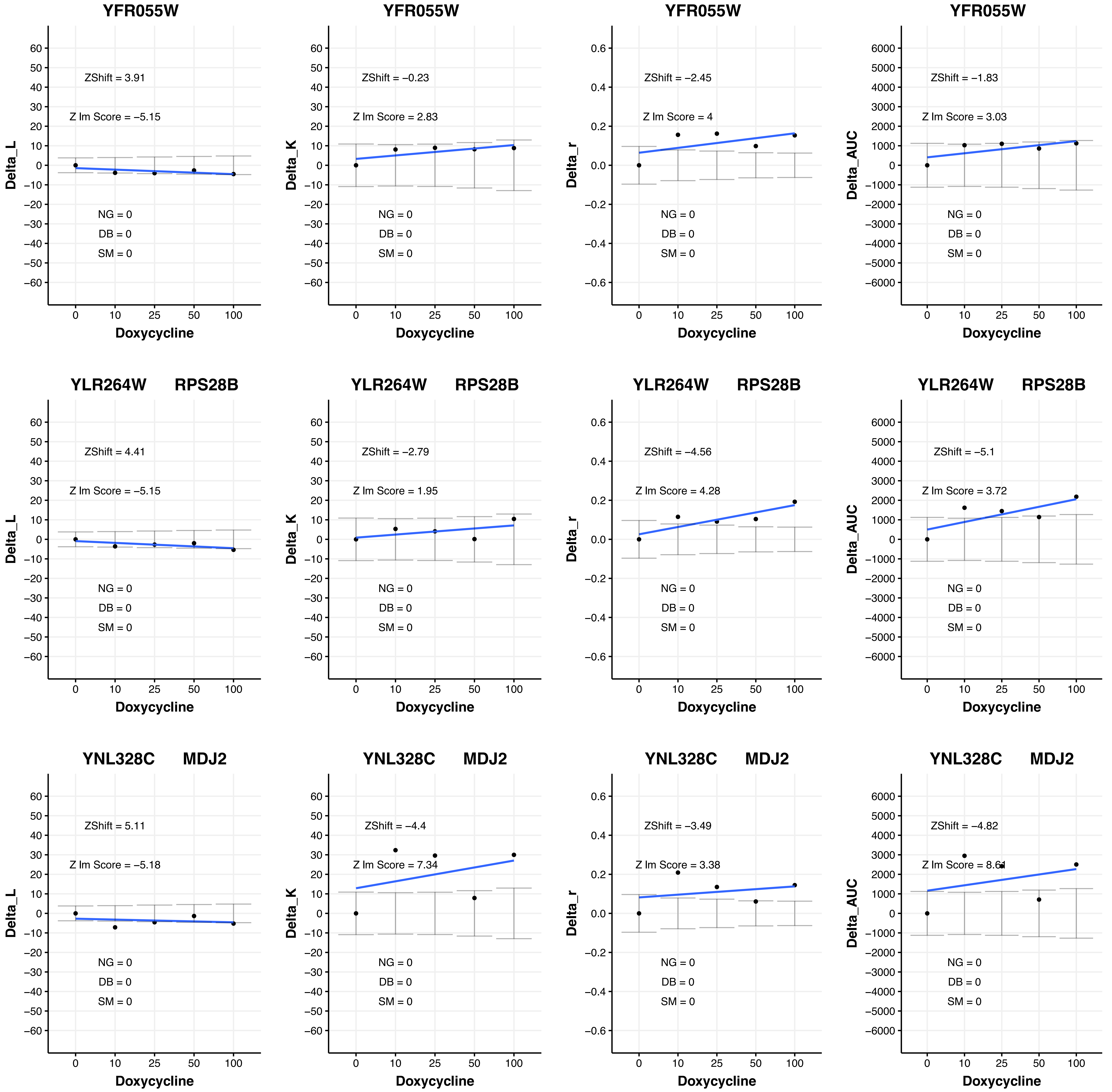
YFR055W
ZShift = 3.91
Z lm Score = −5.15
NG = 0
DB = 0
SM = 0
Delta_L
Doxycycline
YFR055W
ZShift = −0.23
Z lm Score = 2.83
NG = 0
DB = 0
SM = 0
Delta_K
Doxycycline
YFR055W
ZShift = −2.45
Z lm Score = 4
NG = 0
DB = 0
SM = 0
Delta_r
Doxycycline
YFR055W
ZShift = −1.83
Z lm Score = 3.03
NG = 0
DB = 0
SM = 0
Delta_AUC
Doxycycline
YLR264W RPS28B
ZShift = 4.41
Z lm Score = −5.15
NG = 0
DB = 0
SM = 0
Delta_L
Doxycycline
YLR264W RPS28B
ZShift = −2.79
Z lm Score = 1.95
NG = 0
DB = 0
SM = 0
Delta_K
Doxycycline
YLR264W RPS28B
ZShift = −4.56
Z lm Score = 4.28
NG = 0
DB = 0
SM = 0
Delta_r
Doxycycline
YLR264W RPS28B
ZShift = −5.1
Z lm Score = 3.72
NG = 0
DB = 0
SM = 0
Delta_AUC
Doxycycline
YNL328C MDJ2
ZShift = 5.11
Z lm Score = −5.18
NG = 0
DB = 0
SM = 0
Delta_L
Doxycycline
YNL328C MDJ2
ZShift = −4.4
Z lm Score = 7.34
NG = 0
DB = 0
SM = 0
Delta_K
Doxycycline
YNL328C MDJ2
ZShift = −3.49
Z lm Score = 3.38
NG = 0
DB = 0
SM = 0
Delta_r
Doxycycline
YNL328C MDJ2
ZShift = −4.82
Z lm Score = 8.61
NG = 0
DB = 0
SM = 0
Delta_AUC
Doxycycline

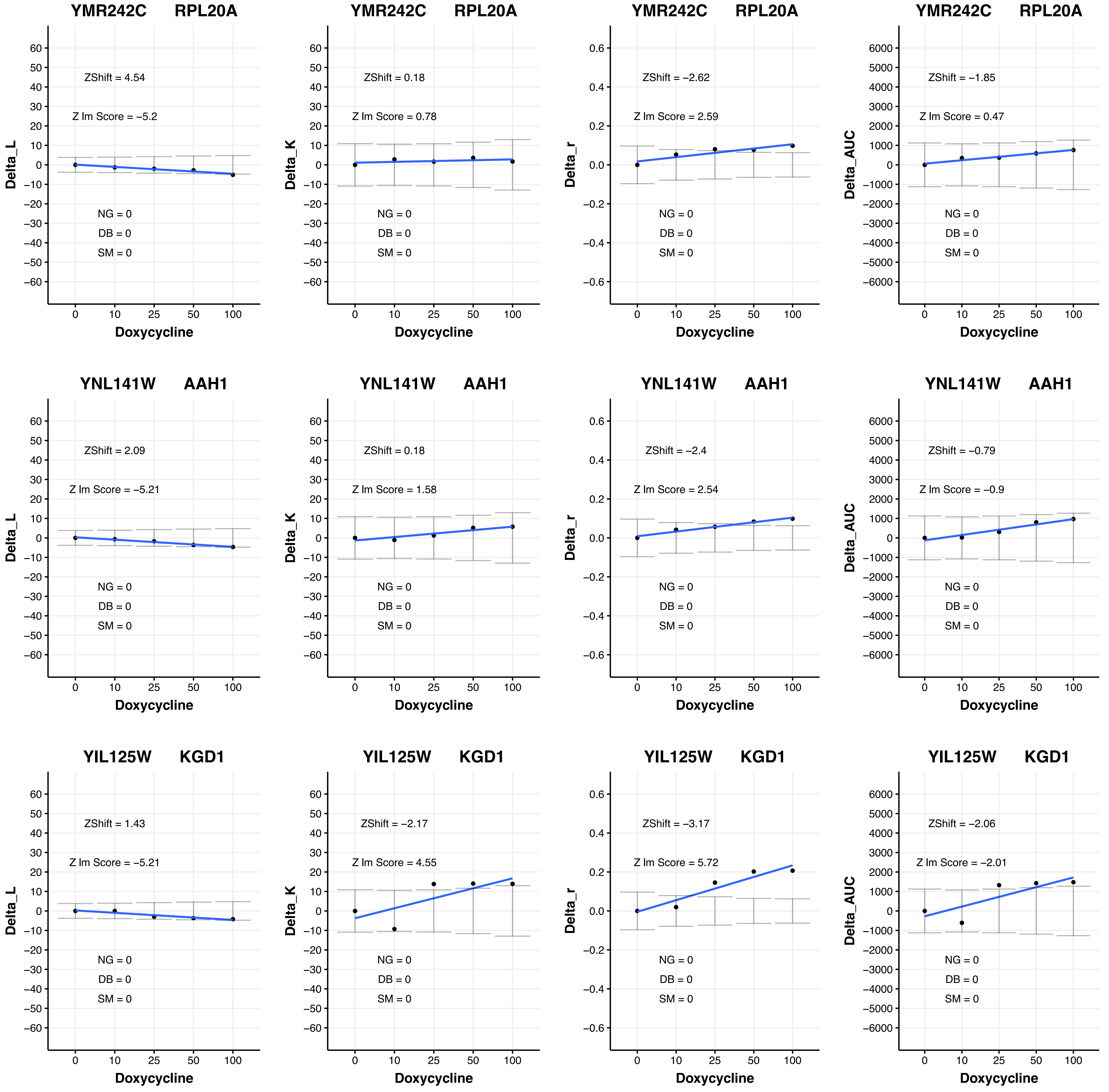
YMR242C RPL20A
ZShift = 4.54
Z lm Score = −5.2
NG = 0
DB = 0
SM = 0
Delta_L
Doxycycline
YMR242C RPL20A
ZShift = 0.18
Z lm Score = 0.78
NG = 0
DB = 0
SM = 0
Delta_K
Doxycycline
YMR242C RPL20A
ZShift = −2.62
Z lm Score = 2.59
NG = 0
DB = 0
SM = 0
Delta_r
Doxycycline
YMR242C RPL20A
ZShift = −1.85
Z lm Score = 0.47
NG = 0
DB = 0
SM = 0
Delta_AUC
Doxycycline
YNL141W AAH1
ZShift = 2.09
Z lm Score = −5.21
NG = 0
DB = 0
SM = 0
Delta_L
Doxycycline
YNL141W AAH1
ZShift = 0.18
Z lm Score = 1.58
NG = 0
DB = 0
SM = 0
Delta_K
Doxycycline
YNL141W AAH1
ZShift = −2.4
Z lm Score = 2.54
NG = 0
DB = 0
SM = 0
Delta_r
Doxycycline
YNL141W AAH1
ZShift = −0.79
Z lm Score = −0.9
NG = 0
DB = 0
SM = 0
Delta_AUC
Doxycycline
YIL125W KGD1
ZShift = 1.43
Z lm Score = −5.21
NG = 0
DB = 0
SM = 0
Delta_L
Doxycycline
YIL125W KGD1
ZShift = −2.17
Z lm Score = 4.55
NG = 0
DB = 0
SM = 0
Delta_K
Doxycycline
YIL125W KGD1
ZShift = −3.17
Z lm Score = 5.72
NG = 0
DB = 0
SM = 0
Delta_r
Doxycycline
YIL125W KGD1
ZShift = −2.06
Z lm Score = −2.01
NG = 0
DB = 0
SM = 0
Delta_AUC
Doxycycline

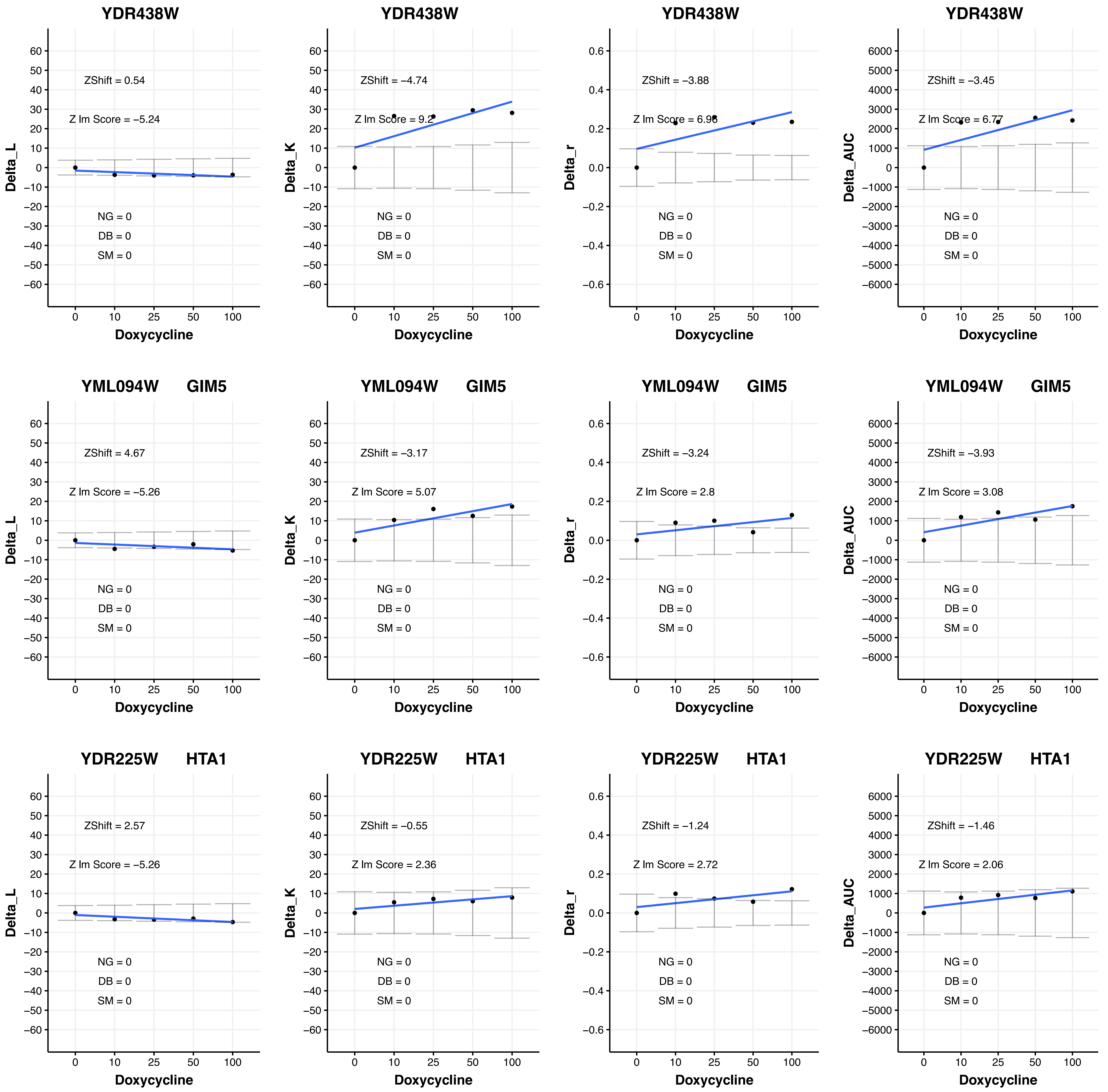

YDR438W
ZShift = 0.54
Z lm Score = −5.24
NG = 0
DB = 0
SM = 0
Delta_L
Doxycycline
YDR438W
ZShift = −4.74
Z lm Score = 9.2
NG = 0
DB = 0
SM = 0
Delta_K
Doxycycline
YDR438W
ZShift = −3.88
Z lm Score = 6.96
NG = 0
DB = 0
SM = 0
Delta_r
Doxycycline
YDR438W
ZShift = −3.45
Z lm Score = 6.77
NG = 0
DB = 0
SM = 0
Delta_AUC
Doxycycline
YML094W GIM5
ZShift = 4.67
Z lm Score = −5.26
NG = 0
DB = 0
SM = 0
Delta_L
Doxycycline
YML094W GIM5
ZShift = −3.17
Z lm Score = 5.07
NG = 0
DB = 0
SM = 0
Delta_K
Doxycycline
YML094W GIM5
ZShift = −3.24
Z lm Score = 2.8
NG = 0
DB = 0
SM = 0
Delta_r
Doxycycline
YML094W GIM5
ZShift = −3.93
Z lm Score = 3.08
NG = 0
DB = 0
SM = 0
Delta_AUC
Doxycycline
YDR225W HTA1
ZShift = 2.57
Z lm Score = −5.26
NG = 0
DB = 0
SM = 0
Delta_L
Doxycycline
YDR225W HTA1
ZShift = −0.55
Z lm Score = 2.36
NG = 0
DB = 0
SM = 0
Delta_K
Doxycycline
YDR225W HTA1
ZShift = −1.24
Z lm Score = 2.72
NG = 0
DB = 0
SM = 0
Delta_r
Doxycycline
YDR225W HTA1
ZShift = −1.46
Z lm Score = 2.06
NG = 0
DB = 0
SM = 0
Delta_AUC
Doxycycline

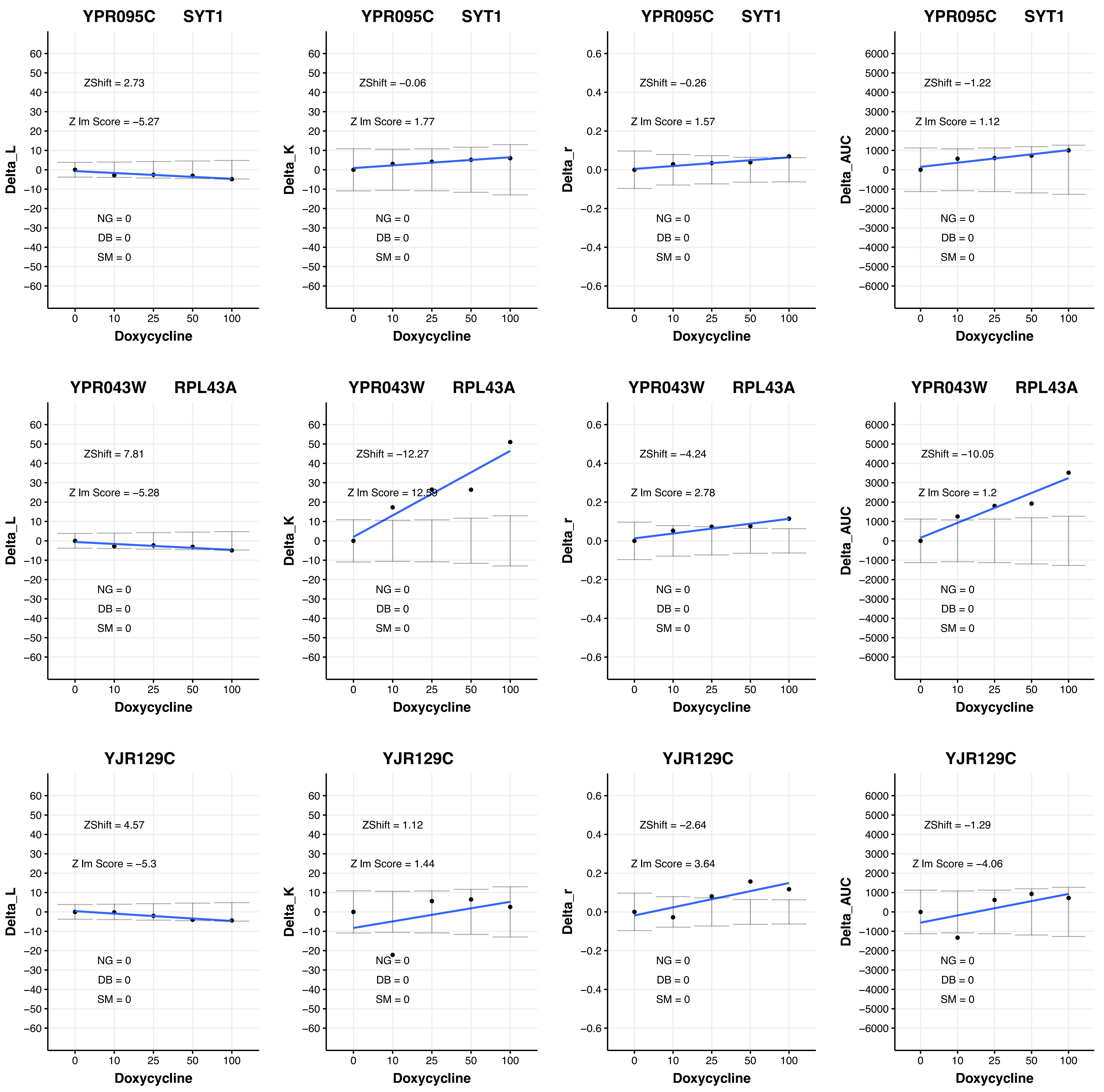

YPR095C SYT1
ZShift = 2.73
Z lm Score = −5.27
NG = 0
DB = 0
SM = 0
Delta_L
Doxycycline
YPR095C SYT1
ZShift = −0.06
Z lm Score = 1.77
NG = 0
DB = 0
SM = 0
Delta_K
Doxycycline
YPR095C SYT1
ZShift = −0.26
Z lm Score = 1.57
NG = 0
DB = 0
SM = 0
Delta_r
Doxycycline
YPR095C SYT1
ZShift = −1.22
Z lm Score = 1.12
NG = 0
DB = 0
SM = 0
Delta_AUC
Doxycycline
YPR043W RPL43A
ZShift = 7.81
Z lm Score = −5.28
NG = 0
DB = 0
SM = 0
Delta_L
Doxycycline
YPR043W RPL43A
ZShift = −12.27
Z lm Score = 12.59
NG = 0
DB = 0
SM = 0
Delta_K
Doxycycline
YPR043W RPL43A
ZShift = −4.24
Z lm Score = 2.78
NG = 0
DB = 0
SM = 0
Delta_r
Doxycycline
YPR043W RPL43A
ZShift = −10.05
Z lm Score = 1.2
NG = 0
DB = 0
SM = 0
Delta_AUC
Doxycycline
YJR129C
ZShift = 4.57
Z lm Score = −5.3
NG = 0
DB = 0
SM = 0
Delta_L
Doxycycline
YJR129C
ZShift = 1.12
Z lm Score = 1.44
NG = 0
DB = 0
SM = 0
Delta_K
Doxycycline
YJR129C
ZShift = −2.64
Z lm Score = 3.64
NG = 0
DB = 0
SM = 0
Delta_r
Doxycycline
YJR129C
ZShift = −1.29
Z lm Score = −4.06
NG = 0
DB = 0
SM = 0
Delta_AUC
Doxycycline

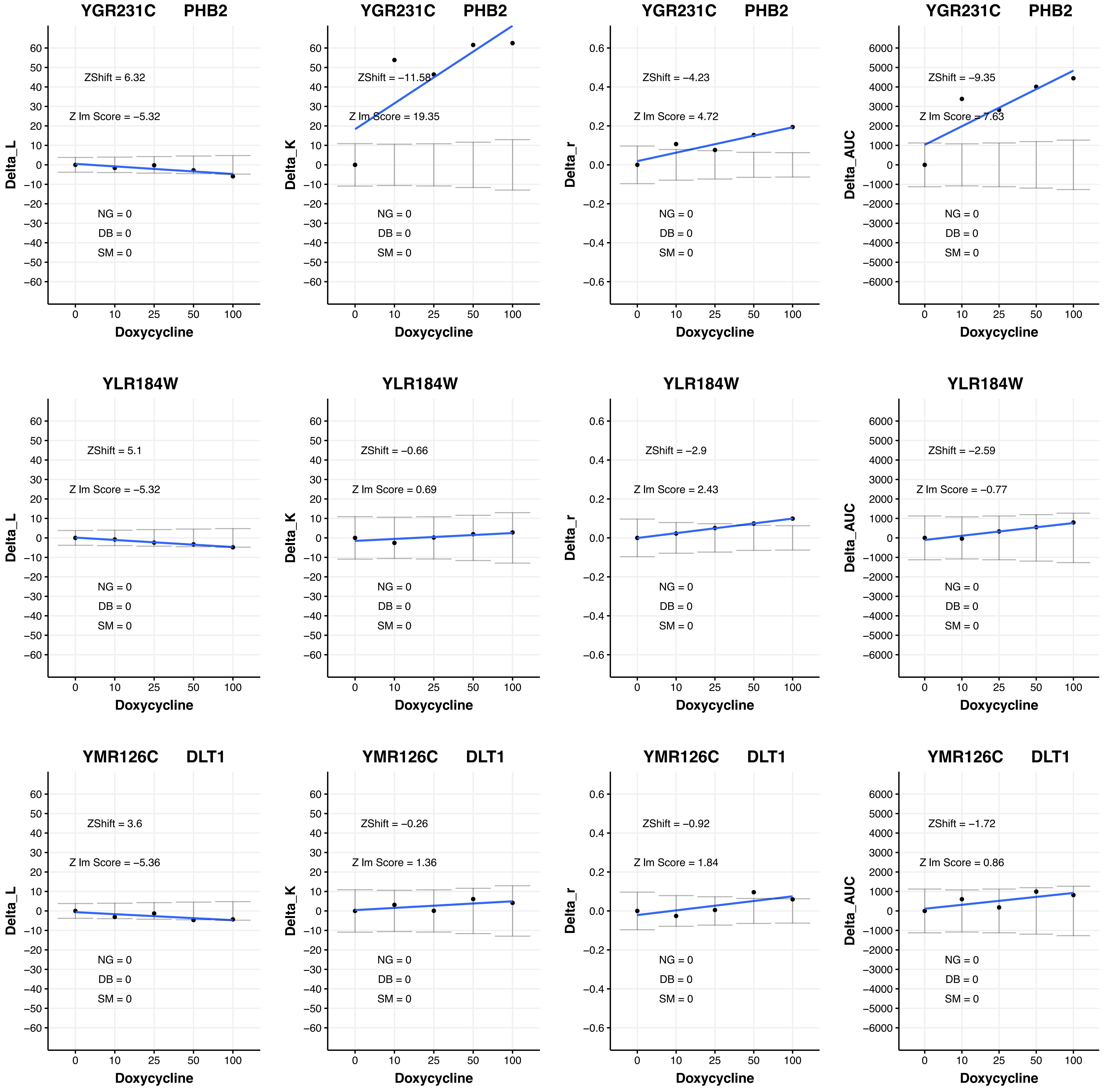

YGR231C PHB2
ZShift = 6.32
Z lm Score = −5.32
NG = 0
DB = 0
SM = 0
Delta_L
Doxycycline
YGR231C PHB2
ZShift = −11.58
Z lm Score = 19.35
NG = 0
DB = 0
SM = 0
Delta_K
Doxycycline
YGR231C PHB2
ZShift = −4.23
Z lm Score = 4.72
NG = 0
DB = 0
SM = 0
Delta_r
Doxycycline
YGR231C PHB2
ZShift = −9.35
Z lm Score = 7.63
NG = 0
DB = 0
SM = 0
Delta_AUC
Doxycycline
YLR184W
ZShift = 5.1
Z lm Score = −5.32
NG = 0
DB = 0
SM = 0
Delta_L
Doxycycline
YLR184W
ZShift = −0.66
Z lm Score = 0.69
NG = 0
DB = 0
SM = 0
Delta_K
Doxycycline
YLR184W
ZShift = −2.9
Z lm Score = 2.43
NG = 0
DB = 0
SM = 0
Delta_r
Doxycycline
YLR184W
ZShift = −2.59
Z lm Score = −0.77
NG = 0
DB = 0
SM = 0
Delta_AUC
Doxycycline
YMR126C DLT1
ZShift = 3.6
Z lm Score = −5.36
NG = 0
DB = 0
SM = 0
Delta_L
Doxycycline
YMR126C DLT1
ZShift = −0.26
Z lm Score = 1.36
NG = 0
DB = 0
SM = 0
Delta_K
Doxycycline
YMR126C DLT1
ZShift = −0.92
Z lm Score = 1.84
NG = 0
DB = 0
SM = 0
Delta_r
Doxycycline
YMR126C DLT1
ZShift = −1.72
Z lm Score = 0.86
NG = 0
DB = 0
SM = 0
Delta_AUC
Doxycycline

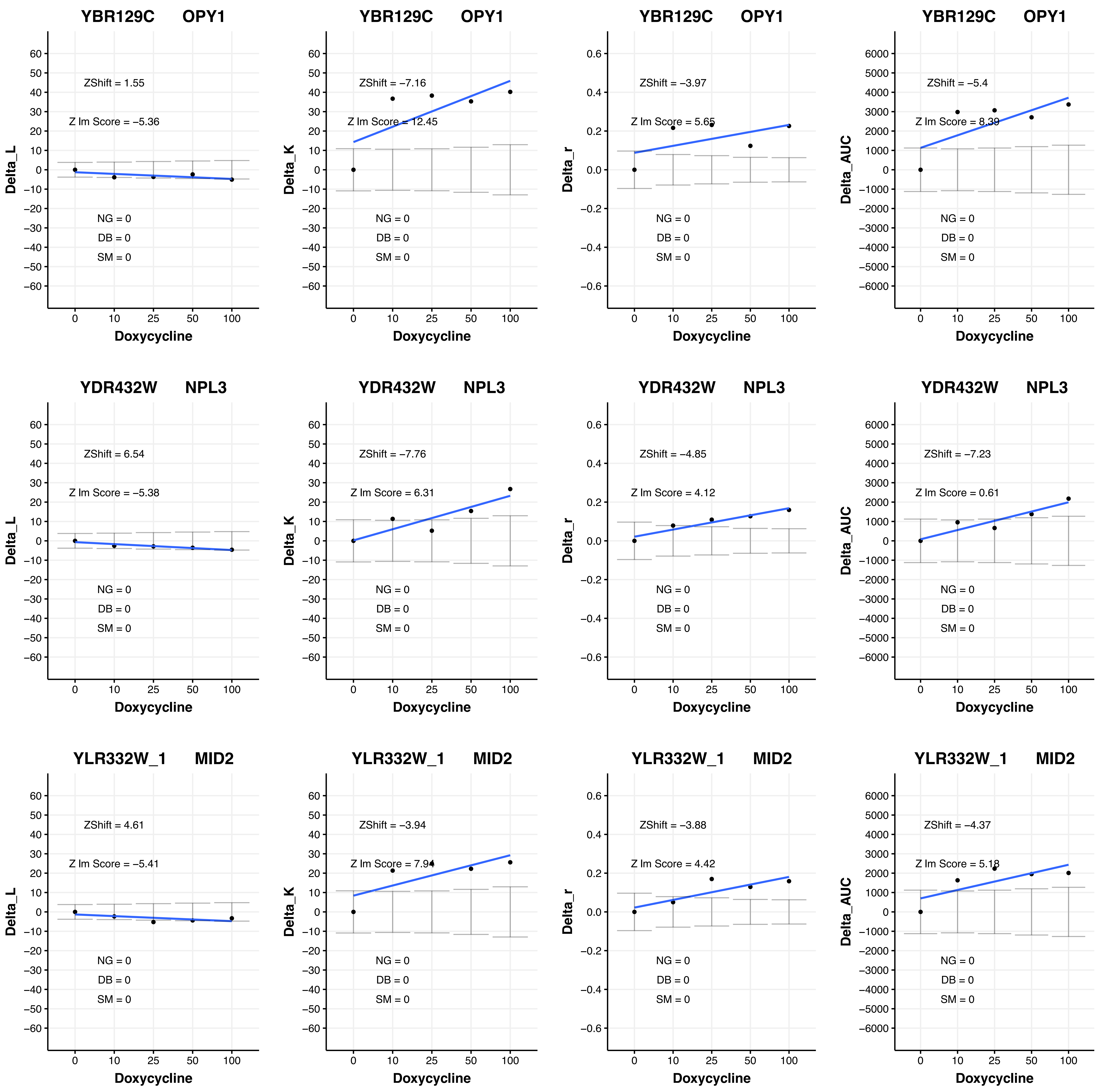

YBR129C OPY1
ZShift = 1.55
Z lm Score = −5.36
NG = 0
DB = 0
SM = 0
Delta_L
Doxycycline
YBR129C OPY1
ZShift = −7.16
Z lm Score = 12.45
NG = 0
DB = 0
SM = 0
Delta_K
Doxycycline
YBR129C OPY1
ZShift = −3.97
Z lm Score = 5.65
NG = 0
DB = 0
SM = 0
Delta_r
Doxycycline
YBR129C OPY1
ZShift = −5.4
Z lm Score = 8.39
NG = 0
DB = 0
SM = 0
Delta_AUC
Doxycycline
YDR432W NPL3
ZShift = 6.54
Z lm Score = −5.38
NG = 0
DB = 0
SM = 0
Delta_L
Doxycycline
YDR432W NPL3
ZShift = −7.76
Z lm Score = 6.31
NG = 0
DB = 0
SM = 0
Delta_K
Doxycycline
YDR432W NPL3
ZShift = −4.85
Z lm Score = 4.12
NG = 0
DB = 0
SM = 0
Delta_r
Doxycycline
YDR432W NPL3
ZShift = −7.23
Z lm Score = 0.61
NG = 0
DB = 0
SM = 0
Delta_AUC
Doxycycline
YLR332W_1 MID2
ZShift = 4.61
Z lm Score = −5.41
NG = 0
DB = 0
SM = 0
Delta_L
Doxycycline
YLR332W_1 MID2
ZShift = −3.94
Z lm Score = 7.94
NG = 0
DB = 0
SM = 0
Delta_K
Doxycycline
YLR332W_1 MID2
ZShift = −3.88
Z lm Score = 4.42
NG = 0
DB = 0
SM = 0
Delta_r
Doxycycline
YLR332W_1 MID2
ZShift = −4.37
Z lm Score = 5.18
NG = 0
DB = 0
SM = 0
Delta_AUC
Doxycycline

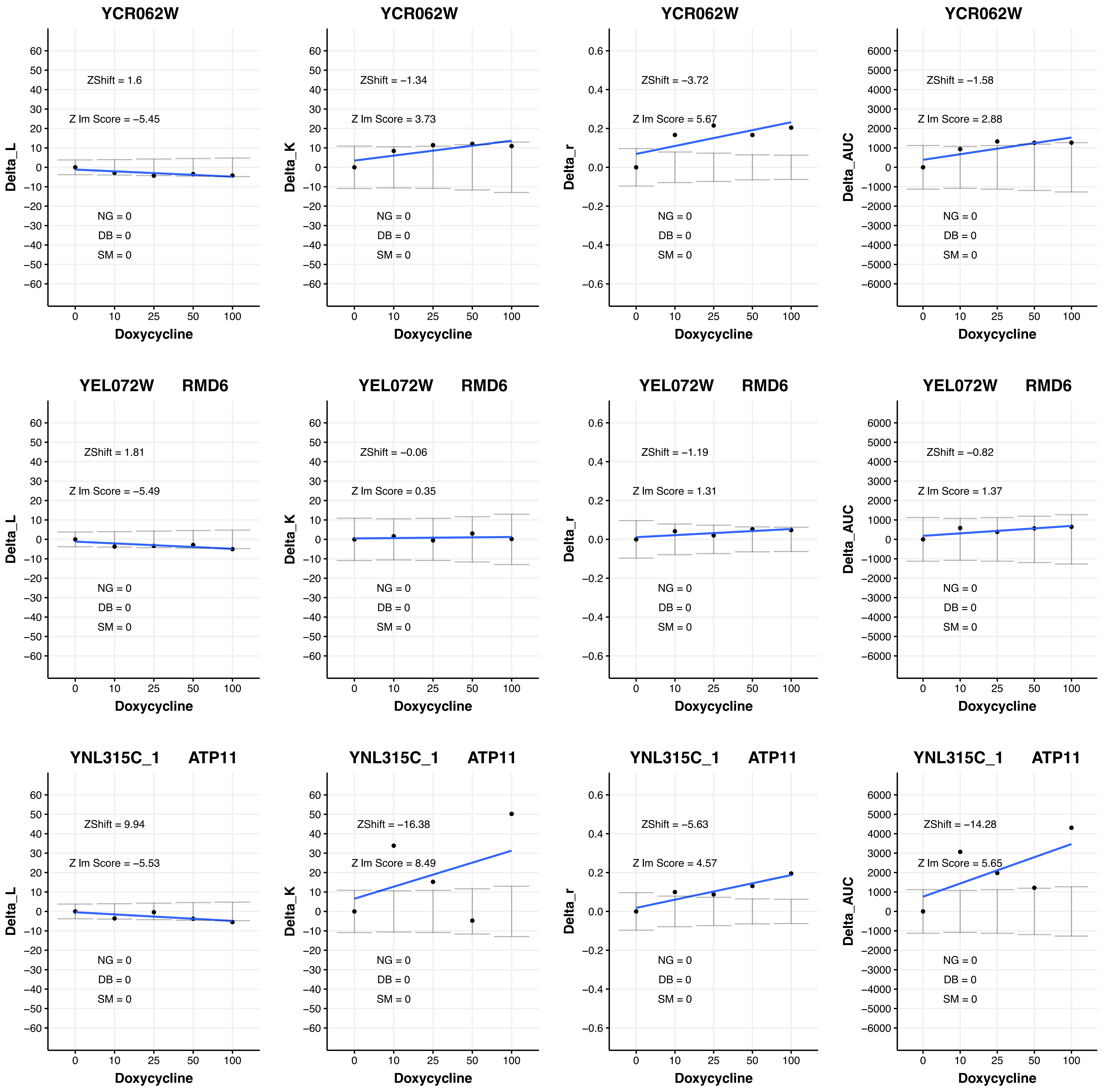
YCR062W
ZShift = 1.6
Z lm Score = −5.45
NG = 0
DB = 0
SM = 0
Delta_L
Doxycycline
YCR062W
ZShift = −1.34
Z lm Score = 3.73
NG = 0
DB = 0
SM = 0
Delta_K
Doxycycline
YCR062W
ZShift = −3.72
Z lm Score = 5.67
NG = 0
DB = 0
SM = 0
Delta_r
Doxycycline
YCR062W
ZShift = −1.58
Z lm Score = 2.88
NG = 0
DB = 0
SM = 0
Delta_AUC
Doxycycline
YEL072W RMD6
ZShift = 1.81
Z lm Score = −5.49
NG = 0
DB = 0
SM = 0
Delta_L
Doxycycline
YEL072W RMD6
ZShift = −0.06
Z lm Score = 0.35
NG = 0
DB = 0
SM = 0
Delta_K
Doxycycline
YEL072W RMD6
ZShift = −1.19
Z lm Score = 1.31
NG = 0
DB = 0
SM = 0
Delta_r
Doxycycline
YEL072W RMD6
ZShift = −0.82
Z lm Score = 1.37
NG = 0
DB = 0
SM = 0
Delta_AUC
Doxycycline
YNL315C_1 ATP11
ZShift = 9.94
Z lm Score = −5.53
NG = 0
DB = 0
SM = 0
Delta_L
Doxycycline
YNL315C_1 ATP11
ZShift = −16.38
Z lm Score = 8.49
NG = 0
DB = 0
SM = 0
Delta_K
Doxycycline
YNL315C_1 ATP11
ZShift = −5.63
Z lm Score = 4.57
NG = 0
DB = 0
SM = 0
Delta_r
Doxycycline
YNL315C_1 ATP11
ZShift = −14.28
Z lm Score = 5.65
NG = 0
DB = 0
SM = 0
Delta_AUC
Doxycycline

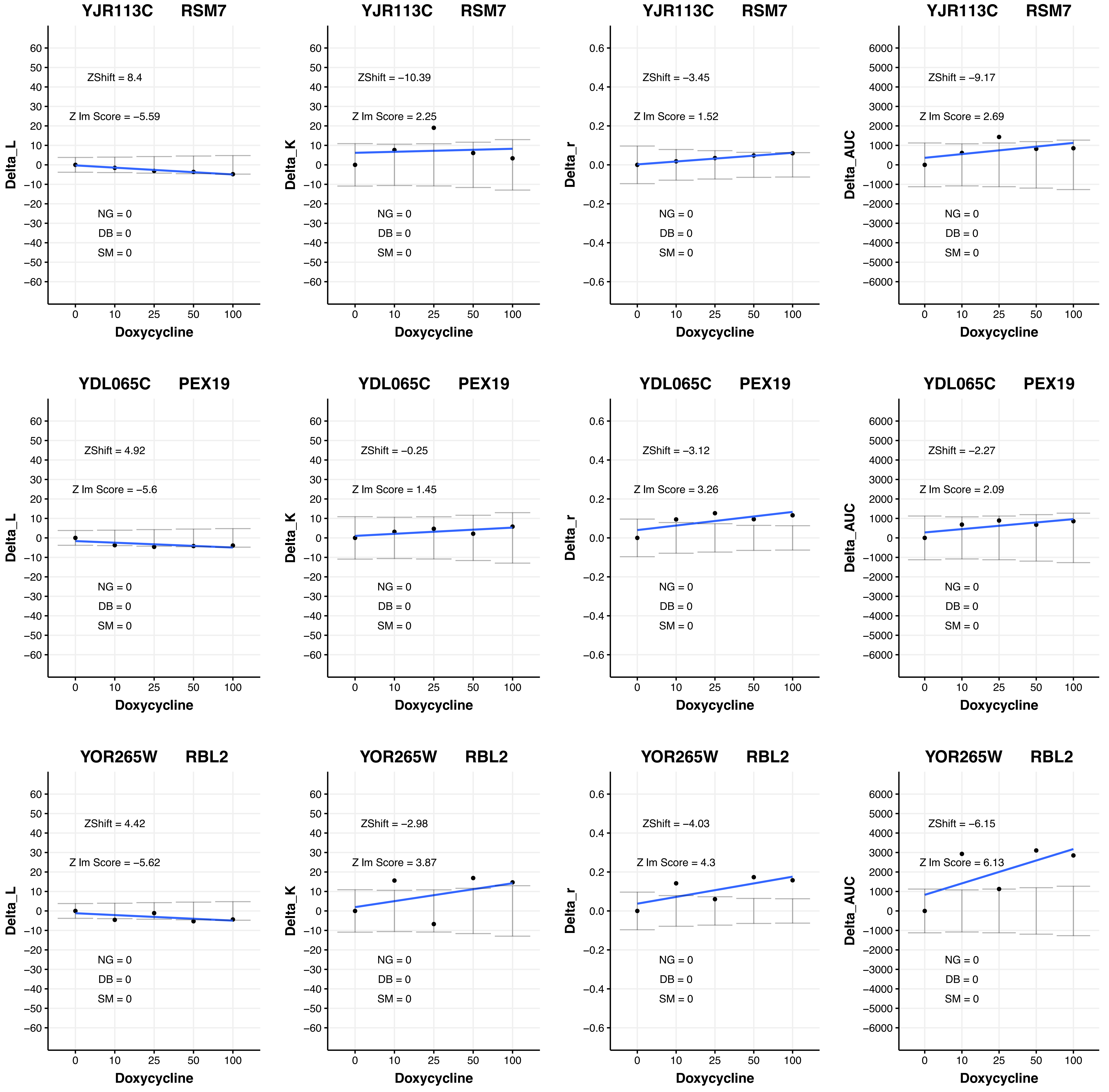

YJR113C RSM7
ZShift = 8.4
Z lm Score = −5.59
NG = 0
DB = 0
SM = 0
Delta_L
Doxycycline
YJR113C RSM7
ZShift = −10.39
Z lm Score = 2.25
NG = 0
DB = 0
SM = 0
Delta_K
Doxycycline
YJR113C RSM7
ZShift = −3.45
Z lm Score = 1.52
NG = 0
DB = 0
SM = 0
Delta_r
Doxycycline
YJR113C RSM7
ZShift = −9.17
Z lm Score = 2.69
NG = 0
DB = 0
SM = 0
Delta_AUC
Doxycycline
YDL065C PEX19
ZShift = 4.92
Z lm Score = −5.6
NG = 0
DB = 0
SM = 0
Delta_L
Doxycycline
YDL065C PEX19
ZShift = −0.25
Z lm Score = 1.45
NG = 0
DB = 0
SM = 0
Delta_K
Doxycycline
YDL065C PEX19
ZShift = −3.12
Z lm Score = 3.26
NG = 0
DB = 0
SM = 0
Delta_r
Doxycycline
YDL065C PEX19
ZShift = −2.27
Z lm Score = 2.09
NG = 0
DB = 0
SM = 0
Delta_AUC
Doxycycline
YOR265W RBL2
ZShift = 4.42
Z lm Score = −5.62
NG = 0
DB = 0
SM = 0
Delta_L
Doxycycline
YOR265W RBL2
ZShift = −2.98
Z lm Score = 3.87
NG = 0
DB = 0
SM = 0
Delta_K
Doxycycline
YOR265W RBL2
ZShift = −4.03
Z lm Score = 4.3
NG = 0
DB = 0
SM = 0
Delta_r
Doxycycline
YOR265W RBL2
ZShift = −6.15
Z lm Score = 6.13
NG = 0
DB = 0
SM = 0
Delta_AUC
Doxycycline

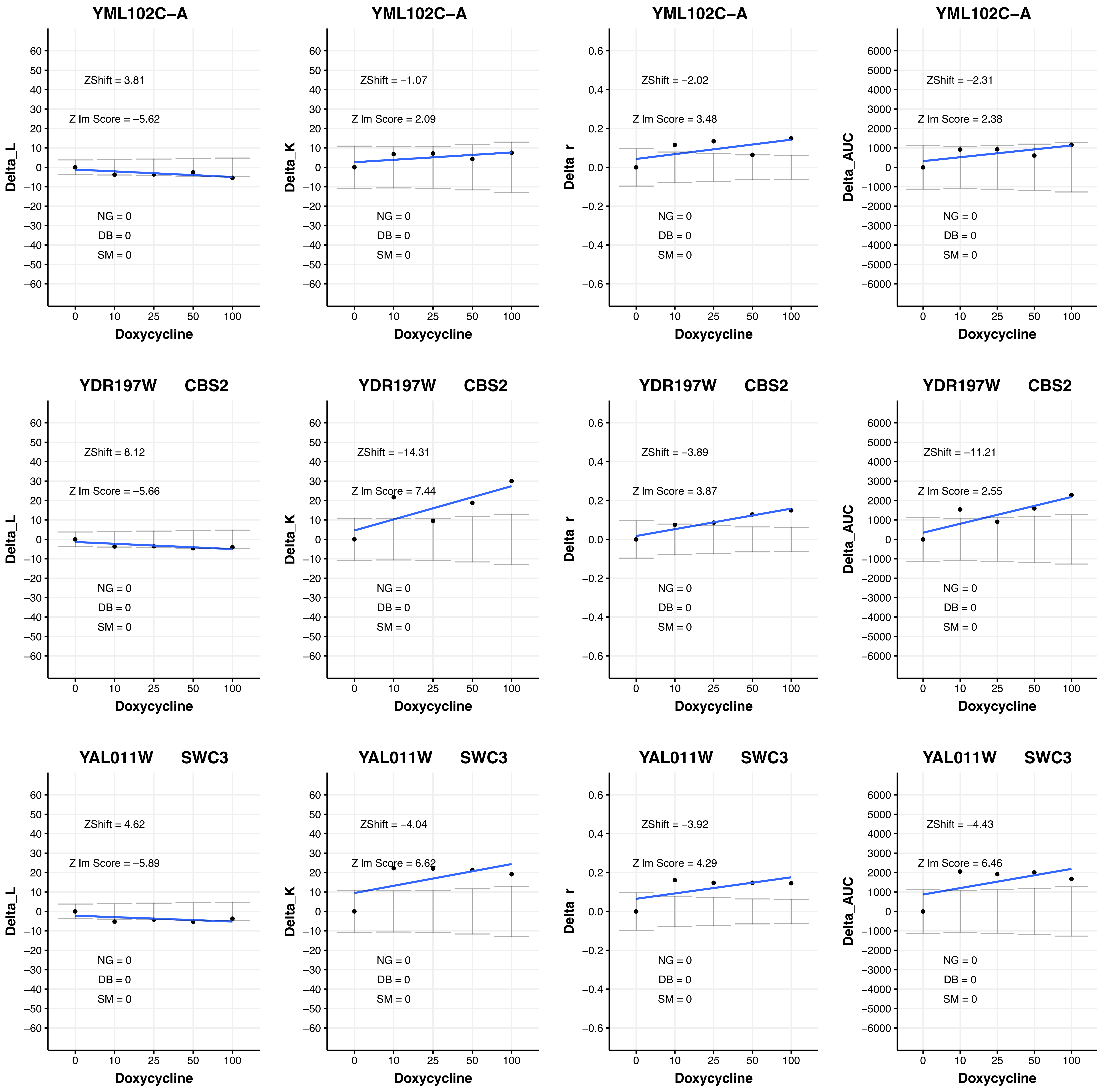
YML102C−A
ZShift = 3.81
Z lm Score = −5.62
NG = 0
DB = 0
SM = 0
Delta_L
Doxycycline
YML102C−A
ZShift = −1.07
Z lm Score = 2.09
Delta_K
YML102C−A
ZShift = −2.02
Z lm Score = 3.48
Delta_r
YML102C−A
ZShift = −2.31
Z lm Score = 2.38
Delta_AUC
YDR197W CBS2
ZShift = 8.12
Z lm Score = −5.66
YDR197W CBS2
ZShift = −14.31
Z lm Score = 7.44
YDR197W CBS2
ZShift = −3.89
Z lm Score = 3.87
YDR197W CBS2
ZShift = −11.21
Z lm Score = 2.55
YAL011W SWC3
ZShift = 4.62
Z lm Score = −5.89
YAL011W SWC3
ZShift = −4.04
Z lm Score = 6.62
YAL011W SWC3
ZShift = −3.92
Z lm Score = 4.29
YAL011W SWC3
ZShift = −4.43
Z lm Score = 6.46

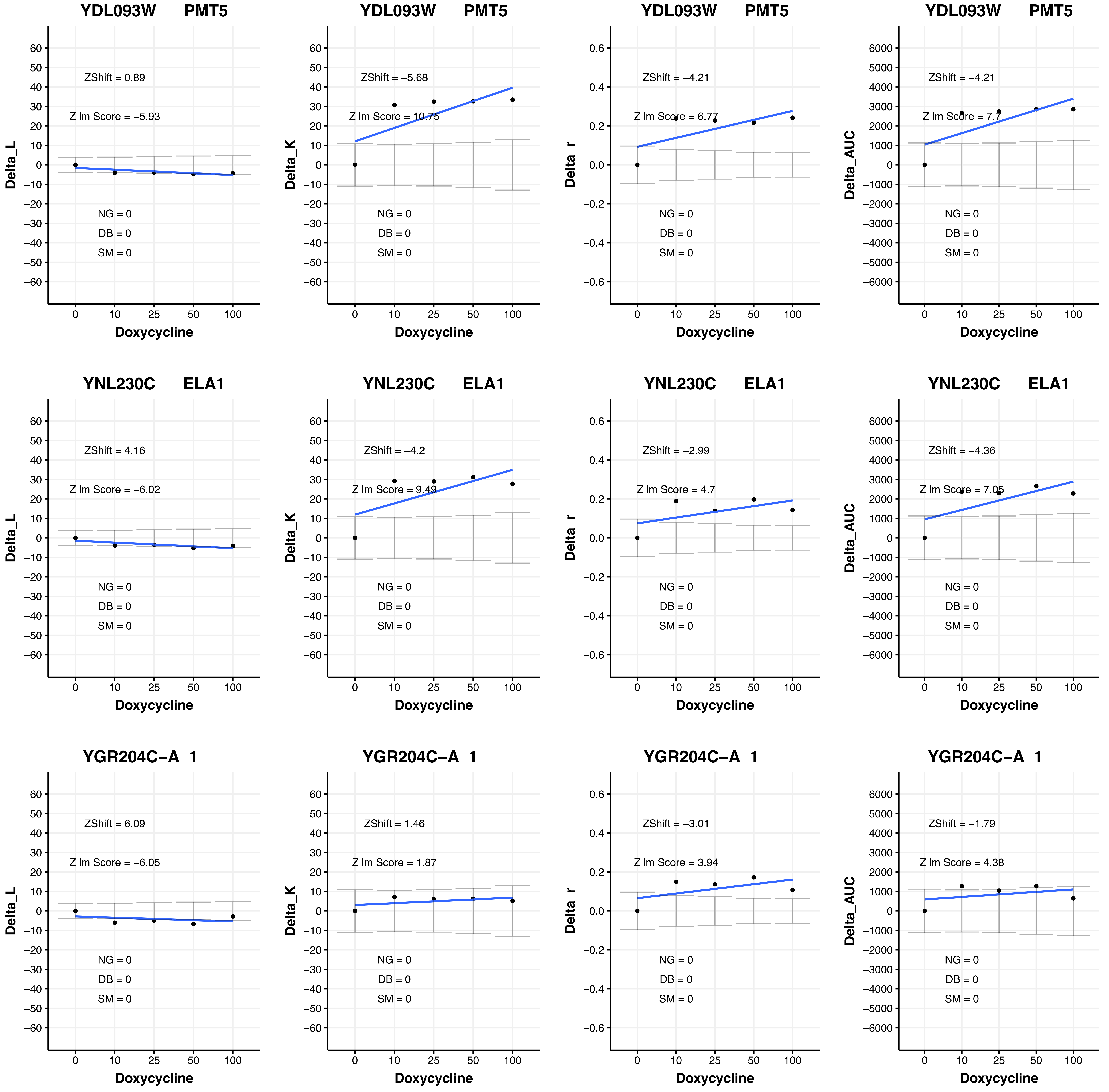

YDL093W PMT5
ZShift = 0.89
Z lm Score = −5.93
NG = 0
DB = 0
SM = 0
Delta_L
Doxycycline
YDL093W PMT5
ZShift = −5.68
Z lm Score = 10.75
NG = 0
DB = 0
SM = 0
Delta_K
Doxycycline
YDL093W PMT5
ZShift = −4.21
Z lm Score = 6.77
NG = 0
DB = 0
SM = 0
Delta_r
Doxycycline
YDL093W PMT5
ZShift = −4.21
Z lm Score = 7.7
NG = 0
DB = 0
SM = 0
Delta_AUC
Doxycycline
YNL230C ELA1
ZShift = 4.16
Z lm Score = −6.02
NG = 0
DB = 0
SM = 0
Delta_L
Doxycycline
YNL230C ELA1
ZShift = −4.2
Z lm Score = 9.49
NG = 0
DB = 0
SM = 0
Delta_K
Doxycycline
YNL230C ELA1
ZShift = −2.99
Z lm Score = 4.7
NG = 0
DB = 0
SM = 0
Delta_r
Doxycycline
YNL230C ELA1
ZShift = −4.36
Z lm Score = 7.05
NG = 0
DB = 0
SM = 0
Delta_AUC
Doxycycline
YGR204C−A_1
ZShift = 6.09
Z lm Score = −6.05
NG = 0
DB = 0
SM = 0
Delta_L
Doxycycline
YGR204C−A_1
ZShift = 1.46
Z lm Score = 1.87
NG = 0
DB = 0
SM = 0
Delta_K
Doxycycline
YGR204C−A_1
ZShift = −3.01
Z lm Score = 3.94
NG = 0
DB = 0
SM = 0
Delta_r
Doxycycline
YGR204C−A_1
ZShift = −1.79
Z lm Score = 4.38
NG = 0
DB = 0
SM = 0
Delta_AUC
Doxycycline

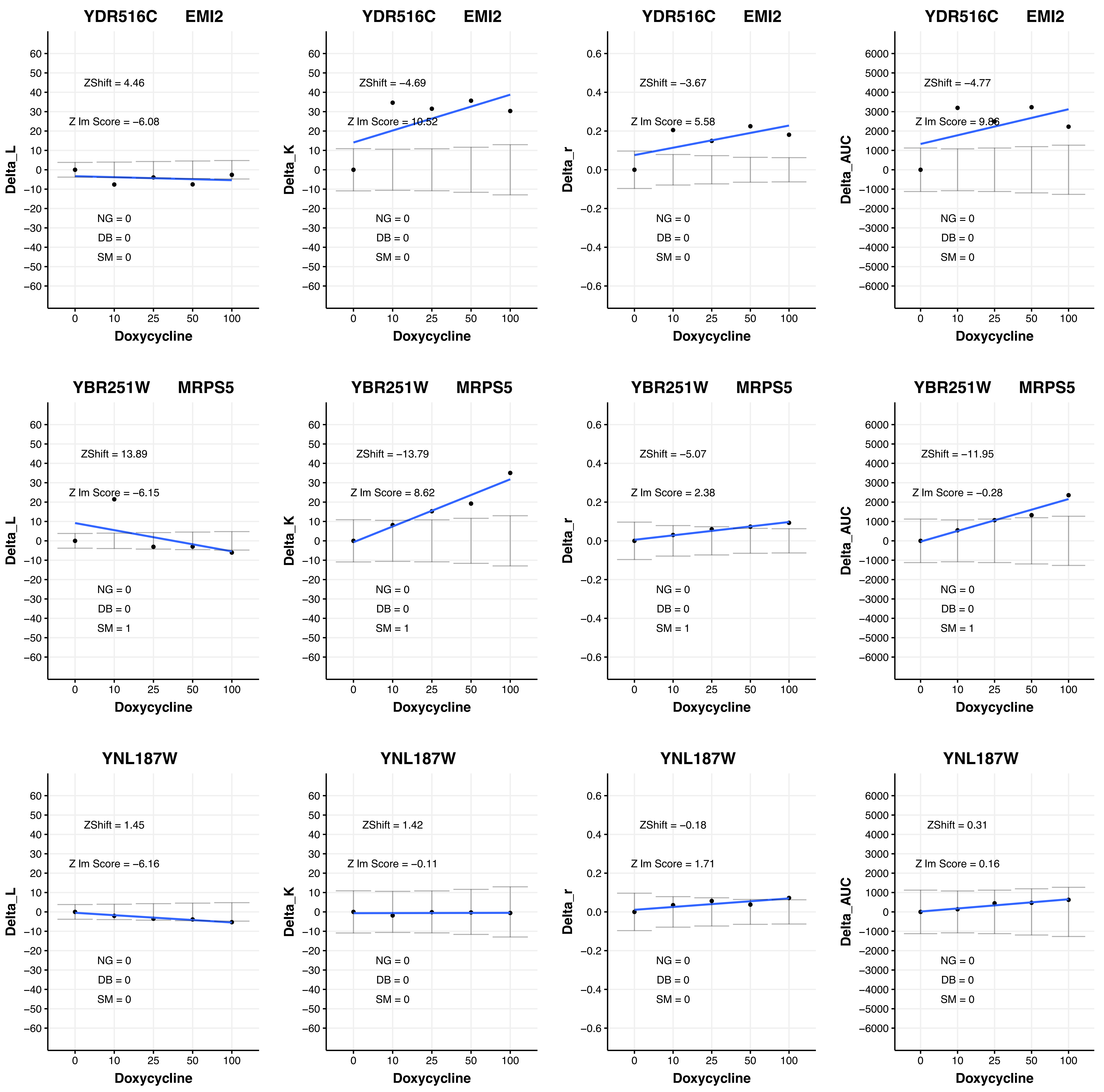

YDR516C EMI2
ZShift = 4.46
Z lm Score = −6.08
NG = 0
DB = 0
SM = 0
Delta_L
Doxycycline
YDR516C EMI2
ZShift = −4.69
Z lm Score = 10.52
NG = 0
DB = 0
SM = 0
Delta_K
Doxycycline
YDR516C EMI2
ZShift = −3.67
Z lm Score = 5.58
NG = 0
DB = 0
SM = 0
Delta_r
Doxycycline
YDR516C EMI2
ZShift = −4.77
Z lm Score = 9.86
NG = 0
DB = 0
SM = 0
Delta_AUC
Doxycycline
YBR251W MRPS5
ZShift = 13.89
Z lm Score = −6.15
NG = 0
DB = 0
SM = 1
Delta_L
Doxycycline
YBR251W MRPS5
ZShift = −13.79
Z lm Score = 8.62
NG = 0
DB = 0
SM = 1
Delta_K
Doxycycline
YBR251W MRPS5
ZShift = −5.07
Z lm Score = 2.38
NG = 0
DB = 0
SM = 1
Delta_r
Doxycycline
YBR251W MRPS5
ZShift = −11.95
Z lm Score = −0.28
NG = 0
DB = 0
SM = 1
Delta_AUC
Doxycycline
YNL187W
ZShift = 1.45
Z lm Score = −6.16
NG = 0
DB = 0
SM = 0
Delta_L
Doxycycline
YNL187W
ZShift = 1.42
Z lm Score = −0.11
NG = 0
DB = 0
SM = 0
Delta_K
Doxycycline
YNL187W
ZShift = −0.18
Z lm Score = 1.71
NG = 0
DB = 0
SM = 0
Delta_r
Doxycycline
YNL187W
ZShift = 0.31
Z lm Score = 0.16
NG = 0
DB = 0
SM = 0
Delta_AUC
Doxycycline

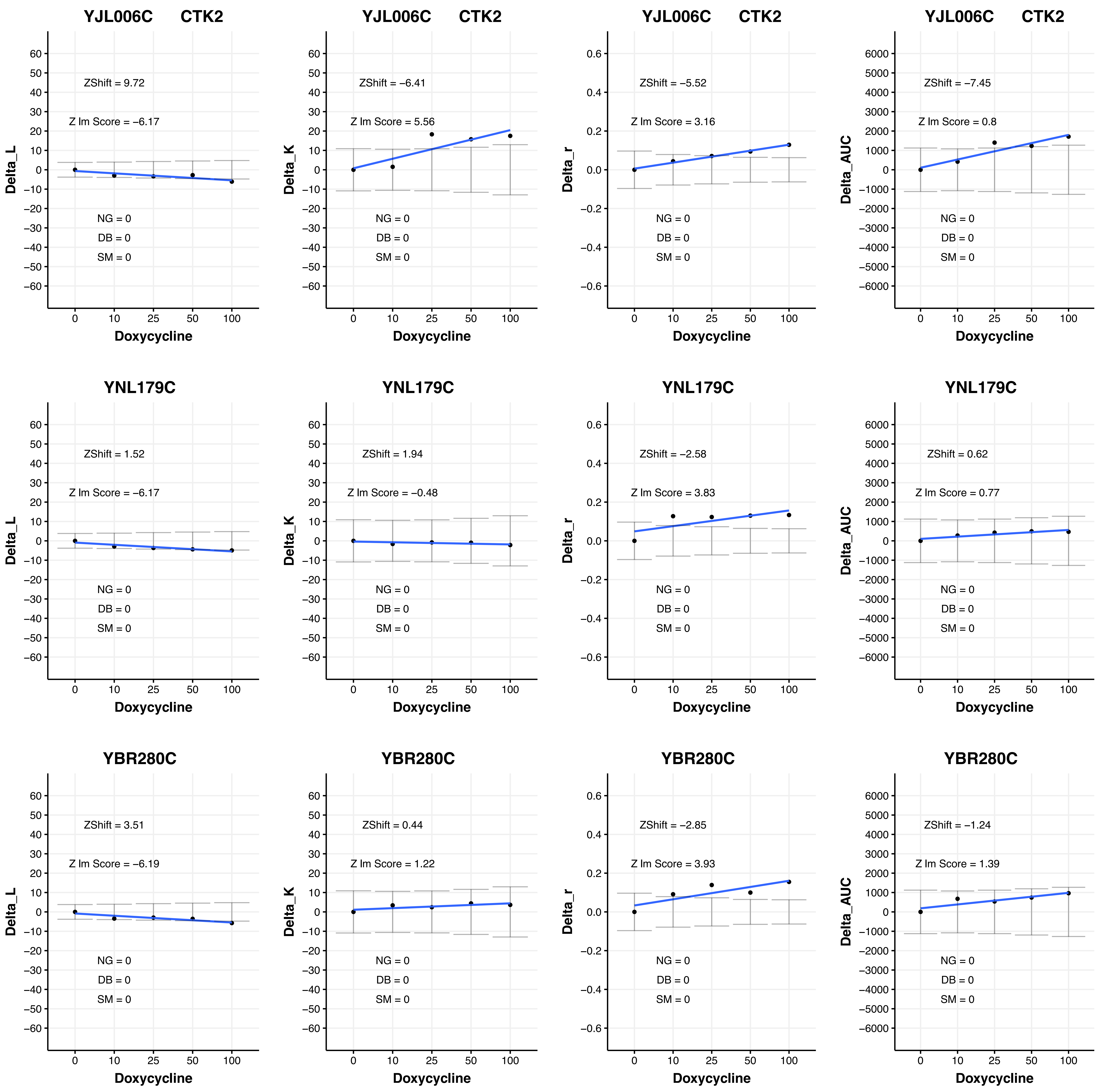

YJL006C CTK2
ZShift = 9.72
Z lm Score = −6.17
NG = 0
DB = 0
SM = 0
Delta_L
Doxycycline
YJL006C CTK2
ZShift = −6.41
Z lm Score = 5.56
NG = 0
DB = 0
SM = 0
Delta_K
Doxycycline
YJL006C CTK2
ZShift = −5.52
Z lm Score = 3.16
NG = 0
DB = 0
SM = 0
Delta_r
Doxycycline
YJL006C CTK2
ZShift = −7.45
Z lm Score = 0.8
NG = 0
DB = 0
SM = 0
Delta_AUC
Doxycycline
YNL179C
ZShift = 1.52
Z lm Score = −6.17
NG = 0
DB = 0
SM = 0
Delta_L
Doxycycline
YNL179C
ZShift = 1.94
Z lm Score = −0.48
NG = 0
DB = 0
SM = 0
Delta_K
Doxycycline
YNL179C
ZShift = −2.58
Z lm Score = 3.83
NG = 0
DB = 0
SM = 0
Delta_r
Doxycycline
YNL179C
ZShift = 0.62
Z lm Score = 0.77
NG = 0
DB = 0
SM = 0
Delta_AUC
Doxycycline
YBR280C
ZShift = 3.51
Z lm Score = −6.19
NG = 0
DB = 0
SM = 0
Delta_L
Doxycycline
YBR280C
ZShift = 0.44
Z lm Score = 1.22
NG = 0
DB = 0
SM = 0
Delta_K
Doxycycline
YBR280C
ZShift = −2.85
Z lm Score = 3.93
NG = 0
DB = 0
SM = 0
Delta_r
Doxycycline
YBR280C
ZShift = −1.24
Z lm Score = 1.39
NG = 0
DB = 0
SM = 0
Delta_AUC
Doxycycline

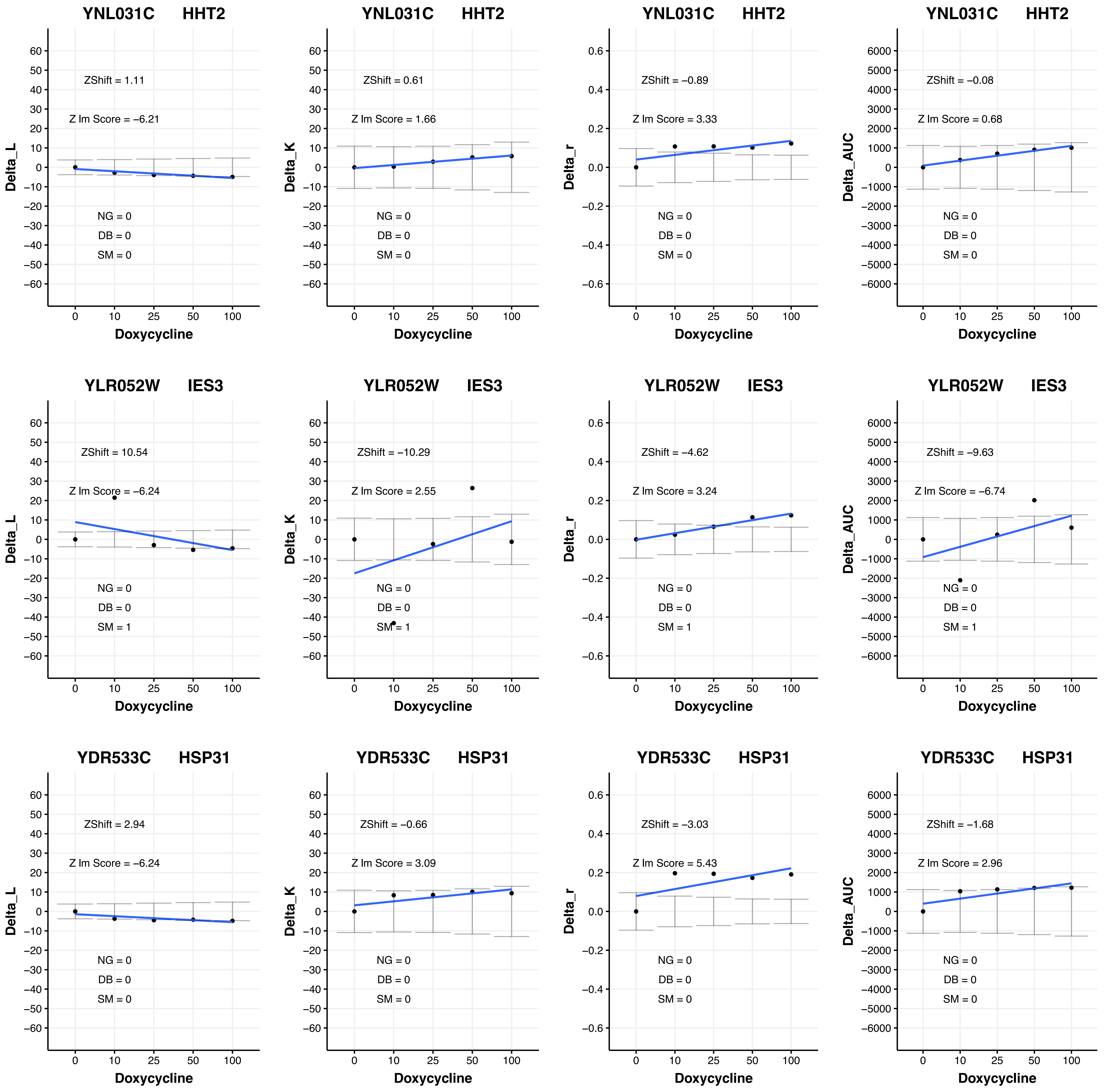

YNL031C HHT2
ZShift = 1.11
Z lm Score = −6.21
NG = 0
DB = 0
SM = 0
Delta_L
Doxycycline
YNL031C HHT2
ZShift = 0.61
Z lm Score = 1.66
NG = 0
DB = 0
SM = 0
Delta_K
Doxycycline
YNL031C HHT2
ZShift = −0.89
Z lm Score = 3.33
NG = 0
DB = 0
SM = 0
Delta_r
Doxycycline
YNL031C HHT2
ZShift = −0.08
Z lm Score = 0.68
NG = 0
DB = 0
SM = 0
Delta_AUC
Doxycycline
YLR052W IES3
ZShift = 10.54
Z lm Score = −6.24
NG = 0
DB = 0
SM = 1
Delta_L
Doxycycline
YLR052W IES3
ZShift = −10.29
Z lm Score = 2.55
NG = 0
DB = 0
SM = 1
Delta_K
Doxycycline
YLR052W IES3
ZShift = −4.62
Z lm Score = 3.24
NG = 0
DB = 0
SM = 1
Delta_r
Doxycycline
YLR052W IES3
ZShift = −9.63
Z lm Score = −6.74
NG = 0
DB = 0
SM = 1
Delta_AUC
Doxycycline
YDR533C HSP31
ZShift = 2.94
Z lm Score = −6.24
NG = 0
DB = 0
SM = 0
Delta_L
Doxycycline
YDR533C HSP31
ZShift = −0.66
Z lm Score = 3.09
NG = 0
DB = 0
SM = 0
Delta_K
Doxycycline
YDR533C HSP31
ZShift = −3.03
Z lm Score = 5.43
NG = 0
DB = 0
SM = 0
Delta_r
Doxycycline
YDR533C HSP31
ZShift = −1.68
Z lm Score = 2.96
NG = 0
DB = 0
SM = 0
Delta_AUC
Doxycycline

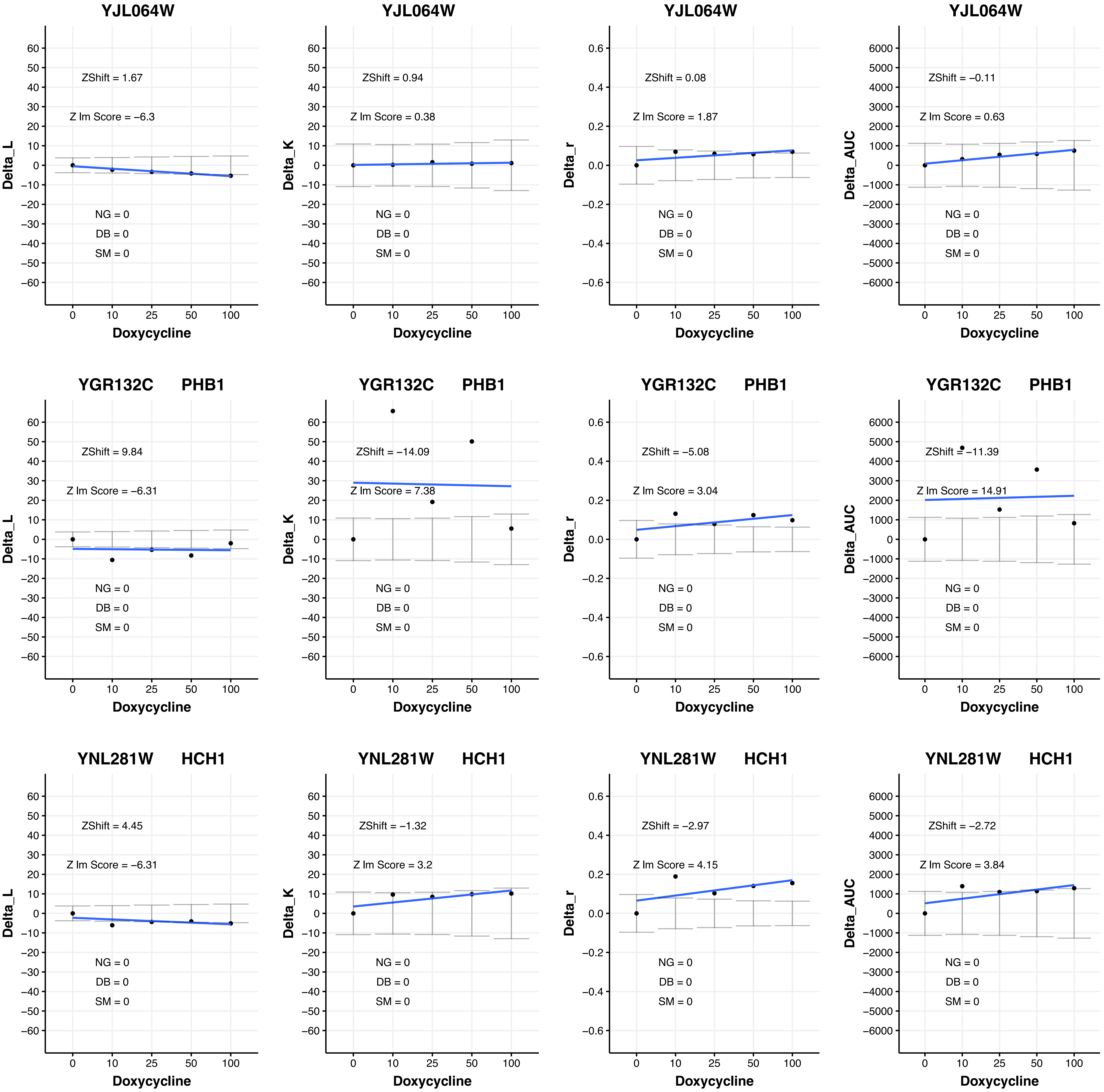
YJL064W
ZShift = 1.67
Z lm Score = −6.3
NG = 0
DB = 0
SM = 0
Delta_L
Doxycycline
YJL064W
ZShift = 0.94
Z lm Score = 0.38
NG = 0
DB = 0
SM = 0
Delta_K
Doxycycline
YJL064W
ZShift = 0.08
Z lm Score = 1.87
NG = 0
DB = 0
SM = 0
Delta_r
Doxycycline
YJL064W
ZShift = −0.11
Z lm Score = 0.63
NG = 0
DB = 0
SM = 0
Delta_AUC
Doxycycline
YGR132C PHB1
ZShift = 9.84
Z lm Score = −6.31
NG = 0
DB = 0
SM = 0
Delta_L
Doxycycline
YGR132C PHB1
ZShift = −14.09
Z lm Score = 7.38
NG = 0
DB = 0
SM = 0
Delta_K
Doxycycline
YGR132C PHB1
ZShift = −5.08
Z lm Score = 3.04
NG = 0
DB = 0
SM = 0
Delta_r
Doxycycline
YGR132C PHB1
ZShift = −11.39
Z lm Score = 14.91
NG = 0
DB = 0
SM = 0
Delta_AUC
Doxycycline
YNL281W HCH1
ZShift = 4.45
Z lm Score = −6.31
NG = 0
DB = 0
SM = 0
Delta_L
Doxycycline
YNL281W HCH1
ZShift = −1.32
Z lm Score = 3.2
NG = 0
DB = 0
SM = 0
Delta_K
Doxycycline
YNL281W HCH1
ZShift = −2.97
Z lm Score = 4.15
NG = 0
DB = 0
SM = 0
Delta_r
Doxycycline
YNL281W HCH1
ZShift = −2.72
Z lm Score = 3.84
NG = 0
DB = 0
SM = 0
Delta_AUC
Doxycycline

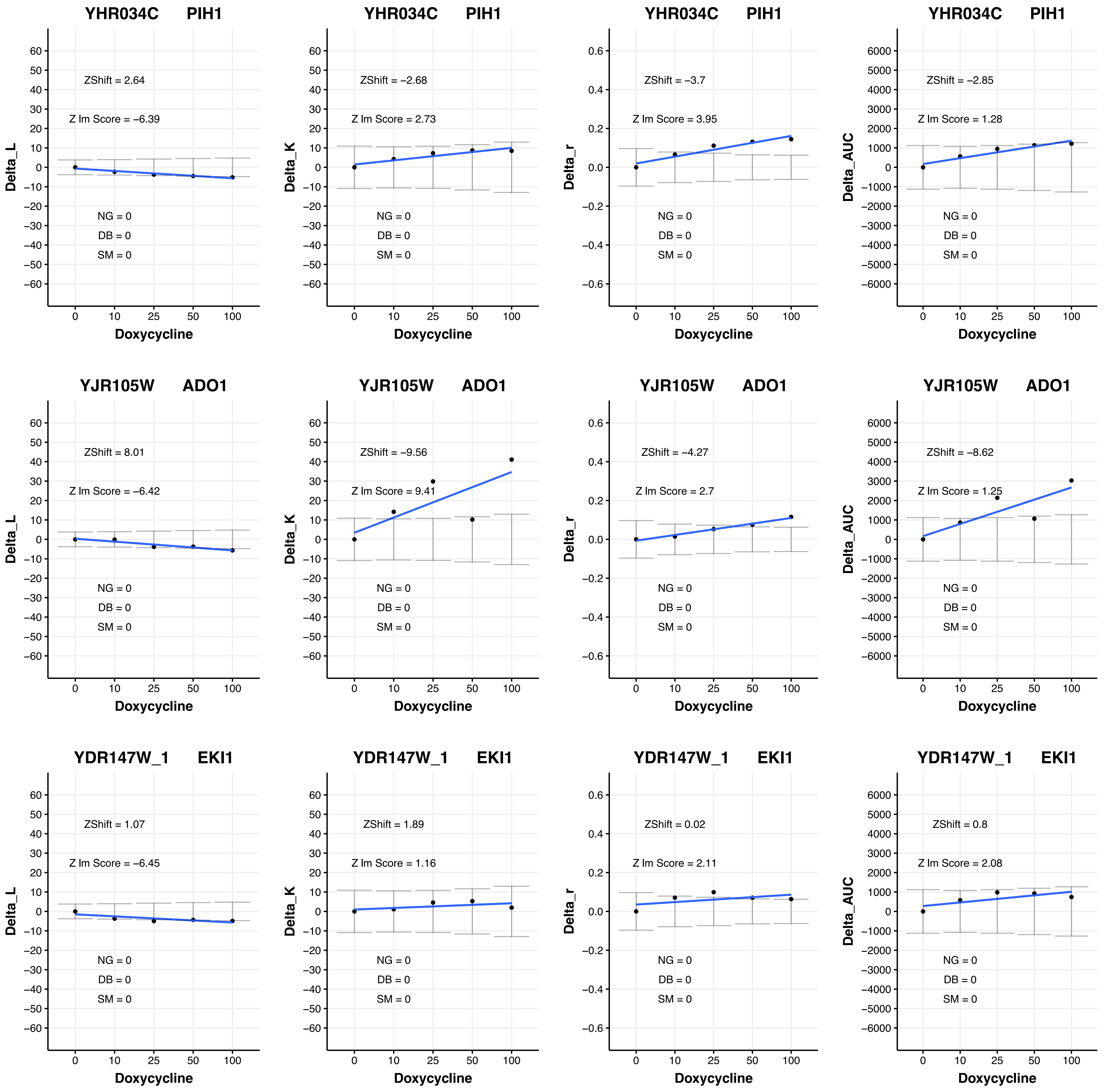
YHR034C PIH1
ZShift = 2.64
Z lm Score = −6.39
NG = 0
DB = 0
SM = 0
Delta_L
Doxycycline
YHR034C PIH1
ZShift = −2.68
Z lm Score = 2.73
NG = 0
DB = 0
SM = 0
Delta_K
Doxycycline
YHR034C PIH1
ZShift = −3.7
Z lm Score = 3.95
NG = 0
DB = 0
SM = 0
Delta_r
Doxycycline
YHR034C PIH1
ZShift = −2.85
Z lm Score = 1.28
NG = 0
DB = 0
SM = 0
Delta_AUC
Doxycycline
YJR105W ADO1
ZShift = 8.01
Z lm Score = −6.42
NG = 0
DB = 0
SM = 0
Delta_L
Doxycycline
YJR105W ADO1
ZShift = −9.56
Z lm Score = 9.41
NG = 0
DB = 0
SM = 0
Delta_K
Doxycycline
YJR105W ADO1
ZShift = −4.27
Z lm Score = 2.7
NG = 0
DB = 0
SM = 0
Delta_r
Doxycycline
YJR105W ADO1
ZShift = −8.62
Z lm Score = 1.25
NG = 0
DB = 0
SM = 0
Delta_AUC
Doxycycline
YDR147W_1 EKI1
ZShift = 1.07
Z lm Score = −6.45
NG = 0
DB = 0
SM = 0
Delta_L
Doxycycline
YDR147W_1 EKI1
ZShift = 1.89
Z lm Score = 1.16
NG = 0
DB = 0
SM = 0
Delta_K
Doxycycline
YDR147W_1 EKI1
ZShift = 0.02
Z lm Score = 2.11
NG = 0
DB = 0
SM = 0
Delta_r
Doxycycline
YDR147W_1 EKI1
ZShift = 0.8
Z lm Score = 2.08
NG = 0
DB = 0
SM = 0
Delta_AUC
Doxycycline

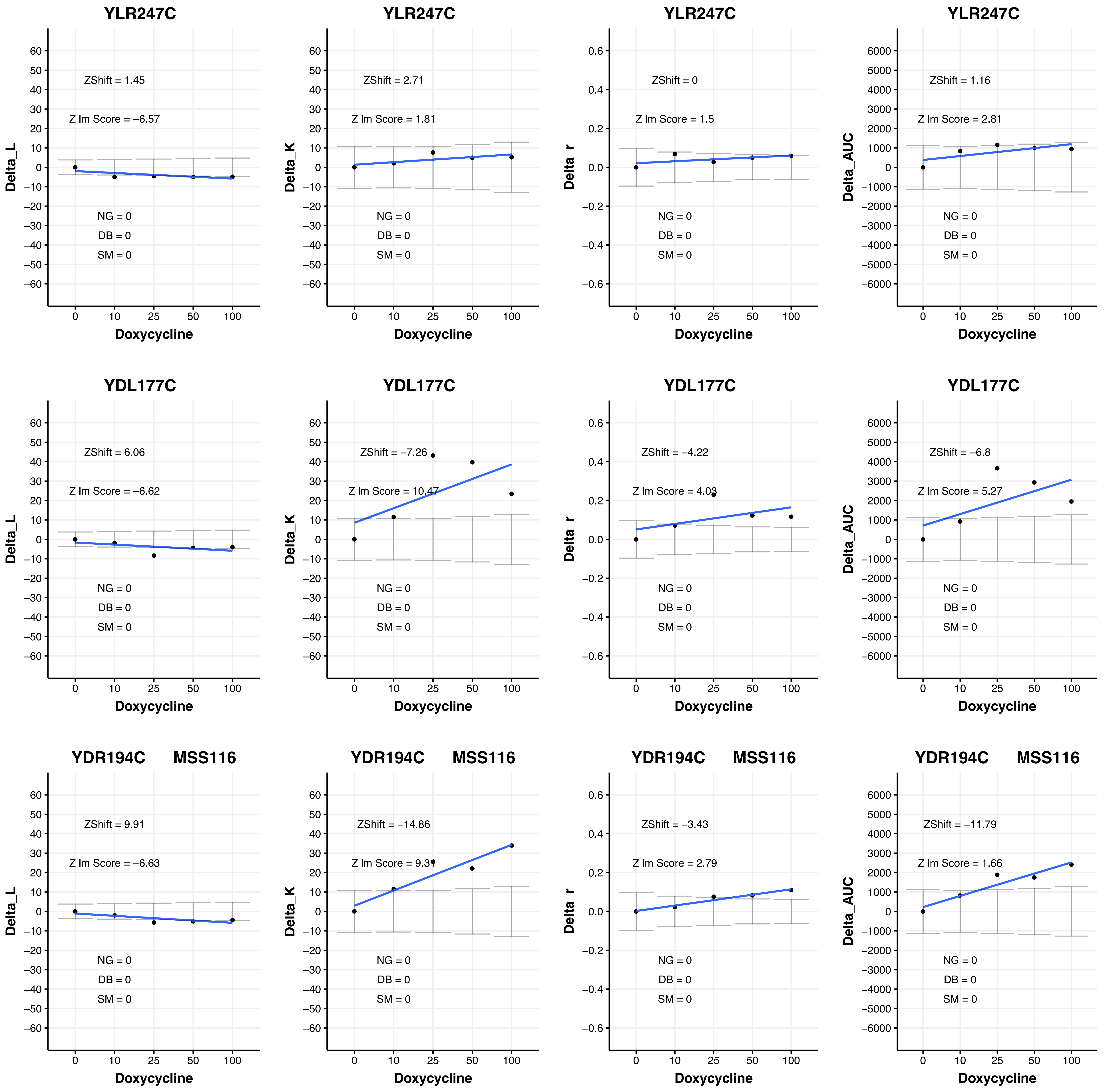
YLR247C
ZShift = 1.45
Z lm Score = −6.57
NG = 0
DB = 0
SM = 0
Delta_L
Doxycycline
YLR247C
ZShift = 2.71
Z lm Score = 1.81
Delta_K
YLR247C
ZShift = 0
Z lm Score = 1.5
Delta_r
YLR247C
ZShift = 1.16
Z lm Score = 2.81
Delta_AUC
YDL177C
ZShift = 6.06
Z lm Score = −6.62
YDL177C
ZShift = −7.26
Z lm Score = 10.47
YDL177C
ZShift = −4.22
Z lm Score = 4.03
YDL177C
ZShift = −6.8
Z lm Score = 5.27
YDR194C MSS116
ZShift = 9.91
Z lm Score = −6.63
YDR194C MSS116
ZShift = −14.86
Z lm Score = 9.37
YDR194C MSS116
ZShift = −3.43
Z lm Score = 2.79
YDR194C MSS116
ZShift = −11.79
Z lm Score = 1.66

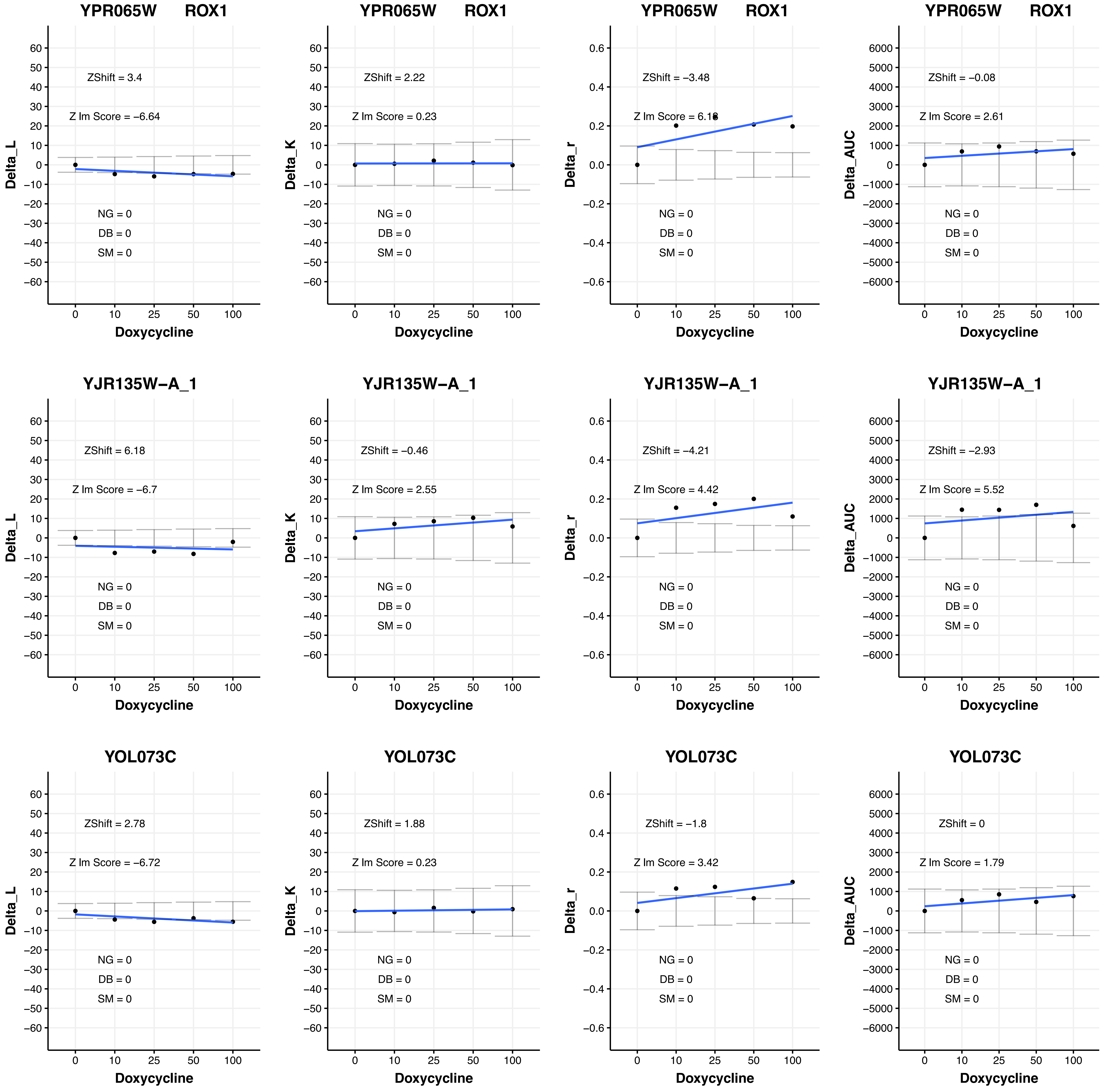

YPR065W ROX1
ZShift = 3.4
Z lm Score = −6.64
NG = 0
DB = 0
SM = 0
Delta_L
Doxycycline
YPR065W ROX1
ZShift = 2.22
Z lm Score = 0.23
NG = 0
DB = 0
SM = 0
Delta_K
Doxycycline
YPR065W ROX1
ZShift = −3.48
Z lm Score = 6.18
NG = 0
DB = 0
SM = 0
Delta_r
Doxycycline
YPR065W ROX1
ZShift = −0.08
Z lm Score = 2.61
NG = 0
DB = 0
SM = 0
Delta_AUC
Doxycycline
YJR135W−A_1
ZShift = 6.18
Z lm Score = −6.7
NG = 0
DB = 0
SM = 0
Delta_L
Doxycycline
YJR135W−A_1
ZShift = −0.46
Z lm Score = 2.55
NG = 0
DB = 0
SM = 0
Delta_K
Doxycycline
YJR135W−A_1
ZShift = −4.21
Z lm Score = 4.42
NG = 0
DB = 0
SM = 0
Delta_r
Doxycycline
YJR135W−A_1
ZShift = −2.93
Z lm Score = 5.52
NG = 0
DB = 0
SM = 0
Delta_AUC
Doxycycline
YOL073C
ZShift = 2.78
Z lm Score = −6.72
NG = 0
DB = 0
SM = 0
Delta_L
Doxycycline
YOL073C
ZShift = 1.88
Z lm Score = 0.23
NG = 0
DB = 0
SM = 0
Delta_K
Doxycycline
YOL073C
ZShift = −1.8
Z lm Score = 3.42
NG = 0
DB = 0
SM = 0
Delta_r
Doxycycline
YOL073C
ZShift = 0
Z lm Score = 1.79
NG = 0
DB = 0
SM = 0
Delta_AUC
Doxycycline

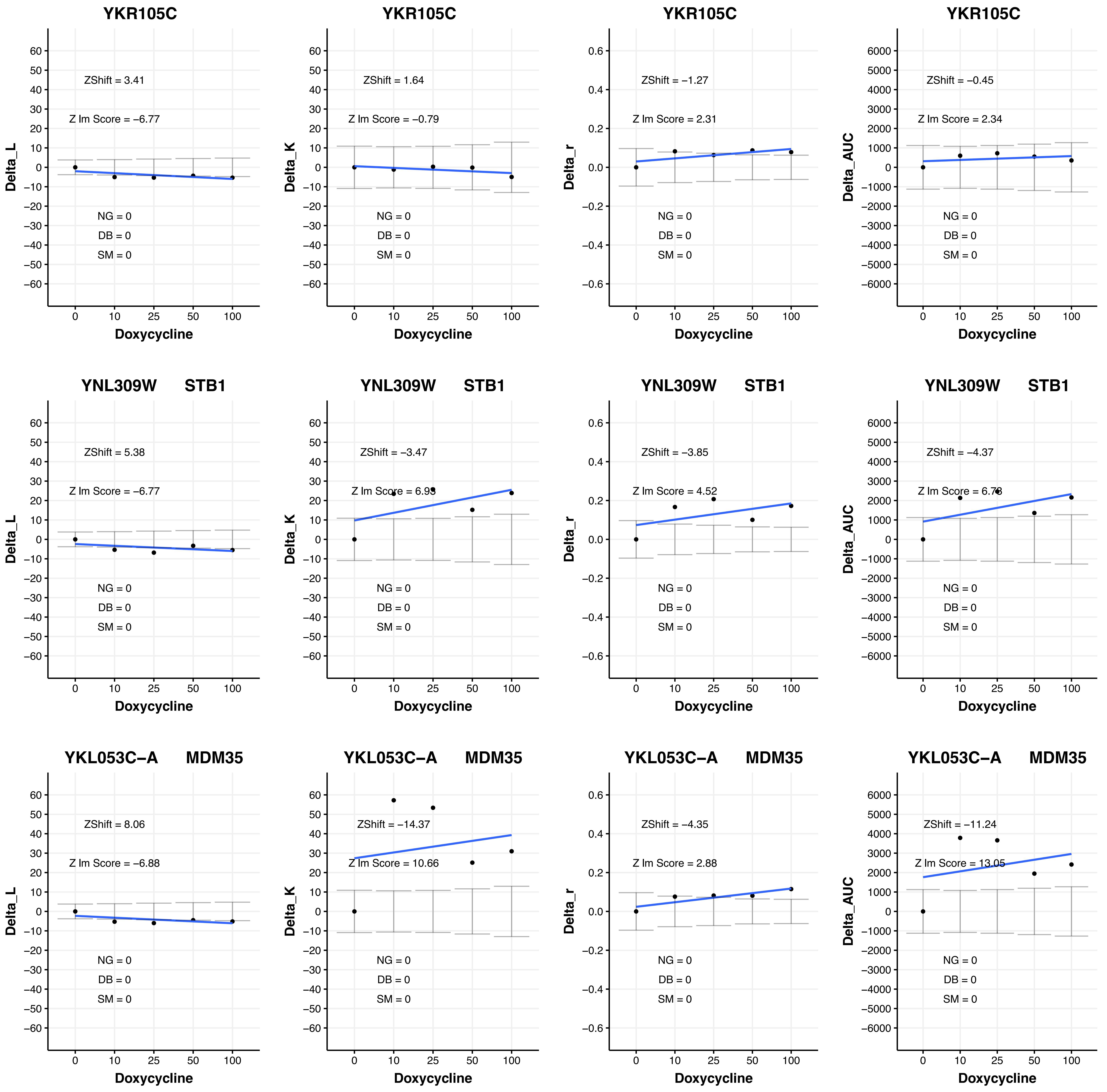

YKR105C
ZShift = 3.41
Z lm Score = −6.77
NG = 0
DB = 0
SM = 0
Delta_L
Doxycycline
YKR105C
ZShift = 1.64
Z lm Score = −0.79
NG = 0
DB = 0
SM = 0
Delta_K
Doxycycline
YKR105C
ZShift = −1.27
Z lm Score = 2.31
NG = 0
DB = 0
SM = 0
Delta_r
Doxycycline
YKR105C
ZShift = −0.45
Z lm Score = 2.34
NG = 0
DB = 0
SM = 0
Delta_AUC
Doxycycline
YNL309W STB1
ZShift = 5.38
Z lm Score = −6.77
NG = 0
DB = 0
SM = 0
Delta_L
Doxycycline
YNL309W STB1
ZShift = −3.47
Z lm Score = 6.93
NG = 0
DB = 0
SM = 0
Delta_K
Doxycycline
YNL309W STB1
ZShift = −3.85
Z lm Score = 4.52
NG = 0
DB = 0
SM = 0
Delta_r
Doxycycline
YNL309W STB1
ZShift = −4.37
Z lm Score = 6.78
NG = 0
DB = 0
SM = 0
Delta_AUC
Doxycycline
YKL053C−A MDM35
ZShift = 8.06
Z lm Score = −6.88
NG = 0
DB = 0
SM = 0
Delta_L
Doxycycline
YKL053C−A MDM35
ZShift = −14.37
Z lm Score = 10.66
NG = 0
DB = 0
SM = 0
Delta_K
Doxycycline
YKL053C−A MDM35
ZShift = −4.35
Z lm Score = 2.88
NG = 0
DB = 0
SM = 0
Delta_r
Doxycycline
YKL053C−A MDM35
ZShift = −11.24
Z lm Score = 13.05
NG = 0
DB = 0
SM = 0
Delta_AUC
Doxycycline

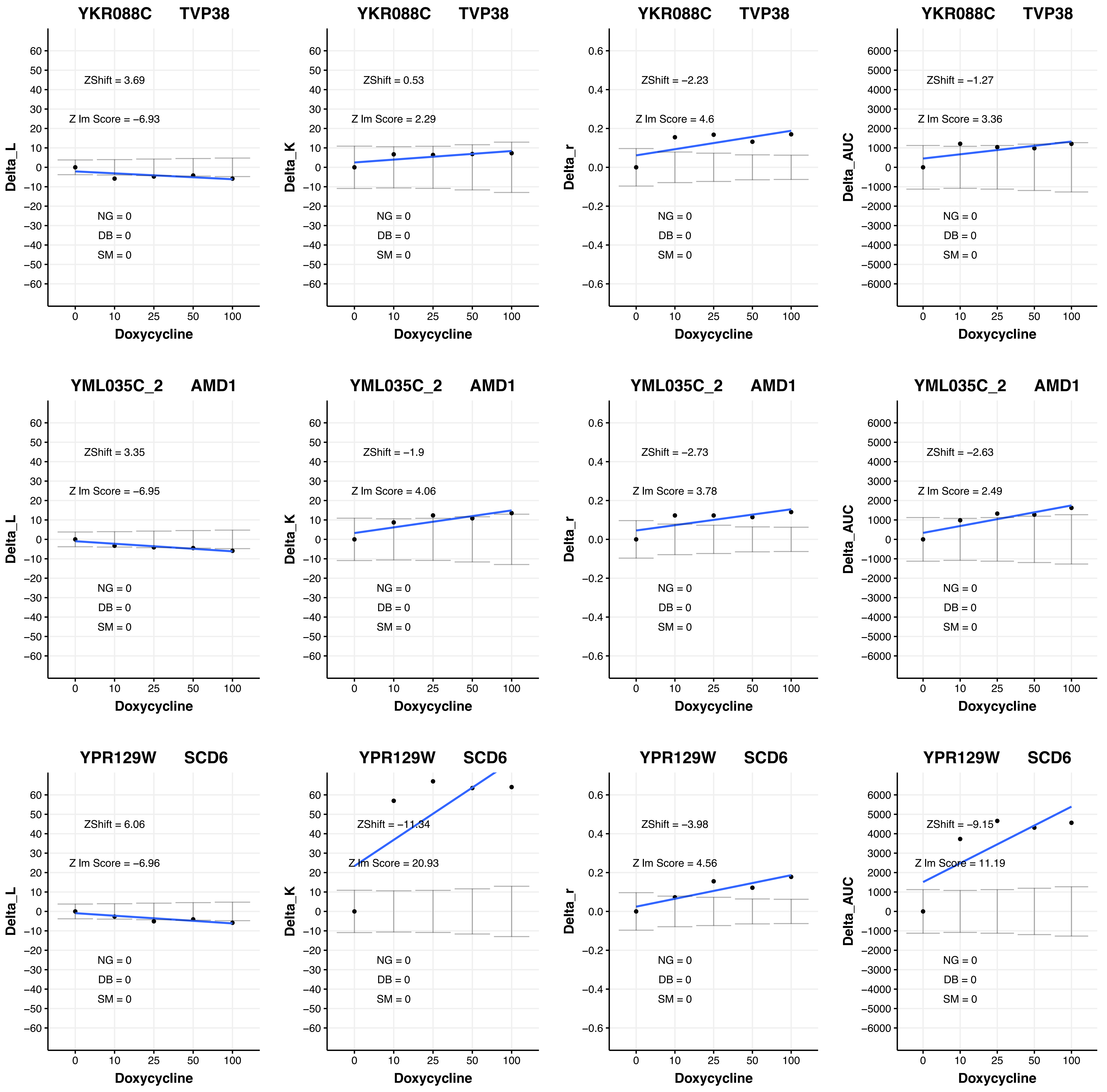
YKR088C TVP38
Delta_L
ZShift = 3.69
Z lm Score = −6.93
NG = 0
DB = 0
SM = 0
Doxycycline
YKR088C TVP38
Delta_K
ZShift = 0.53
Z lm Score = 2.29
NG = 0
DB = 0
SM = 0
Doxycycline
YKR088C TVP38
Delta_r
ZShift = −2.23
Z lm Score = 4.6
NG = 0
DB = 0
SM = 0
Doxycycline
YKR088C TVP38
Delta_AUC
ZShift = −1.27
Z lm Score = 3.36
NG = 0
DB = 0
SM = 0
Doxycycline
YML035C_2 AMD1
Delta_L
ZShift = 3.35
Z lm Score = −6.95
NG = 0
DB = 0
SM = 0
Doxycycline
YML035C_2 AMD1
Delta_K
ZShift = −1.9
Z lm Score = 4.06
NG = 0
DB = 0
SM = 0
Doxycycline
YML035C_2 AMD1
Delta_r
ZShift = −2.73
Z lm Score = 3.78
NG = 0
DB = 0
SM = 0
Doxycycline
YML035C_2 AMD1
Delta_AUC
ZShift = −2.63
Z lm Score = 2.49
NG = 0
DB = 0
SM = 0
Doxycycline
YPR129W SCD6
Delta_L
ZShift = 6.06
Z lm Score = −6.96
NG = 0
DB = 0
SM = 0
Doxycycline
YPR129W SCD6
Delta_K
ZShift = −11.34
Z lm Score = 20.93
NG = 0
DB = 0
SM = 0
Doxycycline
YPR129W SCD6
Delta_r
ZShift = −3.98
Z lm Score = 4.56
NG = 0
DB = 0
SM = 0
Doxycycline
YPR129W SCD6
Delta_AUC
ZShift = −9.15
Z lm Score = 11.19
NG = 0
DB = 0
SM = 0
Doxycycline

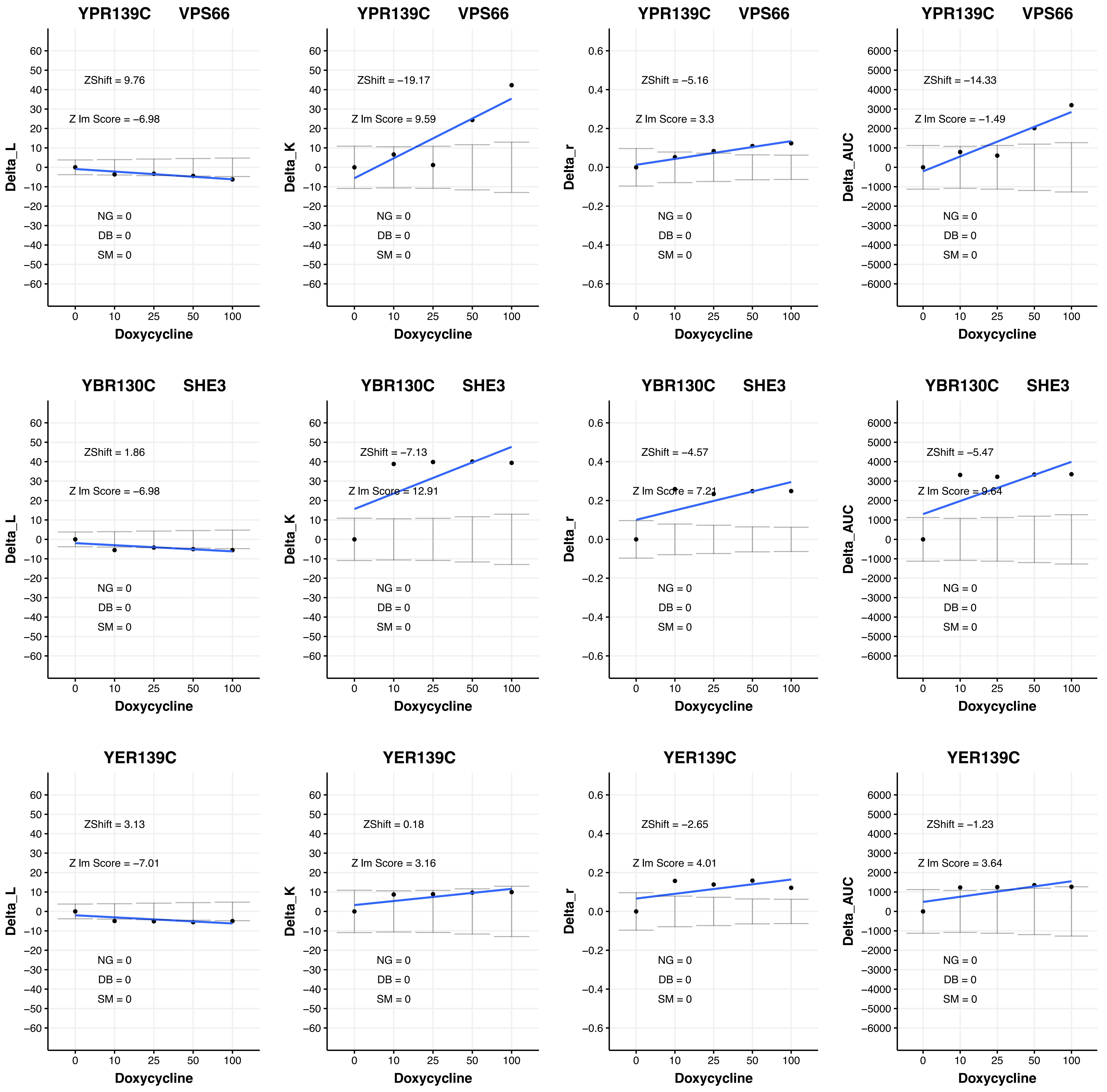
YPR139C VPS66
ZShift = 9.76
Z lm Score = −6.98
NG = 0
DB = 0
SM = 0
Delta_L
Doxycycline
YPR139C VPS66
ZShift = −19.17
Z lm Score = 9.59
NG = 0
DB = 0
SM = 0
Delta_K
Doxycycline
YPR139C VPS66
ZShift = −5.16
Z lm Score = 3.3
NG = 0
DB = 0
SM = 0
Delta_r
Doxycycline
YPR139C VPS66
ZShift = −14.33
Z lm Score = −1.49
NG = 0
DB = 0
SM = 0
Delta_AUC
Doxycycline
YBR130C SHE3
ZShift = 1.86
Z lm Score = −6.98
NG = 0
DB = 0
SM = 0
Delta_L
Doxycycline
YBR130C SHE3
ZShift = −7.13
Z lm Score = 12.91
NG = 0
DB = 0
SM = 0
Delta_K
Doxycycline
YBR130C SHE3
ZShift = −4.57
Z lm Score = 7.21
NG = 0
DB = 0
SM = 0
Delta_r
Doxycycline
YBR130C SHE3
ZShift = −5.47
Z lm Score = 9.64
NG = 0
DB = 0
SM = 0
Delta_AUC
Doxycycline
YER139C
ZShift = 3.13
Z lm Score = −7.01
NG = 0
DB = 0
SM = 0
Delta_L
Doxycycline
YER139C
ZShift = 0.18
Z lm Score = 3.16
NG = 0
DB = 0
SM = 0
Delta_K
Doxycycline
YER139C
ZShift = −2.65
Z lm Score = 4.01
NG = 0
DB = 0
SM = 0
Delta_r
Doxycycline
YER139C
ZShift = −1.23
Z lm Score = 3.64
NG = 0
DB = 0
SM = 0
Delta_AUC
Doxycycline

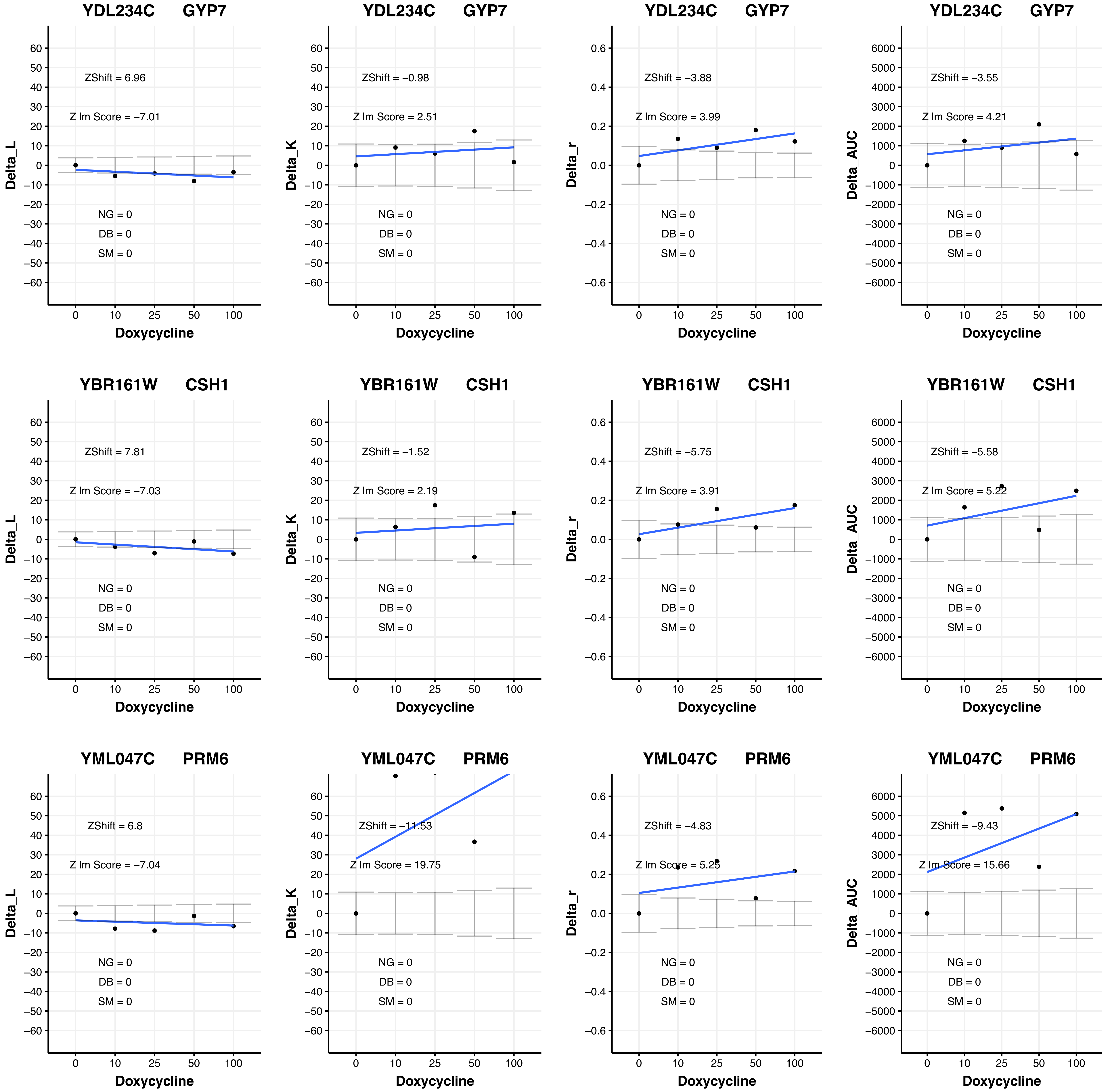

YDL234C GYP7
ZShift = 6.96
Z lm Score = −7.01
NG = 0
DB = 0
SM = 0
Delta_L
Doxycycline
YDL234C GYP7
ZShift = −0.98
Z lm Score = 2.51
NG = 0
DB = 0
SM = 0
Delta_K
Doxycycline
YDL234C GYP7
ZShift = −3.88
Z lm Score = 3.99
NG = 0
DB = 0
SM = 0
Delta_r
Doxycycline
YDL234C GYP7
ZShift = −3.55
Z lm Score = 4.21
NG = 0
DB = 0
SM = 0
Delta_AUC
Doxycycline
YBR161W CSH1
ZShift = 7.81
Z lm Score = −7.03
NG = 0
DB = 0
SM = 0
Delta_L
Doxycycline
YBR161W CSH1
ZShift = −1.52
Z lm Score = 2.19
NG = 0
DB = 0
SM = 0
Delta_K
Doxycycline
YBR161W CSH1
ZShift = −5.75
Z lm Score = 3.91
NG = 0
DB = 0
SM = 0
Delta_r
Doxycycline
YBR161W CSH1
ZShift = −5.58
Z lm Score = 5.22
NG = 0
DB = 0
SM = 0
Delta_AUC
Doxycycline
YML047C PRM6
ZShift = 6.8
Z lm Score = −7.04
NG = 0
DB = 0
SM = 0
Delta_L
Doxycycline
YML047C PRM6
ZShift = −11.53
Z lm Score = 19.75
NG = 0
DB = 0
SM = 0
Delta_K
Doxycycline
YML047C PRM6
ZShift = −4.83
Z lm Score = 5.25
NG = 0
DB = 0
SM = 0
Delta_r
Doxycycline
YML047C PRM6
ZShift = −9.43
Z lm Score = 15.66
NG = 0
DB = 0
SM = 0
Delta_AUC
Doxycycline

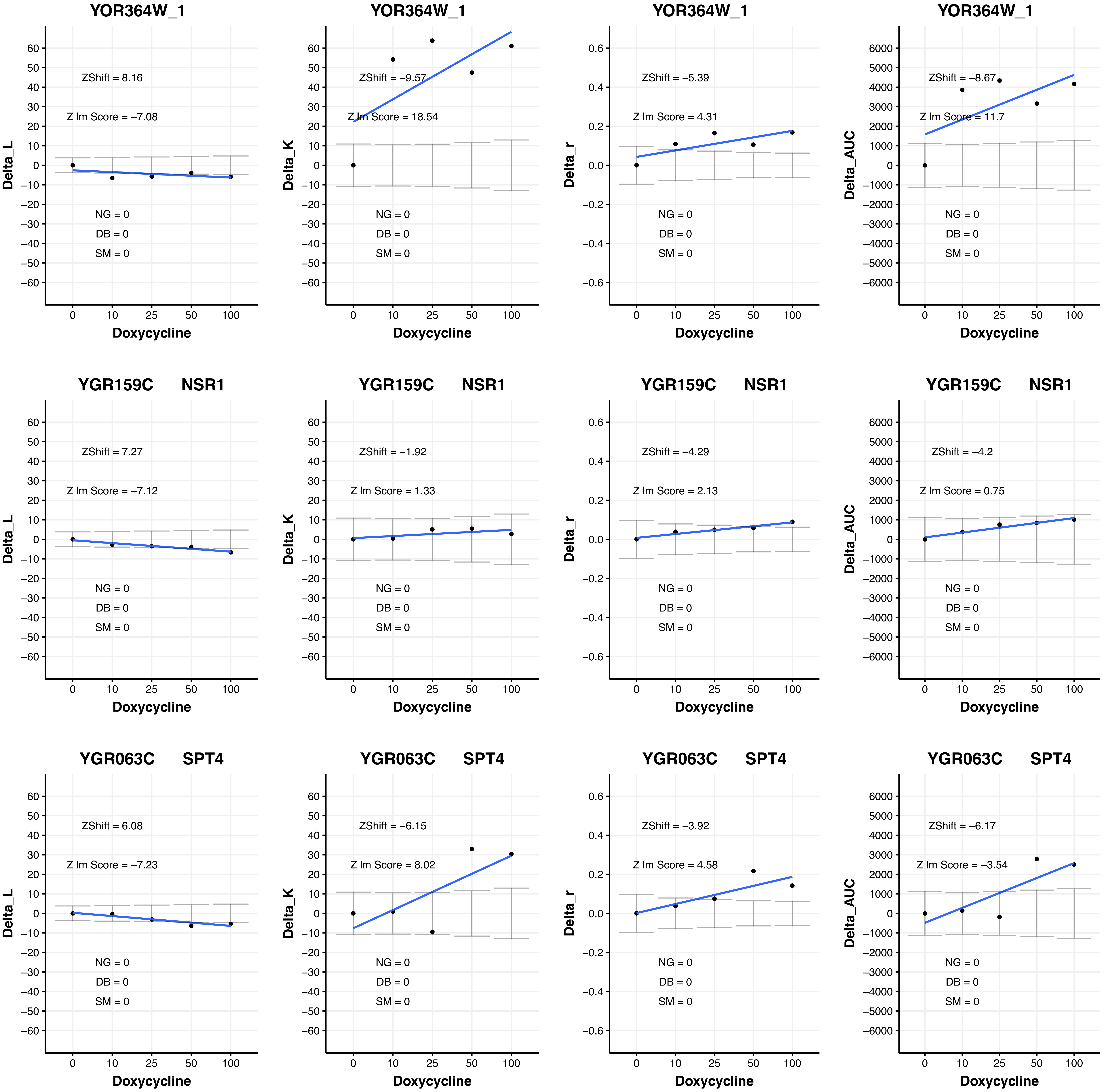
YOR364W_1
ZShift = 8.16
Z lm Score = −7.08
NG = 0
DB = 0
SM = 0
Delta_L
Doxycycline
YOR364W_1
ZShift = −9.57
Z lm Score = 18.54
NG = 0
DB = 0
SM = 0
Delta_K
Doxycycline
YOR364W_1
ZShift = −5.39
Z lm Score = 4.31
NG = 0
DB = 0
SM = 0
Delta_r
Doxycycline
YOR364W_1
ZShift = −8.67
Z lm Score = 11.7
NG = 0
DB = 0
SM = 0
Delta_AUC
Doxycycline
YGR159C NSR1
ZShift = 7.27
Z lm Score = −7.12
NG = 0
DB = 0
SM = 0
Delta_L
Doxycycline
YGR159C NSR1
ZShift = −1.92
Z lm Score = 1.33
NG = 0
DB = 0
SM = 0
Delta_K
Doxycycline
YGR159C NSR1
ZShift = −4.29
Z lm Score = 2.13
NG = 0
DB = 0
SM = 0
Delta_r
Doxycycline
YGR159C NSR1
ZShift = −4.2
Z lm Score = 0.75
NG = 0
DB = 0
SM = 0
Delta_AUC
Doxycycline
YGR063C SPT4
ZShift = 6.08
Z lm Score = −7.23
NG = 0
DB = 0
SM = 0
Delta_L
Doxycycline
YGR063C SPT4
ZShift = −6.15
Z lm Score = 8.02
NG = 0
DB = 0
SM = 0
Delta_K
Doxycycline
YGR063C SPT4
ZShift = −3.92
Z lm Score = 4.58
NG = 0
DB = 0
SM = 0
Delta_r
Doxycycline
YGR063C SPT4
ZShift = −6.17
Z lm Score = −3.54
NG = 0
DB = 0
SM = 0
Delta_AUC
Doxycycline

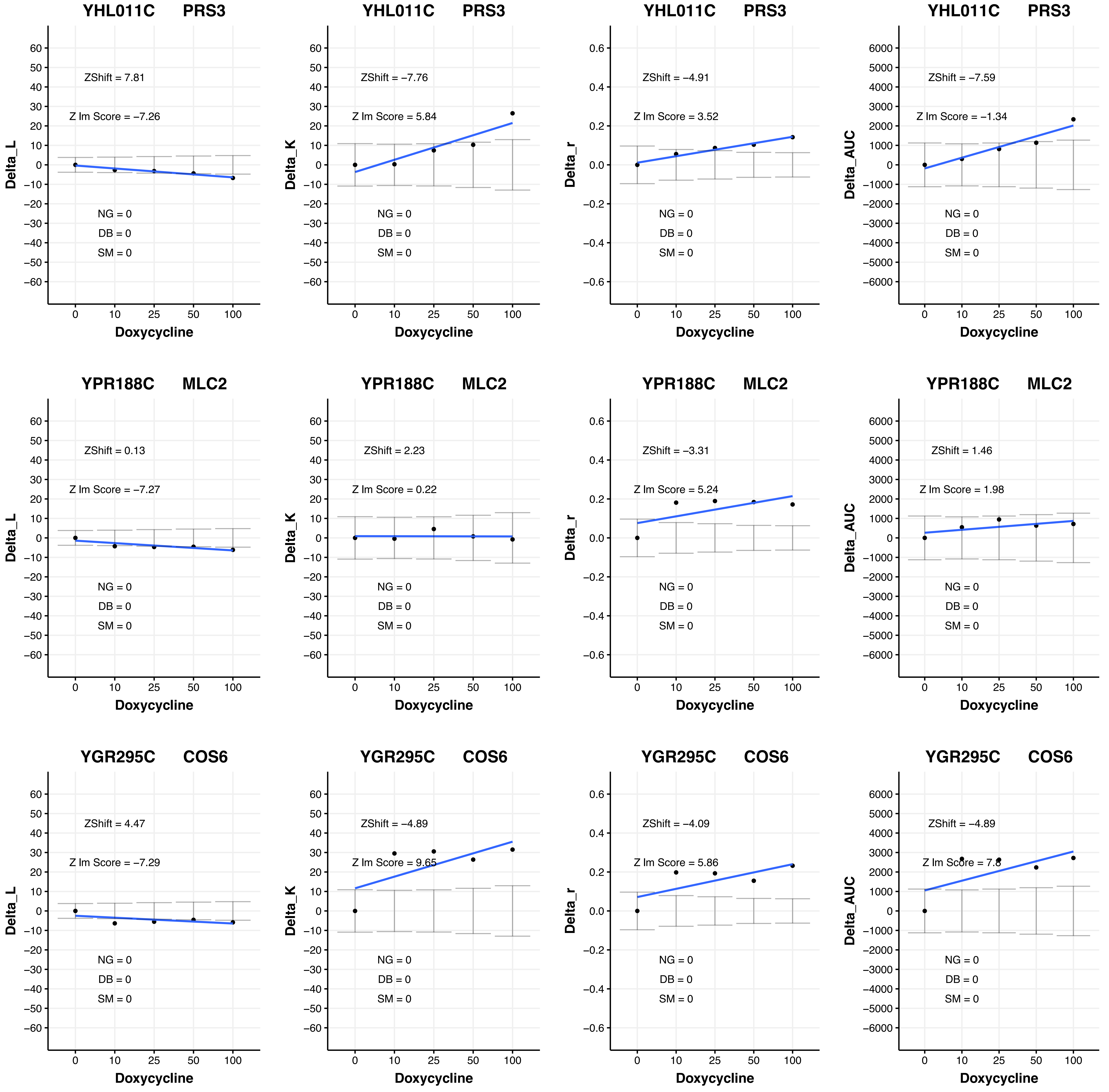

YHL011C PRS3
ZShift = 7.81
Z lm Score = −7.26
NG = 0
DB = 0
SM = 0
Delta_L
Doxycycline
YHL011C PRS3
ZShift = −7.76
Z lm Score = 5.84
NG = 0
DB = 0
SM = 0
Delta_K
Doxycycline
YHL011C PRS3
ZShift = −4.91
Z lm Score = 3.52
NG = 0
DB = 0
SM = 0
Delta_r
Doxycycline
YHL011C PRS3
ZShift = −7.59
Z lm Score = −1.34
NG = 0
DB = 0
SM = 0
Delta_AUC
Doxycycline
YPR188C MLC2
ZShift = 0.13
Z lm Score = −7.27
NG = 0
DB = 0
SM = 0
Delta_L
Doxycycline
YPR188C MLC2
ZShift = 2.23
Z lm Score = 0.22
NG = 0
DB = 0
SM = 0
Delta_K
Doxycycline
YPR188C MLC2
ZShift = −3.31
Z lm Score = 5.24
NG = 0
DB = 0
SM = 0
Delta_r
Doxycycline
YPR188C MLC2
ZShift = 1.46
Z lm Score = 1.98
NG = 0
DB = 0
SM = 0
Delta_AUC
Doxycycline
YGR295C COS6
ZShift = 4.47
Z lm Score = −7.29
NG = 0
DB = 0
SM = 0
Delta_L
Doxycycline
YGR295C COS6
ZShift = −4.89
Z lm Score = 9.65
NG = 0
DB = 0
SM = 0
Delta_K
Doxycycline
YGR295C COS6
ZShift = −4.09
Z lm Score = 5.86
NG = 0
DB = 0
SM = 0
Delta_r
Doxycycline
YGR295C COS6
ZShift = −4.89
Z lm Score = 7.8
NG = 0
DB = 0
SM = 0
Delta_AUC
Doxycycline

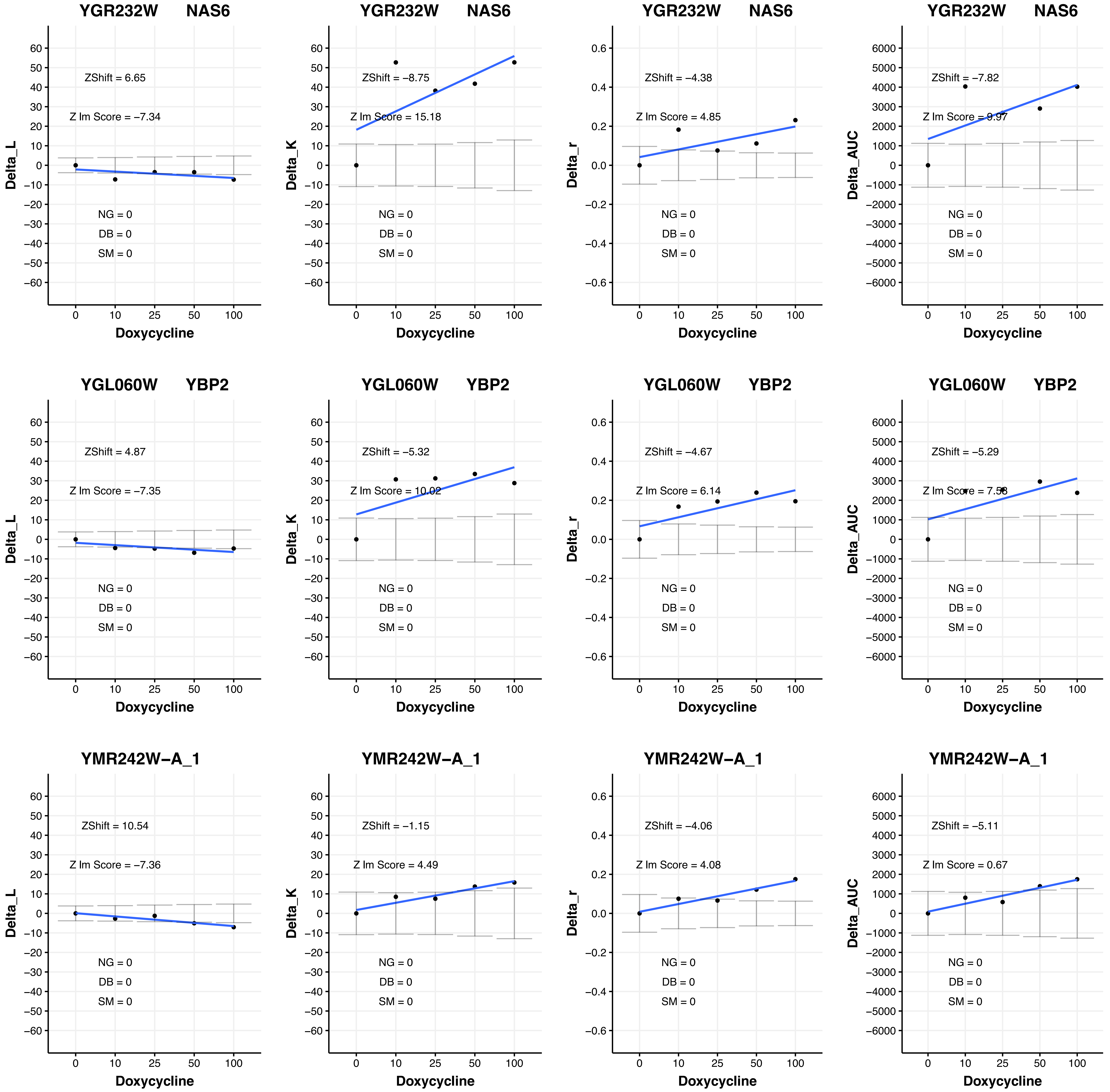

YGR232W NAS6
ZShift = 6.65
Z lm Score = −7.34
NG = 0
DB = 0
SM = 0
Delta_L
Doxycycline
YGR232W NAS6
ZShift = −8.75
Z lm Score = 15.18
NG = 0
DB = 0
SM = 0
Delta_K
Doxycycline
YGR232W NAS6
ZShift = −4.38
Z lm Score = 4.85
NG = 0
DB = 0
SM = 0
Delta_r
Doxycycline
YGR232W NAS6
ZShift = −7.82
Z lm Score = 0.97
NG = 0
DB = 0
SM = 0
Delta_AUC
Doxycycline
YGL060W YBP2
ZShift = 4.87
Z lm Score = −7.35
NG = 0
DB = 0
SM = 0
Delta_L
Doxycycline
YGL060W YBP2
ZShift = −5.32
Z lm Score = 10.02
NG = 0
DB = 0
SM = 0
Delta_K
Doxycycline
YGL060W YBP2
ZShift = −4.67
Z lm Score = 6.14
NG = 0
DB = 0
SM = 0
Delta_r
Doxycycline
YGL060W YBP2
ZShift = −5.29
Z lm Score = 7.58
NG = 0
DB = 0
SM = 0
Delta_AUC
Doxycycline
YMR242W−A_1
ZShift = 10.54
Z lm Score = −7.36
NG = 0
DB = 0
SM = 0
Delta_L
Doxycycline
YMR242W−A_1
ZShift = −1.15
Z lm Score = 4.49
NG = 0
DB = 0
SM = 0
Delta_K
Doxycycline
YMR242W−A_1
ZShift = −4.06
Z lm Score = 4.08
NG = 0
DB = 0
SM = 0
Delta_r
Doxycycline
YMR242W−A_1
ZShift = −5.11
Z lm Score = 0.67
NG = 0
DB = 0
SM = 0
Delta_AUC
Doxycycline

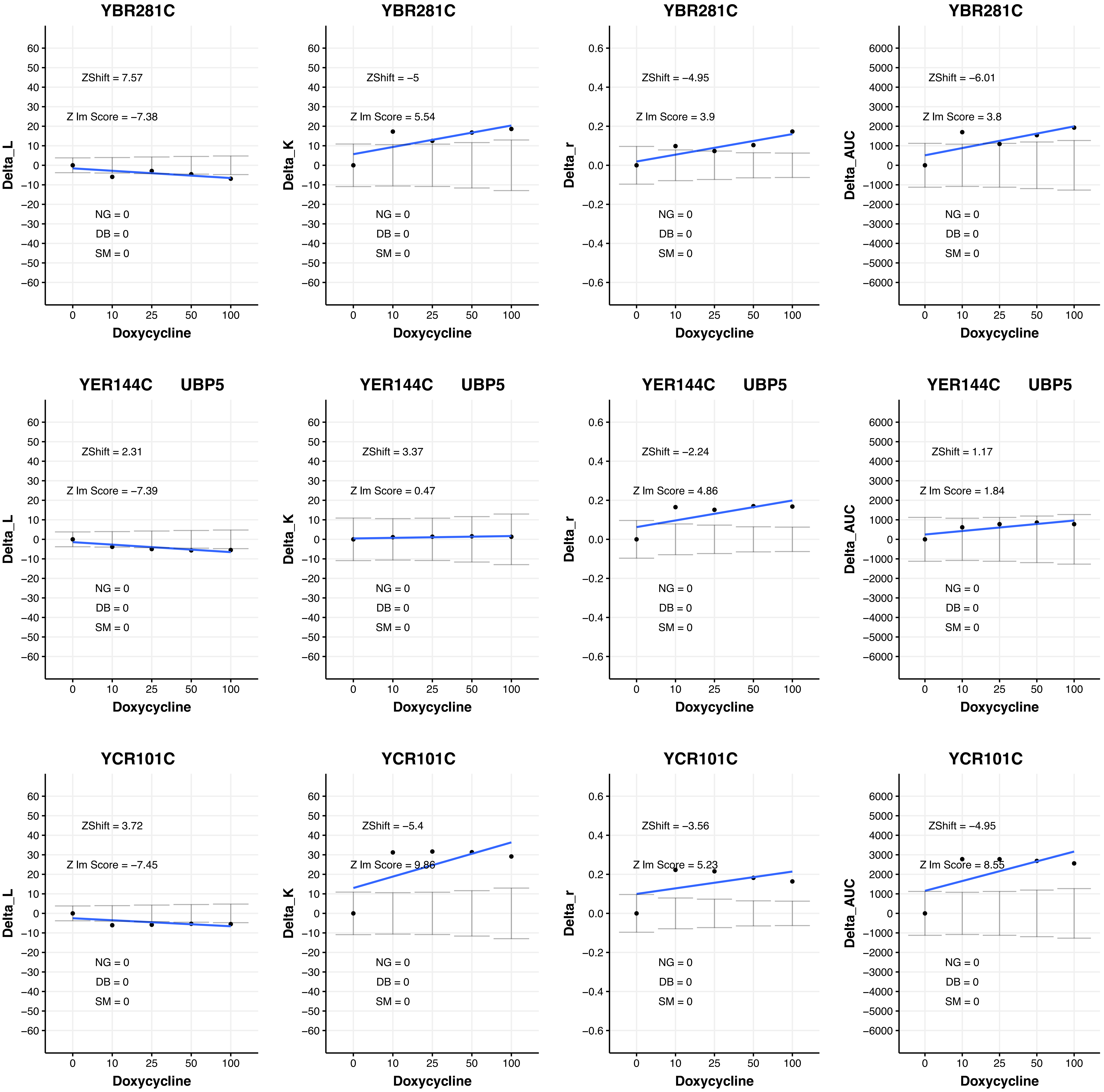
YBR281C
ZShift = 7.57
Z lm Score = −7.38
NG = 0
DB = 0
SM = 0
Delta_L
Doxycycline
YBR281C
ZShift = −5
Z lm Score = 5.54
NG = 0
DB = 0
SM = 0
Delta_K
Doxycycline
YBR281C
ZShift = −4.95
Z lm Score = 3.9
NG = 0
DB = 0
SM = 0
Delta_r
Doxycycline
YBR281C
ZShift = −6.01
Z lm Score = 3.8
NG = 0
DB = 0
SM = 0
Delta_AUC
Doxycycline
YER144C UBP5
ZShift = 2.31
Z lm Score = −7.39
NG = 0
DB = 0
SM = 0
Delta_L
Doxycycline
YER144C UBP5
ZShift = 3.37
Z lm Score = 0.47
NG = 0
DB = 0
SM = 0
Delta_K
Doxycycline
YER144C UBP5
ZShift = −2.24
Z lm Score = 4.86
NG = 0
DB = 0
SM = 0
Delta_r
Doxycycline
YER144C UBP5
ZShift = 1.17
Z lm Score = 1.84
NG = 0
DB = 0
SM = 0
Delta_AUC
Doxycycline
YCR101C
ZShift = 3.72
Z lm Score = −7.45
NG = 0
DB = 0
SM = 0
Delta_L
Doxycycline
YCR101C
ZShift = −5.4
Z lm Score = 9.86
NG = 0
DB = 0
SM = 0
Delta_K
Doxycycline
YCR101C
ZShift = −3.56
Z lm Score = 5.23
NG = 0
DB = 0
SM = 0
Delta_r
Doxycycline
YCR101C
ZShift = −4.95
Z lm Score = 8.55
NG = 0
DB = 0
SM = 0
Delta_AUC
Doxycycline

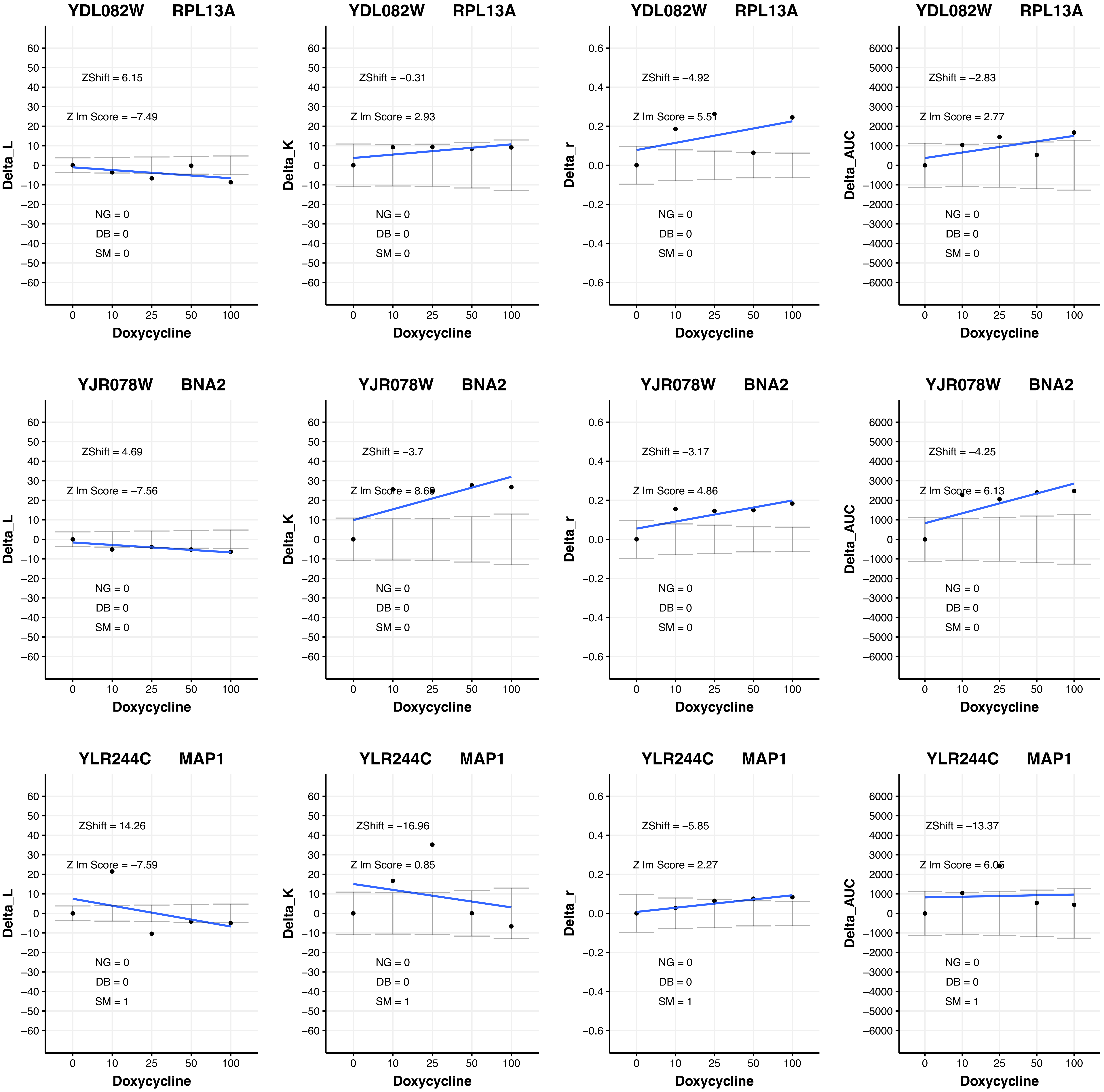
YDL082W RPL13A
ZShift = 6.15
Z lm Score = −7.49
NG = 0
DB = 0
SM = 0
Delta_L
Doxycycline
YDL082W RPL13A
ZShift = −0.31
Z lm Score = 2.93
NG = 0
DB = 0
SM = 0
Delta_K
Doxycycline
YDL082W RPL13A
ZShift = −4.92
Z lm Score = 5.51
NG = 0
DB = 0
SM = 0
Delta_r
Doxycycline
YDL082W RPL13A
ZShift = −2.83
Z lm Score = 2.77
NG = 0
DB = 0
SM = 0
Delta_AUC
Doxycycline
YJR078W BNA2
ZShift = 4.69
Z lm Score = −7.56
NG = 0
DB = 0
SM = 0
Delta_L
Doxycycline
YJR078W BNA2
ZShift = −3.7
Z lm Score = 8.69
NG = 0
DB = 0
SM = 0
Delta_K
Doxycycline
YJR078W BNA2
ZShift = −3.17
Z lm Score = 4.86
NG = 0
DB = 0
SM = 0
Delta_r
Doxycycline
YJR078W BNA2
ZShift = −4.25
Z lm Score = 6.13
NG = 0
DB = 0
SM = 0
Delta_AUC
Doxycycline
YLR244C MAP1
ZShift = 14.26
Z lm Score = −7.59
NG = 0
DB = 0
SM = 1
Delta_L
Doxycycline
YLR244C MAP1
ZShift = −16.96
Z lm Score = 0.85
NG = 0
DB = 0
SM = 1
Delta_K
Doxycycline
YLR244C MAP1
ZShift = −5.85
Z lm Score = 2.27
NG = 0
DB = 0
SM = 1
Delta_r
Doxycycline
YLR244C MAP1
ZShift = −13.37
Z lm Score = 6.05
NG = 0
DB = 0
SM = 1
Delta_AUC
Doxycycline
0 10 25 50 100

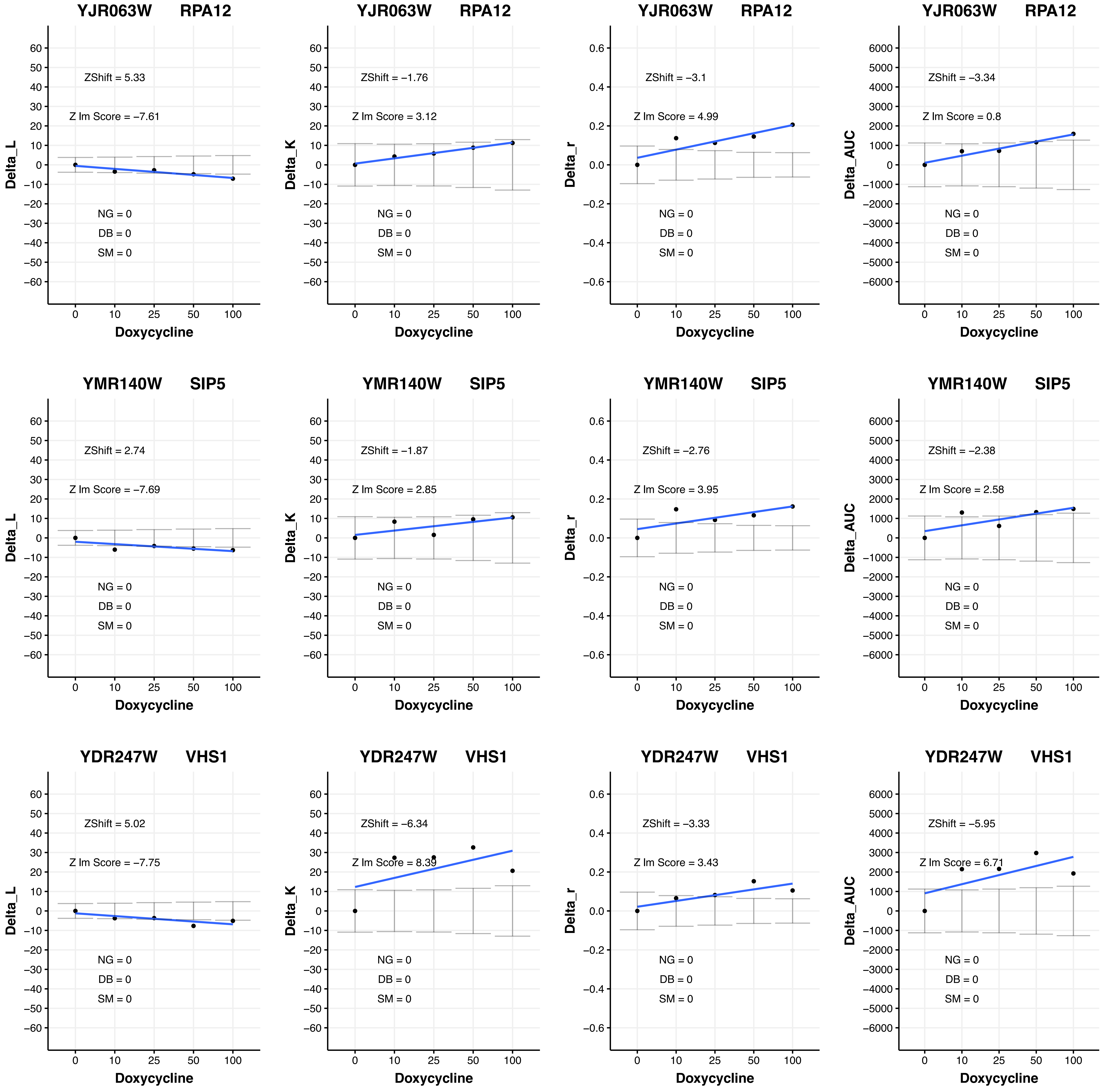

YJR063W RPA12
ZShift = 5.33
Z lm Score = −7.61
NG = 0
DB = 0
SM = 0
Delta_L
Doxycycline
YJR063W RPA12
ZShift = −1.76
Z lm Score = 3.12
NG = 0
DB = 0
SM = 0
Delta_K
Doxycycline
YJR063W RPA12
ZShift = −3.1
Z lm Score = 4.99
NG = 0
DB = 0
SM = 0
Delta_r
Doxycycline
YJR063W RPA12
ZShift = −3.34
Z lm Score = 0.8
NG = 0
DB = 0
SM = 0
Delta_AUC
Doxycycline
YMR140W SIP5
ZShift = 2.74
Z lm Score = −7.69
NG = 0
DB = 0
SM = 0
Delta_L
Doxycycline
YMR140W SIP5
ZShift = −1.87
Z lm Score = 2.85
NG = 0
DB = 0
SM = 0
Delta_K
Doxycycline
YMR140W SIP5
ZShift = −2.76
Z lm Score = 3.95
NG = 0
DB = 0
SM = 0
Delta_r
Doxycycline
YMR140W SIP5
ZShift = −2.38
Z lm Score = 2.58
NG = 0
DB = 0
SM = 0
Delta_AUC
Doxycycline
YDR247W VHS1
ZShift = 5.02
Z lm Score = −7.75
NG = 0
DB = 0
SM = 0
Delta_L
Doxycycline
YDR247W VHS1
ZShift = −6.34
Z lm Score = 8.39
NG = 0
DB = 0
SM = 0
Delta_K
Doxycycline
YDR247W VHS1
ZShift = −3.33
Z lm Score = 3.43
NG = 0
DB = 0
SM = 0
Delta_r
Doxycycline
YDR247W VHS1
ZShift = −5.95
Z lm Score = 6.71
NG = 0
DB = 0
SM = 0
Delta_AUC
Doxycycline

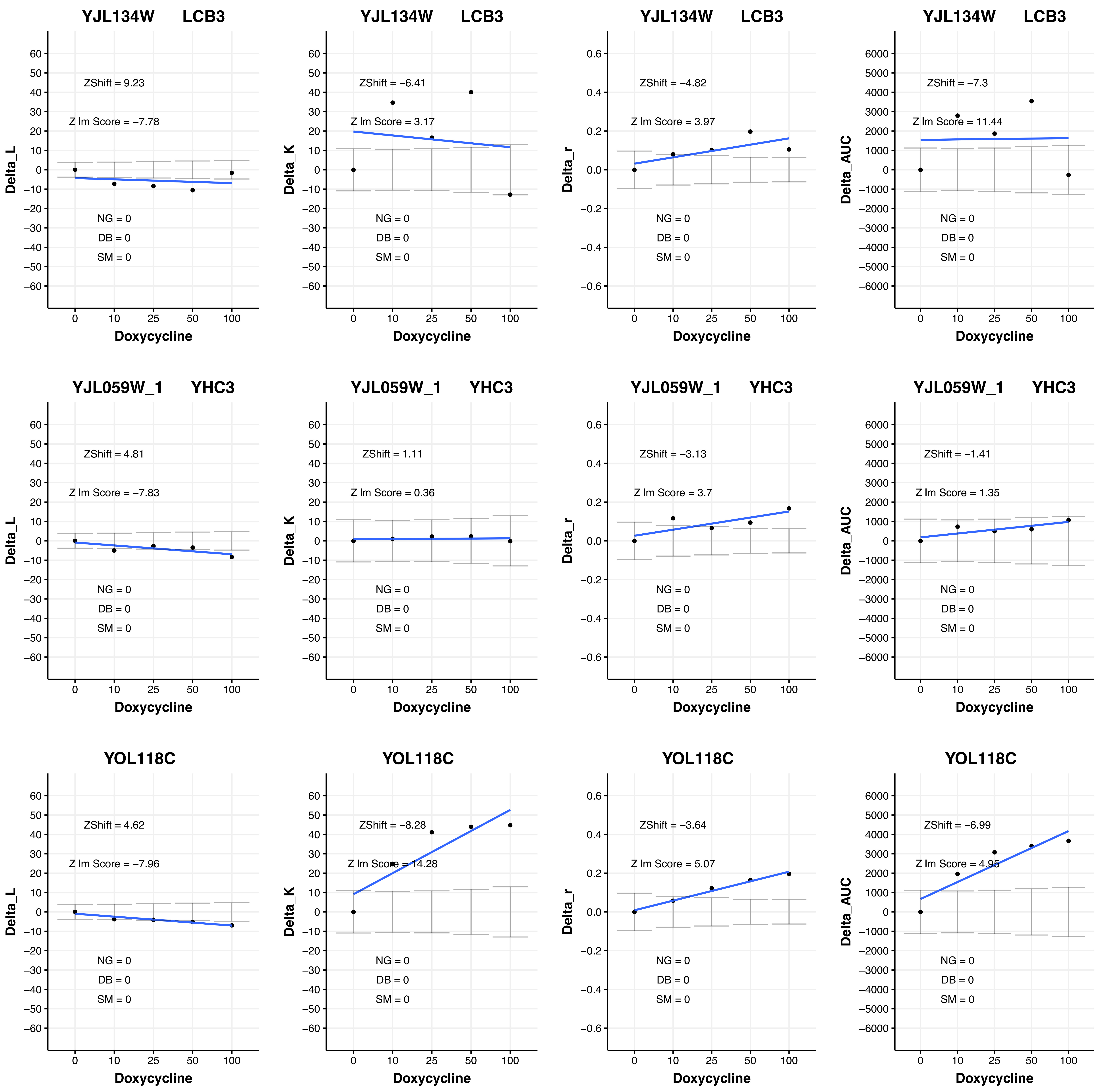

YJL134W LCB3
ZShift = 9.23
Z lm Score = −7.78
NG = 0
DB = 0
SM = 0
Delta_L
Doxycycline
YJL134W LCB3
ZShift = −6.41
Z lm Score = 3.17
NG = 0
DB = 0
SM = 0
Delta_K
Doxycycline
YJL134W LCB3
ZShift = −4.82
Z lm Score = 3.97
NG = 0
DB = 0
SM = 0
Delta_r
Doxycycline
YJL134W LCB3
ZShift = −7.3
Z lm Score = 11.44
NG = 0
DB = 0
SM = 0
Delta_AUC
Doxycycline
YJL059W_1 YHC3
ZShift = 4.81
Z lm Score = −7.83
NG = 0
DB = 0
SM = 0
Delta_L
Doxycycline
YJL059W_1 YHC3
ZShift = 1.11
Z lm Score = 0.36
NG = 0
DB = 0
SM = 0
Delta_K
Doxycycline
YJL059W_1 YHC3
ZShift = −3.13
Z lm Score = 3.7
NG = 0
DB = 0
SM = 0
Delta_r
Doxycycline
YJL059W_1 YHC3
ZShift = −1.41
Z lm Score = 1.35
NG = 0
DB = 0
SM = 0
Delta_AUC
Doxycycline
YOL118C
ZShift = 4.62
Z lm Score = −7.96
NG = 0
DB = 0
SM = 0
Delta_L
Doxycycline
YOL118C
ZShift = −8.28
Z lm Score = 14.28
NG = 0
DB = 0
SM = 0
Delta_K
Doxycycline
YOL118C
ZShift = −3.64
Z lm Score = 5.07
NG = 0
DB = 0
SM = 0
Delta_r
Doxycycline
YOL118C
ZShift = −6.99
Z lm Score = 4.95
NG = 0
DB = 0
SM = 0
Delta_AUC
Doxycycline

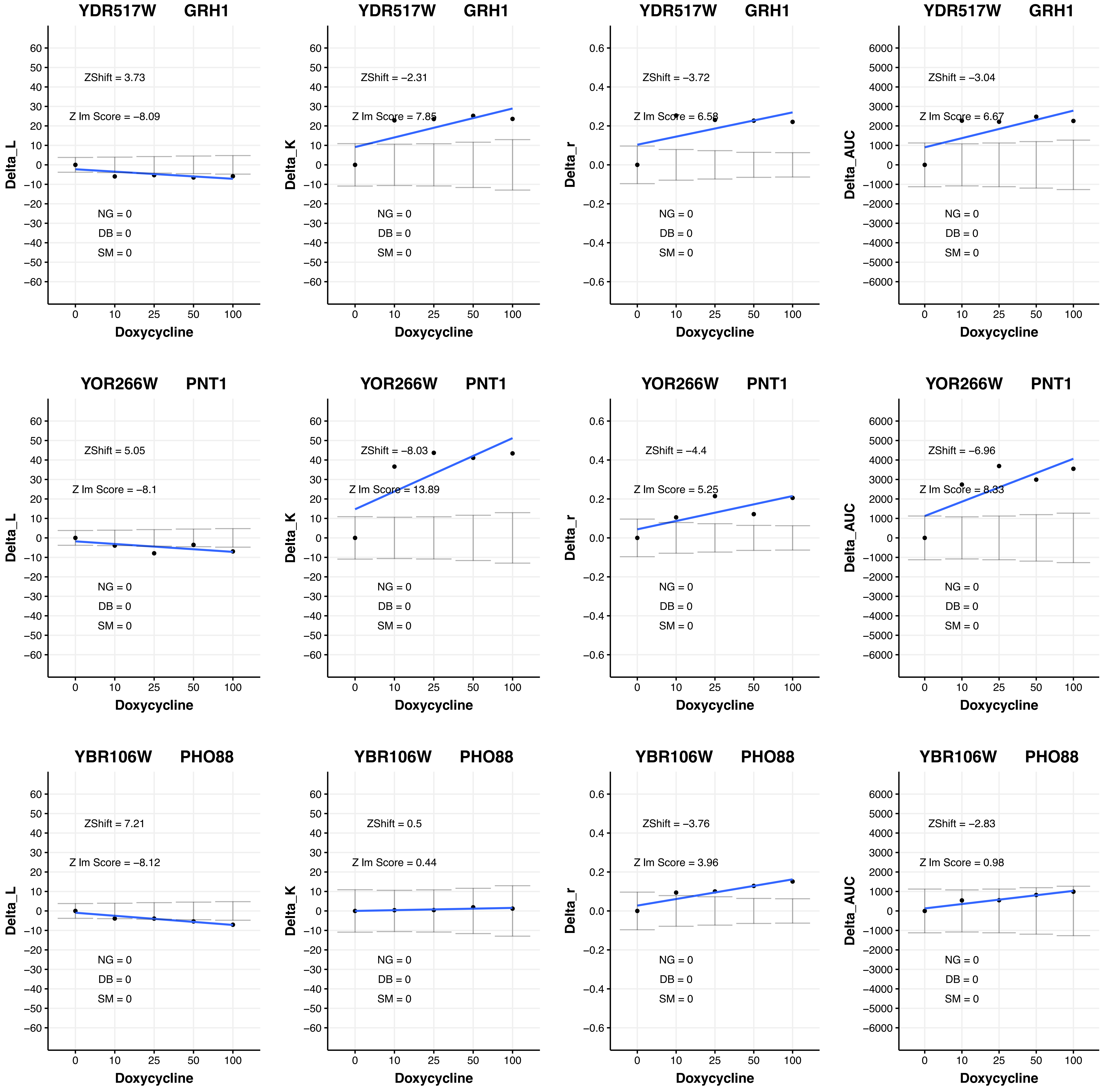

YDR517W GRH1
ZShift = 3.73
Z lm Score = −8.09
NG = 0
DB = 0
SM = 0
Delta_L
Doxycycline
YDR517W GRH1
ZShift = −2.31
Z lm Score = 7.85
NG = 0
DB = 0
SM = 0
Delta_K
Doxycycline
YDR517W GRH1
ZShift = −3.72
Z lm Score = 6.58
NG = 0
DB = 0
SM = 0
Delta_r
Doxycycline
YDR517W GRH1
ZShift = −3.04
Z lm Score = 6.67
NG = 0
DB = 0
SM = 0
Delta_AUC
Doxycycline
YOR266W PNT1
ZShift = 5.05
Z lm Score = −8.1
NG = 0
DB = 0
SM = 0
Delta_L
Doxycycline
YOR266W PNT1
ZShift = −8.03
Z lm Score = 13.89
NG = 0
DB = 0
SM = 0
Delta_K
Doxycycline
YOR266W PNT1
ZShift = −4.4
Z lm Score = 5.25
NG = 0
DB = 0
SM = 0
Delta_r
Doxycycline
YOR266W PNT1
ZShift = −6.96
Z lm Score = 8.33
NG = 0
DB = 0
SM = 0
Delta_AUC
Doxycycline
YBR106W PHO88
ZShift = 7.21
Z lm Score = −8.12
NG = 0
DB = 0
SM = 0
Delta_L
Doxycycline
YBR106W PHO88
ZShift = 0.5
Z lm Score = 0.44
NG = 0
DB = 0
SM = 0
Delta_K
Doxycycline
YBR106W PHO88
ZShift = −3.76
Z lm Score = 3.96
NG = 0
DB = 0
SM = 0
Delta_r
Doxycycline
YBR106W PHO88
ZShift = −2.83
Z lm Score = 0.98
NG = 0
DB = 0
SM = 0
Delta_AUC
Doxycycline

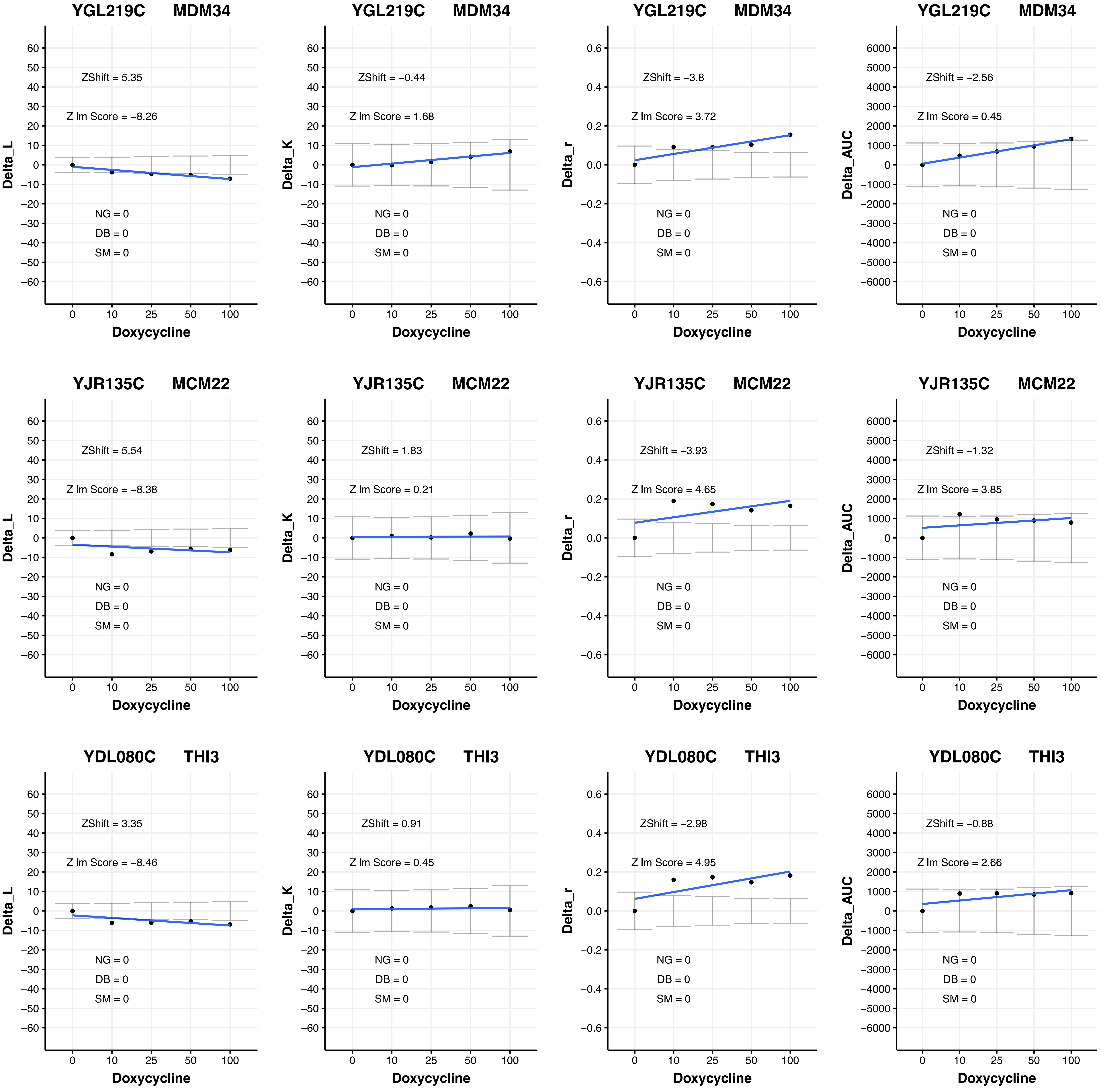
YGL219C MDM34
ZShift = 5.35
Z lm Score = −8.26
NG = 0
DB = 0
SM = 0
Delta_L
Doxycycline
YGL219C MDM34
ZShift = −0.44
Z lm Score = 1.68
NG = 0
DB = 0
SM = 0
Delta_K
Doxycycline
YGL219C MDM34
ZShift = −3.8
Z lm Score = 3.72
NG = 0
DB = 0
SM = 0
Delta_r
Doxycycline
YGL219C MDM34
ZShift = −2.56
Z lm Score = 0.45
NG = 0
DB = 0
SM = 0
Delta_AUC
Doxycycline
YJR135C MCM22
ZShift = 5.54
Z lm Score = −8.38
NG = 0
DB = 0
SM = 0
Delta_L
Doxycycline
YJR135C MCM22
ZShift = 1.83
Z lm Score = 0.21
NG = 0
DB = 0
SM = 0
Delta_K
Doxycycline
YJR135C MCM22
ZShift = −3.93
Z lm Score = 4.65
NG = 0
DB = 0
SM = 0
Delta_r
Doxycycline
YJR135C MCM22
ZShift = −1.32
Z lm Score = 3.85
NG = 0
DB = 0
SM = 0
Delta_AUC
Doxycycline
YDL080C THI3
ZShift = 3.35
Z lm Score = −8.46
NG = 0
DB = 0
SM = 0
Delta_L
Doxycycline
YDL080C THI3
ZShift = 0.91
Z lm Score = 0.45
NG = 0
DB = 0
SM = 0
Delta_K
Doxycycline
YDL080C THI3
ZShift = −2.98
Z lm Score = 4.95
NG = 0
DB = 0
SM = 0
Delta_r
Doxycycline
YDL080C THI3
ZShift = −0.88
Z lm Score = 2.66
NG = 0
DB = 0
SM = 0
Delta_AUC
Doxycycline

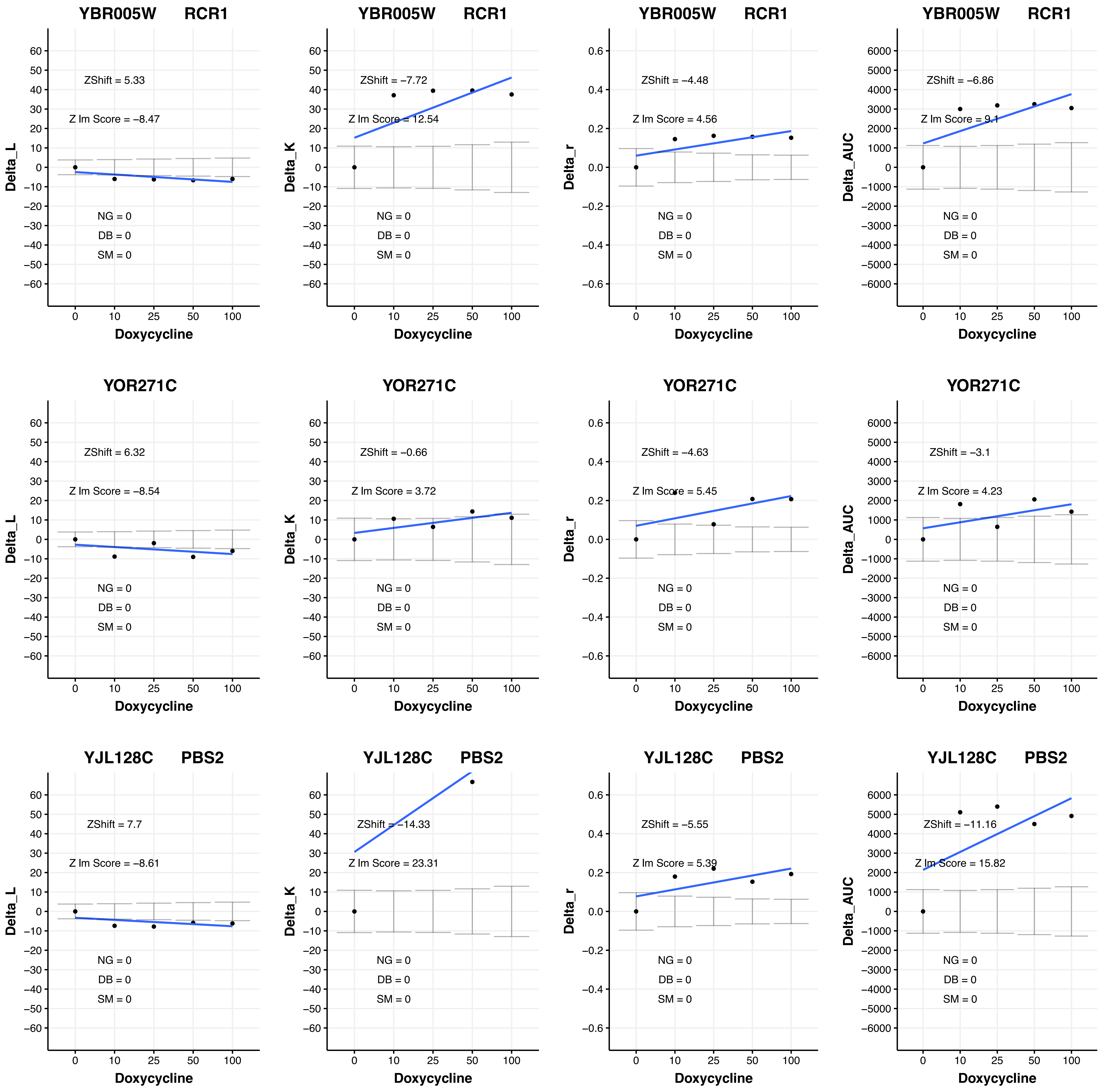
YBR005W RCR1
ZShift = 5.33
Z lm Score = −8.47
NG = 0
DB = 0
SM = 0
Delta_L
Doxycycline
YBR005W RCR1
ZShift = −7.72
Z lm Score = 12.54
NG = 0
DB = 0
SM = 0
Delta_K
Doxycycline
YBR005W RCR1
ZShift = −4.48
Z lm Score = 4.56
NG = 0
DB = 0
SM = 0
Delta_r
Doxycycline
YBR005W RCR1
ZShift = −6.86
Z lm Score = 9.1
NG = 0
DB = 0
SM = 0
Delta_AUC
Doxycycline
YOR271C
ZShift = 6.32
Z lm Score = −8.54
NG = 0
DB = 0
SM = 0
Delta_L
Doxycycline
YOR271C
ZShift = −0.66
Z lm Score = 3.72
NG = 0
DB = 0
SM = 0
Delta_K
Doxycycline
YOR271C
ZShift = −4.63
Z lm Score = 5.45
NG = 0
DB = 0
SM = 0
Delta_r
Doxycycline
YOR271C
ZShift = −3.1
Z lm Score = 4.23
NG = 0
DB = 0
SM = 0
Delta_AUC
Doxycycline
YJL128C PBS2
ZShift = 7.7
Z lm Score = −8.61
NG = 0
DB = 0
SM = 0
Delta_L
Doxycycline
YJL128C PBS2
ZShift = −14.33
Z lm Score = 23.31
NG = 0
DB = 0
SM = 0
Delta_K
Doxycycline
YJL128C PBS2
ZShift = −5.55
Z lm Score = 5.39
NG = 0
DB = 0
SM = 0
Delta_r
Doxycycline
YJL128C PBS2
ZShift = −11.16
Z lm Score = 15.82
NG = 0
DB = 0
SM = 0
Delta_AUC
Doxycycline

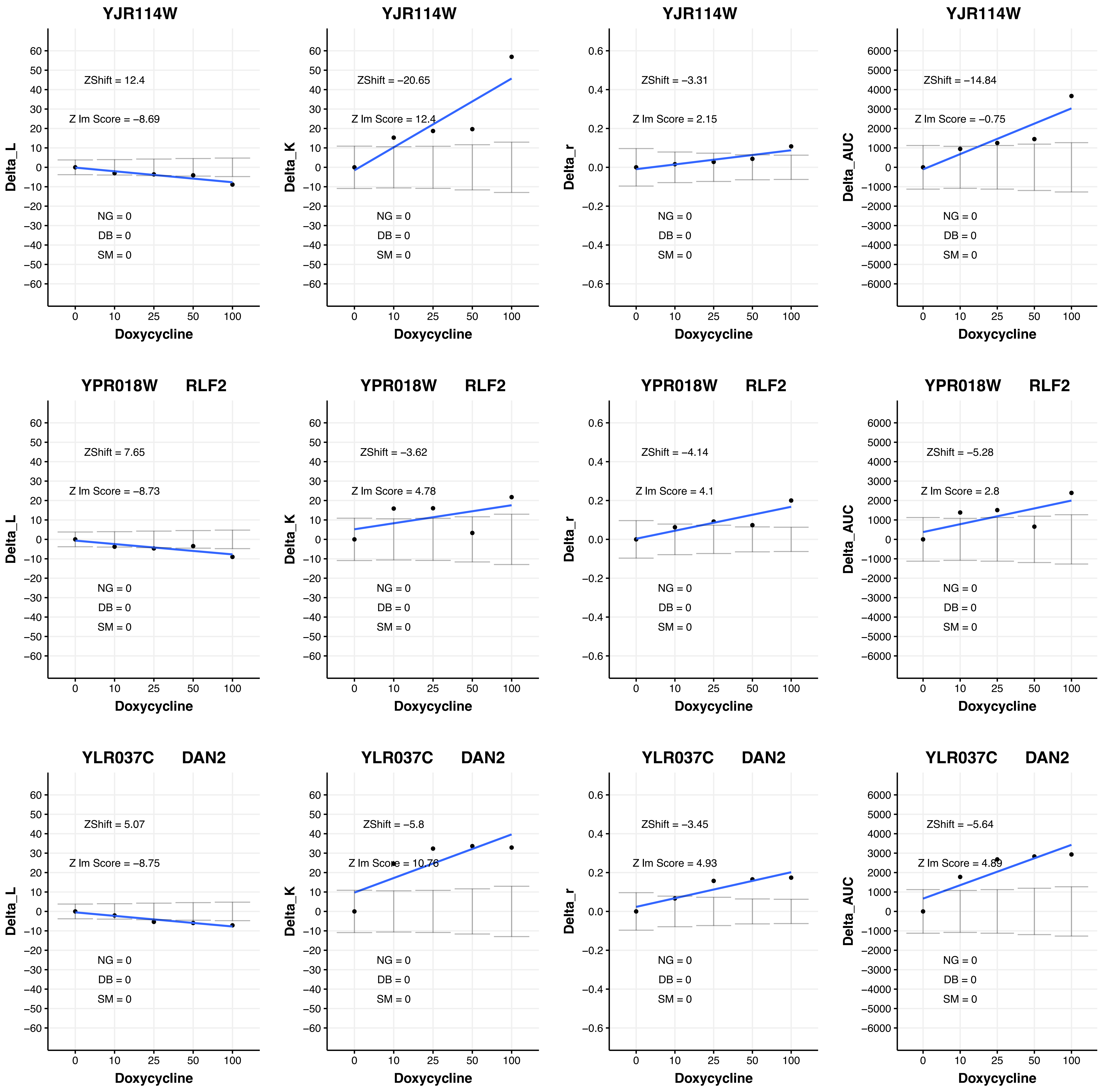
YJR114W
ZShift = 12.4
Z lm Score = −8.69
NG = 0
DB = 0
SM = 0
Delta_L
Doxycycline
YJR114W
ZShift = −20.65
Z lm Score = 12.4
NG = 0
DB = 0
SM = 0
Delta_K
Doxycycline
YJR114W
ZShift = −3.31
Z lm Score = 2.15
NG = 0
DB = 0
SM = 0
Delta_r
Doxycycline
YJR114W
ZShift = −14.84
Z lm Score = −0.75
NG = 0
DB = 0
SM = 0
Delta_AUC
Doxycycline
YPR018W RLF2
ZShift = 7.65
Z lm Score = −8.73
NG = 0
DB = 0
SM = 0
Delta_L
Doxycycline
YPR018W RLF2
ZShift = −3.62
Z lm Score = 4.78
NG = 0
DB = 0
SM = 0
Delta_K
Doxycycline
YPR018W RLF2
ZShift = −4.14
Z lm Score = 4.1
NG = 0
DB = 0
SM = 0
Delta_r
Doxycycline
YPR018W RLF2
ZShift = −5.28
Z lm Score = 2.8
NG = 0
DB = 0
SM = 0
Delta_AUC
Doxycycline
YLR037C DAN2
ZShift = 5.07
Z lm Score = −8.75
NG = 0
DB = 0
SM = 0
Delta_L
Doxycycline
YLR037C DAN2
ZShift = −5.8
Z lm Score = 10.76
NG = 0
DB = 0
SM = 0
Delta_K
Doxycycline
YLR037C DAN2
ZShift = −3.45
Z lm Score = 4.93
NG = 0
DB = 0
SM = 0
Delta_r
Doxycycline
YLR037C DAN2
ZShift = −5.64
Z lm Score = 4.89
NG = 0
DB = 0
SM = 0
Delta_AUC
Doxycycline

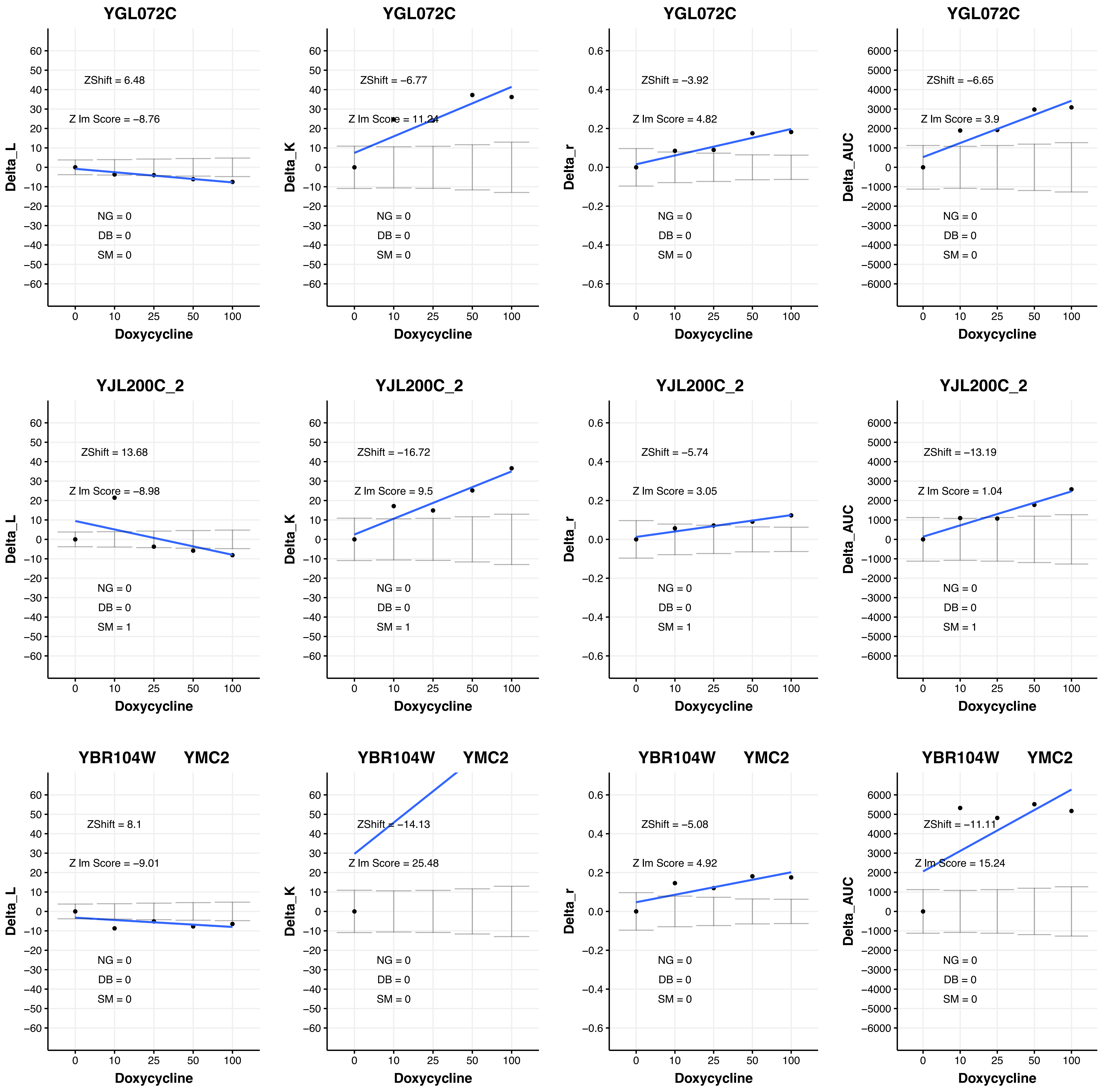
YGL072C
ZShift = 6.48
Z lm Score = −8.76
NG = 0
DB = 0
SM = 0
Delta_L
Doxycycline
YGL072C
ZShift = −6.77
Z lm Score = 11.24
NG = 0
DB = 0
SM = 0
Delta_K
Doxycycline
YGL072C
ZShift = −3.92
Z lm Score = 4.82
NG = 0
DB = 0
SM = 0
Delta_r
Doxycycline
YGL072C
ZShift = −6.65
Z lm Score = 3.9
NG = 0
DB = 0
SM = 0
Delta_AUC
Doxycycline
YJL200C_2
ZShift = 13.68
Z lm Score = −8.98
NG = 0
DB = 0
SM = 1
Delta_L
Doxycycline
YJL200C_2
ZShift = −16.72
Z lm Score = 9.5
NG = 0
DB = 0
SM = 1
Delta_K
Doxycycline
YJL200C_2
ZShift = −5.74
Z lm Score = 3.05
NG = 0
DB = 0
SM = 1
Delta_r
Doxycycline
YJL200C_2
ZShift = −13.19
Z lm Score = 1.04
NG = 0
DB = 0
SM = 1
Delta_AUC
Doxycycline
YBR104W YMC2
ZShift = 8.1
Z lm Score = −9.01
NG = 0
DB = 0
SM = 0
Delta_L
Doxycycline
YBR104W YMC2
ZShift = −14.13
Z lm Score = 25.48
NG = 0
DB = 0
SM = 0
Delta_K
Doxycycline
YBR104W YMC2
ZShift = −5.08
Z lm Score = 4.92
NG = 0
DB = 0
SM = 0
Delta_r
Doxycycline
YBR104W YMC2
ZShift = −11.11
Z lm Score = 15.24
NG = 0
DB = 0
SM = 0
Delta_AUC
Doxycycline
0 10 25 50 100

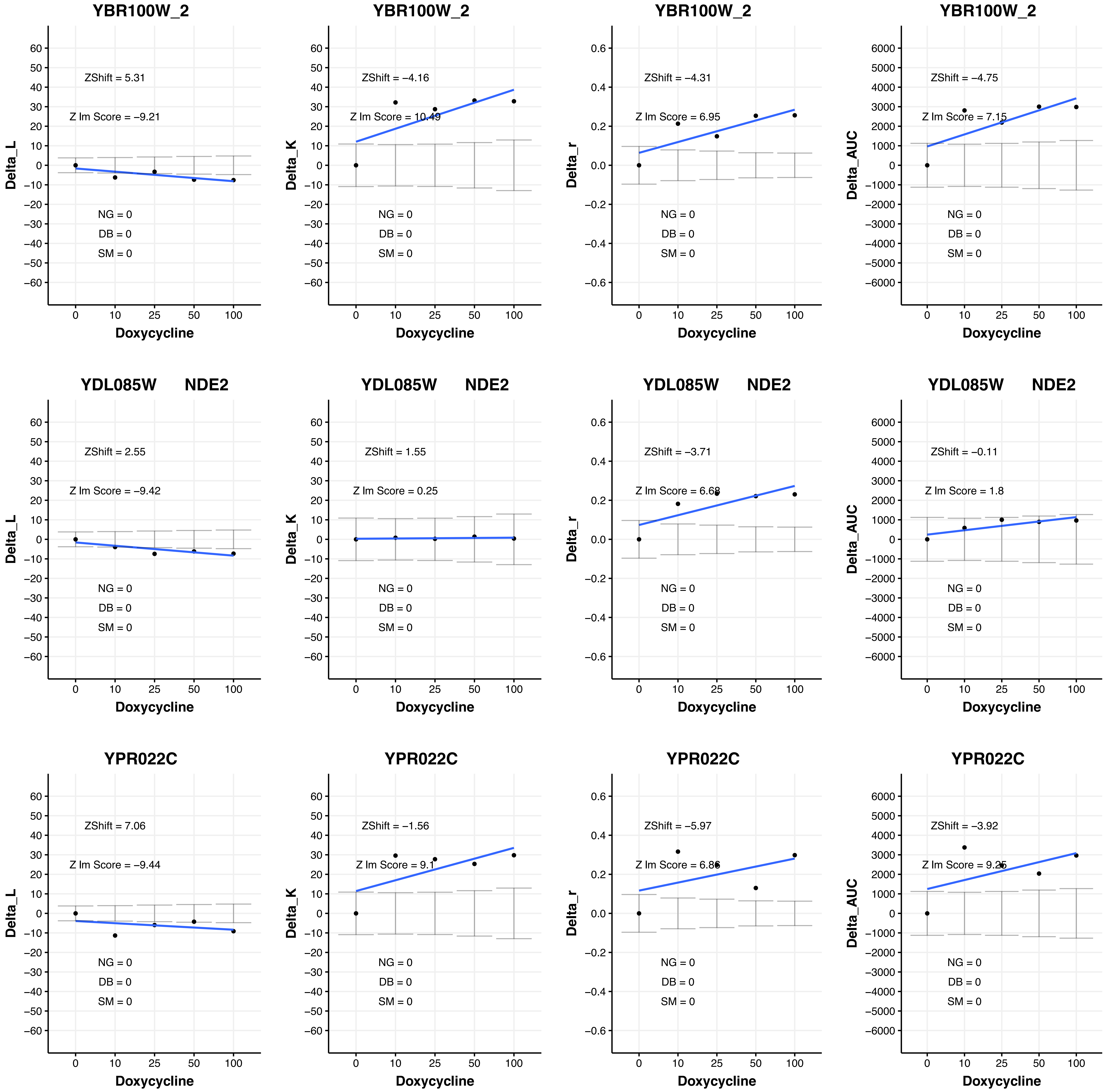
YBR100W_2
ZShift = 5.31
Z lm Score = −9.21
NG = 0
DB = 0
SM = 0
Delta_L
Doxycycline
YBR100W_2
ZShift = −4.16
Z lm Score = 10.49
NG = 0
DB = 0
SM = 0
Delta_K
Doxycycline
YBR100W_2
ZShift = −4.31
Z lm Score = 6.95
NG = 0
DB = 0
SM = 0
Delta_r
Doxycycline
YBR100W_2
ZShift = −4.75
Z lm Score = 7.15
NG = 0
DB = 0
SM = 0
Delta_AUC
Doxycycline
YDL085W NDE2
ZShift = 2.55
Z lm Score = −9.42
NG = 0
DB = 0
SM = 0
Delta_L
Doxycycline
YDL085W NDE2
ZShift = 1.55
Z lm Score = 0.25
NG = 0
DB = 0
SM = 0
Delta_K
Doxycycline
YDL085W NDE2
ZShift = −3.71
Z lm Score = 6.68
NG = 0
DB = 0
SM = 0
Delta_r
Doxycycline
YDL085W NDE2
ZShift = −0.11
Z lm Score = 1.8
NG = 0
DB = 0
SM = 0
Delta_AUC
Doxycycline
YPR022C
ZShift = 7.06
Z lm Score = −9.44
NG = 0
DB = 0
SM = 0
Delta_L
Doxycycline
YPR022C
ZShift = −1.56
Z lm Score = 9.1
NG = 0
DB = 0
SM = 0
Delta_K
Doxycycline
YPR022C
ZShift = −5.97
Z lm Score = 6.86
NG = 0
DB = 0
SM = 0
Delta_r
Doxycycline
YPR022C
ZShift = −3.92
Z lm Score = 9.25
NG = 0
DB = 0
SM = 0
Delta_AUC
Doxycycline

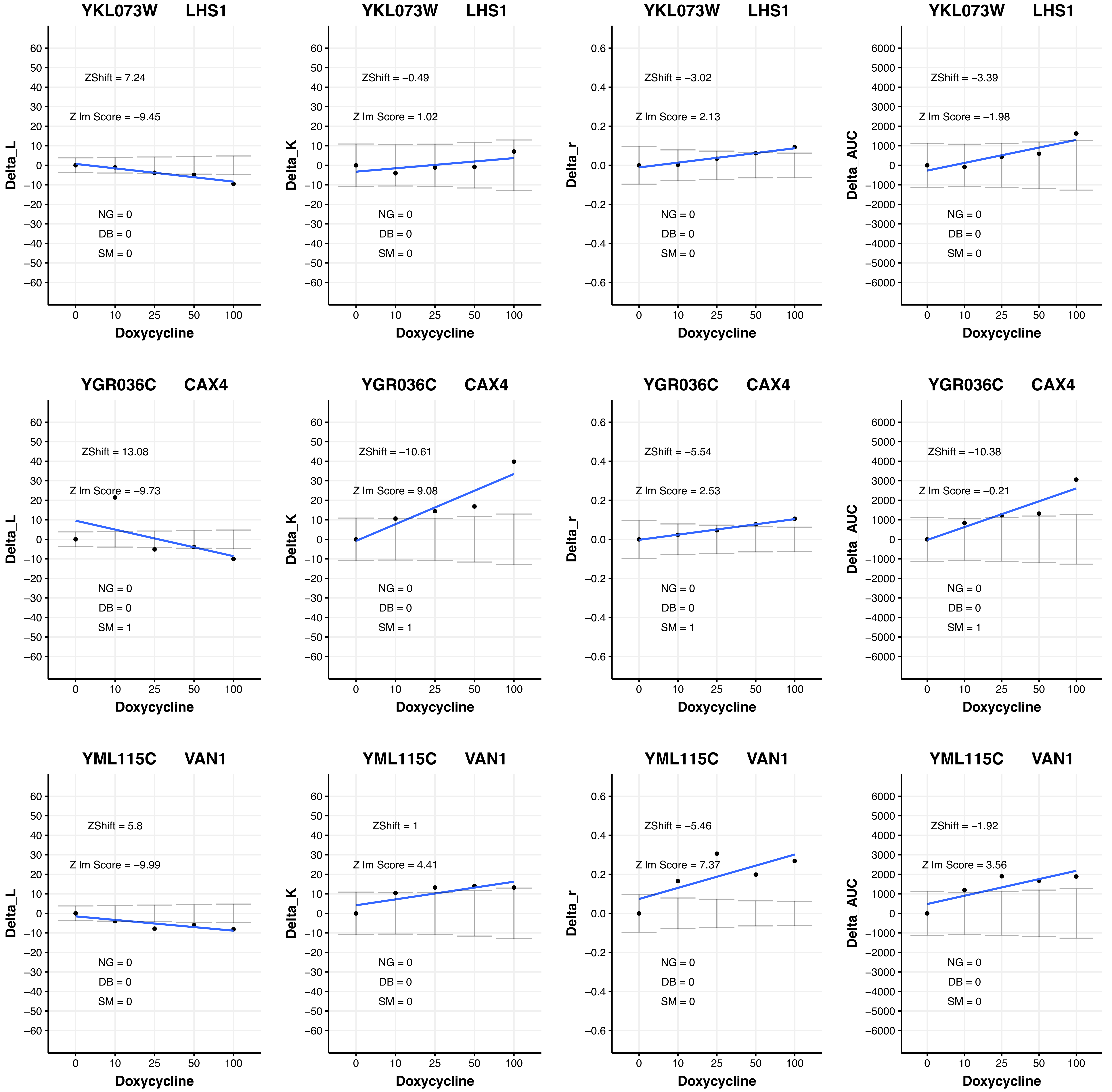
YKL073W LHS1
ZShift = 7.24
Z lm Score = −9.45
NG = 0
DB = 0
SM = 0
Delta_L
Doxycycline
YKL073W LHS1
ZShift = −0.49
Z lm Score = 1.02
NG = 0
DB = 0
SM = 0
Delta_K
Doxycycline
YKL073W LHS1
ZShift = −3.02
Z lm Score = 2.13
NG = 0
DB = 0
SM = 0
Delta_r
Doxycycline
YKL073W LHS1
ZShift = −3.39
Z lm Score = −1.98
NG = 0
DB = 0
SM = 0
Delta_AUC
Doxycycline
YGR036C CAX4
ZShift = 13.08
Z lm Score = −9.73
NG = 0
DB = 0
SM = 1
Delta_L
Doxycycline
YGR036C CAX4
ZShift = −10.61
Z lm Score = 9.08
NG = 0
DB = 0
SM = 1
Delta_K
Doxycycline
YGR036C CAX4
ZShift = −5.54
Z lm Score = 2.53
NG = 0
DB = 0
SM = 1
Delta_r
Doxycycline
YGR036C CAX4
ZShift = −10.38
Z lm Score = −0.21
NG = 0
DB = 0
SM = 1
Delta_AUC
Doxycycline
YML115C VAN1
ZShift = 5.8
Z lm Score = −9.99
NG = 0
DB = 0
SM = 0
Delta_L
Doxycycline
YML115C VAN1
ZShift = 1
Z lm Score = 4.41
NG = 0
DB = 0
SM = 0
Delta_K
Doxycycline
YML115C VAN1
ZShift = −5.46
Z lm Score = 7.37
NG = 0
DB = 0
SM = 0
Delta_r
Doxycycline
YML115C VAN1
ZShift = −1.92
Z lm Score = 3.56
NG = 0
DB = 0
SM = 0
Delta_AUC
Doxycycline

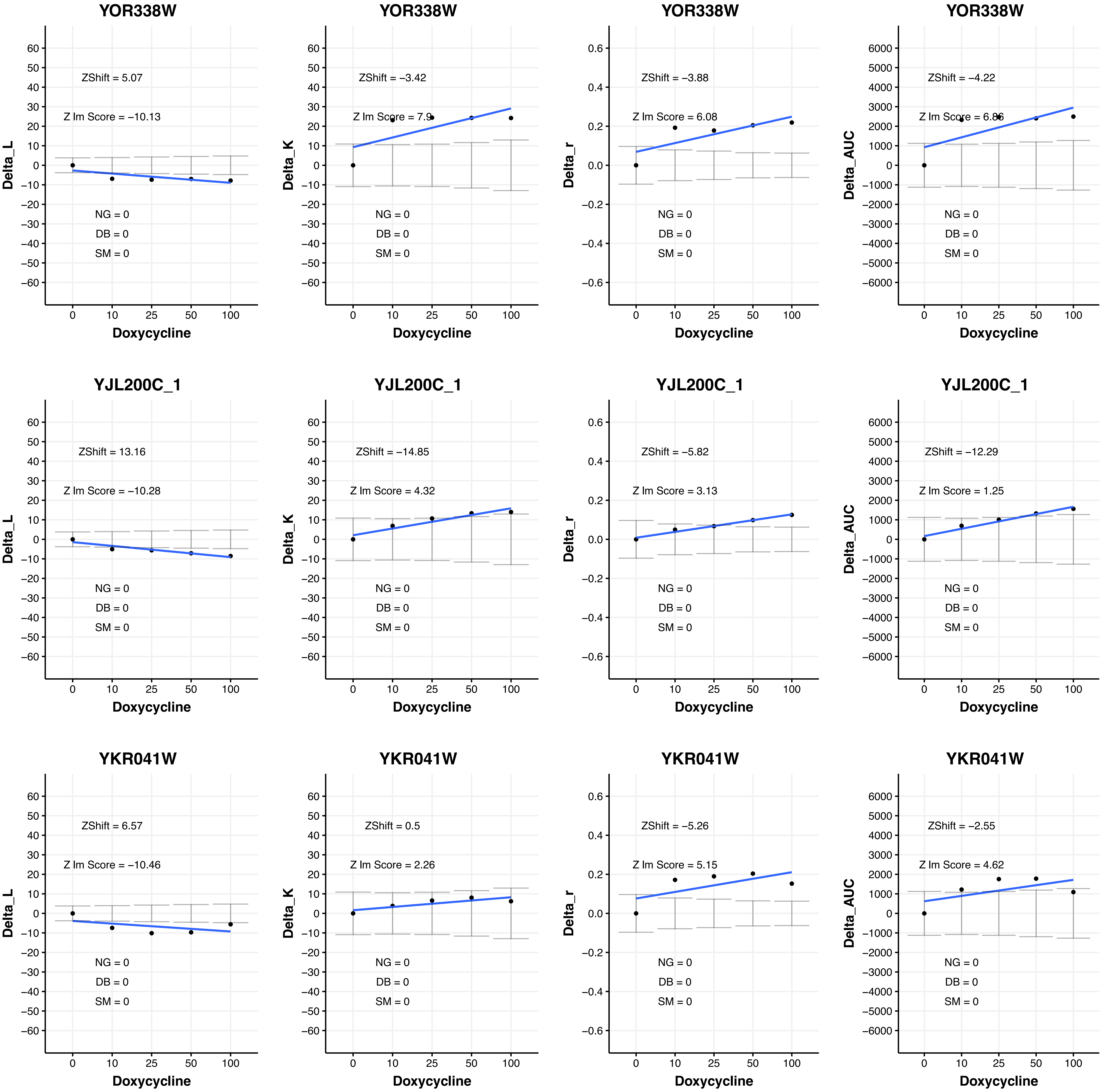
YOR338W
ZShift = 5.07
Z lm Score = −10.13
NG = 0
DB = 0
SM = 0
Delta_L
Doxycycline
YOR338W
ZShift = −3.42
Z lm Score = 7.9
NG = 0
DB = 0
SM = 0
Delta_K
Doxycycline
YOR338W
ZShift = −3.88
Z lm Score = 6.08
NG = 0
DB = 0
SM = 0
Delta_r
Doxycycline
YOR338W
ZShift = −4.22
Z lm Score = 6.86
NG = 0
DB = 0
SM = 0
Delta_AUC
Doxycycline
YJL200C_1
ZShift = 13.16
Z lm Score = −10.28
NG = 0
DB = 0
SM = 0
Delta_L
Doxycycline
YJL200C_1
ZShift = −14.85
Z lm Score = 4.32
NG = 0
DB = 0
SM = 0
Delta_K
Doxycycline
YJL200C_1
ZShift = −5.82
Z lm Score = 3.13
NG = 0
DB = 0
SM = 0
Delta_r
Doxycycline
YJL200C_1
ZShift = −12.29
Z lm Score = 1.25
NG = 0
DB = 0
SM = 0
Delta_AUC
Doxycycline
YKR041W
ZShift = 6.57
Z lm Score = −10.46
NG = 0
DB = 0
SM = 0
Delta_L
Doxycycline
YKR041W
ZShift = 0.5
Z lm Score = 2.26
NG = 0
DB = 0
SM = 0
Delta_K
Doxycycline
YKR041W
ZShift = −5.26
Z lm Score = 5.15
NG = 0
DB = 0
SM = 0
Delta_r
Doxycycline
YKR041W
ZShift = −2.55
Z lm Score = 4.62
NG = 0
DB = 0
SM = 0
Delta_AUC
Doxycycline
0
10
25
50
100

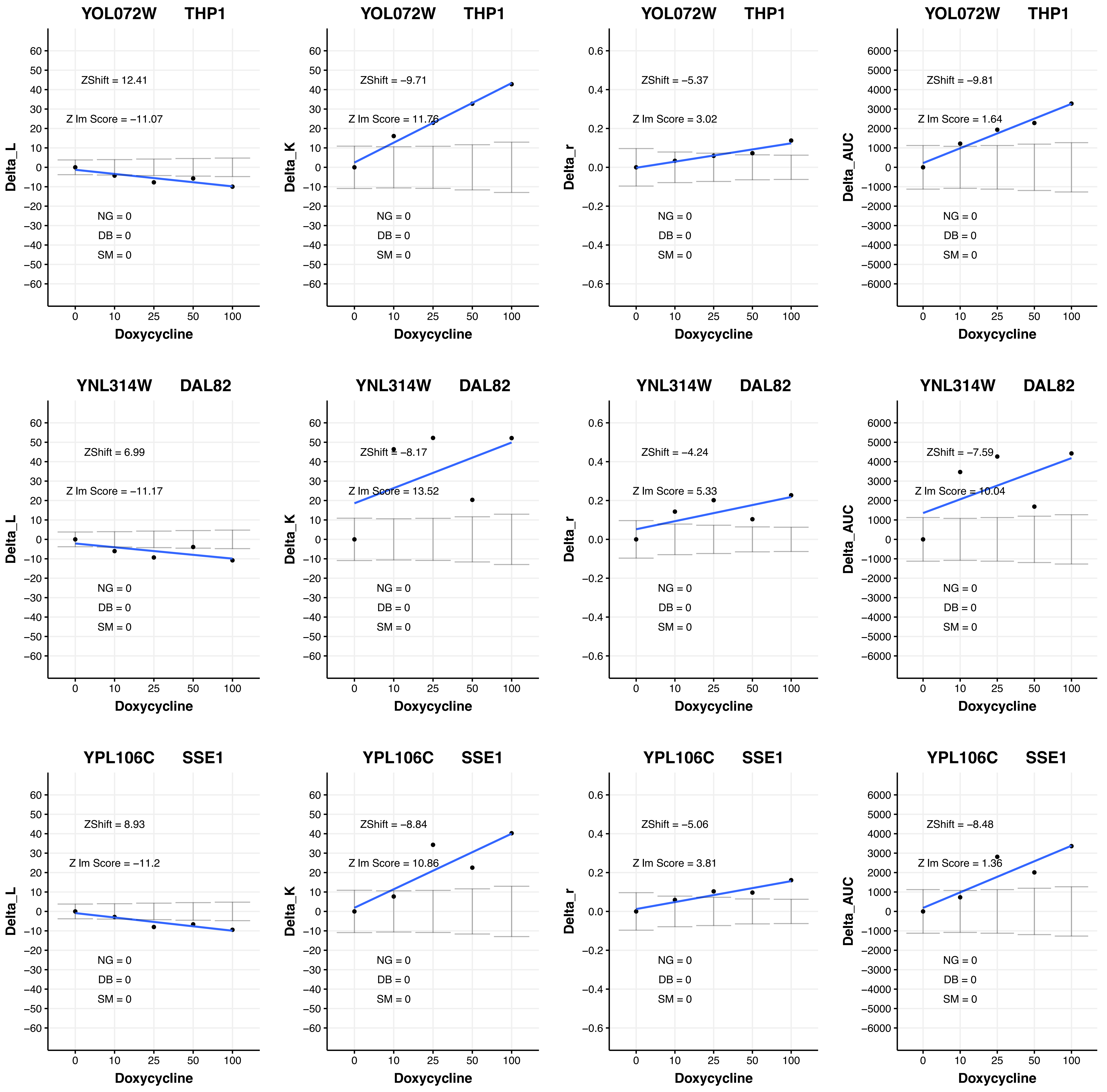

YOL072W THP1
ZShift = 12.41
Z lm Score = −11.07
NG = 0
DB = 0
SM = 0
Delta_L
Doxycycline
YOL072W THP1
ZShift = −9.71
Z lm Score = 11.76
NG = 0
DB = 0
SM = 0
Delta_K
Doxycycline
YOL072W THP1
ZShift = −5.37
Z lm Score = 3.02
NG = 0
DB = 0
SM = 0
Delta_r
Doxycycline
YOL072W THP1
ZShift = −9.81
Z lm Score = 1.64
NG = 0
DB = 0
SM = 0
Delta_AUC
Doxycycline
YNL314W DAL82
ZShift = 6.99
Z lm Score = −11.17
NG = 0
DB = 0
SM = 0
Delta_L
Doxycycline
YNL314W DAL82
ZShift = −8.17
Z lm Score = 13.52
NG = 0
DB = 0
SM = 0
Delta_K
Doxycycline
YNL314W DAL82
ZShift = −4.24
Z lm Score = 5.33
NG = 0
DB = 0
SM = 0
Delta_r
Doxycycline
YNL314W DAL82
ZShift = −7.59
Z lm Score = 10.04
NG = 0
DB = 0
SM = 0
Delta_AUC
Doxycycline
YPL106C SSE1
ZShift = 8.93
Z lm Score = −11.2
NG = 0
DB = 0
SM = 0
Delta_L
Doxycycline
YPL106C SSE1
ZShift = −8.84
Z lm Score = 10.86
NG = 0
DB = 0
SM = 0
Delta_K
Doxycycline
YPL106C SSE1
ZShift = −5.06
Z lm Score = 3.81
NG = 0
DB = 0
SM = 0
Delta_r
Doxycycline
YPL106C SSE1
ZShift = −8.48
Z lm Score = 1.36
NG = 0
DB = 0
SM = 0
Delta_AUC
Doxycycline

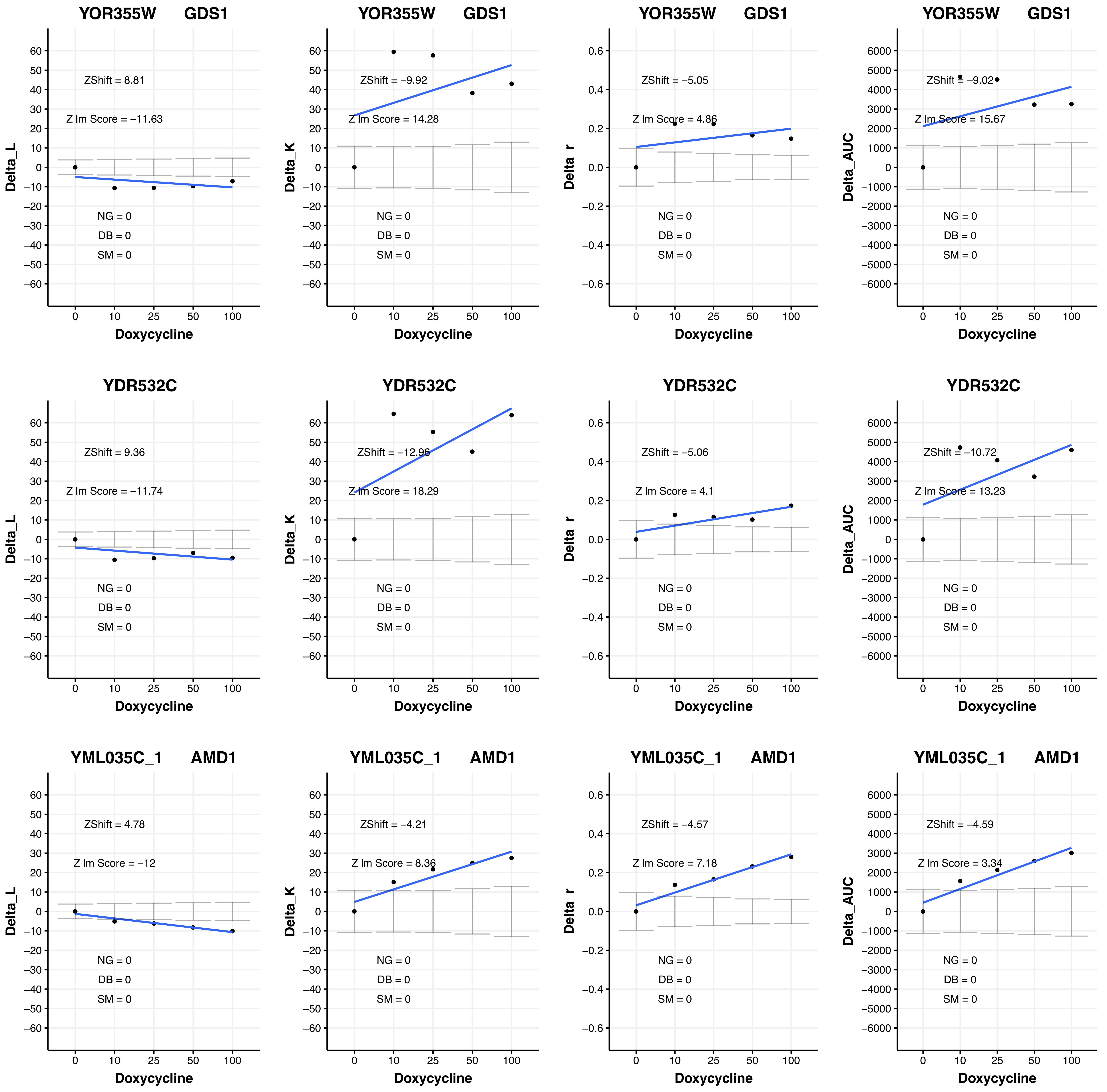
YOR355W GDS1
ZShift = 8.81
Z lm Score = −11.63
NG = 0
DB = 0
SM = 0
Delta_L
Doxycycline
YOR355W GDS1
ZShift = −9.92
Z lm Score = 14.28
NG = 0
DB = 0
SM = 0
Delta_K
Doxycycline
YOR355W GDS1
ZShift = −5.05
Z lm Score = 4.86
NG = 0
DB = 0
SM = 0
Delta_r
Doxycycline
YOR355W GDS1
ZShift = −9.02
Z lm Score = 15.67
NG = 0
DB = 0
SM = 0
Delta_AUC
Doxycycline
YDR532C
ZShift = 9.36
Z lm Score = −11.74
NG = 0
DB = 0
SM = 0
Delta_L
Doxycycline
YDR532C
ZShift = −12.96
Z lm Score = 18.29
NG = 0
DB = 0
SM = 0
Delta_K
Doxycycline
YDR532C
ZShift = −5.06
Z lm Score = 4.1
NG = 0
DB = 0
SM = 0
Delta_r
Doxycycline
YDR532C
ZShift = −10.72
Z lm Score = 13.23
NG = 0
DB = 0
SM = 0
Delta_AUC
Doxycycline
YML035C_1 AMD1
ZShift = 4.78
Z lm Score = −12
NG = 0
DB = 0
SM = 0
Delta_L
Doxycycline
YML035C_1 AMD1
ZShift = −4.21
Z lm Score = 8.36
NG = 0
DB = 0
SM = 0
Delta_K
Doxycycline
YML035C_1 AMD1
ZShift = −4.57
Z lm Score = 7.18
NG = 0
DB = 0
SM = 0
Delta_r
Doxycycline
YML035C_1 AMD1
ZShift = −4.59
Z lm Score = 3.34
NG = 0
DB = 0
SM = 0
Delta_AUC
Doxycycline

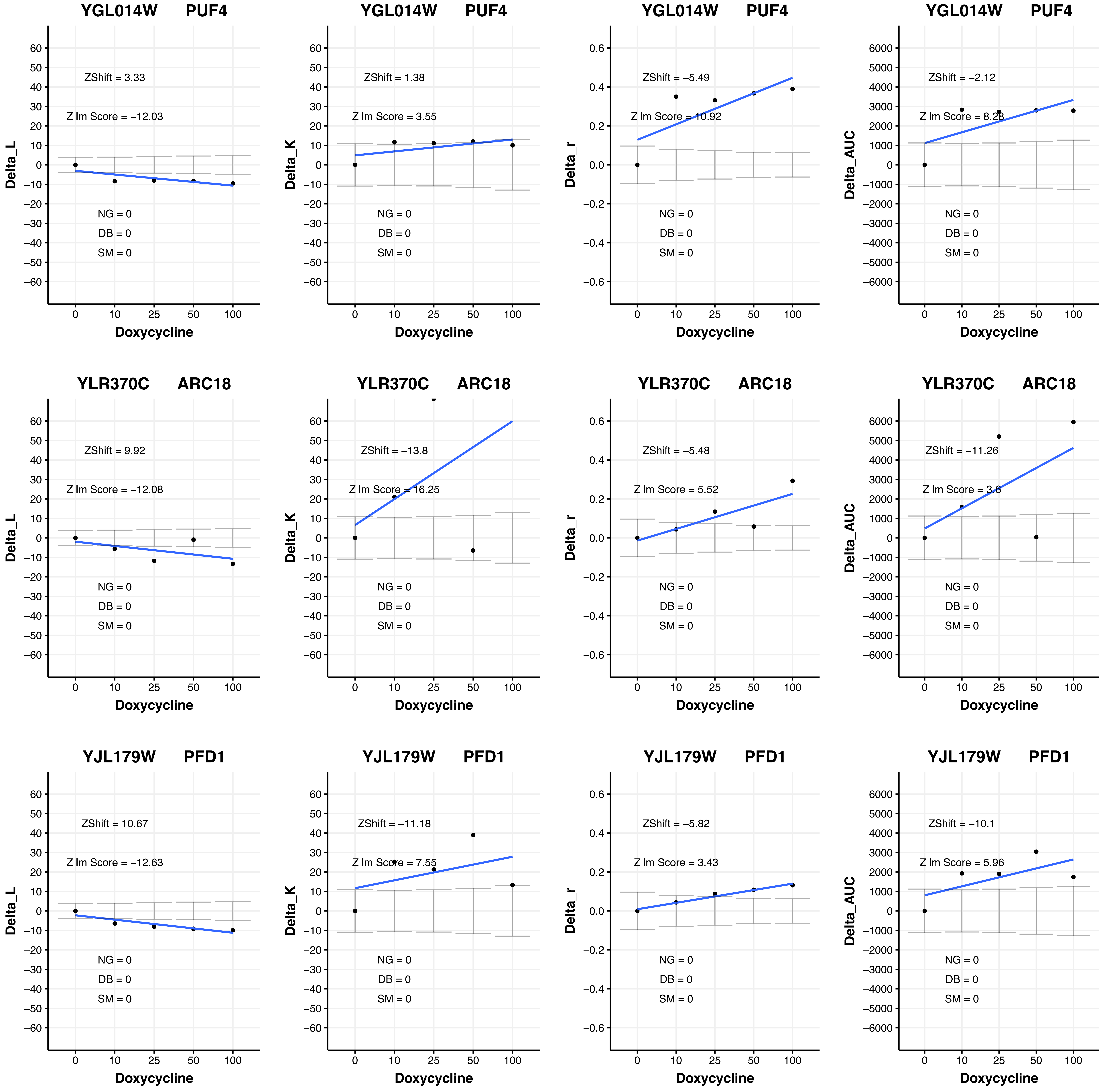

YGL014W PUF4
ZShift = 3.33
Z lm Score = −12.03
NG = 0
DB = 0
SM = 0
Delta_L
Doxycycline
YGL014W PUF4
ZShift = 1.38
Z lm Score = 3.55
NG = 0
DB = 0
SM = 0
Delta_K
Doxycycline
YGL014W PUF4
ZShift = −5.49
Z lm Score = 10.92
NG = 0
DB = 0
SM = 0
Delta_r
Doxycycline
YGL014W PUF4
ZShift = −2.12
Z lm Score = 8.28
NG = 0
DB = 0
SM = 0
Delta_AUC
Doxycycline
YLR370C ARC18
ZShift = 9.92
Z lm Score = −12.08
NG = 0
DB = 0
SM = 0
Delta_L
Doxycycline
YLR370C ARC18
ZShift = −13.8
Z lm Score = 16.25
NG = 0
DB = 0
SM = 0
Delta_K
Doxycycline
YLR370C ARC18
ZShift = −5.48
Z lm Score = 5.52
NG = 0
DB = 0
SM = 0
Delta_r
Doxycycline
YLR370C ARC18
ZShift = −11.26
Z lm Score = 3.6
NG = 0
DB = 0
SM = 0
Delta_AUC
Doxycycline
YJL179W PFD1
ZShift = 10.67
Z lm Score = −12.63
NG = 0
DB = 0
SM = 0
Delta_L
Doxycycline
YJL179W PFD1
ZShift = −11.18
Z lm Score = 7.55
NG = 0
DB = 0
SM = 0
Delta_K
Doxycycline
YJL179W PFD1
ZShift = −5.82
Z lm Score = 3.43
NG = 0
DB = 0
SM = 0
Delta_r
Doxycycline
YJL179W PFD1
ZShift = −10.1
Z lm Score = 5.96
NG = 0
DB = 0
SM = 0
Delta_AUC
Doxycycline

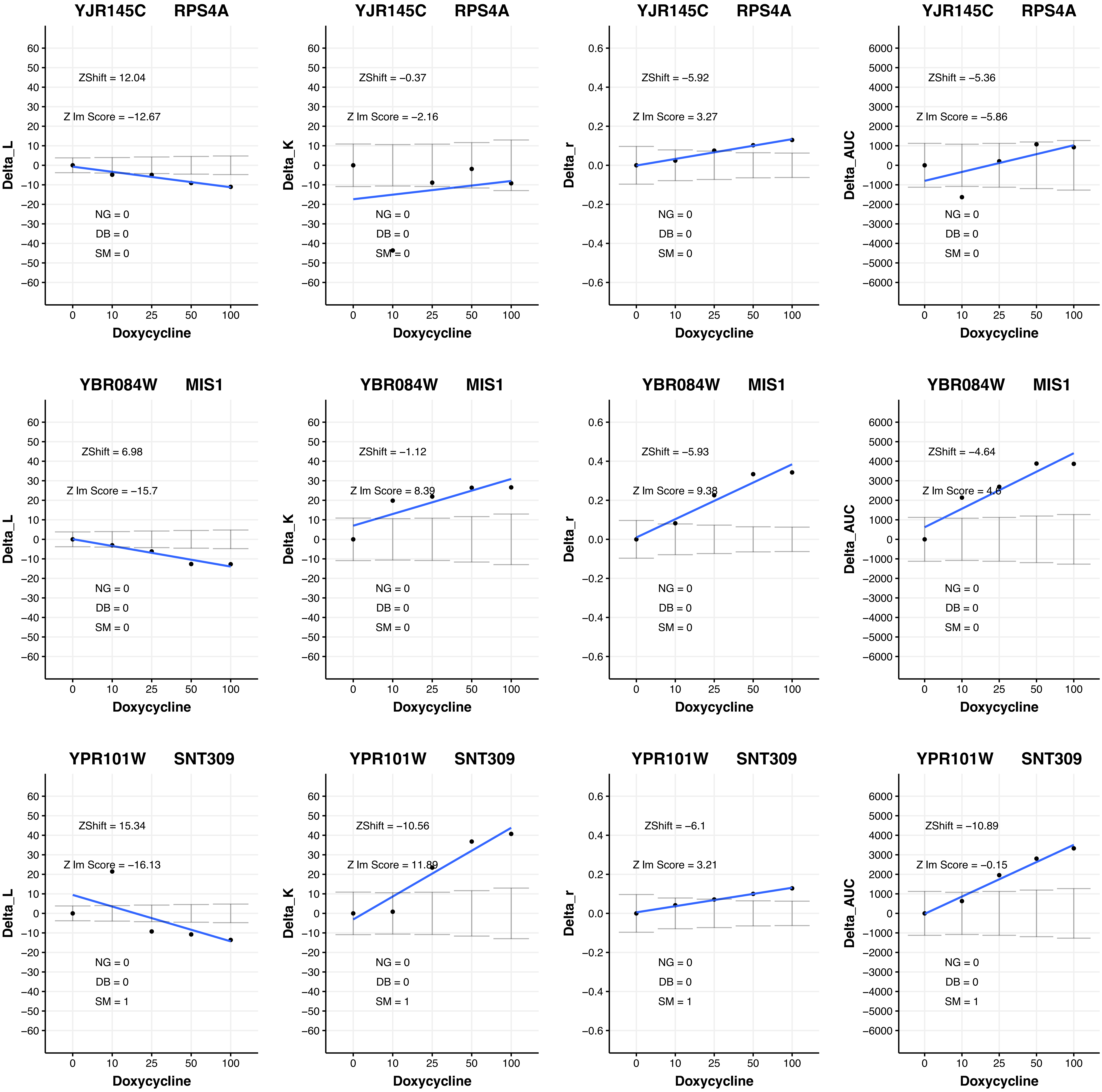
YJR145C RPS4A
ZShift = 12.04
Z lm Score = −12.67
NG = 0
DB = 0
SM = 0
Delta_L
Doxycycline
YJR145C RPS4A
ZShift = −0.37
Z lm Score = −2.16
NG = 0
DB = 0
SM = 0
Delta_K
Doxycycline
YJR145C RPS4A
ZShift = −5.92
Z lm Score = 3.27
NG = 0
DB = 0
SM = 0
Delta_r
Doxycycline
YJR145C RPS4A
ZShift = −5.36
Z lm Score = −5.86
NG = 0
DB = 0
SM = 0
Delta_AUC
Doxycycline
YBR084W MIS1
ZShift = 6.98
Z lm Score = −15.7
NG = 0
DB = 0
SM = 0
Delta_L
Doxycycline
YBR084W MIS1
ZShift = −1.12
Z lm Score = 8.39
NG = 0
DB = 0
SM = 0
Delta_K
Doxycycline
YBR084W MIS1
ZShift = −5.93
Z lm Score = 9.38
NG = 0
DB = 0
SM = 0
Delta_r
Doxycycline
YBR084W MIS1
ZShift = −4.64
Z lm Score = 4.6
NG = 0
DB = 0
SM = 0
Delta_AUC
Doxycycline
YPR101W SNT309
ZShift = 15.34
Z lm Score = −16.13
NG = 0
DB = 0
SM = 1
Delta_L
Doxycycline
YPR101W SNT309
ZShift = −10.56
Z lm Score = 11.89
NG = 0
DB = 0
SM = 1
Delta_K
Doxycycline
YPR101W SNT309
ZShift = −6.1
Z lm Score = 3.21
NG = 0
DB = 0
SM = 1
Delta_r
Doxycycline
YPR101W SNT309
ZShift = −10.89
Z lm Score = −0.15
NG = 0
DB = 0
SM = 1
Delta_AUC
Doxycycline

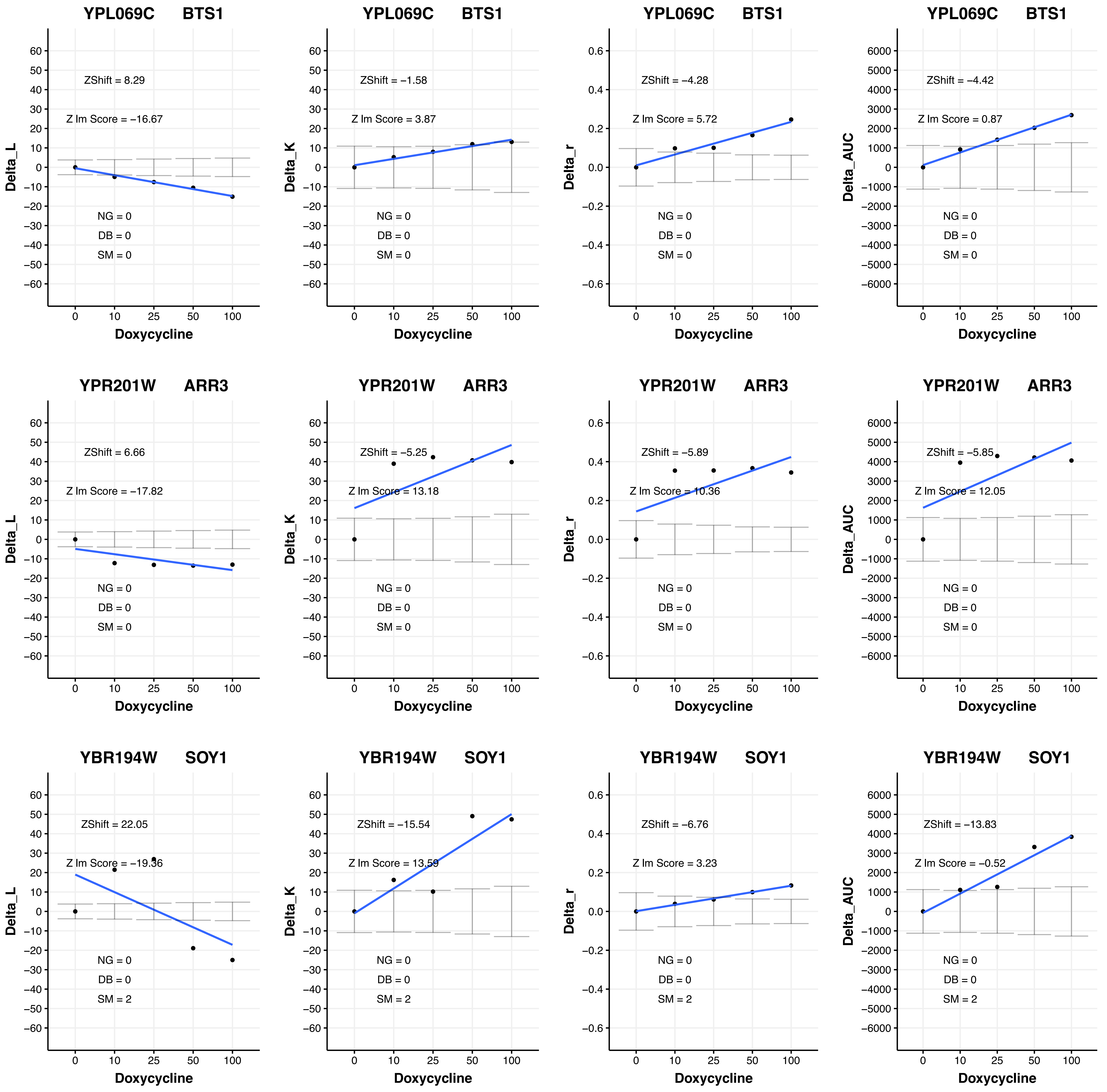

YPL069C BTS1
ZShift = 8.29
Z lm Score = −16.67
NG = 0
DB = 0
SM = 0
Delta_L
Doxycycline
YPL069C BTS1
ZShift = −1.58
Z lm Score = 3.87
NG = 0
DB = 0
SM = 0
Delta_K
Doxycycline
YPL069C BTS1
ZShift = −4.28
Z lm Score = 5.72
NG = 0
DB = 0
SM = 0
Delta_r
Doxycycline
YPL069C BTS1
ZShift = −4.42
Z lm Score = 0.87
NG = 0
DB = 0
SM = 0
Delta_AUC
Doxycycline
YPR201W ARR3
ZShift = 6.66
Z lm Score = −17.82
NG = 0
DB = 0
SM = 0
Delta_L
Doxycycline
YPR201W ARR3
ZShift = −5.25
Z lm Score = 13.18
NG = 0
DB = 0
SM = 0
Delta_K
Doxycycline
YPR201W ARR3
ZShift = −5.89
Z lm Score = 10.36
NG = 0
DB = 0
SM = 0
Delta_r
Doxycycline
YPR201W ARR3
ZShift = −5.85
Z lm Score = 12.05
NG = 0
DB = 0
SM = 0
Delta_AUC
Doxycycline
YBR194W SOY1
ZShift = 22.05
Z lm Score = −19.36
NG = 0
DB = 0
SM = 2
Delta_L
Doxycycline
YBR194W SOY1
ZShift = −15.54
Z lm Score = 13.59
NG = 0
DB = 0
SM = 2
Delta_K
Doxycycline
YBR194W SOY1
ZShift = −6.76
Z lm Score = 3.23
NG = 0
DB = 0
SM = 2
Delta_r
Doxycycline
YBR194W SOY1
ZShift = −13.83
Z lm Score = −0.52
NG = 0
DB = 0
SM = 2
Delta_AUC
Doxycycline

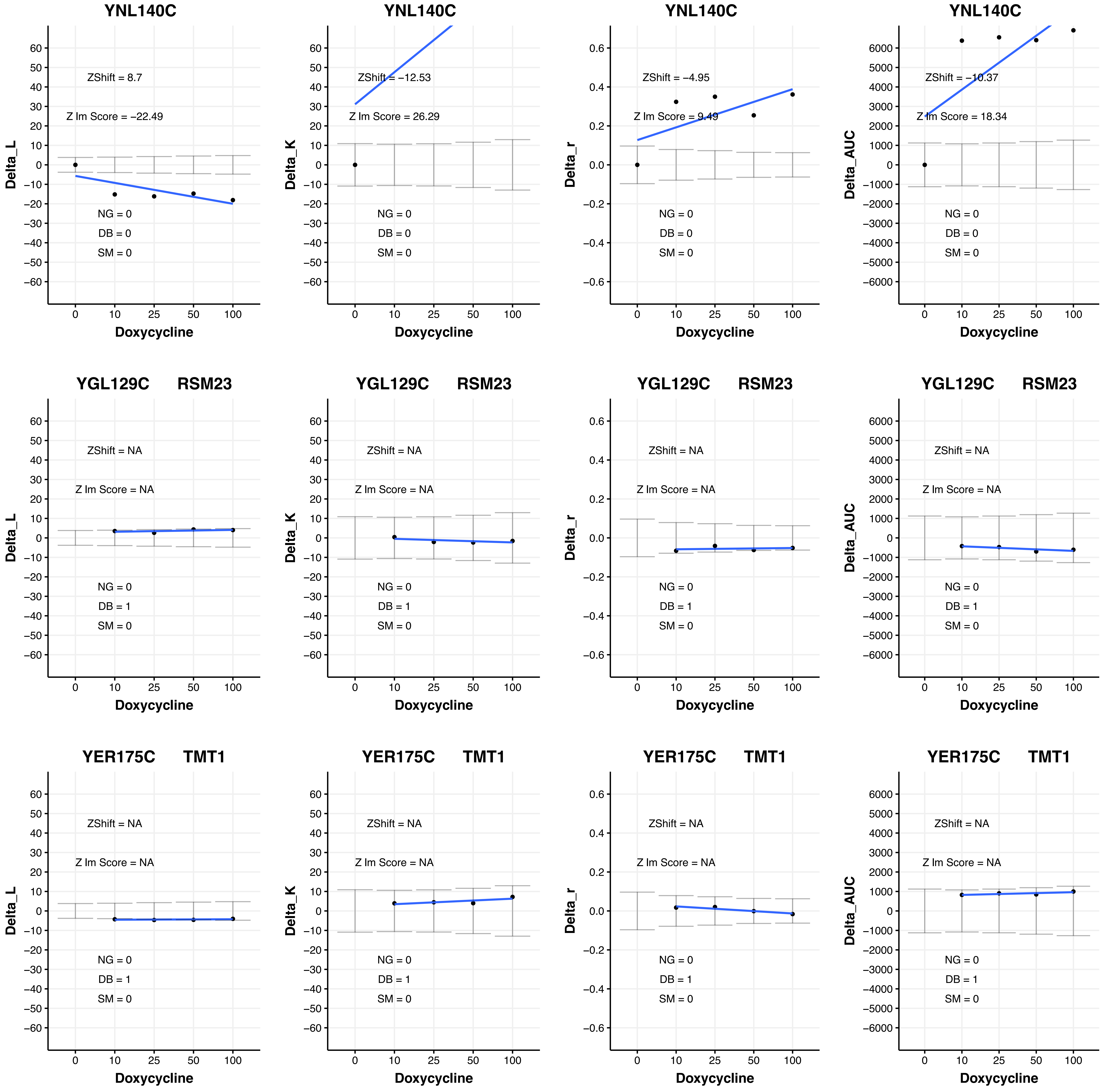
YNL140C
ZShift = 8.7
Z lm Score = −22.49
NG = 0
DB = 0
SM = 0
Delta_L
Doxycycline
YNL140C
ZShift = −12.53
Z lm Score = 26.29
NG = 0
DB = 0
SM = 0
Delta_K
Doxycycline
YNL140C
ZShift = −4.95
Z lm Score = 9.49
NG = 0
DB = 0
SM = 0
Delta_r
Doxycycline
YNL140C
ZShift = −10.37
Z lm Score = 18.34
NG = 0
DB = 0
SM = 0
Delta_AUC
Doxycycline
YGL129C RSM23
ZShift = NA
Z lm Score = NA
NG = 0
DB = 1
SM = 0
Delta_L
Delta_K
Delta_r
Delta_AUC
Doxycycline
YER175C TMT1
ZShift = NA
Z lm Score = NA
NG = 0
DB = 1
SM = 0
Delta_L
Delta_K
Delta_r
Delta_AUC
Doxycycline
