## Additional File 4 for "A humanized yeast phenomic model of deoxycytidine kinase to predict genetic buffering of nucleoside analog cytotoxicity": B - InteractionPlots_DAmPs_Cytarabine.pdf

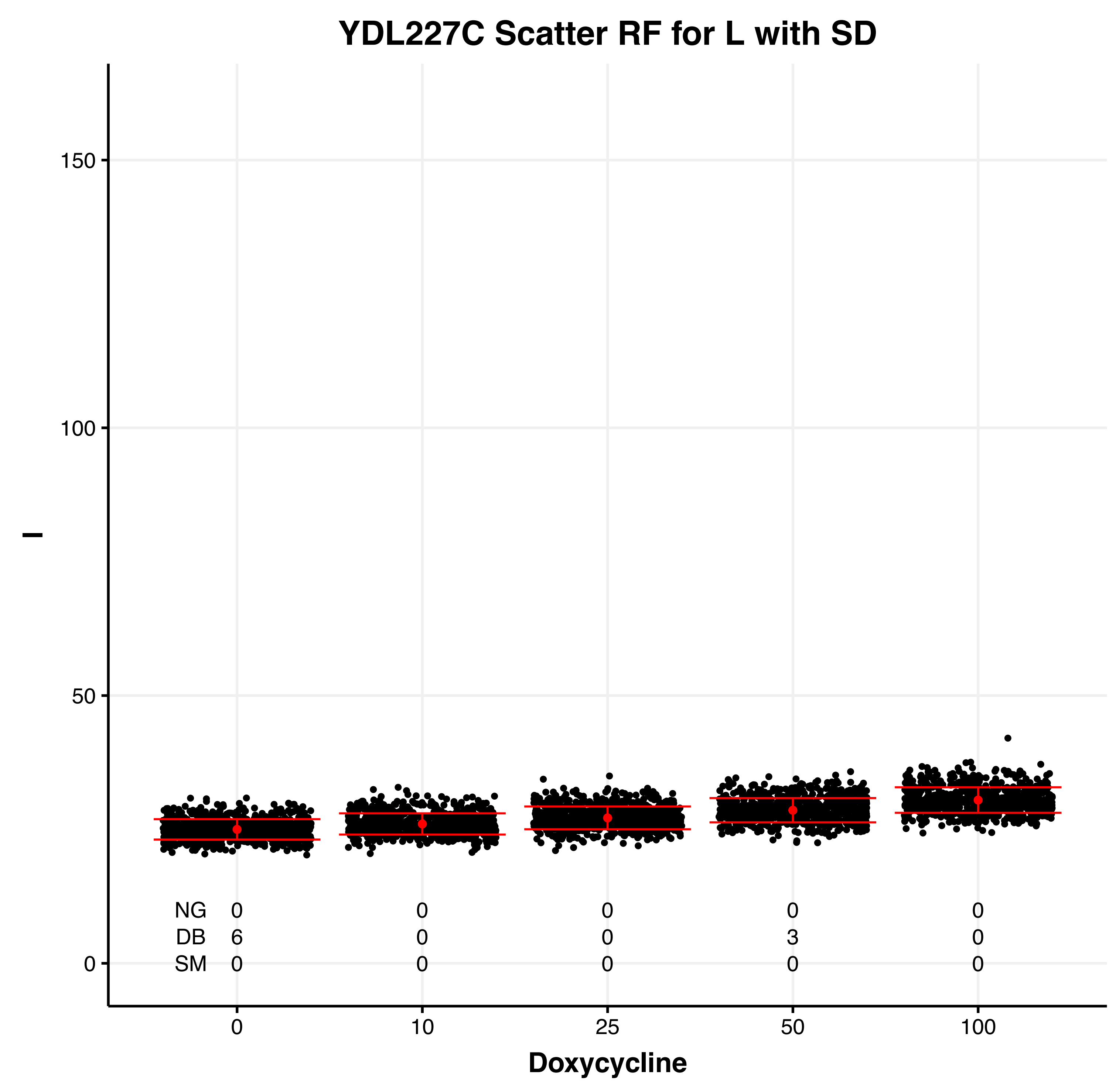

YDL227C Scatter RF for L with SD
l
150
100
50
0
NG 0 0 0 0 0
DB 6 0 0 3 0
SM 0 0 0 0 0
0 10 25 50 100
Doxycycline


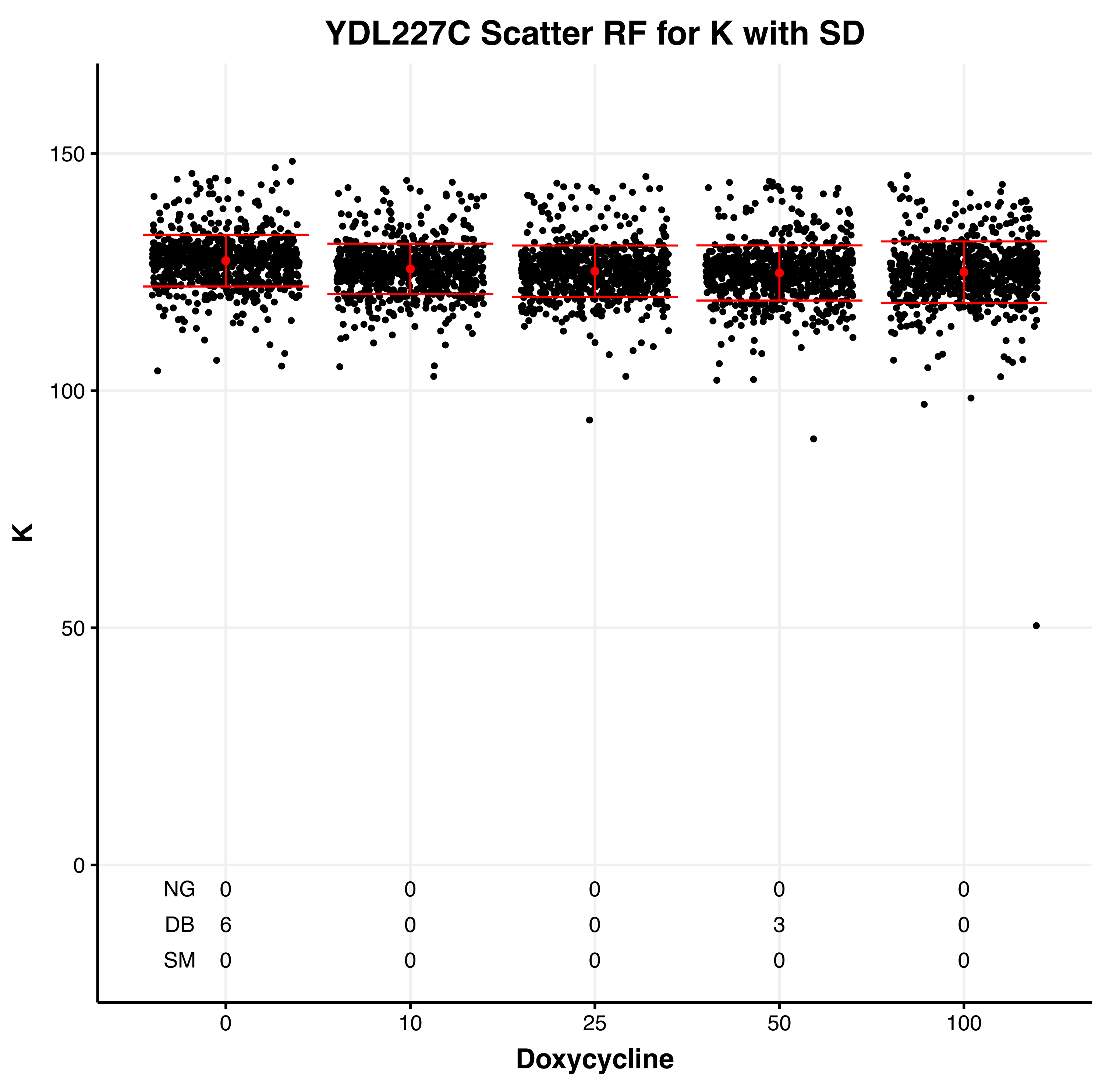

YDL227C Scatter RF for K with SD
K
150
100
50
0
NG 0 0 0 0 0
DB 6 0 0 3 0
SM 0 0 0 0 0
0 10 25 50 100
Doxycycline


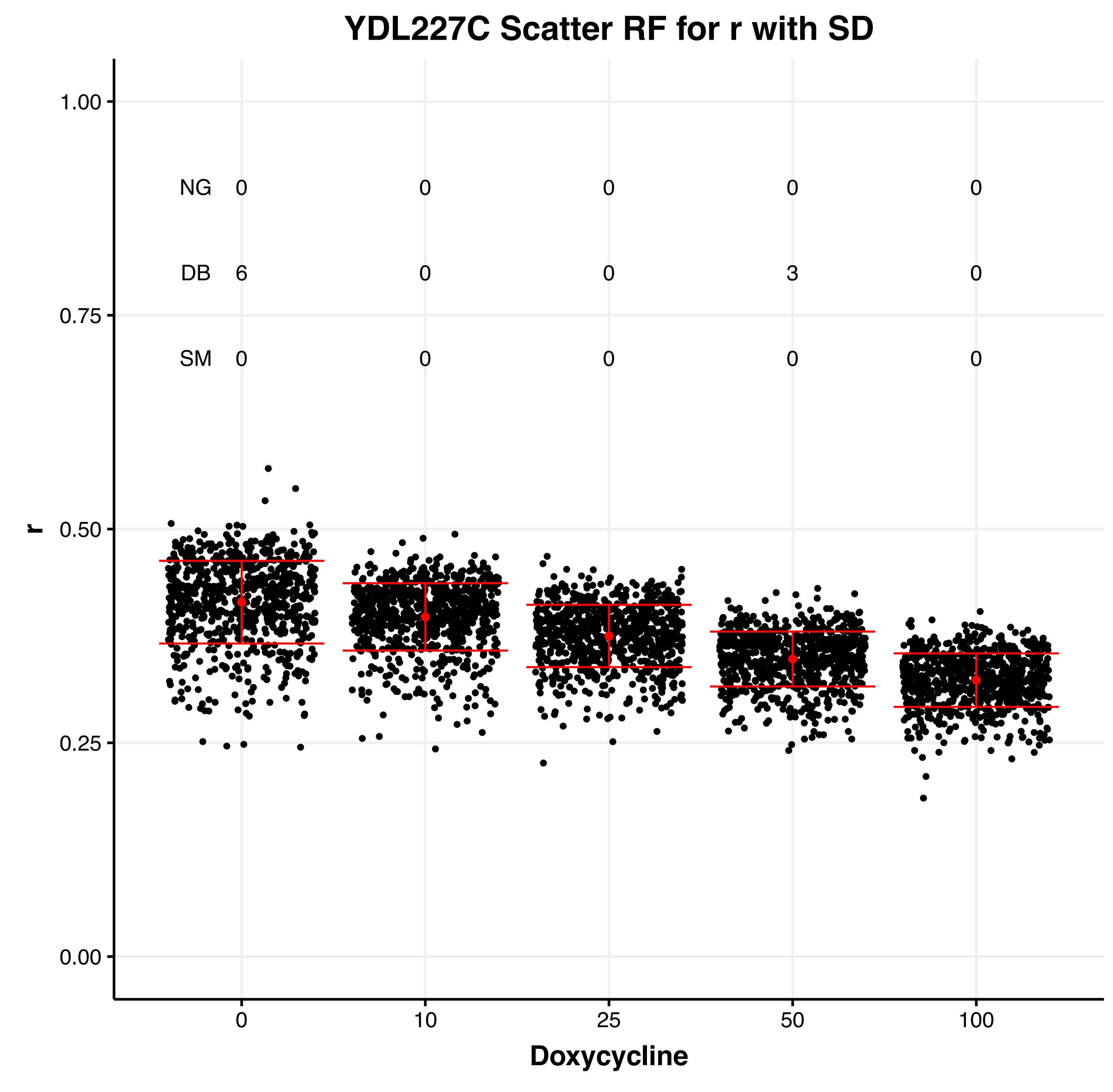

YDL227C Scatter RF for r with SD
r
1.00
0.75
0.50
0.25
0.00
NG 0 0 0 0 0
DB 6 0 0 3 0
SM 0 0 0 0 0
0 10 25 50 100
Doxycycline


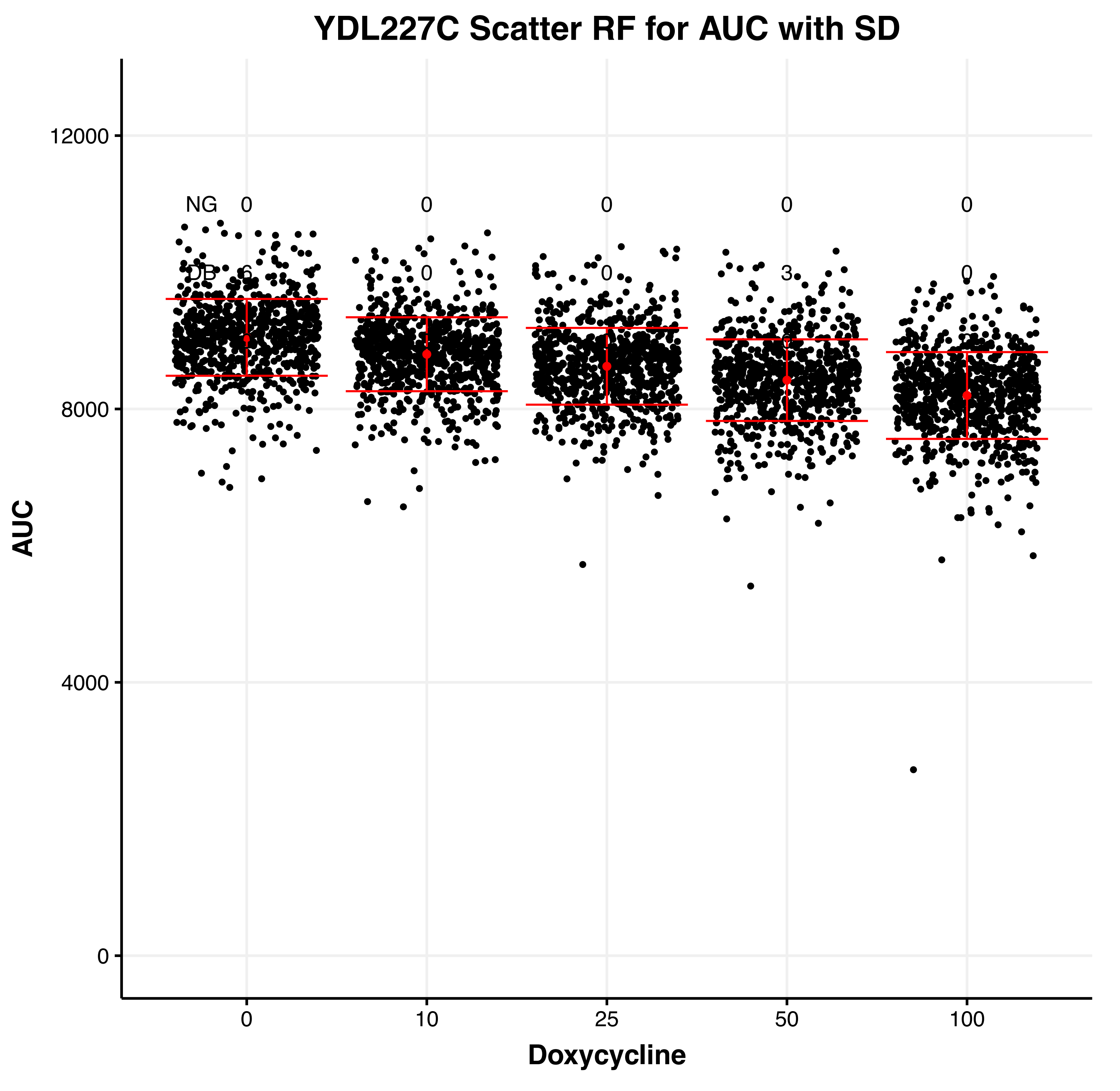

YDL227C Scatter RF for AUC with SD
AUC
12000
8000
4000
0
NG 0 0 0 0 0
DB 6 0 0 3 0
0 10 25 50 100
Doxycycline

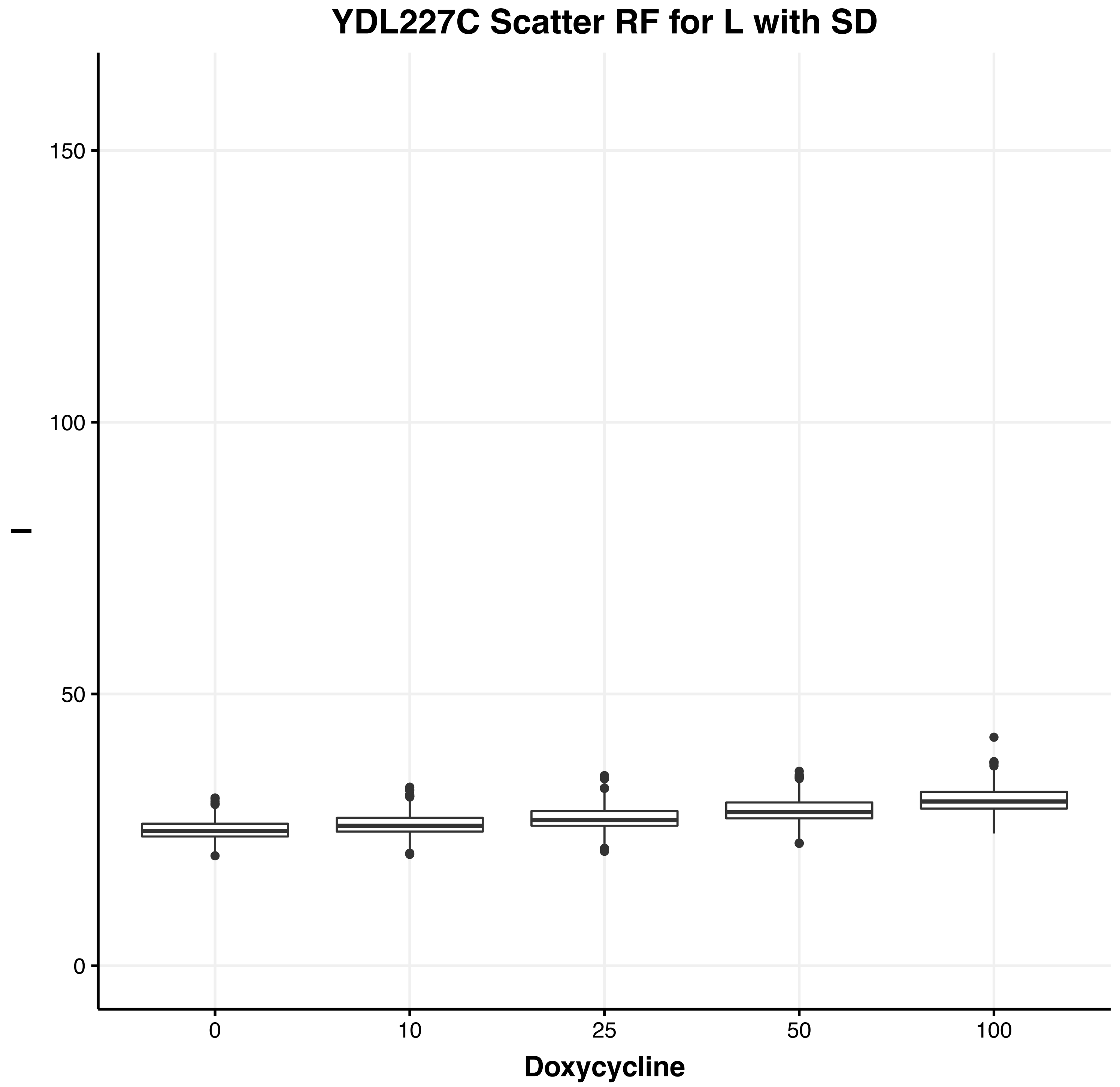
YDL227C Scatter RF for L with SD
l
0
50
100
150
0
10
25
50
100
Doxycycline

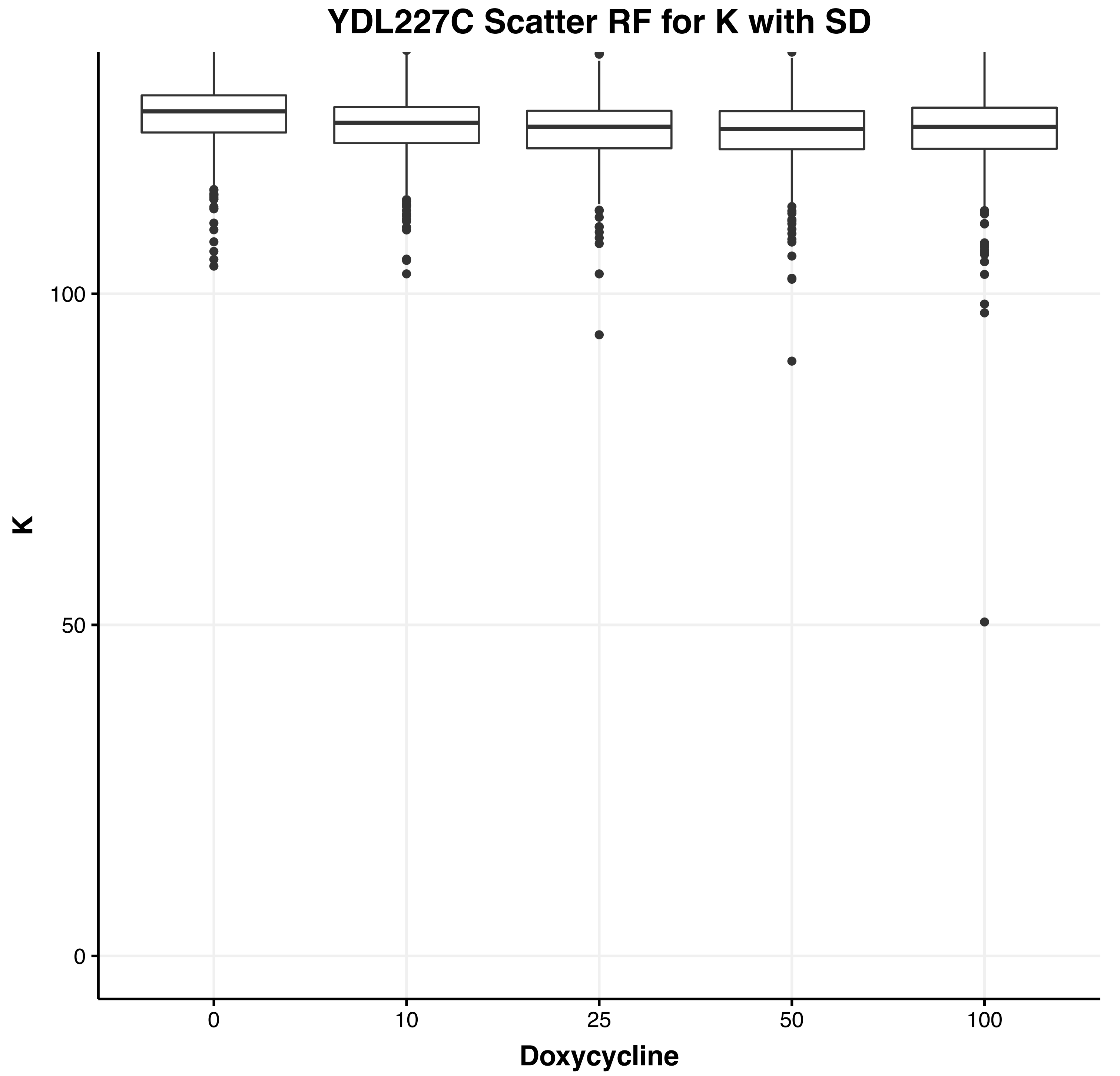
YDL227C Scatter RF for K with SD
K
0
50
100
0
10
25
50
100
Doxycycline

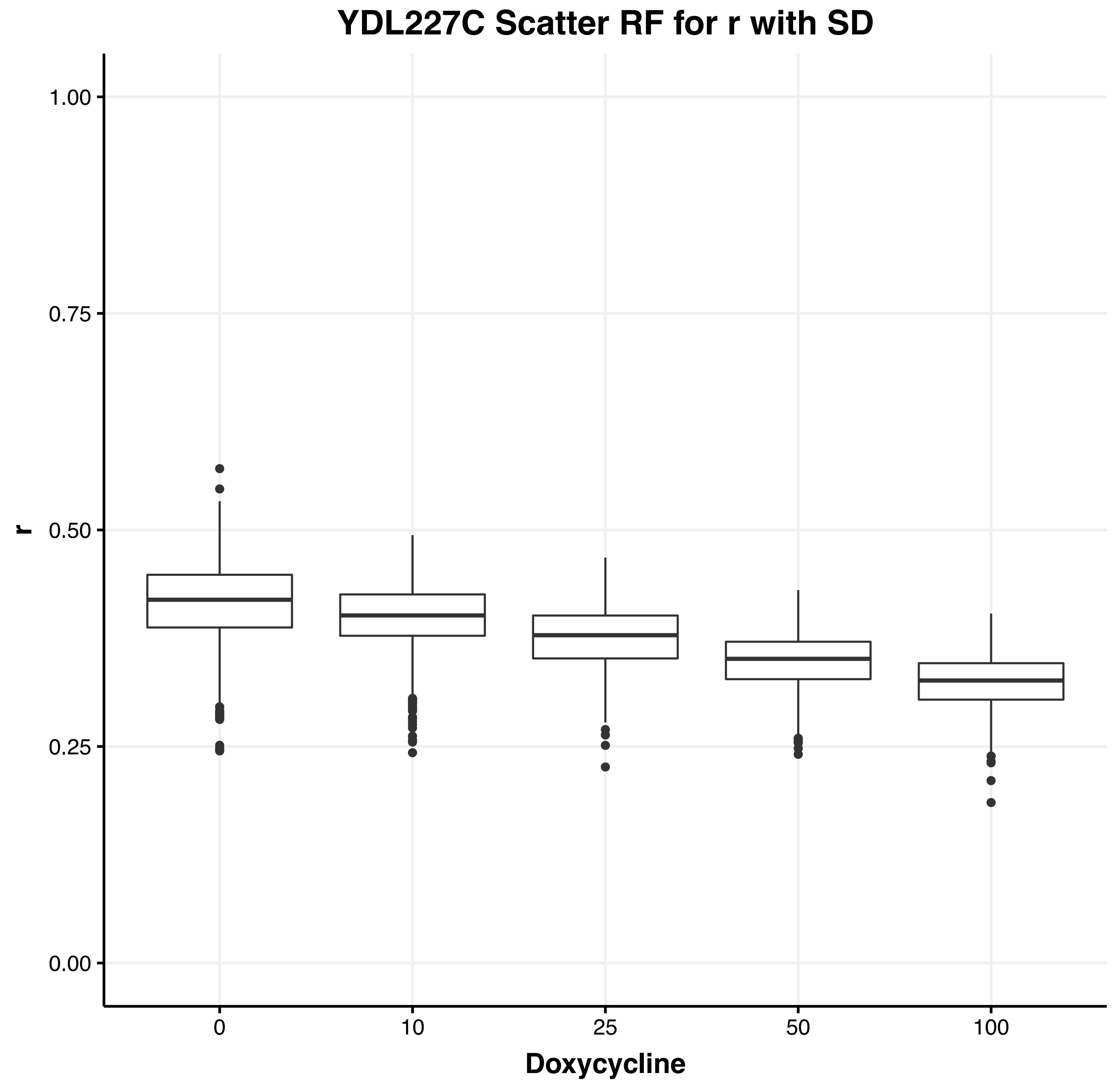
YDL227C Scatter RF for r with SD
r
0.00
0.25
0.50
0.75
1.00
0
10
25
50
100
Doxycycline

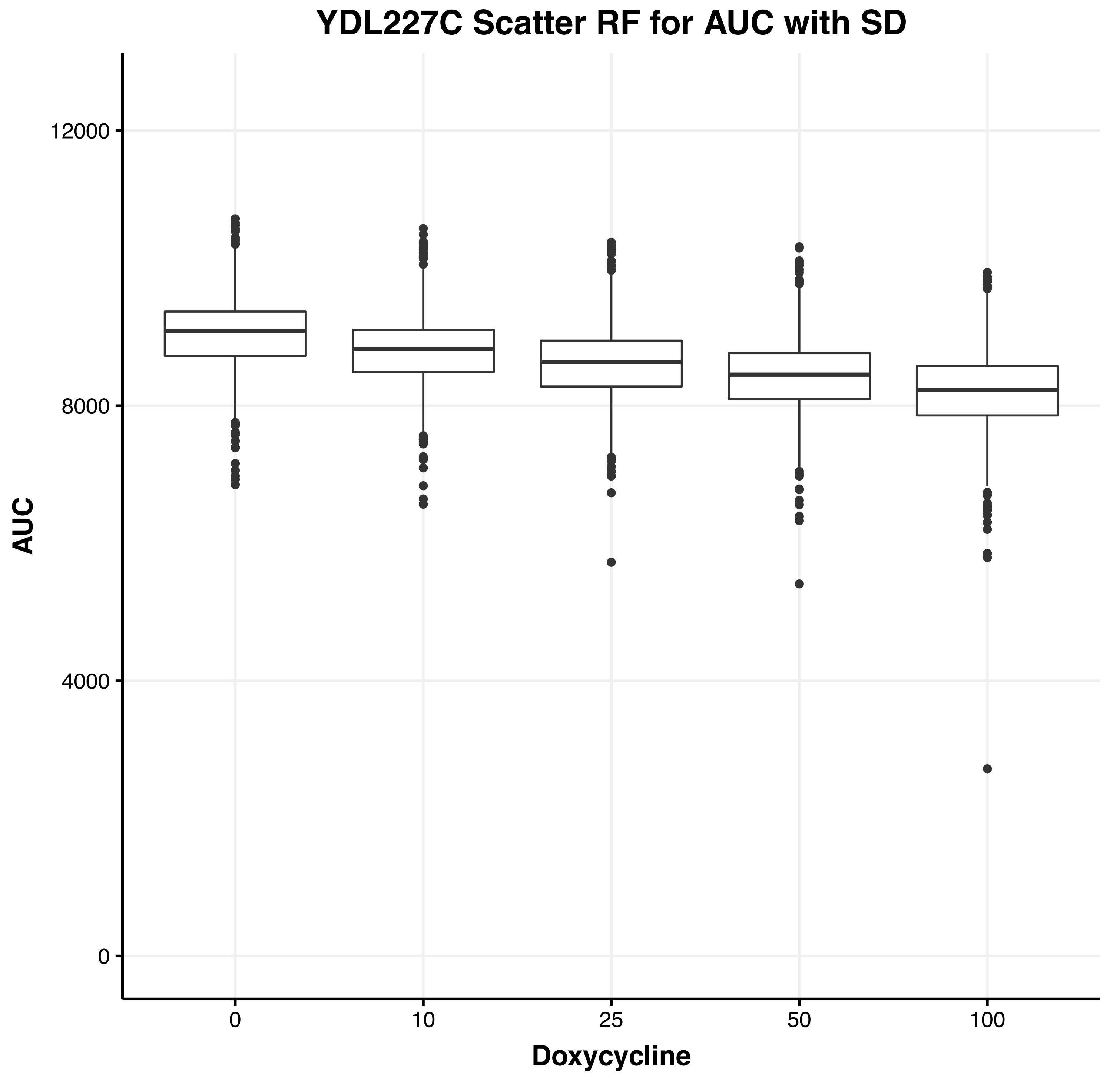
YDL227C Scatter RF for AUC with SD
AUC
0
4000
8000
12000
0
10
25
50
100
Doxycycline

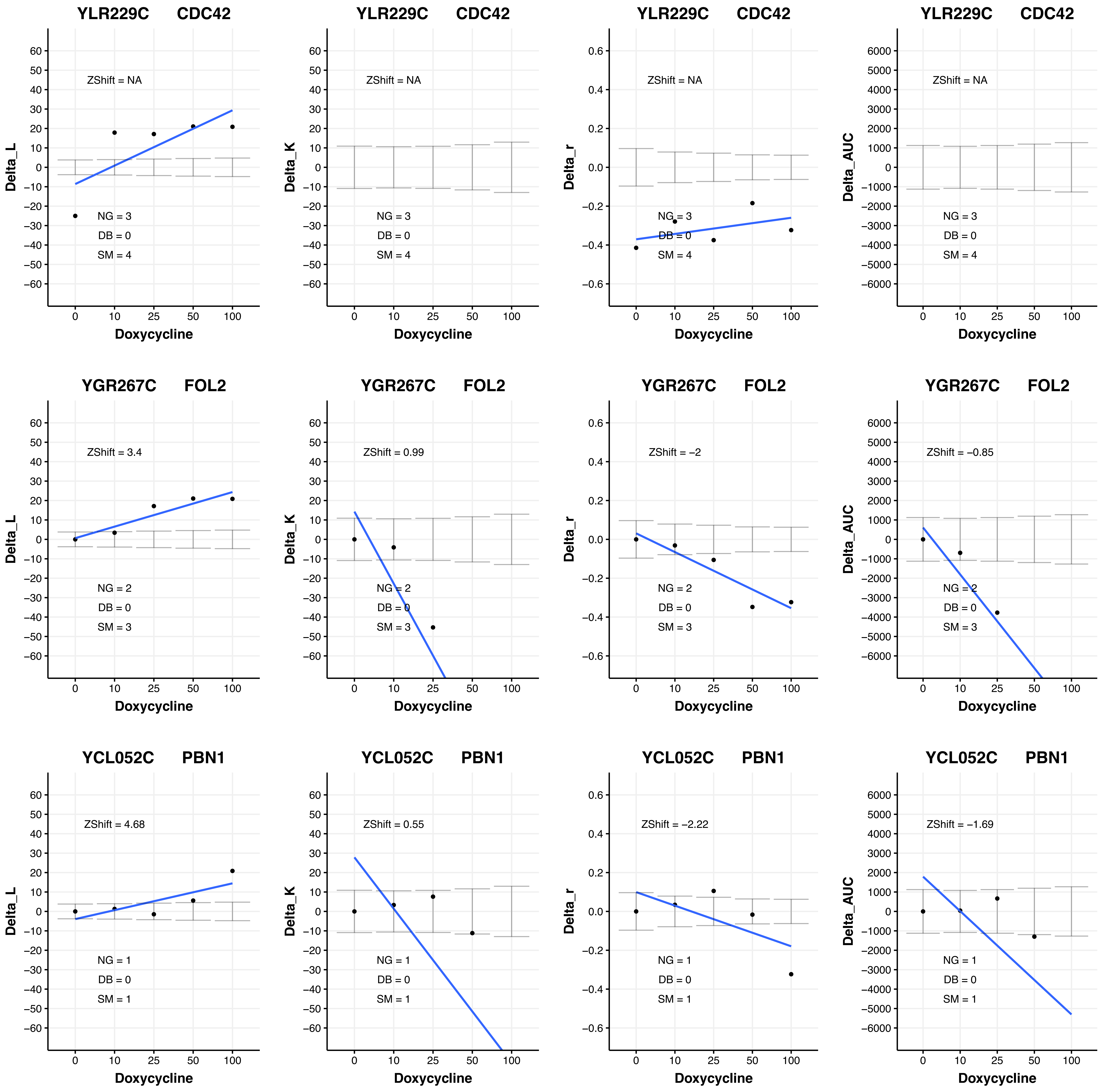

YLR229C CDC42
ZShift = NA
NG = 3
DB = 0
SM = 4
Delta_L
Delta_K
Delta_r
Delta_AUC
Doxycycline
0 10 25 50 100
YGR267C FOL2
ZShift = 3.4
ZShift = 0.99
ZShift = −2
ZShift = −0.85
NG = 2
DB = 0
SM = 3
YCL052C PBN1
ZShift = 4.68
ZShift = 0.55
ZShift = −2.22
ZShift = −1.69
NG = 1
DB = 0
SM = 1

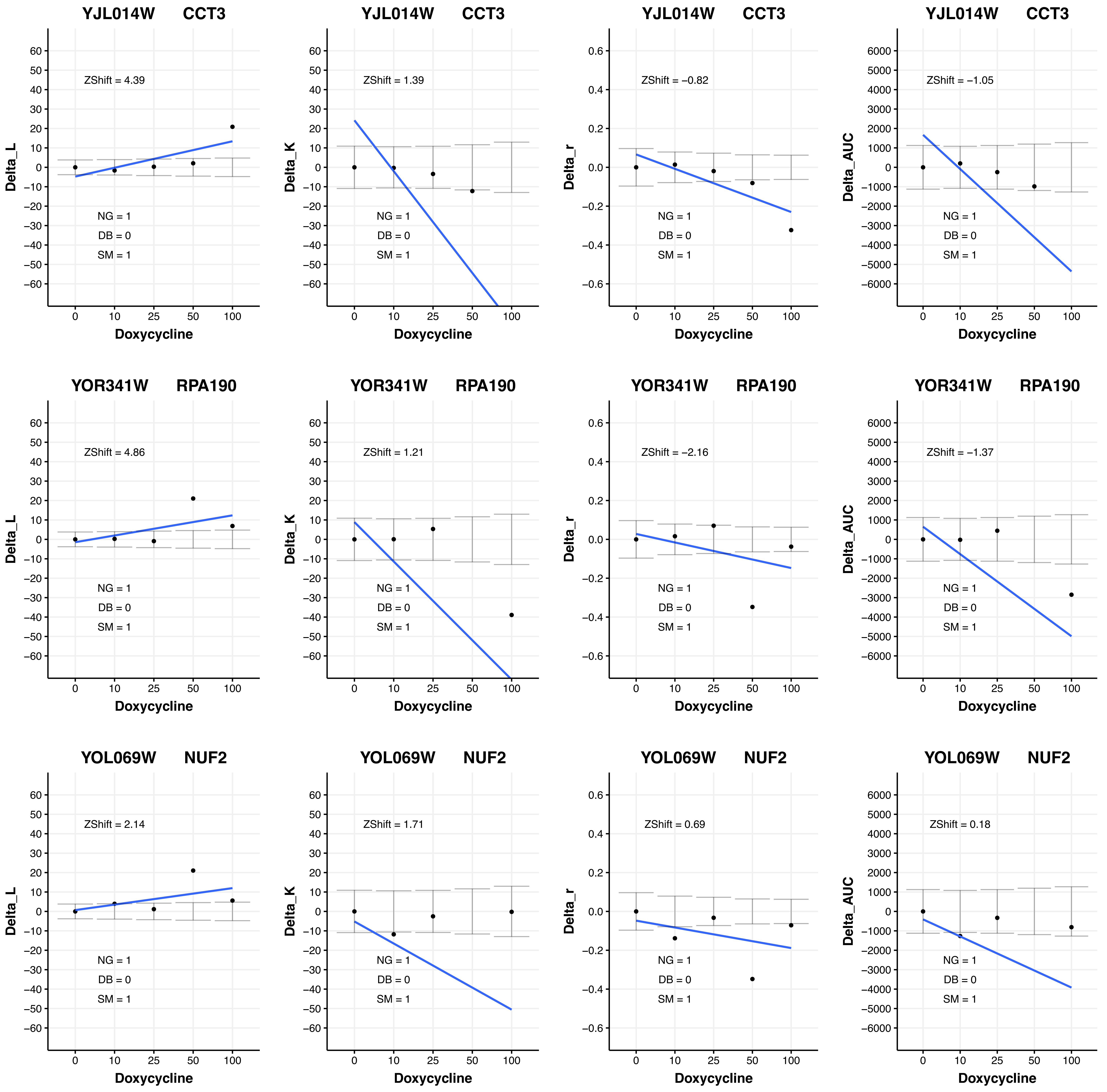

YJL014W CCT3
ZShift = 4.39
NG = 1
DB = 0
SM = 1
Delta_L
Doxycycline
YJL014W CCT3
ZShift = 1.39
NG = 1
DB = 0
SM = 1
Delta_K
Doxycycline
YJL014W CCT3
ZShift = −0.82
NG = 1
DB = 0
SM = 1
Delta_r
Doxycycline
YJL014W CCT3
ZShift = −1.05
NG = 1
DB = 0
SM = 1
Delta_AUC
Doxycycline
YOR341W RPA190
ZShift = 4.86
NG = 1
DB = 0
SM = 1
Delta_L
Doxycycline
YOR341W RPA190
ZShift = 1.21
NG = 1
DB = 0
SM = 1
Delta_K
Doxycycline
YOR341W RPA190
ZShift = −2.16
NG = 1
DB = 0
SM = 1
Delta_r
Doxycycline
YOR341W RPA190
ZShift = −1.37
NG = 1
DB = 0
SM = 1
Delta_AUC
Doxycycline
YOL069W NUF2
ZShift = 2.14
NG = 1
DB = 0
SM = 1
Delta_L
Doxycycline
YOL069W NUF2
ZShift = 1.71
NG = 1
DB = 0
SM = 1
Delta_K
Doxycycline
YOL069W NUF2
ZShift = 0.69
NG = 1
DB = 0
SM = 1
Delta_r
Doxycycline
YOL069W NUF2
ZShift = 0.18
NG = 1
DB = 0
SM = 1
Delta_AUC
Doxycycline

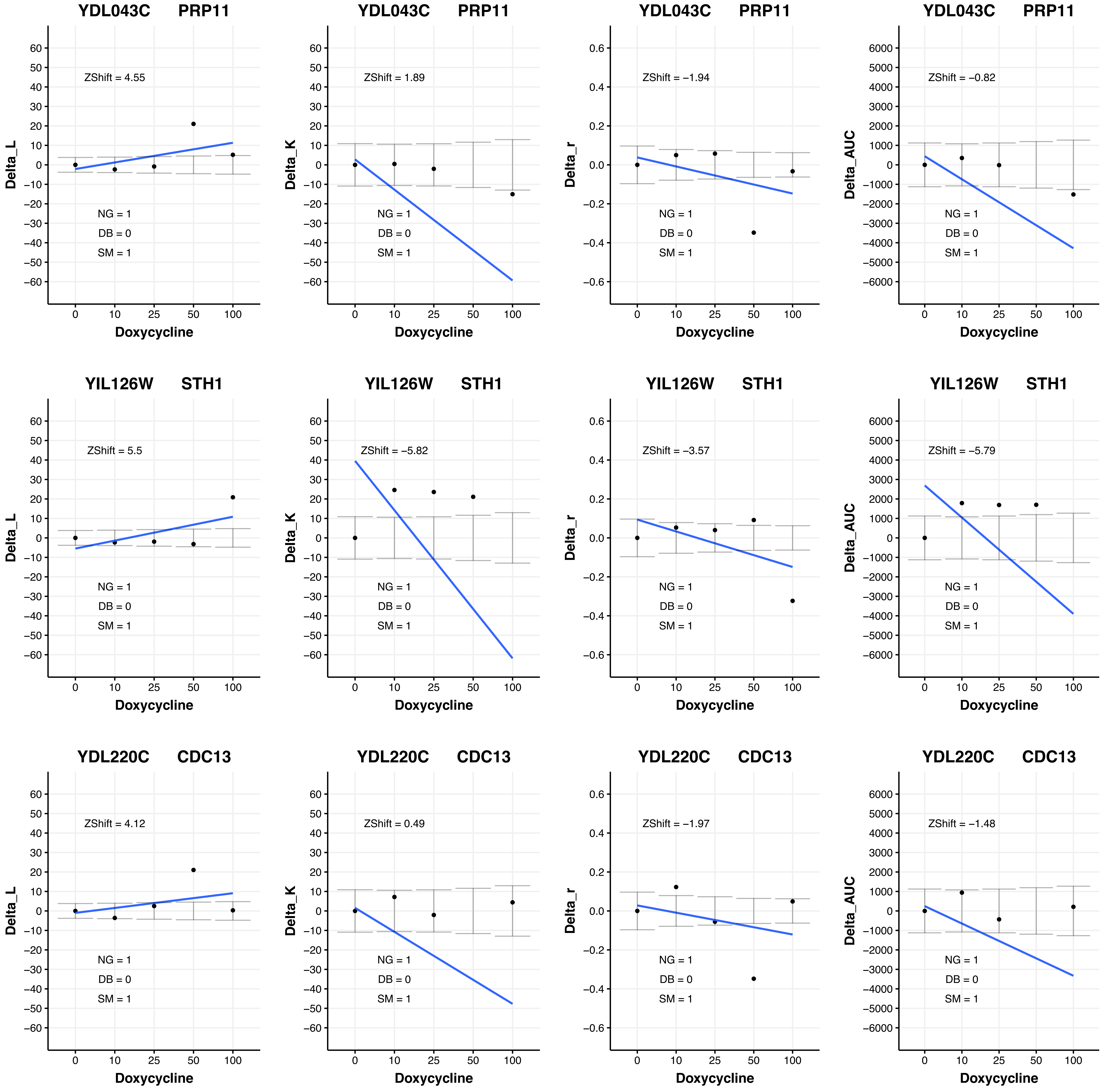

YDL043C PRP11
ZShift = 4.55
NG = 1
DB = 0
SM = 1
Delta_L
Doxycycline
YDL043C PRP11
ZShift = 1.89
NG = 1
DB = 0
SM = 1
Delta_K
Doxycycline
YDL043C PRP11
ZShift = −1.94
NG = 1
DB = 0
SM = 1
Delta_r
Doxycycline
YDL043C PRP11
ZShift = −0.82
NG = 1
DB = 0
SM = 1
Delta_AUC
Doxycycline
YIL126W STH1
ZShift = 5.5
NG = 1
DB = 0
SM = 1
Delta_L
Doxycycline
YIL126W STH1
ZShift = −5.82
NG = 1
DB = 0
SM = 1
Delta_K
Doxycycline
YIL126W STH1
ZShift = −3.57
NG = 1
DB = 0
SM = 1
Delta_r
Doxycycline
YIL126W STH1
ZShift = −5.79
NG = 1
DB = 0
SM = 1
Delta_AUC
Doxycycline
YDL220C CDC13
ZShift = 4.12
NG = 1
DB = 0
SM = 1
Delta_L
Doxycycline
YDL220C CDC13
ZShift = 0.49
NG = 1
DB = 0
SM = 1
Delta_K
Doxycycline
YDL220C CDC13
ZShift = −1.97
NG = 1
DB = 0
SM = 1
Delta_r
Doxycycline
YDL220C CDC13
ZShift = −1.48
NG = 1
DB = 0
SM = 1
Delta_AUC
Doxycycline

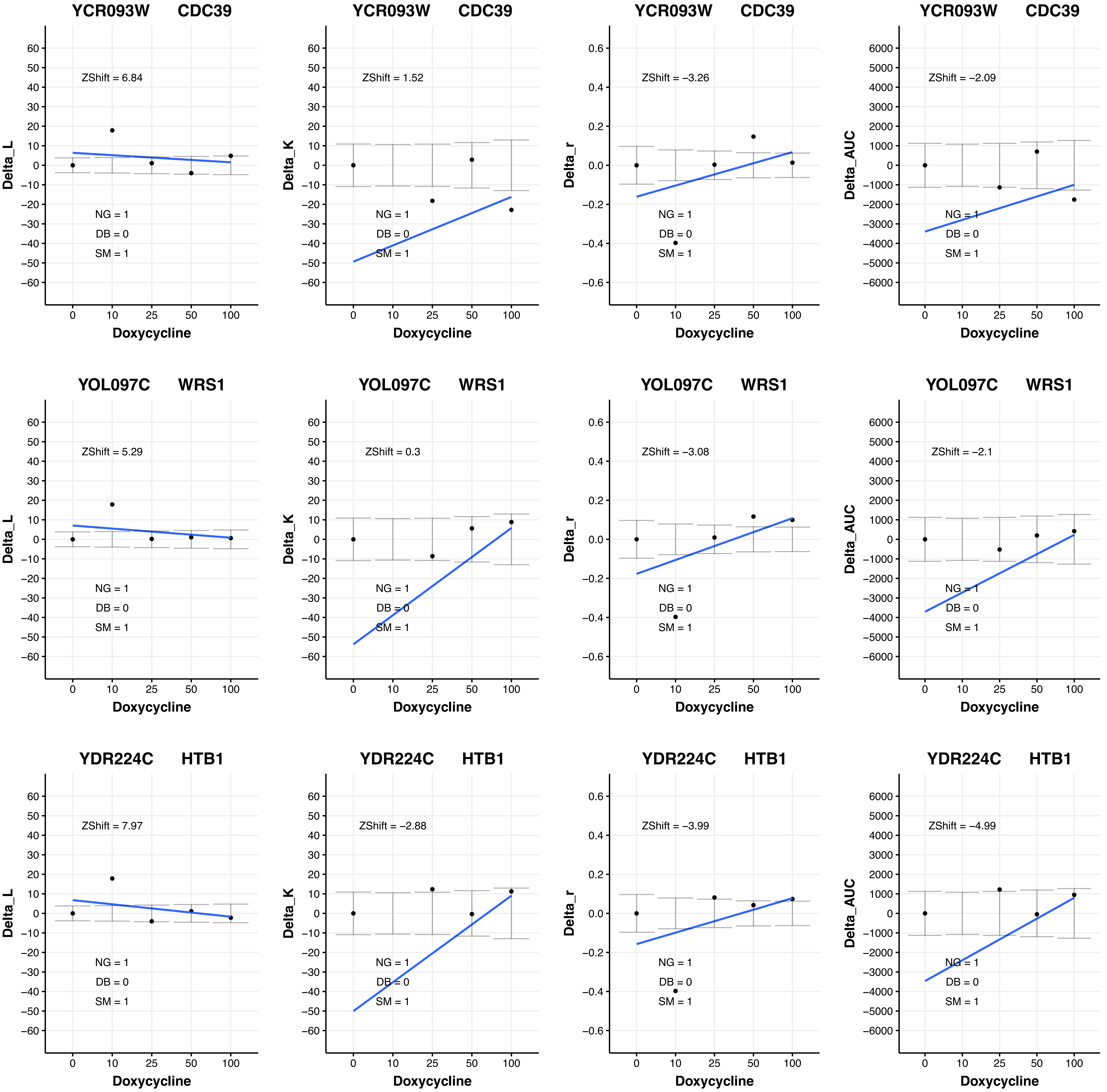

YCR093W CDC39
ZShift = 6.84
NG = 1
DB = 0
SM = 1
Delta_L
Doxycycline
YCR093W CDC39
ZShift = 1.52
NG = 1
DB = 0
SM = 1
Delta_K
Doxycycline
YCR093W CDC39
ZShift = −3.26
NG = 1
DB = 0
SM = 1
Delta_r
Doxycycline
YCR093W CDC39
ZShift = −2.09
NG = 1
DB = 0
SM = 1
Delta_AUC
Doxycycline
YOL097C WRS1
ZShift = 5.29
NG = 1
DB = 0
SM = 1
Delta_L
Doxycycline
YOL097C WRS1
ZShift = 0.3
NG = 1
DB = 0
SM = 1
Delta_K
Doxycycline
YOL097C WRS1
ZShift = −3.08
NG = 1
DB = 0
SM = 1
Delta_r
Doxycycline
YOL097C WRS1
ZShift = −2.1
NG = 1
DB = 0
SM = 1
Delta_AUC
Doxycycline
YDR224C HTB1
ZShift = 7.97
NG = 1
DB = 0
SM = 1
Delta_L
Doxycycline
YDR224C HTB1
ZShift = −2.88
NG = 1
DB = 0
SM = 1
Delta_K
Doxycycline
YDR224C HTB1
ZShift = −3.99
NG = 1
DB = 0
SM = 1
Delta_r
Doxycycline
YDR224C HTB1
ZShift = −4.99
NG = 1
DB = 0
SM = 1
Delta_AUC
Doxycycline

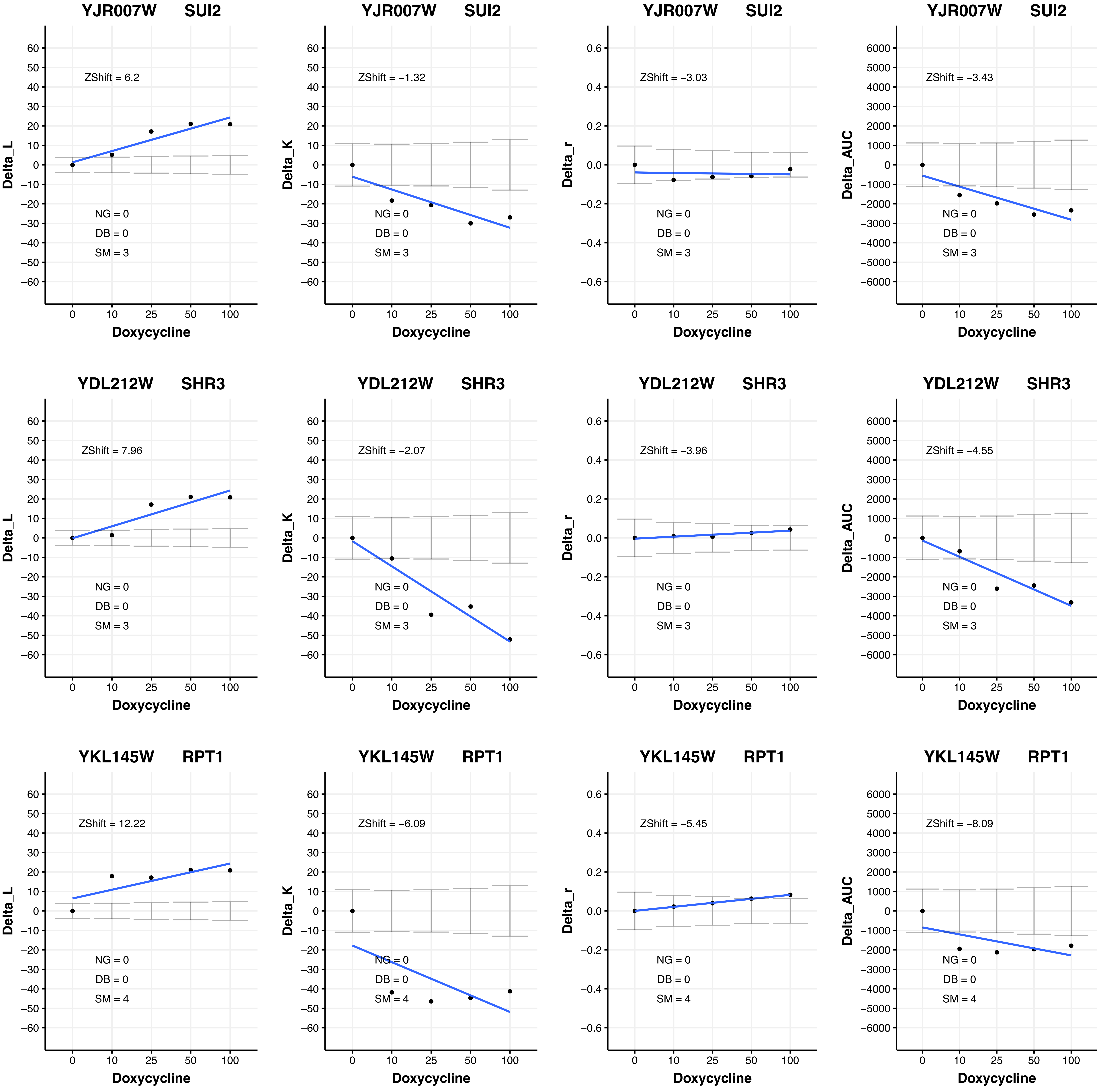
YJR007W SUI2
ZShift = 6.2
NG = 0
DB = 0
SM = 3
Delta_L
Doxycycline
YJR007W SUI2
ZShift = −1.32
NG = 0
DB = 0
SM = 3
Delta_K
Doxycycline
YJR007W SUI2
ZShift = −3.03
NG = 0
DB = 0
SM = 3
Delta_r
Doxycycline
YJR007W SUI2
ZShift = −3.43
NG = 0
DB = 0
SM = 3
Delta_AUC
Doxycycline
YDL212W SHR3
ZShift = 7.96
NG = 0
DB = 0
SM = 3
Delta_L
Doxycycline
YDL212W SHR3
ZShift = −2.07
NG = 0
DB = 0
SM = 3
Delta_K
Doxycycline
YDL212W SHR3
ZShift = −3.96
NG = 0
DB = 0
SM = 3
Delta_r
Doxycycline
YDL212W SHR3
ZShift = −4.55
NG = 0
DB = 0
SM = 3
Delta_AUC
Doxycycline
YKL145W RPT1
ZShift = 12.22
NG = 0
DB = 0
SM = 4
Delta_L
Doxycycline
YKL145W RPT1
ZShift = −6.09
NG = 0
DB = 0
SM = 4
Delta_K
Doxycycline
YKL145W RPT1
ZShift = −5.45
NG = 0
DB = 0
SM = 4
Delta_r
Doxycycline
YKL145W RPT1
ZShift = −8.09
NG = 0
DB = 0
SM = 4
Delta_AUC
Doxycycline

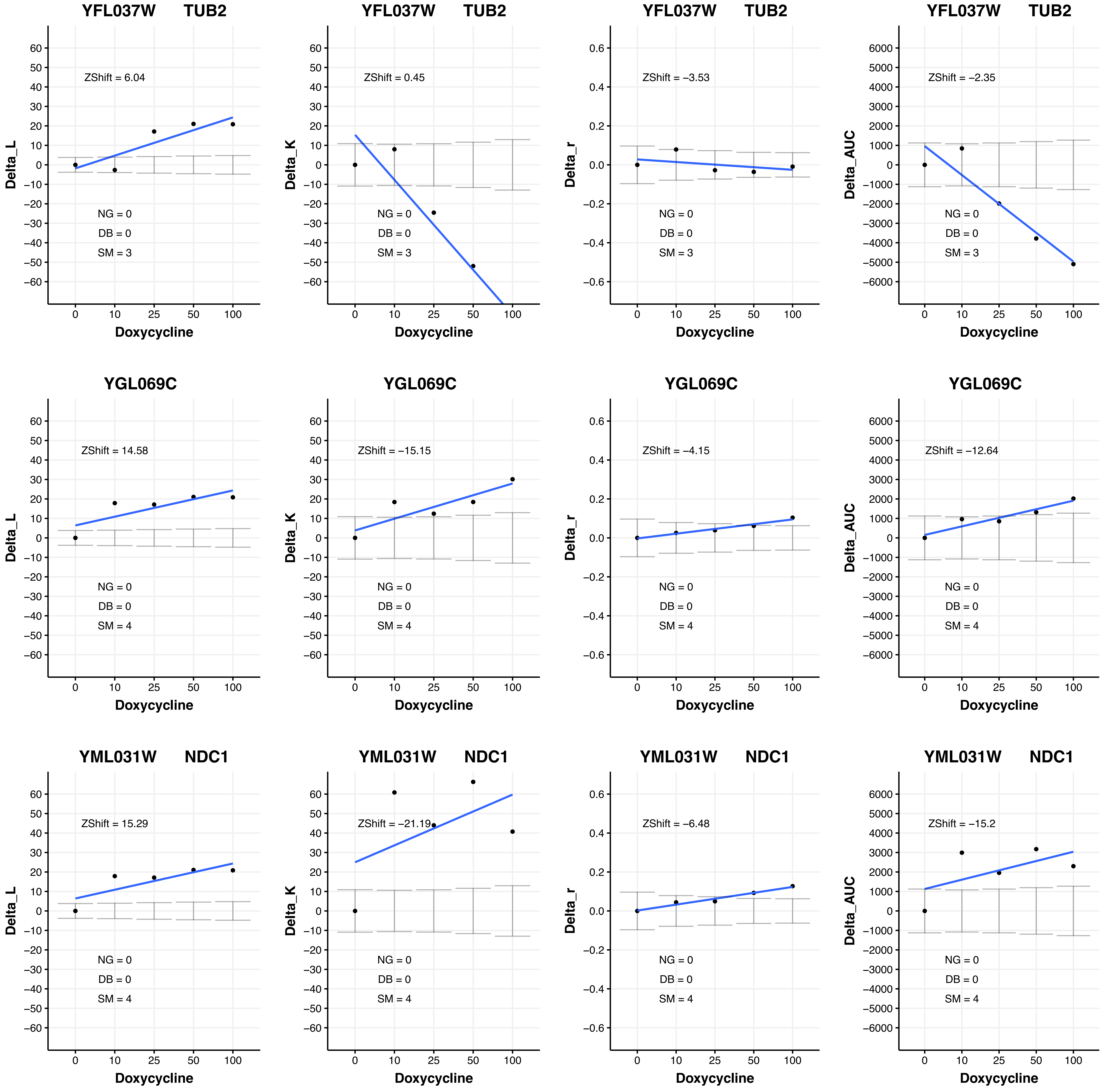
YFL037W TUB2
ZShift = 6.04
NG = 0
DB = 0
SM = 3
Delta_L
Doxycycline
YFL037W TUB2
ZShift = 0.45
NG = 0
DB = 0
SM = 3
Delta_K
Doxycycline
YFL037W TUB2
ZShift = −3.53
NG = 0
DB = 0
SM = 3
Delta_r
Doxycycline
YFL037W TUB2
ZShift = −2.35
NG = 0
DB = 0
SM = 3
Delta_AUC
Doxycycline
YGL069C
ZShift = 14.58
NG = 0
DB = 0
SM = 4
Delta_L
Doxycycline
YGL069C
ZShift = −15.15
NG = 0
DB = 0
SM = 4
Delta_K
Doxycycline
YGL069C
ZShift = −4.15
NG = 0
DB = 0
SM = 4
Delta_r
Doxycycline
YGL069C
ZShift = −12.64
NG = 0
DB = 0
SM = 4
Delta_AUC
Doxycycline
YML031W NDC1
ZShift = 15.29
NG = 0
DB = 0
SM = 4
Delta_L
Doxycycline
YML031W NDC1
ZShift = −21.19
NG = 0
DB = 0
SM = 4
Delta_K
Doxycycline
YML031W NDC1
ZShift = −6.48
NG = 0
DB = 0
SM = 4
Delta_r
Doxycycline
YML031W NDC1
ZShift = −15.2
NG = 0
DB = 0
SM = 4
Delta_AUC
Doxycycline

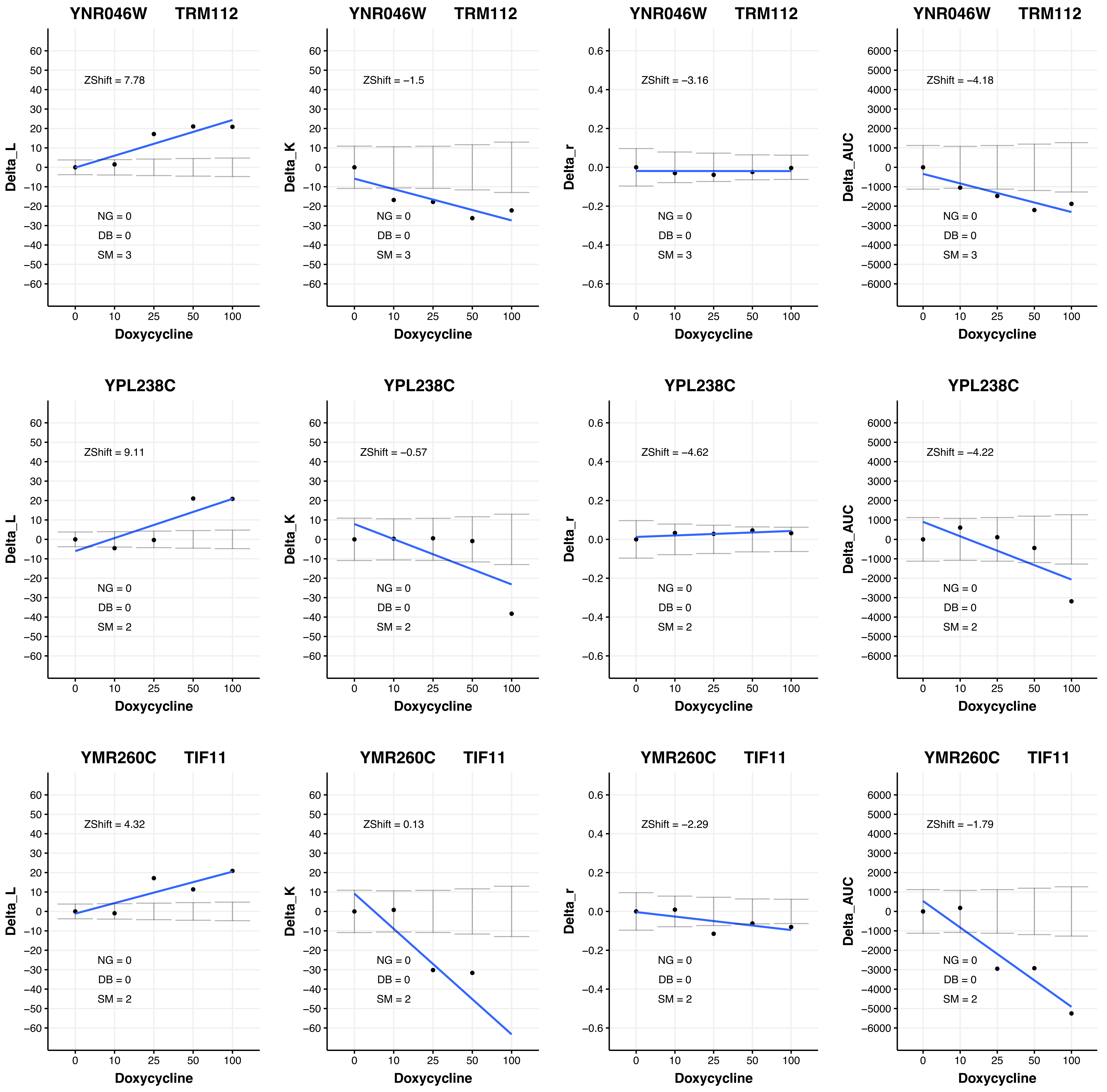

YNR046W TRM112
ZShift = 7.78
NG = 0
DB = 0
SM = 3
Delta_L
Doxycycline
YNR046W TRM112
ZShift = −1.5
NG = 0
DB = 0
SM = 3
Delta_K
Doxycycline
YNR046W TRM112
ZShift = −3.16
NG = 0
DB = 0
SM = 3
Delta_r
Doxycycline
YNR046W TRM112
ZShift = −4.18
NG = 0
DB = 0
SM = 3
Delta_AUC
Doxycycline
YPL238C
ZShift = 9.11
NG = 0
DB = 0
SM = 2
Delta_L
Doxycycline
YPL238C
ZShift = −0.57
NG = 0
DB = 0
SM = 2
Delta_K
Doxycycline
YPL238C
ZShift = −4.62
NG = 0
DB = 0
SM = 2
Delta_r
Doxycycline
YPL238C
ZShift = −4.22
NG = 0
DB = 0
SM = 2
Delta_AUC
Doxycycline
YMR260C TIF11
ZShift = 4.32
NG = 0
DB = 0
SM = 2
Delta_L
Doxycycline
YMR260C TIF11
ZShift = 0.13
NG = 0
DB = 0
SM = 2
Delta_K
Doxycycline
YMR260C TIF11
ZShift = −2.29
NG = 0
DB = 0
SM = 2
Delta_r
Doxycycline
YMR260C TIF11
ZShift = −1.79
NG = 0
DB = 0
SM = 2
Delta_AUC
Doxycycline

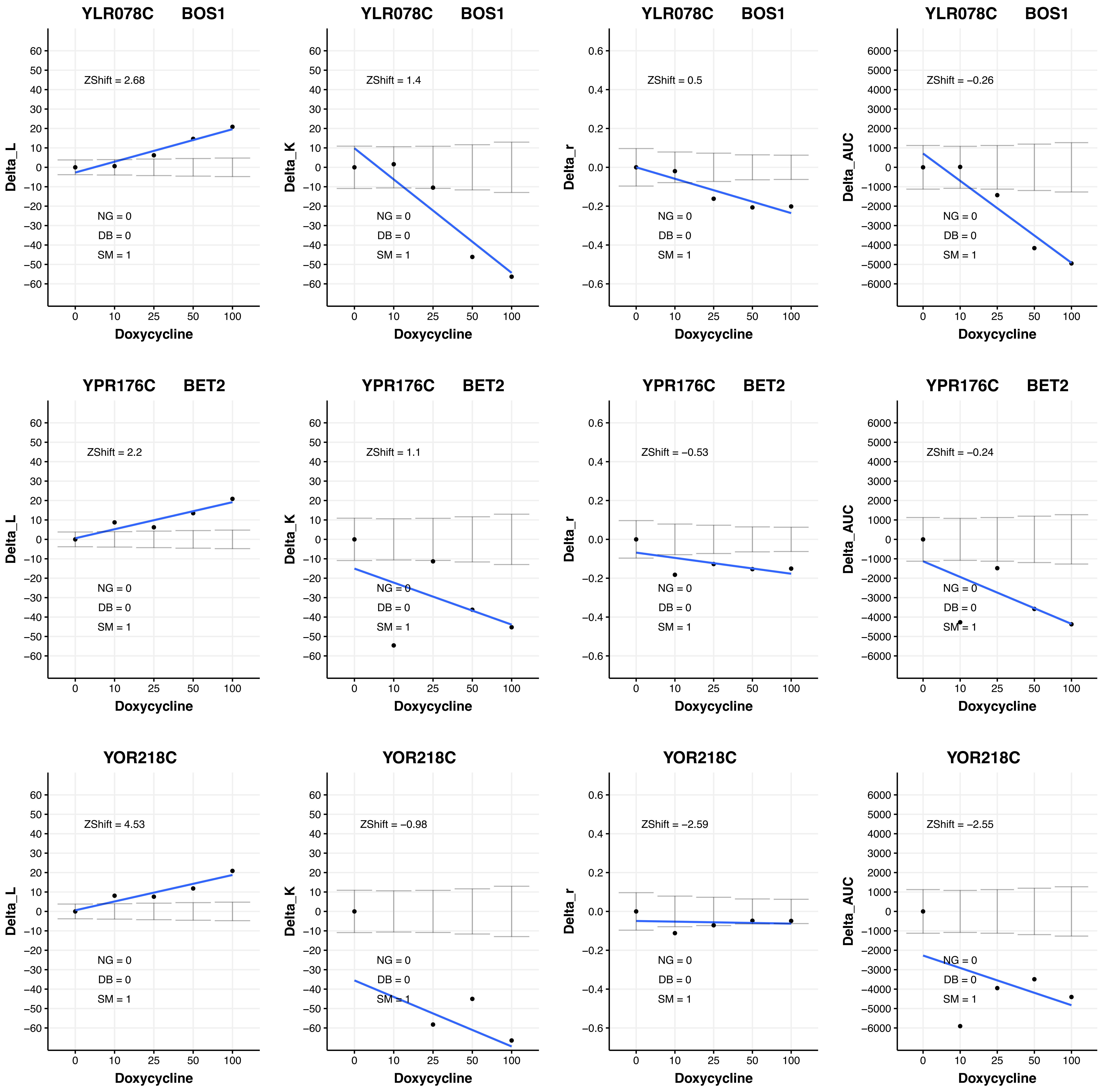
YLR078C BOS1
ZShift = 2.68
NG = 0
DB = 0
SM = 1
Delta_L
Doxycycline
YLR078C BOS1
ZShift = 1.4
NG = 0
DB = 0
SM = 1
Delta_K
Doxycycline
YLR078C BOS1
ZShift = 0.5
NG = 0
DB = 0
SM = 1
Delta_r
Doxycycline
YLR078C BOS1
ZShift = −0.26
NG = 0
DB = 0
SM = 1
Delta_AUC
Doxycycline
YPR176C BET2
ZShift = 2.2
NG = 0
DB = 0
SM = 1
Delta_L
Doxycycline
YPR176C BET2
ZShift = 1.1
NG = 0
DB = 0
SM = 1
Delta_K
Doxycycline
YPR176C BET2
ZShift = −0.53
NG = 0
DB = 0
SM = 1
Delta_r
Doxycycline
YPR176C BET2
ZShift = −0.24
NG = 0
DB = 0
SM = 1
Delta_AUC
Doxycycline
YOR218C
ZShift = 4.53
NG = 0
DB = 0
SM = 1
Delta_L
Doxycycline
YOR218C
ZShift = −0.98
NG = 0
DB = 0
SM = 1
Delta_K
Doxycycline
YOR218C
ZShift = −2.59
NG = 0
DB = 0
SM = 1
Delta_r
Doxycycline
YOR218C
ZShift = −2.55
NG = 0
DB = 0
SM = 1
Delta_AUC
Doxycycline

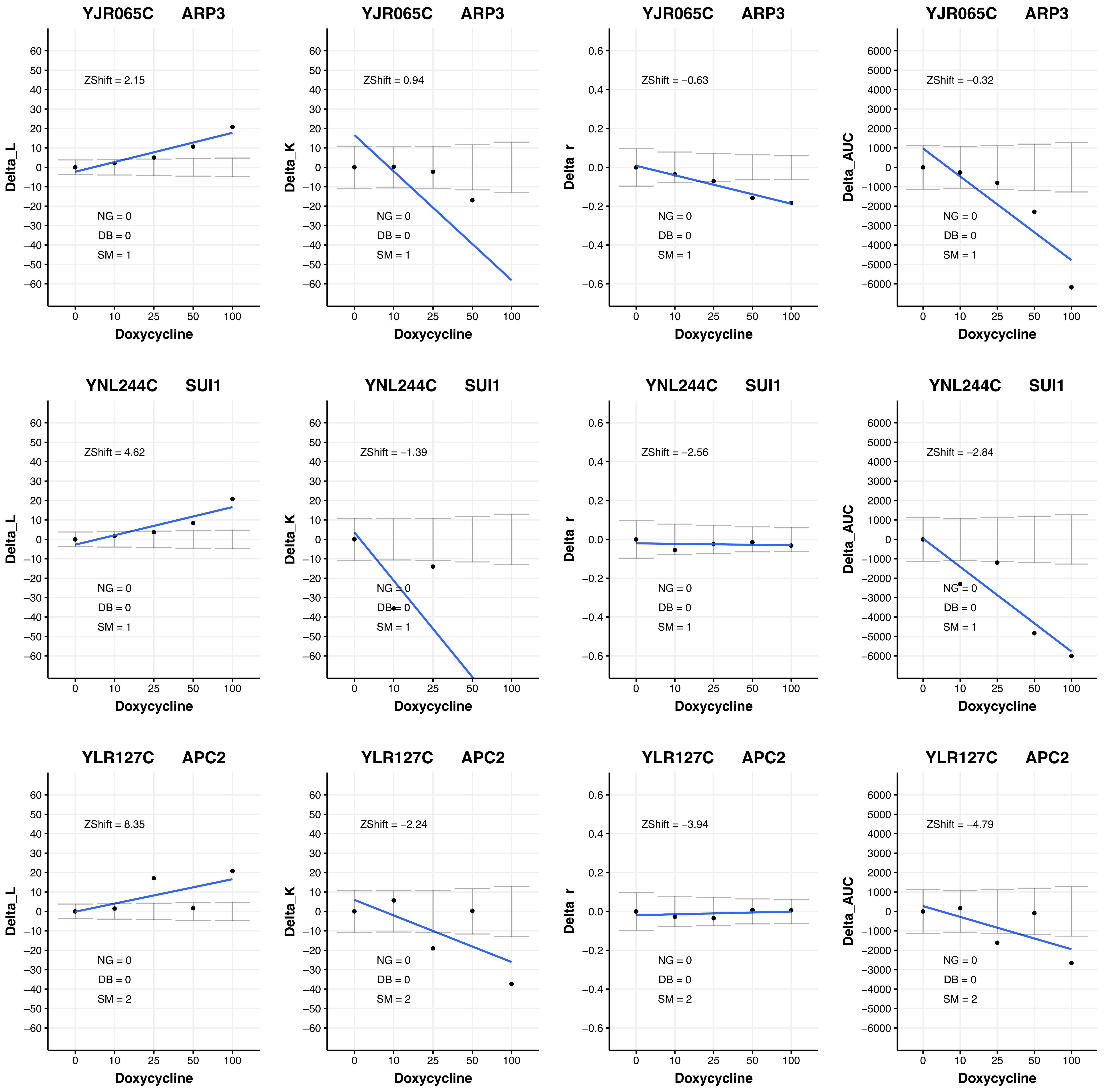

YJR065C ARP3
ZShift = 2.15
NG = 0
DB = 0
SM = 1
Delta_L
Doxycycline
YJR065C ARP3
ZShift = 0.94
NG = 0
DB = 0
SM = 1
Delta_K
Doxycycline
YJR065C ARP3
ZShift = −0.63
NG = 0
DB = 0
SM = 1
Delta_r
Doxycycline
YJR065C ARP3
ZShift = −0.32
NG = 0
DB = 0
SM = 1
Delta_AUC
Doxycycline
YNL244C SUI1
ZShift = 4.62
NG = 0
DB = 0
SM = 1
Delta_L
Doxycycline
YNL244C SUI1
ZShift = −1.39
NG = 0
DB = 0
SM = 1
Delta_K
Doxycycline
YNL244C SUI1
ZShift = −2.56
NG = 0
DB = 0
SM = 1
Delta_r
Doxycycline
YNL244C SUI1
ZShift = −2.84
NG = 0
DB = 0
SM = 1
Delta_AUC
Doxycycline
YLR127C APC2
ZShift = 8.35
NG = 0
DB = 0
SM = 2
Delta_L
Doxycycline
YLR127C APC2
ZShift = −2.24
NG = 0
DB = 0
SM = 2
Delta_K
Doxycycline
YLR127C APC2
ZShift = −3.94
NG = 0
DB = 0
SM = 2
Delta_r
Doxycycline
YLR127C APC2
ZShift = −4.79
NG = 0
DB = 0
SM = 2
Delta_AUC
Doxycycline

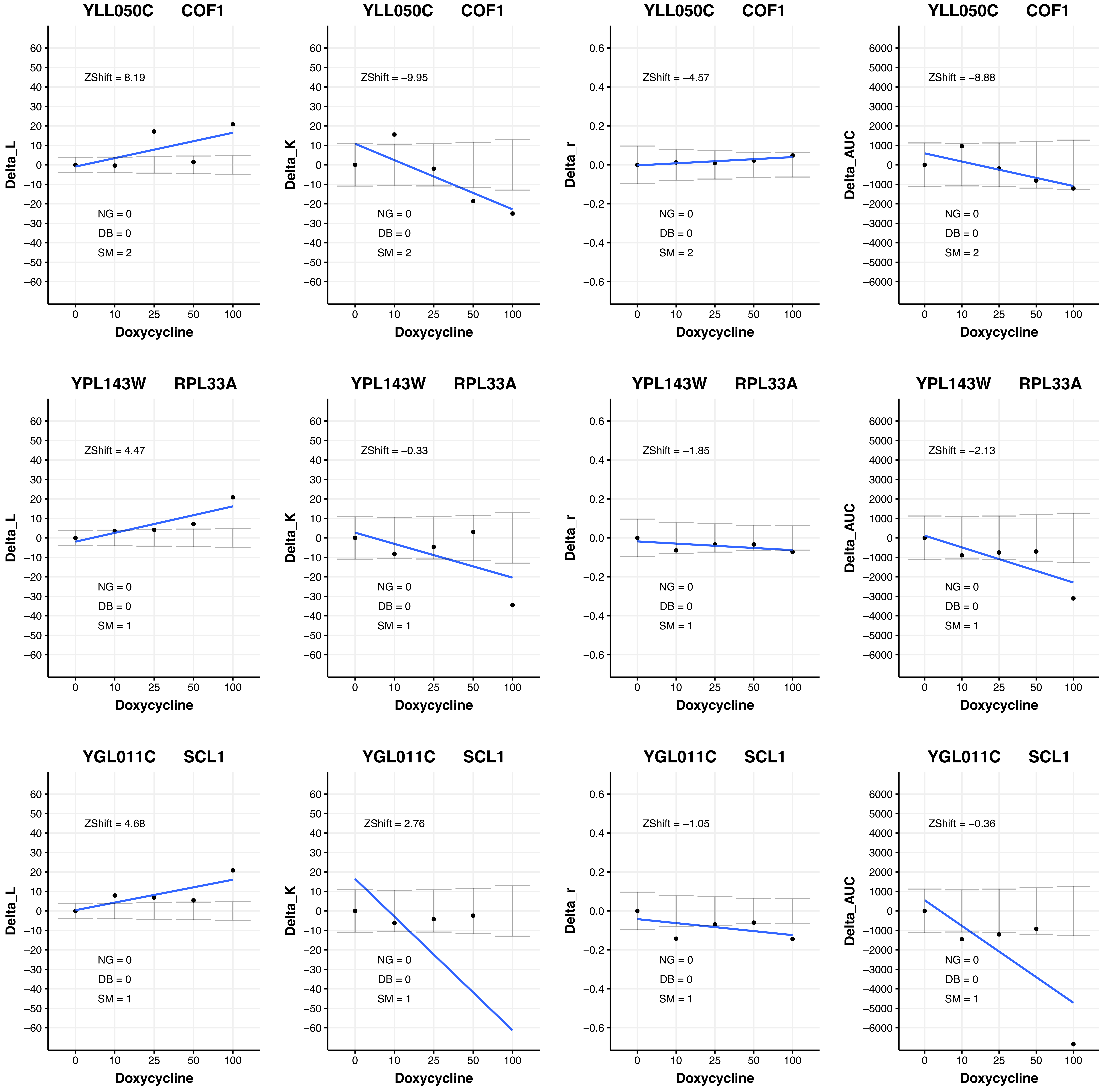

YLL050C COF1
ZShift = 8.19
NG = 0
DB = 0
SM = 2
Delta_L
Doxycycline
YLL050C COF1
ZShift = −9.95
NG = 0
DB = 0
SM = 2
Delta_K
Doxycycline
YLL050C COF1
ZShift = −4.57
NG = 0
DB = 0
SM = 2
Delta_r
Doxycycline
YLL050C COF1
ZShift = −8.88
NG = 0
DB = 0
SM = 2
Delta_AUC
Doxycycline
YPL143W RPL33A
ZShift = 4.47
NG = 0
DB = 0
SM = 1
Delta_L
Doxycycline
YPL143W RPL33A
ZShift = −0.33
NG = 0
DB = 0
SM = 1
Delta_K
Doxycycline
YPL143W RPL33A
ZShift = −1.85
NG = 0
DB = 0
SM = 1
Delta_r
Doxycycline
YPL143W RPL33A
ZShift = −2.13
NG = 0
DB = 0
SM = 1
Delta_AUC
Doxycycline
YGL011C SCL1
ZShift = 4.68
NG = 0
DB = 0
SM = 1
Delta_L
Doxycycline
YGL011C SCL1
ZShift = 2.76
NG = 0
DB = 0
SM = 1
Delta_K
Doxycycline
YGL011C SCL1
ZShift = −1.05
NG = 0
DB = 0
SM = 1
Delta_r
Doxycycline
YGL011C SCL1
ZShift = −0.36
NG = 0
DB = 0
SM = 1
Delta_AUC
Doxycycline

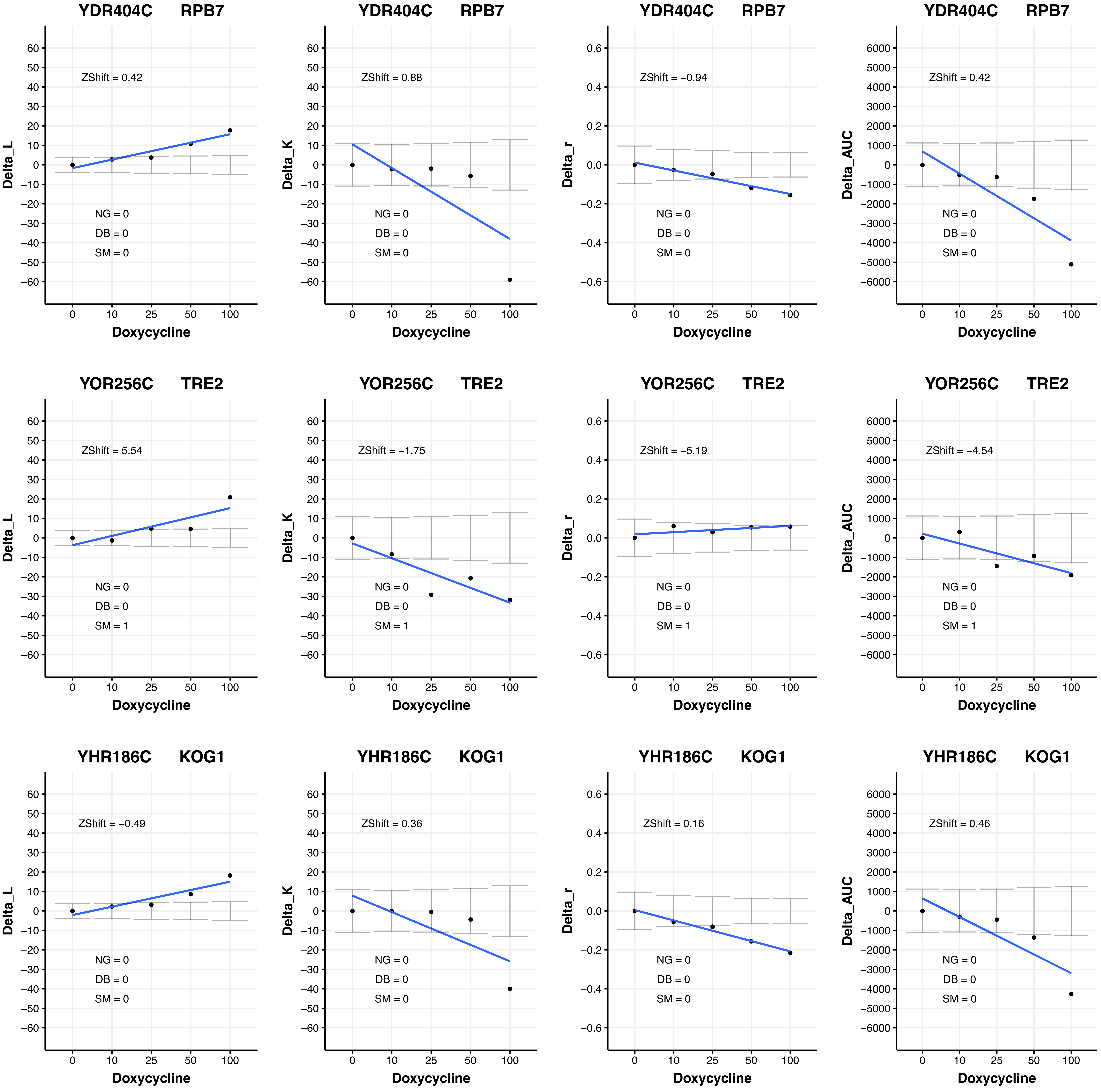
YDR404C RPB7
ZShift = 0.42
NG = 0
DB = 0
SM = 0
Delta_L
Doxycycline
YDR404C RPB7
ZShift = 0.88
NG = 0
DB = 0
SM = 0
Delta_K
Doxycycline
YDR404C RPB7
ZShift = −0.94
NG = 0
DB = 0
SM = 0
Delta_r
Doxycycline
YDR404C RPB7
ZShift = 0.42
NG = 0
DB = 0
SM = 0
Delta_AUC
Doxycycline
YOR256C TRE2
ZShift = 5.54
NG = 0
DB = 0
SM = 1
Delta_L
Doxycycline
YOR256C TRE2
ZShift = −1.75
NG = 0
DB = 0
SM = 1
Delta_K
Doxycycline
YOR256C TRE2
ZShift = −5.19
NG = 0
DB = 0
SM = 1
Delta_r
Doxycycline
YOR256C TRE2
ZShift = −4.54
NG = 0
DB = 0
SM = 1
Delta_AUC
Doxycycline
YHR186C KOG1
ZShift = −0.49
NG = 0
DB = 0
SM = 0
Delta_L
Doxycycline
YHR186C KOG1
ZShift = 0.36
NG = 0
DB = 0
SM = 0
Delta_K
Doxycycline
YHR186C KOG1
ZShift = 0.16
NG = 0
DB = 0
SM = 0
Delta_r
Doxycycline
YHR186C KOG1
ZShift = 0.46
NG = 0
DB = 0
SM = 0
Delta_AUC
Doxycycline

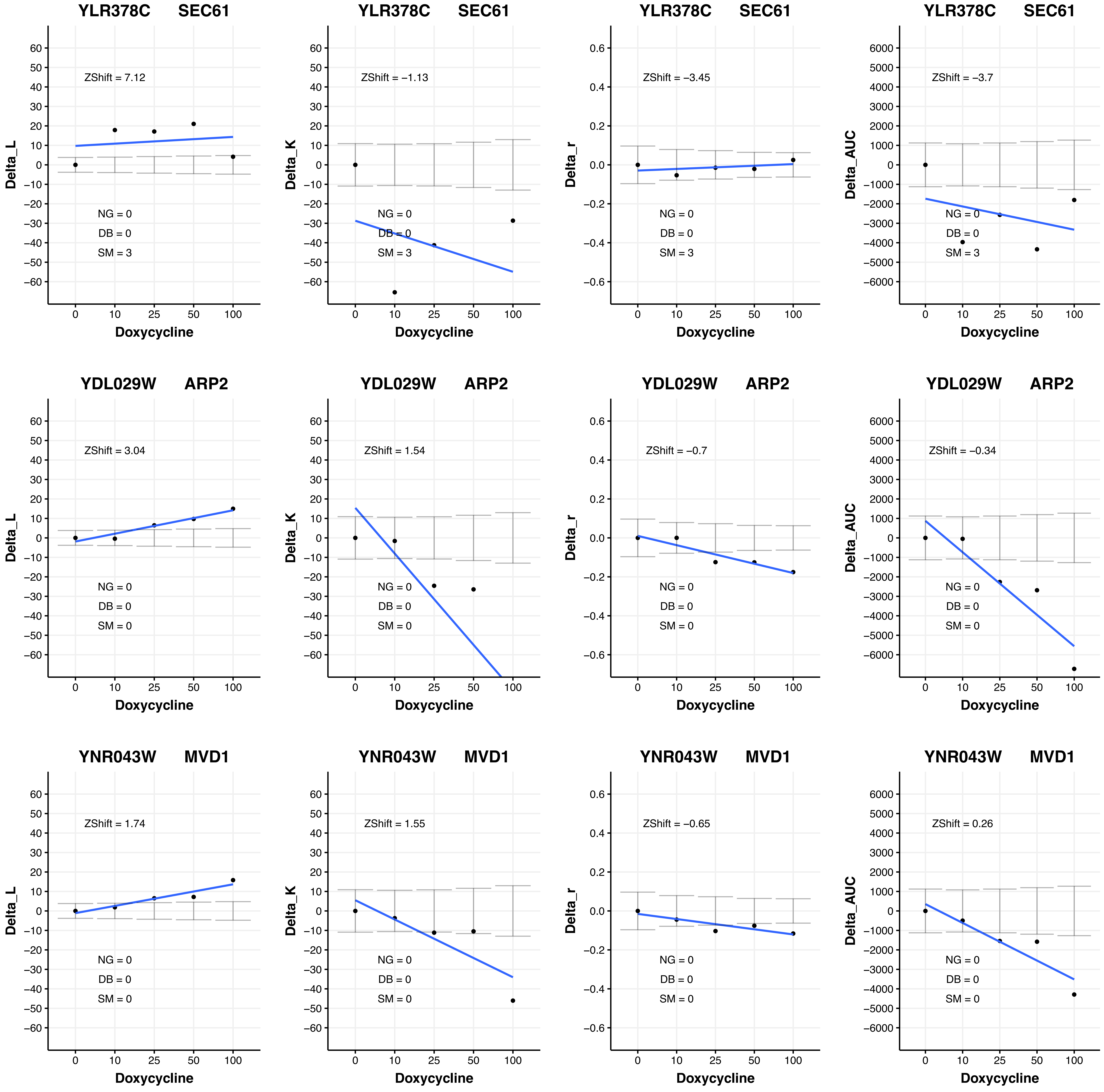

YLR378C SEC61
ZShift = 7.12
NG = 0
DB = 0
SM = 3
Delta_L
Doxycycline
YLR378C SEC61
ZShift = −1.13
NG = 0
DB = 0
SM = 3
Delta_K
Doxycycline
YLR378C SEC61
ZShift = −3.45
NG = 0
DB = 0
SM = 3
Delta_r
Doxycycline
YLR378C SEC61
ZShift = −3.7
NG = 0
DB = 0
SM = 3
Delta_AUC
Doxycycline
YDL029W ARP2
ZShift = 3.04
NG = 0
DB = 0
SM = 0
Delta_L
Doxycycline
YDL029W ARP2
ZShift = 1.54
NG = 0
DB = 0
SM = 0
Delta_K
Doxycycline
YDL029W ARP2
ZShift = −0.7
NG = 0
DB = 0
SM = 0
Delta_r
Doxycycline
YDL029W ARP2
ZShift = −0.34
NG = 0
DB = 0
SM = 0
Delta_AUC
Doxycycline
YNR043W MVD1
ZShift = 1.74
NG = 0
DB = 0
SM = 0
Delta_L
Doxycycline
YNR043W MVD1
ZShift = 1.55
NG = 0
DB = 0
SM = 0
Delta_K
Doxycycline
YNR043W MVD1
ZShift = −0.65
NG = 0
DB = 0
SM = 0
Delta_r
Doxycycline
YNR043W MVD1
ZShift = 0.26
NG = 0
DB = 0
SM = 0
Delta_AUC
Doxycycline

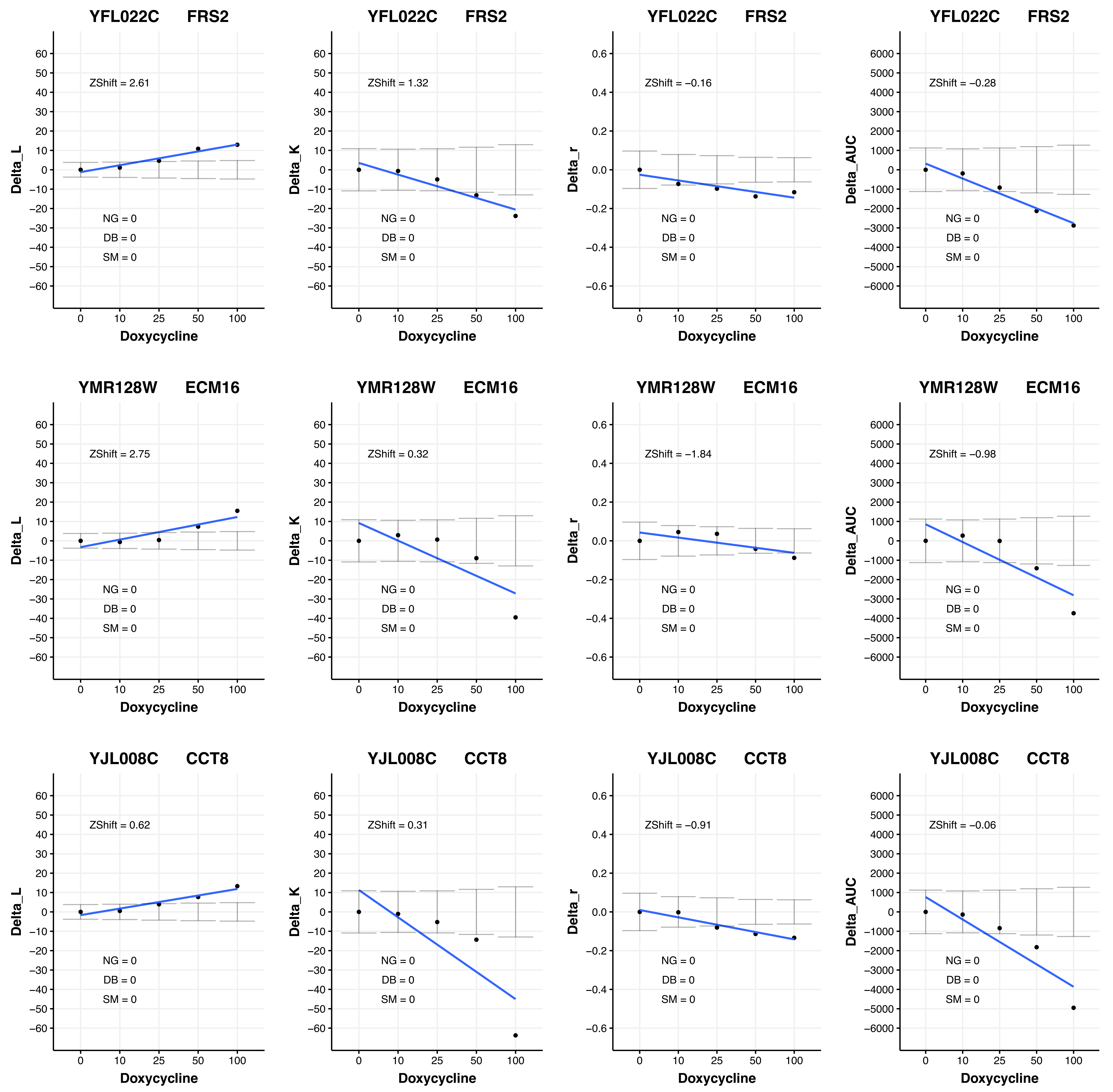
YFL022C FRS2
ZShift = 2.61
NG = 0
DB = 0
SM = 0
Delta_L
Doxycycline
YFL022C FRS2
ZShift = 1.32
NG = 0
DB = 0
SM = 0
Delta_K
Doxycycline
YFL022C FRS2
ZShift = −0.16
NG = 0
DB = 0
SM = 0
Delta_r
Doxycycline
YFL022C FRS2
ZShift = −0.28
NG = 0
DB = 0
SM = 0
Delta_AUC
Doxycycline
YMR128W ECM16
ZShift = 2.75
NG = 0
DB = 0
SM = 0
Delta_L
Doxycycline
YMR128W ECM16
ZShift = 0.32
NG = 0
DB = 0
SM = 0
Delta_K
Doxycycline
YMR128W ECM16
ZShift = −1.84
NG = 0
DB = 0
SM = 0
Delta_r
Doxycycline
YMR128W ECM16
ZShift = −0.98
NG = 0
DB = 0
SM = 0
Delta_AUC
Doxycycline
YJL008C CCT8
ZShift = 0.62
NG = 0
DB = 0
SM = 0
Delta_L
Doxycycline
YJL008C CCT8
ZShift = 0.31
NG = 0
DB = 0
SM = 0
Delta_K
Doxycycline
YJL008C CCT8
ZShift = −0.91
NG = 0
DB = 0
SM = 0
Delta_r
Doxycycline
YJL008C CCT8
ZShift = −0.06
NG = 0
DB = 0
SM = 0
Delta_AUC
Doxycycline
0
10
25
50
100

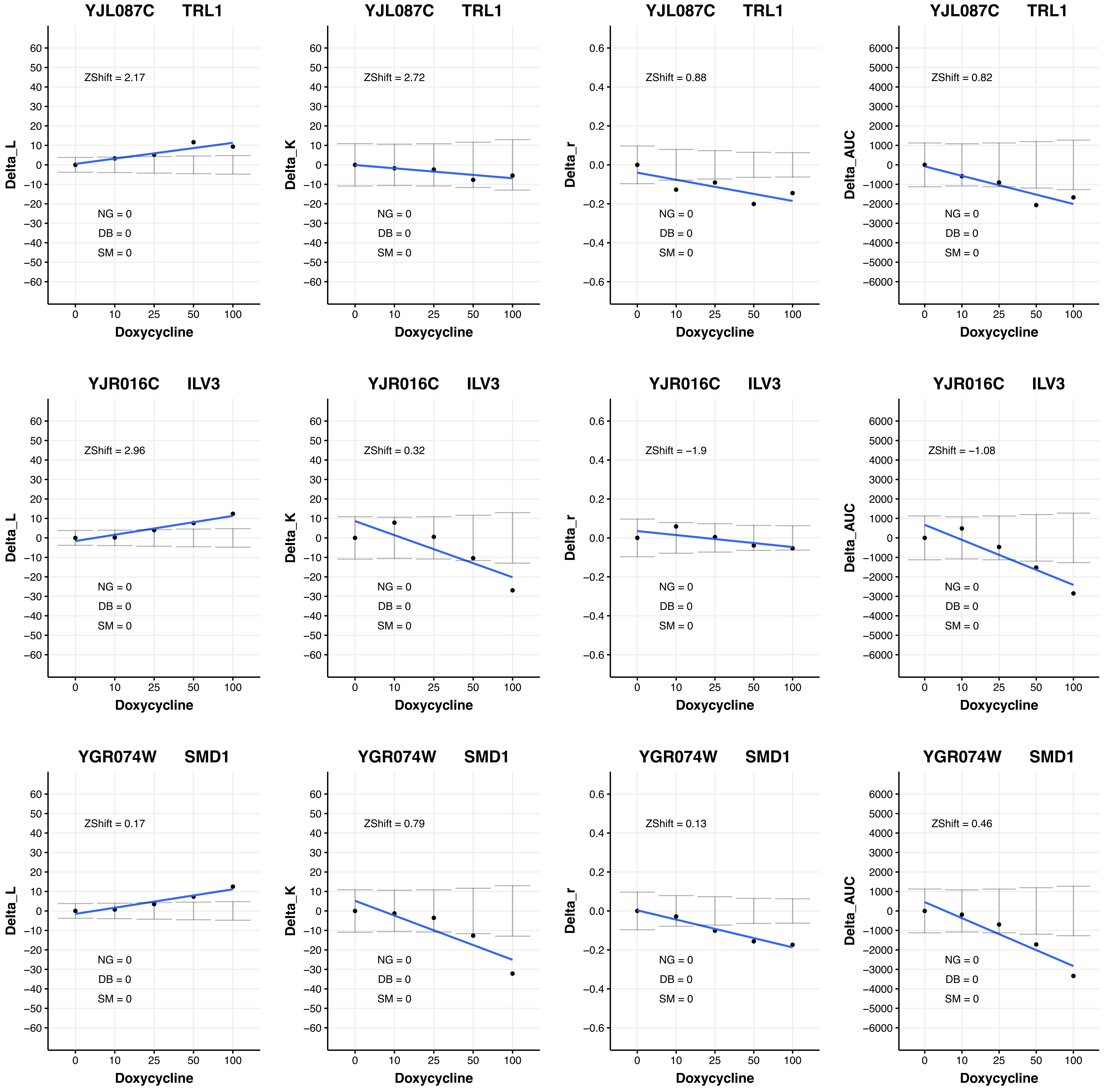

YJL087C TRL1
ZShift = 2.17
NG = 0
DB = 0
SM = 0
Delta_L
Doxycycline
YJL087C TRL1
ZShift = 2.72
NG = 0
DB = 0
SM = 0
Delta_K
Doxycycline
YJL087C TRL1
ZShift = 0.88
NG = 0
DB = 0
SM = 0
Delta_r
Doxycycline
YJL087C TRL1
ZShift = 0.82
NG = 0
DB = 0
SM = 0
Delta_AUC
Doxycycline
YJR016C ILV3
ZShift = 2.96
NG = 0
DB = 0
SM = 0
Delta_L
Doxycycline
YJR016C ILV3
ZShift = 0.32
NG = 0
DB = 0
SM = 0
Delta_K
Doxycycline
YJR016C ILV3
ZShift = −1.9
NG = 0
DB = 0
SM = 0
Delta_r
Doxycycline
YJR016C ILV3
ZShift = −1.08
NG = 0
DB = 0
SM = 0
Delta_AUC
Doxycycline
YGR074W SMD1
ZShift = 0.17
NG = 0
DB = 0
SM = 0
Delta_L
Doxycycline
YGR074W SMD1
ZShift = 0.79
NG = 0
DB = 0
SM = 0
Delta_K
Doxycycline
YGR074W SMD1
ZShift = 0.13
NG = 0
DB = 0
SM = 0
Delta_r
Doxycycline
YGR074W SMD1
ZShift = 0.46
NG = 0
DB = 0
SM = 0
Delta_AUC
Doxycycline

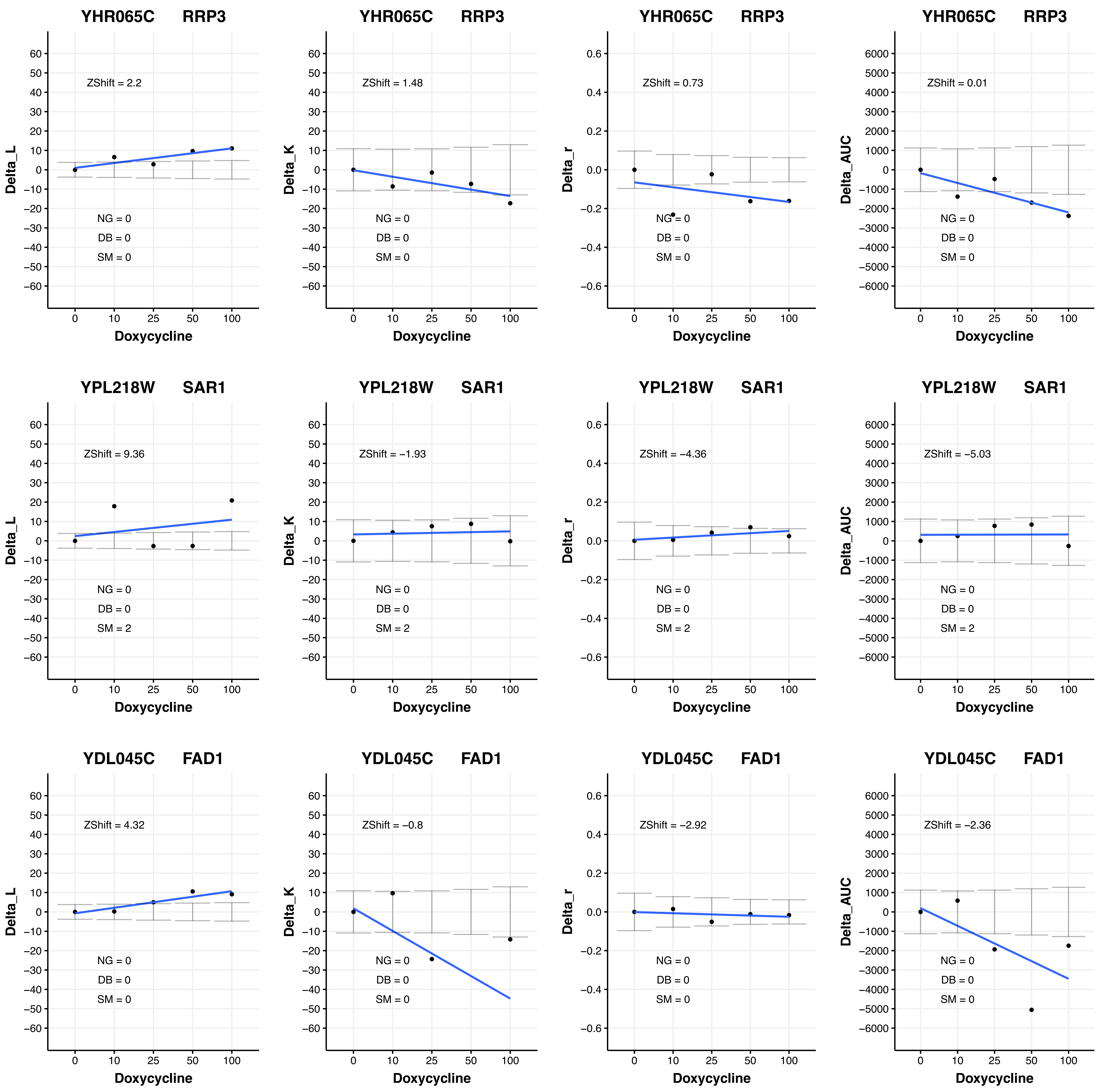
YHR065C RRP3
ZShift = 2.2
NG = 0
DB = 0
SM = 0
Delta_L
Doxycycline
YHR065C RRP3
ZShift = 1.48
NG = 0
DB = 0
SM = 0
Delta_K
Doxycycline
YHR065C RRP3
ZShift = 0.73
NG = 0
DB = 0
SM = 0
Delta_r
Doxycycline
YHR065C RRP3
ZShift = 0.01
NG = 0
DB = 0
SM = 0
Delta_AUC
Doxycycline
YPL218W SAR1
ZShift = 9.36
NG = 0
DB = 0
SM = 2
Delta_L
Doxycycline
YPL218W SAR1
ZShift = −1.93
NG = 0
DB = 0
SM = 2
Delta_K
Doxycycline
YPL218W SAR1
ZShift = −4.36
NG = 0
DB = 0
SM = 2
Delta_r
Doxycycline
YPL218W SAR1
ZShift = −5.03
NG = 0
DB = 0
SM = 2
Delta_AUC
Doxycycline
YDL045C FAD1
ZShift = 4.32
NG = 0
DB = 0
SM = 0
Delta_L
Doxycycline
YDL045C FAD1
ZShift = −0.8
NG = 0
DB = 0
SM = 0
Delta_K
Doxycycline
YDL045C FAD1
ZShift = −2.92
NG = 0
DB = 0
SM = 0
Delta_r
Doxycycline
YDL045C FAD1
ZShift = −2.36
NG = 0
DB = 0
SM = 0
Delta_AUC
Doxycycline

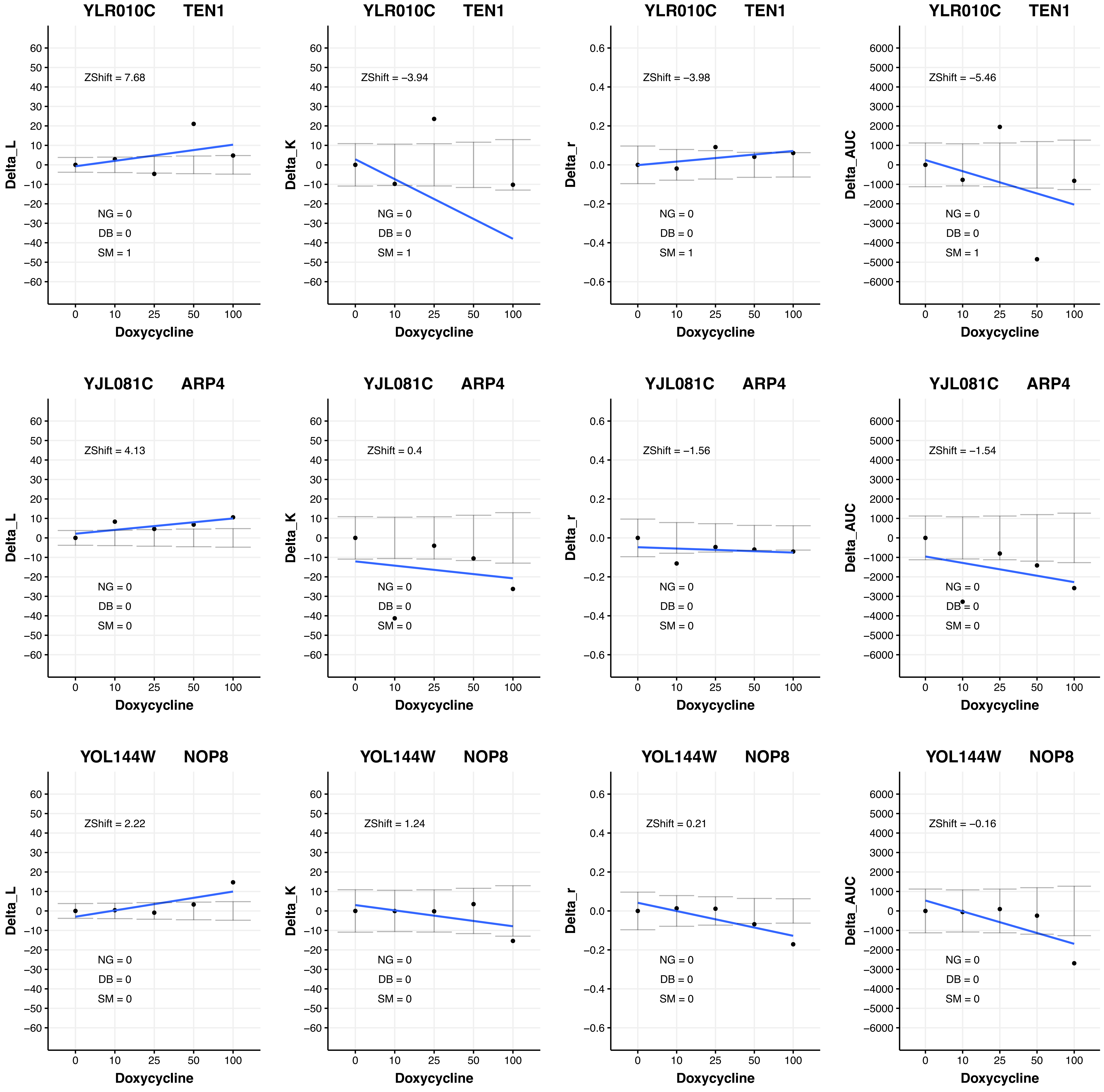
YLR010C TEN1
ZShift = 7.68
NG = 0
DB = 0
SM = 1
Delta_L
Doxycycline
YLR010C TEN1
ZShift = −3.94
NG = 0
DB = 0
SM = 1
Delta_K
Doxycycline
YLR010C TEN1
ZShift = −3.98
NG = 0
DB = 0
SM = 1
Delta_r
Doxycycline
YLR010C TEN1
ZShift = −5.46
NG = 0
DB = 0
SM = 1
Delta_AUC
Doxycycline
YJL081C ARP4
ZShift = 4.13
NG = 0
DB = 0
SM = 0
Delta_L
Doxycycline
YJL081C ARP4
ZShift = 0.4
NG = 0
DB = 0
SM = 0
Delta_K
Doxycycline
YJL081C ARP4
ZShift = −1.56
NG = 0
DB = 0
SM = 0
Delta_r
Doxycycline
YJL081C ARP4
ZShift = −1.54
NG = 0
DB = 0
SM = 0
Delta_AUC
Doxycycline
YOL144W NOP8
ZShift = 2.22
NG = 0
DB = 0
SM = 0
Delta_L
Doxycycline
YOL144W NOP8
ZShift = 1.24
NG = 0
DB = 0
SM = 0
Delta_K
Doxycycline
YOL144W NOP8
ZShift = 0.21
NG = 0
DB = 0
SM = 0
Delta_r
Doxycycline
YOL144W NOP8
ZShift = −0.16
NG = 0
DB = 0
SM = 0
Delta_AUC
Doxycycline

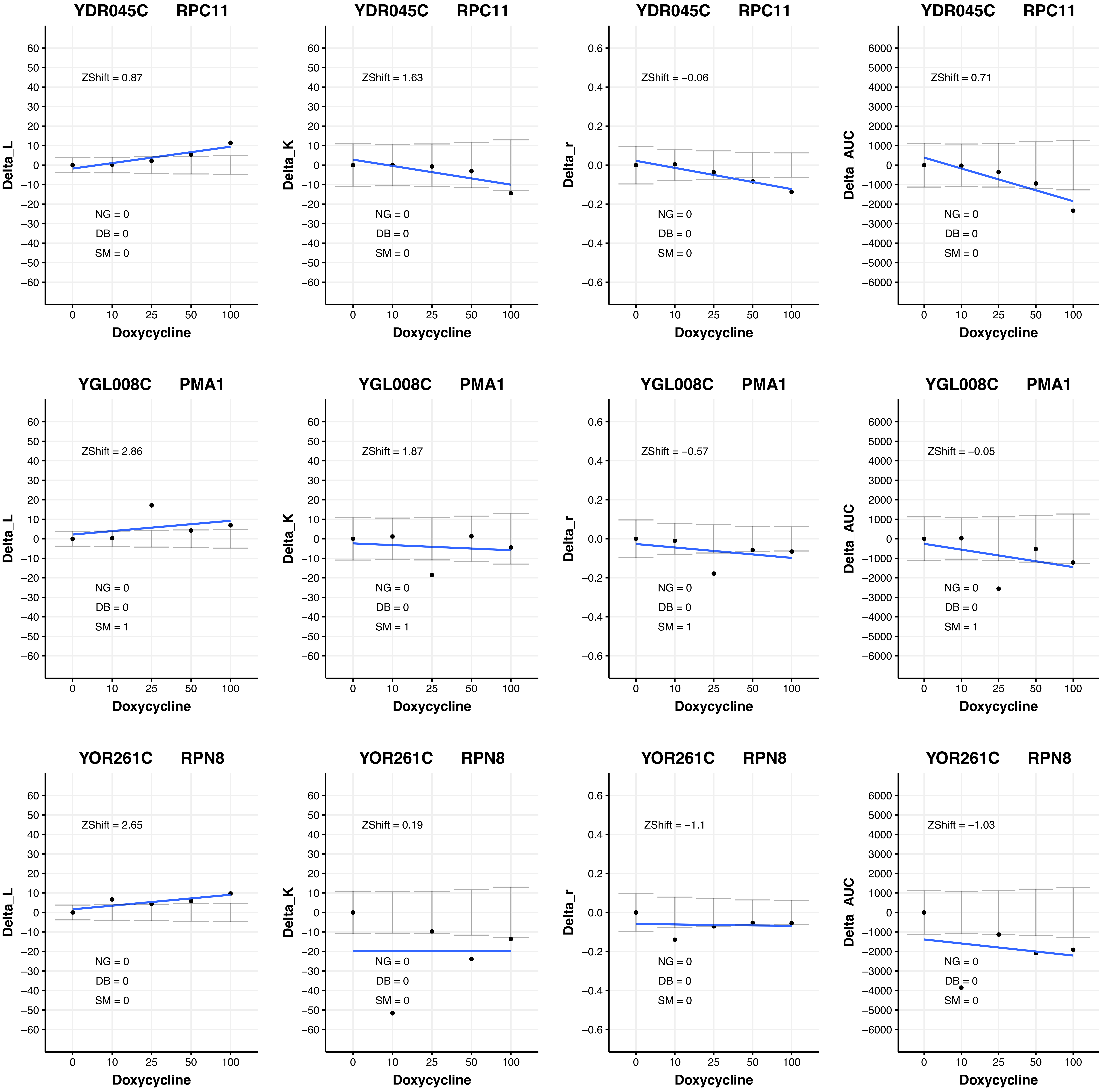

YDR045C RPC11
ZShift = 0.87
NG = 0
DB = 0
SM = 0
Delta_L
Doxycycline
YDR045C RPC11
ZShift = 1.63
NG = 0
DB = 0
SM = 0
Delta_K
Doxycycline
YDR045C RPC11
ZShift = −0.06
NG = 0
DB = 0
SM = 0
Delta_r
Doxycycline
YDR045C RPC11
ZShift = 0.71
NG = 0
DB = 0
SM = 0
Delta_AUC
Doxycycline
YGL008C PMA1
ZShift = 2.86
NG = 0
DB = 0
SM = 1
Delta_L
Doxycycline
YGL008C PMA1
ZShift = 1.87
NG = 0
DB = 0
SM = 1
Delta_K
Doxycycline
YGL008C PMA1
ZShift = −0.57
NG = 0
DB = 0
SM = 1
Delta_r
Doxycycline
YGL008C PMA1
ZShift = −0.05
NG = 0
DB = 0
SM = 1
Delta_AUC
Doxycycline
YOR261C RPN8
ZShift = 2.65
NG = 0
DB = 0
SM = 0
Delta_L
Doxycycline
YOR261C RPN8
ZShift = 0.19
NG = 0
DB = 0
SM = 0
Delta_K
Doxycycline
YOR261C RPN8
ZShift = −1.1
NG = 0
DB = 0
SM = 0
Delta_r
Doxycycline
YOR261C RPN8
ZShift = −1.03
NG = 0
DB = 0
SM = 0
Delta_AUC
Doxycycline
0 10 25 50 100

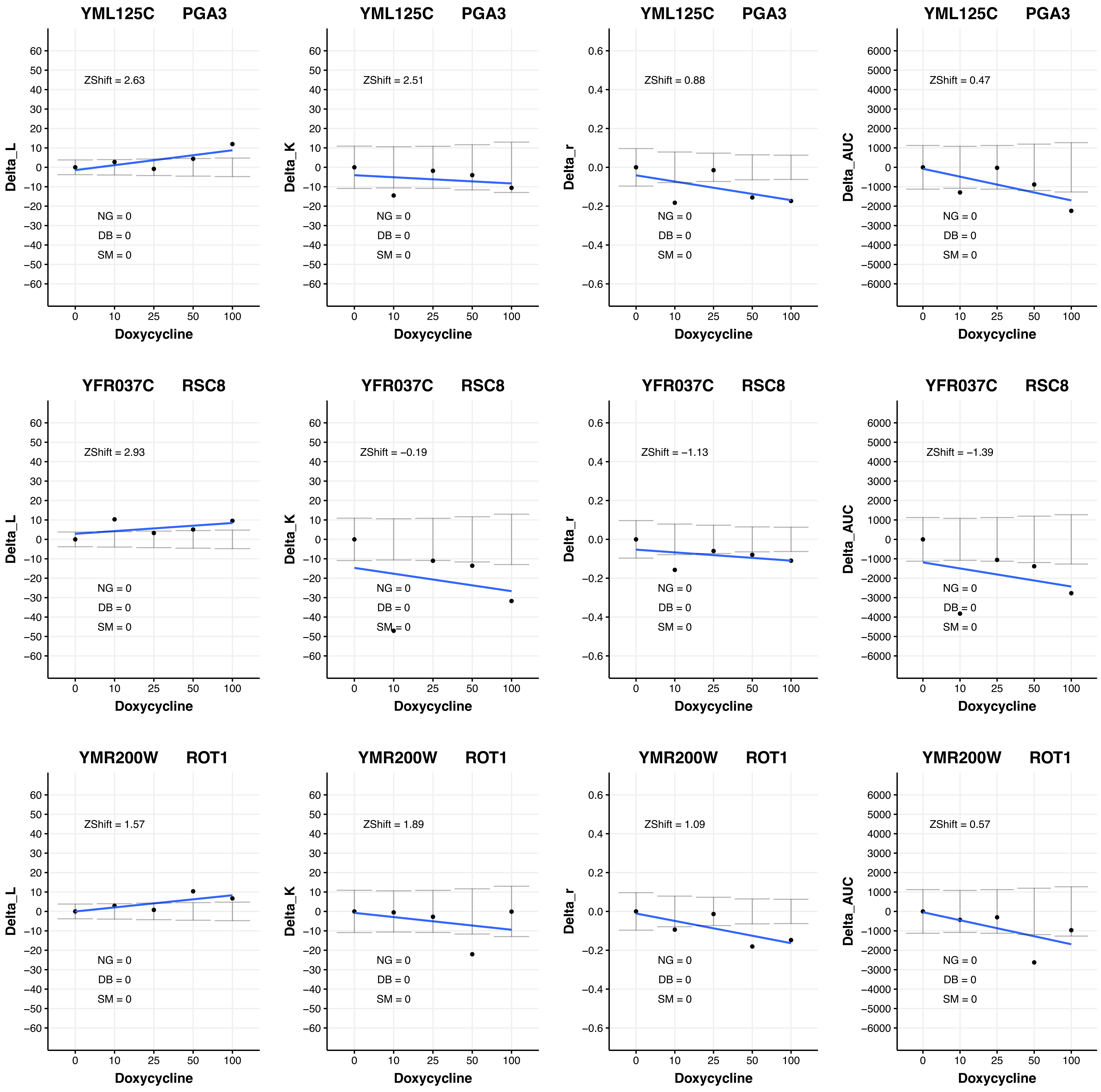

YML125C PGA3
ZShift = 2.63
ZShift = 2.51
ZShift = 0.88
ZShift = 0.47
YFR037C RSC8
ZShift = 2.93
ZShift = −0.19
ZShift = −1.13
ZShift = −1.39
YMR200W ROT1
ZShift = 1.57
ZShift = 1.89
ZShift = 1.09
ZShift = 0.57
NG = 0
DB = 0
SM = 0
Delta_L
Delta_K
Delta_r
Delta_AUC
Doxycycline
0 10 25 50 100

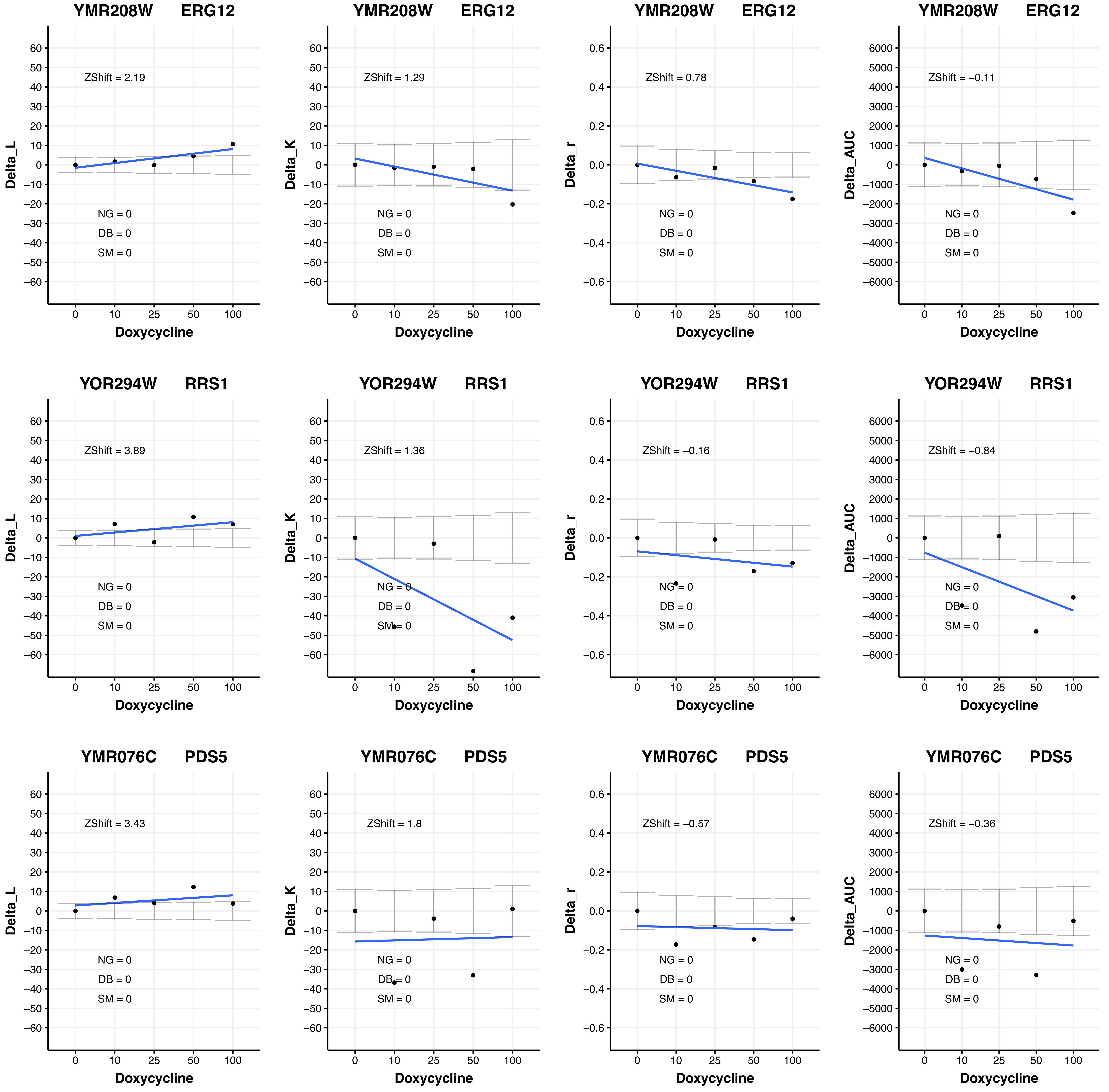
YMR208W ERG12
ZShift = 2.19
NG = 0
DB = 0
SM = 0
Delta_L
Doxycycline
YMR208W ERG12
ZShift = 1.29
NG = 0
DB = 0
SM = 0
Delta_K
Doxycycline
YMR208W ERG12
ZShift = 0.78
NG = 0
DB = 0
SM = 0
Delta_r
Doxycycline
YMR208W ERG12
ZShift = −0.11
NG = 0
DB = 0
SM = 0
Delta_AUC
Doxycycline
YOR294W RRS1
ZShift = 3.89
NG = 0
DB = 0
SM = 0
Delta_L
Doxycycline
YOR294W RRS1
ZShift = 1.36
NG = 0
DB = 0
SM = 0
Delta_K
Doxycycline
YOR294W RRS1
ZShift = −0.16
NG = 0
DB = 0
SM = 0
Delta_r
Doxycycline
YOR294W RRS1
ZShift = −0.84
NG = 0
DB = 0
SM = 0
Delta_AUC
Doxycycline
YMR076C PDS5
ZShift = 3.43
NG = 0
DB = 0
SM = 0
Delta_L
Doxycycline
YMR076C PDS5
ZShift = 1.8
NG = 0
DB = 0
SM = 0
Delta_K
Doxycycline
YMR076C PDS5
ZShift = −0.57
NG = 0
DB = 0
SM = 0
Delta_r
Doxycycline
YMR076C PDS5
ZShift = −0.36
NG = 0
DB = 0
SM = 0
Delta_AUC
Doxycycline

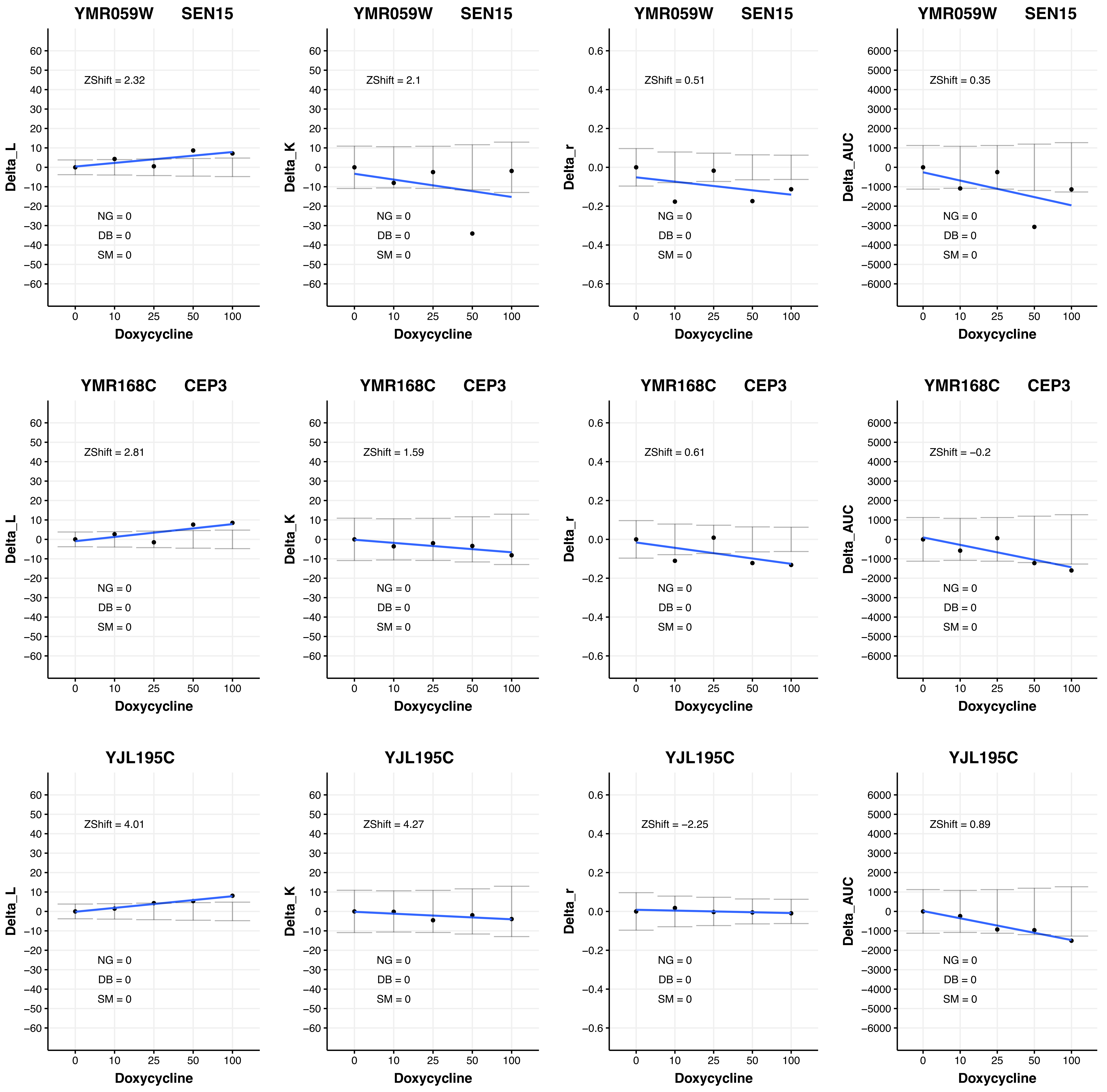
YMR059W SEN15
Delta_L
ZShift = 2.32
NG = 0
DB = 0
SM = 0
Doxycycline
YMR059W SEN15
Delta_K
ZShift = 2.1
NG = 0
DB = 0
SM = 0
Doxycycline
YMR059W SEN15
Delta_r
ZShift = 0.51
NG = 0
DB = 0
SM = 0
Doxycycline
YMR059W SEN15
Delta_AUC
ZShift = 0.35
NG = 0
DB = 0
SM = 0
Doxycycline
YMR168C CEP3
Delta_L
ZShift = 2.81
NG = 0
DB = 0
SM = 0
Doxycycline
YMR168C CEP3
Delta_K
ZShift = 1.59
NG = 0
DB = 0
SM = 0
Doxycycline
YMR168C CEP3
Delta_r
ZShift = 0.61
NG = 0
DB = 0
SM = 0
Doxycycline
YMR168C CEP3
Delta_AUC
ZShift = −0.2
NG = 0
DB = 0
SM = 0
Doxycycline
YJL195C
Delta_L
ZShift = 4.01
NG = 0
DB = 0
SM = 0
Doxycycline
YJL195C
Delta_K
ZShift = 4.27
NG = 0
DB = 0
SM = 0
Doxycycline
YJL195C
Delta_r
ZShift = −2.25
NG = 0
DB = 0
SM = 0
Doxycycline
YJL195C
Delta_AUC
ZShift = 0.89
NG = 0
DB = 0
SM = 0
Doxycycline

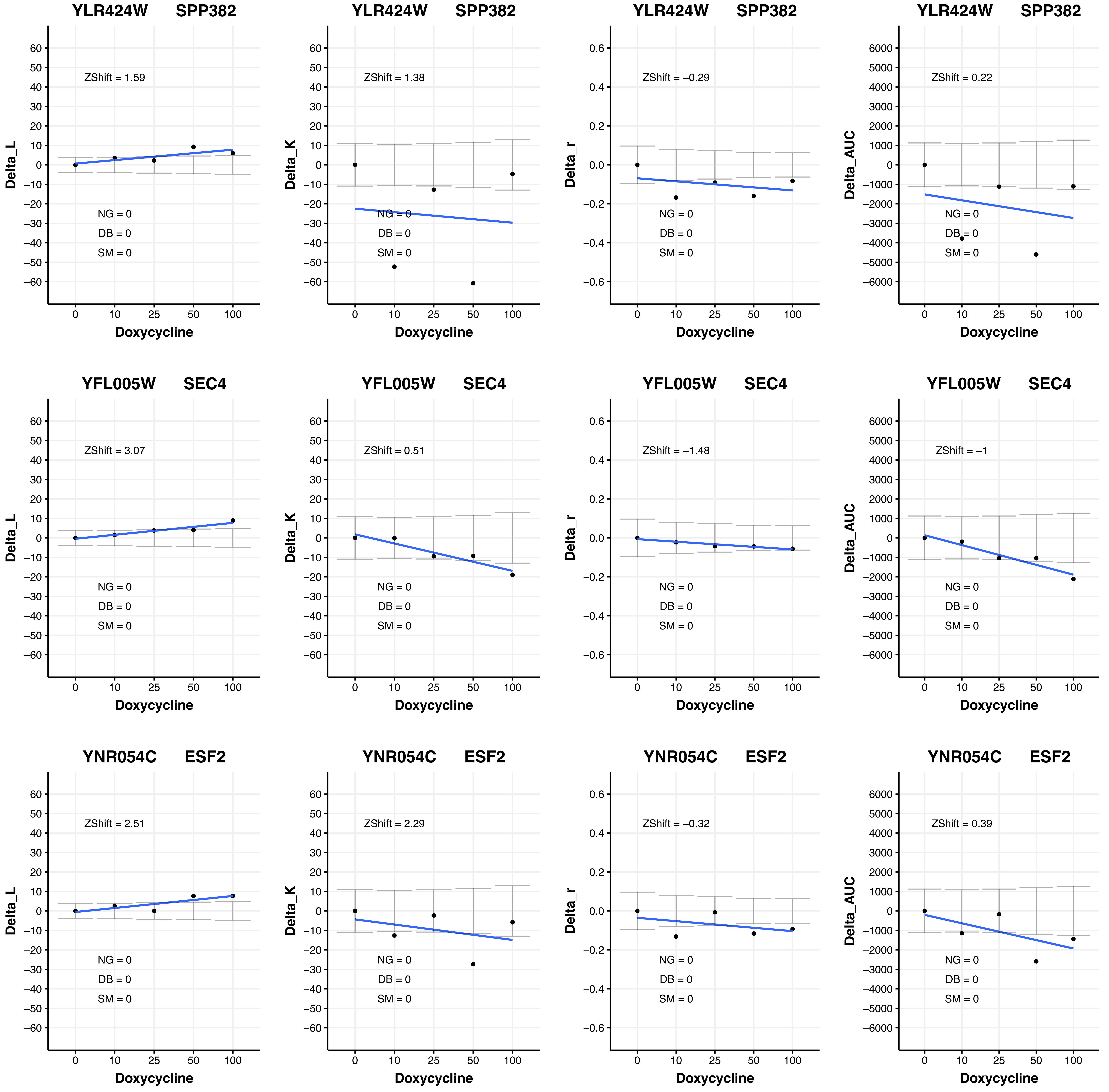

YLR424W SPP382
Delta_L
ZShift = 1.59
NG = 0
DB = 0
SM = 0
Doxycycline
YLR424W SPP382
Delta_K
ZShift = 1.38
NG = 0
DB = 0
SM = 0
Doxycycline
YLR424W SPP382
Delta_r
ZShift = −0.29
NG = 0
DB = 0
SM = 0
Doxycycline
YLR424W SPP382
Delta_AUC
ZShift = 0.22
NG = 0
DB = 0
SM = 0
Doxycycline
YFL005W SEC4
Delta_L
ZShift = 3.07
NG = 0
DB = 0
SM = 0
Doxycycline
YFL005W SEC4
Delta_K
ZShift = 0.51
NG = 0
DB = 0
SM = 0
Doxycycline
YFL005W SEC4
Delta_r
ZShift = −1.48
NG = 0
DB = 0
SM = 0
Doxycycline
YFL005W SEC4
Delta_AUC
ZShift = −1
NG = 0
DB = 0
SM = 0
Doxycycline
YNR054C ESF2
Delta_L
ZShift = 2.51
NG = 0
DB = 0
SM = 0
Doxycycline
YNR054C ESF2
Delta_K
ZShift = 2.29
NG = 0
DB = 0
SM = 0
Doxycycline
YNR054C ESF2
Delta_r
ZShift = −0.32
NG = 0
DB = 0
SM = 0
Doxycycline
YNR054C ESF2
Delta_AUC
ZShift = 0.39
NG = 0
DB = 0
SM = 0
Doxycycline

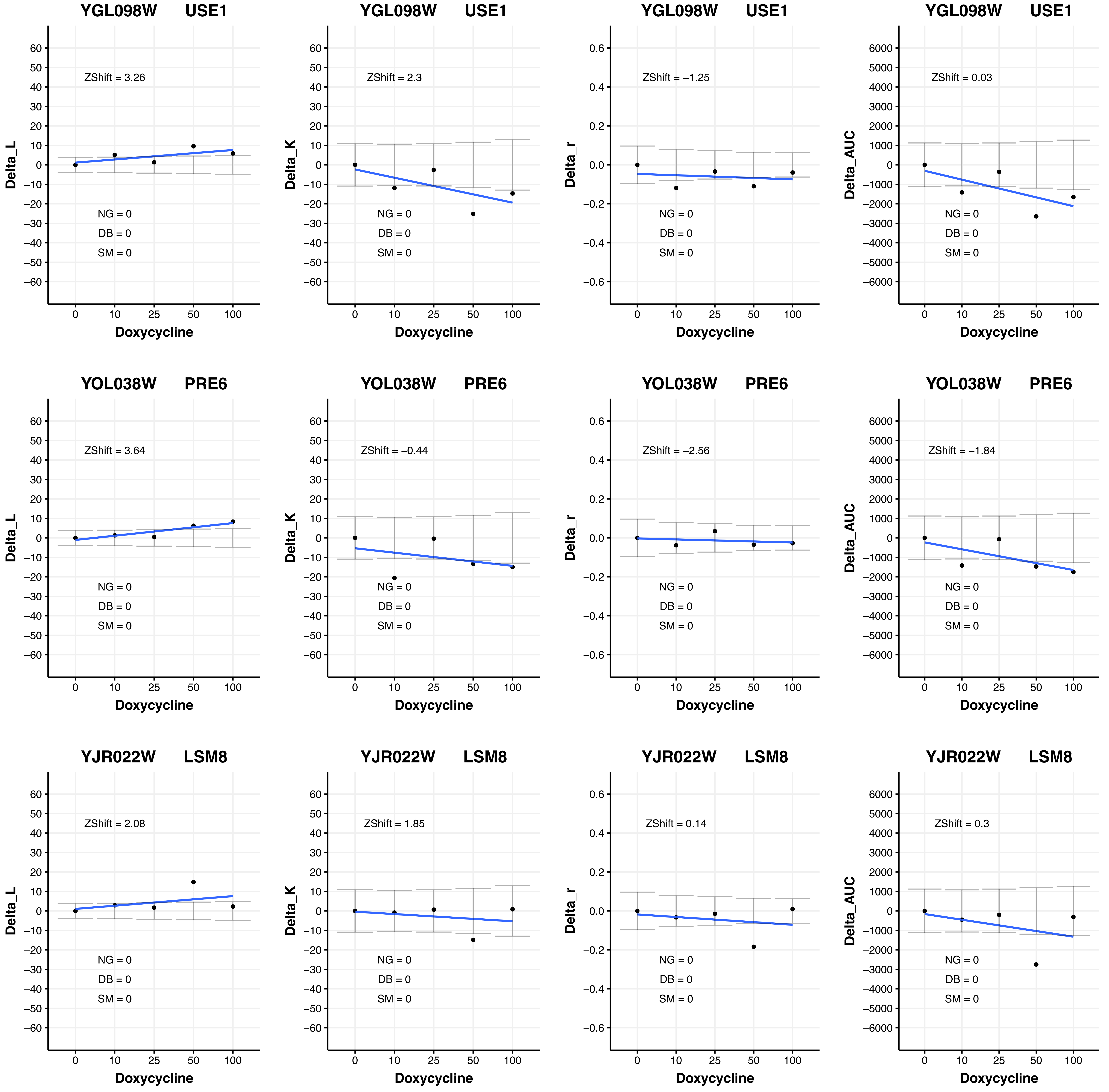
YGL098W USE1
ZShift = 3.26
NG = 0
DB = 0
SM = 0
Delta_L
Doxycycline
YGL098W USE1
ZShift = 2.3
NG = 0
DB = 0
SM = 0
Delta_K
Doxycycline
YGL098W USE1
ZShift = −1.25
NG = 0
DB = 0
SM = 0
Delta_r
Doxycycline
YGL098W USE1
ZShift = 0.03
NG = 0
DB = 0
SM = 0
Delta_AUC
Doxycycline
YOL038W PRE6
ZShift = 3.64
NG = 0
DB = 0
SM = 0
Delta_L
Doxycycline
YOL038W PRE6
ZShift = −0.44
NG = 0
DB = 0
SM = 0
Delta_K
Doxycycline
YOL038W PRE6
ZShift = −2.56
NG = 0
DB = 0
SM = 0
Delta_r
Doxycycline
YOL038W PRE6
ZShift = −1.84
NG = 0
DB = 0
SM = 0
Delta_AUC
Doxycycline
YJR022W LSM8
ZShift = 2.08
NG = 0
DB = 0
SM = 0
Delta_L
Doxycycline
YJR022W LSM8
ZShift = 1.85
NG = 0
DB = 0
SM = 0
Delta_K
Doxycycline
YJR022W LSM8
ZShift = 0.14
NG = 0
DB = 0
SM = 0
Delta_r
Doxycycline
YJR022W LSM8
ZShift = 0.3
NG = 0
DB = 0
SM = 0
Delta_AUC
Doxycycline

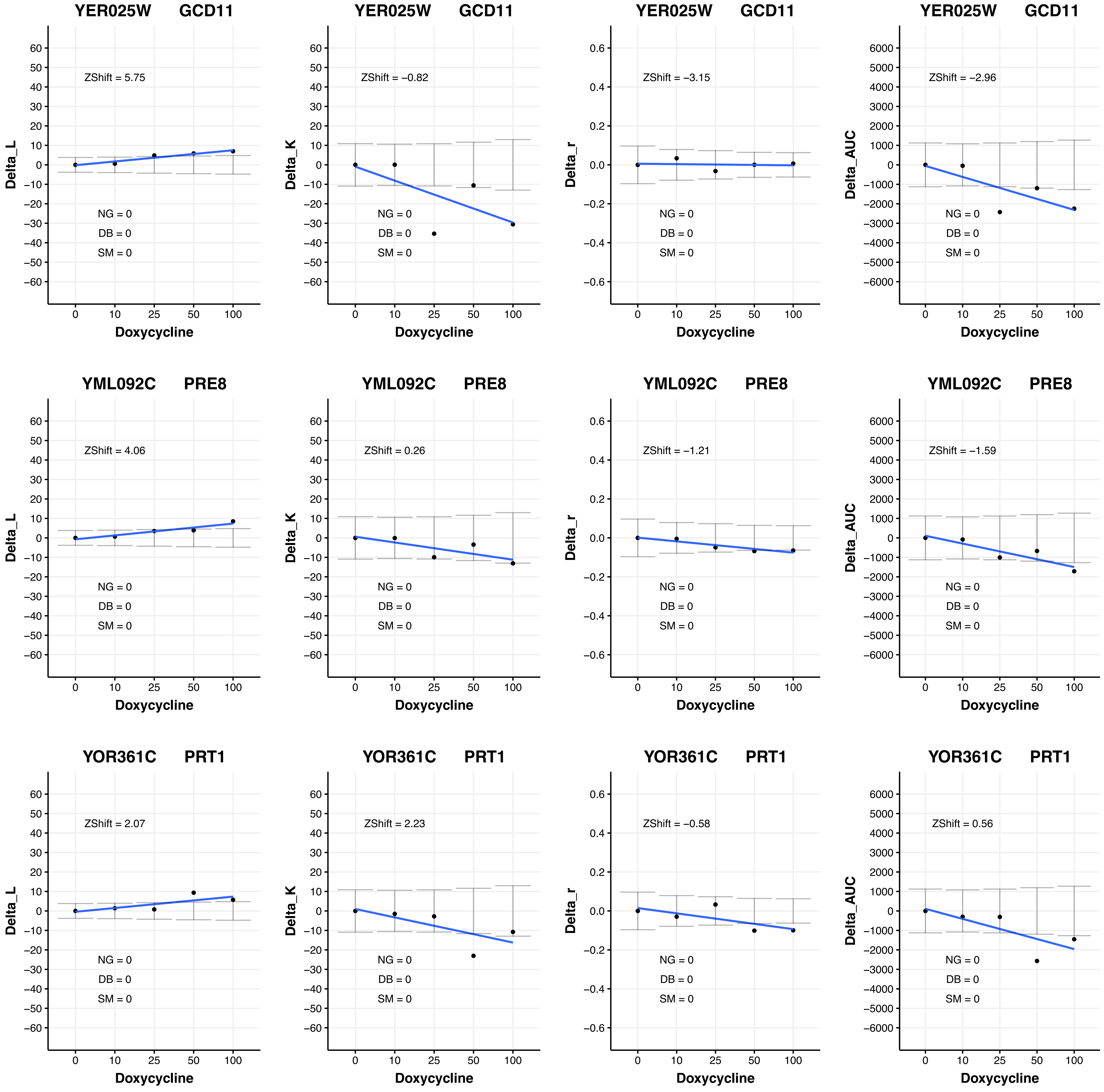

YER025W GCD11
ZShift = 5.75
Delta_L
ZShift = −0.82
Delta_K
ZShift = −3.15
Delta_r
ZShift = −2.96
Delta_AUC
YML092C PRE8
ZShift = 4.06
ZShift = 0.26
ZShift = −1.21
ZShift = −1.59
YOR361C PRT1
ZShift = 2.07
ZShift = 2.23
ZShift = −0.58
ZShift = 0.56
NG = 0
DB = 0
SM = 0
Doxycycline
0
10
25
50
100

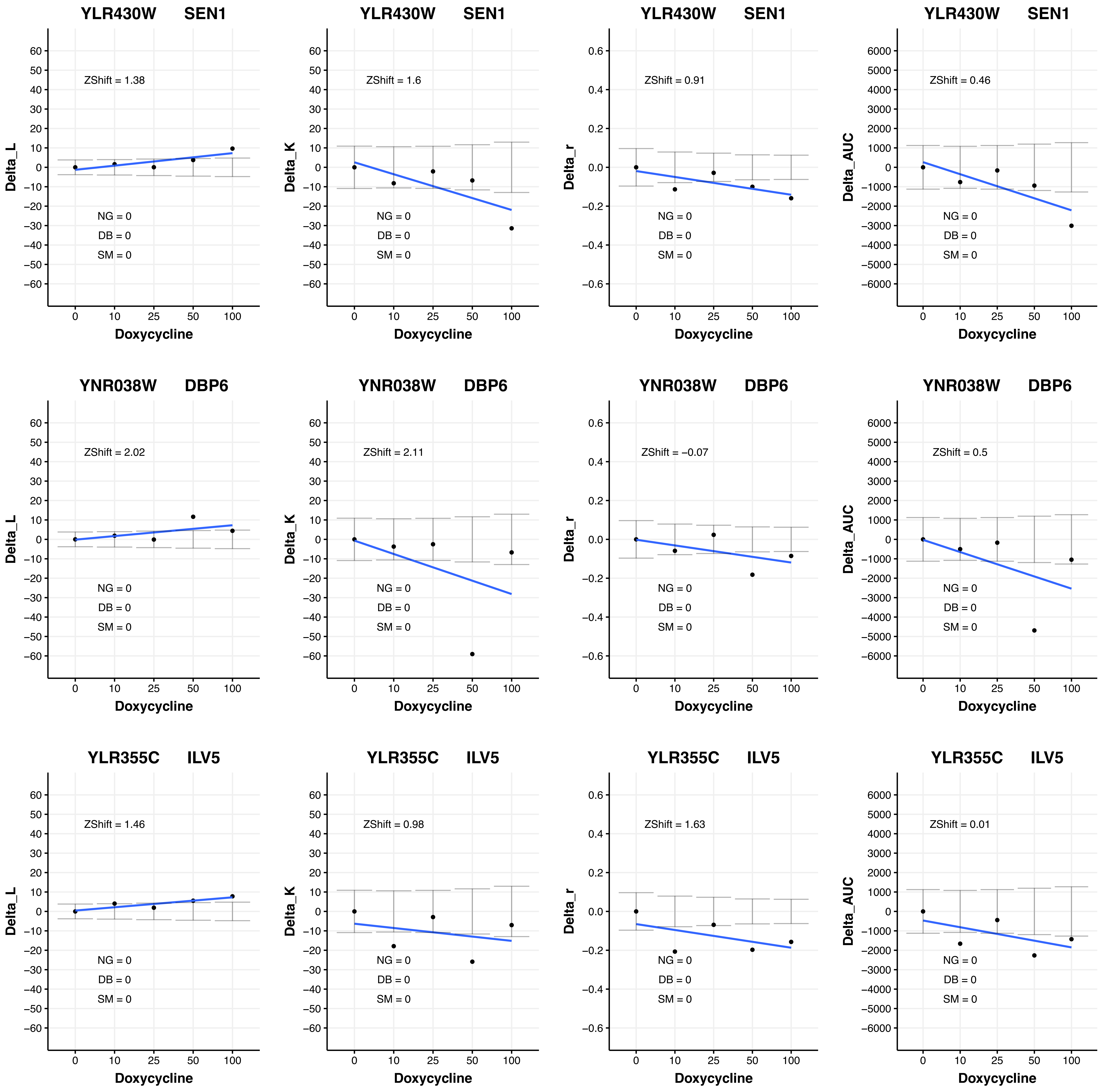

YLR430W SEN1
ZShift = 1.38
NG = 0
DB = 0
SM = 0
Delta_L
Doxycycline
YLR430W SEN1
ZShift = 1.6
NG = 0
DB = 0
SM = 0
Delta_K
Doxycycline
YLR430W SEN1
ZShift = 0.91
NG = 0
DB = 0
SM = 0
Delta_r
Doxycycline
YLR430W SEN1
ZShift = 0.46
NG = 0
DB = 0
SM = 0
Delta_AUC
Doxycycline
YNR038W DBP6
ZShift = 2.02
NG = 0
DB = 0
SM = 0
Delta_L
Doxycycline
YNR038W DBP6
ZShift = 2.11
NG = 0
DB = 0
SM = 0
Delta_K
Doxycycline
YNR038W DBP6
ZShift = −0.07
NG = 0
DB = 0
SM = 0
Delta_r
Doxycycline
YNR038W DBP6
ZShift = 0.5
NG = 0
DB = 0
SM = 0
Delta_AUC
Doxycycline
YLR355C ILV5
ZShift = 1.46
NG = 0
DB = 0
SM = 0
Delta_L
Doxycycline
YLR355C ILV5
ZShift = 0.98
NG = 0
DB = 0
SM = 0
Delta_K
Doxycycline
YLR355C ILV5
ZShift = 1.63
NG = 0
DB = 0
SM = 0
Delta_r
Doxycycline
YLR355C ILV5
ZShift = 0.01
NG = 0
DB = 0
SM = 0
Delta_AUC
Doxycycline

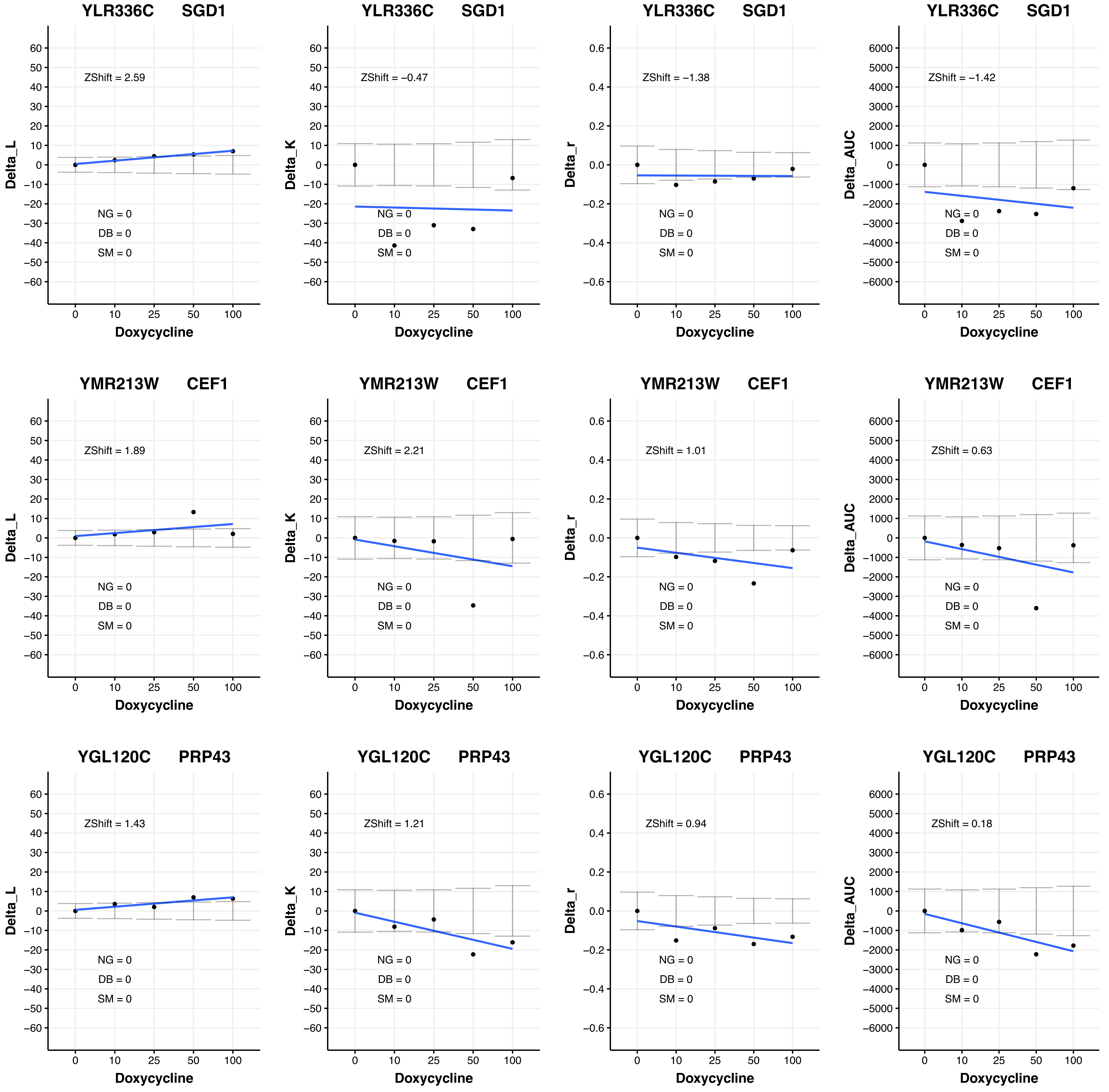
YLR336C SGD1
ZShift = 2.59
NG = 0
DB = 0
SM = 0
Delta_L
Doxycycline
YLR336C SGD1
ZShift = −0.47
NG = 0
DB = 0
SM = 0
Delta_K
Doxycycline
YLR336C SGD1
ZShift = −1.38
NG = 0
DB = 0
SM = 0
Delta_r
Doxycycline
YLR336C SGD1
ZShift = −1.42
NG = 0
DB = 0
SM = 0
Delta_AUC
Doxycycline
YMR213W CEF1
ZShift = 1.89
NG = 0
DB = 0
SM = 0
Delta_L
Doxycycline
YMR213W CEF1
ZShift = 2.21
NG = 0
DB = 0
SM = 0
Delta_K
Doxycycline
YMR213W CEF1
ZShift = 1.01
NG = 0
DB = 0
SM = 0
Delta_r
Doxycycline
YMR213W CEF1
ZShift = 0.63
NG = 0
DB = 0
SM = 0
Delta_AUC
Doxycycline
YGL120C PRP43
ZShift = 1.43
NG = 0
DB = 0
SM = 0
Delta_L
Doxycycline
YGL120C PRP43
ZShift = 1.21
NG = 0
DB = 0
SM = 0
Delta_K
Doxycycline
YGL120C PRP43
ZShift = 0.94
NG = 0
DB = 0
SM = 0
Delta_r
Doxycycline
YGL120C PRP43
ZShift = 0.18
NG = 0
DB = 0
SM = 0
Delta_AUC
Doxycycline

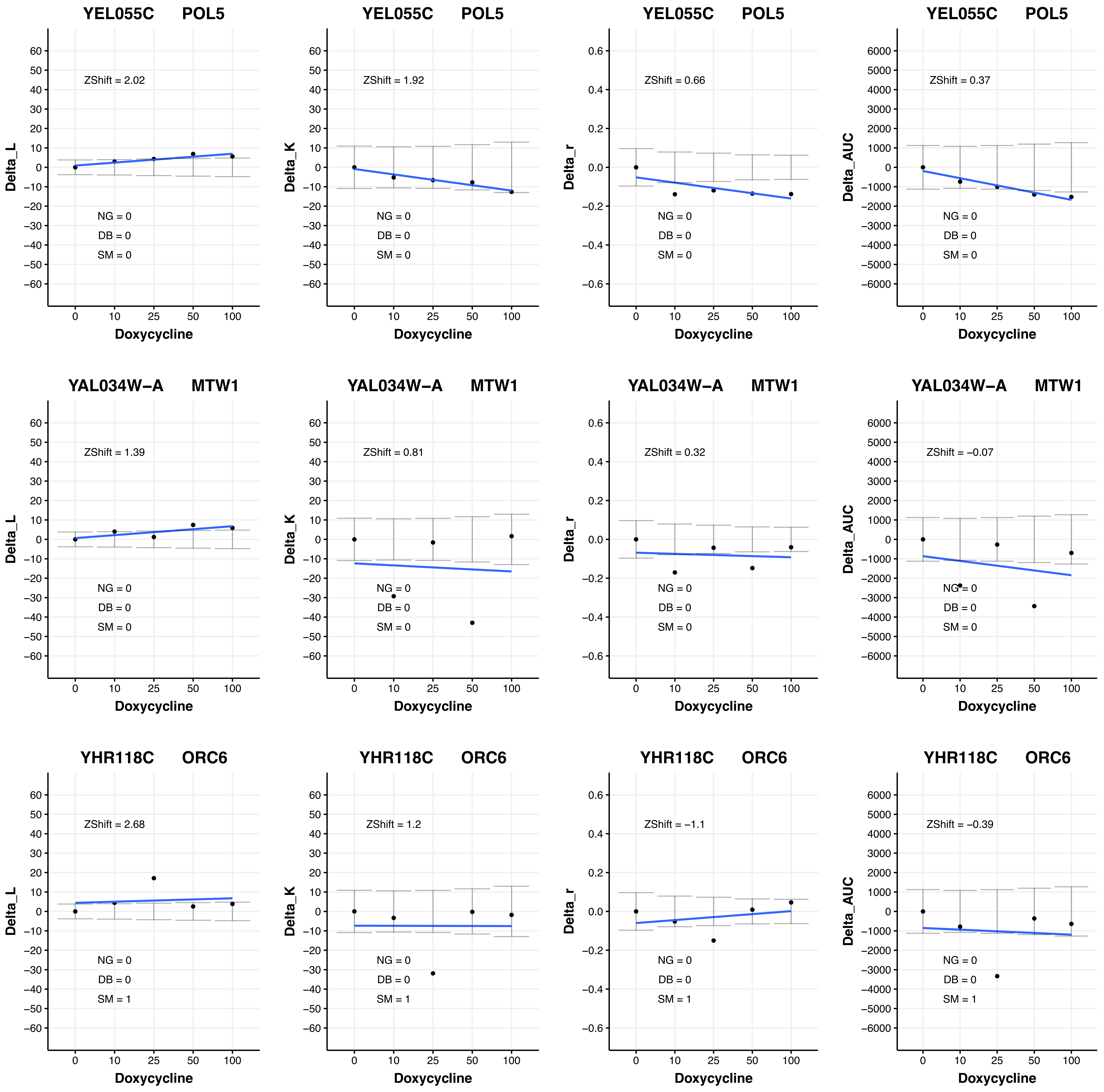

YEL055C POL5
ZShift = 2.02
NG = 0
DB = 0
SM = 0
Delta_L
Doxycycline
YEL055C POL5
ZShift = 1.92
NG = 0
DB = 0
SM = 0
Delta_K
Doxycycline
YEL055C POL5
ZShift = 0.66
NG = 0
DB = 0
SM = 0
Delta_r
Doxycycline
YEL055C POL5
ZShift = 0.37
NG = 0
DB = 0
SM = 0
Delta_AUC
Doxycycline
YAL034W−A MTW1
ZShift = 1.39
NG = 0
DB = 0
SM = 0
Delta_L
Doxycycline
YAL034W−A MTW1
ZShift = 0.81
NG = 0
DB = 0
SM = 0
Delta_K
Doxycycline
YAL034W−A MTW1
ZShift = 0.32
NG = 0
DB = 0
SM = 0
Delta_r
Doxycycline
YAL034W−A MTW1
ZShift = −0.07
NG = 0
DB = 0
SM = 0
Delta_AUC
Doxycycline
YHR118C ORC6
ZShift = 2.68
NG = 0
DB = 0
SM = 1
Delta_L
Doxycycline
YHR118C ORC6
ZShift = 1.2
NG = 0
DB = 0
SM = 1
Delta_K
Doxycycline
YHR118C ORC6
ZShift = −1.1
NG = 0
DB = 0
SM = 1
Delta_r
Doxycycline
YHR118C ORC6
ZShift = −0.39
NG = 0
DB = 0
SM = 1
Delta_AUC
Doxycycline

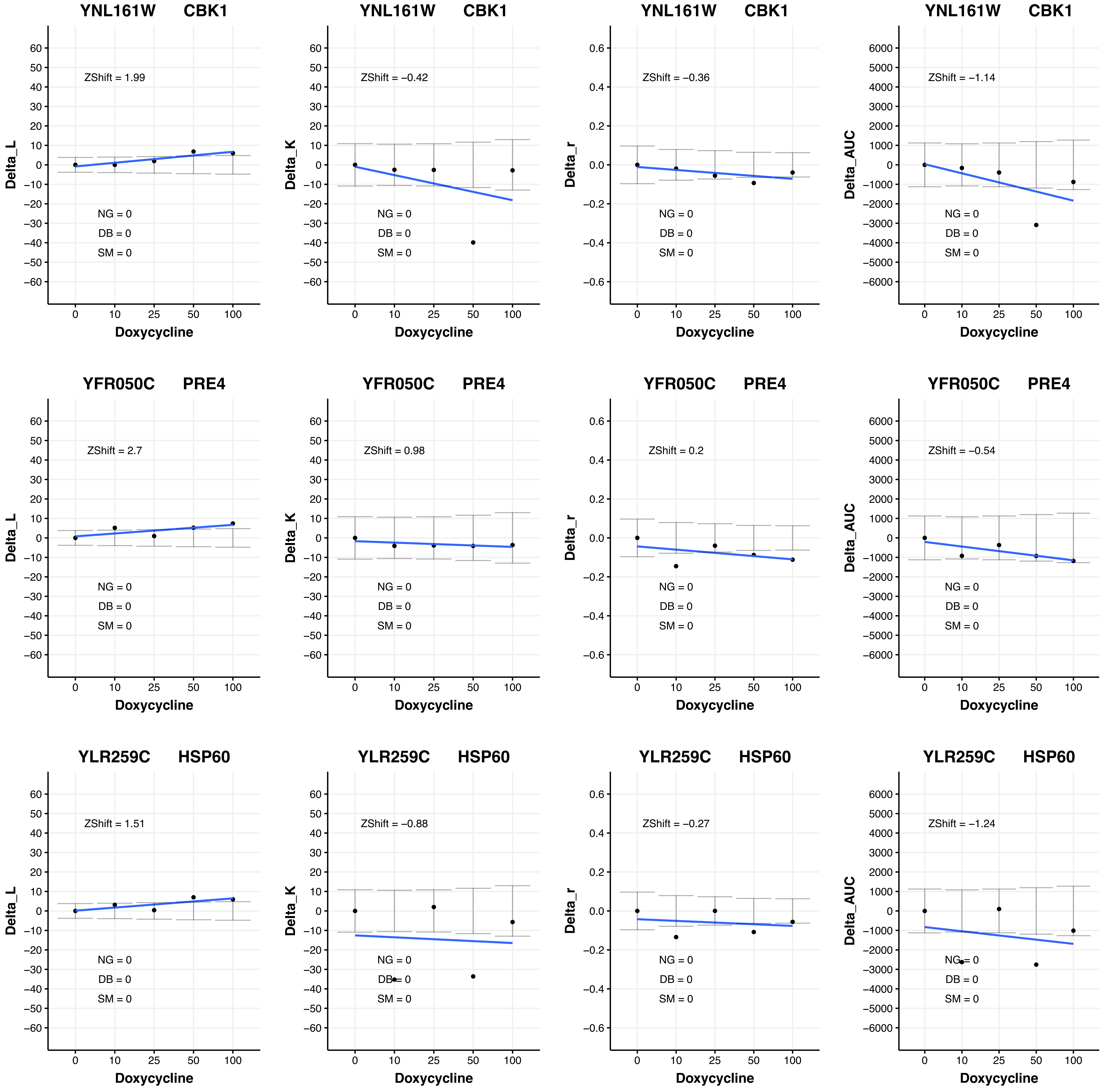
YNL161W CBK1
ZShift = 1.99
ZShift = −0.42
ZShift = −0.36
ZShift = −1.14
YFR050C PRE4
ZShift = 2.7
ZShift = 0.98
ZShift = 0.2
ZShift = −0.54
YLR259C HSP60
ZShift = 1.51
ZShift = −0.88
ZShift = −0.27
ZShift = −1.24
NG = 0
DB = 0
SM = 0
Delta_L
Delta_K
Delta_r
Delta_AUC
Doxycycline

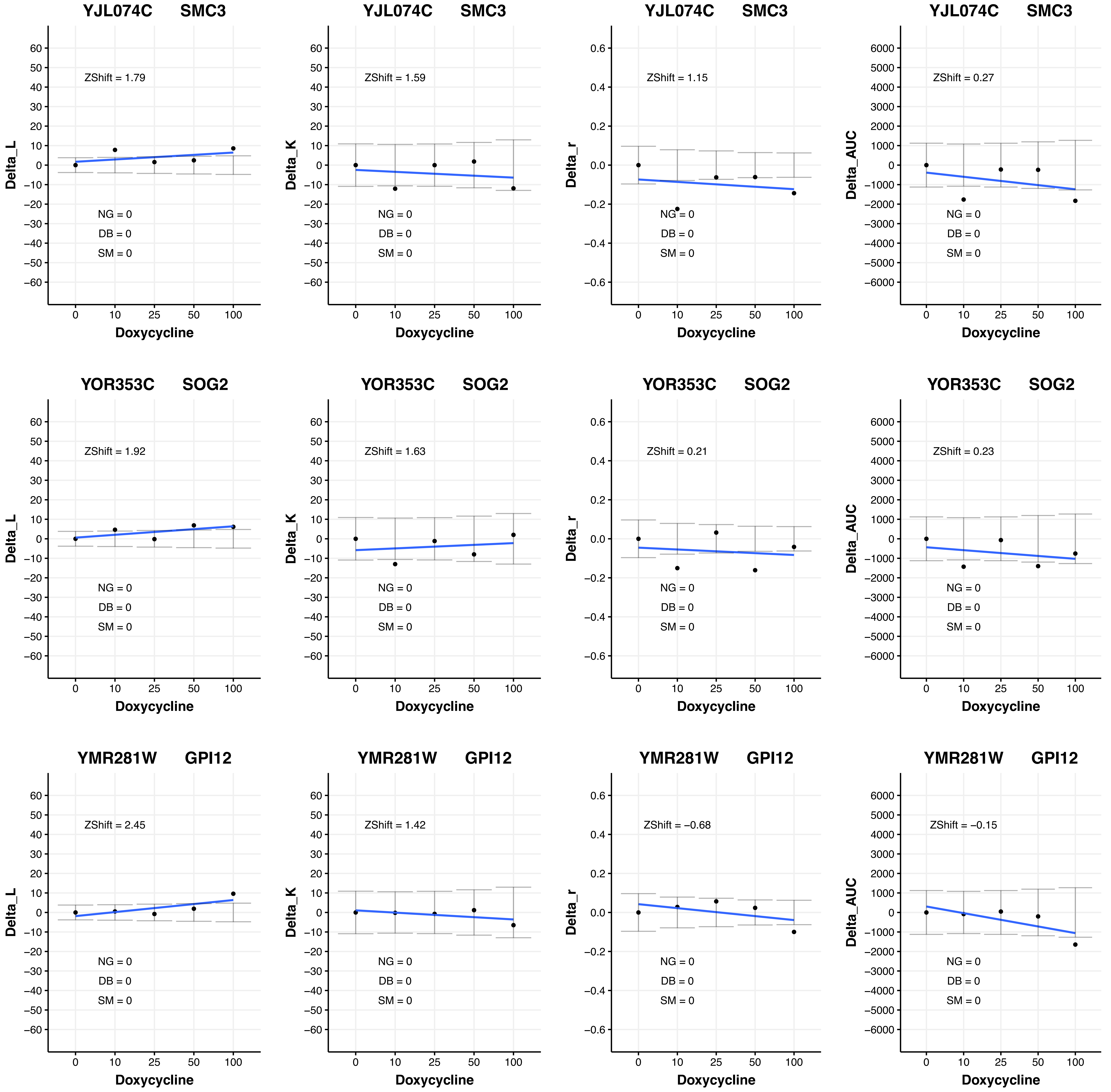

YJL074C SMC3
ZShift = 1.79
NG = 0
DB = 0
SM = 0
Delta_L
Doxycycline
YJL074C SMC3
ZShift = 1.59
NG = 0
DB = 0
SM = 0
Delta_K
Doxycycline
YJL074C SMC3
ZShift = 1.15
NG = 0
DB = 0
SM = 0
Delta_r
Doxycycline
YJL074C SMC3
ZShift = 0.27
NG = 0
DB = 0
SM = 0
Delta_AUC
Doxycycline
YOR353C SOG2
ZShift = 1.92
NG = 0
DB = 0
SM = 0
Delta_L
Doxycycline
YOR353C SOG2
ZShift = 1.63
NG = 0
DB = 0
SM = 0
Delta_K
Doxycycline
YOR353C SOG2
ZShift = 0.21
NG = 0
DB = 0
SM = 0
Delta_r
Doxycycline
YOR353C SOG2
ZShift = 0.23
NG = 0
DB = 0
SM = 0
Delta_AUC
Doxycycline
YMR281W GPI12
ZShift = 2.45
NG = 0
DB = 0
SM = 0
Delta_L
Doxycycline
YMR281W GPI12
ZShift = 1.42
NG = 0
DB = 0
SM = 0
Delta_K
Doxycycline
YMR281W GPI12
ZShift = −0.68
NG = 0
DB = 0
SM = 0
Delta_r
Doxycycline
YMR281W GPI12
ZShift = −0.15
NG = 0
DB = 0
SM = 0
Delta_AUC
Doxycycline

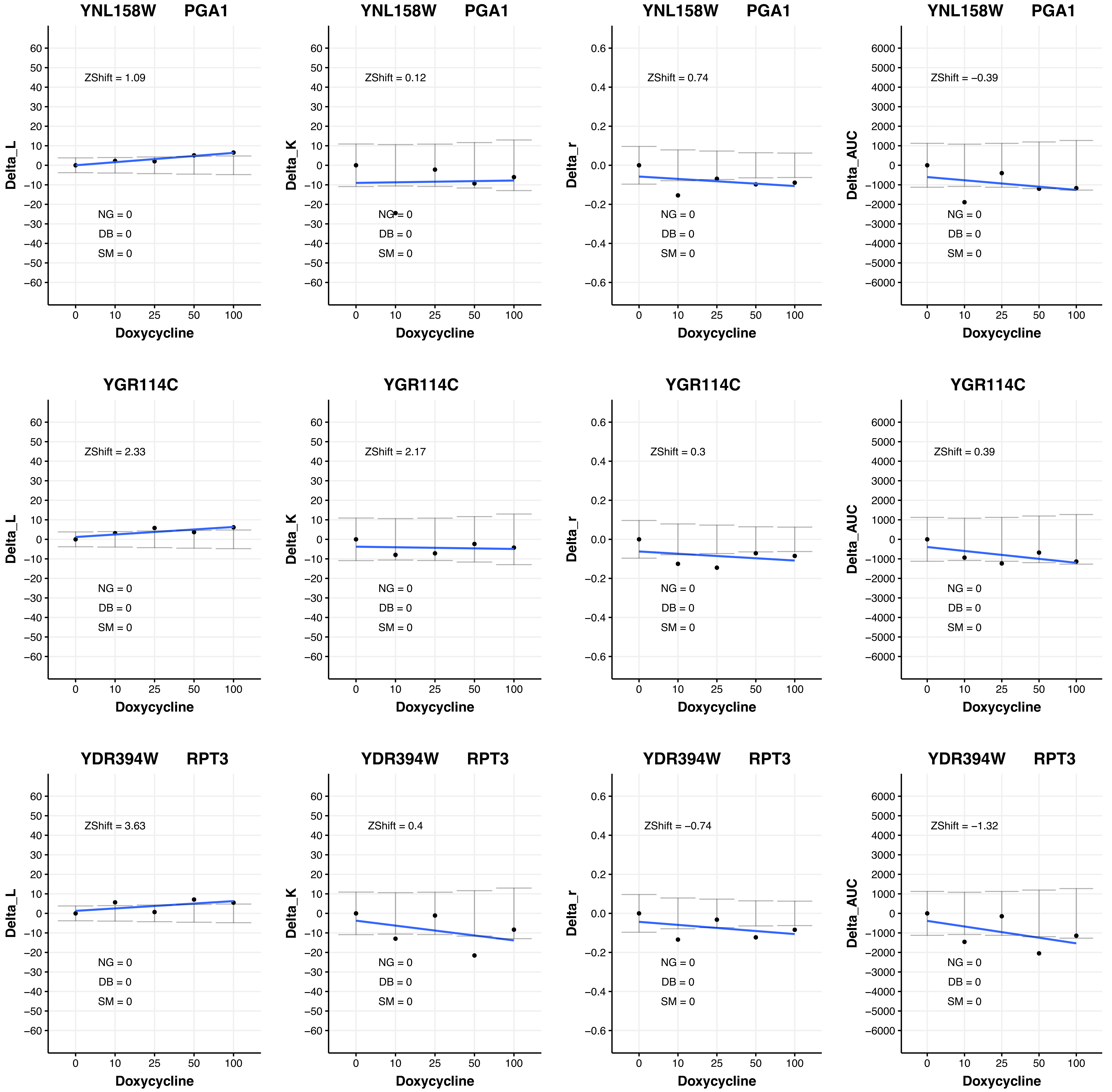

YNL158W PGA1
ZShift = 1.09
ZShift = 0.12
ZShift = 0.74
ZShift = −0.39
YGR114C
ZShift = 2.33
ZShift = 2.17
ZShift = 0.3
ZShift = 0.39
YDR394W RPT3
ZShift = 3.63
ZShift = 0.4
ZShift = −0.74
ZShift = −1.32
NG = 0
DB = 0
SM = 0
Delta_L
Delta_K
Delta_r
Delta_AUC
Doxycycline

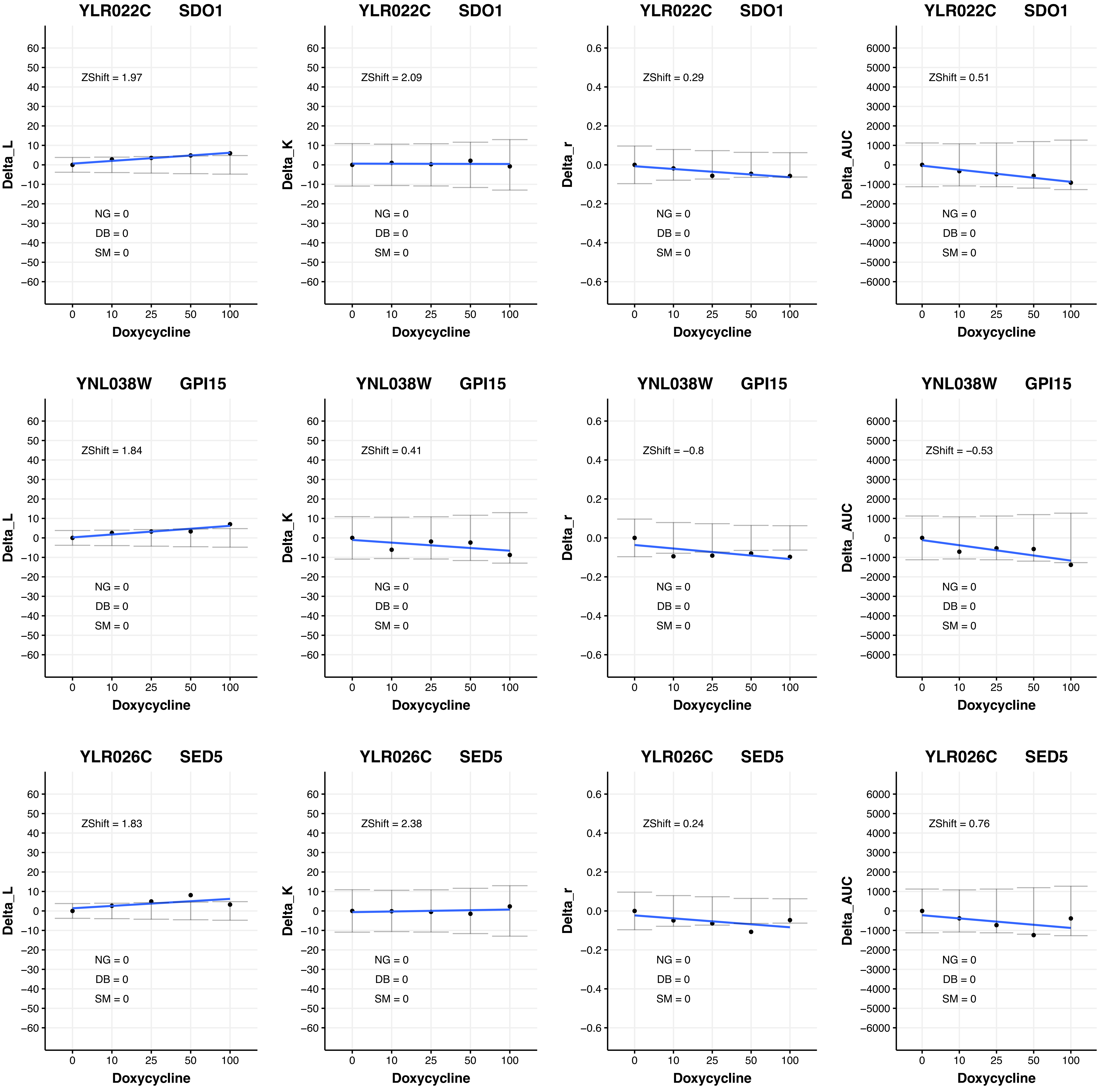
YLR022C SDO1
ZShift = 1.97
NG = 0
DB = 0
SM = 0
Delta_L
Doxycycline
YLR022C SDO1
ZShift = 2.09
NG = 0
DB = 0
SM = 0
Delta_K
Doxycycline
YLR022C SDO1
ZShift = 0.29
NG = 0
DB = 0
SM = 0
Delta_r
Doxycycline
YLR022C SDO1
ZShift = 0.51
NG = 0
DB = 0
SM = 0
Delta_AUC
Doxycycline
YNL038W GPI15
ZShift = 1.84
NG = 0
DB = 0
SM = 0
Delta_L
Doxycycline
YNL038W GPI15
ZShift = 0.41
NG = 0
DB = 0
SM = 0
Delta_K
Doxycycline
YNL038W GPI15
ZShift = −0.8
NG = 0
DB = 0
SM = 0
Delta_r
Doxycycline
YNL038W GPI15
ZShift = −0.53
NG = 0
DB = 0
SM = 0
Delta_AUC
Doxycycline
YLR026C SED5
ZShift = 1.83
NG = 0
DB = 0
SM = 0
Delta_L
Doxycycline
YLR026C SED5
ZShift = 2.38
NG = 0
DB = 0
SM = 0
Delta_K
Doxycycline
YLR026C SED5
ZShift = 0.24
NG = 0
DB = 0
SM = 0
Delta_r
Doxycycline
YLR026C SED5
ZShift = 0.76
NG = 0
DB = 0
SM = 0
Delta_AUC
Doxycycline

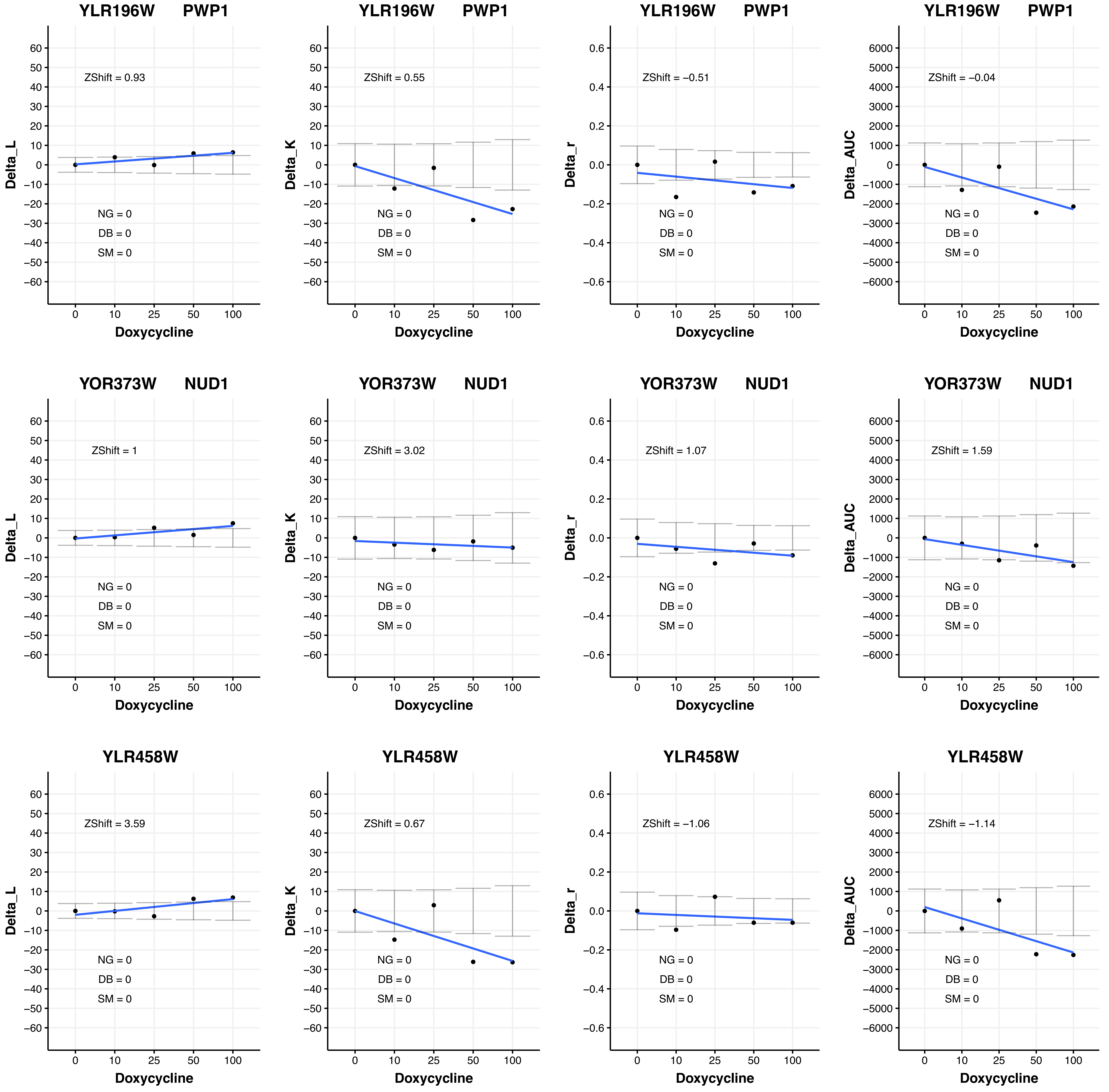
YLR196W PWP1
ZShift = 0.93
Delta_L
NG = 0
DB = 0
SM = 0
Doxycycline
YLR196W PWP1
ZShift = 0.55
Delta_K
NG = 0
DB = 0
SM = 0
Doxycycline
YLR196W PWP1
ZShift = −0.51
Delta_r
NG = 0
DB = 0
SM = 0
Doxycycline
YLR196W PWP1
ZShift = −0.04
Delta_AUC
NG = 0
DB = 0
SM = 0
Doxycycline
YOR373W NUD1
ZShift = 1
Delta_L
NG = 0
DB = 0
SM = 0
Doxycycline
YOR373W NUD1
ZShift = 3.02
Delta_K
NG = 0
DB = 0
SM = 0
Doxycycline
YOR373W NUD1
ZShift = 1.07
Delta_r
NG = 0
DB = 0
SM = 0
Doxycycline
YOR373W NUD1
ZShift = 1.59
Delta_AUC
NG = 0
DB = 0
SM = 0
Doxycycline
YLR458W
ZShift = 3.59
Delta_L
NG = 0
DB = 0
SM = 0
Doxycycline
YLR458W
ZShift = 0.67
Delta_K
NG = 0
DB = 0
SM = 0
Doxycycline
YLR458W
ZShift = −1.06
Delta_r
NG = 0
DB = 0
SM = 0
Doxycycline
YLR458W
ZShift = −1.14
Delta_AUC
NG = 0
DB = 0
SM = 0
Doxycycline

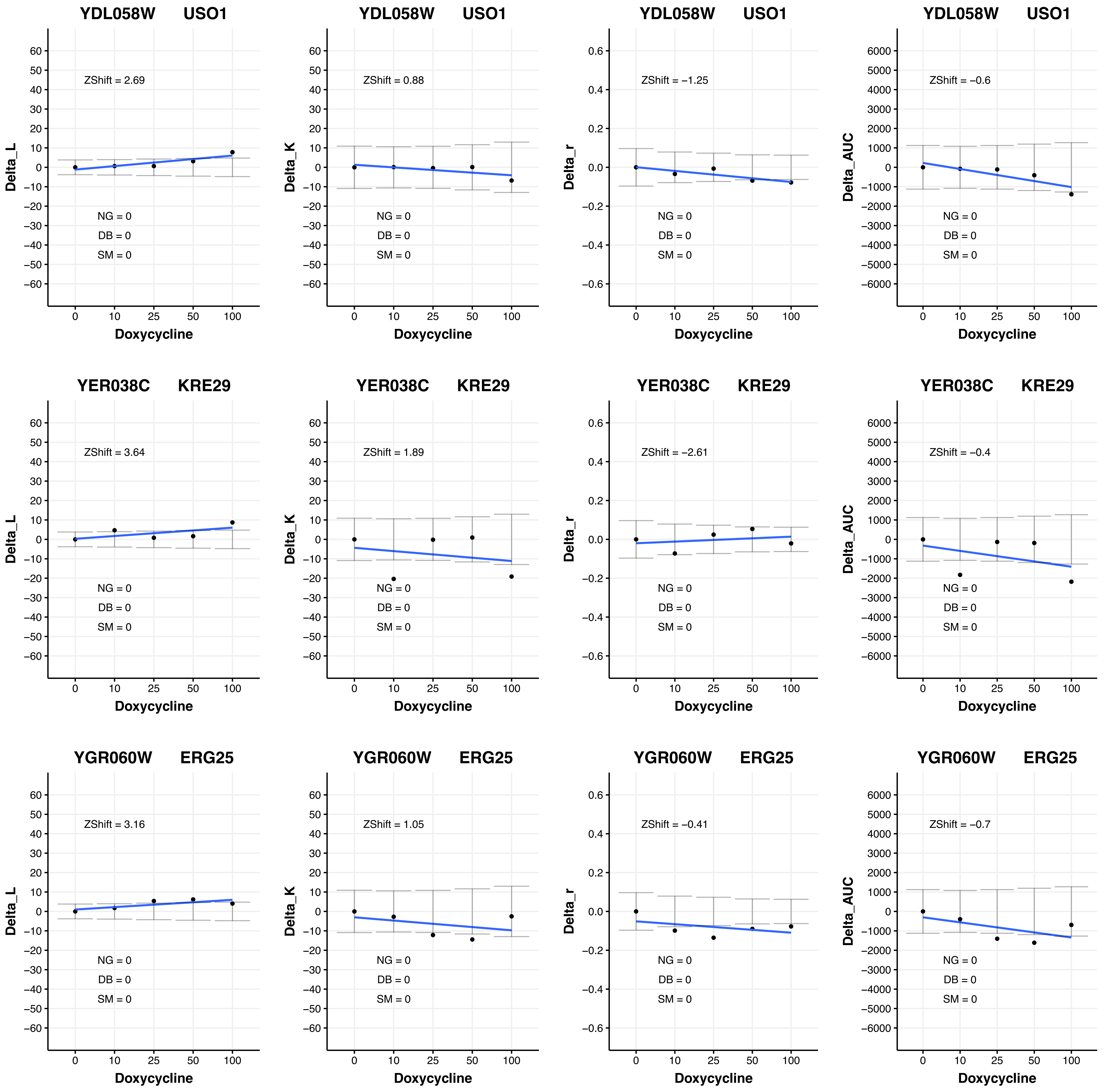
YDL058W USO1
ZShift = 2.69
NG = 0
DB = 0
SM = 0
Delta_L
Doxycycline
YDL058W USO1
ZShift = 0.88
NG = 0
DB = 0
SM = 0
Delta_K
Doxycycline
YDL058W USO1
ZShift = −1.25
NG = 0
DB = 0
SM = 0
Delta_r
Doxycycline
YDL058W USO1
ZShift = −0.6
NG = 0
DB = 0
SM = 0
Delta_AUC
Doxycycline
YER038C KRE29
ZShift = 3.64
NG = 0
DB = 0
SM = 0
Delta_L
Doxycycline
YER038C KRE29
ZShift = 1.89
NG = 0
DB = 0
SM = 0
Delta_K
Doxycycline
YER038C KRE29
ZShift = −2.61
NG = 0
DB = 0
SM = 0
Delta_r
Doxycycline
YER038C KRE29
ZShift = −0.4
NG = 0
DB = 0
SM = 0
Delta_AUC
Doxycycline
YGR060W ERG25
ZShift = 3.16
NG = 0
DB = 0
SM = 0
Delta_L
Doxycycline
YGR060W ERG25
ZShift = 1.05
NG = 0
DB = 0
SM = 0
Delta_K
Doxycycline
YGR060W ERG25
ZShift = −0.41
NG = 0
DB = 0
SM = 0
Delta_r
Doxycycline
YGR060W ERG25
ZShift = −0.7
NG = 0
DB = 0
SM = 0
Delta_AUC
Doxycycline

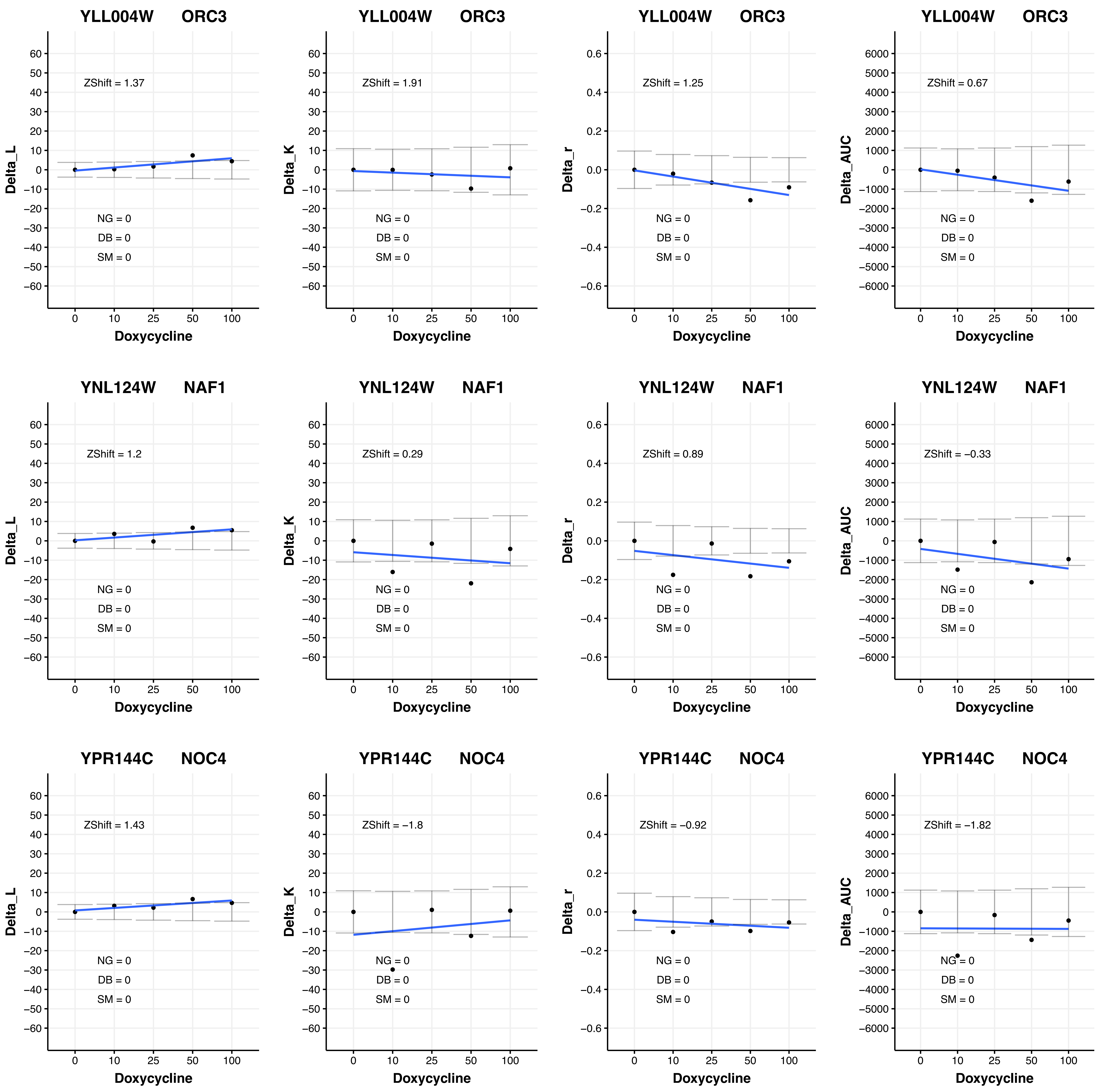

YLL004W ORC3
ZShift = 1.37
ZShift = 1.91
ZShift = 1.25
ZShift = 0.67
YNL124W NAF1
ZShift = 1.2
ZShift = 0.29
ZShift = 0.89
ZShift = −0.33
YPR144C NOC4
ZShift = 1.43
ZShift = −1.8
ZShift = −0.92
ZShift = −1.82
NG = 0
DB = 0
SM = 0
Delta_L
Delta_K
Delta_r
Delta_AUC
Doxycycline

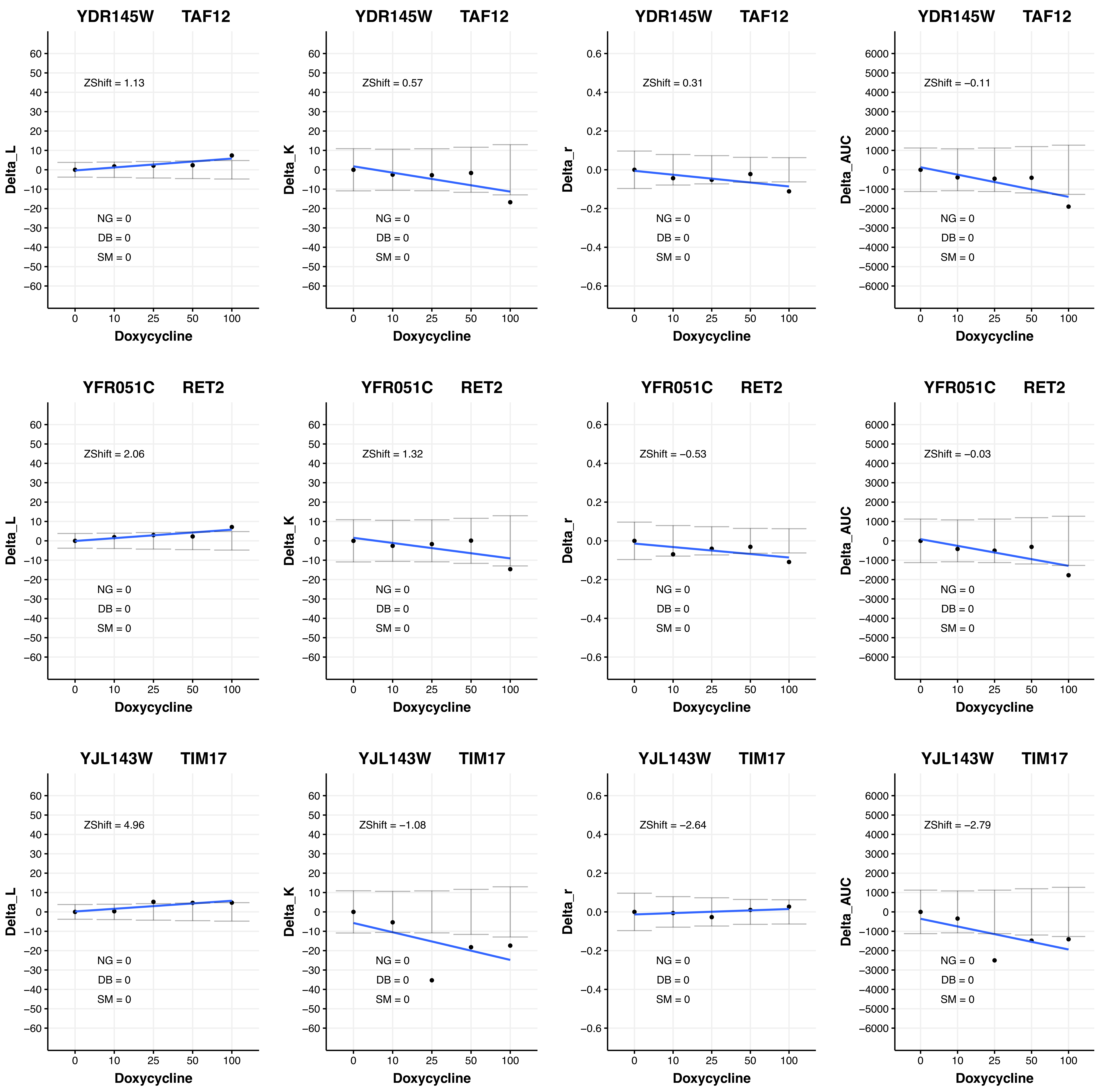

YDR145W TAF12
ZShift = 1.13
NG = 0
DB = 0
SM = 0
Delta_L
Doxycycline
YDR145W TAF12
ZShift = 0.57
NG = 0
DB = 0
SM = 0
Delta_K
Doxycycline
YDR145W TAF12
ZShift = 0.31
NG = 0
DB = 0
SM = 0
Delta_r
Doxycycline
YDR145W TAF12
ZShift = −0.11
NG = 0
DB = 0
SM = 0
Delta_AUC
Doxycycline
YFR051C RET2
ZShift = 2.06
NG = 0
DB = 0
SM = 0
Delta_L
Doxycycline
YFR051C RET2
ZShift = 1.32
NG = 0
DB = 0
SM = 0
Delta_K
Doxycycline
YFR051C RET2
ZShift = −0.53
NG = 0
DB = 0
SM = 0
Delta_r
Doxycycline
YFR051C RET2
ZShift = −0.03
NG = 0
DB = 0
SM = 0
Delta_AUC
Doxycycline
YJL143W TIM17
ZShift = 4.96
NG = 0
DB = 0
SM = 0
Delta_L
Doxycycline
YJL143W TIM17
ZShift = −1.08
NG = 0
DB = 0
SM = 0
Delta_K
Doxycycline
YJL143W TIM17
ZShift = −2.64
NG = 0
DB = 0
SM = 0
Delta_r
Doxycycline
YJL143W TIM17
ZShift = −2.79
NG = 0
DB = 0
SM = 0
Delta_AUC
Doxycycline

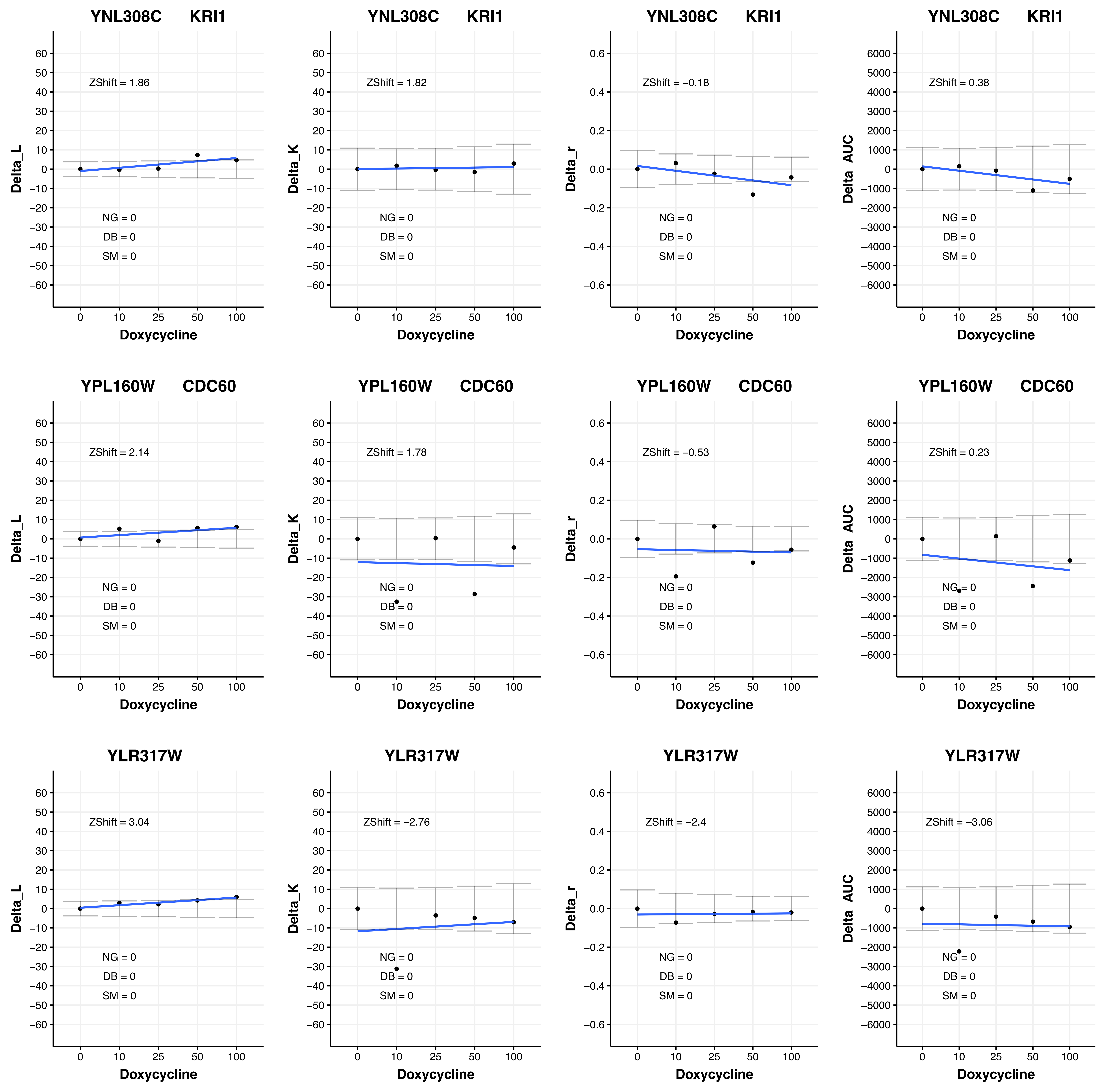
YNL308C KRI1
ZShift = 1.86
NG = 0
DB = 0
SM = 0
Delta_L
Doxycycline
YNL308C KRI1
ZShift = 1.82
NG = 0
DB = 0
SM = 0
Delta_K
Doxycycline
YNL308C KRI1
ZShift = −0.18
NG = 0
DB = 0
SM = 0
Delta_r
Doxycycline
YNL308C KRI1
ZShift = 0.38
NG = 0
DB = 0
SM = 0
Delta_AUC
Doxycycline
YPL160W CDC60
ZShift = 2.14
NG = 0
DB = 0
SM = 0
Delta_L
Doxycycline
YPL160W CDC60
ZShift = 1.78
NG = 0
DB = 0
SM = 0
Delta_K
Doxycycline
YPL160W CDC60
ZShift = −0.53
NG = 0
DB = 0
SM = 0
Delta_r
Doxycycline
YPL160W CDC60
ZShift = 0.23
NG = 0
DB = 0
SM = 0
Delta_AUC
Doxycycline
YLR317W
ZShift = 3.04
NG = 0
DB = 0
SM = 0
Delta_L
Doxycycline
YLR317W
ZShift = −2.76
NG = 0
DB = 0
SM = 0
Delta_K
Doxycycline
YLR317W
ZShift = −2.4
NG = 0
DB = 0
SM = 0
Delta_r
Doxycycline
YLR317W
ZShift = −3.06
NG = 0
DB = 0
SM = 0
Delta_AUC
Doxycycline

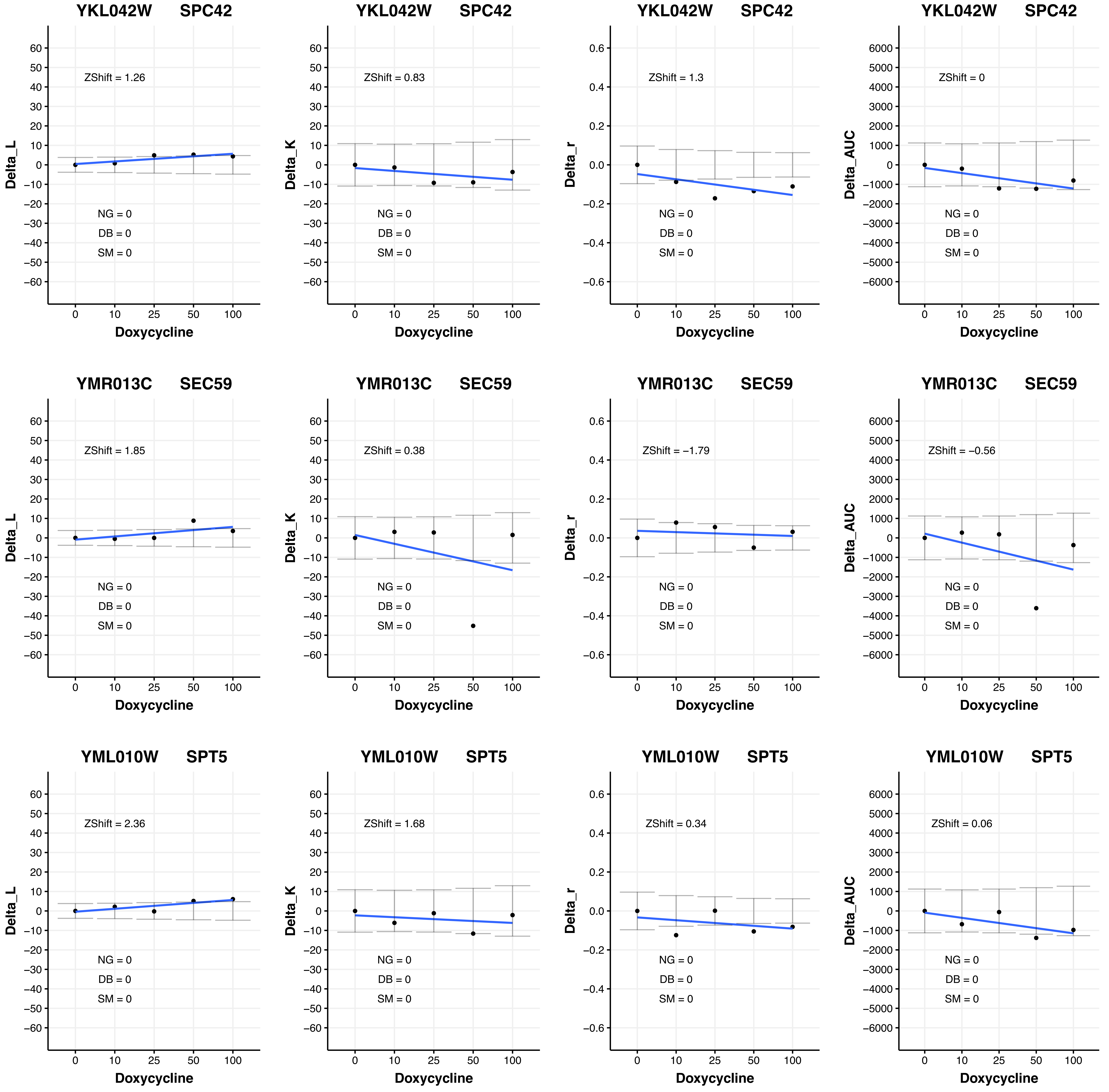
YKL042W SPC42
ZShift = 1.26
NG = 0
DB = 0
SM = 0
Delta_L
Doxycycline
YKL042W SPC42
ZShift = 0.83
NG = 0
DB = 0
SM = 0
Delta_K
Doxycycline
YKL042W SPC42
ZShift = 1.3
NG = 0
DB = 0
SM = 0
Delta_r
Doxycycline
YKL042W SPC42
ZShift = 0
NG = 0
DB = 0
SM = 0
Delta_AUC
Doxycycline
YMR013C SEC59
ZShift = 1.85
NG = 0
DB = 0
SM = 0
Delta_L
Doxycycline
YMR013C SEC59
ZShift = 0.38
NG = 0
DB = 0
SM = 0
Delta_K
Doxycycline
YMR013C SEC59
ZShift = −1.79
NG = 0
DB = 0
SM = 0
Delta_r
Doxycycline
YMR013C SEC59
ZShift = −0.56
NG = 0
DB = 0
SM = 0
Delta_AUC
Doxycycline
YML010W SPT5
ZShift = 2.36
NG = 0
DB = 0
SM = 0
Delta_L
Doxycycline
YML010W SPT5
ZShift = 1.68
NG = 0
DB = 0
SM = 0
Delta_K
Doxycycline
YML010W SPT5
ZShift = 0.34
NG = 0
DB = 0
SM = 0
Delta_r
Doxycycline
YML010W SPT5
ZShift = 0.06
NG = 0
DB = 0
SM = 0
Delta_AUC
Doxycycline

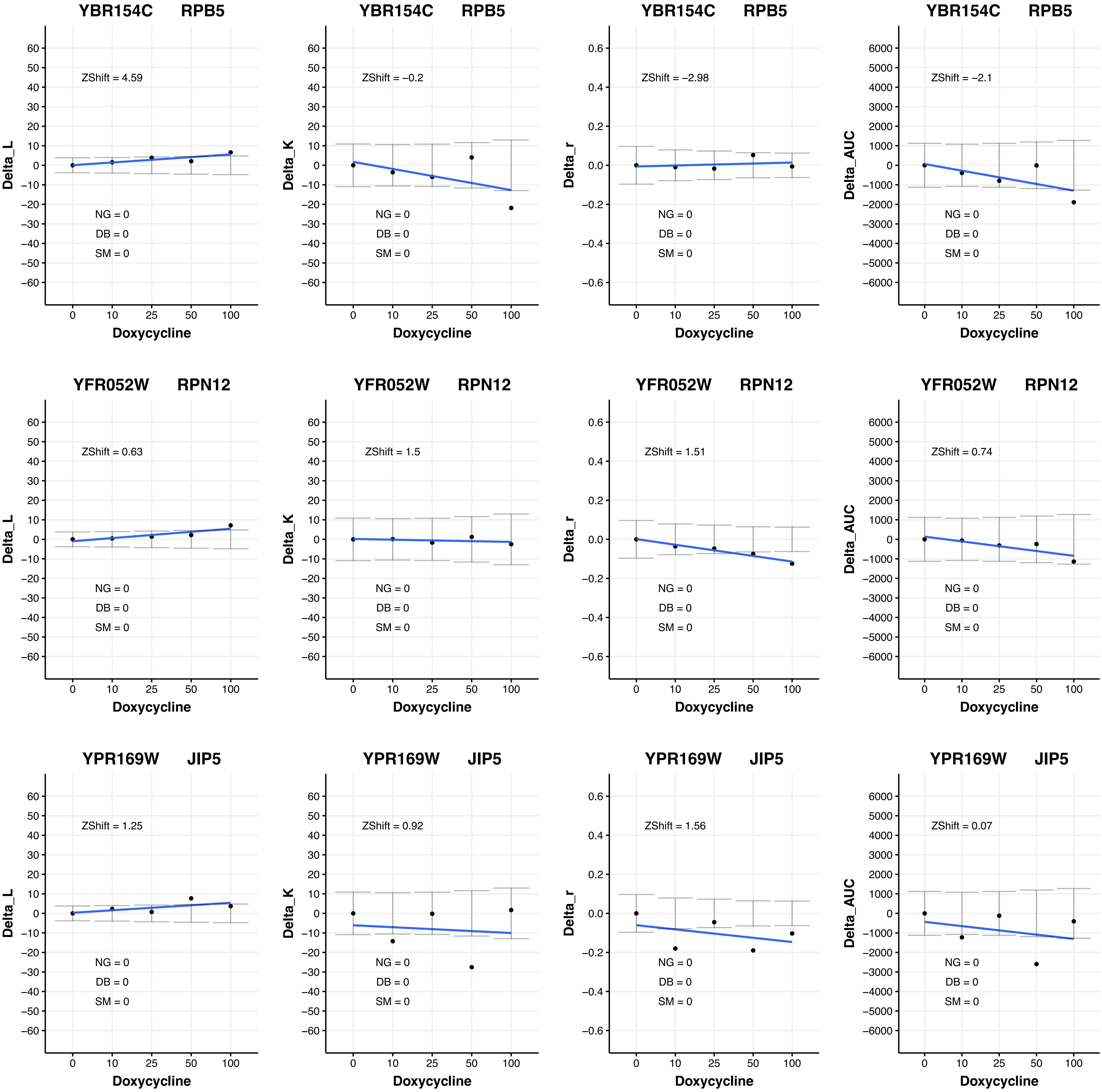

YBR154C RPB5
ZShift = 4.59
NG = 0
DB = 0
SM = 0
Delta_L
Doxycycline
YBR154C RPB5
ZShift = −0.2
NG = 0
DB = 0
SM = 0
Delta_K
Doxycycline
YBR154C RPB5
ZShift = −2.98
NG = 0
DB = 0
SM = 0
Delta_r
Doxycycline
YBR154C RPB5
ZShift = −2.1
NG = 0
DB = 0
SM = 0
Delta_AUC
Doxycycline
YFR052W RPN12
ZShift = 0.63
NG = 0
DB = 0
SM = 0
Delta_L
Doxycycline
YFR052W RPN12
ZShift = 1.5
NG = 0
DB = 0
SM = 0
Delta_K
Doxycycline
YFR052W RPN12
ZShift = 1.51
NG = 0
DB = 0
SM = 0
Delta_r
Doxycycline
YFR052W RPN12
ZShift = 0.74
NG = 0
DB = 0
SM = 0
Delta_AUC
Doxycycline
YPR169W JIP5
ZShift = 1.25
NG = 0
DB = 0
SM = 0
Delta_L
Doxycycline
YPR169W JIP5
ZShift = 0.92
NG = 0
DB = 0
SM = 0
Delta_K
Doxycycline
YPR169W JIP5
ZShift = 1.56
NG = 0
DB = 0
SM = 0
Delta_r
Doxycycline
YPR169W JIP5
ZShift = 0.07
NG = 0
DB = 0
SM = 0
Delta_AUC
Doxycycline

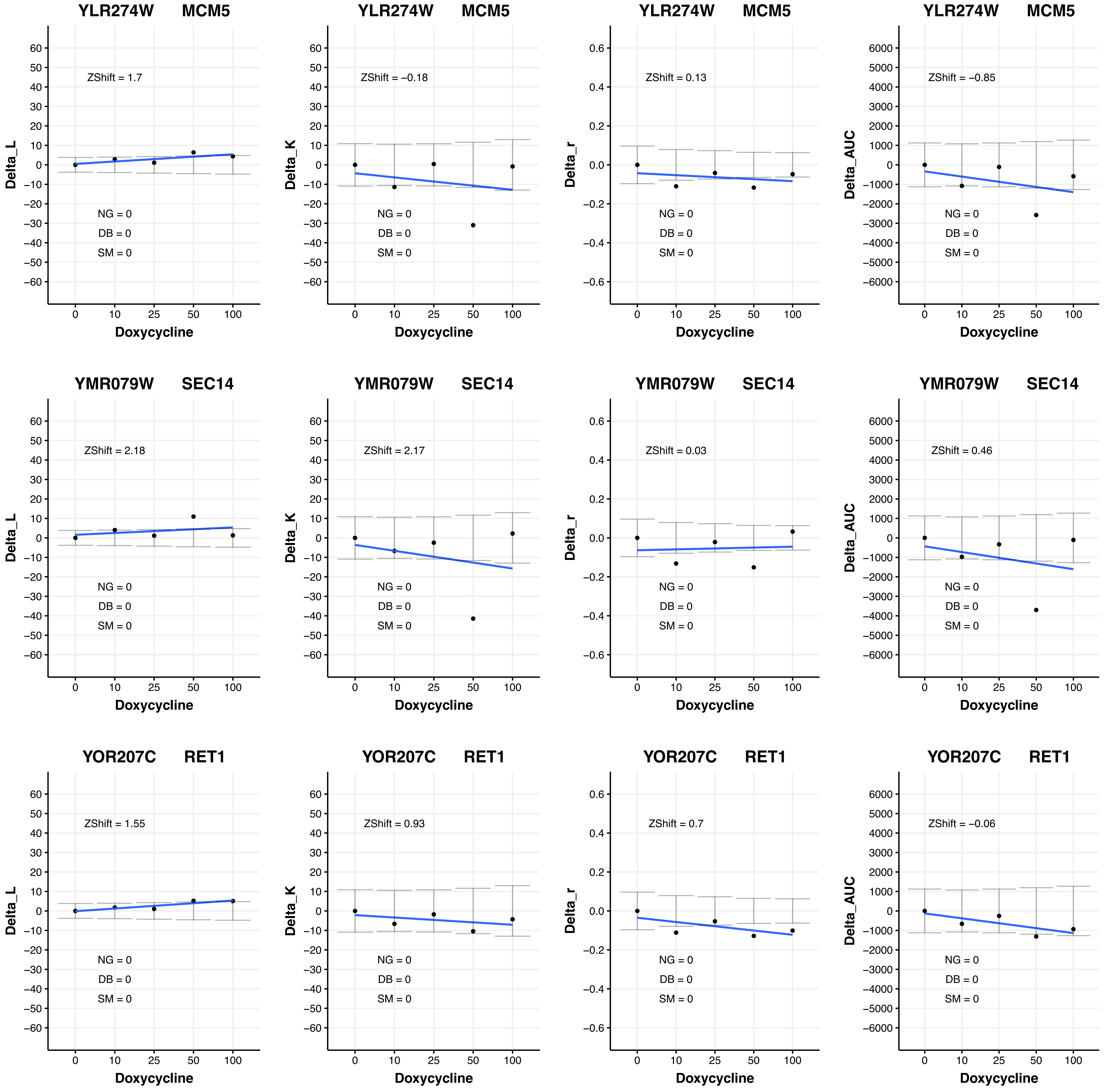

YLR274W MCM5
ZShift = 1.7
Delta_L
NG = 0
DB = 0
SM = 0
Doxycycline
YLR274W MCM5
ZShift = −0.18
Delta_K
NG = 0
DB = 0
SM = 0
Doxycycline
YLR274W MCM5
ZShift = 0.13
Delta_r
NG = 0
DB = 0
SM = 0
Doxycycline
YLR274W MCM5
ZShift = −0.85
Delta_AUC
NG = 0
DB = 0
SM = 0
Doxycycline
YMR079W SEC14
ZShift = 2.18
Delta_L
NG = 0
DB = 0
SM = 0
Doxycycline
YMR079W SEC14
ZShift = 2.17
Delta_K
NG = 0
DB = 0
SM = 0
Doxycycline
YMR079W SEC14
ZShift = 0.03
Delta_r
NG = 0
DB = 0
SM = 0
Doxycycline
YMR079W SEC14
ZShift = 0.46
Delta_AUC
NG = 0
DB = 0
SM = 0
Doxycycline
YOR207C RET1
ZShift = 1.55
Delta_L
NG = 0
DB = 0
SM = 0
Doxycycline
YOR207C RET1
ZShift = 0.93
Delta_K
NG = 0
DB = 0
SM = 0
Doxycycline
YOR207C RET1
ZShift = 0.7
Delta_r
NG = 0
DB = 0
SM = 0
Doxycycline
YOR207C RET1
ZShift = −0.06
Delta_AUC
NG = 0
DB = 0
SM = 0
Doxycycline
0 10 25 50 100

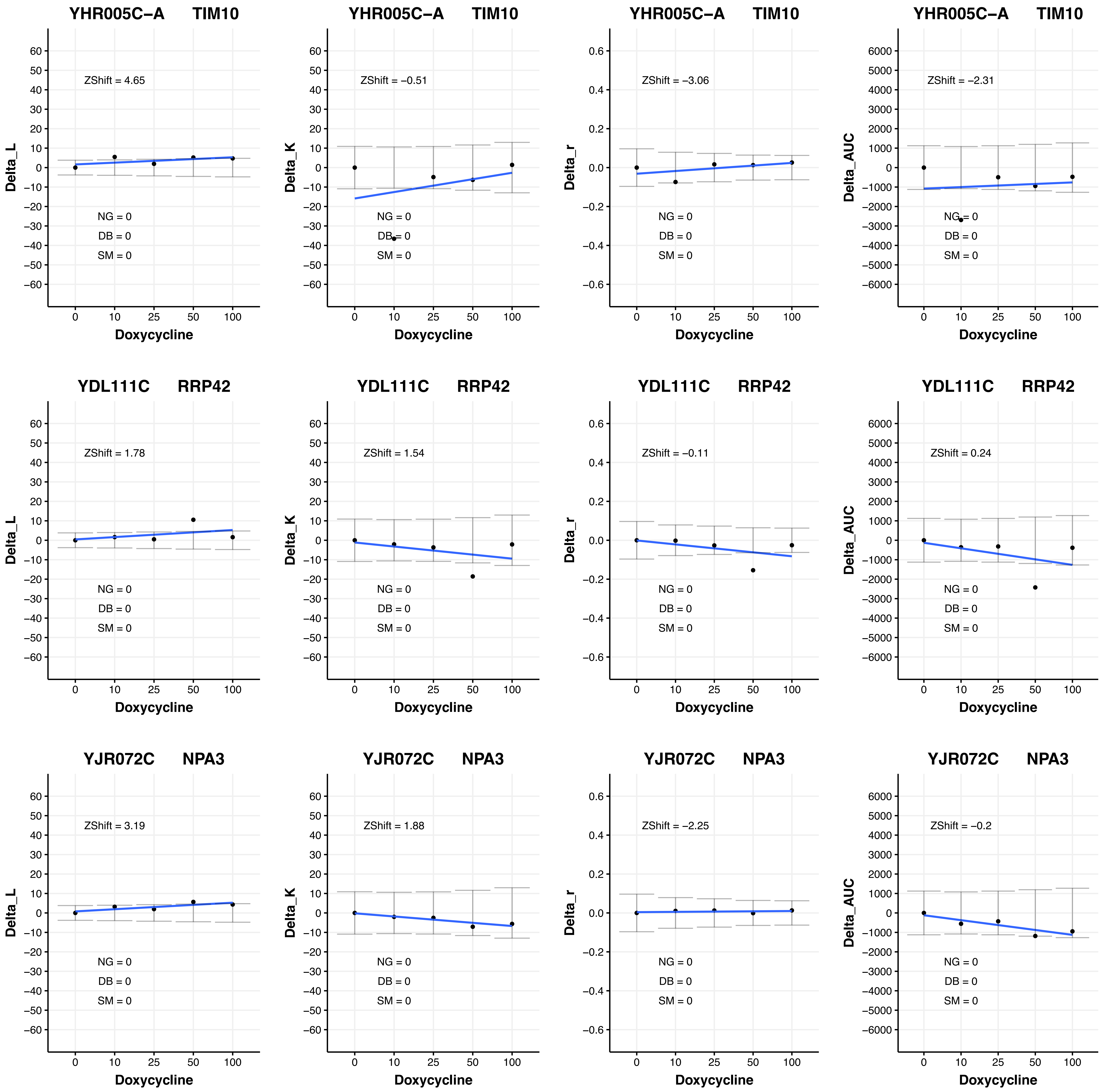

YHR005C−A TIM10
ZShift = 4.65
ZShift = −0.51
ZShift = −3.06
ZShift = −2.31
YDL111C RRP42
ZShift = 1.78
ZShift = 1.54
ZShift = −0.11
ZShift = 0.24
YJR072C NPA3
ZShift = 3.19
ZShift = 1.88
ZShift = −2.25
ZShift = −0.2
NG = 0
DB = 0
SM = 0
Delta_L
Delta_K
Delta_r
Delta_AUC
Doxycycline
0 10 25 50 100

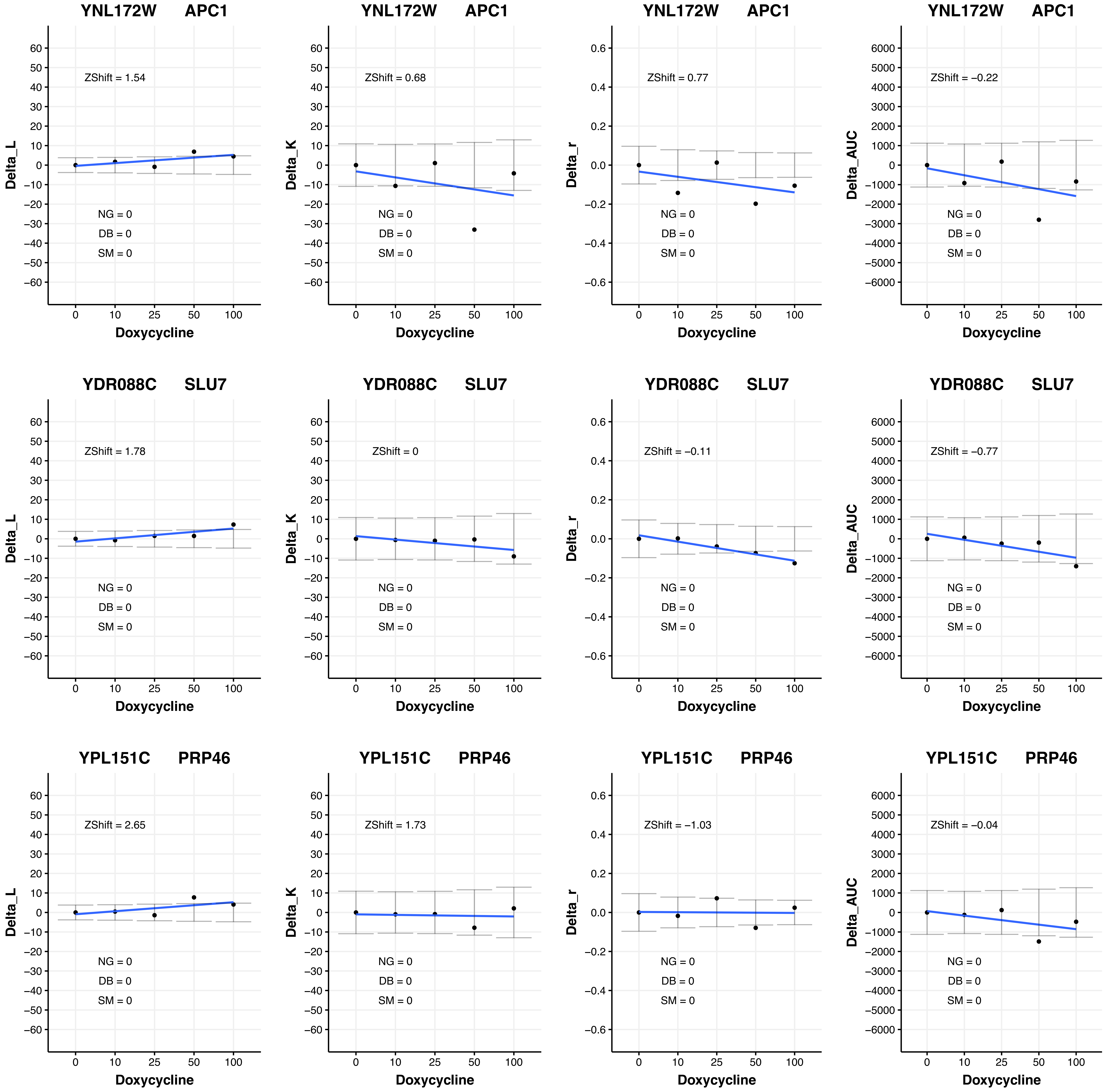
YNL172W APC1
ZShift = 1.54
NG = 0
DB = 0
SM = 0
Delta_L
Doxycycline
YNL172W APC1
ZShift = 0.68
NG = 0
DB = 0
SM = 0
Delta_K
Doxycycline
YNL172W APC1
ZShift = 0.77
NG = 0
DB = 0
SM = 0
Delta_r
Doxycycline
YNL172W APC1
ZShift = −0.22
NG = 0
DB = 0
SM = 0
Delta_AUC
Doxycycline
YDR088C SLU7
ZShift = 1.78
NG = 0
DB = 0
SM = 0
Delta_L
Doxycycline
YDR088C SLU7
ZShift = 0
NG = 0
DB = 0
SM = 0
Delta_K
Doxycycline
YDR088C SLU7
ZShift = −0.11
NG = 0
DB = 0
SM = 0
Delta_r
Doxycycline
YDR088C SLU7
ZShift = −0.77
NG = 0
DB = 0
SM = 0
Delta_AUC
Doxycycline
YPL151C PRP46
ZShift = 2.65
NG = 0
DB = 0
SM = 0
Delta_L
Doxycycline
YPL151C PRP46
ZShift = 1.73
NG = 0
DB = 0
SM = 0
Delta_K
Doxycycline
YPL151C PRP46
ZShift = −1.03
NG = 0
DB = 0
SM = 0
Delta_r
Doxycycline
YPL151C PRP46
ZShift = −0.04
NG = 0
DB = 0
SM = 0
Delta_AUC
Doxycycline

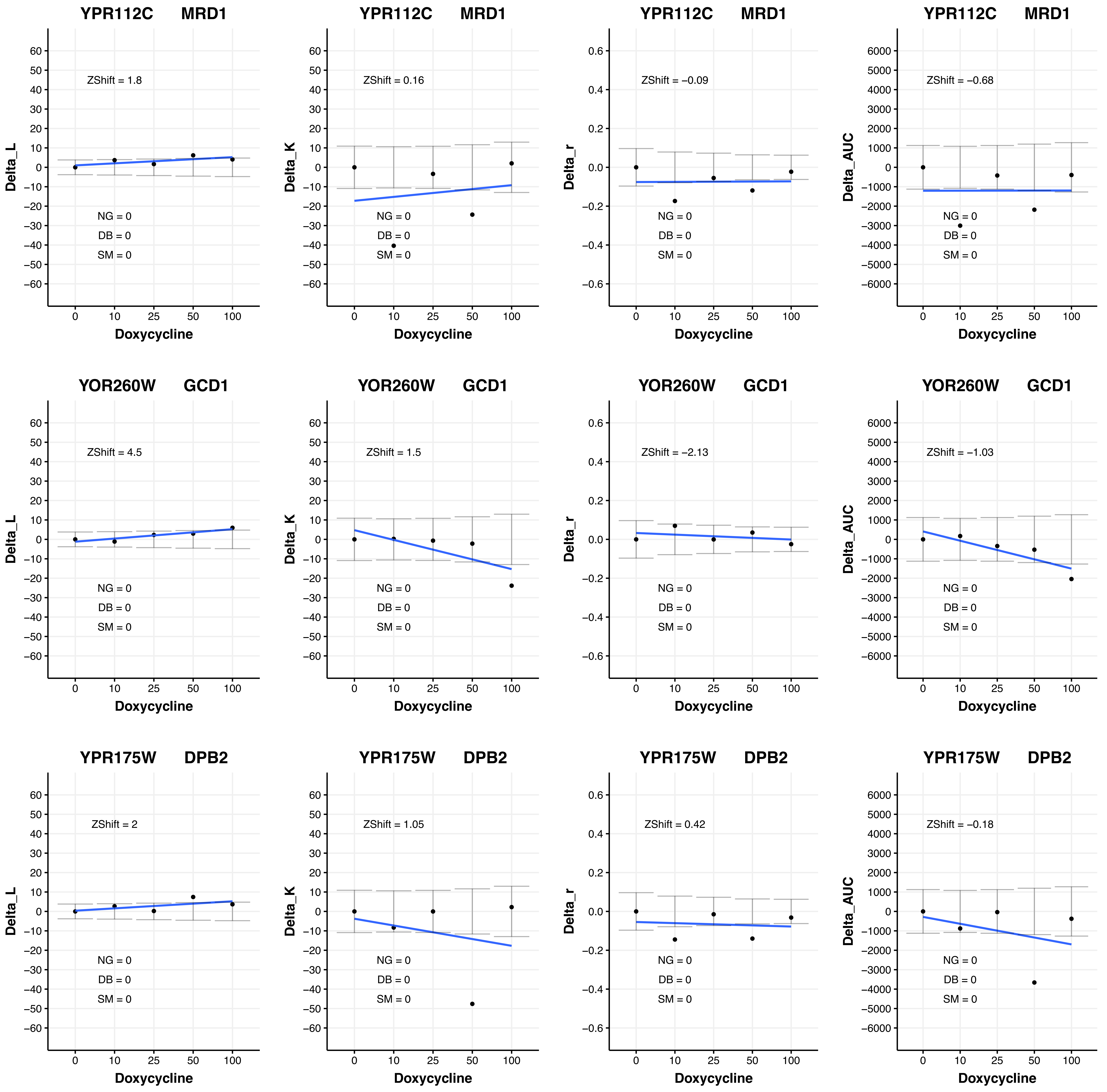
YPR112C MRD1
ZShift = 1.8
NG = 0
DB = 0
SM = 0
Delta_L
Doxycycline
YPR112C MRD1
ZShift = 0.16
NG = 0
DB = 0
SM = 0
Delta_K
Doxycycline
YPR112C MRD1
ZShift = −0.09
NG = 0
DB = 0
SM = 0
Delta_r
Doxycycline
YPR112C MRD1
ZShift = −0.68
NG = 0
DB = 0
SM = 0
Delta_AUC
Doxycycline
YOR260W GCD1
ZShift = 4.5
NG = 0
DB = 0
SM = 0
Delta_L
Doxycycline
YOR260W GCD1
ZShift = 1.5
NG = 0
DB = 0
SM = 0
Delta_K
Doxycycline
YOR260W GCD1
ZShift = −2.13
NG = 0
DB = 0
SM = 0
Delta_r
Doxycycline
YOR260W GCD1
ZShift = −1.03
NG = 0
DB = 0
SM = 0
Delta_AUC
Doxycycline
YPR175W DPB2
ZShift = 2
NG = 0
DB = 0
SM = 0
Delta_L
Doxycycline
YPR175W DPB2
ZShift = 1.05
NG = 0
DB = 0
SM = 0
Delta_K
Doxycycline
YPR175W DPB2
ZShift = 0.42
NG = 0
DB = 0
SM = 0
Delta_r
Doxycycline
YPR175W DPB2
ZShift = −0.18
NG = 0
DB = 0
SM = 0
Delta_AUC
Doxycycline

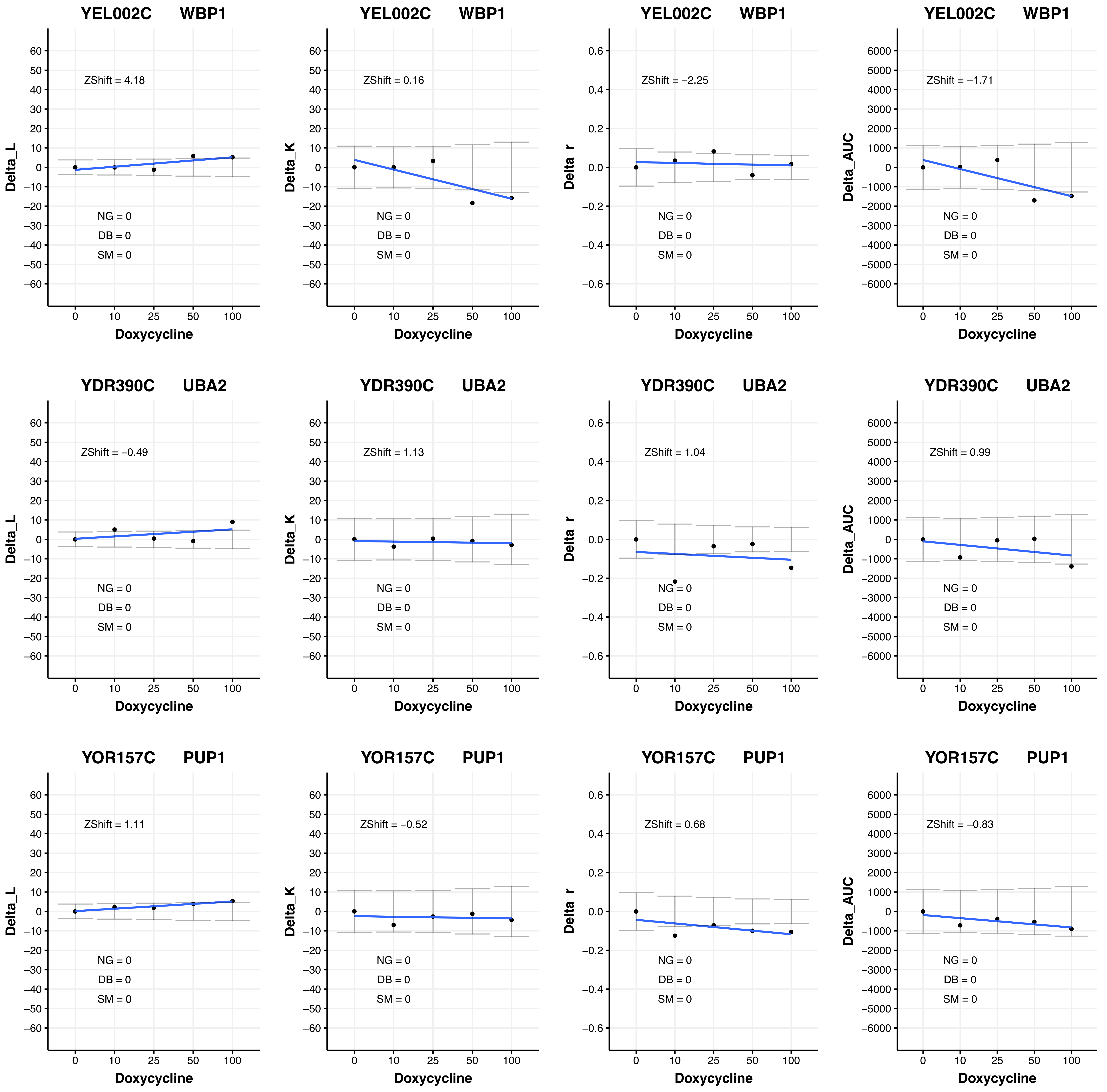
YEL002C WBP1
ZShift = 4.18
ZShift = 0.16
ZShift = −2.25
ZShift = −1.71
YDR390C UBA2
ZShift = −0.49
ZShift = 1.13
ZShift = 1.04
ZShift = 0.99
YOR157C PUP1
ZShift = 1.11
ZShift = −0.52
ZShift = 0.68
ZShift = −0.83
NG = 0
DB = 0
SM = 0
Delta_L
Delta_K
Delta_r
Delta_AUC
Doxycycline
0
10
25
50
100

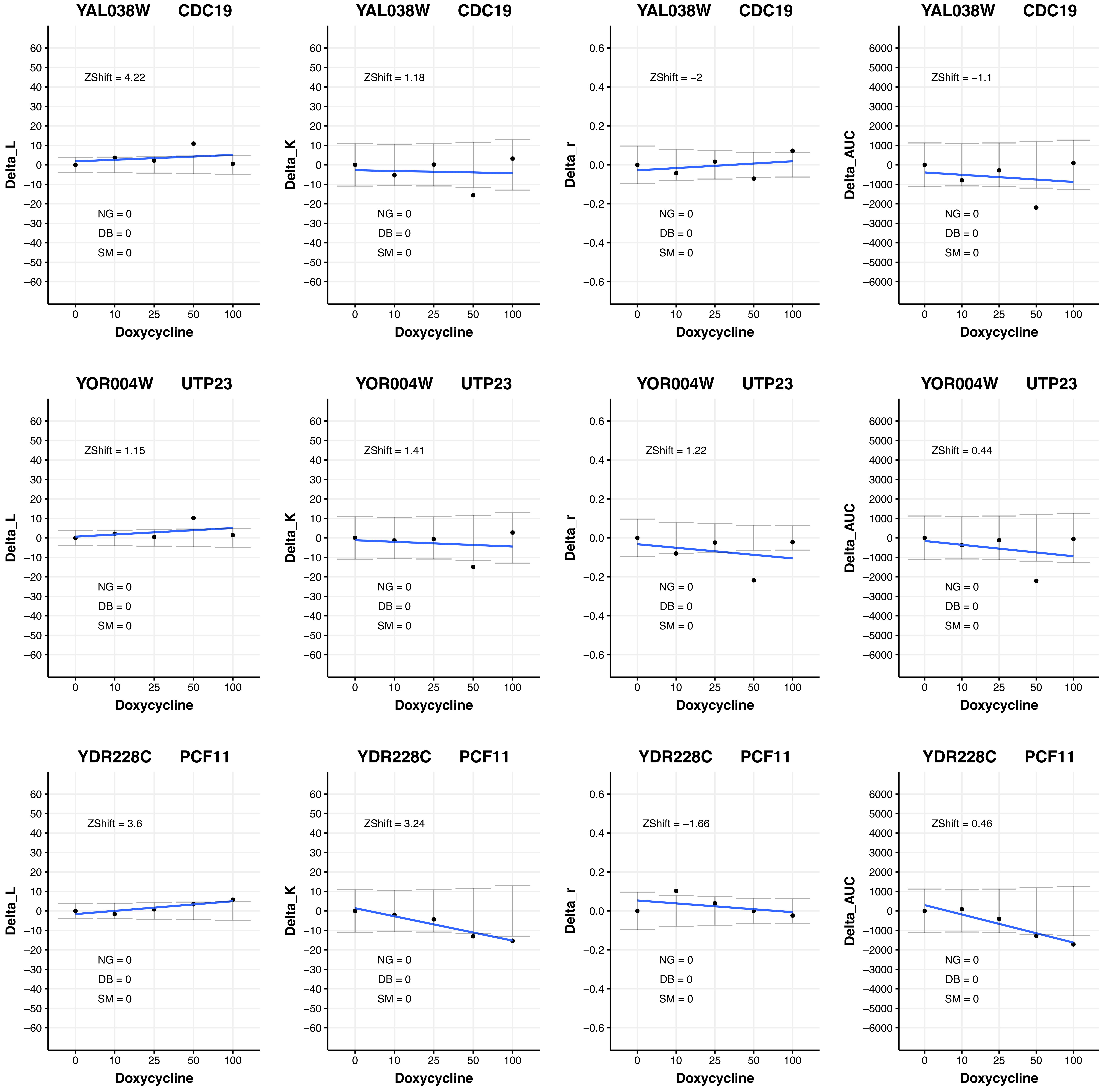

YAL038W CDC19
ZShift = 4.22
Delta_L
NG = 0
DB = 0
SM = 0
Doxycycline
YAL038W CDC19
ZShift = 1.18
Delta_K
NG = 0
DB = 0
SM = 0
Doxycycline
YAL038W CDC19
ZShift = −2
Delta_r
NG = 0
DB = 0
SM = 0
Doxycycline
YAL038W CDC19
ZShift = −1.1
Delta_AUC
NG = 0
DB = 0
SM = 0
Doxycycline
YOR004W UTP23
ZShift = 1.15
Delta_L
NG = 0
DB = 0
SM = 0
Doxycycline
YOR004W UTP23
ZShift = 1.41
Delta_K
NG = 0
DB = 0
SM = 0
Doxycycline
YOR004W UTP23
ZShift = 1.22
Delta_r
NG = 0
DB = 0
SM = 0
Doxycycline
YOR004W UTP23
ZShift = 0.44
Delta_AUC
NG = 0
DB = 0
SM = 0
Doxycycline
YDR228C PCF11
ZShift = 3.6
Delta_L
NG = 0
DB = 0
SM = 0
Doxycycline
YDR228C PCF11
ZShift = 3.24
Delta_K
NG = 0
DB = 0
SM = 0
Doxycycline
YDR228C PCF11
ZShift = −1.66
Delta_r
NG = 0
DB = 0
SM = 0
Doxycycline
YDR228C PCF11
ZShift = 0.46
Delta_AUC
NG = 0
DB = 0
SM = 0
Doxycycline

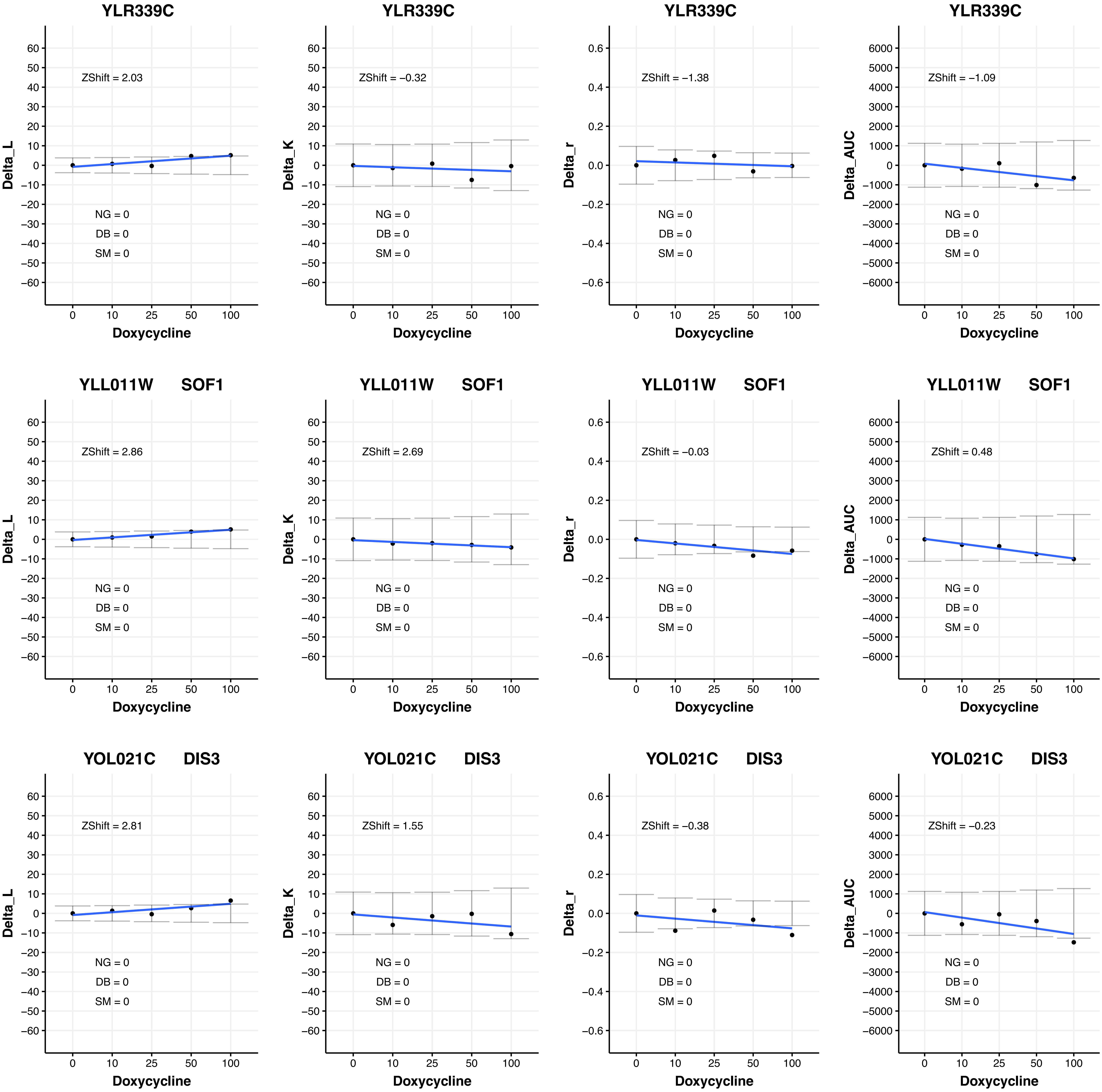

YLR339C
ZShift = 2.03
NG = 0
DB = 0
SM = 0
Delta_L
Doxycycline
YLR339C
ZShift = −0.32
NG = 0
DB = 0
SM = 0
Delta_K
Doxycycline
YLR339C
ZShift = −1.38
NG = 0
DB = 0
SM = 0
Delta_r
Doxycycline
YLR339C
ZShift = −1.09
NG = 0
DB = 0
SM = 0
Delta_AUC
Doxycycline
YLL011W SOF1
ZShift = 2.86
NG = 0
DB = 0
SM = 0
Delta_L
Doxycycline
YLL011W SOF1
ZShift = 2.69
NG = 0
DB = 0
SM = 0
Delta_K
Doxycycline
YLL011W SOF1
ZShift = −0.03
NG = 0
DB = 0
SM = 0
Delta_r
Doxycycline
YLL011W SOF1
ZShift = 0.48
NG = 0
DB = 0
SM = 0
Delta_AUC
Doxycycline
YOL021C DIS3
ZShift = 2.81
NG = 0
DB = 0
SM = 0
Delta_L
Doxycycline
YOL021C DIS3
ZShift = 1.55
NG = 0
DB = 0
SM = 0
Delta_K
Doxycycline
YOL021C DIS3
ZShift = −0.38
NG = 0
DB = 0
SM = 0
Delta_r
Doxycycline
YOL021C DIS3
ZShift = −0.23
NG = 0
DB = 0
SM = 0
Delta_AUC
Doxycycline

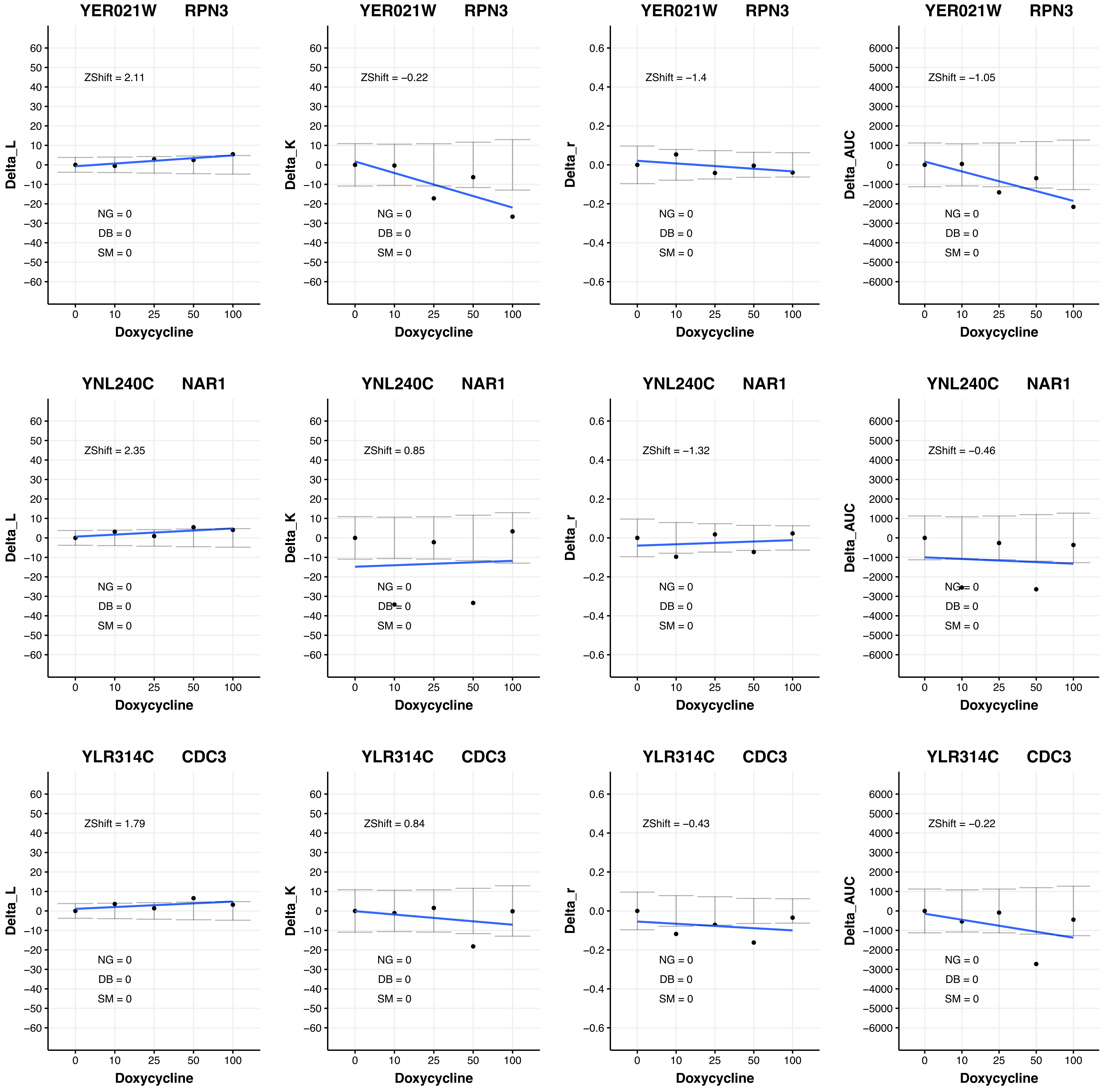

YER021W RPN3
ZShift = 2.11
ZShift = −0.22
ZShift = −1.4
ZShift = −1.05
YNL240C NAR1
ZShift = 2.35
ZShift = 0.85
ZShift = −1.32
ZShift = −0.46
YLR314C CDC3
ZShift = 1.79
ZShift = 0.84
ZShift = −0.43
ZShift = −0.22
NG = 0
DB = 0
SM = 0
Delta_L
Delta_K
Delta_r
Delta_AUC
Doxycycline

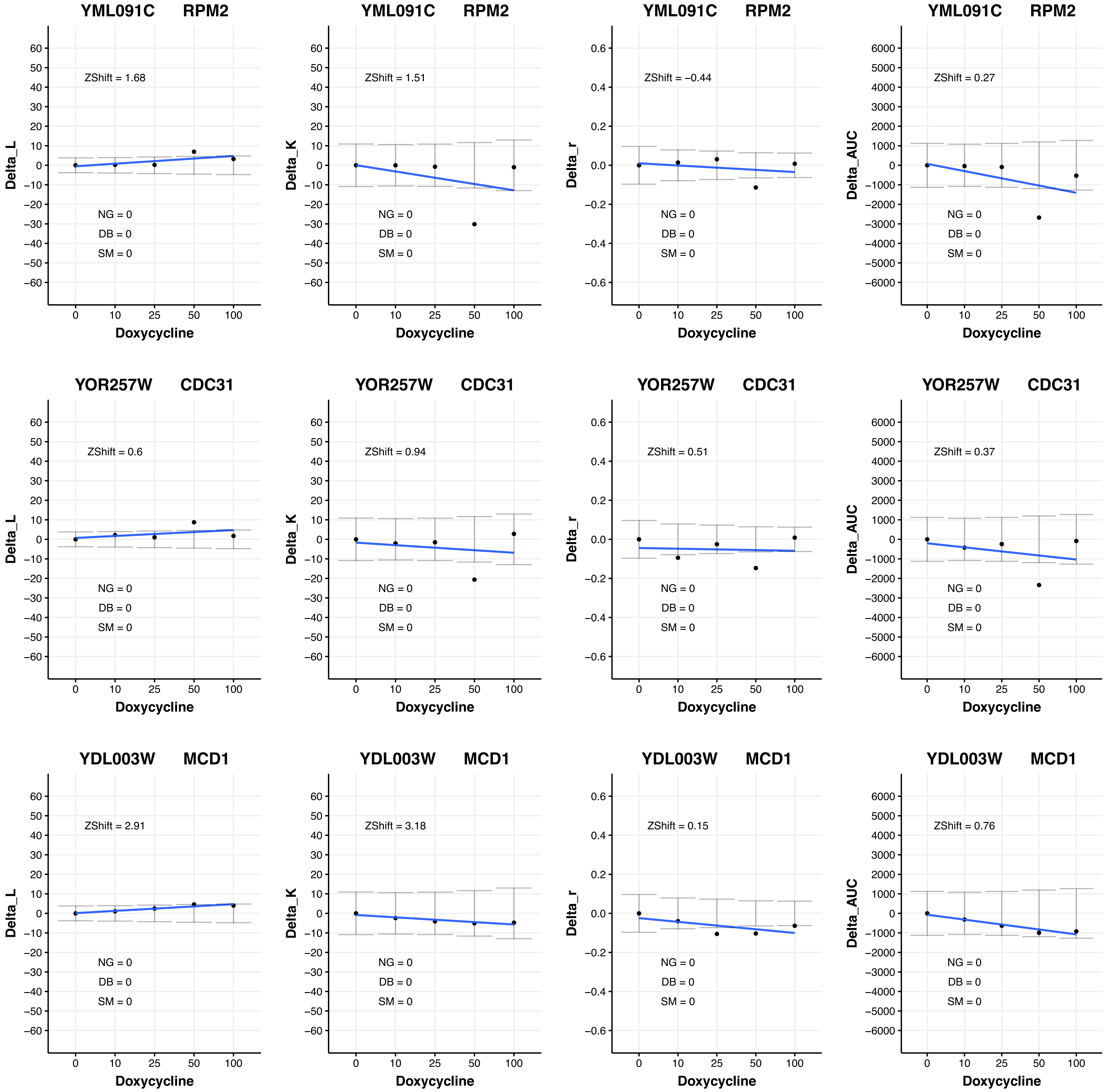
YML091C RPM2
ZShift = 1.68
NG = 0
DB = 0
SM = 0
Delta_L
Doxycycline
YML091C RPM2
ZShift = 1.51
NG = 0
DB = 0
SM = 0
Delta_K
Doxycycline
YML091C RPM2
ZShift = −0.44
NG = 0
DB = 0
SM = 0
Delta_r
Doxycycline
YML091C RPM2
ZShift = 0.27
NG = 0
DB = 0
SM = 0
Delta_AUC
Doxycycline
YOR257W CDC31
ZShift = 0.6
NG = 0
DB = 0
SM = 0
Delta_L
Doxycycline
YOR257W CDC31
ZShift = 0.94
NG = 0
DB = 0
SM = 0
Delta_K
Doxycycline
YOR257W CDC31
ZShift = 0.51
NG = 0
DB = 0
SM = 0
Delta_r
Doxycycline
YOR257W CDC31
ZShift = 0.37
NG = 0
DB = 0
SM = 0
Delta_AUC
Doxycycline
YDL003W MCD1
ZShift = 2.91
NG = 0
DB = 0
SM = 0
Delta_L
Doxycycline
YDL003W MCD1
ZShift = 3.18
NG = 0
DB = 0
SM = 0
Delta_K
Doxycycline
YDL003W MCD1
ZShift = 0.15
NG = 0
DB = 0
SM = 0
Delta_r
Doxycycline
YDL003W MCD1
ZShift = 0.76
NG = 0
DB = 0
SM = 0
Delta_AUC
Doxycycline

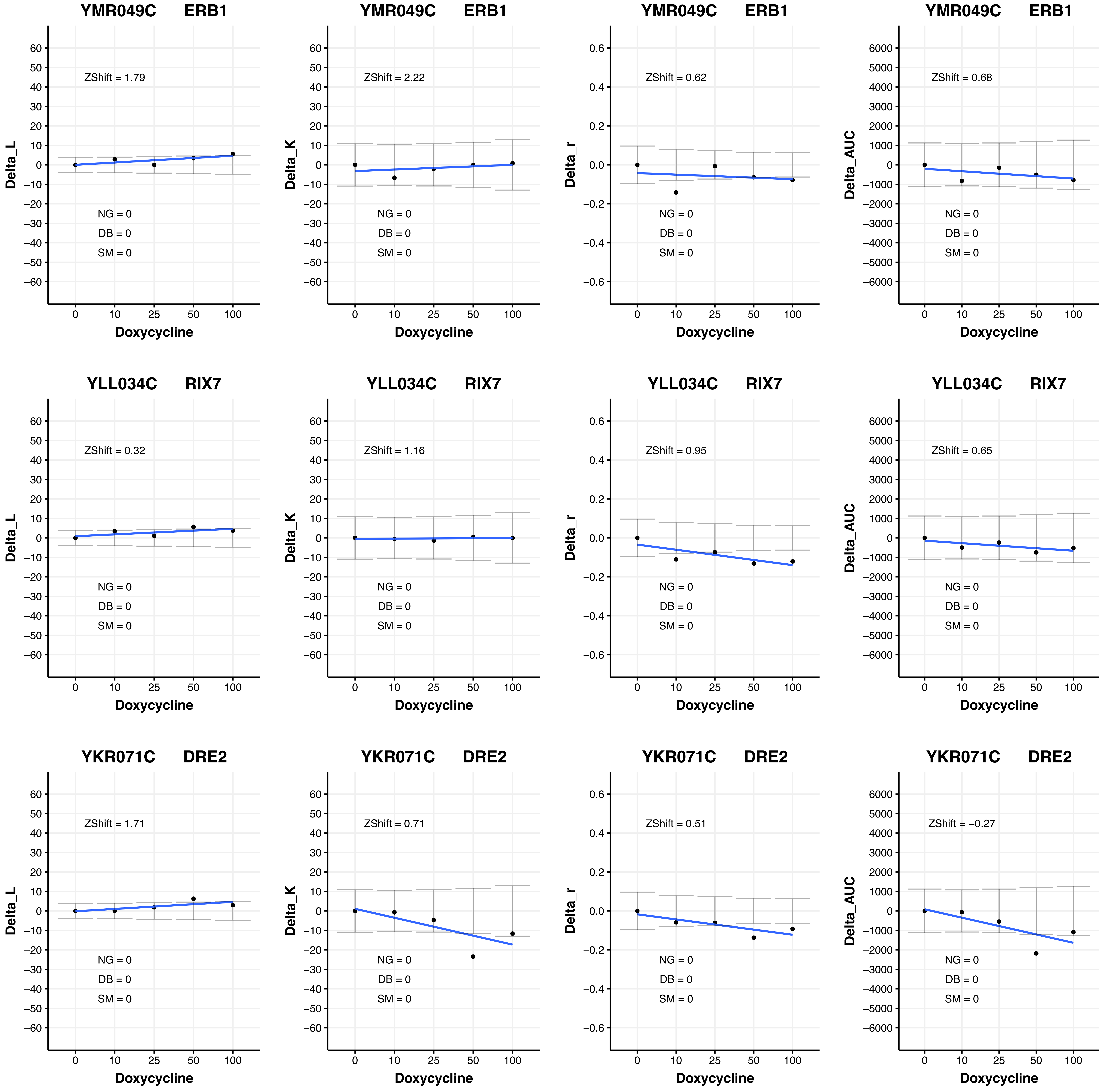
YMR049C ERB1
ZShift = 1.79
NG = 0
DB = 0
SM = 0
Delta_L
Doxycycline
YMR049C ERB1
ZShift = 2.22
NG = 0
DB = 0
SM = 0
Delta_K
Doxycycline
YMR049C ERB1
ZShift = 0.62
NG = 0
DB = 0
SM = 0
Delta_r
Doxycycline
YMR049C ERB1
ZShift = 0.68
NG = 0
DB = 0
SM = 0
Delta_AUC
Doxycycline
YLL034C RIX7
ZShift = 0.32
NG = 0
DB = 0
SM = 0
Delta_L
Doxycycline
YLL034C RIX7
ZShift = 1.16
NG = 0
DB = 0
SM = 0
Delta_K
Doxycycline
YLL034C RIX7
ZShift = 0.95
NG = 0
DB = 0
SM = 0
Delta_r
Doxycycline
YLL034C RIX7
ZShift = 0.65
NG = 0
DB = 0
SM = 0
Delta_AUC
Doxycycline
YKR071C DRE2
ZShift = 1.71
NG = 0
DB = 0
SM = 0
Delta_L
Doxycycline
YKR071C DRE2
ZShift = 0.71
NG = 0
DB = 0
SM = 0
Delta_K
Doxycycline
YKR071C DRE2
ZShift = 0.51
NG = 0
DB = 0
SM = 0
Delta_r
Doxycycline
YKR071C DRE2
ZShift = –0.27
NG = 0
DB = 0
SM = 0
Delta_AUC
Doxycycline

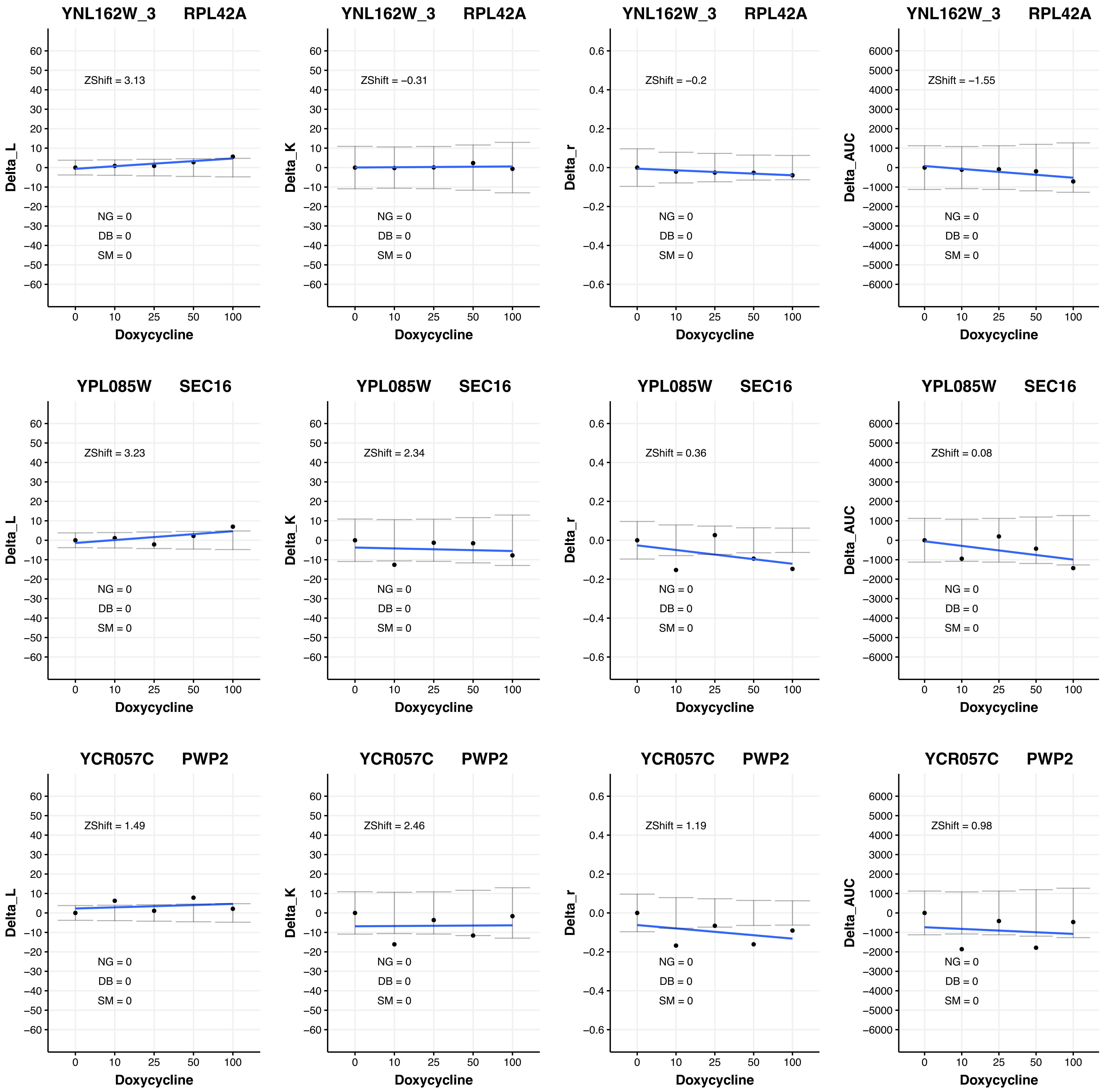
YNL162W_3 RPL42A
ZShift = 3.13
NG = 0
DB = 0
SM = 0
Delta_L
Doxycycline
YNL162W_3 RPL42A
ZShift = −0.31
NG = 0
DB = 0
SM = 0
Delta_K
Doxycycline
YNL162W_3 RPL42A
ZShift = −0.2
NG = 0
DB = 0
SM = 0
Delta_r
Doxycycline
YNL162W_3 RPL42A
ZShift = −1.55
NG = 0
DB = 0
SM = 0
Delta_AUC
Doxycycline
YPL085W SEC16
ZShift = 3.23
NG = 0
DB = 0
SM = 0
Delta_L
Doxycycline
YPL085W SEC16
ZShift = 2.34
NG = 0
DB = 0
SM = 0
Delta_K
Doxycycline
YPL085W SEC16
ZShift = 0.36
NG = 0
DB = 0
SM = 0
Delta_r
Doxycycline
YPL085W SEC16
ZShift = 0.08
NG = 0
DB = 0
SM = 0
Delta_AUC
Doxycycline
YCR057C PWP2
ZShift = 1.49
NG = 0
DB = 0
SM = 0
Delta_L
Doxycycline
YCR057C PWP2
ZShift = 2.46
NG = 0
DB = 0
SM = 0
Delta_K
Doxycycline
YCR057C PWP2
ZShift = 1.19
NG = 0
DB = 0
SM = 0
Delta_r
Doxycycline
YCR057C PWP2
ZShift = 0.98
NG = 0
DB = 0
SM = 0
Delta_AUC
Doxycycline

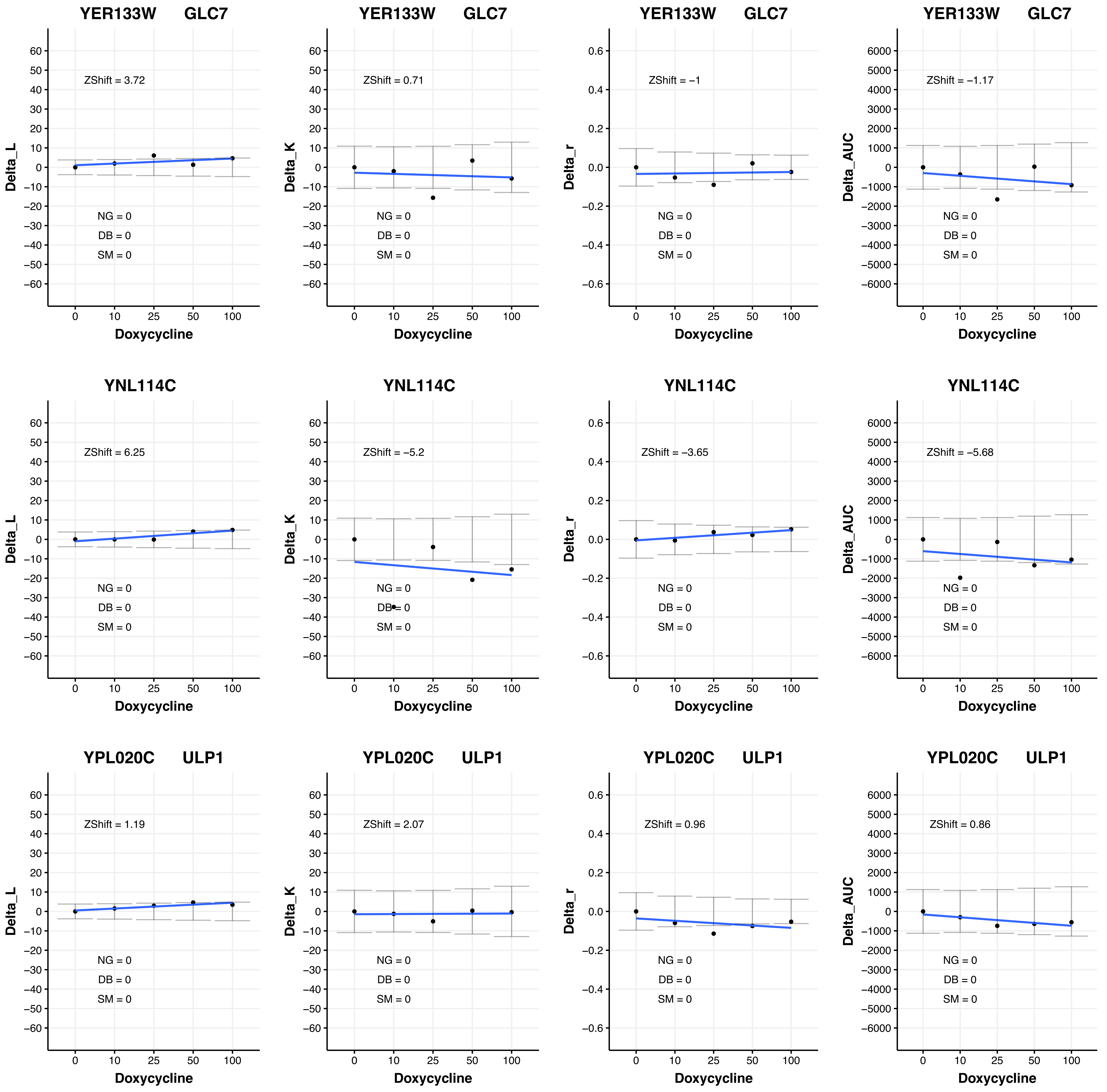
YER133W GLC7
ZShift = 3.72
NG = 0
DB = 0
SM = 0
Delta_L
Doxycycline
YER133W GLC7
ZShift = 0.71
NG = 0
DB = 0
SM = 0
Delta_K
Doxycycline
YER133W GLC7
ZShift = −1
NG = 0
DB = 0
SM = 0
Delta_r
Doxycycline
YER133W GLC7
ZShift = −1.17
NG = 0
DB = 0
SM = 0
Delta_AUC
Doxycycline
YNL114C
ZShift = 6.25
NG = 0
DB = 0
SM = 0
Delta_L
Doxycycline
YNL114C
ZShift = −5.2
NG = 0
DB = 0
SM = 0
Delta_K
Doxycycline
YNL114C
ZShift = −3.65
NG = 0
DB = 0
SM = 0
Delta_r
Doxycycline
YNL114C
ZShift = −5.68
NG = 0
DB = 0
SM = 0
Delta_AUC
Doxycycline
YPL020C ULP1
ZShift = 1.19
NG = 0
DB = 0
SM = 0
Delta_L
Doxycycline
YPL020C ULP1
ZShift = 2.07
NG = 0
DB = 0
SM = 0
Delta_K
Doxycycline
YPL020C ULP1
ZShift = 0.96
NG = 0
DB = 0
SM = 0
Delta_r
Doxycycline
YPL020C ULP1
ZShift = 0.86
NG = 0
DB = 0
SM = 0
Delta_AUC
Doxycycline

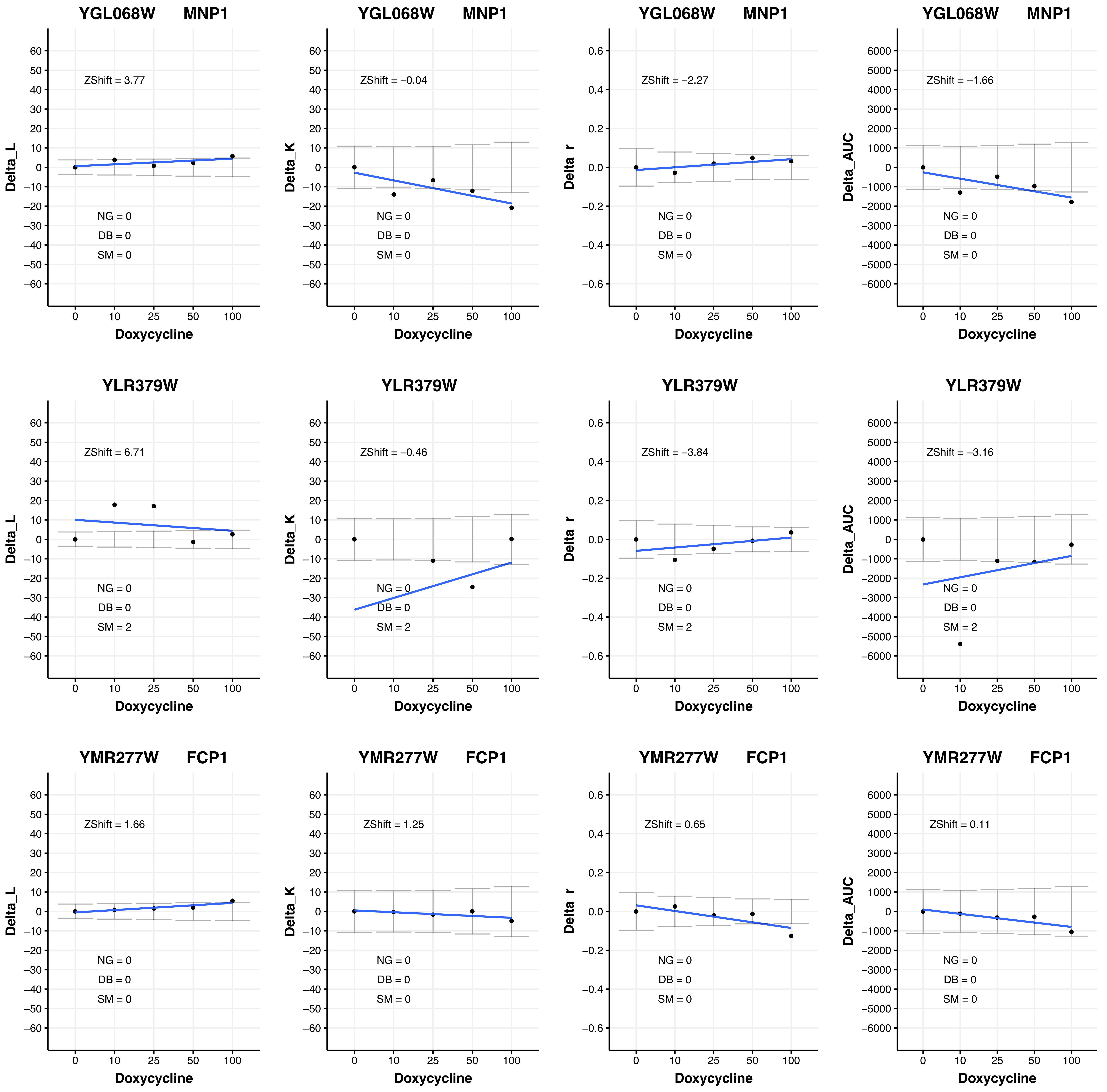

YGL068W MNP1
ZShift = 3.77
NG = 0
DB = 0
SM = 0
Delta_L
Doxycycline
YGL068W MNP1
ZShift = −0.04
NG = 0
DB = 0
SM = 0
Delta_K
Doxycycline
YGL068W MNP1
ZShift = −2.27
NG = 0
DB = 0
SM = 0
Delta_r
Doxycycline
YGL068W MNP1
ZShift = −1.66
NG = 0
DB = 0
SM = 0
Delta_AUC
Doxycycline
YLR379W
ZShift = 6.71
NG = 0
DB = 0
SM = 2
Delta_L
Doxycycline
YLR379W
ZShift = −0.46
NG = 0
DB = 0
SM = 2
Delta_K
Doxycycline
YLR379W
ZShift = −3.84
NG = 0
DB = 0
SM = 2
Delta_r
Doxycycline
YLR379W
ZShift = −3.16
NG = 0
DB = 0
SM = 2
Delta_AUC
Doxycycline
YMR277W FCP1
ZShift = 1.66
NG = 0
DB = 0
SM = 0
Delta_L
Doxycycline
YMR277W FCP1
ZShift = 1.25
NG = 0
DB = 0
SM = 0
Delta_K
Doxycycline
YMR277W FCP1
ZShift = 0.65
NG = 0
DB = 0
SM = 0
Delta_r
Doxycycline
YMR277W FCP1
ZShift = 0.11
NG = 0
DB = 0
SM = 0
Delta_AUC
Doxycycline

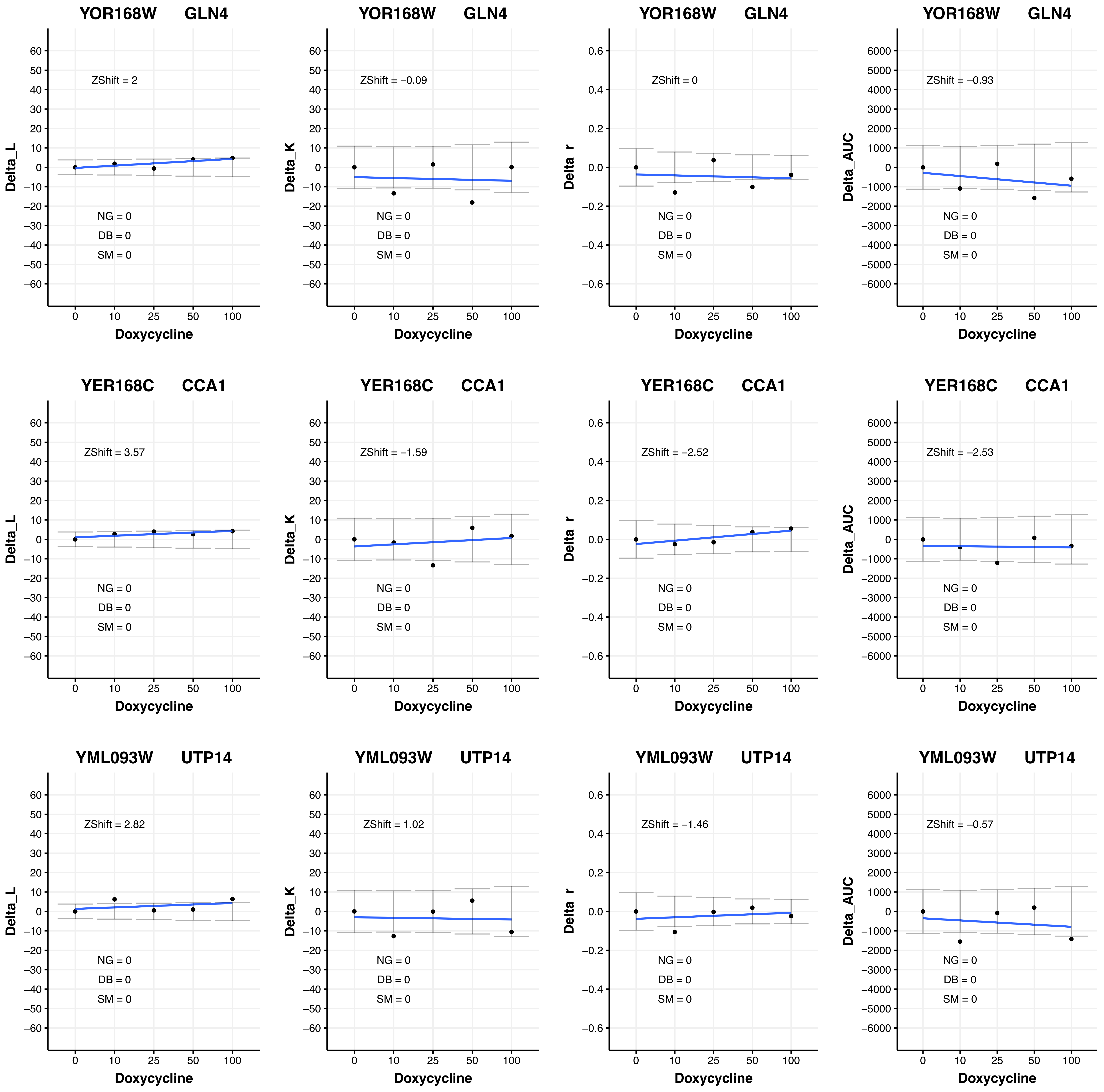
YOR168W GLN4
ZShift = 2
NG = 0
DB = 0
SM = 0
Delta_L
Doxycycline
YOR168W GLN4
ZShift = −0.09
NG = 0
DB = 0
SM = 0
Delta_K
Doxycycline
YOR168W GLN4
ZShift = 0
NG = 0
DB = 0
SM = 0
Delta_r
Doxycycline
YOR168W GLN4
ZShift = −0.93
NG = 0
DB = 0
SM = 0
Delta_AUC
Doxycycline
YER168C CCA1
ZShift = 3.57
NG = 0
DB = 0
SM = 0
Delta_L
Doxycycline
YER168C CCA1
ZShift = −1.59
NG = 0
DB = 0
SM = 0
Delta_K
Doxycycline
YER168C CCA1
ZShift = −2.52
NG = 0
DB = 0
SM = 0
Delta_r
Doxycycline
YER168C CCA1
ZShift = −2.53
NG = 0
DB = 0
SM = 0
Delta_AUC
Doxycycline
YML093W UTP14
ZShift = 2.82
NG = 0
DB = 0
SM = 0
Delta_L
Doxycycline
YML093W UTP14
ZShift = 1.02
NG = 0
DB = 0
SM = 0
Delta_K
Doxycycline
YML093W UTP14
ZShift = −1.46
NG = 0
DB = 0
SM = 0
Delta_r
Doxycycline
YML093W UTP14
ZShift = −0.57
NG = 0
DB = 0
SM = 0
Delta_AUC
Doxycycline

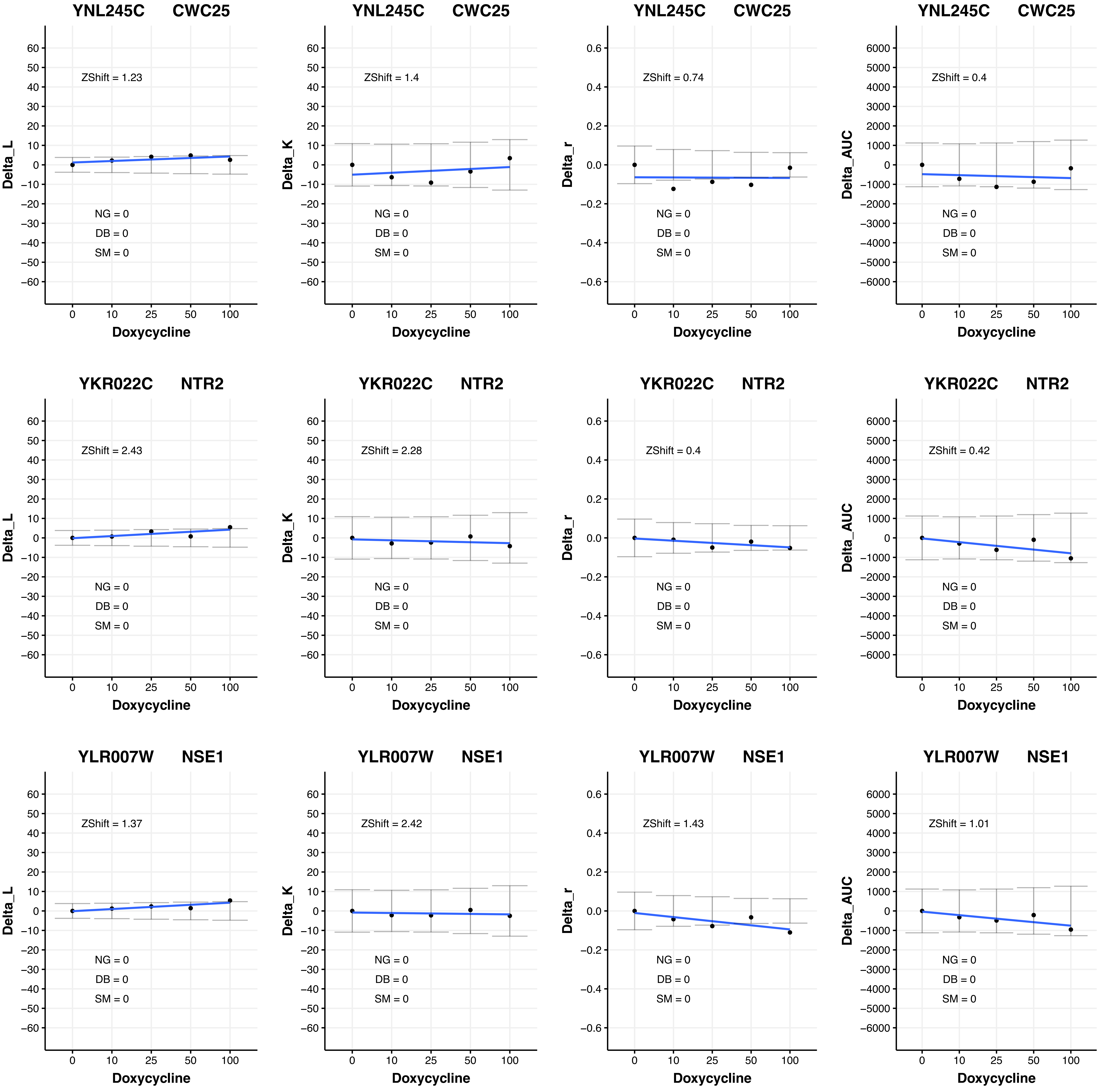

YNL245C CWC25
ZShift = 1.23
Delta_L
NG = 0
DB = 0
SM = 0
Doxycycline
YNL245C CWC25
ZShift = 1.4
Delta_K
NG = 0
DB = 0
SM = 0
Doxycycline
YNL245C CWC25
ZShift = 0.74
Delta_r
NG = 0
DB = 0
SM = 0
Doxycycline
YNL245C CWC25
ZShift = 0.4
Delta_AUC
NG = 0
DB = 0
SM = 0
Doxycycline
YKR022C NTR2
ZShift = 2.43
Delta_L
NG = 0
DB = 0
SM = 0
Doxycycline
YKR022C NTR2
ZShift = 2.28
Delta_K
NG = 0
DB = 0
SM = 0
Doxycycline
YKR022C NTR2
ZShift = 0.4
Delta_r
NG = 0
DB = 0
SM = 0
Doxycycline
YKR022C NTR2
ZShift = 0.42
Delta_AUC
NG = 0
DB = 0
SM = 0
Doxycycline
YLR007W NSE1
ZShift = 1.37
Delta_L
NG = 0
DB = 0
SM = 0
Doxycycline
YLR007W NSE1
ZShift = 2.42
Delta_K
NG = 0
DB = 0
SM = 0
Doxycycline
YLR007W NSE1
ZShift = 1.43
Delta_r
NG = 0
DB = 0
SM = 0
Doxycycline
YLR007W NSE1
ZShift = 1.01
Delta_AUC
NG = 0
DB = 0
SM = 0
Doxycycline

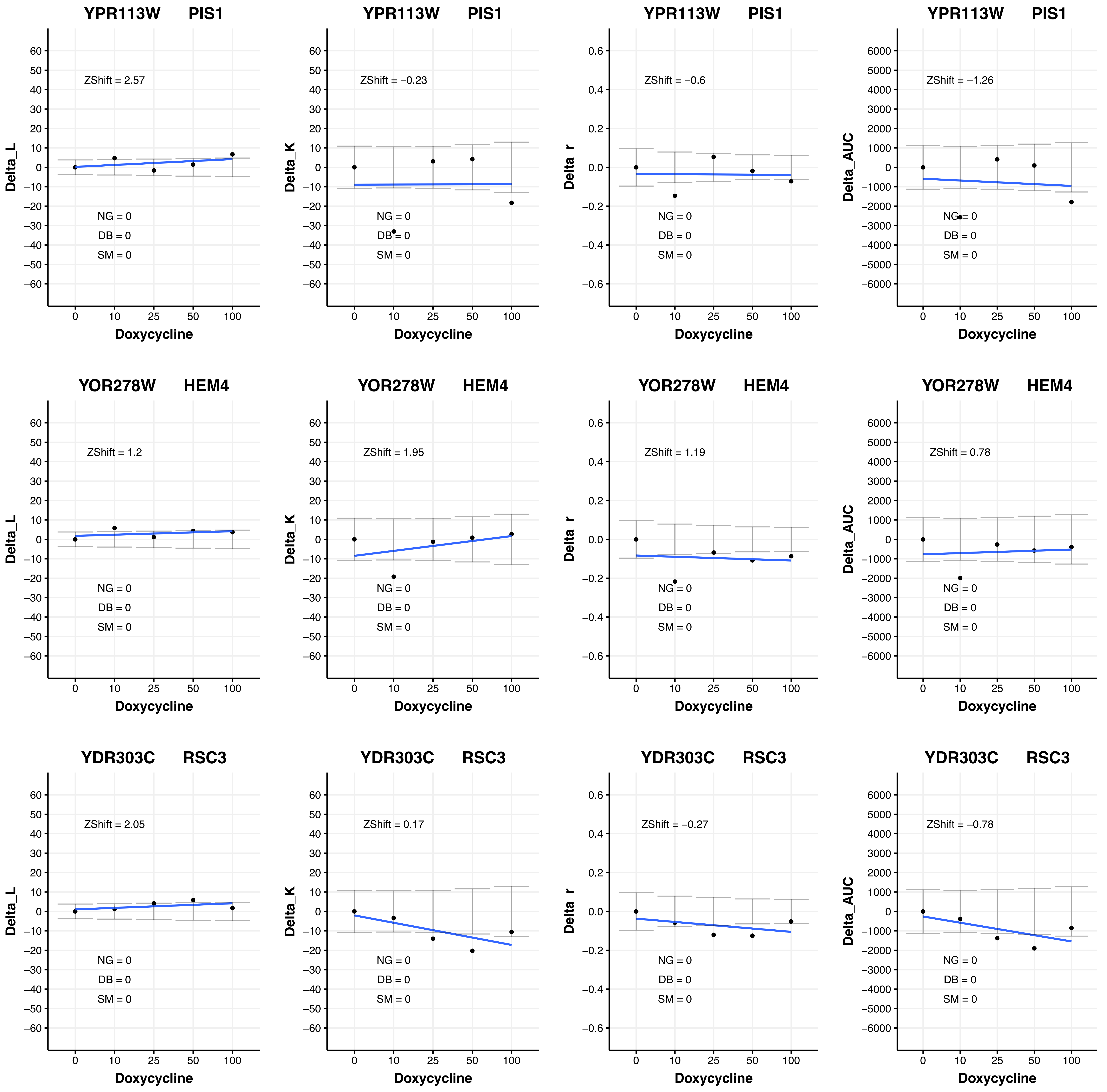
YPR113W PIS1
ZShift = 2.57
NG = 0
DB = 0
SM = 0
Delta_L
Doxycycline
YPR113W PIS1
ZShift = −0.23
NG = 0
DB = 0
SM = 0
Delta_K
Doxycycline
YPR113W PIS1
ZShift = −0.6
NG = 0
DB = 0
SM = 0
Delta_r
Doxycycline
YPR113W PIS1
ZShift = −1.26
NG = 0
DB = 0
SM = 0
Delta_AUC
Doxycycline
YOR278W HEM4
ZShift = 1.2
NG = 0
DB = 0
SM = 0
Delta_L
Doxycycline
YOR278W HEM4
ZShift = 1.95
NG = 0
DB = 0
SM = 0
Delta_K
Doxycycline
YOR278W HEM4
ZShift = 1.19
NG = 0
DB = 0
SM = 0
Delta_r
Doxycycline
YOR278W HEM4
ZShift = 0.78
NG = 0
DB = 0
SM = 0
Delta_AUC
Doxycycline
YDR303C RSC3
ZShift = 2.05
NG = 0
DB = 0
SM = 0
Delta_L
Doxycycline
YDR303C RSC3
ZShift = 0.17
NG = 0
DB = 0
SM = 0
Delta_K
Doxycycline
YDR303C RSC3
ZShift = −0.27
NG = 0
DB = 0
SM = 0
Delta_r
Doxycycline
YDR303C RSC3
ZShift = −0.78
NG = 0
DB = 0
SM = 0
Delta_AUC
Doxycycline

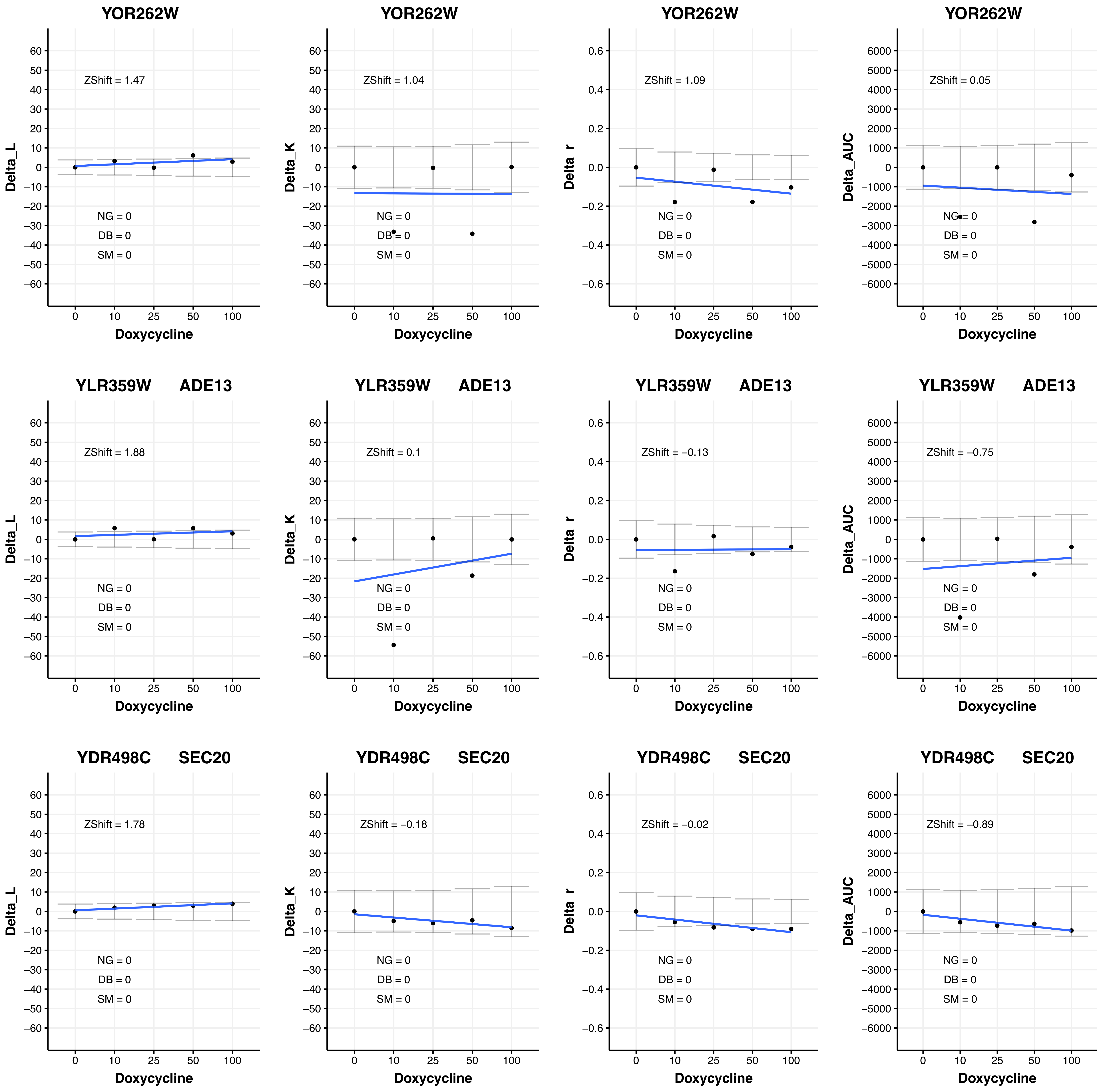
YOR262W
ZShift = 1.47
ZShift = 1.04
ZShift = 1.09
ZShift = 0.05
YLR359W ADE13
ZShift = 1.88
ZShift = 0.1
ZShift = −0.13
ZShift = −0.75
YDR498C SEC20
ZShift = 1.78
ZShift = −0.18
ZShift = −0.02
ZShift = −0.89
NG = 0
DB = 0
SM = 0
Delta_L
Delta_K
Delta_r
Delta_AUC
Doxycycline

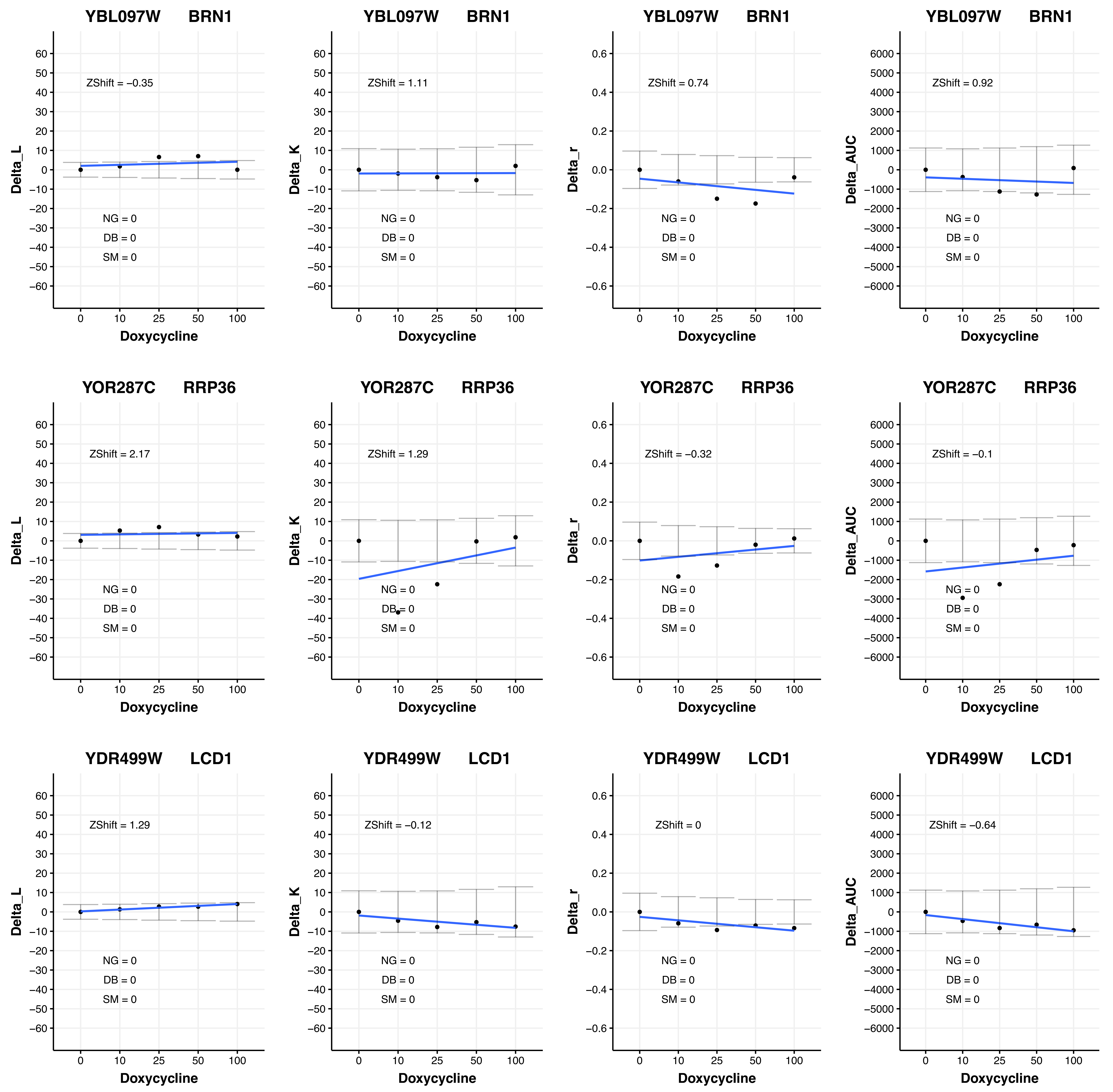
YBL097W BRN1
Delta_L
ZShift = −0.35
NG = 0
DB = 0
SM = 0
Doxycycline
YBL097W BRN1
Delta_K
ZShift = 1.11
NG = 0
DB = 0
SM = 0
Doxycycline
YBL097W BRN1
Delta_r
ZShift = 0.74
NG = 0
DB = 0
SM = 0
Doxycycline
YBL097W BRN1
Delta_AUC
ZShift = 0.92
NG = 0
DB = 0
SM = 0
Doxycycline
YOR287C RRP36
Delta_L
ZShift = 2.17
NG = 0
DB = 0
SM = 0
Doxycycline
YOR287C RRP36
Delta_K
ZShift = 1.29
NG = 0
DB = 0
SM = 0
Doxycycline
YOR287C RRP36
Delta_r
ZShift = −0.32
NG = 0
DB = 0
SM = 0
Doxycycline
YOR287C RRP36
Delta_AUC
ZShift = −0.1
NG = 0
DB = 0
SM = 0
Doxycycline
YDR499W LCD1
Delta_L
ZShift = 1.29
NG = 0
DB = 0
SM = 0
Doxycycline
YDR499W LCD1
Delta_K
ZShift = −0.12
NG = 0
DB = 0
SM = 0
Doxycycline
YDR499W LCD1
Delta_r
ZShift = 0
NG = 0
DB = 0
SM = 0
Doxycycline
YDR499W LCD1
Delta_AUC
ZShift = −0.64
NG = 0
DB = 0
SM = 0
Doxycycline
0
10
25
50
100

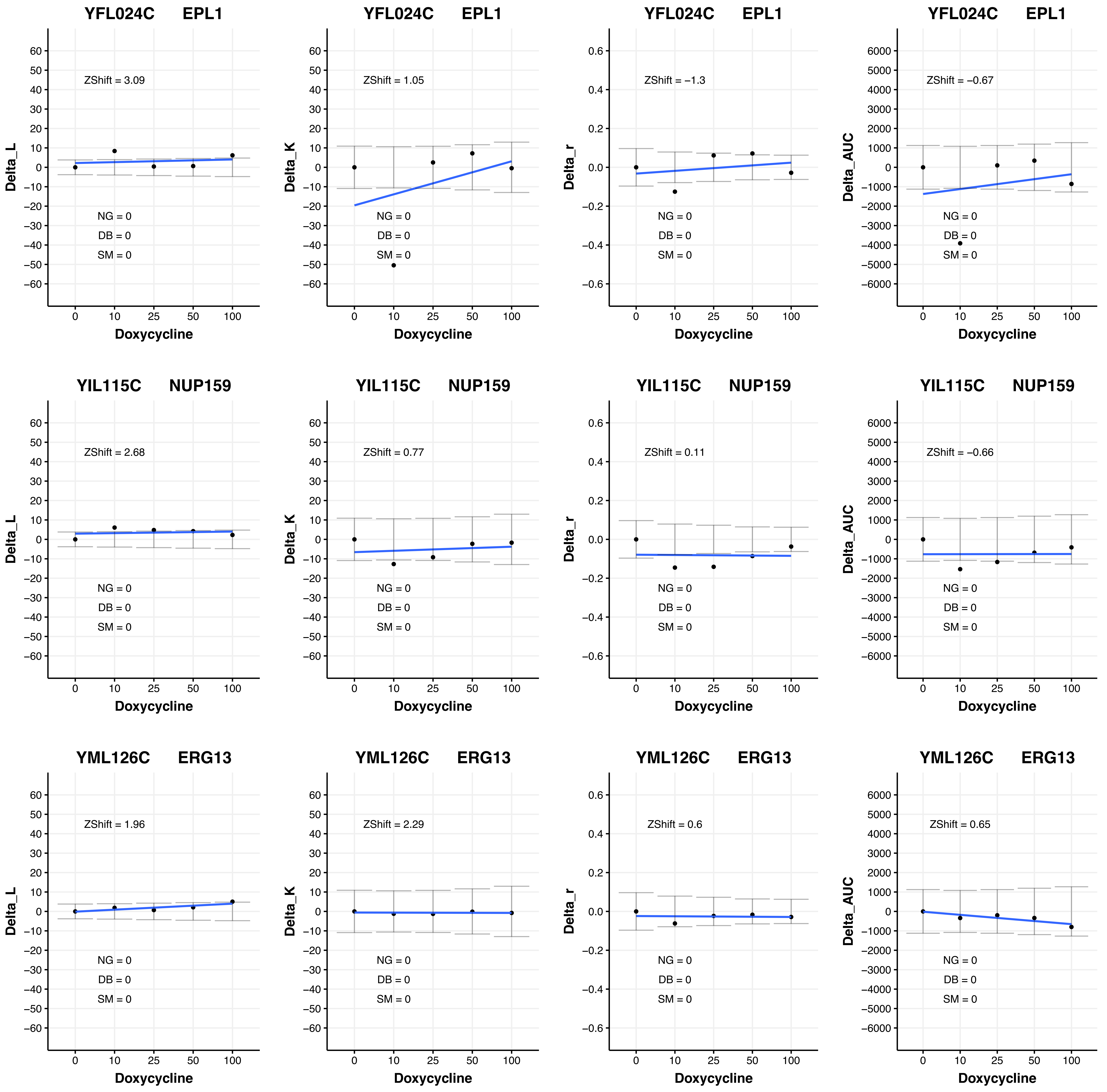
YFL024C EPL1
ZShift = 3.09
NG = 0
DB = 0
SM = 0
Delta_L
Doxycycline
YFL024C EPL1
ZShift = 1.05
NG = 0
DB = 0
SM = 0
Delta_K
Doxycycline
YFL024C EPL1
ZShift = −1.3
NG = 0
DB = 0
SM = 0
Delta_r
Doxycycline
YFL024C EPL1
ZShift = −0.67
NG = 0
DB = 0
SM = 0
Delta_AUC
Doxycycline
YIL115C NUP159
ZShift = 2.68
NG = 0
DB = 0
SM = 0
Delta_L
Doxycycline
YIL115C NUP159
ZShift = 0.77
NG = 0
DB = 0
SM = 0
Delta_K
Doxycycline
YIL115C NUP159
ZShift = 0.11
NG = 0
DB = 0
SM = 0
Delta_r
Doxycycline
YIL115C NUP159
ZShift = −0.66
NG = 0
DB = 0
SM = 0
Delta_AUC
Doxycycline
YML126C ERG13
ZShift = 1.96
NG = 0
DB = 0
SM = 0
Delta_L
Doxycycline
YML126C ERG13
ZShift = 2.29
NG = 0
DB = 0
SM = 0
Delta_K
Doxycycline
YML126C ERG13
ZShift = 0.6
NG = 0
DB = 0
SM = 0
Delta_r
Doxycycline
YML126C ERG13
ZShift = 0.65
NG = 0
DB = 0
SM = 0
Delta_AUC
Doxycycline

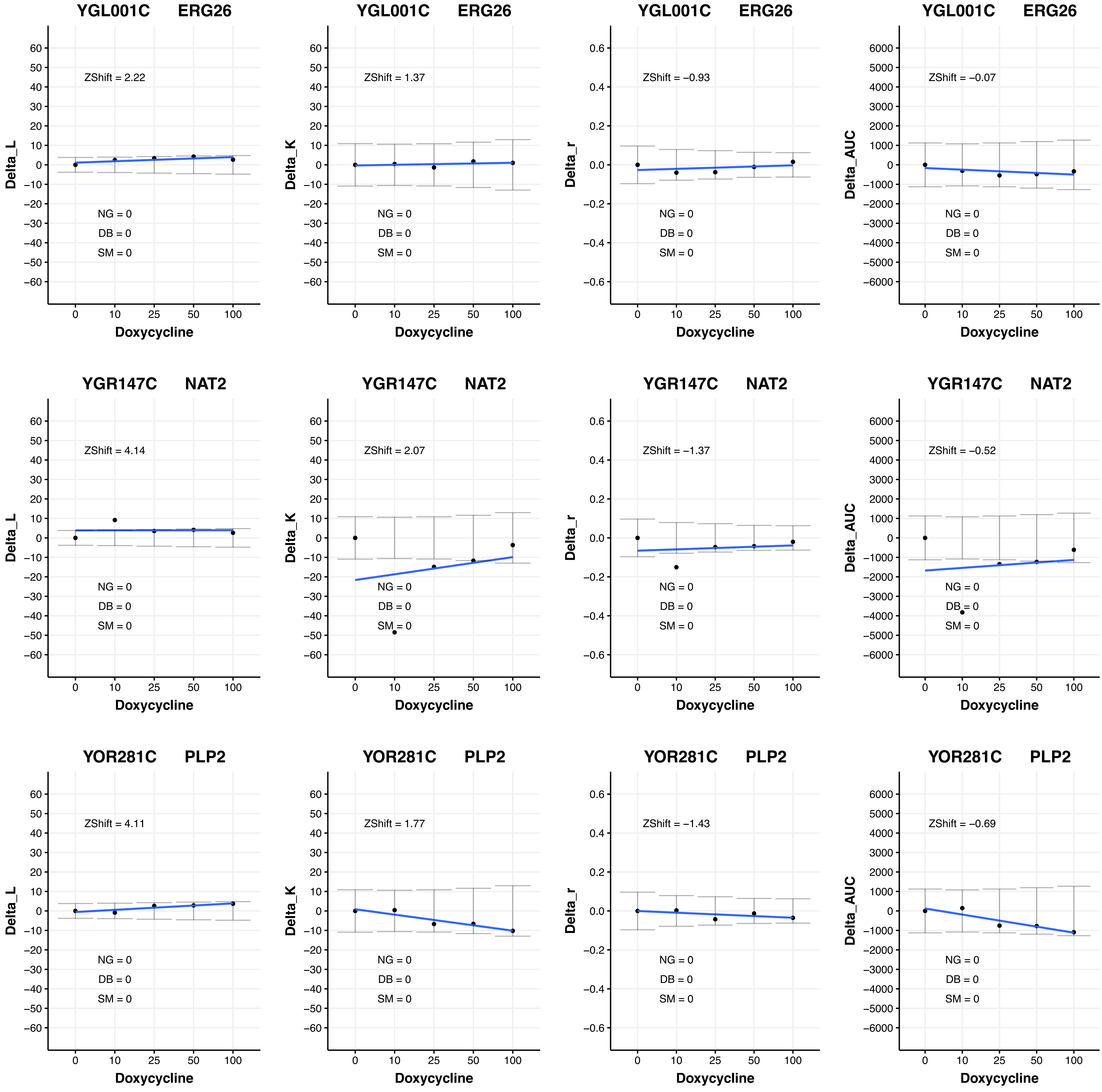
YGL001C ERG26
ZShift = 2.22
NG = 0
DB = 0
SM = 0
Delta_L
Doxycycline
YGL001C ERG26
ZShift = 1.37
NG = 0
DB = 0
SM = 0
Delta_K
Doxycycline
YGL001C ERG26
ZShift = −0.93
NG = 0
DB = 0
SM = 0
Delta_r
Doxycycline
YGL001C ERG26
ZShift = −0.07
NG = 0
DB = 0
SM = 0
Delta_AUC
Doxycycline
YGR147C NAT2
ZShift = 4.14
NG = 0
DB = 0
SM = 0
Delta_L
Doxycycline
YGR147C NAT2
ZShift = 2.07
NG = 0
DB = 0
SM = 0
Delta_K
Doxycycline
YGR147C NAT2
ZShift = −1.37
NG = 0
DB = 0
SM = 0
Delta_r
Doxycycline
YGR147C NAT2
ZShift = −0.52
NG = 0
DB = 0
SM = 0
Delta_AUC
Doxycycline
YOR281C PLP2
ZShift = 4.11
NG = 0
DB = 0
SM = 0
Delta_L
Doxycycline
YOR281C PLP2
ZShift = 1.77
NG = 0
DB = 0
SM = 0
Delta_K
Doxycycline
YOR281C PLP2
ZShift = −1.43
NG = 0
DB = 0
SM = 0
Delta_r
Doxycycline
YOR281C PLP2
ZShift = −0.69
NG = 0
DB = 0
SM = 0
Delta_AUC
Doxycycline

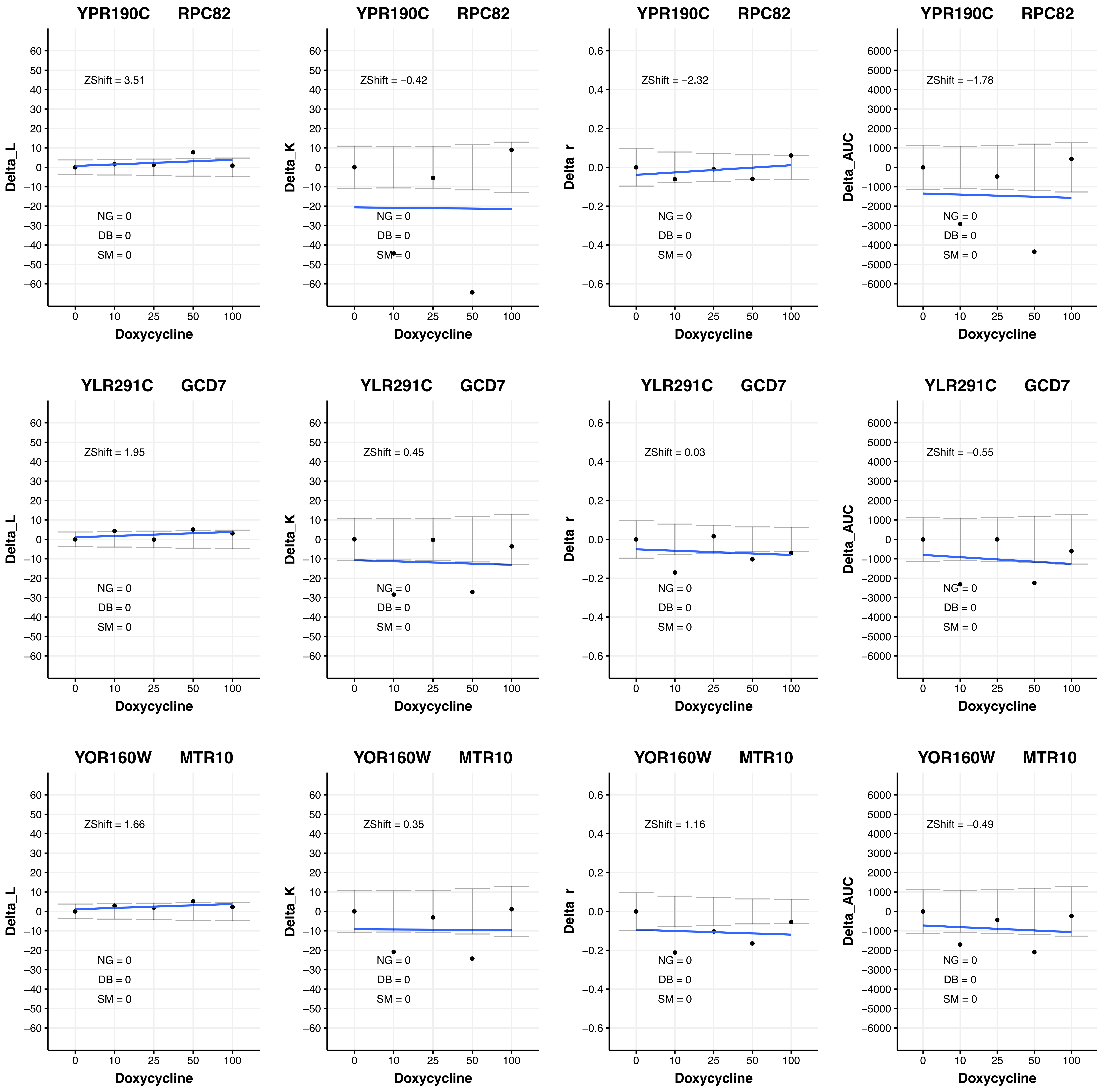

YPR190C RPC82
Delta_L
ZShift = 3.51
NG = 0
DB = 0
SM = 0
Doxycycline
YPR190C RPC82
Delta_K
ZShift = −0.42
NG = 0
DB = 0
SM = 0
Doxycycline
YPR190C RPC82
Delta_r
ZShift = −2.32
NG = 0
DB = 0
SM = 0
Doxycycline
YPR190C RPC82
Delta_AUC
ZShift = −1.78
NG = 0
DB = 0
SM = 0
Doxycycline
YLR291C GCD7
Delta_L
ZShift = 1.95
NG = 0
DB = 0
SM = 0
Doxycycline
YLR291C GCD7
Delta_K
ZShift = 0.45
NG = 0
DB = 0
SM = 0
Doxycycline
YLR291C GCD7
Delta_r
ZShift = 0.03
NG = 0
DB = 0
SM = 0
Doxycycline
YLR291C GCD7
Delta_AUC
ZShift = −0.55
NG = 0
DB = 0
SM = 0
Doxycycline
YOR160W MTR10
Delta_L
ZShift = 1.66
NG = 0
DB = 0
SM = 0
Doxycycline
YOR160W MTR10
Delta_K
ZShift = 0.35
NG = 0
DB = 0
SM = 0
Doxycycline
YOR160W MTR10
Delta_r
ZShift = 1.16
NG = 0
DB = 0
SM = 0
Doxycycline
YOR160W MTR10
Delta_AUC
ZShift = −0.49
NG = 0
DB = 0
SM = 0
Doxycycline

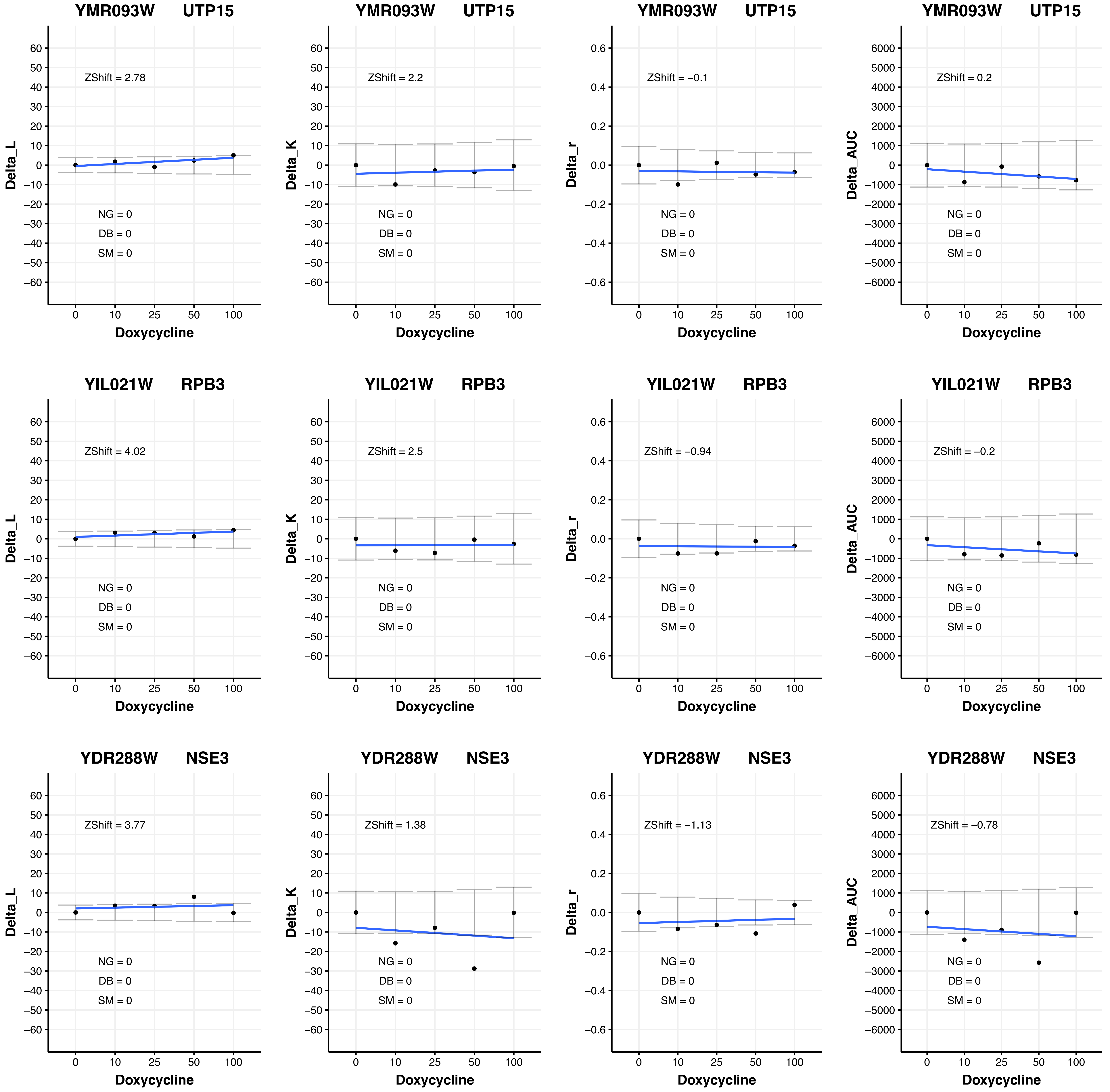

YMR093W UTP15
ZShift = 2.78
NG = 0
DB = 0
SM = 0
Delta_L
Doxycycline
YMR093W UTP15
ZShift = 2.2
NG = 0
DB = 0
SM = 0
Delta_K
Doxycycline
YMR093W UTP15
ZShift = −0.1
NG = 0
DB = 0
SM = 0
Delta_r
Doxycycline
YMR093W UTP15
ZShift = 0.2
NG = 0
DB = 0
SM = 0
Delta_AUC
Doxycycline
YIL021W RPB3
ZShift = 4.02
NG = 0
DB = 0
SM = 0
Delta_L
Doxycycline
YIL021W RPB3
ZShift = 2.5
NG = 0
DB = 0
SM = 0
Delta_K
Doxycycline
YIL021W RPB3
ZShift = −0.94
NG = 0
DB = 0
SM = 0
Delta_r
Doxycycline
YIL021W RPB3
ZShift = −0.2
NG = 0
DB = 0
SM = 0
Delta_AUC
Doxycycline
YDR288W NSE3
ZShift = 3.77
NG = 0
DB = 0
SM = 0
Delta_L
Doxycycline
YDR288W NSE3
ZShift = 1.38
NG = 0
DB = 0
SM = 0
Delta_K
Doxycycline
YDR288W NSE3
ZShift = −1.13
NG = 0
DB = 0
SM = 0
Delta_r
Doxycycline
YDR288W NSE3
ZShift = −0.78
NG = 0
DB = 0
SM = 0
Delta_AUC
Doxycycline

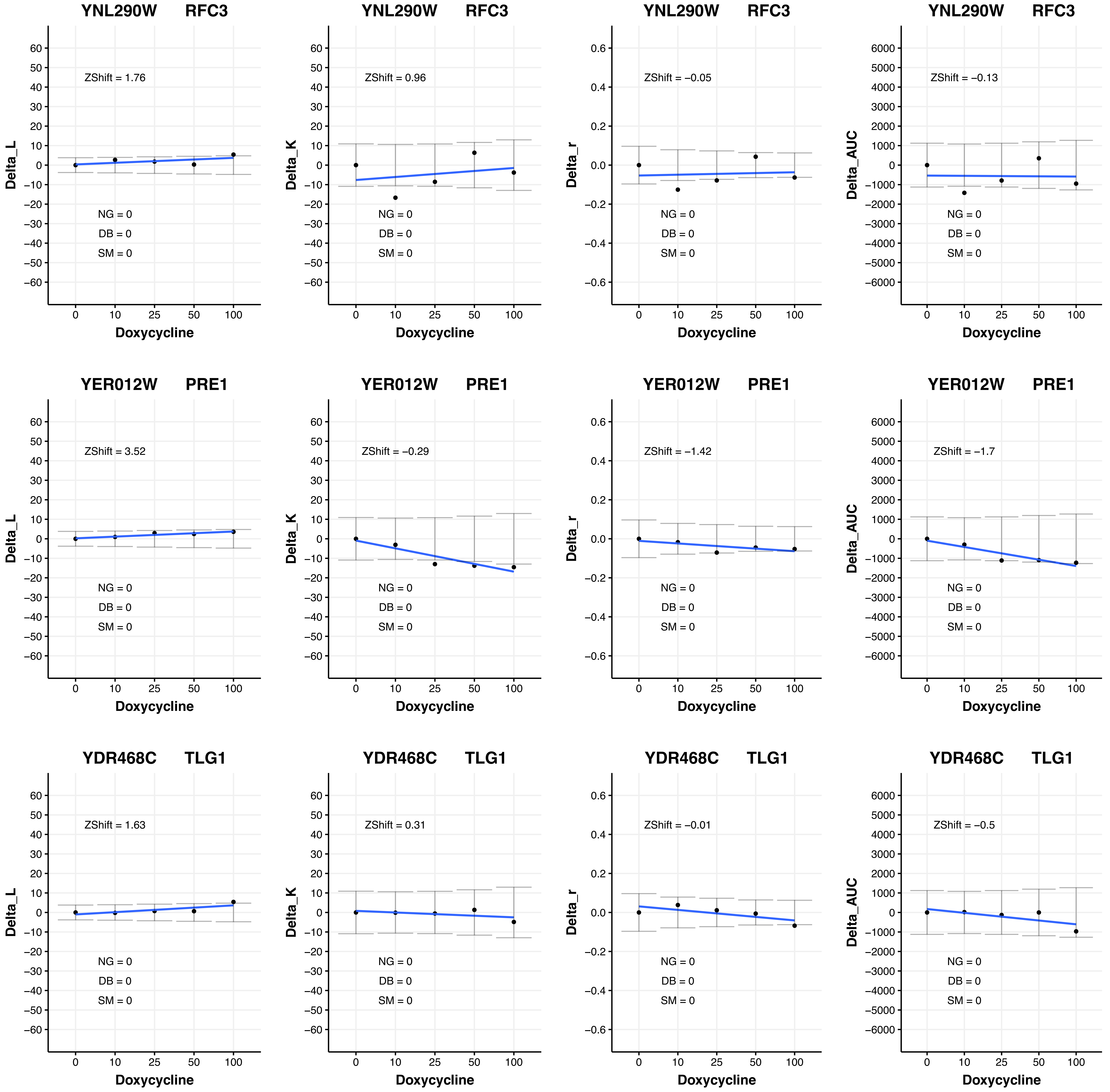
YNL290W RFC3
ZShift = 1.76
Delta_L
NG = 0
DB = 0
SM = 0
Doxycycline
YNL290W RFC3
ZShift = 0.96
Delta_K
NG = 0
DB = 0
SM = 0
Doxycycline
YNL290W RFC3
ZShift = −0.05
Delta_r
NG = 0
DB = 0
SM = 0
Doxycycline
YNL290W RFC3
ZShift = −0.13
Delta_AUC
NG = 0
DB = 0
SM = 0
Doxycycline
YER012W PRE1
ZShift = 3.52
Delta_L
NG = 0
DB = 0
SM = 0
Doxycycline
YER012W PRE1
ZShift = −0.29
Delta_K
NG = 0
DB = 0
SM = 0
Doxycycline
YER012W PRE1
ZShift = −1.42
Delta_r
NG = 0
DB = 0
SM = 0
Doxycycline
YER012W PRE1
ZShift = −1.7
Delta_AUC
NG = 0
DB = 0
SM = 0
Doxycycline
YDR468C TLG1
ZShift = 1.63
Delta_L
NG = 0
DB = 0
SM = 0
Doxycycline
YDR468C TLG1
ZShift = 0.31
Delta_K
NG = 0
DB = 0
SM = 0
Doxycycline
YDR468C TLG1
ZShift = −0.01
Delta_r
NG = 0
DB = 0
SM = 0
Doxycycline
YDR468C TLG1
ZShift = −0.5
Delta_AUC
NG = 0
DB = 0
SM = 0
Doxycycline

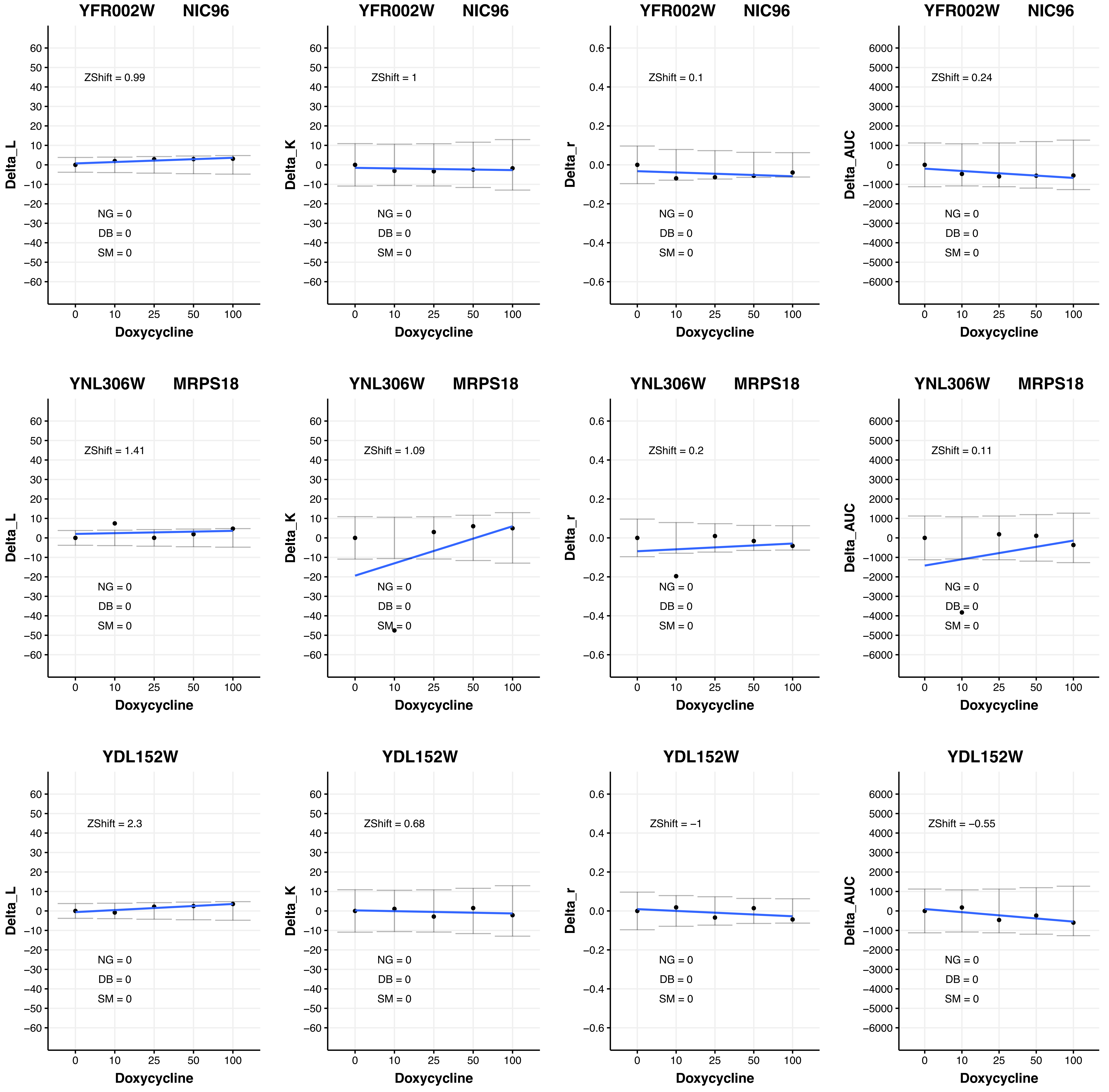

YFR002W NIC96
ZShift = 0.99
Delta_L
NG = 0
DB = 0
SM = 0
Doxycycline
YFR002W NIC96
ZShift = 1
Delta_K
NG = 0
DB = 0
SM = 0
Doxycycline
YFR002W NIC96
ZShift = 0.1
Delta_r
NG = 0
DB = 0
SM = 0
Doxycycline
YFR002W NIC96
ZShift = 0.24
Delta_AUC
NG = 0
DB = 0
SM = 0
Doxycycline
YNL306W MRPS18
ZShift = 1.41
Delta_L
NG = 0
DB = 0
SM = 0
Doxycycline
YNL306W MRPS18
ZShift = 1.09
Delta_K
NG = 0
DB = 0
SM = 0
Doxycycline
YNL306W MRPS18
ZShift = 0.2
Delta_r
NG = 0
DB = 0
SM = 0
Doxycycline
YNL306W MRPS18
ZShift = 0.11
Delta_AUC
NG = 0
DB = 0
SM = 0
Doxycycline
YDL152W
ZShift = 2.3
Delta_L
NG = 0
DB = 0
SM = 0
Doxycycline
YDL152W
ZShift = 0.68
Delta_K
NG = 0
DB = 0
SM = 0
Doxycycline
YDL152W
ZShift = −1
Delta_r
NG = 0
DB = 0
SM = 0
Doxycycline
YDL152W
ZShift = −0.55
Delta_AUC
NG = 0
DB = 0
SM = 0
Doxycycline

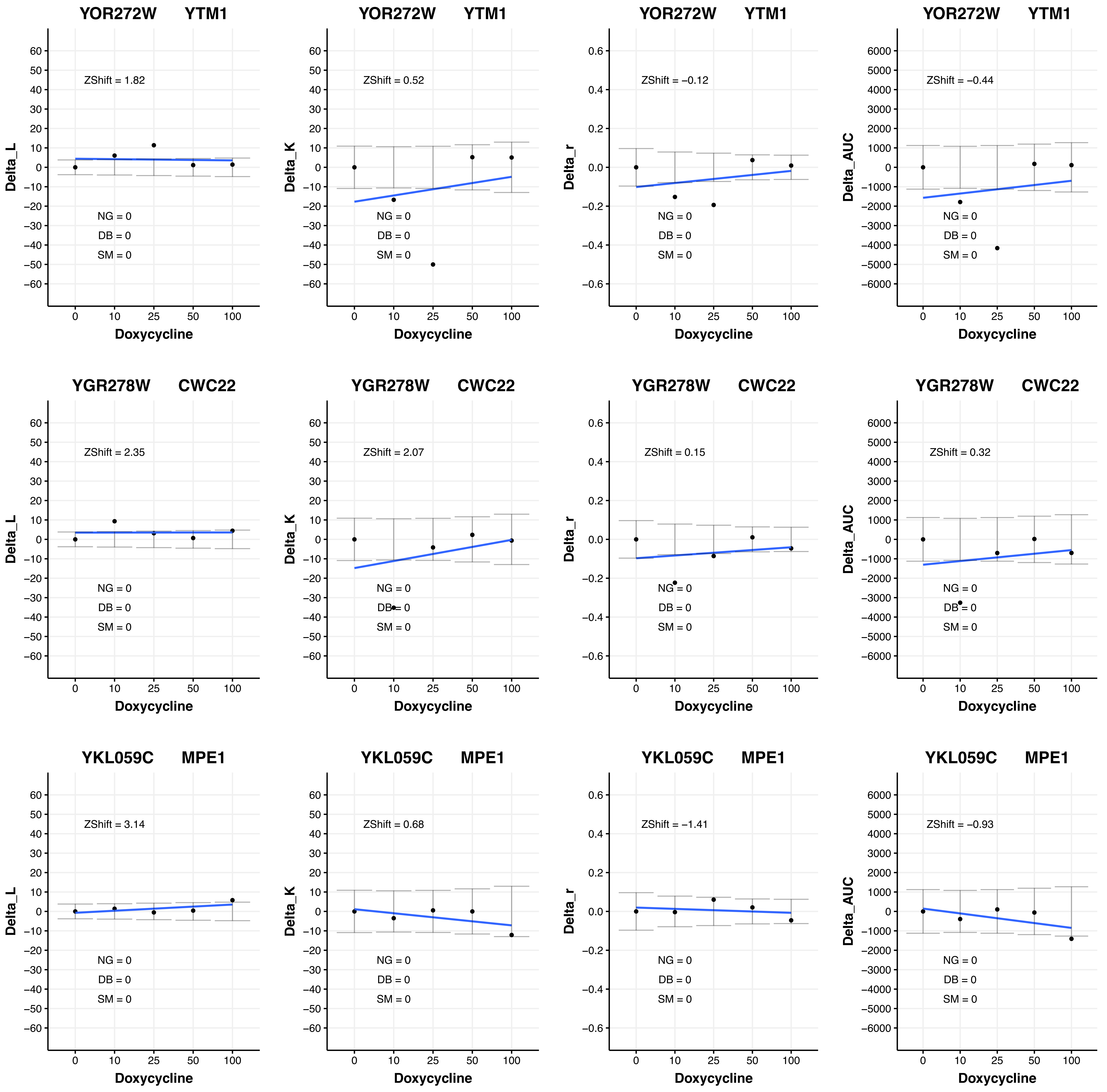

YOR272W YTM1
ZShift = 1.82
Delta_L
NG = 0
DB = 0
SM = 0
Doxycycline
YOR272W YTM1
ZShift = 0.52
Delta_K
NG = 0
DB = 0
SM = 0
Doxycycline
YOR272W YTM1
ZShift = −0.12
Delta_r
NG = 0
DB = 0
SM = 0
Doxycycline
YOR272W YTM1
ZShift = −0.44
Delta_AUC
NG = 0
DB = 0
SM = 0
Doxycycline
YGR278W CWC22
ZShift = 2.35
Delta_L
NG = 0
DB = 0
SM = 0
Doxycycline
YGR278W CWC22
ZShift = 2.07
Delta_K
NG = 0
DB = 0
SM = 0
Doxycycline
YGR278W CWC22
ZShift = 0.15
Delta_r
NG = 0
DB = 0
SM = 0
Doxycycline
YGR278W CWC22
ZShift = 0.32
Delta_AUC
NG = 0
DB = 0
SM = 0
Doxycycline
YKL059C MPE1
ZShift = 3.14
Delta_L
NG = 0
DB = 0
SM = 0
Doxycycline
YKL059C MPE1
ZShift = 0.68
Delta_K
NG = 0
DB = 0
SM = 0
Doxycycline
YKL059C MPE1
ZShift = −1.41
Delta_r
NG = 0
DB = 0
SM = 0
Doxycycline
YKL059C MPE1
ZShift = −0.93
Delta_AUC
NG = 0
DB = 0
SM = 0
Doxycycline

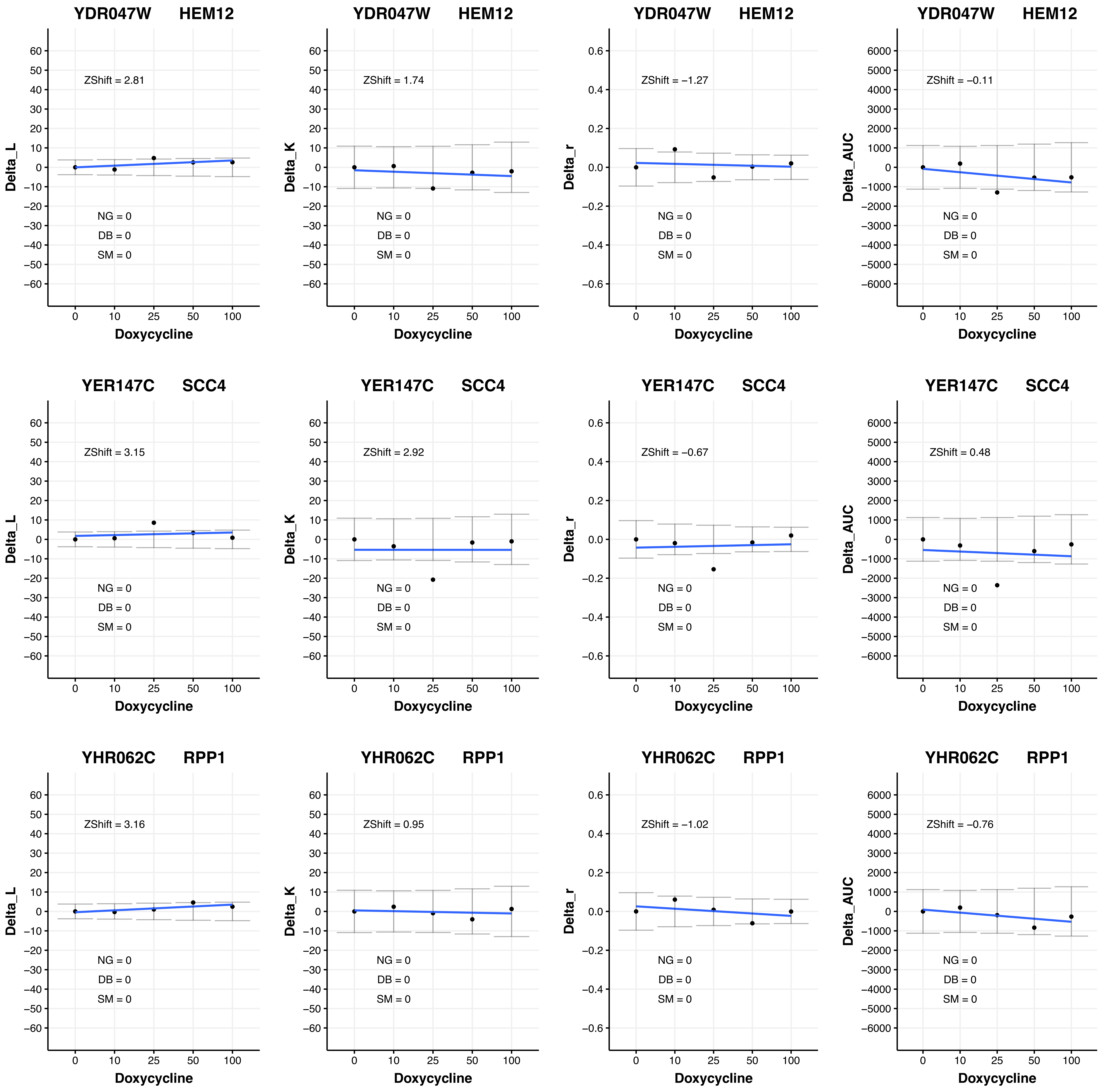
YDR047W HEM12
ZShift = 2.81
NG = 0
DB = 0
SM = 0
Delta_L
Doxycycline
YDR047W HEM12
ZShift = 1.74
NG = 0
DB = 0
SM = 0
Delta_K
Doxycycline
YDR047W HEM12
ZShift = −1.27
NG = 0
DB = 0
SM = 0
Delta_r
Doxycycline
YDR047W HEM12
ZShift = −0.11
NG = 0
DB = 0
SM = 0
Delta_AUC
Doxycycline
YER147C SCC4
ZShift = 3.15
NG = 0
DB = 0
SM = 0
Delta_L
Doxycycline
YER147C SCC4
ZShift = 2.92
NG = 0
DB = 0
SM = 0
Delta_K
Doxycycline
YER147C SCC4
ZShift = −0.67
NG = 0
DB = 0
SM = 0
Delta_r
Doxycycline
YER147C SCC4
ZShift = 0.48
NG = 0
DB = 0
SM = 0
Delta_AUC
Doxycycline
YHR062C RPP1
ZShift = 3.16
NG = 0
DB = 0
SM = 0
Delta_L
Doxycycline
YHR062C RPP1
ZShift = 0.95
NG = 0
DB = 0
SM = 0
Delta_K
Doxycycline
YHR062C RPP1
ZShift = −1.02
NG = 0
DB = 0
SM = 0
Delta_r
Doxycycline
YHR062C RPP1
ZShift = −0.76
NG = 0
DB = 0
SM = 0
Delta_AUC
Doxycycline

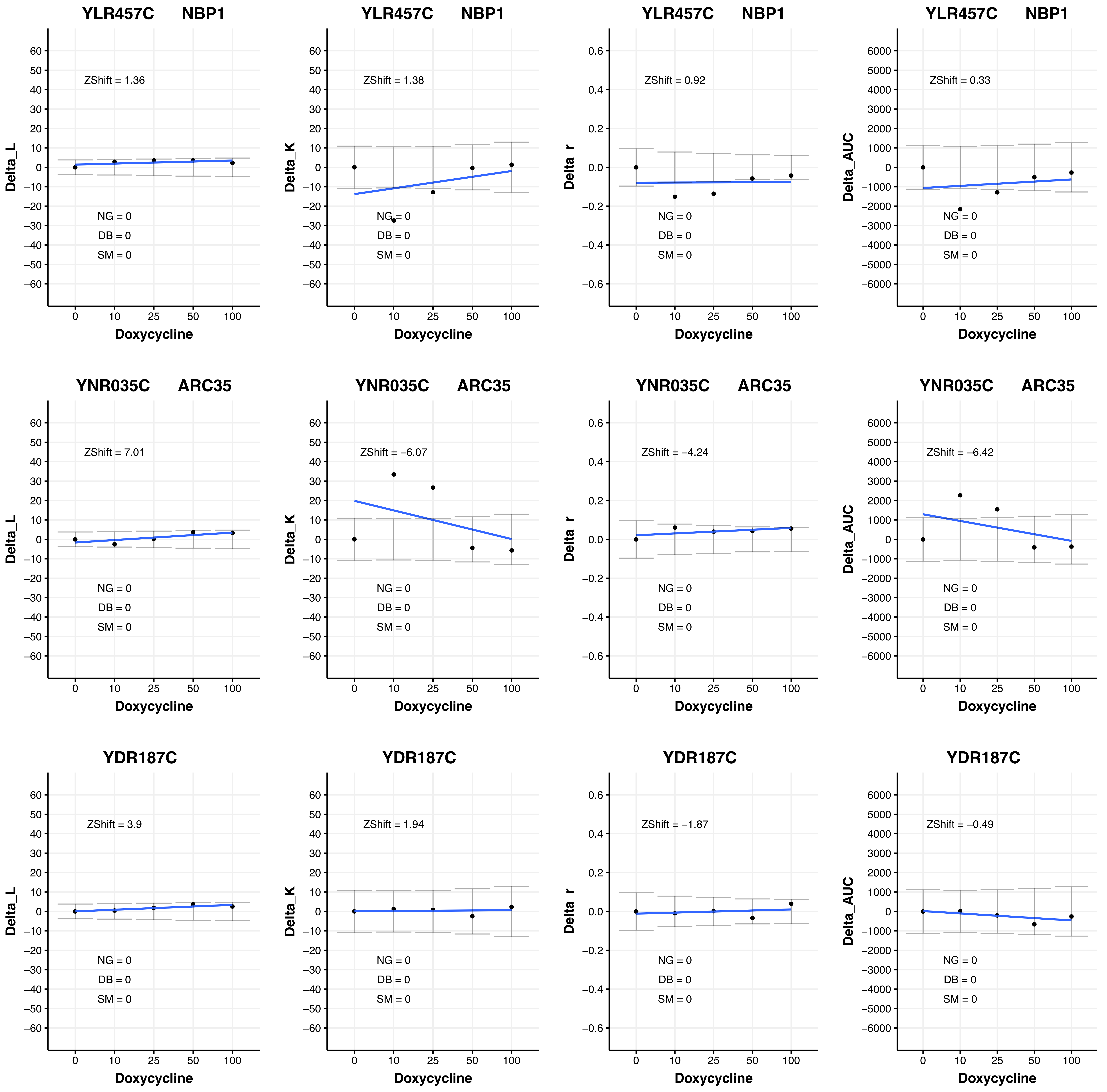

YLR457C NBP1
ZShift = 1.36
NG = 0
DB = 0
SM = 0
Delta_L
Doxycycline
YLR457C NBP1
ZShift = 1.38
NG = 0
DB = 0
SM = 0
Delta_K
Doxycycline
YLR457C NBP1
ZShift = 0.92
NG = 0
DB = 0
SM = 0
Delta_r
Doxycycline
YLR457C NBP1
ZShift = 0.33
NG = 0
DB = 0
SM = 0
Delta_AUC
Doxycycline
YNR035C ARC35
ZShift = 7.01
NG = 0
DB = 0
SM = 0
Delta_L
Doxycycline
YNR035C ARC35
ZShift = −6.07
NG = 0
DB = 0
SM = 0
Delta_K
Doxycycline
YNR035C ARC35
ZShift = −4.24
NG = 0
DB = 0
SM = 0
Delta_r
Doxycycline
YNR035C ARC35
ZShift = −6.42
NG = 0
DB = 0
SM = 0
Delta_AUC
Doxycycline
YDR187C
ZShift = 3.9
NG = 0
DB = 0
SM = 0
Delta_L
Doxycycline
YDR187C
ZShift = 1.94
NG = 0
DB = 0
SM = 0
Delta_K
Doxycycline
YDR187C
ZShift = −1.87
NG = 0
DB = 0
SM = 0
Delta_r
Doxycycline
YDR187C
ZShift = −0.49
NG = 0
DB = 0
SM = 0
Delta_AUC
Doxycycline

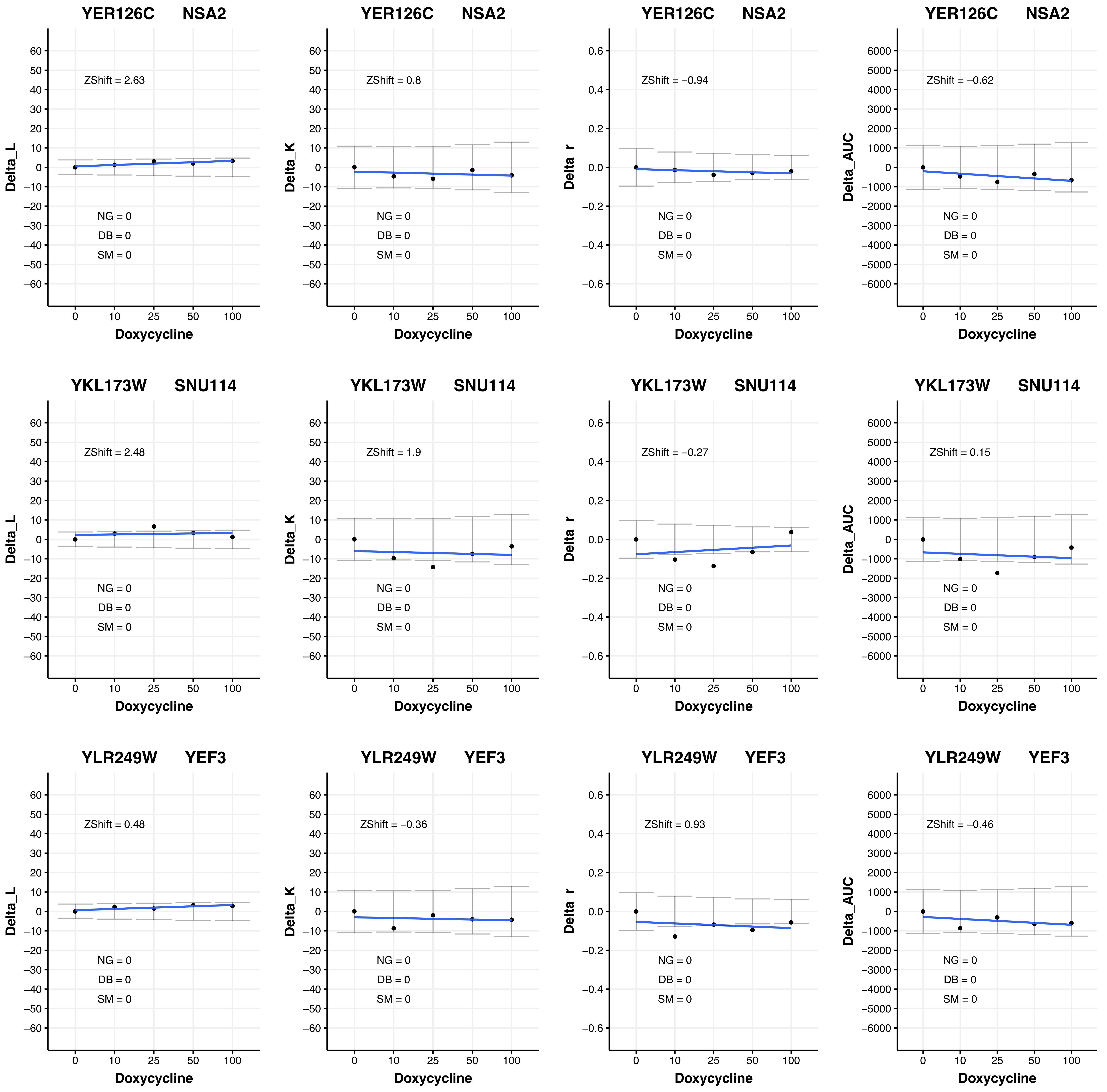

YER126C NSA2
ZShift = 2.63
NG = 0
DB = 0
SM = 0
Delta_L
Doxycycline
YER126C NSA2
ZShift = 0.8
NG = 0
DB = 0
SM = 0
Delta_K
Doxycycline
YER126C NSA2
ZShift = −0.94
NG = 0
DB = 0
SM = 0
Delta_r
Doxycycline
YER126C NSA2
ZShift = −0.62
NG = 0
DB = 0
SM = 0
Delta_AUC
Doxycycline
YKL173W SNU114
ZShift = 2.48
NG = 0
DB = 0
SM = 0
Delta_L
Doxycycline
YKL173W SNU114
ZShift = 1.9
NG = 0
DB = 0
SM = 0
Delta_K
Doxycycline
YKL173W SNU114
ZShift = −0.27
NG = 0
DB = 0
SM = 0
Delta_r
Doxycycline
YKL173W SNU114
ZShift = 0.15
NG = 0
DB = 0
SM = 0
Delta_AUC
Doxycycline
YLR249W YEF3
ZShift = 0.48
NG = 0
DB = 0
SM = 0
Delta_L
Doxycycline
YLR249W YEF3
ZShift = −0.36
NG = 0
DB = 0
SM = 0
Delta_K
Doxycycline
YLR249W YEF3
ZShift = 0.93
NG = 0
DB = 0
SM = 0
Delta_r
Doxycycline
YLR249W YEF3
ZShift = −0.46
NG = 0
DB = 0
SM = 0
Delta_AUC
Doxycycline

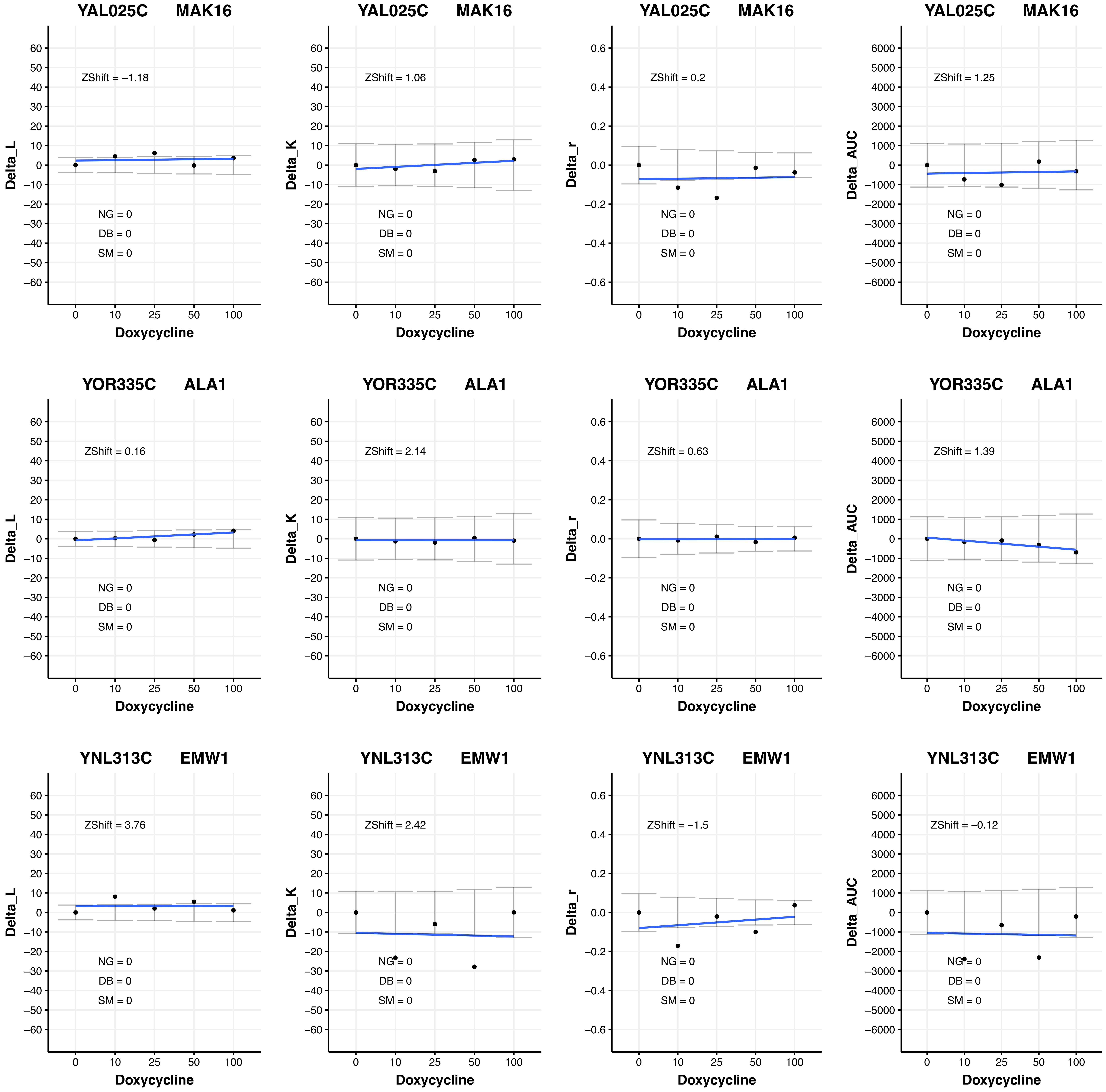
YAL025C MAK16
ZShift = −1.18
Delta_L
NG = 0
DB = 0
SM = 0
Doxycycline
YAL025C MAK16
ZShift = 1.06
Delta_K
NG = 0
DB = 0
SM = 0
Doxycycline
YAL025C MAK16
ZShift = 0.2
Delta_r
NG = 0
DB = 0
SM = 0
Doxycycline
YAL025C MAK16
ZShift = 1.25
Delta_AUC
NG = 0
DB = 0
SM = 0
Doxycycline
YOR335C ALA1
ZShift = 0.16
Delta_L
NG = 0
DB = 0
SM = 0
Doxycycline
YOR335C ALA1
ZShift = 2.14
Delta_K
NG = 0
DB = 0
SM = 0
Doxycycline
YOR335C ALA1
ZShift = 0.63
Delta_r
NG = 0
DB = 0
SM = 0
Doxycycline
YOR335C ALA1
ZShift = 1.39
Delta_AUC
NG = 0
DB = 0
SM = 0
Doxycycline
YNL313C EMW1
ZShift = 3.76
Delta_L
NG = 0
DB = 0
SM = 0
Doxycycline
YNL313C EMW1
ZShift = 2.42
Delta_K
NG = 0
DB = 0
SM = 0
Doxycycline
YNL313C EMW1
ZShift = −1.5
Delta_r
NG = 0
DB = 0
SM = 0
Doxycycline
YNL313C EMW1
ZShift = −0.12
Delta_AUC
NG = 0
DB = 0
SM = 0
Doxycycline

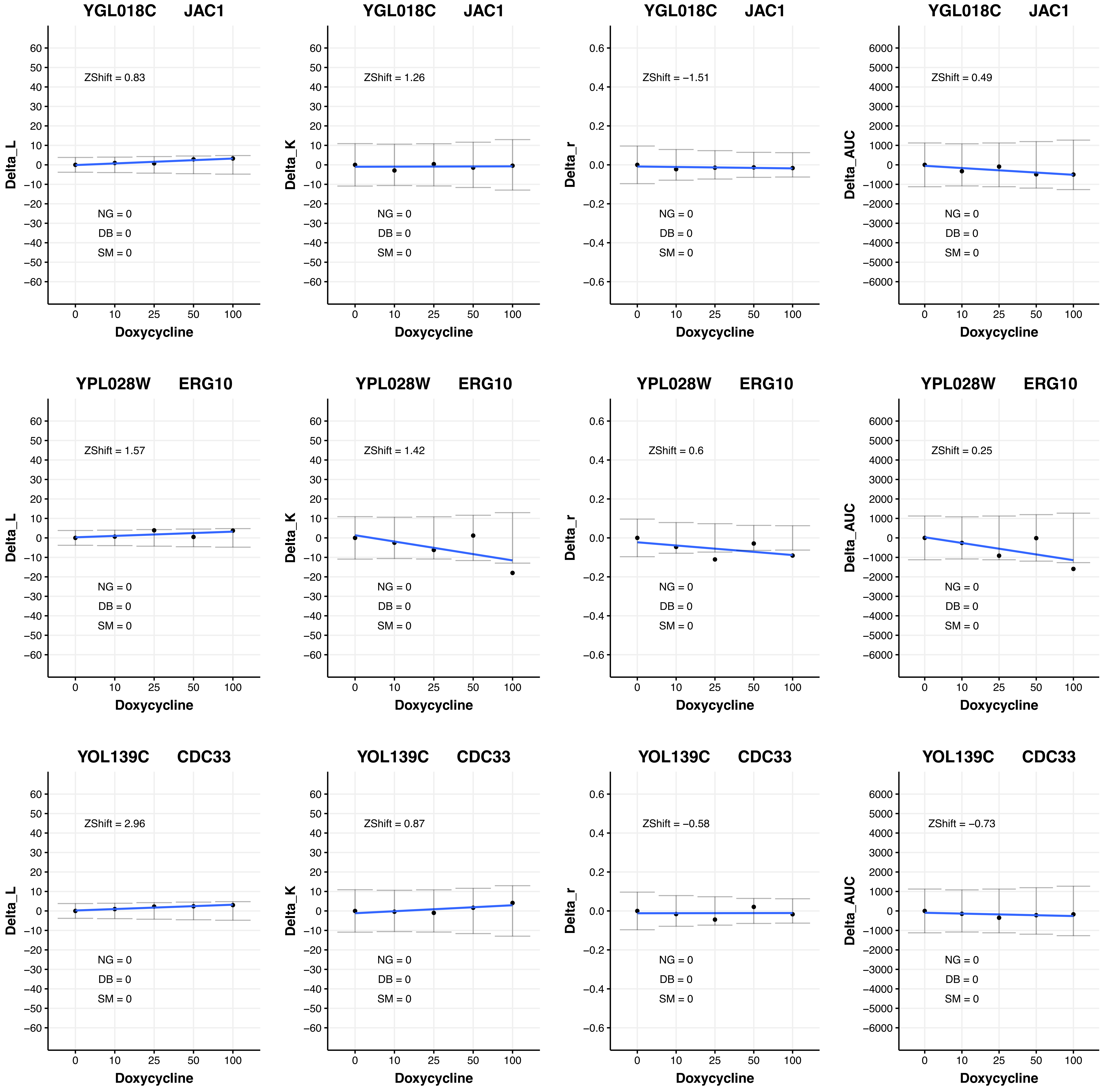

YGL018C JAC1
ZShift = 0.83
NG = 0
DB = 0
SM = 0
Delta_L
Doxycycline
YGL018C JAC1
ZShift = 1.26
NG = 0
DB = 0
SM = 0
Delta_K
Doxycycline
YGL018C JAC1
ZShift = −1.51
NG = 0
DB = 0
SM = 0
Delta_r
Doxycycline
YGL018C JAC1
ZShift = 0.49
NG = 0
DB = 0
SM = 0
Delta_AUC
Doxycycline
YPL028W ERG10
ZShift = 1.57
NG = 0
DB = 0
SM = 0
Delta_L
Doxycycline
YPL028W ERG10
ZShift = 1.42
NG = 0
DB = 0
SM = 0
Delta_K
Doxycycline
YPL028W ERG10
ZShift = 0.6
NG = 0
DB = 0
SM = 0
Delta_r
Doxycycline
YPL028W ERG10
ZShift = 0.25
NG = 0
DB = 0
SM = 0
Delta_AUC
Doxycycline
YOL139C CDC33
ZShift = 2.96
NG = 0
DB = 0
SM = 0
Delta_L
Doxycycline
YOL139C CDC33
ZShift = 0.87
NG = 0
DB = 0
SM = 0
Delta_K
Doxycycline
YOL139C CDC33
ZShift = −0.58
NG = 0
DB = 0
SM = 0
Delta_r
Doxycycline
YOL139C CDC33
ZShift = −0.73
NG = 0
DB = 0
SM = 0
Delta_AUC
Doxycycline

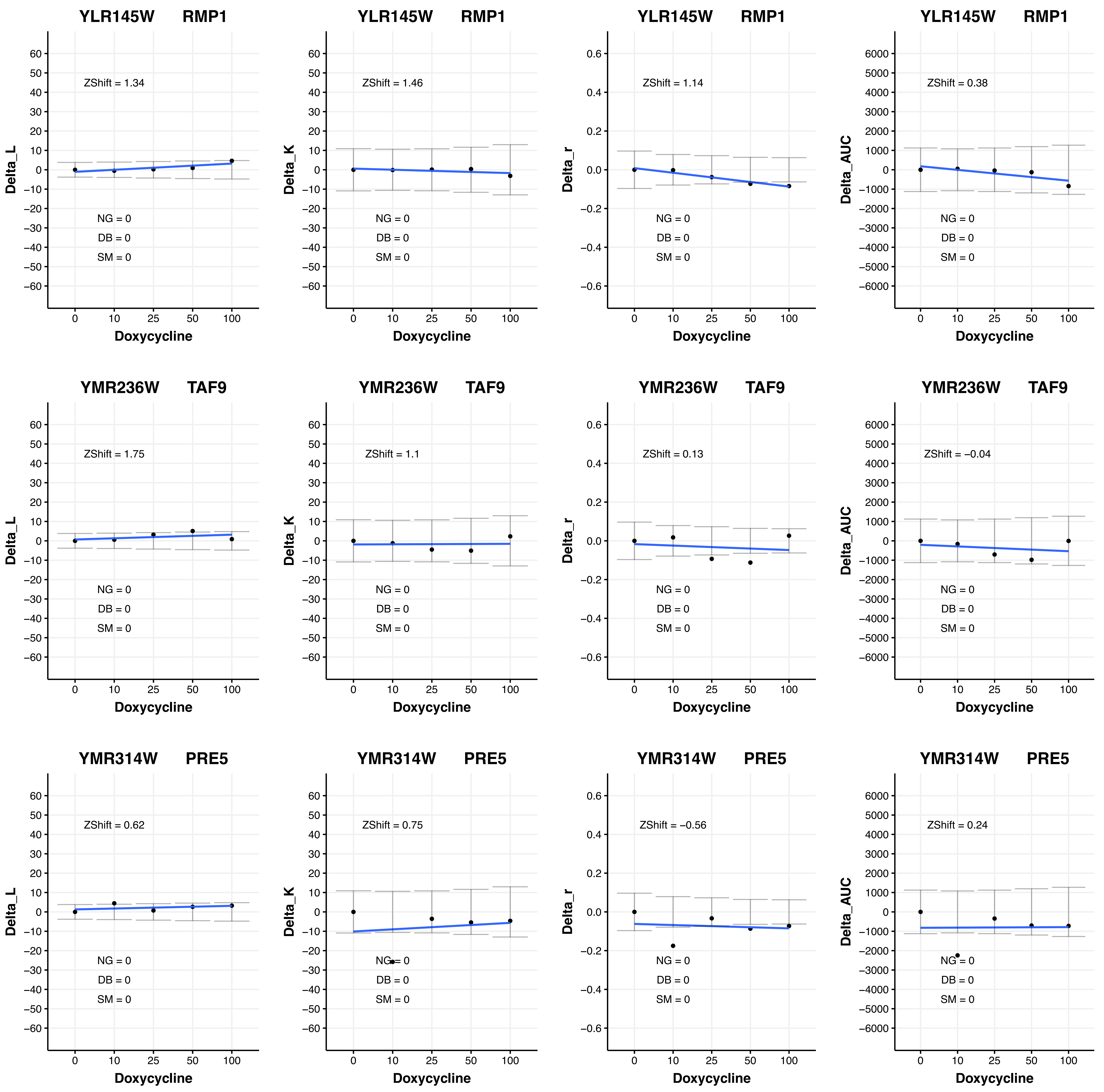
YLR145W RMP1
ZShift = 1.34
NG = 0
DB = 0
SM = 0
Delta_L
Doxycycline
YLR145W RMP1
ZShift = 1.46
NG = 0
DB = 0
SM = 0
Delta_K
Doxycycline
YLR145W RMP1
ZShift = 1.14
NG = 0
DB = 0
SM = 0
Delta_r
Doxycycline
YLR145W RMP1
ZShift = 0.38
NG = 0
DB = 0
SM = 0
Delta_AUC
Doxycycline
YMR236W TAF9
ZShift = 1.75
NG = 0
DB = 0
SM = 0
Delta_L
Doxycycline
YMR236W TAF9
ZShift = 1.1
NG = 0
DB = 0
SM = 0
Delta_K
Doxycycline
YMR236W TAF9
ZShift = 0.13
NG = 0
DB = 0
SM = 0
Delta_r
Doxycycline
YMR236W TAF9
ZShift = −0.04
NG = 0
DB = 0
SM = 0
Delta_AUC
Doxycycline
YMR314W PRE5
ZShift = 0.62
NG = 0
DB = 0
SM = 0
Delta_L
Doxycycline
YMR314W PRE5
ZShift = 0.75
NG = 0
DB = 0
SM = 0
Delta_K
Doxycycline
YMR314W PRE5
ZShift = −0.56
NG = 0
DB = 0
SM = 0
Delta_r
Doxycycline
YMR314W PRE5
ZShift = 0.24
NG = 0
DB = 0
SM = 0
Delta_AUC
Doxycycline

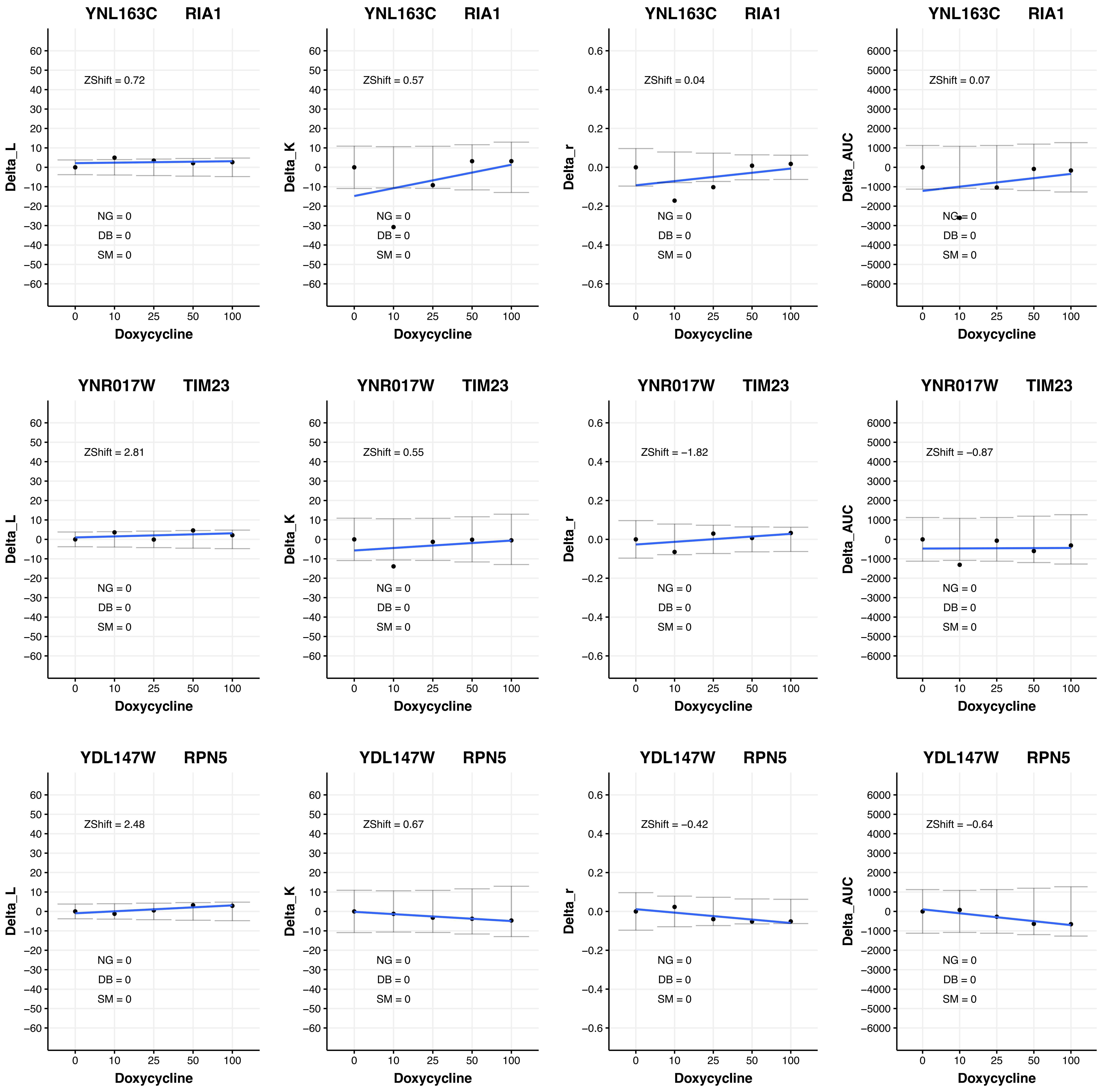
YNL163C RIA1
ZShift = 0.72
NG = 0
DB = 0
SM = 0
Delta_L
Doxycycline
YNL163C RIA1
ZShift = 0.57
NG = 0
DB = 0
SM = 0
Delta_K
Doxycycline
YNL163C RIA1
ZShift = 0.04
NG = 0
DB = 0
SM = 0
Delta_r
Doxycycline
YNL163C RIA1
ZShift = 0.07
NG = 0
DB = 0
SM = 0
Delta_AUC
Doxycycline
YNR017W TIM23
ZShift = 2.81
NG = 0
DB = 0
SM = 0
Delta_L
Doxycycline
YNR017W TIM23
ZShift = 0.55
NG = 0
DB = 0
SM = 0
Delta_K
Doxycycline
YNR017W TIM23
ZShift = −1.82
NG = 0
DB = 0
SM = 0
Delta_r
Doxycycline
YNR017W TIM23
ZShift = −0.87
NG = 0
DB = 0
SM = 0
Delta_AUC
Doxycycline
YDL147W RPN5
ZShift = 2.48
NG = 0
DB = 0
SM = 0
Delta_L
Doxycycline
YDL147W RPN5
ZShift = 0.67
NG = 0
DB = 0
SM = 0
Delta_K
Doxycycline
YDL147W RPN5
ZShift = −0.42
NG = 0
DB = 0
SM = 0
Delta_r
Doxycycline
YDL147W RPN5
ZShift = −0.64
NG = 0
DB = 0
SM = 0
Delta_AUC
Doxycycline

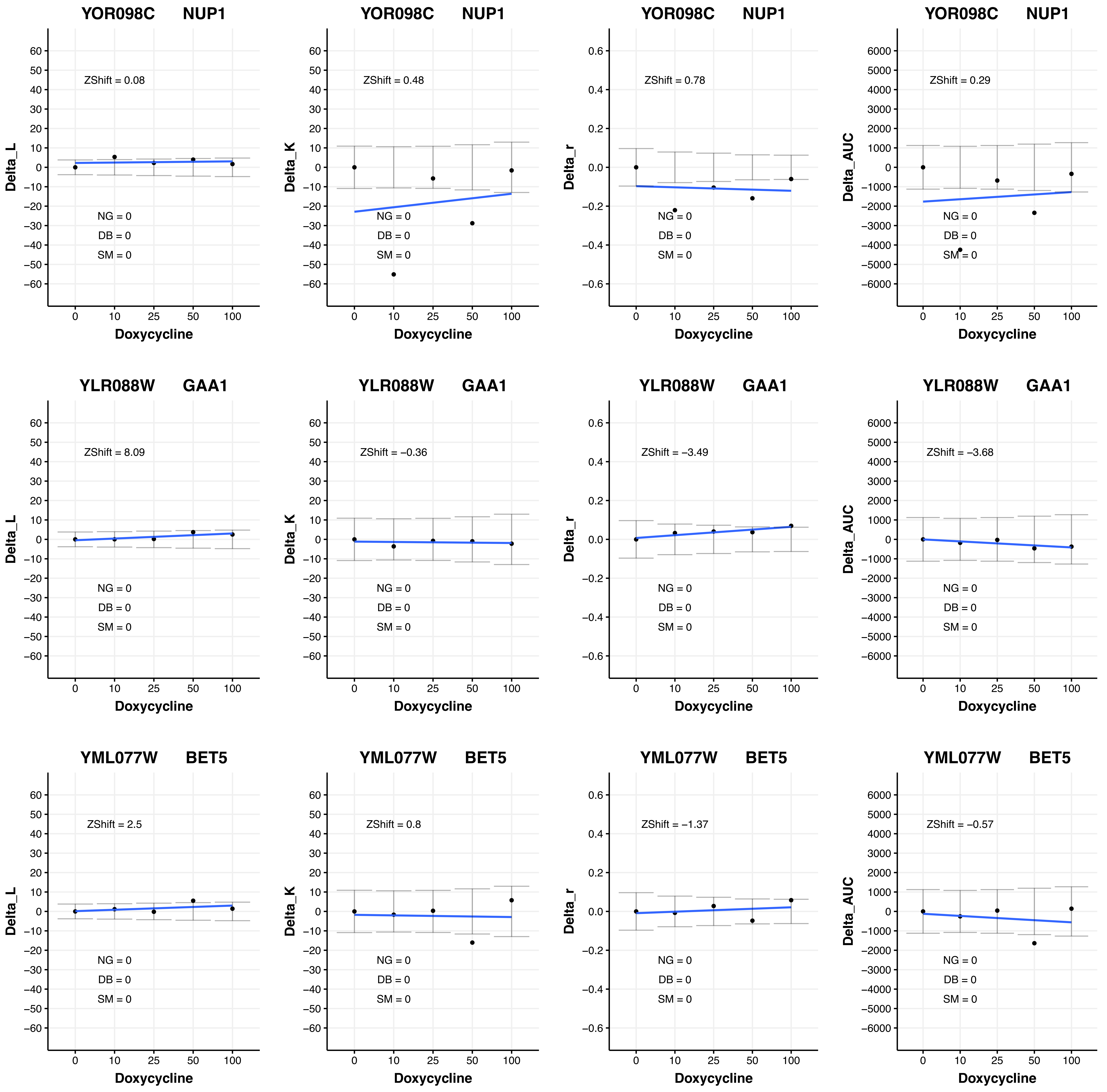
YOR098C NUP1
ZShift = 0.08
NG = 0
DB = 0
SM = 0
Delta_L
Doxycycline
YOR098C NUP1
ZShift = 0.48
NG = 0
DB = 0
SM = 0
Delta_K
Doxycycline
YOR098C NUP1
ZShift = 0.78
NG = 0
DB = 0
SM = 0
Delta_r
Doxycycline
YOR098C NUP1
ZShift = 0.29
NG = 0
DB = 0
SM = 0
Delta_AUC
Doxycycline
YLR088W GAA1
ZShift = 8.09
NG = 0
DB = 0
SM = 0
Delta_L
Doxycycline
YLR088W GAA1
ZShift = −0.36
NG = 0
DB = 0
SM = 0
Delta_K
Doxycycline
YLR088W GAA1
ZShift = −3.49
NG = 0
DB = 0
SM = 0
Delta_r
Doxycycline
YLR088W GAA1
ZShift = −3.68
NG = 0
DB = 0
SM = 0
Delta_AUC
Doxycycline
YML077W BET5
ZShift = 2.5
NG = 0
DB = 0
SM = 0
Delta_L
Doxycycline
YML077W BET5
ZShift = 0.8
NG = 0
DB = 0
SM = 0
Delta_K
Doxycycline
YML077W BET5
ZShift = −1.37
NG = 0
DB = 0
SM = 0
Delta_r
Doxycycline
YML077W BET5
ZShift = −0.57
NG = 0
DB = 0
SM = 0
Delta_AUC
Doxycycline

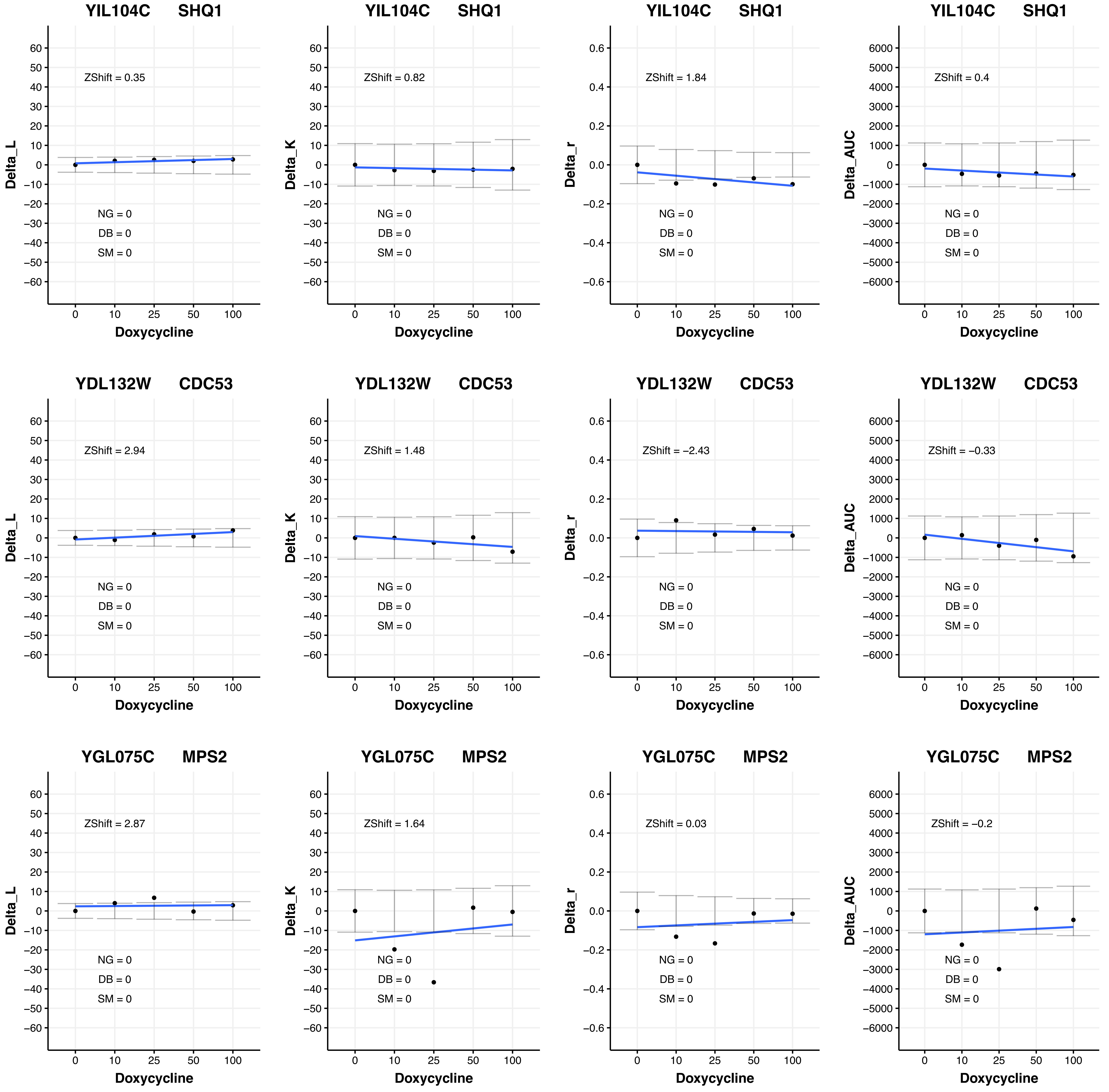
YIL104C SHQ1
ZShift = 0.35
NG = 0
DB = 0
SM = 0
Delta_L
Doxycycline
YIL104C SHQ1
ZShift = 0.82
NG = 0
DB = 0
SM = 0
Delta_K
Doxycycline
YIL104C SHQ1
ZShift = 1.84
NG = 0
DB = 0
SM = 0
Delta_r
Doxycycline
YIL104C SHQ1
ZShift = 0.4
NG = 0
DB = 0
SM = 0
Delta_AUC
Doxycycline
YDL132W CDC53
ZShift = 2.94
NG = 0
DB = 0
SM = 0
Delta_L
Doxycycline
YDL132W CDC53
ZShift = 1.48
NG = 0
DB = 0
SM = 0
Delta_K
Doxycycline
YDL132W CDC53
ZShift = −2.43
NG = 0
DB = 0
SM = 0
Delta_r
Doxycycline
YDL132W CDC53
ZShift = −0.33
NG = 0
DB = 0
SM = 0
Delta_AUC
Doxycycline
YGL075C MPS2
ZShift = 2.87
NG = 0
DB = 0
SM = 0
Delta_L
Doxycycline
YGL075C MPS2
ZShift = 1.64
NG = 0
DB = 0
SM = 0
Delta_K
Doxycycline
YGL075C MPS2
ZShift = 0.03
NG = 0
DB = 0
SM = 0
Delta_r
Doxycycline
YGL075C MPS2
ZShift = −0.2
NG = 0
DB = 0
SM = 0
Delta_AUC
Doxycycline

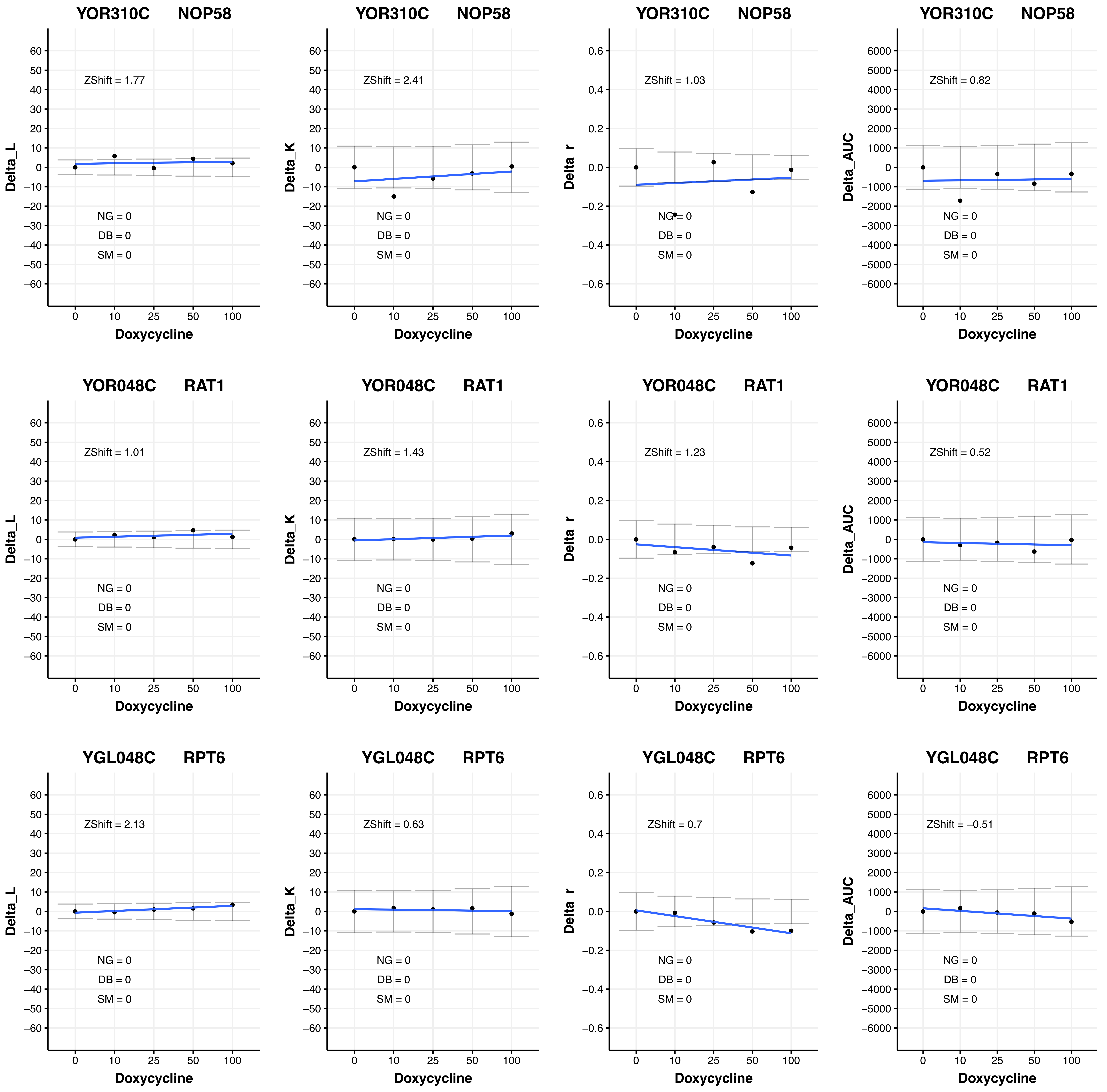
YOR310C NOP58
ZShift = 1.77
Delta_L
NG = 0
DB = 0
SM = 0
Doxycycline
YOR310C NOP58
ZShift = 2.41
Delta_K
NG = 0
DB = 0
SM = 0
Doxycycline
YOR310C NOP58
ZShift = 1.03
Delta_r
NG = 0
DB = 0
SM = 0
Doxycycline
YOR310C NOP58
ZShift = 0.82
Delta_AUC
NG = 0
DB = 0
SM = 0
Doxycycline
YOR048C RAT1
ZShift = 1.01
Delta_L
NG = 0
DB = 0
SM = 0
Doxycycline
YOR048C RAT1
ZShift = 1.43
Delta_K
NG = 0
DB = 0
SM = 0
Doxycycline
YOR048C RAT1
ZShift = 1.23
Delta_r
NG = 0
DB = 0
SM = 0
Doxycycline
YOR048C RAT1
ZShift = 0.52
Delta_AUC
NG = 0
DB = 0
SM = 0
Doxycycline
YGL048C RPT6
ZShift = 2.13
Delta_L
NG = 0
DB = 0
SM = 0
Doxycycline
YGL048C RPT6
ZShift = 0.63
Delta_K
NG = 0
DB = 0
SM = 0
Doxycycline
YGL048C RPT6
ZShift = 0.7
Delta_r
NG = 0
DB = 0
SM = 0
Doxycycline
YGL048C RPT6
ZShift = −0.51
Delta_AUC
NG = 0
DB = 0
SM = 0
Doxycycline

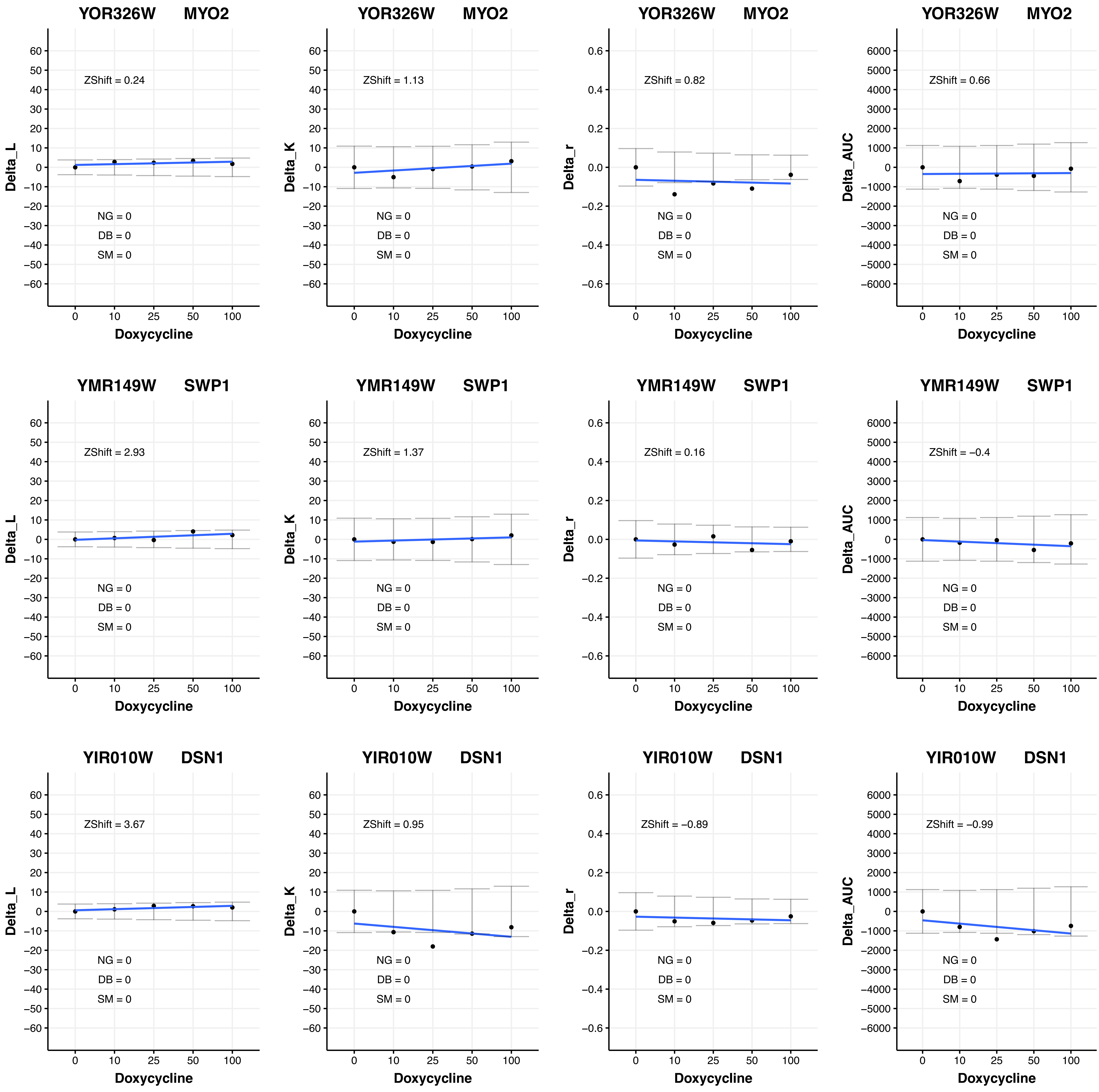
YOR326W MYO2
Delta_L
ZShift = 0.24
NG = 0
DB = 0
SM = 0
Doxycycline
YOR326W MYO2
Delta_K
ZShift = 1.13
NG = 0
DB = 0
SM = 0
Doxycycline
YOR326W MYO2
Delta_r
ZShift = 0.82
NG = 0
DB = 0
SM = 0
Doxycycline
YOR326W MYO2
Delta_AUC
ZShift = 0.66
NG = 0
DB = 0
SM = 0
Doxycycline
YMR149W SWP1
Delta_L
ZShift = 2.93
NG = 0
DB = 0
SM = 0
Doxycycline
YMR149W SWP1
Delta_K
ZShift = 1.37
NG = 0
DB = 0
SM = 0
Doxycycline
YMR149W SWP1
Delta_r
ZShift = 0.16
NG = 0
DB = 0
SM = 0
Doxycycline
YMR149W SWP1
Delta_AUC
ZShift = −0.4
NG = 0
DB = 0
SM = 0
Doxycycline
YIR010W DSN1
Delta_L
ZShift = 3.67
NG = 0
DB = 0
SM = 0
Doxycycline
YIR010W DSN1
Delta_K
ZShift = 0.95
NG = 0
DB = 0
SM = 0
Doxycycline
YIR010W DSN1
Delta_r
ZShift = −0.89
NG = 0
DB = 0
SM = 0
Doxycycline
YIR010W DSN1
Delta_AUC
ZShift = −0.99
NG = 0
DB = 0
SM = 0
Doxycycline

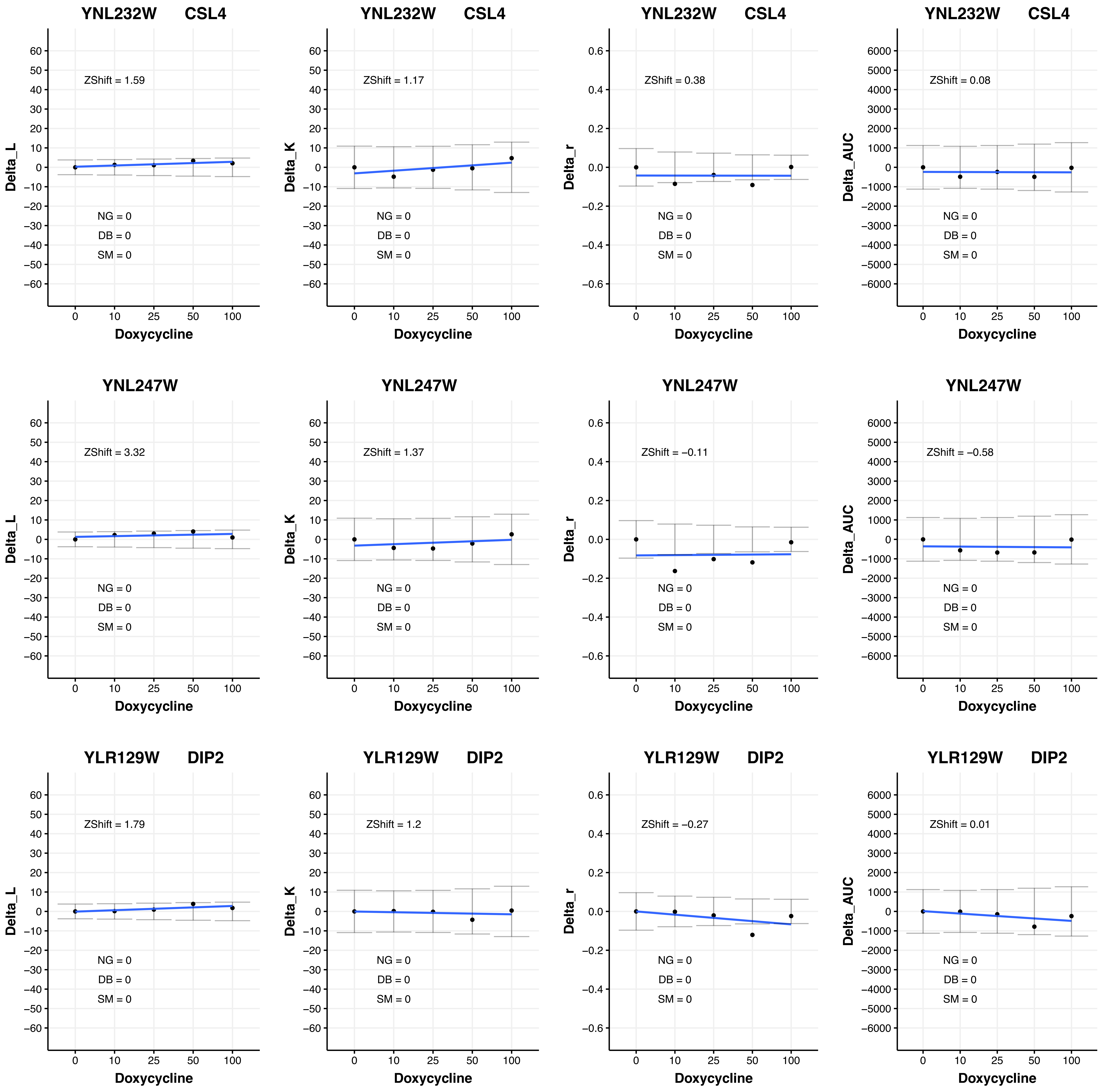
YNL232W CSL4
ZShift = 1.59
Delta_L
NG = 0
DB = 0
SM = 0
Doxycycline
YNL232W CSL4
ZShift = 1.17
Delta_K
NG = 0
DB = 0
SM = 0
Doxycycline
YNL232W CSL4
ZShift = 0.38
Delta_r
NG = 0
DB = 0
SM = 0
Doxycycline
YNL232W CSL4
ZShift = 0.08
Delta_AUC
NG = 0
DB = 0
SM = 0
Doxycycline
YNL247W
ZShift = 3.32
Delta_L
NG = 0
DB = 0
SM = 0
Doxycycline
YNL247W
ZShift = 1.37
Delta_K
NG = 0
DB = 0
SM = 0
Doxycycline
YNL247W
ZShift = −0.11
Delta_r
NG = 0
DB = 0
SM = 0
Doxycycline
YNL247W
ZShift = −0.58
Delta_AUC
NG = 0
DB = 0
SM = 0
Doxycycline
YLR129W DIP2
ZShift = 1.79
Delta_L
NG = 0
DB = 0
SM = 0
Doxycycline
YLR129W DIP2
ZShift = 1.2
Delta_K
NG = 0
DB = 0
SM = 0
Doxycycline
YLR129W DIP2
ZShift = −0.27
Delta_r
NG = 0
DB = 0
SM = 0
Doxycycline
YLR129W DIP2
ZShift = 0.01
Delta_AUC
NG = 0
DB = 0
SM = 0
Doxycycline

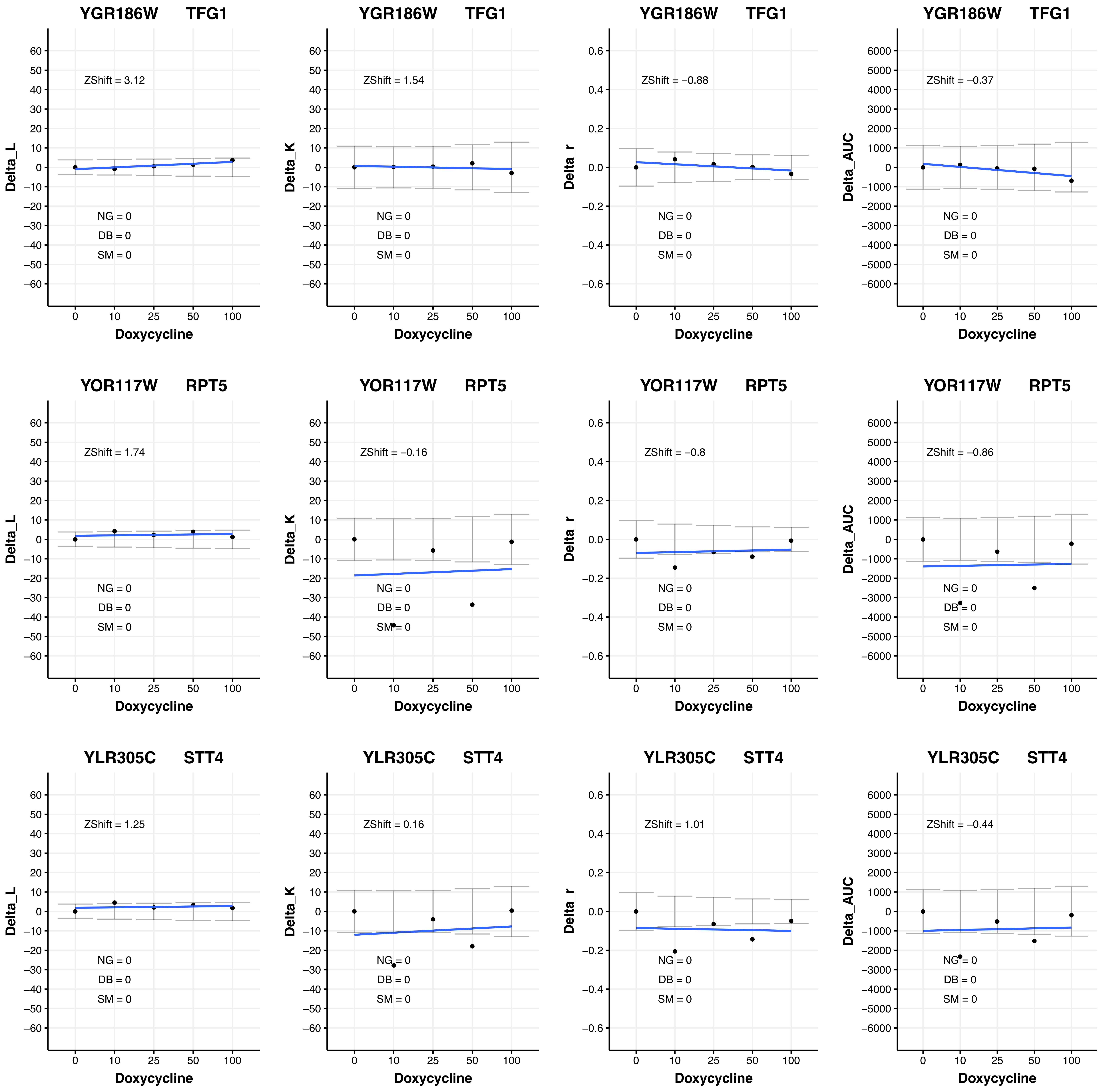
YGR186W TFG1
ZShift = 3.12
ZShift = 1.54
ZShift = −0.88
ZShift = −0.37
YOR117W RPT5
ZShift = 1.74
ZShift = −0.16
ZShift = −0.8
ZShift = −0.86
YLR305C STT4
ZShift = 1.25
ZShift = 0.16
ZShift = 1.01
ZShift = −0.44
NG = 0
DB = 0
SM = 0
Delta_L
Delta_K
Delta_r
Delta_AUC
Doxycycline
0 10 25 50 100

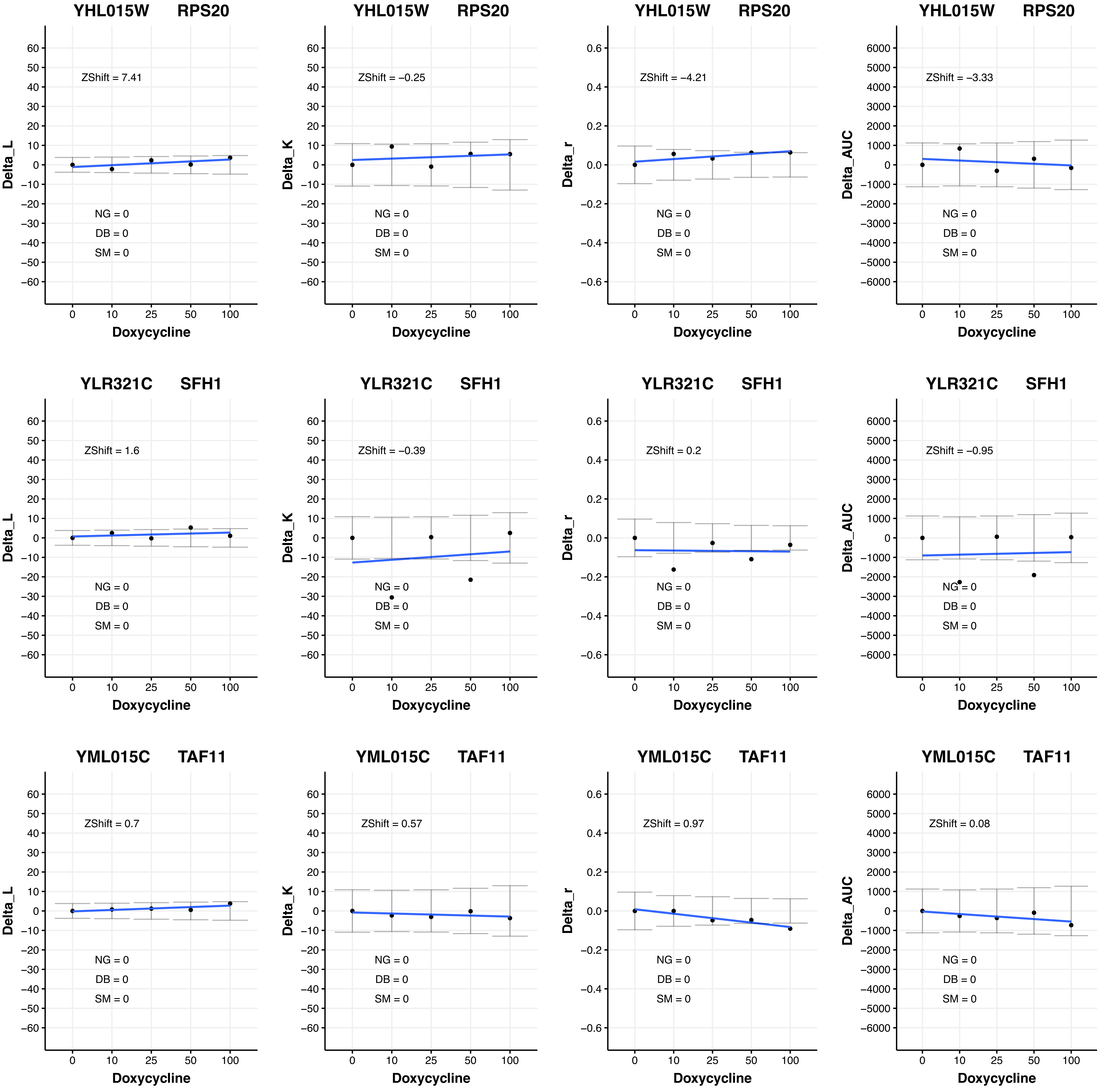
YHL015W RPS20
ZShift = 7.41
ZShift = −0.25
ZShift = −4.21
ZShift = −3.33
YLR321C SFH1
ZShift = 1.6
ZShift = −0.39
ZShift = 0.2
ZShift = −0.95
YML015C TAF11
ZShift = 0.7
ZShift = 0.57
ZShift = 0.97
ZShift = 0.08
NG = 0
DB = 0
SM = 0
Delta_L
Delta_K
Delta_r
Delta_AUC
Doxycycline
0
10
25
50
100

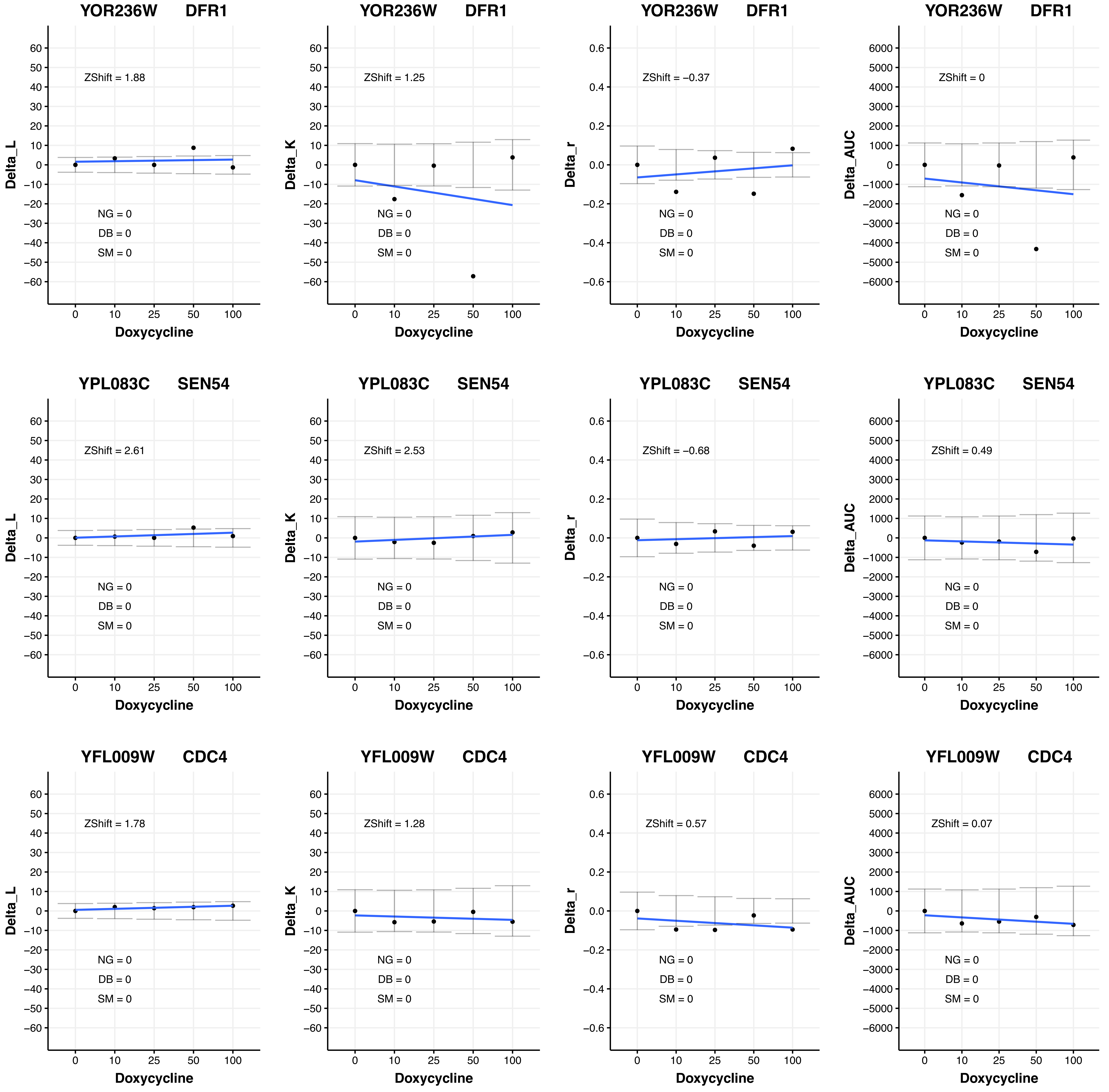
YOR236W DFR1
ZShift = 1.88
NG = 0
DB = 0
SM = 0
Delta_L
Doxycycline
YOR236W DFR1
ZShift = 1.25
NG = 0
DB = 0
SM = 0
Delta_K
Doxycycline
YOR236W DFR1
ZShift = −0.37
NG = 0
DB = 0
SM = 0
Delta_r
Doxycycline
YOR236W DFR1
ZShift = 0
NG = 0
DB = 0
SM = 0
Delta_AUC
Doxycycline
YPL083C SEN54
ZShift = 2.61
NG = 0
DB = 0
SM = 0
Delta_L
Doxycycline
YPL083C SEN54
ZShift = 2.53
NG = 0
DB = 0
SM = 0
Delta_K
Doxycycline
YPL083C SEN54
ZShift = −0.68
NG = 0
DB = 0
SM = 0
Delta_r
Doxycycline
YPL083C SEN54
ZShift = 0.49
NG = 0
DB = 0
SM = 0
Delta_AUC
Doxycycline
YFL009W CDC4
ZShift = 1.78
NG = 0
DB = 0
SM = 0
Delta_L
Doxycycline
YFL009W CDC4
ZShift = 1.28
NG = 0
DB = 0
SM = 0
Delta_K
Doxycycline
YFL009W CDC4
ZShift = 0.57
NG = 0
DB = 0
SM = 0
Delta_r
Doxycycline
YFL009W CDC4
ZShift = 0.07
NG = 0
DB = 0
SM = 0
Delta_AUC
Doxycycline

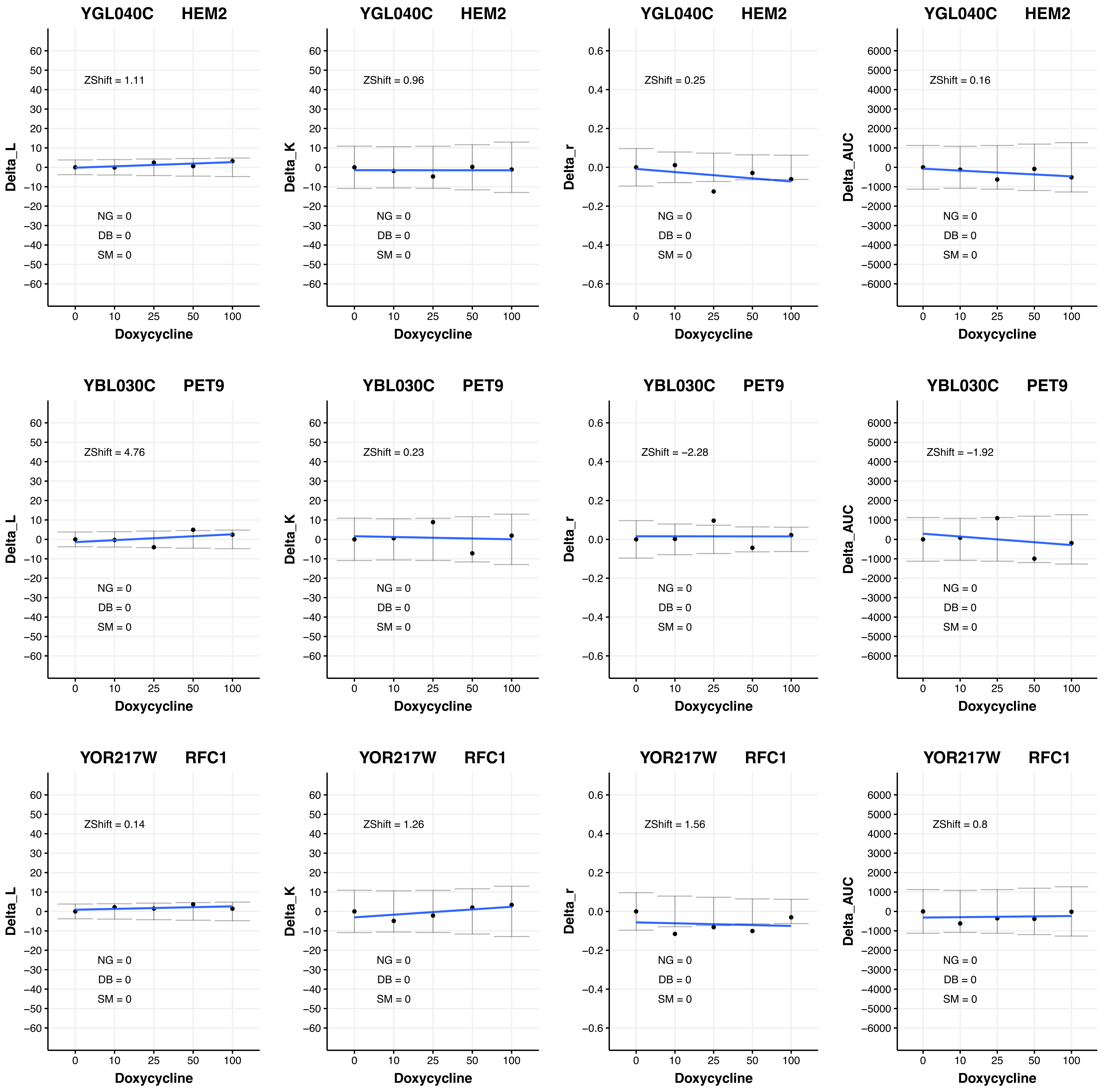
YGL040C HEM2
Delta_L
ZShift = 1.11
NG = 0
DB = 0
SM = 0
Doxycycline
YGL040C HEM2
Delta_K
ZShift = 0.96
NG = 0
DB = 0
SM = 0
Doxycycline
YGL040C HEM2
Delta_r
ZShift = 0.25
NG = 0
DB = 0
SM = 0
Doxycycline
YGL040C HEM2
Delta_AUC
ZShift = 0.16
NG = 0
DB = 0
SM = 0
Doxycycline
YBL030C PET9
Delta_L
ZShift = 4.76
NG = 0
DB = 0
SM = 0
Doxycycline
YBL030C PET9
Delta_K
ZShift = 0.23
NG = 0
DB = 0
SM = 0
Doxycycline
YBL030C PET9
Delta_r
ZShift = −2.28
NG = 0
DB = 0
SM = 0
Doxycycline
YBL030C PET9
Delta_AUC
ZShift = −1.92
NG = 0
DB = 0
SM = 0
Doxycycline
YOR217W RFC1
Delta_L
ZShift = 0.14
NG = 0
DB = 0
SM = 0
Doxycycline
YOR217W RFC1
Delta_K
ZShift = 1.26
NG = 0
DB = 0
SM = 0
Doxycycline
YOR217W RFC1
Delta_r
ZShift = 1.56
NG = 0
DB = 0
SM = 0
Doxycycline
YOR217W RFC1
Delta_AUC
ZShift = 0.8
NG = 0
DB = 0
SM = 0
Doxycycline

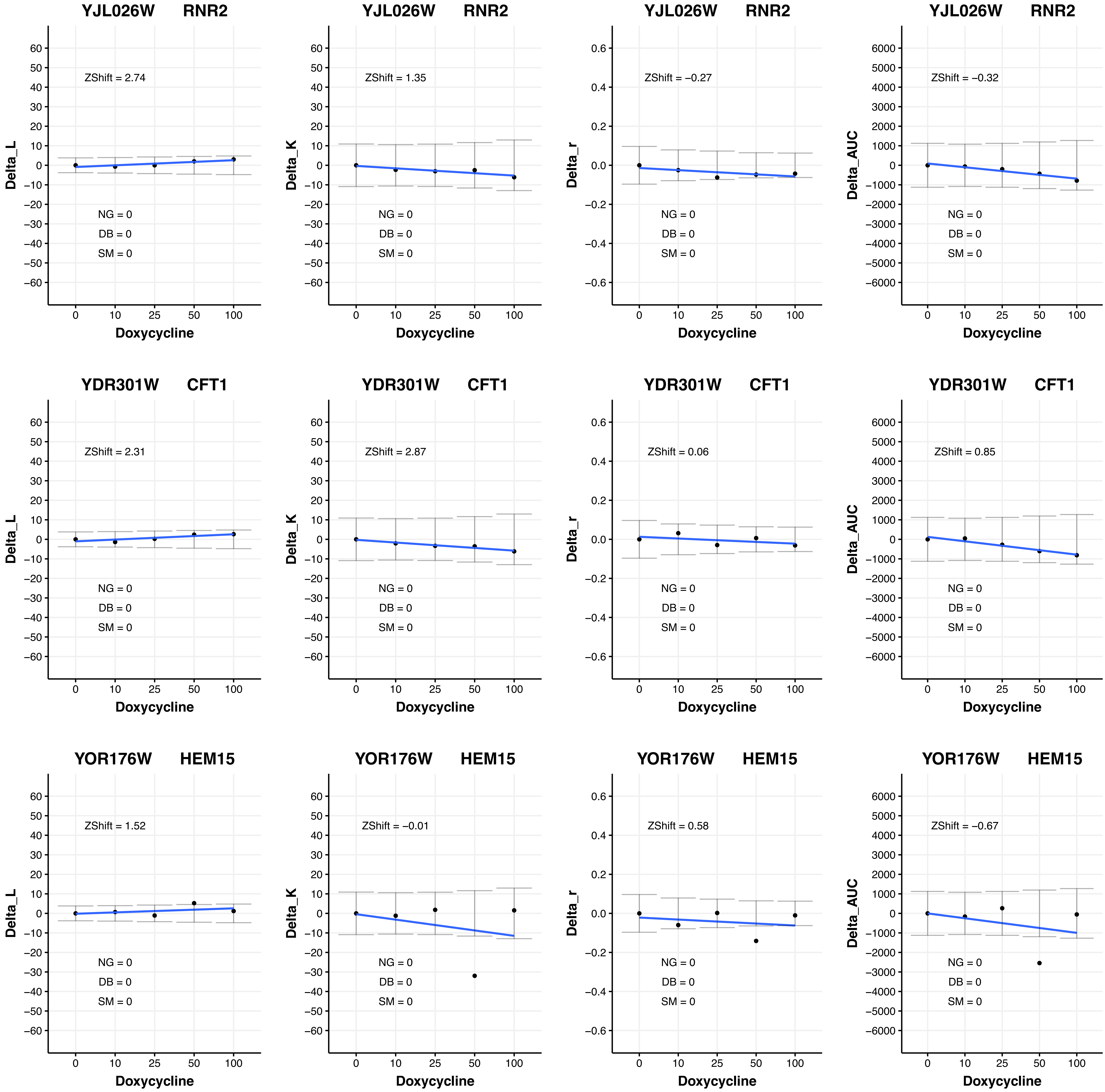
YJL026W RNR2
Delta_L
ZShift = 2.74
NG = 0
DB = 0
SM = 0
Doxycycline
YJL026W RNR2
Delta_K
ZShift = 1.35
NG = 0
DB = 0
SM = 0
Doxycycline
YJL026W RNR2
Delta_r
ZShift = −0.27
NG = 0
DB = 0
SM = 0
Doxycycline
YJL026W RNR2
Delta_AUC
ZShift = −0.32
NG = 0
DB = 0
SM = 0
Doxycycline
YDR301W CFT1
Delta_L
ZShift = 2.31
NG = 0
DB = 0
SM = 0
Doxycycline
YDR301W CFT1
Delta_K
ZShift = 2.87
NG = 0
DB = 0
SM = 0
Doxycycline
YDR301W CFT1
Delta_r
ZShift = 0.06
NG = 0
DB = 0
SM = 0
Doxycycline
YDR301W CFT1
Delta_AUC
ZShift = 0.85
NG = 0
DB = 0
SM = 0
Doxycycline
YOR176W HEM15
Delta_L
ZShift = 1.52
NG = 0
DB = 0
SM = 0
Doxycycline
YOR176W HEM15
Delta_K
ZShift = −0.01
NG = 0
DB = 0
SM = 0
Doxycycline
YOR176W HEM15
Delta_r
ZShift = 0.58
NG = 0
DB = 0
SM = 0
Doxycycline
YOR176W HEM15
Delta_AUC
ZShift = −0.67
NG = 0
DB = 0
SM = 0
Doxycycline

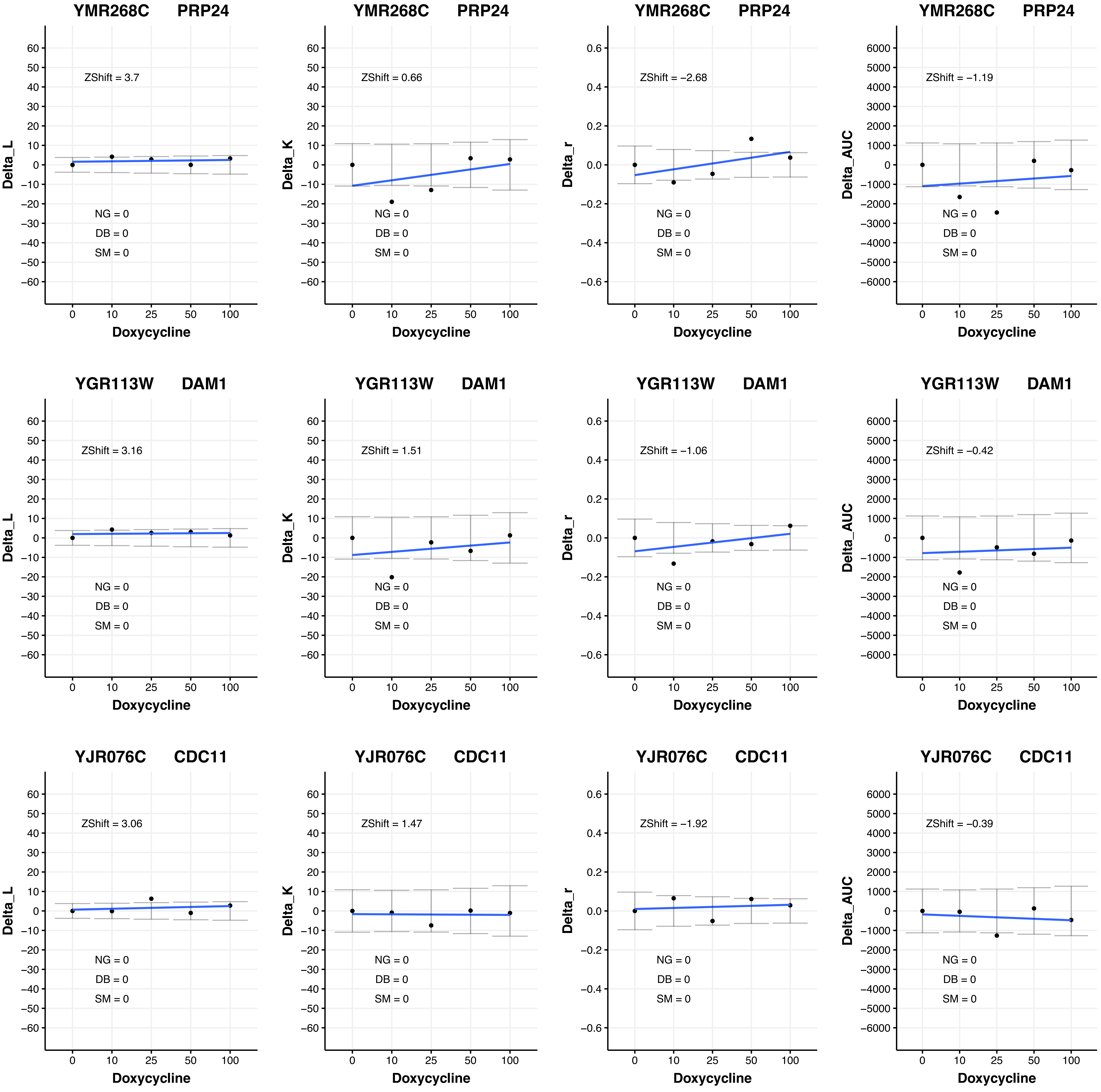
YMR268C PRP24
ZShift = 3.7
NG = 0
DB = 0
SM = 0
Delta_L
Doxycycline
YMR268C PRP24
ZShift = 0.66
NG = 0
DB = 0
SM = 0
Delta_K
Doxycycline
YMR268C PRP24
ZShift = −2.68
NG = 0
DB = 0
SM = 0
Delta_r
Doxycycline
YMR268C PRP24
ZShift = −1.19
NG = 0
DB = 0
SM = 0
Delta_AUC
Doxycycline
YGR113W DAM1
ZShift = 3.16
NG = 0
DB = 0
SM = 0
Delta_L
Doxycycline
YGR113W DAM1
ZShift = 1.51
NG = 0
DB = 0
SM = 0
Delta_K
Doxycycline
YGR113W DAM1
ZShift = −1.06
NG = 0
DB = 0
SM = 0
Delta_r
Doxycycline
YGR113W DAM1
ZShift = −0.42
NG = 0
DB = 0
SM = 0
Delta_AUC
Doxycycline
YJR076C CDC11
ZShift = 3.06
NG = 0
DB = 0
SM = 0
Delta_L
Doxycycline
YJR076C CDC11
ZShift = 1.47
NG = 0
DB = 0
SM = 0
Delta_K
Doxycycline
YJR076C CDC11
ZShift = −1.92
NG = 0
DB = 0
SM = 0
Delta_r
Doxycycline
YJR076C CDC11
ZShift = −0.39
NG = 0
DB = 0
SM = 0
Delta_AUC
Doxycycline

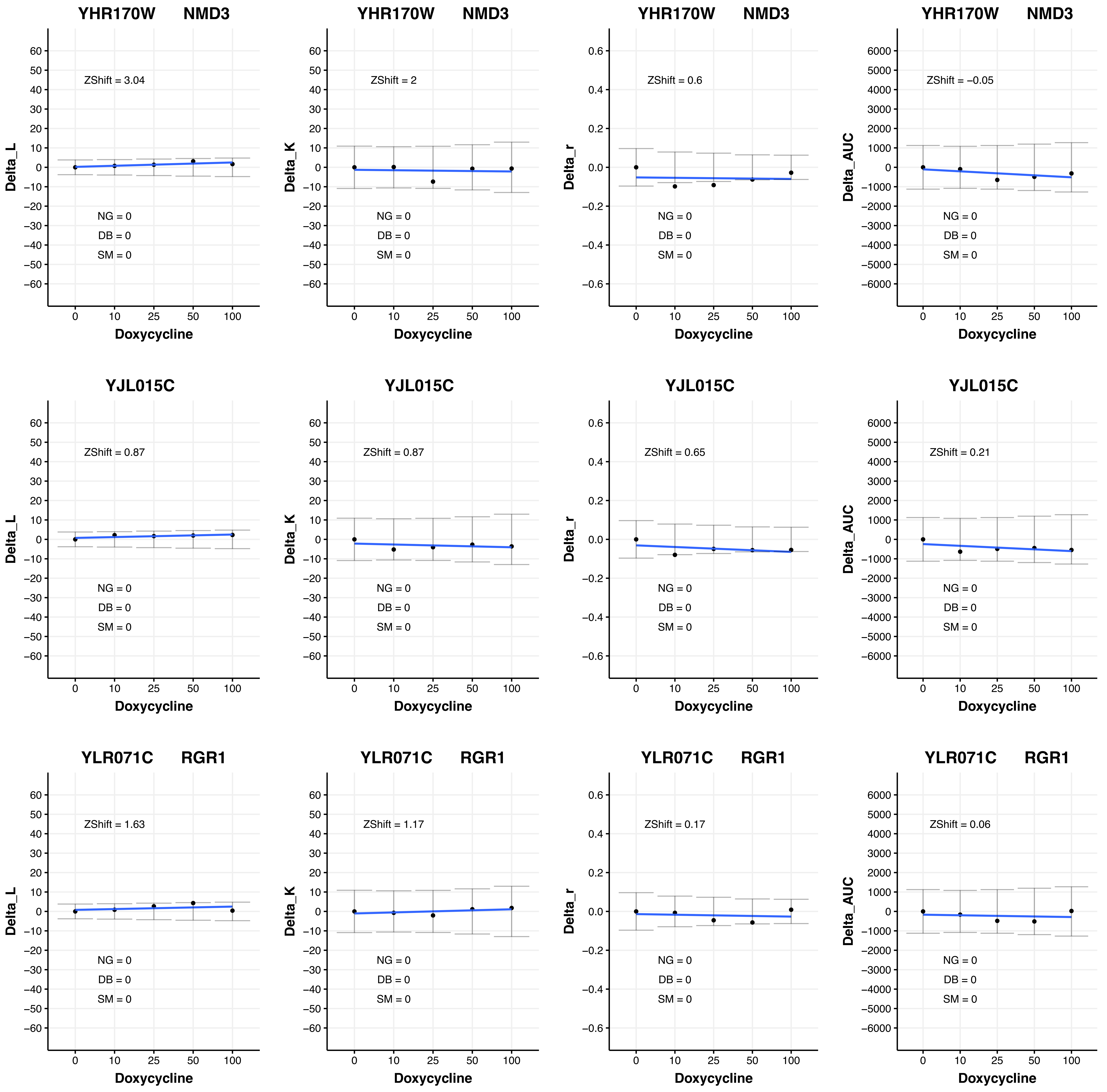
YHR170W NMD3
ZShift = 3.04
ZShift = 2
ZShift = 0.6
ZShift = −0.05
YJL015C
ZShift = 0.87
ZShift = 0.87
ZShift = 0.65
ZShift = 0.21
YLR071C RGR1
ZShift = 1.63
ZShift = 1.17
ZShift = 0.17
ZShift = 0.06
NG = 0
DB = 0
SM = 0
Delta_L
Delta_K
Delta_r
Delta_AUC
Doxycycline
0
10
25
50
100

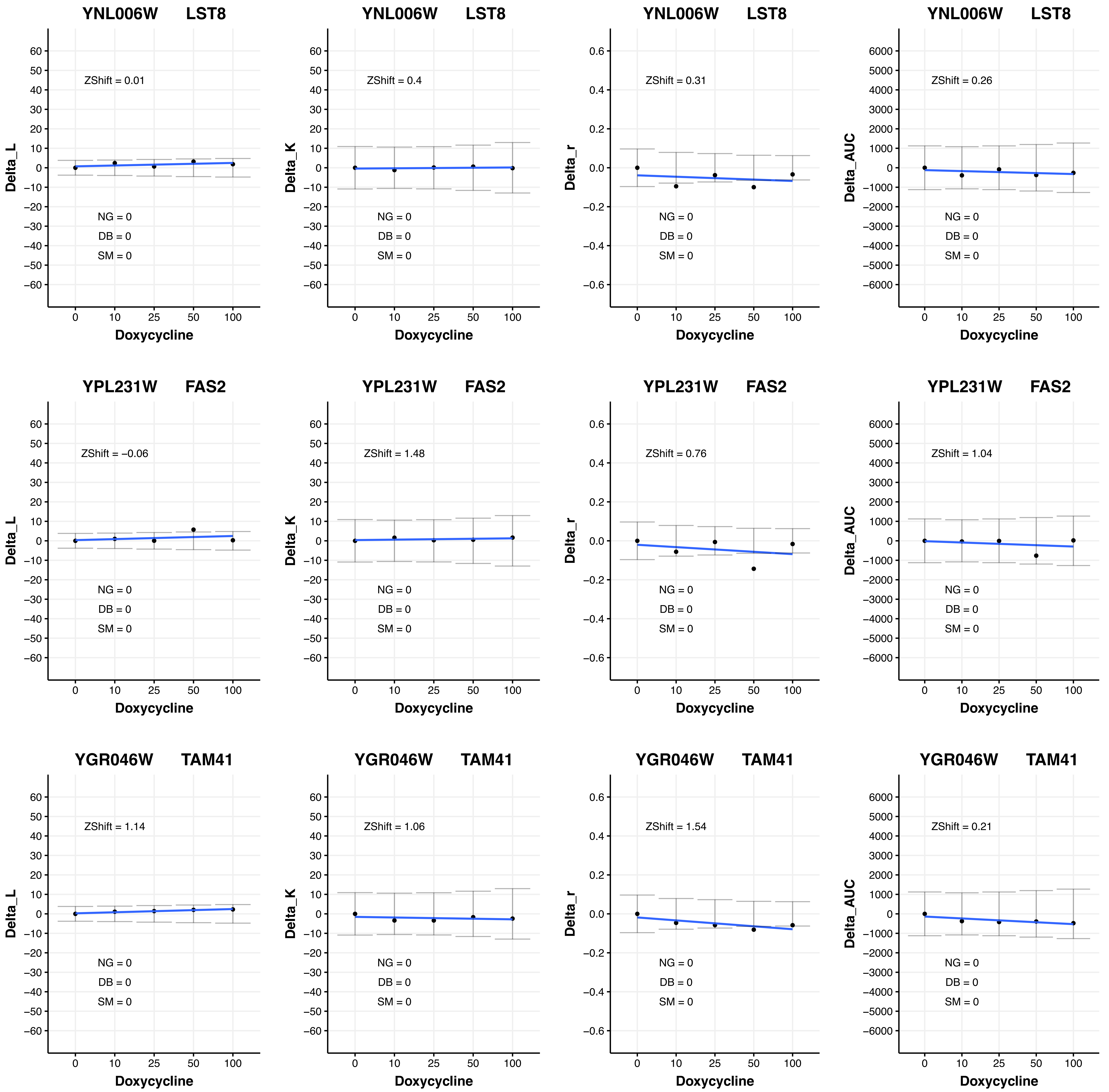

YNL006W LST8
ZShift = 0.01
Delta_L
NG = 0
DB = 0
SM = 0
Doxycycline
YNL006W LST8
ZShift = 0.4
Delta_K
NG = 0
DB = 0
SM = 0
Doxycycline
YNL006W LST8
ZShift = 0.31
Delta_r
NG = 0
DB = 0
SM = 0
Doxycycline
YNL006W LST8
ZShift = 0.26
Delta_AUC
NG = 0
DB = 0
SM = 0
Doxycycline
YPL231W FAS2
ZShift = −0.06
Delta_L
NG = 0
DB = 0
SM = 0
Doxycycline
YPL231W FAS2
ZShift = 1.48
Delta_K
NG = 0
DB = 0
SM = 0
Doxycycline
YPL231W FAS2
ZShift = 0.76
Delta_r
NG = 0
DB = 0
SM = 0
Doxycycline
YPL231W FAS2
ZShift = 1.04
Delta_AUC
NG = 0
DB = 0
SM = 0
Doxycycline
YGR046W TAM41
ZShift = 1.14
Delta_L
NG = 0
DB = 0
SM = 0
Doxycycline
YGR046W TAM41
ZShift = 1.06
Delta_K
NG = 0
DB = 0
SM = 0
Doxycycline
YGR046W TAM41
ZShift = 1.54
Delta_r
NG = 0
DB = 0
SM = 0
Doxycycline
YGR046W TAM41
ZShift = 0.21
Delta_AUC
NG = 0
DB = 0
SM = 0
Doxycycline

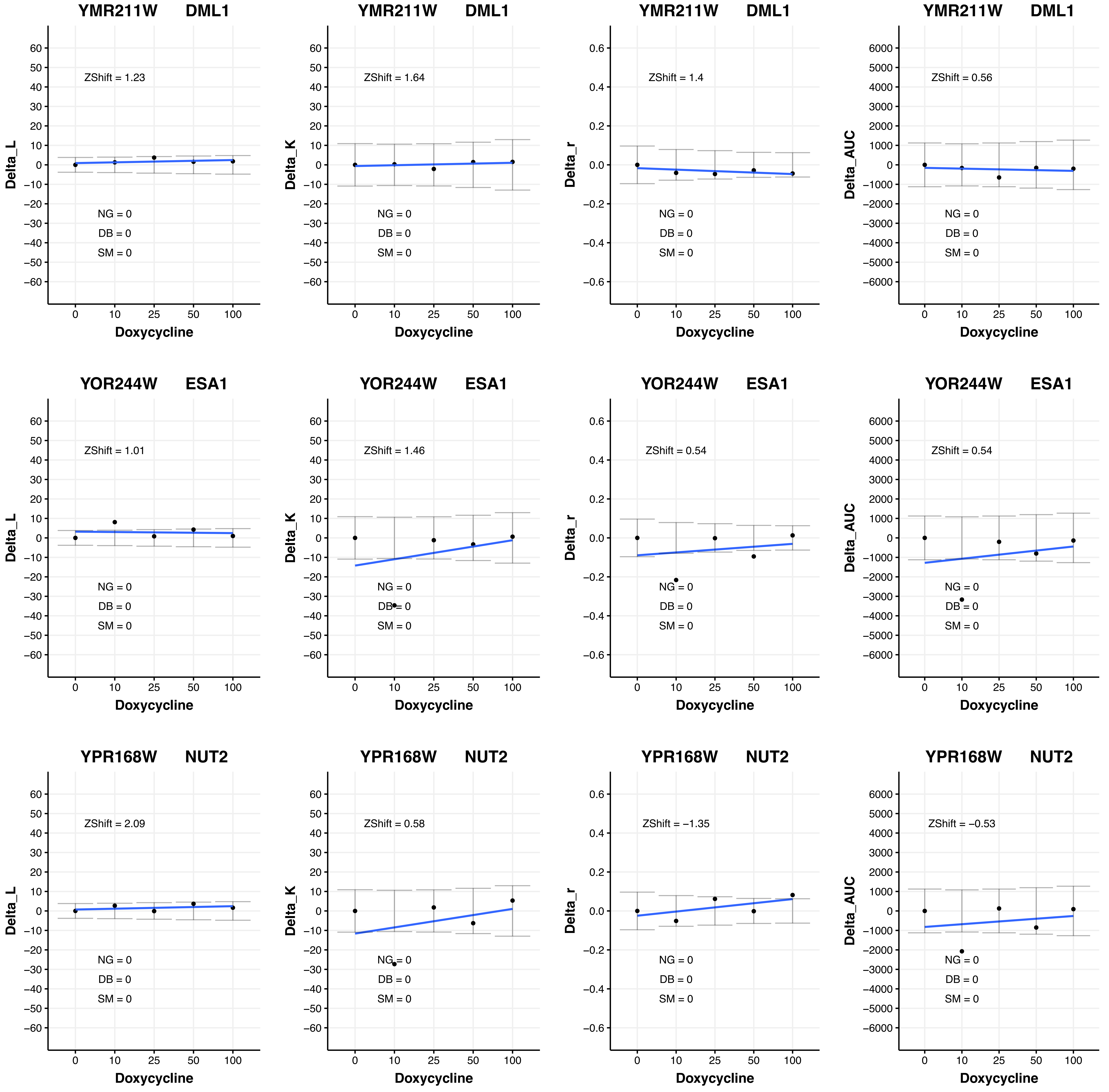

YMR211W DML1
ZShift = 1.23
NG = 0
DB = 0
SM = 0
Delta_L
Doxycycline
YMR211W DML1
ZShift = 1.64
NG = 0
DB = 0
SM = 0
Delta_K
Doxycycline
YMR211W DML1
ZShift = 1.4
NG = 0
DB = 0
SM = 0
Delta_r
Doxycycline
YMR211W DML1
ZShift = 0.56
NG = 0
DB = 0
SM = 0
Delta_AUC
Doxycycline
YOR244W ESA1
ZShift = 1.01
NG = 0
DB = 0
SM = 0
Delta_L
Doxycycline
YOR244W ESA1
ZShift = 1.46
NG = 0
DB = 0
SM = 0
Delta_K
Doxycycline
YOR244W ESA1
ZShift = 0.54
NG = 0
DB = 0
SM = 0
Delta_r
Doxycycline
YOR244W ESA1
ZShift = 0.54
NG = 0
DB = 0
SM = 0
Delta_AUC
Doxycycline
YPR168W NUT2
ZShift = 2.09
NG = 0
DB = 0
SM = 0
Delta_L
Doxycycline
YPR168W NUT2
ZShift = 0.58
NG = 0
DB = 0
SM = 0
Delta_K
Doxycycline
YPR168W NUT2
ZShift = −1.35
NG = 0
DB = 0
SM = 0
Delta_r
Doxycycline
YPR168W NUT2
ZShift = −0.53
NG = 0
DB = 0
SM = 0
Delta_AUC
Doxycycline

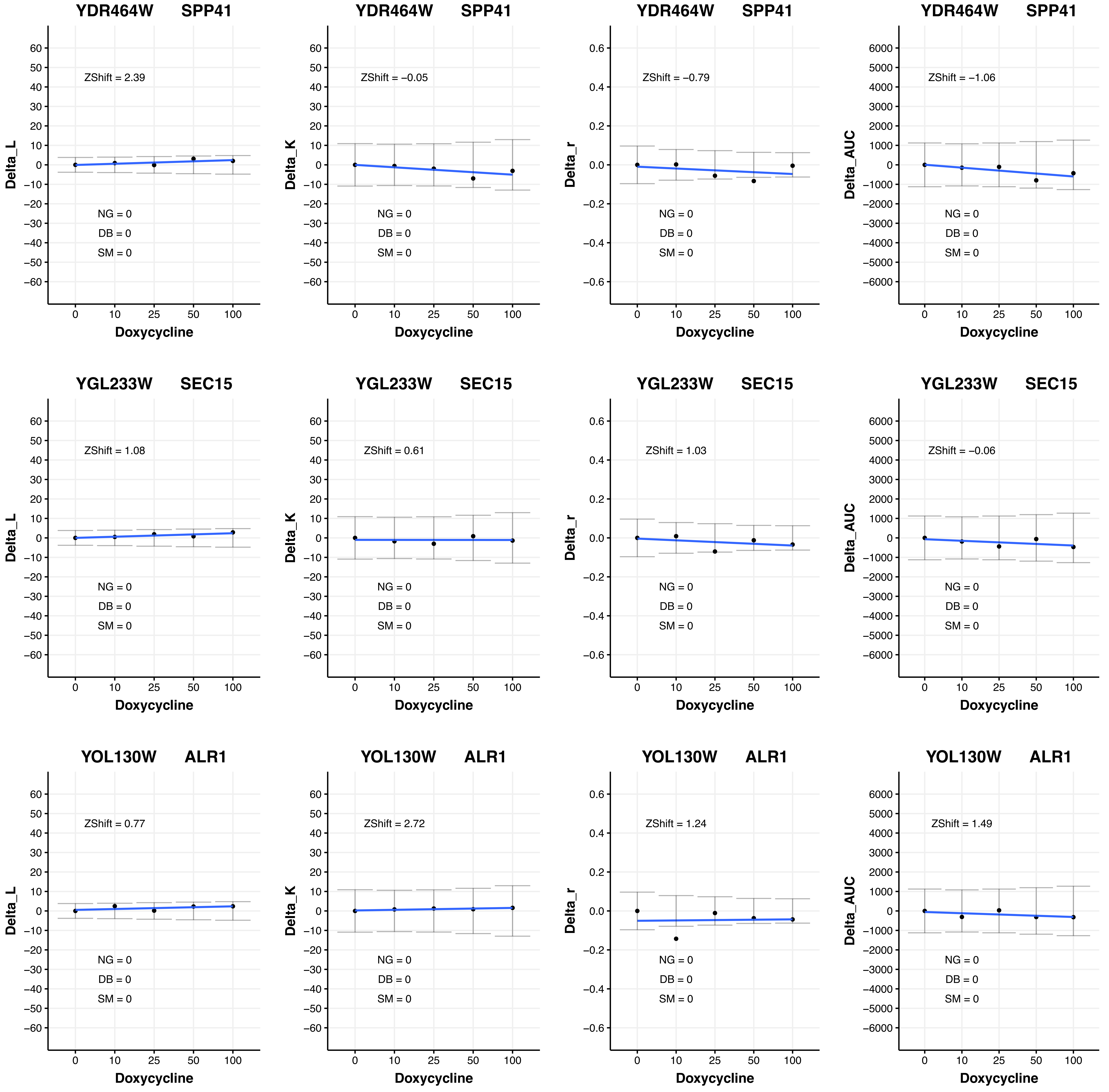

YDR464W SPP41
ZShift = 2.39
ZShift = −0.05
ZShift = −0.79
ZShift = −1.06
YGL233W SEC15
ZShift = 1.08
ZShift = 0.61
ZShift = 1.03
ZShift = −0.06
YOL130W ALR1
ZShift = 0.77
ZShift = 2.72
ZShift = 1.24
ZShift = 1.49
NG = 0
DB = 0
SM = 0
Delta_L
Delta_K
Delta_r
Delta_AUC
Doxycycline
0
10
25
50
100

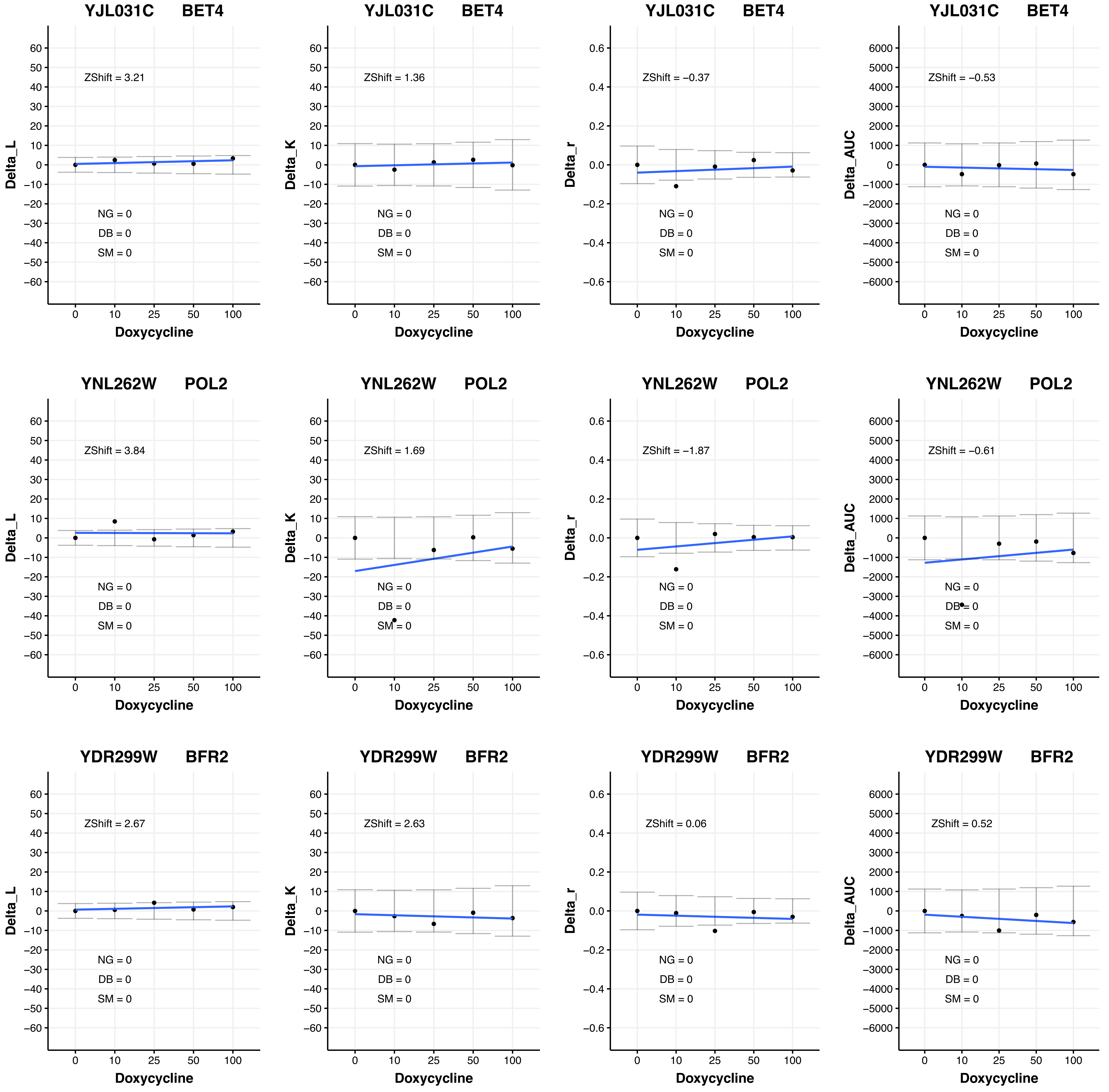

YJL031C BET4
ZShift = 3.21
NG = 0
DB = 0
SM = 0
Delta_L
Doxycycline
YJL031C BET4
ZShift = 1.36
NG = 0
DB = 0
SM = 0
Delta_K
Doxycycline
YJL031C BET4
ZShift = −0.37
NG = 0
DB = 0
SM = 0
Delta_r
Doxycycline
YJL031C BET4
ZShift = −0.53
NG = 0
DB = 0
SM = 0
Delta_AUC
Doxycycline
YNL262W POL2
ZShift = 3.84
NG = 0
DB = 0
SM = 0
Delta_L
Doxycycline
YNL262W POL2
ZShift = 1.69
NG = 0
DB = 0
SM = 0
Delta_K
Doxycycline
YNL262W POL2
ZShift = −1.87
NG = 0
DB = 0
SM = 0
Delta_r
Doxycycline
YNL262W POL2
ZShift = −0.61
NG = 0
DB = 0
SM = 0
Delta_AUC
Doxycycline
YDR299W BFR2
ZShift = 2.67
NG = 0
DB = 0
SM = 0
Delta_L
Doxycycline
YDR299W BFR2
ZShift = 2.63
NG = 0
DB = 0
SM = 0
Delta_K
Doxycycline
YDR299W BFR2
ZShift = 0.06
NG = 0
DB = 0
SM = 0
Delta_r
Doxycycline
YDR299W BFR2
ZShift = 0.52
NG = 0
DB = 0
SM = 0
Delta_AUC
Doxycycline

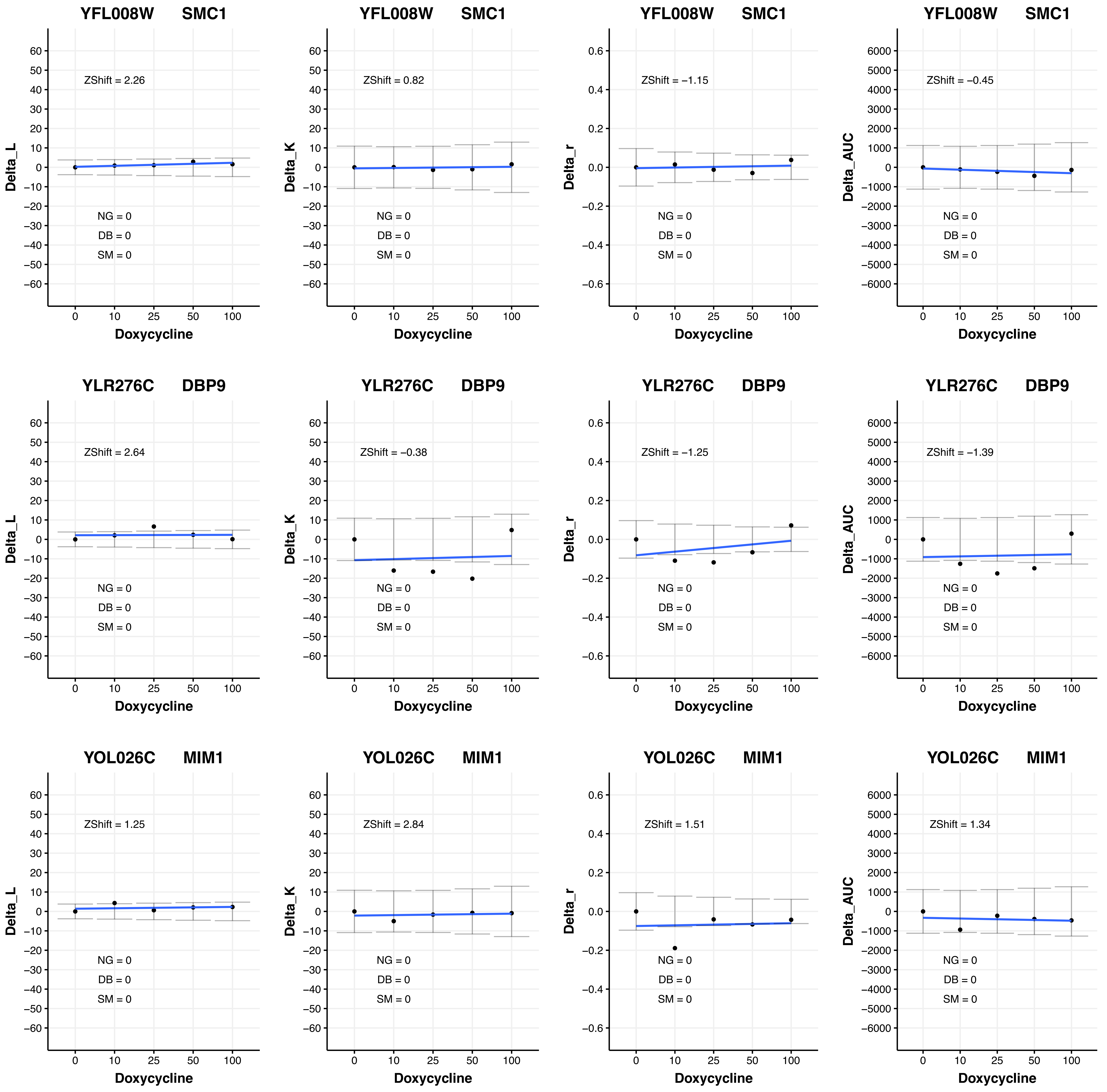

YFL008W SMC1
ZShift = 2.26
NG = 0
DB = 0
SM = 0
Delta_L
Doxycycline
YFL008W SMC1
ZShift = 0.82
NG = 0
DB = 0
SM = 0
Delta_K
Doxycycline
YFL008W SMC1
ZShift = −1.15
NG = 0
DB = 0
SM = 0
Delta_r
Doxycycline
YFL008W SMC1
ZShift = −0.45
NG = 0
DB = 0
SM = 0
Delta_AUC
Doxycycline
YLR276C DBP9
ZShift = 2.64
NG = 0
DB = 0
SM = 0
Delta_L
Doxycycline
YLR276C DBP9
ZShift = −0.38
NG = 0
DB = 0
SM = 0
Delta_K
Doxycycline
YLR276C DBP9
ZShift = −1.25
NG = 0
DB = 0
SM = 0
Delta_r
Doxycycline
YLR276C DBP9
ZShift = −1.39
NG = 0
DB = 0
SM = 0
Delta_AUC
Doxycycline
YOL026C MIM1
ZShift = 1.25
NG = 0
DB = 0
SM = 0
Delta_L
Doxycycline
YOL026C MIM1
ZShift = 2.84
NG = 0
DB = 0
SM = 0
Delta_K
Doxycycline
YOL026C MIM1
ZShift = 1.51
NG = 0
DB = 0
SM = 0
Delta_r
Doxycycline
YOL026C MIM1
ZShift = 1.34
NG = 0
DB = 0
SM = 0
Delta_AUC
Doxycycline
0 10 25 50 100

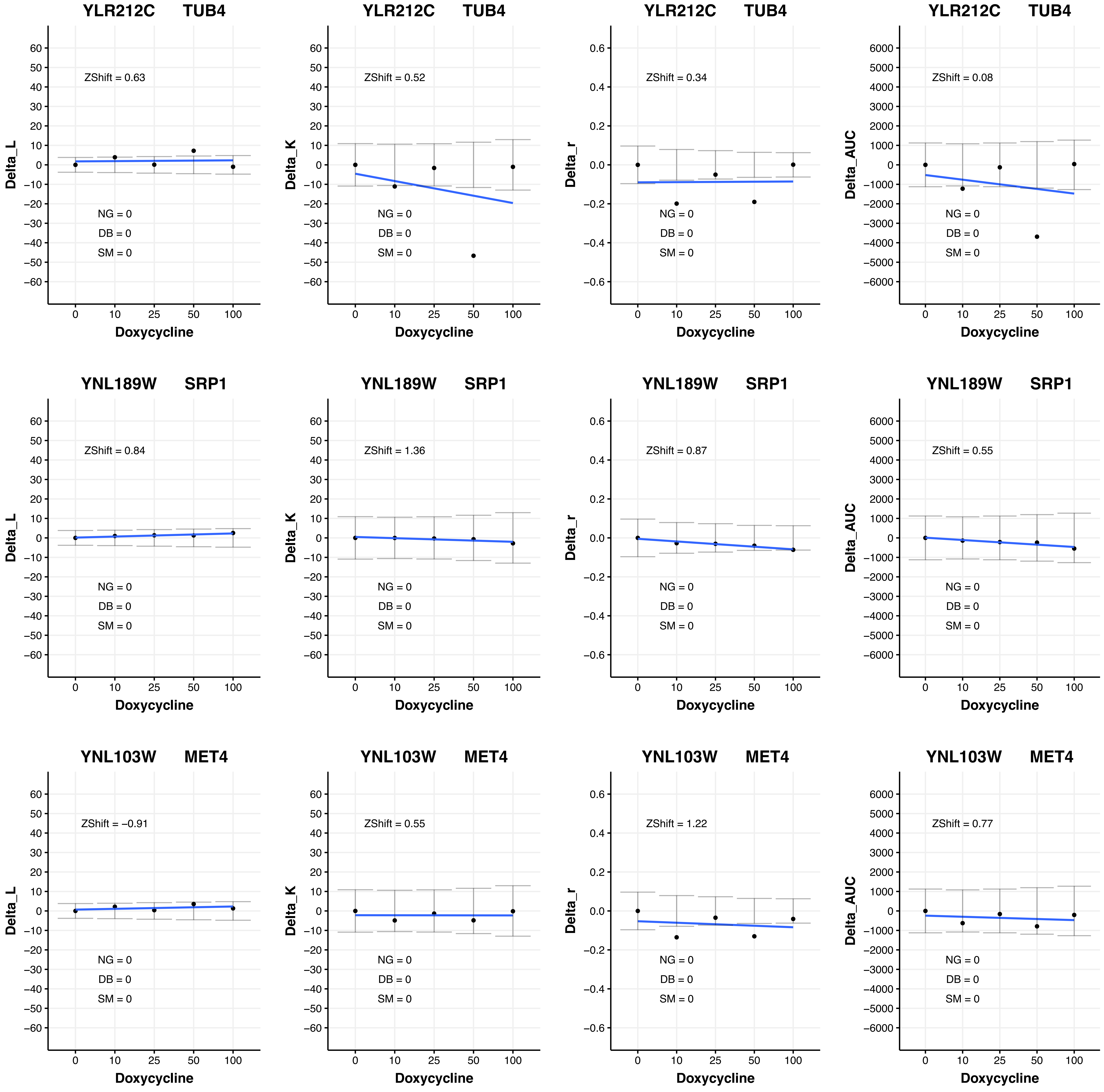

YLR212C TUB4
ZShift = 0.63
NG = 0
DB = 0
SM = 0
Delta_L
Doxycycline
YLR212C TUB4
ZShift = 0.52
NG = 0
DB = 0
SM = 0
Delta_K
Doxycycline
YLR212C TUB4
ZShift = 0.34
NG = 0
DB = 0
SM = 0
Delta_r
Doxycycline
YLR212C TUB4
ZShift = 0.08
NG = 0
DB = 0
SM = 0
Delta_AUC
Doxycycline
YNL189W SRP1
ZShift = 0.84
NG = 0
DB = 0
SM = 0
Delta_L
Doxycycline
YNL189W SRP1
ZShift = 1.36
NG = 0
DB = 0
SM = 0
Delta_K
Doxycycline
YNL189W SRP1
ZShift = 0.87
NG = 0
DB = 0
SM = 0
Delta_r
Doxycycline
YNL189W SRP1
ZShift = 0.55
NG = 0
DB = 0
SM = 0
Delta_AUC
Doxycycline
YNL103W MET4
ZShift = −0.91
NG = 0
DB = 0
SM = 0
Delta_L
Doxycycline
YNL103W MET4
ZShift = 0.55
NG = 0
DB = 0
SM = 0
Delta_K
Doxycycline
YNL103W MET4
ZShift = 1.22
NG = 0
DB = 0
SM = 0
Delta_r
Doxycycline
YNL103W MET4
ZShift = 0.77
NG = 0
DB = 0
SM = 0
Delta_AUC
Doxycycline

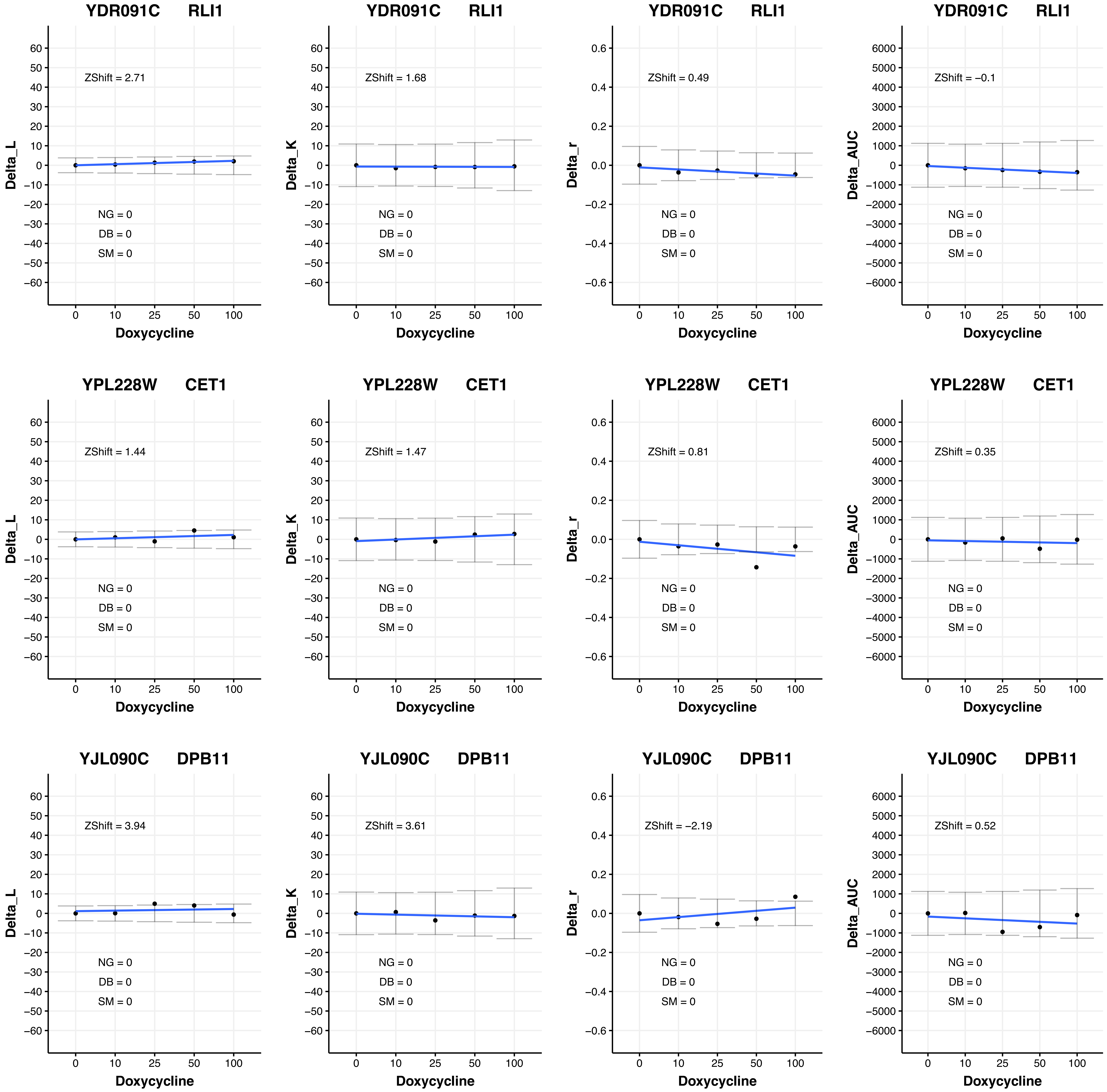
YDR091C RLI1
ZShift = 2.71
NG = 0
DB = 0
SM = 0
Delta_L
Doxycycline
YDR091C RLI1
ZShift = 1.68
NG = 0
DB = 0
SM = 0
Delta_K
Doxycycline
YDR091C RLI1
ZShift = 0.49
NG = 0
DB = 0
SM = 0
Delta_r
Doxycycline
YDR091C RLI1
ZShift = −0.1
NG = 0
DB = 0
SM = 0
Delta_AUC
Doxycycline
YPL228W CET1
ZShift = 1.44
NG = 0
DB = 0
SM = 0
Delta_L
Doxycycline
YPL228W CET1
ZShift = 1.47
NG = 0
DB = 0
SM = 0
Delta_K
Doxycycline
YPL228W CET1
ZShift = 0.81
NG = 0
DB = 0
SM = 0
Delta_r
Doxycycline
YPL228W CET1
ZShift = 0.35
NG = 0
DB = 0
SM = 0
Delta_AUC
Doxycycline
YJL090C DPB11
ZShift = 3.94
NG = 0
DB = 0
SM = 0
Delta_L
Doxycycline
YJL090C DPB11
ZShift = 3.61
NG = 0
DB = 0
SM = 0
Delta_K
Doxycycline
YJL090C DPB11
ZShift = −2.19
NG = 0
DB = 0
SM = 0
Delta_r
Doxycycline
YJL090C DPB11
ZShift = 0.52
NG = 0
DB = 0
SM = 0
Delta_AUC
Doxycycline

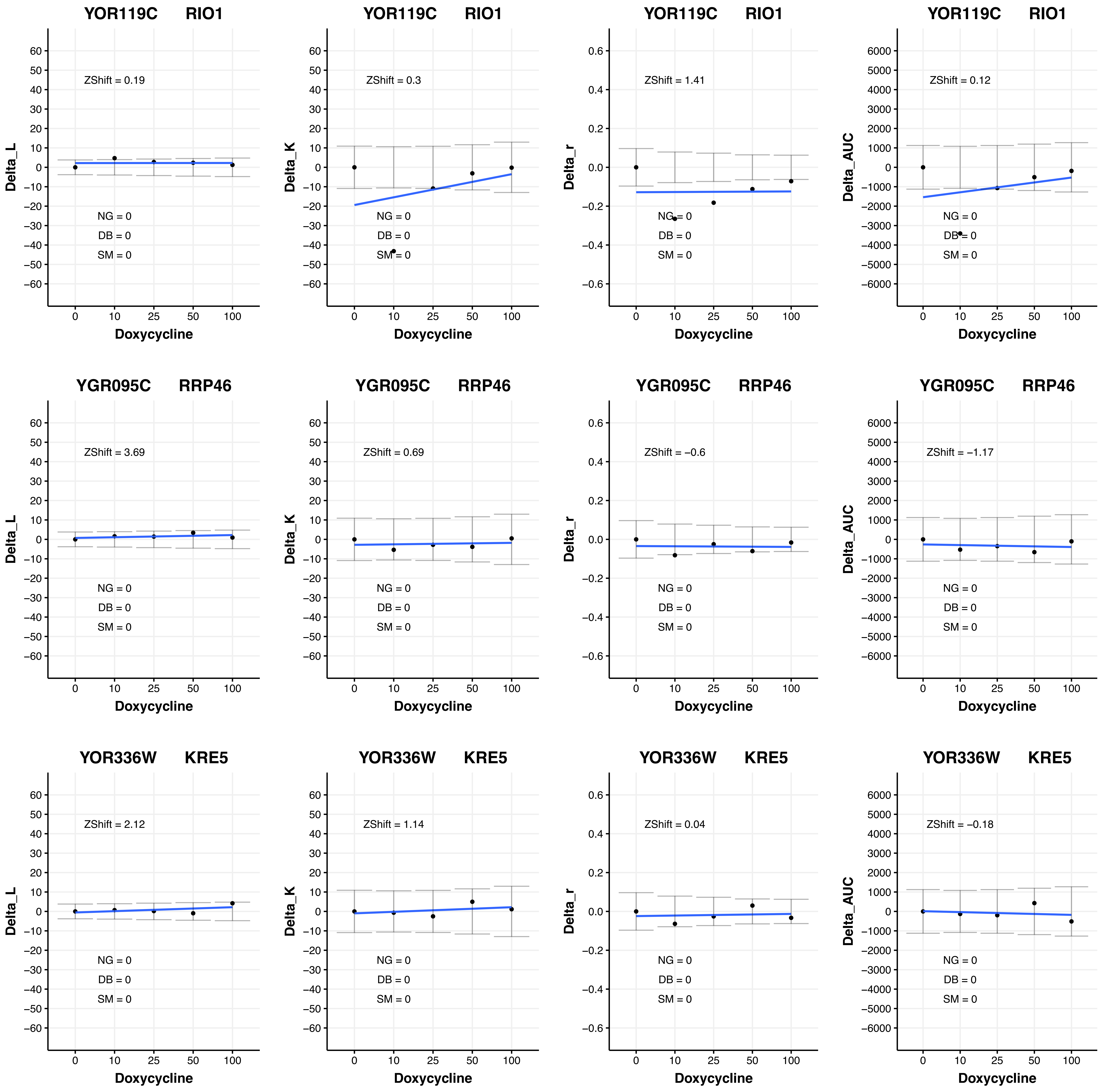

YOR119C RIO1
ZShift = 0.19
NG = 0
DB = 0
SM = 0
Delta_L
Doxycycline
YOR119C RIO1
ZShift = 0.3
NG = 0
DB = 0
SM = 0
Delta_K
Doxycycline
YOR119C RIO1
ZShift = 1.41
NG = 0
DB = 0
SM = 0
Delta_r
Doxycycline
YOR119C RIO1
ZShift = 0.12
NG = 0
DB = 0
SM = 0
Delta_AUC
Doxycycline
YGR095C RRP46
ZShift = 3.69
NG = 0
DB = 0
SM = 0
Delta_L
Doxycycline
YGR095C RRP46
ZShift = 0.69
NG = 0
DB = 0
SM = 0
Delta_K
Doxycycline
YGR095C RRP46
ZShift = −0.6
NG = 0
DB = 0
SM = 0
Delta_r
Doxycycline
YGR095C RRP46
ZShift = −1.17
NG = 0
DB = 0
SM = 0
Delta_AUC
Doxycycline
YOR336W KRE5
ZShift = 2.12
NG = 0
DB = 0
SM = 0
Delta_L
Doxycycline
YOR336W KRE5
ZShift = 1.14
NG = 0
DB = 0
SM = 0
Delta_K
Doxycycline
YOR336W KRE5
ZShift = 0.04
NG = 0
DB = 0
SM = 0
Delta_r
Doxycycline
YOR336W KRE5
ZShift = −0.18
NG = 0
DB = 0
SM = 0
Delta_AUC
Doxycycline

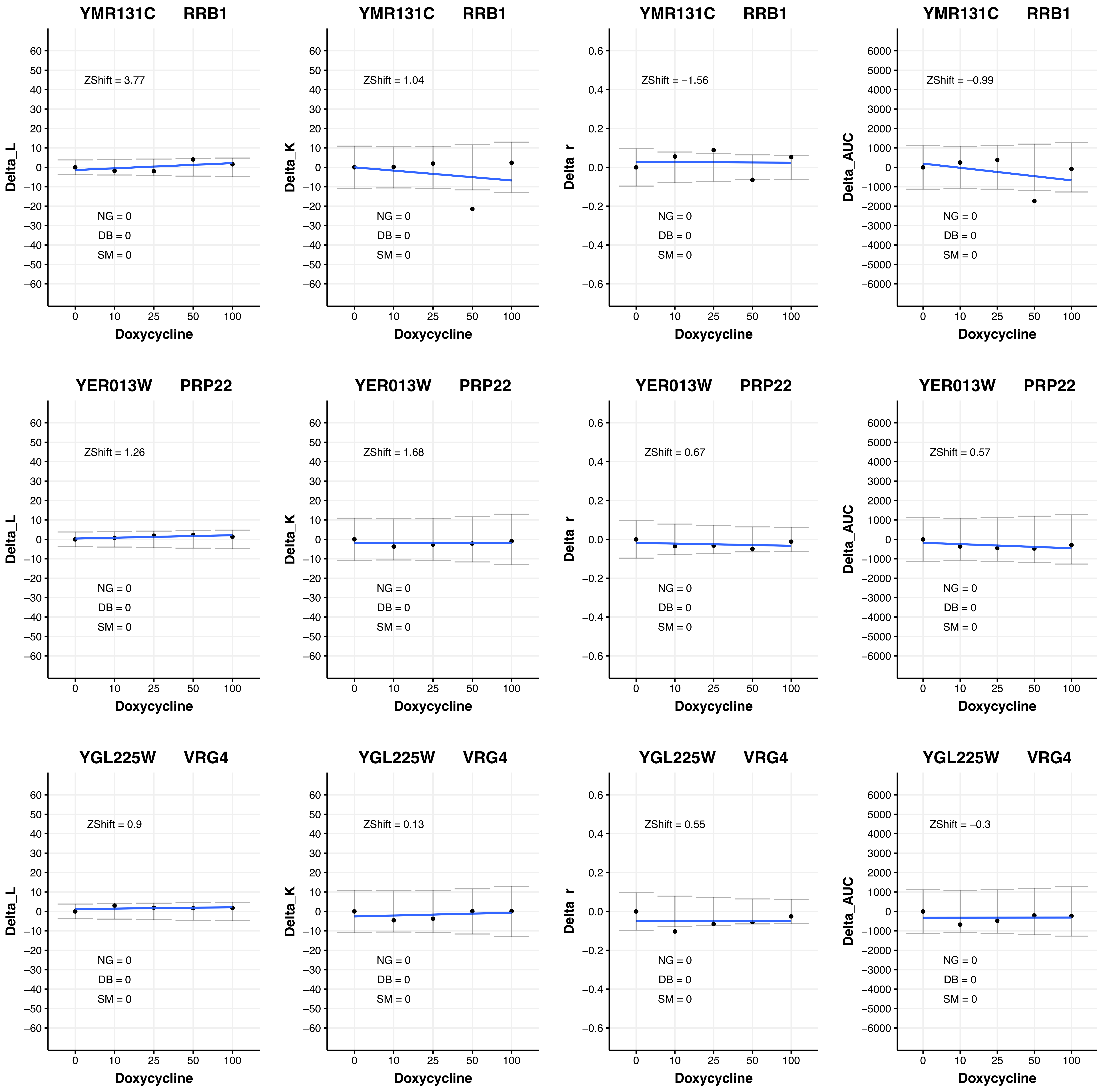

YMR131C RRB1
ZShift = 3.77
NG = 0
DB = 0
SM = 0
Delta_L
Doxycycline
YMR131C RRB1
ZShift = 1.04
NG = 0
DB = 0
SM = 0
Delta_K
Doxycycline
YMR131C RRB1
ZShift = −1.56
NG = 0
DB = 0
SM = 0
Delta_r
Doxycycline
YMR131C RRB1
ZShift = −0.99
NG = 0
DB = 0
SM = 0
Delta_AUC
Doxycycline
YER013W PRP22
ZShift = 1.26
NG = 0
DB = 0
SM = 0
Delta_L
Doxycycline
YER013W PRP22
ZShift = 1.68
NG = 0
DB = 0
SM = 0
Delta_K
Doxycycline
YER013W PRP22
ZShift = 0.67
NG = 0
DB = 0
SM = 0
Delta_r
Doxycycline
YER013W PRP22
ZShift = 0.57
NG = 0
DB = 0
SM = 0
Delta_AUC
Doxycycline
YGL225W VRG4
ZShift = 0.9
NG = 0
DB = 0
SM = 0
Delta_L
Doxycycline
YGL225W VRG4
ZShift = 0.13
NG = 0
DB = 0
SM = 0
Delta_K
Doxycycline
YGL225W VRG4
ZShift = 0.55
NG = 0
DB = 0
SM = 0
Delta_r
Doxycycline
YGL225W VRG4
ZShift = −0.3
NG = 0
DB = 0
SM = 0
Delta_AUC
Doxycycline

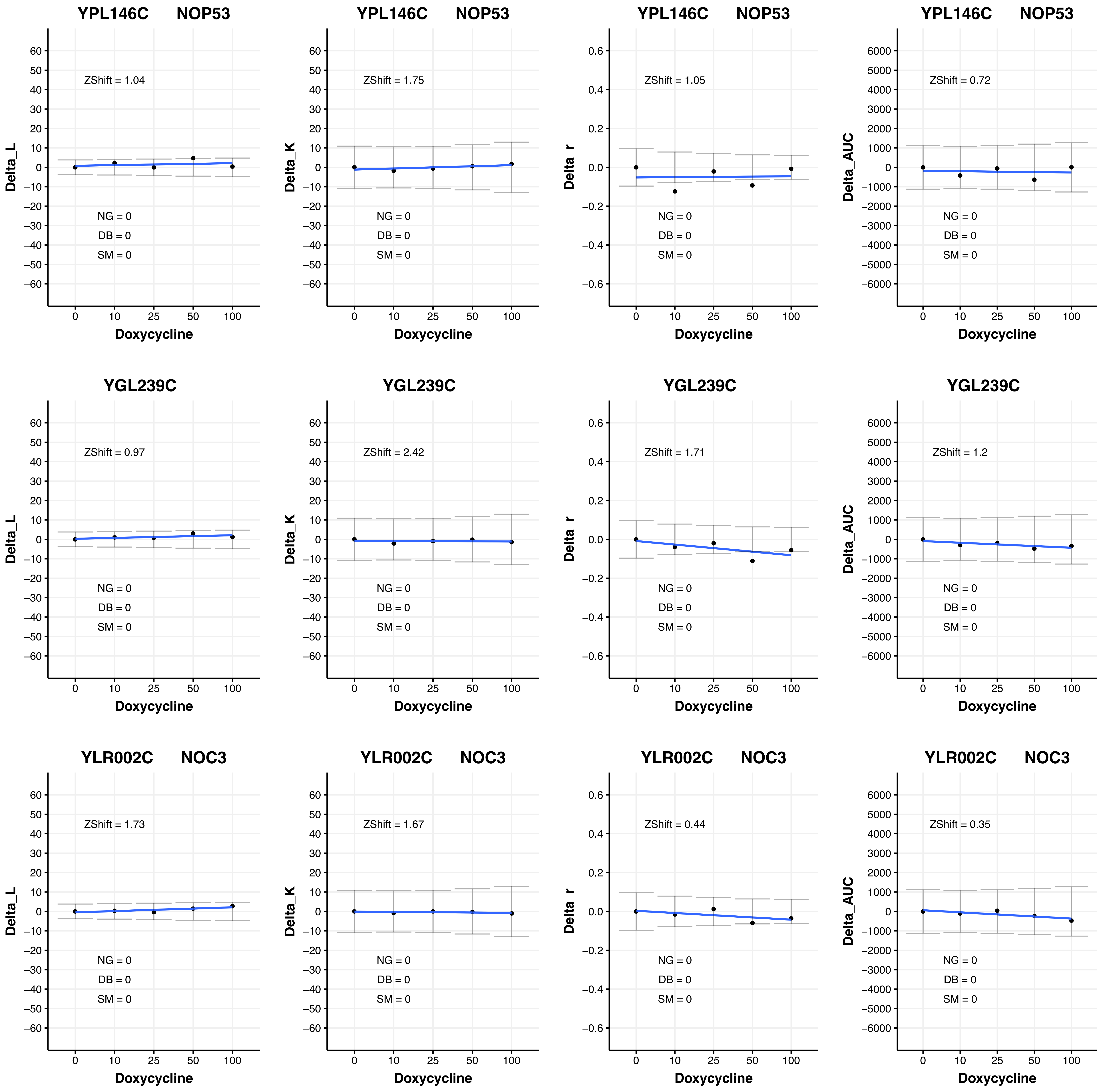

YPL146C NOP53
ZShift = 1.04
NG = 0
DB = 0
SM = 0
Delta_L
Doxycycline
YPL146C NOP53
ZShift = 1.75
NG = 0
DB = 0
SM = 0
Delta_K
Doxycycline
YPL146C NOP53
ZShift = 1.05
NG = 0
DB = 0
SM = 0
Delta_r
Doxycycline
YPL146C NOP53
ZShift = 0.72
NG = 0
DB = 0
SM = 0
Delta_AUC
Doxycycline
YGL239C
ZShift = 0.97
NG = 0
DB = 0
SM = 0
Delta_L
Doxycycline
YGL239C
ZShift = 2.42
NG = 0
DB = 0
SM = 0
Delta_K
Doxycycline
YGL239C
ZShift = 1.71
NG = 0
DB = 0
SM = 0
Delta_r
Doxycycline
YGL239C
ZShift = 1.2
NG = 0
DB = 0
SM = 0
Delta_AUC
Doxycycline
YLR002C NOC3
ZShift = 1.73
NG = 0
DB = 0
SM = 0
Delta_L
Doxycycline
YLR002C NOC3
ZShift = 1.67
NG = 0
DB = 0
SM = 0
Delta_K
Doxycycline
YLR002C NOC3
ZShift = 0.44
NG = 0
DB = 0
SM = 0
Delta_r
Doxycycline
YLR002C NOC3
ZShift = 0.35
NG = 0
DB = 0
SM = 0
Delta_AUC
Doxycycline

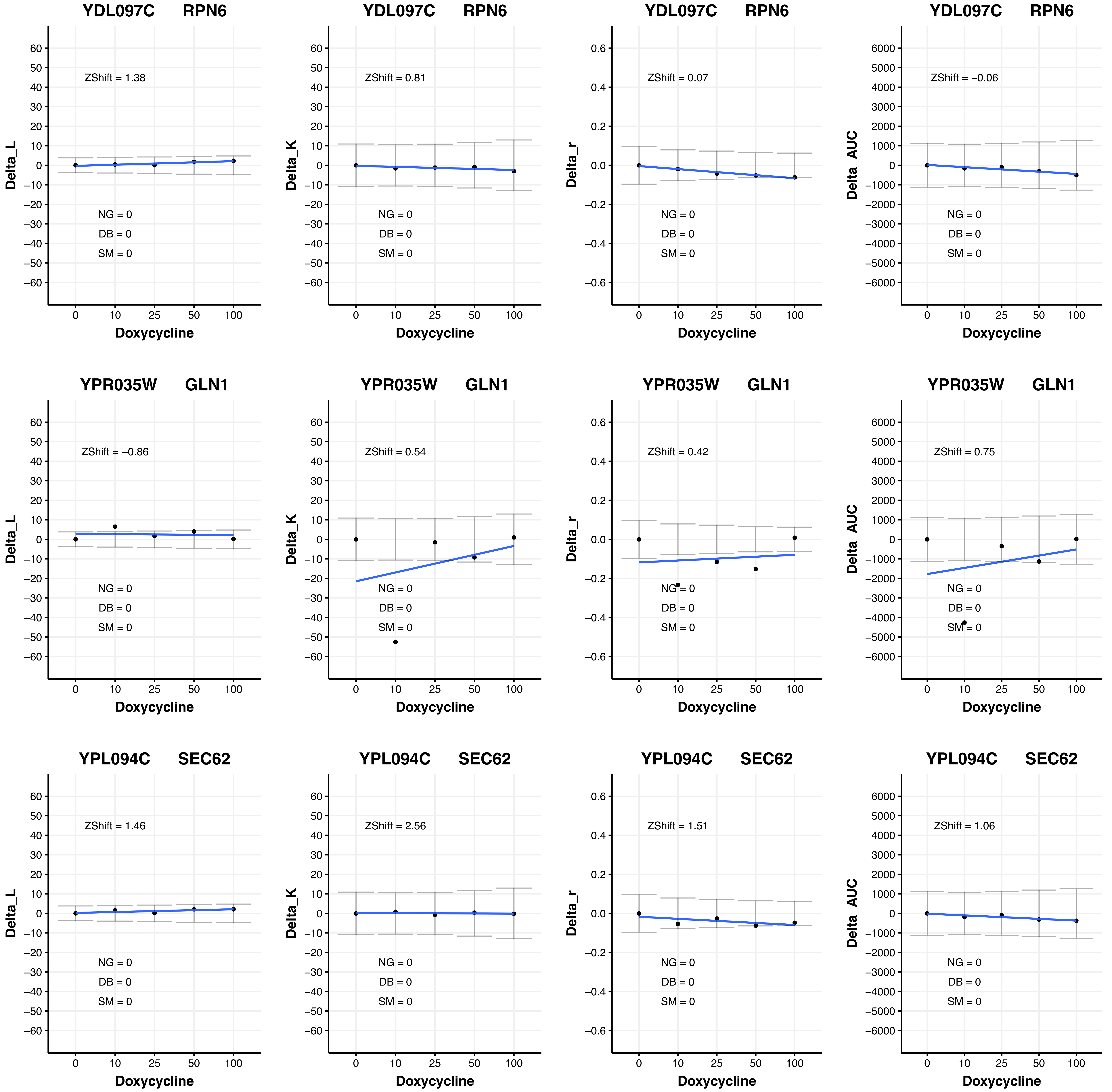

YDL097C RPN6
ZShift = 1.38
NG = 0
DB = 0
SM = 0
Delta_L
Doxycycline
YDL097C RPN6
ZShift = 0.81
NG = 0
DB = 0
SM = 0
Delta_K
Doxycycline
YDL097C RPN6
ZShift = 0.07
NG = 0
DB = 0
SM = 0
Delta_r
Doxycycline
YDL097C RPN6
ZShift = −0.06
NG = 0
DB = 0
SM = 0
Delta_AUC
Doxycycline
YPR035W GLN1
ZShift = −0.86
NG = 0
DB = 0
SM = 0
Delta_L
Doxycycline
YPR035W GLN1
ZShift = 0.54
NG = 0
DB = 0
SM = 0
Delta_K
Doxycycline
YPR035W GLN1
ZShift = 0.42
NG = 0
DB = 0
SM = 0
Delta_r
Doxycycline
YPR035W GLN1
ZShift = 0.75
NG = 0
DB = 0
SM = 0
Delta_AUC
Doxycycline
YPL094C SEC62
ZShift = 1.46
NG = 0
DB = 0
SM = 0
Delta_L
Doxycycline
YPL094C SEC62
ZShift = 2.56
NG = 0
DB = 0
SM = 0
Delta_K
Doxycycline
YPL094C SEC62
ZShift = 1.51
NG = 0
DB = 0
SM = 0
Delta_r
Doxycycline
YPL094C SEC62
ZShift = 1.06
NG = 0
DB = 0
SM = 0
Delta_AUC
Doxycycline

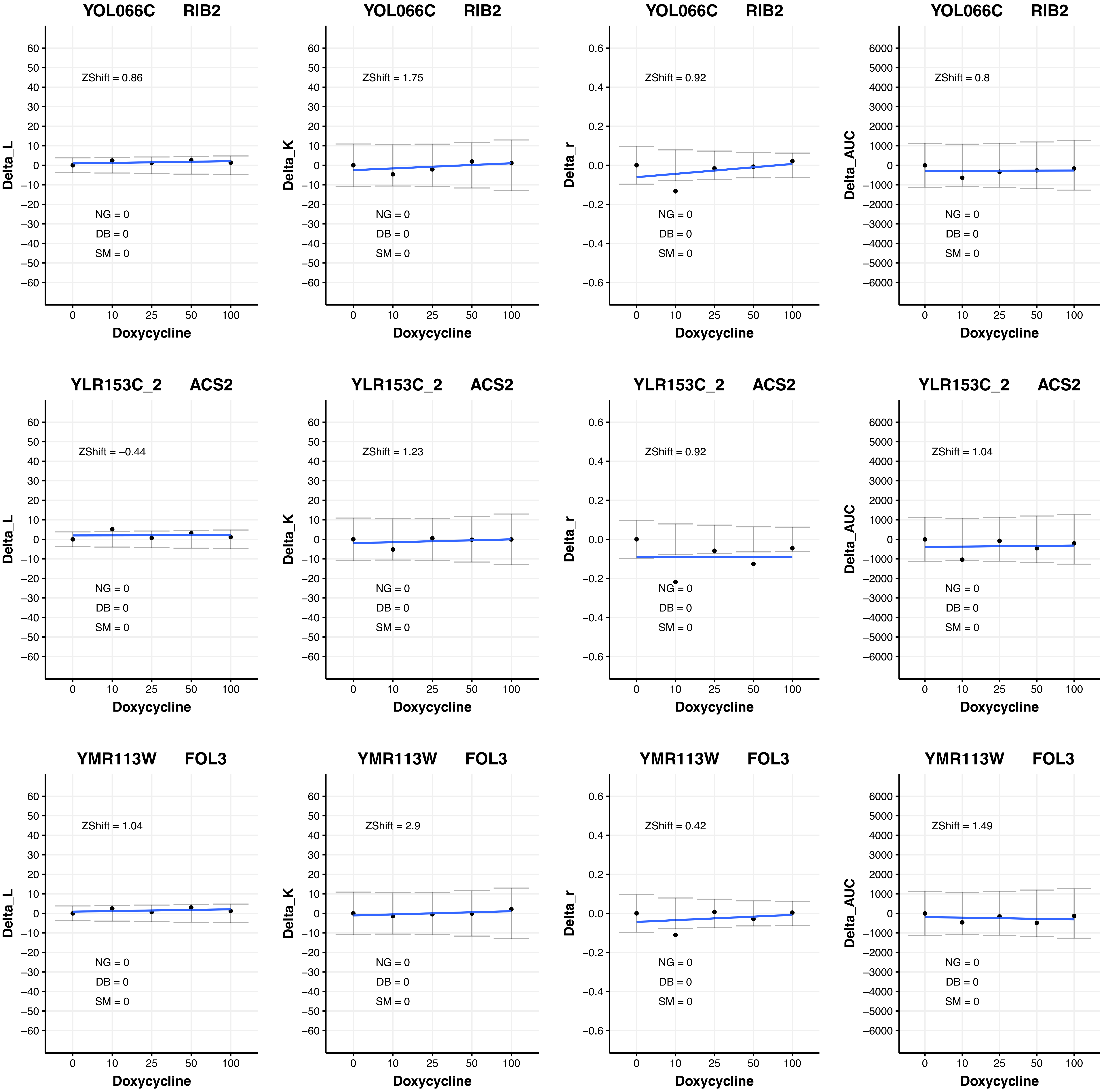

YOL066C RIB2
ZShift = 0.86
Delta_L
NG = 0
DB = 0
SM = 0
Doxycycline
YOL066C RIB2
ZShift = 1.75
Delta_K
NG = 0
DB = 0
SM = 0
Doxycycline
YOL066C RIB2
ZShift = 0.92
Delta_r
NG = 0
DB = 0
SM = 0
Doxycycline
YOL066C RIB2
ZShift = 0.8
Delta_AUC
NG = 0
DB = 0
SM = 0
Doxycycline
YLR153C_2 ACS2
ZShift = −0.44
Delta_L
NG = 0
DB = 0
SM = 0
Doxycycline
YLR153C_2 ACS2
ZShift = 1.23
Delta_K
NG = 0
DB = 0
SM = 0
Doxycycline
YLR153C_2 ACS2
ZShift = 0.92
Delta_r
NG = 0
DB = 0
SM = 0
Doxycycline
YLR153C_2 ACS2
ZShift = 1.04
Delta_AUC
NG = 0
DB = 0
SM = 0
Doxycycline
YMR113W FOL3
ZShift = 1.04
Delta_L
NG = 0
DB = 0
SM = 0
Doxycycline
YMR113W FOL3
ZShift = 2.9
Delta_K
NG = 0
DB = 0
SM = 0
Doxycycline
YMR113W FOL3
ZShift = 0.42
Delta_r
NG = 0
DB = 0
SM = 0
Doxycycline
YMR113W FOL3
ZShift = 1.49
Delta_AUC
NG = 0
DB = 0
SM = 0
Doxycycline

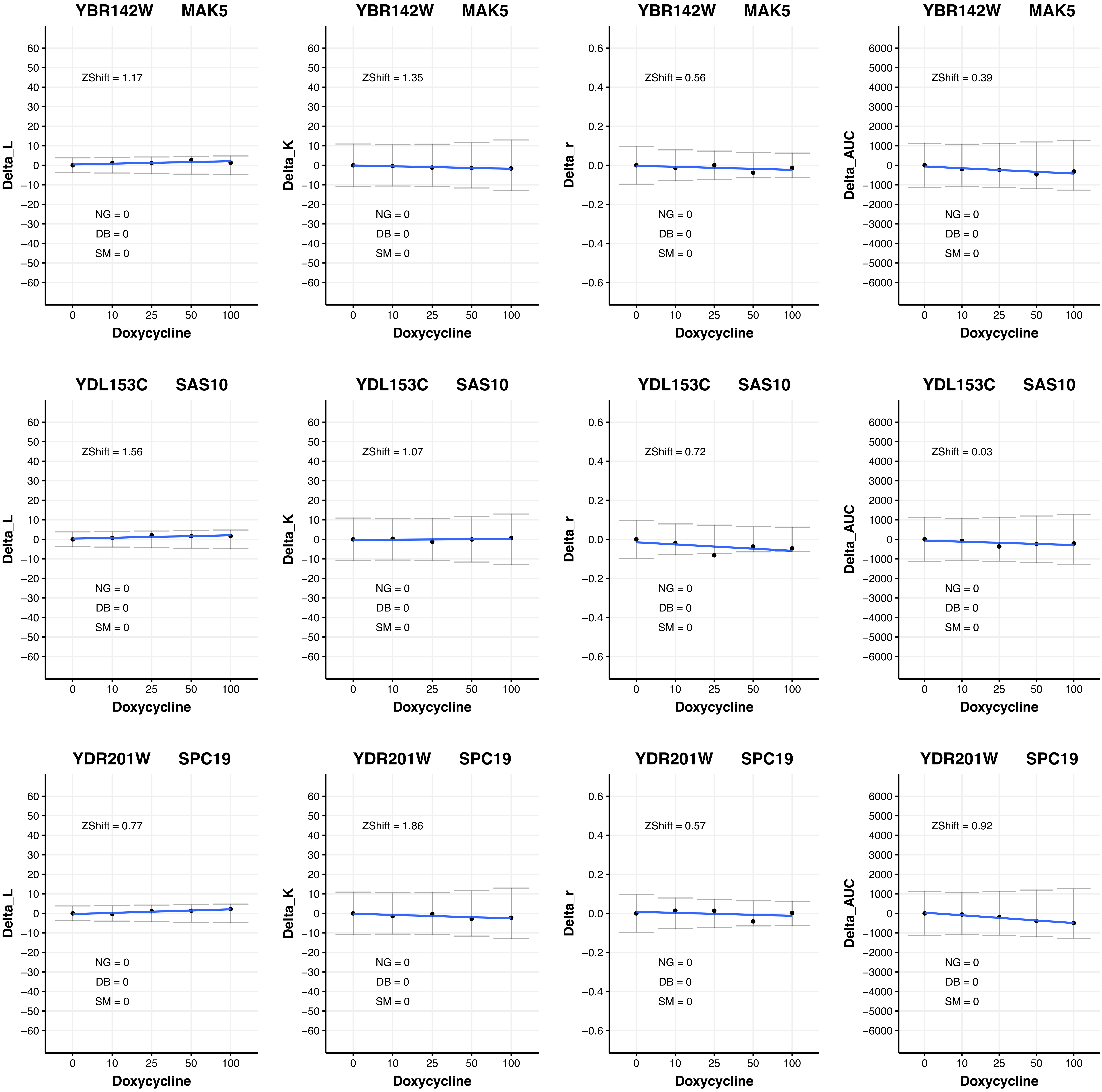
YBR142W MAK5
ZShift = 1.17
NG = 0
DB = 0
SM = 0
Delta_L
YBR142W MAK5
ZShift = 1.35
NG = 0
DB = 0
SM = 0
Delta_K
YBR142W MAK5
ZShift = 0.56
NG = 0
DB = 0
SM = 0
Delta_r
YBR142W MAK5
ZShift = 0.39
NG = 0
DB = 0
SM = 0
Delta_AUC
YDL153C SAS10
ZShift = 1.56
NG = 0
DB = 0
SM = 0
Delta_L
YDL153C SAS10
ZShift = 1.07
NG = 0
DB = 0
SM = 0
Delta_K
YDL153C SAS10
ZShift = 0.72
NG = 0
DB = 0
SM = 0
Delta_r
YDL153C SAS10
ZShift = 0.03
NG = 0
DB = 0
SM = 0
Delta_AUC
YDR201W SPC19
ZShift = 0.77
NG = 0
DB = 0
SM = 0
Delta_L
YDR201W SPC19
ZShift = 1.86
NG = 0
DB = 0
SM = 0
Delta_K
YDR201W SPC19
ZShift = 0.57
NG = 0
DB = 0
SM = 0
Delta_r
YDR201W SPC19
ZShift = 0.92
NG = 0
DB = 0
SM = 0
Delta_AUC
Doxycycline
0 10 25 50 100

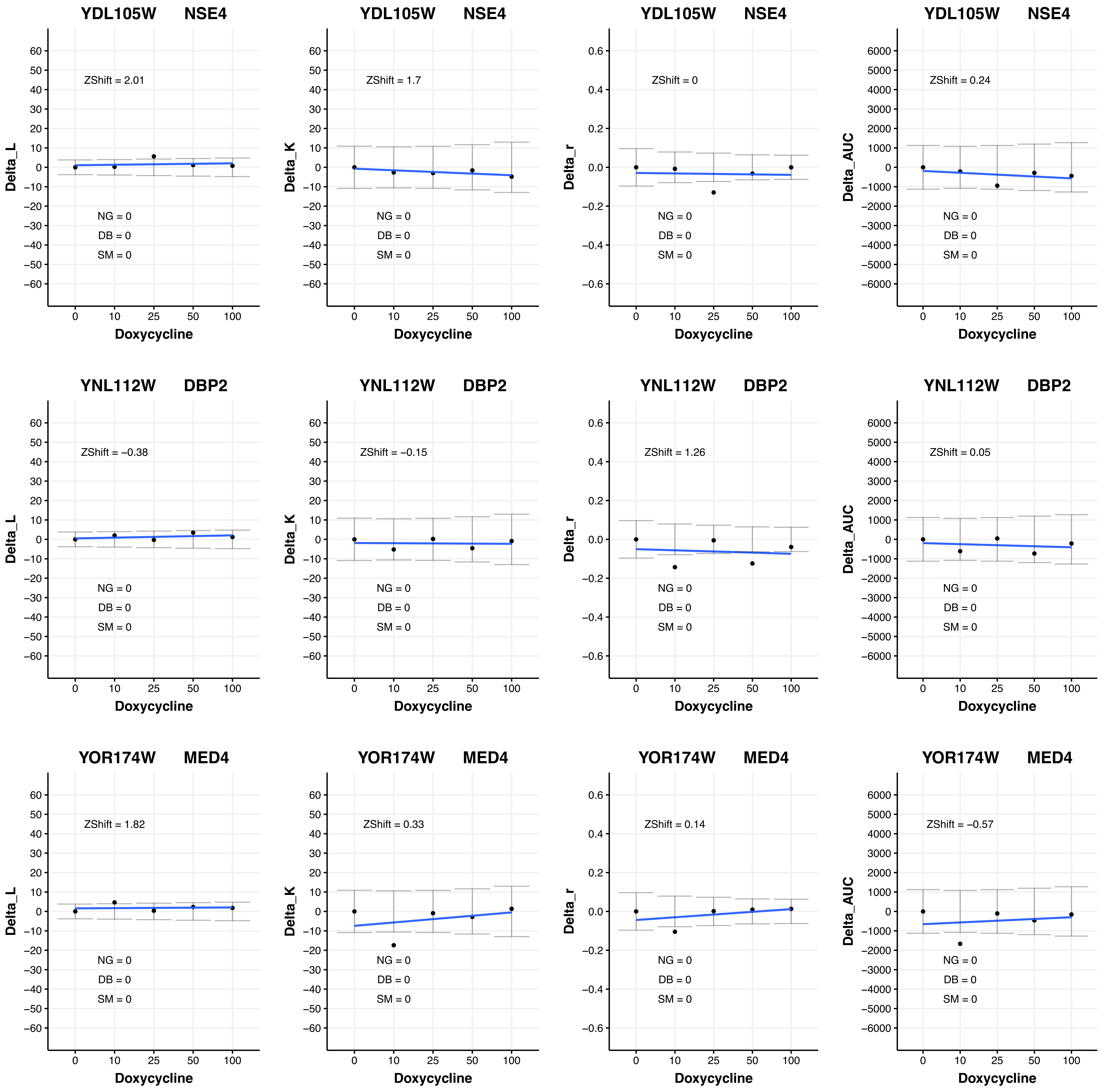
YDL105W NSE4
ZShift = 2.01
NG = 0
DB = 0
SM = 0
Delta_L
Doxycycline
YDL105W NSE4
ZShift = 1.7
NG = 0
DB = 0
SM = 0
Delta_K
Doxycycline
YDL105W NSE4
ZShift = 0
NG = 0
DB = 0
SM = 0
Delta_r
Doxycycline
YDL105W NSE4
ZShift = 0.24
NG = 0
DB = 0
SM = 0
Delta_AUC
Doxycycline
YNL112W DBP2
ZShift = −0.38
NG = 0
DB = 0
SM = 0
Delta_L
Doxycycline
YNL112W DBP2
ZShift = −0.15
NG = 0
DB = 0
SM = 0
Delta_K
Doxycycline
YNL112W DBP2
ZShift = 1.26
NG = 0
DB = 0
SM = 0
Delta_r
Doxycycline
YNL112W DBP2
ZShift = 0.05
NG = 0
DB = 0
SM = 0
Delta_AUC
Doxycycline
YOR174W MED4
ZShift = 1.82
NG = 0
DB = 0
SM = 0
Delta_L
Doxycycline
YOR174W MED4
ZShift = 0.33
NG = 0
DB = 0
SM = 0
Delta_K
Doxycycline
YOR174W MED4
ZShift = 0.14
NG = 0
DB = 0
SM = 0
Delta_r
Doxycycline
YOR174W MED4
ZShift = −0.57
NG = 0
DB = 0
SM = 0
Delta_AUC
Doxycycline

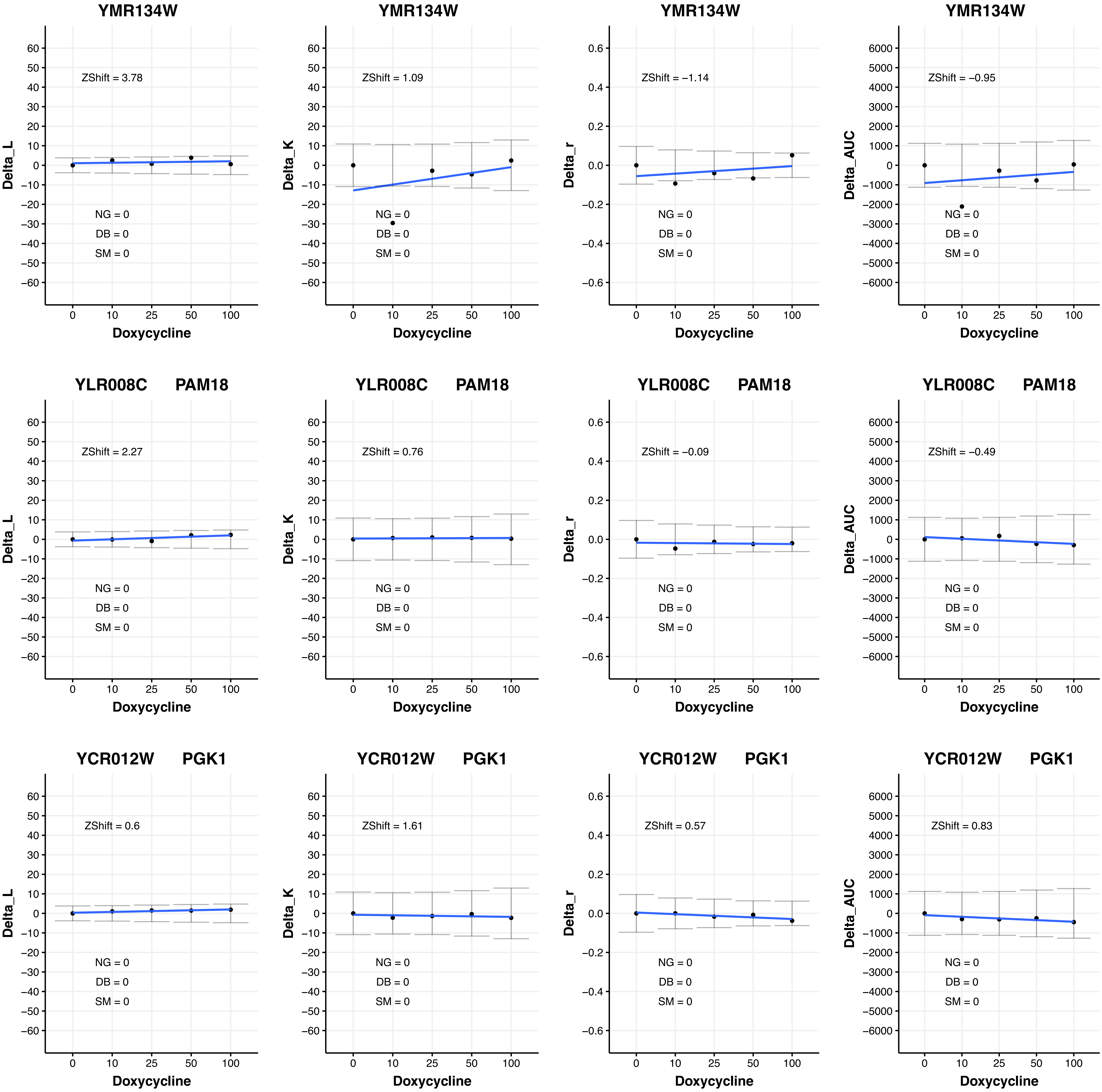
YMR134W
ZShift = 3.78
NG = 0
DB = 0
SM = 0
Delta_L
Doxycycline
YMR134W
ZShift = 1.09
NG = 0
DB = 0
SM = 0
Delta_K
Doxycycline
YMR134W
ZShift = −1.14
NG = 0
DB = 0
SM = 0
Delta_r
Doxycycline
YMR134W
ZShift = −0.95
NG = 0
DB = 0
SM = 0
Delta_AUC
Doxycycline
YLR008C PAM18
ZShift = 2.27
NG = 0
DB = 0
SM = 0
Delta_L
Doxycycline
YLR008C PAM18
ZShift = 0.76
NG = 0
DB = 0
SM = 0
Delta_K
Doxycycline
YLR008C PAM18
ZShift = −0.09
NG = 0
DB = 0
SM = 0
Delta_r
Doxycycline
YLR008C PAM18
ZShift = −0.49
NG = 0
DB = 0
SM = 0
Delta_AUC
Doxycycline
YCR012W PGK1
ZShift = 0.6
NG = 0
DB = 0
SM = 0
Delta_L
Doxycycline
YCR012W PGK1
ZShift = 1.61
NG = 0
DB = 0
SM = 0
Delta_K
Doxycycline
YCR012W PGK1
ZShift = 0.57
NG = 0
DB = 0
SM = 0
Delta_r
Doxycycline
YCR012W PGK1
ZShift = 0.83
NG = 0
DB = 0
SM = 0
Delta_AUC
Doxycycline

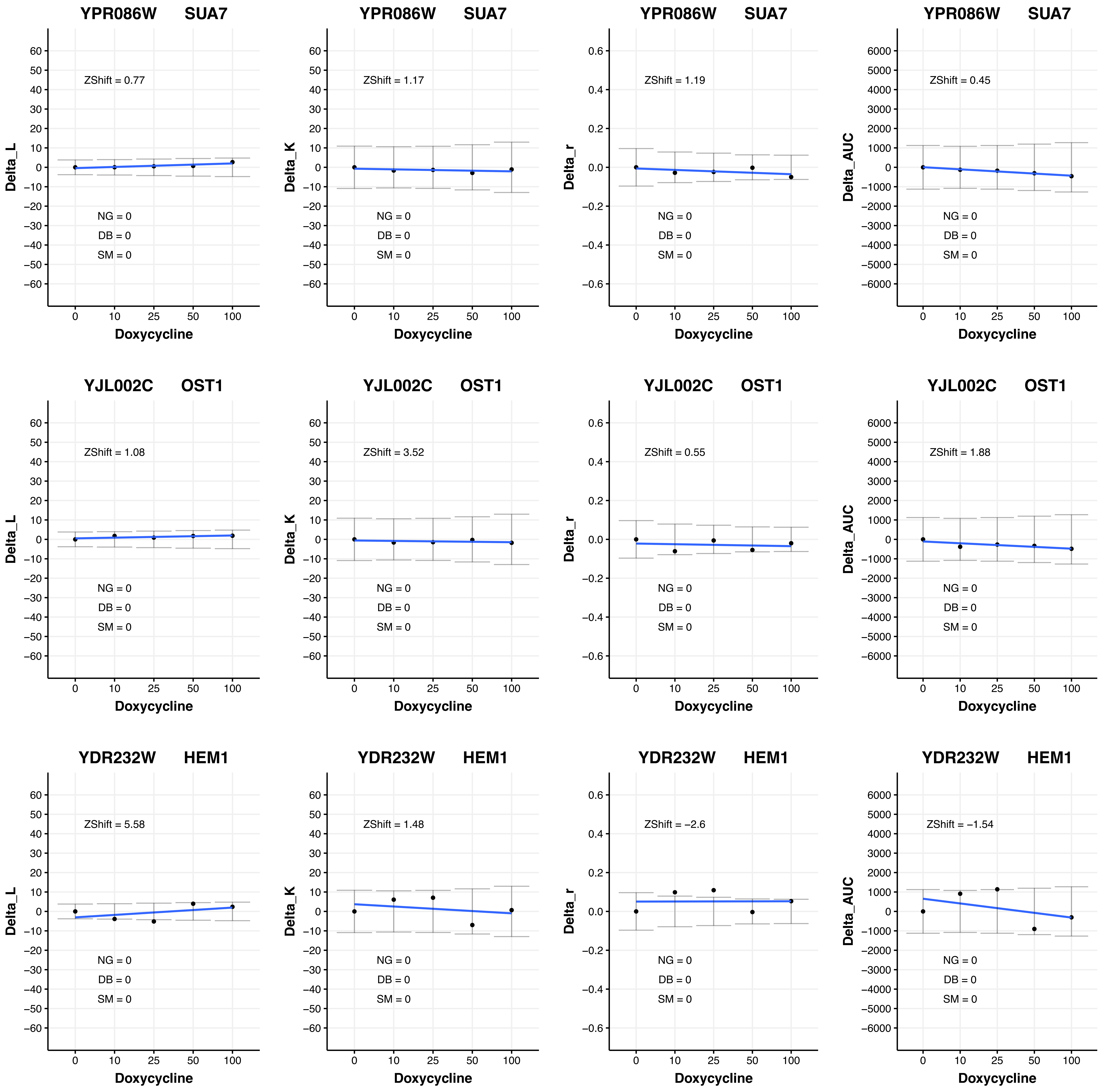
YPR086W SUA7
ZShift = 0.77
NG = 0
DB = 0
SM = 0
Delta_L
Doxycycline
YPR086W SUA7
ZShift = 1.17
NG = 0
DB = 0
SM = 0
Delta_K
Doxycycline
YPR086W SUA7
ZShift = 1.19
NG = 0
DB = 0
SM = 0
Delta_r
Doxycycline
YPR086W SUA7
ZShift = 0.45
NG = 0
DB = 0
SM = 0
Delta_AUC
Doxycycline
YJL002C OST1
ZShift = 1.08
NG = 0
DB = 0
SM = 0
Delta_L
Doxycycline
YJL002C OST1
ZShift = 3.52
NG = 0
DB = 0
SM = 0
Delta_K
Doxycycline
YJL002C OST1
ZShift = 0.55
NG = 0
DB = 0
SM = 0
Delta_r
Doxycycline
YJL002C OST1
ZShift = 1.88
NG = 0
DB = 0
SM = 0
Delta_AUC
Doxycycline
YDR232W HEM1
ZShift = 5.58
NG = 0
DB = 0
SM = 0
Delta_L
Doxycycline
YDR232W HEM1
ZShift = 1.48
NG = 0
DB = 0
SM = 0
Delta_K
Doxycycline
YDR232W HEM1
ZShift = −2.6
NG = 0
DB = 0
SM = 0
Delta_r
Doxycycline
YDR232W HEM1
ZShift = −1.54
NG = 0
DB = 0
SM = 0
Delta_AUC
Doxycycline

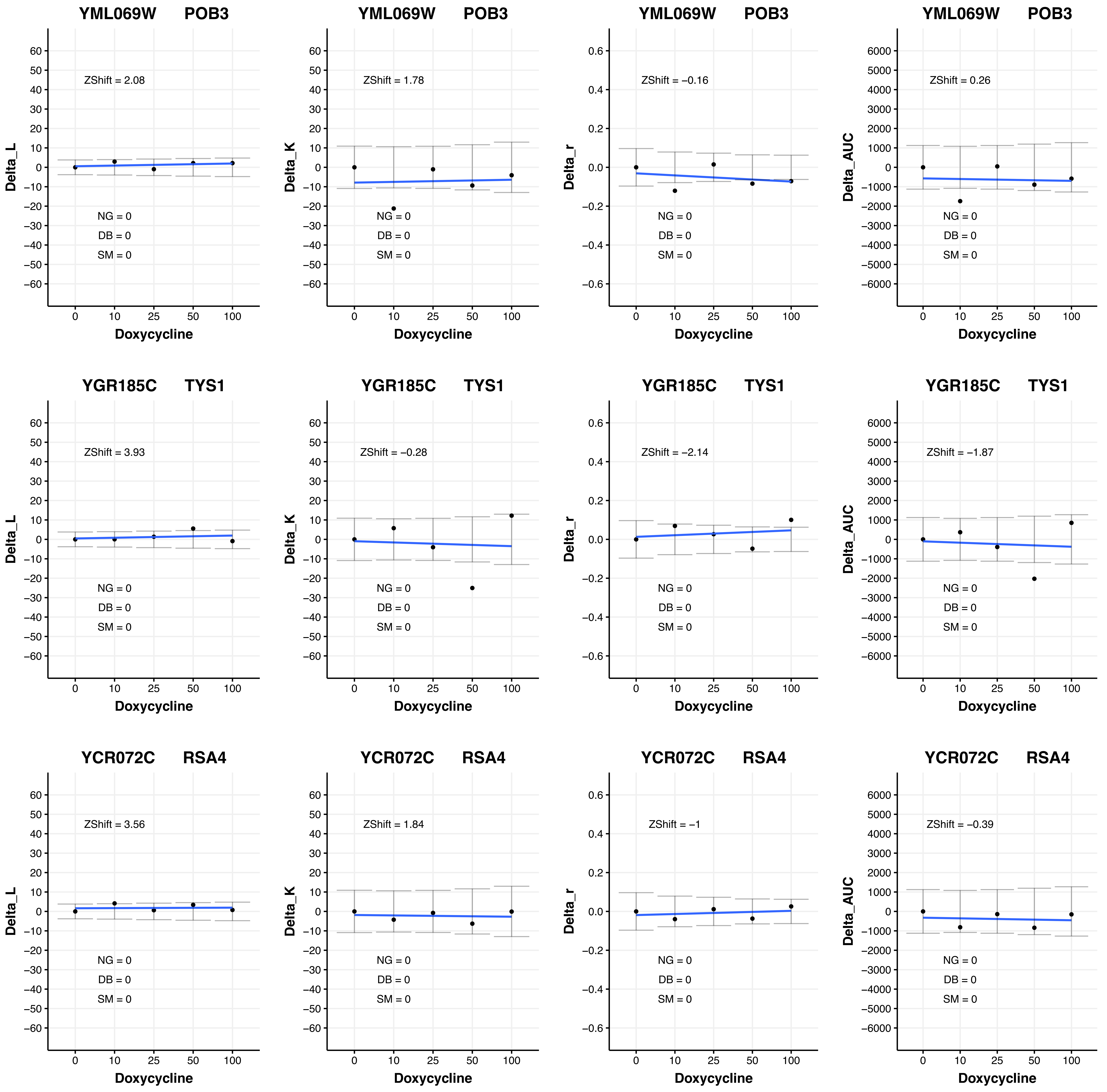
YML069W POB3
ZShift = 2.08
ZShift = 1.78
ZShift = −0.16
ZShift = 0.26
YGR185C TYS1
ZShift = 3.93
ZShift = −0.28
ZShift = −2.14
ZShift = −1.87
YCR072C RSA4
ZShift = 3.56
ZShift = 1.84
ZShift = −1
ZShift = −0.39
NG = 0
DB = 0
SM = 0
Delta_L
Delta_K
Delta_r
Delta_AUC
Doxycycline
0
10
25
50
100

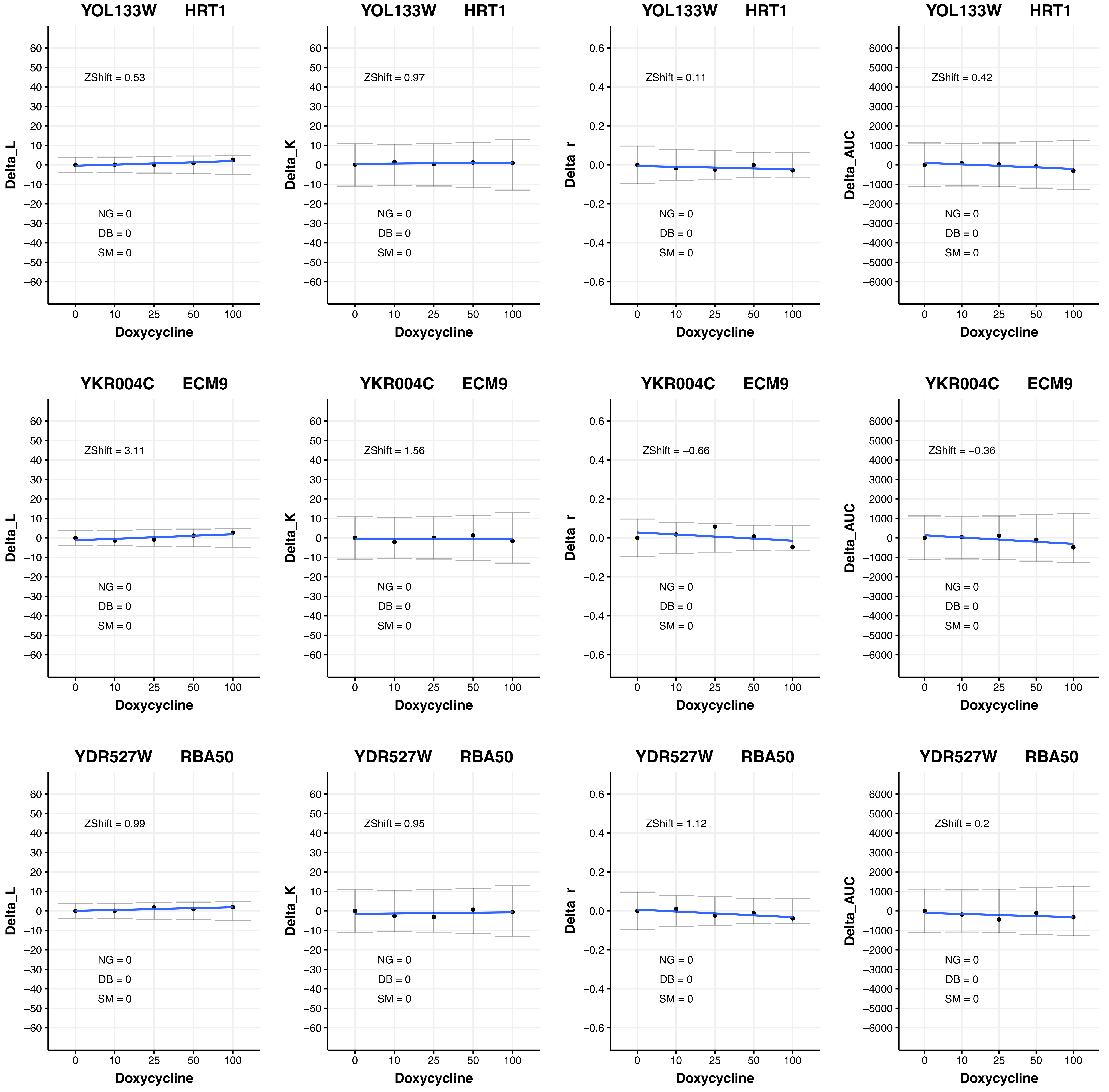
YOL133W HRT1
Delta_L
ZShift = 0.53
NG = 0
DB = 0
SM = 0
Doxycycline
YOL133W HRT1
Delta_K
ZShift = 0.97
NG = 0
DB = 0
SM = 0
Doxycycline
YOL133W HRT1
Delta_r
ZShift = 0.11
NG = 0
DB = 0
SM = 0
Doxycycline
YOL133W HRT1
Delta_AUC
ZShift = 0.42
NG = 0
DB = 0
SM = 0
Doxycycline
YKR004C ECM9
Delta_L
ZShift = 3.11
NG = 0
DB = 0
SM = 0
Doxycycline
YKR004C ECM9
Delta_K
ZShift = 1.56
NG = 0
DB = 0
SM = 0
Doxycycline
YKR004C ECM9
Delta_r
ZShift = −0.66
NG = 0
DB = 0
SM = 0
Doxycycline
YKR004C ECM9
Delta_AUC
ZShift = −0.36
NG = 0
DB = 0
SM = 0
Doxycycline
YDR527W RBA50
Delta_L
ZShift = 0.99
NG = 0
DB = 0
SM = 0
Doxycycline
YDR527W RBA50
Delta_K
ZShift = 0.95
NG = 0
DB = 0
SM = 0
Doxycycline
YDR527W RBA50
Delta_r
ZShift = 1.12
NG = 0
DB = 0
SM = 0
Doxycycline
YDR527W RBA50
Delta_AUC
ZShift = 0.2
NG = 0
DB = 0
SM = 0
Doxycycline

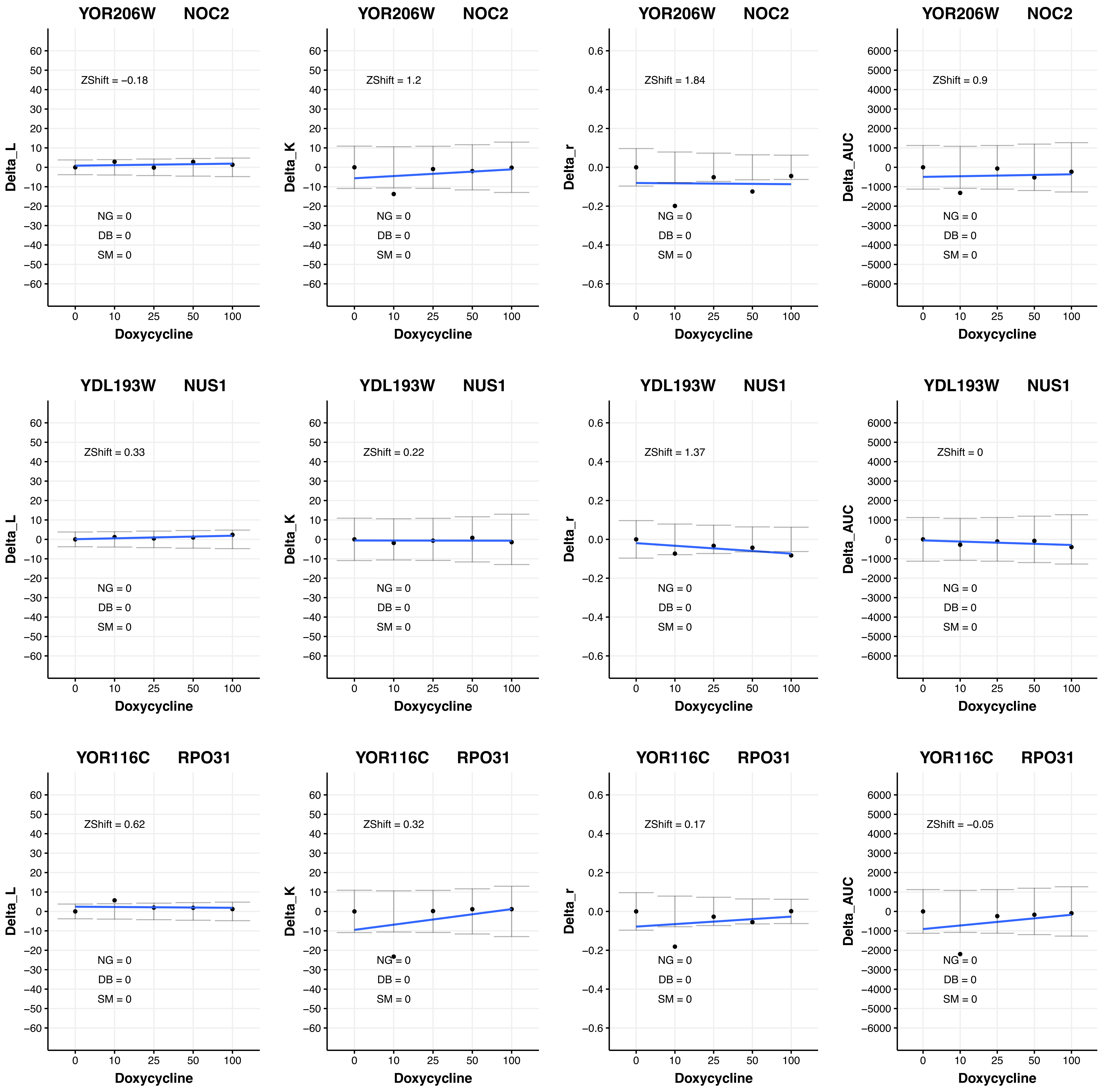
YOR206W NOC2
ZShift = −0.18
ZShift = 1.2
ZShift = 1.84
ZShift = 0.9
YDL193W NUS1
ZShift = 0.33
ZShift = 0.22
ZShift = 1.37
ZShift = 0
YOR116C RPO31
ZShift = 0.62
ZShift = 0.32
ZShift = 0.17
ZShift = −0.05
NG = 0
DB = 0
SM = 0
Delta_L
Delta_K
Delta_r
Delta_AUC
Doxycycline
0
10
25
50
100

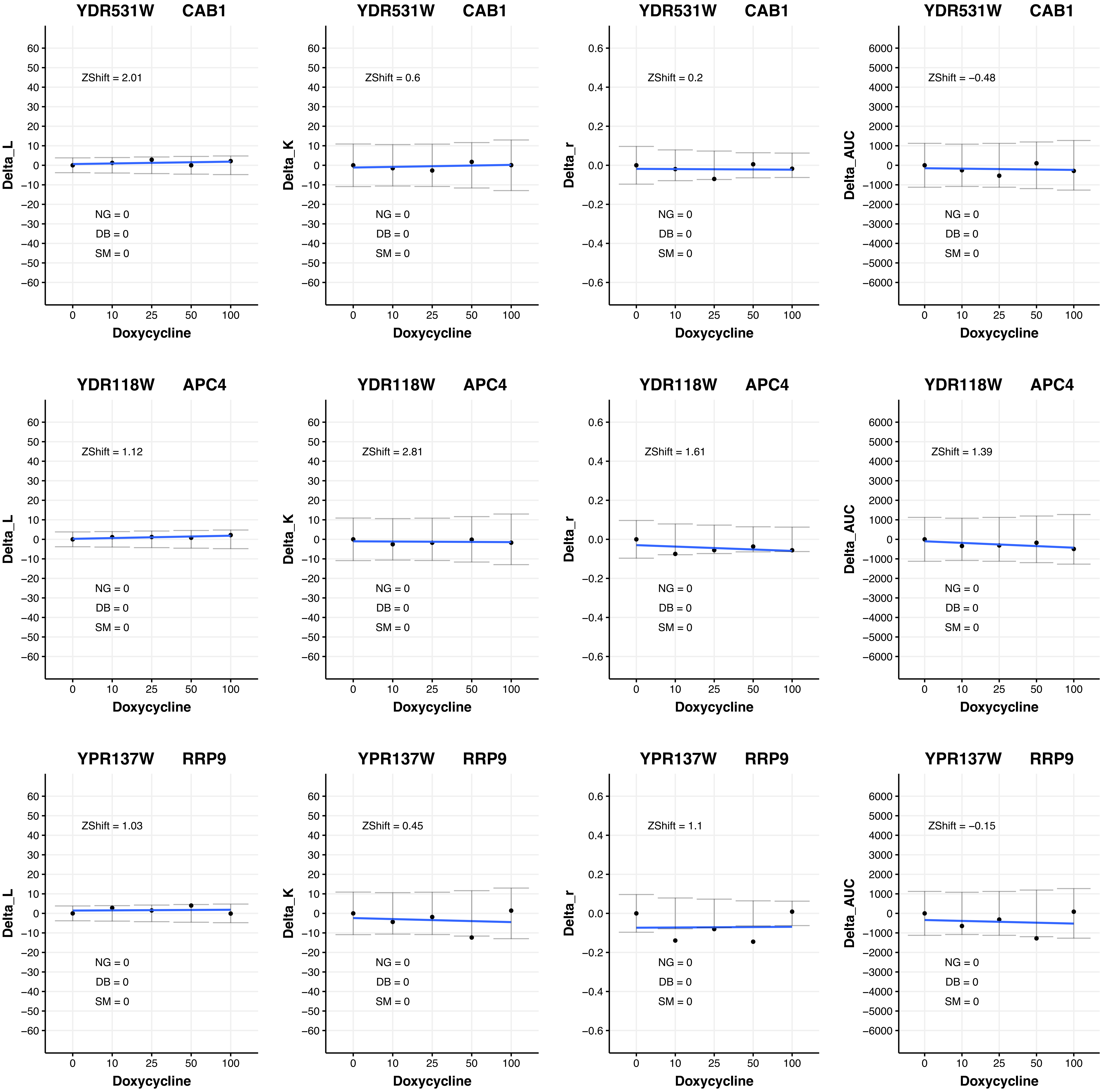
YDR531W CAB1
ZShift = 2.01
NG = 0
DB = 0
SM = 0
Delta_L
Doxycycline
YDR531W CAB1
ZShift = 0.6
NG = 0
DB = 0
SM = 0
Delta_K
Doxycycline
YDR531W CAB1
ZShift = 0.2
NG = 0
DB = 0
SM = 0
Delta_r
Doxycycline
YDR531W CAB1
ZShift = −0.48
NG = 0
DB = 0
SM = 0
Delta_AUC
Doxycycline
YDR118W APC4
ZShift = 1.12
NG = 0
DB = 0
SM = 0
Delta_L
Doxycycline
YDR118W APC4
ZShift = 2.81
NG = 0
DB = 0
SM = 0
Delta_K
Doxycycline
YDR118W APC4
ZShift = 1.61
NG = 0
DB = 0
SM = 0
Delta_r
Doxycycline
YDR118W APC4
ZShift = 1.39
NG = 0
DB = 0
SM = 0
Delta_AUC
Doxycycline
YPR137W RRP9
ZShift = 1.03
NG = 0
DB = 0
SM = 0
Delta_L
Doxycycline
YPR137W RRP9
ZShift = 0.45
NG = 0
DB = 0
SM = 0
Delta_K
Doxycycline
YPR137W RRP9
ZShift = 1.1
NG = 0
DB = 0
SM = 0
Delta_r
Doxycycline
YPR137W RRP9
ZShift = −0.15
NG = 0
DB = 0
SM = 0
Delta_AUC
Doxycycline

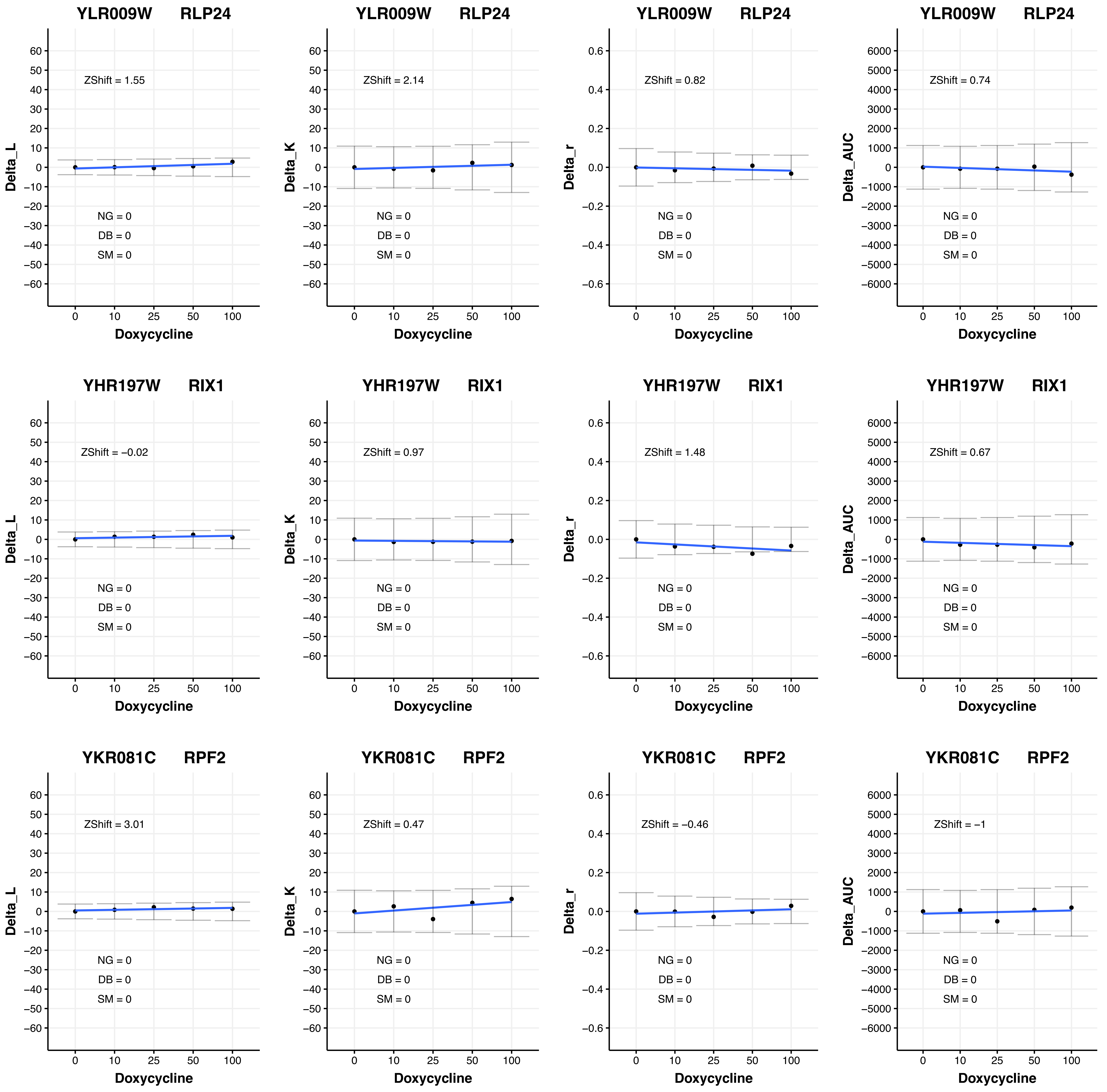
YLR009W RLP24
Delta_L
ZShift = 1.55
NG = 0
DB = 0
SM = 0
Doxycycline
YLR009W RLP24
Delta_K
ZShift = 2.14
NG = 0
DB = 0
SM = 0
Doxycycline
YLR009W RLP24
Delta_r
ZShift = 0.82
NG = 0
DB = 0
SM = 0
Doxycycline
YLR009W RLP24
Delta_AUC
ZShift = 0.74
NG = 0
DB = 0
SM = 0
Doxycycline
YHR197W RIX1
Delta_L
ZShift = −0.02
NG = 0
DB = 0
SM = 0
Doxycycline
YHR197W RIX1
Delta_K
ZShift = 0.97
NG = 0
DB = 0
SM = 0
Doxycycline
YHR197W RIX1
Delta_r
ZShift = 1.48
NG = 0
DB = 0
SM = 0
Doxycycline
YHR197W RIX1
Delta_AUC
ZShift = 0.67
NG = 0
DB = 0
SM = 0
Doxycycline
YKR081C RPF2
Delta_L
ZShift = 3.01
NG = 0
DB = 0
SM = 0
Doxycycline
YKR081C RPF2
Delta_K
ZShift = 0.47
NG = 0
DB = 0
SM = 0
Doxycycline
YKR081C RPF2
Delta_r
ZShift = −0.46
NG = 0
DB = 0
SM = 0
Doxycycline
YKR081C RPF2
Delta_AUC
ZShift = −1
NG = 0
DB = 0
SM = 0
Doxycycline

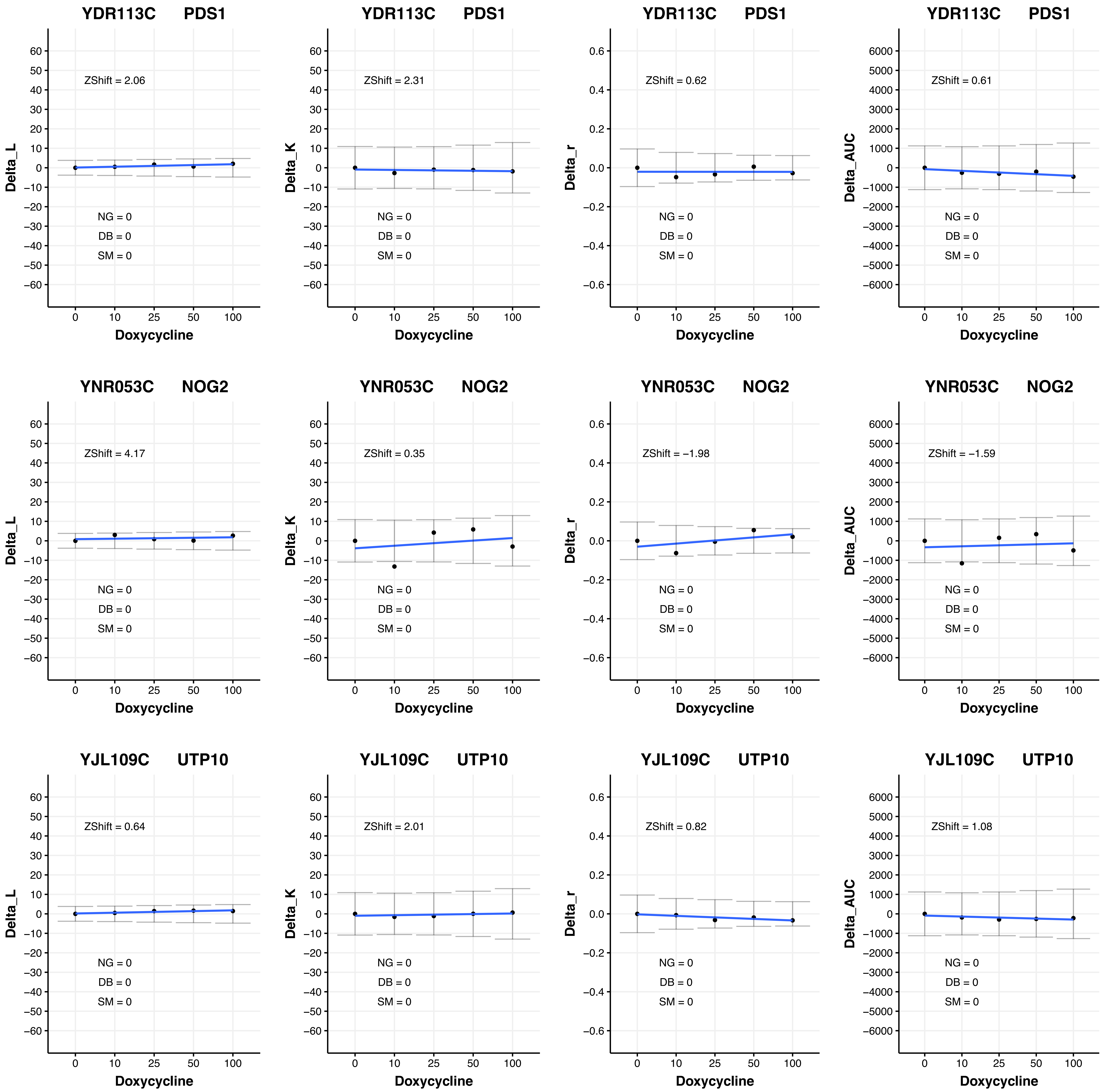

YDR113C PDS1
ZShift = 2.06
ZShift = 2.31
ZShift = 0.62
ZShift = 0.61
YNR053C NOG2
ZShift = 4.17
ZShift = 0.35
ZShift = −1.98
ZShift = −1.59
YJL109C UTP10
ZShift = 0.64
ZShift = 2.01
ZShift = 0.82
ZShift = 1.08
NG = 0
DB = 0
SM = 0
Delta_L
Delta_K
Delta_r
Delta_AUC
Doxycycline
0
10
25
50
100

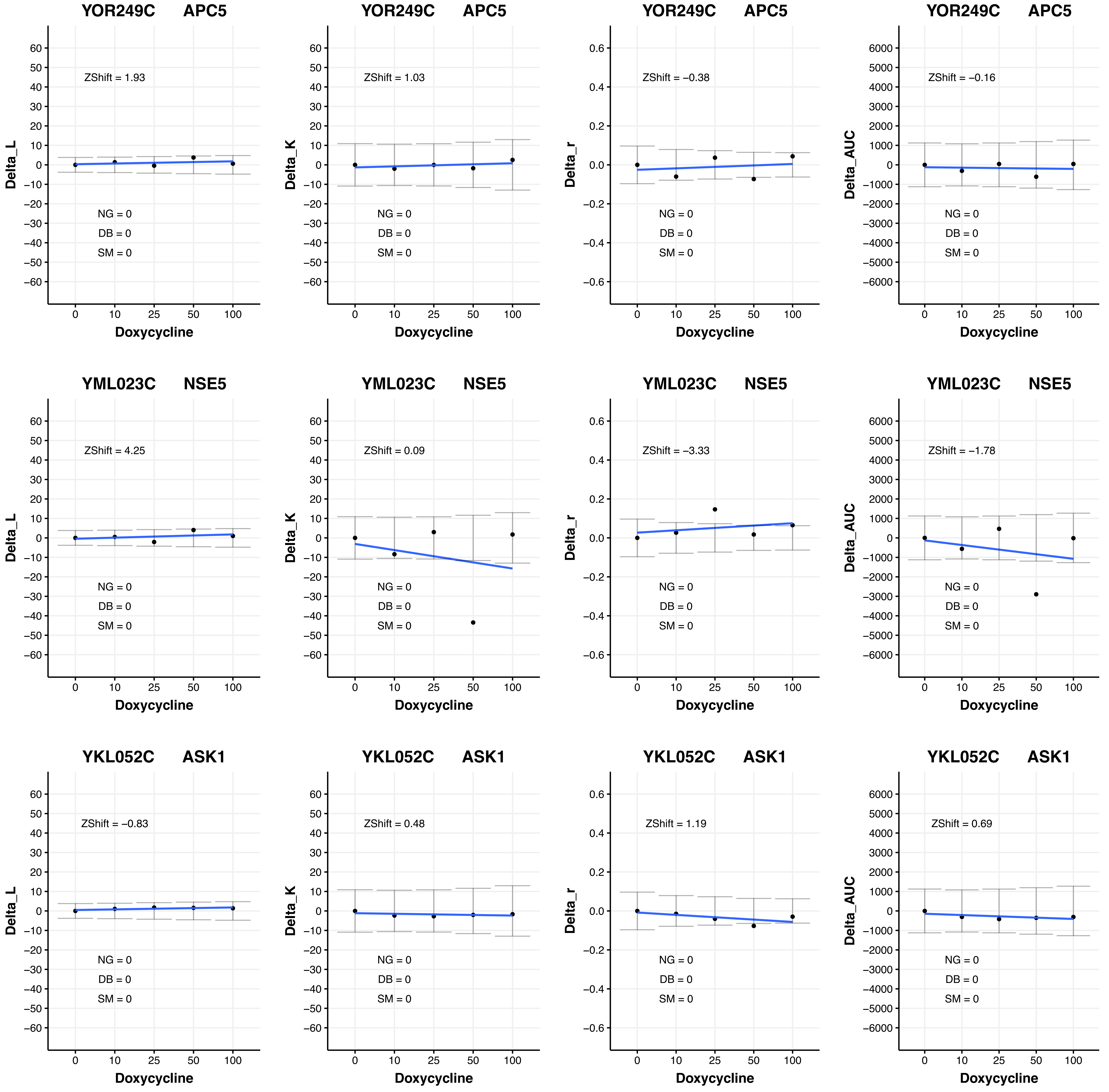

YOR249C APC5
ZShift = 1.93
NG = 0
DB = 0
SM = 0
Delta_L
Doxycycline
YOR249C APC5
ZShift = 1.03
NG = 0
DB = 0
SM = 0
Delta_K
Doxycycline
YOR249C APC5
ZShift = −0.38
NG = 0
DB = 0
SM = 0
Delta_r
Doxycycline
YOR249C APC5
ZShift = −0.16
NG = 0
DB = 0
SM = 0
Delta_AUC
Doxycycline
YML023C NSE5
ZShift = 4.25
NG = 0
DB = 0
SM = 0
Delta_L
Doxycycline
YML023C NSE5
ZShift = 0.09
NG = 0
DB = 0
SM = 0
Delta_K
Doxycycline
YML023C NSE5
ZShift = −3.33
NG = 0
DB = 0
SM = 0
Delta_r
Doxycycline
YML023C NSE5
ZShift = −1.78
NG = 0
DB = 0
SM = 0
Delta_AUC
Doxycycline
YKL052C ASK1
ZShift = −0.83
NG = 0
DB = 0
SM = 0
Delta_L
Doxycycline
YKL052C ASK1
ZShift = 0.48
NG = 0
DB = 0
SM = 0
Delta_K
Doxycycline
YKL052C ASK1
ZShift = 1.19
NG = 0
DB = 0
SM = 0
Delta_r
Doxycycline
YKL052C ASK1
ZShift = 0.69
NG = 0
DB = 0
SM = 0
Delta_AUC
Doxycycline

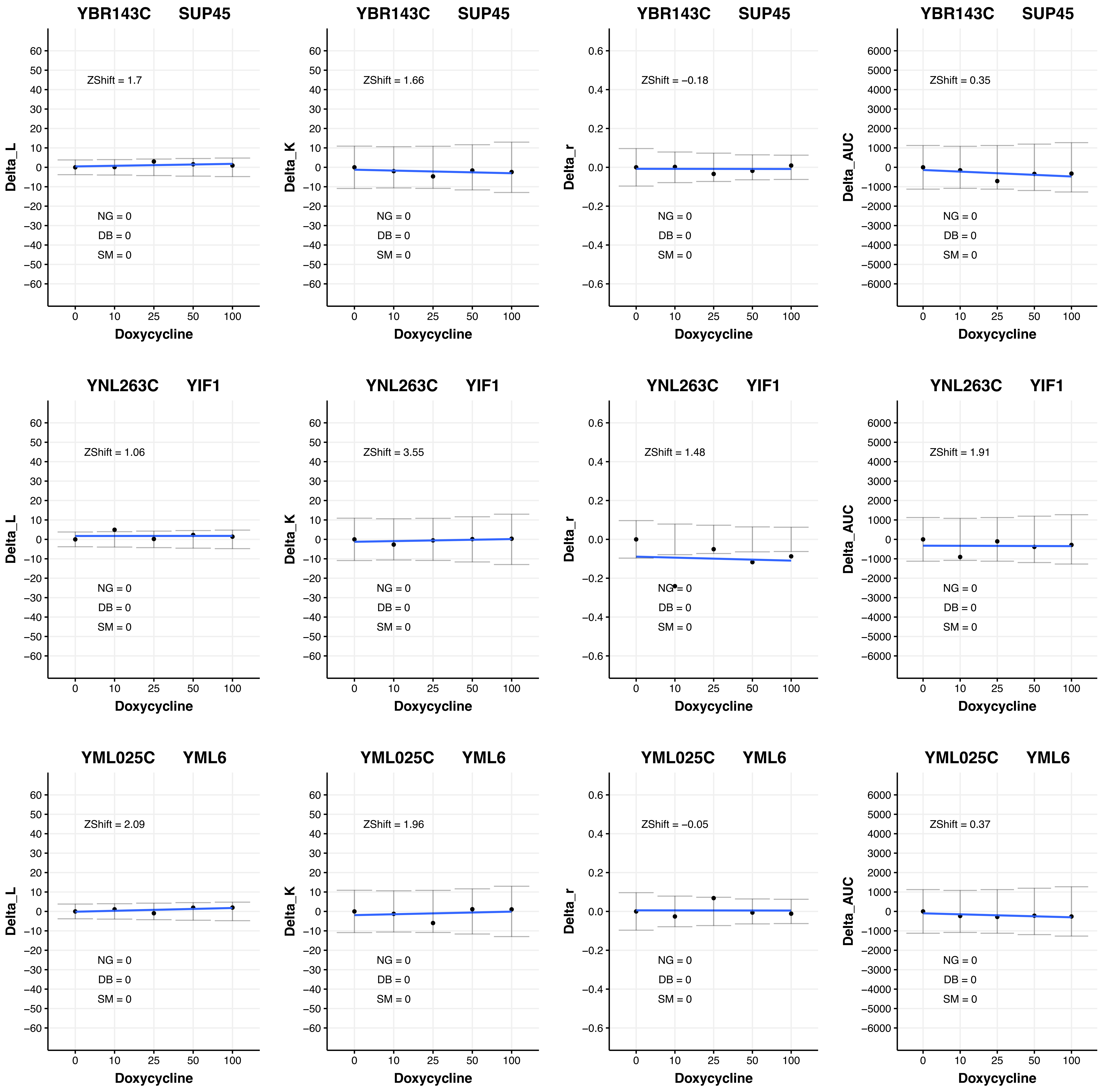

YBR143C SUP45
ZShift = 1.7
NG = 0
DB = 0
SM = 0
Delta_L
Doxycycline
YBR143C SUP45
ZShift = 1.66
NG = 0
DB = 0
SM = 0
Delta_K
Doxycycline
YBR143C SUP45
ZShift = −0.18
NG = 0
DB = 0
SM = 0
Delta_r
Doxycycline
YBR143C SUP45
ZShift = 0.35
NG = 0
DB = 0
SM = 0
Delta_AUC
Doxycycline
YNL263C YIF1
ZShift = 1.06
NG = 0
DB = 0
SM = 0
Delta_L
Doxycycline
YNL263C YIF1
ZShift = 3.55
NG = 0
DB = 0
SM = 0
Delta_K
Doxycycline
YNL263C YIF1
ZShift = 1.48
NG = 0
DB = 0
SM = 0
Delta_r
Doxycycline
YNL263C YIF1
ZShift = 1.91
NG = 0
DB = 0
SM = 0
Delta_AUC
Doxycycline
YML025C YML6
ZShift = 2.09
NG = 0
DB = 0
SM = 0
Delta_L
Doxycycline
YML025C YML6
ZShift = 1.96
NG = 0
DB = 0
SM = 0
Delta_K
Doxycycline
YML025C YML6
ZShift = −0.05
NG = 0
DB = 0
SM = 0
Delta_r
Doxycycline
YML025C YML6
ZShift = 0.37
NG = 0
DB = 0
SM = 0
Delta_AUC
Doxycycline
0 10 25 50 100

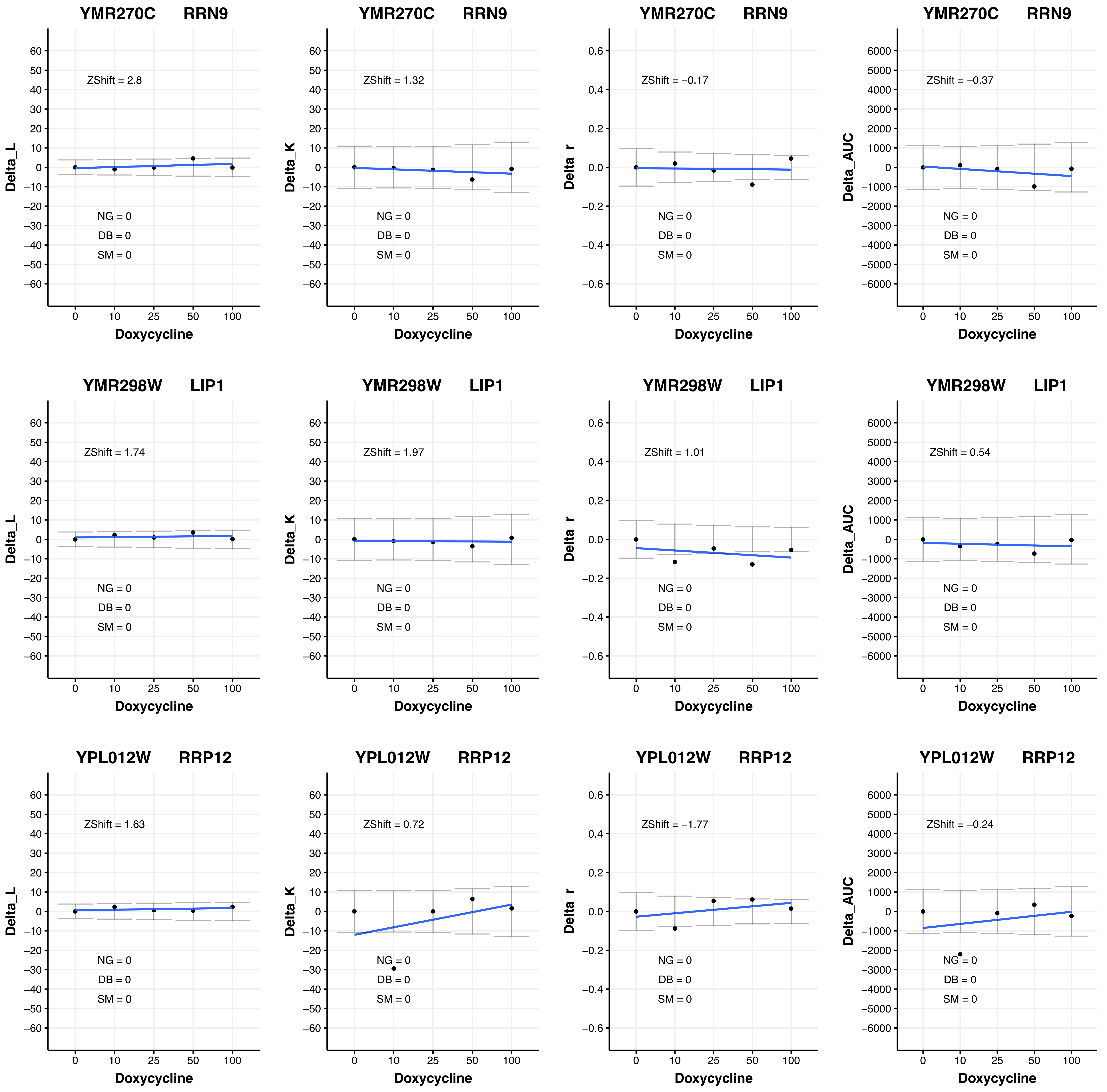

YMR270C RRN9
ZShift = 2.8
NG = 0
DB = 0
SM = 0
Delta_L
Doxycycline
YMR270C RRN9
ZShift = 1.32
NG = 0
DB = 0
SM = 0
Delta_K
Doxycycline
YMR270C RRN9
ZShift = −0.17
NG = 0
DB = 0
SM = 0
Delta_r
Doxycycline
YMR270C RRN9
ZShift = −0.37
NG = 0
DB = 0
SM = 0
Delta_AUC
Doxycycline
YMR298W LIP1
ZShift = 1.74
NG = 0
DB = 0
SM = 0
Delta_L
Doxycycline
YMR298W LIP1
ZShift = 1.97
NG = 0
DB = 0
SM = 0
Delta_K
Doxycycline
YMR298W LIP1
ZShift = 1.01
NG = 0
DB = 0
SM = 0
Delta_r
Doxycycline
YMR298W LIP1
ZShift = 0.54
NG = 0
DB = 0
SM = 0
Delta_AUC
Doxycycline
YPL012W RRP12
ZShift = 1.63
NG = 0
DB = 0
SM = 0
Delta_L
Doxycycline
YPL012W RRP12
ZShift = 0.72
NG = 0
DB = 0
SM = 0
Delta_K
Doxycycline
YPL012W RRP12
ZShift = −1.77
NG = 0
DB = 0
SM = 0
Delta_r
Doxycycline
YPL012W RRP12
ZShift = −0.24
NG = 0
DB = 0
SM = 0
Delta_AUC
Doxycycline

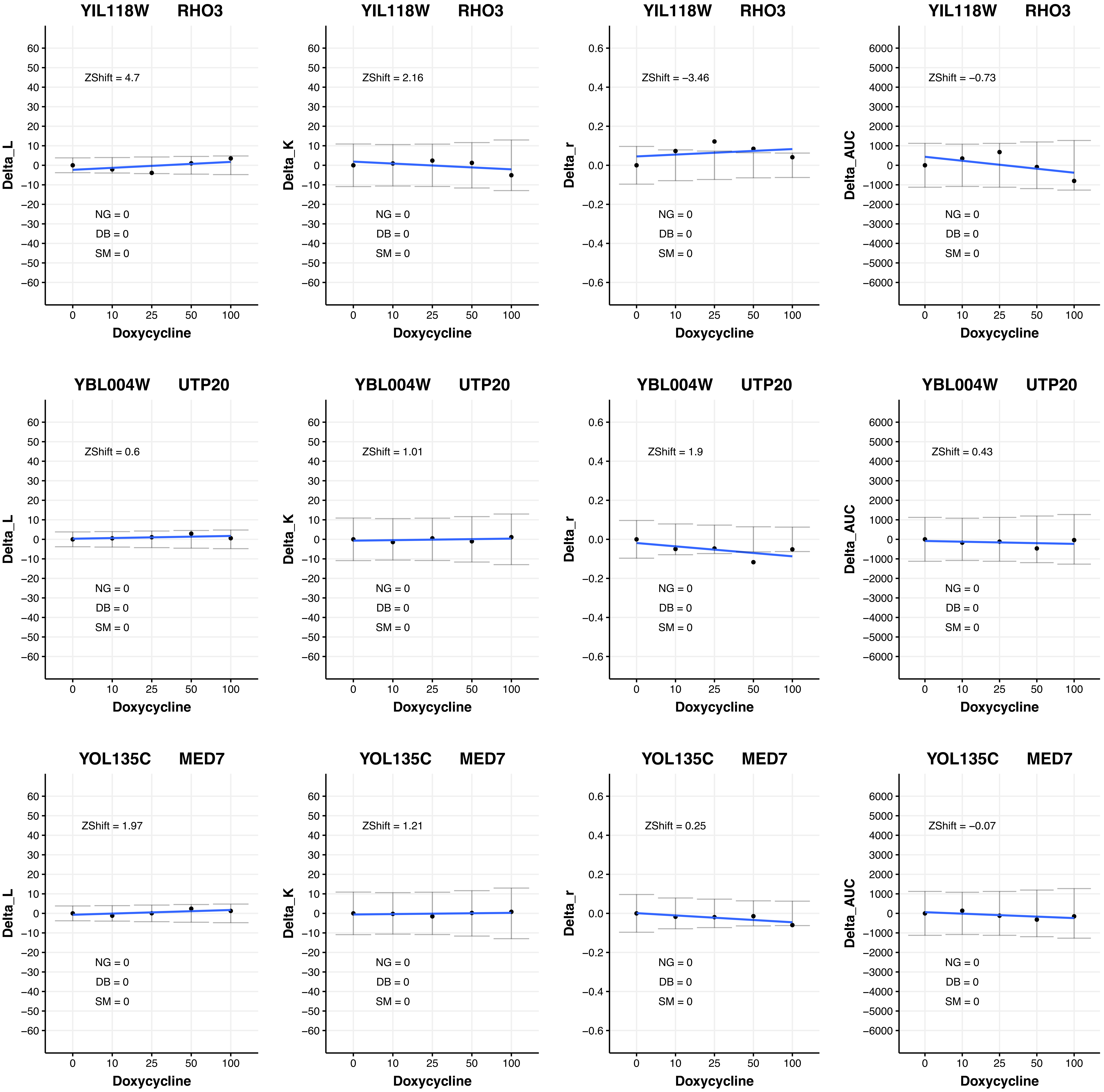
YIL118W RHO3
ZShift = 4.7
NG = 0
DB = 0
SM = 0
Delta_L
Doxycycline
YIL118W RHO3
ZShift = 2.16
NG = 0
DB = 0
SM = 0
Delta_K
Doxycycline
YIL118W RHO3
ZShift = −3.46
NG = 0
DB = 0
SM = 0
Delta_r
Doxycycline
YIL118W RHO3
ZShift = −0.73
NG = 0
DB = 0
SM = 0
Delta_AUC
Doxycycline
YBL004W UTP20
ZShift = 0.6
NG = 0
DB = 0
SM = 0
Delta_L
Doxycycline
YBL004W UTP20
ZShift = 1.01
NG = 0
DB = 0
SM = 0
Delta_K
Doxycycline
YBL004W UTP20
ZShift = 1.9
NG = 0
DB = 0
SM = 0
Delta_r
Doxycycline
YBL004W UTP20
ZShift = 0.43
NG = 0
DB = 0
SM = 0
Delta_AUC
Doxycycline
YOL135C MED7
ZShift = 1.97
NG = 0
DB = 0
SM = 0
Delta_L
Doxycycline
YOL135C MED7
ZShift = 1.21
NG = 0
DB = 0
SM = 0
Delta_K
Doxycycline
YOL135C MED7
ZShift = 0.25
NG = 0
DB = 0
SM = 0
Delta_r
Doxycycline
YOL135C MED7
ZShift = −0.07
NG = 0
DB = 0
SM = 0
Delta_AUC
Doxycycline

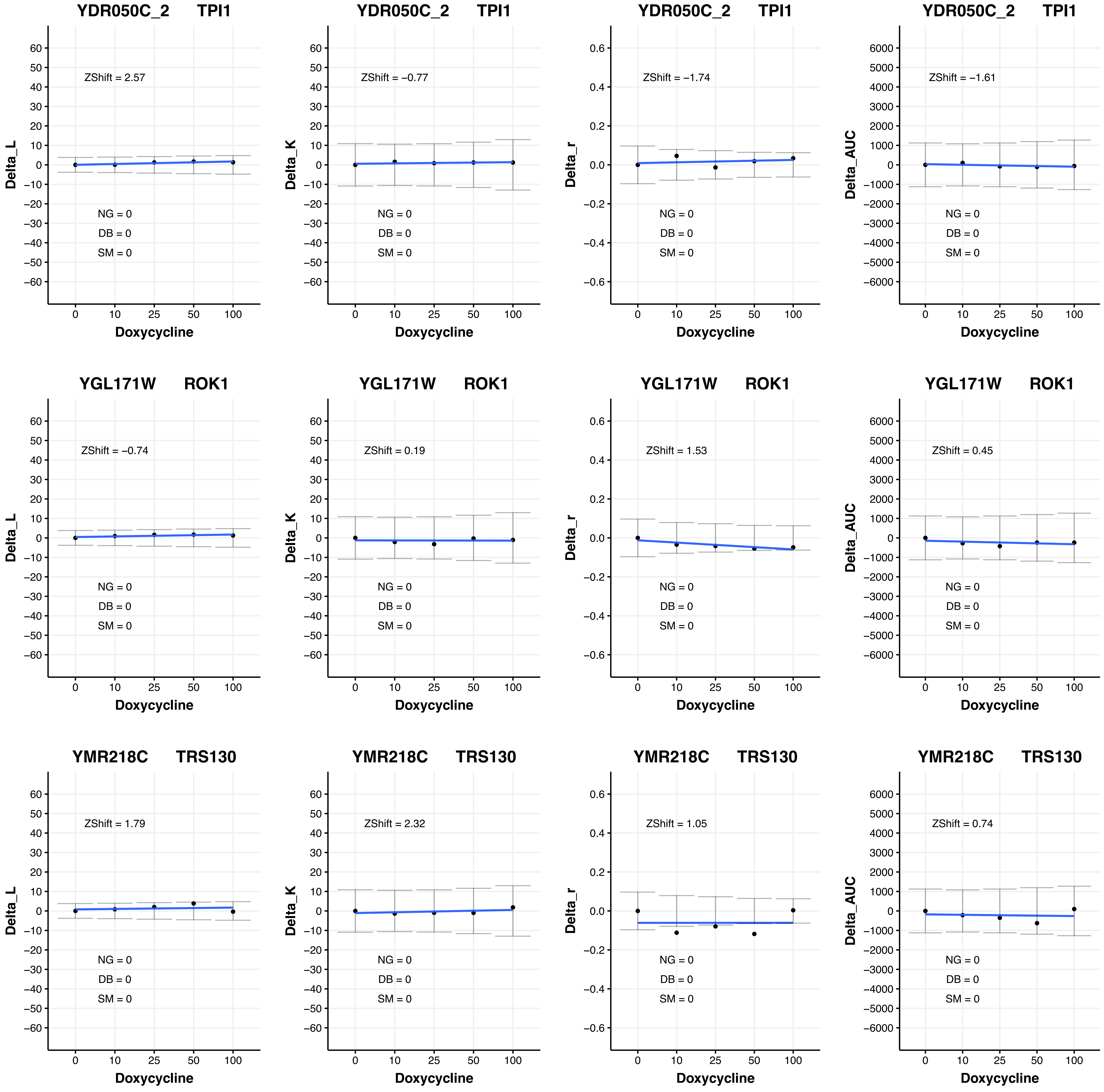
YDR050C_2 TPI1
ZShift = 2.57
NG = 0
DB = 0
SM = 0
Delta_L
Doxycycline
YDR050C_2 TPI1
ZShift = −0.77
NG = 0
DB = 0
SM = 0
Delta_K
Doxycycline
YDR050C_2 TPI1
ZShift = −1.74
NG = 0
DB = 0
SM = 0
Delta_r
Doxycycline
YDR050C_2 TPI1
ZShift = −1.61
NG = 0
DB = 0
SM = 0
Delta_AUC
Doxycycline
YGL171W ROK1
ZShift = −0.74
NG = 0
DB = 0
SM = 0
Delta_L
Doxycycline
YGL171W ROK1
ZShift = 0.19
NG = 0
DB = 0
SM = 0
Delta_K
Doxycycline
YGL171W ROK1
ZShift = 1.53
NG = 0
DB = 0
SM = 0
Delta_r
Doxycycline
YGL171W ROK1
ZShift = 0.45
NG = 0
DB = 0
SM = 0
Delta_AUC
Doxycycline
YMR218C TRS130
ZShift = 1.79
NG = 0
DB = 0
SM = 0
Delta_L
Doxycycline
YMR218C TRS130
ZShift = 2.32
NG = 0
DB = 0
SM = 0
Delta_K
Doxycycline
YMR218C TRS130
ZShift = 1.05
NG = 0
DB = 0
SM = 0
Delta_r
Doxycycline
YMR218C TRS130
ZShift = 0.74
NG = 0
DB = 0
SM = 0
Delta_AUC
Doxycycline

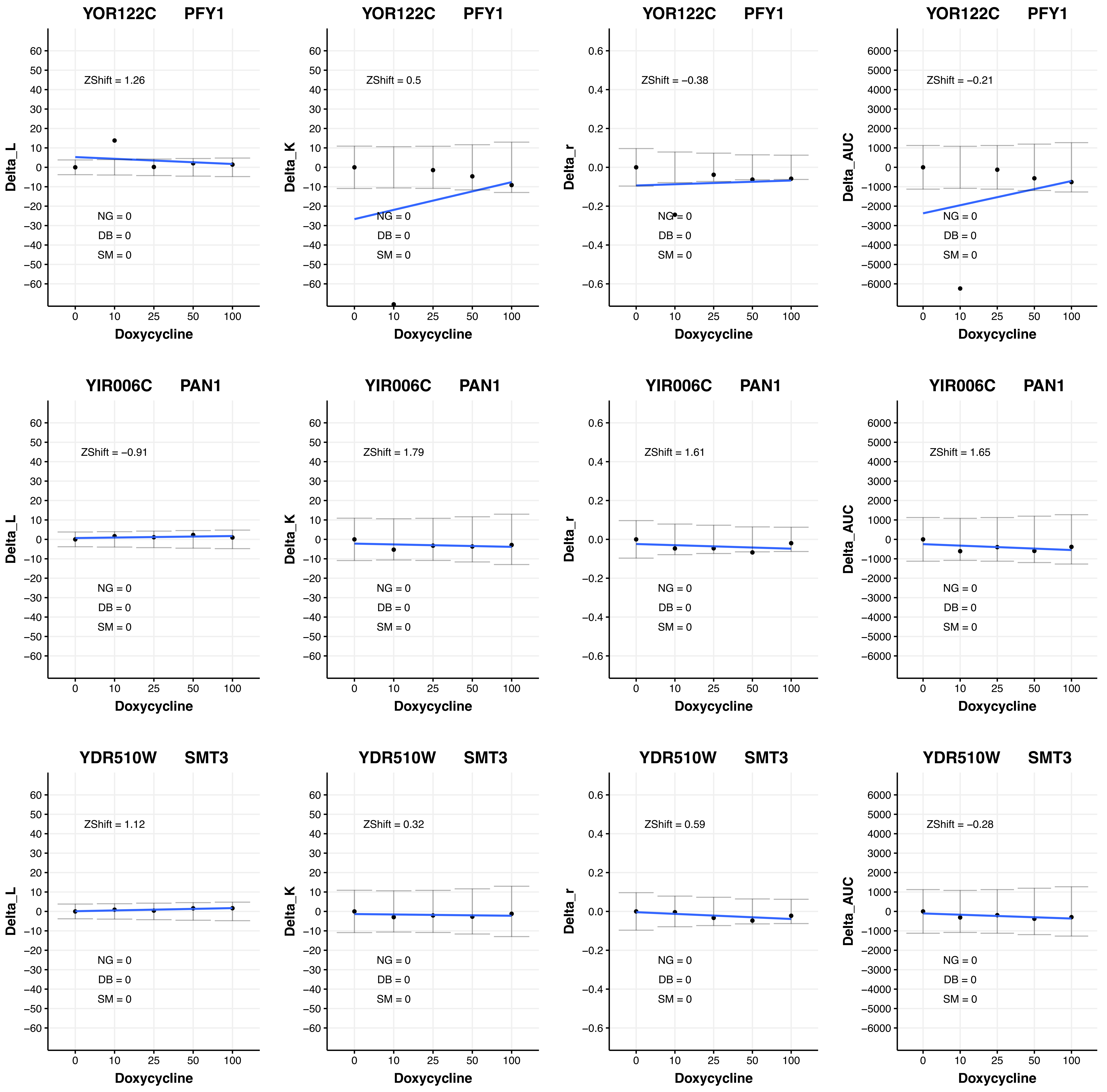

YOR122C PFY1
ZShift = 1.26
ZShift = 0.5
ZShift = −0.38
ZShift = −0.21
YIR006C PAN1
ZShift = −0.91
ZShift = 1.79
ZShift = 1.61
ZShift = 1.65
YDR510W SMT3
ZShift = 1.12
ZShift = 0.32
ZShift = 0.59
ZShift = −0.28
NG = 0
DB = 0
SM = 0
Delta_L
Delta_K
Delta_r
Delta_AUC
Doxycycline
0
10
25
50
100

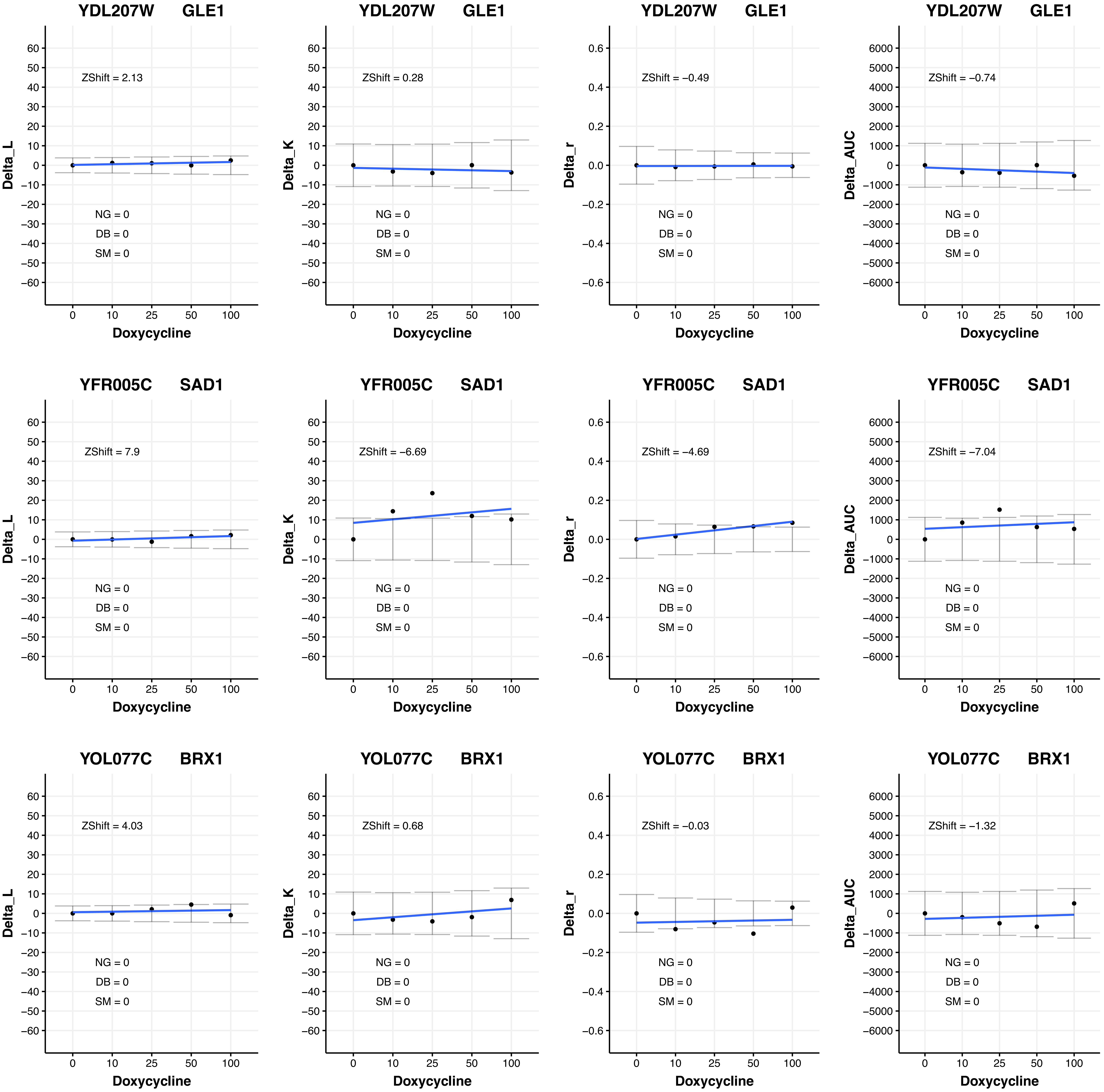

YDL207W GLE1
ZShift = 2.13
Delta_L
ZShift = 0.28
Delta_K
ZShift = −0.49
Delta_r
ZShift = −0.74
Delta_AUC
YFR005C SAD1
ZShift = 7.9
ZShift = −6.69
ZShift = −4.69
ZShift = −7.04
YOL077C BRX1
ZShift = 4.03
ZShift = 0.68
ZShift = −0.03
ZShift = −1.32
NG = 0
DB = 0
SM = 0
Doxycycline
0
10
25
50
100

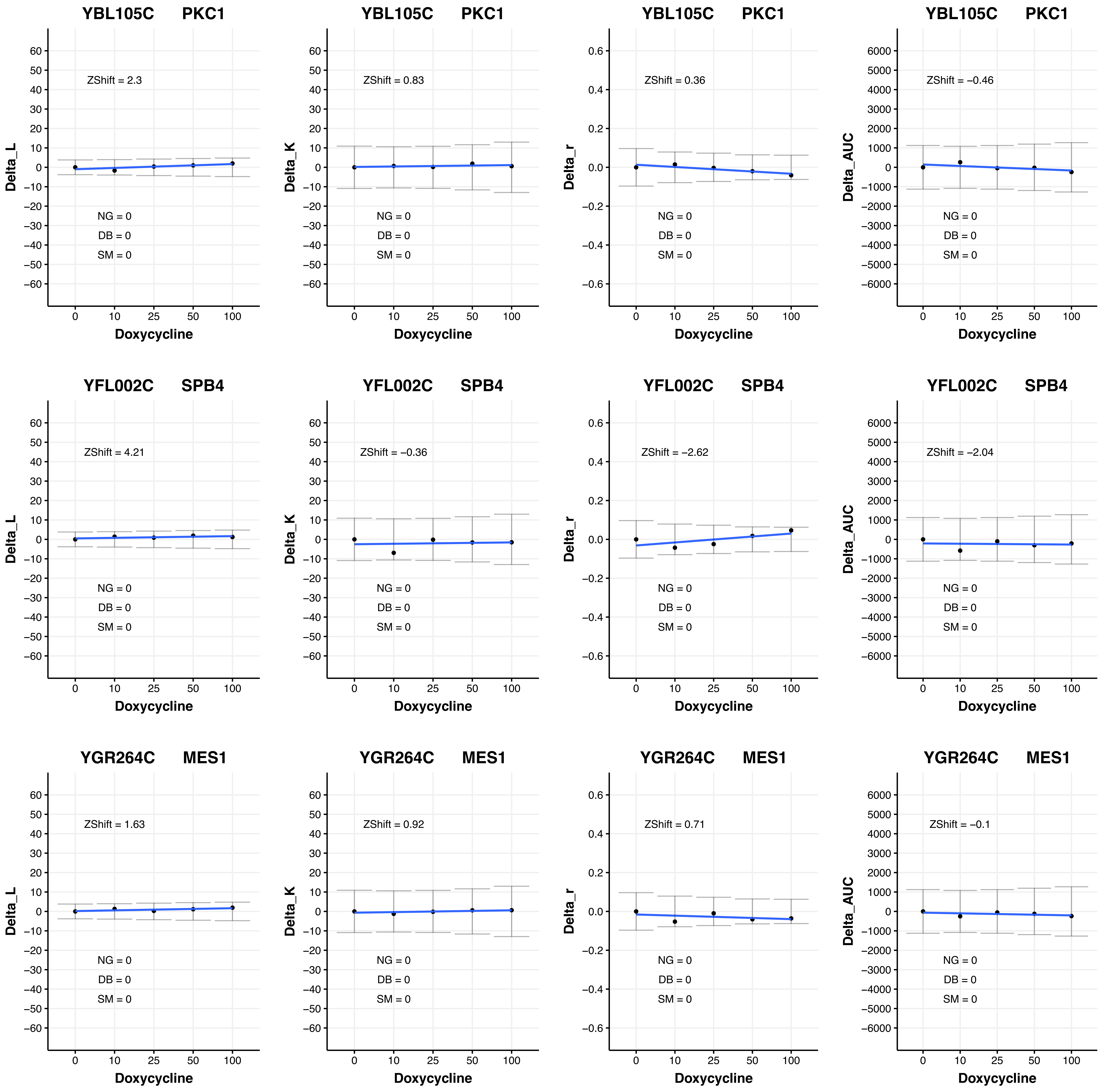

YBL105C PKC1
ZShift = 2.3
Delta_L
NG = 0
DB = 0
SM = 0
Doxycycline
YBL105C PKC1
ZShift = 0.83
Delta_K
NG = 0
DB = 0
SM = 0
Doxycycline
YBL105C PKC1
ZShift = 0.36
Delta_r
NG = 0
DB = 0
SM = 0
Doxycycline
YBL105C PKC1
ZShift = −0.46
Delta_AUC
NG = 0
DB = 0
SM = 0
Doxycycline
YFL002C SPB4
ZShift = 4.21
Delta_L
NG = 0
DB = 0
SM = 0
Doxycycline
YFL002C SPB4
ZShift = −0.36
Delta_K
NG = 0
DB = 0
SM = 0
Doxycycline
YFL002C SPB4
ZShift = −2.62
Delta_r
NG = 0
DB = 0
SM = 0
Doxycycline
YFL002C SPB4
ZShift = −2.04
Delta_AUC
NG = 0
DB = 0
SM = 0
Doxycycline
YGR264C MES1
ZShift = 1.63
Delta_L
NG = 0
DB = 0
SM = 0
Doxycycline
YGR264C MES1
ZShift = 0.92
Delta_K
NG = 0
DB = 0
SM = 0
Doxycycline
YGR264C MES1
ZShift = 0.71
Delta_r
NG = 0
DB = 0
SM = 0
Doxycycline
YGR264C MES1
ZShift = −0.1
Delta_AUC
NG = 0
DB = 0
SM = 0
Doxycycline

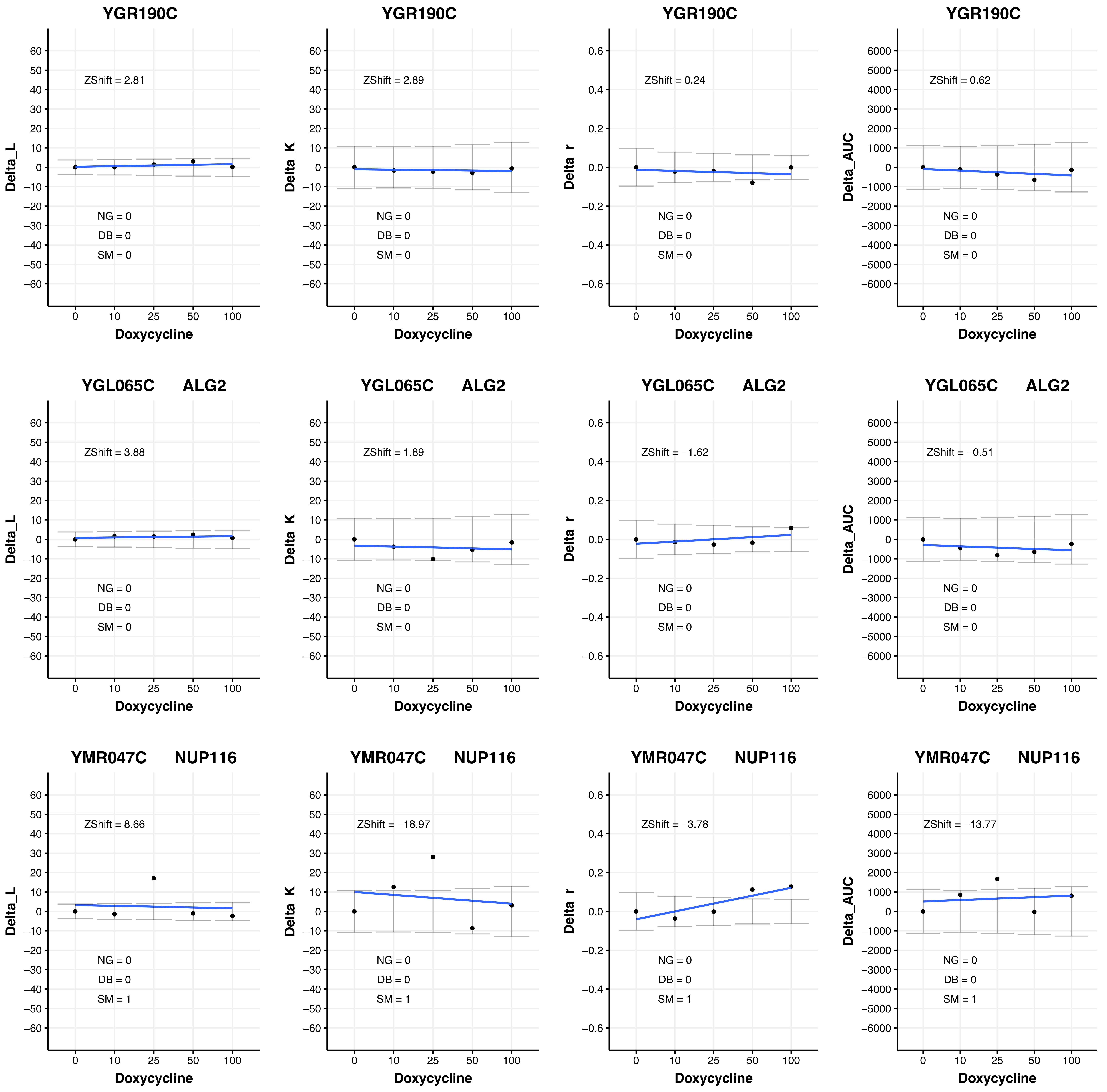
YGR190C
ZShift = 2.81
Delta_L
NG = 0
DB = 0
SM = 0
Doxycycline
YGR190C
ZShift = 2.89
Delta_K
NG = 0
DB = 0
SM = 0
Doxycycline
YGR190C
ZShift = 0.24
Delta_r
NG = 0
DB = 0
SM = 0
Doxycycline
YGR190C
ZShift = 0.62
Delta_AUC
NG = 0
DB = 0
SM = 0
Doxycycline
YGL065C ALG2
ZShift = 3.88
Delta_L
NG = 0
DB = 0
SM = 0
Doxycycline
YGL065C ALG2
ZShift = 1.89
Delta_K
NG = 0
DB = 0
SM = 0
Doxycycline
YGL065C ALG2
ZShift = −1.62
Delta_r
NG = 0
DB = 0
SM = 0
Doxycycline
YGL065C ALG2
ZShift = −0.51
Delta_AUC
NG = 0
DB = 0
SM = 0
Doxycycline
YMR047C NUP116
ZShift = 8.66
Delta_L
NG = 0
DB = 0
SM = 1
Doxycycline
YMR047C NUP116
ZShift = −18.97
Delta_K
NG = 0
DB = 0
SM = 1
Doxycycline
YMR047C NUP116
ZShift = −3.78
Delta_r
NG = 0
DB = 0
SM = 1
Doxycycline
YMR047C NUP116
ZShift = −13.77
Delta_AUC
NG = 0
DB = 0
SM = 1
Doxycycline

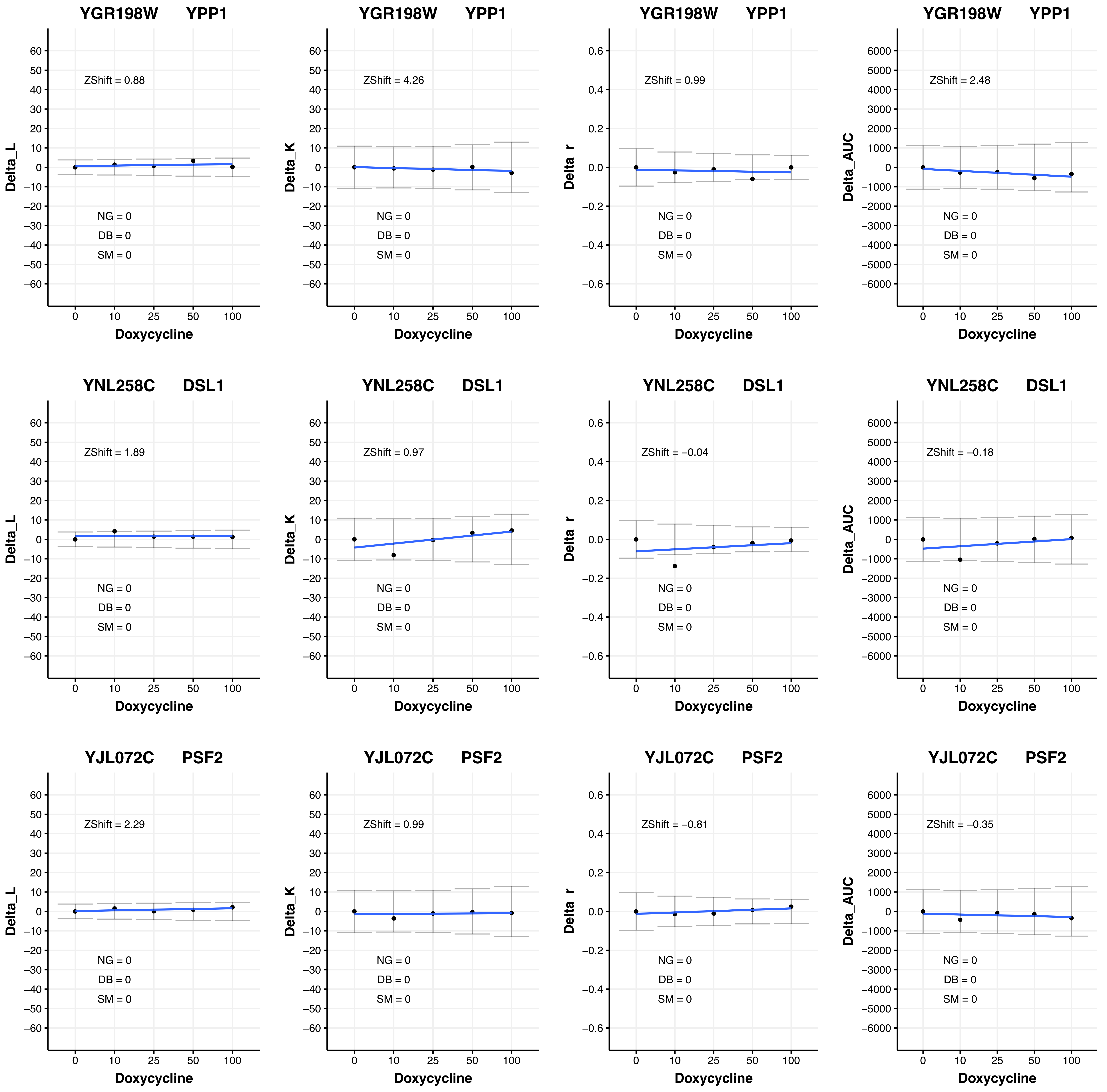
YGR198W YPP1
ZShift = 0.88
NG = 0
DB = 0
SM = 0
Delta_L
Doxycycline
YGR198W YPP1
ZShift = 4.26
NG = 0
DB = 0
SM = 0
Delta_K
Doxycycline
YGR198W YPP1
ZShift = 0.99
NG = 0
DB = 0
SM = 0
Delta_r
Doxycycline
YGR198W YPP1
ZShift = 2.48
NG = 0
DB = 0
SM = 0
Delta_AUC
Doxycycline
YNL258C DSL1
ZShift = 1.89
NG = 0
DB = 0
SM = 0
Delta_L
Doxycycline
YNL258C DSL1
ZShift = 0.97
NG = 0
DB = 0
SM = 0
Delta_K
Doxycycline
YNL258C DSL1
ZShift = −0.04
NG = 0
DB = 0
SM = 0
Delta_r
Doxycycline
YNL258C DSL1
ZShift = −0.18
NG = 0
DB = 0
SM = 0
Delta_AUC
Doxycycline
YJL072C PSF2
ZShift = 2.29
NG = 0
DB = 0
SM = 0
Delta_L
Doxycycline
YJL072C PSF2
ZShift = 0.99
NG = 0
DB = 0
SM = 0
Delta_K
Doxycycline
YJL072C PSF2
ZShift = −0.81
NG = 0
DB = 0
SM = 0
Delta_r
Doxycycline
YJL072C PSF2
ZShift = −0.35
NG = 0
DB = 0
SM = 0
Delta_AUC
Doxycycline

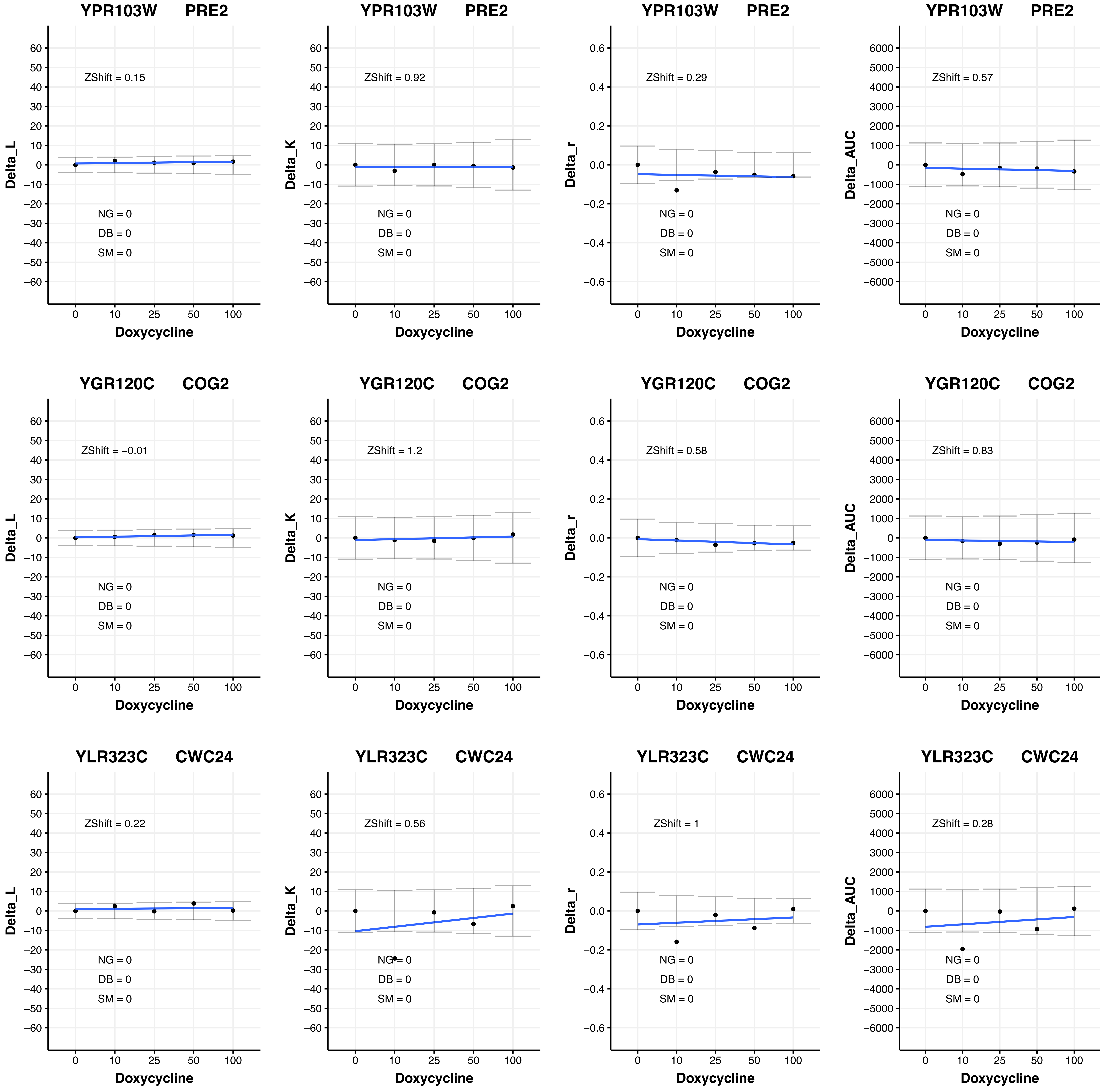

YPR103W PRE2
Delta_L
ZShift = 0.15
NG = 0
DB = 0
SM = 0
Doxycycline
YPR103W PRE2
Delta_K
ZShift = 0.92
NG = 0
DB = 0
SM = 0
Doxycycline
YPR103W PRE2
Delta_r
ZShift = 0.29
NG = 0
DB = 0
SM = 0
Doxycycline
YPR103W PRE2
Delta_AUC
ZShift = 0.57
NG = 0
DB = 0
SM = 0
Doxycycline
YGR120C COG2
Delta_L
ZShift = −0.01
NG = 0
DB = 0
SM = 0
Doxycycline
YGR120C COG2
Delta_K
ZShift = 1.2
NG = 0
DB = 0
SM = 0
Doxycycline
YGR120C COG2
Delta_r
ZShift = 0.58
NG = 0
DB = 0
SM = 0
Doxycycline
YGR120C COG2
Delta_AUC
ZShift = 0.83
NG = 0
DB = 0
SM = 0
Doxycycline
YLR323C CWC24
Delta_L
ZShift = 0.22
NG = 0
DB = 0
SM = 0
Doxycycline
YLR323C CWC24
Delta_K
ZShift = 0.56
NG = 0
DB = 0
SM = 0
Doxycycline
YLR323C CWC24
Delta_r
ZShift = 1
NG = 0
DB = 0
SM = 0
Doxycycline
YLR323C CWC24
Delta_AUC
ZShift = 0.28
NG = 0
DB = 0
SM = 0
Doxycycline

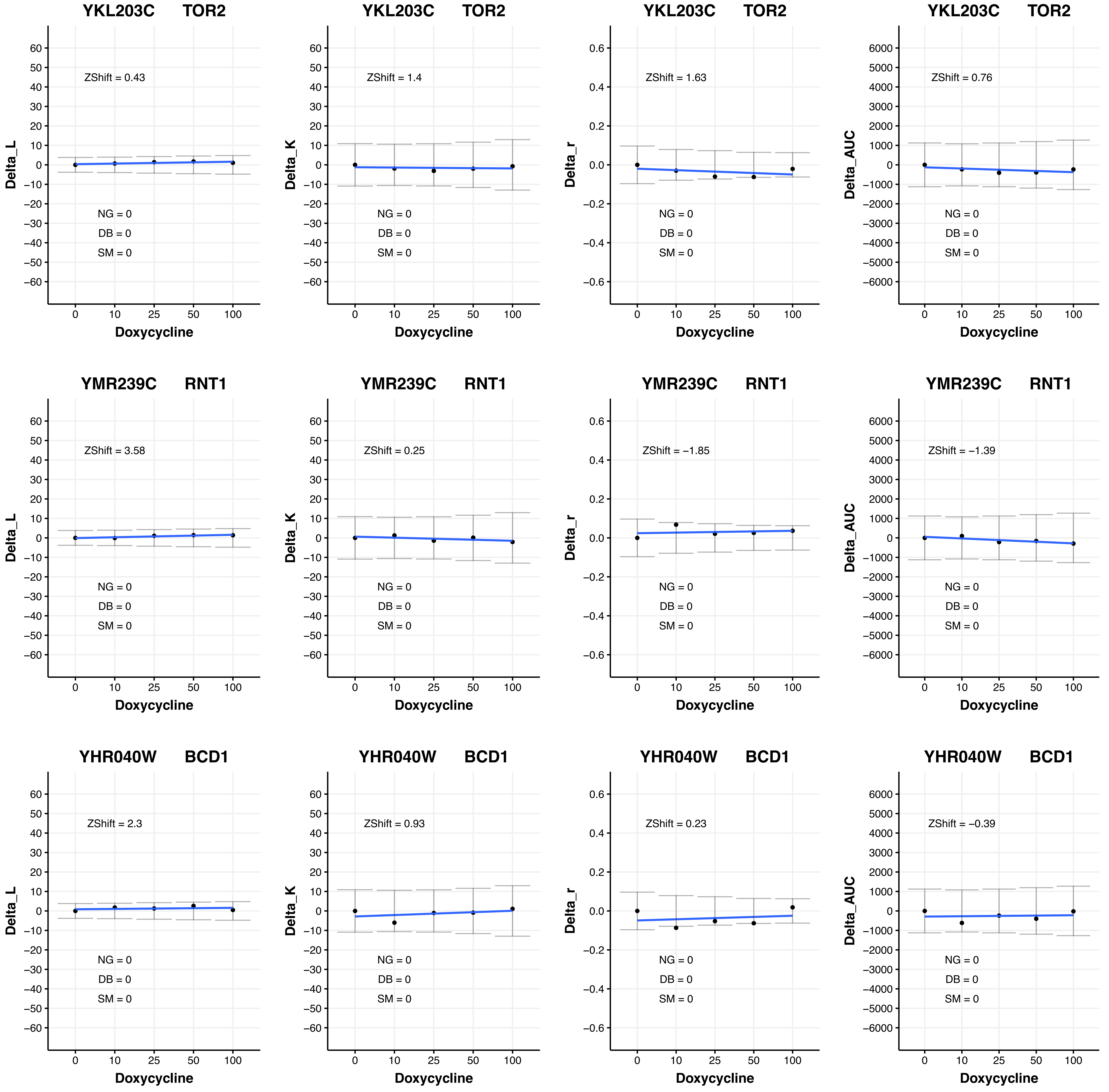

YKL203C TOR2
ZShift = 0.43
NG = 0
DB = 0
SM = 0
Delta_L
Doxycycline
YKL203C TOR2
ZShift = 1.4
NG = 0
DB = 0
SM = 0
Delta_K
Doxycycline
YKL203C TOR2
ZShift = 1.63
NG = 0
DB = 0
SM = 0
Delta_r
Doxycycline
YKL203C TOR2
ZShift = 0.76
NG = 0
DB = 0
SM = 0
Delta_AUC
Doxycycline
YMR239C RNT1
ZShift = 3.58
NG = 0
DB = 0
SM = 0
Delta_L
Doxycycline
YMR239C RNT1
ZShift = 0.25
NG = 0
DB = 0
SM = 0
Delta_K
Doxycycline
YMR239C RNT1
ZShift = −1.85
NG = 0
DB = 0
SM = 0
Delta_r
Doxycycline
YMR239C RNT1
ZShift = −1.39
NG = 0
DB = 0
SM = 0
Delta_AUC
Doxycycline
YHR040W BCD1
ZShift = 2.3
NG = 0
DB = 0
SM = 0
Delta_L
Doxycycline
YHR040W BCD1
ZShift = 0.93
NG = 0
DB = 0
SM = 0
Delta_K
Doxycycline
YHR040W BCD1
ZShift = 0.23
NG = 0
DB = 0
SM = 0
Delta_r
Doxycycline
YHR040W BCD1
ZShift = −0.39
NG = 0
DB = 0
SM = 0
Delta_AUC
Doxycycline

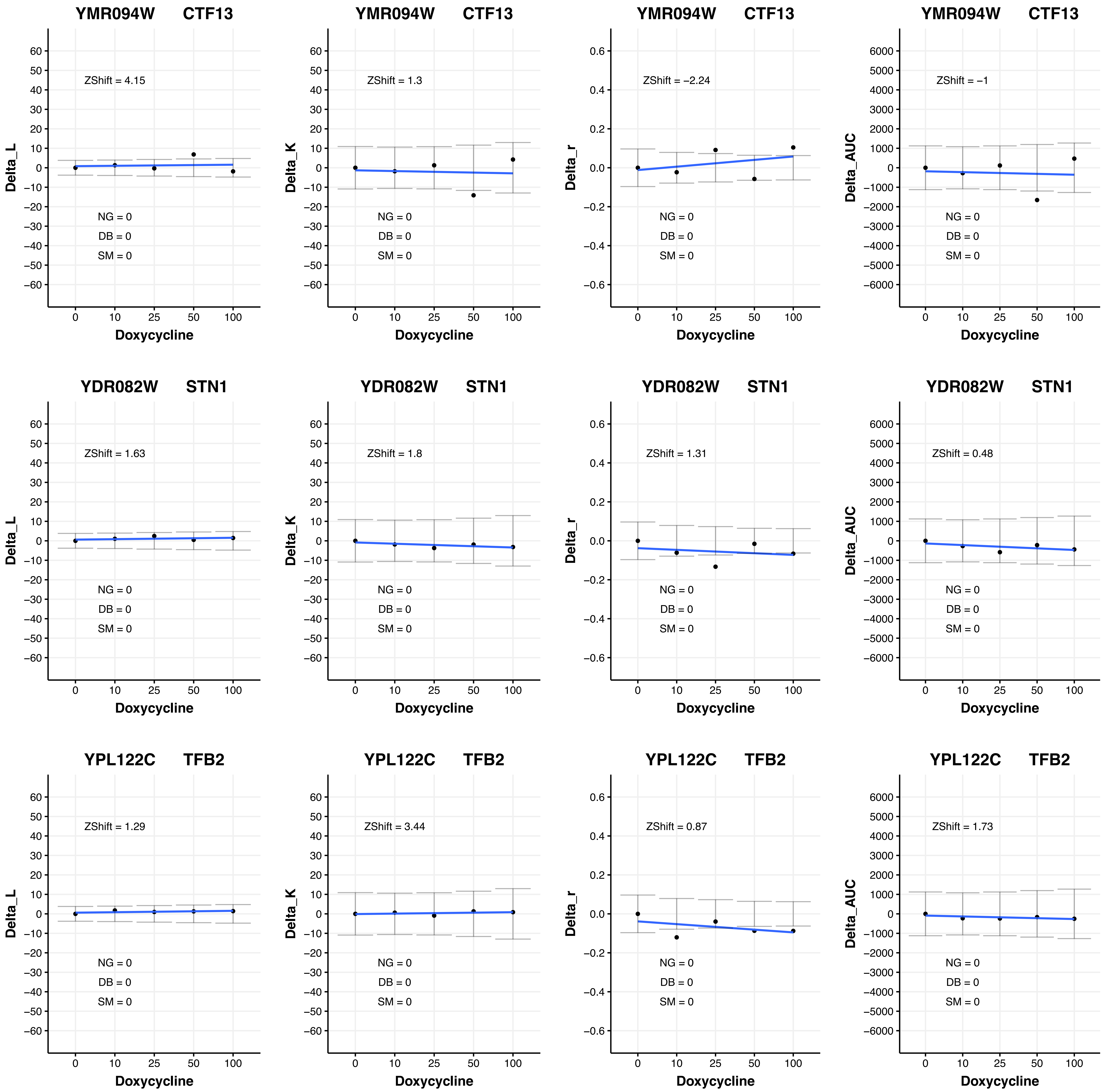
YMR094W CTF13
Delta_L
ZShift = 4.15
NG = 0
DB = 0
SM = 0
Doxycycline
YMR094W CTF13
Delta_K
ZShift = 1.3
NG = 0
DB = 0
SM = 0
Doxycycline
YMR094W CTF13
Delta_r
ZShift = −2.24
NG = 0
DB = 0
SM = 0
Doxycycline
YMR094W CTF13
Delta_AUC
ZShift = −1
NG = 0
DB = 0
SM = 0
Doxycycline
YDR082W STN1
Delta_L
ZShift = 1.63
NG = 0
DB = 0
SM = 0
Doxycycline
YDR082W STN1
Delta_K
ZShift = 1.8
NG = 0
DB = 0
SM = 0
Doxycycline
YDR082W STN1
Delta_r
ZShift = 1.31
NG = 0
DB = 0
SM = 0
Doxycycline
YDR082W STN1
Delta_AUC
ZShift = 0.48
NG = 0
DB = 0
SM = 0
Doxycycline
YPL122C TFB2
Delta_L
ZShift = 1.29
NG = 0
DB = 0
SM = 0
Doxycycline
YPL122C TFB2
Delta_K
ZShift = 3.44
NG = 0
DB = 0
SM = 0
Doxycycline
YPL122C TFB2
Delta_r
ZShift = 0.87
NG = 0
DB = 0
SM = 0
Doxycycline
YPL122C TFB2
Delta_AUC
ZShift = 1.73
NG = 0
DB = 0
SM = 0
Doxycycline

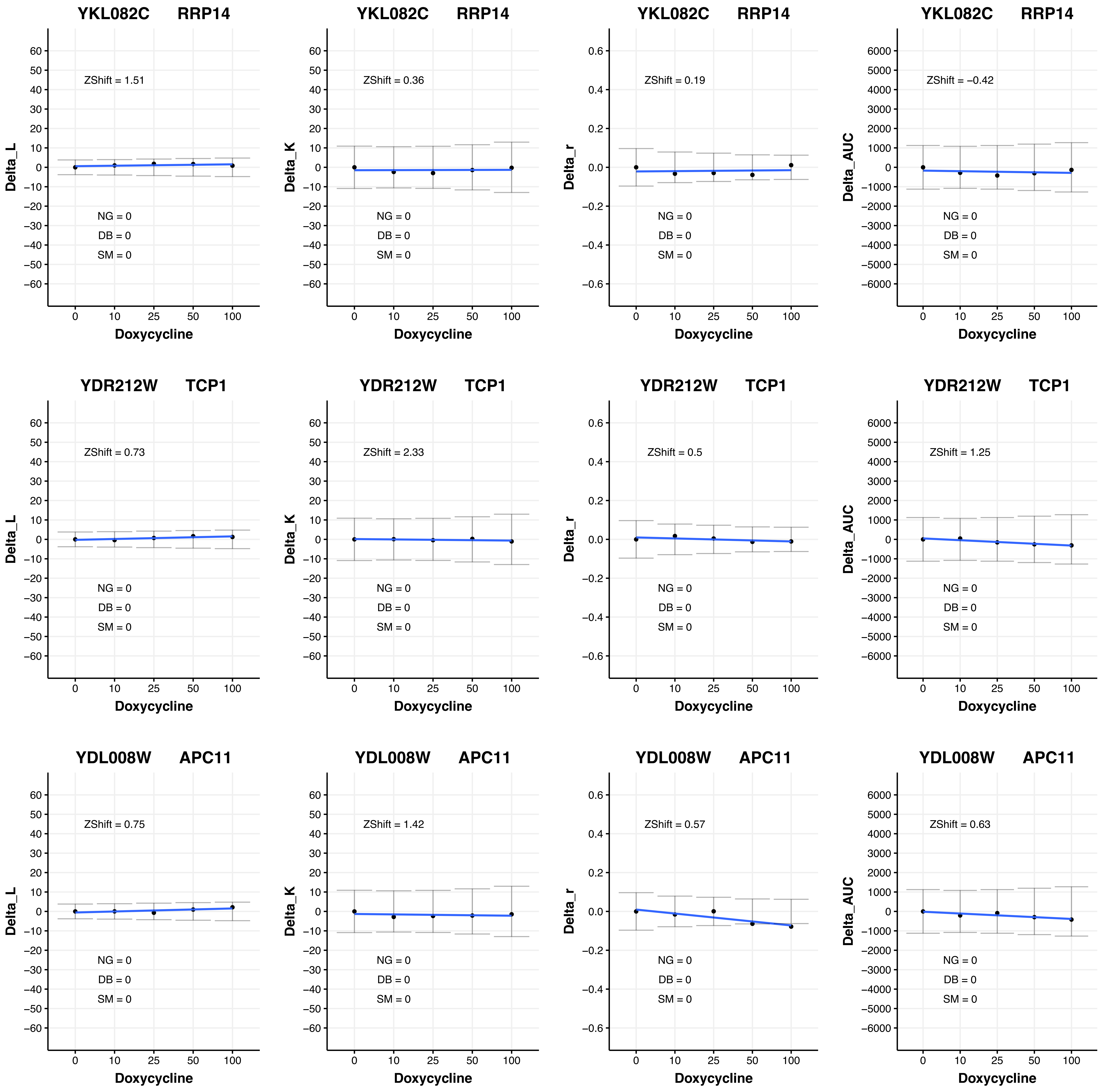

YKL082C RRP14
ZShift = 1.51
Delta_L
NG = 0
DB = 0
SM = 0
Doxycycline
YKL082C RRP14
ZShift = 0.36
Delta_K
NG = 0
DB = 0
SM = 0
Doxycycline
YKL082C RRP14
ZShift = 0.19
Delta_r
NG = 0
DB = 0
SM = 0
Doxycycline
YKL082C RRP14
ZShift = −0.42
Delta_AUC
NG = 0
DB = 0
SM = 0
Doxycycline
YDR212W TCP1
ZShift = 0.73
Delta_L
NG = 0
DB = 0
SM = 0
Doxycycline
YDR212W TCP1
ZShift = 2.33
Delta_K
NG = 0
DB = 0
SM = 0
Doxycycline
YDR212W TCP1
ZShift = 0.5
Delta_r
NG = 0
DB = 0
SM = 0
Doxycycline
YDR212W TCP1
ZShift = 1.25
Delta_AUC
NG = 0
DB = 0
SM = 0
Doxycycline
YDL008W APC11
ZShift = 0.75
Delta_L
NG = 0
DB = 0
SM = 0
Doxycycline
YDL008W APC11
ZShift = 1.42
Delta_K
NG = 0
DB = 0
SM = 0
Doxycycline
YDL008W APC11
ZShift = 0.57
Delta_r
NG = 0
DB = 0
SM = 0
Doxycycline
YDL008W APC11
ZShift = 0.63
Delta_AUC
NG = 0
DB = 0
SM = 0
Doxycycline

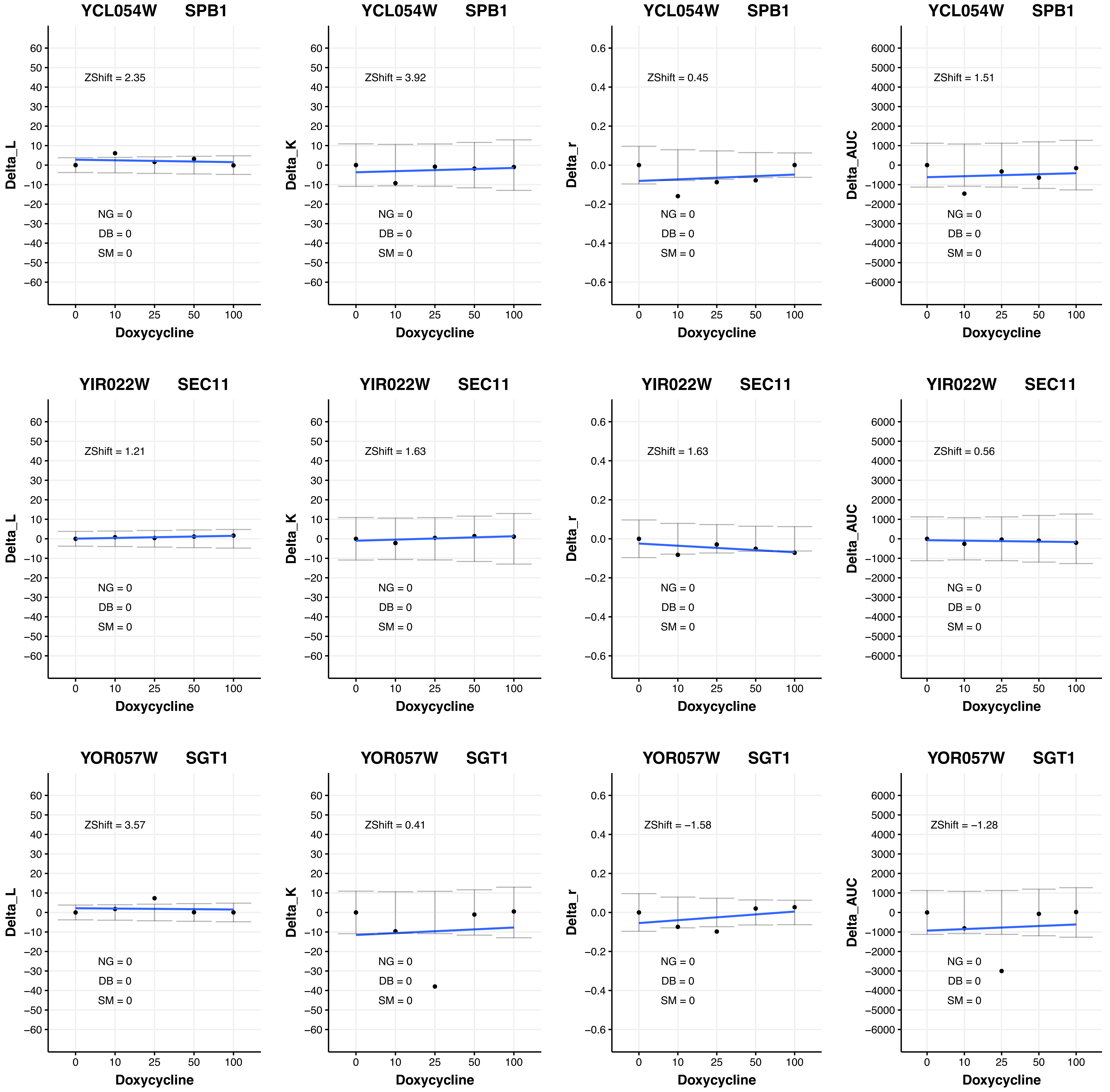

YCL054W SPB1
ZShift = 2.35
NG = 0
DB = 0
SM = 0
Delta_L
Doxycycline
YCL054W SPB1
ZShift = 3.92
NG = 0
DB = 0
SM = 0
Delta_K
Doxycycline
YCL054W SPB1
ZShift = 0.45
NG = 0
DB = 0
SM = 0
Delta_r
Doxycycline
YCL054W SPB1
ZShift = 1.51
NG = 0
DB = 0
SM = 0
Delta_AUC
Doxycycline
YIR022W SEC11
ZShift = 1.21
NG = 0
DB = 0
SM = 0
Delta_L
Doxycycline
YIR022W SEC11
ZShift = 1.63
NG = 0
DB = 0
SM = 0
Delta_K
Doxycycline
YIR022W SEC11
ZShift = 1.63
NG = 0
DB = 0
SM = 0
Delta_r
Doxycycline
YIR022W SEC11
ZShift = 0.56
NG = 0
DB = 0
SM = 0
Delta_AUC
Doxycycline
YOR057W SGT1
ZShift = 3.57
NG = 0
DB = 0
SM = 0
Delta_L
Doxycycline
YOR057W SGT1
ZShift = 0.41
NG = 0
DB = 0
SM = 0
Delta_K
Doxycycline
YOR057W SGT1
ZShift = −1.58
NG = 0
DB = 0
SM = 0
Delta_r
Doxycycline
YOR057W SGT1
ZShift = −1.28
NG = 0
DB = 0
SM = 0
Delta_AUC
Doxycycline

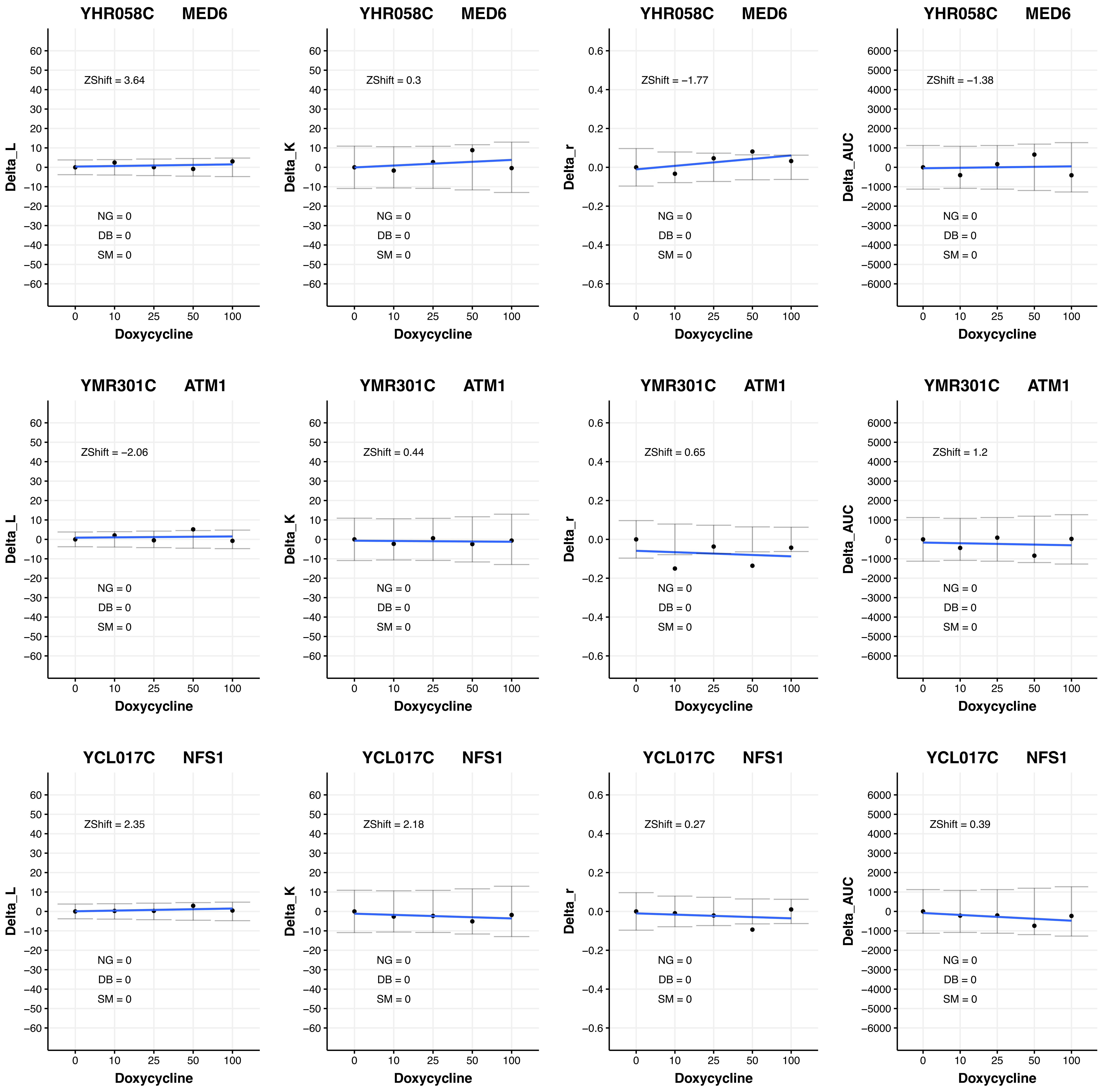
YHR058C MED6
ZShift = 3.64
Delta_L
ZShift = 0.3
Delta_K
ZShift = −1.77
Delta_r
ZShift = −1.38
Delta_AUC
YMR301C ATM1
ZShift = −2.06
ZShift = 0.44
ZShift = 0.65
ZShift = 1.2
YCL017C NFS1
ZShift = 2.35
ZShift = 2.18
ZShift = 0.27
ZShift = 0.39
NG = 0
DB = 0
SM = 0
Doxycycline
0
10
25
50
100

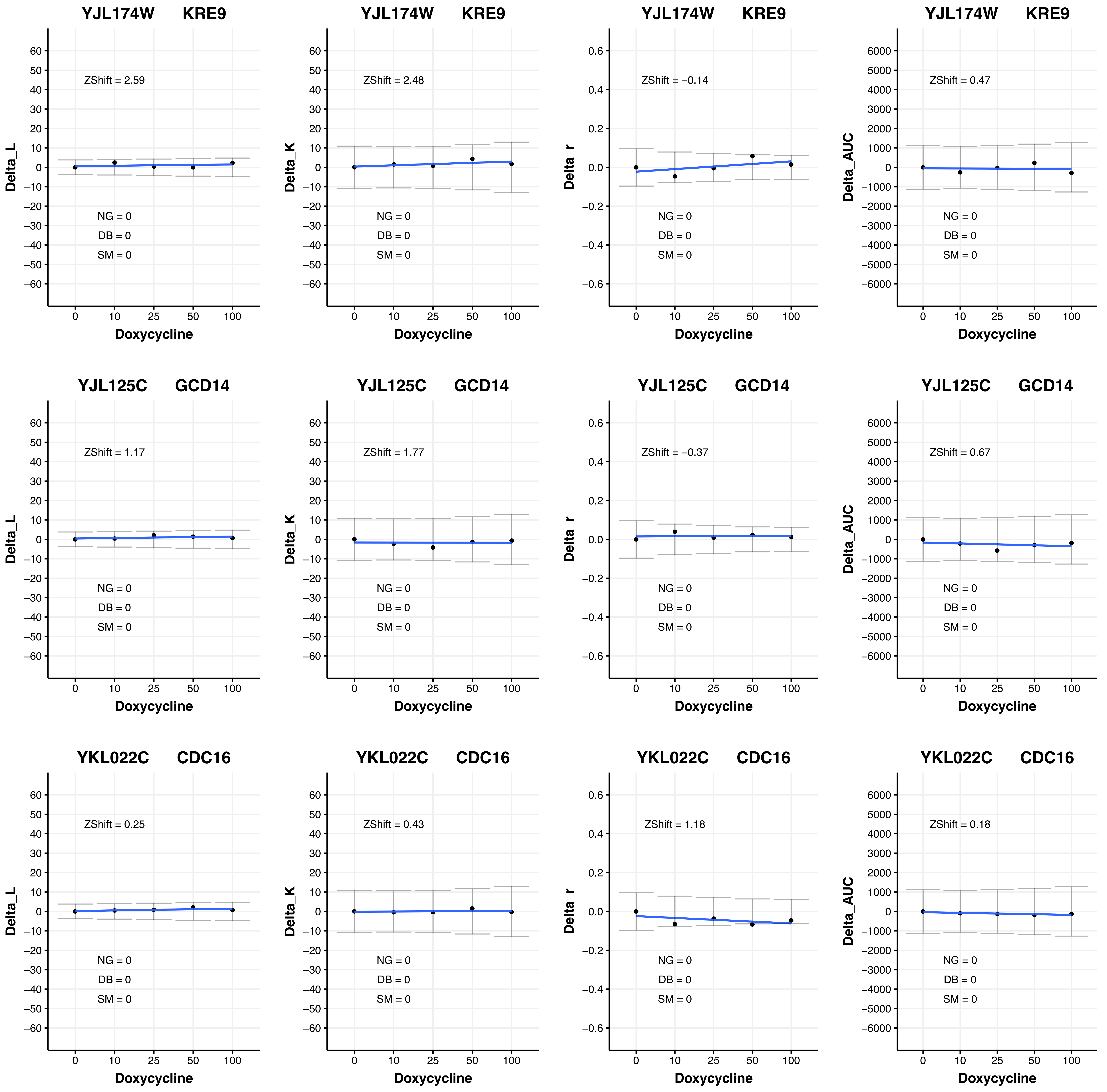
YJL174W KRE9
ZShift = 2.59
NG = 0
DB = 0
SM = 0
Delta_L
Doxycycline
YJL174W KRE9
ZShift = 2.48
NG = 0
DB = 0
SM = 0
Delta_K
Doxycycline
YJL174W KRE9
ZShift = −0.14
NG = 0
DB = 0
SM = 0
Delta_r
Doxycycline
YJL174W KRE9
ZShift = 0.47
NG = 0
DB = 0
SM = 0
Delta_AUC
Doxycycline
YJL125C GCD14
ZShift = 1.17
NG = 0
DB = 0
SM = 0
Delta_L
Doxycycline
YJL125C GCD14
ZShift = 1.77
NG = 0
DB = 0
SM = 0
Delta_K
Doxycycline
YJL125C GCD14
ZShift = −0.37
NG = 0
DB = 0
SM = 0
Delta_r
Doxycycline
YJL125C GCD14
ZShift = 0.67
NG = 0
DB = 0
SM = 0
Delta_AUC
Doxycycline
YKL022C CDC16
ZShift = 0.25
NG = 0
DB = 0
SM = 0
Delta_L
Doxycycline
YKL022C CDC16
ZShift = 0.43
NG = 0
DB = 0
SM = 0
Delta_K
Doxycycline
YKL022C CDC16
ZShift = 1.18
NG = 0
DB = 0
SM = 0
Delta_r
Doxycycline
YKL022C CDC16
ZShift = 0.18
NG = 0
DB = 0
SM = 0
Delta_AUC
Doxycycline

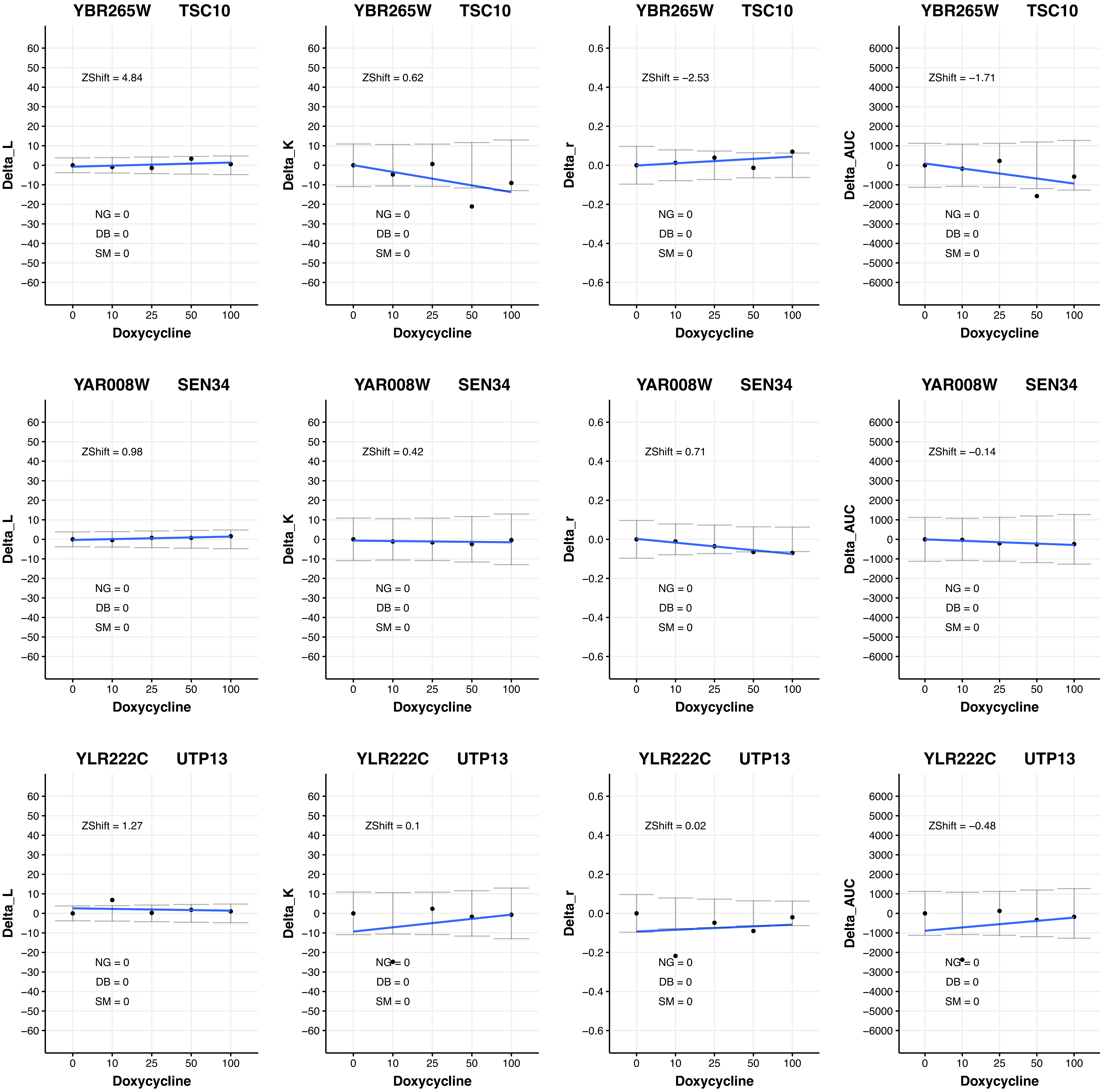

YBR265W TSC10
ZShift = 4.84
Delta_L
NG = 0
DB = 0
SM = 0
Doxycycline
YBR265W TSC10
ZShift = 0.62
Delta_K
NG = 0
DB = 0
SM = 0
Doxycycline
YBR265W TSC10
ZShift = −2.53
Delta_r
NG = 0
DB = 0
SM = 0
Doxycycline
YBR265W TSC10
ZShift = −1.71
Delta_AUC
NG = 0
DB = 0
SM = 0
Doxycycline
YAR008W SEN34
ZShift = 0.98
Delta_L
NG = 0
DB = 0
SM = 0
Doxycycline
YAR008W SEN34
ZShift = 0.42
Delta_K
NG = 0
DB = 0
SM = 0
Doxycycline
YAR008W SEN34
ZShift = 0.71
Delta_r
NG = 0
DB = 0
SM = 0
Doxycycline
YAR008W SEN34
ZShift = −0.14
Delta_AUC
NG = 0
DB = 0
SM = 0
Doxycycline
YLR222C UTP13
ZShift = 1.27
Delta_L
NG = 0
DB = 0
SM = 0
Doxycycline
YLR222C UTP13
ZShift = 0.1
Delta_K
NG = 0
DB = 0
SM = 0
Doxycycline
YLR222C UTP13
ZShift = 0.02
Delta_r
NG = 0
DB = 0
SM = 0
Doxycycline
YLR222C UTP13
ZShift = −0.48
Delta_AUC
NG = 0
DB = 0
SM = 0
Doxycycline

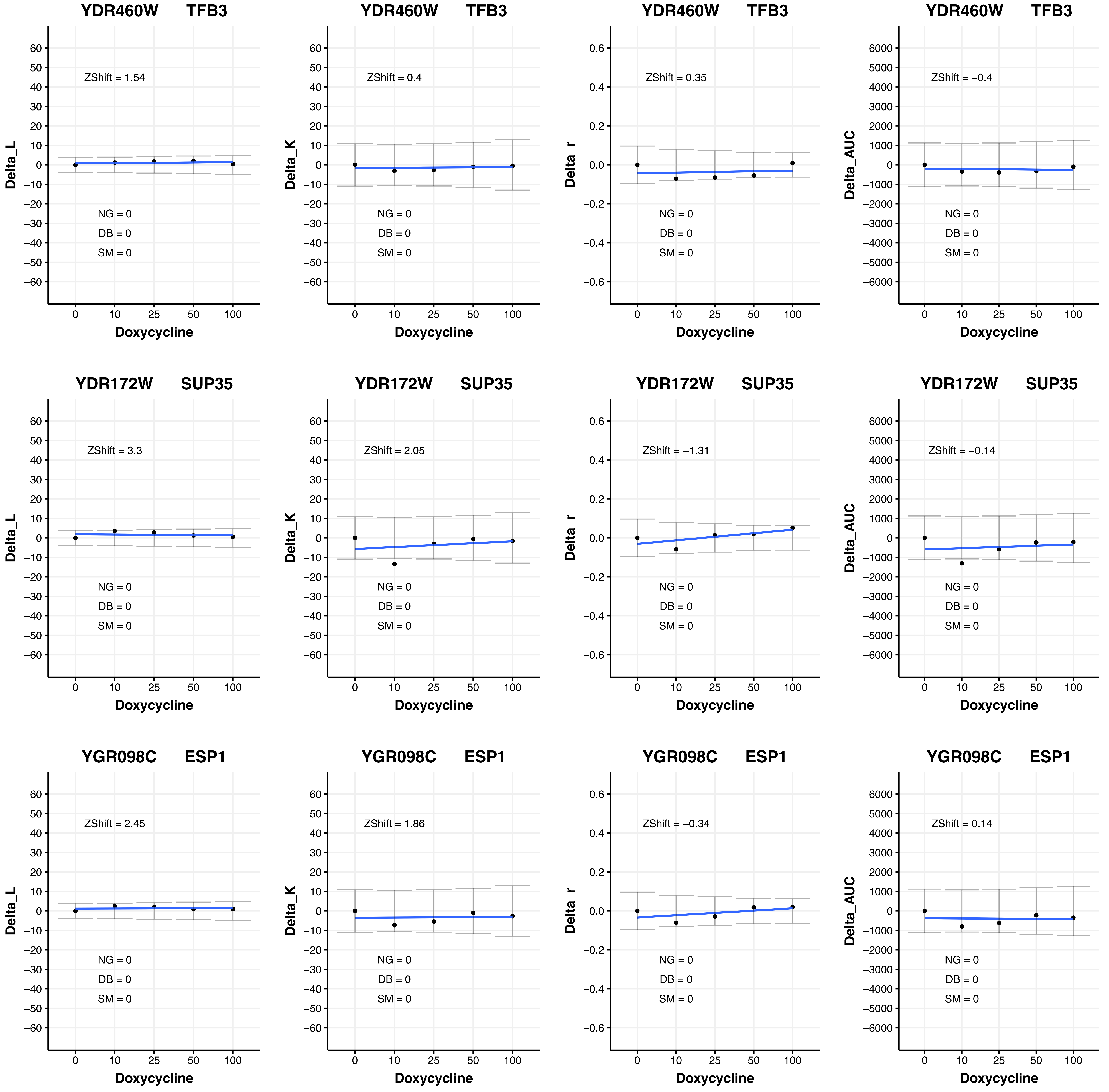

YDR460W TFB3
ZShift = 1.54
NG = 0
DB = 0
SM = 0
Delta_L
Doxycycline
YDR460W TFB3
ZShift = 0.4
NG = 0
DB = 0
SM = 0
Delta_K
Doxycycline
YDR460W TFB3
ZShift = 0.35
NG = 0
DB = 0
SM = 0
Delta_r
Doxycycline
YDR460W TFB3
ZShift = −0.4
NG = 0
DB = 0
SM = 0
Delta_AUC
Doxycycline
YDR172W SUP35
ZShift = 3.3
NG = 0
DB = 0
SM = 0
Delta_L
Doxycycline
YDR172W SUP35
ZShift = 2.05
NG = 0
DB = 0
SM = 0
Delta_K
Doxycycline
YDR172W SUP35
ZShift = −1.31
NG = 0
DB = 0
SM = 0
Delta_r
Doxycycline
YDR172W SUP35
ZShift = −0.14
NG = 0
DB = 0
SM = 0
Delta_AUC
Doxycycline
YGR098C ESP1
ZShift = 2.45
NG = 0
DB = 0
SM = 0
Delta_L
Doxycycline
YGR098C ESP1
ZShift = 1.86
NG = 0
DB = 0
SM = 0
Delta_K
Doxycycline
YGR098C ESP1
ZShift = −0.34
NG = 0
DB = 0
SM = 0
Delta_r
Doxycycline
YGR098C ESP1
ZShift = 0.14
NG = 0
DB = 0
SM = 0
Delta_AUC
Doxycycline

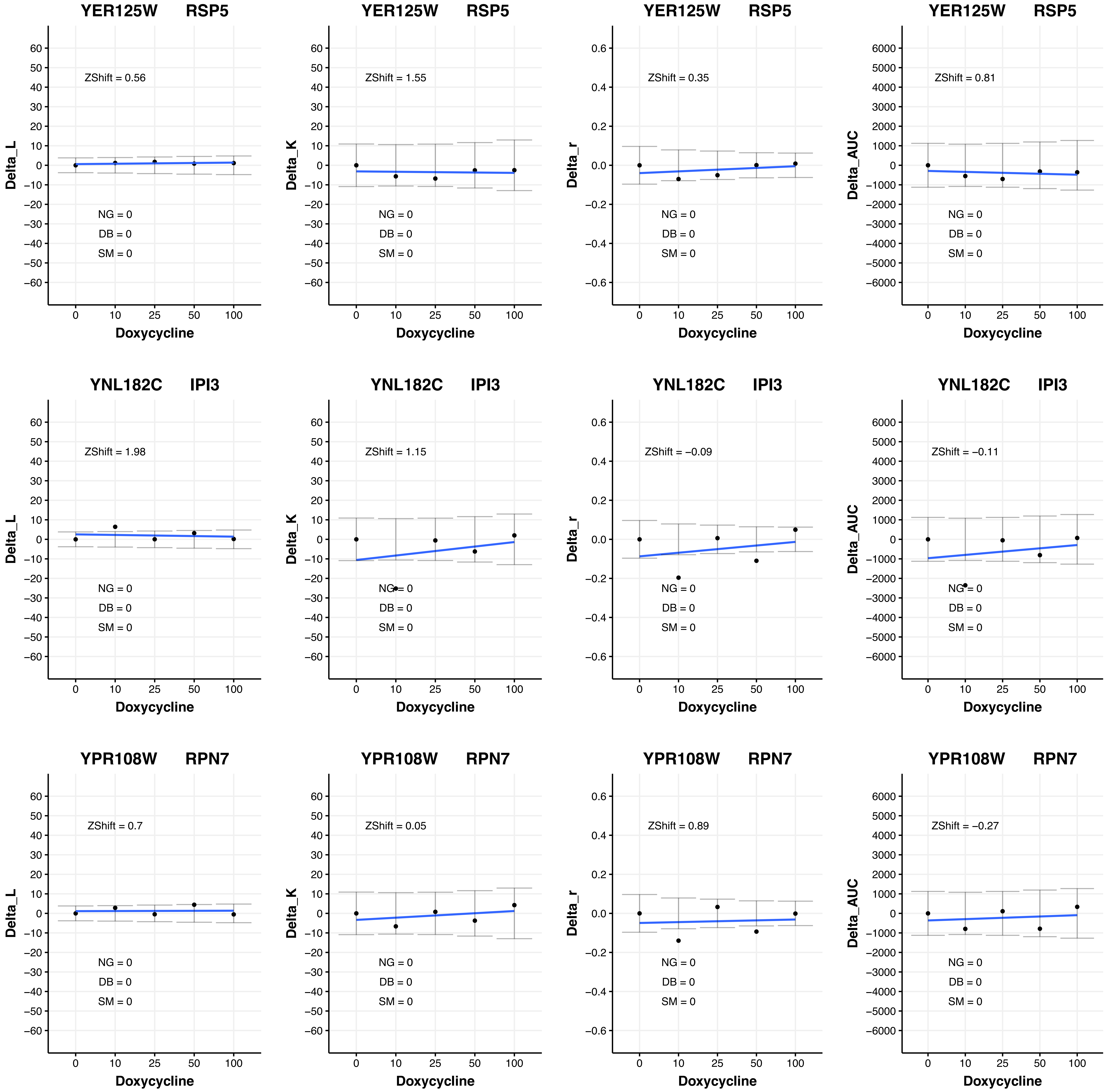
YER125W RSP5
ZShift = 0.56
ZShift = 1.55
ZShift = 0.35
ZShift = 0.81
YNL182C IPI3
ZShift = 1.98
ZShift = 1.15
ZShift = −0.09
ZShift = −0.11
YPR108W RPN7
ZShift = 0.7
ZShift = 0.05
ZShift = 0.89
ZShift = −0.27
NG = 0
DB = 0
SM = 0
Delta_L
Delta_K
Delta_r
Delta_AUC
Doxycycline
0
10
25
50
100

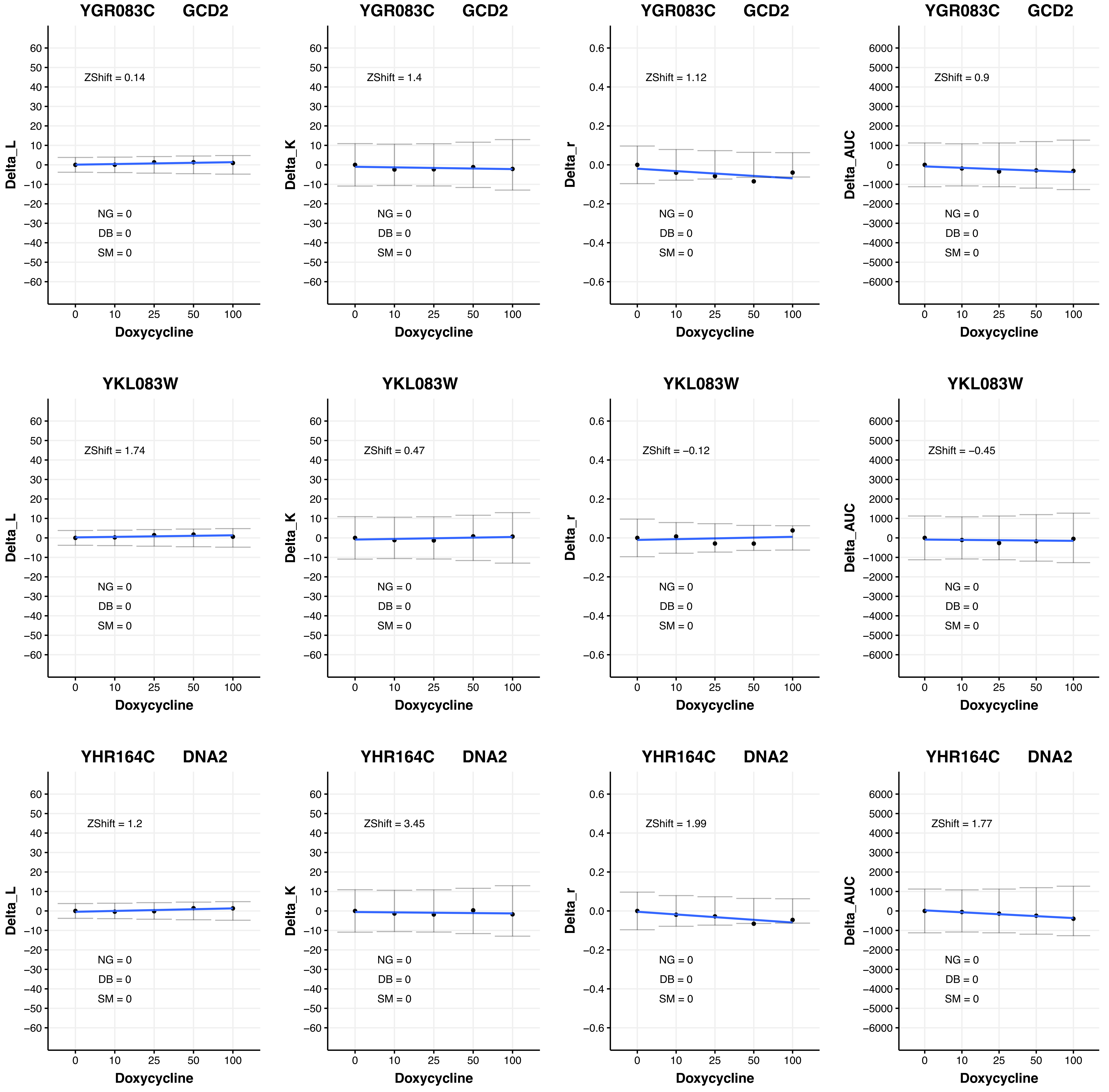

YGR083C GCD2
ZShift = 0.14
Delta_L
NG = 0
DB = 0
SM = 0
Doxycycline
YGR083C GCD2
ZShift = 1.4
Delta_K
NG = 0
DB = 0
SM = 0
Doxycycline
YGR083C GCD2
ZShift = 1.12
Delta_r
NG = 0
DB = 0
SM = 0
Doxycycline
YGR083C GCD2
ZShift = 0.9
Delta_AUC
NG = 0
DB = 0
SM = 0
Doxycycline
YKL083W
ZShift = 1.74
Delta_L
NG = 0
DB = 0
SM = 0
Doxycycline
YKL083W
ZShift = 0.47
Delta_K
NG = 0
DB = 0
SM = 0
Doxycycline
YKL083W
ZShift = −0.12
Delta_r
NG = 0
DB = 0
SM = 0
Doxycycline
YKL083W
ZShift = −0.45
Delta_AUC
NG = 0
DB = 0
SM = 0
Doxycycline
YHR164C DNA2
ZShift = 1.2
Delta_L
NG = 0
DB = 0
SM = 0
Doxycycline
YHR164C DNA2
ZShift = 3.45
Delta_K
NG = 0
DB = 0
SM = 0
Doxycycline
YHR164C DNA2
ZShift = 1.99
Delta_r
NG = 0
DB = 0
SM = 0
Doxycycline
YHR164C DNA2
ZShift = 1.77
Delta_AUC
NG = 0
DB = 0
SM = 0
Doxycycline

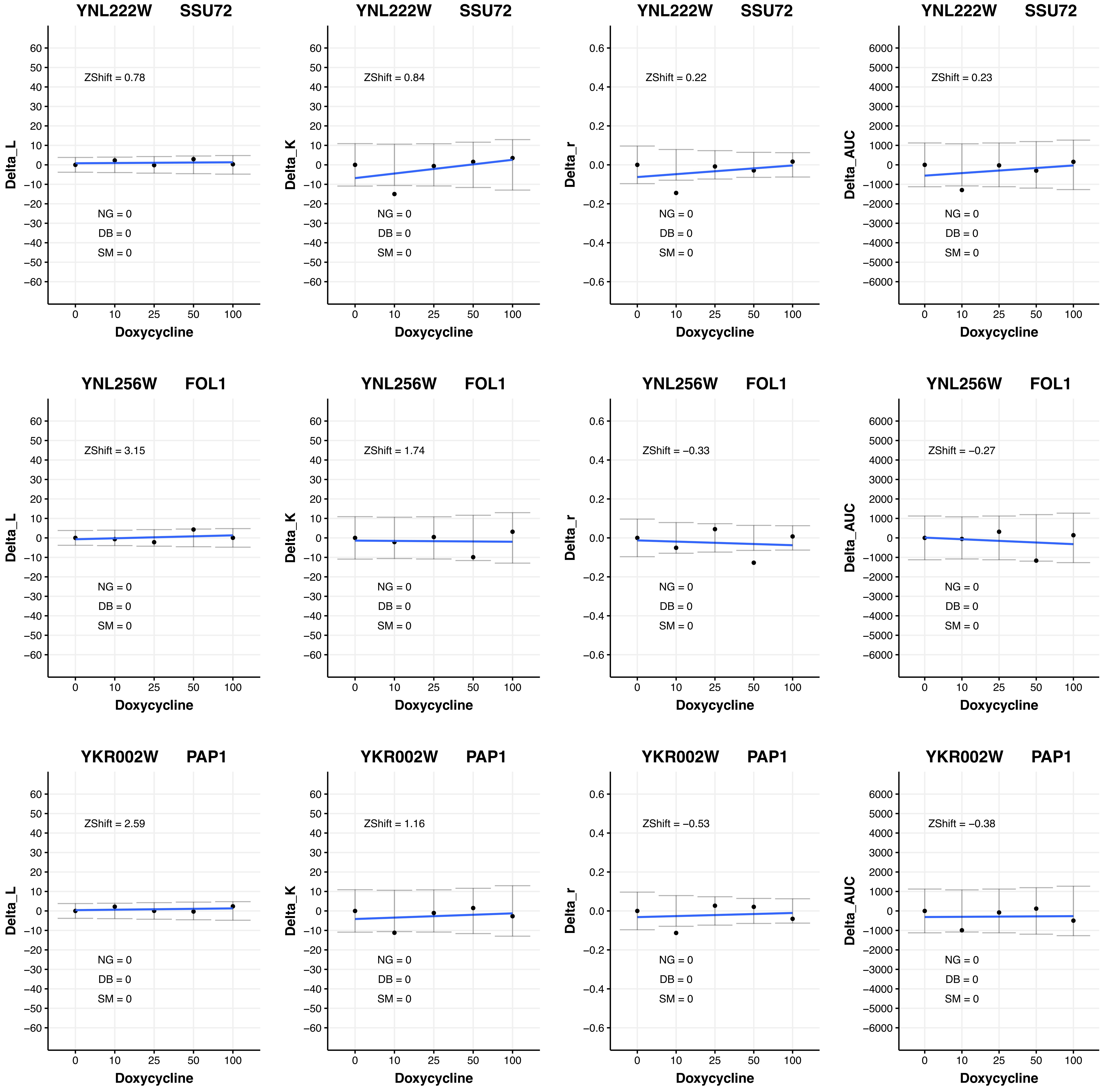

YNL222W SSU72
ZShift = 0.78
NG = 0
DB = 0
SM = 0
Delta_L
Doxycycline
YNL222W SSU72
ZShift = 0.84
NG = 0
DB = 0
SM = 0
Delta_K
Doxycycline
YNL222W SSU72
ZShift = 0.22
NG = 0
DB = 0
SM = 0
Delta_r
Doxycycline
YNL222W SSU72
ZShift = 0.23
NG = 0
DB = 0
SM = 0
Delta_AUC
Doxycycline
YNL256W FOL1
ZShift = 3.15
NG = 0
DB = 0
SM = 0
Delta_L
Doxycycline
YNL256W FOL1
ZShift = 1.74
NG = 0
DB = 0
SM = 0
Delta_K
Doxycycline
YNL256W FOL1
ZShift = −0.33
NG = 0
DB = 0
SM = 0
Delta_r
Doxycycline
YNL256W FOL1
ZShift = −0.27
NG = 0
DB = 0
SM = 0
Delta_AUC
Doxycycline
YKR002W PAP1
ZShift = 2.59
NG = 0
DB = 0
SM = 0
Delta_L
Doxycycline
YKR002W PAP1
ZShift = 1.16
NG = 0
DB = 0
SM = 0
Delta_K
Doxycycline
YKR002W PAP1
ZShift = −0.53
NG = 0
DB = 0
SM = 0
Delta_r
Doxycycline
YKR002W PAP1
ZShift = −0.38
NG = 0
DB = 0
SM = 0
Delta_AUC
Doxycycline

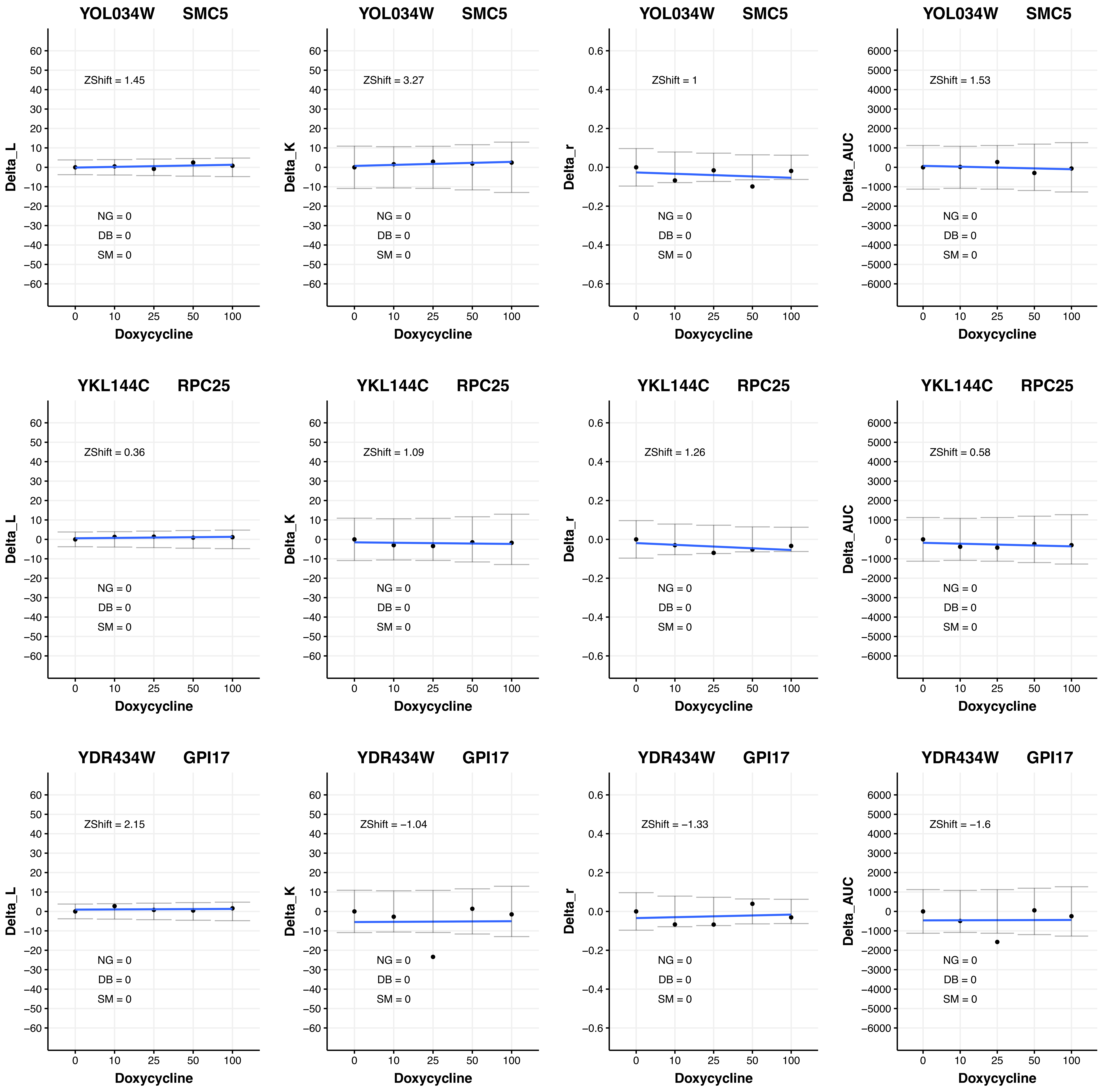

YOL034W SMC5
ZShift = 1.45
NG = 0
DB = 0
SM = 0
Delta_L
Doxycycline
YOL034W SMC5
ZShift = 3.27
NG = 0
DB = 0
SM = 0
Delta_K
Doxycycline
YOL034W SMC5
ZShift = 1
NG = 0
DB = 0
SM = 0
Delta_r
Doxycycline
YOL034W SMC5
ZShift = 1.53
NG = 0
DB = 0
SM = 0
Delta_AUC
Doxycycline
YKL144C RPC25
ZShift = 0.36
NG = 0
DB = 0
SM = 0
Delta_L
Doxycycline
YKL144C RPC25
ZShift = 1.09
NG = 0
DB = 0
SM = 0
Delta_K
Doxycycline
YKL144C RPC25
ZShift = 1.26
NG = 0
DB = 0
SM = 0
Delta_r
Doxycycline
YKL144C RPC25
ZShift = 0.58
NG = 0
DB = 0
SM = 0
Delta_AUC
Doxycycline
YDR434W GPI17
ZShift = 2.15
NG = 0
DB = 0
SM = 0
Delta_L
Doxycycline
YDR434W GPI17
ZShift = −1.04
NG = 0
DB = 0
SM = 0
Delta_K
Doxycycline
YDR434W GPI17
ZShift = −1.33
NG = 0
DB = 0
SM = 0
Delta_r
Doxycycline
YDR434W GPI17
ZShift = −1.6
NG = 0
DB = 0
SM = 0
Delta_AUC
Doxycycline

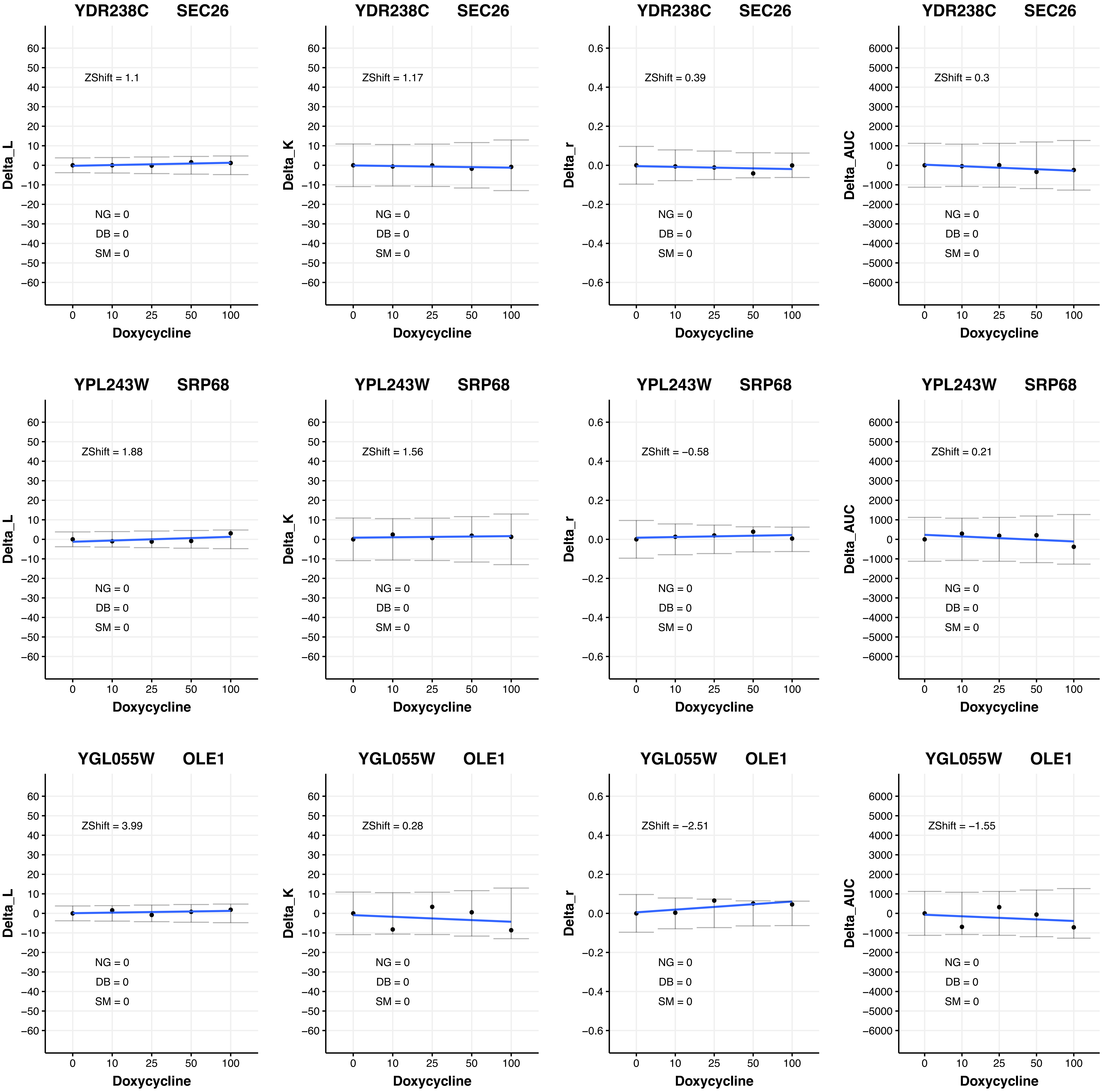

YDR238C SEC26
ZShift = 1.1
Delta_L
NG = 0
DB = 0
SM = 0
Doxycycline
YDR238C SEC26
ZShift = 1.17
Delta_K
NG = 0
DB = 0
SM = 0
Doxycycline
YDR238C SEC26
ZShift = 0.39
Delta_r
NG = 0
DB = 0
SM = 0
Doxycycline
YDR238C SEC26
ZShift = 0.3
Delta_AUC
NG = 0
DB = 0
SM = 0
Doxycycline
YPL243W SRP68
ZShift = 1.88
Delta_L
NG = 0
DB = 0
SM = 0
Doxycycline
YPL243W SRP68
ZShift = 1.56
Delta_K
NG = 0
DB = 0
SM = 0
Doxycycline
YPL243W SRP68
ZShift = −0.58
Delta_r
NG = 0
DB = 0
SM = 0
Doxycycline
YPL243W SRP68
ZShift = 0.21
Delta_AUC
NG = 0
DB = 0
SM = 0
Doxycycline
YGL055W OLE1
ZShift = 3.99
Delta_L
NG = 0
DB = 0
SM = 0
Doxycycline
YGL055W OLE1
ZShift = 0.28
Delta_K
NG = 0
DB = 0
SM = 0
Doxycycline
YGL055W OLE1
ZShift = −2.51
Delta_r
NG = 0
DB = 0
SM = 0
Doxycycline
YGL055W OLE1
ZShift = −1.55
Delta_AUC
NG = 0
DB = 0
SM = 0
Doxycycline

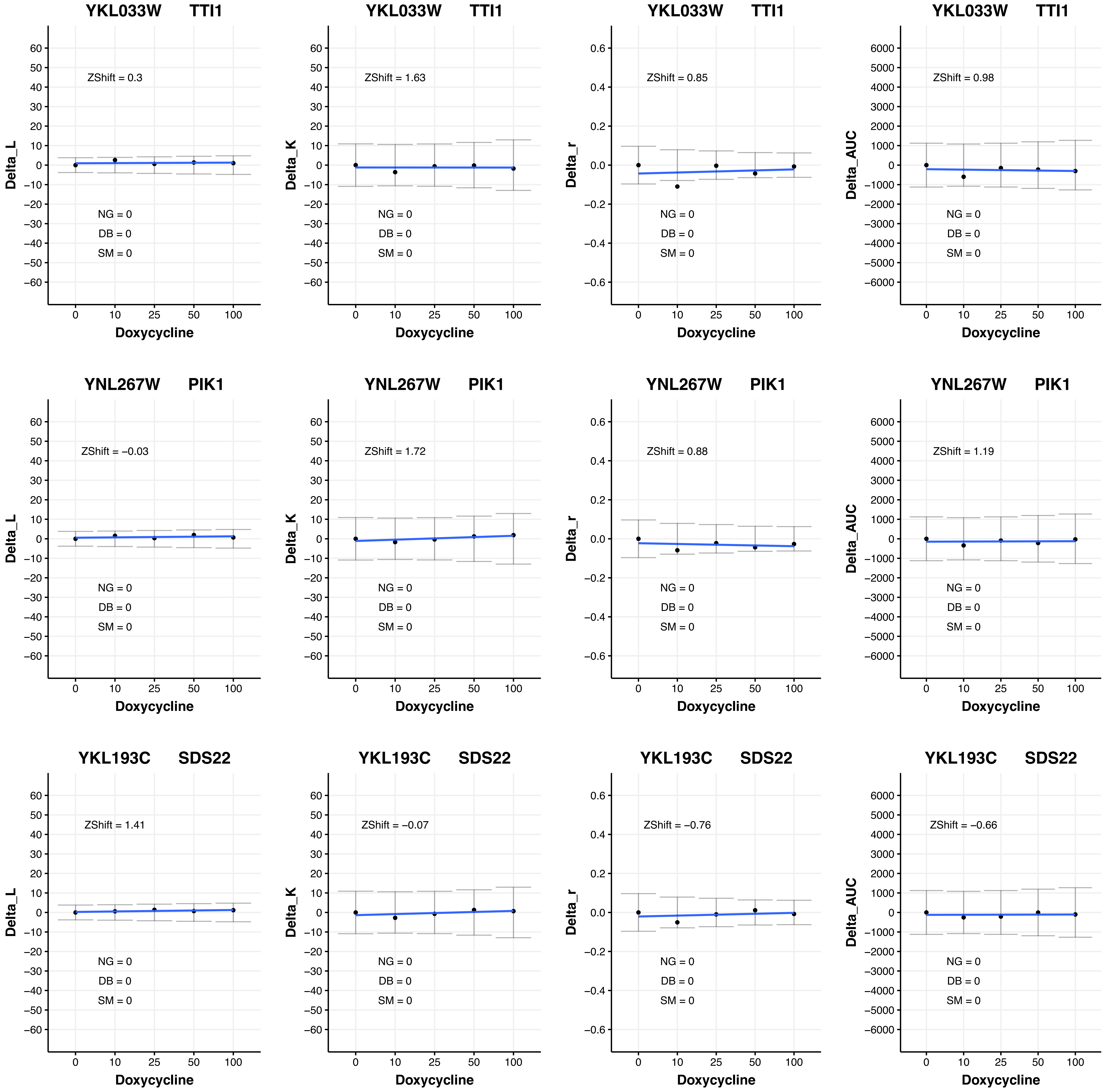

YKL033W TTI1
ZShift = 0.3
ZShift = 1.63
ZShift = 0.85
ZShift = 0.98
YNL267W PIK1
ZShift = −0.03
ZShift = 1.72
ZShift = 0.88
ZShift = 1.19
YKL193C SDS22
ZShift = 1.41
ZShift = −0.07
ZShift = −0.76
ZShift = −0.66
NG = 0
DB = 0
SM = 0
Delta_L
Delta_K
Delta_r
Delta_AUC
Doxycycline
0
10
25
50
100

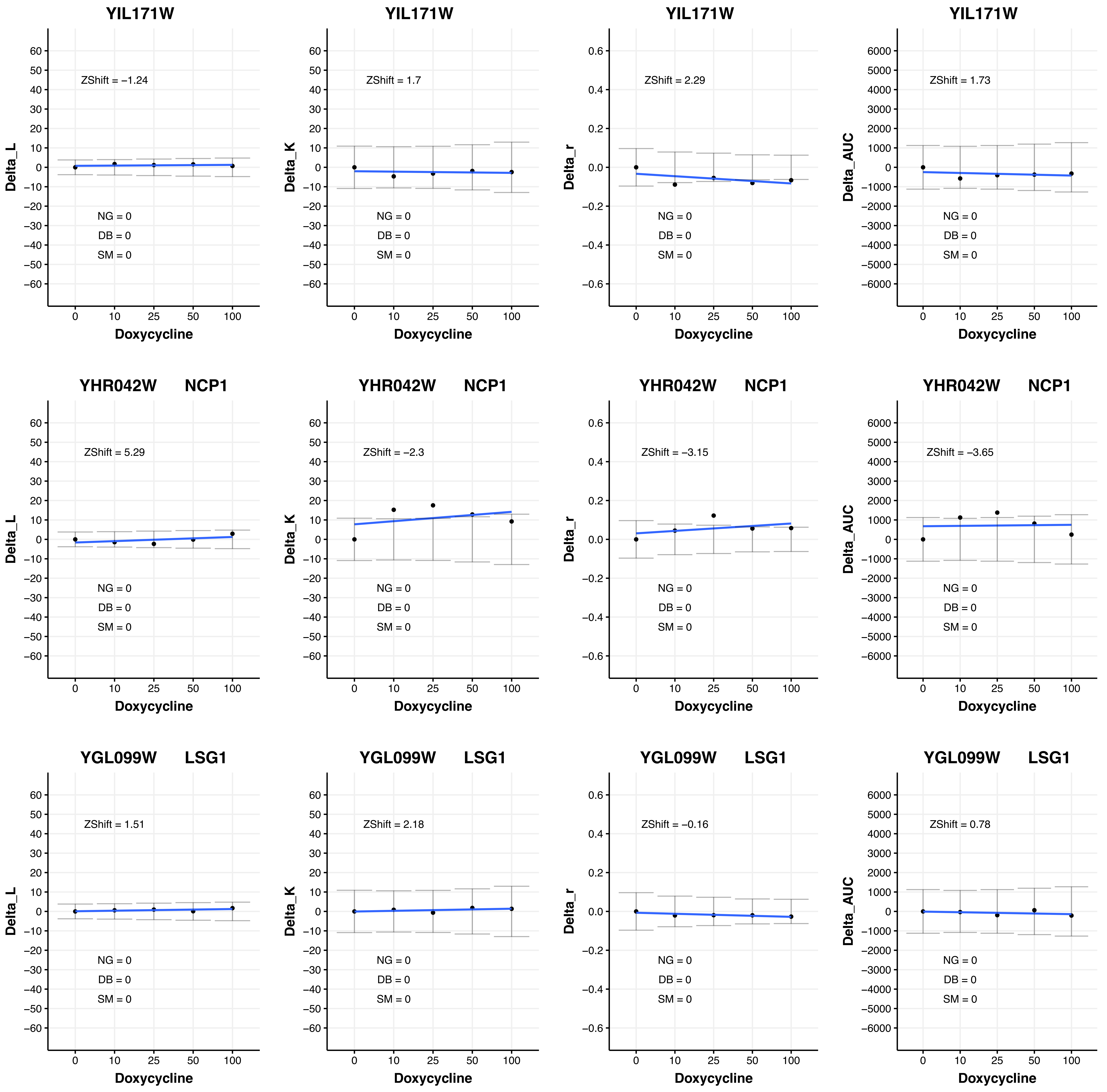

YIL171W
ZShift = −1.24
NG = 0
DB = 0
SM = 0
Delta_L
Doxycycline
YIL171W
ZShift = 1.7
NG = 0
DB = 0
SM = 0
Delta_K
Doxycycline
YIL171W
ZShift = 2.29
NG = 0
DB = 0
SM = 0
Delta_r
Doxycycline
YIL171W
ZShift = 1.73
NG = 0
DB = 0
SM = 0
Delta_AUC
Doxycycline
YHR042W NCP1
ZShift = 5.29
NG = 0
DB = 0
SM = 0
Delta_L
Doxycycline
YHR042W NCP1
ZShift = −2.3
NG = 0
DB = 0
SM = 0
Delta_K
Doxycycline
YHR042W NCP1
ZShift = −3.15
NG = 0
DB = 0
SM = 0
Delta_r
Doxycycline
YHR042W NCP1
ZShift = −3.65
NG = 0
DB = 0
SM = 0
Delta_AUC
Doxycycline
YGL099W LSG1
ZShift = 1.51
NG = 0
DB = 0
SM = 0
Delta_L
Doxycycline
YGL099W LSG1
ZShift = 2.18
NG = 0
DB = 0
SM = 0
Delta_K
Doxycycline
YGL099W LSG1
ZShift = −0.16
NG = 0
DB = 0
SM = 0
Delta_r
Doxycycline
YGL099W LSG1
ZShift = 0.78
NG = 0
DB = 0
SM = 0
Delta_AUC
Doxycycline
0 10 25 50 100

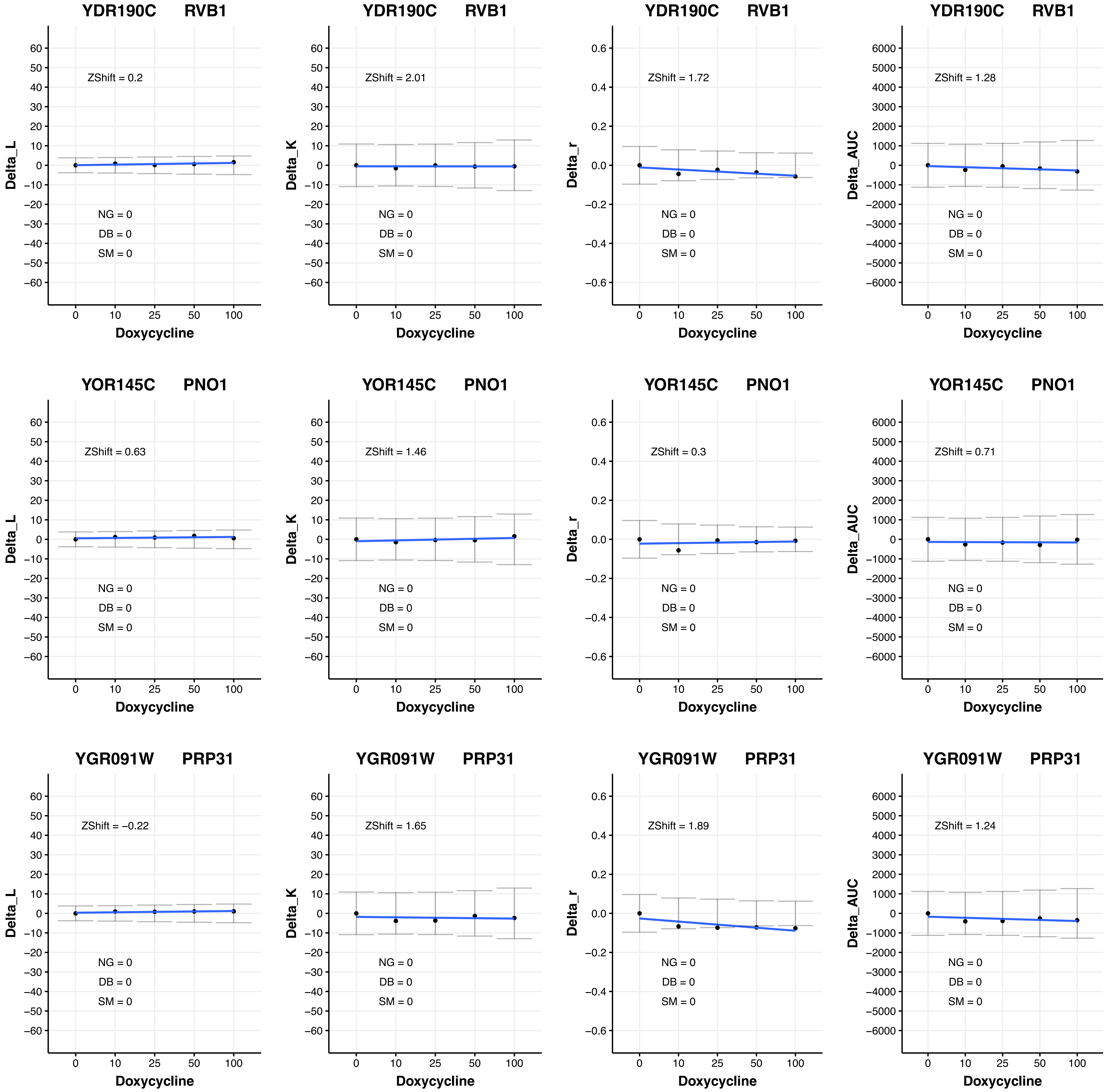

YDR190C RVB1
ZShift = 0.2
ZShift = 2.01
ZShift = 1.72
ZShift = 1.28
YOR145C PNO1
ZShift = 0.63
ZShift = 1.46
ZShift = 0.3
ZShift = 0.71
YGR091W PRP31
ZShift = −0.22
ZShift = 1.65
ZShift = 1.89
ZShift = 1.24
NG = 0
DB = 0
SM = 0
Delta_L
Delta_K
Delta_r
Delta_AUC
Doxycycline

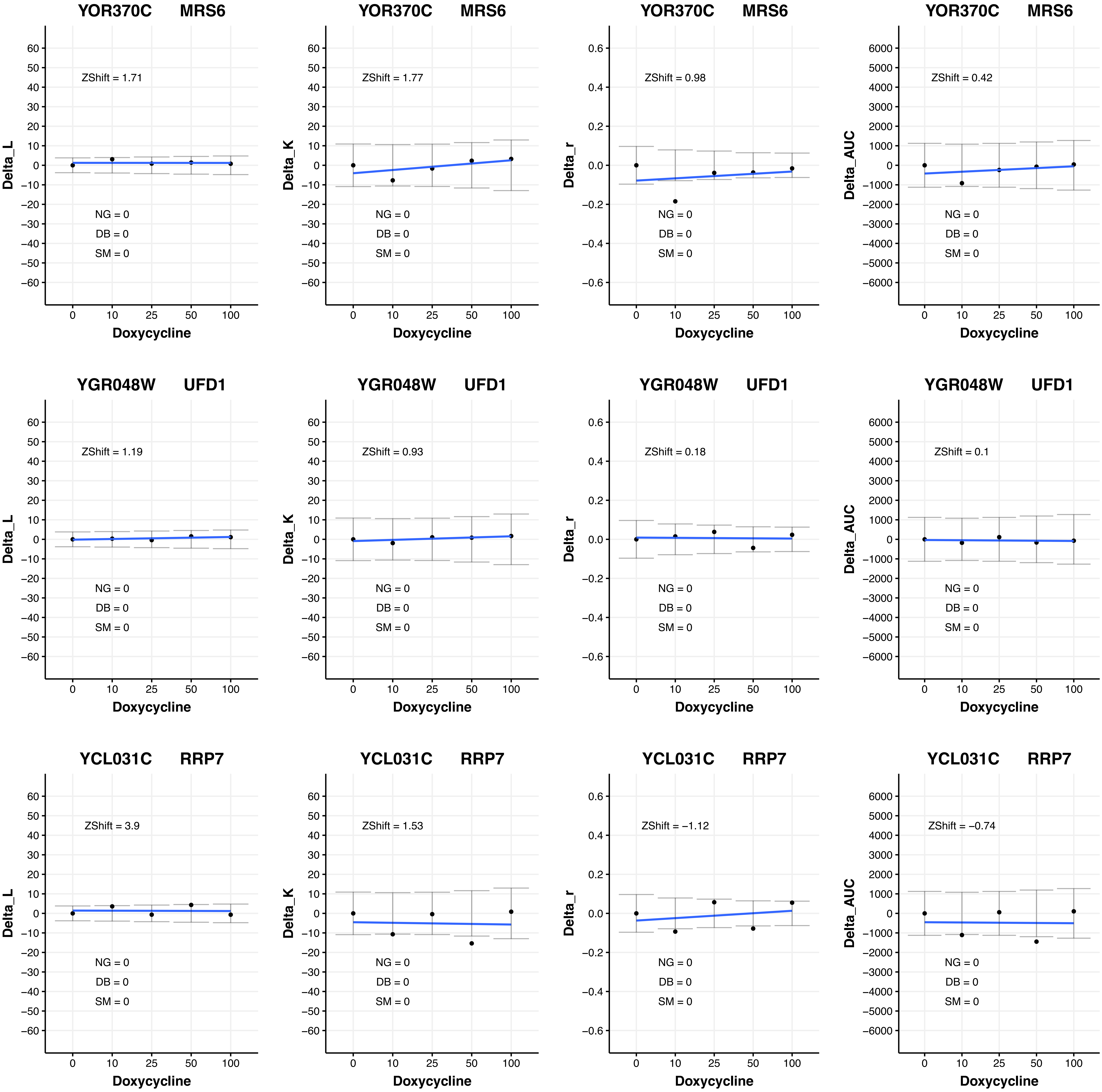

YOR370C MRS6
ZShift = 1.71
NG = 0
DB = 0
SM = 0
Delta_L
Doxycycline
YOR370C MRS6
ZShift = 1.77
NG = 0
DB = 0
SM = 0
Delta_K
Doxycycline
YOR370C MRS6
ZShift = 0.98
NG = 0
DB = 0
SM = 0
Delta_r
Doxycycline
YOR370C MRS6
ZShift = 0.42
NG = 0
DB = 0
SM = 0
Delta_AUC
Doxycycline
YGR048W UFD1
ZShift = 1.19
NG = 0
DB = 0
SM = 0
Delta_L
Doxycycline
YGR048W UFD1
ZShift = 0.93
NG = 0
DB = 0
SM = 0
Delta_K
Doxycycline
YGR048W UFD1
ZShift = 0.18
NG = 0
DB = 0
SM = 0
Delta_r
Doxycycline
YGR048W UFD1
ZShift = 0.1
NG = 0
DB = 0
SM = 0
Delta_AUC
Doxycycline
YCL031C RRP7
ZShift = 3.9
NG = 0
DB = 0
SM = 0
Delta_L
Doxycycline
YCL031C RRP7
ZShift = 1.53
NG = 0
DB = 0
SM = 0
Delta_K
Doxycycline
YCL031C RRP7
ZShift = −1.12
NG = 0
DB = 0
SM = 0
Delta_r
Doxycycline
YCL031C RRP7
ZShift = −0.74
NG = 0
DB = 0
SM = 0
Delta_AUC
Doxycycline

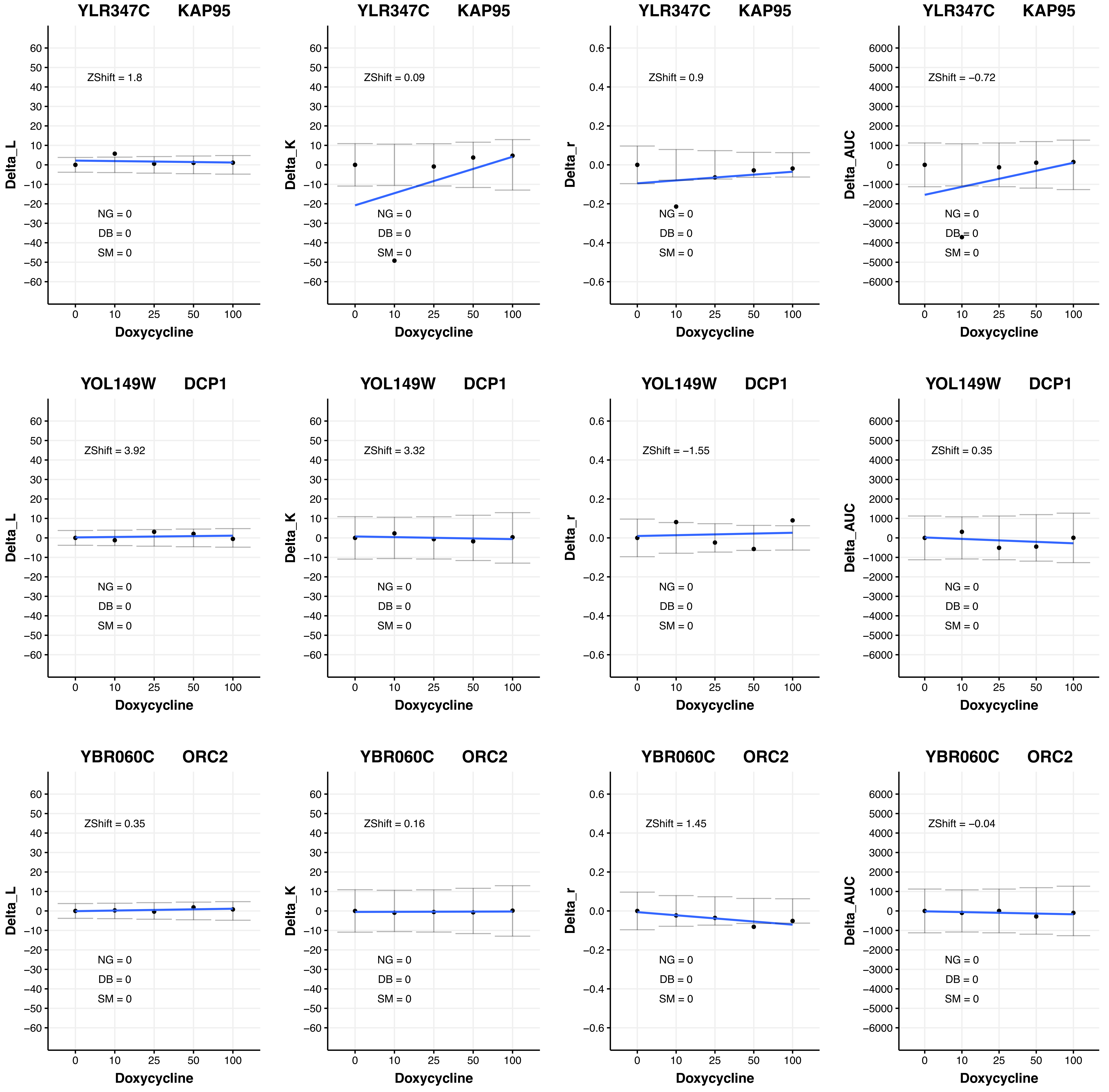
YLR347C KAP95
ZShift = 1.8
Delta_L
ZShift = 0.09
Delta_K
ZShift = 0.9
Delta_r
ZShift = −0.72
Delta_AUC
YOL149W DCP1
ZShift = 3.92
ZShift = 3.32
ZShift = −1.55
ZShift = 0.35
YBR060C ORC2
ZShift = 0.35
ZShift = 0.16
ZShift = 1.45
ZShift = −0.04
NG = 0
DB = 0
SM = 0
Doxycycline
0
10
25
50
100

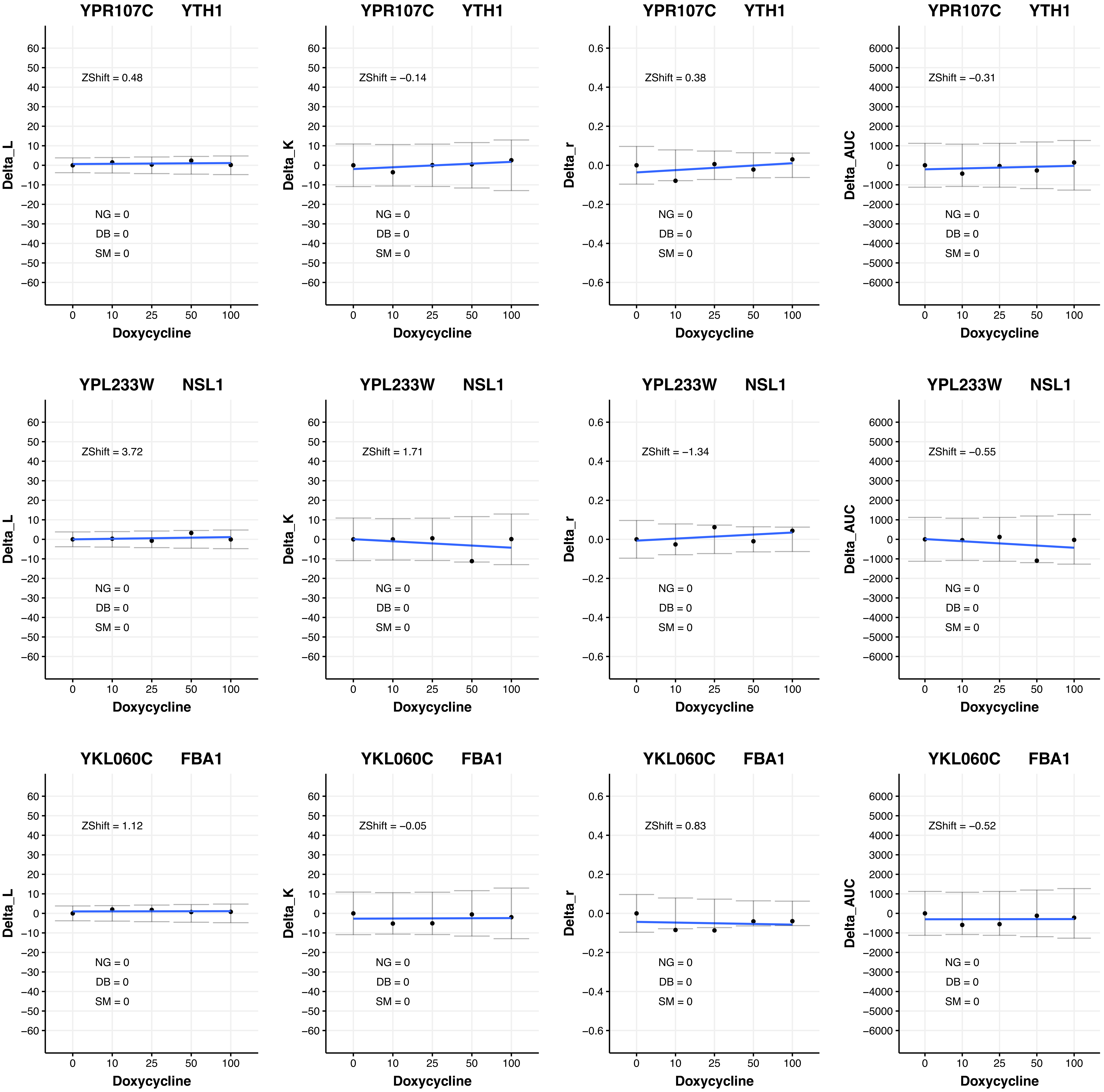
YPR107C YTH1
ZShift = 0.48
NG = 0
DB = 0
SM = 0
Delta_L
Doxycycline
YPR107C YTH1
ZShift = −0.14
NG = 0
DB = 0
SM = 0
Delta_K
Doxycycline
YPR107C YTH1
ZShift = 0.38
NG = 0
DB = 0
SM = 0
Delta_r
Doxycycline
YPR107C YTH1
ZShift = −0.31
NG = 0
DB = 0
SM = 0
Delta_AUC
Doxycycline
YPL233W NSL1
ZShift = 3.72
NG = 0
DB = 0
SM = 0
Delta_L
Doxycycline
YPL233W NSL1
ZShift = 1.71
NG = 0
DB = 0
SM = 0
Delta_K
Doxycycline
YPL233W NSL1
ZShift = −1.34
NG = 0
DB = 0
SM = 0
Delta_r
Doxycycline
YPL233W NSL1
ZShift = −0.55
NG = 0
DB = 0
SM = 0
Delta_AUC
Doxycycline
YKL060C FBA1
ZShift = 1.12
NG = 0
DB = 0
SM = 0
Delta_L
Doxycycline
YKL060C FBA1
ZShift = −0.05
NG = 0
DB = 0
SM = 0
Delta_K
Doxycycline
YKL060C FBA1
ZShift = 0.83
NG = 0
DB = 0
SM = 0
Delta_r
Doxycycline
YKL060C FBA1
ZShift = −0.52
NG = 0
DB = 0
SM = 0
Delta_AUC
Doxycycline

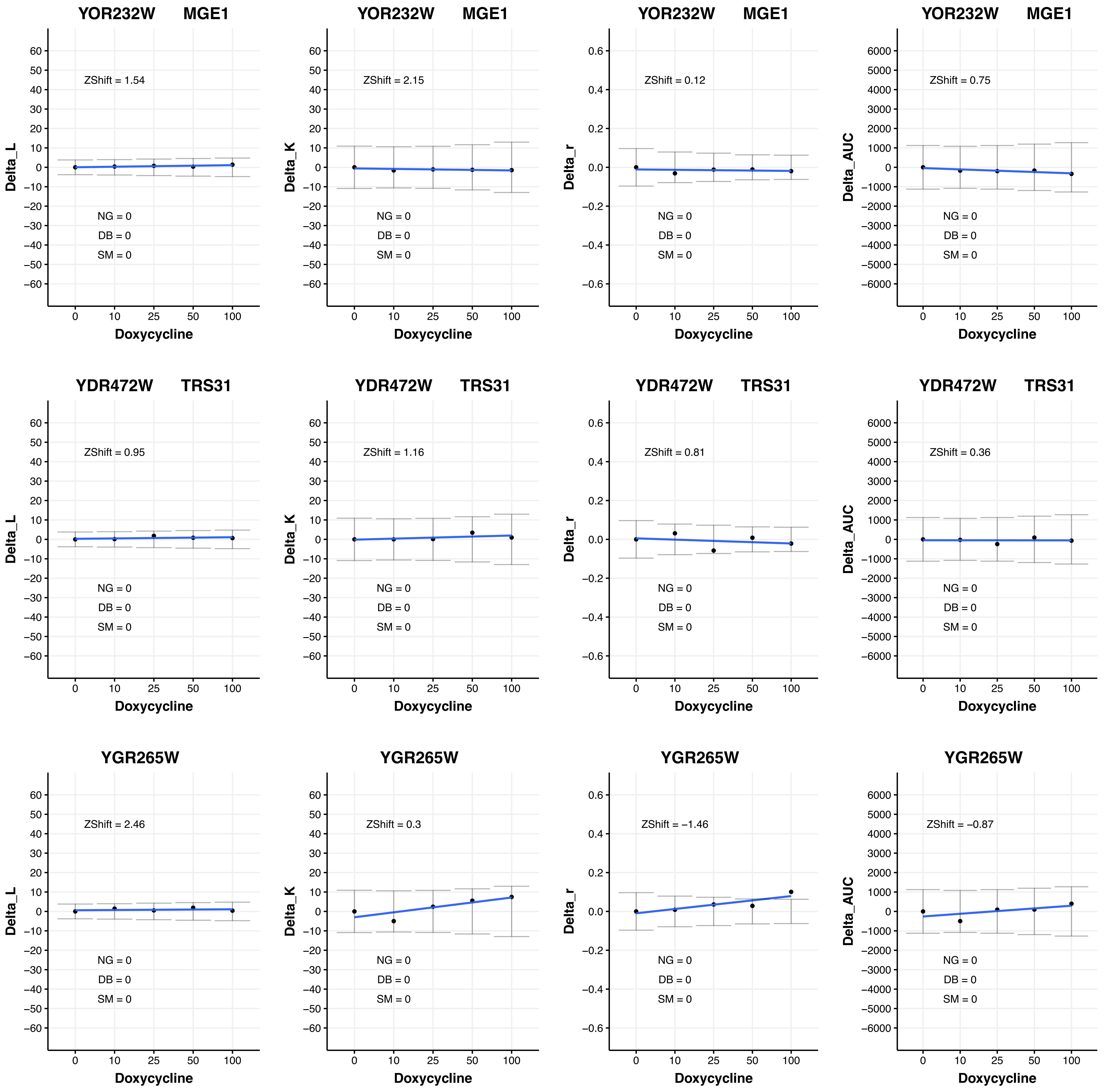

YOR232W MGE1
Delta_L
ZShift = 1.54
NG = 0
DB = 0
SM = 0
Doxycycline
YOR232W MGE1
Delta_K
ZShift = 2.15
NG = 0
DB = 0
SM = 0
Doxycycline
YOR232W MGE1
Delta_r
ZShift = 0.12
NG = 0
DB = 0
SM = 0
Doxycycline
YOR232W MGE1
Delta_AUC
ZShift = 0.75
NG = 0
DB = 0
SM = 0
Doxycycline
YDR472W TRS31
Delta_L
ZShift = 0.95
NG = 0
DB = 0
SM = 0
Doxycycline
YDR472W TRS31
Delta_K
ZShift = 1.16
NG = 0
DB = 0
SM = 0
Doxycycline
YDR472W TRS31
Delta_r
ZShift = 0.81
NG = 0
DB = 0
SM = 0
Doxycycline
YDR472W TRS31
Delta_AUC
ZShift = 0.36
NG = 0
DB = 0
SM = 0
Doxycycline
YGR265W
Delta_L
ZShift = 2.46
NG = 0
DB = 0
SM = 0
Doxycycline
YGR265W
Delta_K
ZShift = 0.3
NG = 0
DB = 0
SM = 0
Doxycycline
YGR265W
Delta_r
ZShift = −1.46
NG = 0
DB = 0
SM = 0
Doxycycline
YGR265W
Delta_AUC
ZShift = −0.87
NG = 0
DB = 0
SM = 0
Doxycycline

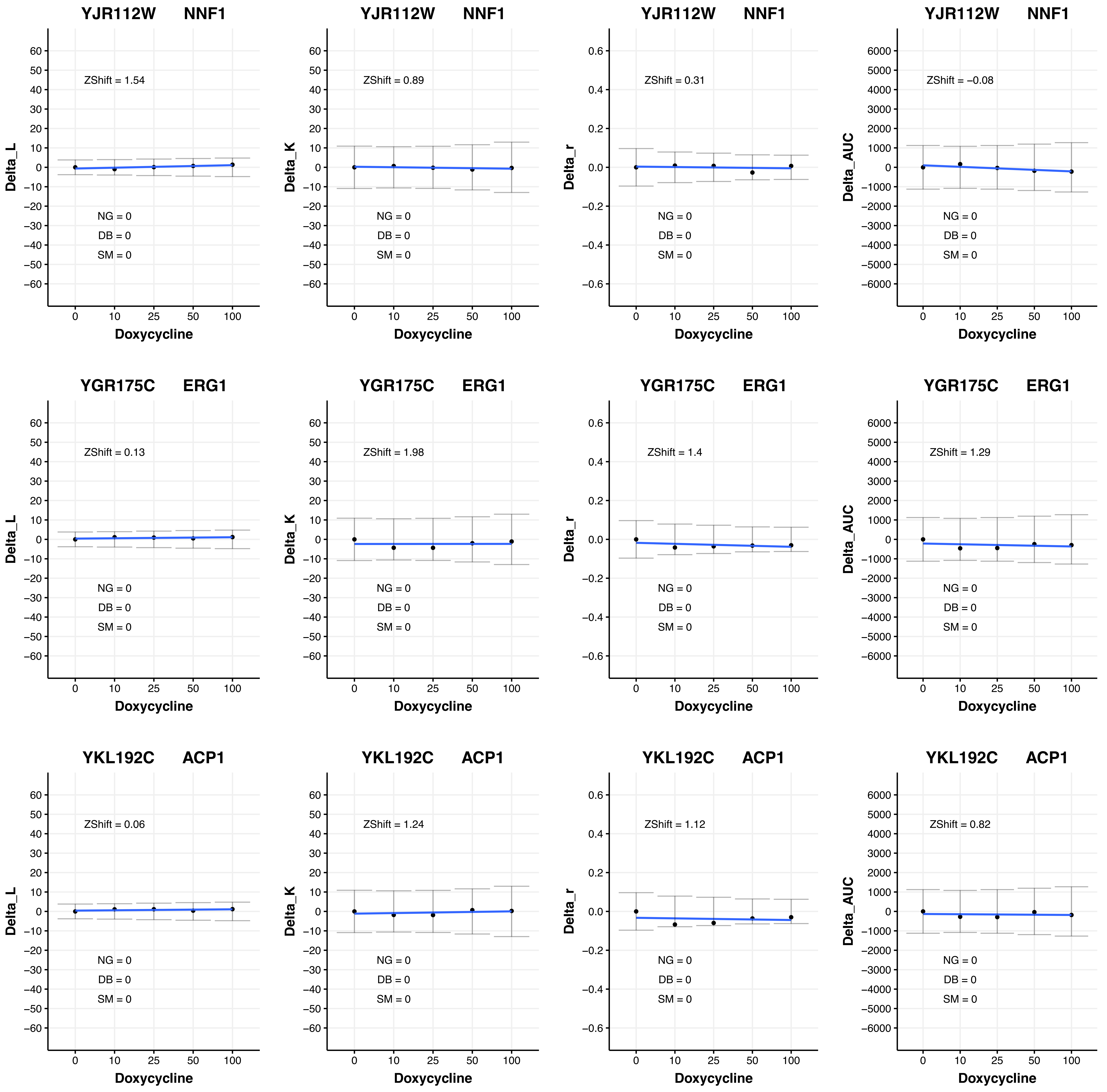
YJR112W NNF1
ZShift = 1.54
Delta_L
NG = 0
DB = 0
SM = 0
Doxycycline
YJR112W NNF1
ZShift = 0.89
Delta_K
NG = 0
DB = 0
SM = 0
Doxycycline
YJR112W NNF1
ZShift = 0.31
Delta_r
NG = 0
DB = 0
SM = 0
Doxycycline
YJR112W NNF1
ZShift = −0.08
Delta_AUC
NG = 0
DB = 0
SM = 0
Doxycycline
YGR175C ERG1
ZShift = 0.13
Delta_L
NG = 0
DB = 0
SM = 0
Doxycycline
YGR175C ERG1
ZShift = 1.98
Delta_K
NG = 0
DB = 0
SM = 0
Doxycycline
YGR175C ERG1
ZShift = 1.4
Delta_r
NG = 0
DB = 0
SM = 0
Doxycycline
YGR175C ERG1
ZShift = 1.29
Delta_AUC
NG = 0
DB = 0
SM = 0
Doxycycline
YKL192C ACP1
ZShift = 0.06
Delta_L
NG = 0
DB = 0
SM = 0
Doxycycline
YKL192C ACP1
ZShift = 1.24
Delta_K
NG = 0
DB = 0
SM = 0
Doxycycline
YKL192C ACP1
ZShift = 1.12
Delta_r
NG = 0
DB = 0
SM = 0
Doxycycline
YKL192C ACP1
ZShift = 0.82
Delta_AUC
NG = 0
DB = 0
SM = 0
Doxycycline

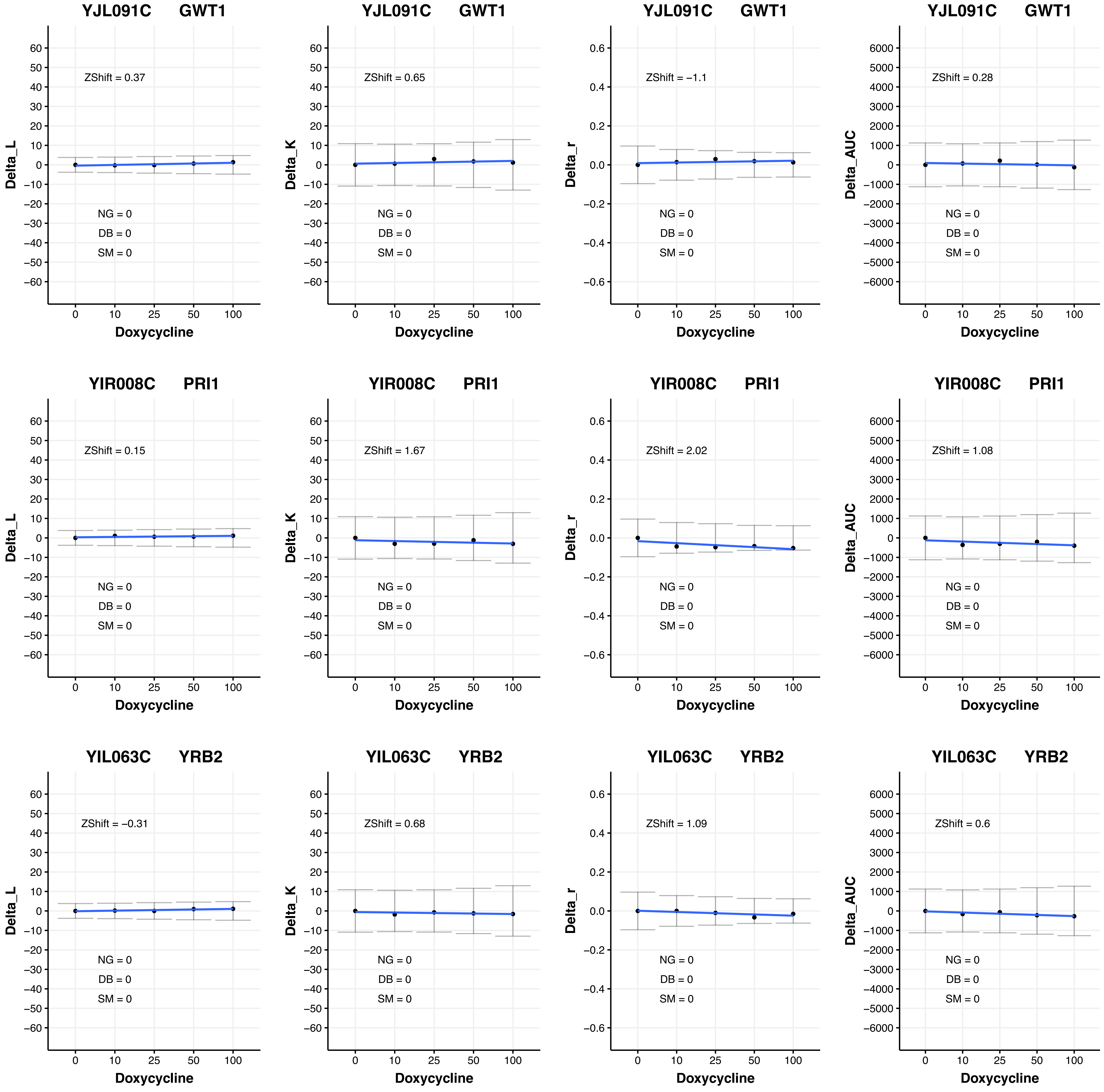
YJL091C GWT1
ZShift = 0.37
NG = 0
DB = 0
SM = 0
Delta_L
Doxycycline
YJL091C GWT1
ZShift = 0.65
NG = 0
DB = 0
SM = 0
Delta_K
Doxycycline
YJL091C GWT1
ZShift = −1.1
NG = 0
DB = 0
SM = 0
Delta_r
Doxycycline
YJL091C GWT1
ZShift = 0.28
NG = 0
DB = 0
SM = 0
Delta_AUC
Doxycycline
YIR008C PRI1
ZShift = 0.15
NG = 0
DB = 0
SM = 0
Delta_L
Doxycycline
YIR008C PRI1
ZShift = 1.67
NG = 0
DB = 0
SM = 0
Delta_K
Doxycycline
YIR008C PRI1
ZShift = 2.02
NG = 0
DB = 0
SM = 0
Delta_r
Doxycycline
YIR008C PRI1
ZShift = 1.08
NG = 0
DB = 0
SM = 0
Delta_AUC
Doxycycline
YIL063C YRB2
ZShift = −0.31
NG = 0
DB = 0
SM = 0
Delta_L
Doxycycline
YIL063C YRB2
ZShift = 0.68
NG = 0
DB = 0
SM = 0
Delta_K
Doxycycline
YIL063C YRB2
ZShift = 1.09
NG = 0
DB = 0
SM = 0
Delta_r
Doxycycline
YIL063C YRB2
ZShift = 0.6
NG = 0
DB = 0
SM = 0
Delta_AUC
Doxycycline

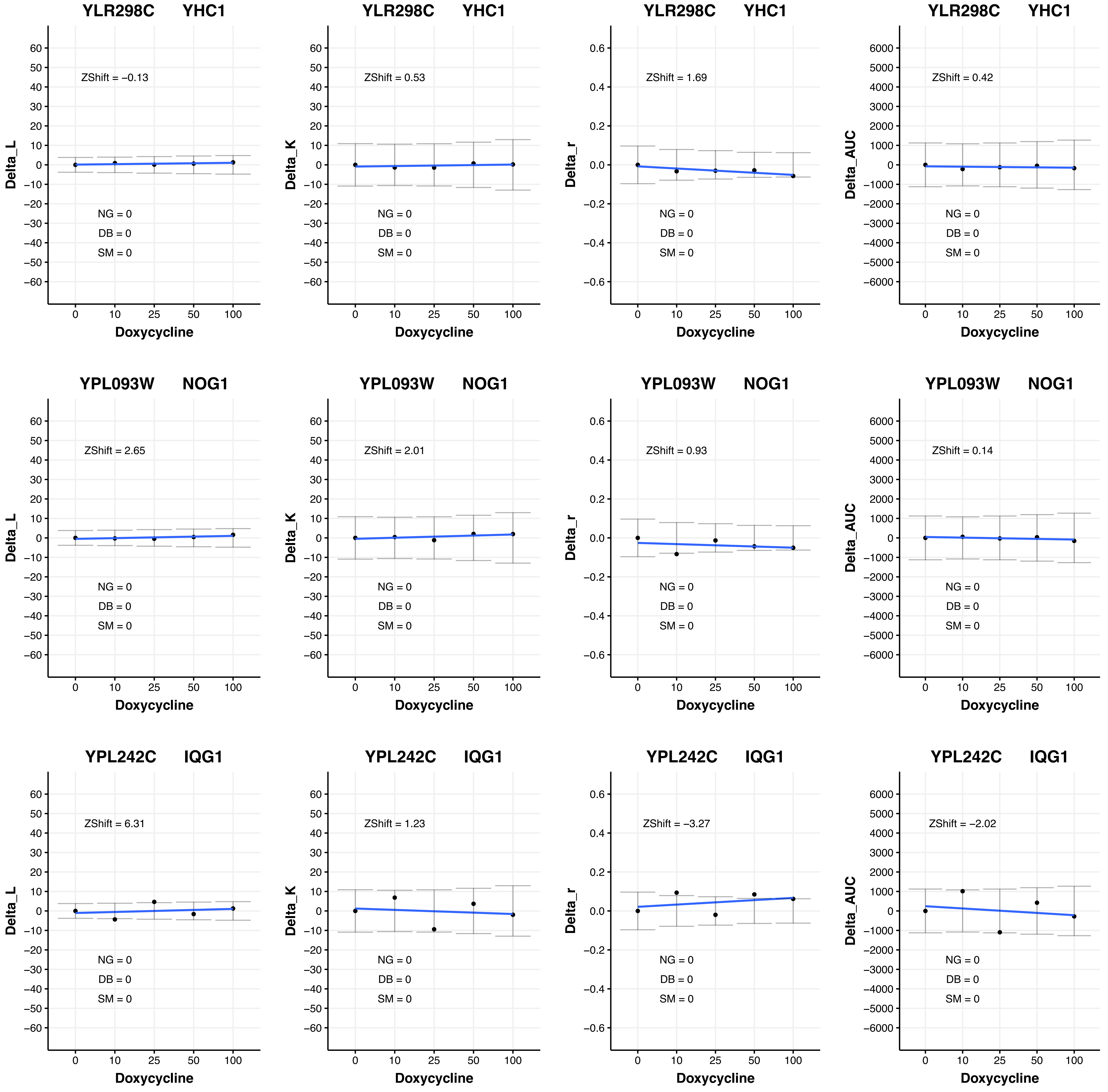

YLR298C YHC1
ZShift = −0.13
NG = 0
DB = 0
SM = 0
Delta_L
Doxycycline
YLR298C YHC1
ZShift = 0.53
NG = 0
DB = 0
SM = 0
Delta_K
Doxycycline
YLR298C YHC1
ZShift = 1.69
NG = 0
DB = 0
SM = 0
Delta_r
Doxycycline
YLR298C YHC1
ZShift = 0.42
NG = 0
DB = 0
SM = 0
Delta_AUC
Doxycycline
YPL093W NOG1
ZShift = 2.65
NG = 0
DB = 0
SM = 0
Delta_L
Doxycycline
YPL093W NOG1
ZShift = 2.01
NG = 0
DB = 0
SM = 0
Delta_K
Doxycycline
YPL093W NOG1
ZShift = 0.93
NG = 0
DB = 0
SM = 0
Delta_r
Doxycycline
YPL093W NOG1
ZShift = 0.14
NG = 0
DB = 0
SM = 0
Delta_AUC
Doxycycline
YPL242C IQG1
ZShift = 6.31
NG = 0
DB = 0
SM = 0
Delta_L
Doxycycline
YPL242C IQG1
ZShift = 1.23
NG = 0
DB = 0
SM = 0
Delta_K
Doxycycline
YPL242C IQG1
ZShift = −3.27
NG = 0
DB = 0
SM = 0
Delta_r
Doxycycline
YPL242C IQG1
ZShift = −2.02
NG = 0
DB = 0
SM = 0
Delta_AUC
Doxycycline

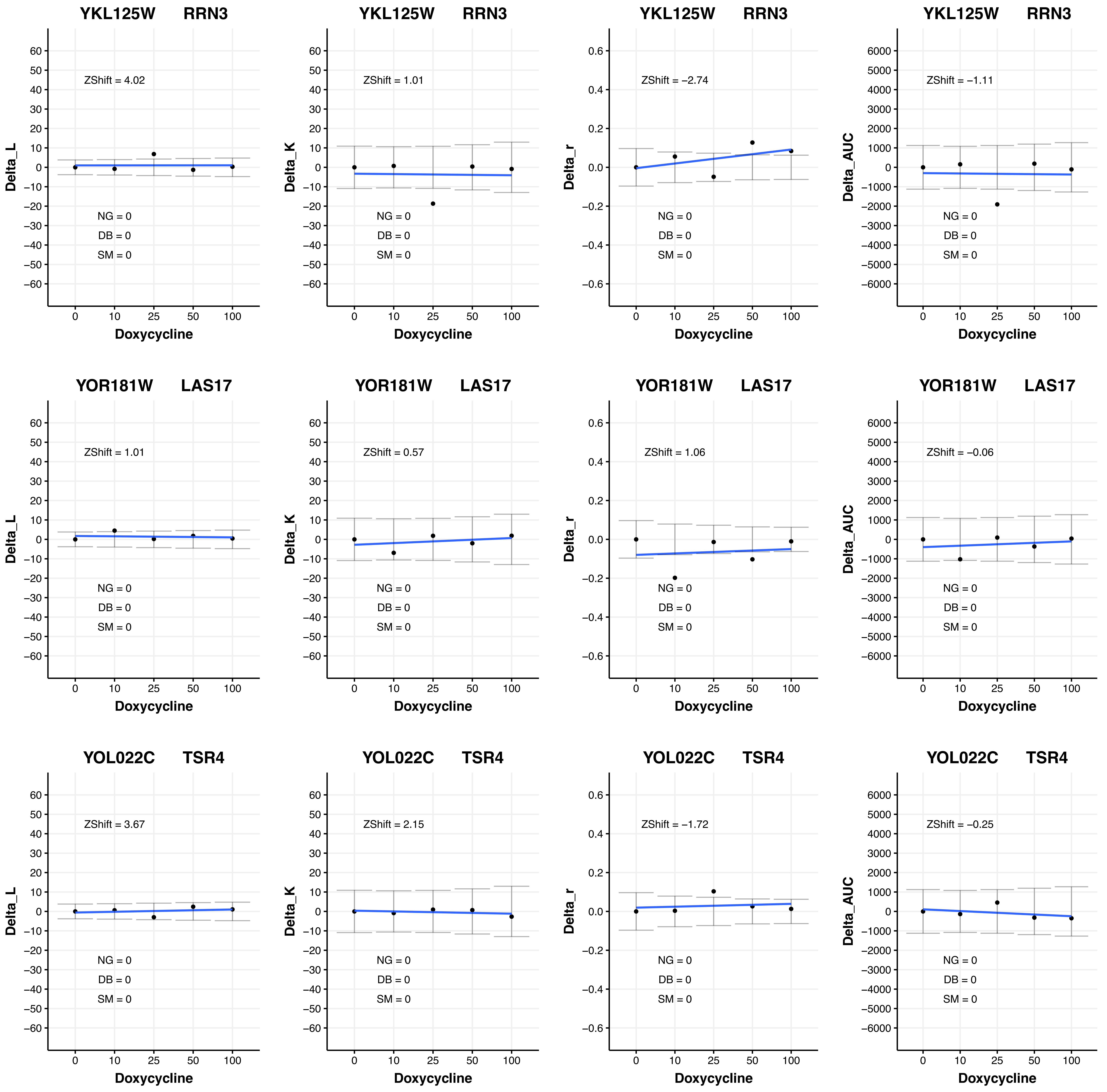
YKL125W RRN3
ZShift = 4.02
ZShift = 1.01
ZShift = −2.74
ZShift = −1.11
YOR181W LAS17
ZShift = 1.01
ZShift = 0.57
ZShift = 1.06
ZShift = −0.06
YOL022C TSR4
ZShift = 3.67
ZShift = 2.15
ZShift = −1.72
ZShift = −0.25
NG = 0
DB = 0
SM = 0
Delta_L
Delta_K
Delta_r
Delta_AUC
Doxycycline

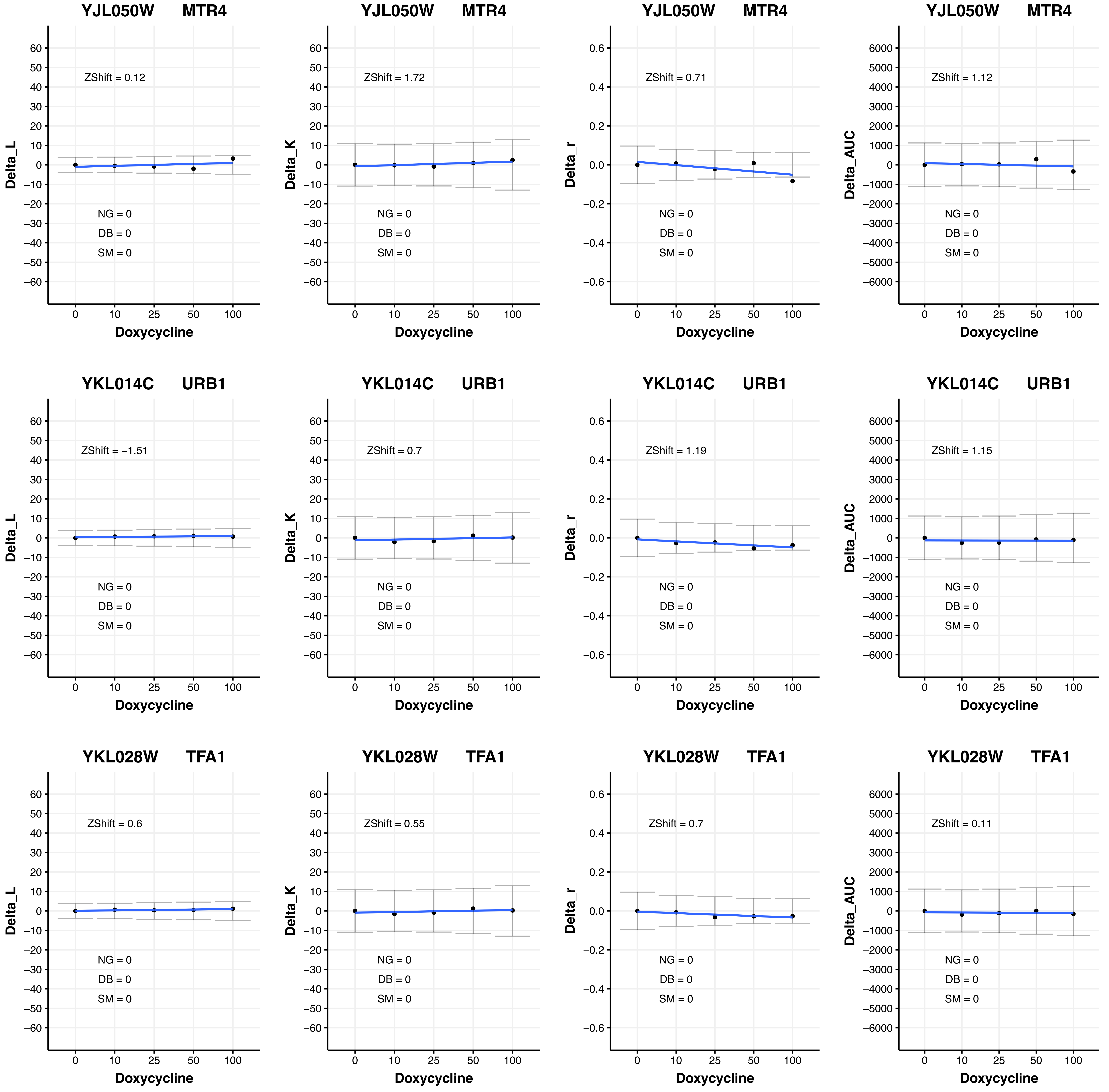

YJL050W MTR4
ZShift = 0.12
ZShift = 1.72
ZShift = 0.71
ZShift = 1.12
YKL014C URB1
ZShift = −1.51
ZShift = 0.7
ZShift = 1.19
ZShift = 1.15
YKL028W TFA1
ZShift = 0.6
ZShift = 0.55
ZShift = 0.7
ZShift = 0.11
NG = 0
DB = 0
SM = 0
Delta_L
Delta_K
Delta_r
Delta_AUC
Doxycycline

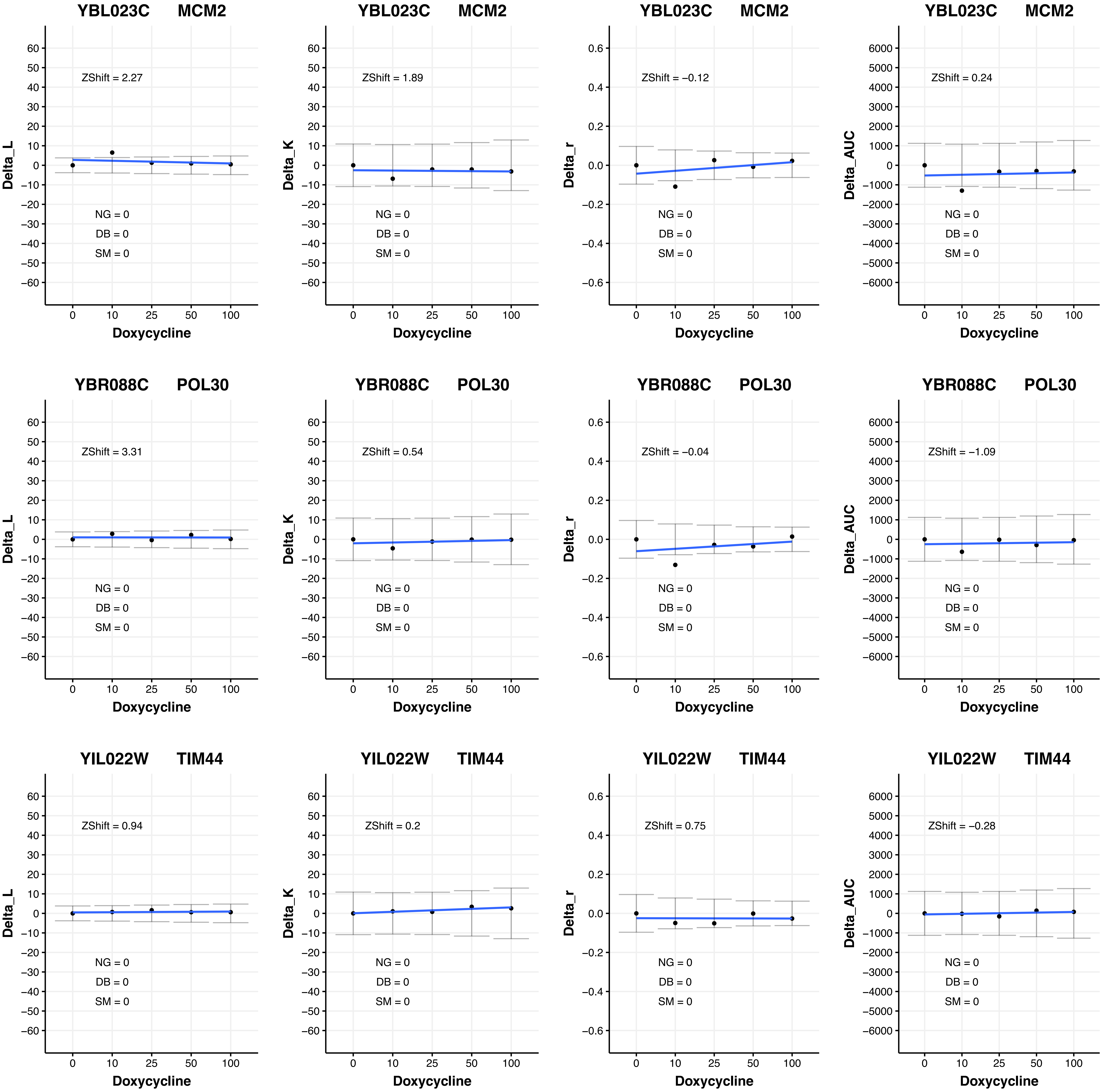

YBL023C MCM2
Delta_L
ZShift = 2.27
NG = 0
DB = 0
SM = 0
Doxycycline
YBL023C MCM2
Delta_K
ZShift = 1.89
NG = 0
DB = 0
SM = 0
Doxycycline
YBL023C MCM2
Delta_r
ZShift = −0.12
NG = 0
DB = 0
SM = 0
Doxycycline
YBL023C MCM2
Delta_AUC
ZShift = 0.24
NG = 0
DB = 0
SM = 0
Doxycycline
YBR088C POL30
Delta_L
ZShift = 3.31
NG = 0
DB = 0
SM = 0
Doxycycline
YBR088C POL30
Delta_K
ZShift = 0.54
NG = 0
DB = 0
SM = 0
Doxycycline
YBR088C POL30
Delta_r
ZShift = −0.04
NG = 0
DB = 0
SM = 0
Doxycycline
YBR088C POL30
Delta_AUC
ZShift = −1.09
NG = 0
DB = 0
SM = 0
Doxycycline
YIL022W TIM44
Delta_L
ZShift = 0.94
NG = 0
DB = 0
SM = 0
Doxycycline
YIL022W TIM44
Delta_K
ZShift = 0.2
NG = 0
DB = 0
SM = 0
Doxycycline
YIL022W TIM44
Delta_r
ZShift = 0.75
NG = 0
DB = 0
SM = 0
Doxycycline
YIL022W TIM44
Delta_AUC
ZShift = −0.28
NG = 0
DB = 0
SM = 0
Doxycycline

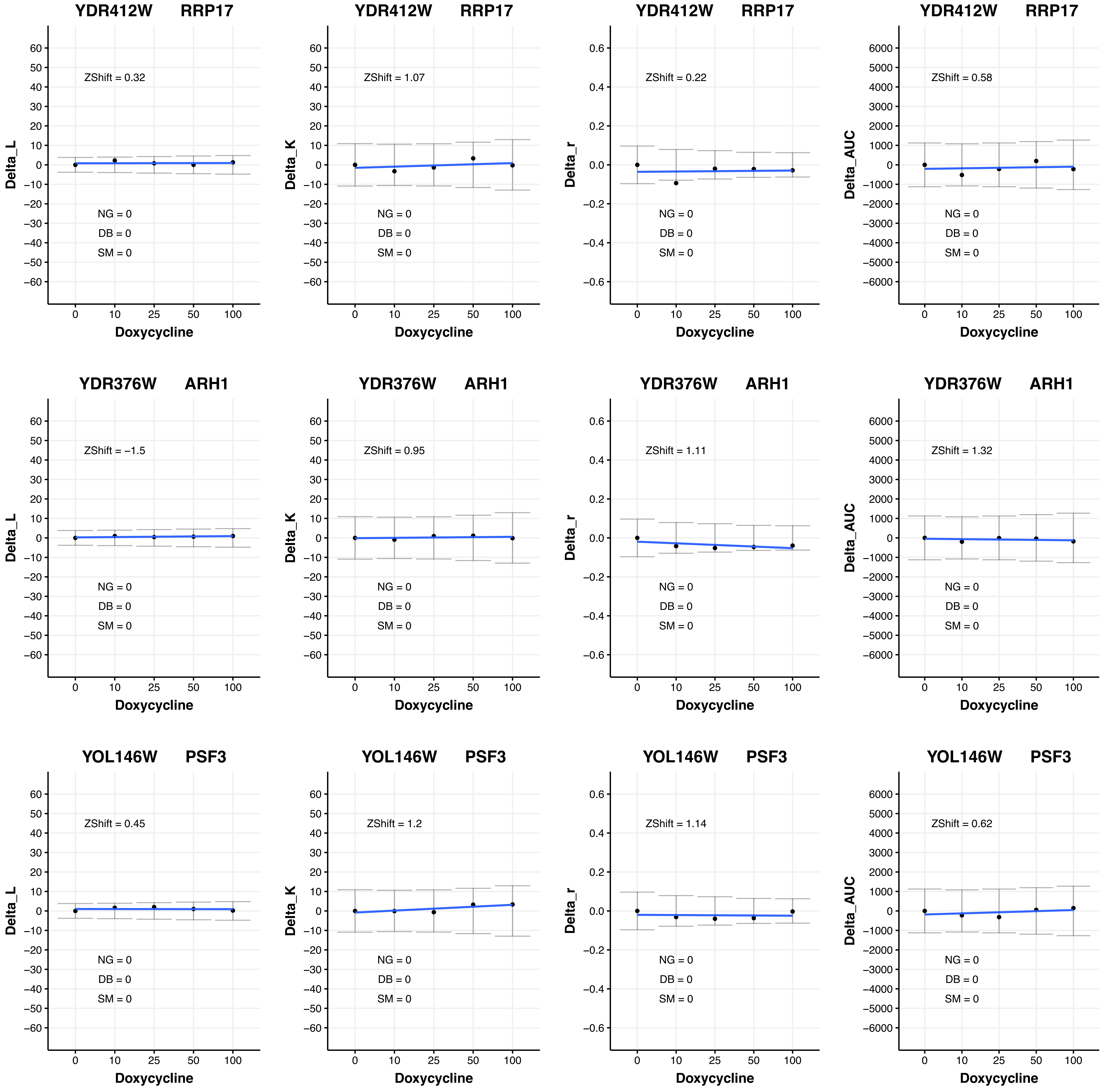
YDR412W RRP17
ZShift = 0.32
ZShift = 1.07
ZShift = 0.22
ZShift = 0.58
YDR376W ARH1
ZShift = −1.5
ZShift = 0.95
ZShift = 1.11
ZShift = 1.32
YOL146W PSF3
ZShift = 0.45
ZShift = 1.2
ZShift = 1.14
ZShift = 0.62
NG = 0
DB = 0
SM = 0
Delta_L
Delta_K
Delta_r
Delta_AUC
Doxycycline
0
10
25
50
100

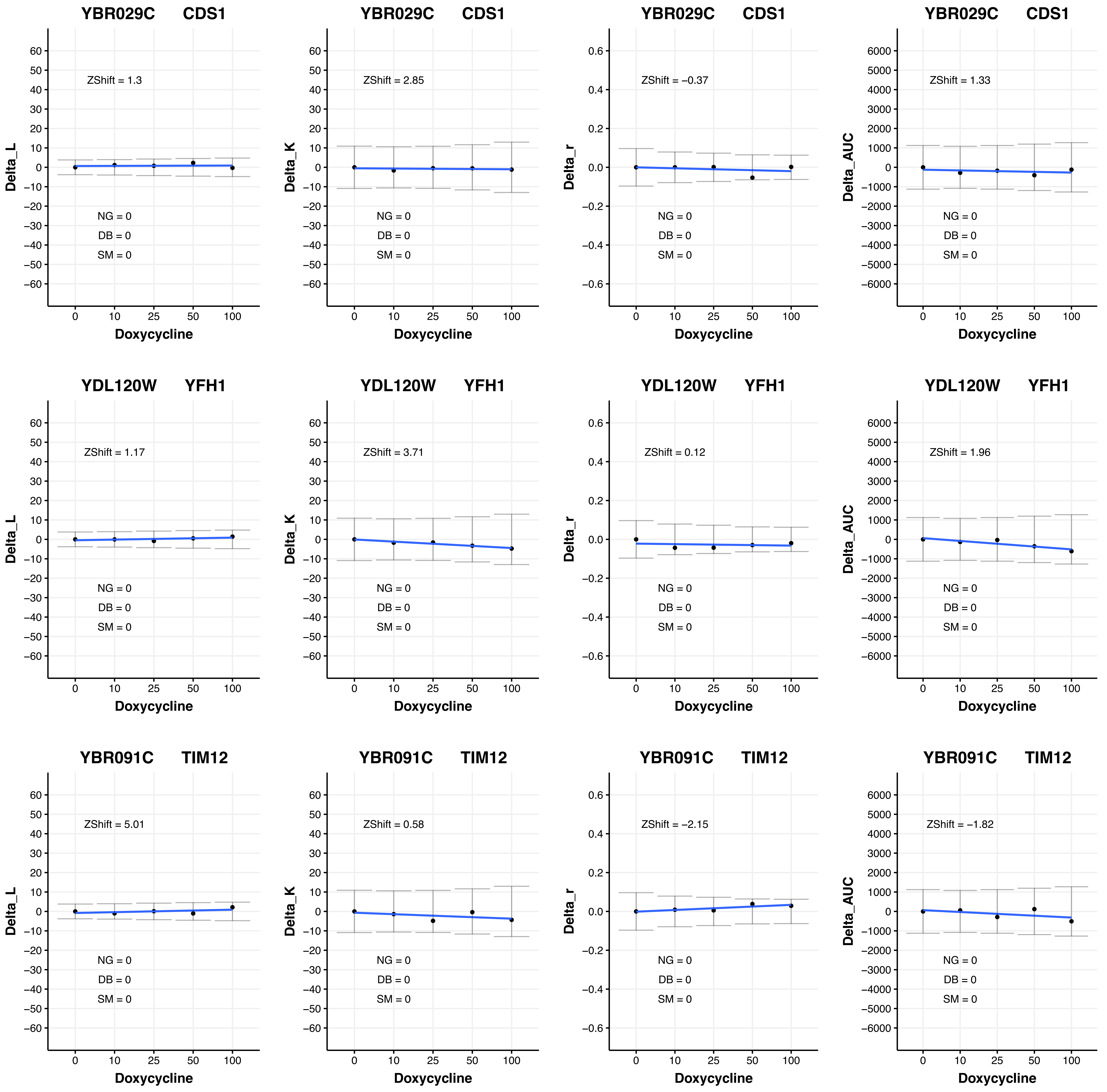
YBR029C CDS1
Delta_L
ZShift = 1.3
Delta_K
ZShift = 2.85
Delta_r
ZShift = −0.37
Delta_AUC
ZShift = 1.33
YDL120W YFH1
ZShift = 1.17
ZShift = 3.71
ZShift = 0.12
ZShift = 1.96
YBR091C TIM12
ZShift = 5.01
ZShift = 0.58
ZShift = −2.15
ZShift = −1.82
NG = 0
DB = 0
SM = 0
Doxycycline
0
10
25
50
100

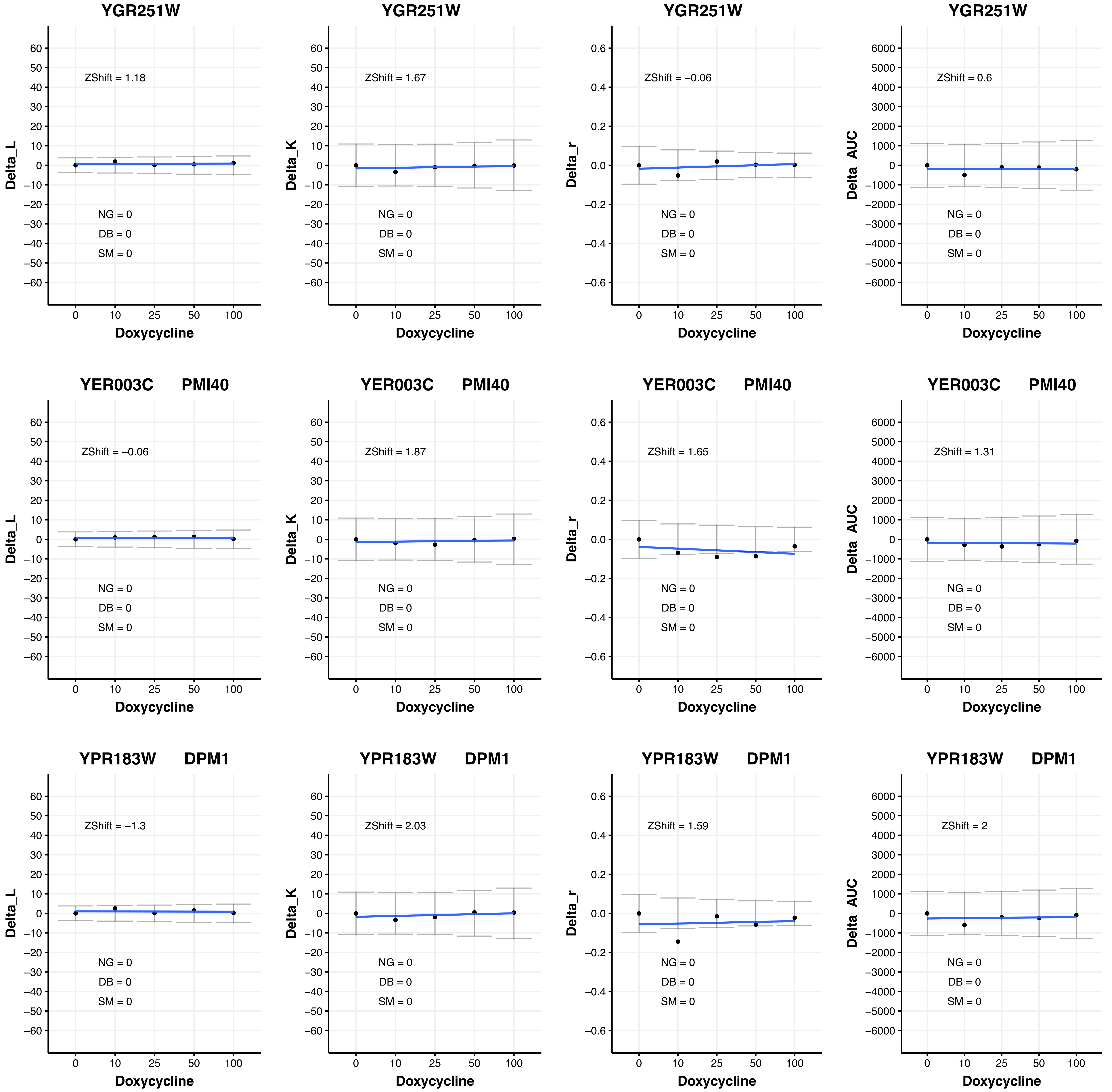
YGR251W
ZShift = 1.18
NG = 0
DB = 0
SM = 0
Delta_L
Doxycycline
YGR251W
ZShift = 1.67
Delta_K
YGR251W
ZShift = −0.06
Delta_r
YGR251W
ZShift = 0.6
Delta_AUC
YER003C PMI40
ZShift = −0.06
YER003C PMI40
ZShift = 1.87
YER003C PMI40
ZShift = 1.65
YER003C PMI40
ZShift = 1.31
YPR183W DPM1
ZShift = −1.3
YPR183W DPM1
ZShift = 2.03
YPR183W DPM1
ZShift = 1.59
YPR183W DPM1
ZShift = 2

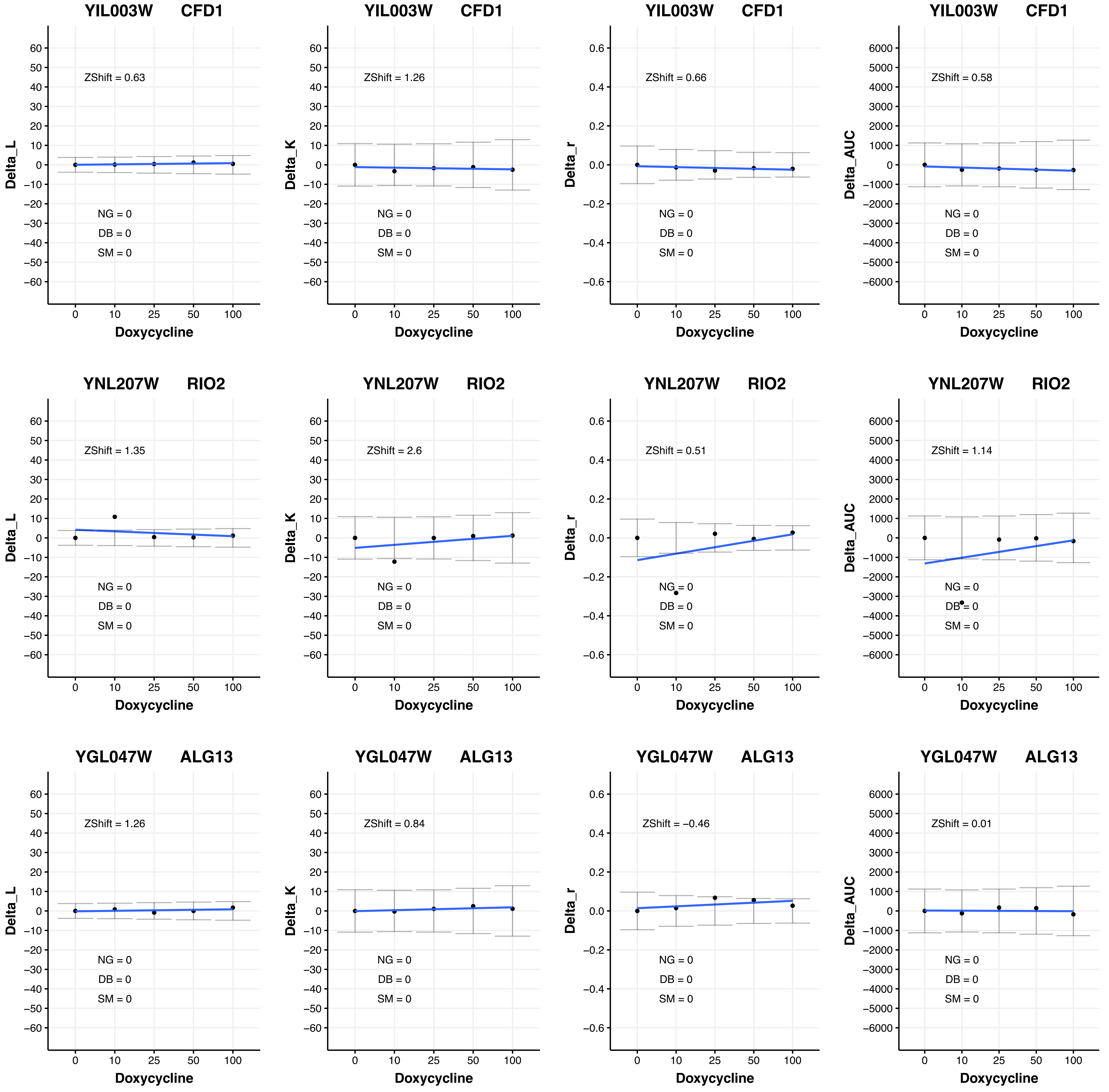
YIL003W CFD1
ZShift = 0.63
NG = 0
DB = 0
SM = 0
Delta_L
Doxycycline
YIL003W CFD1
ZShift = 1.26
NG = 0
DB = 0
SM = 0
Delta_K
Doxycycline
YIL003W CFD1
ZShift = 0.66
NG = 0
DB = 0
SM = 0
Delta_r
Doxycycline
YIL003W CFD1
ZShift = 0.58
NG = 0
DB = 0
SM = 0
Delta_AUC
Doxycycline
YNL207W RIO2
ZShift = 1.35
NG = 0
DB = 0
SM = 0
Delta_L
Doxycycline
YNL207W RIO2
ZShift = 2.6
NG = 0
DB = 0
SM = 0
Delta_K
Doxycycline
YNL207W RIO2
ZShift = 0.51
NG = 0
DB = 0
SM = 0
Delta_r
Doxycycline
YNL207W RIO2
ZShift = 1.14
NG = 0
DB = 0
SM = 0
Delta_AUC
Doxycycline
YGL047W ALG13
ZShift = 1.26
NG = 0
DB = 0
SM = 0
Delta_L
Doxycycline
YGL047W ALG13
ZShift = 0.84
NG = 0
DB = 0
SM = 0
Delta_K
Doxycycline
YGL047W ALG13
ZShift = −0.46
NG = 0
DB = 0
SM = 0
Delta_r
Doxycycline
YGL047W ALG13
ZShift = 0.01
NG = 0
DB = 0
SM = 0
Delta_AUC
Doxycycline

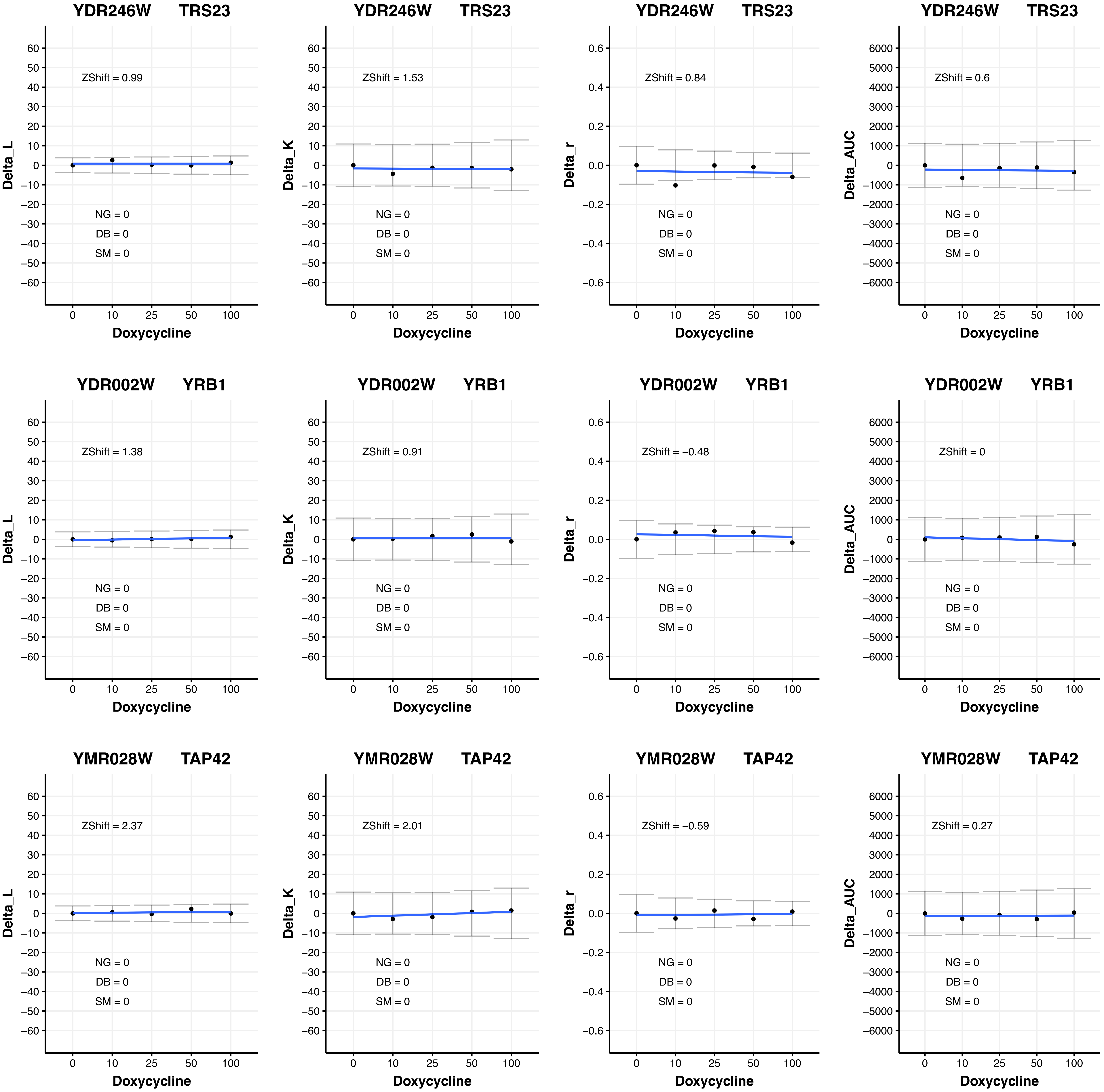
YDR246W TRS23
ZShift = 0.99
NG = 0
DB = 0
SM = 0
Delta_L
Doxycycline
0 10 25 50 100
YDR246W TRS23
ZShift = 1.53
NG = 0
DB = 0
SM = 0
Delta_K
Doxycycline
YDR246W TRS23
ZShift = 0.84
NG = 0
DB = 0
SM = 0
Delta_r
Doxycycline
YDR246W TRS23
ZShift = 0.6
NG = 0
DB = 0
SM = 0
Delta_AUC
Doxycycline
YDR002W YRB1
ZShift = 1.38
NG = 0
DB = 0
SM = 0
Delta_L
Doxycycline
YDR002W YRB1
ZShift = 0.91
NG = 0
DB = 0
SM = 0
Delta_K
Doxycycline
YDR002W YRB1
ZShift = −0.48
NG = 0
DB = 0
SM = 0
Delta_r
Doxycycline
YDR002W YRB1
ZShift = 0
NG = 0
DB = 0
SM = 0
Delta_AUC
Doxycycline
YMR028W TAP42
ZShift = 2.37
NG = 0
DB = 0
SM = 0
Delta_L
Doxycycline
YMR028W TAP42
ZShift = 2.01
NG = 0
DB = 0
SM = 0
Delta_K
Doxycycline
YMR028W TAP42
ZShift = −0.59
NG = 0
DB = 0
SM = 0
Delta_r
Doxycycline
YMR028W TAP42
ZShift = 0.27
NG = 0
DB = 0
SM = 0
Delta_AUC
Doxycycline

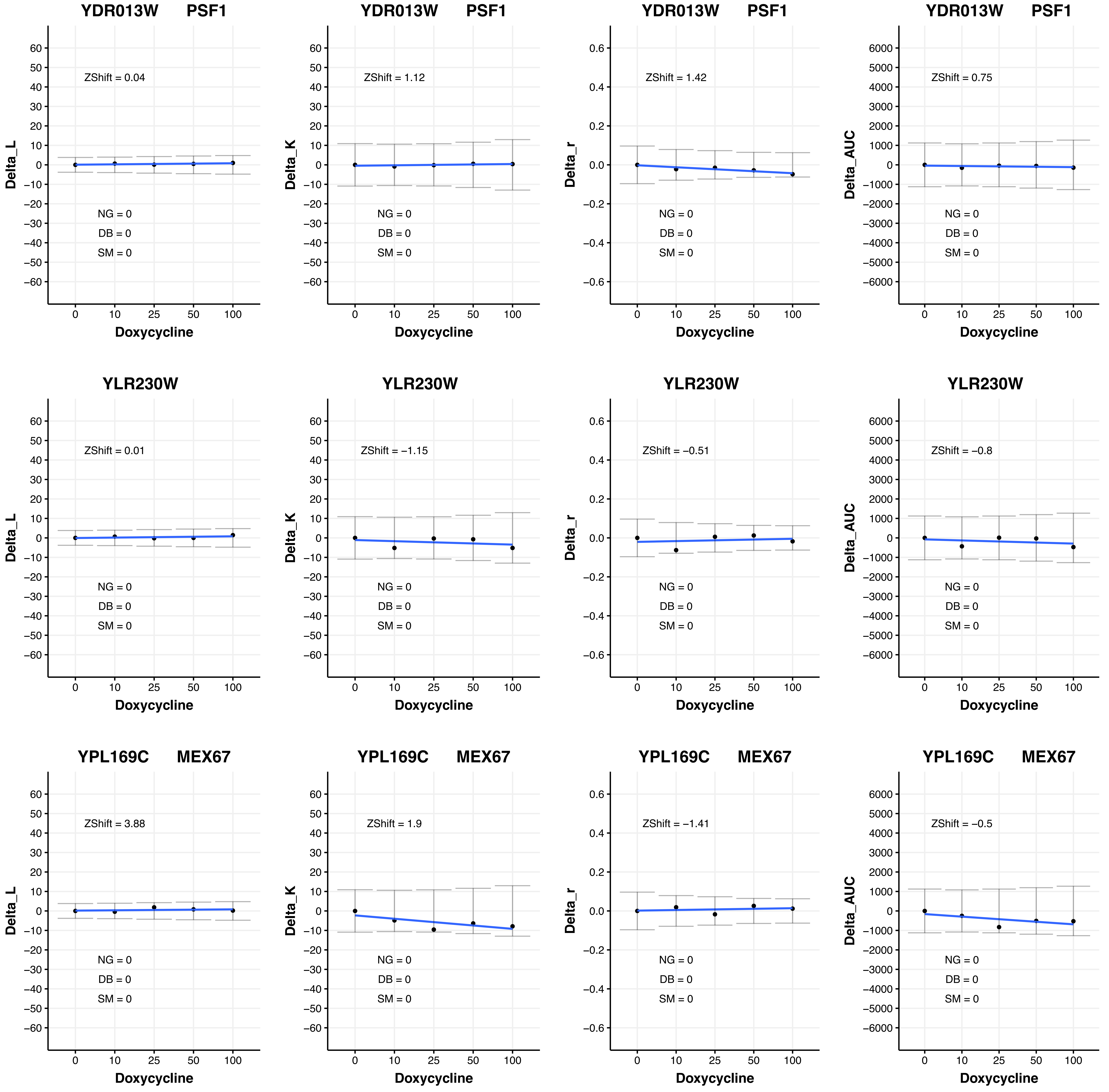

YDR013W PSF1
Delta_L
ZShift = 0.04
NG = 0
DB = 0
SM = 0
Doxycycline
YDR013W PSF1
Delta_K
ZShift = 1.12
NG = 0
DB = 0
SM = 0
Doxycycline
YDR013W PSF1
Delta_r
ZShift = 1.42
NG = 0
DB = 0
SM = 0
Doxycycline
YDR013W PSF1
Delta_AUC
ZShift = 0.75
NG = 0
DB = 0
SM = 0
Doxycycline
YLR230W
Delta_L
ZShift = 0.01
NG = 0
DB = 0
SM = 0
Doxycycline
YLR230W
Delta_K
ZShift = −1.15
NG = 0
DB = 0
SM = 0
Doxycycline
YLR230W
Delta_r
ZShift = −0.51
NG = 0
DB = 0
SM = 0
Doxycycline
YLR230W
Delta_AUC
ZShift = −0.8
NG = 0
DB = 0
SM = 0
Doxycycline
YPL169C MEX67
Delta_L
ZShift = 3.88
NG = 0
DB = 0
SM = 0
Doxycycline
YPL169C MEX67
Delta_K
ZShift = 1.9
NG = 0
DB = 0
SM = 0
Doxycycline
YPL169C MEX67
Delta_r
ZShift = −1.41
NG = 0
DB = 0
SM = 0
Doxycycline
YPL169C MEX67
Delta_AUC
ZShift = −0.5
NG = 0
DB = 0
SM = 0
Doxycycline

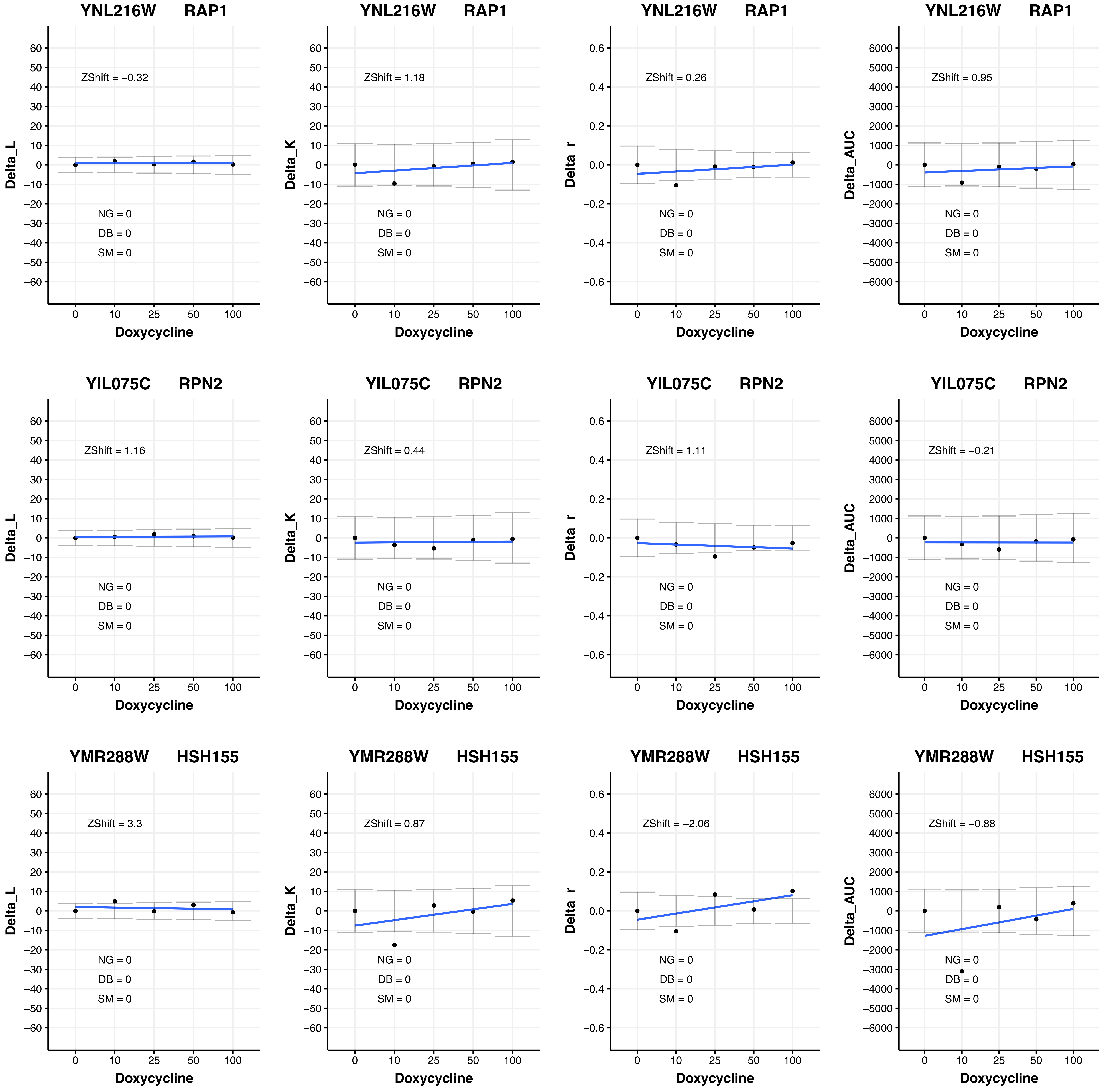
YNL216W RAP1
ZShift = −0.32
NG = 0
DB = 0
SM = 0
Delta_L
Doxycycline
YNL216W RAP1
ZShift = 1.18
NG = 0
DB = 0
SM = 0
Delta_K
Doxycycline
YNL216W RAP1
ZShift = 0.26
NG = 0
DB = 0
SM = 0
Delta_r
Doxycycline
YNL216W RAP1
ZShift = 0.95
NG = 0
DB = 0
SM = 0
Delta_AUC
Doxycycline
YIL075C RPN2
ZShift = 1.16
NG = 0
DB = 0
SM = 0
Delta_L
Doxycycline
YIL075C RPN2
ZShift = 0.44
NG = 0
DB = 0
SM = 0
Delta_K
Doxycycline
YIL075C RPN2
ZShift = 1.11
NG = 0
DB = 0
SM = 0
Delta_r
Doxycycline
YIL075C RPN2
ZShift = −0.21
NG = 0
DB = 0
SM = 0
Delta_AUC
Doxycycline
YMR288W HSH155
ZShift = 3.3
NG = 0
DB = 0
SM = 0
Delta_L
Doxycycline
YMR288W HSH155
ZShift = 0.87
NG = 0
DB = 0
SM = 0
Delta_K
Doxycycline
YMR288W HSH155
ZShift = −2.06
NG = 0
DB = 0
SM = 0
Delta_r
Doxycycline
YMR288W HSH155
ZShift = −0.88
NG = 0
DB = 0
SM = 0
Delta_AUC
Doxycycline

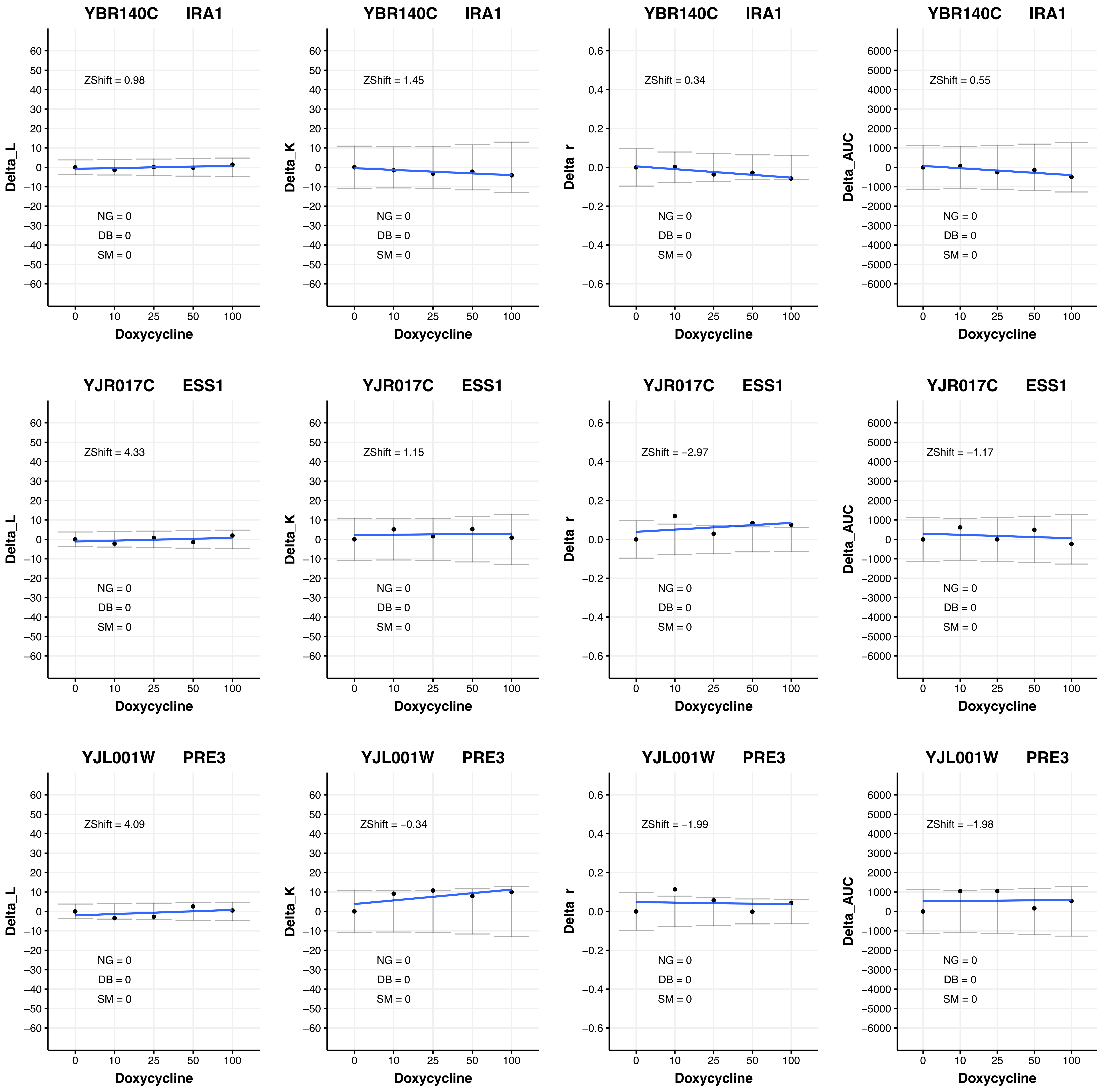

YBR140C IRA1
Delta_L
ZShift = 0.98
NG = 0
DB = 0
SM = 0
Doxycycline
YBR140C IRA1
Delta_K
ZShift = 1.45
NG = 0
DB = 0
SM = 0
Doxycycline
YBR140C IRA1
Delta_r
ZShift = 0.34
NG = 0
DB = 0
SM = 0
Doxycycline
YBR140C IRA1
Delta_AUC
ZShift = 0.55
NG = 0
DB = 0
SM = 0
Doxycycline
YJR017C ESS1
Delta_L
ZShift = 4.33
NG = 0
DB = 0
SM = 0
Doxycycline
YJR017C ESS1
Delta_K
ZShift = 1.15
NG = 0
DB = 0
SM = 0
Doxycycline
YJR017C ESS1
Delta_r
ZShift = −2.97
NG = 0
DB = 0
SM = 0
Doxycycline
YJR017C ESS1
Delta_AUC
ZShift = −1.17
NG = 0
DB = 0
SM = 0
Doxycycline
YJL001W PRE3
Delta_L
ZShift = 4.09
NG = 0
DB = 0
SM = 0
Doxycycline
YJL001W PRE3
Delta_K
ZShift = −0.34
NG = 0
DB = 0
SM = 0
Doxycycline
YJL001W PRE3
Delta_r
ZShift = −1.99
NG = 0
DB = 0
SM = 0
Doxycycline
YJL001W PRE3
Delta_AUC
ZShift = −1.98
NG = 0
DB = 0
SM = 0
Doxycycline

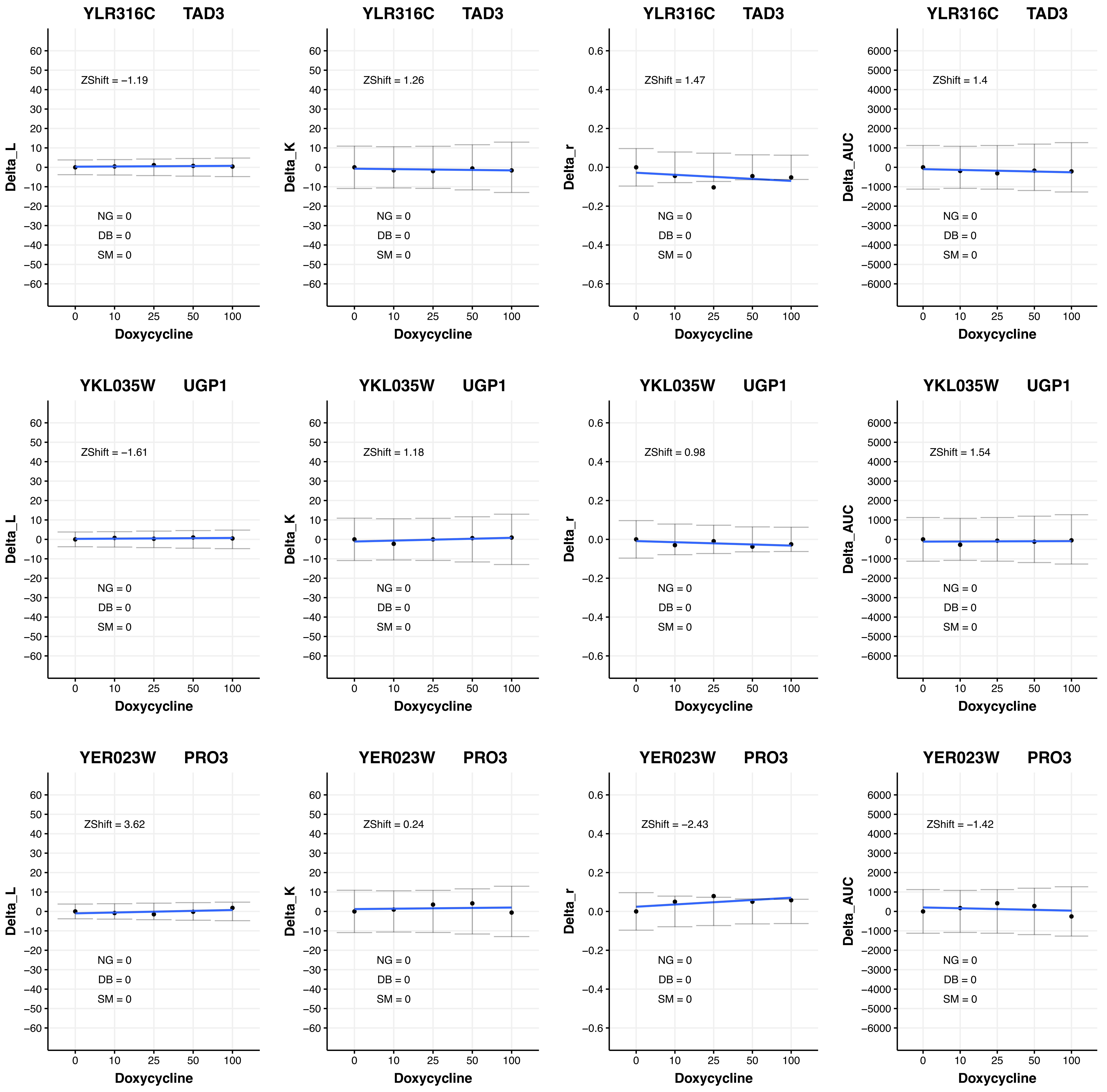

YLR316C TAD3
ZShift = −1.19
Delta_L
NG = 0
DB = 0
SM = 0
Doxycycline
YLR316C TAD3
ZShift = 1.26
Delta_K
NG = 0
DB = 0
SM = 0
Doxycycline
YLR316C TAD3
ZShift = 1.47
Delta_r
NG = 0
DB = 0
SM = 0
Doxycycline
YLR316C TAD3
ZShift = 1.4
Delta_AUC
NG = 0
DB = 0
SM = 0
Doxycycline
YKL035W UGP1
ZShift = −1.61
Delta_L
NG = 0
DB = 0
SM = 0
Doxycycline
YKL035W UGP1
ZShift = 1.18
Delta_K
NG = 0
DB = 0
SM = 0
Doxycycline
YKL035W UGP1
ZShift = 0.98
Delta_r
NG = 0
DB = 0
SM = 0
Doxycycline
YKL035W UGP1
ZShift = 1.54
Delta_AUC
NG = 0
DB = 0
SM = 0
Doxycycline
YER023W PRO3
ZShift = 3.62
Delta_L
NG = 0
DB = 0
SM = 0
Doxycycline
YER023W PRO3
ZShift = 0.24
Delta_K
NG = 0
DB = 0
SM = 0
Doxycycline
YER023W PRO3
ZShift = −2.43
Delta_r
NG = 0
DB = 0
SM = 0
Doxycycline
YER023W PRO3
ZShift = −1.42
Delta_AUC
NG = 0
DB = 0
SM = 0
Doxycycline

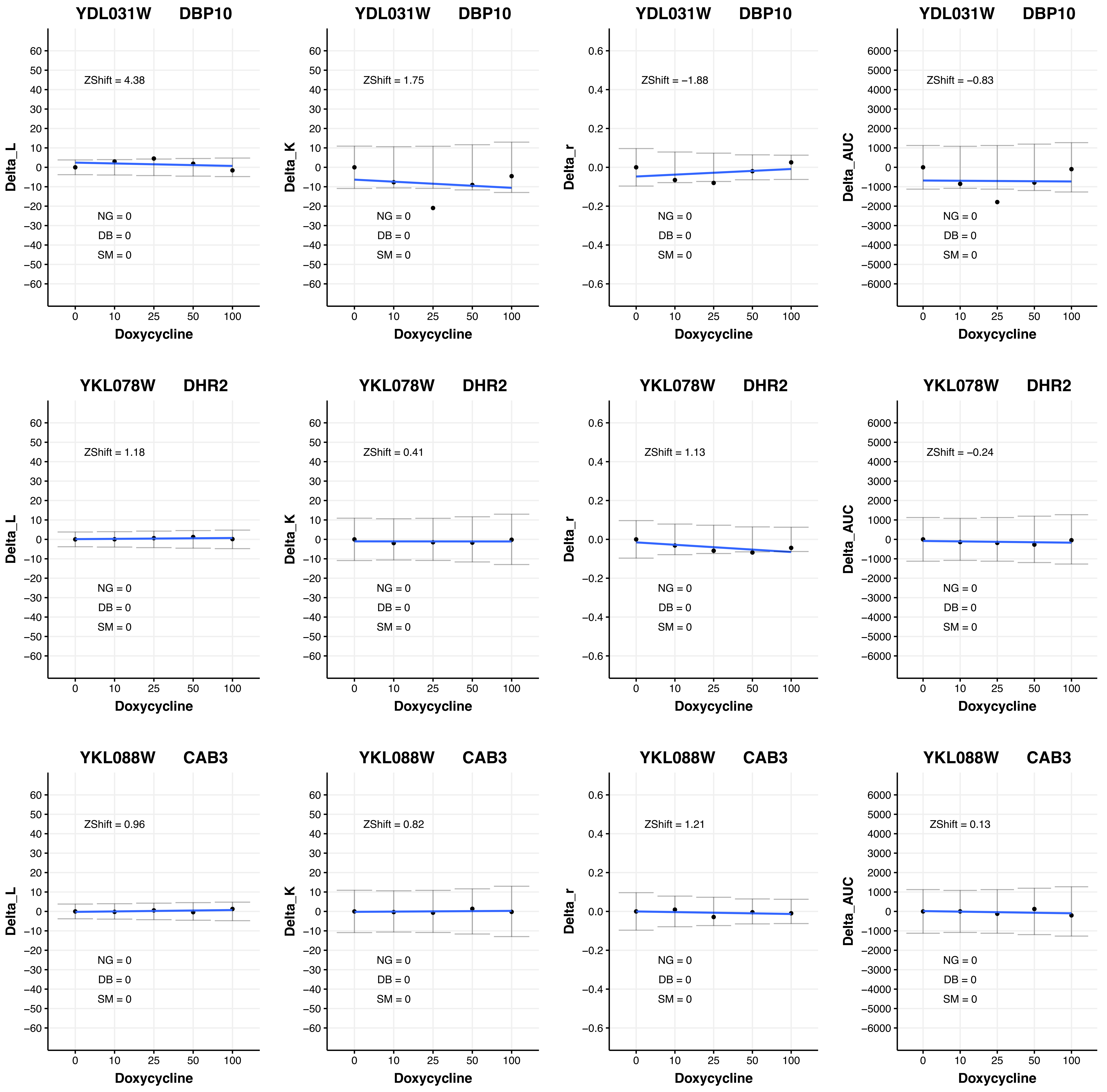
YDL031W DBP10
ZShift = 4.38
NG = 0
DB = 0
SM = 0
Delta_L
Doxycycline
YDL031W DBP10
ZShift = 1.75
NG = 0
DB = 0
SM = 0
Delta_K
Doxycycline
YDL031W DBP10
ZShift = −1.88
NG = 0
DB = 0
SM = 0
Delta_r
Doxycycline
YDL031W DBP10
ZShift = −0.83
NG = 0
DB = 0
SM = 0
Delta_AUC
Doxycycline
YKL078W DHR2
ZShift = 1.18
NG = 0
DB = 0
SM = 0
Delta_L
Doxycycline
YKL078W DHR2
ZShift = 0.41
NG = 0
DB = 0
SM = 0
Delta_K
Doxycycline
YKL078W DHR2
ZShift = 1.13
NG = 0
DB = 0
SM = 0
Delta_r
Doxycycline
YKL078W DHR2
ZShift = −0.24
NG = 0
DB = 0
SM = 0
Delta_AUC
Doxycycline
YKL088W CAB3
ZShift = 0.96
NG = 0
DB = 0
SM = 0
Delta_L
Doxycycline
YKL088W CAB3
ZShift = 0.82
NG = 0
DB = 0
SM = 0
Delta_K
Doxycycline
YKL088W CAB3
ZShift = 1.21
NG = 0
DB = 0
SM = 0
Delta_r
Doxycycline
YKL088W CAB3
ZShift = 0.13
NG = 0
DB = 0
SM = 0
Delta_AUC
Doxycycline

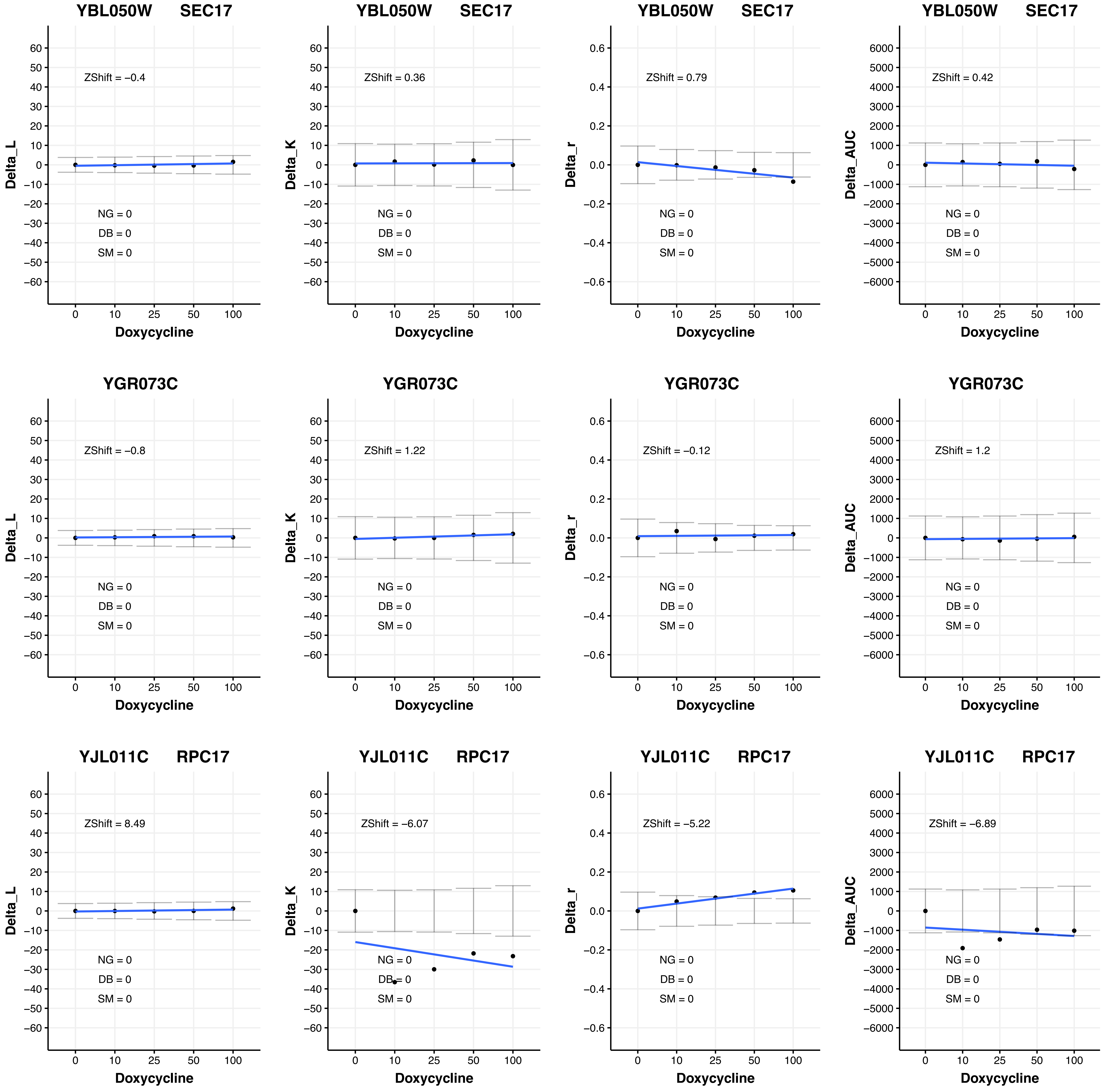

YBL050W SEC17
ZShift = −0.4
ZShift = 0.36
ZShift = 0.79
ZShift = 0.42
YGR073C
ZShift = −0.8
ZShift = 1.22
ZShift = −0.12
ZShift = 1.2
YJL011C RPC17
ZShift = 8.49
ZShift = −6.07
ZShift = −5.22
ZShift = −6.89
NG = 0
DB = 0
SM = 0
Delta_L
Delta_K
Delta_r
Delta_AUC
Doxycycline
0 10 25 50 100

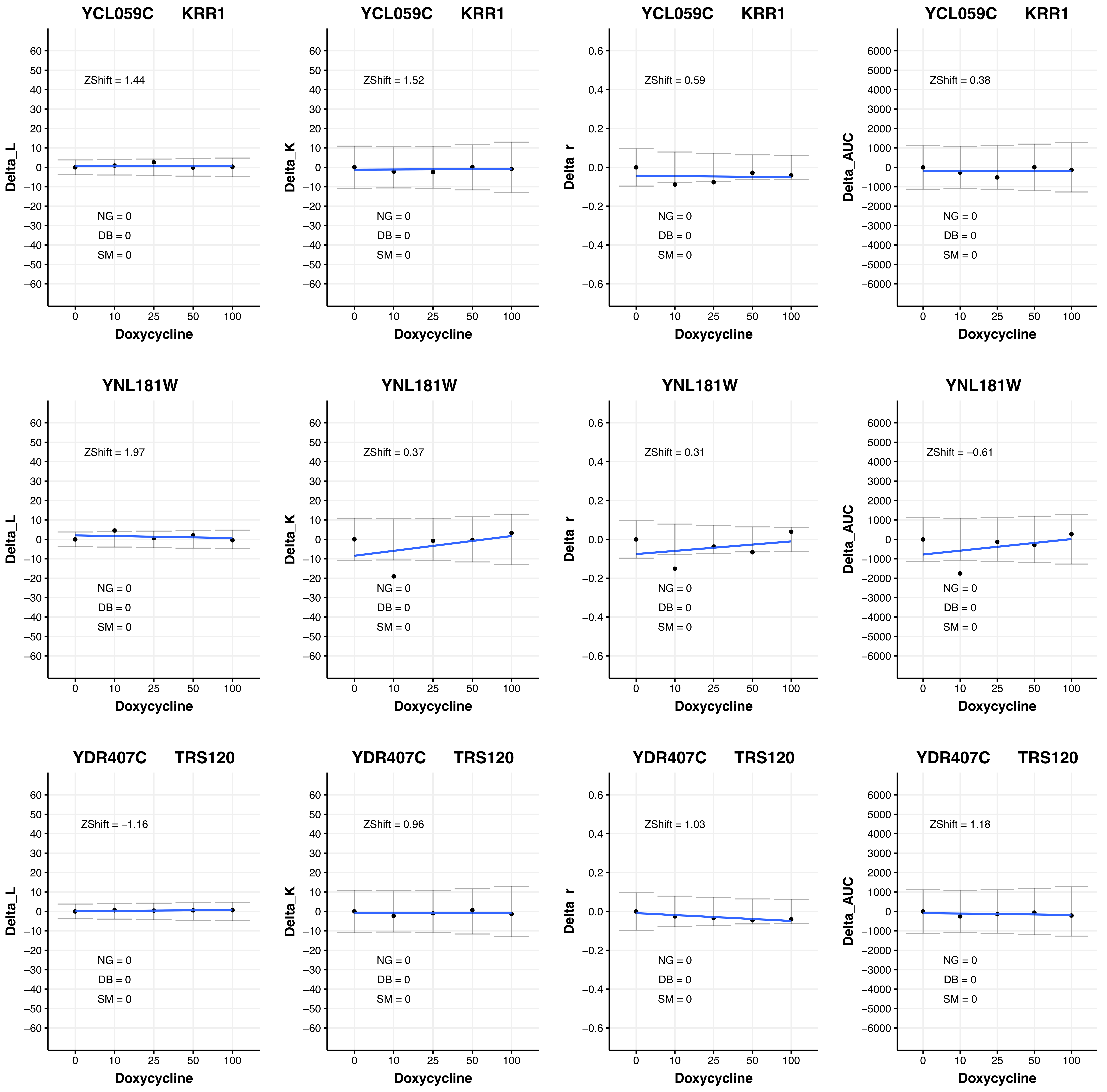

YCL059C KRR1
ZShift = 1.44
NG = 0
DB = 0
SM = 0
Delta_L
Doxycycline
YCL059C KRR1
ZShift = 1.52
NG = 0
DB = 0
SM = 0
Delta_K
Doxycycline
YCL059C KRR1
ZShift = 0.59
NG = 0
DB = 0
SM = 0
Delta_r
Doxycycline
YCL059C KRR1
ZShift = 0.38
NG = 0
DB = 0
SM = 0
Delta_AUC
Doxycycline
YNL181W
ZShift = 1.97
NG = 0
DB = 0
SM = 0
Delta_L
Doxycycline
YNL181W
ZShift = 0.37
NG = 0
DB = 0
SM = 0
Delta_K
Doxycycline
YNL181W
ZShift = 0.31
NG = 0
DB = 0
SM = 0
Delta_r
Doxycycline
YNL181W
ZShift = −0.61
NG = 0
DB = 0
SM = 0
Delta_AUC
Doxycycline
YDR407C TRS120
ZShift = −1.16
NG = 0
DB = 0
SM = 0
Delta_L
Doxycycline
YDR407C TRS120
ZShift = 0.96
NG = 0
DB = 0
SM = 0
Delta_K
Doxycycline
YDR407C TRS120
ZShift = 1.03
NG = 0
DB = 0
SM = 0
Delta_r
Doxycycline
YDR407C TRS120
ZShift = 1.18
NG = 0
DB = 0
SM = 0
Delta_AUC
Doxycycline

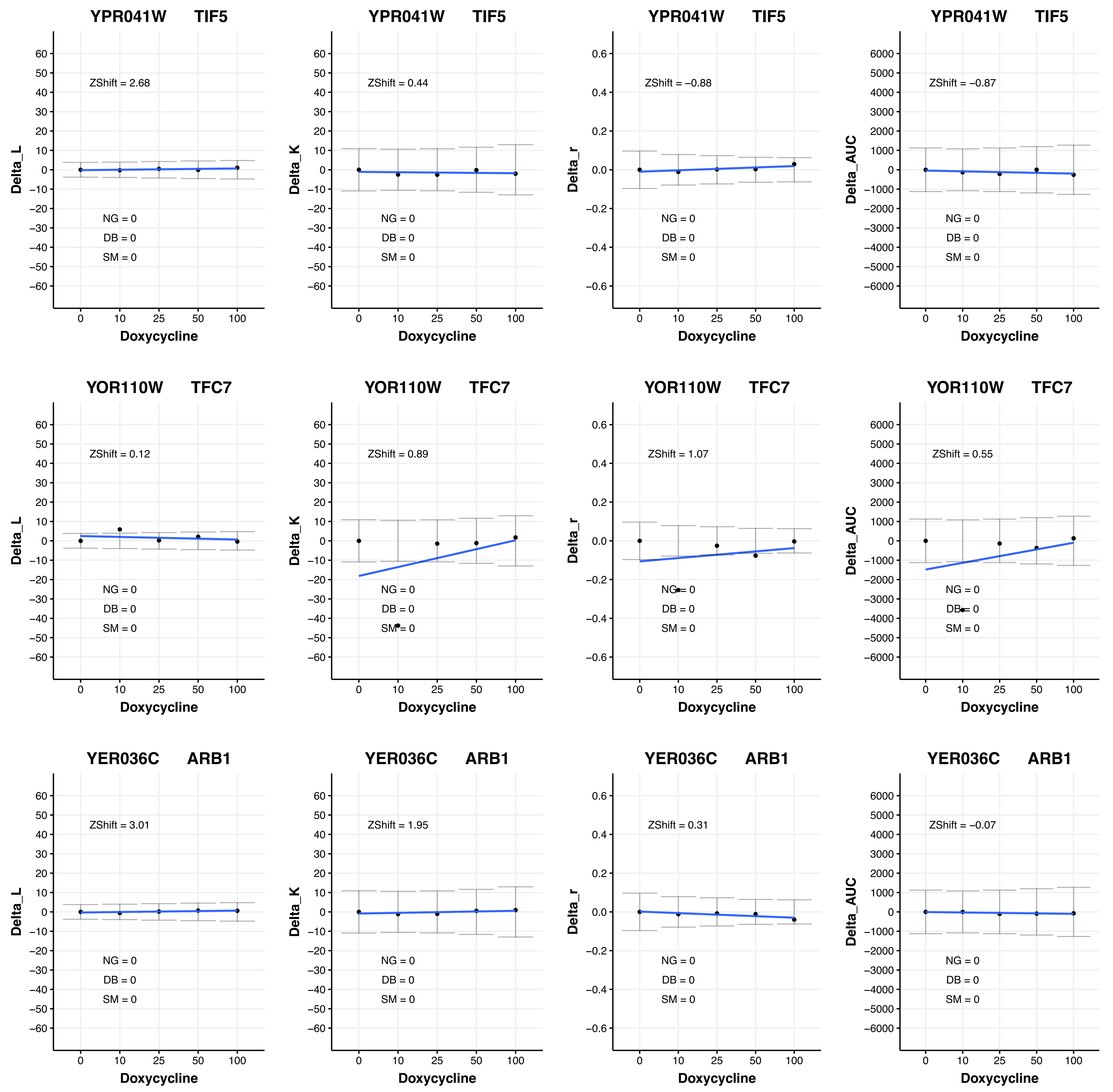
YPR041W TIF5
ZShift = 2.68
NG = 0
DB = 0
SM = 0
Delta_L
Doxycycline
YPR041W TIF5
ZShift = 0.44
NG = 0
DB = 0
SM = 0
Delta_K
Doxycycline
YPR041W TIF5
ZShift = −0.88
NG = 0
DB = 0
SM = 0
Delta_r
Doxycycline
YPR041W TIF5
ZShift = −0.87
NG = 0
DB = 0
SM = 0
Delta_AUC
Doxycycline
YOR110W TFC7
ZShift = 0.12
NG = 0
DB = 0
SM = 0
Delta_L
Doxycycline
YOR110W TFC7
ZShift = 0.89
NG = 0
DB = 0
SM = 0
Delta_K
Doxycycline
YOR110W TFC7
ZShift = 1.07
NG = 0
DB = 0
SM = 0
Delta_r
Doxycycline
YOR110W TFC7
ZShift = 0.55
NG = 0
DB = 0
SM = 0
Delta_AUC
Doxycycline
YER036C ARB1
ZShift = 3.01
NG = 0
DB = 0
SM = 0
Delta_L
Doxycycline
YER036C ARB1
ZShift = 1.95
NG = 0
DB = 0
SM = 0
Delta_K
Doxycycline
YER036C ARB1
ZShift = 0.31
NG = 0
DB = 0
SM = 0
Delta_r
Doxycycline
YER036C ARB1
ZShift = −0.07
NG = 0
DB = 0
SM = 0
Delta_AUC
Doxycycline

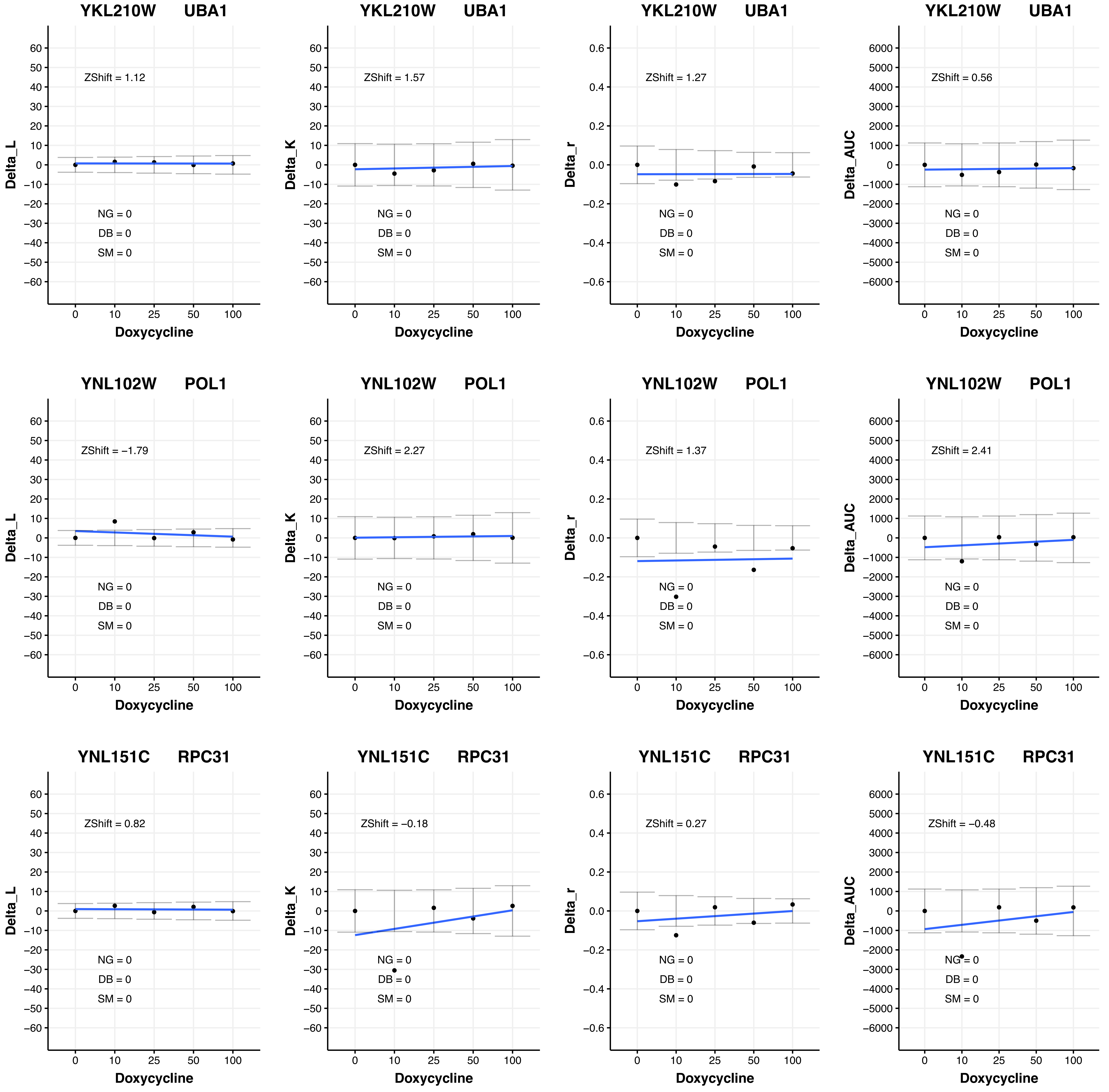

YKL210W UBA1
ZShift = 1.12
NG = 0
DB = 0
SM = 0
Delta_L
Doxycycline
YKL210W UBA1
ZShift = 1.57
NG = 0
DB = 0
SM = 0
Delta_K
Doxycycline
YKL210W UBA1
ZShift = 1.27
NG = 0
DB = 0
SM = 0
Delta_r
Doxycycline
YKL210W UBA1
ZShift = 0.56
NG = 0
DB = 0
SM = 0
Delta_AUC
Doxycycline
YNL102W POL1
ZShift = −1.79
NG = 0
DB = 0
SM = 0
Delta_L
Doxycycline
YNL102W POL1
ZShift = 2.27
NG = 0
DB = 0
SM = 0
Delta_K
Doxycycline
YNL102W POL1
ZShift = 1.37
NG = 0
DB = 0
SM = 0
Delta_r
Doxycycline
YNL102W POL1
ZShift = 2.41
NG = 0
DB = 0
SM = 0
Delta_AUC
Doxycycline
YNL151C RPC31
ZShift = 0.82
NG = 0
DB = 0
SM = 0
Delta_L
Doxycycline
YNL151C RPC31
ZShift = −0.18
NG = 0
DB = 0
SM = 0
Delta_K
Doxycycline
YNL151C RPC31
ZShift = 0.27
NG = 0
DB = 0
SM = 0
Delta_r
Doxycycline
YNL151C RPC31
ZShift = −0.48
NG = 0
DB = 0
SM = 0
Delta_AUC
Doxycycline

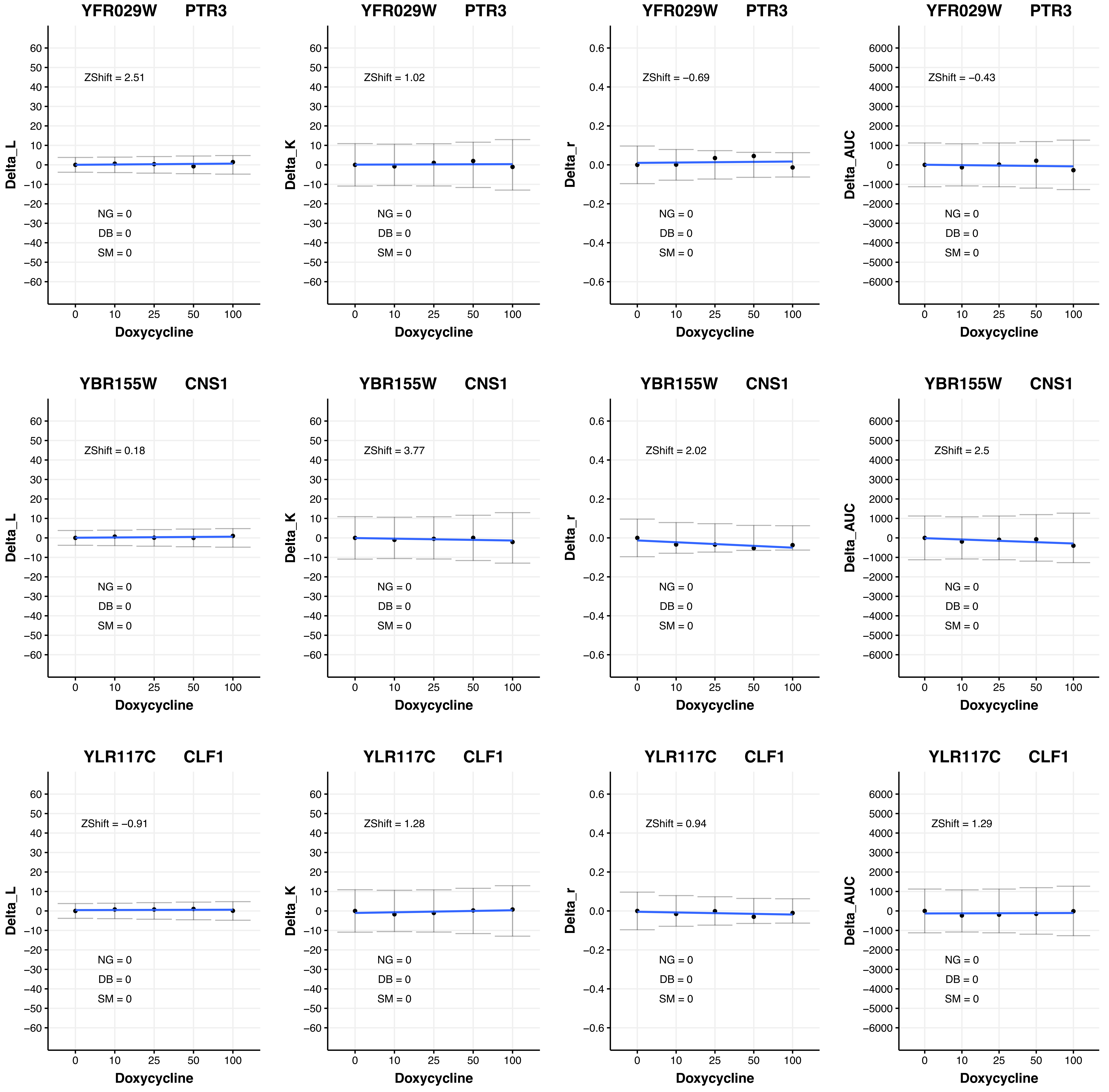
YFR029W PTR3
ZShift = 2.51
ZShift = 1.02
ZShift = −0.69
ZShift = −0.43
YBR155W CNS1
ZShift = 0.18
ZShift = 3.77
ZShift = 2.02
ZShift = 2.5
YLR117C CLF1
ZShift = −0.91
ZShift = 1.28
ZShift = 0.94
ZShift = 1.29
NG = 0
DB = 0
SM = 0
Delta_L
Delta_K
Delta_r
Delta_AUC
Doxycycline

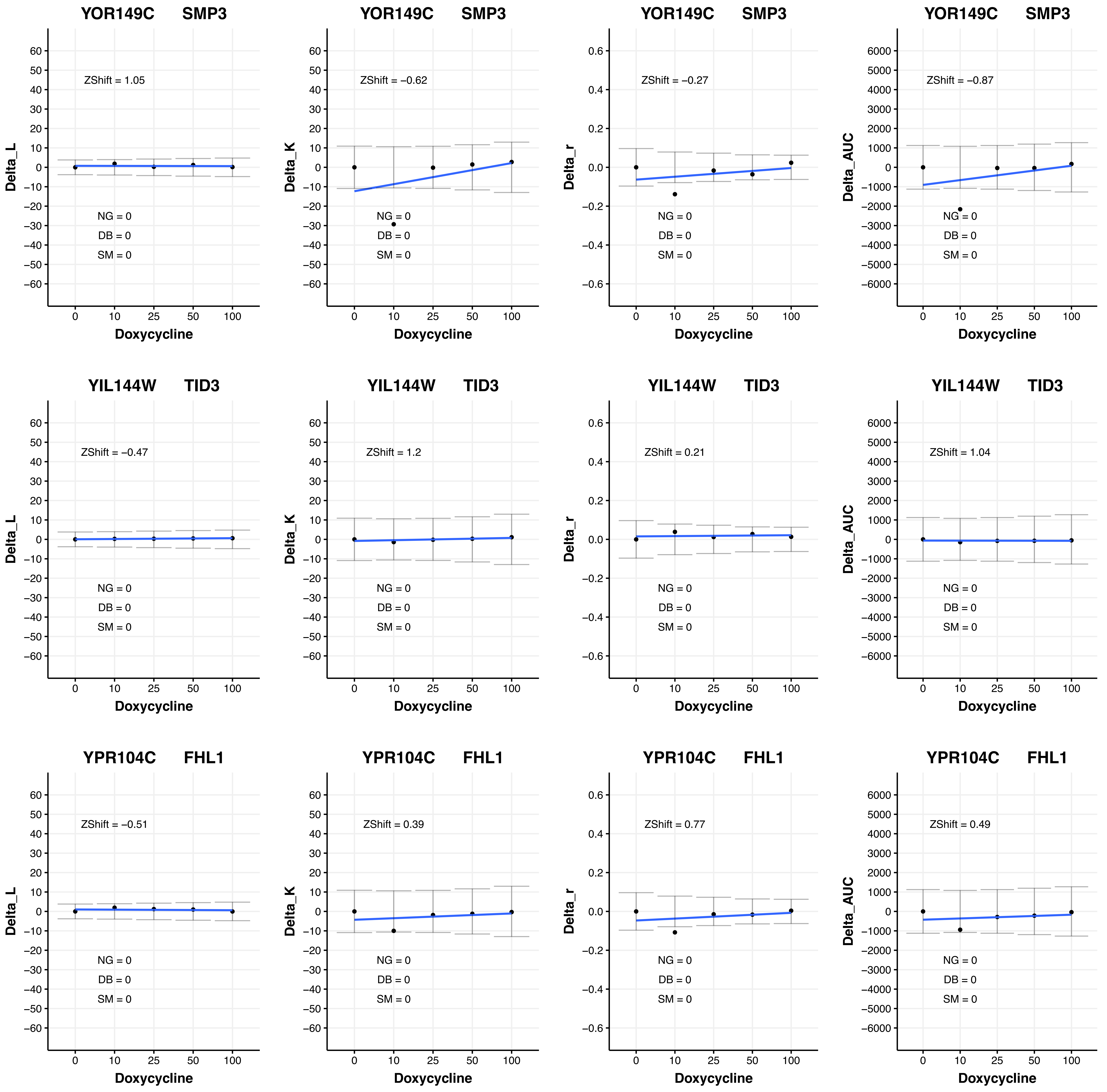
YOR149C SMP3
ZShift = 1.05
NG = 0
DB = 0
SM = 0
Delta_L
Doxycycline
YOR149C SMP3
ZShift = −0.62
NG = 0
DB = 0
SM = 0
Delta_K
Doxycycline
YOR149C SMP3
ZShift = −0.27
NG = 0
DB = 0
SM = 0
Delta_r
Doxycycline
YOR149C SMP3
ZShift = −0.87
NG = 0
DB = 0
SM = 0
Delta_AUC
Doxycycline
YIL144W TID3
ZShift = −0.47
NG = 0
DB = 0
SM = 0
Delta_L
Doxycycline
YIL144W TID3
ZShift = 1.2
NG = 0
DB = 0
SM = 0
Delta_K
Doxycycline
YIL144W TID3
ZShift = 0.21
NG = 0
DB = 0
SM = 0
Delta_r
Doxycycline
YIL144W TID3
ZShift = 1.04
NG = 0
DB = 0
SM = 0
Delta_AUC
Doxycycline
YPR104C FHL1
ZShift = −0.51
NG = 0
DB = 0
SM = 0
Delta_L
Doxycycline
YPR104C FHL1
ZShift = 0.39
NG = 0
DB = 0
SM = 0
Delta_K
Doxycycline
YPR104C FHL1
ZShift = 0.77
NG = 0
DB = 0
SM = 0
Delta_r
Doxycycline
YPR104C FHL1
ZShift = 0.49
NG = 0
DB = 0
SM = 0
Delta_AUC
Doxycycline

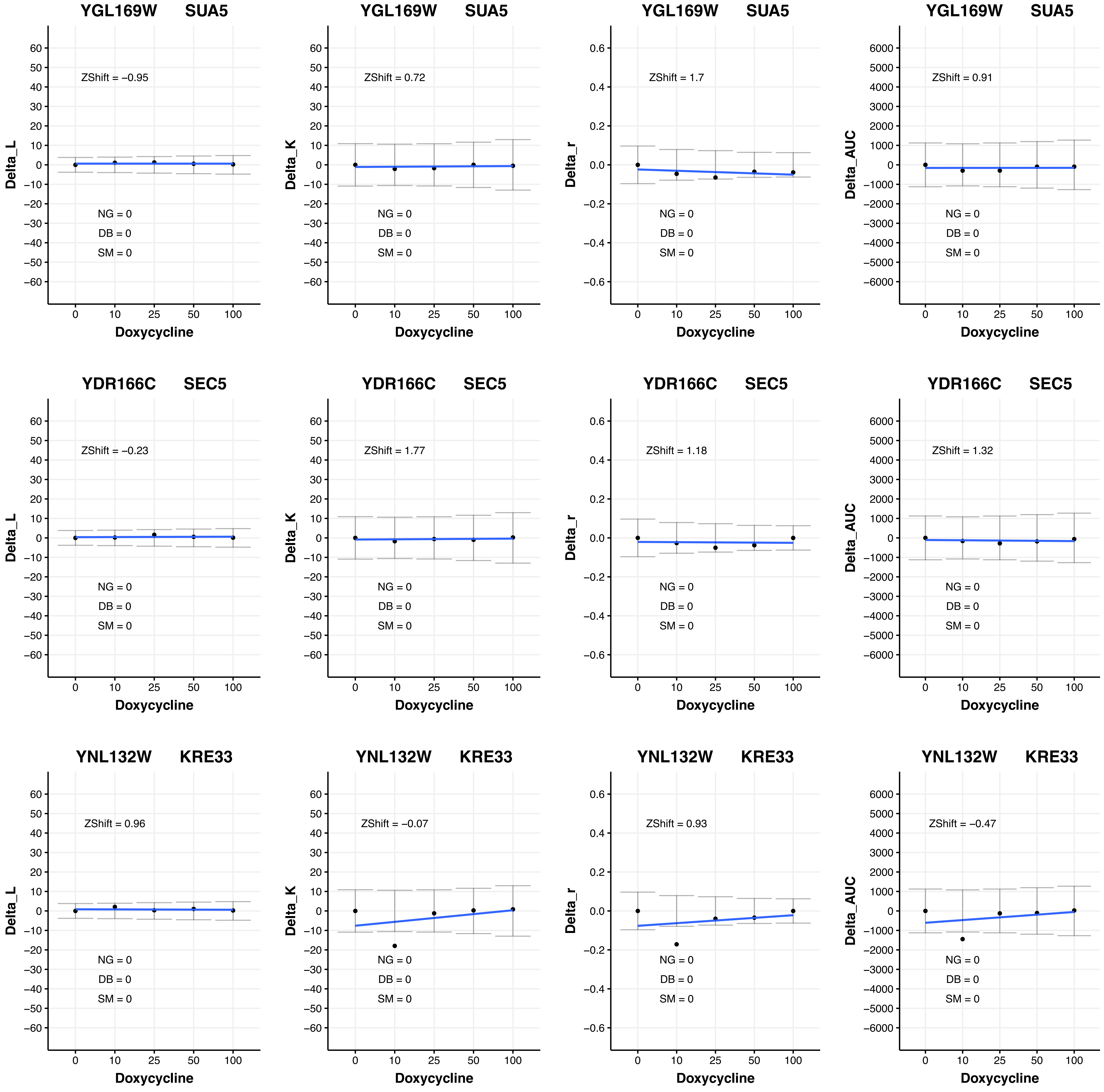

YGL169W SUA5
Delta_L
ZShift = −0.95
NG = 0
DB = 0
SM = 0
Doxycycline
YGL169W SUA5
Delta_K
ZShift = 0.72
NG = 0
DB = 0
SM = 0
Doxycycline
YGL169W SUA5
Delta_r
ZShift = 1.7
NG = 0
DB = 0
SM = 0
Doxycycline
YGL169W SUA5
Delta_AUC
ZShift = 0.91
NG = 0
DB = 0
SM = 0
Doxycycline
YDR166C SEC5
Delta_L
ZShift = −0.23
NG = 0
DB = 0
SM = 0
Doxycycline
YDR166C SEC5
Delta_K
ZShift = 1.77
NG = 0
DB = 0
SM = 0
Doxycycline
YDR166C SEC5
Delta_r
ZShift = 1.18
NG = 0
DB = 0
SM = 0
Doxycycline
YDR166C SEC5
Delta_AUC
ZShift = 1.32
NG = 0
DB = 0
SM = 0
Doxycycline
YNL132W KRE33
Delta_L
ZShift = 0.96
NG = 0
DB = 0
SM = 0
Doxycycline
YNL132W KRE33
Delta_K
ZShift = −0.07
NG = 0
DB = 0
SM = 0
Doxycycline
YNL132W KRE33
Delta_r
ZShift = 0.93
NG = 0
DB = 0
SM = 0
Doxycycline
YNL132W KRE33
Delta_AUC
ZShift = −0.47
NG = 0
DB = 0
SM = 0
Doxycycline

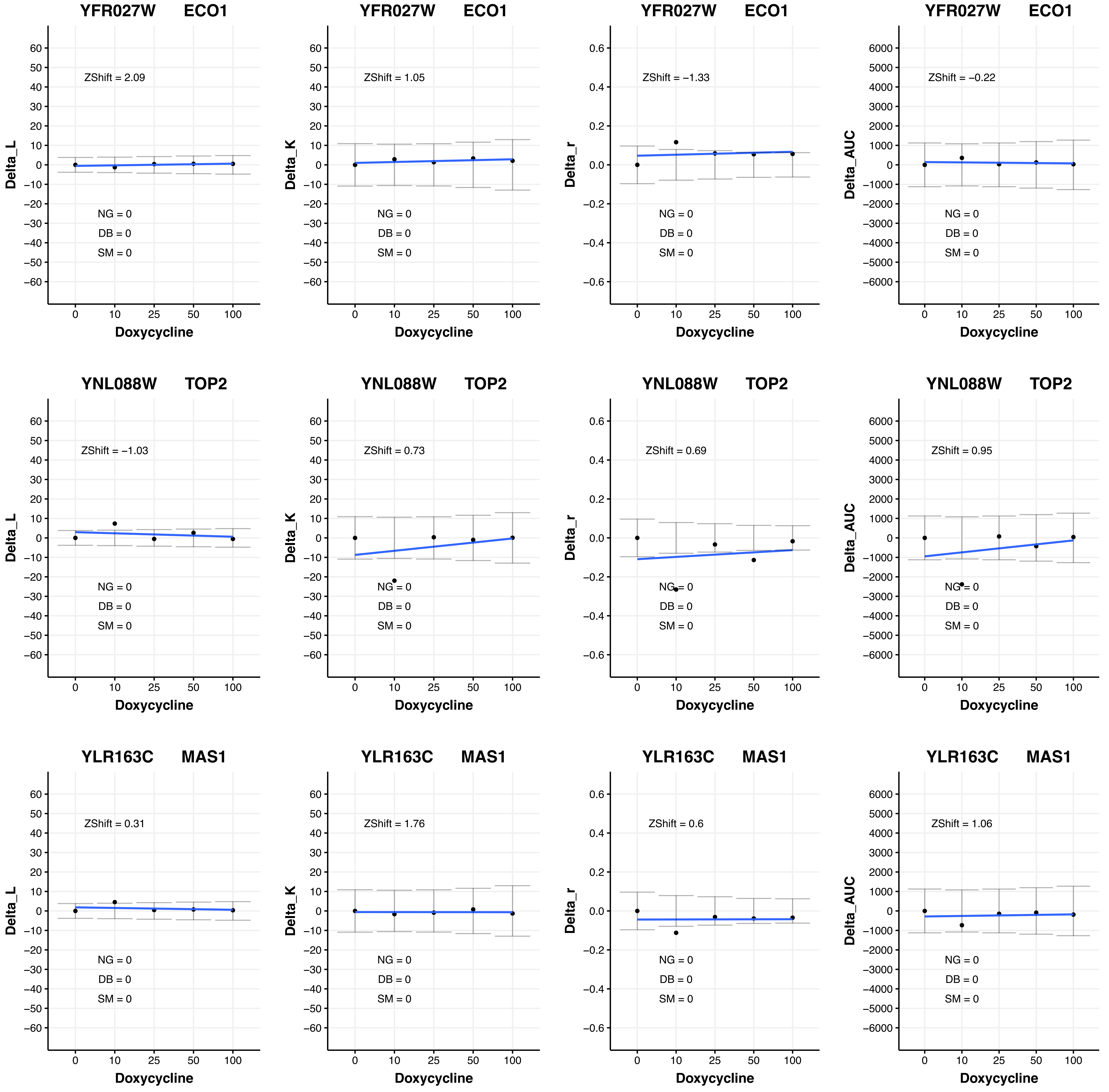
YFR027W ECO1
ZShift = 2.09
NG = 0
DB = 0
SM = 0
Delta_L
Doxycycline
YFR027W ECO1
ZShift = 1.05
NG = 0
DB = 0
SM = 0
Delta_K
Doxycycline
YFR027W ECO1
ZShift = −1.33
NG = 0
DB = 0
SM = 0
Delta_r
Doxycycline
YFR027W ECO1
ZShift = −0.22
NG = 0
DB = 0
SM = 0
Delta_AUC
Doxycycline
YNL088W TOP2
ZShift = −1.03
NG = 0
DB = 0
SM = 0
Delta_L
Doxycycline
YNL088W TOP2
ZShift = 0.73
NG = 0
DB = 0
SM = 0
Delta_K
Doxycycline
YNL088W TOP2
ZShift = 0.69
NG = 0
DB = 0
SM = 0
Delta_r
Doxycycline
YNL088W TOP2
ZShift = 0.95
NG = 0
DB = 0
SM = 0
Delta_AUC
Doxycycline
YLR163C MAS1
ZShift = 0.31
NG = 0
DB = 0
SM = 0
Delta_L
Doxycycline
YLR163C MAS1
ZShift = 1.76
NG = 0
DB = 0
SM = 0
Delta_K
Doxycycline
YLR163C MAS1
ZShift = 0.6
NG = 0
DB = 0
SM = 0
Delta_r
Doxycycline
YLR163C MAS1
ZShift = 1.06
NG = 0
DB = 0
SM = 0
Delta_AUC
Doxycycline

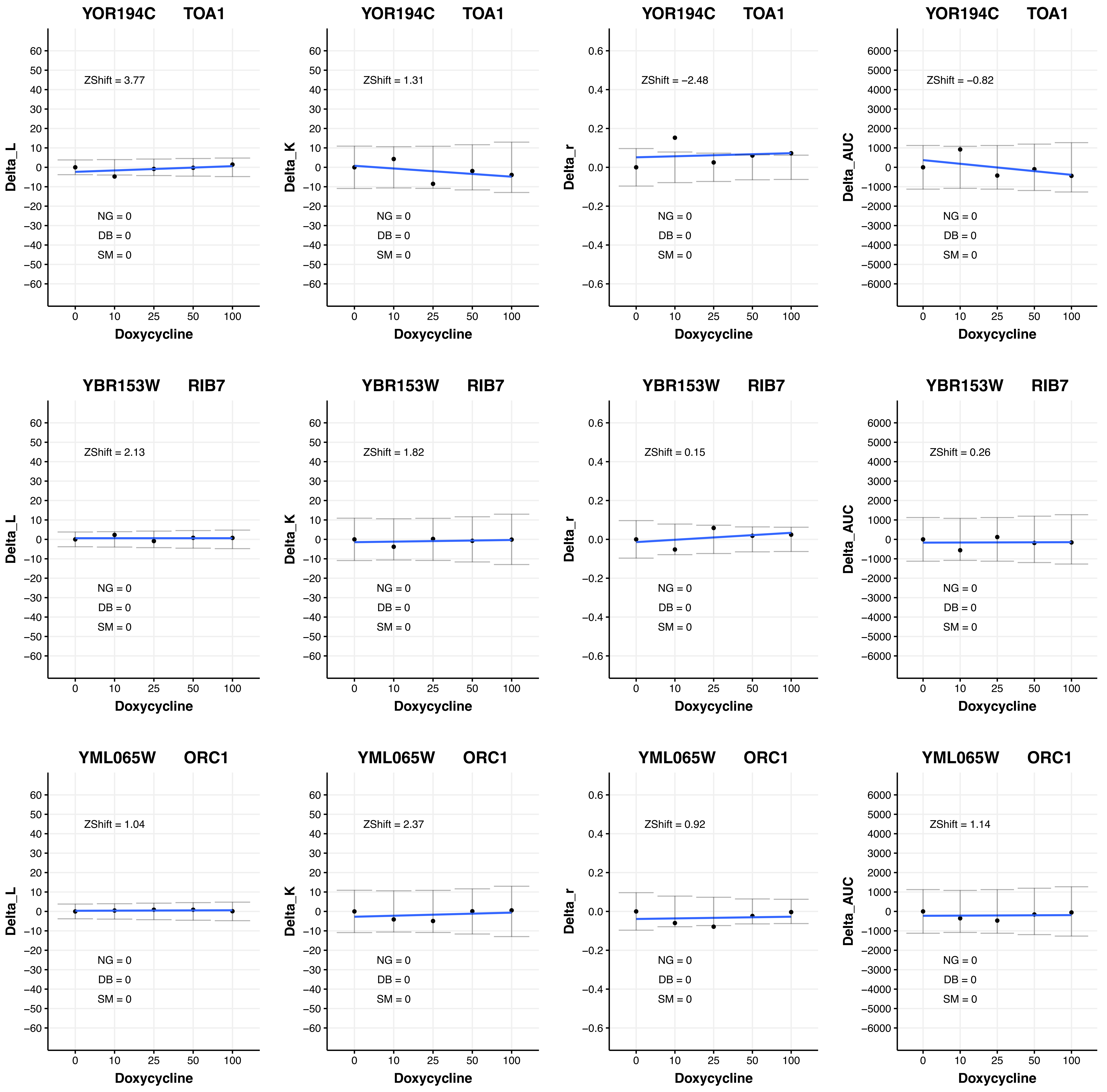

YOR194C TOA1
ZShift = 3.77
NG = 0
DB = 0
SM = 0
Delta_L
Doxycycline
YOR194C TOA1
ZShift = 1.31
NG = 0
DB = 0
SM = 0
Delta_K
Doxycycline
YOR194C TOA1
ZShift = −2.48
NG = 0
DB = 0
SM = 0
Delta_r
Doxycycline
YOR194C TOA1
ZShift = −0.82
NG = 0
DB = 0
SM = 0
Delta_AUC
Doxycycline
YBR153W RIB7
ZShift = 2.13
NG = 0
DB = 0
SM = 0
Delta_L
Doxycycline
YBR153W RIB7
ZShift = 1.82
NG = 0
DB = 0
SM = 0
Delta_K
Doxycycline
YBR153W RIB7
ZShift = 0.15
NG = 0
DB = 0
SM = 0
Delta_r
Doxycycline
YBR153W RIB7
ZShift = 0.26
NG = 0
DB = 0
SM = 0
Delta_AUC
Doxycycline
YML065W ORC1
ZShift = 1.04
NG = 0
DB = 0
SM = 0
Delta_L
Doxycycline
YML065W ORC1
ZShift = 2.37
NG = 0
DB = 0
SM = 0
Delta_K
Doxycycline
YML065W ORC1
ZShift = 0.92
NG = 0
DB = 0
SM = 0
Delta_r
Doxycycline
YML065W ORC1
ZShift = 1.14
NG = 0
DB = 0
SM = 0
Delta_AUC
Doxycycline

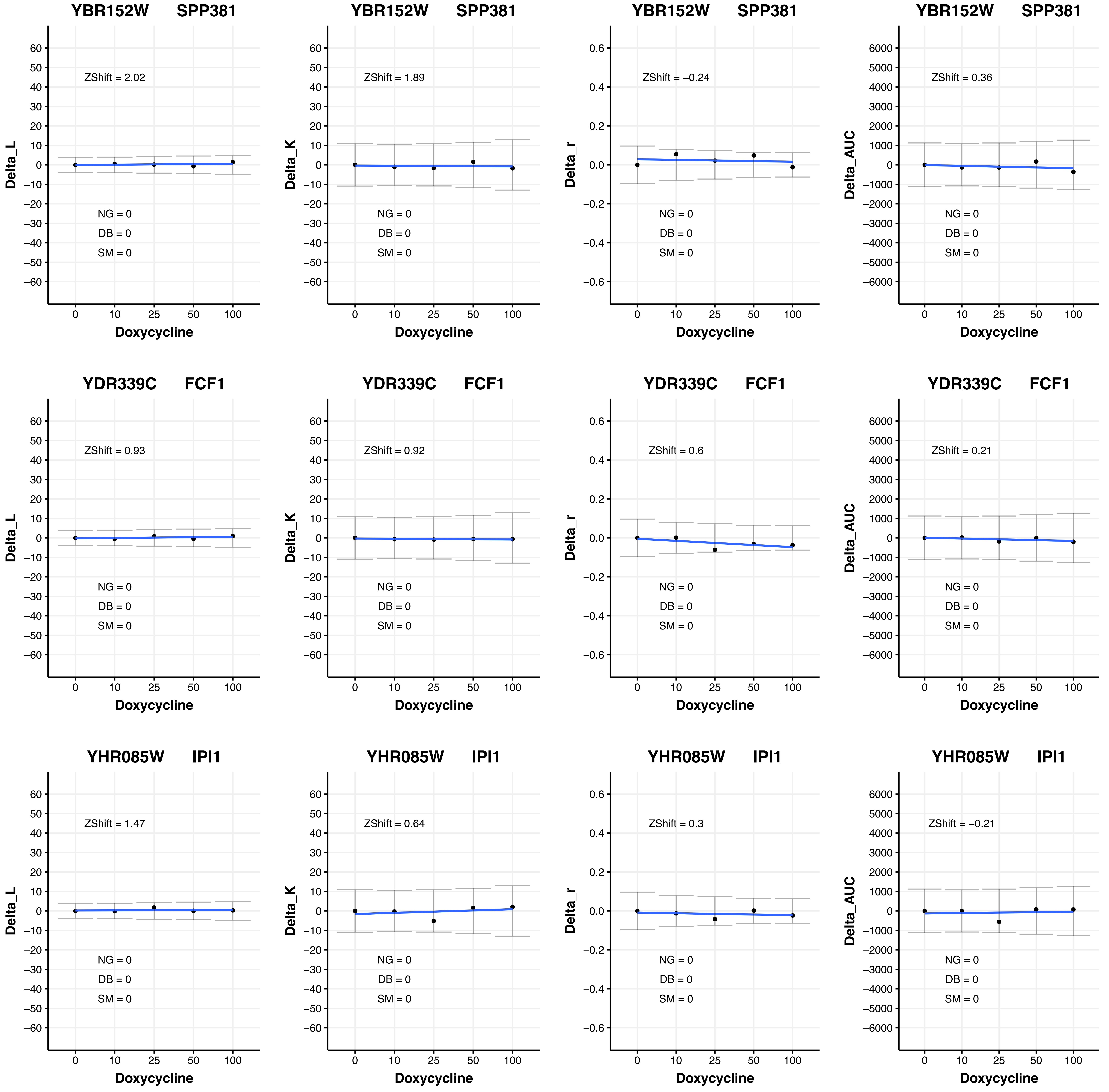

YBR152W SPP381
ZShift = 2.02
ZShift = 1.89
ZShift = −0.24
ZShift = 0.36
YDR339C FCF1
ZShift = 0.93
ZShift = 0.92
ZShift = 0.6
ZShift = 0.21
YHR085W IPI1
ZShift = 1.47
ZShift = 0.64
ZShift = 0.3
ZShift = −0.21
NG = 0
DB = 0
SM = 0
Delta_L
Delta_K
Delta_r
Delta_AUC
Doxycycline
0 10 25 50 100

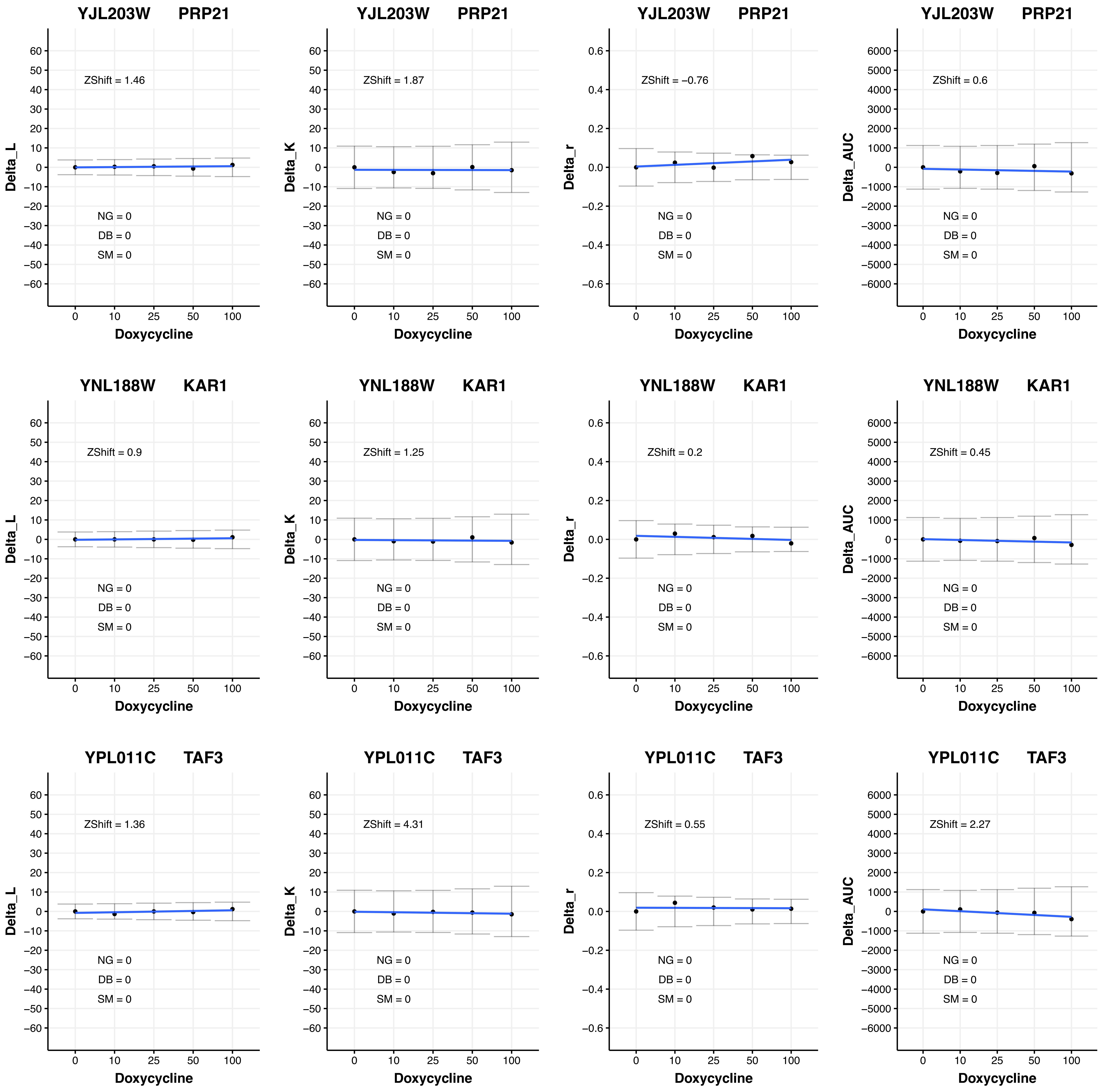

YJL203W PRP21
ZShift = 1.46
NG = 0
DB = 0
SM = 0
Delta_L
Doxycycline
YJL203W PRP21
ZShift = 1.87
NG = 0
DB = 0
SM = 0
Delta_K
Doxycycline
YJL203W PRP21
ZShift = −0.76
NG = 0
DB = 0
SM = 0
Delta_r
Doxycycline
YJL203W PRP21
ZShift = 0.6
NG = 0
DB = 0
SM = 0
Delta_AUC
Doxycycline
YNL188W KAR1
ZShift = 0.9
NG = 0
DB = 0
SM = 0
Delta_L
Doxycycline
YNL188W KAR1
ZShift = 1.25
NG = 0
DB = 0
SM = 0
Delta_K
Doxycycline
YNL188W KAR1
ZShift = 0.2
NG = 0
DB = 0
SM = 0
Delta_r
Doxycycline
YNL188W KAR1
ZShift = 0.45
NG = 0
DB = 0
SM = 0
Delta_AUC
Doxycycline
YPL011C TAF3
ZShift = 1.36
NG = 0
DB = 0
SM = 0
Delta_L
Doxycycline
YPL011C TAF3
ZShift = 4.31
NG = 0
DB = 0
SM = 0
Delta_K
Doxycycline
YPL011C TAF3
ZShift = 0.55
NG = 0
DB = 0
SM = 0
Delta_r
Doxycycline
YPL011C TAF3
ZShift = 2.27
NG = 0
DB = 0
SM = 0
Delta_AUC
Doxycycline

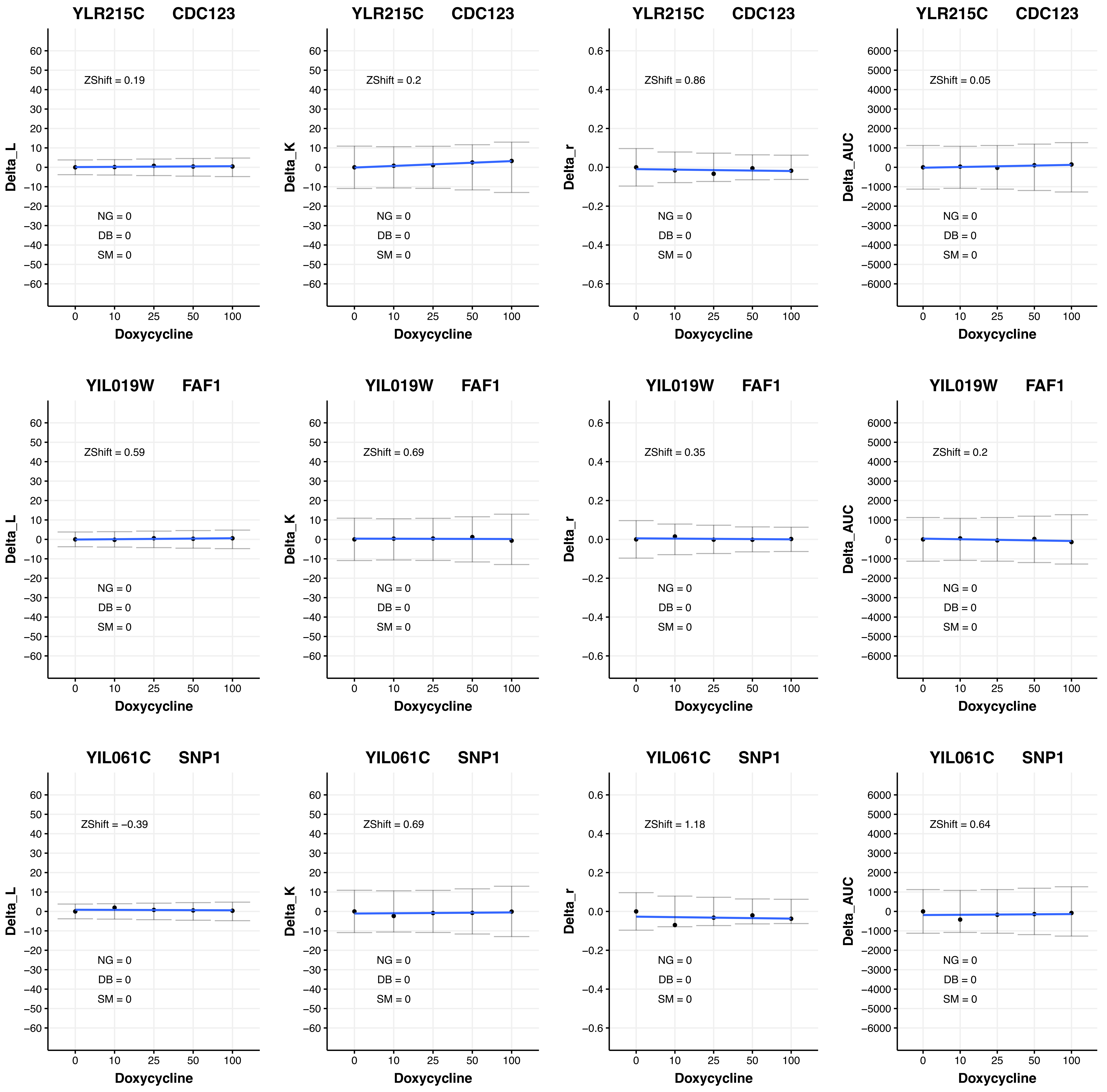

YLR215C CDC123
ZShift = 0.19
NG = 0
DB = 0
SM = 0
Delta_L
Doxycycline
YLR215C CDC123
ZShift = 0.2
NG = 0
DB = 0
SM = 0
Delta_K
Doxycycline
YLR215C CDC123
ZShift = 0.86
NG = 0
DB = 0
SM = 0
Delta_r
Doxycycline
YLR215C CDC123
ZShift = 0.05
NG = 0
DB = 0
SM = 0
Delta_AUC
Doxycycline
YIL019W FAF1
ZShift = 0.59
NG = 0
DB = 0
SM = 0
Delta_L
Doxycycline
YIL019W FAF1
ZShift = 0.69
NG = 0
DB = 0
SM = 0
Delta_K
Doxycycline
YIL019W FAF1
ZShift = 0.35
NG = 0
DB = 0
SM = 0
Delta_r
Doxycycline
YIL019W FAF1
ZShift = 0.2
NG = 0
DB = 0
SM = 0
Delta_AUC
Doxycycline
YIL061C SNP1
ZShift = −0.39
NG = 0
DB = 0
SM = 0
Delta_L
Doxycycline
YIL061C SNP1
ZShift = 0.69
NG = 0
DB = 0
SM = 0
Delta_K
Doxycycline
YIL061C SNP1
ZShift = 1.18
NG = 0
DB = 0
SM = 0
Delta_r
Doxycycline
YIL061C SNP1
ZShift = 0.64
NG = 0
DB = 0
SM = 0
Delta_AUC
Doxycycline
0
10
25
50
100

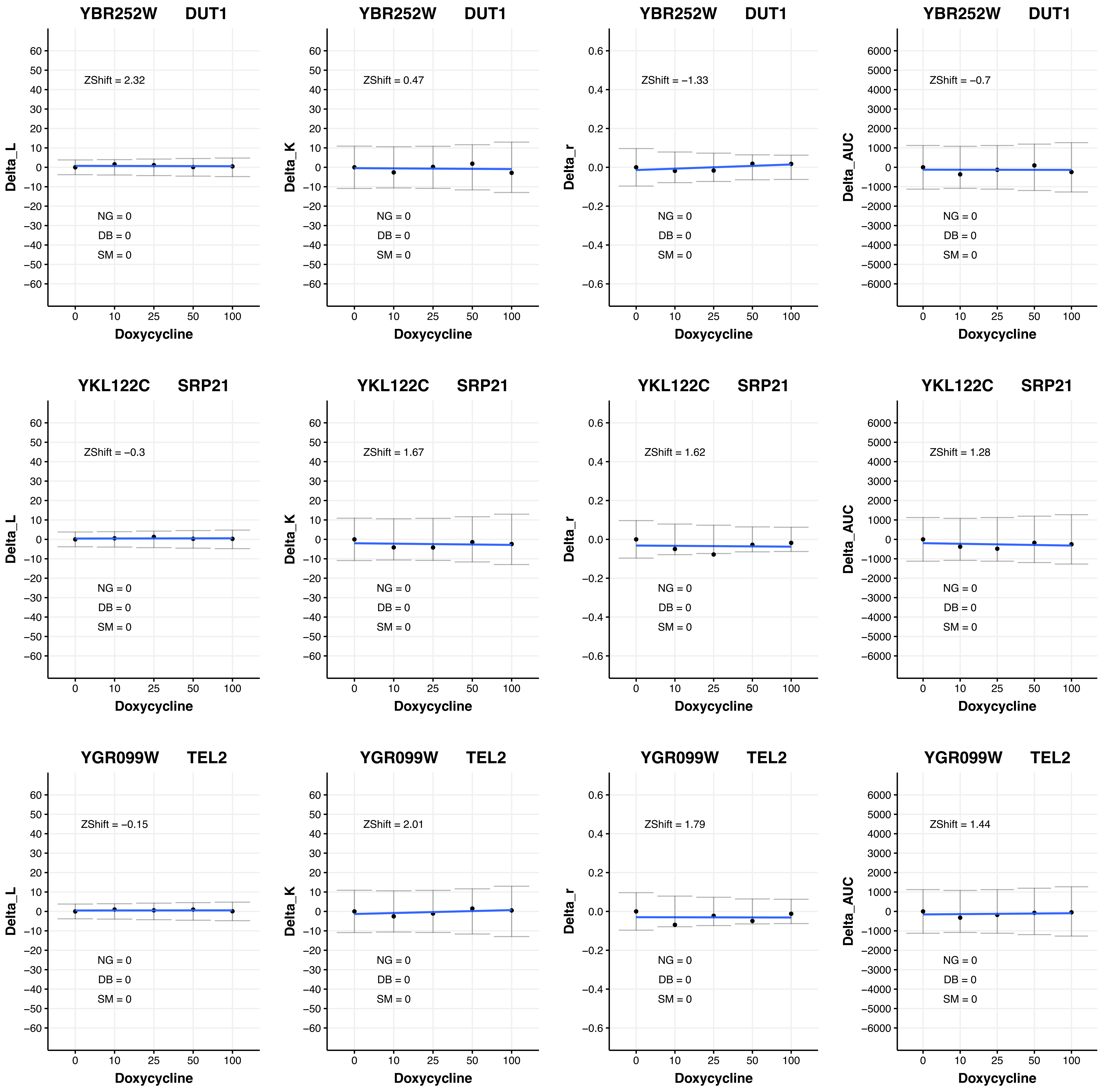

YBR252W DUT1
ZShift = 2.32
Delta_L
NG = 0
DB = 0
SM = 0
Doxycycline
YBR252W DUT1
ZShift = 0.47
Delta_K
NG = 0
DB = 0
SM = 0
Doxycycline
YBR252W DUT1
ZShift = −1.33
Delta_r
NG = 0
DB = 0
SM = 0
Doxycycline
YBR252W DUT1
ZShift = −0.7
Delta_AUC
NG = 0
DB = 0
SM = 0
Doxycycline
YKL122C SRP21
ZShift = −0.3
Delta_L
NG = 0
DB = 0
SM = 0
Doxycycline
YKL122C SRP21
ZShift = 1.67
Delta_K
NG = 0
DB = 0
SM = 0
Doxycycline
YKL122C SRP21
ZShift = 1.62
Delta_r
NG = 0
DB = 0
SM = 0
Doxycycline
YKL122C SRP21
ZShift = 1.28
Delta_AUC
NG = 0
DB = 0
SM = 0
Doxycycline
YGR099W TEL2
ZShift = −0.15
Delta_L
NG = 0
DB = 0
SM = 0
Doxycycline
YGR099W TEL2
ZShift = 2.01
Delta_K
NG = 0
DB = 0
SM = 0
Doxycycline
YGR099W TEL2
ZShift = 1.79
Delta_r
NG = 0
DB = 0
SM = 0
Doxycycline
YGR099W TEL2
ZShift = 1.44
Delta_AUC
NG = 0
DB = 0
SM = 0
Doxycycline

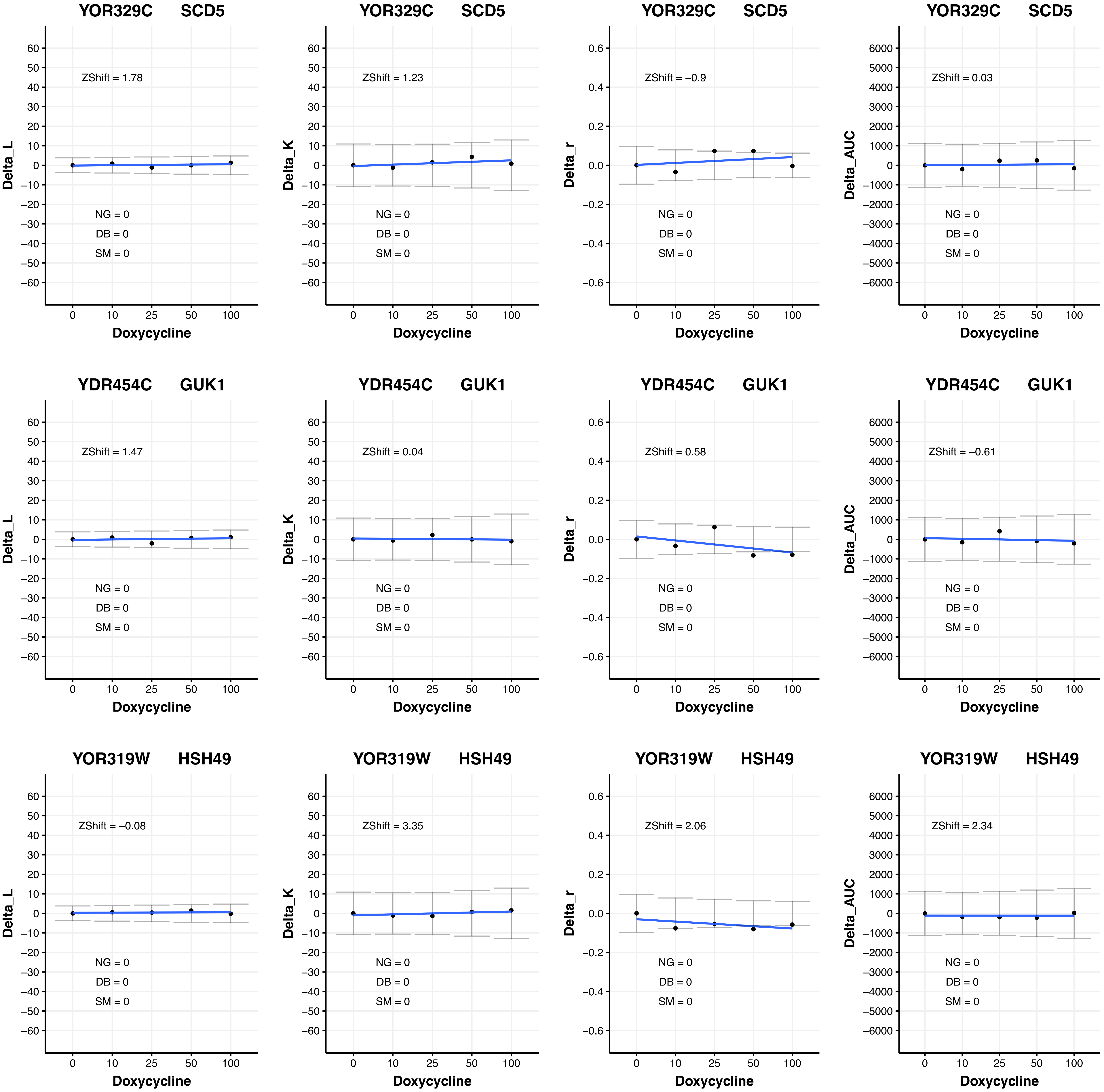
YOR329C SCD5
Delta_L
ZShift = 1.78
NG = 0
DB = 0
SM = 0
Doxycycline
YOR329C SCD5
Delta_K
ZShift = 1.23
NG = 0
DB = 0
SM = 0
Doxycycline
YOR329C SCD5
Delta_r
ZShift = −0.9
NG = 0
DB = 0
SM = 0
Doxycycline
YOR329C SCD5
Delta_AUC
ZShift = 0.03
NG = 0
DB = 0
SM = 0
Doxycycline
YDR454C GUK1
Delta_L
ZShift = 1.47
NG = 0
DB = 0
SM = 0
Doxycycline
YDR454C GUK1
Delta_K
ZShift = 0.04
NG = 0
DB = 0
SM = 0
Doxycycline
YDR454C GUK1
Delta_r
ZShift = 0.58
NG = 0
DB = 0
SM = 0
Doxycycline
YDR454C GUK1
Delta_AUC
ZShift = −0.61
NG = 0
DB = 0
SM = 0
Doxycycline
YOR319W HSH49
Delta_L
ZShift = −0.08
NG = 0
DB = 0
SM = 0
Doxycycline
YOR319W HSH49
Delta_K
ZShift = 3.35
NG = 0
DB = 0
SM = 0
Doxycycline
YOR319W HSH49
Delta_r
ZShift = 2.06
NG = 0
DB = 0
SM = 0
Doxycycline
YOR319W HSH49
Delta_AUC
ZShift = 2.34
NG = 0
DB = 0
SM = 0
Doxycycline
0
10
25
50
100

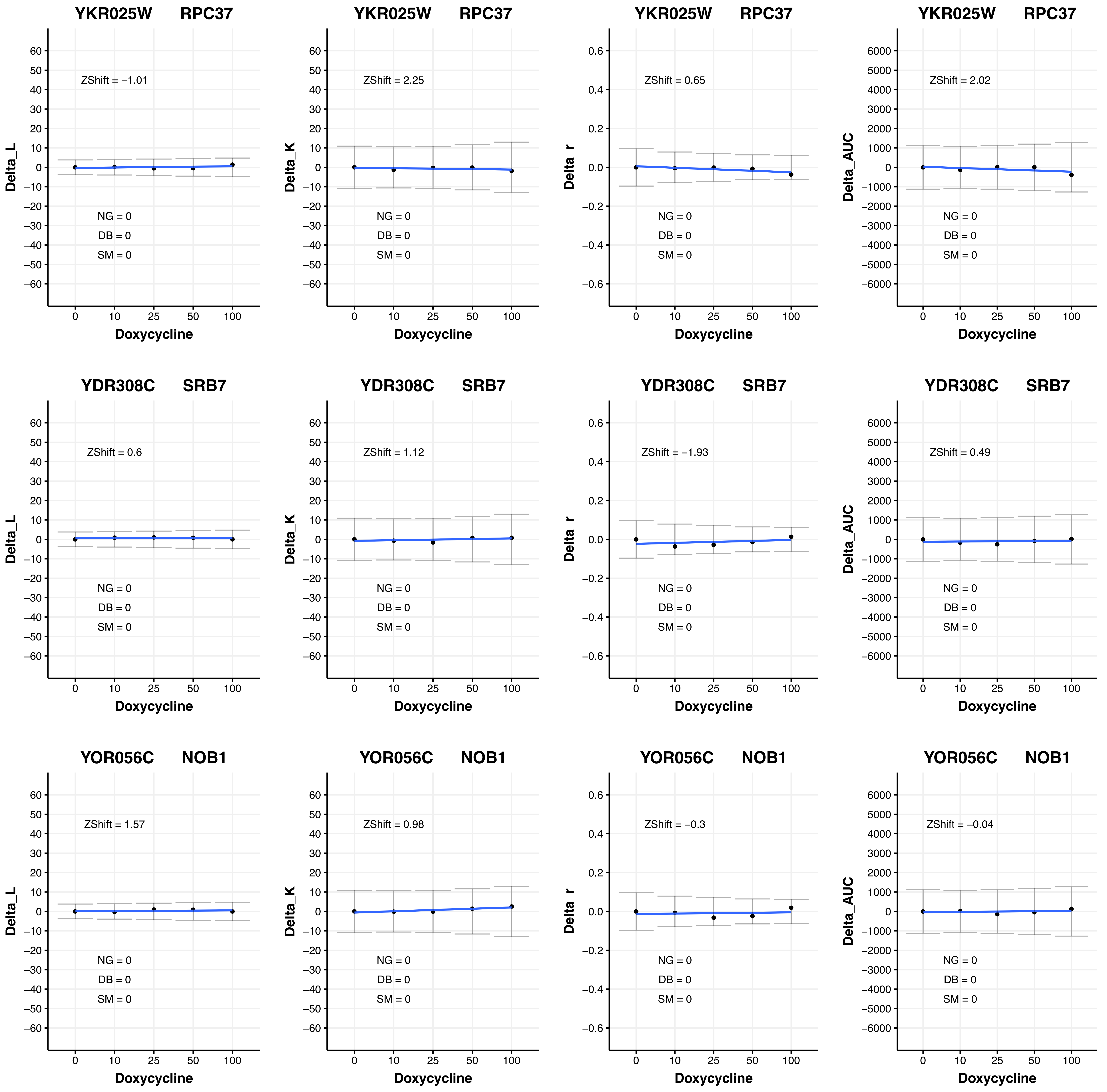

YKR025W RPC37
ZShift = −1.01
NG = 0
DB = 0
SM = 0
Delta_L
Doxycycline
YKR025W RPC37
ZShift = 2.25
NG = 0
DB = 0
SM = 0
Delta_K
Doxycycline
YKR025W RPC37
ZShift = 0.65
NG = 0
DB = 0
SM = 0
Delta_r
Doxycycline
YKR025W RPC37
ZShift = 2.02
NG = 0
DB = 0
SM = 0
Delta_AUC
Doxycycline
YDR308C SRB7
ZShift = 0.6
NG = 0
DB = 0
SM = 0
Delta_L
Doxycycline
YDR308C SRB7
ZShift = 1.12
NG = 0
DB = 0
SM = 0
Delta_K
Doxycycline
YDR308C SRB7
ZShift = −1.93
NG = 0
DB = 0
SM = 0
Delta_r
Doxycycline
YDR308C SRB7
ZShift = 0.49
NG = 0
DB = 0
SM = 0
Delta_AUC
Doxycycline
YOR056C NOB1
ZShift = 1.57
NG = 0
DB = 0
SM = 0
Delta_L
Doxycycline
YOR056C NOB1
ZShift = 0.98
NG = 0
DB = 0
SM = 0
Delta_K
Doxycycline
YOR056C NOB1
ZShift = −0.3
NG = 0
DB = 0
SM = 0
Delta_r
Doxycycline
YOR056C NOB1
ZShift = −0.04
NG = 0
DB = 0
SM = 0
Delta_AUC
Doxycycline

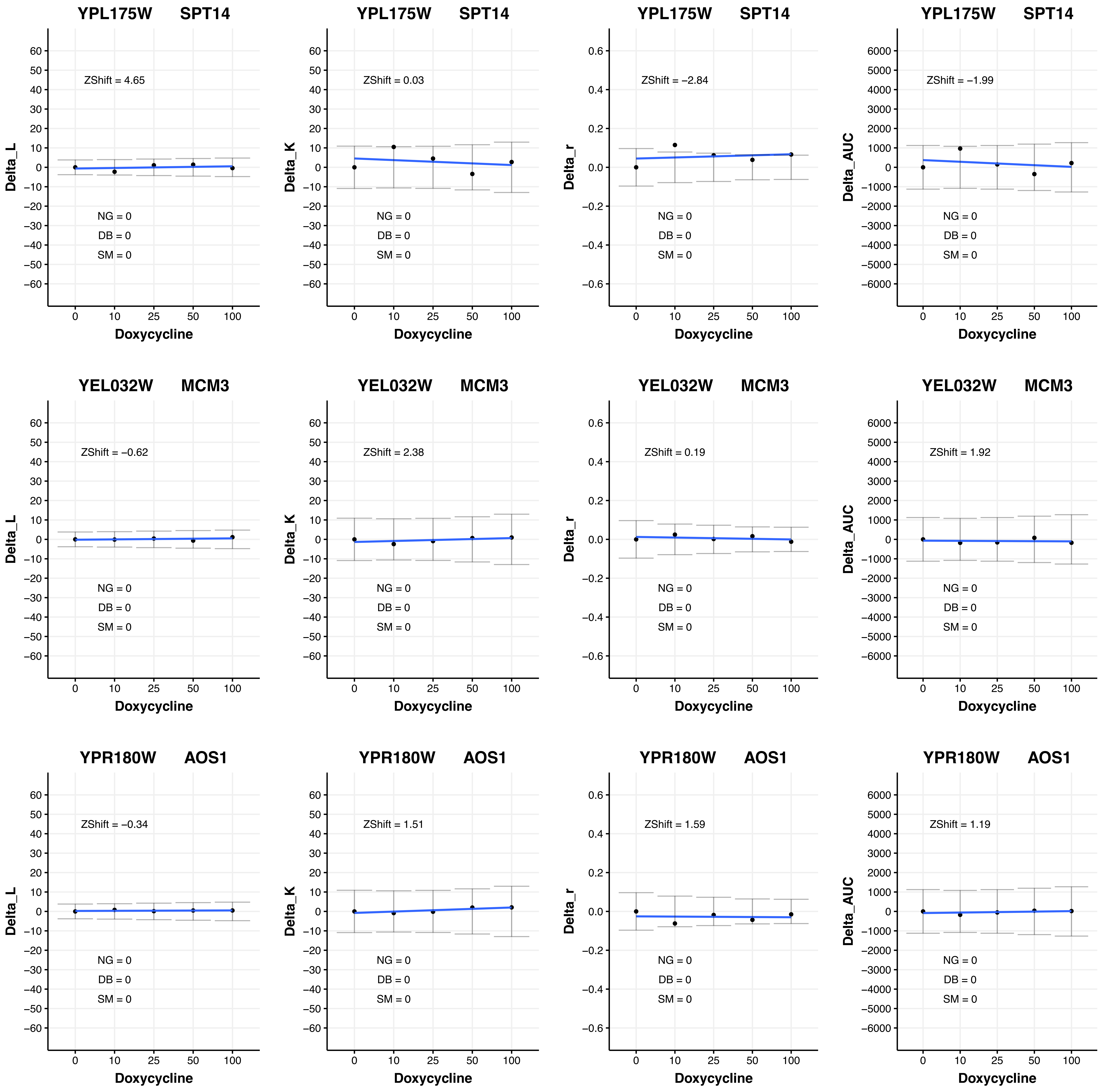

YPL175W SPT14
ZShift = 4.65
Delta_L
NG = 0
DB = 0
SM = 0
Doxycycline
0 10 25 50 100
YPL175W SPT14
ZShift = 0.03
Delta_K
NG = 0
DB = 0
SM = 0
Doxycycline
YPL175W SPT14
ZShift = −2.84
Delta_r
NG = 0
DB = 0
SM = 0
Doxycycline
YPL175W SPT14
ZShift = −1.99
Delta_AUC
NG = 0
DB = 0
SM = 0
Doxycycline
YEL032W MCM3
ZShift = −0.62
Delta_L
NG = 0
DB = 0
SM = 0
Doxycycline
YEL032W MCM3
ZShift = 2.38
Delta_K
NG = 0
DB = 0
SM = 0
Doxycycline
YEL032W MCM3
ZShift = 0.19
Delta_r
NG = 0
DB = 0
SM = 0
Doxycycline
YEL032W MCM3
ZShift = 1.92
Delta_AUC
NG = 0
DB = 0
SM = 0
Doxycycline
YPR180W AOS1
ZShift = −0.34
Delta_L
NG = 0
DB = 0
SM = 0
Doxycycline
YPR180W AOS1
ZShift = 1.51
Delta_K
NG = 0
DB = 0
SM = 0
Doxycycline
YPR180W AOS1
ZShift = 1.59
Delta_r
NG = 0
DB = 0
SM = 0
Doxycycline
YPR180W AOS1
ZShift = 1.19
Delta_AUC
NG = 0
DB = 0
SM = 0
Doxycycline

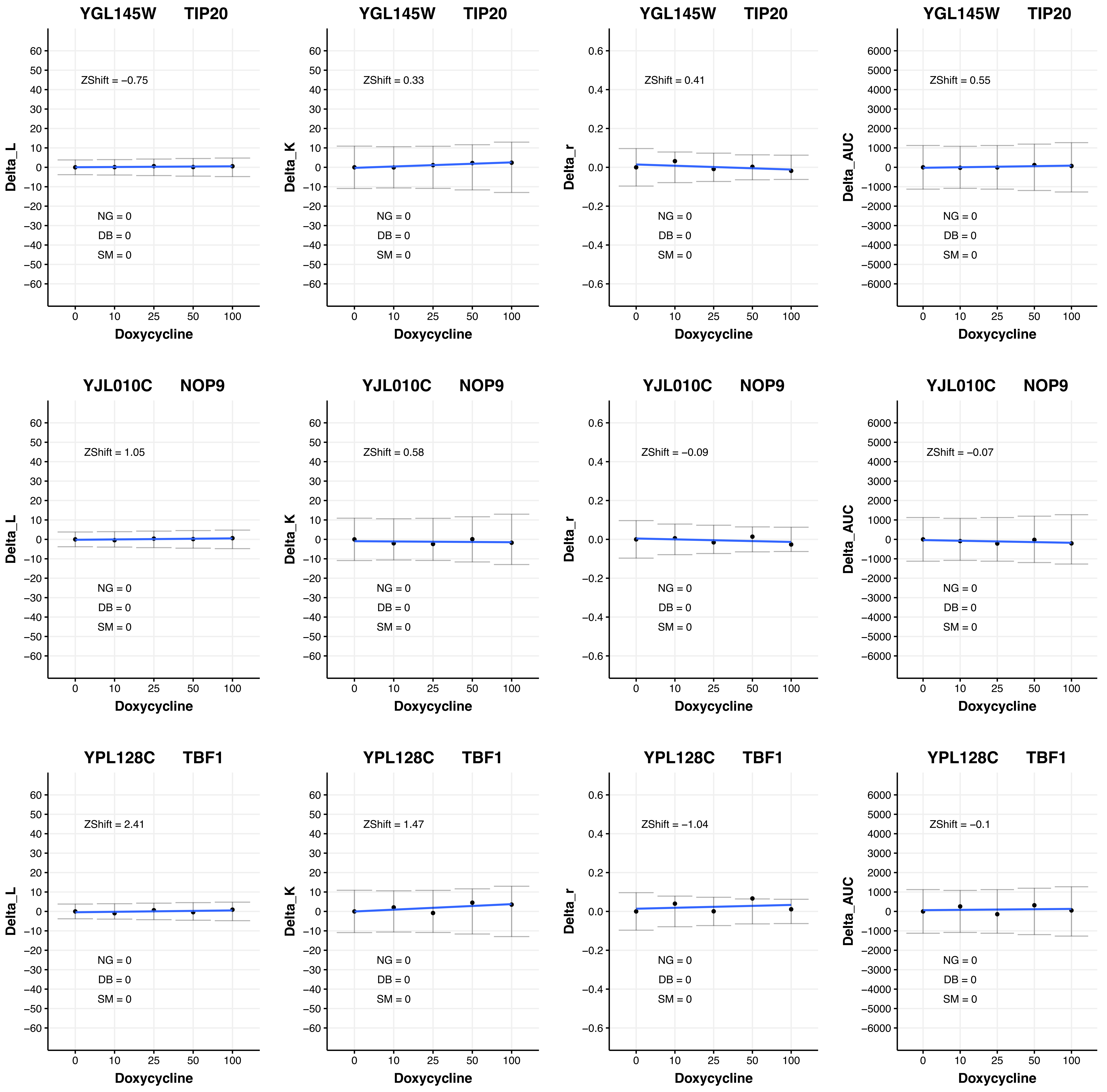

YGL145W TIP20
ZShift = −0.75
ZShift = 0.33
ZShift = 0.41
ZShift = 0.55
YJL010C NOP9
ZShift = 1.05
ZShift = 0.58
ZShift = −0.09
ZShift = −0.07
YPL128C TBF1
ZShift = 2.41
ZShift = 1.47
ZShift = −1.04
ZShift = −0.1
NG = 0
DB = 0
SM = 0
Delta_L
Delta_K
Delta_r
Delta_AUC
Doxycycline
0
10
25
50
100

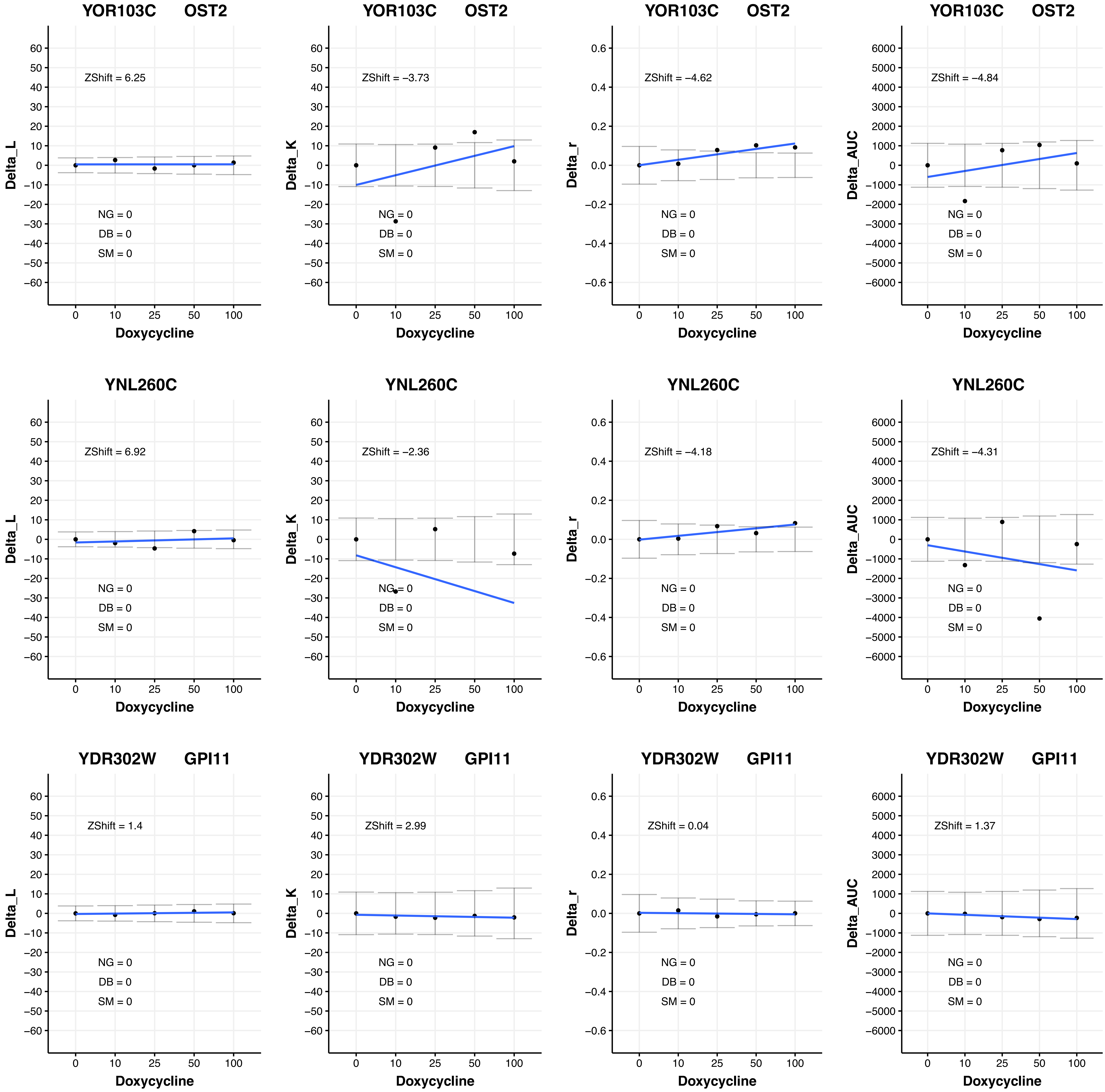
YOR103C OST2
ZShift = 6.25
ZShift = −3.73
ZShift = −4.62
ZShift = −4.84
YNL260C
ZShift = 6.92
ZShift = −2.36
ZShift = −4.18
ZShift = −4.31
YDR302W GPI11
ZShift = 1.4
ZShift = 2.99
ZShift = 0.04
ZShift = 1.37
NG = 0
DB = 0
SM = 0
Delta_L
Delta_K
Delta_r
Delta_AUC
Doxycycline
0
10
25
50
100

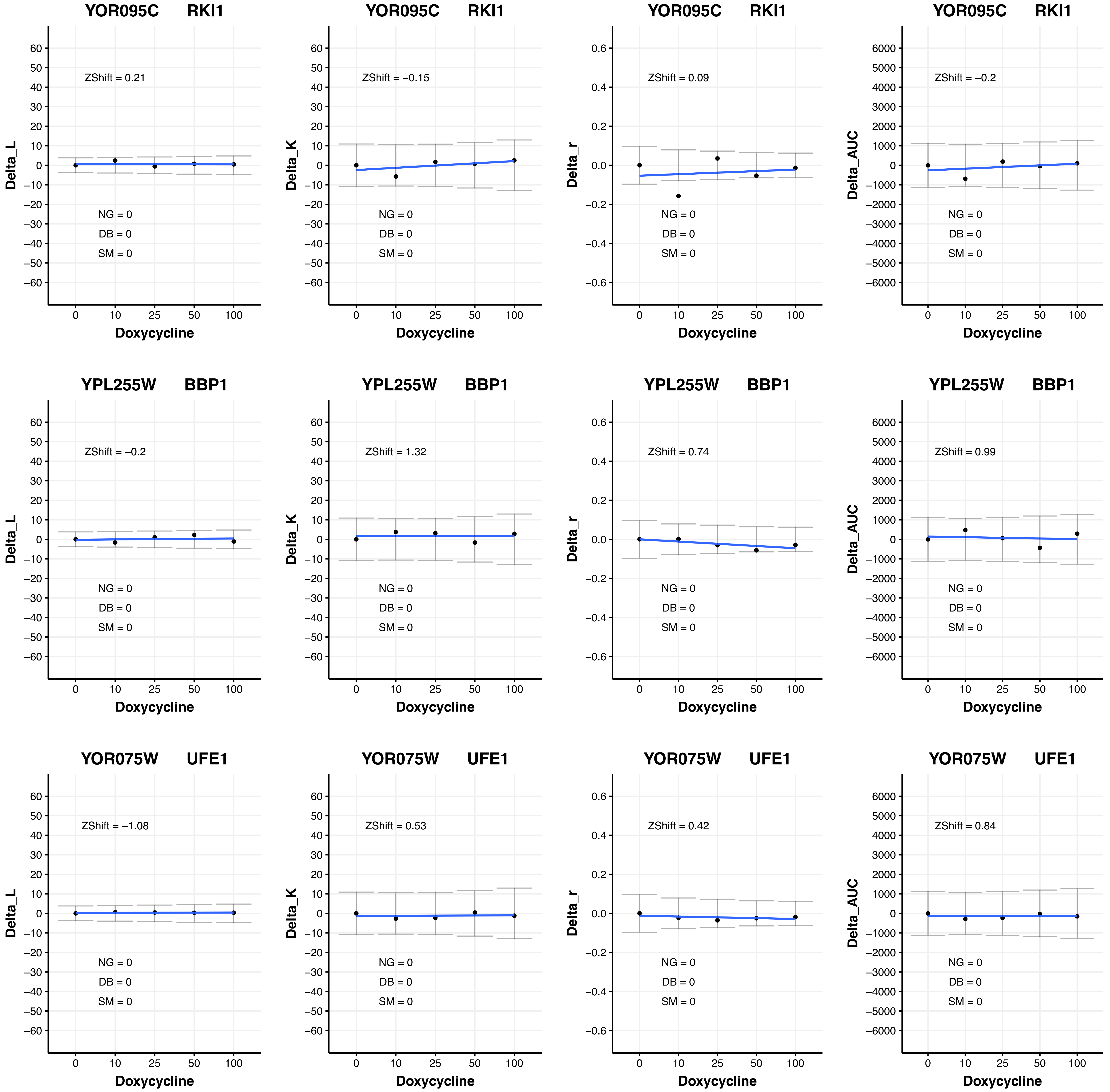

YOR095C RKI1
ZShift = 0.21
ZShift = −0.15
ZShift = 0.09
ZShift = −0.2
YPL255W BBP1
ZShift = −0.2
ZShift = 1.32
ZShift = 0.74
ZShift = 0.99
YOR075W UFE1
ZShift = −1.08
ZShift = 0.53
ZShift = 0.42
ZShift = 0.84
NG = 0
DB = 0
SM = 0
Delta_L
Delta_K
Delta_r
Delta_AUC
Doxycycline
0
10
25
50
100

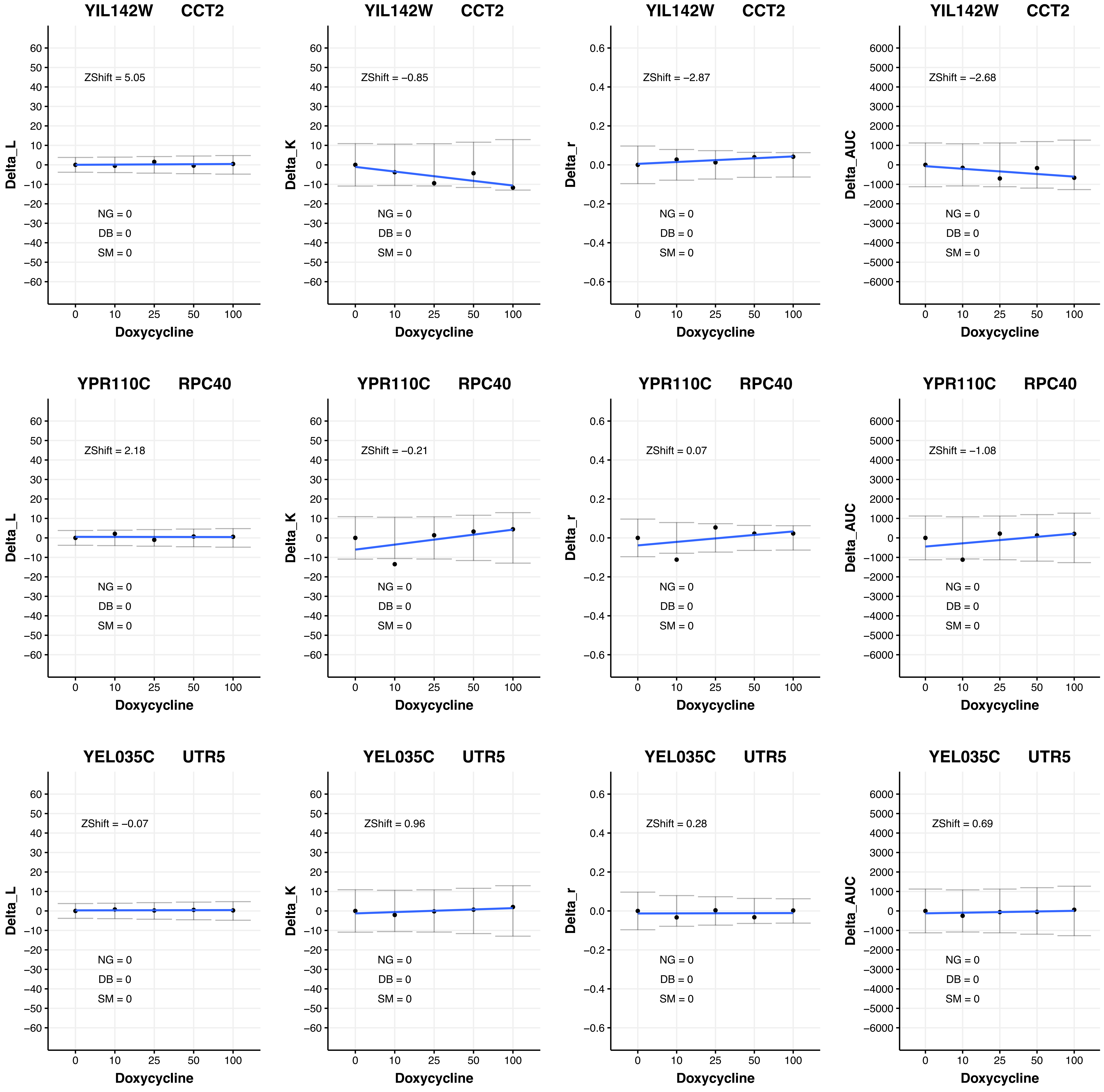
YIL142W CCT2
Delta_L
ZShift = 5.05
NG = 0
DB = 0
SM = 0
Doxycycline
YIL142W CCT2
Delta_K
ZShift = −0.85
NG = 0
DB = 0
SM = 0
Doxycycline
YIL142W CCT2
Delta_r
ZShift = −2.87
NG = 0
DB = 0
SM = 0
Doxycycline
YIL142W CCT2
Delta_AUC
ZShift = −2.68
NG = 0
DB = 0
SM = 0
Doxycycline
YPR110C RPC40
Delta_L
ZShift = 2.18
NG = 0
DB = 0
SM = 0
Doxycycline
YPR110C RPC40
Delta_K
ZShift = −0.21
NG = 0
DB = 0
SM = 0
Doxycycline
YPR110C RPC40
Delta_r
ZShift = 0.07
NG = 0
DB = 0
SM = 0
Doxycycline
YPR110C RPC40
Delta_AUC
ZShift = −1.08
NG = 0
DB = 0
SM = 0
Doxycycline
YEL035C UTR5
Delta_L
ZShift = −0.07
NG = 0
DB = 0
SM = 0
Doxycycline
YEL035C UTR5
Delta_K
ZShift = 0.96
NG = 0
DB = 0
SM = 0
Doxycycline
YEL035C UTR5
Delta_r
ZShift = 0.28
NG = 0
DB = 0
SM = 0
Doxycycline
YEL035C UTR5
Delta_AUC
ZShift = 0.69
NG = 0
DB = 0
SM = 0
Doxycycline

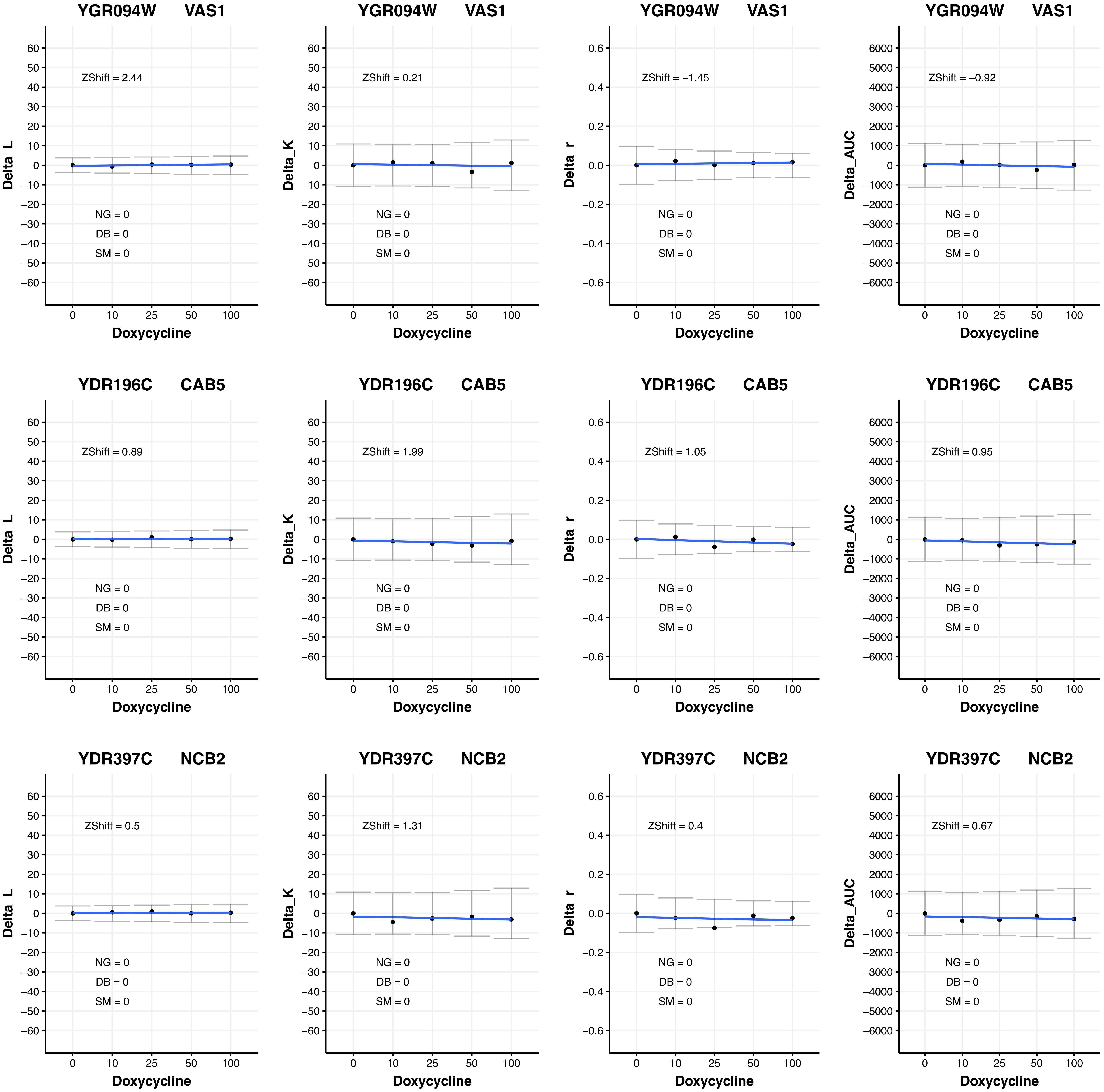
YGR094W VAS1
ZShift = 2.44
NG = 0
DB = 0
SM = 0
Delta_L
Doxycycline
YGR094W VAS1
ZShift = 0.21
NG = 0
DB = 0
SM = 0
Delta_K
Doxycycline
YGR094W VAS1
ZShift = −1.45
NG = 0
DB = 0
SM = 0
Delta_r
Doxycycline
YGR094W VAS1
ZShift = −0.92
NG = 0
DB = 0
SM = 0
Delta_AUC
Doxycycline
YDR196C CAB5
ZShift = 0.89
NG = 0
DB = 0
SM = 0
Delta_L
Doxycycline
YDR196C CAB5
ZShift = 1.99
NG = 0
DB = 0
SM = 0
Delta_K
Doxycycline
YDR196C CAB5
ZShift = 1.05
NG = 0
DB = 0
SM = 0
Delta_r
Doxycycline
YDR196C CAB5
ZShift = 0.95
NG = 0
DB = 0
SM = 0
Delta_AUC
Doxycycline
YDR397C NCB2
ZShift = 0.5
NG = 0
DB = 0
SM = 0
Delta_L
Doxycycline
YDR397C NCB2
ZShift = 1.31
NG = 0
DB = 0
SM = 0
Delta_K
Doxycycline
YDR397C NCB2
ZShift = 0.4
NG = 0
DB = 0
SM = 0
Delta_r
Doxycycline
YDR397C NCB2
ZShift = 0.67
NG = 0
DB = 0
SM = 0
Delta_AUC
Doxycycline

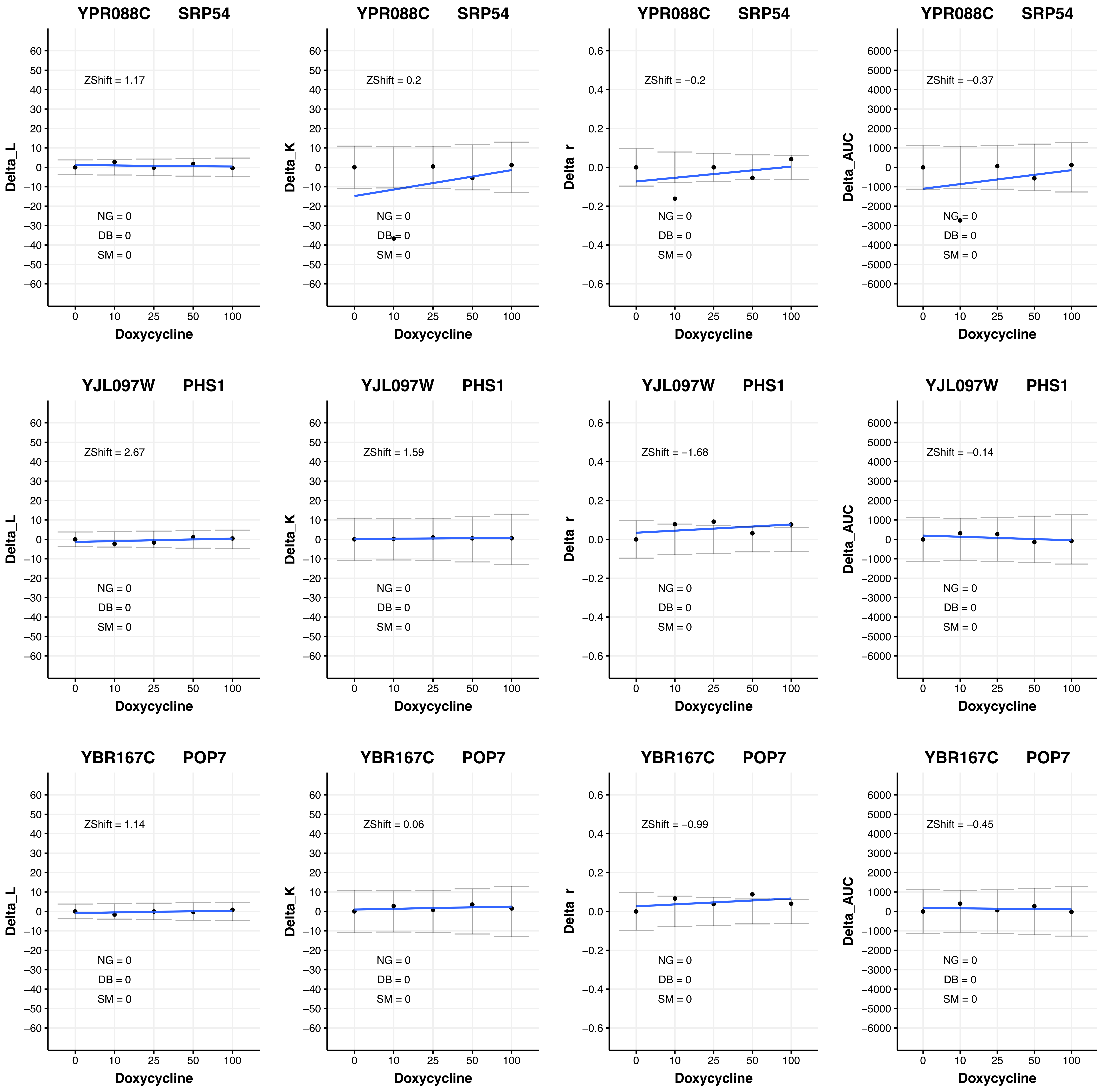

YPR088C SRP54
ZShift = 1.17
Delta_L
NG = 0
DB = 0
SM = 0
Doxycycline
YPR088C SRP54
ZShift = 0.2
Delta_K
NG = 0
DB = 0
SM = 0
Doxycycline
YPR088C SRP54
ZShift = −0.2
Delta_r
NG = 0
DB = 0
SM = 0
Doxycycline
YPR088C SRP54
ZShift = −0.37
Delta_AUC
NG = 0
DB = 0
SM = 0
Doxycycline
YJL097W PHS1
ZShift = 2.67
Delta_L
NG = 0
DB = 0
SM = 0
Doxycycline
YJL097W PHS1
ZShift = 1.59
Delta_K
NG = 0
DB = 0
SM = 0
Doxycycline
YJL097W PHS1
ZShift = −1.68
Delta_r
NG = 0
DB = 0
SM = 0
Doxycycline
YJL097W PHS1
ZShift = −0.14
Delta_AUC
NG = 0
DB = 0
SM = 0
Doxycycline
YBR167C POP7
ZShift = 1.14
Delta_L
NG = 0
DB = 0
SM = 0
Doxycycline
YBR167C POP7
ZShift = 0.06
Delta_K
NG = 0
DB = 0
SM = 0
Doxycycline
YBR167C POP7
ZShift = −0.99
Delta_r
NG = 0
DB = 0
SM = 0
Doxycycline
YBR167C POP7
ZShift = −0.45
Delta_AUC
NG = 0
DB = 0
SM = 0
Doxycycline

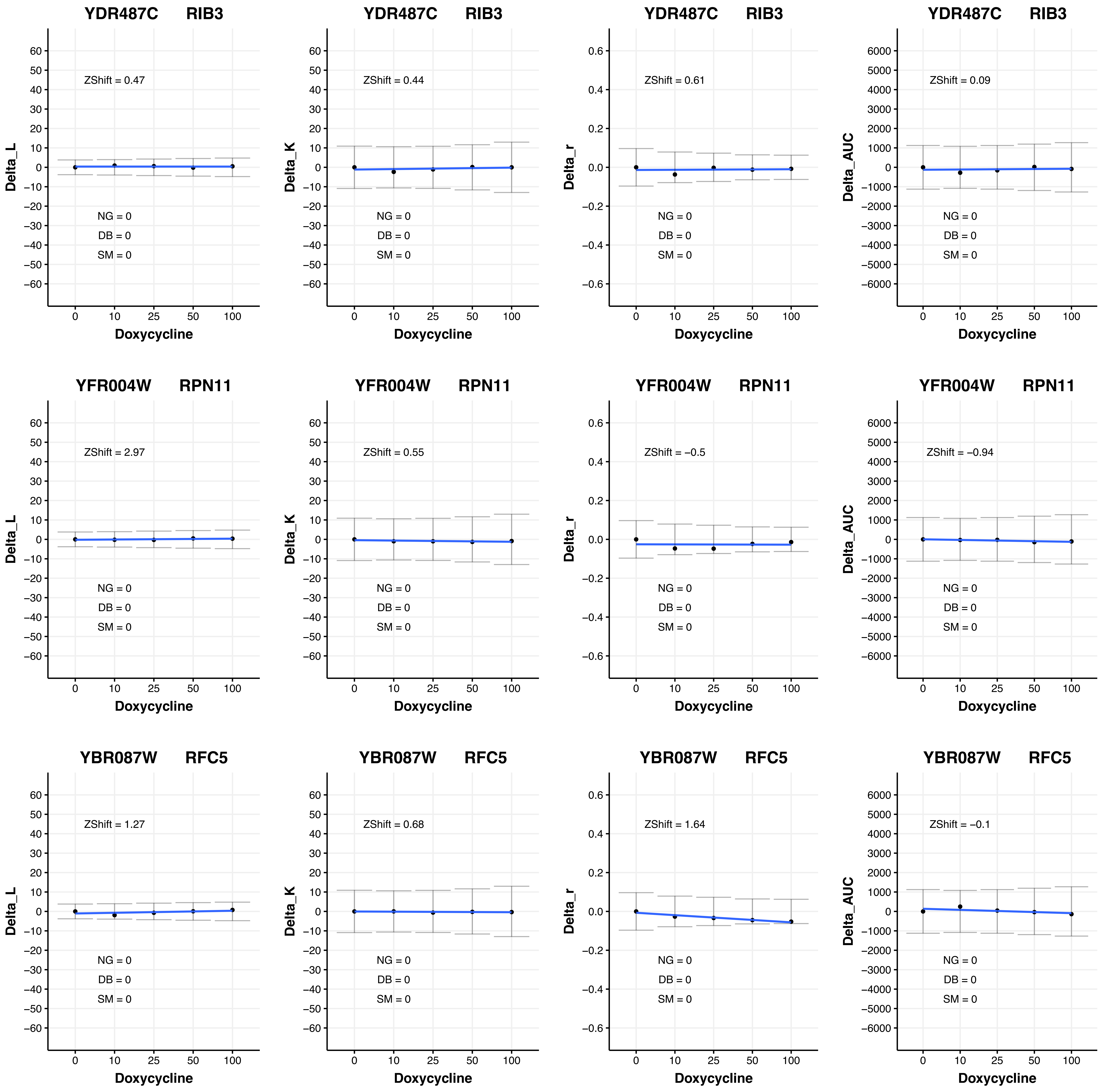

YDR487C RIB3
ZShift = 0.47
Delta_L
NG = 0
DB = 0
SM = 0
Doxycycline
YDR487C RIB3
ZShift = 0.44
Delta_K
NG = 0
DB = 0
SM = 0
Doxycycline
YDR487C RIB3
ZShift = 0.61
Delta_r
NG = 0
DB = 0
SM = 0
Doxycycline
YDR487C RIB3
ZShift = 0.09
Delta_AUC
NG = 0
DB = 0
SM = 0
Doxycycline
YFR004W RPN11
ZShift = 2.97
Delta_L
NG = 0
DB = 0
SM = 0
Doxycycline
YFR004W RPN11
ZShift = 0.55
Delta_K
NG = 0
DB = 0
SM = 0
Doxycycline
YFR004W RPN11
ZShift = −0.5
Delta_r
NG = 0
DB = 0
SM = 0
Doxycycline
YFR004W RPN11
ZShift = −0.94
Delta_AUC
NG = 0
DB = 0
SM = 0
Doxycycline
YBR087W RFC5
ZShift = 1.27
Delta_L
NG = 0
DB = 0
SM = 0
Doxycycline
YBR087W RFC5
ZShift = 0.68
Delta_K
NG = 0
DB = 0
SM = 0
Doxycycline
YBR087W RFC5
ZShift = 1.64
Delta_r
NG = 0
DB = 0
SM = 0
Doxycycline
YBR087W RFC5
ZShift = −0.1
Delta_AUC
NG = 0
DB = 0
SM = 0
Doxycycline

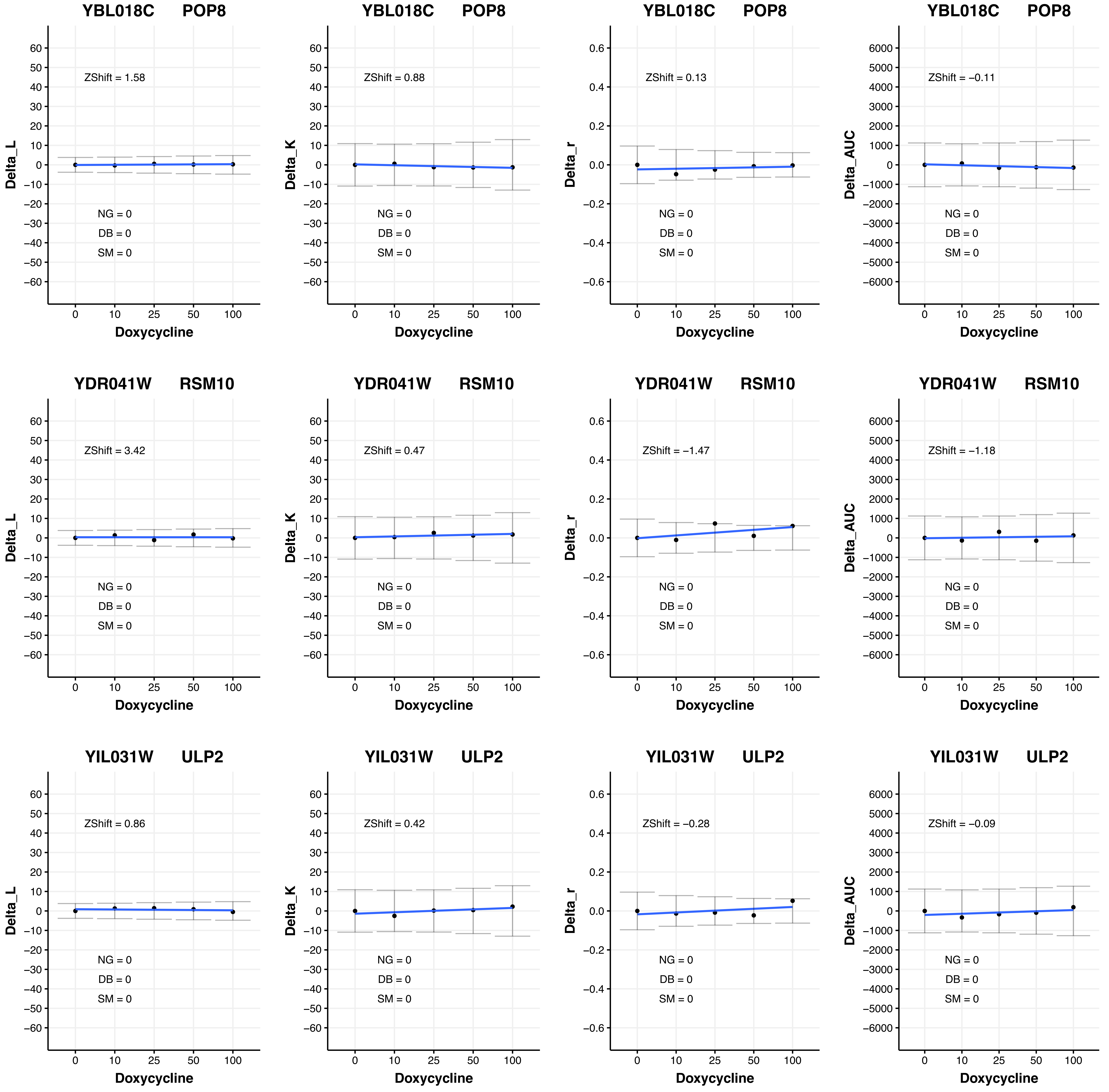

YBL018C POP8
ZShift = 1.58
NG = 0
DB = 0
SM = 0
Delta_L
Doxycycline
YBL018C POP8
ZShift = 0.88
NG = 0
DB = 0
SM = 0
Delta_K
Doxycycline
YBL018C POP8
ZShift = 0.13
NG = 0
DB = 0
SM = 0
Delta_r
Doxycycline
YBL018C POP8
ZShift = −0.11
NG = 0
DB = 0
SM = 0
Delta_AUC
Doxycycline
YDR041W RSM10
ZShift = 3.42
NG = 0
DB = 0
SM = 0
Delta_L
Doxycycline
YDR041W RSM10
ZShift = 0.47
NG = 0
DB = 0
SM = 0
Delta_K
Doxycycline
YDR041W RSM10
ZShift = −1.47
NG = 0
DB = 0
SM = 0
Delta_r
Doxycycline
YDR041W RSM10
ZShift = −1.18
NG = 0
DB = 0
SM = 0
Delta_AUC
Doxycycline
YIL031W ULP2
ZShift = 0.86
NG = 0
DB = 0
SM = 0
Delta_L
Doxycycline
YIL031W ULP2
ZShift = 0.42
NG = 0
DB = 0
SM = 0
Delta_K
Doxycycline
YIL031W ULP2
ZShift = −0.28
NG = 0
DB = 0
SM = 0
Delta_r
Doxycycline
YIL031W ULP2
ZShift = −0.09
NG = 0
DB = 0
SM = 0
Delta_AUC
Doxycycline

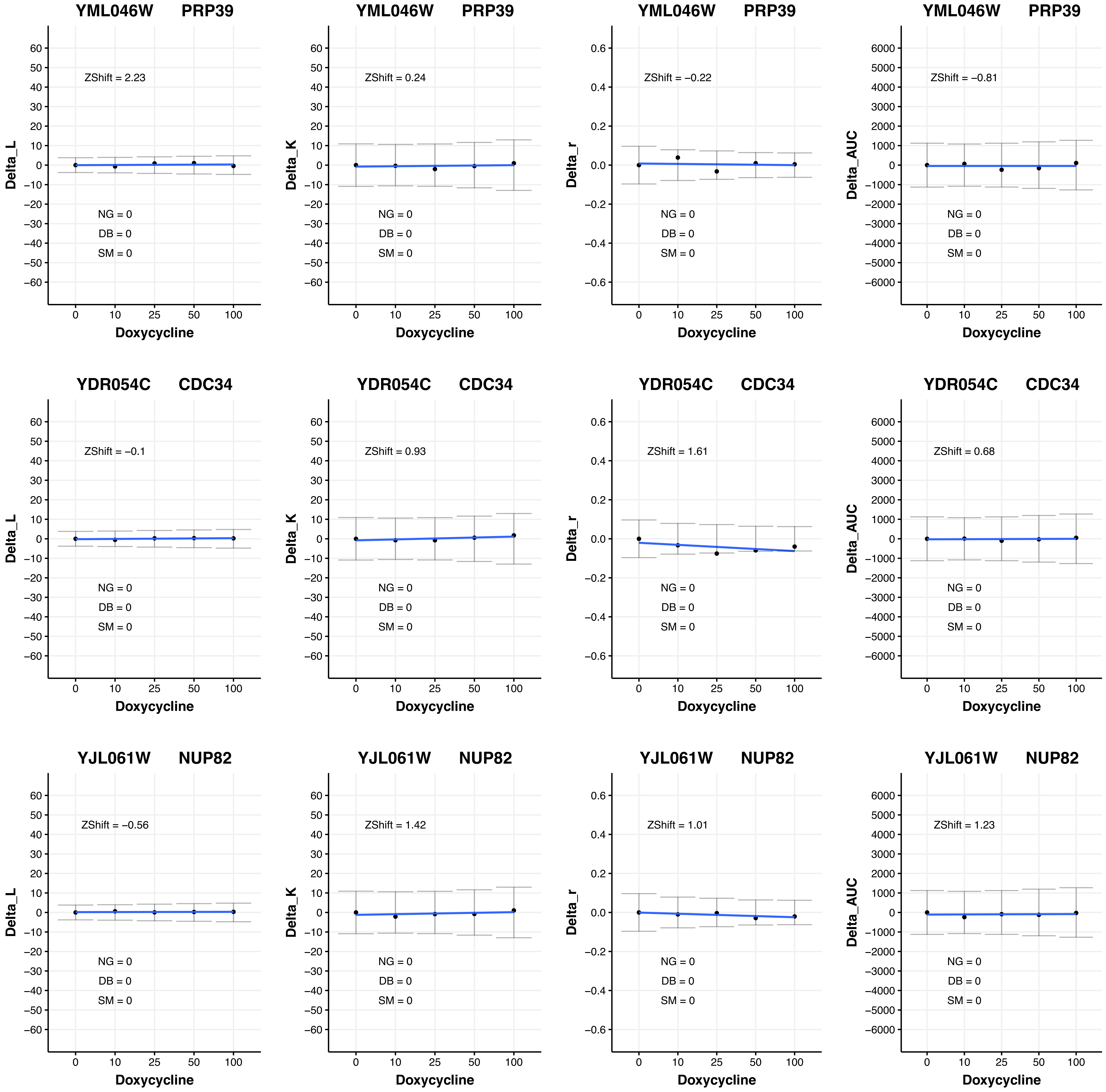
YML046W PRP39
Delta_L
ZShift = 2.23
NG = 0
DB = 0
SM = 0
Doxycycline
YML046W PRP39
Delta_K
ZShift = 0.24
NG = 0
DB = 0
SM = 0
Doxycycline
YML046W PRP39
Delta_r
ZShift = −0.22
NG = 0
DB = 0
SM = 0
Doxycycline
YML046W PRP39
Delta_AUC
ZShift = −0.81
NG = 0
DB = 0
SM = 0
Doxycycline
YDR054C CDC34
Delta_L
ZShift = −0.1
NG = 0
DB = 0
SM = 0
Doxycycline
YDR054C CDC34
Delta_K
ZShift = 0.93
NG = 0
DB = 0
SM = 0
Doxycycline
YDR054C CDC34
Delta_r
ZShift = 1.61
NG = 0
DB = 0
SM = 0
Doxycycline
YDR054C CDC34
Delta_AUC
ZShift = 0.68
NG = 0
DB = 0
SM = 0
Doxycycline
YJL061W NUP82
Delta_L
ZShift = −0.56
NG = 0
DB = 0
SM = 0
Doxycycline
YJL061W NUP82
Delta_K
ZShift = 1.42
NG = 0
DB = 0
SM = 0
Doxycycline
YJL061W NUP82
Delta_r
ZShift = 1.01
NG = 0
DB = 0
SM = 0
Doxycycline
YJL061W NUP82
Delta_AUC
ZShift = 1.23
NG = 0
DB = 0
SM = 0
Doxycycline

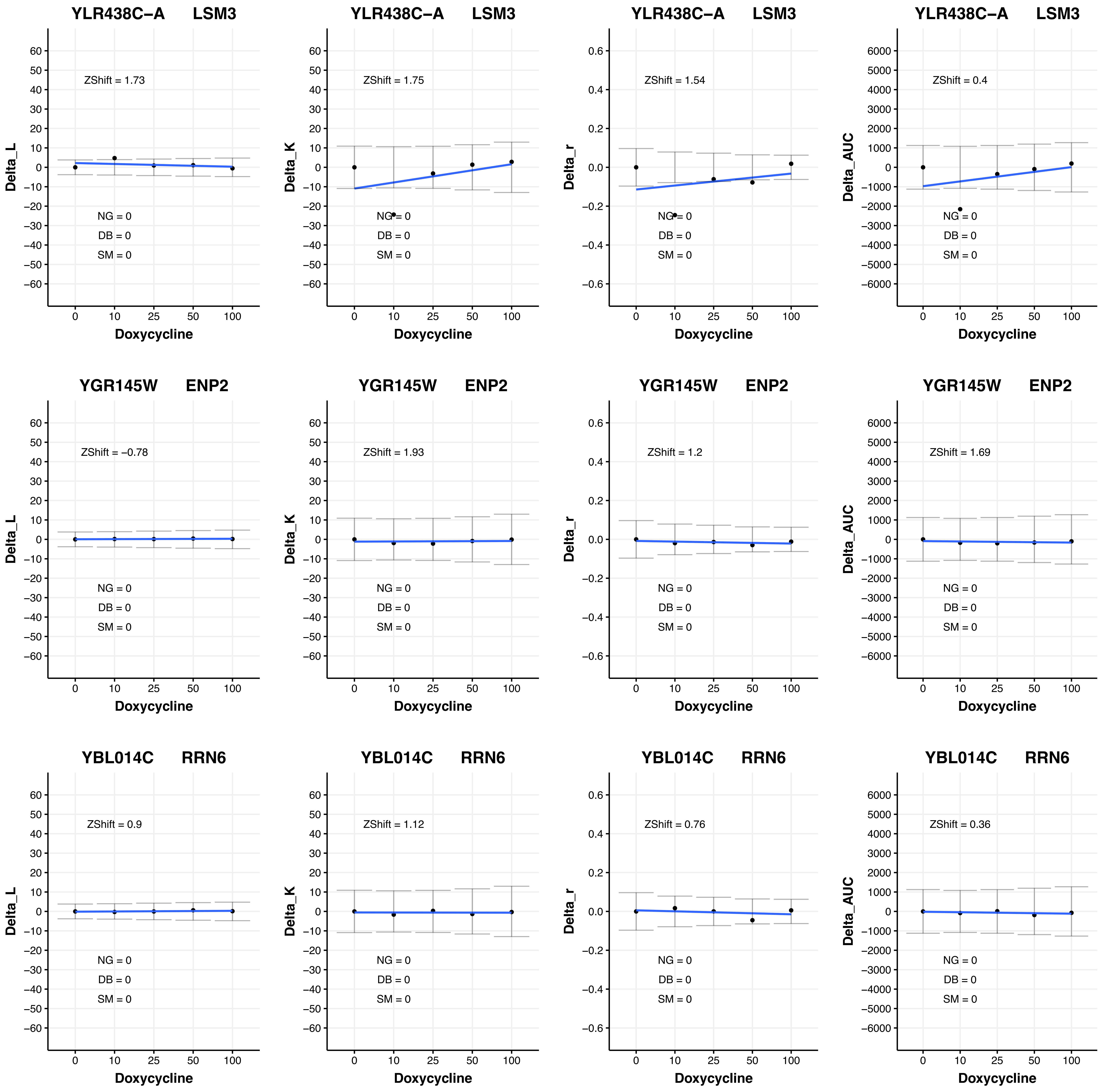

YLR438C−A LSM3
ZShift = 1.73
NG = 0
DB = 0
SM = 0
Delta_L
Doxycycline
YLR438C−A LSM3
ZShift = 1.75
NG = 0
DB = 0
SM = 0
Delta_K
Doxycycline
YLR438C−A LSM3
ZShift = 1.54
NG = 0
DB = 0
SM = 0
Delta_r
Doxycycline
YLR438C−A LSM3
ZShift = 0.4
NG = 0
DB = 0
SM = 0
Delta_AUC
Doxycycline
YGR145W ENP2
ZShift = −0.78
NG = 0
DB = 0
SM = 0
Delta_L
Doxycycline
YGR145W ENP2
ZShift = 1.93
NG = 0
DB = 0
SM = 0
Delta_K
Doxycycline
YGR145W ENP2
ZShift = 1.2
NG = 0
DB = 0
SM = 0
Delta_r
Doxycycline
YGR145W ENP2
ZShift = 1.69
NG = 0
DB = 0
SM = 0
Delta_AUC
Doxycycline
YBL014C RRN6
ZShift = 0.9
NG = 0
DB = 0
SM = 0
Delta_L
Doxycycline
YBL014C RRN6
ZShift = 1.12
NG = 0
DB = 0
SM = 0
Delta_K
Doxycycline
YBL014C RRN6
ZShift = 0.76
NG = 0
DB = 0
SM = 0
Delta_r
Doxycycline
YBL014C RRN6
ZShift = 0.36
NG = 0
DB = 0
SM = 0
Delta_AUC
Doxycycline

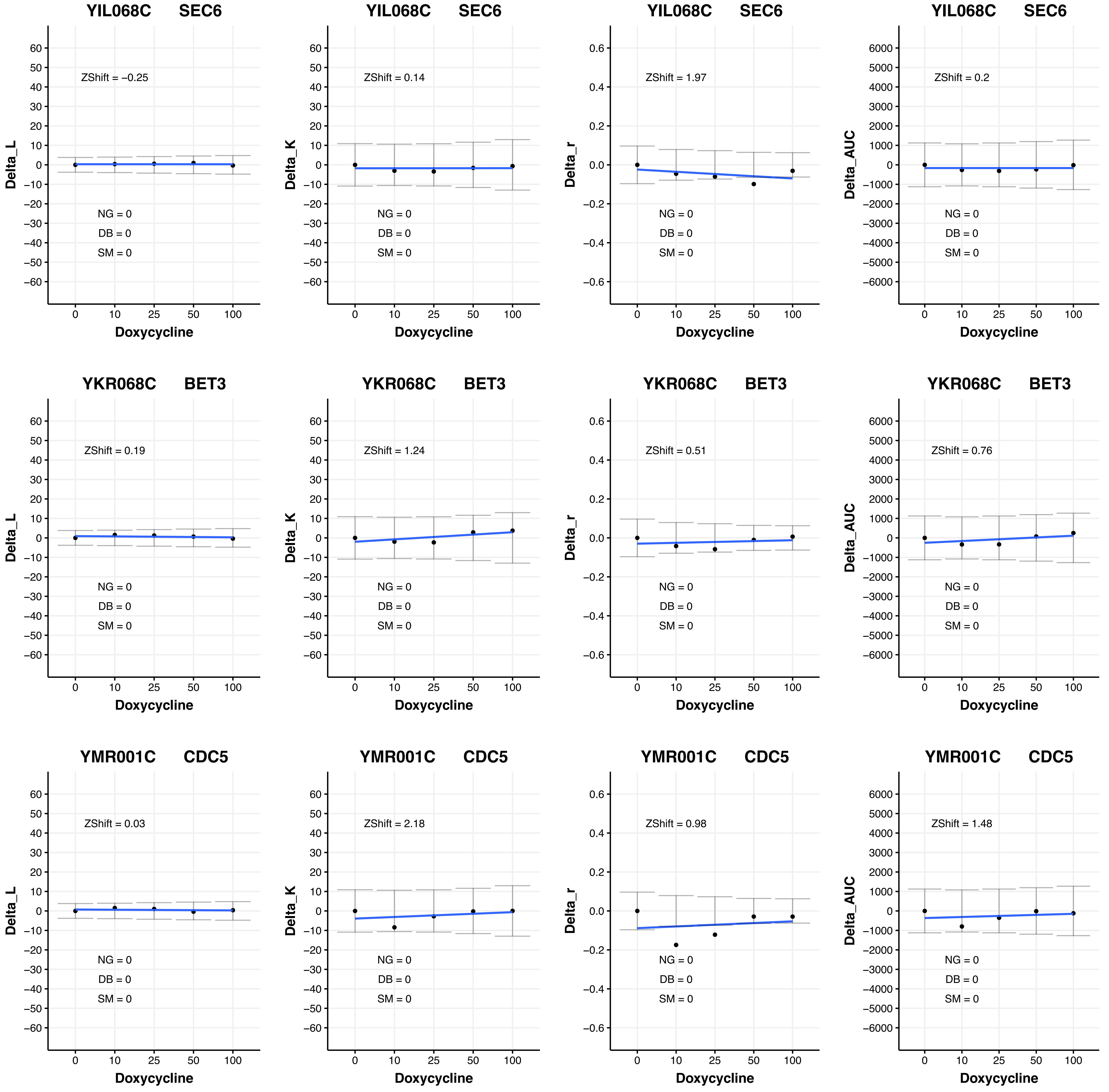
YIL068C SEC6
ZShift = −0.25
NG = 0
DB = 0
SM = 0
Delta_L
Doxycycline
YIL068C SEC6
ZShift = 0.14
NG = 0
DB = 0
SM = 0
Delta_K
Doxycycline
YIL068C SEC6
ZShift = 1.97
NG = 0
DB = 0
SM = 0
Delta_r
Doxycycline
YIL068C SEC6
ZShift = 0.2
NG = 0
DB = 0
SM = 0
Delta_AUC
Doxycycline
YKR068C BET3
ZShift = 0.19
NG = 0
DB = 0
SM = 0
Delta_L
Doxycycline
YKR068C BET3
ZShift = 1.24
NG = 0
DB = 0
SM = 0
Delta_K
Doxycycline
YKR068C BET3
ZShift = 0.51
NG = 0
DB = 0
SM = 0
Delta_r
Doxycycline
YKR068C BET3
ZShift = 0.76
NG = 0
DB = 0
SM = 0
Delta_AUC
Doxycycline
YMR001C CDC5
ZShift = 0.03
NG = 0
DB = 0
SM = 0
Delta_L
Doxycycline
YMR001C CDC5
ZShift = 2.18
NG = 0
DB = 0
SM = 0
Delta_K
Doxycycline
YMR001C CDC5
ZShift = 0.98
NG = 0
DB = 0
SM = 0
Delta_r
Doxycycline
YMR001C CDC5
ZShift = 1.48
NG = 0
DB = 0
SM = 0
Delta_AUC
Doxycycline
0
10
25
50
100

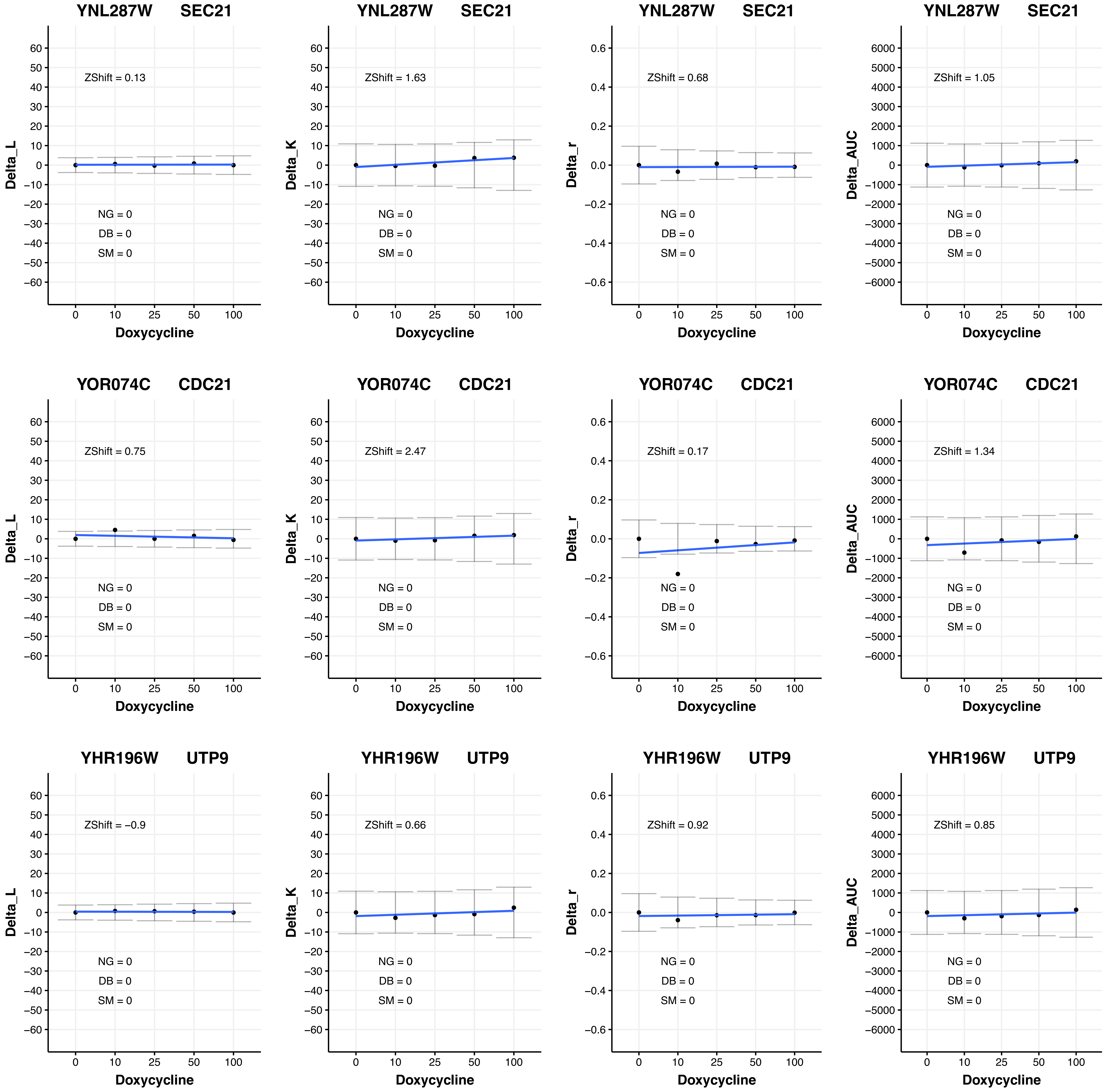

YNL287W SEC21
ZShift = 0.13
NG = 0
DB = 0
SM = 0
Delta_L
Doxycycline
YNL287W SEC21
ZShift = 1.63
NG = 0
DB = 0
SM = 0
Delta_K
Doxycycline
YNL287W SEC21
ZShift = 0.68
NG = 0
DB = 0
SM = 0
Delta_r
Doxycycline
YNL287W SEC21
ZShift = 1.05
NG = 0
DB = 0
SM = 0
Delta_AUC
Doxycycline
YOR074C CDC21
ZShift = 0.75
NG = 0
DB = 0
SM = 0
Delta_L
Doxycycline
YOR074C CDC21
ZShift = 2.47
NG = 0
DB = 0
SM = 0
Delta_K
Doxycycline
YOR074C CDC21
ZShift = 0.17
NG = 0
DB = 0
SM = 0
Delta_r
Doxycycline
YOR074C CDC21
ZShift = 1.34
NG = 0
DB = 0
SM = 0
Delta_AUC
Doxycycline
YHR196W UTP9
ZShift = −0.9
NG = 0
DB = 0
SM = 0
Delta_L
Doxycycline
YHR196W UTP9
ZShift = 0.66
NG = 0
DB = 0
SM = 0
Delta_K
Doxycycline
YHR196W UTP9
ZShift = 0.92
NG = 0
DB = 0
SM = 0
Delta_r
Doxycycline
YHR196W UTP9
ZShift = 0.85
NG = 0
DB = 0
SM = 0
Delta_AUC
Doxycycline

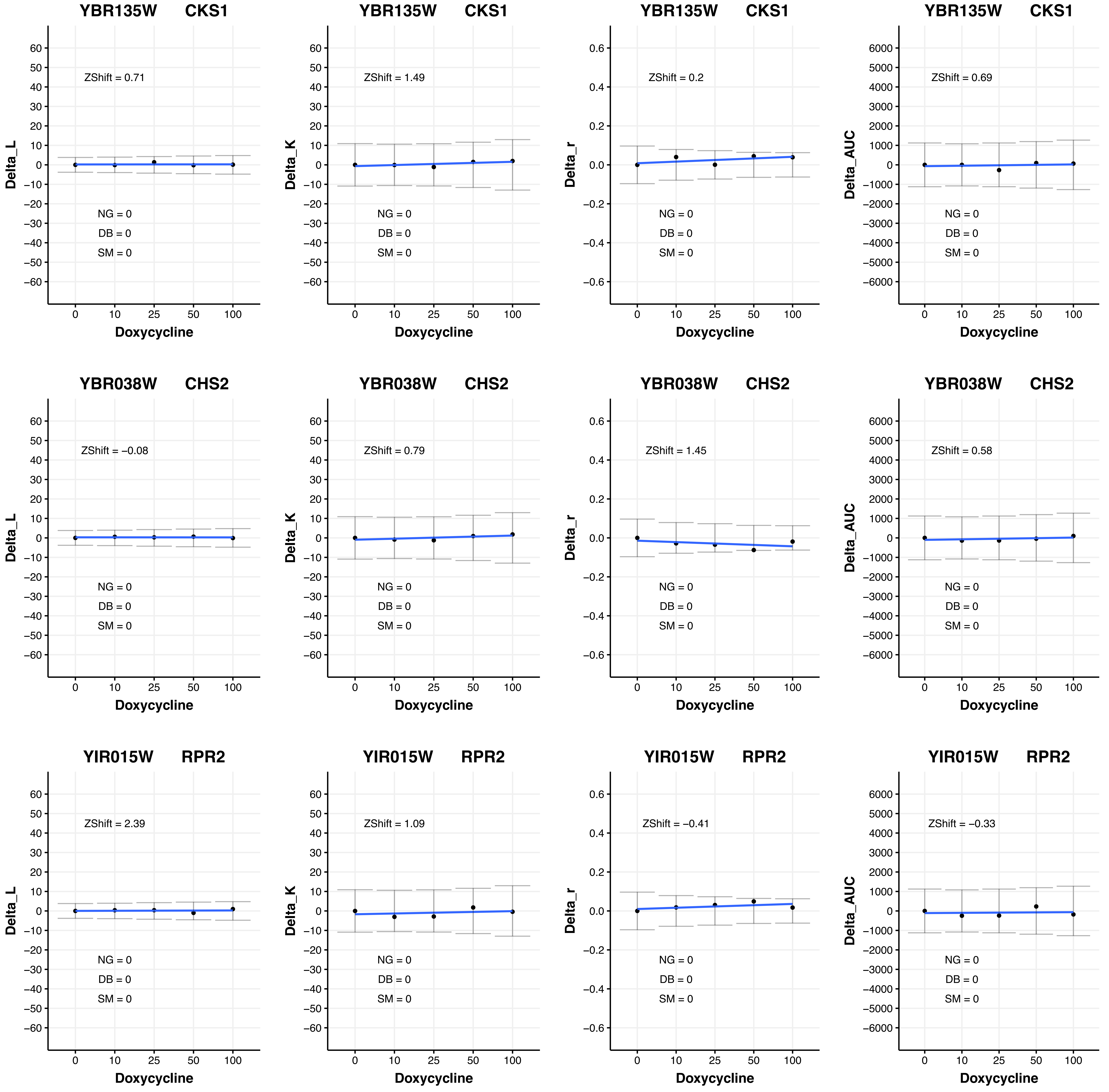

YBR135W CKS1
ZShift = 0.71
NG = 0
DB = 0
SM = 0
Delta_L
Doxycycline
YBR135W CKS1
ZShift = 1.49
NG = 0
DB = 0
SM = 0
Delta_K
Doxycycline
YBR135W CKS1
ZShift = 0.2
NG = 0
DB = 0
SM = 0
Delta_r
Doxycycline
YBR135W CKS1
ZShift = 0.69
NG = 0
DB = 0
SM = 0
Delta_AUC
Doxycycline
YBR038W CHS2
ZShift = −0.08
NG = 0
DB = 0
SM = 0
Delta_L
Doxycycline
YBR038W CHS2
ZShift = 0.79
NG = 0
DB = 0
SM = 0
Delta_K
Doxycycline
YBR038W CHS2
ZShift = 1.45
NG = 0
DB = 0
SM = 0
Delta_r
Doxycycline
YBR038W CHS2
ZShift = 0.58
NG = 0
DB = 0
SM = 0
Delta_AUC
Doxycycline
YIR015W RPR2
ZShift = 2.39
NG = 0
DB = 0
SM = 0
Delta_L
Doxycycline
YIR015W RPR2
ZShift = 1.09
NG = 0
DB = 0
SM = 0
Delta_K
Doxycycline
YIR015W RPR2
ZShift = −0.41
NG = 0
DB = 0
SM = 0
Delta_r
Doxycycline
YIR015W RPR2
ZShift = −0.33
NG = 0
DB = 0
SM = 0
Delta_AUC
Doxycycline

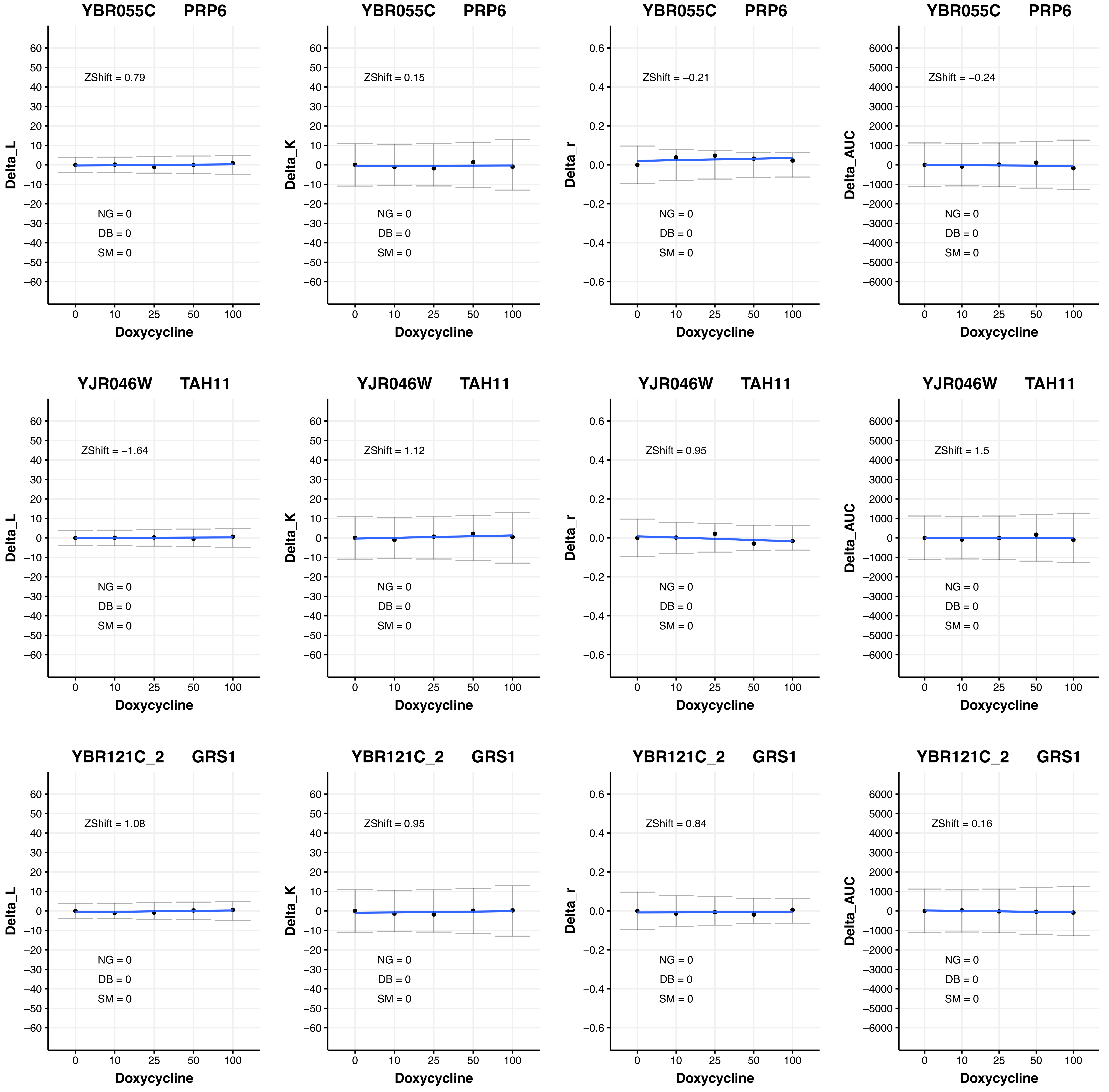
YBR055C PRP6
Delta_L
ZShift = 0.79
NG = 0
DB = 0
SM = 0
Doxycycline
YBR055C PRP6
Delta_K
ZShift = 0.15
NG = 0
DB = 0
SM = 0
Doxycycline
YBR055C PRP6
Delta_r
ZShift = −0.21
NG = 0
DB = 0
SM = 0
Doxycycline
YBR055C PRP6
Delta_AUC
ZShift = −0.24
NG = 0
DB = 0
SM = 0
Doxycycline
YJR046W TAH11
Delta_L
ZShift = −1.64
NG = 0
DB = 0
SM = 0
Doxycycline
YJR046W TAH11
Delta_K
ZShift = 1.12
NG = 0
DB = 0
SM = 0
Doxycycline
YJR046W TAH11
Delta_r
ZShift = 0.95
NG = 0
DB = 0
SM = 0
Doxycycline
YJR046W TAH11
Delta_AUC
ZShift = 1.5
NG = 0
DB = 0
SM = 0
Doxycycline
YBR121C_2 GRS1
Delta_L
ZShift = 1.08
NG = 0
DB = 0
SM = 0
Doxycycline
YBR121C_2 GRS1
Delta_K
ZShift = 0.95
NG = 0
DB = 0
SM = 0
Doxycycline
YBR121C_2 GRS1
Delta_r
ZShift = 0.84
NG = 0
DB = 0
SM = 0
Doxycycline
YBR121C_2 GRS1
Delta_AUC
ZShift = 0.16
NG = 0
DB = 0
SM = 0
Doxycycline

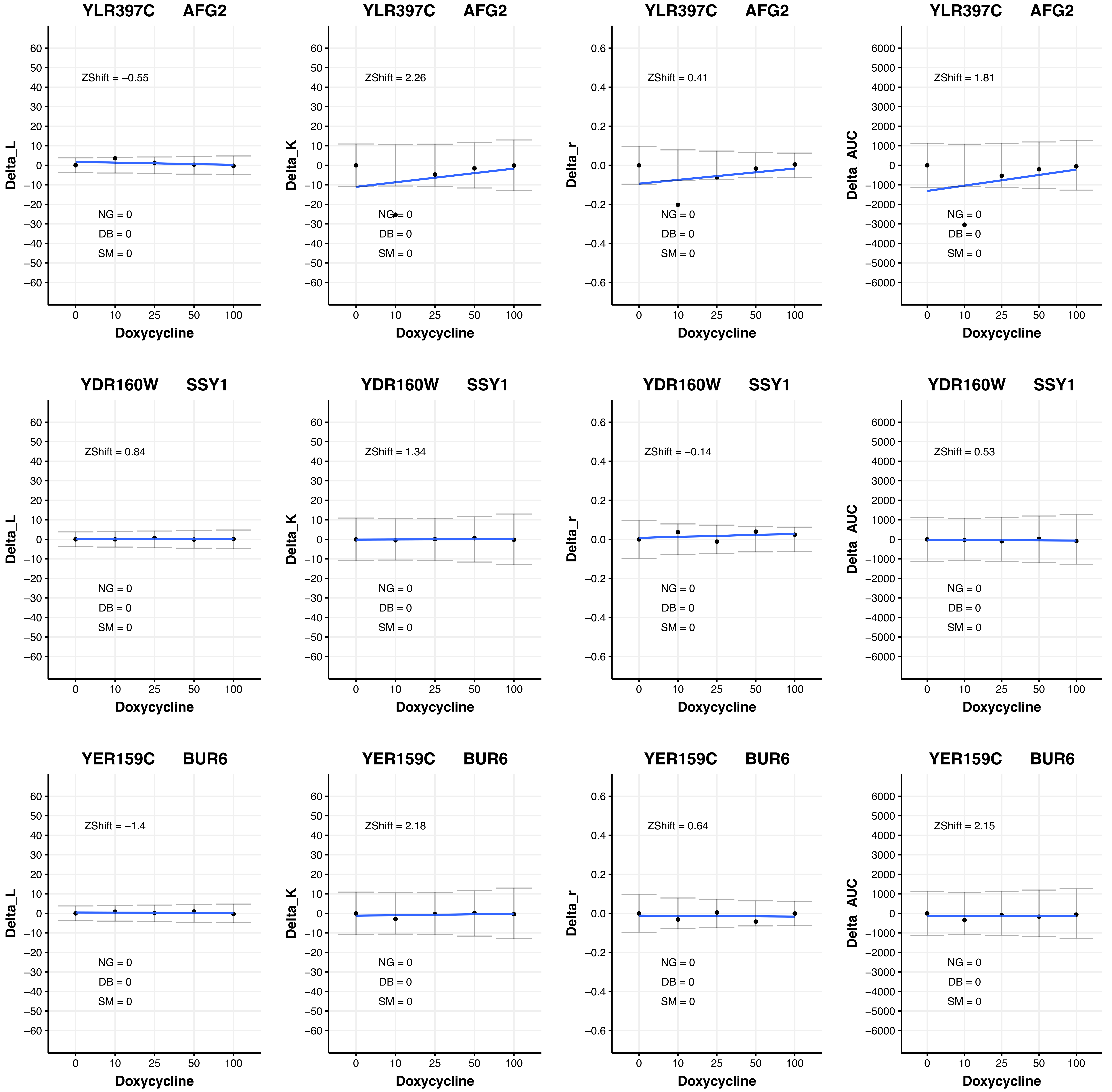
YLR397C AFG2
ZShift = −0.55
NG = 0
DB = 0
SM = 0
Delta_L
Doxycycline
YLR397C AFG2
ZShift = 2.26
NG = 0
DB = 0
SM = 0
Delta_K
Doxycycline
YLR397C AFG2
ZShift = 0.41
NG = 0
DB = 0
SM = 0
Delta_r
Doxycycline
YLR397C AFG2
ZShift = 1.81
NG = 0
DB = 0
SM = 0
Delta_AUC
Doxycycline
YDR160W SSY1
ZShift = 0.84
NG = 0
DB = 0
SM = 0
Delta_L
Doxycycline
YDR160W SSY1
ZShift = 1.34
NG = 0
DB = 0
SM = 0
Delta_K
Doxycycline
YDR160W SSY1
ZShift = −0.14
NG = 0
DB = 0
SM = 0
Delta_r
Doxycycline
YDR160W SSY1
ZShift = 0.53
NG = 0
DB = 0
SM = 0
Delta_AUC
Doxycycline
YER159C BUR6
ZShift = −1.4
NG = 0
DB = 0
SM = 0
Delta_L
Doxycycline
YER159C BUR6
ZShift = 2.18
NG = 0
DB = 0
SM = 0
Delta_K
Doxycycline
YER159C BUR6
ZShift = 0.64
NG = 0
DB = 0
SM = 0
Delta_r
Doxycycline
YER159C BUR6
ZShift = 2.15
NG = 0
DB = 0
SM = 0
Delta_AUC
Doxycycline

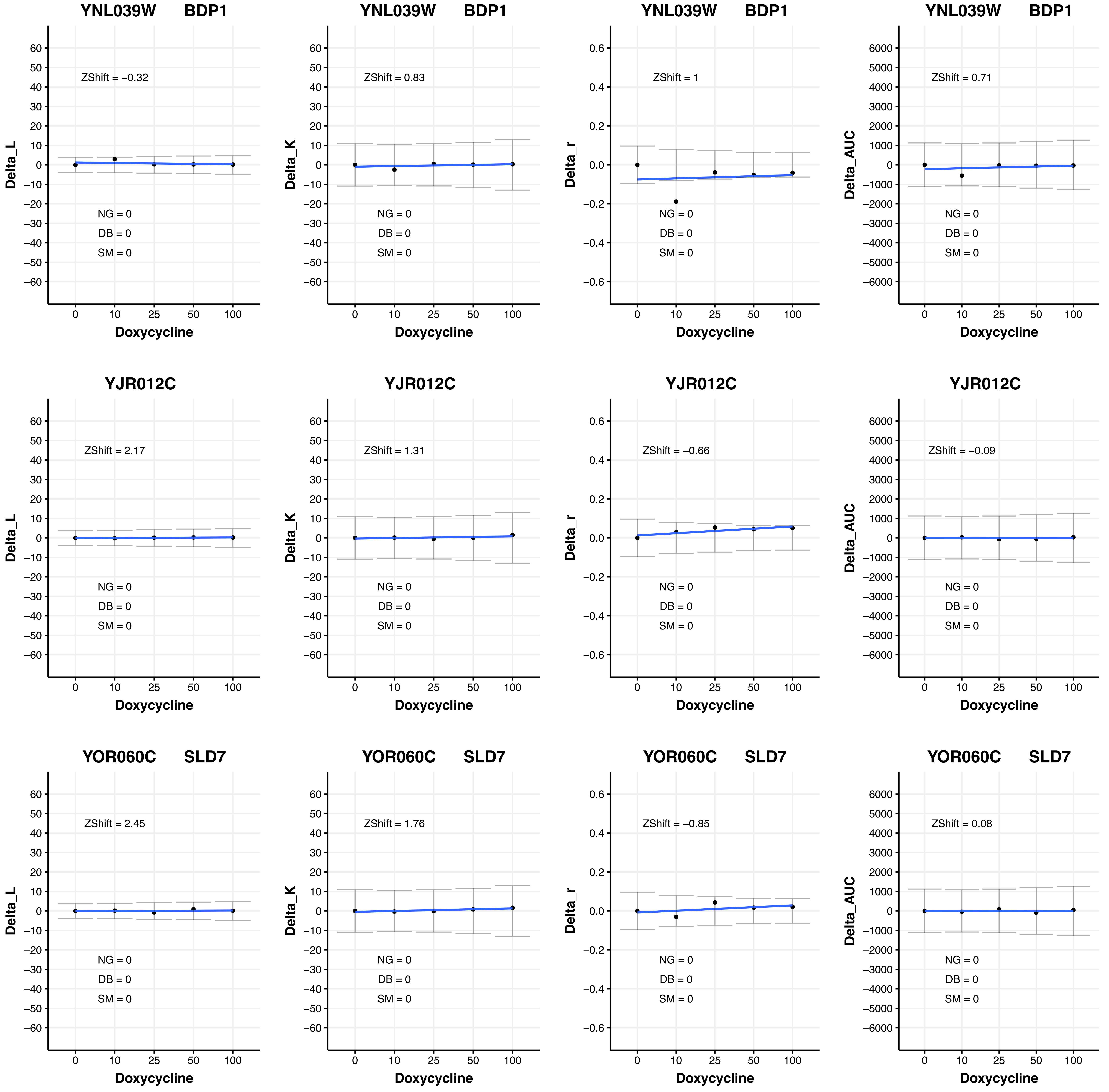
YNL039W BDP1
ZShift = −0.32
ZShift = 0.83
ZShift = 1
ZShift = 0.71
YJR012C
ZShift = 2.17
ZShift = 1.31
ZShift = −0.66
ZShift = −0.09
YOR060C SLD7
ZShift = 2.45
ZShift = 1.76
ZShift = −0.85
ZShift = 0.08
NG = 0
DB = 0
SM = 0
Delta_L
Delta_K
Delta_r
Delta_AUC
Doxycycline

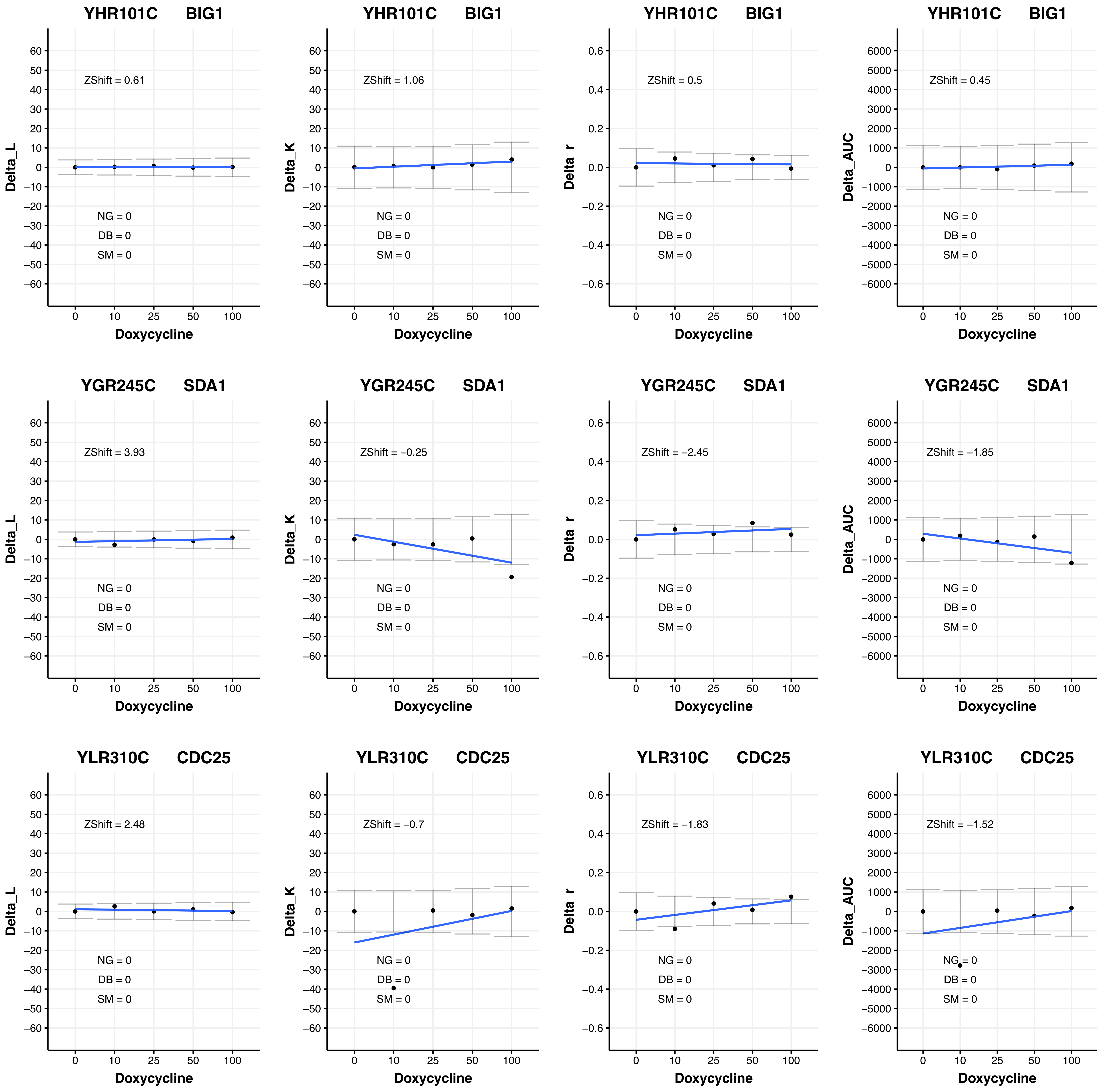
YHR101C BIG1
ZShift = 0.61
NG = 0
DB = 0
SM = 0
Delta_L
Doxycycline
YHR101C BIG1
ZShift = 1.06
NG = 0
DB = 0
SM = 0
Delta_K
Doxycycline
YHR101C BIG1
ZShift = 0.5
NG = 0
DB = 0
SM = 0
Delta_r
Doxycycline
YHR101C BIG1
ZShift = 0.45
NG = 0
DB = 0
SM = 0
Delta_AUC
Doxycycline
YGR245C SDA1
ZShift = 3.93
NG = 0
DB = 0
SM = 0
Delta_L
Doxycycline
YGR245C SDA1
ZShift = −0.25
NG = 0
DB = 0
SM = 0
Delta_K
Doxycycline
YGR245C SDA1
ZShift = −2.45
NG = 0
DB = 0
SM = 0
Delta_r
Doxycycline
YGR245C SDA1
ZShift = −1.85
NG = 0
DB = 0
SM = 0
Delta_AUC
Doxycycline
YLR310C CDC25
ZShift = 2.48
NG = 0
DB = 0
SM = 0
Delta_L
Doxycycline
YLR310C CDC25
ZShift = −0.7
NG = 0
DB = 0
SM = 0
Delta_K
Doxycycline
YLR310C CDC25
ZShift = −1.83
NG = 0
DB = 0
SM = 0
Delta_r
Doxycycline
YLR310C CDC25
ZShift = −1.52
NG = 0
DB = 0
SM = 0
Delta_AUC
Doxycycline

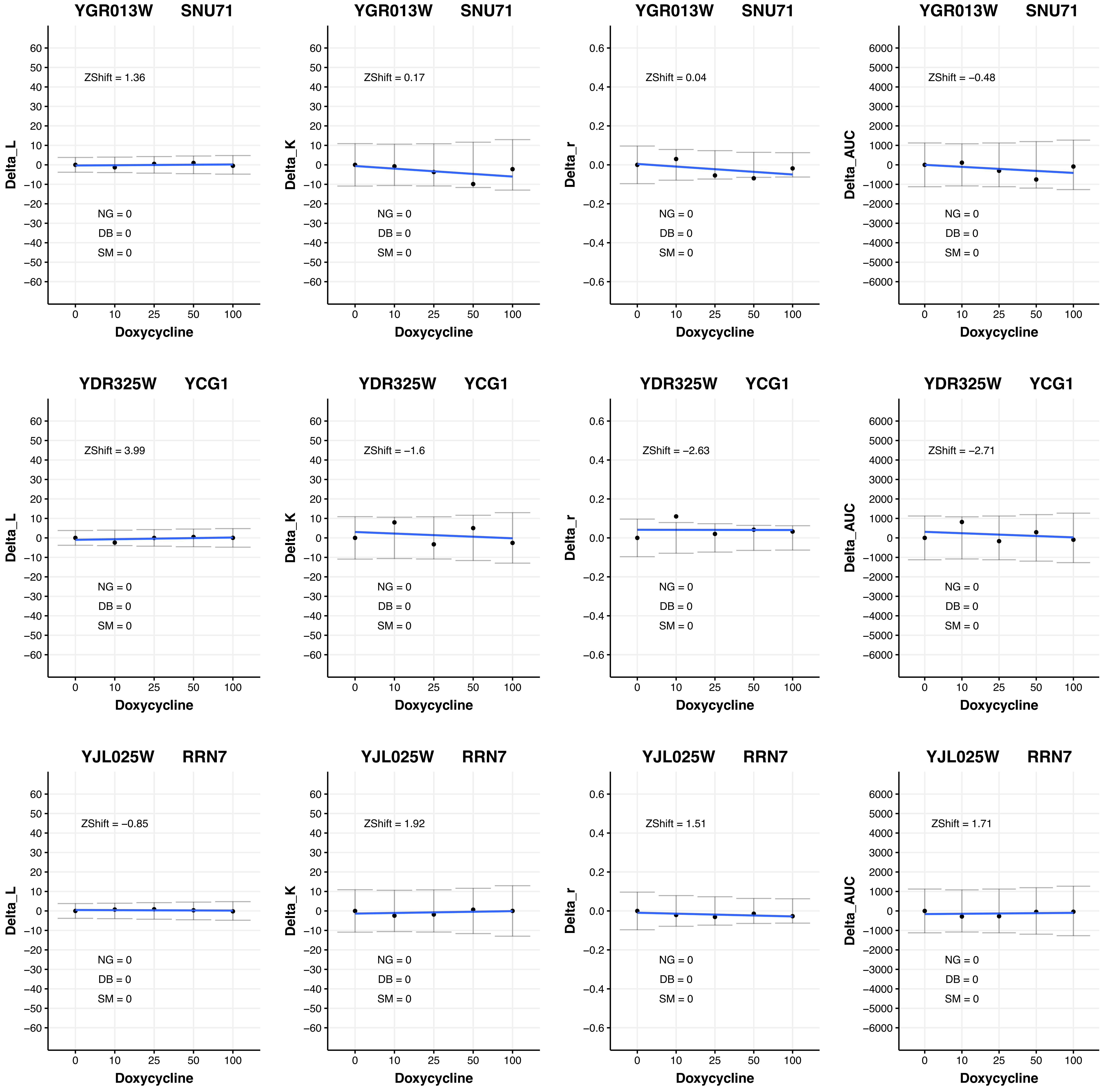
YGR013W SNU71
ZShift = 1.36
Delta_L
NG = 0
DB = 0
SM = 0
Doxycycline
YGR013W SNU71
ZShift = 0.17
Delta_K
NG = 0
DB = 0
SM = 0
Doxycycline
YGR013W SNU71
ZShift = 0.04
Delta_r
NG = 0
DB = 0
SM = 0
Doxycycline
YGR013W SNU71
ZShift = −0.48
Delta_AUC
NG = 0
DB = 0
SM = 0
Doxycycline
YDR325W YCG1
ZShift = 3.99
Delta_L
NG = 0
DB = 0
SM = 0
Doxycycline
YDR325W YCG1
ZShift = −1.6
Delta_K
NG = 0
DB = 0
SM = 0
Doxycycline
YDR325W YCG1
ZShift = −2.63
Delta_r
NG = 0
DB = 0
SM = 0
Doxycycline
YDR325W YCG1
ZShift = −2.71
Delta_AUC
NG = 0
DB = 0
SM = 0
Doxycycline
YJL025W RRN7
ZShift = −0.85
Delta_L
NG = 0
DB = 0
SM = 0
Doxycycline
YJL025W RRN7
ZShift = 1.92
Delta_K
NG = 0
DB = 0
SM = 0
Doxycycline
YJL025W RRN7
ZShift = 1.51
Delta_r
NG = 0
DB = 0
SM = 0
Doxycycline
YJL025W RRN7
ZShift = 1.71
Delta_AUC
NG = 0
DB = 0
SM = 0
Doxycycline

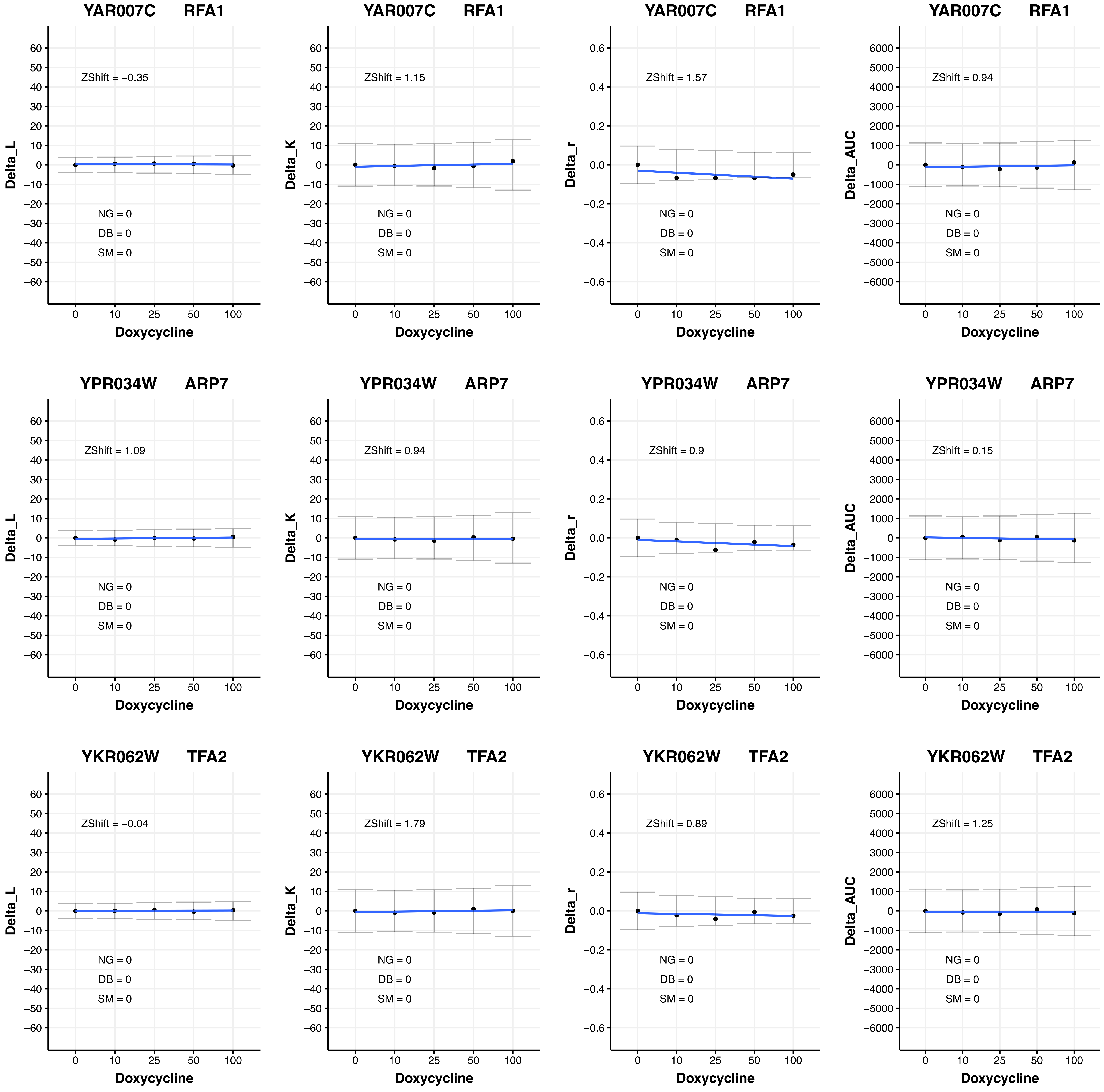
YAR007C RFA1
ZShift = −0.35
Delta_L
NG = 0
DB = 0
SM = 0
Doxycycline
YAR007C RFA1
ZShift = 1.15
Delta_K
NG = 0
DB = 0
SM = 0
Doxycycline
YAR007C RFA1
ZShift = 1.57
Delta_r
NG = 0
DB = 0
SM = 0
Doxycycline
YAR007C RFA1
ZShift = 0.94
Delta_AUC
NG = 0
DB = 0
SM = 0
Doxycycline
YPR034W ARP7
ZShift = 1.09
Delta_L
NG = 0
DB = 0
SM = 0
Doxycycline
YPR034W ARP7
ZShift = 0.94
Delta_K
NG = 0
DB = 0
SM = 0
Doxycycline
YPR034W ARP7
ZShift = 0.9
Delta_r
NG = 0
DB = 0
SM = 0
Doxycycline
YPR034W ARP7
ZShift = 0.15
Delta_AUC
NG = 0
DB = 0
SM = 0
Doxycycline
YKR062W TFA2
ZShift = −0.04
Delta_L
NG = 0
DB = 0
SM = 0
Doxycycline
YKR062W TFA2
ZShift = 1.79
Delta_K
NG = 0
DB = 0
SM = 0
Doxycycline
YKR062W TFA2
ZShift = 0.89
Delta_r
NG = 0
DB = 0
SM = 0
Doxycycline
YKR062W TFA2
ZShift = 1.25
Delta_AUC
NG = 0
DB = 0
SM = 0
Doxycycline

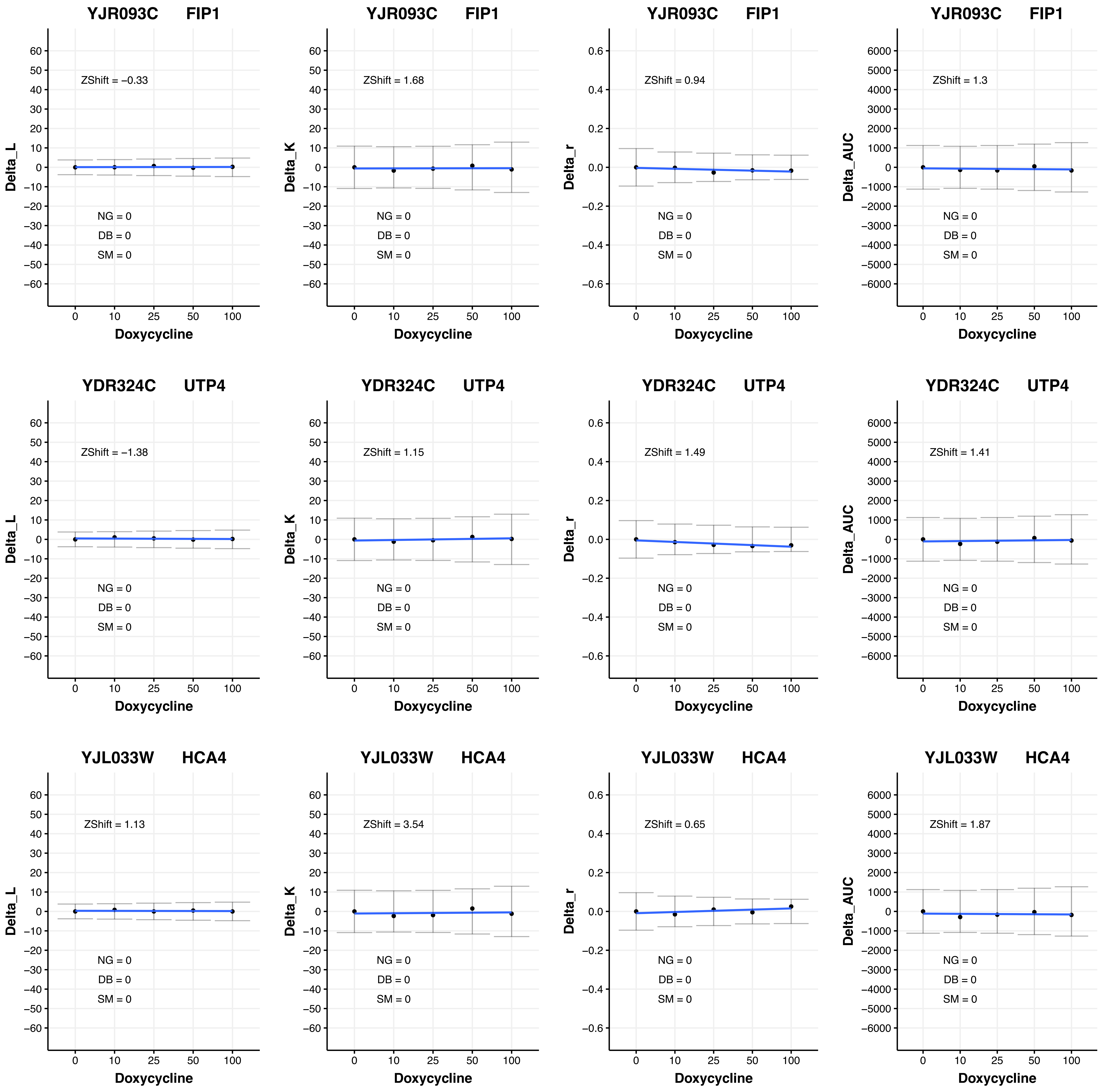
YJR093C FIP1
Delta_L
ZShift = −0.33
NG = 0
DB = 0
SM = 0
Doxycycline
YJR093C FIP1
Delta_K
ZShift = 1.68
NG = 0
DB = 0
SM = 0
Doxycycline
YJR093C FIP1
Delta_r
ZShift = 0.94
NG = 0
DB = 0
SM = 0
Doxycycline
YJR093C FIP1
Delta_AUC
ZShift = 1.3
NG = 0
DB = 0
SM = 0
Doxycycline
YDR324C UTP4
Delta_L
ZShift = −1.38
NG = 0
DB = 0
SM = 0
Doxycycline
YDR324C UTP4
Delta_K
ZShift = 1.15
NG = 0
DB = 0
SM = 0
Doxycycline
YDR324C UTP4
Delta_r
ZShift = 1.49
NG = 0
DB = 0
SM = 0
Doxycycline
YDR324C UTP4
Delta_AUC
ZShift = 1.41
NG = 0
DB = 0
SM = 0
Doxycycline
YJL033W HCA4
Delta_L
ZShift = 1.13
NG = 0
DB = 0
SM = 0
Doxycycline
YJL033W HCA4
Delta_K
ZShift = 3.54
NG = 0
DB = 0
SM = 0
Doxycycline
YJL033W HCA4
Delta_r
ZShift = 0.65
NG = 0
DB = 0
SM = 0
Doxycycline
YJL033W HCA4
Delta_AUC
ZShift = 1.87
NG = 0
DB = 0
SM = 0
Doxycycline

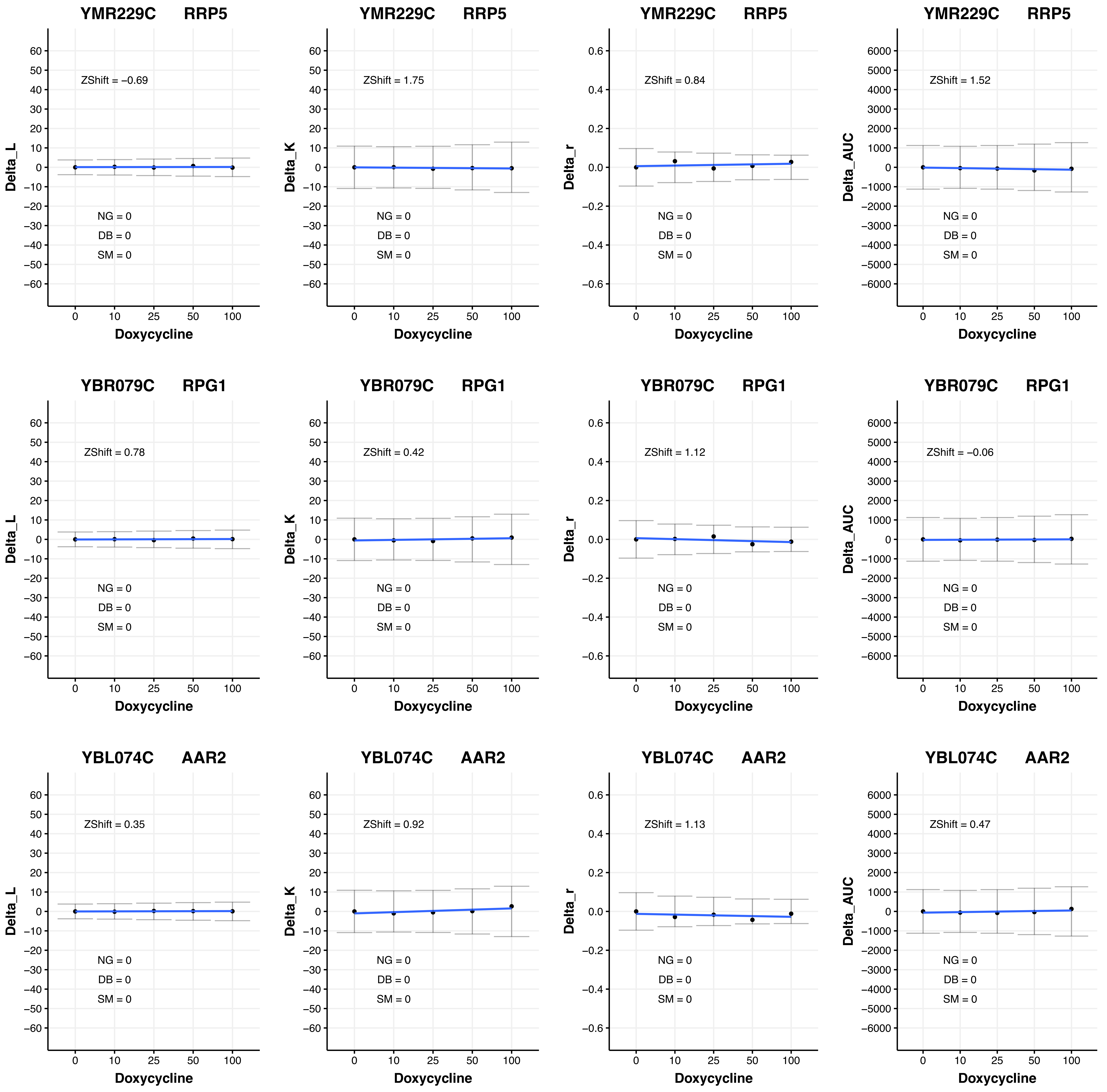
YMR229C RRP5
ZShift = −0.69
ZShift = 1.75
ZShift = 0.84
ZShift = 1.52
YBR079C RPG1
ZShift = 0.78
ZShift = 0.42
ZShift = 1.12
ZShift = −0.06
YBL074C AAR2
ZShift = 0.35
ZShift = 0.92
ZShift = 1.13
ZShift = 0.47
NG = 0
DB = 0
SM = 0
Delta_L
Delta_K
Delta_r
Delta_AUC
Doxycycline
0 10 25 50 100

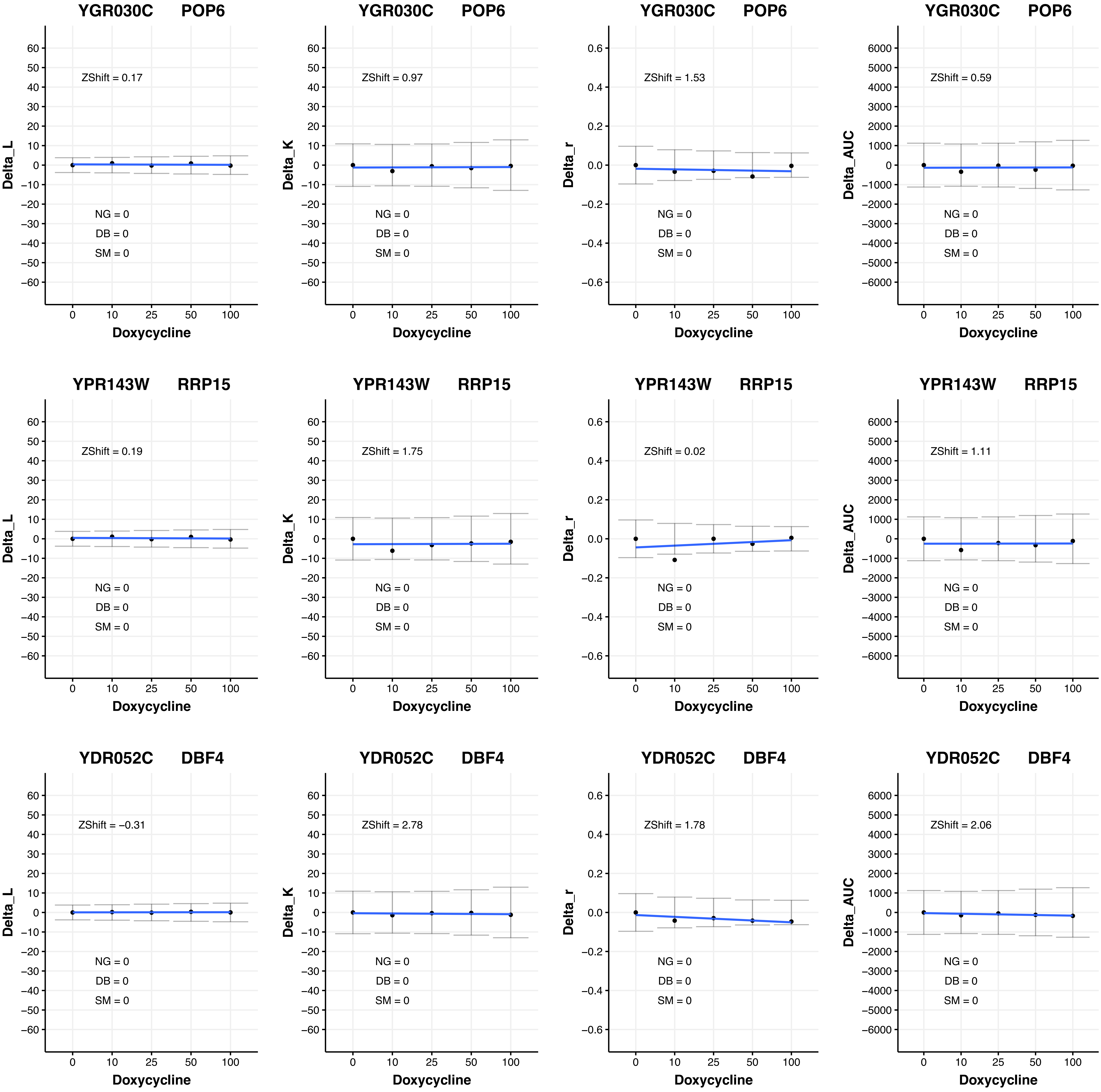
YGR030C POP6
ZShift = 0.17
NG = 0
DB = 0
SM = 0
Delta_L
Doxycycline
YGR030C POP6
ZShift = 0.97
NG = 0
DB = 0
SM = 0
Delta_K
Doxycycline
YGR030C POP6
ZShift = 1.53
NG = 0
DB = 0
SM = 0
Delta_r
Doxycycline
YGR030C POP6
ZShift = 0.59
NG = 0
DB = 0
SM = 0
Delta_AUC
Doxycycline
YPR143W RRP15
ZShift = 0.19
NG = 0
DB = 0
SM = 0
Delta_L
Doxycycline
YPR143W RRP15
ZShift = 1.75
NG = 0
DB = 0
SM = 0
Delta_K
Doxycycline
YPR143W RRP15
ZShift = 0.02
NG = 0
DB = 0
SM = 0
Delta_r
Doxycycline
YPR143W RRP15
ZShift = 1.11
NG = 0
DB = 0
SM = 0
Delta_AUC
Doxycycline
YDR052C DBF4
ZShift = −0.31
NG = 0
DB = 0
SM = 0
Delta_L
Doxycycline
YDR052C DBF4
ZShift = 2.78
NG = 0
DB = 0
SM = 0
Delta_K
Doxycycline
YDR052C DBF4
ZShift = 1.78
NG = 0
DB = 0
SM = 0
Delta_r
Doxycycline
YDR052C DBF4
ZShift = 2.06
NG = 0
DB = 0
SM = 0
Delta_AUC
Doxycycline
0
10
25
50
100

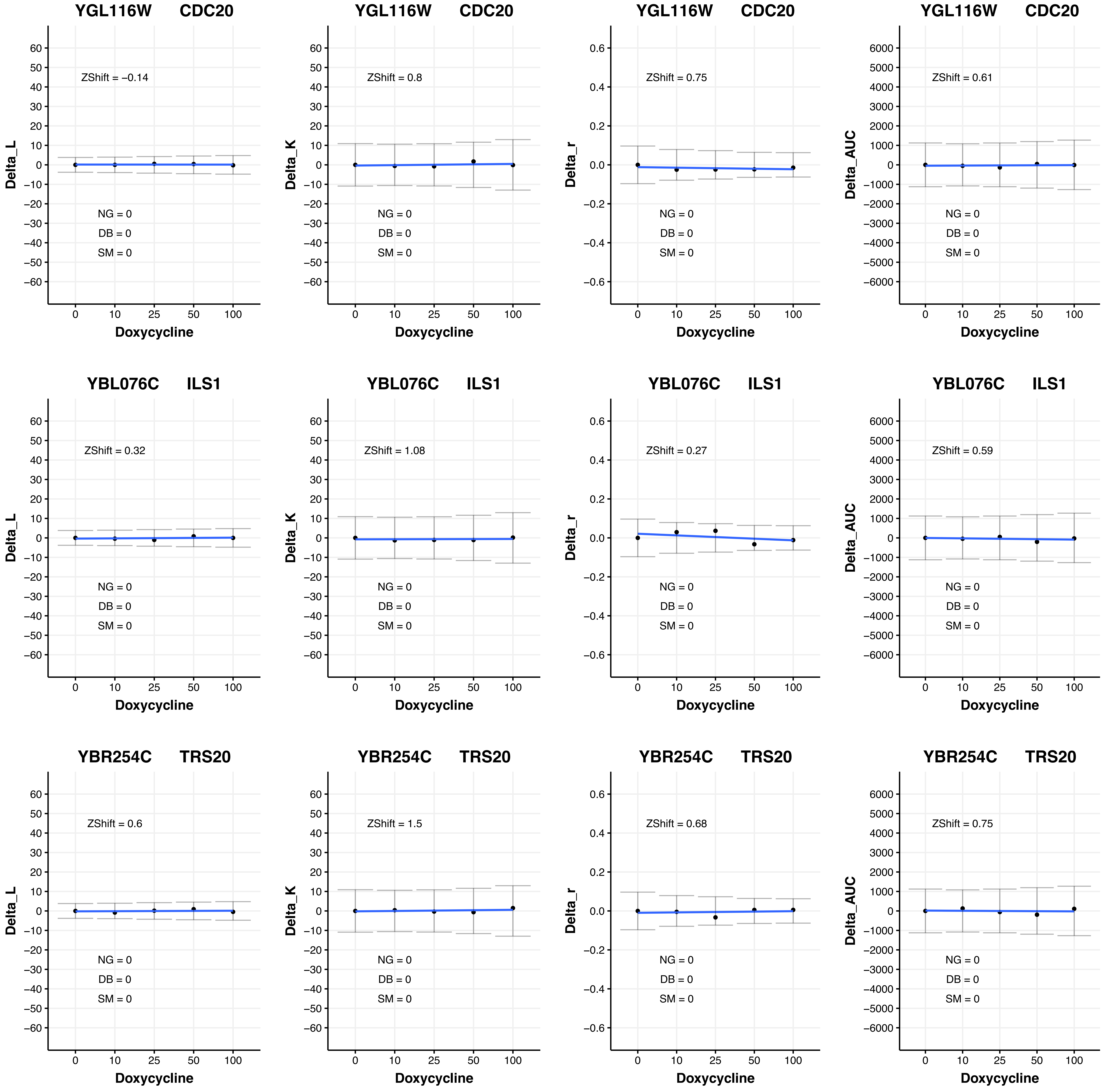
YGL116W CDC20
ZShift = −0.14
NG = 0
DB = 0
SM = 0
Delta_L
Doxycycline
YGL116W CDC20
ZShift = 0.8
NG = 0
DB = 0
SM = 0
Delta_K
Doxycycline
YGL116W CDC20
ZShift = 0.75
NG = 0
DB = 0
SM = 0
Delta_r
Doxycycline
YGL116W CDC20
ZShift = 0.61
NG = 0
DB = 0
SM = 0
Delta_AUC
Doxycycline
YBL076C ILS1
ZShift = 0.32
NG = 0
DB = 0
SM = 0
Delta_L
Doxycycline
YBL076C ILS1
ZShift = 1.08
NG = 0
DB = 0
SM = 0
Delta_K
Doxycycline
YBL076C ILS1
ZShift = 0.27
NG = 0
DB = 0
SM = 0
Delta_r
Doxycycline
YBL076C ILS1
ZShift = 0.59
NG = 0
DB = 0
SM = 0
Delta_AUC
Doxycycline
YBR254C TRS20
ZShift = 0.6
NG = 0
DB = 0
SM = 0
Delta_L
Doxycycline
YBR254C TRS20
ZShift = 1.5
NG = 0
DB = 0
SM = 0
Delta_K
Doxycycline
YBR254C TRS20
ZShift = 0.68
NG = 0
DB = 0
SM = 0
Delta_r
Doxycycline
YBR254C TRS20
ZShift = 0.75
NG = 0
DB = 0
SM = 0
Delta_AUC
Doxycycline
0 10 25 50 100

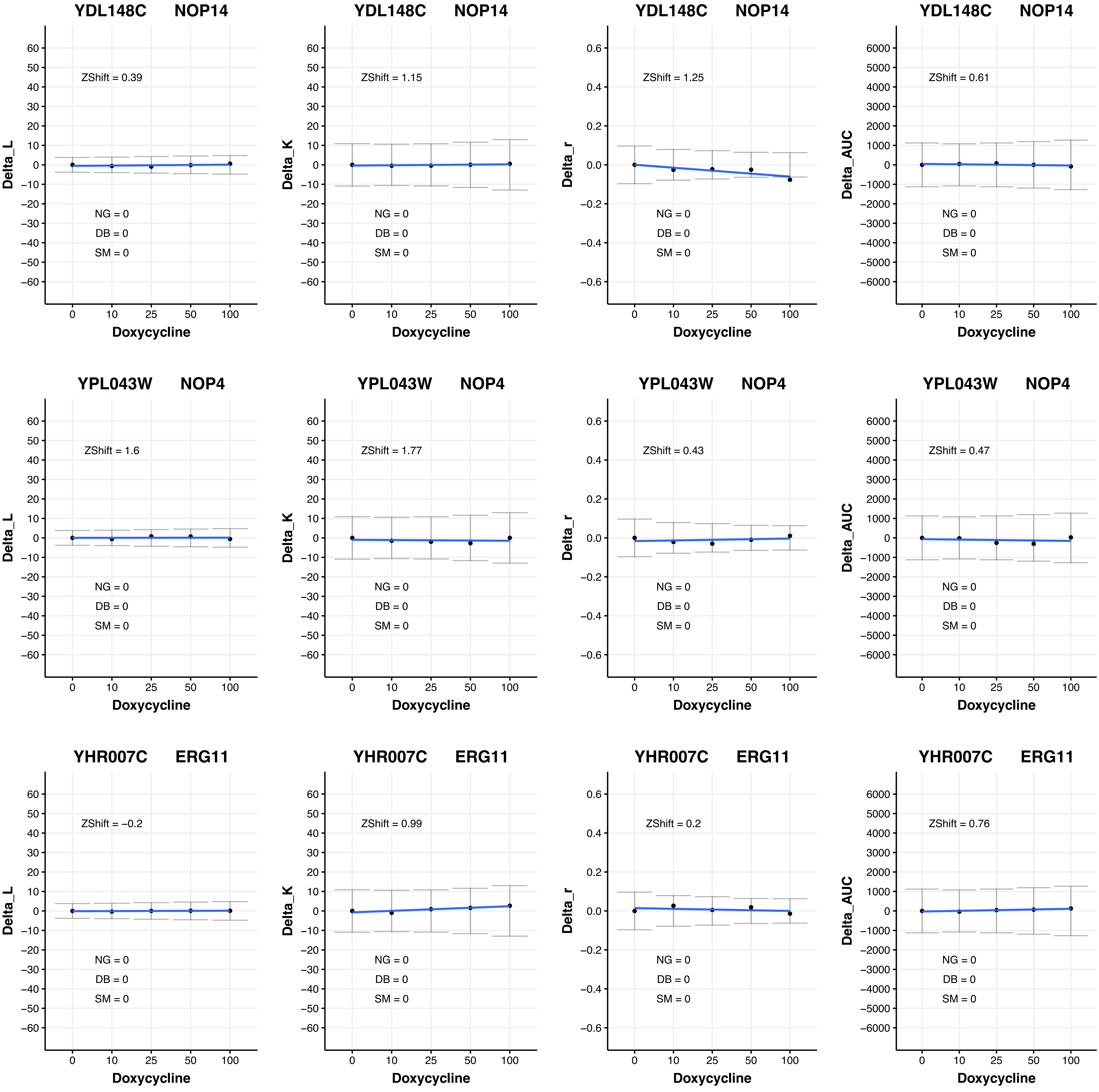
YDL148C NOP14
Delta_L
ZShift = 0.39
NG = 0
DB = 0
SM = 0
Doxycycline
YDL148C NOP14
Delta_K
ZShift = 1.15
NG = 0
DB = 0
SM = 0
Doxycycline
YDL148C NOP14
Delta_r
ZShift = 1.25
NG = 0
DB = 0
SM = 0
Doxycycline
YDL148C NOP14
Delta_AUC
ZShift = 0.61
NG = 0
DB = 0
SM = 0
Doxycycline
YPL043W NOP4
Delta_L
ZShift = 1.6
NG = 0
DB = 0
SM = 0
Doxycycline
YPL043W NOP4
Delta_K
ZShift = 1.77
NG = 0
DB = 0
SM = 0
Doxycycline
YPL043W NOP4
Delta_r
ZShift = 0.43
NG = 0
DB = 0
SM = 0
Doxycycline
YPL043W NOP4
Delta_AUC
ZShift = 0.47
NG = 0
DB = 0
SM = 0
Doxycycline
YHR007C ERG11
Delta_L
ZShift = −0.2
NG = 0
DB = 0
SM = 0
Doxycycline
YHR007C ERG11
Delta_K
ZShift = 0.99
NG = 0
DB = 0
SM = 0
Doxycycline
YHR007C ERG11
Delta_r
ZShift = 0.2
NG = 0
DB = 0
SM = 0
Doxycycline
YHR007C ERG11
Delta_AUC
ZShift = 0.76
NG = 0
DB = 0
SM = 0
Doxycycline
0 10 25 50 100

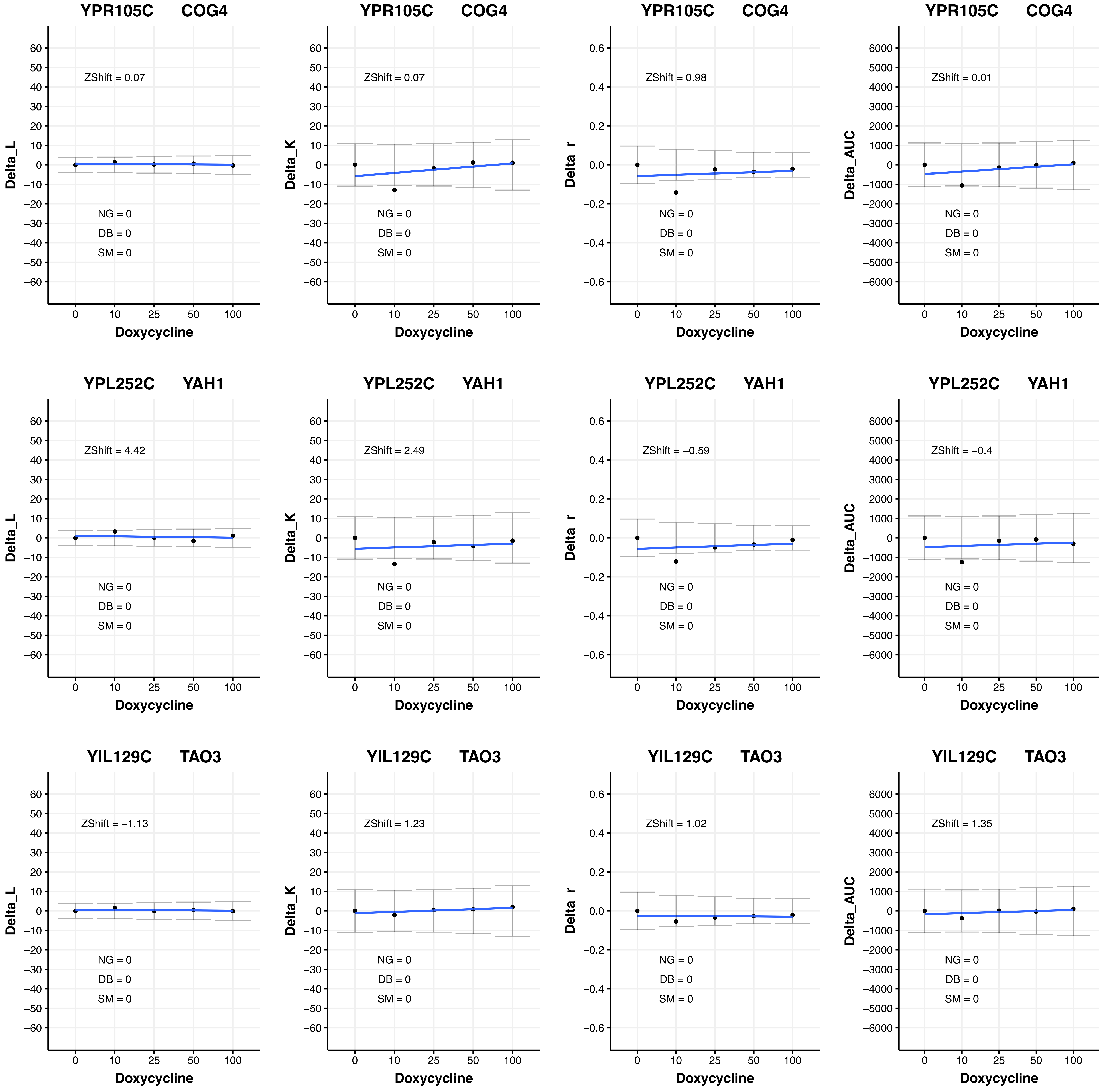

YPR105C COG4
ZShift = 0.07
ZShift = 0.07
ZShift = 0.98
ZShift = 0.01
YPL252C YAH1
ZShift = 4.42
ZShift = 2.49
ZShift = −0.59
ZShift = −0.4
YIL129C TAO3
ZShift = −1.13
ZShift = 1.23
ZShift = 1.02
ZShift = 1.35
NG = 0
DB = 0
SM = 0
Delta_L
Delta_K
Delta_r
Delta_AUC
Doxycycline

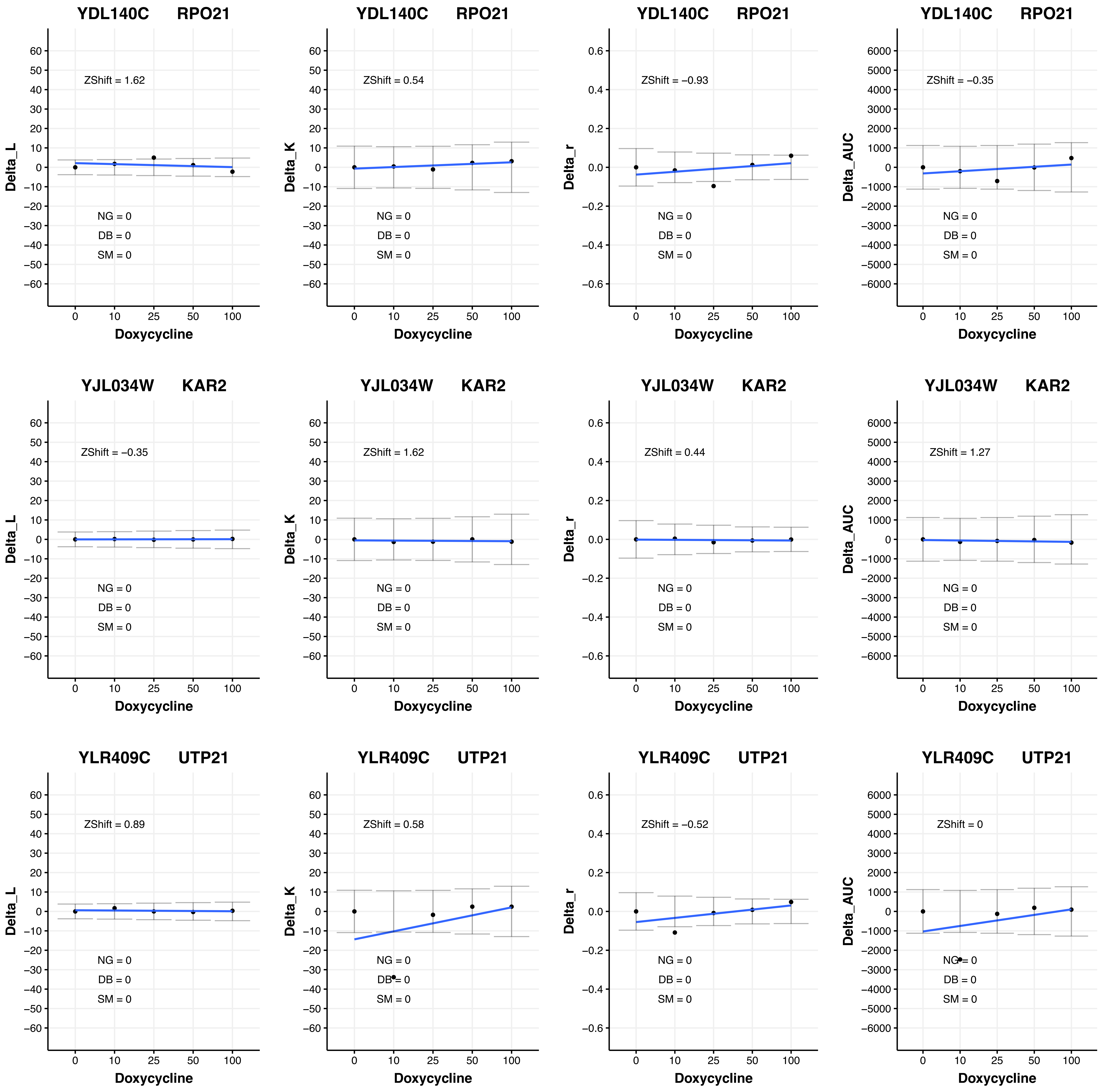
YDL140C RPO21
ZShift = 1.62
ZShift = 0.54
ZShift = −0.93
ZShift = −0.35
YJL034W KAR2
ZShift = −0.35
ZShift = 1.62
ZShift = 0.44
ZShift = 1.27
YLR409C UTP21
ZShift = 0.89
ZShift = 0.58
ZShift = −0.52
ZShift = 0
NG = 0
DB = 0
SM = 0
Delta_L
Delta_K
Delta_r
Delta_AUC
Doxycycline

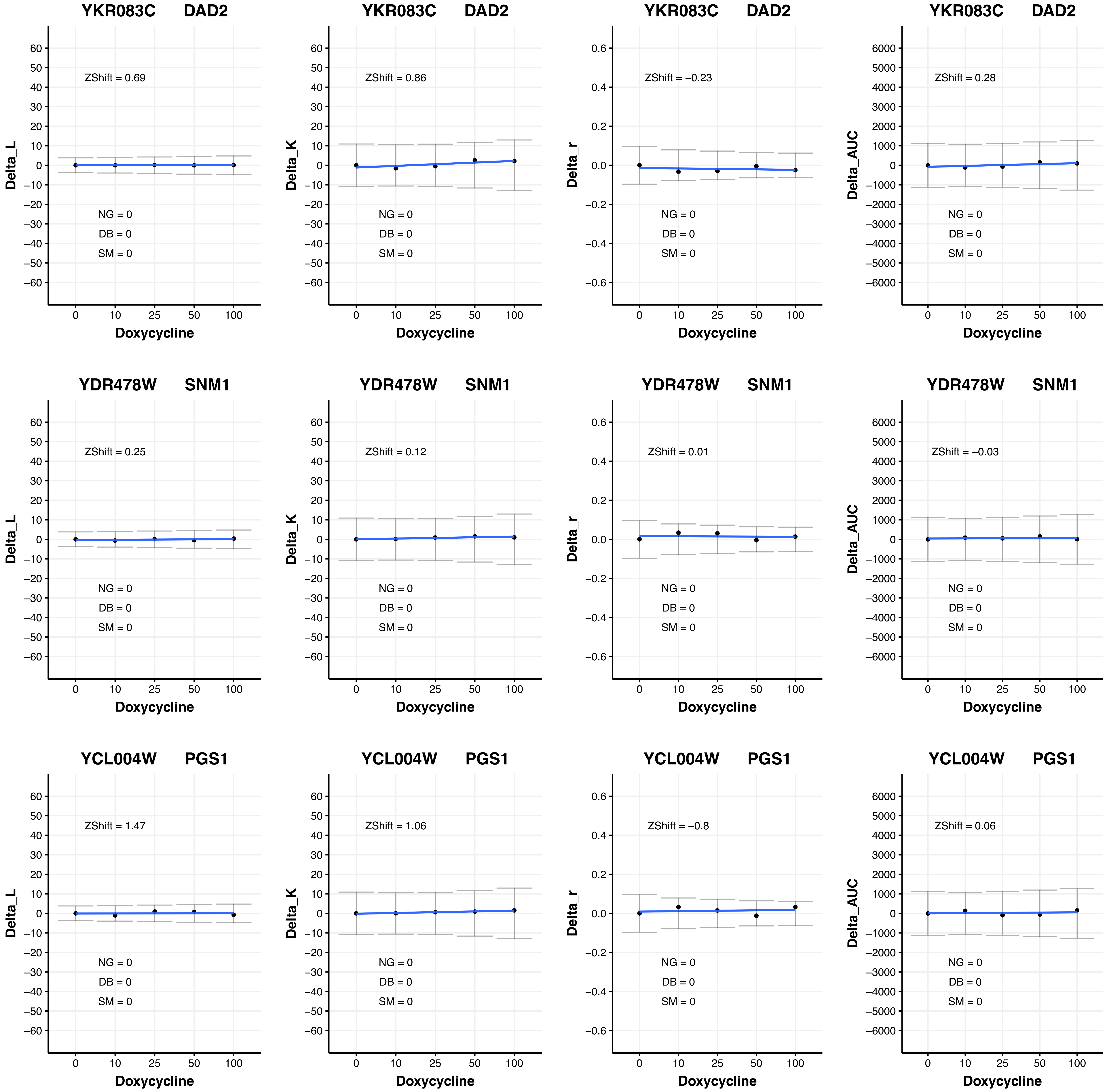

YKR083C DAD2
ZShift = 0.69
NG = 0
DB = 0
SM = 0
Delta_L
Doxycycline
YKR083C DAD2
ZShift = 0.86
NG = 0
DB = 0
SM = 0
Delta_K
Doxycycline
YKR083C DAD2
ZShift = −0.23
NG = 0
DB = 0
SM = 0
Delta_r
Doxycycline
YKR083C DAD2
ZShift = 0.28
NG = 0
DB = 0
SM = 0
Delta_AUC
Doxycycline
YDR478W SNM1
ZShift = 0.25
NG = 0
DB = 0
SM = 0
Delta_L
Doxycycline
YDR478W SNM1
ZShift = 0.12
NG = 0
DB = 0
SM = 0
Delta_K
Doxycycline
YDR478W SNM1
ZShift = 0.01
NG = 0
DB = 0
SM = 0
Delta_r
Doxycycline
YDR478W SNM1
ZShift = −0.03
NG = 0
DB = 0
SM = 0
Delta_AUC
Doxycycline
YCL004W PGS1
ZShift = 1.47
NG = 0
DB = 0
SM = 0
Delta_L
Doxycycline
YCL004W PGS1
ZShift = 1.06
NG = 0
DB = 0
SM = 0
Delta_K
Doxycycline
YCL004W PGS1
ZShift = −0.8
NG = 0
DB = 0
SM = 0
Delta_r
Doxycycline
YCL004W PGS1
ZShift = 0.06
NG = 0
DB = 0
SM = 0
Delta_AUC
Doxycycline

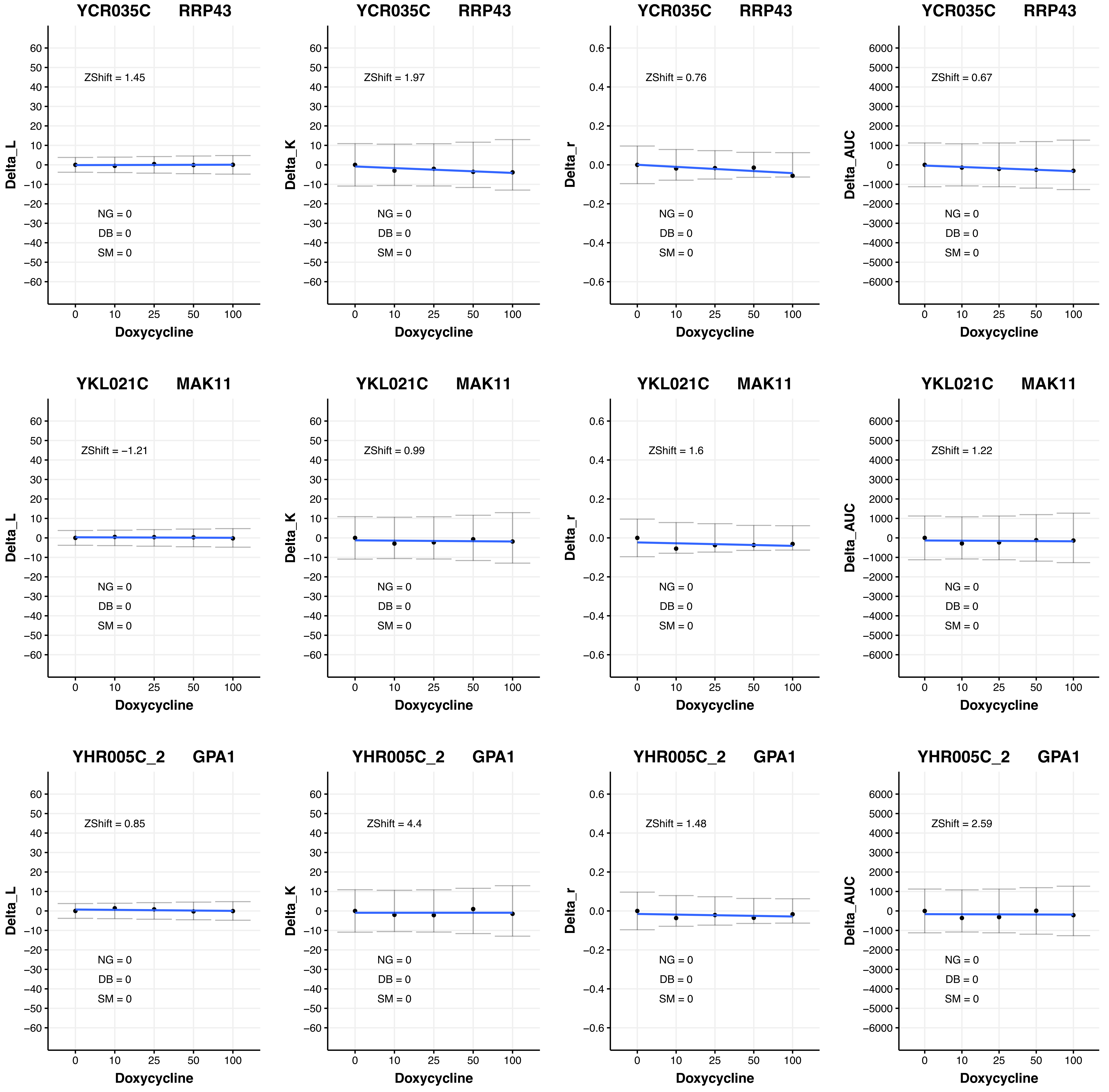
YCR035C RRP43
ZShift = 1.45
Delta_L
NG = 0
DB = 0
SM = 0
Doxycycline
YCR035C RRP43
ZShift = 1.97
Delta_K
NG = 0
DB = 0
SM = 0
Doxycycline
YCR035C RRP43
ZShift = 0.76
Delta_r
NG = 0
DB = 0
SM = 0
Doxycycline
YCR035C RRP43
ZShift = 0.67
Delta_AUC
NG = 0
DB = 0
SM = 0
Doxycycline
YKL021C MAK11
ZShift = −1.21
Delta_L
NG = 0
DB = 0
SM = 0
Doxycycline
YKL021C MAK11
ZShift = 0.99
Delta_K
NG = 0
DB = 0
SM = 0
Doxycycline
YKL021C MAK11
ZShift = 1.6
Delta_r
NG = 0
DB = 0
SM = 0
Doxycycline
YKL021C MAK11
ZShift = 1.22
Delta_AUC
NG = 0
DB = 0
SM = 0
Doxycycline
YHR005C_2 GPA1
ZShift = 0.85
Delta_L
NG = 0
DB = 0
SM = 0
Doxycycline
YHR005C_2 GPA1
ZShift = 4.4
Delta_K
NG = 0
DB = 0
SM = 0
Doxycycline
YHR005C_2 GPA1
ZShift = 1.48
Delta_r
NG = 0
DB = 0
SM = 0
Doxycycline
YHR005C_2 GPA1
ZShift = 2.59
Delta_AUC
NG = 0
DB = 0
SM = 0
Doxycycline

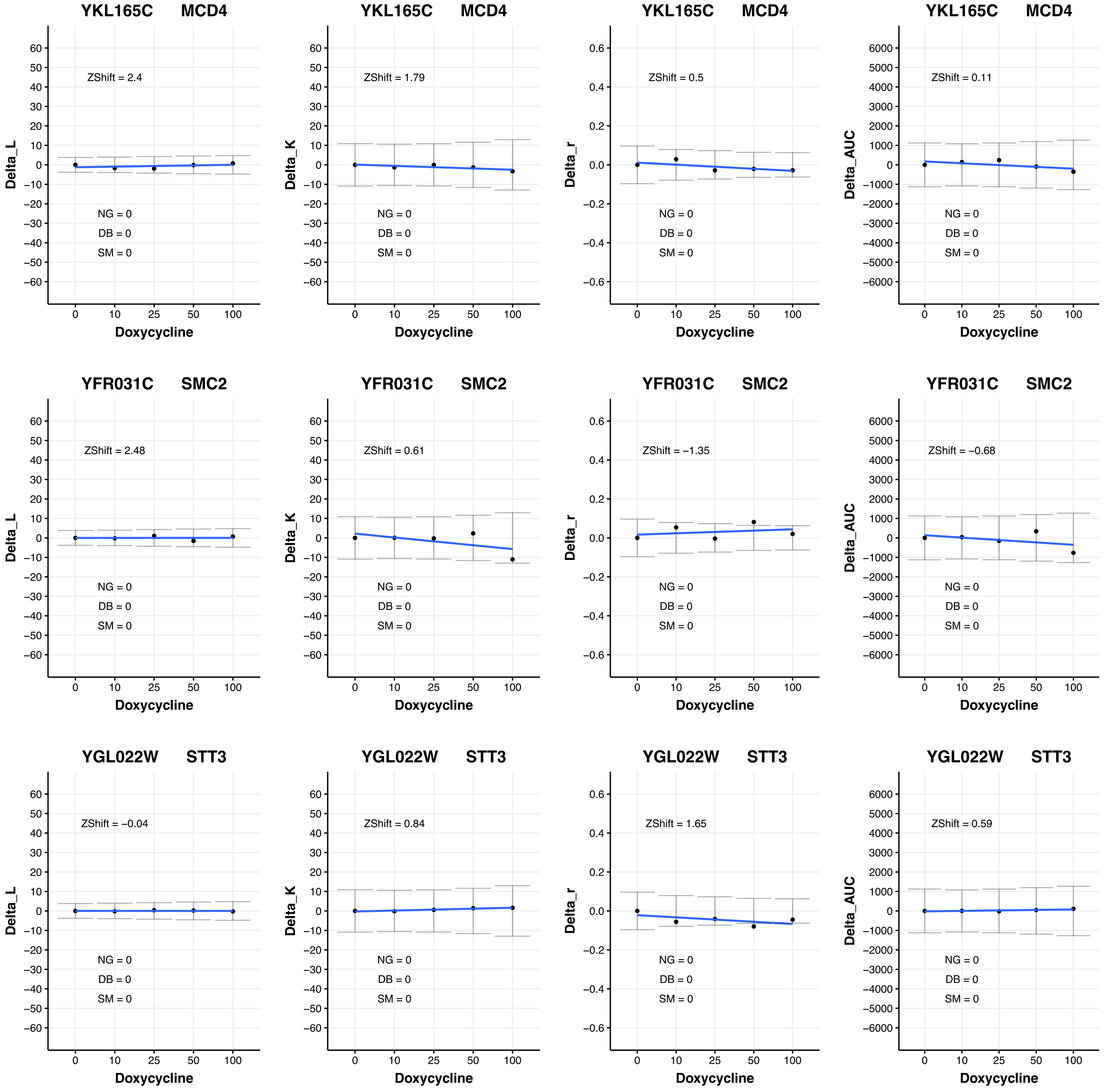
YKL165C MCD4
ZShift = 2.4
NG = 0
DB = 0
SM = 0
Delta_L
Doxycycline
YKL165C MCD4
ZShift = 1.79
NG = 0
DB = 0
SM = 0
Delta_K
Doxycycline
YKL165C MCD4
ZShift = 0.5
NG = 0
DB = 0
SM = 0
Delta_r
Doxycycline
YKL165C MCD4
ZShift = 0.11
NG = 0
DB = 0
SM = 0
Delta_AUC
Doxycycline
YFR031C SMC2
ZShift = 2.48
NG = 0
DB = 0
SM = 0
Delta_L
Doxycycline
YFR031C SMC2
ZShift = 0.61
NG = 0
DB = 0
SM = 0
Delta_K
Doxycycline
YFR031C SMC2
ZShift = −1.35
NG = 0
DB = 0
SM = 0
Delta_r
Doxycycline
YFR031C SMC2
ZShift = −0.68
NG = 0
DB = 0
SM = 0
Delta_AUC
Doxycycline
YGL022W STT3
ZShift = −0.04
NG = 0
DB = 0
SM = 0
Delta_L
Doxycycline
YGL022W STT3
ZShift = 0.84
NG = 0
DB = 0
SM = 0
Delta_K
Doxycycline
YGL022W STT3
ZShift = 1.65
NG = 0
DB = 0
SM = 0
Delta_r
Doxycycline
YGL022W STT3
ZShift = 0.59
NG = 0
DB = 0
SM = 0
Delta_AUC
Doxycycline
0 10 25 50 100

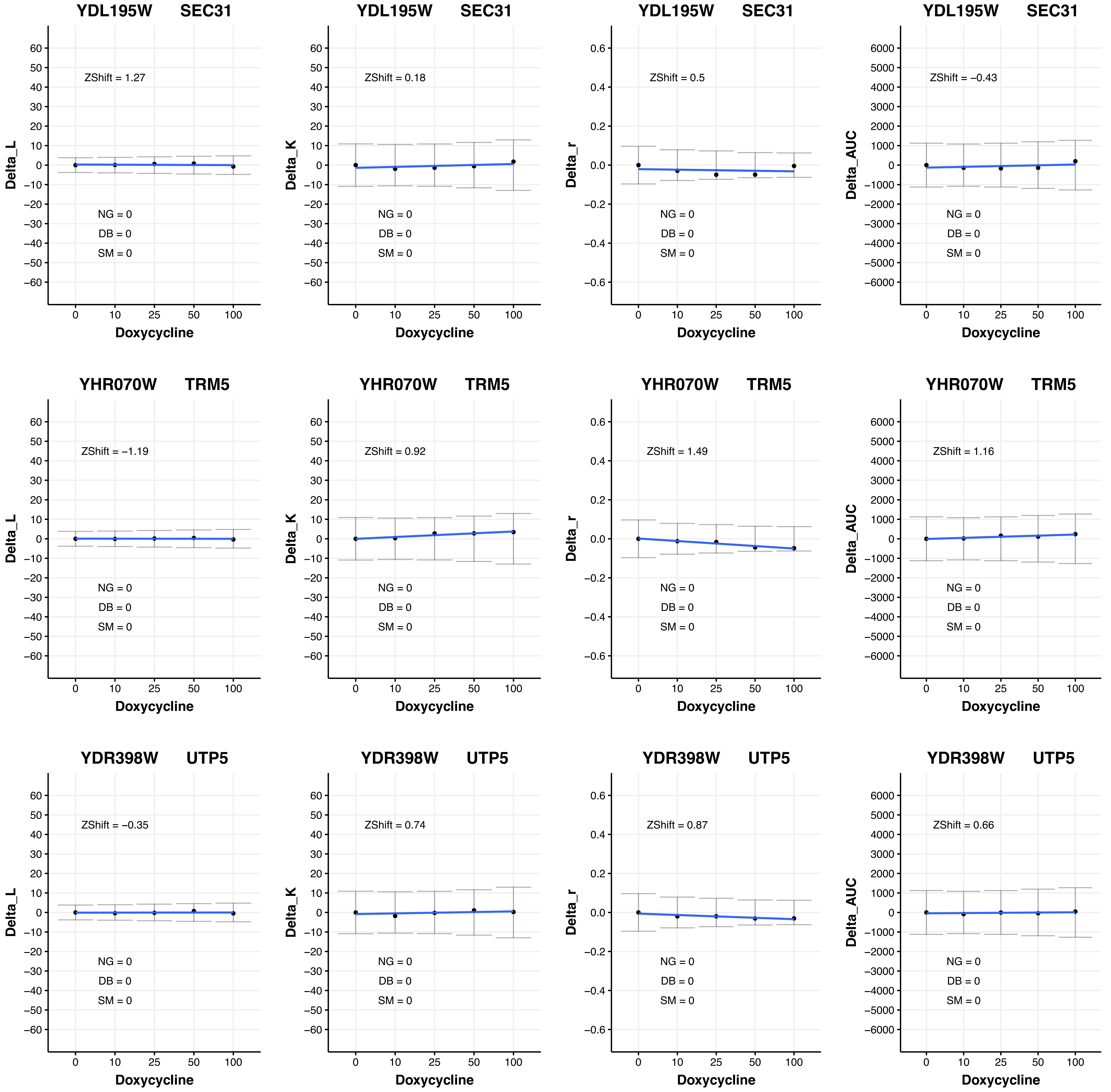
YDL195W SEC31
Delta_L
ZShift = 1.27
NG = 0
DB = 0
SM = 0
Doxycycline
YDL195W SEC31
Delta_K
ZShift = 0.18
NG = 0
DB = 0
SM = 0
Doxycycline
YDL195W SEC31
Delta_r
ZShift = 0.5
NG = 0
DB = 0
SM = 0
Doxycycline
YDL195W SEC31
Delta_AUC
ZShift = −0.43
NG = 0
DB = 0
SM = 0
Doxycycline
YHR070W TRM5
Delta_L
ZShift = −1.19
NG = 0
DB = 0
SM = 0
Doxycycline
YHR070W TRM5
Delta_K
ZShift = 0.92
NG = 0
DB = 0
SM = 0
Doxycycline
YHR070W TRM5
Delta_r
ZShift = 1.49
NG = 0
DB = 0
SM = 0
Doxycycline
YHR070W TRM5
Delta_AUC
ZShift = 1.16
NG = 0
DB = 0
SM = 0
Doxycycline
YDR398W UTP5
Delta_L
ZShift = −0.35
NG = 0
DB = 0
SM = 0
Doxycycline
YDR398W UTP5
Delta_K
ZShift = 0.74
NG = 0
DB = 0
SM = 0
Doxycycline
YDR398W UTP5
Delta_r
ZShift = 0.87
NG = 0
DB = 0
SM = 0
Doxycycline
YDR398W UTP5
Delta_AUC
ZShift = 0.66
NG = 0
DB = 0
SM = 0
Doxycycline

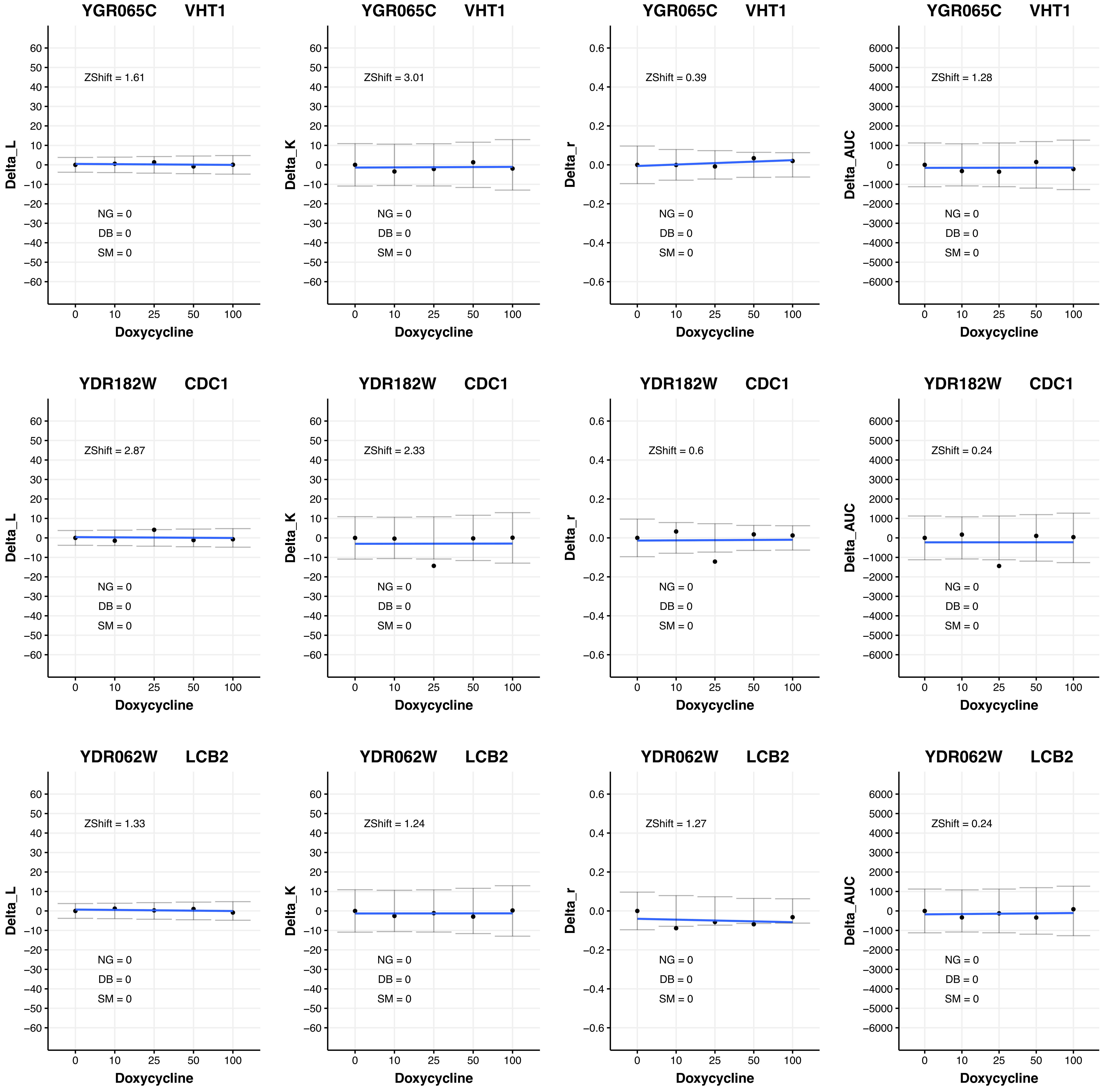

YGR065C VHT1
ZShift = 1.61
ZShift = 3.01
ZShift = 0.39
ZShift = 1.28
YDR182W CDC1
ZShift = 2.87
ZShift = 2.33
ZShift = 0.6
ZShift = 0.24
YDR062W LCB2
ZShift = 1.33
ZShift = 1.24
ZShift = 1.27
ZShift = 0.24
NG = 0
DB = 0
SM = 0
Delta_L
Delta_K
Delta_r
Delta_AUC
Doxycycline
0
10
25
50
100

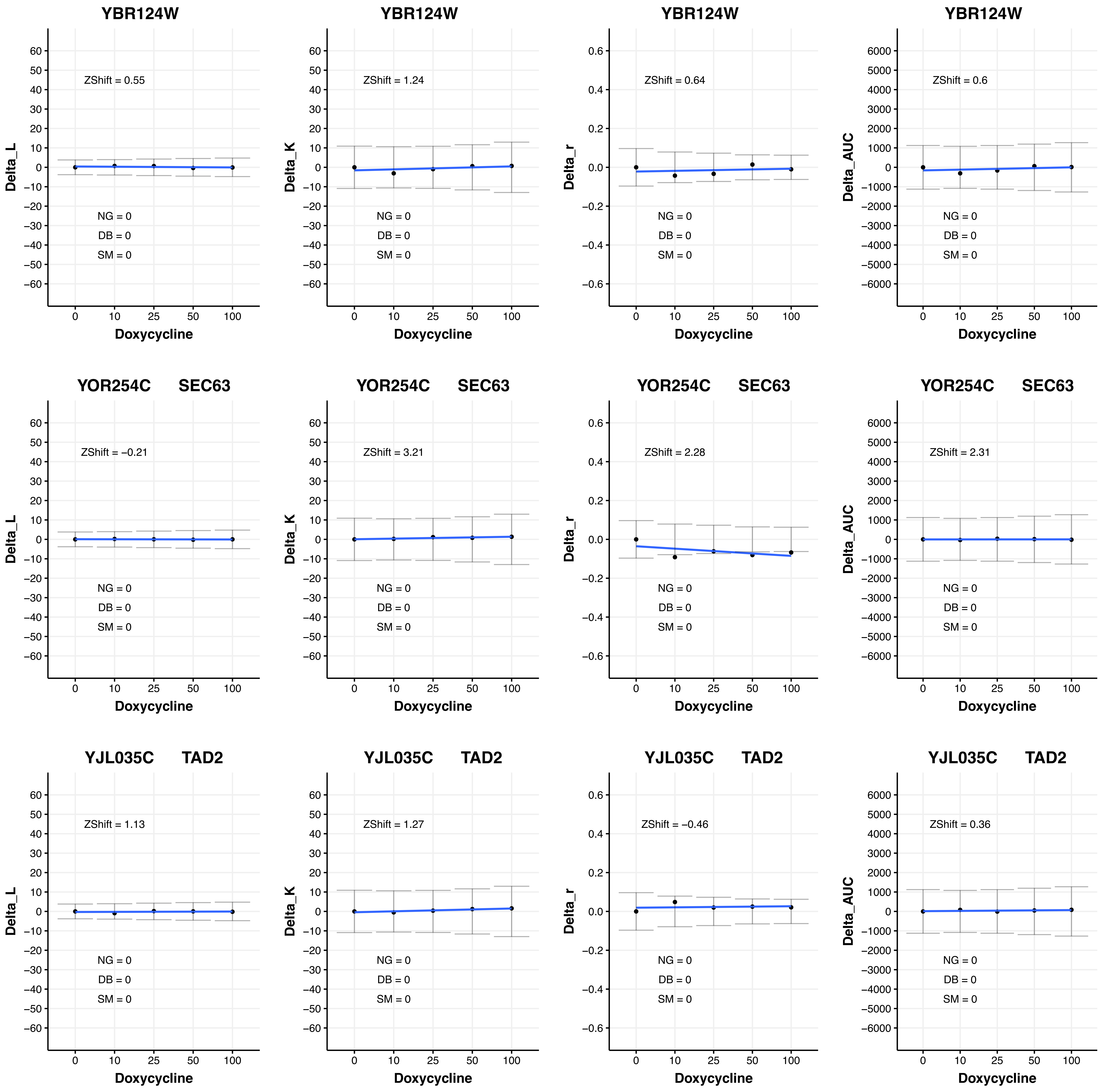
YBR124W
ZShift = 0.55
NG = 0
DB = 0
SM = 0
Delta_L
Doxycycline
YBR124W
ZShift = 1.24
NG = 0
DB = 0
SM = 0
Delta_K
Doxycycline
YBR124W
ZShift = 0.64
NG = 0
DB = 0
SM = 0
Delta_r
Doxycycline
YBR124W
ZShift = 0.6
NG = 0
DB = 0
SM = 0
Delta_AUC
Doxycycline
YOR254C SEC63
ZShift = −0.21
NG = 0
DB = 0
SM = 0
Delta_L
Doxycycline
YOR254C SEC63
ZShift = 3.21
NG = 0
DB = 0
SM = 0
Delta_K
Doxycycline
YOR254C SEC63
ZShift = 2.28
NG = 0
DB = 0
SM = 0
Delta_r
Doxycycline
YOR254C SEC63
ZShift = 2.31
NG = 0
DB = 0
SM = 0
Delta_AUC
Doxycycline
YJL035C TAD2
ZShift = 1.13
NG = 0
DB = 0
SM = 0
Delta_L
Doxycycline
YJL035C TAD2
ZShift = 1.27
NG = 0
DB = 0
SM = 0
Delta_K
Doxycycline
YJL035C TAD2
ZShift = −0.46
NG = 0
DB = 0
SM = 0
Delta_r
Doxycycline
YJL035C TAD2
ZShift = 0.36
NG = 0
DB = 0
SM = 0
Delta_AUC
Doxycycline

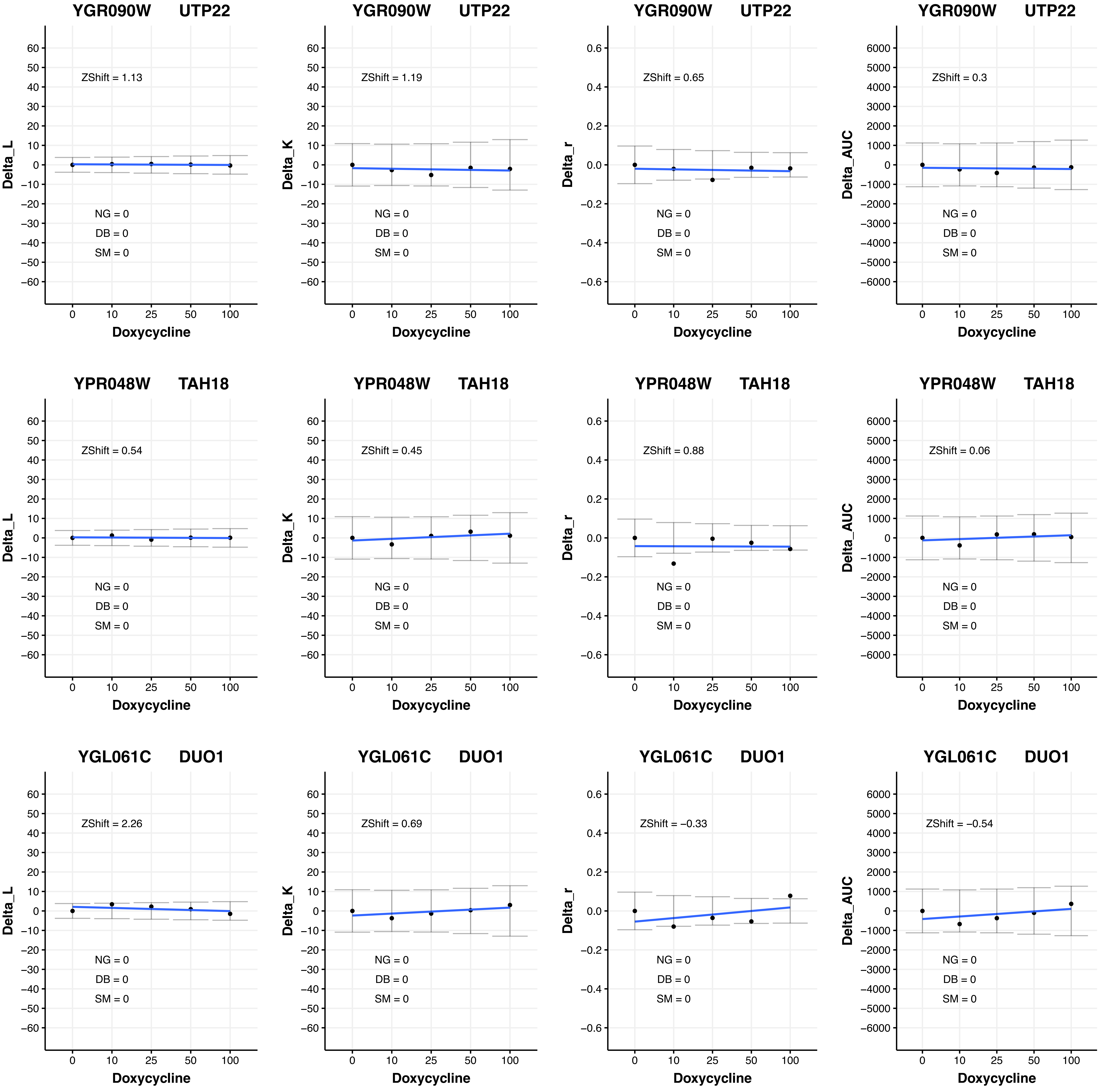

YGR090W UTP22
Delta_L
ZShift = 1.13
NG = 0
DB = 0
SM = 0
Doxycycline
YGR090W UTP22
Delta_K
ZShift = 1.19
NG = 0
DB = 0
SM = 0
Doxycycline
YGR090W UTP22
Delta_r
ZShift = 0.65
NG = 0
DB = 0
SM = 0
Doxycycline
YGR090W UTP22
Delta_AUC
ZShift = 0.3
NG = 0
DB = 0
SM = 0
Doxycycline
YPR048W TAH18
Delta_L
ZShift = 0.54
NG = 0
DB = 0
SM = 0
Doxycycline
YPR048W TAH18
Delta_K
ZShift = 0.45
NG = 0
DB = 0
SM = 0
Doxycycline
YPR048W TAH18
Delta_r
ZShift = 0.88
NG = 0
DB = 0
SM = 0
Doxycycline
YPR048W TAH18
Delta_AUC
ZShift = 0.06
NG = 0
DB = 0
SM = 0
Doxycycline
YGL061C DUO1
Delta_L
ZShift = 2.26
NG = 0
DB = 0
SM = 0
Doxycycline
YGL061C DUO1
Delta_K
ZShift = 0.69
NG = 0
DB = 0
SM = 0
Doxycycline
YGL061C DUO1
Delta_r
ZShift = −0.33
NG = 0
DB = 0
SM = 0
Doxycycline
YGL061C DUO1
Delta_AUC
ZShift = −0.54
NG = 0
DB = 0
SM = 0
Doxycycline

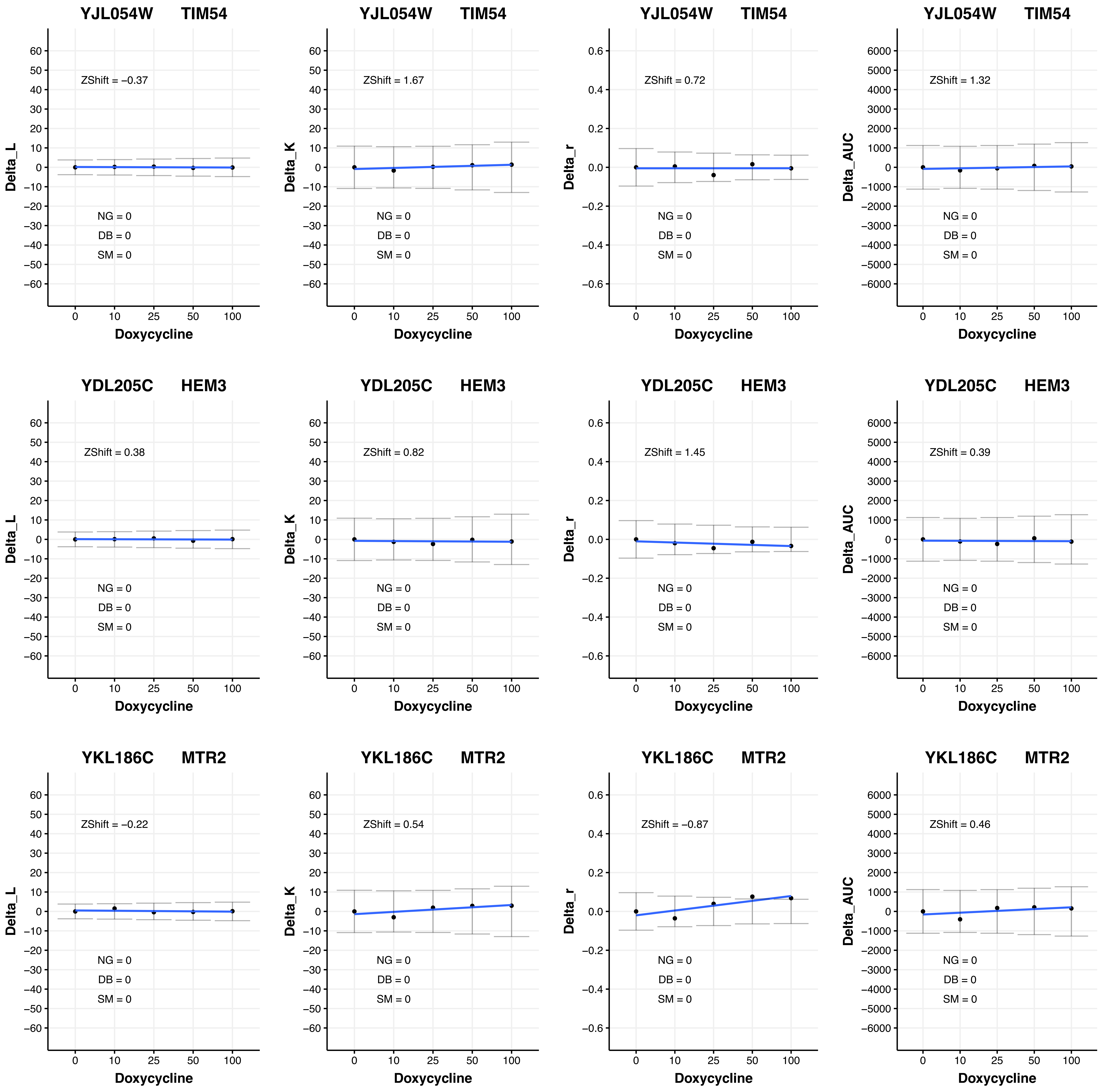

YJL054W TIM54
Delta_L
ZShift = −0.37
NG = 0
DB = 0
SM = 0
Doxycycline
YJL054W TIM54
Delta_K
ZShift = 1.67
NG = 0
DB = 0
SM = 0
Doxycycline
YJL054W TIM54
Delta_r
ZShift = 0.72
NG = 0
DB = 0
SM = 0
Doxycycline
YJL054W TIM54
Delta_AUC
ZShift = 1.32
NG = 0
DB = 0
SM = 0
Doxycycline
YDL205C HEM3
Delta_L
ZShift = 0.38
NG = 0
DB = 0
SM = 0
Doxycycline
YDL205C HEM3
Delta_K
ZShift = 0.82
NG = 0
DB = 0
SM = 0
Doxycycline
YDL205C HEM3
Delta_r
ZShift = 1.45
NG = 0
DB = 0
SM = 0
Doxycycline
YDL205C HEM3
Delta_AUC
ZShift = 0.39
NG = 0
DB = 0
SM = 0
Doxycycline
YKL186C MTR2
Delta_L
ZShift = −0.22
NG = 0
DB = 0
SM = 0
Doxycycline
YKL186C MTR2
Delta_K
ZShift = 0.54
NG = 0
DB = 0
SM = 0
Doxycycline
YKL186C MTR2
Delta_r
ZShift = −0.87
NG = 0
DB = 0
SM = 0
Doxycycline
YKL186C MTR2
Delta_AUC
ZShift = 0.46
NG = 0
DB = 0
SM = 0
Doxycycline

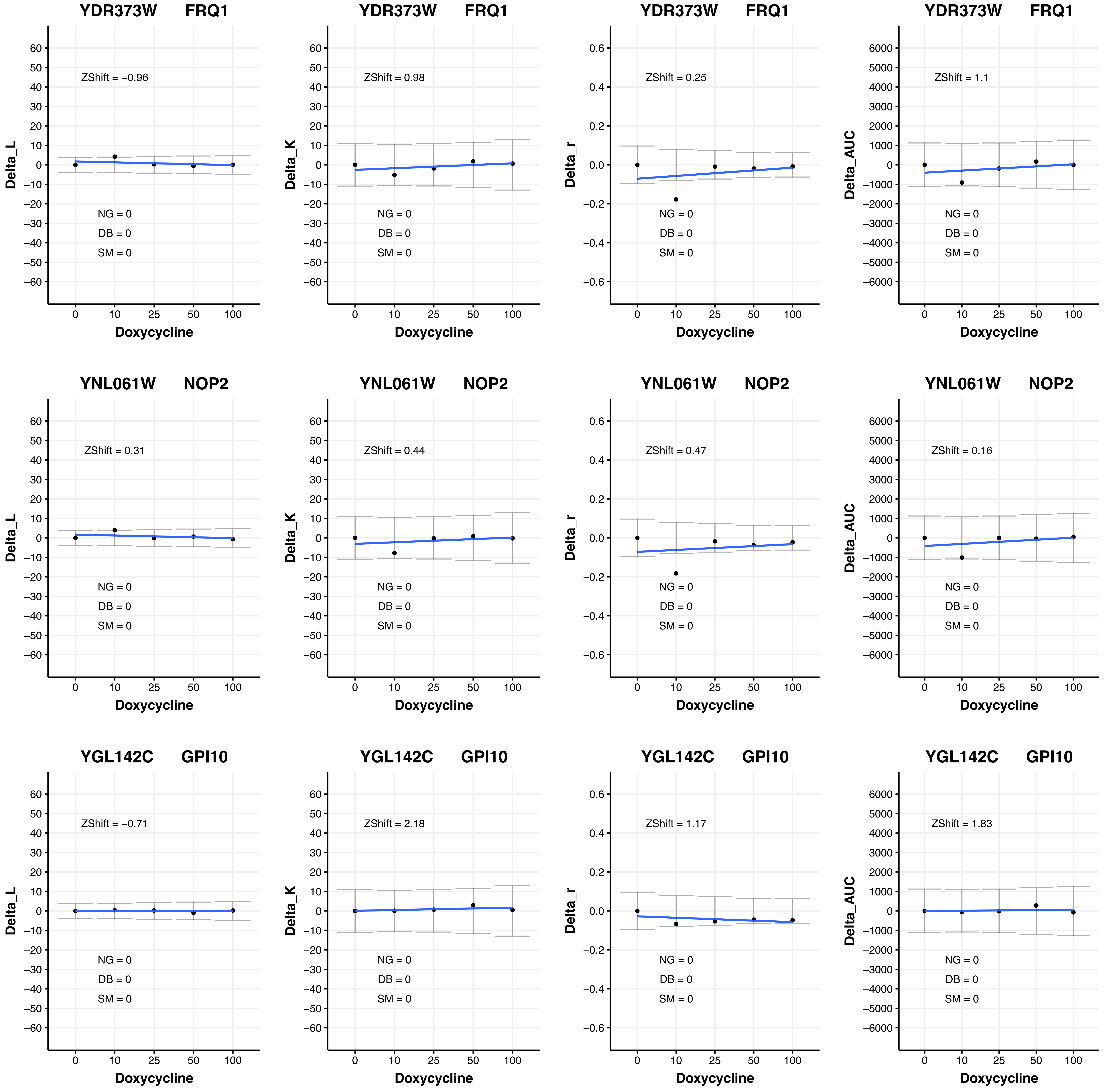
YDR373W FRQ1
Delta_L
ZShift = −0.96
NG = 0
DB = 0
SM = 0
Doxycycline
YDR373W FRQ1
Delta_K
ZShift = 0.98
NG = 0
DB = 0
SM = 0
Doxycycline
YDR373W FRQ1
Delta_r
ZShift = 0.25
NG = 0
DB = 0
SM = 0
Doxycycline
YDR373W FRQ1
Delta_AUC
ZShift = 1.1
NG = 0
DB = 0
SM = 0
Doxycycline
YNL061W NOP2
Delta_L
ZShift = 0.31
NG = 0
DB = 0
SM = 0
Doxycycline
YNL061W NOP2
Delta_K
ZShift = 0.44
NG = 0
DB = 0
SM = 0
Doxycycline
YNL061W NOP2
Delta_r
ZShift = 0.47
NG = 0
DB = 0
SM = 0
Doxycycline
YNL061W NOP2
Delta_AUC
ZShift = 0.16
NG = 0
DB = 0
SM = 0
Doxycycline
YGL142C GPI10
Delta_L
ZShift = −0.71
NG = 0
DB = 0
SM = 0
Doxycycline
YGL142C GPI10
Delta_K
ZShift = 2.18
NG = 0
DB = 0
SM = 0
Doxycycline
YGL142C GPI10
Delta_r
ZShift = 1.17
NG = 0
DB = 0
SM = 0
Doxycycline
YGL142C GPI10
Delta_AUC
ZShift = 1.83
NG = 0
DB = 0
SM = 0
Doxycycline

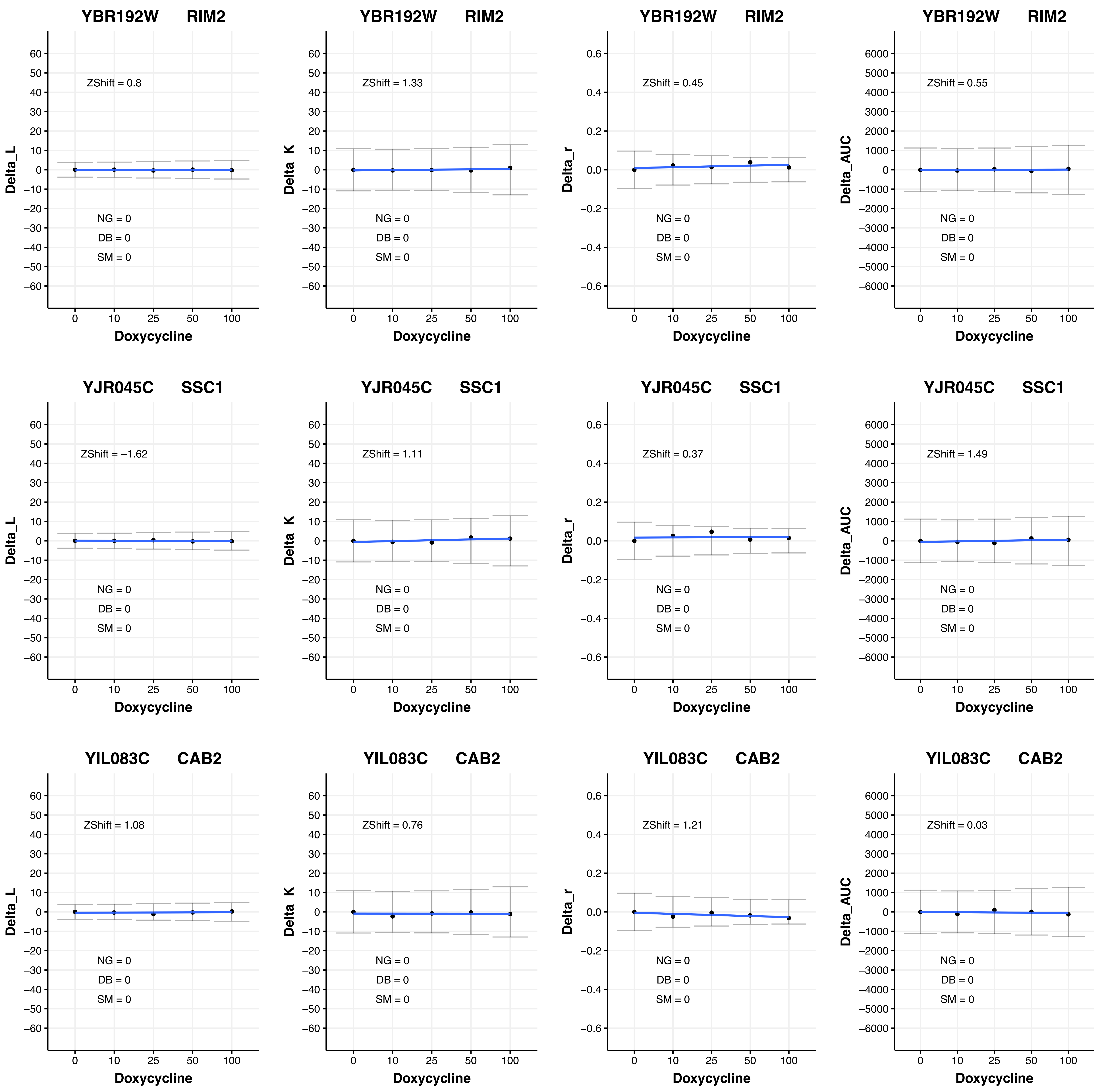

YBR192W RIM2
ZShift = 0.8
ZShift = 1.33
ZShift = 0.45
ZShift = 0.55
YJR045C SSC1
ZShift = −1.62
ZShift = 1.11
ZShift = 0.37
ZShift = 1.49
YIL083C CAB2
ZShift = 1.08
ZShift = 0.76
ZShift = 1.21
ZShift = 0.03
NG = 0
DB = 0
SM = 0
Delta_L
Delta_K
Delta_r
Delta_AUC
Doxycycline

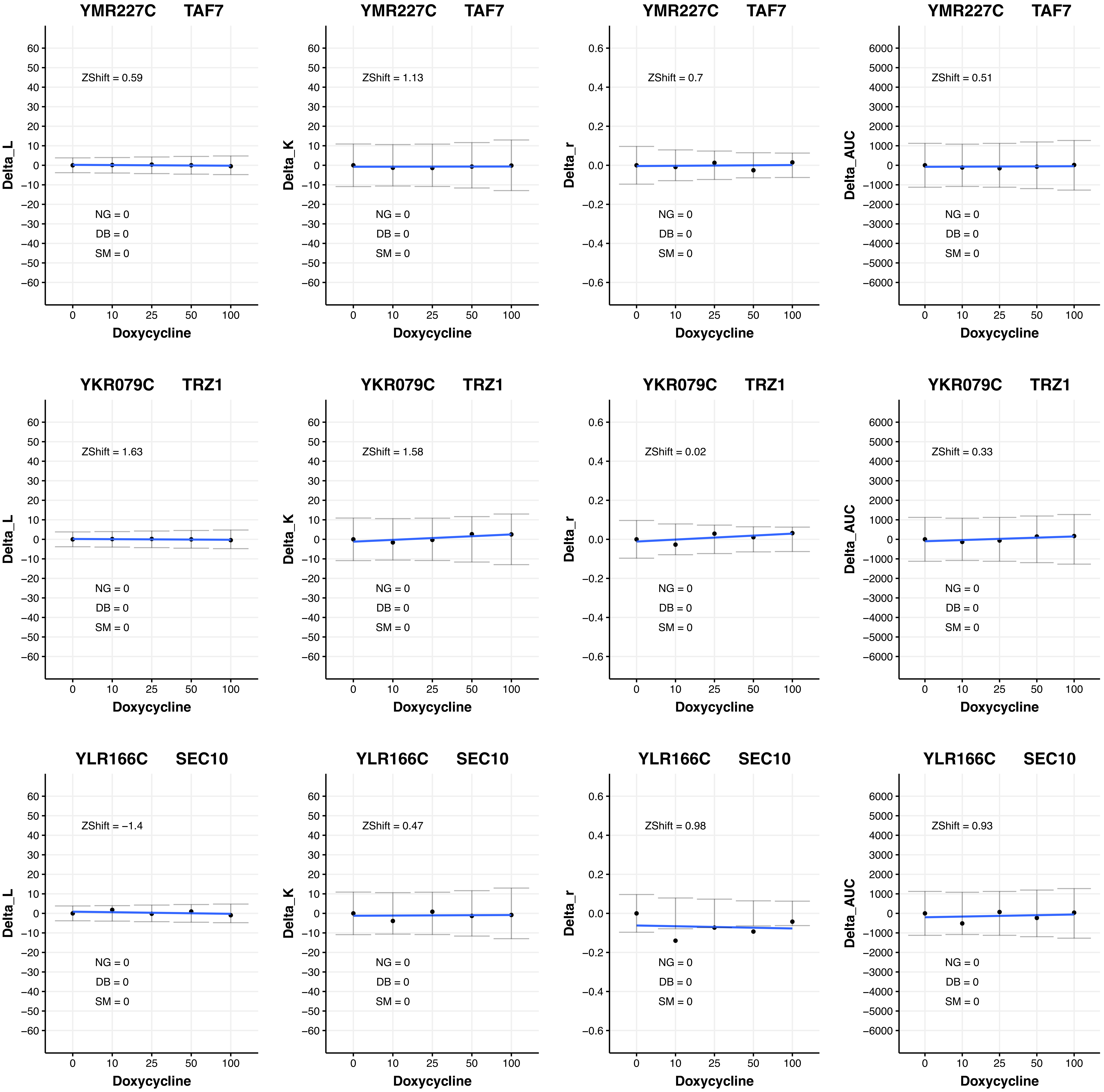
YMR227C TAF7
Delta_L
ZShift = 0.59
NG = 0
DB = 0
SM = 0
Doxycycline
YMR227C TAF7
Delta_K
ZShift = 1.13
NG = 0
DB = 0
SM = 0
Doxycycline
YMR227C TAF7
Delta_r
ZShift = 0.7
NG = 0
DB = 0
SM = 0
Doxycycline
YMR227C TAF7
Delta_AUC
ZShift = 0.51
NG = 0
DB = 0
SM = 0
Doxycycline
YKR079C TRZ1
Delta_L
ZShift = 1.63
NG = 0
DB = 0
SM = 0
Doxycycline
YKR079C TRZ1
Delta_K
ZShift = 1.58
NG = 0
DB = 0
SM = 0
Doxycycline
YKR079C TRZ1
Delta_r
ZShift = 0.02
NG = 0
DB = 0
SM = 0
Doxycycline
YKR079C TRZ1
Delta_AUC
ZShift = 0.33
NG = 0
DB = 0
SM = 0
Doxycycline
YLR166C SEC10
Delta_L
ZShift = −1.4
NG = 0
DB = 0
SM = 0
Doxycycline
YLR166C SEC10
Delta_K
ZShift = 0.47
NG = 0
DB = 0
SM = 0
Doxycycline
YLR166C SEC10
Delta_r
ZShift = 0.98
NG = 0
DB = 0
SM = 0
Doxycycline
YLR166C SEC10
Delta_AUC
ZShift = 0.93
NG = 0
DB = 0
SM = 0
Doxycycline

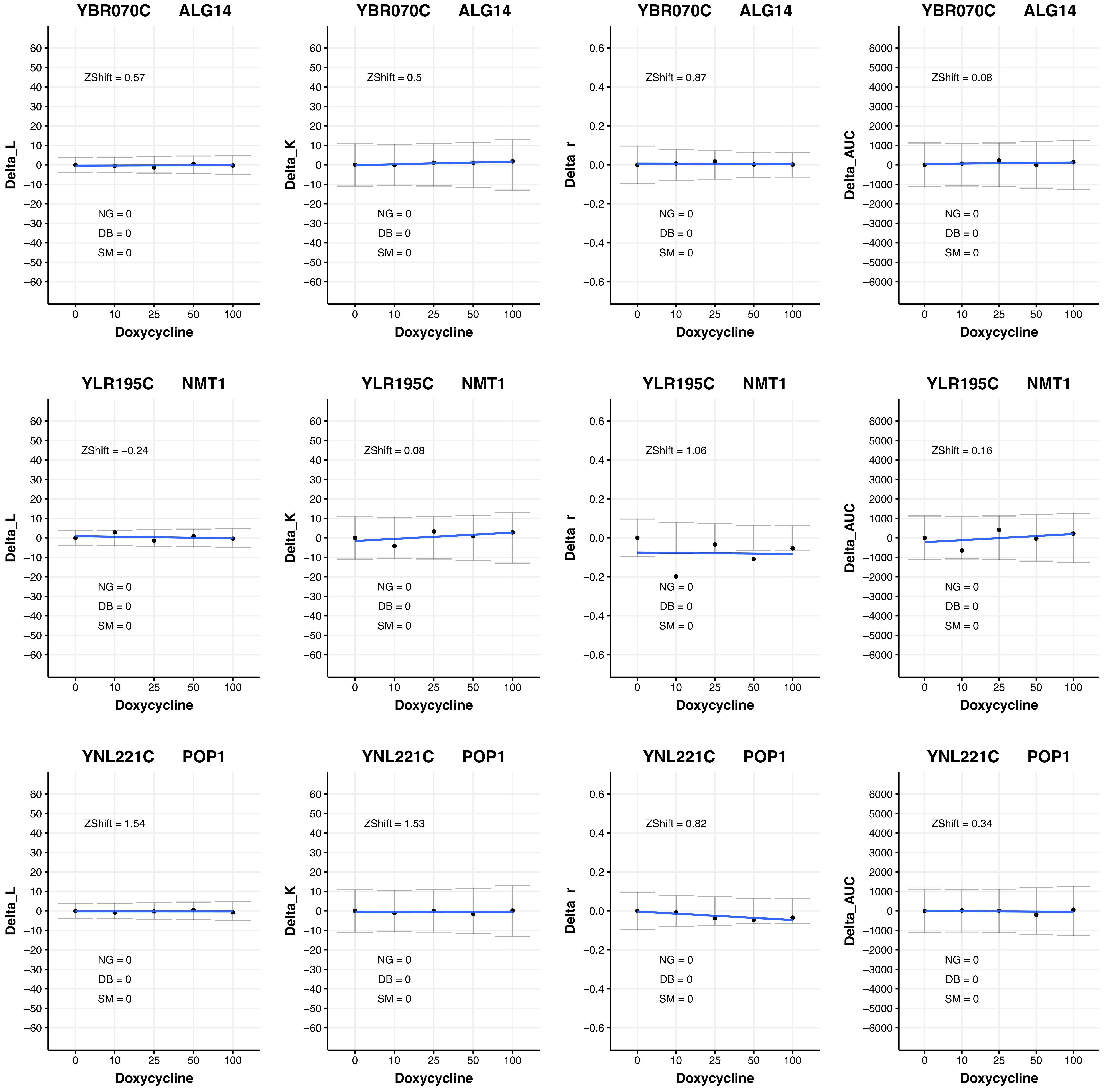

YBR070C ALG14
ZShift = 0.57
NG = 0
DB = 0
SM = 0
Delta_L
Doxycycline
YBR070C ALG14
ZShift = 0.5
NG = 0
DB = 0
SM = 0
Delta_K
Doxycycline
YBR070C ALG14
ZShift = 0.87
NG = 0
DB = 0
SM = 0
Delta_r
Doxycycline
YBR070C ALG14
ZShift = 0.08
NG = 0
DB = 0
SM = 0
Delta_AUC
Doxycycline
YLR195C NMT1
ZShift = −0.24
NG = 0
DB = 0
SM = 0
Delta_L
Doxycycline
YLR195C NMT1
ZShift = 0.08
NG = 0
DB = 0
SM = 0
Delta_K
Doxycycline
YLR195C NMT1
ZShift = 1.06
NG = 0
DB = 0
SM = 0
Delta_r
Doxycycline
YLR195C NMT1
ZShift = 0.16
NG = 0
DB = 0
SM = 0
Delta_AUC
Doxycycline
YNL221C POP1
ZShift = 1.54
NG = 0
DB = 0
SM = 0
Delta_L
Doxycycline
YNL221C POP1
ZShift = 1.53
NG = 0
DB = 0
SM = 0
Delta_K
Doxycycline
YNL221C POP1
ZShift = 0.82
NG = 0
DB = 0
SM = 0
Delta_r
Doxycycline
YNL221C POP1
ZShift = 0.34
NG = 0
DB = 0
SM = 0
Delta_AUC
Doxycycline

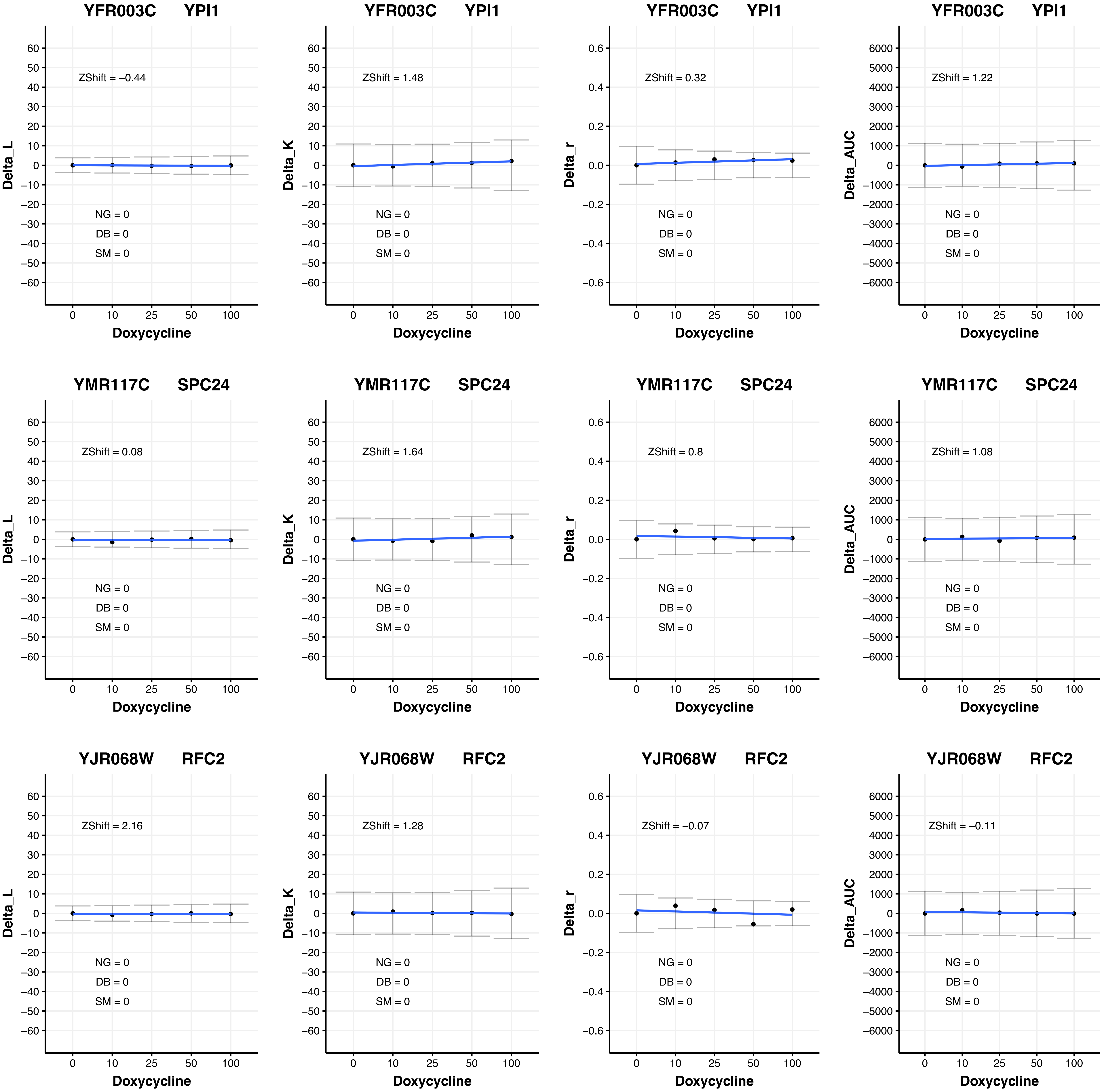

YFR003C YPI1
ZShift = −0.44
NG = 0
DB = 0
SM = 0
Delta_L
Doxycycline
YFR003C YPI1
ZShift = 1.48
NG = 0
DB = 0
SM = 0
Delta_K
Doxycycline
YFR003C YPI1
ZShift = 0.32
NG = 0
DB = 0
SM = 0
Delta_r
Doxycycline
YFR003C YPI1
ZShift = 1.22
NG = 0
DB = 0
SM = 0
Delta_AUC
Doxycycline
YMR117C SPC24
ZShift = 0.08
NG = 0
DB = 0
SM = 0
Delta_L
Doxycycline
YMR117C SPC24
ZShift = 1.64
NG = 0
DB = 0
SM = 0
Delta_K
Doxycycline
YMR117C SPC24
ZShift = 0.8
NG = 0
DB = 0
SM = 0
Delta_r
Doxycycline
YMR117C SPC24
ZShift = 1.08
NG = 0
DB = 0
SM = 0
Delta_AUC
Doxycycline
YJR068W RFC2
ZShift = 2.16
NG = 0
DB = 0
SM = 0
Delta_L
Doxycycline
YJR068W RFC2
ZShift = 1.28
NG = 0
DB = 0
SM = 0
Delta_K
Doxycycline
YJR068W RFC2
ZShift = −0.07
NG = 0
DB = 0
SM = 0
Delta_r
Doxycycline
YJR068W RFC2
ZShift = −0.11
NG = 0
DB = 0
SM = 0
Delta_AUC
Doxycycline

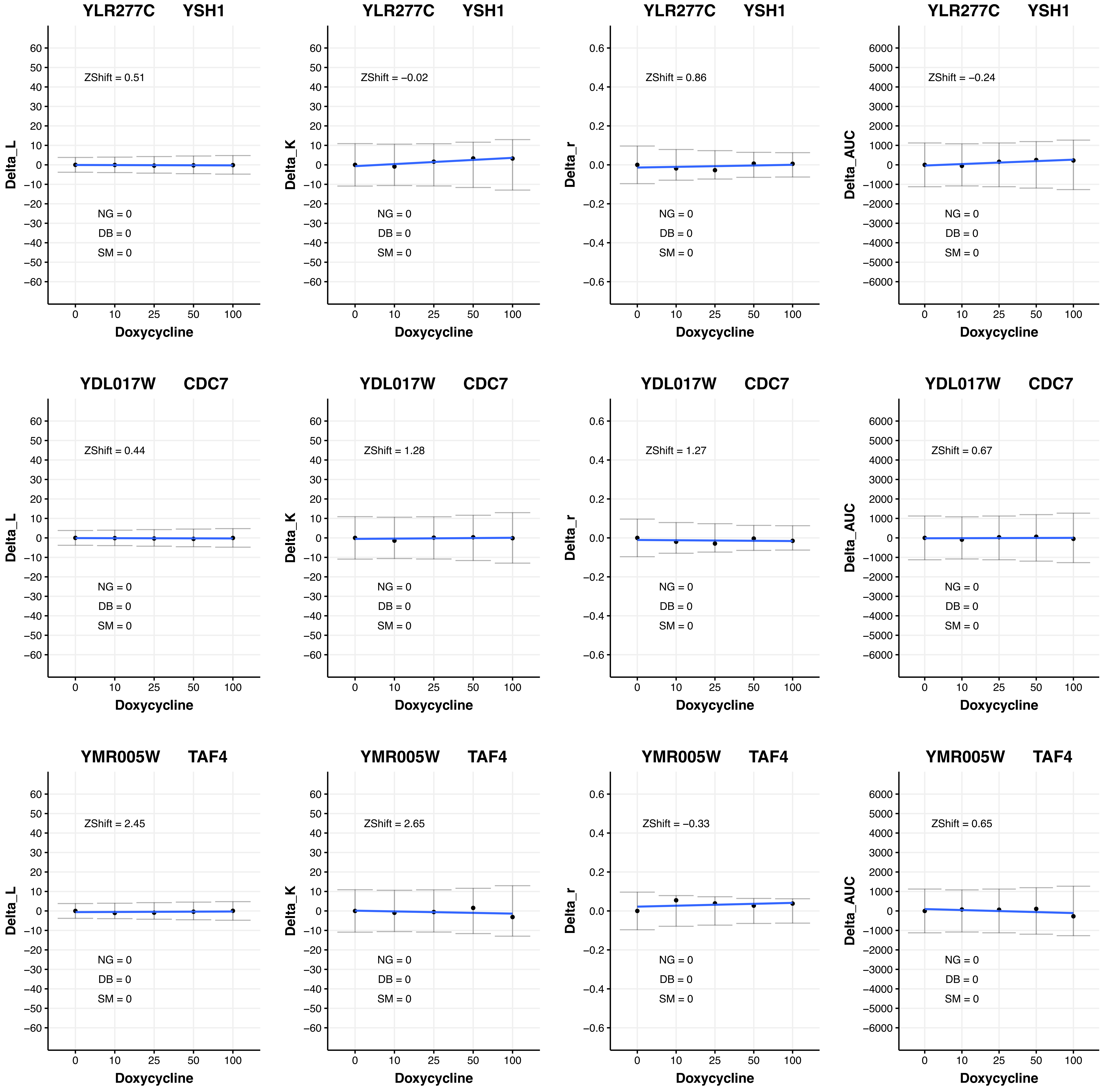

YLR277C YSH1
ZShift = 0.51
NG = 0
DB = 0
SM = 0
Delta_L
Doxycycline
YLR277C YSH1
ZShift = −0.02
NG = 0
DB = 0
SM = 0
Delta_K
Doxycycline
YLR277C YSH1
ZShift = 0.86
NG = 0
DB = 0
SM = 0
Delta_r
Doxycycline
YLR277C YSH1
ZShift = −0.24
NG = 0
DB = 0
SM = 0
Delta_AUC
Doxycycline
YDL017W CDC7
ZShift = 0.44
NG = 0
DB = 0
SM = 0
Delta_L
Doxycycline
YDL017W CDC7
ZShift = 1.28
NG = 0
DB = 0
SM = 0
Delta_K
Doxycycline
YDL017W CDC7
ZShift = 1.27
NG = 0
DB = 0
SM = 0
Delta_r
Doxycycline
YDL017W CDC7
ZShift = 0.67
NG = 0
DB = 0
SM = 0
Delta_AUC
Doxycycline
YMR005W TAF4
ZShift = 2.45
NG = 0
DB = 0
SM = 0
Delta_L
Doxycycline
YMR005W TAF4
ZShift = 2.65
NG = 0
DB = 0
SM = 0
Delta_K
Doxycycline
YMR005W TAF4
ZShift = −0.33
NG = 0
DB = 0
SM = 0
Delta_r
Doxycycline
YMR005W TAF4
ZShift = 0.65
NG = 0
DB = 0
SM = 0
Delta_AUC
Doxycycline

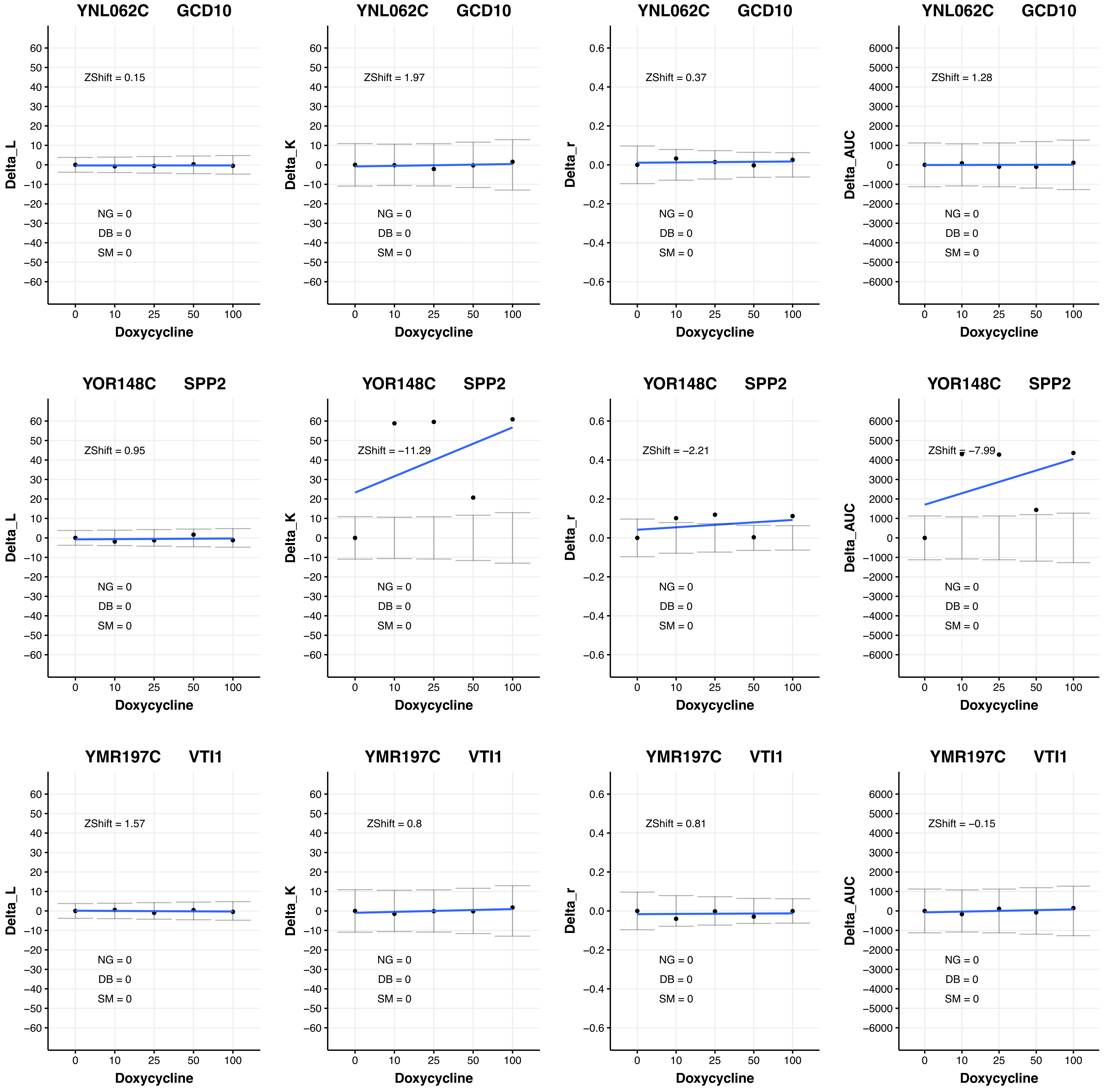

YNL062C GCD10
ZShift = 0.15
NG = 0
DB = 0
SM = 0
Delta_L
Doxycycline
YNL062C GCD10
ZShift = 1.97
NG = 0
DB = 0
SM = 0
Delta_K
Doxycycline
YNL062C GCD10
ZShift = 0.37
NG = 0
DB = 0
SM = 0
Delta_r
Doxycycline
YNL062C GCD10
ZShift = 1.28
NG = 0
DB = 0
SM = 0
Delta_AUC
Doxycycline
YOR148C SPP2
ZShift = 0.95
NG = 0
DB = 0
SM = 0
Delta_L
Doxycycline
YOR148C SPP2
ZShift = −11.29
NG = 0
DB = 0
SM = 0
Delta_K
Doxycycline
YOR148C SPP2
ZShift = −2.21
NG = 0
DB = 0
SM = 0
Delta_r
Doxycycline
YOR148C SPP2
ZShift = −7.99
NG = 0
DB = 0
SM = 0
Delta_AUC
Doxycycline
YMR197C VTI1
ZShift = 1.57
NG = 0
DB = 0
SM = 0
Delta_L
Doxycycline
YMR197C VTI1
ZShift = 0.8
NG = 0
DB = 0
SM = 0
Delta_K
Doxycycline
YMR197C VTI1
ZShift = 0.81
NG = 0
DB = 0
SM = 0
Delta_r
Doxycycline
YMR197C VTI1
ZShift = −0.15
NG = 0
DB = 0
SM = 0
Delta_AUC
Doxycycline

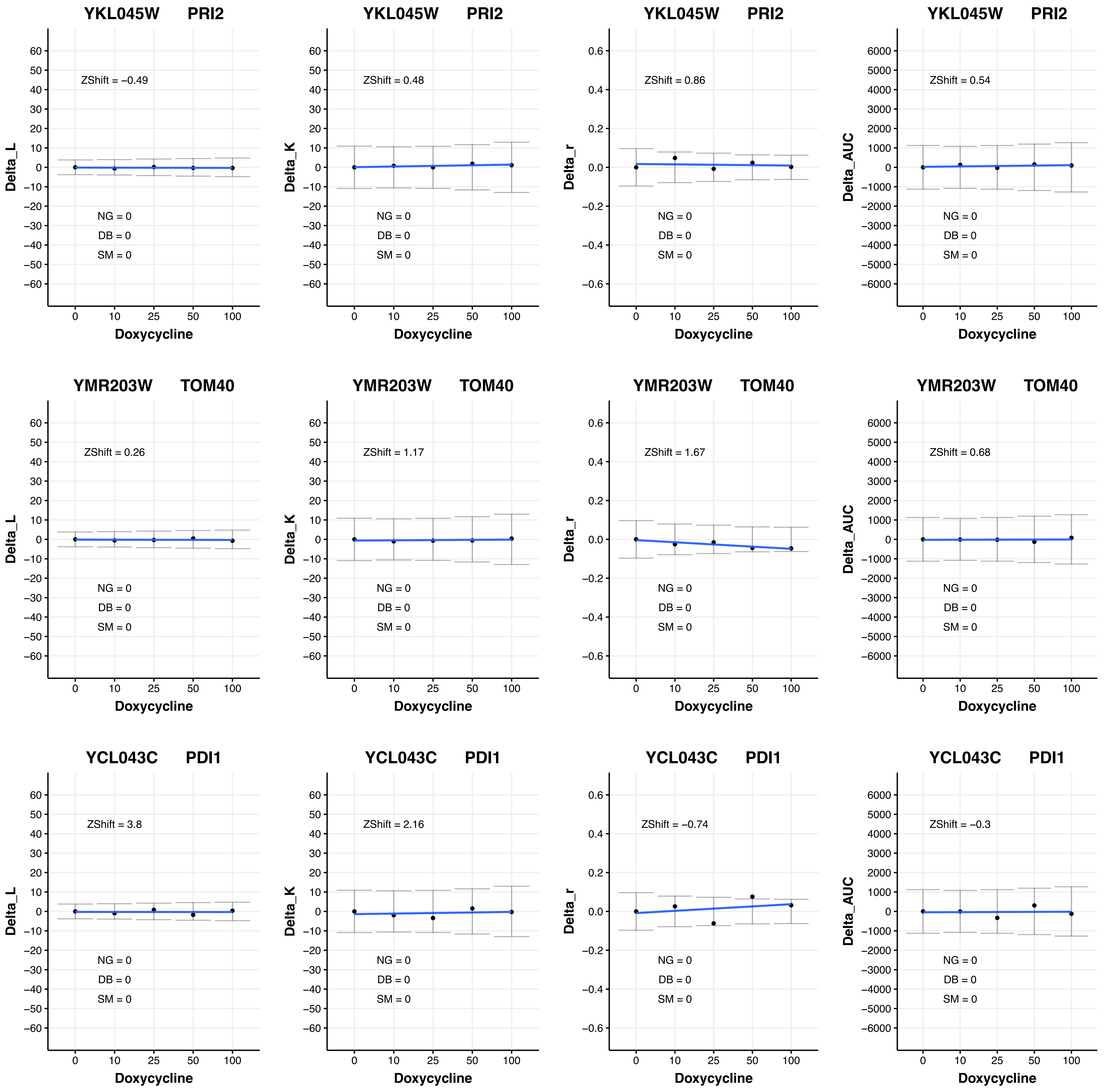
YKL045W PRI2
ZShift = −0.49
NG = 0
DB = 0
SM = 0
Delta_L
Doxycycline
YKL045W PRI2
ZShift = 0.48
NG = 0
DB = 0
SM = 0
Delta_K
Doxycycline
YKL045W PRI2
ZShift = 0.86
NG = 0
DB = 0
SM = 0
Delta_r
Doxycycline
YKL045W PRI2
ZShift = 0.54
NG = 0
DB = 0
SM = 0
Delta_AUC
Doxycycline
YMR203W TOM40
ZShift = 0.26
NG = 0
DB = 0
SM = 0
Delta_L
Doxycycline
YMR203W TOM40
ZShift = 1.17
NG = 0
DB = 0
SM = 0
Delta_K
Doxycycline
YMR203W TOM40
ZShift = 1.67
NG = 0
DB = 0
SM = 0
Delta_r
Doxycycline
YMR203W TOM40
ZShift = 0.68
NG = 0
DB = 0
SM = 0
Delta_AUC
Doxycycline
YCL043C PDI1
ZShift = 3.8
NG = 0
DB = 0
SM = 0
Delta_L
Doxycycline
YCL043C PDI1
ZShift = 2.16
NG = 0
DB = 0
SM = 0
Delta_K
Doxycycline
YCL043C PDI1
ZShift = −0.74
NG = 0
DB = 0
SM = 0
Delta_r
Doxycycline
YCL043C PDI1
ZShift = −0.3
NG = 0
DB = 0
SM = 0
Delta_AUC
Doxycycline

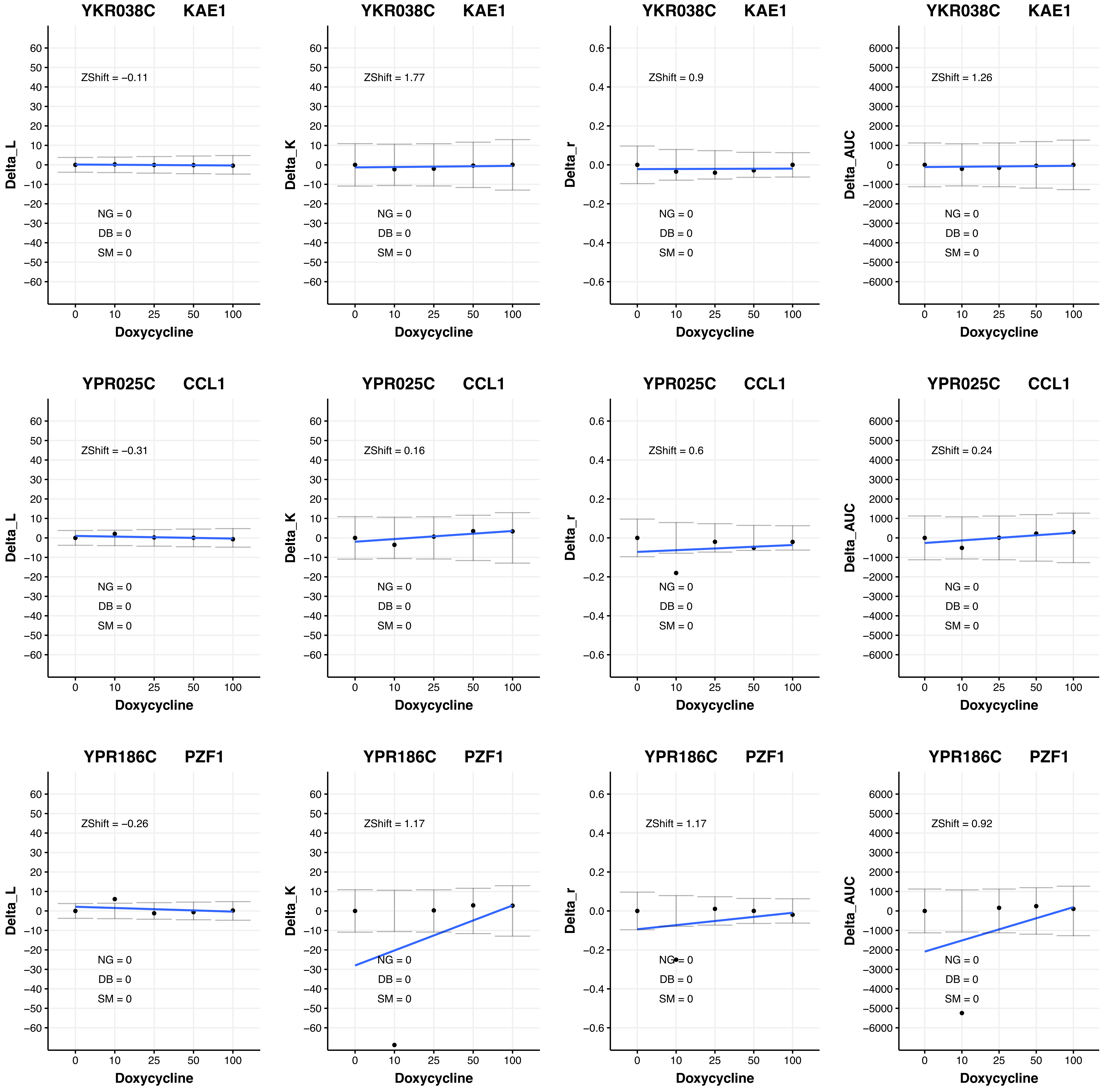
YKR038C KAE1
ZShift = −0.11
Delta_L
NG = 0
DB = 0
SM = 0
Doxycycline
YKR038C KAE1
ZShift = 1.77
Delta_K
NG = 0
DB = 0
SM = 0
Doxycycline
YKR038C KAE1
ZShift = 0.9
Delta_r
NG = 0
DB = 0
SM = 0
Doxycycline
YKR038C KAE1
ZShift = 1.26
Delta_AUC
NG = 0
DB = 0
SM = 0
Doxycycline
YPR025C CCL1
ZShift = −0.31
Delta_L
NG = 0
DB = 0
SM = 0
Doxycycline
YPR025C CCL1
ZShift = 0.16
Delta_K
NG = 0
DB = 0
SM = 0
Doxycycline
YPR025C CCL1
ZShift = 0.6
Delta_r
NG = 0
DB = 0
SM = 0
Doxycycline
YPR025C CCL1
ZShift = 0.24
Delta_AUC
NG = 0
DB = 0
SM = 0
Doxycycline
YPR186C PZF1
ZShift = −0.26
Delta_L
NG = 0
DB = 0
SM = 0
Doxycycline
YPR186C PZF1
ZShift = 1.17
Delta_K
NG = 0
DB = 0
SM = 0
Doxycycline
YPR186C PZF1
ZShift = 1.17
Delta_r
NG = 0
DB = 0
SM = 0
Doxycycline
YPR186C PZF1
ZShift = 0.92
Delta_AUC
NG = 0
DB = 0
SM = 0
Doxycycline

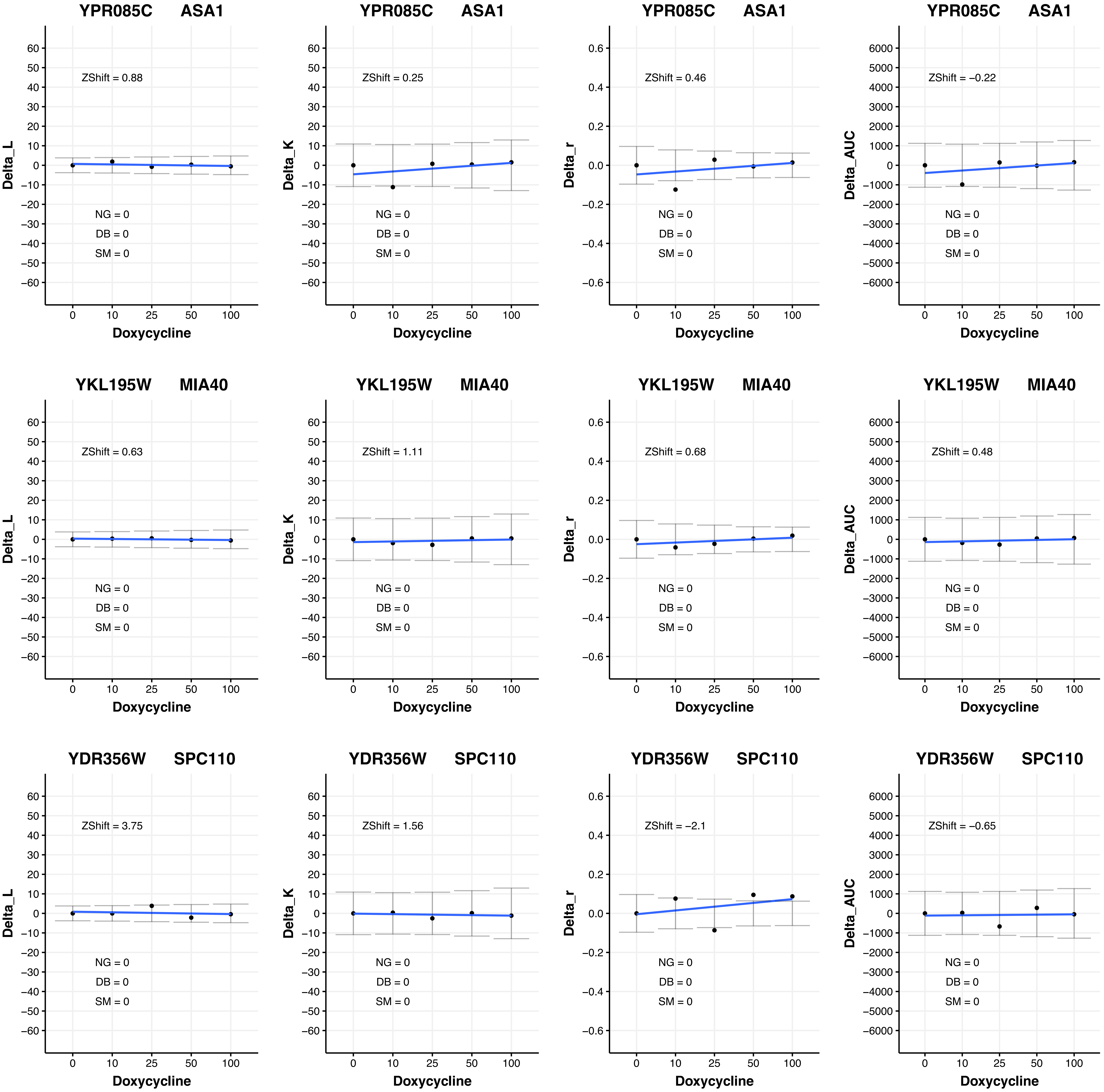
YPR085C ASA1
ZShift = 0.88
NG = 0
DB = 0
SM = 0
Delta_L
Doxycycline
YPR085C ASA1
ZShift = 0.25
NG = 0
DB = 0
SM = 0
Delta_K
Doxycycline
YPR085C ASA1
ZShift = 0.46
NG = 0
DB = 0
SM = 0
Delta_r
Doxycycline
YPR085C ASA1
ZShift = −0.22
NG = 0
DB = 0
SM = 0
Delta_AUC
Doxycycline
YKL195W MIA40
ZShift = 0.63
NG = 0
DB = 0
SM = 0
Delta_L
Doxycycline
YKL195W MIA40
ZShift = 1.11
NG = 0
DB = 0
SM = 0
Delta_K
Doxycycline
YKL195W MIA40
ZShift = 0.68
NG = 0
DB = 0
SM = 0
Delta_r
Doxycycline
YKL195W MIA40
ZShift = 0.48
NG = 0
DB = 0
SM = 0
Delta_AUC
Doxycycline
YDR356W SPC110
ZShift = 3.75
NG = 0
DB = 0
SM = 0
Delta_L
Doxycycline
YDR356W SPC110
ZShift = 1.56
NG = 0
DB = 0
SM = 0
Delta_K
Doxycycline
YDR356W SPC110
ZShift = −2.1
NG = 0
DB = 0
SM = 0
Delta_r
Doxycycline
YDR356W SPC110
ZShift = −0.65
NG = 0
DB = 0
SM = 0
Delta_AUC
Doxycycline

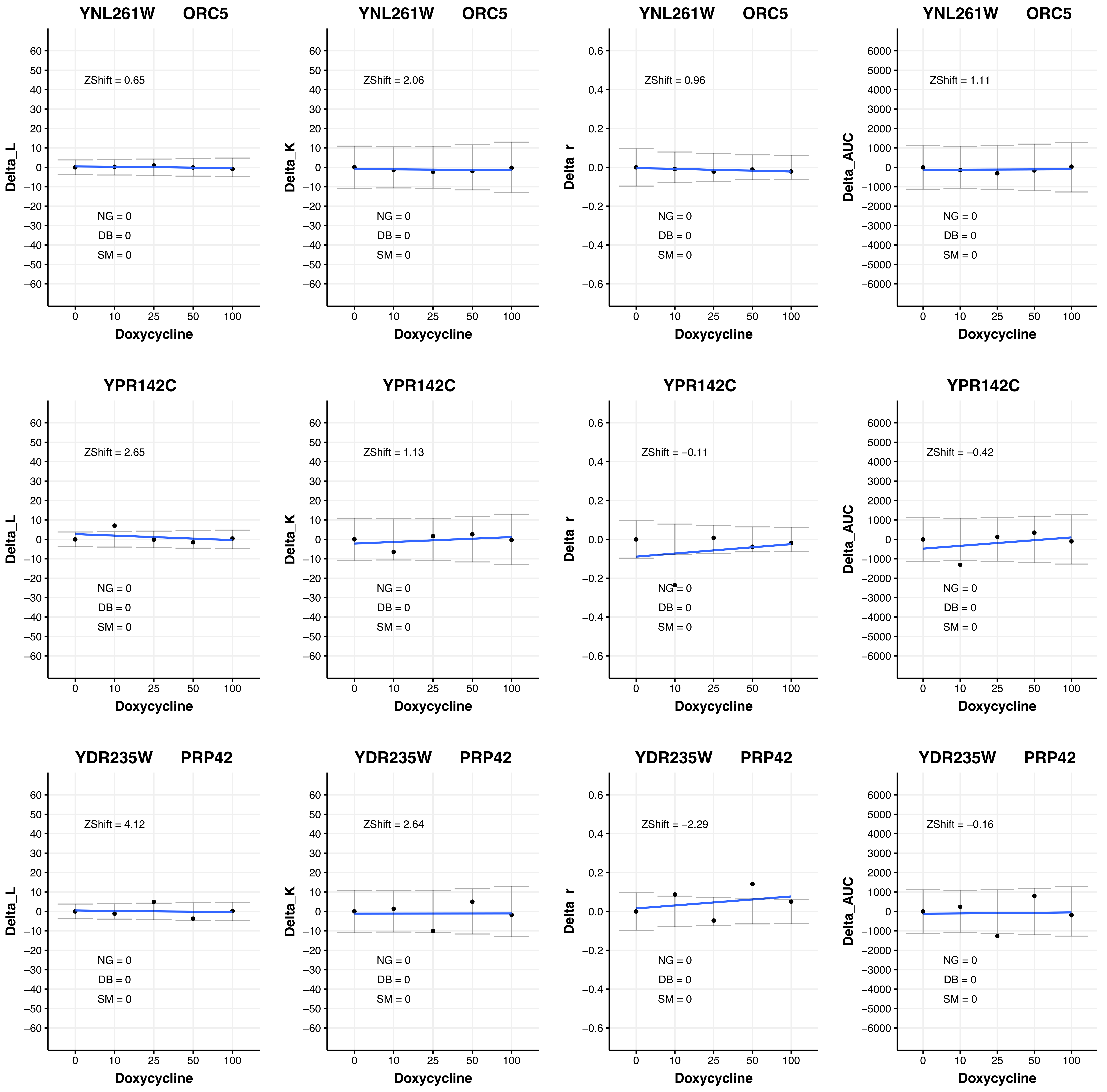

YNL261W ORC5
ZShift = 0.65
Delta_L
NG = 0
DB = 0
SM = 0
Doxycycline
0 10 25 50 100
YNL261W ORC5
ZShift = 2.06
Delta_K
NG = 0
DB = 0
SM = 0
Doxycycline
YNL261W ORC5
ZShift = 0.96
Delta_r
NG = 0
DB = 0
SM = 0
Doxycycline
YNL261W ORC5
ZShift = 1.11
Delta_AUC
NG = 0
DB = 0
SM = 0
Doxycycline
YPR142C
ZShift = 2.65
Delta_L
NG = 0
DB = 0
SM = 0
Doxycycline
YPR142C
ZShift = 1.13
Delta_K
NG = 0
DB = 0
SM = 0
Doxycycline
YPR142C
ZShift = −0.11
Delta_r
NG = 0
DB = 0
SM = 0
Doxycycline
YPR142C
ZShift = −0.42
Delta_AUC
NG = 0
DB = 0
SM = 0
Doxycycline
YDR235W PRP42
ZShift = 4.12
Delta_L
NG = 0
DB = 0
SM = 0
Doxycycline
YDR235W PRP42
ZShift = 2.64
Delta_K
NG = 0
DB = 0
SM = 0
Doxycycline
YDR235W PRP42
ZShift = −2.29
Delta_r
NG = 0
DB = 0
SM = 0
Doxycycline
YDR235W PRP42
ZShift = −0.16
Delta_AUC
NG = 0
DB = 0
SM = 0
Doxycycline

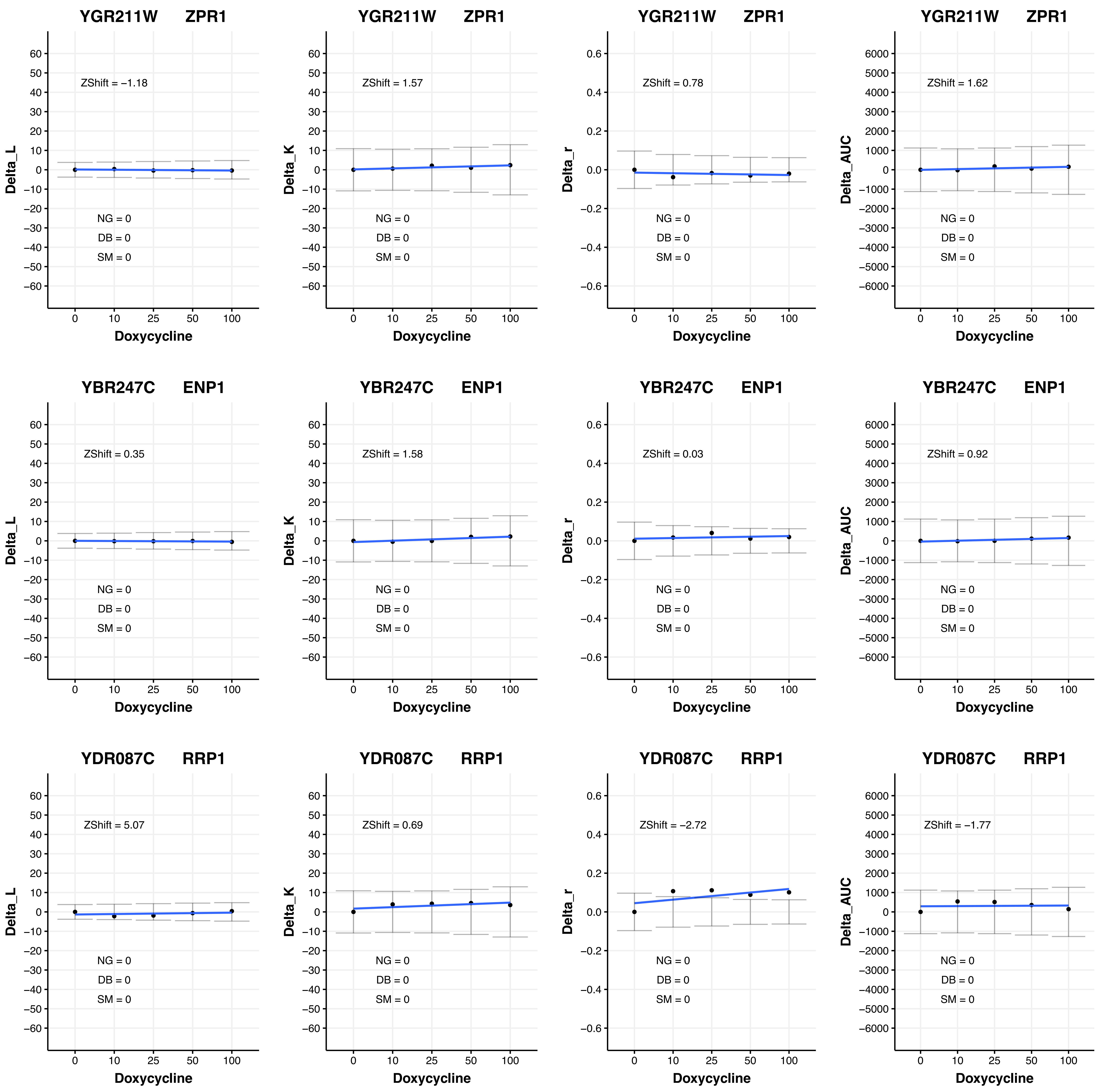
YGR211W ZPR1
ZShift = −1.18
NG = 0
DB = 0
SM = 0
Delta_L
Doxycycline
YGR211W ZPR1
ZShift = 1.57
NG = 0
DB = 0
SM = 0
Delta_K
Doxycycline
YGR211W ZPR1
ZShift = 0.78
NG = 0
DB = 0
SM = 0
Delta_r
Doxycycline
YGR211W ZPR1
ZShift = 1.62
NG = 0
DB = 0
SM = 0
Delta_AUC
Doxycycline
YBR247C ENP1
ZShift = 0.35
NG = 0
DB = 0
SM = 0
Delta_L
Doxycycline
YBR247C ENP1
ZShift = 1.58
NG = 0
DB = 0
SM = 0
Delta_K
Doxycycline
YBR247C ENP1
ZShift = 0.03
NG = 0
DB = 0
SM = 0
Delta_r
Doxycycline
YBR247C ENP1
ZShift = 0.92
NG = 0
DB = 0
SM = 0
Delta_AUC
Doxycycline
YDR087C RRP1
ZShift = 5.07
NG = 0
DB = 0
SM = 0
Delta_L
Doxycycline
YDR087C RRP1
ZShift = 0.69
NG = 0
DB = 0
SM = 0
Delta_K
Doxycycline
YDR087C RRP1
ZShift = −2.72
NG = 0
DB = 0
SM = 0
Delta_r
Doxycycline
YDR087C RRP1
ZShift = −1.77
NG = 0
DB = 0
SM = 0
Delta_AUC
Doxycycline

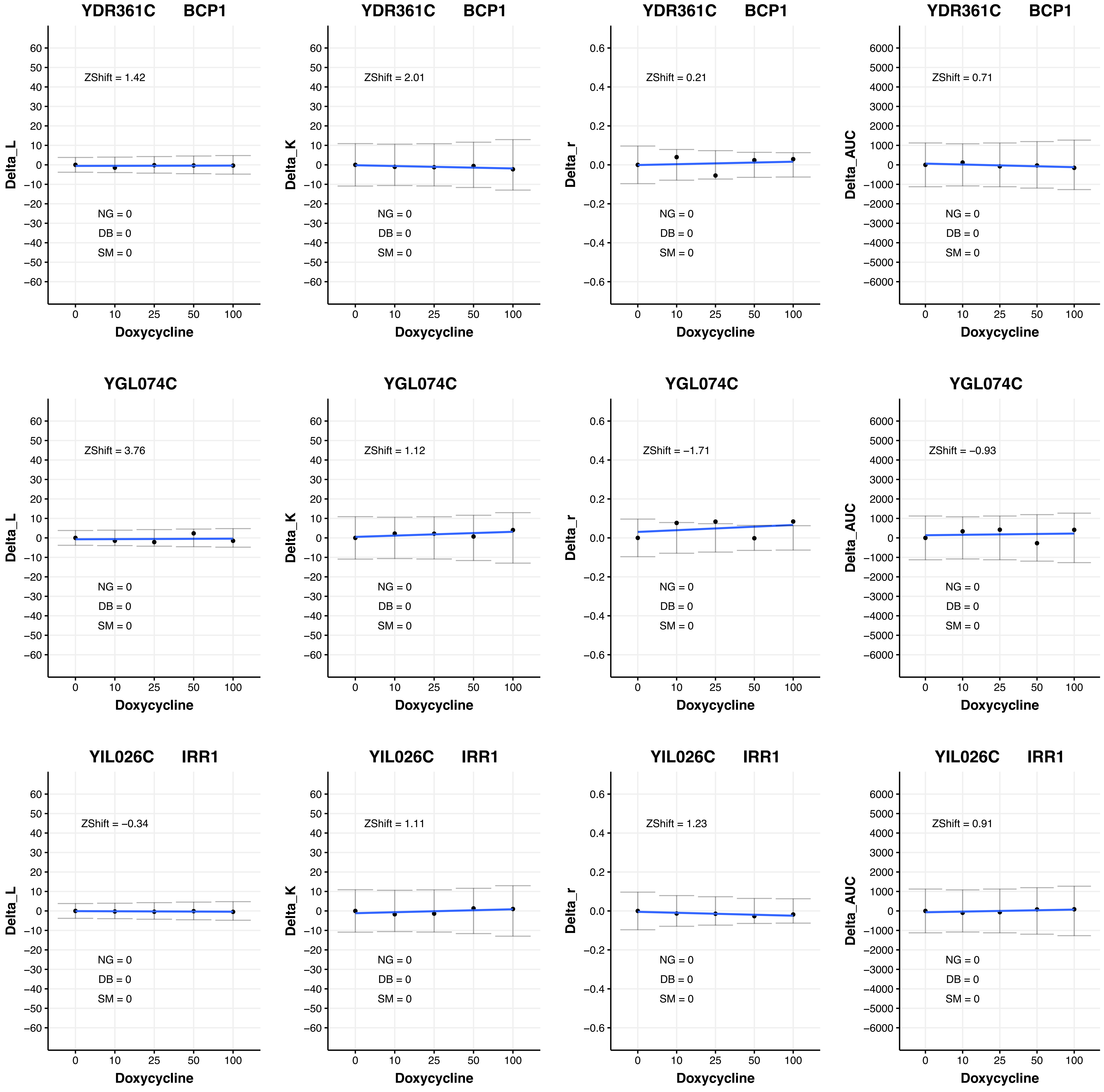
YDR361C BCP1
ZShift = 1.42
NG = 0
DB = 0
SM = 0
Delta_L
Doxycycline
YDR361C BCP1
ZShift = 2.01
NG = 0
DB = 0
SM = 0
Delta_K
Doxycycline
YDR361C BCP1
ZShift = 0.21
NG = 0
DB = 0
SM = 0
Delta_r
Doxycycline
YDR361C BCP1
ZShift = 0.71
NG = 0
DB = 0
SM = 0
Delta_AUC
Doxycycline
YGL074C
ZShift = 3.76
NG = 0
DB = 0
SM = 0
Delta_L
Doxycycline
YGL074C
ZShift = 1.12
NG = 0
DB = 0
SM = 0
Delta_K
Doxycycline
YGL074C
ZShift = −1.71
NG = 0
DB = 0
SM = 0
Delta_r
Doxycycline
YGL074C
ZShift = −0.93
NG = 0
DB = 0
SM = 0
Delta_AUC
Doxycycline
YIL026C IRR1
ZShift = −0.34
NG = 0
DB = 0
SM = 0
Delta_L
Doxycycline
YIL026C IRR1
ZShift = 1.11
NG = 0
DB = 0
SM = 0
Delta_K
Doxycycline
YIL026C IRR1
ZShift = 1.23
NG = 0
DB = 0
SM = 0
Delta_r
Doxycycline
YIL026C IRR1
ZShift = 0.91
NG = 0
DB = 0
SM = 0
Delta_AUC
Doxycycline

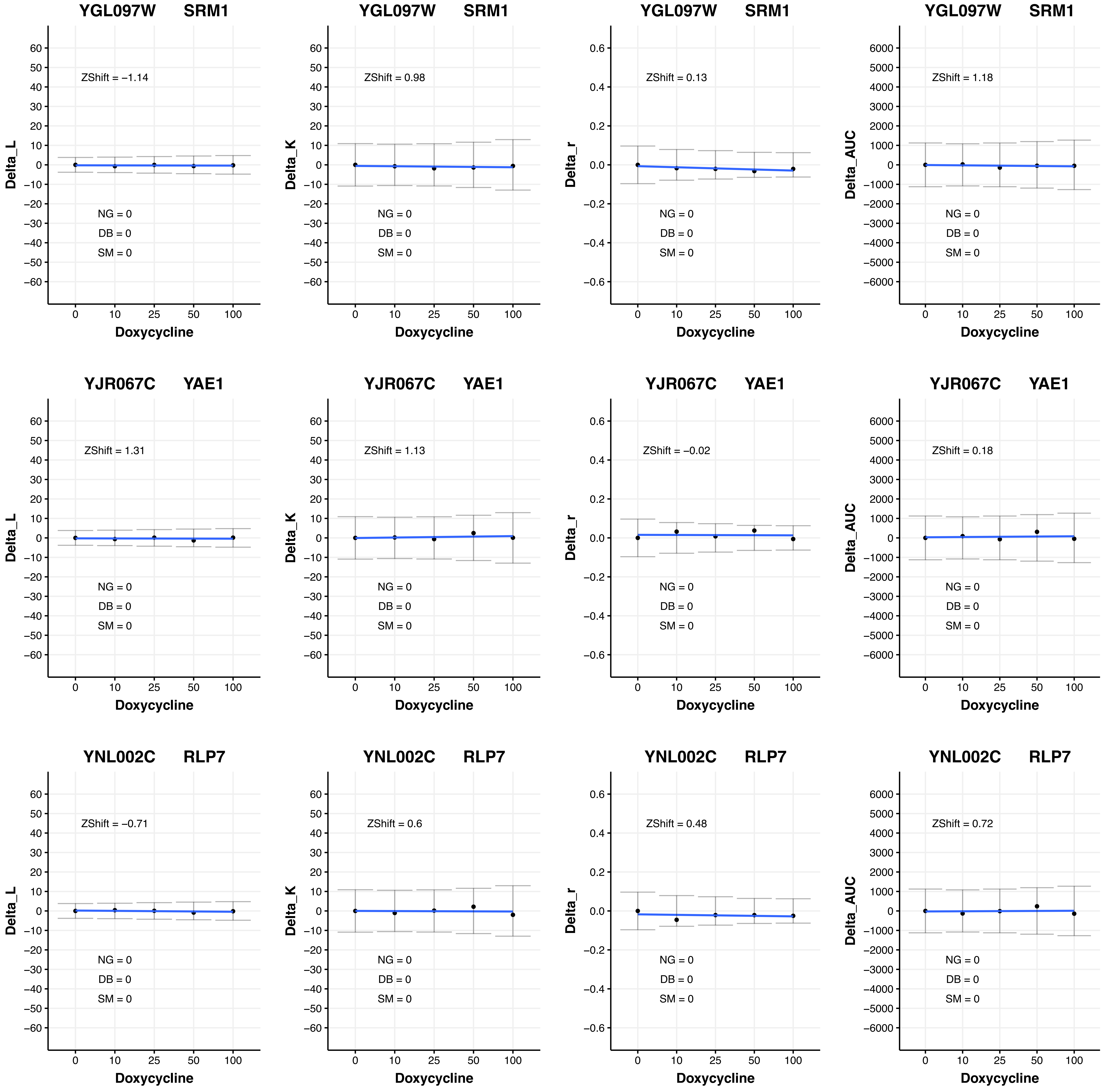
YGL097W SRM1
ZShift = −1.14
NG = 0
DB = 0
SM = 0
Delta_L
Doxycycline
YGL097W SRM1
ZShift = 0.98
NG = 0
DB = 0
SM = 0
Delta_K
Doxycycline
YGL097W SRM1
ZShift = 0.13
NG = 0
DB = 0
SM = 0
Delta_r
Doxycycline
YGL097W SRM1
ZShift = 1.18
NG = 0
DB = 0
SM = 0
Delta_AUC
Doxycycline
YJR067C YAE1
ZShift = 1.31
NG = 0
DB = 0
SM = 0
Delta_L
Doxycycline
YJR067C YAE1
ZShift = 1.13
NG = 0
DB = 0
SM = 0
Delta_K
Doxycycline
YJR067C YAE1
ZShift = −0.02
NG = 0
DB = 0
SM = 0
Delta_r
Doxycycline
YJR067C YAE1
ZShift = 0.18
NG = 0
DB = 0
SM = 0
Delta_AUC
Doxycycline
YNL002C RLP7
ZShift = −0.71
NG = 0
DB = 0
SM = 0
Delta_L
Doxycycline
YNL002C RLP7
ZShift = 0.6
NG = 0
DB = 0
SM = 0
Delta_K
Doxycycline
YNL002C RLP7
ZShift = 0.48
NG = 0
DB = 0
SM = 0
Delta_r
Doxycycline
YNL002C RLP7
ZShift = 0.72
NG = 0
DB = 0
SM = 0
Delta_AUC
Doxycycline

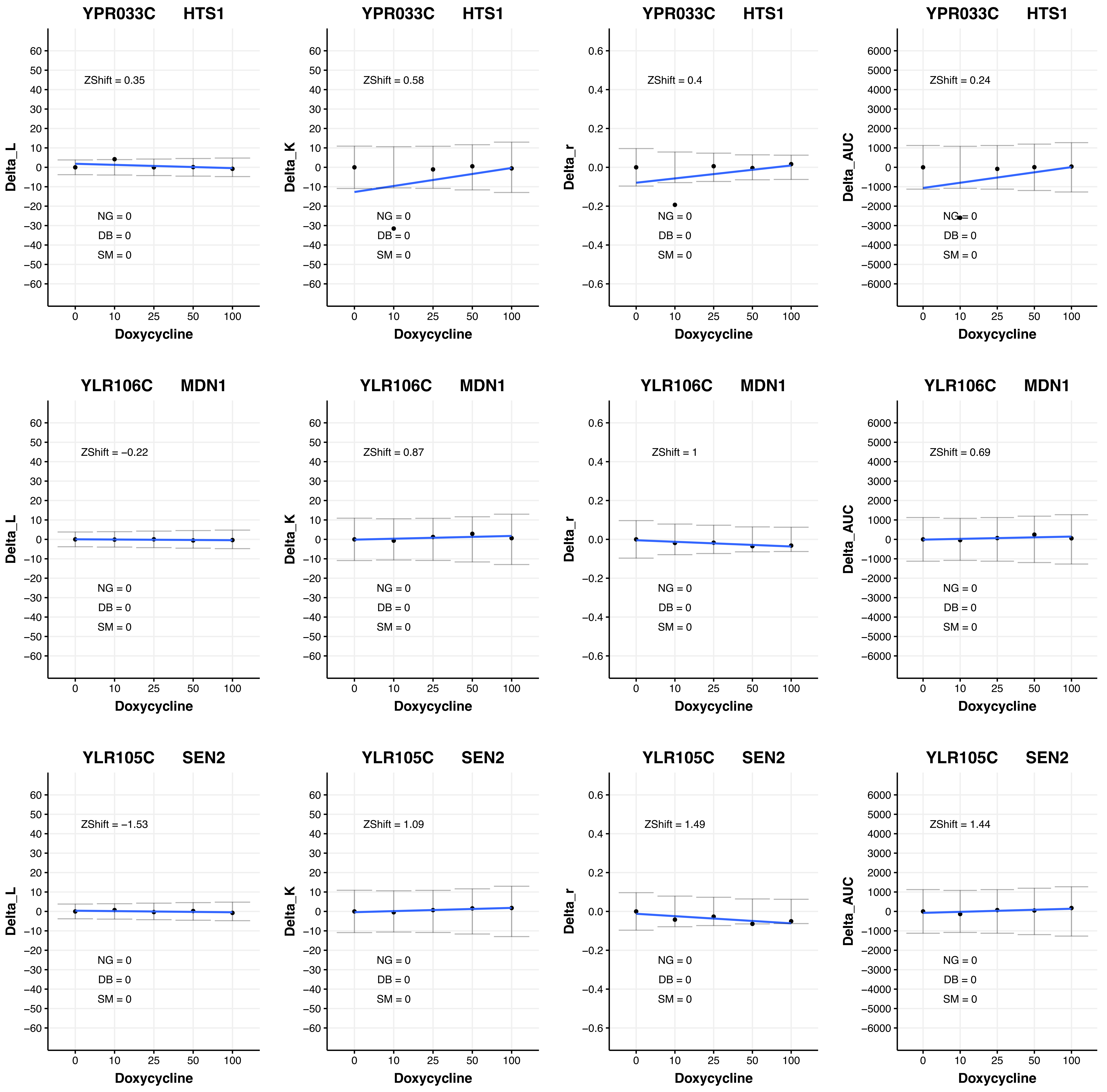
YPR033C HTS1
Delta_L
ZShift = 0.35
NG = 0
DB = 0
SM = 0
Doxycycline
YPR033C HTS1
Delta_K
ZShift = 0.58
NG = 0
DB = 0
SM = 0
Doxycycline
YPR033C HTS1
Delta_r
ZShift = 0.4
NG = 0
DB = 0
SM = 0
Doxycycline
YPR033C HTS1
Delta_AUC
ZShift = 0.24
NG = 0
DB = 0
SM = 0
Doxycycline
YLR106C MDN1
Delta_L
ZShift = −0.22
NG = 0
DB = 0
SM = 0
Doxycycline
YLR106C MDN1
Delta_K
ZShift = 0.87
NG = 0
DB = 0
SM = 0
Doxycycline
YLR106C MDN1
Delta_r
ZShift = 1
NG = 0
DB = 0
SM = 0
Doxycycline
YLR106C MDN1
Delta_AUC
ZShift = 0.69
NG = 0
DB = 0
SM = 0
Doxycycline
YLR105C SEN2
Delta_L
ZShift = −1.53
NG = 0
DB = 0
SM = 0
Doxycycline
YLR105C SEN2
Delta_K
ZShift = 1.09
NG = 0
DB = 0
SM = 0
Doxycycline
YLR105C SEN2
Delta_r
ZShift = 1.49
NG = 0
DB = 0
SM = 0
Doxycycline
YLR105C SEN2
Delta_AUC
ZShift = 1.44
NG = 0
DB = 0
SM = 0
Doxycycline

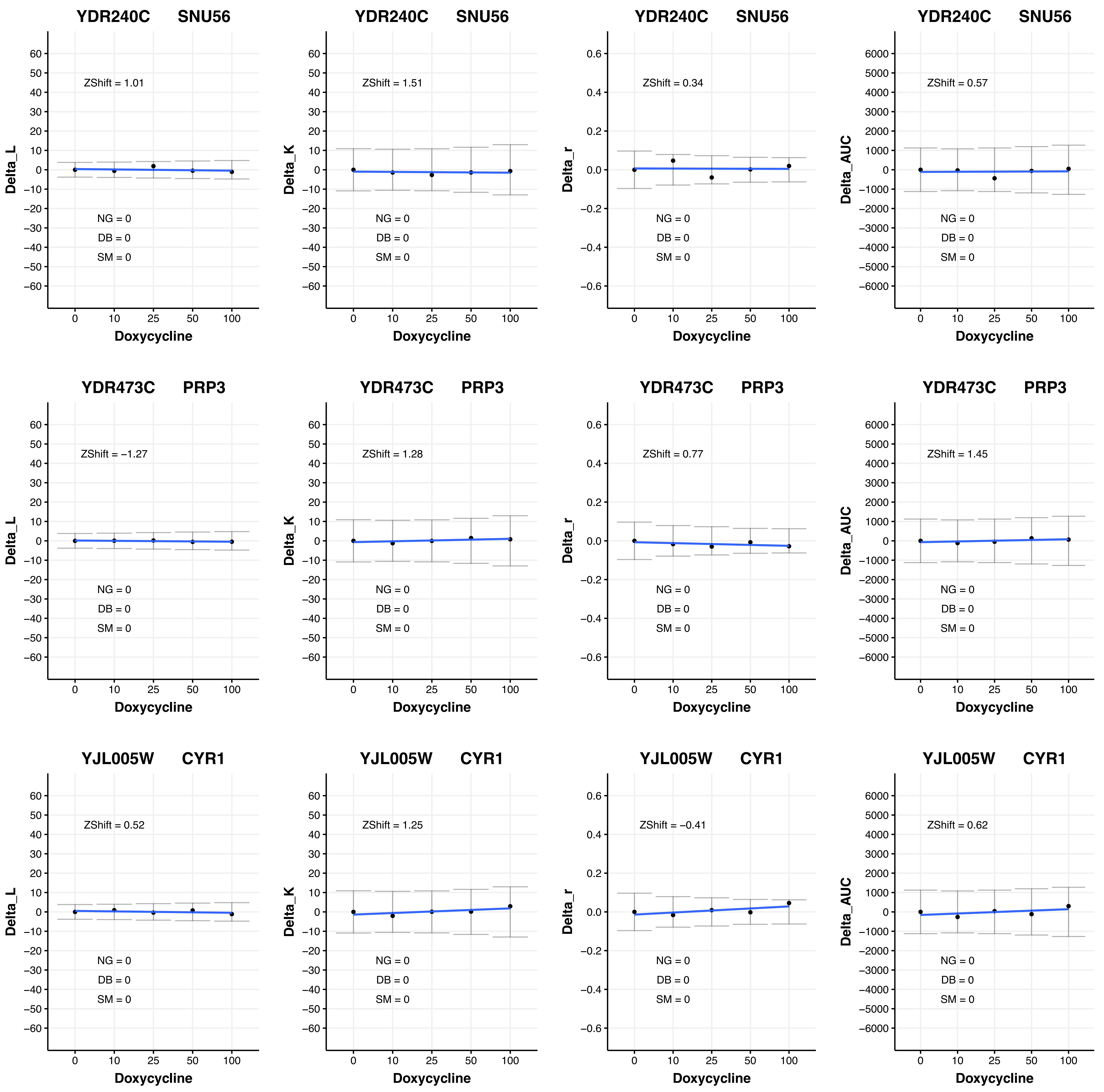

YDR240C SNU56
ZShift = 1.01
NG = 0
DB = 0
SM = 0
Delta_L
Doxycycline
YDR240C SNU56
ZShift = 1.51
NG = 0
DB = 0
SM = 0
Delta_K
Doxycycline
YDR240C SNU56
ZShift = 0.34
NG = 0
DB = 0
SM = 0
Delta_r
Doxycycline
YDR240C SNU56
ZShift = 0.57
NG = 0
DB = 0
SM = 0
Delta_AUC
Doxycycline
YDR473C PRP3
ZShift = −1.27
NG = 0
DB = 0
SM = 0
Delta_L
Doxycycline
YDR473C PRP3
ZShift = 1.28
NG = 0
DB = 0
SM = 0
Delta_K
Doxycycline
YDR473C PRP3
ZShift = 0.77
NG = 0
DB = 0
SM = 0
Delta_r
Doxycycline
YDR473C PRP3
ZShift = 1.45
NG = 0
DB = 0
SM = 0
Delta_AUC
Doxycycline
YJL005W CYR1
ZShift = 0.52
NG = 0
DB = 0
SM = 0
Delta_L
Doxycycline
YJL005W CYR1
ZShift = 1.25
NG = 0
DB = 0
SM = 0
Delta_K
Doxycycline
YJL005W CYR1
ZShift = −0.41
NG = 0
DB = 0
SM = 0
Delta_r
Doxycycline
YJL005W CYR1
ZShift = 0.62
NG = 0
DB = 0
SM = 0
Delta_AUC
Doxycycline
0 10 25 50 100

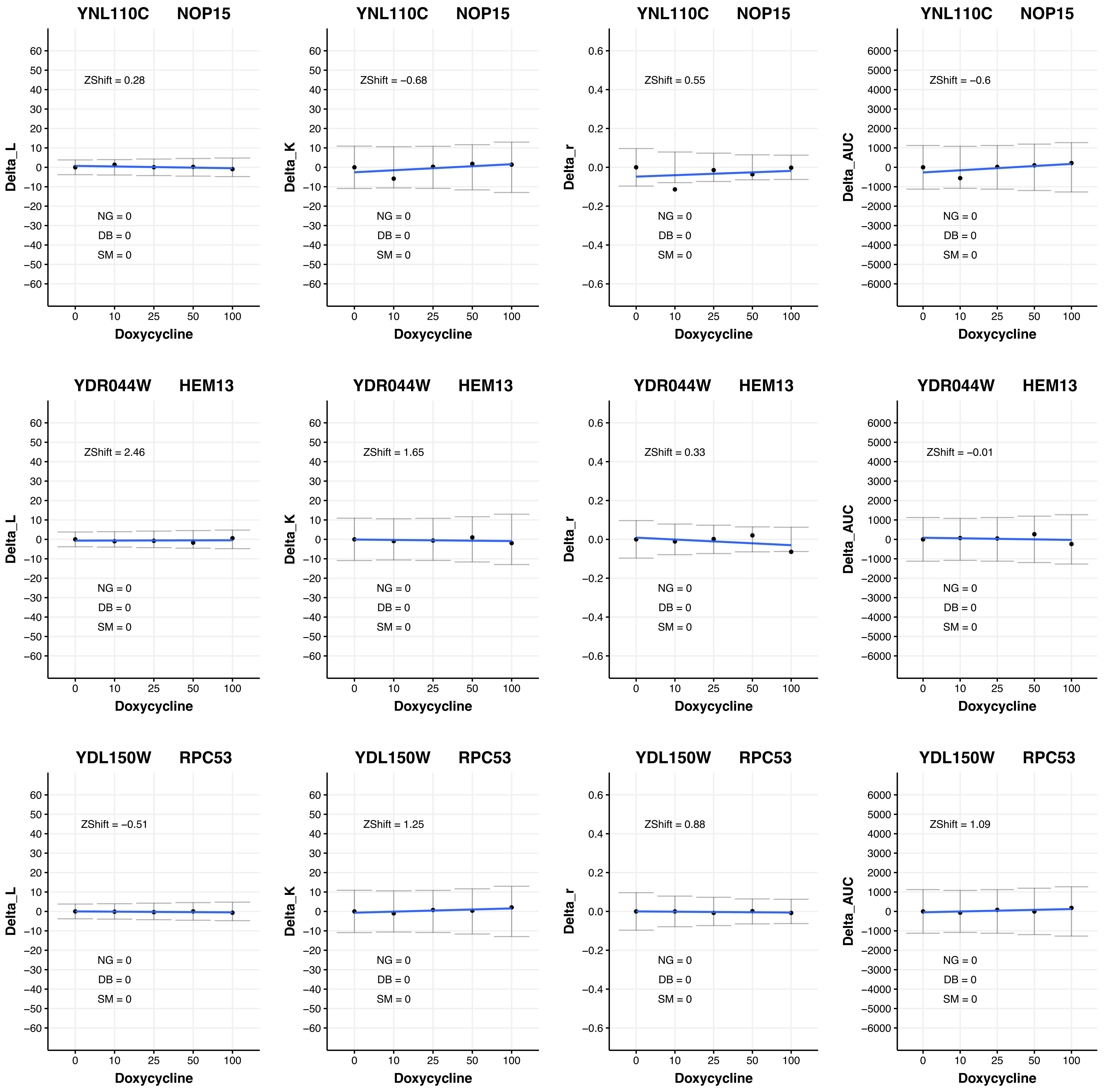
YNL110C NOP15
ZShift = 0.28
NG = 0
DB = 0
SM = 0
Delta_L
Doxycycline
YNL110C NOP15
ZShift = −0.68
NG = 0
DB = 0
SM = 0
Delta_K
Doxycycline
YNL110C NOP15
ZShift = 0.55
NG = 0
DB = 0
SM = 0
Delta_r
Doxycycline
YNL110C NOP15
ZShift = −0.6
NG = 0
DB = 0
SM = 0
Delta_AUC
Doxycycline
YDR044W HEM13
ZShift = 2.46
NG = 0
DB = 0
SM = 0
Delta_L
Doxycycline
YDR044W HEM13
ZShift = 1.65
NG = 0
DB = 0
SM = 0
Delta_K
Doxycycline
YDR044W HEM13
ZShift = 0.33
NG = 0
DB = 0
SM = 0
Delta_r
Doxycycline
YDR044W HEM13
ZShift = −0.01
NG = 0
DB = 0
SM = 0
Delta_AUC
Doxycycline
YDL150W RPC53
ZShift = −0.51
NG = 0
DB = 0
SM = 0
Delta_L
Doxycycline
YDL150W RPC53
ZShift = 1.25
NG = 0
DB = 0
SM = 0
Delta_K
Doxycycline
YDL150W RPC53
ZShift = 0.88
NG = 0
DB = 0
SM = 0
Delta_r
Doxycycline
YDL150W RPC53
ZShift = 1.09
NG = 0
DB = 0
SM = 0
Delta_AUC
Doxycycline

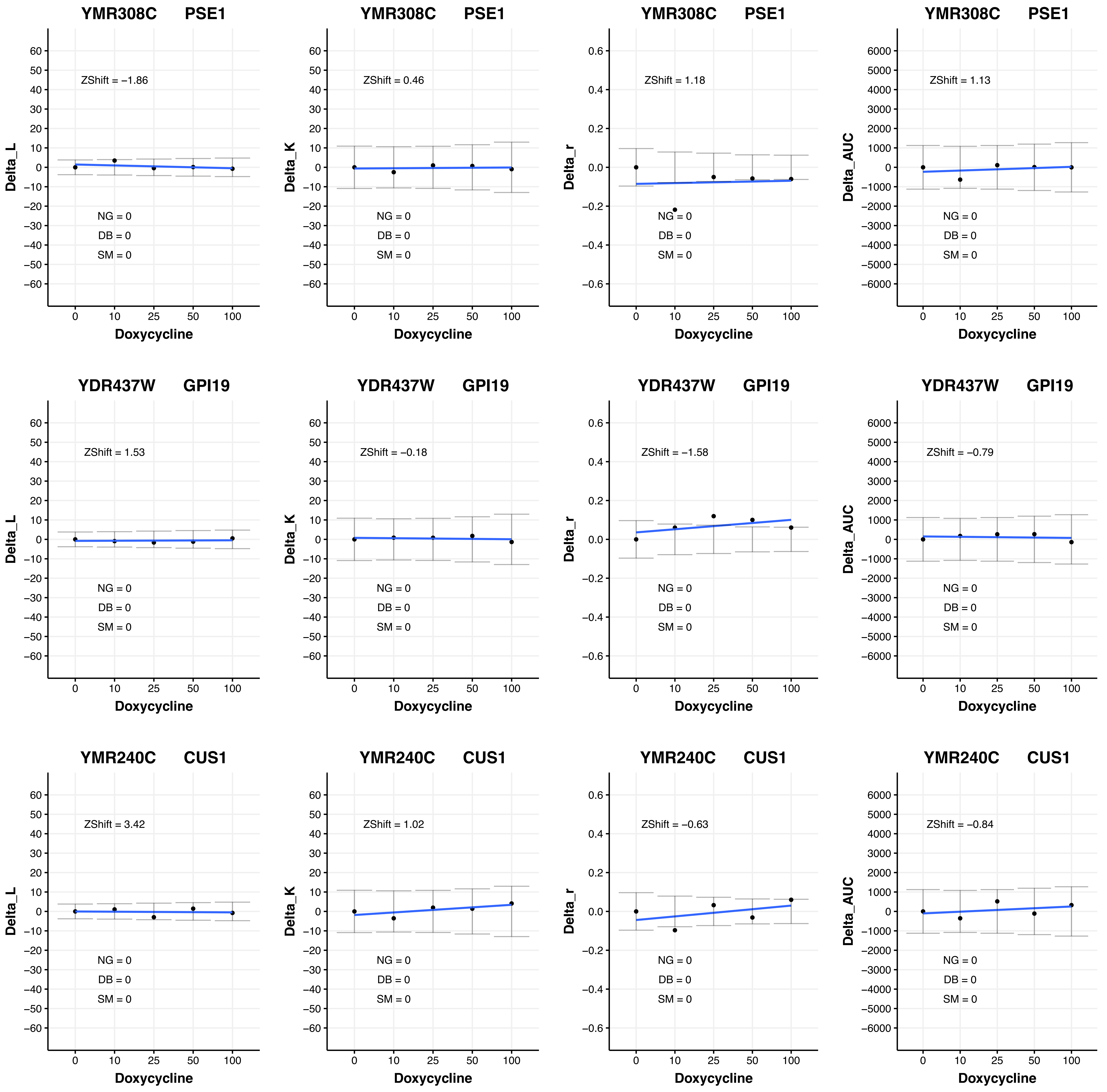

YMR308C PSE1
ZShift = −1.86
NG = 0
DB = 0
SM = 0
Delta_L
Doxycycline
YMR308C PSE1
ZShift = 0.46
NG = 0
DB = 0
SM = 0
Delta_K
Doxycycline
YMR308C PSE1
ZShift = 1.18
NG = 0
DB = 0
SM = 0
Delta_r
Doxycycline
YMR308C PSE1
ZShift = 1.13
NG = 0
DB = 0
SM = 0
Delta_AUC
Doxycycline
YDR437W GPI19
ZShift = 1.53
NG = 0
DB = 0
SM = 0
Delta_L
Doxycycline
YDR437W GPI19
ZShift = −0.18
NG = 0
DB = 0
SM = 0
Delta_K
Doxycycline
YDR437W GPI19
ZShift = −1.58
NG = 0
DB = 0
SM = 0
Delta_r
Doxycycline
YDR437W GPI19
ZShift = −0.79
NG = 0
DB = 0
SM = 0
Delta_AUC
Doxycycline
YMR240C CUS1
ZShift = 3.42
NG = 0
DB = 0
SM = 0
Delta_L
Doxycycline
YMR240C CUS1
ZShift = 1.02
NG = 0
DB = 0
SM = 0
Delta_K
Doxycycline
YMR240C CUS1
ZShift = −0.63
NG = 0
DB = 0
SM = 0
Delta_r
Doxycycline
YMR240C CUS1
ZShift = −0.84
NG = 0
DB = 0
SM = 0
Delta_AUC
Doxycycline

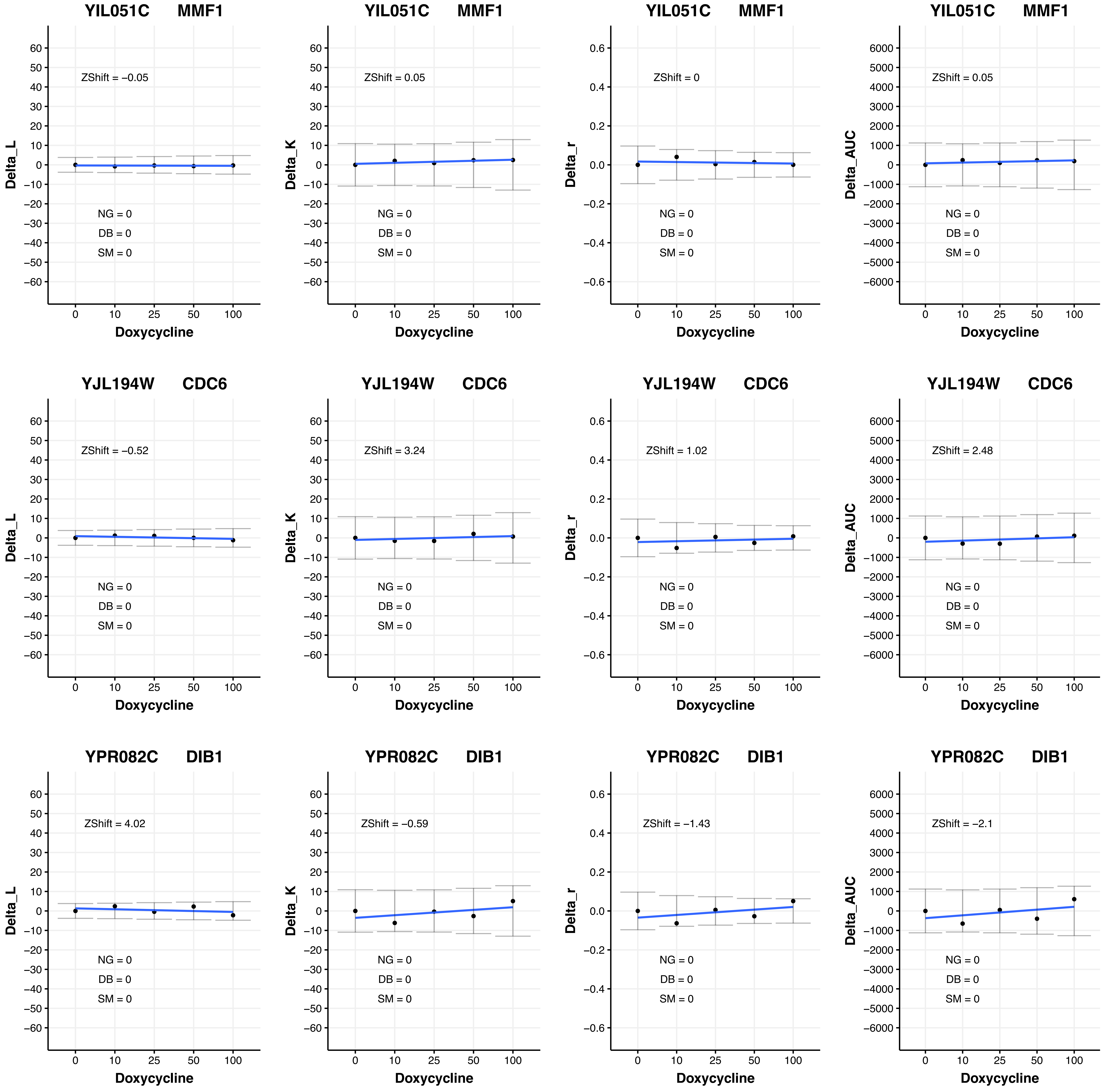
YIL051C MMF1
ZShift = −0.05
ZShift = 0.05
ZShift = 0
ZShift = 0.05
YJL194W CDC6
ZShift = −0.52
ZShift = 3.24
ZShift = 1.02
ZShift = 2.48
YPR082C DIB1
ZShift = 4.02
ZShift = −0.59
ZShift = −1.43
ZShift = −2.1
NG = 0
DB = 0
SM = 0
Delta_L
Delta_K
Delta_r
Delta_AUC
Doxycycline
0
10
25
50
100

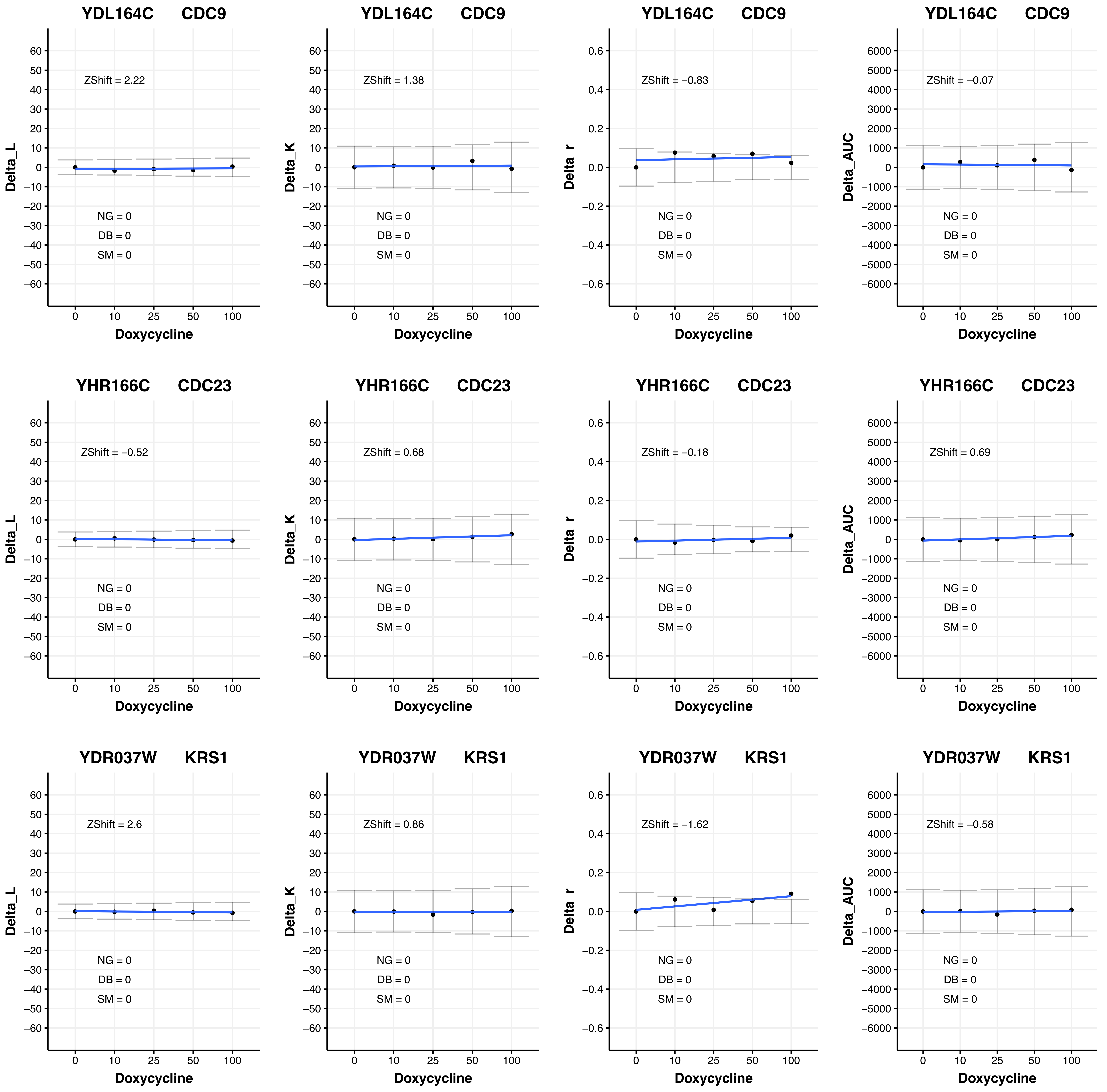

YDL164C CDC9
ZShift = 2.22
NG = 0
DB = 0
SM = 0
Delta_L
Doxycycline
YDL164C CDC9
ZShift = 1.38
NG = 0
DB = 0
SM = 0
Delta_K
Doxycycline
YDL164C CDC9
ZShift = −0.83
NG = 0
DB = 0
SM = 0
Delta_r
Doxycycline
YDL164C CDC9
ZShift = −0.07
NG = 0
DB = 0
SM = 0
Delta_AUC
Doxycycline
YHR166C CDC23
ZShift = −0.52
NG = 0
DB = 0
SM = 0
Delta_L
Doxycycline
YHR166C CDC23
ZShift = 0.68
NG = 0
DB = 0
SM = 0
Delta_K
Doxycycline
YHR166C CDC23
ZShift = −0.18
NG = 0
DB = 0
SM = 0
Delta_r
Doxycycline
YHR166C CDC23
ZShift = 0.69
NG = 0
DB = 0
SM = 0
Delta_AUC
Doxycycline
YDR037W KRS1
ZShift = 2.6
NG = 0
DB = 0
SM = 0
Delta_L
Doxycycline
YDR037W KRS1
ZShift = 0.86
NG = 0
DB = 0
SM = 0
Delta_K
Doxycycline
YDR037W KRS1
ZShift = −1.62
NG = 0
DB = 0
SM = 0
Delta_r
Doxycycline
YDR037W KRS1
ZShift = −0.58
NG = 0
DB = 0
SM = 0
Delta_AUC
Doxycycline

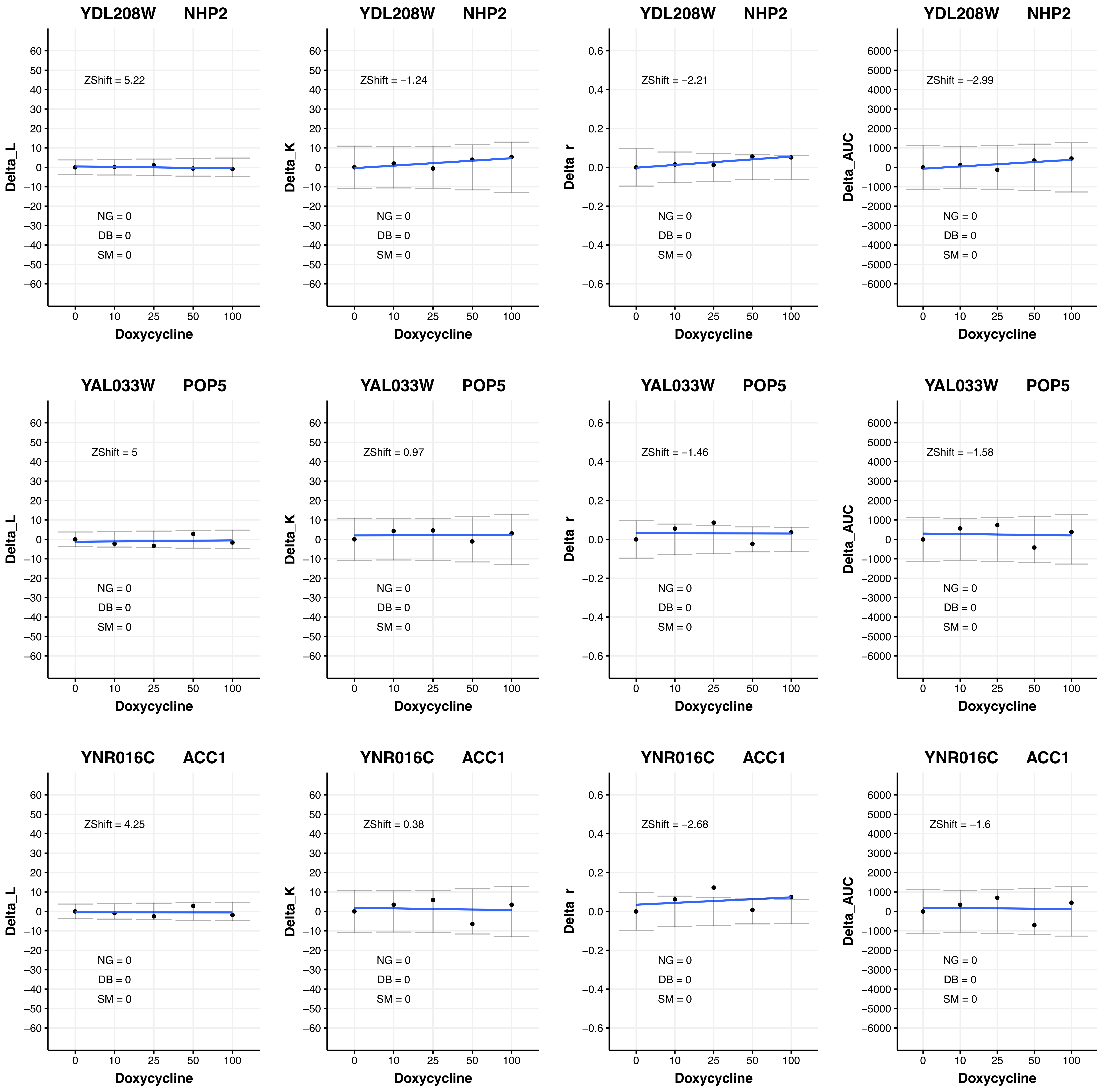
YDL208W NHP2
ZShift = 5.22
Delta_L
NG = 0
DB = 0
SM = 0
Doxycycline
YDL208W NHP2
ZShift = −1.24
Delta_K
NG = 0
DB = 0
SM = 0
Doxycycline
YDL208W NHP2
ZShift = −2.21
Delta_r
NG = 0
DB = 0
SM = 0
Doxycycline
YDL208W NHP2
ZShift = −2.99
Delta_AUC
NG = 0
DB = 0
SM = 0
Doxycycline
YAL033W POP5
ZShift = 5
Delta_L
NG = 0
DB = 0
SM = 0
Doxycycline
YAL033W POP5
ZShift = 0.97
Delta_K
NG = 0
DB = 0
SM = 0
Doxycycline
YAL033W POP5
ZShift = −1.46
Delta_r
NG = 0
DB = 0
SM = 0
Doxycycline
YAL033W POP5
ZShift = −1.58
Delta_AUC
NG = 0
DB = 0
SM = 0
Doxycycline
YNR016C ACC1
ZShift = 4.25
Delta_L
NG = 0
DB = 0
SM = 0
Doxycycline
YNR016C ACC1
ZShift = 0.38
Delta_K
NG = 0
DB = 0
SM = 0
Doxycycline
YNR016C ACC1
ZShift = −2.68
Delta_r
NG = 0
DB = 0
SM = 0
Doxycycline
YNR016C ACC1
ZShift = −1.6
Delta_AUC
NG = 0
DB = 0
SM = 0
Doxycycline

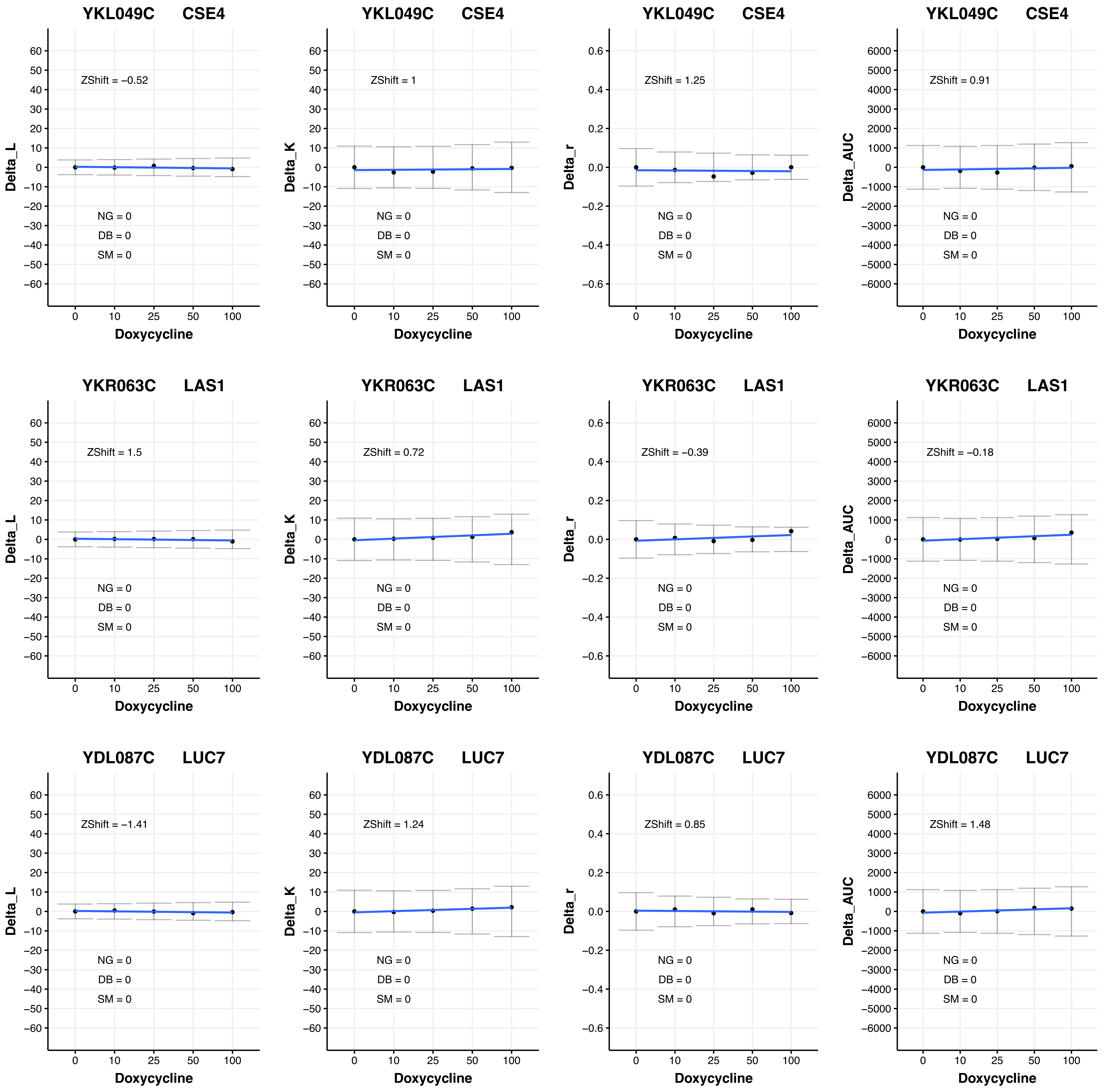

YKL049C CSE4
ZShift = −0.52
NG = 0
DB = 0
SM = 0
Delta_L
Doxycycline
YKL049C CSE4
ZShift = 1
NG = 0
DB = 0
SM = 0
Delta_K
Doxycycline
YKL049C CSE4
ZShift = 1.25
NG = 0
DB = 0
SM = 0
Delta_r
Doxycycline
YKL049C CSE4
ZShift = 0.91
NG = 0
DB = 0
SM = 0
Delta_AUC
Doxycycline
YKR063C LAS1
ZShift = 1.5
NG = 0
DB = 0
SM = 0
Delta_L
Doxycycline
YKR063C LAS1
ZShift = 0.72
NG = 0
DB = 0
SM = 0
Delta_K
Doxycycline
YKR063C LAS1
ZShift = −0.39
NG = 0
DB = 0
SM = 0
Delta_r
Doxycycline
YKR063C LAS1
ZShift = −0.18
NG = 0
DB = 0
SM = 0
Delta_AUC
Doxycycline
YDL087C LUC7
ZShift = −1.41
NG = 0
DB = 0
SM = 0
Delta_L
Doxycycline
YDL087C LUC7
ZShift = 1.24
NG = 0
DB = 0
SM = 0
Delta_K
Doxycycline
YDL087C LUC7
ZShift = 0.85
NG = 0
DB = 0
SM = 0
Delta_r
Doxycycline
YDL087C LUC7
ZShift = 1.48
NG = 0
DB = 0
SM = 0
Delta_AUC
Doxycycline
0
10
25
50
100

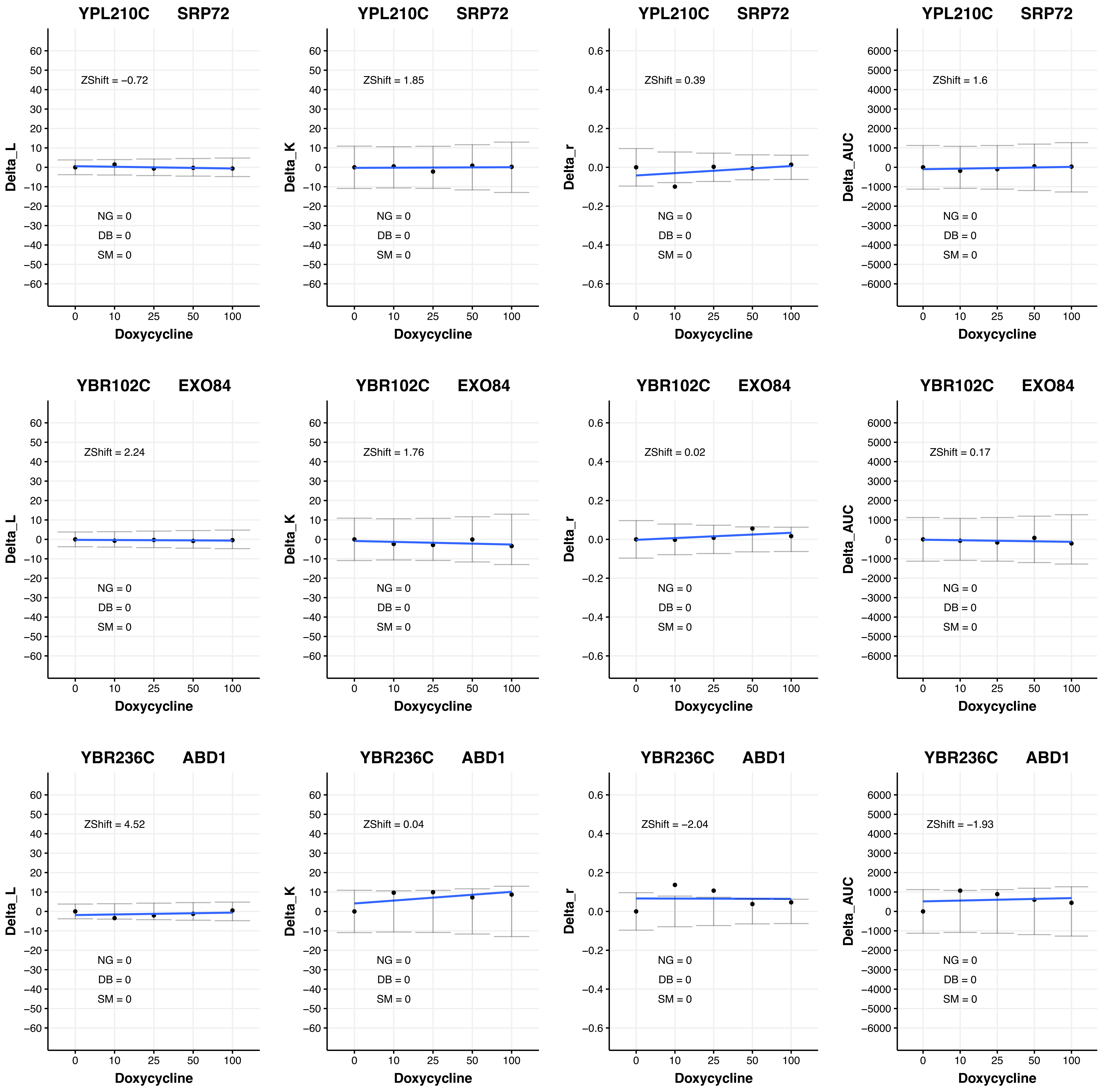
YPL210C SRP72
ZShift = −0.72
NG = 0
DB = 0
SM = 0
Delta_L
Doxycycline
YPL210C SRP72
ZShift = 1.85
NG = 0
DB = 0
SM = 0
Delta_K
Doxycycline
YPL210C SRP72
ZShift = 0.39
NG = 0
DB = 0
SM = 0
Delta_r
Doxycycline
YPL210C SRP72
ZShift = 1.6
NG = 0
DB = 0
SM = 0
Delta_AUC
Doxycycline
YBR102C EXO84
ZShift = 2.24
NG = 0
DB = 0
SM = 0
Delta_L
Doxycycline
YBR102C EXO84
ZShift = 1.76
NG = 0
DB = 0
SM = 0
Delta_K
Doxycycline
YBR102C EXO84
ZShift = 0.02
NG = 0
DB = 0
SM = 0
Delta_r
Doxycycline
YBR102C EXO84
ZShift = 0.17
NG = 0
DB = 0
SM = 0
Delta_AUC
Doxycycline
YBR236C ABD1
ZShift = 4.52
NG = 0
DB = 0
SM = 0
Delta_L
Doxycycline
YBR236C ABD1
ZShift = 0.04
NG = 0
DB = 0
SM = 0
Delta_K
Doxycycline
YBR236C ABD1
ZShift = −2.04
NG = 0
DB = 0
SM = 0
Delta_r
Doxycycline
YBR236C ABD1
ZShift = −1.93
NG = 0
DB = 0
SM = 0
Delta_AUC
Doxycycline

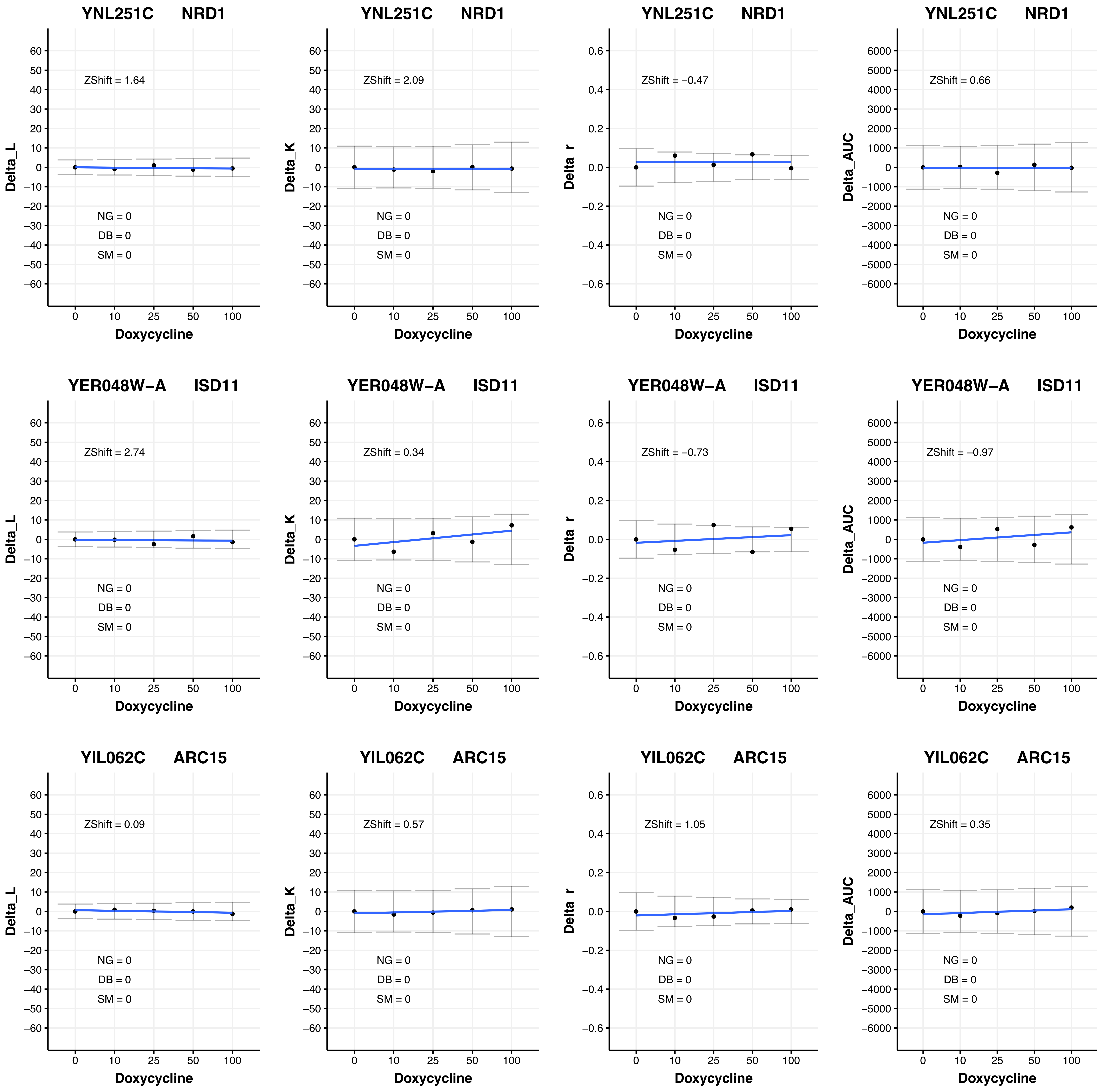
YNL251C NRD1
ZShift = 1.64
NG = 0
DB = 0
SM = 0
Delta_L
Doxycycline
YNL251C NRD1
ZShift = 2.09
NG = 0
DB = 0
SM = 0
Delta_K
Doxycycline
YNL251C NRD1
ZShift = −0.47
NG = 0
DB = 0
SM = 0
Delta_r
Doxycycline
YNL251C NRD1
ZShift = 0.66
NG = 0
DB = 0
SM = 0
Delta_AUC
Doxycycline
YER048W−A ISD11
ZShift = 2.74
NG = 0
DB = 0
SM = 0
Delta_L
Doxycycline
YER048W−A ISD11
ZShift = 0.34
NG = 0
DB = 0
SM = 0
Delta_K
Doxycycline
YER048W−A ISD11
ZShift = −0.73
NG = 0
DB = 0
SM = 0
Delta_r
Doxycycline
YER048W−A ISD11
ZShift = −0.97
NG = 0
DB = 0
SM = 0
Delta_AUC
Doxycycline
YIL062C ARC15
ZShift = 0.09
NG = 0
DB = 0
SM = 0
Delta_L
Doxycycline
YIL062C ARC15
ZShift = 0.57
NG = 0
DB = 0
SM = 0
Delta_K
Doxycycline
YIL062C ARC15
ZShift = 1.05
NG = 0
DB = 0
SM = 0
Delta_r
Doxycycline
YIL062C ARC15
ZShift = 0.35
NG = 0
DB = 0
SM = 0
Delta_AUC
Doxycycline

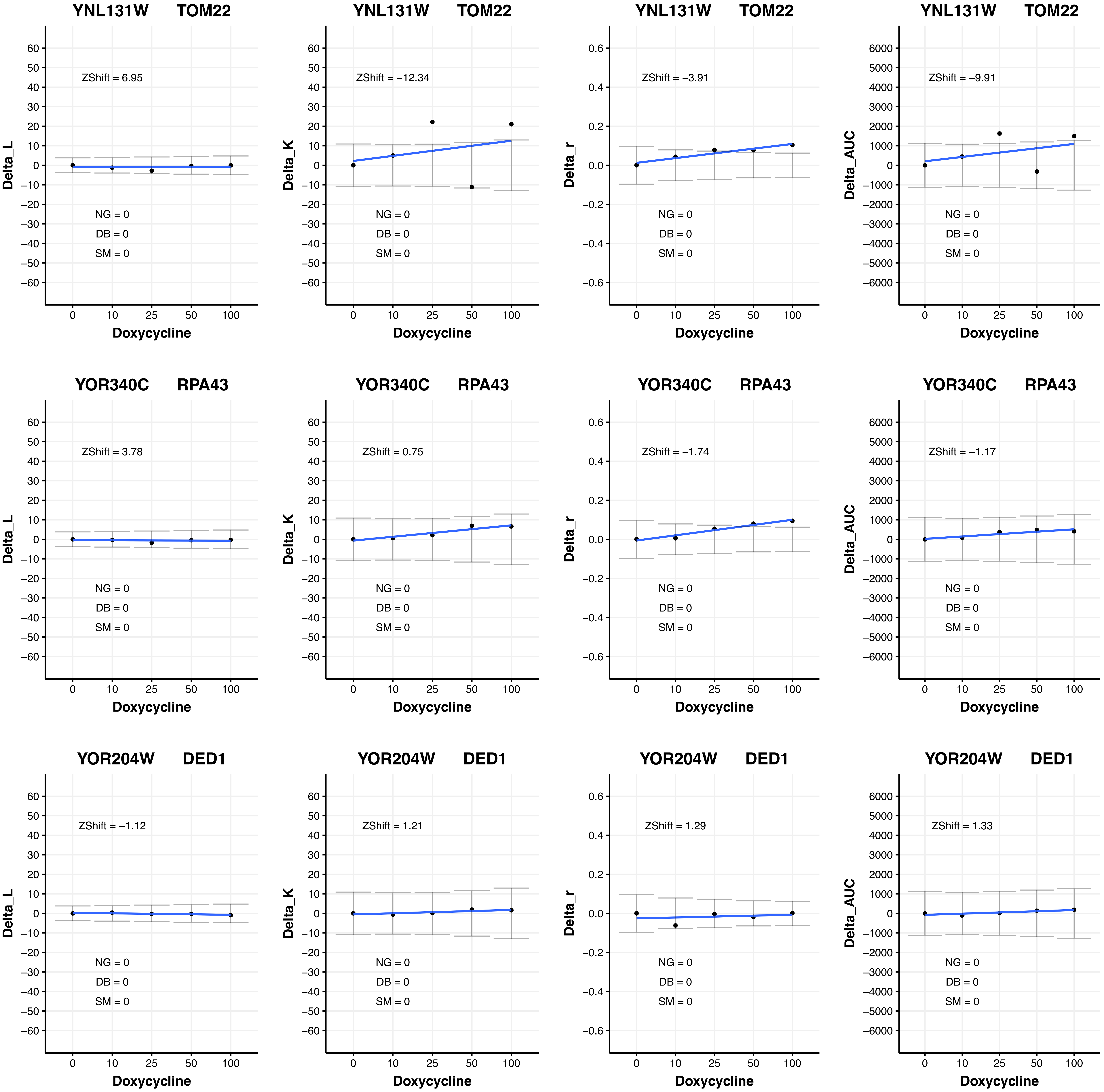

YNL131W TOM22
ZShift = 6.95
Delta_L
NG = 0
DB = 0
SM = 0
Doxycycline
YNL131W TOM22
ZShift = −12.34
Delta_K
NG = 0
DB = 0
SM = 0
Doxycycline
YNL131W TOM22
ZShift = −3.91
Delta_r
NG = 0
DB = 0
SM = 0
Doxycycline
YNL131W TOM22
ZShift = −9.91
Delta_AUC
NG = 0
DB = 0
SM = 0
Doxycycline
YOR340C RPA43
ZShift = 3.78
Delta_L
NG = 0
DB = 0
SM = 0
Doxycycline
YOR340C RPA43
ZShift = 0.75
Delta_K
NG = 0
DB = 0
SM = 0
Doxycycline
YOR340C RPA43
ZShift = −1.74
Delta_r
NG = 0
DB = 0
SM = 0
Doxycycline
YOR340C RPA43
ZShift = −1.17
Delta_AUC
NG = 0
DB = 0
SM = 0
Doxycycline
YOR204W DED1
ZShift = −1.12
Delta_L
NG = 0
DB = 0
SM = 0
Doxycycline
YOR204W DED1
ZShift = 1.21
Delta_K
NG = 0
DB = 0
SM = 0
Doxycycline
YOR204W DED1
ZShift = 1.29
Delta_r
NG = 0
DB = 0
SM = 0
Doxycycline
YOR204W DED1
ZShift = 1.33
Delta_AUC
NG = 0
DB = 0
SM = 0
Doxycycline

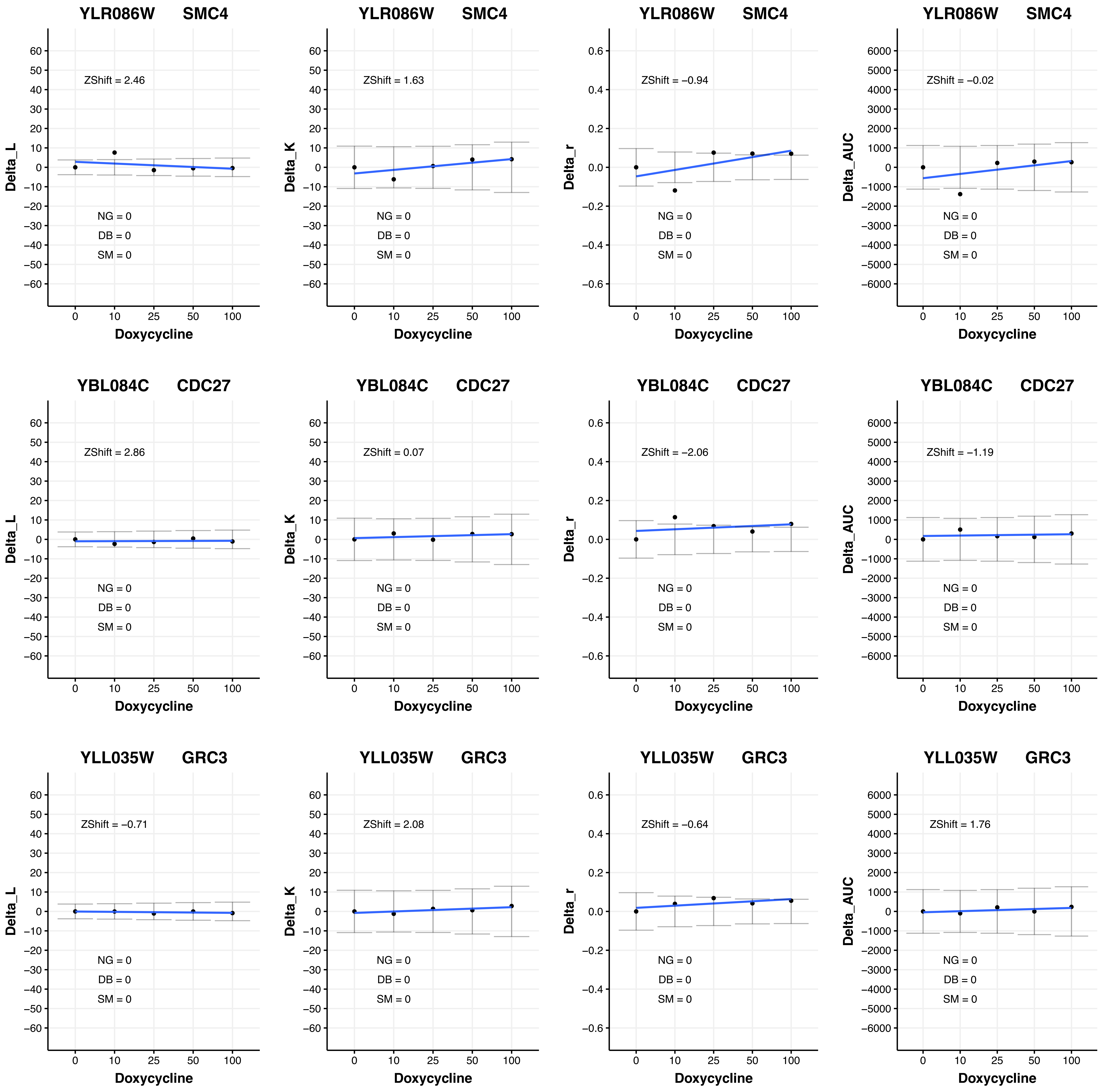
YLR086W SMC4
ZShift = 2.46
Delta_L
NG = 0
DB = 0
SM = 0
Doxycycline
YLR086W SMC4
ZShift = 1.63
Delta_K
NG = 0
DB = 0
SM = 0
Doxycycline
YLR086W SMC4
ZShift = −0.94
Delta_r
NG = 0
DB = 0
SM = 0
Doxycycline
YLR086W SMC4
ZShift = −0.02
Delta_AUC
NG = 0
DB = 0
SM = 0
Doxycycline
YBL084C CDC27
ZShift = 2.86
Delta_L
NG = 0
DB = 0
SM = 0
Doxycycline
YBL084C CDC27
ZShift = 0.07
Delta_K
NG = 0
DB = 0
SM = 0
Doxycycline
YBL084C CDC27
ZShift = −2.06
Delta_r
NG = 0
DB = 0
SM = 0
Doxycycline
YBL084C CDC27
ZShift = −1.19
Delta_AUC
NG = 0
DB = 0
SM = 0
Doxycycline
YLL035W GRC3
ZShift = −0.71
Delta_L
NG = 0
DB = 0
SM = 0
Doxycycline
YLL035W GRC3
ZShift = 2.08
Delta_K
NG = 0
DB = 0
SM = 0
Doxycycline
YLL035W GRC3
ZShift = −0.64
Delta_r
NG = 0
DB = 0
SM = 0
Doxycycline
YLL035W GRC3
ZShift = 1.76
Delta_AUC
NG = 0
DB = 0
SM = 0
Doxycycline

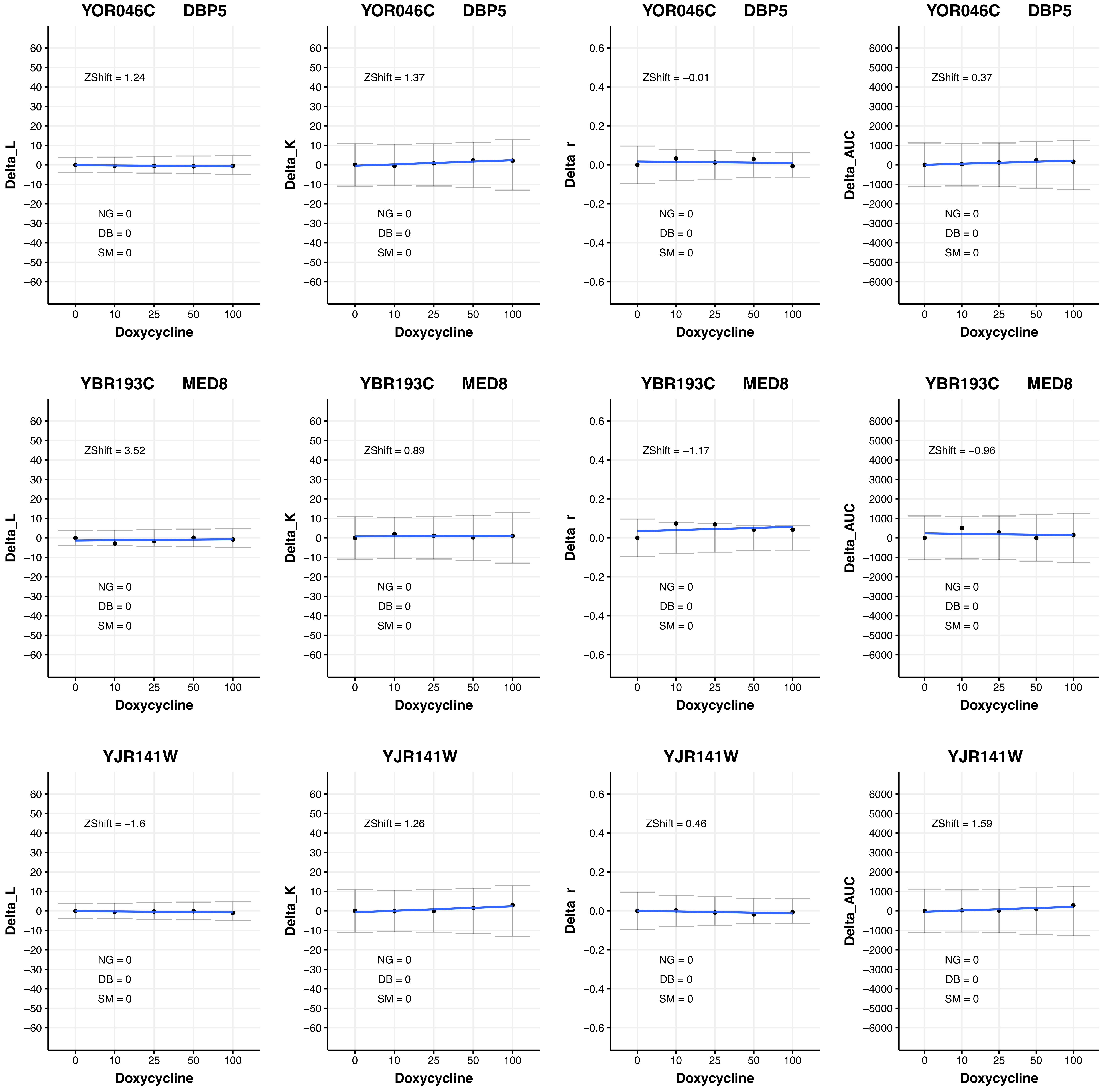

YOR046C DBP5
ZShift = 1.24
Delta_L
NG = 0
DB = 0
SM = 0
Doxycycline
YOR046C DBP5
ZShift = 1.37
Delta_K
NG = 0
DB = 0
SM = 0
Doxycycline
YOR046C DBP5
ZShift = −0.01
Delta_r
NG = 0
DB = 0
SM = 0
Doxycycline
YOR046C DBP5
ZShift = 0.37
Delta_AUC
NG = 0
DB = 0
SM = 0
Doxycycline
YBR193C MED8
ZShift = 3.52
Delta_L
NG = 0
DB = 0
SM = 0
Doxycycline
YBR193C MED8
ZShift = 0.89
Delta_K
NG = 0
DB = 0
SM = 0
Doxycycline
YBR193C MED8
ZShift = −1.17
Delta_r
NG = 0
DB = 0
SM = 0
Doxycycline
YBR193C MED8
ZShift = −0.96
Delta_AUC
NG = 0
DB = 0
SM = 0
Doxycycline
YJR141W
ZShift = −1.6
Delta_L
NG = 0
DB = 0
SM = 0
Doxycycline
YJR141W
ZShift = 1.26
Delta_K
NG = 0
DB = 0
SM = 0
Doxycycline
YJR141W
ZShift = 0.46
Delta_r
NG = 0
DB = 0
SM = 0
Doxycycline
YJR141W
ZShift = 1.59
Delta_AUC
NG = 0
DB = 0
SM = 0
Doxycycline

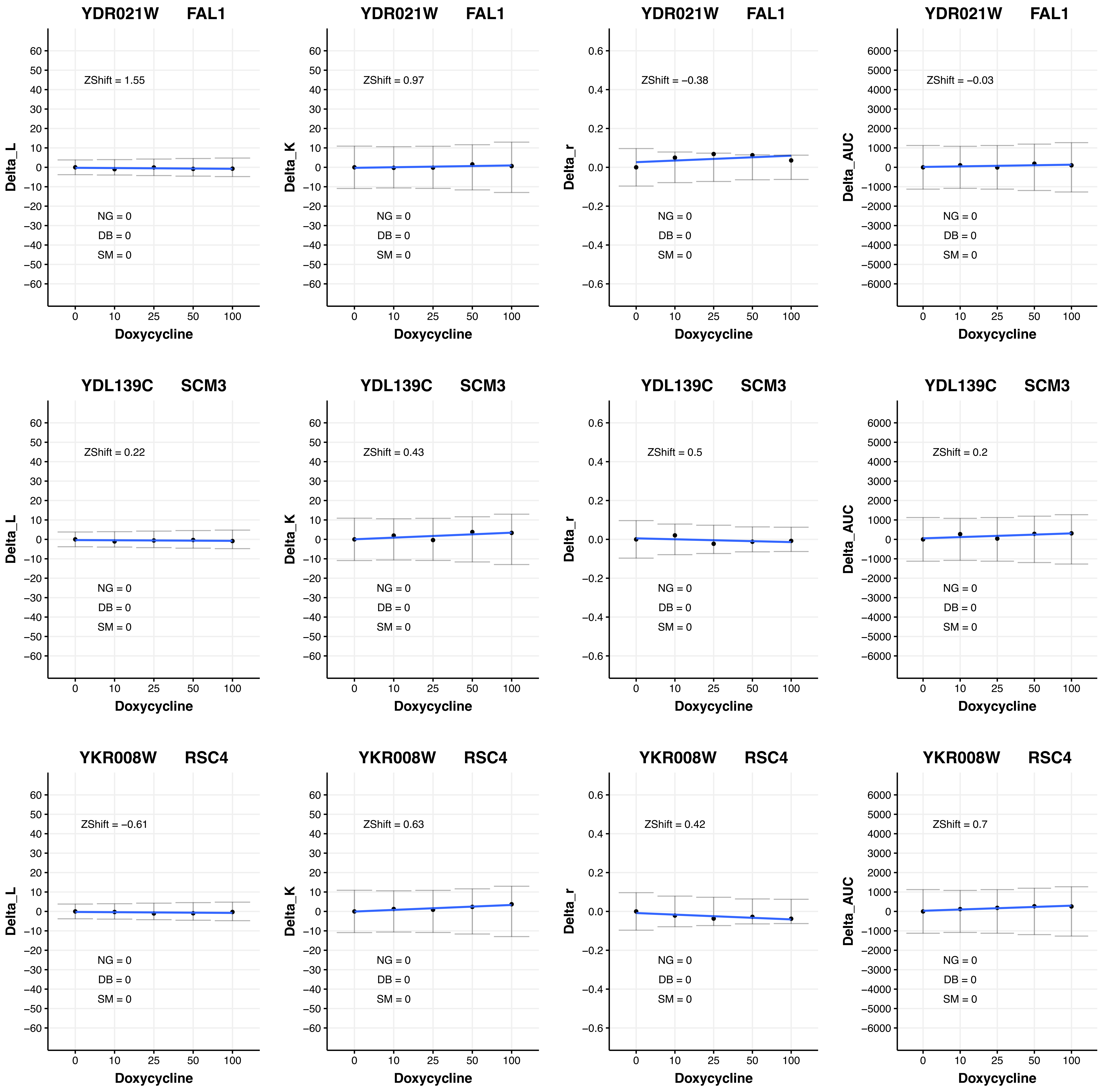
YDR021W FAL1
ZShift = 1.55
Delta_L
NG = 0
DB = 0
SM = 0
Doxycycline
YDR021W FAL1
ZShift = 0.97
Delta_K
NG = 0
DB = 0
SM = 0
Doxycycline
YDR021W FAL1
ZShift = −0.38
Delta_r
NG = 0
DB = 0
SM = 0
Doxycycline
YDR021W FAL1
ZShift = −0.03
Delta_AUC
NG = 0
DB = 0
SM = 0
Doxycycline
YDL139C SCM3
ZShift = 0.22
Delta_L
NG = 0
DB = 0
SM = 0
Doxycycline
YDL139C SCM3
ZShift = 0.43
Delta_K
NG = 0
DB = 0
SM = 0
Doxycycline
YDL139C SCM3
ZShift = 0.5
Delta_r
NG = 0
DB = 0
SM = 0
Doxycycline
YDL139C SCM3
ZShift = 0.2
Delta_AUC
NG = 0
DB = 0
SM = 0
Doxycycline
YKR008W RSC4
ZShift = −0.61
Delta_L
NG = 0
DB = 0
SM = 0
Doxycycline
YKR008W RSC4
ZShift = 0.63
Delta_K
NG = 0
DB = 0
SM = 0
Doxycycline
YKR008W RSC4
ZShift = 0.42
Delta_r
NG = 0
DB = 0
SM = 0
Doxycycline
YKR008W RSC4
ZShift = 0.7
Delta_AUC
NG = 0
DB = 0
SM = 0
Doxycycline

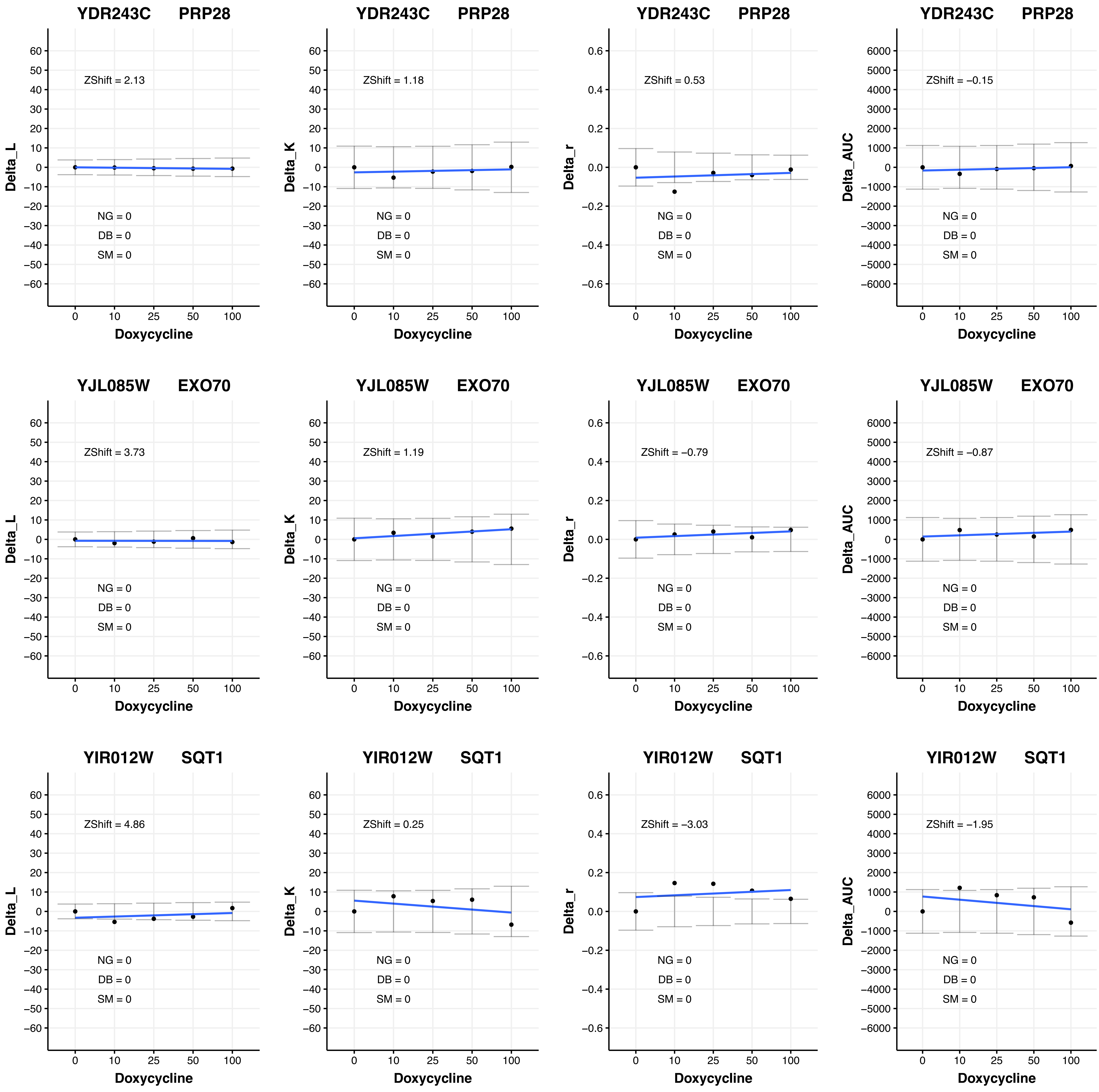

YDR243C PRP28
ZShift = 2.13
NG = 0
DB = 0
SM = 0
Delta_L
Doxycycline
YDR243C PRP28
ZShift = 1.18
NG = 0
DB = 0
SM = 0
Delta_K
Doxycycline
YDR243C PRP28
ZShift = 0.53
NG = 0
DB = 0
SM = 0
Delta_r
Doxycycline
YDR243C PRP28
ZShift = −0.15
NG = 0
DB = 0
SM = 0
Delta_AUC
Doxycycline
YJL085W EXO70
ZShift = 3.73
NG = 0
DB = 0
SM = 0
Delta_L
Doxycycline
YJL085W EXO70
ZShift = 1.19
NG = 0
DB = 0
SM = 0
Delta_K
Doxycycline
YJL085W EXO70
ZShift = −0.79
NG = 0
DB = 0
SM = 0
Delta_r
Doxycycline
YJL085W EXO70
ZShift = −0.87
NG = 0
DB = 0
SM = 0
Delta_AUC
Doxycycline
YIR012W SQT1
ZShift = 4.86
NG = 0
DB = 0
SM = 0
Delta_L
Doxycycline
YIR012W SQT1
ZShift = 0.25
NG = 0
DB = 0
SM = 0
Delta_K
Doxycycline
YIR012W SQT1
ZShift = −3.03
NG = 0
DB = 0
SM = 0
Delta_r
Doxycycline
YIR012W SQT1
ZShift = −1.95
NG = 0
DB = 0
SM = 0
Delta_AUC
Doxycycline

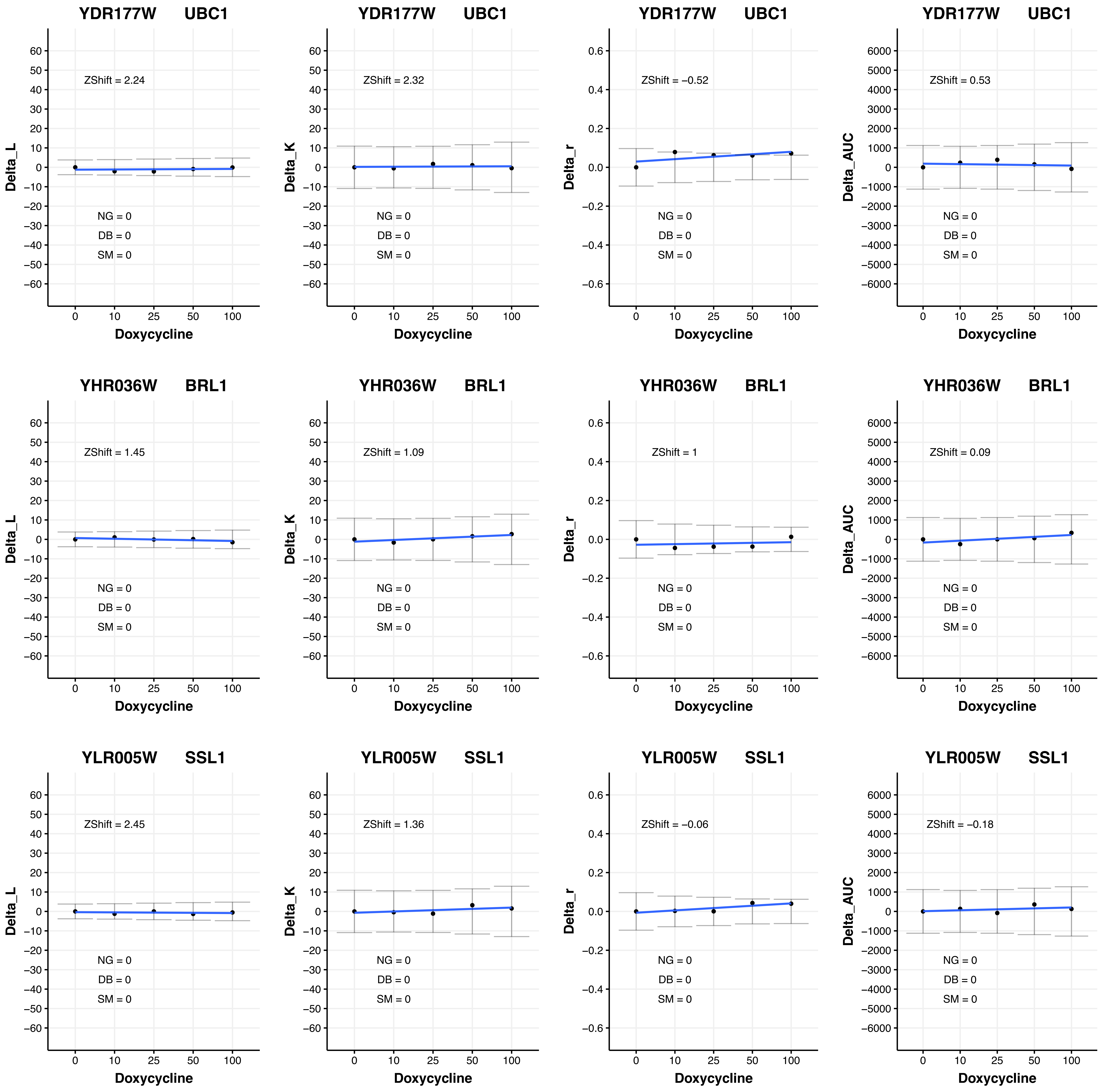
YDR177W UBC1
ZShift = 2.24
NG = 0
DB = 0
SM = 0
Delta_L
Doxycycline
YDR177W UBC1
ZShift = 2.32
NG = 0
DB = 0
SM = 0
Delta_K
Doxycycline
YDR177W UBC1
ZShift = −0.52
NG = 0
DB = 0
SM = 0
Delta_r
Doxycycline
YDR177W UBC1
ZShift = 0.53
NG = 0
DB = 0
SM = 0
Delta_AUC
Doxycycline
YHR036W BRL1
ZShift = 1.45
NG = 0
DB = 0
SM = 0
Delta_L
Doxycycline
YHR036W BRL1
ZShift = 1.09
NG = 0
DB = 0
SM = 0
Delta_K
Doxycycline
YHR036W BRL1
ZShift = 1
NG = 0
DB = 0
SM = 0
Delta_r
Doxycycline
YHR036W BRL1
ZShift = 0.09
NG = 0
DB = 0
SM = 0
Delta_AUC
Doxycycline
YLR005W SSL1
ZShift = 2.45
NG = 0
DB = 0
SM = 0
Delta_L
Doxycycline
YLR005W SSL1
ZShift = 1.36
NG = 0
DB = 0
SM = 0
Delta_K
Doxycycline
YLR005W SSL1
ZShift = −0.06
NG = 0
DB = 0
SM = 0
Delta_r
Doxycycline
YLR005W SSL1
ZShift = −0.18
NG = 0
DB = 0
SM = 0
Delta_AUC
Doxycycline

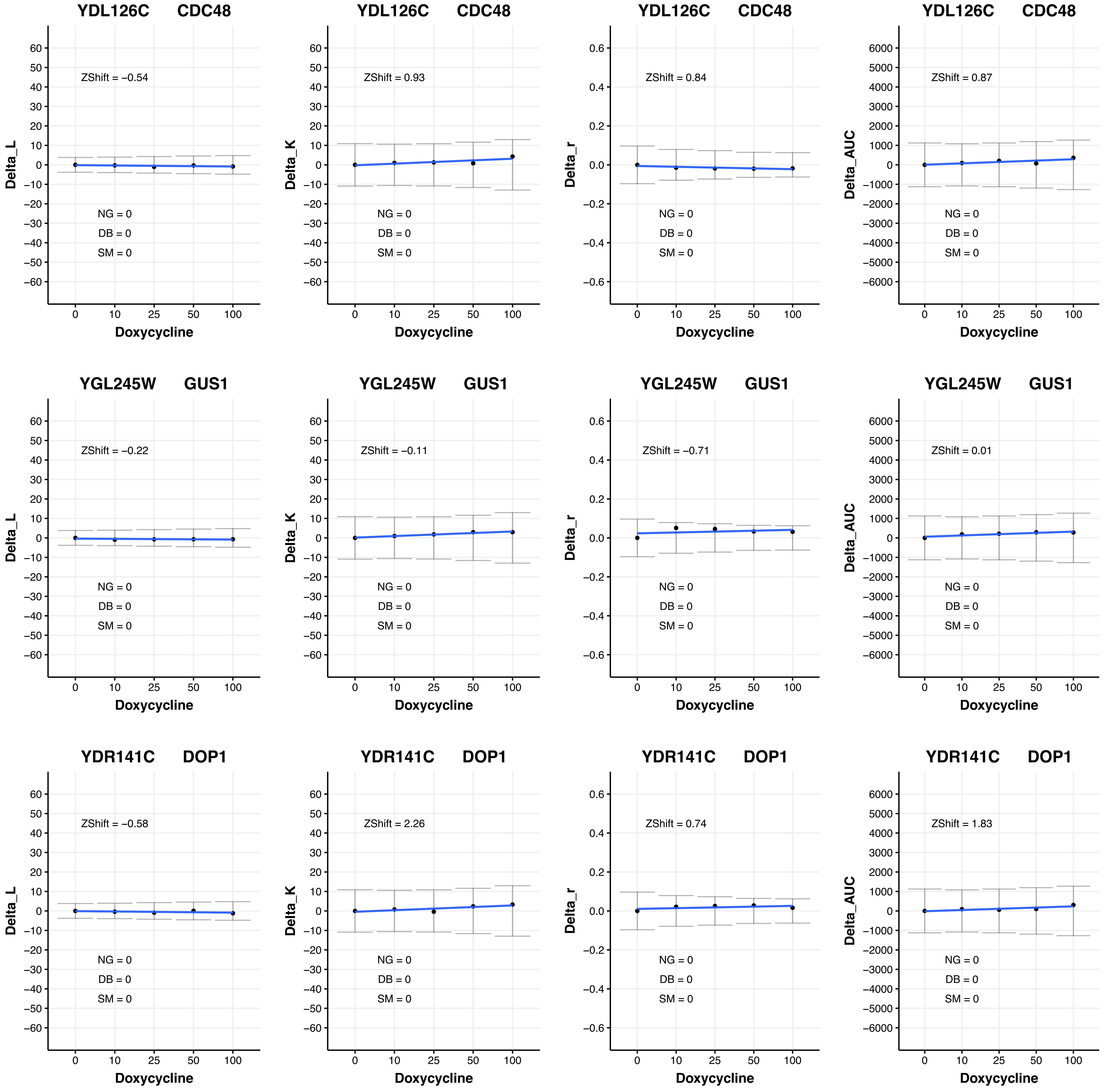

YDL126C CDC48
Delta_L
ZShift = −0.54
NG = 0
DB = 0
SM = 0
Doxycycline
YDL126C CDC48
Delta_K
ZShift = 0.93
NG = 0
DB = 0
SM = 0
Doxycycline
YDL126C CDC48
Delta_r
ZShift = 0.84
NG = 0
DB = 0
SM = 0
Doxycycline
YDL126C CDC48
Delta_AUC
ZShift = 0.87
NG = 0
DB = 0
SM = 0
Doxycycline
YGL245W GUS1
Delta_L
ZShift = −0.22
NG = 0
DB = 0
SM = 0
Doxycycline
YGL245W GUS1
Delta_K
ZShift = −0.11
NG = 0
DB = 0
SM = 0
Doxycycline
YGL245W GUS1
Delta_r
ZShift = −0.71
NG = 0
DB = 0
SM = 0
Doxycycline
YGL245W GUS1
Delta_AUC
ZShift = 0.01
NG = 0
DB = 0
SM = 0
Doxycycline
YDR141C DOP1
Delta_L
ZShift = −0.58
NG = 0
DB = 0
SM = 0
Doxycycline
YDR141C DOP1
Delta_K
ZShift = 2.26
NG = 0
DB = 0
SM = 0
Doxycycline
YDR141C DOP1
Delta_r
ZShift = 0.74
NG = 0
DB = 0
SM = 0
Doxycycline
YDR141C DOP1
Delta_AUC
ZShift = 1.83
NG = 0
DB = 0
SM = 0
Doxycycline

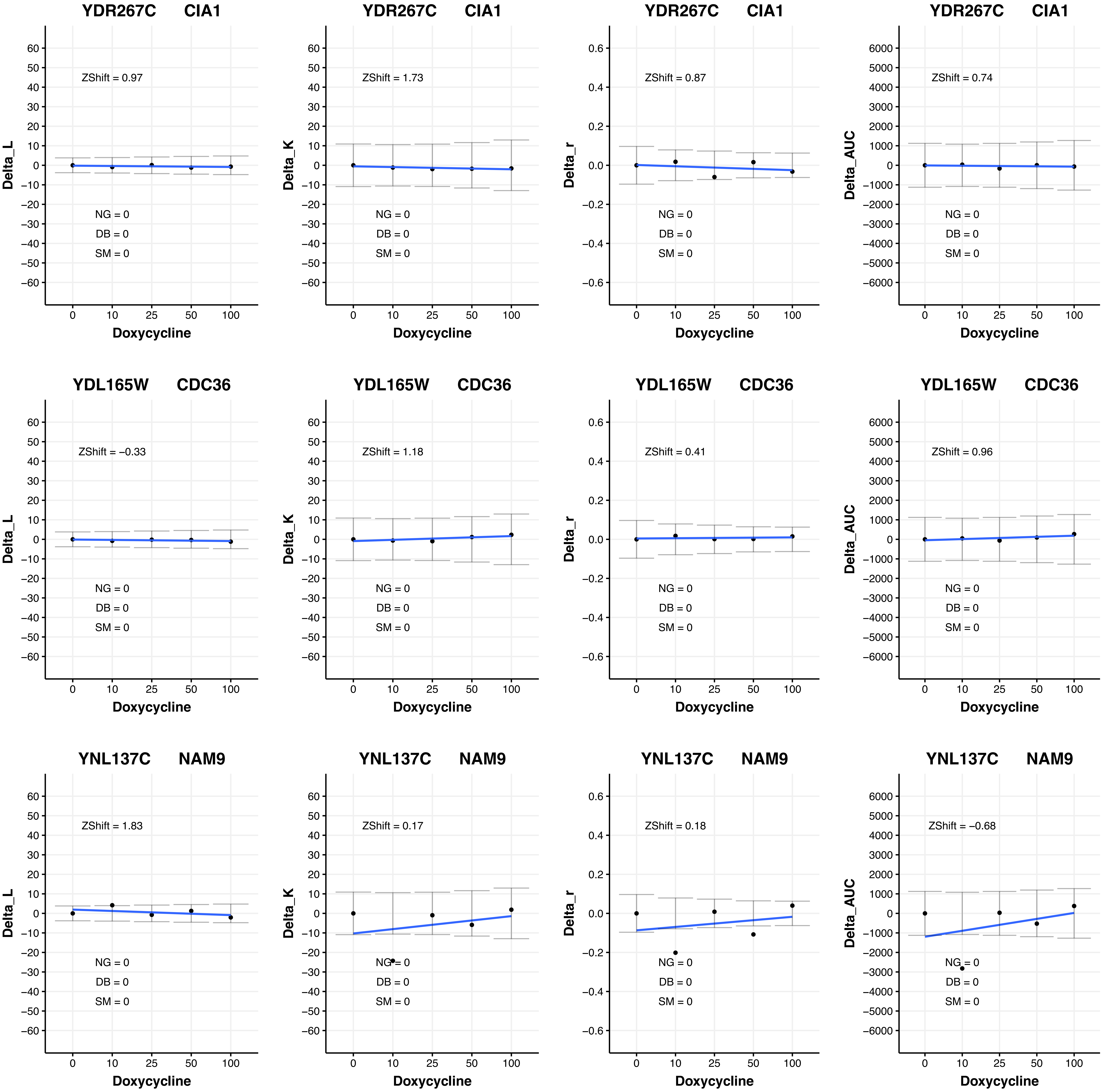

YDR267C CIA1
ZShift = 0.97
NG = 0
DB = 0
SM = 0
Delta_L
Doxycycline
YDR267C CIA1
ZShift = 1.73
NG = 0
DB = 0
SM = 0
Delta_K
Doxycycline
YDR267C CIA1
ZShift = 0.87
NG = 0
DB = 0
SM = 0
Delta_r
Doxycycline
YDR267C CIA1
ZShift = 0.74
NG = 0
DB = 0
SM = 0
Delta_AUC
Doxycycline
YDL165W CDC36
ZShift = −0.33
NG = 0
DB = 0
SM = 0
Delta_L
Doxycycline
YDL165W CDC36
ZShift = 1.18
NG = 0
DB = 0
SM = 0
Delta_K
Doxycycline
YDL165W CDC36
ZShift = 0.41
NG = 0
DB = 0
SM = 0
Delta_r
Doxycycline
YDL165W CDC36
ZShift = 0.96
NG = 0
DB = 0
SM = 0
Delta_AUC
Doxycycline
YNL137C NAM9
ZShift = 1.83
NG = 0
DB = 0
SM = 0
Delta_L
Doxycycline
YNL137C NAM9
ZShift = 0.17
NG = 0
DB = 0
SM = 0
Delta_K
Doxycycline
YNL137C NAM9
ZShift = 0.18
NG = 0
DB = 0
SM = 0
Delta_r
Doxycycline
YNL137C NAM9
ZShift = −0.68
NG = 0
DB = 0
SM = 0
Delta_AUC
Doxycycline

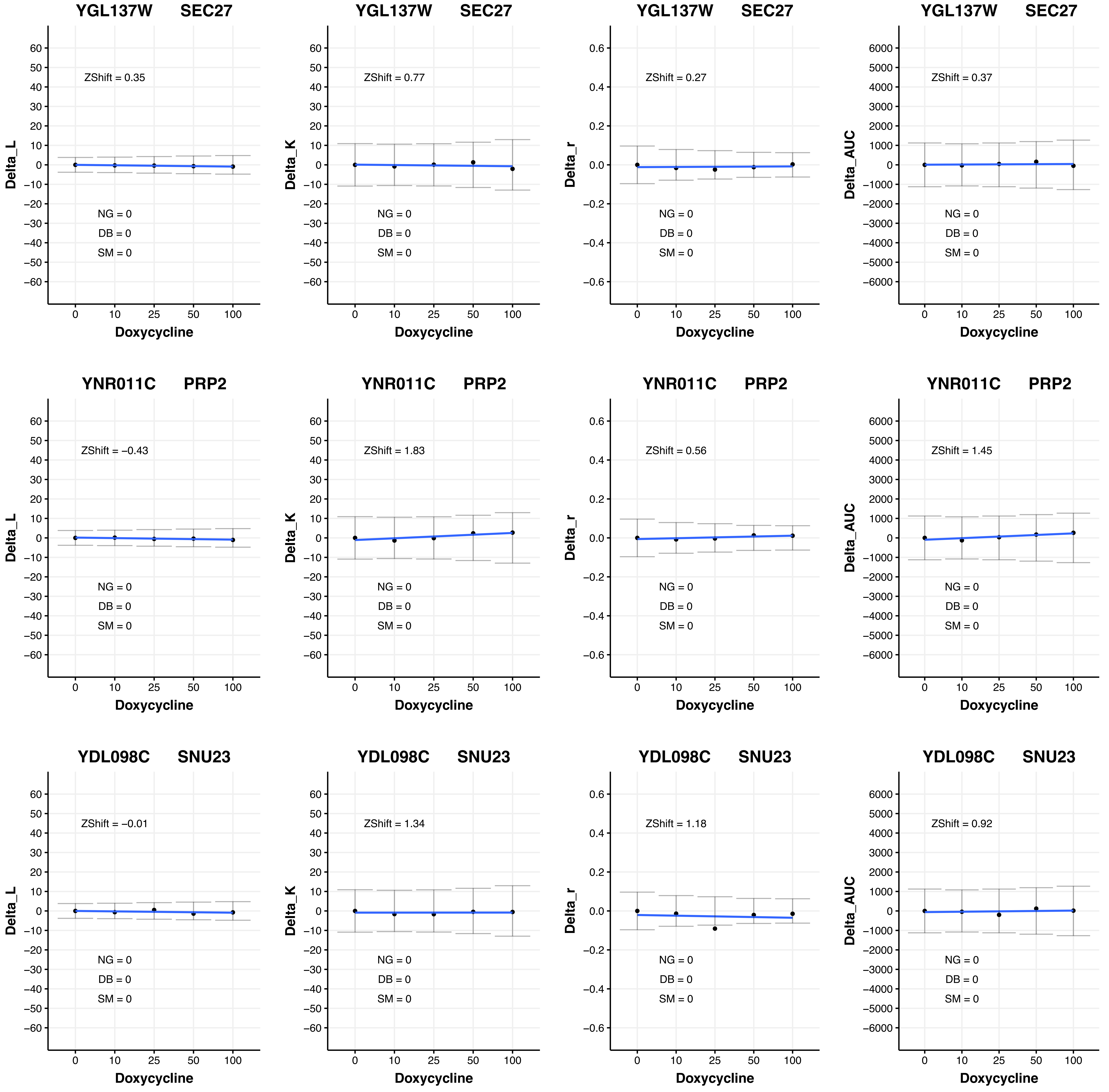

YGL137W SEC27
ZShift = 0.35
Delta_L
NG = 0
DB = 0
SM = 0
Doxycycline
YGL137W SEC27
ZShift = 0.77
Delta_K
NG = 0
DB = 0
SM = 0
Doxycycline
YGL137W SEC27
ZShift = 0.27
Delta_r
NG = 0
DB = 0
SM = 0
Doxycycline
YGL137W SEC27
ZShift = 0.37
Delta_AUC
NG = 0
DB = 0
SM = 0
Doxycycline
YNR011C PRP2
ZShift = −0.43
Delta_L
NG = 0
DB = 0
SM = 0
Doxycycline
YNR011C PRP2
ZShift = 1.83
Delta_K
NG = 0
DB = 0
SM = 0
Doxycycline
YNR011C PRP2
ZShift = 0.56
Delta_r
NG = 0
DB = 0
SM = 0
Doxycycline
YNR011C PRP2
ZShift = 1.45
Delta_AUC
NG = 0
DB = 0
SM = 0
Doxycycline
YDL098C SNU23
ZShift = −0.01
Delta_L
NG = 0
DB = 0
SM = 0
Doxycycline
YDL098C SNU23
ZShift = 1.34
Delta_K
NG = 0
DB = 0
SM = 0
Doxycycline
YDL098C SNU23
ZShift = 1.18
Delta_r
NG = 0
DB = 0
SM = 0
Doxycycline
YDL098C SNU23
ZShift = 0.92
Delta_AUC
NG = 0
DB = 0
SM = 0
Doxycycline

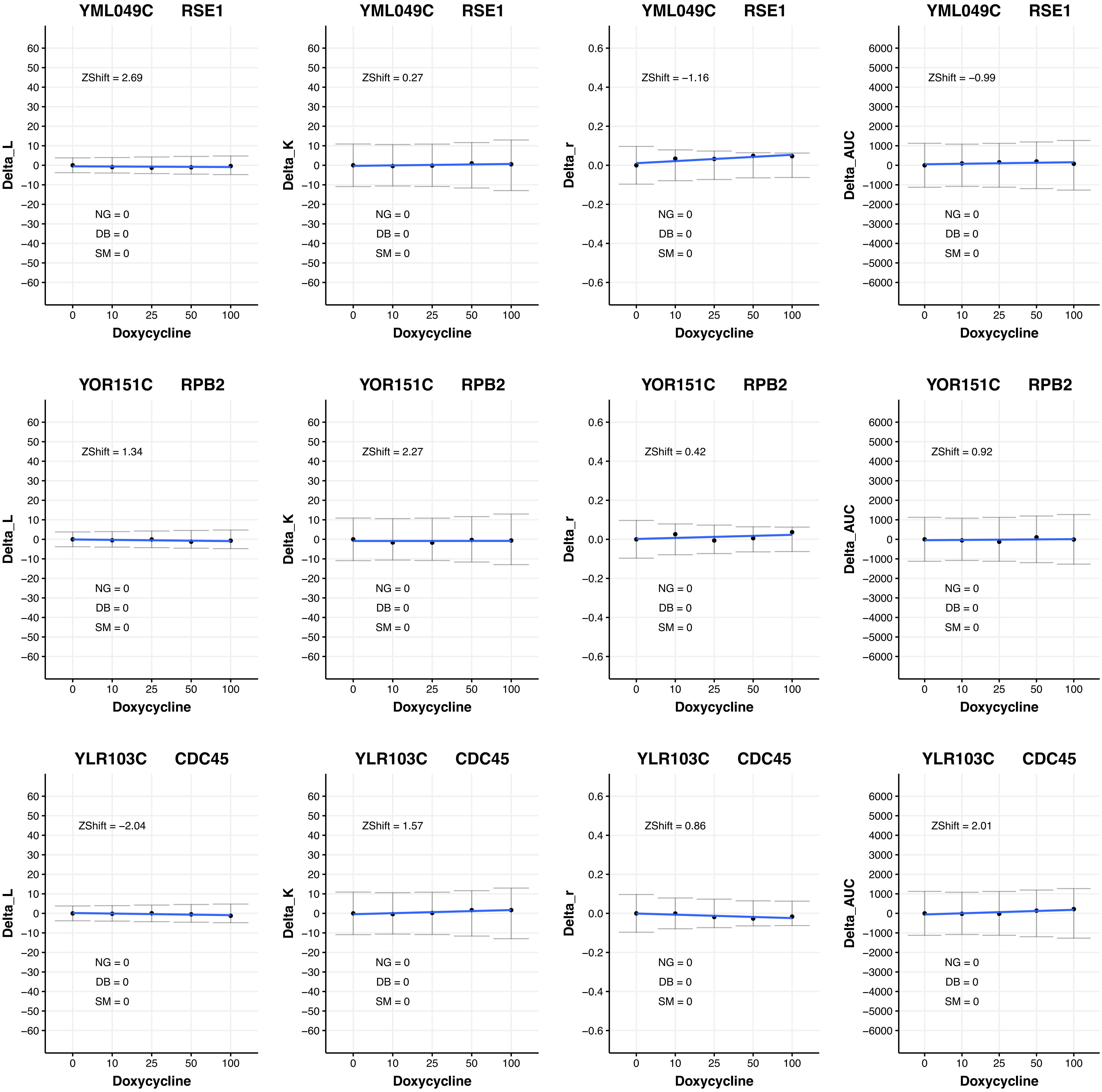

YML049C RSE1
ZShift = 2.69
NG = 0
DB = 0
SM = 0
Delta_L
Doxycycline
YML049C RSE1
ZShift = 0.27
NG = 0
DB = 0
SM = 0
Delta_K
Doxycycline
YML049C RSE1
ZShift = −1.16
NG = 0
DB = 0
SM = 0
Delta_r
Doxycycline
YML049C RSE1
ZShift = −0.99
NG = 0
DB = 0
SM = 0
Delta_AUC
Doxycycline
YOR151C RPB2
ZShift = 1.34
NG = 0
DB = 0
SM = 0
Delta_L
Doxycycline
YOR151C RPB2
ZShift = 2.27
NG = 0
DB = 0
SM = 0
Delta_K
Doxycycline
YOR151C RPB2
ZShift = 0.42
NG = 0
DB = 0
SM = 0
Delta_r
Doxycycline
YOR151C RPB2
ZShift = 0.92
NG = 0
DB = 0
SM = 0
Delta_AUC
Doxycycline
YLR103C CDC45
ZShift = −2.04
NG = 0
DB = 0
SM = 0
Delta_L
Doxycycline
YLR103C CDC45
ZShift = 1.57
NG = 0
DB = 0
SM = 0
Delta_K
Doxycycline
YLR103C CDC45
ZShift = 0.86
NG = 0
DB = 0
SM = 0
Delta_r
Doxycycline
YLR103C CDC45
ZShift = 2.01
NG = 0
DB = 0
SM = 0
Delta_AUC
Doxycycline

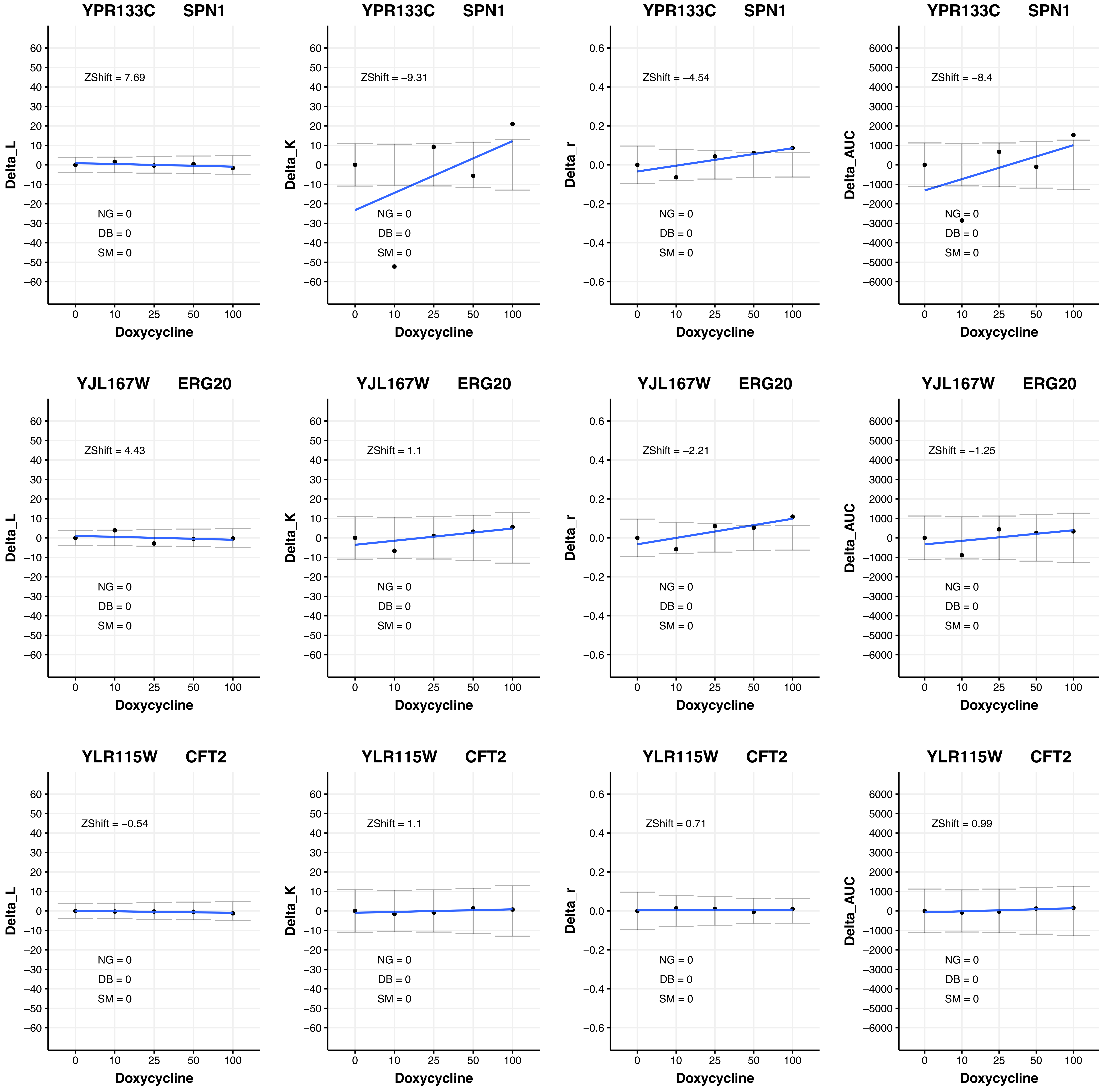

YPR133C SPN1
ZShift = 7.69
NG = 0
DB = 0
SM = 0
Delta_L
Doxycycline
YPR133C SPN1
ZShift = −9.31
NG = 0
DB = 0
SM = 0
Delta_K
Doxycycline
YPR133C SPN1
ZShift = −4.54
NG = 0
DB = 0
SM = 0
Delta_r
Doxycycline
YPR133C SPN1
ZShift = −8.4
NG = 0
DB = 0
SM = 0
Delta_AUC
Doxycycline
YJL167W ERG20
ZShift = 4.43
NG = 0
DB = 0
SM = 0
Delta_L
Doxycycline
YJL167W ERG20
ZShift = 1.1
NG = 0
DB = 0
SM = 0
Delta_K
Doxycycline
YJL167W ERG20
ZShift = −2.21
NG = 0
DB = 0
SM = 0
Delta_r
Doxycycline
YJL167W ERG20
ZShift = −1.25
NG = 0
DB = 0
SM = 0
Delta_AUC
Doxycycline
YLR115W CFT2
ZShift = −0.54
NG = 0
DB = 0
SM = 0
Delta_L
Doxycycline
YLR115W CFT2
ZShift = 1.1
NG = 0
DB = 0
SM = 0
Delta_K
Doxycycline
YLR115W CFT2
ZShift = 0.71
NG = 0
DB = 0
SM = 0
Delta_r
Doxycycline
YLR115W CFT2
ZShift = 0.99
NG = 0
DB = 0
SM = 0
Delta_AUC
Doxycycline

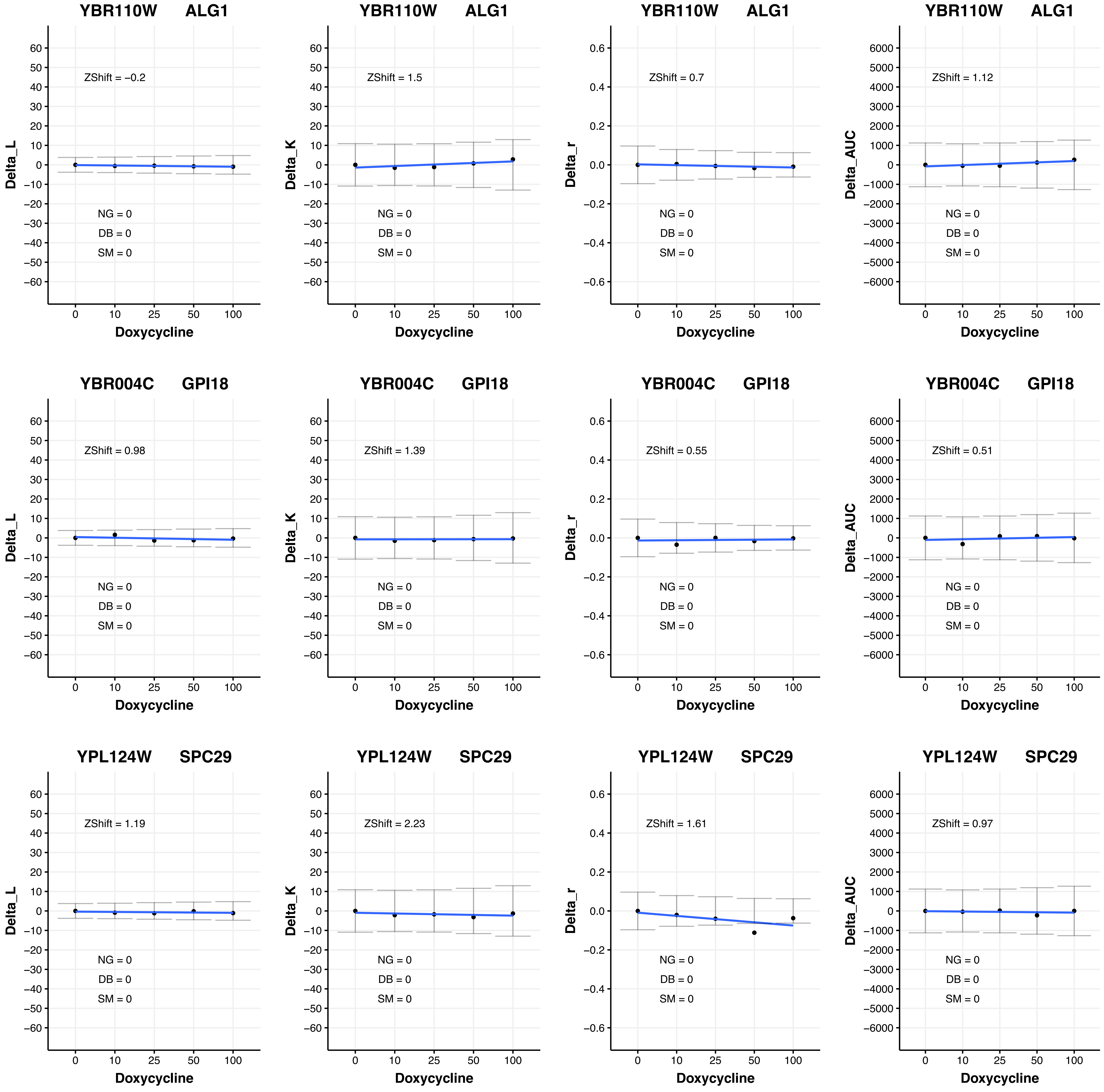
YBR110W ALG1
ZShift = –0.2
NG = 0
DB = 0
SM = 0
Delta_L
Doxycycline
YBR110W ALG1
ZShift = 1.5
NG = 0
DB = 0
SM = 0
Delta_K
Doxycycline
YBR110W ALG1
ZShift = 0.7
NG = 0
DB = 0
SM = 0
Delta_r
Doxycycline
YBR110W ALG1
ZShift = 1.12
NG = 0
DB = 0
SM = 0
Delta_AUC
Doxycycline
YBR004C GPI18
ZShift = 0.98
NG = 0
DB = 0
SM = 0
Delta_L
Doxycycline
YBR004C GPI18
ZShift = 1.39
NG = 0
DB = 0
SM = 0
Delta_K
Doxycycline
YBR004C GPI18
ZShift = 0.55
NG = 0
DB = 0
SM = 0
Delta_r
Doxycycline
YBR004C GPI18
ZShift = 0.51
NG = 0
DB = 0
SM = 0
Delta_AUC
Doxycycline
YPL124W SPC29
ZShift = 1.19
NG = 0
DB = 0
SM = 0
Delta_L
Doxycycline
YPL124W SPC29
ZShift = 2.23
NG = 0
DB = 0
SM = 0
Delta_K
Doxycycline
YPL124W SPC29
ZShift = 1.61
NG = 0
DB = 0
SM = 0
Delta_r
Doxycycline
YPL124W SPC29
ZShift = 0.97
NG = 0
DB = 0
SM = 0
Delta_AUC
Doxycycline

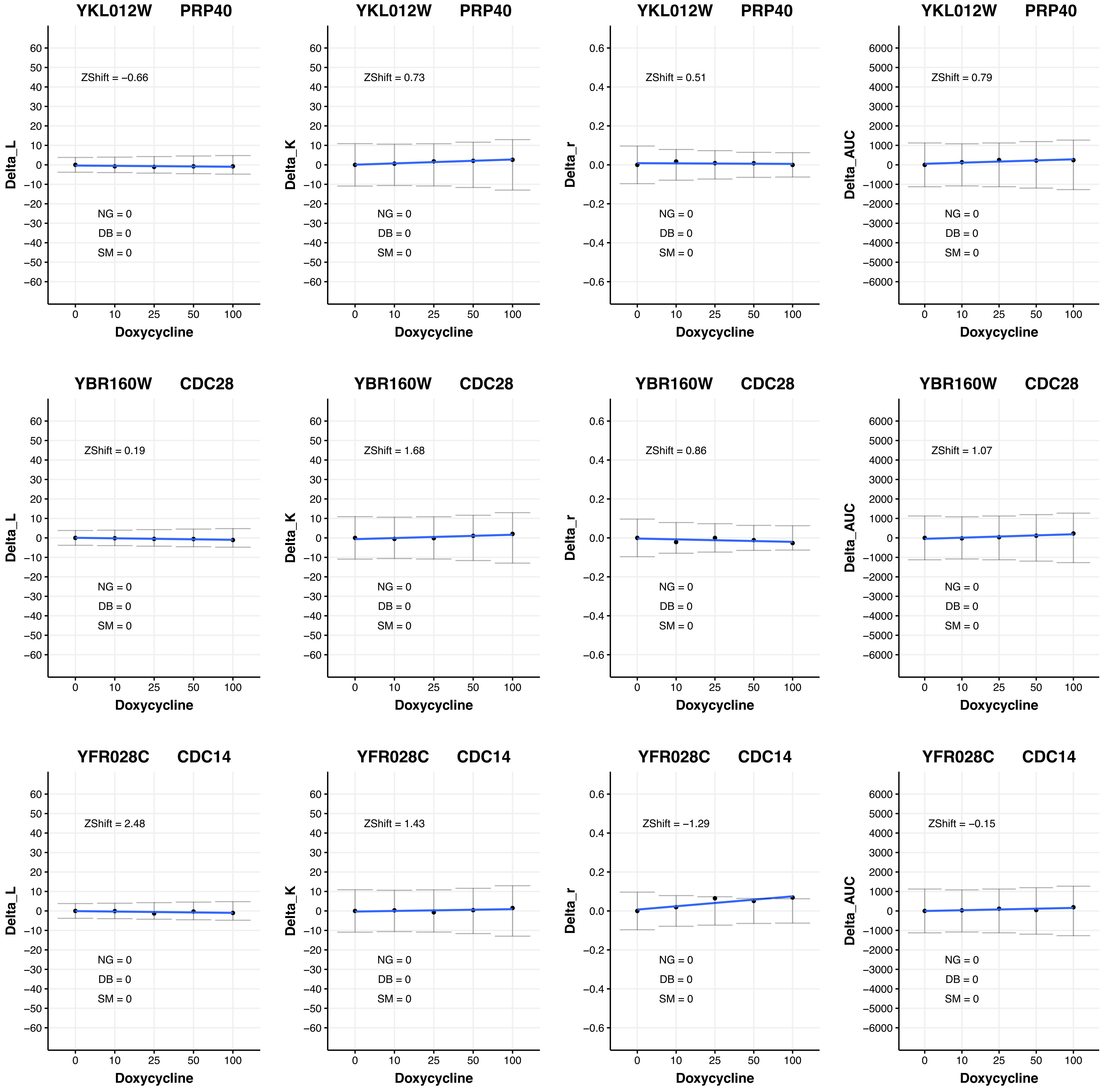
YKL012W PRP40
ZShift = −0.66
ZShift = 0.73
ZShift = 0.51
ZShift = 0.79
YBR160W CDC28
ZShift = 0.19
ZShift = 1.68
ZShift = 0.86
ZShift = 1.07
YFR028C CDC14
ZShift = 2.48
ZShift = 1.43
ZShift = −1.29
ZShift = −0.15
NG = 0
DB = 0
SM = 0
Delta_L
Delta_K
Delta_r
Delta_AUC
Doxycycline
0
10
25
50
100

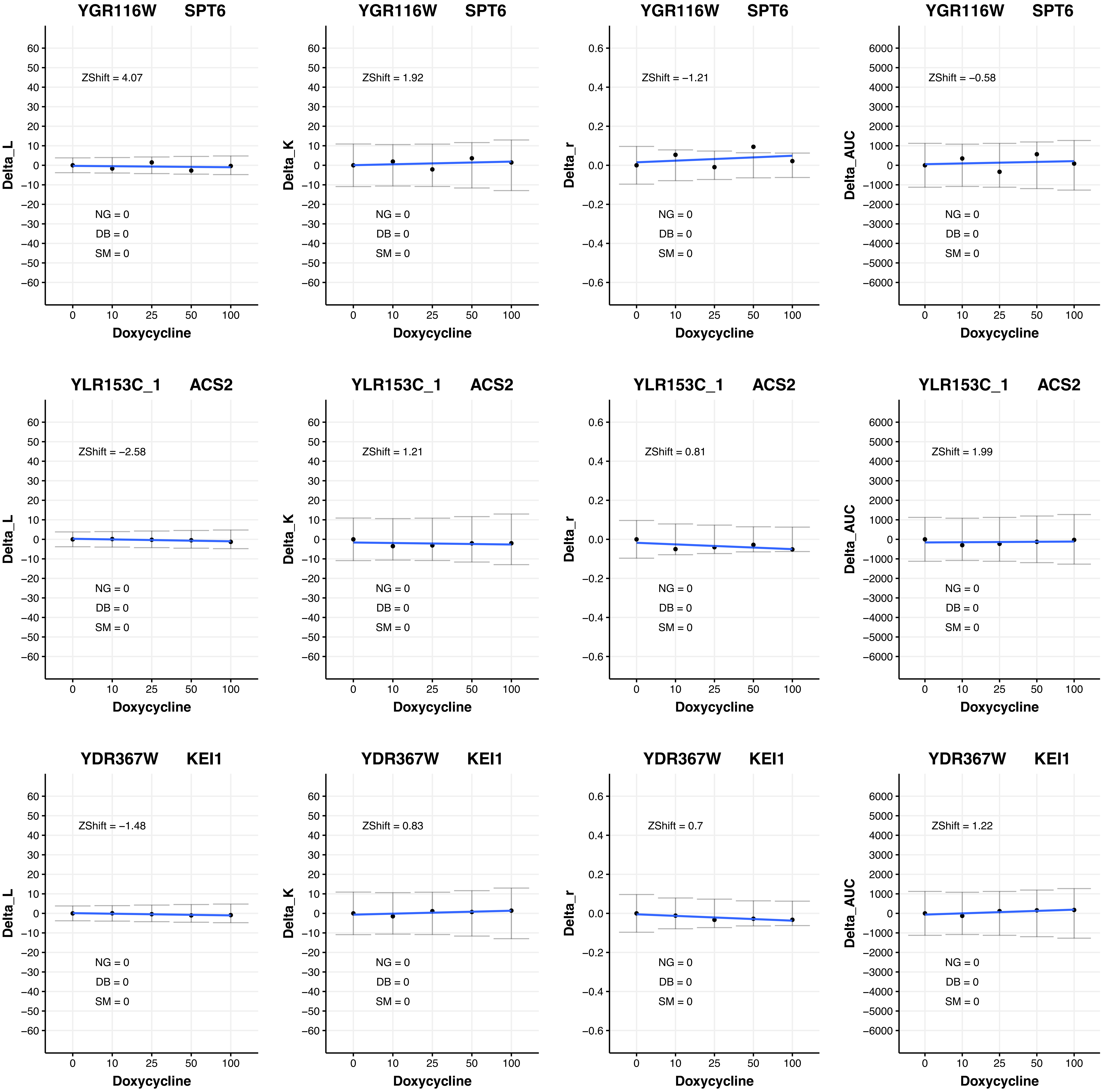
YGR116W SPT6
ZShift = 4.07
Delta_L
NG = 0
DB = 0
SM = 0
Doxycycline
0 10 25 50 100
YGR116W SPT6
ZShift = 1.92
Delta_K
NG = 0
DB = 0
SM = 0
Doxycycline
YGR116W SPT6
ZShift = −1.21
Delta_r
NG = 0
DB = 0
SM = 0
Doxycycline
YGR116W SPT6
ZShift = −0.58
Delta_AUC
NG = 0
DB = 0
SM = 0
Doxycycline
YLR153C_1 ACS2
ZShift = −2.58
Delta_L
NG = 0
DB = 0
SM = 0
Doxycycline
YLR153C_1 ACS2
ZShift = 1.21
Delta_K
NG = 0
DB = 0
SM = 0
Doxycycline
YLR153C_1 ACS2
ZShift = 0.81
Delta_r
NG = 0
DB = 0
SM = 0
Doxycycline
YLR153C_1 ACS2
ZShift = 1.99
Delta_AUC
NG = 0
DB = 0
SM = 0
Doxycycline
YDR367W KEI1
ZShift = −1.48
Delta_L
NG = 0
DB = 0
SM = 0
Doxycycline
YDR367W KEI1
ZShift = 0.83
Delta_K
NG = 0
DB = 0
SM = 0
Doxycycline
YDR367W KEI1
ZShift = 0.7
Delta_r
NG = 0
DB = 0
SM = 0
Doxycycline
YDR367W KEI1
ZShift = 1.22
Delta_AUC
NG = 0
DB = 0
SM = 0
Doxycycline

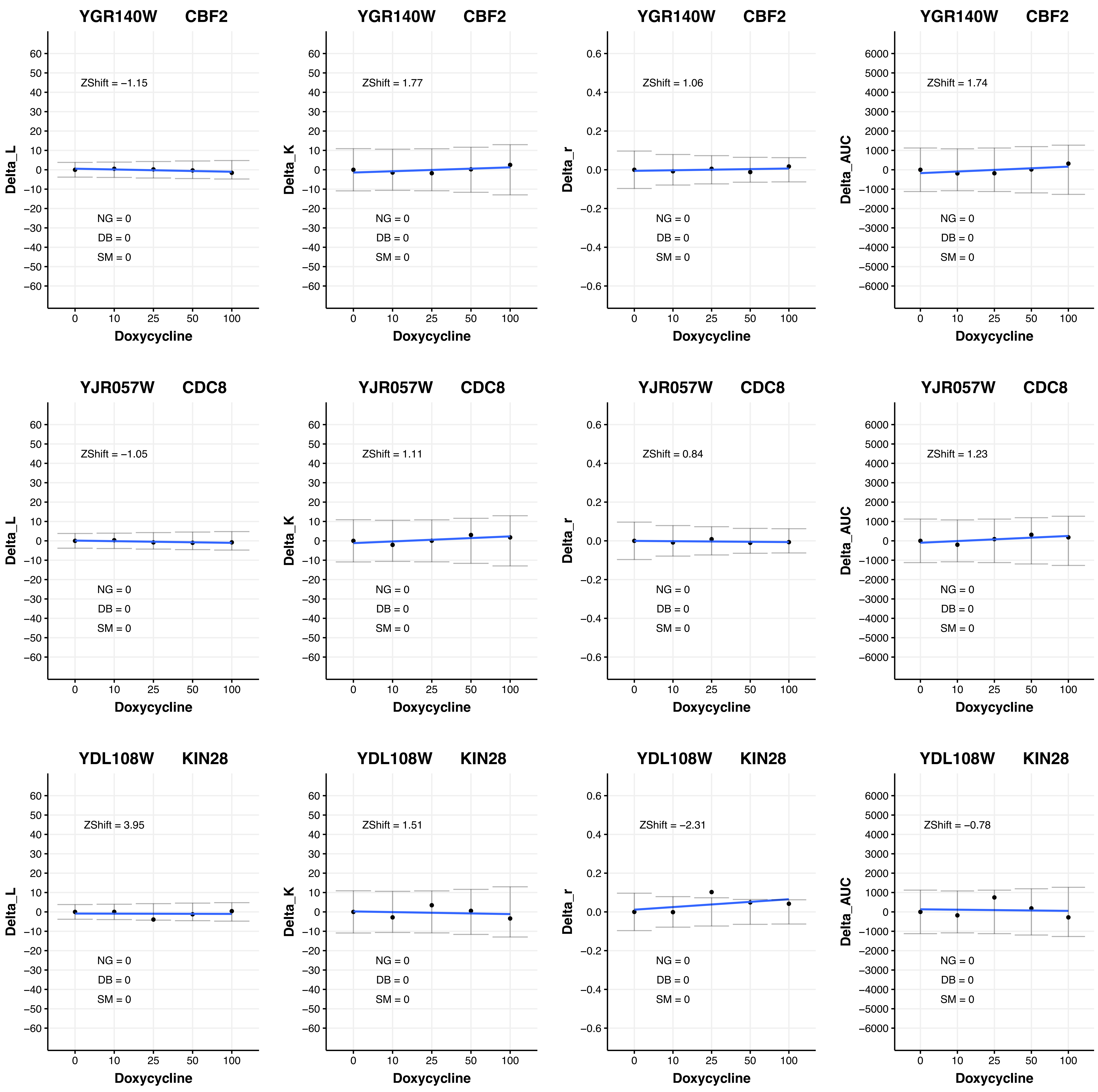
YGR140W CBF2
Delta_L
ZShift = −1.15
NG = 0
DB = 0
SM = 0
Doxycycline
YGR140W CBF2
Delta_K
ZShift = 1.77
NG = 0
DB = 0
SM = 0
Doxycycline
YGR140W CBF2
Delta_r
ZShift = 1.06
NG = 0
DB = 0
SM = 0
Doxycycline
YGR140W CBF2
Delta_AUC
ZShift = 1.74
NG = 0
DB = 0
SM = 0
Doxycycline
YJR057W CDC8
Delta_L
ZShift = −1.05
NG = 0
DB = 0
SM = 0
Doxycycline
YJR057W CDC8
Delta_K
ZShift = 1.11
NG = 0
DB = 0
SM = 0
Doxycycline
YJR057W CDC8
Delta_r
ZShift = 0.84
NG = 0
DB = 0
SM = 0
Doxycycline
YJR057W CDC8
Delta_AUC
ZShift = 1.23
NG = 0
DB = 0
SM = 0
Doxycycline
YDL108W KIN28
Delta_L
ZShift = 3.95
NG = 0
DB = 0
SM = 0
Doxycycline
YDL108W KIN28
Delta_K
ZShift = 1.51
NG = 0
DB = 0
SM = 0
Doxycycline
YDL108W KIN28
Delta_r
ZShift = −2.31
NG = 0
DB = 0
SM = 0
Doxycycline
YDL108W KIN28
Delta_AUC
ZShift = −0.78
NG = 0
DB = 0
SM = 0
Doxycycline

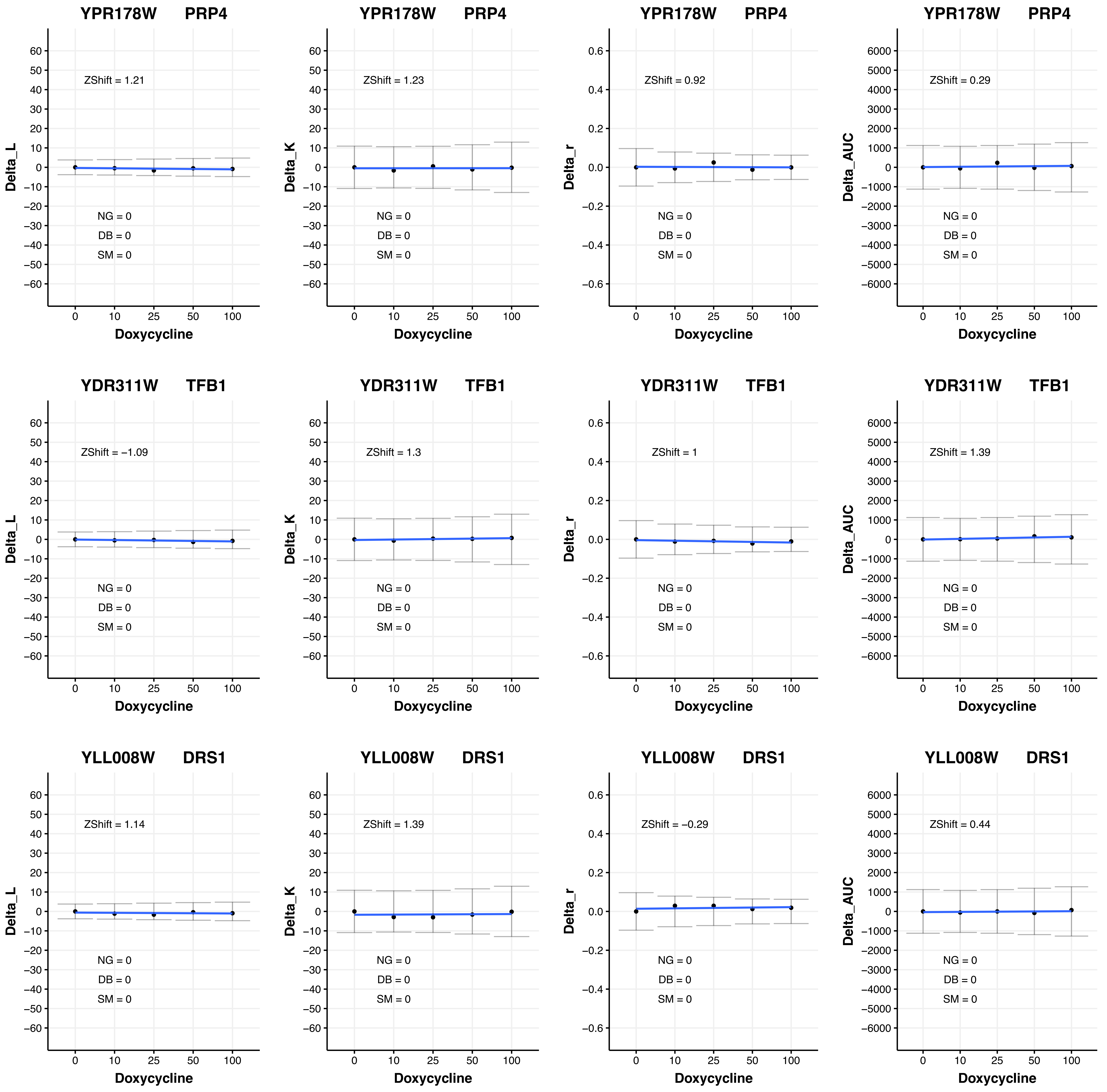

YPR178W PRP4
ZShift = 1.21
Delta_L
NG = 0
DB = 0
SM = 0
Doxycycline
YPR178W PRP4
ZShift = 1.23
Delta_K
NG = 0
DB = 0
SM = 0
Doxycycline
YPR178W PRP4
ZShift = 0.92
Delta_r
NG = 0
DB = 0
SM = 0
Doxycycline
YPR178W PRP4
ZShift = 0.29
Delta_AUC
NG = 0
DB = 0
SM = 0
Doxycycline
YDR311W TFB1
ZShift = −1.09
Delta_L
NG = 0
DB = 0
SM = 0
Doxycycline
YDR311W TFB1
ZShift = 1.3
Delta_K
NG = 0
DB = 0
SM = 0
Doxycycline
YDR311W TFB1
ZShift = 1
Delta_r
NG = 0
DB = 0
SM = 0
Doxycycline
YDR311W TFB1
ZShift = 1.39
Delta_AUC
NG = 0
DB = 0
SM = 0
Doxycycline
YLL008W DRS1
ZShift = 1.14
Delta_L
NG = 0
DB = 0
SM = 0
Doxycycline
YLL008W DRS1
ZShift = 1.39
Delta_K
NG = 0
DB = 0
SM = 0
Doxycycline
YLL008W DRS1
ZShift = −0.29
Delta_r
NG = 0
DB = 0
SM = 0
Doxycycline
YLL008W DRS1
ZShift = 0.44
Delta_AUC
NG = 0
DB = 0
SM = 0
Doxycycline

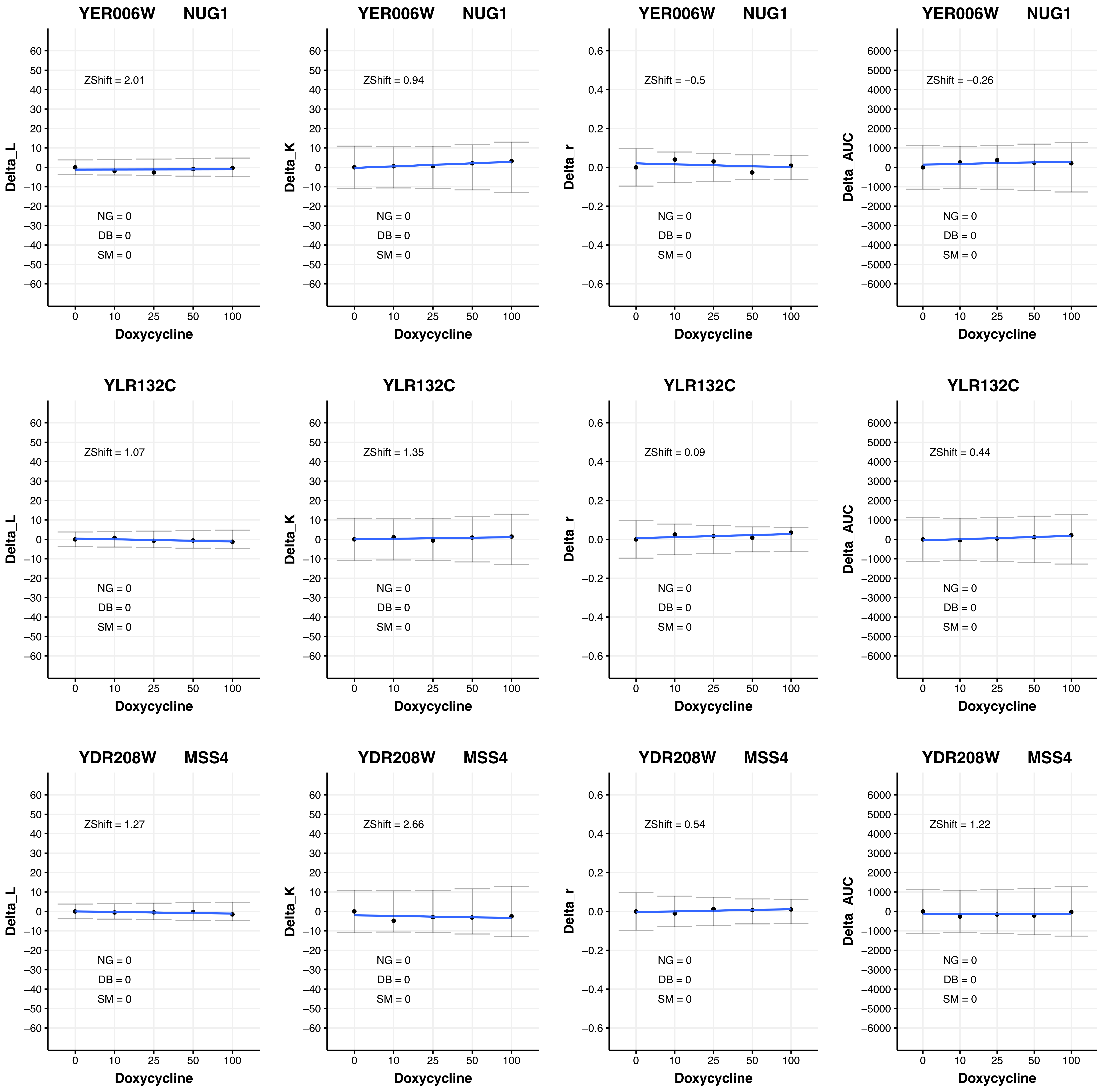
YER006W NUG1
ZShift = 2.01
NG = 0
DB = 0
SM = 0
Delta_L
Doxycycline
YER006W NUG1
ZShift = 0.94
NG = 0
DB = 0
SM = 0
Delta_K
Doxycycline
YER006W NUG1
ZShift = −0.5
NG = 0
DB = 0
SM = 0
Delta_r
Doxycycline
YER006W NUG1
ZShift = −0.26
NG = 0
DB = 0
SM = 0
Delta_AUC
Doxycycline
YLR132C
ZShift = 1.07
NG = 0
DB = 0
SM = 0
Delta_L
Doxycycline
YLR132C
ZShift = 1.35
NG = 0
DB = 0
SM = 0
Delta_K
Doxycycline
YLR132C
ZShift = 0.09
NG = 0
DB = 0
SM = 0
Delta_r
Doxycycline
YLR132C
ZShift = 0.44
NG = 0
DB = 0
SM = 0
Delta_AUC
Doxycycline
YDR208W MSS4
ZShift = 1.27
NG = 0
DB = 0
SM = 0
Delta_L
Doxycycline
YDR208W MSS4
ZShift = 2.66
NG = 0
DB = 0
SM = 0
Delta_K
Doxycycline
YDR208W MSS4
ZShift = 0.54
NG = 0
DB = 0
SM = 0
Delta_r
Doxycycline
YDR208W MSS4
ZShift = 1.22
NG = 0
DB = 0
SM = 0
Delta_AUC
Doxycycline

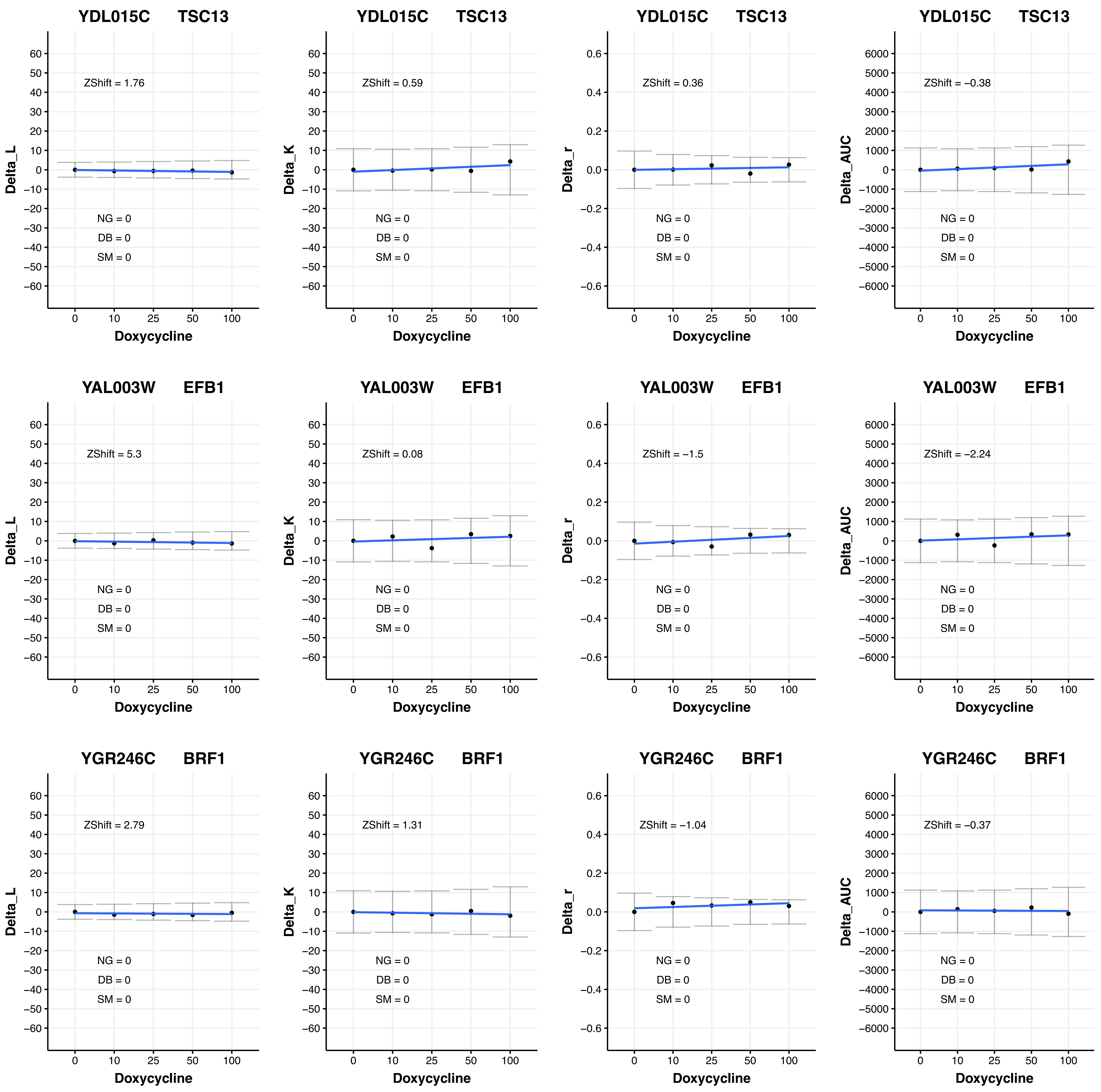

YDL015C TSC13
Delta_L
ZShift = 1.76
NG = 0
DB = 0
SM = 0
Doxycycline
YDL015C TSC13
Delta_K
ZShift = 0.59
NG = 0
DB = 0
SM = 0
Doxycycline
YDL015C TSC13
Delta_r
ZShift = 0.36
NG = 0
DB = 0
SM = 0
Doxycycline
YDL015C TSC13
Delta_AUC
ZShift = −0.38
NG = 0
DB = 0
SM = 0
Doxycycline
YAL003W EFB1
Delta_L
ZShift = 5.3
NG = 0
DB = 0
SM = 0
Doxycycline
YAL003W EFB1
Delta_K
ZShift = 0.08
NG = 0
DB = 0
SM = 0
Doxycycline
YAL003W EFB1
Delta_r
ZShift = −1.5
NG = 0
DB = 0
SM = 0
Doxycycline
YAL003W EFB1
Delta_AUC
ZShift = −2.24
NG = 0
DB = 0
SM = 0
Doxycycline
YGR246C BRF1
Delta_L
ZShift = 2.79
NG = 0
DB = 0
SM = 0
Doxycycline
YGR246C BRF1
Delta_K
ZShift = 1.31
NG = 0
DB = 0
SM = 0
Doxycycline
YGR246C BRF1
Delta_r
ZShift = −1.04
NG = 0
DB = 0
SM = 0
Doxycycline
YGR246C BRF1
Delta_AUC
ZShift = −0.37
NG = 0
DB = 0
SM = 0
Doxycycline

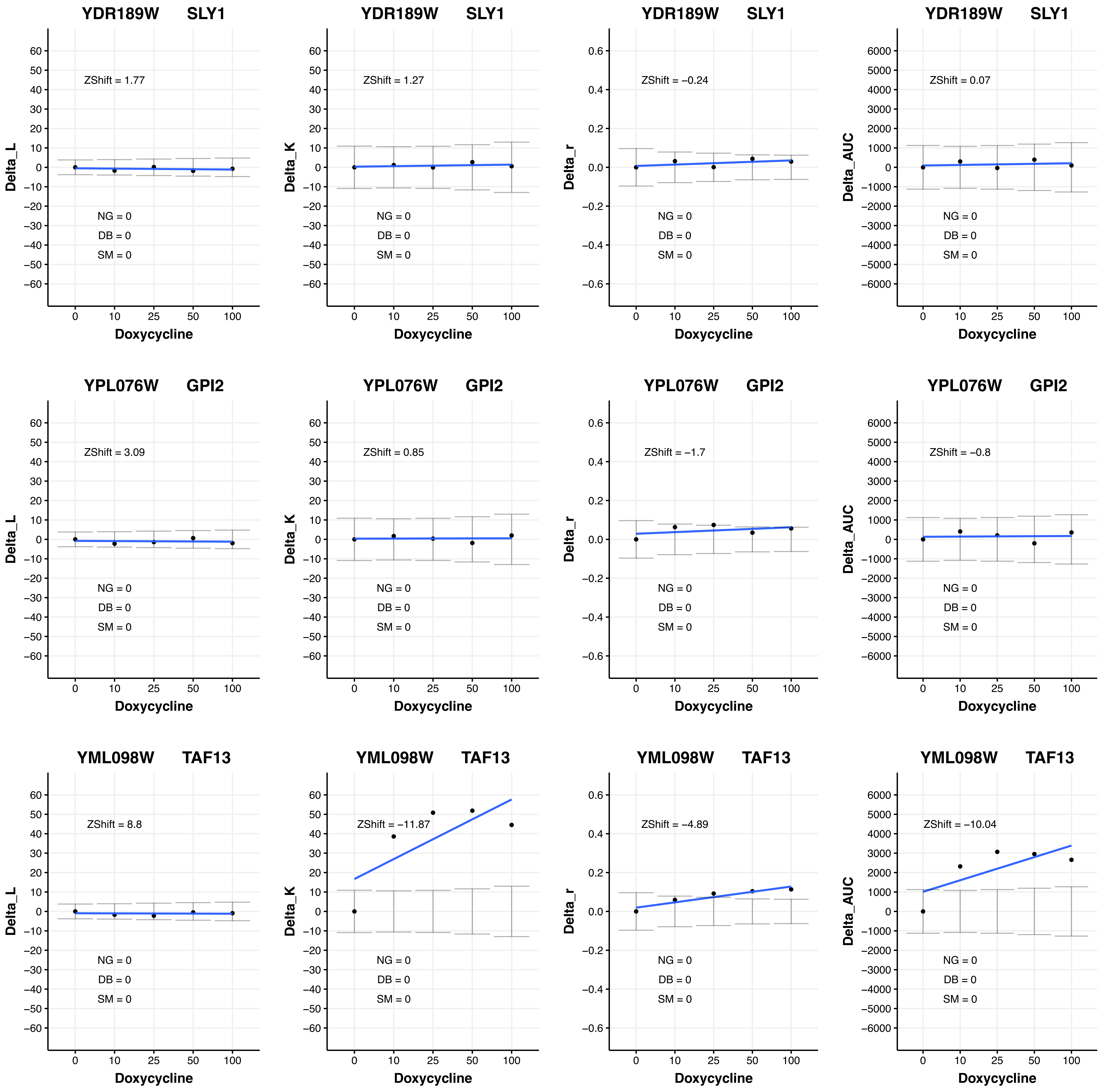
YDR189W SLY1
ZShift = 1.77
NG = 0
DB = 0
SM = 0
Delta_L
Doxycycline
YDR189W SLY1
ZShift = 1.27
NG = 0
DB = 0
SM = 0
Delta_K
Doxycycline
YDR189W SLY1
ZShift = −0.24
NG = 0
DB = 0
SM = 0
Delta_r
Doxycycline
YDR189W SLY1
ZShift = 0.07
NG = 0
DB = 0
SM = 0
Delta_AUC
Doxycycline
YPL076W GPI2
ZShift = 3.09
NG = 0
DB = 0
SM = 0
Delta_L
Doxycycline
YPL076W GPI2
ZShift = 0.85
NG = 0
DB = 0
SM = 0
Delta_K
Doxycycline
YPL076W GPI2
ZShift = −1.7
NG = 0
DB = 0
SM = 0
Delta_r
Doxycycline
YPL076W GPI2
ZShift = −0.8
NG = 0
DB = 0
SM = 0
Delta_AUC
Doxycycline
YML098W TAF13
ZShift = 8.8
NG = 0
DB = 0
SM = 0
Delta_L
Doxycycline
YML098W TAF13
ZShift = −11.87
NG = 0
DB = 0
SM = 0
Delta_K
Doxycycline
YML098W TAF13
ZShift = −4.89
NG = 0
DB = 0
SM = 0
Delta_r
Doxycycline
YML098W TAF13
ZShift = −10.04
NG = 0
DB = 0
SM = 0
Delta_AUC
Doxycycline

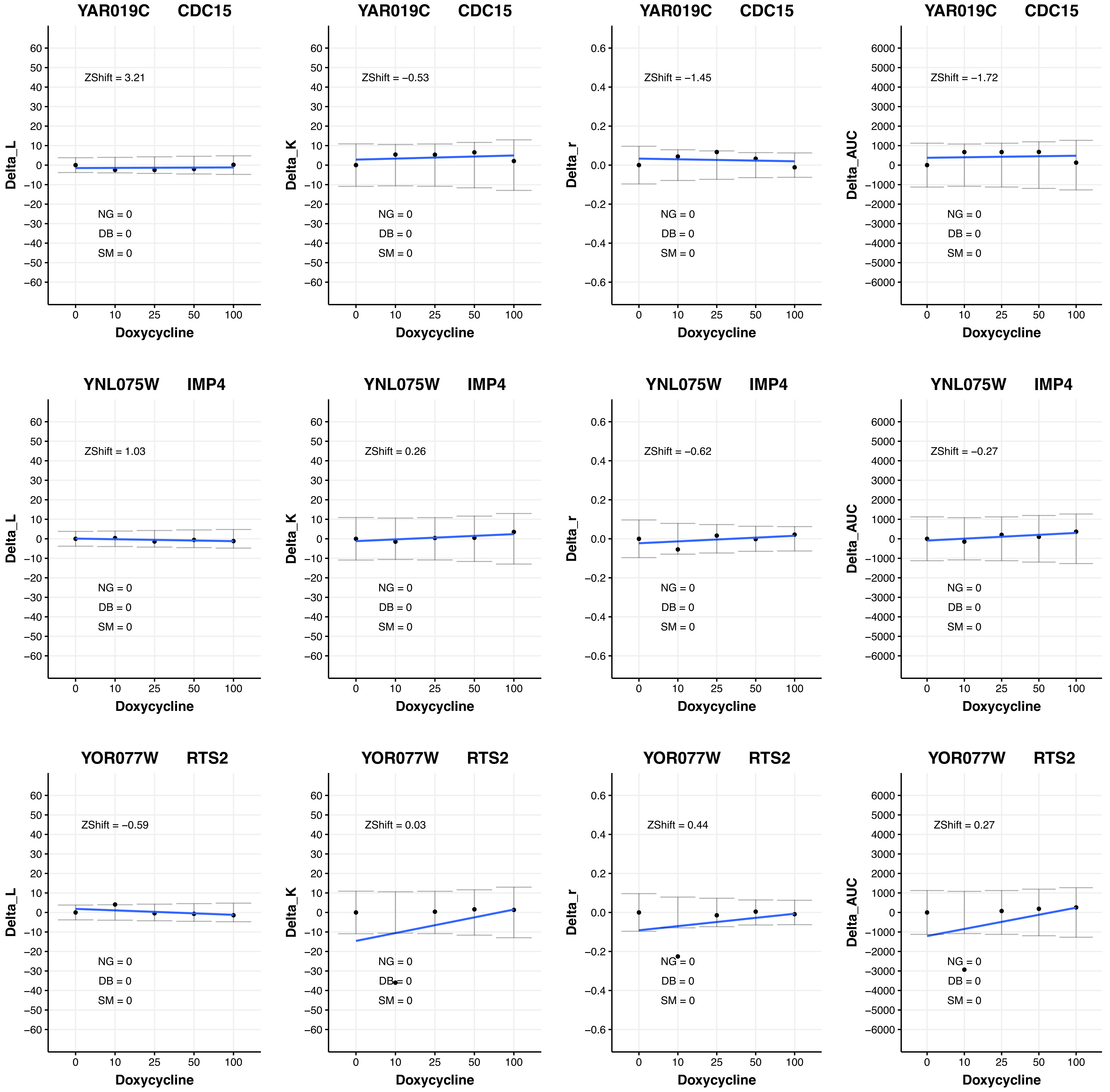

YAR019C CDC15
ZShift = 3.21
NG = 0
DB = 0
SM = 0
Delta_L
Doxycycline
YAR019C CDC15
ZShift = −0.53
NG = 0
DB = 0
SM = 0
Delta_K
Doxycycline
YAR019C CDC15
ZShift = −1.45
NG = 0
DB = 0
SM = 0
Delta_r
Doxycycline
YAR019C CDC15
ZShift = −1.72
NG = 0
DB = 0
SM = 0
Delta_AUC
Doxycycline
YNL075W IMP4
ZShift = 1.03
NG = 0
DB = 0
SM = 0
Delta_L
Doxycycline
YNL075W IMP4
ZShift = 0.26
NG = 0
DB = 0
SM = 0
Delta_K
Doxycycline
YNL075W IMP4
ZShift = −0.62
NG = 0
DB = 0
SM = 0
Delta_r
Doxycycline
YNL075W IMP4
ZShift = −0.27
NG = 0
DB = 0
SM = 0
Delta_AUC
Doxycycline
YOR077W RTS2
ZShift = −0.59
NG = 0
DB = 0
SM = 0
Delta_L
Doxycycline
YOR077W RTS2
ZShift = 0.03
NG = 0
DB = 0
SM = 0
Delta_K
Doxycycline
YOR077W RTS2
ZShift = 0.44
NG = 0
DB = 0
SM = 0
Delta_r
Doxycycline
YOR077W RTS2
ZShift = 0.27
NG = 0
DB = 0
SM = 0
Delta_AUC
Doxycycline

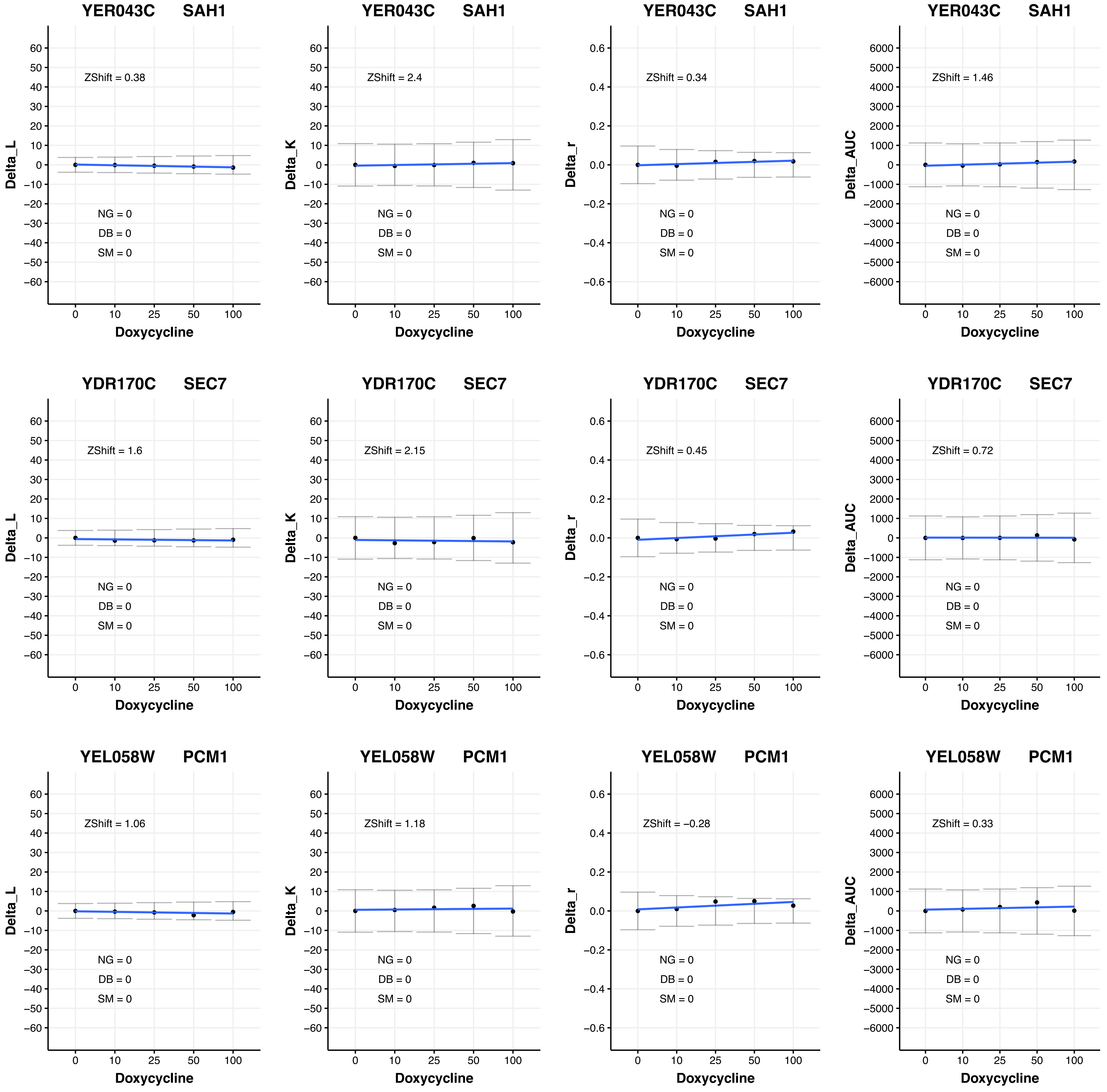

YER043C SAH1
ZShift = 0.38
NG = 0
DB = 0
SM = 0
Delta_L
Doxycycline
YER043C SAH1
ZShift = 2.4
NG = 0
DB = 0
SM = 0
Delta_K
Doxycycline
YER043C SAH1
ZShift = 0.34
NG = 0
DB = 0
SM = 0
Delta_r
Doxycycline
YER043C SAH1
ZShift = 1.46
NG = 0
DB = 0
SM = 0
Delta_AUC
Doxycycline
YDR170C SEC7
ZShift = 1.6
NG = 0
DB = 0
SM = 0
Delta_L
Doxycycline
YDR170C SEC7
ZShift = 2.15
NG = 0
DB = 0
SM = 0
Delta_K
Doxycycline
YDR170C SEC7
ZShift = 0.45
NG = 0
DB = 0
SM = 0
Delta_r
Doxycycline
YDR170C SEC7
ZShift = 0.72
NG = 0
DB = 0
SM = 0
Delta_AUC
Doxycycline
YEL058W PCM1
ZShift = 1.06
NG = 0
DB = 0
SM = 0
Delta_L
Doxycycline
YEL058W PCM1
ZShift = 1.18
NG = 0
DB = 0
SM = 0
Delta_K
Doxycycline
YEL058W PCM1
ZShift = −0.28
NG = 0
DB = 0
SM = 0
Delta_r
Doxycycline
YEL058W PCM1
ZShift = 0.33
NG = 0
DB = 0
SM = 0
Delta_AUC
Doxycycline

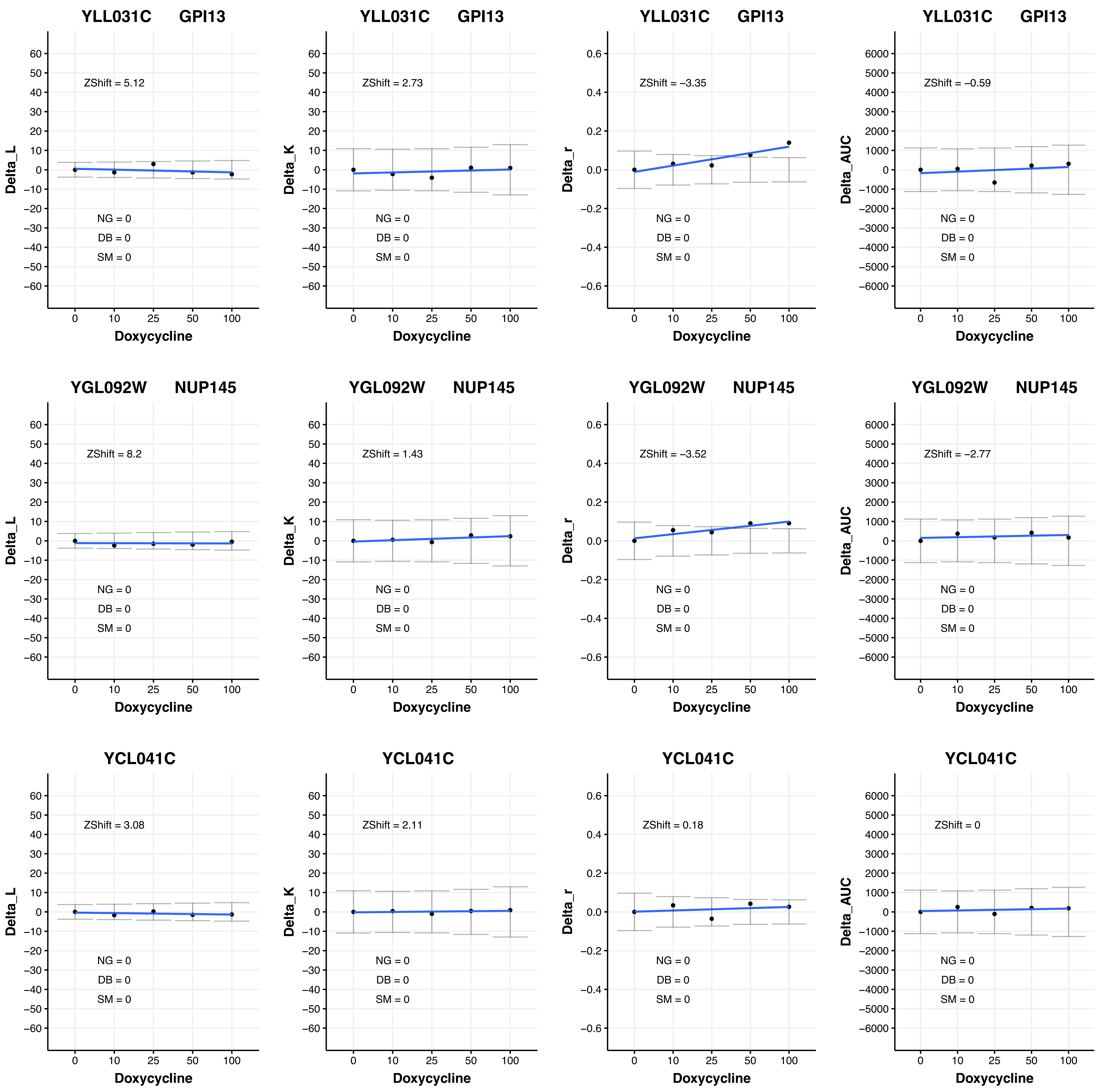

YLL031C GPI13
ZShift = 5.12
ZShift = 2.73
ZShift = −3.35
ZShift = −0.59
YGL092W NUP145
ZShift = 8.2
ZShift = 1.43
ZShift = −3.52
ZShift = −2.77
YCL041C
ZShift = 3.08
ZShift = 2.11
ZShift = 0.18
ZShift = 0
NG = 0
DB = 0
SM = 0
Delta_L
Delta_K
Delta_r
Delta_AUC
Doxycycline
0
10
25
50
100

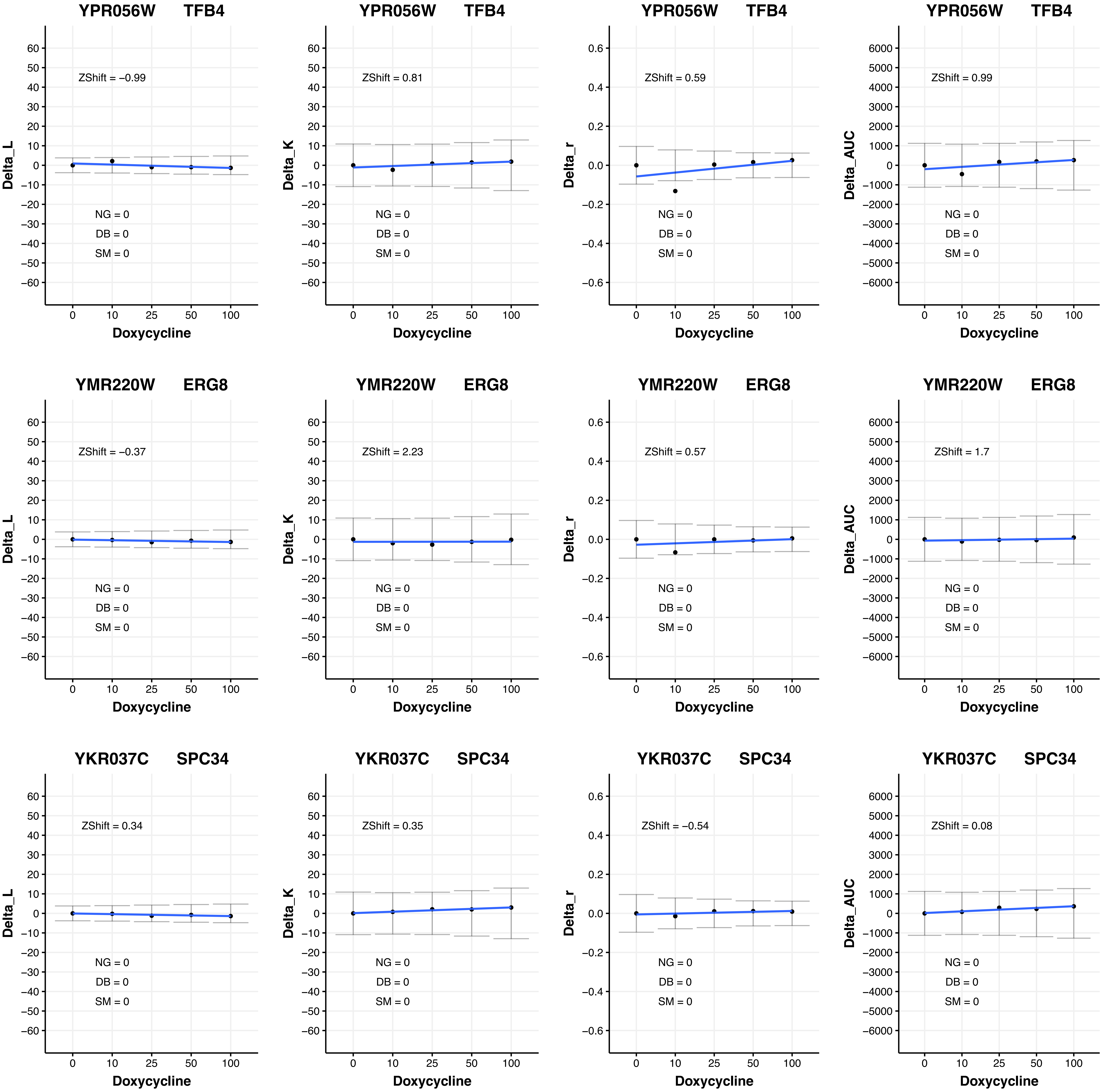

YPR056W TFB4
Delta_L
ZShift = −0.99
NG = 0
DB = 0
SM = 0
Doxycycline
YPR056W TFB4
Delta_K
ZShift = 0.81
NG = 0
DB = 0
SM = 0
Doxycycline
YPR056W TFB4
Delta_r
ZShift = 0.59
NG = 0
DB = 0
SM = 0
Doxycycline
YPR056W TFB4
Delta_AUC
ZShift = 0.99
NG = 0
DB = 0
SM = 0
Doxycycline
YMR220W ERG8
Delta_L
ZShift = −0.37
NG = 0
DB = 0
SM = 0
Doxycycline
YMR220W ERG8
Delta_K
ZShift = 2.23
NG = 0
DB = 0
SM = 0
Doxycycline
YMR220W ERG8
Delta_r
ZShift = 0.57
NG = 0
DB = 0
SM = 0
Doxycycline
YMR220W ERG8
Delta_AUC
ZShift = 1.7
NG = 0
DB = 0
SM = 0
Doxycycline
YKR037C SPC34
Delta_L
ZShift = 0.34
NG = 0
DB = 0
SM = 0
Doxycycline
YKR037C SPC34
Delta_K
ZShift = 0.35
NG = 0
DB = 0
SM = 0
Doxycycline
YKR037C SPC34
Delta_r
ZShift = −0.54
NG = 0
DB = 0
SM = 0
Doxycycline
YKR037C SPC34
Delta_AUC
ZShift = 0.08
NG = 0
DB = 0
SM = 0
Doxycycline

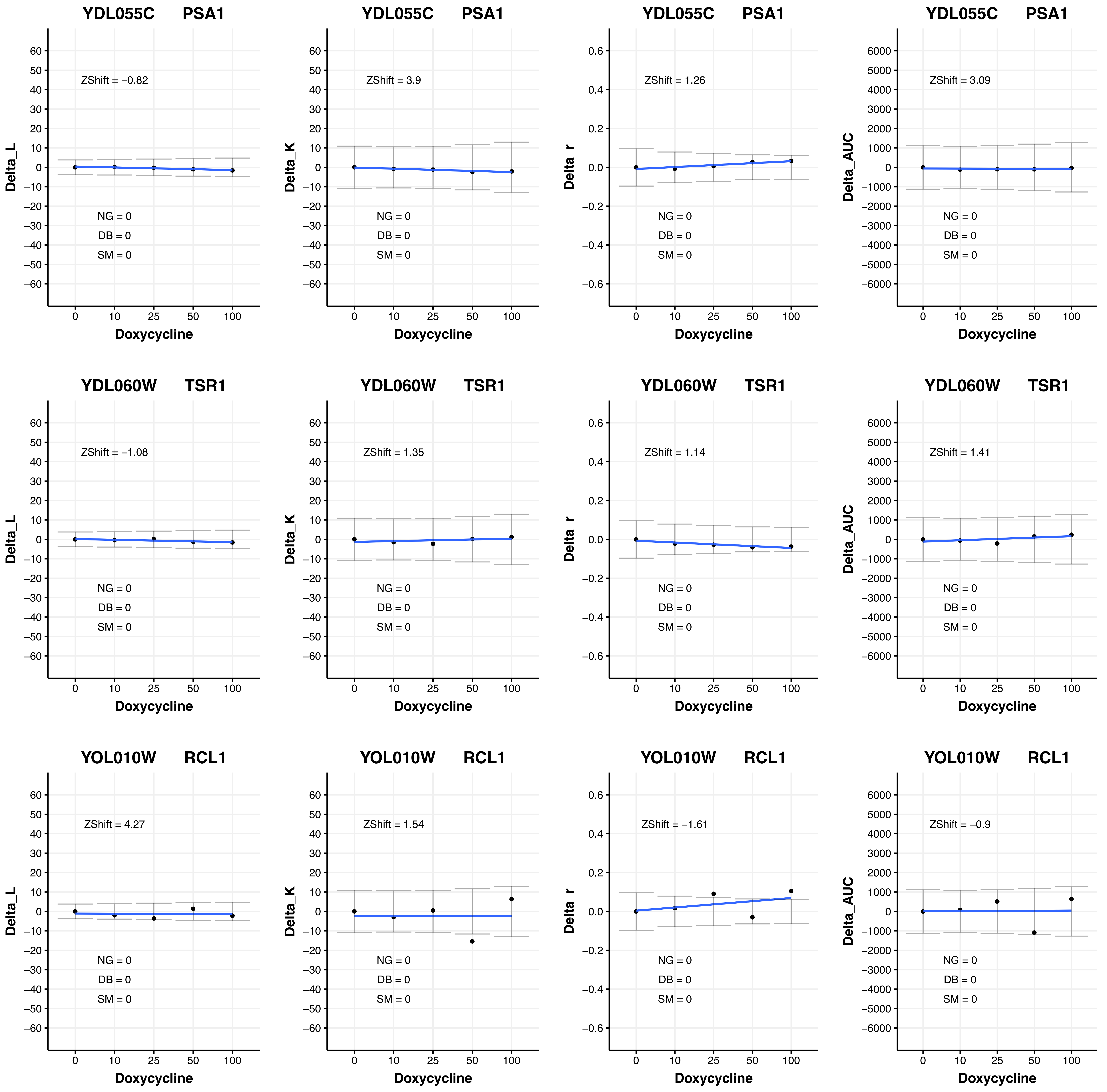

YDL055C PSA1
ZShift = −0.82
NG = 0
DB = 0
SM = 0
Delta_L
Doxycycline
YDL055C PSA1
ZShift = 3.9
NG = 0
DB = 0
SM = 0
Delta_K
Doxycycline
YDL055C PSA1
ZShift = 1.26
NG = 0
DB = 0
SM = 0
Delta_r
Doxycycline
YDL055C PSA1
ZShift = 3.09
NG = 0
DB = 0
SM = 0
Delta_AUC
Doxycycline
YDL060W TSR1
ZShift = −1.08
NG = 0
DB = 0
SM = 0
Delta_L
Doxycycline
YDL060W TSR1
ZShift = 1.35
NG = 0
DB = 0
SM = 0
Delta_K
Doxycycline
YDL060W TSR1
ZShift = 1.14
NG = 0
DB = 0
SM = 0
Delta_r
Doxycycline
YDL060W TSR1
ZShift = 1.41
NG = 0
DB = 0
SM = 0
Delta_AUC
Doxycycline
YOL010W RCL1
ZShift = 4.27
NG = 0
DB = 0
SM = 0
Delta_L
Doxycycline
YOL010W RCL1
ZShift = 1.54
NG = 0
DB = 0
SM = 0
Delta_K
Doxycycline
YOL010W RCL1
ZShift = −1.61
NG = 0
DB = 0
SM = 0
Delta_r
Doxycycline
YOL010W RCL1
ZShift = −0.9
NG = 0
DB = 0
SM = 0
Delta_AUC
Doxycycline

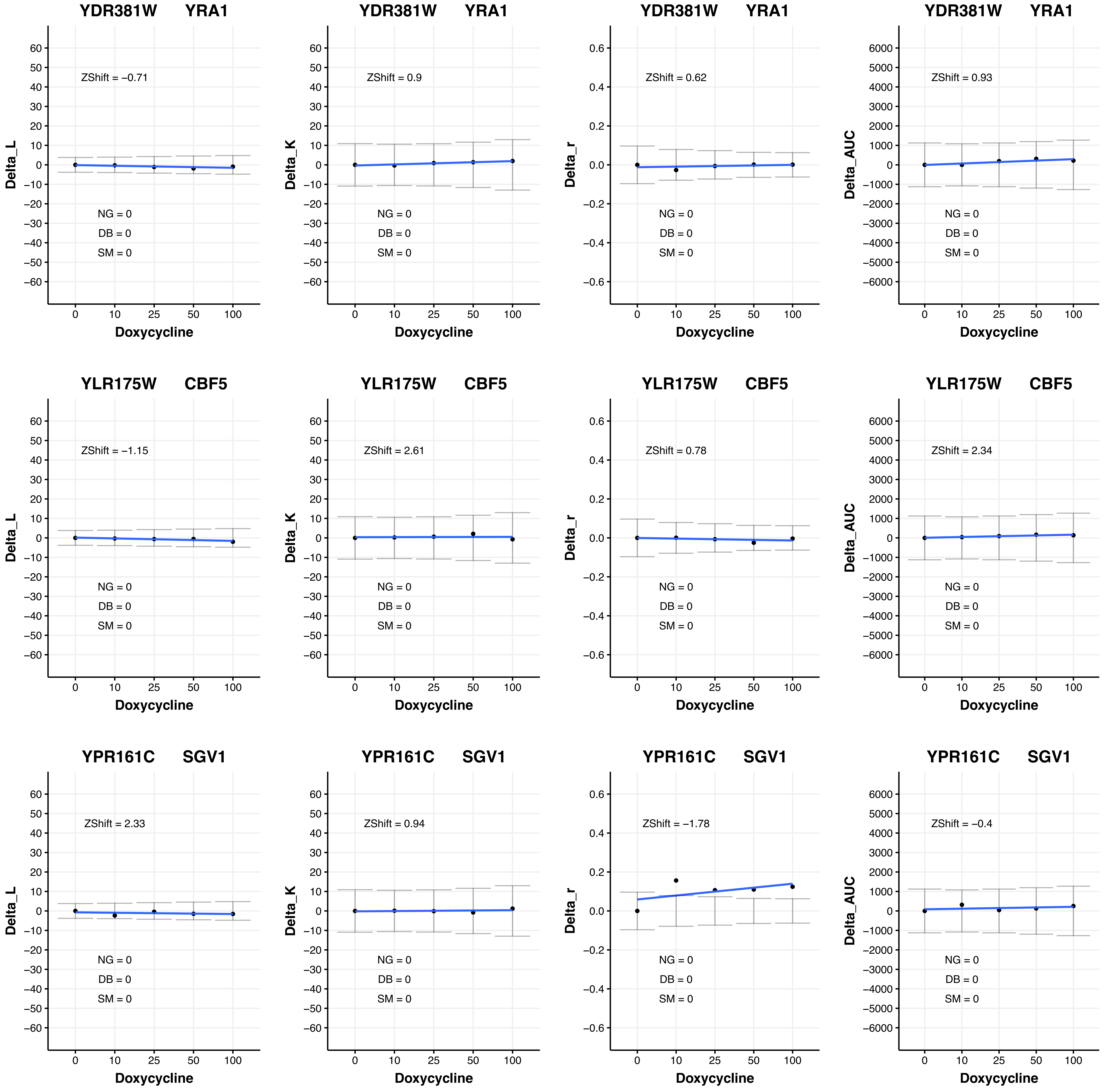
YDR381W YRA1
ZShift = −0.71
NG = 0
DB = 0
SM = 0
Delta_L
Doxycycline
YDR381W YRA1
ZShift = 0.9
NG = 0
DB = 0
SM = 0
Delta_K
Doxycycline
YDR381W YRA1
ZShift = 0.62
NG = 0
DB = 0
SM = 0
Delta_r
Doxycycline
YDR381W YRA1
ZShift = 0.93
NG = 0
DB = 0
SM = 0
Delta_AUC
Doxycycline
YLR175W CBF5
ZShift = −1.15
NG = 0
DB = 0
SM = 0
Delta_L
Doxycycline
YLR175W CBF5
ZShift = 2.61
NG = 0
DB = 0
SM = 0
Delta_K
Doxycycline
YLR175W CBF5
ZShift = 0.78
NG = 0
DB = 0
SM = 0
Delta_r
Doxycycline
YLR175W CBF5
ZShift = 2.34
NG = 0
DB = 0
SM = 0
Delta_AUC
Doxycycline
YPR161C SGV1
ZShift = 2.33
NG = 0
DB = 0
SM = 0
Delta_L
Doxycycline
YPR161C SGV1
ZShift = 0.94
NG = 0
DB = 0
SM = 0
Delta_K
Doxycycline
YPR161C SGV1
ZShift = −1.78
NG = 0
DB = 0
SM = 0
Delta_r
Doxycycline
YPR161C SGV1
ZShift = −0.4
NG = 0
DB = 0
SM = 0
Delta_AUC
Doxycycline

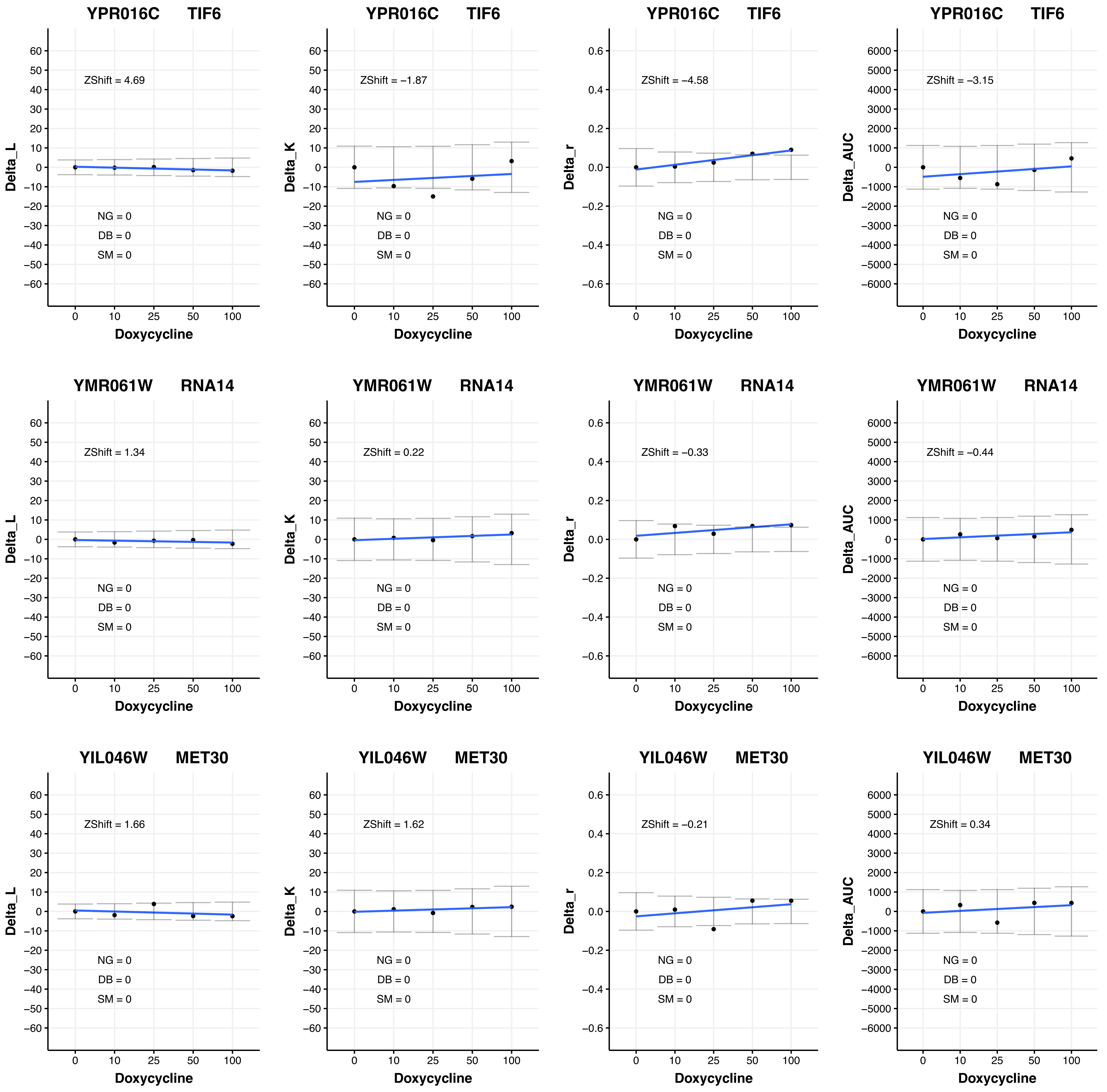

YPR016C TIF6
ZShift = 4.69
Delta_L
NG = 0
DB = 0
SM = 0
Doxycycline
YPR016C TIF6
ZShift = −1.87
Delta_K
NG = 0
DB = 0
SM = 0
Doxycycline
YPR016C TIF6
ZShift = −4.58
Delta_r
NG = 0
DB = 0
SM = 0
Doxycycline
YPR016C TIF6
ZShift = −3.15
Delta_AUC
NG = 0
DB = 0
SM = 0
Doxycycline
YMR061W RNA14
ZShift = 1.34
Delta_L
NG = 0
DB = 0
SM = 0
Doxycycline
YMR061W RNA14
ZShift = 0.22
Delta_K
NG = 0
DB = 0
SM = 0
Doxycycline
YMR061W RNA14
ZShift = −0.33
Delta_r
NG = 0
DB = 0
SM = 0
Doxycycline
YMR061W RNA14
ZShift = −0.44
Delta_AUC
NG = 0
DB = 0
SM = 0
Doxycycline
YIL046W MET30
ZShift = 1.66
Delta_L
NG = 0
DB = 0
SM = 0
Doxycycline
YIL046W MET30
ZShift = 1.62
Delta_K
NG = 0
DB = 0
SM = 0
Doxycycline
YIL046W MET30
ZShift = −0.21
Delta_r
NG = 0
DB = 0
SM = 0
Doxycycline
YIL046W MET30
ZShift = 0.34
Delta_AUC
NG = 0
DB = 0
SM = 0
Doxycycline

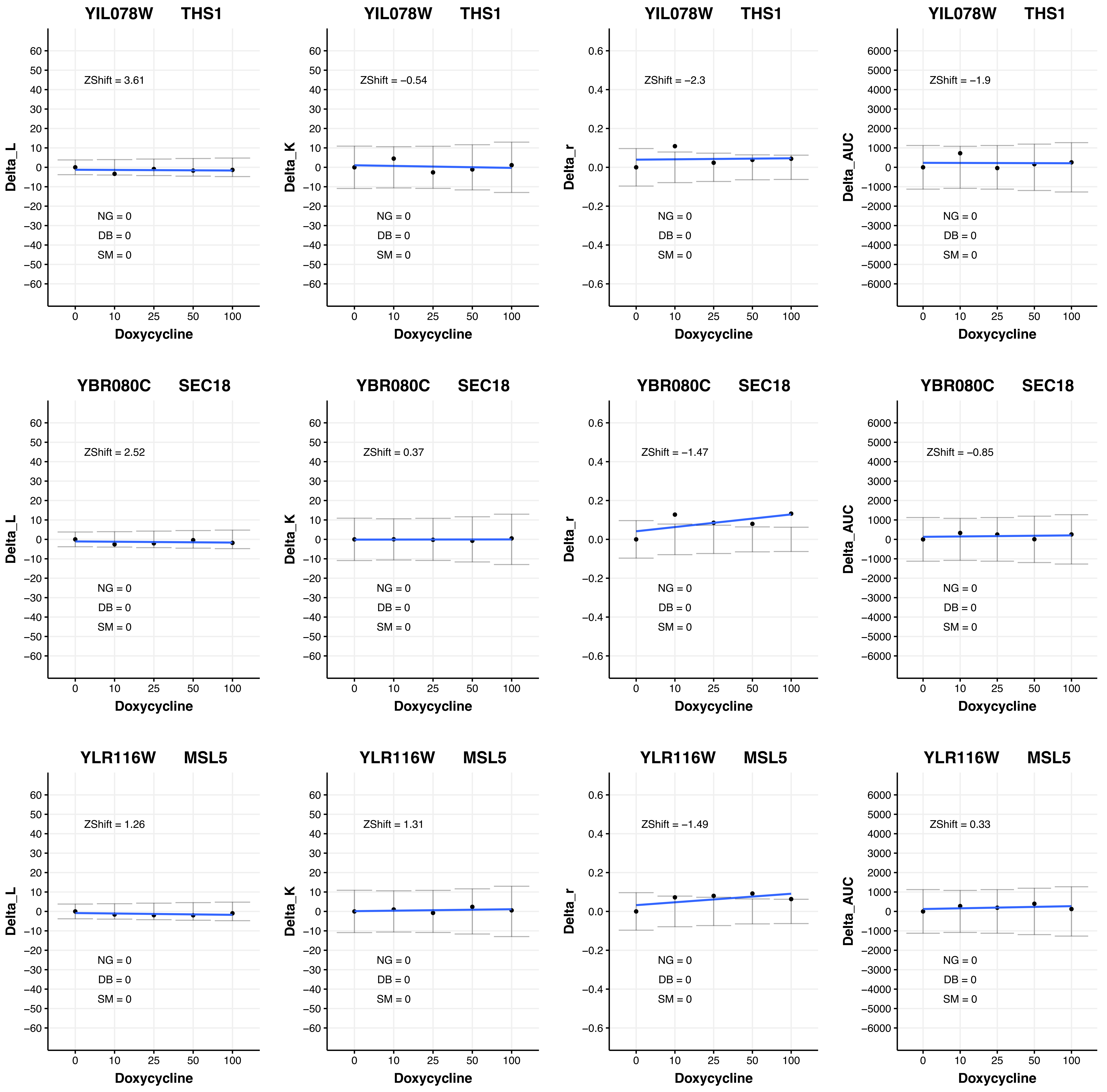
YIL078W THS1
Delta_L
ZShift = 3.61
NG = 0
DB = 0
SM = 0
Doxycycline
YIL078W THS1
Delta_K
ZShift = −0.54
NG = 0
DB = 0
SM = 0
Doxycycline
YIL078W THS1
Delta_r
ZShift = −2.3
NG = 0
DB = 0
SM = 0
Doxycycline
YIL078W THS1
Delta_AUC
ZShift = −1.9
NG = 0
DB = 0
SM = 0
Doxycycline
YBR080C SEC18
Delta_L
ZShift = 2.52
NG = 0
DB = 0
SM = 0
Doxycycline
YBR080C SEC18
Delta_K
ZShift = 0.37
NG = 0
DB = 0
SM = 0
Doxycycline
YBR080C SEC18
Delta_r
ZShift = −1.47
NG = 0
DB = 0
SM = 0
Doxycycline
YBR080C SEC18
Delta_AUC
ZShift = −0.85
NG = 0
DB = 0
SM = 0
Doxycycline
YLR116W MSL5
Delta_L
ZShift = 1.26
NG = 0
DB = 0
SM = 0
Doxycycline
YLR116W MSL5
Delta_K
ZShift = 1.31
NG = 0
DB = 0
SM = 0
Doxycycline
YLR116W MSL5
Delta_r
ZShift = −1.49
NG = 0
DB = 0
SM = 0
Doxycycline
YLR116W MSL5
Delta_AUC
ZShift = 0.33
NG = 0
DB = 0
SM = 0
Doxycycline

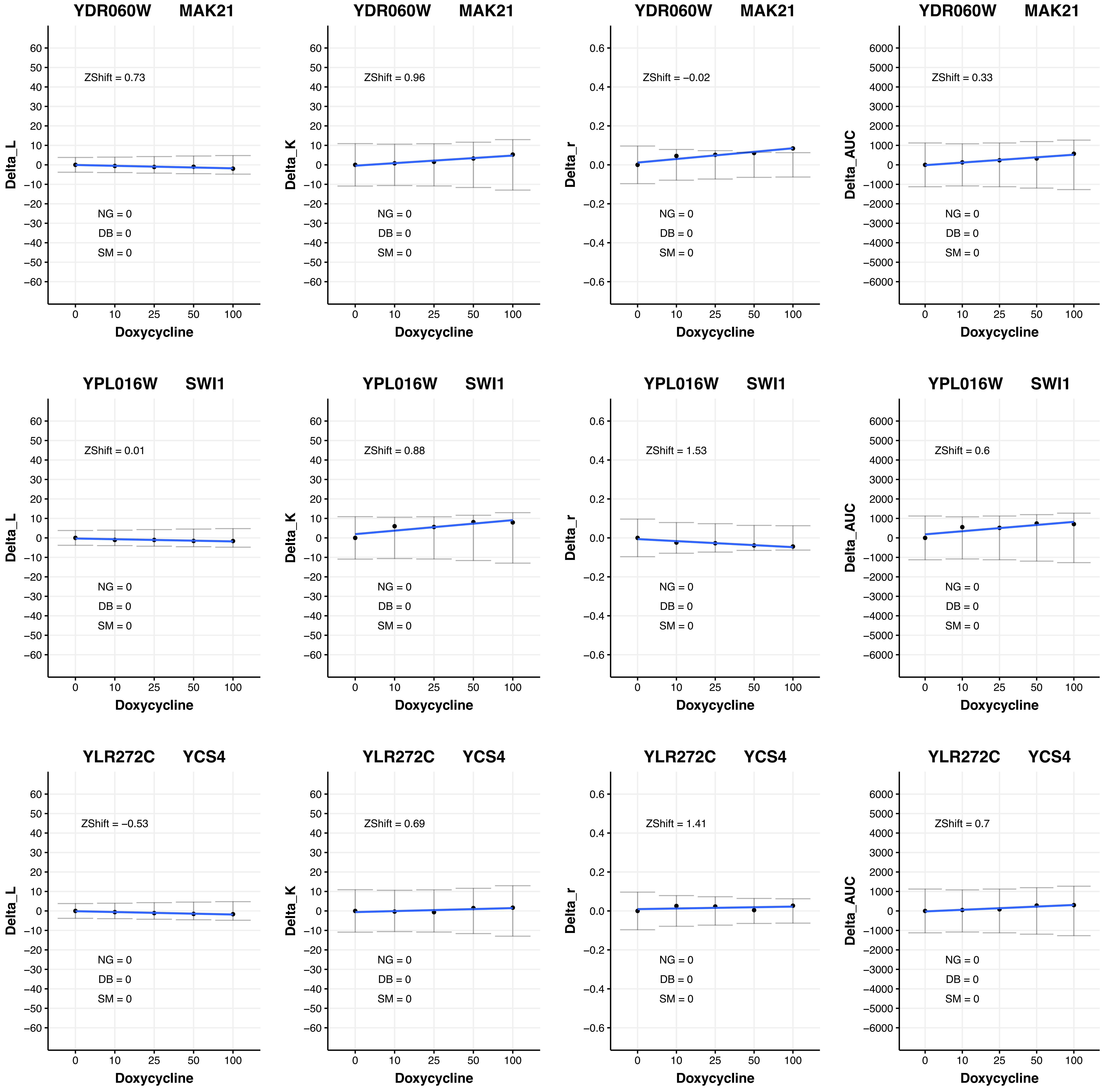

YDR060W MAK21
ZShift = 0.73
NG = 0
DB = 0
SM = 0
Delta_L
Doxycycline
YDR060W MAK21
ZShift = 0.96
NG = 0
DB = 0
SM = 0
Delta_K
Doxycycline
YDR060W MAK21
ZShift = −0.02
NG = 0
DB = 0
SM = 0
Delta_r
Doxycycline
YDR060W MAK21
ZShift = 0.33
NG = 0
DB = 0
SM = 0
Delta_AUC
Doxycycline
YPL016W SWI1
ZShift = 0.01
NG = 0
DB = 0
SM = 0
Delta_L
Doxycycline
YPL016W SWI1
ZShift = 0.88
NG = 0
DB = 0
SM = 0
Delta_K
Doxycycline
YPL016W SWI1
ZShift = 1.53
NG = 0
DB = 0
SM = 0
Delta_r
Doxycycline
YPL016W SWI1
ZShift = 0.6
NG = 0
DB = 0
SM = 0
Delta_AUC
Doxycycline
YLR272C YCS4
ZShift = −0.53
NG = 0
DB = 0
SM = 0
Delta_L
Doxycycline
YLR272C YCS4
ZShift = 0.69
NG = 0
DB = 0
SM = 0
Delta_K
Doxycycline
YLR272C YCS4
ZShift = 1.41
NG = 0
DB = 0
SM = 0
Delta_r
Doxycycline
YLR272C YCS4
ZShift = 0.7
NG = 0
DB = 0
SM = 0
Delta_AUC
Doxycycline

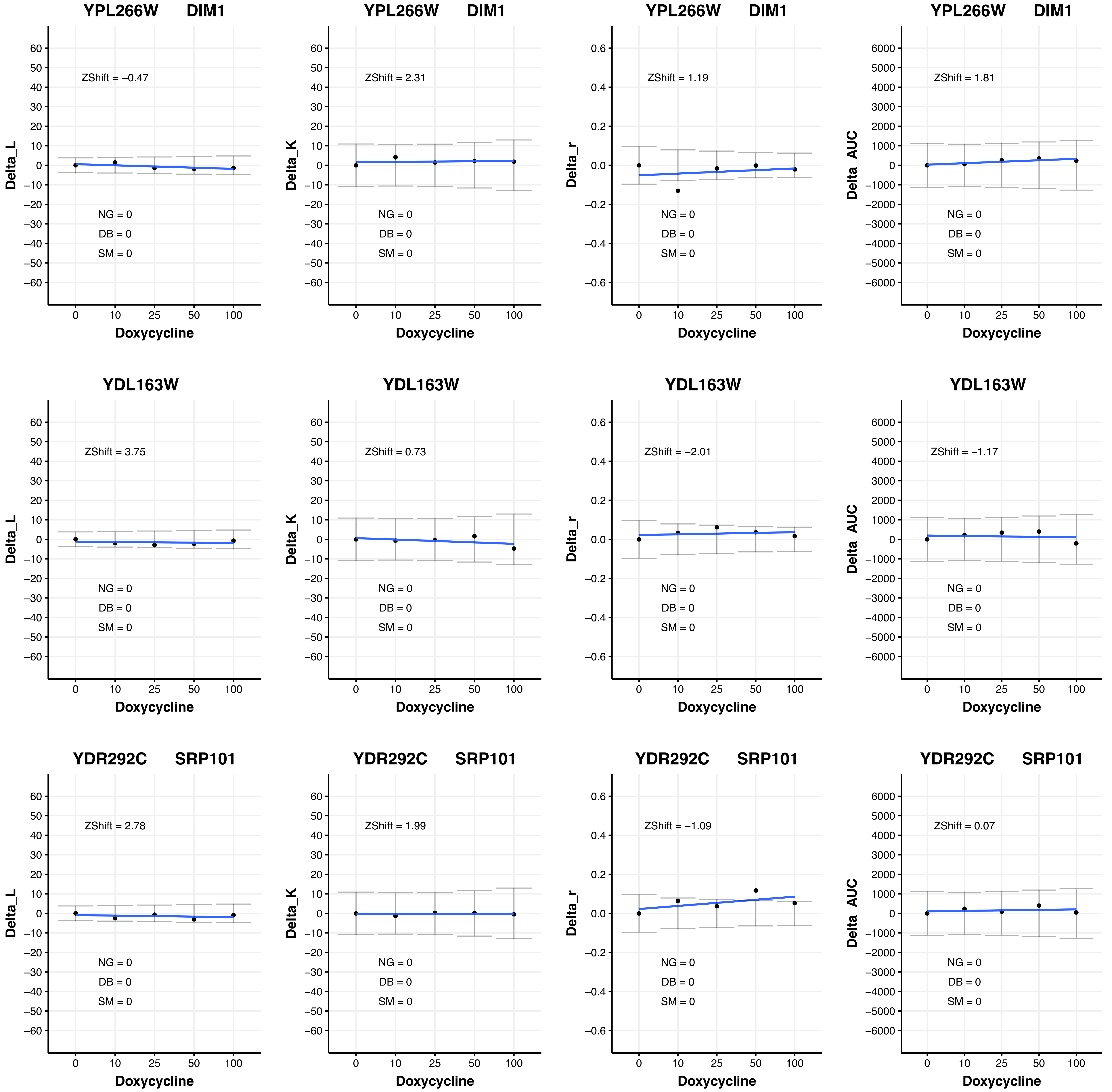

YPL266W DIM1
ZShift = −0.47
ZShift = 2.31
ZShift = 1.19
ZShift = 1.81
YDL163W
ZShift = 3.75
ZShift = 0.73
ZShift = −2.01
ZShift = −1.17
YDR292C SRP101
ZShift = 2.78
ZShift = 1.99
ZShift = −1.09
ZShift = 0.07
NG = 0
DB = 0
SM = 0
Delta_L
Delta_K
Delta_r
Delta_AUC
Doxycycline

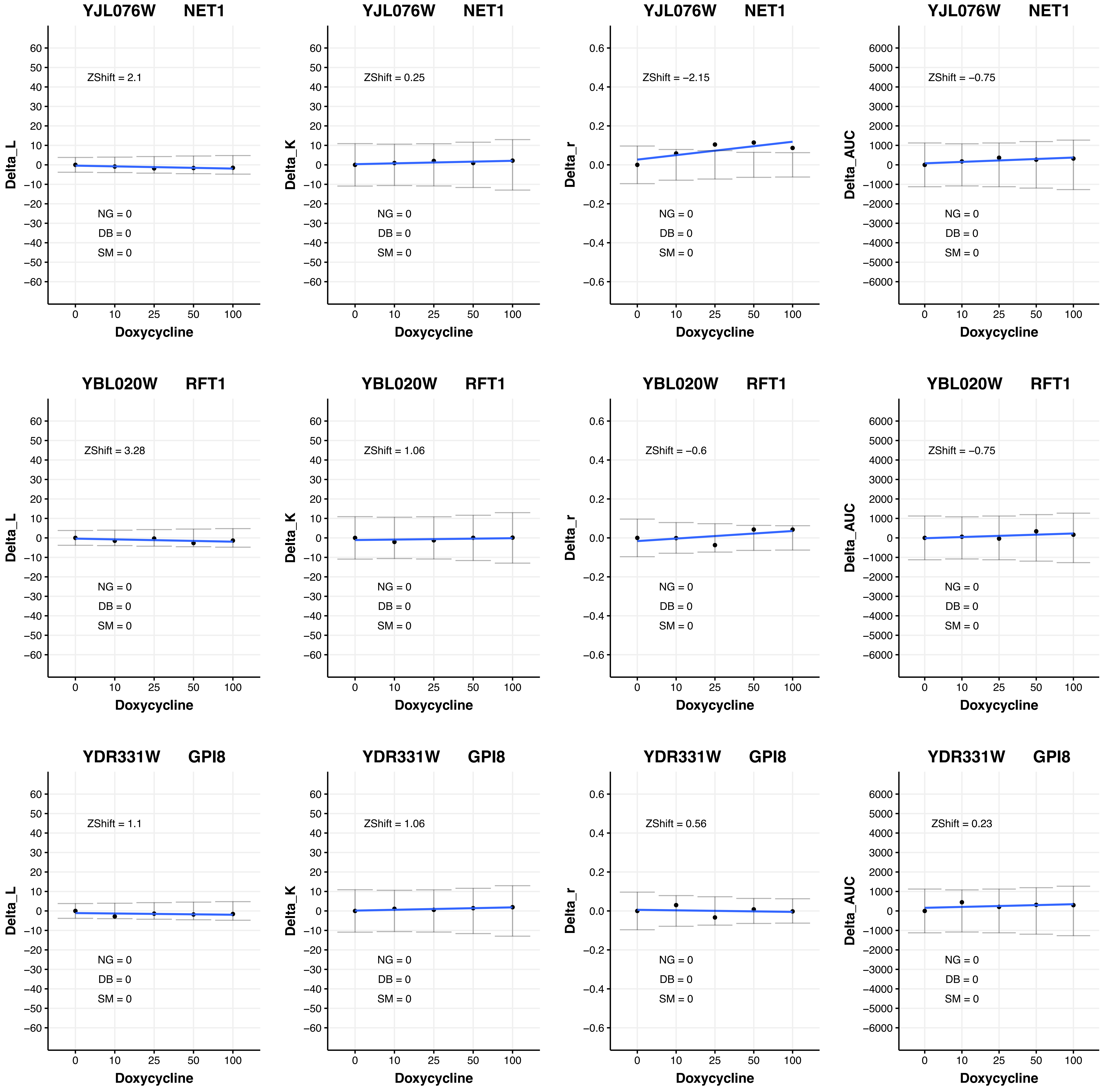
YJL076W NET1
ZShift = 2.1
NG = 0
DB = 0
SM = 0
Delta_L
Doxycycline
YJL076W NET1
ZShift = 0.25
NG = 0
DB = 0
SM = 0
Delta_K
Doxycycline
YJL076W NET1
ZShift = −2.15
NG = 0
DB = 0
SM = 0
Delta_r
Doxycycline
YJL076W NET1
ZShift = −0.75
NG = 0
DB = 0
SM = 0
Delta_AUC
Doxycycline
YBL020W RFT1
ZShift = 3.28
NG = 0
DB = 0
SM = 0
Delta_L
Doxycycline
YBL020W RFT1
ZShift = 1.06
NG = 0
DB = 0
SM = 0
Delta_K
Doxycycline
YBL020W RFT1
ZShift = −0.6
NG = 0
DB = 0
SM = 0
Delta_r
Doxycycline
YBL020W RFT1
ZShift = −0.75
NG = 0
DB = 0
SM = 0
Delta_AUC
Doxycycline
YDR331W GPI8
ZShift = 1.1
NG = 0
DB = 0
SM = 0
Delta_L
Doxycycline
YDR331W GPI8
ZShift = 1.06
NG = 0
DB = 0
SM = 0
Delta_K
Doxycycline
YDR331W GPI8
ZShift = 0.56
NG = 0
DB = 0
SM = 0
Delta_r
Doxycycline
YDR331W GPI8
ZShift = 0.23
NG = 0
DB = 0
SM = 0
Delta_AUC
Doxycycline

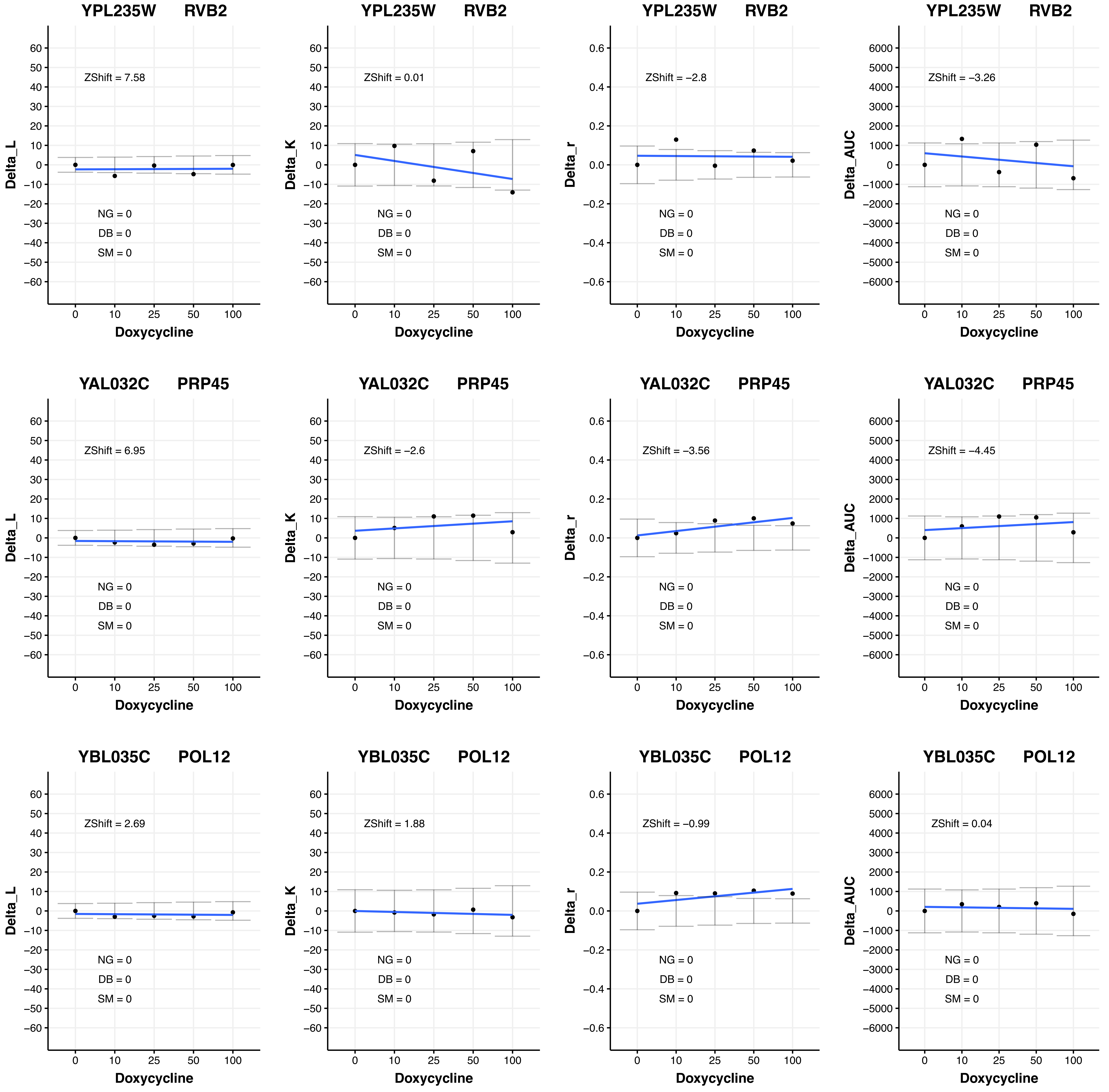
YPL235W RVB2
ZShift = 7.58
NG = 0
DB = 0
SM = 0
Delta_L
Doxycycline
YPL235W RVB2
ZShift = 0.01
NG = 0
DB = 0
SM = 0
Delta_K
Doxycycline
YPL235W RVB2
ZShift = −2.8
NG = 0
DB = 0
SM = 0
Delta_r
Doxycycline
YPL235W RVB2
ZShift = −3.26
NG = 0
DB = 0
SM = 0
Delta_AUC
Doxycycline
YAL032C PRP45
ZShift = 6.95
NG = 0
DB = 0
SM = 0
Delta_L
Doxycycline
YAL032C PRP45
ZShift = −2.6
NG = 0
DB = 0
SM = 0
Delta_K
Doxycycline
YAL032C PRP45
ZShift = −3.56
NG = 0
DB = 0
SM = 0
Delta_r
Doxycycline
YAL032C PRP45
ZShift = −4.45
NG = 0
DB = 0
SM = 0
Delta_AUC
Doxycycline
YBL035C POL12
ZShift = 2.69
NG = 0
DB = 0
SM = 0
Delta_L
Doxycycline
YBL035C POL12
ZShift = 1.88
NG = 0
DB = 0
SM = 0
Delta_K
Doxycycline
YBL035C POL12
ZShift = −0.99
NG = 0
DB = 0
SM = 0
Delta_r
Doxycycline
YBL035C POL12
ZShift = 0.04
NG = 0
DB = 0
SM = 0
Delta_AUC
Doxycycline

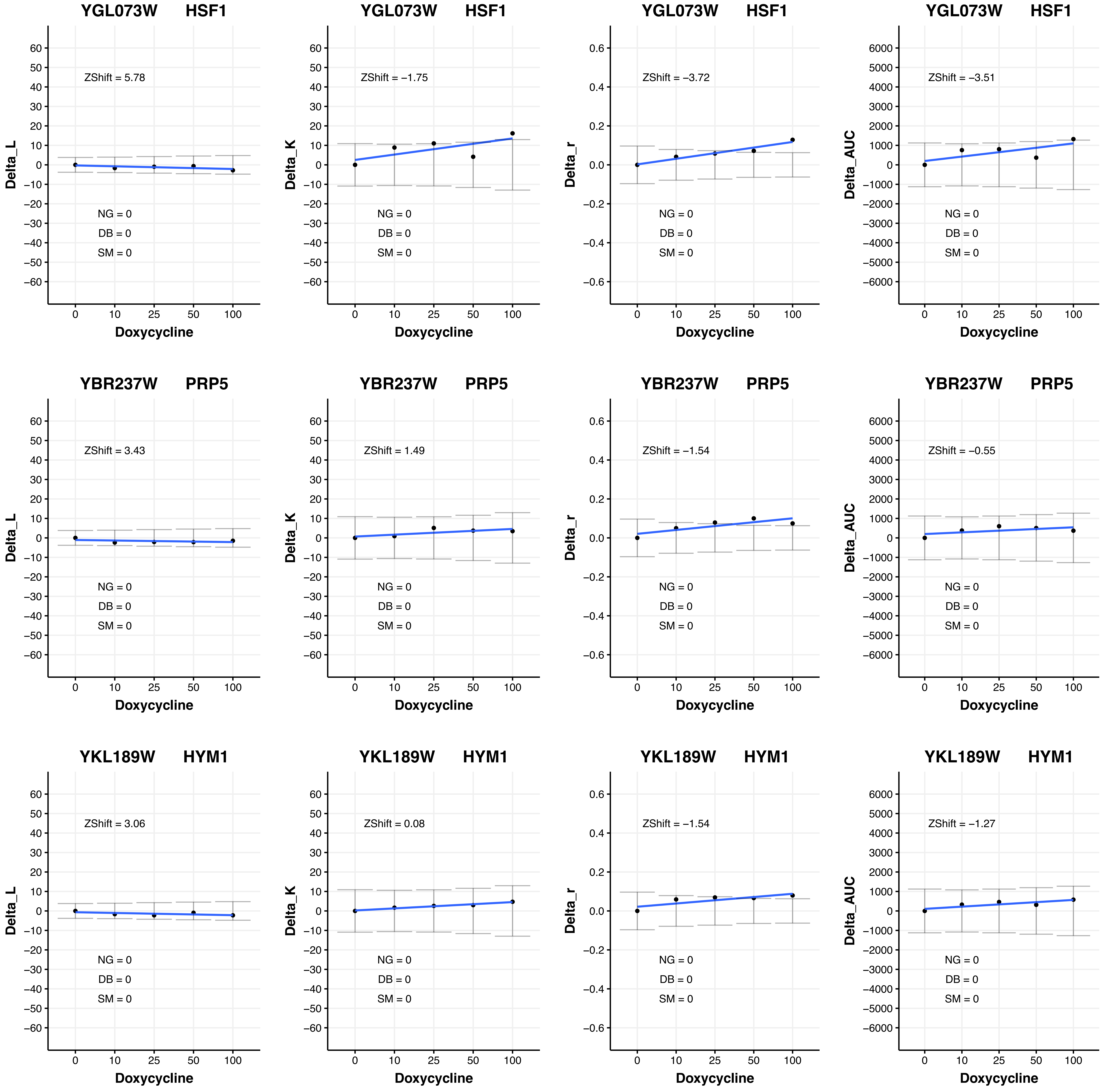

YGL073W HSF1
ZShift = 5.78
NG = 0
DB = 0
SM = 0
Delta_L
Doxycycline
YGL073W HSF1
ZShift = −1.75
NG = 0
DB = 0
SM = 0
Delta_K
Doxycycline
YGL073W HSF1
ZShift = −3.72
NG = 0
DB = 0
SM = 0
Delta_r
Doxycycline
YGL073W HSF1
ZShift = −3.51
NG = 0
DB = 0
SM = 0
Delta_AUC
Doxycycline
YBR237W PRP5
ZShift = 3.43
NG = 0
DB = 0
SM = 0
Delta_L
Doxycycline
YBR237W PRP5
ZShift = 1.49
NG = 0
DB = 0
SM = 0
Delta_K
Doxycycline
YBR237W PRP5
ZShift = −1.54
NG = 0
DB = 0
SM = 0
Delta_r
Doxycycline
YBR237W PRP5
ZShift = −0.55
NG = 0
DB = 0
SM = 0
Delta_AUC
Doxycycline
YKL189W HYM1
ZShift = 3.06
NG = 0
DB = 0
SM = 0
Delta_L
Doxycycline
YKL189W HYM1
ZShift = 0.08
NG = 0
DB = 0
SM = 0
Delta_K
Doxycycline
YKL189W HYM1
ZShift = −1.54
NG = 0
DB = 0
SM = 0
Delta_r
Doxycycline
YKL189W HYM1
ZShift = −1.27
NG = 0
DB = 0
SM = 0
Delta_AUC
Doxycycline

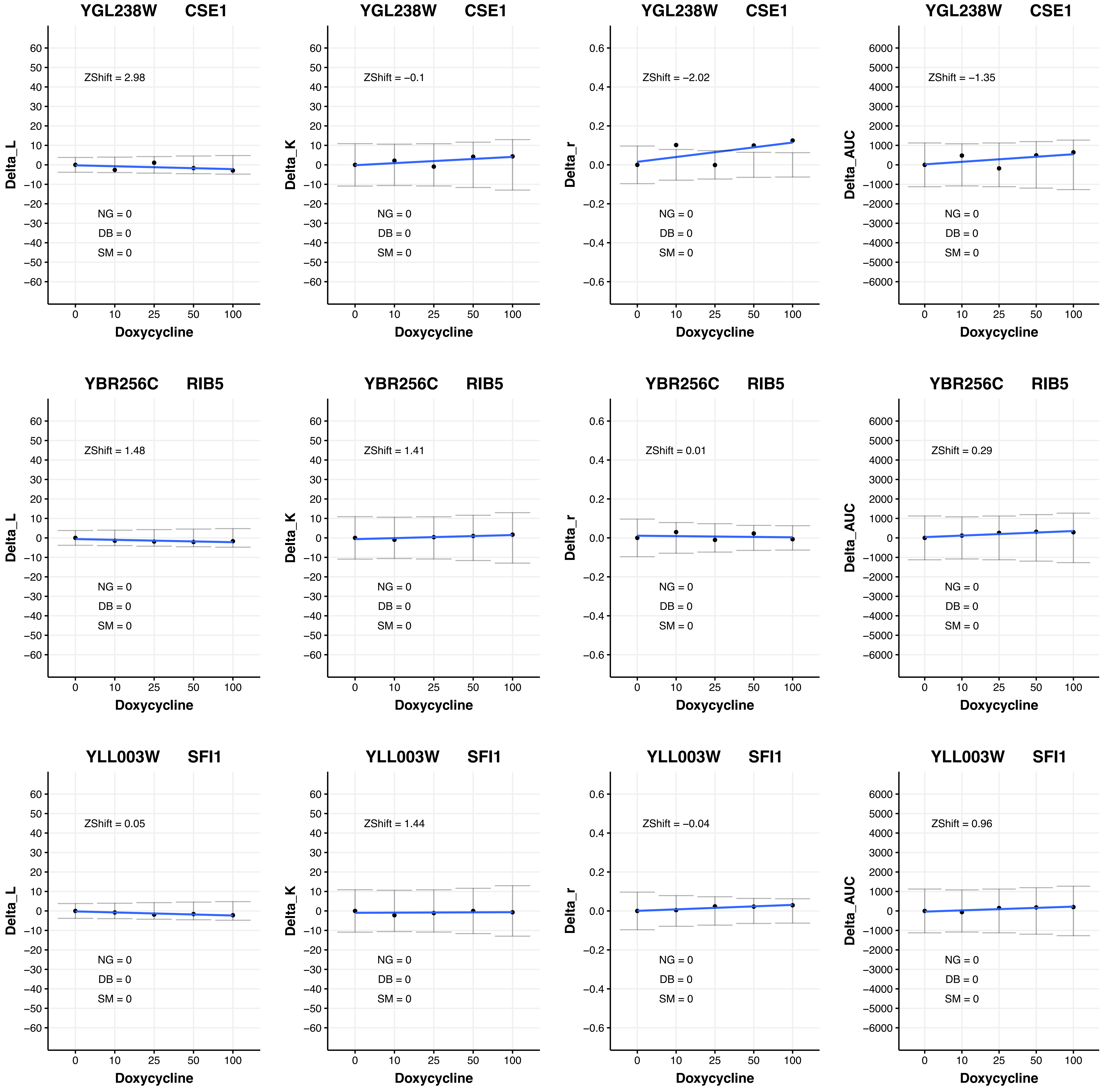
YGL238W CSE1
Delta_L
ZShift = 2.98
NG = 0
DB = 0
SM = 0
Doxycycline
0 10 25 50 100
YGL238W CSE1
Delta_K
ZShift = −0.1
NG = 0
DB = 0
SM = 0
Doxycycline
YGL238W CSE1
Delta_r
ZShift = −2.02
NG = 0
DB = 0
SM = 0
Doxycycline
YGL238W CSE1
Delta_AUC
ZShift = −1.35
NG = 0
DB = 0
SM = 0
Doxycycline
YBR256C RIB5
Delta_L
ZShift = 1.48
NG = 0
DB = 0
SM = 0
Doxycycline
YBR256C RIB5
Delta_K
ZShift = 1.41
NG = 0
DB = 0
SM = 0
Doxycycline
YBR256C RIB5
Delta_r
ZShift = 0.01
NG = 0
DB = 0
SM = 0
Doxycycline
YBR256C RIB5
Delta_AUC
ZShift = 0.29
NG = 0
DB = 0
SM = 0
Doxycycline
YLL003W SFI1
Delta_L
ZShift = 0.05
NG = 0
DB = 0
SM = 0
Doxycycline
YLL003W SFI1
Delta_K
ZShift = 1.44
NG = 0
DB = 0
SM = 0
Doxycycline
YLL003W SFI1
Delta_r
ZShift = −0.04
NG = 0
DB = 0
SM = 0
Doxycycline
YLL003W SFI1
Delta_AUC
ZShift = 0.96
NG = 0
DB = 0
SM = 0
Doxycycline

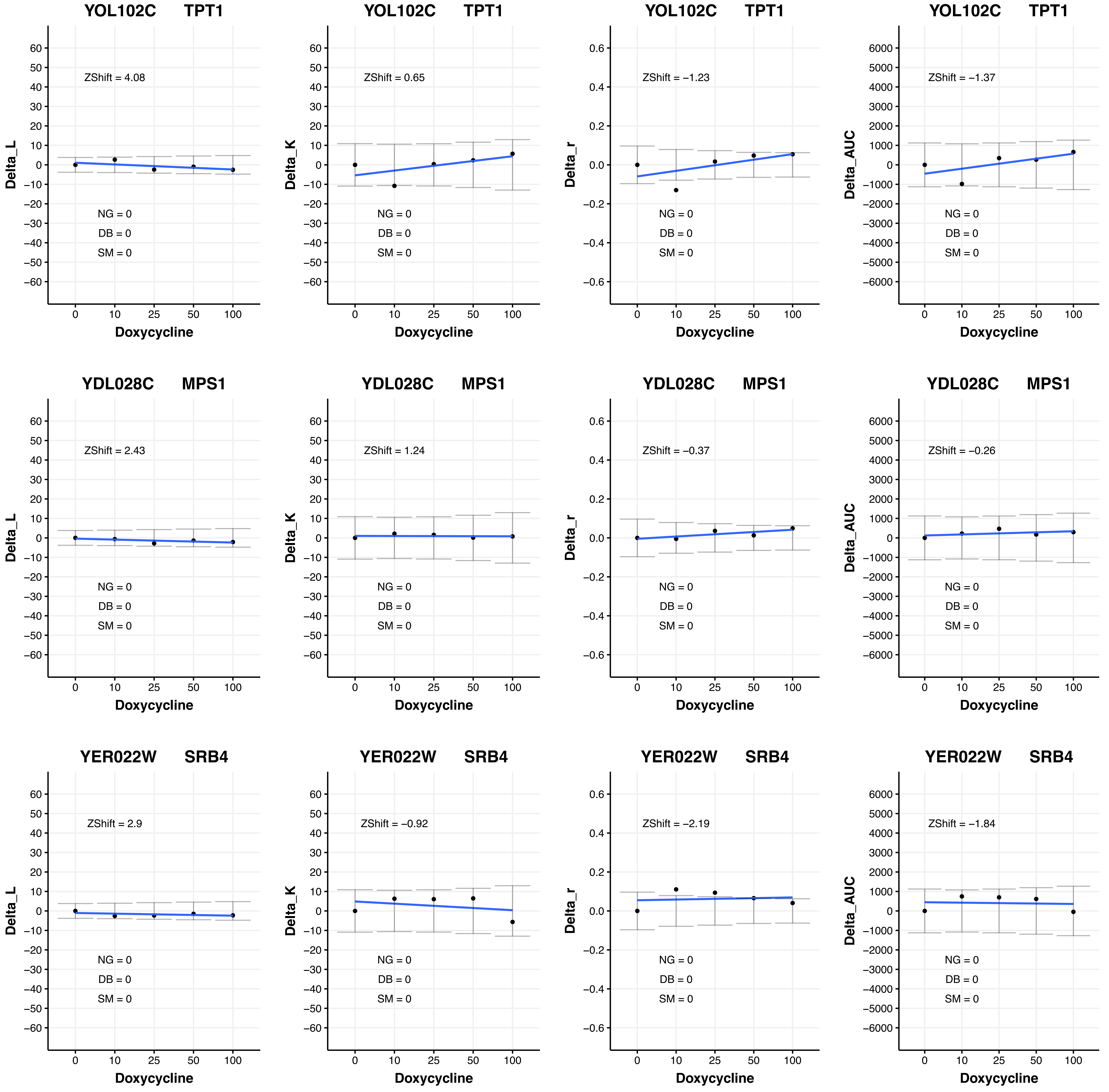
YOL102C TPT1
ZShift = 4.08
ZShift = 0.65
ZShift = −1.23
ZShift = −1.37
YDL028C MPS1
ZShift = 2.43
ZShift = 1.24
ZShift = −0.37
ZShift = −0.26
YER022W SRB4
ZShift = 2.9
ZShift = −0.92
ZShift = −2.19
ZShift = −1.84
NG = 0
DB = 0
SM = 0
Delta_L
Delta_K
Delta_r
Delta_AUC
Doxycycline

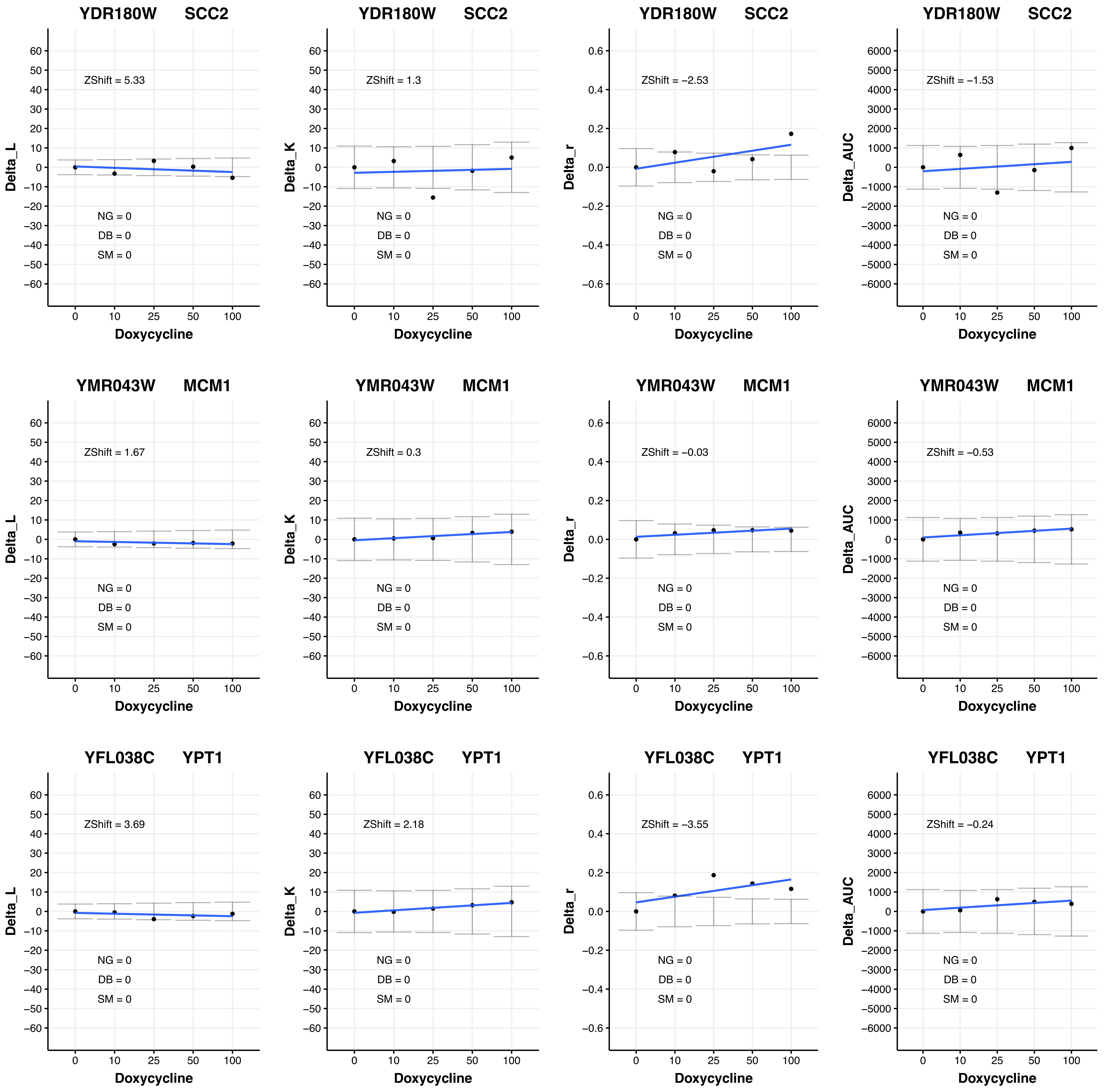
YDR180W SCC2
Delta_L
ZShift = 5.33
NG = 0
DB = 0
SM = 0
Doxycycline
YDR180W SCC2
Delta_K
ZShift = 1.3
NG = 0
DB = 0
SM = 0
Doxycycline
YDR180W SCC2
Delta_r
ZShift = −2.53
NG = 0
DB = 0
SM = 0
Doxycycline
YDR180W SCC2
Delta_AUC
ZShift = −1.53
NG = 0
DB = 0
SM = 0
Doxycycline
YMR043W MCM1
Delta_L
ZShift = 1.67
NG = 0
DB = 0
SM = 0
Doxycycline
YMR043W MCM1
Delta_K
ZShift = 0.3
NG = 0
DB = 0
SM = 0
Doxycycline
YMR043W MCM1
Delta_r
ZShift = −0.03
NG = 0
DB = 0
SM = 0
Doxycycline
YMR043W MCM1
Delta_AUC
ZShift = −0.53
NG = 0
DB = 0
SM = 0
Doxycycline
YFL038C YPT1
Delta_L
ZShift = 3.69
NG = 0
DB = 0
SM = 0
Doxycycline
YFL038C YPT1
Delta_K
ZShift = 2.18
NG = 0
DB = 0
SM = 0
Doxycycline
YFL038C YPT1
Delta_r
ZShift = −3.55
NG = 0
DB = 0
SM = 0
Doxycycline
YFL038C YPT1
Delta_AUC
ZShift = −0.24
NG = 0
DB = 0
SM = 0
Doxycycline

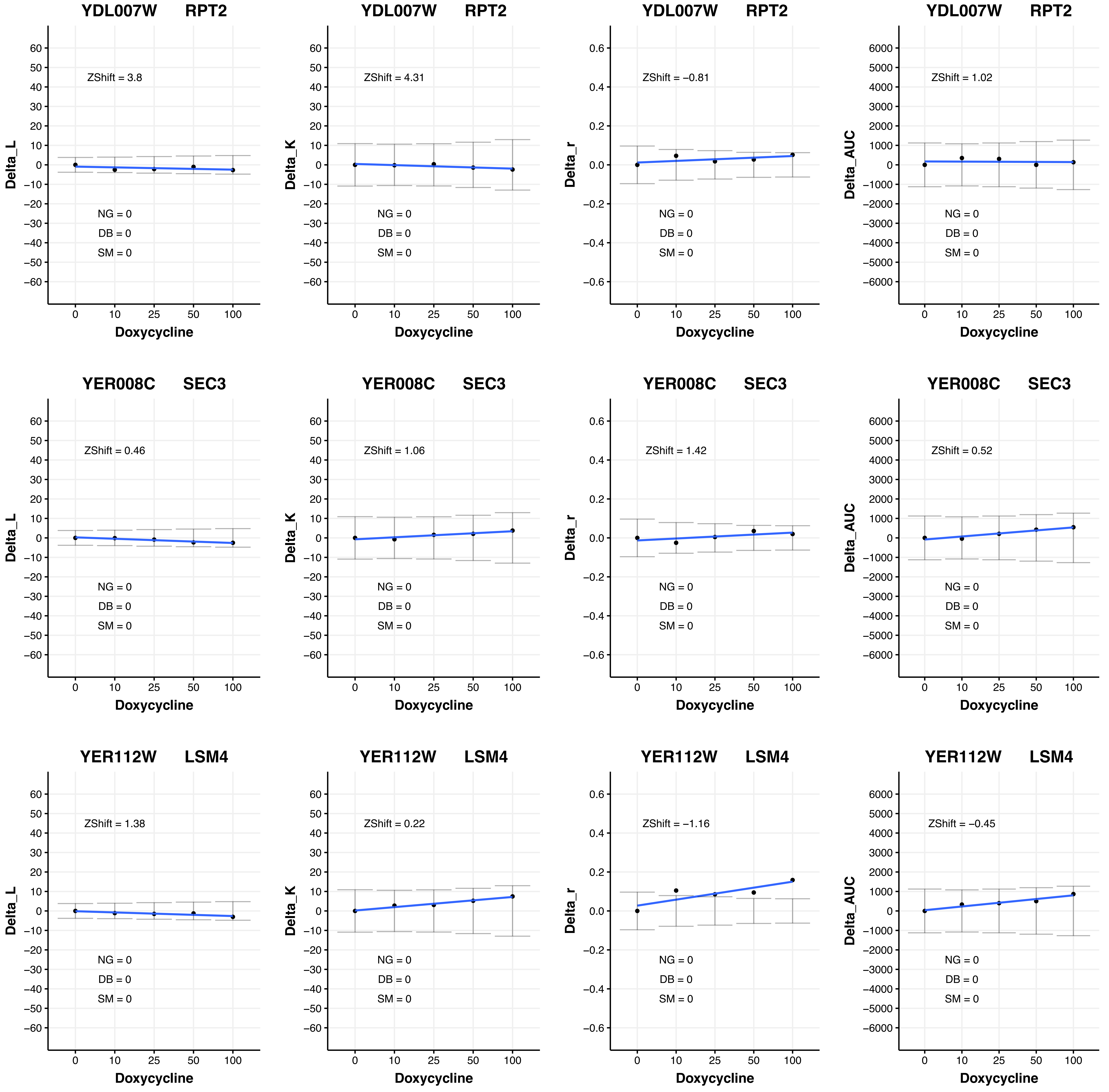
YDL007W RPT2
ZShift = 3.8
NG = 0
DB = 0
SM = 0
Delta_L
YDL007W RPT2
ZShift = 4.31
NG = 0
DB = 0
SM = 0
Delta_K
YDL007W RPT2
ZShift = −0.81
NG = 0
DB = 0
SM = 0
Delta_r
YDL007W RPT2
ZShift = 1.02
NG = 0
DB = 0
SM = 0
Delta_AUC
YER008C SEC3
ZShift = 0.46
NG = 0
DB = 0
SM = 0
Delta_L
YER008C SEC3
ZShift = 1.06
NG = 0
DB = 0
SM = 0
Delta_K
YER008C SEC3
ZShift = 1.42
NG = 0
DB = 0
SM = 0
Delta_r
YER008C SEC3
ZShift = 0.52
NG = 0
DB = 0
SM = 0
Delta_AUC
YER112W LSM4
ZShift = 1.38
NG = 0
DB = 0
SM = 0
Delta_L
YER112W LSM4
ZShift = 0.22
NG = 0
DB = 0
SM = 0
Delta_K
YER112W LSM4
ZShift = −1.16
NG = 0
DB = 0
SM = 0
Delta_r
YER112W LSM4
ZShift = −0.45
NG = 0
DB = 0
SM = 0
Delta_AUC
Doxycycline
0
10
25
50
100

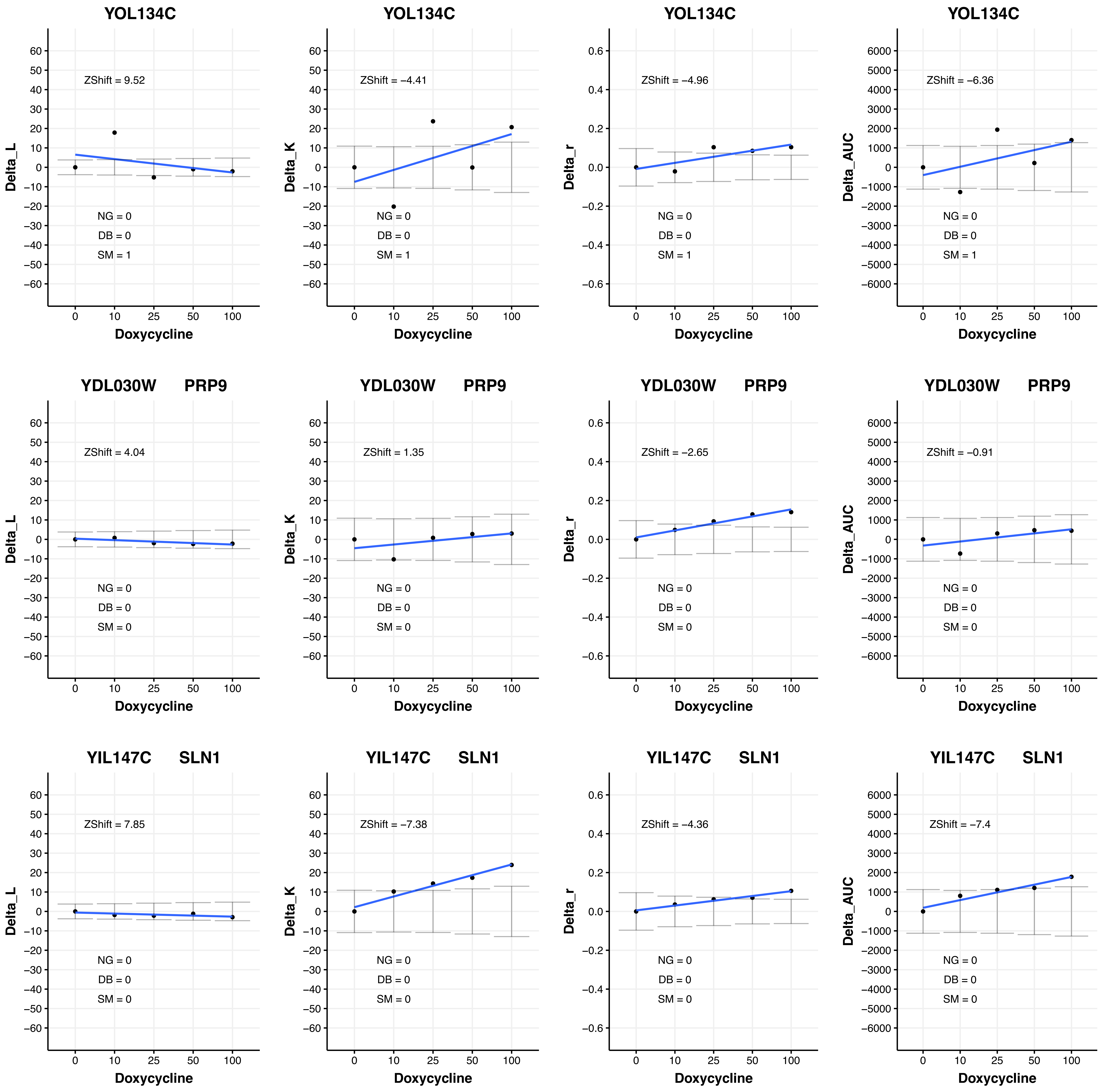

YOL134C
ZShift = 9.52
NG = 0
DB = 0
SM = 1
Delta_L
Doxycycline
YOL134C
ZShift = −4.41
NG = 0
DB = 0
SM = 1
Delta_K
Doxycycline
YOL134C
ZShift = −4.96
NG = 0
DB = 0
SM = 1
Delta_r
Doxycycline
YOL134C
ZShift = −6.36
NG = 0
DB = 0
SM = 1
Delta_AUC
Doxycycline
YDL030W PRP9
ZShift = 4.04
NG = 0
DB = 0
SM = 0
Delta_L
Doxycycline
YDL030W PRP9
ZShift = 1.35
NG = 0
DB = 0
SM = 0
Delta_K
Doxycycline
YDL030W PRP9
ZShift = −2.65
NG = 0
DB = 0
SM = 0
Delta_r
Doxycycline
YDL030W PRP9
ZShift = −0.91
NG = 0
DB = 0
SM = 0
Delta_AUC
Doxycycline
YIL147C SLN1
ZShift = 7.85
NG = 0
DB = 0
SM = 0
Delta_L
Doxycycline
YIL147C SLN1
ZShift = −7.38
NG = 0
DB = 0
SM = 0
Delta_K
Doxycycline
YIL147C SLN1
ZShift = −4.36
NG = 0
DB = 0
SM = 0
Delta_r
Doxycycline
YIL147C SLN1
ZShift = −7.4
NG = 0
DB = 0
SM = 0
Delta_AUC
Doxycycline

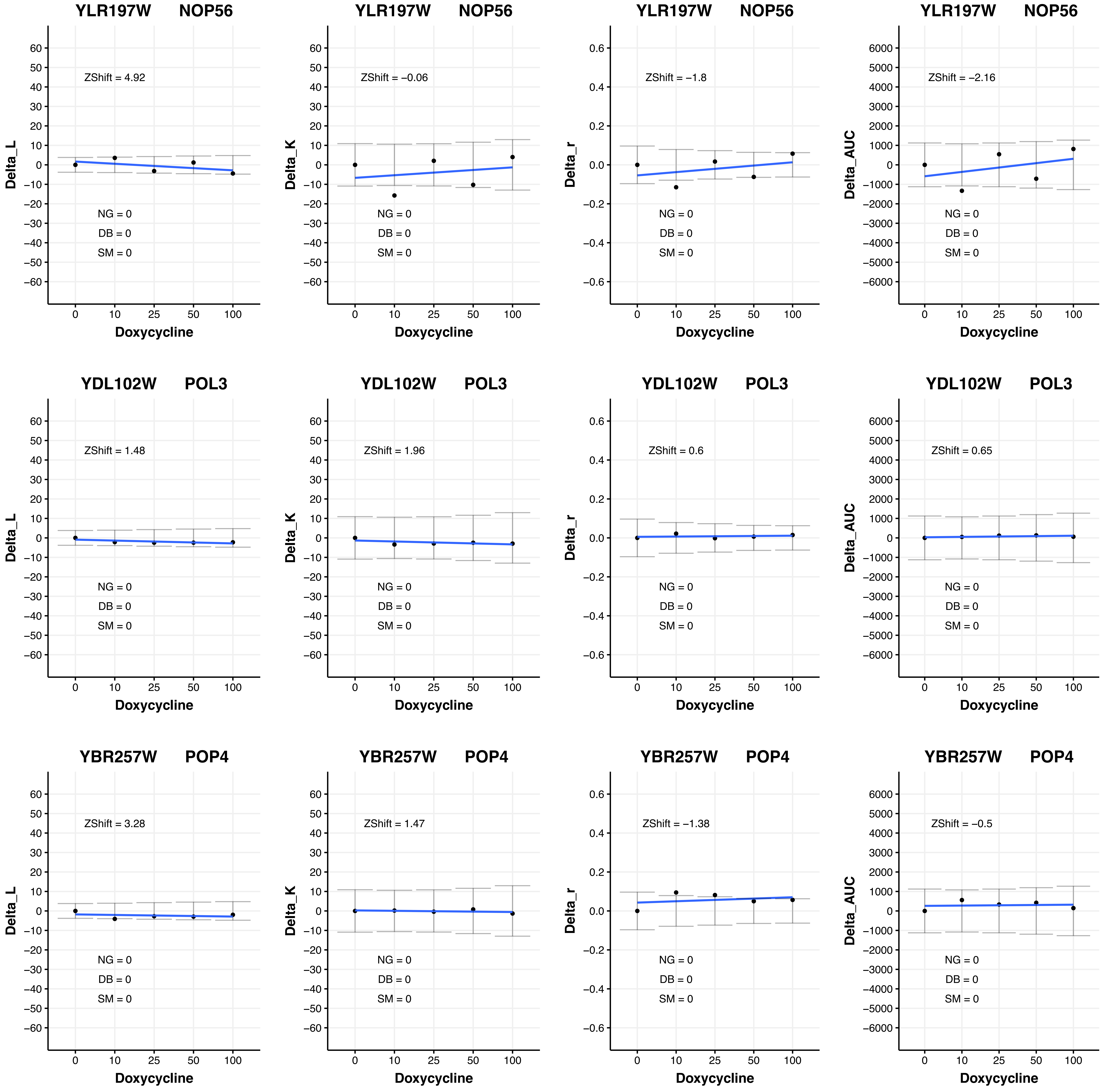
YLR197W NOP56
ZShift = 4.92
NG = 0
DB = 0
SM = 0
Delta_L
Doxycycline
YLR197W NOP56
ZShift = −0.06
NG = 0
DB = 0
SM = 0
Delta_K
Doxycycline
YLR197W NOP56
ZShift = −1.8
NG = 0
DB = 0
SM = 0
Delta_r
Doxycycline
YLR197W NOP56
ZShift = −2.16
NG = 0
DB = 0
SM = 0
Delta_AUC
Doxycycline
YDL102W POL3
ZShift = 1.48
NG = 0
DB = 0
SM = 0
Delta_L
Doxycycline
YDL102W POL3
ZShift = 1.96
NG = 0
DB = 0
SM = 0
Delta_K
Doxycycline
YDL102W POL3
ZShift = 0.6
NG = 0
DB = 0
SM = 0
Delta_r
Doxycycline
YDL102W POL3
ZShift = 0.65
NG = 0
DB = 0
SM = 0
Delta_AUC
Doxycycline
YBR257W POP4
ZShift = 3.28
NG = 0
DB = 0
SM = 0
Delta_L
Doxycycline
YBR257W POP4
ZShift = 1.47
NG = 0
DB = 0
SM = 0
Delta_K
Doxycycline
YBR257W POP4
ZShift = −1.38
NG = 0
DB = 0
SM = 0
Delta_r
Doxycycline
YBR257W POP4
ZShift = −0.5
NG = 0
DB = 0
SM = 0
Delta_AUC
Doxycycline

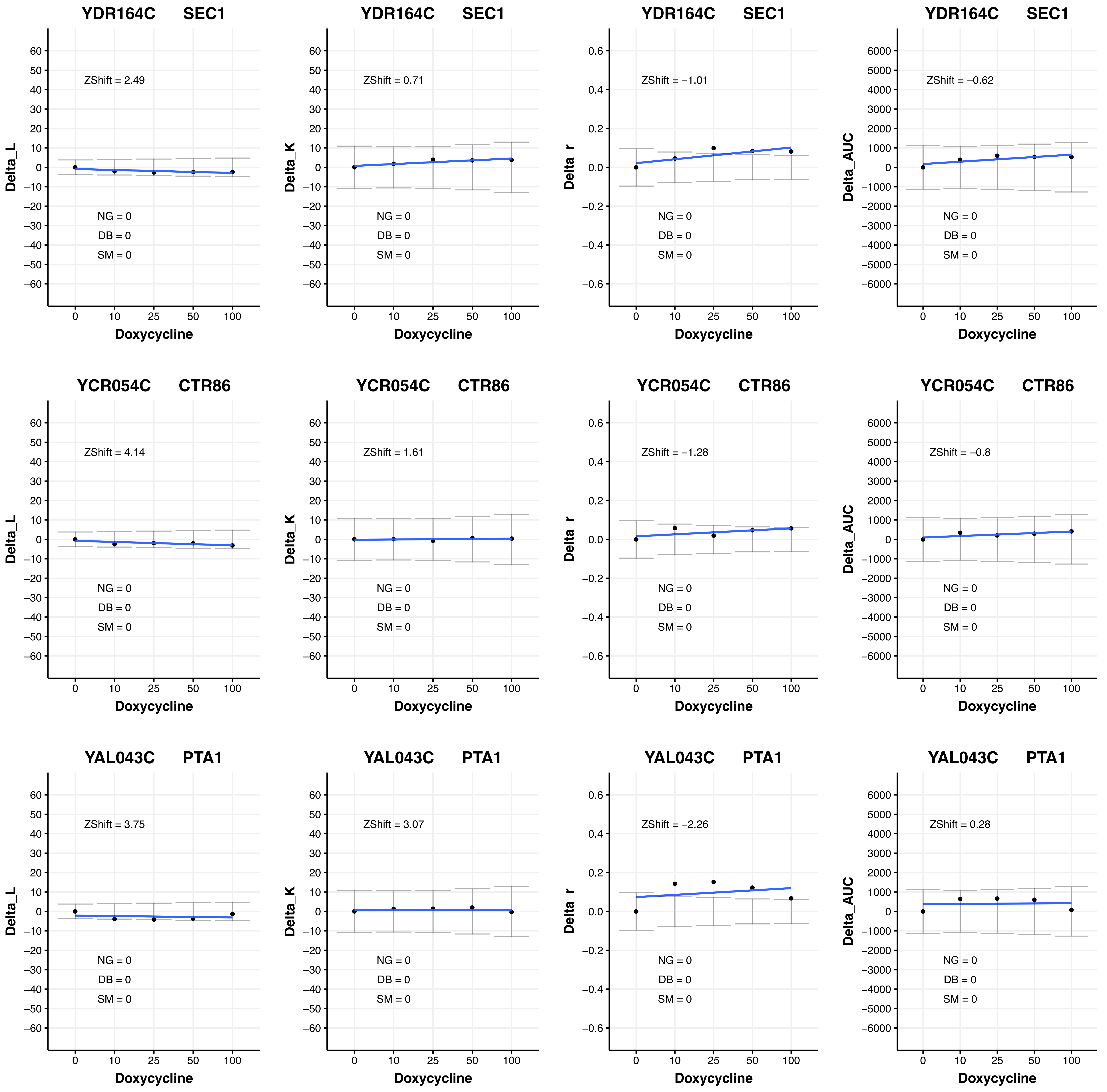

YDR164C SEC1
ZShift = 2.49
NG = 0
DB = 0
SM = 0
Delta_L
Doxycycline
YDR164C SEC1
ZShift = 0.71
NG = 0
DB = 0
SM = 0
Delta_K
Doxycycline
YDR164C SEC1
ZShift = −1.01
NG = 0
DB = 0
SM = 0
Delta_r
Doxycycline
YDR164C SEC1
ZShift = −0.62
NG = 0
DB = 0
SM = 0
Delta_AUC
Doxycycline
YCR054C CTR86
ZShift = 4.14
NG = 0
DB = 0
SM = 0
Delta_L
Doxycycline
YCR054C CTR86
ZShift = 1.61
NG = 0
DB = 0
SM = 0
Delta_K
Doxycycline
YCR054C CTR86
ZShift = −1.28
NG = 0
DB = 0
SM = 0
Delta_r
Doxycycline
YCR054C CTR86
ZShift = −0.8
NG = 0
DB = 0
SM = 0
Delta_AUC
Doxycycline
YAL043C PTA1
ZShift = 3.75
NG = 0
DB = 0
SM = 0
Delta_L
Doxycycline
YAL043C PTA1
ZShift = 3.07
NG = 0
DB = 0
SM = 0
Delta_K
Doxycycline
YAL043C PTA1
ZShift = −2.26
NG = 0
DB = 0
SM = 0
Delta_r
Doxycycline
YAL043C PTA1
ZShift = 0.28
NG = 0
DB = 0
SM = 0
Delta_AUC
Doxycycline

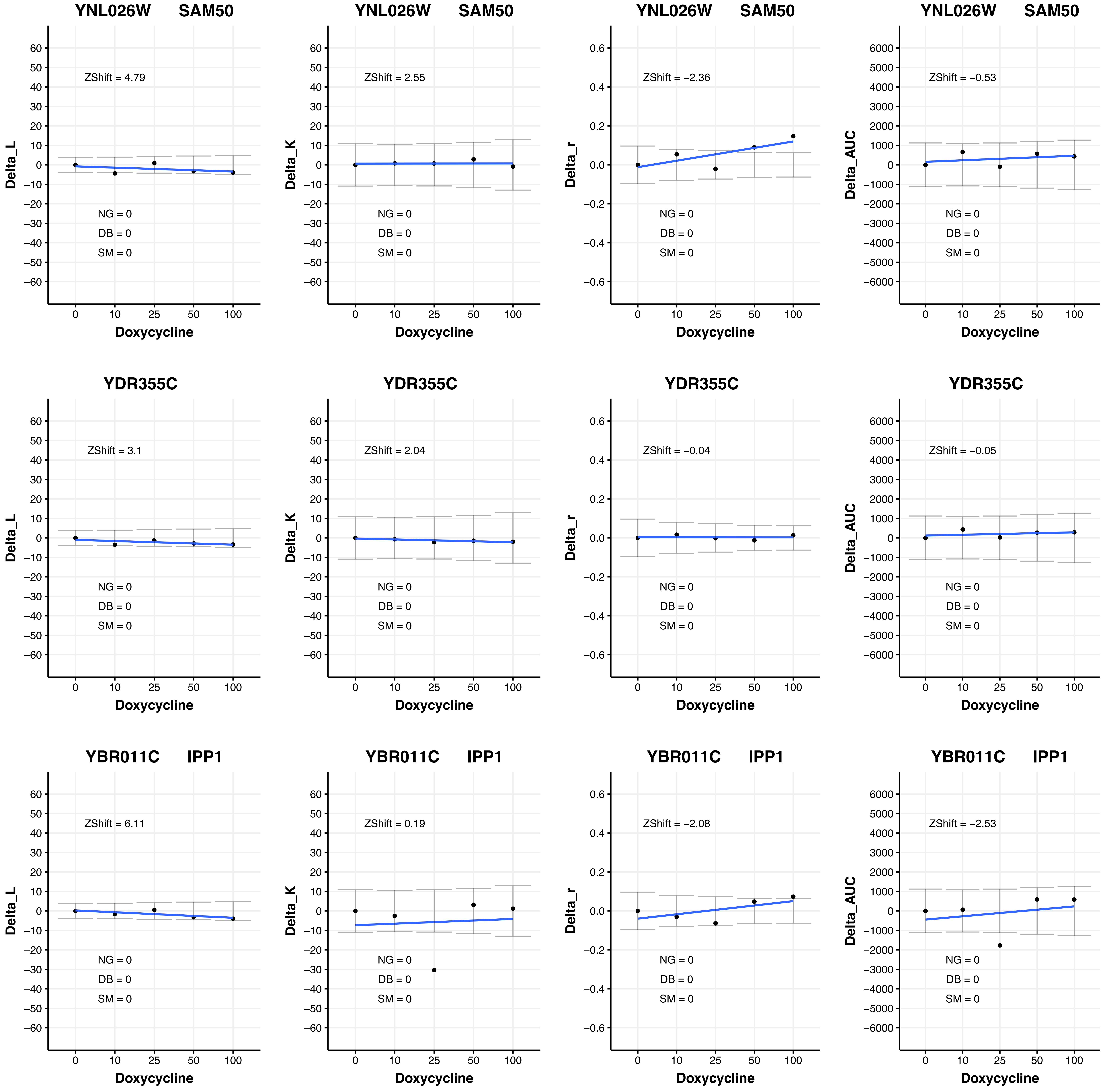

YNL026W SAM50
ZShift = 4.79
ZShift = 2.55
ZShift = −2.36
ZShift = −0.53
YDR355C
ZShift = 3.1
ZShift = 2.04
ZShift = −0.04
ZShift = −0.05
YBR011C IPP1
ZShift = 6.11
ZShift = 0.19
ZShift = −2.08
ZShift = −2.53
NG = 0
DB = 0
SM = 0
Delta_L
Delta_K
Delta_r
Delta_AUC
Doxycycline

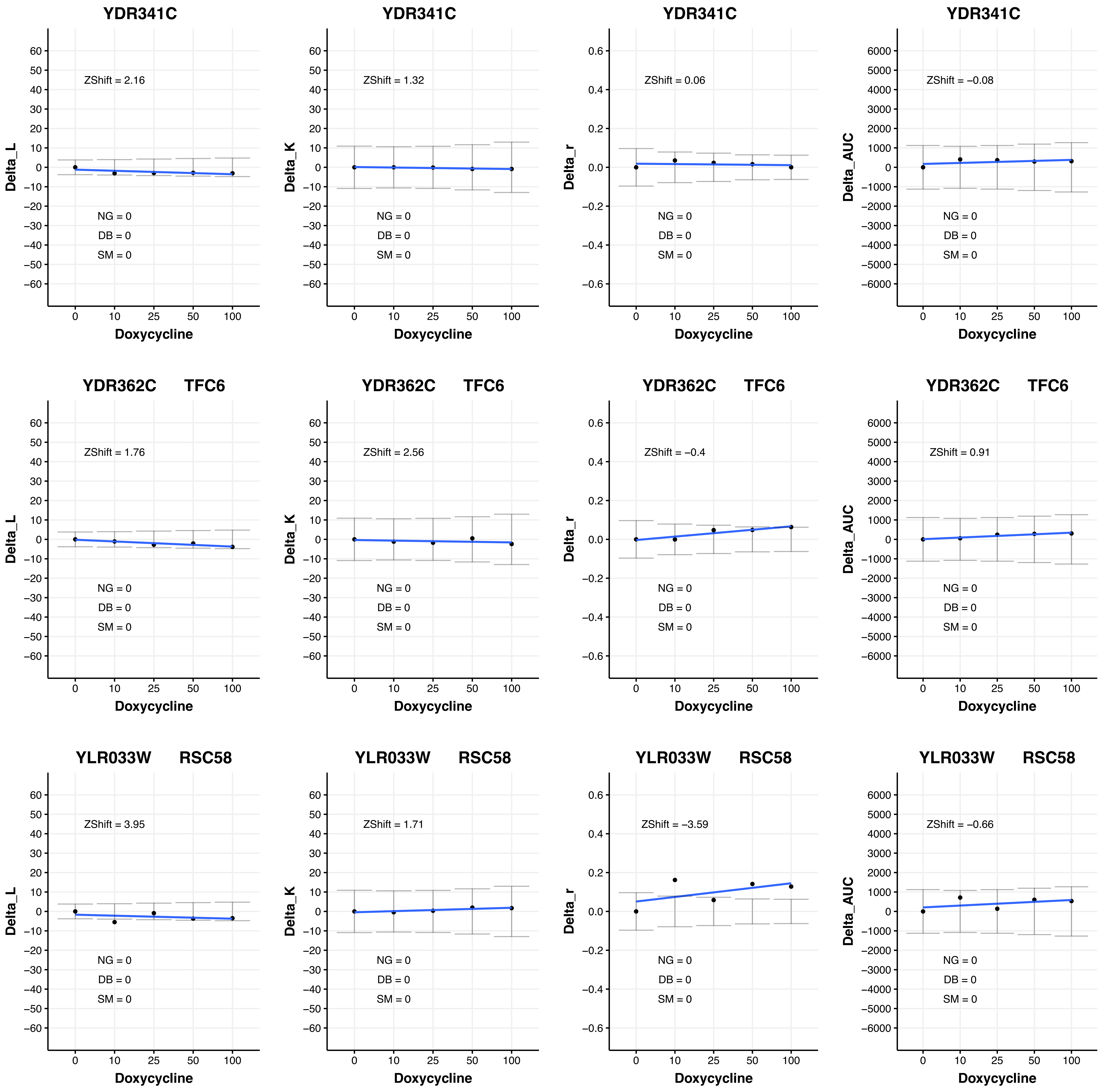
YDR341C
ZShift = 2.16
NG = 0
DB = 0
SM = 0
Delta_L
Doxycycline
0 10 25 50 100
YDR341C
ZShift = 1.32
NG = 0
DB = 0
SM = 0
Delta_K
Doxycycline
YDR341C
ZShift = 0.06
NG = 0
DB = 0
SM = 0
Delta_r
Doxycycline
YDR341C
ZShift = −0.08
NG = 0
DB = 0
SM = 0
Delta_AUC
Doxycycline
YDR362C TFC6
ZShift = 1.76
NG = 0
DB = 0
SM = 0
Delta_L
Doxycycline
YDR362C TFC6
ZShift = 2.56
NG = 0
DB = 0
SM = 0
Delta_K
Doxycycline
YDR362C TFC6
ZShift = −0.4
NG = 0
DB = 0
SM = 0
Delta_r
Doxycycline
YDR362C TFC6
ZShift = 0.91
NG = 0
DB = 0
SM = 0
Delta_AUC
Doxycycline
YLR033W RSC58
ZShift = 3.95
NG = 0
DB = 0
SM = 0
Delta_L
Doxycycline
YLR033W RSC58
ZShift = 1.71
NG = 0
DB = 0
SM = 0
Delta_K
Doxycycline
YLR033W RSC58
ZShift = −3.59
NG = 0
DB = 0
SM = 0
Delta_r
Doxycycline
YLR033W RSC58
ZShift = −0.66
NG = 0
DB = 0
SM = 0
Delta_AUC
Doxycycline

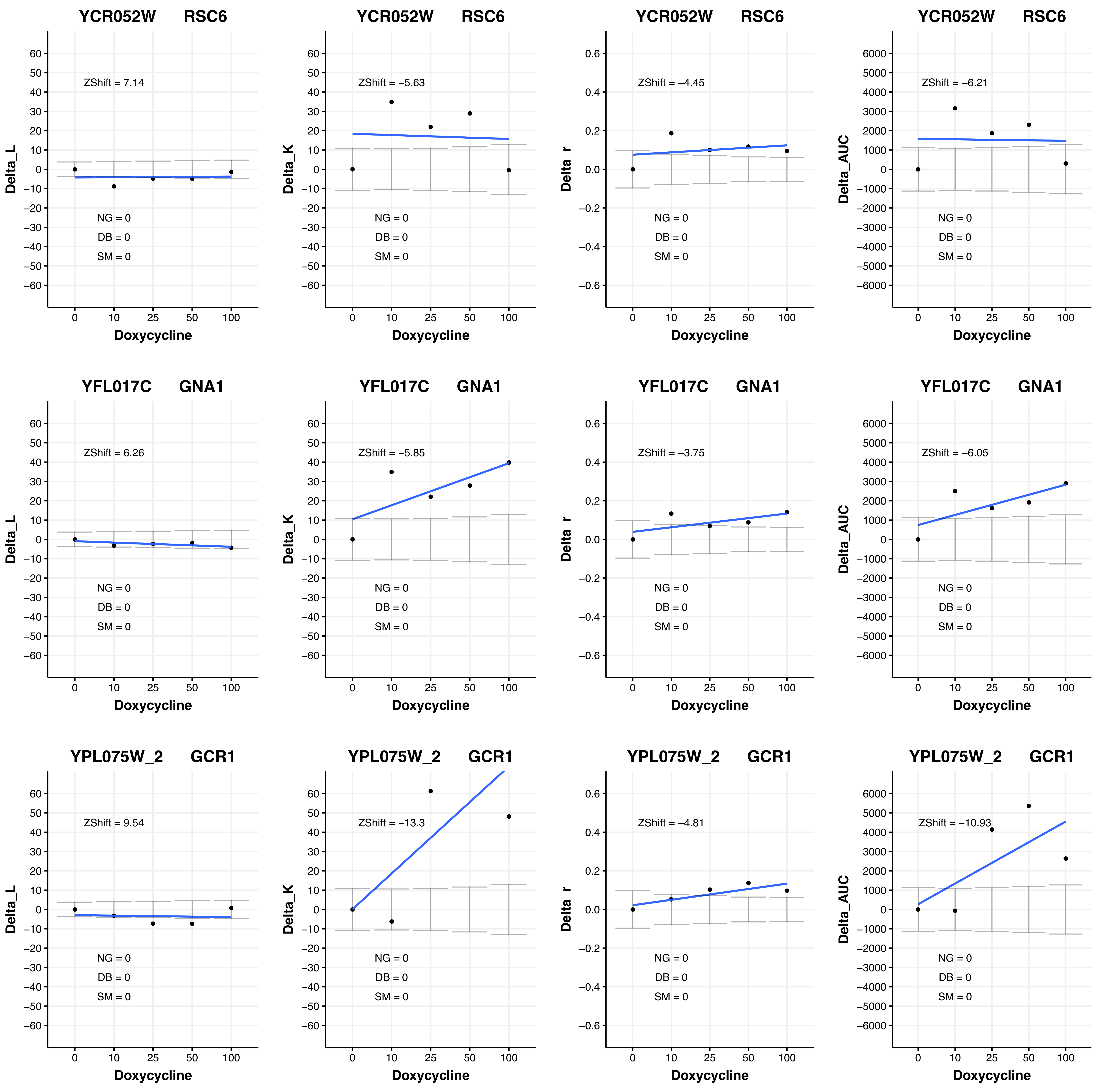
YCR052W RSC6
Delta_L
ZShift = 7.14
NG = 0
DB = 0
SM = 0
Doxycycline
YCR052W RSC6
Delta_K
ZShift = −5.63
NG = 0
DB = 0
SM = 0
Doxycycline
YCR052W RSC6
Delta_r
ZShift = −4.45
NG = 0
DB = 0
SM = 0
Doxycycline
YCR052W RSC6
Delta_AUC
ZShift = −6.21
NG = 0
DB = 0
SM = 0
Doxycycline
YFL017C GNA1
Delta_L
ZShift = 6.26
NG = 0
DB = 0
SM = 0
Doxycycline
YFL017C GNA1
Delta_K
ZShift = −5.85
NG = 0
DB = 0
SM = 0
Doxycycline
YFL017C GNA1
Delta_r
ZShift = −3.75
NG = 0
DB = 0
SM = 0
Doxycycline
YFL017C GNA1
Delta_AUC
ZShift = −6.05
NG = 0
DB = 0
SM = 0
Doxycycline
YPL075W_2 GCR1
Delta_L
ZShift = 9.54
NG = 0
DB = 0
SM = 0
Doxycycline
YPL075W_2 GCR1
Delta_K
ZShift = −13.3
NG = 0
DB = 0
SM = 0
Doxycycline
YPL075W_2 GCR1
Delta_r
ZShift = −4.81
NG = 0
DB = 0
SM = 0
Doxycycline
YPL075W_2 GCR1
Delta_AUC
ZShift = −10.93
NG = 0
DB = 0
SM = 0
Doxycycline

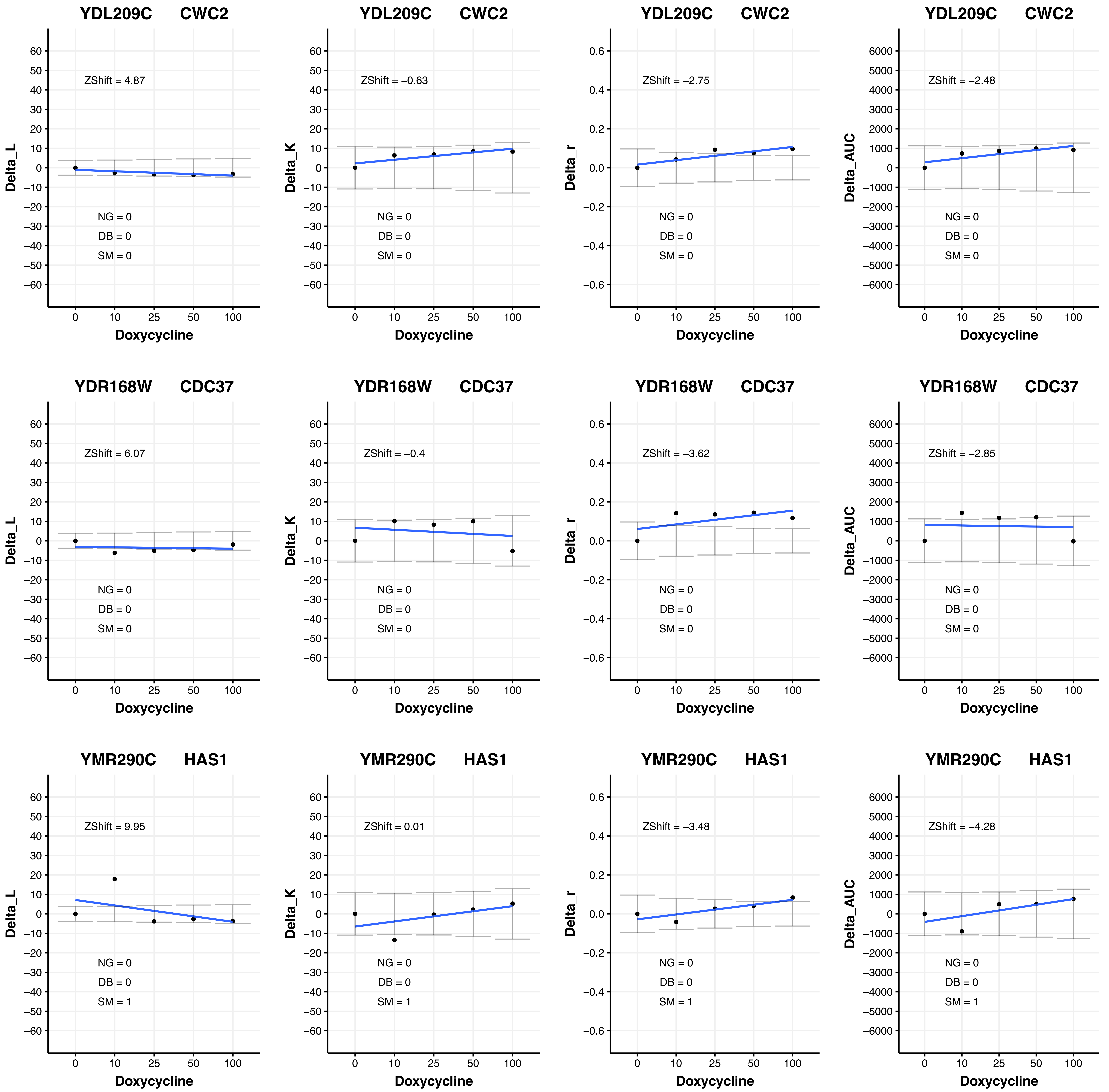
YDL209C CWC2
ZShift = 4.87
NG = 0
DB = 0
SM = 0
Delta_L
Doxycycline
YDL209C CWC2
ZShift = −0.63
NG = 0
DB = 0
SM = 0
Delta_K
Doxycycline
YDL209C CWC2
ZShift = −2.75
NG = 0
DB = 0
SM = 0
Delta_r
Doxycycline
YDL209C CWC2
ZShift = −2.48
NG = 0
DB = 0
SM = 0
Delta_AUC
Doxycycline
YDR168W CDC37
ZShift = 6.07
NG = 0
DB = 0
SM = 0
Delta_L
Doxycycline
YDR168W CDC37
ZShift = −0.4
NG = 0
DB = 0
SM = 0
Delta_K
Doxycycline
YDR168W CDC37
ZShift = −3.62
NG = 0
DB = 0
SM = 0
Delta_r
Doxycycline
YDR168W CDC37
ZShift = −2.85
NG = 0
DB = 0
SM = 0
Delta_AUC
Doxycycline
YMR290C HAS1
ZShift = 9.95
NG = 0
DB = 0
SM = 1
Delta_L
Doxycycline
YMR290C HAS1
ZShift = 0.01
NG = 0
DB = 0
SM = 1
Delta_K
Doxycycline
YMR290C HAS1
ZShift = −3.48
NG = 0
DB = 0
SM = 1
Delta_r
Doxycycline
YMR290C HAS1
ZShift = −4.28
NG = 0
DB = 0
SM = 1
Delta_AUC
Doxycycline

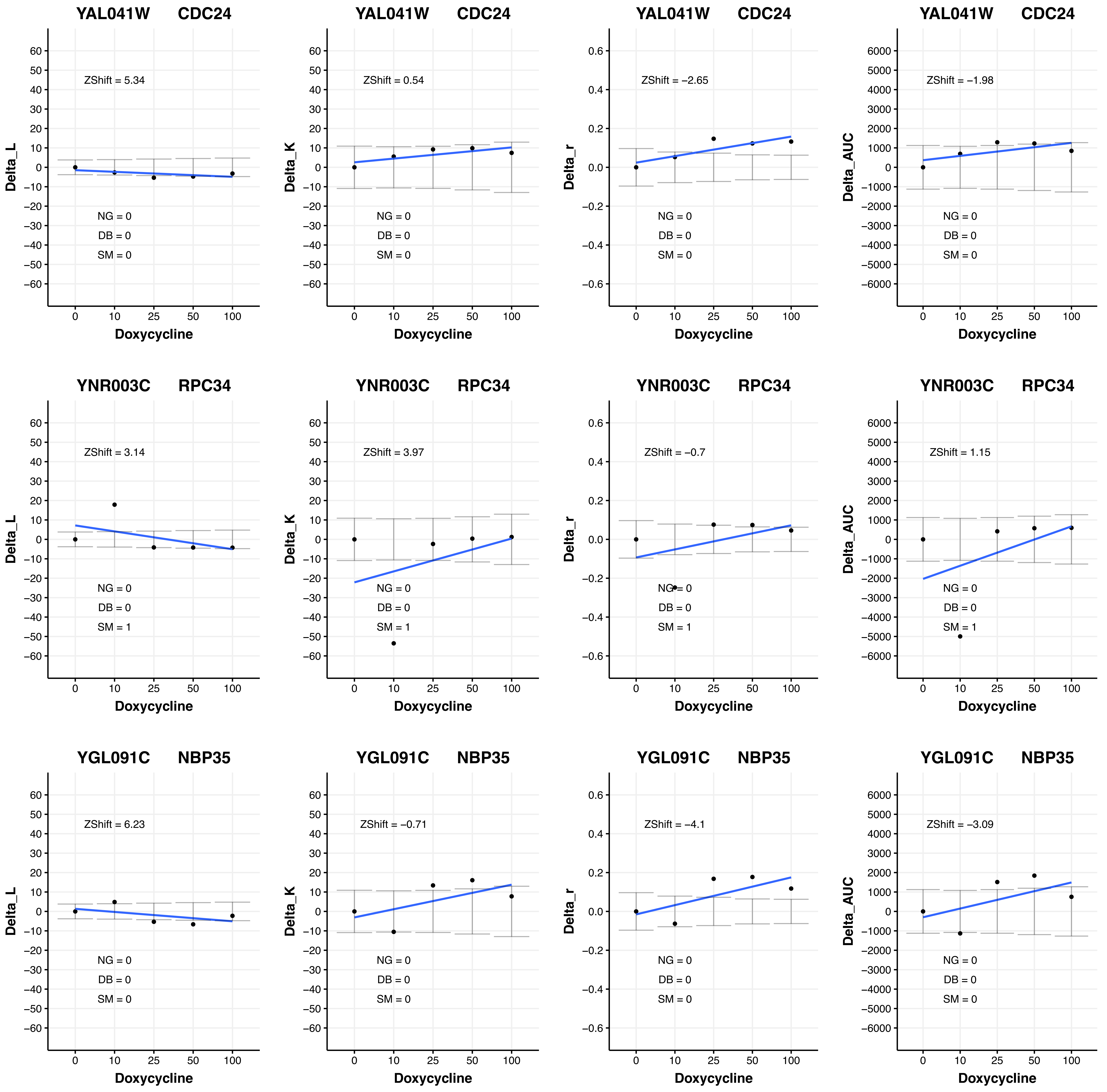

YAL041W CDC24
ZShift = 5.34
NG = 0
DB = 0
SM = 0
Delta_L
Doxycycline
YAL041W CDC24
ZShift = 0.54
NG = 0
DB = 0
SM = 0
Delta_K
Doxycycline
YAL041W CDC24
ZShift = −2.65
NG = 0
DB = 0
SM = 0
Delta_r
Doxycycline
YAL041W CDC24
ZShift = −1.98
NG = 0
DB = 0
SM = 0
Delta_AUC
Doxycycline
YNR003C RPC34
ZShift = 3.14
NG = 0
DB = 0
SM = 1
Delta_L
Doxycycline
YNR003C RPC34
ZShift = 3.97
NG = 0
DB = 0
SM = 1
Delta_K
Doxycycline
YNR003C RPC34
ZShift = −0.7
NG = 0
DB = 0
SM = 1
Delta_r
Doxycycline
YNR003C RPC34
ZShift = 1.15
NG = 0
DB = 0
SM = 1
Delta_AUC
Doxycycline
YGL091C NBP35
ZShift = 6.23
NG = 0
DB = 0
SM = 0
Delta_L
Doxycycline
YGL091C NBP35
ZShift = −0.71
NG = 0
DB = 0
SM = 0
Delta_K
Doxycycline
YGL091C NBP35
ZShift = −4.1
NG = 0
DB = 0
SM = 0
Delta_r
Doxycycline
YGL091C NBP35
ZShift = −3.09
NG = 0
DB = 0
SM = 0
Delta_AUC
Doxycycline

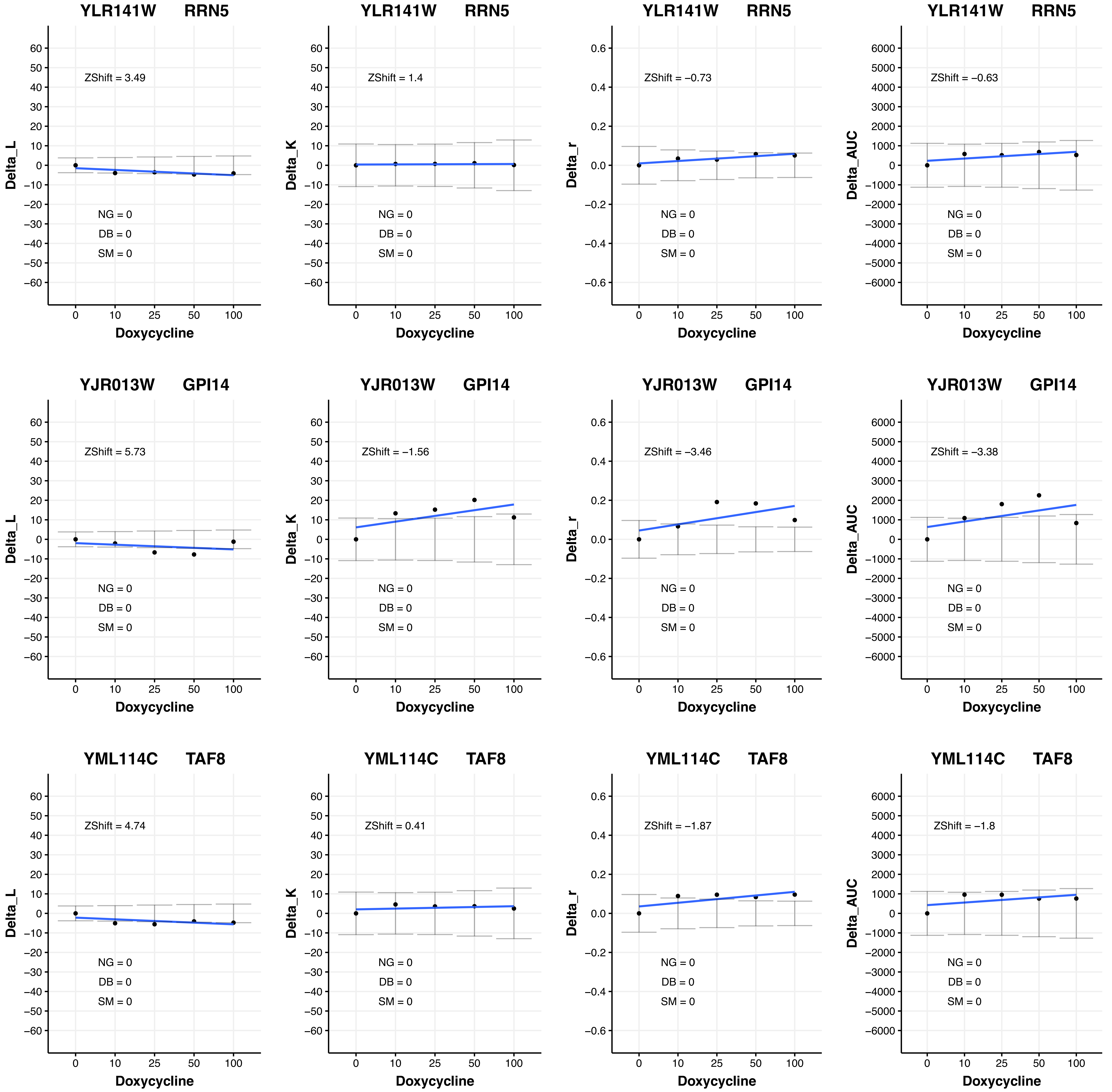
YLR141W RRN5
ZShift = 3.49
NG = 0
DB = 0
SM = 0
Delta_L
Doxycycline
YLR141W RRN5
ZShift = 1.4
NG = 0
DB = 0
SM = 0
Delta_K
Doxycycline
YLR141W RRN5
ZShift = −0.73
NG = 0
DB = 0
SM = 0
Delta_r
Doxycycline
YLR141W RRN5
ZShift = −0.63
NG = 0
DB = 0
SM = 0
Delta_AUC
Doxycycline
YJR013W GPI14
ZShift = 5.73
NG = 0
DB = 0
SM = 0
Delta_L
Doxycycline
YJR013W GPI14
ZShift = −1.56
NG = 0
DB = 0
SM = 0
Delta_K
Doxycycline
YJR013W GPI14
ZShift = −3.46
NG = 0
DB = 0
SM = 0
Delta_r
Doxycycline
YJR013W GPI14
ZShift = −3.38
NG = 0
DB = 0
SM = 0
Delta_AUC
Doxycycline
YML114C TAF8
ZShift = 4.74
NG = 0
DB = 0
SM = 0
Delta_L
Doxycycline
YML114C TAF8
ZShift = 0.41
NG = 0
DB = 0
SM = 0
Delta_K
Doxycycline
YML114C TAF8
ZShift = −1.87
NG = 0
DB = 0
SM = 0
Delta_r
Doxycycline
YML114C TAF8
ZShift = −1.8
NG = 0
DB = 0
SM = 0
Delta_AUC
Doxycycline

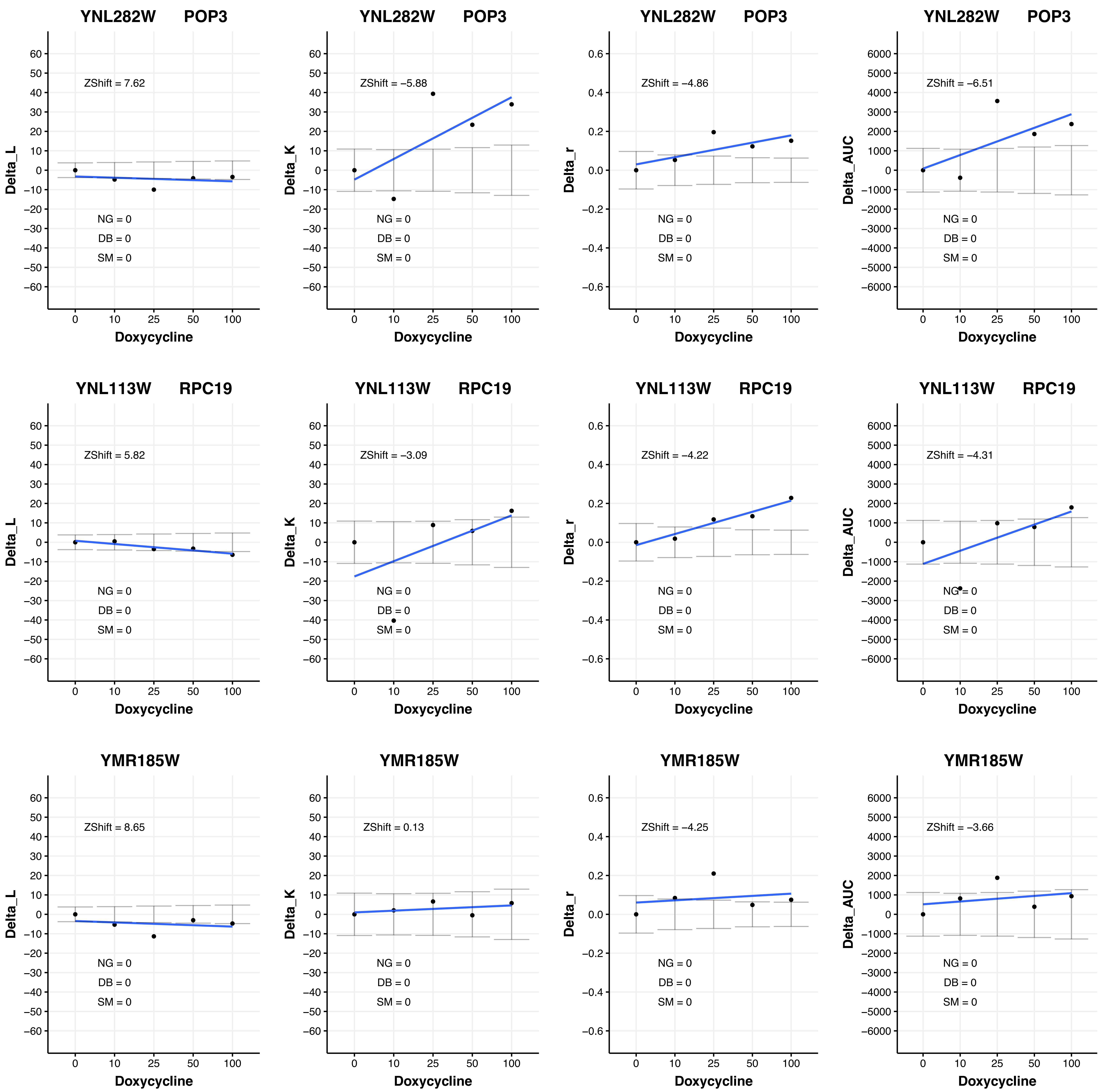
YNL282W POP3
ZShift = 7.62
Delta_L
NG = 0
DB = 0
SM = 0
Doxycycline
YNL282W POP3
ZShift = −5.88
Delta_K
NG = 0
DB = 0
SM = 0
Doxycycline
YNL282W POP3
ZShift = −4.86
Delta_r
NG = 0
DB = 0
SM = 0
Doxycycline
YNL282W POP3
ZShift = −6.51
Delta_AUC
NG = 0
DB = 0
SM = 0
Doxycycline
YNL113W RPC19
ZShift = 5.82
Delta_L
NG = 0
DB = 0
SM = 0
Doxycycline
YNL113W RPC19
ZShift = −3.09
Delta_K
NG = 0
DB = 0
SM = 0
Doxycycline
YNL113W RPC19
ZShift = −4.22
Delta_r
NG = 0
DB = 0
SM = 0
Doxycycline
YNL113W RPC19
ZShift = −4.31
Delta_AUC
NG = 0
DB = 0
SM = 0
Doxycycline
YMR185W
ZShift = 8.65
Delta_L
NG = 0
DB = 0
SM = 0
Doxycycline
YMR185W
ZShift = 0.13
Delta_K
NG = 0
DB = 0
SM = 0
Doxycycline
YMR185W
ZShift = −4.25
Delta_r
NG = 0
DB = 0
SM = 0
Doxycycline
YMR185W
ZShift = −3.66
Delta_AUC
NG = 0
DB = 0
SM = 0
Doxycycline

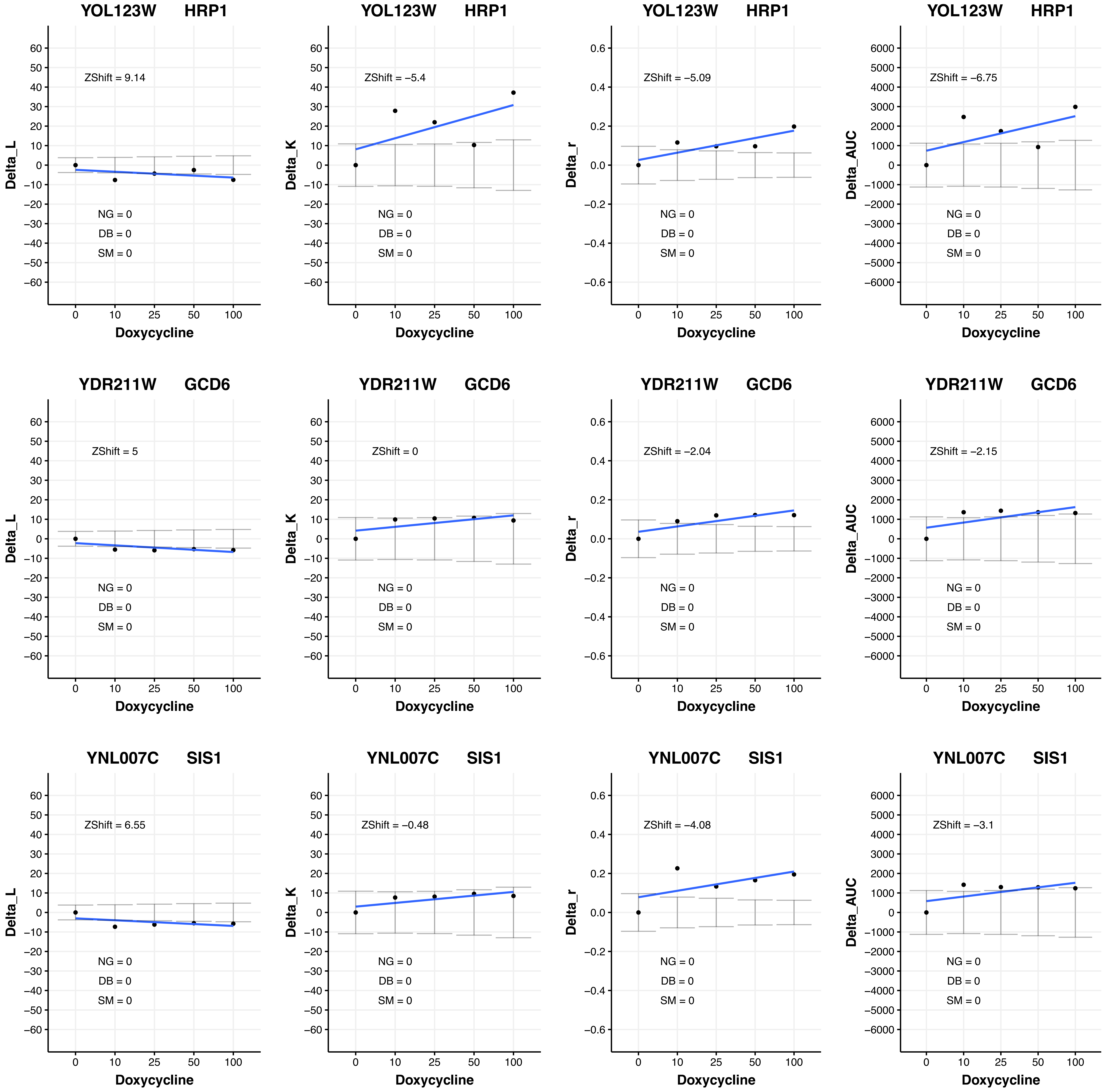
YOL123W HRP1
Delta_L
ZShift = 9.14
NG = 0
DB = 0
SM = 0
Doxycycline
YOL123W HRP1
Delta_K
ZShift = −5.4
NG = 0
DB = 0
SM = 0
Doxycycline
YOL123W HRP1
Delta_r
ZShift = −5.09
NG = 0
DB = 0
SM = 0
Doxycycline
YOL123W HRP1
Delta_AUC
ZShift = −6.75
NG = 0
DB = 0
SM = 0
Doxycycline
YDR211W GCD6
Delta_L
ZShift = 5
NG = 0
DB = 0
SM = 0
Doxycycline
YDR211W GCD6
Delta_K
ZShift = 0
NG = 0
DB = 0
SM = 0
Doxycycline
YDR211W GCD6
Delta_r
ZShift = −2.04
NG = 0
DB = 0
SM = 0
Doxycycline
YDR211W GCD6
Delta_AUC
ZShift = −2.15
NG = 0
DB = 0
SM = 0
Doxycycline
YNL007C SIS1
Delta_L
ZShift = 6.55
NG = 0
DB = 0
SM = 0
Doxycycline
YNL007C SIS1
Delta_K
ZShift = −0.48
NG = 0
DB = 0
SM = 0
Doxycycline
YNL007C SIS1
Delta_r
ZShift = −4.08
NG = 0
DB = 0
SM = 0
Doxycycline
YNL007C SIS1
Delta_AUC
ZShift = −3.1
NG = 0
DB = 0
SM = 0
Doxycycline

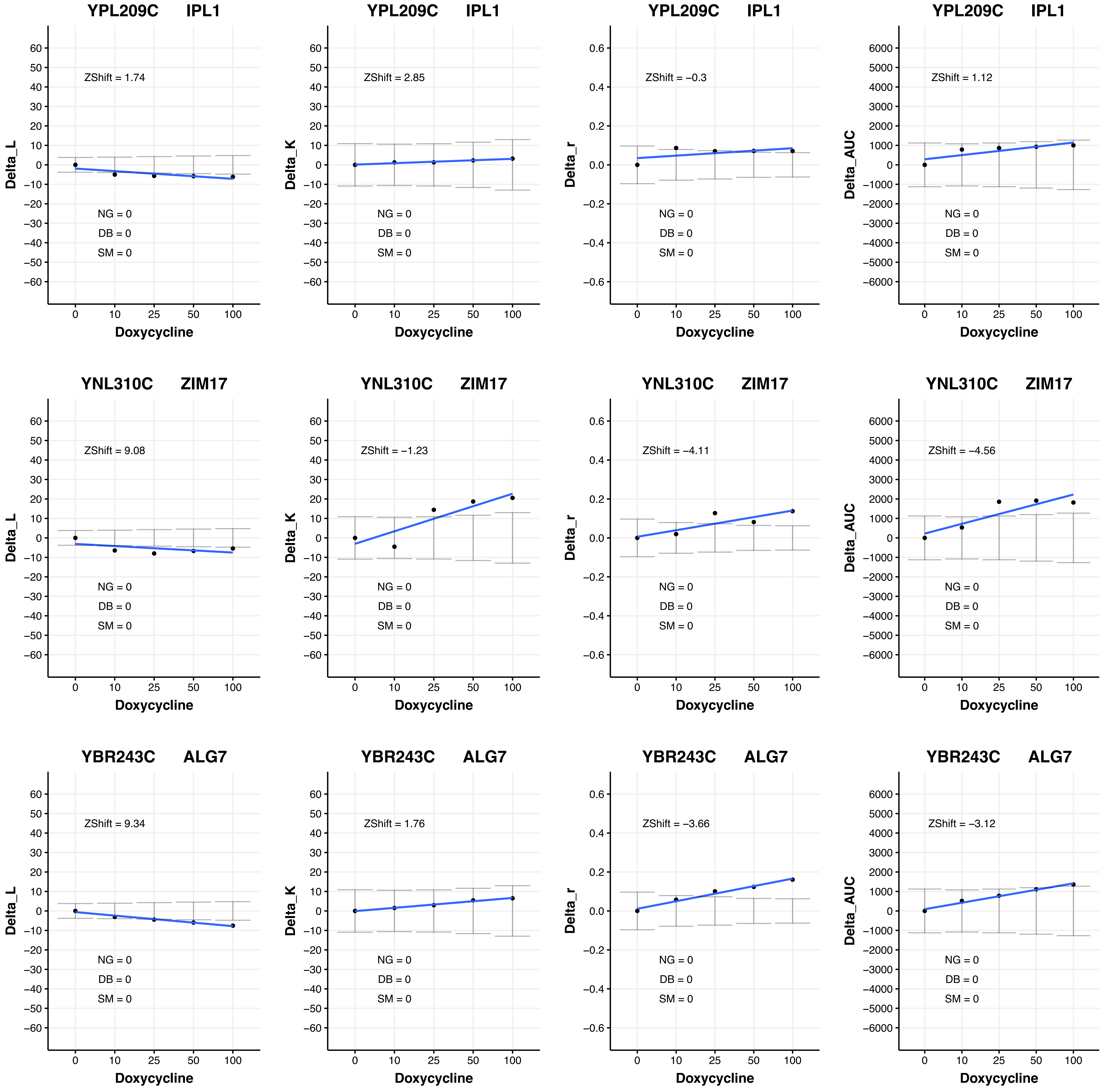
YPL209C IPL1
ZShift = 1.74
NG = 0
DB = 0
SM = 0
Delta_L
Doxycycline
YPL209C IPL1
ZShift = 2.85
NG = 0
DB = 0
SM = 0
Delta_K
Doxycycline
YPL209C IPL1
ZShift = −0.3
NG = 0
DB = 0
SM = 0
Delta_r
Doxycycline
YPL209C IPL1
ZShift = 1.12
NG = 0
DB = 0
SM = 0
Delta_AUC
Doxycycline
YNL310C ZIM17
ZShift = 9.08
NG = 0
DB = 0
SM = 0
Delta_L
Doxycycline
YNL310C ZIM17
ZShift = −1.23
NG = 0
DB = 0
SM = 0
Delta_K
Doxycycline
YNL310C ZIM17
ZShift = −4.11
NG = 0
DB = 0
SM = 0
Delta_r
Doxycycline
YNL310C ZIM17
ZShift = −4.56
NG = 0
DB = 0
SM = 0
Delta_AUC
Doxycycline
YBR243C ALG7
ZShift = 9.34
NG = 0
DB = 0
SM = 0
Delta_L
Doxycycline
YBR243C ALG7
ZShift = 1.76
NG = 0
DB = 0
SM = 0
Delta_K
Doxycycline
YBR243C ALG7
ZShift = −3.66
NG = 0
DB = 0
SM = 0
Delta_r
Doxycycline
YBR243C ALG7
ZShift = −3.12
NG = 0
DB = 0
SM = 0
Delta_AUC
Doxycycline

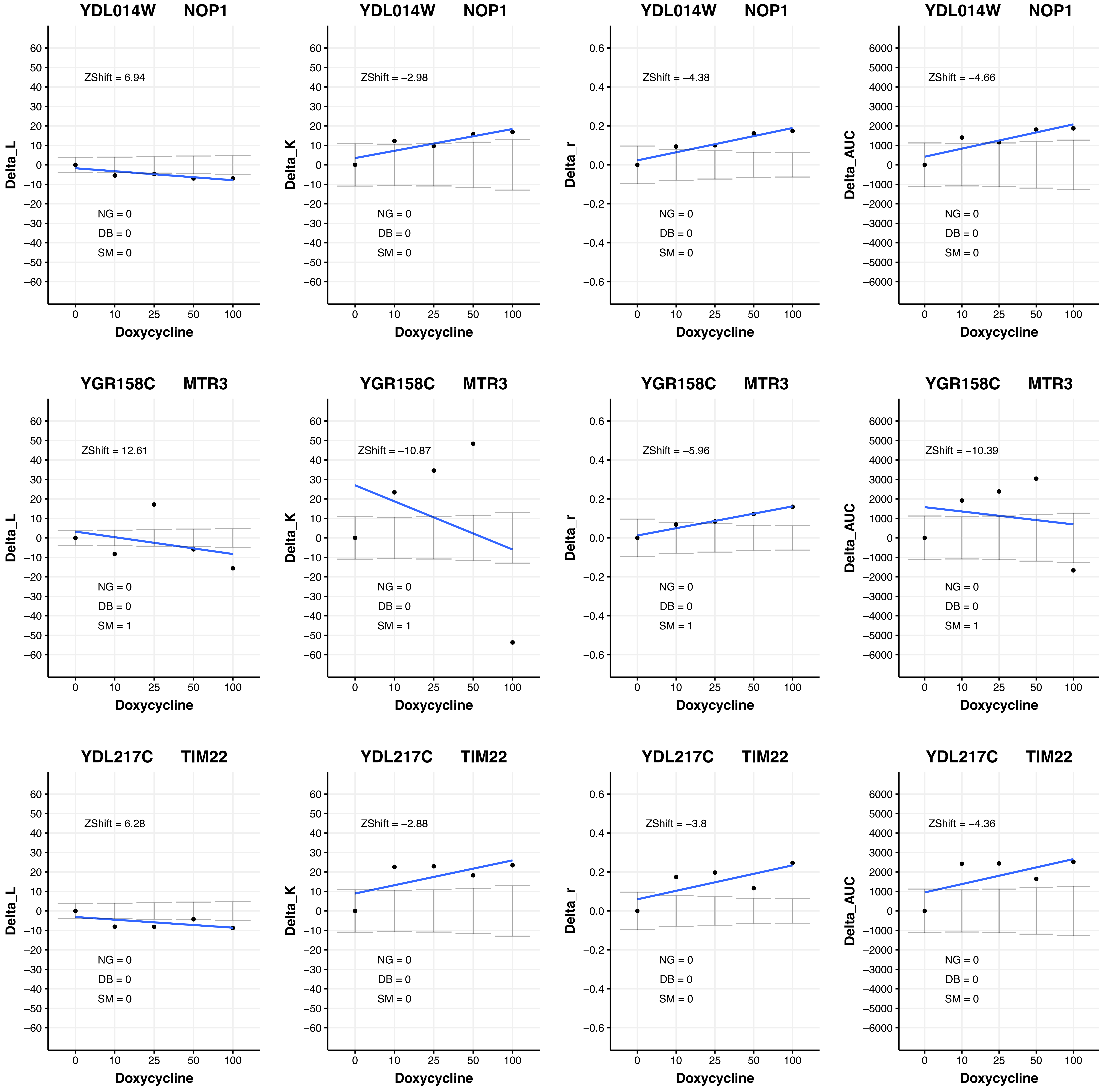

YDL014W NOP1
ZShift = 6.94
NG = 0
DB = 0
SM = 0
Delta_L
Doxycycline
YDL014W NOP1
ZShift = −2.98
NG = 0
DB = 0
SM = 0
Delta_K
Doxycycline
YDL014W NOP1
ZShift = −4.38
NG = 0
DB = 0
SM = 0
Delta_r
Doxycycline
YDL014W NOP1
ZShift = −4.66
NG = 0
DB = 0
SM = 0
Delta_AUC
Doxycycline
YGR158C MTR3
ZShift = 12.61
NG = 0
DB = 0
SM = 1
Delta_L
Doxycycline
YGR158C MTR3
ZShift = −10.87
NG = 0
DB = 0
SM = 1
Delta_K
Doxycycline
YGR158C MTR3
ZShift = −5.96
NG = 0
DB = 0
SM = 1
Delta_r
Doxycycline
YGR158C MTR3
ZShift = −10.39
NG = 0
DB = 0
SM = 1
Delta_AUC
Doxycycline
YDL217C TIM22
ZShift = 6.28
NG = 0
DB = 0
SM = 0
Delta_L
Doxycycline
YDL217C TIM22
ZShift = −2.88
NG = 0
DB = 0
SM = 0
Delta_K
Doxycycline
YDL217C TIM22
ZShift = −3.8
NG = 0
DB = 0
SM = 0
Delta_r
Doxycycline
YDL217C TIM22
ZShift = −4.36
NG = 0
DB = 0
SM = 0
Delta_AUC
Doxycycline

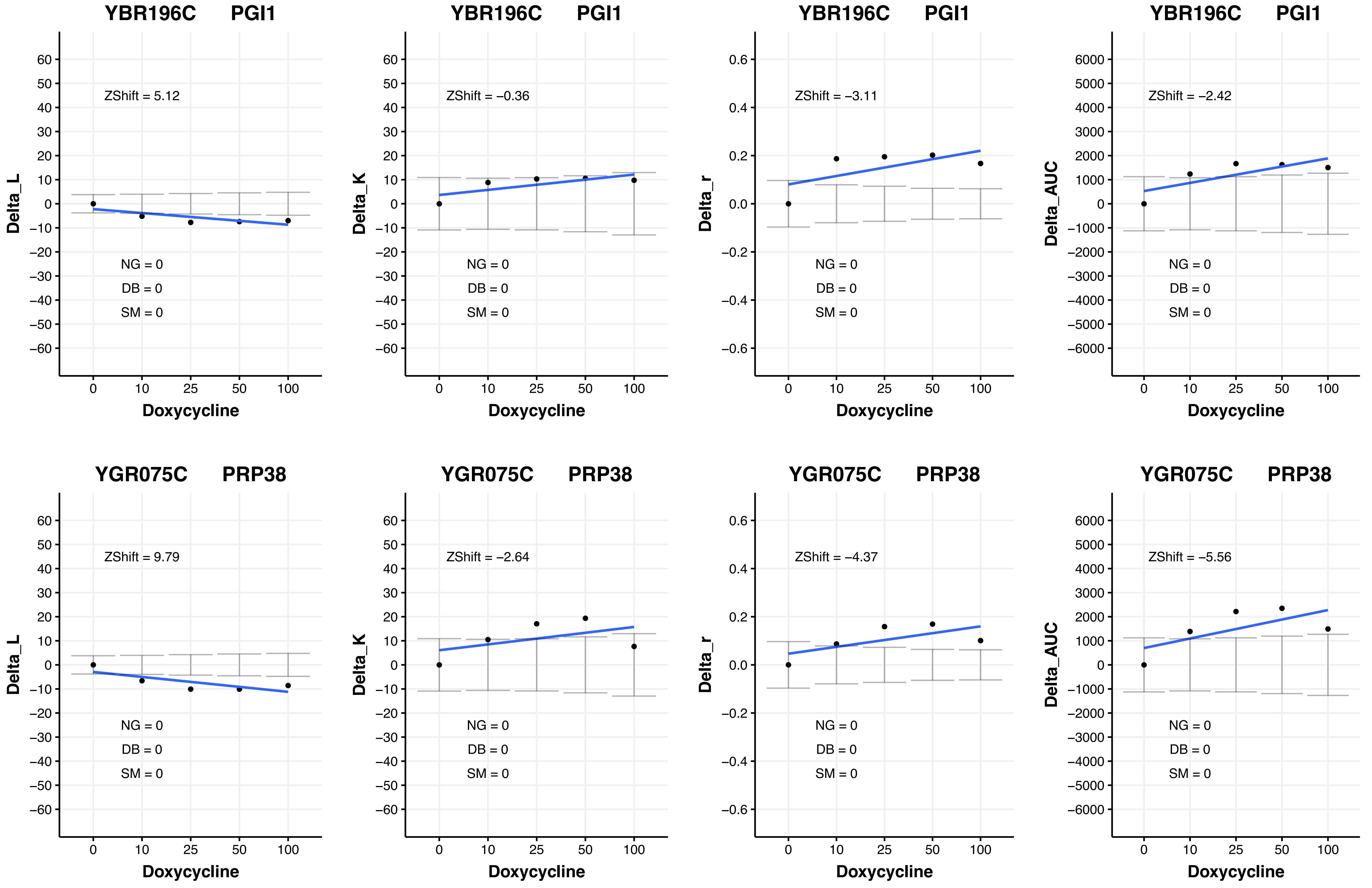
YBR196C PGI1
ZShift = 5.12
NG = 0
DB = 0
SM = 0
Delta_L
Doxycycline
YBR196C PGI1
ZShift = −0.36
NG = 0
DB = 0
SM = 0
Delta_K
Doxycycline
YBR196C PGI1
ZShift = −3.11
NG = 0
DB = 0
SM = 0
Delta_r
Doxycycline
YBR196C PGI1
ZShift = −2.42
NG = 0
DB = 0
SM = 0
Delta_AUC
Doxycycline
YGR075C PRP38
ZShift = 9.79
NG = 0
DB = 0
SM = 0
Delta_L
Doxycycline
YGR075C PRP38
ZShift = −2.64
NG = 0
DB = 0
SM = 0
Delta_K
Doxycycline
YGR075C PRP38
ZShift = −4.37
NG = 0
DB = 0
SM = 0
Delta_r
Doxycycline
YGR075C PRP38
ZShift = −5.56
NG = 0
DB = 0
SM = 0
Delta_AUC
Doxycycline
