## Additional File 5 for "A humanized yeast phenomic model of deoxycytidine kinase to predict genetic buffering of nucleoside analog cytotoxicity": B - Heatmaps.pdf

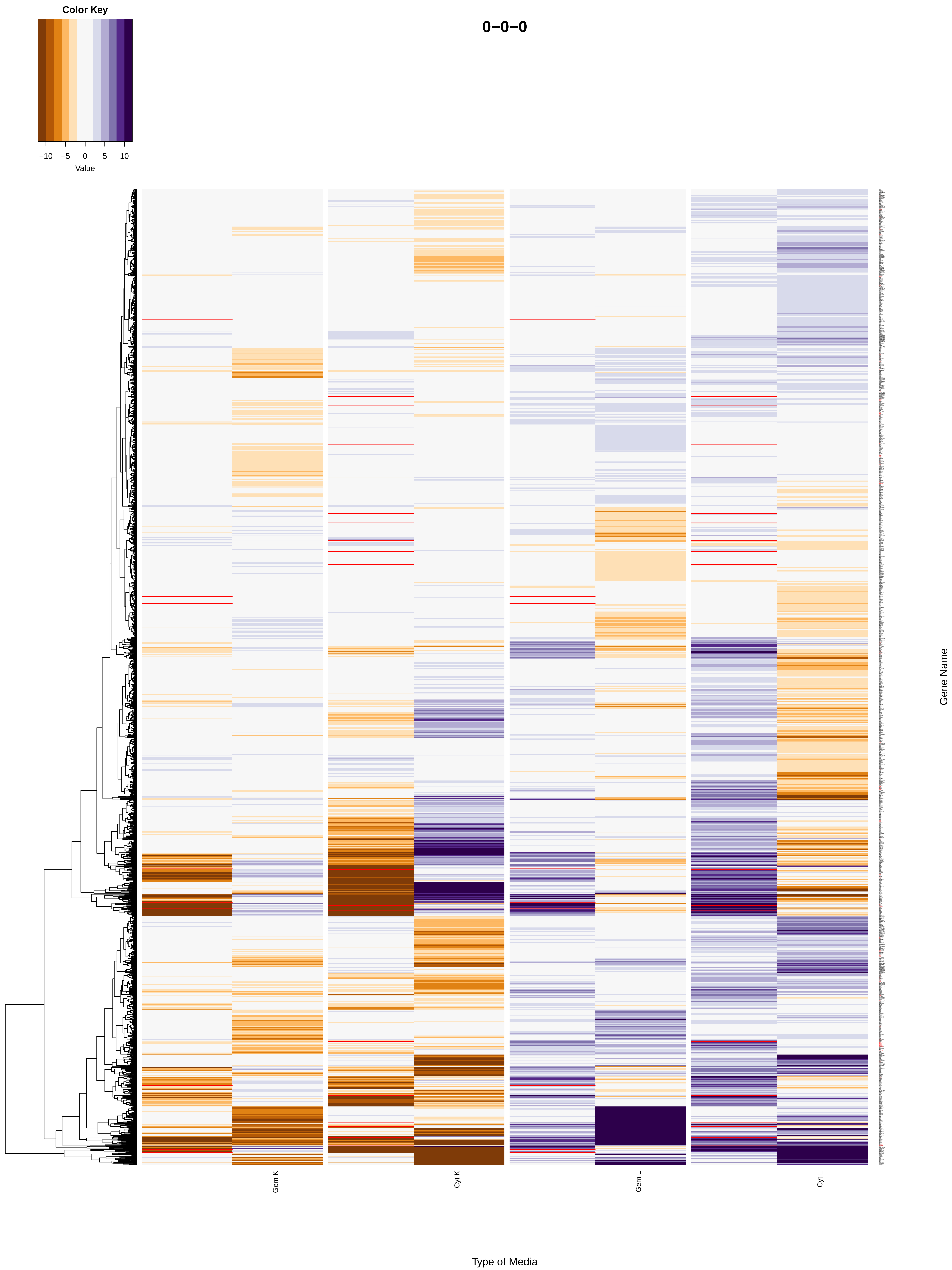
Color Key
−10
−5
0
5
10
Value
0−0−0
Gem K
Cyt K
Gem L
Cyt L
Gene Name
Type of Media

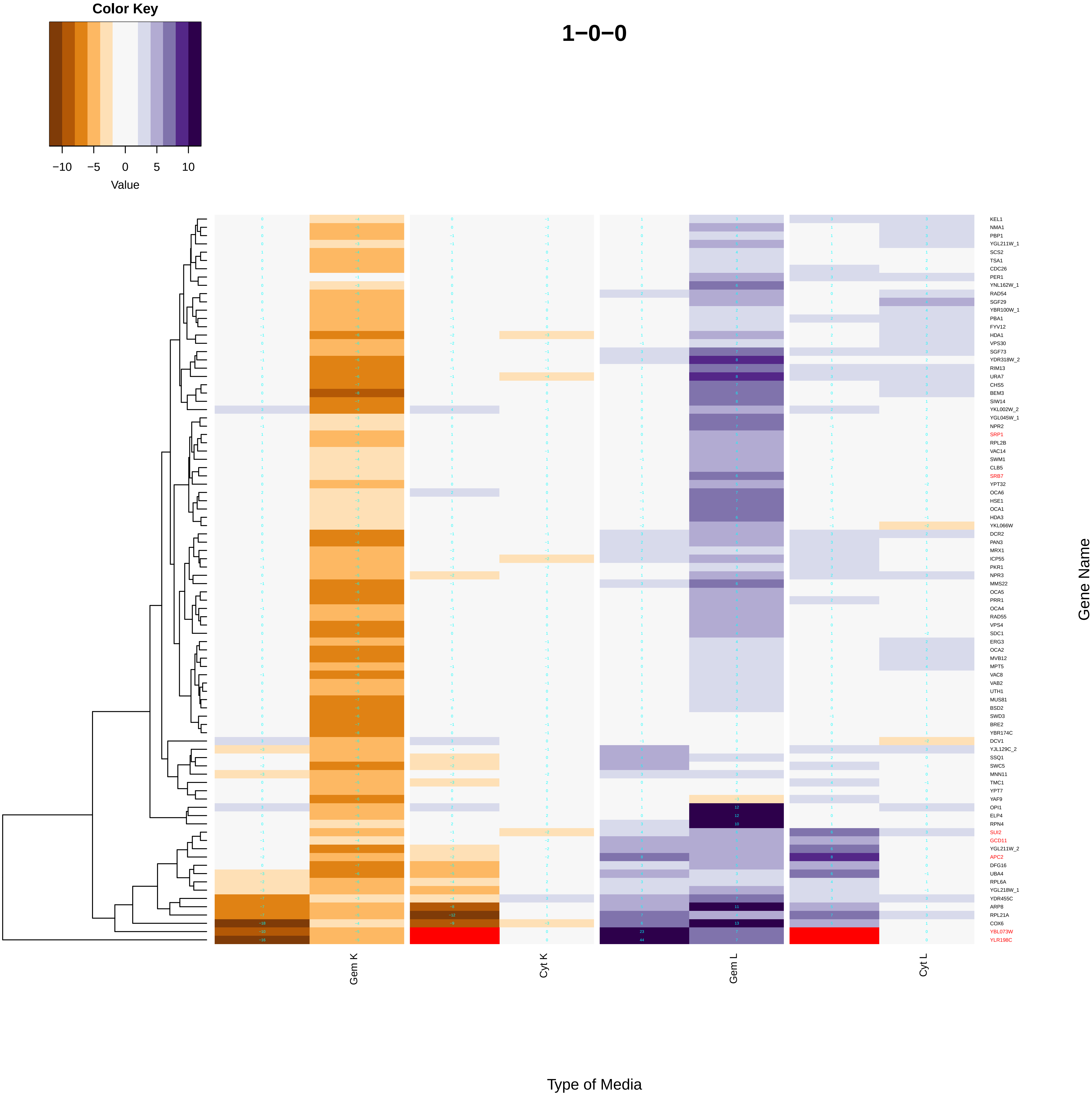
Color Key
−10
−5
0
5
10
Value
1−0−0
KEL1
NMA1
PBP1
YGL211W_1
SCS2
TSA1
CDC26
PER1
YNL162W_1
RAD54
SGF29
YBR100W_1
PBA1
FYV12
HDA1
VPS30
SGF73
YDR318W_2
RIM13
URA7
CHS5
BEM3
SIW14
YKL002W_2
YGL045W_1
NPR2
SRP1
RPL2B
VAC14
SWM1
CLB5
SRB7
YPT32
OCA6
HSE1
OCA1
HDA3
YKL066W
DCR2
PAN3
MRX1
ICP55
PKR1
NPR3
MMS22
OCA5
PRR1
OCA4
RAD55
VPS4
SDC1
ERG3
OCA2
MVB12
MPT5
VAC8
VAB2
UTH1
MUS81
BSD2
SWD3
BRE2
YBR174C
DCV1
YJL129C_2
SSQ1
SWC5
MNN11
TMC1
YPT7
YAF9
OPI1
ELP4
RPN4
SUI2
GCD11
YGL211W_2
APC2
DFG16
UBA4
RPL6A
YGL218W_1
YDR455C
ARP8
RPL21A
COX6
YBL073W
YLR198C
Gene Name
Gem K
Cyt K
Gem L
Cyt L
Type of Media

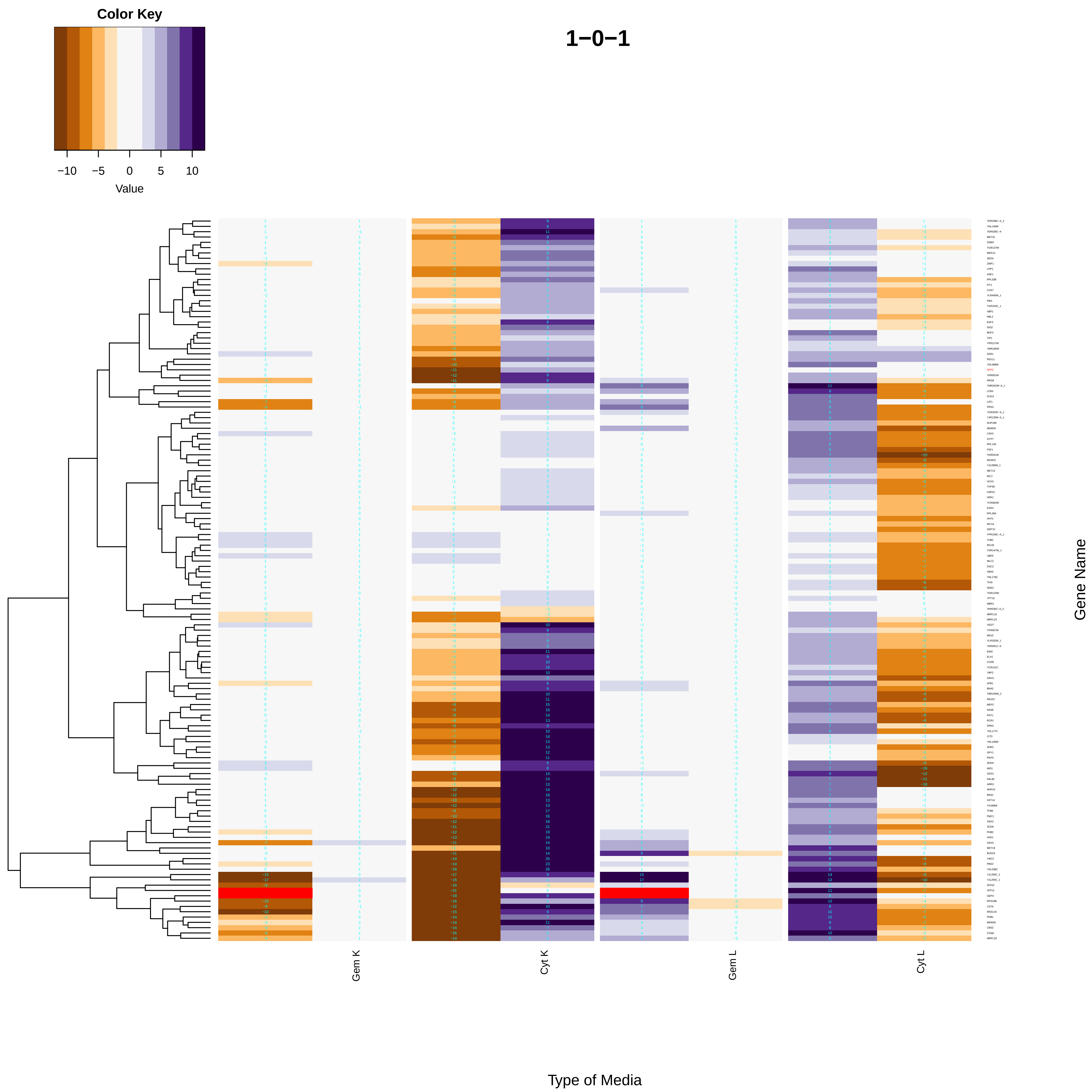
1−0−1
Color Key
Value
−10
−5
0
5
10
Gem K
Cyt K
Gem L
Cyt L
Type of Media
Gene Name

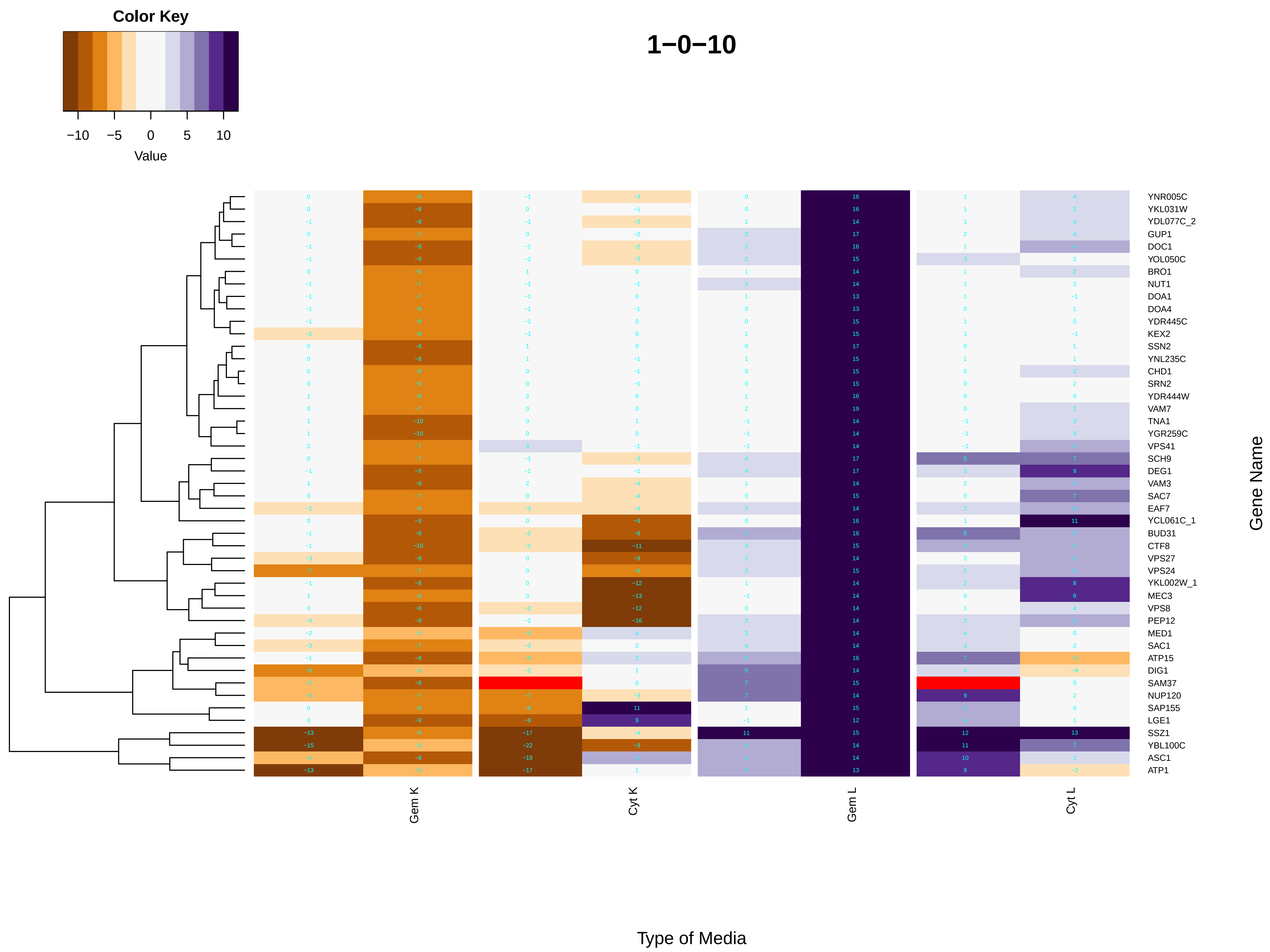

Color Key
−10
−5
0
5
10
Value
1−0−10
Gem K
Cyt K
Gem L
Cyt L
Type of Media
Gene Name
YNR005C
YKL031W
YDL077C_2
GUP1
DOC1
YOL050C
BRO1
NUT1
DOA1
DOA4
YDR445C
KEX2
SSN2
YNL235C
CHD1
SRN2
YDR444W
VAM7
TNA1
YGR259C
VPS41
SCH9
DEG1
VAM3
SAC7
EAF7
YCL061C_1
BUD31
CTF8
VPS27
VPS24
YKL002W_1
MEC3
VPS8
PEP12
MED1
SAC1
ATP15
DIG1
SAM37
NUP120
SAP155
LGE1
SSZ1
YBL100C
ASC1
ATP1

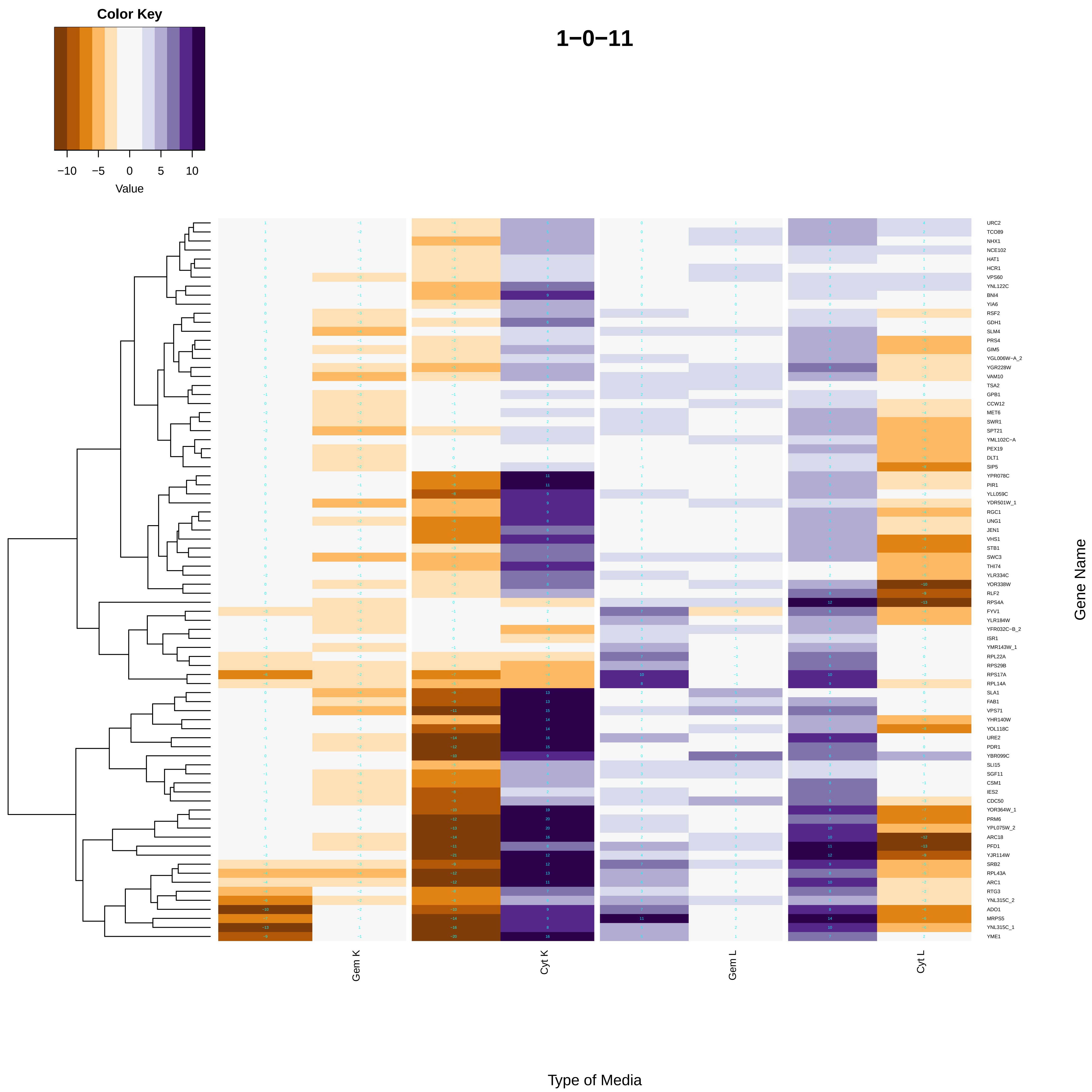
Color Key
−10
−5
0
5
10
Value
1−0−11
URC2
TCO89
NHX1
NCE102
HAT1
HCR1
VPS60
YNL122C
BNI4
YIA6
RSF2
GDH1
SLM4
PRS4
GIM5
YGL006W−A_2
YGR228W
VAM10
TSA2
GPB1
CCW12
MET6
SWR1
SPT21
YML102C−A
PEX19
DLT1
SIP5
YPR078C
PIR1
YLL059C
YDR501W_1
RGC1
UNG1
JEN1
VHS1
STB1
SWC3
THI74
YLR334C
YOR338W
RLF2
RPS4A
FYV1
YLR184W
YFR032C−B_2
ISR1
YMR143W_1
RPL22A
RPS29B
RPS17A
RPL14A
SLA1
FAB1
VPS71
YHR140W
YOL118C
URE2
PDR1
YBR099C
SLI15
SGF11
CSM1
IES2
CDC50
YOR364W_1
PRM6
YPL075W_2
ARC18
PFD1
YJR114W
SRB2
RPL43A
ARC1
RTG3
YNL315C_2
ADO1
MRPS5
YNL315C_1
YME1
Gene Name
Gem K
Cyt K
Gem L
Cyt L
Type of Media

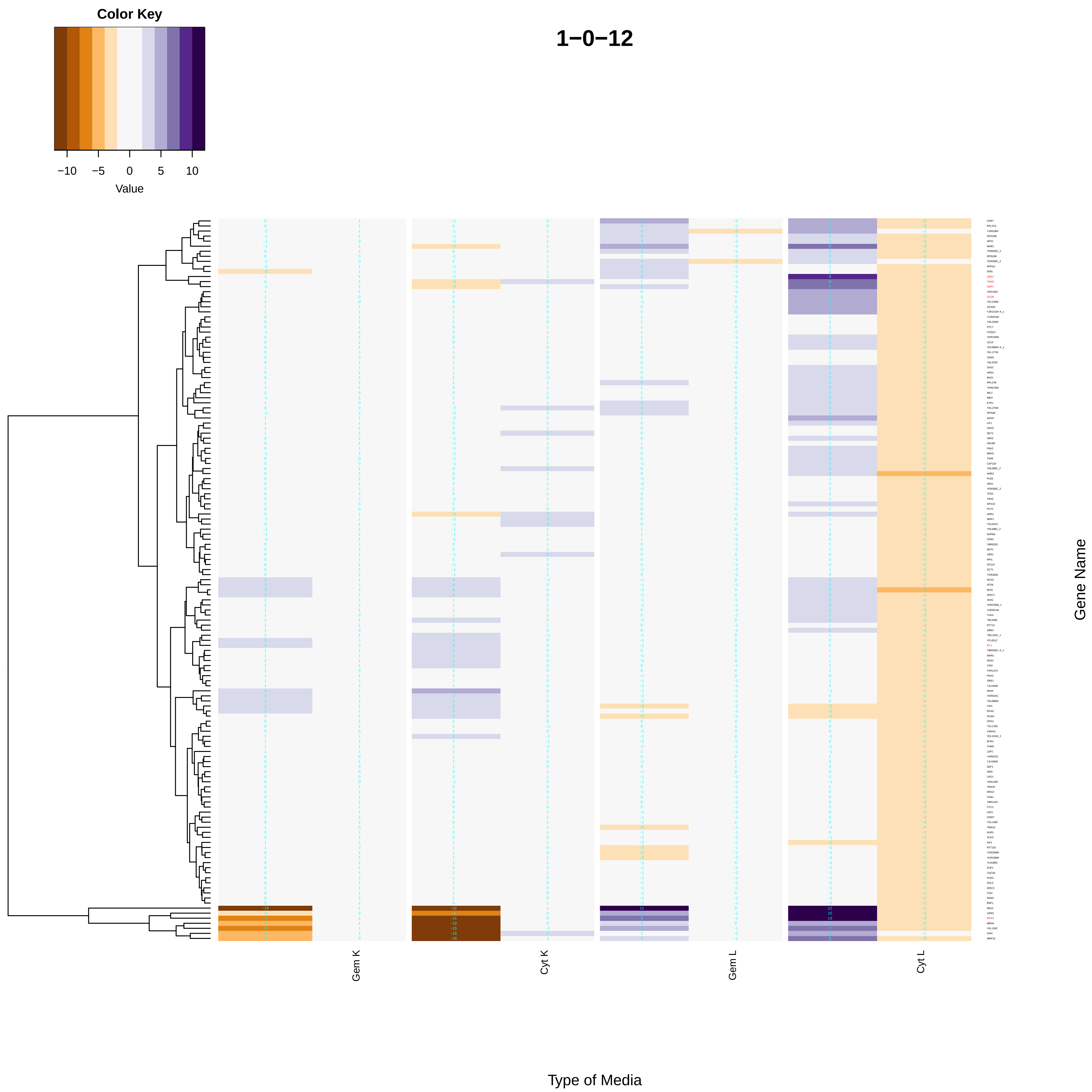
Color Key
−10
−5
0
5
10
Value
1−0−12
Gem K
Cyt K
Gem L
Cyt L
Gene Name
Type of Media

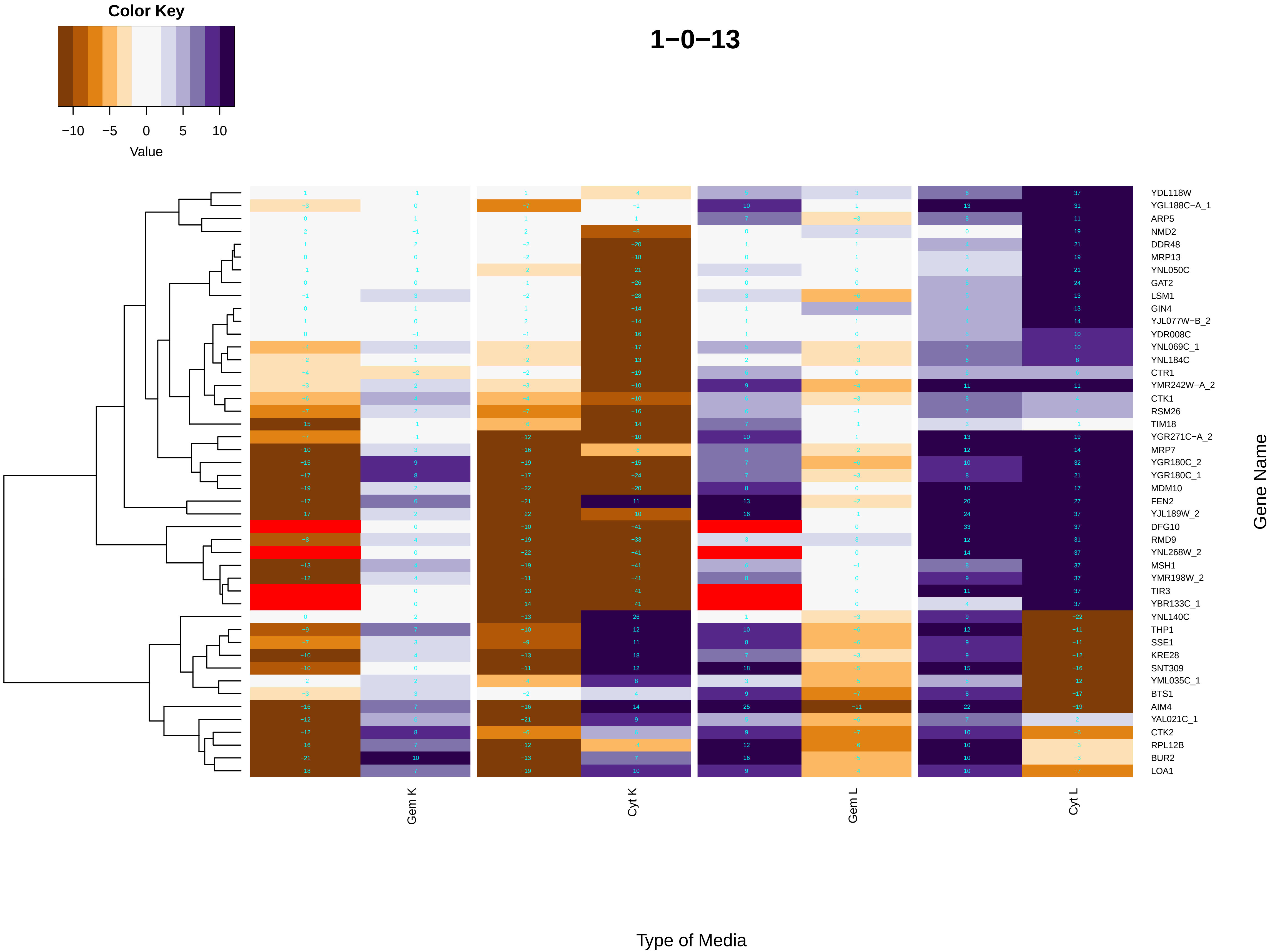

Color Key
−10 −5 0 5 10
Value
1−0−13
YDL118W
YGL188C−A_1
ARP5
NMD2
DDR48
MRP13
YNL050C
GAT2
LSM1
GIN4
YJL077W−B_2
YDR008C
YNL069C_1
YNL184C
CTR1
YMR242W−A_2
CTK1
RSM26
TIM18
YGR271C−A_2
MRP7
YGR180C_2
YGR180C_1
MDM10
FEN2
YJL189W_2
DFG10
RMD9
YNL268W_2
MSH1
YMR198W_2
TIR3
YBR133C_1
YNL140C
THP1
SSE1
KRE28
SNT309
YML035C_1
BTS1
AIM4
YAL021C_1
CTK2
RPL12B
BUR2
LOA1
Gene Name
Gem K
Cyt K
Gem L
Cyt L
Type of Media

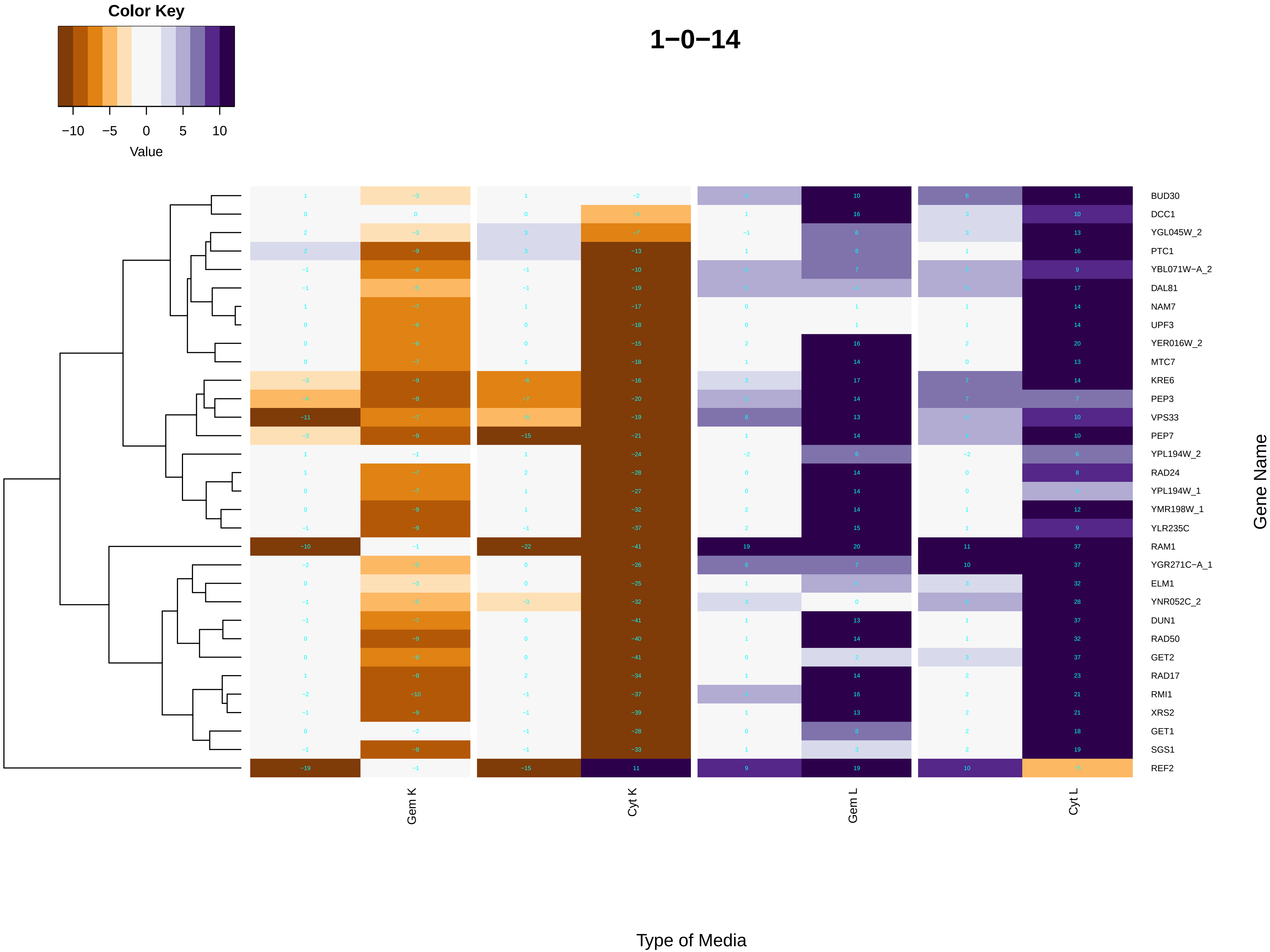
Color Key
−10 −5 0 5 10
Value
1−0−14
BUD30
DCC1
YGL045W_2
PTC1
YBL071W−A_2
DAL81
NAM7
UPF3
YER016W_2
MTC7
KRE6
PEP3
VPS33
PEP7
YPL194W_2
RAD24
YPL194W_1
YMR198W_1
YLR235C
RAM1
YGR271C−A_1
ELM1
YNR052C_2
DUN1
RAD50
GET2
RAD17
RMI1
XRS2
GET1
SGS1
REF2
Gene Name
Gem K
Cyt K
Gem L
Cyt L
Type of Media

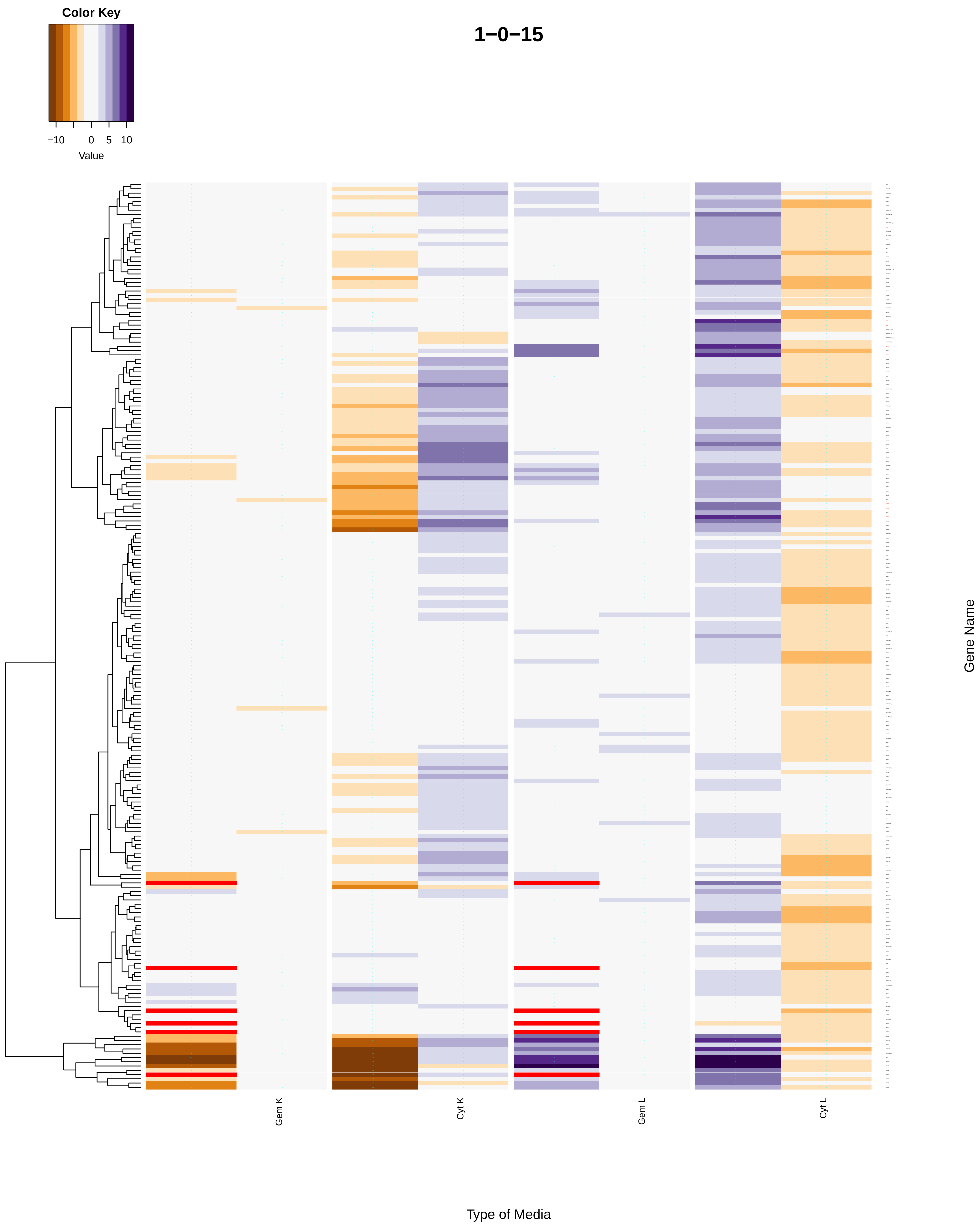

Color Key
−10
0
5
10
Value
1−0−15
Gene Name
Gem K
Cyt K
Gem L
Cyt L
Type of Media

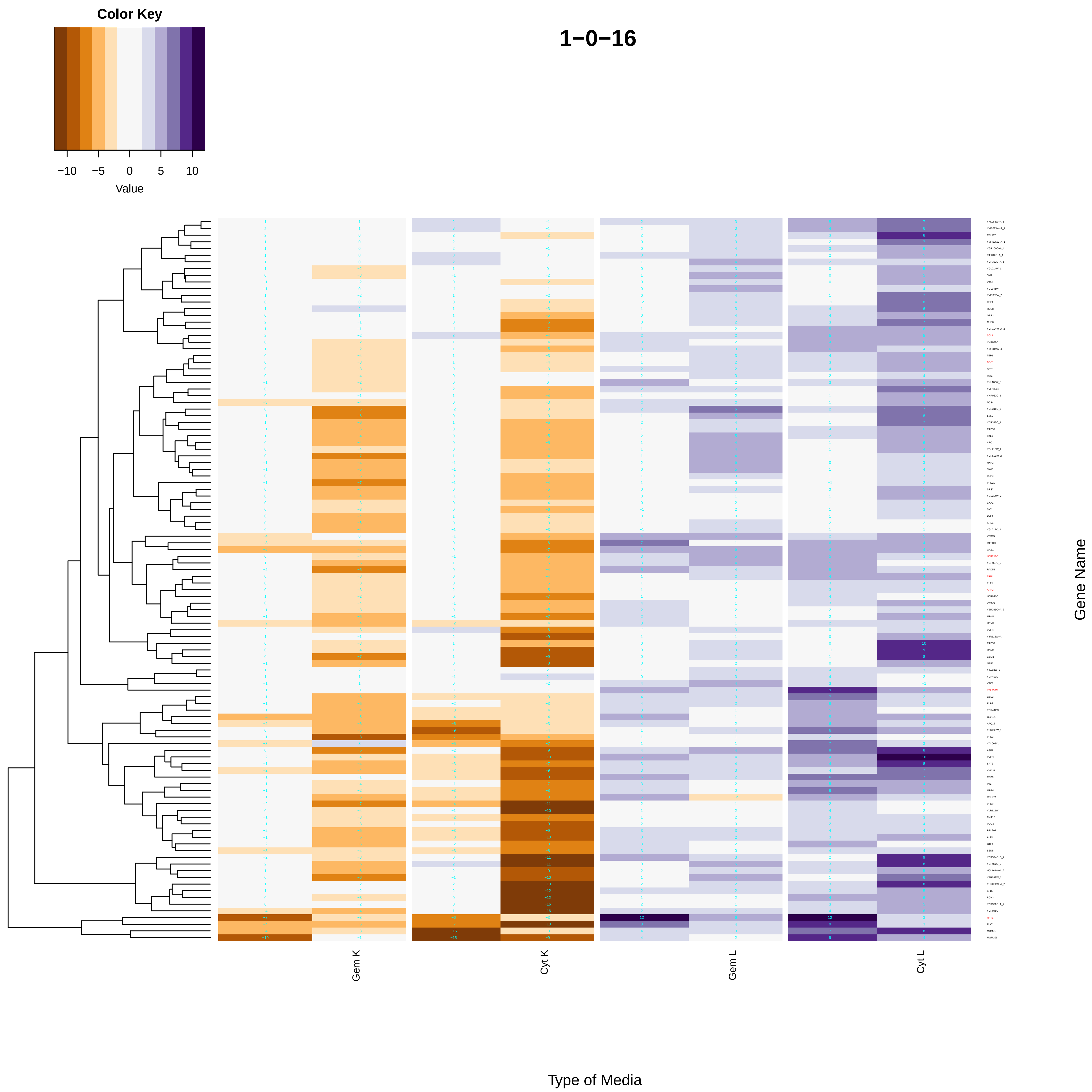

Color Key
−10
−5
0
5
10
Value
1−0−16
Gem K
Cyt K
Gem L
Cyt L
Gene Name
Type of Media

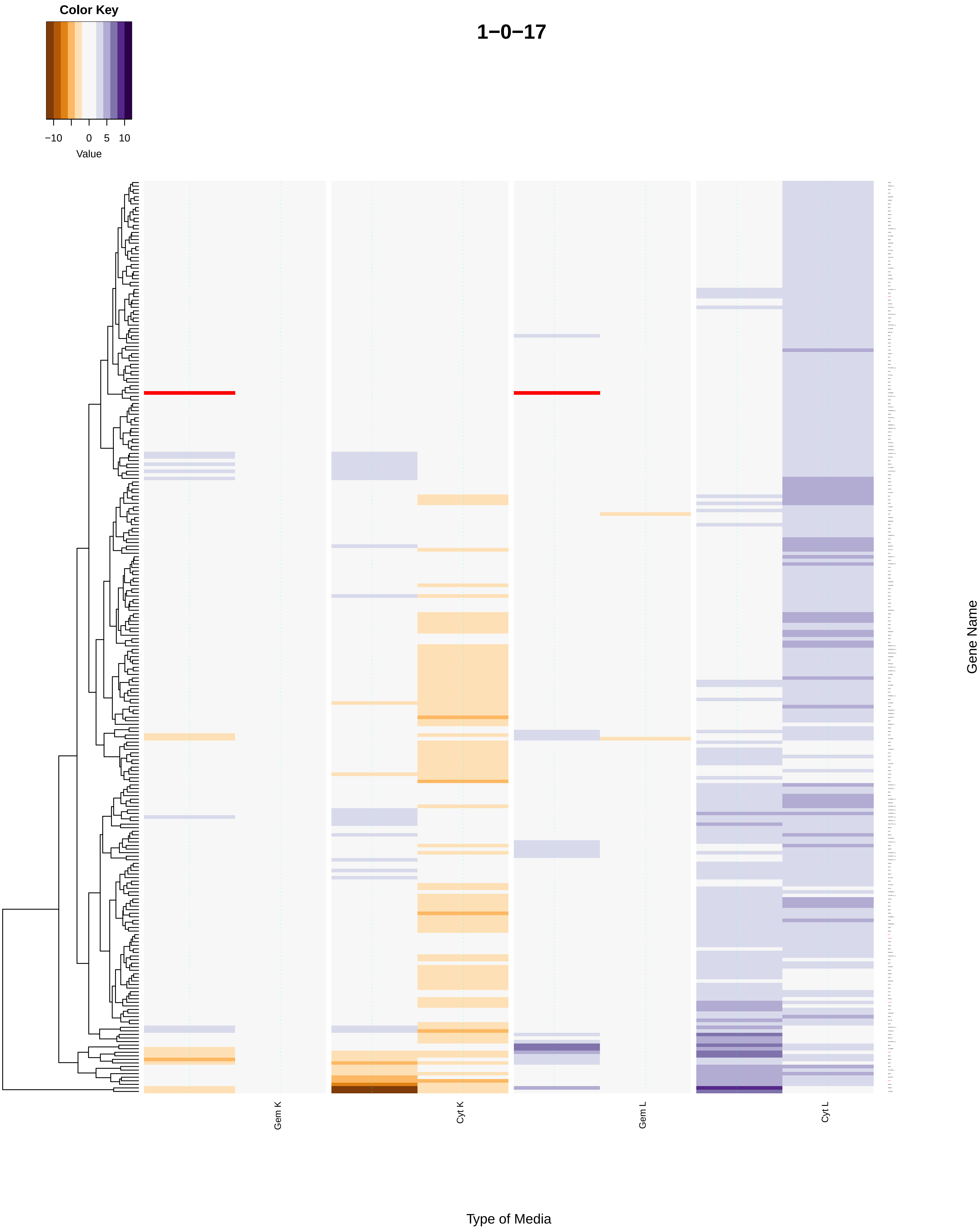
Color Key
−10
0
5
10
Value
1−0−17
Gem K
Cyt K
Gem L
Cyt L
Gene Name
Type of Media

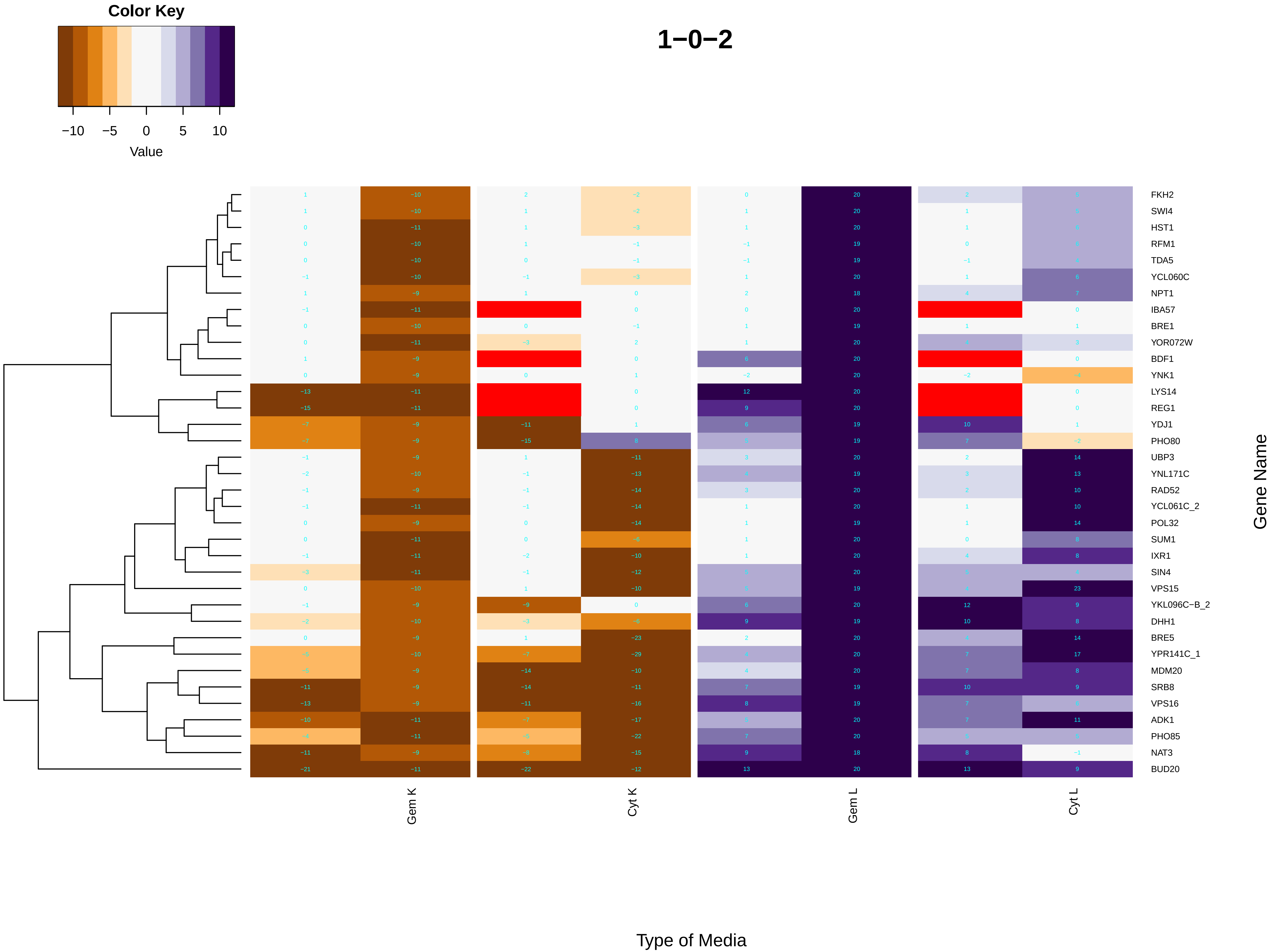
Color Key
−10 −5 0 5 10
Value
1−0−2
FKH2
SWI4
HST1
RFM1
TDA5
YCL060C
NPT1
IBA57
BRE1
YOR072W
BDF1
YNK1
LYS14
REG1
YDJ1
PHO80
UBP3
YNL171C
RAD52
YCL061C_2
POL32
SUM1
IXR1
SIN4
VPS15
YKL096C−B_2
DHH1
BRE5
YPR141C_1
MDM20
SRB8
VPS16
ADK1
PHO85
NAT3
BUD20
Gene Name
Gem K
Cyt K
Gem L
Cyt L
Type of Media

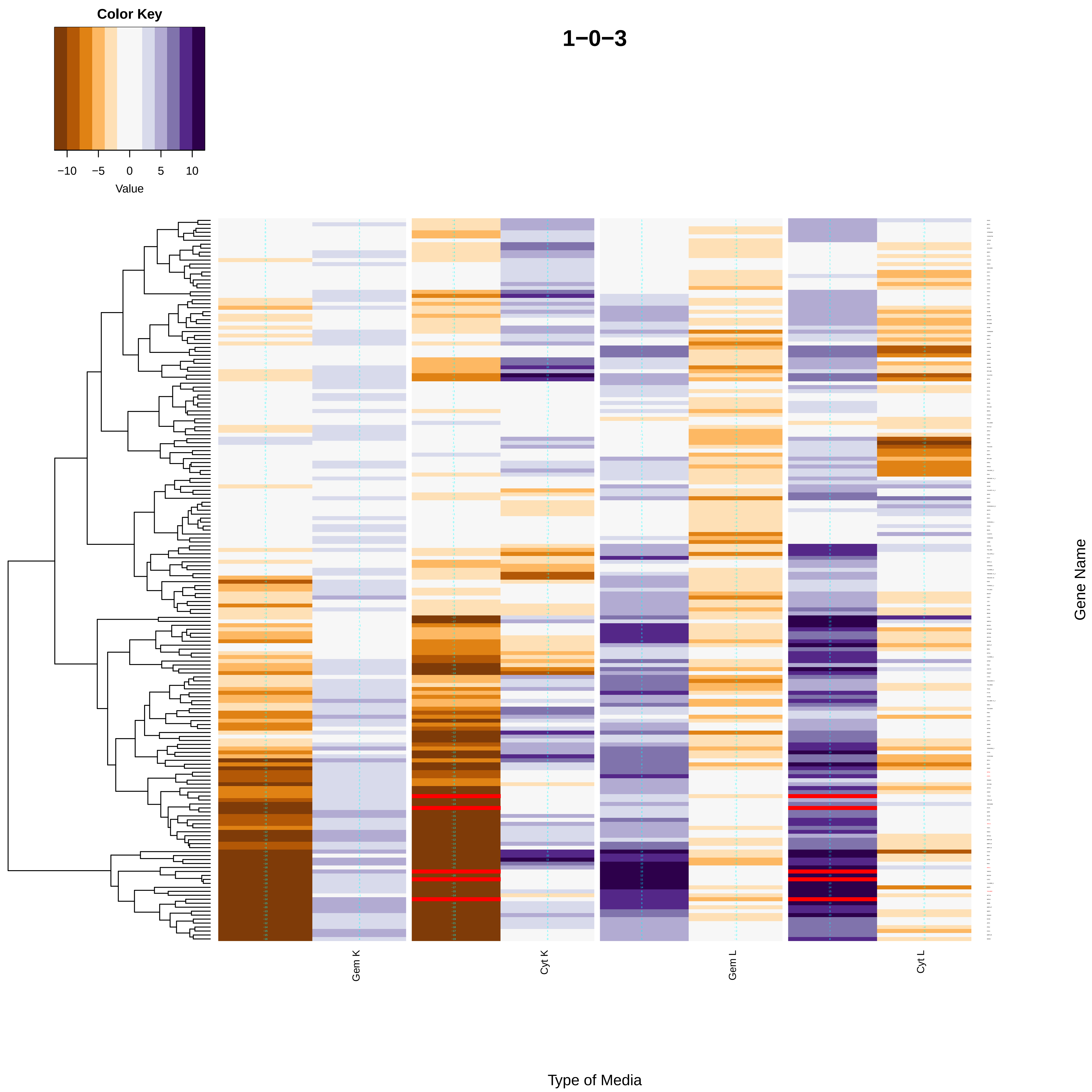
Color Key
−10
−5
0
5
10
Value
1−0−3
Gem K
Cyt K
Gem L
Cyt L
Gene Name
Type of Media

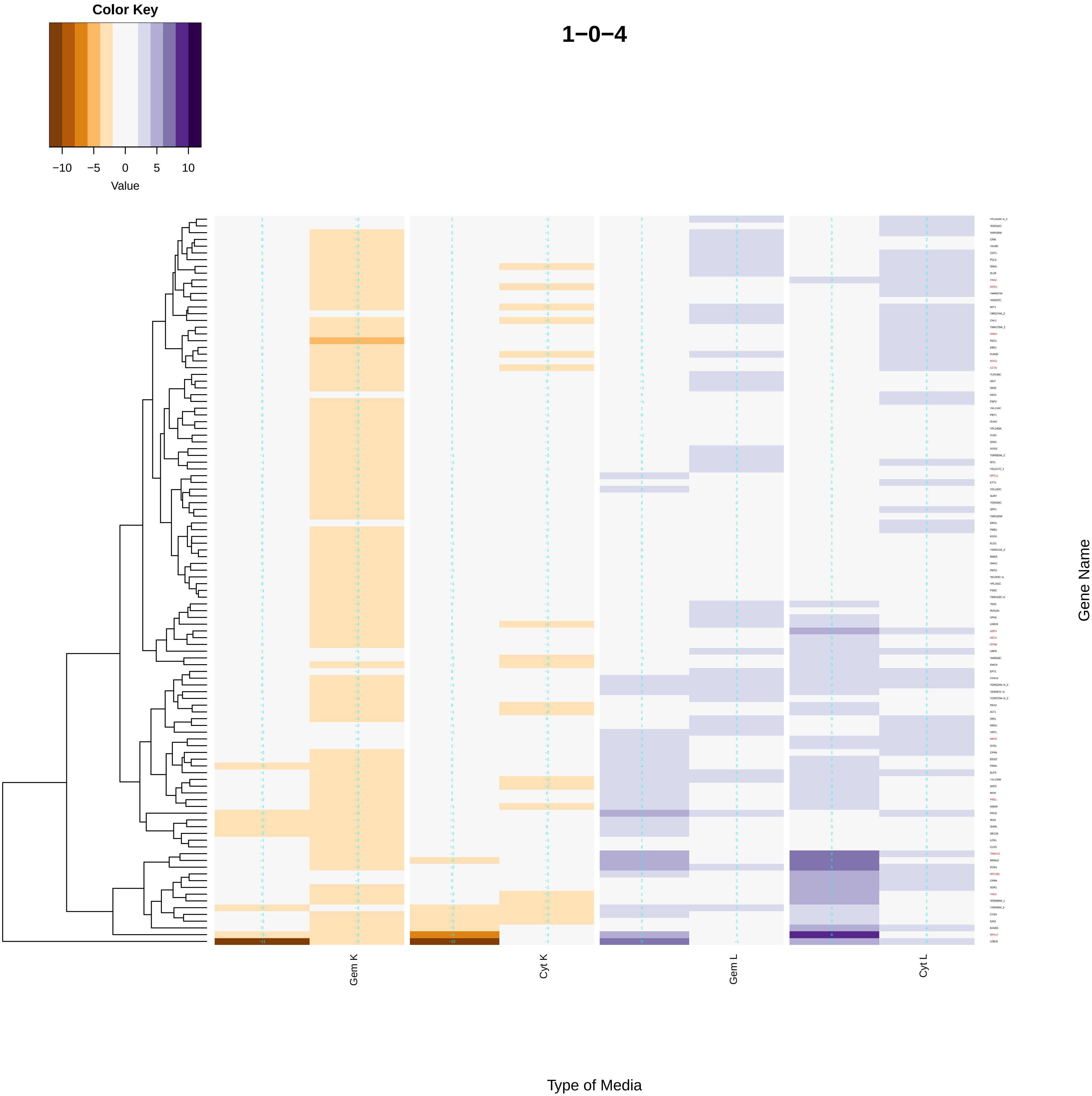
Color Key
−10
−5
0
5
10
Value
1−0−4
Gem K
Cyt K
Gem L
Cyt L
Type of Media
Gene Name

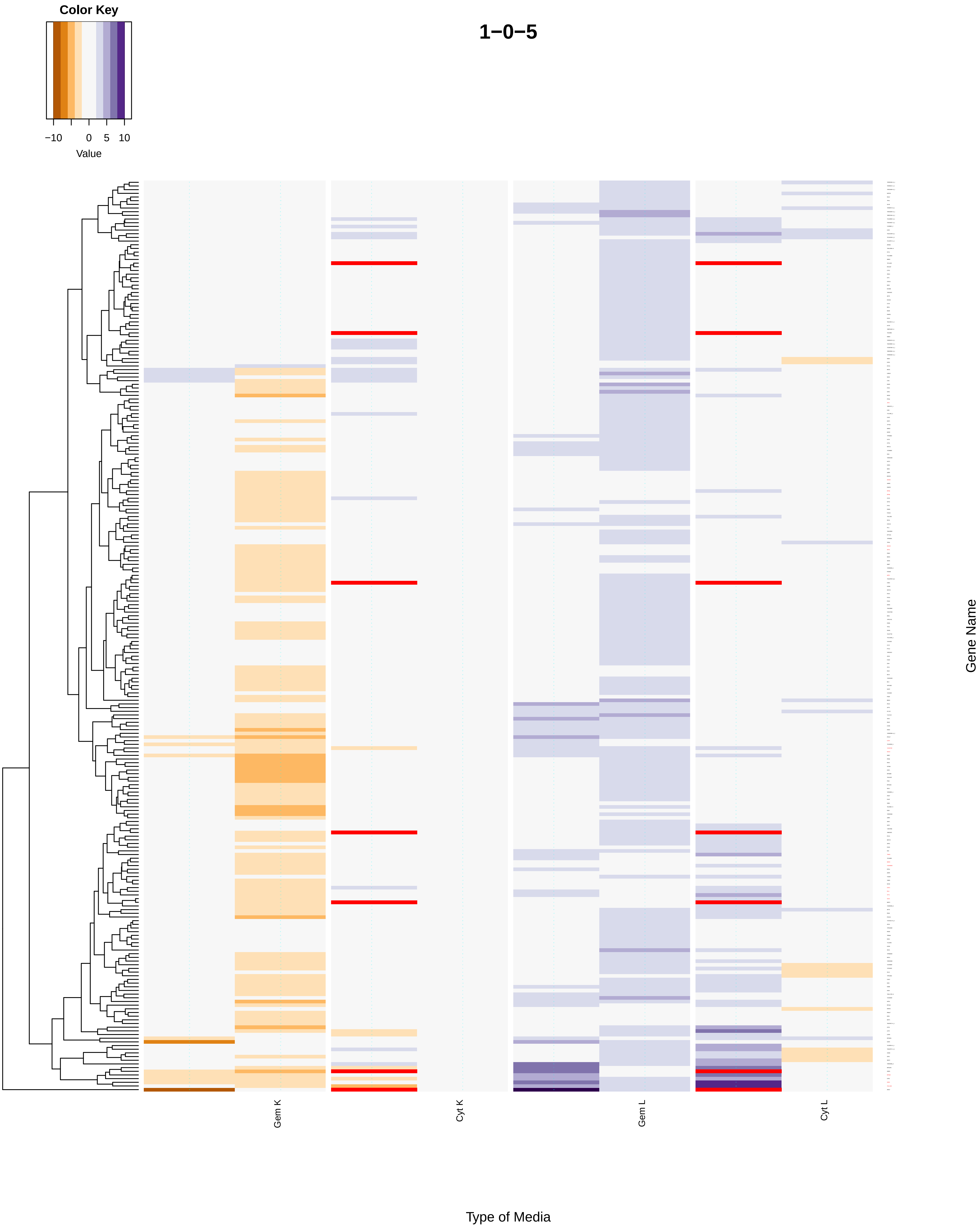
Color Key
−10
0
5
10
Value
1−0−5
Gem K
Cyt K
Gem L
Cyt L
Type of Media
Gene Name

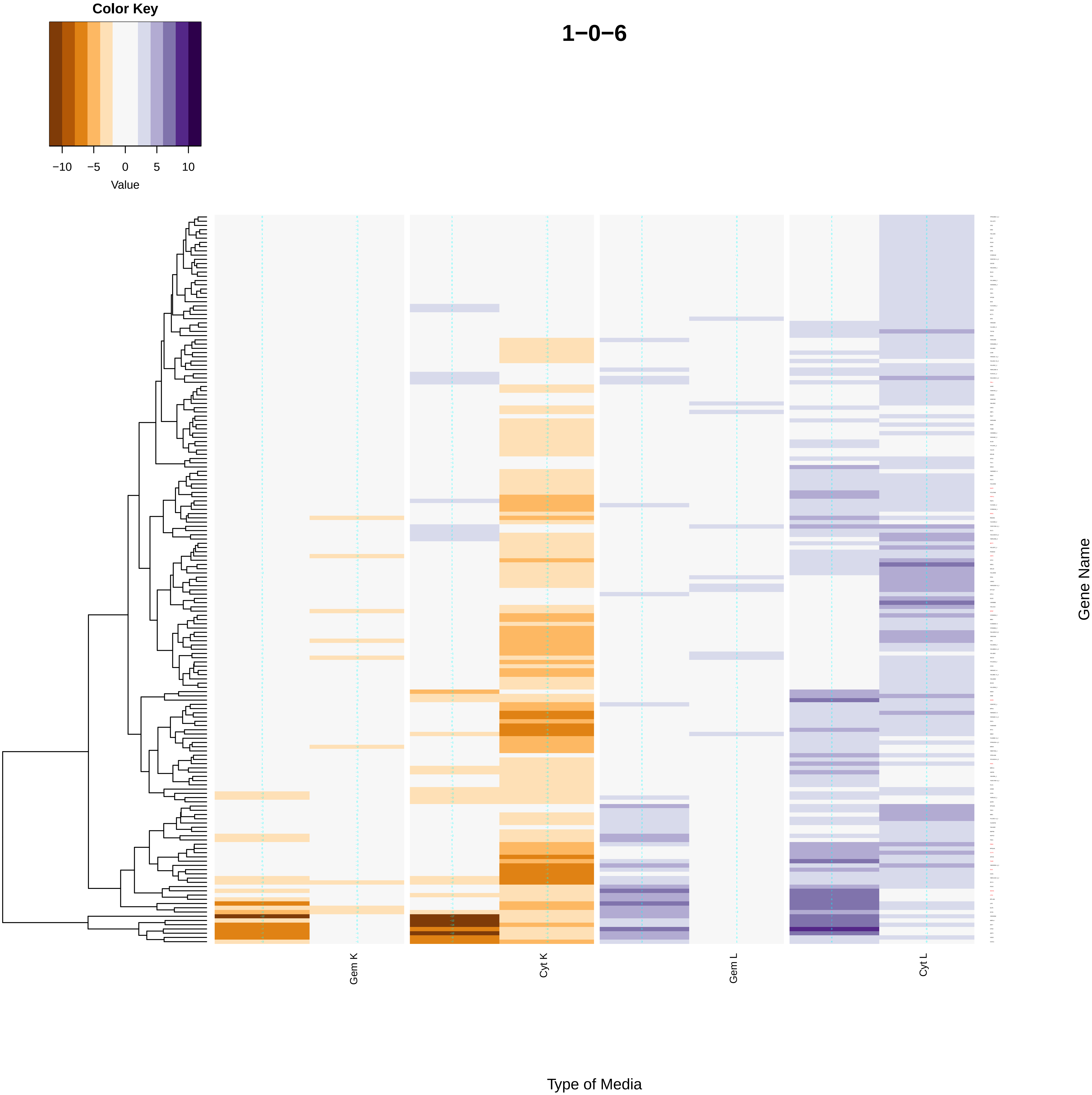
Color Key
−10
−5
0
5
10
Value
1−0−6
Gem K
Cyt K
Gem L
Cyt L
Gene Name
Type of Media

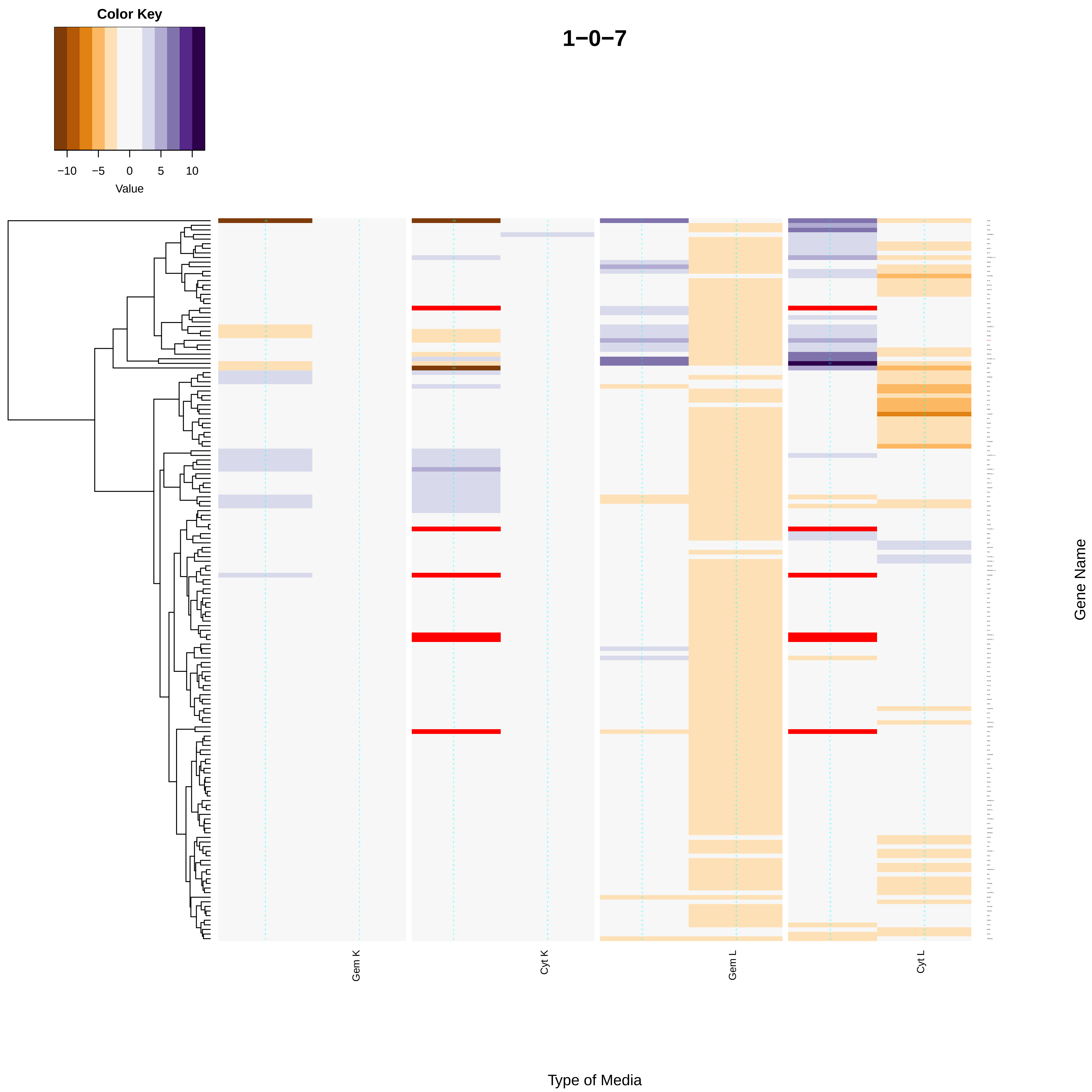

Color Key
−10
−5
0
5
10
Value
1−0−7
Gem K
Cyt K
Gem L
Cyt L
Gene Name
Type of Media

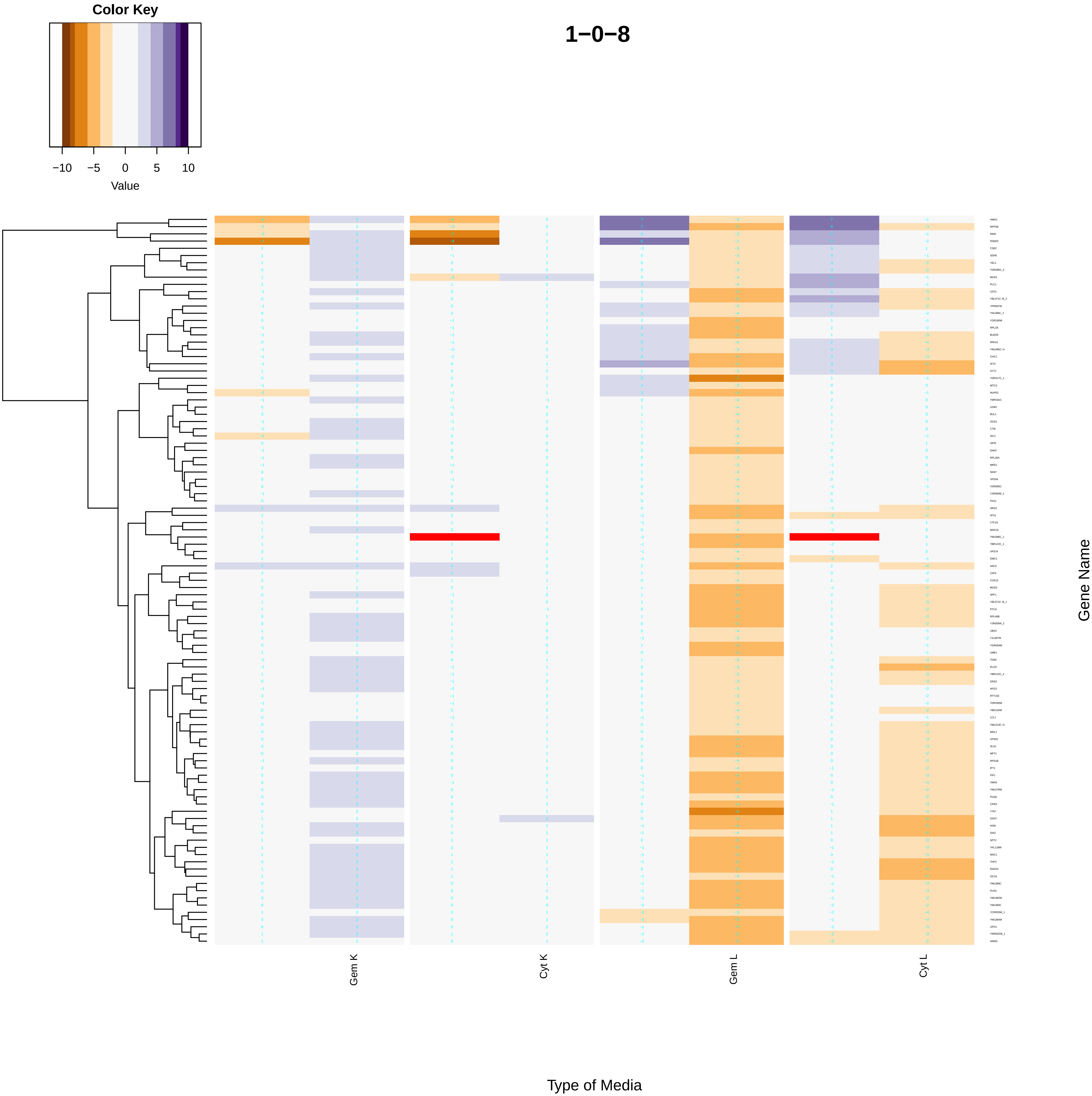
Color Key
−10 −5 0 5 10
Value
1−0−8
Gem K
Cyt K
Gem L
Cyt L
Gene Name
Type of Media

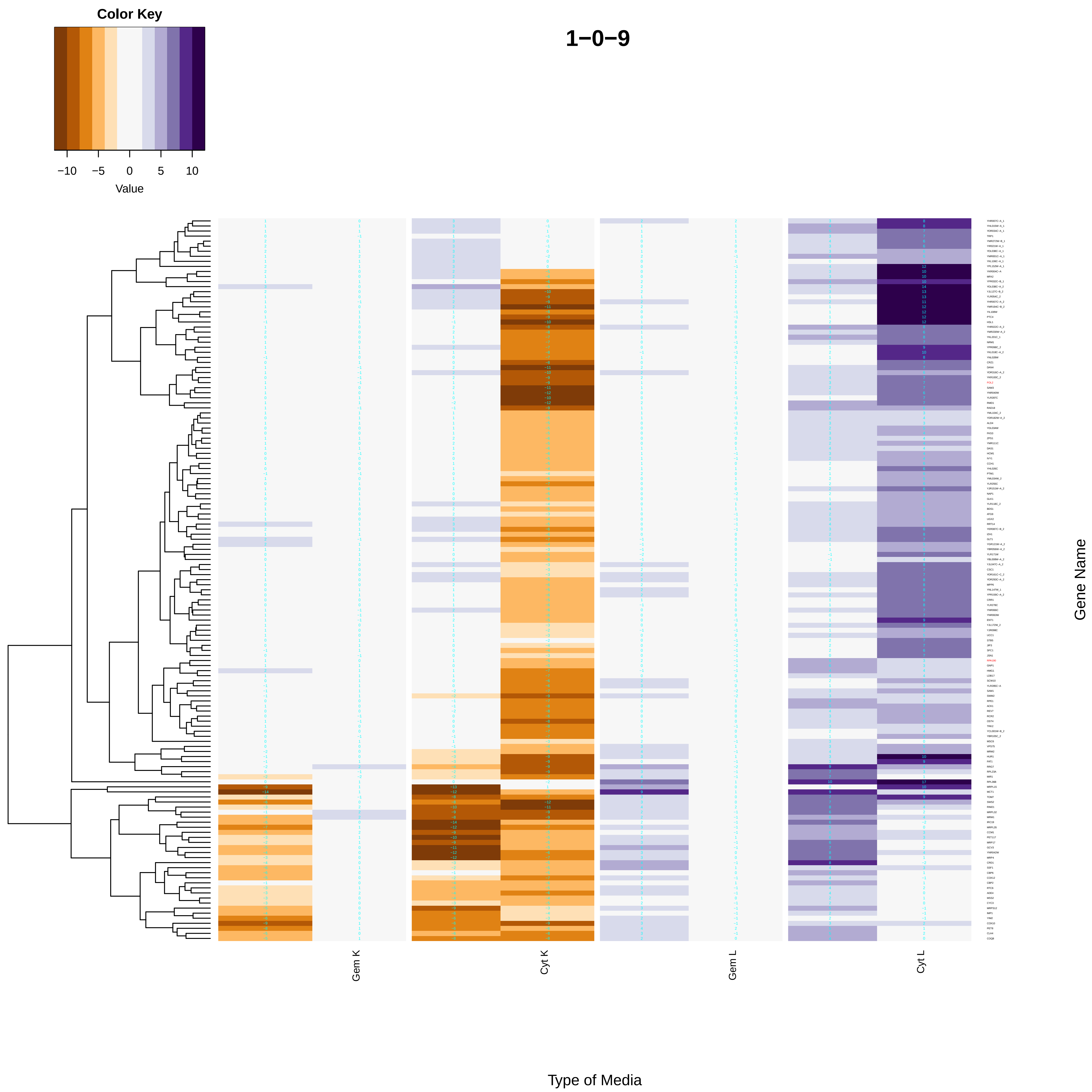

Color Key
−10
−5
0
5
10
Value
1−0−9
Gem K
Cyt K
Gem L
Cyt L
Type of Media
Gene Name

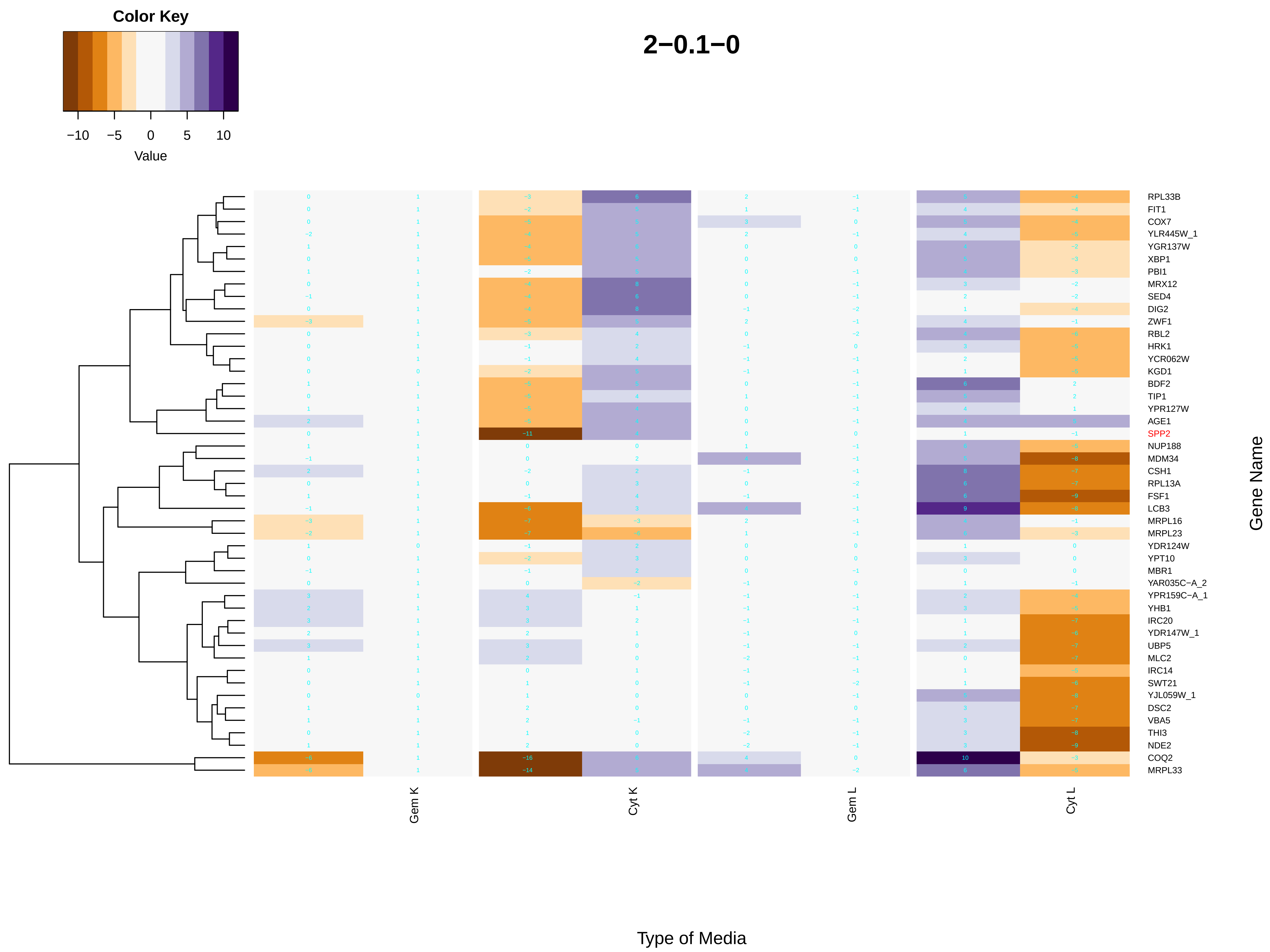
Color Key
−10
−5
0
5
10
Value
2−0.1−0
RPL33B
FIT1
COX7
YLR445W_1
YGR137W
XBP1
PBI1
MRX12
SED4
DIG2
ZWF1
RBL2
HRK1
YCR062W
KGD1
BDF2
TIP1
YPR127W
AGE1
SPP2
NUP188
MDM34
CSH1
RPL13A
FSF1
LCB3
MRPL16
MRPL23
YDR124W
YPT10
MBR1
YAR035C−A_2
YPR159C−A_1
YHB1
IRC20
YDR147W_1
UBP5
MLC2
IRC14
SWT21
YJL059W_1
DSC2
VBA5
THI3
NDE2
COQ2
MRPL33
Gene Name
Gem K
Cyt K
Gem L
Cyt L
Type of Media

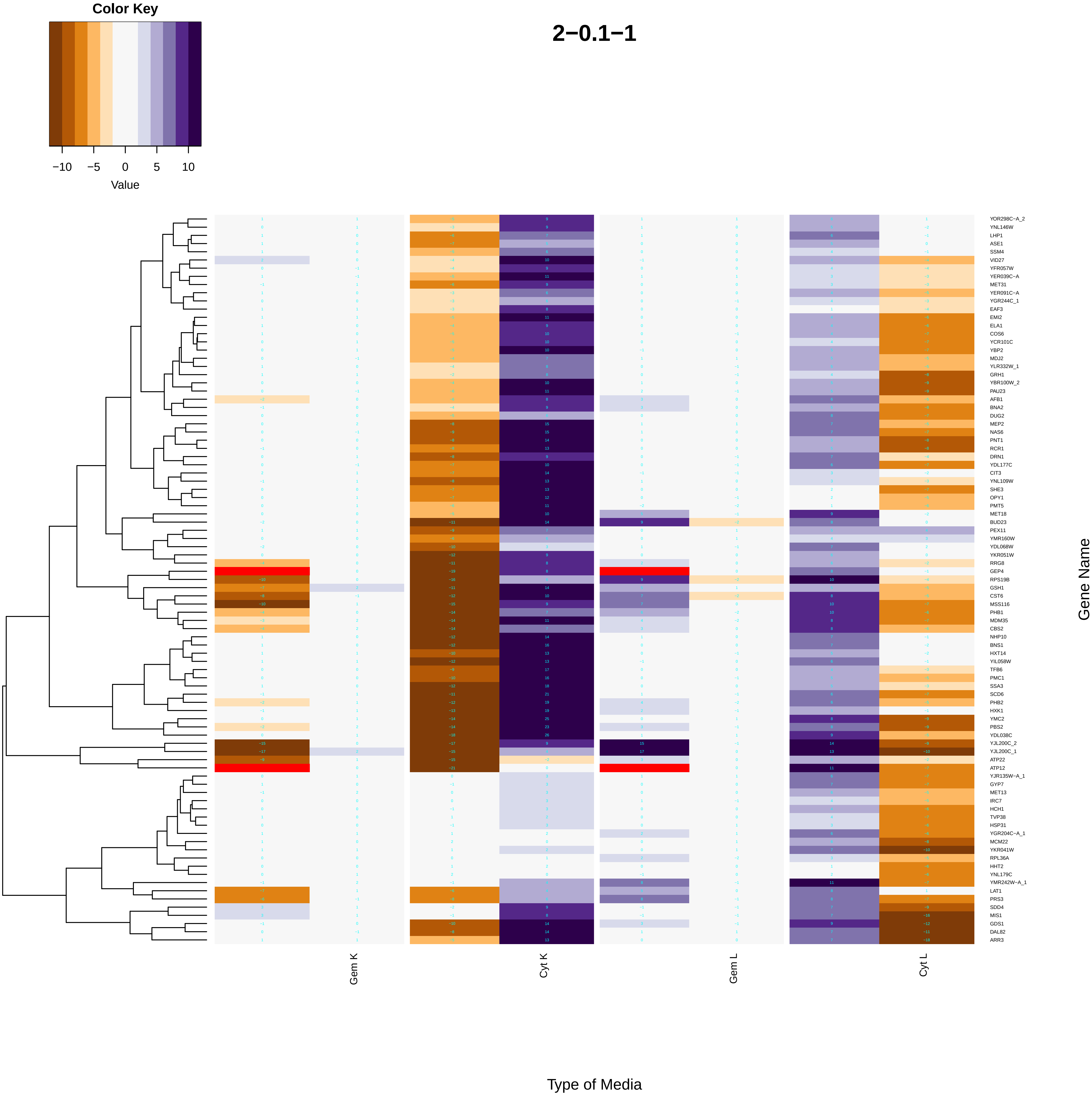

Color Key
−10
−5
0
5
10
Value
2−0.1−1
YOR298C−A_2
YNL146W
LHP1
ASE1
SSM4
VID27
YFR057W
YER039C−A
MET31
YER091C−A
YGR244C_1
EAF3
EMI2
ELA1
COS6
YCR101C
YBP2
MDJ2
YLR332W_1
GRH1
YBR100W_2
PAU23
AFB1
BNA2
DUG2
MEP2
NAS6
PNT1
RCR1
DRN1
YDL177C
CIT3
YNL109W
SHE3
OPY1
PMT5
MET18
BUD23
PEX11
YMR160W
YDL068W
YKR051W
RRG8
GEP4
RPS19B
GSH1
CST6
MSS116
PHB1
MDM35
CBS2
NHP10
BNS1
HXT14
YIL058W
TFB6
PMC1
SSA3
SCD6
PHB2
HXK1
YMC2
PBS2
YDL038C
YJL200C_2
YJL200C_1
ATP22
ATP12
YJR135W−A_1
GYP7
MET13
IRC7
HCH1
TVP38
HSP31
YGR204C−A_1
MCM22
YKR041W
RPL36A
HHT2
YNL179C
YMR242W−A_1
LAT1
PRS3
SDD4
MIS1
GDS1
DAL82
ARR3
Gene Name
Gem K
Cyt K
Gem L
Cyt L
Type of Media

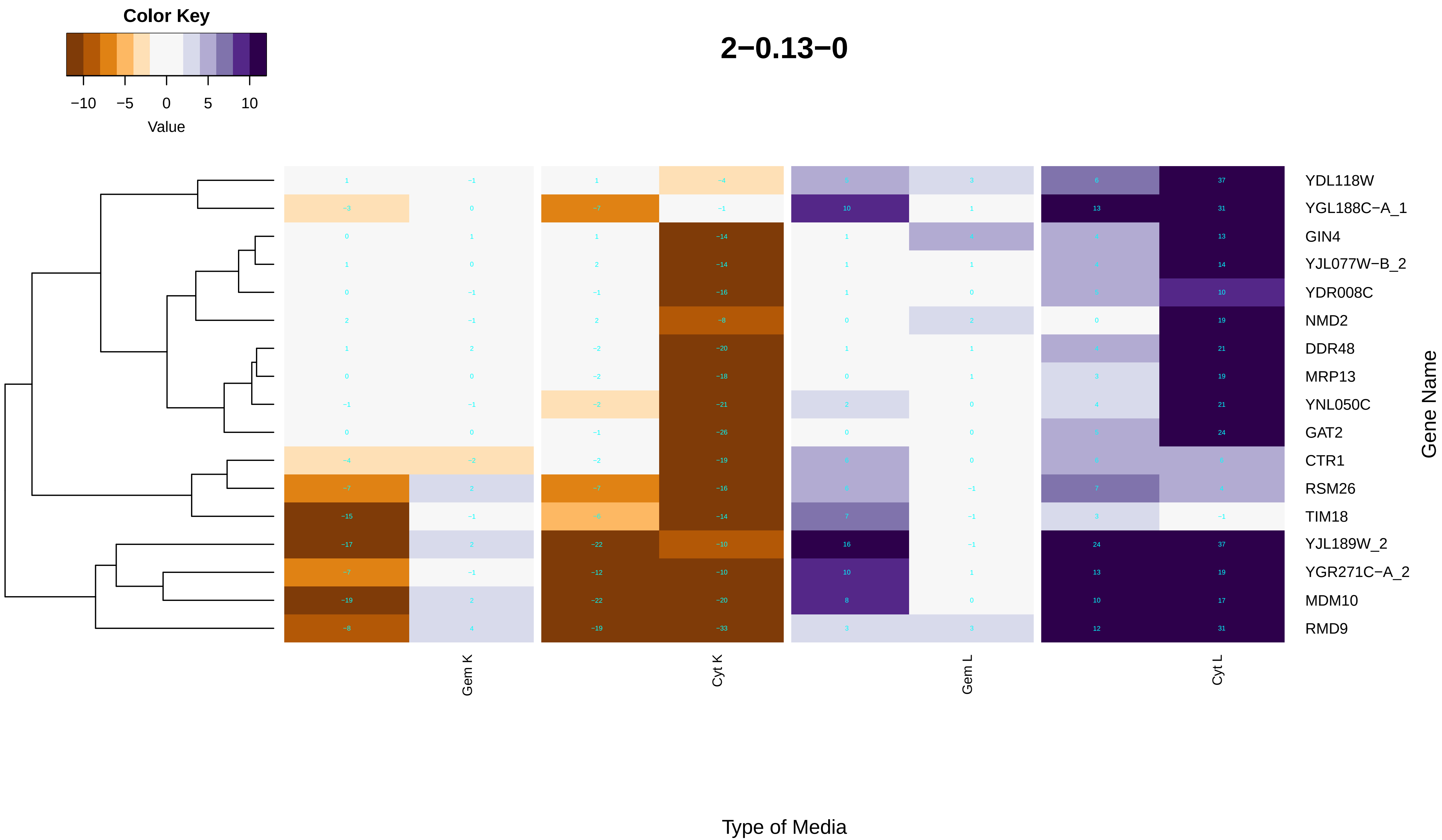

Color Key
−10
−5
0
5
10
Value
2−0.13−0
YDL118W
YGL188C−A_1
GIN4
YJL077W−B_2
YDR008C
NMD2
DDR48
MRP13
YNL050C
GAT2
CTR1
RSM26
TIM18
YJL189W_2
YGR271C−A_2
MDM10
RMD9
Gene Name
Gem K
Cyt K
Gem L
Cyt L
Type of Media

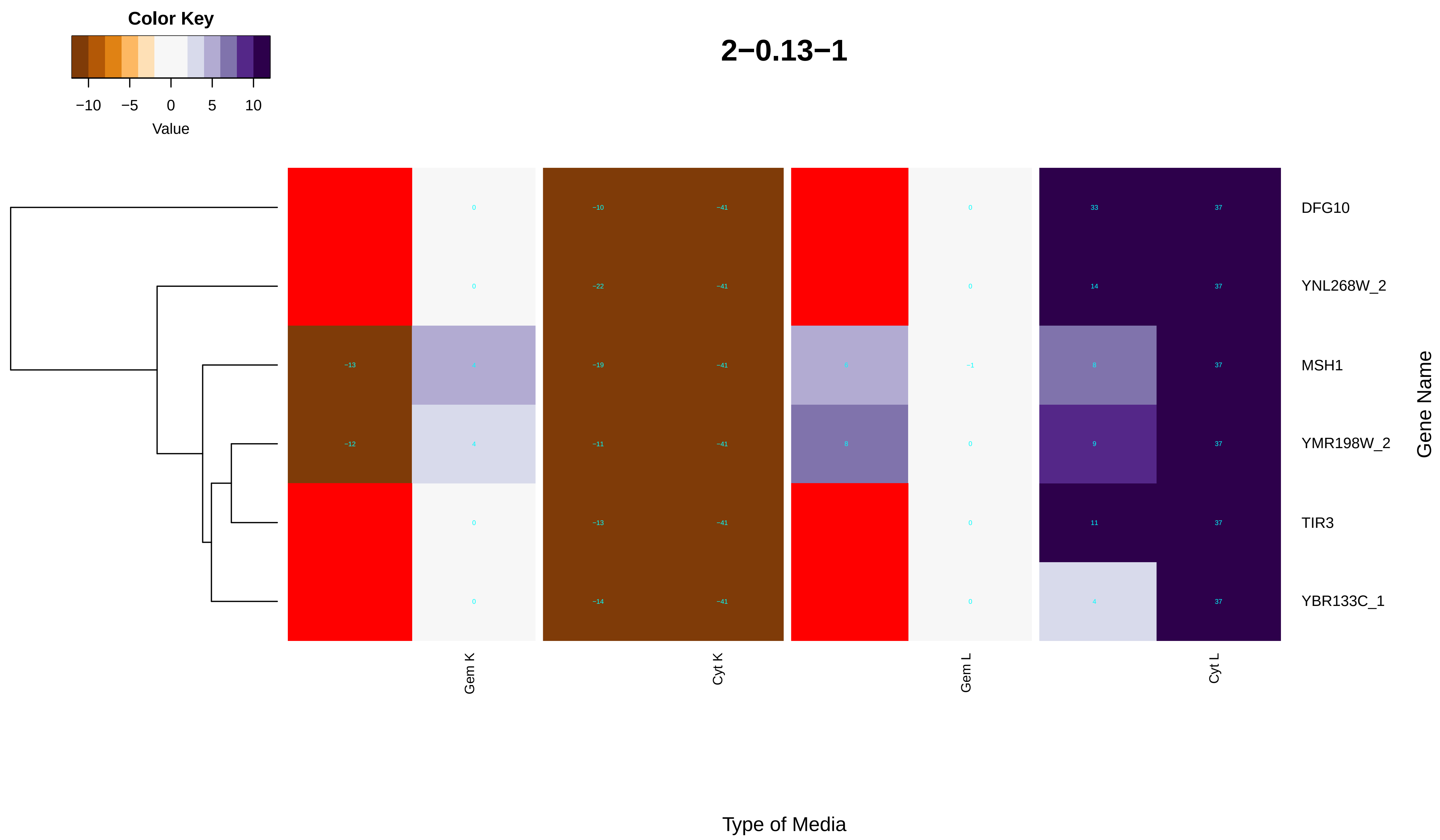

Color Key
−10
−5
0
5
10
Value
2−0.13−1
DFG10
YNL268W_2
MSH1
YMR198W_2
TIR3
YBR133C_1
Gene Name
Gem K
Cyt K
Gem L
Cyt L
Type of Media
0 −10 −41 0 33 37
0 −22 −41 0 14 37
−13 −19 −41 −1 8 37
−12 4 −11 −41 8 0 9 37
0 −13 −41 0 11 37
0 −14 −41 0 4 37

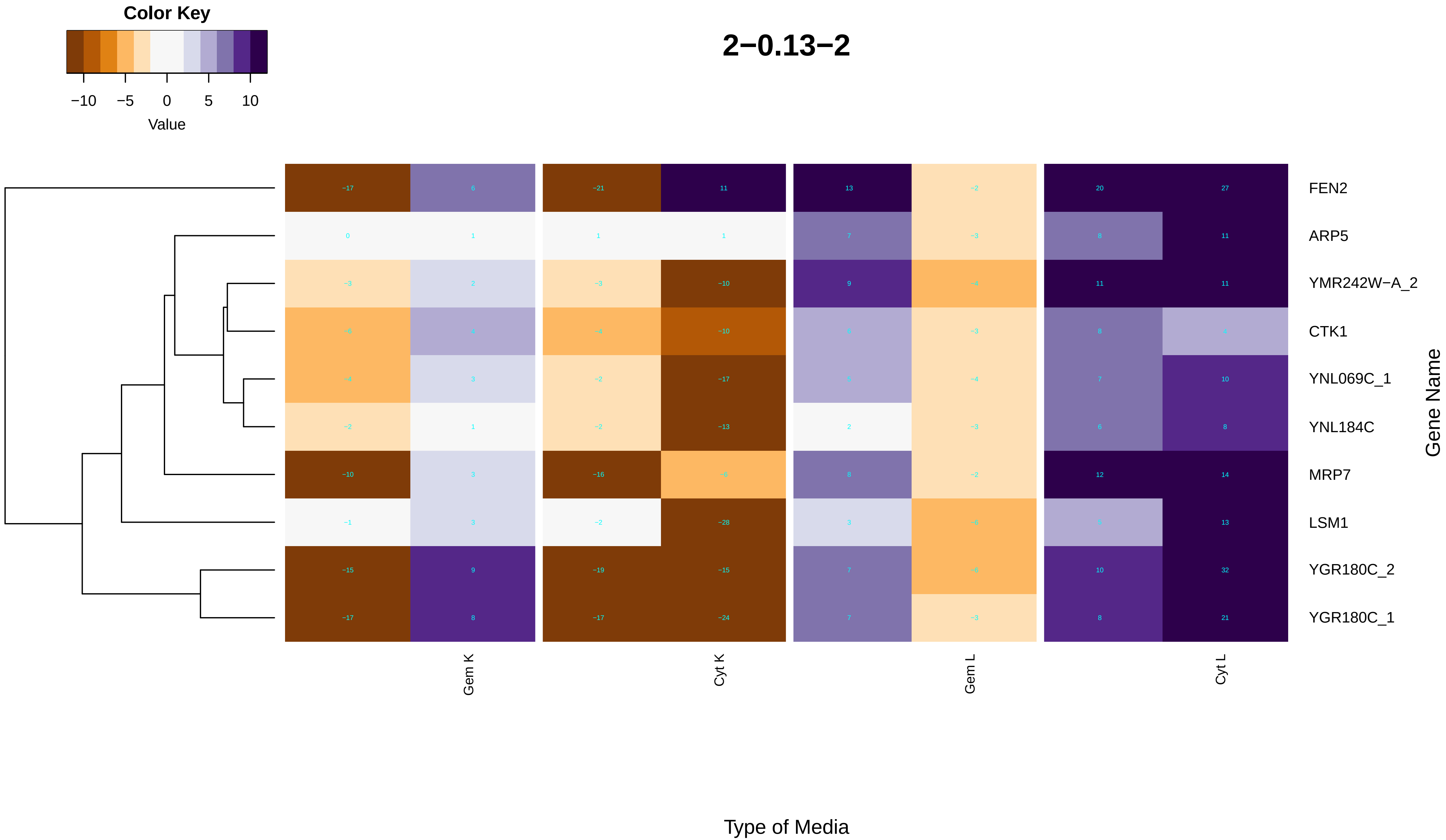
Color Key
−10
−5
0
5
10
Value
2−0.13−2
FEN2
ARP5
YMR242W−A_2
CTK1
YNL069C_1
YNL184C
MRP7
LSM1
YGR180C_2
YGR180C_1
Gene Name
Gem K
Cyt K
Gem L
Cyt L
Type of Media

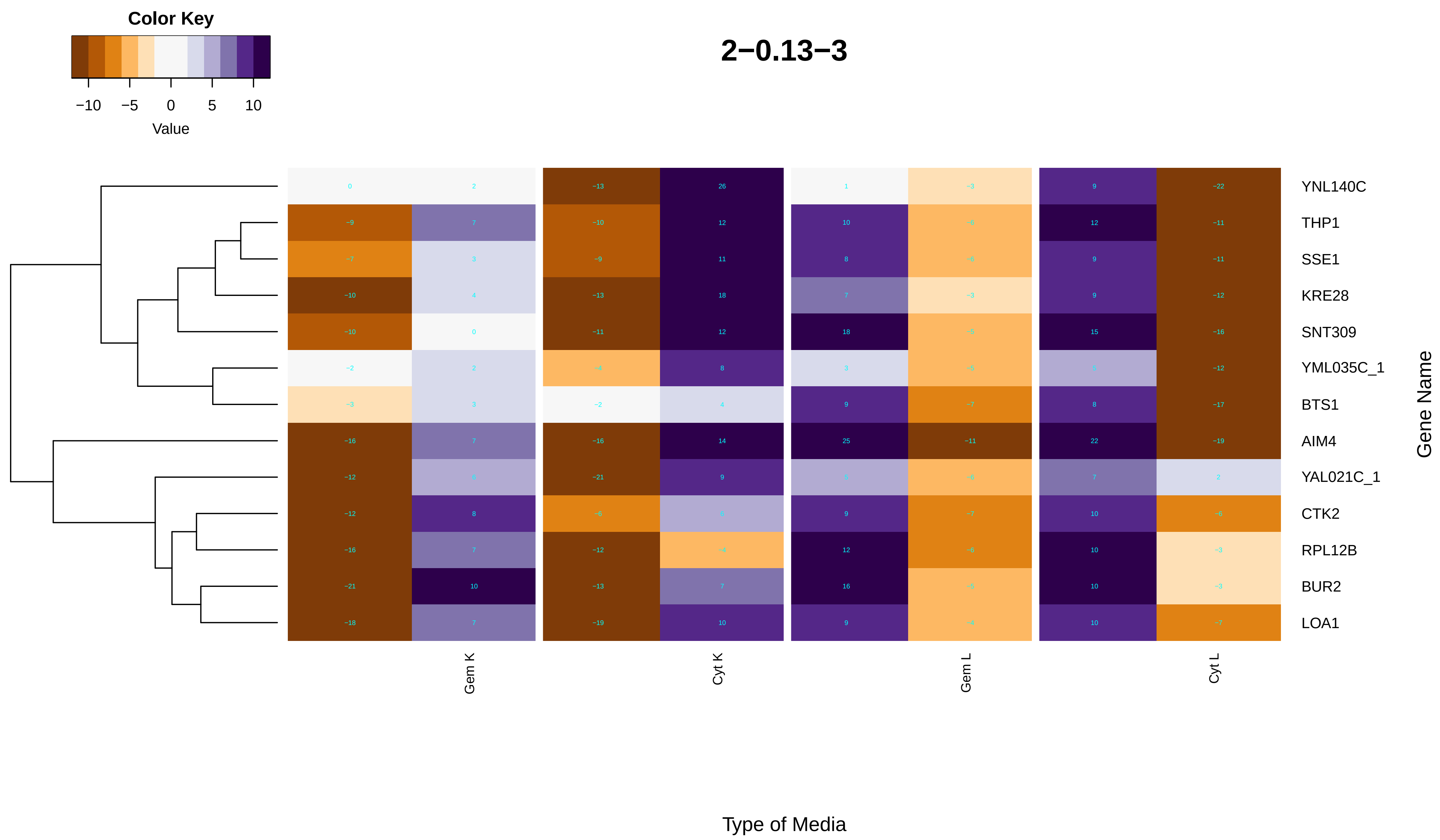
Color Key
−10 −5 0 5 10
Value
2−0.13−3
YNL140C
THP1
SSE1
KRE28
SNT309
YML035C_1
BTS1
AIM4
YAL021C_1
CTK2
RPL12B
BUR2
LOA1
Gene Name
Gem K
Cyt K
Gem L
Cyt L
Type of Media

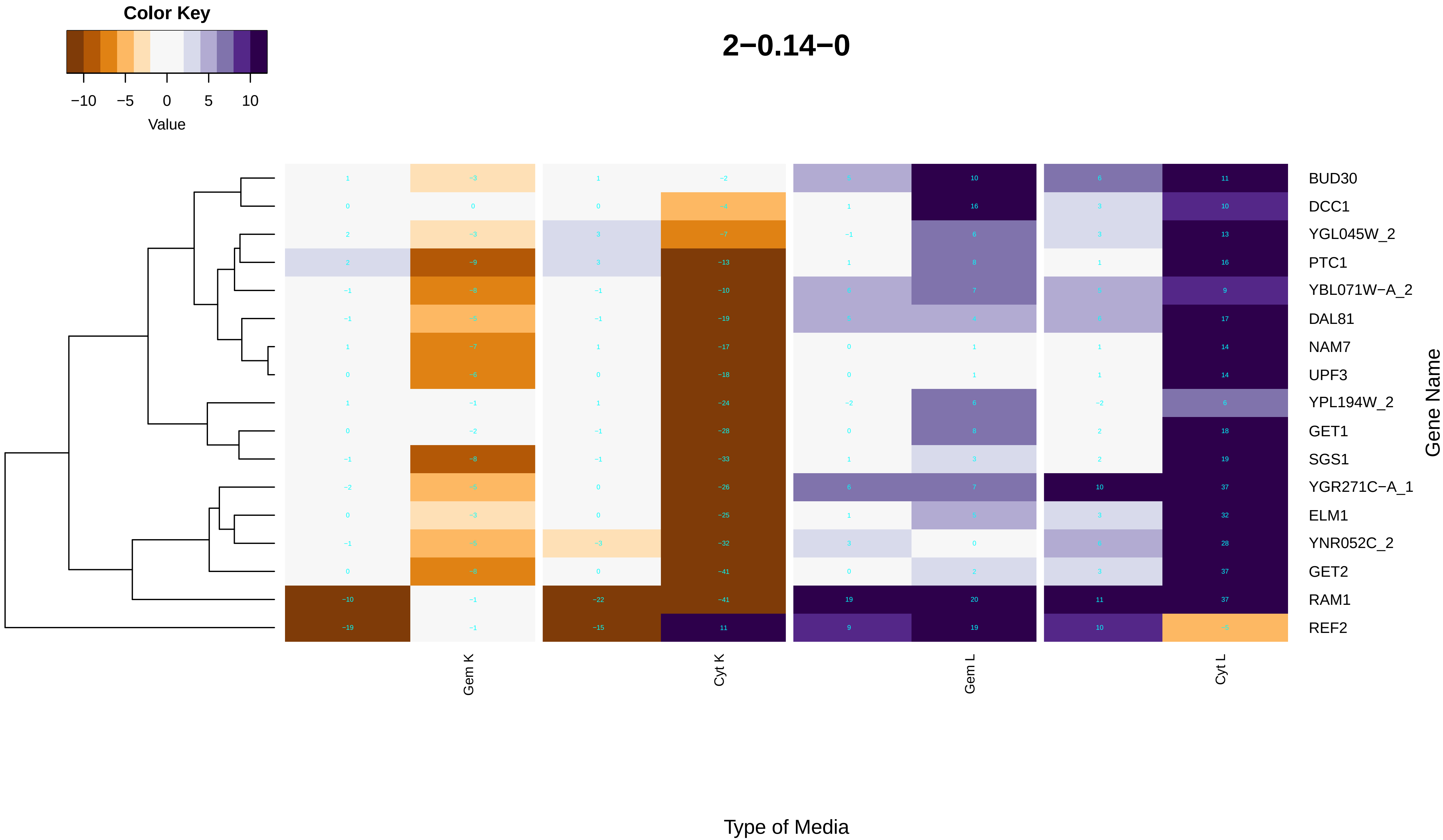

Color Key
−10
−5
0
5
10
Value
2−0.14−0
BUD30
DCC1
YGL045W_2
PTC1
YBL071W−A_2
DAL81
NAM7
UPF3
YPL194W_2
GET1
SGS1
YGR271C−A_1
ELM1
YNR052C_2
GET2
RAM1
REF2
Gene Name
Gem K
Cyt K
Gem L
Cyt L
Type of Media

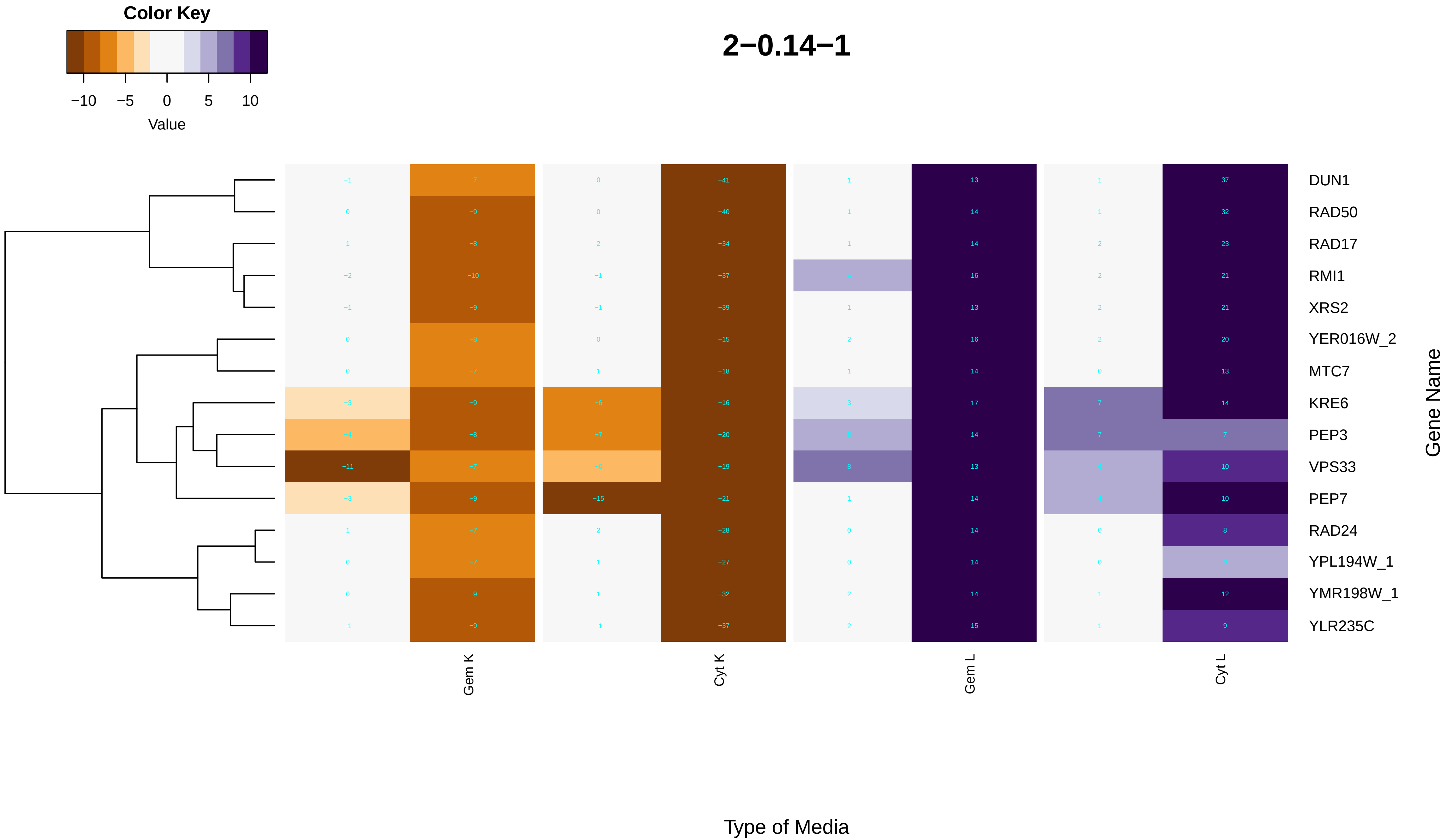
Color Key
−10 −5 0 5 10
Value
2−0.14−1
DUN1
RAD50
RAD17
RMI1
XRS2
YER016W_2
MTC7
KRE6
PEP3
VPS33
PEP7
RAD24
YPL194W_1
YMR198W_1
YLR235C
Gene Name
Gem K
Cyt K
Gem L
Cyt L
Type of Media

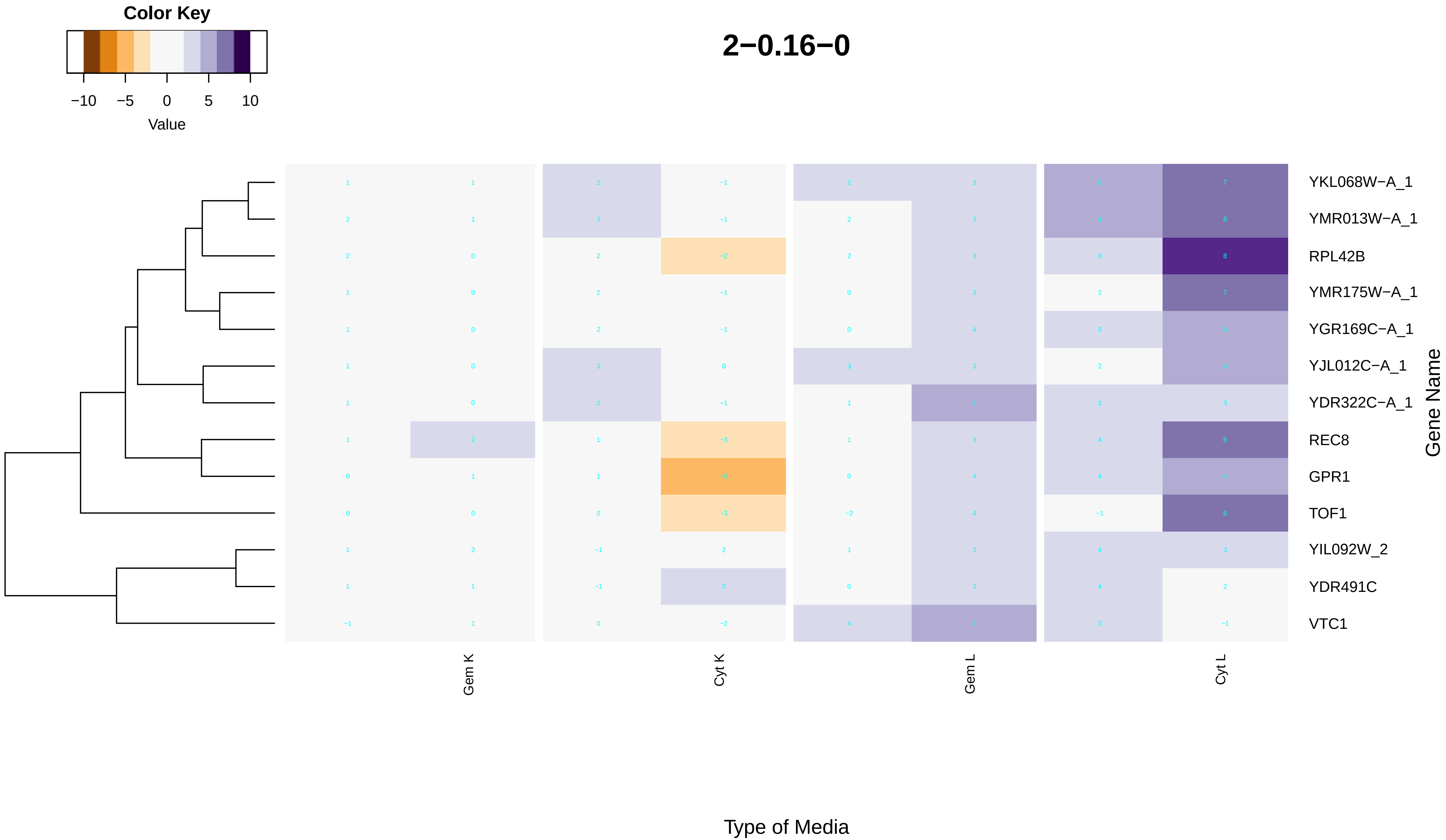

2−0.16−0
Color Key
−10
−5
0
5
10
Value
YKL068W−A_1
YMR013W−A_1
RPL42B
YMR175W−A_1
YGR169C−A_1
YJL012C−A_1
YDR322C−A_1
REC8
GPR1
TOF1
YIL092W_2
YDR491C
VTC1
Gene Name
Gem K
Cyt K
Gem L
Cyt L
Type of Media

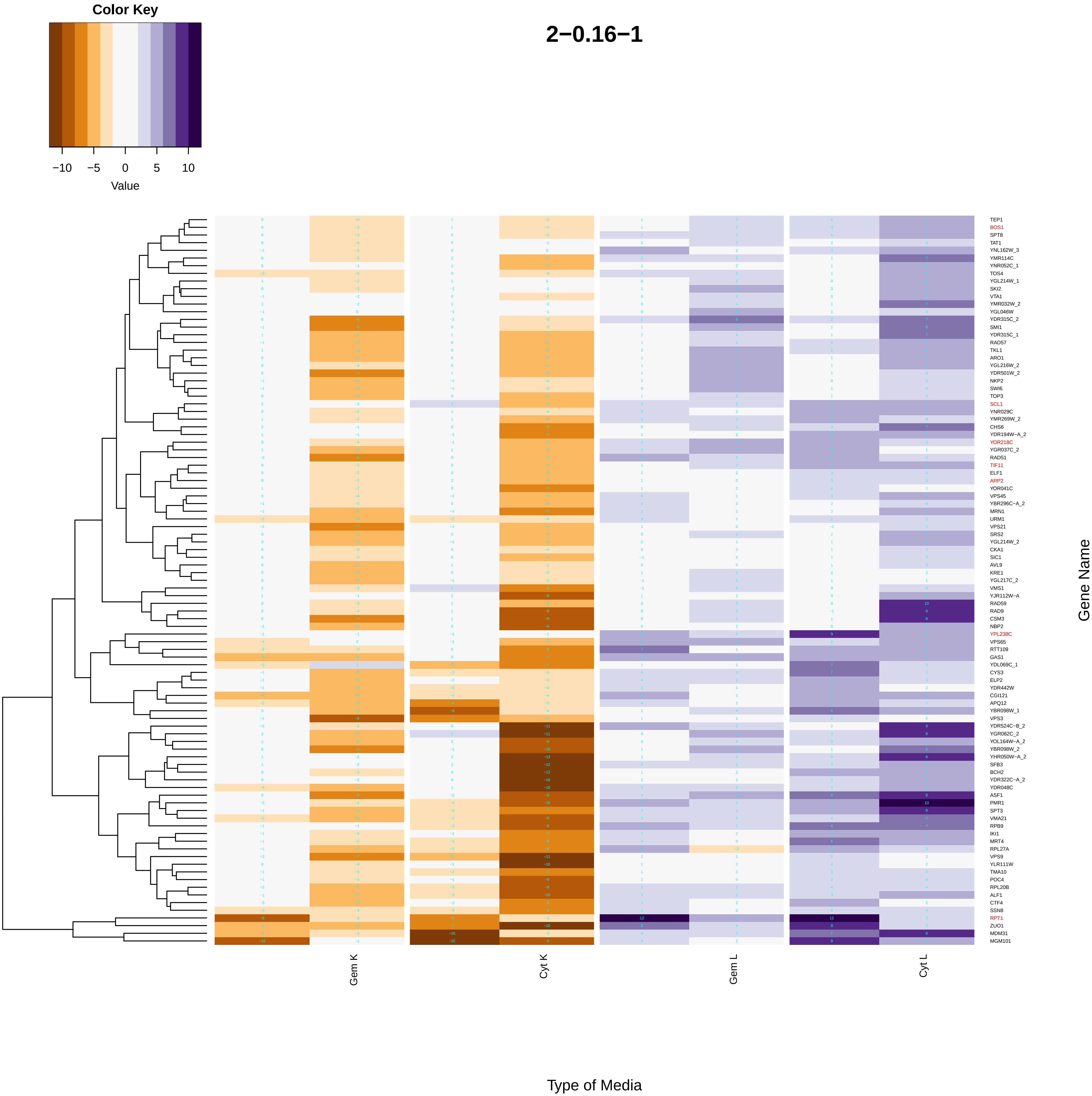
Color Key
−10
−5
0
5
10
Value
2−0.16−1
TEP1
BOS1
SPT8
TAT1
YNL162W_3
YMR114C
YNR052C_1
TOS4
YGL214W_1
SKI2
VTA1
YMR032W_2
YGL046W
YDR315C_2
SMI1
YDR315C_1
RAD57
TKL1
ARO1
YGL216W_2
YDR501W_2
NKP2
SWI6
TOP3
SCL1
YNR029C
YMR269W_2
CHS6
YDR194W−A_2
YOR218C
YGR037C_2
RAD51
TIF11
ELF1
ARP2
YOR041C
VPS45
YBR296C−A_2
MRN1
URM1
VPS21
SRS2
YGL214W_2
CKA1
SIC1
AVL9
KRE1
YGL217C_2
VMS1
YJR112W−A
RAD59
RAD9
CSM3
NBP2
YPL238C
VPS65
RTT109
GAS1
YDL069C_1
CYS3
ELP2
YDR442W
CGI121
APQ12
YBR098W_1
VPS3
YDR524C−B_2
YGR062C_2
YOL164W−A_2
YBR098W_2
YHR050W−A_2
SFB3
BCH2
YDR322C−A_2
YDR048C
ASF1
PMR1
SPT3
VMA21
RPB9
IKI1
MRT4
RPL27A
VPS9
YLR111W
TMA10
POC4
RPL20B
ALF1
CTF4
SSN8
RPT1
ZUO1
MDM31
MGM101
Gem K
Cyt K
Gem L
Cyt L
Gene Name
Type of Media

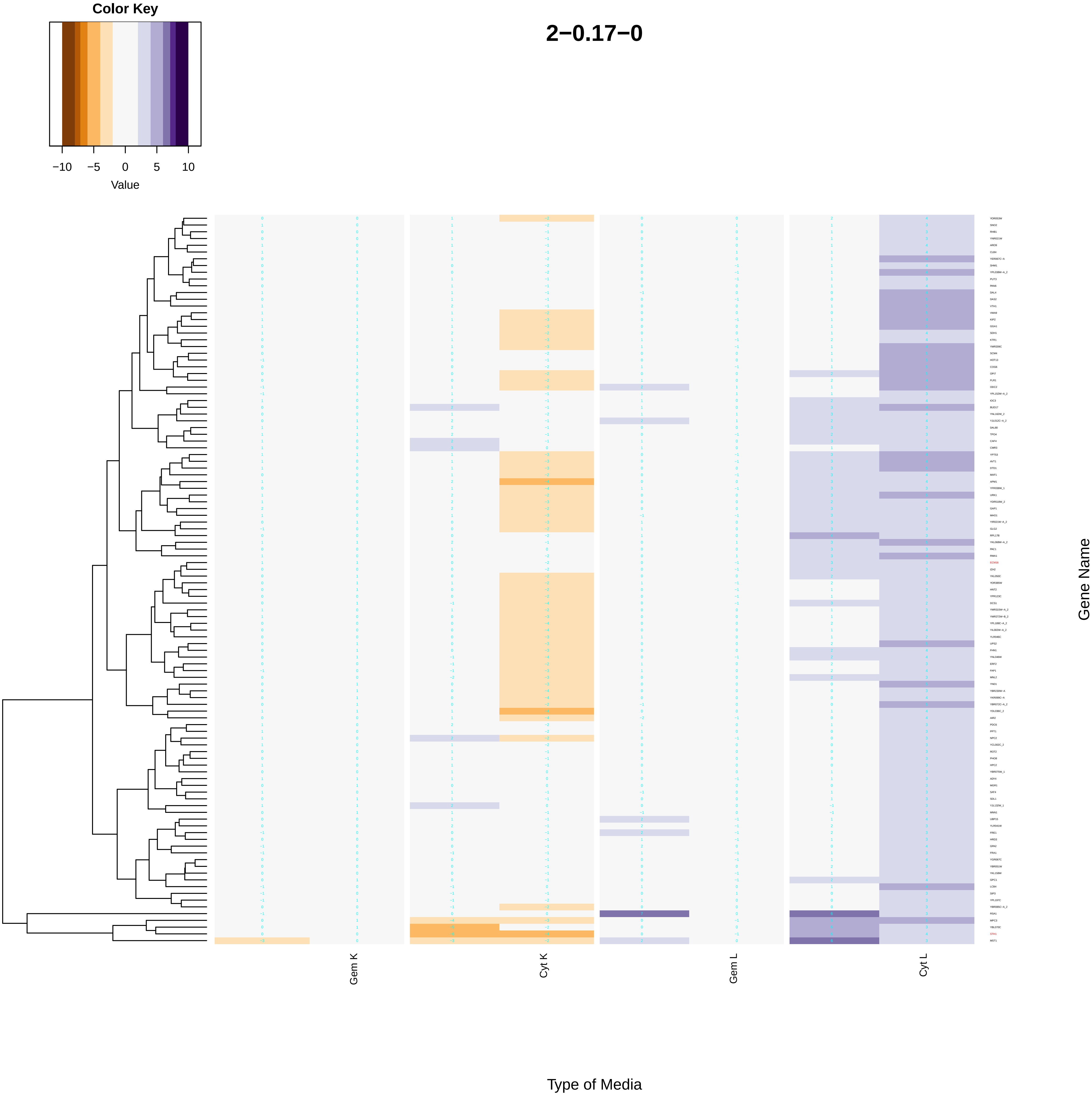
Color Key
−10 −5 0 5 10
Value
2−0.17−0
Gem K
Cyt K
Gem L
Cyt L
Gene Name
Type of Media

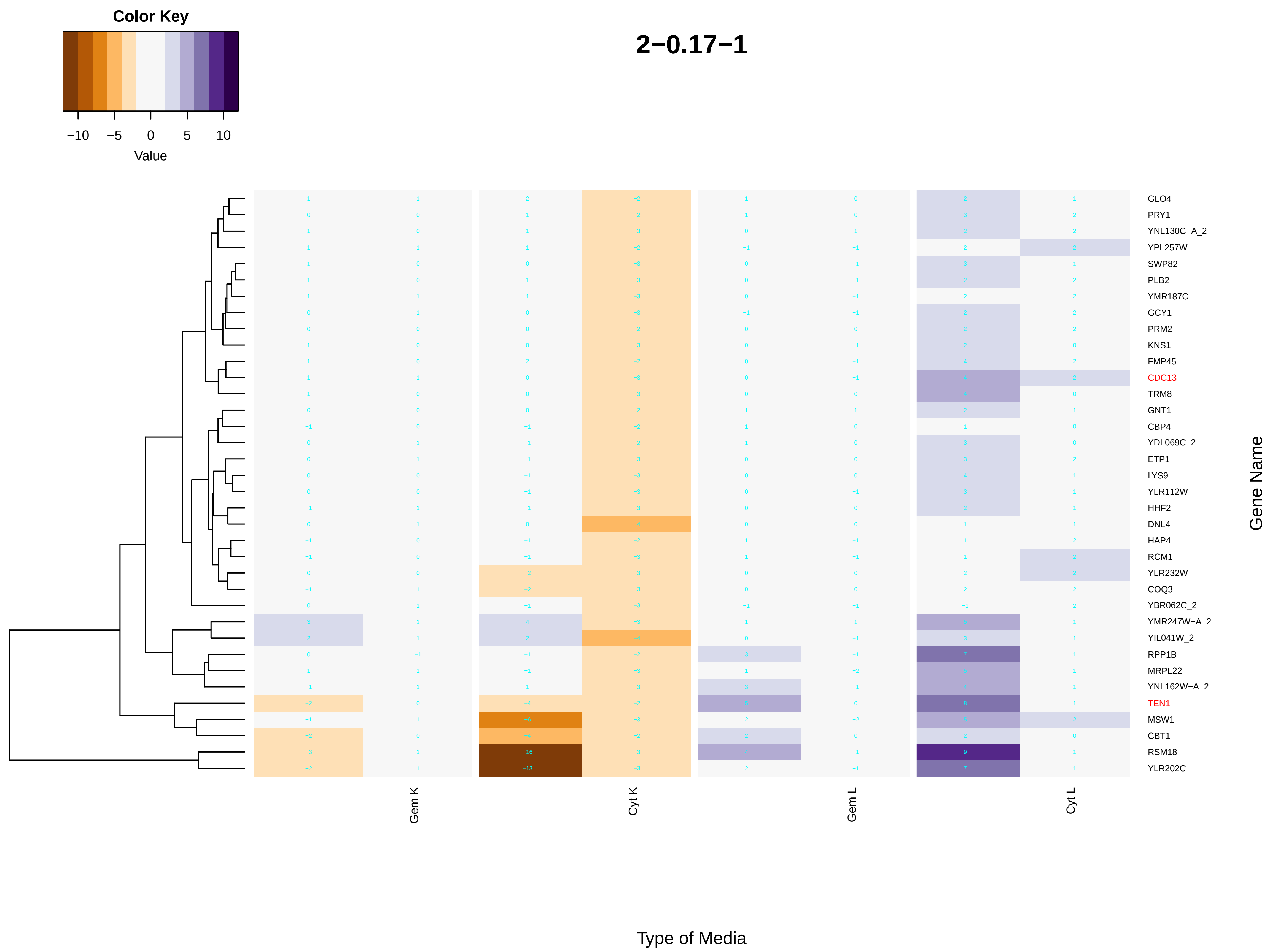

Color Key
−10
−5
0
5
10
Value
2−0.17−1
GLO4
PRY1
YNL130C−A_2
YPL257W
SWP82
PLB2
YMR187C
GCY1
PRM2
KNS1
FMP45
CDC13
TRM8
GNT1
CBP4
YDL069C_2
ETP1
LYS9
YLR112W
HHF2
DNL4
HAP4
RCM1
YLR232W
COQ3
YBR062C_2
YMR247W−A_2
YIL041W_2
RPP1B
MRPL22
YNL162W−A_2
TEN1
MSW1
CBT1
RSM18
YLR202C
Gene Name
Gem K
Cyt K
Gem L
Cyt L
Type of Media

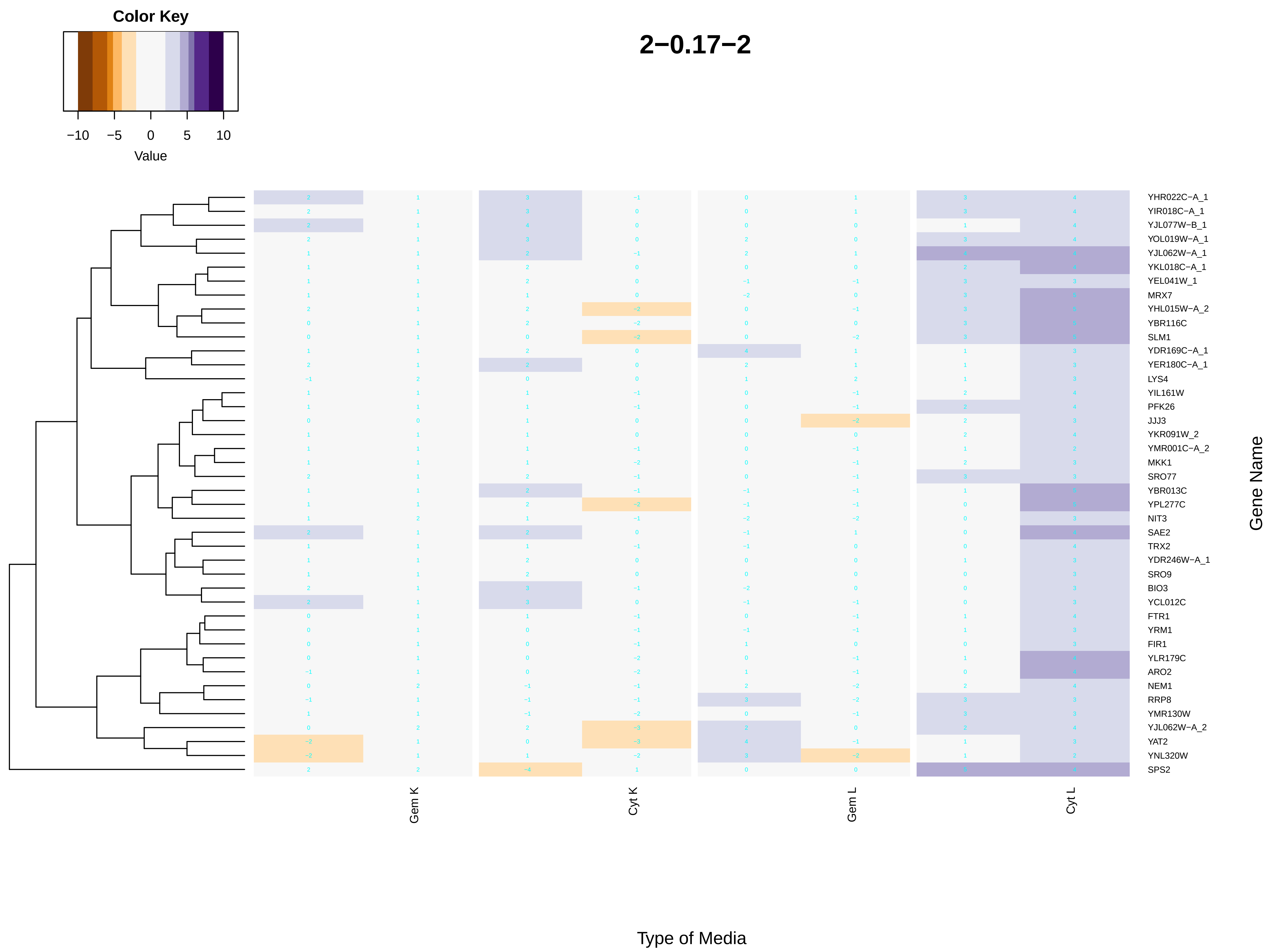
Color Key
−10
−5
0
5
10
Value
2−0.17−2
YHR022C−A_1
YIR018C−A_1
YJL077W−B_1
YOL019W−A_1
YJL062W−A_1
YKL018C−A_1
YEL041W_1
MRX7
YHL015W−A_2
YBR116C
SLM1
YDR169C−A_1
YER180C−A_1
LYS4
YIL161W
PFK26
JJJ3
YKR091W_2
YMR001C−A_2
MKK1
SRO77
YBR013C
YPL277C
NIT3
SAE2
TRX2
YDR246W−A_1
SRO9
BIO3
YCL012C
FTR1
YRM1
FIR1
YLR179C
ARO2
NEM1
RRP8
YMR130W
YJL062W−A_2
YAT2
YNL320W
SPS2
Gene Name
Gem K
Cyt K
Gem L
Cyt L
Type of Media

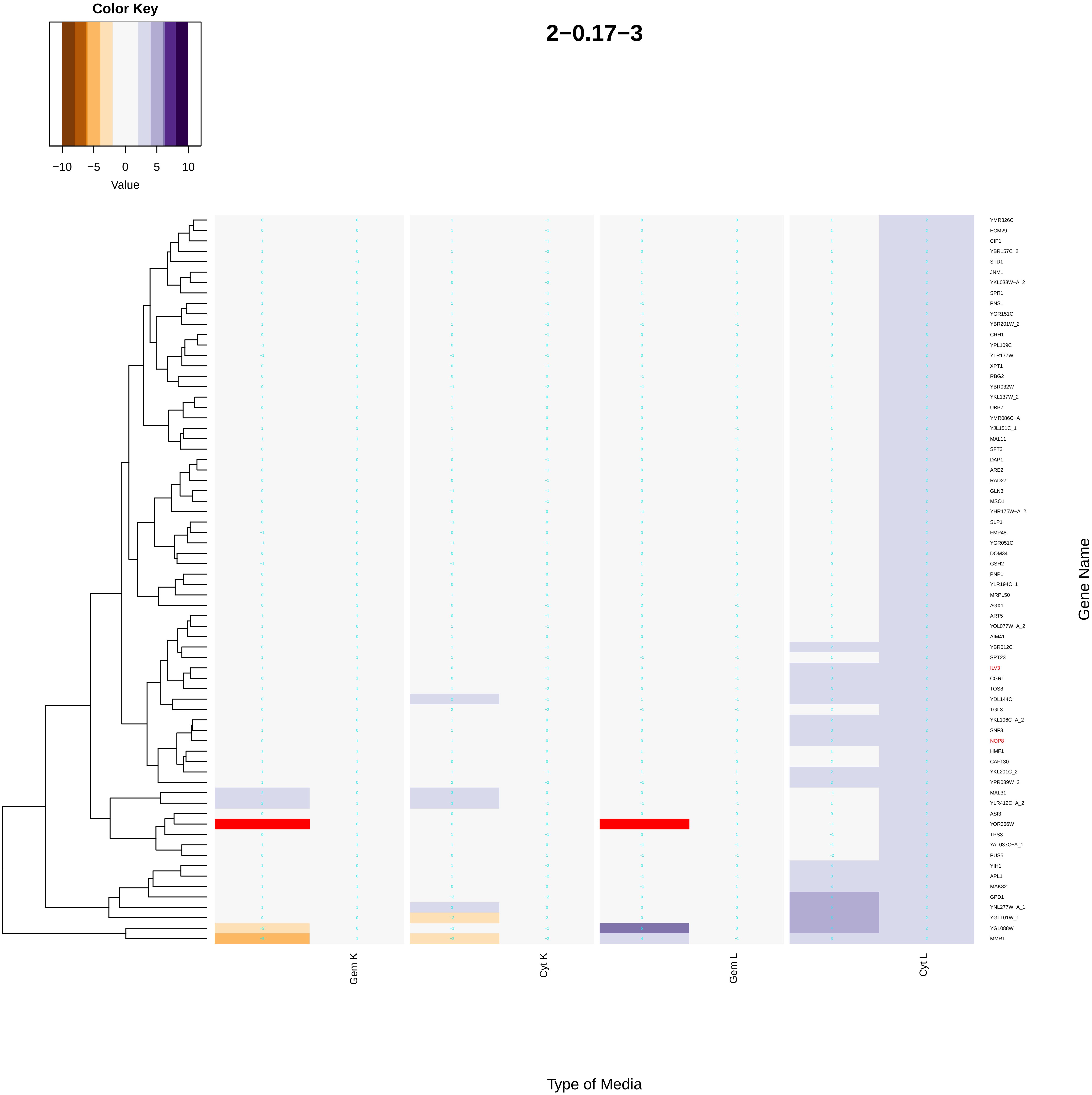

Color Key
−10
−5
0
5
10
Value
2−0.17−3
YMR326C
ECM29
CIP1
YBR157C_2
STD1
JNM1
YKL033W−A_2
SPR1
PNS1
YGR151C
YBR201W_2
CRH1
YPL109C
YLR177W
XPT1
RBG2
YBR032W
YKL137W_2
UBP7
YMR086C−A
YJL151C_1
MAL11
SFT2
DAP1
ARE2
RAD27
GLN3
MSO1
YHR175W−A_2
SLP1
FMP48
YGR051C
DOM34
GSH2
PNP1
YLR194C_1
MRPL50
AGX1
ART5
YOL077W−A_2
AIM41
YBR012C
SPT23
ILV3
CGR1
TOS8
YDL144C
TGL3
YKL106C−A_2
SNF3
NOP8
HMF1
CAF130
YKL201C_2
YPR089W_2
MAL31
YLR412C−A_2
ASI3
YOR366W
TPS3
YAL037C−A_1
PUS5
YIH1
APL1
MAK32
GPD1
YNL277W−A_1
YGL101W_1
YGL088W
MMR1
Gene Name
Gem K
Cyt K
Gem L
Cyt L
Type of Media

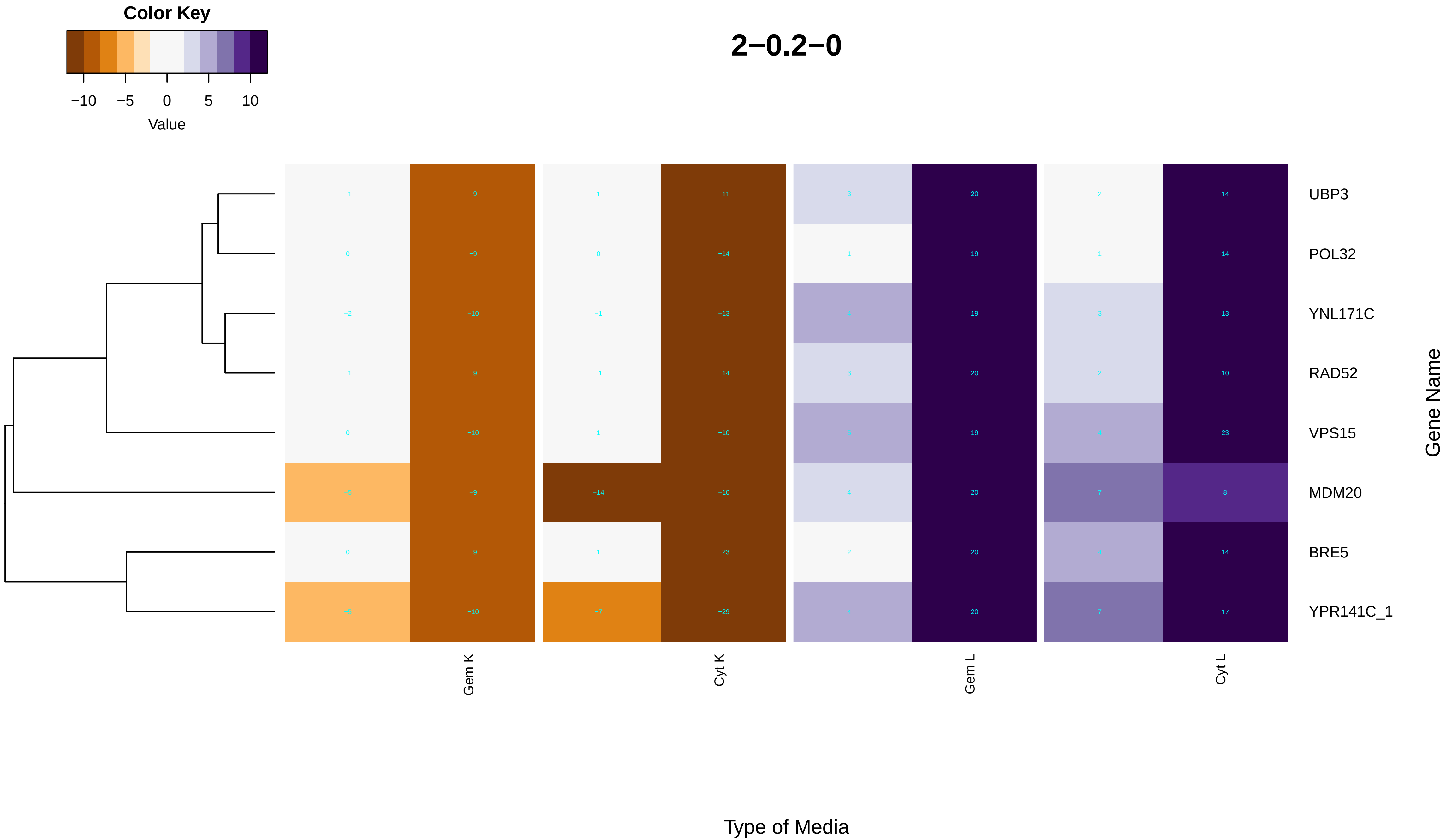

Color Key
−10
−5
0
5
10
Value
2−0.2−0
UBP3
POL32
YNL171C
RAD52
VPS15
MDM20
BRE5
YPR141C_1
Gene Name
Gem K
Cyt K
Gem L
Cyt L
Type of Media

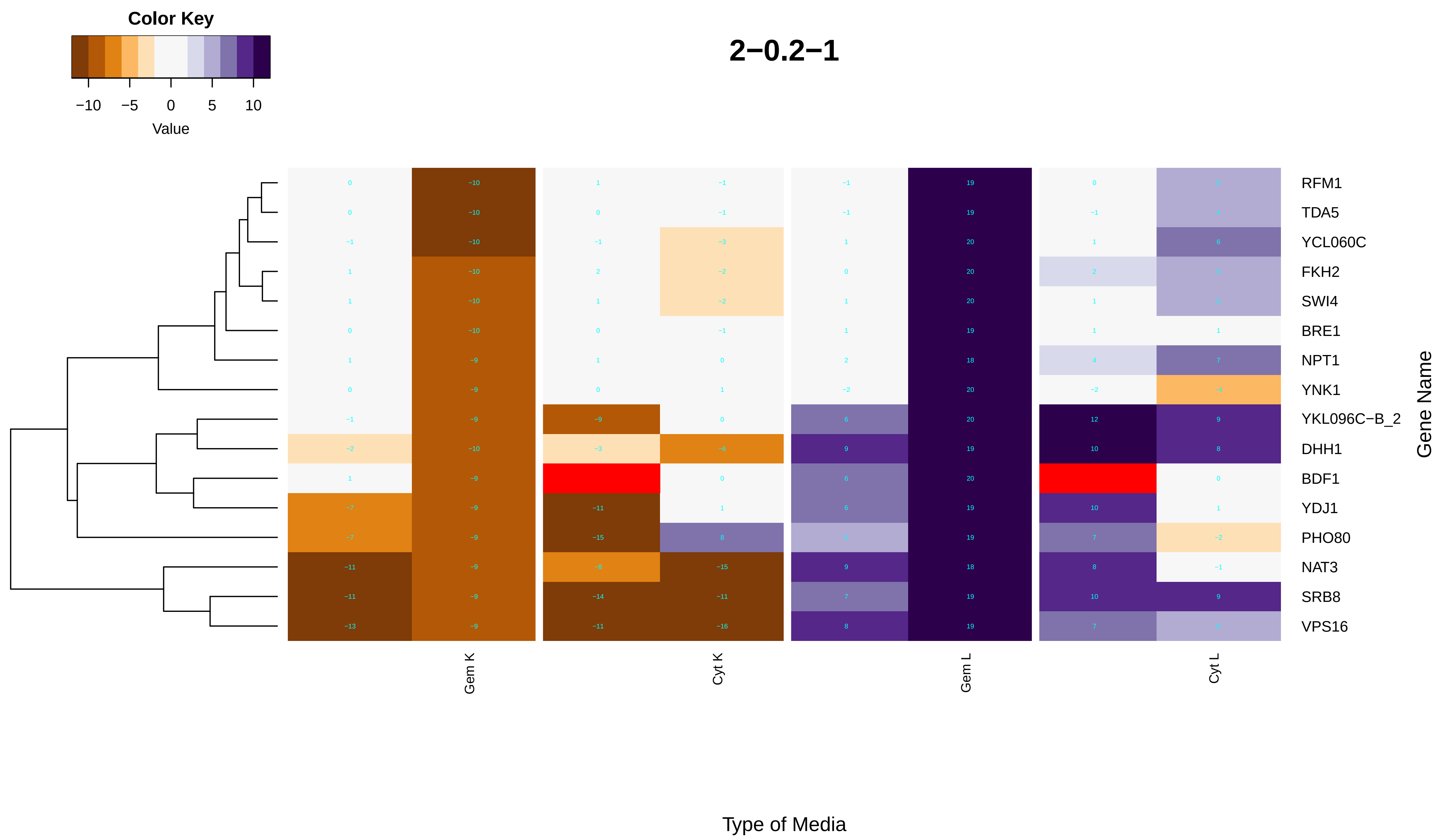
Color Key
−10 −5 0 5 10
Value
2−0.2−1
RFM1
TDA5
YCL060C
FKH2
SWI4
BRE1
NPT1
YNK1
YKL096C−B_2
DHH1
BDF1
YDJ1
PHO80
NAT3
SRB8
VPS16
Gene Name
Gem K
Cyt K
Gem L
Cyt L
Type of Media

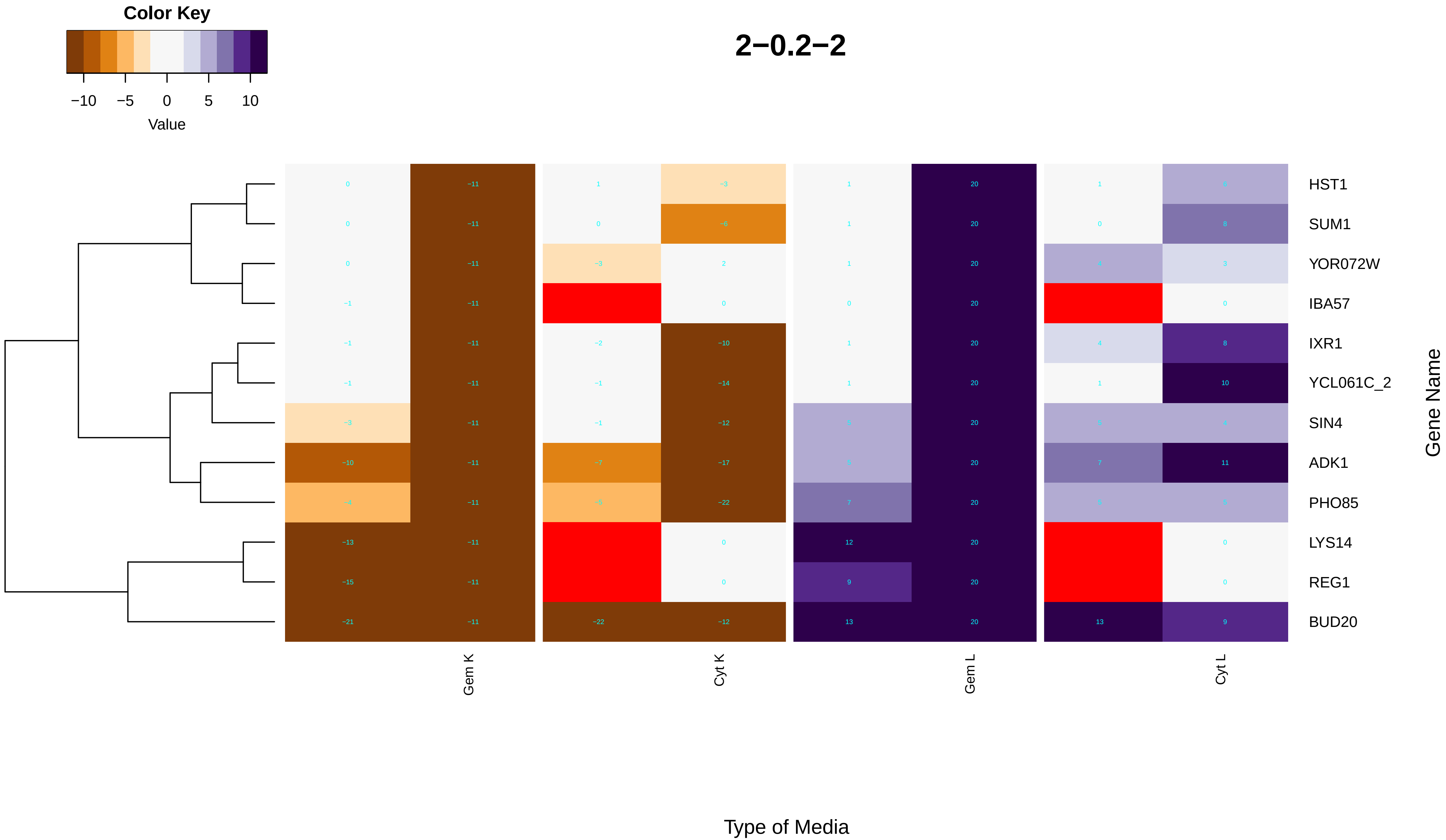

Color Key
−10
−5
0
5
10
Value
2−0.2−2
HST1
SUM1
YOR072W
IBA57
IXR1
YCL061C_2
SIN4
ADK1
PHO85
LYS14
REG1
BUD20
Gene Name
Gem K
Cyt K
Gem L
Cyt L
Type of Media

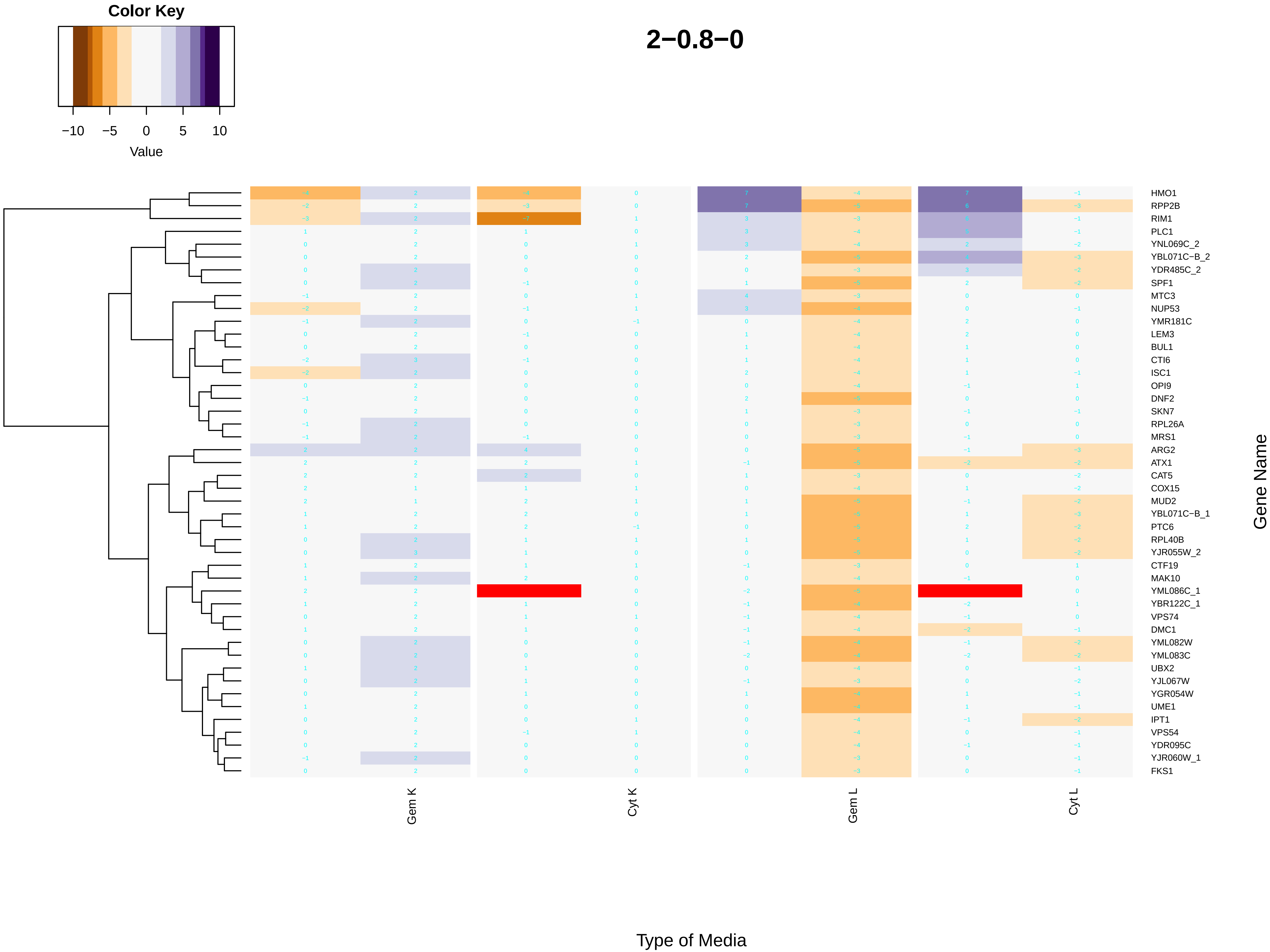
Color Key
−10
−5
0
5
10
Value
2−0.8−0
HMO1
RPP2B
RIM1
PLC1
YNL069C_2
YBL071C−B_2
YDR485C_2
SPF1
MTC3
NUP53
YMR181C
LEM3
BUL1
CTI6
ISC1
OPI9
DNF2
SKN7
RPL26A
MRS1
ARG2
ATX1
CAT5
COX15
MUD2
YBL071C−B_1
PTC6
RPL40B
YJR055W_2
CTF19
MAK10
YML086C_1
YBR122C_1
VPS74
DMC1
YML082W
YML083C
UBX2
YJL067W
YGR054W
UME1
IPT1
VPS54
YDR095C
YJR060W_1
FKS1
Gene Name
Gem K
Cyt K
Gem L
Cyt L
Type of Media

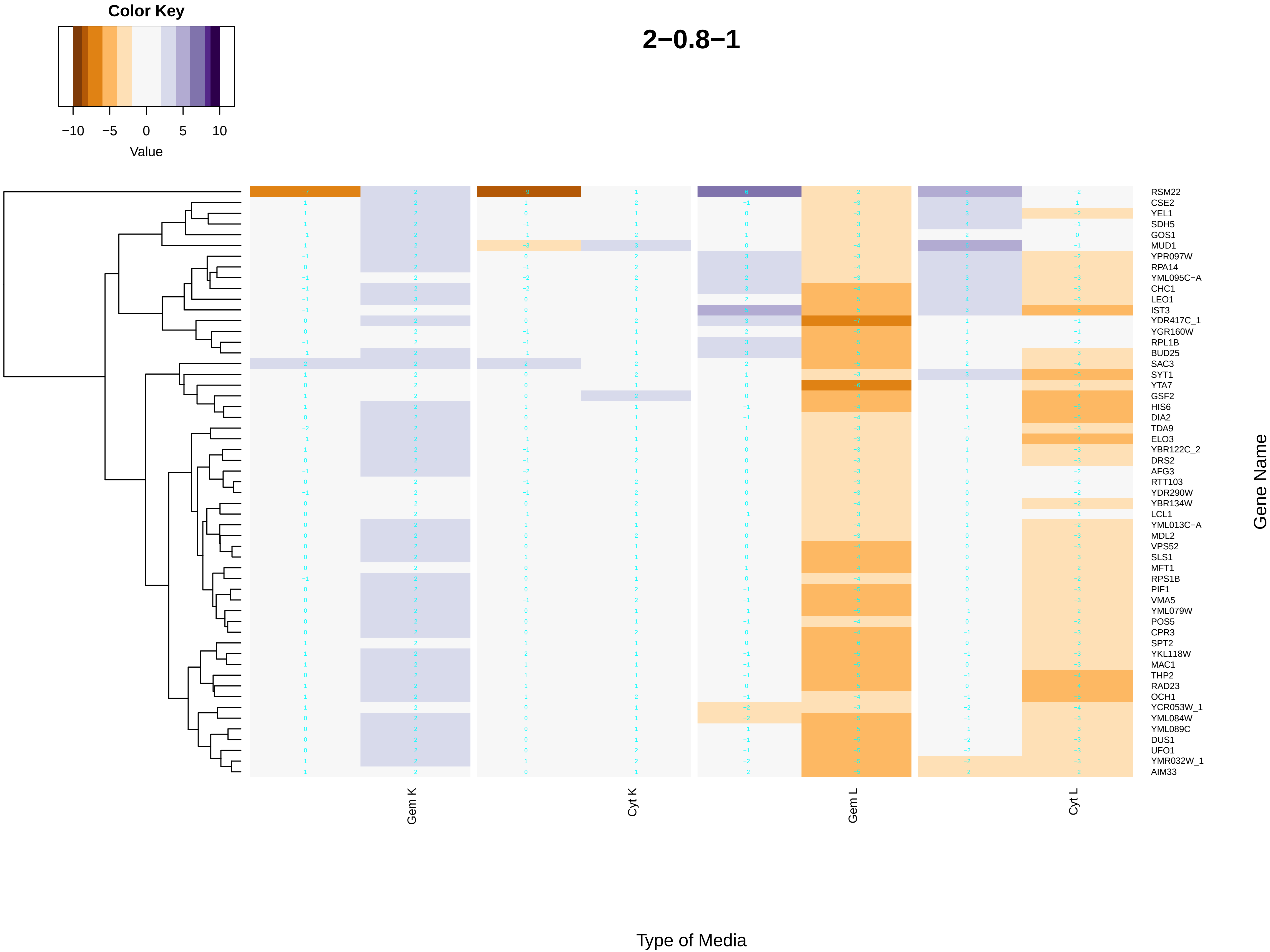
Color Key
−10
−5
0
5
10
Value
2−0.8−1
RSM22
CSE2
YEL1
SDH5
GOS1
MUD1
YPR097W
RPA14
YML095C−A
CHC1
LEO1
IST3
YDR417C_1
YGR160W
RPL1B
BUD25
SAC3
SYT1
YTA7
GSF2
HIS6
DIA2
TDA9
ELO3
YBR122C_2
DRS2
AFG3
RTT103
YDR290W
YBR134W
LCL1
YML013C−A
MDL2
VPS52
SLS1
MFT1
RPS1B
PIF1
VMA5
YML079W
POS5
CPR3
SPT2
YKL118W
MAC1
THP2
RAD23
OCH1
YCR053W_1
YML084W
YML089C
DUS1
UFO1
YMR032W_1
AIM33
Gene Name
Gem K
Cyt K
Gem L
Cyt L
Type of Media

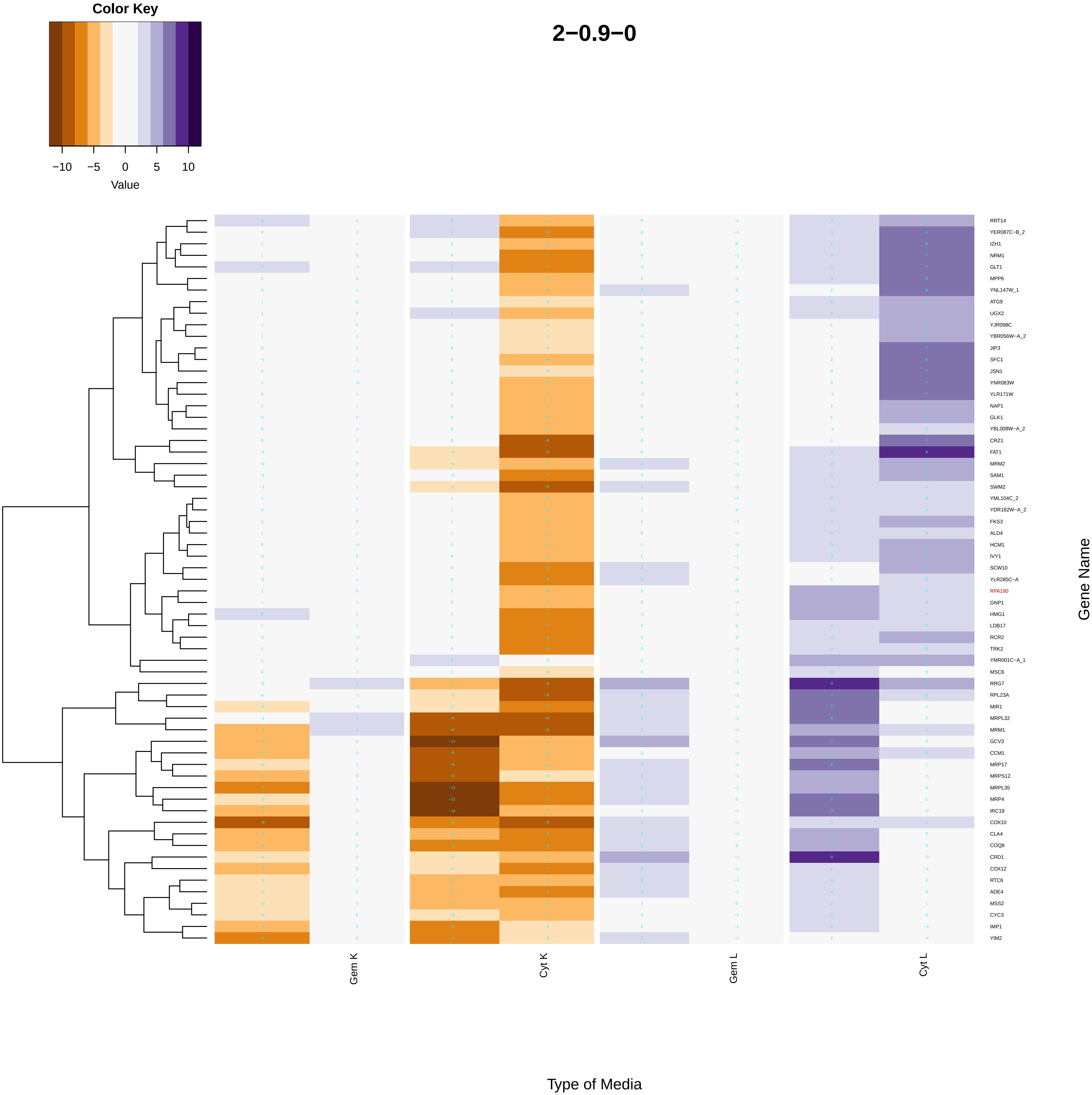

Color Key
−10
−5
0
5
10
Value
2−0.9−0
RRT14
YER087C−B_2
IZH1
NRM1
GLT1
MPP6
YNL147W_1
ATG9
UGX2
YJR098C
YBR056W−A_2
JIP3
SFC1
JSN1
YNR063W
YLR171W
NAP1
GLK1
YBL008W−A_2
CRZ1
FAT1
MRM2
SAM1
SWM2
YML104C_2
YDR182W−A_2
FKS3
ALD4
HCM1
IVY1
SCW10
YLR285C−A
RPA190
GNP1
HMG1
LDB17
RCR2
TRK2
YMR001C−A_1
MSC6
RRG7
RPL23A
MIR1
MRPL32
MRM1
GCV3
CCM1
MRP17
MRPS12
MRPL35
MRP4
IRC19
COX10
CLA4
COQ8
CRD1
COX12
RTC6
ADE4
MSS2
CYC3
IMP1
YIM2
Gene Name
Gem K
Cyt K
Gem L
Cyt L
Type of Media

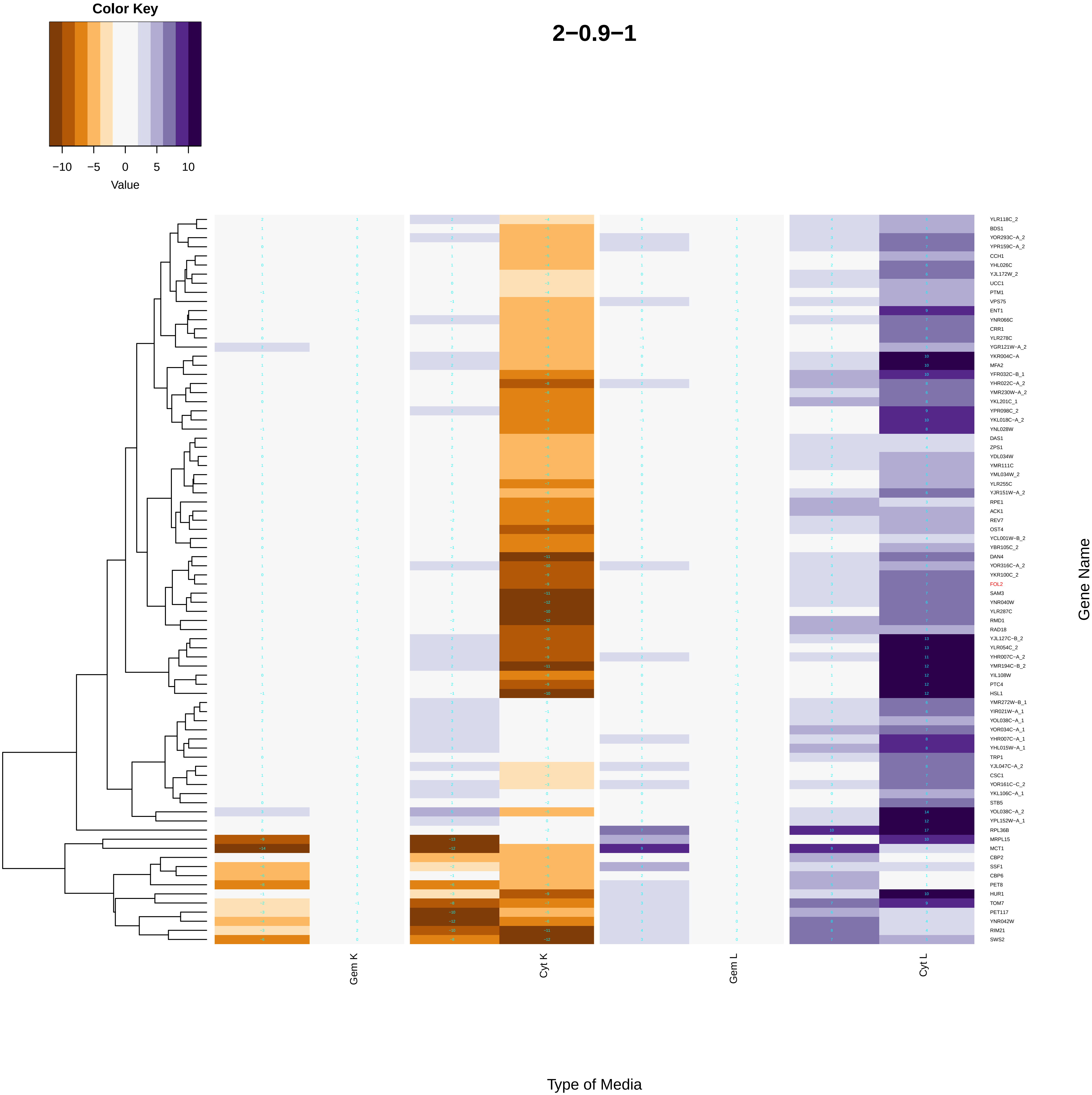

Color Key
−10
−5
0
5
10
Value
2−0.9−1
YLR118C_2
BDS1
YOR293C−A_2
YPR159C−A_2
CCH1
YHL026C
YJL172W_2
UCC1
PTM1
VPS75
ENT1
YNR066C
CRR1
YLR278C
YGR121W−A_2
YKR004C−A
MFA2
YFR032C−B_1
YHR022C−A_2
YMR230W−A_2
YKL201C_1
YPR098C_2
YKL018C−A_2
YNL028W
DAS1
ZPS1
YDL034W
YMR111C
YML034W_2
YLR255C
YJR151W−A_2
RPE1
ACK1
REV7
OST4
YCL001W−B_2
YBR105C_2
DAN4
YOR316C−A_2
YKR100C_2
FOL2
SAM3
YNR040W
YLR287C
RMD1
RAD18
YJL127C−B_2
YLR054C_2
YHR007C−A_2
YMR194C−B_2
YIL108W
PTC4
HSL1
YMR272W−B_1
YIR021W−A_1
YOL038C−A_1
YOR034C−A_1
YHR007C−A_1
YHL015W−A_1
TRP1
YJL047C−A_2
CSC1
YOR161C−C_2
YKL106C−A_1
STB5
YOL038C−A_2
YPL152W−A_1
RPL36B
MRPL15
MCT1
CBP2
SSF1
CBP6
PET8
HUR1
TOM7
PET117
YNR042W
RIM21
SWS2
Gene Name
Gem K
Cyt K
Gem L
Cyt L
Type of Media

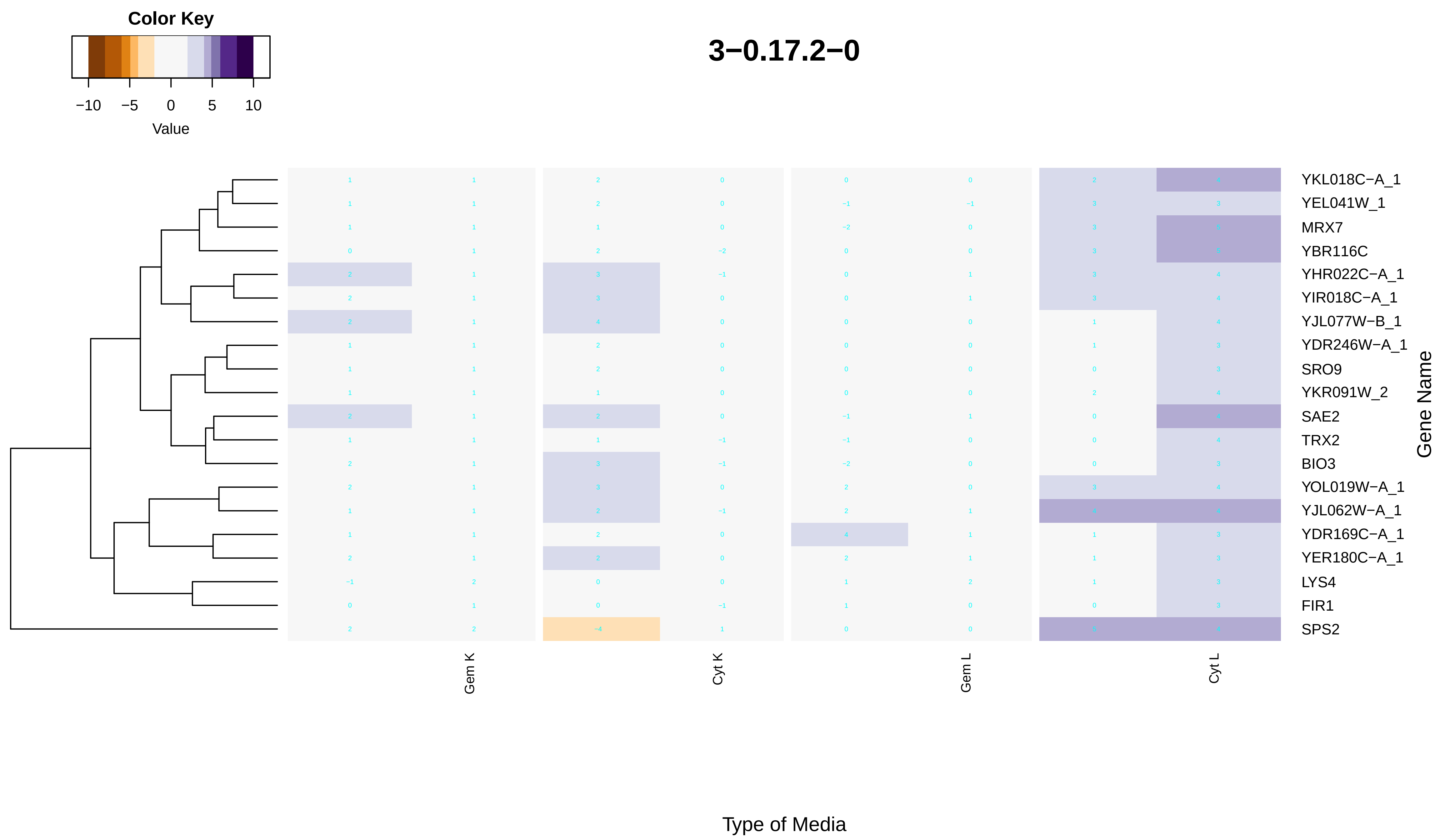
Color Key
−10 −5 0 5 10
Value
3−0.17.2−0
YKL018C−A_1
YEL041W_1
MRX7
YBR116C
YHR022C−A_1
YIR018C−A_1
YJL077W−B_1
YDR246W−A_1
SRO9
YKR091W_2
SAE2
TRX2
BIO3
YOL019W−A_1
YJL062W−A_1
YDR169C−A_1
YER180C−A_1
LYS4
FIR1
SPS2
Gene Name
Gem K
Cyt K
Gem L
Cyt L
Type of Media

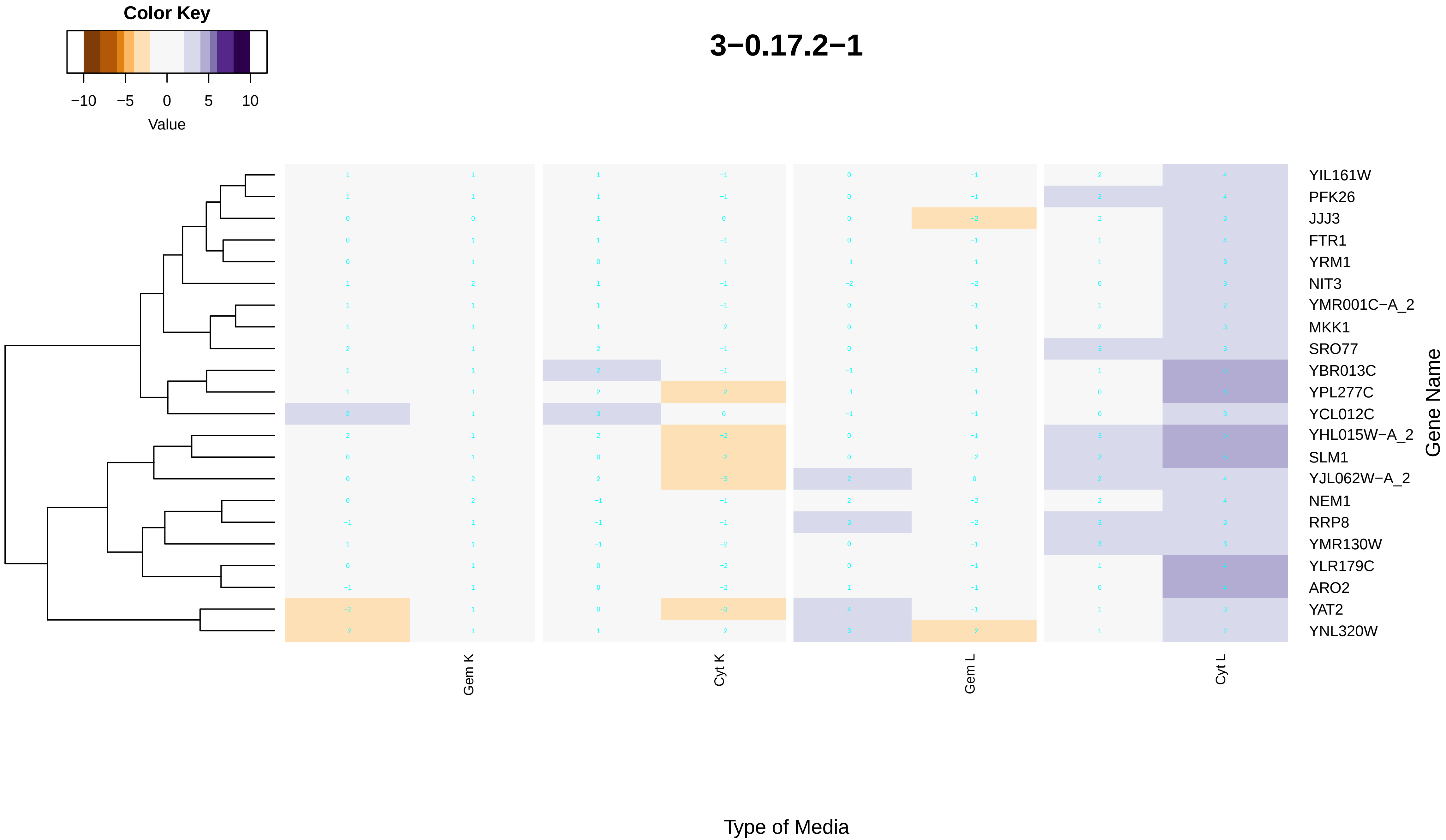

Color Key
−10
−5
0
5
10
Value
3−0.17.2−1
YIL161W
PFK26
JJJ3
FTR1
YRM1
NIT3
YMR001C−A_2
MKK1
SRO77
YBR013C
YPL277C
YCL012C
YHL015W−A_2
SLM1
YJL062W−A_2
NEM1
RRP8
YMR130W
YLR179C
ARO2
YAT2
YNL320W
Gene Name
Gem K
Cyt K
Gem L
Cyt L
Type of Media

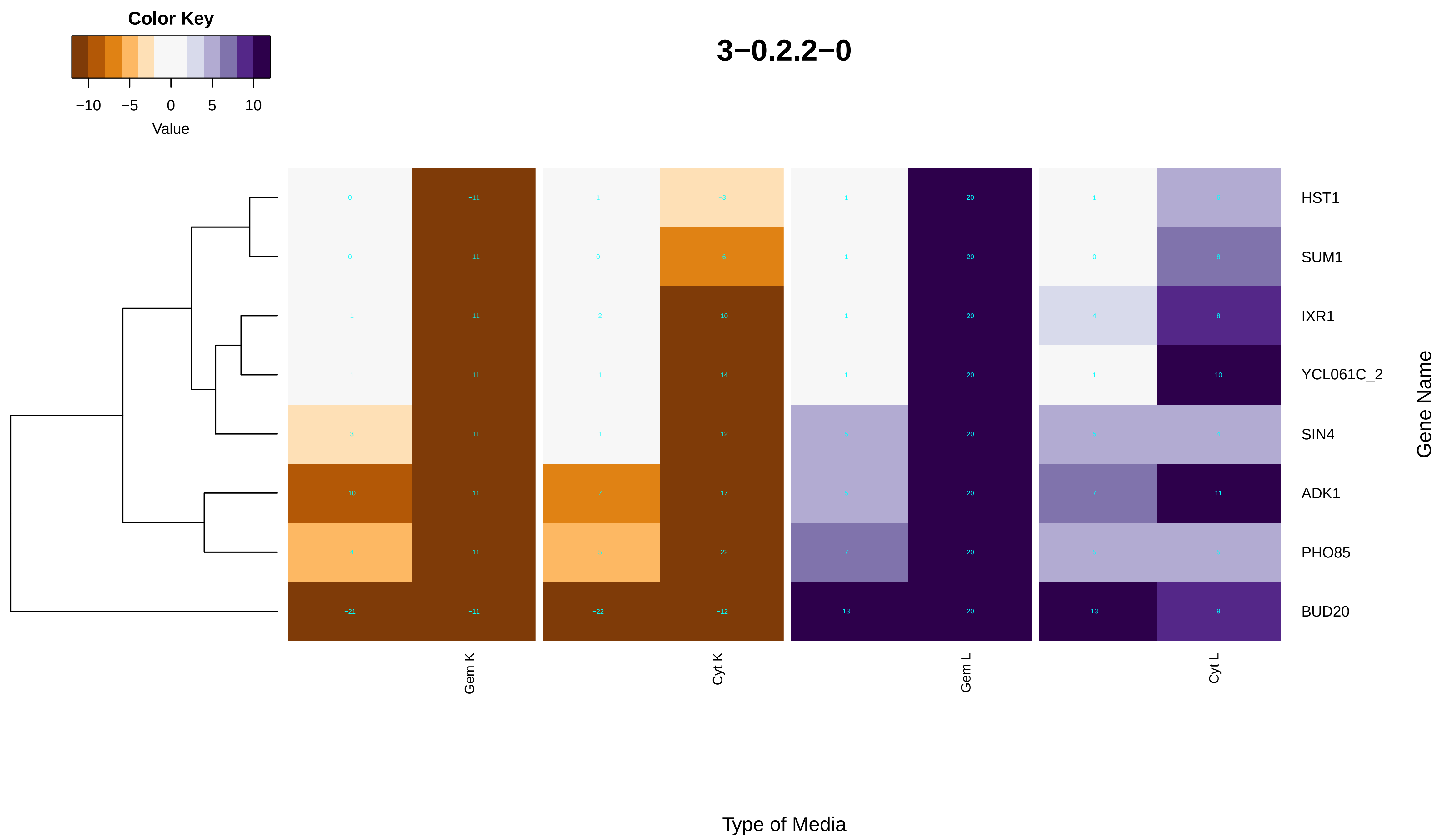

Color Key
−10 −5 0 5 10
Value
3−0.2.2−0
HST1
SUM1
IXR1
YCL061C_2
SIN4
ADK1
PHO85
BUD20
Gene Name
Gem K
Cyt K
Gem L
Cyt L
Type of Media

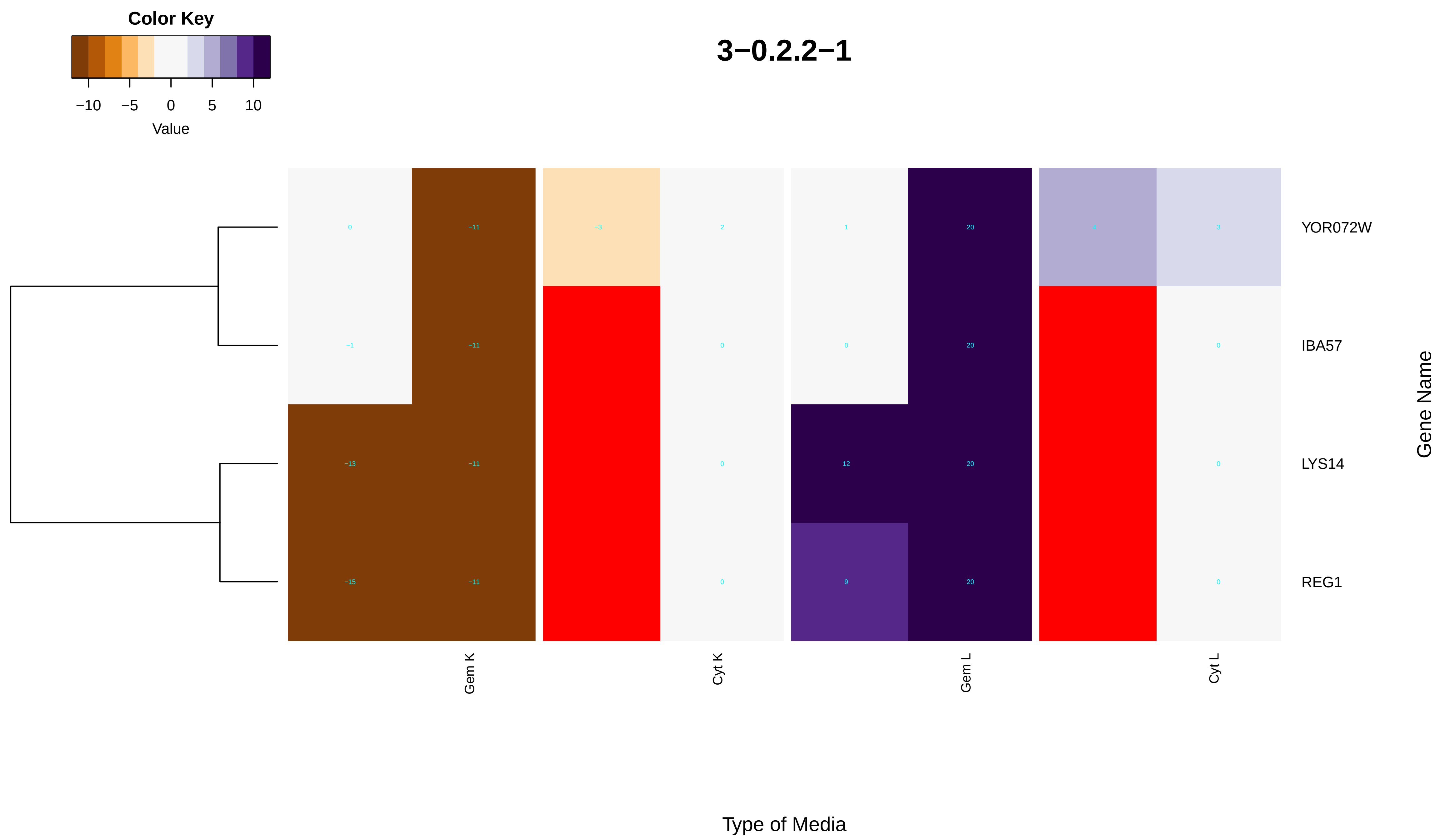

Color Key
−10
−5
0
5
10
Value
3−0.2.2−1
0
−11
−3
2
1
20
3
YOR072W
−1
−11
0
0
20
0
IBA57
−13
−11
0
12
20
0
LYS14
−15
−11
0
9
20
0
REG1
Gene Name
Gem K
Cyt K
Gem L
Cyt L
Type of Media

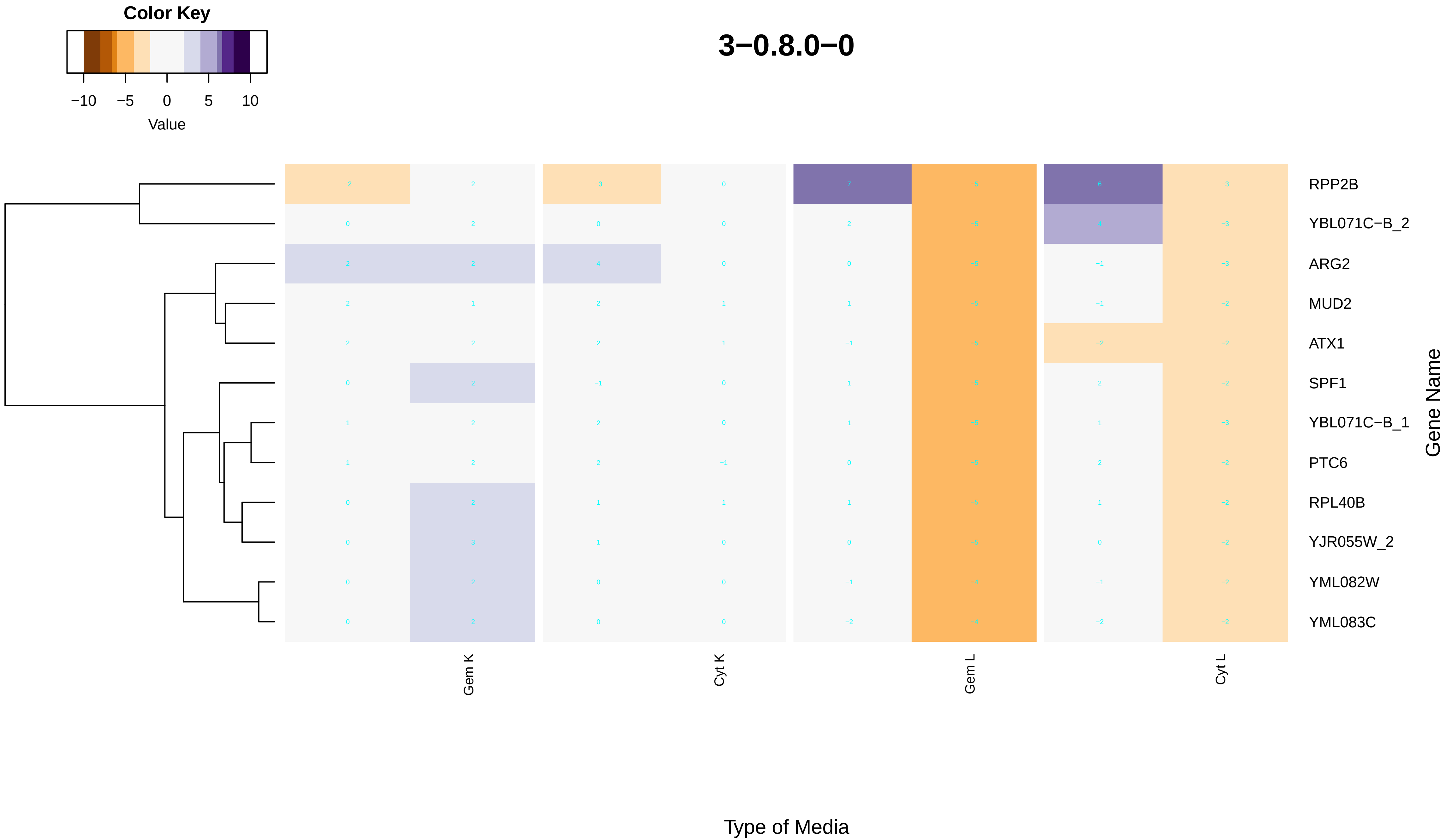
Color Key
−10
−5
0
5
10
Value
3−0.8.0−0
RPP2B
YBL071C−B_2
ARG2
MUD2
ATX1
SPF1
YBL071C−B_1
PTC6
RPL40B
YJR055W_2
YML082W
YML083C
Gene Name
Gem K
Cyt K
Gem L
Cyt L
Type of Media

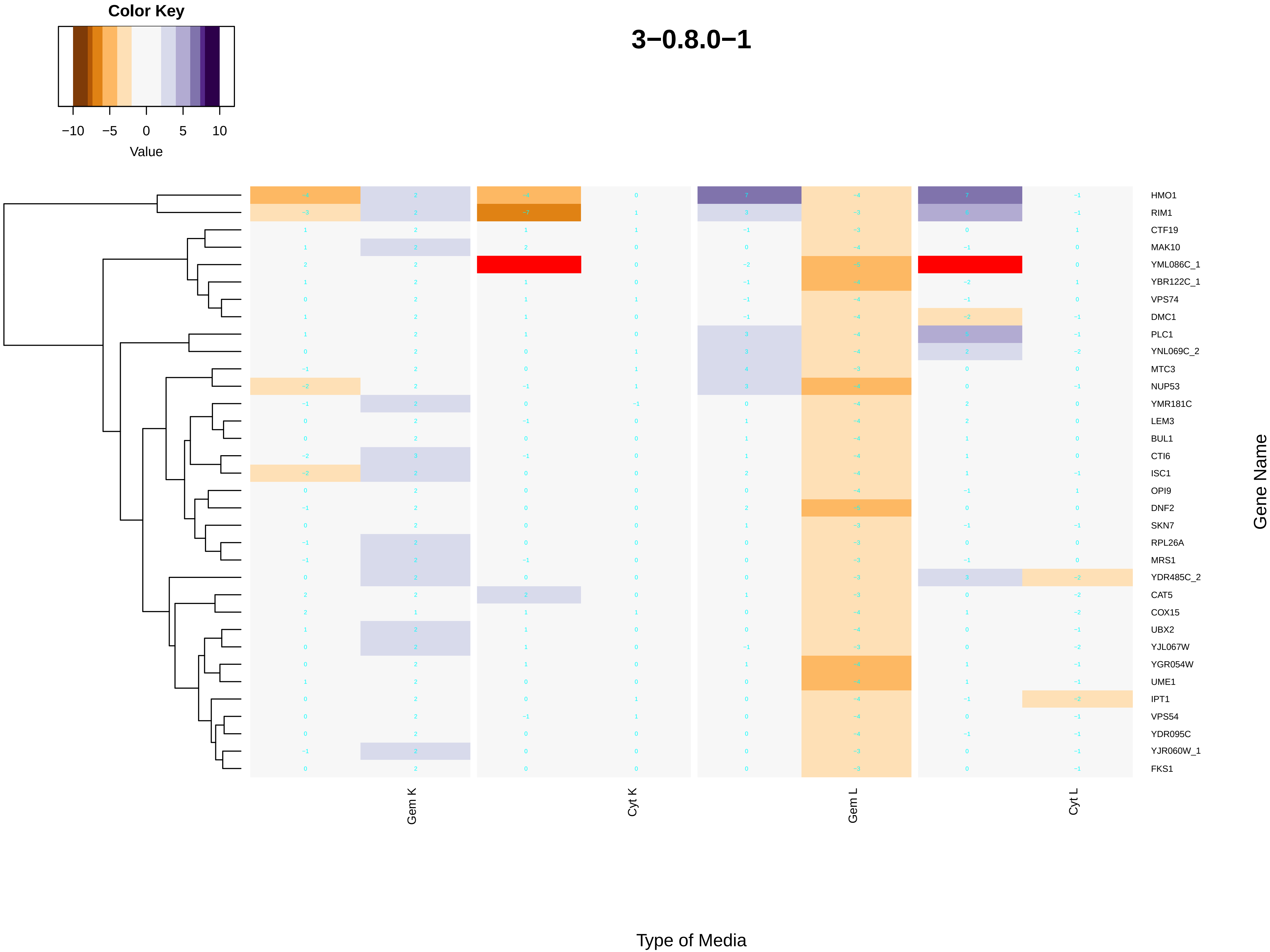
Color Key
−10 −5 0 5 10
Value
3−0.8.0−1
HMO1
RIM1
CTF19
MAK10
YML086C_1
YBR122C_1
VPS74
DMC1
PLC1
YNL069C_2
MTC3
NUP53
YMR181C
LEM3
BUL1
CTI6
ISC1
OPI9
DNF2
SKN7
RPL26A
MRS1
YDR485C_2
CAT5
COX15
UBX2
YJL067W
YGR054W
UME1
IPT1
VPS54
YDR095C
YJR060W_1
FKS1
Gene Name
Gem K
Cyt K
Gem L
Cyt L
Type of Media
