## Additional File 7 for "A humanized yeast phenomic model of deoxycytidine kinase to predict genetic buffering of nucleoside analog cytotoxicity": anion_transport.pdf

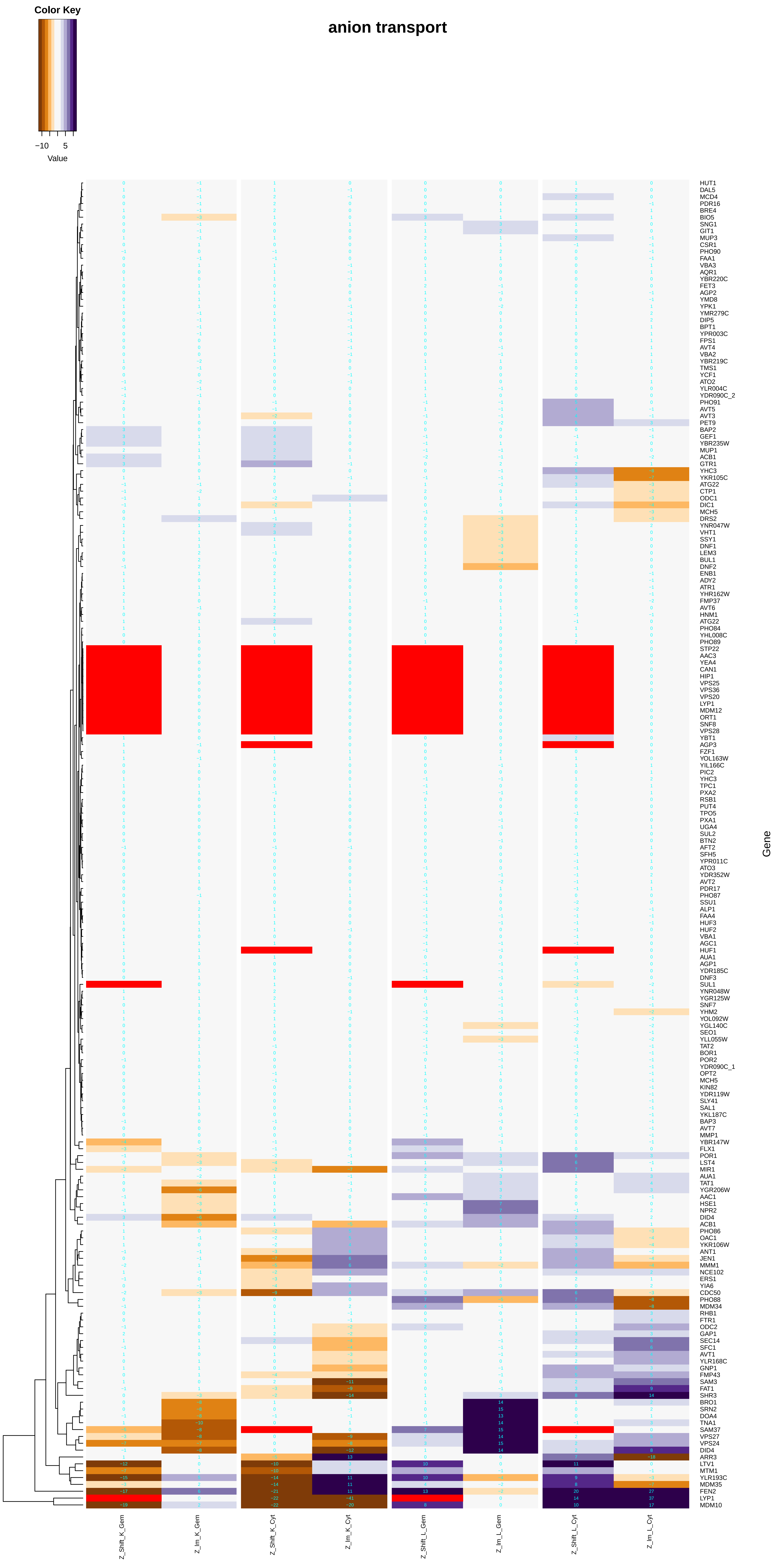
anion transport
Color Key
−10
5
Value
Gene
Z_Shift_K_Gem
Z_Im_K_Gem
Z_Shift_K_Cyt
Z_Im_K_Cyt
Z_Shift_L_Gem
Z_Im_L_Gem
Z_Shift_L_Cyt
Z_Im_L_Cyt
HUT1
DAL5
MCD4
PDR16
BRE4
BIO5
SNG1
GIT1
MUP3
CSR1
PHO90
FAA1
VBA3
AQR1
YBR220C
FET3
AGP2
YMD8
YPK1
YMR279C
DIP5
BPT1
YPR003C
FPS1
AVT4
VBA2
YBR219C
TMS1
YCF1
ATO2
YLR004C
YDR090C_2
PHO91
AVT5
AVT3
PET9
BAP2
GEF1
YBR235W
MUP1
ACB1
GTR1
YHC3
YKR105C
ATG22
CTP1
ODC1
DIC1
MCH5
DRS2
YNR047W
VHT1
SSY1
DNF1
LEM3
BUL1
DNF2
ENB1
ADY2
ATR1
YHR162W
FMP37
AVT6
HNM1
ATG22
PHO84
YHL008C
PHO89
STP22
AAC3
YEA4
CAN1
HIP1
VPS25
VPS36
VPS20
LYP1
MDM12
ORT1
SNF8
VPS28
YBT1
AGP3
FZF1
YOL163W
YIL166C
PIC2
YHC3
TPC1
PXA2
RSB1
PUT4
TPO5
PXA1
UGA4
SUL2
BTN2
AFT2
SFH5
YPR011C
ATO3
YDR352W
AVT2
PDR17
PHO87
SSU1
ALP1
FAA4
HUF3
HUF2
VBA1
AGC1
HUF1
AUA1
AGP1
YDR185C
DNF3
SUL1
YNR048W
YGR125W
SNF7
YHM2
YOL092W
YGL140C
SEO1
YLL055W
TAT2
BOR1
POR2
YDR090C_1
OPT2
MCH5
KIN82
YDR119W
SLY41
SAL1
YKL187C
BAP3
AVT7
MMP1
YBR147W
FLX1
POR1
LST4
MIR1
AUA1
TAT1
YGR206W
AAC1
HSE1
NPR2
DID4
ACB1
PHO86
OAC1
YKR106W
ANT1
JEN1
MMM1
NCE102
ERS1
YIA6
CDC50
PHO88
MDM34
RHB1
FTR1
ODC2
GAP1
SEC14
SFC1
AVT1
YLR168C
GNP1
FMP43
SAM3
FAT1
SHR3
BRO1
SRN2
DOA4
TNA1
SAM37
VPS27
VPS24
DID4
ARR3
LTV1
MTM1
YLR193C
MDM35
FEN2
LYP1
MDM10

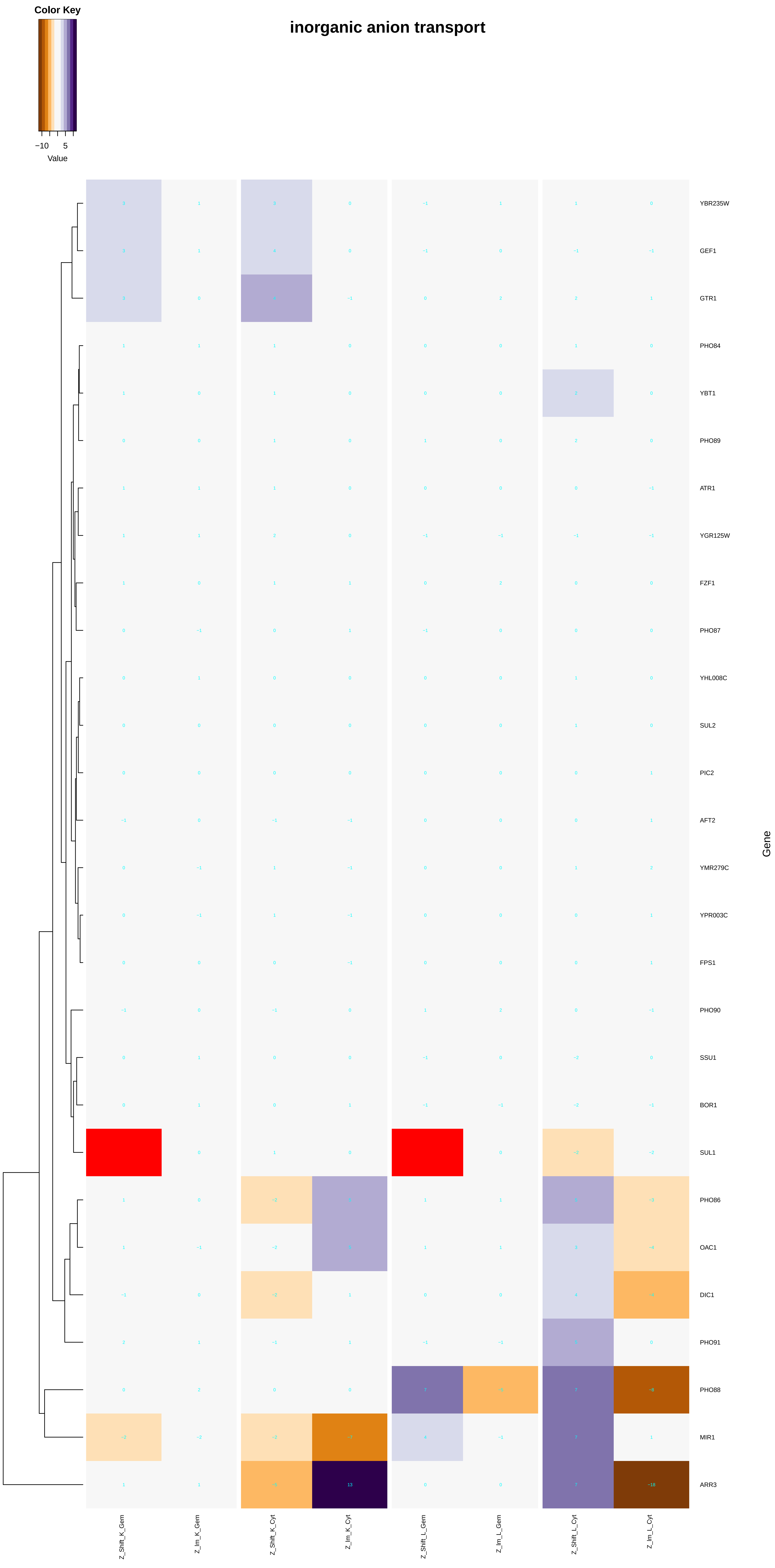
inorganic anion transport
Color Key
−10
5
Value
YBR235W
GEF1
GTR1
PHO84
YBT1
PHO89
ATR1
YGR125W
FZF1
PHO87
YHL008C
SUL2
PIC2
AFT2
YMR279C
YPR003C
FPS1
PHO90
SSU1
BOR1
SUL1
PHO86
OAC1
DIC1
PHO91
PHO88
MIR1
ARR3
Gene
Z_Shift_K_Gem
Z_lm_K_Gem
Z_Shift_K_Cyt
Z_lm_K_Cyt
Z_Shift_L_Gem
Z_lm_L_Gem
Z_Shift_L_Cyt
Z_lm_L_Cyt

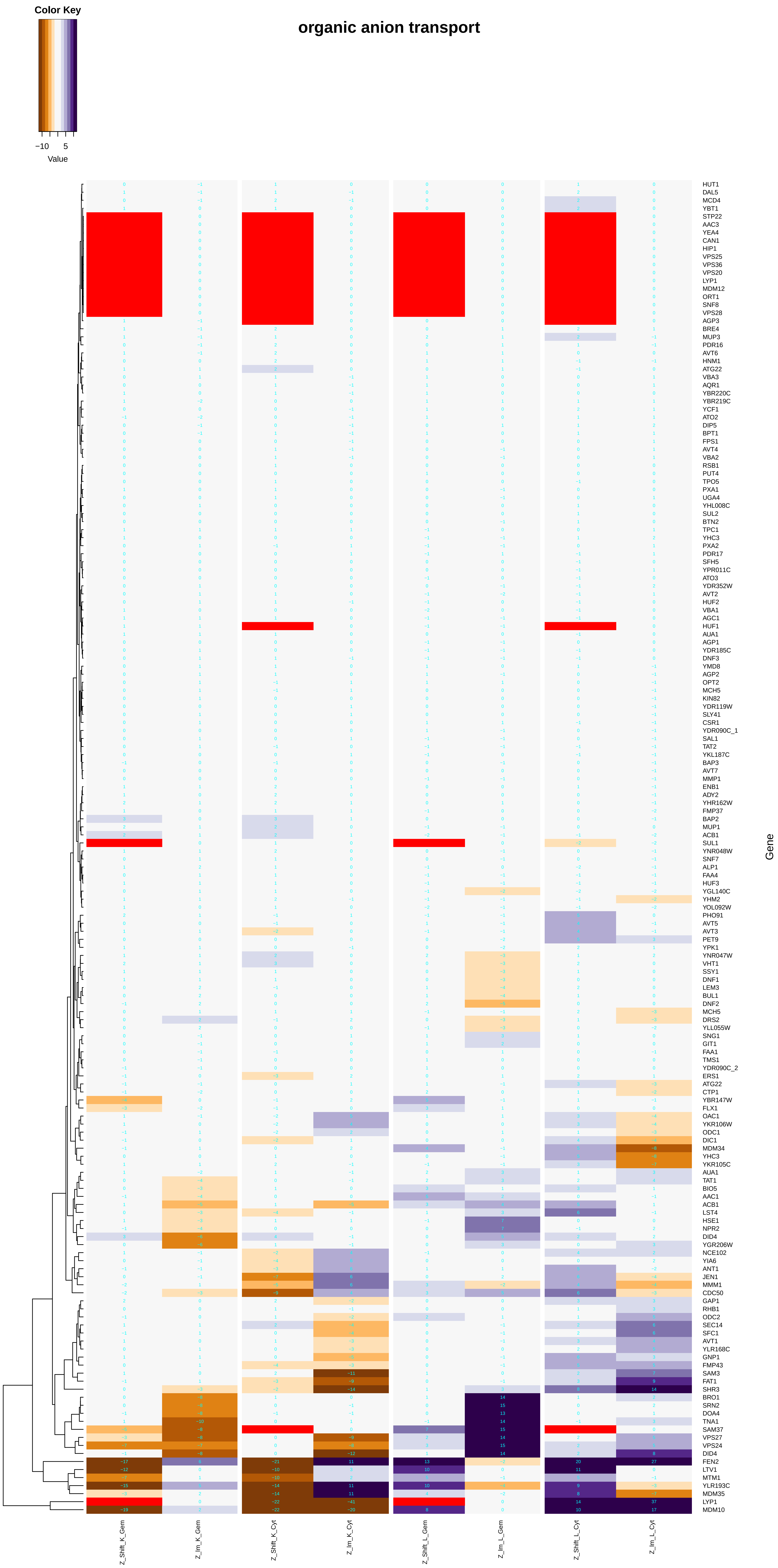
organic anion transport
Color Key
−10
5
Value
Gene
HUT1
DAL5
MCD4
YBT1
STP22
AAC3
YEA4
CAN1
HIP1
VPS25
VPS36
VPS20
LYP1
MDM12
ORT1
SNF8
VPS28
AGP3
BRE4
MUP3
PDR16
AVT6
HNM1
ATG22
VBA3
AQR1
YBR220C
YBR219C
YCF1
ATO2
DIP5
BPT1
FPS1
AVT4
VBA2
RSB1
PUT4
TPO5
PXA1
UGA4
YHL008C
SUL2
BTN2
TPC1
YHC3
PXA2
PDR17
SFH5
YPR011C
ATO3
YDR352W
AVT2
HUF2
VBA1
AGC1
HUF1
AUA1
AGP1
YDR185C
DNF3
YMD8
AGP2
OPT2
MCH5
KIN82
YDR119W
SLY41
CSR1
YDR090C_1
SAL1
TAT2
YKL187C
BAP3
AVT7
MMP1
ENB1
ADY2
YHR162W
FMP37
BAP2
MUP1
ACB1
SUL1
YNR048W
SNF7
ALP1
FAA4
HUF3
YGL140C
YHM2
YOL092W
PHO91
AVT5
AVT3
PET9
YPK1
YNR047W
VHT1
SSY1
DNF1
LEM3
BUL1
DNF2
MCH5
DRS2
YLL055W
SNG1
GIT1
FAA1
TMS1
YDR090C_2
ERS1
ATG22
CTP1
YBR147W
FLX1
OAC1
YKR106W
ODC1
DIC1
MDM34
YHC3
YKR105C
AUA1
TAT1
BIO5
AAC1
ACB1
LST4
HSE1
NPR2
DID4
YGR206W
NCE102
YIA6
ANT1
JEN1
MMM1
CDC50
GAP1
RHB1
ODC2
SEC14
SFC1
AVT1
YLR168C
GNP1
FMP43
SAM3
FAT1
SHR3
BRO1
SRN2
DOA4
TNA1
SAM37
VPS27
VPS24
DID4
FEN2
LTV1
MTM1
YLR193C
MDM35
LYP1
MDM10
Z_Shift_K_Gem
Z_Im_K_Gem
Z_Shift_K_Cyt
Z_Im_K_Cyt
Z_Shift_L_Gem
Z_Im_L_Gem
Z_Shift_L_Cyt
Z_Im_L_Cyt

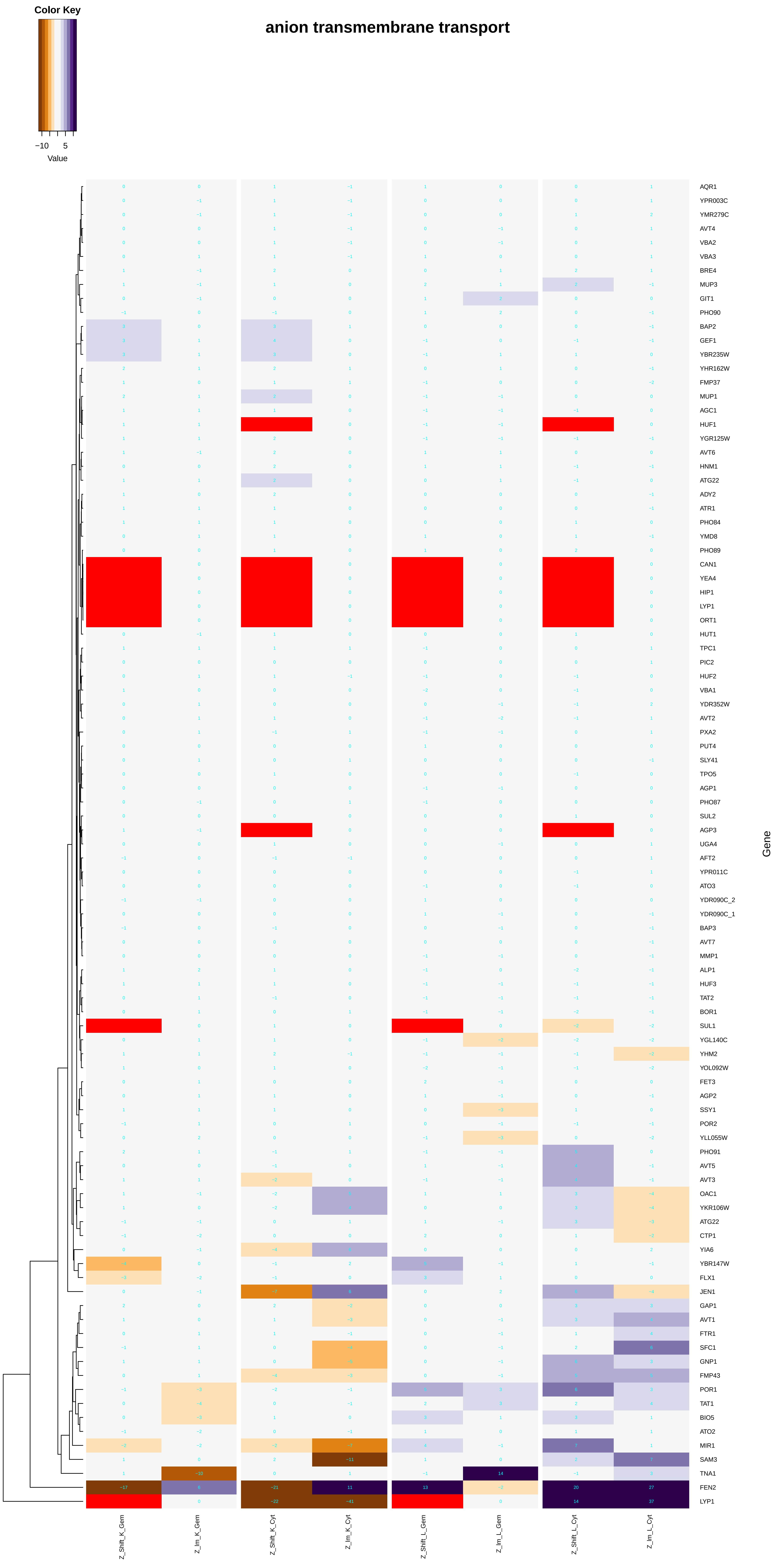

Color Key
−10
5
Value
anion transmembrane transport
AQR1
YPR003C
YMR279C
AVT4
VBA2
VBA3
BRE4
MUP3
GIT1
PHO90
BAP2
GEF1
YBR235W
YHR162W
FMP37
MUP1
AGC1
HUF1
YGR125W
AVT6
HNM1
ATG22
ADY2
ATR1
PHO84
YMD8
PHO89
CAN1
YEA4
HIP1
LYP1
ORT1
HUT1
TPC1
PIC2
HUF2
VBA1
YDR352W
AVT2
PXA2
PUT4
SLY41
TPO5
AGP1
PHO87
SUL2
AGP3
UGA4
AFT2
YPR011C
ATO3
YDR090C_2
YDR090C_1
BAP3
AVT7
MMP1
ALP1
HUF3
TAT2
BOR1
SUL1
YGL140C
YHM2
YOL092W
FET3
AGP2
SSY1
POR2
YLL055W
PHO91
AVT5
AVT3
OAC1
YKR106W
ATG22
CTP1
YIA6
YBR147W
FLX1
JEN1
GAP1
AVT1
FTR1
SFC1
GNP1
FMP43
POR1
TAT1
BIO5
ATO2
MIR1
SAM3
TNA1
FEN2
LYP1
Gene
Z_Shift_K_Gem
Z_Im_K_Gem
Z_Shift_K_Cyt
Z_Im_K_Cyt
Z_Shift_L_Gem
Z_Im_L_Gem
Z_Shift_L_Cyt
Z_Im_L_Cyt

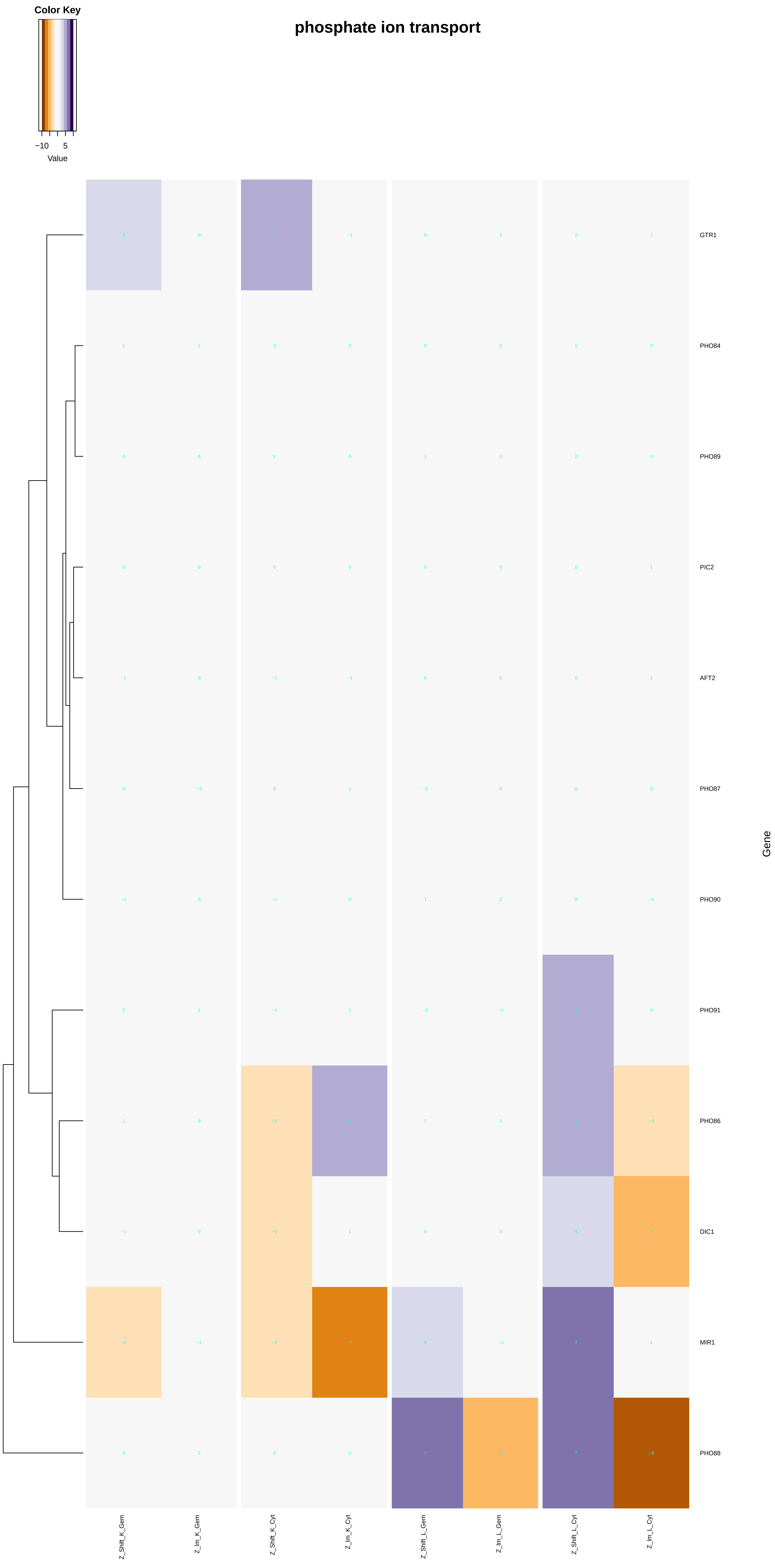
phosphate ion transport
Color Key
−10
5
Value
GTR1
PHO84
PHO89
PIC2
AFT2
PHO87
PHO90
PHO91
PHO86
DIC1
MIR1
PHO88
Gene
Z_Shift_K_Gem
Z_Im_K_Gem
Z_Shift_K_Cyt
Z_Im_K_Cyt
Z_Shift_L_Gem
Z_Im_L_Gem
Z_Shift_L_Cyt
Z_Im_L_Cyt

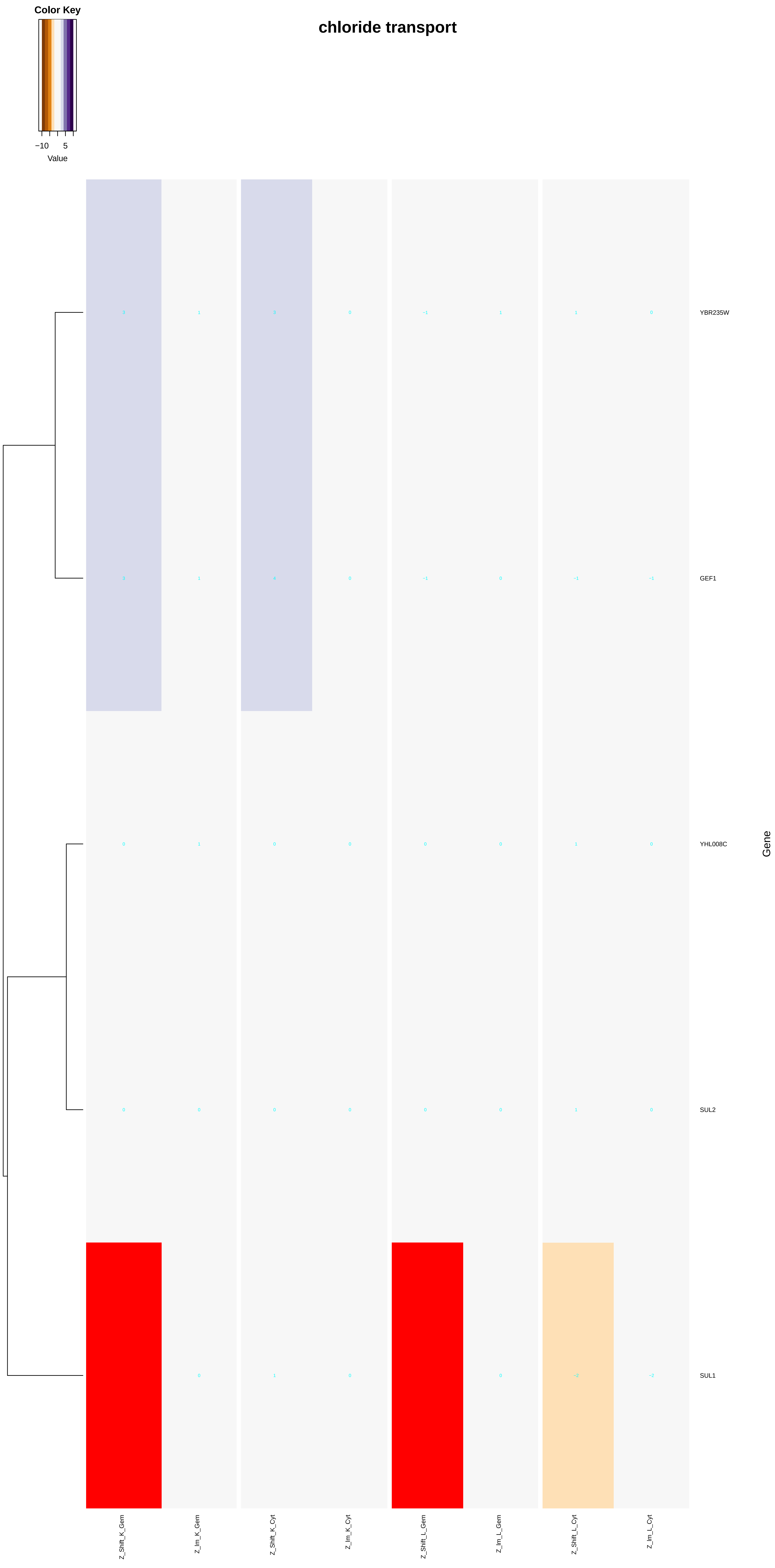

Color Key
−10
5
Value
chloride transport
YBR235W
GEF1
YHL008C
SUL2
SUL1
Gene
Z_Shift_K_Gem
Z_Im_K_Gem
Z_Shift_K_Cyt
Z_Im_K_Cyt
Z_Shift_L_Gem
Z_Im_L_Gem
Z_Shift_L_Cyt
Z_Im_L_Cyt

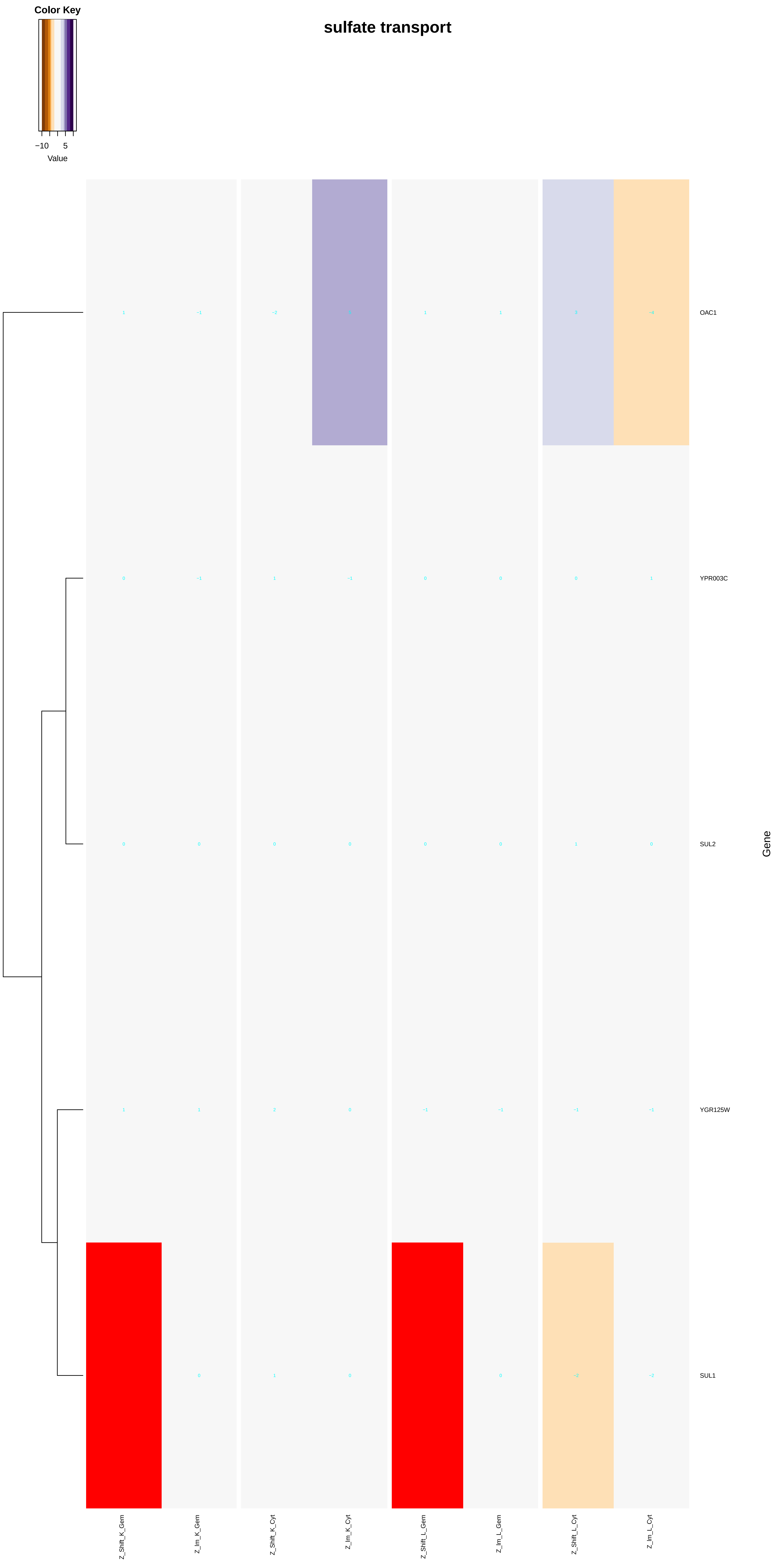
Color Key
−10
5
Value
sulfate transport
OAC1
YPR003C
SUL2
YGR125W
SUL1
Gene
Z_Shift_K_Gem
Z_Im_K_Gem
Z_Shift_K_Cyt
Z_Im_K_Cyt
Z_Shift_L_Gem
Z_Im_L_Gem
Z_Shift_L_Cyt
Z_Im_L_Cyt

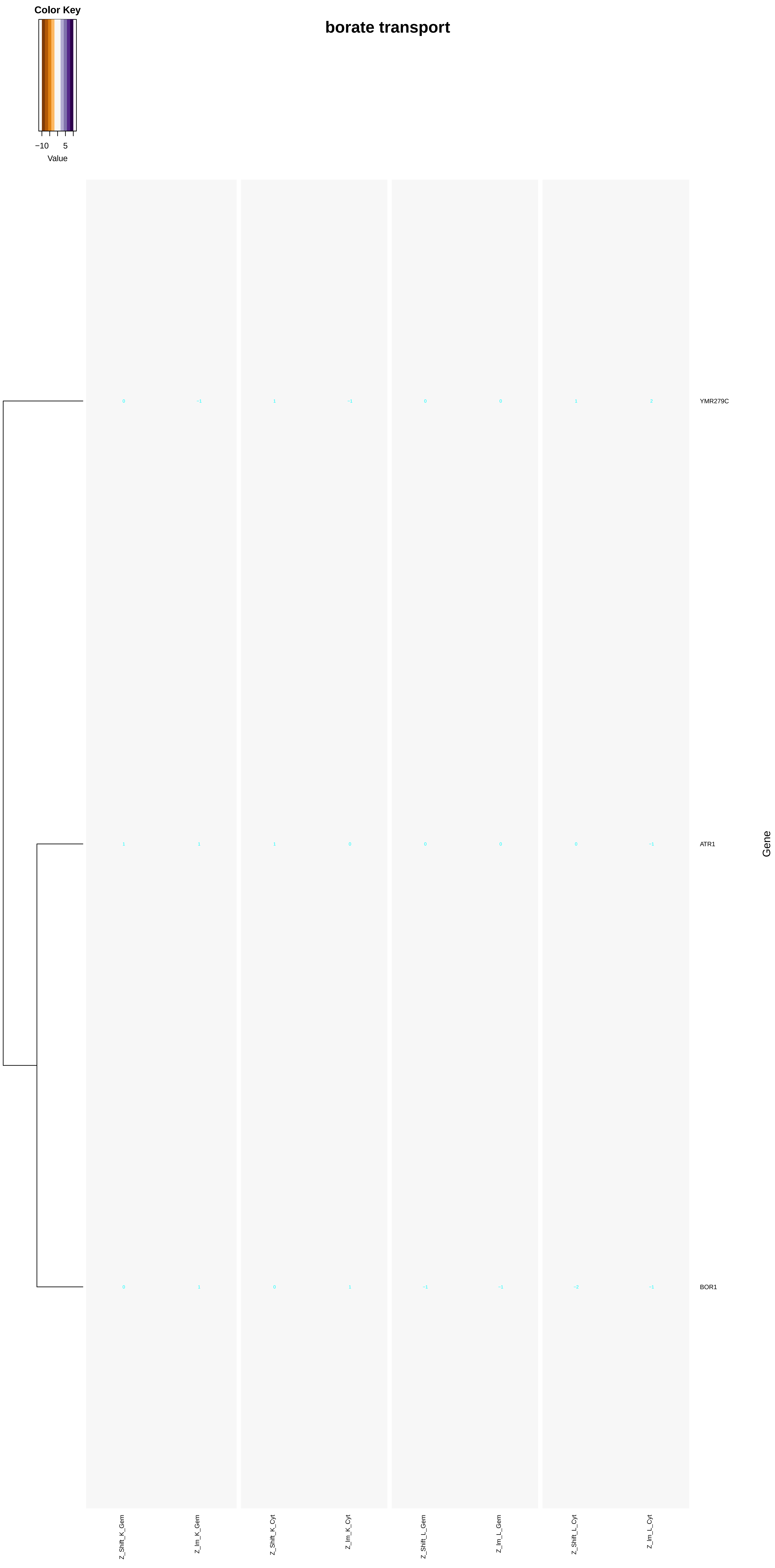

borate transport
Color Key
−10
5
Value
YMR279C
ATR1
BOR1
Gene
Z_Shift_K_Gem
Z_Im_K_Gem
Z_Shift_K_Cyt
Z_Im_K_Cyt
Z_Shift_L_Gem
Z_Im_L_Gem
Z_Shift_L_Cyt
Z_Im_L_Cyt

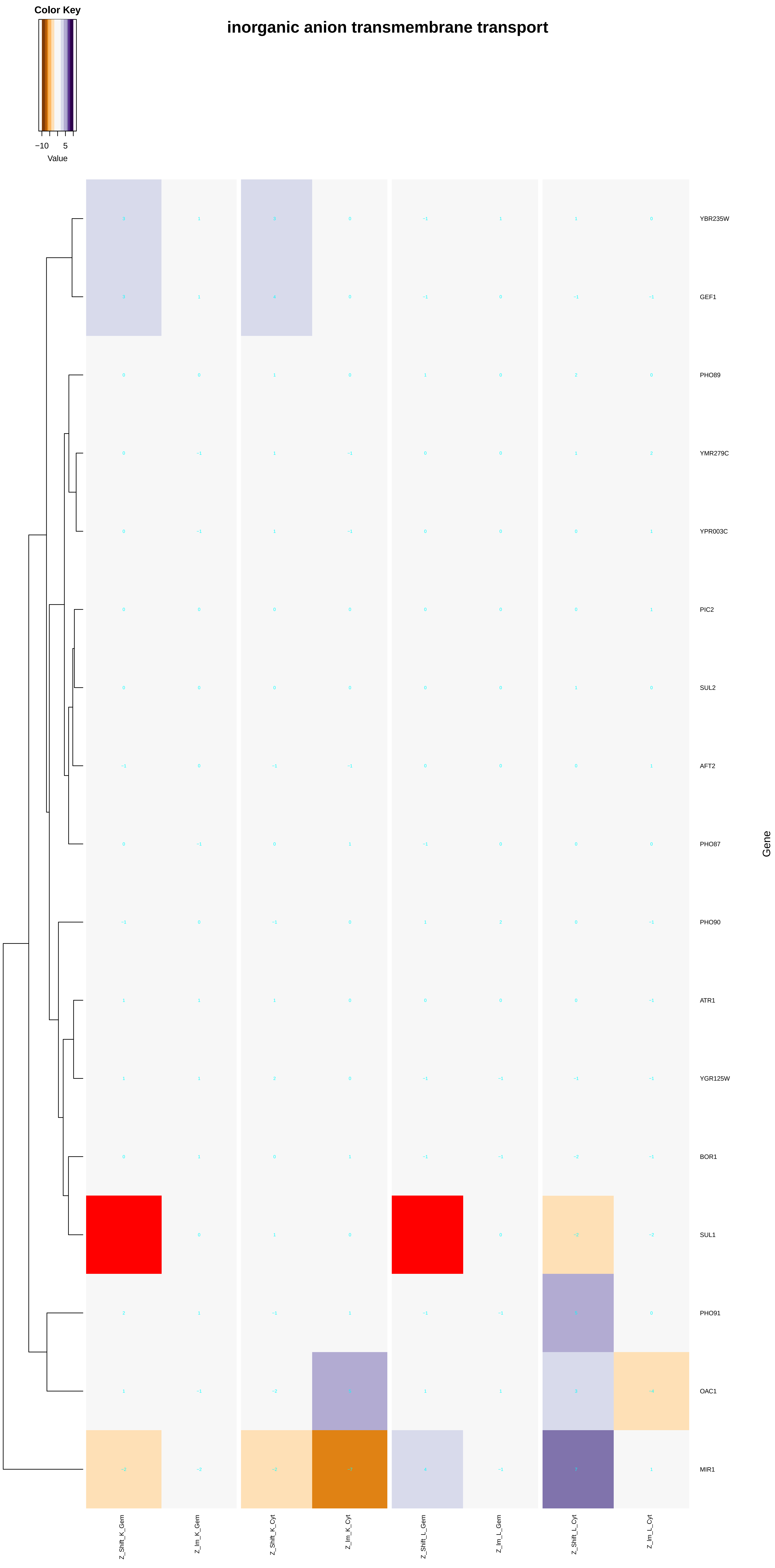

Color Key
−10
5
Value
inorganic anion transmembrane transport
YBR235W
GEF1
PHO89
YMR279C
YPR003C
PIC2
SUL2
AFT2
PHO87
PHO90
ATR1
YGR125W
BOR1
SUL1
PHO91
OAC1
MIR1
Gene
Z_Shift_K_Gem
Z_Im_K_Gem
Z_Shift_K_Cyt
Z_Im_K_Cyt
Z_Shift_L_Gem
Z_Im_L_Gem
Z_Shift_L_Cyt
Z_Im_L_Cyt

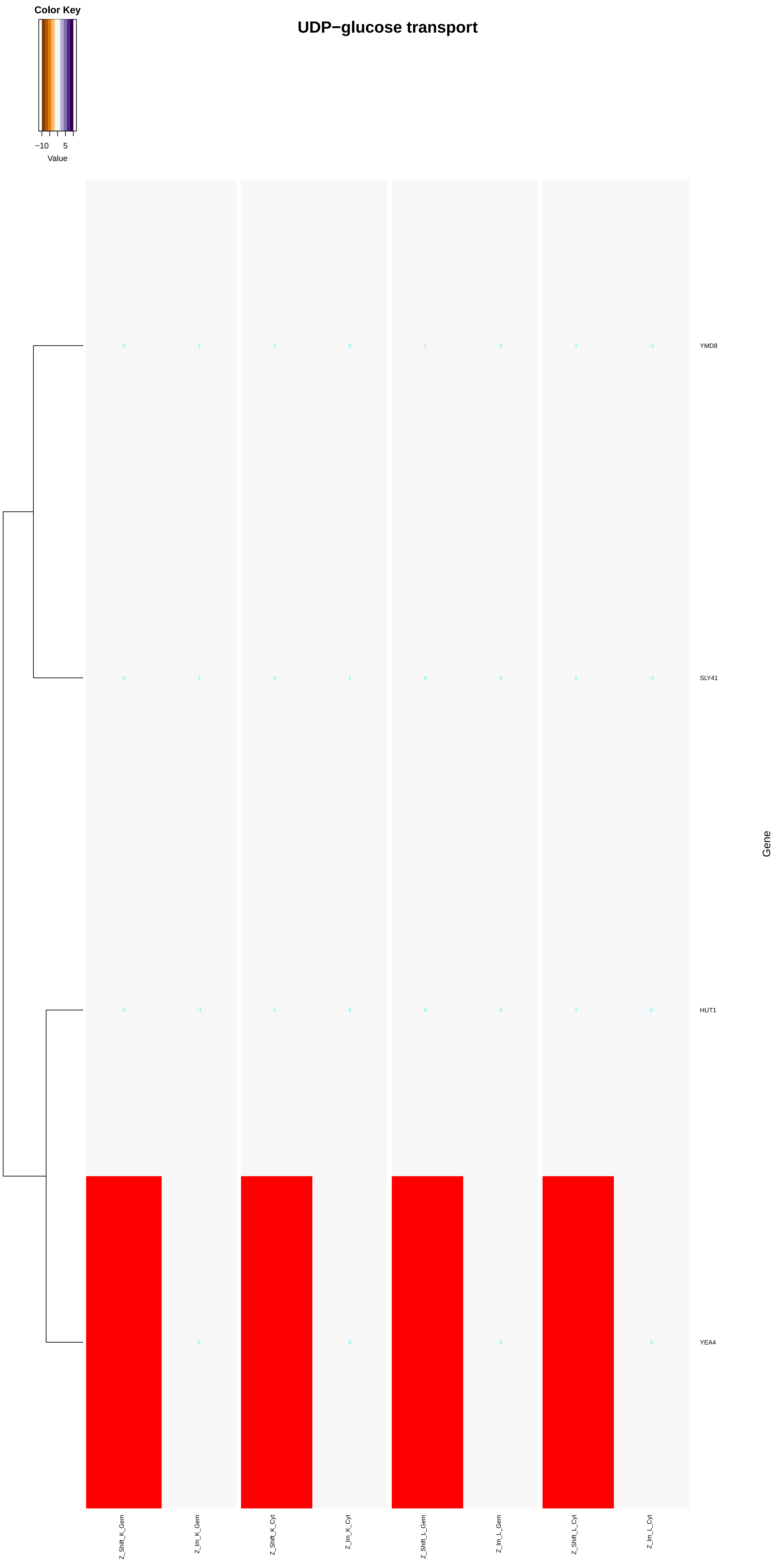
Color Key
−10 5
Value
UDP−glucose transport
YMD8
SLY41
HUT1
YEA4
Gene
Z_Shift_K_Gem
Z_Im_K_Gem
Z_Shift_K_Cyt
Z_Im_K_Cyt
Z_Shift_L_Gem
Z_Im_L_Gem
Z_Shift_L_Cyt
Z_Im_L_Cyt

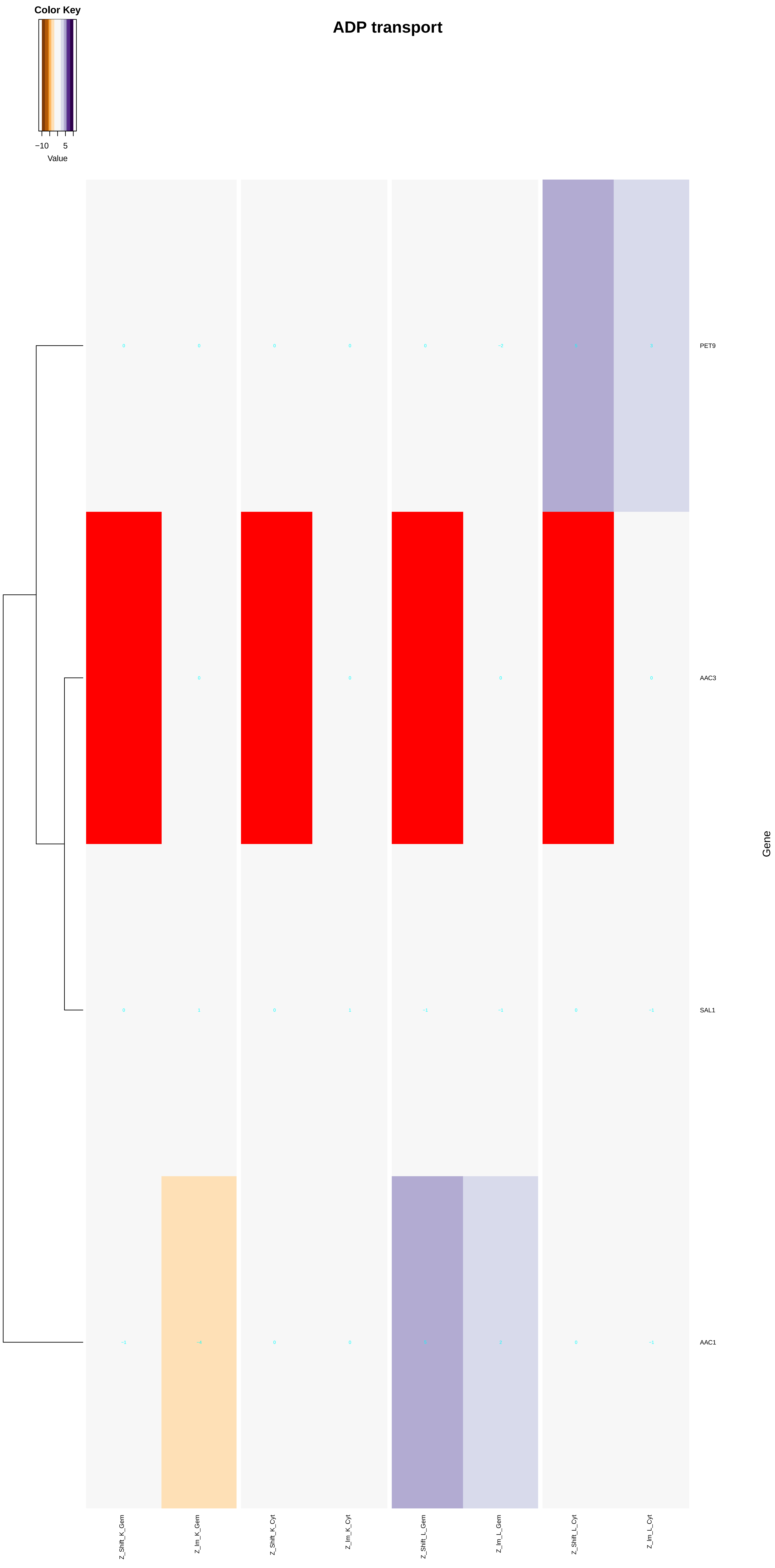
Color Key
−10
5
Value
ADP transport
PET9
AAC3
SAL1
AAC1
Gene
Z_Shift_K_Gem
Z_Im_K_Gem
Z_Shift_K_Cyt
Z_Im_K_Cyt
Z_Shift_L_Gem
Z_Im_L_Gem
Z_Shift_L_Cyt
Z_Im_L_Cyt

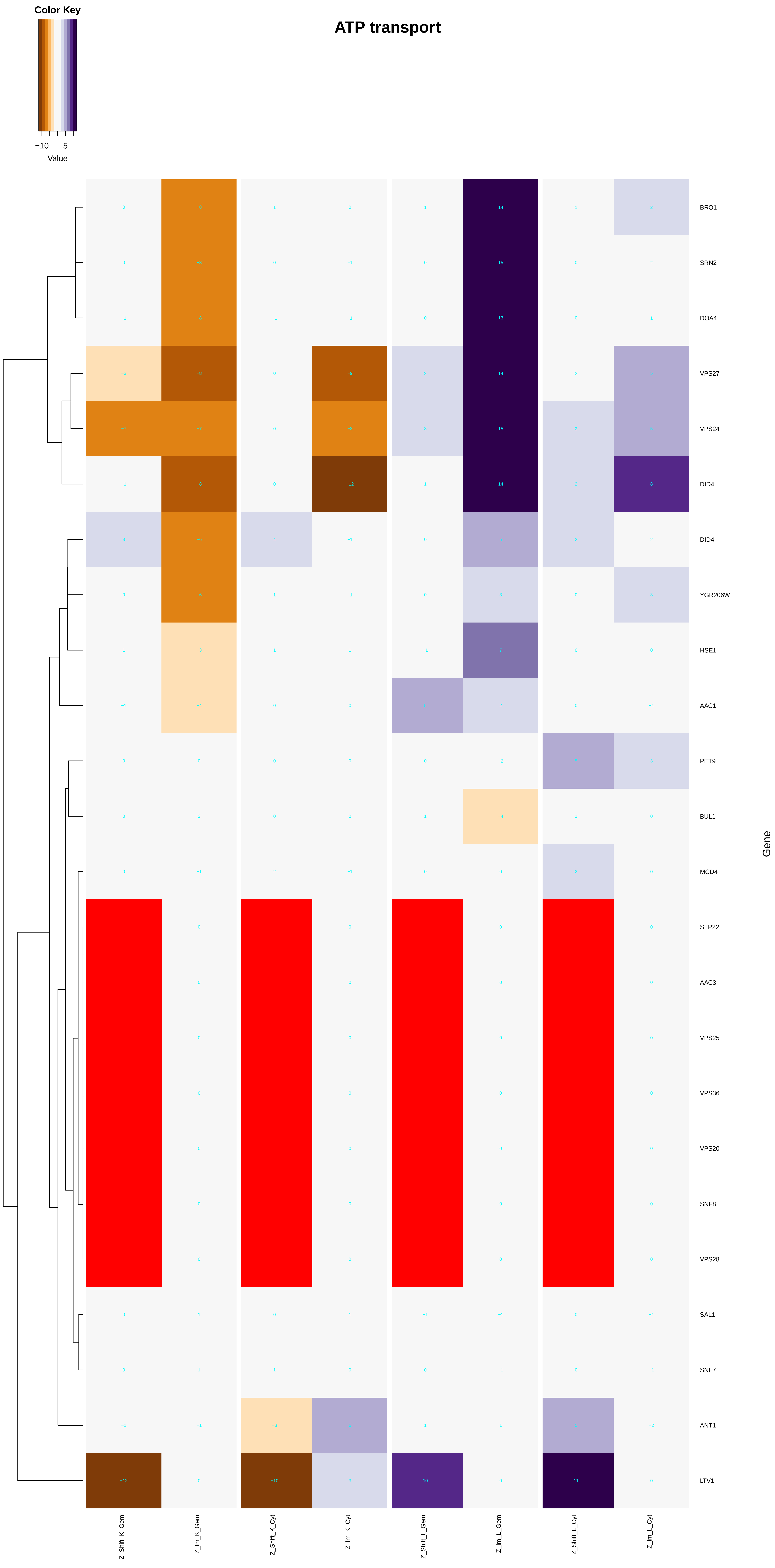

Color Key
−10
5
Value
ATP transport
BRO1
SRN2
DOA4
VPS27
VPS24
DID4
DID4
YGR206W
HSE1
AAC1
PET9
BUL1
MCD4
STP22
AAC3
VPS25
VPS36
VPS20
SNF8
VPS28
SAL1
SNF7
ANT1
LTV1
Gene
Z_Shift_K_Gem
Z_Im_K_Gem
Z_Shift_K_Cyt
Z_Im_K_Cyt
Z_Shift_L_Gem
Z_Im_L_Gem
Z_Shift_L_Cyt
Z_Im_L_Cyt

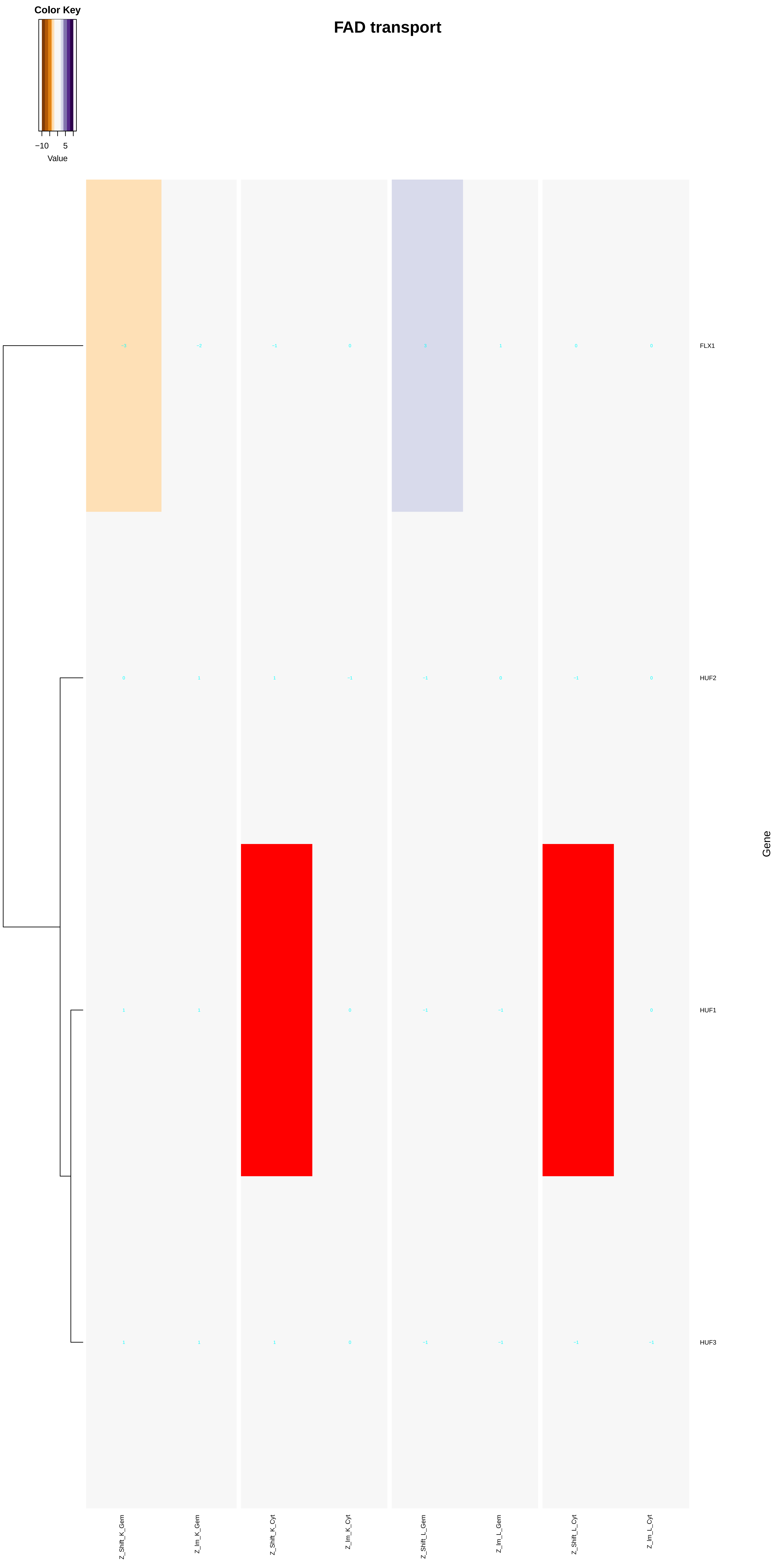
FAD transport
Color Key
−10
5
Value
FLX1
HUF2
HUF1
HUF3
Gene
Z_Shift_K_Gem
Z_Im_K_Gem
Z_Shift_K_Cyt
Z_Im_K_Cyt
Z_Shift_L_Gem
Z_Im_L_Gem
Z_Shift_L_Cyt
Z_Im_L_Cyt

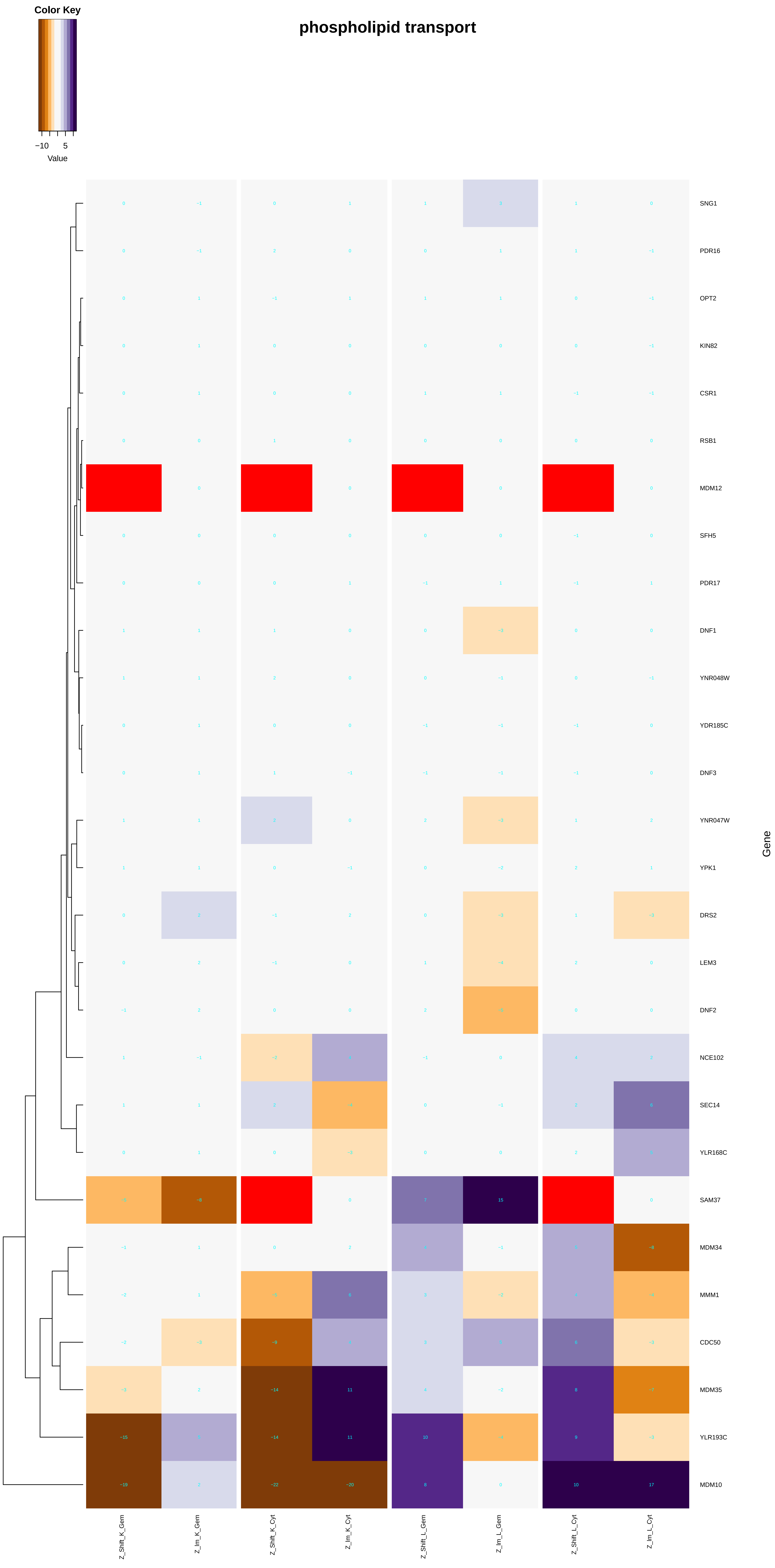
Color Key
−10
5
Value
phospholipid transport
Gene
SNG1
PDR16
OPT2
KIN82
CSR1
RSB1
MDM12
SFH5
PDR17
DNF1
YNR048W
YDR185C
DNF3
YNR047W
YPK1
DRS2
LEM3
DNF2
NCE102
SEC14
YLR168C
SAM37
MDM34
MMM1
CDC50
MDM35
YLR193C
MDM10
Z_Shift_K_Gem
Z_Im_K_Gem
Z_Shift_K_Cyt
Z_Im_K_Cyt
Z_Shift_L_Gem
Z_Im_L_Gem
Z_Shift_L_Cyt
Z_Im_L_Cyt

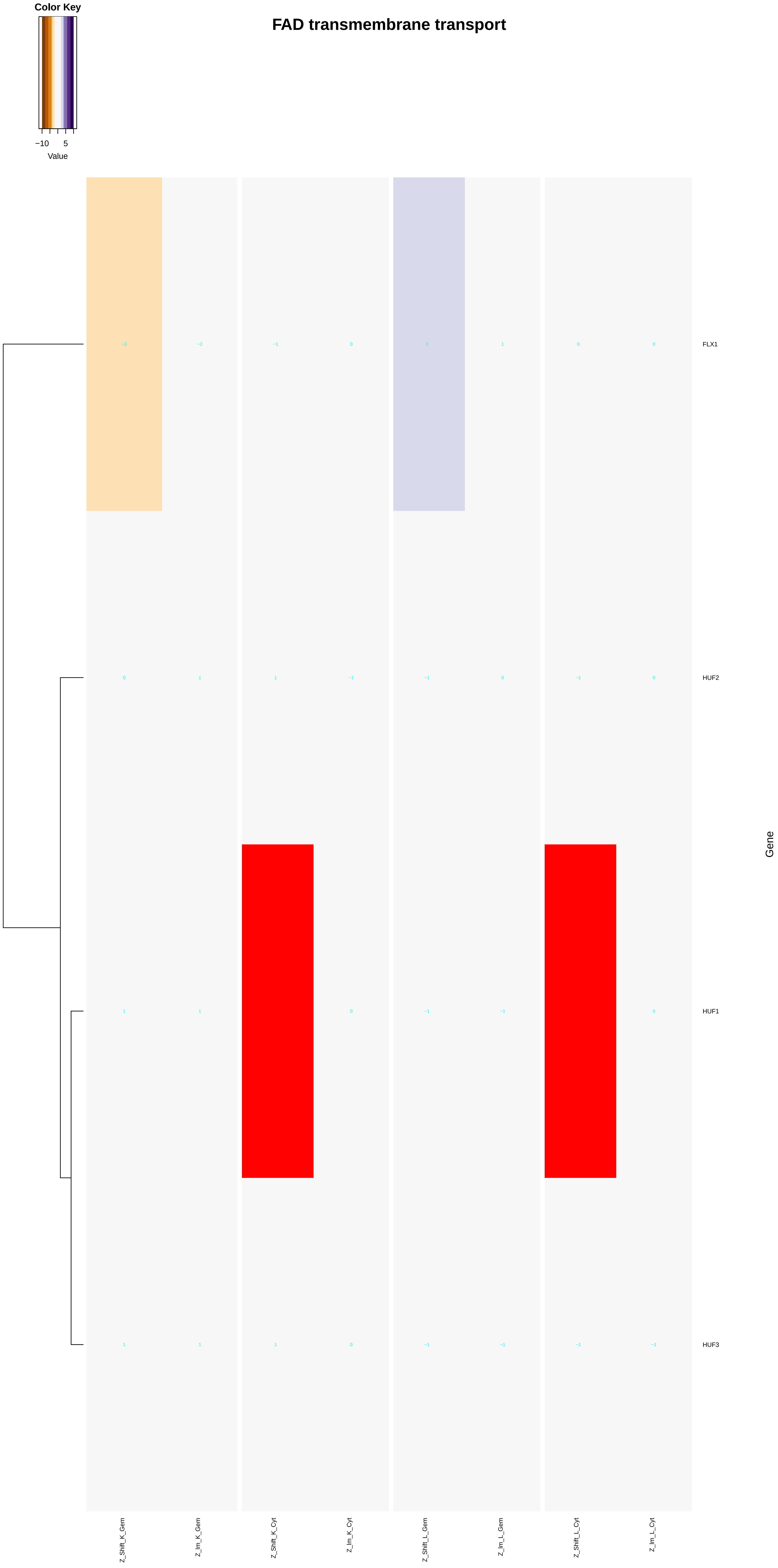
FAD transmembrane transport
Color Key
−10
5
Value
FLX1
HUF2
HUF1
HUF3
Gene
Z_Shift_K_Gem
Z_Im_K_Gem
Z_Shift_K_Cyt
Z_Im_K_Cyt
Z_Shift_L_Gem
Z_Im_L_Gem
Z_Shift_L_Cyt
Z_Im_L_Cyt

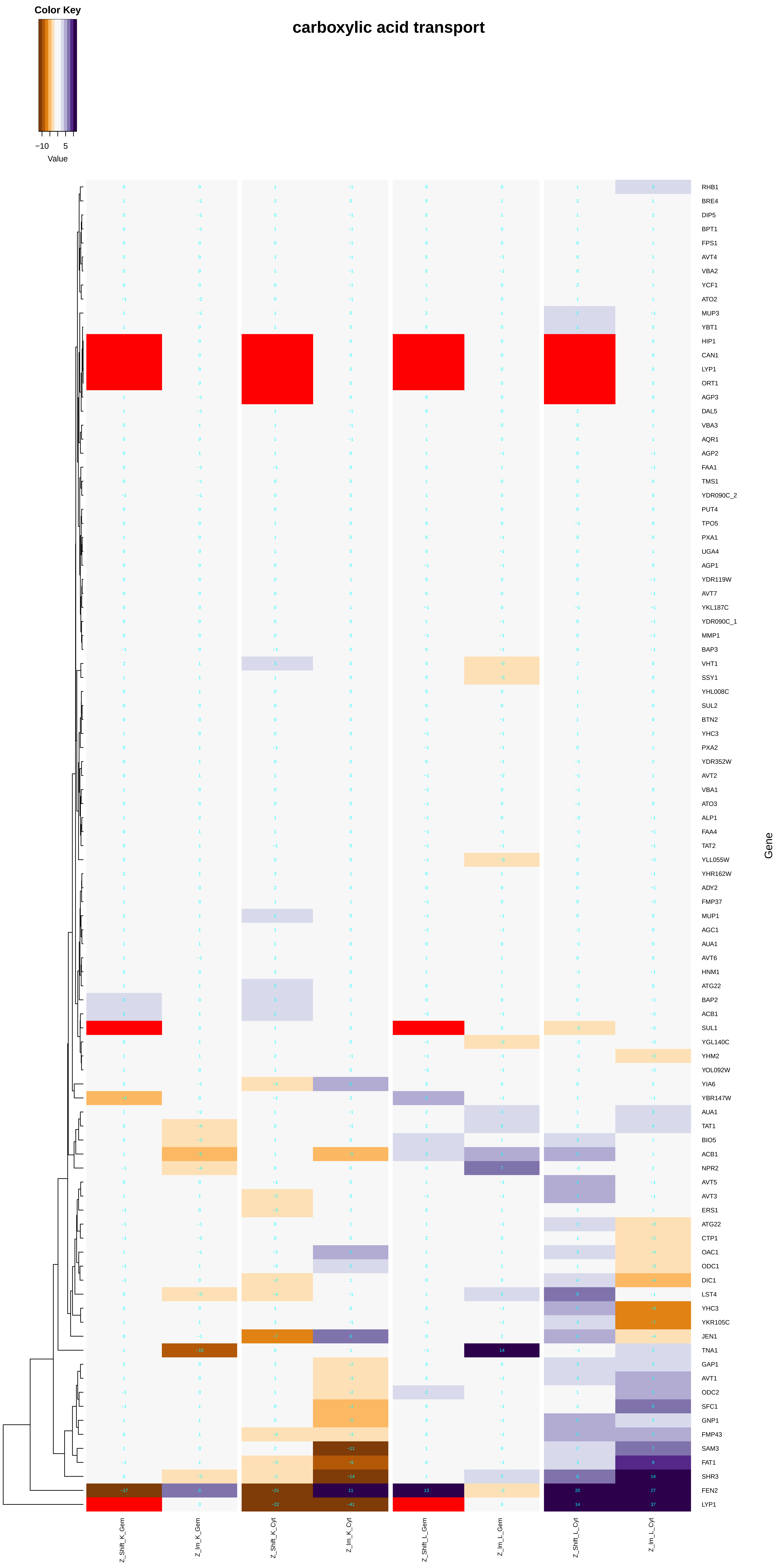
carboxylic acid transport
Color Key
−10
5
Value
Gene
RHB1
BRE4
DIP5
BPT1
FPS1
AVT4
VBA2
YCF1
ATO2
MUP3
YBT1
HIP1
CAN1
LYP1
ORT1
AGP3
DAL5
VBA3
AQR1
AGP2
FAA1
TMS1
YDR090C_2
PUT4
TPO5
PXA1
UGA4
AGP1
YDR119W
AVT7
YKL187C
YDR090C_1
MMP1
BAP3
VHT1
SSY1
YHL008C
SUL2
BTN2
YHC3
PXA2
YDR352W
AVT2
VBA1
ATO3
ALP1
FAA4
TAT2
YLL055W
YHR162W
ADY2
FMP37
MUP1
AGC1
AUA1
AVT6
HNM1
ATG22
BAP2
ACB1
SUL1
YGL140C
YHM2
YOL092W
YIA6
YBR147W
AUA1
TAT1
BIO5
ACB1
NPR2
AVT5
AVT3
ERS1
ATG22
CTP1
OAC1
ODC1
DIC1
LST4
YHC3
YKR105C
JEN1
TNA1
GAP1
AVT1
ODC2
SFC1
GNP1
FMP43
SAM3
FAT1
SHR3
FEN2
LYP1
Z_Shift_K_Gem
Z_Im_K_Gem
Z_Shift_K_Cyt
Z_Im_K_Cyt
Z_Shift_L_Gem
Z_Im_L_Gem
Z_Shift_L_Cyt
Z_Im_L_Cyt

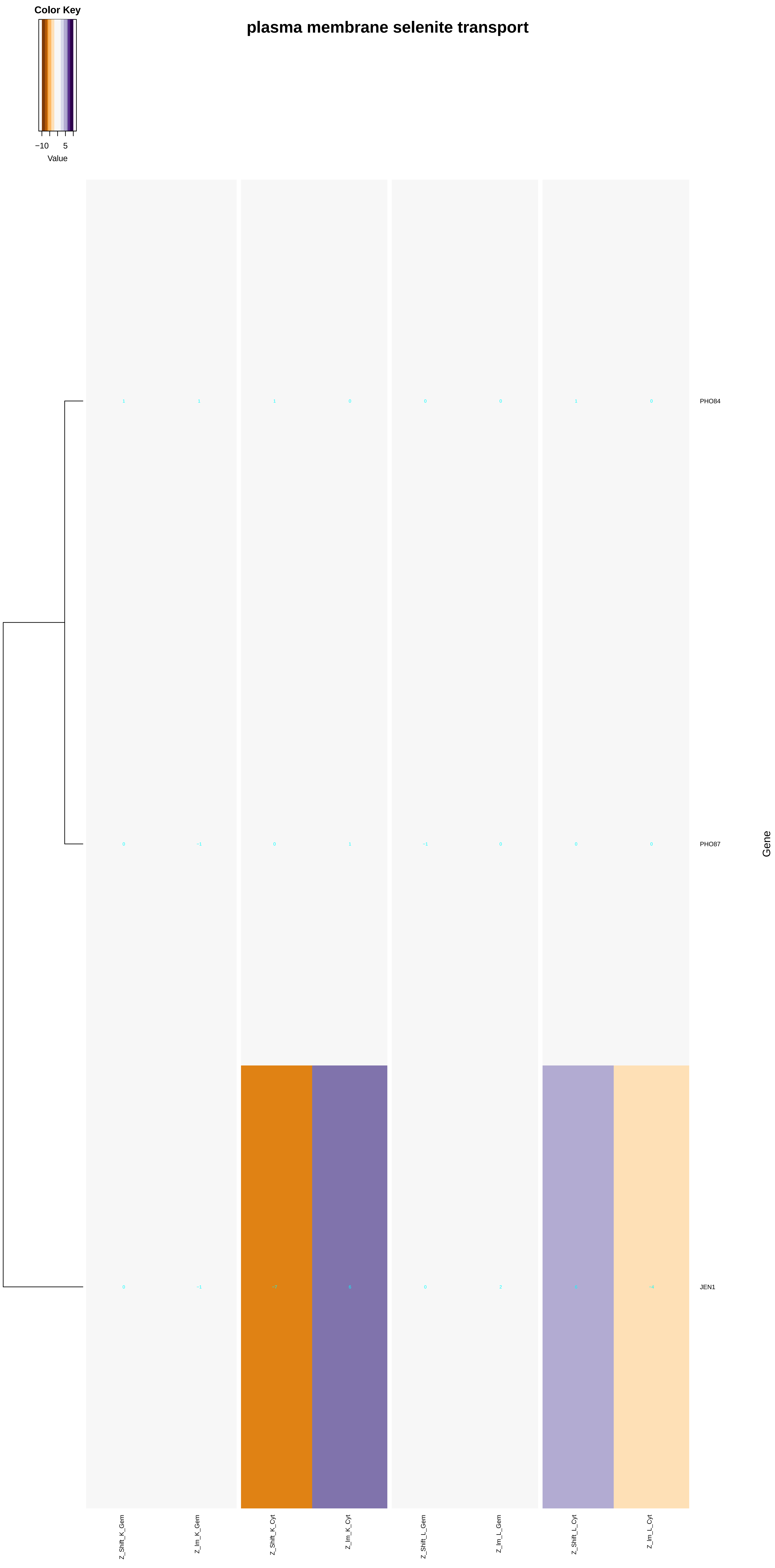
Color Key
−10
5
Value
plasma membrane selenite transport
PHO84
PHO87
JEN1
Gene
Z_Shift_K_Gem
Z_Im_K_Gem
Z_Shift_K_Cyt
Z_Im_K_Cyt
Z_Shift_L_Gem
Z_Im_L_Gem
Z_Shift_L_Cyt
Z_Im_L_Cyt

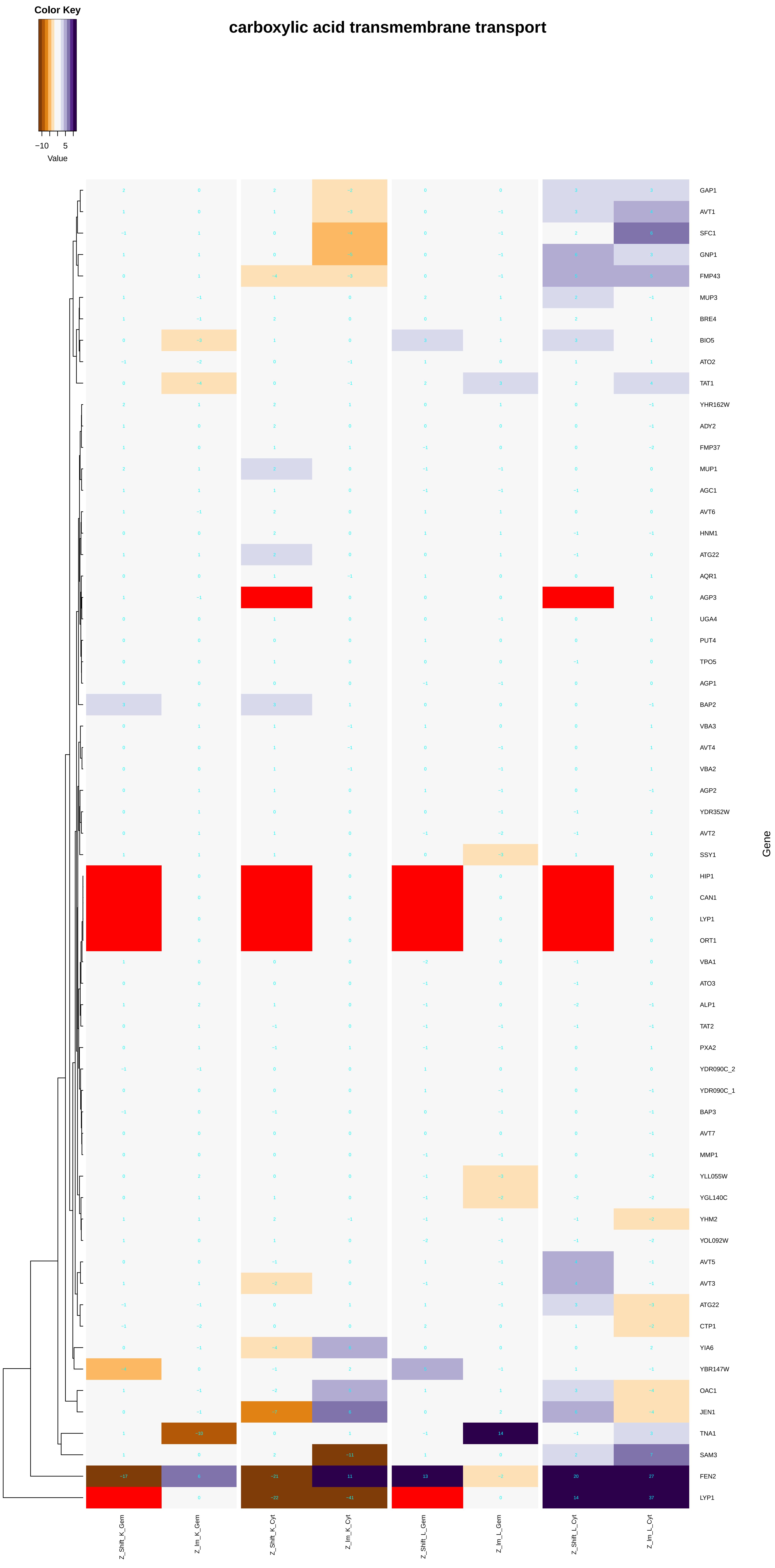

Color Key
−10
5
Value
carboxylic acid transmembrane transport
GAP1
AVT1
SFC1
GNP1
FMP43
MUP3
BRE4
BIO5
ATO2
TAT1
YHR162W
ADY2
FMP37
MUP1
AGC1
AVT6
HNM1
ATG22
AQR1
AGP3
UGA4
PUT4
TPO5
AGP1
BAP2
VBA3
AVT4
VBA2
AGP2
YDR352W
AVT2
SSY1
HIP1
CAN1
LYP1
ORT1
VBA1
ATO3
ALP1
TAT2
PXA2
YDR090C_2
YDR090C_1
BAP3
AVT7
MMP1
YLL055W
YGL140C
YHM2
YOL092W
AVT5
AVT3
ATG22
CTP1
YIA6
YBR147W
OAC1
JEN1
TNA1
SAM3
FEN2
LYP1
Gene
Z_Shift_K_Gem
Z_Im_K_Gem
Z_Shift_K_Cyt
Z_Im_K_Cyt
Z_Shift_L_Gem
Z_Im_L_Gem
Z_Shift_L_Cyt
Z_Im_L_Cyt

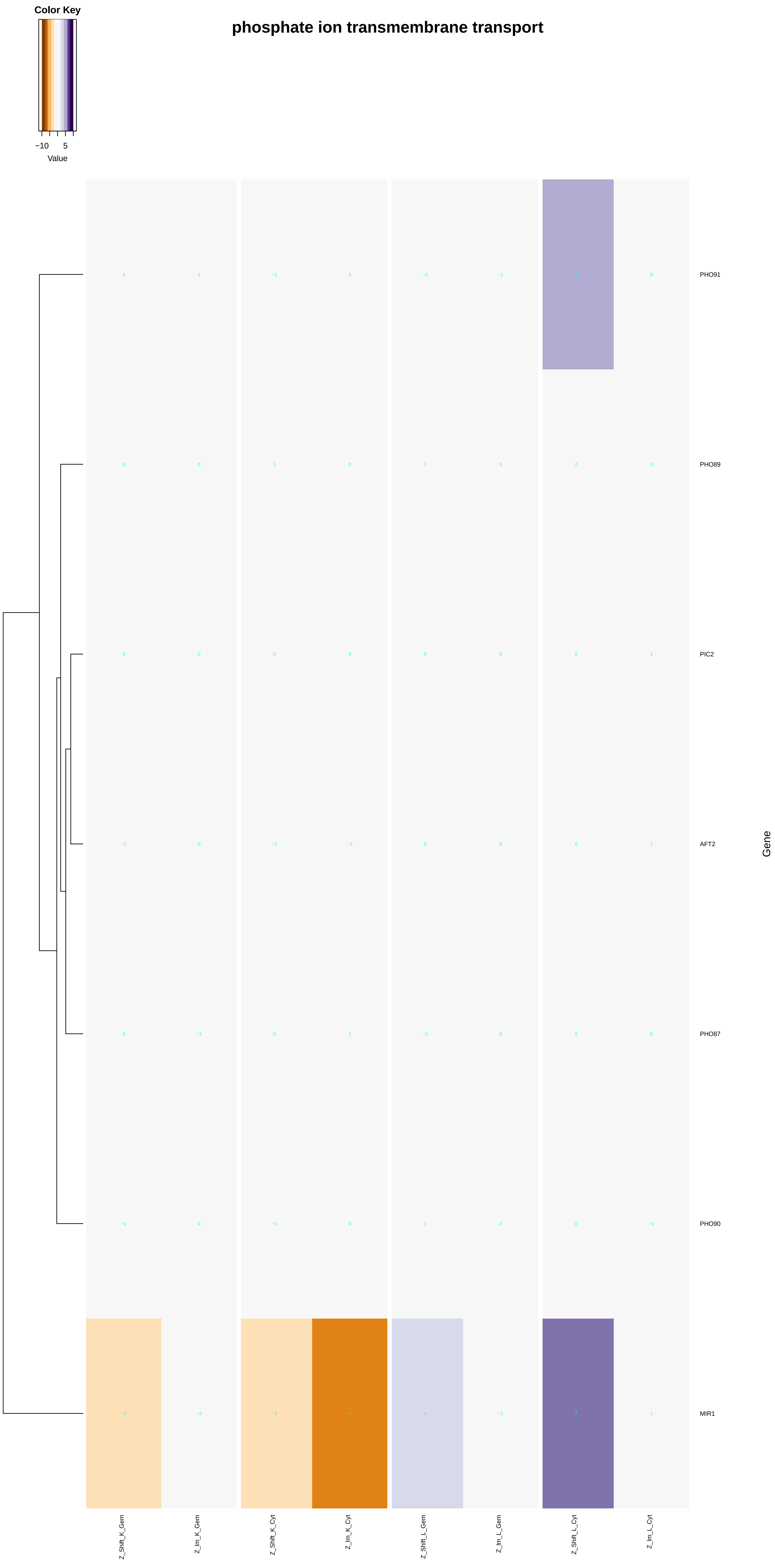
Color Key
−10
5
Value
phosphate ion transmembrane transport
PHO91
PHO89
PIC2
AFT2
PHO87
PHO90
MIR1
Gene
Z_Shift_K_Gem
Z_Im_K_Gem
Z_Shift_K_Cyt
Z_Im_K_Cyt
Z_Shift_L_Gem
Z_Im_L_Gem
Z_Shift_L_Cyt
Z_Im_L_Cyt

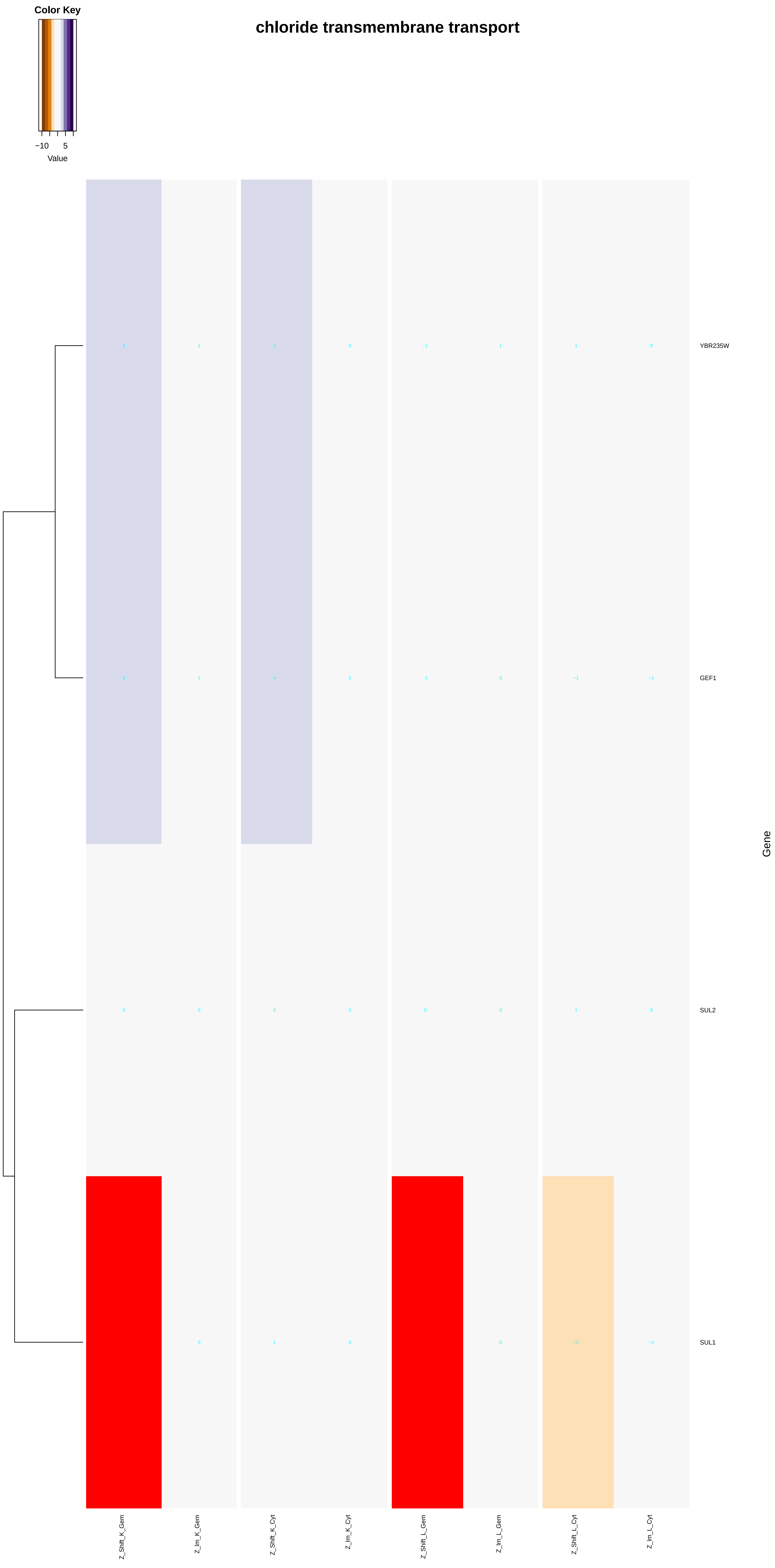
Color Key
−10
5
Value
chloride transmembrane transport
3
1
3
0
−1
1
1
0
YBR235W
3
1
4
0
−1
0
−1
−1
GEF1
0
0
0
0
0
0
1
0
SUL2
0
1
0
0
−2
−2
SUL1
Gene
Z_Shift_K_Gem
Z_Im_K_Gem
Z_Shift_K_Cyt
Z_Im_K_Cyt
Z_Shift_L_Gem
Z_Im_L_Gem
Z_Shift_L_Cyt
Z_Im_L_Cyt

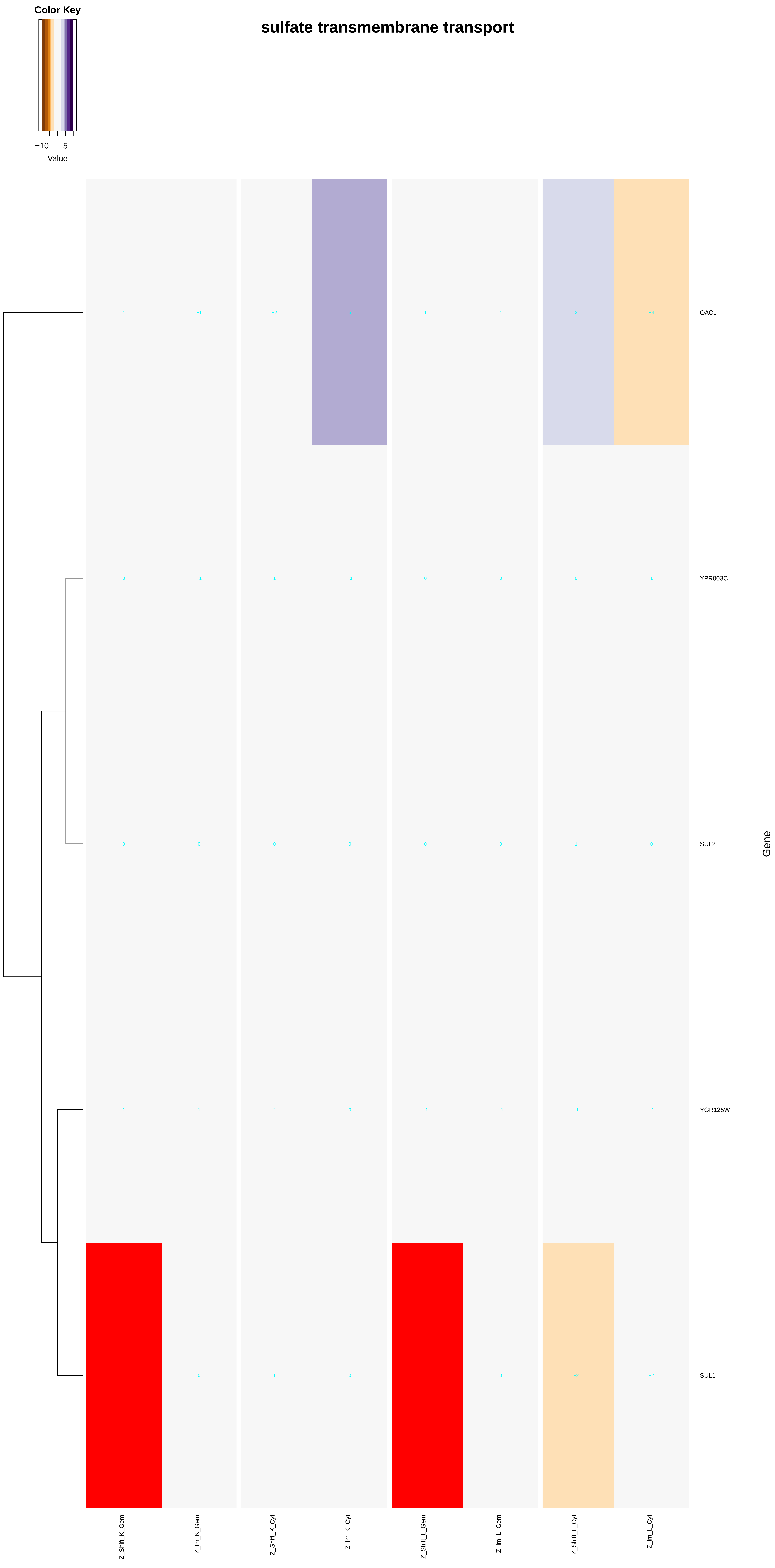

Color Key
−10
5
Value
sulfate transmembrane transport
OAC1
YPR003C
SUL2
YGR125W
SUL1
Gene
Z_Shift_K_Gem
Z_Im_K_Gem
Z_Shift_K_Cyt
Z_Im_K_Cyt
Z_Shift_L_Gem
Z_Im_L_Gem
Z_Shift_L_Cyt
Z_Im_L_Cyt

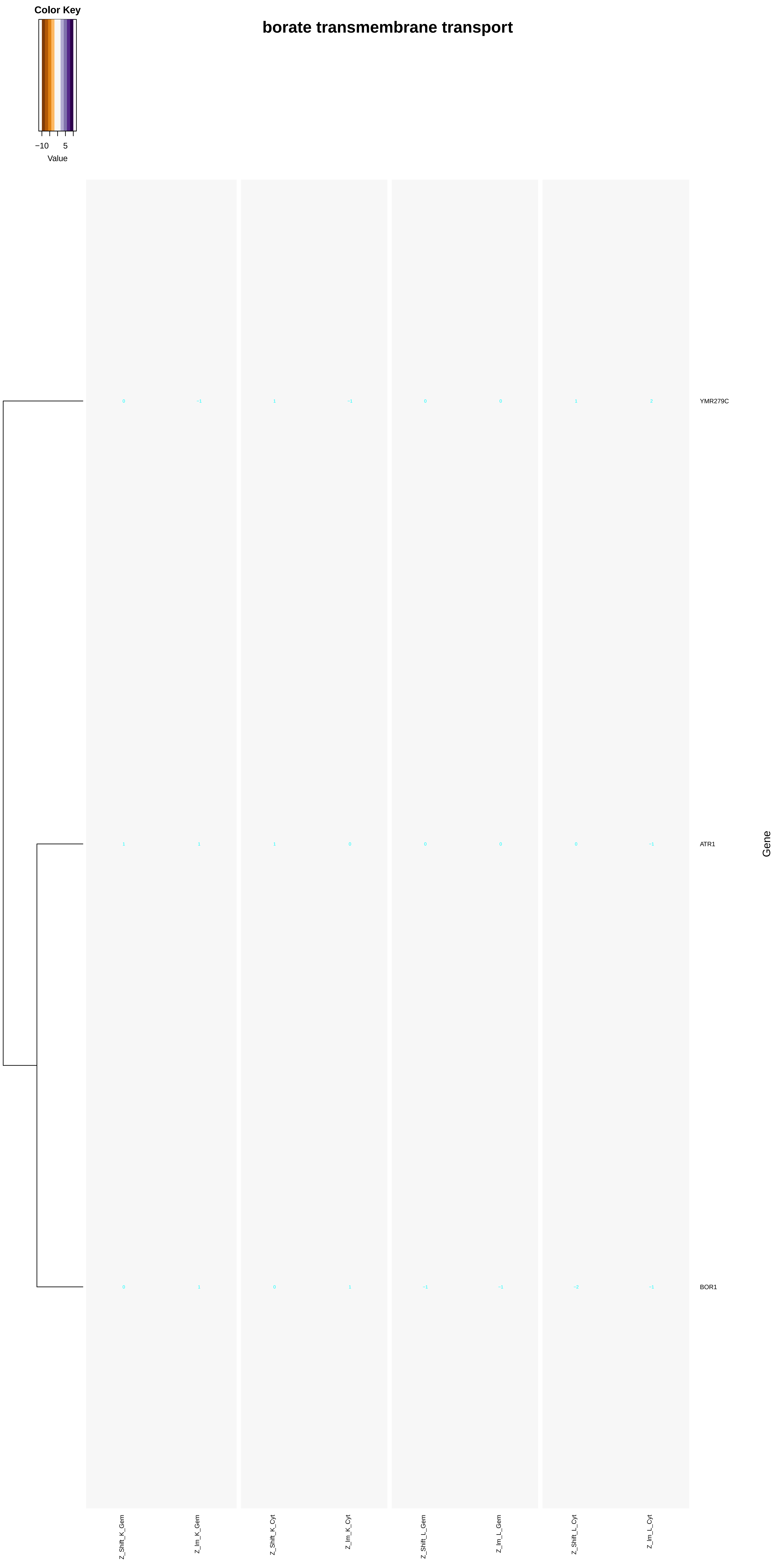
Color Key
−10
5
Value
borate transmembrane transport
0
−1
1
−1
0
0
1
2
YMR279C
1
1
1
0
0
0
0
−1
ATR1
0
1
0
1
−1
−1
−2
−1
BOR1
Gene
Z_Shift_K_Gem
Z_Im_K_Gem
Z_Shift_K_Cyt
Z_Im_K_Cyt
Z_Shift_L_Gem
Z_Im_L_Gem
Z_Shift_L_Cyt
Z_Im_L_Cyt

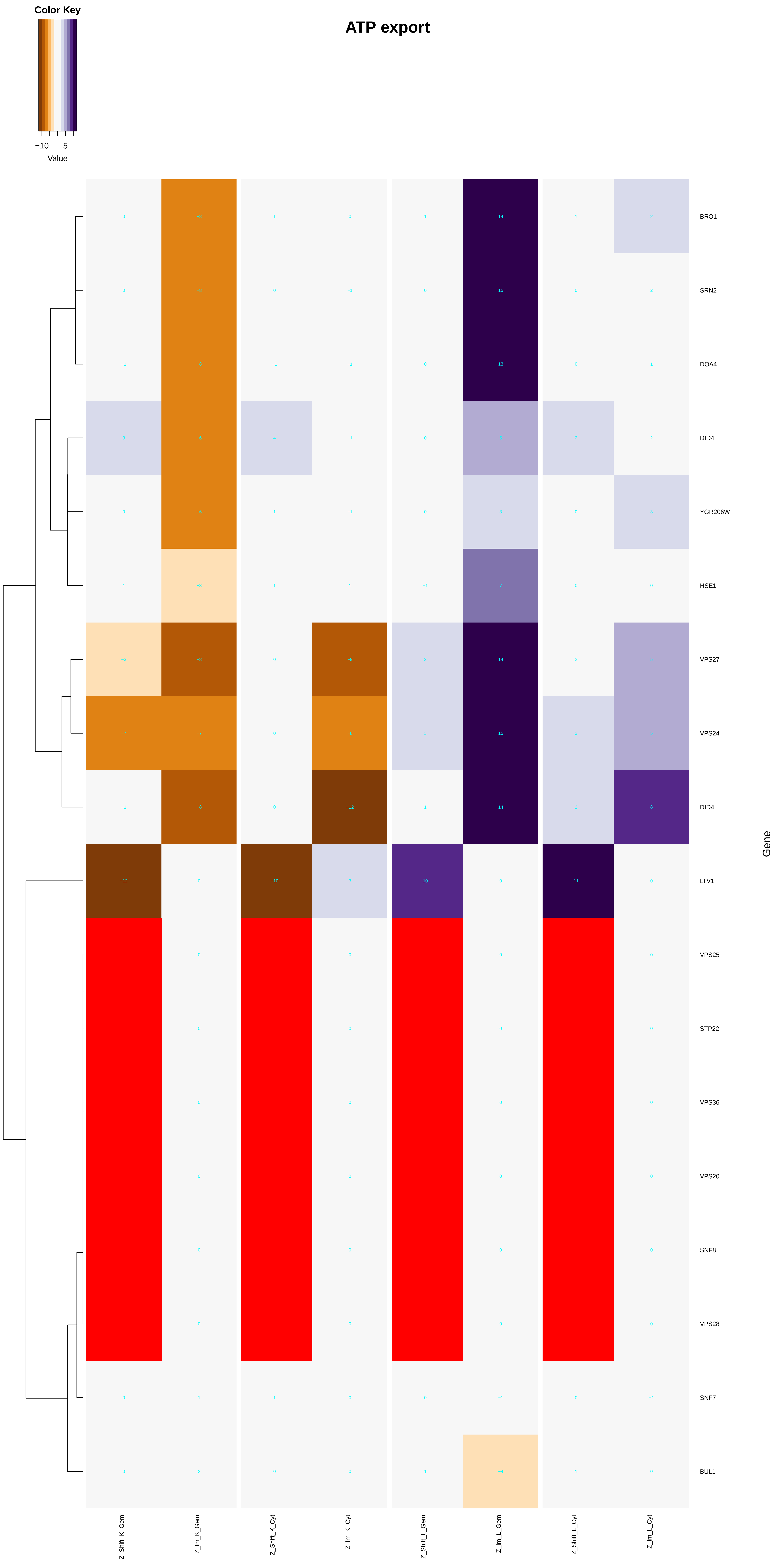
Color Key
−10 5
Value
ATP export
BRO1
SRN2
DOA4
DID4
YGR206W
HSE1
VPS27
VPS24
DID4
LTV1
VPS25
STP22
VPS36
VPS20
SNF8
VPS28
SNF7
BUL1
Gene
Z_Shift_K_Gem
Z_Im_K_Gem
Z_Shift_K_Cyt
Z_Im_K_Cyt
Z_Shift_L_Gem
Z_Im_L_Gem
Z_Shift_L_Cyt
Z_Im_L_Cyt

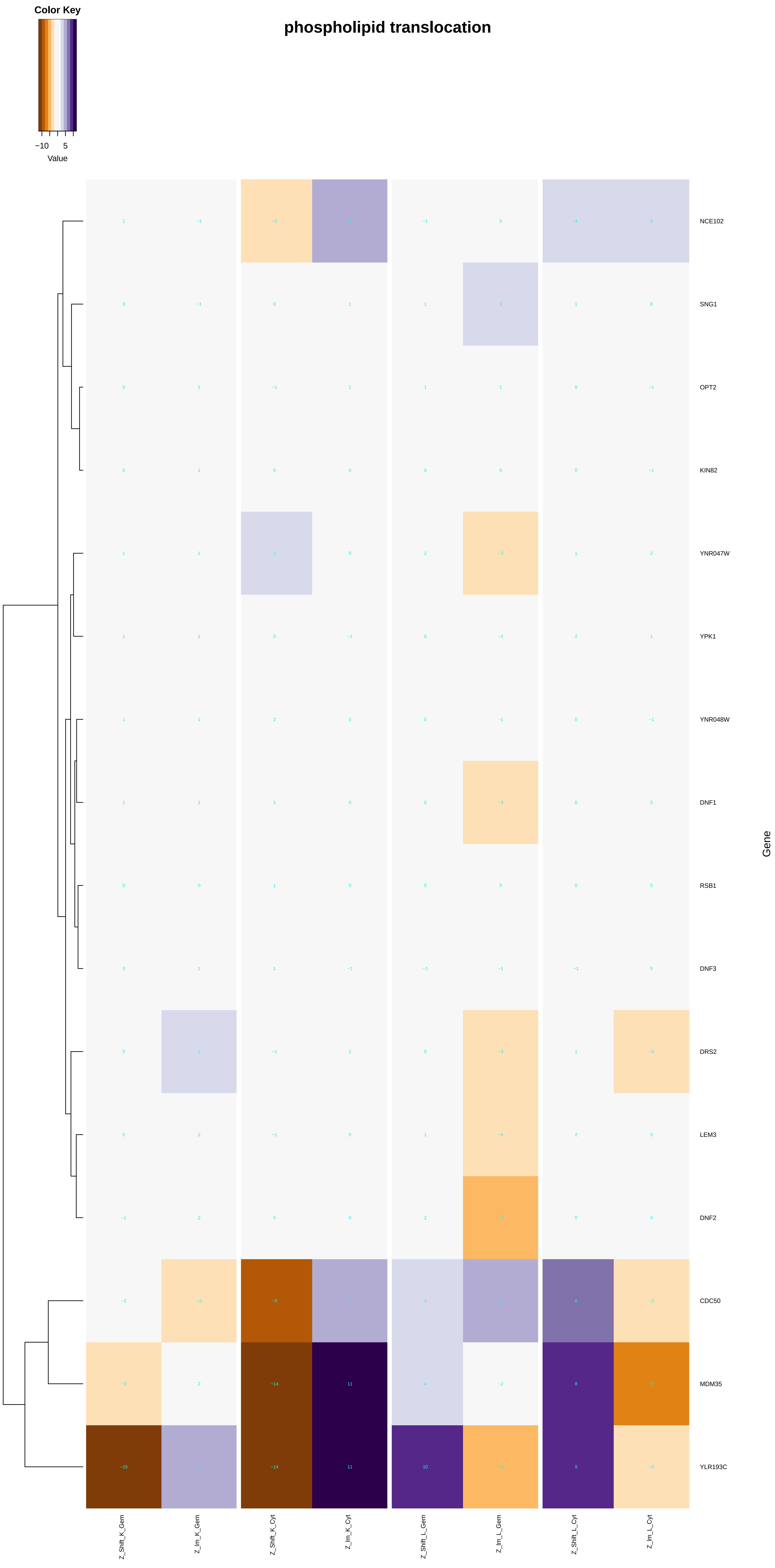
Color Key
−10
5
Value
phospholipid translocation
NCE102
SNG1
OPT2
KIN82
YNR047W
YPK1
YNR048W
DNF1
RSB1
DNF3
DRS2
LEM3
DNF2
CDC50
MDM35
YLR193C
Gene
Z_Shift_K_Gem
Z_Im_K_Gem
Z_Shift_K_Cyt
Z_Im_K_Cyt
Z_Shift_L_Gem
Z_Im_L_Gem
Z_Shift_L_Cyt
Z_Im_L_Cyt

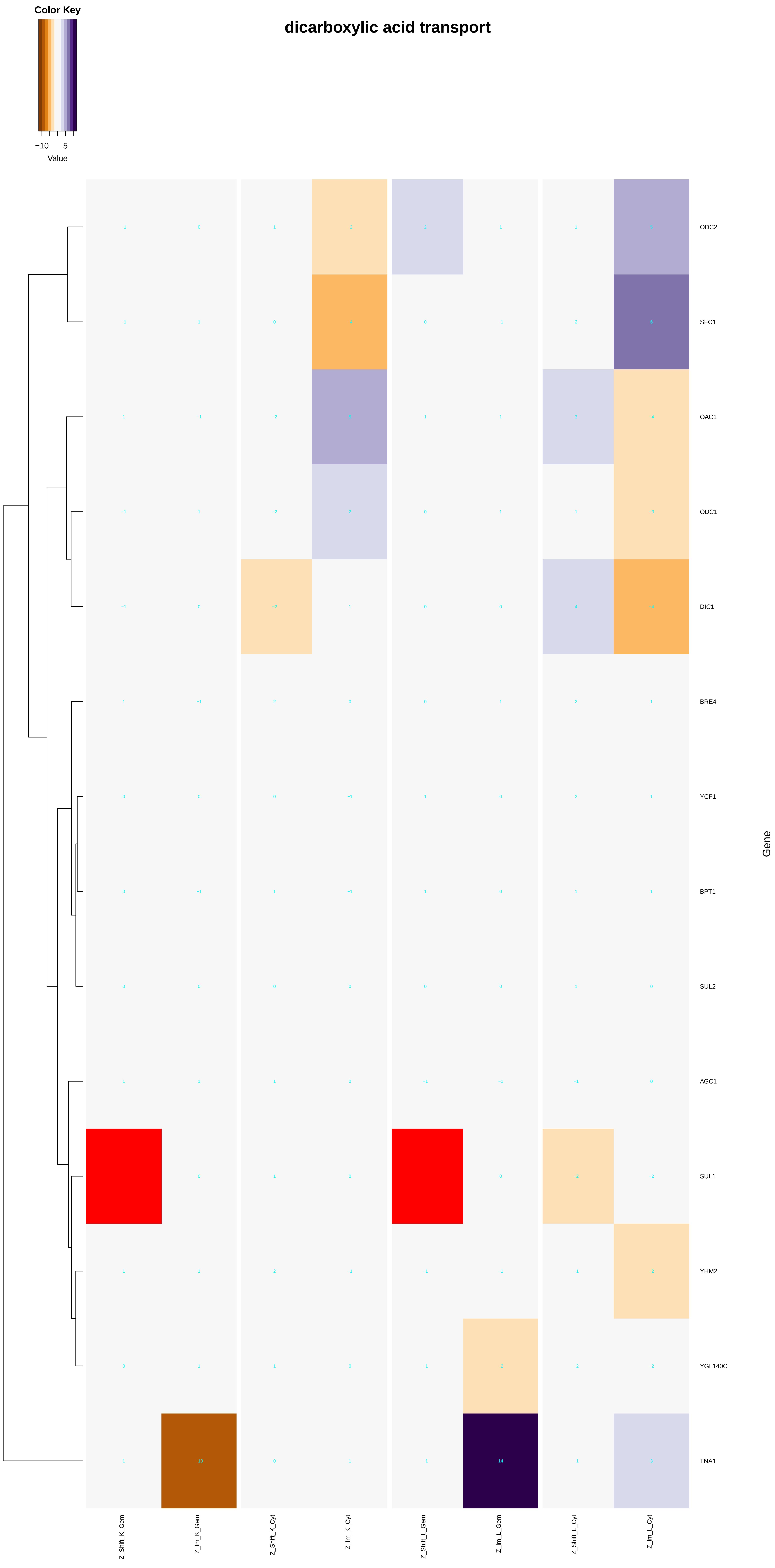
dicarboxylic acid transport
Color Key
−10
5
Value
ODC2
SFC1
OAC1
ODC1
DIC1
BRE4
YCF1
BPT1
SUL2
AGC1
SUL1
YHM2
YGL140C
TNA1
Gene
Z_Shift_K_Gem
Z_Im_K_Gem
Z_Shift_K_Cyt
Z_Im_K_Cyt
Z_Shift_L_Gem
Z_Im_L_Gem
Z_Shift_L_Cyt
Z_Im_L_Cyt

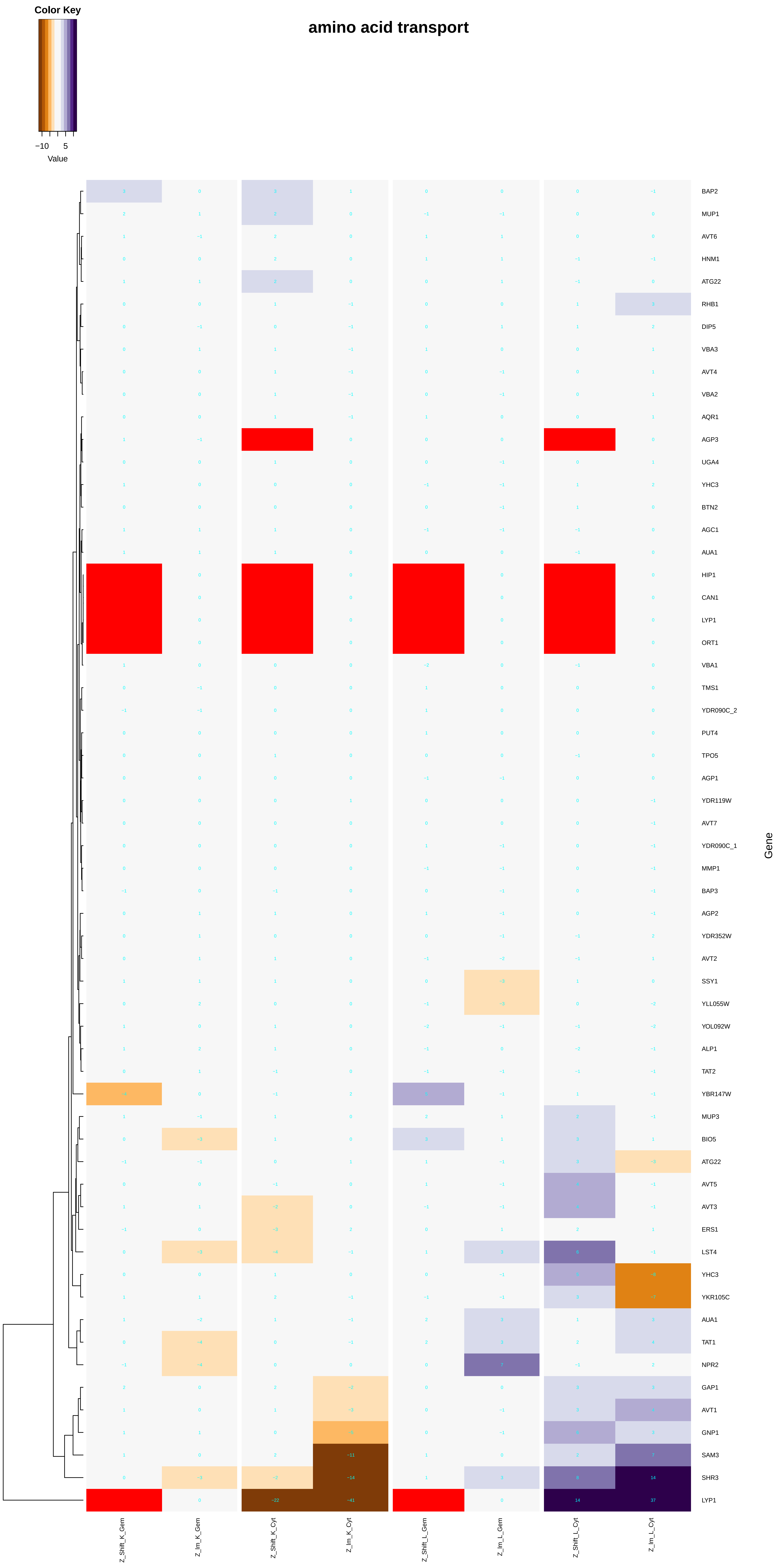

Color Key
−10
5
Value
amino acid transport
BAP2
MUP1
AVT6
HNM1
ATG22
RHB1
DIP5
VBA3
AVT4
VBA2
AQR1
AGP3
UGA4
YHC3
BTN2
AGC1
AUA1
HIP1
CAN1
LYP1
ORT1
VBA1
TMS1
YDR090C_2
PUT4
TPO5
AGP1
YDR119W
AVT7
YDR090C_1
MMP1
BAP3
AGP2
YDR352W
AVT2
SSY1
YLL055W
YOL092W
ALP1
TAT2
YBR147W
MUP3
BIO5
ATG22
AVT5
AVT3
ERS1
LST4
YHC3
YKR105C
AUA1
TAT1
NPR2
GAP1
AVT1
GNP1
SAM3
SHR3
LYP1
Gene
Z_Shift_K_Gem
Z_lm_K_Gem
Z_Shift_K_Cyt
Z_lm_K_Cyt
Z_Shift_L_Gem
Z_lm_L_Gem
Z_Shift_L_Cyt
Z_lm_L_Cyt

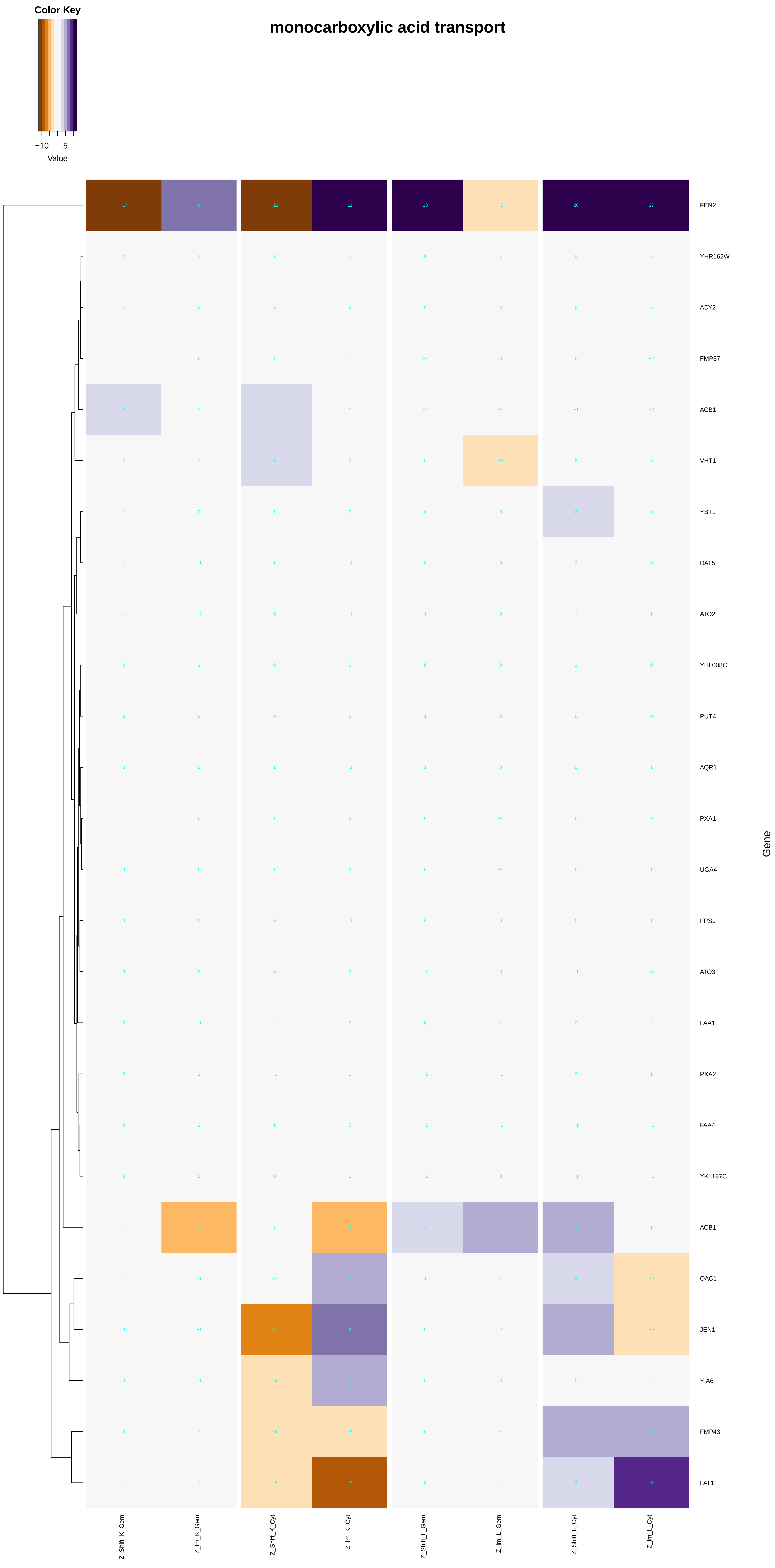

Color Key
−10
5
Value
monocarboxylic acid transport
FEN2
YHR162W
ADY2
FMP37
ACB1
VHT1
YBT1
DAL5
ATO2
YHL008C
PUT4
AQR1
PXA1
UGA4
FPS1
ATO3
FAA1
PXA2
FAA4
YKL187C
ACB1
OAC1
JEN1
YIA6
FMP43
FAT1
Gene
Z_Shift_K_Gem
Z_Im_K_Gem
Z_Shift_K_Cyt
Z_Im_K_Cyt
Z_Shift_L_Gem
Z_Im_L_Gem
Z_Shift_L_Cyt
Z_Im_L_Cyt

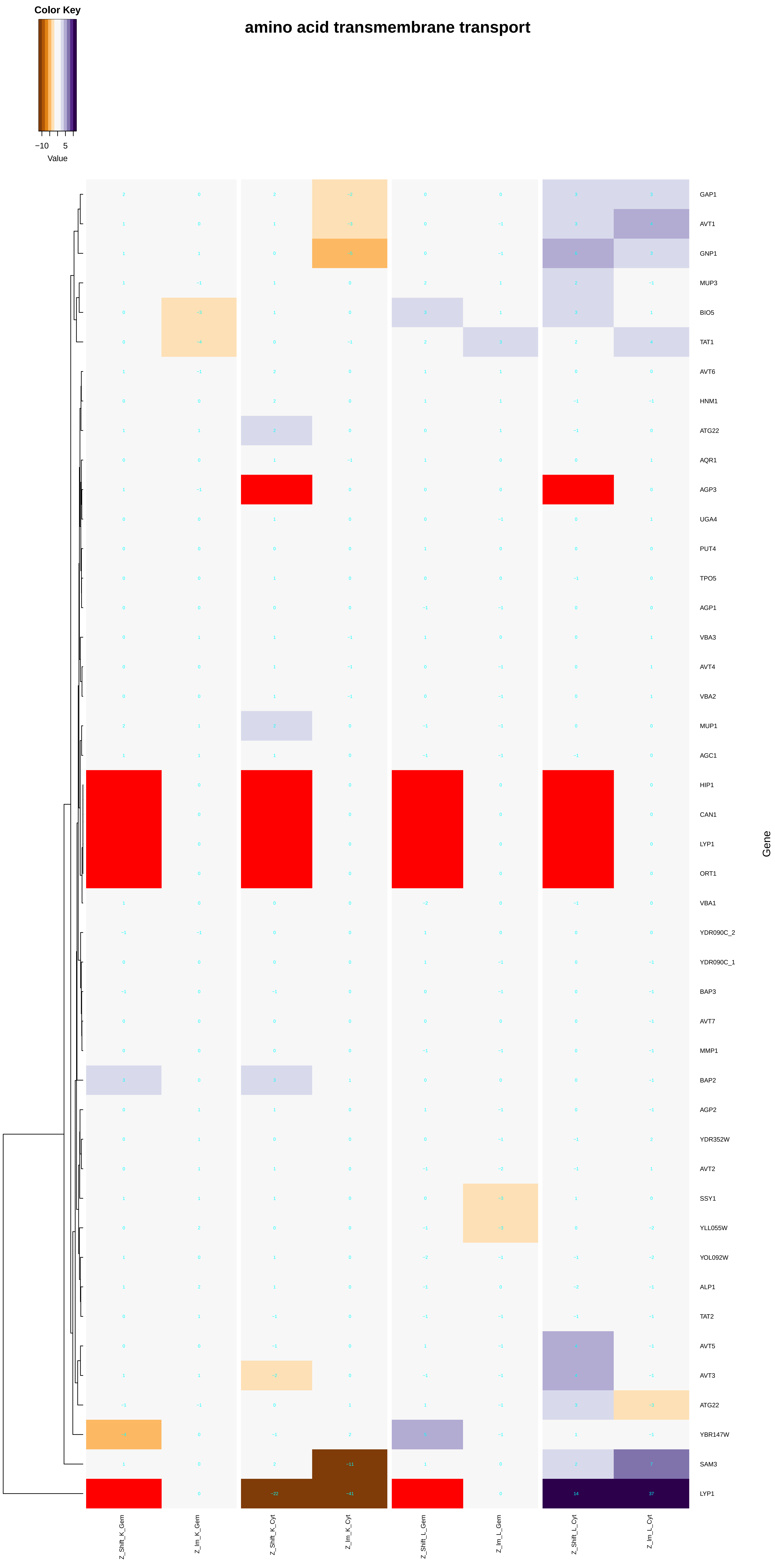
Color Key
−10
5
Value
amino acid transmembrane transport
GAP1
AVT1
GNP1
MUP3
BIO5
TAT1
AVT6
HNM1
ATG22
AQR1
AGP3
UGA4
PUT4
TPO5
AGP1
VBA3
AVT4
VBA2
MUP1
AGC1
HIP1
CAN1
LYP1
ORT1
VBA1
YDR090C_2
YDR090C_1
BAP3
AVT7
MMP1
BAP2
AGP2
YDR352W
AVT2
SSY1
YLL055W
YOL092W
ALP1
TAT2
AVT5
AVT3
ATG22
YBR147W
SAM3
LYP1
Gene
Z_Shift_K_Gem
Z_Im_K_Gem
Z_Shift_K_Cyt
Z_Im_K_Cyt
Z_Shift_L_Gem
Z_Im_L_Gem
Z_Shift_L_Cyt
Z_Im_L_Cyt

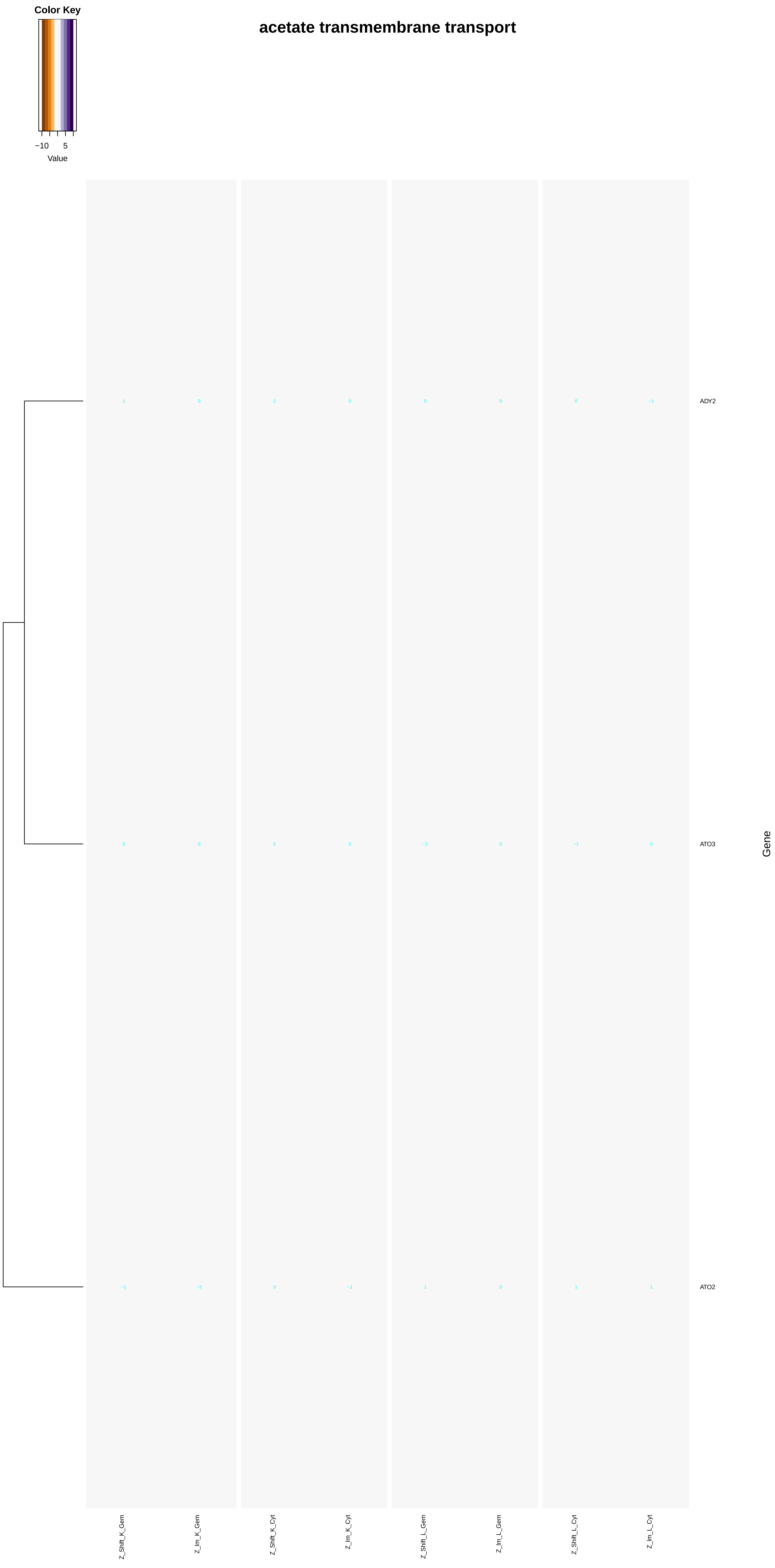
Color Key
−10
5
Value
acetate transmembrane transport
ADY2
ATO3
ATO2
Gene
Z_Shift_K_Gem
Z_Im_K_Gem
Z_Shift_K_Cyt
Z_Im_K_Cyt
Z_Shift_L_Gem
Z_Im_L_Gem
Z_Shift_L_Cyt
Z_Im_L_Cyt

### pyruvate transmembrane transport

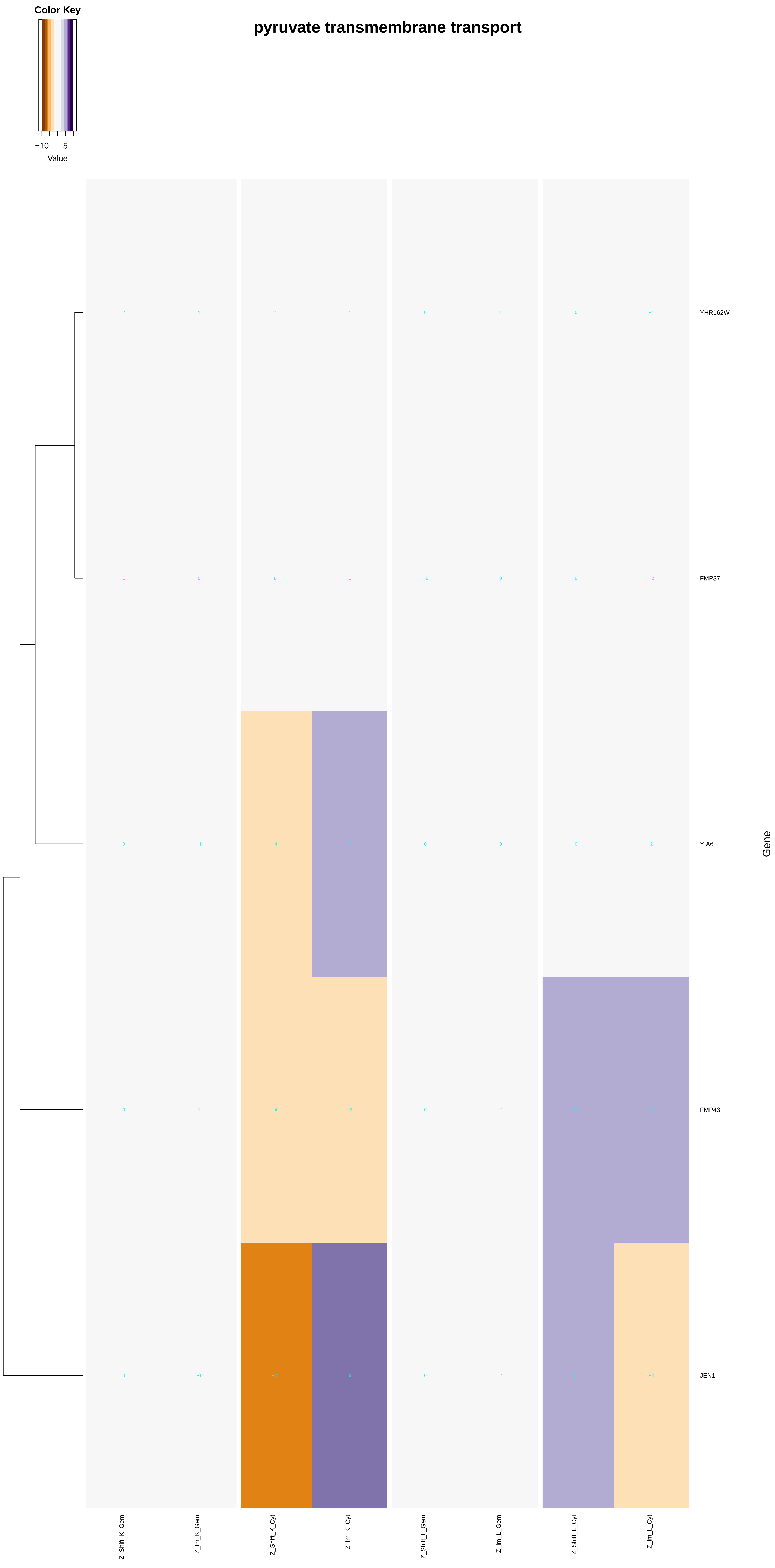

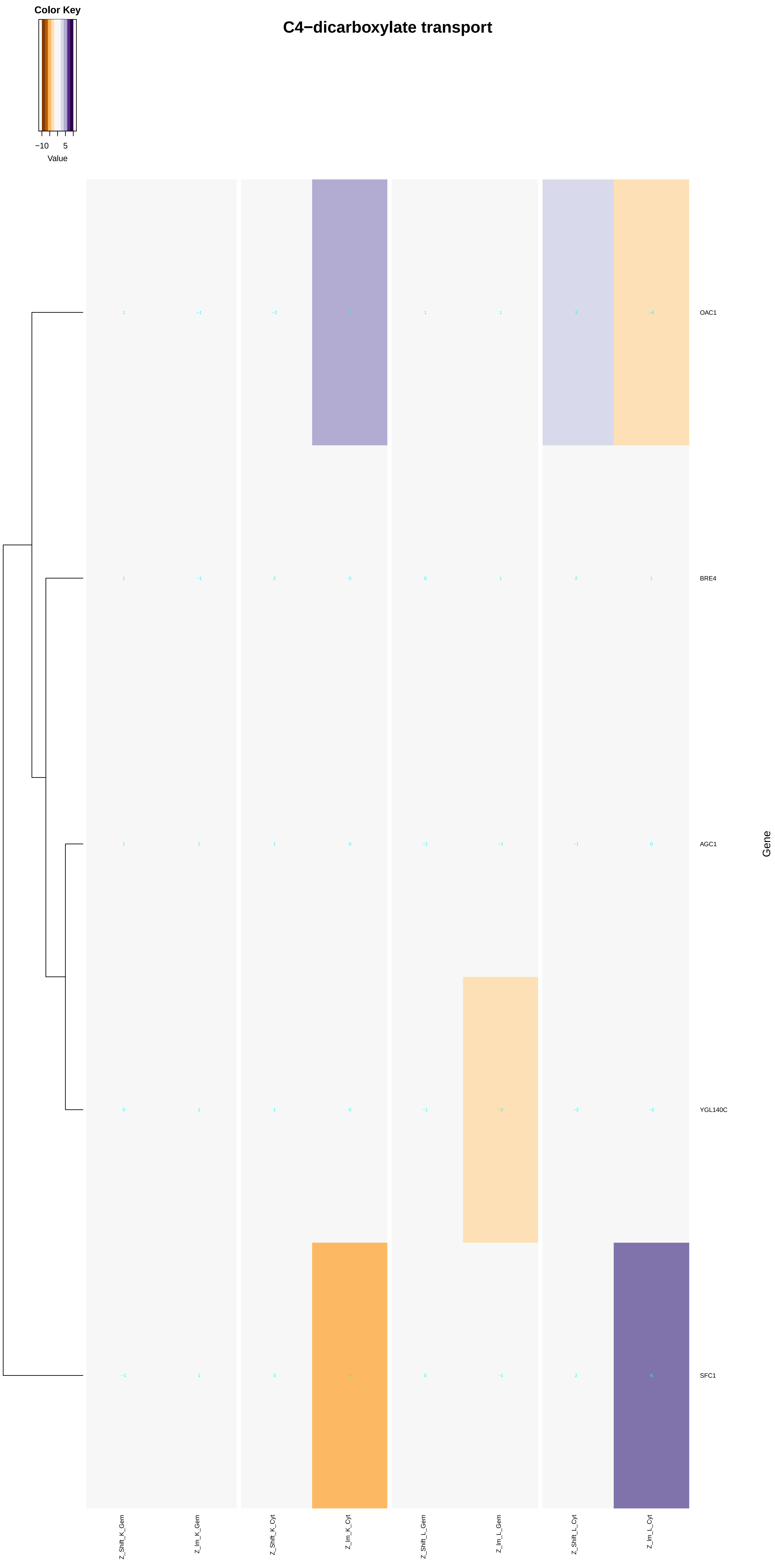
C4−dicarboxylate transport
Color Key
−10
5
Value
OAC1
BRE4
AGC1
YGL140C
SFC1
Gene
Z_Shift_K_Gem
Z_Im_K_Gem
Z_Shift_K_Cyt
Z_Im_K_Cyt
Z_Shift_L_Gem
Z_Im_L_Gem
Z_Shift_L_Cyt
Z_Im_L_Cyt

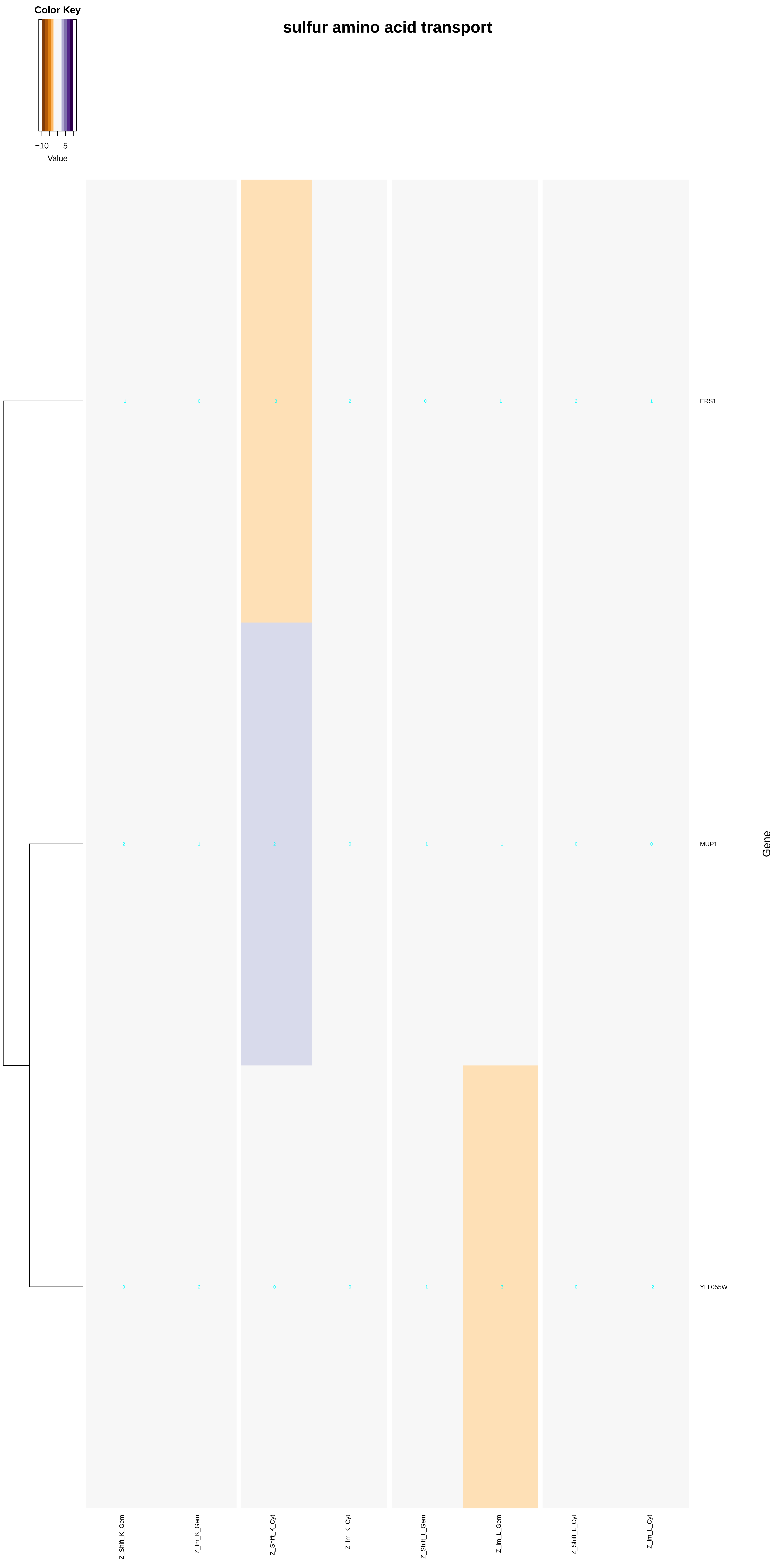

Color Key
−10
5
Value
sulfur amino acid transport
ERS1
MUP1
YLL055W
Gene
-1
0
-3
2
0
1
2
1
2
1
2
0
-1
-1
0
0
0
2
0
0
-1
-3
0
-2
Z_Shift_K_Gem
Z_Im_K_Gem
Z_Shift_K_Cyt
Z_Im_K_Cyt
Z_Shift_L_Gem
Z_Im_L_Gem
Z_Shift_L_Cyt
Z_Im_L_Cyt

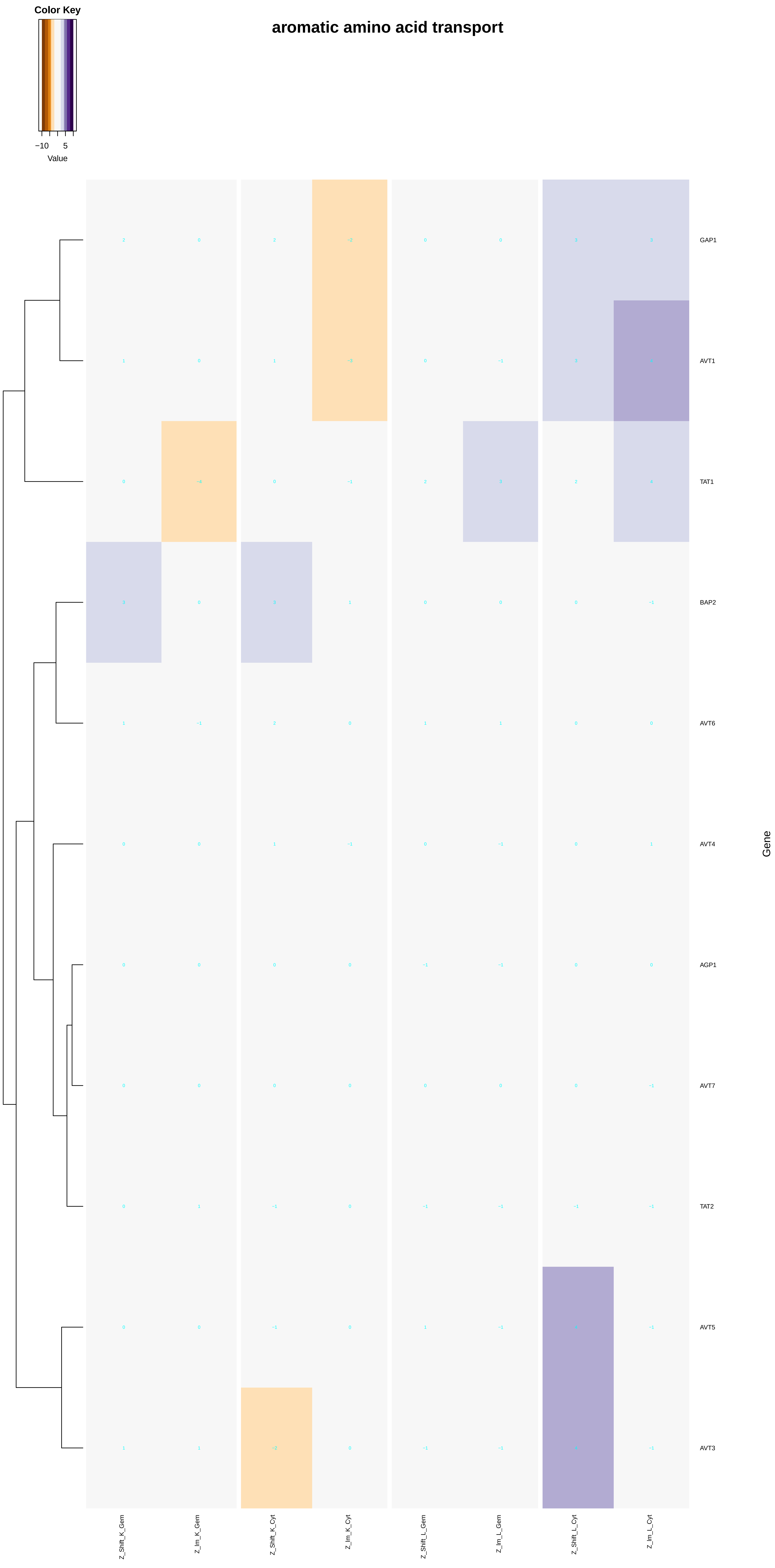
aromatic amino acid transport
Color Key
−10
5
Value
GAP1
AVT1
TAT1
BAP2
AVT6
AVT4
AGP1
AVT7
TAT2
AVT5
AVT3
Gene
Z_Shift_K_Gem
Z_Im_K_Gem
Z_Shift_K_Cyt
Z_Im_K_Cyt
Z_Shift_L_Gem
Z_Im_L_Gem
Z_Shift_L_Cyt
Z_Im_L_Cyt

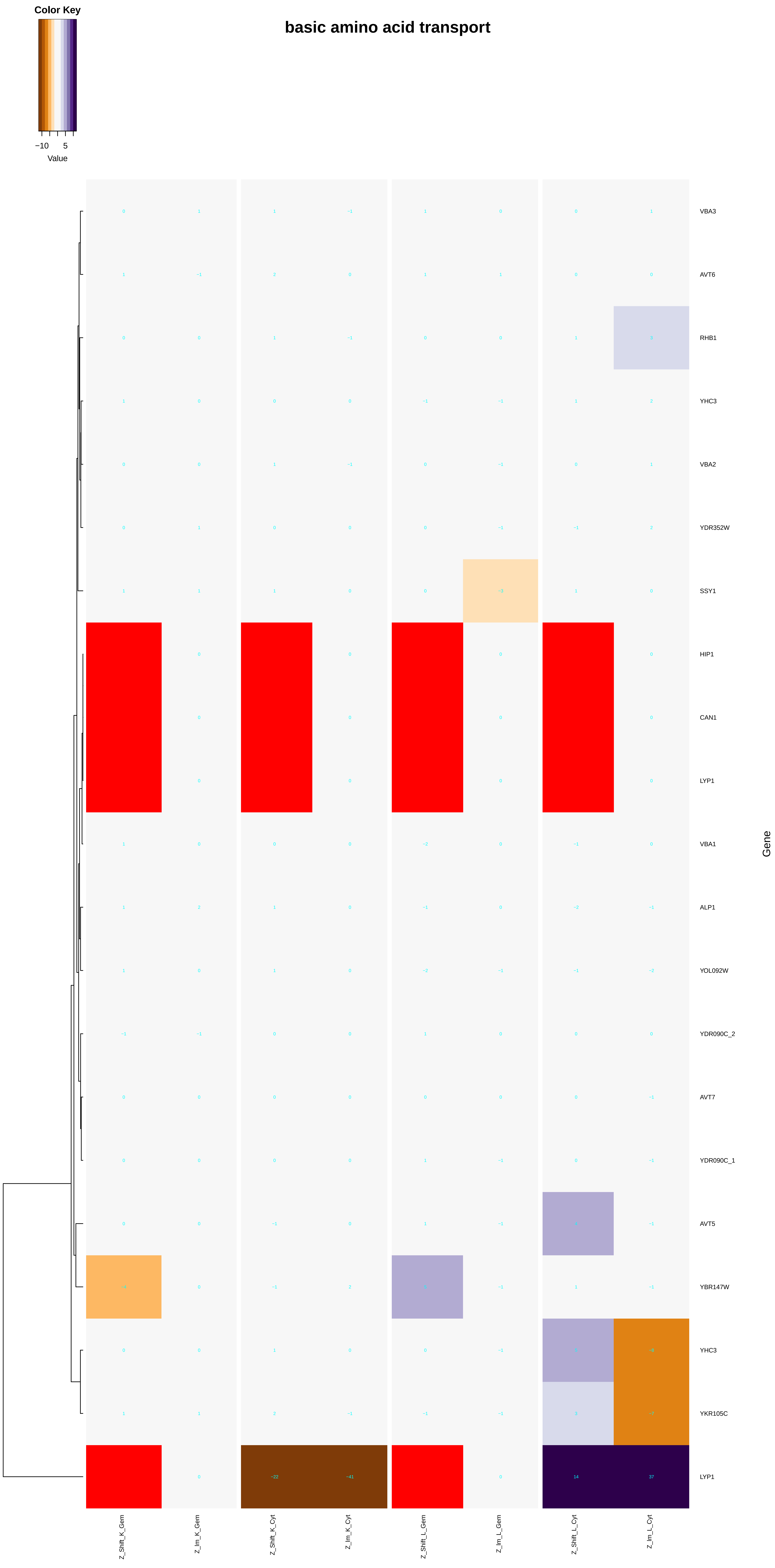
basic amino acid transport
Color Key
−10
5
Value
VBA3
AVT6
RHB1
YHC3
VBA2
YDR352W
SSY1
HIP1
CAN1
LYP1
VBA1
ALP1
YOL092W
YDR090C_2
AVT7
YDR090C_1
AVT5
YBR147W
YHC3
YKR105C
LYP1
Gene
Z_Shift_K_Gem
Z_Im_K_Gem
Z_Shift_K_Cyt
Z_Im_K_Cyt
Z_Shift_L_Gem
Z_Im_L_Gem
Z_Shift_L_Cyt
Z_Im_L_Cyt

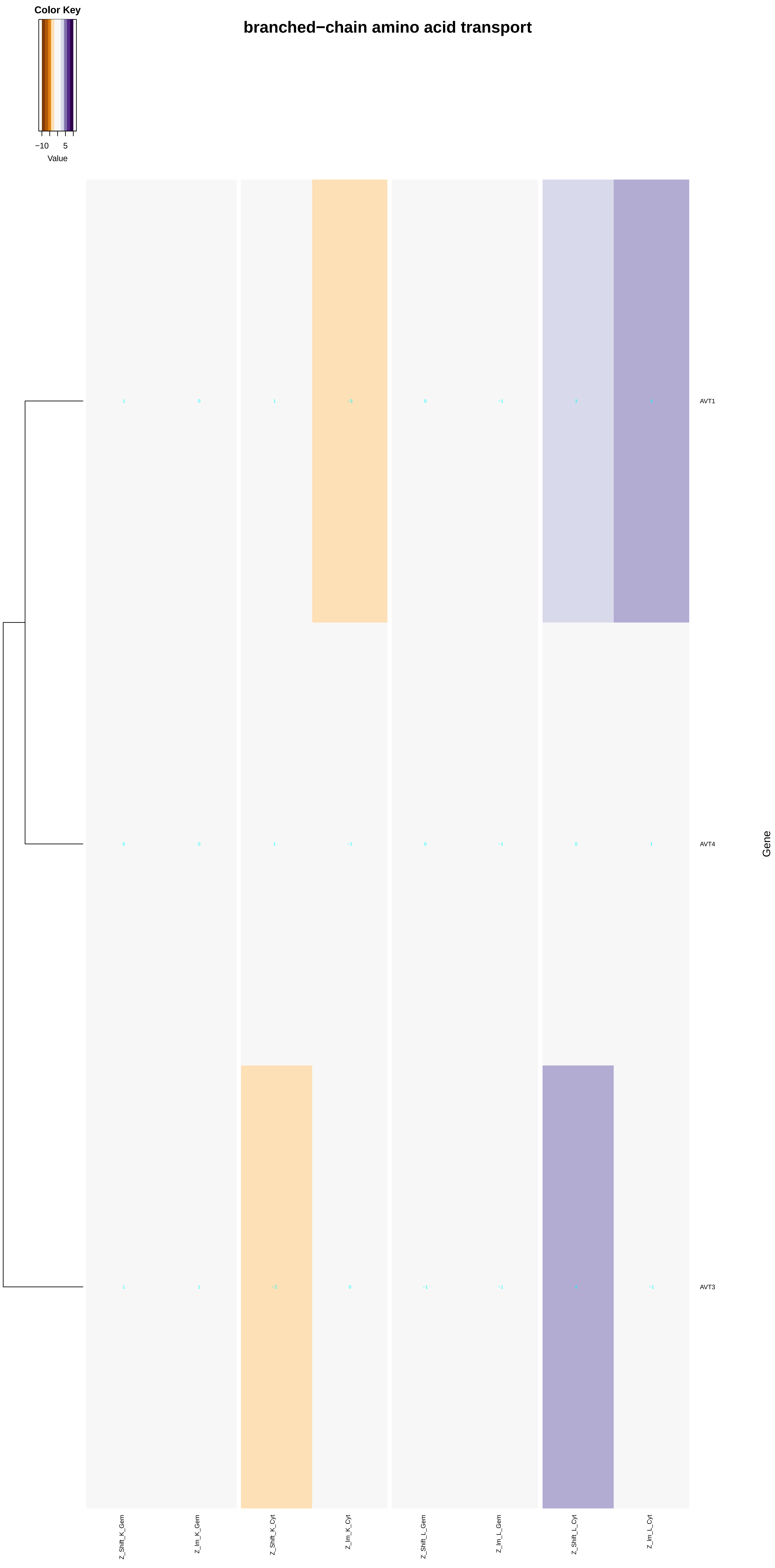
branched−chain amino acid transport
Color Key
−10
5
Value
AVT1
AVT4
AVT3
Gene
Z_Shift_K_Gem
Z_Im_K_Gem
Z_Shift_K_Cyt
Z_Im_K_Cyt
Z_Shift_L_Gem
Z_Im_L_Gem
Z_Shift_L_Cyt
Z_Im_L_Cyt

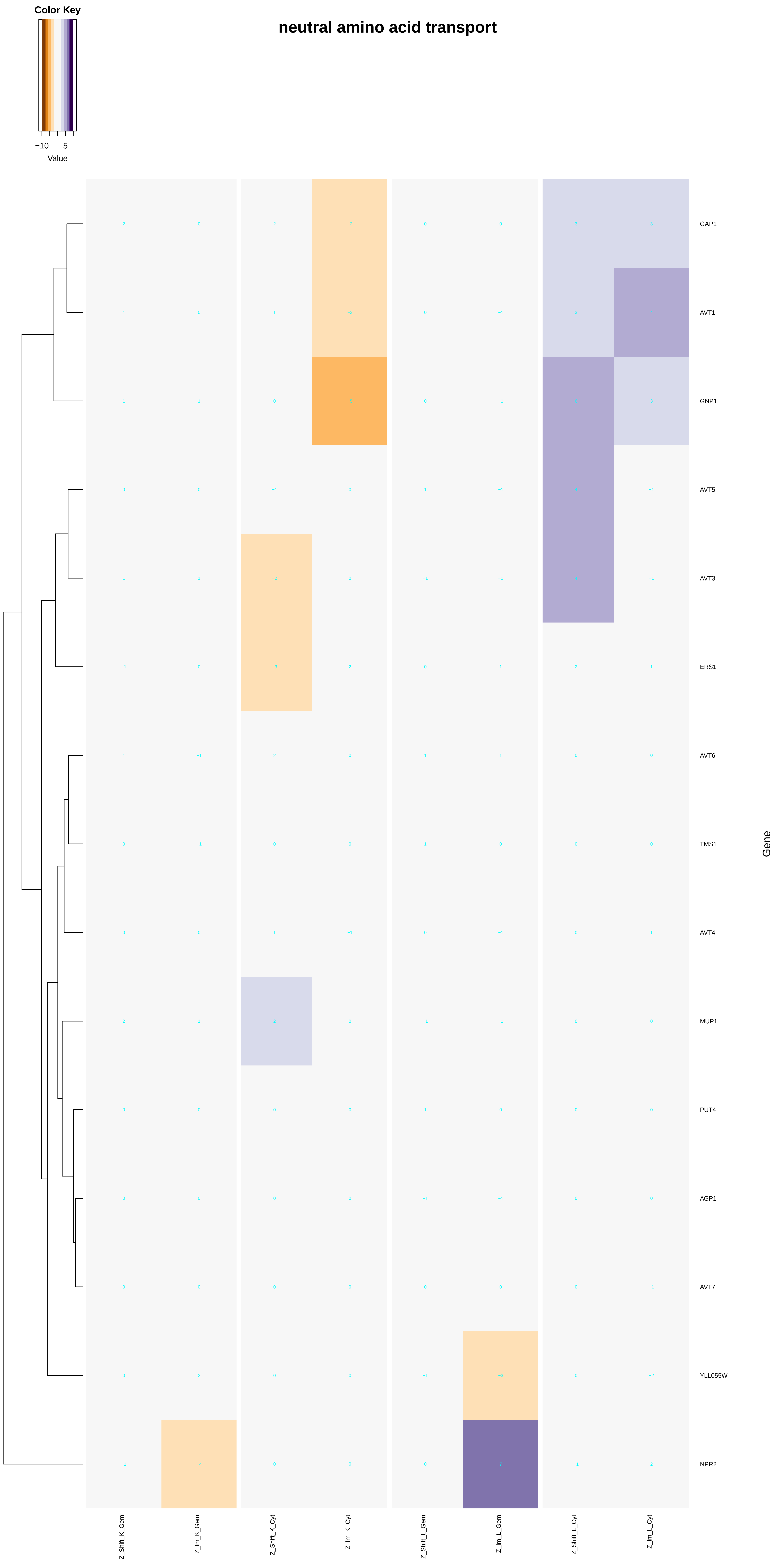
neutral amino acid transport
Color Key
−10
5
Value
GAP1
AVT1
GNP1
AVT5
AVT3
ERS1
AVT6
TMS1
AVT4
MUP1
PUT4
AGP1
AVT7
YLL055W
NPR2
Gene
Z_Shift_K_Gem
Z_Im_K_Gem
Z_Shift_K_Cyt
Z_Im_K_Cyt
Z_Shift_L_Gem
Z_Im_L_Gem
Z_Shift_L_Cyt
Z_Im_L_Cyt

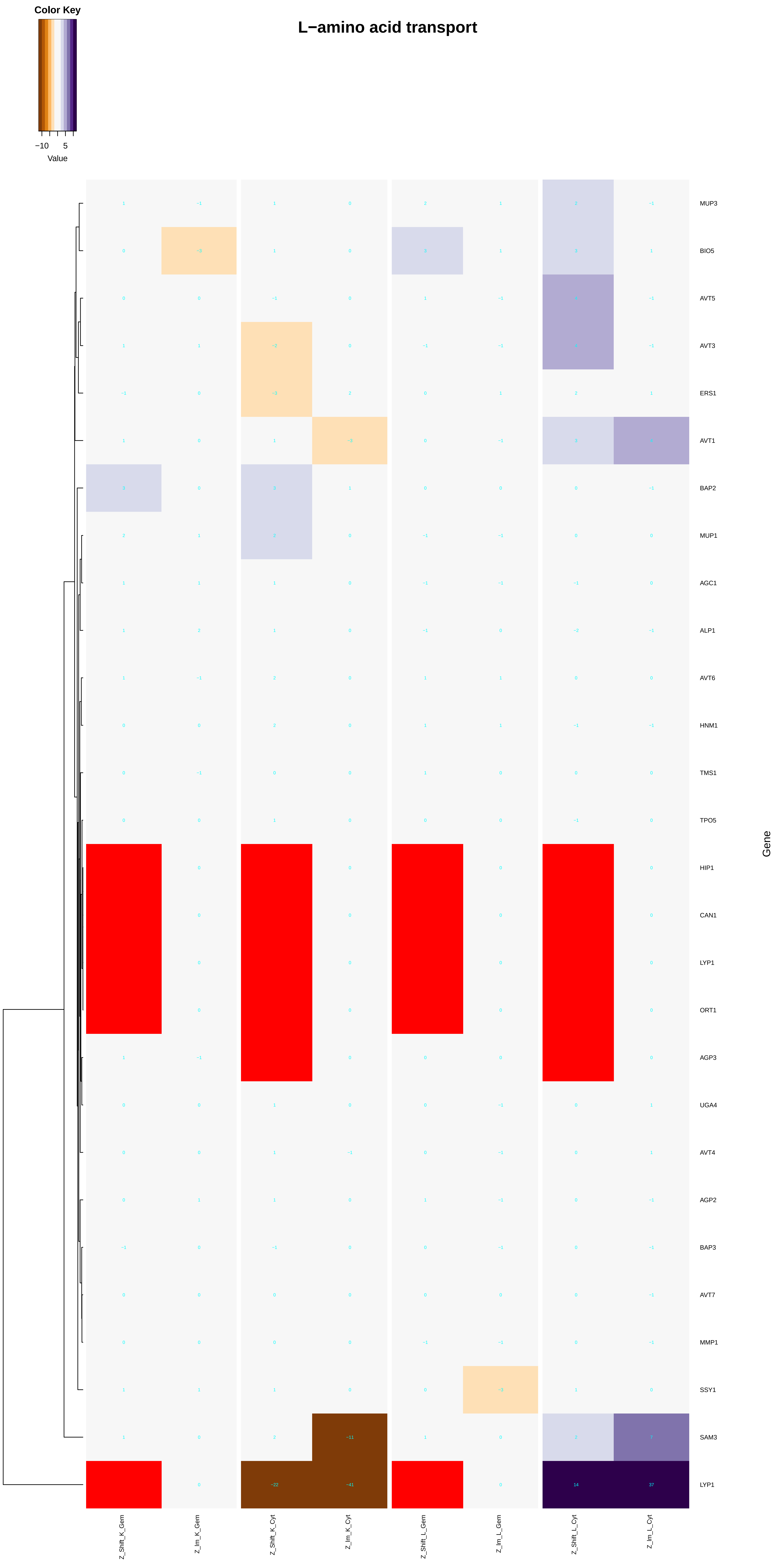

Color Key
−10
5
Value
L−amino acid transport
MUP3
BIO5
AVT5
AVT3
ERS1
AVT1
BAP2
MUP1
AGC1
ALP1
AVT6
HNM1
TMS1
TPO5
HIP1
CAN1
LYP1
ORT1
AGP3
UGA4
AVT4
AGP2
BAP3
AVT7
MMP1
SSY1
SAM3
LYP1
Gene
Z_Shift_K_Gem
Z_Im_K_Gem
Z_Shift_K_Cyt
Z_Im_K_Cyt
Z_Shift_L_Gem
Z_Im_L_Gem
Z_Shift_L_Cyt
Z_Im_L_Cyt

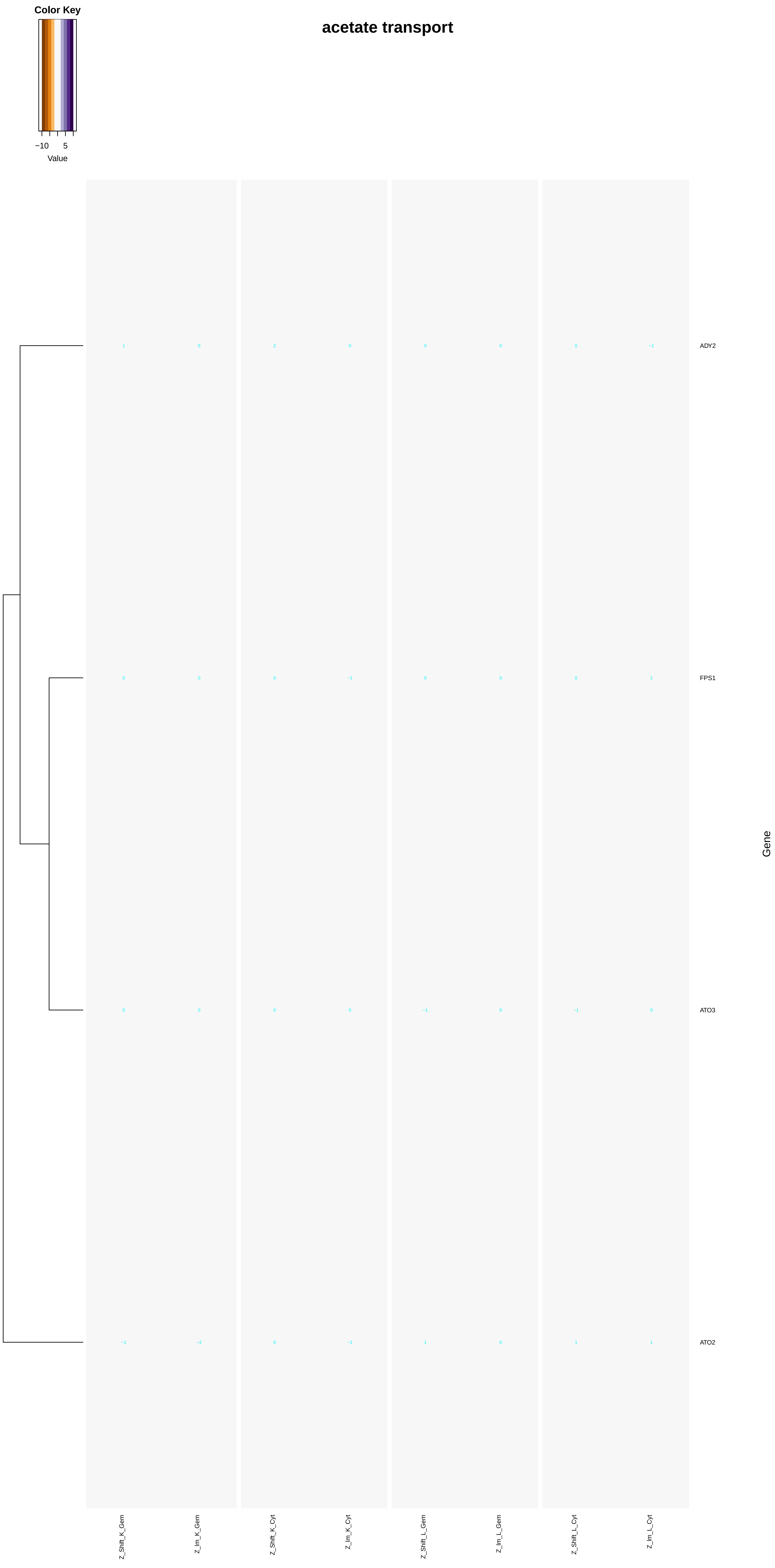
Color Key
−10
5
Value
acetate transport
ADY2
FPS1
ATO3
ATO2
Gene
Z_Shift_K_Gem
Z_Im_K_Gem
Z_Shift_K_Cyt
Z_Im_K_Cyt
Z_Shift_L_Gem
Z_Im_L_Gem
Z_Shift_L_Cyt
Z_Im_L_Cyt

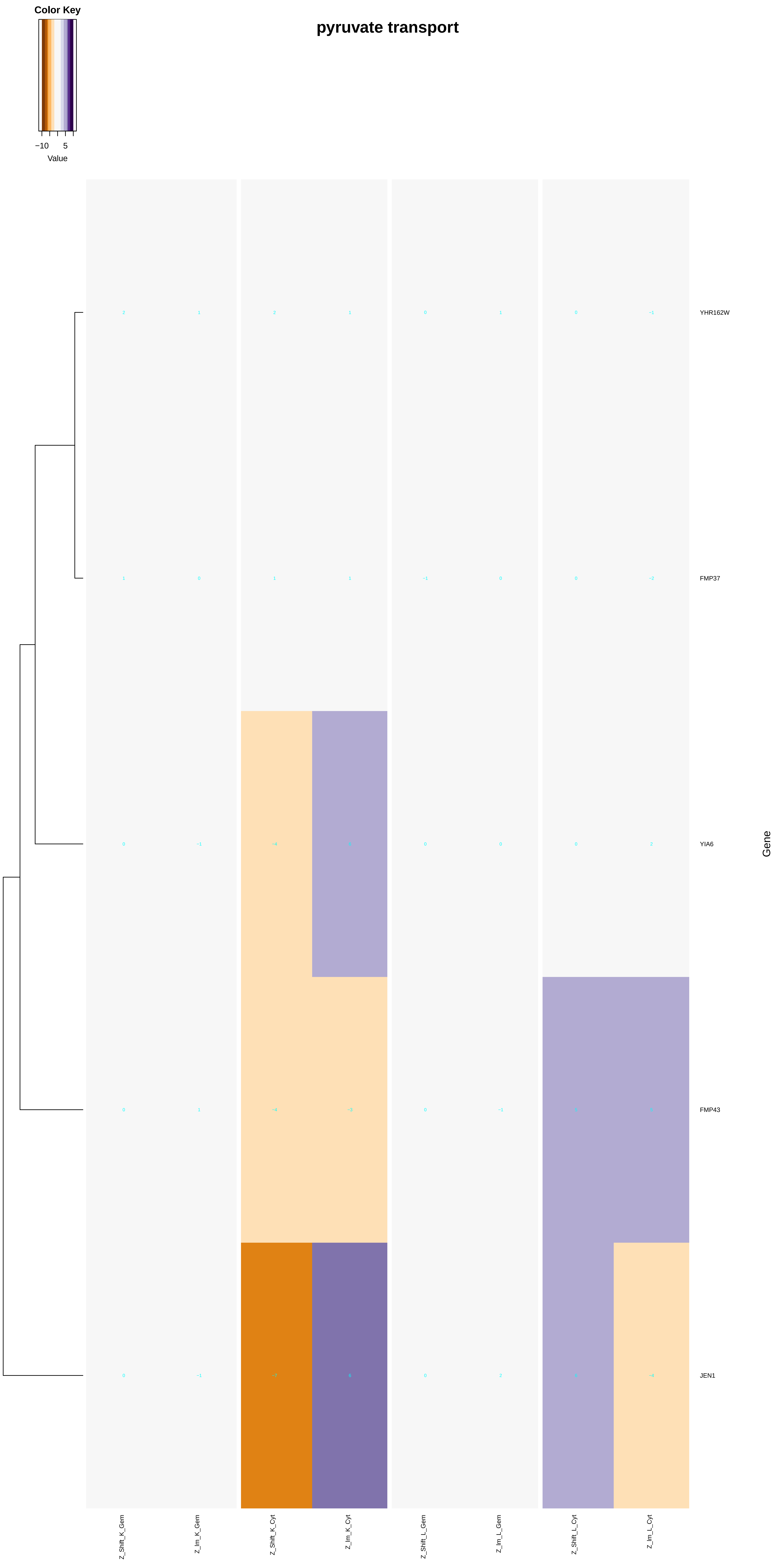
Color Key
−10
5
Value
pyruvate transport
YHR162W
FMP37
YIA6
FMP43
JEN1
Gene
Z_Shift_K_Gem
Z_Im_K_Gem
Z_Shift_K_Cyt
Z_Im_K_Cyt
Z_Shift_L_Gem
Z_Im_L_Gem
Z_Shift_L_Cyt
Z_Im_L_Cyt

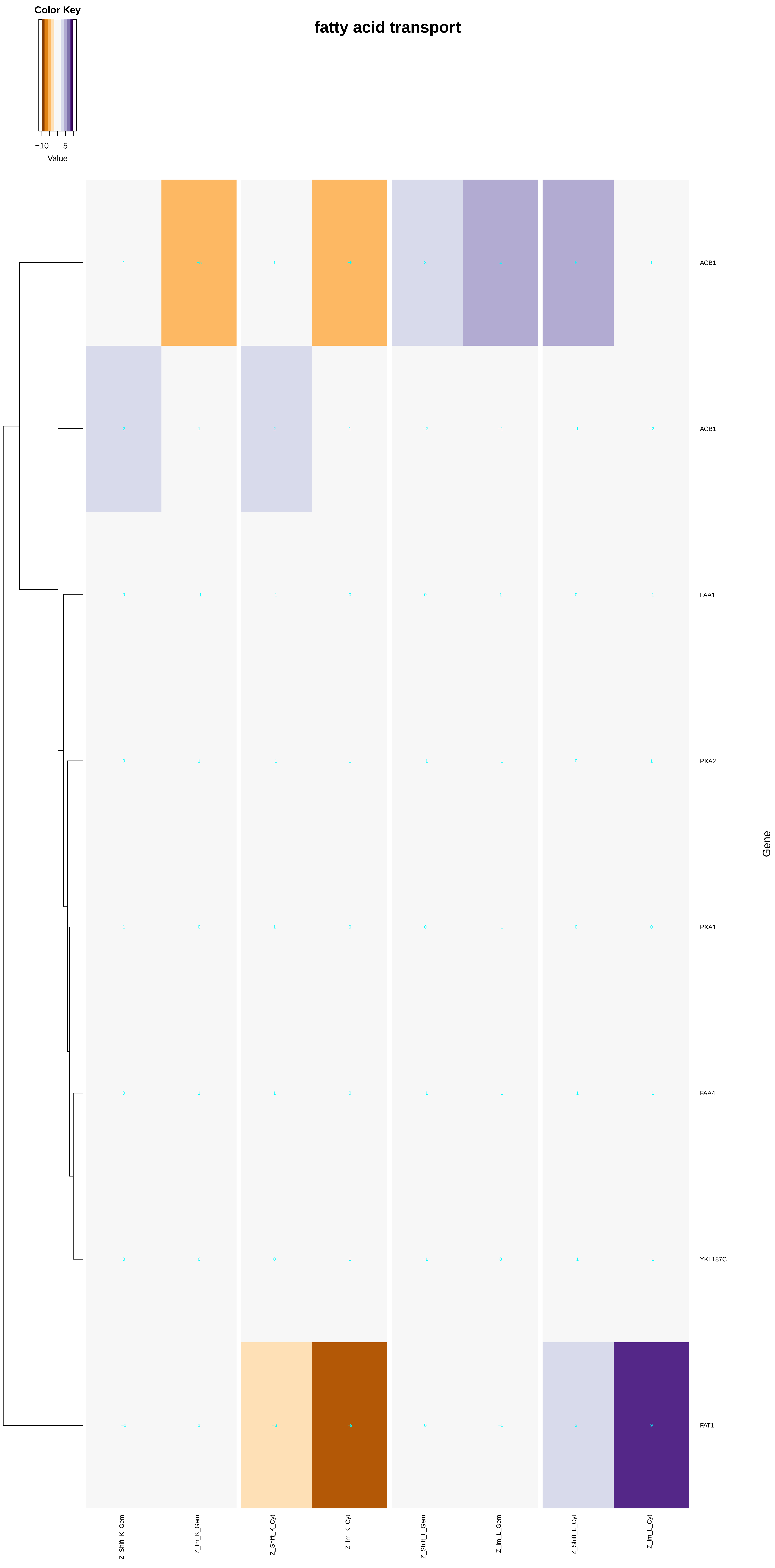

Color Key
−10
5
Value
fatty acid transport
ACB1
ACB1
FAA1
PXA2
PXA1
FAA4
YKL187C
FAT1
Gene
Z_Shift_K_Gem
Z_Im_K_Gem
Z_Shift_K_Cyt
Z_Im_K_Cyt
Z_Shift_L_Gem
Z_Im_L_Gem
Z_Shift_L_Cyt
Z_Im_L_Cyt

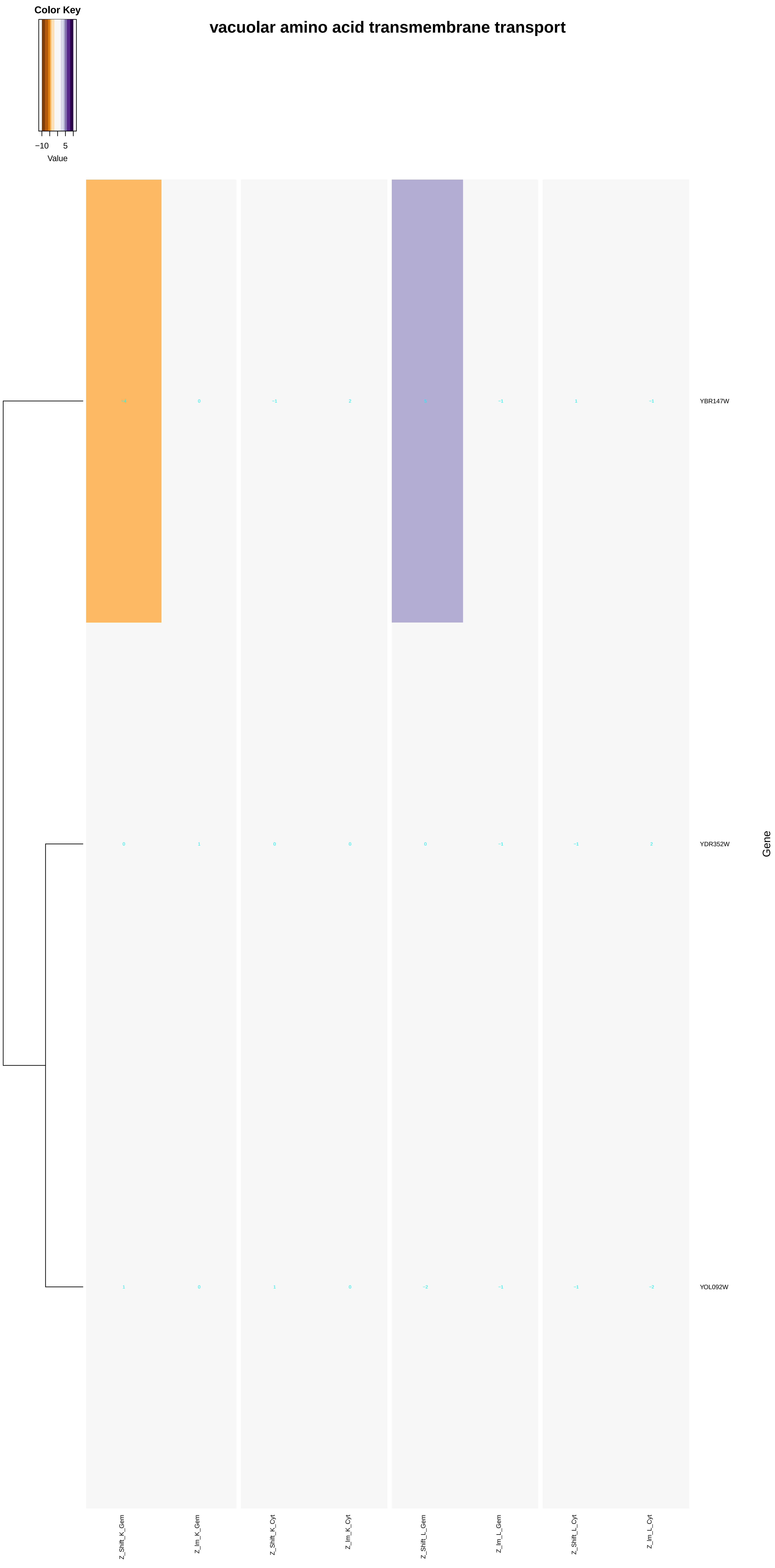

Color Key
−10
5
Value
vacuolar amino acid transmembrane transport
YBR147W
YDR352W
YOL092W
Gene
Z_Shift_K_Gem
Z_Im_K_Gem
Z_Shift_K_Cyt
Z_Im_K_Cyt
Z_Shift_L_Gem
Z_Im_L_Gem
Z_Shift_L_Cyt
Z_Im_L_Cyt

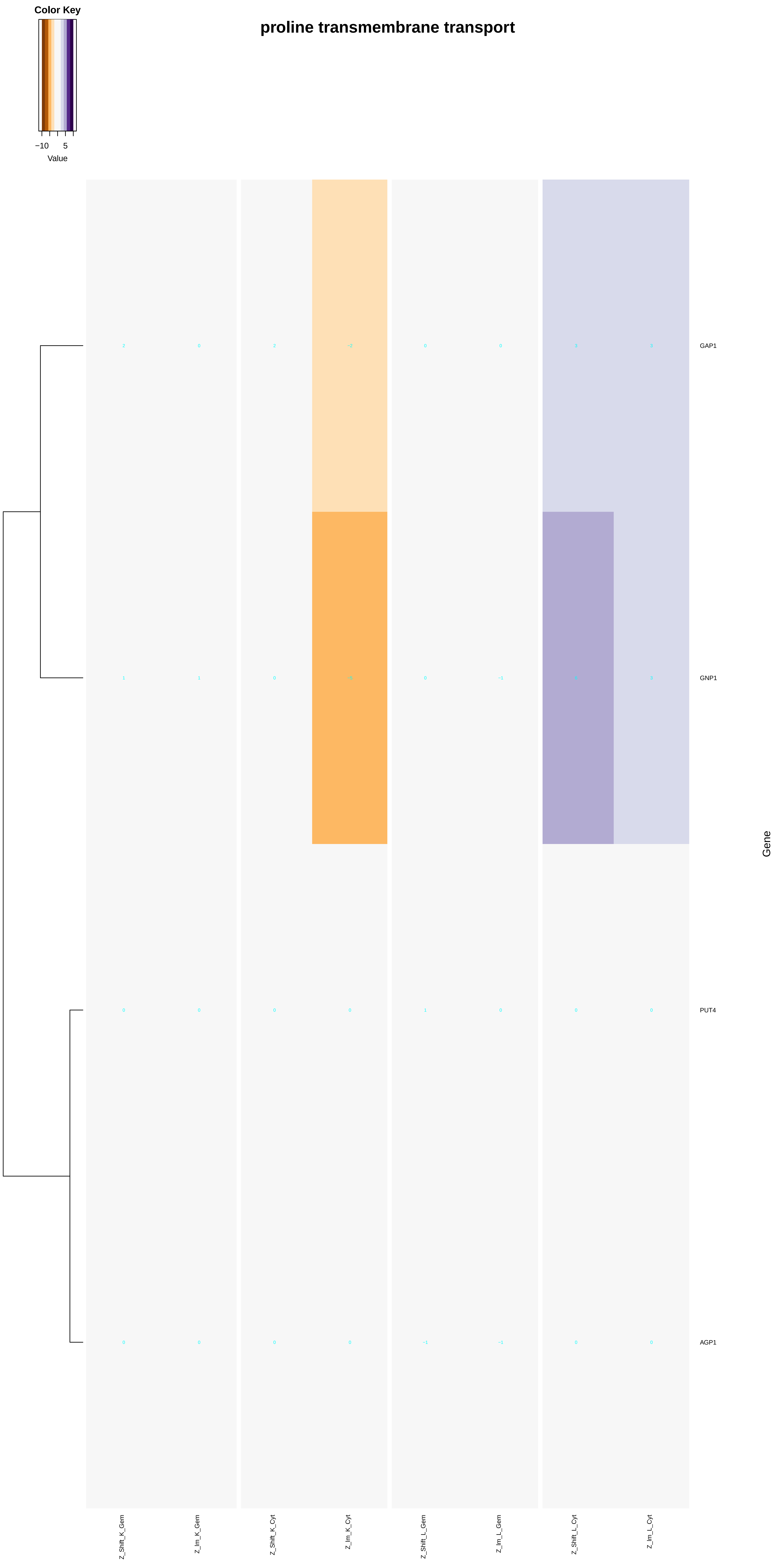
Color Key
−10
5
Value
proline transmembrane transport
GAP1
GNP1
PUT4
AGP1
Gene
Z_Shift_K_Gem
Z_Im_K_Gem
Z_Shift_K_Cyt
Z_Im_K_Cyt
Z_Shift_L_Gem
Z_Im_L_Gem
Z_Shift_L_Cyt
Z_Im_L_Cyt

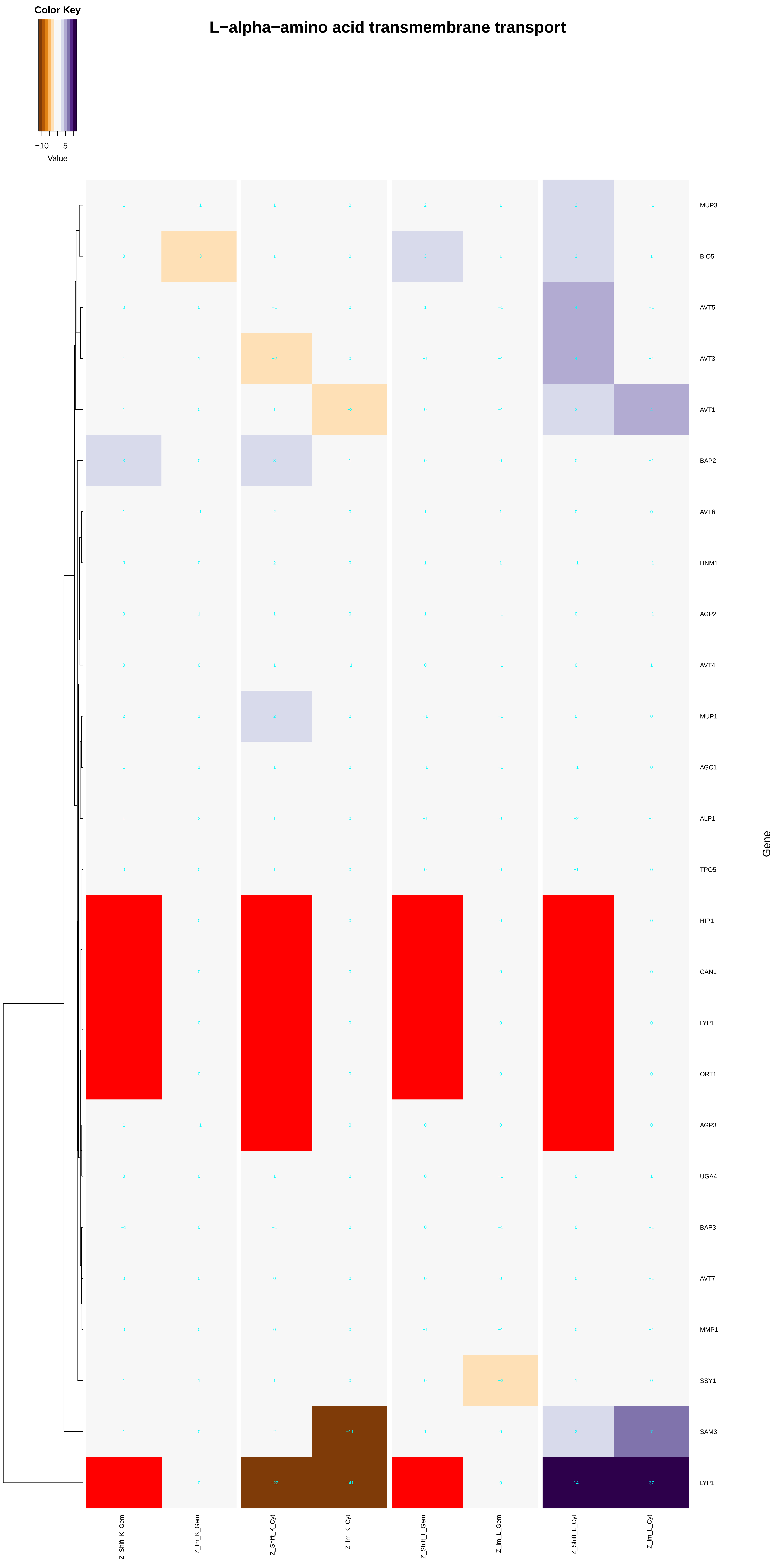

Color Key
−10
5
Value
L−alpha−amino acid transmembrane transport
MUP3
BIO5
AVT5
AVT3
AVT1
BAP2
AVT6
HNM1
AGP2
AVT4
MUP1
AGC1
ALP1
TPO5
HIP1
CAN1
LYP1
ORT1
AGP3
UGA4
BAP3
AVT7
MMP1
SSY1
SAM3
LYP1
Gene
Z_Shift_K_Gem
Z_Im_K_Gem
Z_Shift_K_Cyt
Z_Im_K_Cyt
Z_Shift_L_Gem
Z_Im_L_Gem
Z_Shift_L_Cyt
Z_Im_L_Cyt

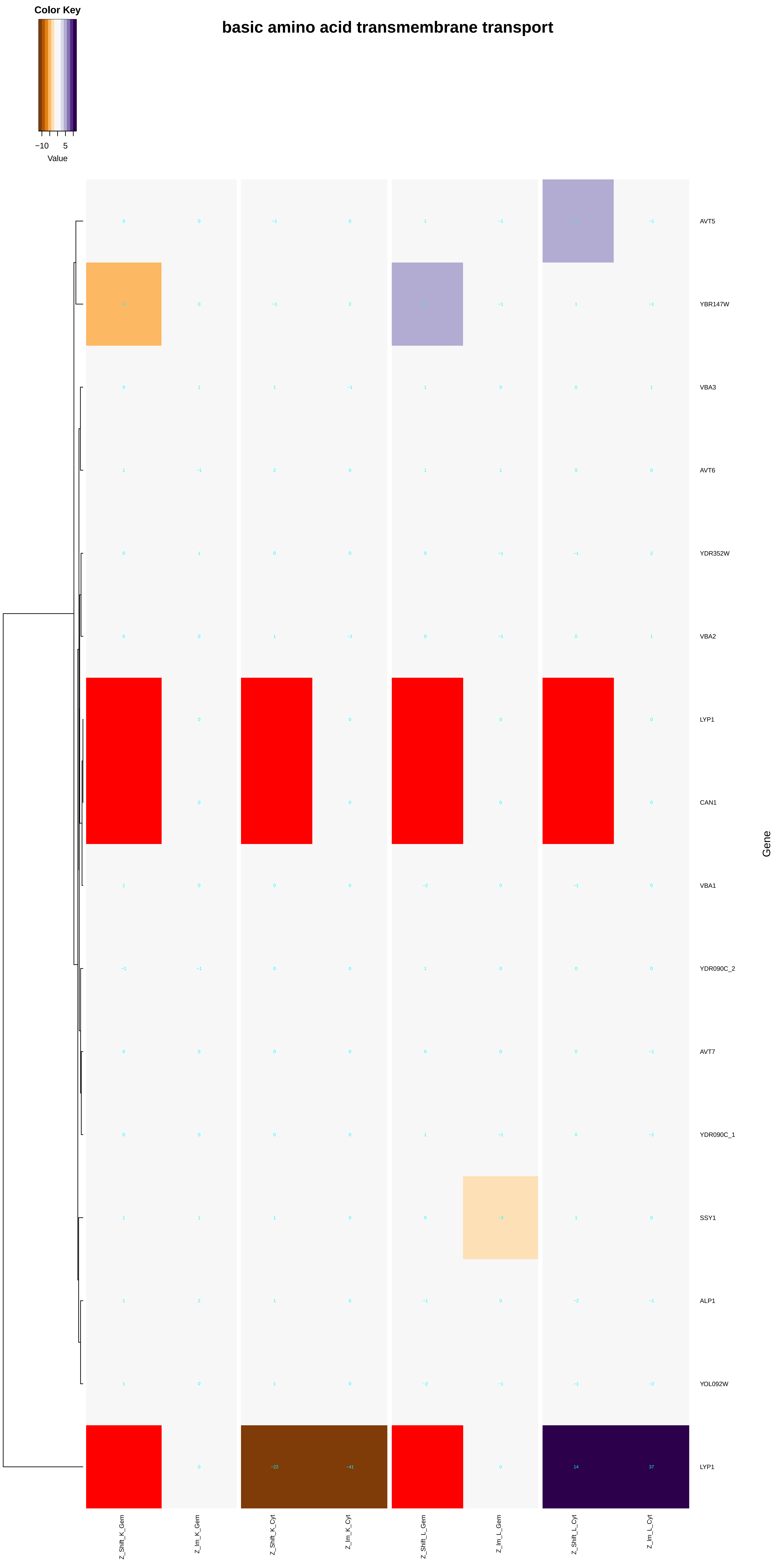
Color Key
−10
5
Value
basic amino acid transmembrane transport
AVT5
YBR147W
VBA3
AVT6
YDR352W
VBA2
LYP1
CAN1
VBA1
YDR090C_2
AVT7
YDR090C_1
SSY1
ALP1
YOL092W
LYP1
Gene
Z_Shift_K_Gem
Z_Im_K_Gem
Z_Shift_K_Cyt
Z_Im_K_Cyt
Z_Shift_L_Gem
Z_Im_L_Gem
Z_Shift_L_Cyt
Z_Im_L_Cyt

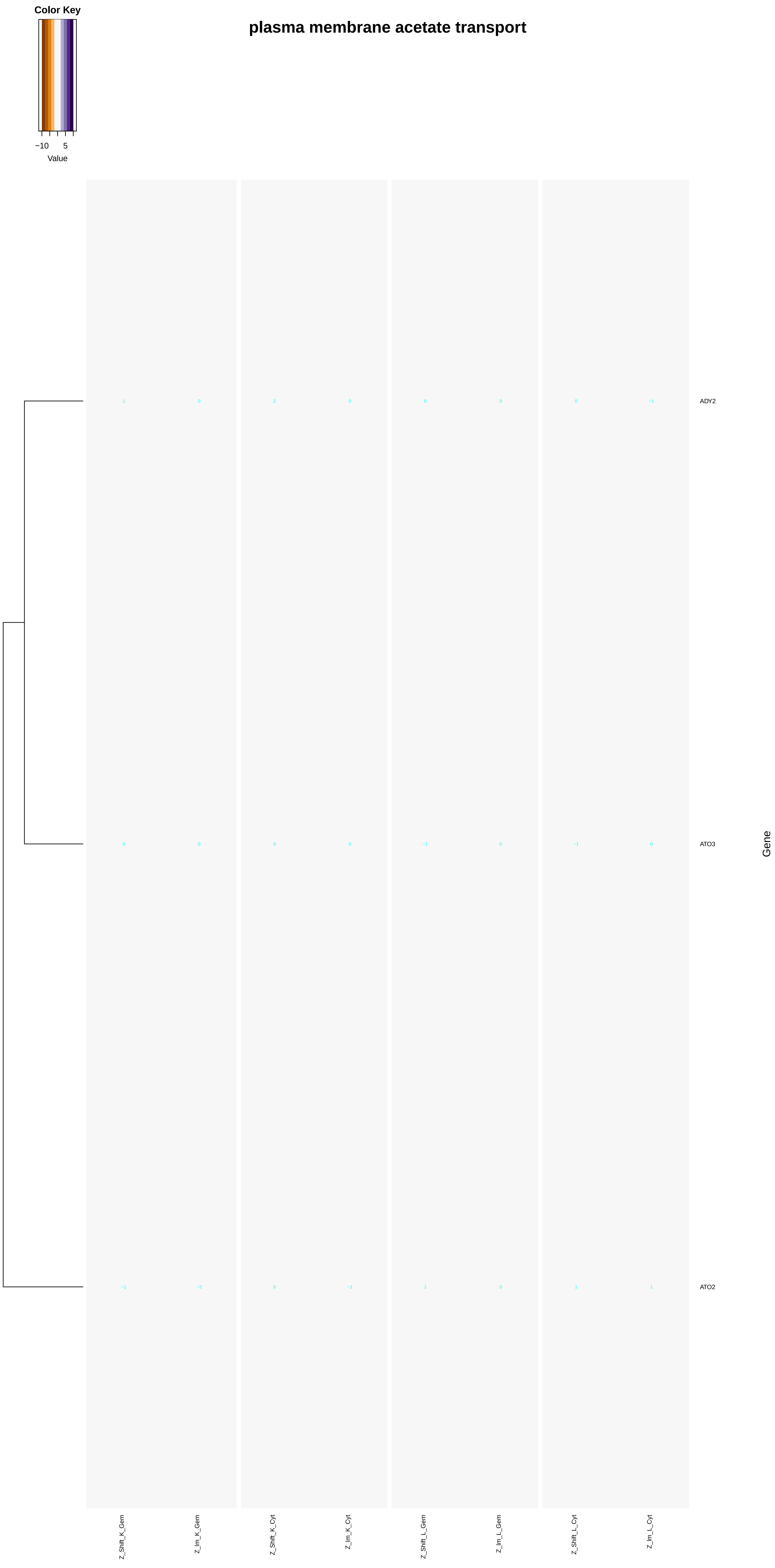
Color Key
−10
5
Value
plasma membrane acetate transport
1
0
2
0
0
0
0
−1
ADY2
0
0
0
0
−1
0
−1
0
ATO3
−1
−2
0
−1
1
0
1
1
ATO2
Gene
Z_Shift_K_Gem
Z_Im_K_Gem
Z_Shift_K_Cyt
Z_Im_K_Cyt
Z_Shift_L_Gem
Z_Im_L_Gem
Z_Shift_L_Cyt
Z_Im_L_Cyt

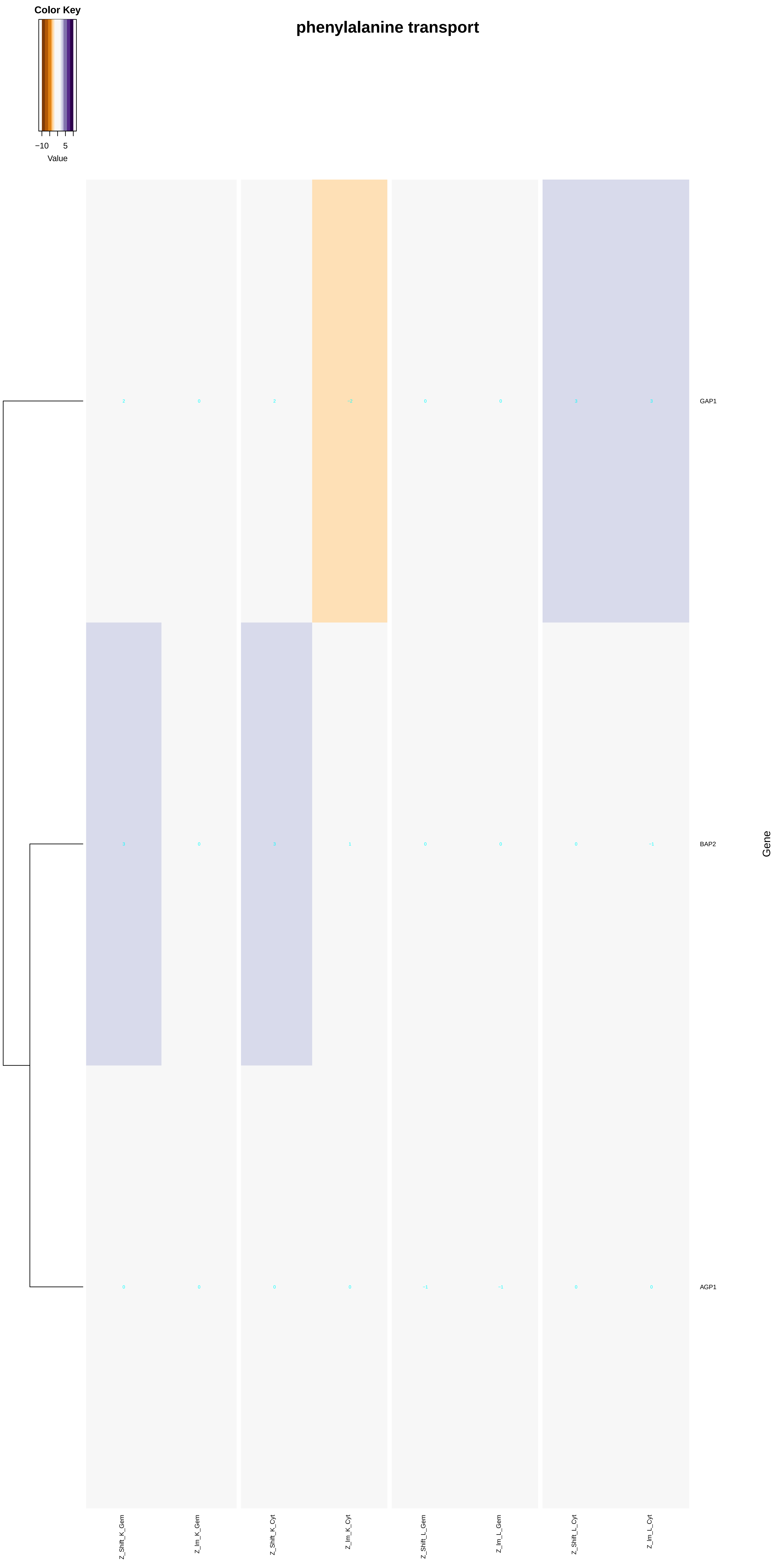

Color Key
−10
5
Value
phenylalanine transport
GAP1
BAP2
AGP1
Gene
Z_Shift_K_Gem
Z_Im_K_Gem
Z_Shift_K_Cyt
Z_Im_K_Cyt
Z_Shift_L_Gem
Z_Im_L_Gem
Z_Shift_L_Cyt
Z_Im_L_Cyt

### tyrosine transport

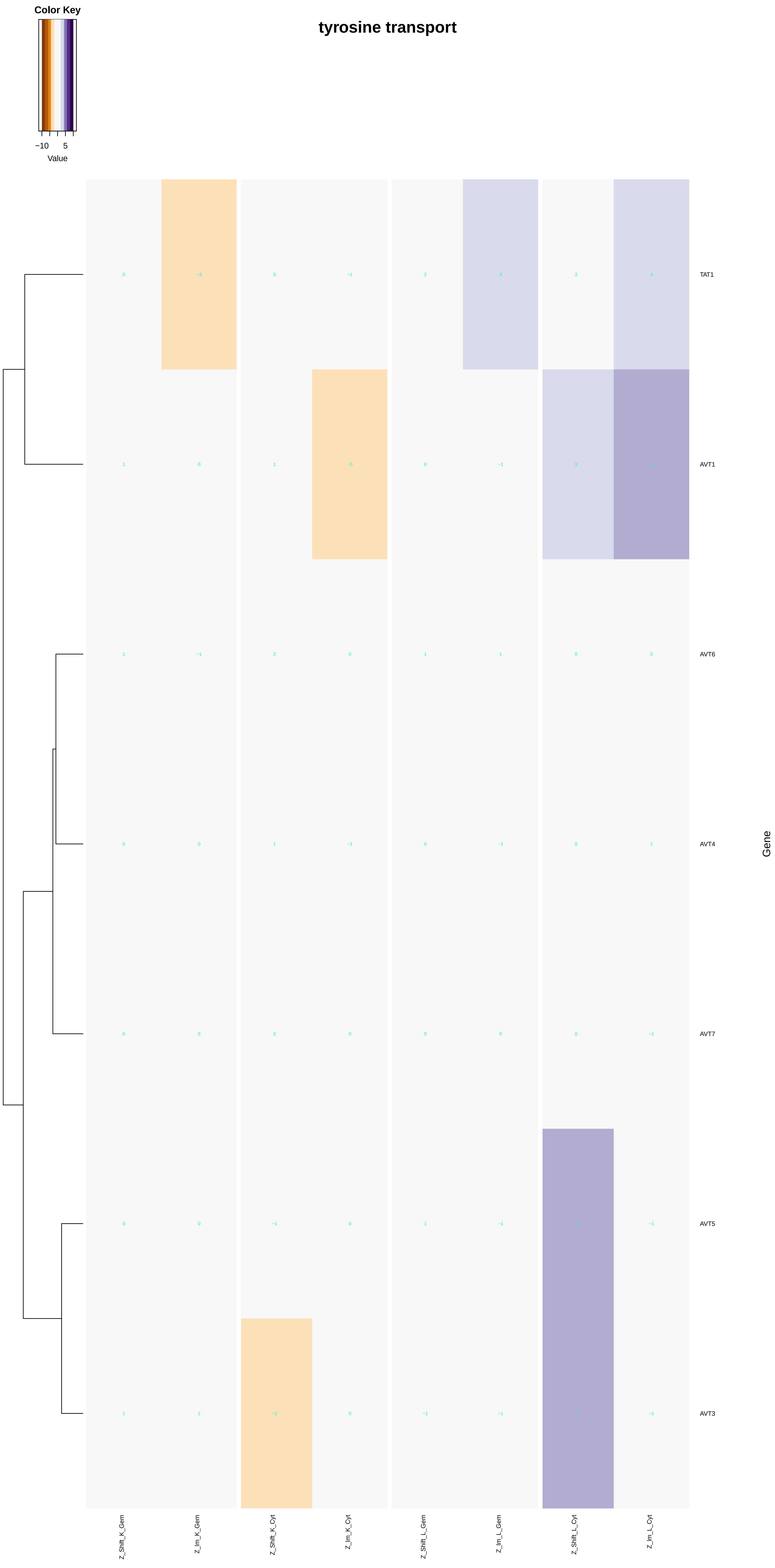

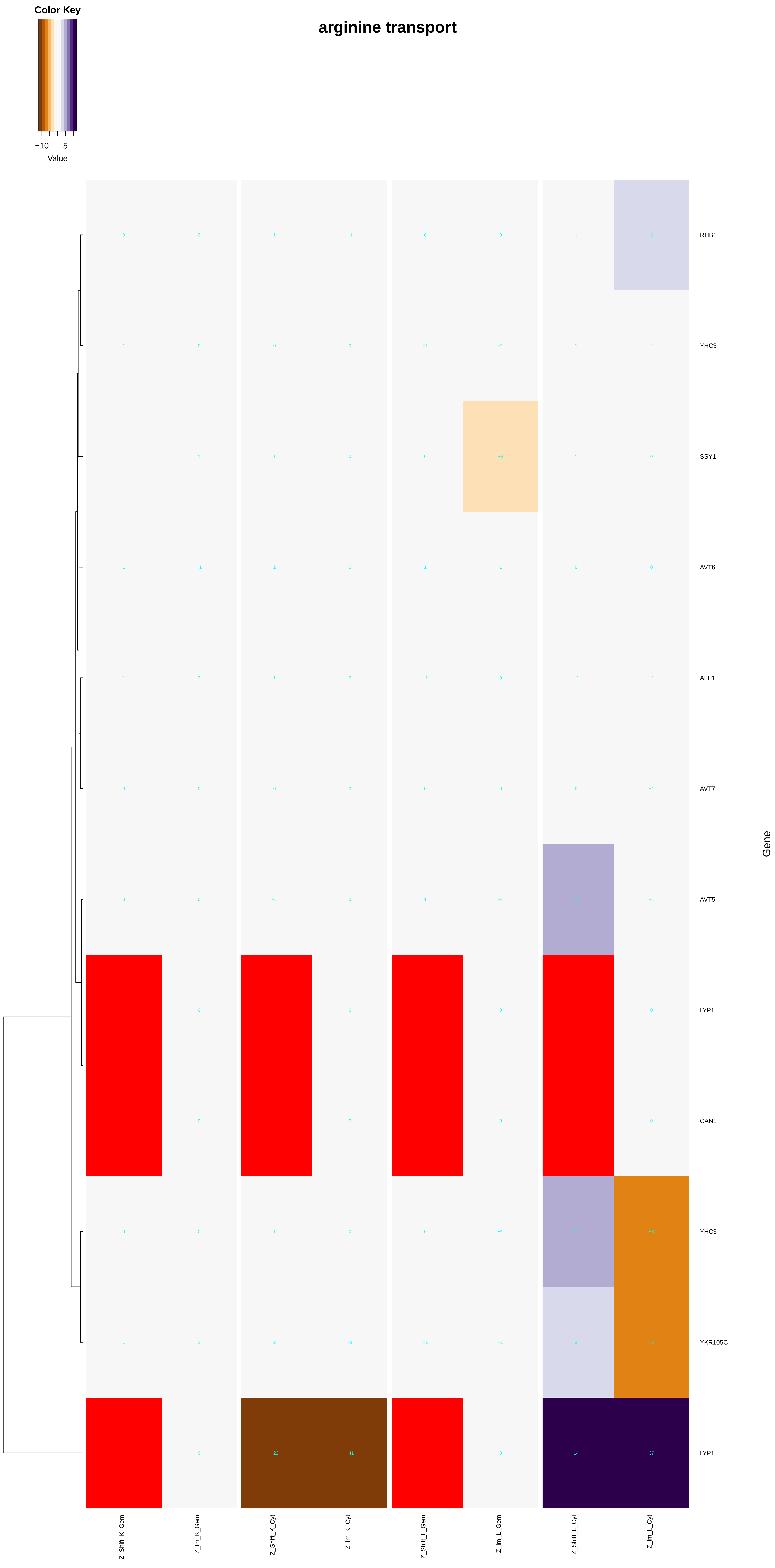
Color Key
−10
5
Value
arginine transport
RHB1
YHC3
SSY1
AVT6
ALP1
AVT7
AVT5
LYP1
CAN1
YHC3
YKR105C
LYP1
Gene
Z_Shift_K_Gem
Z_Im_K_Gem
Z_Shift_K_Cyt
Z_Im_K_Cyt
Z_Shift_L_Gem
Z_Im_L_Gem
Z_Shift_L_Cyt
Z_Im_L_Cyt

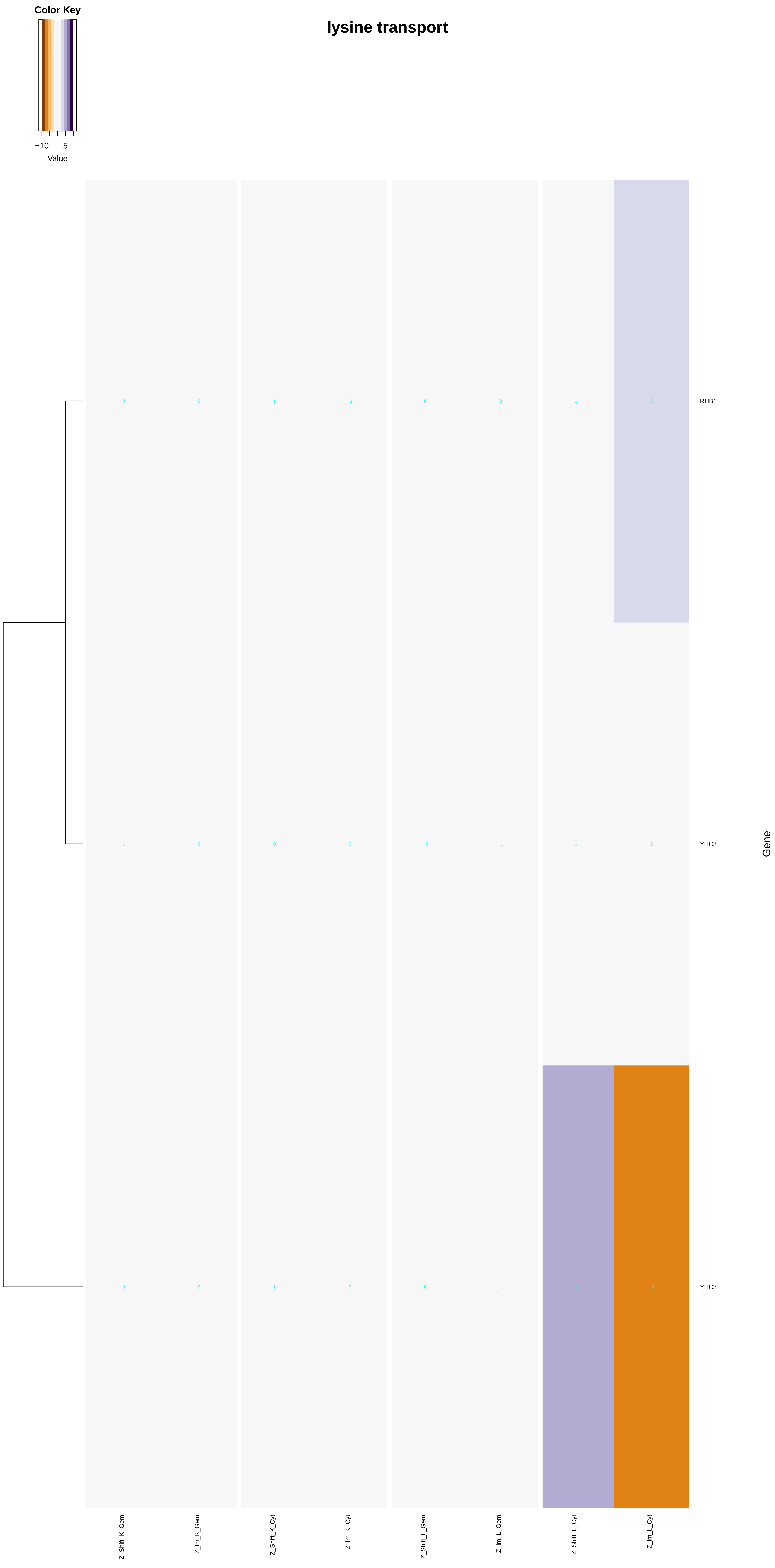
Color Key
−10
5
Value
lysine transport
RHB1
YHC3
YHC3
Gene
Z_Shift_K_Gem
Z_Im_K_Gem
Z_Shift_K_Cyt
Z_Im_K_Cyt
Z_Shift_L_Gem
Z_Im_L_Gem
Z_Shift_L_Cyt
Z_Im_L_Cyt

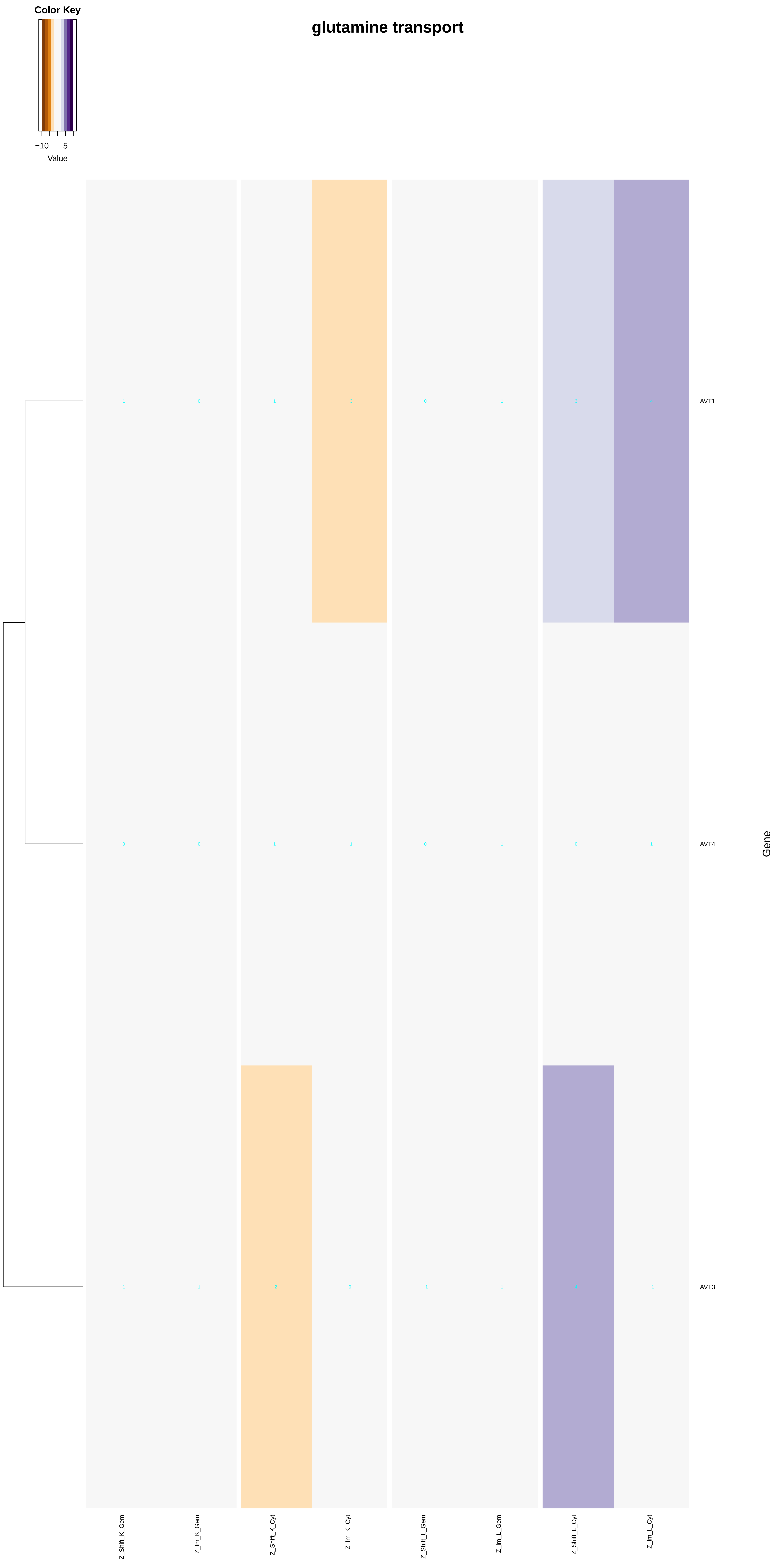
Color Key
−10
5
Value
glutamine transport
AVT1
AVT4
AVT3
Gene
Z_Shift_K_Gem
Z_Im_K_Gem
Z_Shift_K_Cyt
Z_Im_K_Cyt
Z_Shift_L_Gem
Z_Im_L_Gem
Z_Shift_L_Cyt
Z_Im_L_Cyt

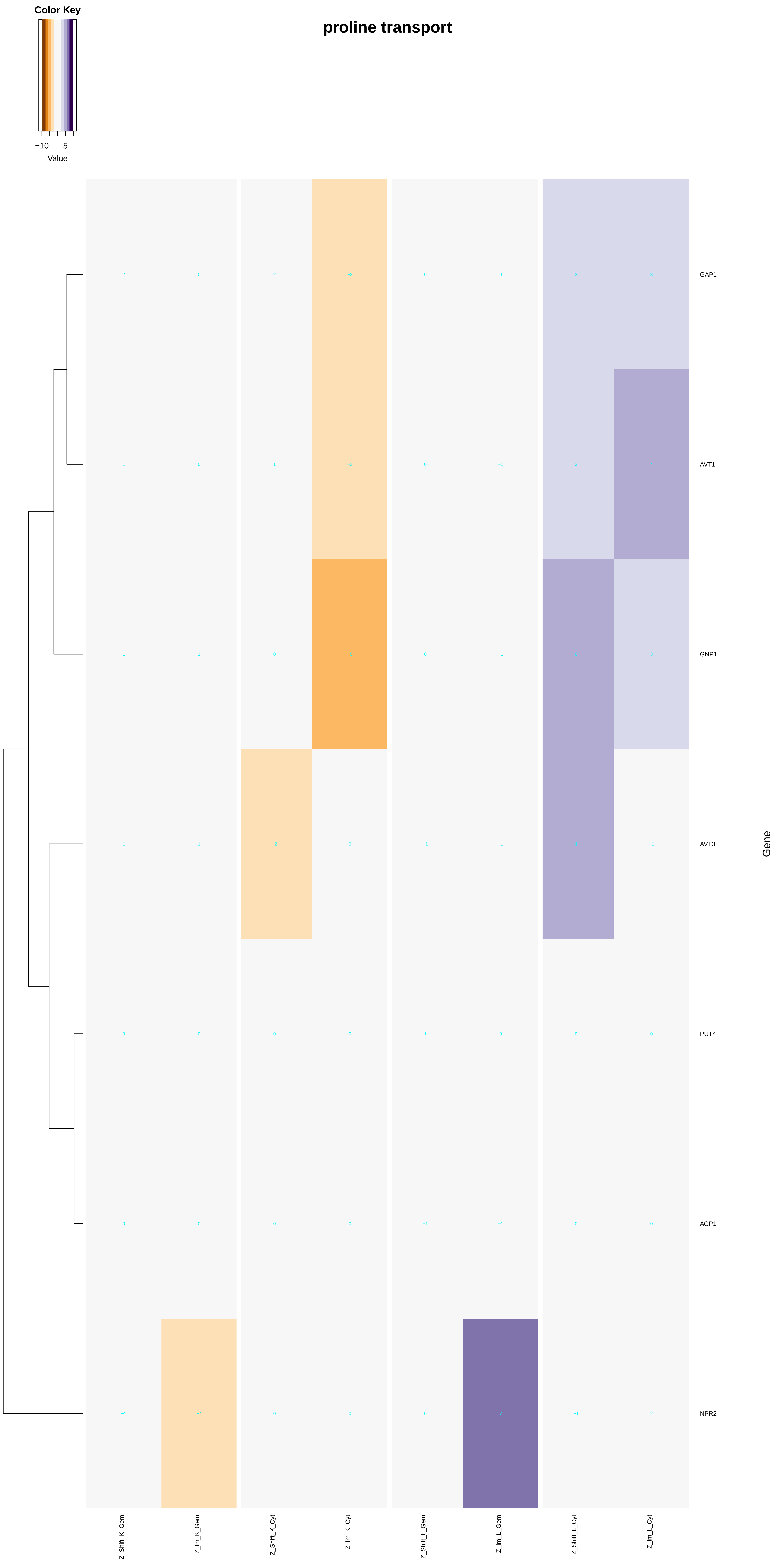
Color Key
−10
5
Value
proline transport
GAP1
AVT1
GNP1
AVT3
PUT4
AGP1
NPR2
Gene
Z_Shift_K_Gem
Z_Im_K_Gem
Z_Shift_K_Cyt
Z_Im_K_Cyt
Z_Shift_L_Gem
Z_Im_L_Gem
Z_Shift_L_Cyt
Z_Im_L_Cyt

#### serine transport

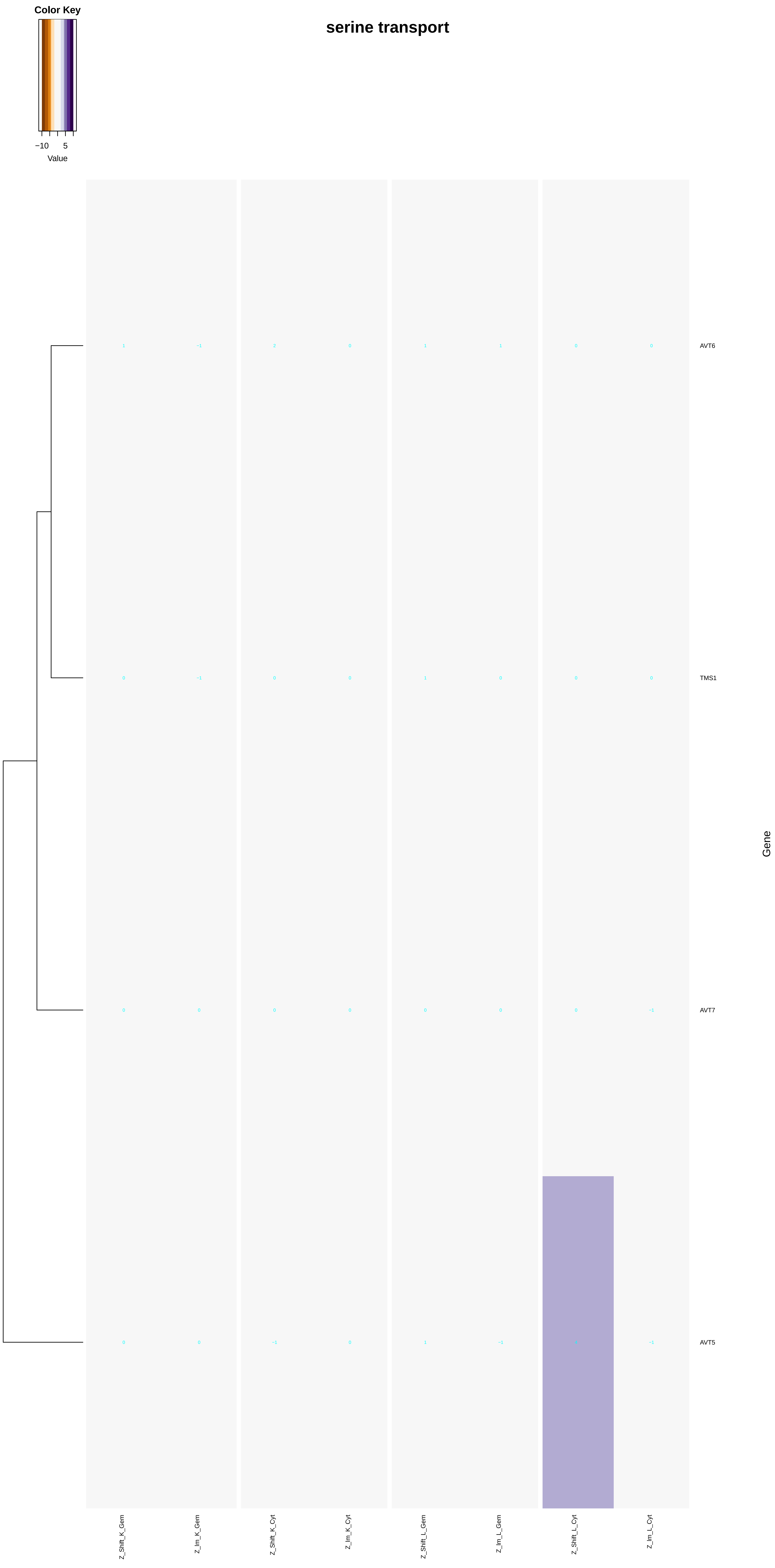

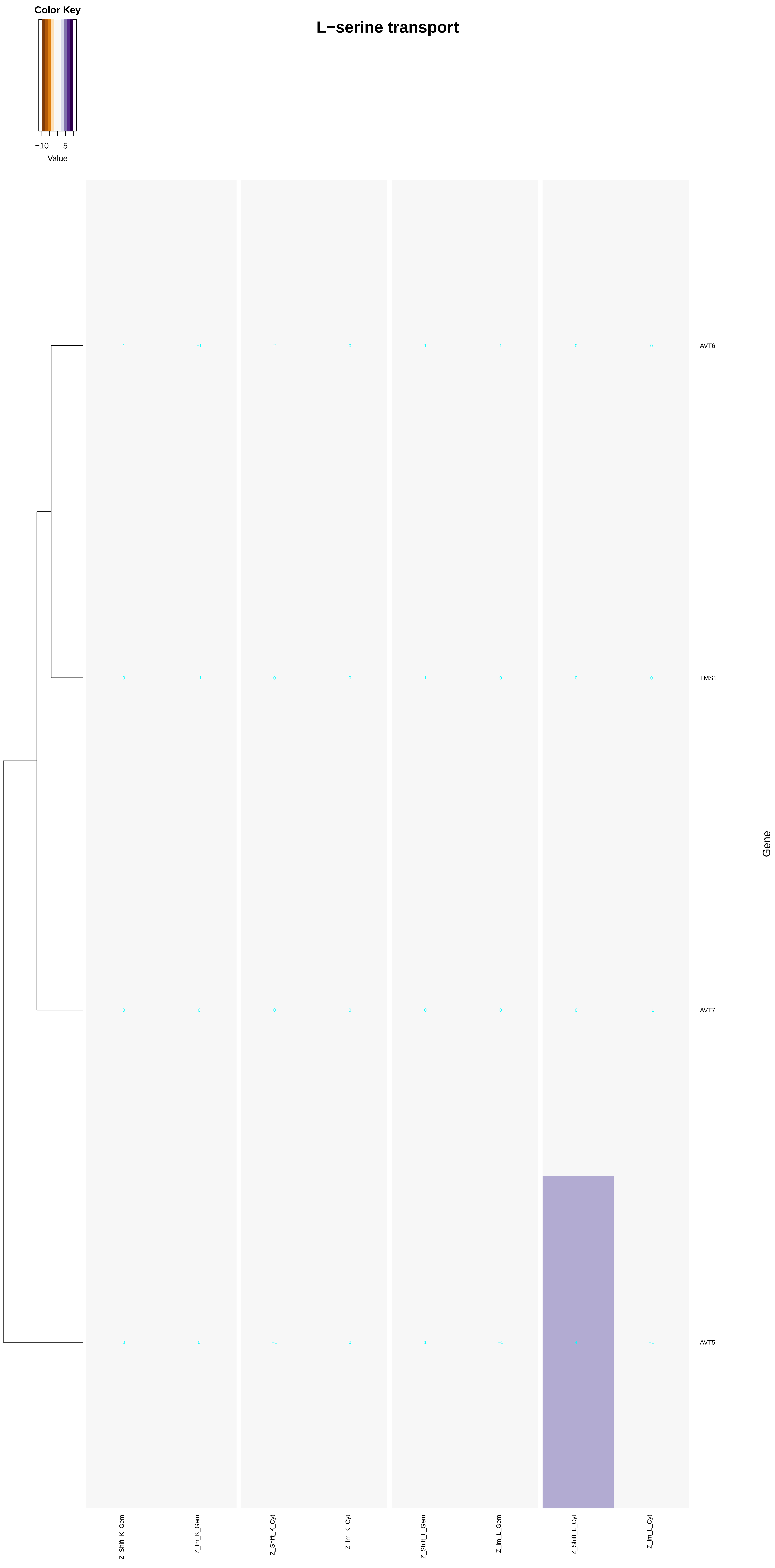
Color Key
−10
5
Value
L−serine transport
AVT6
TMS1
AVT7
AVT5
Gene
Z_Shift_K_Gem
Z_Im_K_Gem
Z_Shift_K_Cyt
Z_Im_K_Cyt
Z_Shift_L_Gem
Z_Im_L_Gem
Z_Shift_L_Cyt
Z_Im_L_Cyt

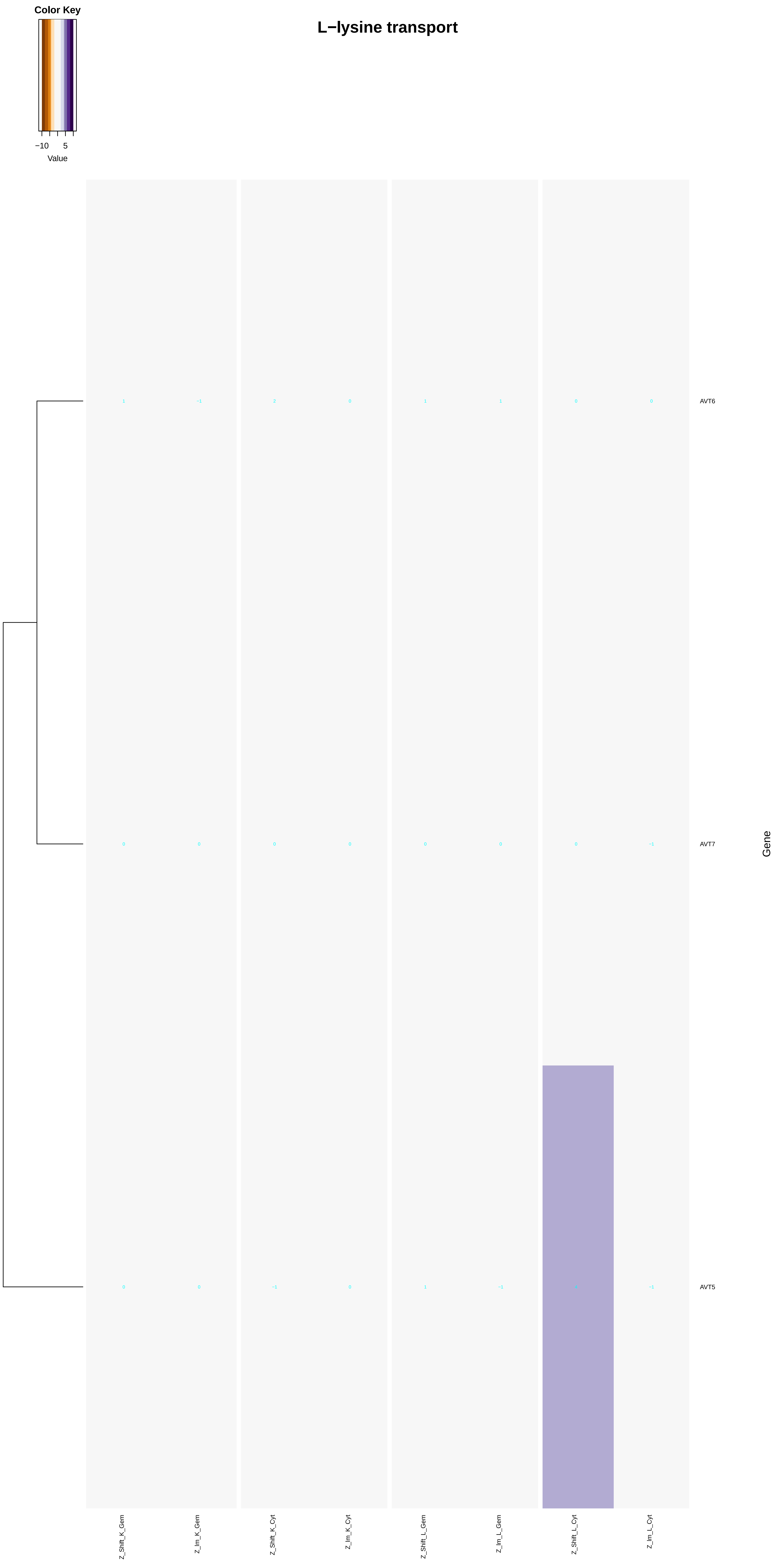
Color Key
−10
5
Value
L−lysine transport
AVT6
AVT7
AVT5
Gene
Z_Shift_K_Gem
Z_Im_K_Gem
Z_Shift_K_Cyt
Z_Im_K_Cyt
Z_Shift_L_Gem
Z_Im_L_Gem
Z_Shift_L_Cyt
Z_Im_L_Cyt

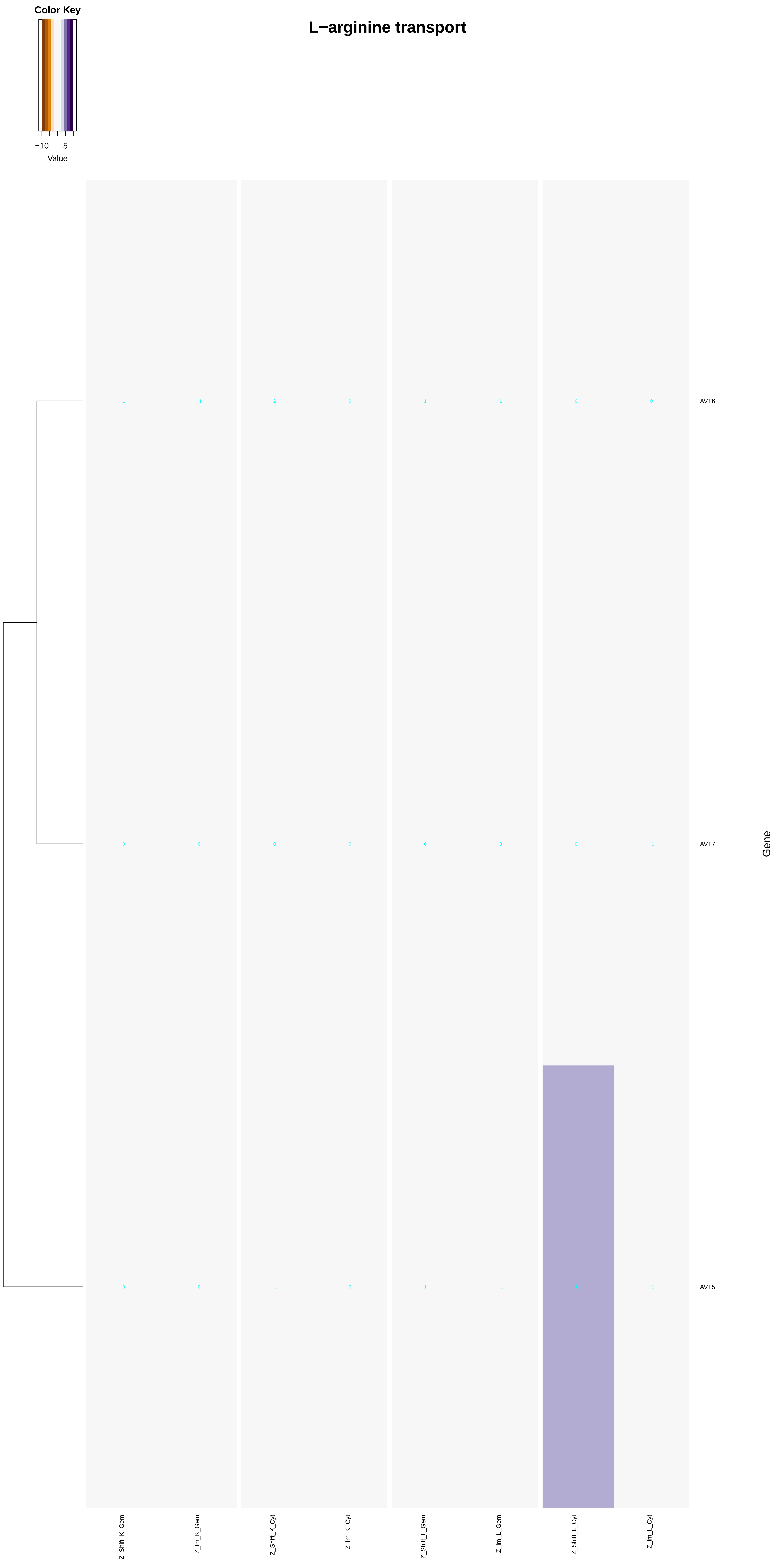

Color Key
−10
5
Value
L−arginine transport
AVT6
AVT7
AVT5
Gene
Z_Shift_K_Gem
Z_Im_K_Gem
Z_Shift_K_Cyt
Z_Im_K_Cyt
Z_Shift_L_Gem
Z_Im_L_Gem
Z_Shift_L_Cyt
Z_Im_L_Cyt

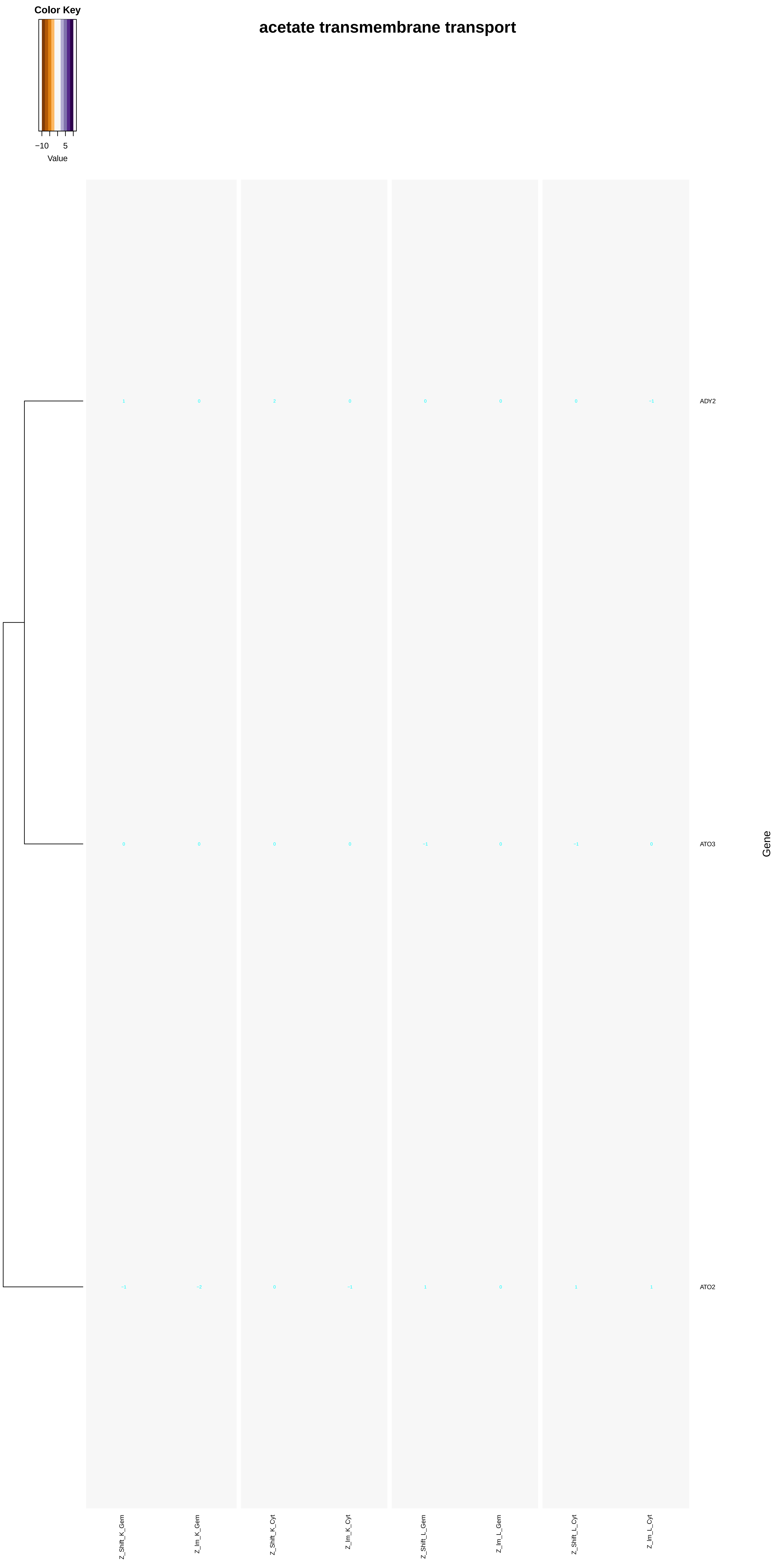
Color Key
−10
5
Value
acetate transmembrane transport
ADY2
ATO3
ATO2
Gene
Z_Shift_K_Gem
Z_Im_K_Gem
Z_Shift_K_Cyt
Z_Im_K_Cyt
Z_Shift_L_Gem
Z_Im_L_Gem
Z_Shift_L_Cyt
Z_Im_L_Cyt

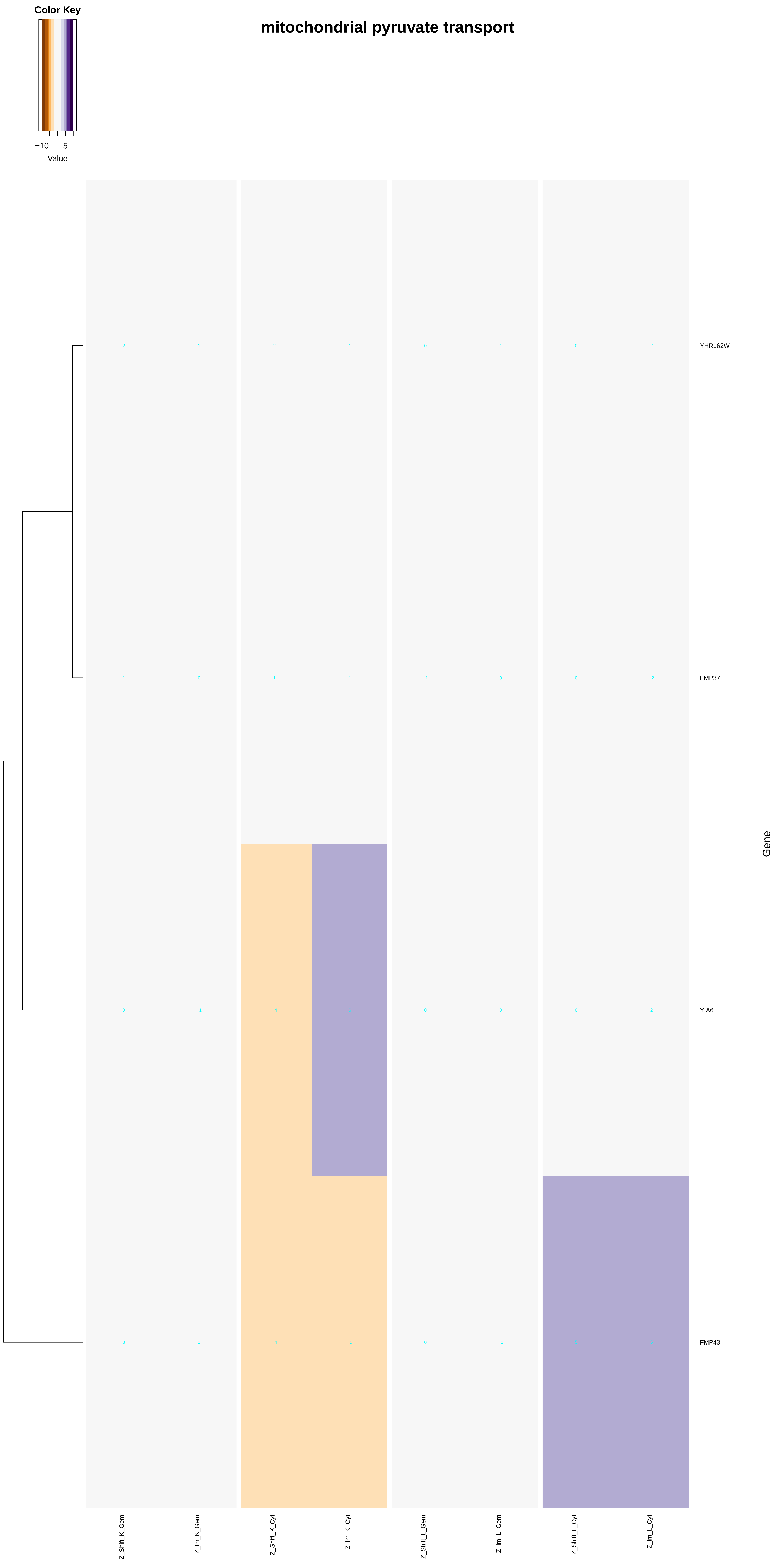
Color Key
mitochondrial pyruvate transport
−10
5
Value
YHR162W
FMP37
YIA6
FMP43
Gene
Z_Shift_K_Gem
Z_Im_K_Gem
Z_Shift_K_Cyt
Z_Im_K_Cyt
Z_Shift_L_Gem
Z_Im_L_Gem
Z_Shift_L_Cyt
Z_Im_L_Cyt

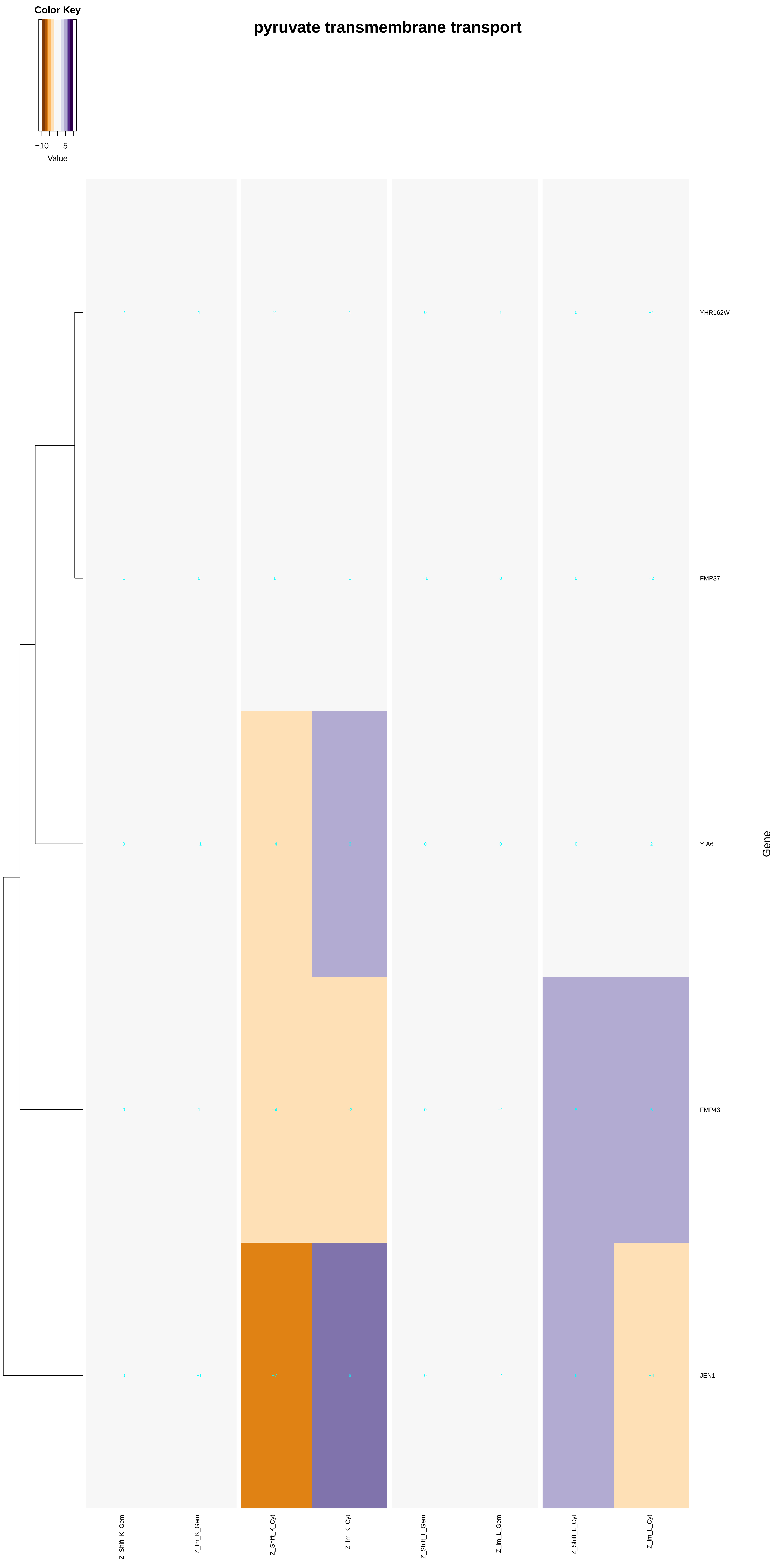

Color Key
−10
5
Value
pyruvate transmembrane transport
YHR162W
FMP37
YIA6
FMP43
JEN1
Gene
Z_Shift_K_Gem
Z_lm_K_Gem
Z_Shift_K_Cyt
Z_lm_K_Cyt
Z_Shift_L_Gem
Z_lm_L_Gem
Z_Shift_L_Cyt
Z_lm_L_Cyt

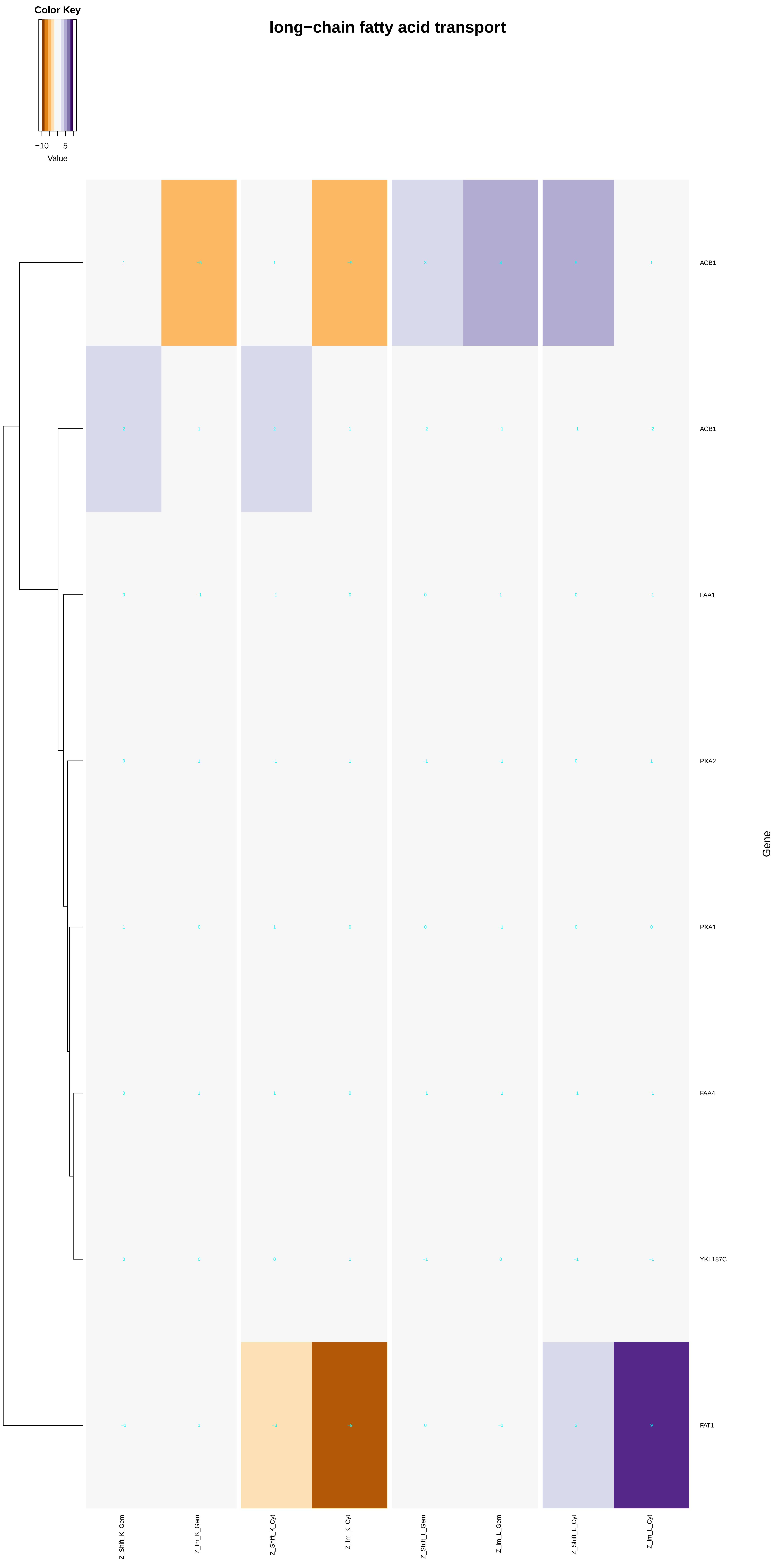
Color Key
−10
5
Value
long−chain fatty acid transport
ACB1
ACB1
FAA1
PXA2
PXA1
FAA4
YKL187C
FAT1
Gene
Z_Shift_K_Gem
Z_Im_K_Gem
Z_Shift_K_Cyt
Z_Im_K_Cyt
Z_Shift_L_Gem
Z_Im_L_Gem
Z_Shift_L_Cyt
Z_Im_L_Cyt

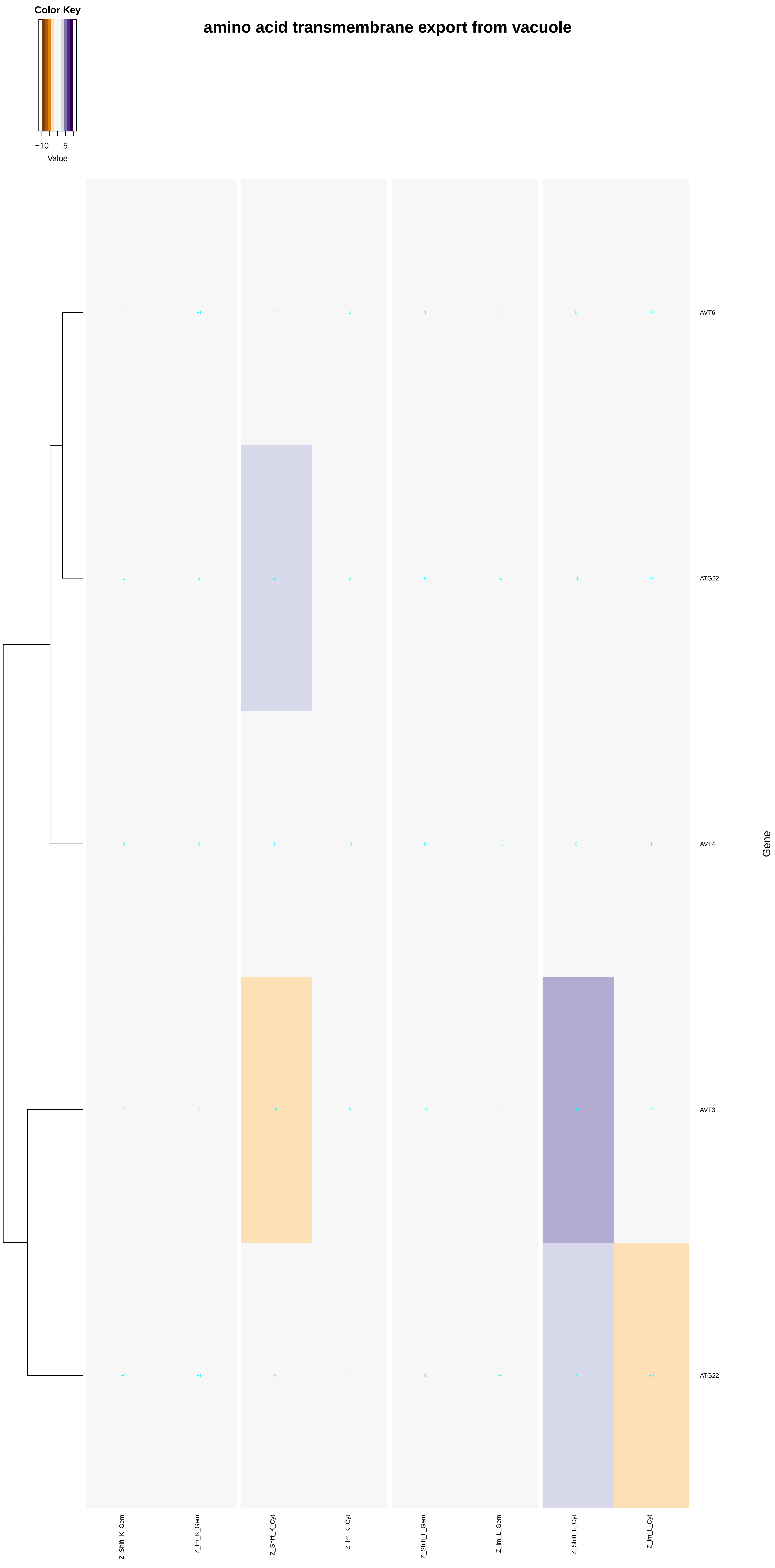
amino acid transmembrane export from vacuole
Color Key
−10
5
Value
AVT6
ATG22
AVT4
AVT3
ATG22
Gene
Z_Shift_K_Gem
Z_Im_K_Gem
Z_Shift_K_Cyt
Z_Im_K_Cyt
Z_Shift_L_Gem
Z_Im_L_Gem
Z_Shift_L_Cyt
Z_Im_L_Cyt

### L−histidine transmembrane transport

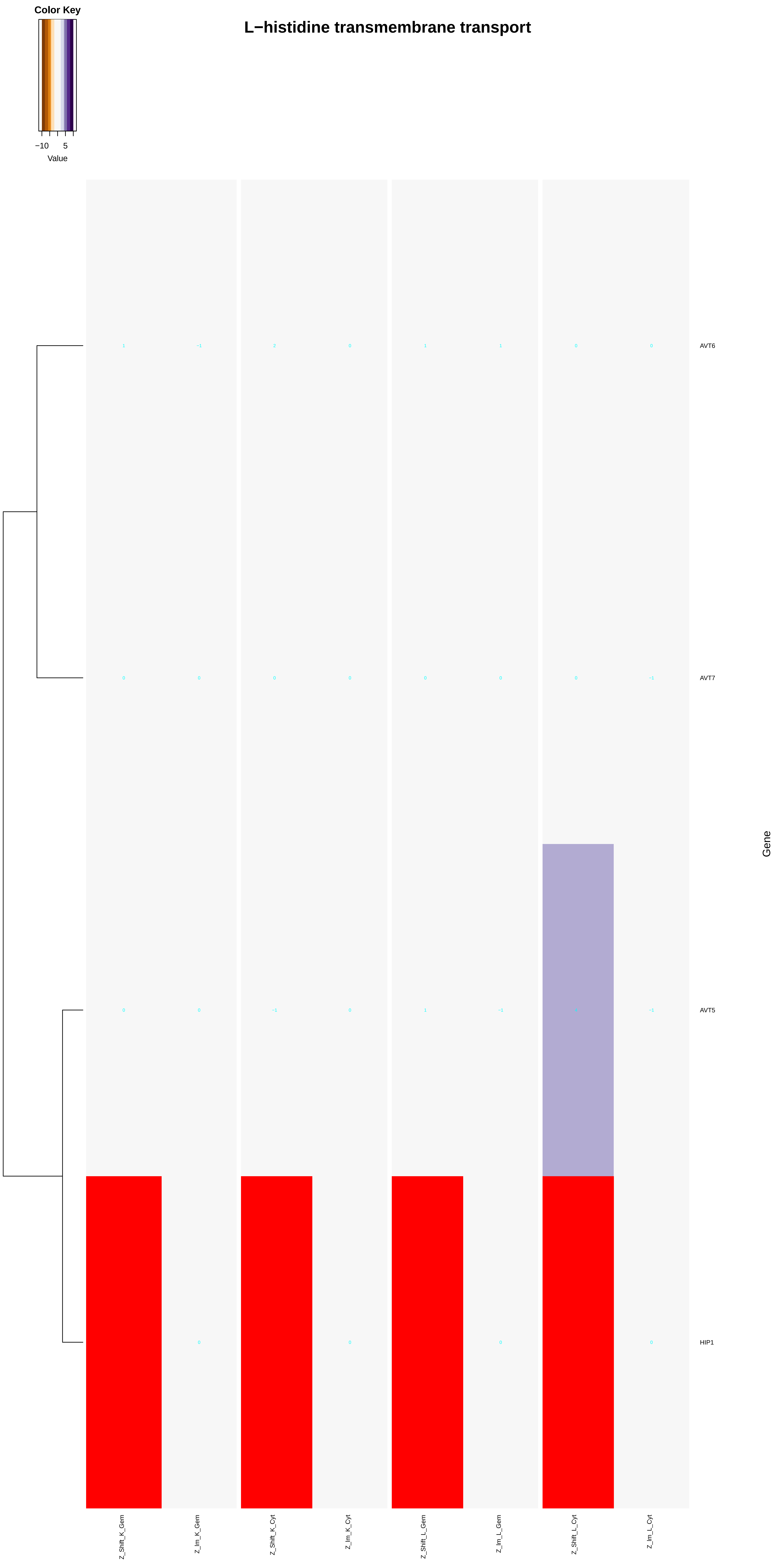

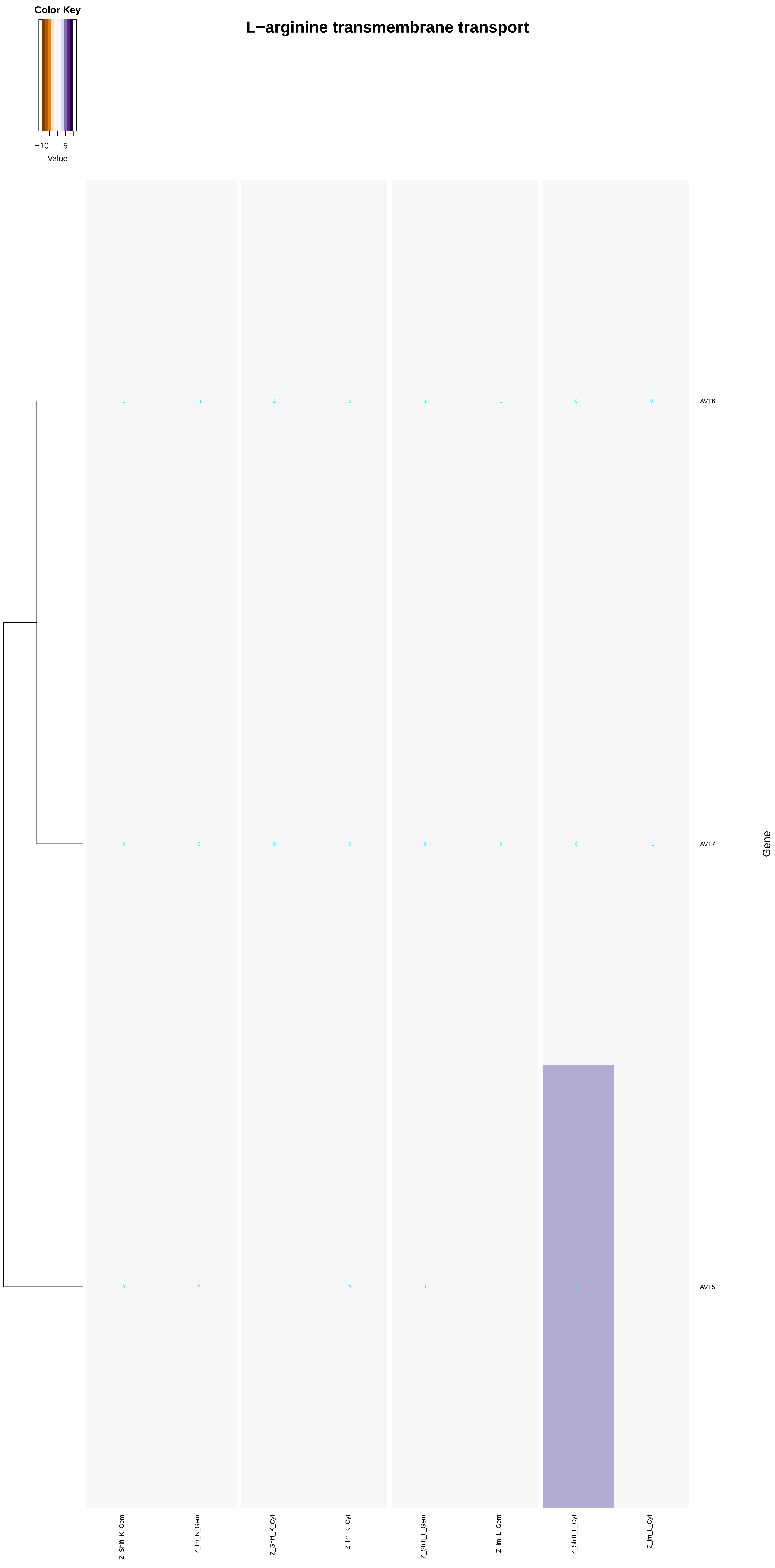
Color Key
−10
5
Value
L−arginine transmembrane transport
1
−1
2
0
1
1
0
0
AVT6
0
0
0
0
0
0
0
−1
AVT7
0
0
−1
0
1
−1
4
−1
AVT5
Gene
Z_Shift_K_Gem
Z_Im_K_Gem
Z_Shift_K_Cyt
Z_Im_K_Cyt
Z_Shift_L_Gem
Z_Im_L_Gem
Z_Shift_L_Cyt
Z_Im_L_Cyt

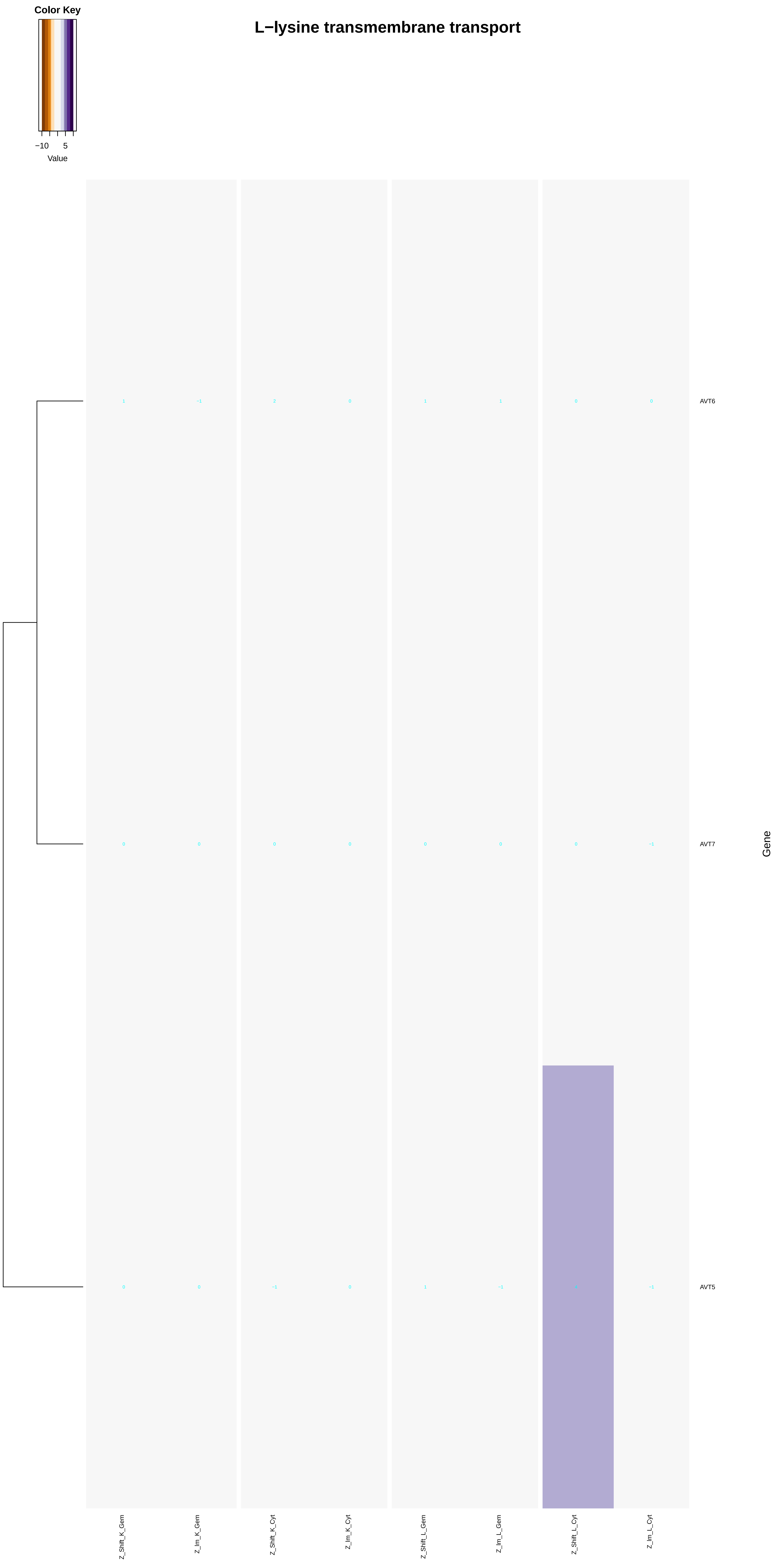

L−lysine transmembrane transport
Color Key
−10
5
Value
AVT6
AVT7
AVT5
Gene
Z_Shift_K_Gem
Z_Im_K_Gem
Z_Shift_K_Cyt
Z_Im_K_Cyt
Z_Shift_L_Gem
Z_Im_L_Gem
Z_Shift_L_Cyt
Z_Im_L_Cyt
1 −1 2 0 1 1 0 0
0 0 0 0 0 0 0 −1
0 0 −1 0 1 −1 −1

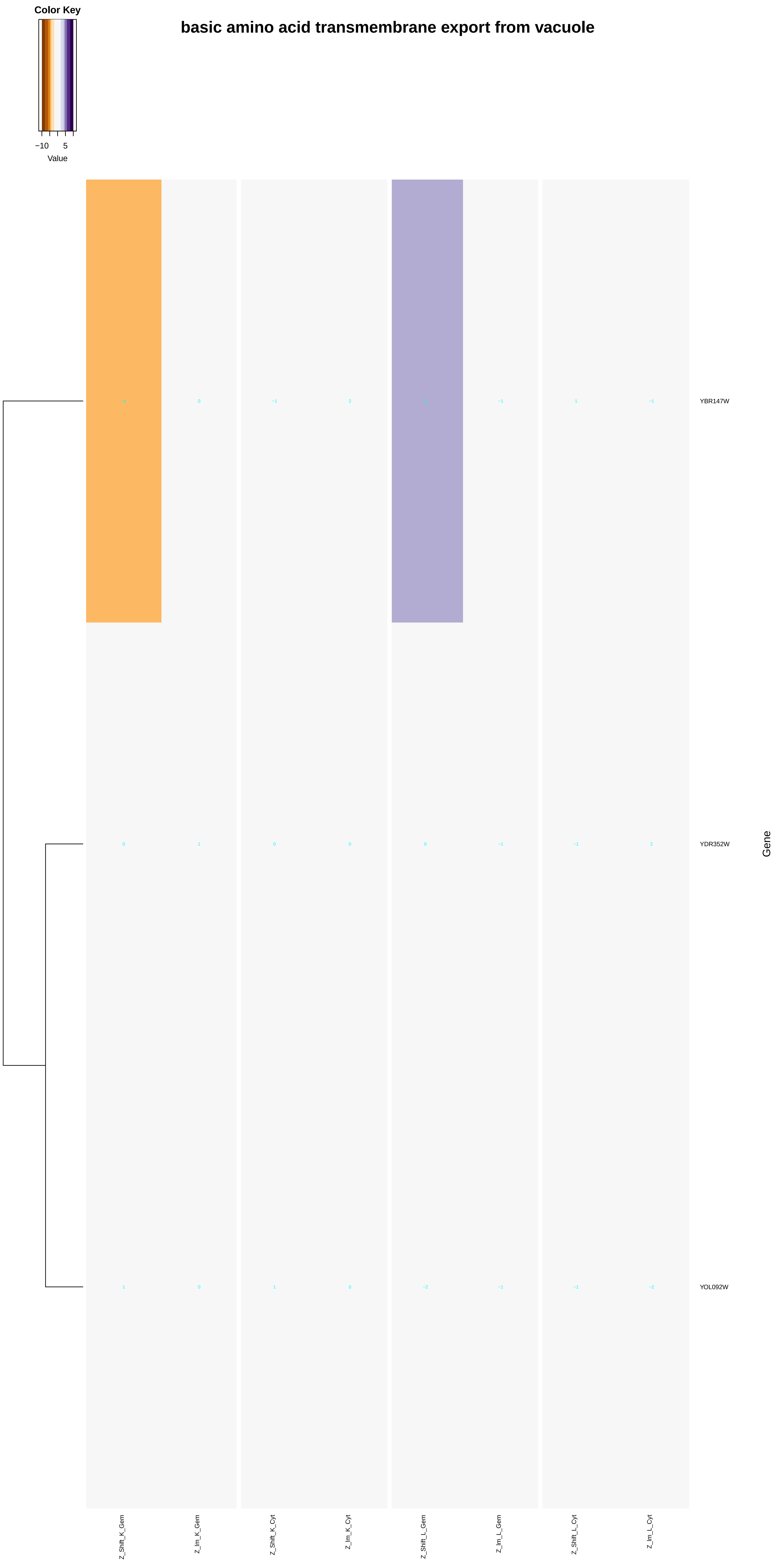
basic amino acid transmembrane export from vacuole
Color Key
−10
5
Value
YBR147W
YDR352W
YOL092W
Gene
Z_Shift_K_Gem
Z_Im_K_Gem
Z_Shift_K_Cyt
Z_Im_K_Cyt
Z_Shift_L_Gem
Z_Im_L_Gem
Z_Shift_L_Cyt
Z_Im_L_Cyt

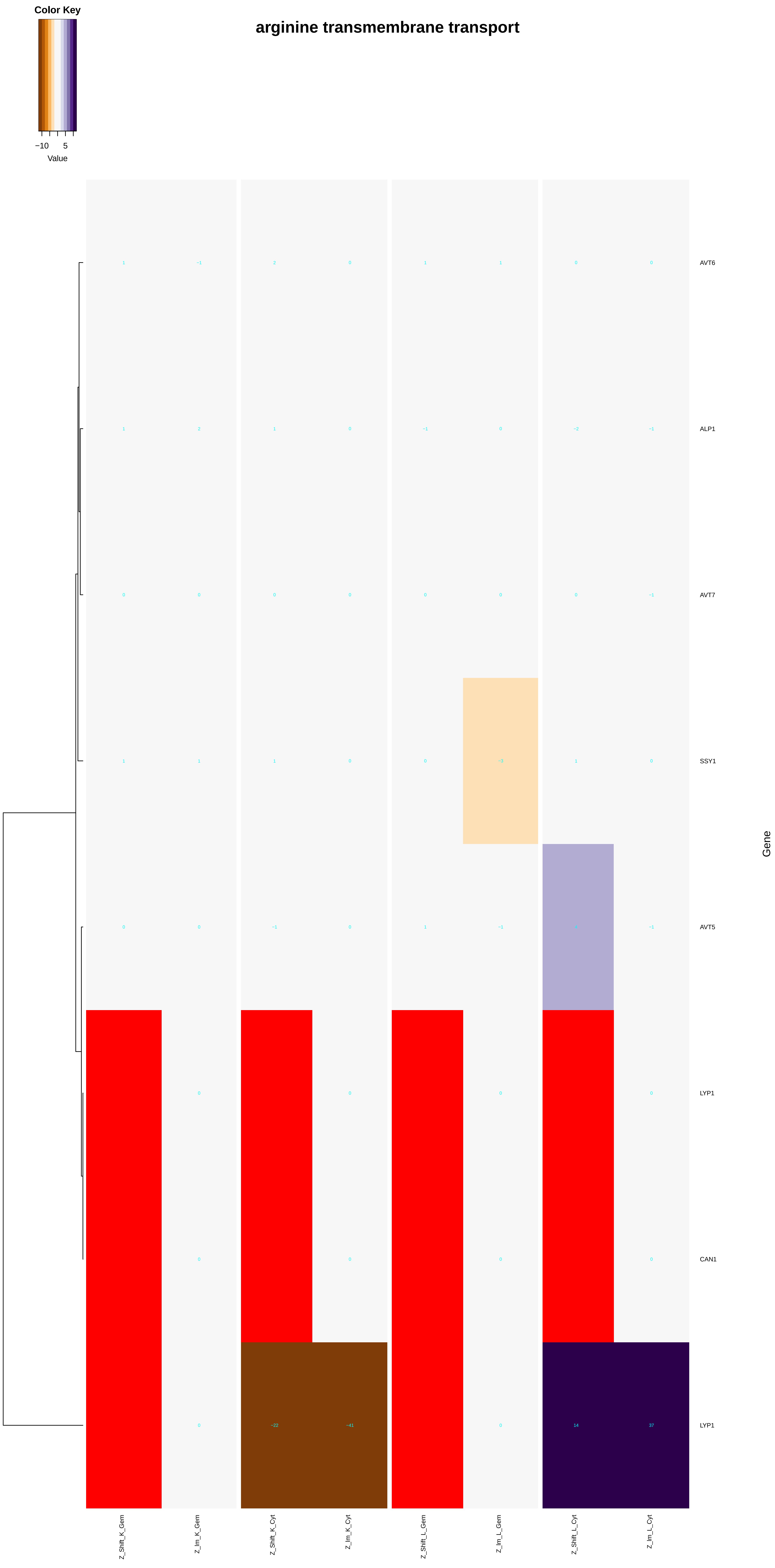

arginine transmembrane transport
Color Key
−10
5
Value
AVT6
ALP1
AVT7
SSY1
AVT5
LYP1
CAN1
LYP1
Gene
Z_Shift_K_Gem
Z_Im_K_Gem
Z_Shift_K_Cyt
Z_Im_K_Cyt
Z_Shift_L_Gem
Z_Im_L_Gem
Z_Shift_L_Cyt
Z_Im_L_Cyt

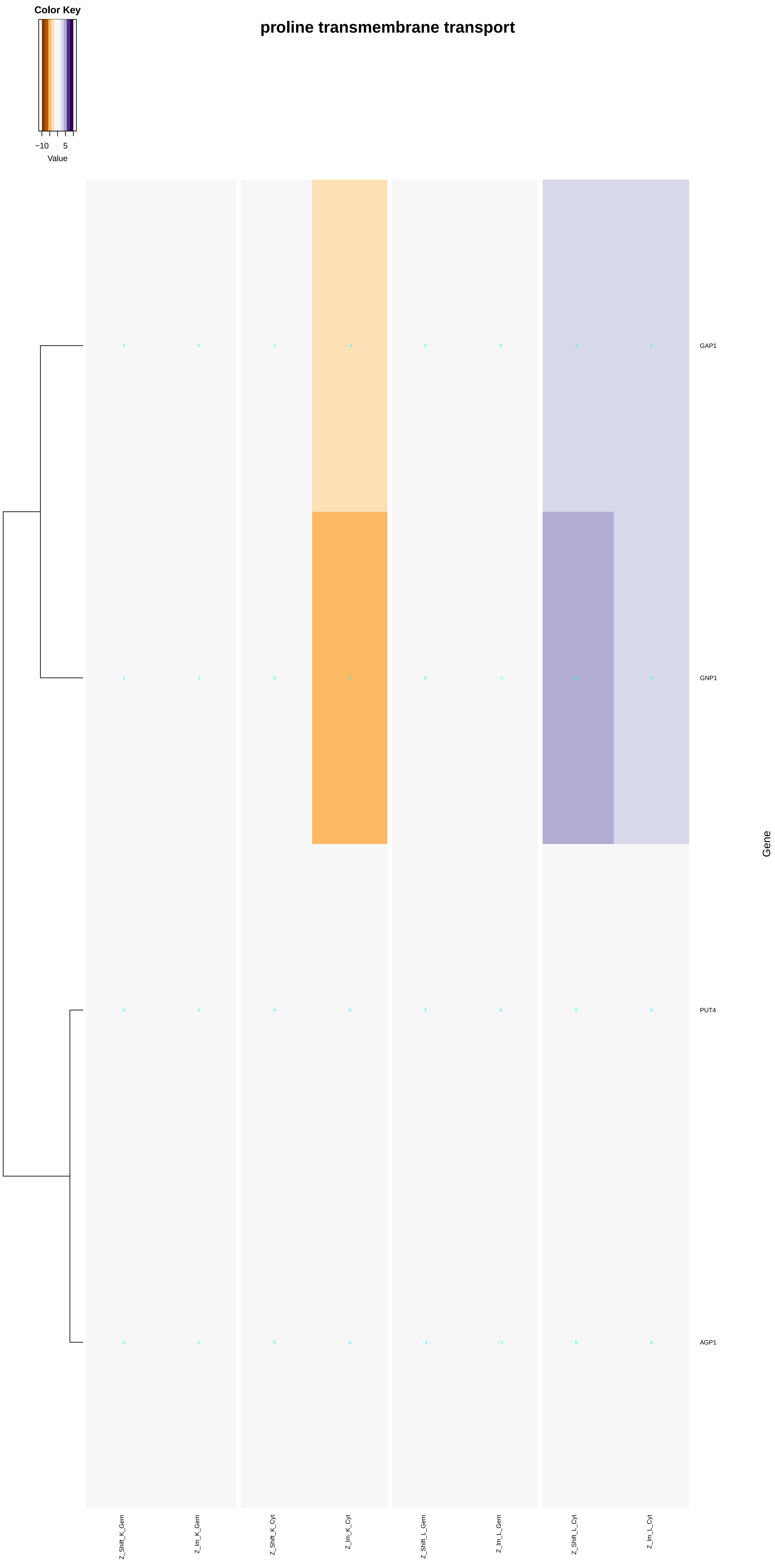
proline transmembrane transport
Color Key
−10
5
Value
GAP1
GNP1
PUT4
AGP1
Gene
Z_Shift_K_Gem
Z_Im_K_Gem
Z_Shift_K_Cyt
Z_Im_K_Cyt
Z_Shift_L_Gem
Z_Im_L_Gem
Z_Shift_L_Cyt
Z_Im_L_Cyt

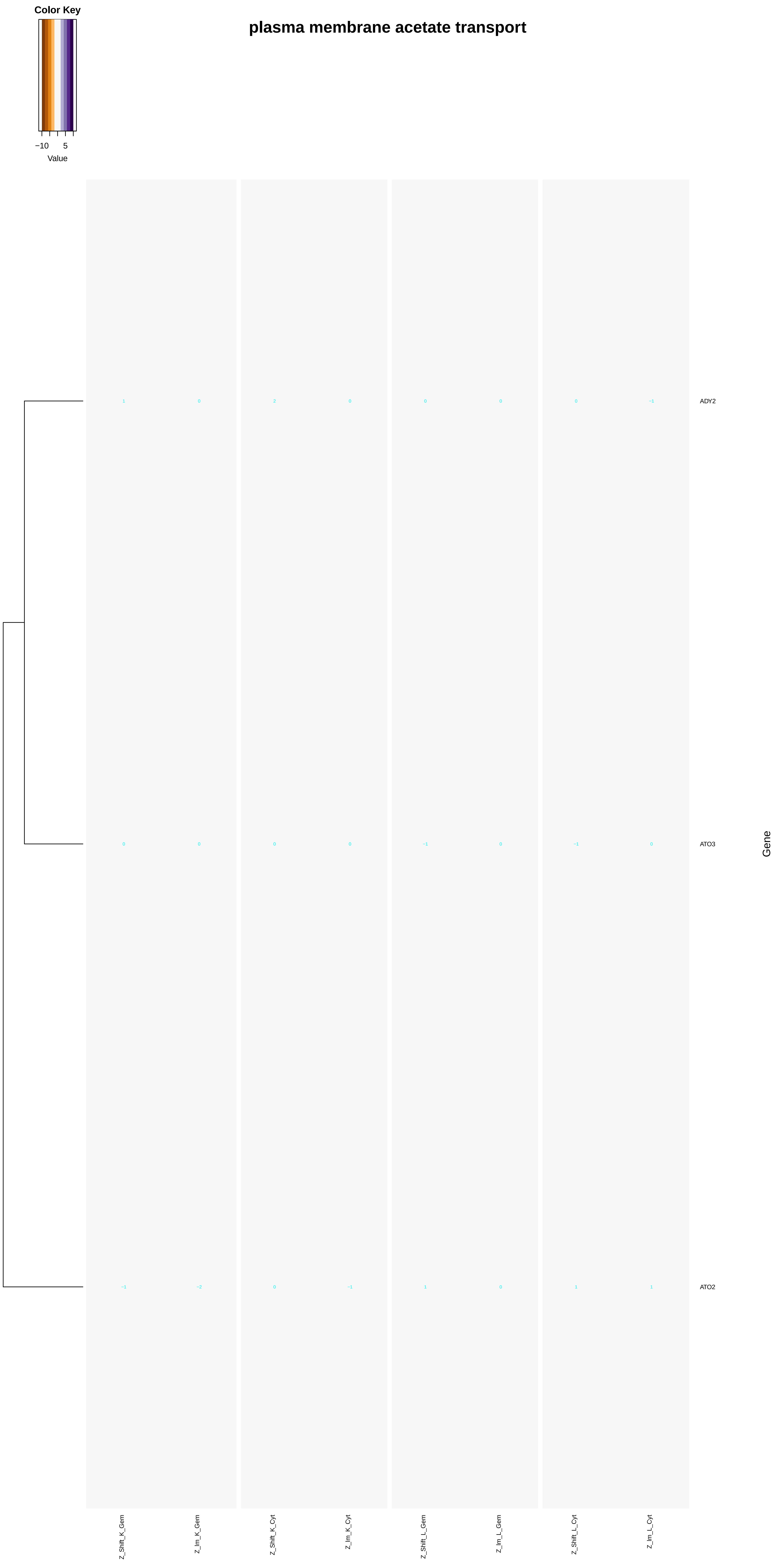

Color Key
−10
5
Value
plasma membrane acetate transport
1
0
2
0
0
0
0
−1
ADY2
0
0
0
0
−1
0
−1
0
ATO3
−1
−2
0
−1
1
0
1
1
ATO2
Gene
Z_Shift_K_Gem
Z_Im_K_Gem
Z_Shift_K_Cyt
Z_Im_K_Cyt
Z_Shift_L_Gem
Z_Im_L_Gem
Z_Shift_L_Cyt
Z_Im_L_Cyt

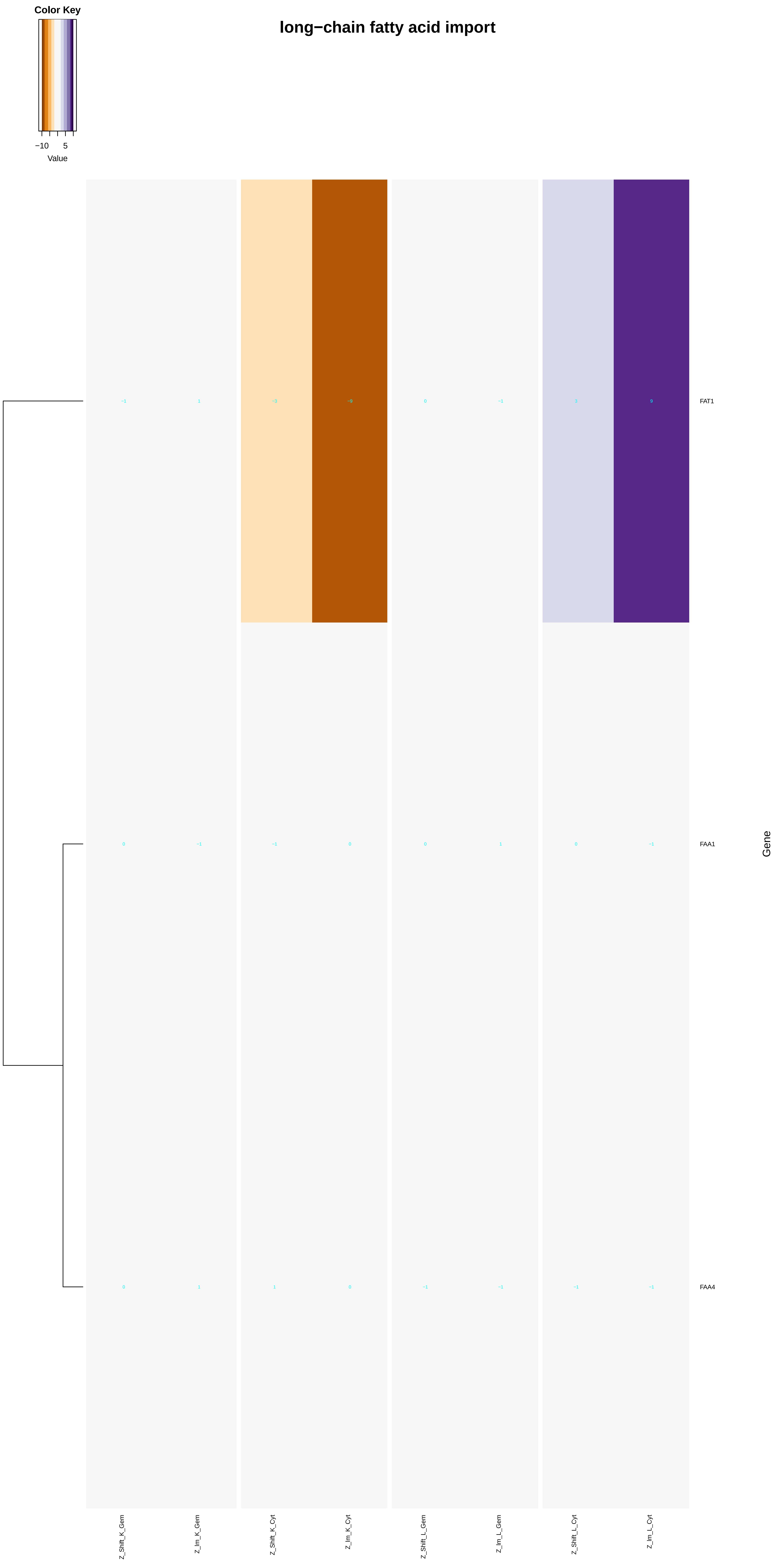
Color Key
−10
5
Value
long−chain fatty acid import
FAT1
FAA1
FAA4
Gene
Z_Shift_K_Gem
Z_Im_K_Gem
Z_Shift_K_Cyt
Z_Im_K_Cyt
Z_Shift_L_Gem
Z_Im_L_Gem
Z_Shift_L_Cyt
Z_Im_L_Cyt
