## Additional File 7 for "A humanized yeast phenomic model of deoxycytidine kinase to predict genetic buffering of nucleoside analog cytotoxicity": aromatic_compound_catabolic_process.pdf

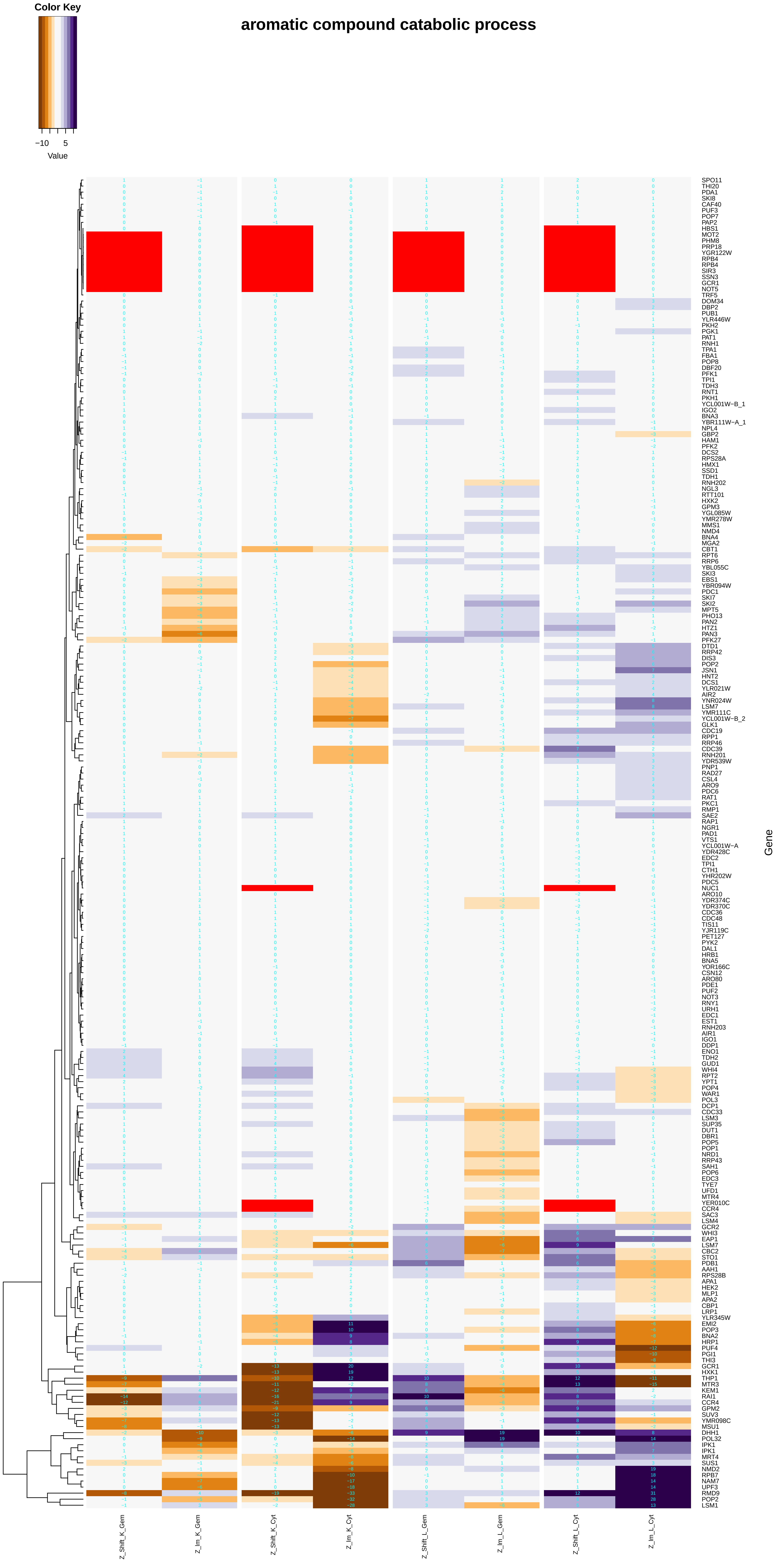

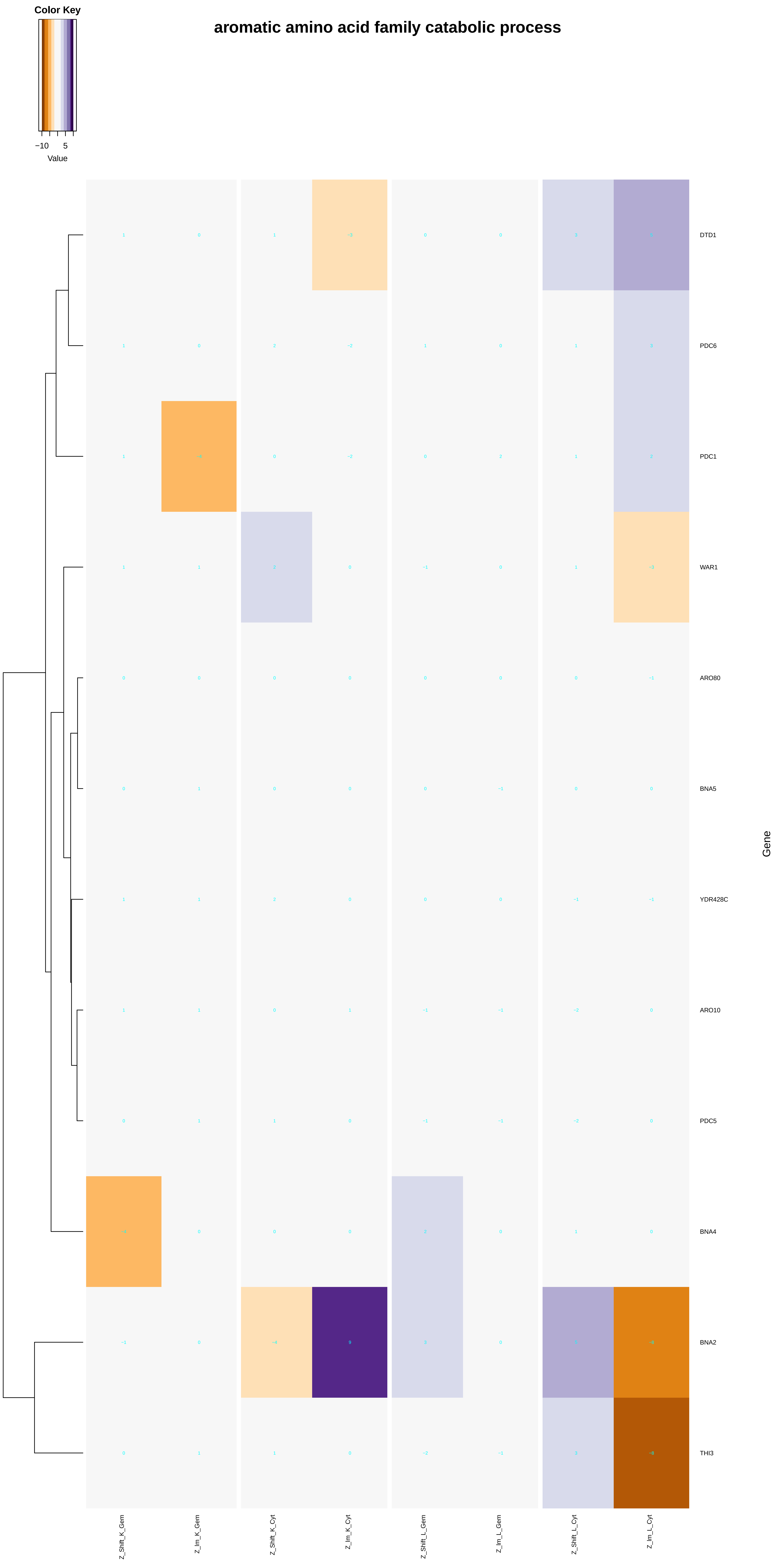
Color Key
−10 5
Value
aromatic amino acid family catabolic process
DTD1
PDC6
PDC1
WAR1
ARO80
BNA5
YDR428C
ARO10
PDC5
BNA4
BNA2
THI3
Gene
Z_Shift_K_Gem
Z_Im_K_Gem
Z_Shift_K_Cyt
Z_Im_K_Cyt
Z_Shift_L_Gem
Z_Im_L_Gem
Z_Shift_L_Cyt
Z_Im_L_Cyt

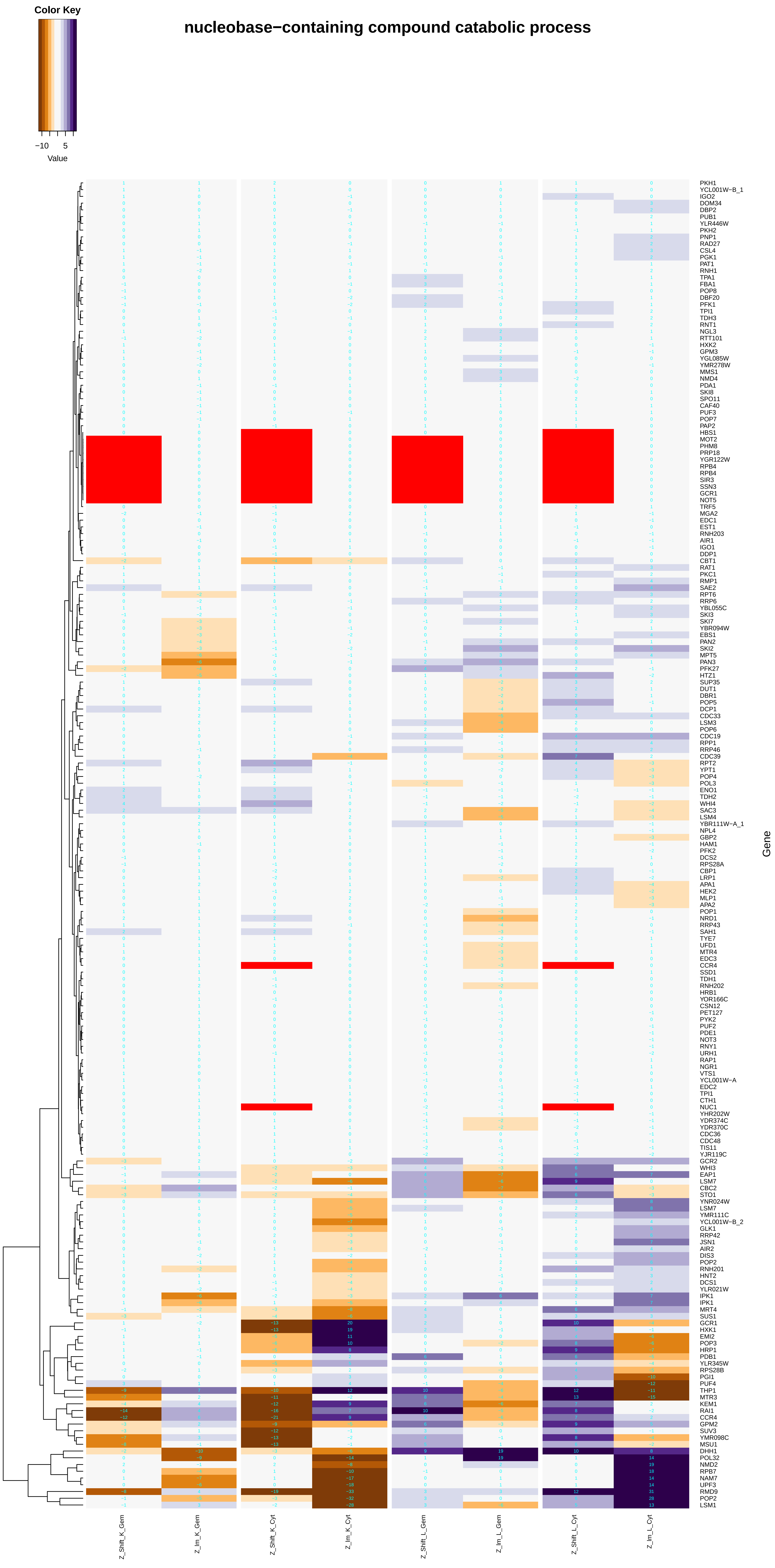

Color Key
−10 5
Value
nucleobase−containing compound catabolic process
PKH1
YCL001W−B_1
IGO2
DOM34
DBP2
PUB1
YLR446W
PKH2
PNP1
RAD27
CSL4
PGK1
PAT1
RNH1
TPA1
FBA1
POP8
DBF20
PFK1
TPI1
TDH3
RNT1
NGL3
RTT101
HXK2
GPM3
YGL085W
YMR278W
MMS1
NMD4
PDA1
SKI8
SPO11
CAF40
PUF3
POP7
PAP2
HBS1
MOT2
PHM8
PRP18
YGR122W
RPB4
RPB4
SIR3
SSN3
GCR1
NOT5
TRF5
MGA2
EDC1
EST1
RNH203
AIR1
IGO1
DDP1
CBT1
RAT1
PKC1
RMP1
SAE2
RPT6
RRP6
YBL055C
SKI3
SKI7
YBR094W
EBS1
PAN2
SKI2
MPT5
PAN3
PFK27
HTZ1
SUP35
DUT1
DBR1
POP5
DCP1
CDC33
LSM3
POP6
CDC19
RPP1
RRP46
CDC39
RPT2
YPT1
POP4
POL3
ENO1
TDH2
WHI4
SAC3
LSM4
YBR111W−A_1
NPL4
GBP2
HAM1
PFK2
DCS2
RPS28A
CBP1
LRP1
APA1
HEK2
MLP1
APA2
POP1
NRD1
RRP43
SAH1
TYE7
UFD1
MTR4
EDC3
CCR4
SSD1
TDH1
RNH202
HRB1
YOR166C
CSN12
PET127
PYK2
PUF2
PDE1
NOT3
RNY1
URH1
RAP1
NGR1
VTS1
YCL001W−A
EDC2
TPI1
CTH1
NUC1
YHR202W
YDR374C
YDR370C
CDC36
CDC48
TIS11
YJR119C
GCR2
WHI3
EAP1
LSM7
CBC2
STO1
YNR024W
LSM7
YMR111C
YCL001W−B_2
GLK1
RRP42
JSN1
AIR2
DIS3
POP2
RNH201
HNT2
DCS1
YLR021W
IPK1
IPK1
MRT4
SUS1
GCR1
HXK1
EMI2
POP3
HRP1
PDB1
YLR345W
RPS28B
PGI1
PUF4
THP1
MTR3
KEM1
RAI1
CCR4
GPM2
SUV3
YMR098C
MSU1
DHH1
POL32
NMD2
RPB7
NAM7
UPF3
RMD9
POP2
LSM1
Gene
Z_Shift_K_Gem
Z_lm_K_Gem
Z_Shift_K_Cyt
Z_lm_K_Cyt
Z_Shift_L_Gem
Z_lm_L_Gem
Z_Shift_L_Cyt
Z_lm_L_Cyt

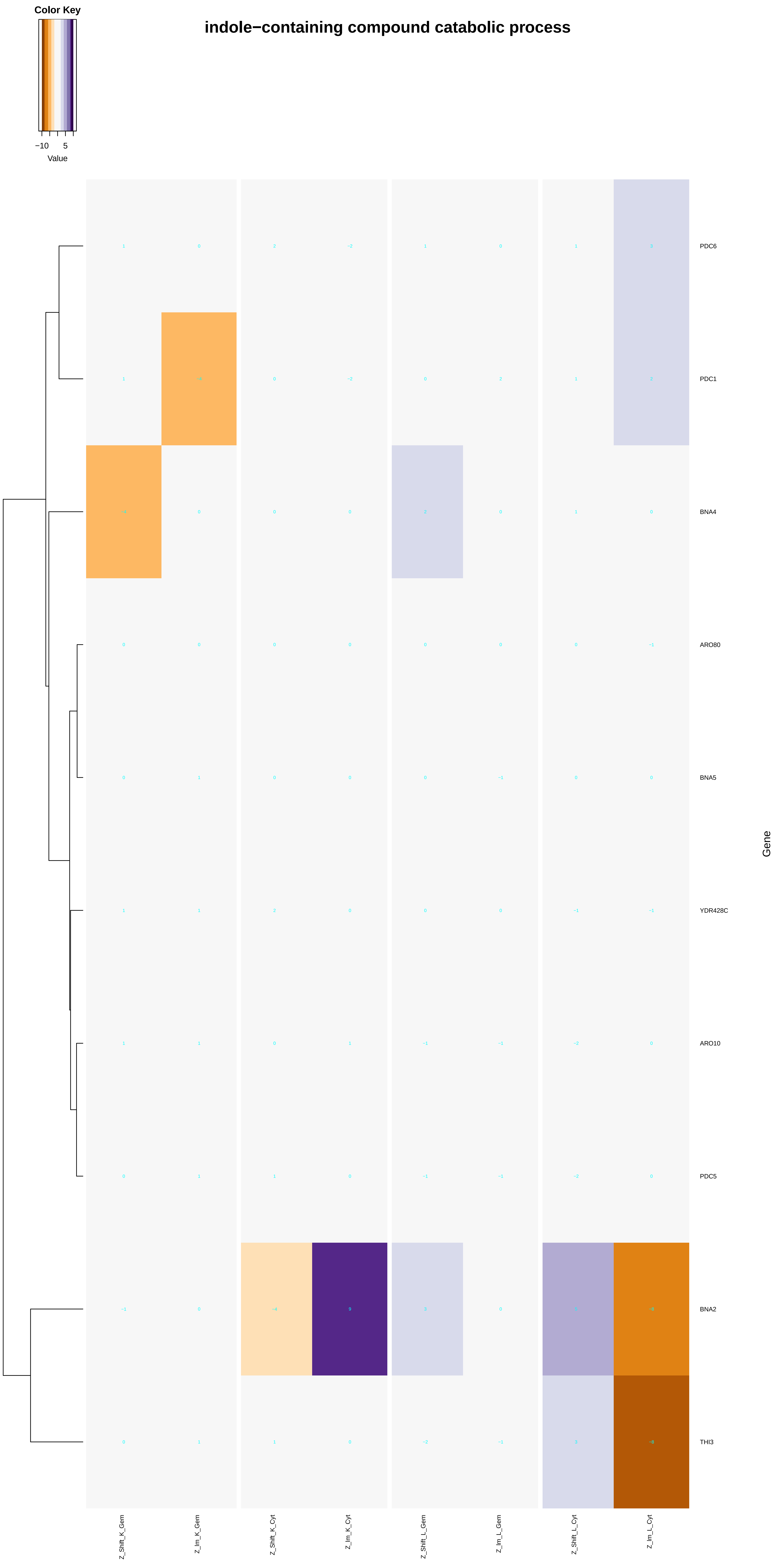

Color Key
−10
5
Value
indole−containing compound catabolic process
PDC6
PDC1
BNA4
ARO80
BNA5
YDR428C
ARO10
PDC5
BNA2
THI3
Gene
Z_Shift_K_Gem
Z_Im_K_Gem
Z_Shift_K_Cyt
Z_Im_K_Cyt
Z_Shift_L_Gem
Z_Im_L_Gem
Z_Shift_L_Cyt
Z_Im_L_Cyt

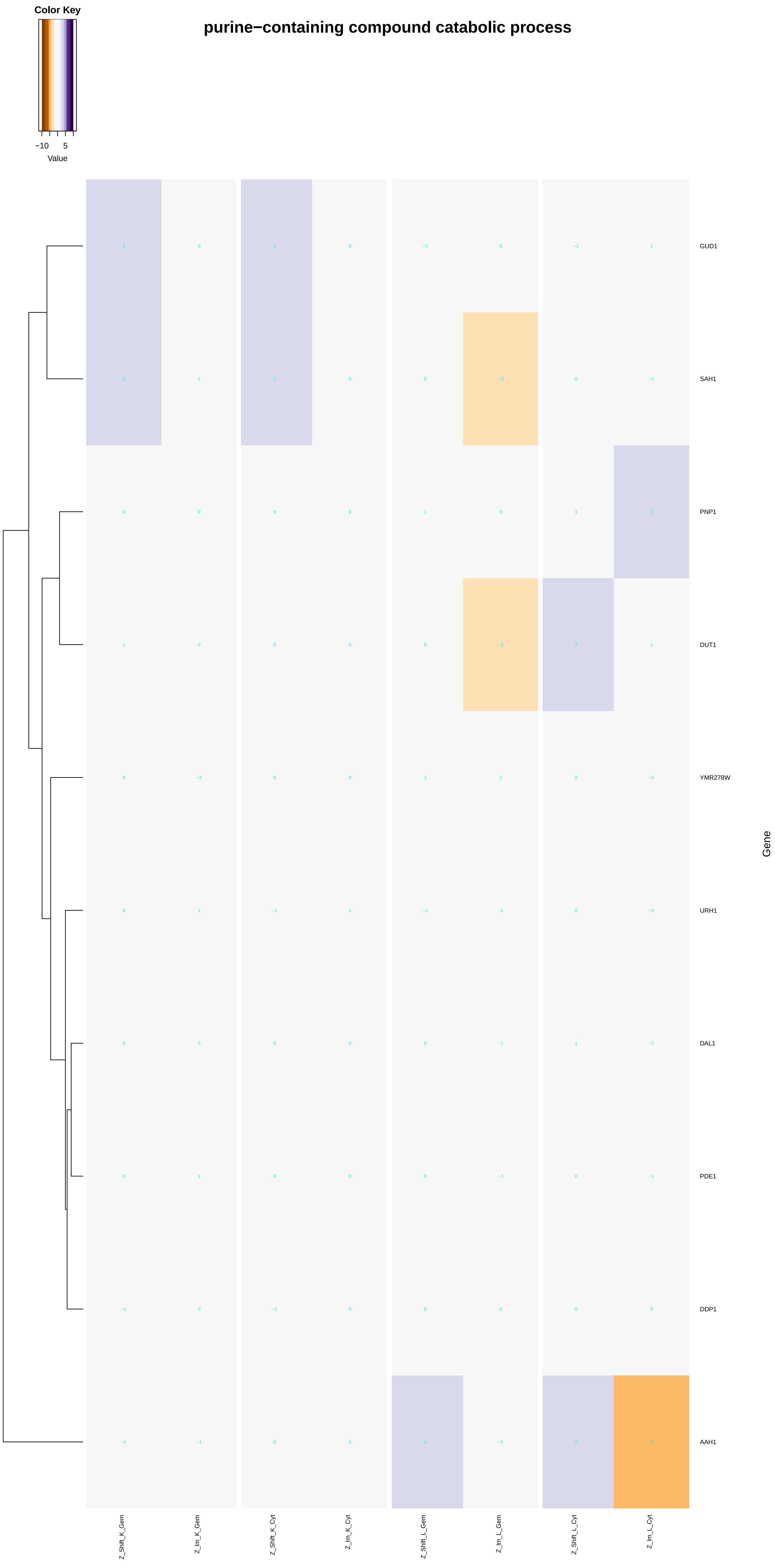
Color Key
−10
5
Value
purine−containing compound catabolic process
GUD1
SAH1
PNP1
DUT1
YMR278W
URH1
DAL1
PDE1
DDP1
AAH1
Gene
Z_Shift_K_Gem
Z_Im_K_Gem
Z_Shift_K_Cyt
Z_Im_K_Cyt
Z_Shift_L_Gem
Z_Im_L_Gem
Z_Shift_L_Cyt
Z_Im_L_Cyt

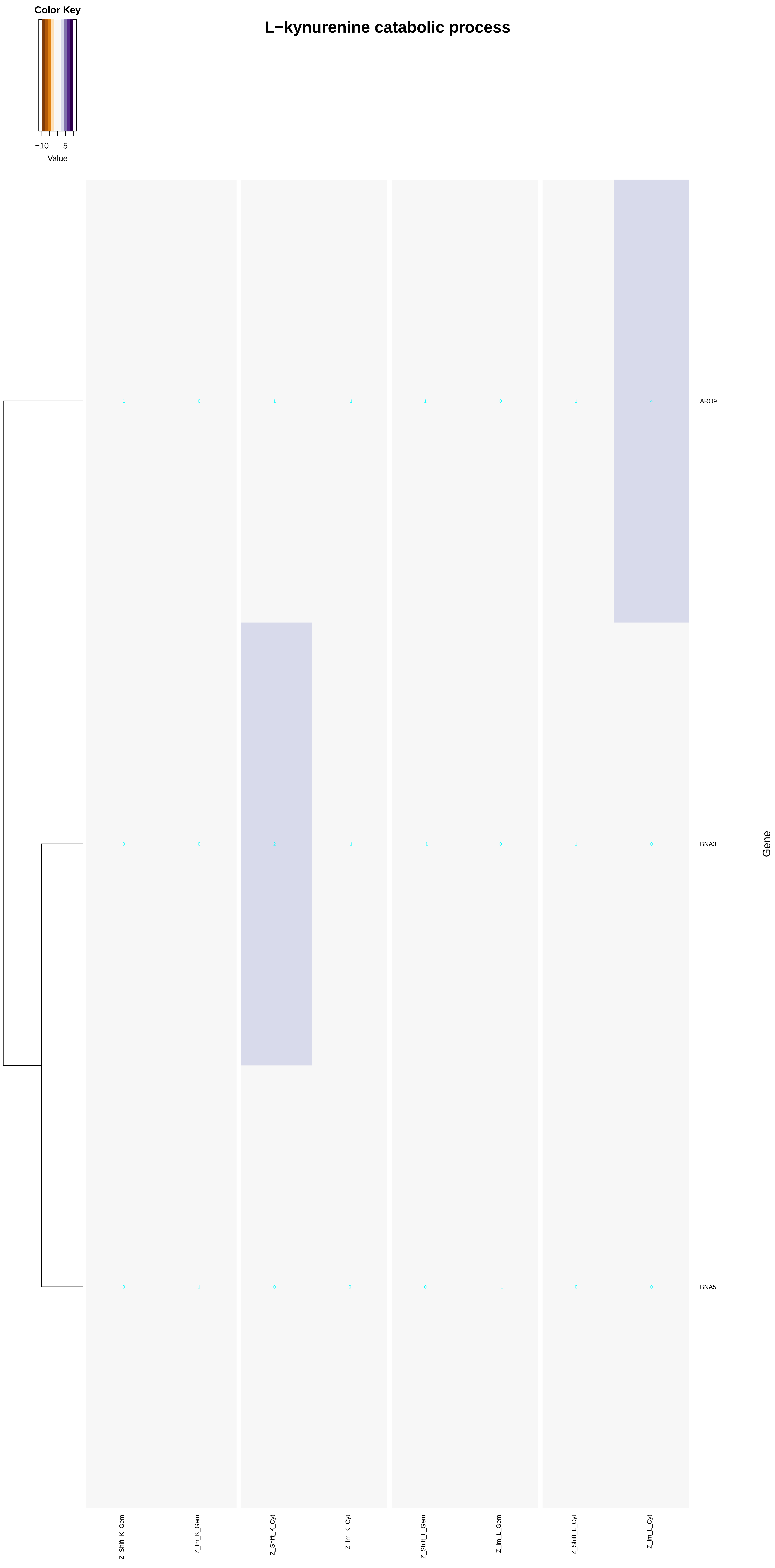
Color Key
−10
5
Value
L−kynurenine catabolic process
ARO9
BNA3
BNA5
Gene
Z_Shift_K_Gem
Z_Im_K_Gem
Z_Shift_K_Cyt
Z_Im_K_Cyt
Z_Shift_L_Gem
Z_Im_L_Gem
Z_Shift_L_Cyt
Z_Im_L_Cyt

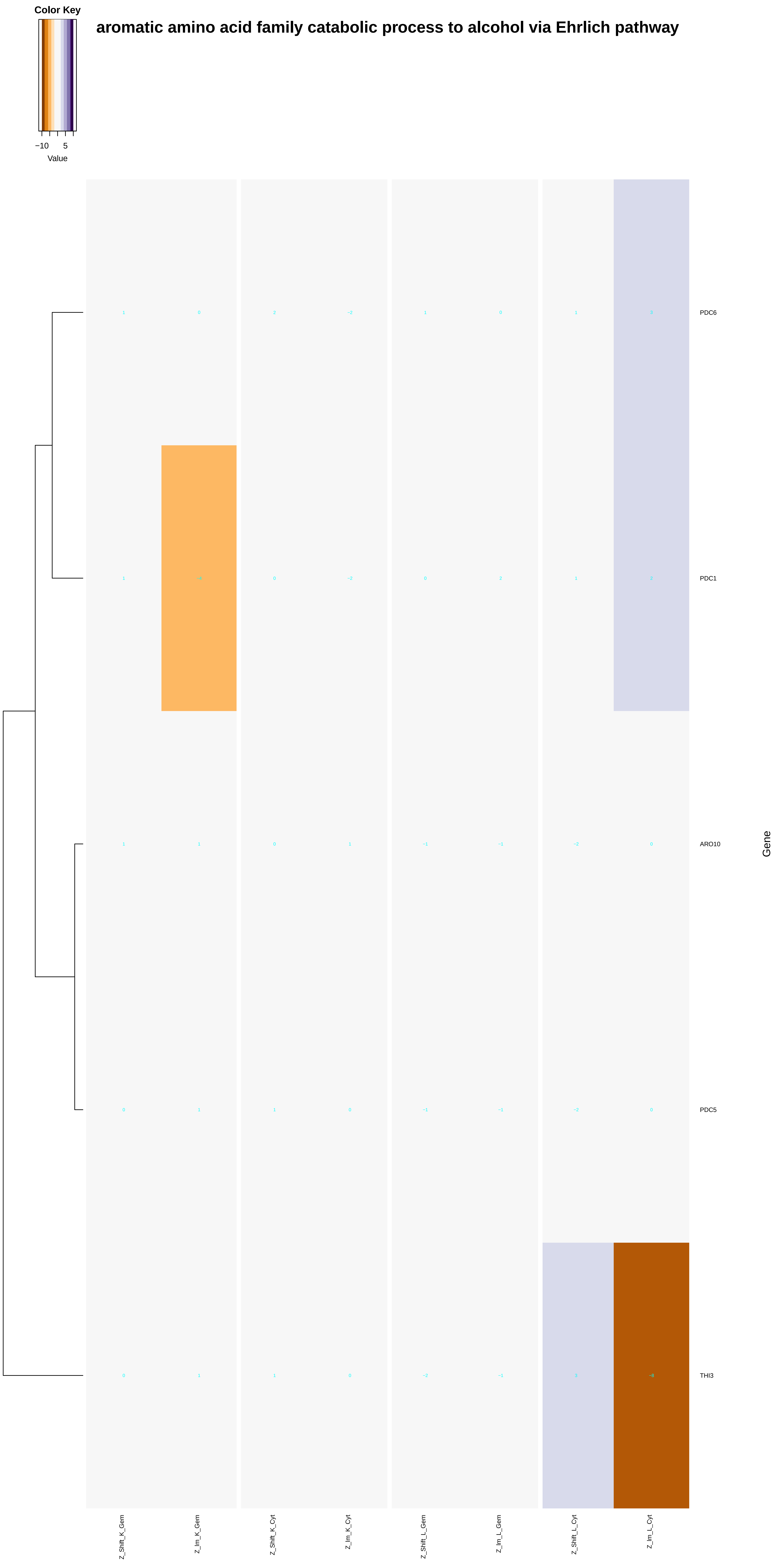

Color Key
−10
5
Value
aromatic amino acid family catabolic process to alcohol via Ehrlich pathway
PDC6
PDC1
ARO10
PDC5
THI3
Gene
Z_Shift_K_Gem
Z_Im_K_Gem
Z_Shift_K_Cyt
Z_Im_K_Cyt
Z_Shift_L_Gem
Z_Im_L_Gem
Z_Shift_L_Cyt
Z_Im_L_Cyt

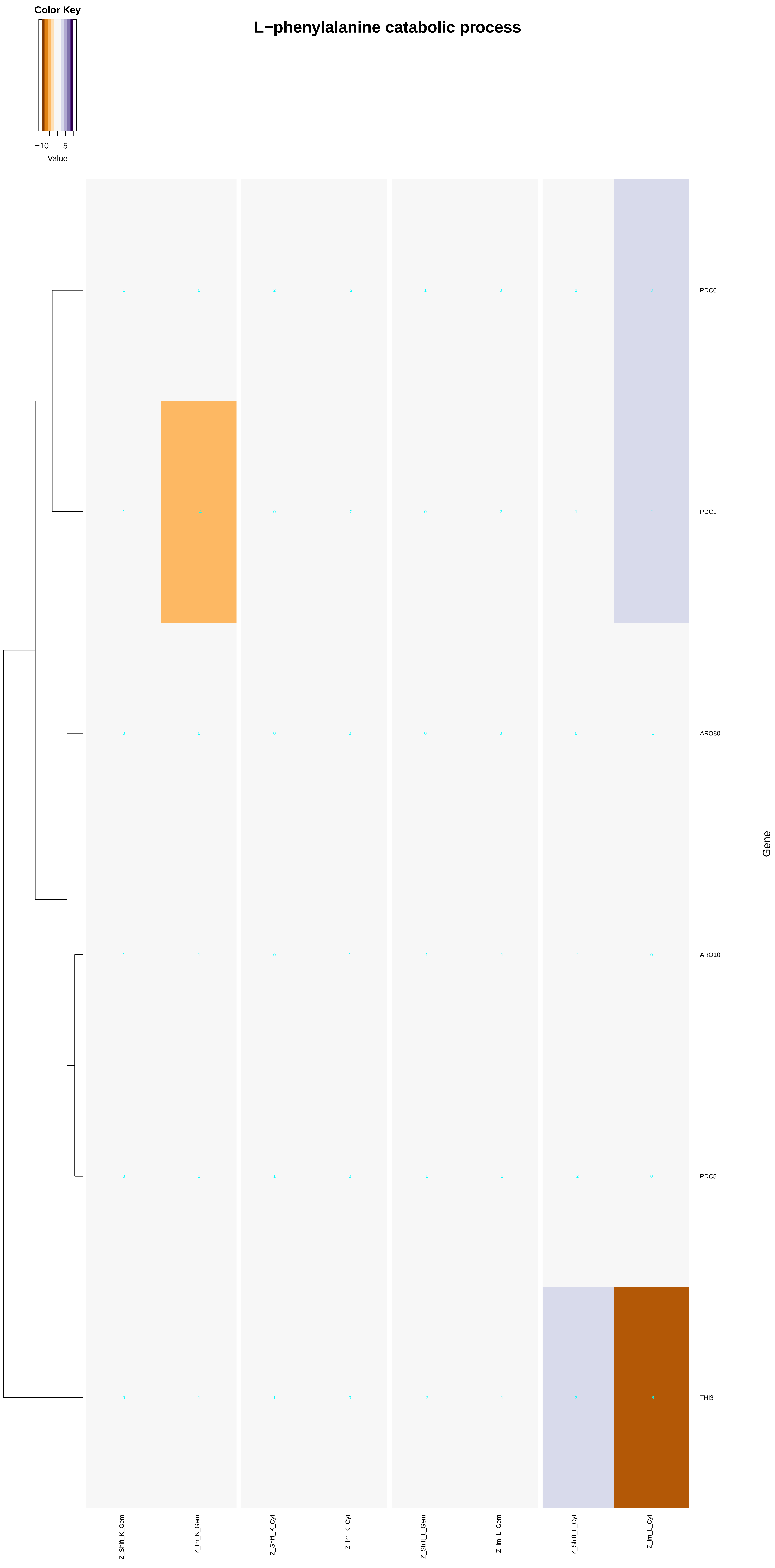
L−phenylalanine catabolic process
Color Key
−10
5
Value
PDC6
PDC1
ARO80
ARO10
PDC5
THI3
Gene
Z_Shift_K_Gem
Z_Im_K_Gem
Z_Shift_K_Cyt
Z_Im_K_Cyt
Z_Shift_L_Gem
Z_Im_L_Gem
Z_Shift_L_Cyt
Z_Im_L_Cyt

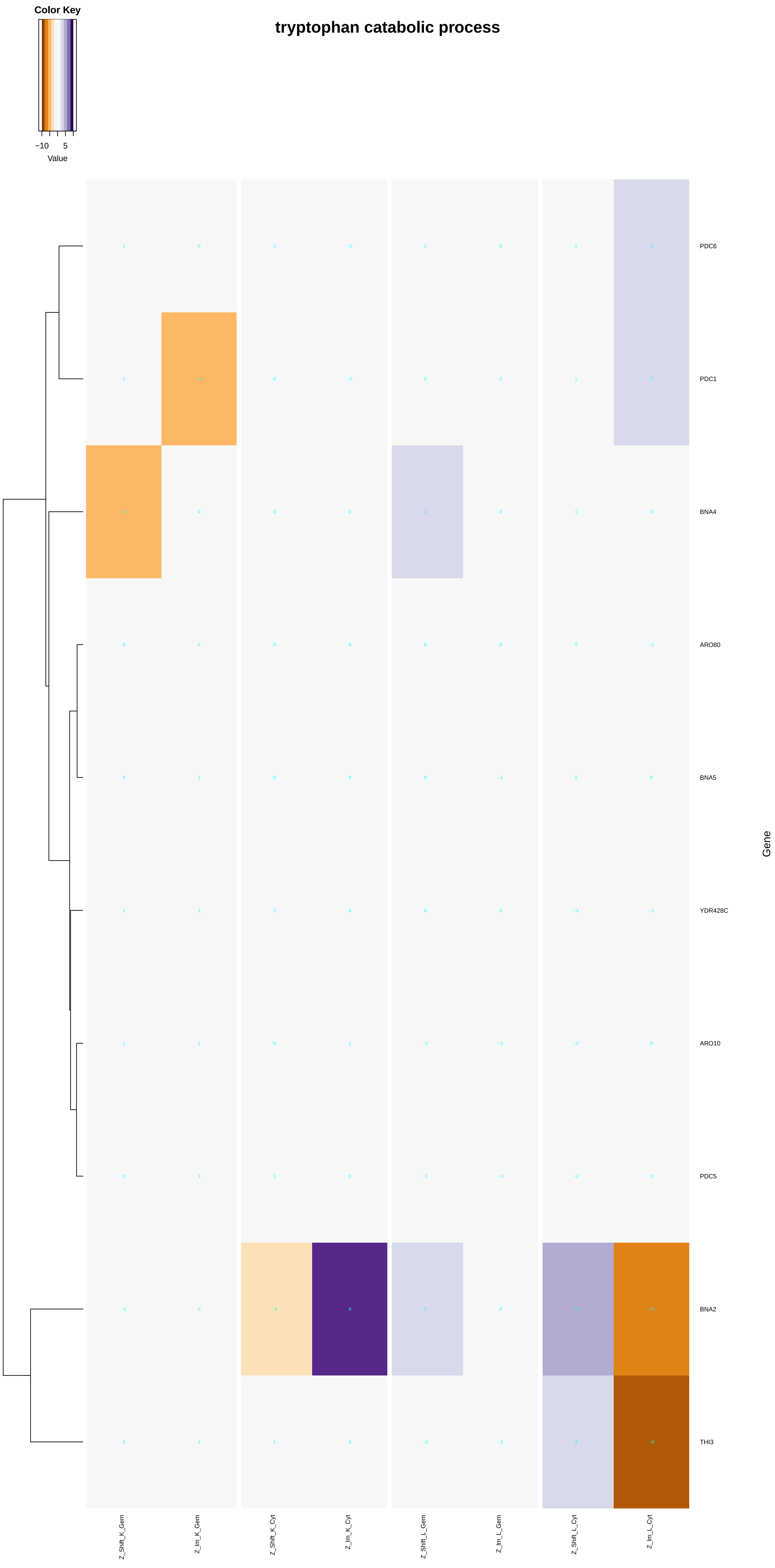

Color Key
−10
5
Value
tryptophan catabolic process
PDC6
PDC1
BNA4
ARO80
BNA5
YDR428C
ARO10
PDC5
BNA2
THI3
Gene
Z_Shift_K_Gem
Z_Im_K_Gem
Z_Shift_K_Cyt
Z_Im_K_Cyt
Z_Shift_L_Gem
Z_Im_L_Gem
Z_Shift_L_Cyt
Z_Im_L_Cyt

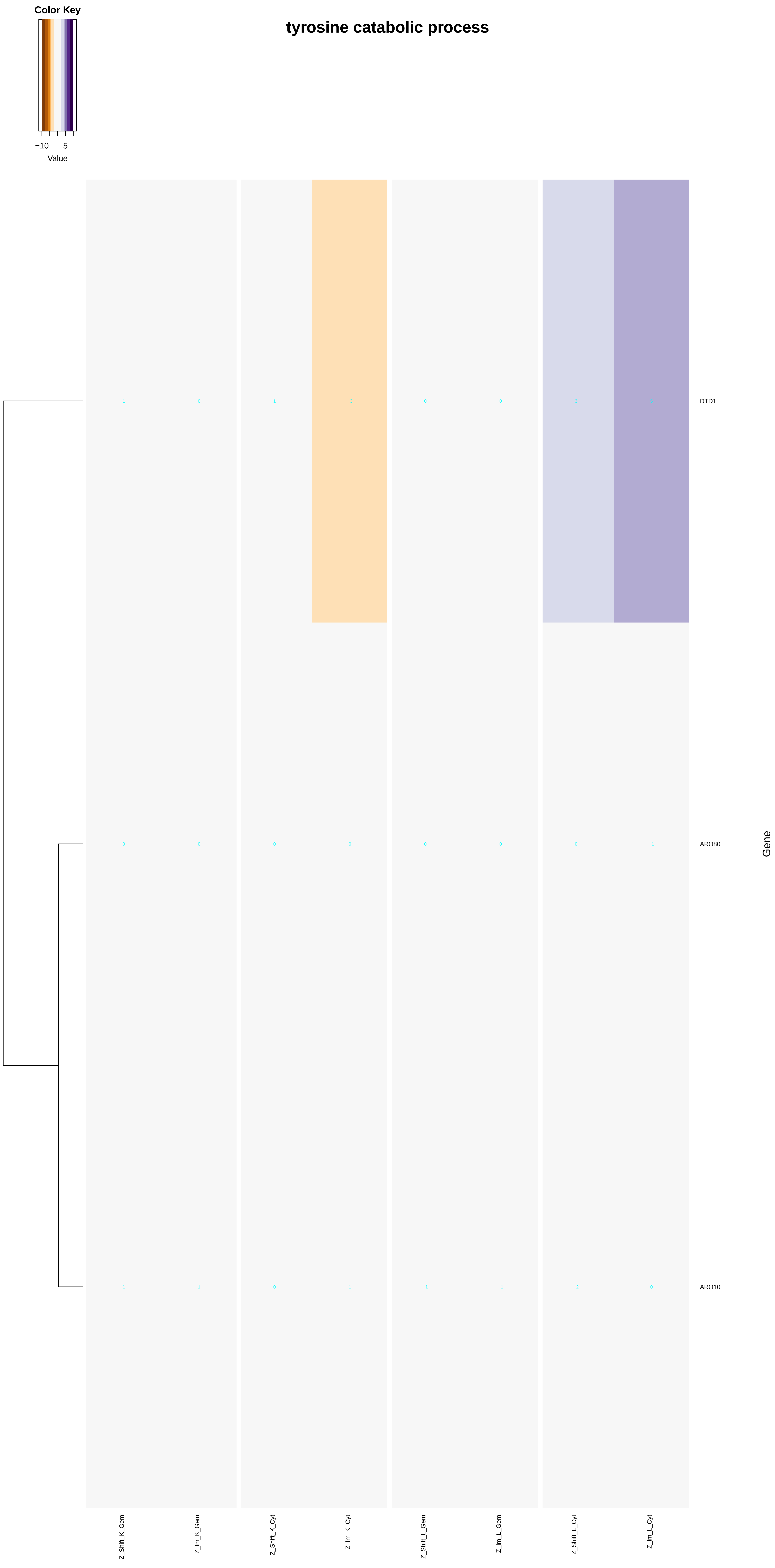
tyrosine catabolic process
Color Key
−10
5
Value
DTD1
ARO80
ARO10
Gene
Z_Shift_K_Gem
Z_Im_K_Gem
Z_Shift_K_Cyt
Z_Im_K_Cyt
Z_Shift_L_Gem
Z_Im_L_Gem
Z_Shift_L_Cyt
Z_Im_L_Cyt

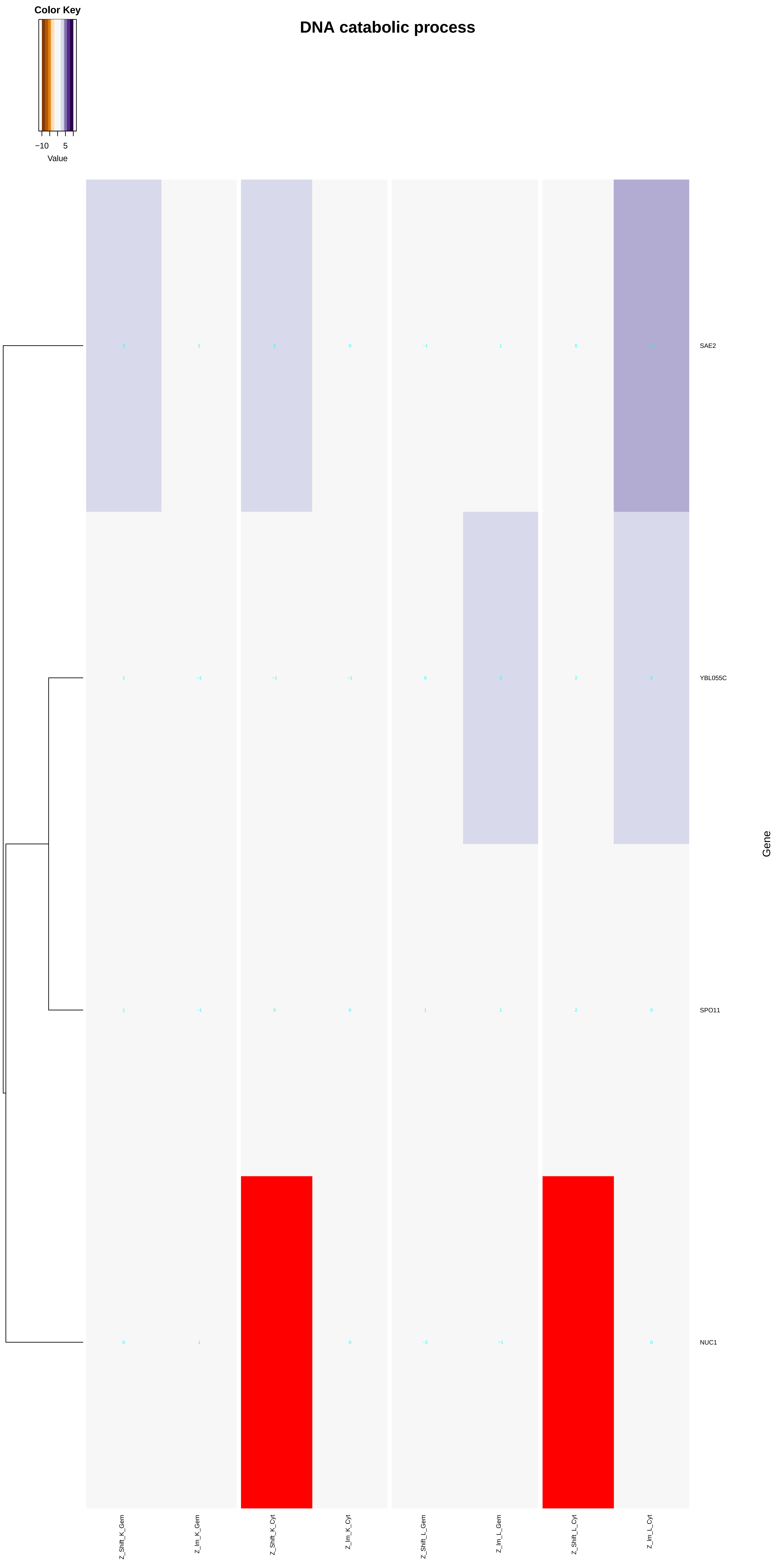
DNA catabolic process
Color Key
−10
5
Value
SAE2
YBL055C
SPO11
NUC1
Gene
Z_Shift_K_Gem
Z_Im_K_Gem
Z_Shift_K_Cyt
Z_Im_K_Cyt
Z_Shift_L_Gem
Z_Im_L_Gem
Z_Shift_L_Cyt
Z_Im_L_Cyt

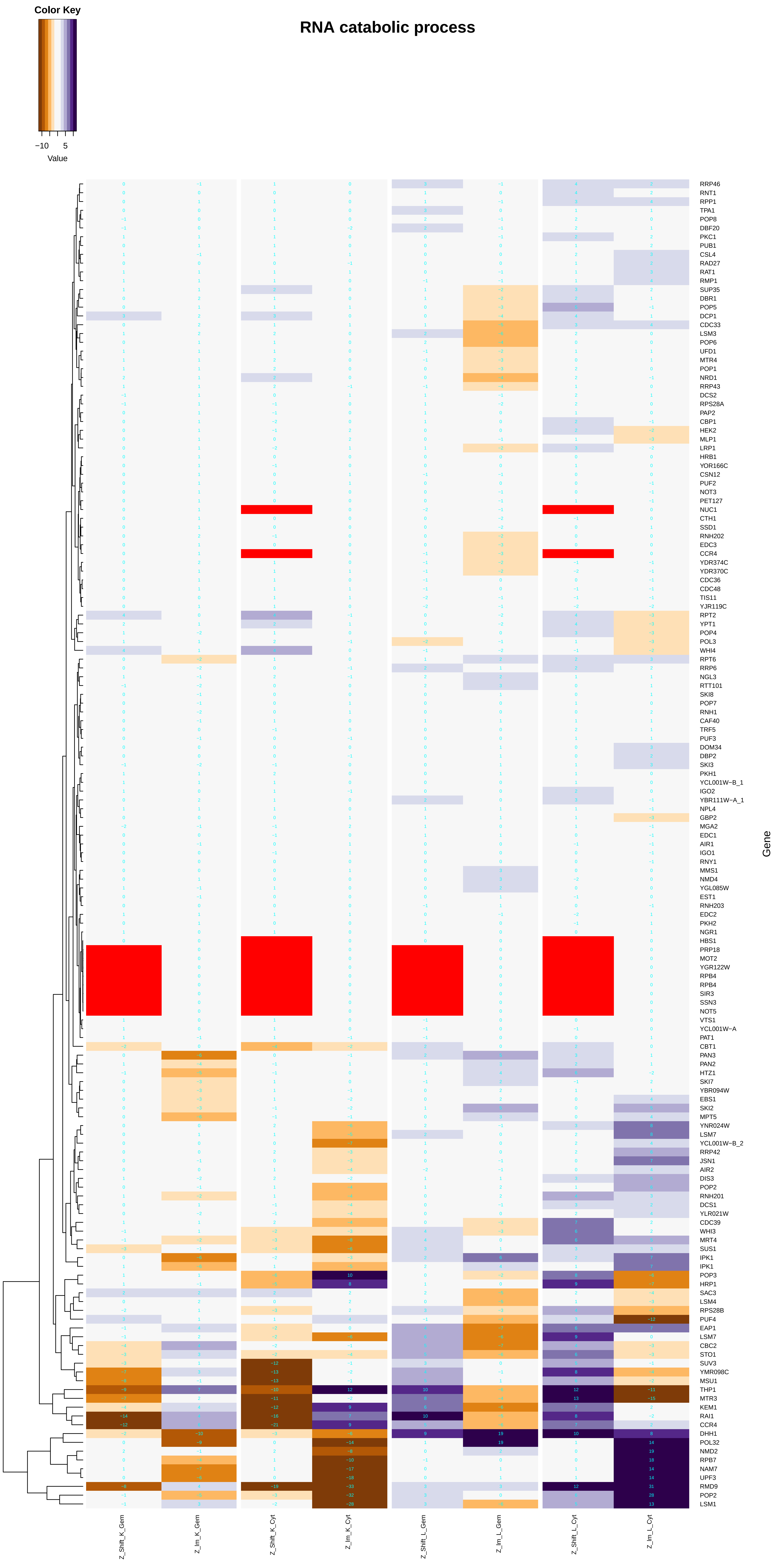

RNA catabolic process
Color Key
−10
5
Value
Gene
Z_Shift_K_Gem
Z_Im_K_Gem
Z_Shift_K_Cyt
Z_Im_K_Cyt
Z_Shift_L_Gem
Z_Im_L_Gem
Z_Shift_L_Cyt
Z_Im_L_Cyt
RRP46
RNT1
RPP1
TPA1
POP8
DBF20
PKC1
PUB1
CSL4
RAD27
RAT1
RMP1
SUP35
DBR1
POP5
DCP1
CDC33
LSM3
POP6
UFD1
MTR4
POP1
NRD1
RRP43
DCS2
RPS28A
PAP2
CBP1
HEK2
MLP1
LRP1
HRB1
YOR166C
CSN12
PUF2
NOT3
PET127
NUC1
CTH1
SSD1
RNH202
EDC3
CCR4
YDR374C
YDR370C
CDC36
CDC48
TIS11
YJR119C
RPT2
YPT1
POP4
POL3
WHI4
RPT6
RRP6
NGL3
RTT101
SKI8
POP7
RNH1
CAF40
TRF5
PUF3
DOM34
DBP2
SKI3
PKH1
YCL001W−B_1
IGO2
YBR111W−A_1
NPL4
GBP2
MGA2
EDC1
AIR1
IGO1
RNY1
MMS1
NMD4
YGL085W
EST1
RNH203
EDC2
PKH2
NGR1
HBS1
PRP18
MOT2
YGR122W
RPB4
RPB4
SIR3
SSN3
NOT5
VTS1
YCL001W−A
PAT1
CBT1
PAN3
PAN2
HTZ1
SKI7
YBR094W
EBS1
SKI2
MPT5
YNR024W
LSM7
YCL001W−B_2
RRP42
JSN1
AIR2
DIS3
POP2
RNH201
DCS1
YLR021W
CDC39
WHI3
MRT4
SUS1
IPK1
IPK1
POP3
HRP1
SAC3
LSM4
RPS28B
PUF4
EAP1
LSM7
CBC2
STO1
SUV3
YMR098C
MSU1
THP1
MTR3
KEM1
RAI1
CCR4
DHH1
POL32
NMD2
RPB7
NAM7
UPF3
RMD9
POP2
LSM1

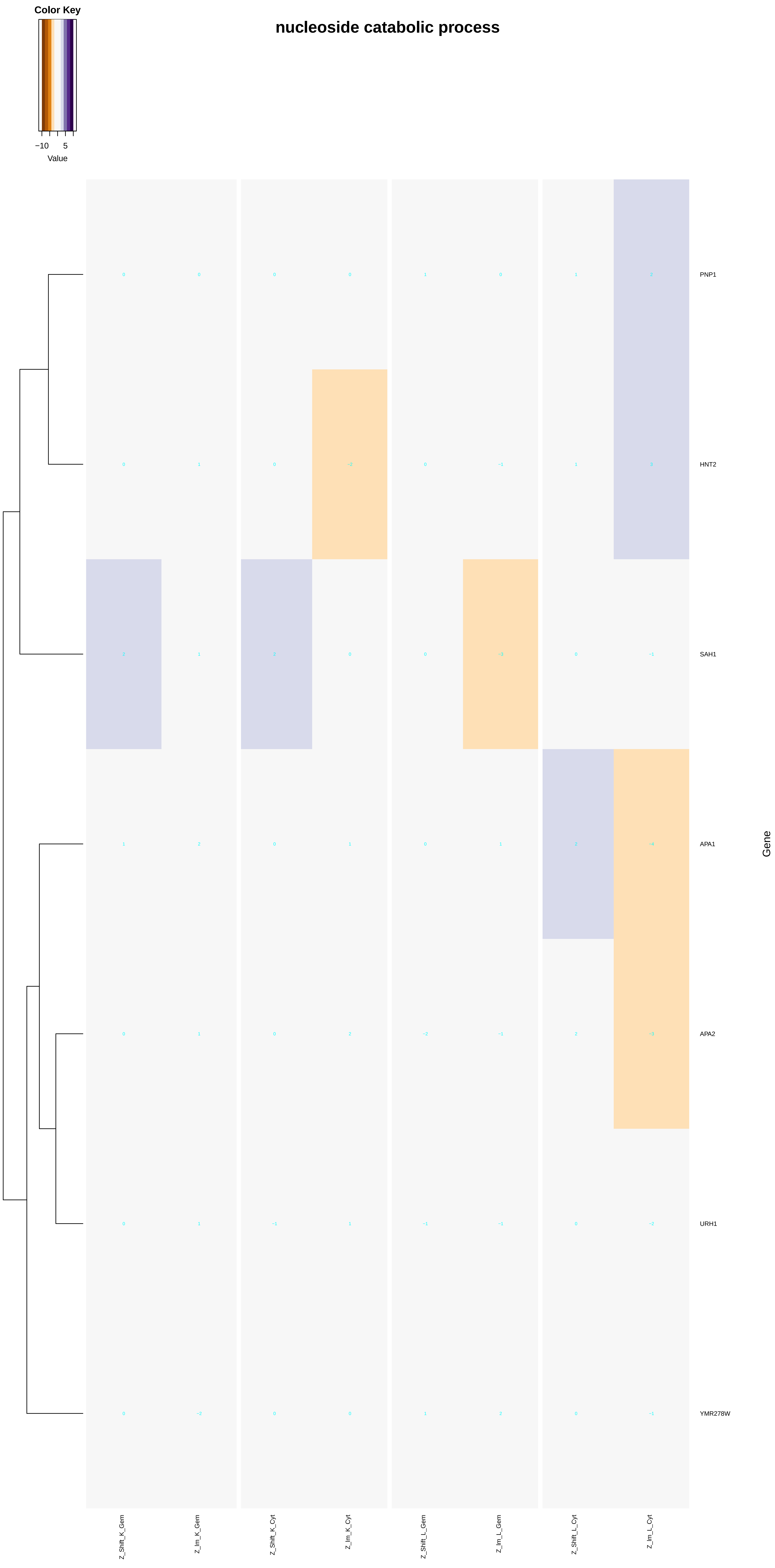
nucleoside catabolic process
Color Key
−10
5
Value
PNP1
HNT2
SAH1
APA1
APA2
URH1
YMR278W
Gene
Z_Shift_K_Gem
Z_Im_K_Gem
Z_Shift_K_Cyt
Z_Im_K_Cyt
Z_Shift_L_Gem
Z_Im_L_Gem
Z_Shift_L_Cyt
Z_Im_L_Cyt

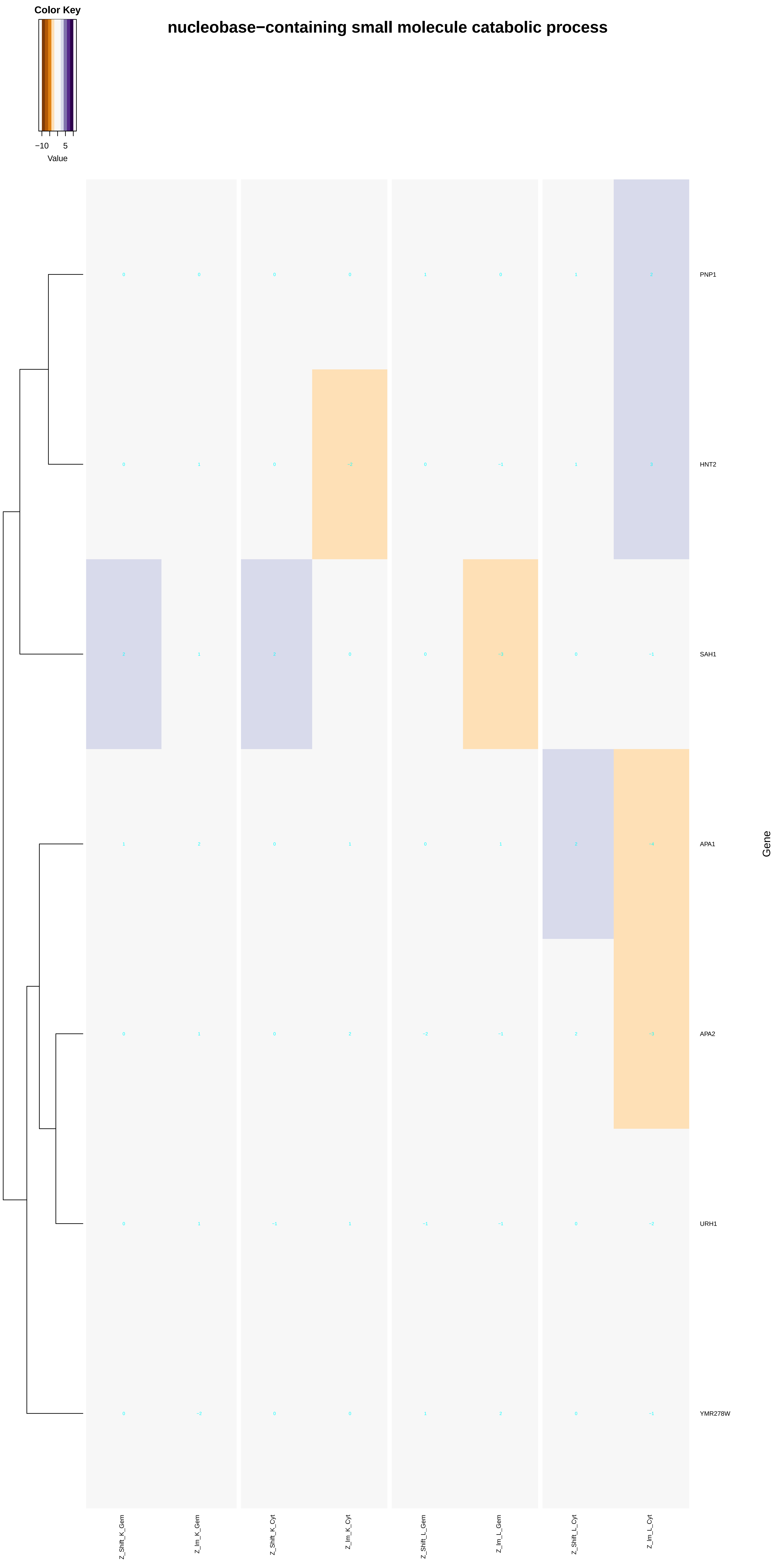

Color Key
−10
5
Value
nucleobase−containing small molecule catabolic process
PNP1
HNT2
SAH1
APA1
APA2
URH1
YMR278W
Gene
Z_Shift_K_Gem
Z_Im_K_Gem
Z_Shift_K_Cyt
Z_Im_K_Cyt
Z_Shift_L_Gem
Z_Im_L_Gem
Z_Shift_L_Cyt
Z_Im_L_Cyt

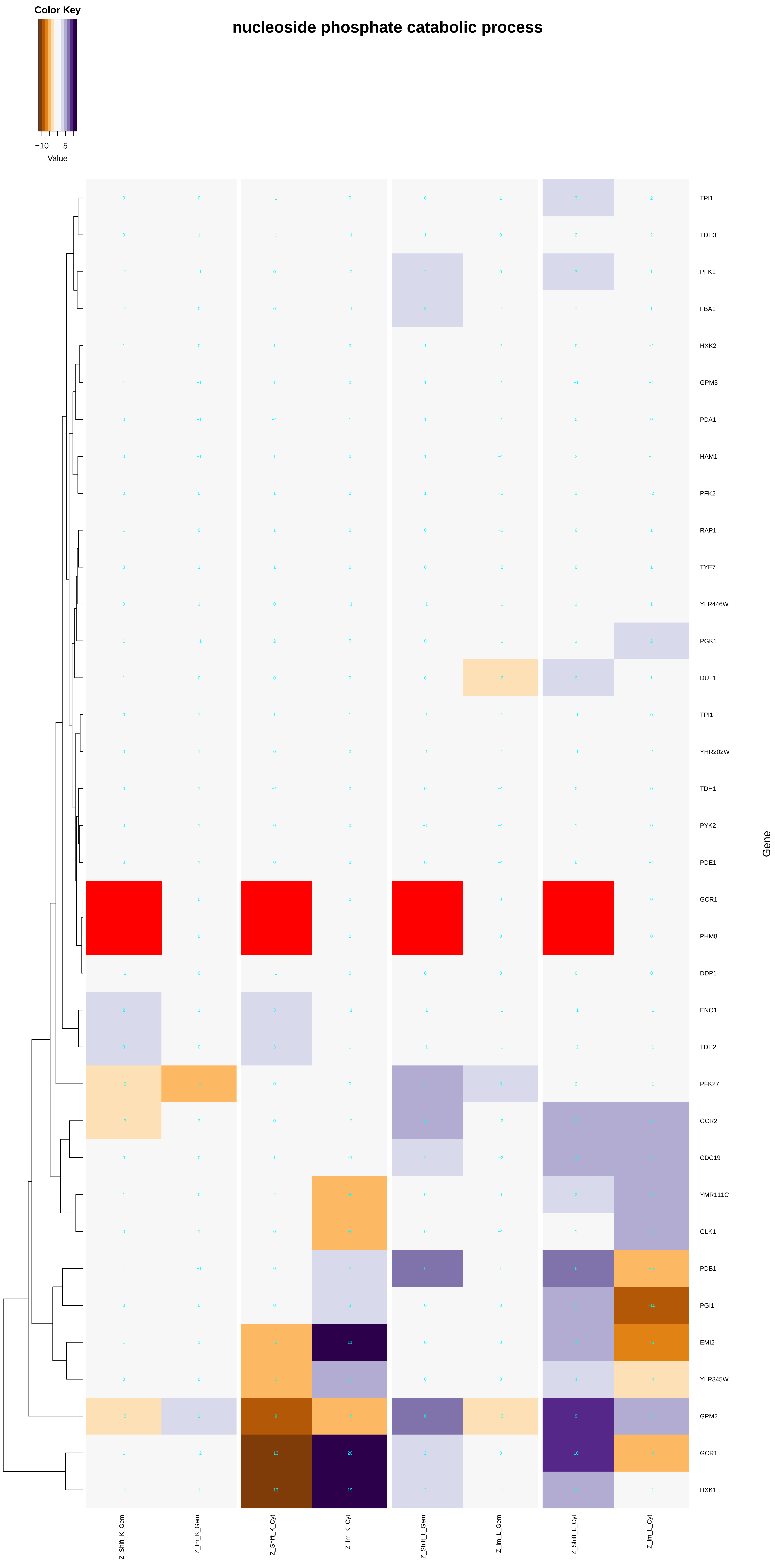

Color Key
−10
5
Value
nucleoside phosphate catabolic process
TPI1
TDH3
PFK1
FBA1
HXK2
GPM3
PDA1
HAM1
PFK2
RAP1
TYE7
YLR446W
PGK1
DUT1
TPI1
YHR202W
TDH1
PYK2
PDE1
GCR1
PHM8
DDP1
ENO1
TDH2
PFK27
GCR2
CDC19
YMR111C
GLK1
PDB1
PGI1
EMI2
YLR345W
GPM2
GCR1
HXK1
Gene
Z_Shift_K_Gem
Z_Im_K_Gem
Z_Shift_K_Cyt
Z_Im_K_Cyt
Z_Shift_L_Gem
Z_Im_L_Gem
Z_Shift_L_Cyt
Z_Im_L_Cyt

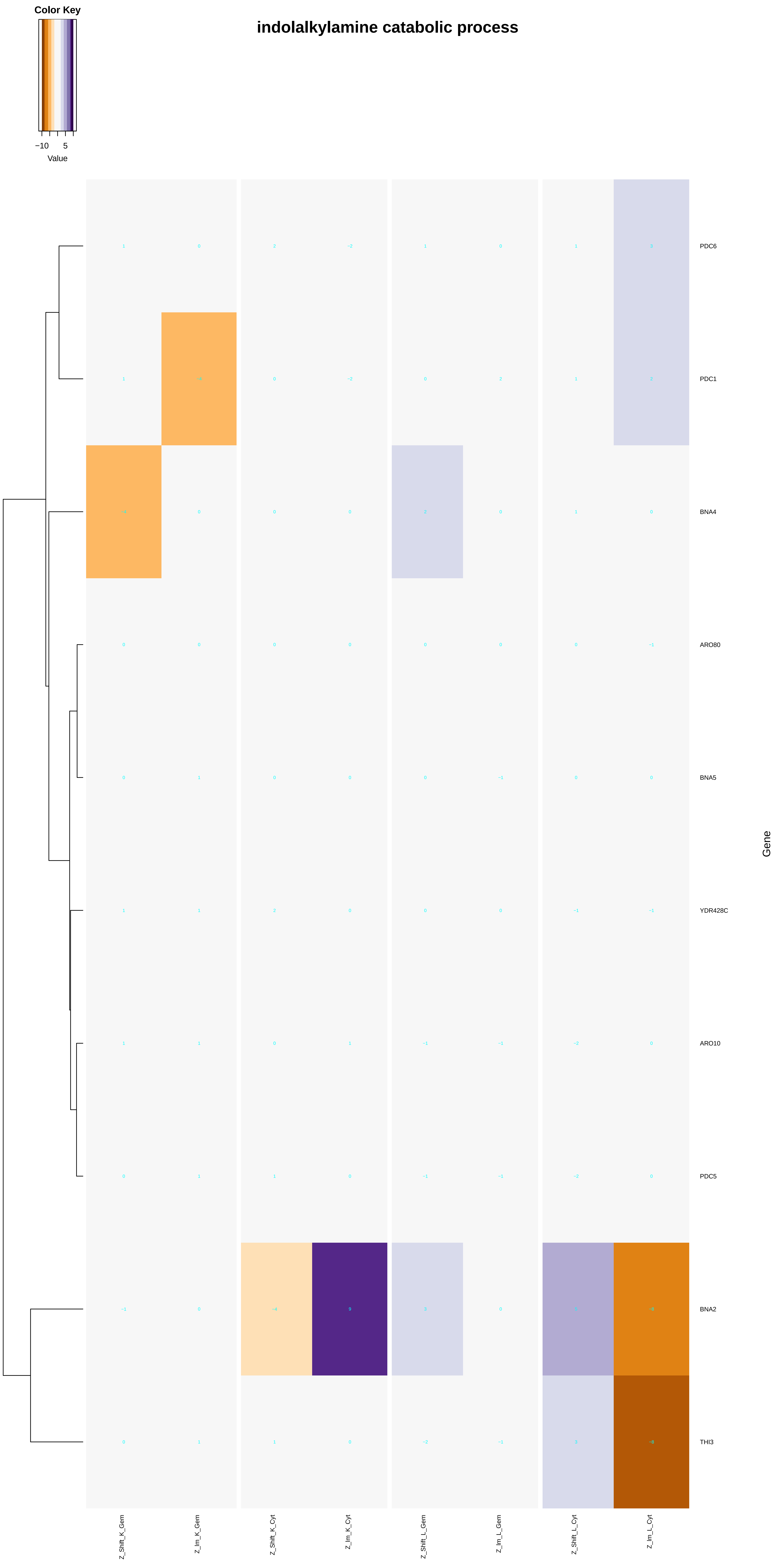
Color Key
−10
5
Value
indolalkylamine catabolic process
PDC6
PDC1
BNA4
ARO80
BNA5
YDR428C
ARO10
PDC5
BNA2
THI3
Gene
Z_Shift_K_Gem
Z_Im_K_Gem
Z_Shift_K_Cyt
Z_Im_K_Cyt
Z_Shift_L_Gem
Z_Im_L_Gem
Z_Shift_L_Cyt
Z_Im_L_Cyt

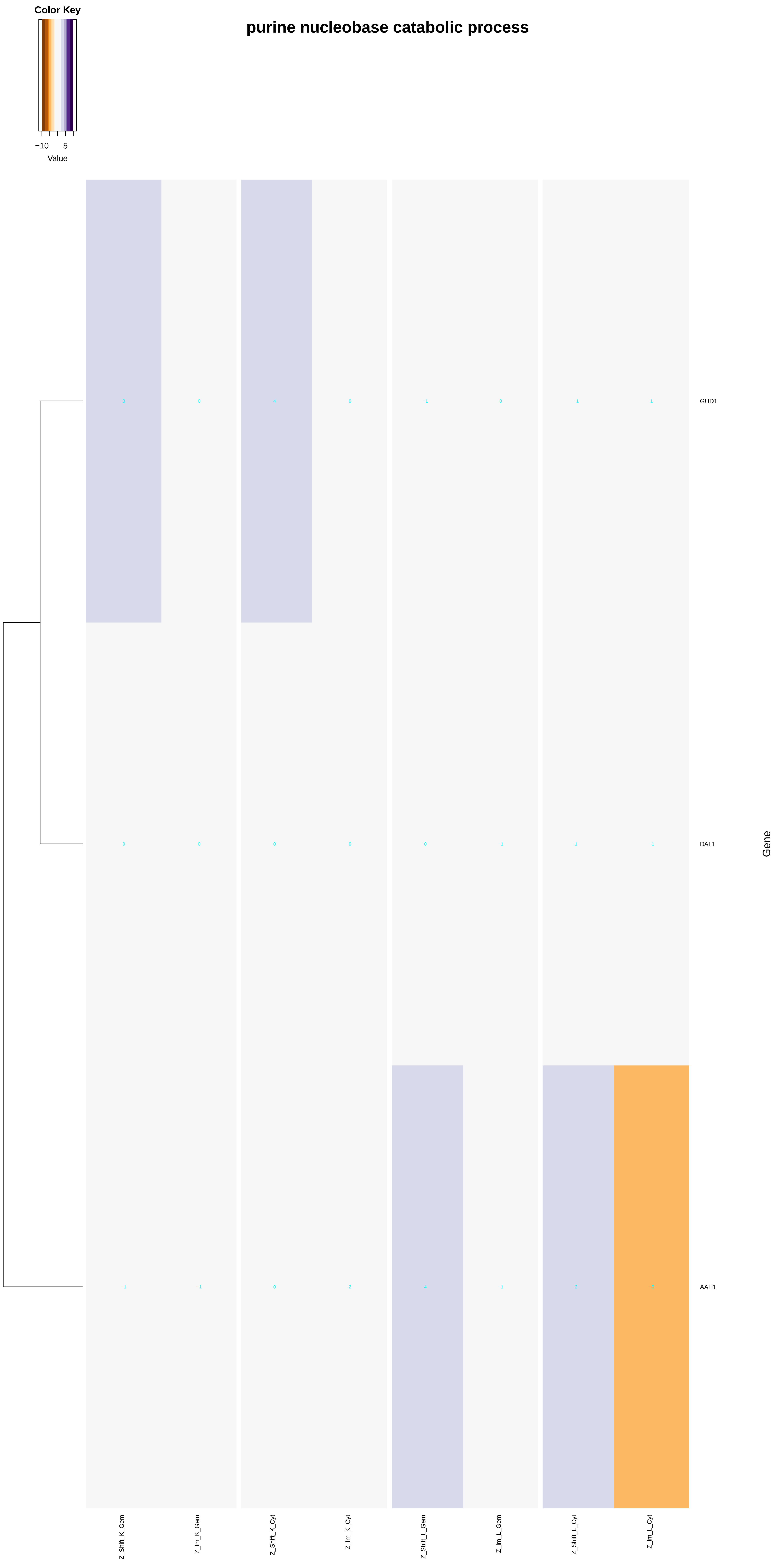

Color Key
−10
5
Value
purine nucleobase catabolic process
3
0
4
0
−1
0
−1
1
GUD1
0
0
0
0
0
−1
1
−1
DAL1
−1
−1
0
2
4
−1
2
−5
AAH1
Gene
Z_Shift_K_Gem
Z_Im_K_Gem
Z_Shift_K_Cyt
Z_Im_K_Cyt
Z_Shift_L_Gem
Z_Im_L_Gem
Z_Shift_L_Cyt
Z_Im_L_Cyt

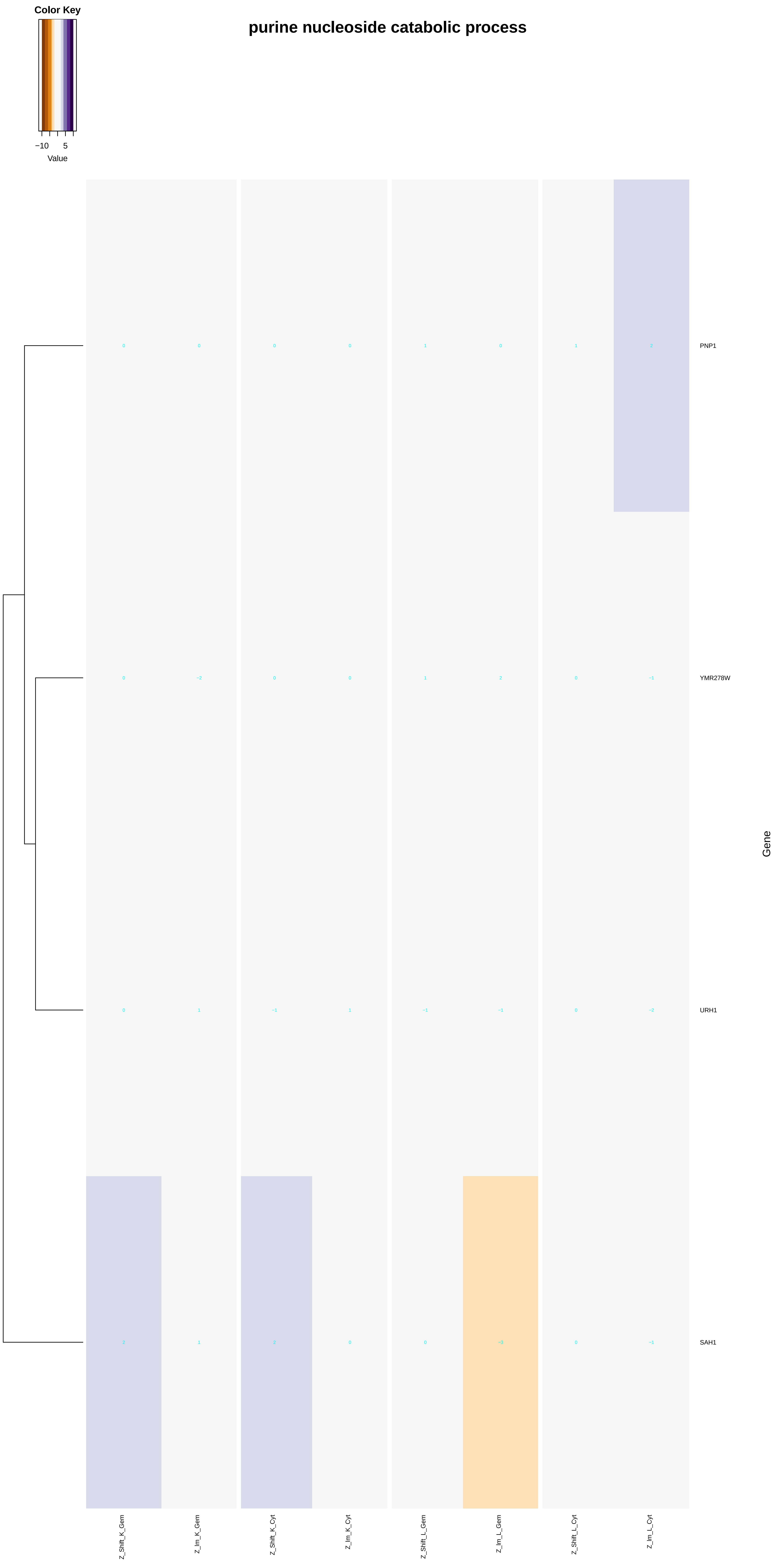

Color Key
−10
5
Value
purine nucleoside catabolic process
PNP1
YMR278W
URH1
SAH1
Gene
Z_Shift_K_Gem
Z_Im_K_Gem
Z_Shift_K_Cyt
Z_Im_K_Cyt
Z_Shift_L_Gem
Z_Im_L_Gem
Z_Shift_L_Cyt
Z_Im_L_Cyt

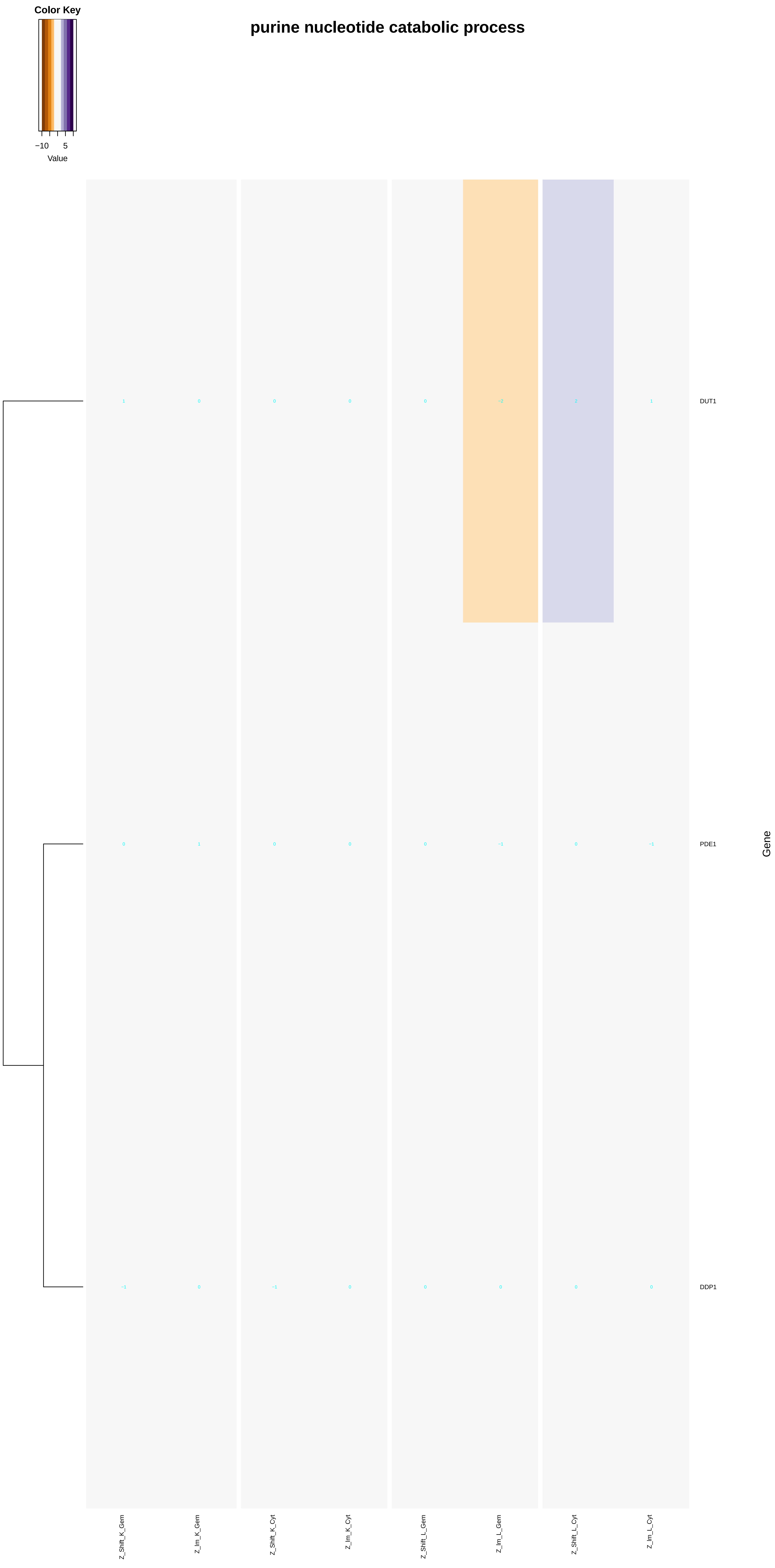

Color Key
−10
5
Value
purine nucleotide catabolic process
DUT1
PDE1
DDP1
Gene
Z_Shift_K_Gem
Z_lm_K_Gem
Z_Shift_K_Cyt
Z_lm_K_Cyt
Z_Shift_L_Gem
Z_lm_L_Gem
Z_Shift_L_Cyt
Z_lm_L_Cyt

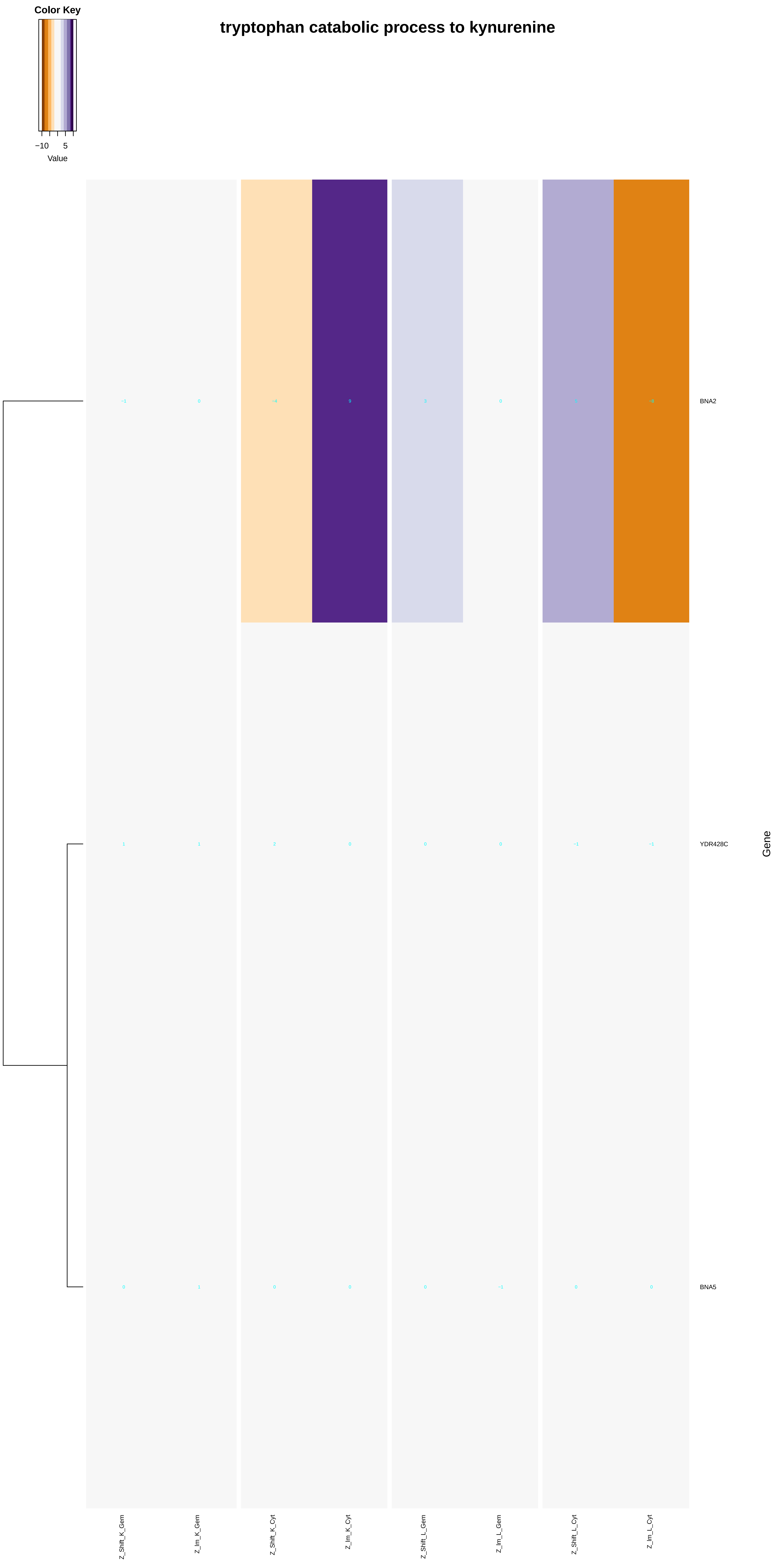

Color Key
−10
5
Value
tryptophan catabolic process to kynurenine
BNA2
YDR428C
BNA5
Gene
Z_Shift_K_Gem
Z_Im_K_Gem
Z_Shift_K_Cyt
Z_Im_K_Cyt
Z_Shift_L_Gem
Z_Im_L_Gem
Z_Shift_L_Cyt
Z_Im_L_Cyt

### DNA catabolic process, endonucleolytic

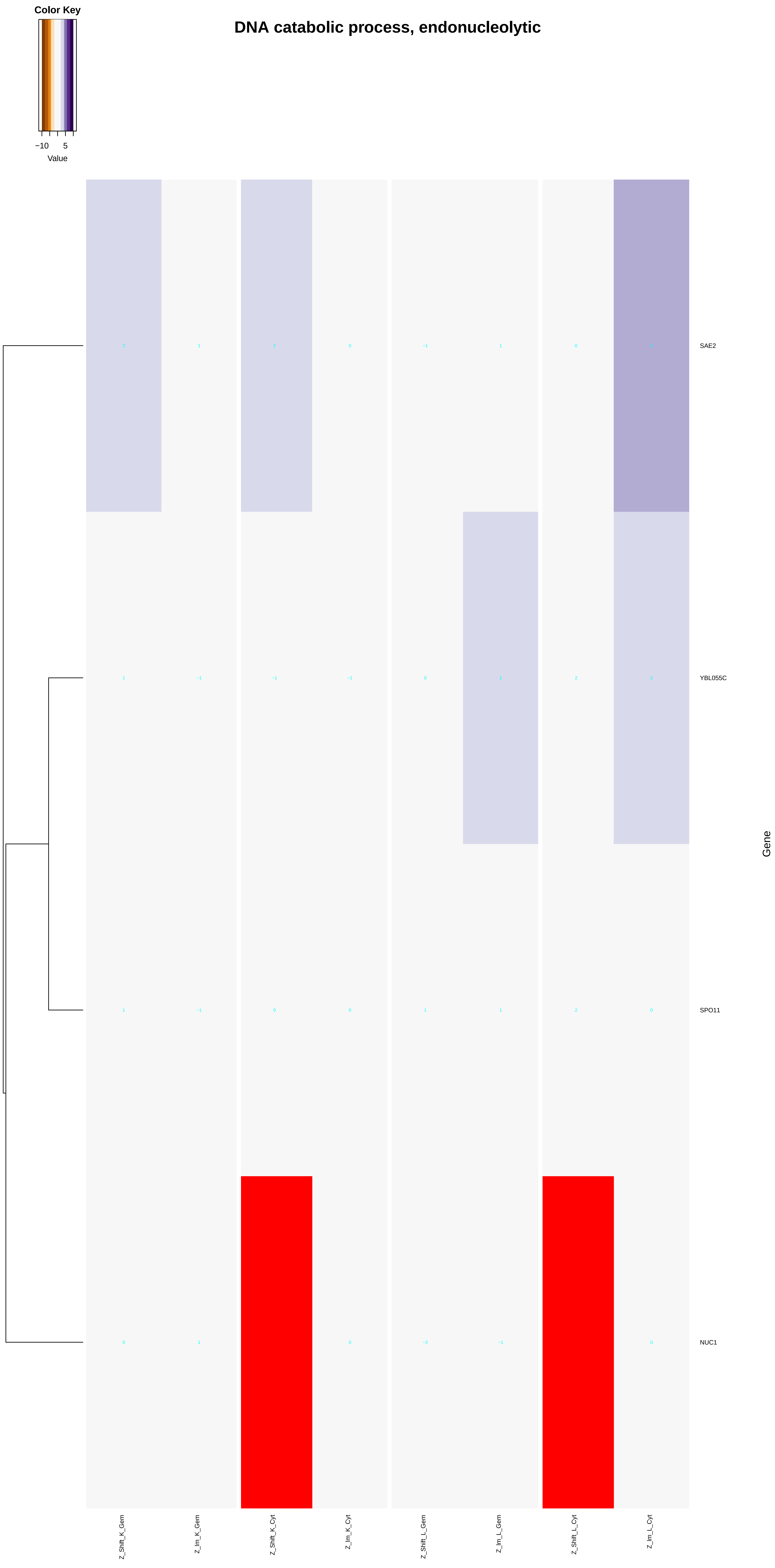

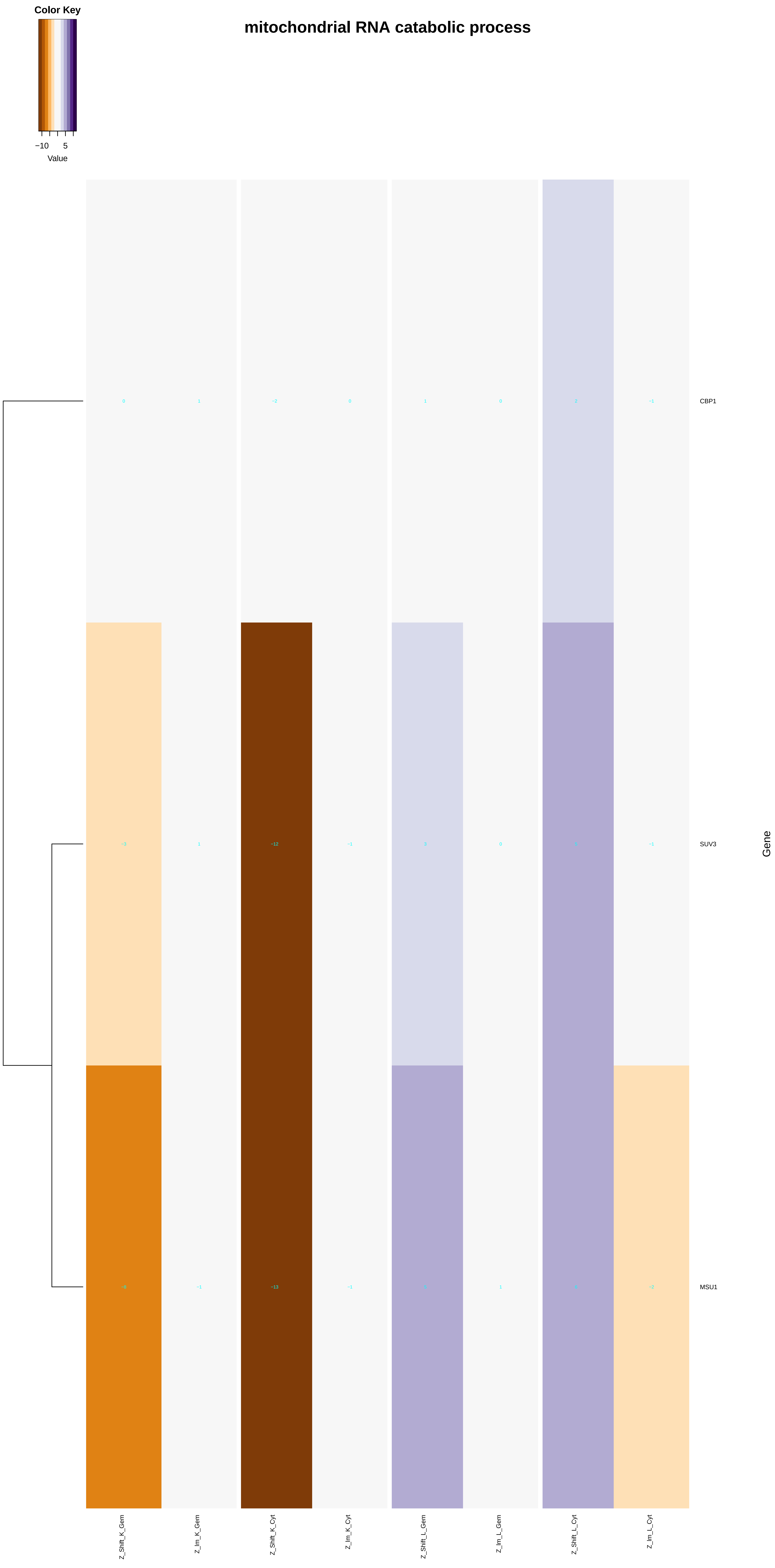

Color Key
−10
5
Value
mitochondrial RNA catabolic process
0
1
−2
0
1
0
2
−1
CBP1
−3
1
−12
−1
3
0
5
−1
SUV3
−8
−1
−13
−1
5
1
6
−2
MSU1
Gene
Z_Shift_K_Gem
Z_Im_K_Gem
Z_Shift_K_Cyt
Z_Im_K_Cyt
Z_Shift_L_Gem
Z_Im_L_Gem
Z_Shift_L_Cyt
Z_Im_L_Cyt

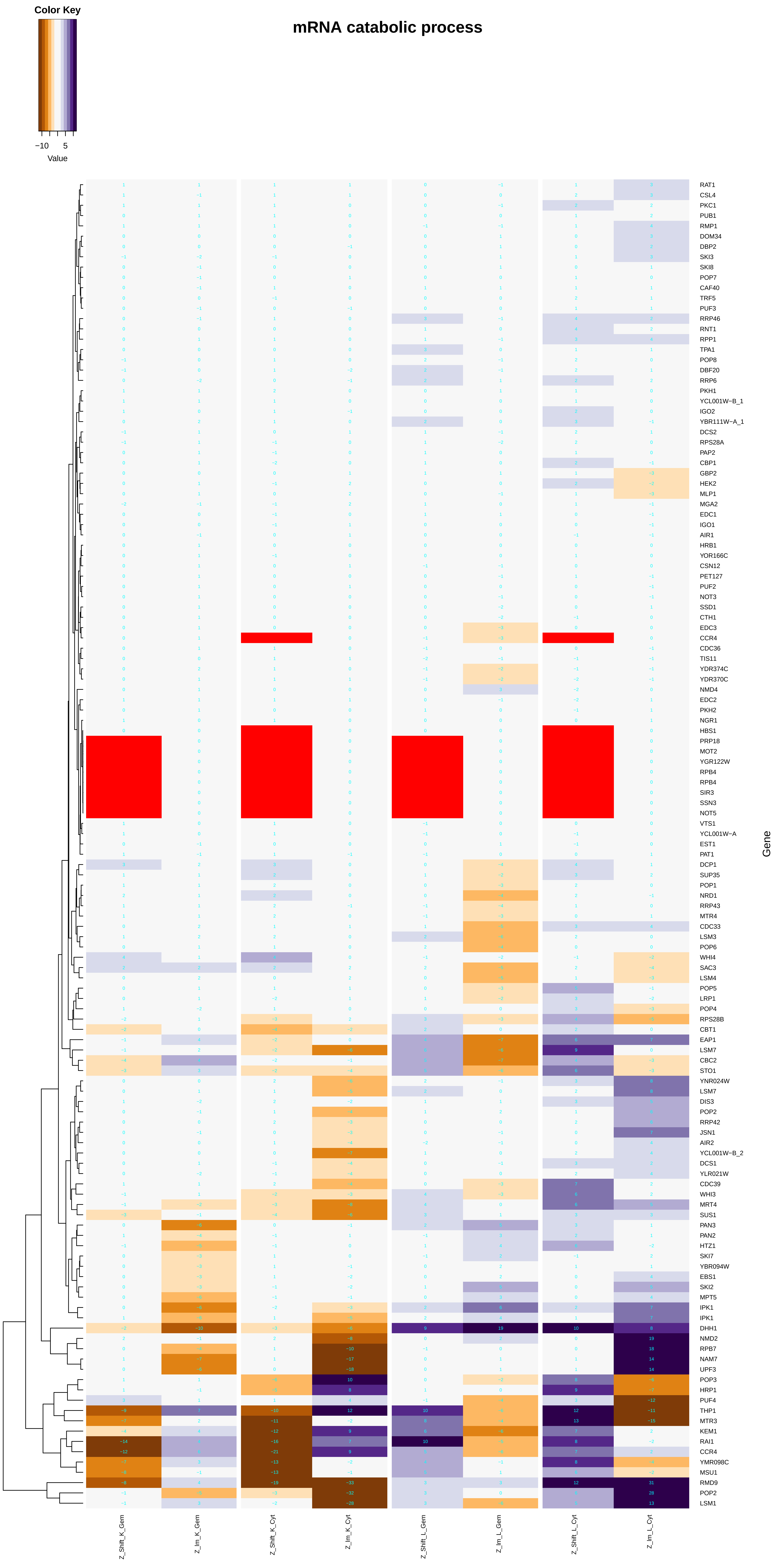
mRNA catabolic process
Color Key
−10
5
Value
Gene
RAT1
CSL4
PKC1
PUB1
RMP1
DOM34
DBP2
SKI3
SKI8
POP7
CAF40
TRF5
PUF3
RRP46
RNT1
RPP1
TPA1
POP8
DBF20
RRP6
PKH1
YCL001W−B_1
IGO2
YBR111W−A_1
DCS2
RPS28A
PAP2
CBP1
GBP2
HEK2
MLP1
MGA2
EDC1
IGO1
AIR1
HRB1
YOR166C
CSN12
PET127
PUF2
NOT3
SSD1
CTH1
EDC3
CCR4
CDC36
TIS11
YDR374C
YDR370C
NMD4
EDC2
PKH2
NGR1
HBS1
PRP18
MOT2
YGR122W
RPB4
RPB4
SIR3
SSN3
NOT5
VTS1
YCL001W−A
EST1
PAT1
DCP1
SUP35
POP1
NRD1
RRP43
MTR4
CDC33
LSM3
POP6
WHI4
SAC3
LSM4
POP5
LRP1
POP4
RPS28B
CBT1
EAP1
LSM7
CBC2
STO1
YNR024W
LSM7
DIS3
POP2
RRP42
JSN1
AIR2
YCL001W−B_2
DCS1
YLR021W
CDC39
WHI3
MRT4
SUS1
PAN3
PAN2
HTZ1
SKI7
YBR094W
EBS1
SKI2
MPT5
IPK1
IPK1
DHH1
NMD2
RPB7
NAM7
UPF3
POP3
HRP1
PUF4
THP1
MTR3
KEM1
RAI1
CCR4
YMR098C
MSU1
RMD9
POP2
LSM1
Z_Shift_K_Gem
Z_Im_K_Gem
Z_Shift_K_Cyt
Z_Im_K_Cyt
Z_Shift_L_Gem
Z_Im_L_Gem
Z_Shift_L_Cyt
Z_Im_L_Cyt

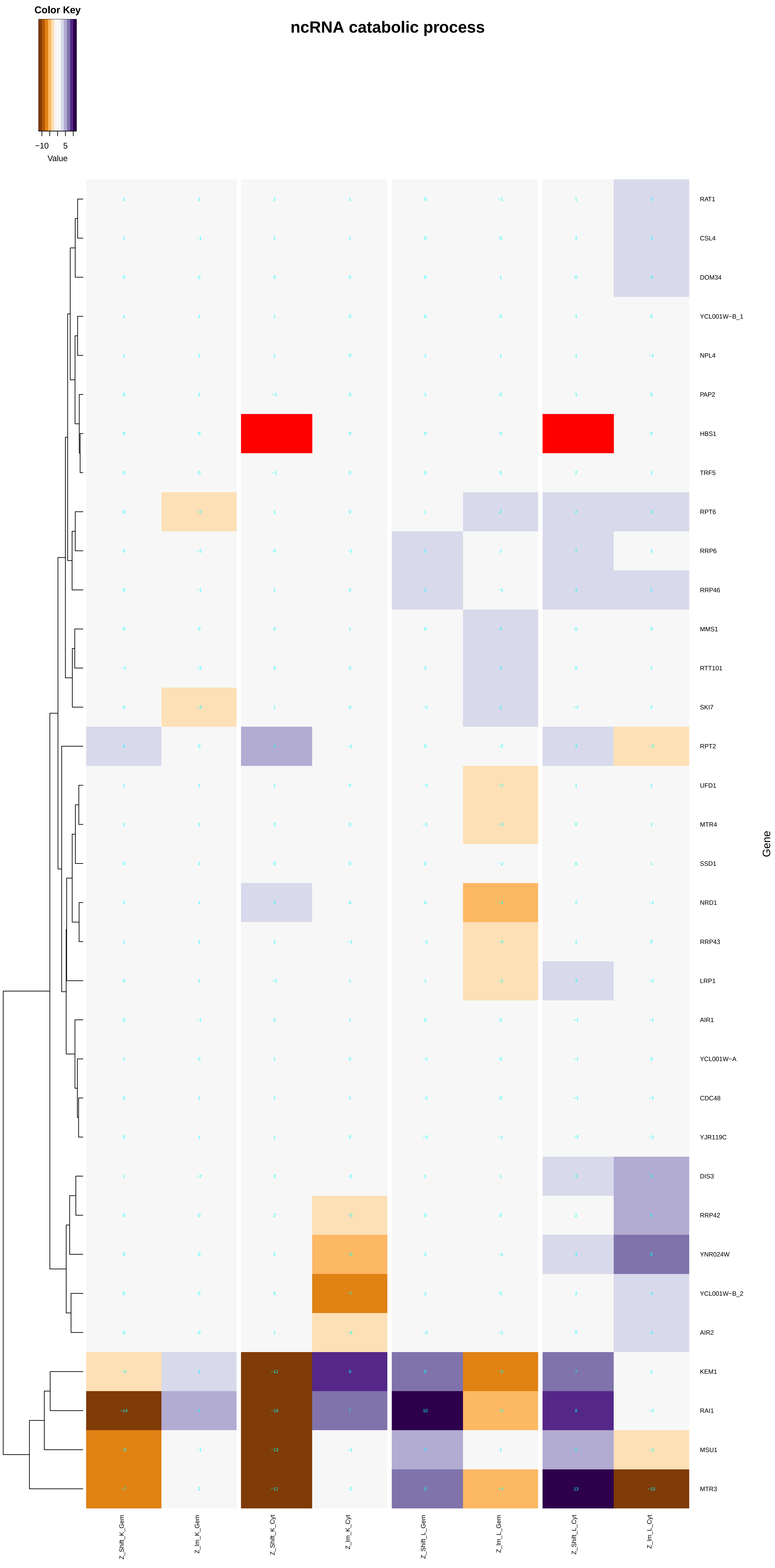

ncRNA catabolic process
Color Key
−10
5
Value
RAT1
CSL4
DOM34
YCL001W−B_1
NPL4
PAP2
HBS1
TRF5
RPT6
RRP6
RRP46
MMS1
RTT101
SKI7
RPT2
UFD1
MTR4
SSD1
NRD1
RRP43
LRP1
AIR1
YCL001W−A
CDC48
YJR119C
DIS3
RRP42
YNR024W
YCL001W−B_2
AIR2
KEM1
RAI1
MSU1
MTR3
Gene
Z_Shift_K_Gem
Z_lm_K_Gem
Z_Shift_K_Cyt
Z_lm_K_Cyt
Z_Shift_L_Gem
Z_lm_L_Gem
Z_Shift_L_Cyt
Z_lm_L_Cyt

### DNA replication, removal of RNA primer

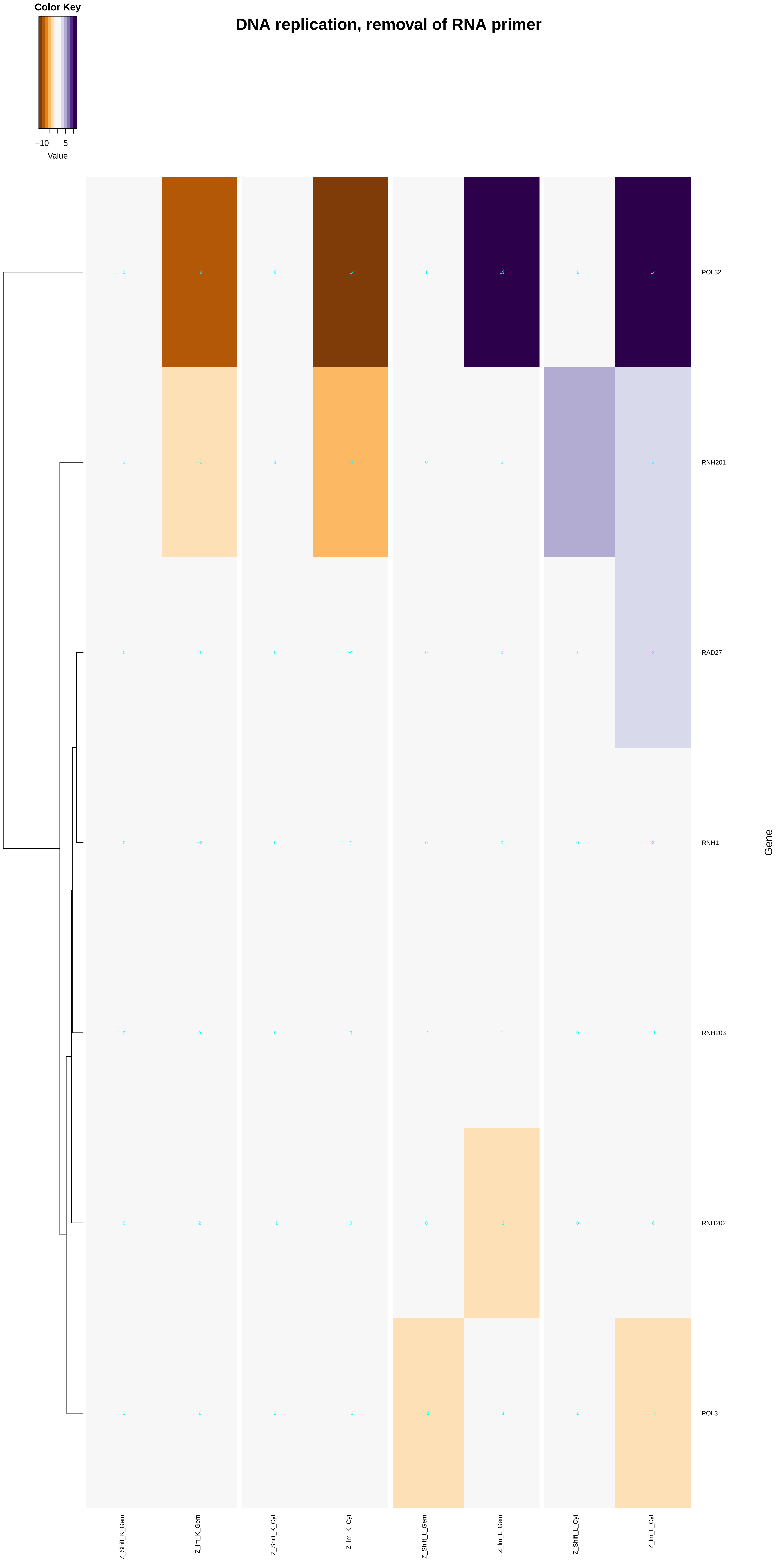

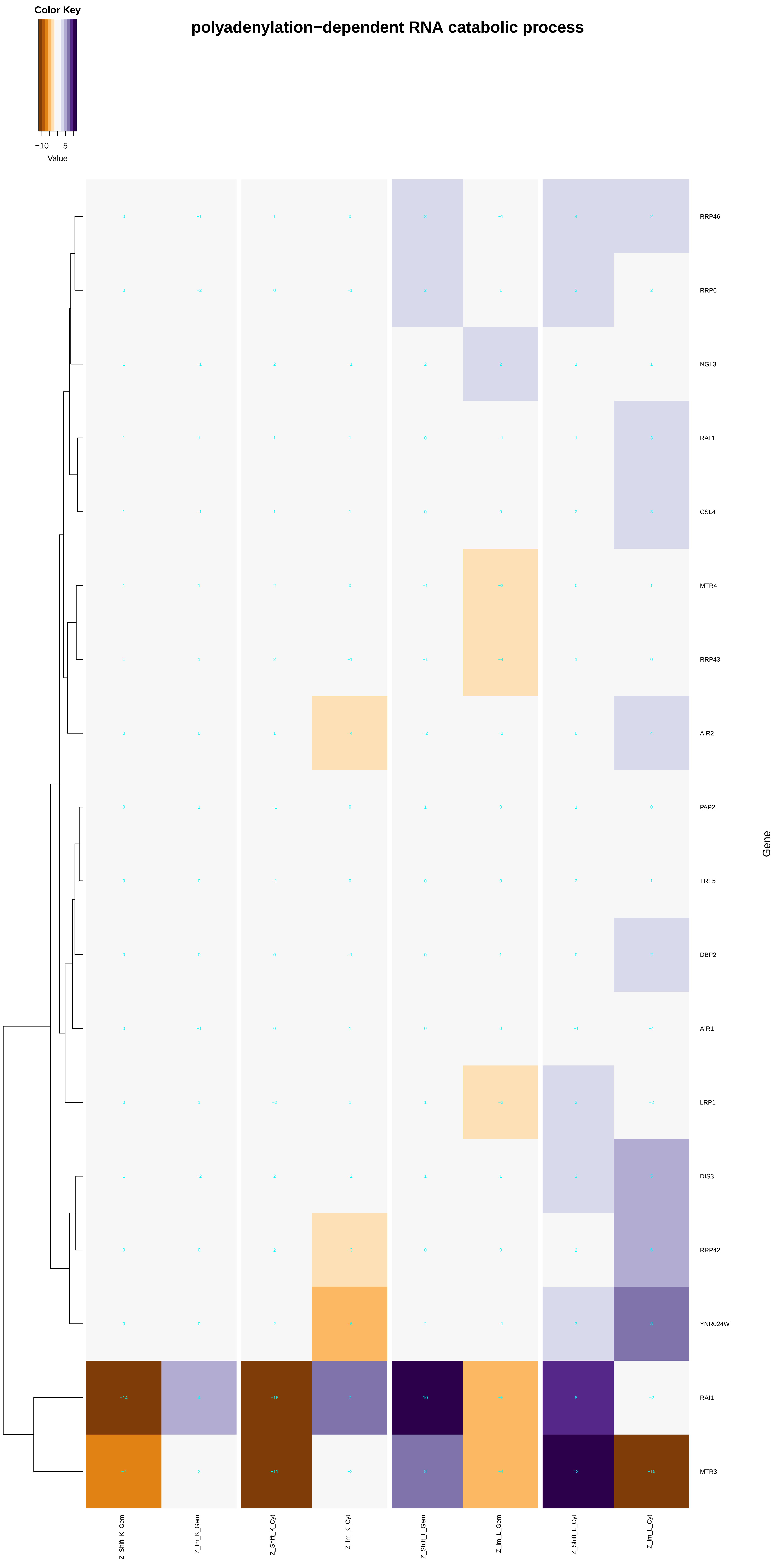
Color Key
−10
5
Value
polyadenylation−dependent RNA catabolic process
RRP46
RRP6
NGL3
RAT1
CSL4
MTR4
RRP43
AIR2
PAP2
TRF5
DBP2
AIR1
LRP1
DIS3
RRP42
YNR024W
RAI1
MTR3
Gene
Z_Shift_K_Gem
Z_Im_K_Gem
Z_Shift_K_Cyt
Z_Im_K_Cyt
Z_Shift_L_Gem
Z_Im_L_Gem
Z_Shift_L_Cyt
Z_Im_L_Cyt

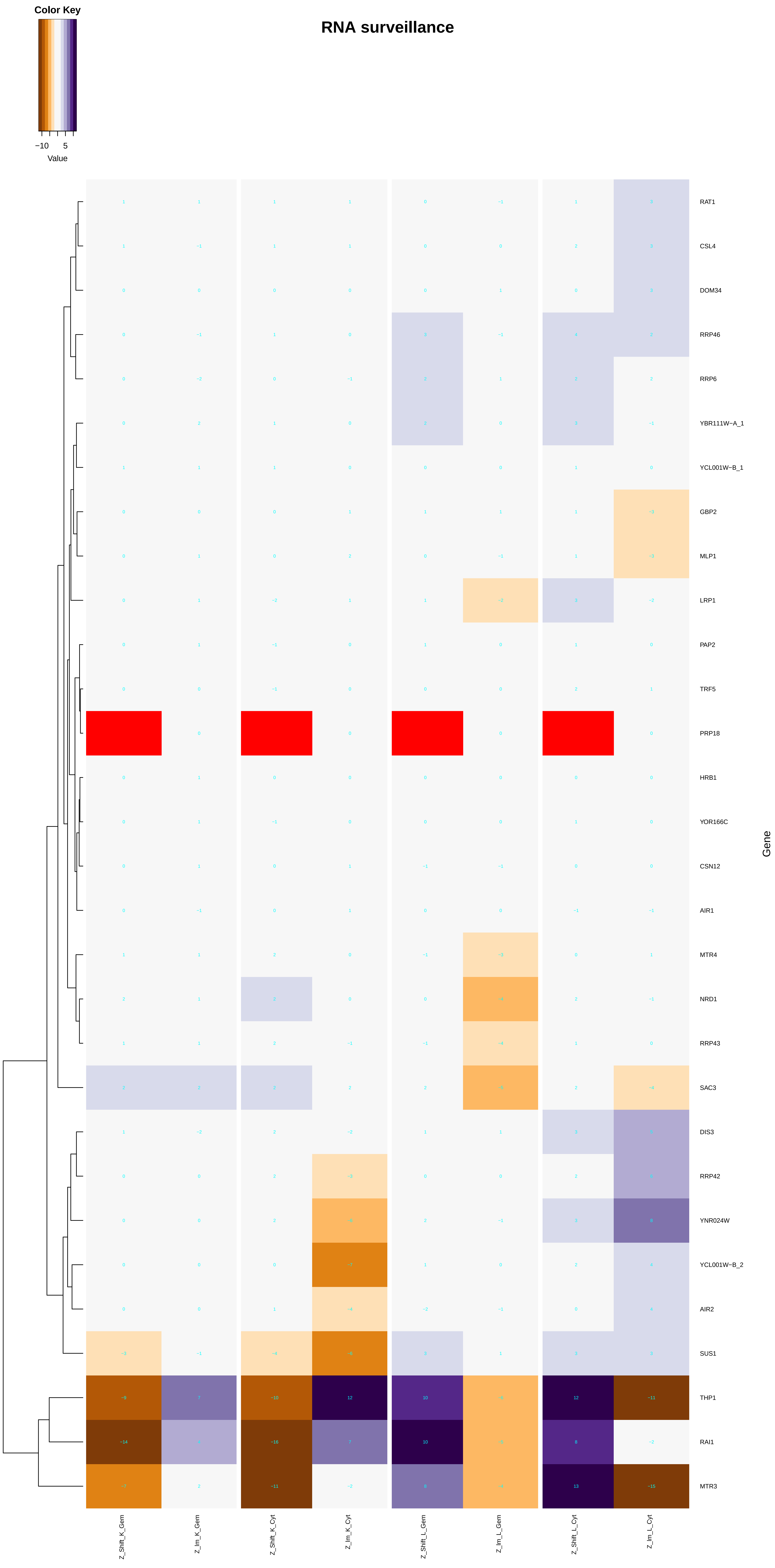

Color Key
−10
5
Value
RNA surveillance
RAT1
CSL4
DOM34
RRP46
RRP6
YBR111W−A_1
YCL001W−B_1
GBP2
MLP1
LRP1
PAP2
TRF5
PRP18
HRB1
YOR166C
CSN12
AIR1
MTR4
NRD1
RRP43
SAC3
DIS3
RRP42
YNR024W
YCL001W−B_2
AIR2
SUS1
THP1
RAI1
MTR3
Gene
Z_Shift_K_Gem
Z_Im_K_Gem
Z_Shift_K_Cyt
Z_Im_K_Cyt
Z_Shift_L_Gem
Z_Im_L_Gem
Z_Shift_L_Cyt
Z_Im_L_Cyt

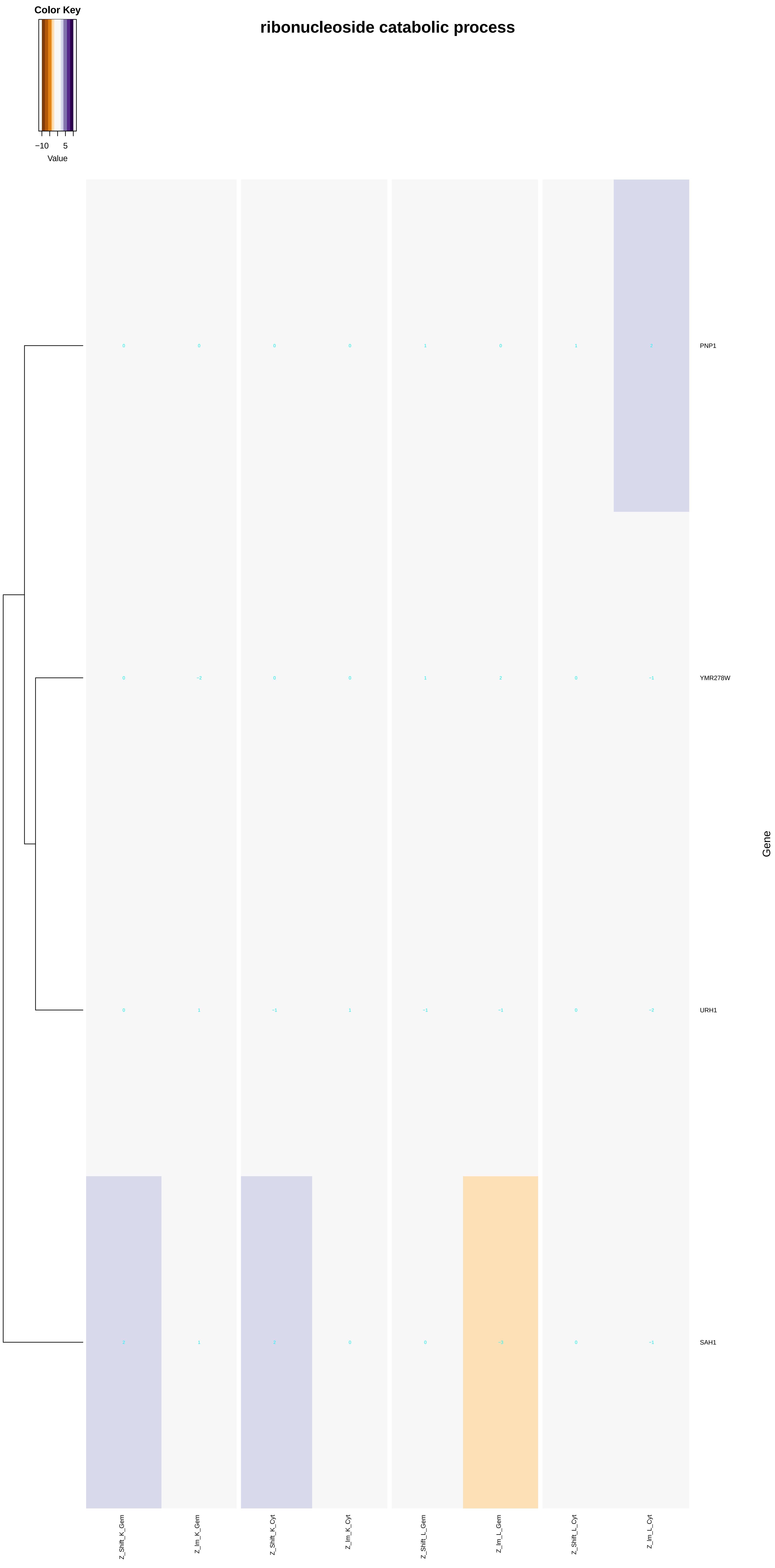
ribonucleoside catabolic process
Color Key
−10
5
Value
PNP1
YMR278W
URH1
SAH1
Gene
Z_Shift_K_Gem
Z_Im_K_Gem
Z_Shift_K_Cyt
Z_Im_K_Cyt
Z_Shift_L_Gem
Z_Im_L_Gem
Z_Shift_L_Cyt
Z_Im_L_Cyt

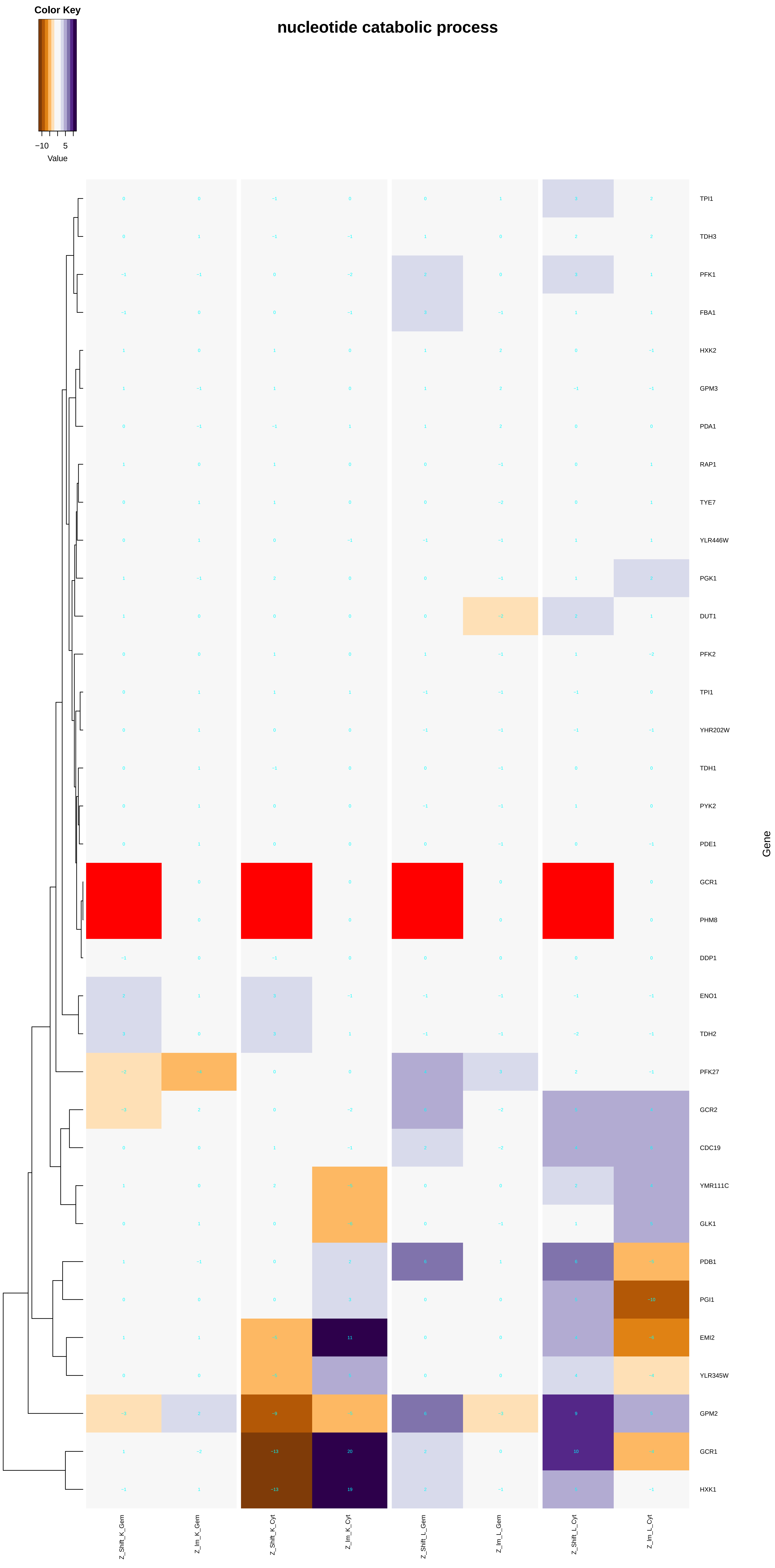
Color Key
−10
5
Value
nucleotide catabolic process
TPI1
TDH3
PFK1
FBA1
HXK2
GPM3
PDA1
RAP1
TYE7
YLR446W
PGK1
DUT1
PFK2
TPI1
YHR202W
TDH1
PYK2
PDE1
GCR1
PHM8
DDP1
ENO1
TDH2
PFK27
GCR2
CDC19
YMR111C
GLK1
PDB1
PGI1
EMI2
YLR345W
GPM2
GCR1
HXK1
Gene
Z_Shift_K_Gem
Z_lm_K_Gem
Z_Shift_K_Cyt
Z_lm_K_Cyt
Z_Shift_L_Gem
Z_lm_L_Gem
Z_Shift_L_Cyt
Z_lm_L_Cyt

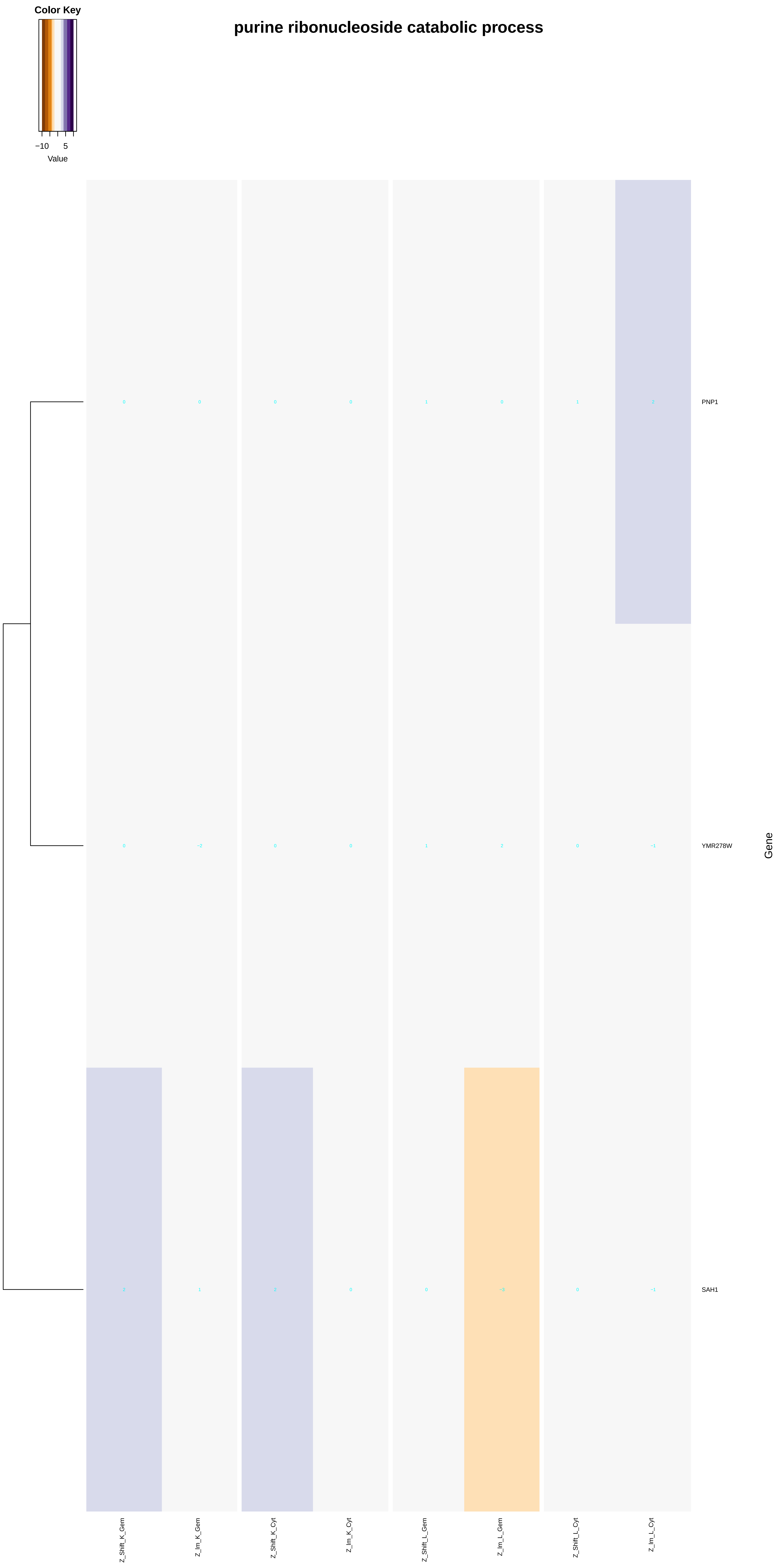
purine ribonucleoside catabolic process
Color Key
−10
5
Value
PNP1
YMR278W
SAH1
Gene
Z_Shift_K_Gem
Z_Im_K_Gem
Z_Shift_K_Cyt
Z_Im_K_Cyt
Z_Shift_L_Gem
Z_Im_L_Gem
Z_Shift_L_Cyt
Z_Im_L_Cyt
0 0 0 0 1 0 1 2
0 −2 0 0 1 2 0 −1
2 1 2 0 0 −3 0 −1

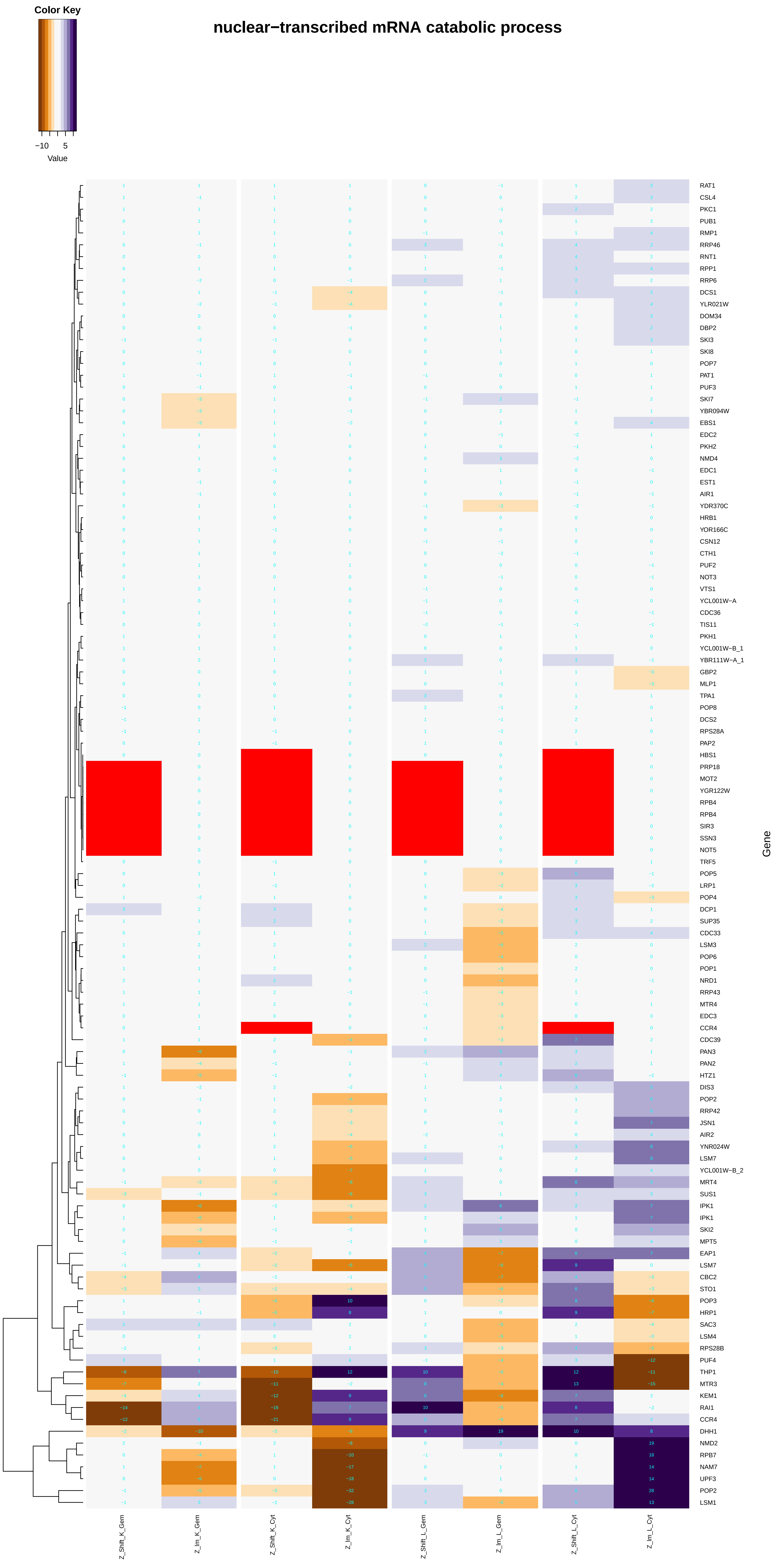
Color Key
−10
5
Value
nuclear−transcribed mRNA catabolic process
RAT1
CSL4
PKC1
PUB1
RMP1
RRP46
RNT1
RPP1
RRP6
DCS1
YLR021W
DOM34
DBP2
SKI3
SKI8
POP7
PAT1
PUF3
SKI7
YBR094W
EBS1
EDC2
PKH2
NMD4
EDC1
EST1
AIR1
YDR370C
HRB1
YOR166C
CSN12
CTH1
PUF2
NOT3
VTS1
YCL001W−A
CDC36
TIS11
PKH1
YCL001W−B_1
YBR111W−A_1
GBP2
MLP1
TPA1
POP8
DCS2
RPS28A
PAP2
HBS1
PRP18
MOT2
YGR122W
RPB4
RPB4
SIR3
SSN3
NOT5
TRF5
POP5
LRP1
POP4
DCP1
SUP35
CDC33
LSM3
POP6
POP1
NRD1
RRP43
MTR4
EDC3
CCR4
CDC39
PAN3
PAN2
HTZ1
DIS3
POP2
RRP42
JSN1
AIR2
YNR024W
LSM7
YCL001W−B_2
MRT4
SUS1
IPK1
IPK1
SKI2
MPT5
EAP1
LSM7
CBC2
STO1
POP3
HRP1
SAC3
LSM4
RPS28B
PUF4
THP1
MTR3
KEM1
RAI1
CCR4
DHH1
NMD2
RPB7
NAM7
UPF3
POP2
LSM1
Gene
Z_Shift_K_Gem
Z_Im_K_Gem
Z_Shift_K_Cyt
Z_Im_K_Cyt
Z_Shift_L_Gem
Z_Im_L_Gem
Z_Shift_L_Cyt
Z_Im_L_Cyt

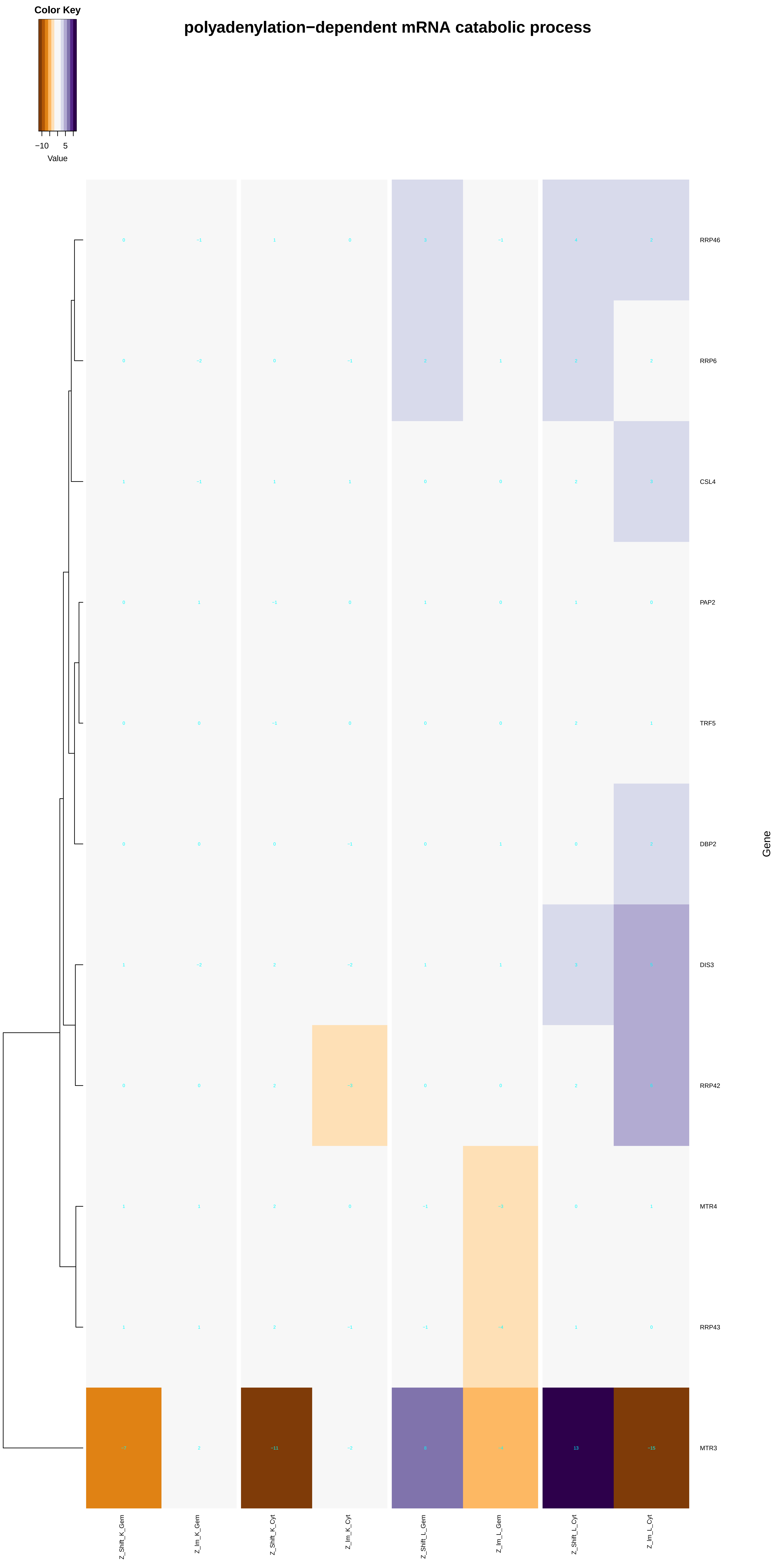
polyadenylation−dependent mRNA catabolic process
Color Key
−10
5
Value
RRP46
RRP6
CSL4
PAP2
TRF5
DBP2
DIS3
RRP42
MTR4
RRP43
MTR3
Gene
Z_Shift_K_Gem
Z_Im_K_Gem
Z_Shift_K_Cyt
Z_Im_K_Cyt
Z_Shift_L_Gem
Z_Im_L_Gem
Z_Shift_L_Cyt
Z_Im_L_Cyt

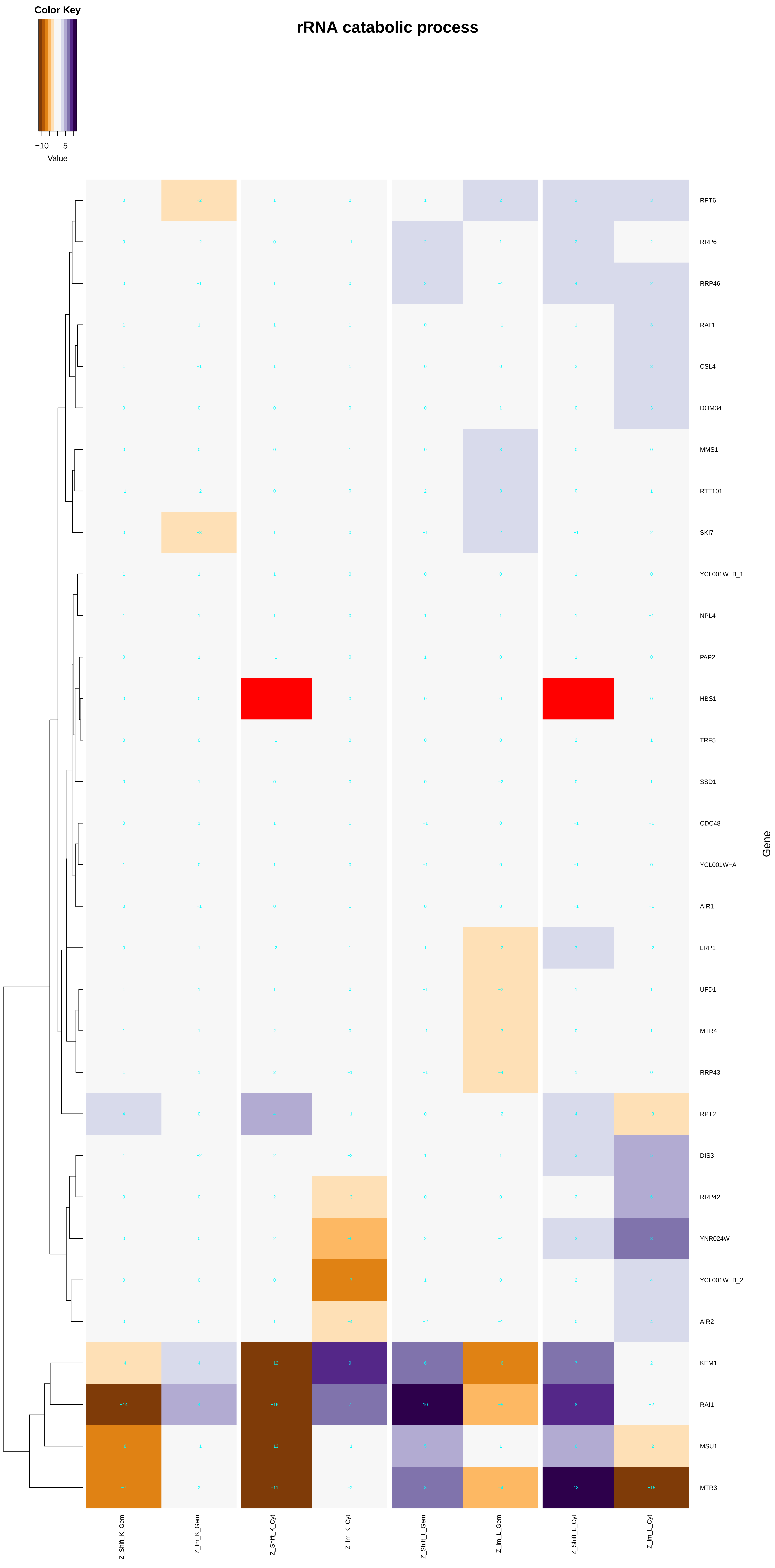
rRNA catabolic process
Color Key
−10
5
Value
RPT6
RRP6
RRP46
RAT1
CSL4
DOM34
MMS1
RTT101
SKI7
YCL001W−B_1
NPL4
PAP2
HBS1
TRF5
SSD1
CDC48
YCL001W−A
AIR1
LRP1
UFD1
MTR4
RRP43
RPT2
DIS3
RRP42
YNR024W
YCL001W−B_2
AIR2
KEM1
RAI1
MSU1
MTR3
Gene
Z_Shift_K_Gem
Z_Im_K_Gem
Z_Shift_K_Cyt
Z_Im_K_Cyt
Z_Shift_L_Gem
Z_Im_L_Gem
Z_Shift_L_Cyt
Z_Im_L_Cyt

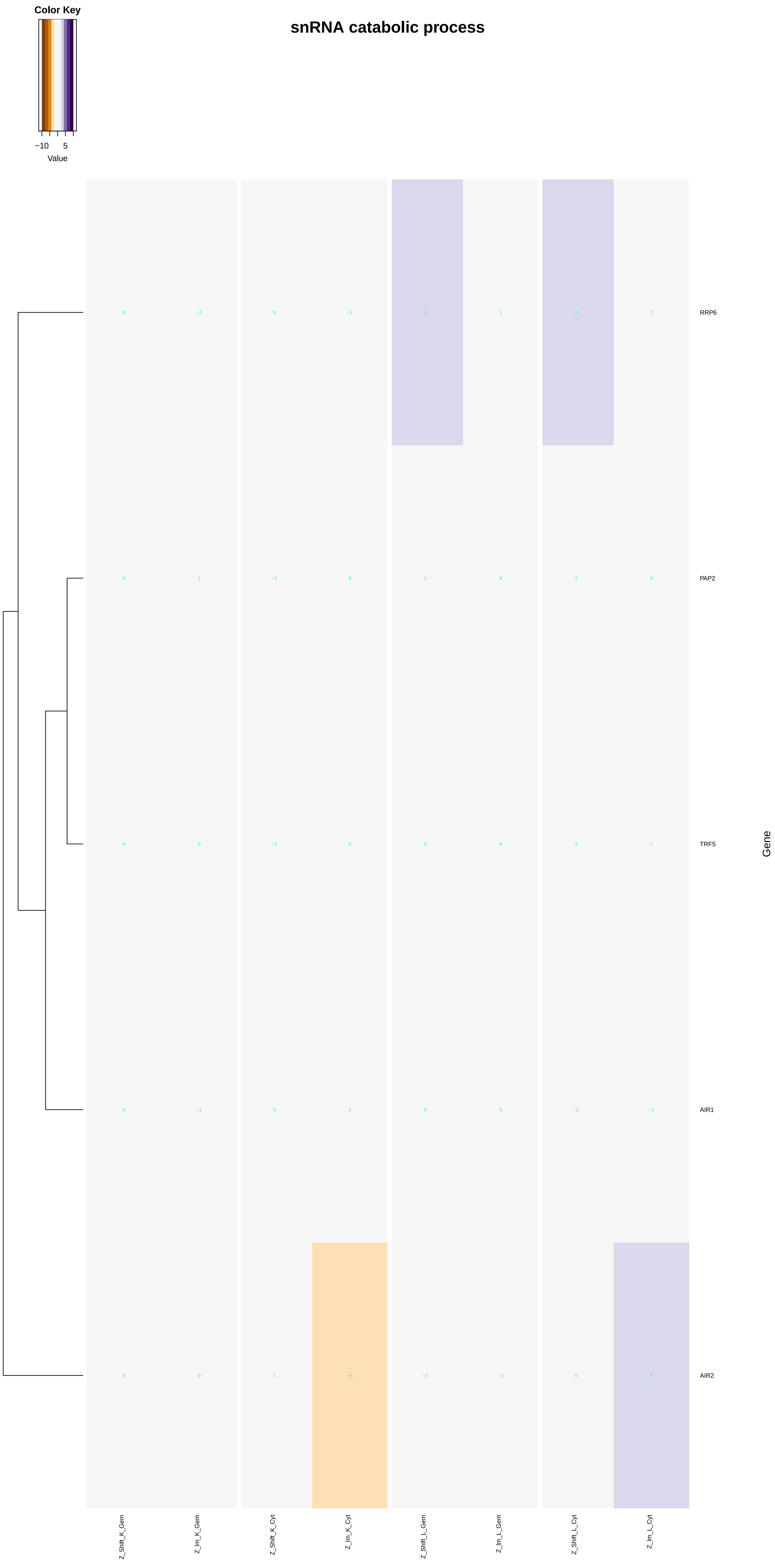
Color Key
−10
5
Value
snRNA catabolic process
RRP6
PAP2
TRF5
AIR1
AIR2
Gene
Z_Shift_K_Gem
Z_Im_K_Gem
Z_Shift_K_Cyt
Z_Im_K_Cyt
Z_Shift_L_Gem
Z_Im_L_Gem
Z_Shift_L_Cyt
Z_Im_L_Cyt

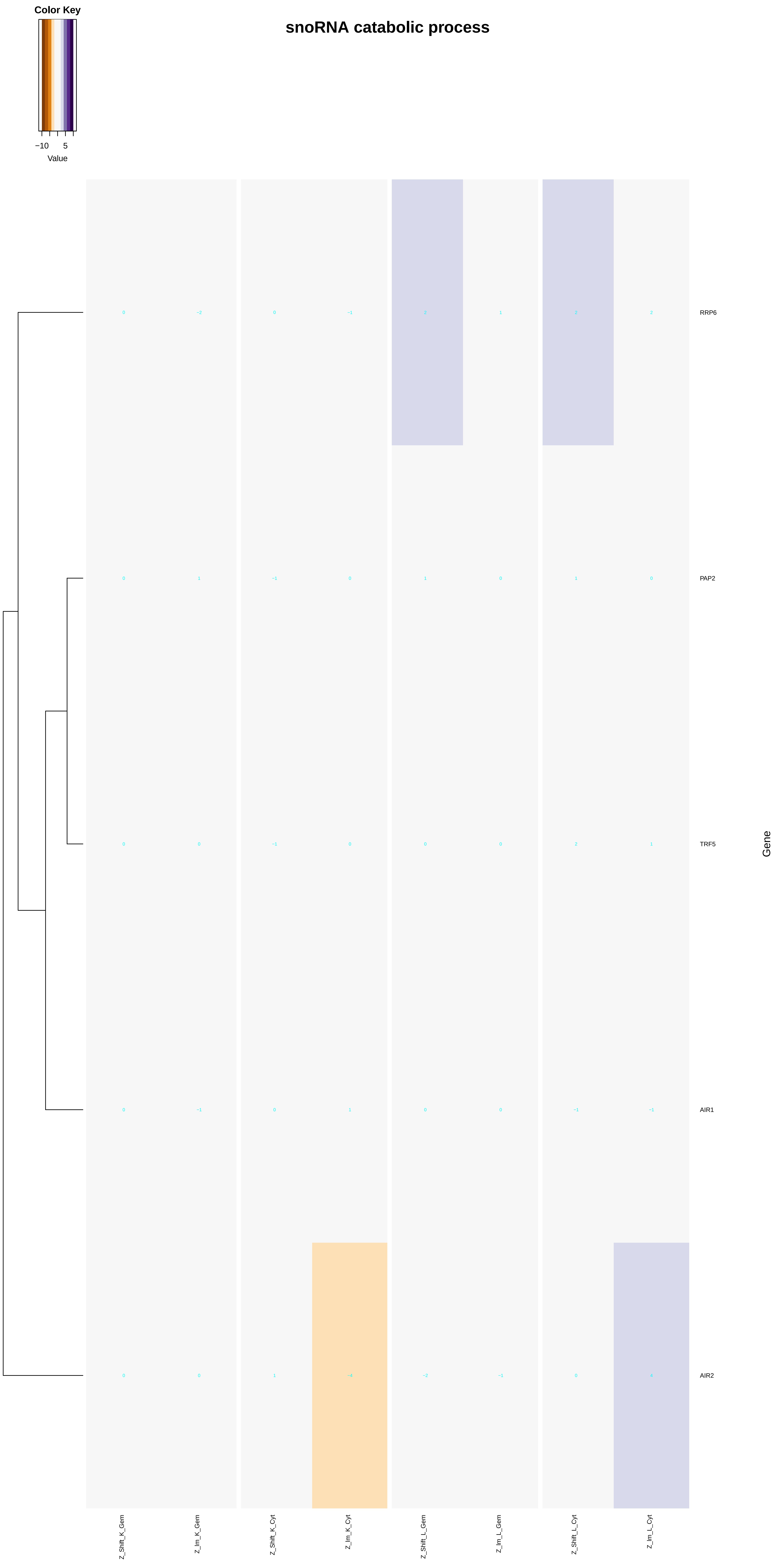
Color Key
−10
5
Value
snoRNA catabolic process
RRP6
PAP2
TRF5
AIR1
AIR2
Gene
Z_Shift_K_Gem
Z_Im_K_Gem
Z_Shift_K_Cyt
Z_Im_K_Cyt
Z_Shift_L_Gem
Z_Im_L_Gem
Z_Shift_L_Cyt
Z_Im_L_Cyt

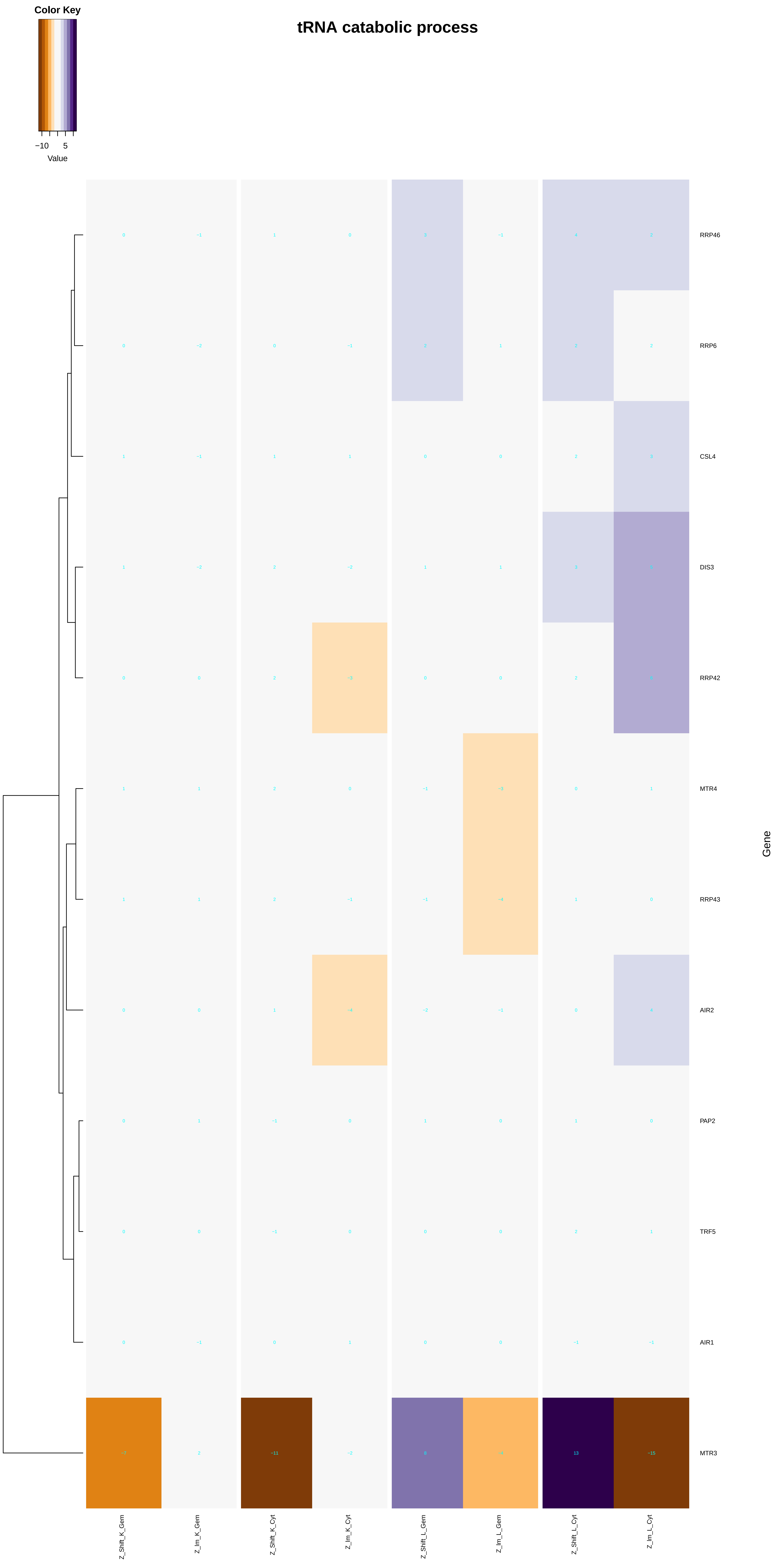
tRNA catabolic process
Color Key
−10
5
Value
RRP46
RRP6
CSL4
DIS3
RRP42
MTR4
RRP43
AIR2
PAP2
TRF5
AIR1
MTR3
Gene
Z_Shift_K_Gem
Z_Im_K_Gem
Z_Shift_K_Cyt
Z_Im_K_Cyt
Z_Shift_L_Gem
Z_Im_L_Gem
Z_Shift_L_Cyt
Z_Im_L_Cyt

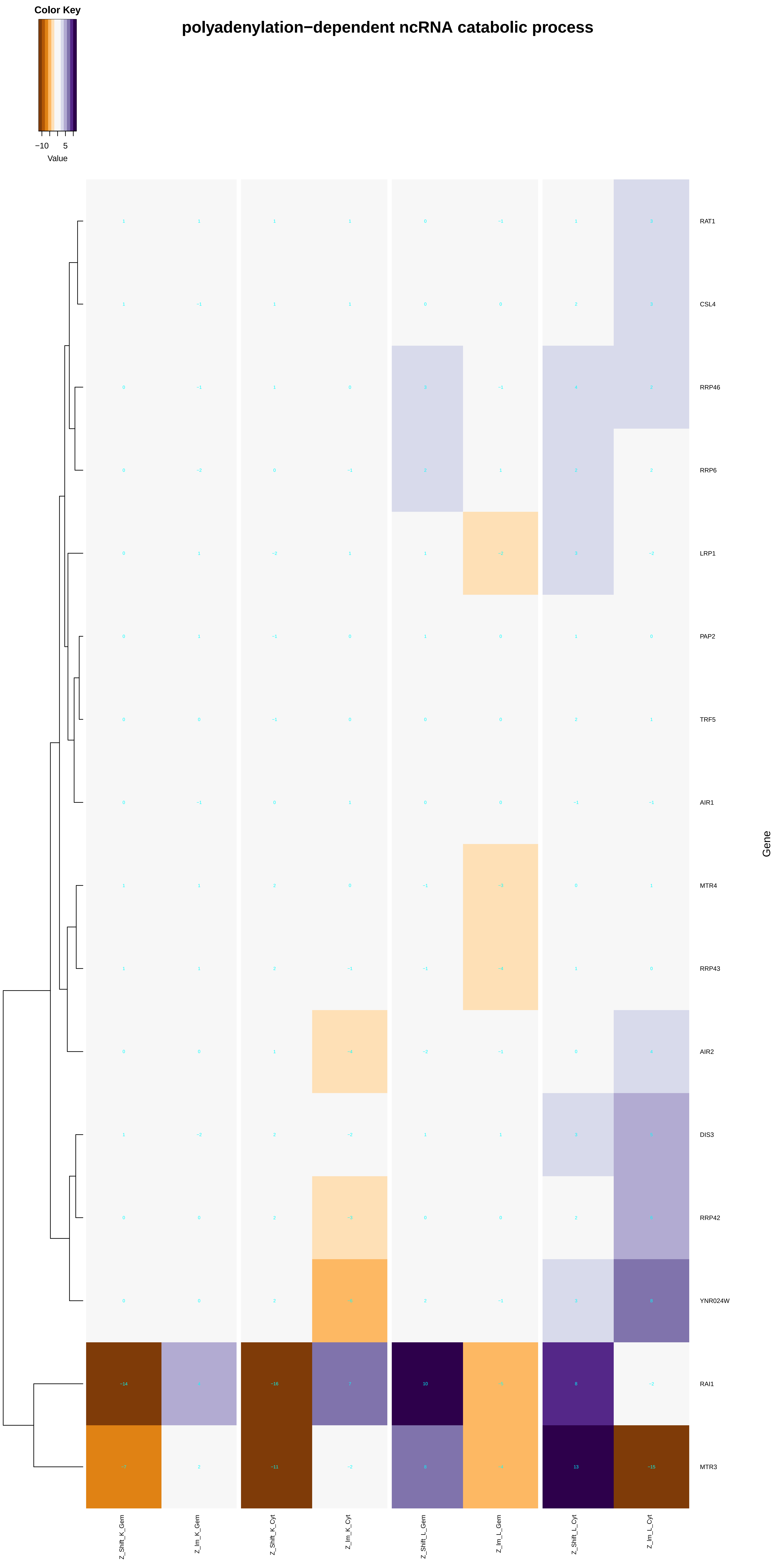
Color Key
−10
5
Value
polyadenylation−dependent ncRNA catabolic process
RAT1
CSL4
RRP46
RRP6
LRP1
PAP2
TRF5
AIR1
MTR4
RRP43
AIR2
DIS3
RRP42
YNR024W
RAI1
MTR3
Gene
Z_Shift_K_Gem
Z_Im_K_Gem
Z_Shift_K_Cyt
Z_Im_K_Cyt
Z_Shift_L_Gem
Z_Im_L_Gem
Z_Shift_L_Cyt
Z_Im_L_Cyt

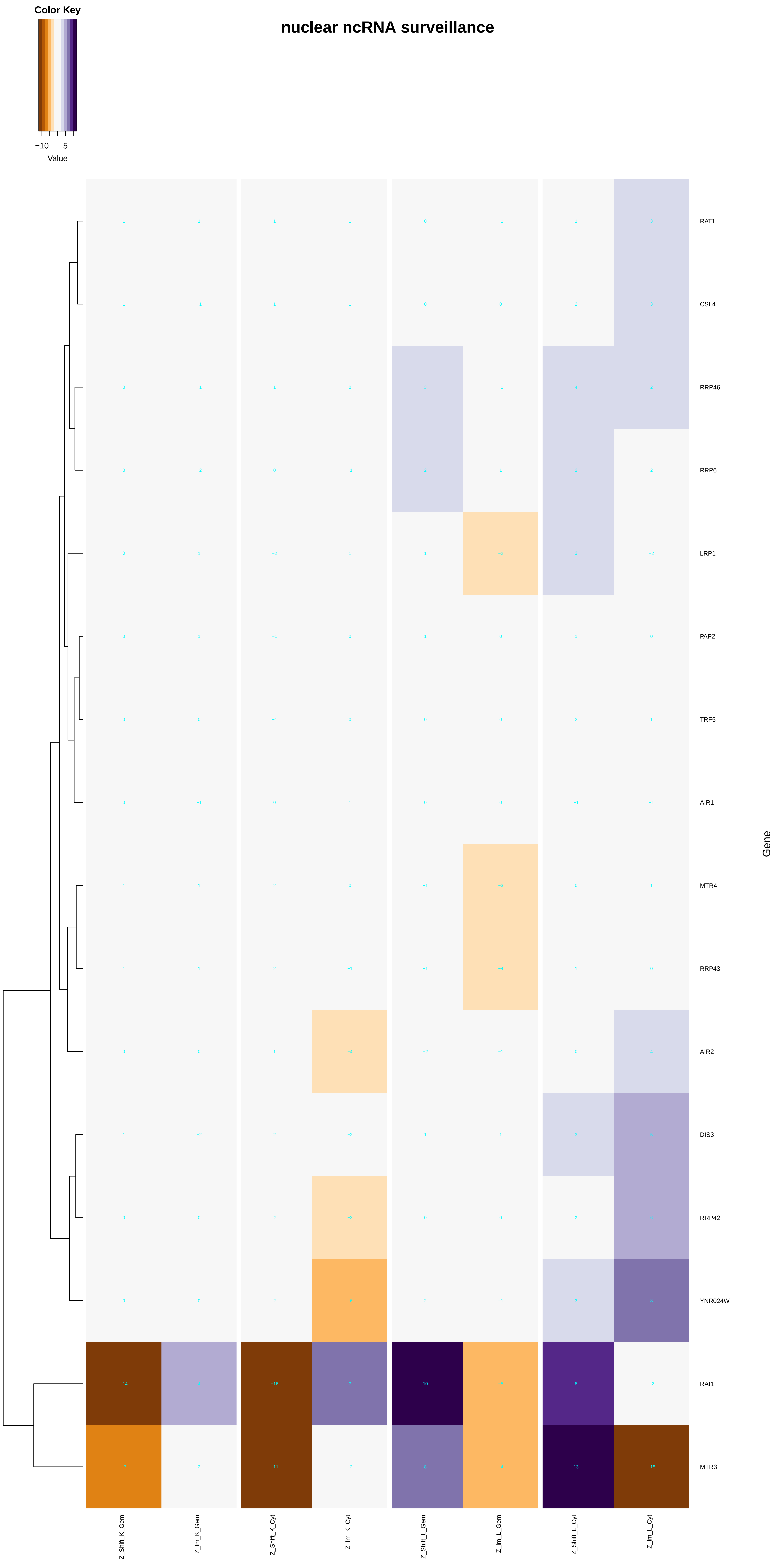
Color Key
−10
5
Value
nuclear ncRNA surveillance
RAT1
CSL4
RRP46
RRP6
LRP1
PAP2
TRF5
AIR1
MTR4
RRP43
AIR2
DIS3
RRP42
YNR024W
RAI1
MTR3
Gene
Z_Shift_K_Gem
Z_Im_K_Gem
Z_Shift_K_Cyt
Z_Im_K_Cyt
Z_Shift_L_Gem
Z_Im_L_Gem
Z_Shift_L_Cyt
Z_Im_L_Cyt

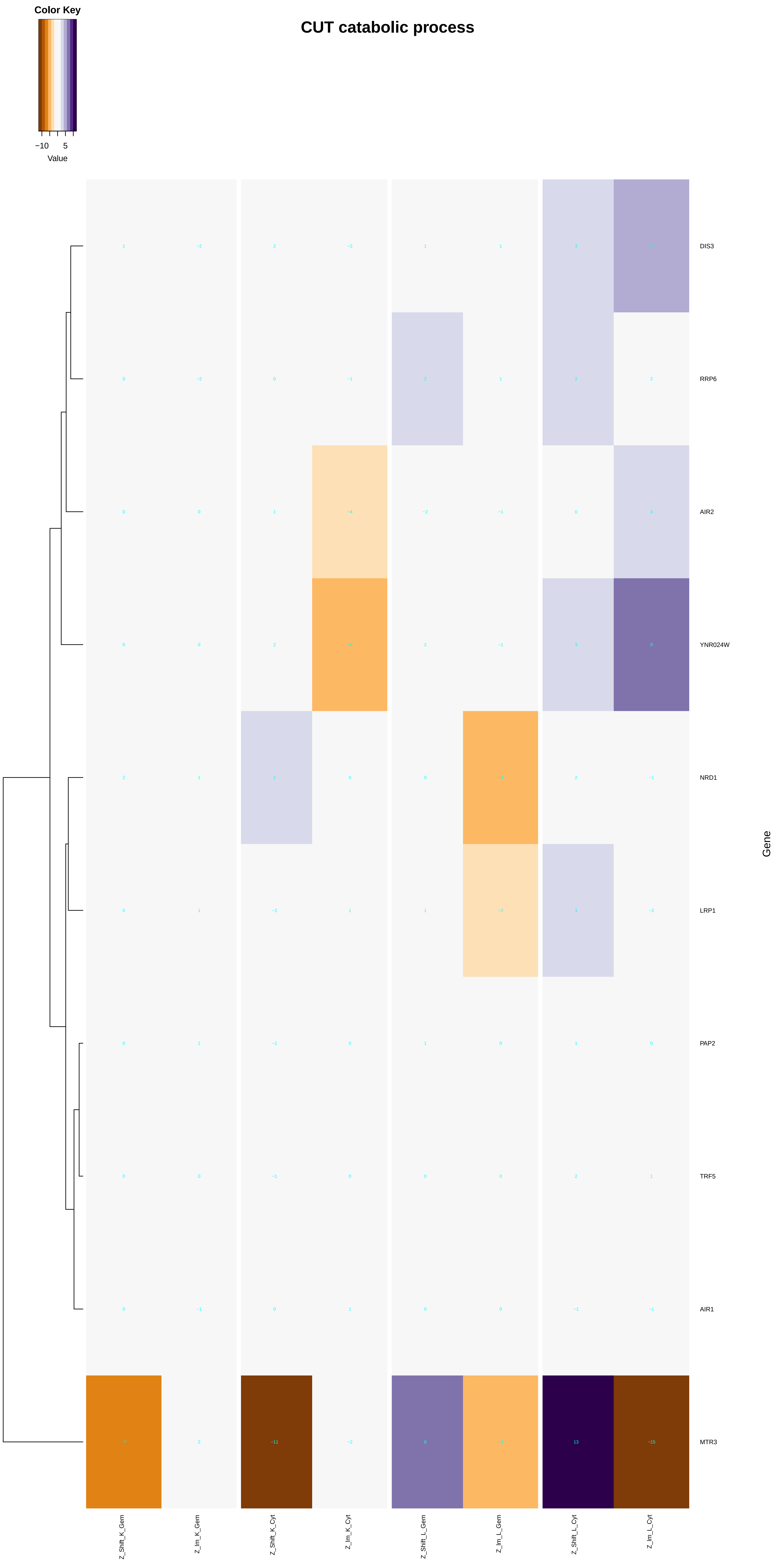

Color Key
−10
5
Value
CUT catabolic process
DIS3
RRP6
AIR2
YNR024W
NRD1
LRP1
PAP2
TRF5
AIR1
MTR3
Gene
Z_Shift_K_Gem
Z_Im_K_Gem
Z_Shift_K_Cyt
Z_Im_K_Cyt
Z_Shift_L_Gem
Z_Im_L_Gem
Z_Shift_L_Cyt
Z_Im_L_Cyt

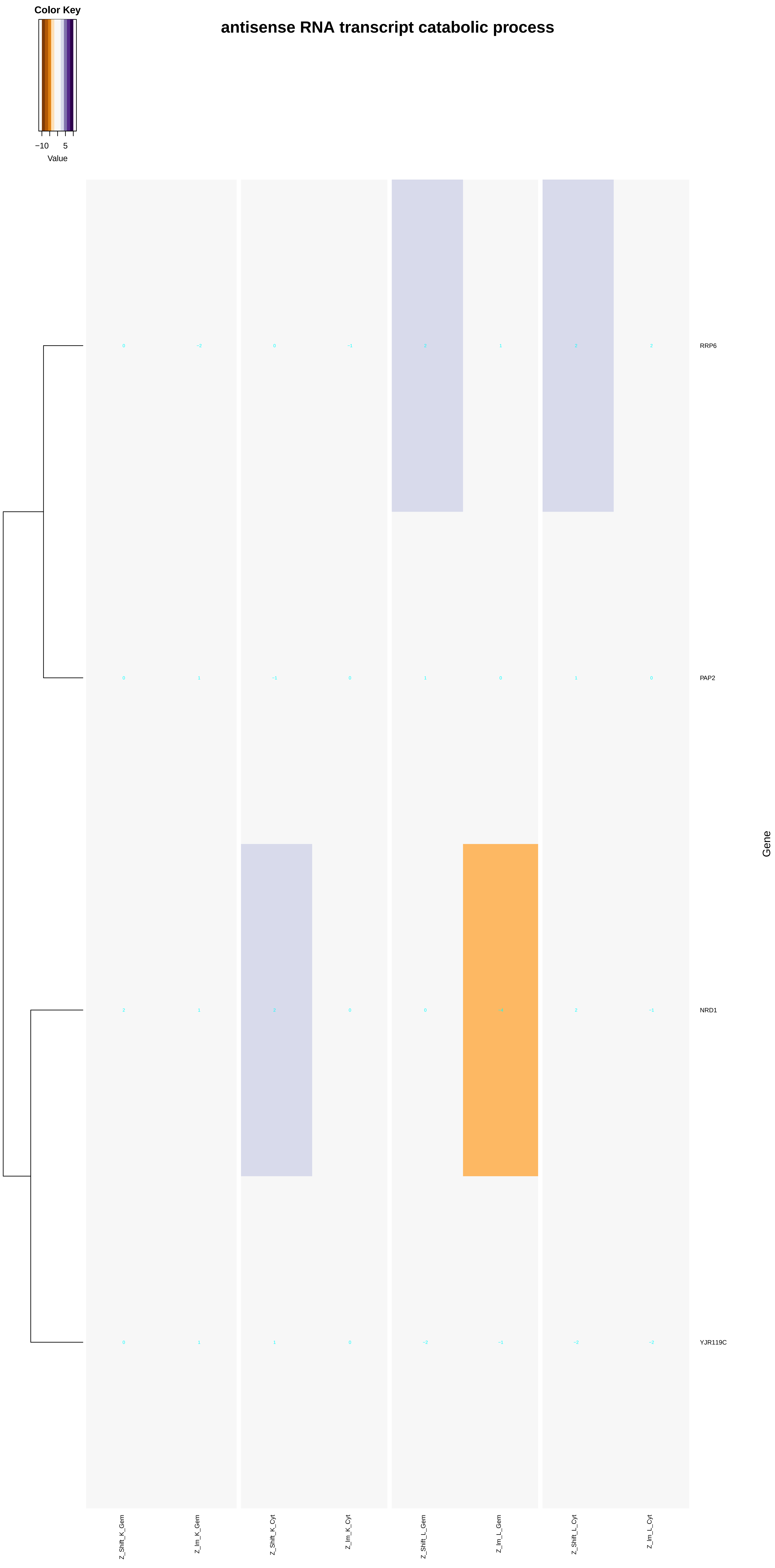
Color Key
−10
5
Value
antisense RNA transcript catabolic process
RRP6
PAP2
NRD1
YJR119C
Gene
Z_Shift_K_Gem
Z_Im_K_Gem
Z_Shift_K_Cyt
Z_Im_K_Cyt
Z_Shift_L_Gem
Z_Im_L_Gem
Z_Shift_L_Cyt
Z_Im_L_Cyt

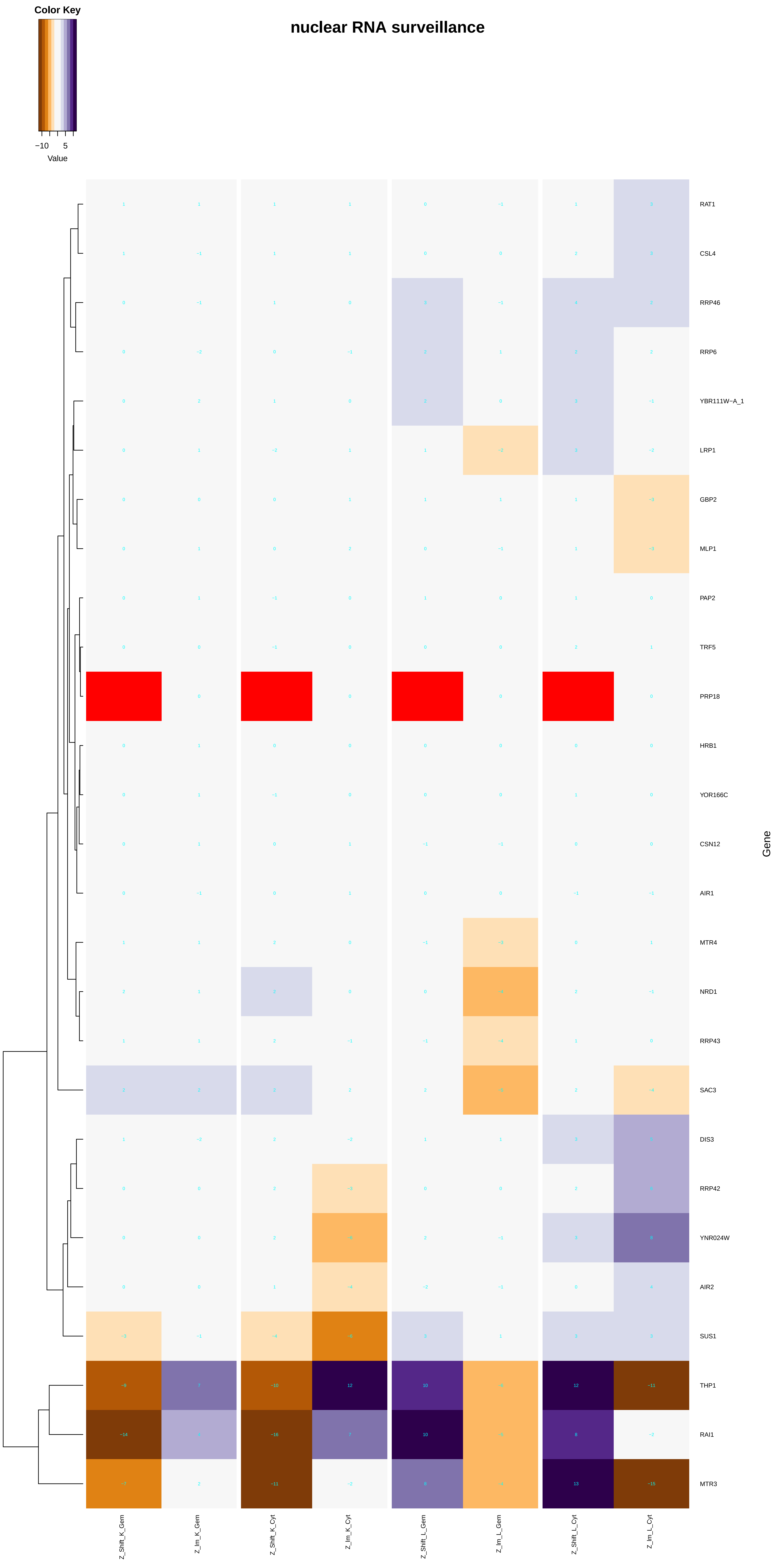

Color Key
−10
5
Value
nuclear RNA surveillance
RAT1
CSL4
RRP46
RRP6
YBR111W−A_1
LRP1
GBP2
MLP1
PAP2
TRF5
PRP18
HRB1
YOR166C
CSN12
AIR1
MTR4
NRD1
RRP43
SAC3
DIS3
RRP42
YNR024W
AIR2
SUS1
THP1
RAI1
MTR3
Gene
Z_Shift_K_Gem
Z_Im_K_Gem
Z_Shift_K_Cyt
Z_Im_K_Cyt
Z_Shift_L_Gem
Z_Im_L_Gem
Z_Shift_L_Cyt
Z_Im_L_Cyt

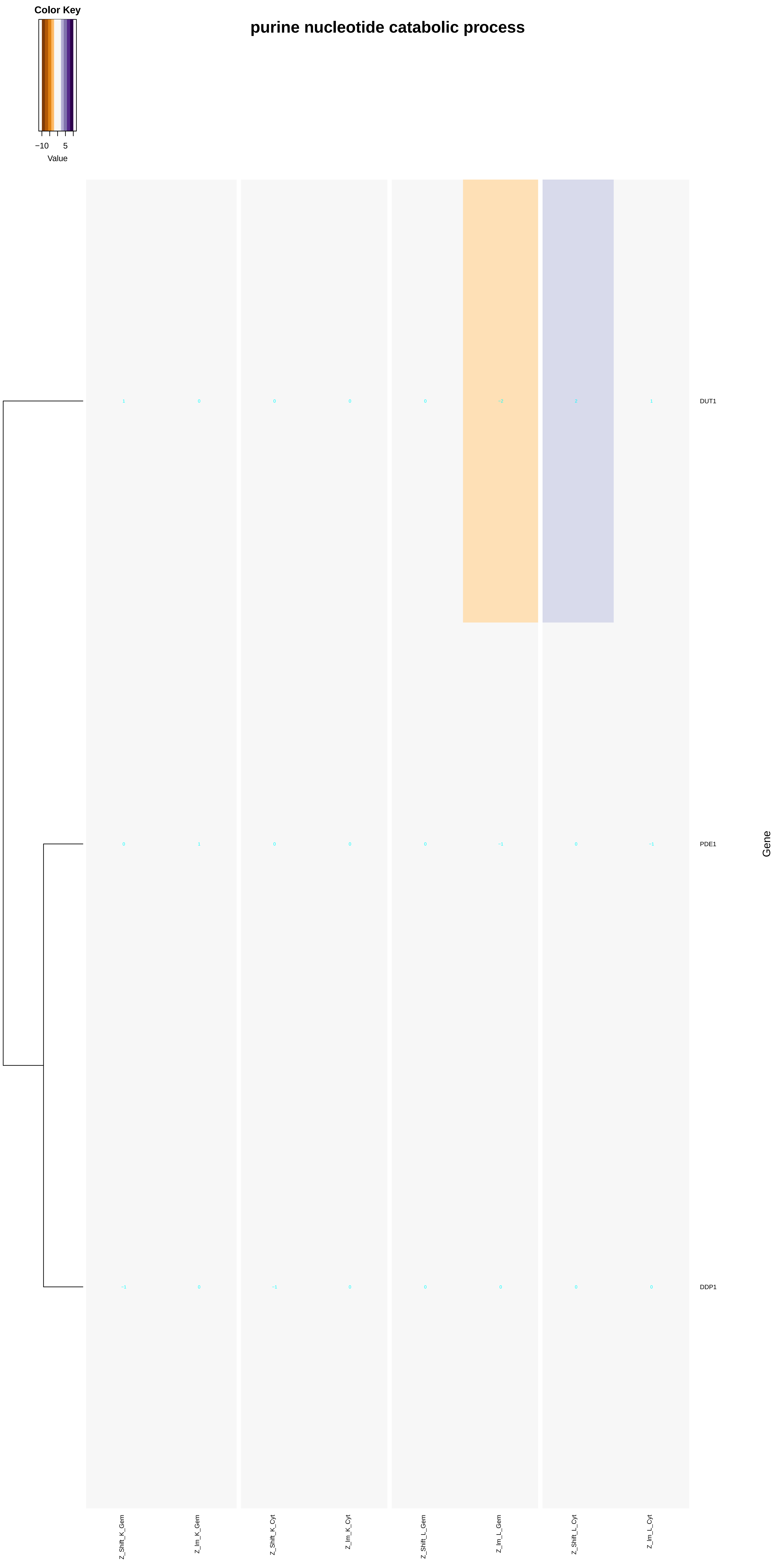

Color Key
−10
5
Value
purine nucleotide catabolic process
DUT1
PDE1
DDP1
Gene
Z_Shift_K_Gem
Z_Im_K_Gem
Z_Shift_K_Cyt
Z_Im_K_Cyt
Z_Shift_L_Gem
Z_Im_L_Gem
Z_Shift_L_Cyt
Z_Im_L_Cyt

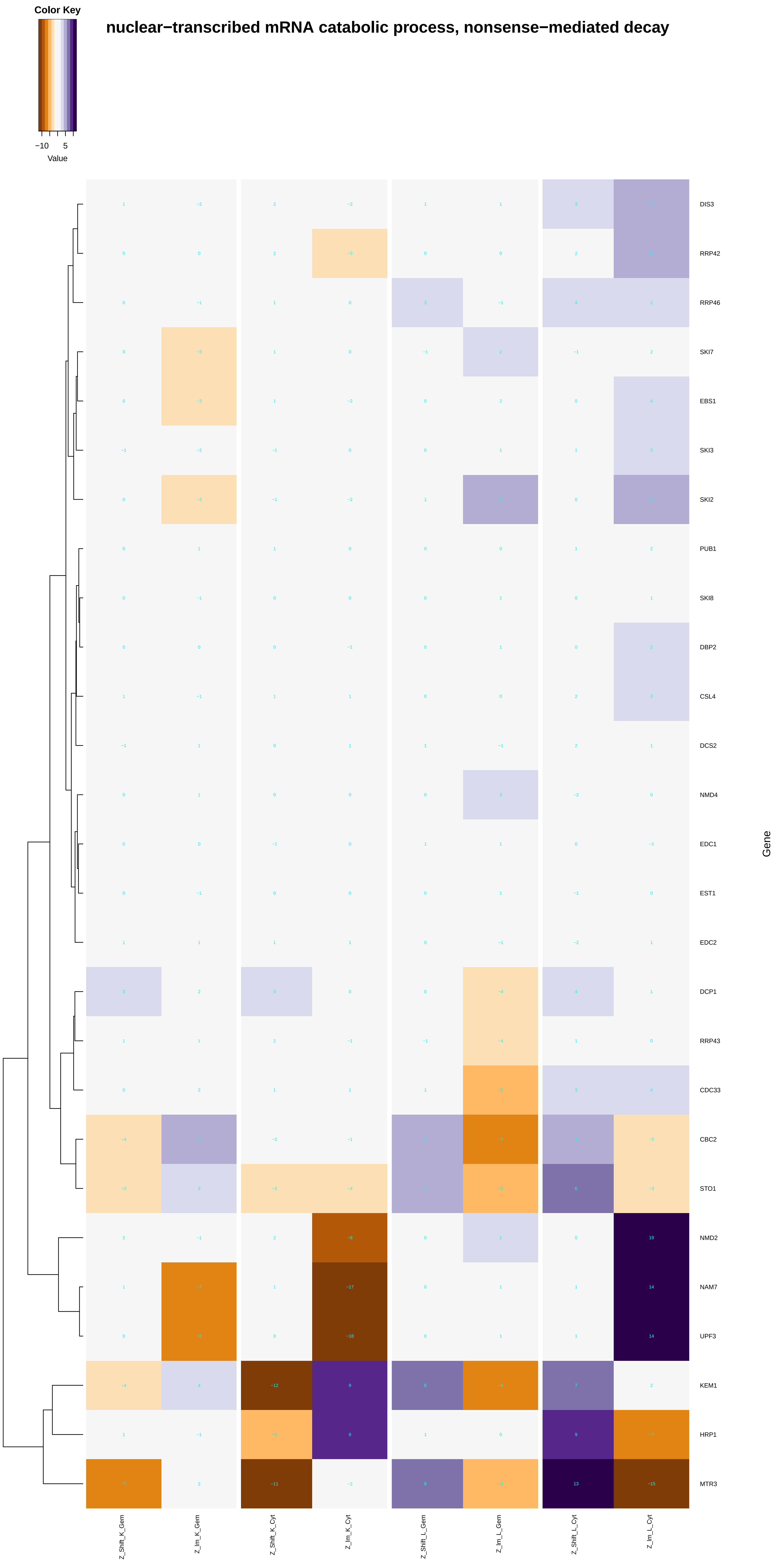
Color Key
−10
5
Value
nuclear−transcribed mRNA catabolic process, nonsense−mediated decay
DIS3
RRP42
RRP46
SKI7
EBS1
SKI3
SKI2
PUB1
SKI8
DBP2
CSL4
DCS2
NMD4
EDC1
EST1
EDC2
DCP1
RRP43
CDC33
CBC2
STO1
NMD2
NAM7
UPF3
KEM1
HRP1
MTR3
Gene
Z_Shift_K_Gem
Z_Im_K_Gem
Z_Shift_K_Cyt
Z_Im_K_Cyt
Z_Shift_L_Gem
Z_Im_L_Gem
Z_Shift_L_Cyt
Z_Im_L_Cyt

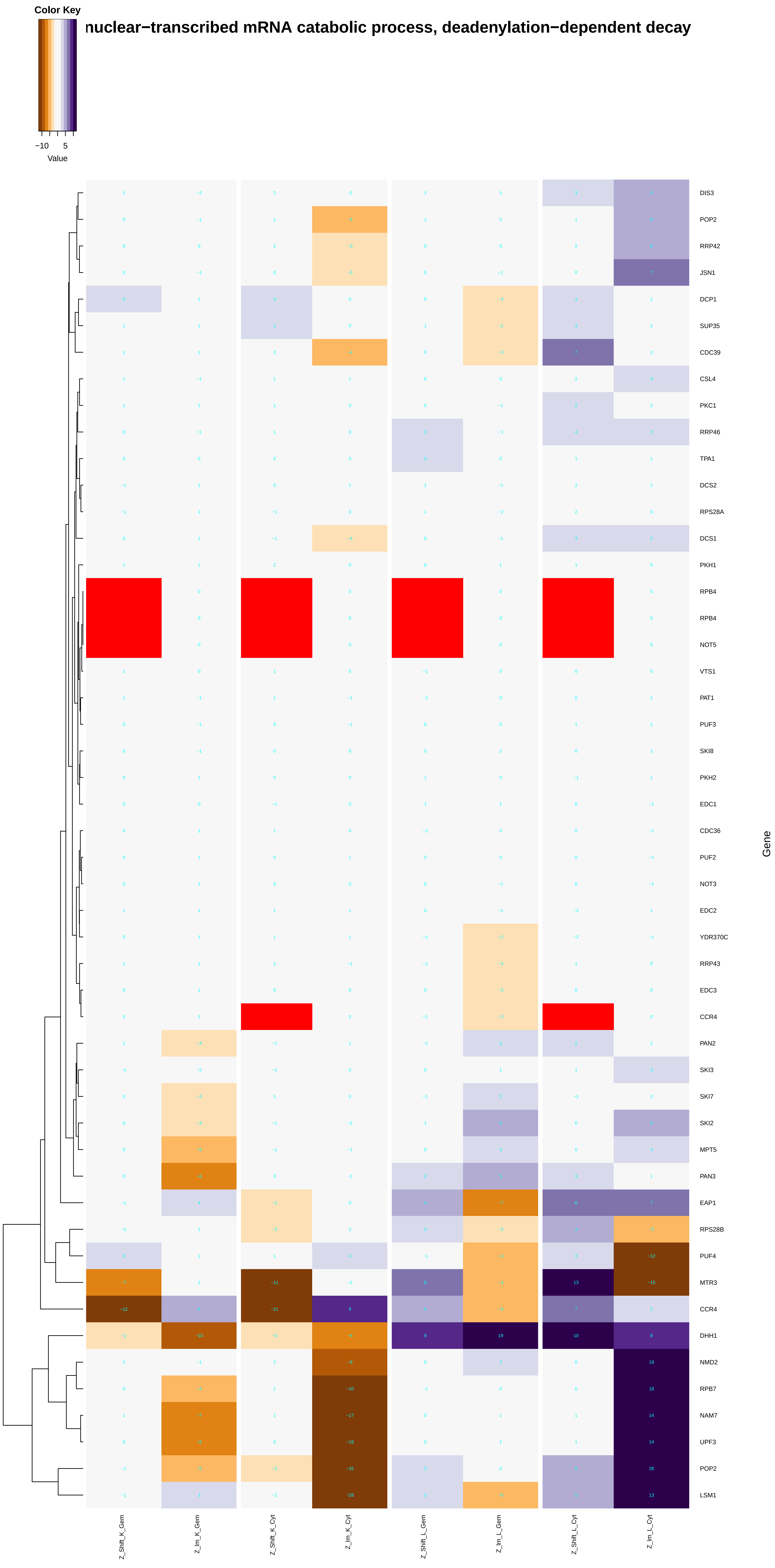

Color Key
−10
5
Value
nuclear−transcribed mRNA catabolic process, deadenylation−dependent decay
DIS3
POP2
RRP42
JSN1
DCP1
SUP35
CDC39
CSL4
PKC1
RRP46
TPA1
DCS2
RPS28A
DCS1
PKH1
RPB4
RPB4
NOT5
VTS1
PAT1
PUF3
SKI8
PKH2
EDC1
CDC36
PUF2
NOT3
EDC2
YDR370C
RRP43
EDC3
CCR4
PAN2
SKI3
SKI7
SKI2
MPT5
PAN3
EAP1
RPS28B
PUF4
MTR3
CCR4
DHH1
NMD2
RPB7
NAM7
UPF3
POP2
LSM1
Gene
Z_Shift_K_Gem
Z_Im_K_Gem
Z_Shift_K_Cyt
Z_Im_K_Cyt
Z_Shift_L_Gem
Z_Im_L_Gem
Z_Shift_L_Cyt
Z_Im_L_Cyt

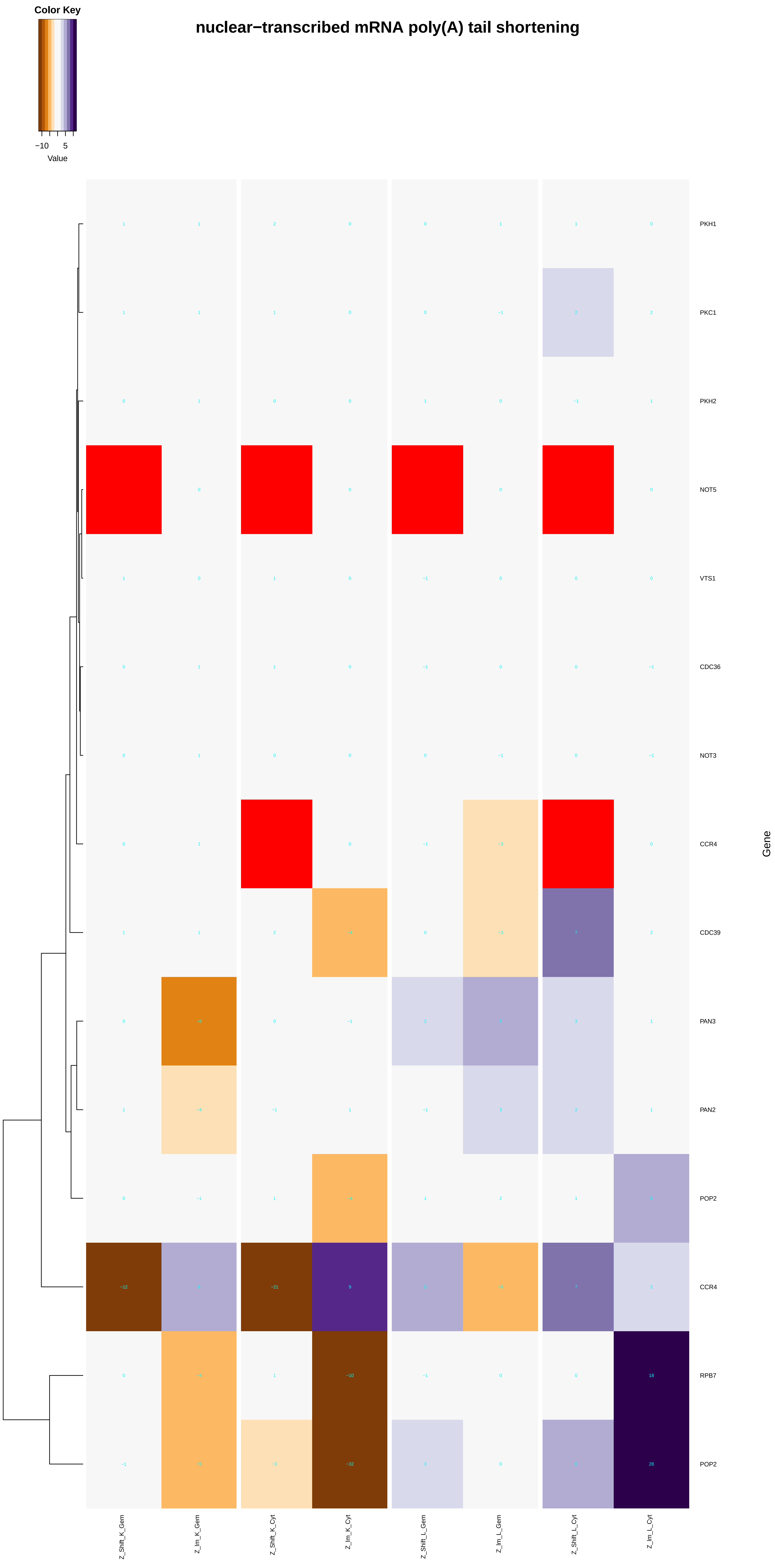
nuclear−transcribed mRNA poly(A) tail shortening
Color Key
−10
5
Value
PKH1
PKC1
PKH2
NOT5
VTS1
CDC36
NOT3
CCR4
CDC39
PAN3
PAN2
POP2
CCR4
RPB7
POP2
Gene
Z_Shift_K_Gem
Z_Im_K_Gem
Z_Shift_K_Cyt
Z_Im_K_Cyt
Z_Shift_L_Gem
Z_Im_L_Gem
Z_Shift_L_Cyt
Z_Im_L_Cyt

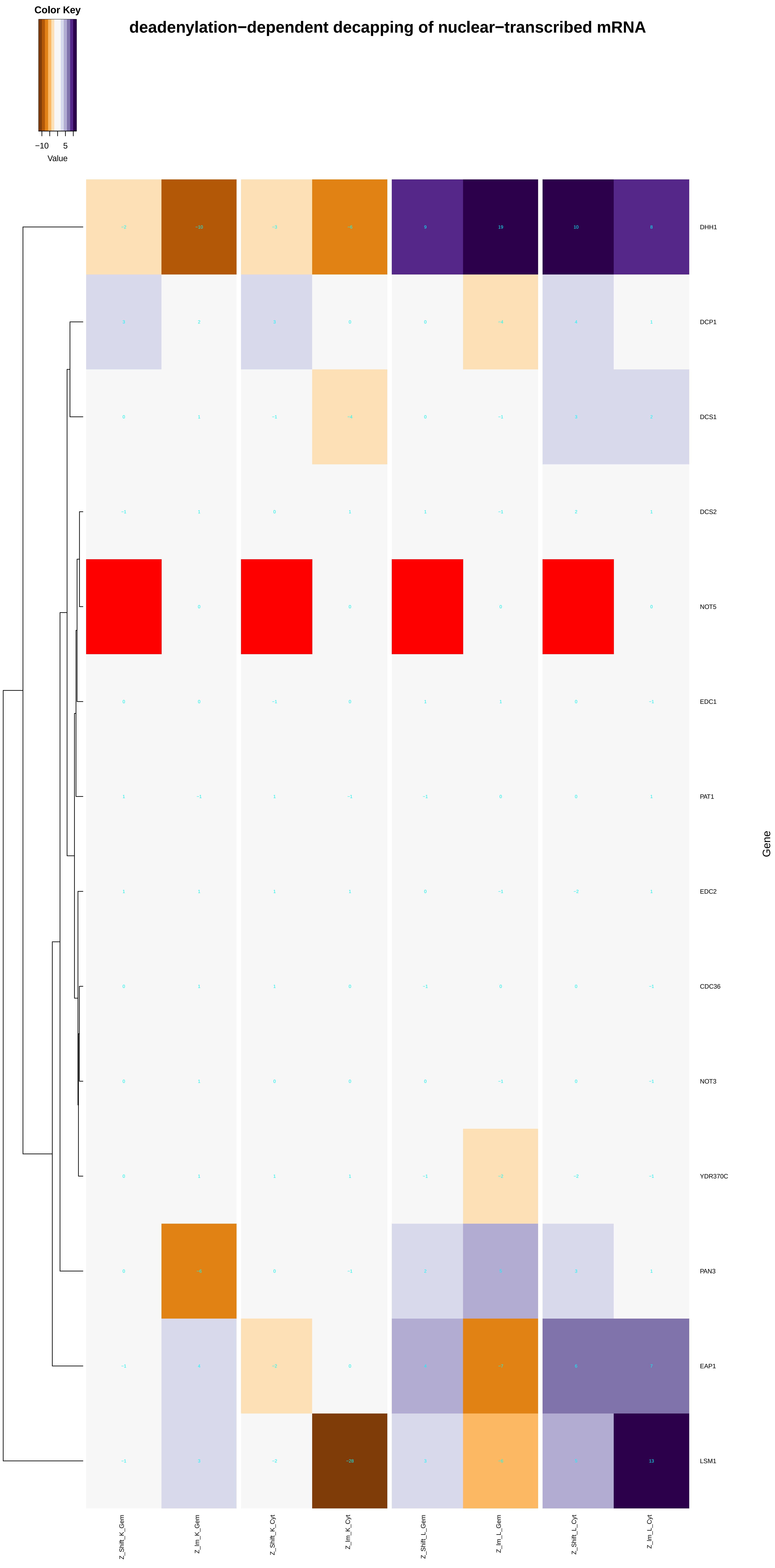
Color Key
−10 5
Value
deadenylation−dependent decapping of nuclear−transcribed mRNA
DHH1
DCP1
DCS1
DCS2
NOT5
EDC1
PAT1
EDC2
CDC36
NOT3
YDR370C
PAN3
EAP1
LSM1
Gene
Z_Shift_K_Gem
Z_Im_K_Gem
Z_Shift_K_Cyt
Z_Im_K_Cyt
Z_Shift_L_Gem
Z_Im_L_Gem
Z_Shift_L_Cyt
Z_Im_L_Cyt

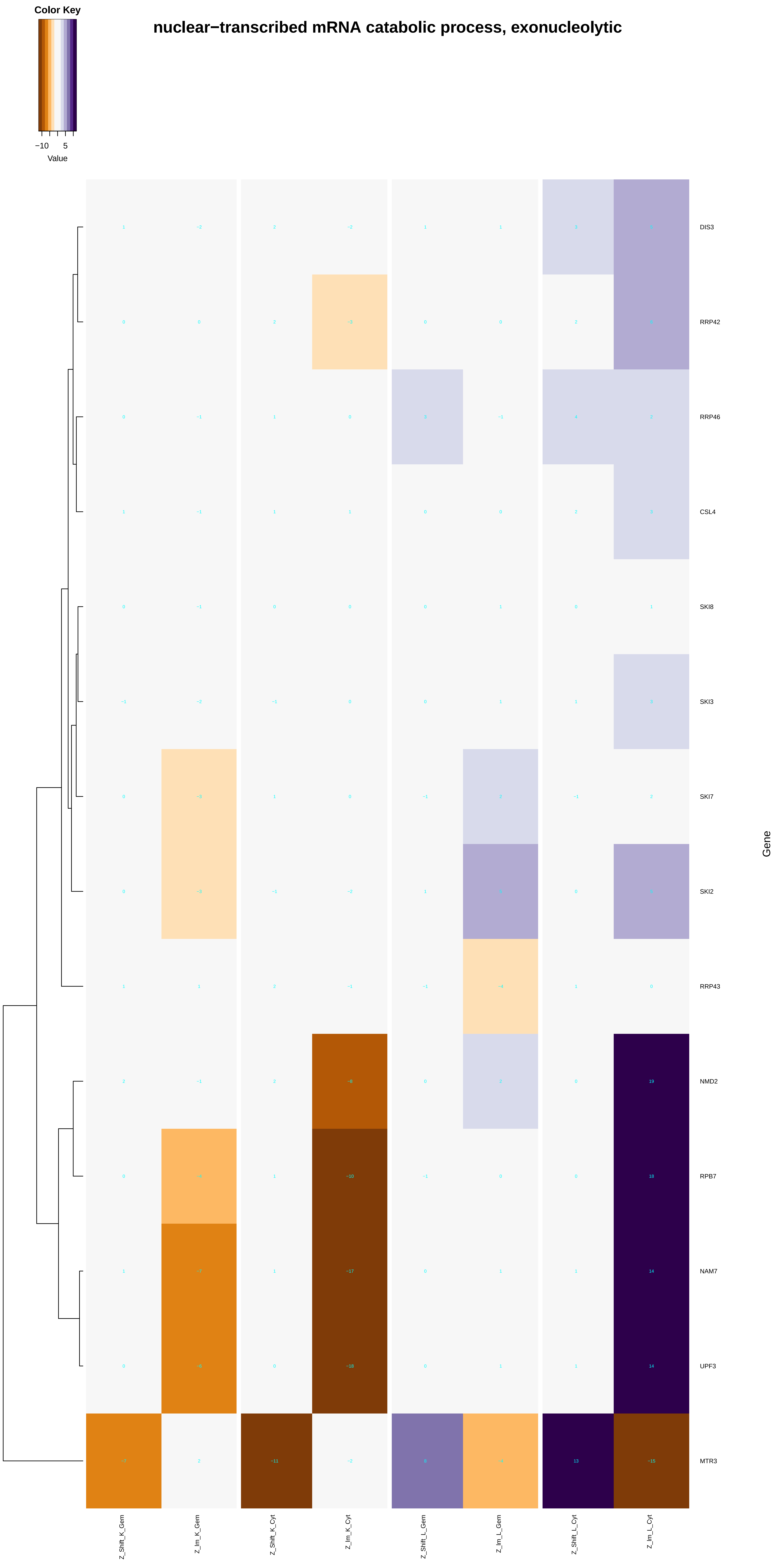

Color Key
−10
5
Value
nuclear−transcribed mRNA catabolic process, exonucleolytic
DIS3
RRP42
RRP46
CSL4
SKI8
SKI3
SKI7
SKI2
RRP43
NMD2
RPB7
NAM7
UPF3
MTR3
Gene
Z_Shift_K_Gem
Z_lm_K_Gem
Z_Shift_K_Cyt
Z_lm_K_Cyt
Z_Shift_L_Gem
Z_lm_L_Gem
Z_Shift_L_Cyt
Z_lm_L_Cyt

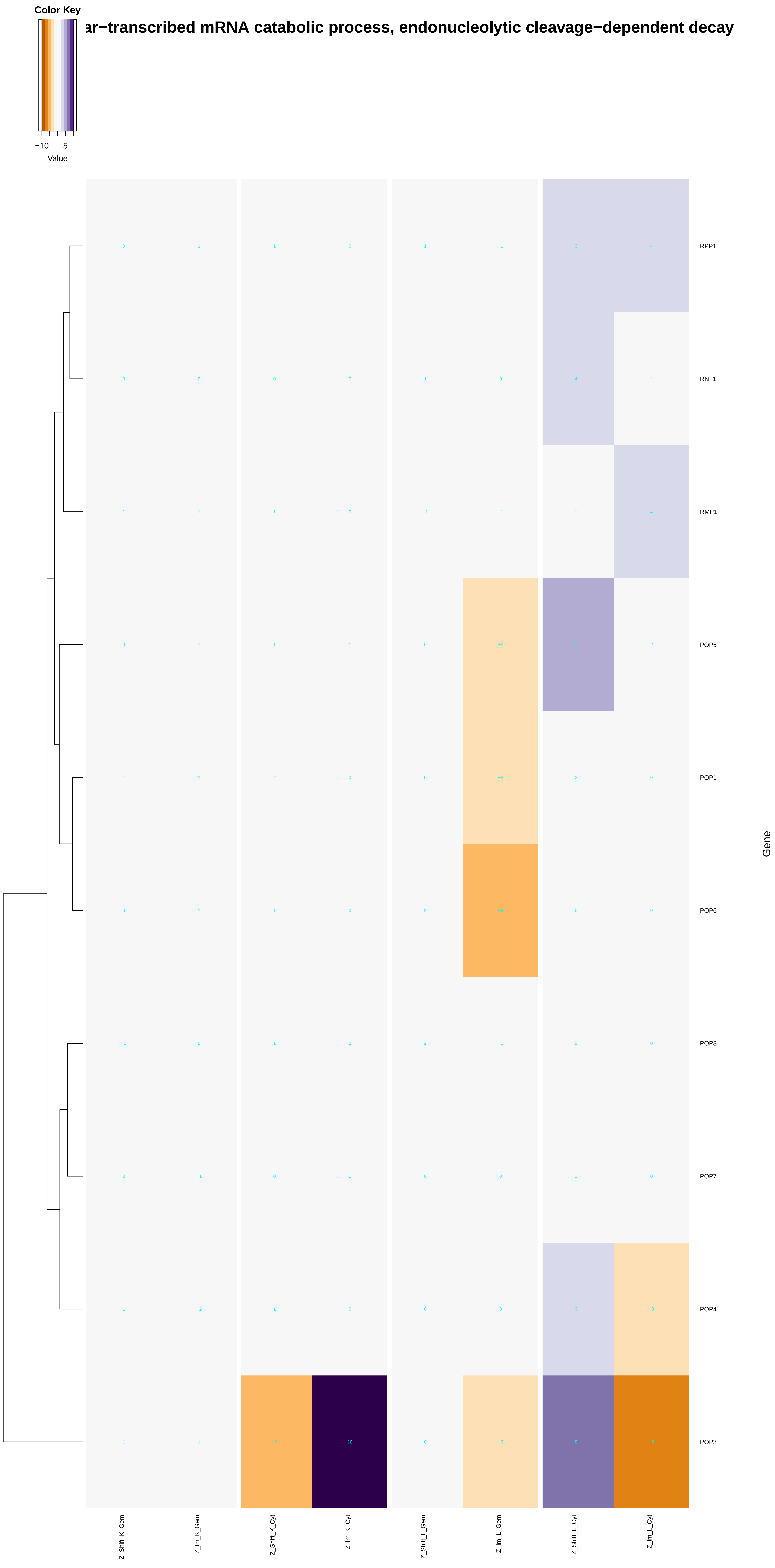

Color Key
−10
5
Value
ar−transcribed mRNA catabolic process, endonucleolytic cleavage−dependent decay
RPP1
RNT1
RMP1
POP5
POP1
POP6
POP8
POP7
POP4
POP3
Gene
Z_Shift_K_Gem
Z_Im_K_Gem
Z_Shift_K_Cyt
Z_Im_K_Cyt
Z_Shift_L_Gem
Z_Im_L_Gem
Z_Shift_L_Cyt
Z_Im_L_Cyt

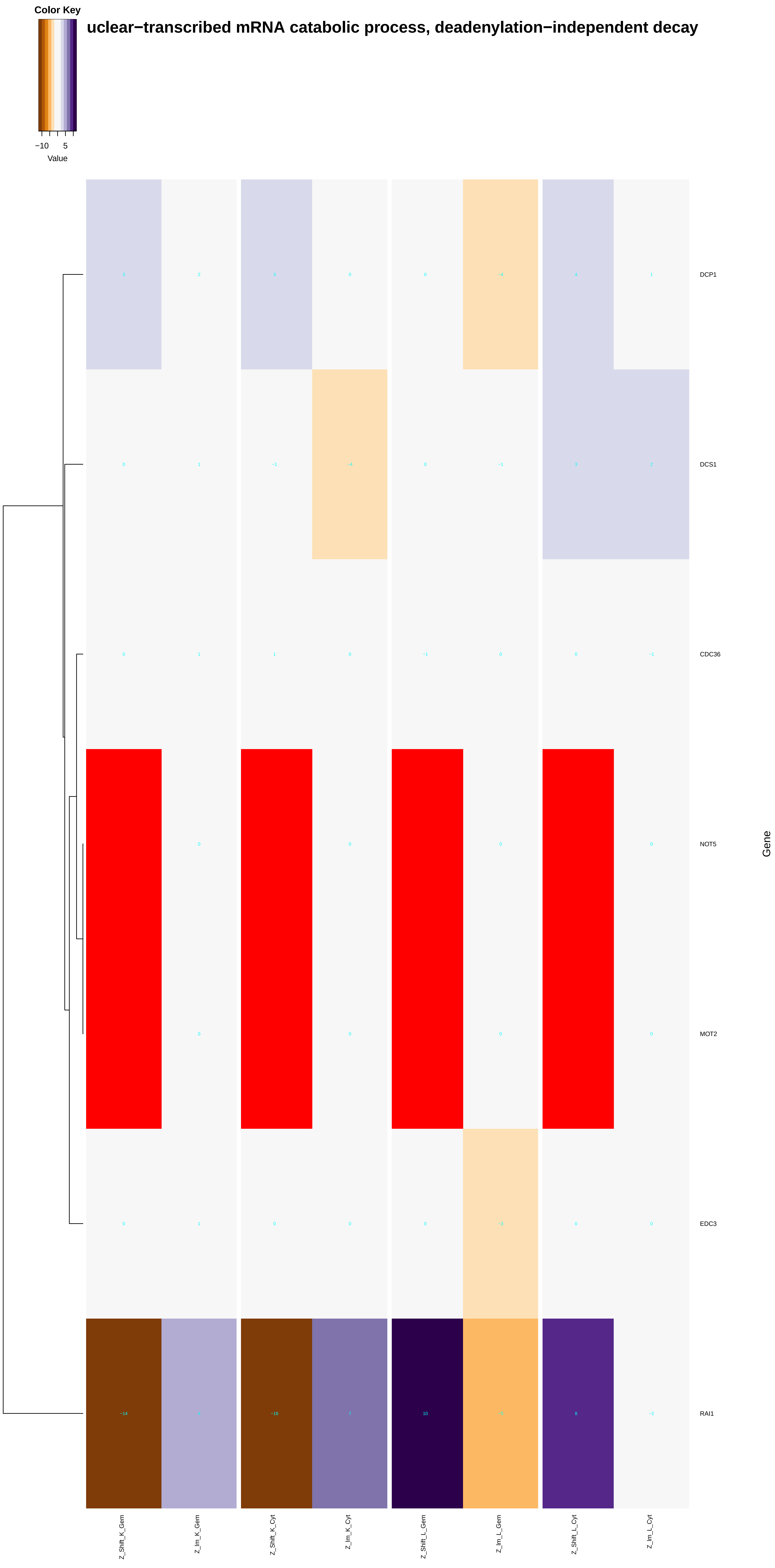
Color Key
−10
5
Value
uclear−transcribed mRNA catabolic process, deadenylation−independent decay
DCP1
DCS1
CDC36
NOT5
MOT2
EDC3
RAI1
Gene
Z_Shift_K_Gem
Z_lm_K_Gem
Z_Shift_K_Cyt
Z_lm_K_Cyt
Z_Shift_L_Gem
Z_lm_L_Gem
Z_Shift_L_Cyt
Z_lm_L_Cyt

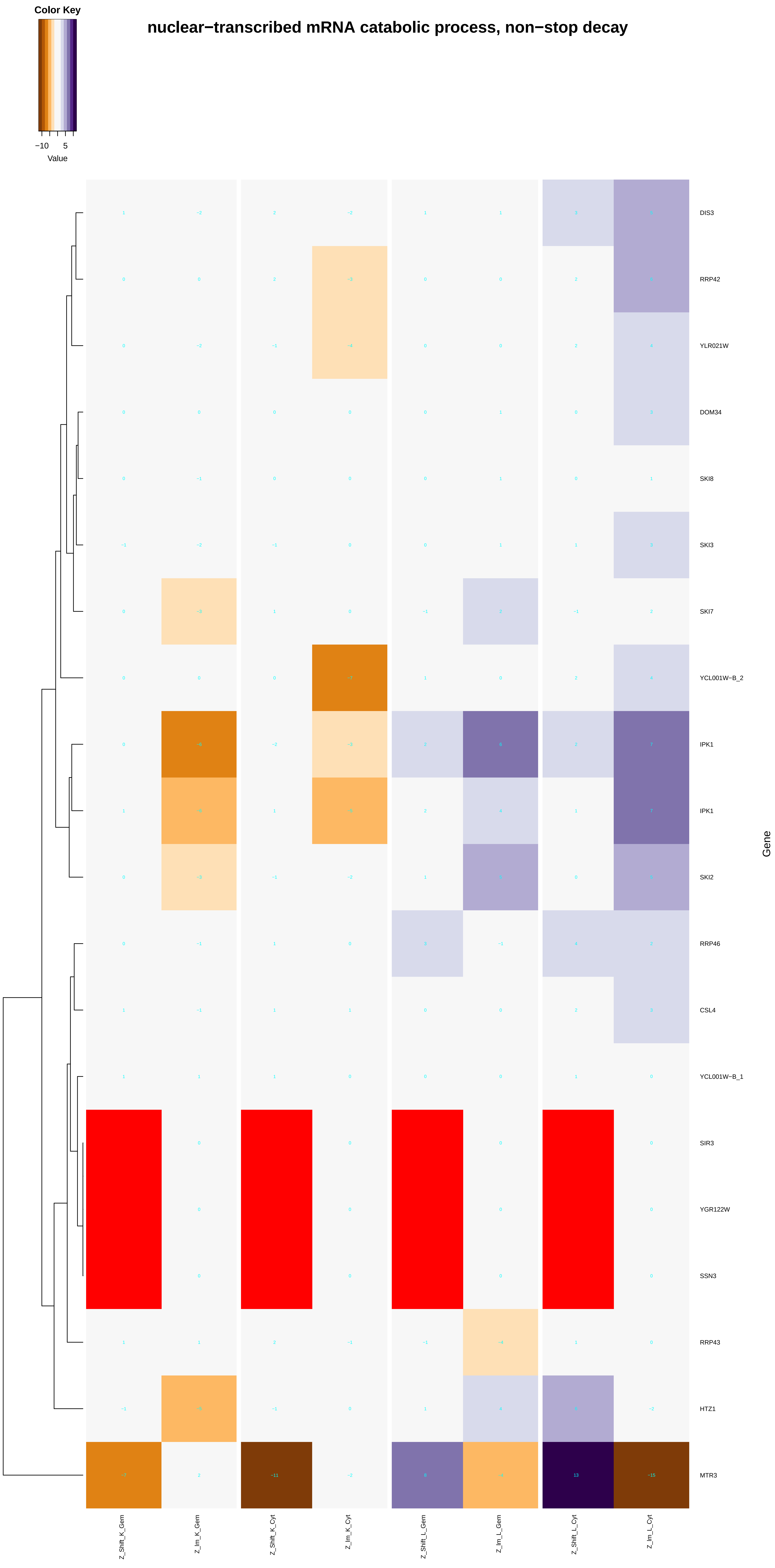

Color Key
−10
5
Value
nuclear−transcribed mRNA catabolic process, non−stop decay
DIS3
RRP42
YLR021W
DOM34
SKI8
SKI3
SKI7
YCL001W−B_2
IPK1
IPK1
SKI2
RRP46
CSL4
YCL001W−B_1
SIR3
YGR122W
SSN3
RRP43
HTZ1
MTR3
Gene
Z_Shift_K_Gem
Z_Im_K_Gem
Z_Shift_K_Cyt
Z_Im_K_Cyt
Z_Shift_L_Gem
Z_Im_L_Gem
Z_Shift_L_Cyt
Z_Im_L_Cyt

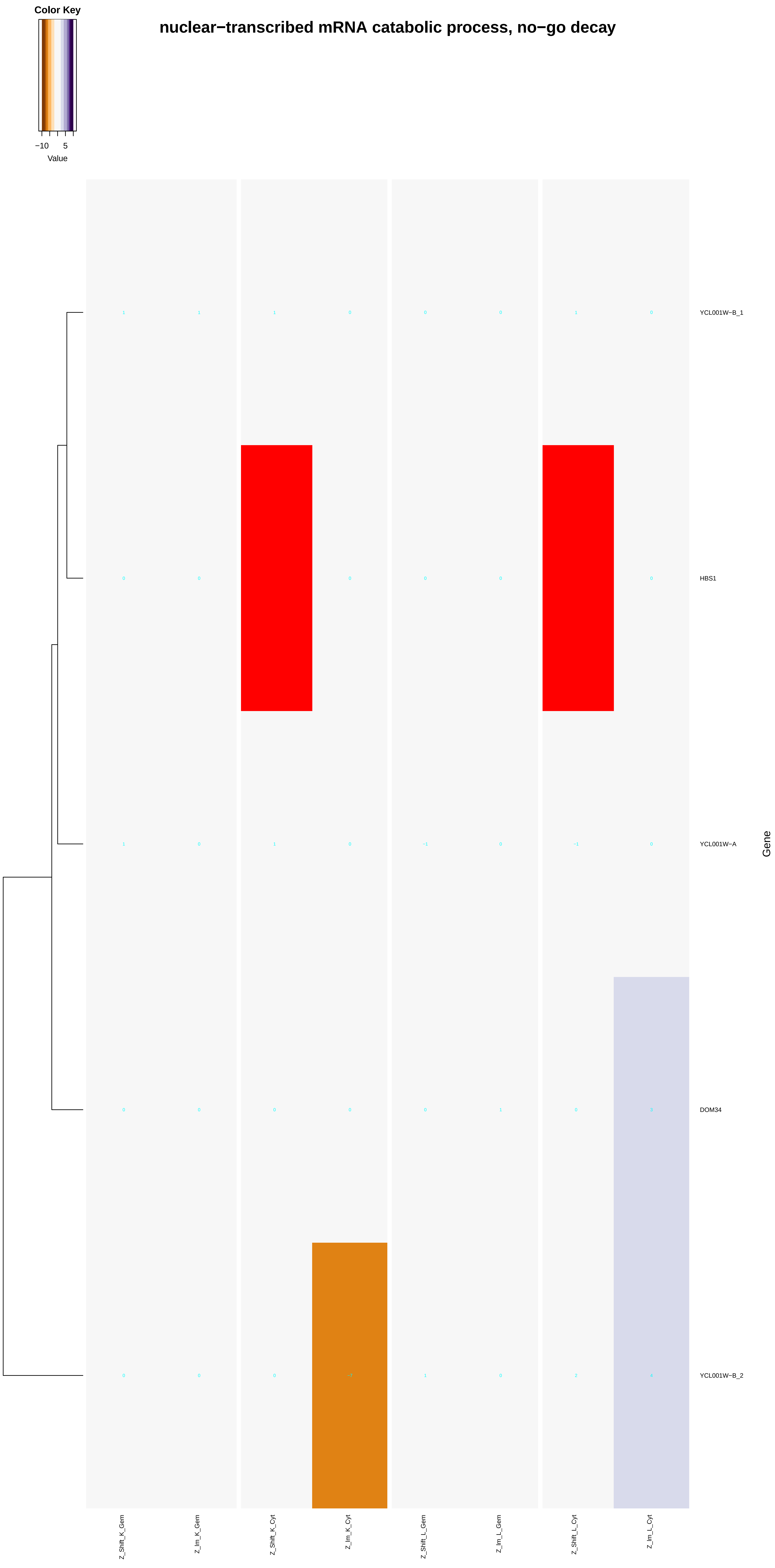

Color Key
−10
5
Value
nuclear−transcribed mRNA catabolic process, no−go decay
YCL001W−B_1
HBS1
YCL001W−A
DOM34
YCL001W−B_2
Gene
Z_Shift_K_Gem
Z_Im_K_Gem
Z_Shift_K_Cyt
Z_Im_K_Cyt
Z_Shift_L_Gem
Z_Im_L_Gem
Z_Shift_L_Cyt
Z_Im_L_Cyt

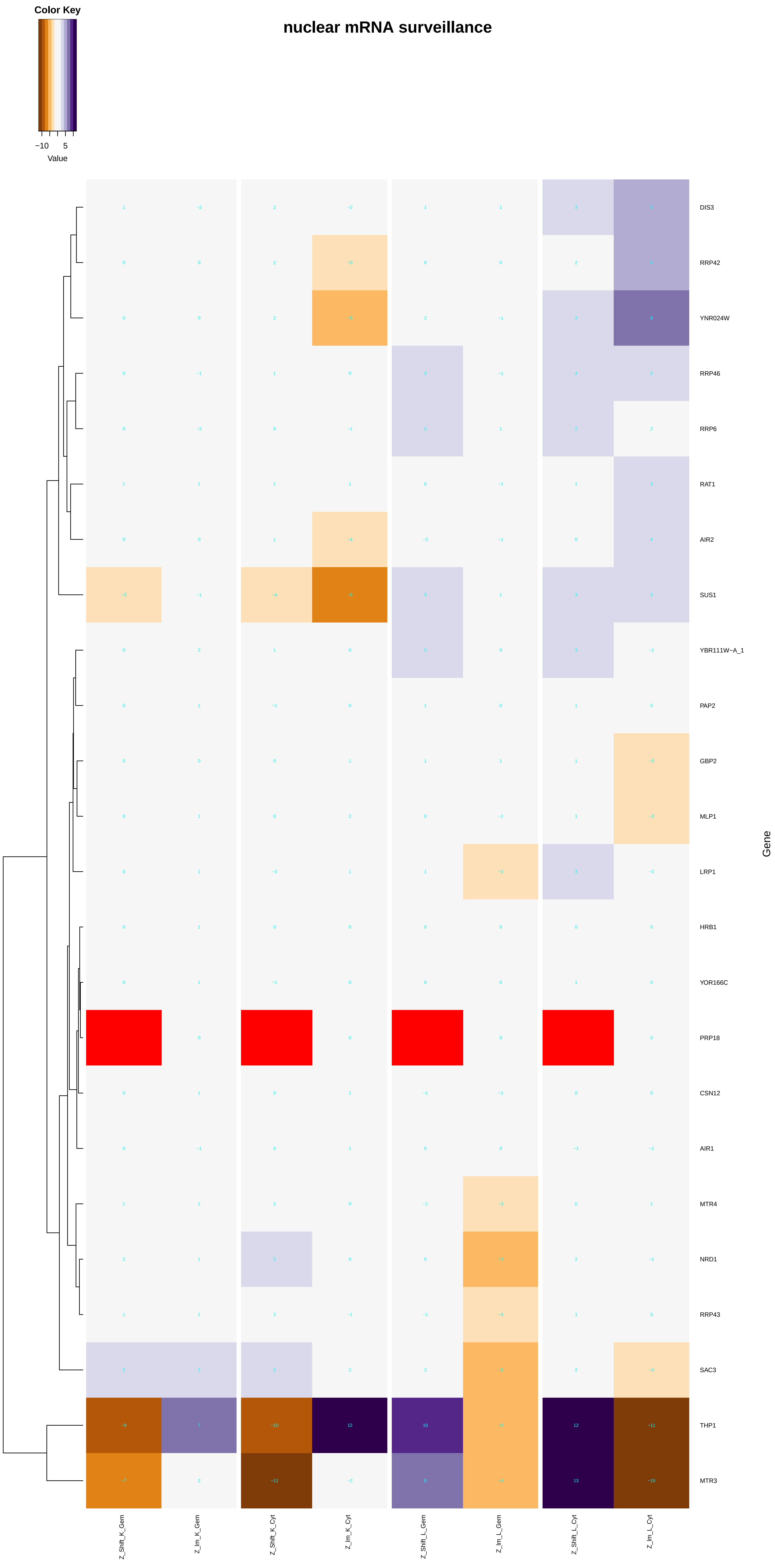
Color Key
−10
5
Value
nuclear mRNA surveillance
DIS3
RRP42
YNR024W
RRP46
RRP6
RAT1
AIR2
SUS1
YBR111W−A_1
PAP2
GBP2
MLP1
LRP1
HRB1
YOR166C
PRP18
CSN12
AIR1
MTR4
NRD1
RRP43
SAC3
THP1
MTR3
Gene
Z_Shift_K_Gem
Z_Im_K_Gem
Z_Shift_K_Cyt
Z_Im_K_Cyt
Z_Shift_L_Gem
Z_Im_L_Gem
Z_Shift_L_Cyt
Z_Im_L_Cyt

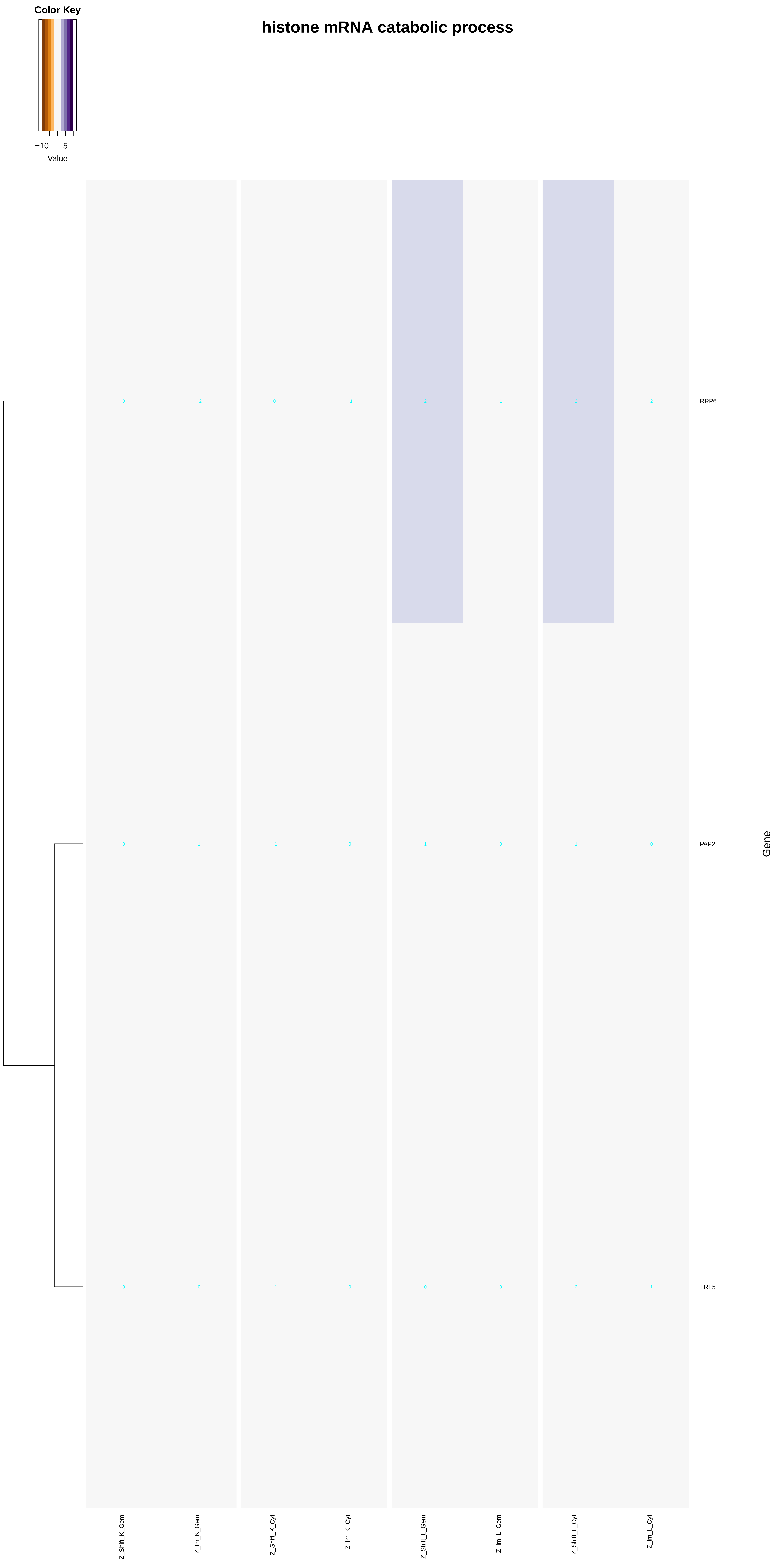

Color Key
−10
5
Value
histone mRNA catabolic process
RRP6
PAP2
TRF5
Gene
Z_Shift_K_Gem
Z_Im_K_Gem
Z_Shift_K_Cyt
Z_Im_K_Cyt
Z_Shift_L_Gem
Z_Im_L_Gem
Z_Shift_L_Cyt
Z_Im_L_Cyt

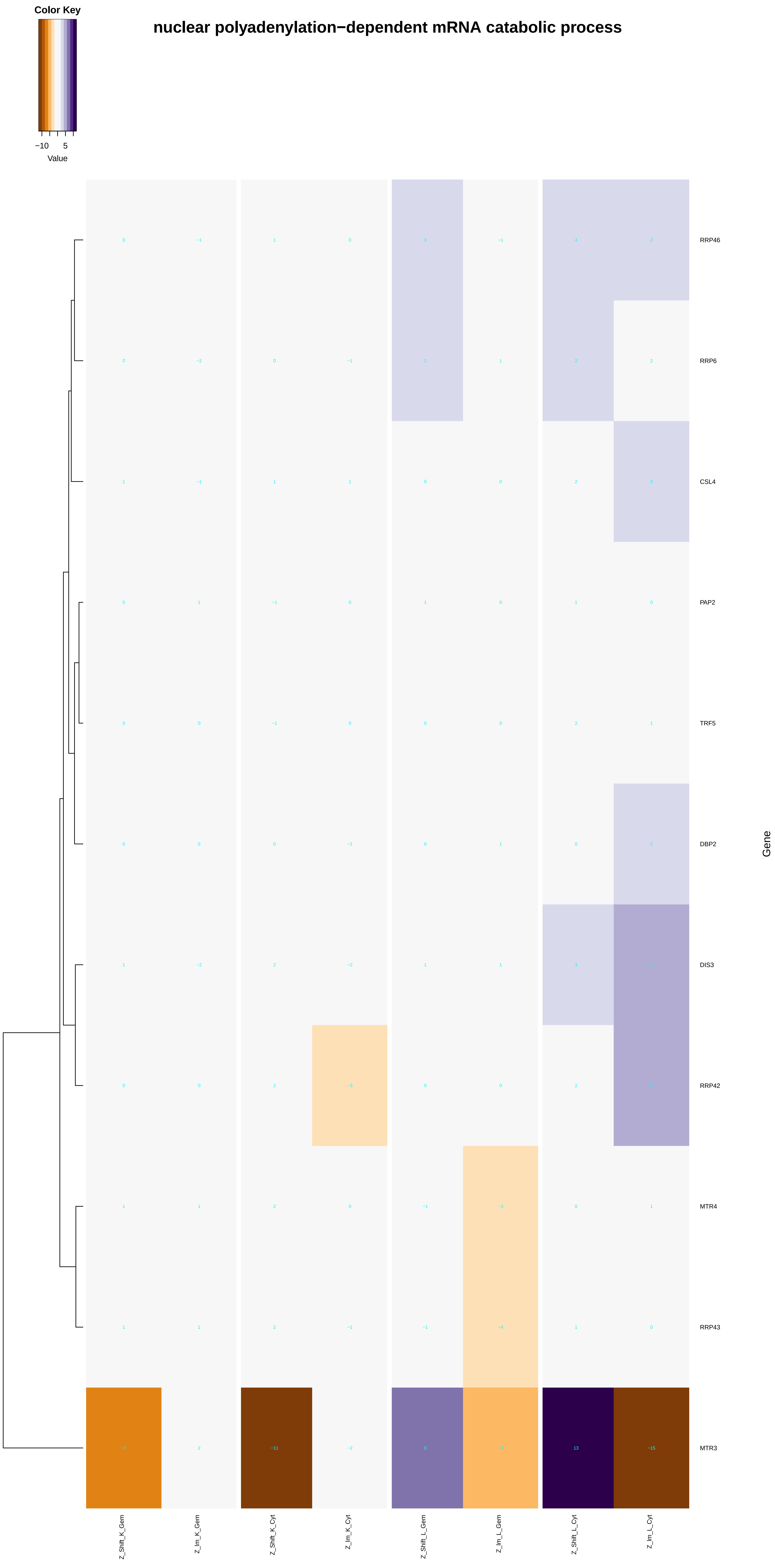
Color Key
−10
5
Value
nuclear polyadenylation−dependent mRNA catabolic process
RRP46
RRP6
CSL4
PAP2
TRF5
DBP2
DIS3
RRP42
MTR4
RRP43
MTR3
Gene
Z_Shift_K_Gem
Z_Im_K_Gem
Z_Shift_K_Cyt
Z_Im_K_Cyt
Z_Shift_L_Gem
Z_Im_L_Gem
Z_Shift_L_Cyt
Z_Im_L_Cyt

### nonfunctional rRNA decay

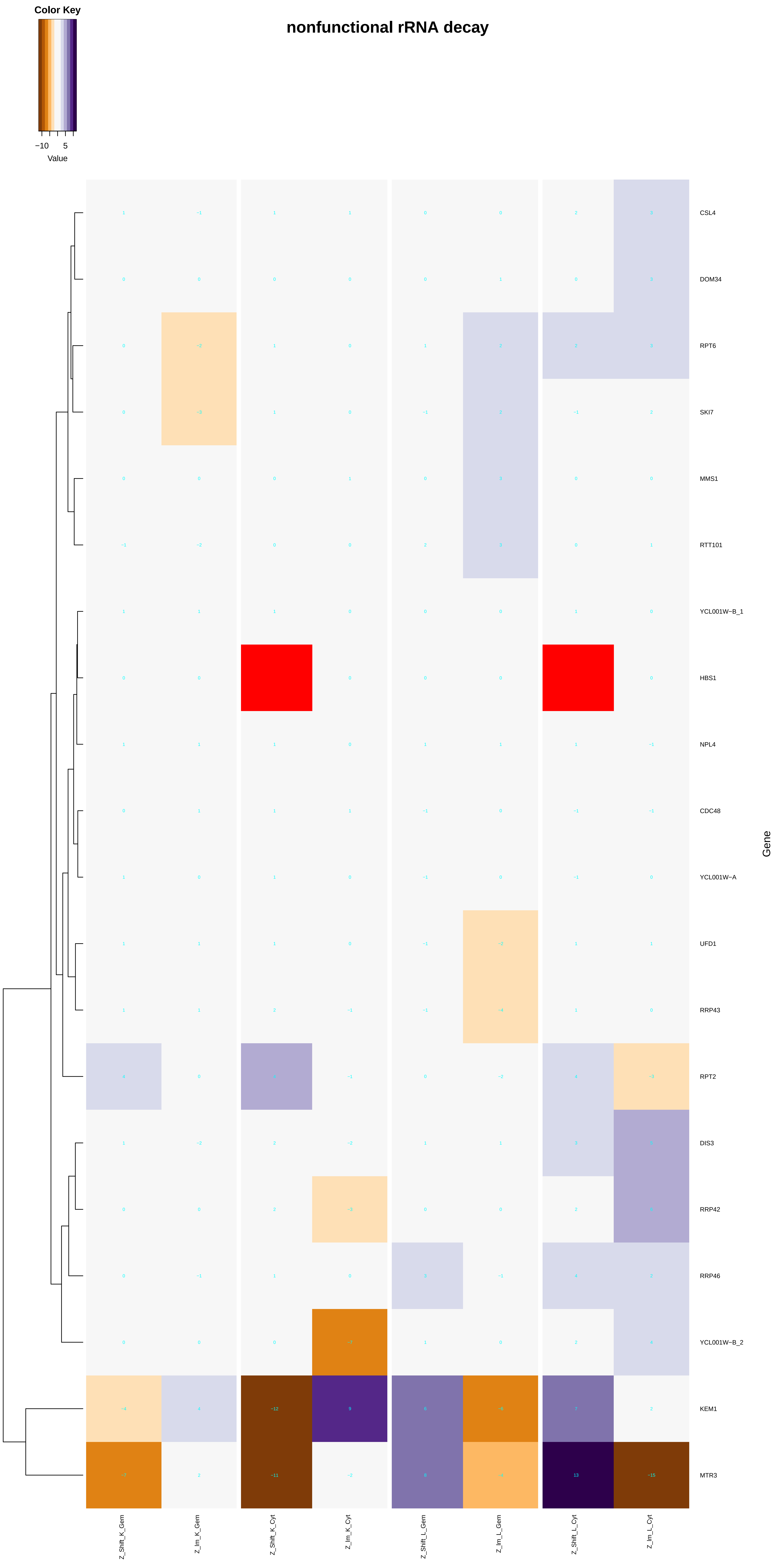

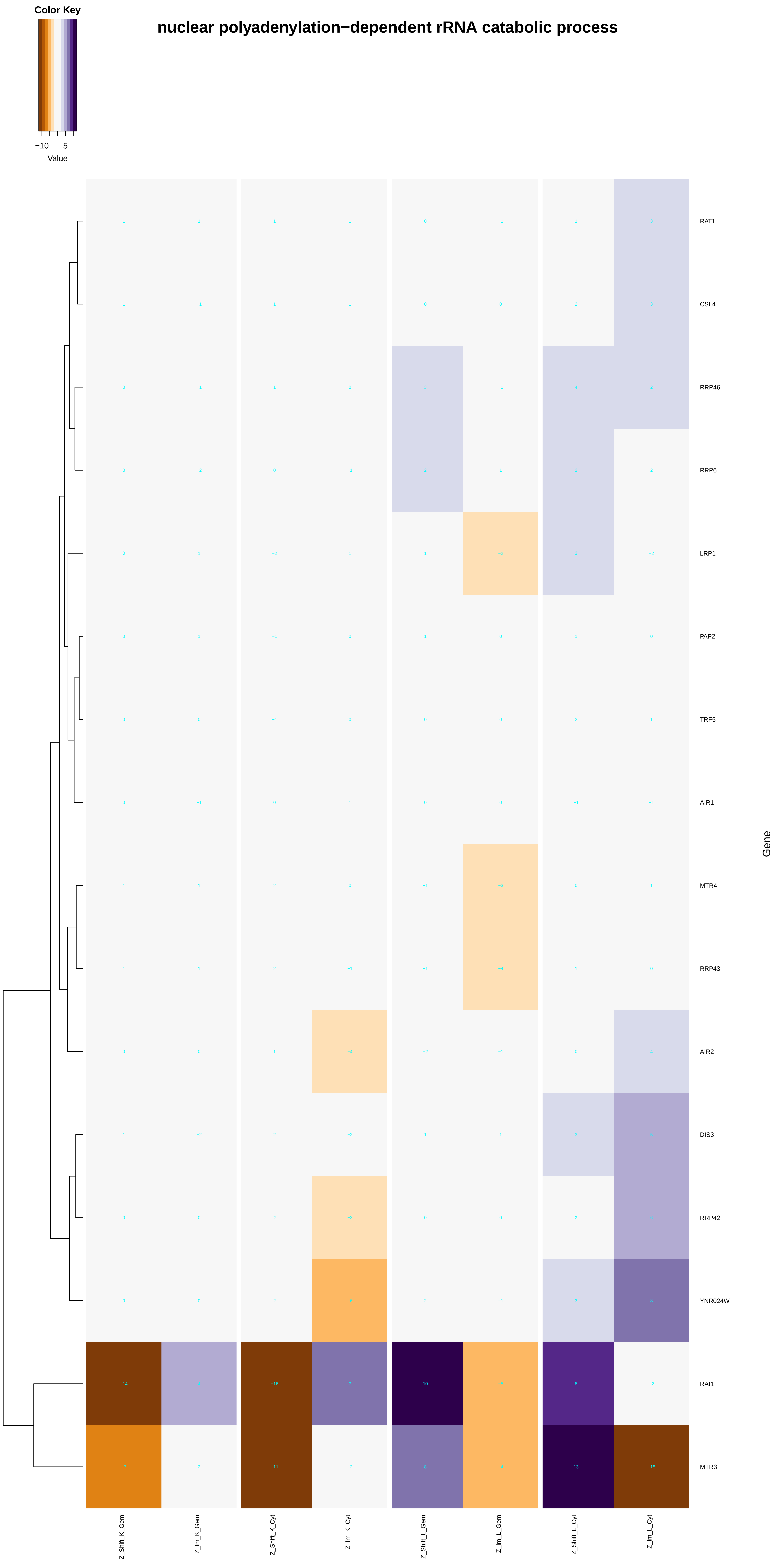
Color Key
−10
5
Value
nuclear polyadenylation−dependent rRNA catabolic process
RAT1
CSL4
RRP46
RRP6
LRP1
PAP2
TRF5
AIR1
MTR4
RRP43
AIR2
DIS3
RRP42
YNR024W
RAI1
MTR3
Gene
Z_Shift_K_Gem
Z_Im_K_Gem
Z_Shift_K_Cyt
Z_Im_K_Cyt
Z_Shift_L_Gem
Z_Im_L_Gem
Z_Shift_L_Cyt
Z_Im_L_Cyt

### nuclear polyadenylation−dependent snRNA catabolic process

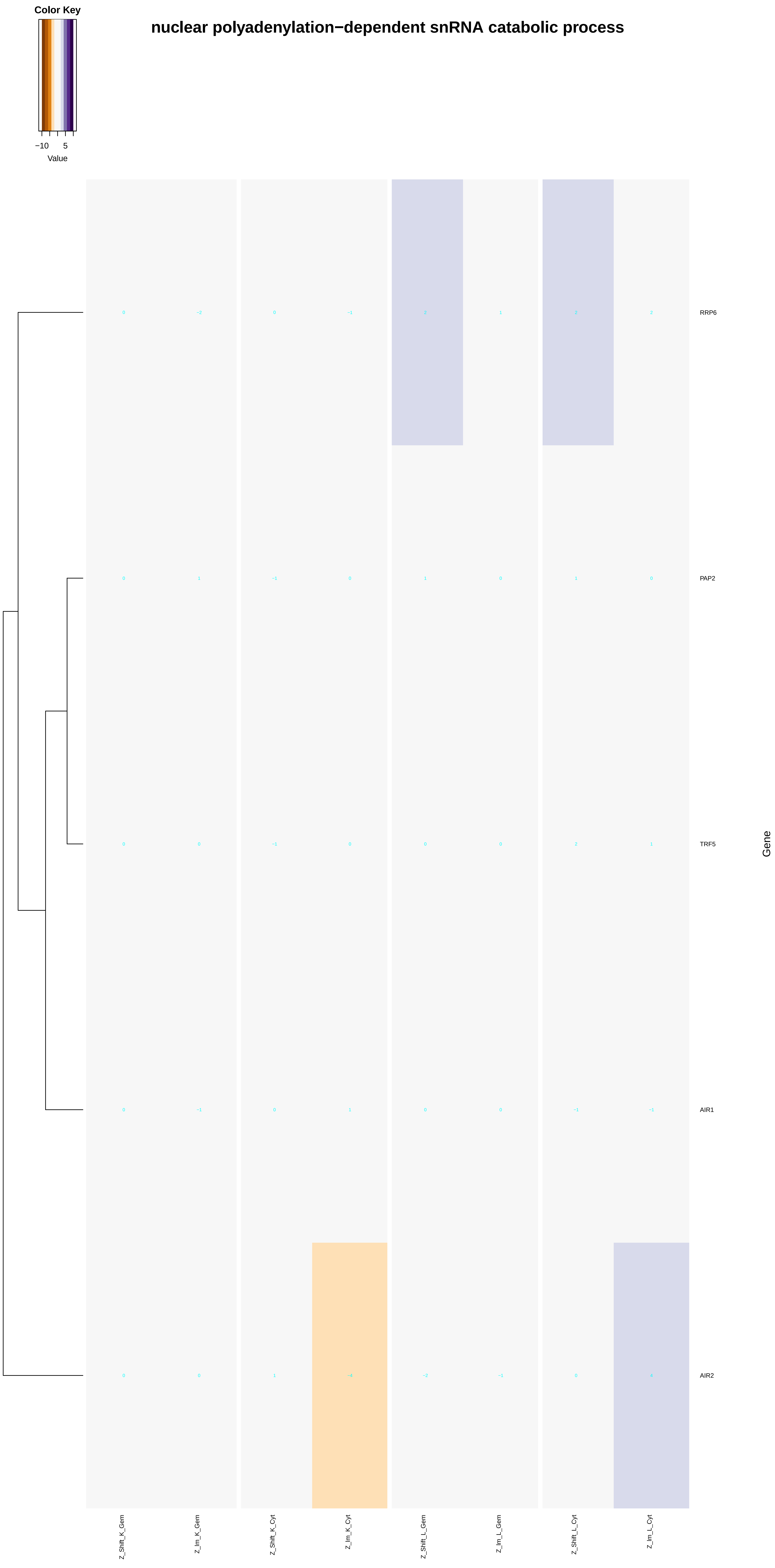

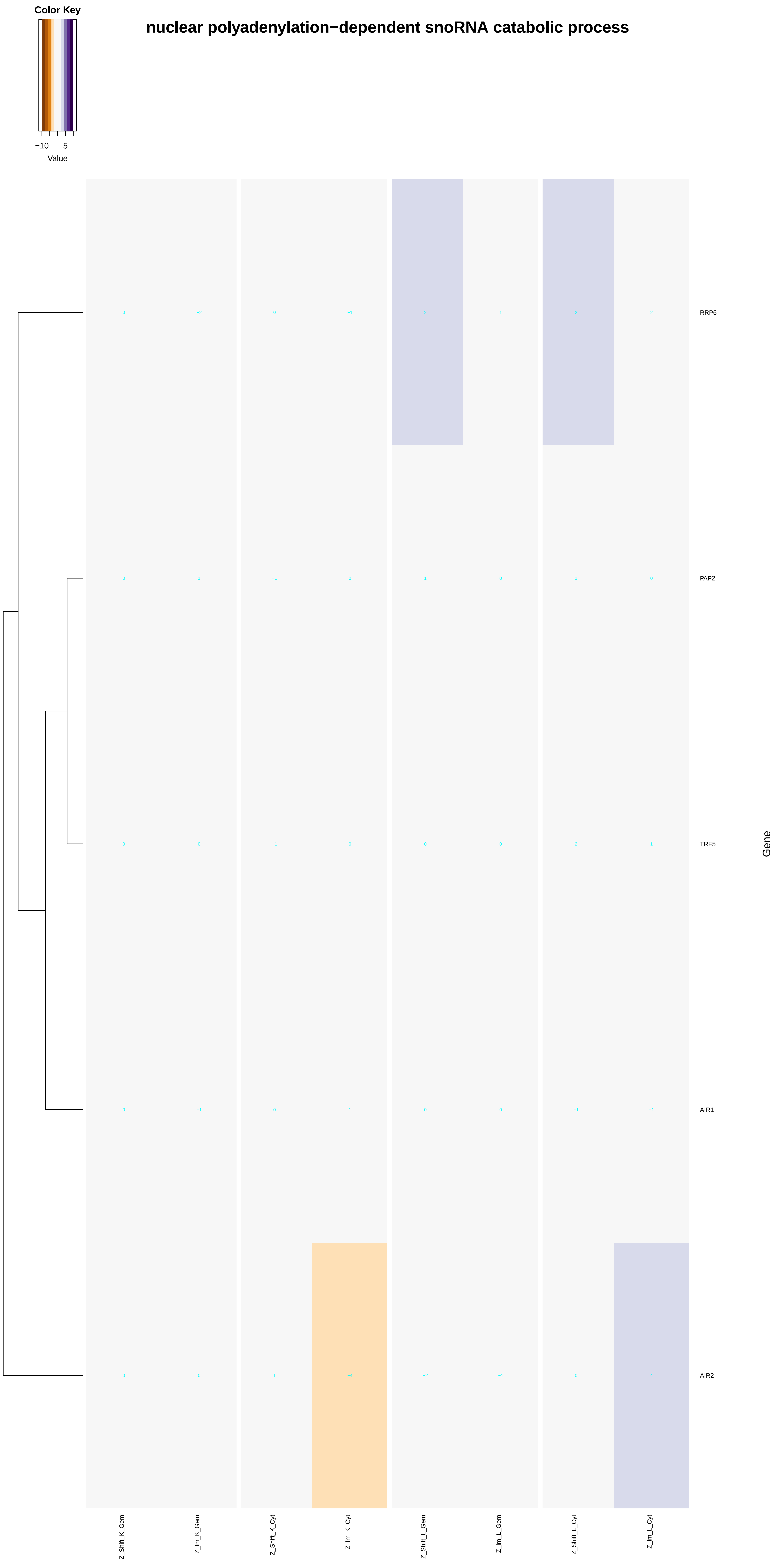

nuclear polyadenylation−dependent snoRNA catabolic process
Color Key
−10
5
Value
RRP6
PAP2
TRF5
AIR1
AIR2
Gene
Z_Shift_K_Gem
Z_Im_K_Gem
Z_Shift_K_Cyt
Z_Im_K_Cyt
Z_Shift_L_Gem
Z_Im_L_Gem
Z_Shift_L_Cyt
Z_Im_L_Cyt

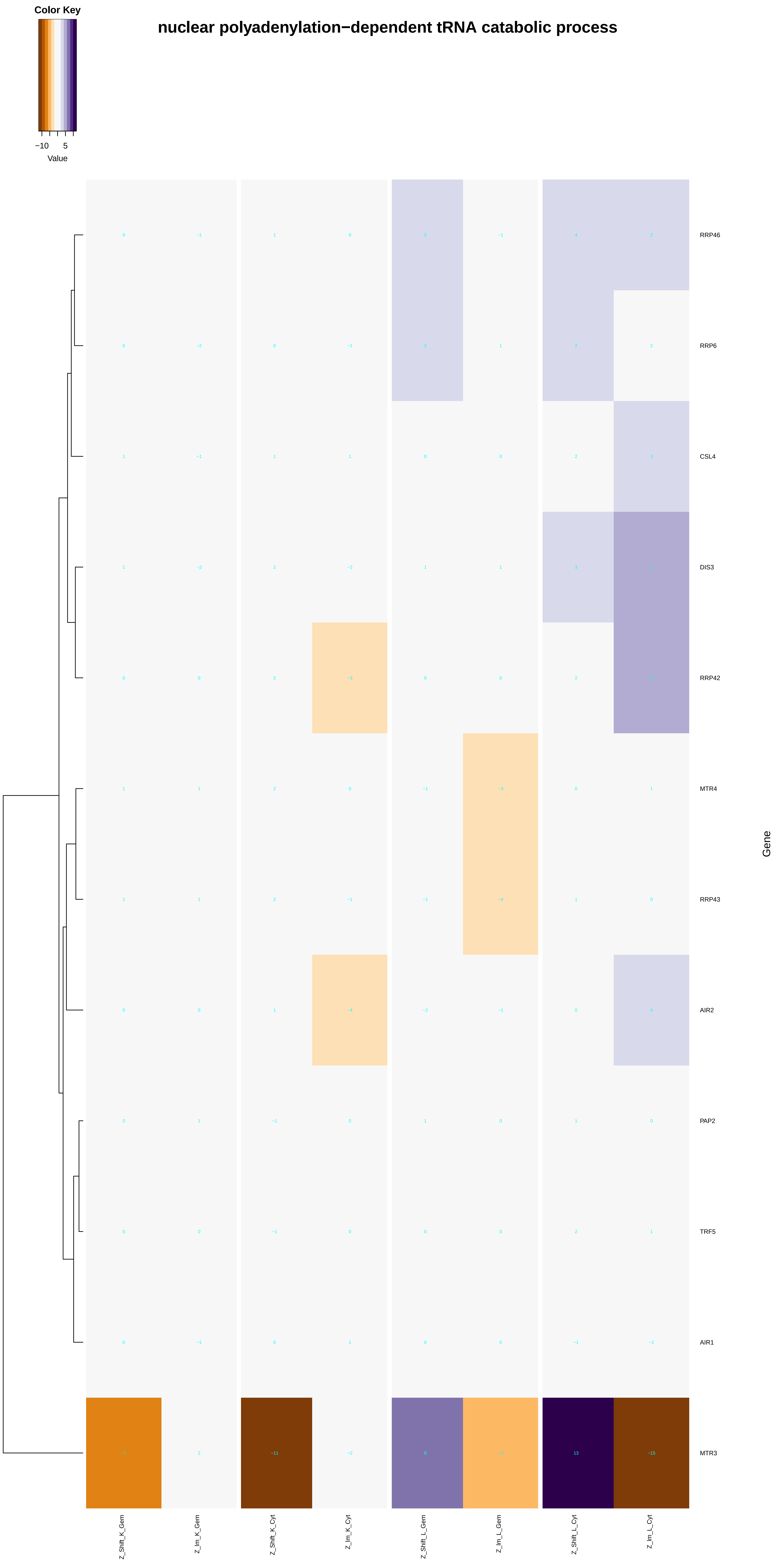

Color Key
−10
5
Value
nuclear polyadenylation−dependent tRNA catabolic process
RRP46
RRP6
CSL4
DIS3
RRP42
MTR4
RRP43
AIR2
PAP2
TRF5
AIR1
MTR3
Gene
Z_Shift_K_Gem
Z_Im_K_Gem
Z_Shift_K_Cyt
Z_Im_K_Cyt
Z_Shift_L_Gem
Z_Im_L_Gem
Z_Shift_L_Cyt
Z_Im_L_Cyt

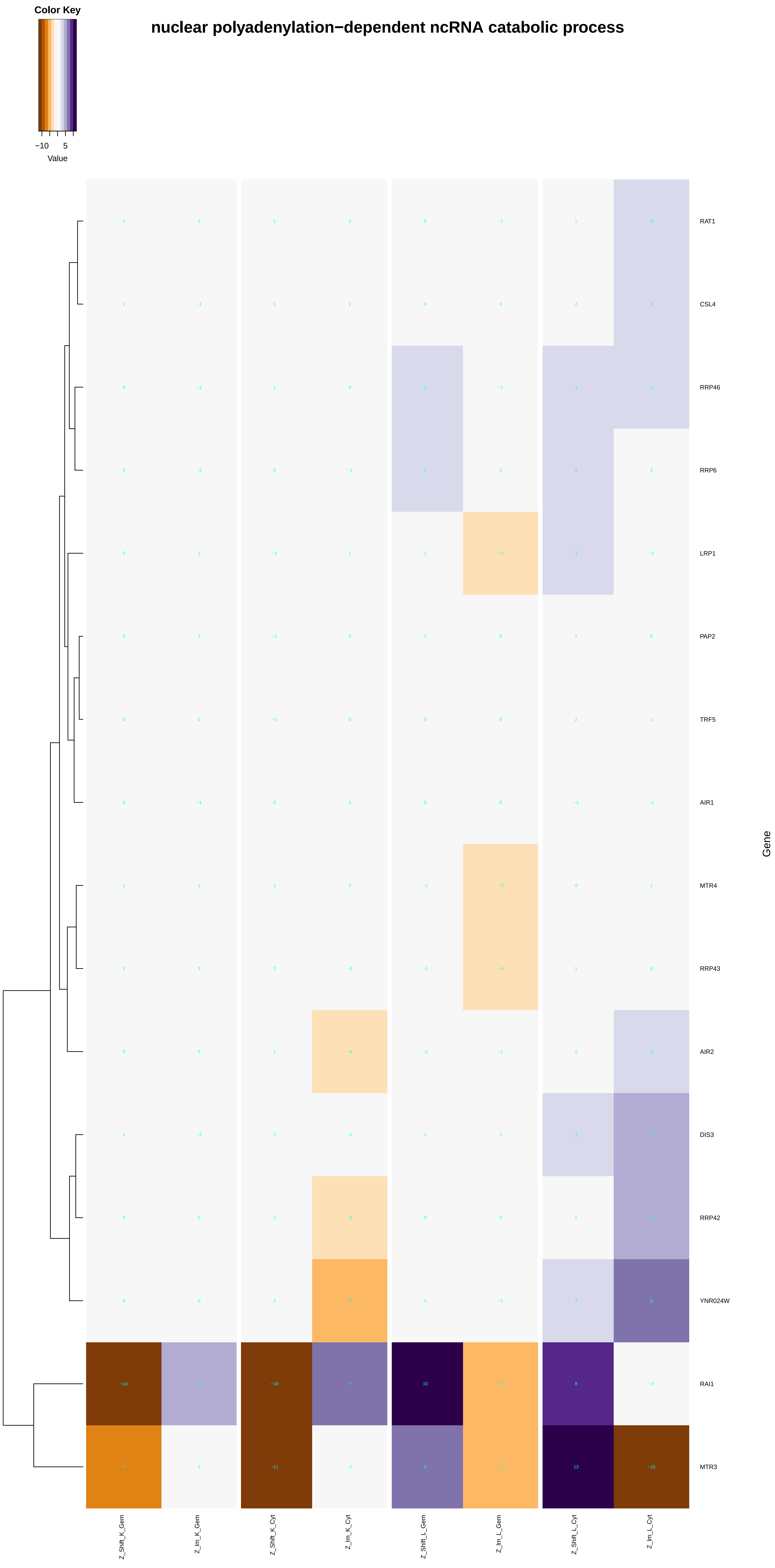
Color Key
−10
5
Value
nuclear polyadenylation−dependent ncRNA catabolic process
RAT1
CSL4
RRP46
RRP6
LRP1
PAP2
TRF5
AIR1
MTR4
RRP43
AIR2
DIS3
RRP42
YNR024W
RAI1
MTR3
Gene
Z_Shift_K_Gem
Z_Im_K_Gem
Z_Shift_K_Cyt
Z_Im_K_Cyt
Z_Shift_L_Gem
Z_Im_L_Gem
Z_Shift_L_Cyt
Z_Im_L_Cyt

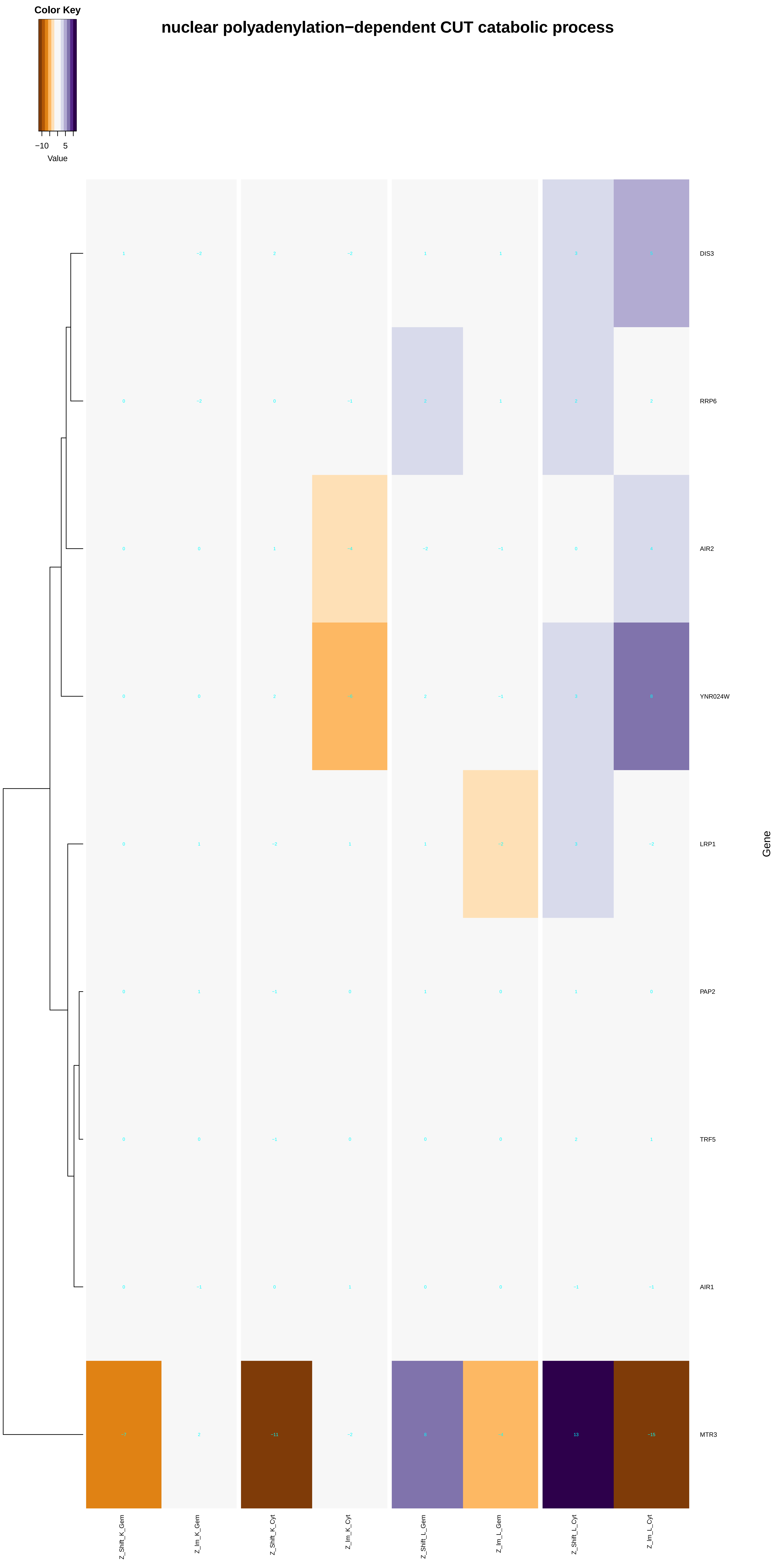
Color Key
−10
5
Value
nuclear polyadenylation−dependent CUT catabolic process
DIS3
RRP6
AIR2
YNR024W
LRP1
PAP2
TRF5
AIR1
MTR3
Gene
Z_Shift_K_Gem
Z_Im_K_Gem
Z_Shift_K_Cyt
Z_Im_K_Cyt
Z_Shift_L_Gem
Z_Im_L_Gem
Z_Shift_L_Cyt
Z_Im_L_Cyt

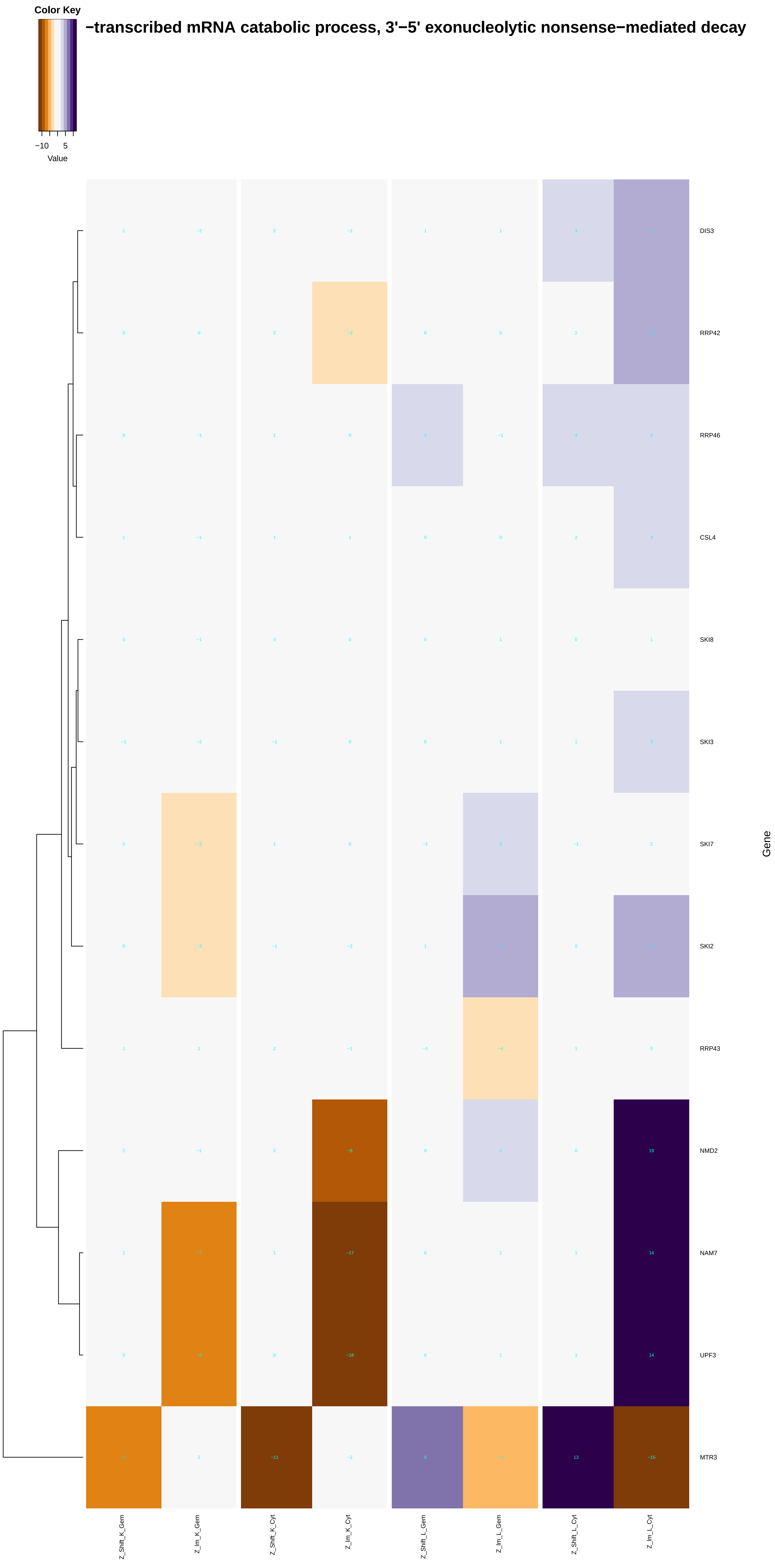
Color Key
−10 5
Value
−transcribed mRNA catabolic process, 3'−5' exonucleolytic nonsense−mediated decay
DIS3
RRP42
RRP46
CSL4
SKI8
SKI3
SKI7
SKI2
RRP43
NMD2
NAM7
UPF3
MTR3
Gene
Z_Shift_K_Gem
Z_Im_K_Gem
Z_Shift_K_Cyt
Z_Im_K_Cyt
Z_Shift_L_Gem
Z_Im_L_Gem
Z_Shift_L_Cyt
Z_Im_L_Cyt

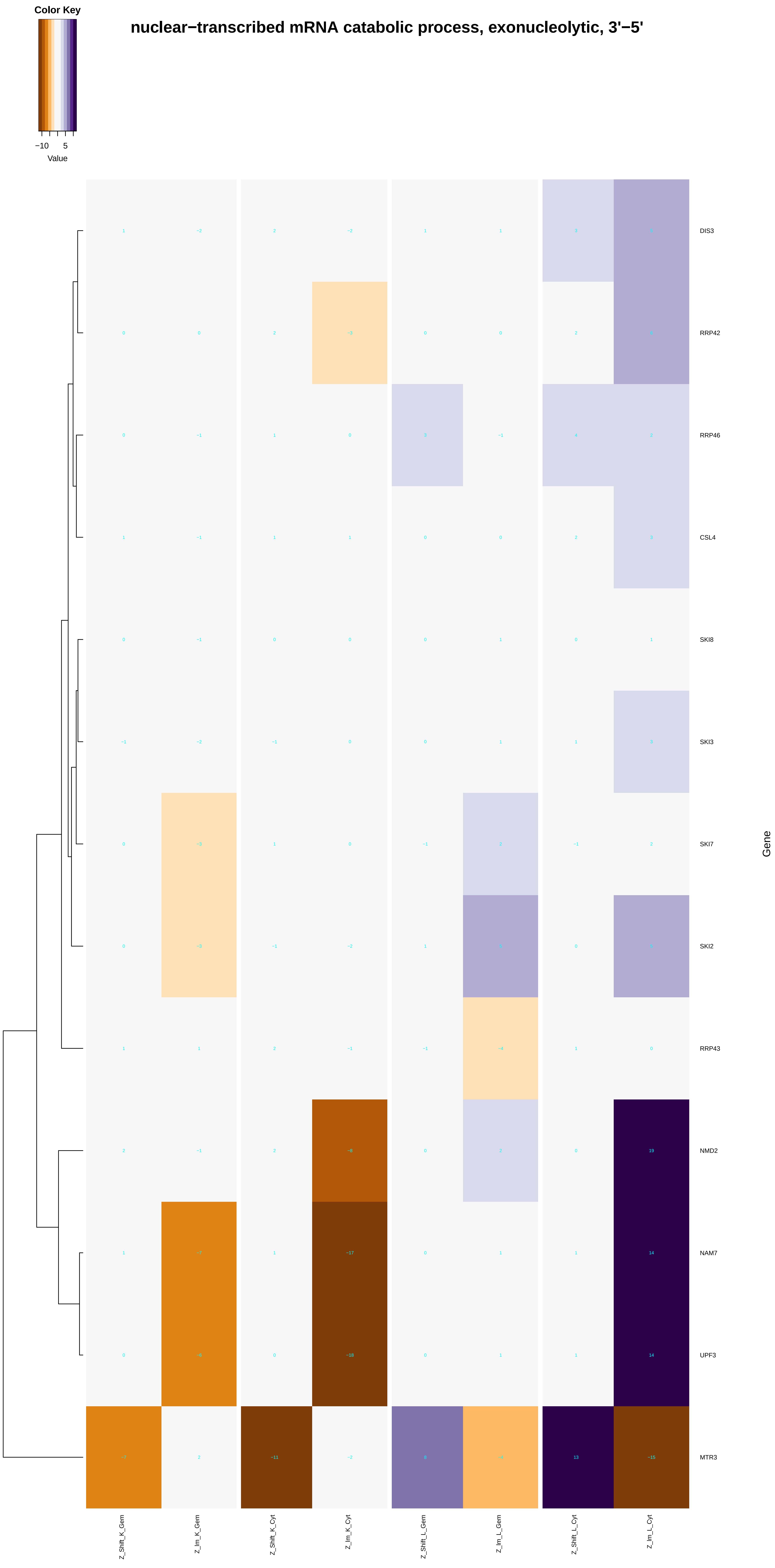
nuclear−transcribed mRNA catabolic process, exonucleolytic, 3'−5'
Color Key
−10
5
Value
DIS3
RRP42
RRP46
CSL4
SKI8
SKI3
SKI7
SKI2
RRP43
NMD2
NAM7
UPF3
MTR3
Gene
Z_Shift_K_Gem
Z_Im_K_Gem
Z_Shift_K_Cyt
Z_Im_K_Cyt
Z_Shift_L_Gem
Z_Im_L_Gem
Z_Shift_L_Cyt
Z_Im_L_Cyt

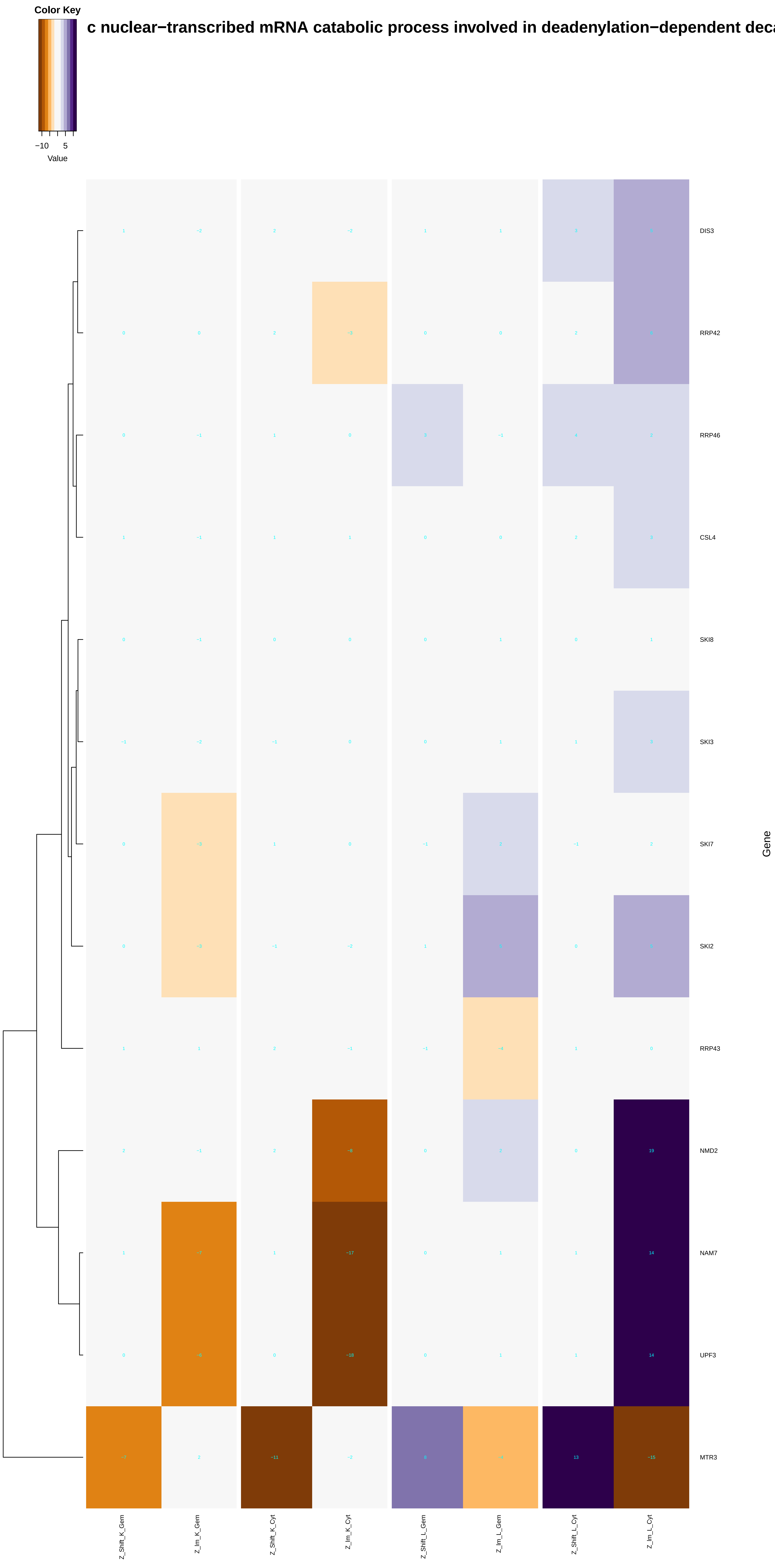

Color Key
−10
5
Value
c nuclear−transcribed mRNA catabolic process involved in deadenylation−dependent deca
DIS3
RRP42
RRP46
CSL4
SKI8
SKI3
SKI7
SKI2
RRP43
NMD2
NAM7
UPF3
MTR3
Gene
Z_Shift_K_Gem
Z_Im_K_Gem
Z_Shift_K_Cyt
Z_Im_K_Cyt
Z_Shift_L_Gem
Z_Im_L_Gem
Z_Shift_L_Cyt
Z_Im_L_Cyt

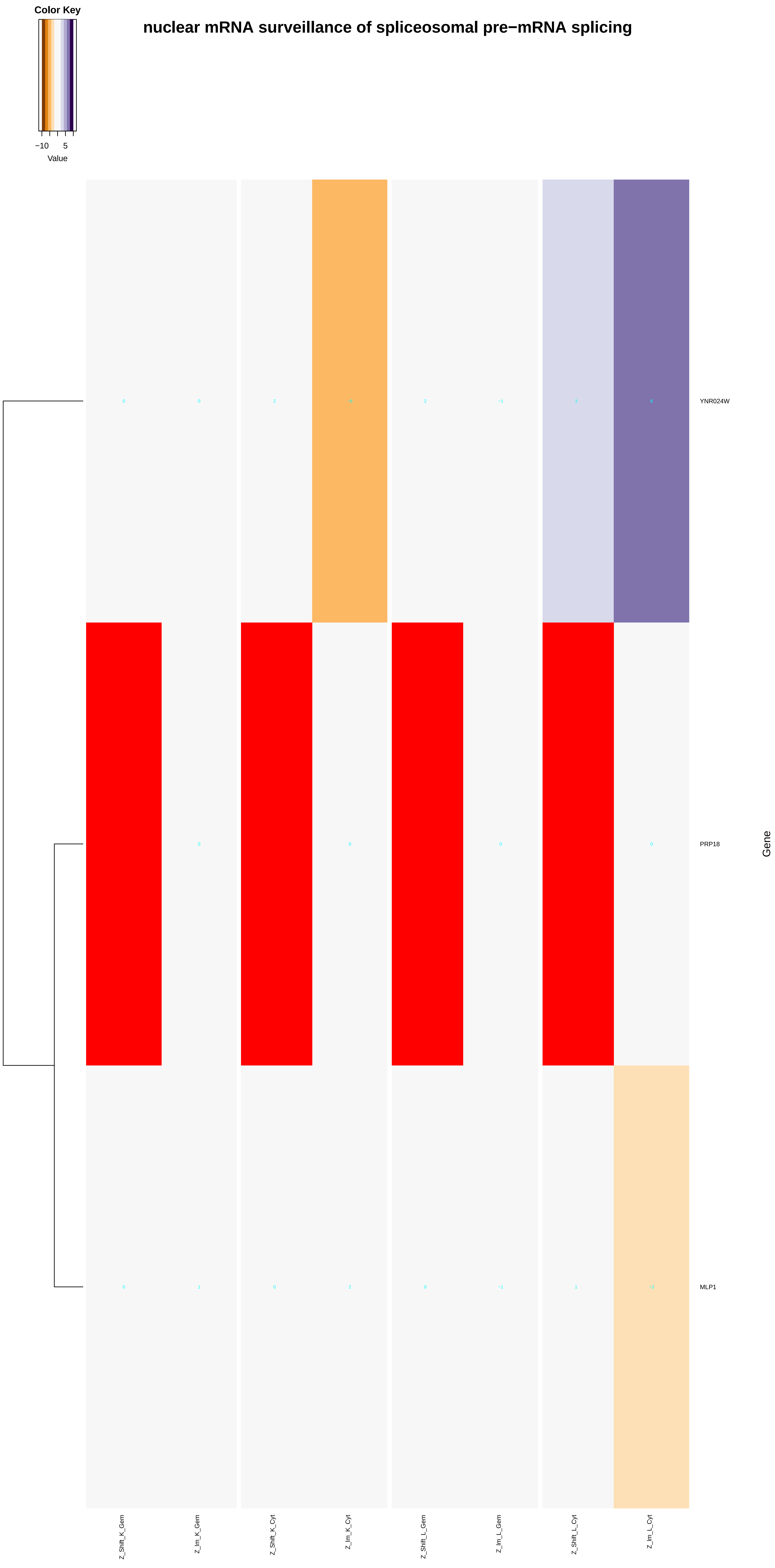
Color Key
−10
5
Value
nuclear mRNA surveillance of spliceosomal pre−mRNA splicing
YNR024W
PRP18
MLP1
Gene
Z_Shift_K_Gem
Z_Im_K_Gem
Z_Shift_K_Cyt
Z_Im_K_Cyt
Z_Shift_L_Gem
Z_Im_L_Gem
Z_Shift_L_Cyt
Z_Im_L_Cyt

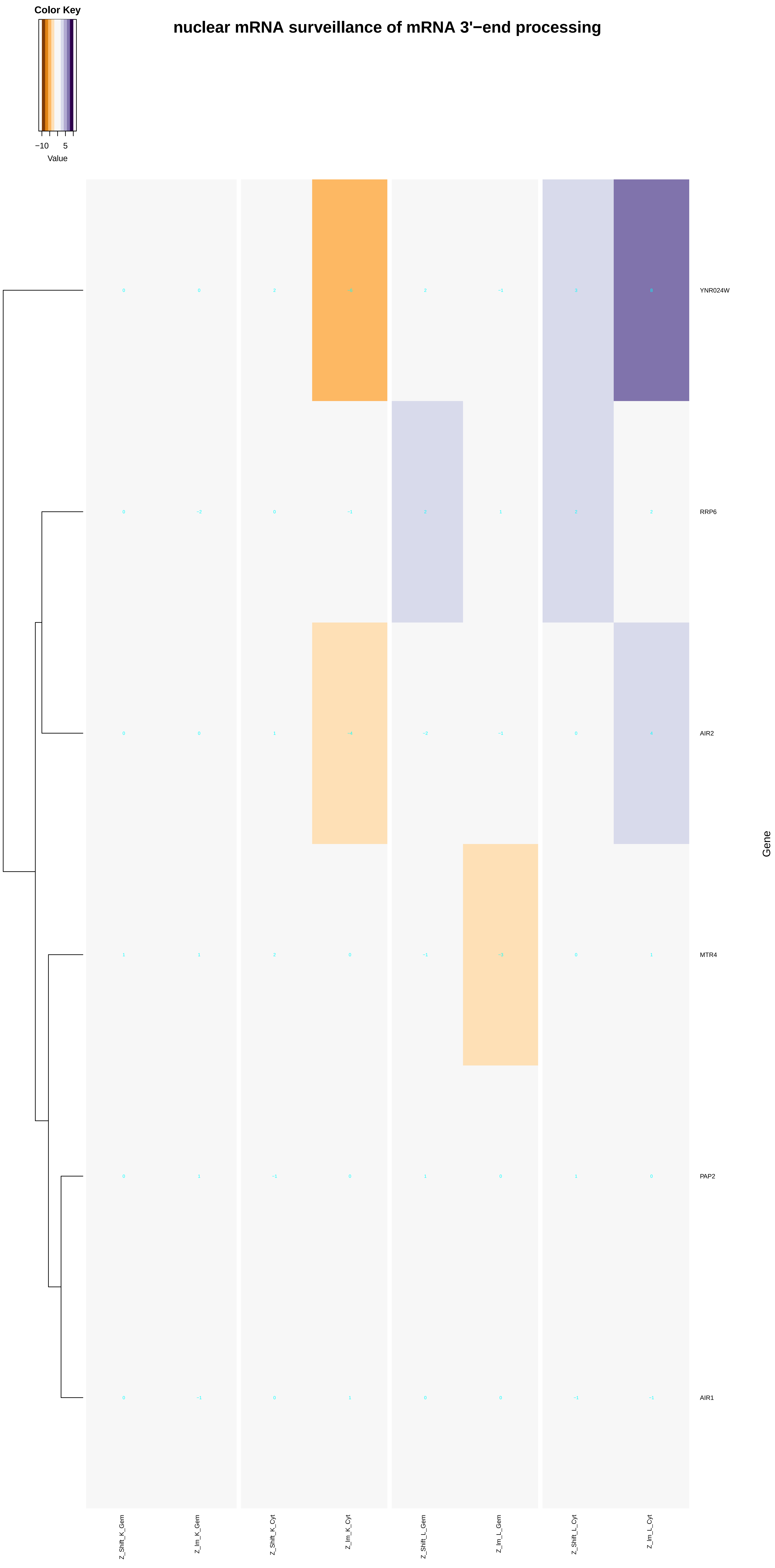
Color Key
−10 5
Value
nuclear mRNA surveillance of mRNA 3'−end processing
YNR024W
RRP6
AIR2
MTR4
PAP2
AIR1
Gene
Z_Shift_K_Gem
Z_Im_K_Gem
Z_Shift_K_Cyt
Z_Im_K_Cyt
Z_Shift_L_Gem
Z_Im_L_Gem
Z_Shift_L_Cyt
Z_Im_L_Cyt
