## Additional File 7 for "A humanized yeast phenomic model of deoxycytidine kinase to predict genetic buffering of nucleoside analog cytotoxicity": ATPase_activity.pdf

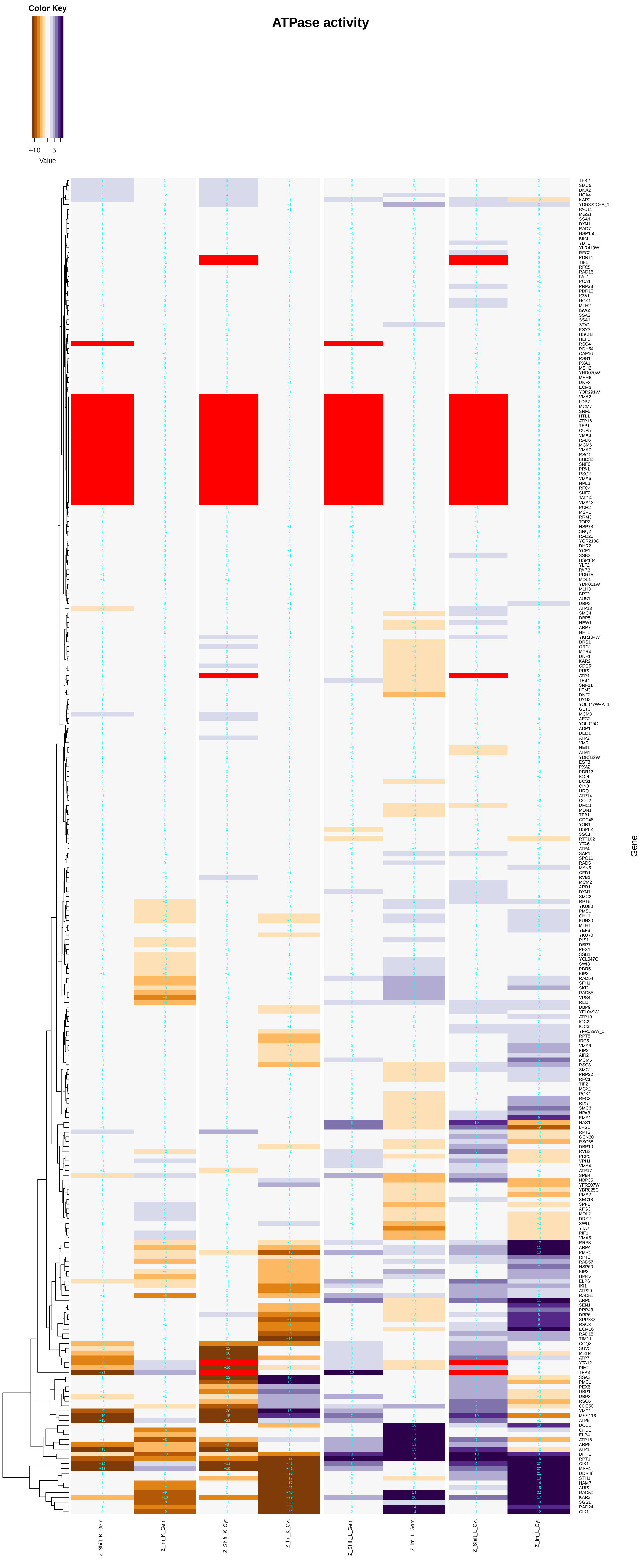
Color Key
−10 5
Value
ATPase activity
Gene
Z_Shift_K_Gem
Z_Im_K_Gem
Z_Shift_K_Cyt
Z_Im_K_Cyt
Z_Shift_L_Gem
Z_Im_L_Gem
Z_Shift_L_Cyt
Z_Im_L_Cyt

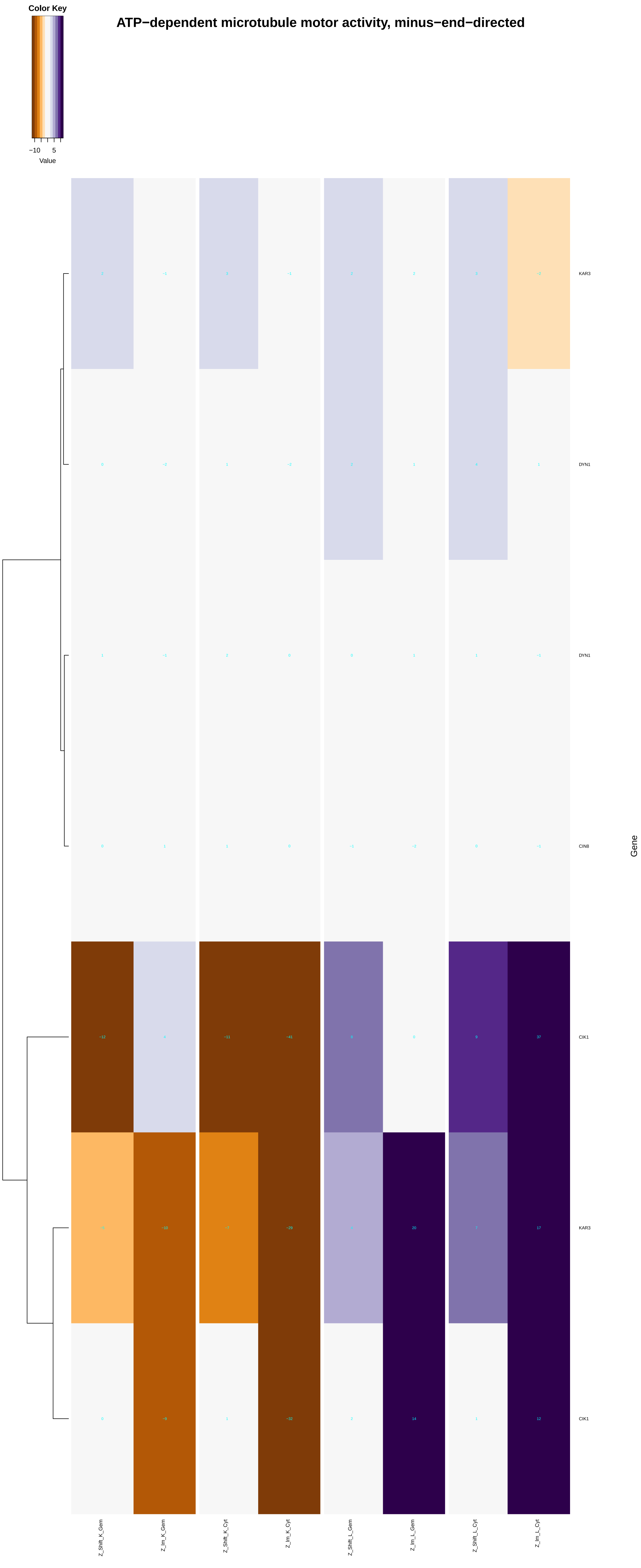
ATP−dependent microtubule motor activity, minus−end−directed
Color Key
−10
5
Value
KAR3
DYN1
DYN1
CIN8
CIK1
KAR3
CIK1
Gene
Z_Shift_K_Gem
Z_Im_K_Gem
Z_Shift_K_Cyt
Z_Im_K_Cyt
Z_Shift_L_Gem
Z_Im_L_Gem
Z_Shift_L_Cyt
Z_Im_L_Cyt

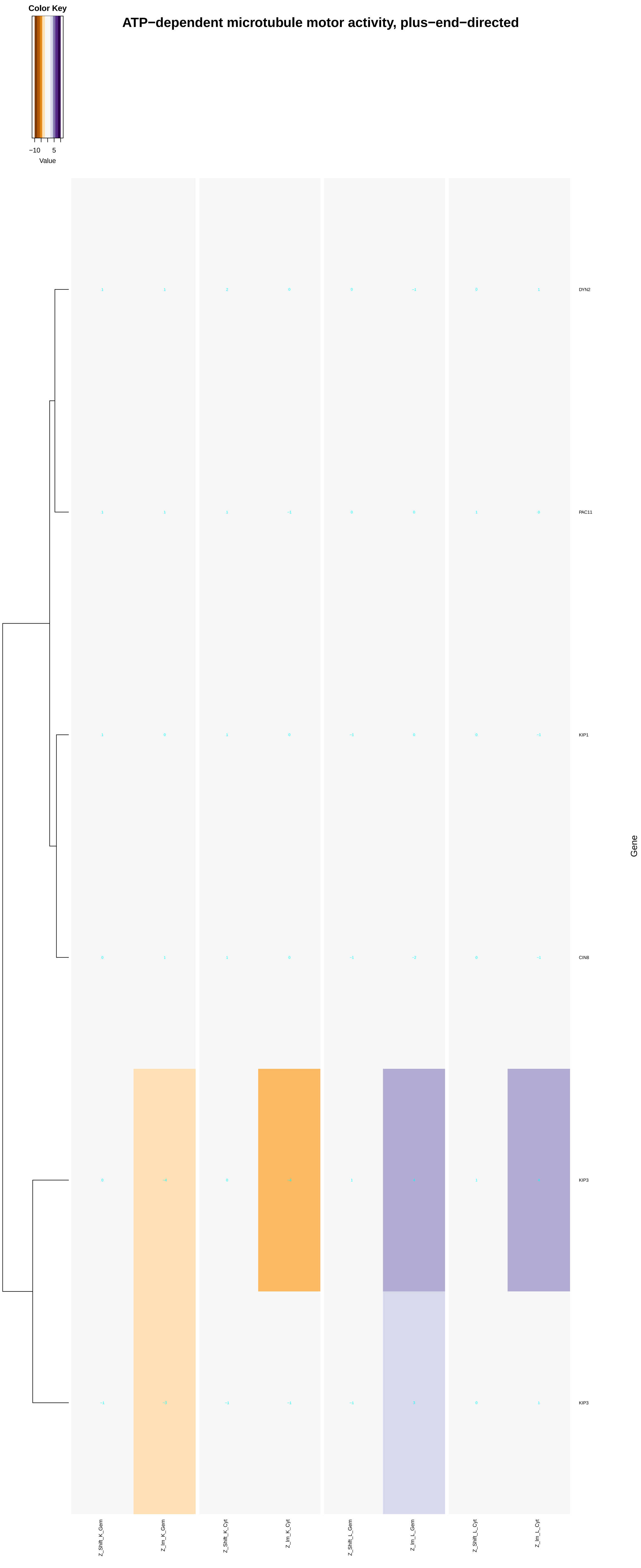
ATP−dependent microtubule motor activity, plus−end−directed
Color Key
−10
5
Value
DYN2
PAC11
KIP1
CIN8
KIP3
KIP3
Gene
Z_Shift_K_Gem
Z_Im_K_Gem
Z_Shift_K_Cyt
Z_Im_K_Cyt
Z_Shift_L_Gem
Z_Im_L_Gem
Z_Shift_L_Cyt
Z_Im_L_Cyt

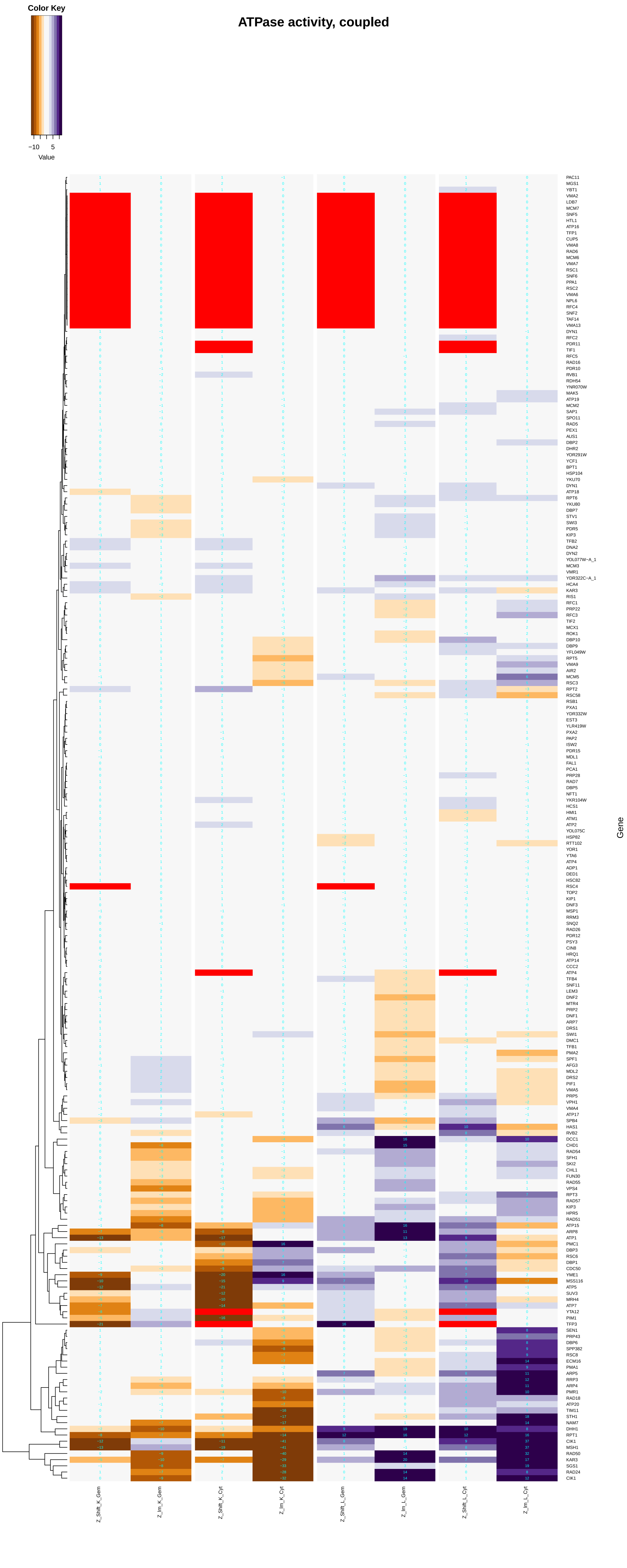

Color Key
−10
5
Value
ATPase activity, coupled
Gene
Z_Shift_K_Gem
Z_Im_K_Gem
Z_Shift_K_Cyt
Z_Im_K_Cyt
Z_Shift_L_Gem
Z_Im_L_Gem
Z_Shift_L_Cyt
Z_Im_L_Cyt

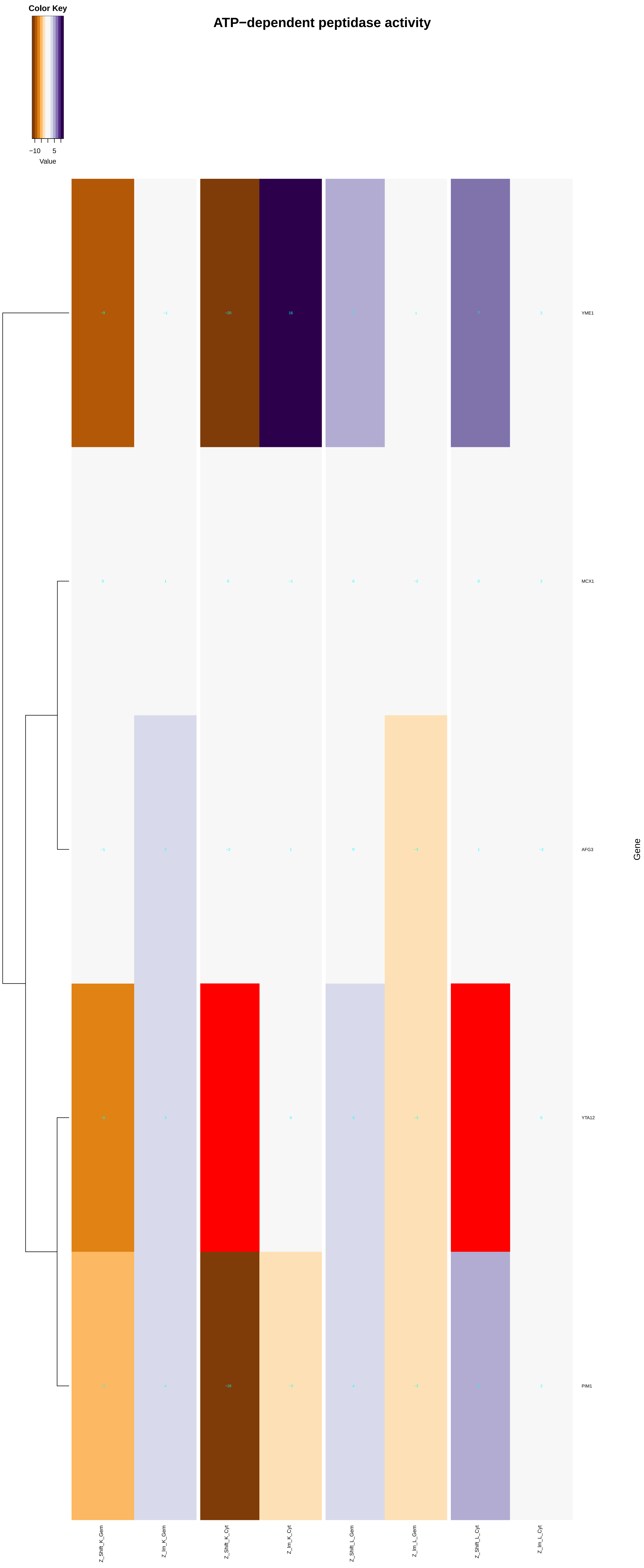

Color Key
−10
5
Value
ATP−dependent peptidase activity
YME1
MCX1
AFG3
YTA12
PIM1
Gene
Z_Shift_K_Gem
Z_Im_K_Gem
Z_Shift_K_Cyt
Z_Im_K_Cyt
Z_Shift_L_Gem
Z_Im_L_Gem
Z_Shift_L_Cyt
Z_Im_L_Cyt

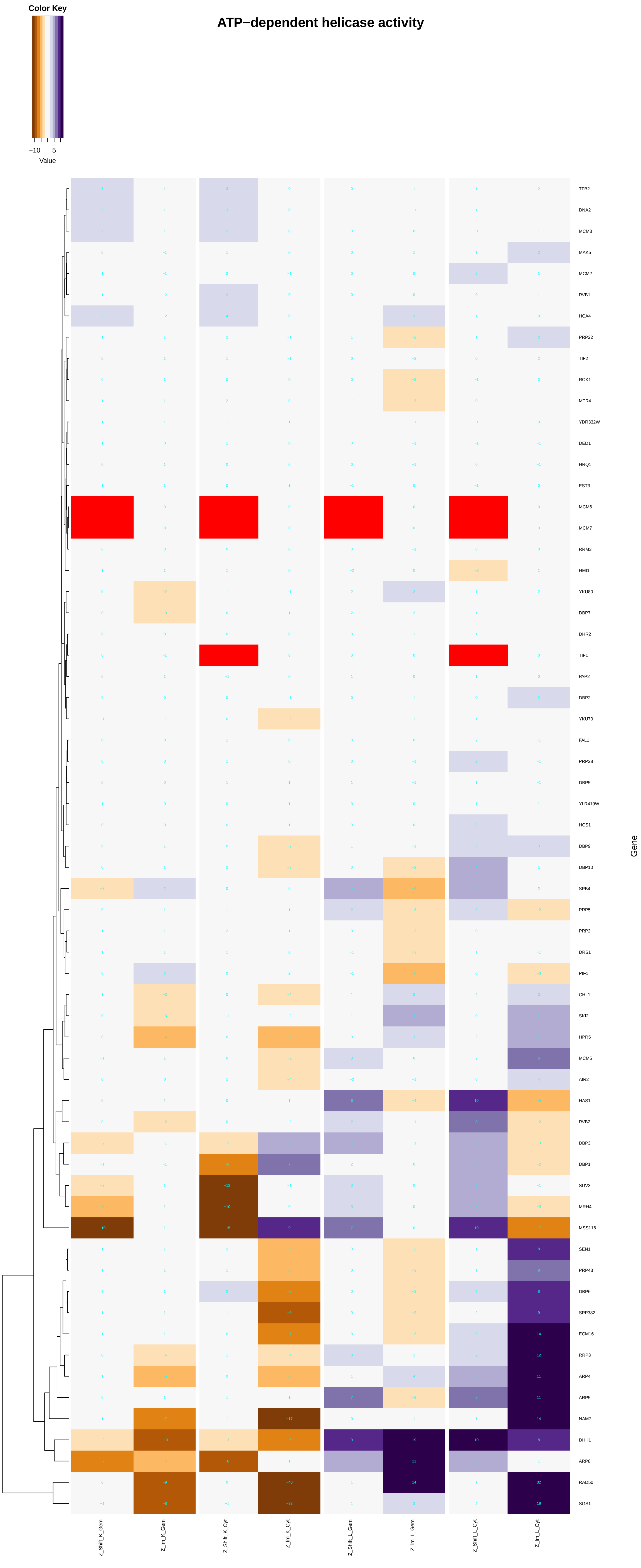
ATP−dependent helicase activity
Color Key
−10
5
Value
TFB2
DNA2
MCM3
MAK5
MCM2
RVB1
HCA4
PRP22
TIF2
ROK1
MTR4
YDR332W
DED1
HRQ1
EST3
MCM6
MCM7
RRM3
HMI1
YKU80
DBP7
DHR2
TIF1
PAP2
DBP2
YKU70
FAL1
PRP28
DBP5
YLR419W
HCS1
DBP9
DBP10
SPB4
PRP5
PRP2
DRS1
PIF1
CHL1
SKI2
HPR5
MCM5
AIR2
HAS1
RVB2
DBP3
DBP1
SUV3
MRH4
MSS116
SEN1
PRP43
DBP6
SPP382
ECM16
RRP3
ARP4
ARP5
NAM7
DHH1
ARP8
RAD50
SGS1
Gene
Z_Shift_K_Gem
Z_Im_K_Gem
Z_Shift_K_Cyt
Z_Im_K_Cyt
Z_Shift_L_Gem
Z_Im_L_Gem
Z_Shift_L_Cyt
Z_Im_L_Cyt

### DNA−dependent ATPase activity

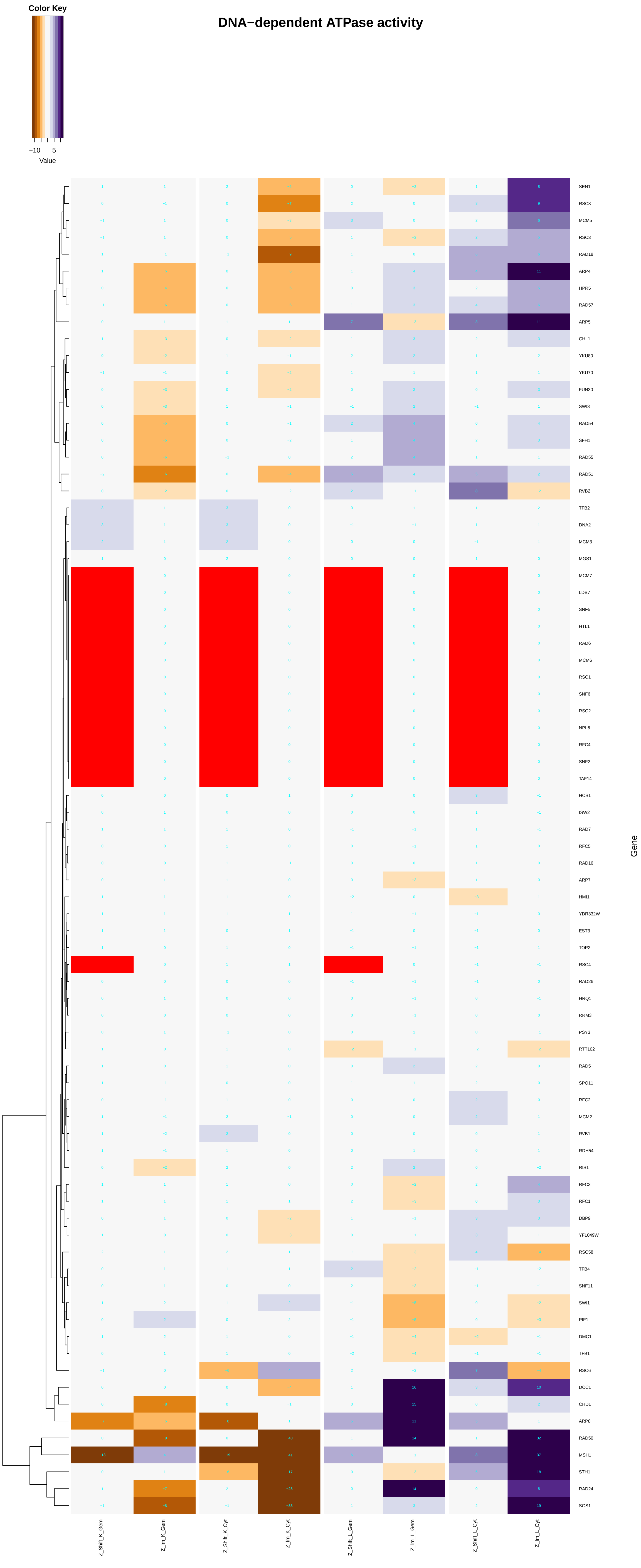

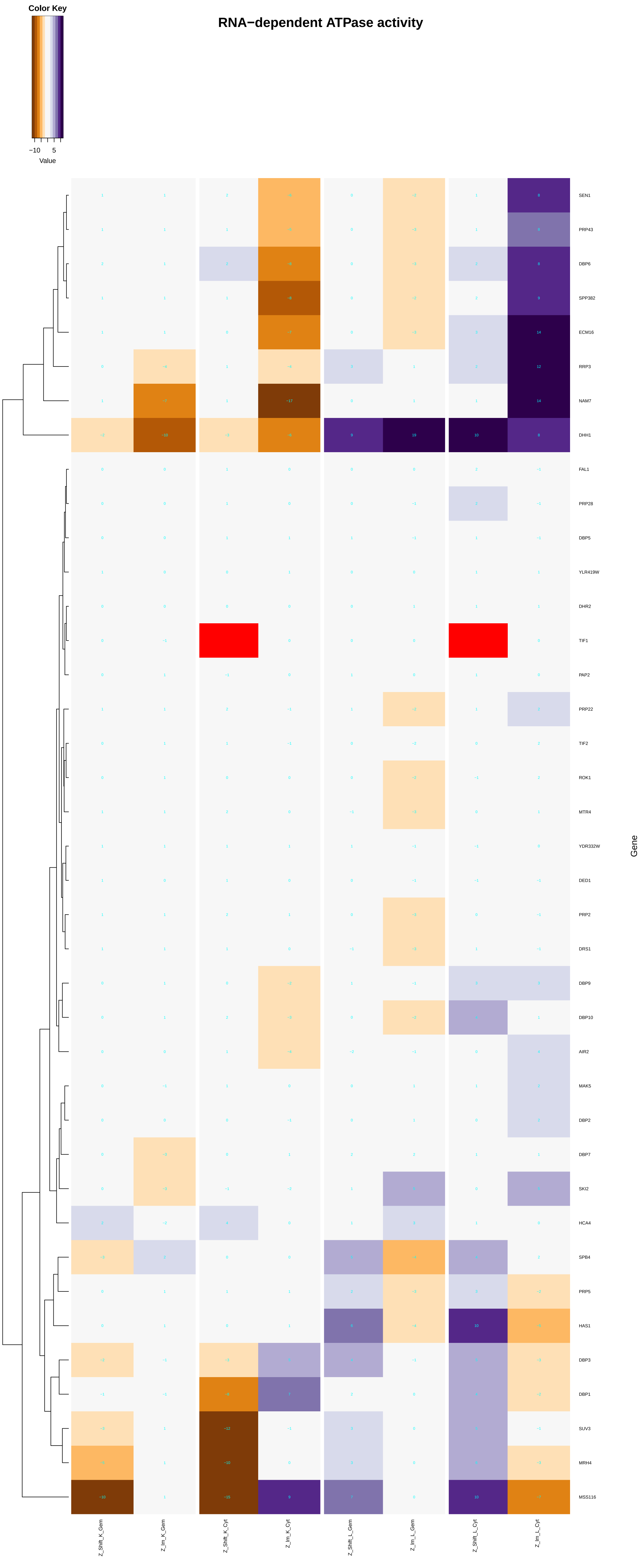

Color Key
−10
5
Value
RNA−dependent ATPase activity
SEN1
PRP43
DBP6
SPP382
ECM16
RRP3
NAM7
DHH1
FAL1
PRP28
DBP5
YLR419W
DHR2
TIF1
PAP2
PRP22
TIF2
ROK1
MTR4
YDR332W
DED1
PRP2
DRS1
DBP9
DBP10
AIR2
MAK5
DBP2
DBP7
SKI2
HCA4
SPB4
PRP5
HAS1
DBP3
DBP1
SUV3
MRH4
MSS116
Gene
Z_Shift_K_Gem
Z_Im_K_Gem
Z_Shift_K_Cyt
Z_Im_K_Cyt
Z_Shift_L_Gem
Z_Im_L_Gem
Z_Shift_L_Cyt
Z_Im_L_Cyt

#### microtubule−severing ATPase activity

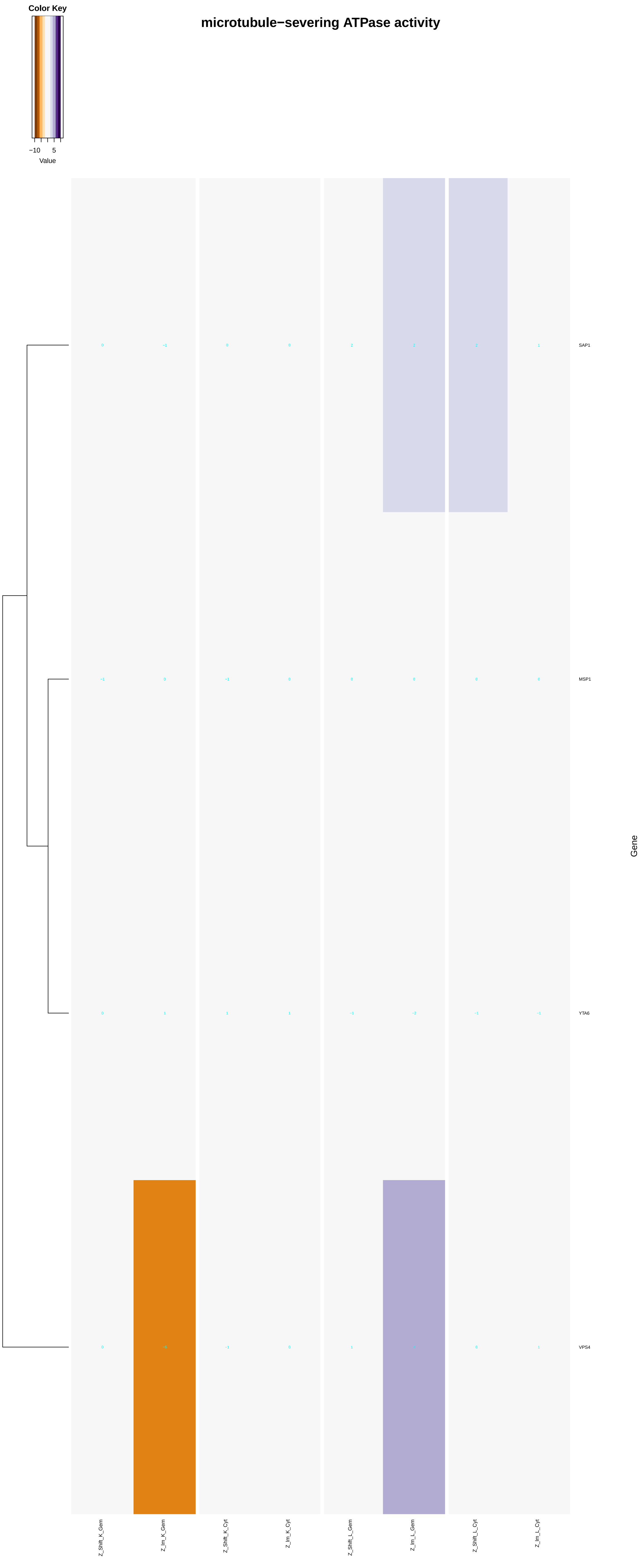

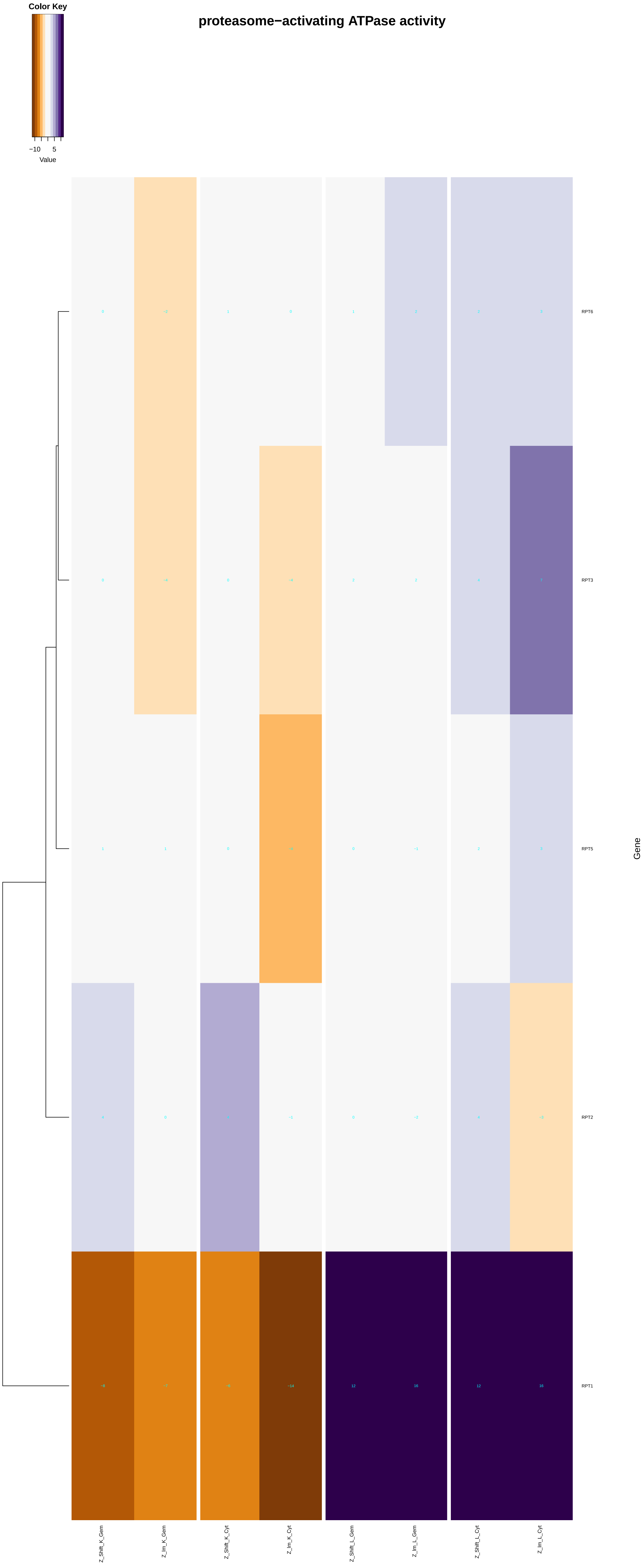
proteasome−activating ATPase activity
Color Key
−10
5
Value
RPT6
RPT3
RPT5
RPT2
RPT1
Gene
Z_Shift_K_Gem
Z_Im_K_Gem
Z_Shift_K_Cyt
Z_Im_K_Cyt
Z_Shift_L_Gem
Z_Im_L_Gem
Z_Shift_L_Cyt
Z_Im_L_Cyt

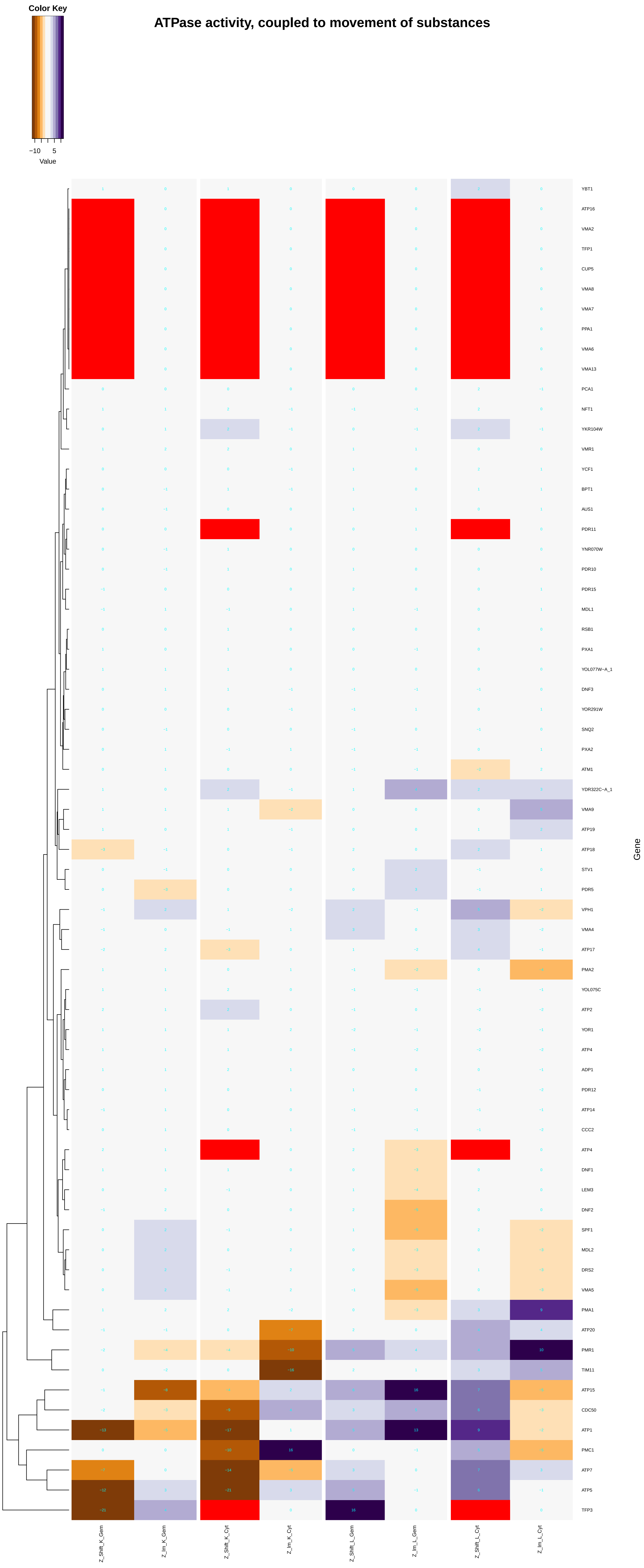
ATPase activity, coupled to movement of substances
Color Key
−10
5
Value
YBT1
ATP16
VMA2
TFP1
CUP5
VMA8
VMA7
PPA1
VMA6
VMA13
PCA1
NFT1
YKR104W
VMR1
YCF1
BPT1
AUS1
PDR11
YNR070W
PDR10
PDR15
MDL1
RSB1
PXA1
YOL077W−A_1
DNF3
YOR291W
SNQ2
PXA2
ATM1
YDR322C−A_1
VMA9
ATP19
ATP18
STV1
PDR5
VPH1
VMA4
ATP17
PMA2
YOL075C
ATP2
YOR1
ATP4
ADP1
PDR12
ATP14
CCC2
ATP4
DNF1
LEM3
DNF2
SPF1
MDL2
DRS2
VMA5
PMA1
ATP20
PMR1
TIM11
ATP15
CDC50
ATP1
PMC1
ATP7
ATP5
TFP3
Gene
Z_Shift_K_Gem
Z_Im_K_Gem
Z_Shift_K_Cyt
Z_Im_K_Cyt
Z_Shift_L_Gem
Z_Im_L_Gem
Z_Shift_L_Cyt
Z_Im_L_Cyt

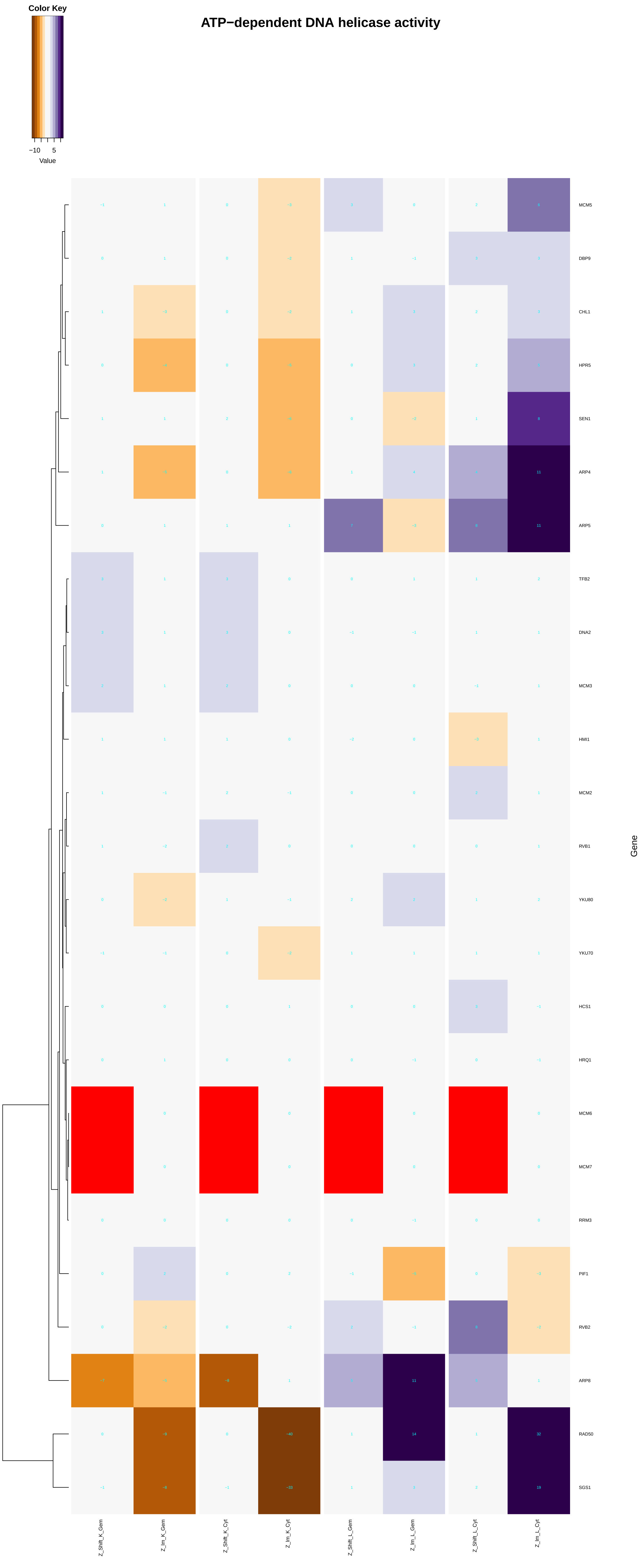
ATP−dependent DNA helicase activity
Color Key
−10
5
Value
MCM5
DBP9
CHL1
HPR5
SEN1
ARP4
ARP5
TFB2
DNA2
MCM3
HMI1
MCM2
RVB1
YKU80
YKU70
HCS1
HRQ1
MCM6
MCM7
RRM3
PIF1
RVB2
ARP8
RAD50
SGS1
Gene
Z_Shift_K_Gem
Z_Im_K_Gem
Z_Shift_K_Cyt
Z_Im_K_Cyt
Z_Shift_L_Gem
Z_Im_L_Gem
Z_Shift_L_Cyt
Z_Im_L_Cyt

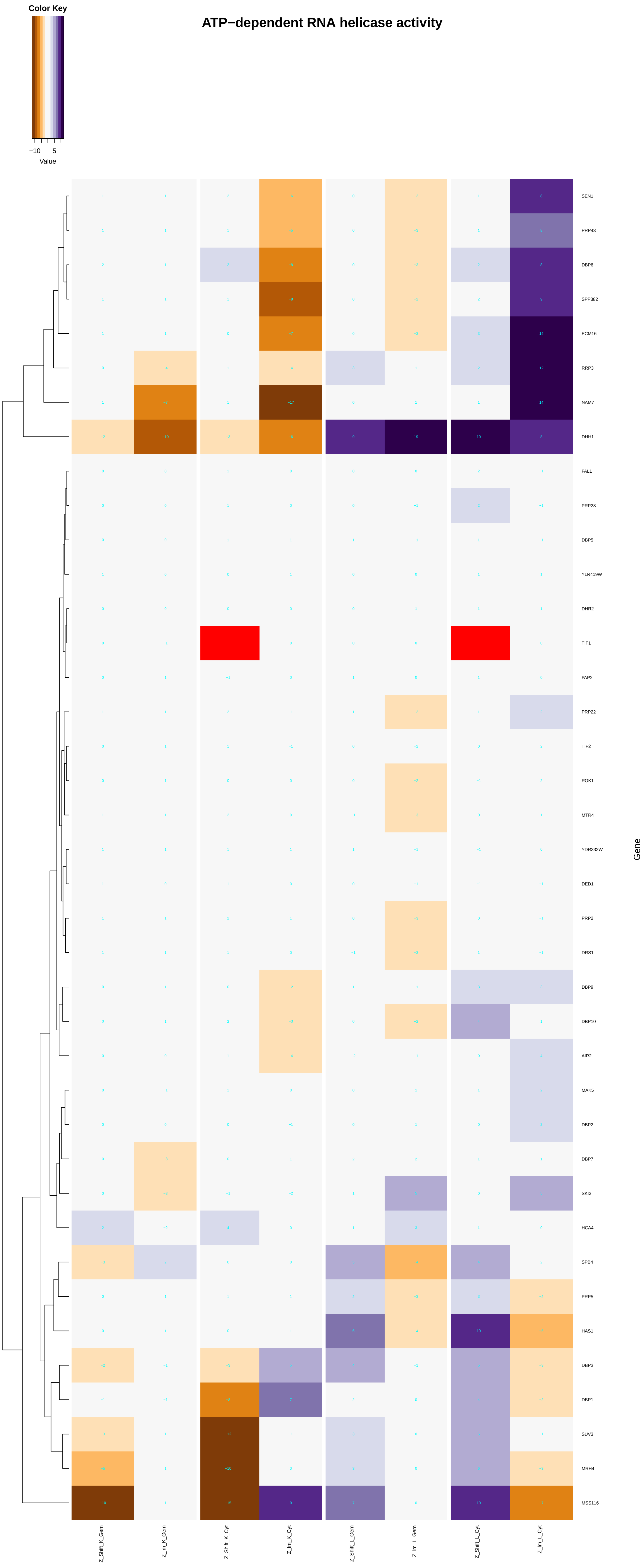

Color Key
−10
5
Value
ATP−dependent RNA helicase activity
SEN1
PRP43
DBP6
SPP382
ECM16
RRP3
NAM7
DHH1
FAL1
PRP28
DBP5
YLR419W
DHR2
TIF1
PAP2
PRP22
TIF2
ROK1
MTR4
YDR332W
DED1
PRP2
DRS1
DBP9
DBP10
AIR2
MAK5
DBP2
DBP7
SKI2
HCA4
SPB4
PRP5
HAS1
DBP3
DBP1
SUV3
MRH4
MSS116
Gene
Z_Shift_K_Gem
Z_Im_K_Gem
Z_Shift_K_Cyt
Z_Im_K_Cyt
Z_Shift_L_Gem
Z_Im_L_Gem
Z_Shift_L_Cyt
Z_Im_L_Cyt

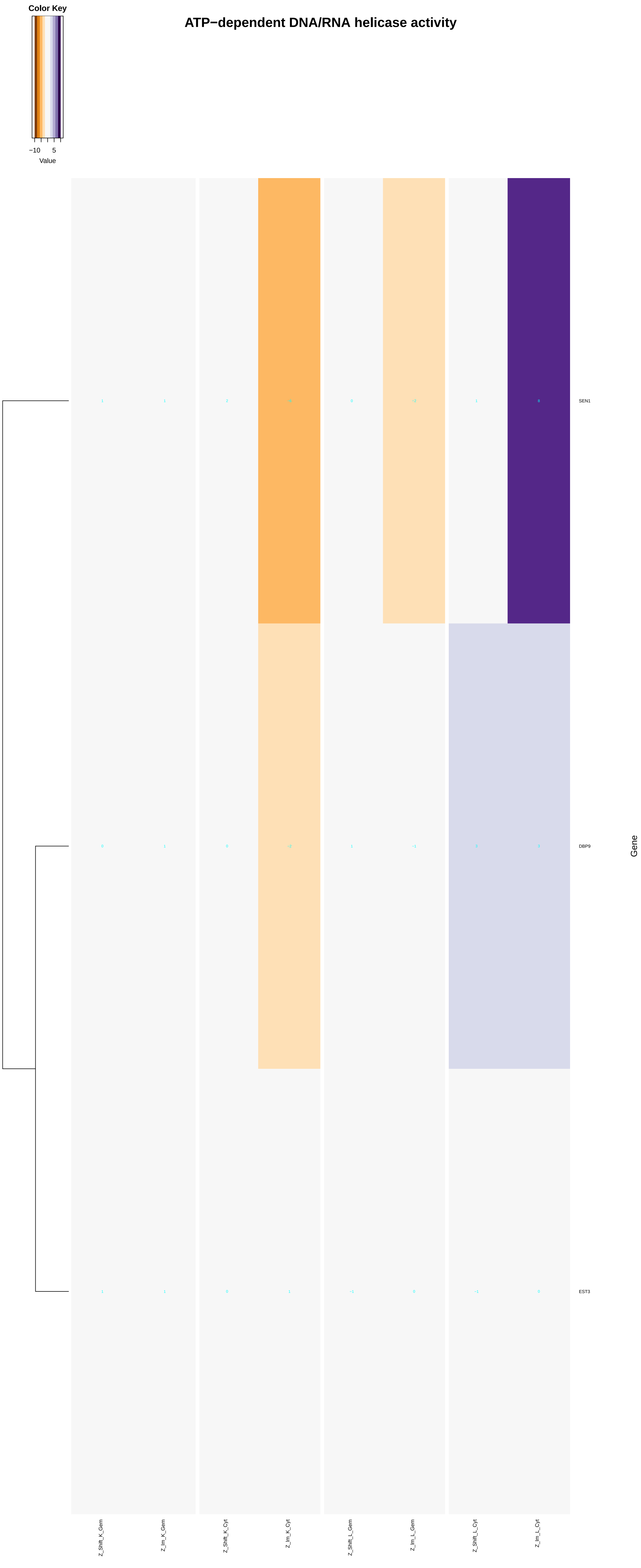
ATP−dependent DNA/RNA helicase activity
Color Key
−10
5
Value
SEN1
DBP9
EST3
Gene
Z_Shift_K_Gem
Z_Im_K_Gem
Z_Shift_K_Cyt
Z_Im_K_Cyt
Z_Shift_L_Gem
Z_Im_L_Gem
Z_Shift_L_Cyt
Z_Im_L_Cyt

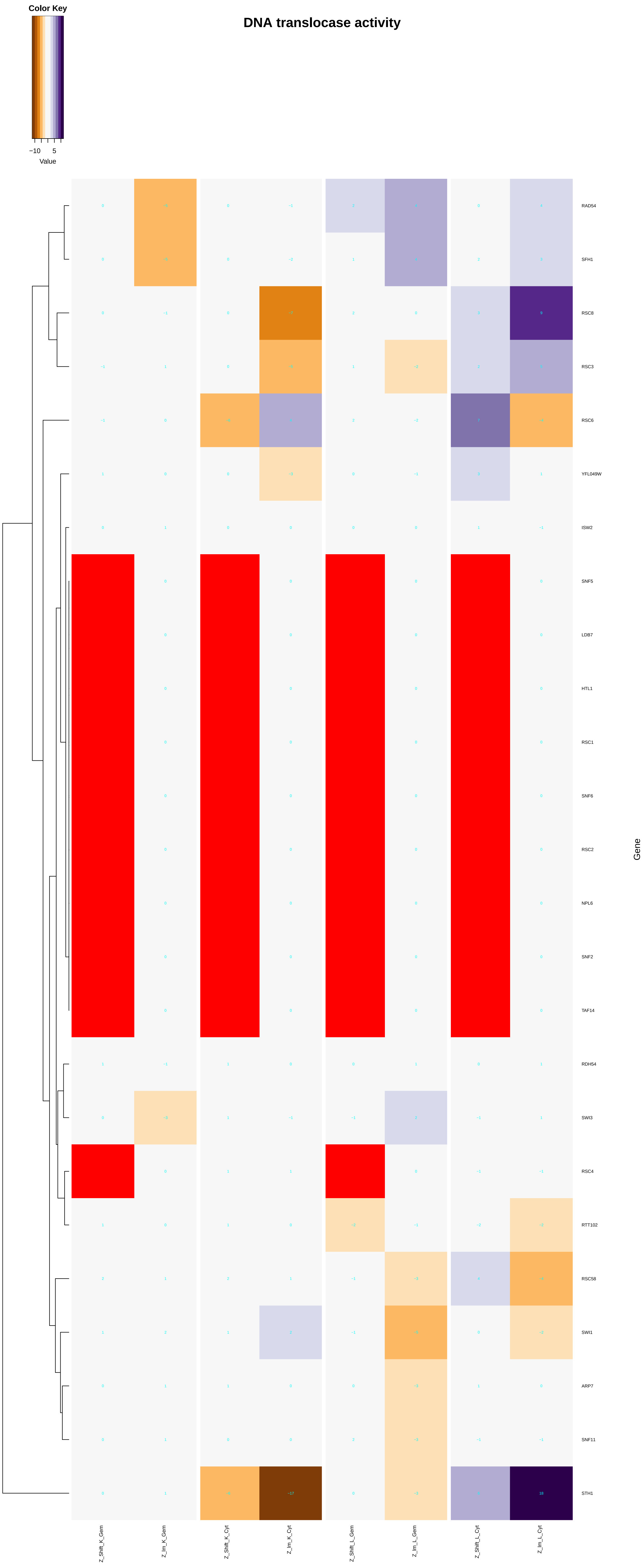
DNA translocase activity
Color Key
−10
5
Value
RAD54
SFH1
RSC8
RSC3
RSC6
YFL049W
ISW2
SNF5
LDB7
HTL1
RSC1
SNF6
RSC2
NPL6
SNF2
TAF14
RDH54
SWI3
RSC4
RTT102
RSC58
SWI1
ARP7
SNF11
STH1
Gene
Z_Shift_K_Gem
Z_lm_K_Gem
Z_Shift_K_Cyt
Z_lm_K_Cyt
Z_Shift_L_Gem
Z_lm_L_Gem
Z_Shift_L_Cyt
Z_lm_L_Cyt

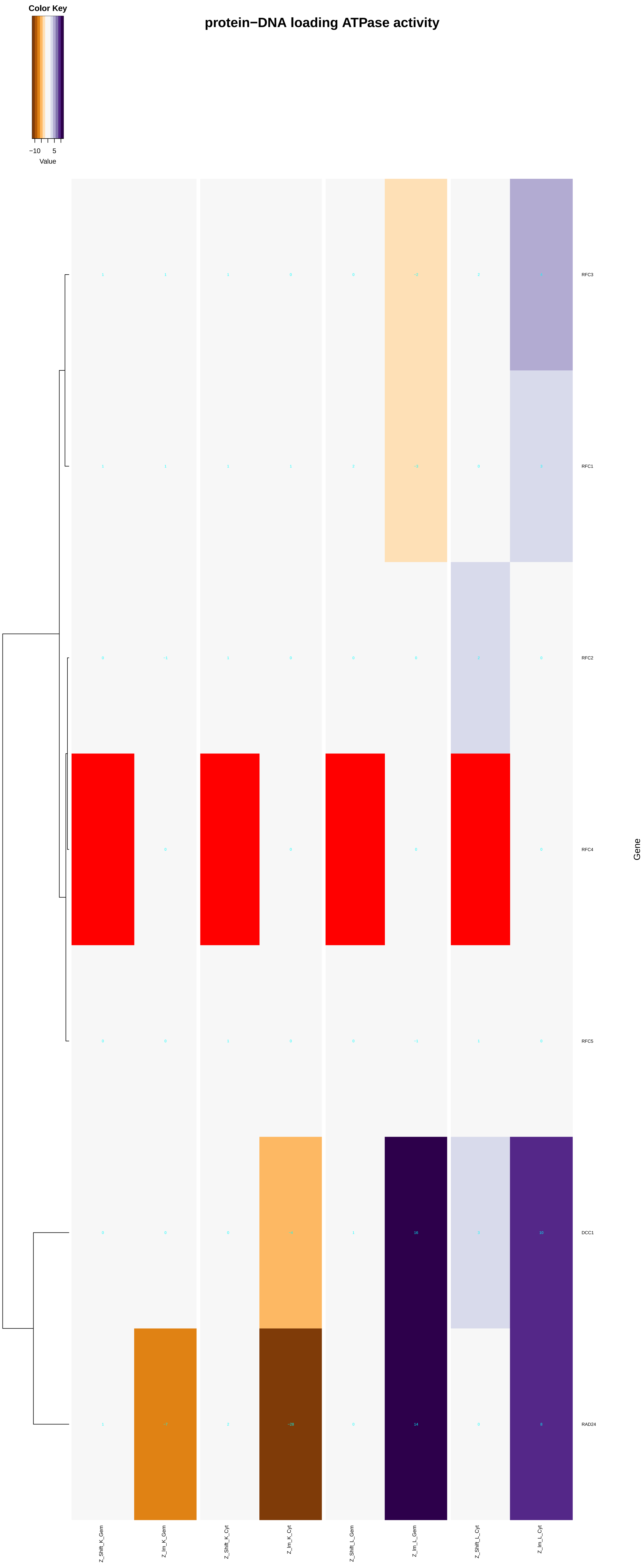

protein−DNA loading ATPase activity
Color Key
−10
5
Value
RFC3
RFC1
RFC2
RFC4
RFC5
DCC1
RAD24
Gene
Z_Shift_K_Gem
Z_Im_K_Gem
Z_Shift_K_Cyt
Z_Im_K_Cyt
Z_Shift_L_Gem
Z_Im_L_Gem
Z_Shift_L_Cyt
Z_Im_L_Cyt

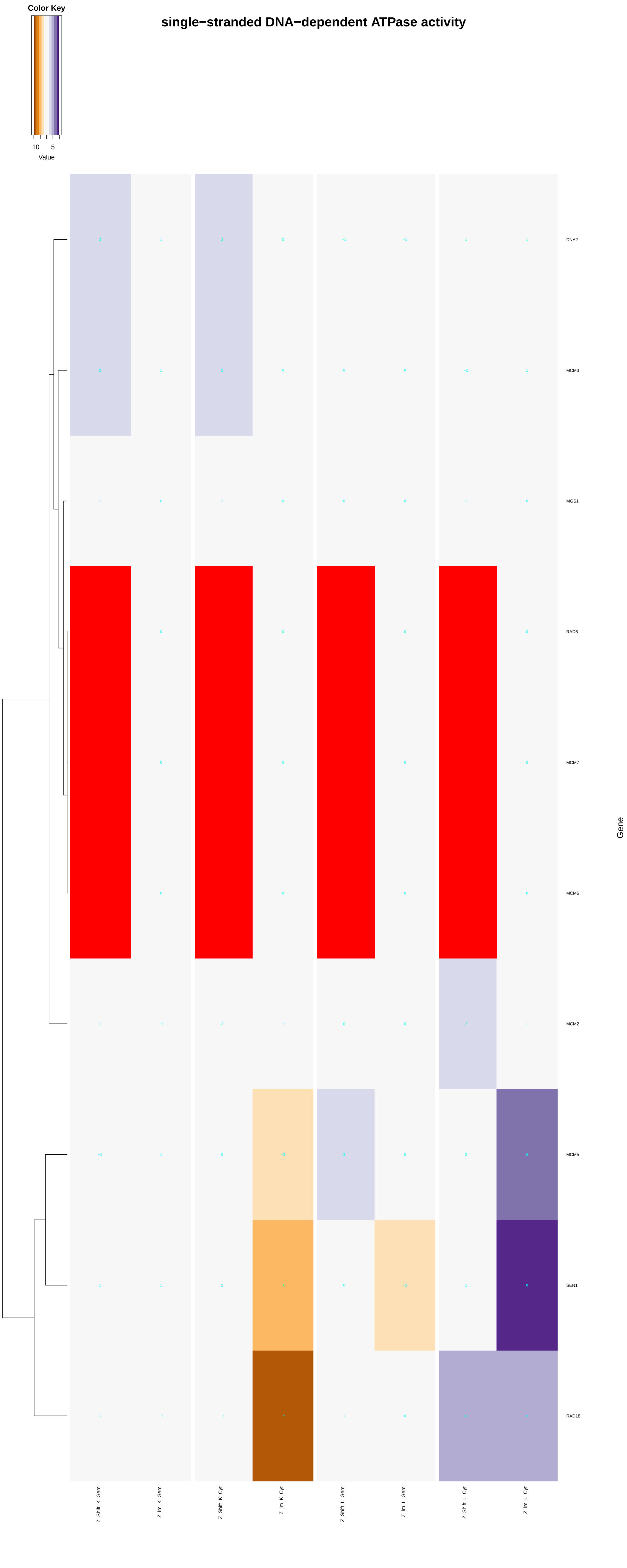
single−stranded DNA−dependent ATPase activity
Color Key
−10
5
Value
DNA2
MCM3
MGS1
RAD6
MCM7
MCM6
MCM2
MCM5
SEN1
RAD18
Gene
Z_Shift_K_Gem
Z_Im_K_Gem
Z_Shift_K_Cyt
Z_Im_K_Cyt
Z_Shift_L_Gem
Z_Im_L_Gem
Z_Shift_L_Cyt
Z_Im_L_Cyt

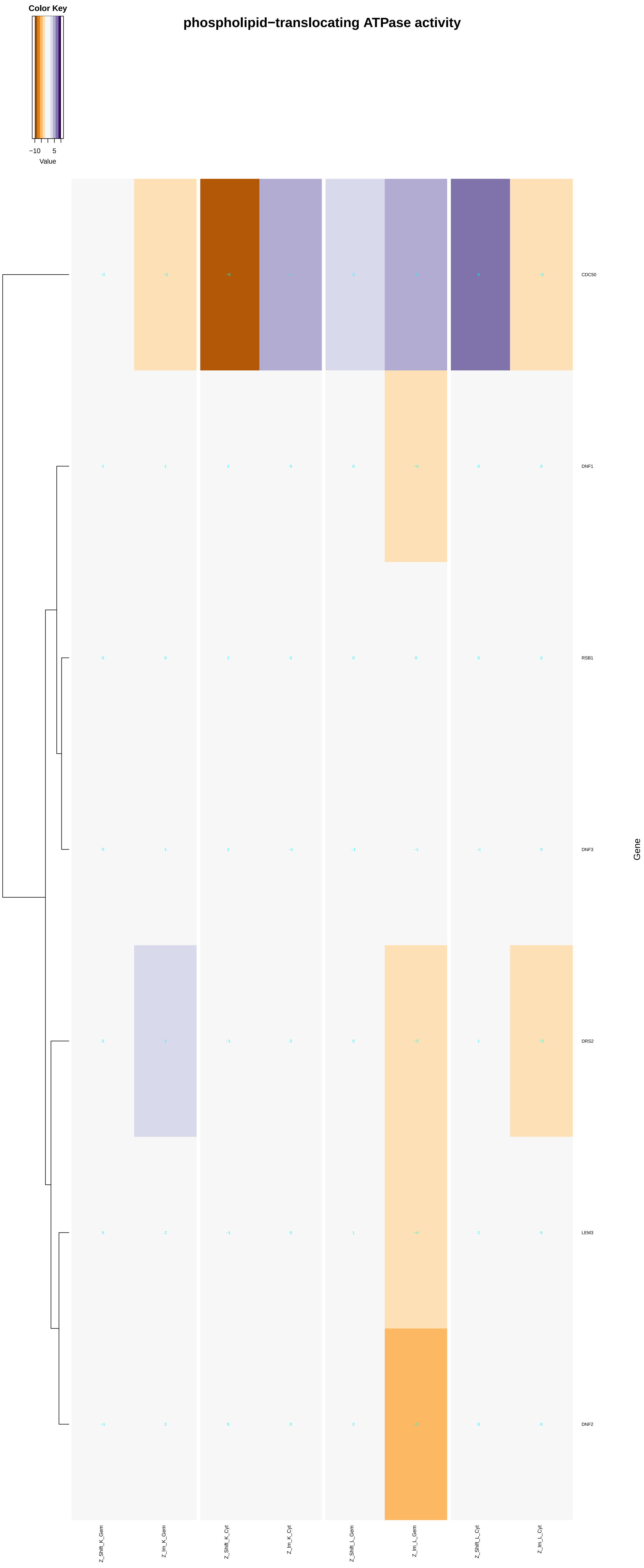

Color Key
−10
5
Value
phospholipid−translocating ATPase activity
CDC50
DNF1
RSB1
DNF3
DRS2
LEM3
DNF2
Gene
Z_Shift_K_Gem
Z_Im_K_Gem
Z_Shift_K_Cyt
Z_Im_K_Cyt
Z_Shift_L_Gem
Z_Im_L_Gem
Z_Shift_L_Cyt
Z_Im_L_Cyt

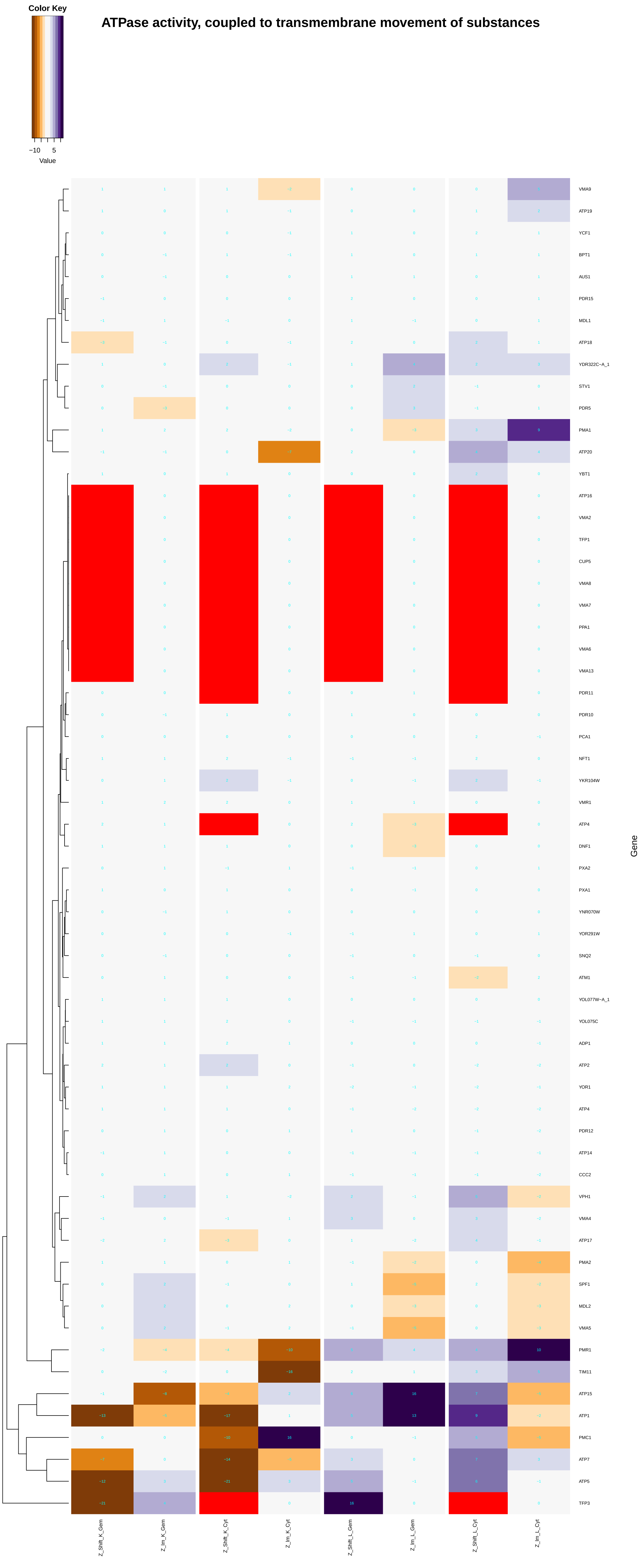
Color Key
−10
5
Value
ATPase activity, coupled to transmembrane movement of substances
VMA9
ATP19
YCF1
BPT1
AUS1
PDR15
MDL1
ATP18
YDR322C−A_1
STV1
PDR5
PMA1
ATP20
YBT1
ATP16
VMA2
TFP1
CUP5
VMA8
VMA7
PPA1
VMA6
VMA13
PDR11
PDR10
PCA1
NFT1
YKR104W
VMR1
ATP4
DNF1
PXA2
PXA1
YNR070W
YOR291W
SNQ2
ATM1
YOL077W−A_1
YOL075C
ADP1
ATP2
YOR1
ATP4
PDR12
ATP14
CCC2
VPH1
VMA4
ATP17
PMA2
SPF1
MDL2
VMA5
PMR1
TIM11
ATP15
ATP1
PMC1
ATP7
ATP5
TFP3
Gene
Z_Shift_K_Gem
Z_Im_K_Gem
Z_Shift_K_Cyt
Z_Im_K_Cyt
Z_Shift_L_Gem
Z_Im_L_Gem
Z_Shift_L_Cyt
Z_Im_L_Cyt

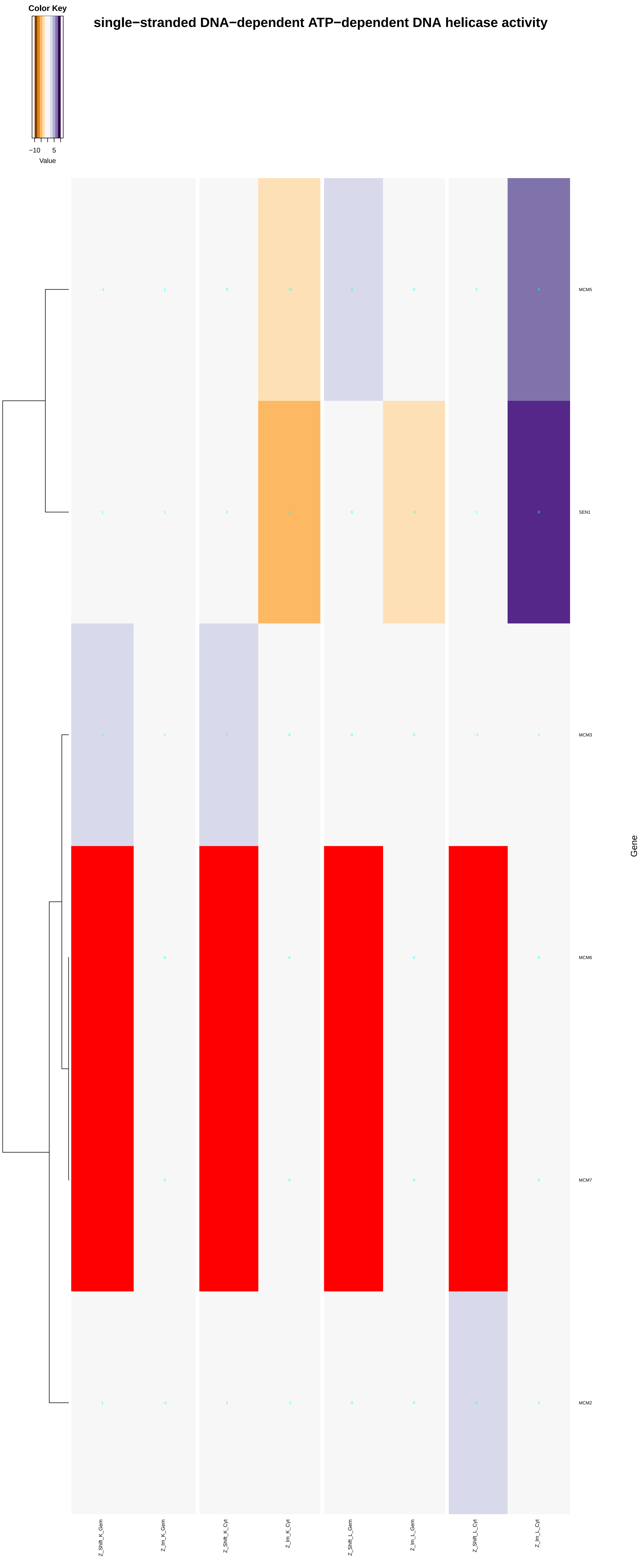
single−stranded DNA−dependent ATP−dependent DNA helicase activity
Color Key
−10
5
Value
MCM5
SEN1
MCM3
MCM6
MCM7
MCM2
Gene
Z_Shift_K_Gem
Z_Im_K_Gem
Z_Shift_K_Cyt
Z_Im_K_Cyt
Z_Shift_L_Gem
Z_Im_L_Gem
Z_Shift_L_Cyt
Z_Im_L_Cyt

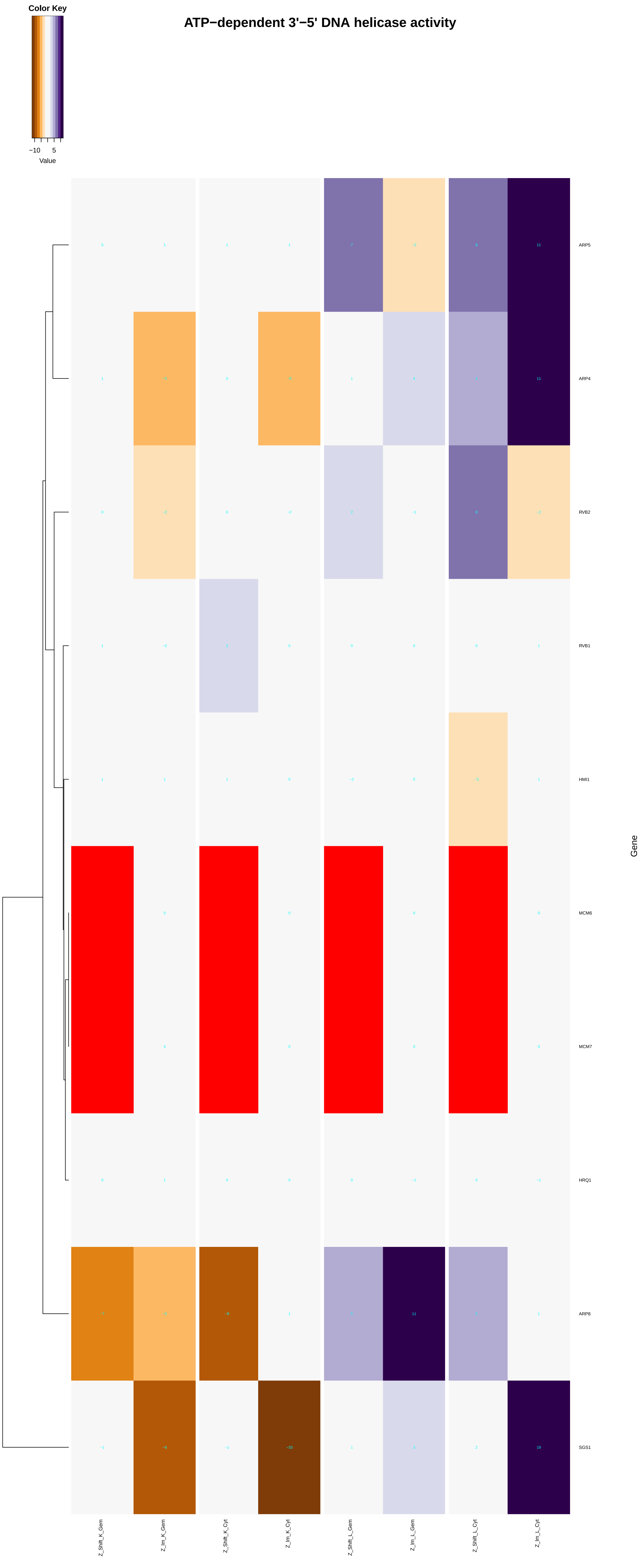

Color Key
−10
5
Value
ATP−dependent 3'−5' DNA helicase activity
ARP5
ARP4
RVB2
RVB1
HMI1
MCM6
MCM7
HRQ1
ARP8
SGS1
Gene
Z_Shift_K_Gem
Z_Im_K_Gem
Z_Shift_K_Cyt
Z_Im_K_Cyt
Z_Shift_L_Gem
Z_Im_L_Gem
Z_Shift_L_Cyt
Z_Im_L_Cyt

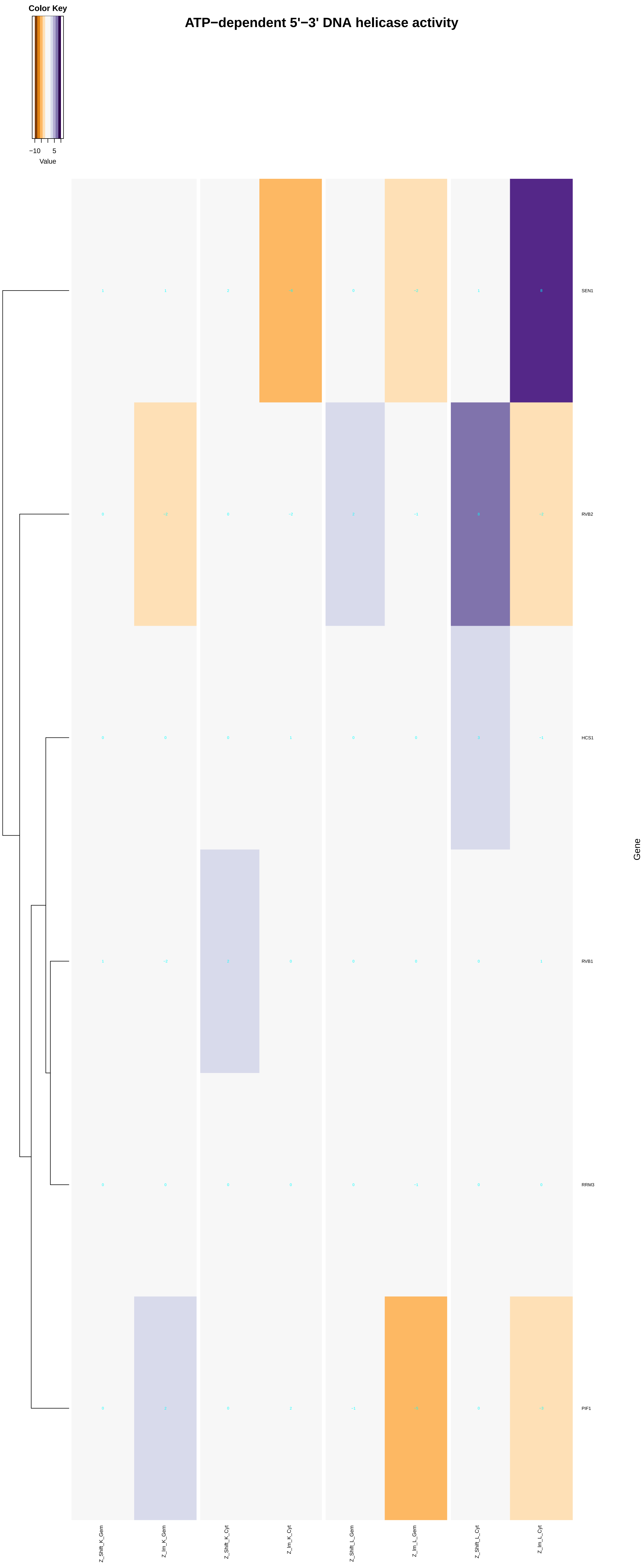
ATP−dependent 5'−3' DNA helicase activity
Color Key
−10
5
Value
SEN1
RVB2
HCS1
RVB1
RRM3
PIF1
Gene
Z_Shift_K_Gem
Z_lm_K_Gem
Z_Shift_K_Cyt
Z_lm_K_Cyt
Z_Shift_L_Gem
Z_lm_L_Gem
Z_Shift_L_Cyt
Z_lm_L_Cyt

### ATP−dependent 3'−5' RNA helicase activity

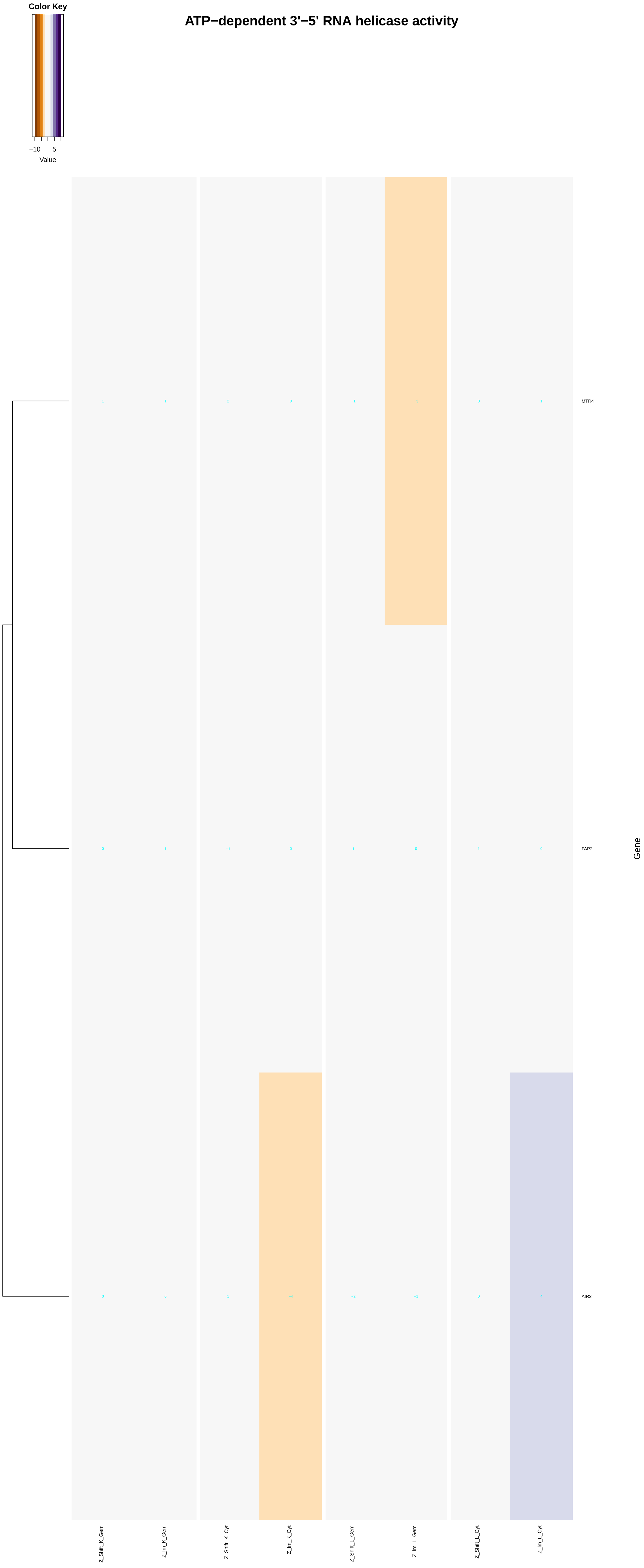

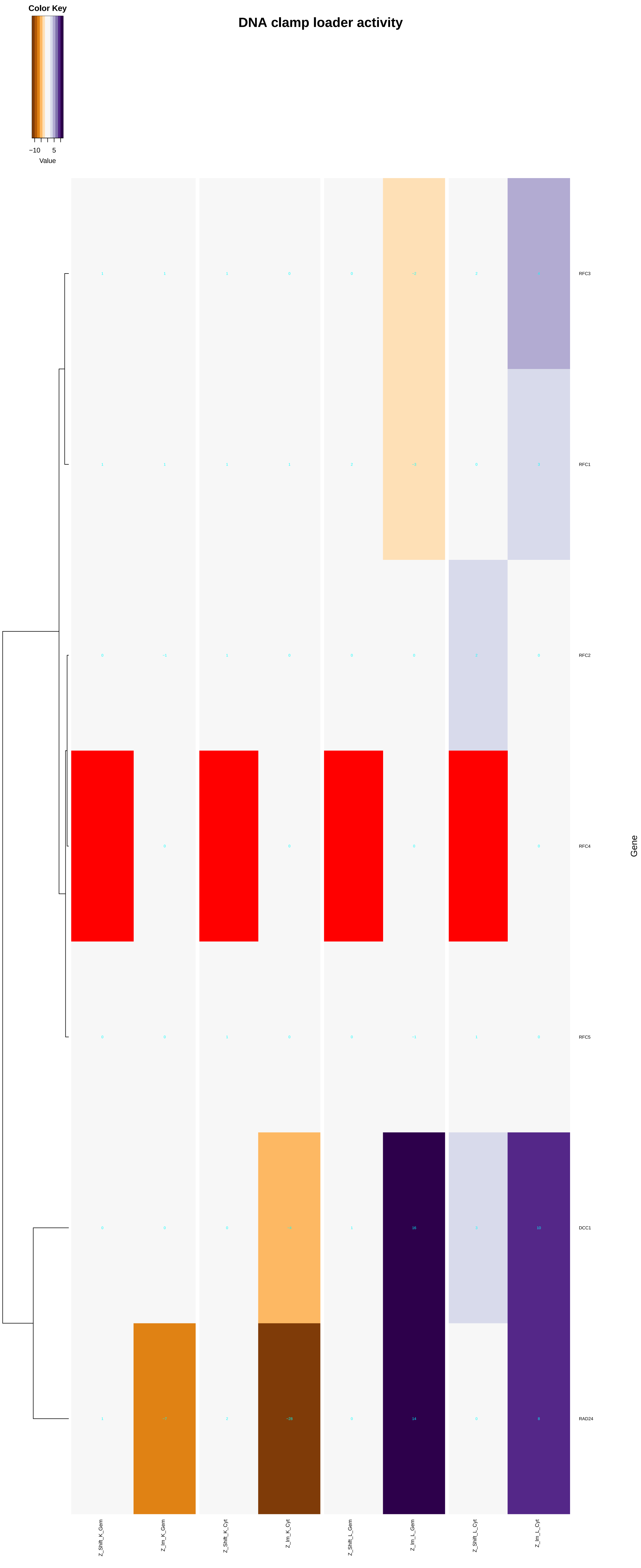
DNA clamp loader activity
Color Key
−10
5
Value
RFC3
RFC1
RFC2
RFC4
RFC5
DCC1
RAD24
Gene
Z_Shift_K_Gem
Z_Im_K_Gem
Z_Shift_K_Cyt
Z_Im_K_Cyt
Z_Shift_L_Gem
Z_Im_L_Gem
Z_Shift_L_Cyt
Z_Im_L_Cyt

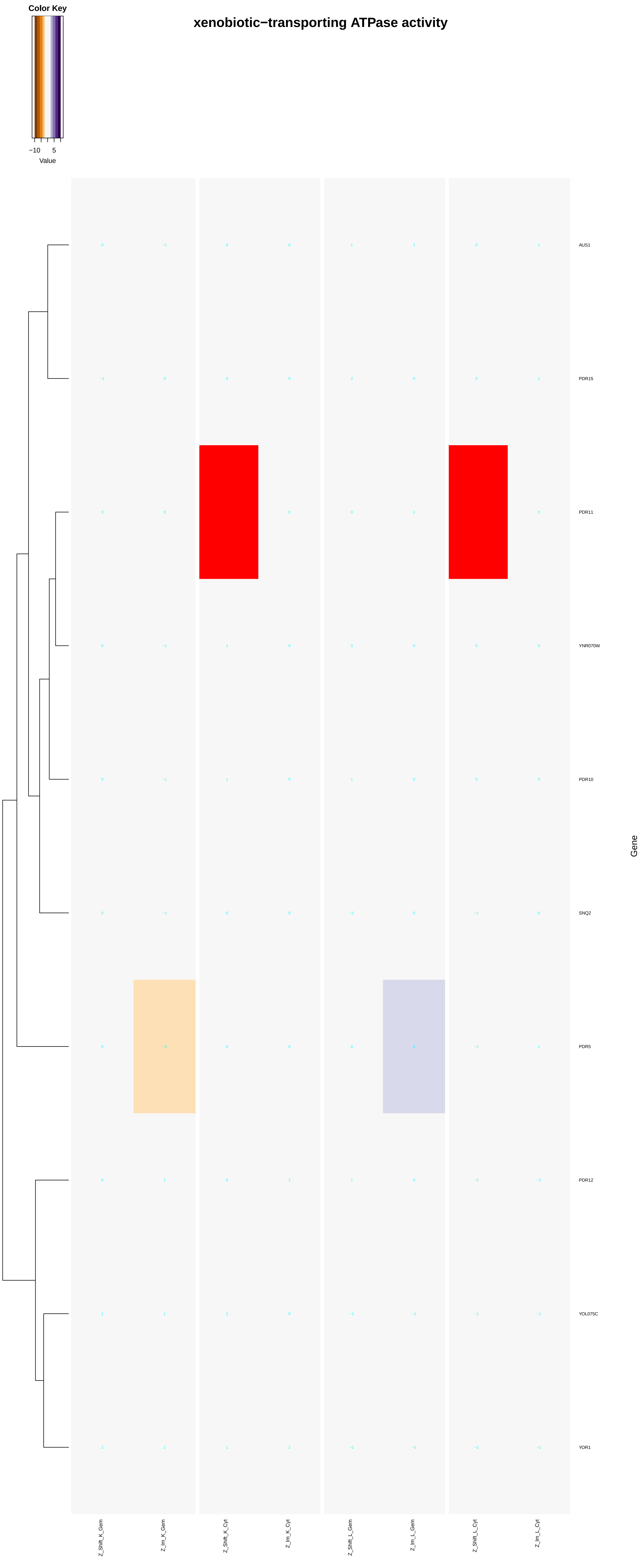
xenobiotic−transporting ATPase activity
Color Key
−10
5
Value
AUS1
PDR15
PDR11
YNR070W
PDR10
SNQ2
PDR5
PDR12
YOL075C
YOR1
Gene
Z_Shift_K_Gem
Z_Im_K_Gem
Z_Shift_K_Cyt
Z_Im_K_Cyt
Z_Shift_L_Gem
Z_Im_L_Gem
Z_Shift_L_Cyt
Z_Im_L_Cyt

### ATPase coupled ion transmembrane transporter activity

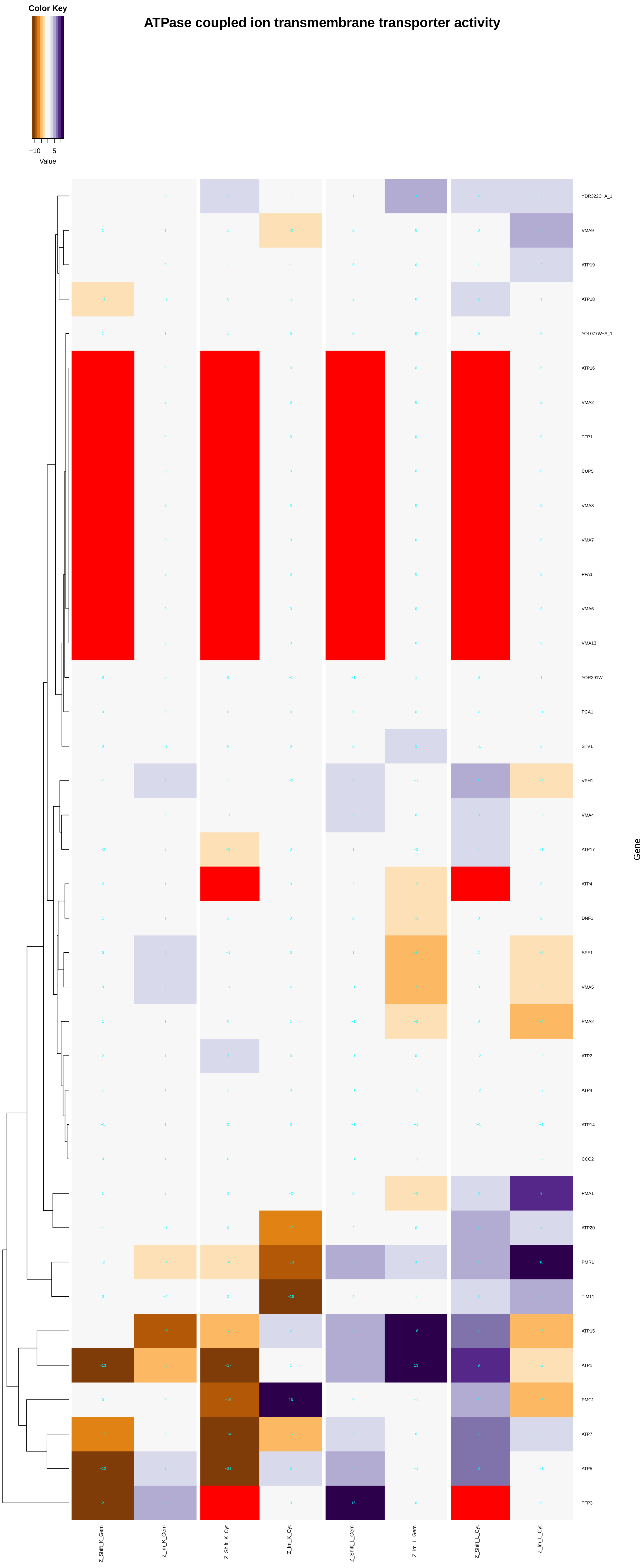

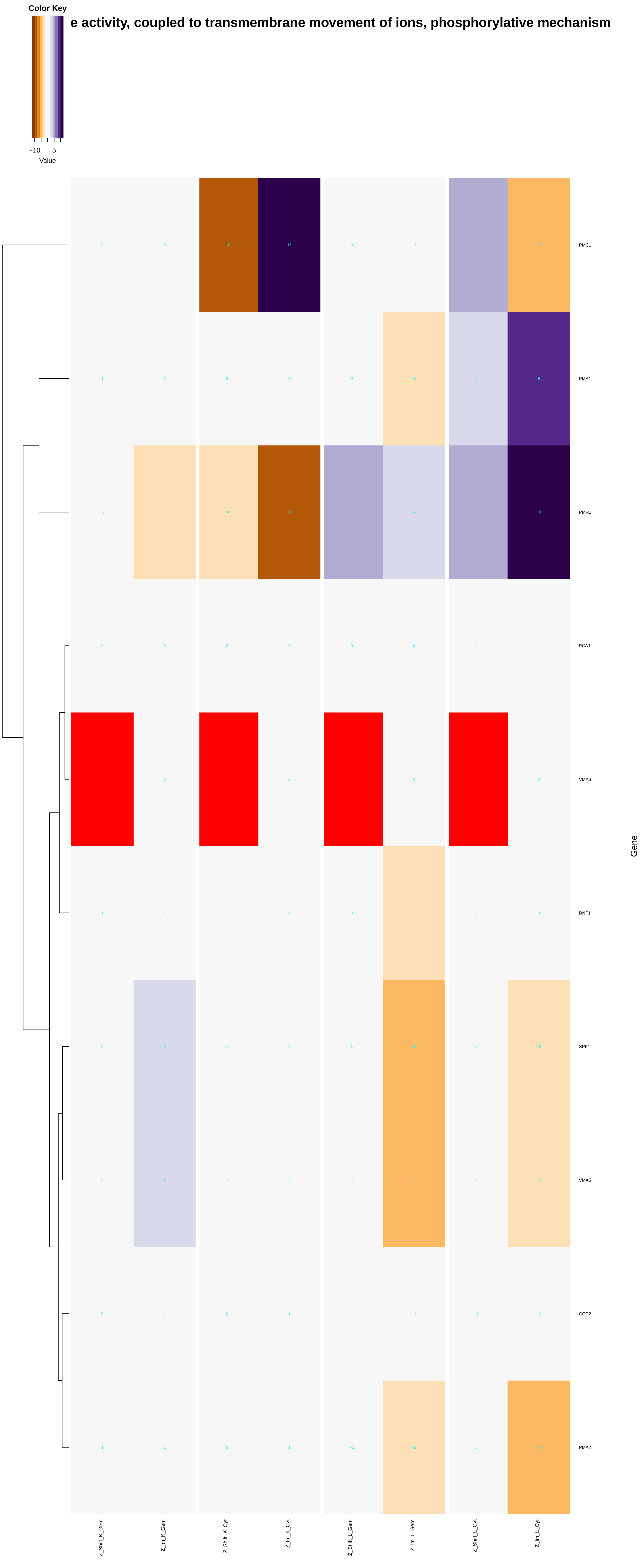
e activity, coupled to transmembrane movement of ions, phosphorylative mechanism
Color Key
−10
5
Value
PMC1
PMA1
PMR1
PCA1
VMA6
DNF1
SPF1
VMA5
CCC2
PMA2
Gene
Z_Shift_K_Gem
Z_Im_K_Gem
Z_Shift_K_Cyt
Z_Im_K_Cyt
Z_Shift_L_Gem
Z_Im_L_Gem
Z_Shift_L_Cyt
Z_Im_L_Cyt

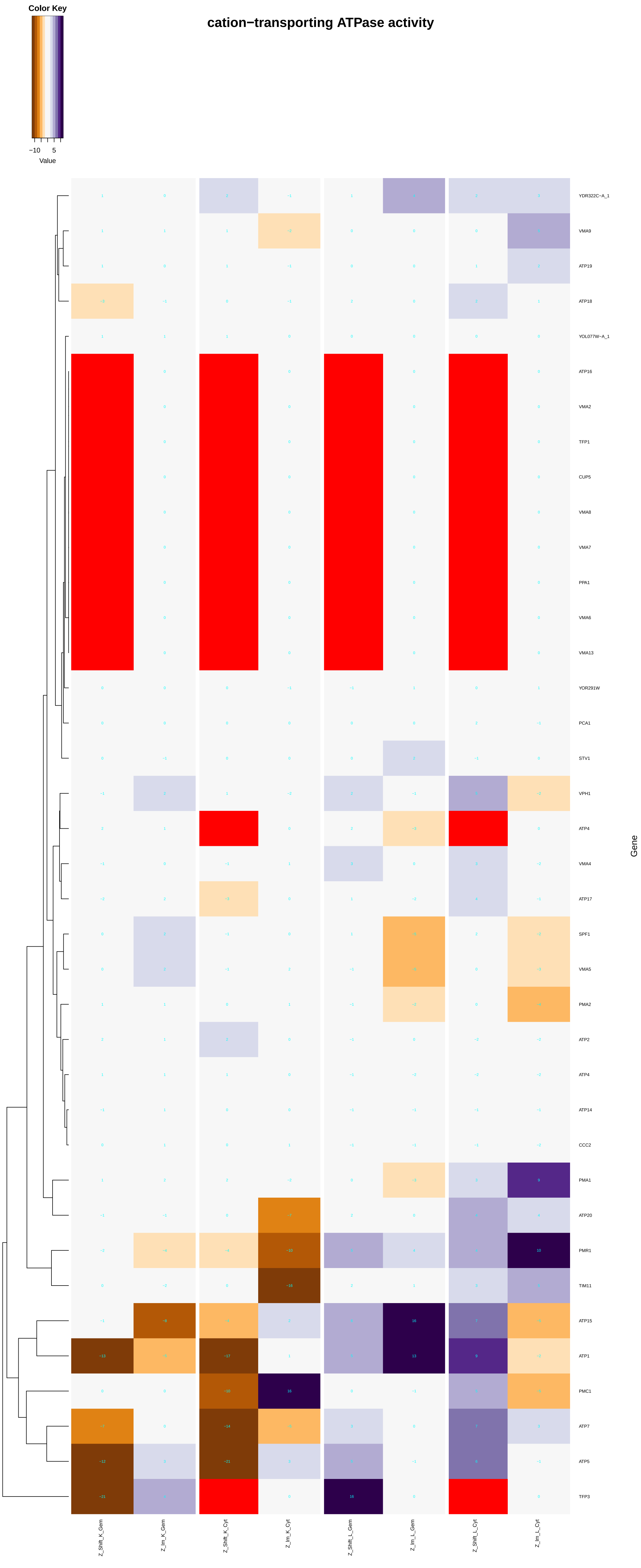
cation−transporting ATPase activity
Color Key
−10
5
Value
YDR322C−A_1
VMA9
ATP19
ATP18
YOL077W−A_1
ATP16
VMA2
TFP1
CUP5
VMA8
VMA7
PPA1
VMA6
VMA13
YOR291W
PCA1
STV1
VPH1
ATP4
VMA4
ATP17
SPF1
VMA5
PMA2
ATP2
ATP4
ATP14
CCC2
PMA1
ATP20
PMR1
TIM11
ATP15
ATP1
PMC1
ATP7
ATP5
TFP3
Gene
Z_Shift_K_Gem
Z_Im_K_Gem
Z_Shift_K_Cyt
Z_Im_K_Cyt
Z_Shift_L_Gem
Z_Im_L_Gem
Z_Shift_L_Cyt
Z_Im_L_Cyt

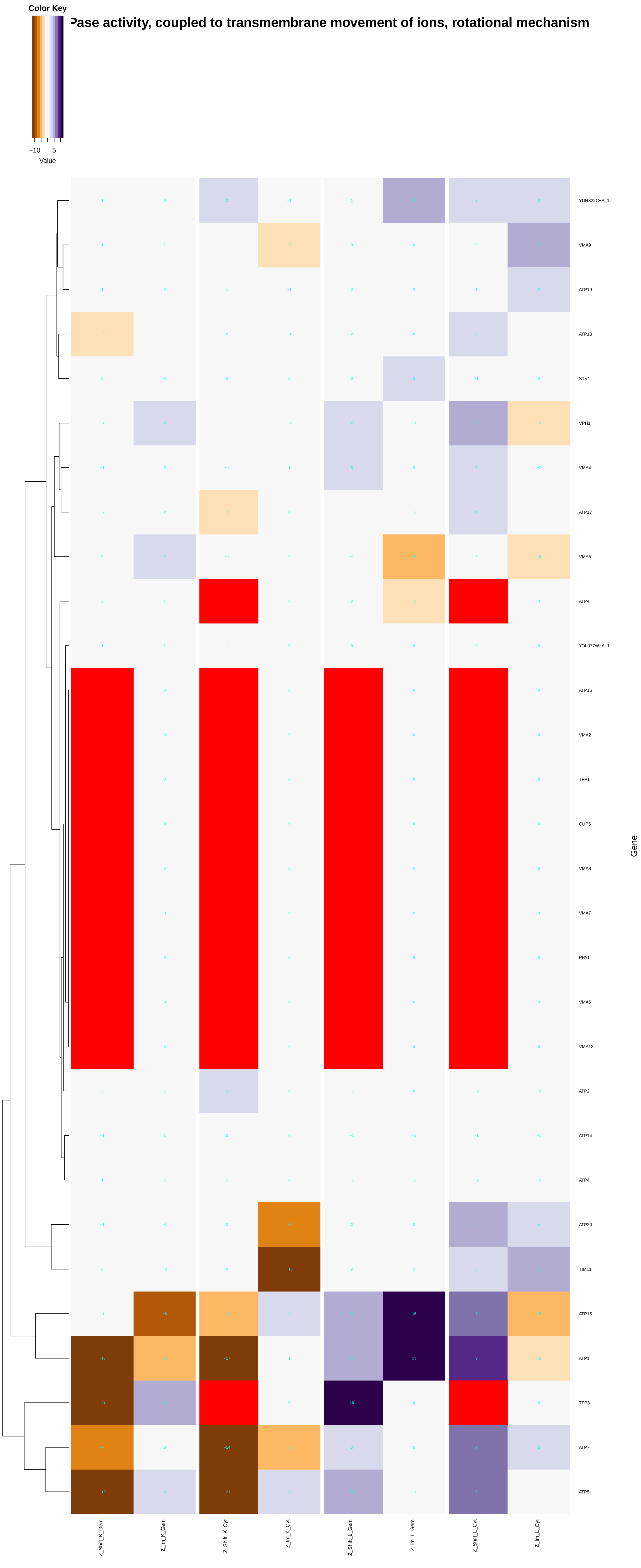
Color Key
−10 5
Value
Pase activity, coupled to transmembrane movement of ions, rotational mechanism
YDR322C−A_1
VMA9
ATP19
ATP18
STV1
VPH1
VMA4
ATP17
VMA5
ATP4
YOL077W−A_1
ATP16
VMA2
TFP1
CUP5
VMA8
VMA7
PPA1
VMA6
VMA13
ATP2
ATP14
ATP4
ATP20
TIM11
ATP15
ATP1
TFP3
ATP7
ATP5
Gene
Z_Shift_K_Gem
Z_Im_K_Gem
Z_Shift_K_Cyt
Z_Im_K_Cyt
Z_Shift_L_Gem
Z_Im_L_Gem
Z_Shift_L_Cyt
Z_Im_L_Cyt

#### hydrogen−exporting ATPase activity, phosphorylative mechanism

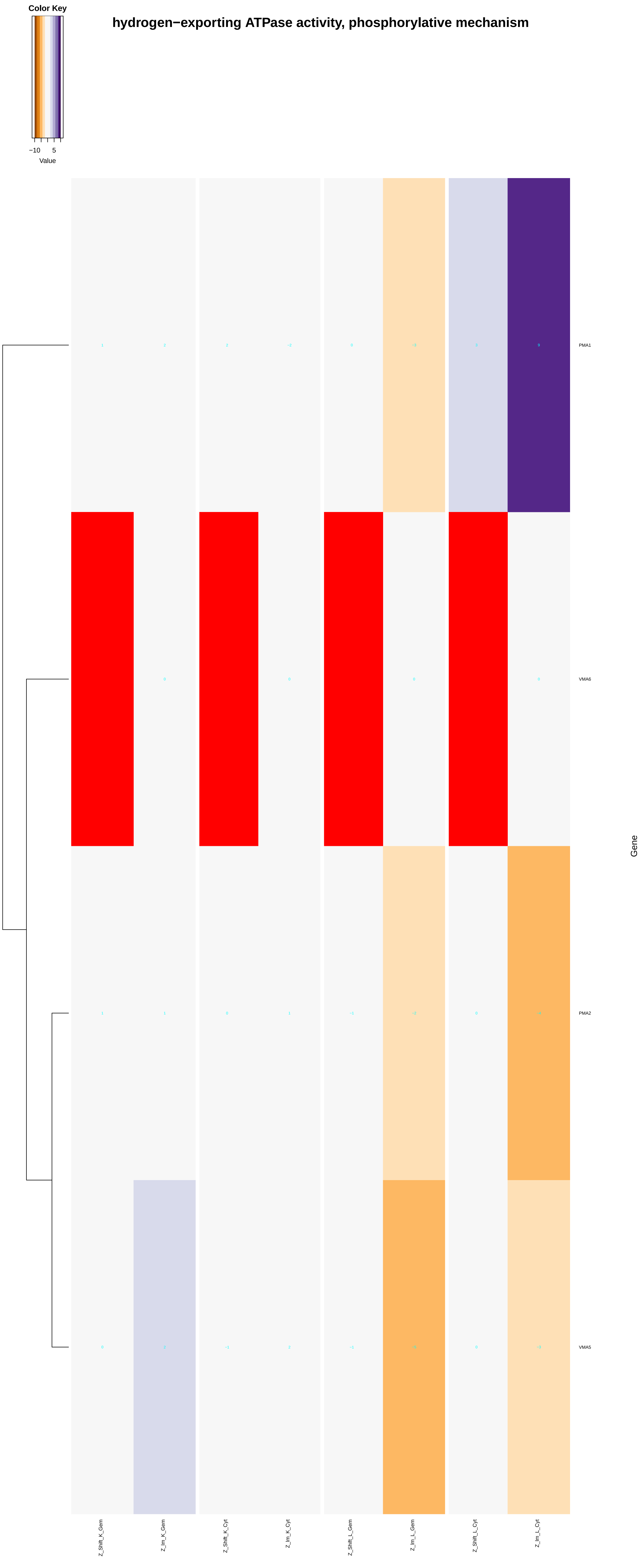

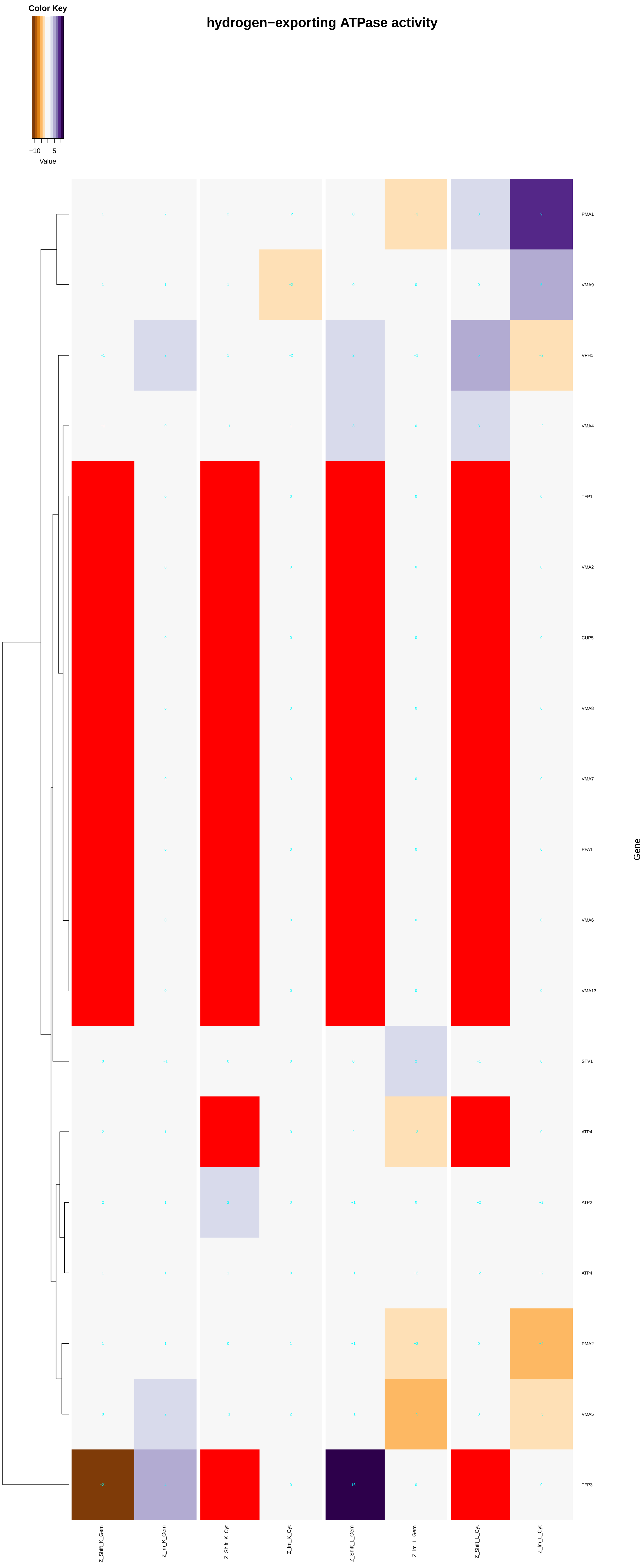

Color Key
−10
5
Value
hydrogen−exporting ATPase activity
PMA1
VMA9
VPH1
VMA4
TFP1
VMA2
CUP5
VMA8
VMA7
PPA1
VMA6
VMA13
STV1
ATP4
ATP2
ATP4
PMA2
VMA5
TFP3
Gene
Z_Shift_K_Gem
Z_Im_K_Gem
Z_Shift_K_Cyt
Z_Im_K_Cyt
Z_Shift_L_Gem
Z_Im_L_Gem
Z_Shift_L_Cyt
Z_Im_L_Cyt

### proton−transporting ATP synthase activity, rotational mechanism

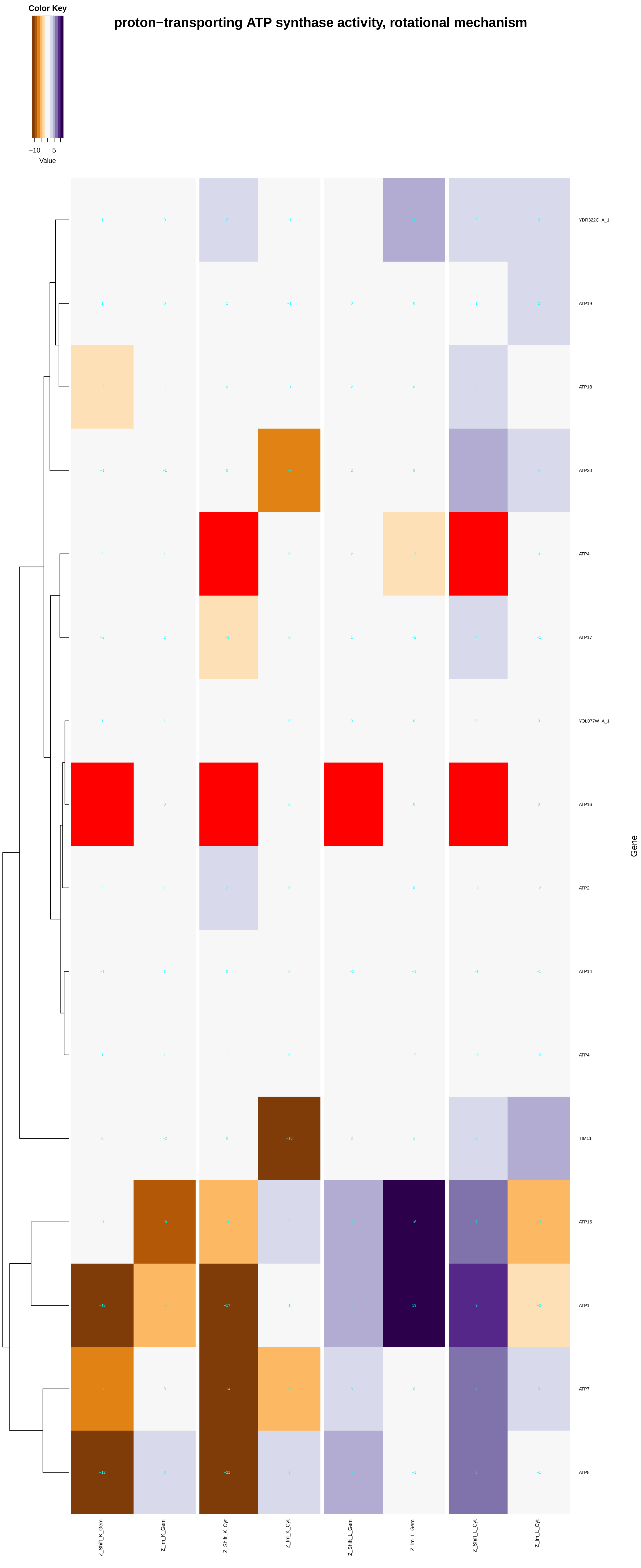

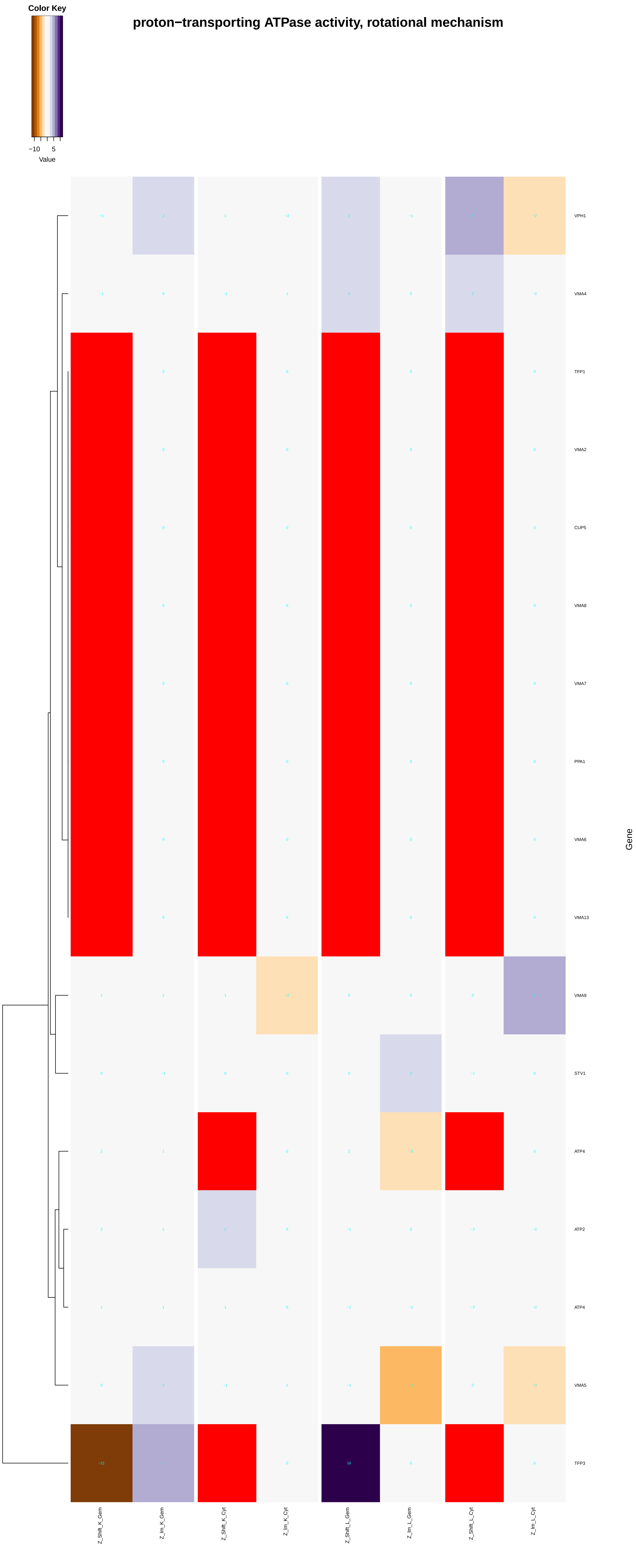
proton−transporting ATPase activity, rotational mechanism
Color Key
−10
5
Value
VPH1
VMA4
TFP1
VMA2
CUP5
VMA8
VMA7
PPA1
VMA6
VMA13
VMA9
STV1
ATP4
ATP2
ATP4
VMA5
TFP3
Gene
Z_Shift_K_Gem
Z_Im_K_Gem
Z_Shift_K_Cyt
Z_Im_K_Cyt
Z_Shift_L_Gem
Z_Im_L_Gem
Z_Shift_L_Cyt
Z_Im_L_Cyt
