## Additional File 7 for "A humanized yeast phenomic model of deoxycytidine kinase to predict genetic buffering of nucleoside analog cytotoxicity": autophagy.pdf

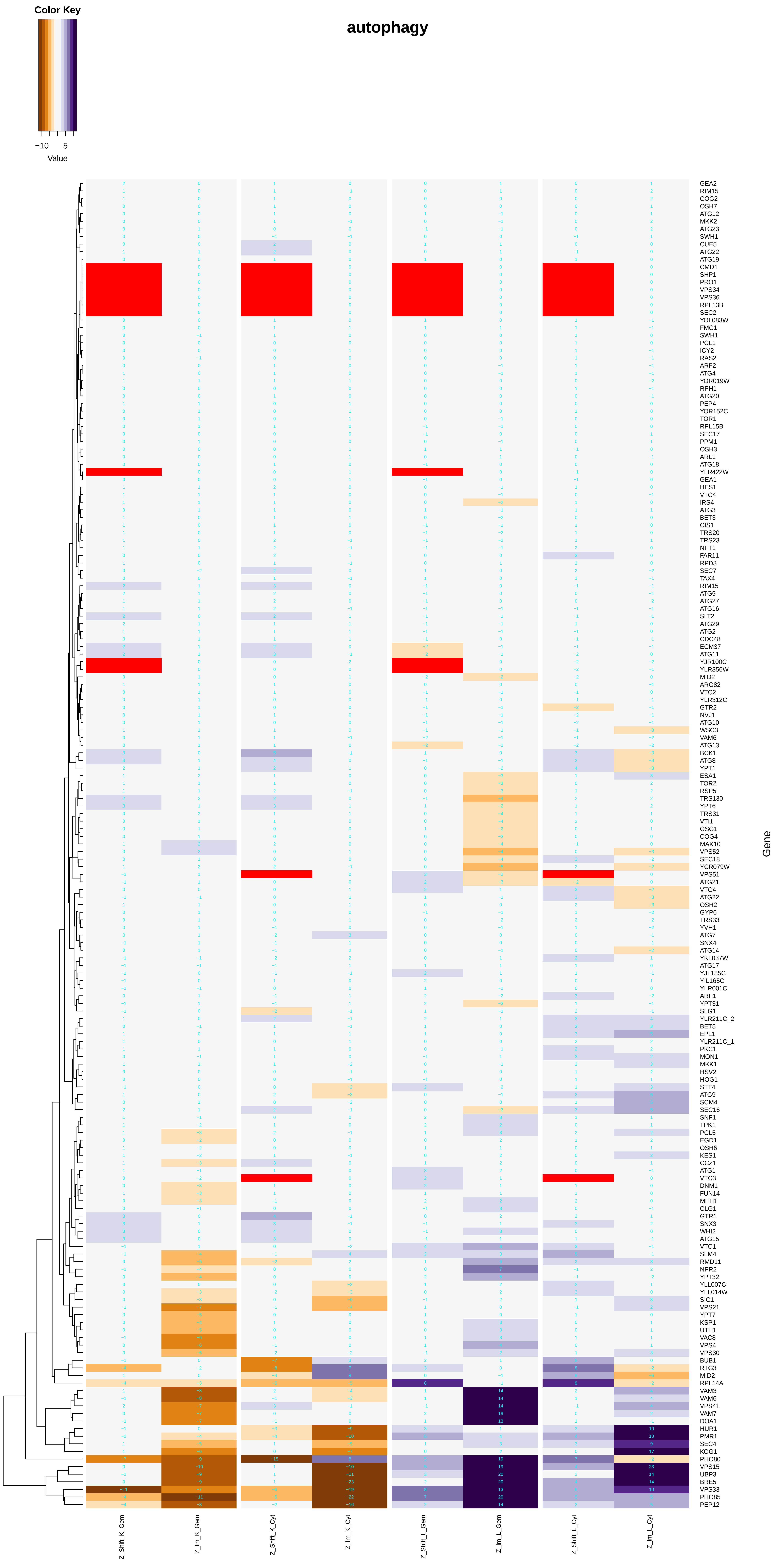

Color Key
−10
5
Value
autophagy
Gene
Z_Shift_K_Gem
Z_Im_K_Gem
Z_Shift_K_Cyt
Z_Im_K_Cyt
Z_Shift_L_Gem
Z_Im_L_Gem
Z_Shift_L_Cyt
Z_Im_L_Cyt
GEA2
RIM15
COG2
OSH7
ATG12
MKK2
ATG23
SWH1
CUE5
ATG22
ATG19
CMD1
SHP1
PRO1
VPS34
VPS36
RPL13B
SEC2
YOL083W
FMC1
SWH1
PCL1
ICY2
RAS2
ARF2
ATG4
YOR019W
RPH1
ATG20
PEP4
YOR152C
TOR1
RPL15B
SEC17
PPM1
OSH3
ARL1
ATG18
YLR422W
GEA1
HES1
VTC4
IRS4
ATG3
BET3
CIS1
TRS20
TRS23
NFT1
FAR11
RPD3
SEC7
TAX4
RIM15
ATG5
ATG27
ATG16
SLT2
ATG29
ATG2
CDC48
ECM37
ATG11
YJR100C
YLR356W
MID2
ARG82
VTC2
YLR312C
GTR2
NVJ1
ATG10
WSC3
VAM6
ATG13
BCK1
ATG8
YPT1
ESA1
TOR2
RSP5
TRS130
YPT6
TRS31
VTI1
GSG1
COG4
MAK10
VPS52
SEC18
YCR079W
VPS51
ATG21
VTC4
ATG22
OSH2
GYP6
TRS33
YVH1
ATG7
SNX4
ATG14
YKL037W
ATG17
YJL185C
YIL165C
YLR001C
ARF1
YPT31
SLG1
YLR211C_2
BET5
EPL1
YLR211C_1
PKC1
MON1
MKK1
HSV2
HOG1
STT4
ATG9
SCM4
SEC16
SNF1
TPK1
PCL5
EGD1
OSH6
KES1
CCZ1
ATG1
VTC3
DNM1
FUN14
MEH1
CLG1
GTR1
SNX3
WHI2
ATG15
VTC1
SLM4
RMD11
NPR2
YPT32
YLL007C
YLL014W
SIC1
VPS21
YPT7
KSP1
UTH1
VAC8
VPS4
VPS30
BUB1
RTG3
MID2
RPL14A
VAM3
VAM6
VPS41
VAM7
DOA1
HUR1
PMR1
SEC4
KOG1
PHO80
VPS15
UBP3
BRE5
VPS33
PHO85
PEP12

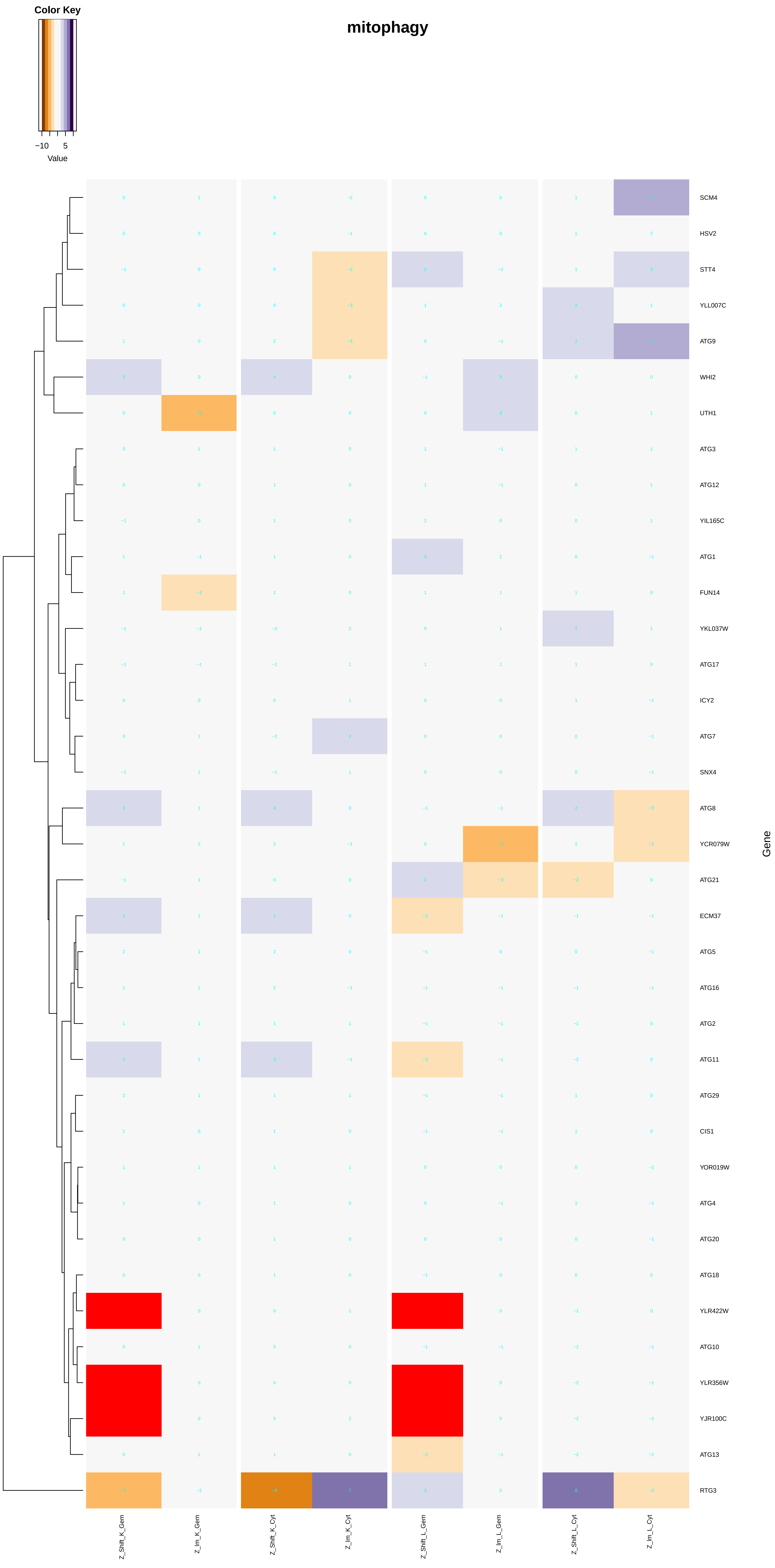

Color Key
−10
5
Value
mitophagy
SCM4
HSV2
STT4
YLL007C
ATG9
WHI2
UTH1
ATG3
ATG12
YIL165C
ATG1
FUN14
YKL037W
ATG17
ICY2
ATG7
SNX4
ATG8
YCR079W
ATG21
ECM37
ATG5
ATG16
ATG2
ATG11
ATG29
CIS1
YOR019W
ATG4
ATG20
ATG18
YLR422W
ATG10
YLR356W
YJR100C
ATG13
RTG3
Gene
Z_Shift_K_Gem
Z_Im_K_Gem
Z_Shift_K_Cyt
Z_Im_K_Cyt
Z_Shift_L_Gem
Z_Im_L_Gem
Z_Shift_L_Cyt
Z_Im_L_Cyt

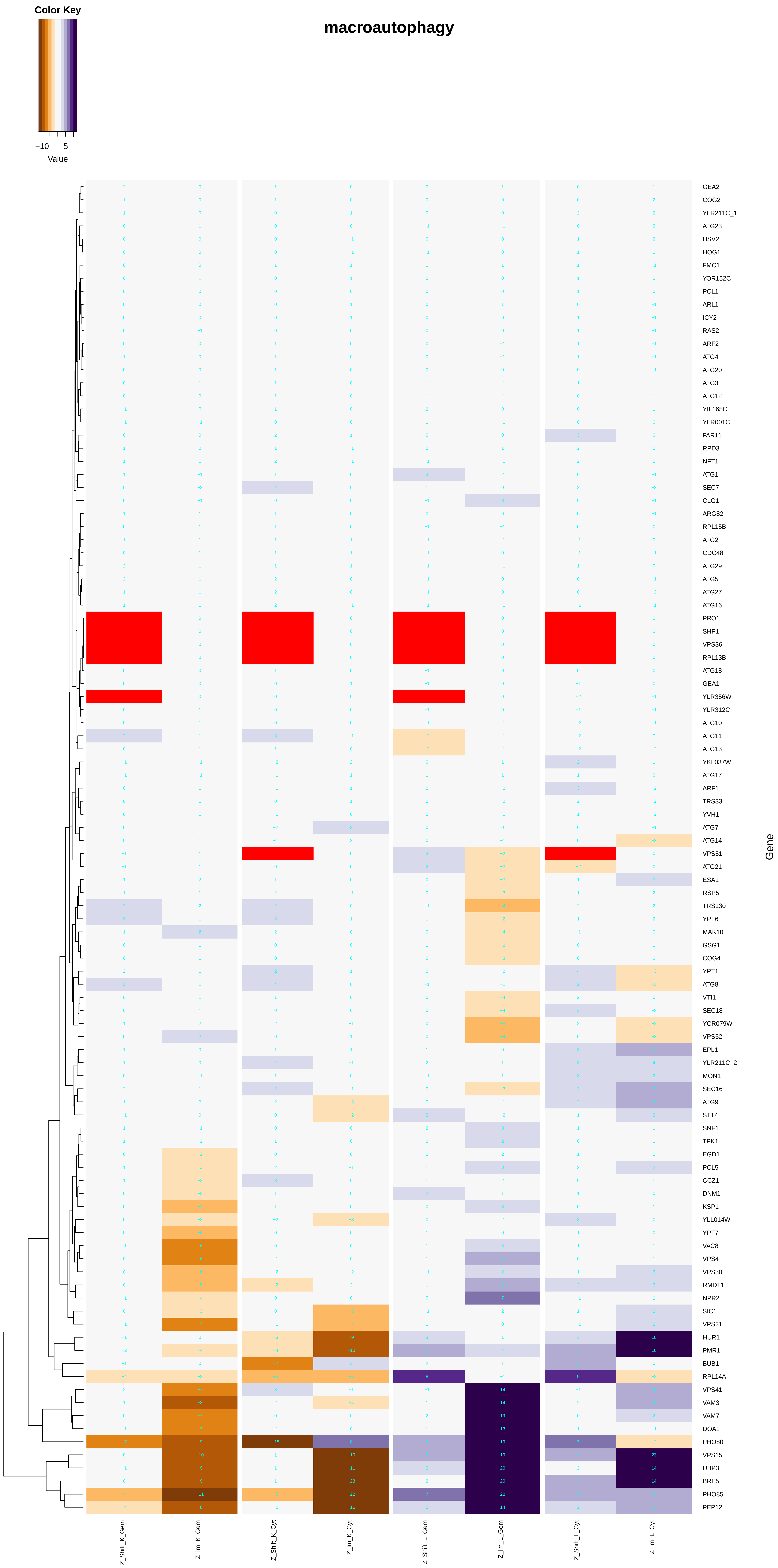
macroautophagy
Color Key
−10
5
Value
Gene
GEA2
COG2
YLR211C_1
ATG23
HSV2
HOG1
FMC1
YOR152C
PCL1
ARL1
ICY2
RAS2
ARF2
ATG4
ATG20
ATG3
ATG12
YIL165C
YLR001C
FAR11
RPD3
NFT1
ATG1
SEC7
CLG1
ARG82
RPL15B
ATG2
CDC48
ATG29
ATG5
ATG27
ATG16
PRO1
SHP1
VPS36
RPL13B
ATG18
GEA1
YLR356W
YLR312C
ATG10
ATG11
ATG13
YKL037W
ATG17
ARF1
TRS33
YVH1
ATG7
ATG14
VPS51
ATG21
ESA1
RSP5
TRS130
YPT6
MAK10
GSG1
COG4
YPT1
ATG8
VTI1
SEC18
YCR079W
VPS52
EPL1
YLR211C_2
MON1
SEC16
ATG9
STT4
SNF1
TPK1
EGD1
PCL5
CCZ1
DNM1
KSP1
YLL014W
YPT7
VAC8
VPS4
VPS30
RMD11
NPR2
SIC1
VPS21
HUR1
PMR1
BUB1
RPL14A
VPS41
VAM3
VAM7
DOA1
PHO80
VPS15
UBP3
BRE5
PHO85
PEP12
Z_Shift_K_Gem
Z_Im_K_Gem
Z_Shift_K_Cyt
Z_Im_K_Cyt
Z_Shift_L_Gem
Z_Im_L_Gem
Z_Shift_L_Cyt
Z_Im_L_Cyt

### lysosomal microautophagy

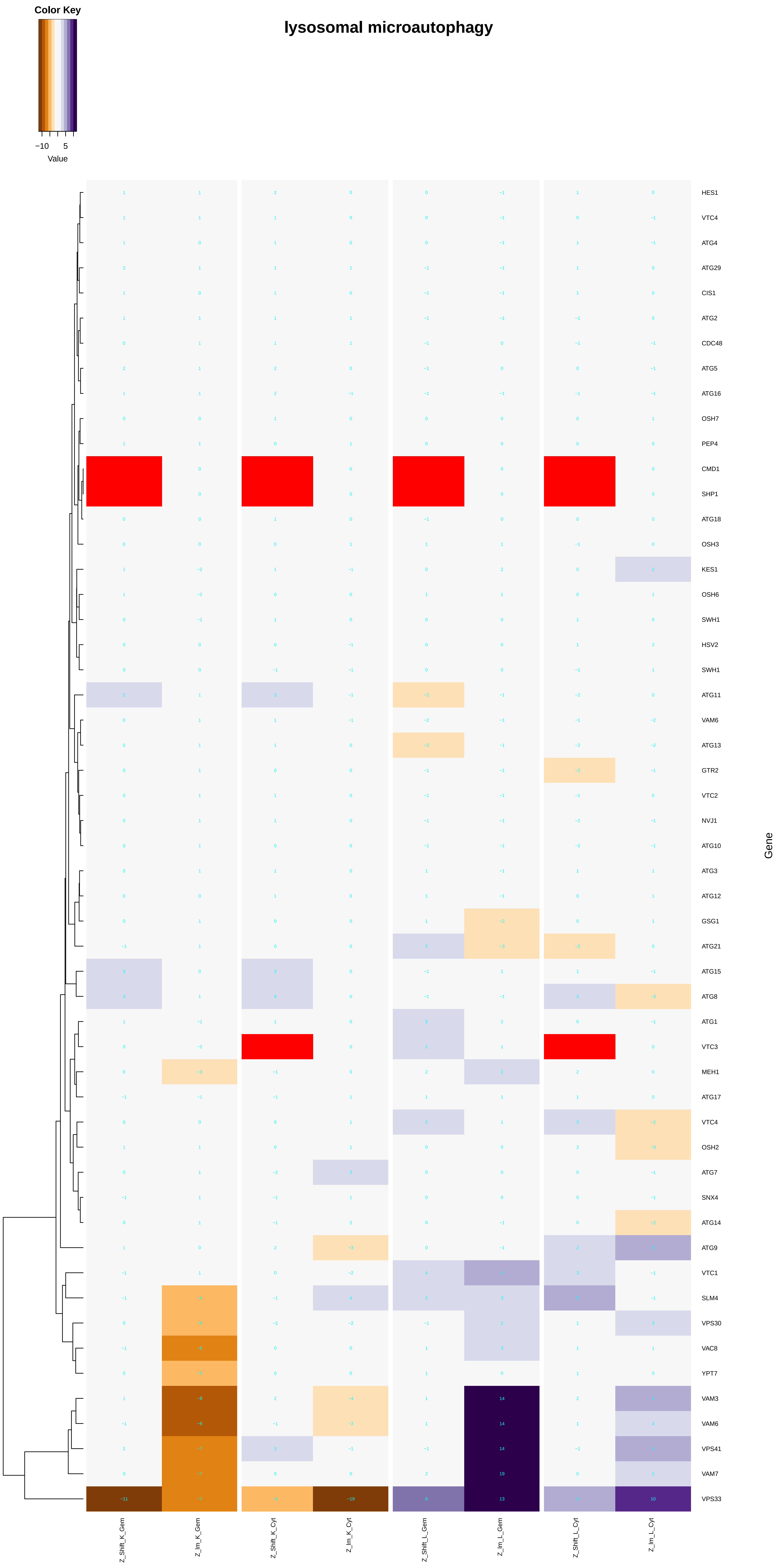

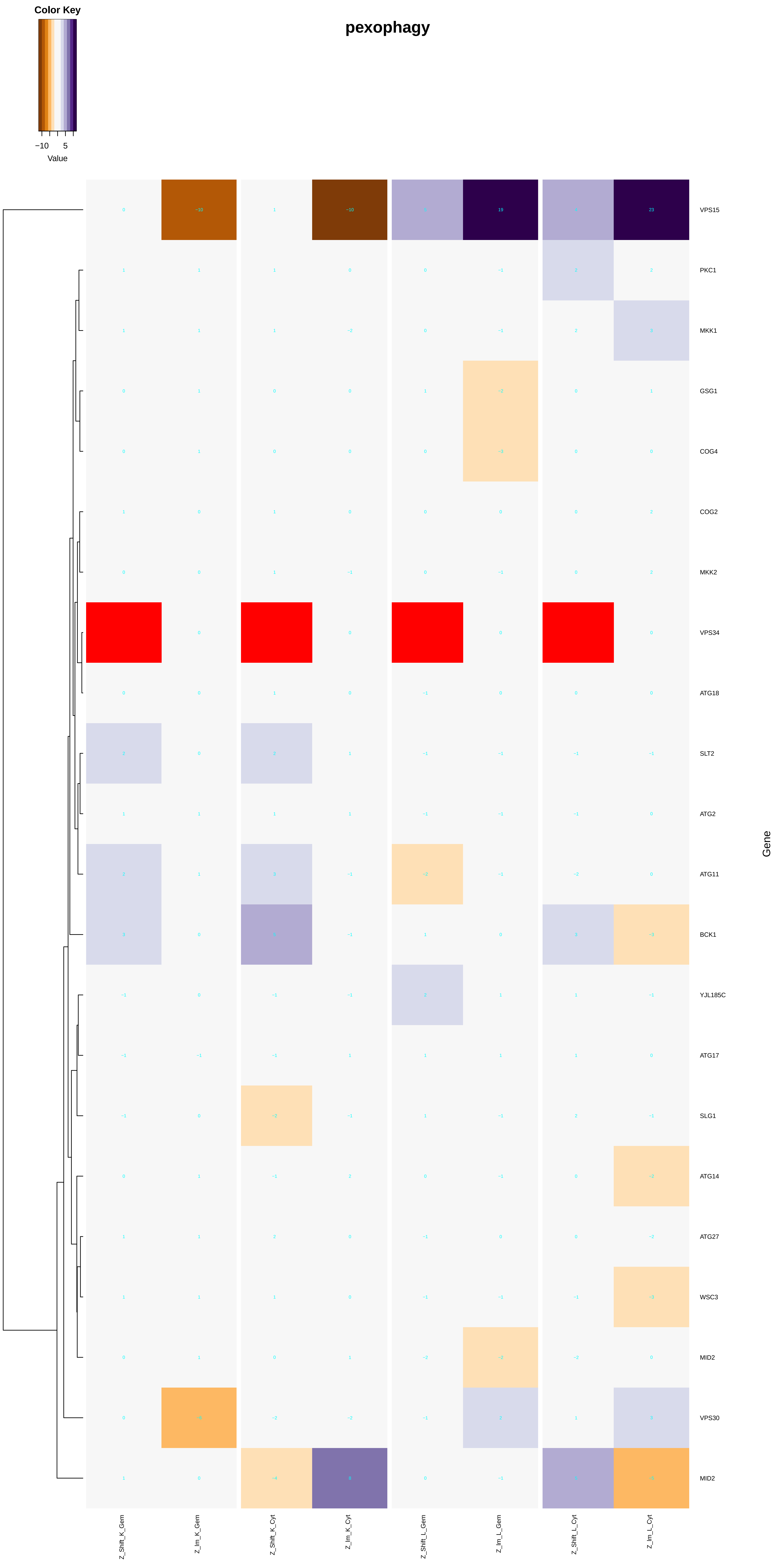
pexophagy
Color Key
−10
5
Value
VPS15
PKC1
MKK1
GSG1
COG4
COG2
MKK2
VPS34
ATG18
SLT2
ATG2
ATG11
BCK1
YJL185C
ATG17
SLG1
ATG14
ATG27
WSC3
MID2
VPS30
MID2
Gene
Z_Shift_K_Gem
Z_Im_K_Gem
Z_Shift_K_Cyt
Z_Im_K_Cyt
Z_Shift_L_Gem
Z_Im_L_Gem
Z_Shift_L_Cyt
Z_Im_L_Cyt

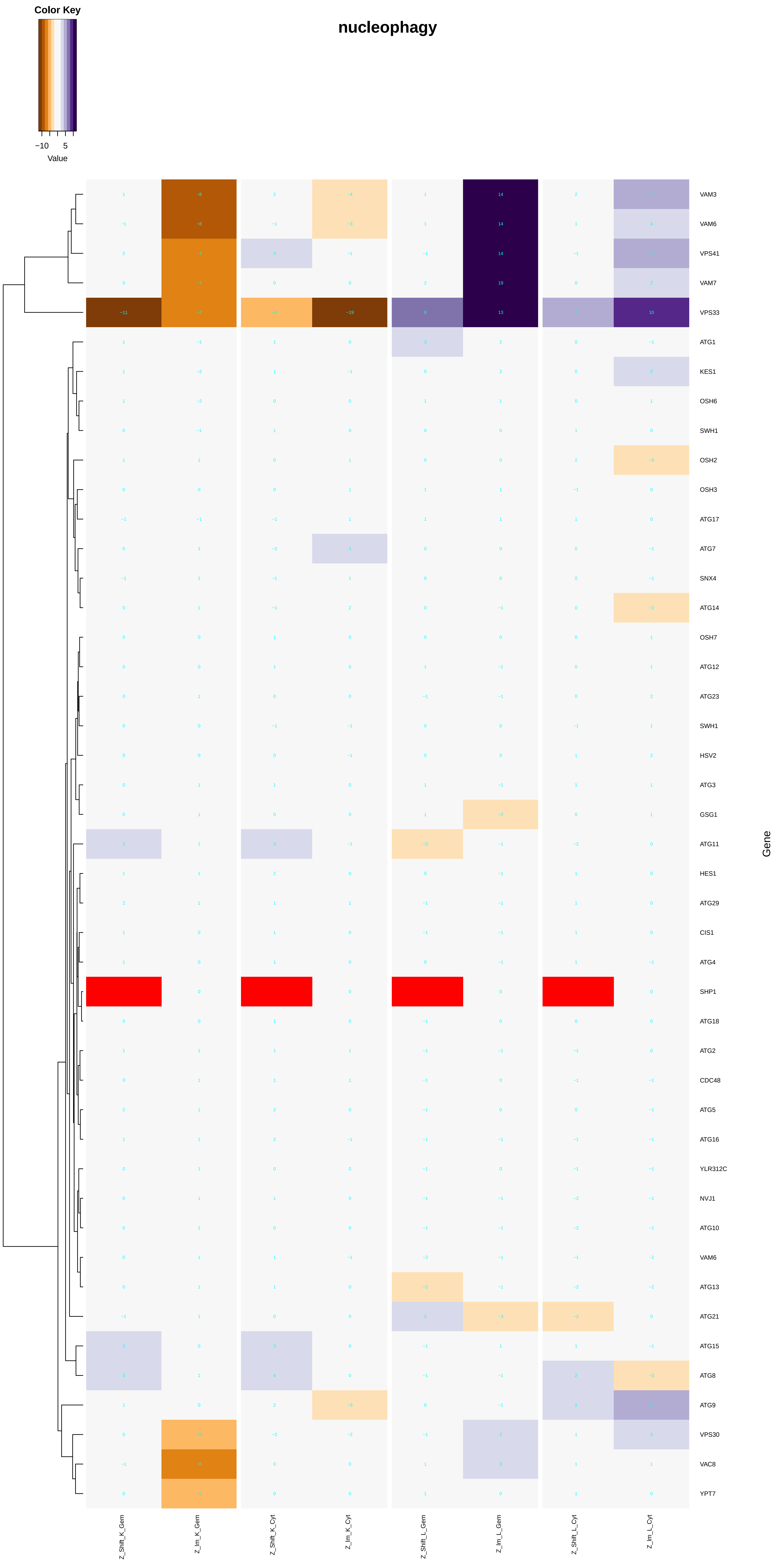
nucleophagy
Color Key
−10
5
Value
VAM3
VAM6
VPS41
VAM7
VPS33
ATG1
KES1
OSH6
SWH1
OSH2
OSH3
ATG17
ATG7
SNX4
ATG14
OSH7
ATG12
ATG23
SWH1
HSV2
ATG3
GSG1
ATG11
HES1
ATG29
CIS1
ATG4
SHP1
ATG18
ATG2
CDC48
ATG5
ATG16
YLR312C
NVJ1
ATG10
VAM6
ATG13
ATG21
ATG15
ATG8
ATG9
VPS30
VAC8
YPT7
Gene
Z_Shift_K_Gem
Z_Im_K_Gem
Z_Shift_K_Cyt
Z_Im_K_Cyt
Z_Shift_L_Gem
Z_Im_L_Gem
Z_Shift_L_Cyt
Z_Im_L_Cyt

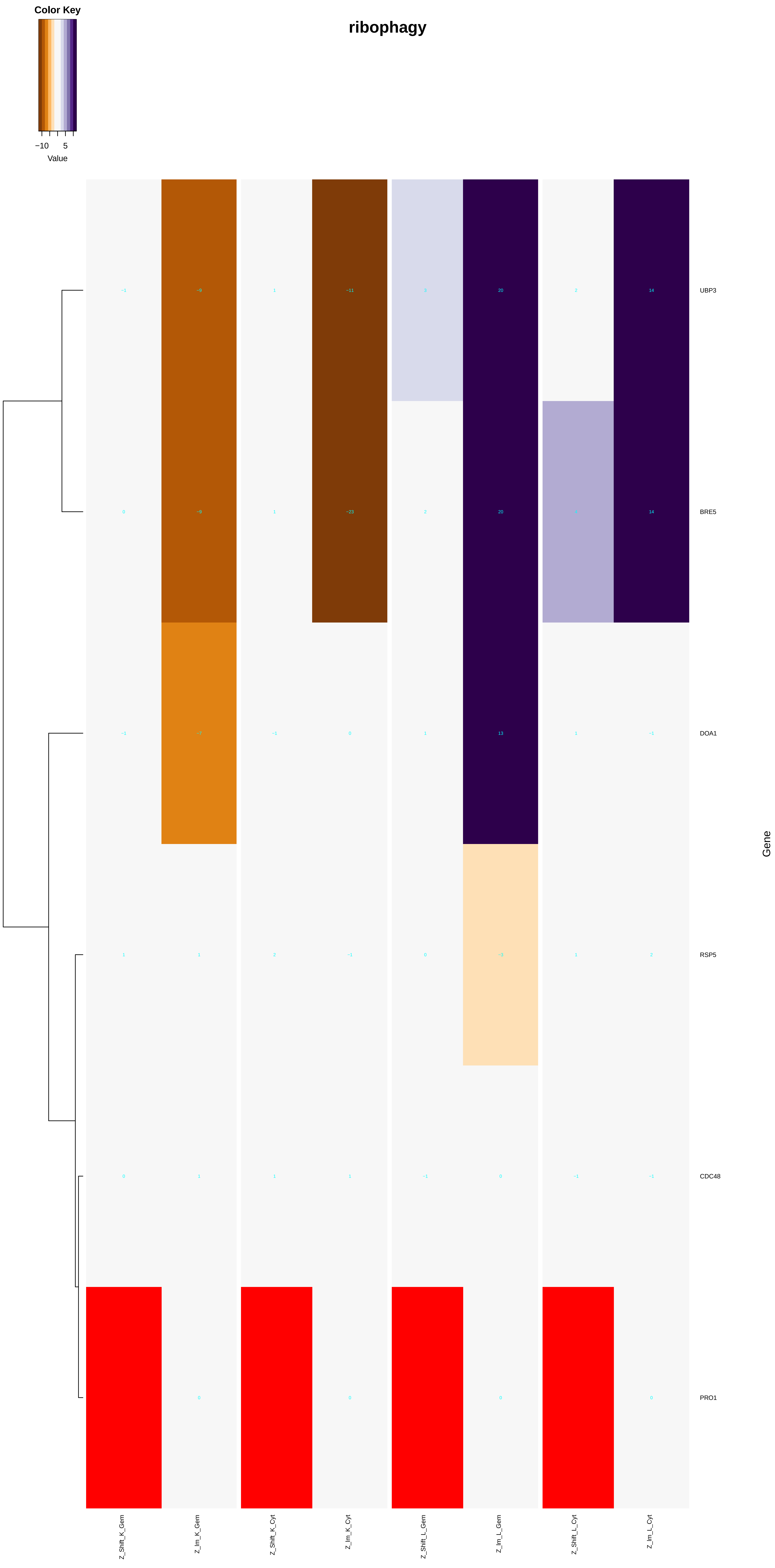
Color Key
−10
5
Value
ribophagy
UBP3
BRE5
DOA1
RSP5
CDC48
PRO1
Gene
Z_Shift_K_Gem
Z_Im_K_Gem
Z_Shift_K_Cyt
Z_Im_K_Cyt
Z_Shift_L_Gem
Z_Im_L_Gem
Z_Shift_L_Cyt
Z_Im_L_Cyt

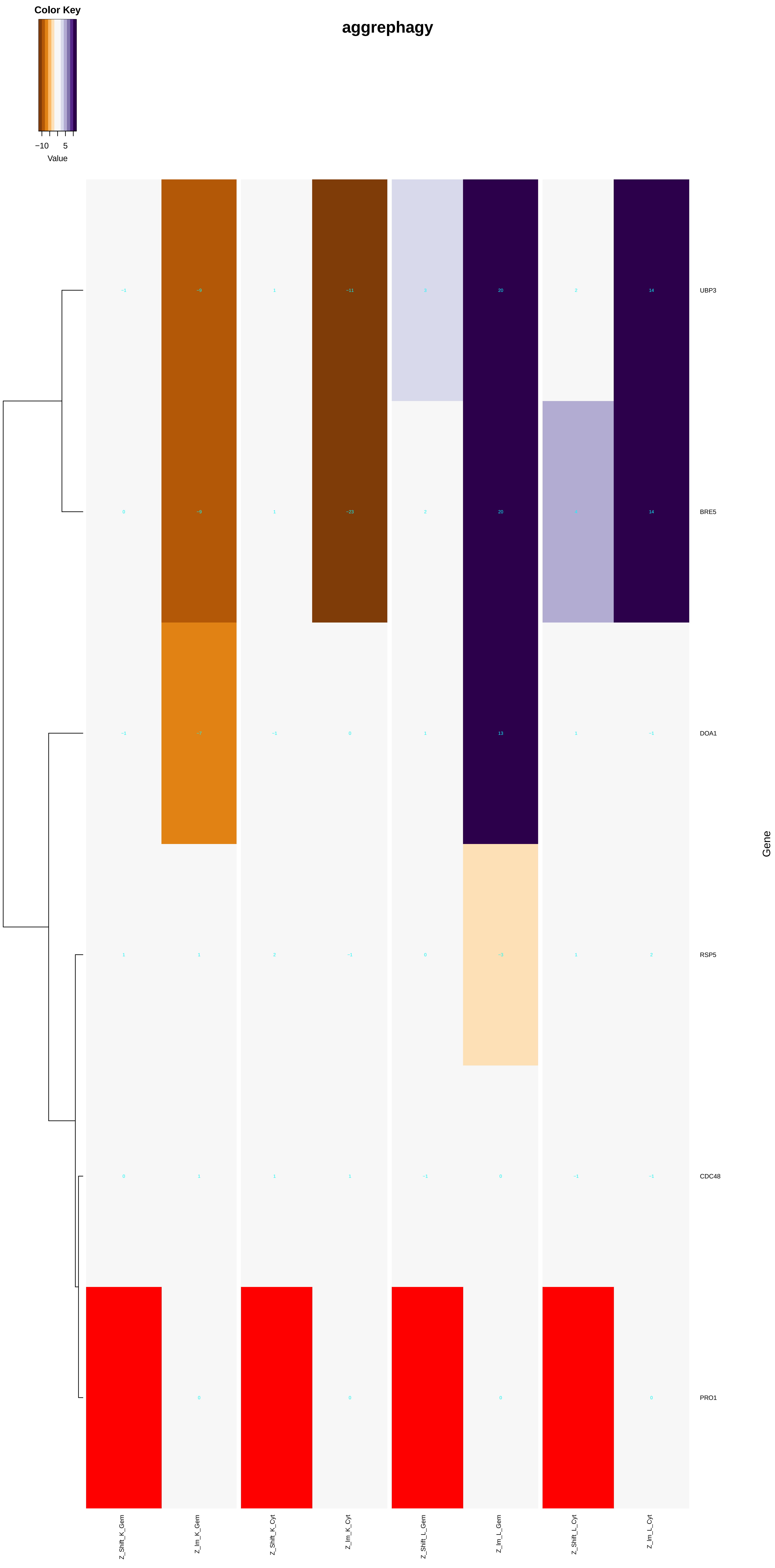

Color Key
−10
5
Value
aggrephagy
UBP3
BRE5
DOA1
RSP5
CDC48
PRO1
Gene
Z_Shift_K_Gem
Z_Im_K_Gem
Z_Shift_K_Cyt
Z_Im_K_Cyt
Z_Shift_L_Gem
Z_Im_L_Gem
Z_Shift_L_Cyt
Z_Im_L_Cyt

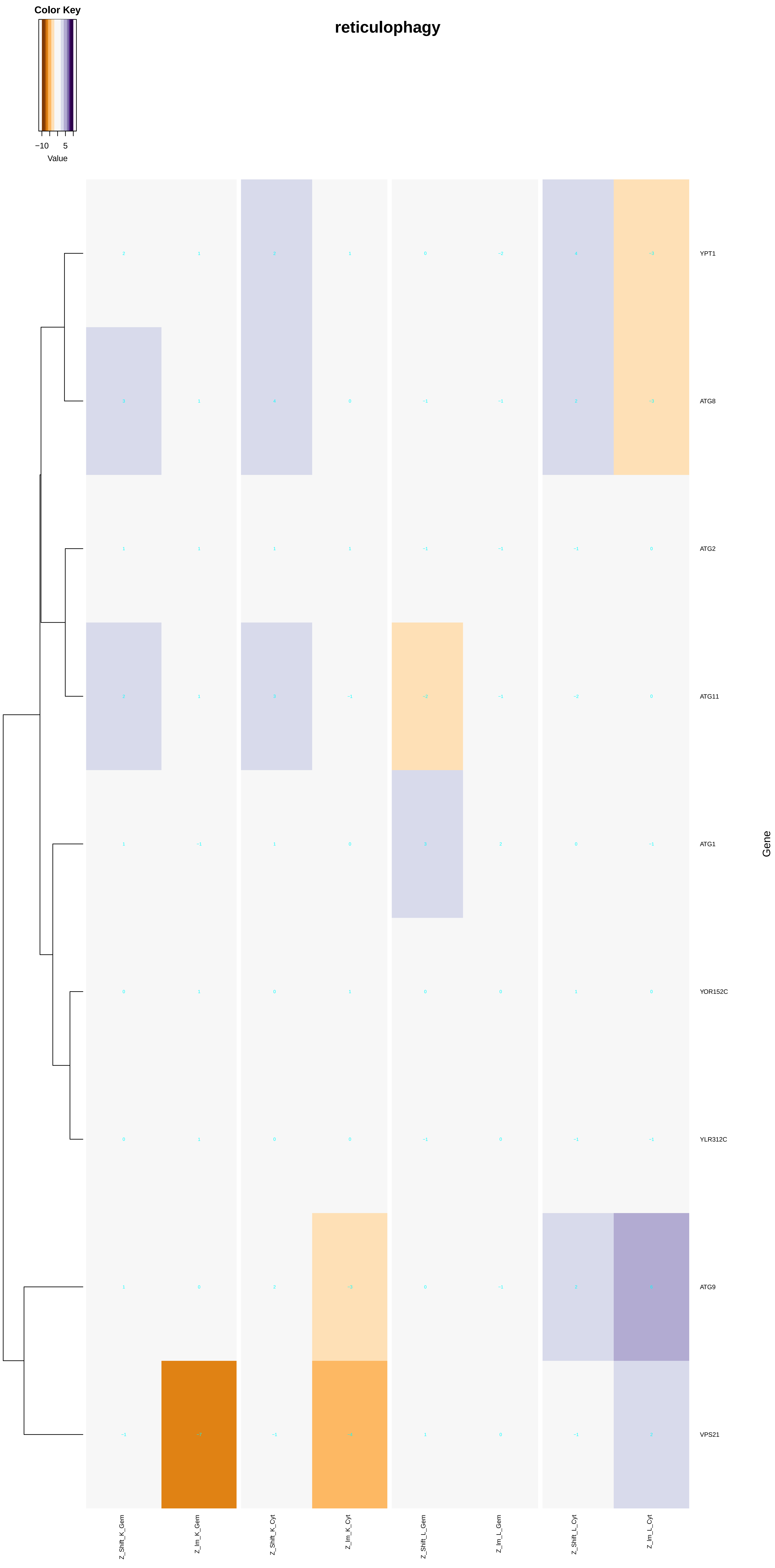

Color Key
−10
5
Value
reticulophagy
YPT1
ATG8
ATG2
ATG11
ATG1
YOR152C
YLR312C
ATG9
VPS21
Gene
Z_Shift_K_Gem
Z_Im_K_Gem
Z_Shift_K_Cyt
Z_Im_K_Cyt
Z_Shift_L_Gem
Z_Im_L_Gem
Z_Shift_L_Cyt
Z_Im_L_Cyt

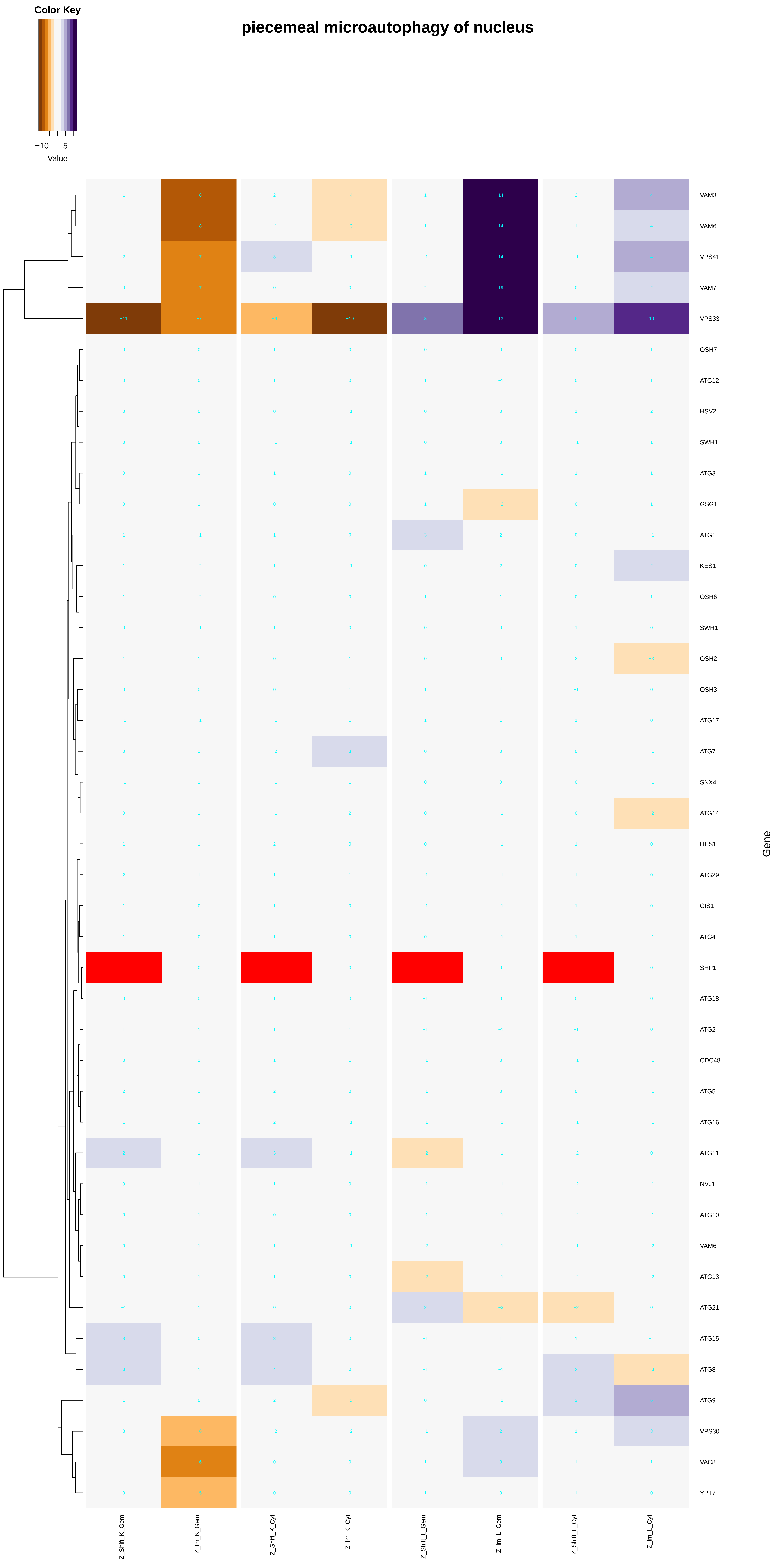

Color Key
−10
5
Value
piecemeal microautophagy of nucleus
VAM3
VAM6
VPS41
VAM7
VPS33
OSH7
ATG12
HSV2
SWH1
ATG3
GSG1
ATG1
KES1
OSH6
SWH1
OSH2
OSH3
ATG17
ATG7
SNX4
ATG14
HES1
ATG29
CIS1
ATG4
SHP1
ATG18
ATG2
CDC48
ATG5
ATG16
ATG11
NVJ1
ATG10
VAM6
ATG13
ATG21
ATG15
ATG8
ATG9
VPS30
VAC8
YPT7
Gene
Z_Shift_K_Gem
Z_lm_K_Gem
Z_Shift_K_Cyt
Z_lm_K_Cyt
Z_Shift_L_Gem
Z_lm_L_Gem
Z_Shift_L_Cyt
Z_lm_L_Cyt

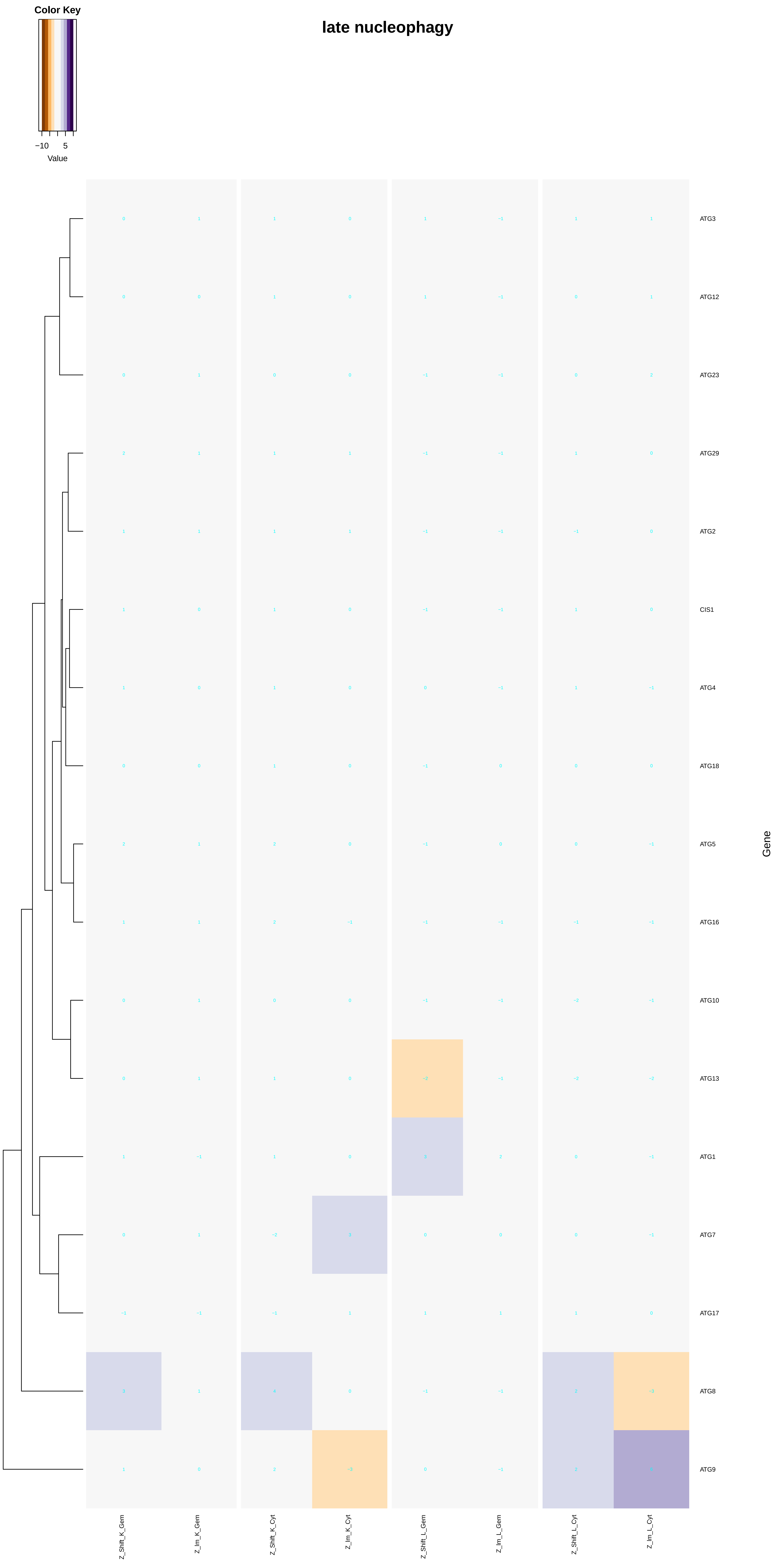
Color Key
−10
5
Value
late nucleophagy
ATG3
ATG12
ATG23
ATG29
ATG2
CIS1
ATG4
ATG18
ATG5
ATG16
ATG10
ATG13
ATG1
ATG7
ATG17
ATG8
ATG9
Gene
Z_Shift_K_Gem
Z_Im_K_Gem
Z_Shift_K_Cyt
Z_Im_K_Cyt
Z_Shift_L_Gem
Z_Im_L_Gem
Z_Shift_L_Cyt
Z_Im_L_Cyt
