## Additional File 7 for "A humanized yeast phenomic model of deoxycytidine kinase to predict genetic buffering of nucleoside analog cytotoxicity": carbohydrate_derivative_binding.pdf

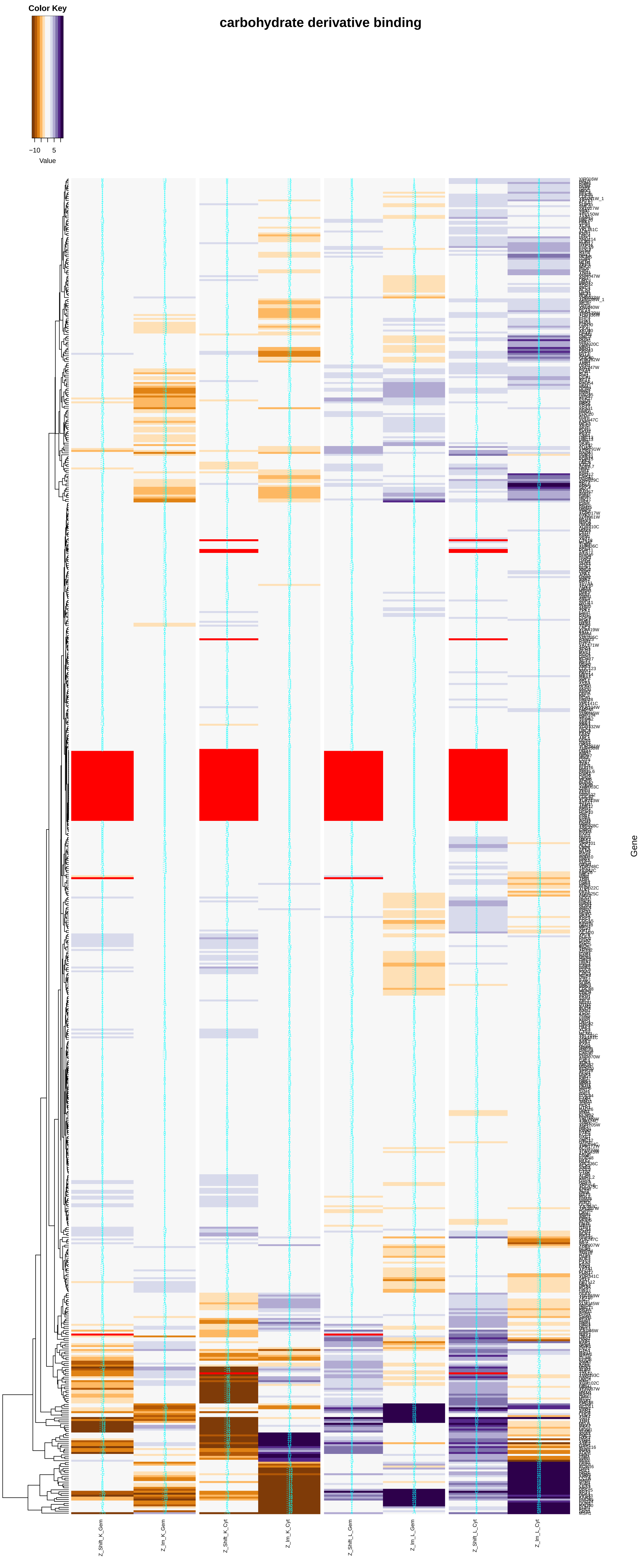

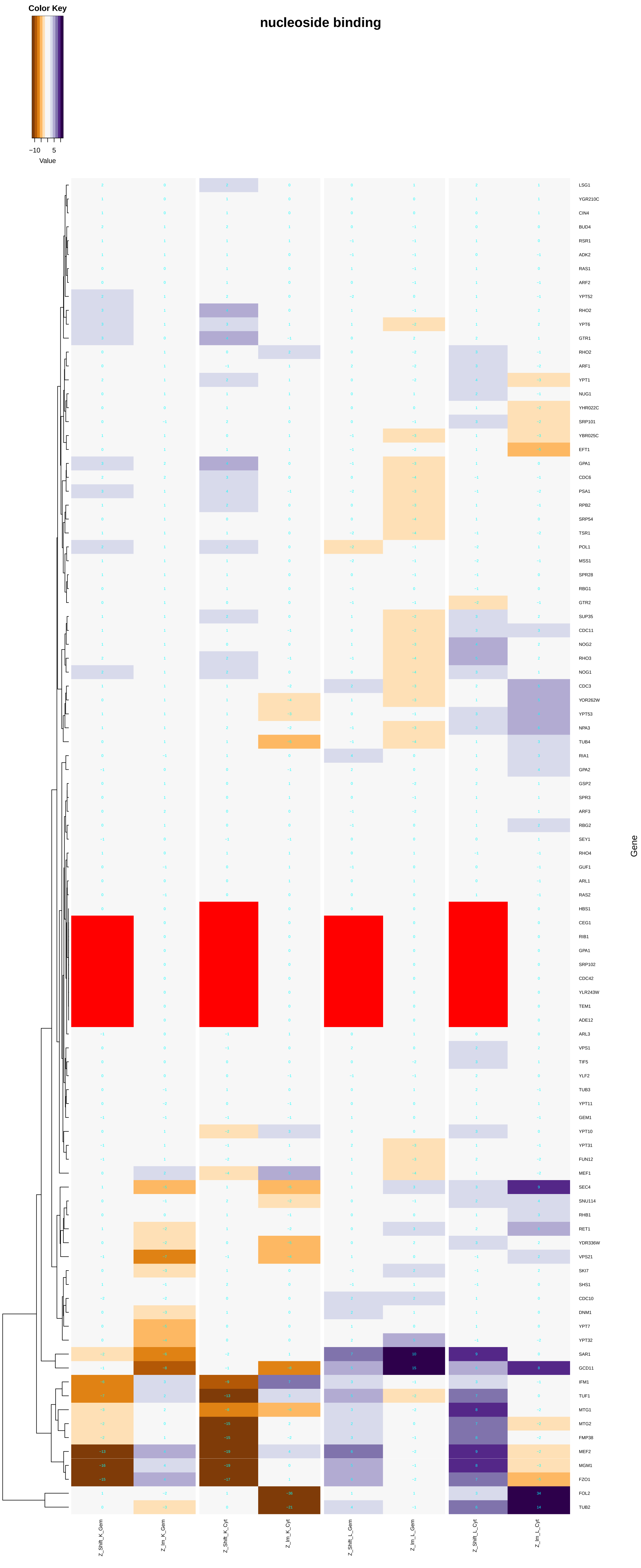
nucleoside binding
Color Key
−10
5
Value
Gene
LSG1
YGR210C
CIN4
BUD4
RSR1
ADK2
RAS1
ARF2
YPT52
RHO2
YPT6
GTR1
RHO2
ARF1
YPT1
NUG1
YHR022C
SRP101
YBR025C
EFT1
GPA1
CDC6
PSA1
RPB2
SRP54
TSR1
POL1
MSS1
SPR28
RBG1
GTR2
SUP35
CDC11
NOG2
RHO3
NOG1
CDC3
YOR262W
YPT53
NPA3
TUB4
RIA1
GPA2
GSP2
SPR3
ARF3
RBG2
SEY1
RHO4
GUF1
ARL1
RAS2
HBS1
CEG1
RIB1
GPA1
SRP102
CDC42
YLR243W
TEM1
ADE12
ARL3
VPS1
TIF5
YLF2
TUB3
YPT11
GEM1
YPT10
YPT31
FUN12
MEF1
SEC4
SNU114
RHB1
RET1
YDR336W
VPS21
SKI7
SHS1
CDC10
DNM1
YPT7
YPT32
SAR1
GCD11
IFM1
TUF1
MTG1
MTG2
FMP38
MEF2
MGM1
FZO1
FOL2
TUB2
Z_Shift_K_Gem
Z_Im_K_Gem
Z_Shift_K_Cyt
Z_Im_K_Cyt
Z_Shift_L_Gem
Z_Im_L_Gem
Z_Shift_L_Cyt
Z_Im_L_Cyt

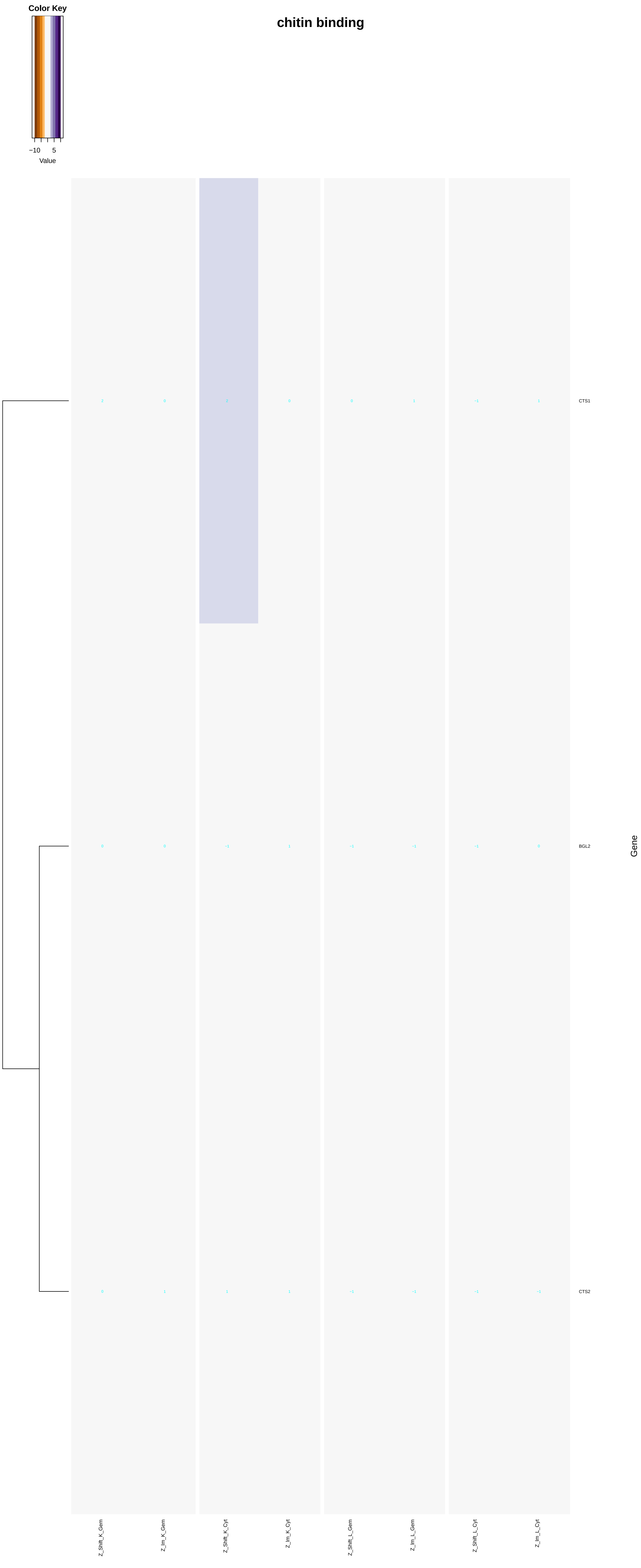

chitin binding
Color Key
−10
5
Value
CTS1
BGL2
CTS2
Gene
Z_Shift_K_Gem
Z_Im_K_Gem
Z_Shift_K_Cyt
Z_Im_K_Cyt
Z_Shift_L_Gem
Z_Im_L_Gem
Z_Shift_L_Cyt
Z_Im_L_Cyt

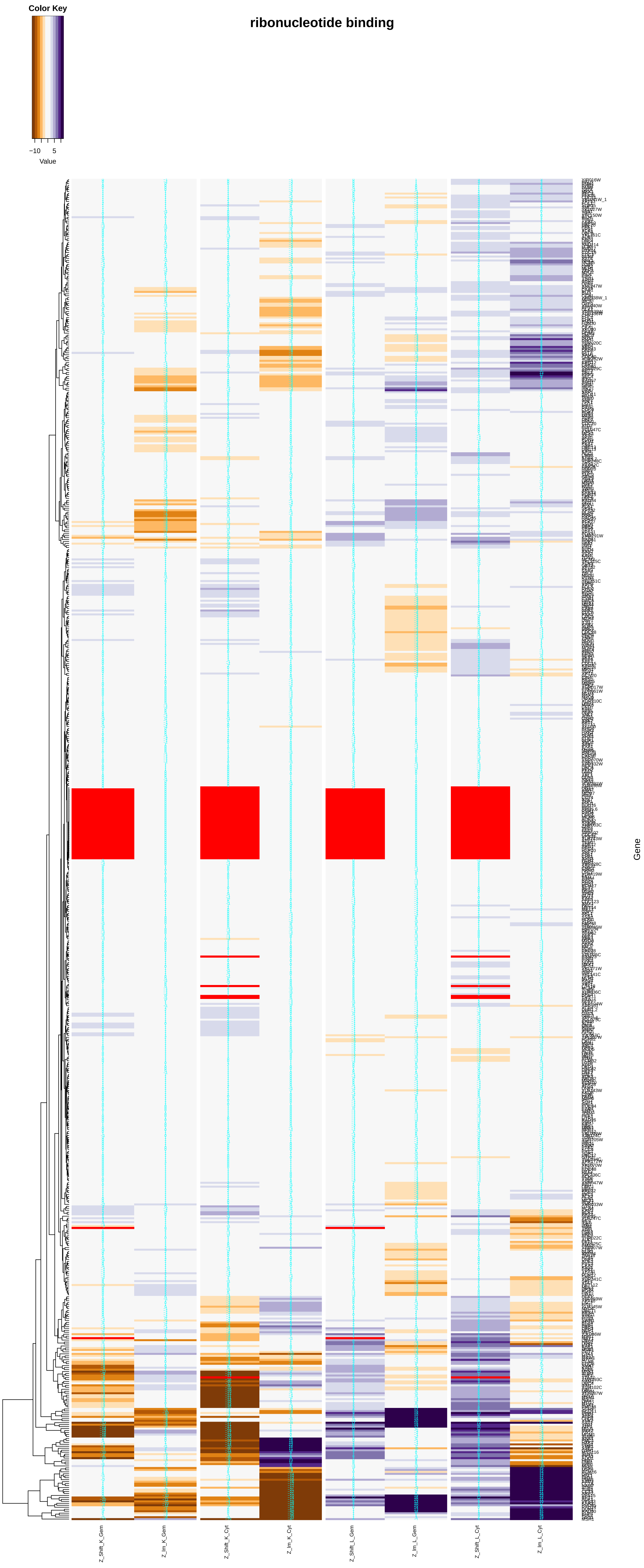
Color Key
−10
5
Value
ribonucleotide binding
Gene
Z_Shift_K_Gem
Z_Im_K_Gem
Z_Shift_K_Cyt
Z_Im_K_Cyt
Z_Shift_L_Gem
Z_Im_L_Gem
Z_Shift_L_Cyt
Z_Im_L_Cyt

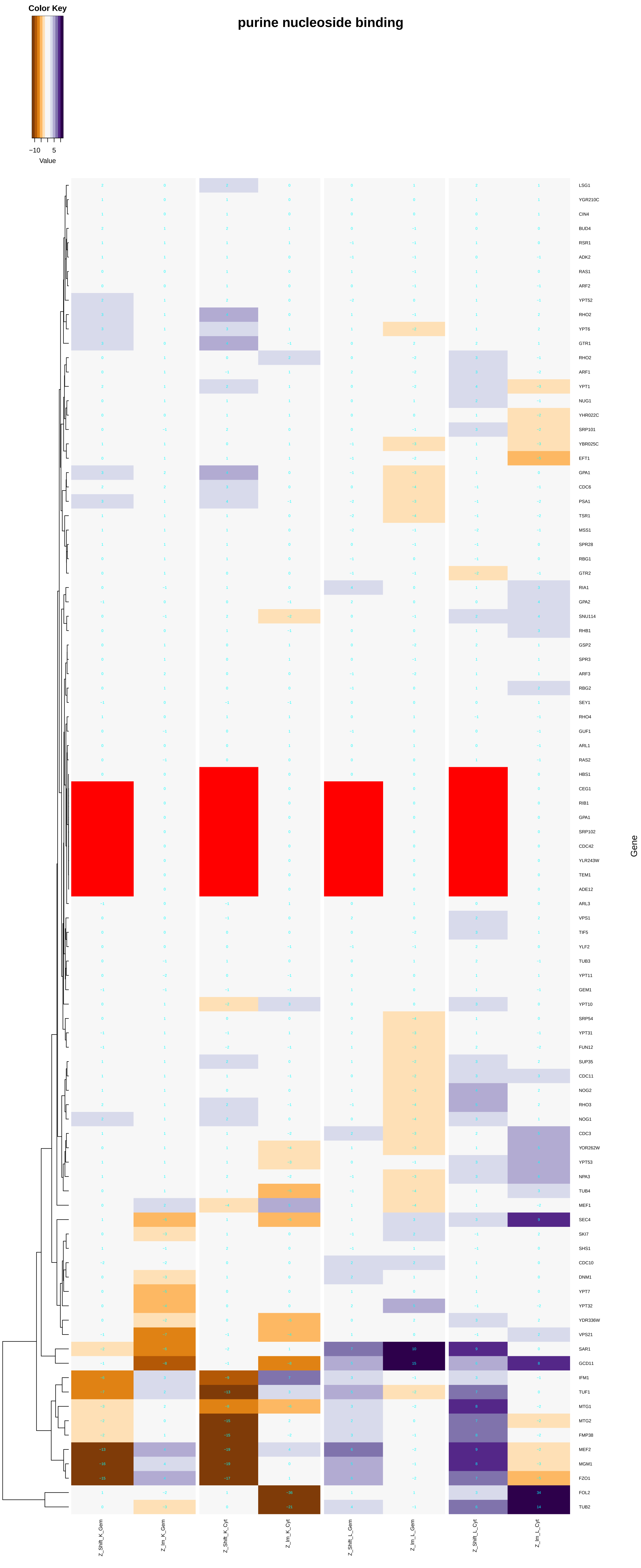
purine nucleoside binding
Color Key
−10
5
Value
LSG1
YGR210C
CIN4
BUD4
RSR1
ADK2
RAS1
ARF2
YPT52
RHO2
YPT6
GTR1
RHO2
ARF1
YPT1
NUG1
YHR022C
SRP101
YBR025C
EFT1
GPA1
CDC6
PSA1
TSR1
MSS1
SPR28
RBG1
GTR2
RIA1
GPA2
SNU114
RHB1
GSP2
SPR3
ARF3
RBG2
SEY1
RHO4
GUF1
ARL1
RAS2
HBS1
CEG1
RIB1
GPA1
SRP102
CDC42
YLR243W
TEM1
ADE12
ARL3
VPS1
TIF5
YLF2
TUB3
YPT11
GEM1
YPT10
SRP54
YPT31
FUN12
SUP35
CDC11
NOG2
RHO3
NOG1
CDC3
YOR262W
YPT53
NPA3
TUB4
MEF1
SEC4
SKI7
SHS1
CDC10
DNM1
YPT7
YPT32
YDR336W
VPS21
SAR1
GCD11
IFM1
TUF1
MTG1
MTG2
FMP38
MEF2
MGM1
FZO1
FOL2
TUB2
Gene
Z_Shift_K_Gem
Z_Im_K_Gem
Z_Shift_K_Cyt
Z_Im_K_Cyt
Z_Shift_L_Gem
Z_Im_L_Gem
Z_Shift_L_Cyt
Z_Im_L_Cyt

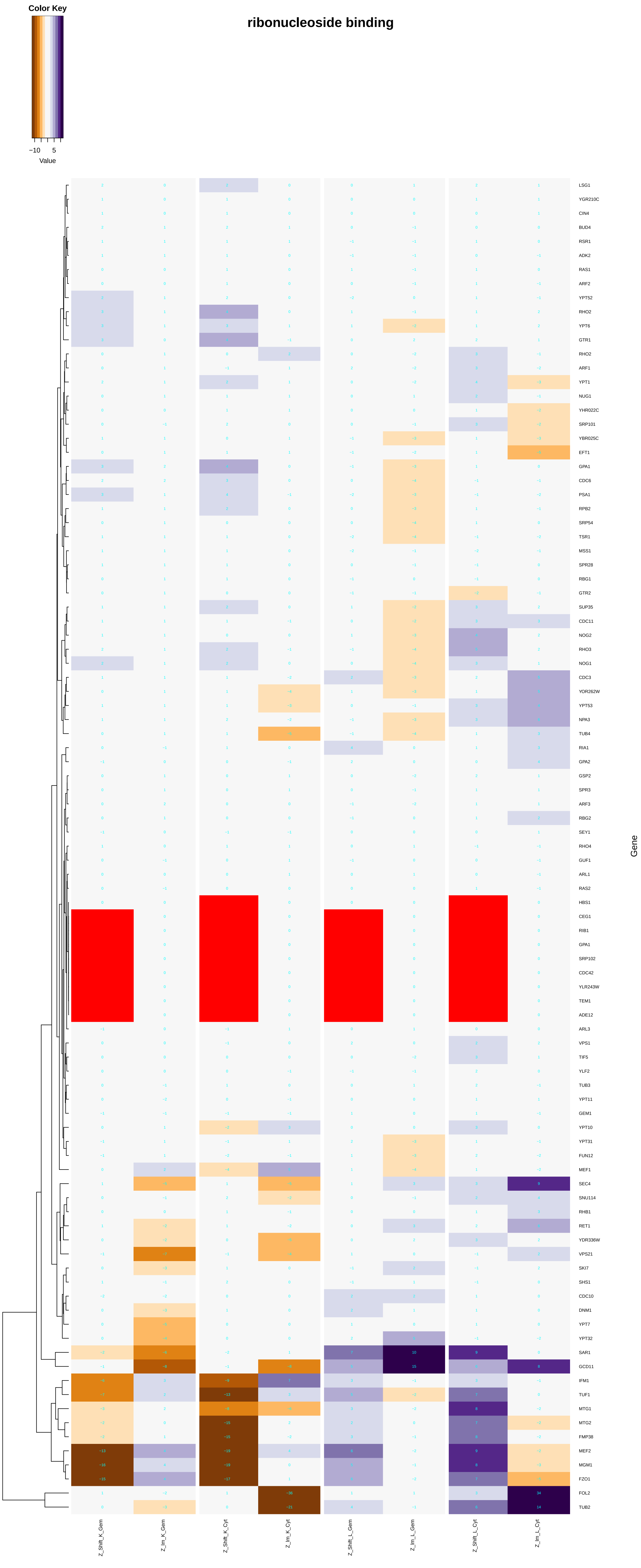
Color Key
−10
5
Value
ribonucleoside binding
LSG1
YGR210C
CIN4
BUD4
RSR1
ADK2
RAS1
ARF2
YPT52
RHO2
YPT6
GTR1
RHO2
ARF1
YPT1
NUG1
YHR022C
SRP101
YBR025C
EFT1
GPA1
CDC6
PSA1
RPB2
SRP54
TSR1
MSS1
SPR28
RBG1
GTR2
SUP35
CDC11
NOG2
RHO3
NOG1
CDC3
YOR262W
YPT53
NPA3
TUB4
RIA1
GPA2
GSP2
SPR3
ARF3
RBG2
SEY1
RHO4
GUF1
ARL1
RAS2
HBS1
CEG1
RIB1
GPA1
SRP102
CDC42
YLR243W
TEM1
ADE12
ARL3
VPS1
TIF5
YLF2
TUB3
YPT11
GEM1
YPT10
YPT31
FUN12
MEF1
SEC4
SNU114
RHB1
RET1
YDR336W
VPS21
SKI7
SHS1
CDC10
DNM1
YPT7
YPT32
SAR1
GCD11
IFM1
TUF1
MTG1
MTG2
FMP38
MEF2
MGM1
FZO1
FOL2
TUB2
Gene
Z_Shift_K_Gem
Z_Im_K_Gem
Z_Shift_K_Cyt
Z_Im_K_Cyt
Z_Shift_L_Gem
Z_Im_L_Gem
Z_Shift_L_Cyt
Z_Im_L_Cyt

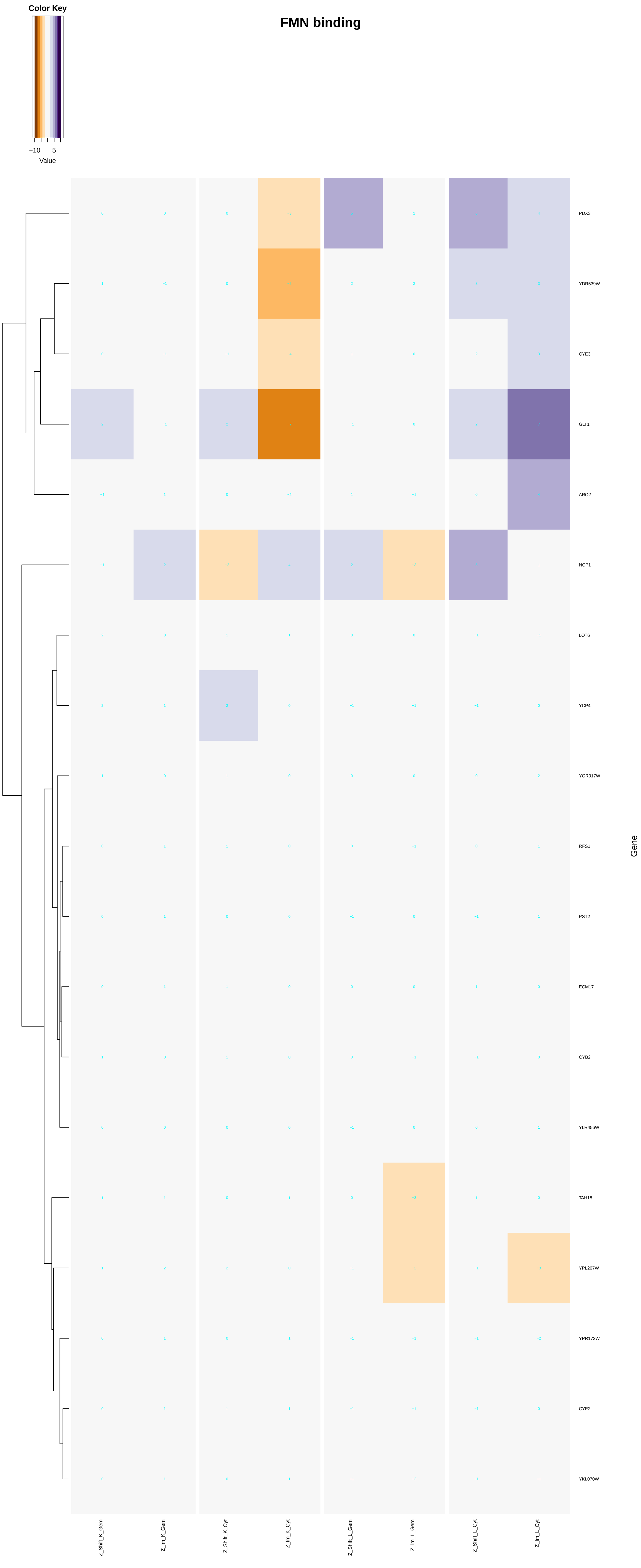

FMN binding
Color Key
−10
5
Value
PDX3
YDR539W
OYE3
GLT1
ARO2
NCP1
LOT6
YCP4
YGR017W
RFS1
PST2
ECM17
CYB2
YLR456W
TAH18
YPL207W
YPR172W
OYE2
YKL070W
Gene
Z_Shift_K_Gem
Z_Im_K_Gem
Z_Shift_K_Cyt
Z_Im_K_Cyt
Z_Shift_L_Gem
Z_Im_L_Gem
Z_Shift_L_Cyt
Z_Im_L_Cyt

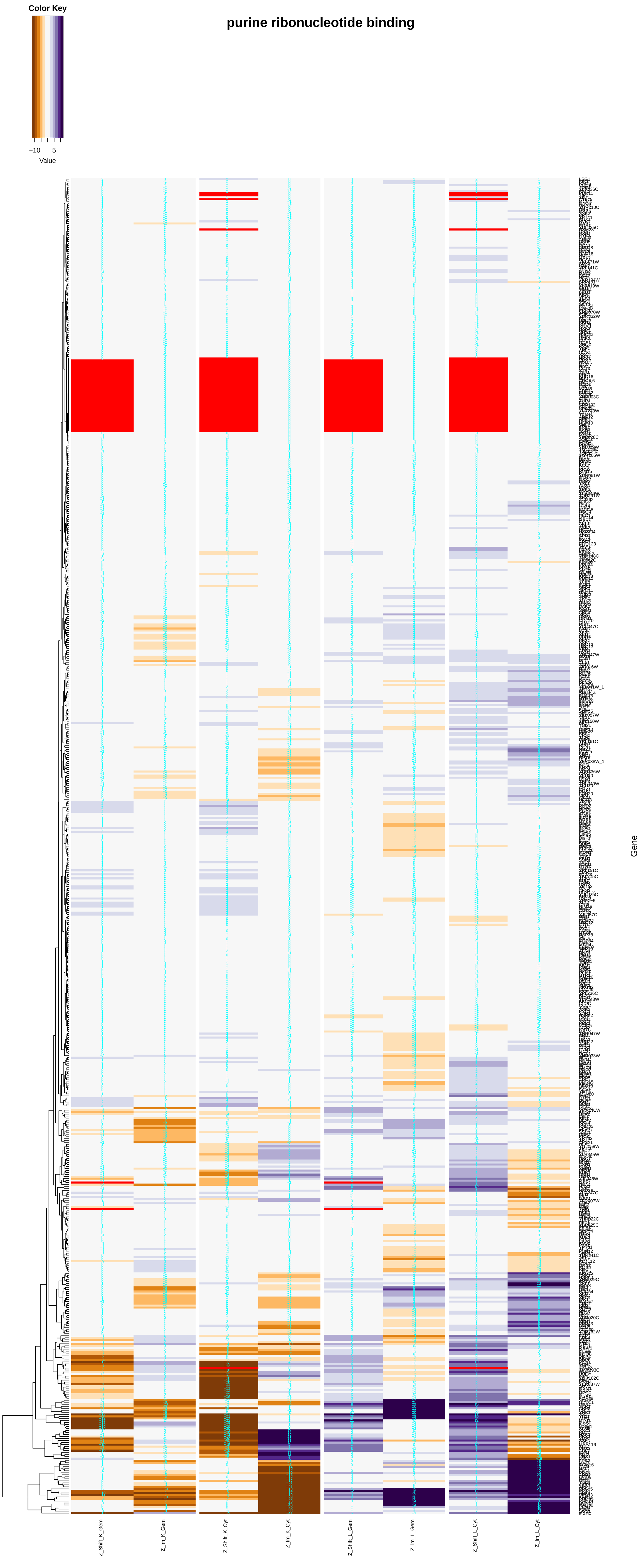
purine ribonucleotide binding
Color Key
−10
5
Value
Gene
Z_Shift_K_Gem
Z_Im_K_Gem
Z_Shift_K_Cyt
Z_Im_K_Cyt
Z_Shift_L_Gem
Z_Im_L_Gem
Z_Shift_L_Cyt
Z_Im_L_Cyt

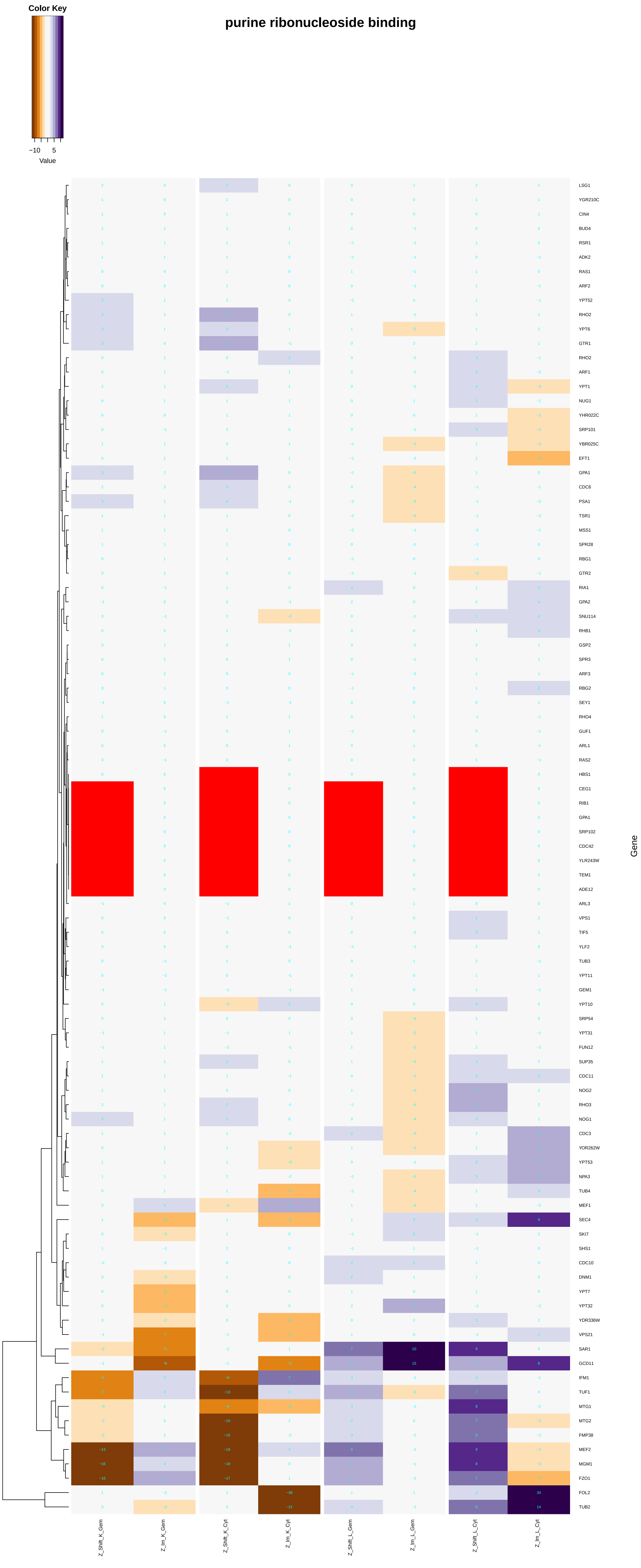
Color Key
−10
5
Value
purine ribonucleoside binding
Gene
LSG1
YGR210C
CIN4
BUD4
RSR1
ADK2
RAS1
ARF2
YPT52
RHO2
YPT6
GTR1
RHO2
ARF1
YPT1
NUG1
YHR022C
SRP101
YBR025C
EFT1
GPA1
CDC6
PSA1
TSR1
MSS1
SPR28
RBG1
GTR2
RIA1
GPA2
SNU114
RHB1
GSP2
SPR3
ARF3
RBG2
SEY1
RHO4
GUF1
ARL1
RAS2
HBS1
CEG1
RIB1
GPA1
SRP102
CDC42
YLR243W
TEM1
ADE12
ARL3
VPS1
TIF5
YLF2
TUB3
YPT11
GEM1
YPT10
SRP54
YPT31
FUN12
SUP35
CDC11
NOG2
RHO3
NOG1
CDC3
YOR262W
YPT53
NPA3
TUB4
MEF1
SEC4
SKI7
SHS1
CDC10
DNM1
YPT7
YPT32
YDR336W
VPS21
SAR1
GCD11
IFM1
TUF1
MTG1
MTG2
FMP38
MEF2
MGM1
FZO1
FOL2
TUB2
Z_Shift_K_Gem
Z_Im_K_Gem
Z_Shift_K_Cyt
Z_Im_K_Cyt
Z_Shift_L_Gem
Z_Im_L_Gem
Z_Shift_L_Cyt
Z_Im_L_Cyt

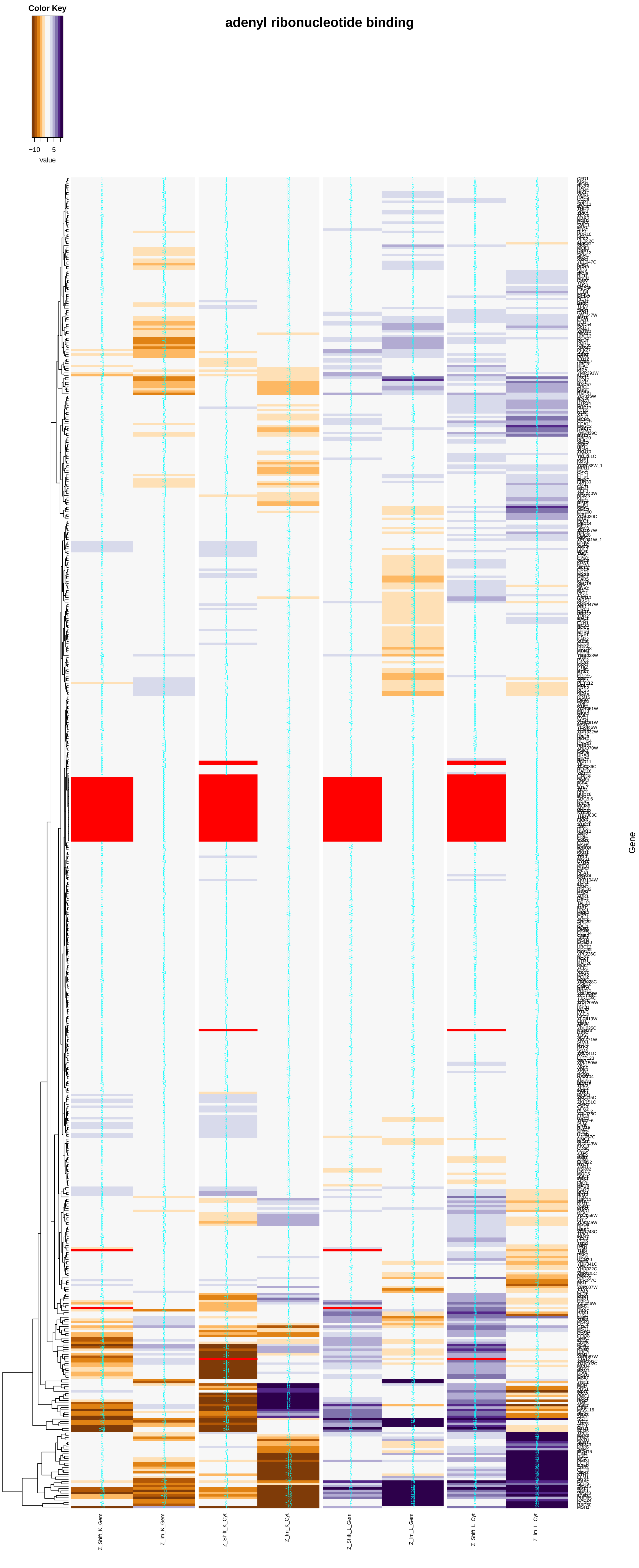
adenyl ribonucleotide binding
Color Key
−10
5
Value
Gene
Z_Shift_K_Gem
Z_Im_K_Gem
Z_Shift_K_Cyt
Z_Im_K_Cyt
Z_Shift_L_Gem
Z_Im_L_Gem
Z_Shift_L_Cyt
Z_Im_L_Cyt

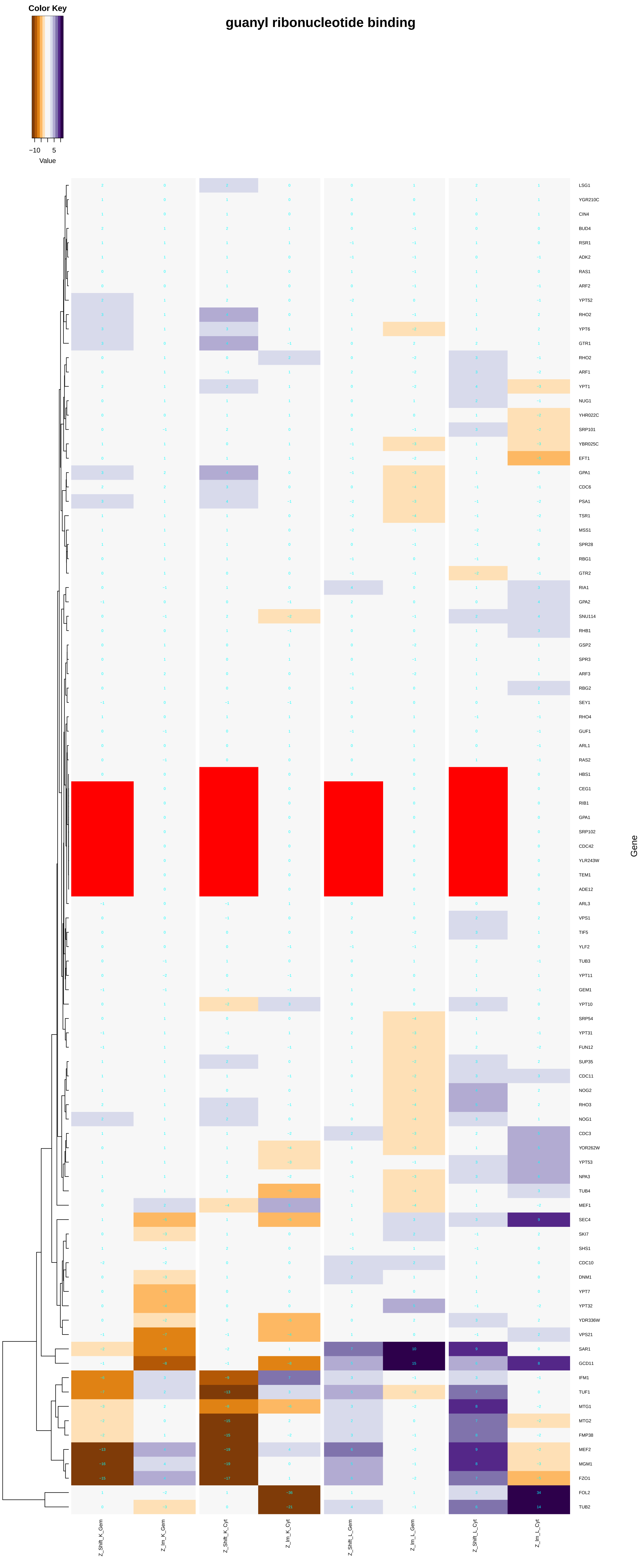
guanyl ribonucleotide binding
Color Key
−10
5
Value
LSG1
YGR210C
CIN4
BUD4
RSR1
ADK2
RAS1
ARF2
YPT52
RHO2
YPT6
GTR1
RHO2
ARF1
YPT1
NUG1
YHR022C
SRP101
YBR025C
EFT1
GPA1
CDC6
PSA1
TSR1
MSS1
SPR28
RBG1
GTR2
RIA1
GPA2
SNU114
RHB1
GSP2
SPR3
ARF3
RBG2
SEY1
RHO4
GUF1
ARL1
RAS2
HBS1
CEG1
RIB1
GPA1
SRP102
CDC42
YLR243W
TEM1
ADE12
ARL3
VPS1
TIF5
YLF2
TUB3
YPT11
GEM1
YPT10
SRP54
YPT31
FUN12
SUP35
CDC11
NOG2
RHO3
NOG1
CDC3
YOR262W
YPT53
NPA3
TUB4
MEF1
SEC4
SKI7
SHS1
CDC10
DNM1
YPT7
YPT32
YDR336W
VPS21
SAR1
GCD11
IFM1
TUF1
MTG1
MTG2
FMP38
MEF2
MGM1
FZO1
FOL2
TUB2
Gene
Z_Shift_K_Gem
Z_Im_K_Gem
Z_Shift_K_Cyt
Z_Im_K_Cyt
Z_Shift_L_Gem
Z_Im_L_Gem
Z_Shift_L_Cyt
Z_Im_L_Cyt

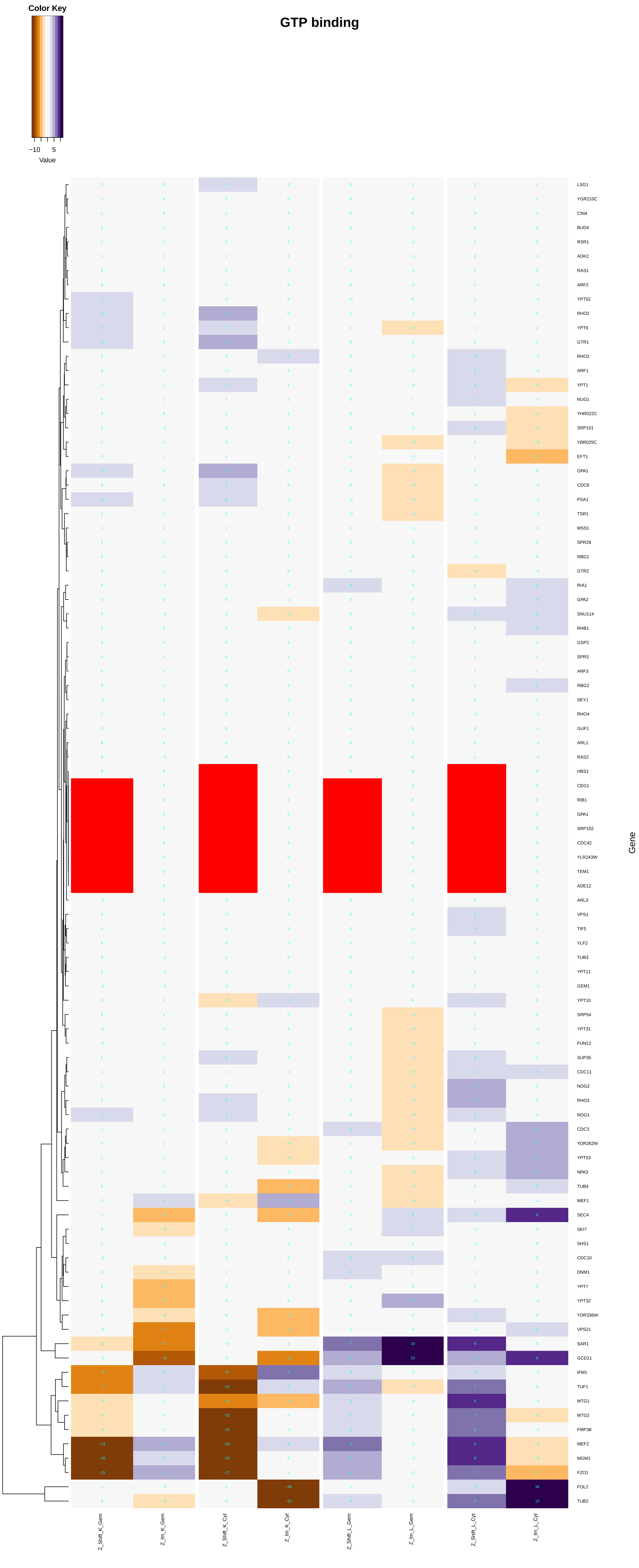
Color Key
−10 5
Value
GTP binding
Gene
Z_Shift_K_Gem
Z_Im_K_Gem
Z_Shift_K_Cyt
Z_Im_K_Cyt
Z_Shift_L_Gem
Z_Im_L_Gem
Z_Shift_L_Cyt
Z_Im_L_Cyt
LSG1
YGR210C
CIN4
BUD4
RSR1
ADK2
RAS1
ARF2
YPT52
RHO2
YPT6
GTR1
RHO2
ARF1
YPT1
NUG1
YHR022C
SRP101
YBR025C
EFT1
GPA1
CDC6
PSA1
TSR1
MSS1
SPR28
RBG1
GTR2
RIA1
GPA2
SNU114
RHB1
GSP2
SPR3
ARF3
RBG2
SEY1
RHO4
GUF1
ARL1
RAS2
HBS1
CEG1
RIB1
GPA1
SRP102
CDC42
YLR243W
TEM1
ADE12
ARL3
VPS1
TIF5
YLF2
TUB3
YPT11
GEM1
YPT10
SRP54
YPT31
FUN12
SUP35
CDC11
NOG2
RHO3
NOG1
CDC3
YOR262W
YPT53
NPA3
TUB4
MEF1
SEC4
SKI7
SHS1
CDC10
DNM1
YPT7
YPT32
YDR336W
VPS21
SAR1
GCD11
IFM1
TUF1
MTG1
MTG2
FMP38
MEF2
MGM1
FZO1
FOL2
TUB2

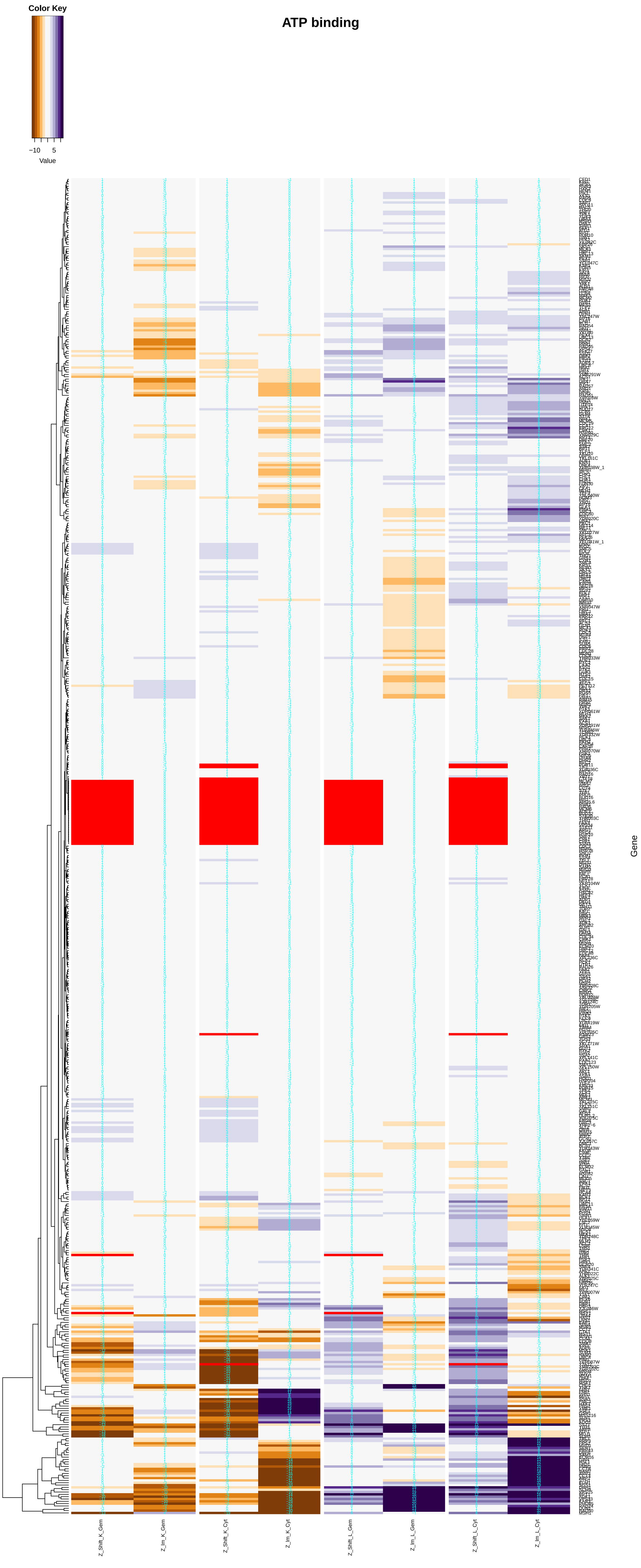
ATP binding
Color Key
−10
5
Value
Gene
Z_Shift_K_Gem
Z_Im_K_Gem
Z_Shift_K_Cyt
Z_Im_K_Cyt
Z_Shift_L_Gem
Z_Im_L_Gem
Z_Shift_L_Cyt
Z_Im_L_Cyt

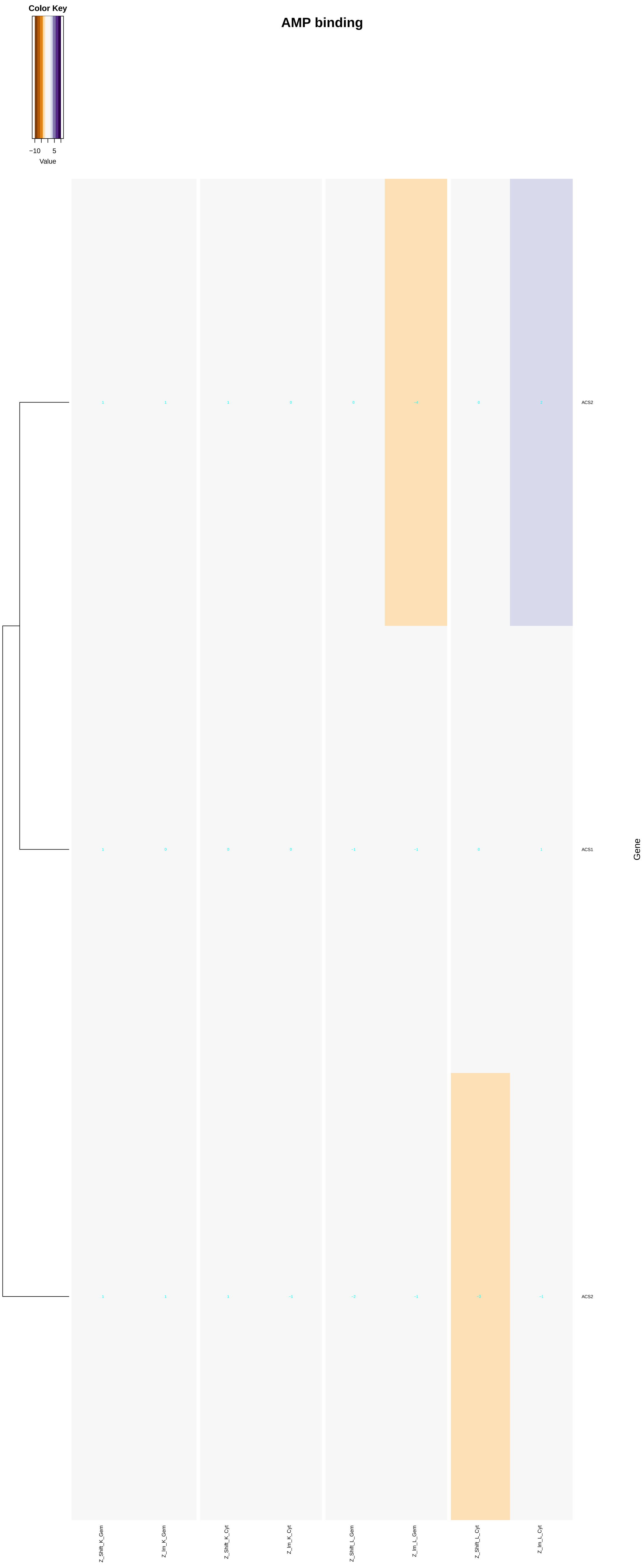
AMP binding
Color Key
−10
5
Value
ACS2
ACS1
ACS2
Gene
Z_Shift_K_Gem
Z_Im_K_Gem
Z_Shift_K_Cyt
Z_Im_K_Cyt
Z_Shift_L_Gem
Z_Im_L_Gem
Z_Shift_L_Cyt
Z_Im_L_Cyt
