## Additional File 7 for "A humanized yeast phenomic model of deoxycytidine kinase to predict genetic buffering of nucleoside analog cytotoxicity": cell_cycle_checkpoint.pdf

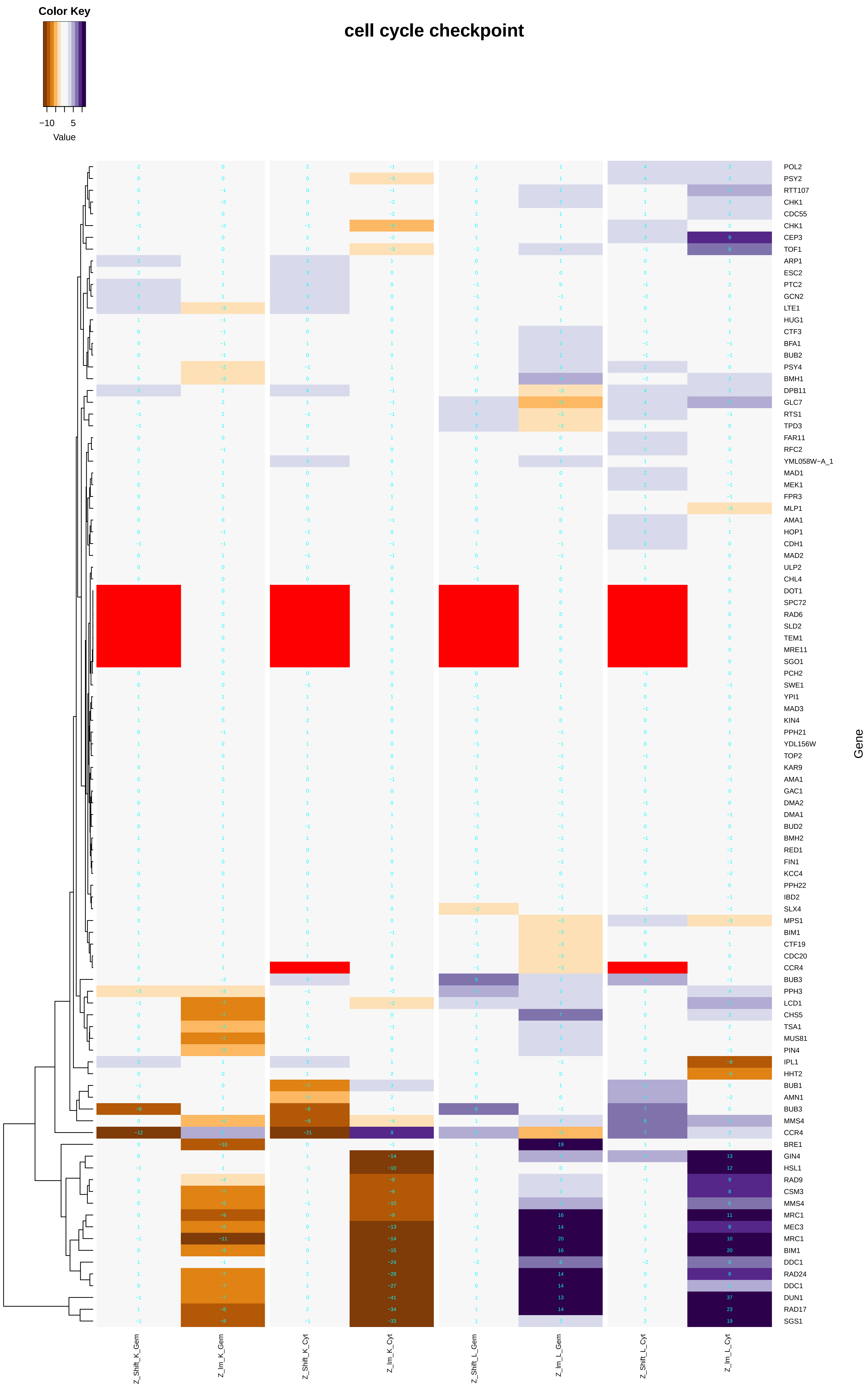
cell cycle checkpoint
Color Key
−10
5
Value
POL2
PSY2
RTT107
CHK1
CDC55
CHK1
CEP3
TOF1
ARP1
ESC2
PTC2
GCN2
LTE1
HUG1
CTF3
BFA1
BUB2
PSY4
BMH1
DPB11
GLC7
RTS1
TPD3
FAR11
RFC2
YML058W−A_1
MAD1
MEK1
FPR3
MLP1
AMA1
HOP1
CDH1
MAD2
ULP2
CHL4
DOT1
SPC72
RAD6
SLD2
TEM1
MRE11
SGO1
PCH2
SWE1
YPI1
MAD3
KIN4
PPH21
YDL156W
TOP2
KAR9
AMA1
GAC1
DMA2
DMA1
BUD2
BMH2
RED1
FIN1
KCC4
PPH22
IBD2
SLX4
MPS1
BIM1
CTF19
CDC20
CCR4
BUB3
PPH3
LCD1
CHS5
TSA1
MUS81
PIN4
IPL1
HHT2
BUB1
AMN1
BUB3
MMS4
CCR4
BRE1
GIN4
HSL1
RAD9
CSM3
MMS4
MRC1
MEC3
MRC1
BIM1
DDC1
RAD24
DDC1
DUN1
RAD17
SGS1
Gene
Z_Shift_K_Gem
Z_Im_K_Gem
Z_Shift_K_Cyt
Z_Im_K_Cyt
Z_Shift_L_Gem
Z_Im_L_Gem
Z_Shift_L_Cyt
Z_Im_L_Cyt

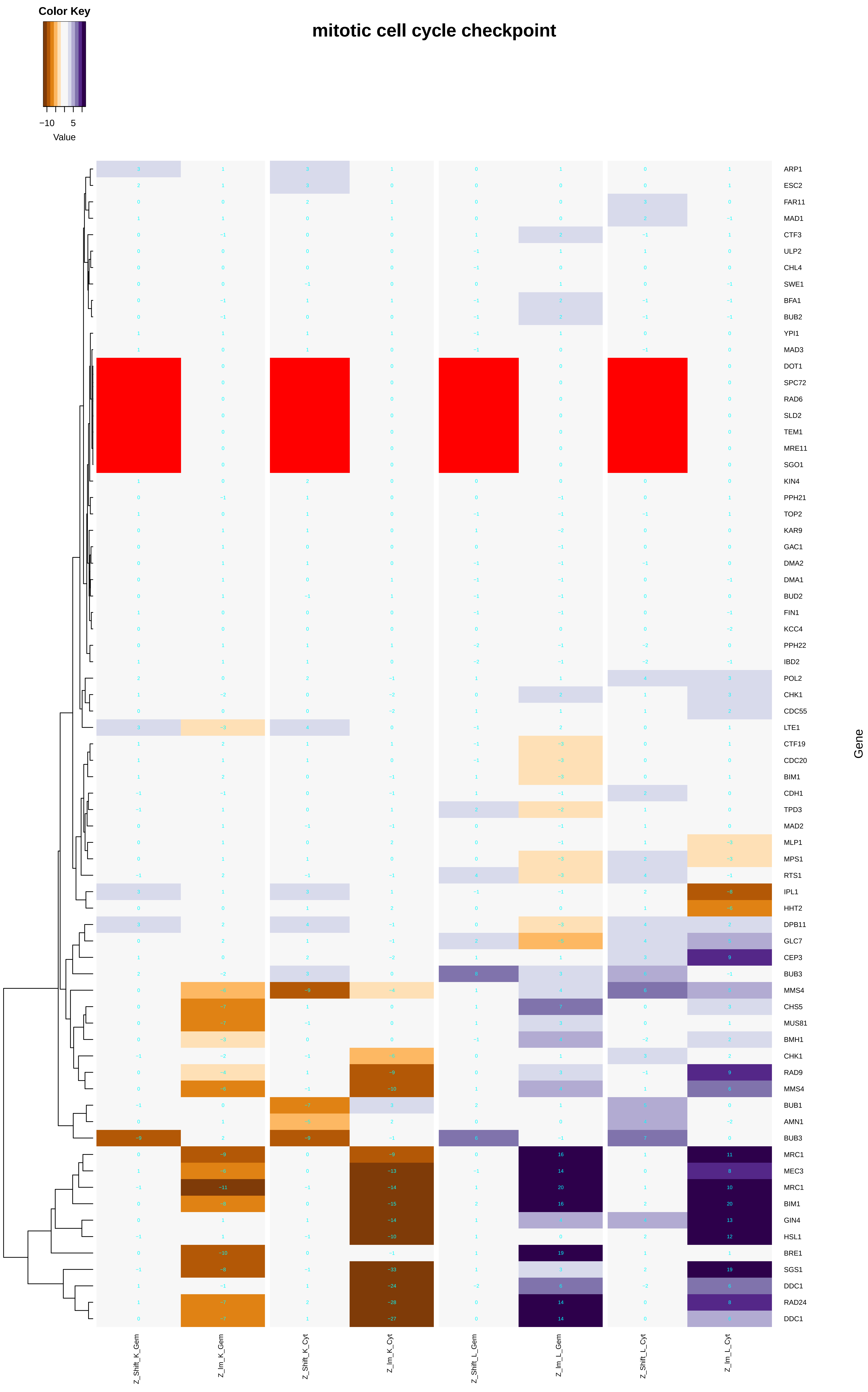
Color Key
−10
5
Value
mitotic cell cycle checkpoint
ARP1
ESC2
FAR11
MAD1
CTF3
ULP2
CHL4
SWE1
BFA1
BUB2
YPI1
MAD3
DOT1
SPC72
RAD6
SLD2
TEM1
MRE11
SGO1
KIN4
PPH21
TOP2
KAR9
GAC1
DMA2
DMA1
BUD2
FIN1
KCC4
PPH22
IBD2
POL2
CHK1
CDC55
LTE1
CTF19
CDC20
BIM1
CDH1
TPD3
MAD2
MLP1
MPS1
RTS1
IPL1
HHT2
DPB11
GLC7
CEP3
BUB3
MMS4
CHS5
MUS81
BMH1
CHK1
RAD9
MMS4
BUB1
AMN1
BUB3
MRC1
MEC3
MRC1
BIM1
GIN4
HSL1
BRE1
SGS1
DDC1
RAD24
DDC1
Gene
Z_Shift_K_Gem
Z_Im_K_Gem
Z_Shift_K_Cyt
Z_Im_K_Cyt
Z_Shift_L_Gem
Z_Im_L_Gem
Z_Shift_L_Cyt
Z_Im_L_Cyt

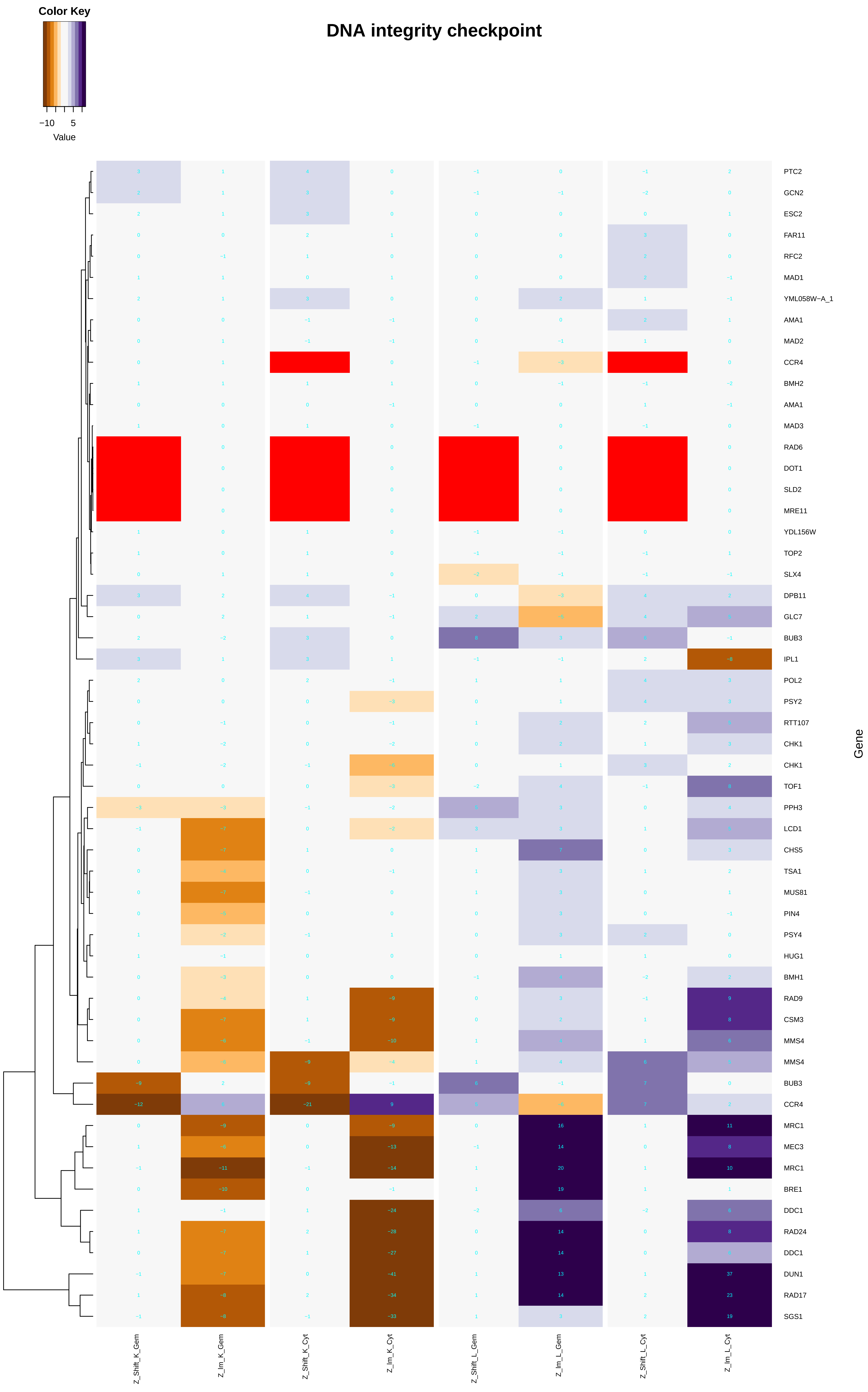

Color Key
−10
5
Value
DNA integrity checkpoint
PTC2
GCN2
ESC2
FAR11
RFC2
MAD1
YML058W−A_1
AMA1
MAD2
CCR4
BMH2
AMA1
MAD3
RAD6
DOT1
SLD2
MRE11
YDL156W
TOP2
SLX4
DPB11
GLC7
BUB3
IPL1
POL2
PSY2
RTT107
CHK1
CHK1
TOF1
PPH3
LCD1
CHS5
TSA1
MUS81
PIN4
PSY4
HUG1
BMH1
RAD9
CSM3
MMS4
MMS4
BUB3
CCR4
MRC1
MEC3
MRC1
BRE1
DDC1
RAD24
DDC1
DUN1
RAD17
SGS1
Gene
Z_Shift_K_Gem
Z_Im_K_Gem
Z_Shift_K_Cyt
Z_Im_K_Cyt
Z_Shift_L_Gem
Z_Im_L_Gem
Z_Shift_L_Cyt
Z_Im_L_Cyt

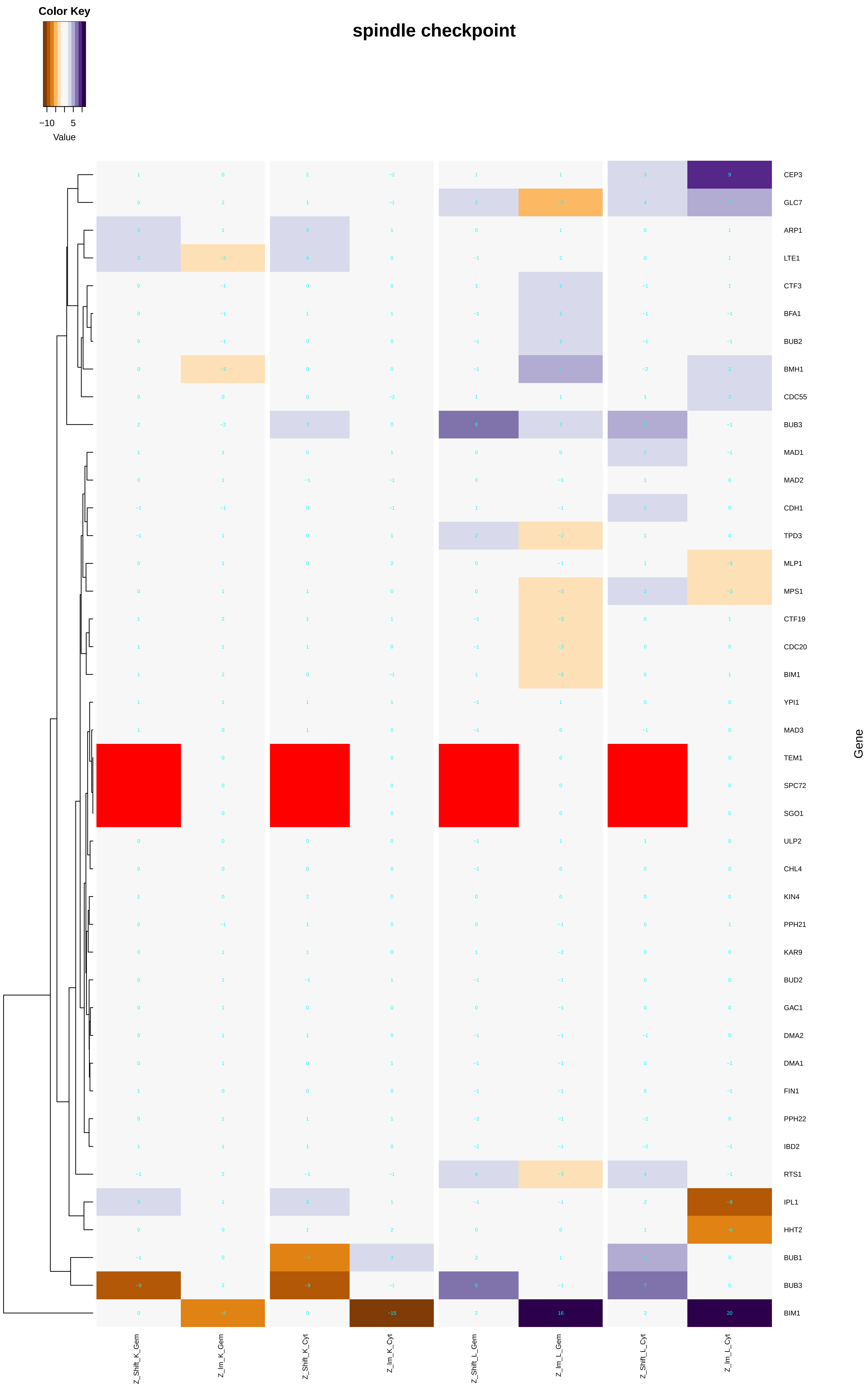
spindle checkpoint
Color Key
−10
5
Value
CEP3
GLC7
ARP1
LTE1
CTF3
BFA1
BUB2
BMH1
CDC55
BUB3
MAD1
MAD2
CDH1
TPD3
MLP1
MPS1
CTF19
CDC20
BIM1
YPI1
MAD3
TEM1
SPC72
SGO1
ULP2
CHL4
KIN4
PPH21
KAR9
BUD2
GAC1
DMA2
DMA1
FIN1
PPH22
IBD2
RTS1
IPL1
HHT2
BUB1
BUB3
BIM1
Gene
Z_Shift_K_Gem
Z_Im_K_Gem
Z_Shift_K_Cyt
Z_Im_K_Cyt
Z_Shift_L_Gem
Z_Im_L_Gem
Z_Shift_L_Cyt
Z_Im_L_Cyt

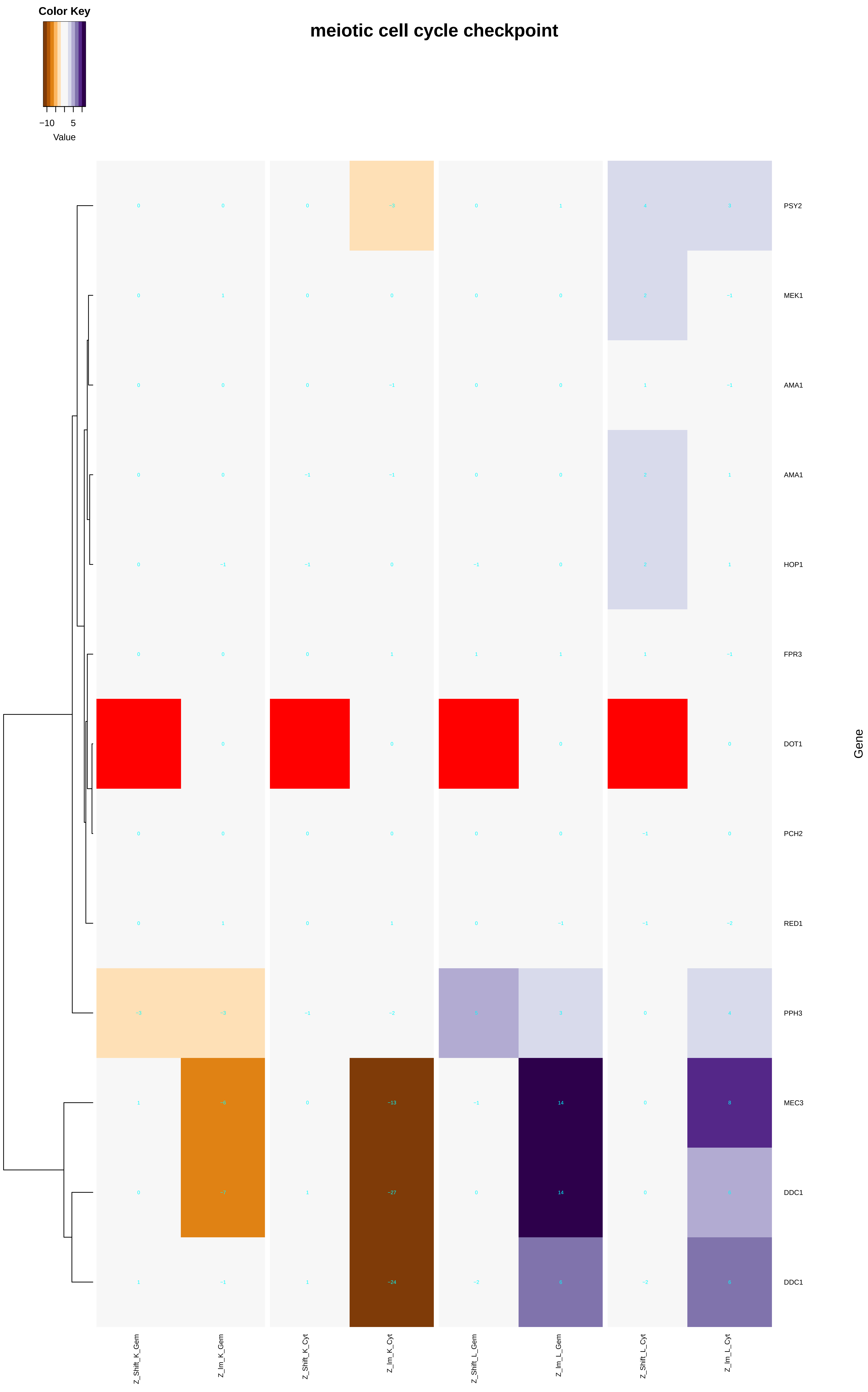

Color Key
−10
5
Value
meiotic cell cycle checkpoint
PSY2
MEK1
AMA1
AMA1
HOP1
FPR3
DOT1
PCH2
RED1
PPH3
MEC3
DDC1
DDC1
Gene
Z_Shift_K_Gem
Z_Im_K_Gem
Z_Shift_K_Cyt
Z_Im_K_Cyt
Z_Shift_L_Gem
Z_Im_L_Gem
Z_Shift_L_Cyt
Z_Im_L_Cyt

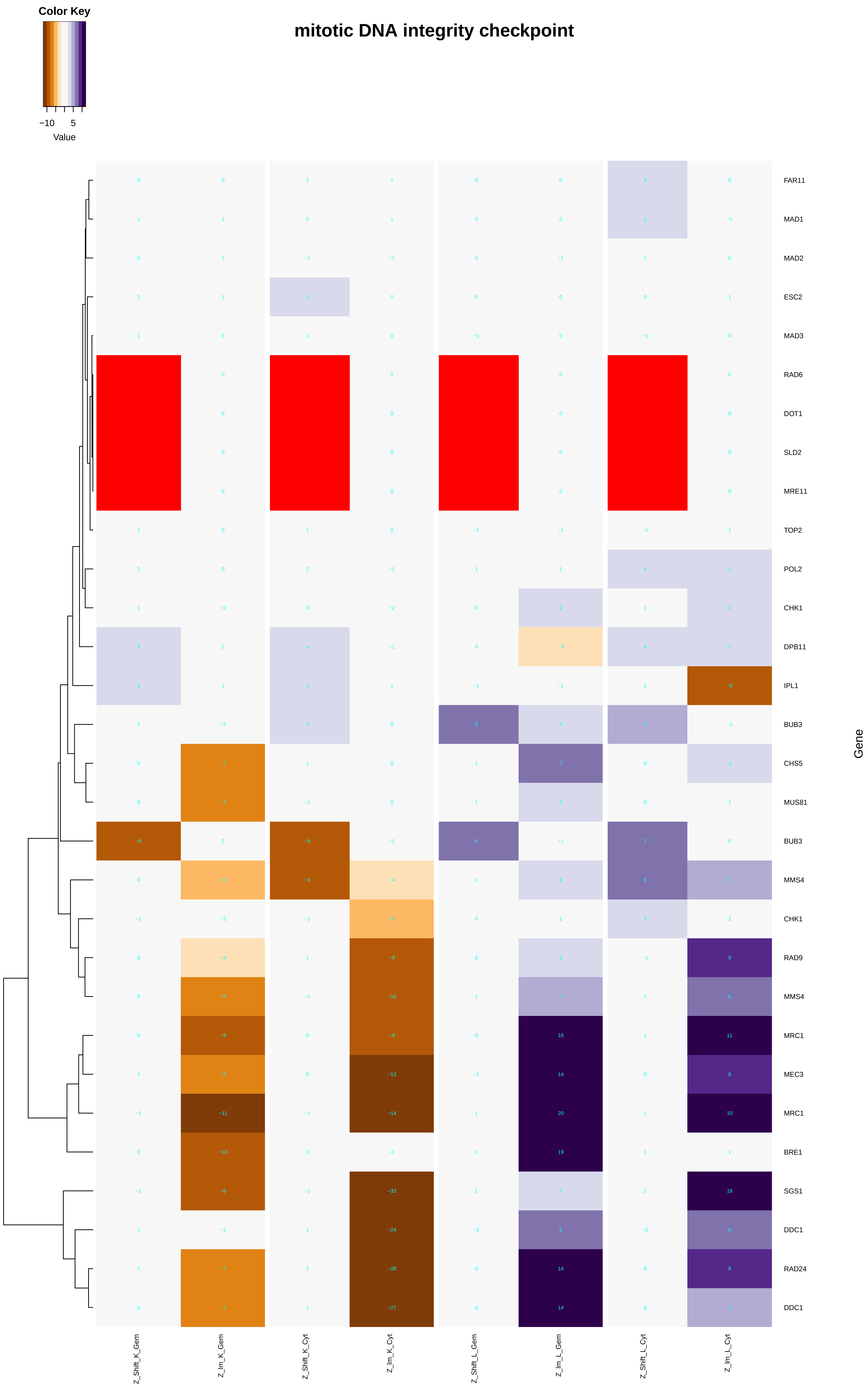

Color Key
−10 5
Value
mitotic DNA integrity checkpoint
FAR11
MAD1
MAD2
ESC2
MAD3
RAD6
DOT1
SLD2
MRE11
TOP2
POL2
CHK1
DPB11
IPL1
BUB3
CHS5
MUS81
BUB3
MMS4
CHK1
RAD9
MMS4
MRC1
MEC3
MRC1
BRE1
SGS1
DDC1
RAD24
DDC1
Gene
Z_Shift_K_Gem
Z_Im_K_Gem
Z_Shift_K_Cyt
Z_Im_K_Cyt
Z_Shift_L_Gem
Z_Im_L_Gem
Z_Shift_L_Cyt
Z_Im_L_Cyt

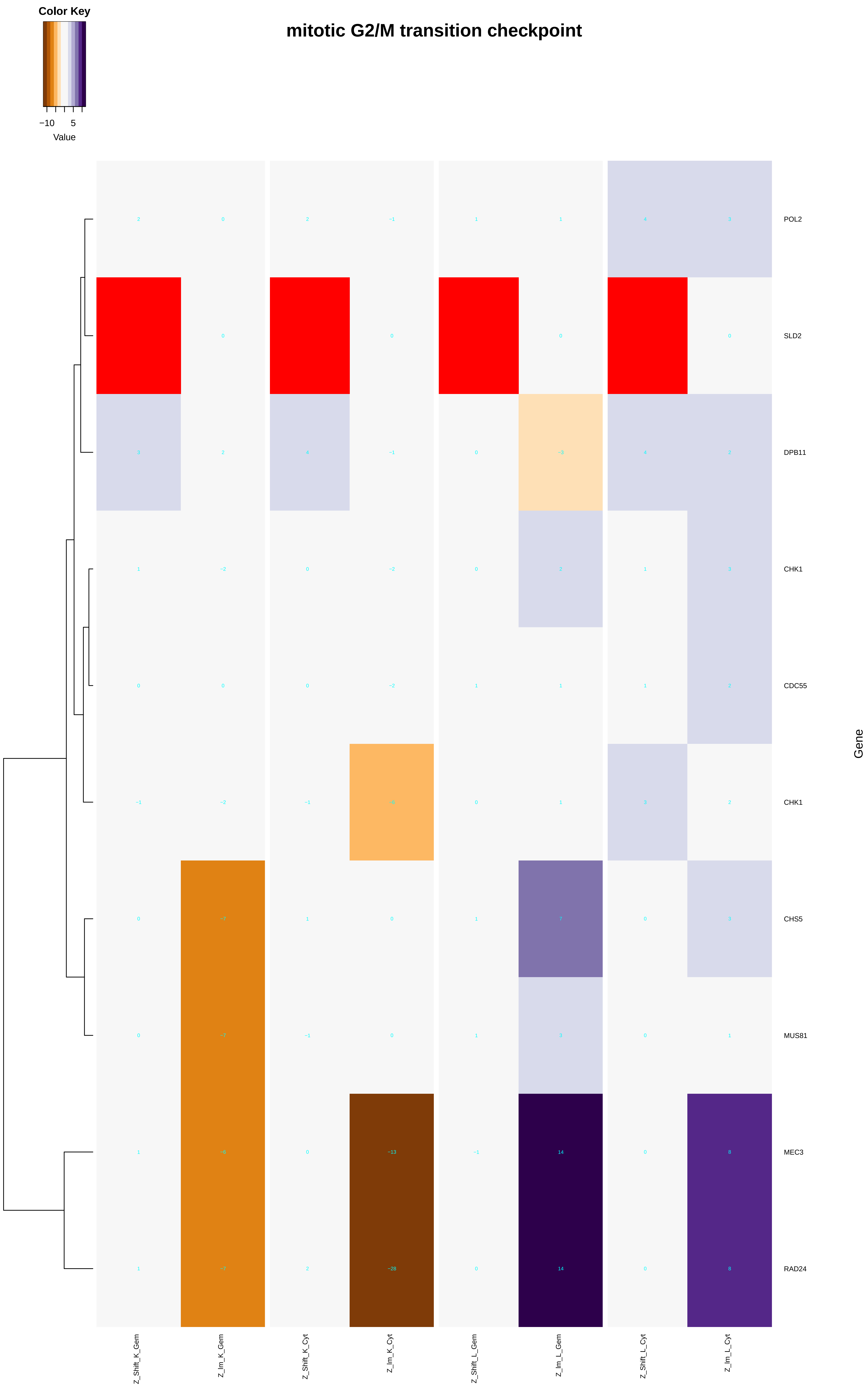
Color Key
−10
5
Value
mitotic G2/M transition checkpoint
POL2
SLD2
DPB11
CHK1
CDC55
CHK1
CHS5
MUS81
MEC3
RAD24
Gene
Z_Shift_K_Gem
Z_lm_K_Gem
Z_Shift_K_Cyt
Z_lm_K_Cyt
Z_Shift_L_Gem
Z_lm_L_Gem
Z_Shift_L_Cyt
Z_lm_L_Cyt

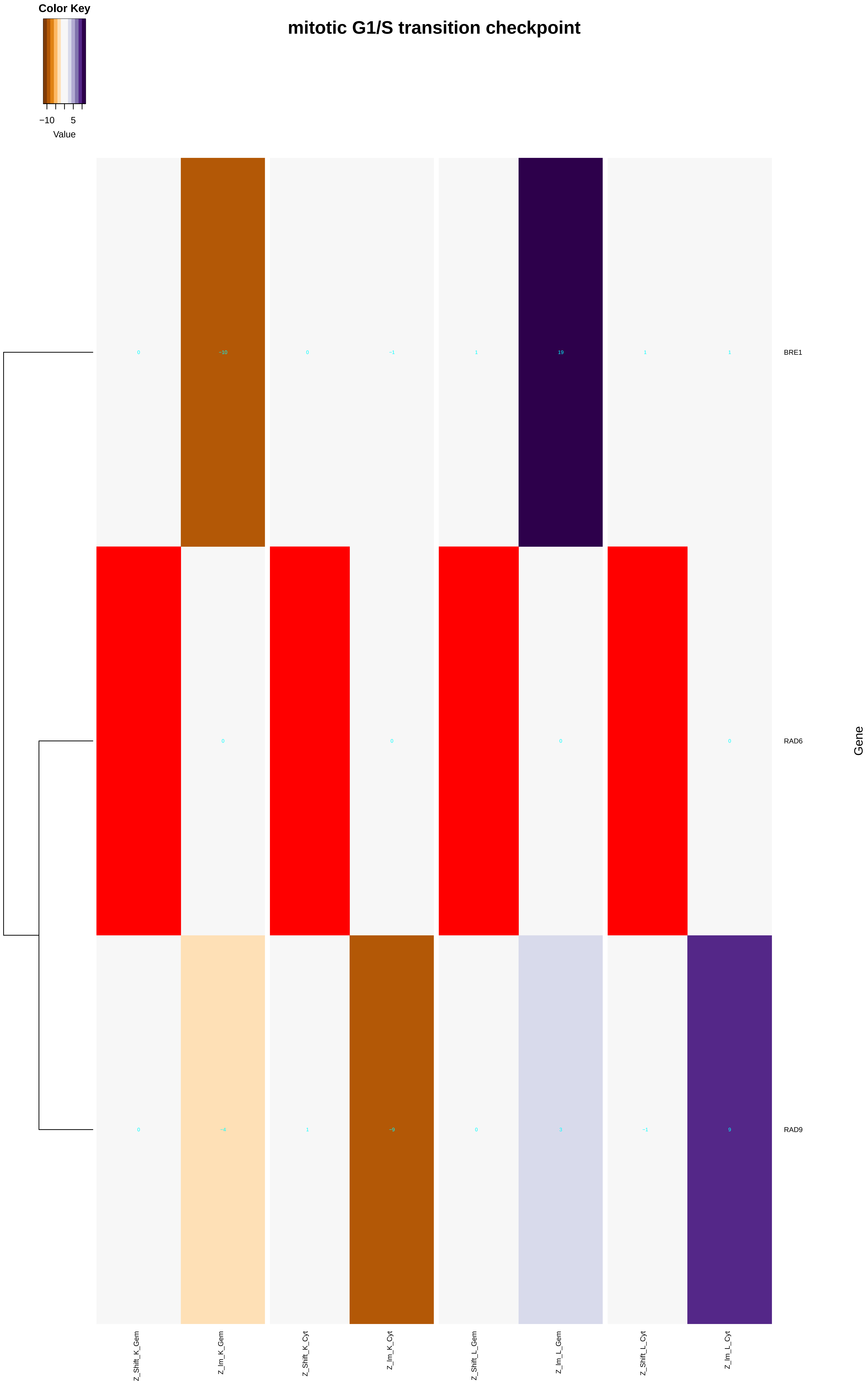

Color Key
−10
5
Value
mitotic G1/S transition checkpoint
BRE1
RAD6
RAD9
Gene
Z_Shift_K_Gem
Z_Im_K_Gem
Z_Shift_K_Cyt
Z_Im_K_Cyt
Z_Shift_L_Gem
Z_Im_L_Gem
Z_Shift_L_Cyt
Z_Im_L_Cyt

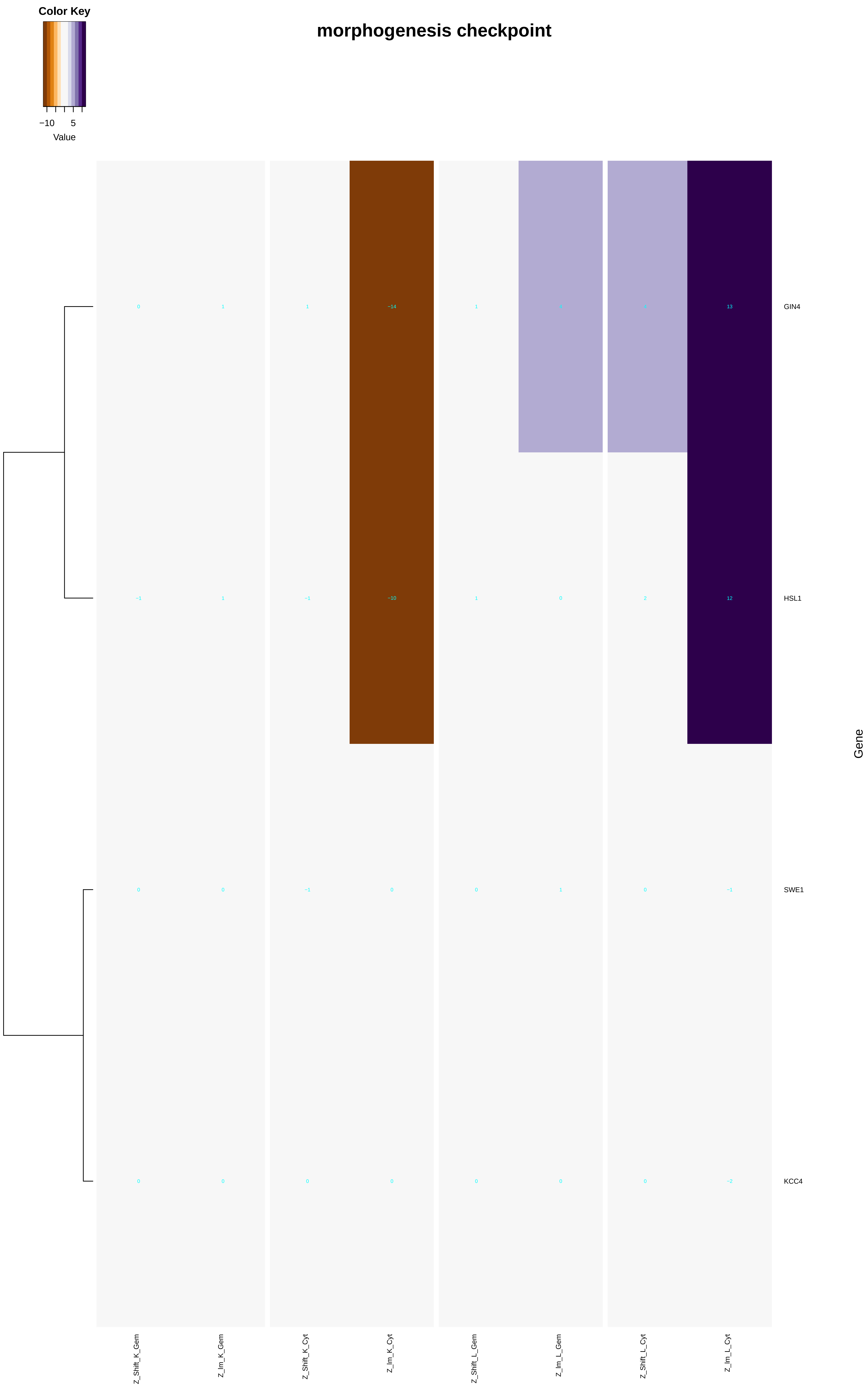
Color Key
−10
5
Value
morphogenesis checkpoint
GIN4
HSL1
SWE1
KCC4
Gene
Z_Shift_K_Gem
Z_Im_K_Gem
Z_Shift_K_Cyt
Z_Im_K_Cyt
Z_Shift_L_Gem
Z_Im_L_Gem
Z_Shift_L_Cyt
Z_Im_L_Cyt
0
1
1
−14
1
4
4
13
−1
1
−1
−10
1
0
2
12
0
0
−1
0
0
1
0
−1
0
0
0
0
0
0
0
−2

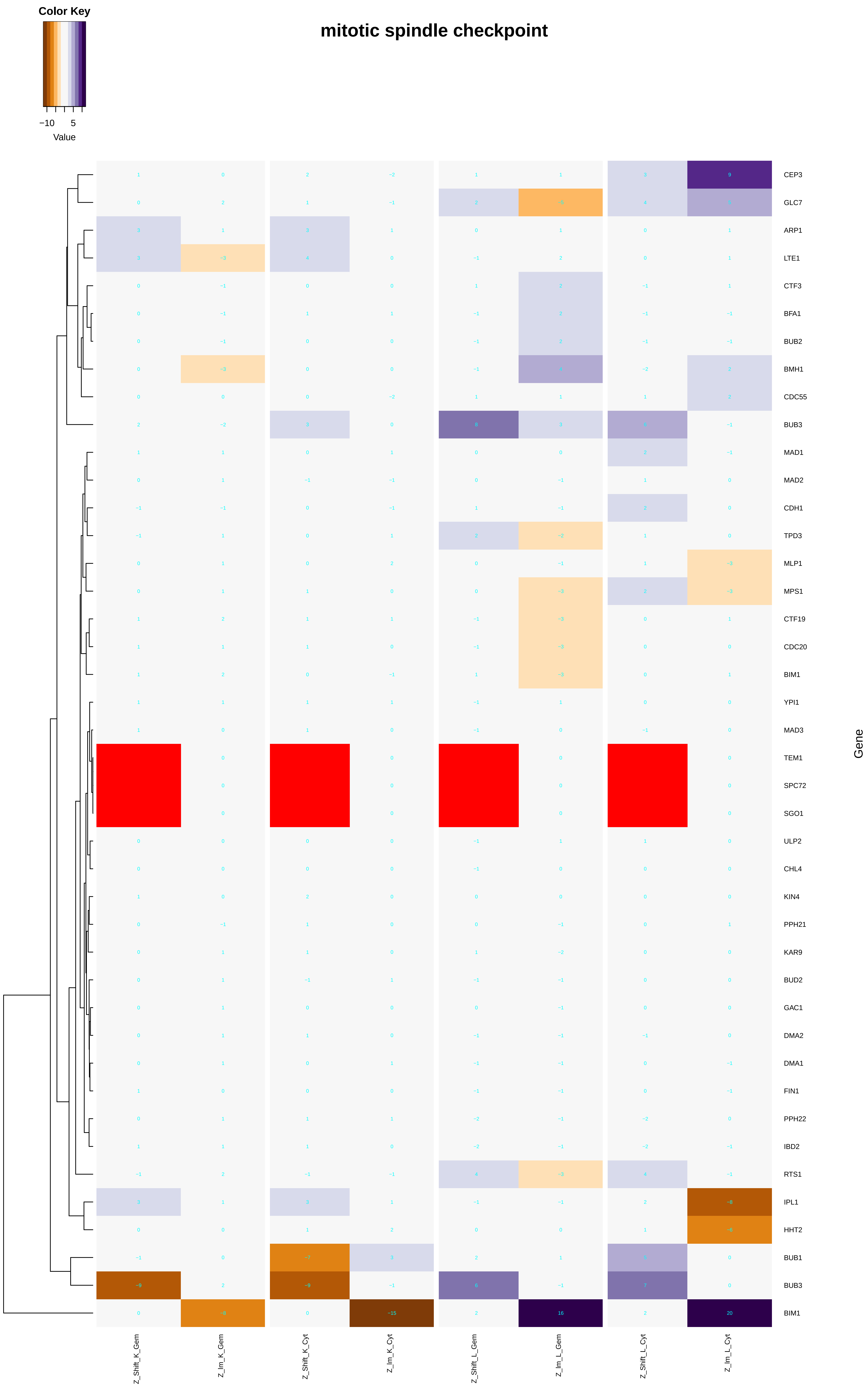

mitotic spindle checkpoint
Color Key
−10 5
Value
Gene
CEP3
GLC7
ARP1
LTE1
CTF3
BFA1
BUB2
BMH1
CDC55
BUB3
MAD1
MAD2
CDH1
TPD3
MLP1
MPS1
CTF19
CDC20
BIM1
YPI1
MAD3
TEM1
SPC72
SGO1
ULP2
CHL4
KIN4
PPH21
KAR9
BUD2
GAC1
DMA2
DMA1
FIN1
PPH22
IBD2
RTS1
IPL1
HHT2
BUB1
BUB3
BIM1
Z_Shift_K_Gem
Z_Im_K_Gem
Z_Shift_K_Cyt
Z_Im_K_Cyt
Z_Shift_L_Gem
Z_Im_L_Gem
Z_Shift_L_Cyt
Z_Im_L_Cyt

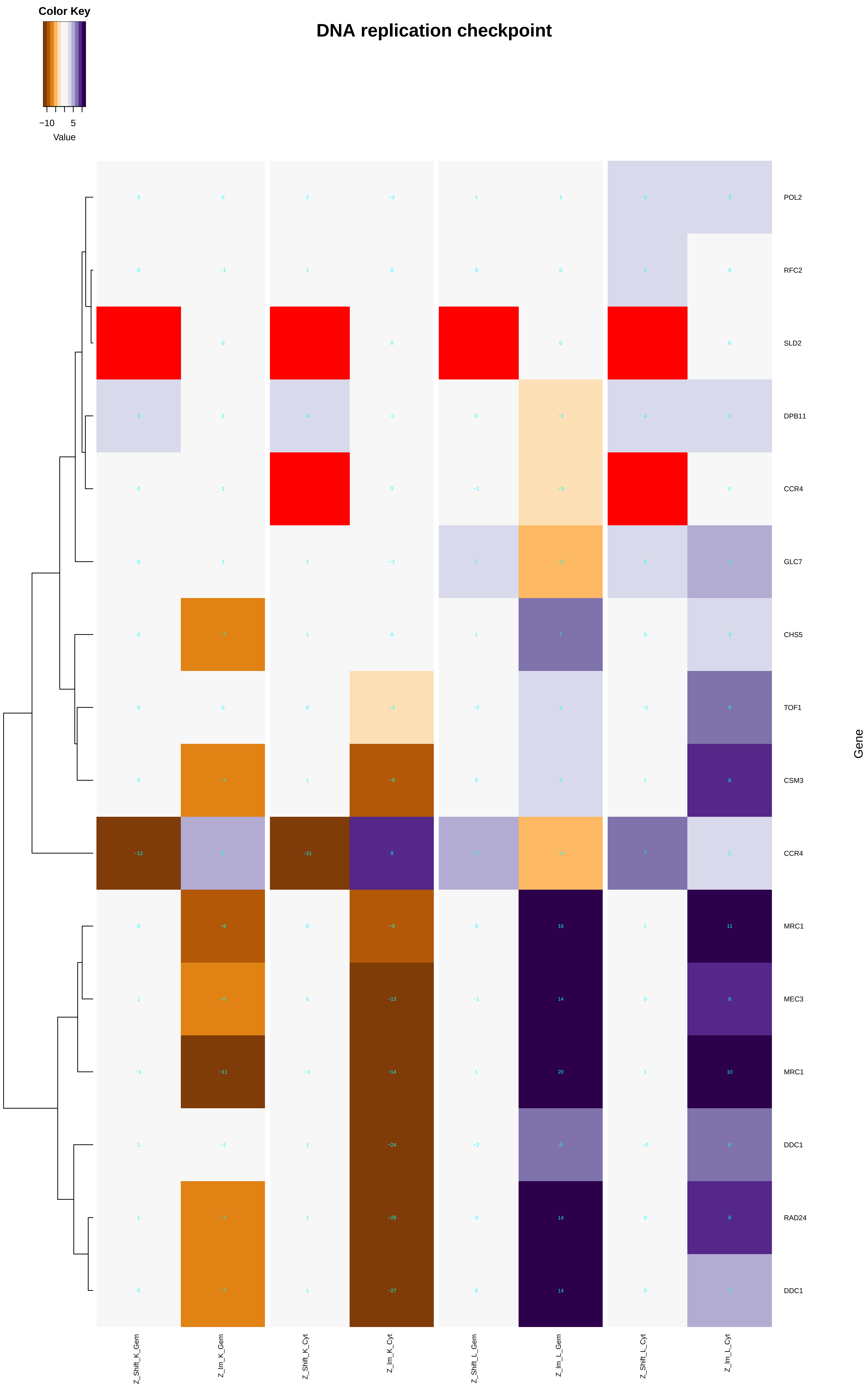
Color Key
−10 5
Value
DNA replication checkpoint
POL2
RFC2
SLD2
DPB11
CCR4
GLC7
CHS5
TOF1
CSM3
CCR4
MRC1
MEC3
MRC1
DDC1
RAD24
DDC1
Gene
Z_Shift_K_Gem
Z_Im_K_Gem
Z_Shift_K_Cyt
Z_Im_K_Cyt
Z_Shift_L_Gem
Z_Im_L_Gem
Z_Shift_L_Cyt
Z_Im_L_Cyt

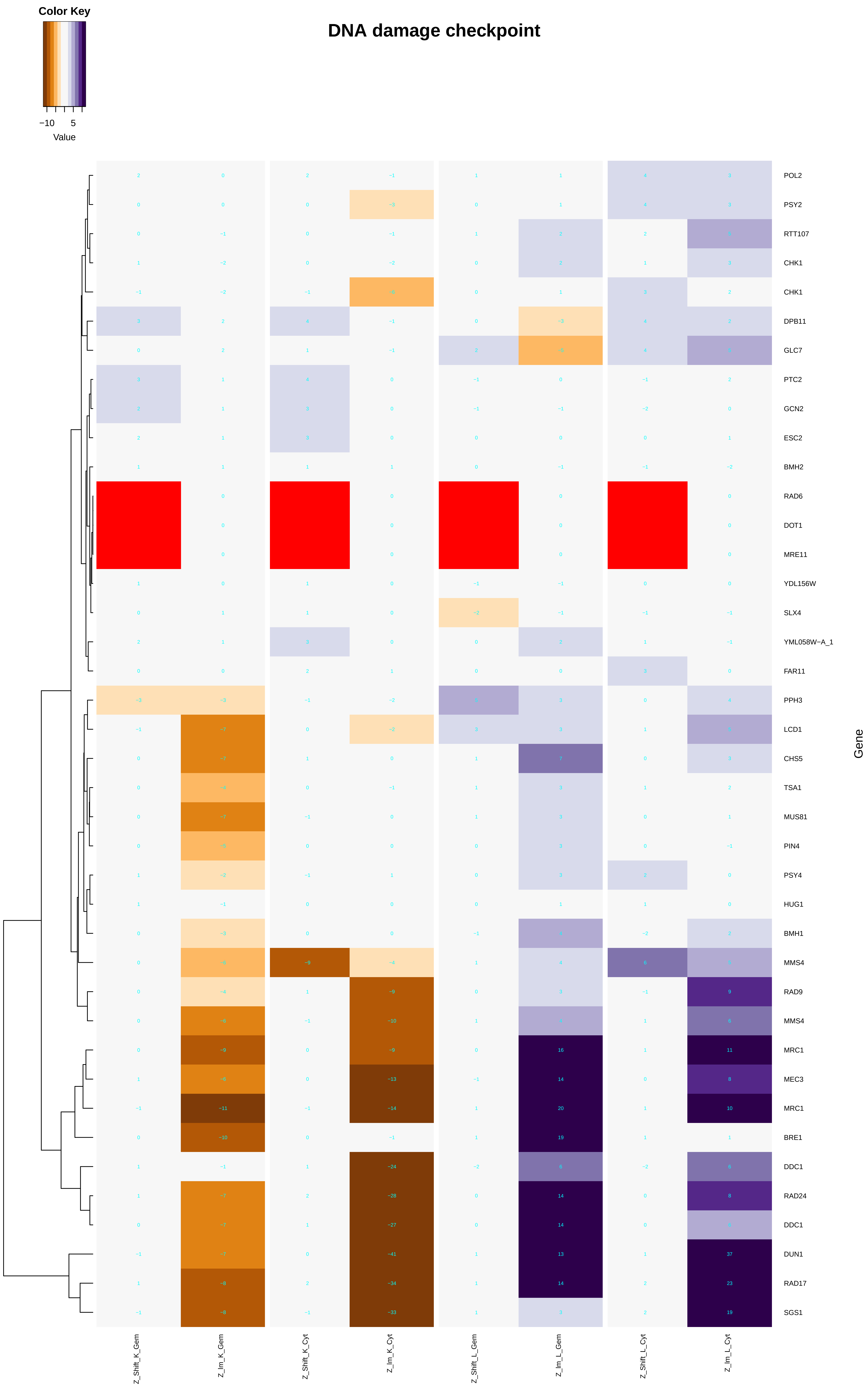
Color Key
−10
5
Value
DNA damage checkpoint
POL2
PSY2
RTT107
CHK1
CHK1
DPB11
GLC7
PTC2
GCN2
ESC2
BMH2
RAD6
DOT1
MRE11
YDL156W
SLX4
YML058W−A_1
FAR11
PPH3
LCD1
CHS5
TSA1
MUS81
PIN4
PSY4
HUG1
BMH1
MMS4
RAD9
MMS4
MRC1
MEC3
MRC1
BRE1
DDC1
RAD24
DDC1
DUN1
RAD17
SGS1
Gene
Z_Shift_K_Gem
Z_Im_K_Gem
Z_Shift_K_Cyt
Z_Im_K_Cyt
Z_Shift_L_Gem
Z_Im_L_Gem
Z_Shift_L_Cyt
Z_Im_L_Cyt

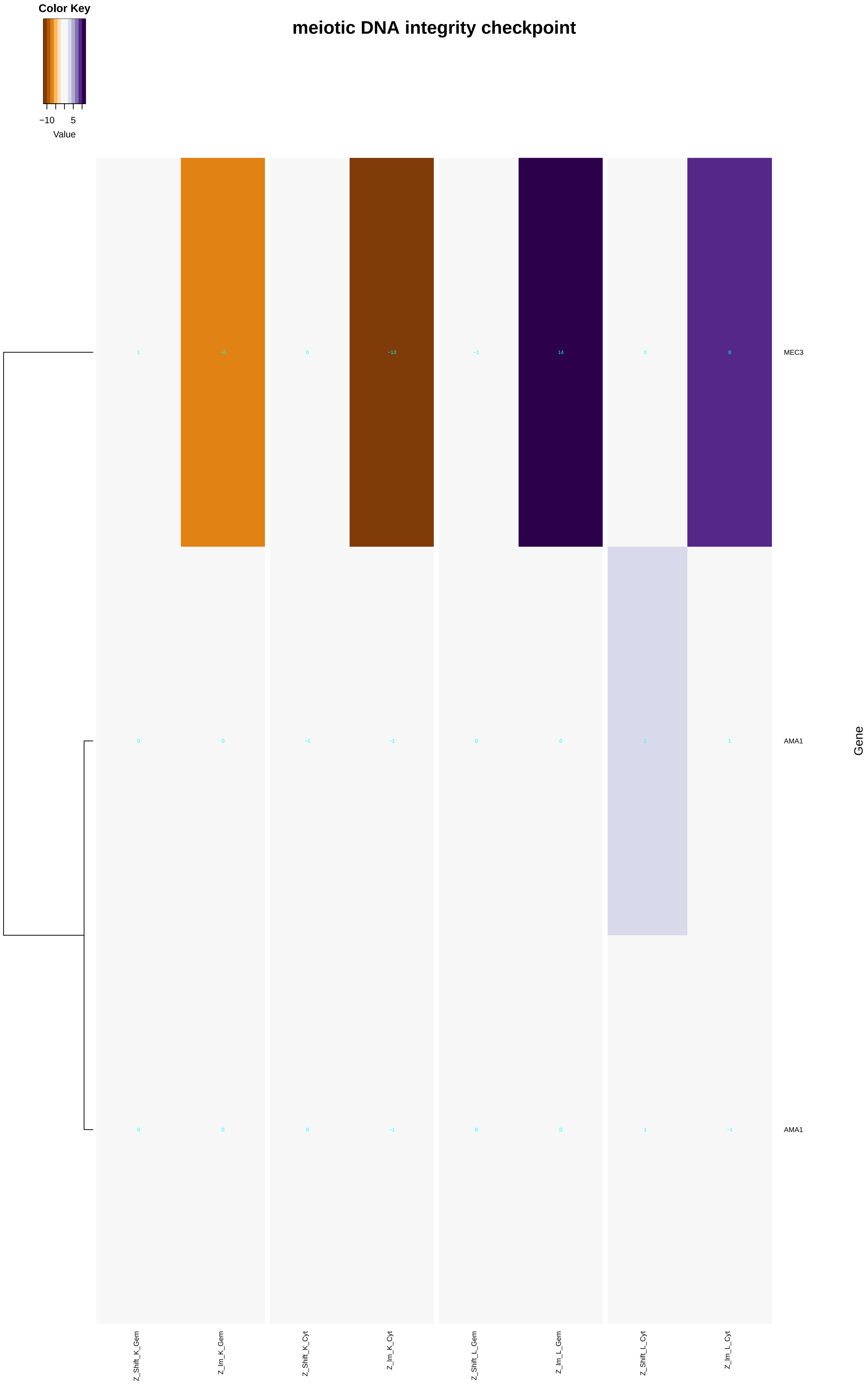

meiotic DNA integrity checkpoint
Color Key
−10
5
Value
MEC3
AMA1
AMA1
Gene
Z_Shift_K_Gem
Z_Im_K_Gem
Z_Shift_K_Cyt
Z_Im_K_Cyt
Z_Shift_L_Gem
Z_Im_L_Gem
Z_Shift_L_Cyt
Z_Im_L_Cyt

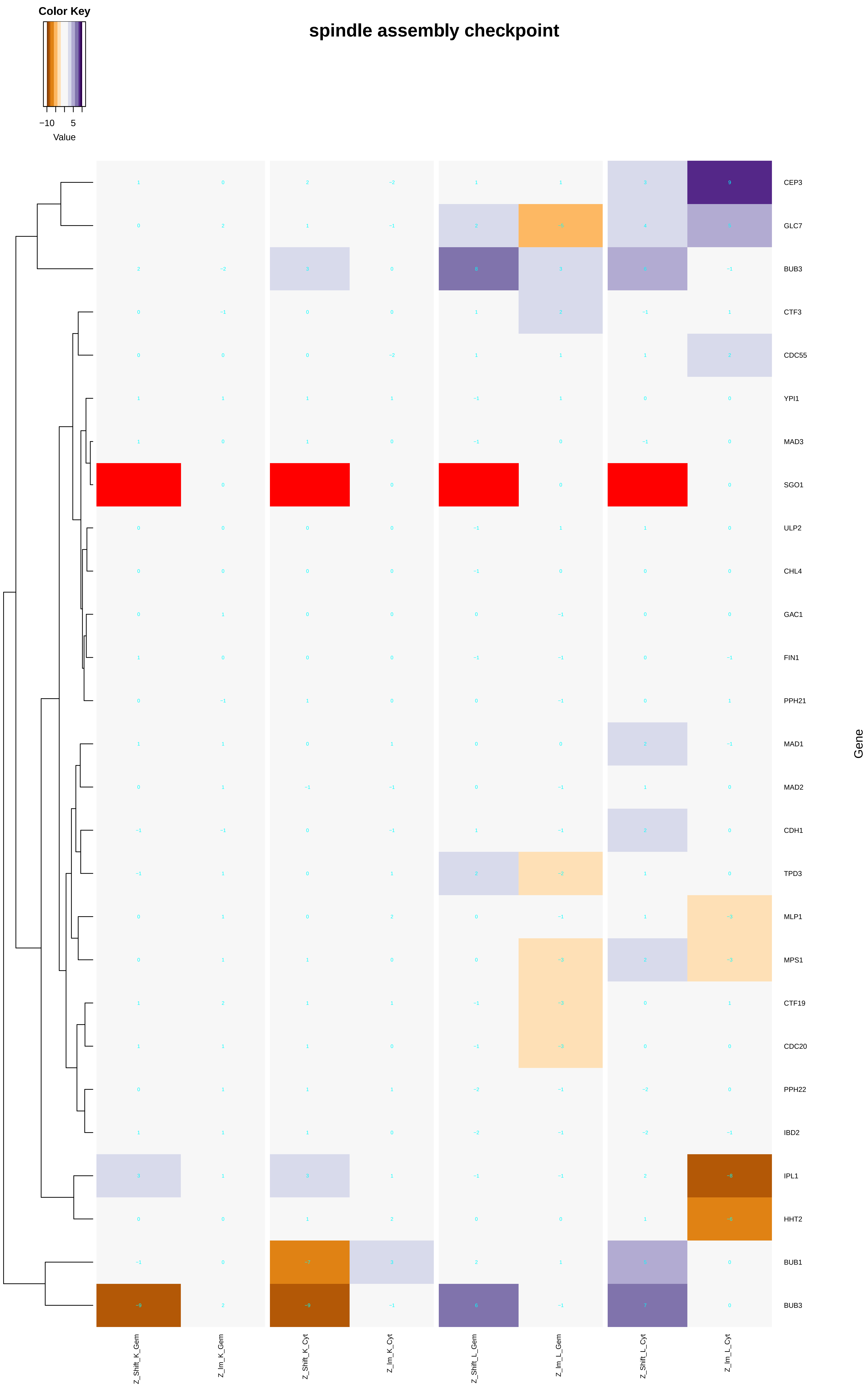

Color Key
−10
5
Value
spindle assembly checkpoint
CEP3
GLC7
BUB3
CTF3
CDC55
YPI1
MAD3
SGO1
ULP2
CHL4
GAC1
FIN1
PPH21
MAD1
MAD2
CDH1
TPD3
MLP1
MPS1
CTF19
CDC20
PPH22
IBD2
IPL1
HHT2
BUB1
BUB3
Gene
Z_Shift_K_Gem
Z_Im_K_Gem
Z_Shift_K_Cyt
Z_Im_K_Cyt
Z_Shift_L_Gem
Z_Im_L_Gem
Z_Shift_L_Cyt
Z_Im_L_Cyt

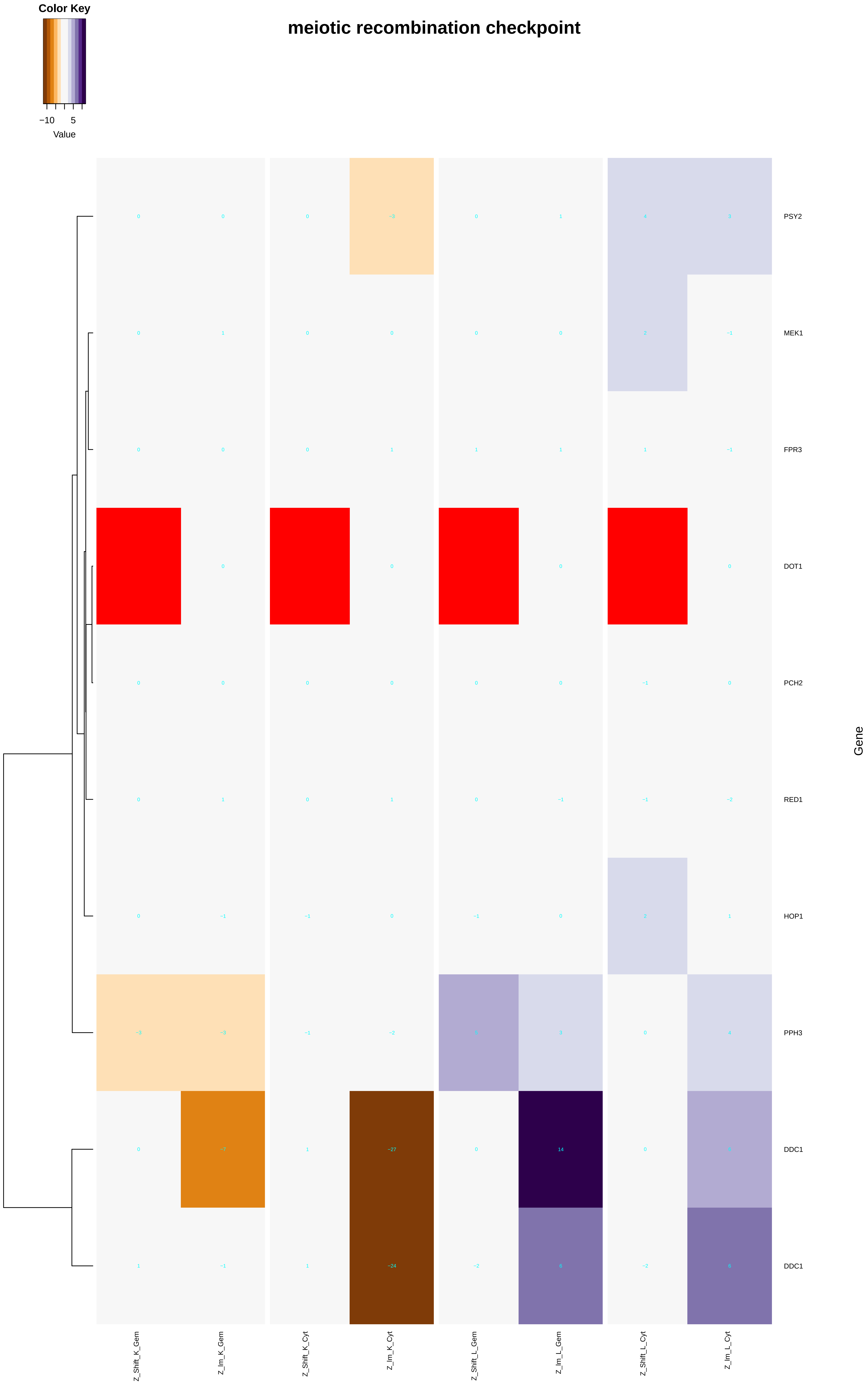

Color Key
−10
5
Value
meiotic recombination checkpoint
PSY2
MEK1
FPR3
DOT1
PCH2
RED1
HOP1
PPH3
DDC1
DDC1
Gene
Z_Shift_K_Gem
Z_Im_K_Gem
Z_Shift_K_Cyt
Z_Im_K_Cyt
Z_Shift_L_Gem
Z_Im_L_Gem
Z_Shift_L_Cyt
Z_Im_L_Cyt

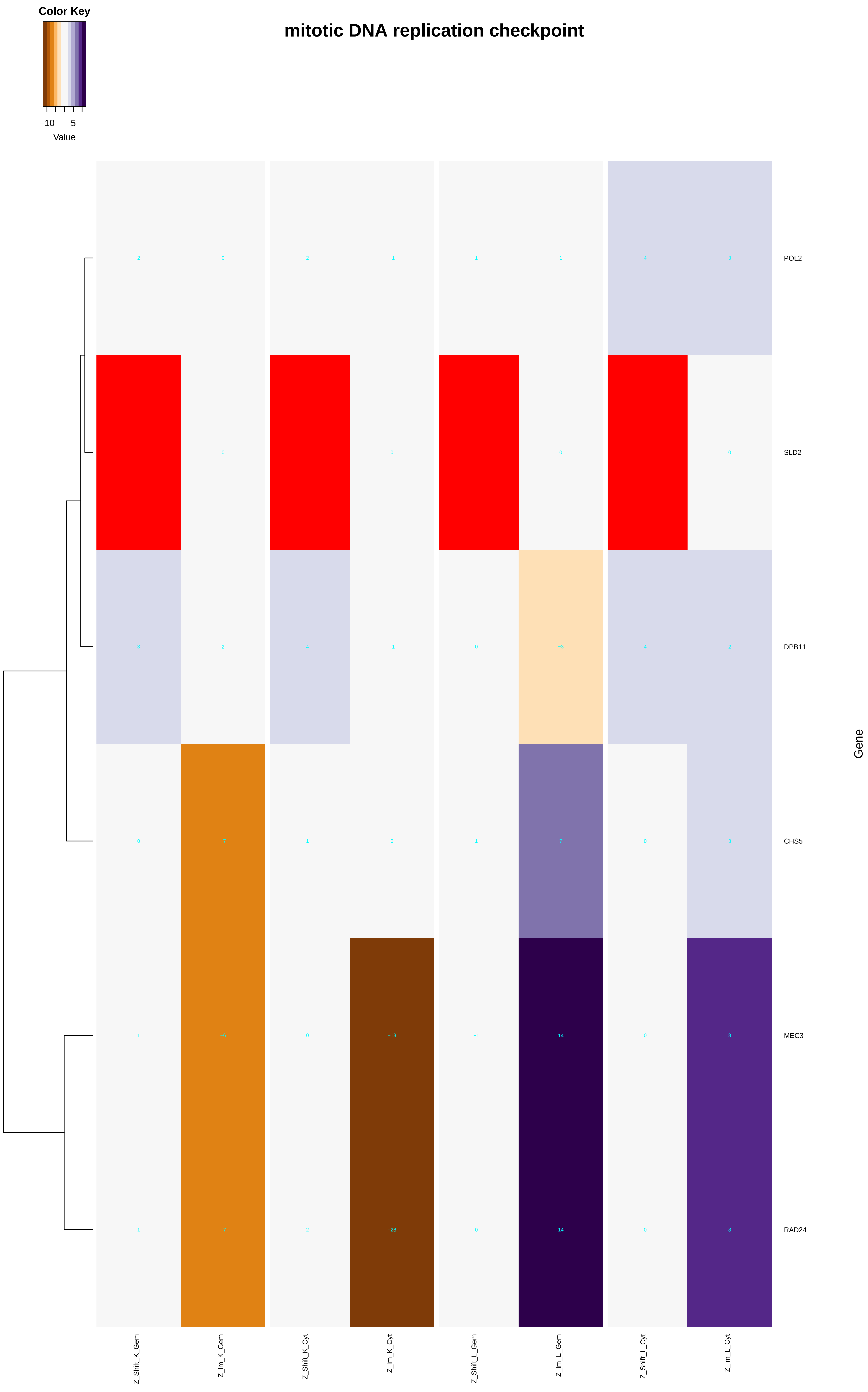
Color Key
−10
5
Value
mitotic DNA replication checkpoint
2 0 2 −1 1 1 4 3
0 0 0 0
3 2 4 −1 0 −3 4 2
0 −7 1 0 1 7 0 3
1 −6 0 −13 −1 14 0 8
1 −7 2 −28 0 14 0 8
POL2
SLD2
DPB11
CHS5
MEC3
RAD24
Gene
Z_Shift_K_Gem
Z_Im_K_Gem
Z_Shift_K_Cyt
Z_Im_K_Cyt
Z_Shift_L_Gem
Z_Im_L_Gem
Z_Shift_L_Cyt
Z_Im_L_Cyt

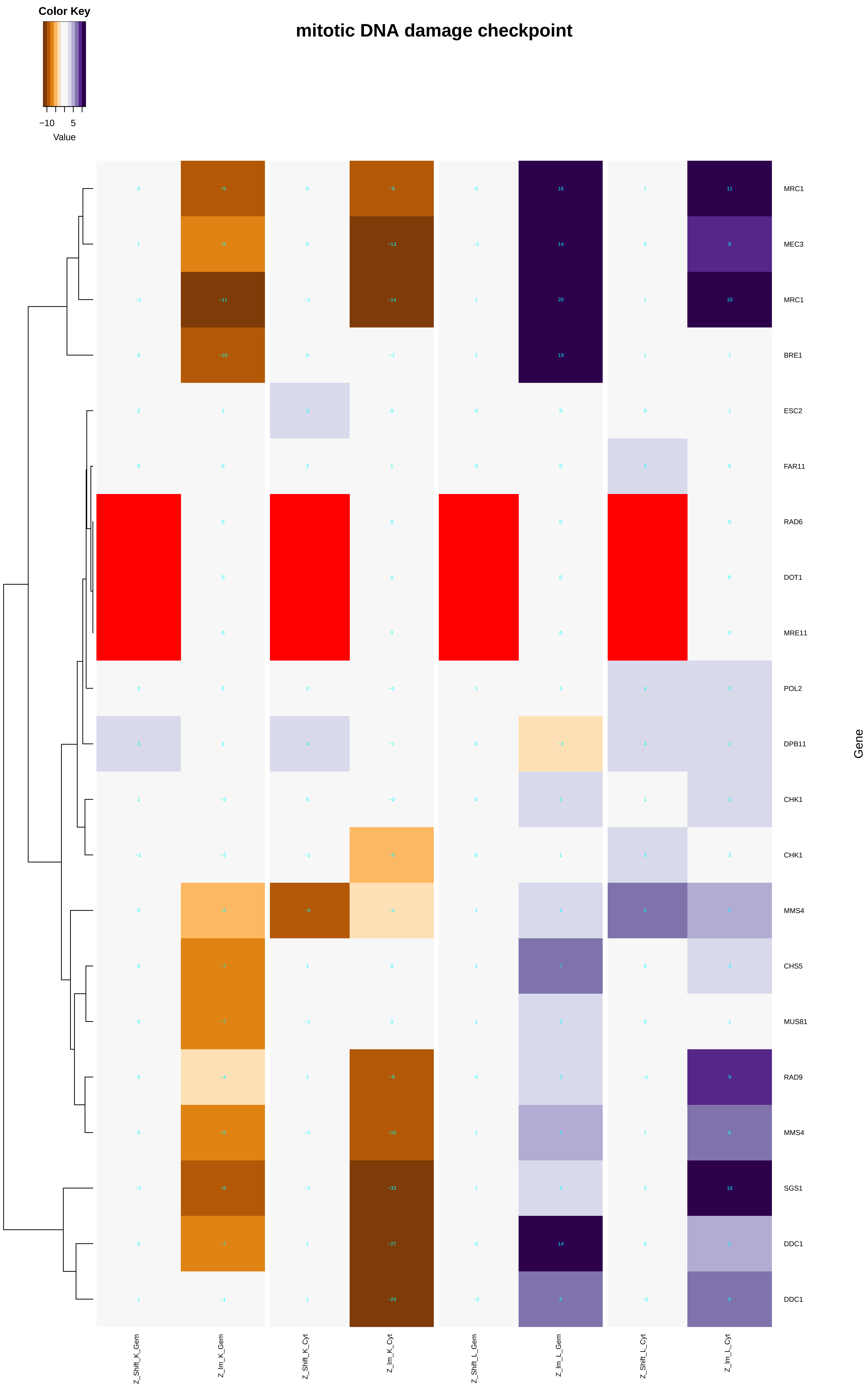

Color Key
−10
5
Value
mitotic DNA damage checkpoint
MRC1
MEC3
MRC1
BRE1
ESC2
FAR11
RAD6
DOT1
MRE11
POL2
DPB11
CHK1
CHK1
MMS4
CHS5
MUS81
RAD9
MMS4
SGS1
DDC1
DDC1
Gene
Z_Shift_K_Gem
Z_Im_K_Gem
Z_Shift_K_Cyt
Z_Im_K_Cyt
Z_Shift_L_Gem
Z_Im_L_Gem
Z_Shift_L_Cyt
Z_Im_L_Cyt

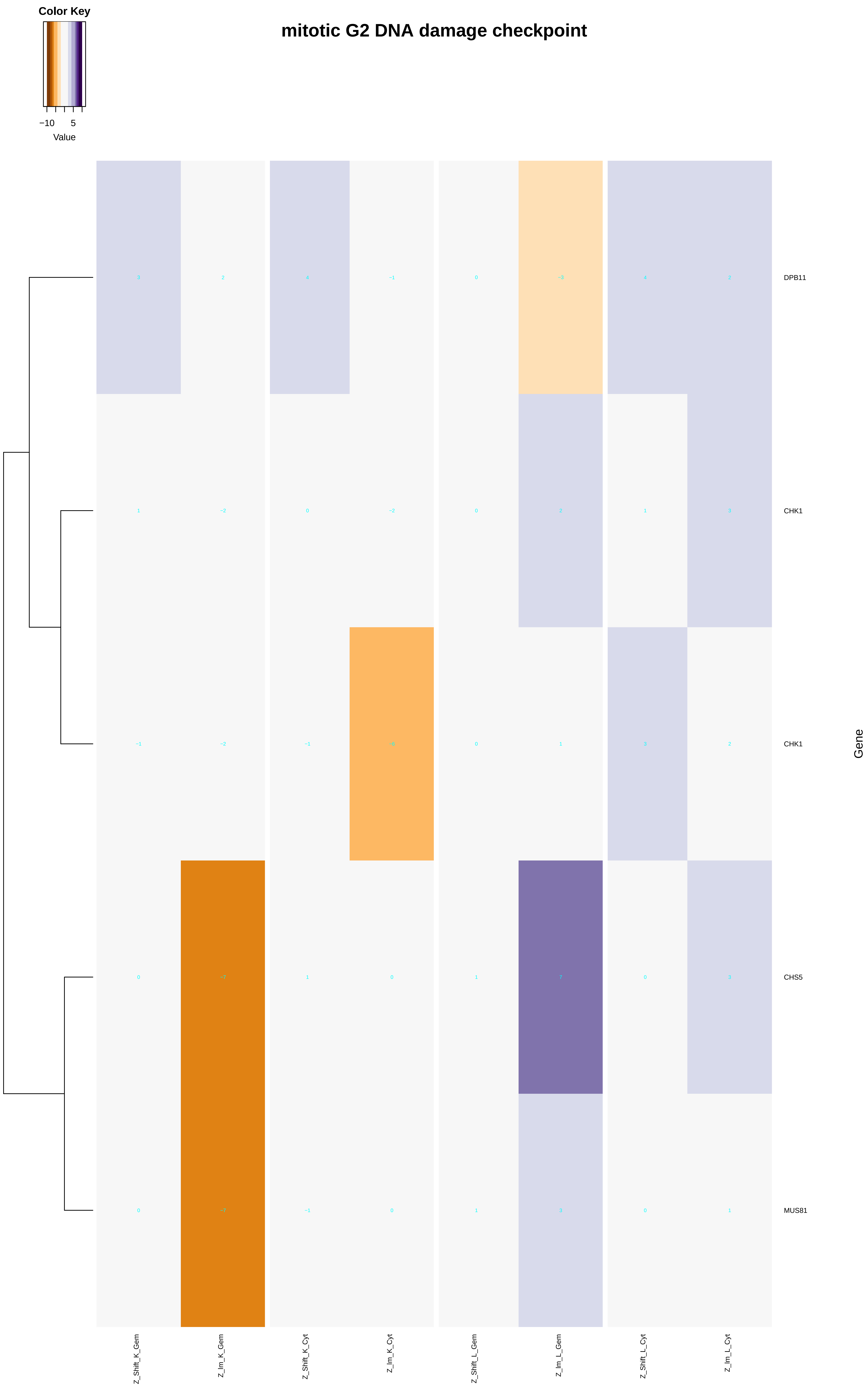

Color Key
−10
5
Value
mitotic G2 DNA damage checkpoint
DPB11
CHK1
CHK1
CHS5
MUS81
Gene
Z_Shift_K_Gem
Z_Im_K_Gem
Z_Shift_K_Cyt
Z_Im_K_Cyt
Z_Shift_L_Gem
Z_Im_L_Gem
Z_Shift_L_Cyt
Z_Im_L_Cyt

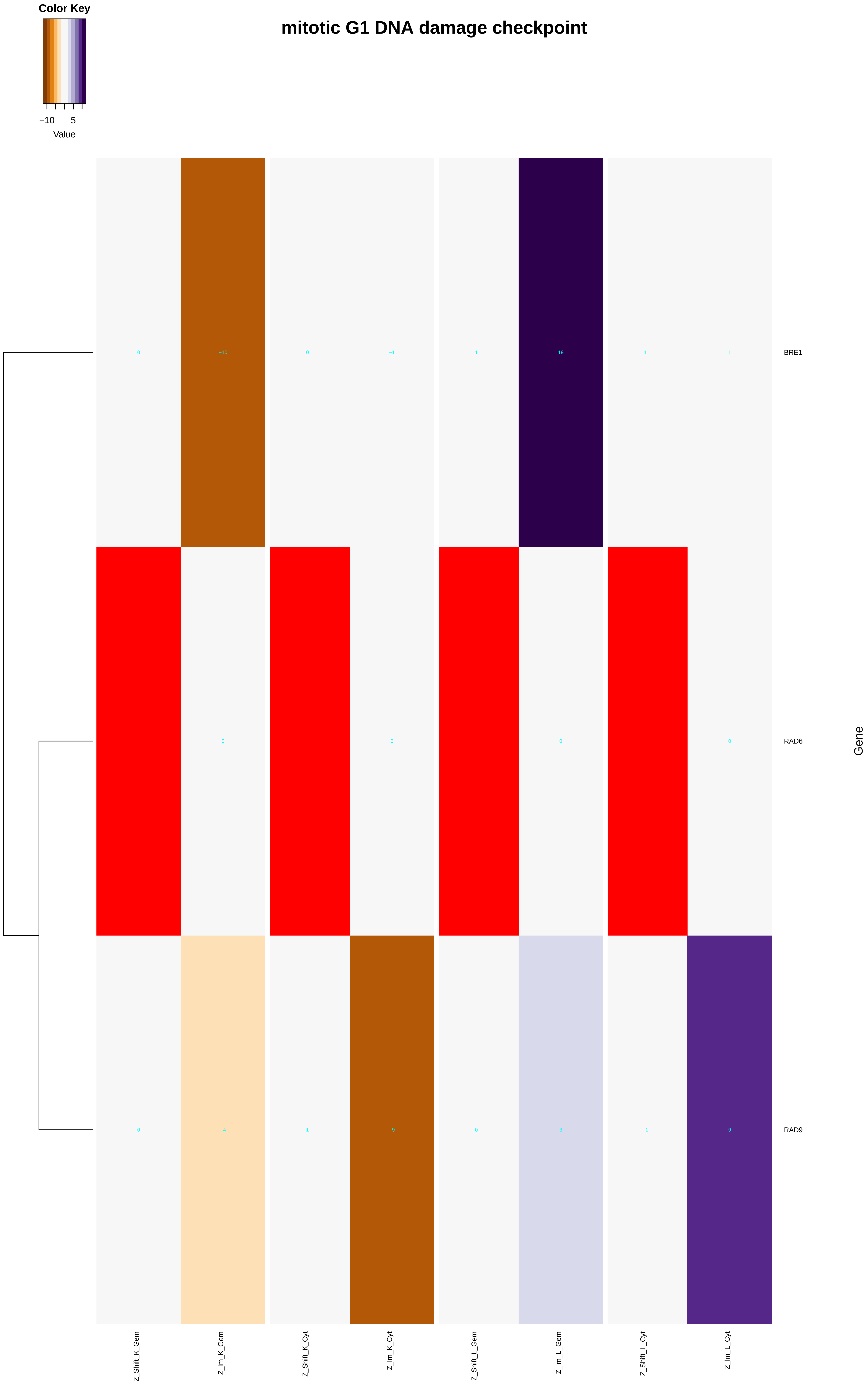
Color Key
−10
5
Value
mitotic G1 DNA damage checkpoint
0
−10
0
−1
1
19
1
1
BRE1
0
0
0
0
RAD6
Gene
0
−4
1
−9
0
3
−1
9
RAD9
Z_Shift_K_Gem
Z_Im_K_Gem
Z_Shift_K_Cyt
Z_Im_K_Cyt
Z_Shift_L_Gem
Z_Im_L_Gem
Z_Shift_L_Cyt
Z_Im_L_Cyt

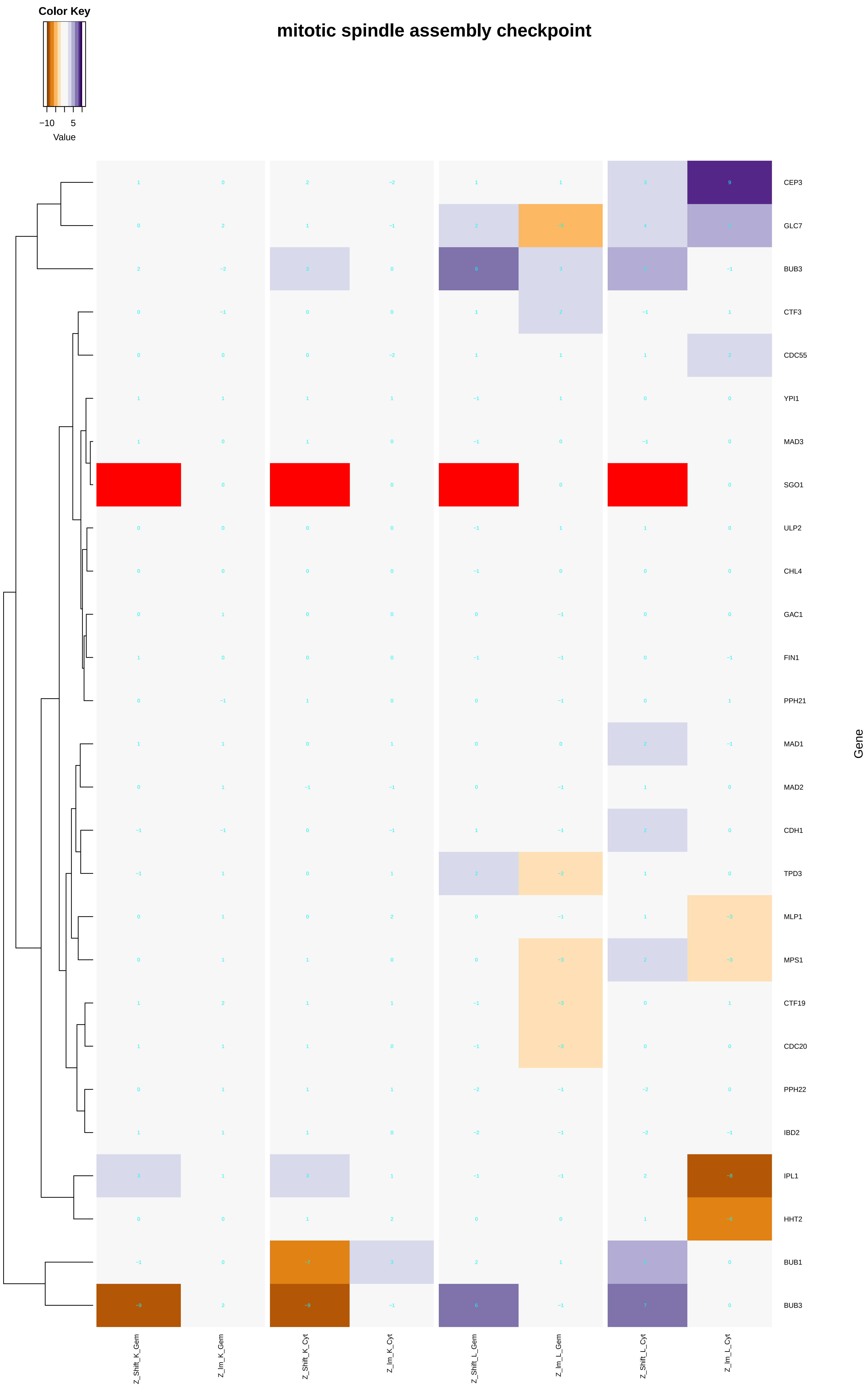

Color Key
−10
5
Value
mitotic spindle assembly checkpoint
CEP3
GLC7
BUB3
CTF3
CDC55
YPI1
MAD3
SGO1
ULP2
CHL4
GAC1
FIN1
PPH21
MAD1
MAD2
CDH1
TPD3
MLP1
MPS1
CTF19
CDC20
PPH22
IBD2
IPL1
HHT2
BUB1
BUB3
Gene
Z_Shift_K_Gem
Z_Im_K_Gem
Z_Shift_K_Cyt
Z_Im_K_Cyt
Z_Shift_L_Gem
Z_Im_L_Gem
Z_Shift_L_Cyt
Z_Im_L_Cyt

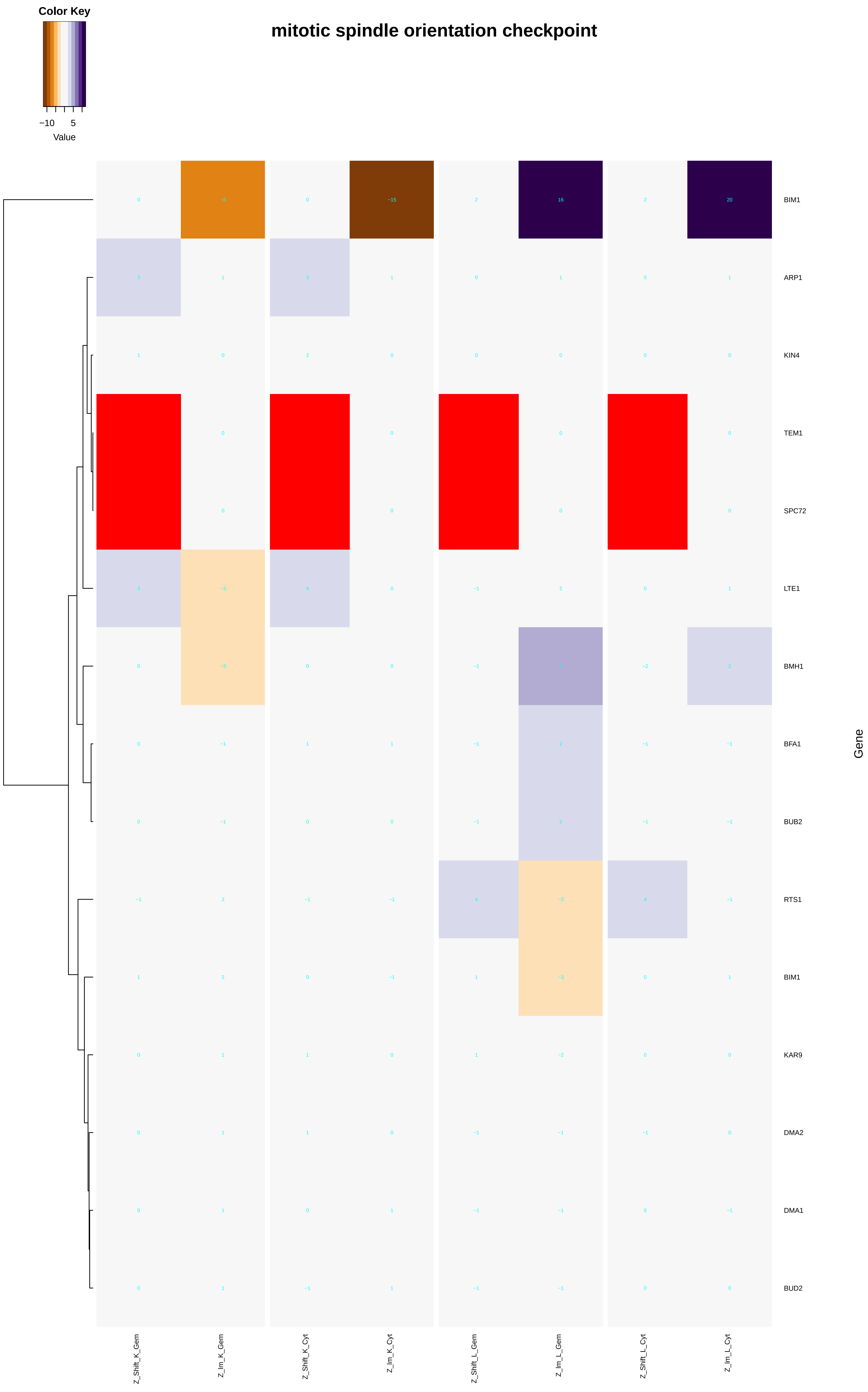

mitotic spindle orientation checkpoint
Color Key
−10
5
Value
BIM1
ARP1
KIN4
TEM1
SPC72
LTE1
BMH1
BFA1
BUB2
RTS1
BIM1
KAR9
DMA2
DMA1
BUD2
Gene
Z_Shift_K_Gem
Z_Im_K_Gem
Z_Shift_K_Cyt
Z_Im_K_Cyt
Z_Shift_L_Gem
Z_Im_L_Gem
Z_Shift_L_Cyt
Z_Im_L_Cyt

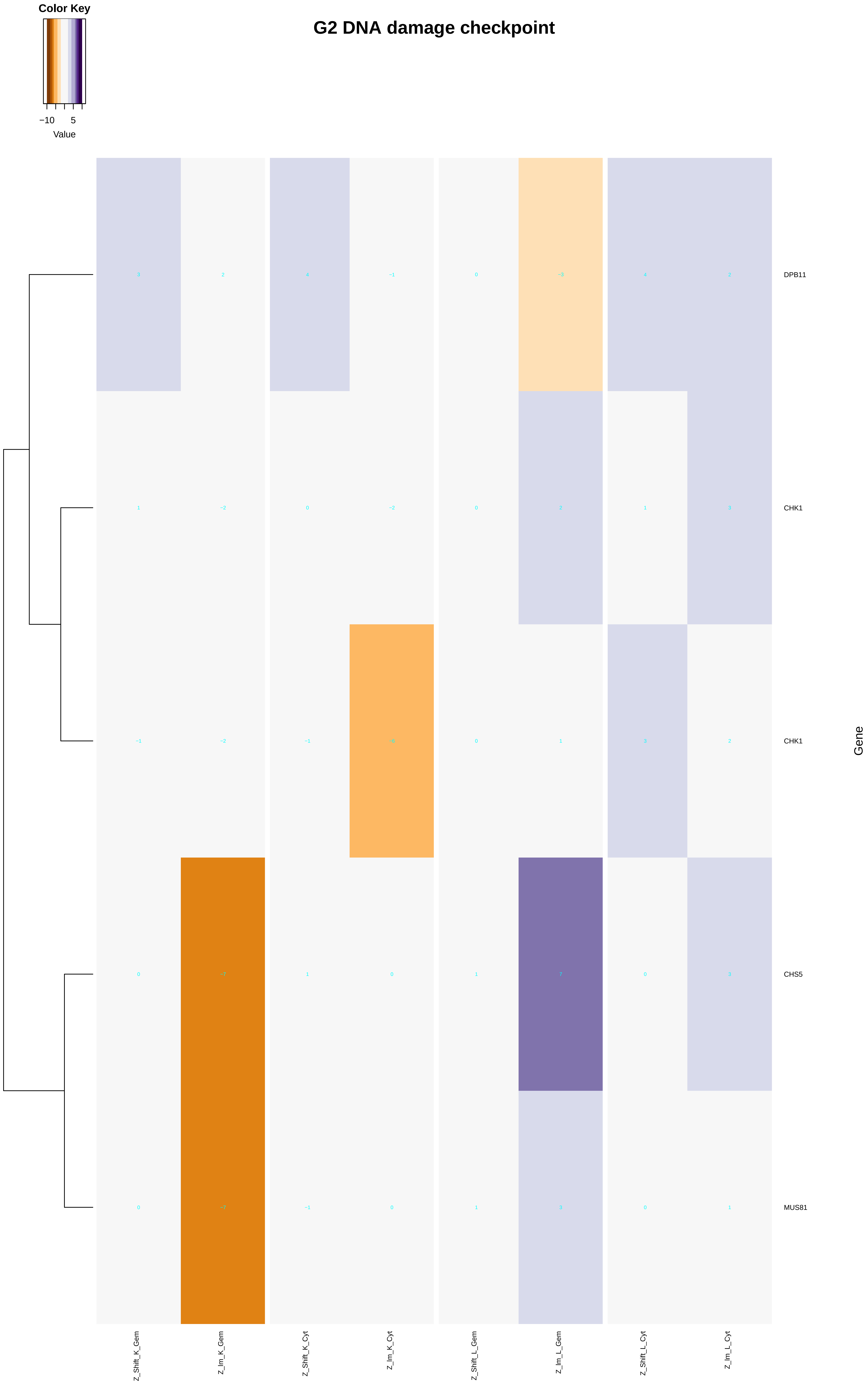
Color Key
−10
5
Value
G2 DNA damage checkpoint
DPB11
CHK1
CHK1
CHS5
MUS81
Gene
Z_Shift_K_Gem
Z_Im_K_Gem
Z_Shift_K_Cyt
Z_Im_K_Cyt
Z_Shift_L_Gem
Z_Im_L_Gem
Z_Shift_L_Cyt
Z_Im_L_Cyt

### G1 DNA damage checkpoint

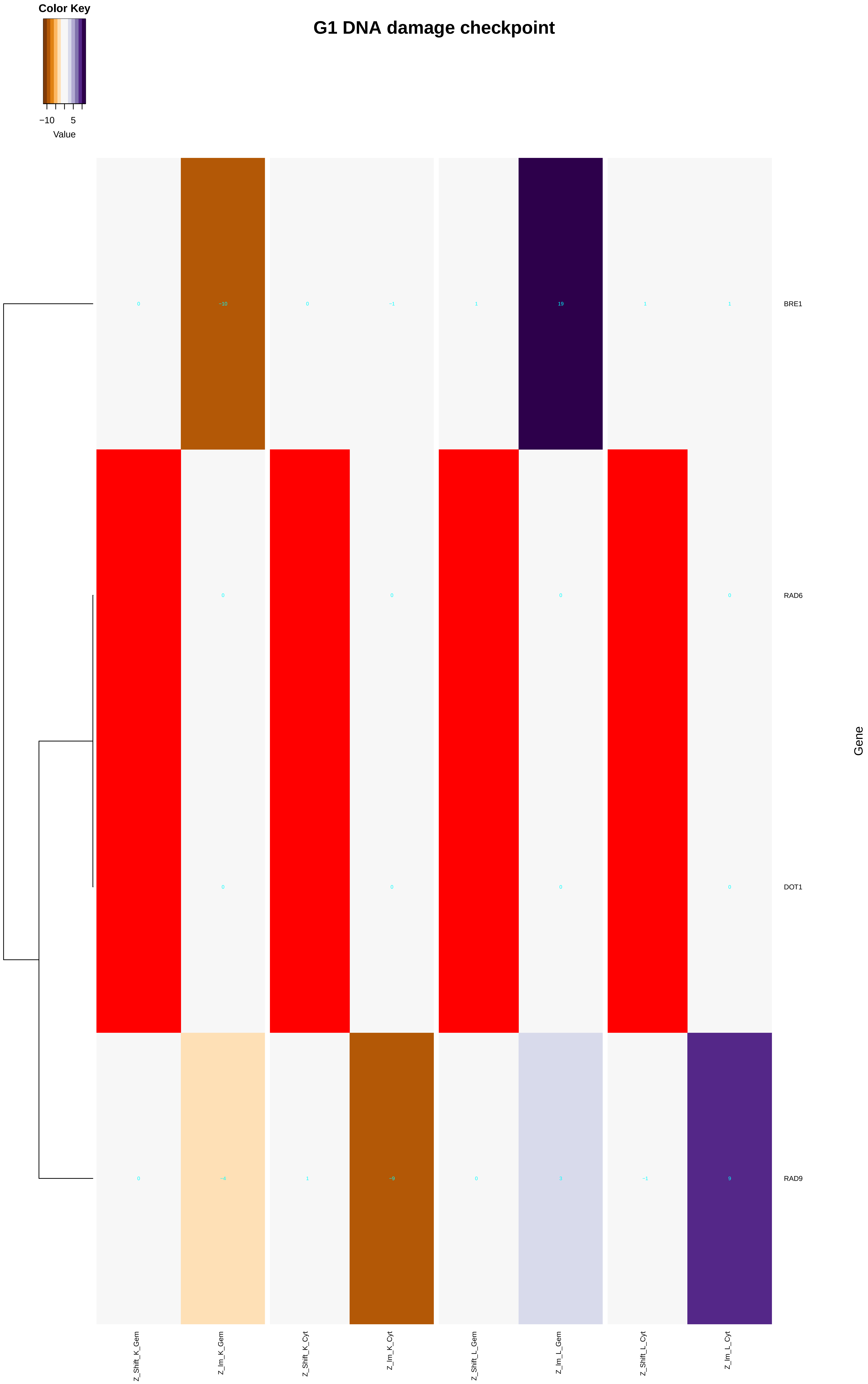

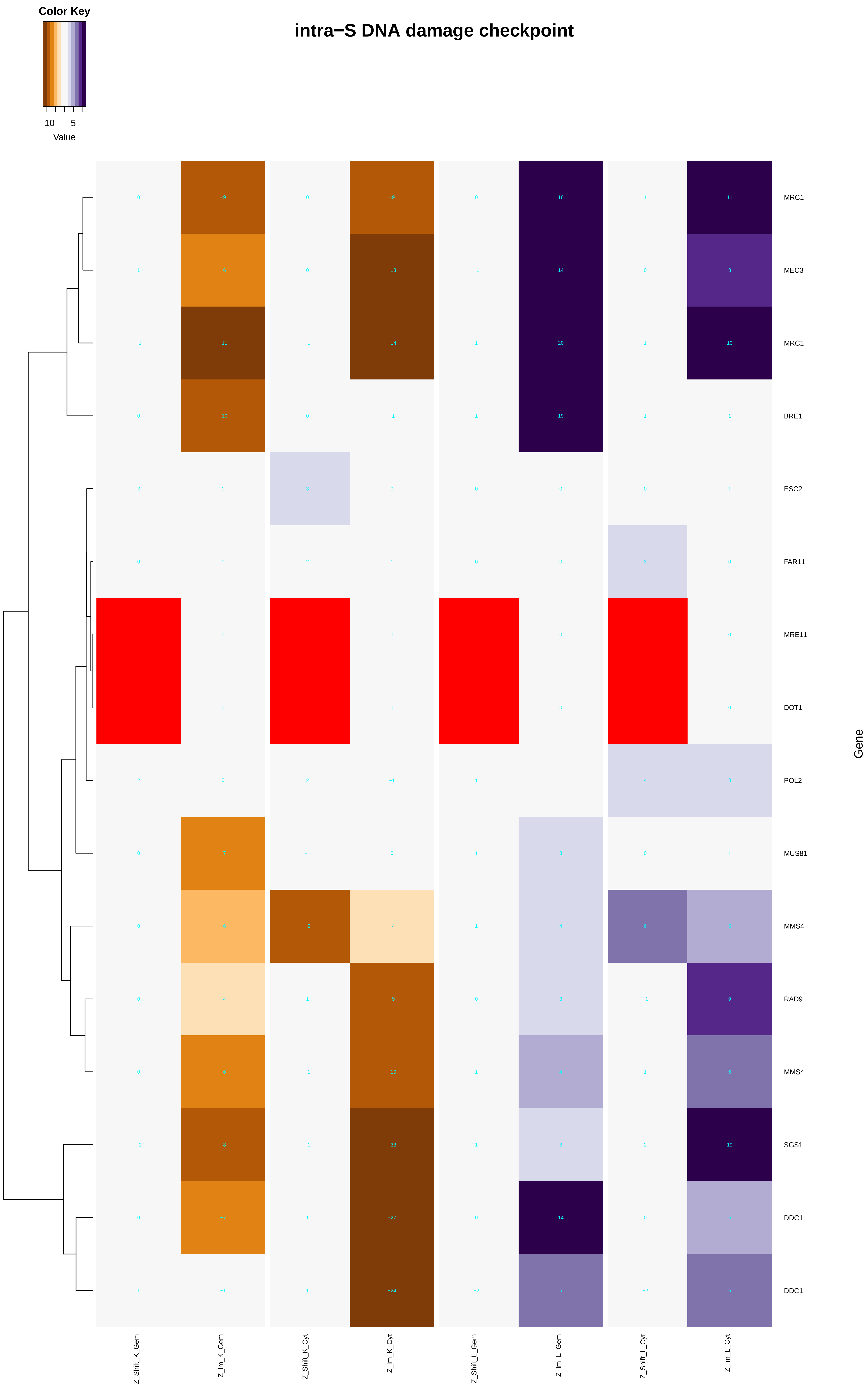

Color Key
−10
5
Value
intra−S DNA damage checkpoint
MRC1
MEC3
MRC1
BRE1
ESC2
FAR11
MRE11
DOT1
POL2
MUS81
MMS4
RAD9
MMS4
SGS1
DDC1
DDC1
Gene
Z_Shift_K_Gem
Z_Im_K_Gem
Z_Shift_K_Cyt
Z_Im_K_Cyt
Z_Shift_L_Gem
Z_Im_L_Gem
Z_Shift_L_Cyt
Z_Im_L_Cyt
