## Additional File 7 for "A humanized yeast phenomic model of deoxycytidine kinase to predict genetic buffering of nucleoside analog cytotoxicity": cellular_amide_metabolic_process.pdf

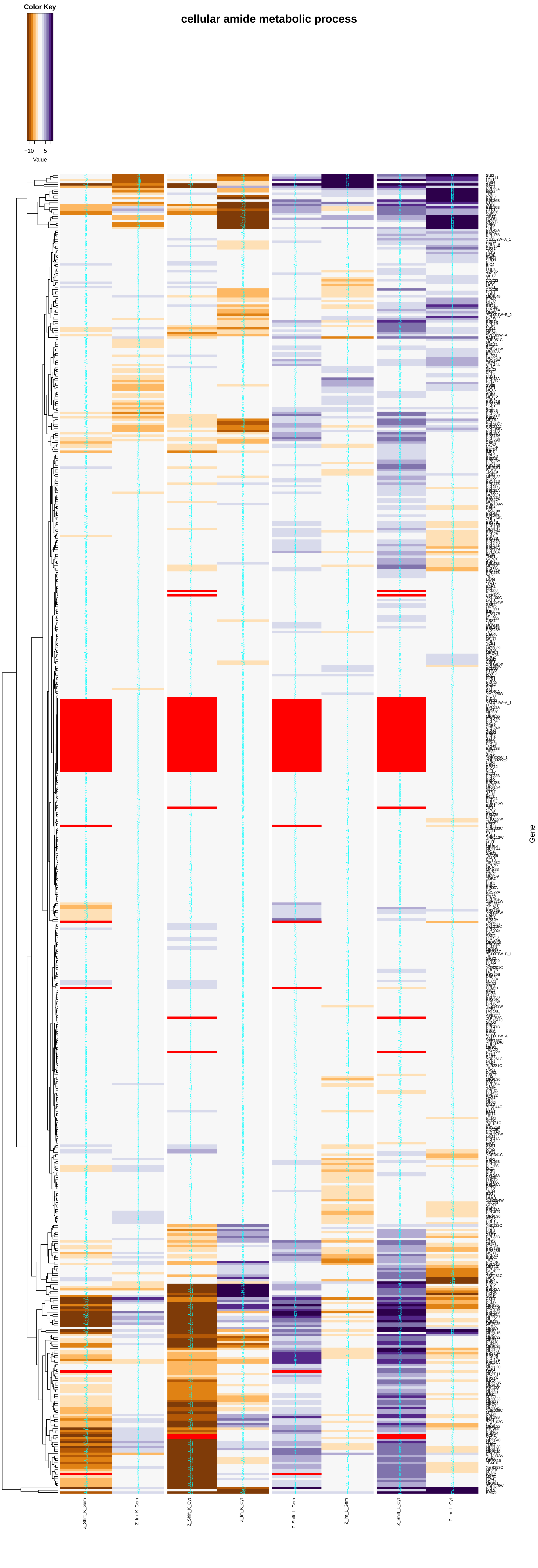

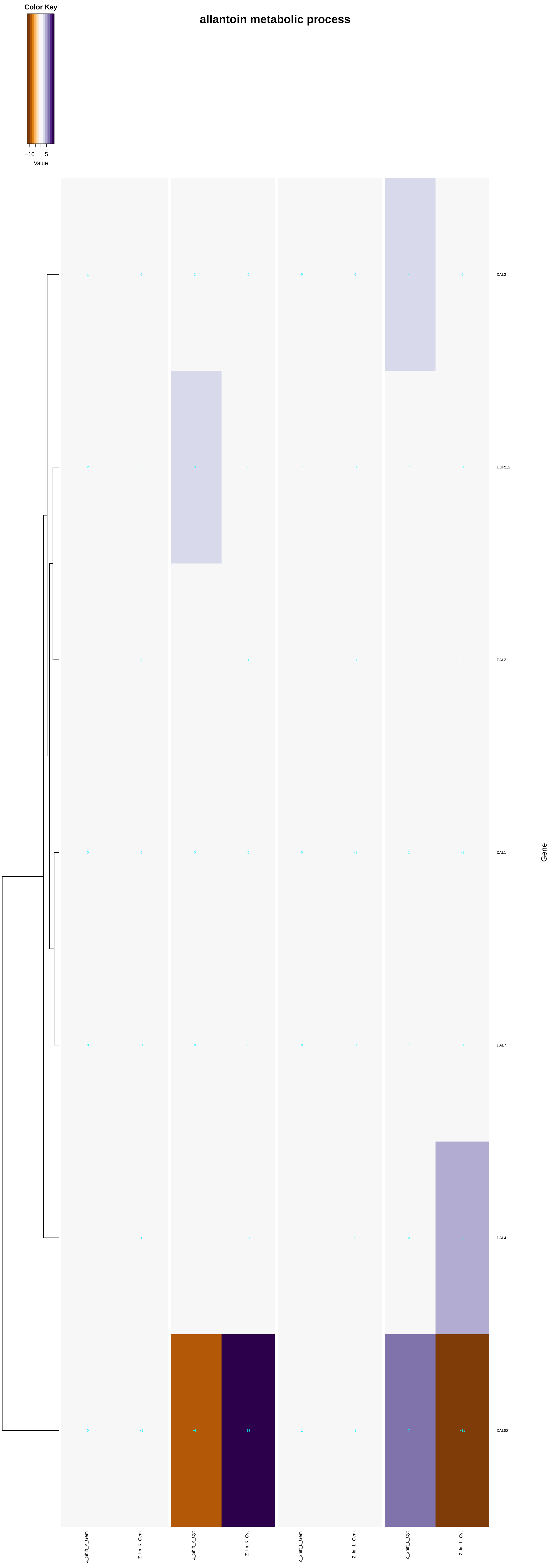
allantoin metabolic process
Color Key
−10
5
Value
DAL3
DUR1,2
DAL2
DAL1
DAL7
DAL4
DAL82
Gene
Z_Shift_K_Gem
Z_Im_K_Gem
Z_Shift_K_Cyt
Z_Im_K_Cyt
Z_Shift_L_Gem
Z_Im_L_Gem
Z_Shift_L_Cyt
Z_Im_L_Cyt

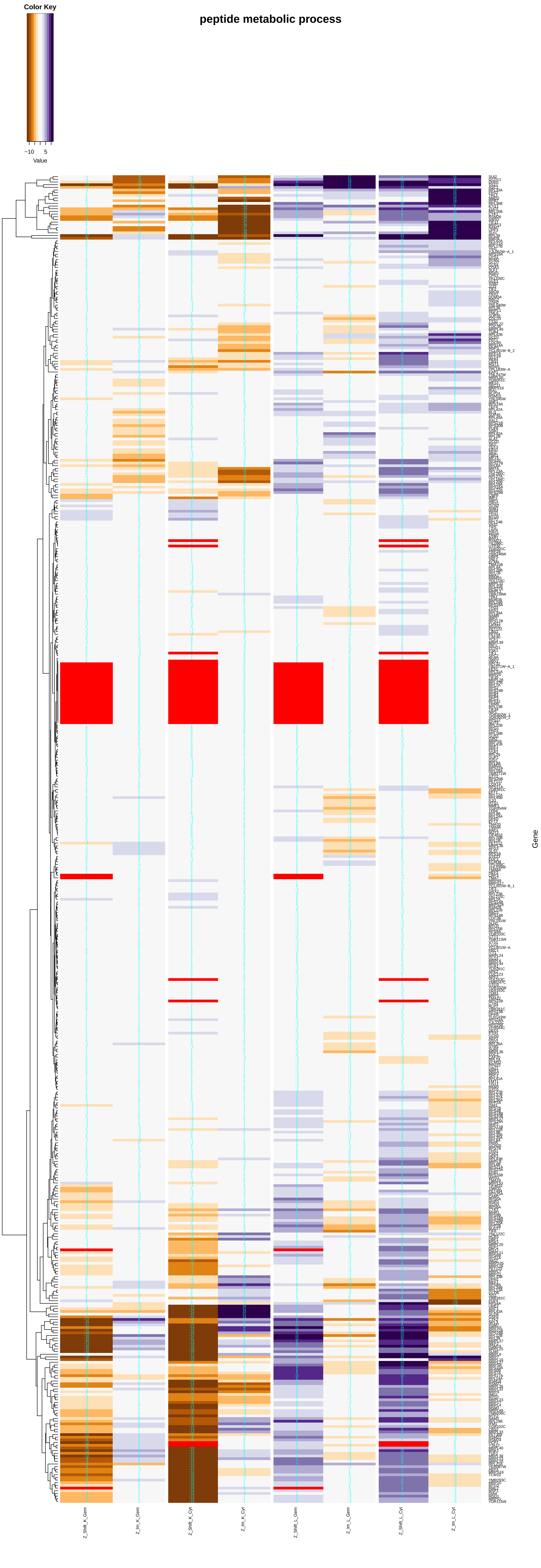
peptide metabolic process
Color Key
−10
5
Value
Z_Shift_K_Gem
Z_Im_K_Gem
Z_Shift_K_Cyt
Z_Im_K_Cyt
Z_Shift_L_Gem
Z_Im_L_Gem
Z_Shift_L_Cyt
Z_Im_L_Cyt
Gene

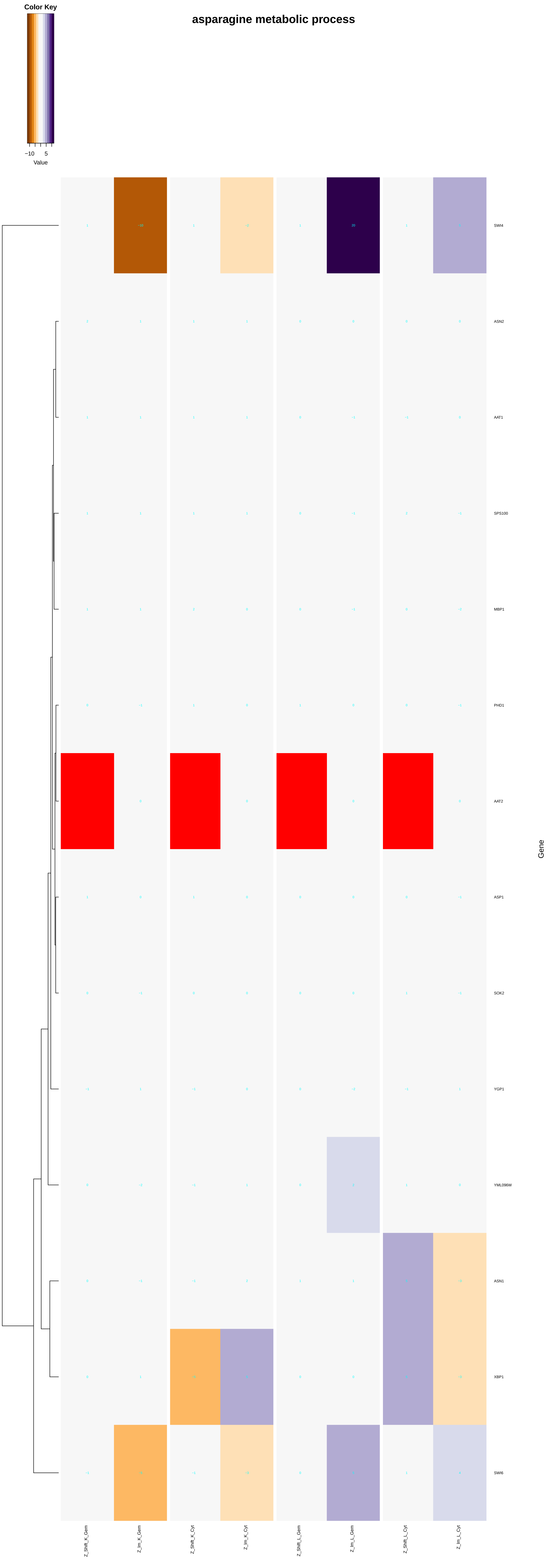
asparagine metabolic process
Color Key
−10
5
Value
SWI4
ASN2
AAT1
SPS100
MBP1
PHD1
AAT2
ASP1
SOK2
YGP1
YML096W
ASN1
XBP1
SWI6
Gene
Z_Shift_K_Gem
Z_Im_K_Gem
Z_Shift_K_Cyt
Z_Im_K_Cyt
Z_Shift_L_Gem
Z_Im_L_Gem
Z_Shift_L_Cyt
Z_Im_L_Cyt

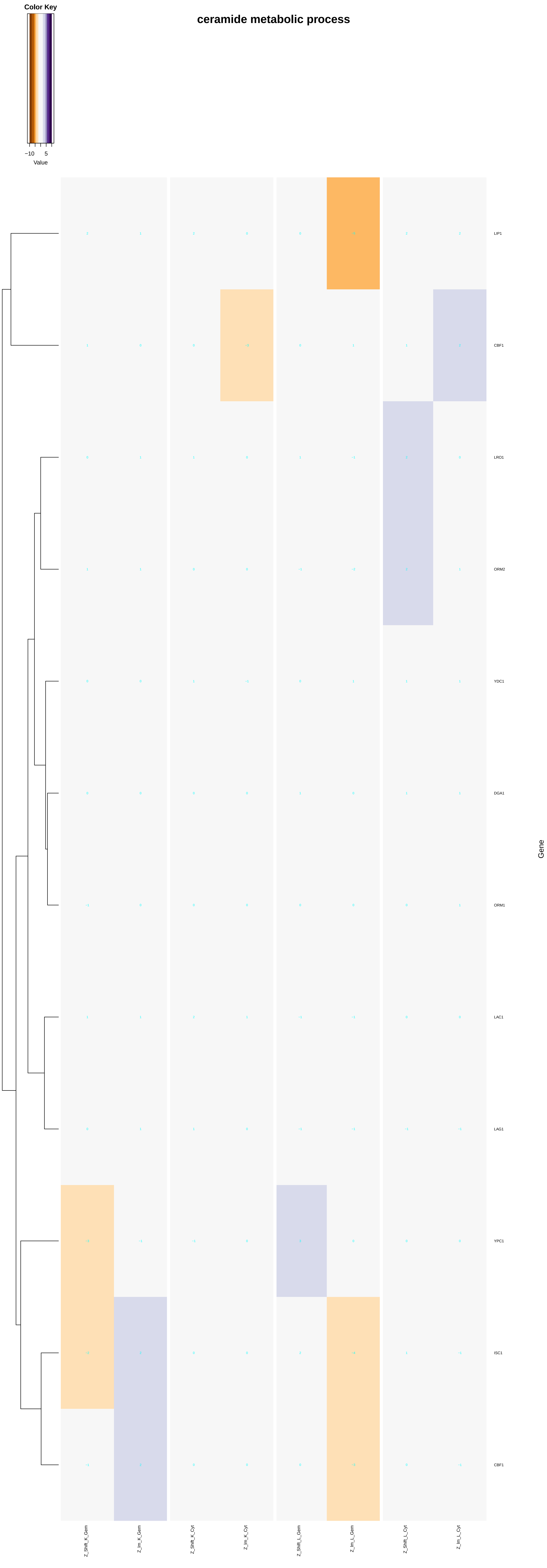
ceramide metabolic process
Color Key
−10
5
Value
LIP1
CBF1
LRO1
ORM2
YDC1
DGA1
ORM1
LAC1
LAG1
YPC1
ISC1
CBF1
Gene
Z_Shift_K_Gem
Z_Im_K_Gem
Z_Shift_K_Cyt
Z_Im_K_Cyt
Z_Shift_L_Gem
Z_Im_L_Gem
Z_Shift_L_Cyt
Z_Im_L_Cyt

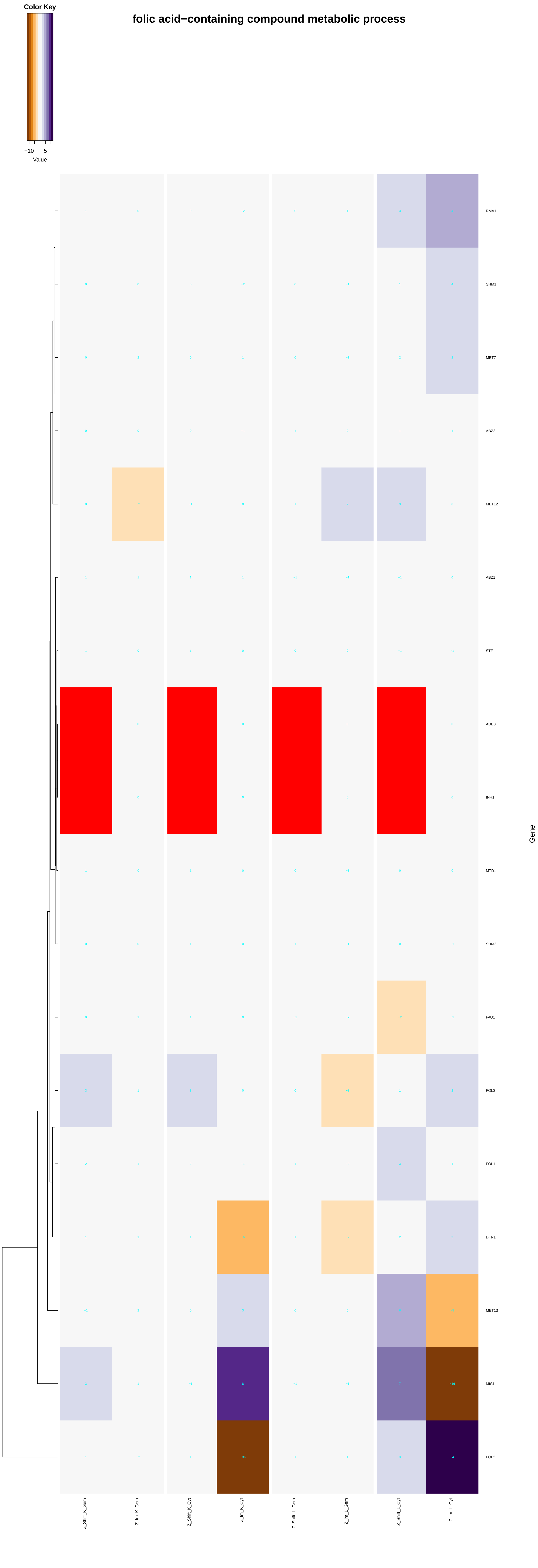
folic acid−containing compound metabolic process
Color Key
−10
5
Value
RMA1
SHM1
MET7
ABZ2
MET12
ABZ1
STF1
ADE3
INH1
MTD1
SHM2
FAU1
FOL3
FOL1
DFR1
MET13
MIS1
FOL2
Gene
Z_Shift_K_Gem
Z_Im_K_Gem
Z_Shift_K_Cyt
Z_Im_K_Cyt
Z_Shift_L_Gem
Z_Im_L_Gem
Z_Shift_L_Cyt
Z_Im_L_Cyt

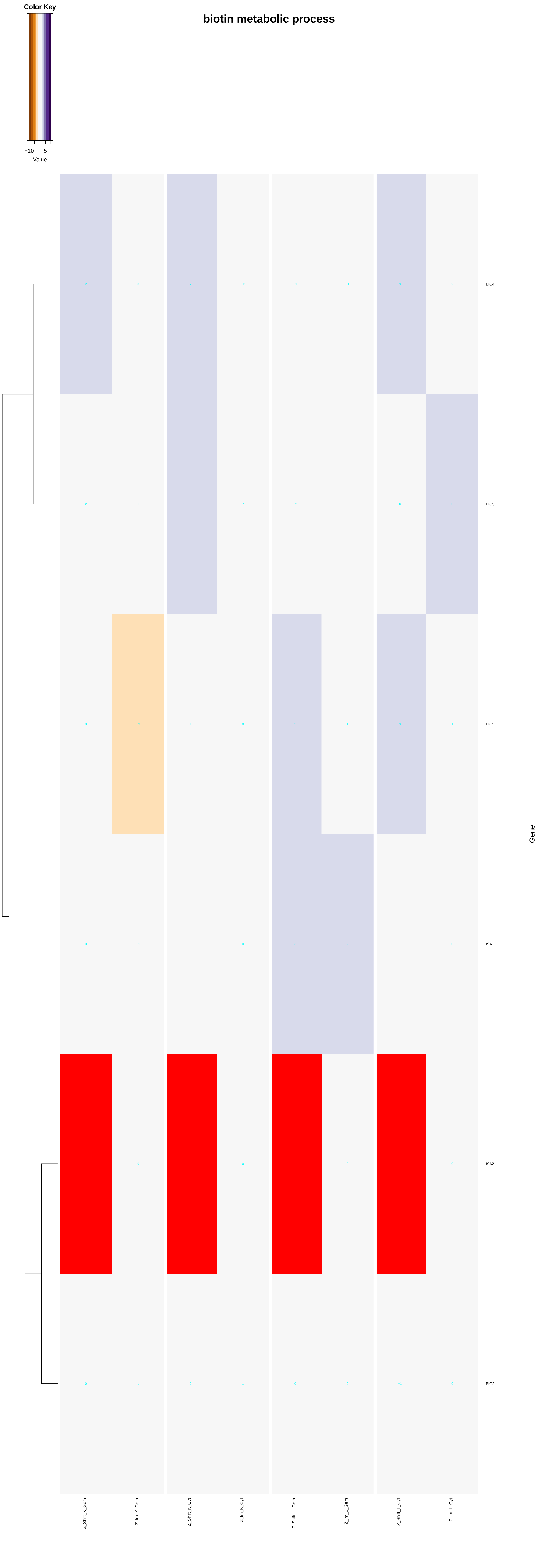
Color Key
−10
5
Value
biotin metabolic process
BIO4
BIO3
BIO5
ISA1
ISA2
BIO2
Gene
Z_Shift_K_Gem
Z_Im_K_Gem
Z_Shift_K_Cyt
Z_Im_K_Cyt
Z_Shift_L_Gem
Z_Im_L_Gem
Z_Shift_L_Cyt
Z_Im_L_Cyt

### pantothenate metabolic process

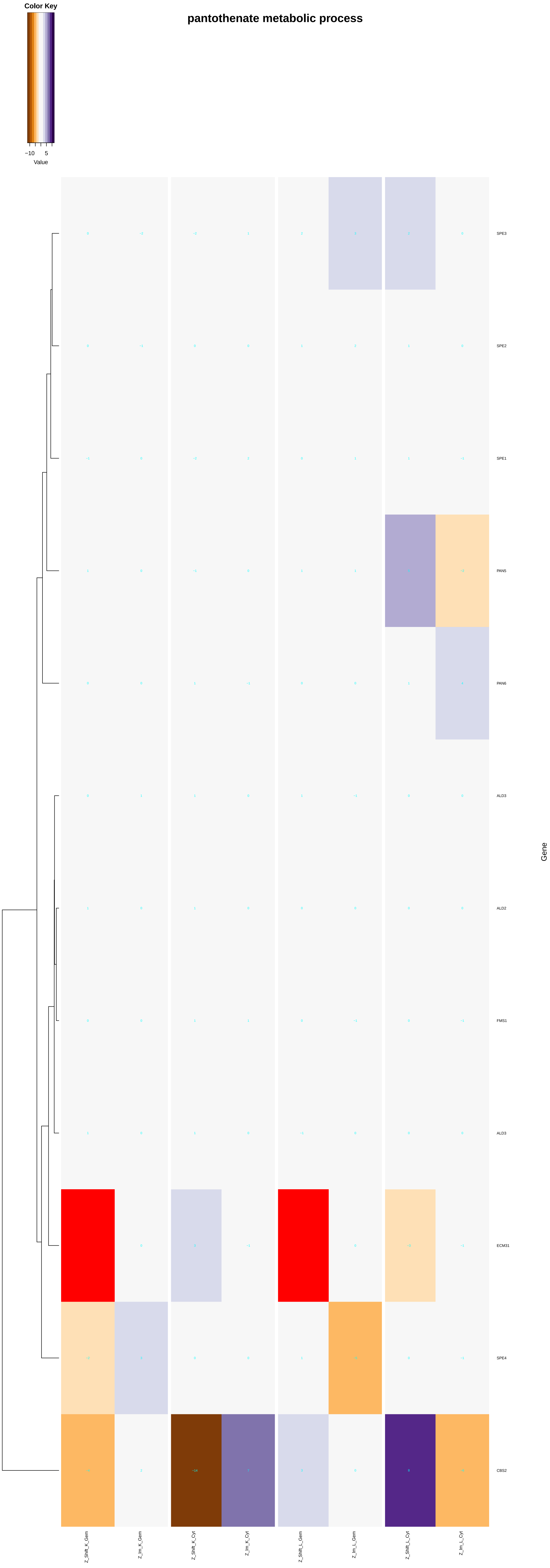

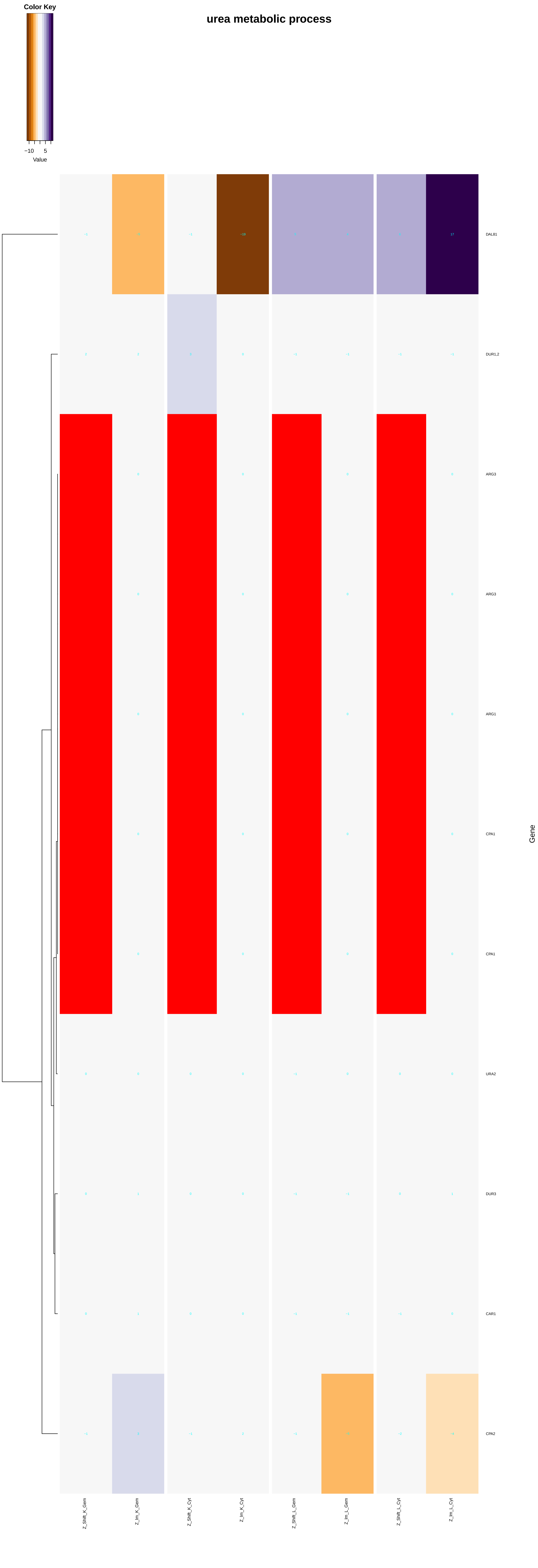

Color Key
−10
5
Value
urea metabolic process
DAL81
DUR1,2
ARG3
ARG3
ARG1
CPA1
CPA1
URA2
DUR3
CAR1
CPA2
Gene
Z_Shift_K_Gem
Z_Im_K_Gem
Z_Shift_K_Cyt
Z_Im_K_Cyt
Z_Shift_L_Gem
Z_Im_L_Gem
Z_Shift_L_Cyt
Z_Im_L_Cyt

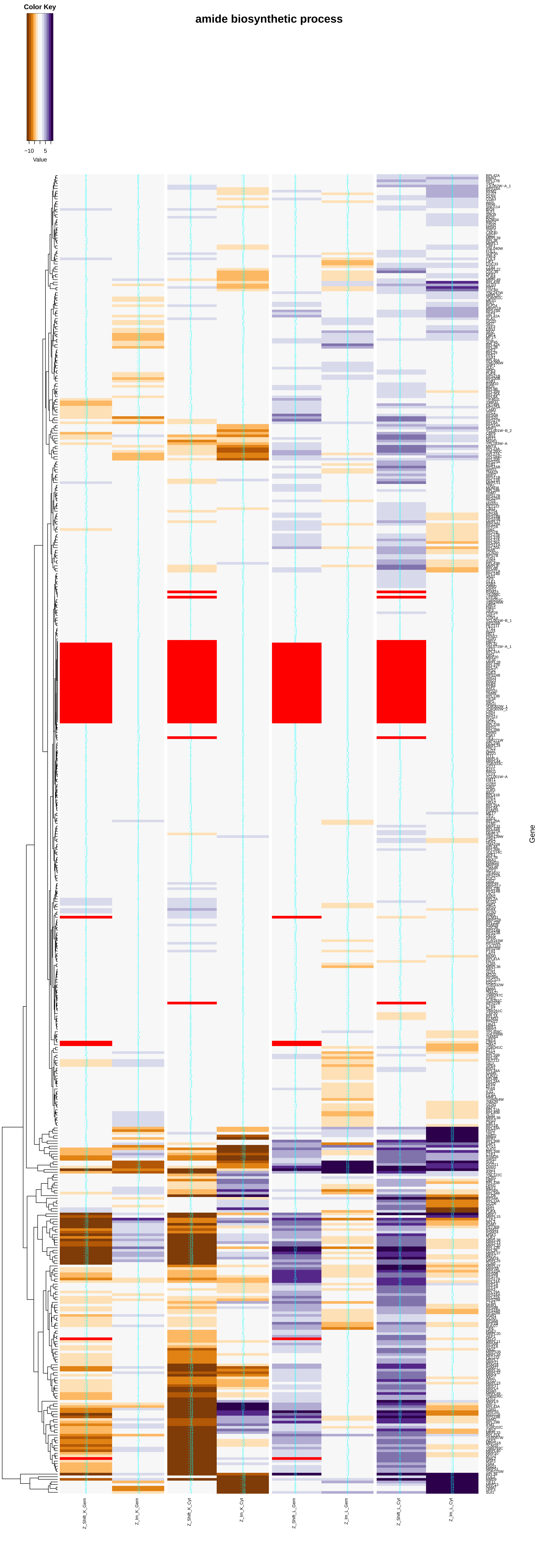
amide biosynthetic process
Color Key
−10
5
Value
Gene
Z_Shift_K_Gem
Z_Im_K_Gem
Z_Shift_K_Cyt
Z_Im_K_Cyt
Z_Shift_L_Gem
Z_Im_L_Gem
Z_Shift_L_Cyt
Z_Im_L_Cyt

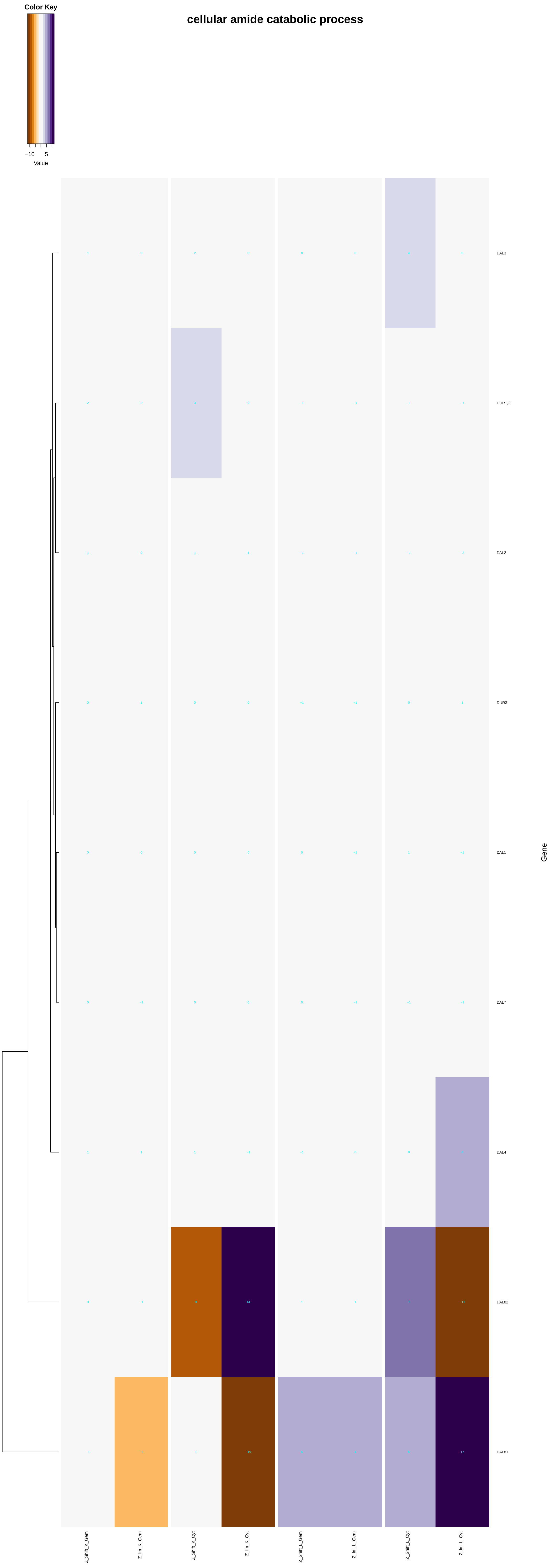
cellular amide catabolic process
Color Key
−10
5
Value
DAL3
DUR1,2
DAL2
DUR3
DAL1
DAL7
DAL4
DAL82
DAL81
Gene
Z_Shift_K_Gem
Z_Im_K_Gem
Z_Shift_K_Cyt
Z_Im_K_Cyt
Z_Shift_L_Gem
Z_Im_L_Gem
Z_Shift_L_Cyt
Z_Im_L_Cyt

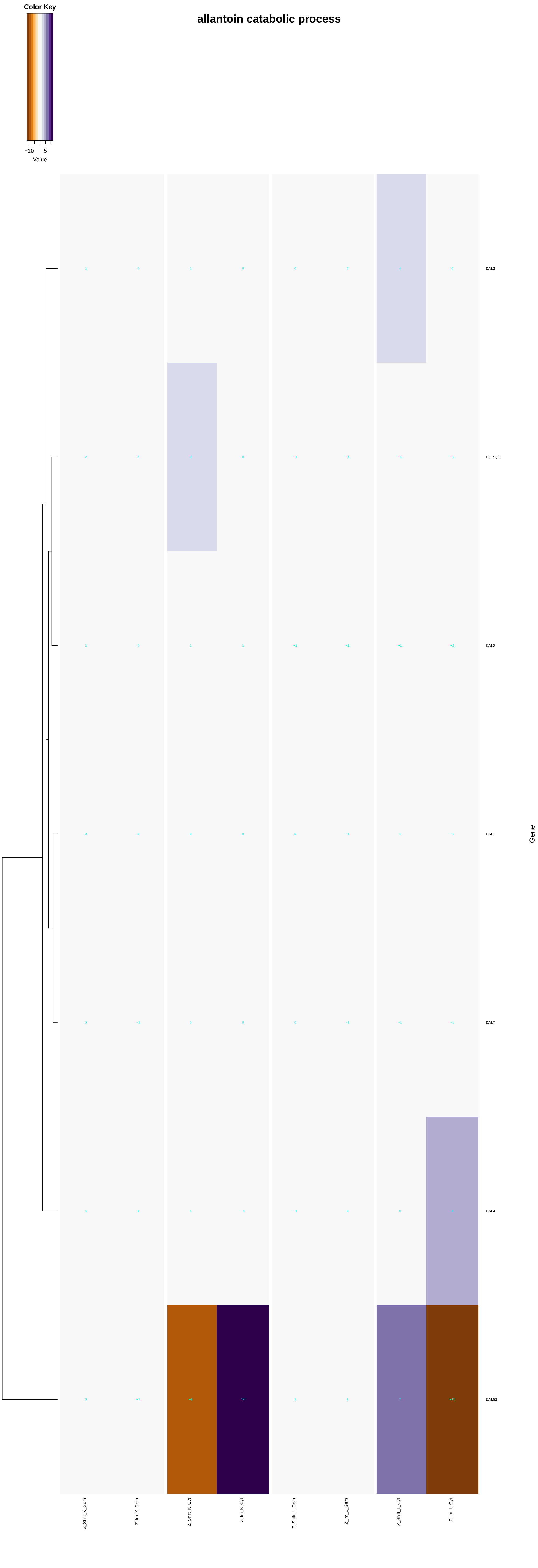

allantoin catabolic process
Color Key
−10
5
Value
DAL3
DUR1,2
DAL2
DAL1
DAL7
DAL4
DAL82
Gene
Z_Shift_K_Gem
Z_Im_K_Gem
Z_Shift_K_Cyt
Z_Im_K_Cyt
Z_Shift_L_Gem
Z_Im_L_Gem
Z_Shift_L_Cyt
Z_Im_L_Cyt

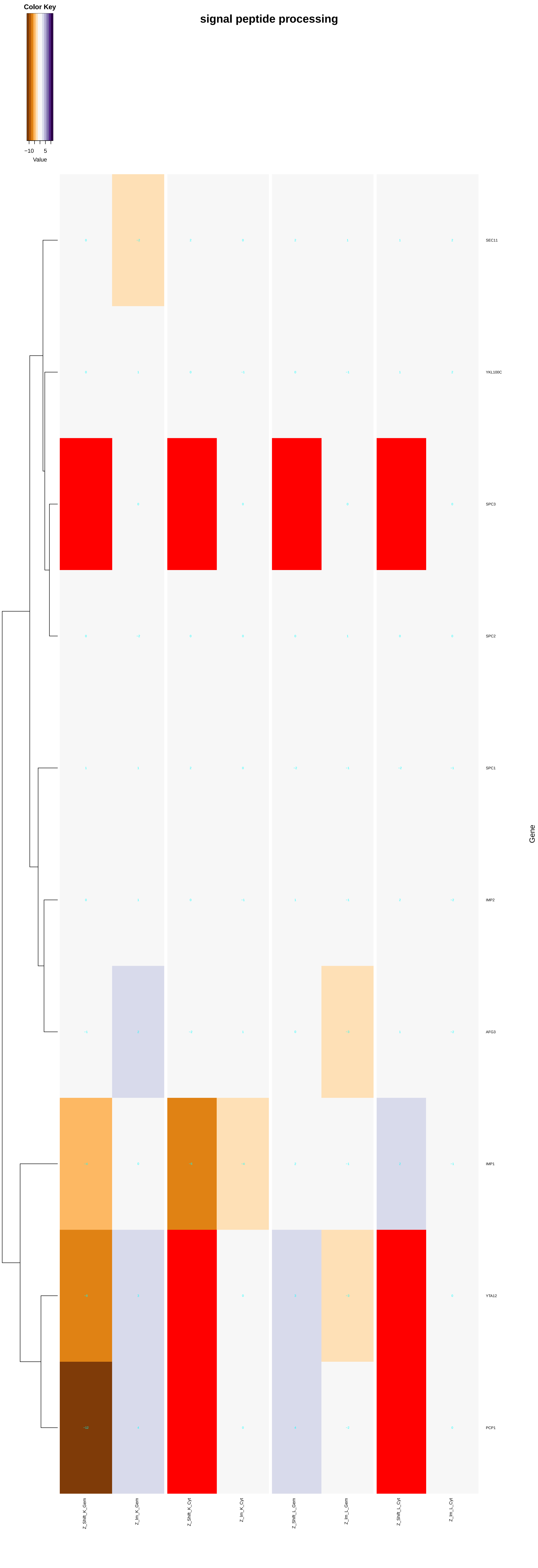

signal peptide processing
Color Key
−10
5
Value
SEC11
YKL100C
SPC3
SPC2
SPC1
IMP2
AFG3
IMP1
YTA12
PCP1
Gene
Z_Shift_K_Gem
Z_Im_K_Gem
Z_Shift_K_Cyt
Z_Im_K_Cyt
Z_Shift_L_Gem
Z_Im_L_Gem
Z_Shift_L_Cyt
Z_Im_L_Cyt

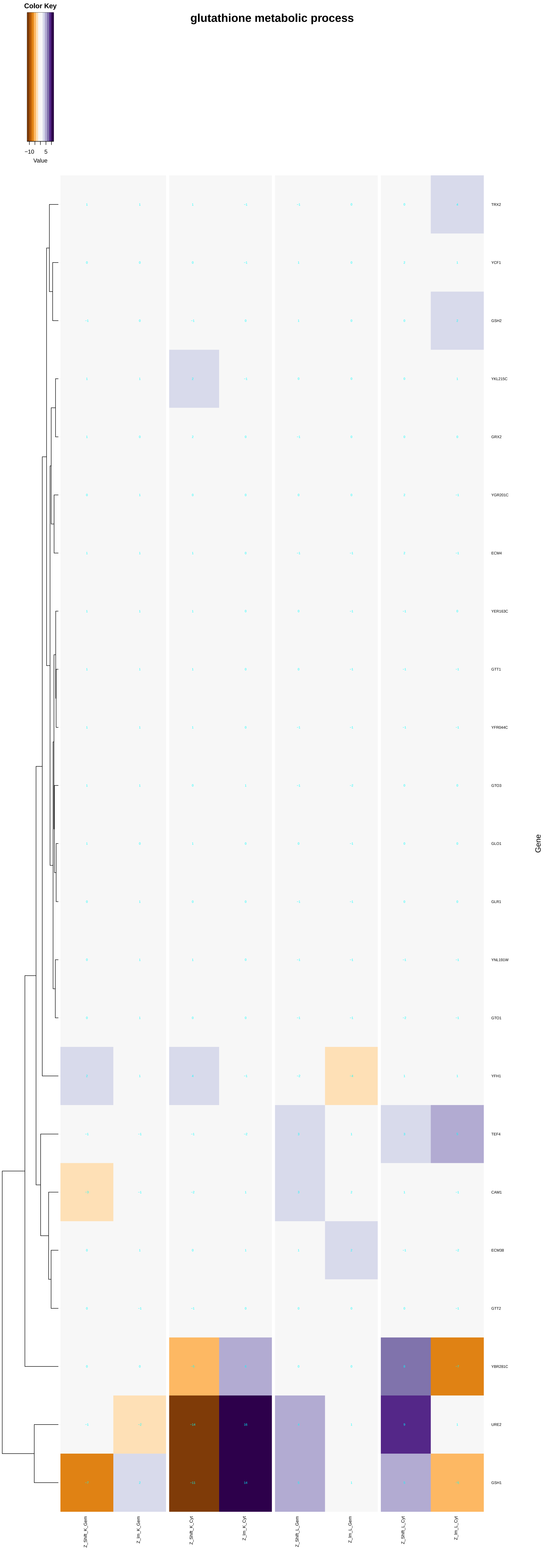
glutathione metabolic process
Color Key
−10
5
Value
TRX2
YCF1
GSH2
YKL215C
GRX2
YGR201C
ECM4
YER163C
GTT1
YFR044C
GTO3
GLO1
GLR1
YNL191W
GTO1
YFH1
TEF4
CAM1
ECM38
GTT2
YBR281C
URE2
GSH1
Gene
Z_Shift_K_Gem
Z_Im_K_Gem
Z_Shift_K_Cyt
Z_Im_K_Cyt
Z_Shift_L_Gem
Z_Im_L_Gem
Z_Shift_L_Cyt
Z_Im_L_Cyt

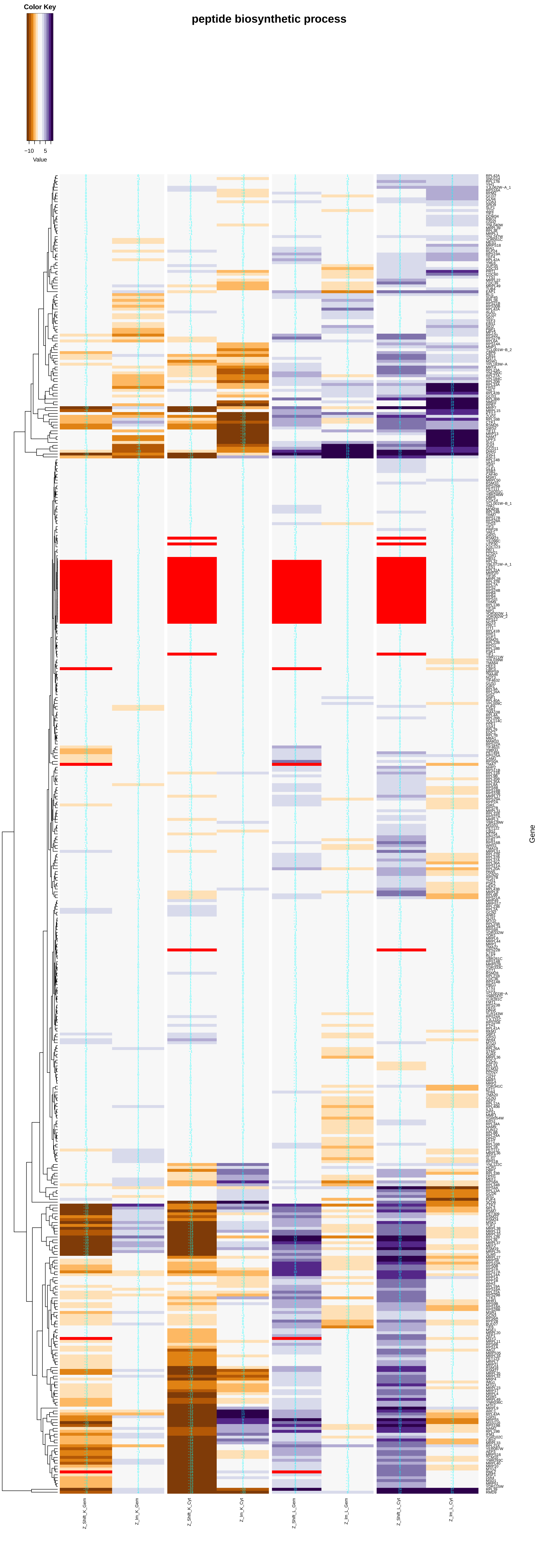
peptide biosynthetic process
Color Key
−10
5
Value
Gene
Z_Shift_K_Gem
Z_Im_K_Gem
Z_Shift_K_Cyt
Z_Im_K_Cyt
Z_Shift_L_Gem
Z_Im_L_Gem
Z_Shift_L_Cyt
Z_Im_L_Cyt

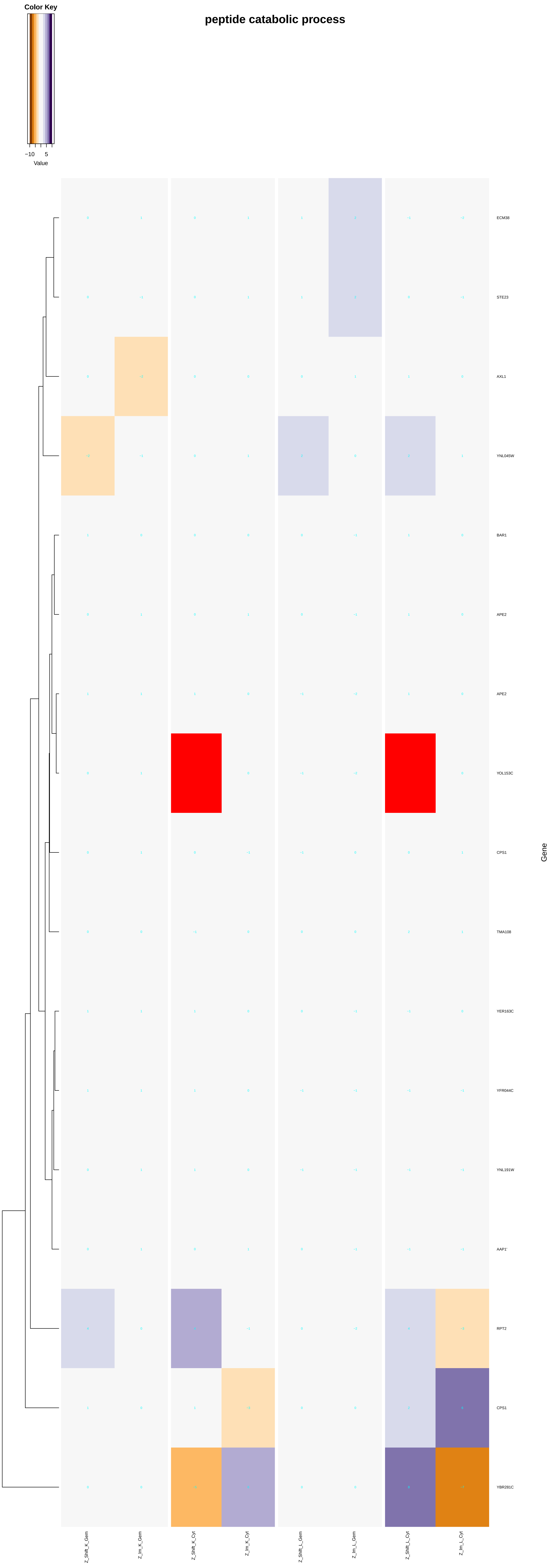

Color Key
−10
5
Value
peptide catabolic process
ECM38
STE23
AXL1
YNL045W
BAR1
APE2
APE2
YOL153C
CPS1
TMA108
YER163C
YFR044C
YNL191W
AAP1'
RPT2
CPS1
YBR281C
Gene
Z_Shift_K_Gem
Z_Im_K_Gem
Z_Shift_K_Cyt
Z_Im_K_Cyt
Z_Shift_L_Gem
Z_Im_L_Gem
Z_Shift_L_Cyt
Z_Im_L_Cyt

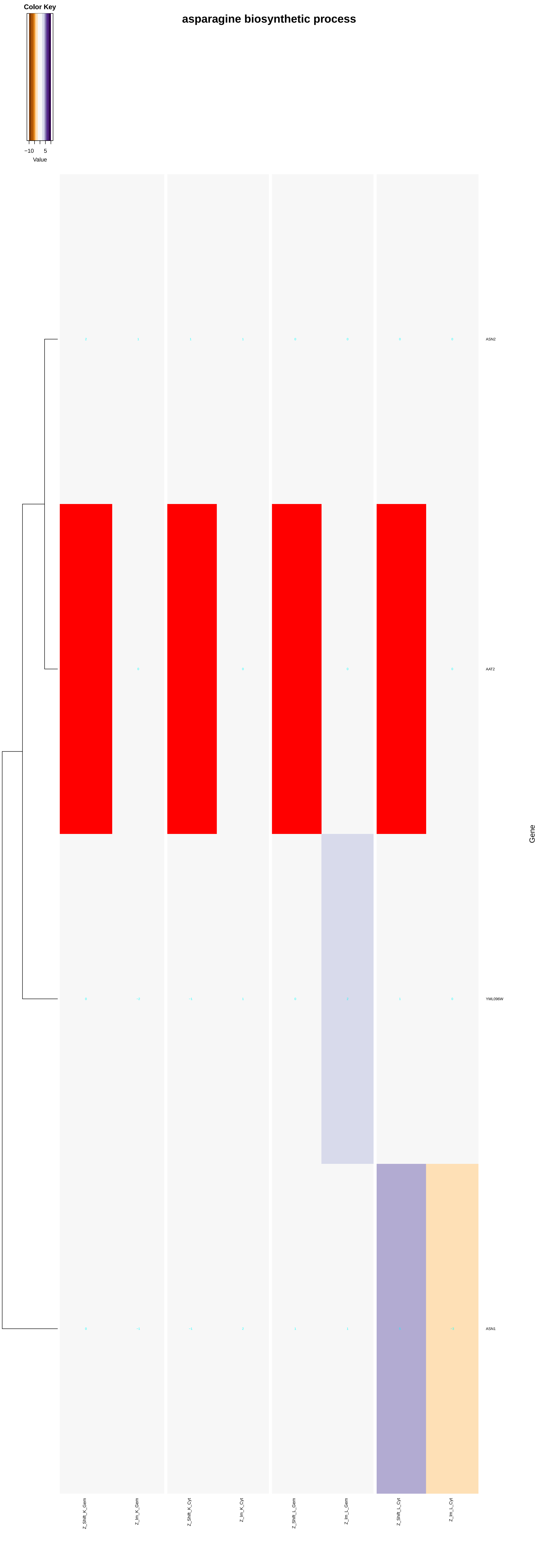
asparagine biosynthetic process
Color Key
−10
5
Value
ASN2
AAT2
YML096W
ASN1
Gene
Z_Shift_K_Gem
Z_lm_K_Gem
Z_Shift_K_Cyt
Z_lm_K_Cyt
Z_Shift_L_Gem
Z_lm_L_Gem
Z_Shift_L_Cyt
Z_lm_L_Cyt

### asparagine catabolic process

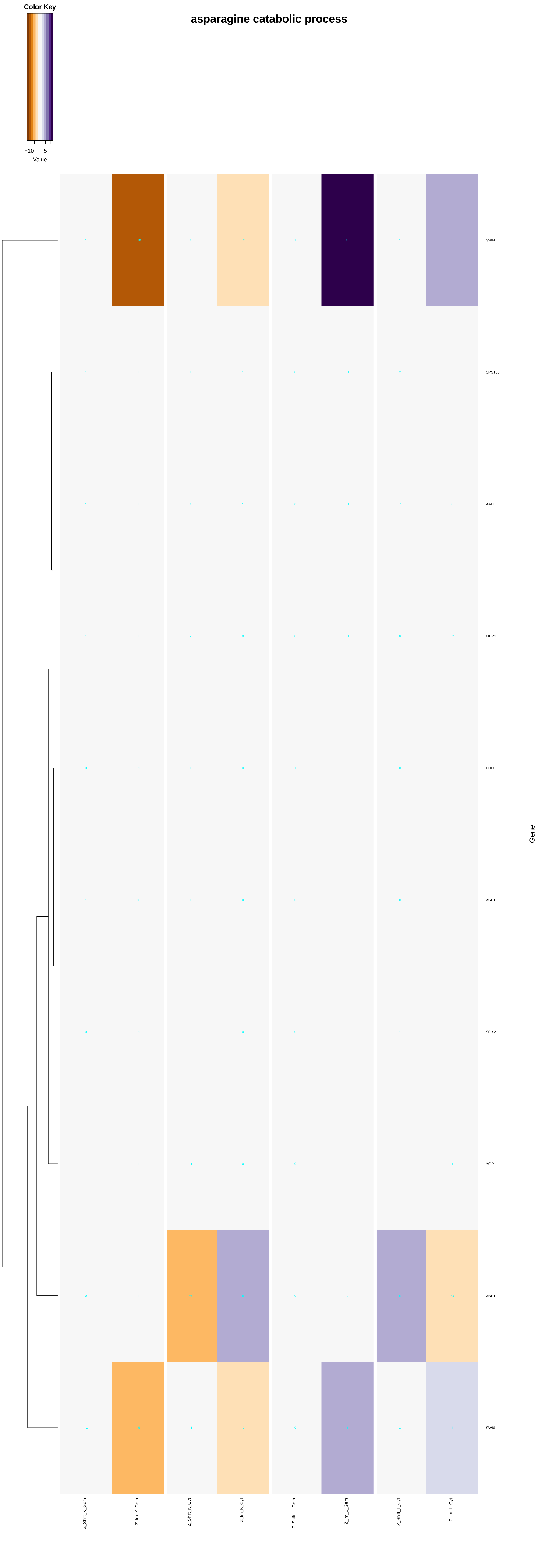

#### ceramide biosynthetic process

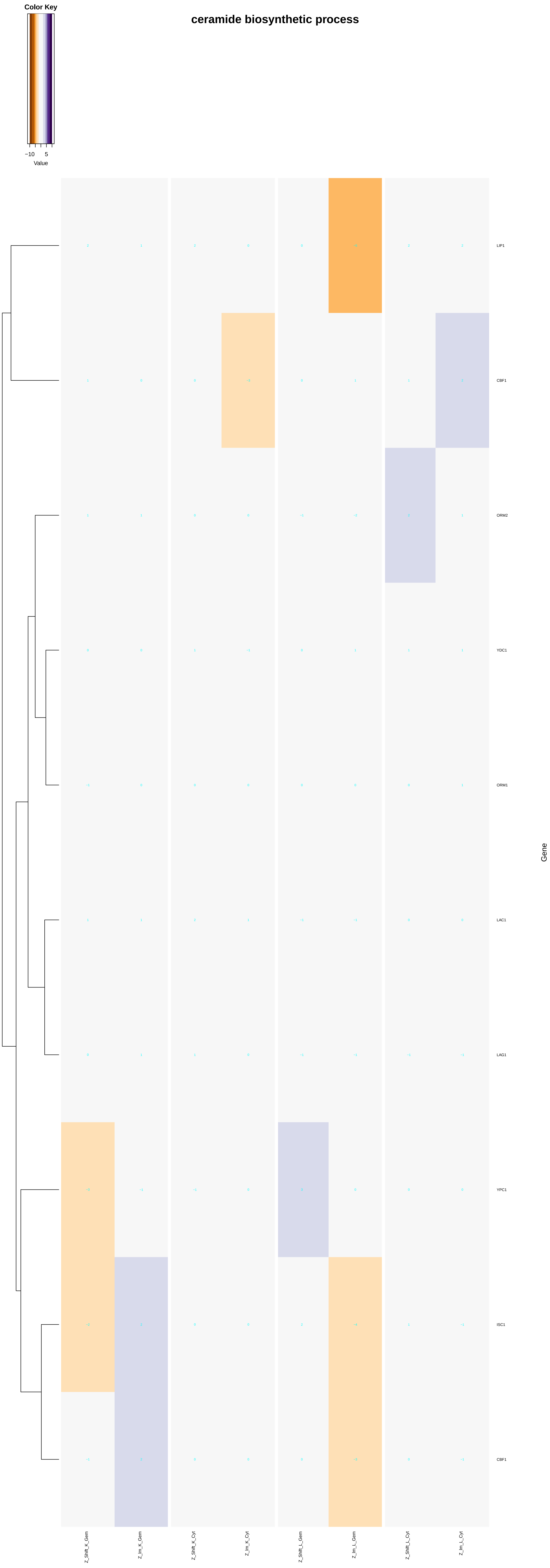

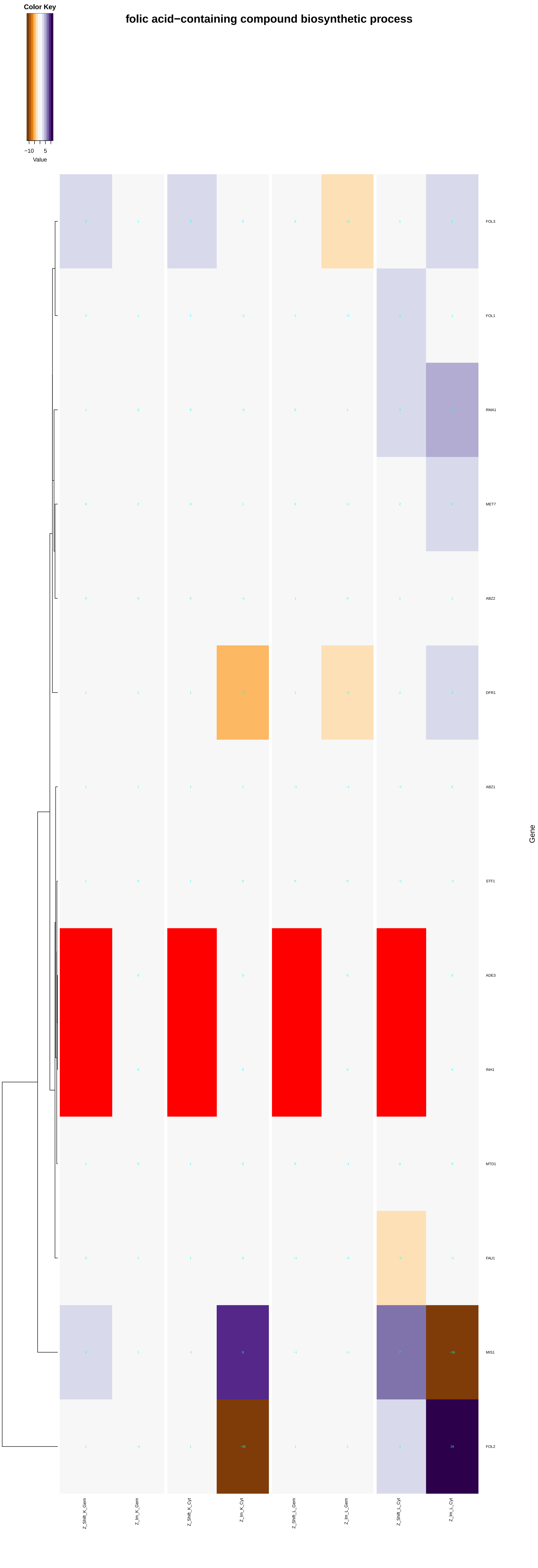
folic acid−containing compound biosynthetic process
Color Key
−10
5
Value
FOL3
FOL1
RMA1
MET7
ABZ2
DFR1
ABZ1
STF1
ADE3
INH1
MTD1
FAU1
MIS1
FOL2
Gene
Z_Shift_K_Gem
Z_Im_K_Gem
Z_Shift_K_Cyt
Z_Im_K_Cyt
Z_Shift_L_Gem
Z_Im_L_Gem
Z_Shift_L_Cyt
Z_Im_L_Cyt

### dihydrofolate metabolic process

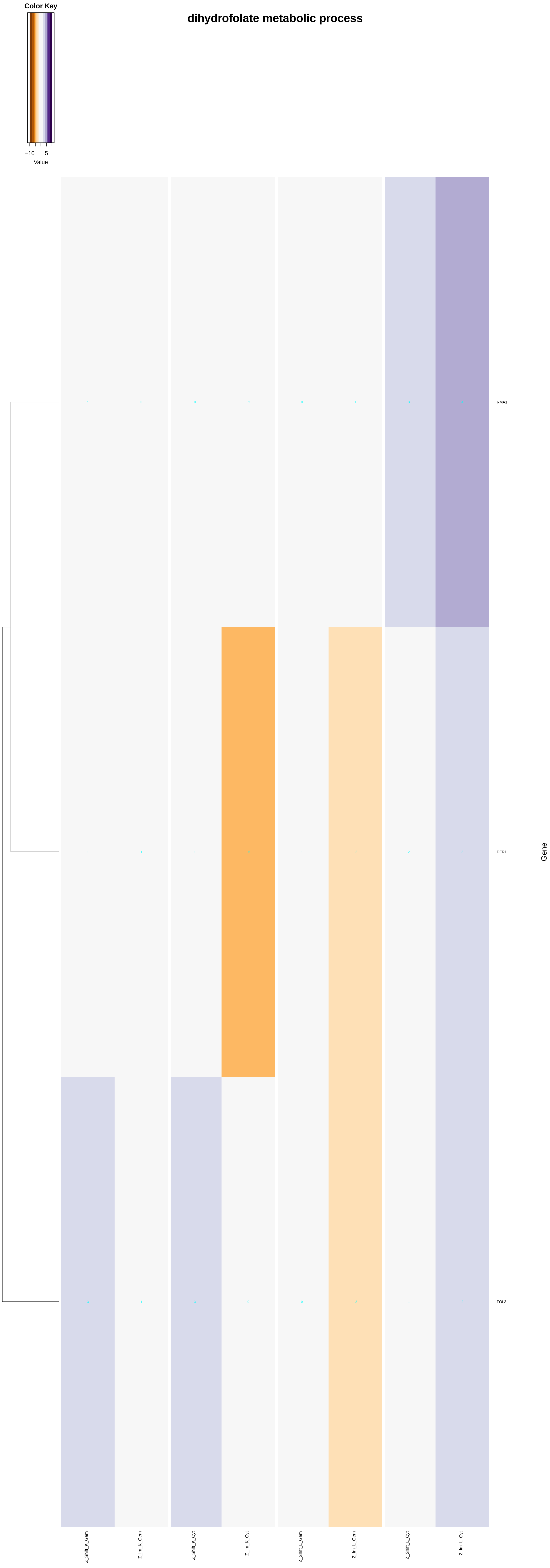

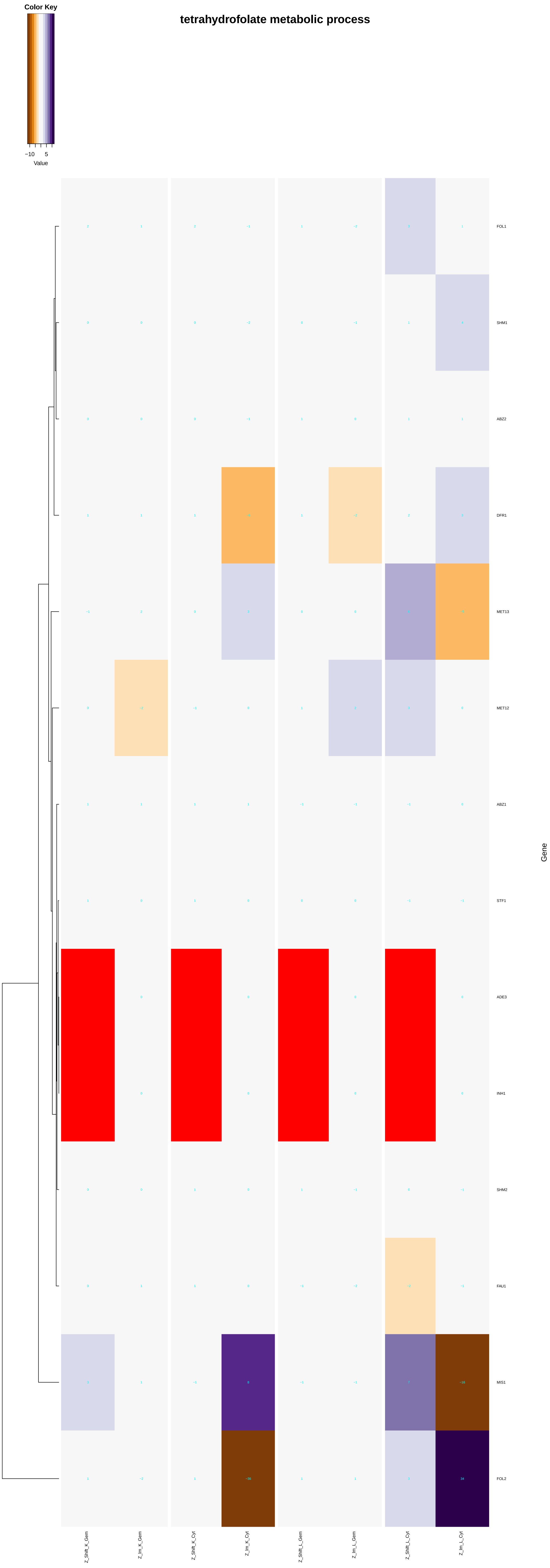

tetrahydrofolate metabolic process
Color Key
−10
5
Value
FOL1
SHM1
ABZ2
DFR1
MET13
MET12
ABZ1
STF1
ADE3
INH1
SHM2
FAU1
MIS1
FOL2
Gene
Z_Shift_K_Gem
Z_Im_K_Gem
Z_Shift_K_Cyt
Z_Im_K_Cyt
Z_Shift_L_Gem
Z_Im_L_Gem
Z_Shift_L_Cyt
Z_Im_L_Cyt

#### folic acid metabolic process

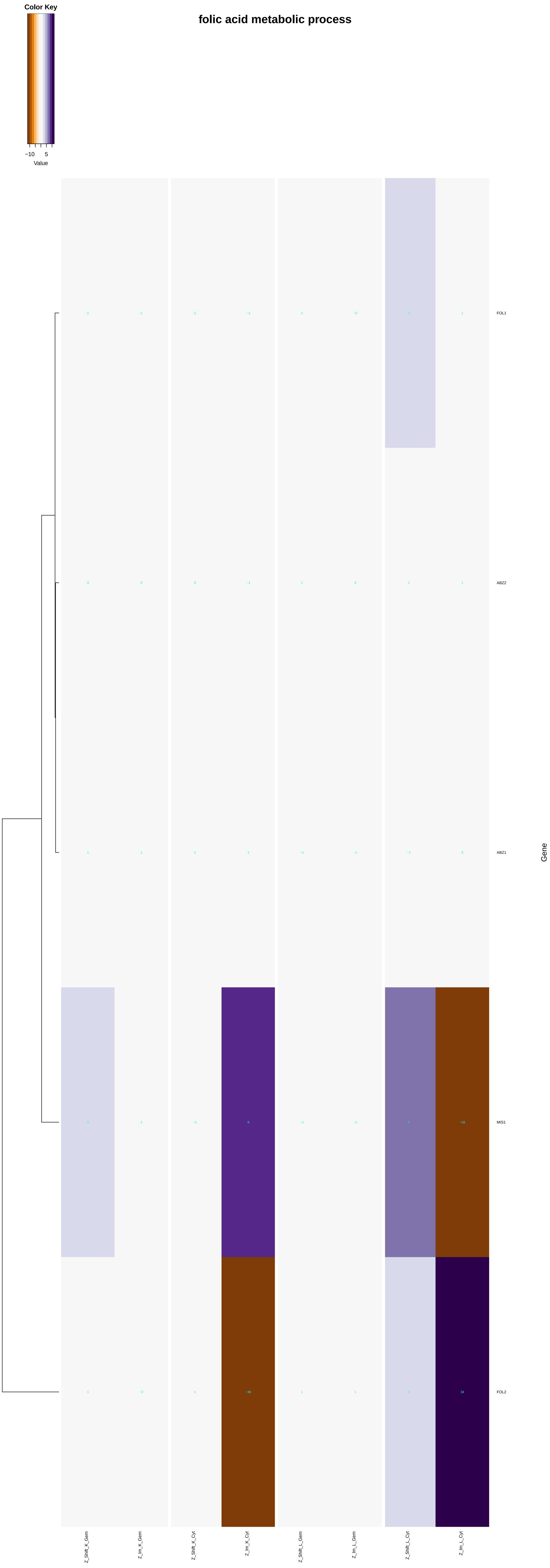

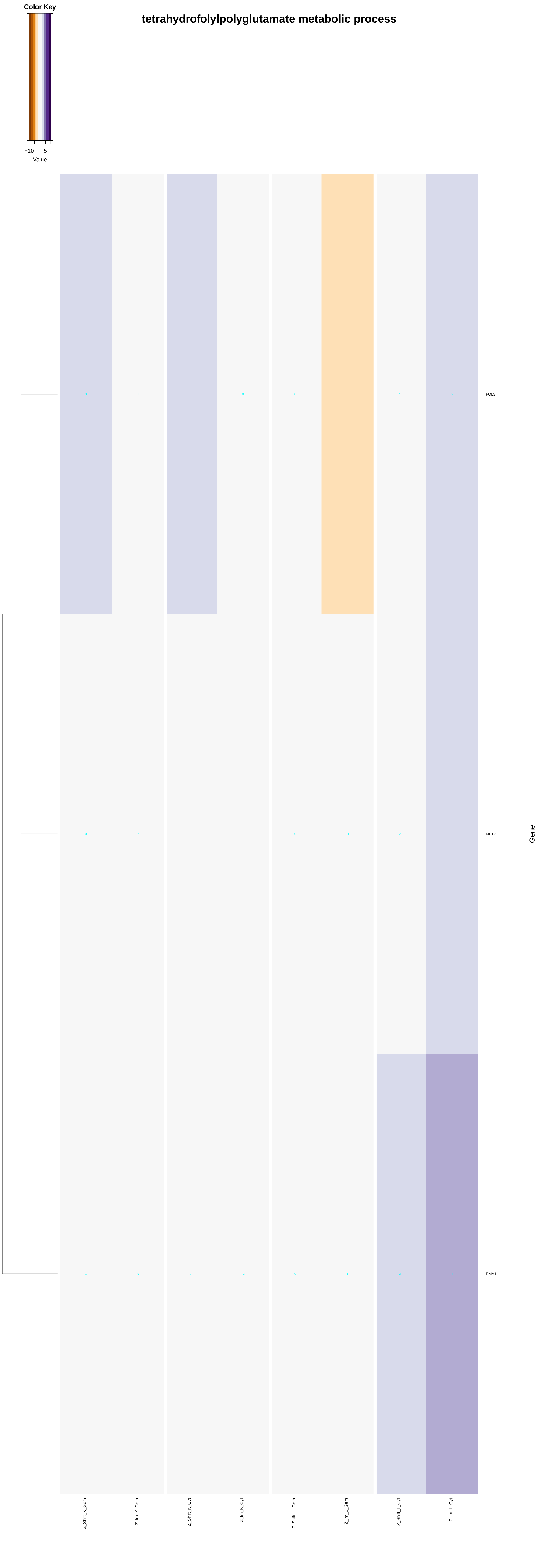
tetrahydrofolylpolyglutamate metabolic process
Color Key
−10
5
Value
FOL3
MET7
RMA1
Gene
Z_Shift_K_Gem
Z_Im_K_Gem
Z_Shift_K_Cyt
Z_Im_K_Cyt
Z_Shift_L_Gem
Z_Im_L_Gem
Z_Shift_L_Cyt
Z_Im_L_Cyt

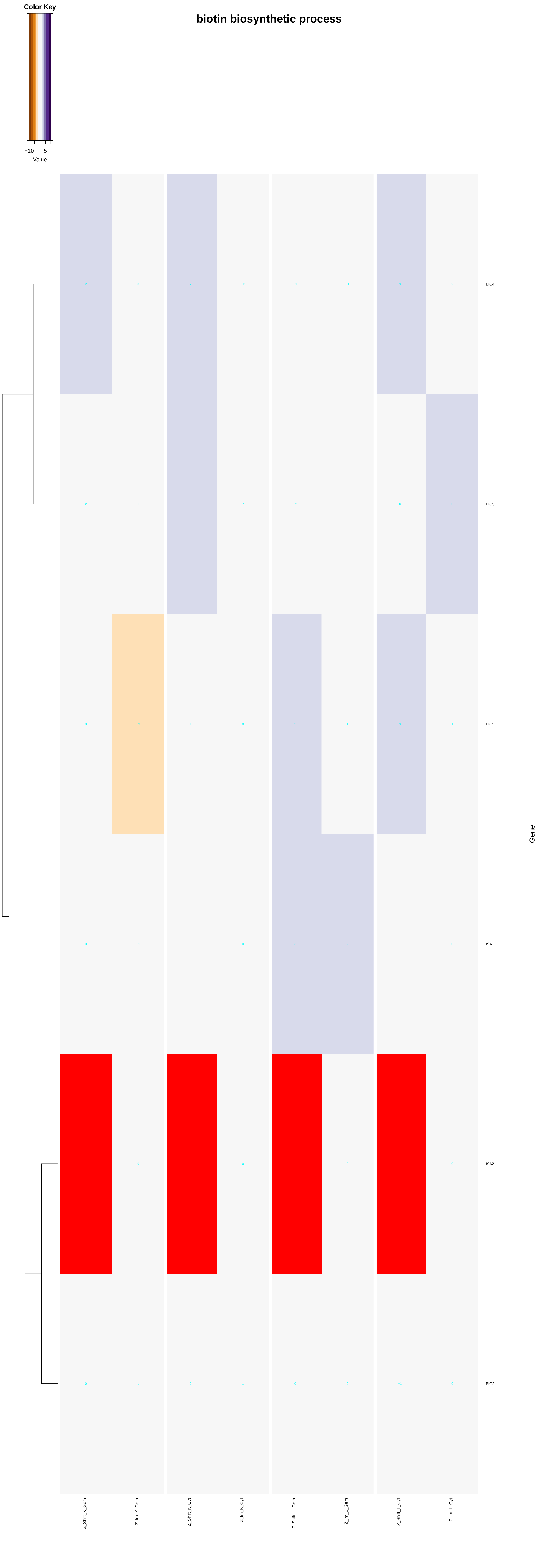

biotin biosynthetic process
Color Key
−10
5
Value
BIO4
BIO3
BIO5
ISA1
ISA2
BIO2
Gene
Z_Shift_K_Gem
Z_Im_K_Gem
Z_Shift_K_Cyt
Z_Im_K_Cyt
Z_Shift_L_Gem
Z_Im_L_Gem
Z_Shift_L_Cyt
Z_Im_L_Cyt

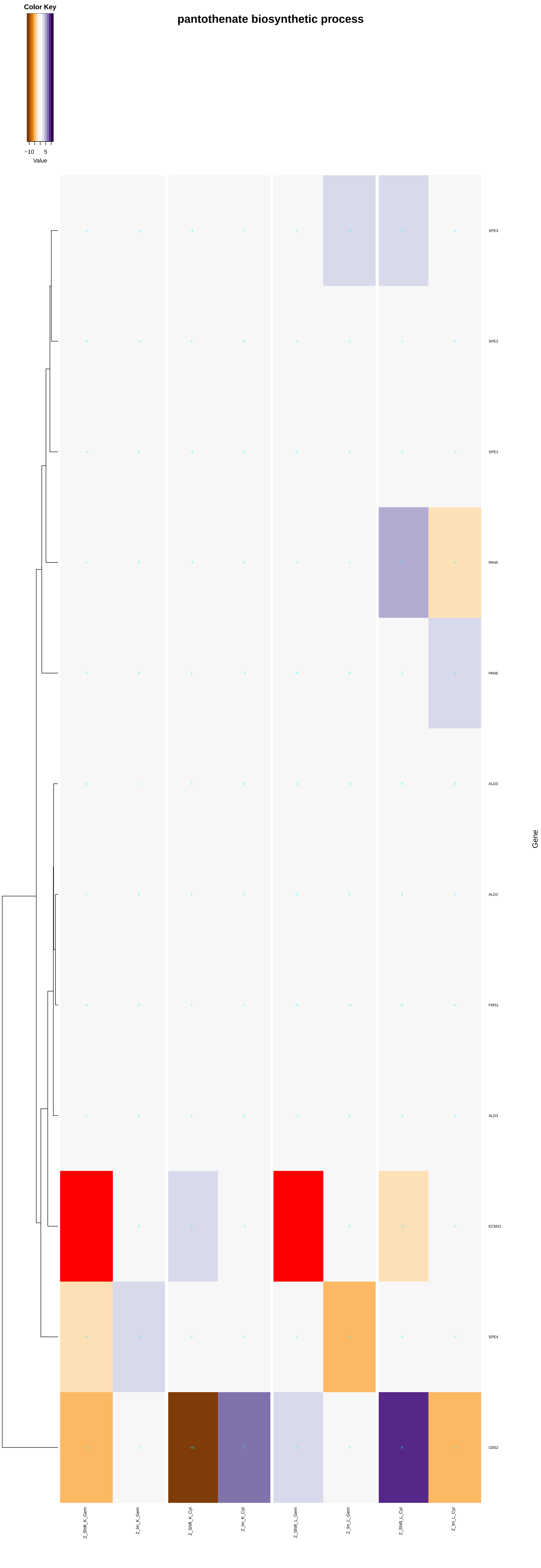

Color Key
−10
5
Value
pantothenate biosynthetic process
SPE3
SPE2
SPE1
PAN5
PAN6
ALD3
ALD2
FMS1
ALD3
ECM31
SPE4
CBS2
Gene
Z_Shift_K_Gem
Z_Im_K_Gem
Z_Shift_K_Cyt
Z_Im_K_Cyt
Z_Shift_L_Gem
Z_Im_L_Gem
Z_Shift_L_Cyt
Z_Im_L_Cyt

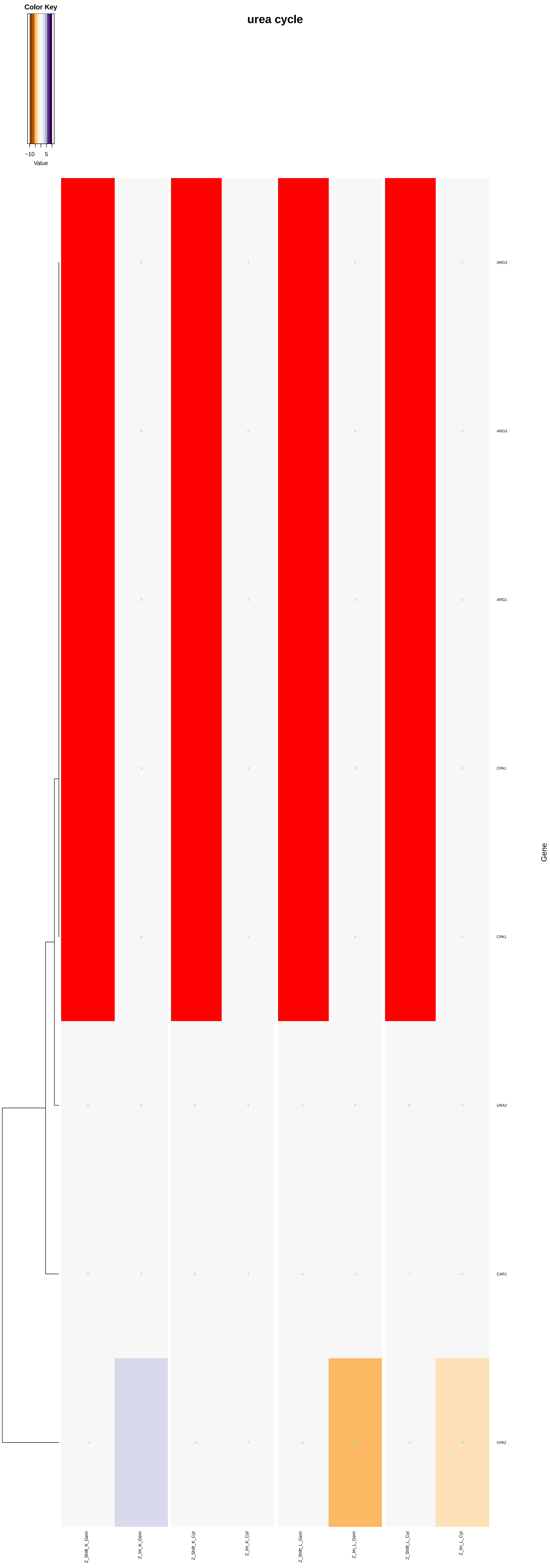

Color Key
−10
5
Value
urea cycle
ARG3
ARG3
ARG1
CPA1
CPA1
URA2
CAR1
CPA2
Gene
Z_Shift_K_Gem
Z_Im_K_Gem
Z_Shift_K_Cyt
Z_Im_K_Cyt
Z_Shift_L_Gem
Z_Im_L_Gem
Z_Shift_L_Cyt
Z_Im_L_Cyt

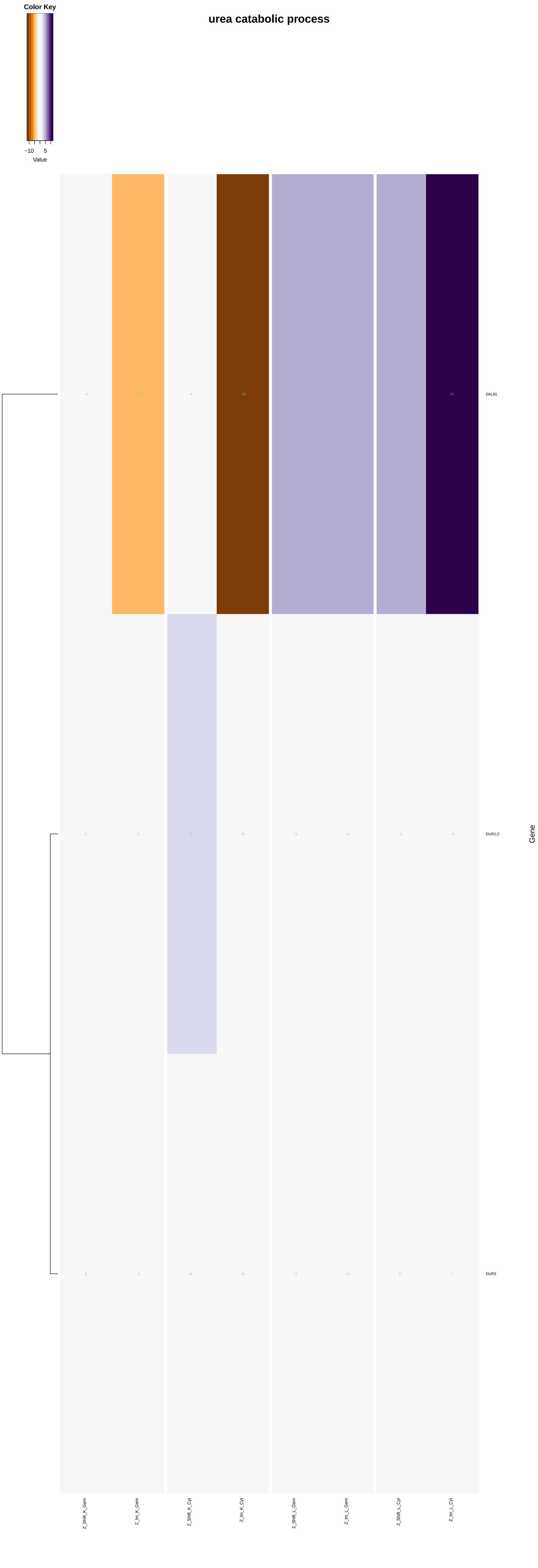

urea catabolic process
Color Key
−10
5
Value
−1
−5
−1
−19
5
4
6
17
DAL81
2
2
3
0
−1
−1
−1
−1
DUR1,2
0
1
0
0
−1
−1
0
1
DUR3
Gene
Z_Shift_K_Gem
Z_Im_K_Gem
Z_Shift_K_Cyt
Z_Im_K_Cyt
Z_Shift_L_Gem
Z_Im_L_Gem
Z_Shift_L_Cyt
Z_Im_L_Cyt

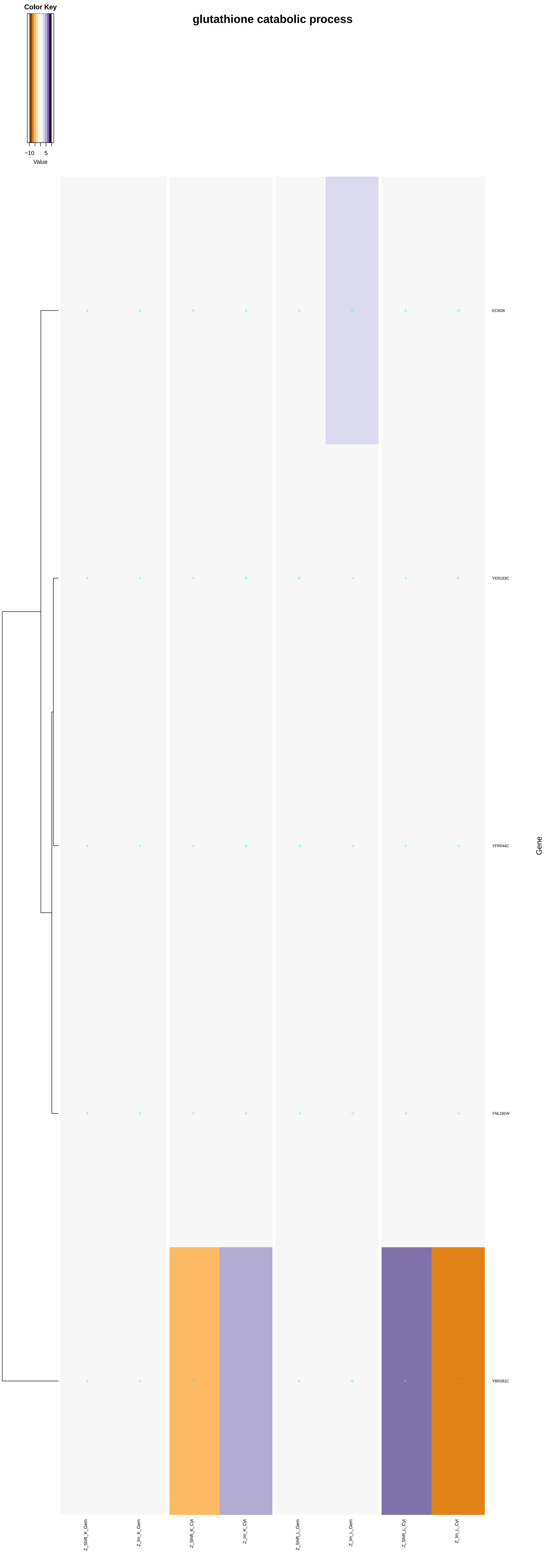
glutathione catabolic process
Color Key
−10
5
Value
ECM38
YER163C
YFR044C
YNL191W
YBR281C
Gene
Z_Shift_K_Gem
Z_Im_K_Gem
Z_Shift_K_Cyt
Z_Im_K_Cyt
Z_Shift_L_Gem
Z_Im_L_Gem
Z_Shift_L_Cyt
Z_Im_L_Cyt

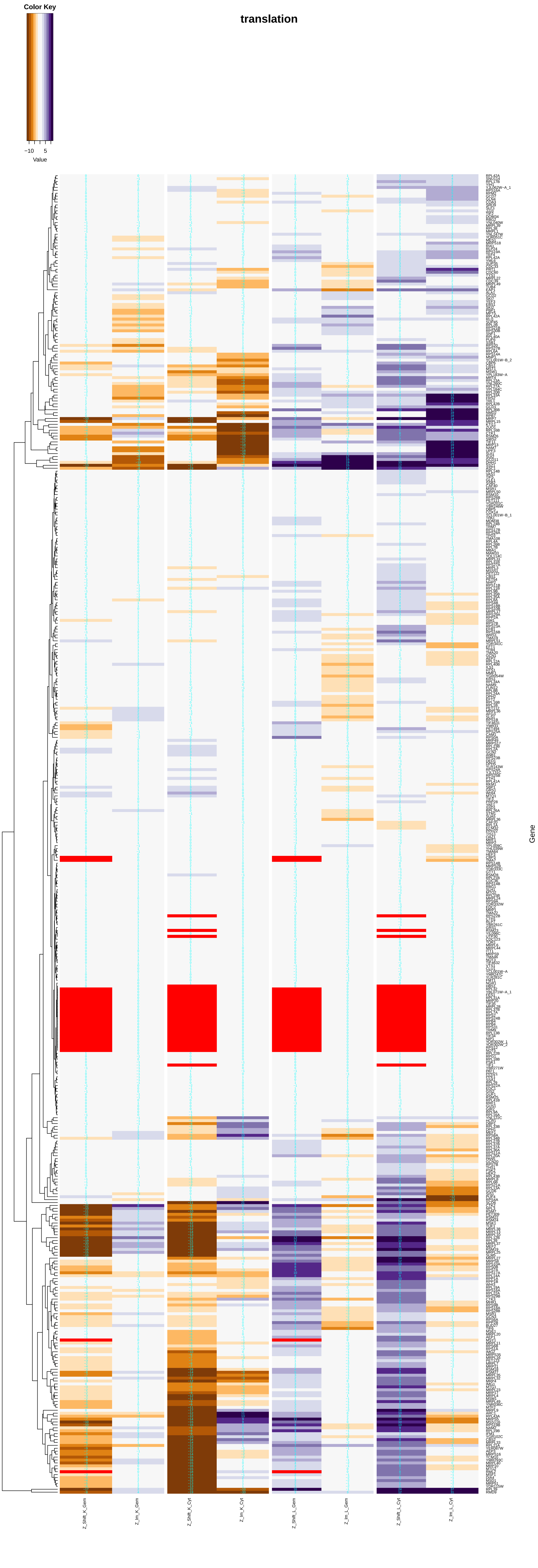
Color Key
−10
5
Value
translation
Gene
Z_Shift_K_Gem
Z_Im_K_Gem
Z_Shift_K_Cyt
Z_Im_K_Cyt
Z_Shift_L_Gem
Z_Im_L_Gem
Z_Shift_L_Cyt
Z_Im_L_Cyt

#### tetrahydrofolate biosynthetic process

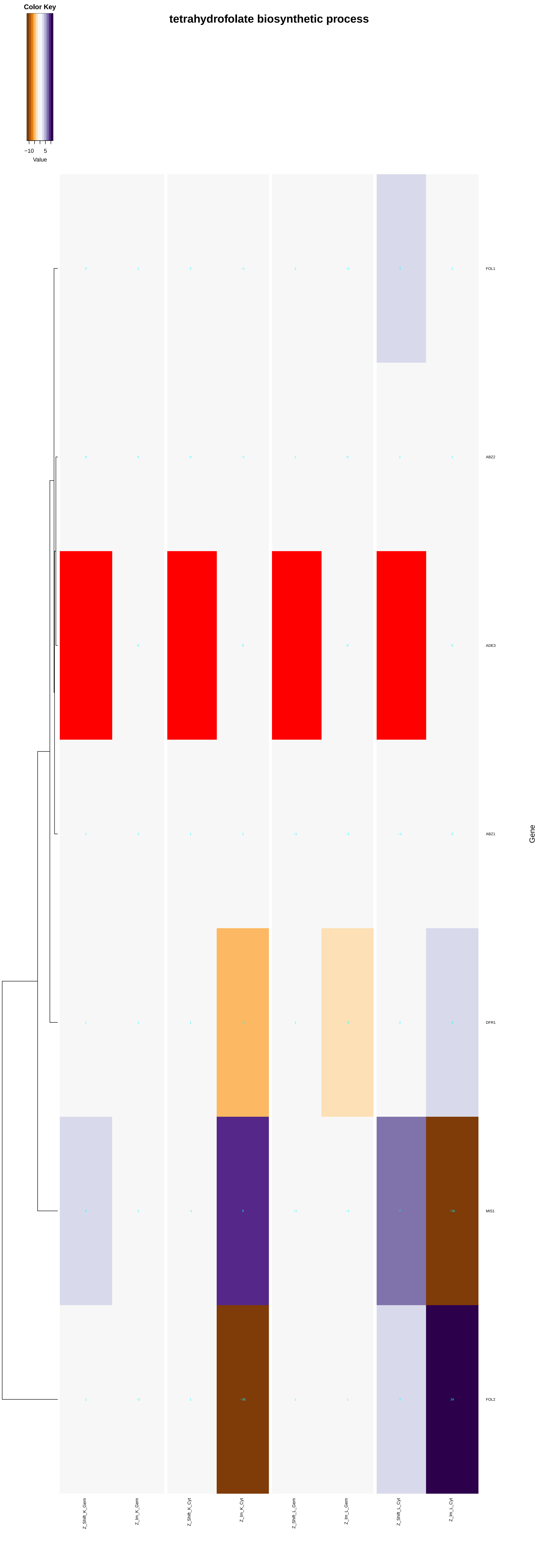

### folic acid biosynthetic process

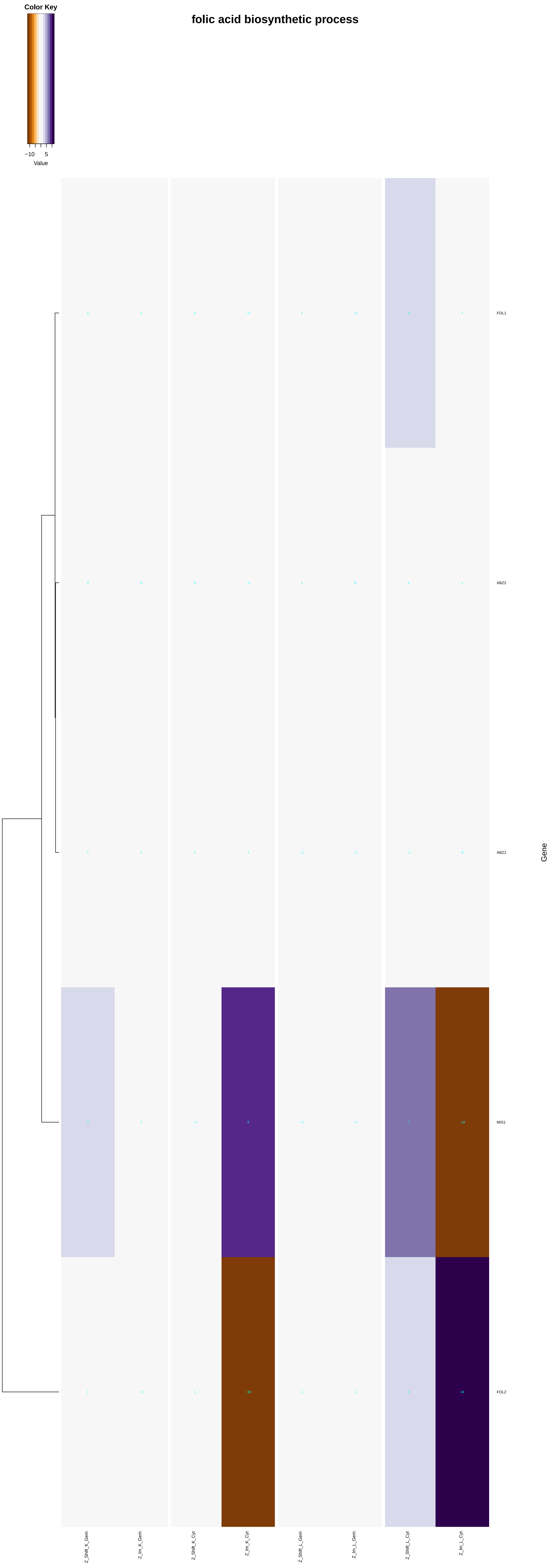

### tetrahydrofolylpolyglutamate biosynthetic process

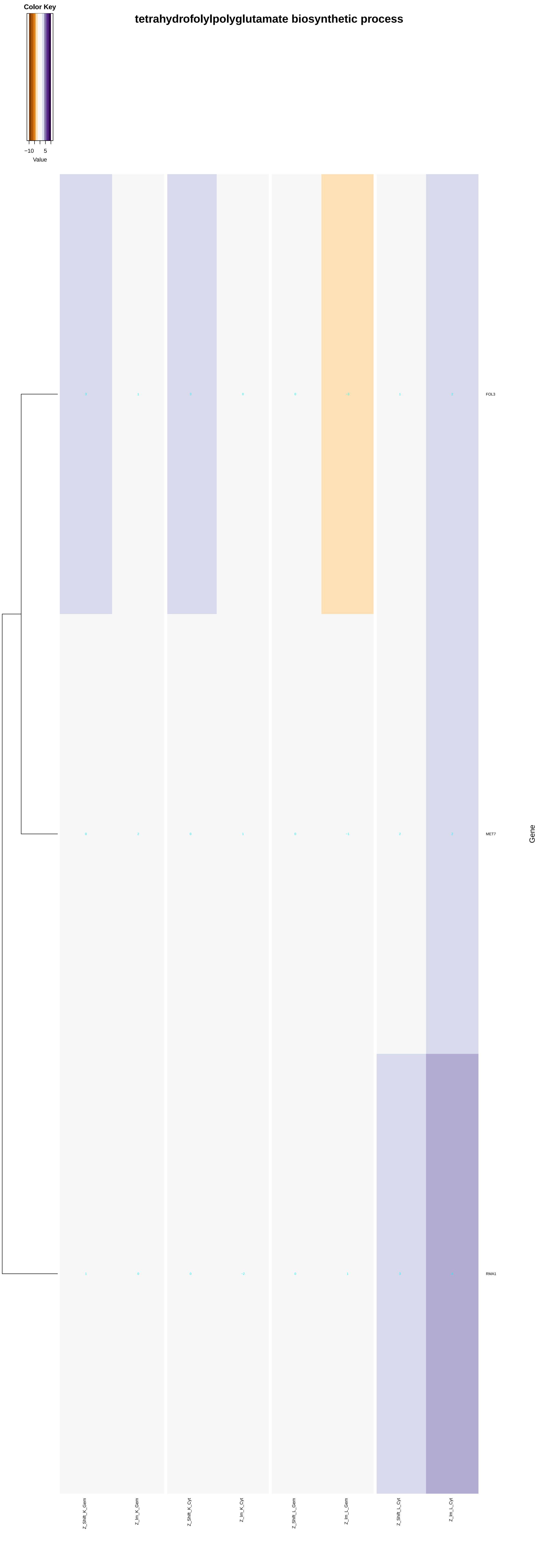

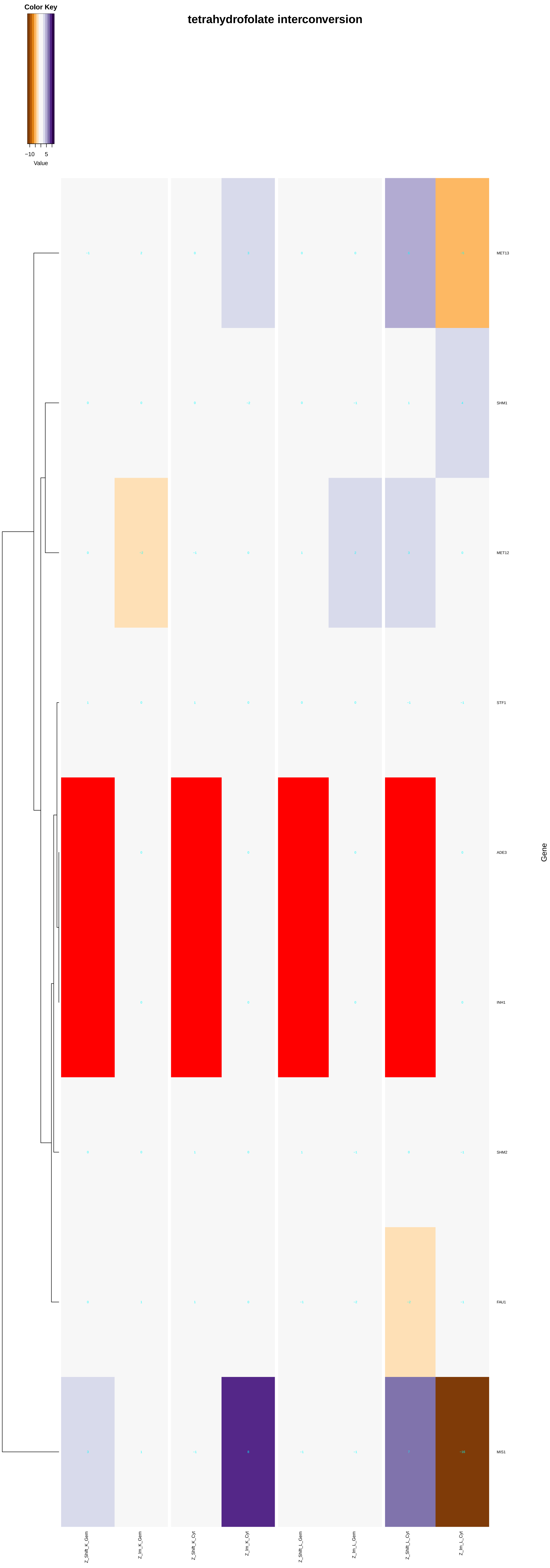
tetrahydrofolate interconversion
Color Key
−10
5
Value
MET13
SHM1
MET12
STF1
ADE3
INH1
SHM2
FAU1
MIS1
Gene
Z_Shift_K_Gem
Z_Im_K_Gem
Z_Shift_K_Cyt
Z_Im_K_Cyt
Z_Shift_L_Gem
Z_Im_L_Gem
Z_Shift_L_Cyt
Z_Im_L_Cyt

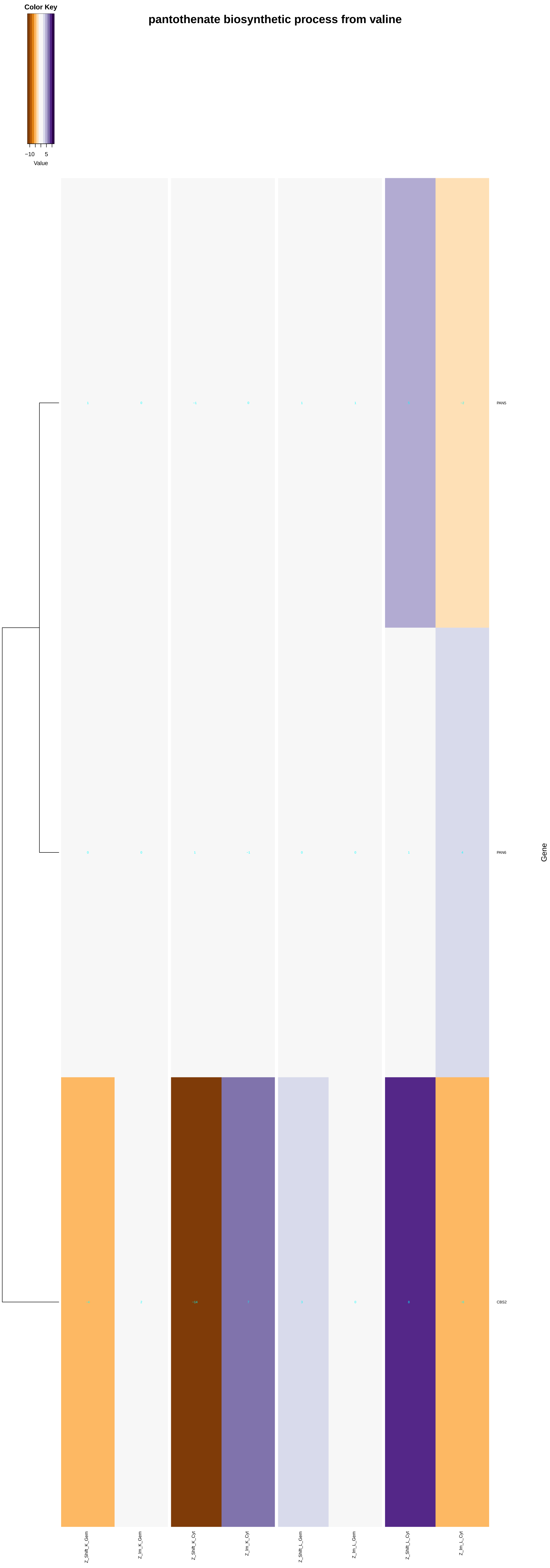
pantothenate biosynthetic process from valine
Color Key
−10
5
Value
PAN5
PAN6
CBS2
Gene
Z_Shift_K_Gem
Z_Im_K_Gem
Z_Shift_K_Cyt
Z_Im_K_Cyt
Z_Shift_L_Gem
Z_Im_L_Gem
Z_Shift_L_Cyt
Z_Im_L_Cyt

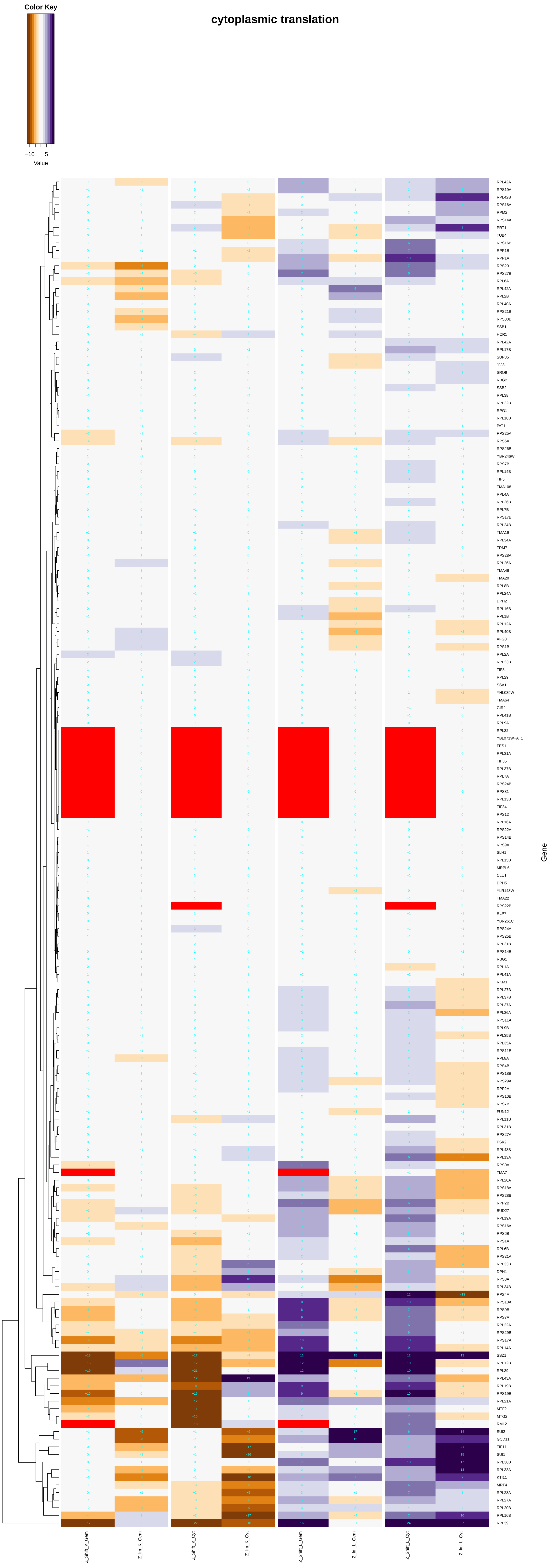
cytoplasmic translation
Color Key
−10
5
Value
RPL42A
RPS19A
RPL42B
RPS16A
RPM2
RPS14A
PRT1
TUB4
RPS16B
RPP1B
RPP1A
RPS20
RPS27B
RPL6A
RPL42A
RPL2B
RPL40A
RPS21B
RPS30B
SSB1
HCR1
RPL42A
RPL17B
SUP35
JJJ3
SRO9
RBG2
SSB2
RPL38
RPL22B
RPG1
RPL18B
PAT1
RPS25A
RPS6A
RPS26B
YBR246W
RPS7B
RPL14B
TIF5
TMA108
RPL4A
RPL26B
RPL7B
RPS17B
RPL24B
TMA19
RPL34A
TRM7
RPS28A
RPL26A
TMA46
TMA20
RPL8B
RPL24A
DPH2
RPL16B
RPL1B
RPL12A
RPL40B
AFG3
RPS1B
RPL2A
RPL23B
TIF3
RPL29
SSA1
YHL039W
TMA64
GIR2
RPL41B
RPL9A
RPL32
YBL071W−A_1
FES1
RPL31A
TIF35
RPL37B
RPL7A
RPS24B
RPS31
RPL13B
TIF34
RPS12
RPL16A
RPS22A
RPS14B
RPS9A
SLH1
RPL15B
MRPL6
CLU1
DPH5
YLR143W
TMA22
RPS22B
RLP7
YBR261C
RPS24A
RPS25B
RPL21B
RPS14B
RBG1
RPL1A
RPL41A
RKM1
RPL27B
RPL37B
RPL37A
RPL36A
RPS11A
RPL9B
RPL35B
RPL35A
RPS11B
RPL8A
RPS4B
RPS18B
RPS29A
RPP2A
RPS10B
RPS7B
FUN12
RPL11B
RPL31B
RPS27A
PSK2
RPL43B
RPL13A
RPS0A
TMA7
RPL20A
RPS18A
RPS28B
RPP2B
BUD27
RPL19A
RPS16A
RPS6B
RPS1A
RPL6B
RPS21A
RPL33B
DPH1
RPS8A
RPL34B
RPS4A
RPS10A
RPS0B
RPS7A
RPL22A
RPS29B
RPS17A
RPL14A
SSZ1
RPL12B
RPL39
RPL43A
RPL19B
RPS19B
RPL21A
MTF2
MTG2
RML2
SUI2
GCD11
TIF11
SUI1
RPL36B
RPL33A
KTI11
MRT4
RPL23A
RPL27A
RPL20B
RPL16B
RPL39
Gene
Z_Shift_K_Gem
Z_Im_K_Gem
Z_Shift_K_Cyt
Z_Im_K_Cyt
Z_Shift_L_Gem
Z_Im_L_Gem
Z_Shift_L_Cyt
Z_Im_L_Cyt

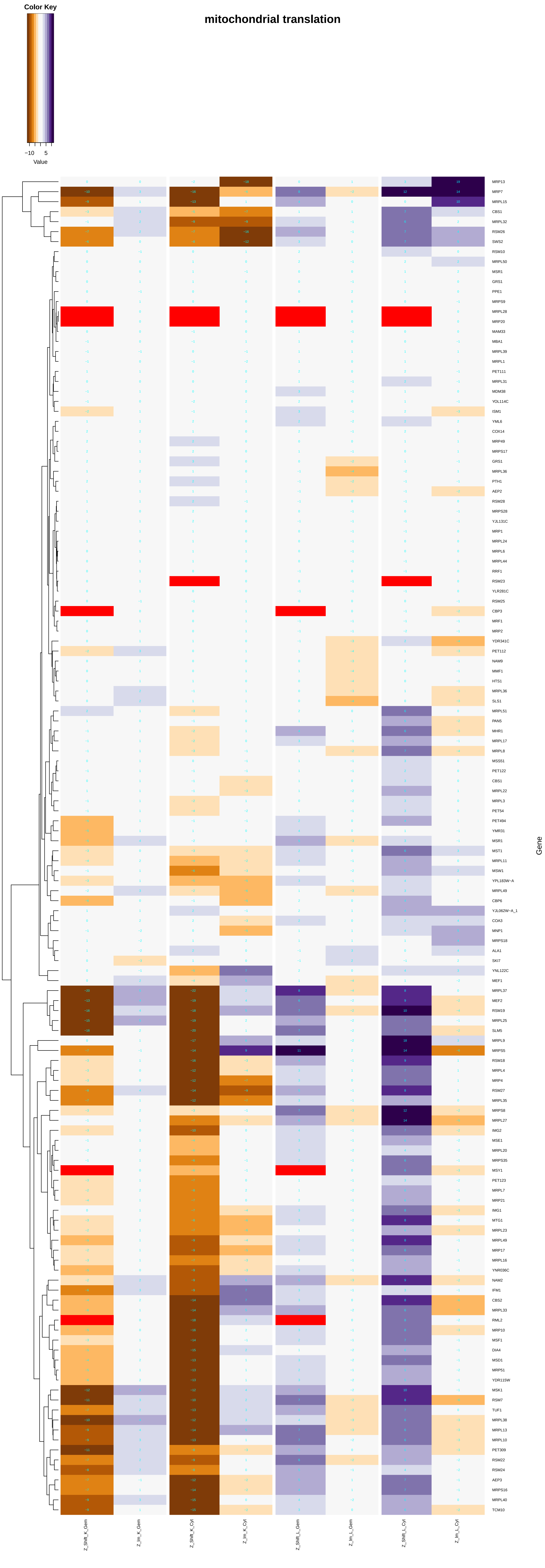
Color Key
−10
5
Value
mitochondrial translation
Gene
Z_Shift_K_Gem
Z_Im_K_Gem
Z_Shift_K_Cyt
Z_Im_K_Cyt
Z_Shift_L_Gem
Z_Im_L_Gem
Z_Shift_L_Cyt
Z_Im_L_Cyt
