## Additional File 7 for "A humanized yeast phenomic model of deoxycytidine kinase to predict genetic buffering of nucleoside analog cytotoxicity": cellular_protein_localization.pdf

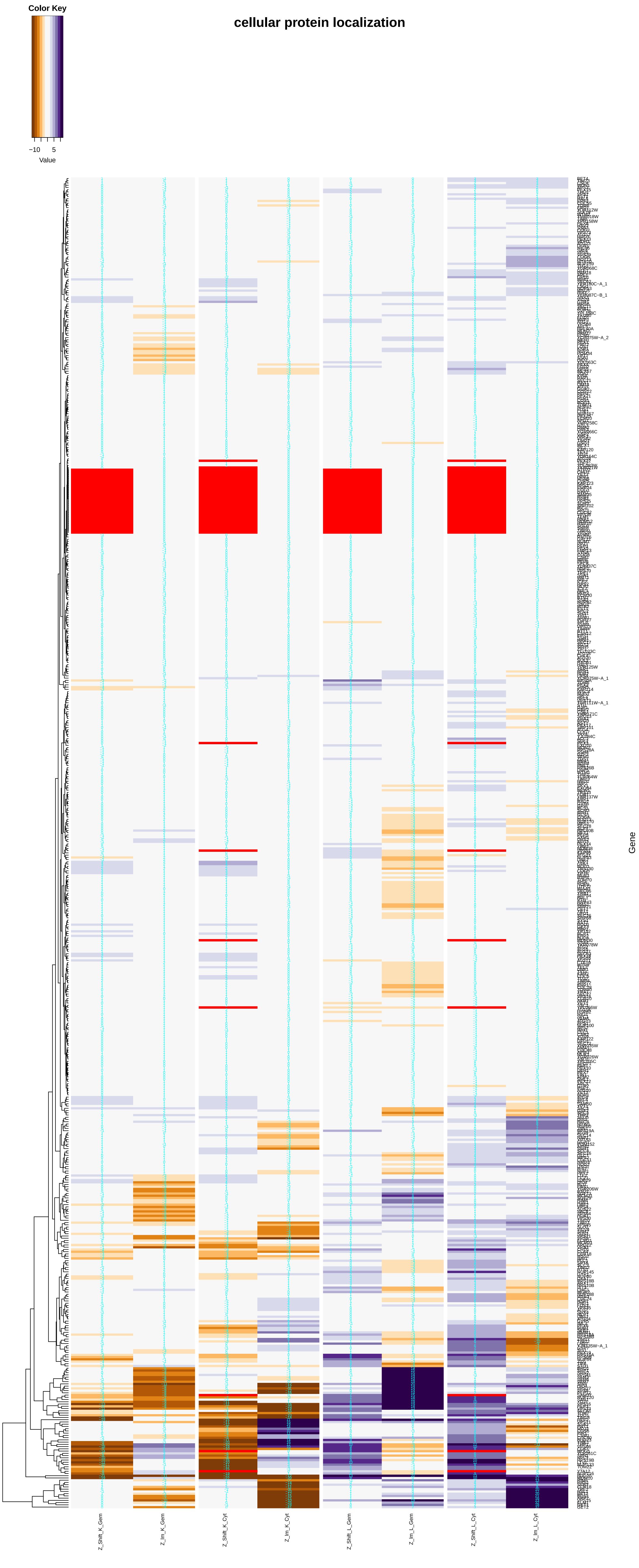
Color Key
−10
5
Value
cellular protein localization
Z_Shift_K_Gem
Z_Im_K_Gem
Z_Shift_K_Cyt
Z_Im_K_Cyt
Z_Shift_L_Gem
Z_Im_L_Gem
Z_Shift_L_Cyt
Z_Im_L_Cyt
Gene

### protein localization to organelle

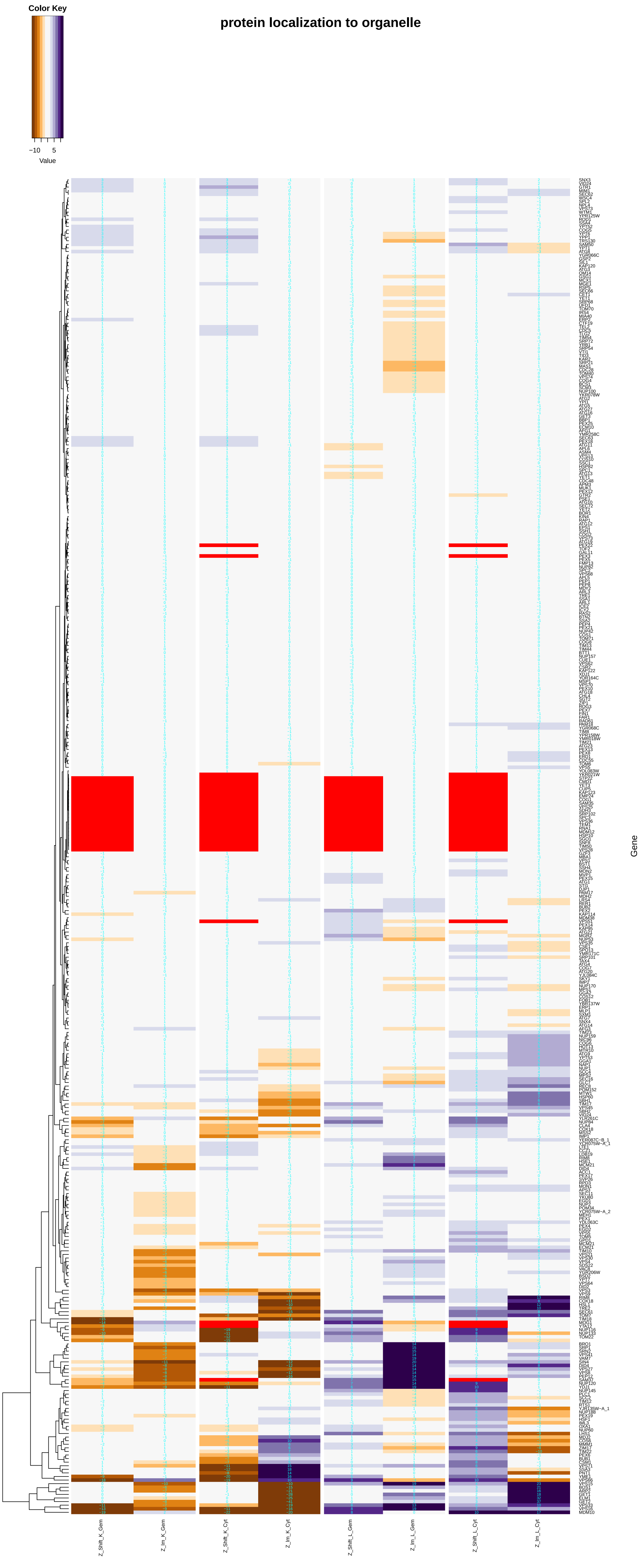

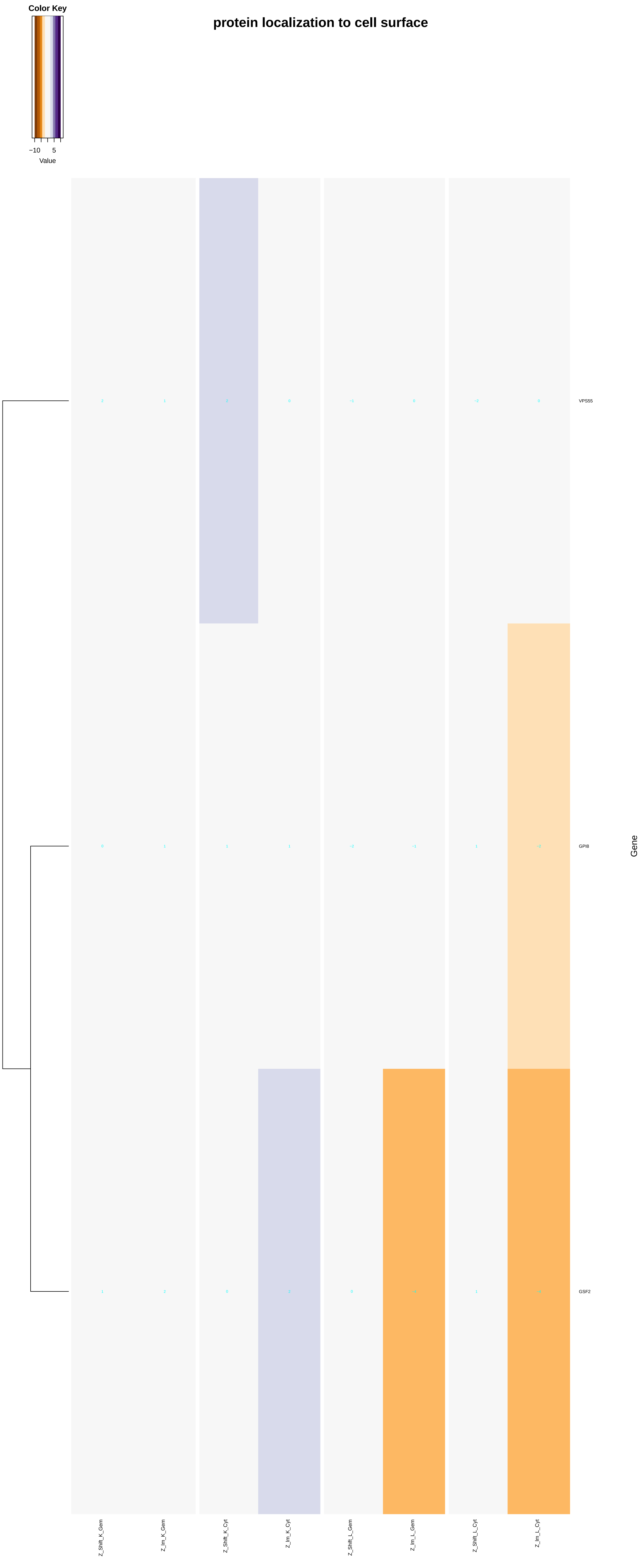
protein localization to cell surface
Color Key
−10
5
Value
VPS55
GPI8
GSF2
Gene
Z_Shift_K_Gem
Z_Im_K_Gem
Z_Shift_K_Cyt
Z_Im_K_Cyt
Z_Shift_L_Gem
Z_Im_L_Gem
Z_Shift_L_Cyt
Z_Im_L_Cyt

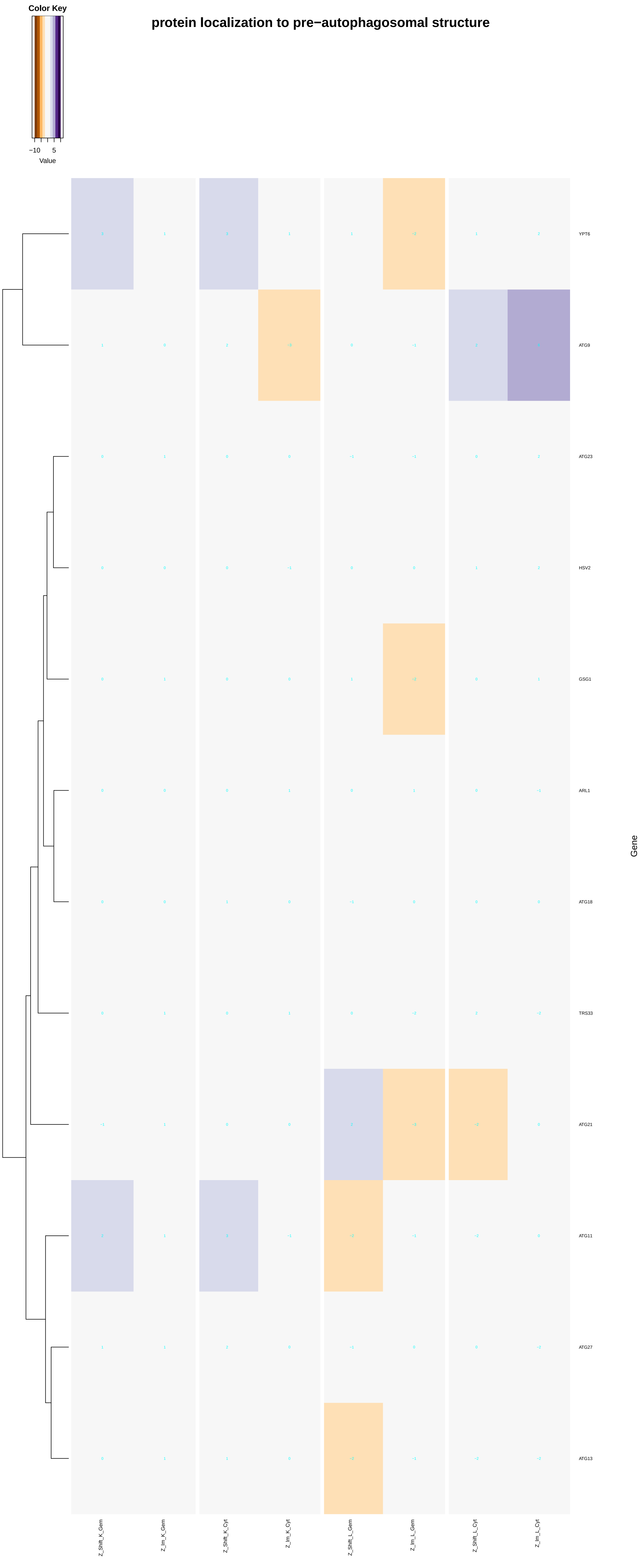

Color Key
−10
5
Value
protein localization to pre−autophagosomal structure
YPT6
ATG9
ATG23
HSV2
GSG1
ARL1
ATG18
TRS33
ATG21
ATG11
ATG27
ATG13
Gene
Z_Shift_K_Gem
Z_Im_K_Gem
Z_Shift_K_Cyt
Z_Im_K_Cyt
Z_Shift_L_Gem
Z_Im_L_Gem
Z_Shift_L_Cyt
Z_Im_L_Cyt

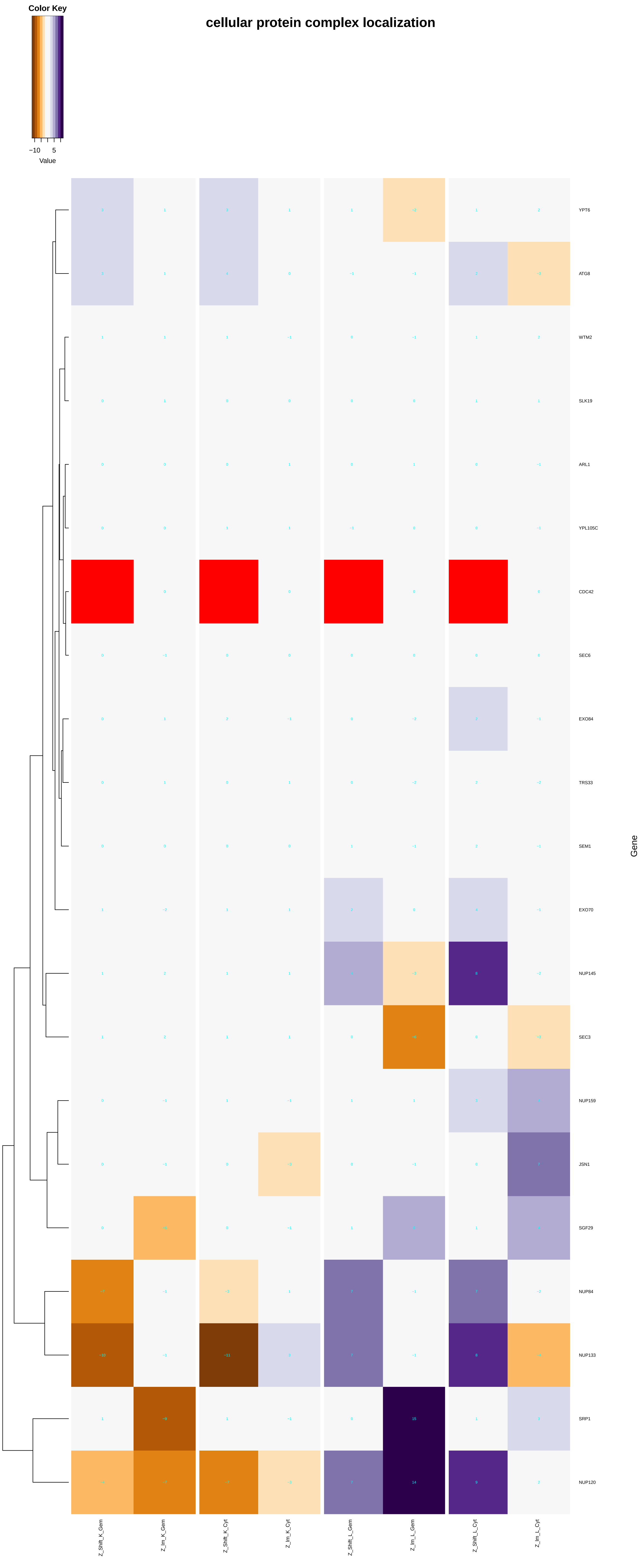

Color Key
−10
5
Value
cellular protein complex localization
YPT6
ATG8
WTM2
SLK19
ARL1
YPL105C
CDC42
SEC6
EXO84
TRS33
SEM1
EXO70
NUP145
SEC3
NUP159
JSN1
SGF29
NUP84
NUP133
SRP1
NUP120
Gene
Z_Shift_K_Gem
Z_lm_K_Gem
Z_Shift_K_Cyt
Z_lm_K_Cyt
Z_Shift_L_Gem
Z_lm_L_Gem
Z_Shift_L_Cyt
Z_lm_L_Cyt

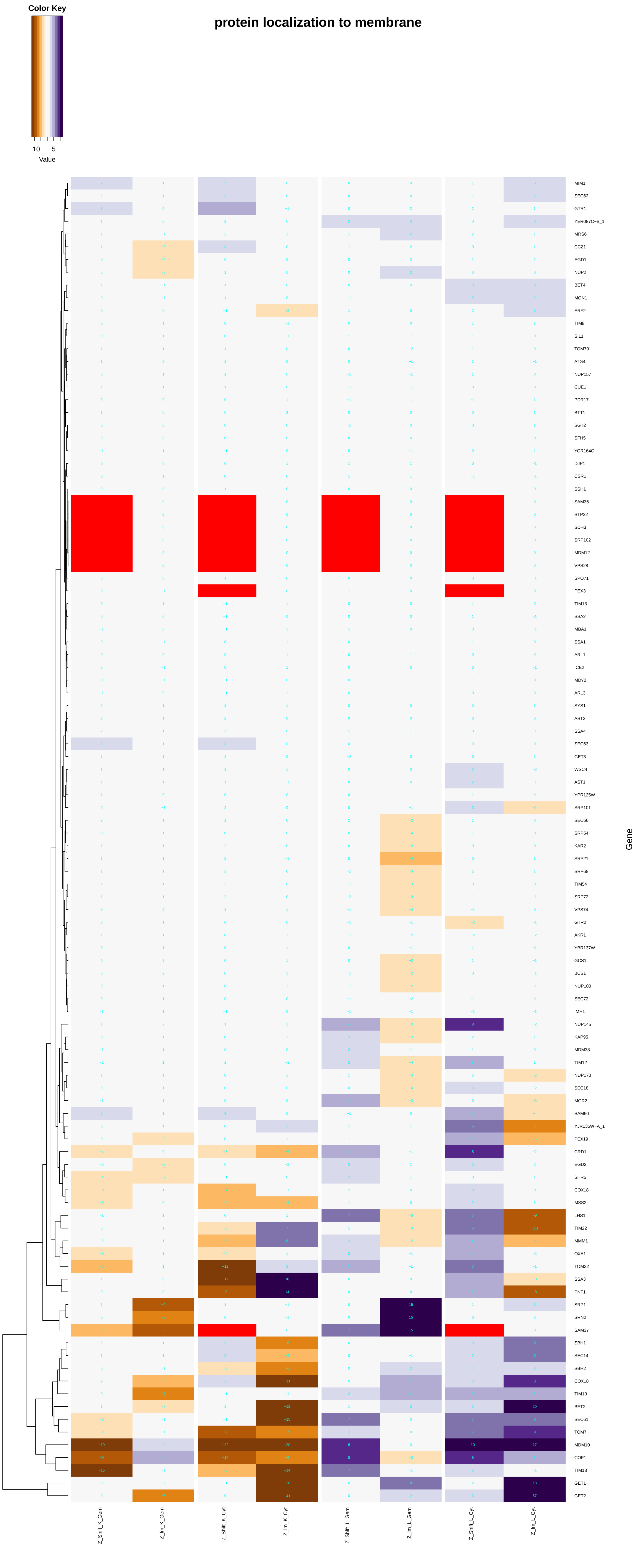

protein localization to membrane
Color Key
−10
5
Value
MIM1
SEC62
GTR1
YER087C−B_1
MRS6
CCZ1
EGD1
NUP2
BET4
MON1
ERF2
TIM8
SIL1
TOM70
ATG4
NUP157
CUE1
PDR17
BTT1
SGT2
SFH5
YOR164C
DJP1
CSR1
SSH1
SAM35
STP22
SDH3
SRP102
MDM12
VPS28
SPO71
PEX3
TIM13
SSA2
MBA1
SSA1
ARL1
ICE2
MDY2
ARL3
SYS1
AST2
SSA4
SEC63
GET3
WSC4
AST1
YPR125W
SRP101
SEC66
SRP54
KAR2
SRP21
SRP68
TIM54
SRP72
VPS74
GTR2
AKR1
YBR137W
GCS1
BCS1
NUP100
SEC72
IMH1
NUP145
KAP95
MDM38
TIM12
NUP170
SEC18
MGR2
SAM50
YJR135W−A_1
PEX19
CRD1
EGD2
SHR5
COX18
MSS2
LHS1
TIM22
MMM1
OXA1
TOM22
SSA3
PNT1
SRP1
SRN2
SAM37
SBH1
SEC14
SBH2
COX18
TIM10
BET2
SEC61
TOM7
MDM10
COF1
TIM18
GET1
GET2
Gene
Z_Shift_K_Gem
Z_Im_K_Gem
Z_Shift_K_Cyt
Z_Im_K_Cyt
Z_Shift_L_Gem
Z_Im_L_Gem
Z_Shift_L_Cyt
Z_Im_L_Cyt

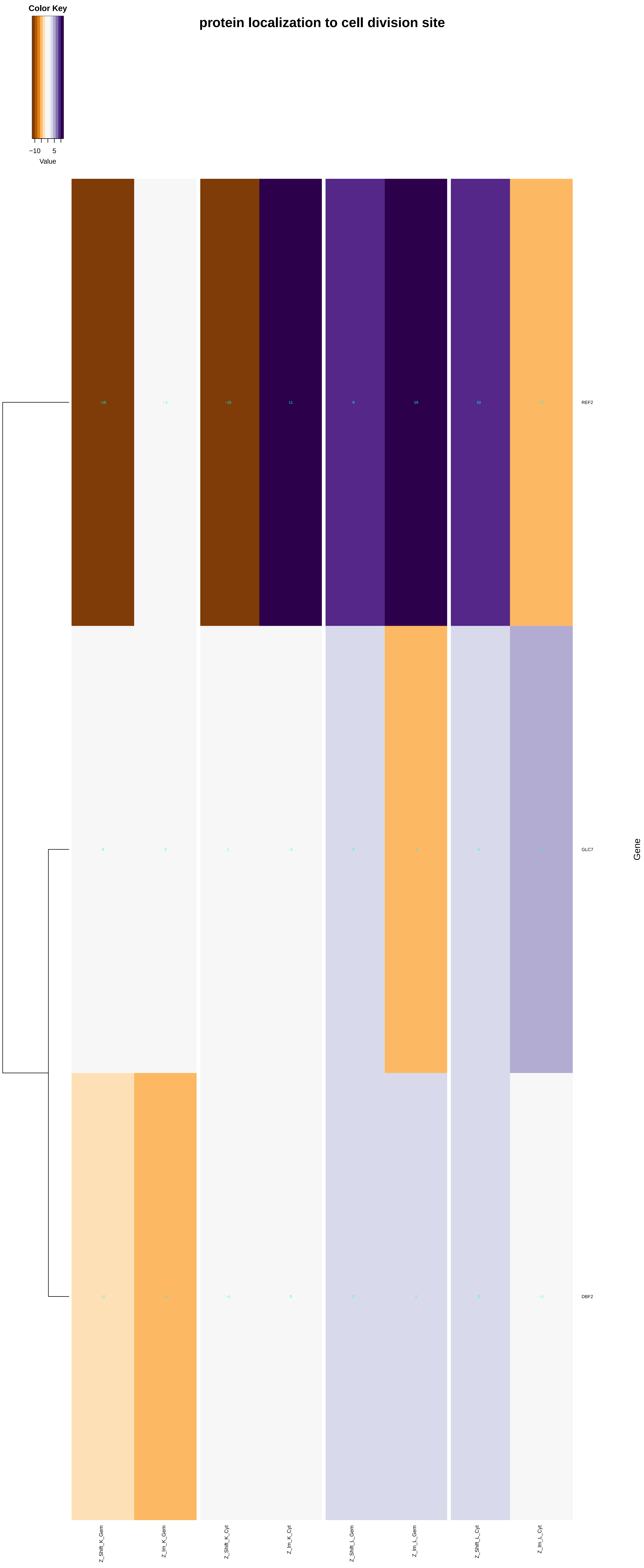

Color Key
−10
5
Value
protein localization to cell division site
−19
−1
−15
11
9
19
10
−5
REF2
0
2
1
−1
2
−5
4
5
GLC7
−2
−5
−1
0
2
2
3
−1
DBF2
Gene
Z_Shift_K_Gem
Z_Im_K_Gem
Z_Shift_K_Cyt
Z_Im_K_Cyt
Z_Shift_L_Gem
Z_Im_L_Gem
Z_Shift_L_Cyt
Z_Im_L_Cyt

#### protein localization to bud neck

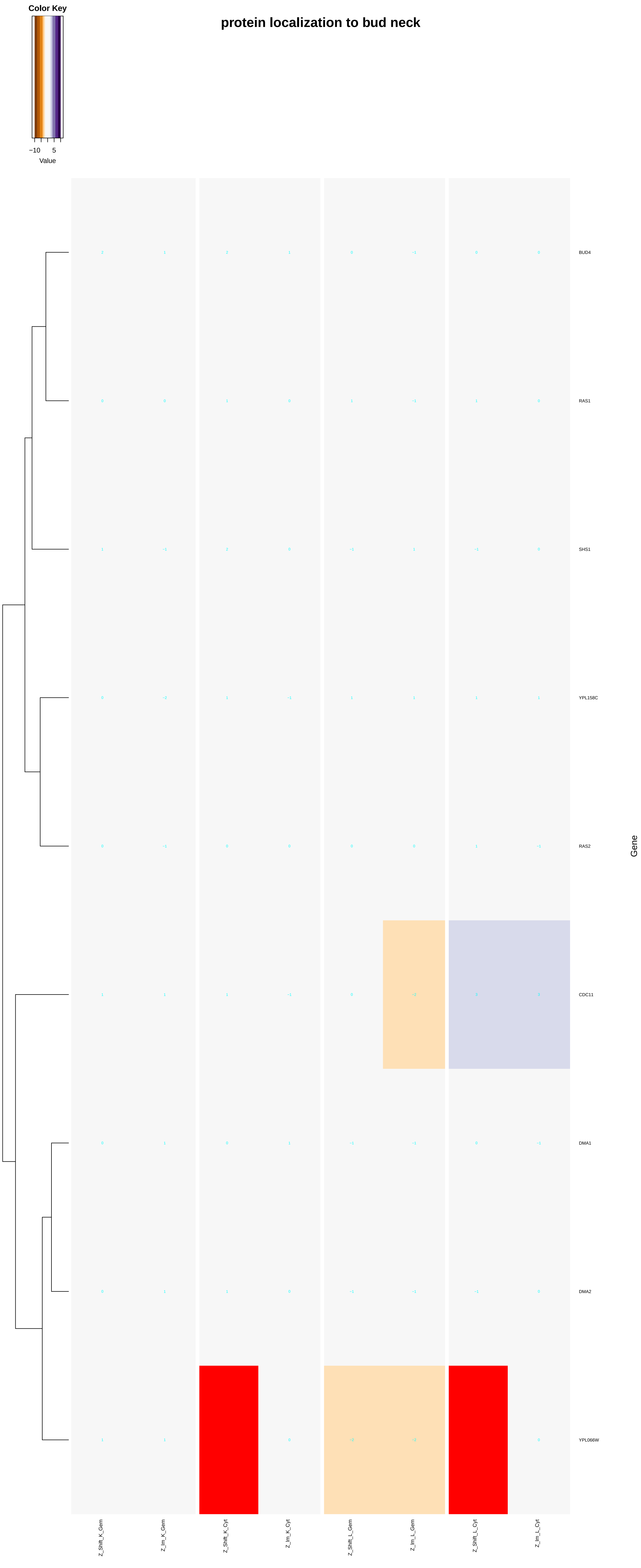

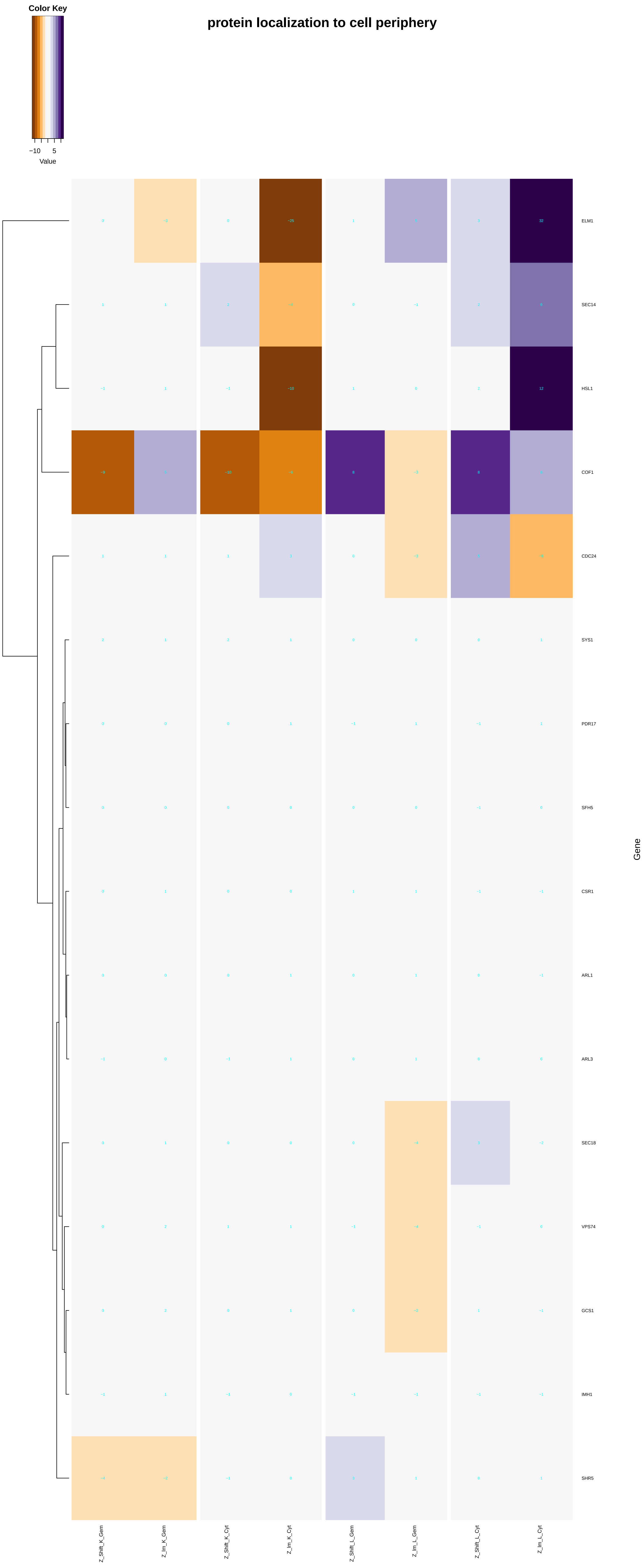

protein localization to cell periphery
Color Key
−10
5
Value
ELM1
SEC14
HSL1
COF1
CDC24
SYS1
PDR17
SFH5
CSR1
ARL1
ARL3
SEC18
VPS74
GCS1
IMH1
SHR5
Gene
Z_Shift_K_Gem
Z_Im_K_Gem
Z_Shift_K_Cyt
Z_Im_K_Cyt
Z_Shift_L_Gem
Z_Im_L_Gem
Z_Shift_L_Cyt
Z_Im_L_Cyt

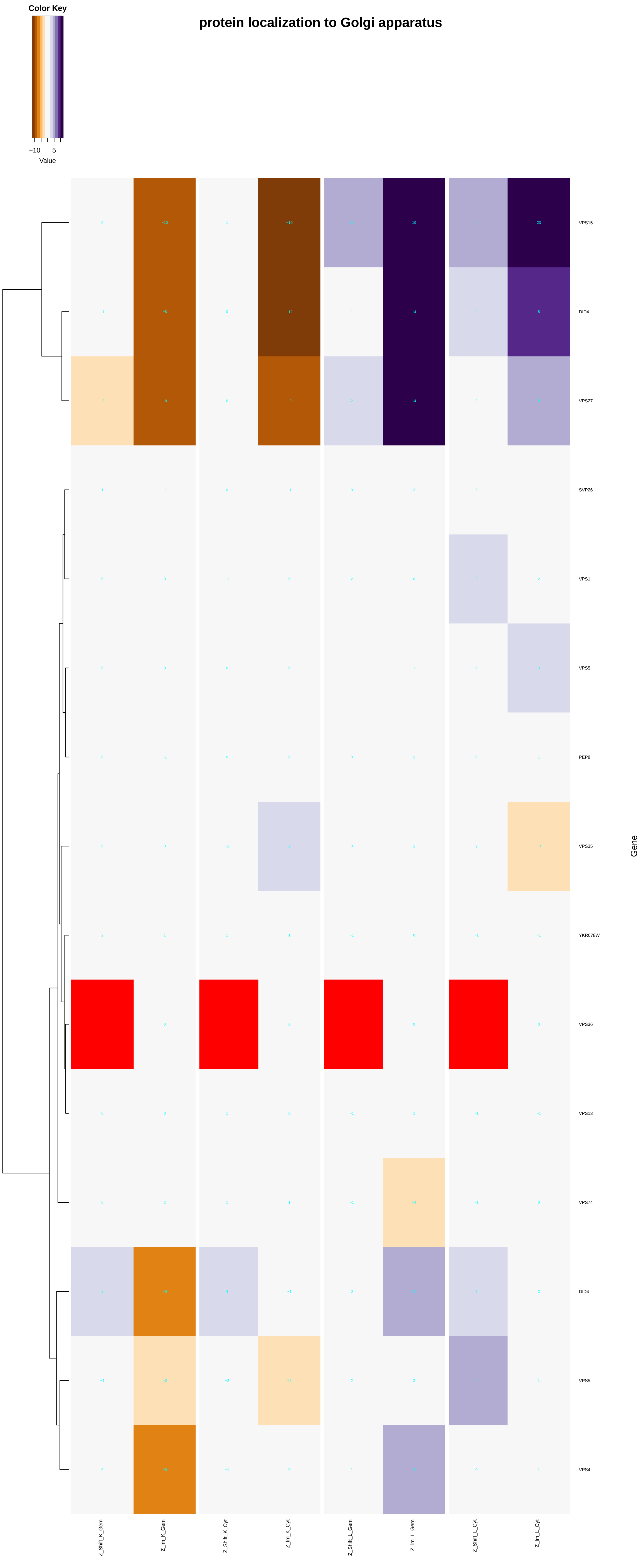

Color Key
−10 5
Value
protein localization to Golgi apparatus
VPS15
DID4
VPS27
SVP26
VPS1
VPS5
PEP8
VPS35
YKR078W
VPS36
VPS13
VPS74
DID4
VPS5
VPS4
Gene
Z_Shift_K_Gem
Z_Im_K_Gem
Z_Shift_K_Cyt
Z_Im_K_Cyt
Z_Shift_L_Gem
Z_Im_L_Gem
Z_Shift_L_Cyt
Z_Im_L_Cyt

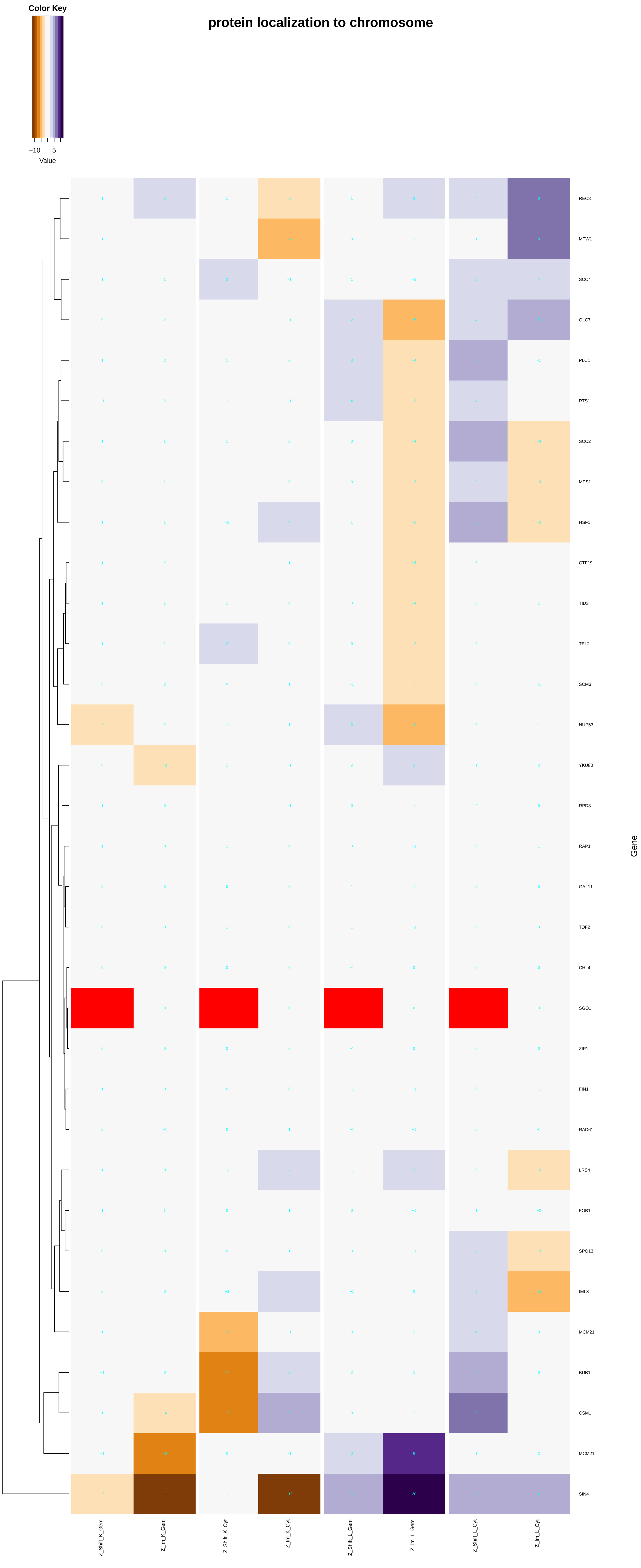
protein localization to chromosome
Color Key
−10
5
Value
REC8
MTW1
SCC4
GLC7
PLC1
RTS1
SCC2
MPS1
HSF1
CTF19
TID3
TEL2
SCM3
NUP53
YKU80
RPD3
RAP1
GAL11
TOF2
CHL4
SGO1
ZIP1
FIN1
RAD61
LRS4
FOB1
SPO13
IML3
MCM21
BUB1
CSM1
MCM21
SIN4
Gene
Z_Shift_K_Gem
Z_lm_K_Gem
Z_Shift_K_Cyt
Z_lm_K_Cyt
Z_Shift_L_Gem
Z_lm_L_Gem
Z_Shift_L_Cyt
Z_lm_L_Cyt

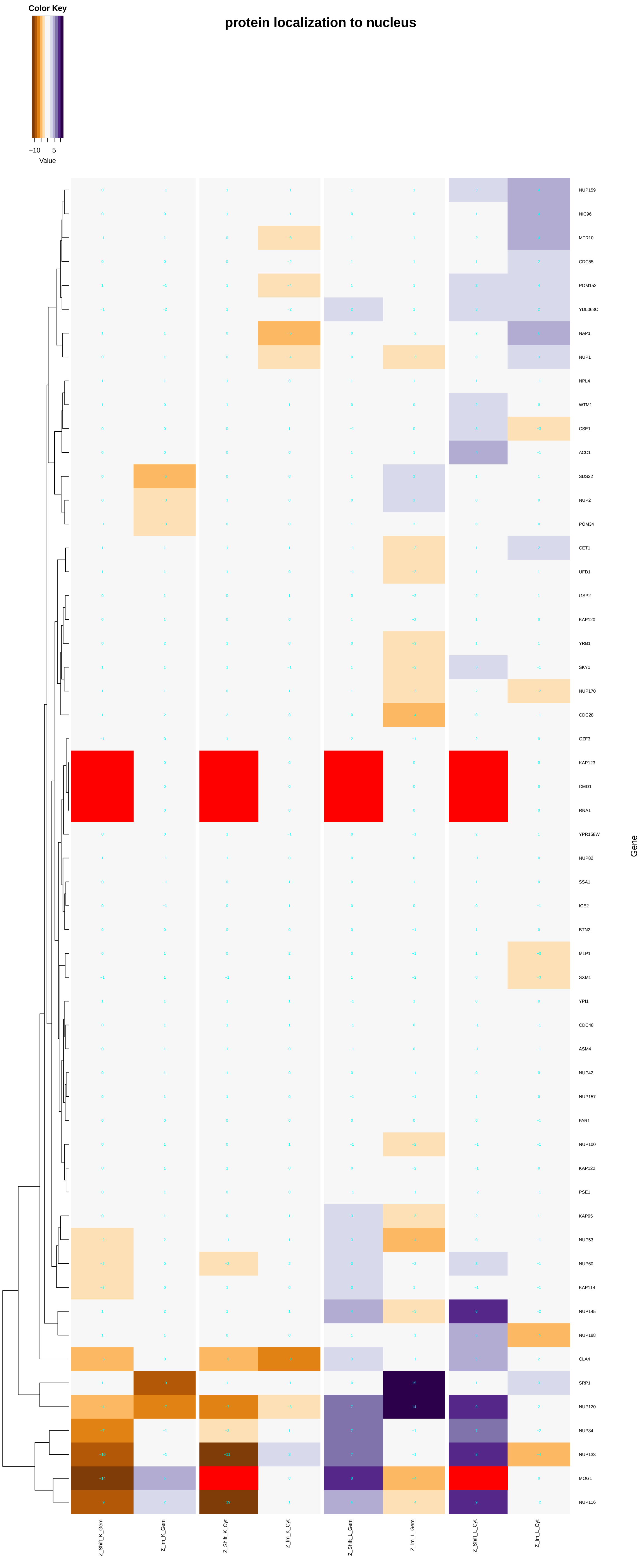

Color Key
−10
5
Value
protein localization to nucleus
NUP159
NIC96
MTR10
CDC55
POM152
YDL063C
NAP1
NUP1
NPL4
WTM1
CSE1
ACC1
SDS22
NUP2
POM34
CET1
UFD1
GSP2
KAP120
YRB1
SKY1
NUP170
CDC28
GZF3
KAP123
CMD1
RNA1
YPR158W
NUP82
SSA1
ICE2
BTN2
MLP1
SXM1
YPI1
CDC48
ASM4
NUP42
NUP157
FAR1
NUP100
KAP122
PSE1
KAP95
NUP53
NUP60
KAP114
NUP145
NUP188
CLA4
SRP1
NUP120
NUP84
NUP133
MOG1
NUP116
Gene
Z_Shift_K_Gem
Z_Im_K_Gem
Z_Shift_K_Cyt
Z_Im_K_Cyt
Z_Shift_L_Gem
Z_Im_L_Gem
Z_Shift_L_Cyt
Z_Im_L_Cyt

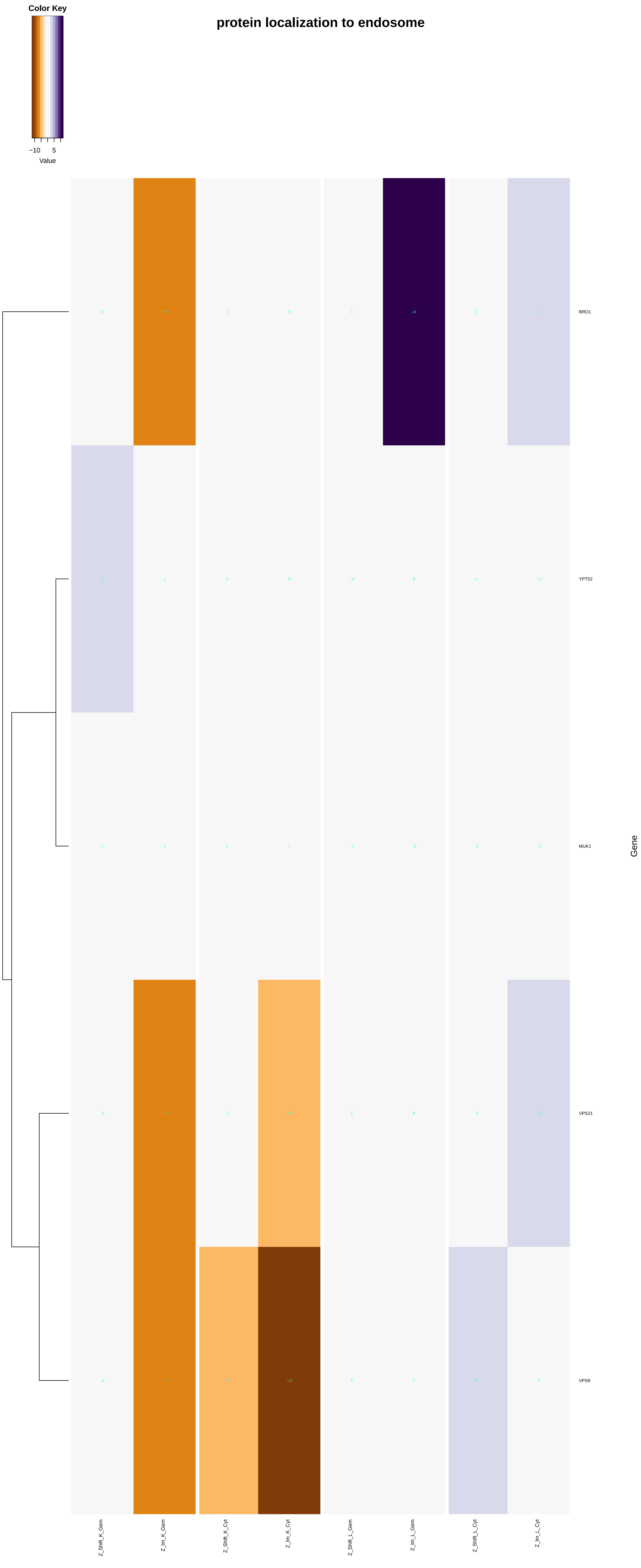
protein localization to endosome
Color Key
−10
5
Value
BRO1
YPT52
MUK1
VPS21
VPS9
Gene
Z_Shift_K_Gem
Z_Im_K_Gem
Z_Shift_K_Cyt
Z_Im_K_Cyt
Z_Shift_L_Gem
Z_Im_L_Gem
Z_Shift_L_Cyt
Z_Im_L_Cyt

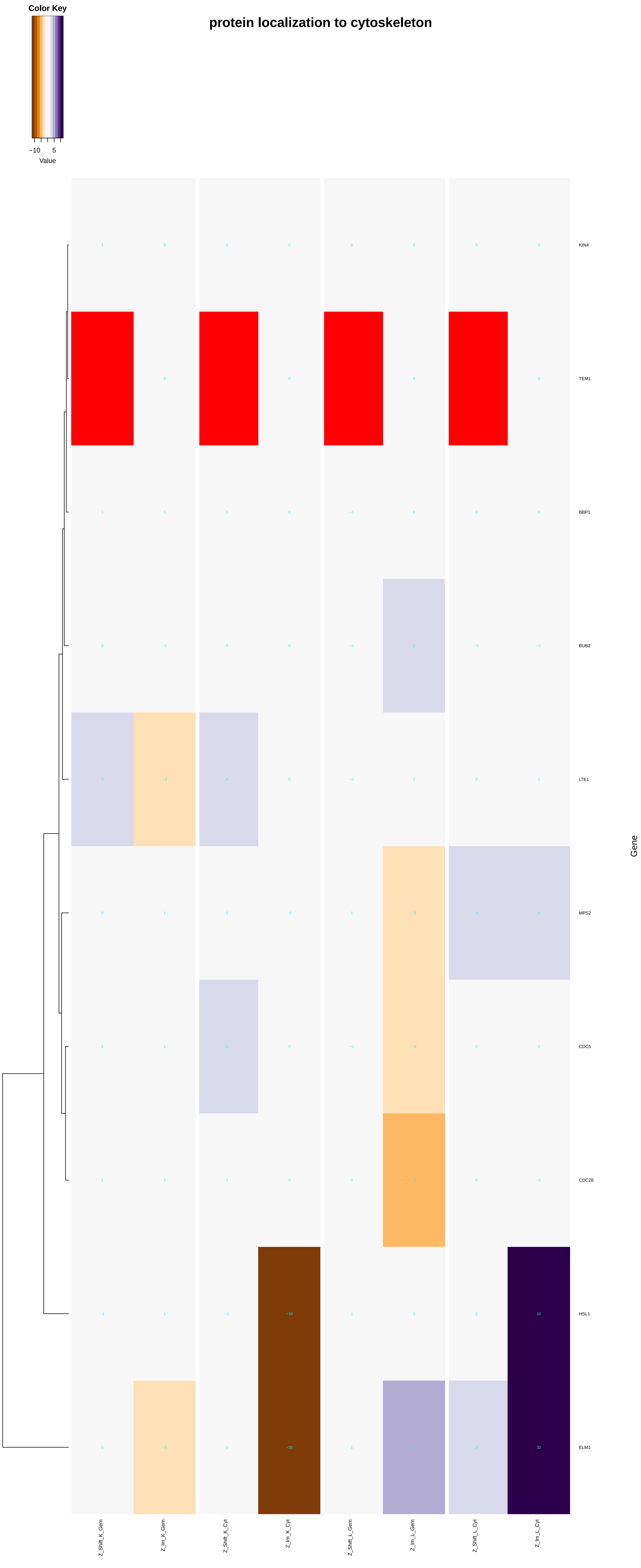

protein localization to cytoskeleton
Color Key
−10
5
Value
KIN4
TEM1
BBP1
BUB2
LTE1
MPS2
CDC5
CDC28
HSL1
ELM1
Gene
Z_Shift_K_Gem
Z_Im_K_Gem
Z_Shift_K_Cyt
Z_Im_K_Cyt
Z_Shift_L_Gem
Z_Im_L_Gem
Z_Shift_L_Cyt
Z_Im_L_Cyt

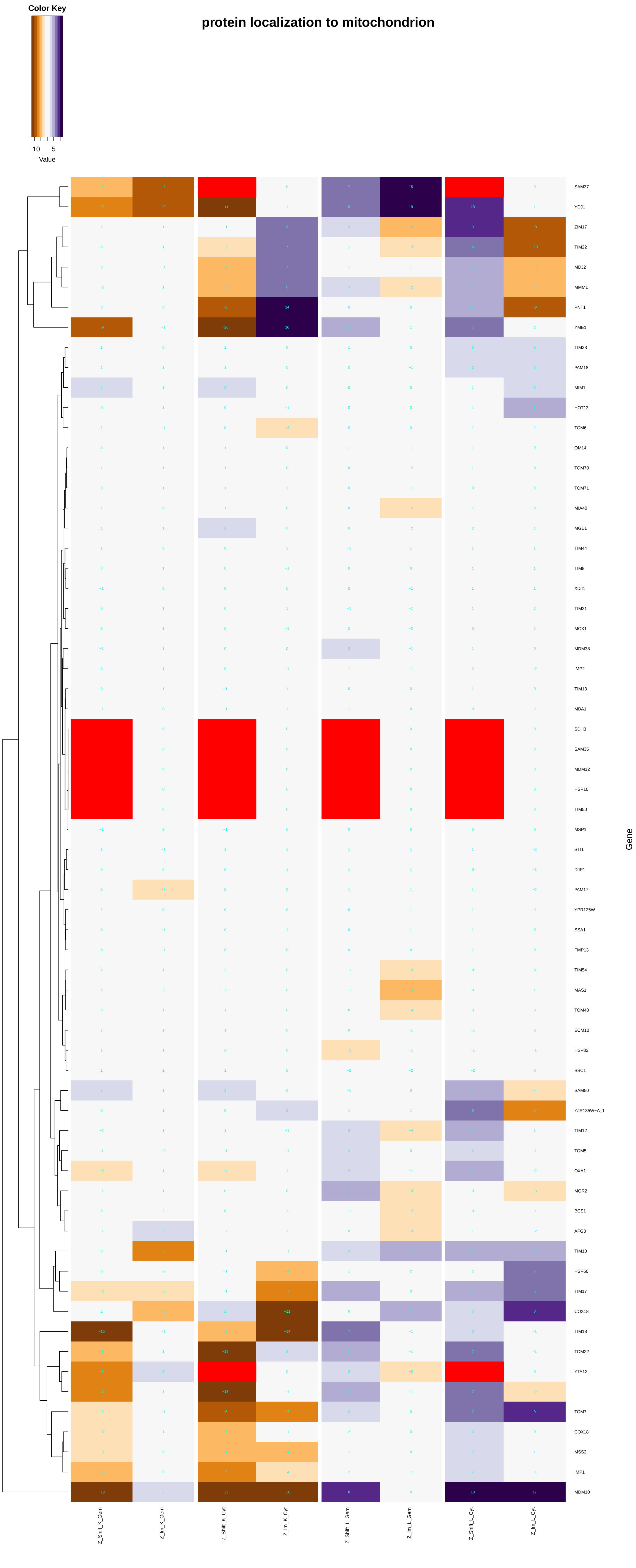
Color Key
−10
5
Value
protein localization to mitochondrion
SAM37
YDJ1
ZIM17
TIM22
MDJ2
MMM1
PNT1
YME1
TIM23
PAM18
MIM1
HOT13
TOM6
OM14
TOM70
TOM71
MIA40
MGE1
TIM44
TIM8
XDJ1
TIM21
MCX1
MDM38
IMP2
TIM13
MBA1
SDH3
SAM35
MDM12
HSP10
TIM50
MSP1
STI1
DJP1
PAM17
YPR125W
SSA1
FMP13
TIM54
MAS1
TOM40
ECM10
HSP82
SSC1
SAM50
YJR135W−A_1
TIM12
TOM5
OXA1
MGR2
BCS1
AFG3
TIM10
HSP60
TIM17
COX18
TIM18
TOM22
YTA12
TOM7
COX18
MSS2
IMP1
MDM10
Gene
Z_Shift_K_Gem
Z_Im_K_Gem
Z_Shift_K_Cyt
Z_Im_K_Cyt
Z_Shift_L_Gem
Z_Im_L_Gem
Z_Shift_L_Cyt
Z_Im_L_Cyt

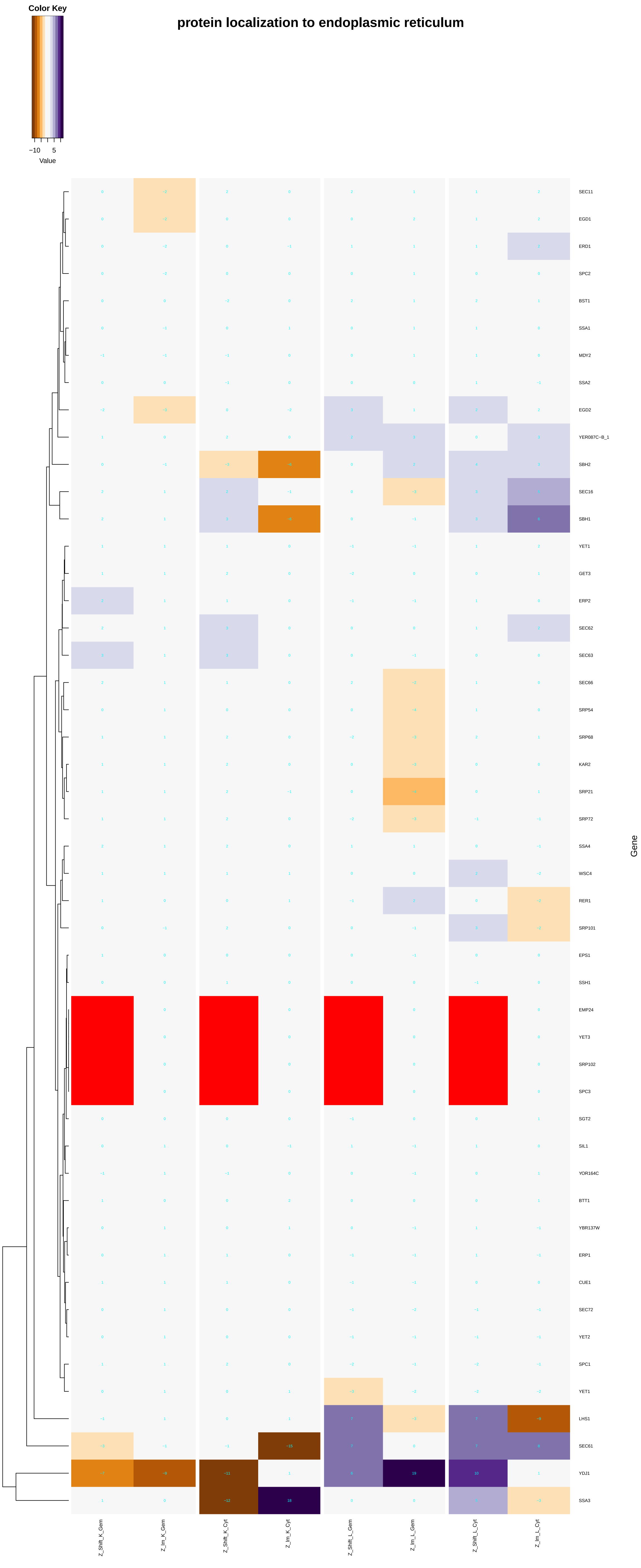

protein localization to endoplasmic reticulum
Color Key
−10
5
Value
SEC11
EGD1
ERD1
SPC2
BST1
SSA1
MDY2
SSA2
EGD2
YER087C−B_1
SBH2
SEC16
SBH1
YET1
GET3
ERP2
SEC62
SEC63
SEC66
SRP54
SRP68
KAR2
SRP21
SRP72
SSA4
WSC4
RER1
SRP101
EPS1
SSH1
EMP24
YET3
SRP102
SPC3
SGT2
SIL1
YOR164C
BTT1
YBR137W
ERP1
CUE1
SEC72
YET2
SPC1
YET1
LHS1
SEC61
YDJ1
SSA3
Gene
Z_Shift_K_Gem
Z_Im_K_Gem
Z_Shift_K_Cyt
Z_Im_K_Cyt
Z_Shift_L_Gem
Z_Im_L_Gem
Z_Shift_L_Cyt
Z_Im_L_Cyt

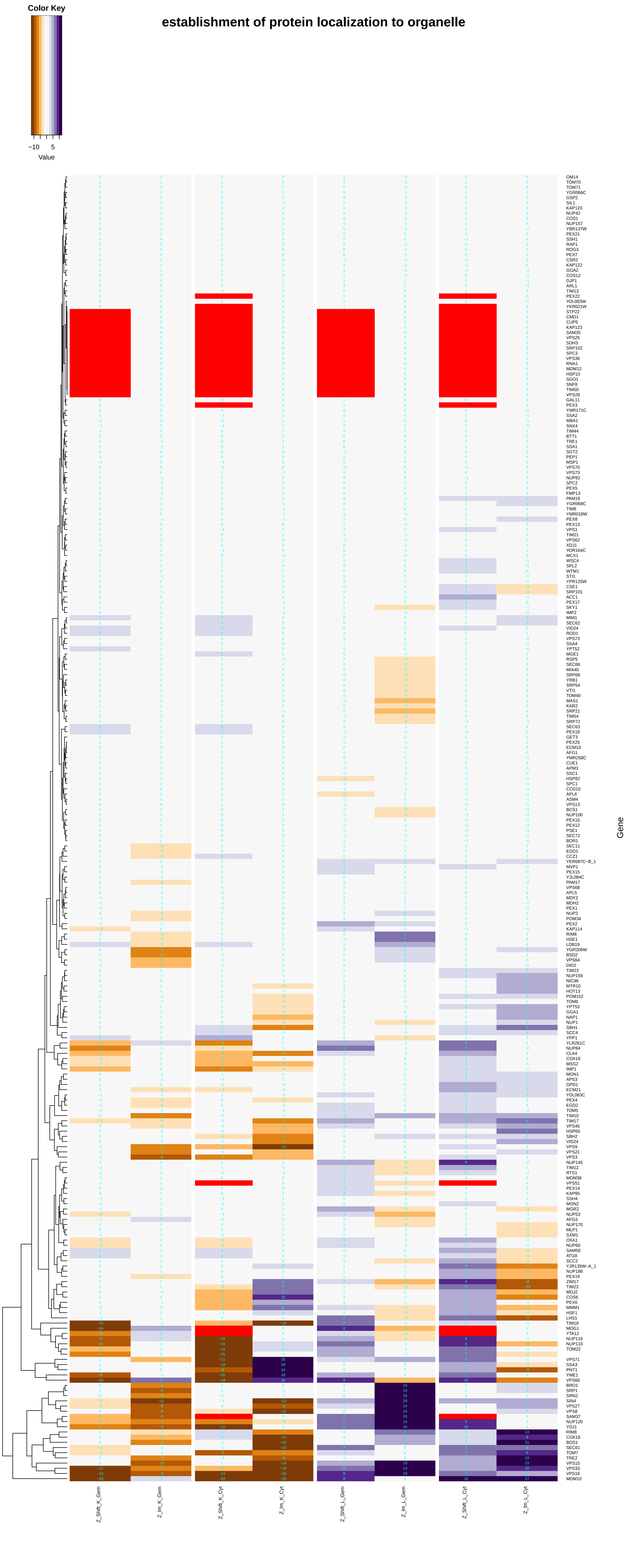
establishment of protein localization to organelle
Color Key
−10
5
Value
Gene
Z_Shift_K_Gem
Z_Im_K_Gem
Z_Shift_K_Cyt
Z_Im_K_Cyt
Z_Shift_L_Gem
Z_Im_L_Gem
Z_Shift_L_Cyt
Z_Im_L_Cyt
OM14
TOM70
TOM71
YGR066C
GSP2
SIL1
KAP120
NUP42
COS1
NUP157
YBR137W
PEX21
SSH1
RAP1
ROG3
PEX7
CSR2
KAP122
GGA2
COS12
DJP1
ARL1
TIM13
PEX22
YOL083W
YKR021W
STP22
CMD1
CUP5
KAP123
SAM35
VPS25
SDH3
SRP102
SPC3
VPS36
RNA1
MDM12
HSP10
SGO1
SNF8
TIM50
VPS28
GAL11
PEX3
YMR171C
SSA2
MBA1
SNX4
TIM44
BTT1
TRE1
SSA1
SGT2
PEP1
MSP1
VPS70
VPS73
NUP82
SPC2
PEX5
FMP13
PAM18
YGR068C
TIM8
YMR018W
PEX8
PEX13
VPS1
TIM21
VPS62
XDJ1
YOR164C
MCX1
WSC4
SPL2
WTM1
STI1
YPR125W
CSE1
SRP101
ACC1
PEX17
SKY1
IMP2
MIM1
SEC62
VID24
ROD1
VPS73
SSA4
YPT52
MGE1
RSP5
SEC66
MIA40
SRP68
YRB1
SRP54
VTI1
TOM40
MAS1
KAR2
SRP21
TIM54
SRP72
SEC63
PEX18
GET3
PEX25
ECM10
AFG1
YMR258C
CUE1
APM3
SSC1
HSP82
SPC1
COS10
APL6
ASM4
VPS13
BCS1
NUP100
PEX10
PEX12
PSE1
SEC72
BOR1
SEC11
EGD1
CCZ1
YER087C−B_1
MVP1
PEX15
YJL084C
PAM17
VPS68
APL5
MDY2
MDH2
PEX1
NUP2
POM34
PEX2
KAP114
RIM8
HSE1
LDB19
YGR206W
BSD2
VPS64
DID2
TIM23
NUP159
NIC96
MTR10
HOT13
POM152
TOM6
YPT53
GGA1
NAP1
NUP1
SBH1
SCC4
YPP1
YLR261C
NUP84
CLA4
COX18
MSS2
IMP1
MON1
APS3
GPD1
ECM21
YDL063C
PEX4
EGD2
TOM5
TIM10
TIM17
VPS45
HSP60
SBH2
VID24
VPS9
VPS21
VPS3
NUP145
TIM12
RTS1
MDM38
VPS51
PEX14
KAP95
SSH4
MON2
MGR2
NUP53
APG3
NUP170
MLP1
SXM1
OXA1
NUP60
SAM50
ATG8
SCC2
YJR135W−A_1
NUP188
PEX19
ZIM17
TIM22
MDJ2
COS6
PEX6
MMM1
HSF1
LHS1
TIM18
MOG1
YTA12
NUP116
NUP133
TOM22
VPS71
SSA3
PNT1
YME1
VPS66
BRO1
SRP1
SRN2
SIN4
VPS27
VPS8
SAM37
NUP120
YDJ1
RIM8
COX18
BOS1
SEC61
TOM7
TRE2
VPS15
VPS33
VPS16
MDM10

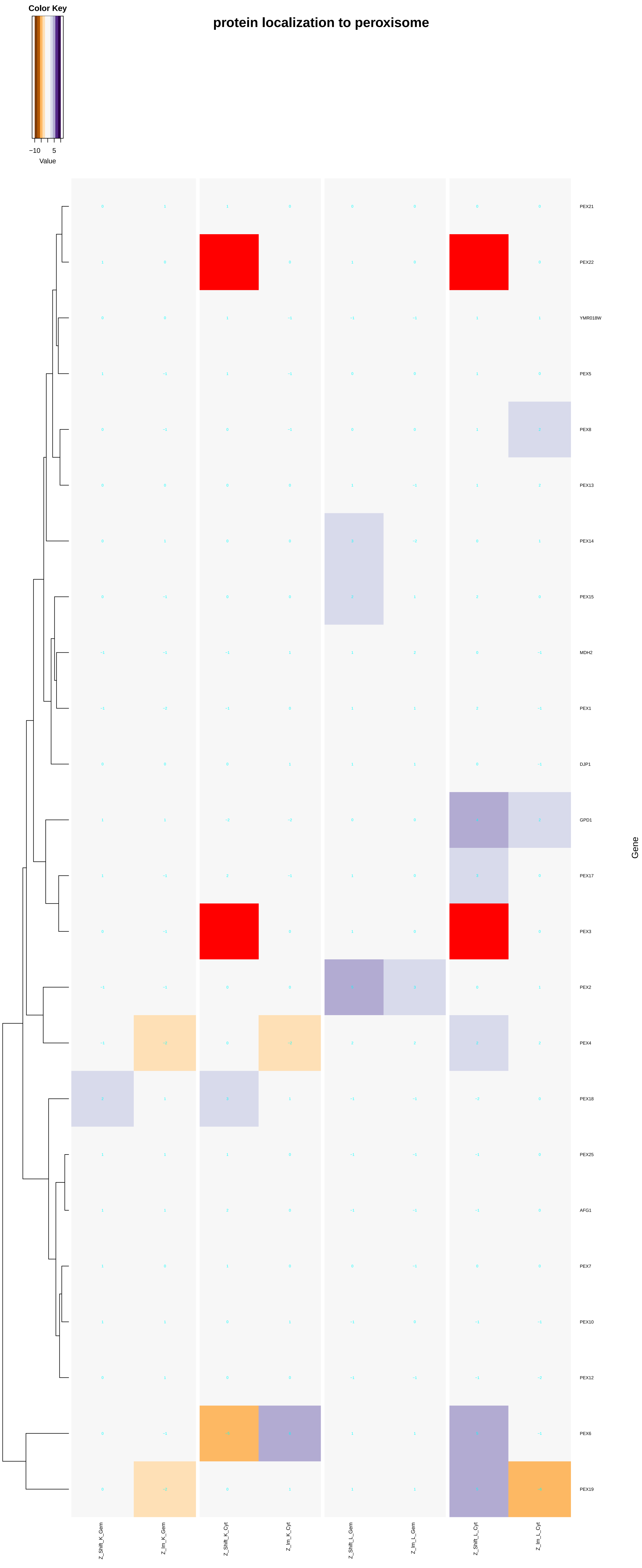

protein localization to peroxisome
Color Key
−10
5
Value
PEX21
PEX22
YMR018W
PEX5
PEX8
PEX13
PEX14
PEX15
MDH2
PEX1
DJP1
GPD1
PEX17
PEX3
PEX2
PEX4
PEX18
PEX25
AFG1
PEX7
PEX10
PEX12
PEX6
PEX19
Gene
Z_Shift_K_Gem
Z_lm_K_Gem
Z_Shift_K_Cyt
Z_lm_K_Cyt
Z_Shift_L_Gem
Z_lm_L_Gem
Z_Shift_L_Cyt
Z_lm_L_Cyt

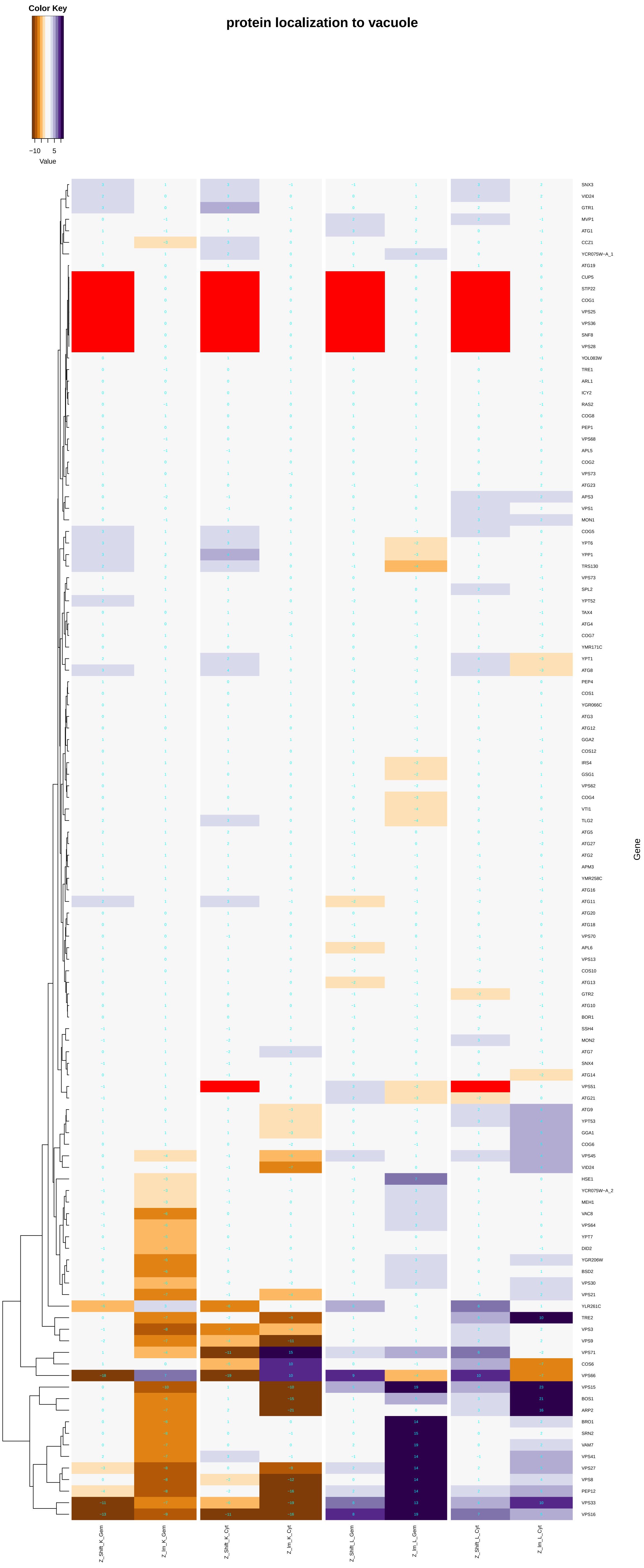
protein localization to vacuole
Color Key
−10
5
Value
Gene
Z_Shift_K_Gem
Z_Im_K_Gem
Z_Shift_K_Cyt
Z_Im_K_Cyt
Z_Shift_L_Gem
Z_Im_L_Gem
Z_Shift_L_Cyt
Z_Im_L_Cyt
SNX3
VID24
GTR1
MVP1
ATG1
CCZ1
YCR075W−A_1
ATG19
CUP5
STP22
COG1
VPS25
VPS36
SNF8
VPS28
YOL083W
TRE1
ARL1
ICY2
RAS2
COG8
PEP1
VPS68
APL5
COG2
VPS73
ATG23
APS3
VPS1
MON1
COG5
YPT6
YPP1
TRS130
VPS73
SPL2
YPT52
TAX4
ATG4
COG7
YMR171C
YPT1
ATG8
PEP4
COS1
YGR066C
ATG3
ATG12
GGA2
COS12
IRS4
GSG1
VPS62
COG4
VTI1
TLG2
ATG5
ATG27
ATG2
APM3
YMR258C
ATG16
ATG11
ATG20
ATG18
VPS70
APL6
VPS13
COS10
ATG13
GTR2
ATG10
BOR1
SSH4
MON2
ATG7
SNX4
ATG14
VPS51
ATG21
ATG9
YPT53
GGA1
COG6
VPS45
VID24
HSE1
YCR075W−A_2
MEH1
VAC8
VPS64
YPT7
DID2
YGR206W
BSD2
VPS30
VPS21
YLR261C
TRE2
VPS3
VPS9
VPS71
COS6
VPS66
VPS15
BOS1
ARP2
BRO1
SRN2
VAM7
VPS41
VPS27
VPS8
PEP12
VPS33
VPS16

### RNA polymerase II complex import to nucleus

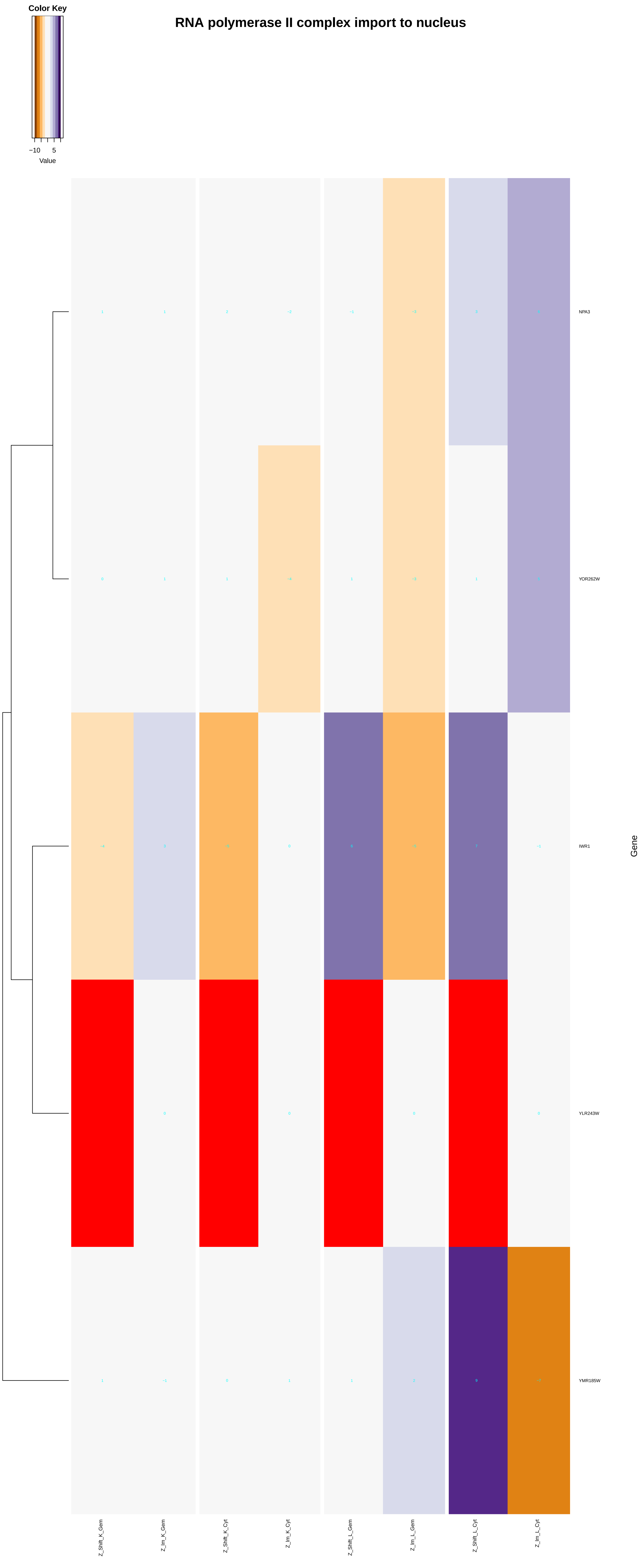

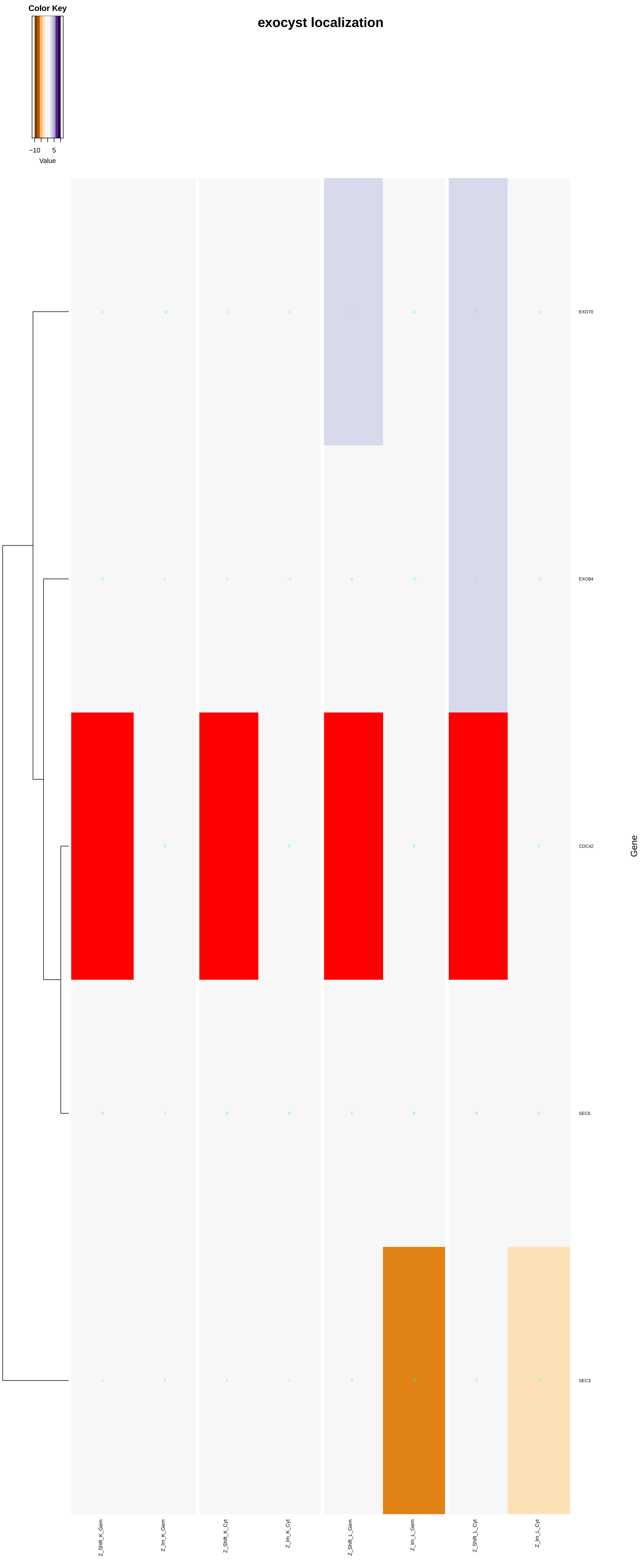
exocyst localization
Color Key
−10
5
Value
EXO70
EXO84
CDC42
SEC6
SEC3
Gene
Z_Shift_K_Gem
Z_Im_K_Gem
Z_Shift_K_Cyt
Z_Im_K_Cyt
Z_Shift_L_Gem
Z_Im_L_Gem
Z_Shift_L_Cyt
Z_Im_L_Cyt

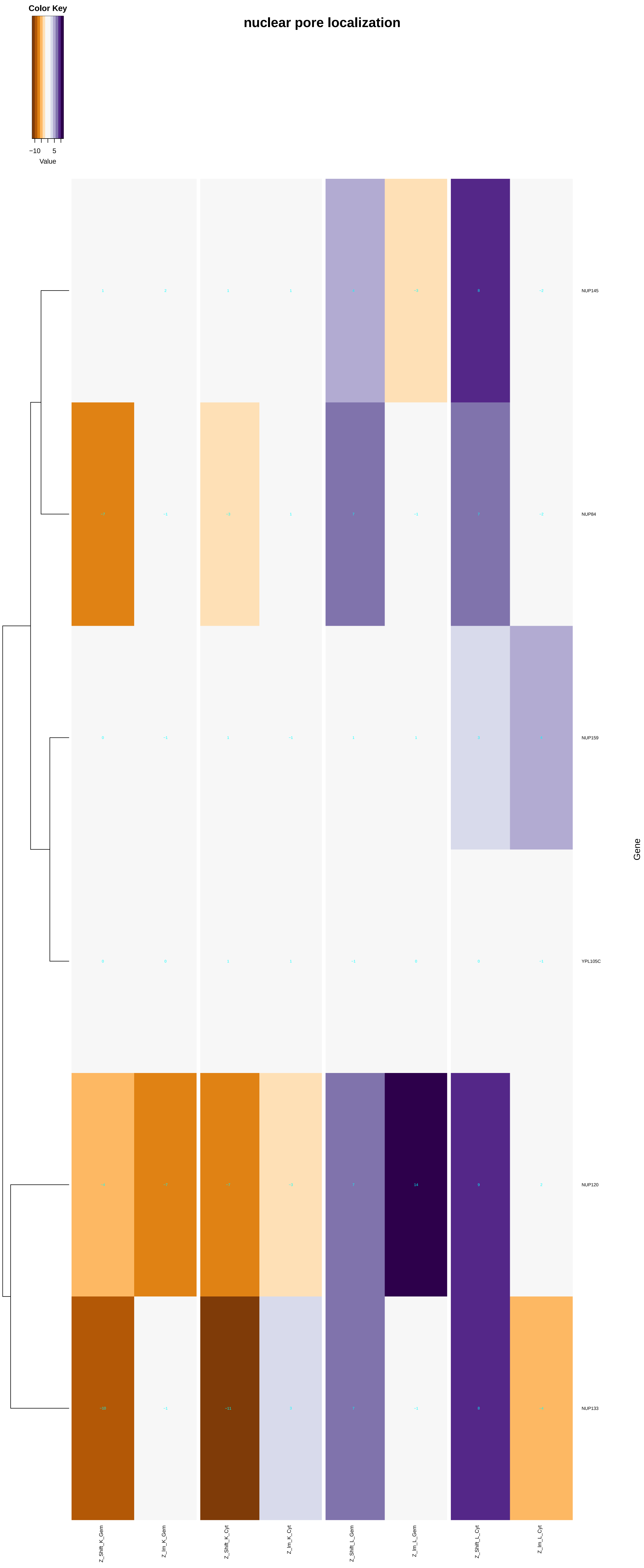

Color Key
−10 5
Value
nuclear pore localization
NUP145
NUP84
NUP159
YPL105C
NUP120
NUP133
Gene
Z_Shift_K_Gem
Z_Im_K_Gem
Z_Shift_K_Cyt
Z_Im_K_Cyt
Z_Shift_L_Gem
Z_Im_L_Gem
Z_Shift_L_Cyt
Z_Im_L_Cyt

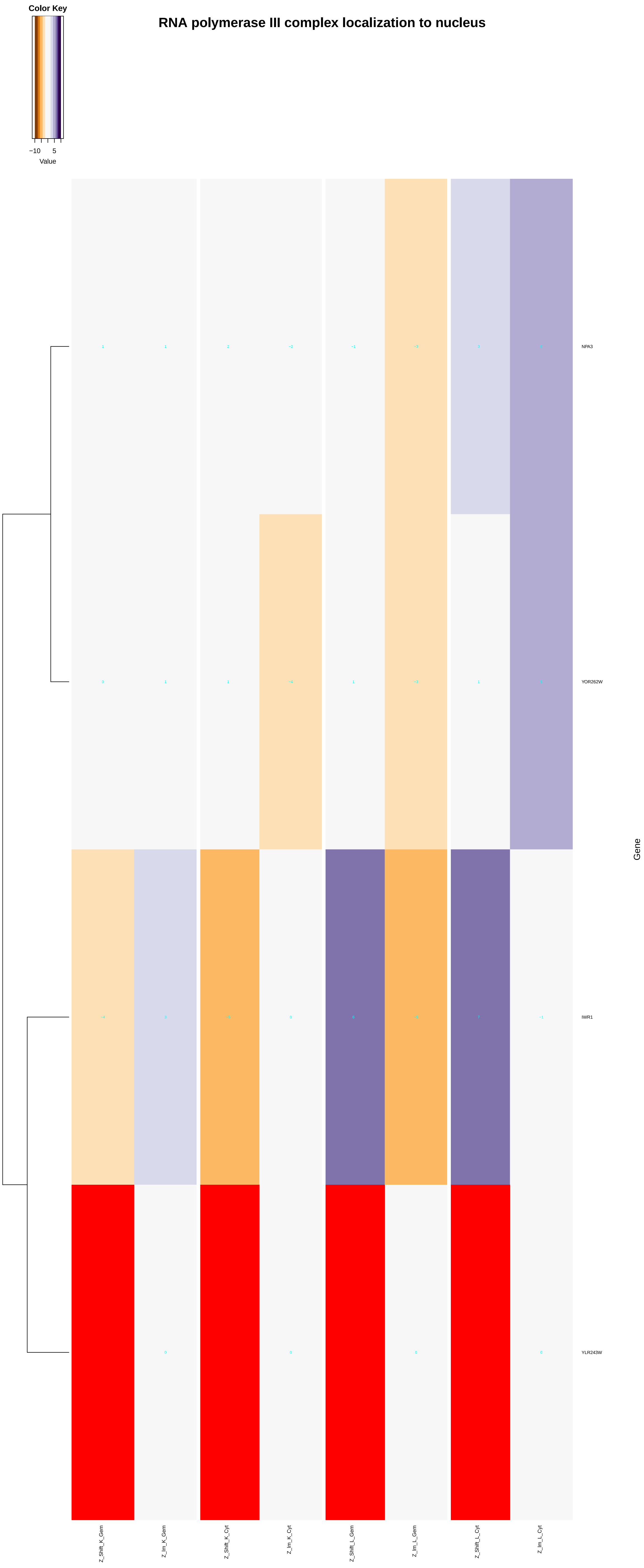
Color Key
−10
5
Value
RNA polymerase III complex localization to nucleus
NPA3
YOR262W
IWR1
YLR243W
Gene
Z_Shift_K_Gem
Z_Im_K_Gem
Z_Shift_K_Cyt
Z_Im_K_Cyt
Z_Shift_L_Gem
Z_Im_L_Gem
Z_Shift_L_Cyt
Z_Im_L_Cyt

#### protein localization to plasma membrane

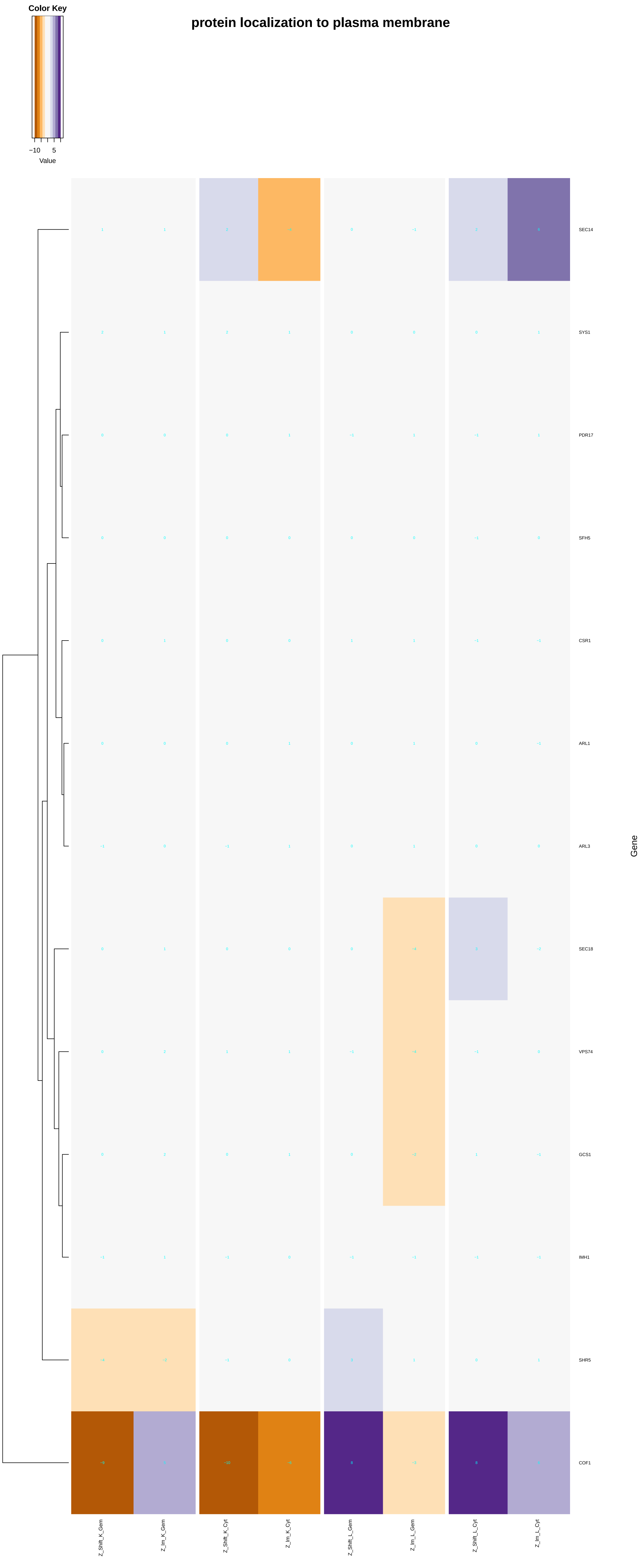

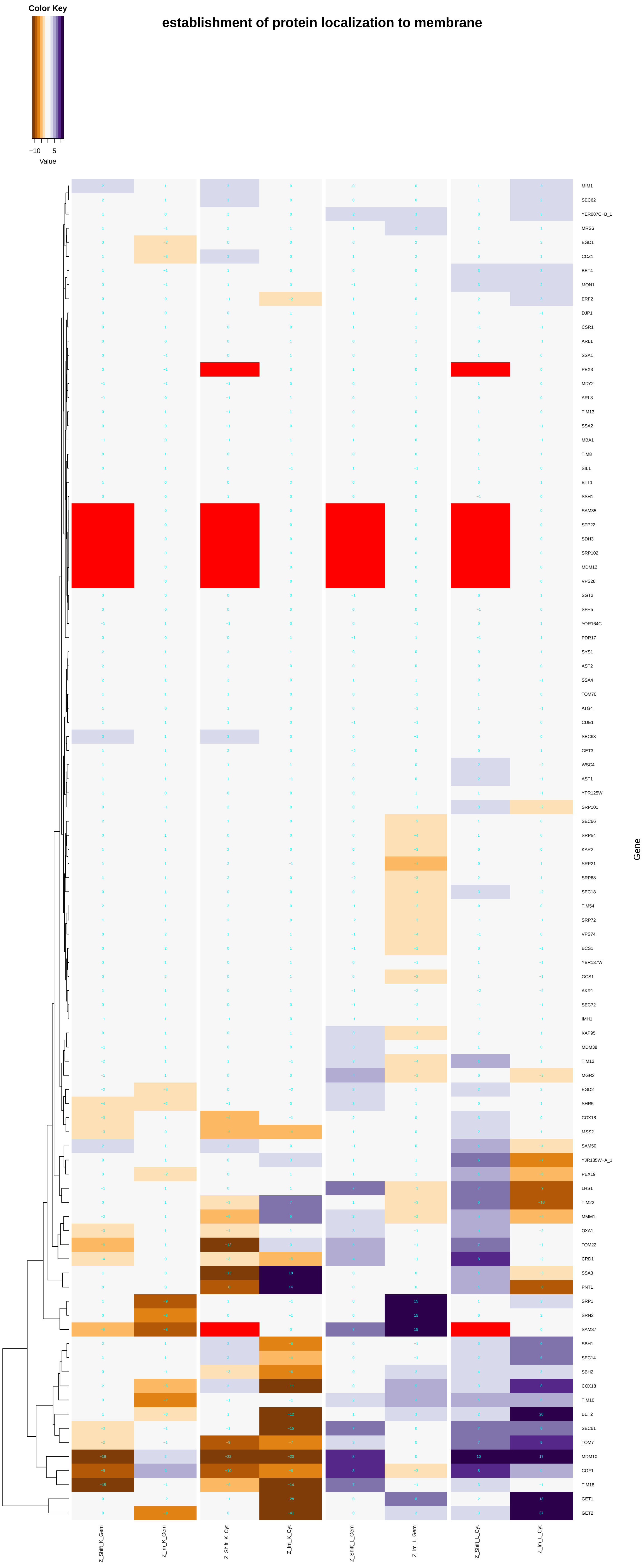

establishment of protein localization to membrane
Color Key
−10
5
Value
MIM1
SEC62
YER087C−B_1
MRS6
EGD1
CCZ1
BET4
MON1
ERF2
DJP1
CSR1
ARL1
SSA1
PEX3
MDY2
ARL3
TIM13
SSA2
MBA1
TIM8
SIL1
BTT1
SSH1
SAM35
STP22
SDH3
SRP102
MDM12
VPS28
SGT2
SFH5
YOR164C
PDR17
SYS1
AST2
SSA4
TOM70
ATG4
CUE1
SEC63
GET3
WSC4
AST1
YPR125W
SRP101
SEC66
SRP54
KAR2
SRP21
SRP68
SEC18
TIM54
SRP72
VPS74
BCS1
YBR137W
GCS1
AKR1
SEC72
IMH1
KAP95
MDM38
TIM12
MGR2
EGD2
SHR5
COX18
MSS2
SAM50
YJR135W−A_1
PEX19
LHS1
TIM22
MMM1
OXA1
TOM22
CRD1
SSA3
PNT1
SRP1
SRN2
SAM37
SBH1
SEC14
SBH2
COX18
TIM10
BET2
SEC61
TOM7
MDM10
COF1
TIM18
GET1
GET2
Gene
Z_Shift_K_Gem
Z_Im_K_Gem
Z_Shift_K_Cyt
Z_Im_K_Cyt
Z_Shift_L_Gem
Z_Im_L_Gem
Z_Shift_L_Cyt
Z_Im_L_Cyt

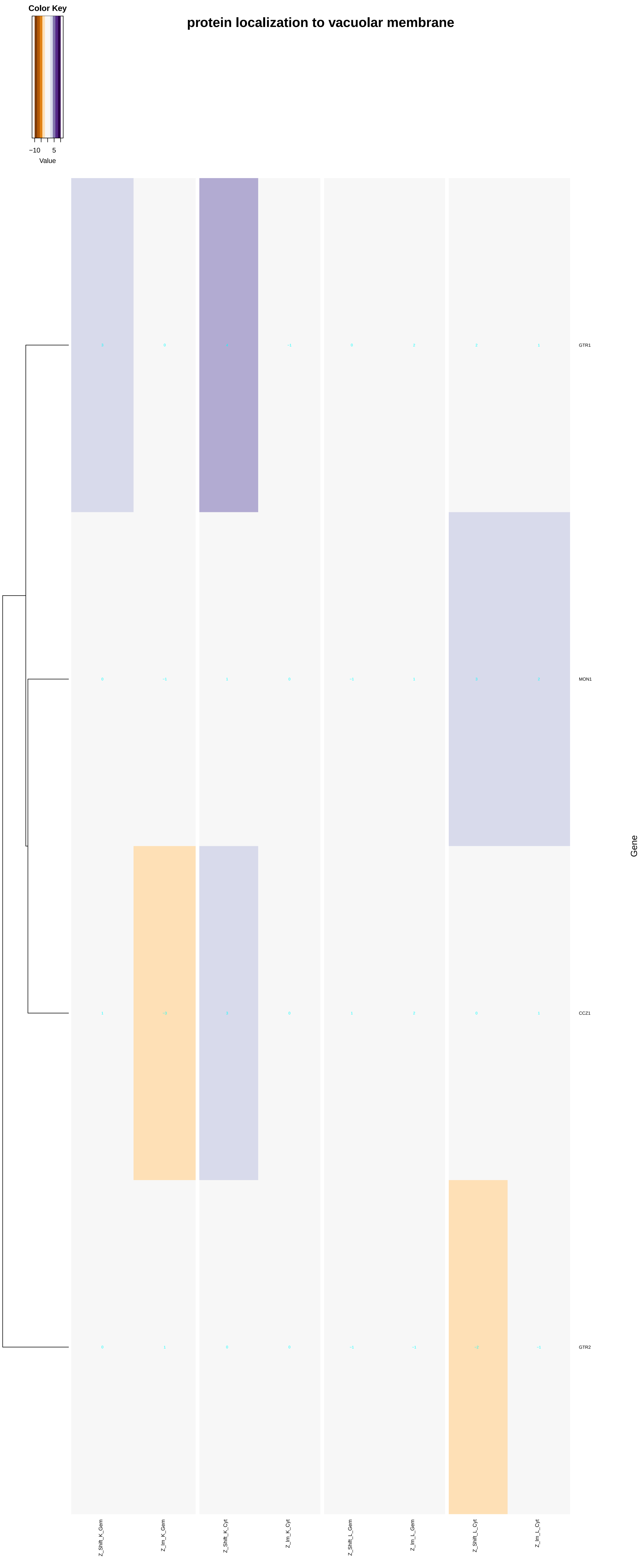

Color Key
−10
5
Value
protein localization to vacuolar membrane
GTR1
MON1
CCZ1
GTR2
Gene
Z_Shift_K_Gem
Z_Im_K_Gem
Z_Shift_K_Cyt
Z_Im_K_Cyt
Z_Shift_L_Gem
Z_Im_L_Gem
Z_Shift_L_Cyt
Z_Im_L_Cyt

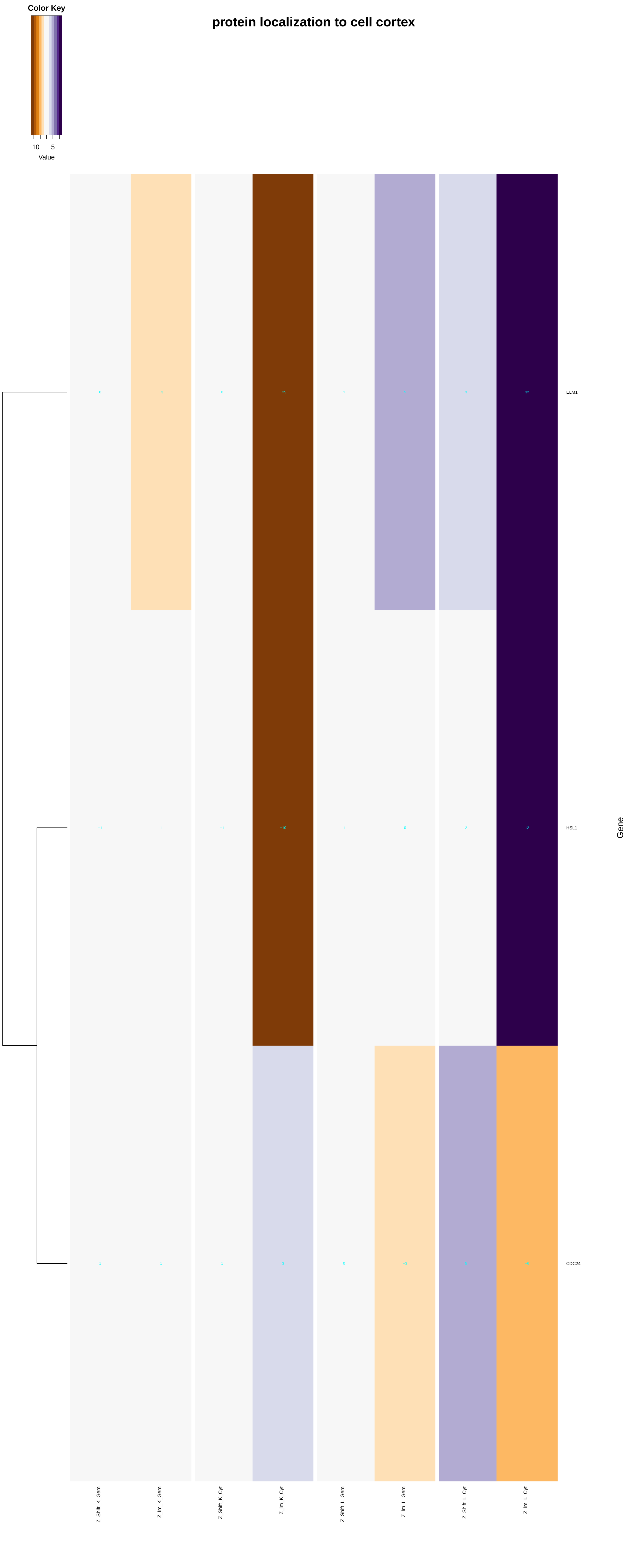

Color Key
−10
5
Value
protein localization to cell cortex
ELM1
HSL1
CDC24
Gene
Z_Shift_K_Gem
Z_Im_K_Gem
Z_Shift_K_Cyt
Z_Im_K_Cyt
Z_Shift_L_Gem
Z_Im_L_Gem
Z_Shift_L_Cyt
Z_Im_L_Cyt

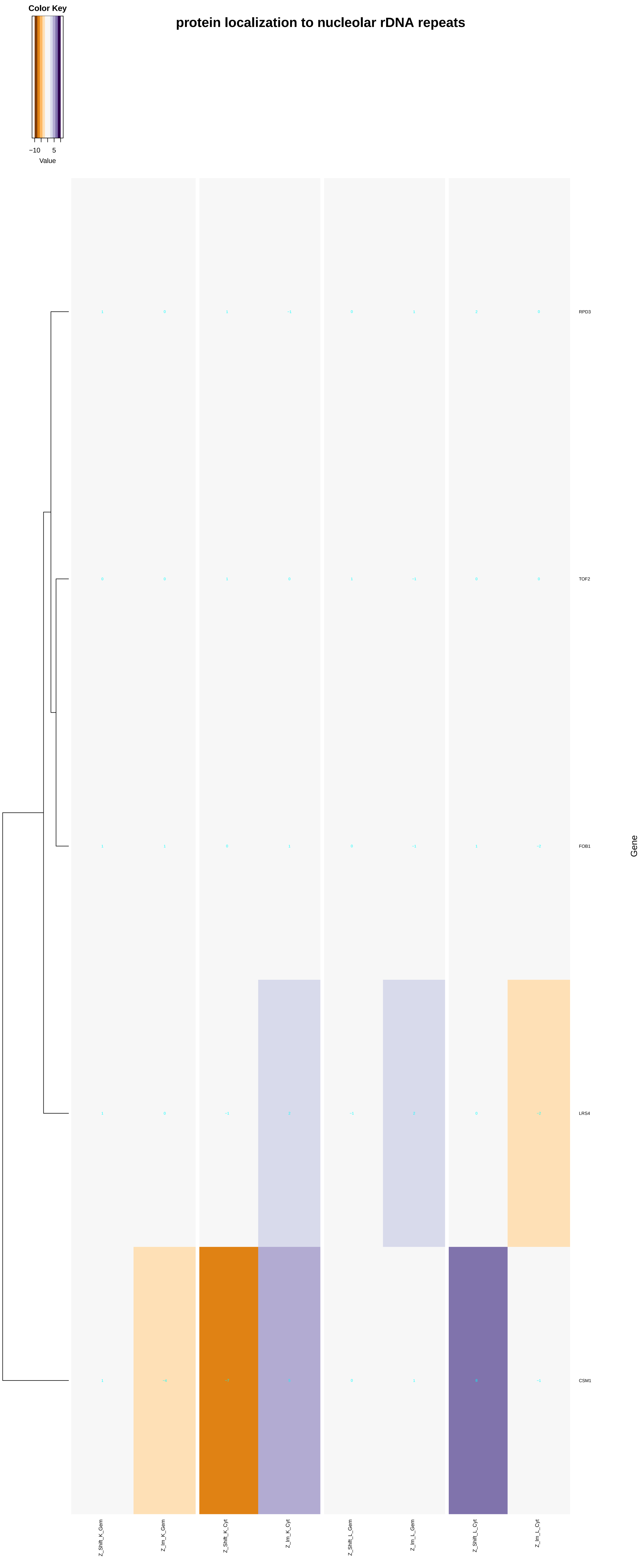

protein localization to nucleolar rDNA repeats
Color Key
−10
5
Value
RPD3
TOF2
FOB1
LRS4
CSM1
Gene
Z_Shift_K_Gem
Z_lm_K_Gem
Z_Shift_K_Cyt
Z_lm_K_Cyt
Z_Shift_L_Gem
Z_lm_L_Gem
Z_Shift_L_Cyt
Z_lm_L_Cyt

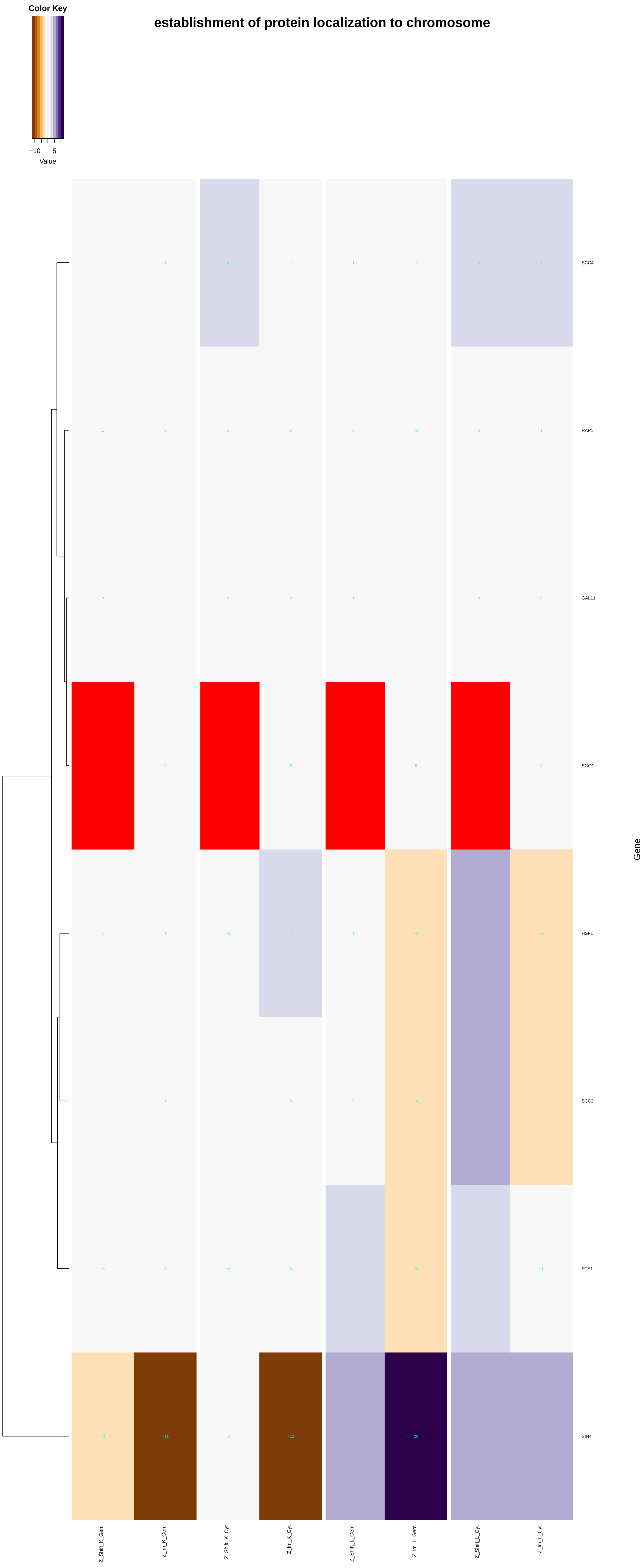

Color Key
−10
5
Value
establishment of protein localization to chromosome
SCC4
RAP1
GAL11
SGO1
HSF1
SCC2
RTS1
SIN4
Gene
Z_Shift_K_Gem
Z_Int_K_Gem
Z_Shift_K_Cyt
Z_Int_K_Cyt
Z_Shift_L_Gem
Z_Int_L_Gem
Z_Shift_L_Cyt
Z_Int_L_Cyt

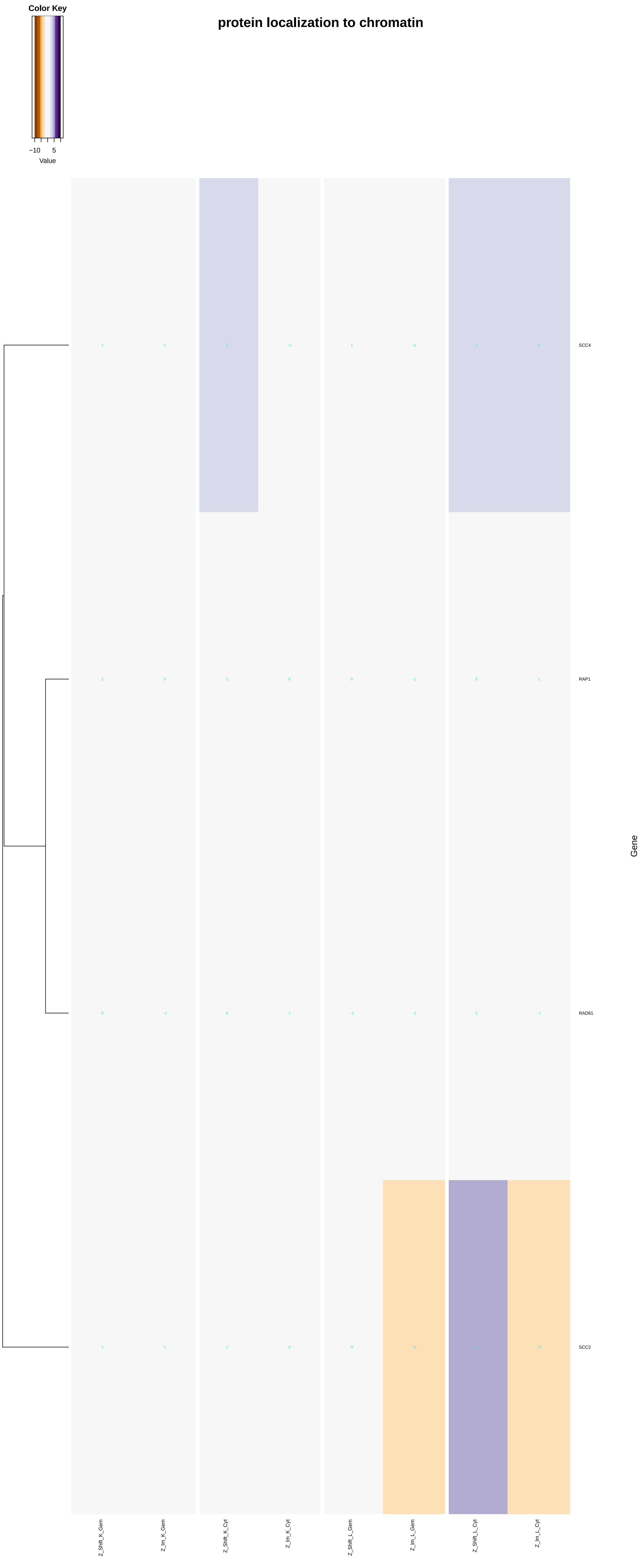

protein localization to chromatin
Color Key
−10
5
Value
SCC4
RAP1
RAD61
SCC2
Gene
Z_Shift_K_Gem
Z_Im_K_Gem
Z_Shift_K_Cyt
Z_Im_K_Cyt
Z_Shift_L_Gem
Z_Im_L_Gem
Z_Shift_L_Cyt
Z_Im_L_Cyt

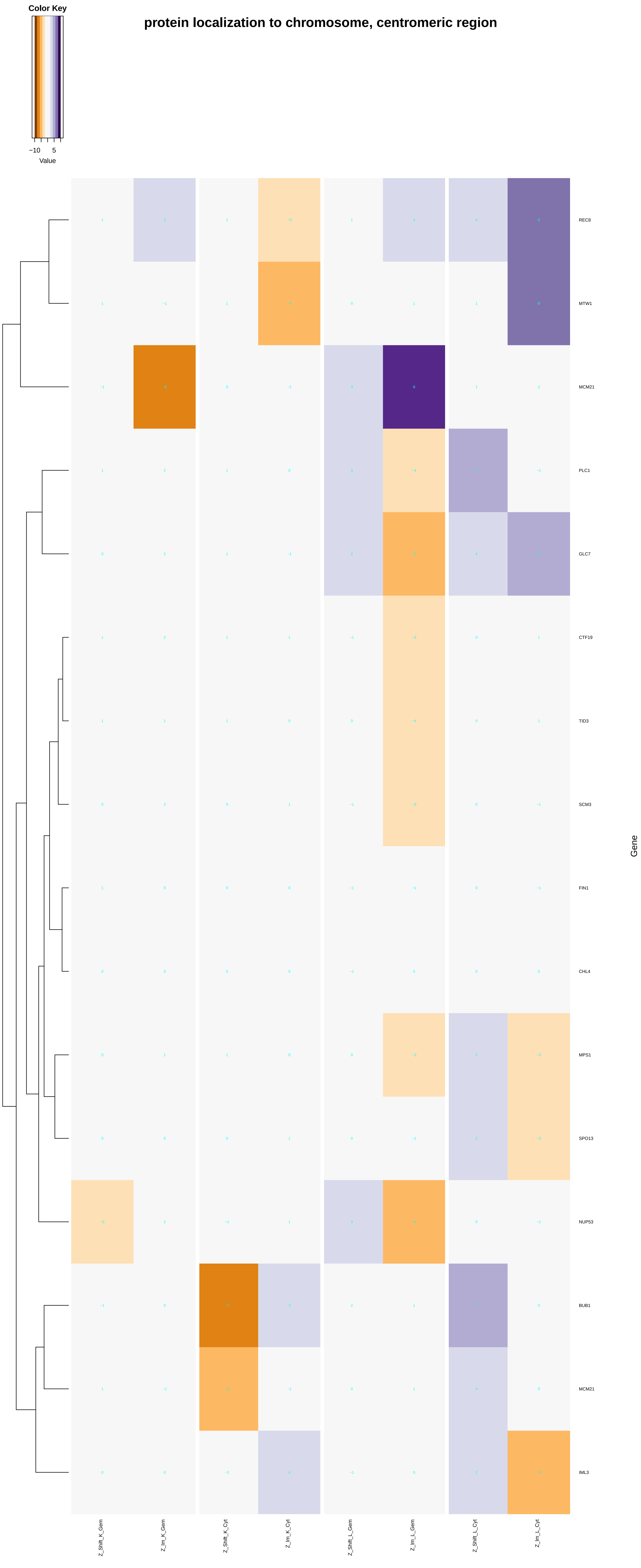
Color Key
−10
5
Value
protein localization to chromosome, centromeric region
REC8
MTW1
MCM21
PLC1
GLC7
CTF19
TID3
SCM3
FIN1
CHL4
MPS1
SPO13
NUP53
BUB1
MCM21
IML3
Gene
Z_Shift_K_Gem
Z_lm_K_Gem
Z_Shift_K_Cyt
Z_lm_K_Cyt
Z_Shift_L_Gem
Z_lm_L_Gem
Z_Shift_L_Cyt
Z_lm_L_Cyt

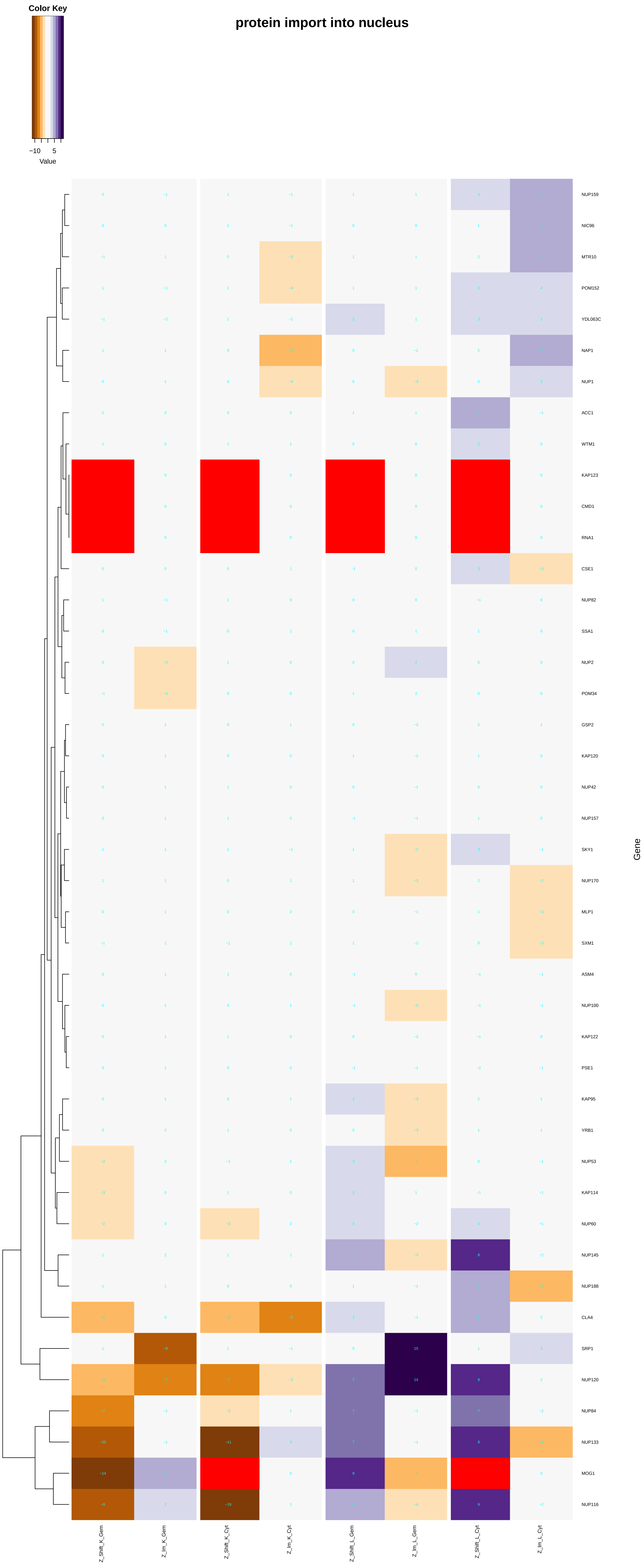

Color Key
−10
5
Value
protein import into nucleus
NUP159
NIC96
MTR10
POM152
YDL063C
NAP1
NUP1
ACC1
WTM1
KAP123
CMD1
RNA1
CSE1
NUP82
SSA1
NUP2
POM34
GSP2
KAP120
NUP42
NUP157
SKY1
NUP170
MLP1
SXM1
ASM4
NUP100
KAP122
PSE1
KAP95
YRB1
NUP53
KAP114
NUP60
NUP145
NUP188
CLA4
SRP1
NUP120
NUP84
NUP133
MOG1
NUP116
Gene
Z_Shift_K_Gem
Z_Im_K_Gem
Z_Shift_K_Cyt
Z_Im_K_Cyt
Z_Shift_L_Gem
Z_Im_L_Gem
Z_Shift_L_Cyt
Z_Im_L_Cyt

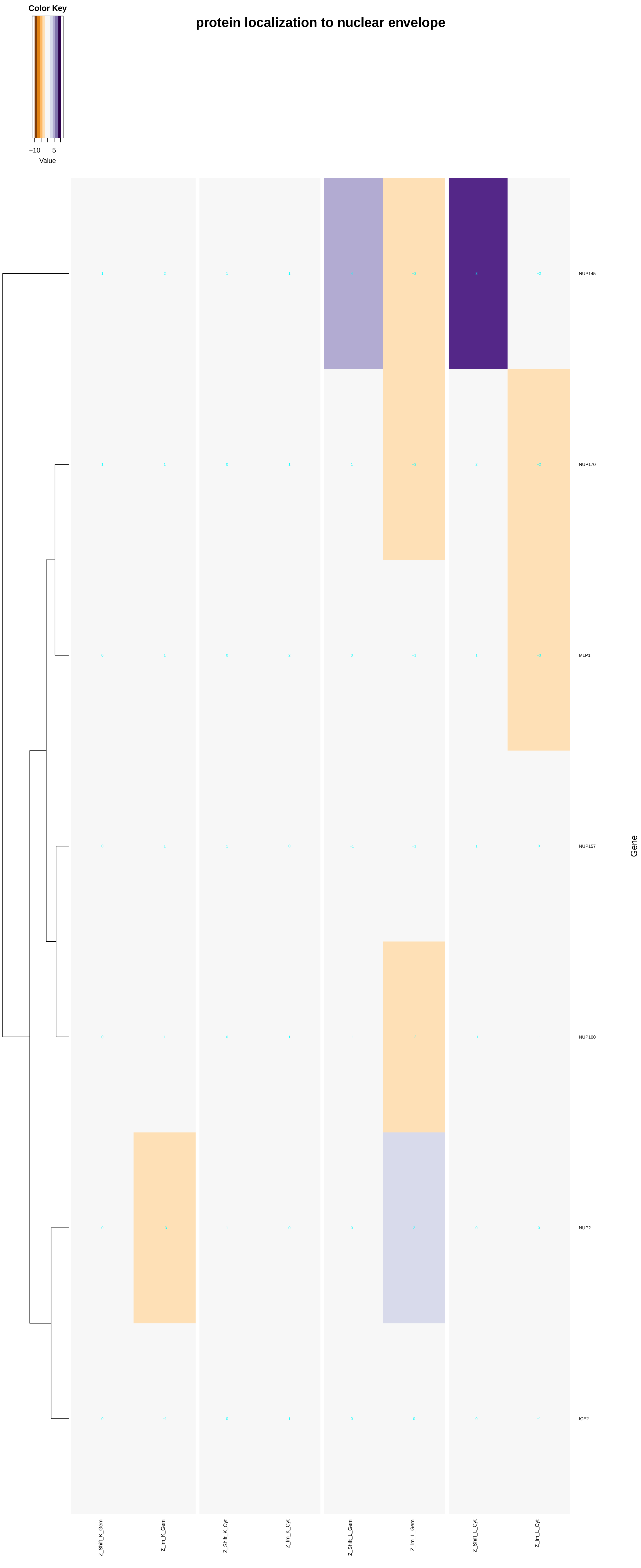

protein localization to nuclear envelope
Color Key
−10
5
Value
NUP145
NUP170
MLP1
NUP157
NUP100
NUP2
ICE2
Gene
Z_Shift_K_Gem
Z_Im_K_Gem
Z_Shift_K_Cyt
Z_Im_K_Cyt
Z_Shift_L_Gem
Z_Im_L_Gem
Z_Shift_L_Cyt
Z_Im_L_Cyt

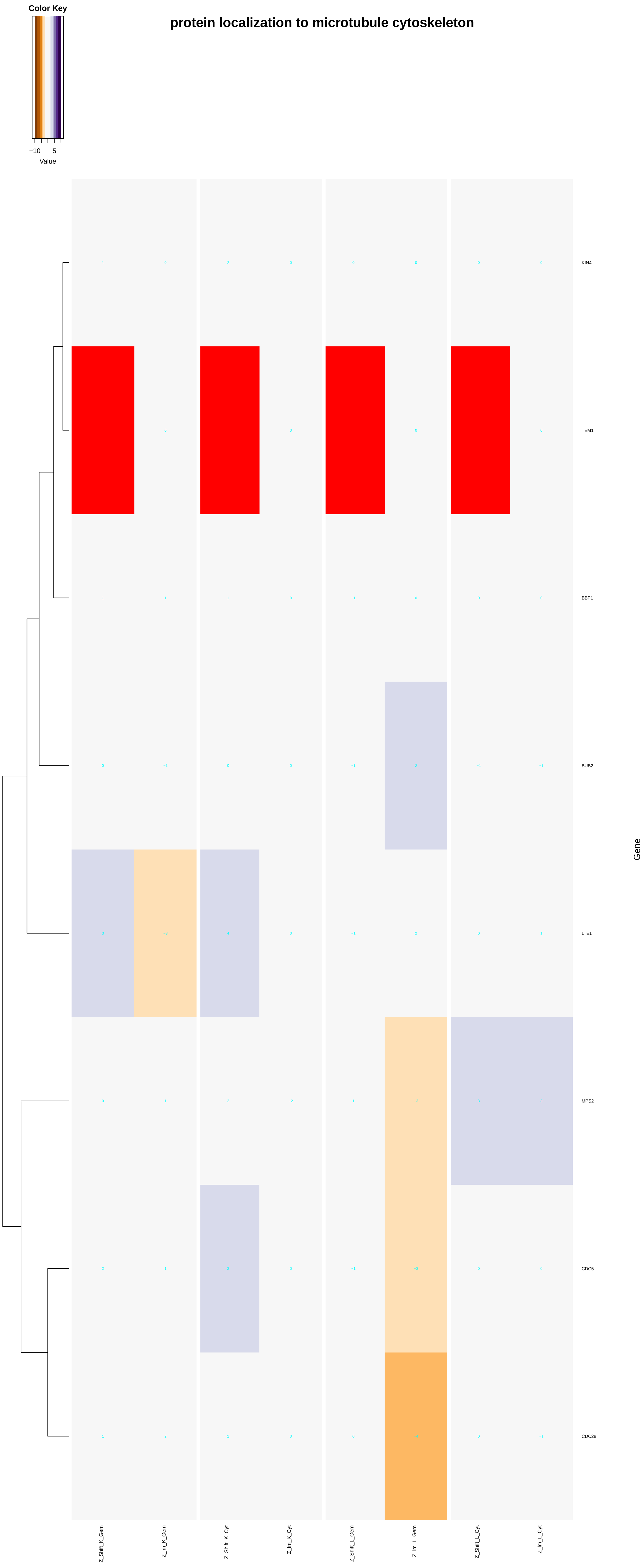

protein localization to microtubule cytoskeleton
Color Key
−10
5
Value
KIN4
TEM1
BBP1
BUB2
LTE1
MPS2
CDC5
CDC28
Gene
Z_Shift_K_Gem
Z_Im_K_Gem
Z_Shift_K_Cyt
Z_Im_K_Cyt
Z_Shift_L_Gem
Z_Im_L_Gem
Z_Shift_L_Cyt
Z_Im_L_Cyt

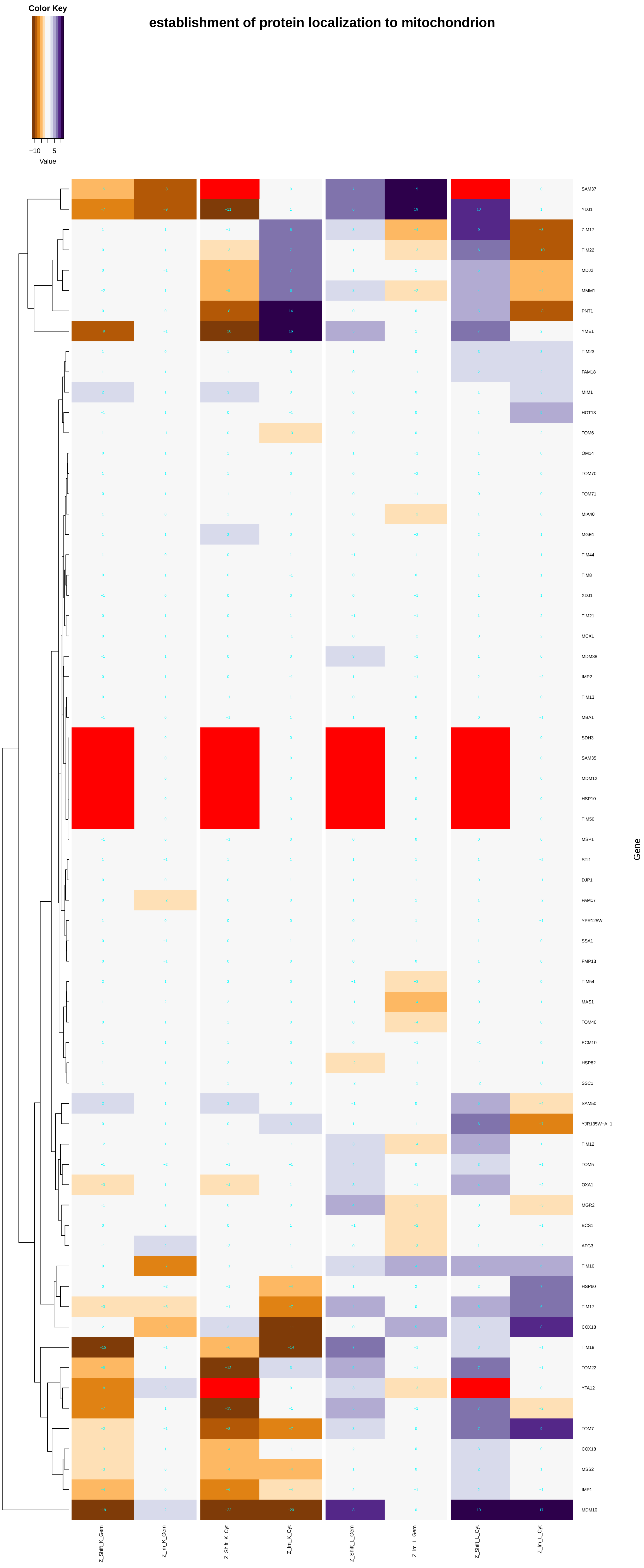

Color Key
−10
5
Value
establishment of protein localization to mitochondrion
SAM37
YDJ1
ZIM17
TIM22
MDJ2
MMM1
PNT1
YME1
TIM23
PAM18
MIM1
HOT13
TOM6
OM14
TOM70
TOM71
MIA40
MGE1
TIM44
TIM8
XDJ1
TIM21
MCX1
MDM38
IMP2
TIM13
MBA1
SDH3
SAM35
MDM12
HSP10
TIM50
MSP1
STI1
DJP1
PAM17
YPR125W
SSA1
FMP13
TIM54
MAS1
TOM40
ECM10
HSP82
SSC1
SAM50
YJR135W−A_1
TIM12
TOM5
OXA1
MGR2
BCS1
AFG3
TIM10
HSP60
TIM17
COX18
TIM18
TOM22
YTA12
TOM7
COX18
MSS2
IMP1
MDM10
Gene
Z_Shift_K_Gem
Z_Im_K_Gem
Z_Shift_K_Cyt
Z_Im_K_Cyt
Z_Shift_L_Gem
Z_Im_L_Gem
Z_Shift_L_Cyt
Z_Im_L_Cyt

### protein localization to endoplasmic reticulum exit site

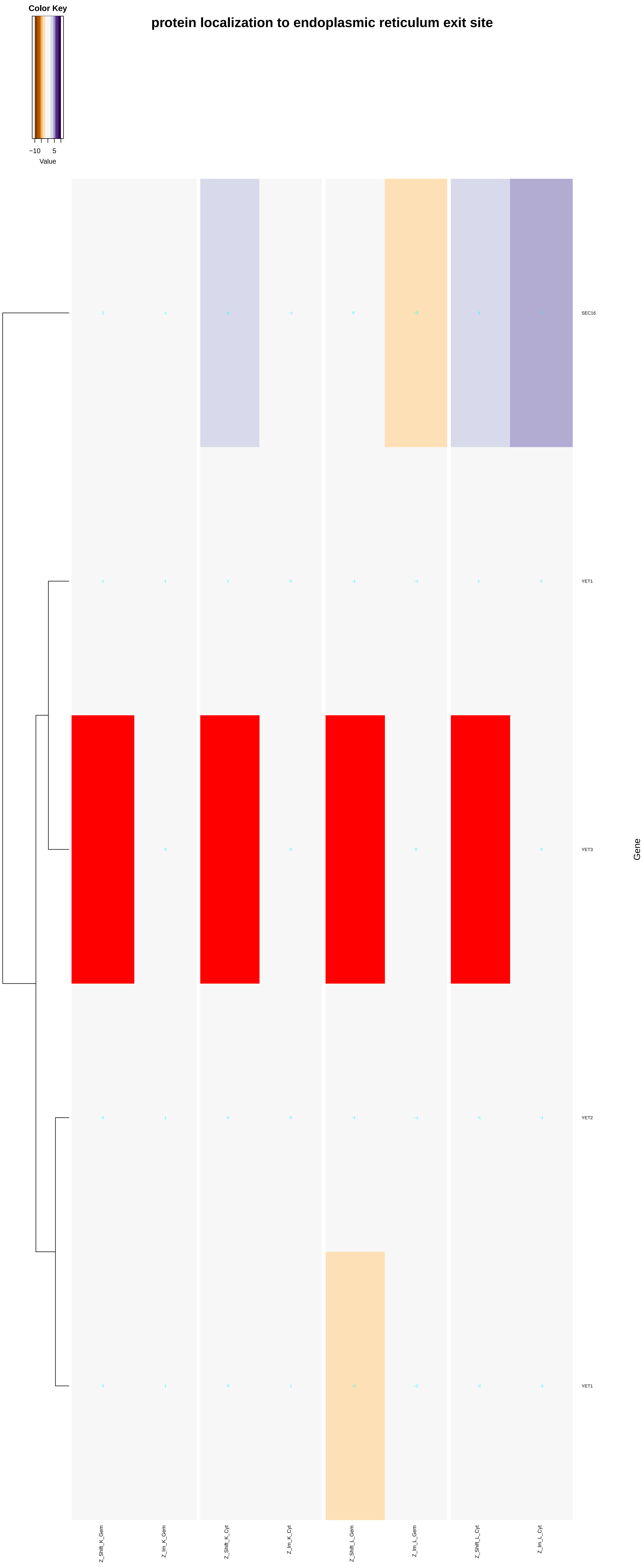

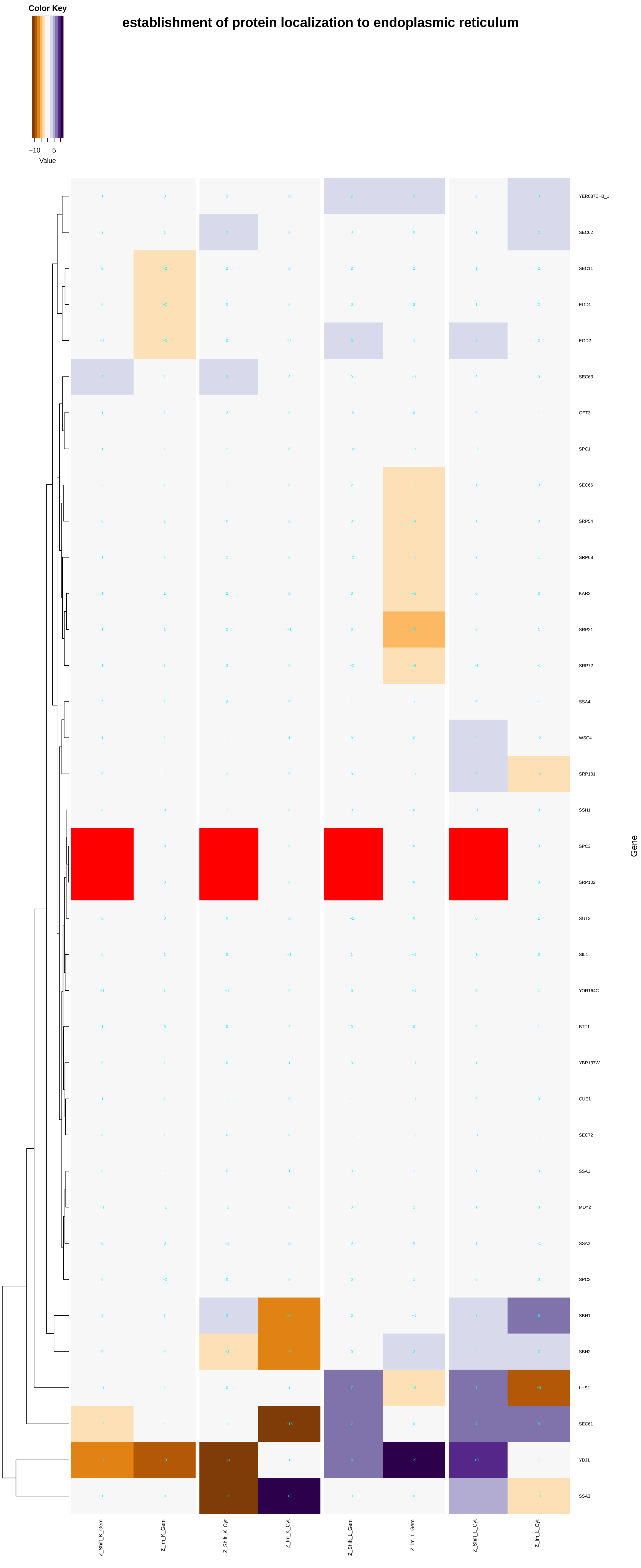
establishment of protein localization to endoplasmic reticulum
Color Key
−10
5
Value
YER087C−B_1
SEC62
SEC11
EGD1
EGD2
SEC63
GET3
SPC1
SEC66
SRP54
SRP68
KAR2
SRP21
SRP72
SSA4
WSC4
SRP101
SSH1
SPC3
SRP102
SGT2
SIL1
YOR164C
BTT1
YBR137W
CUE1
SEC72
SSA1
MDY2
SSA2
SPC2
SBH1
SBH2
LHS1
SEC61
YDJ1
SSA3
Gene
Z_Shift_K_Gem
Z_lm_K_Gem
Z_Shift_K_Cyt
Z_lm_K_Cyt
Z_Shift_L_Gem
Z_lm_L_Gem
Z_Shift_L_Cyt
Z_lm_L_Cyt

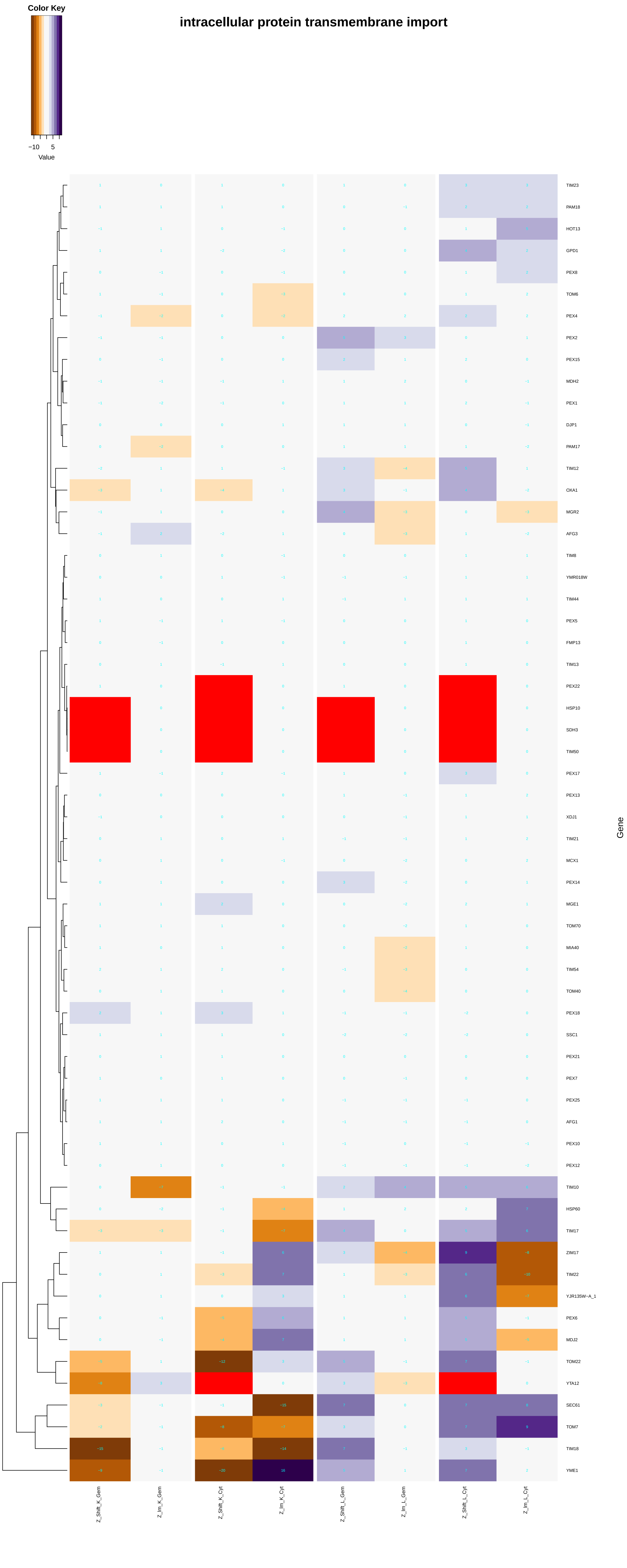

intracellular protein transmembrane import
Color Key
−10
5
Value
Gene
TIM23
PAM18
HOT13
GPD1
PEX8
TOM6
PEX4
PEX2
PEX15
MDH2
PEX1
DJP1
PAM17
TIM12
OXA1
MGR2
AFG3
TIM8
YMR018W
TIM44
PEX5
FMP13
TIM13
PEX22
HSP10
SDH3
TIM50
PEX17
PEX13
XDJ1
TIM21
MCX1
PEX14
MGE1
TOM70
MIA40
TIM54
TOM40
PEX18
SSC1
PEX21
PEX7
PEX25
AFG1
PEX10
PEX12
TIM10
HSP60
TIM17
ZIM17
TIM22
YJR135W−A_1
PEX6
MDJ2
TOM22
YTA12
SEC61
TOM7
TIM18
YME1
Z_Shift_K_Gem
Z_Im_K_Gem
Z_Shift_K_Cyt
Z_Im_K_Cyt
Z_Shift_L_Gem
Z_Im_L_Gem
Z_Shift_L_Cyt
Z_Im_L_Cyt

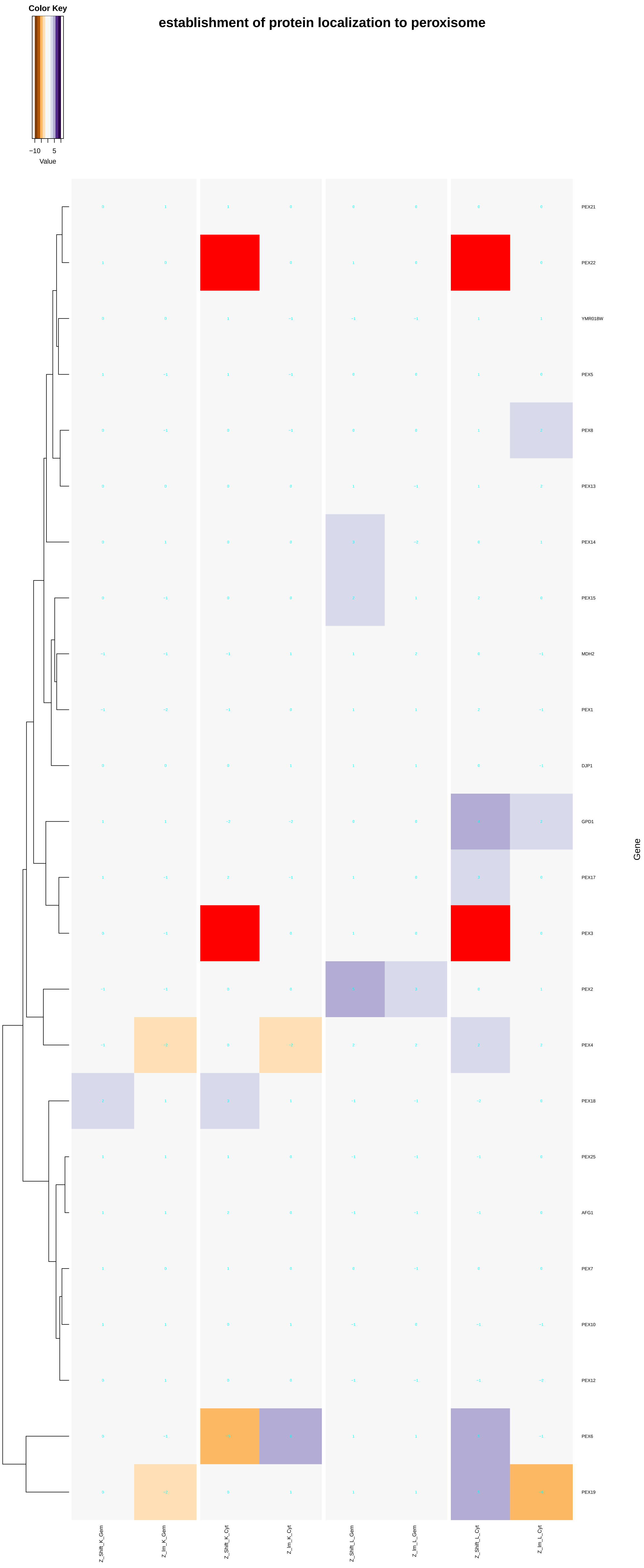

Color Key
−10 5
Value
establishment of protein localization to peroxisome
PEX21
PEX22
YMR018W
PEX5
PEX8
PEX13
PEX14
PEX15
MDH2
PEX1
DJP1
GPD1
PEX17
PEX3
PEX2
PEX4
PEX18
PEX25
AFG1
PEX7
PEX10
PEX12
PEX6
PEX19
Gene
Z_Shift_K_Gem
Z_Im_K_Gem
Z_Shift_K_Cyt
Z_Im_K_Cyt
Z_Shift_L_Gem
Z_Im_L_Gem
Z_Shift_L_Cyt
Z_Im_L_Cyt

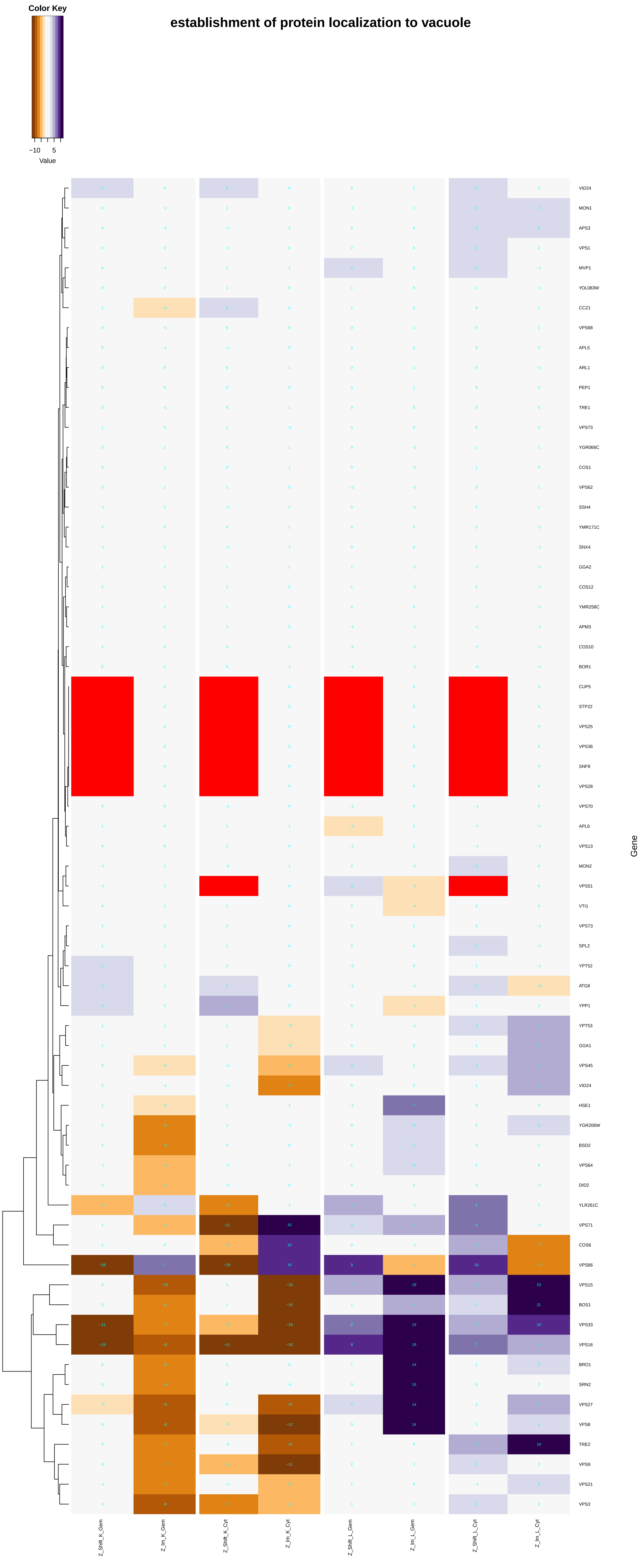
establishment of protein localization to vacuole
Color Key
−10
5
Value
Gene
VID24
MON1
APS3
VPS1
MVP1
YOL083W
CCZ1
VPS68
APL5
ARL1
PEP1
TRE1
VPS73
YGR066C
COS1
VPS62
SSH4
YMR171C
SNX4
GGA2
COS12
YMR258C
APM3
COS10
BOR1
CUP5
STP22
VPS25
VPS36
SNF8
VPS28
VPS70
APL6
VPS13
MON2
VPS51
VTI1
VPS73
SPL2
YPT52
ATG8
YPP1
YPT53
GGA1
VPS45
VID24
HSE1
YGR206W
BSD2
VPS64
DID2
YLR261C
VPS71
COS6
VPS66
VPS15
BOS1
VPS33
VPS16
BRO1
SRN2
VPS27
VPS8
TRE2
VPS9
VPS21
VPS3
Z_Shift_K_Gem
Z_lm_K_Gem
Z_Shift_K_Cyt
Z_lm_K_Cyt
Z_Shift_L_Gem
Z_lm_L_Gem
Z_Shift_L_Cyt
Z_lm_L_Cyt

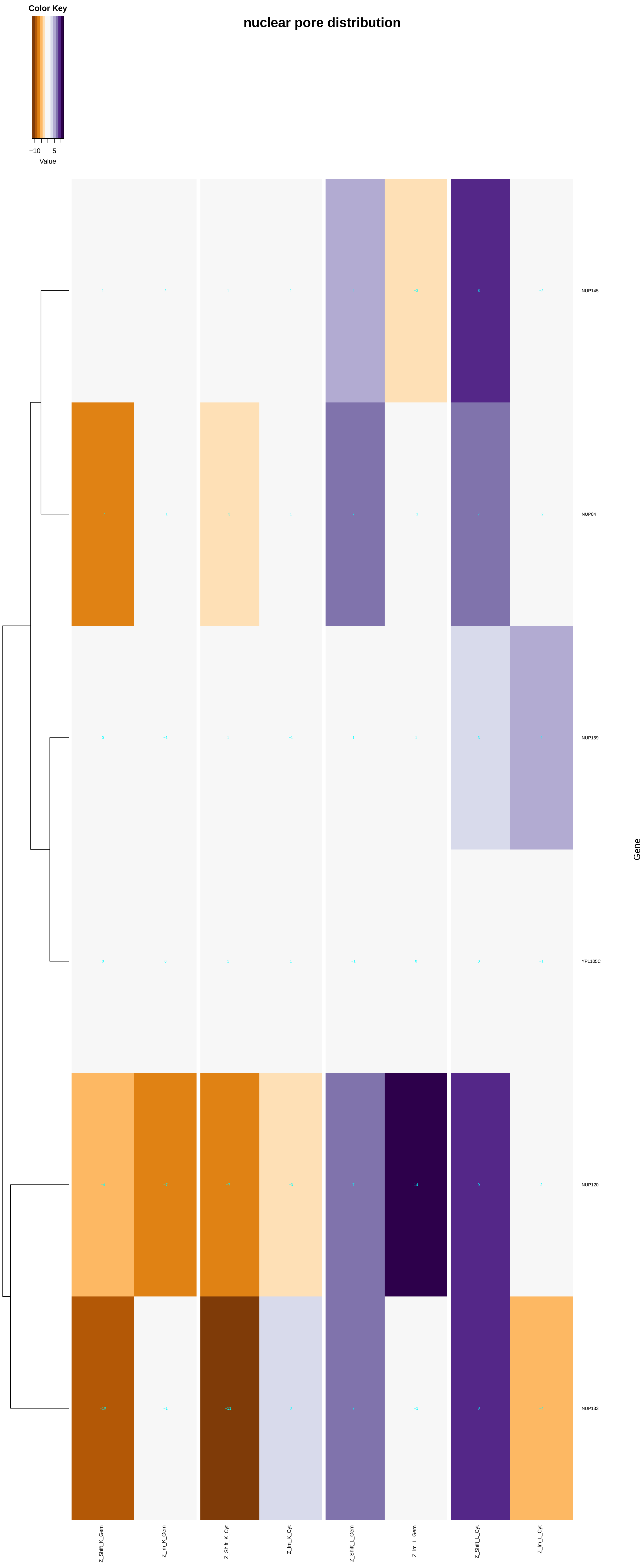
Color Key
−10
5
Value
nuclear pore distribution
1
2
1
1
4
−3
8
−2
NUP145
−7
−1
−3
1
7
−1
7
−2
NUP84
0
−1
1
−1
1
1
3
4
NUP159
0
0
1
1
−1
0
0
−1
YPL105C
−4
−7
−7
−3
7
14
9
2
NUP120
−10
−1
−11
3
7
−1
8
−4
NUP133
Gene
Z_Shift_K_Gem
Z_Im_K_Gem
Z_Shift_K_Cyt
Z_Im_K_Cyt
Z_Shift_L_Gem
Z_Im_L_Gem
Z_Shift_L_Cyt
Z_Im_L_Cyt

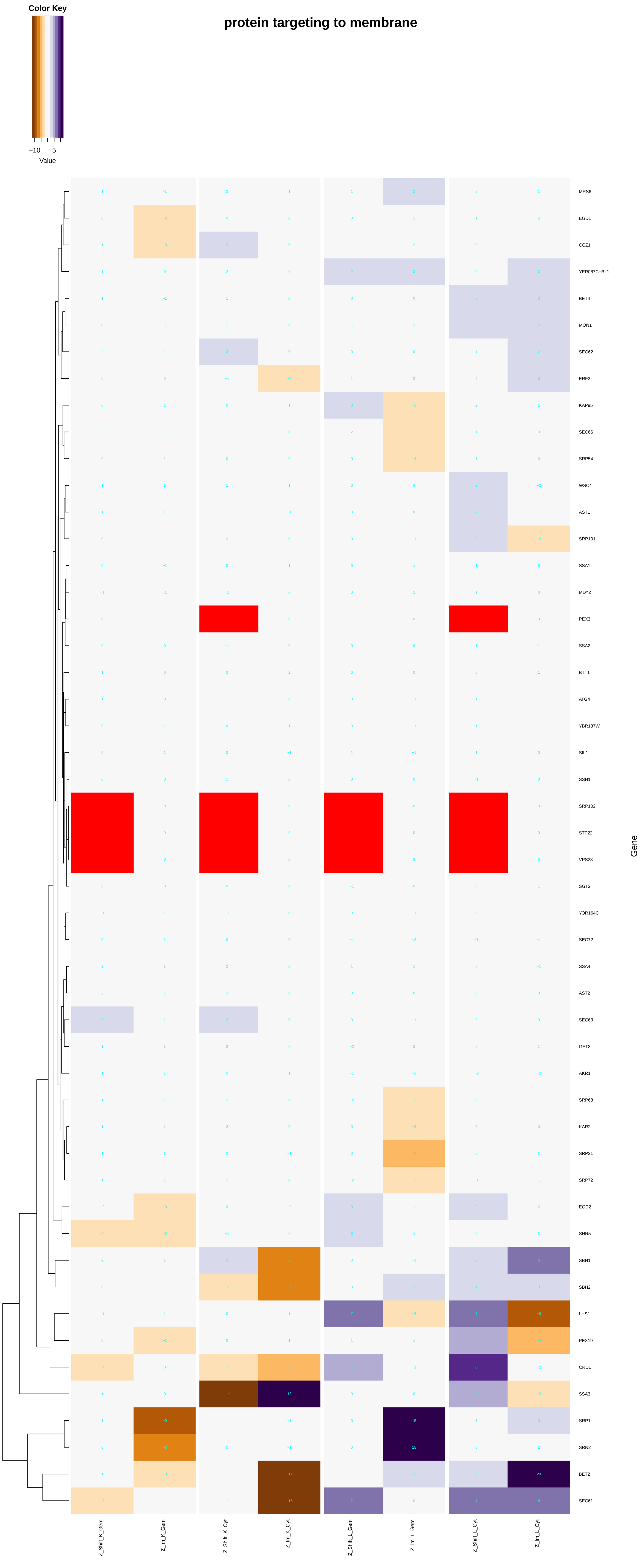

protein targeting to membrane
Color Key
−10
5
Value
Gene
MRS6
EGD1
CCZ1
YER087C−B_1
BET4
MON1
SEC62
ERF2
KAP95
SEC66
SRP54
WSC4
AST1
SRP101
SSA1
MDY2
PEX3
SSA2
BTT1
ATG4
YBR137W
SIL1
SSH1
SRP102
STP22
VPS28
SGT2
YOR164C
SEC72
SSA4
AST2
SEC63
GET3
AKR1
SRP68
KAR2
SRP21
SRP72
EGD2
SHR5
SBH1
SBH2
LHS1
PEX19
CRD1
SSA3
SRP1
SRN2
BET2
SEC61
Z_Shift_K_Gem
Z_Im_K_Gem
Z_Shift_K_Cyt
Z_Im_K_Cyt
Z_Shift_L_Gem
Z_Im_L_Gem
Z_Shift_L_Cyt
Z_Im_L_Cyt

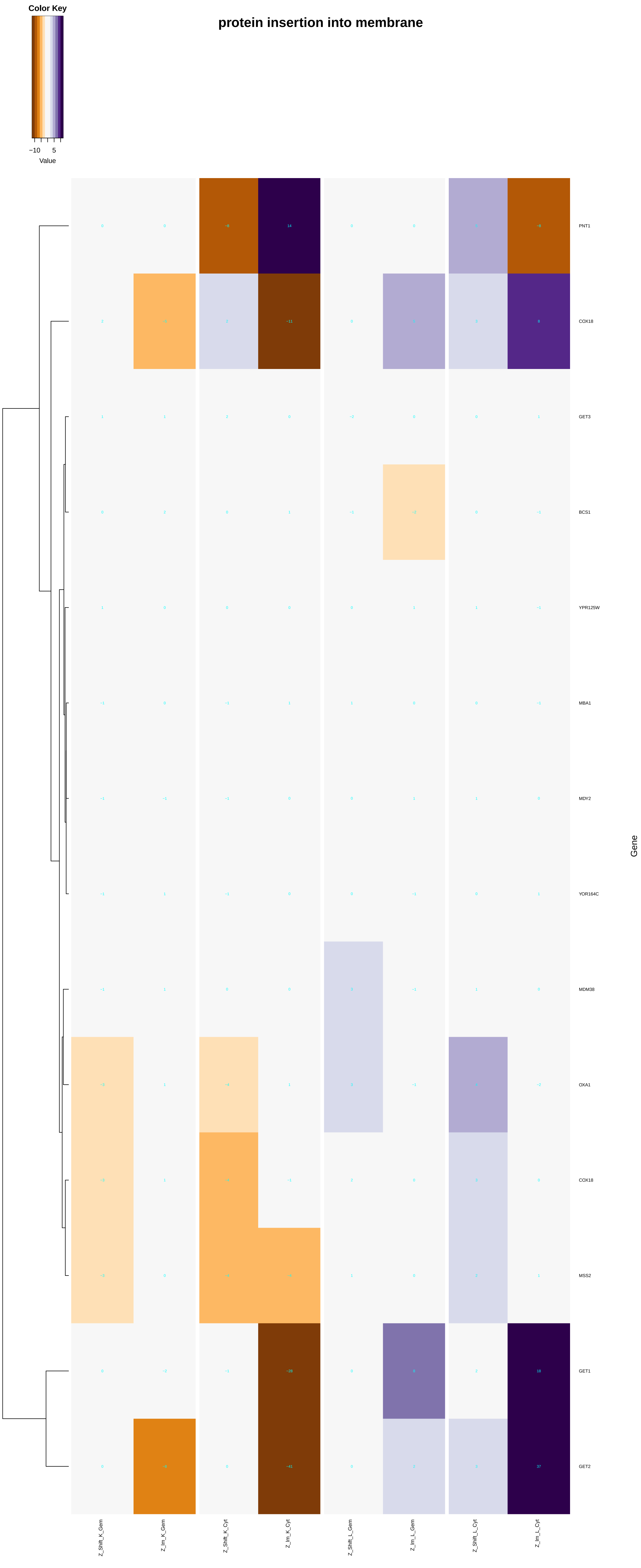

Color Key
−10
5
Value
protein insertion into membrane
PNT1
COX18
GET3
BCS1
YPR125W
MBA1
MDY2
YOR164C
MDM38
OXA1
COX18
MSS2
GET1
GET2
Gene
Z_Shift_K_Gem
Z_Im_K_Gem
Z_Shift_K_Cyt
Z_Im_K_Cyt
Z_Shift_L_Gem
Z_Im_L_Gem
Z_Shift_L_Cyt
Z_Im_L_Cyt

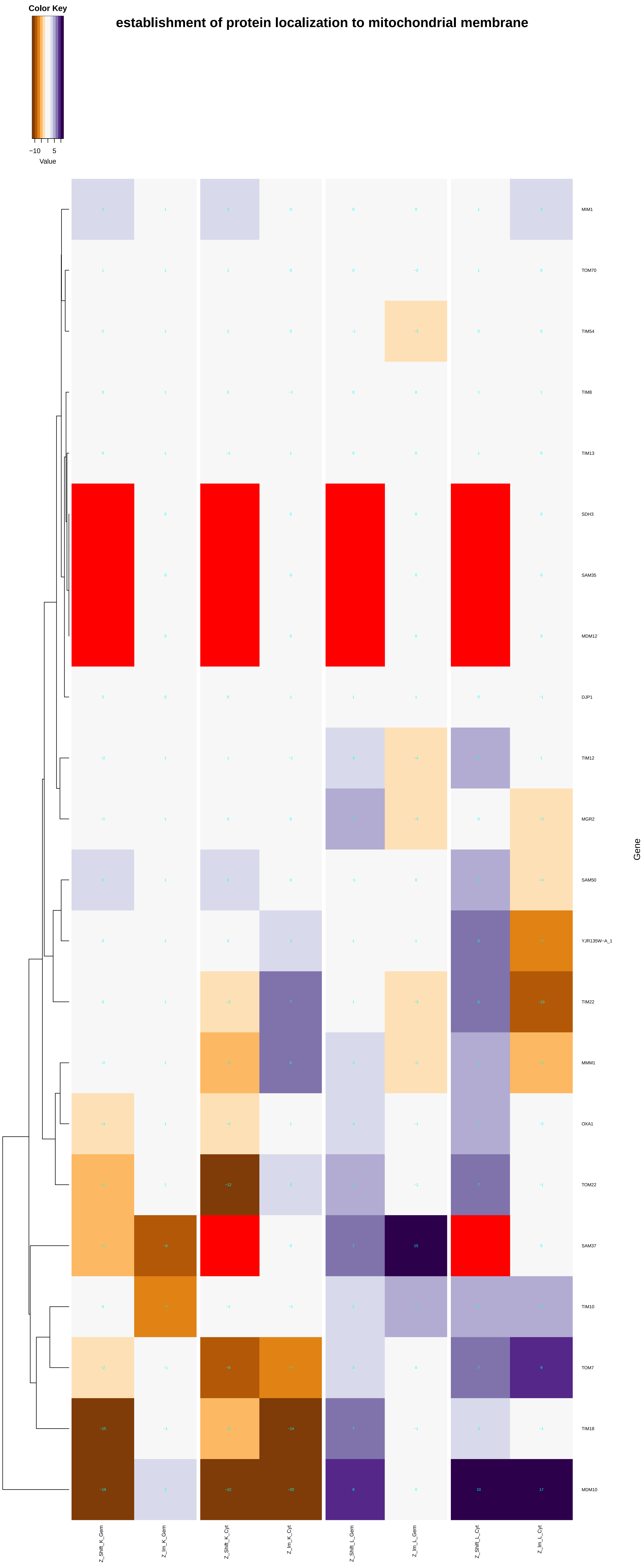
establishment of protein localization to mitochondrial membrane
Color Key
−10
5
Value
MIM1
TOM70
TIM54
TIM8
TIM13
SDH3
SAM35
MDM12
DJP1
TIM12
MGR2
SAM50
YJR135W−A_1
TIM22
MMM1
OXA1
TOM22
SAM37
TIM10
TOM7
TIM18
MDM10
Gene
Z_Shift_K_Gem
Z_Im_K_Gem
Z_Shift_K_Cyt
Z_Im_K_Cyt
Z_Shift_L_Gem
Z_Im_L_Gem
Z_Shift_L_Cyt
Z_Im_L_Cyt

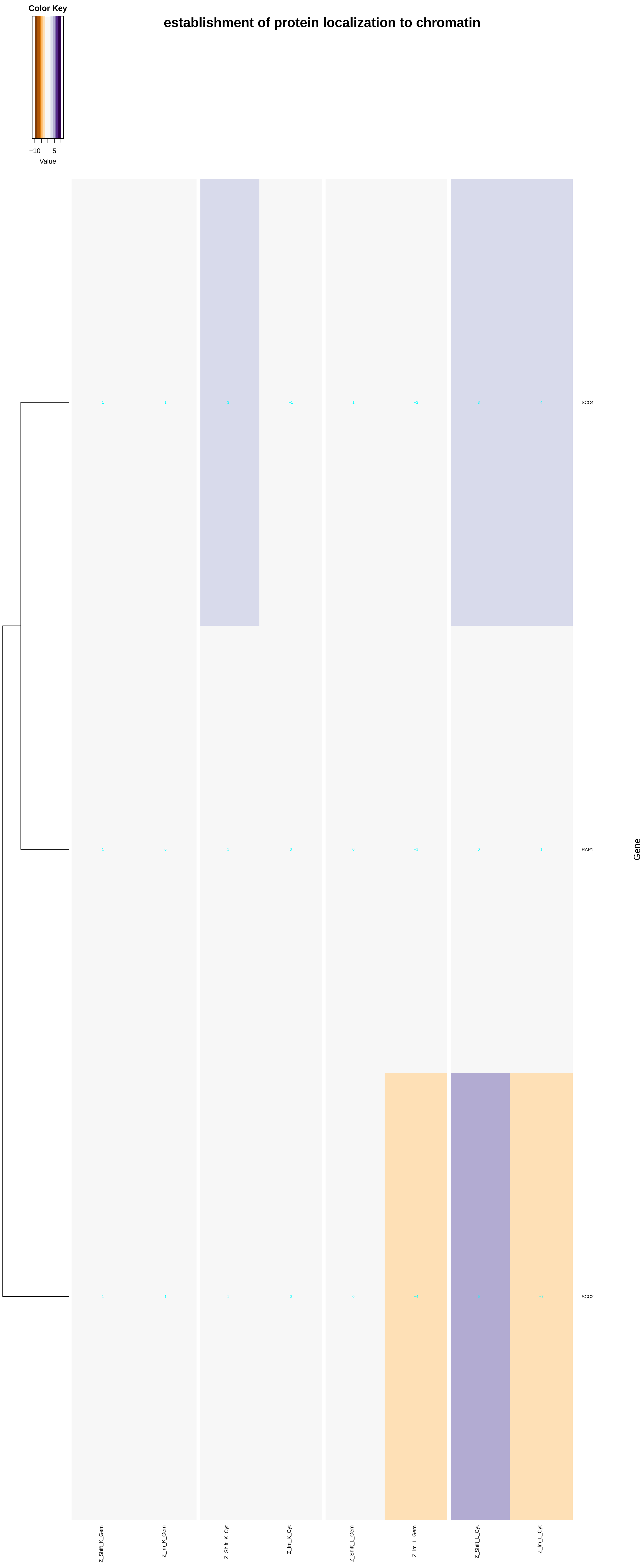

establishment of protein localization to chromatin
Color Key
−10
5
Value
SCC4
RAP1
SCC2
Gene
Z_Shift_K_Gem
Z_Im_K_Gem
Z_Shift_K_Cyt
Z_Im_K_Cyt
Z_Shift_L_Gem
Z_Im_L_Gem
Z_Shift_L_Cyt
Z_Im_L_Cyt

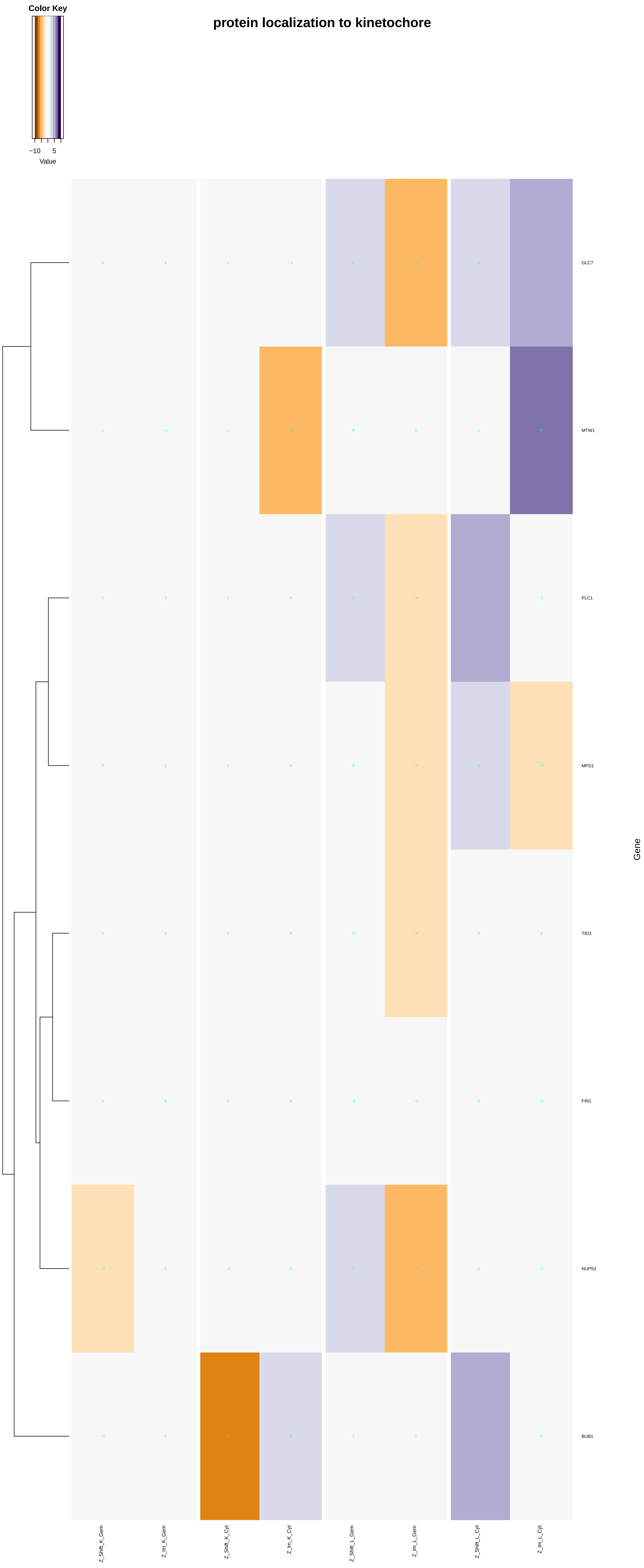
protein localization to kinetochore
Color Key
−10
5
Value
GLC7
MTW1
PLC1
MPS1
TID3
FIN1
NUP53
BUB1
Gene
Z_Shift_K_Gem
Z_Im_K_Gem
Z_Shift_K_Cyt
Z_Im_K_Cyt
Z_Shift_L_Gem
Z_Im_L_Gem
Z_Shift_L_Cyt
Z_Im_L_Cyt

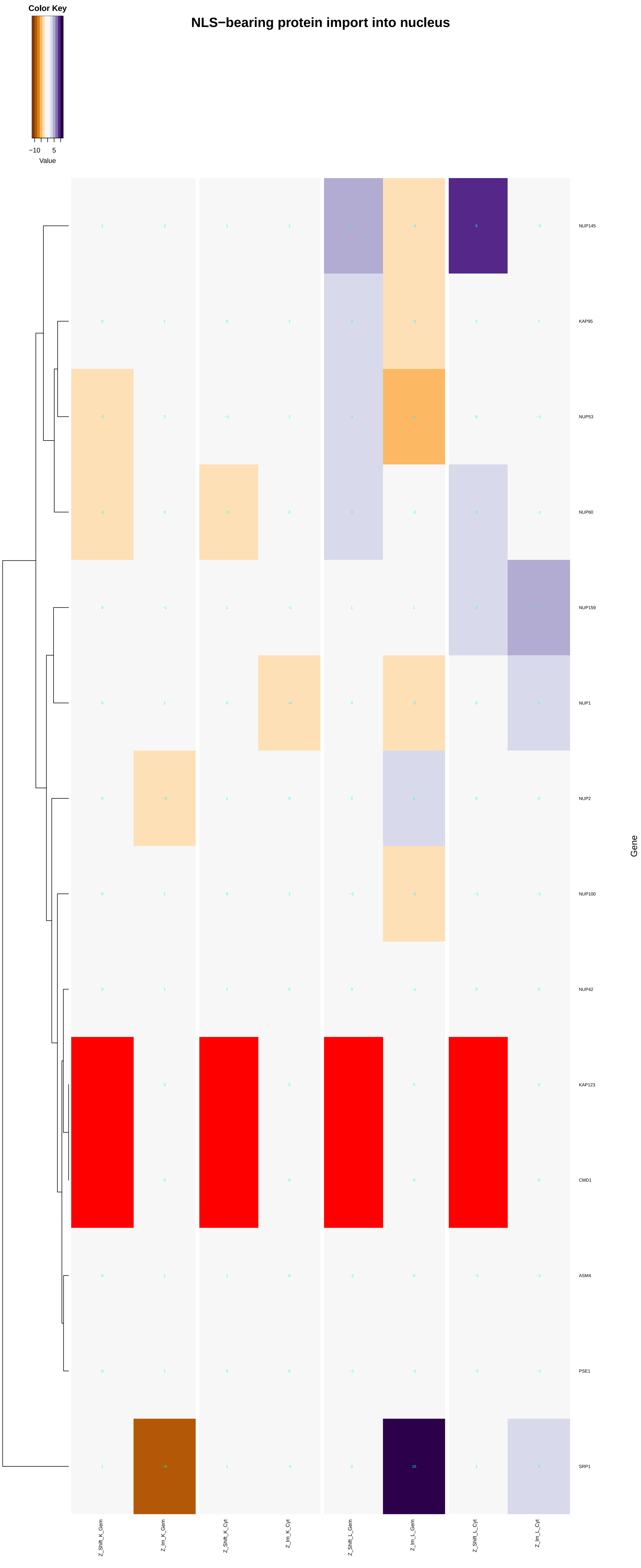

Color Key
−10
5
Value
NLS−bearing protein import into nucleus
NUP145
KAP95
NUP53
NUP60
NUP159
NUP1
NUP2
NUP100
NUP42
KAP123
CMD1
ASM4
PSE1
SRP1
Gene
Z_Shift_K_Gem
Z_Im_K_Gem
Z_Shift_K_Cyt
Z_Im_K_Cyt
Z_Shift_L_Gem
Z_Im_L_Gem
Z_Shift_L_Cyt
Z_Im_L_Cyt

### ribosomal protein import into nucleus

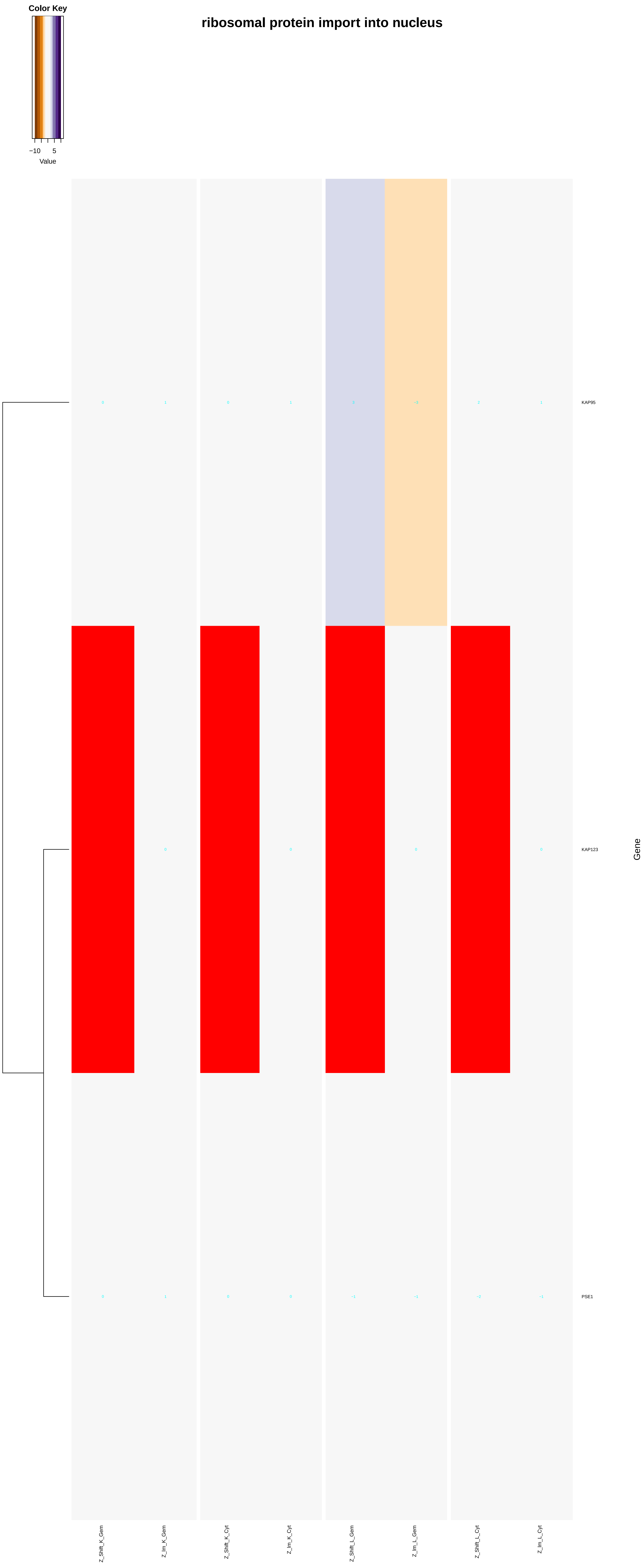

### transcription factor import into nucleus

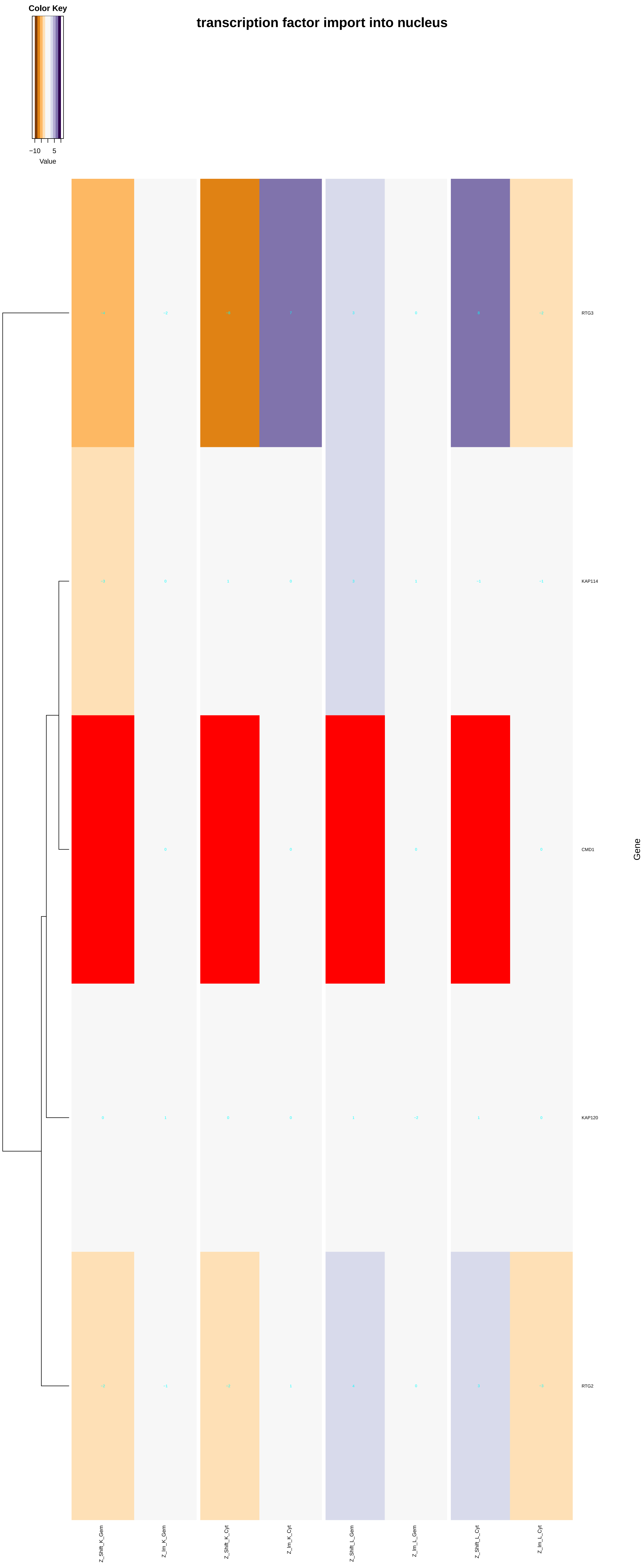

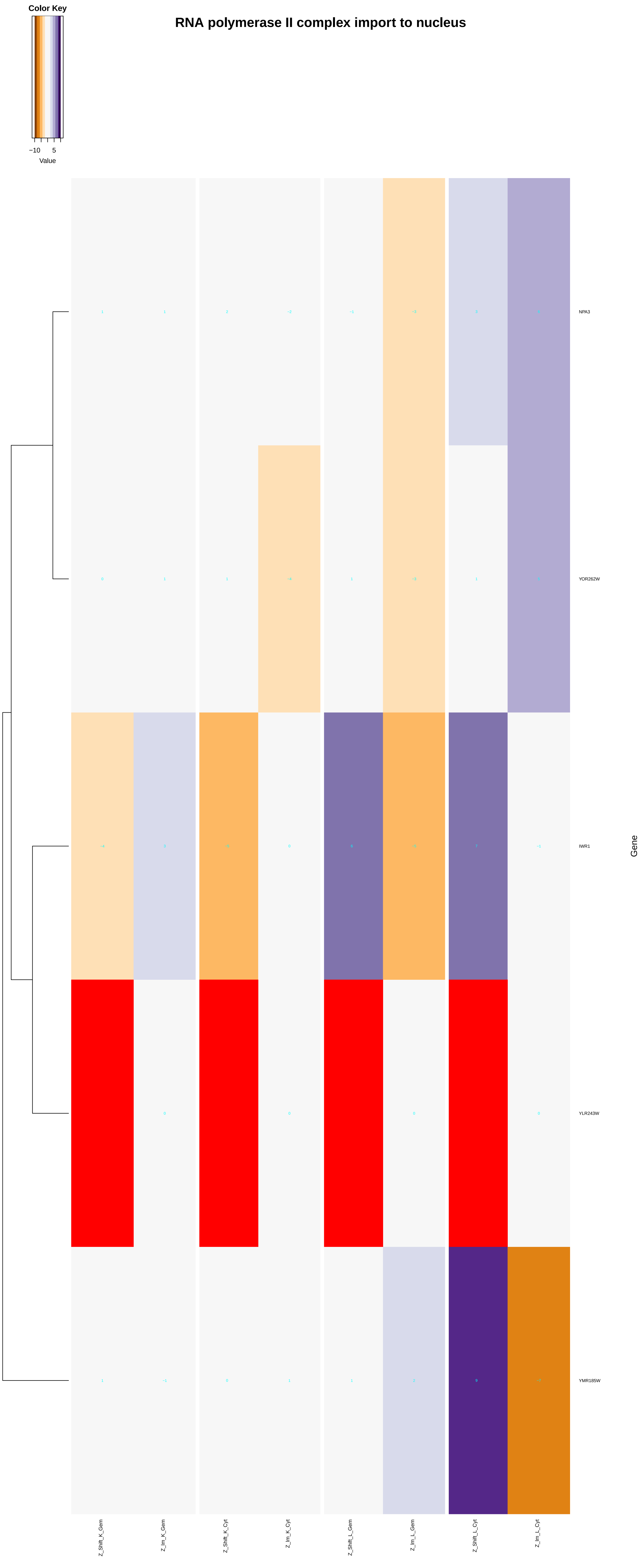
Color Key
−10
5
Value
RNA polymerase II complex import to nucleus
NPA3
YOR262W
IWR1
YLR243W
YMR185W
Gene
Z_Shift_K_Gem
Z_Im_K_Gem
Z_Shift_K_Cyt
Z_Im_K_Cyt
Z_Shift_L_Gem
Z_Im_L_Gem
Z_Shift_L_Cyt
Z_Im_L_Cyt

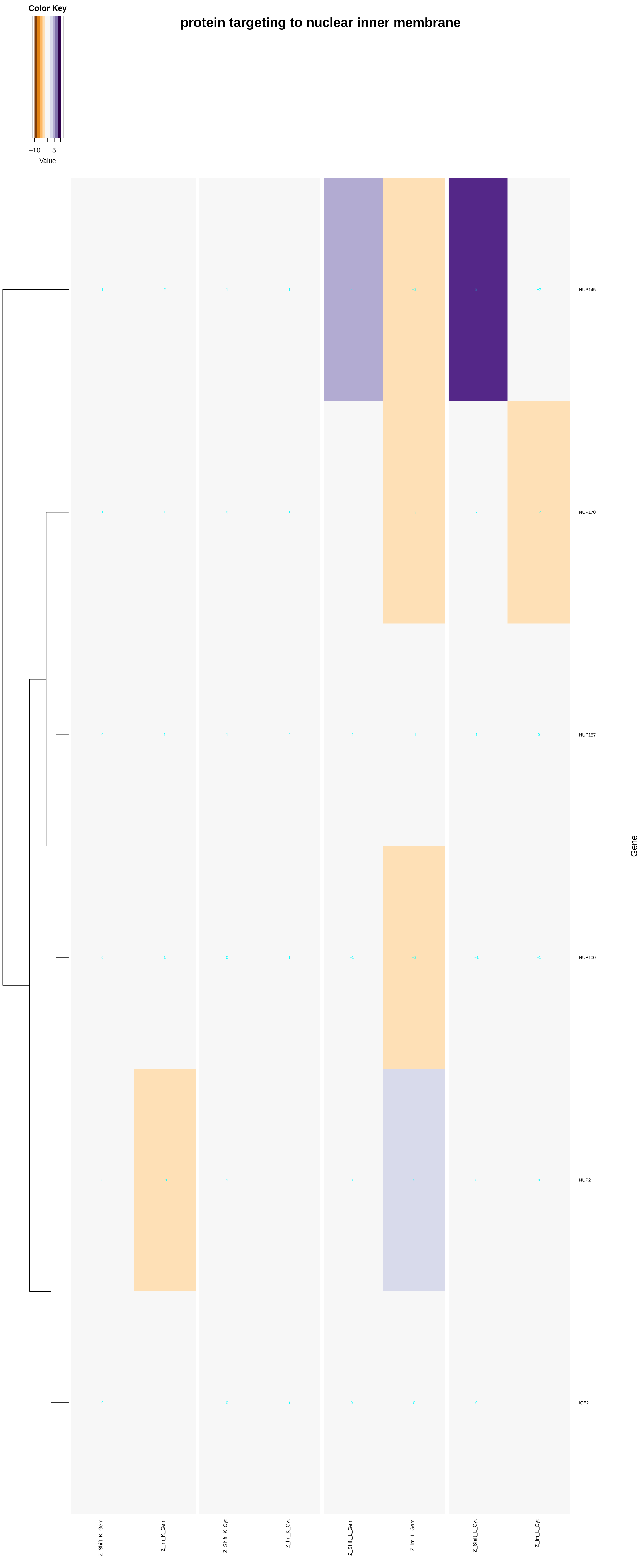
protein targeting to nuclear inner membrane
Color Key
−10
5
Value
NUP145
NUP170
NUP157
NUP100
NUP2
ICE2
Gene
Z_Shift_K_Gem
Z_Im_K_Gem
Z_Shift_K_Cyt
Z_Im_K_Cyt
Z_Shift_L_Gem
Z_Im_L_Gem
Z_Shift_L_Cyt
Z_Im_L_Cyt

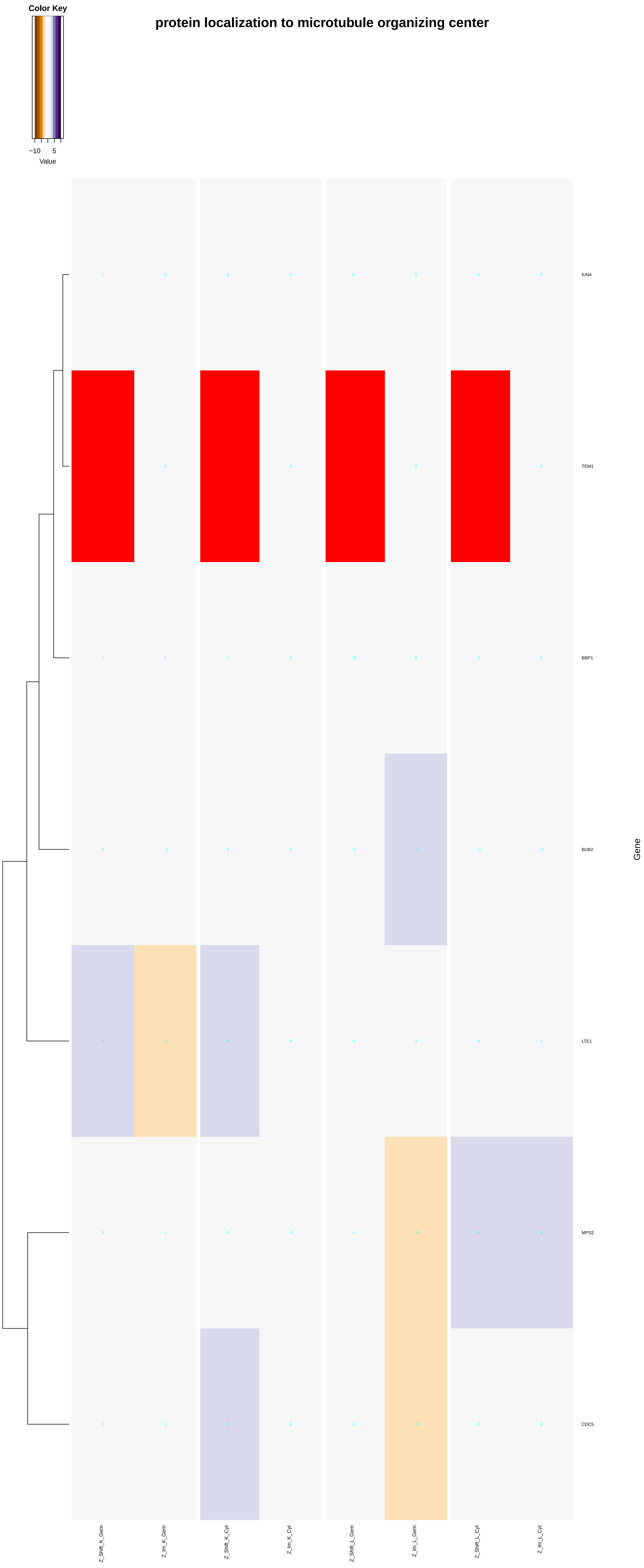

Color Key
−10
5
Value
protein localization to microtubule organizing center
KIN4
TEM1
BBP1
BUB2
LTE1
MPS2
CDC5
Gene
Z_Shift_K_Gem
Z_lm_K_Gem
Z_Shift_K_Cyt
Z_lm_K_Cyt
Z_Shift_L_Gem
Z_lm_L_Gem
Z_Shift_L_Cyt
Z_lm_L_Cyt

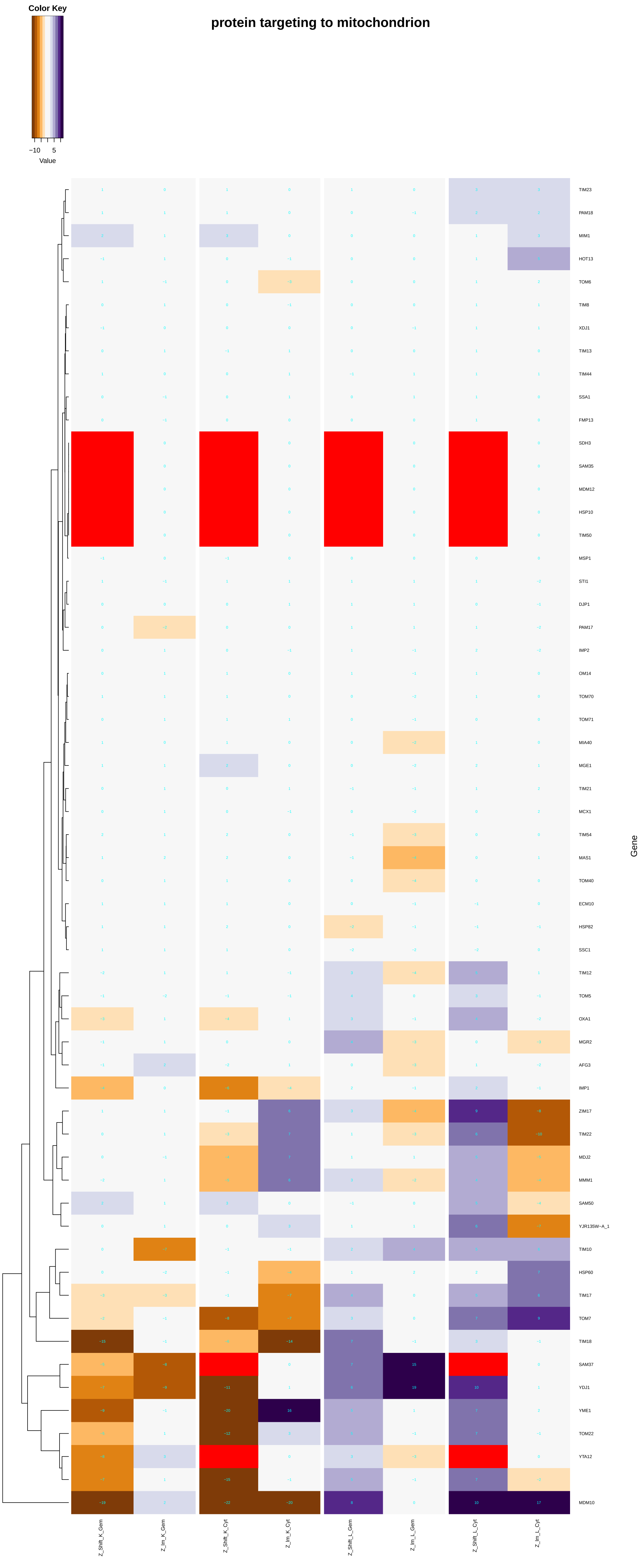

protein targeting to mitochondrion
Color Key
−10
5
Value
TIM23
PAM18
MIM1
HOT13
TOM6
TIM8
XDJ1
TIM13
TIM44
SSA1
FMP13
SDH3
SAM35
MDM12
HSP10
TIM50
MSP1
STI1
DJP1
PAM17
IMP2
OM14
TOM70
TOM71
MIA40
MGE1
TIM21
MCX1
TIM54
MAS1
TOM40
ECM10
HSP82
SSC1
TIM12
TOM5
OXA1
MGR2
AFG3
IMP1
ZIM17
TIM22
MDJ2
MMM1
SAM50
YJR135W−A_1
TIM10
HSP60
TIM17
TOM7
TIM18
SAM37
YDJ1
YME1
TOM22
YTA12
MDM10
Gene
Z_Shift_K_Gem
Z_lm_K_Gem
Z_Shift_K_Cyt
Z_lm_K_Cyt
Z_Shift_L_Gem
Z_lm_L_Gem
Z_Shift_L_Cyt
Z_lm_L_Cyt

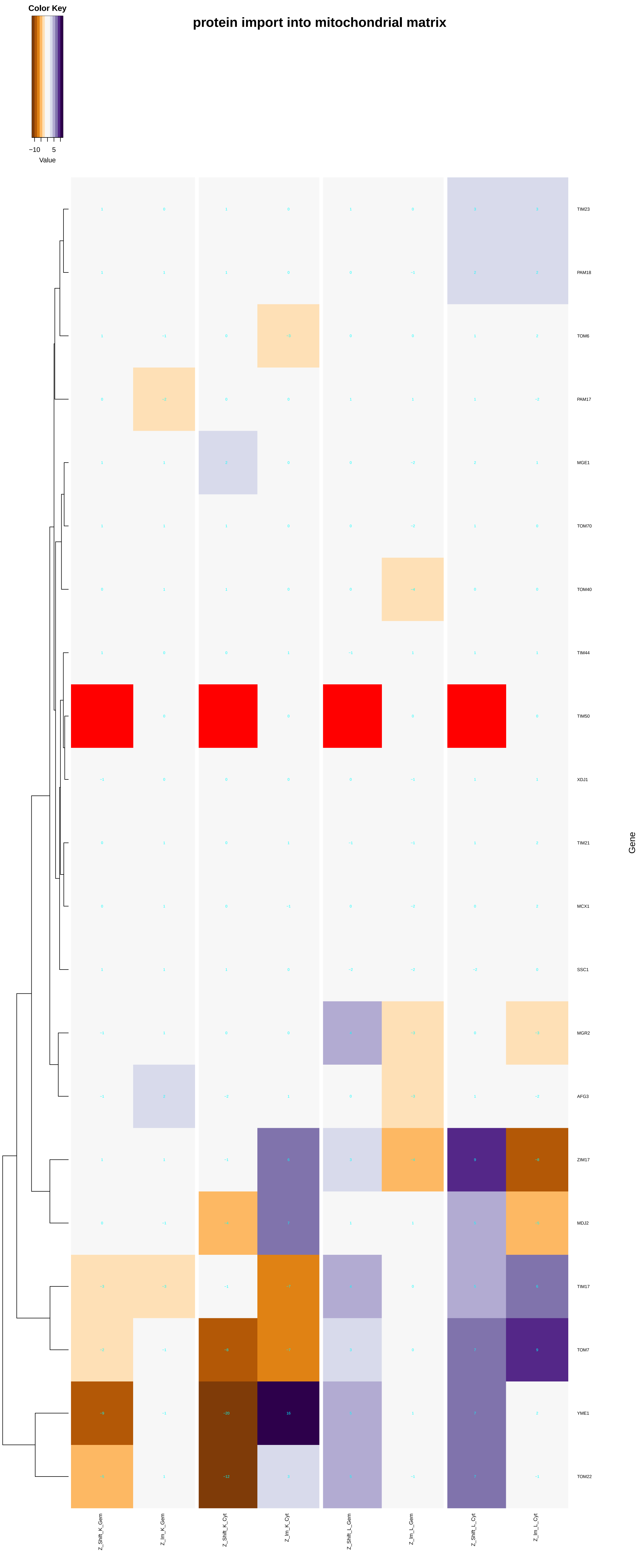

protein import into mitochondrial matrix
Color Key
−10
5
Value
TIM23
PAM18
TOM6
PAM17
MGE1
TOM70
TOM40
TIM44
TIM50
XDJ1
TIM21
MCX1
SSC1
MGR2
AFG3
ZIM17
MDJ2
TIM17
TOM7
YME1
TOM22
Gene
Z_Shift_K_Gem
Z_Im_K_Gem
Z_Shift_K_Cyt
Z_Im_K_Cyt
Z_Shift_L_Gem
Z_Im_L_Gem
Z_Shift_L_Cyt
Z_Im_L_Cyt

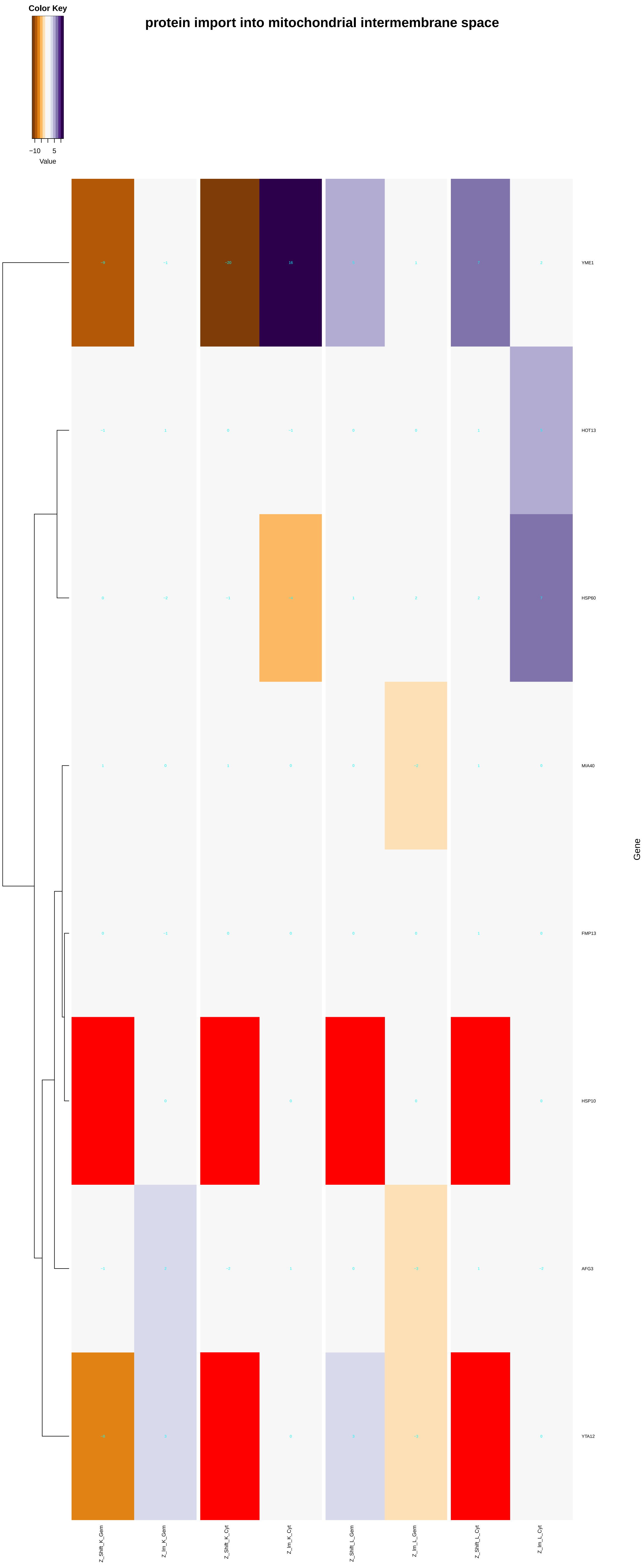
protein import into mitochondrial intermembrane space
Color Key
−10
5
Value
YME1
HOT13
HSP60
MIA40
FMP13
HSP10
AFG3
YTA12
Gene
Z_Shift_K_Gem
Z_Im_K_Gem
Z_Shift_K_Cyt
Z_Im_K_Cyt
Z_Shift_L_Gem
Z_Im_L_Gem
Z_Shift_L_Cyt
Z_Im_L_Cyt

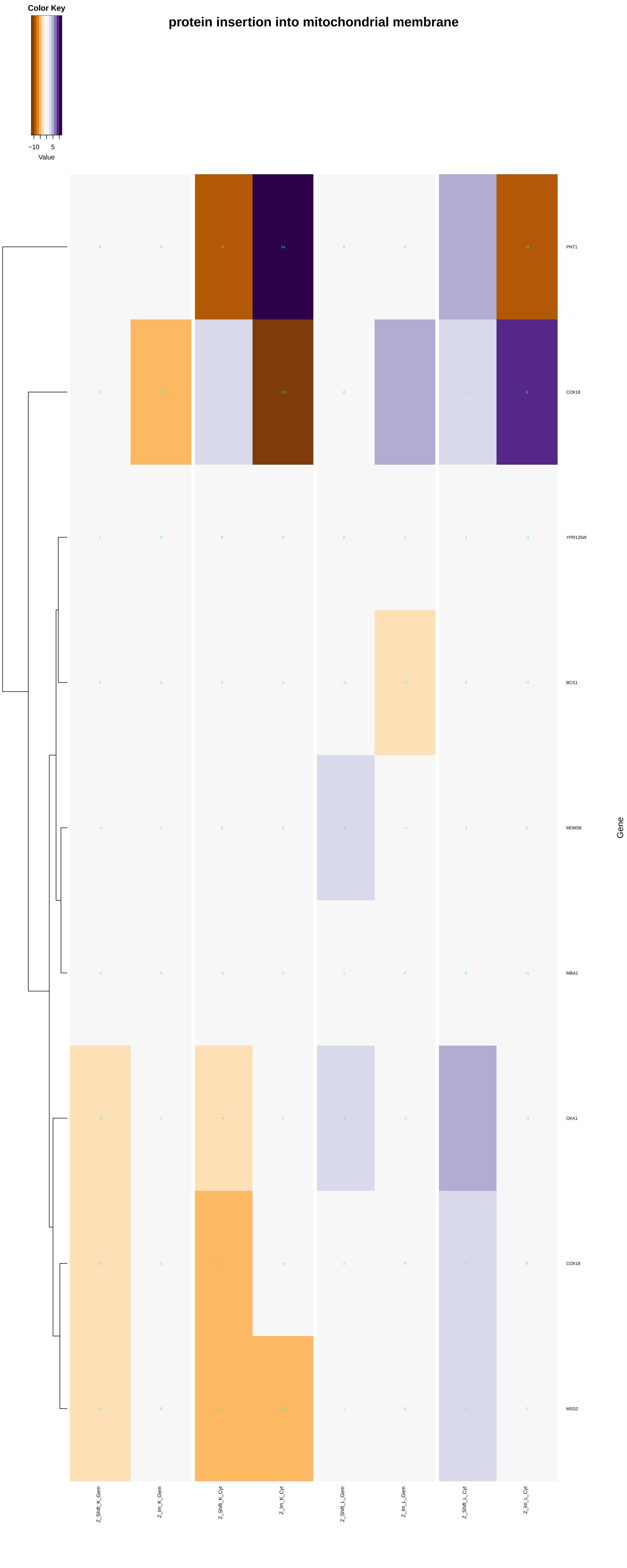
Color Key
−10
5
Value
protein insertion into mitochondrial membrane
PNT1
COX18
YPR125W
BCS1
MDM38
MBA1
OXA1
COX18
MSS2
Gene
Z_Shift_K_Gem
Z_Im_K_Gem
Z_Shift_K_Cyt
Z_Im_K_Cyt
Z_Shift_L_Gem
Z_Im_L_Gem
Z_Shift_L_Cyt
Z_Im_L_Cyt

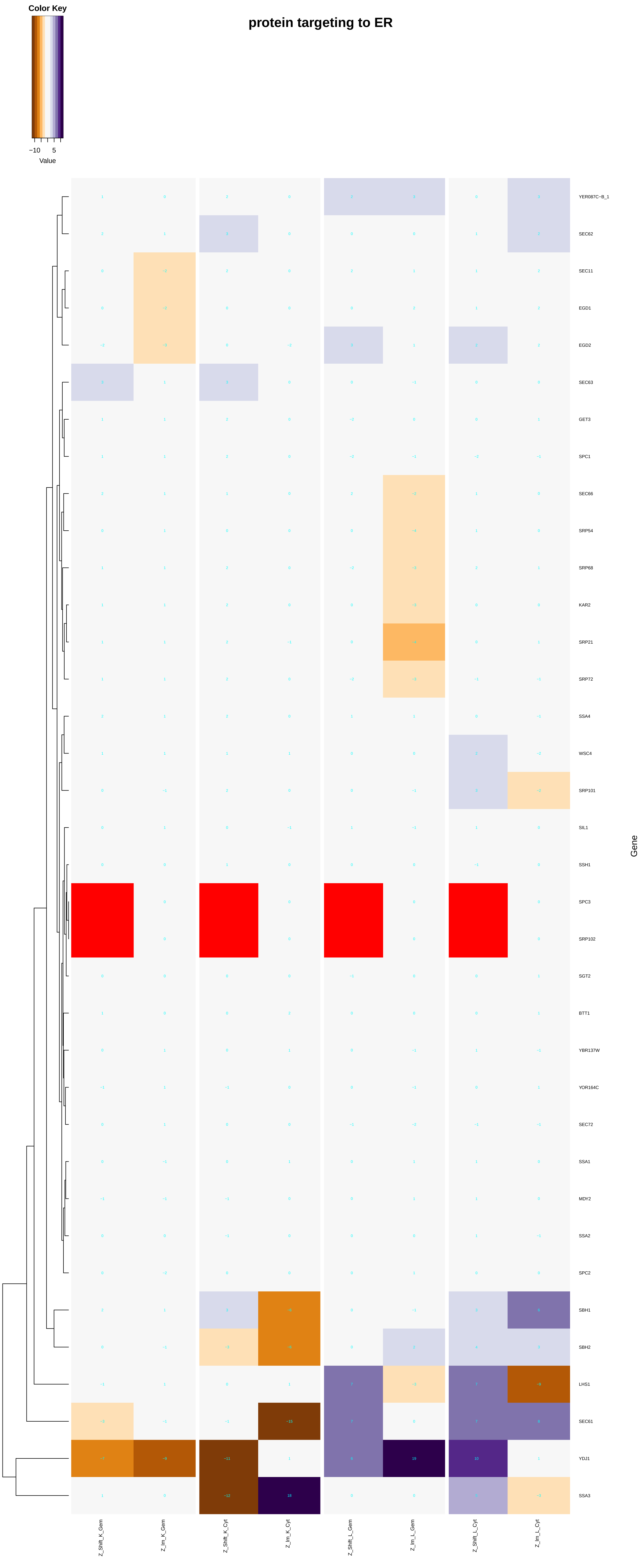
Color Key
−10
5
Value
protein targeting to ER
Gene
YER087C−B_1
SEC62
SEC11
EGD1
EGD2
SEC63
GET3
SPC1
SEC66
SRP54
SRP68
KAR2
SRP21
SRP72
SSA4
WSC4
SRP101
SIL1
SSH1
SPC3
SRP102
SGT2
BTT1
YBR137W
YOR164C
SEC72
SSA1
MDY2
SSA2
SPC2
SBH1
SBH2
LHS1
SEC61
YDJ1
SSA3
Z_Shift_K_Gem
Z_Im_K_Gem
Z_Shift_K_Cyt
Z_Im_K_Cyt
Z_Shift_L_Gem
Z_Im_L_Gem
Z_Shift_L_Cyt
Z_Im_L_Cyt

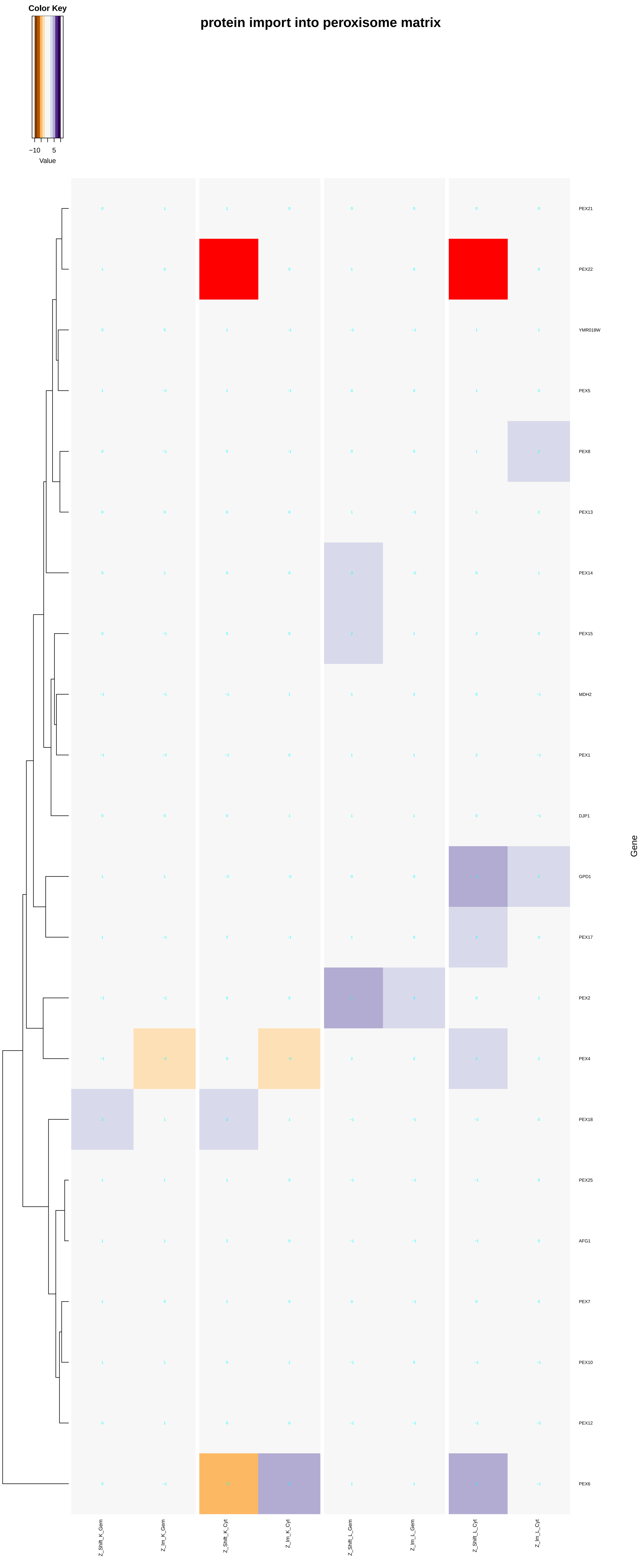

Color Key
−10
5
Value
protein import into peroxisome matrix
PEX21
PEX22
YMR018W
PEX5
PEX8
PEX13
PEX14
PEX15
MDH2
PEX1
DJP1
GPD1
PEX17
PEX2
PEX4
PEX18
PEX25
AFG1
PEX7
PEX10
PEX12
PEX6
Gene
Z_Shift_K_Gem
Z_Im_K_Gem
Z_Shift_K_Cyt
Z_Im_K_Cyt
Z_Shift_L_Gem
Z_Im_L_Gem
Z_Shift_L_Cyt
Z_Im_L_Cyt

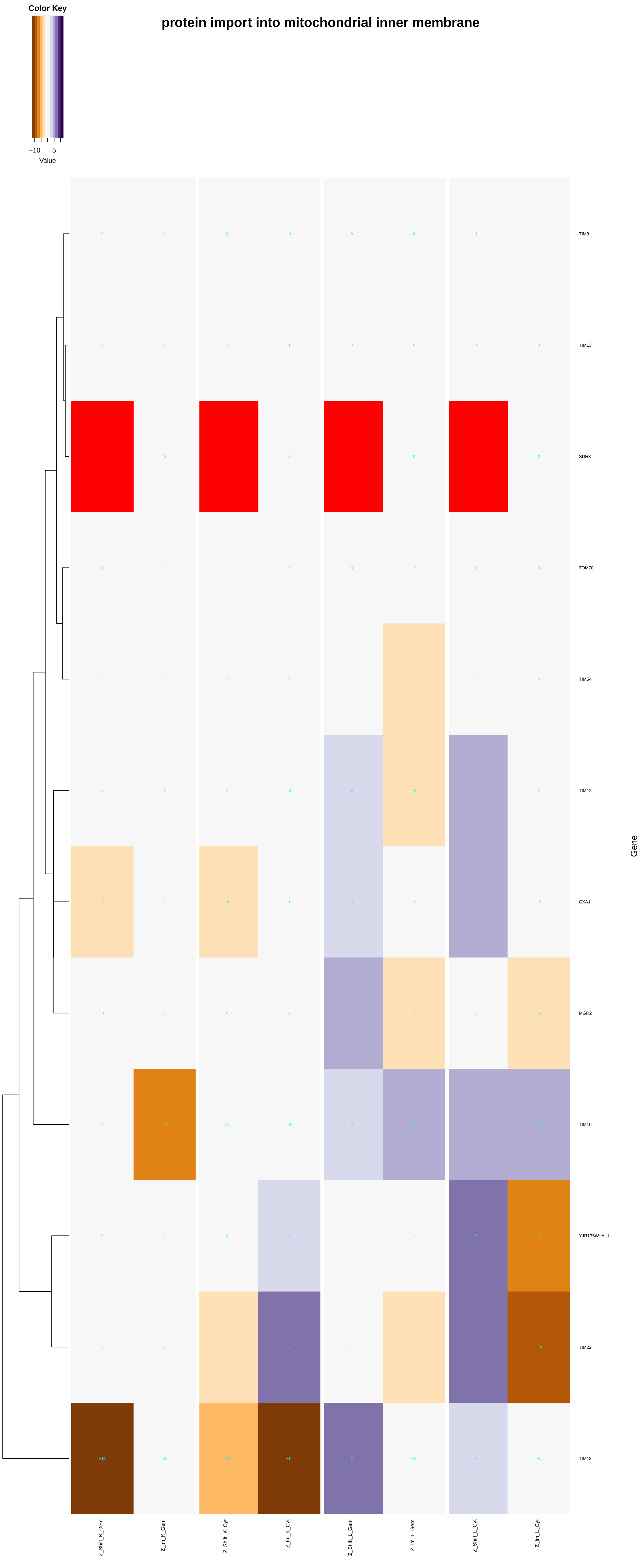
Color Key
−10
5
Value
protein import into mitochondrial inner membrane
TIM8
TIM13
SDH3
TOM70
TIM54
TIM12
OXA1
MGR2
TIM10
YJR135W−A_1
TIM22
TIM18
Gene
Z_Shift_K_Gem
Z_Im_K_Gem
Z_Shift_K_Cyt
Z_Im_K_Cyt
Z_Shift_L_Gem
Z_Im_L_Gem
Z_Shift_L_Cyt
Z_Im_L_Cyt

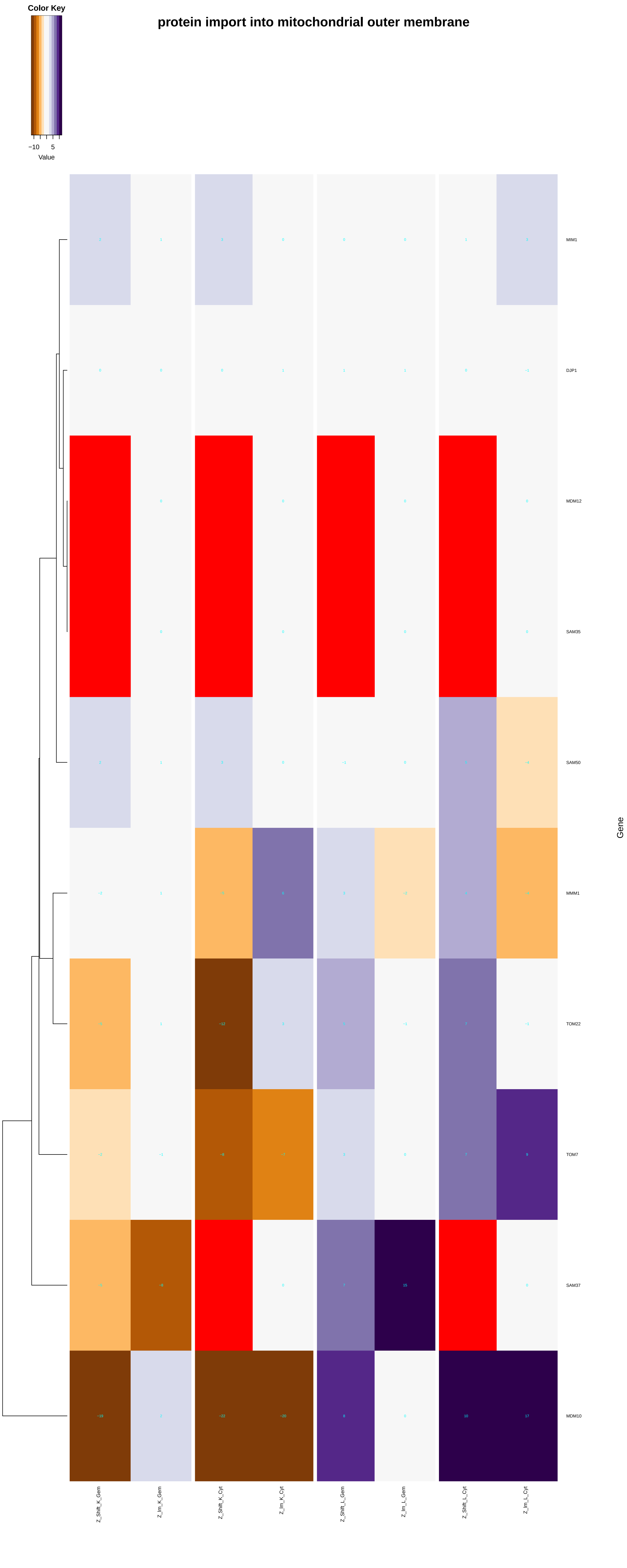
Color Key
−10
5
Value
protein import into mitochondrial outer membrane
MIM1
DJP1
MDM12
SAM35
SAM50
MMM1
TOM22
TOM7
SAM37
MDM10
Gene
Z_Shift_K_Gem
Z_Im_K_Gem
Z_Shift_K_Cyt
Z_Im_K_Cyt
Z_Shift_L_Gem
Z_Im_L_Gem
Z_Shift_L_Cyt
Z_Im_L_Cyt

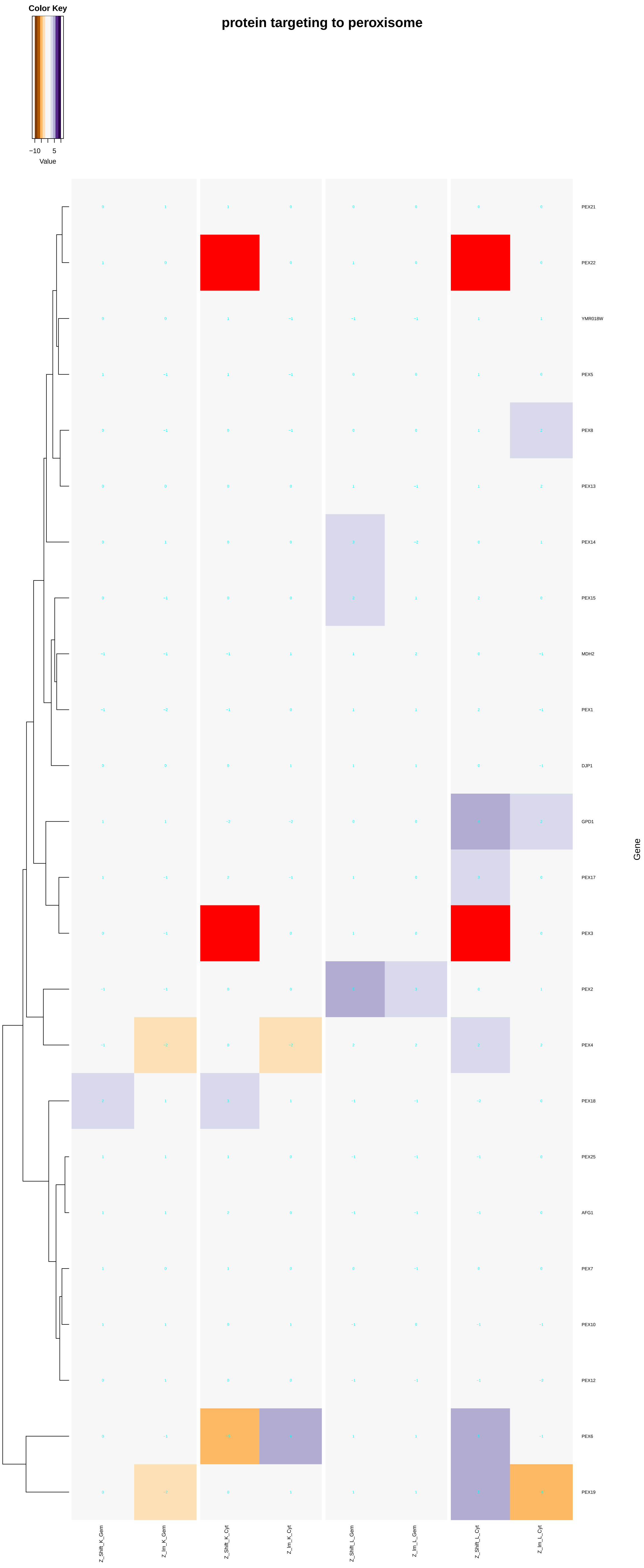

Color Key
−10
5
Value
protein targeting to peroxisome
PEX21
PEX22
YMR018W
PEX5
PEX8
PEX13
PEX14
PEX15
MDH2
PEX1
DJP1
GPD1
PEX17
PEX3
PEX2
PEX4
PEX18
PEX25
AFG1
PEX7
PEX10
PEX12
PEX6
PEX19
Gene
Z_Shift_K_Gem
Z_Im_K_Gem
Z_Shift_K_Cyt
Z_Im_K_Cyt
Z_Shift_L_Gem
Z_Im_L_Gem
Z_Shift_L_Cyt
Z_Im_L_Cyt

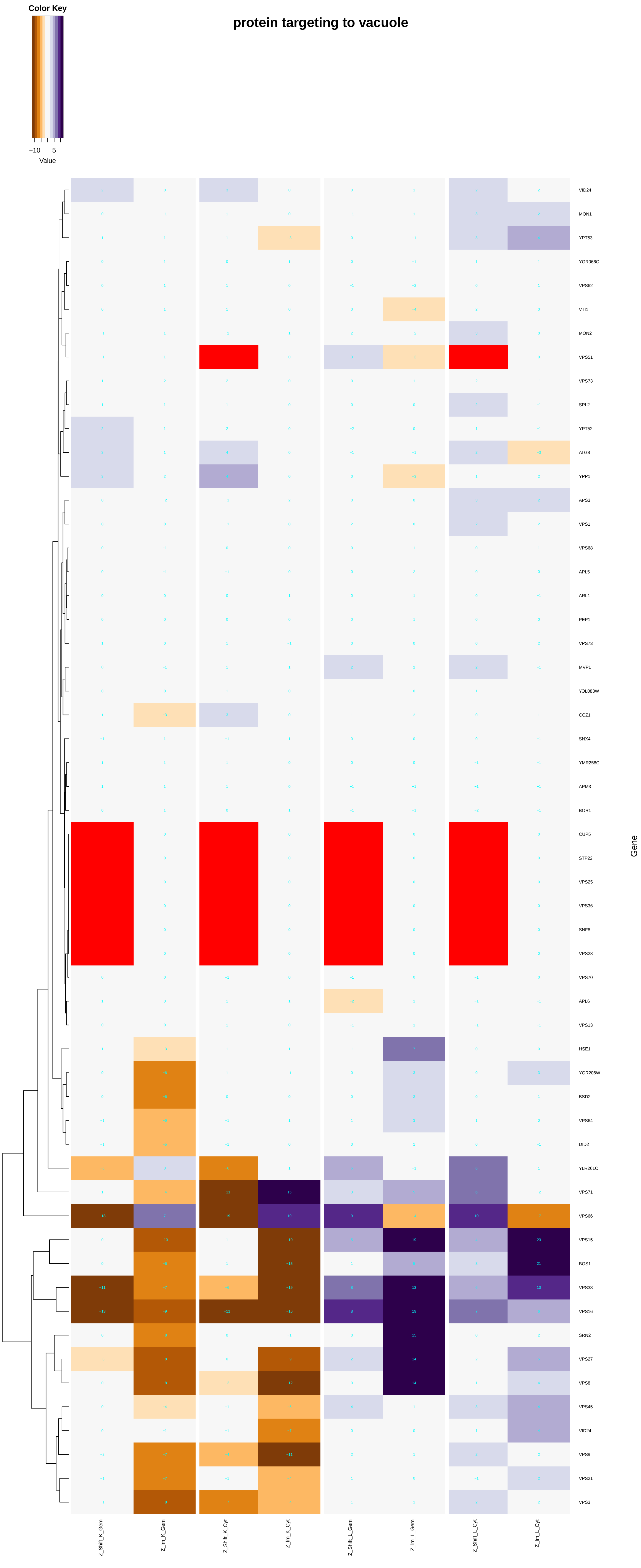

Color Key
−10
5
Value
protein targeting to vacuole
VID24
MON1
YPT53
YGR066C
VPS62
VTI1
MON2
VPS51
VPS73
SPL2
YPT52
ATG8
YPP1
APS3
VPS1
VPS68
APL5
ARL1
PEP1
VPS73
MVP1
YOL083W
CCZ1
SNX4
YMR258C
APM3
BOR1
CUP5
STP22
VPS25
VPS36
SNF8
VPS28
VPS70
APL6
VPS13
HSE1
YGR206W
BSD2
VPS64
DID2
YLR261C
VPS71
VPS66
VPS15
BOS1
VPS33
VPS16
SRN2
VPS27
VPS8
VPS45
VID24
VPS9
VPS21
VPS3
Gene
Z_Shift_K_Gem
Z_Im_K_Gem
Z_Shift_K_Cyt
Z_Im_K_Cyt
Z_Shift_L_Gem
Z_Im_L_Gem
Z_Shift_L_Cyt
Z_Im_L_Cyt

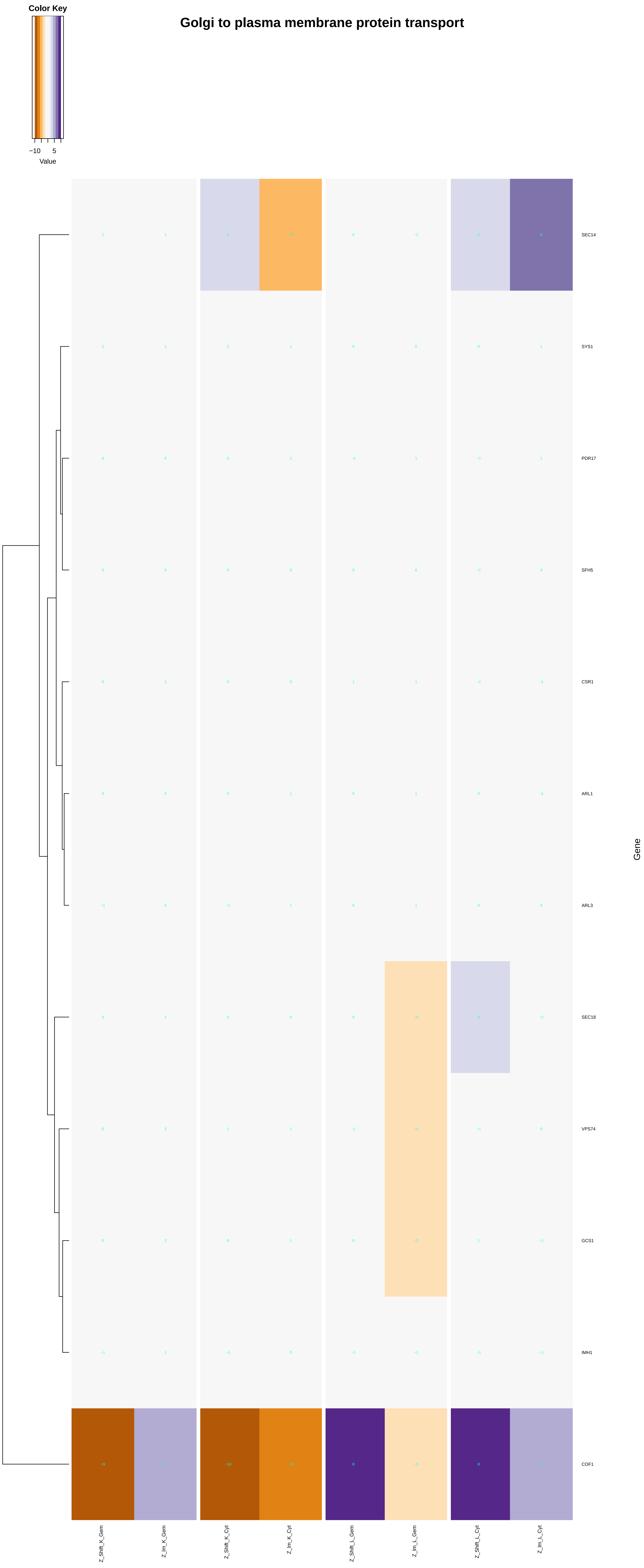

Golgi to plasma membrane protein transport
Color Key
−10
5
Value
SEC14
SYS1
PDR17
SFH5
CSR1
ARL1
ARL3
SEC18
VPS74
GCS1
IMH1
COF1
Gene
Z_Shift_K_Gem
Z_Im_K_Gem
Z_Shift_K_Cyt
Z_Im_K_Cyt
Z_Shift_L_Gem
Z_Im_L_Gem
Z_Shift_L_Cyt
Z_Im_L_Cyt

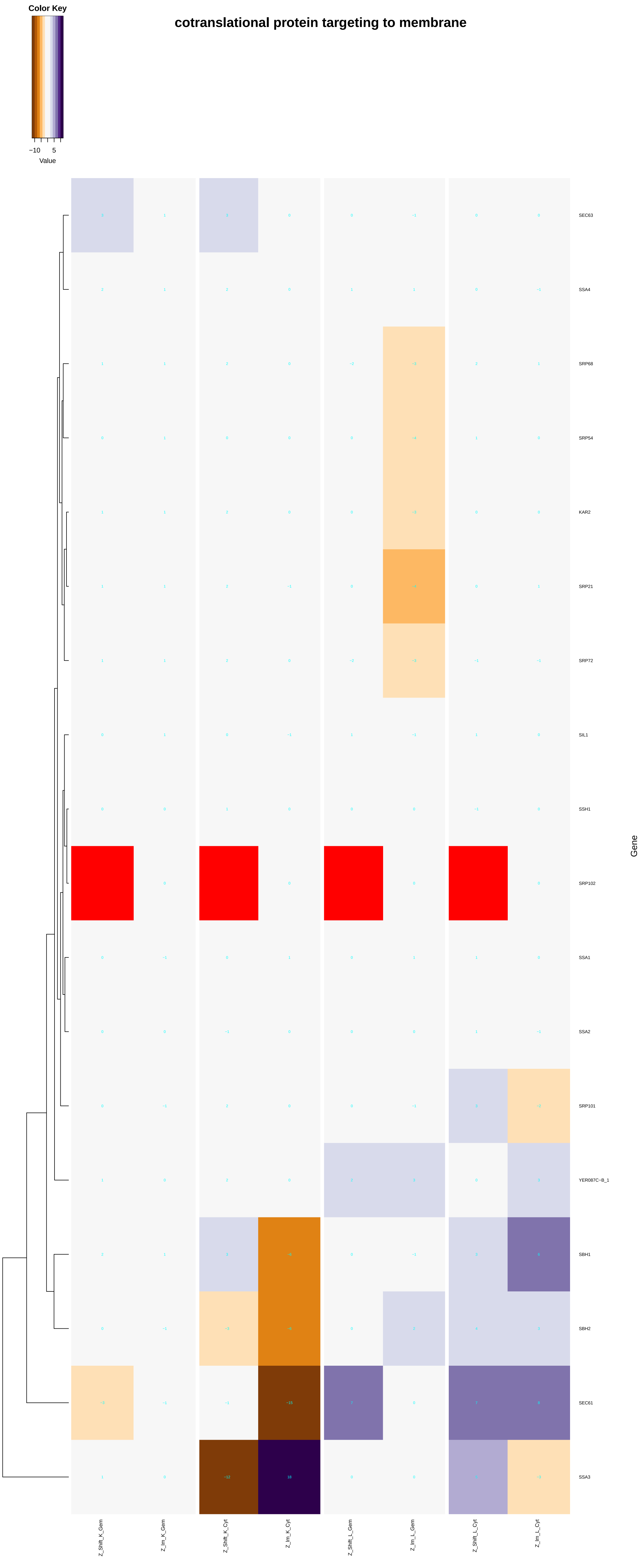

Color Key
−10
5
Value
cotranslational protein targeting to membrane
SEC63
SSA4
SRP68
SRP54
KAR2
SRP21
SRP72
SIL1
SSH1
SRP102
SSA1
SSA2
SRP101
YER087C−B_1
SBH1
SBH2
SEC61
SSA3
Gene
Z_Shift_K_Gem
Z_Im_K_Gem
Z_Shift_K_Cyt
Z_Im_K_Cyt
Z_Shift_L_Gem
Z_Im_L_Gem
Z_Shift_L_Cyt
Z_Im_L_Cyt

### posttranslational protein targeting to membrane

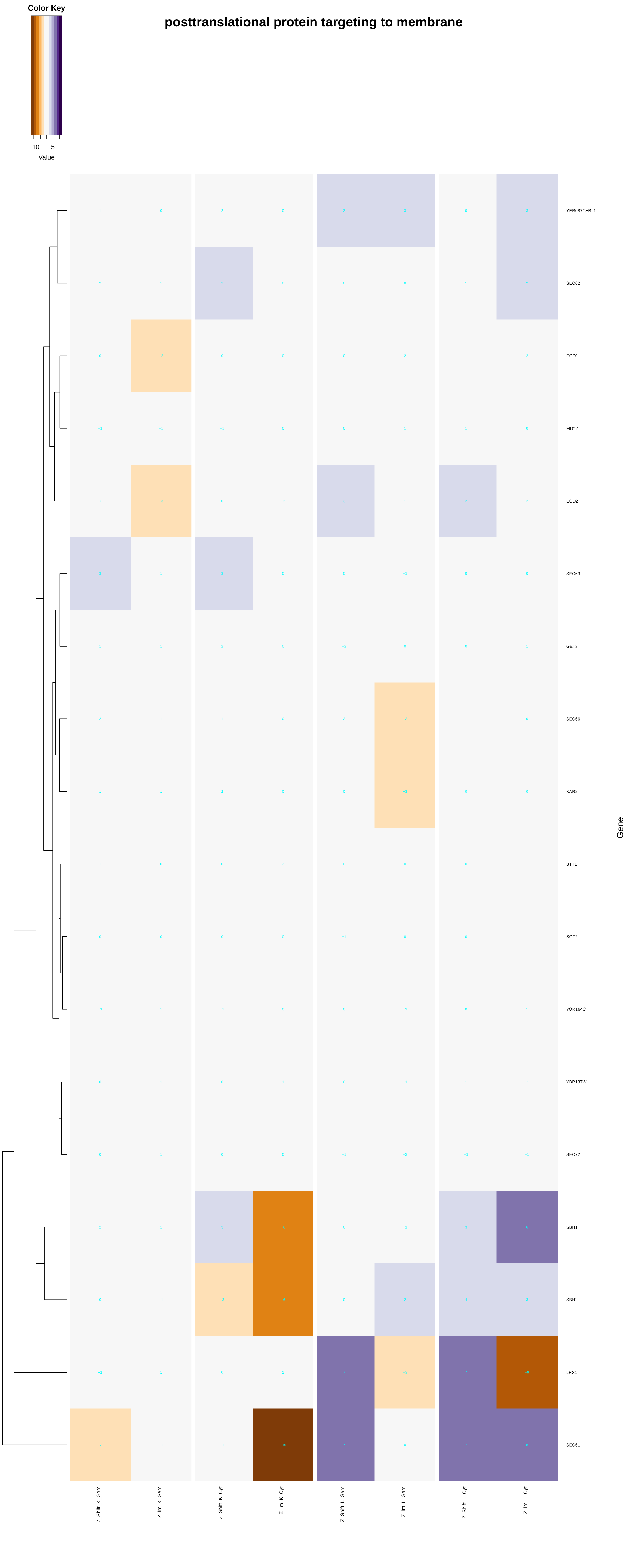

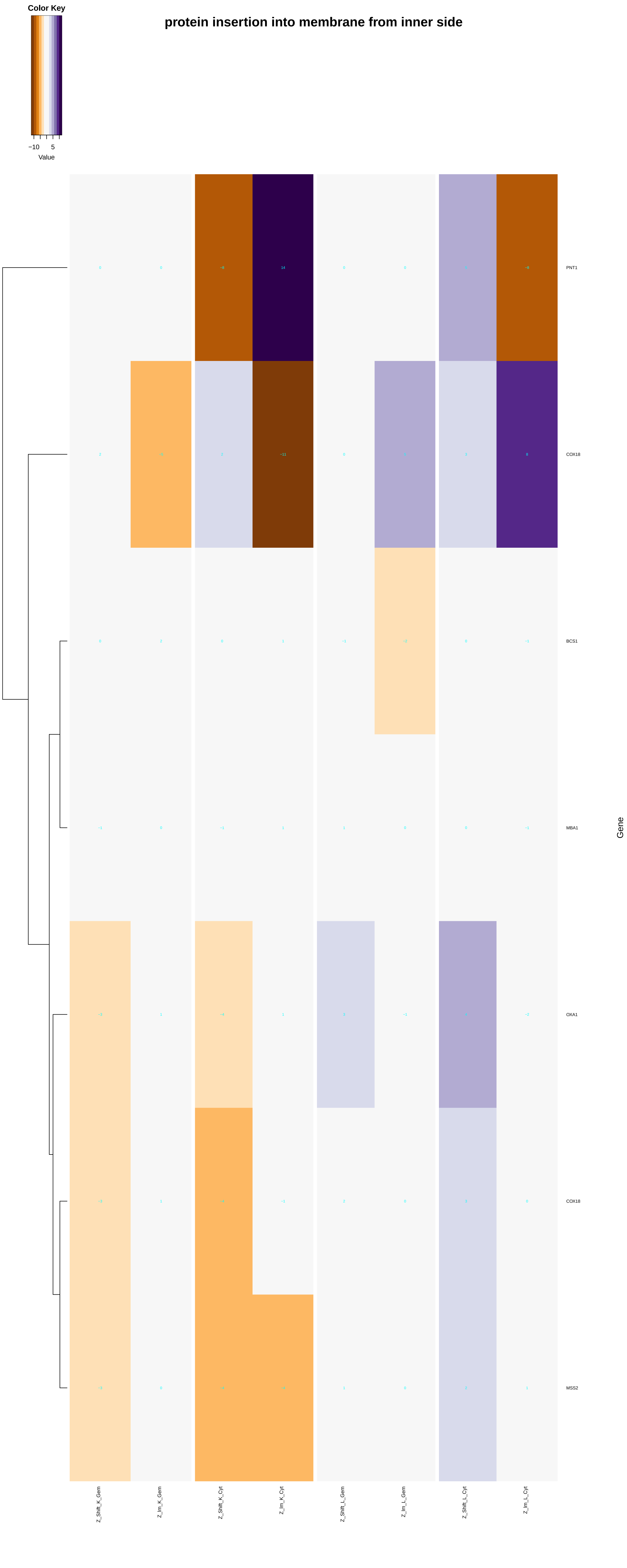

protein insertion into membrane from inner side
Color Key
−10
5
Value
PNT1
COX18
BCS1
MBA1
OXA1
COX18
MSS2
Gene
Z_Shift_K_Gem
Z_Im_K_Gem
Z_Shift_K_Cyt
Z_Im_K_Cyt
Z_Shift_L_Gem
Z_Im_L_Gem
Z_Shift_L_Cyt
Z_Im_L_Cyt

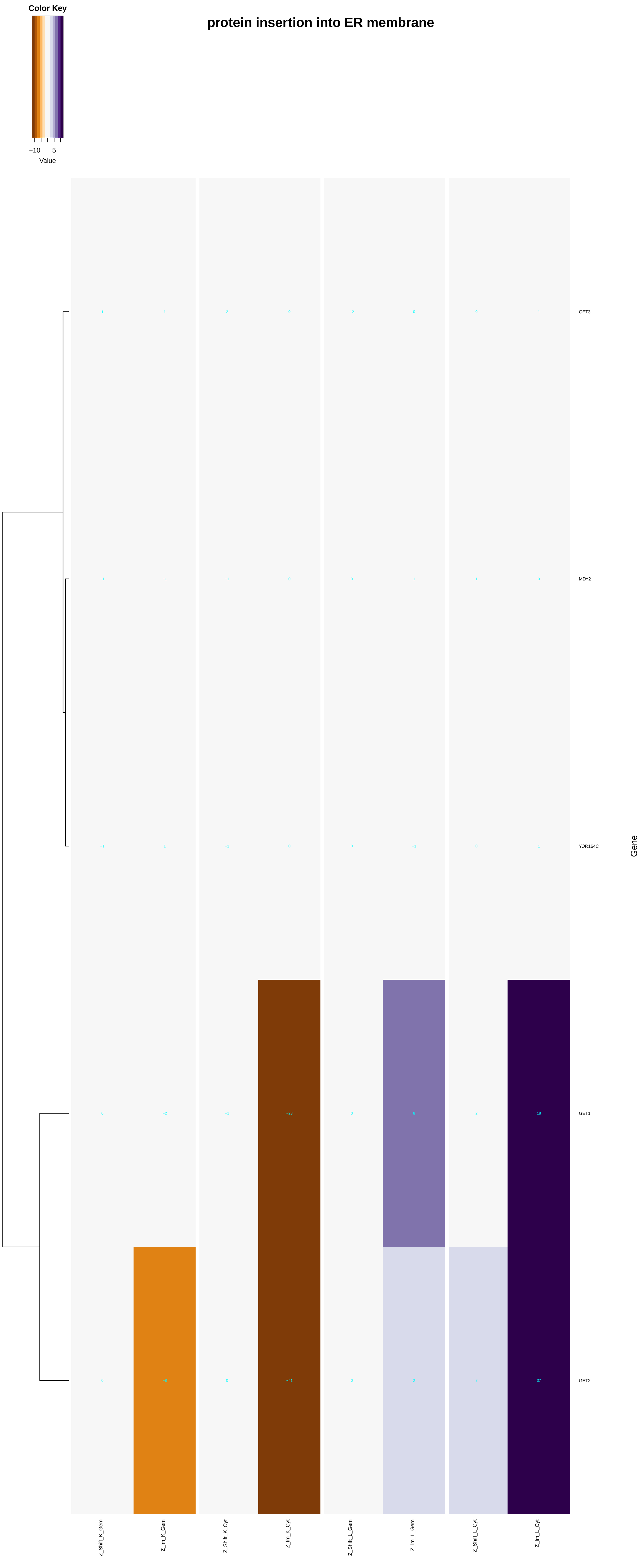
protein insertion into ER membrane
Color Key
−10
5
Value
GET3
MDY2
YOR164C
GET1
GET2
Gene
Z_Shift_K_Gem
Z_Im_K_Gem
Z_Shift_K_Cyt
Z_Im_K_Cyt
Z_Shift_L_Gem
Z_Im_L_Gem
Z_Shift_L_Cyt
Z_Im_L_Cyt

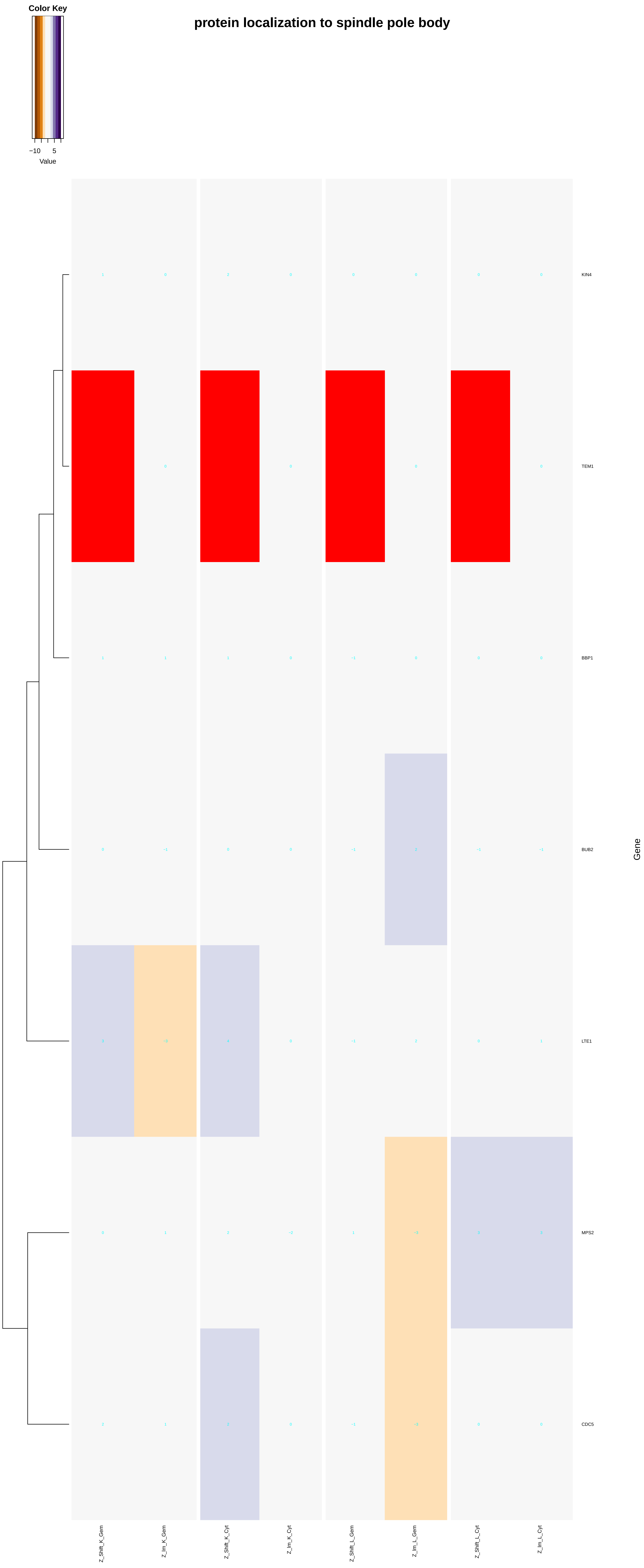

Color Key
−10
5
Value
protein localization to spindle pole body
KIN4
TEM1
BBP1
BUB2
LTE1
MPS2
CDC5
Gene
Z_Shift_K_Gem
Z_Im_K_Gem
Z_Shift_K_Cyt
Z_Im_K_Cyt
Z_Shift_L_Gem
Z_Im_L_Gem
Z_Shift_L_Cyt
Z_Im_L_Cyt

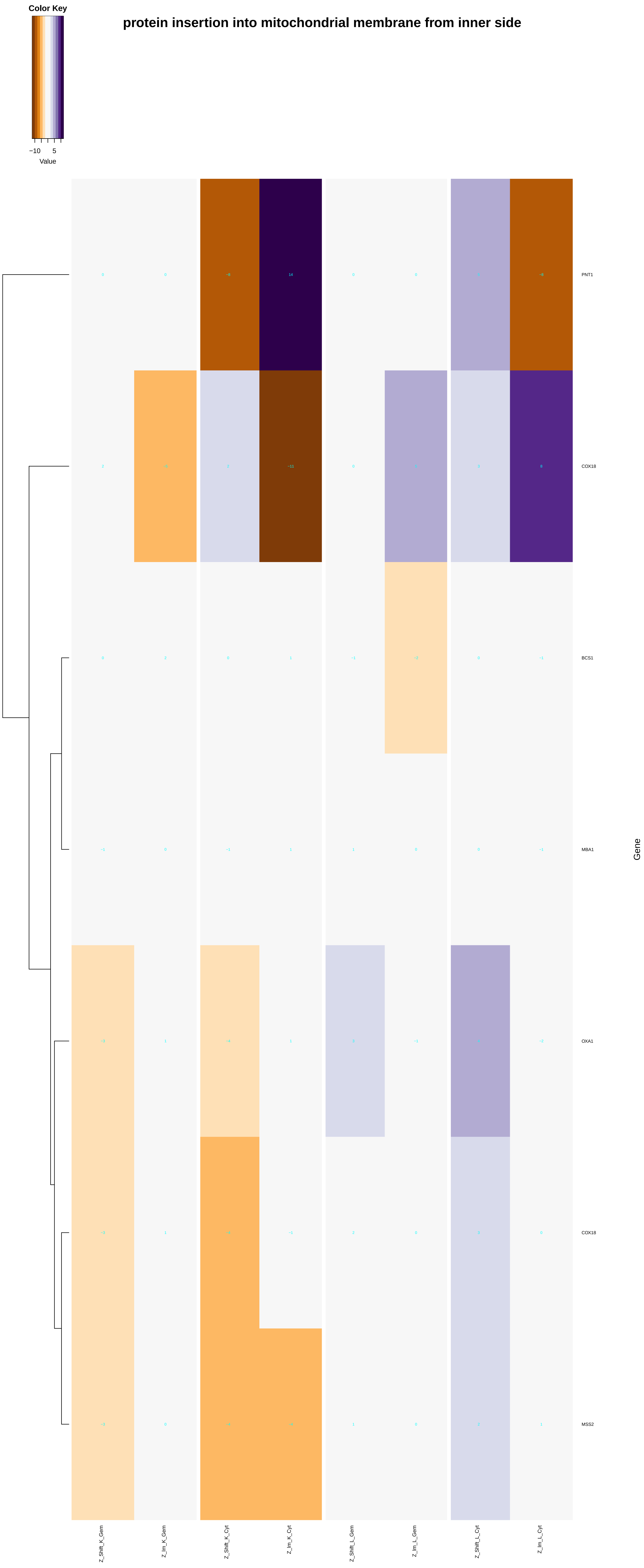
Color Key
−10
5
Value
protein insertion into mitochondrial membrane from inner side
PNT1
COX18
BCS1
MBA1
OXA1
COX18
MSS2
Gene
Z_Shift_K_Gem
Z_Im_K_Gem
Z_Shift_K_Cyt
Z_Im_K_Cyt
Z_Shift_L_Gem
Z_Im_L_Gem
Z_Shift_L_Cyt
Z_Im_L_Cyt

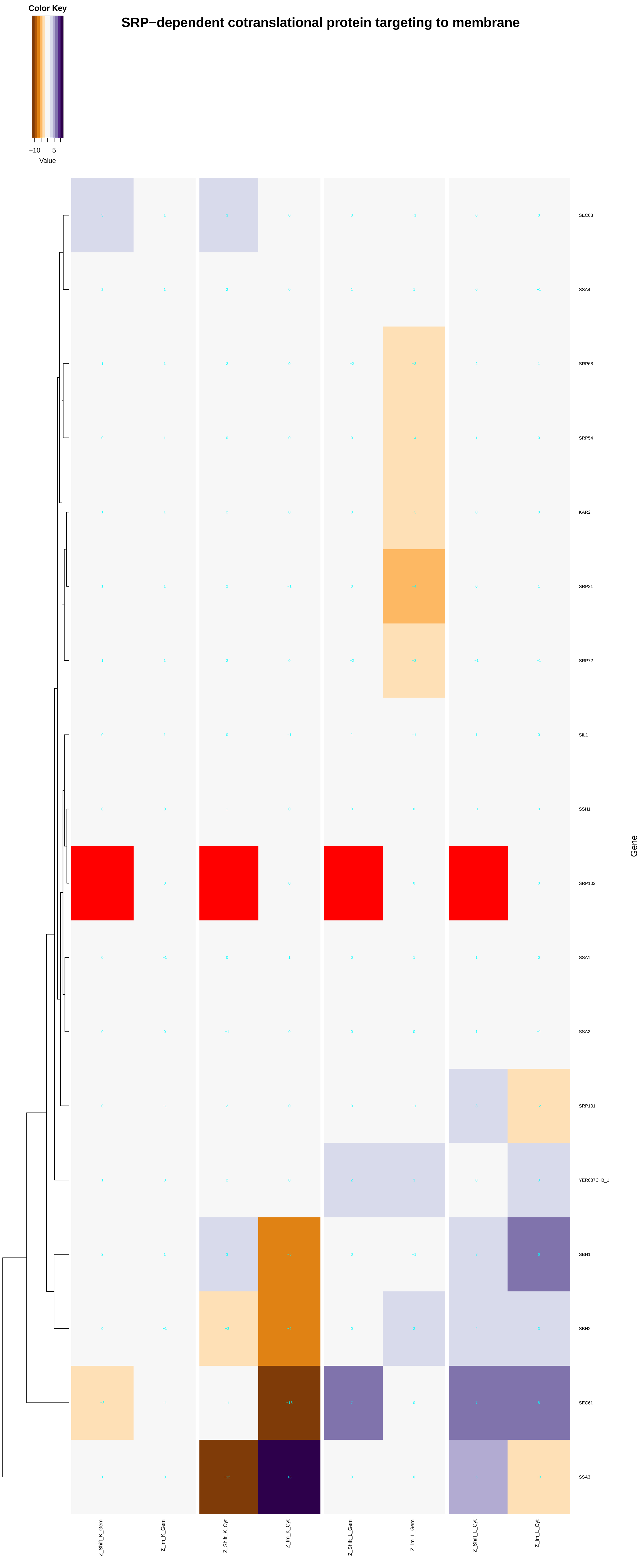

Color Key
−10 5
Value
SRP−dependent cotranslational protein targeting to membrane
SEC63
SSA4
SRP68
SRP54
KAR2
SRP21
SRP72
SIL1
SSH1
SRP102
SSA1
SSA2
SRP101
YER087C−B_1
SBH1
SBH2
SEC61
SSA3
Gene
Z_Shift_K_Gem
Z_Im_K_Gem
Z_Shift_K_Cyt
Z_Im_K_Cyt
Z_Shift_L_Gem
Z_Im_L_Gem
Z_Shift_L_Cyt
Z_Im_L_Cyt

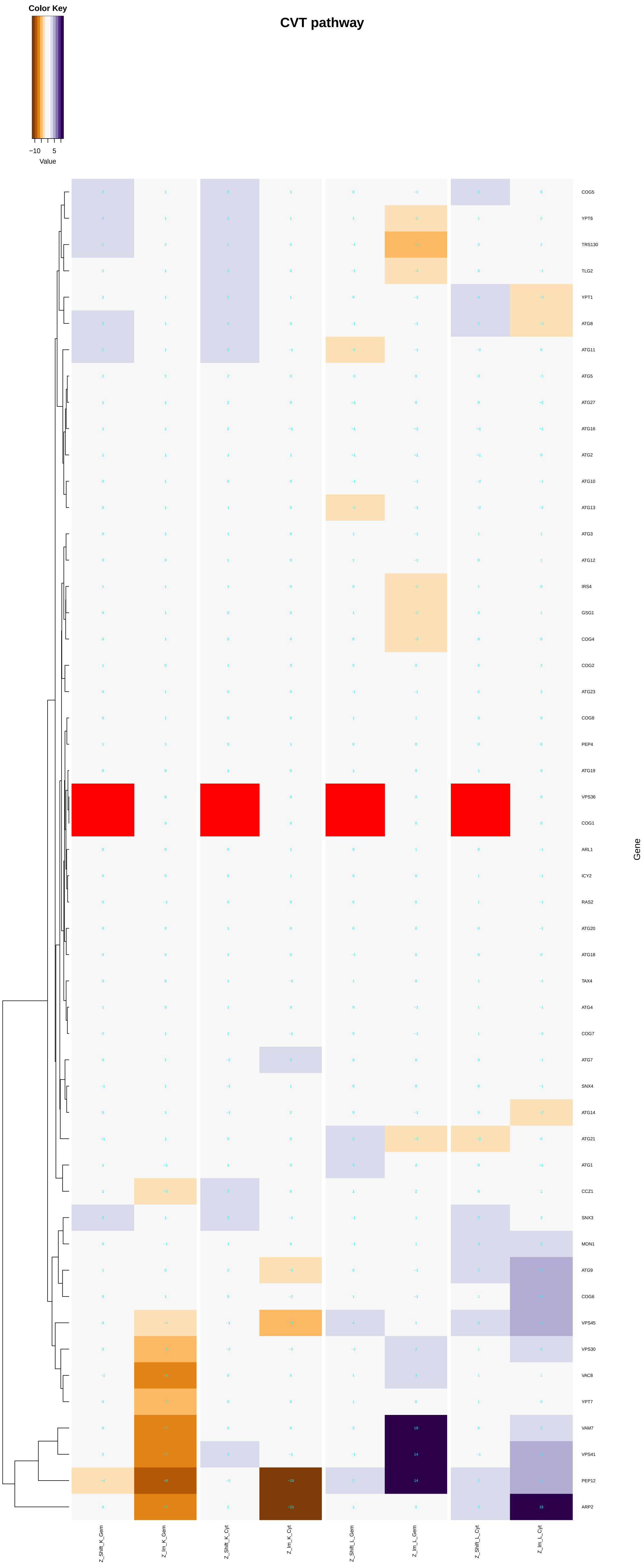
CVT pathway
Color Key
−10
5
Value
COG5
YPT6
TRS130
TLG2
YPT1
ATG8
ATG11
ATG5
ATG27
ATG16
ATG2
ATG10
ATG13
ATG3
ATG12
IRS4
GSG1
COG4
COG2
ATG23
COG8
PEP4
ATG19
VPS36
COG1
ARL1
ICY2
RAS2
ATG20
ATG18
TAX4
ATG4
COG7
ATG7
SNX4
ATG14
ATG21
ATG1
CCZ1
SNX3
MON1
ATG9
COG6
VPS45
VPS30
VAC8
YPT7
VAM7
VPS41
PEP12
ARP2
Gene
Z_Shift_K_Gem
Z_Im_K_Gem
Z_Shift_K_Cyt
Z_Im_K_Cyt
Z_Shift_L_Gem
Z_Im_L_Gem
Z_Shift_L_Cyt
Z_Im_L_Cyt

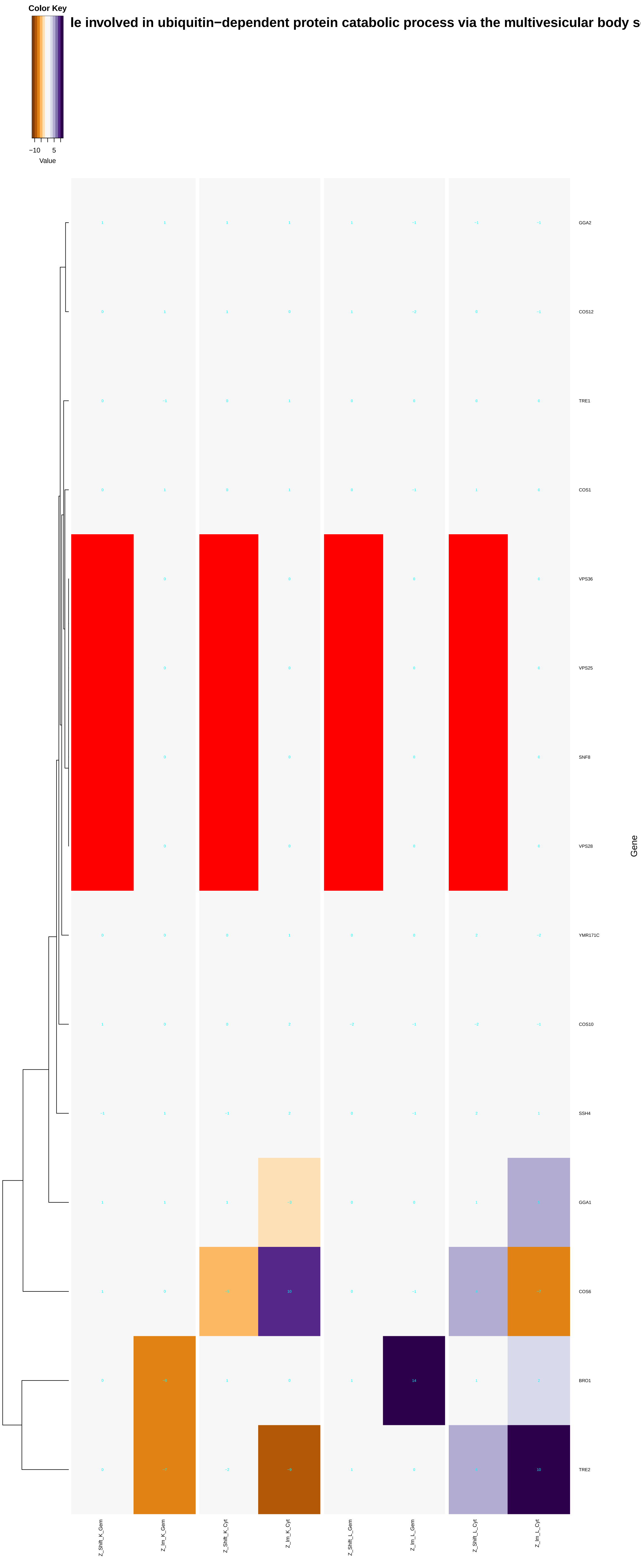

Color Key
−10 5
Value
le involved in ubiquitin−dependent protein catabolic process via the multivesicular body s
GGA2
COS12
TRE1
COS1
VPS36
VPS25
SNF8
VPS28
YMR171C
COS10
SSH4
GGA1
COS6
BRO1
TRE2
Gene
Z_Shift_K_Gem
Z_lm_K_Gem
Z_Shift_K_Cyt
Z_lm_K_Cyt
Z_Shift_L_Gem
Z_lm_L_Gem
Z_Shift_L_Cyt
Z_lm_L_Cyt

### protein localization to mitotic spindle pole body

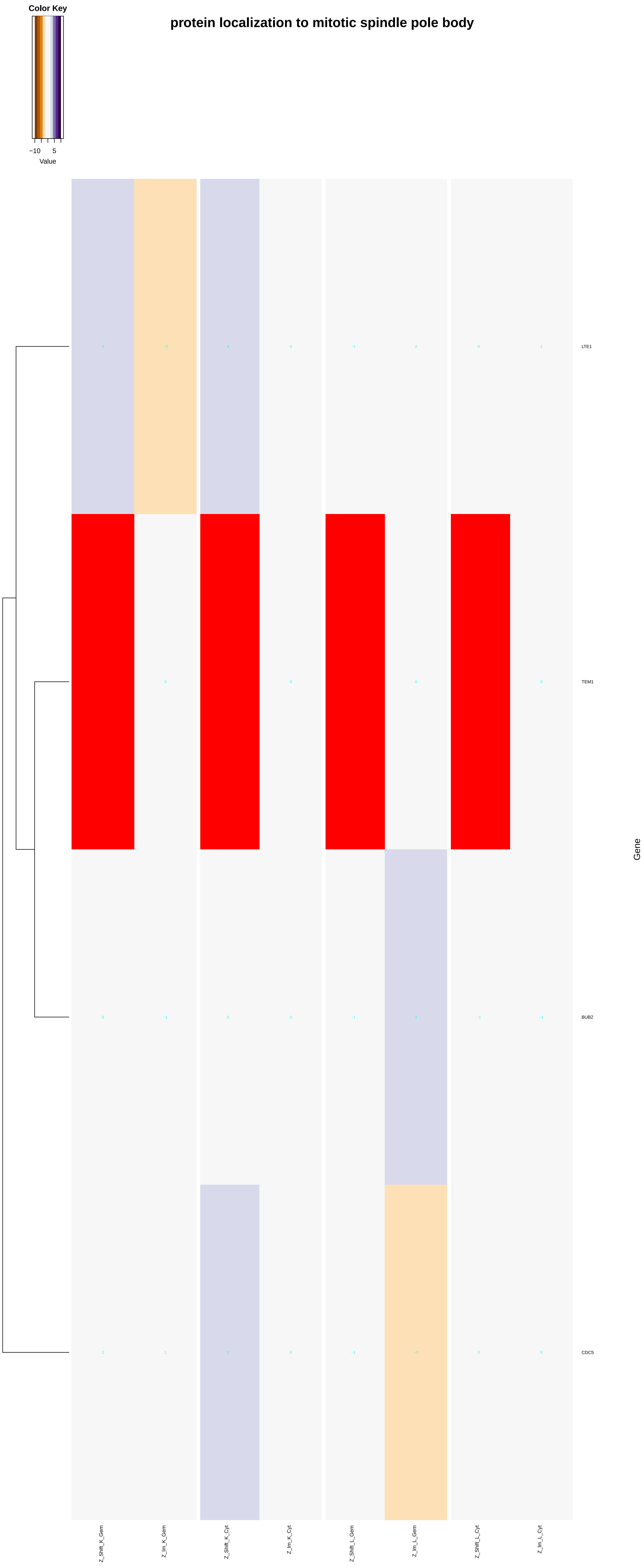
