## Additional File 7 for "A humanized yeast phenomic model of deoxycytidine kinase to predict genetic buffering of nucleoside analog cytotoxicity": cellular_response_to_DNA_damage_stimulus.pdf

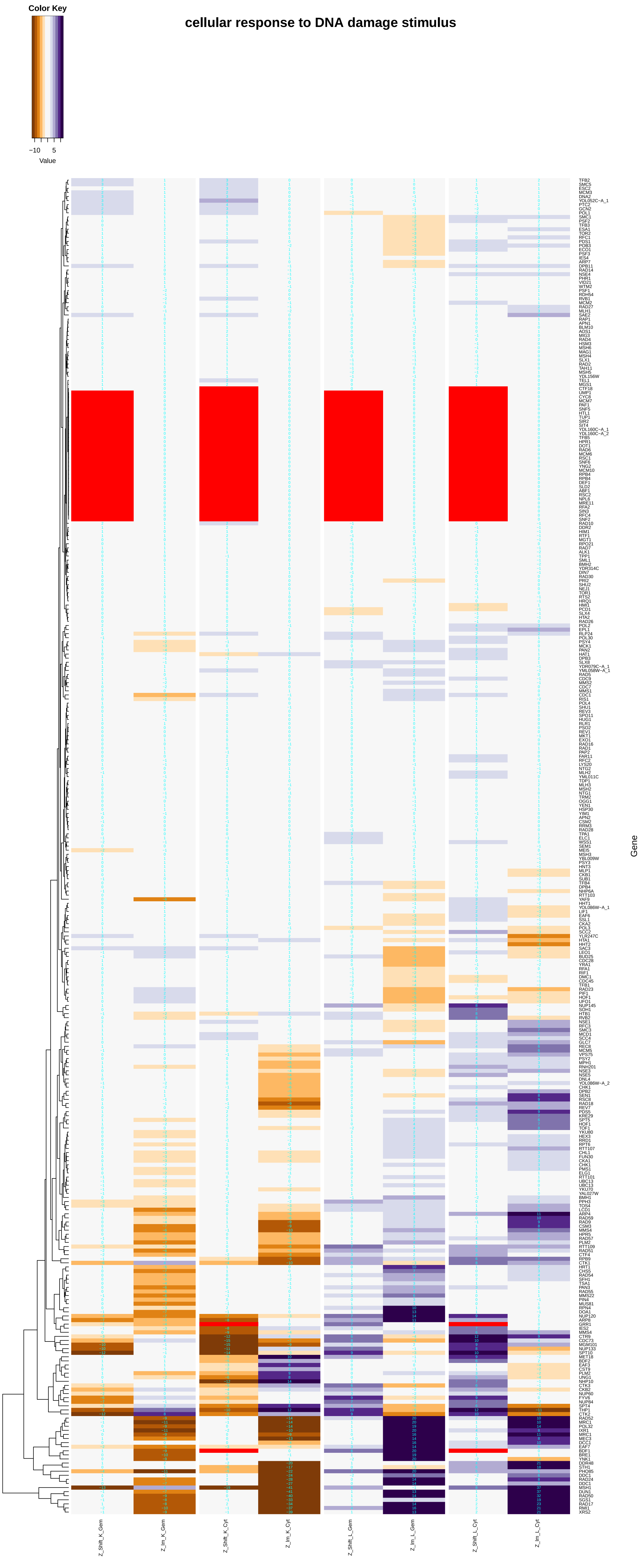
cellular response to DNA damage stimulus
Color Key
−10 5
Value
Gene
Z_Shift_K_Gem
Z_Im_K_Gem
Z_Shift_K_Cyt
Z_Im_K_Cyt
Z_Shift_L_Gem
Z_Im_L_Gem
Z_Shift_L_Cyt
Z_Im_L_Cyt

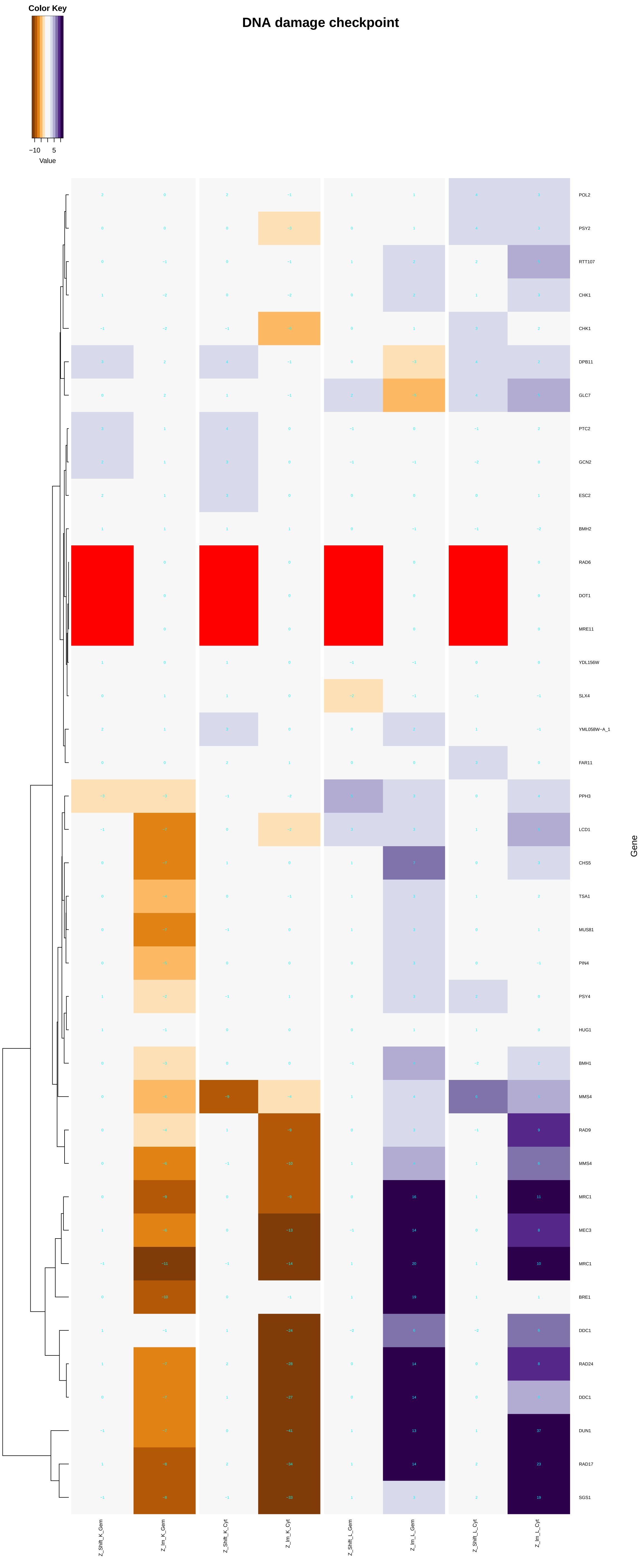

Color Key
−10 5
Value
DNA damage checkpoint
POL2
PSY2
RTT107
CHK1
CHK1
DPB11
GLC7
PTC2
GCN2
ESC2
BMH2
RAD6
DOT1
MRE11
YDL156W
SLX4
YML058W−A_1
FAR11
PPH3
LCD1
CHS5
TSA1
MUS81
PIN4
PSY4
HUG1
BMH1
MMS4
RAD9
MMS4
MRC1
MEC3
MRC1
BRE1
DDC1
RAD24
DDC1
DUN1
RAD17
SGS1
Gene
Z_Shift_K_Gem
Z_lm_K_Gem
Z_Shift_K_Cyt
Z_lm_K_Cyt
Z_Shift_L_Gem
Z_lm_L_Gem
Z_Shift_L_Cyt
Z_lm_L_Cyt

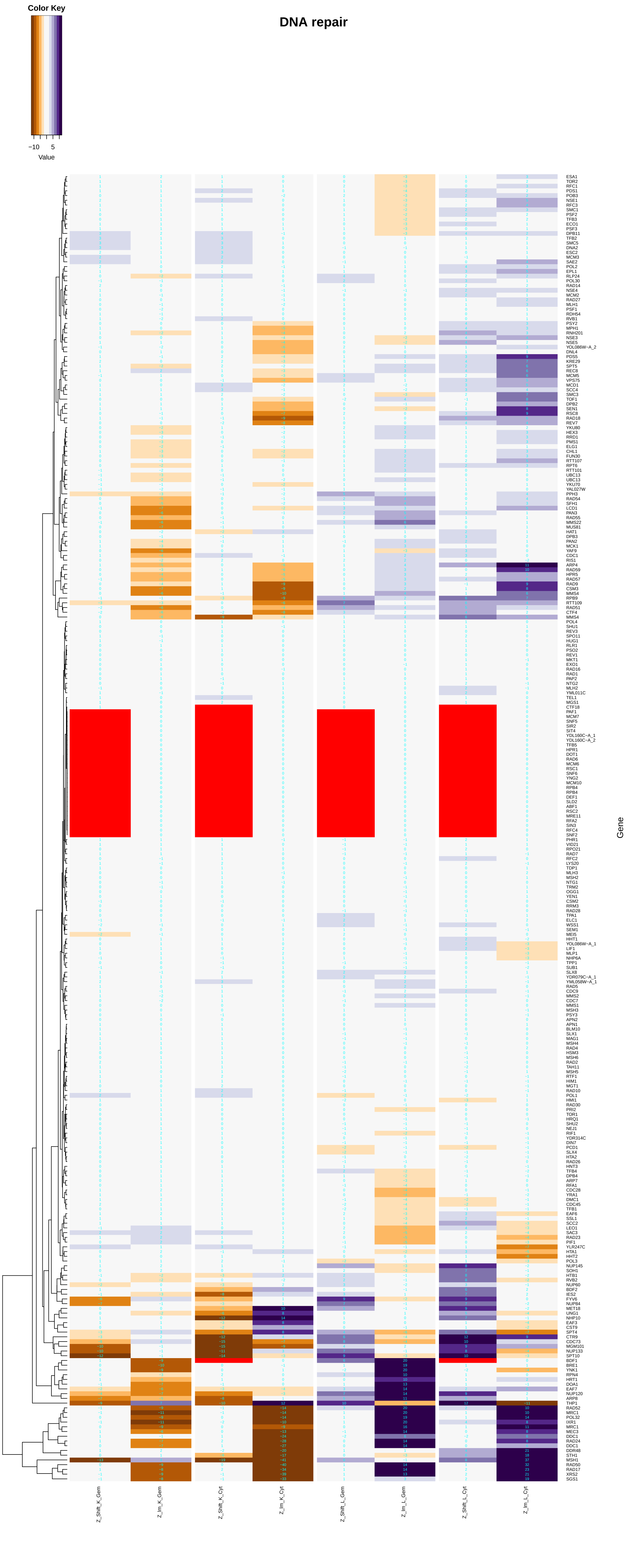

Color Key
−10
5
Value
DNA repair
Gene
Z_Shift_K_Gem
Z_Im_K_Gem
Z_Shift_K_Cyt
Z_Im_K_Cyt
Z_Shift_L_Gem
Z_Im_L_Gem
Z_Shift_L_Cyt
Z_Im_L_Cyt

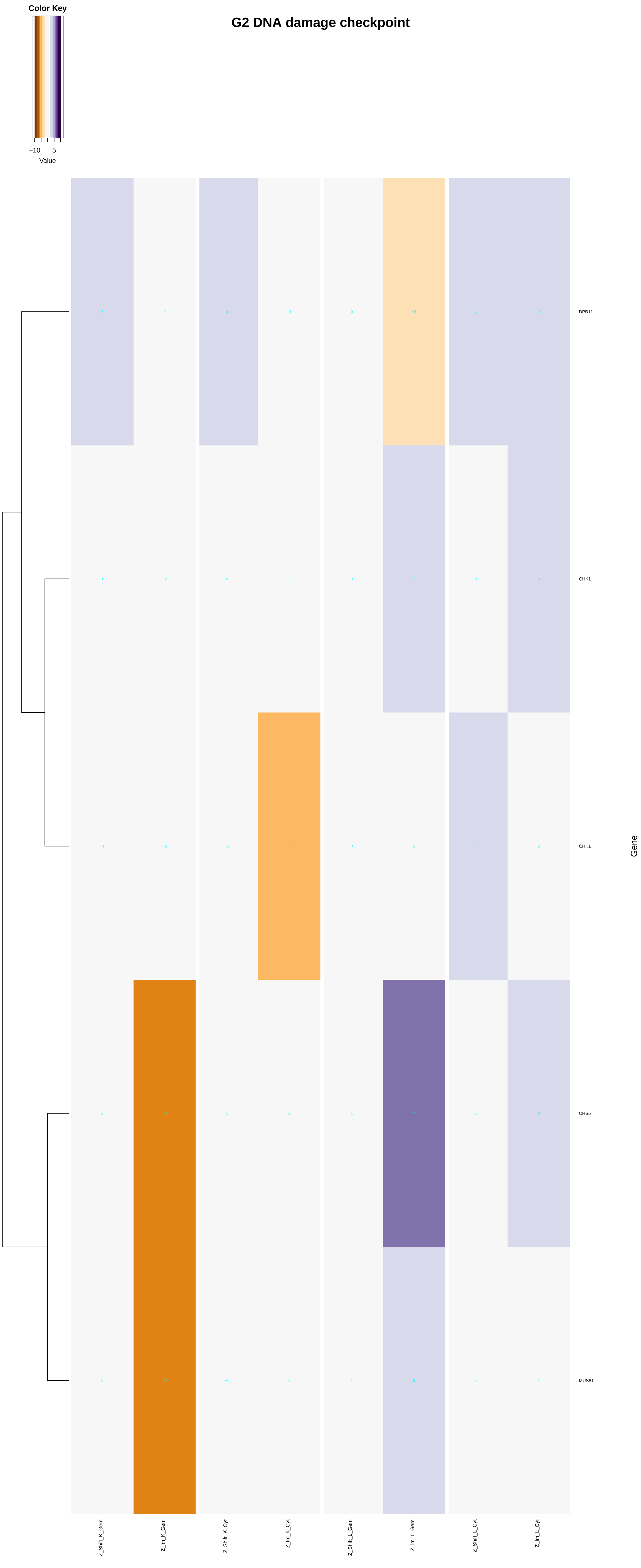

Color Key
−10
5
Value
G2 DNA damage checkpoint
DPB11
CHK1
CHK1
CHS5
MUS81
Gene
Z_Shift_K_Gem
Z_Im_K_Gem
Z_Shift_K_Cyt
Z_Im_K_Cyt
Z_Shift_L_Gem
Z_Im_L_Gem
Z_Shift_L_Cyt
Z_Im_L_Cyt

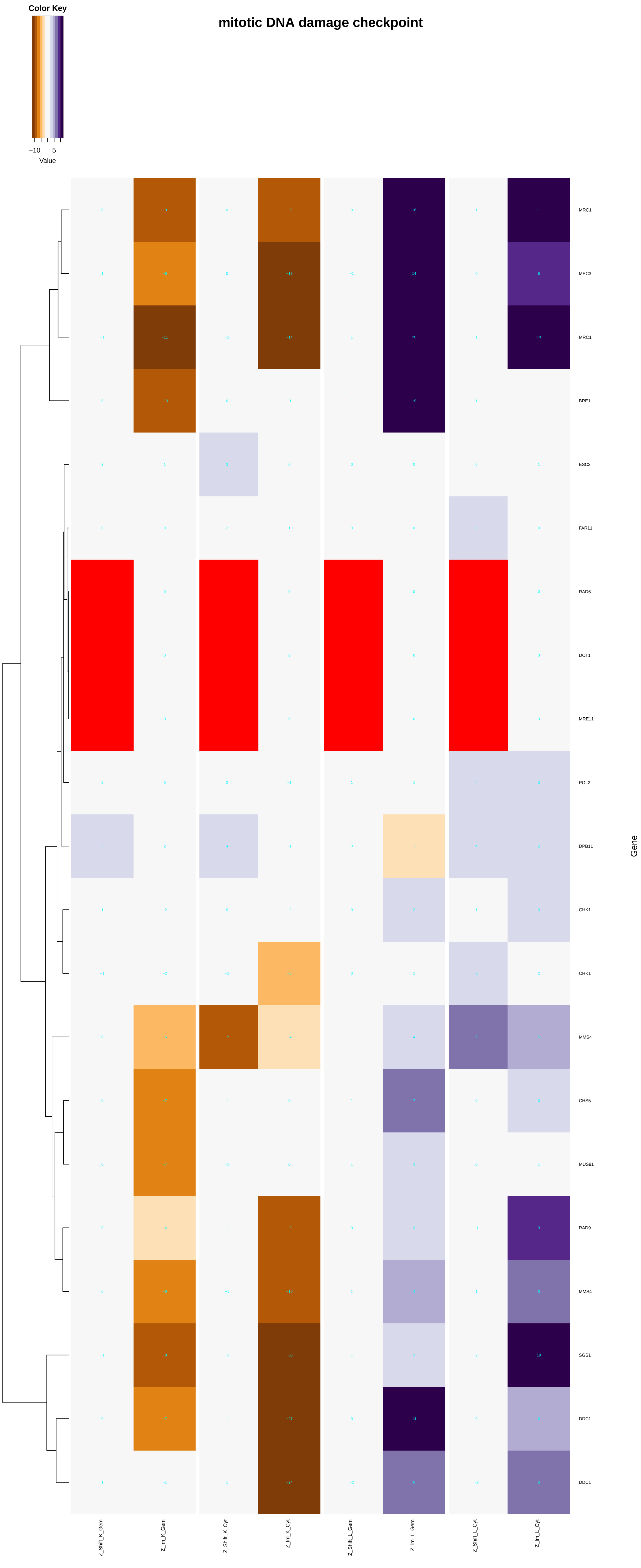

Color Key
−10 5
Value
mitotic DNA damage checkpoint
MRC1
MEC3
MRC1
BRE1
ESC2
FAR11
RAD6
DOT1
MRE11
POL2
DPB11
CHK1
CHK1
MMS4
CHS5
MUS81
RAD9
MMS4
SGS1
DDC1
DDC1
Gene
Z_Shift_K_Gem
Z_Im_K_Gem
Z_Shift_K_Cyt
Z_Im_K_Cyt
Z_Shift_L_Gem
Z_Im_L_Gem
Z_Shift_L_Cyt
Z_Im_L_Cyt

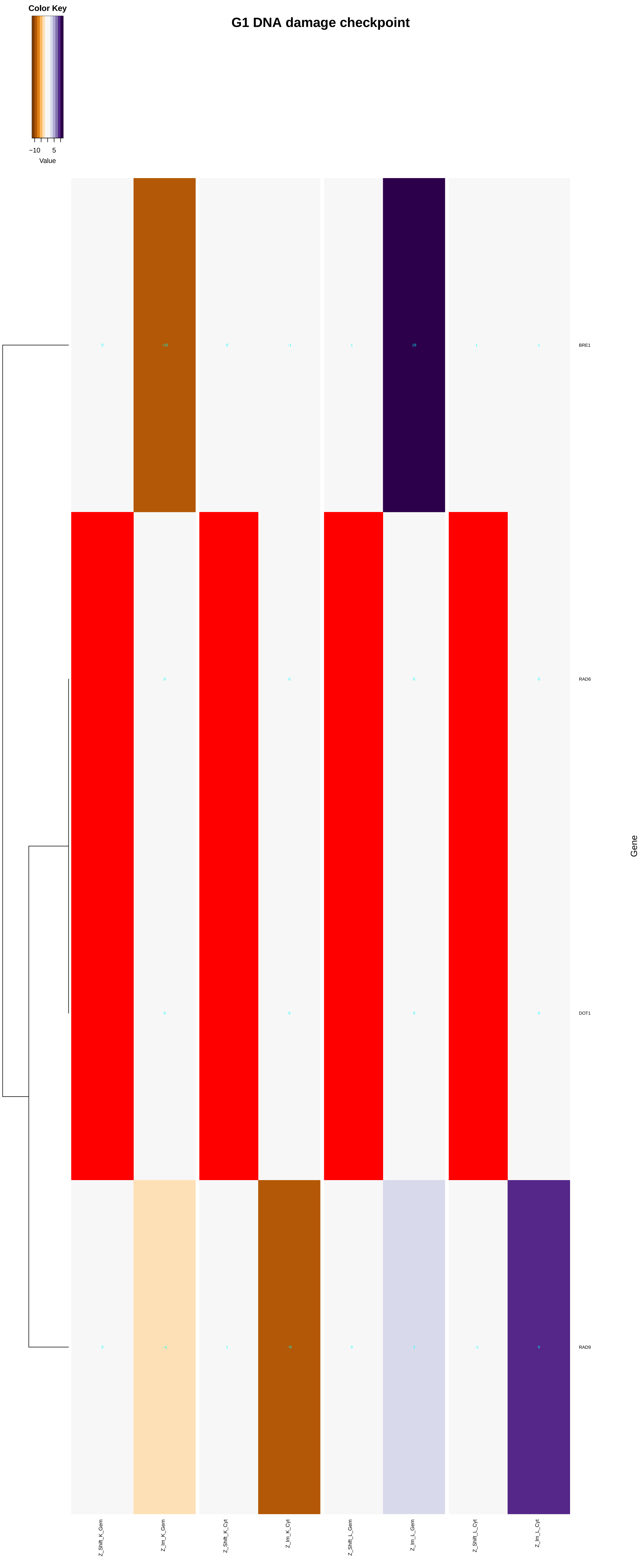
Color Key
−10
5
Value
G1 DNA damage checkpoint
BRE1
RAD6
DOT1
RAD9
Gene
Z_Shift_K_Gem
Z_Im_K_Gem
Z_Shift_K_Cyt
Z_Im_K_Cyt
Z_Shift_L_Gem
Z_Im_L_Gem
Z_Shift_L_Cyt
Z_Im_L_Cyt

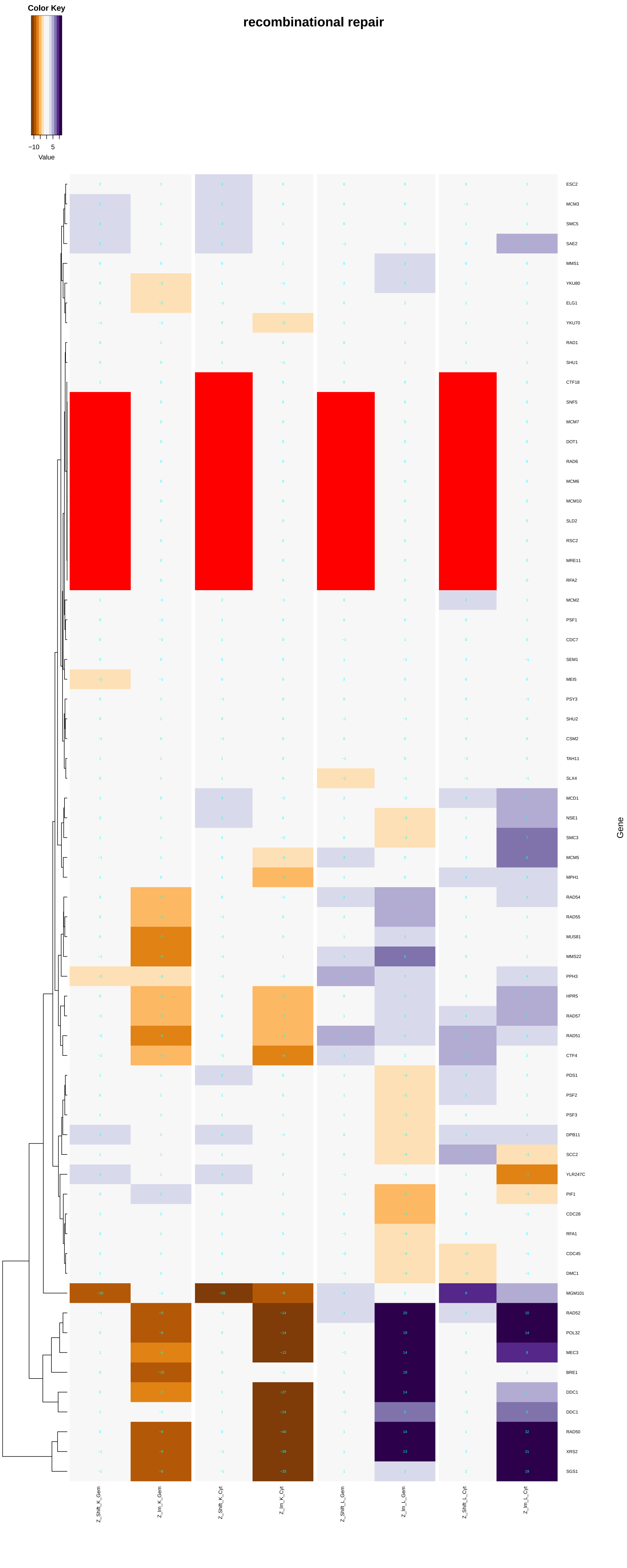

recombinational repair
Color Key
−10
5
Value
ESC2
MCM3
SMC5
SAE2
MMS1
YKU80
ELG1
YKU70
RAD1
SHU1
CTF18
SNF5
MCM7
DOT1
RAD6
MCM6
MCM10
SLD2
RSC2
MRE11
RFA2
MCM2
PSF1
CDC7
SEM1
MEI5
PSY3
SHU2
CSM2
TAH11
SLX4
MCD1
NSE1
SMC3
MCM5
MPH1
RAD54
RAD55
MUS81
MMS22
PPH3
HPR5
RAD57
RAD51
CTF4
PDS1
PSF2
PSF3
DPB11
SCC2
YLR247C
PIF1
CDC28
RFA1
CDC45
DMC1
MGM101
RAD52
POL32
MEC3
BRE1
DDC1
DDC1
RAD50
XRS2
SGS1
Gene
Z_Shift_K_Gem
Z_Im_K_Gem
Z_Shift_K_Cyt
Z_Im_K_Cyt
Z_Shift_L_Gem
Z_Im_L_Gem
Z_Shift_L_Cyt
Z_Im_L_Cyt

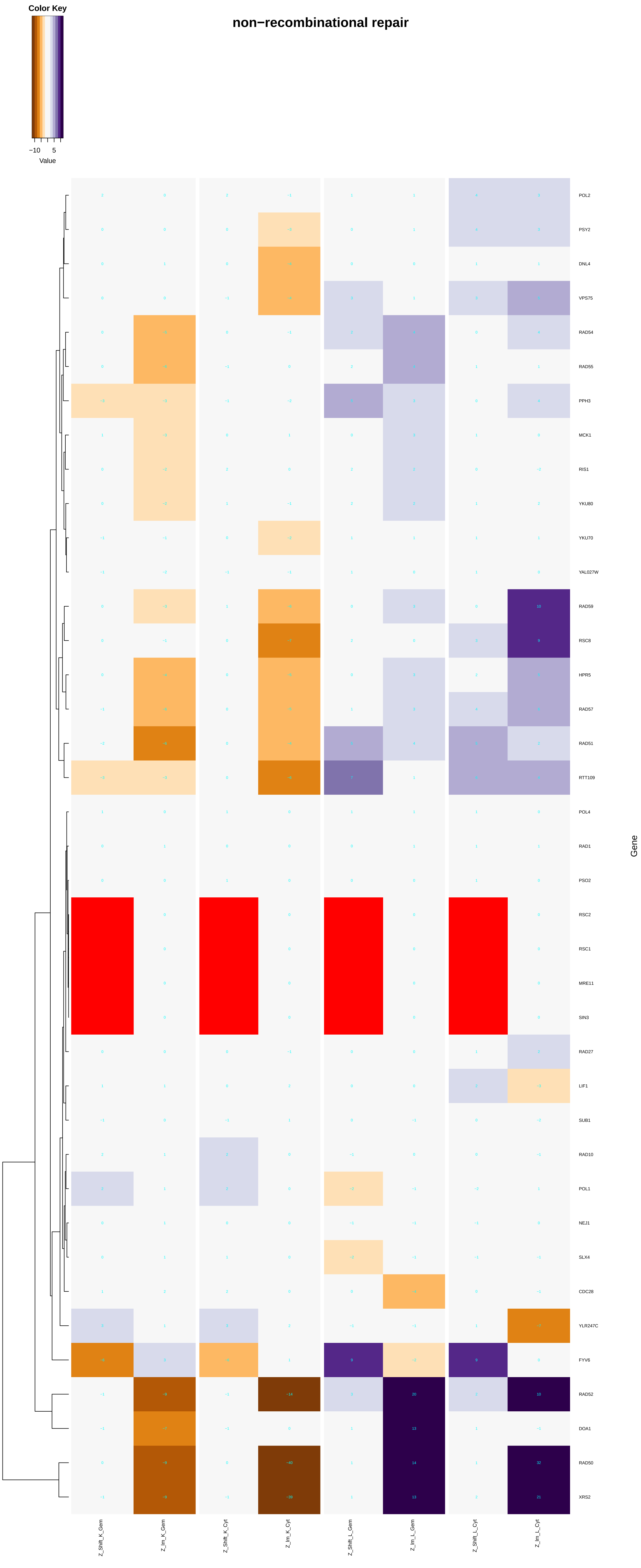

Color Key
−10 5
Value
non−recombinational repair
POL2
PSY2
DNL4
VPS75
RAD54
RAD55
PPH3
MCK1
RIS1
YKU80
YKU70
YAL027W
RAD59
RSC8
HPR5
RAD57
RAD51
RTT109
POL4
RAD1
PSO2
RSC2
RSC1
MRE11
SIN3
RAD27
LIF1
SUB1
RAD10
POL1
NEJ1
SLX4
CDC28
YLR247C
FYV6
RAD52
DOA1
RAD50
XRS2
Gene
Z_Shift_K_Gem
Z_Im_K_Gem
Z_Shift_K_Cyt
Z_Im_K_Cyt
Z_Shift_L_Gem
Z_Im_L_Gem
Z_Shift_L_Cyt
Z_Im_L_Cyt

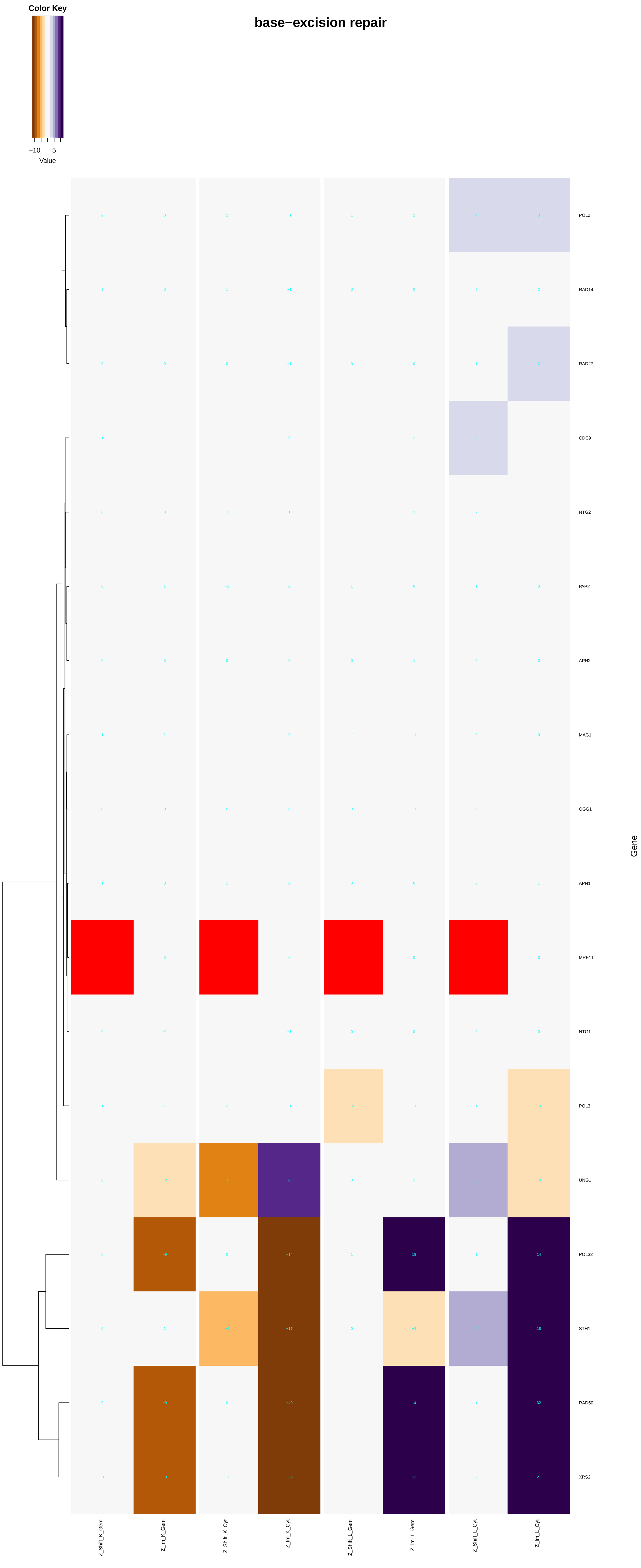
base−excision repair
Color Key
−10
5
Value
POL2
RAD14
RAD27
CDC9
NTG2
PAP2
APN2
MAG1
OGG1
APN1
MRE11
NTG1
POL3
UNG1
POL32
STH1
RAD50
XRS2
Gene
Z_Shift_K_Gem
Z_Im_K_Gem
Z_Shift_K_Cyt
Z_Im_K_Cyt
Z_Shift_L_Gem
Z_Im_L_Gem
Z_Shift_L_Cyt
Z_Im_L_Cyt

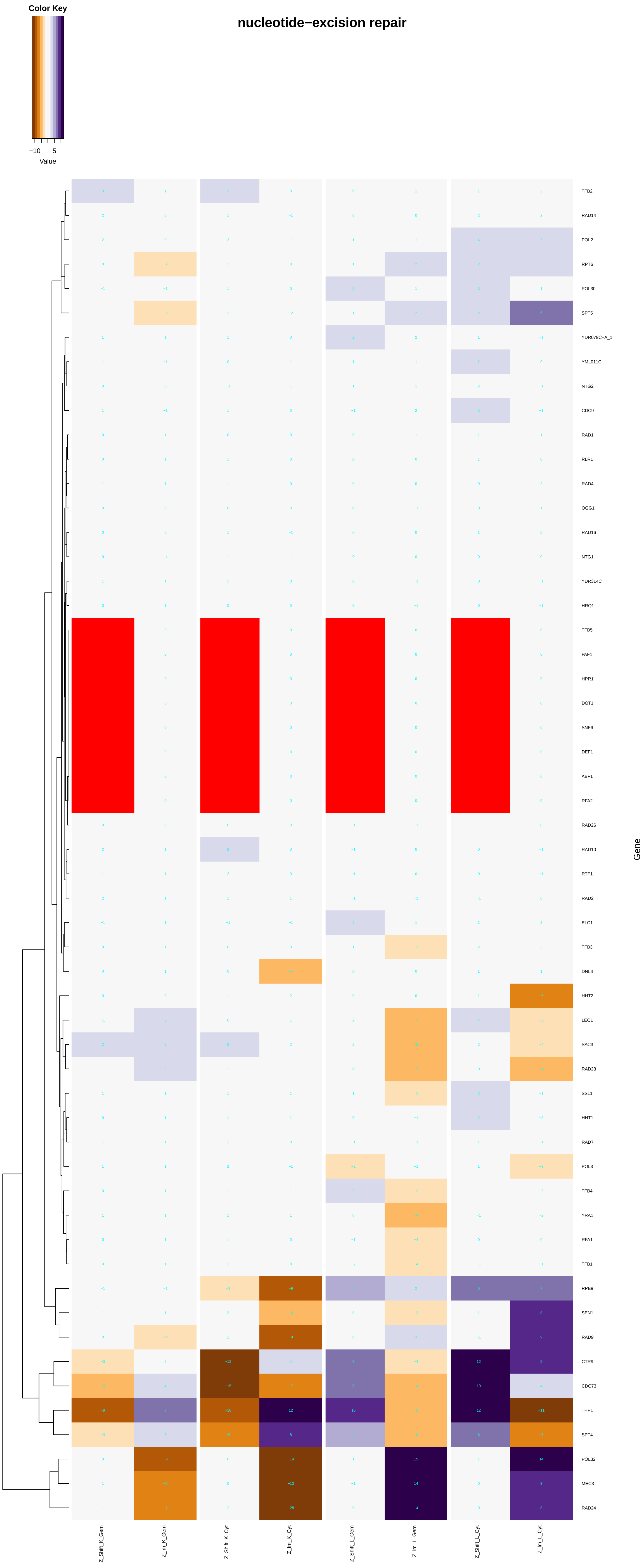

Color Key
−10
5
Value
nucleotide−excision repair
Gene
TFB2
RAD14
POL2
RPT6
POL30
SPT5
YDR079C−A_1
YML011C
NTG2
CDC9
RAD1
RLR1
RAD4
OGG1
RAD16
NTG1
YDR314C
HRQ1
TFB5
PAF1
HPR1
DOT1
SNF6
DEF1
ABF1
RFA2
RAD26
RAD10
RTF1
RAD2
ELC1
TFB3
DNL4
HHT2
LEO1
SAC3
RAD23
SSL1
HHT1
RAD7
POL3
TFB4
YRA1
RFA1
TFB1
RPB9
SEN1
RAD9
CTR9
CDC73
THP1
SPT4
POL32
MEC3
RAD24
Z_Shift_K_Gem
Z_Im_K_Gem
Z_Shift_K_Cyt
Z_Im_K_Cyt
Z_Shift_L_Gem
Z_Im_L_Gem
Z_Shift_L_Cyt
Z_Im_L_Cyt

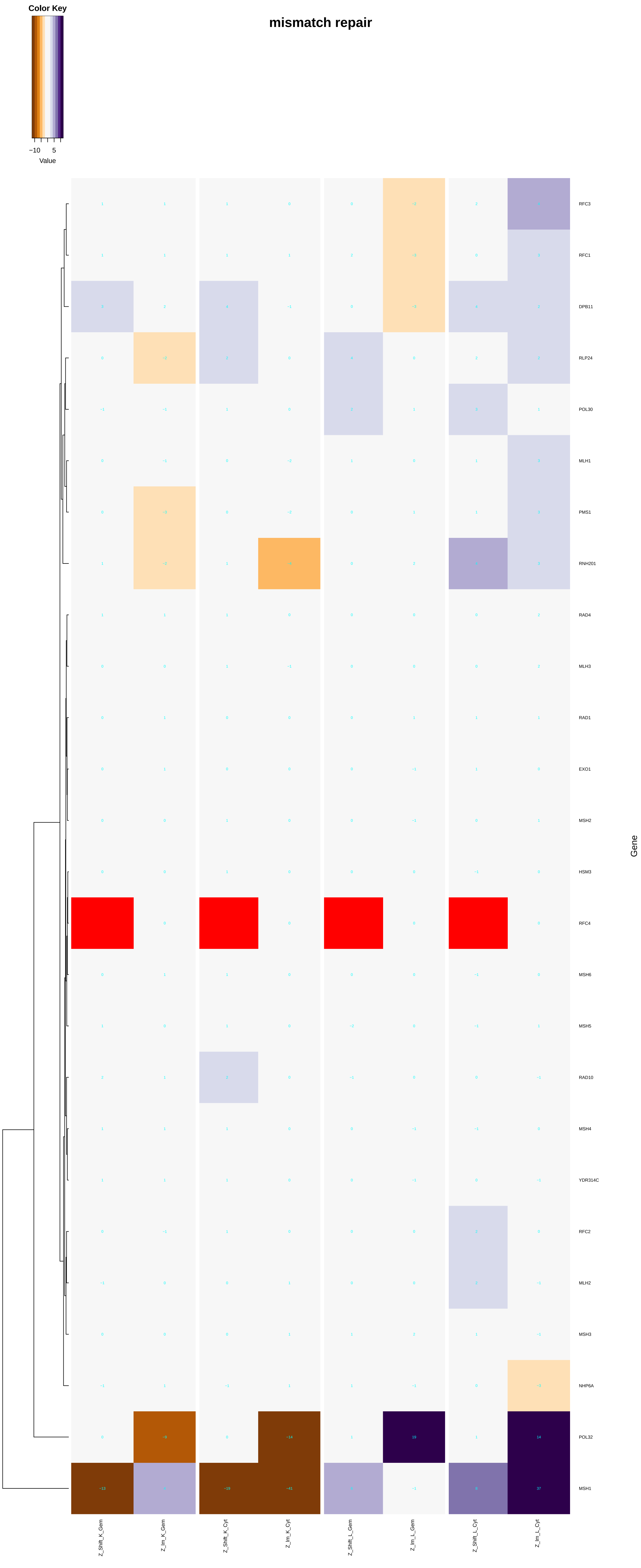

Color Key
−10
5
Value
mismatch repair
RFC3
RFC1
DPB11
RLP24
POL30
MLH1
PMS1
RNH201
RAD4
MLH3
RAD1
EXO1
MSH2
HSM3
RFC4
MSH6
MSH5
RAD10
MSH4
YDR314C
RFC2
MLH2
MSH3
NHP6A
POL32
MSH1
Gene
Z_Shift_K_Gem
Z_Im_K_Gem
Z_Shift_K_Cyt
Z_Im_K_Cyt
Z_Shift_L_Gem
Z_Im_L_Gem
Z_Shift_L_Cyt
Z_Im_L_Cyt

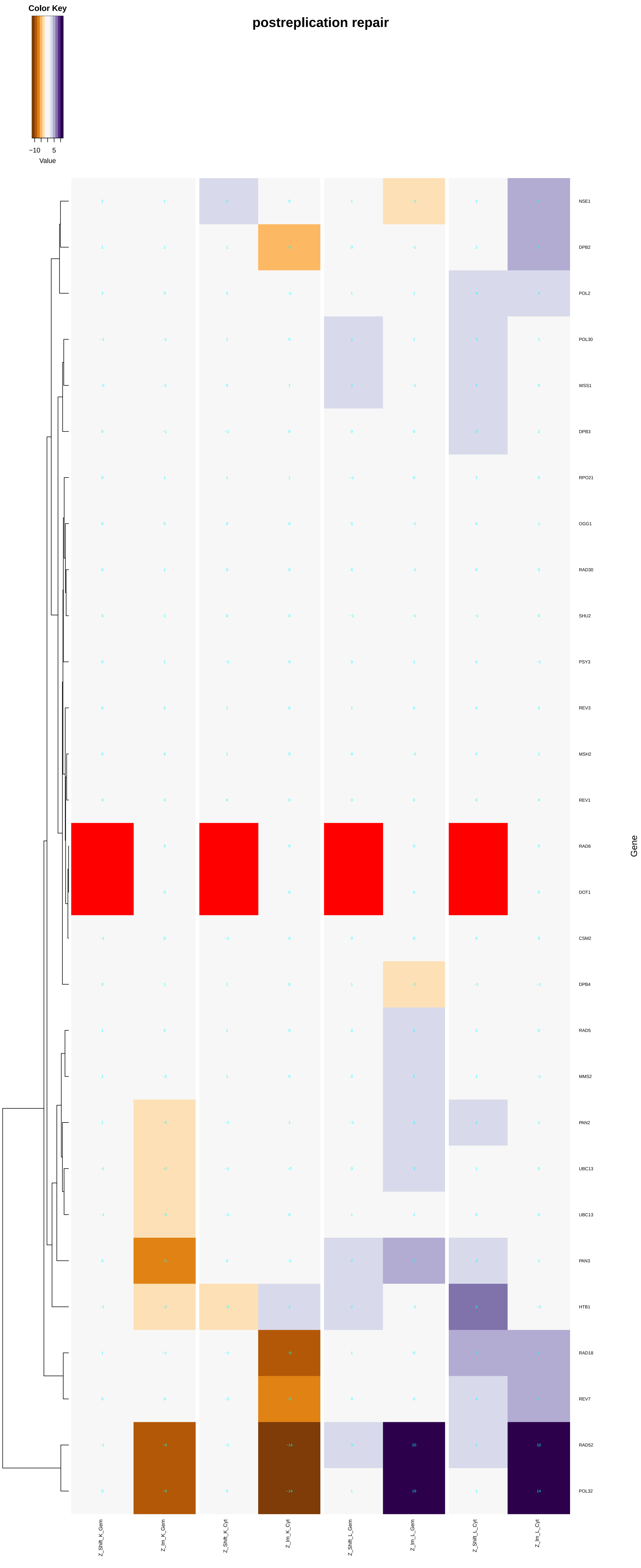

Color Key
−10
5
Value
postreplication repair
NSE1
DPB2
POL2
POL30
WSS1
DPB3
RPO21
OGG1
RAD30
SHU2
PSY3
REV3
MSH2
REV1
RAD6
DOT1
CSM2
DPB4
RAD5
MMS2
PAN2
UBC13
UBC13
PAN3
HTB1
RAD18
REV7
RAD52
POL32
Gene
Z_Shift_K_Gem
Z_Im_K_Gem
Z_Shift_K_Cyt
Z_Im_K_Cyt
Z_Shift_L_Gem
Z_Im_L_Gem
Z_Shift_L_Cyt
Z_Im_L_Cyt

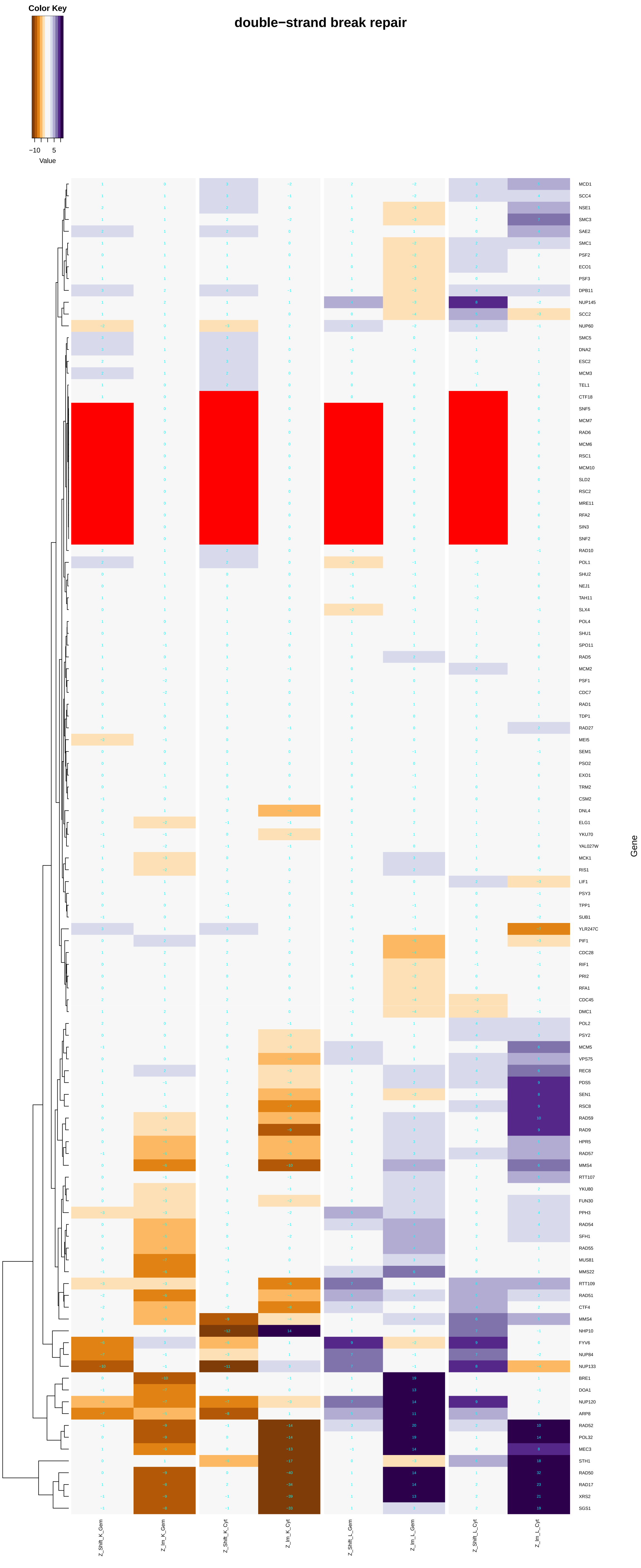
double−strand break repair
Color Key
−10
5
Value
MCD1
SCC4
NSE1
SMC3
SAE2
SMC1
PSF2
ECO1
PSF3
DPB11
NUP145
SCC2
NUP60
SMC5
DNA2
ESC2
MCM3
TEL1
CTF18
SNF5
MCM7
RAD6
MCM6
RSC1
MCM10
SLD2
RSC2
MRE11
RFA2
SIN3
SNF2
RAD10
POL1
SHU2
NEJ1
TAH11
SLX4
POL4
SHU1
SPO11
RAD5
MCM2
PSF1
CDC7
RAD1
TDP1
RAD27
MEI5
SEM1
PSO2
EXO1
TRM2
CSM2
DNL4
ELG1
YKU70
YAL027W
MCK1
RIS1
LIF1
PSY3
TPP1
SUB1
YLR247C
PIF1
CDC28
RIF1
PRI2
RFA1
CDC45
DMC1
POL2
PSY2
MCM5
VPS75
REC8
PDS5
SEN1
RSC8
RAD59
RAD9
HPR5
RAD57
MMS4
RTT107
YKU80
FUN30
PPH3
RAD54
SFH1
RAD55
MUS81
MMS22
RTT109
RAD51
CTF4
MMS4
NHP10
FYV6
NUP84
NUP133
BRE1
DOA1
NUP120
ARP8
RAD52
POL32
MEC3
STH1
RAD50
RAD17
XRS2
SGS1
Gene
Z_Shift_K_Gem
Z_Im_K_Gem
Z_Shift_K_Cyt
Z_Im_K_Cyt
Z_Shift_L_Gem
Z_Im_L_Gem
Z_Shift_L_Cyt
Z_Im_L_Cyt

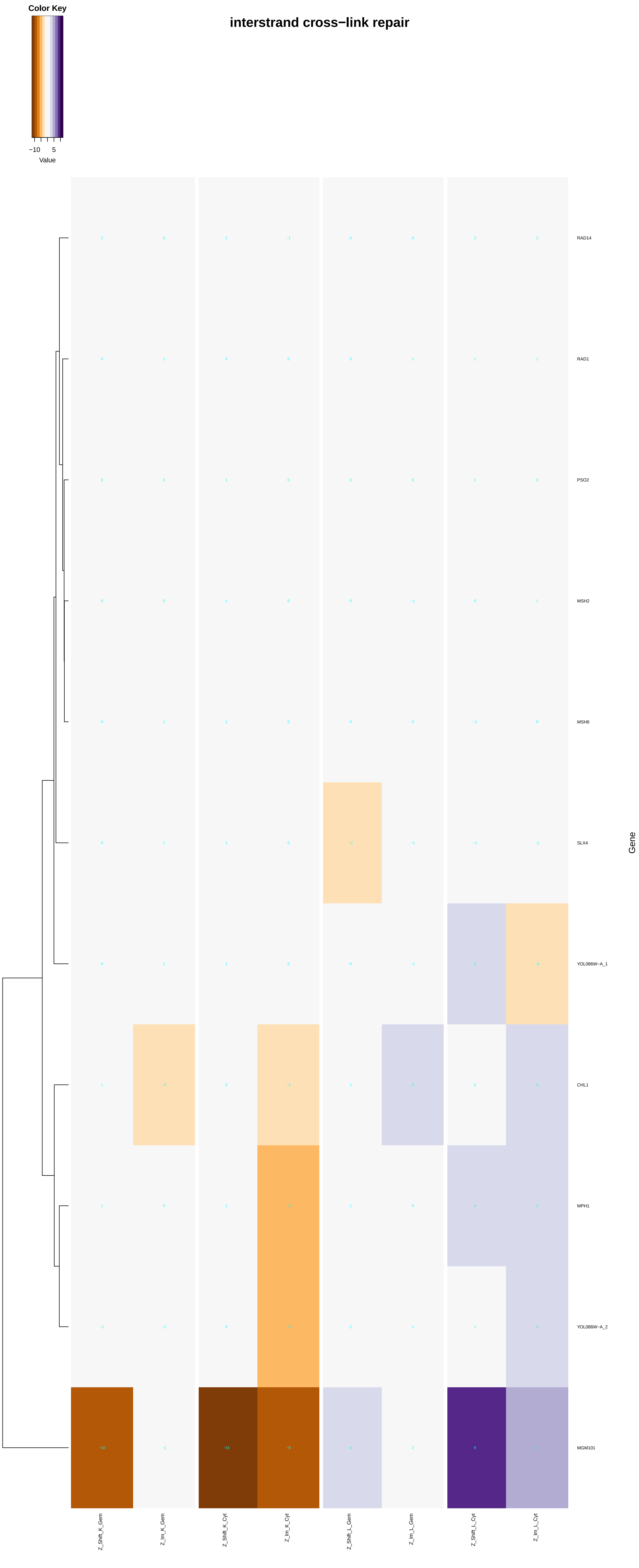
interstrand cross−link repair
Color Key
−10
5
Value
RAD14
RAD1
PSO2
MSH2
MSH6
SLX4
YOL086W−A_1
CHL1
MPH1
YOL086W−A_2
MGM101
Gene
Z_Shift_K_Gem
Z_Im_K_Gem
Z_Shift_K_Cyt
Z_Im_K_Cyt
Z_Shift_L_Gem
Z_Im_L_Gem
Z_Shift_L_Cyt
Z_Im_L_Cyt

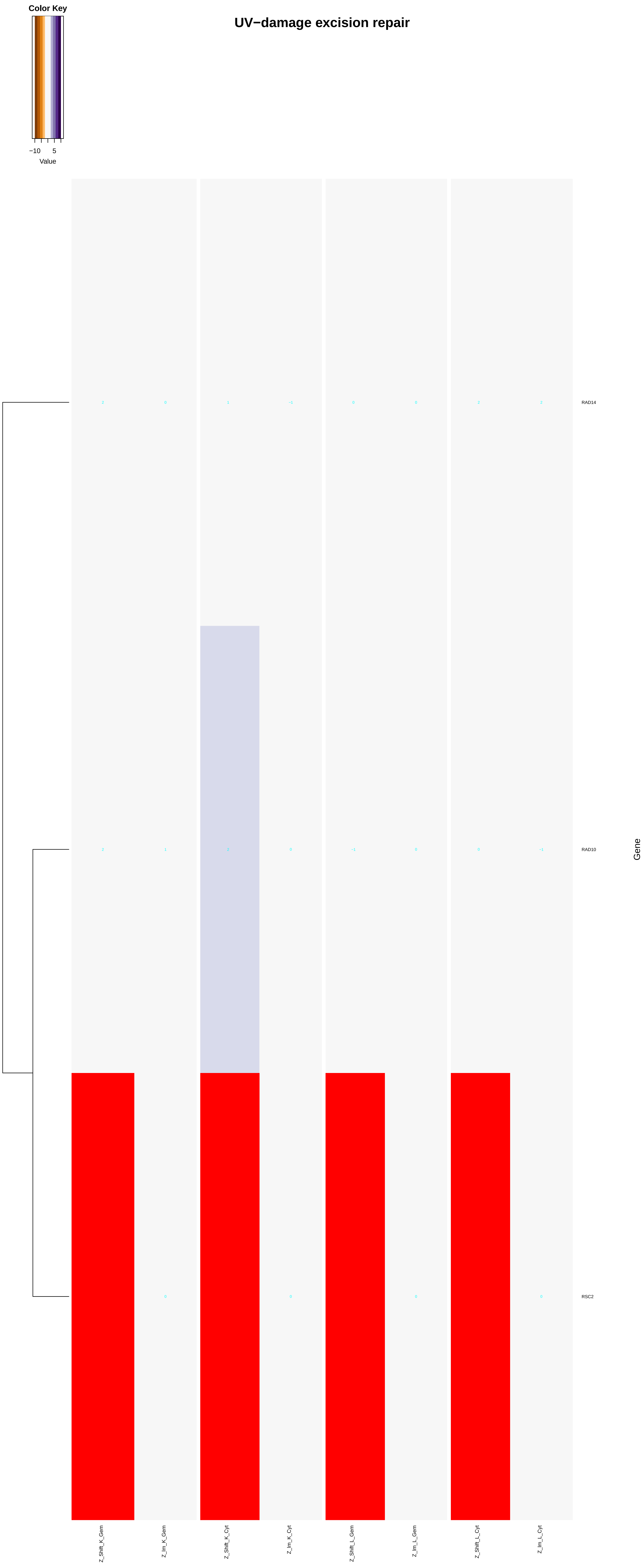
Color Key
−10
5
Value
UV−damage excision repair
2
0
1
−1
0
0
2
2
RAD14
2
1
2
0
−1
0
0
−1
RAD10
0
0
0
0
RSC2
Gene
Z_Shift_K_Gem
Z_Im_K_Gem
Z_Shift_K_Cyt
Z_Im_K_Cyt
Z_Shift_L_Gem
Z_Im_L_Gem
Z_Shift_L_Cyt
Z_Im_L_Cyt

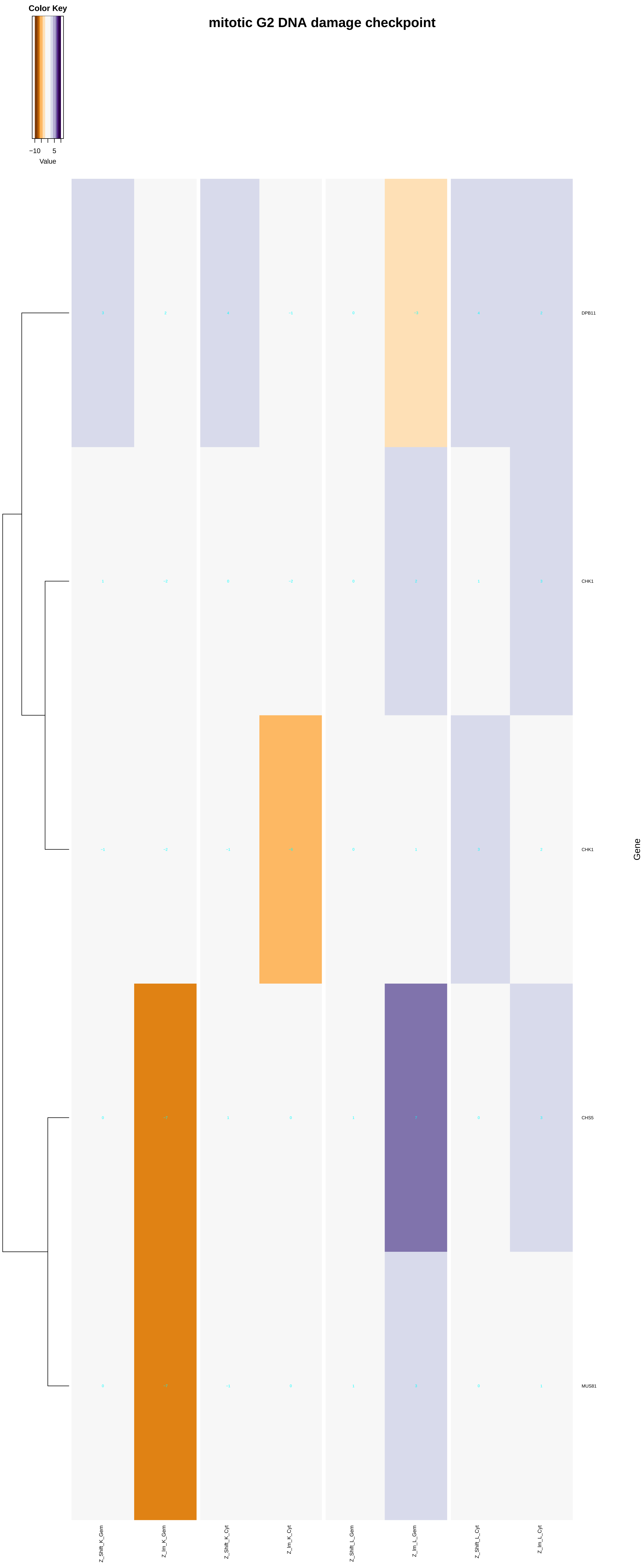
mitotic G2 DNA damage checkpoint
Color Key
−10
5
Value
DPB11
CHK1
CHK1
CHS5
MUS81
Gene
Z_Shift_K_Gem
Z_Im_K_Gem
Z_Shift_K_Cyt
Z_Im_K_Cyt
Z_Shift_L_Gem
Z_Im_L_Gem
Z_Shift_L_Cyt
Z_Im_L_Cyt

### mitotic G1 DNA damage checkpoint

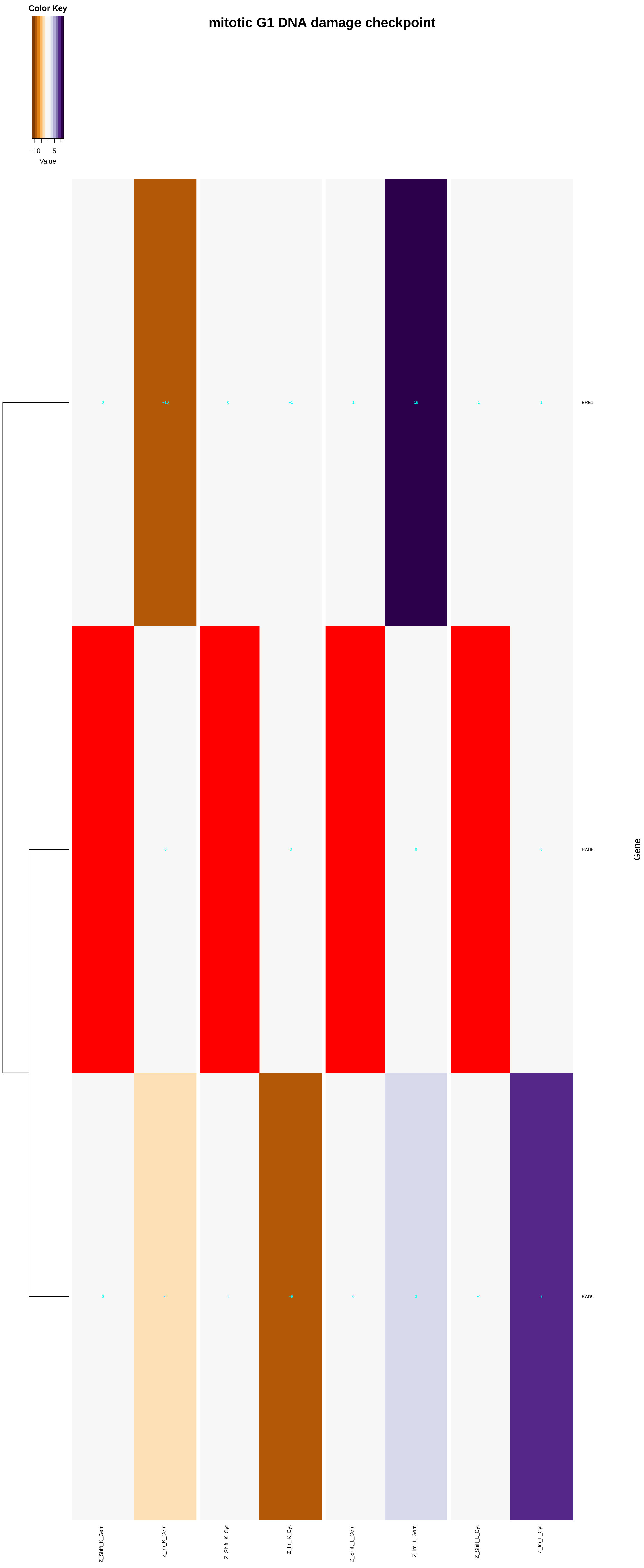

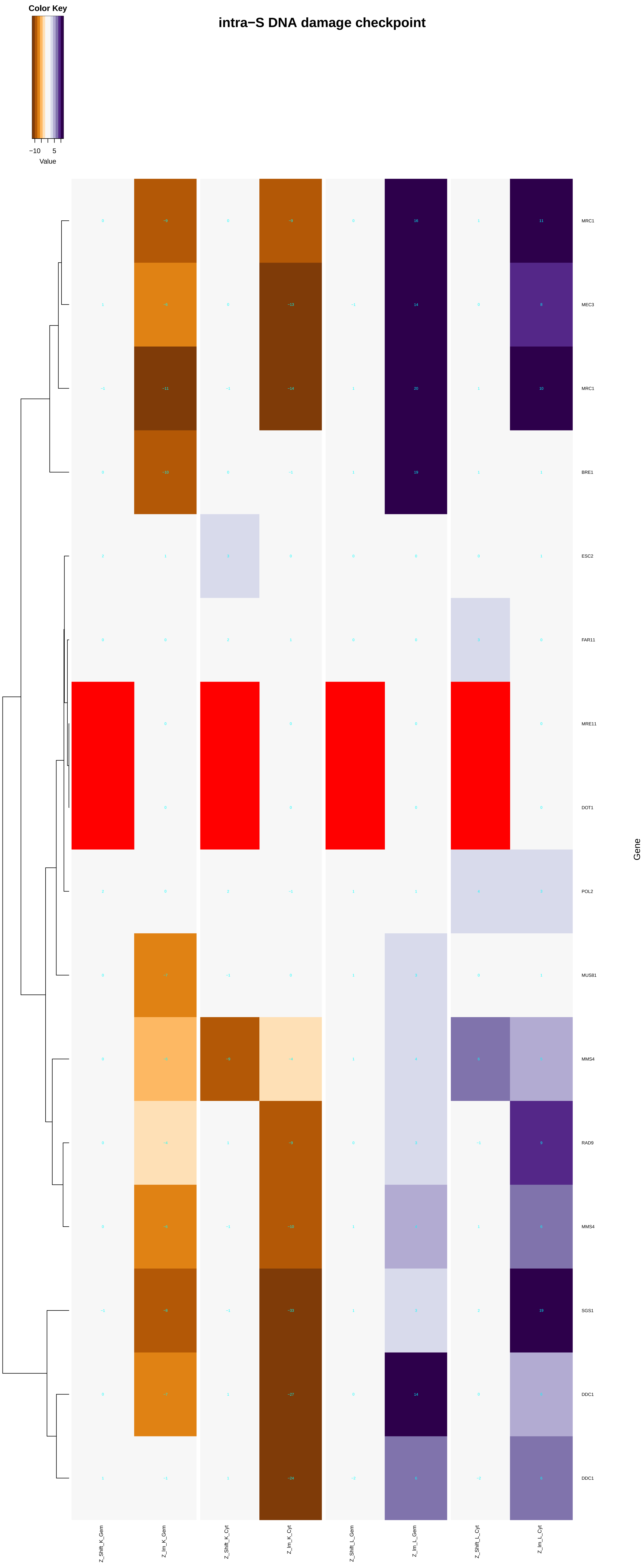

Color Key
−10
5
Value
intra−S DNA damage checkpoint
MRC1
MEC3
MRC1
BRE1
ESC2
FAR11
MRE11
DOT1
POL2
MUS81
MMS4
RAD9
MMS4
SGS1
DDC1
DDC1
Gene
Z_Shift_K_Gem
Z_Im_K_Gem
Z_Shift_K_Cyt
Z_Im_K_Cyt
Z_Shift_L_Gem
Z_Im_L_Gem
Z_Shift_L_Cyt
Z_Im_L_Cyt

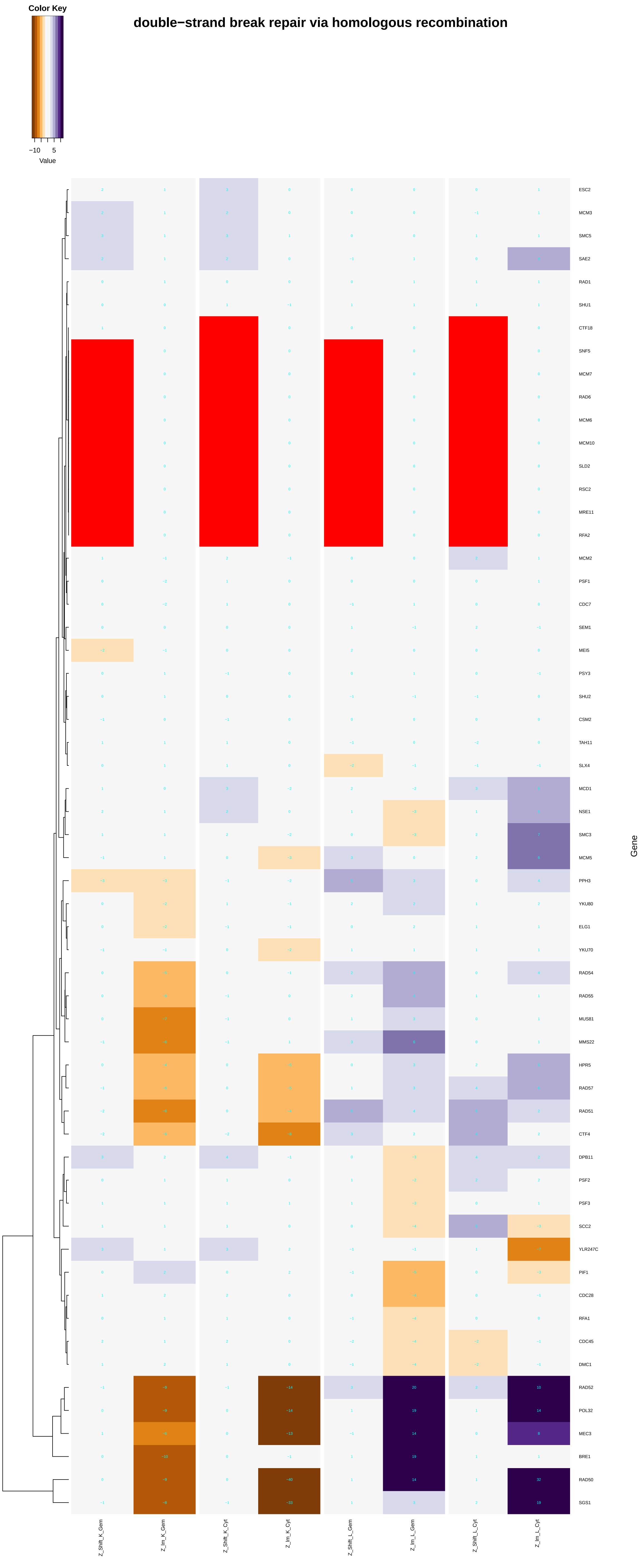
double−strand break repair via homologous recombination
Color Key
−10
5
Value
ESC2
MCM3
SMC5
SAE2
RAD1
SHU1
CTF18
SNF5
MCM7
RAD6
MCM6
MCM10
SLD2
RSC2
MRE11
RFA2
MCM2
PSF1
CDC7
SEM1
MEI5
PSY3
SHU2
CSM2
TAH11
SLX4
MCD1
NSE1
SMC3
MCM5
PPH3
YKU80
ELG1
YKU70
RAD54
RAD55
MUS81
MMS22
HPR5
RAD57
RAD51
CTF4
DPB11
PSF2
PSF3
SCC2
YLR247C
PIF1
CDC28
RFA1
CDC45
DMC1
RAD52
POL32
MEC3
BRE1
RAD50
SGS1
Gene
Z_Shift_K_Gem
Z_Im_K_Gem
Z_Shift_K_Cyt
Z_Im_K_Cyt
Z_Shift_L_Gem
Z_Im_L_Gem
Z_Shift_L_Cyt
Z_Im_L_Cyt

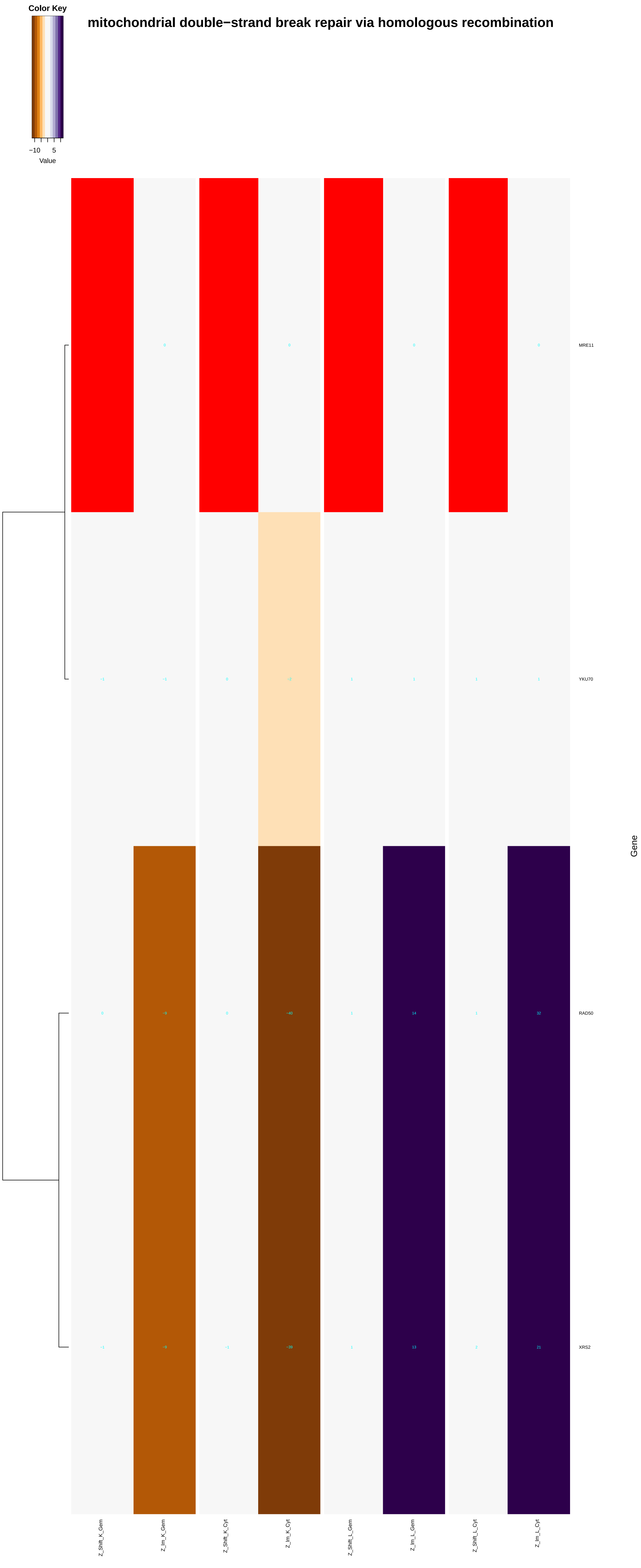
mitochondrial double−strand break repair via homologous recombination
Color Key
−10
5
Value
MRE11
YKU70
RAD50
XRS2
Gene
Z_Shift_K_Gem
Z_Im_K_Gem
Z_Shift_K_Cyt
Z_Im_K_Cyt
Z_Shift_L_Gem
Z_Im_L_Gem
Z_Shift_L_Cyt
Z_Im_L_Cyt

### double−strand break repair via nonhomologous end joining

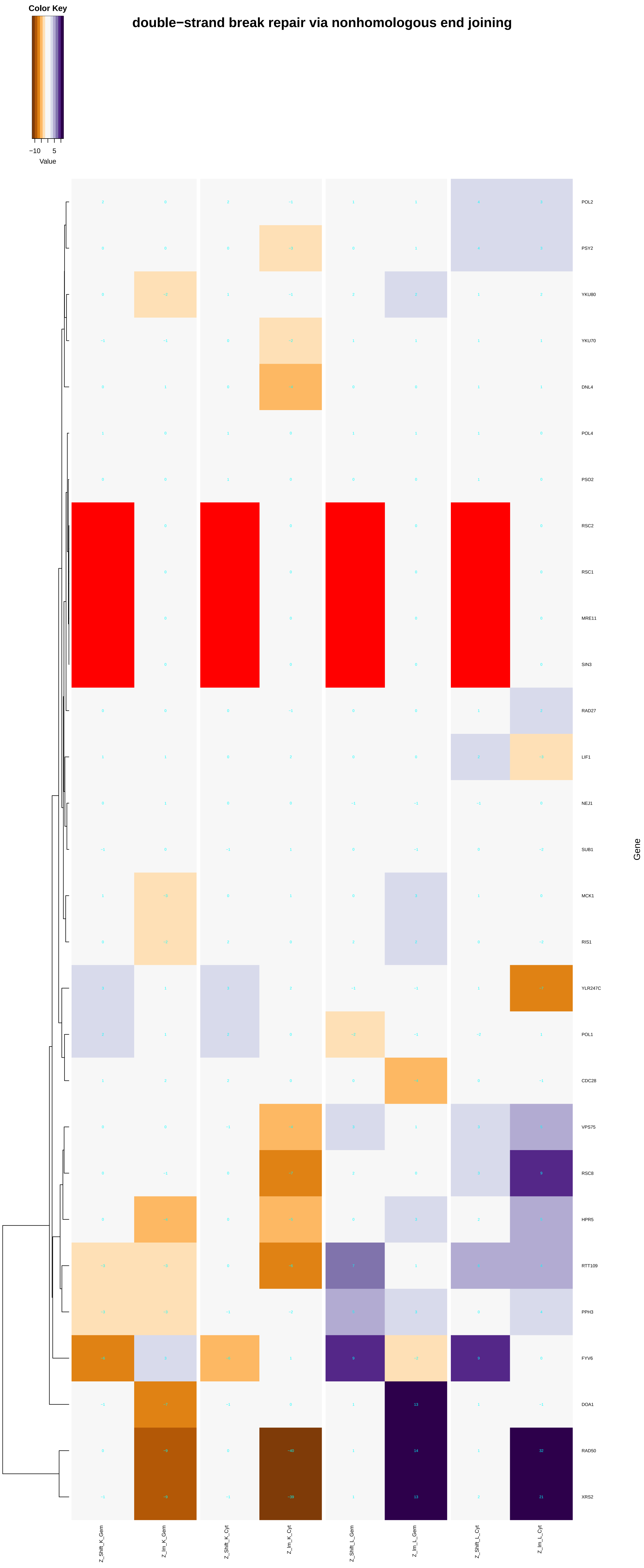

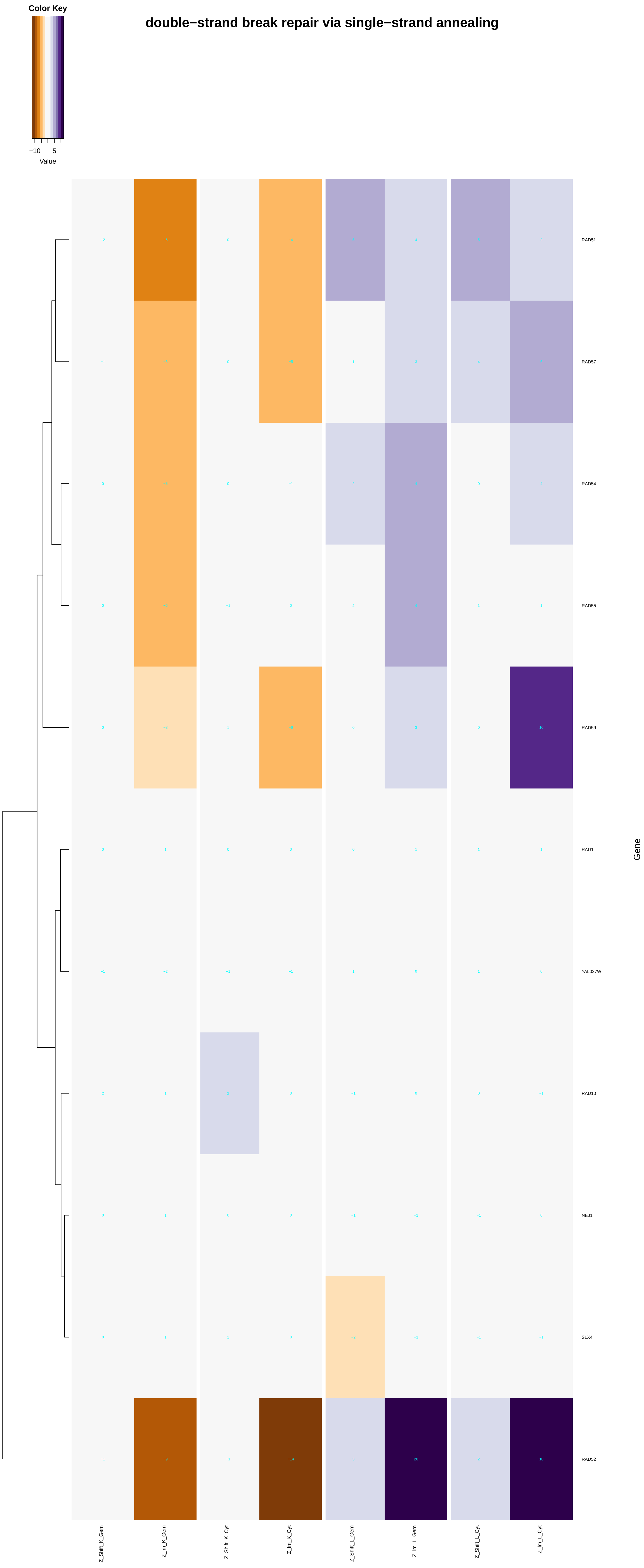

Color Key
−10
5
Value
double−strand break repair via single−strand annealing
RAD51
RAD57
RAD54
RAD55
RAD59
RAD1
YAL027W
RAD10
NEJ1
SLX4
RAD52
Gene
Z_Shift_K_Gem
Z_Im_K_Gem
Z_Shift_K_Cyt
Z_Im_K_Cyt
Z_Shift_L_Gem
Z_Im_L_Gem
Z_Shift_L_Cyt
Z_Im_L_Cyt

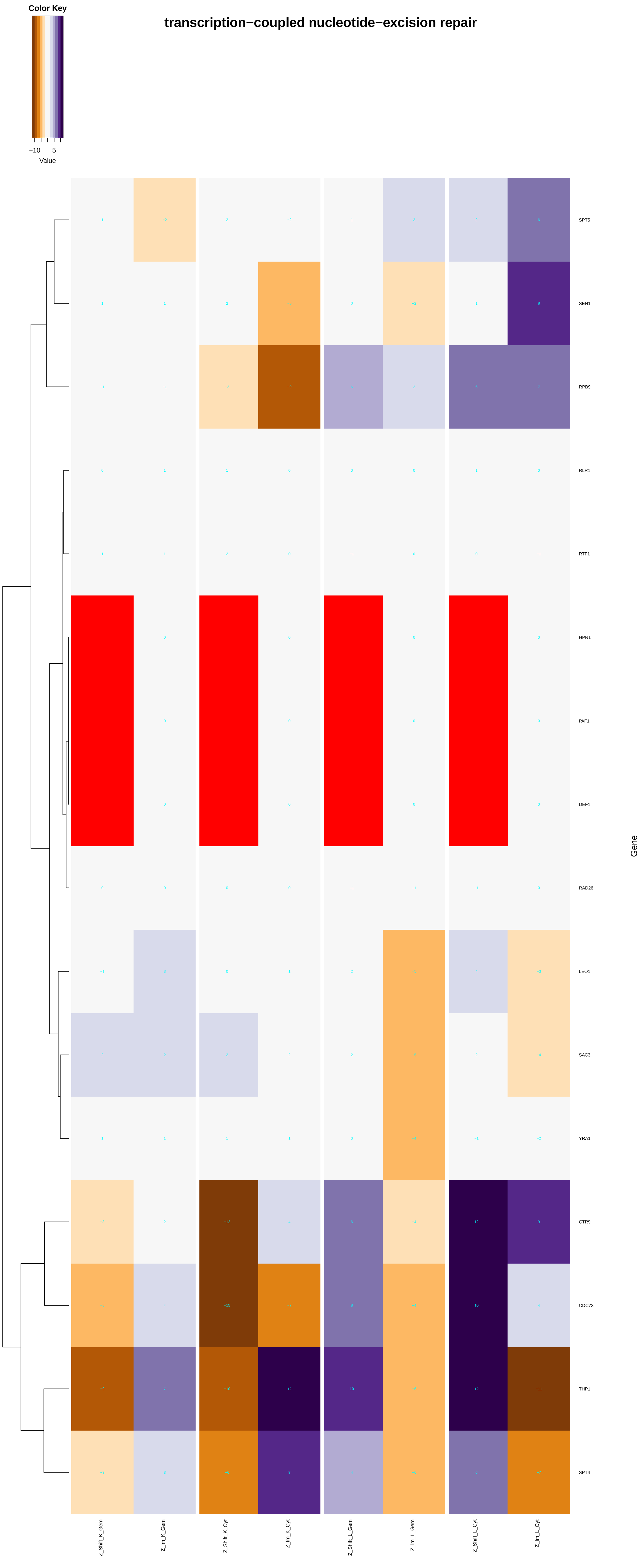
transcription−coupled nucleotide−excision repair
Color Key
−10
5
Value
SPT5
SEN1
RPB9
RLR1
RTF1
HPR1
PAF1
DEF1
RAD26
LEO1
SAC3
YRA1
CTR9
CDC73
THP1
SPT4
Gene
Z_Shift_K_Gem
Z_Im_K_Gem
Z_Shift_K_Cyt
Z_Im_K_Cyt
Z_Shift_L_Gem
Z_Im_L_Gem
Z_Shift_L_Cyt
Z_Im_L_Cyt

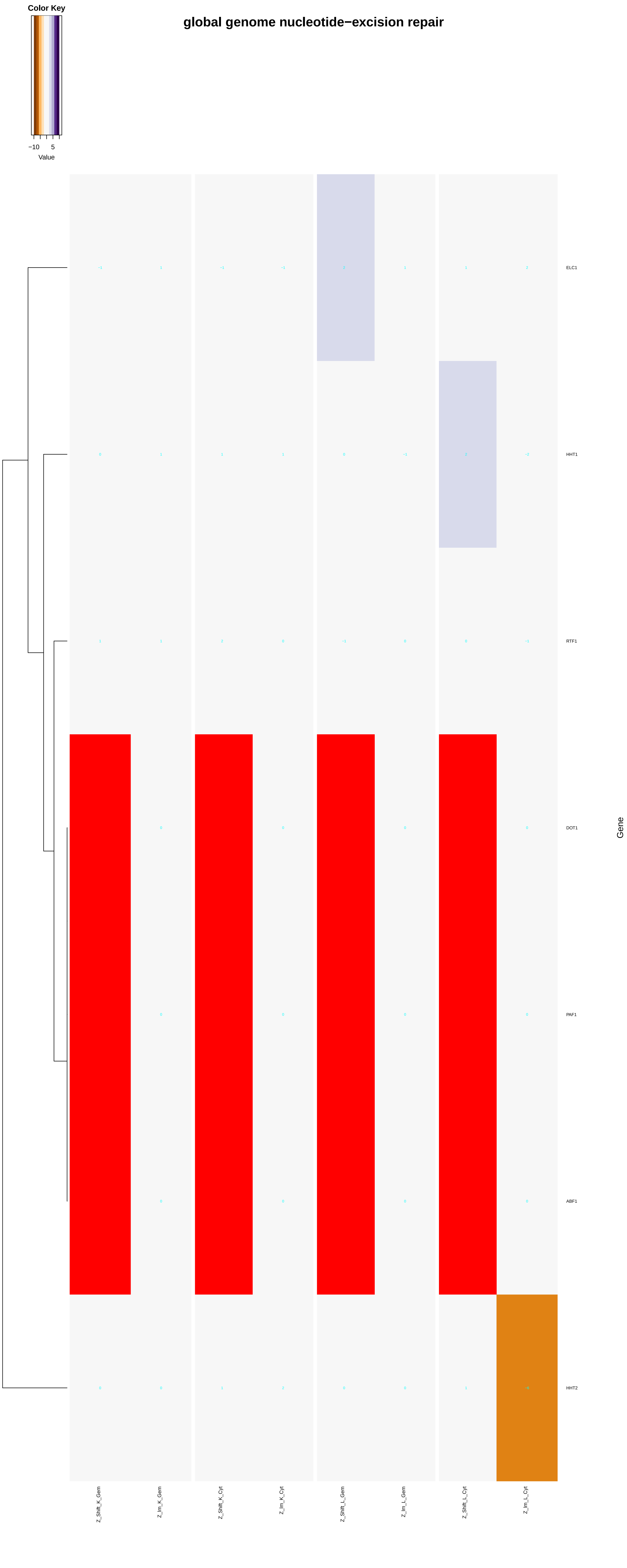

Color Key
−10
5
Value
global genome nucleotide−excision repair
ELC1
HHT1
RTF1
DOT1
PAF1
ABF1
HHT2
Gene
Z_Shift_K_Gem
Z_Im_K_Gem
Z_Shift_K_Cyt
Z_Im_K_Cyt
Z_Shift_L_Gem
Z_Im_L_Gem
Z_Shift_L_Cyt
Z_Im_L_Cyt

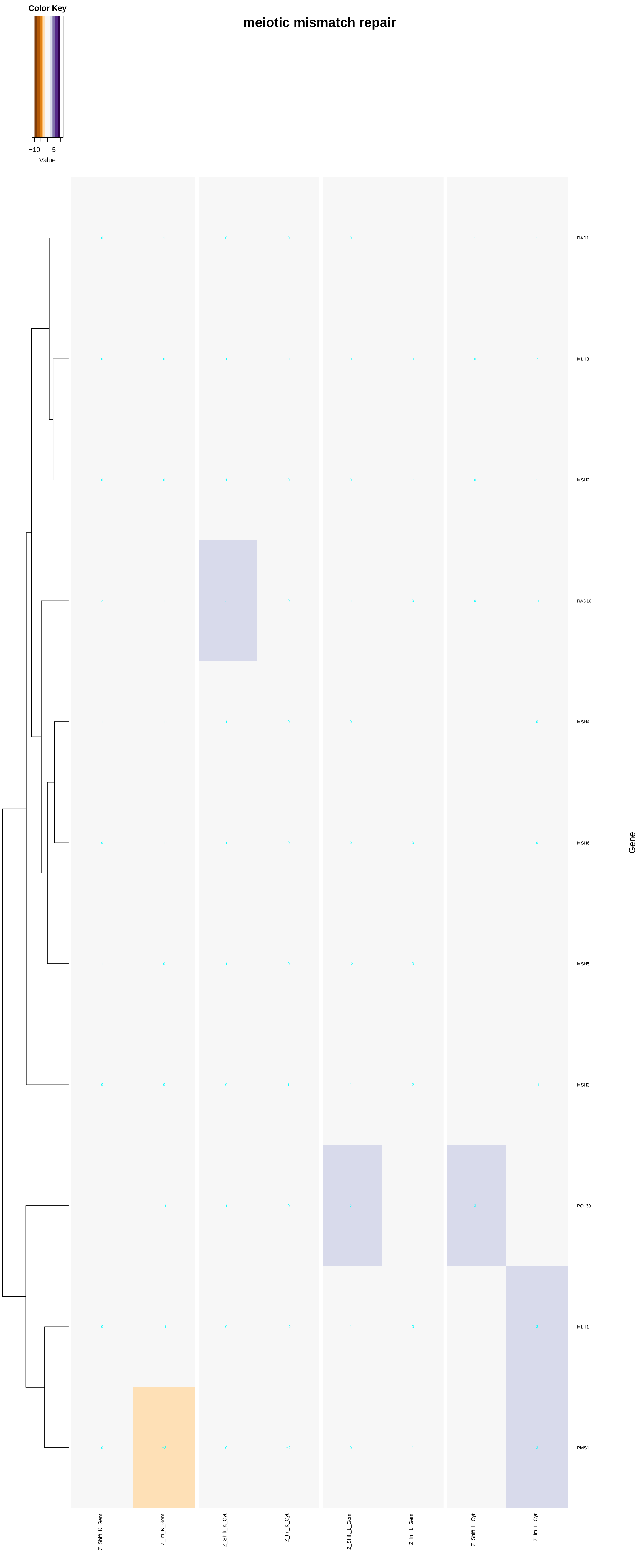
meiotic mismatch repair
Color Key
−10
5
Value
RAD1
MLH3
MSH2
RAD10
MSH4
MSH6
MSH5
MSH3
POL30
MLH1
PMS1
Gene
Z_Shift_K_Gem
Z_Im_K_Gem
Z_Shift_K_Cyt
Z_Im_K_Cyt
Z_Shift_L_Gem
Z_Im_L_Gem
Z_Shift_L_Cyt
Z_Im_L_Cyt

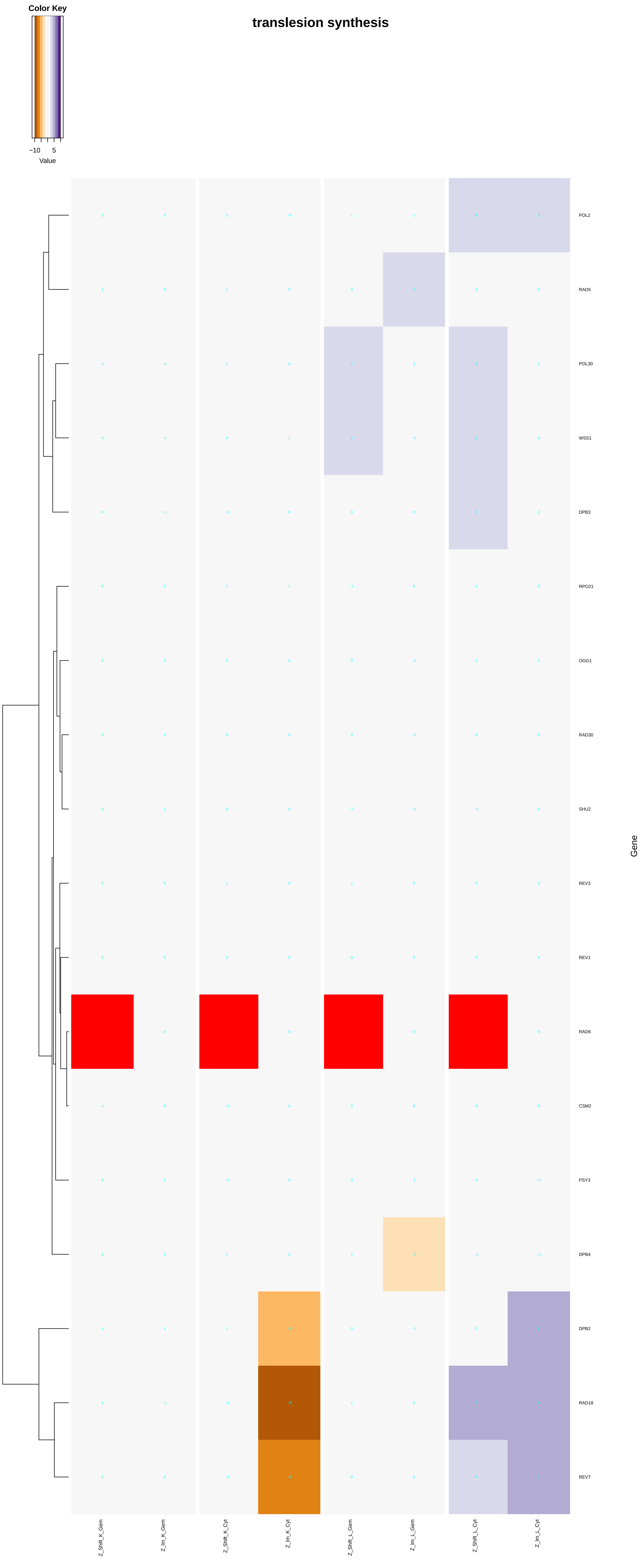
Color Key
−10
5
Value
translesion synthesis
POL2
RAD5
POL30
WSS1
DPB3
RPO21
OGG1
RAD30
SHU2
REV3
REV1
RAD6
CSM2
PSY3
DPB4
DPB2
RAD18
REV7
Gene
Z_Shift_K_Gem
Z_Im_K_Gem
Z_Shift_K_Cyt
Z_Im_K_Cyt
Z_Shift_L_Gem
Z_Im_L_Gem
Z_Shift_L_Cyt
Z_Im_L_Cyt

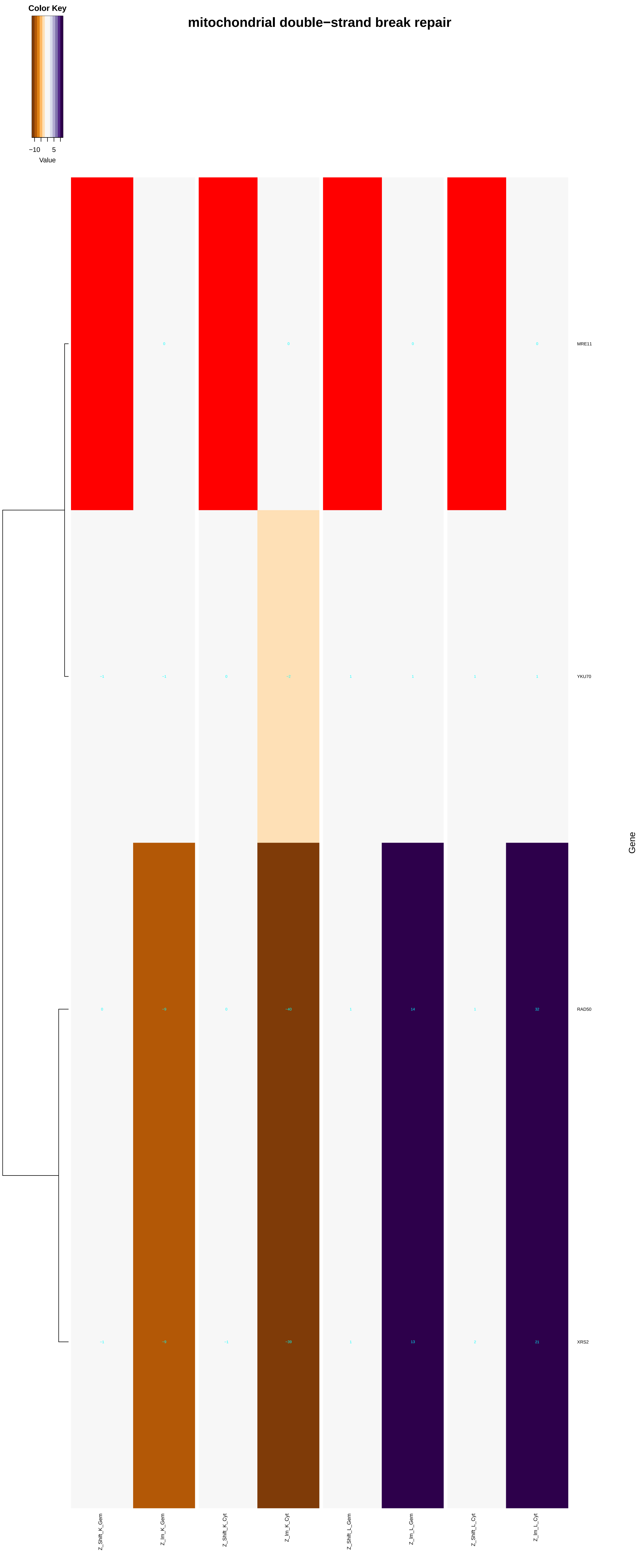

mitochondrial double−strand break repair
Color Key
−10
5
Value
MRE11
YKU70
RAD50
XRS2
Gene
Z_Shift_K_Gem
Z_Im_K_Gem
Z_Shift_K_Cyt
Z_Im_K_Cyt
Z_Shift_L_Gem
Z_Im_L_Gem
Z_Shift_L_Cyt
Z_Im_L_Cyt

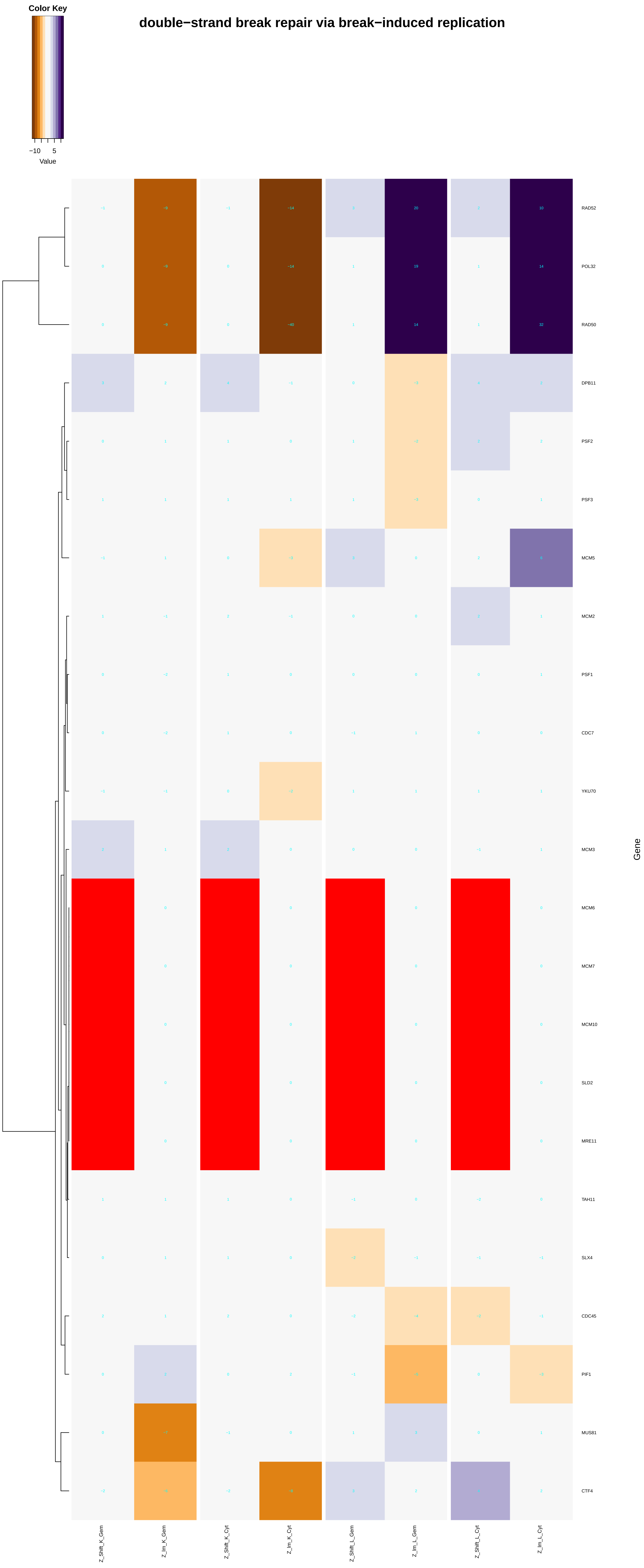

double−strand break repair via break−induced replication
Color Key
−10
5
Value
RAD52
POL32
RAD50
DPB11
PSF2
PSF3
MCM5
MCM2
PSF1
CDC7
YKU70
MCM3
MCM6
MCM7
MCM10
SLD2
MRE11
TAH11
SLX4
CDC45
PIF1
MUS81
CTF4
Gene
Z_Shift_K_Gem
Z_Im_K_Gem
Z_Shift_K_Cyt
Z_Im_K_Cyt
Z_Shift_L_Gem
Z_Im_L_Gem
Z_Shift_L_Cyt
Z_Im_L_Cyt

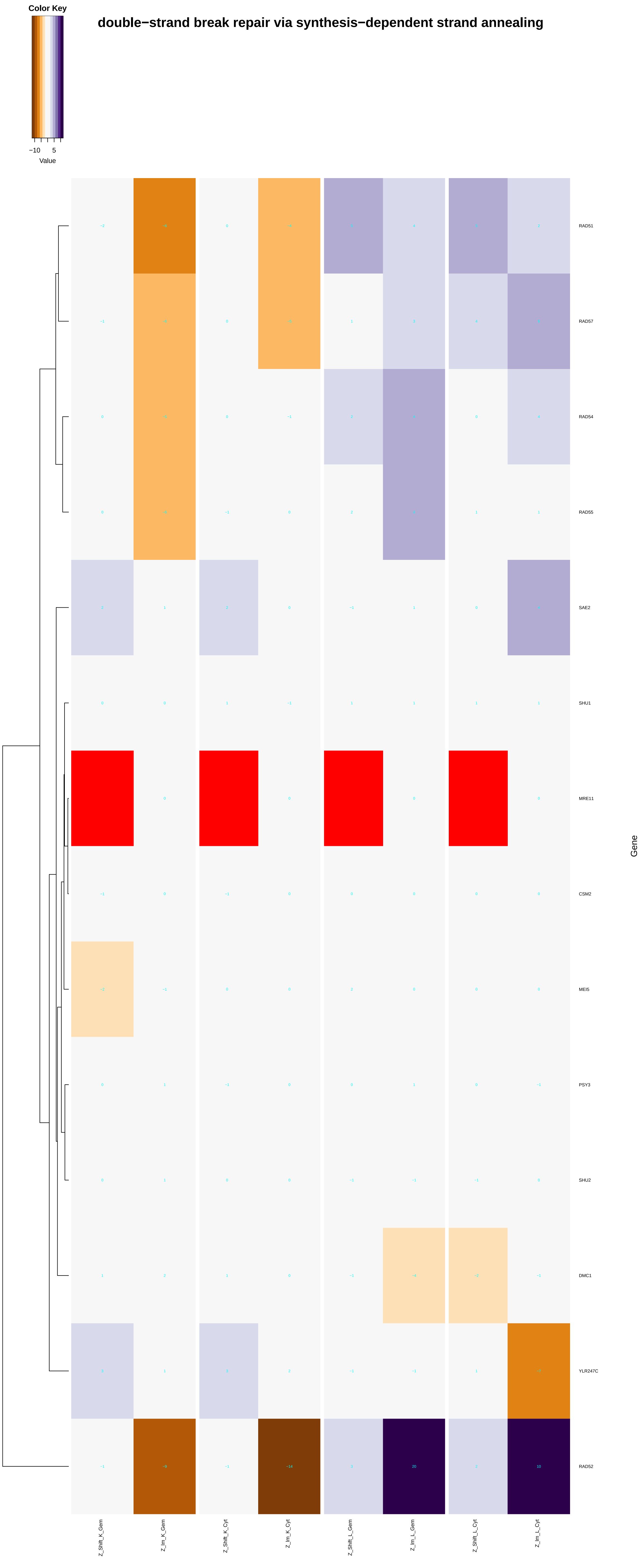

Color Key
−10
5
Value
double−strand break repair via synthesis−dependent strand annealing
RAD51
RAD57
RAD54
RAD55
SAE2
SHU1
MRE11
CSM2
MEI5
PSY3
SHU2
DMC1
YLR247C
RAD52
Gene
Z_Shift_K_Gem
Z_Im_K_Gem
Z_Shift_K_Cyt
Z_Im_K_Cyt
Z_Shift_L_Gem
Z_Im_L_Gem
Z_Shift_L_Cyt
Z_Im_L_Cyt

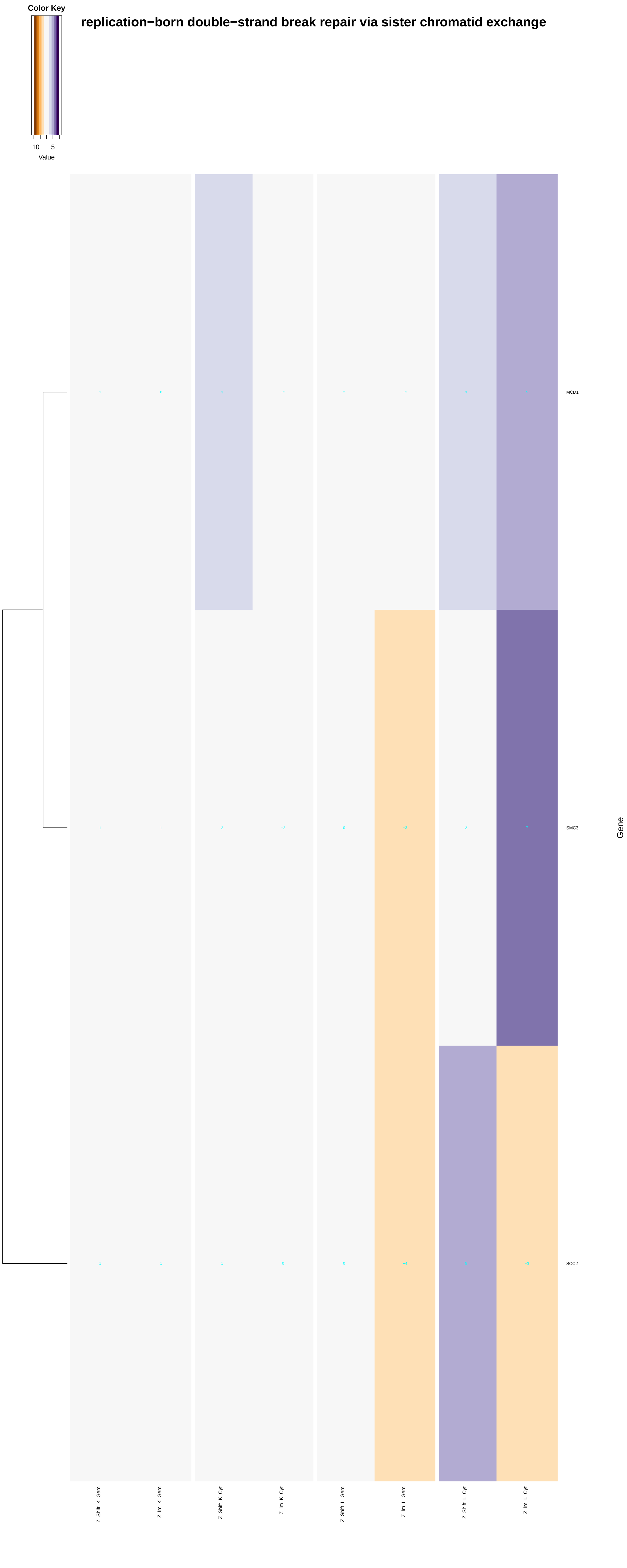

Color Key
−10
5
Value
replication−born double−strand break repair via sister chromatid exchange
MCD1
SMC3
SCC2
Gene
Z_Shift_K_Gem
Z_lm_K_Gem
Z_Shift_K_Cyt
Z_lm_K_Cyt
Z_Shift_L_Gem
Z_lm_L_Gem
Z_Shift_L_Cyt
Z_lm_L_Cyt

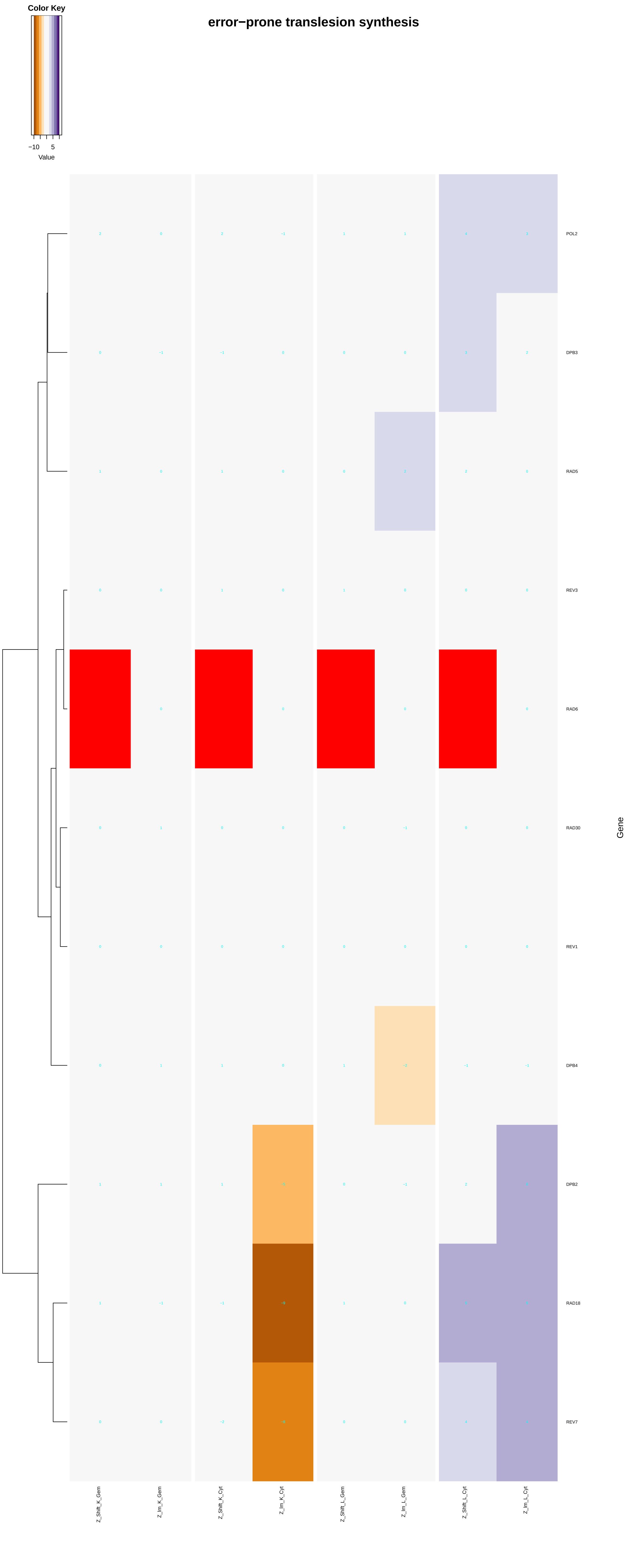
error−prone translesion synthesis
Color Key
−10
5
Value
POL2
DPB3
RAD5
REV3
RAD6
RAD30
REV1
DPB4
DPB2
RAD18
REV7
Gene
Z_Shift_K_Gem
Z_Im_K_Gem
Z_Shift_K_Cyt
Z_Im_K_Cyt
Z_Shift_L_Gem
Z_Im_L_Gem
Z_Shift_L_Cyt
Z_Im_L_Cyt

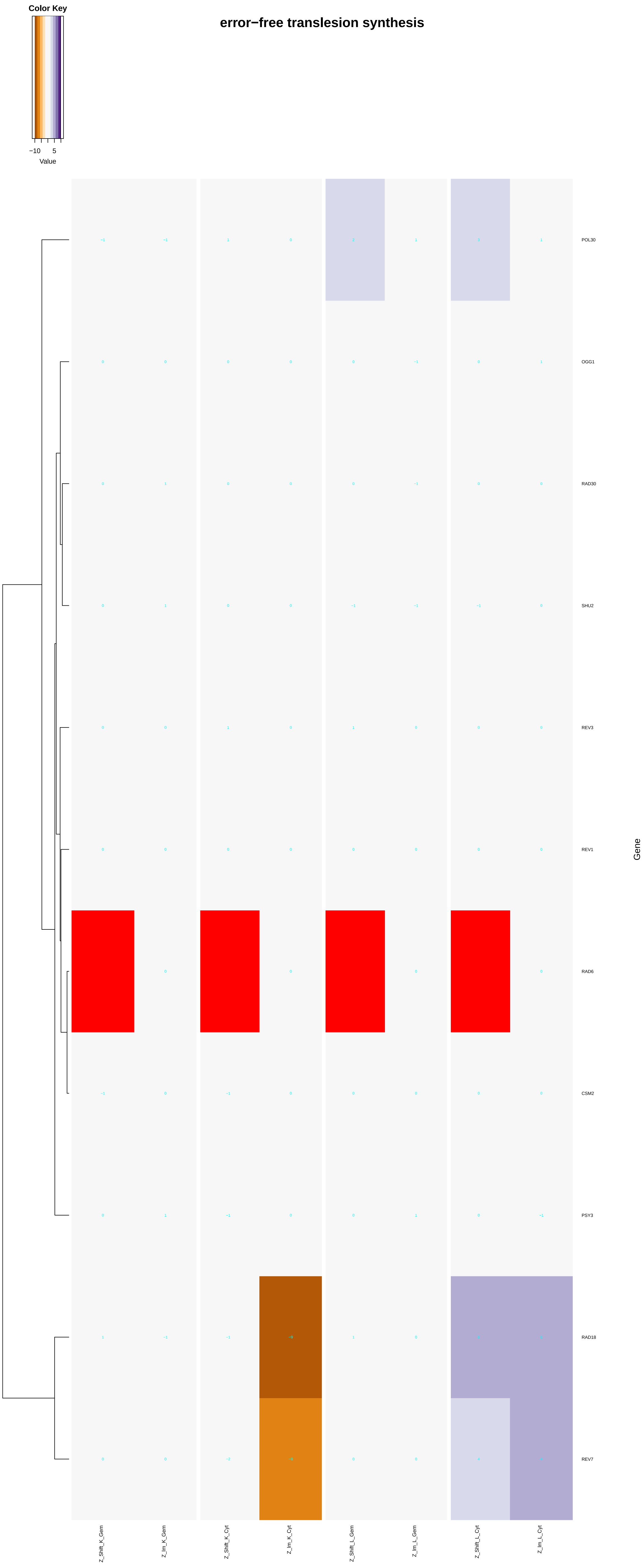

Color Key
−10
5
Value
error−free translesion synthesis
POL30
OGG1
RAD30
SHU2
REV3
REV1
RAD6
CSM2
PSY3
RAD18
REV7
Gene
Z_Shift_K_Gem
Z_Im_K_Gem
Z_Shift_K_Cyt
Z_Im_K_Cyt
Z_Shift_L_Gem
Z_Im_L_Gem
Z_Shift_L_Cyt
Z_Im_L_Cyt
