## Additional File 7 for "A humanized yeast phenomic model of deoxycytidine kinase to predict genetic buffering of nucleoside analog cytotoxicity": cellular_response_to_stress.pdf

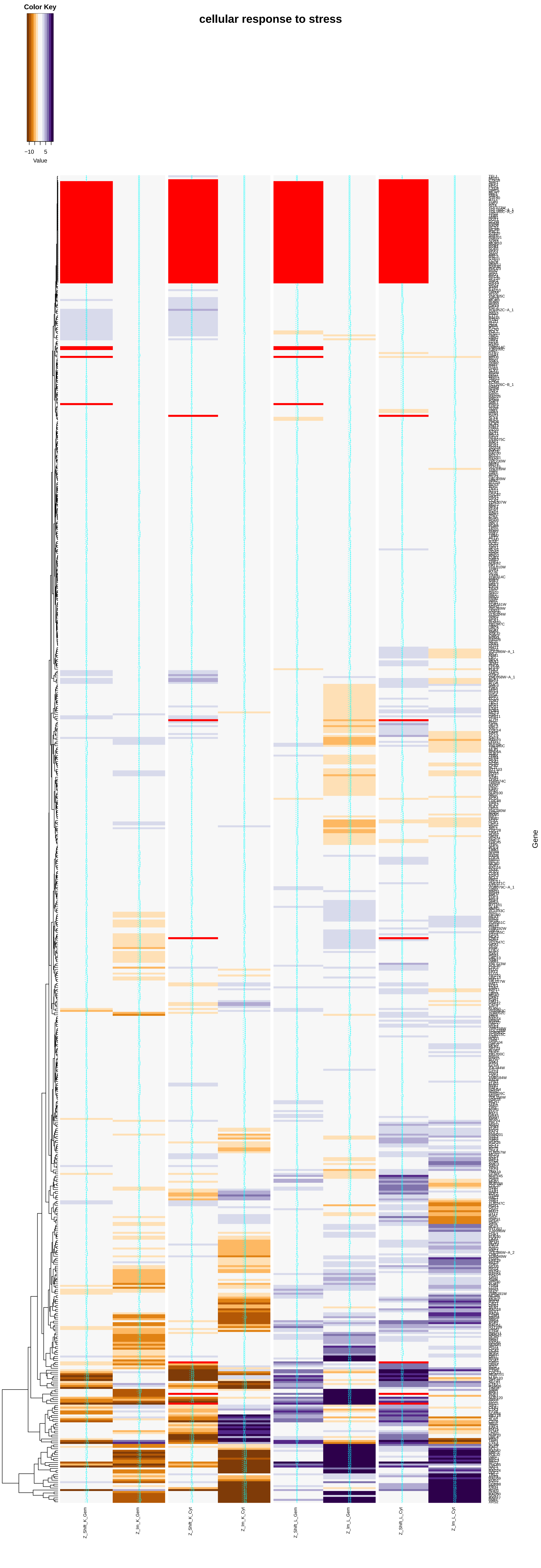
Color Key
−10
5
Value
cellular response to stress
Gene
Z_Shift_K_Gem
Z_Im_K_Gem
Z_Shift_K_Cyt
Z_Im_K_Cyt
Z_Shift_L_Gem
Z_Im_L_Gem
Z_Shift_L_Cyt
Z_Im_L_Cyt

### cellular response to DNA damage stimulus

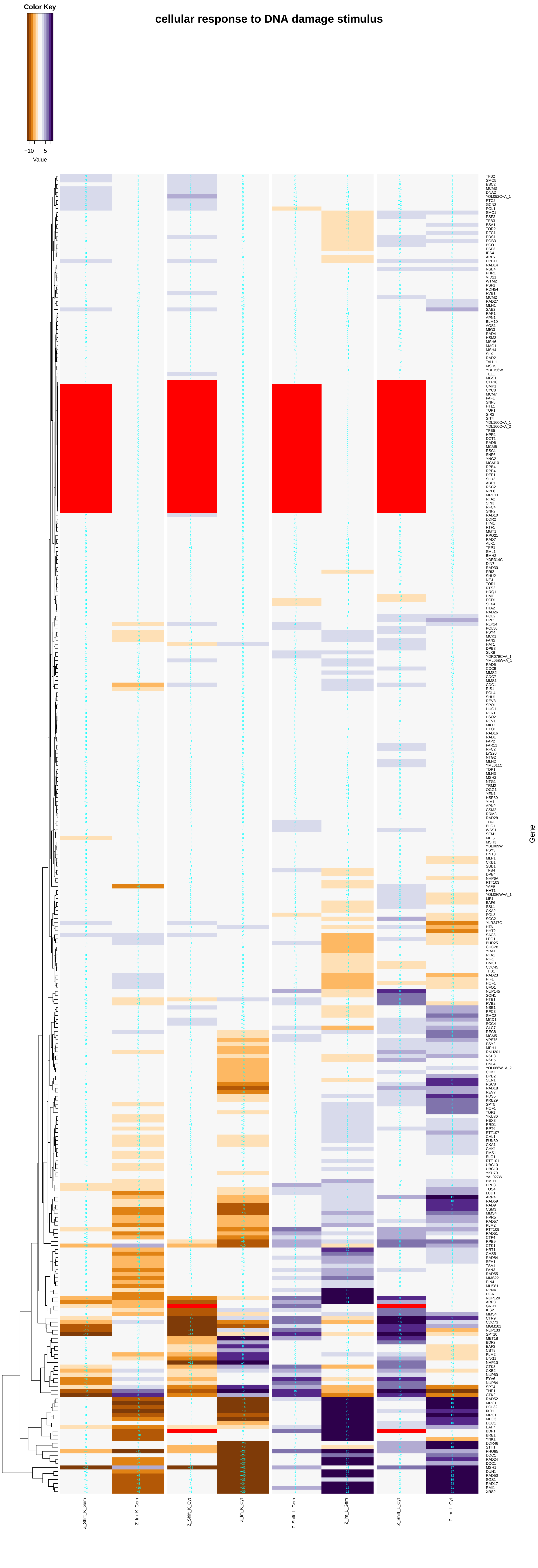

### cellular response to starvation

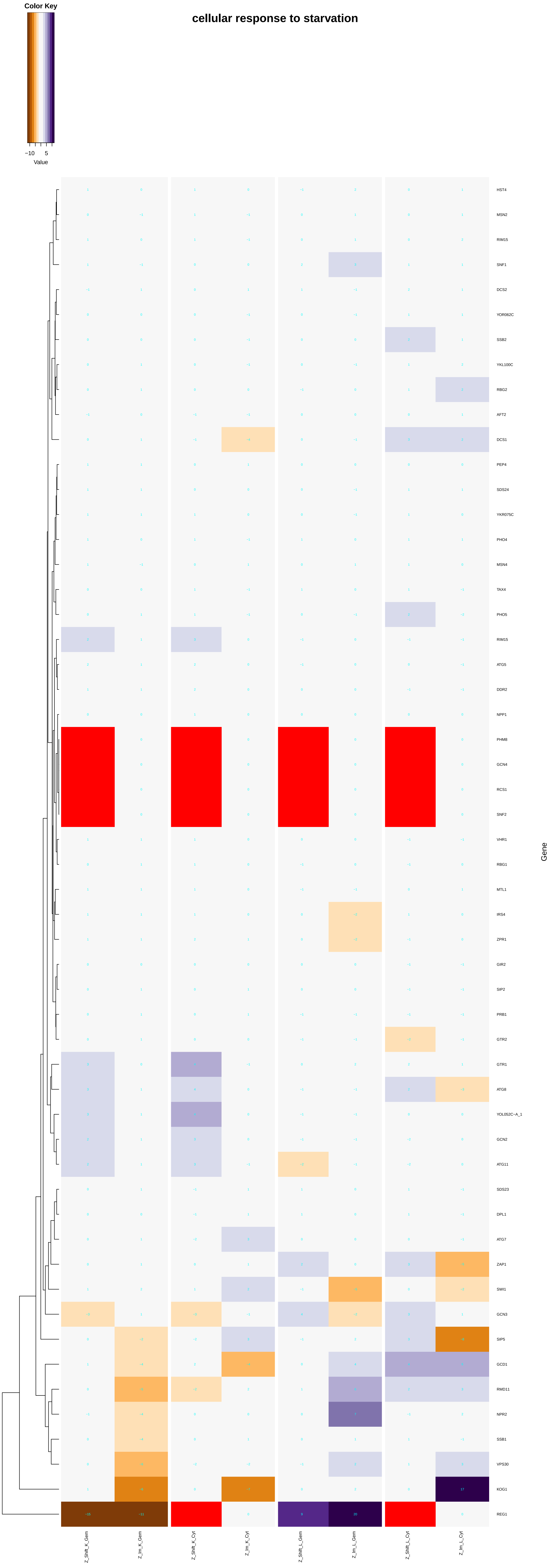

### stress−activated protein kinase signaling cascade

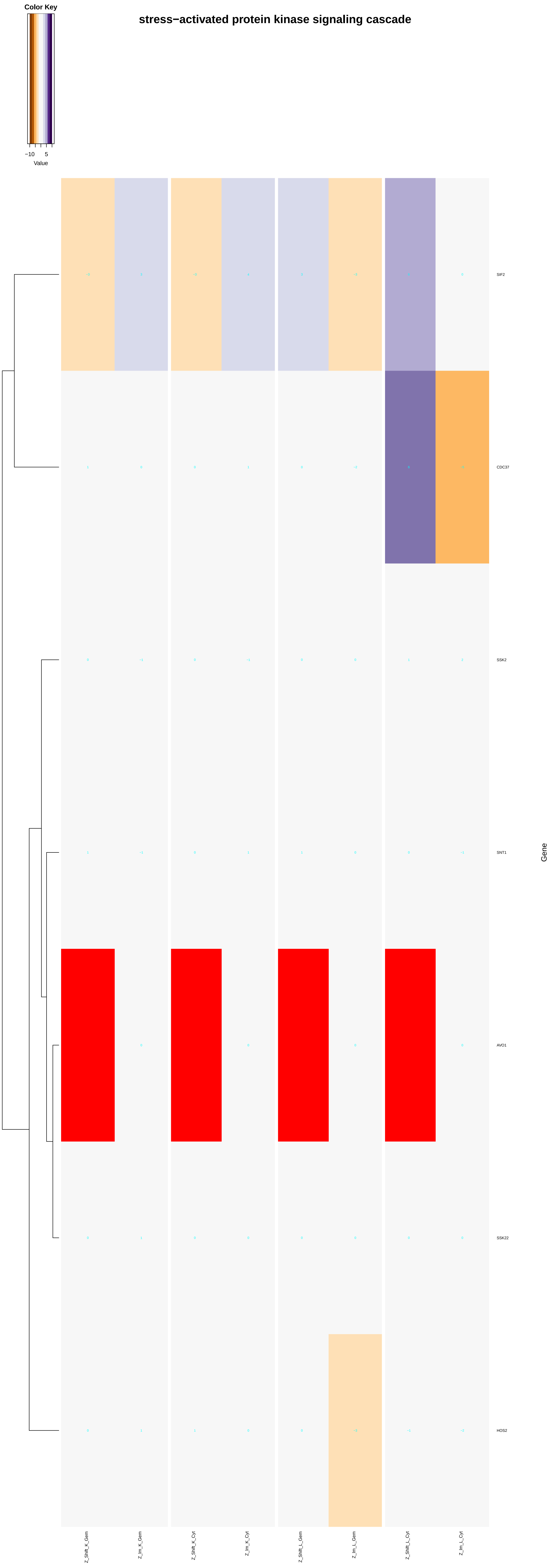

### cellular response to oxidative stress

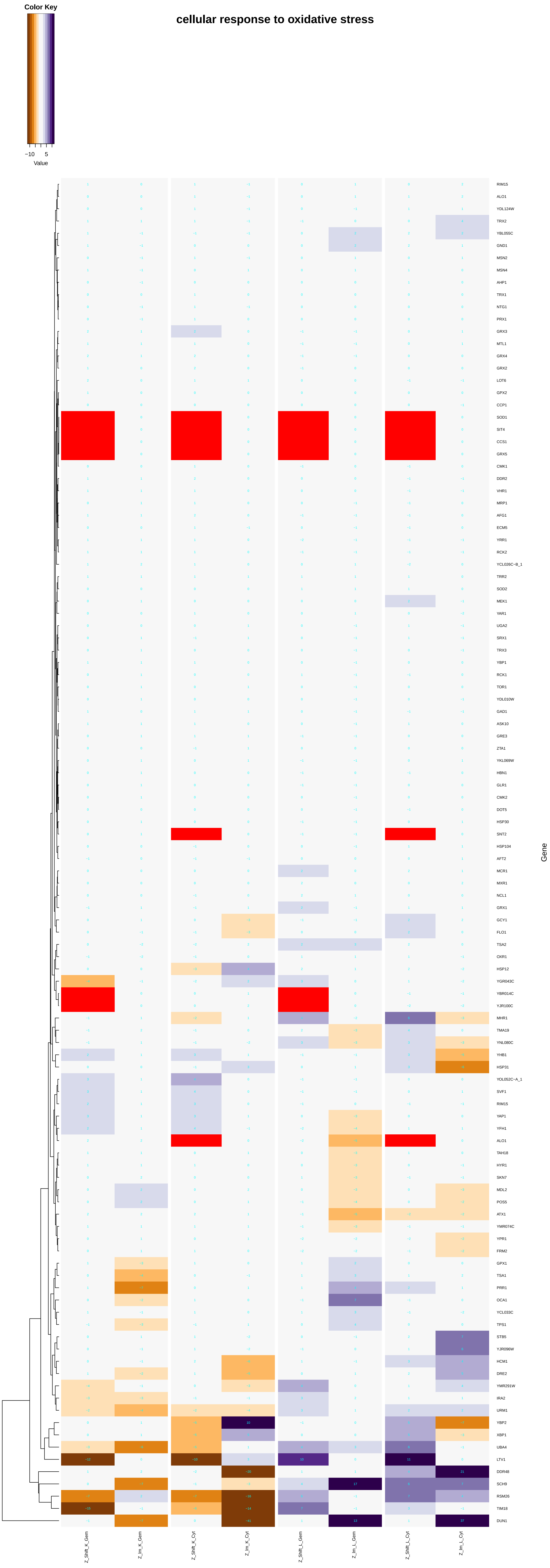

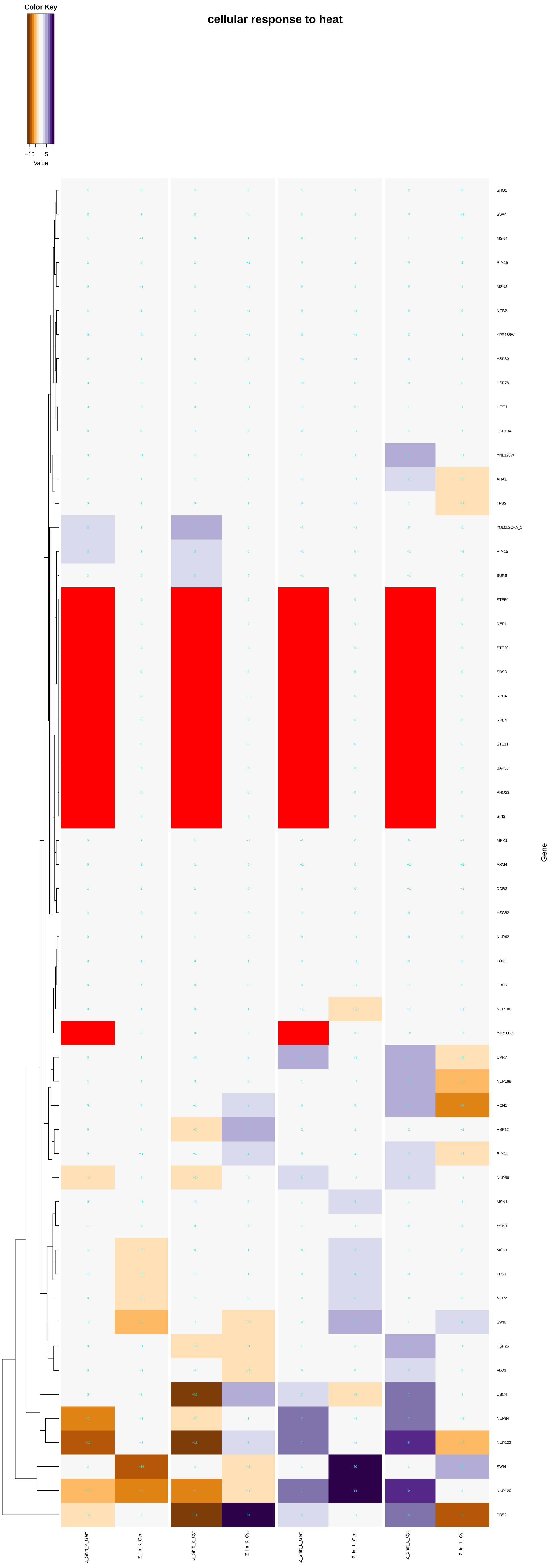
cellular response to heat
Color Key
−10 5
Value
SHO1
SSA4
MSN4
RIM15
MSN2
NCB2
YPR158W
HSP30
HSP78
HOG1
HSP104
YNL123W
AHA1
TPS2
YOL052C−A_1
RIM15
BUR6
STE50
DEP1
STE20
SDS3
RPB4
RPB4
STE11
SAP30
PHO23
SIN3
MRK1
ASM4
DDR2
HSC82
NUP42
TOR1
UBC5
NUP100
YJR100C
CPR7
NUP188
HCH1
HSP12
RIM11
NUP60
MSN1
YGK3
MCK1
TPS1
NUP2
SWI6
HSP26
FLO1
UBC4
NUP84
NUP133
SWI4
NUP120
PBS2
Gene
Z_Shift_K_Gem
Z_Im_K_Gem
Z_Shift_K_Cyt
Z_Im_K_Cyt
Z_Shift_L_Gem
Z_Im_L_Gem
Z_Shift_L_Cyt
Z_Im_L_Cyt

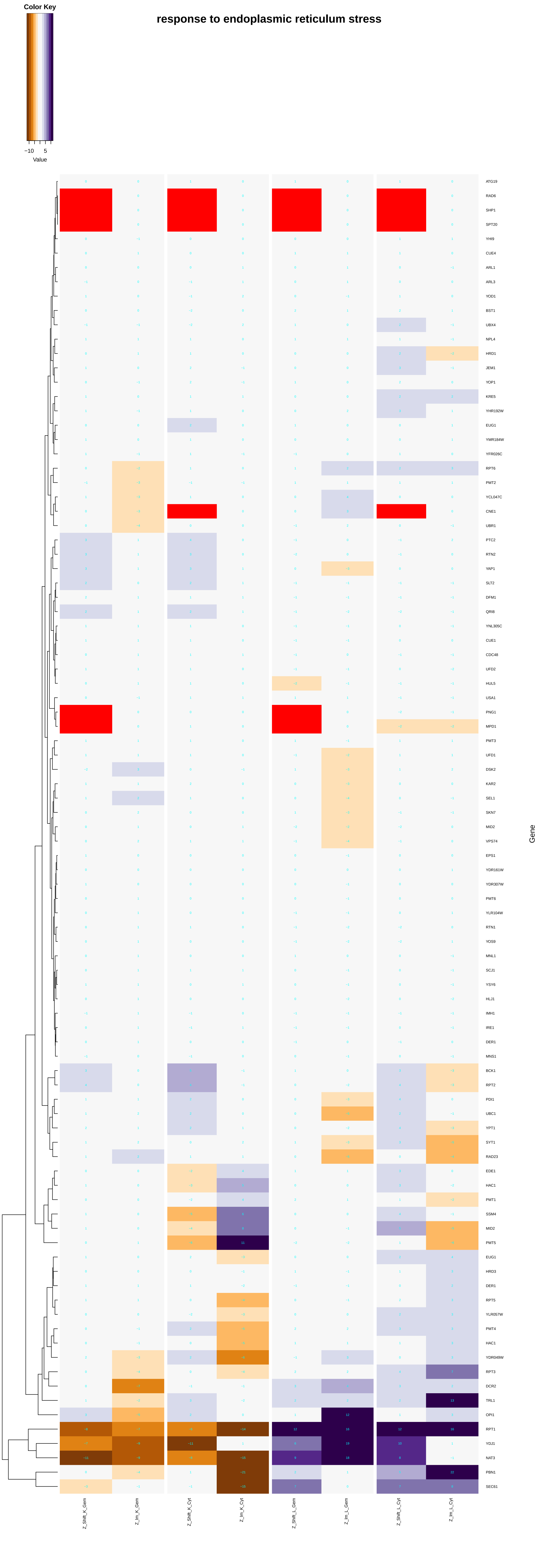
response to endoplasmic reticulum stress
Color Key
−10
5
Value
ATG19
RAD6
SHP1
SPT20
YHI9
CUE4
ARL1
ARL3
YOD1
BST1
UBX4
NPL4
HRD1
JEM1
YOP1
KRE5
YHR192W
EUG1
YMR184W
YFR026C
RPT6
PMT2
YCL047C
CNE1
UBR1
PTC2
RTN2
YAP1
SLT2
DFM1
QRI8
YNL305C
CUE1
CDC48
UFD2
HUL5
USA1
PNG1
MPD1
PMT3
UFD1
DSK2
KAR2
SEL1
SKN7
MID2
VPS74
EPS1
YDR161W
YDR307W
PMT6
YLR104W
RTN1
YOS9
MNL1
SCJ1
YSY6
HLJ1
IMH1
IRE1
DER1
MNS1
BCK1
RPT2
PDI1
UBC1
YPT1
SYT1
RAD23
EDE1
HAC1
PMT1
SSM4
MID2
PMT5
EUG1
HRD3
DER1
RPT5
YLR057W
PMT4
HAC1
YDR049W
RPT3
DCR2
TRL1
OPI1
RPT1
YDJ1
NAT3
PBN1
SEC61
Gene
Z_Shift_K_Gem
Z_Im_K_Gem
Z_Shift_K_Cyt
Z_Im_K_Cyt
Z_Shift_L_Gem
Z_Im_L_Gem
Z_Shift_L_Cyt
Z_Im_L_Cyt

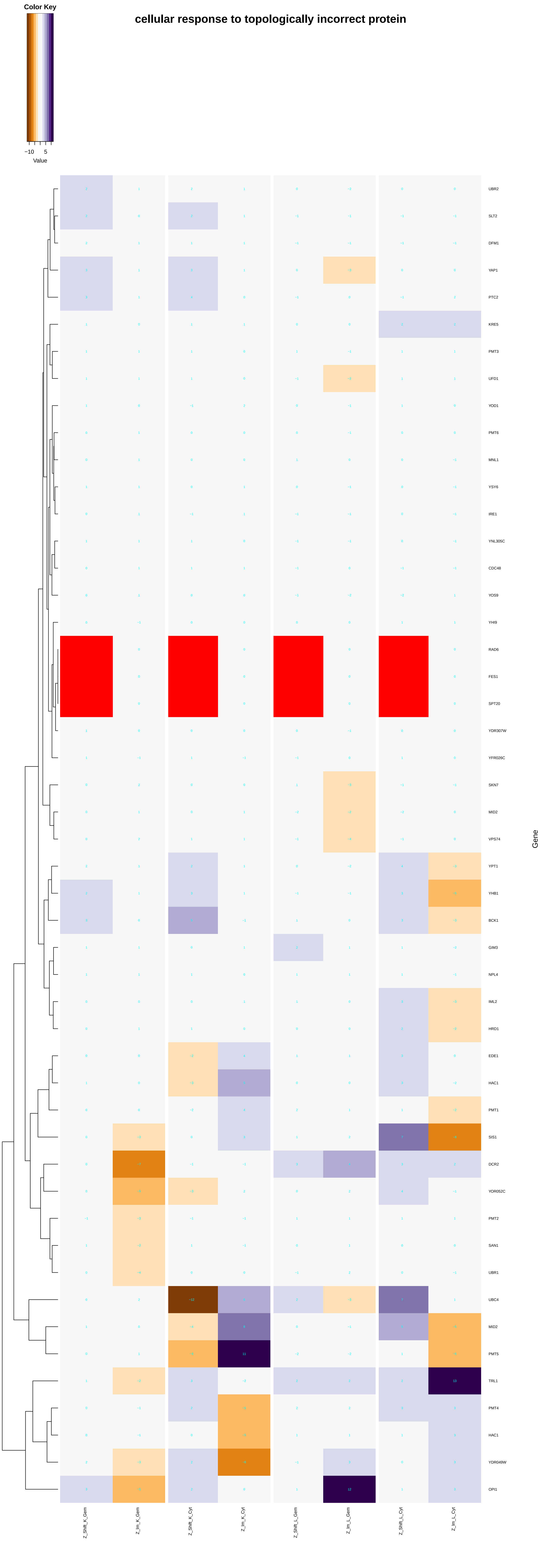

cellular response to topologically incorrect protein
Color Key
−10
5
Value
UBR2
SLT2
DFM1
YAP1
PTC2
KRE5
PMT3
UFD1
YOD1
PMT6
MNL1
YSY6
IRE1
YNL305C
CDC48
YOS9
YHI9
RAD6
FES1
SPT20
YDR307W
YFR026C
SKN7
MID2
VPS74
YPT1
YHB1
BCK1
GIM3
NPL4
IML2
HRD1
EDE1
HAC1
PMT1
SIS1
DCR2
YOR052C
PMT2
SAN1
UBR1
UBC4
MID2
PMT5
TRL1
PMT4
HAC1
YDR049W
OPI1
Gene
Z_Shift_K_Gem
Z_Im_K_Gem
Z_Shift_K_Cyt
Z_Im_K_Cyt
Z_Shift_L_Gem
Z_Im_L_Gem
Z_Shift_L_Cyt
Z_Im_L_Cyt

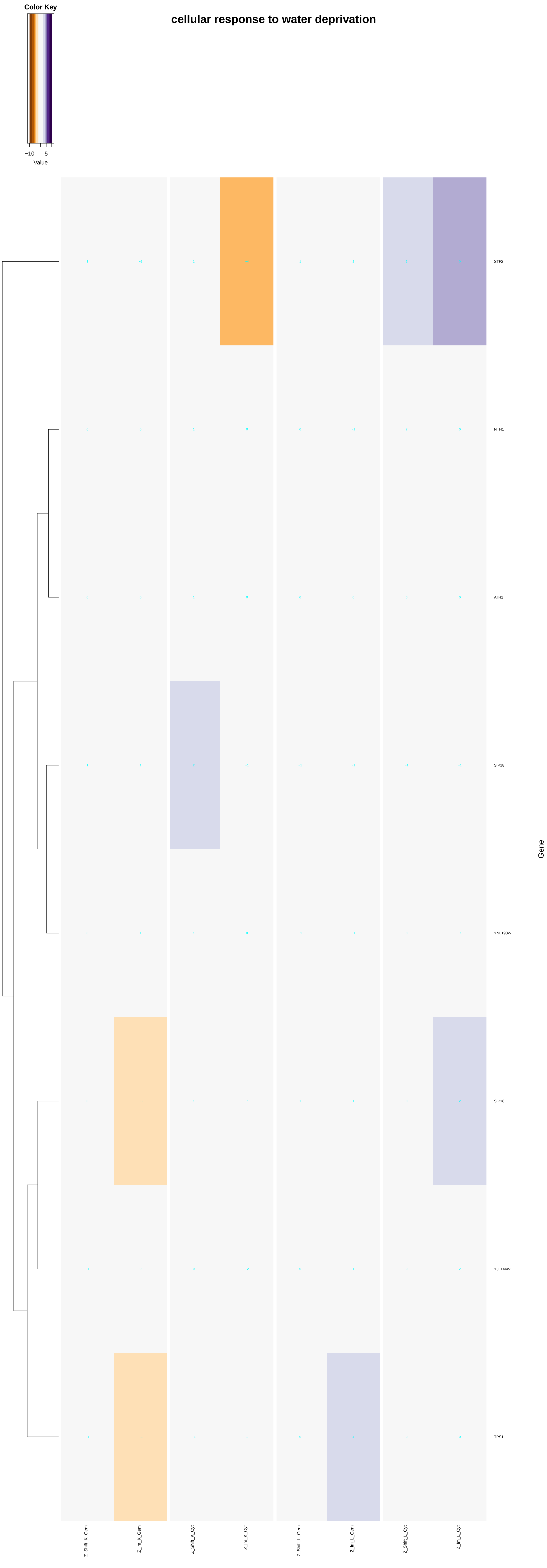

Color Key
−10
5
Value
cellular response to water deprivation
STF2
NTH1
ATH1
SIP18
YNL190W
SIP18
YJL144W
TPS1
Gene
Z_Shift_K_Gem
Z_lm_K_Gem
Z_Shift_K_Cyt
Z_lm_K_Cyt
Z_Shift_L_Gem
Z_lm_L_Gem
Z_Shift_L_Cyt
Z_lm_L_Cyt

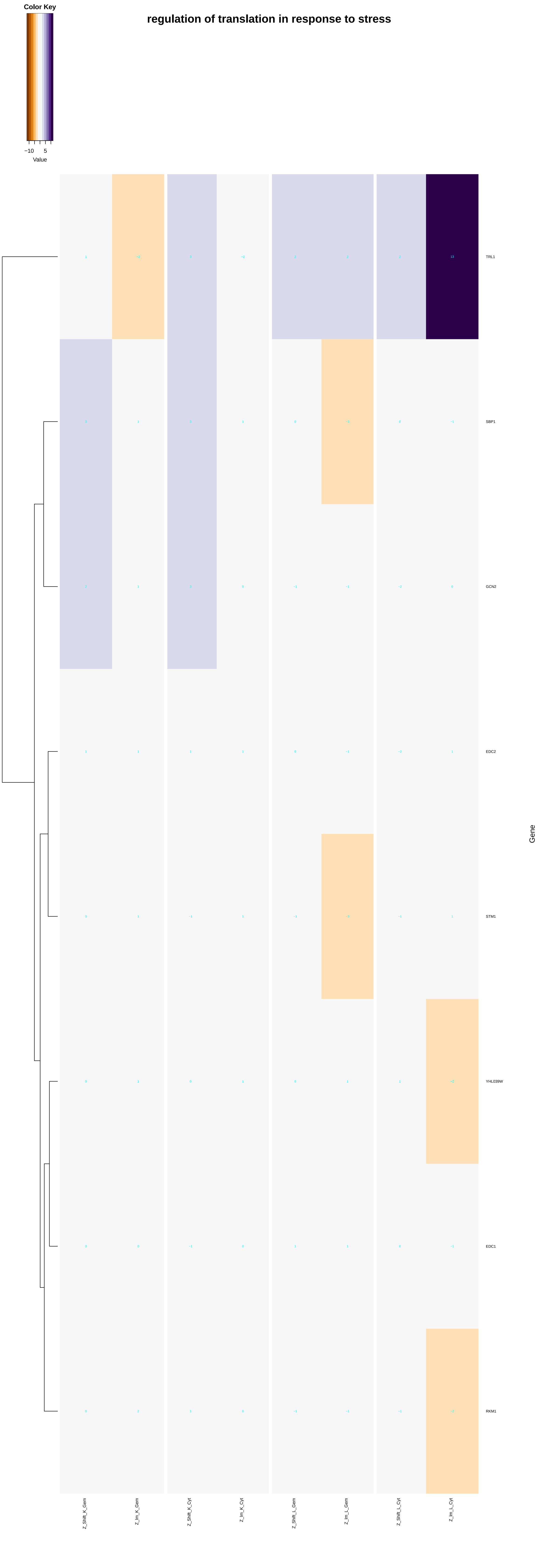
regulation of translation in response to stress
Color Key
−10
5
Value
TRL1
SBP1
GCN2
EDC2
STM1
YHL039W
EDC1
RKM1
Gene
Z_Shift_K_Gem
Z_Im_K_Gem
Z_Shift_K_Cyt
Z_Im_K_Cyt
Z_Shift_L_Gem
Z_Im_L_Gem
Z_Shift_L_Cyt
Z_Im_L_Cyt

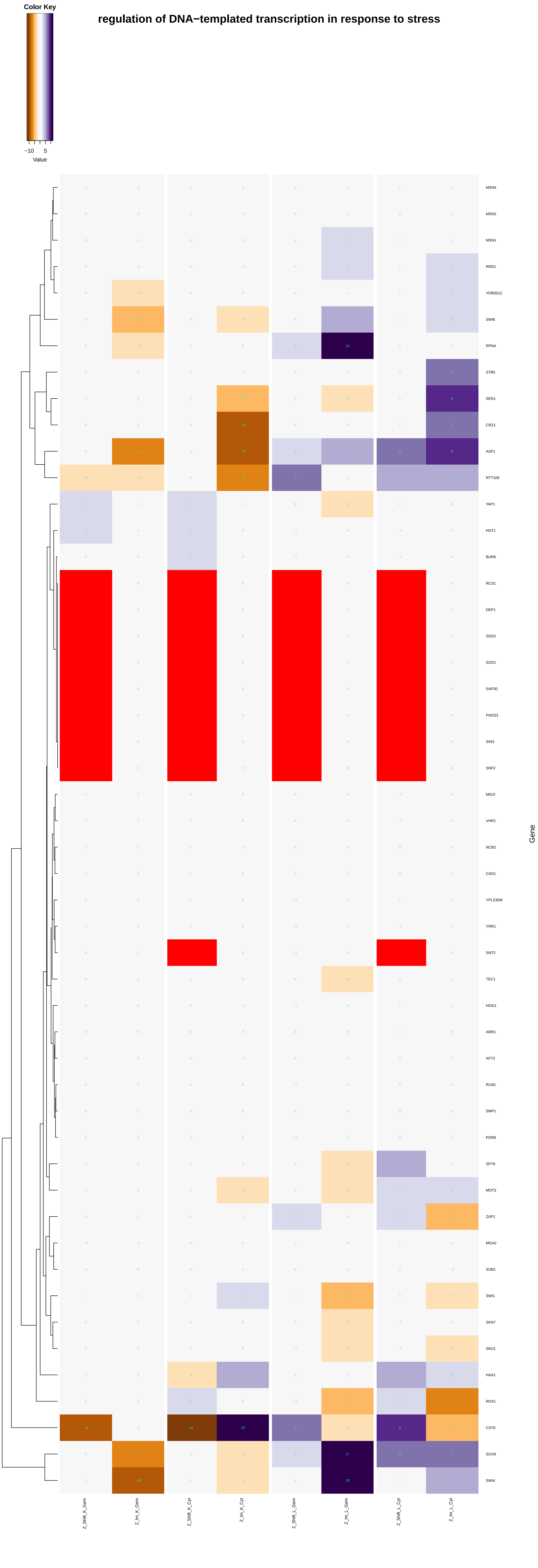
regulation of DNA−templated transcription in response to stress
Color Key
−10
5
Value
MSN4
MSN2
MSN1
RRD1
YOR051C
SWI6
RPN4
STB5
SEN1
CRZ1
ASF1
RTT109
YAP1
HOT1
BUR6
RCS1
DEP1
SDS3
SOD1
SAP30
PHO23
SIN3
SNF2
MIG3
VHR1
NCB2
CAD1
YPL230W
YRR1
SNT2
TEC1
HOG1
ARR1
AFT2
RLM1
SMP1
PDR8
SPT6
MOT3
ZAP1
MGA2
SUB1
SWI1
SKN7
SKO1
HAA1
ROX1
CST6
SCH9
SWI4
Gene
Z_Shift_K_Gem
Z_Im_K_Gem
Z_Shift_K_Cyt
Z_Im_K_Cyt
Z_Shift_L_Gem
Z_Im_L_Gem
Z_Shift_L_Cyt
Z_Im_L_Cyt

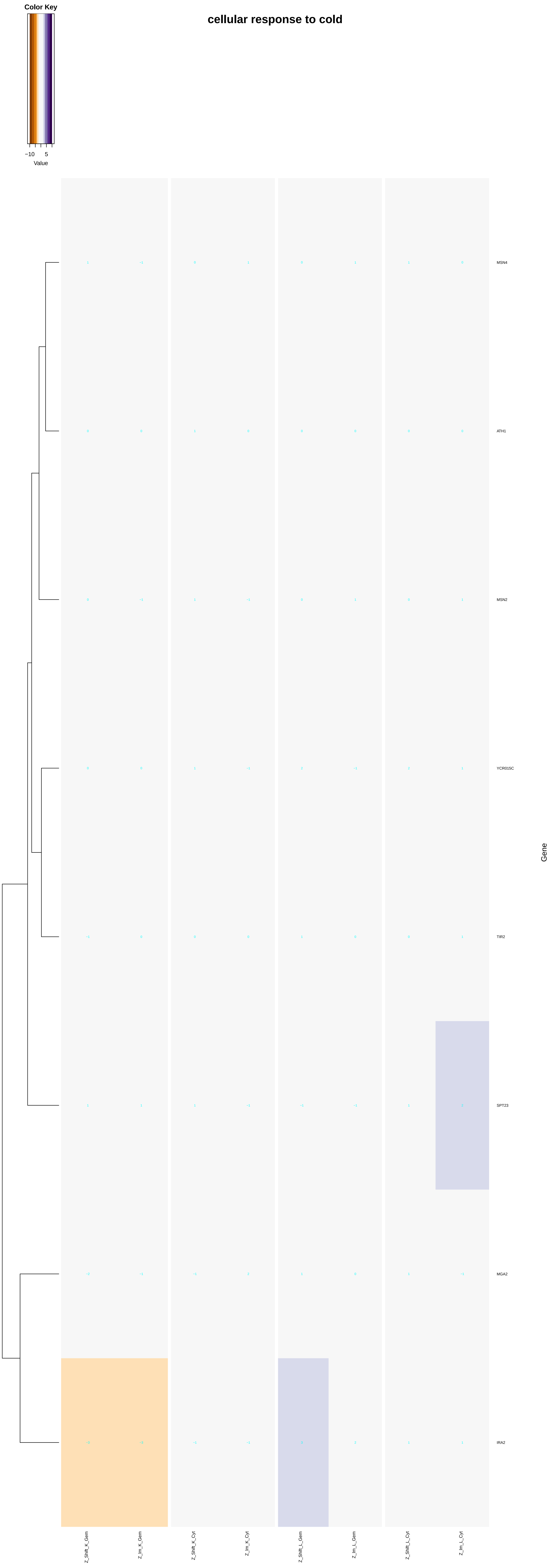
Color Key
−10
5
Value
cellular response to cold
MSN4
ATH1
MSN2
YCR015C
TIR2
SPT23
MGA2
IRA2
Gene
Z_Shift_K_Gem
Z_Im_K_Gem
Z_Shift_K_Cyt
Z_Im_K_Cyt
Z_Shift_L_Gem
Z_Im_L_Gem
Z_Shift_L_Cyt
Z_Im_L_Cyt

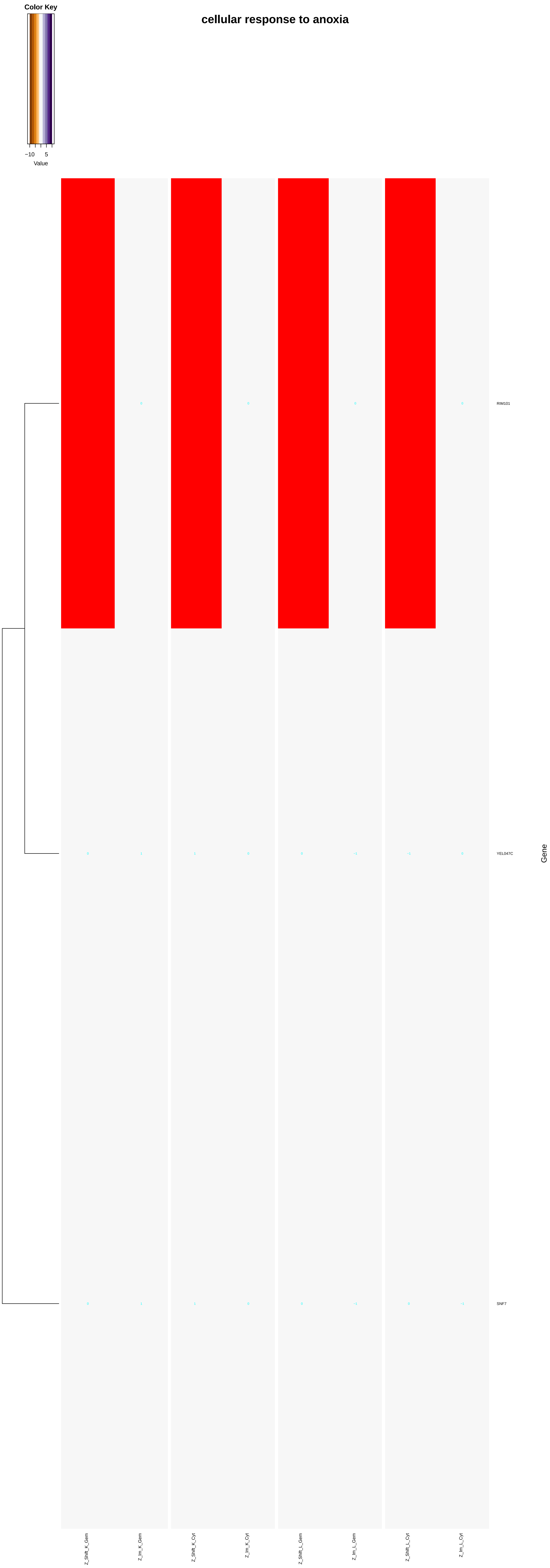
Color Key
−10
5
Value
cellular response to anoxia
RIM101
YEL047C
SNF7
Gene
Z_Shift_K_Gem
Z_lm_K_Gem
Z_Shift_K_Cyt
Z_lm_K_Cyt
Z_Shift_L_Gem
Z_lm_L_Gem
Z_Shift_L_Cyt
Z_lm_L_Cyt

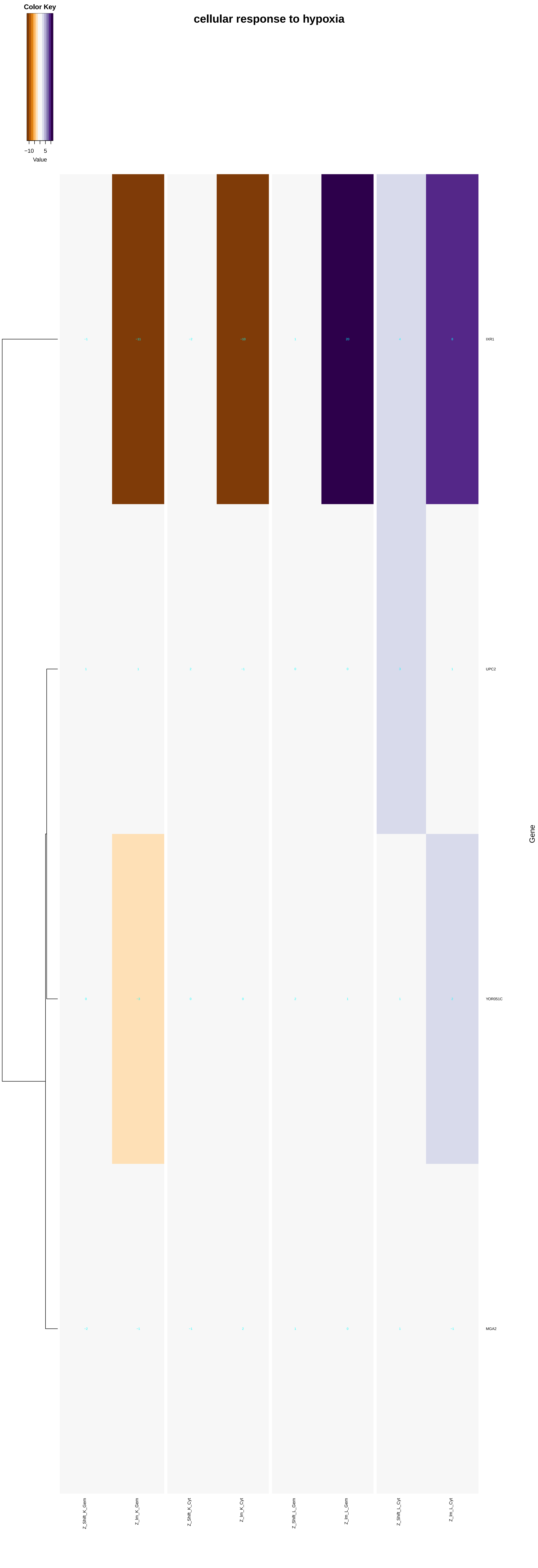
cellular response to hypoxia
Color Key
−10
5
Value
IXR1
UPC2
YOR051C
MGA2
Gene
Z_Shift_K_Gem
Z_Im_K_Gem
Z_Shift_K_Cyt
Z_Im_K_Cyt
Z_Shift_L_Gem
Z_Im_L_Gem
Z_Shift_L_Cyt
Z_Im_L_Cyt

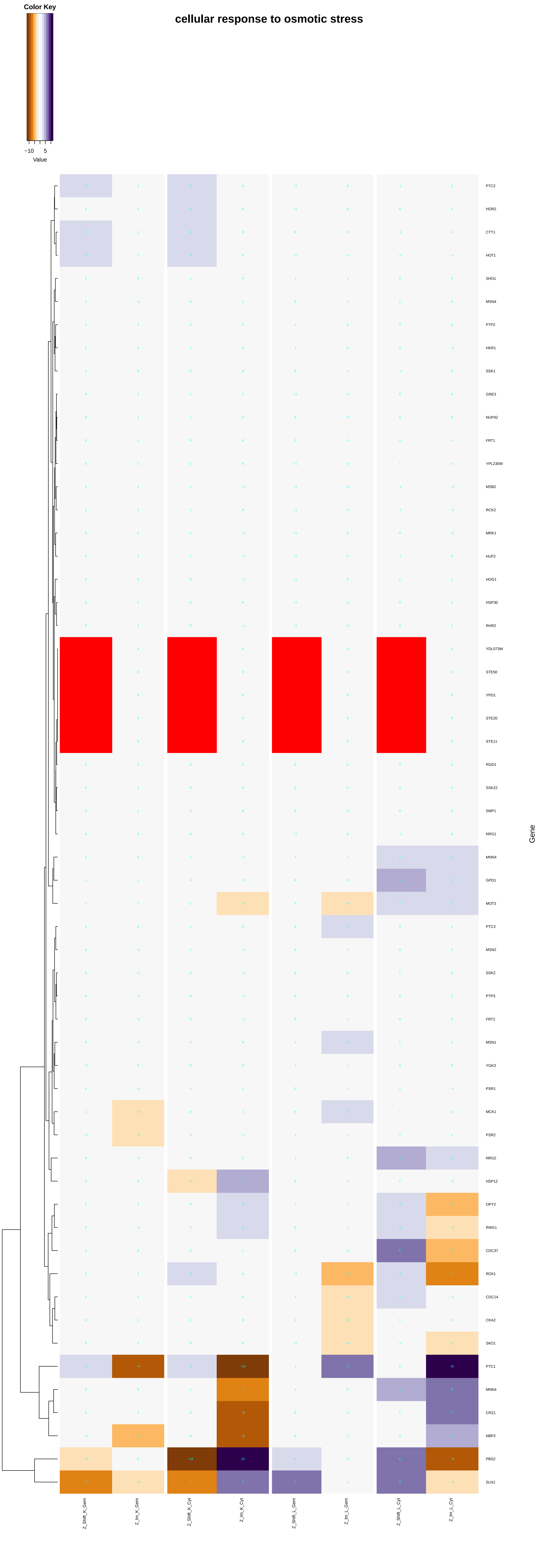
cellular response to osmotic stress
Color Key
−10
5
Value
PTC2
HOR2
CTT1
HOT1
SHO1
MSN4
PTP2
HKR1
SSK1
GRE3
NUP42
FRT1
YPL230W
MSB2
RCK2
MRK1
HUF2
HOG1
HSP30
RHR2
YDL073W
STE50
YPD1
STE20
STE11
RGD1
SSK22
SMP1
NRG1
MNN4
GPD1
MOT3
PTC3
MSN2
SSK2
PTP3
FRT2
MSN1
YGK3
PSR1
MCK1
PSR2
NRG2
HSP12
OPY2
RIM11
CDC37
ROX1
CDC14
CKA2
SKO1
PTC1
MNN4
CRZ1
NBP2
PBS2
SLN1
Gene
Z_Shift_K_Gen
Z_Im_K_Gen
Z_Shift_K_Cyt
Z_Im_K_Cyt
Z_Shift_L_Gen
Z_Im_L_Gen
Z_Shift_L_Cyt
Z_Im_L_Cyt

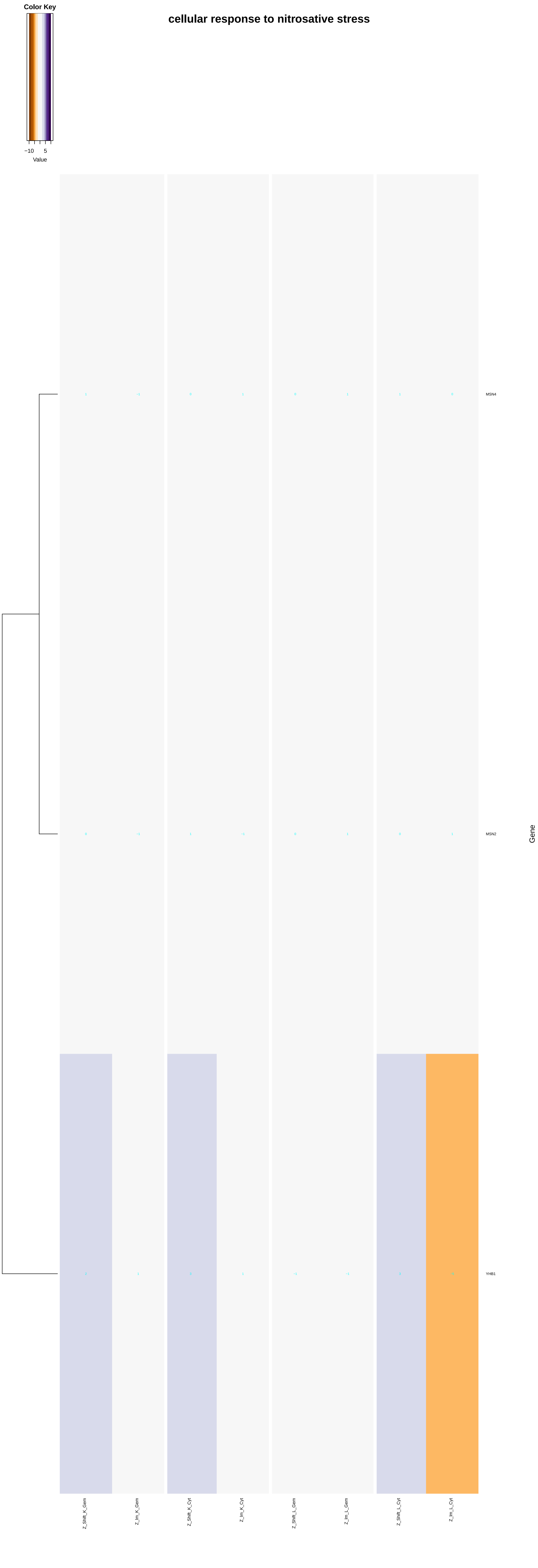

cellular response to nitrosative stress
Color Key
−10
5
Value
MSN4
MSN2
YHB1
Gene
Z_Shift_K_Gem
Z_Im_K_Gem
Z_Shift_K_Cyt
Z_Im_K_Cyt
Z_Shift_L_Gem
Z_Im_L_Gem
Z_Shift_L_Cyt
Z_Im_L_Cyt

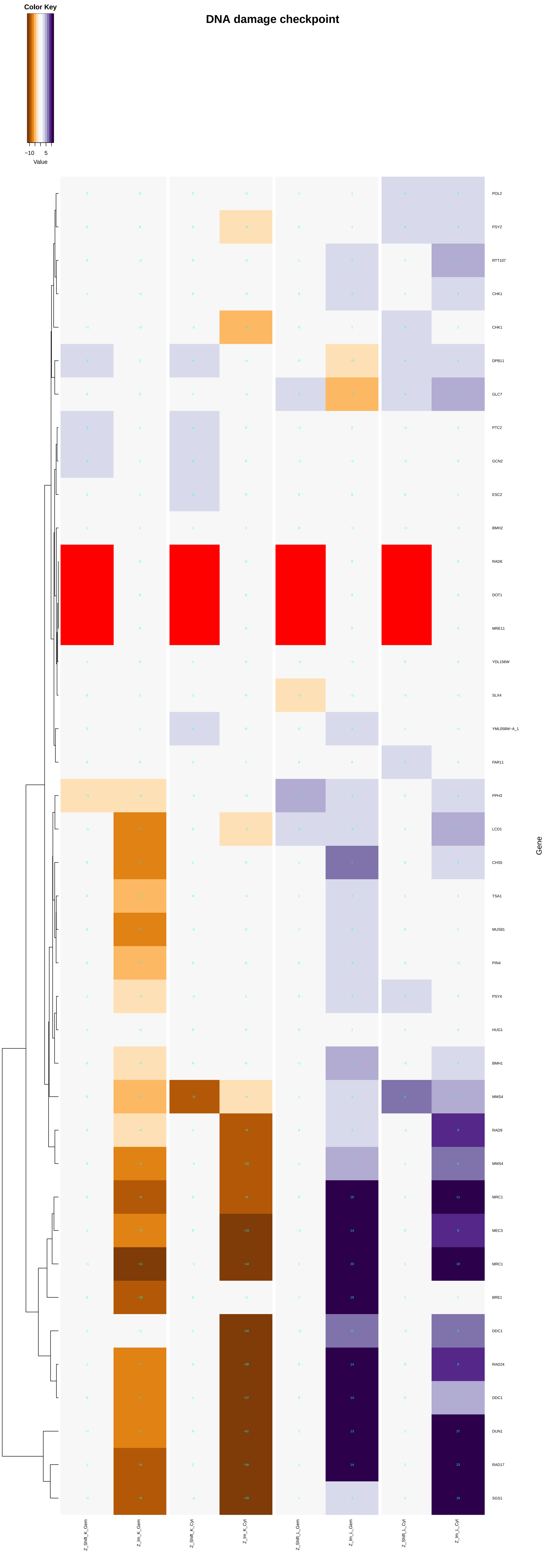
DNA damage checkpoint
Color Key
−10
5
Value
POL2
PSY2
RTT107
CHK1
CHK1
DPB11
GLC7
PTC2
GCN2
ESC2
BMH2
RAD6
DOT1
MRE11
YDL156W
SLX4
YML058W−A_1
FAR11
PPH3
LCD1
CHS5
TSA1
MUS81
PIN4
PSY4
HUG1
BMH1
MMS4
RAD9
MMS4
MRC1
MEC3
MRC1
BRE1
DDC1
RAD24
DDC1
DUN1
RAD17
SGS1
Gene
Z_Shift_K_Gem
Z_Im_K_Gem
Z_Shift_K_Cyt
Z_Im_K_Cyt
Z_Shift_L_Gem
Z_Im_L_Gem
Z_Shift_L_Cyt
Z_Im_L_Cyt

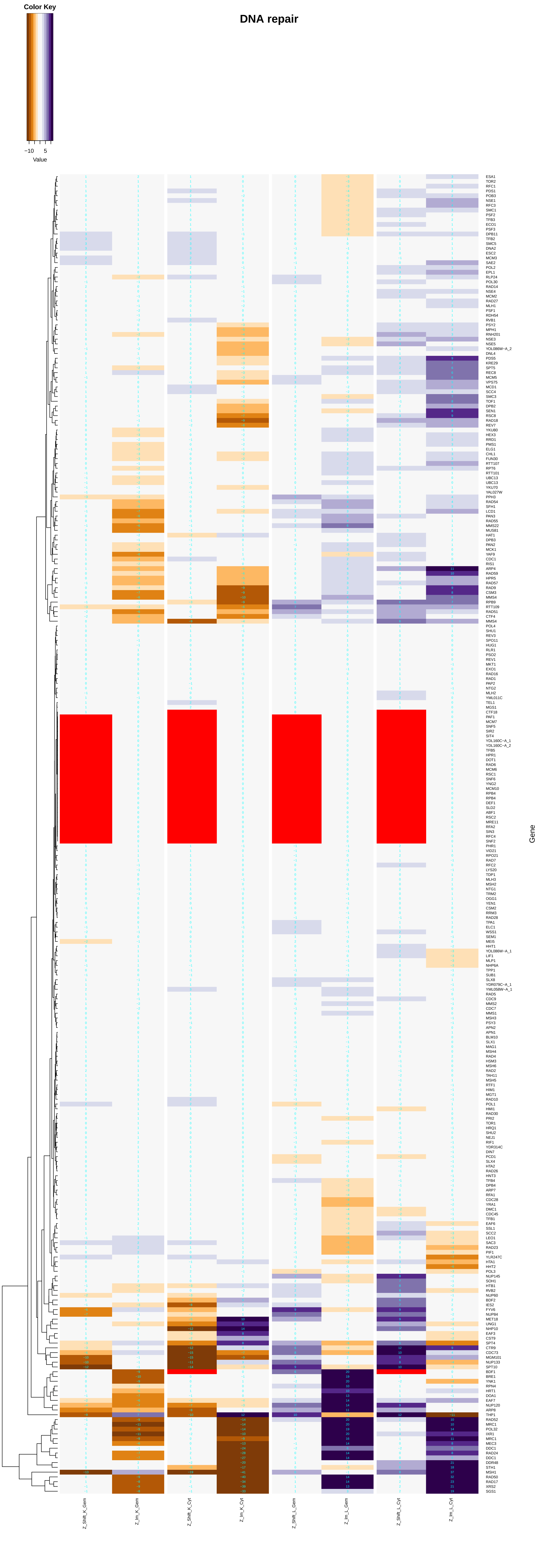
DNA repair
Color Key
−10
5
Value
Gene
Z_Shift_K_Gem
Z_Im_K_Gem
Z_Shift_K_Cyt
Z_Im_K_Cyt
Z_Shift_L_Gem
Z_Im_L_Gem
Z_Shift_L_Cyt
Z_Im_L_Cyt

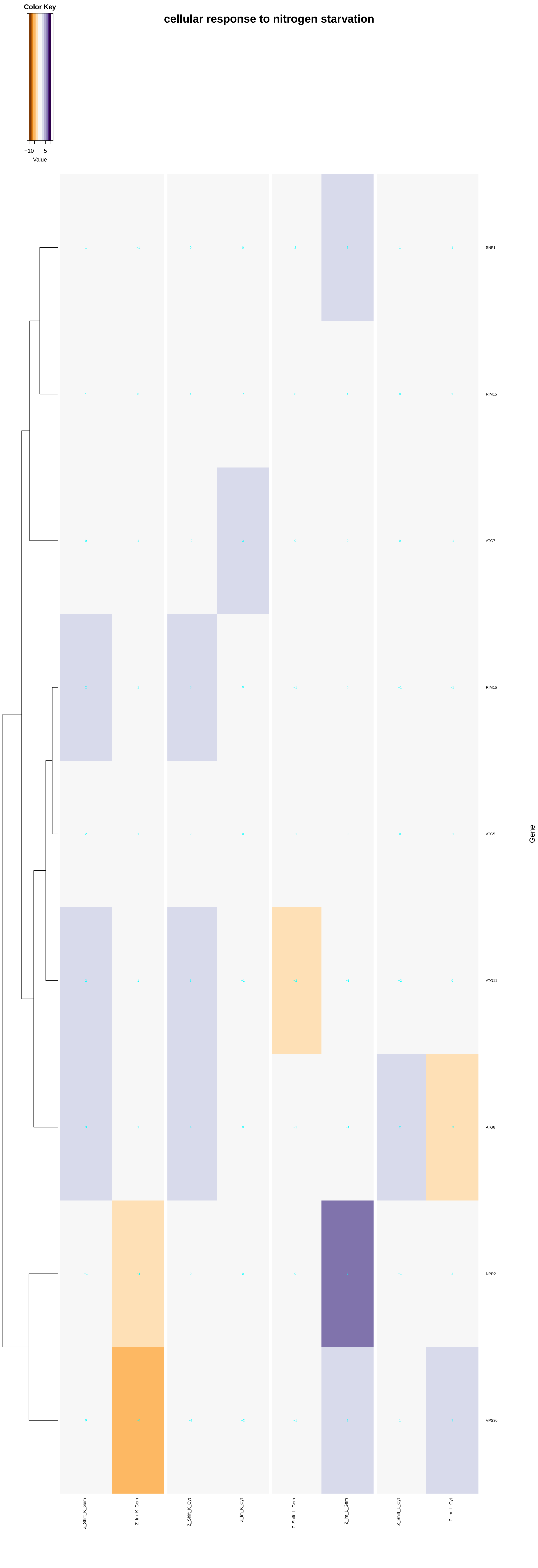
cellular response to nitrogen starvation
Color Key
−10
5
Value
SNF1
RIM15
ATG7
RIM15
ATG5
ATG11
ATG8
NPR2
VPS30
Gene
Z_Shift_K_Gem
Z_Im_K_Gem
Z_Shift_K_Cyt
Z_Im_K_Cyt
Z_Shift_L_Gem
Z_Im_L_Gem
Z_Shift_L_Cyt
Z_Im_L_Cyt

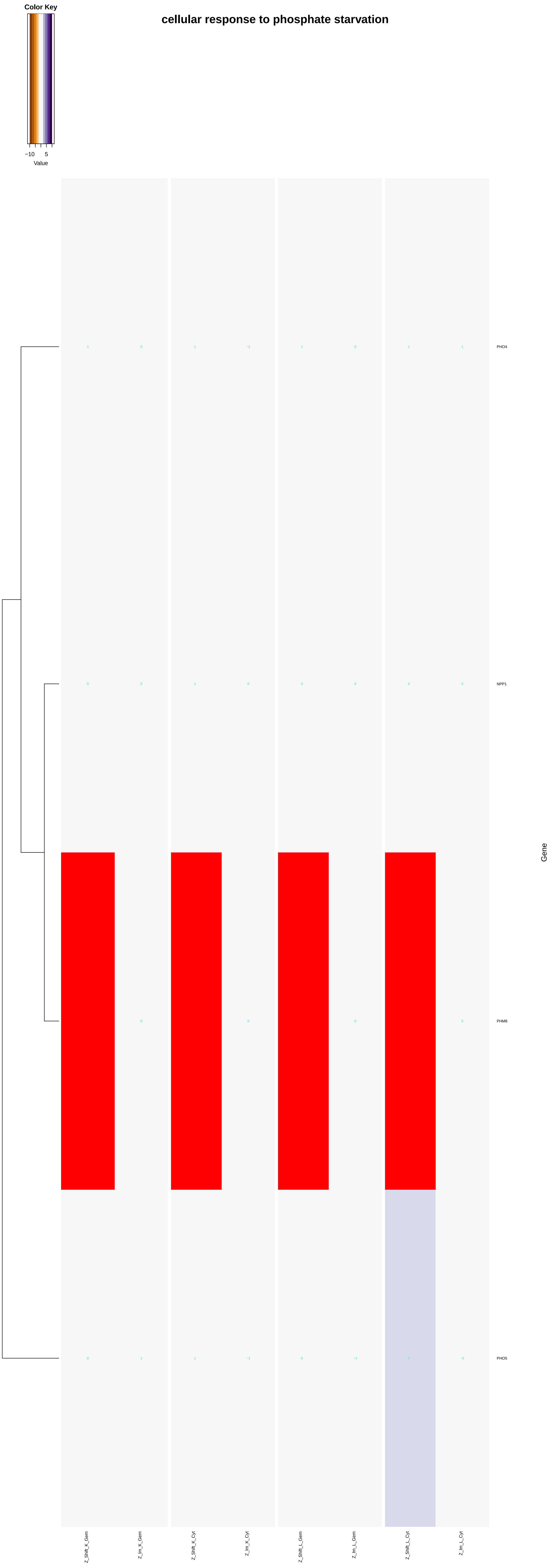

cellular response to phosphate starvation
Color Key
−10
5
Value
PHO4
NPP1
PHM8
PHO5
Gene
Z_Shift_K_Gem
Z_Im_K_Gem
Z_Shift_K_Cyt
Z_Im_K_Cyt
Z_Shift_L_Gem
Z_Im_L_Gem
Z_Shift_L_Cyt
Z_Im_L_Cyt

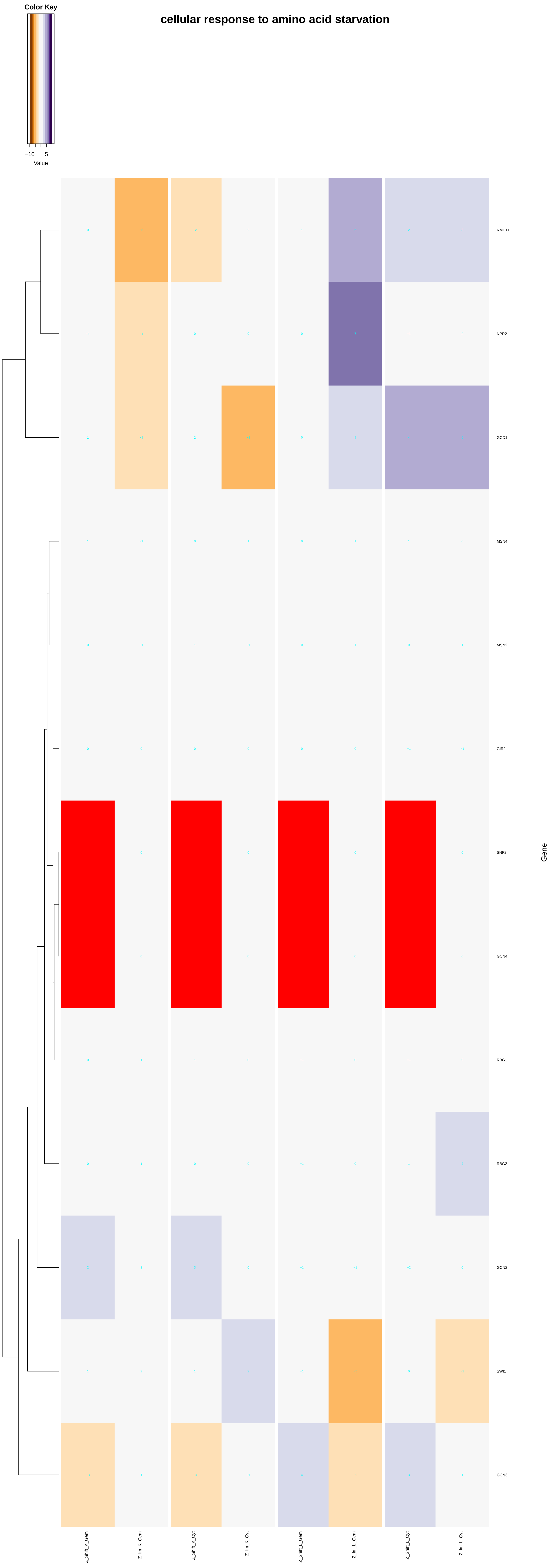

Color Key
−10
5
Value
cellular response to amino acid starvation
RMD11
NPR2
GCD1
MSN4
MSN2
GIR2
SNF2
GCN4
RBG1
RBG2
GCN2
SWI1
GCN3
Gene
Z_Shift_K_Gem
Z_Im_K_Gem
Z_Shift_K_Cyt
Z_Im_K_Cyt
Z_Shift_L_Gem
Z_Im_L_Gem
Z_Shift_L_Cyt
Z_Im_L_Cyt

### cellular response to zinc ion starvation

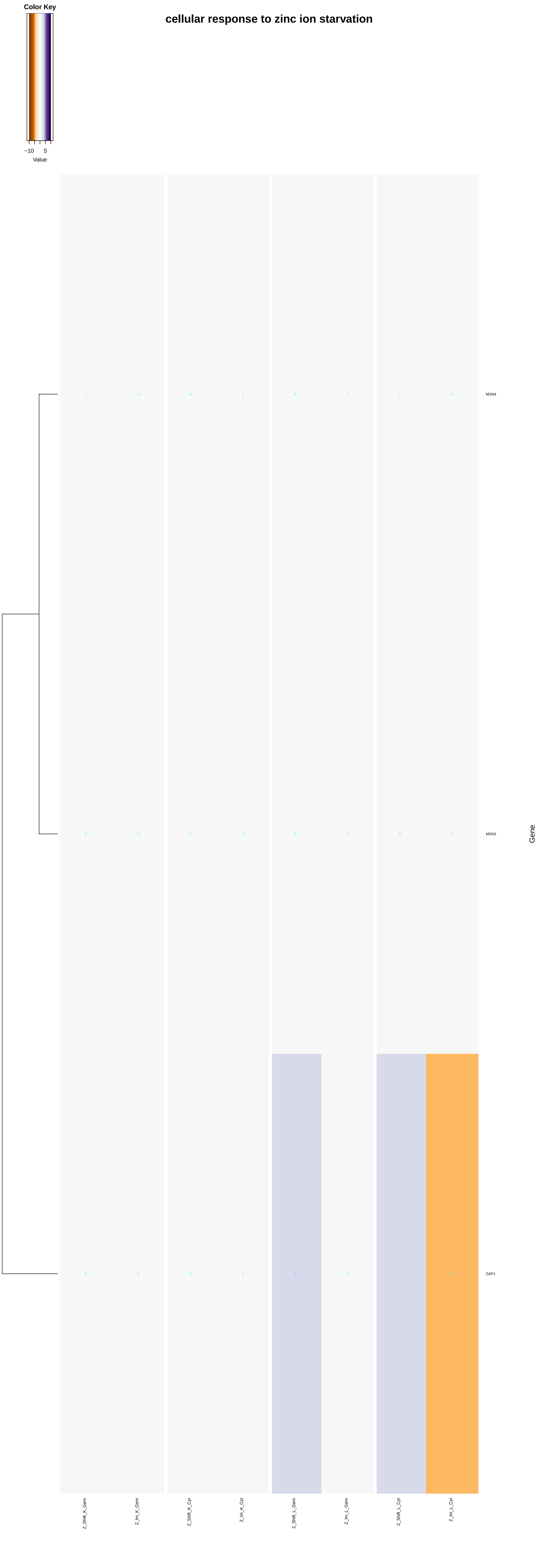

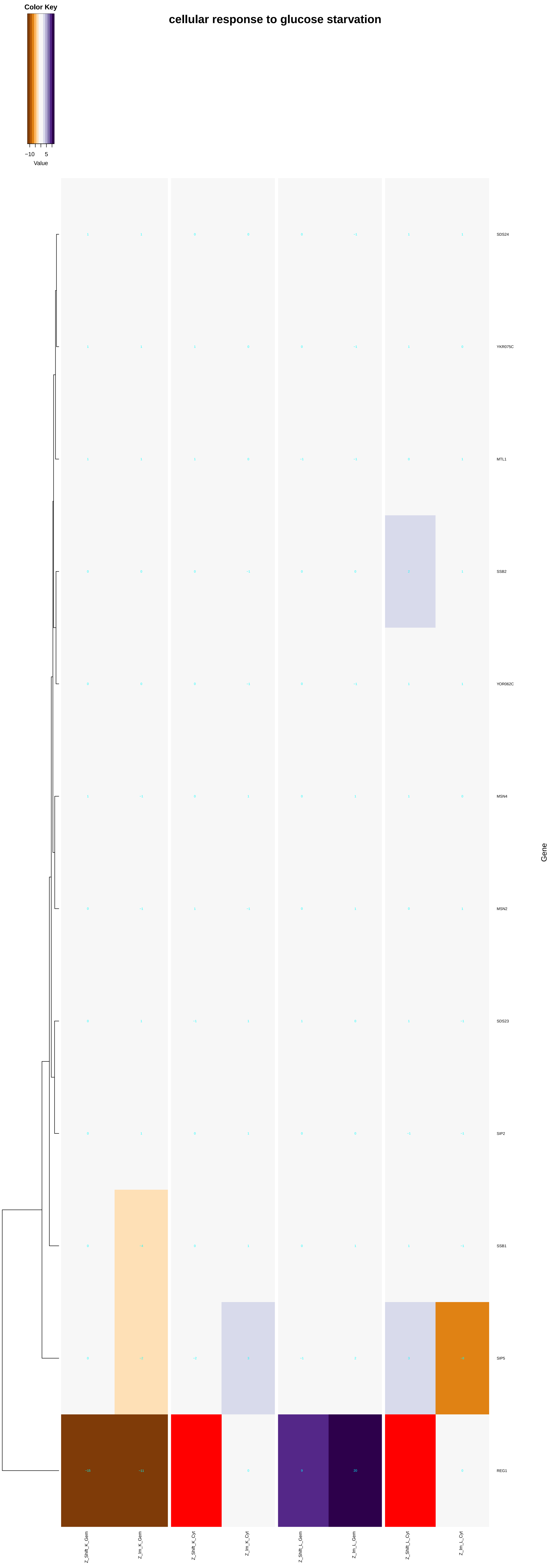

Color Key
−10
5
Value
cellular response to glucose starvation
SDS24
YKR075C
MTL1
SSB2
YOR062C
MSN4
MSN2
SDS23
SIP2
SSB1
SIP5
REG1
Gene
Z_Shift_K_Gem
Z_Im_K_Gem
Z_Shift_K_Cyt
Z_Im_K_Cyt
Z_Shift_L_Gem
Z_Im_L_Gem
Z_Shift_L_Cyt
Z_Im_L_Cyt

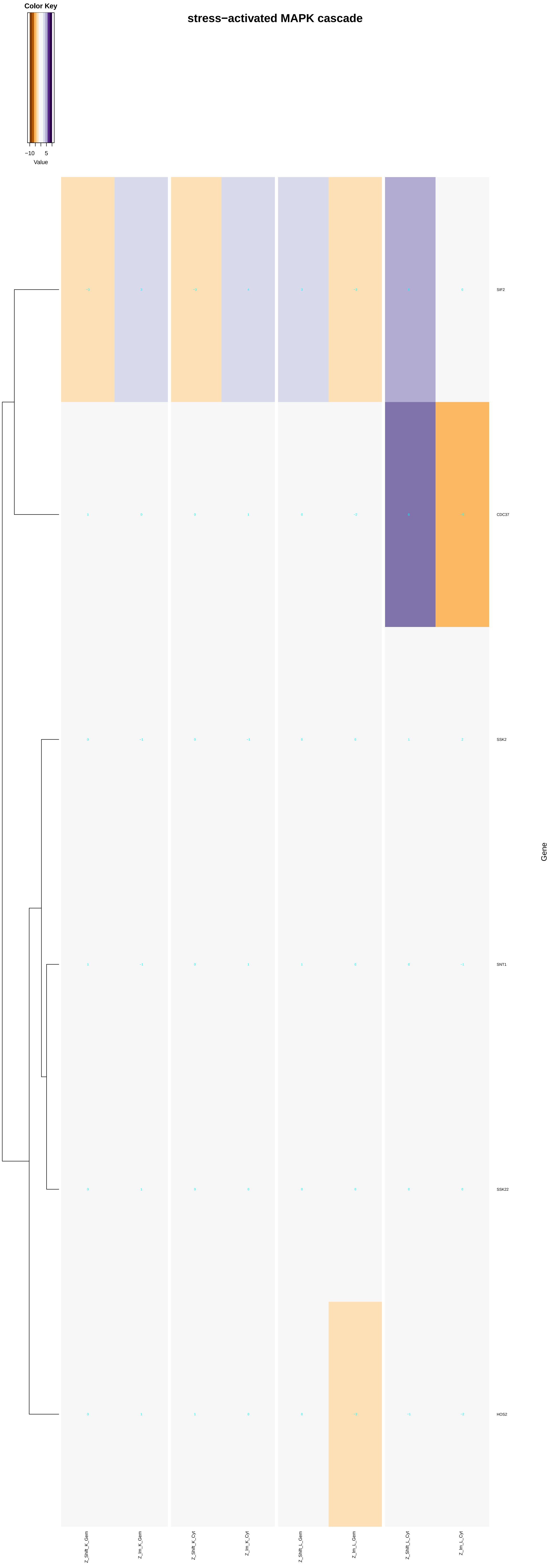
stress−activated MAPK cascade
Color Key
−10
5
Value
SIF2
CDC37
SSK2
SNT1
SSK22
HOS2
Gene
Z_Shift_K_Gem
Z_Im_K_Gem
Z_Shift_K_Cyt
Z_Im_K_Cyt
Z_Shift_L_Gem
Z_Im_L_Gem
Z_Shift_L_Cyt
Z_Im_L_Cyt

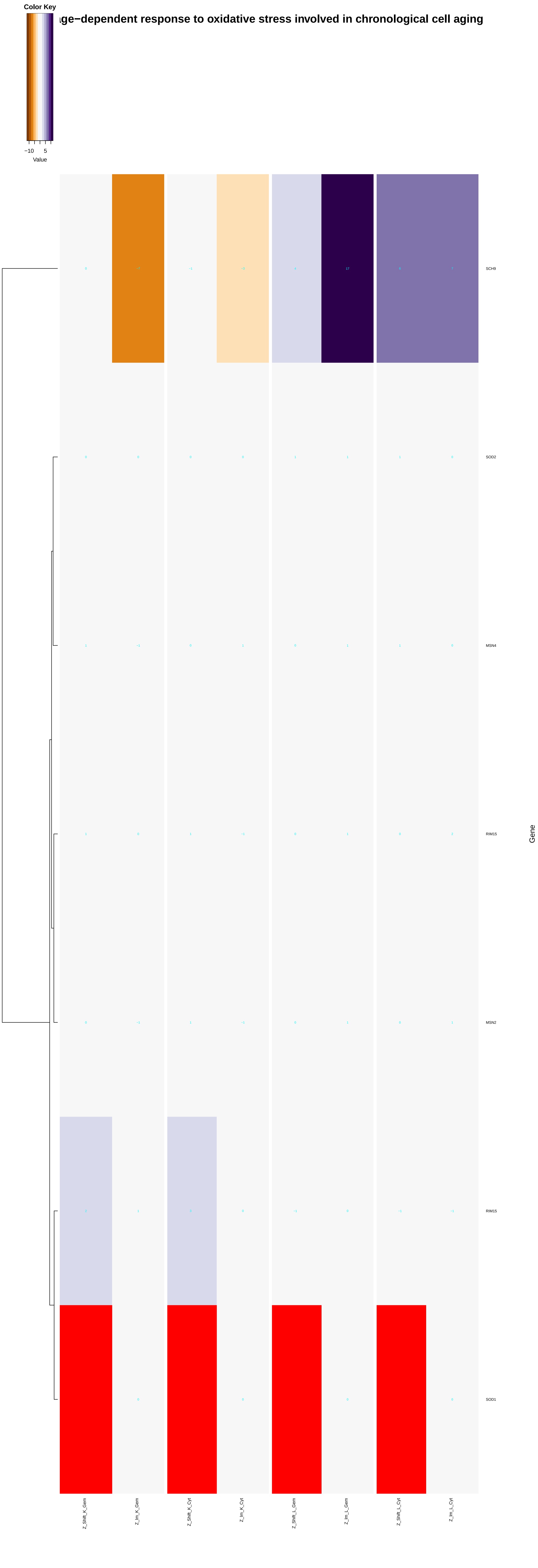

ge−dependent response to oxidative stress involved in chronological cell aging
Color Key
−10
5
Value
SCH9
SOD2
MSN4
RIM15
MSN2
RIM15
SOD1
Gene
Z_Shift_K_Gem
Z_Im_K_Gem
Z_Shift_K_Cyt
Z_Im_K_Cyt
Z_Shift_L_Gem
Z_Im_L_Gem
Z_Shift_L_Cyt
Z_Im_L_Cyt

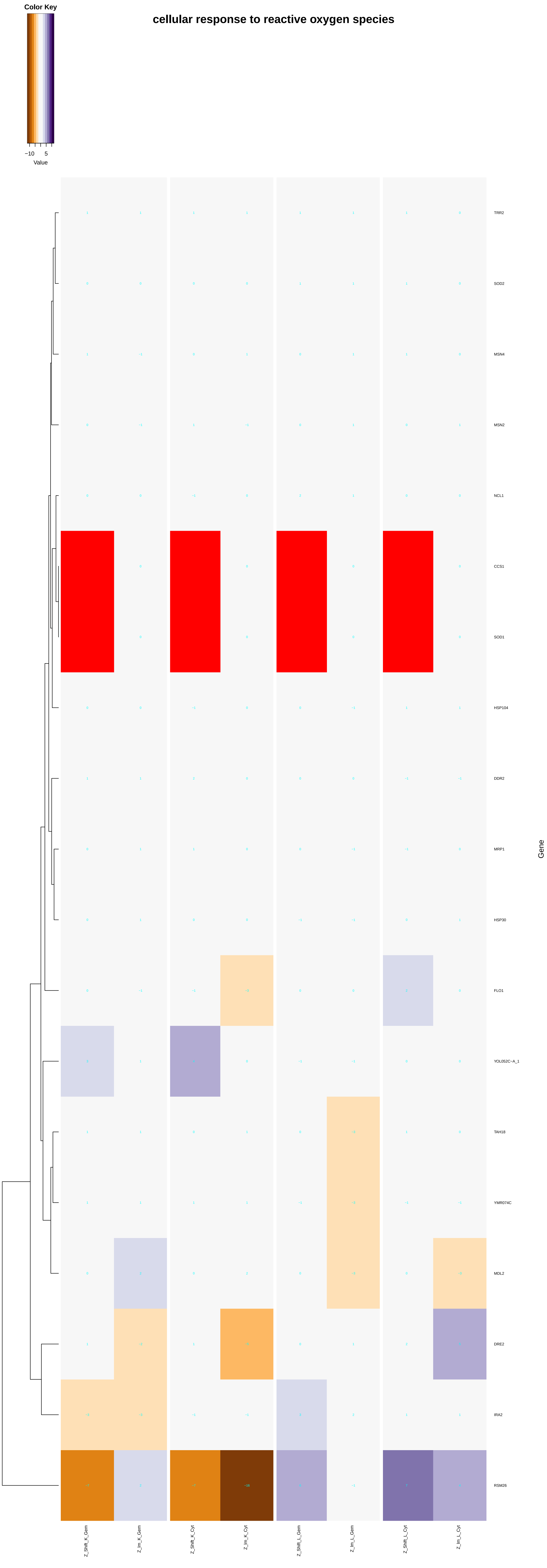

cellular response to reactive oxygen species
Color Key
−10
5
Value
TRR2
SOD2
MSN4
MSN2
NCL1
CCS1
SOD1
HSP104
DDR2
MRP1
HSP30
FLO1
YOL052C−A_1
TAH18
YMR074C
MDL2
DRE2
IRA2
RSM26
Gene
Z_Shift_K_Gem
Z_Im_K_Gem
Z_Shift_K_Cyt
Z_Im_K_Cyt
Z_Shift_L_Gem
Z_Im_L_Gem
Z_Shift_L_Cyt
Z_Im_L_Cyt

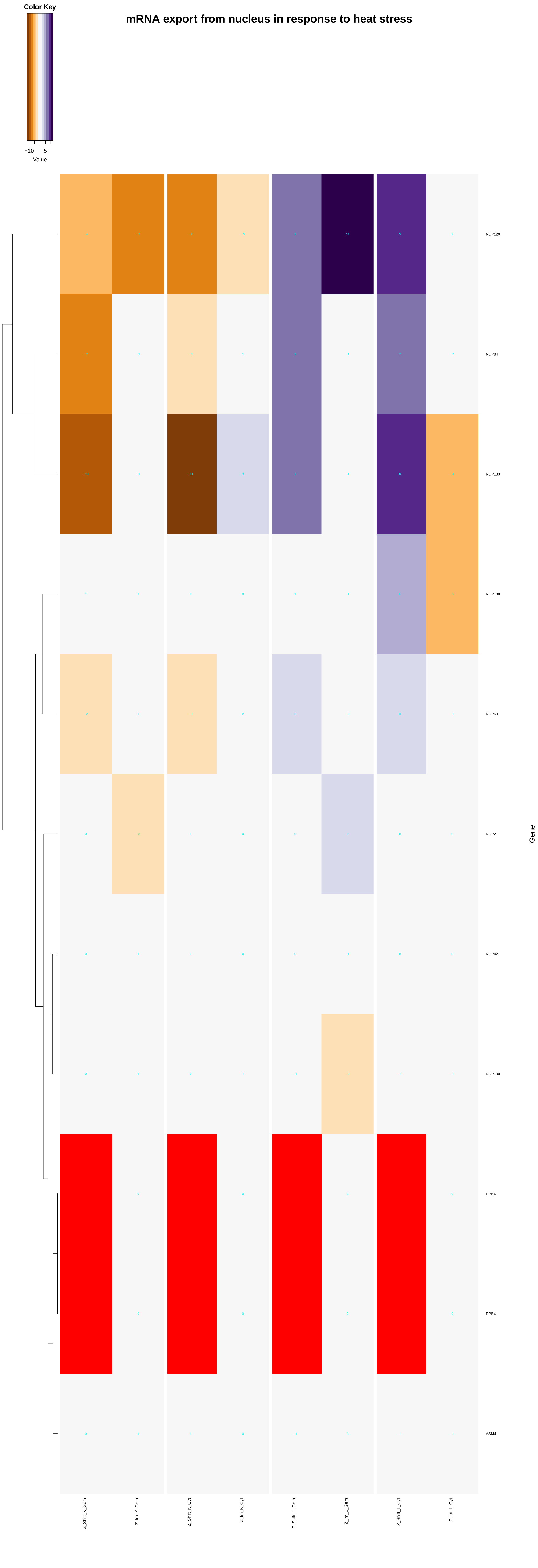
mRNA export from nucleus in response to heat stress
Color Key
−10
5
Value
NUP120
NUP84
NUP133
NUP188
NUP60
NUP2
NUP42
NUP100
RPB4
RPB4
ASM4
Gene
Z_Shift_K_Gem
Z_Im_K_Gem
Z_Shift_K_Cyt
Z_Im_K_Cyt
Z_Shift_L_Gem
Z_Im_L_Gem
Z_Shift_L_Cyt
Z_Im_L_Cyt

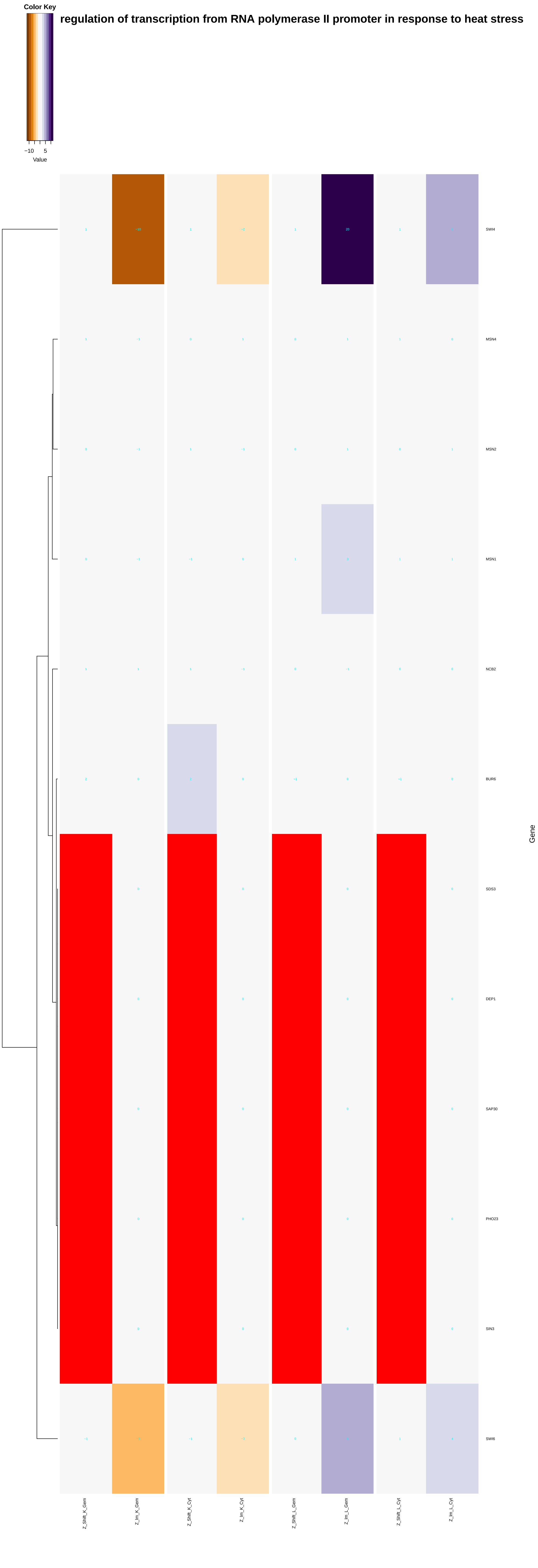

Color Key
−10 5
Value
regulation of transcription from RNA polymerase II promoter in response to heat stress
SWI4
MSN4
MSN2
MSN1
NCB2
BUR6
SDS3
DEP1
SAP30
PHO23
SIN3
SWI6
Gene
Z_Shift_K_Gem
Z_Im_K_Gem
Z_Shift_K_Cyt
Z_Im_K_Cyt
Z_Shift_L_Gem
Z_Im_L_Gem
Z_Shift_L_Cyt
Z_Im_L_Cyt

### endoplasmic reticulum unfolded protein response

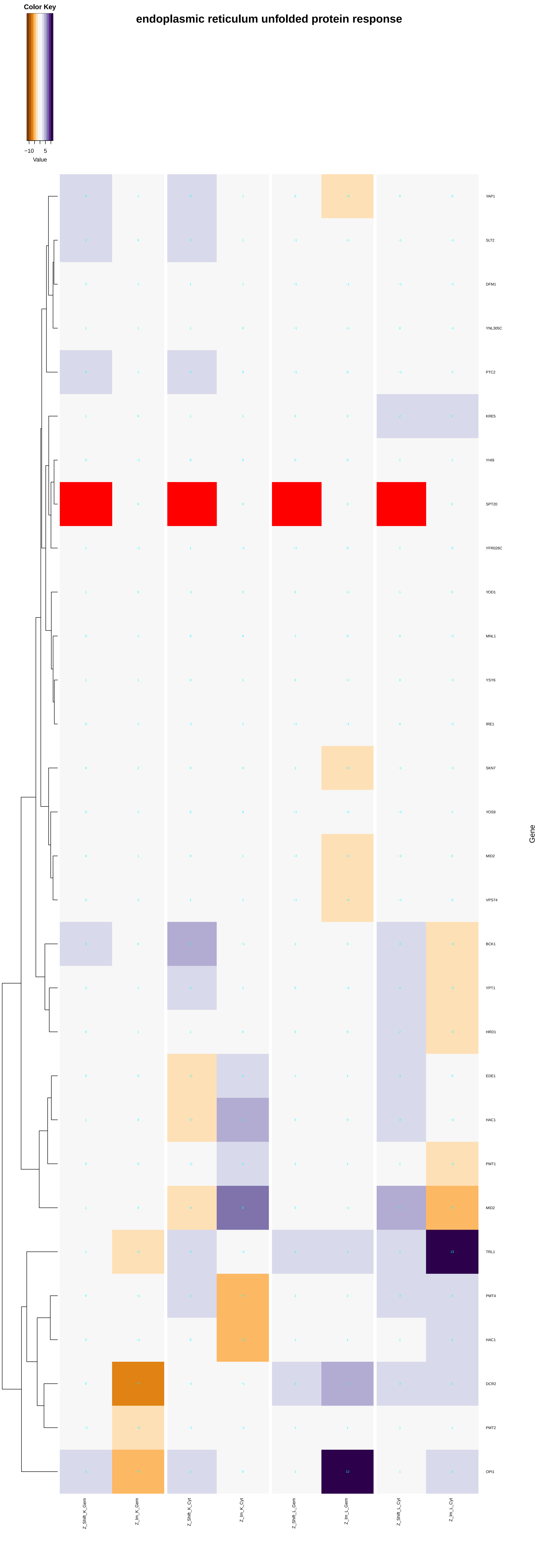

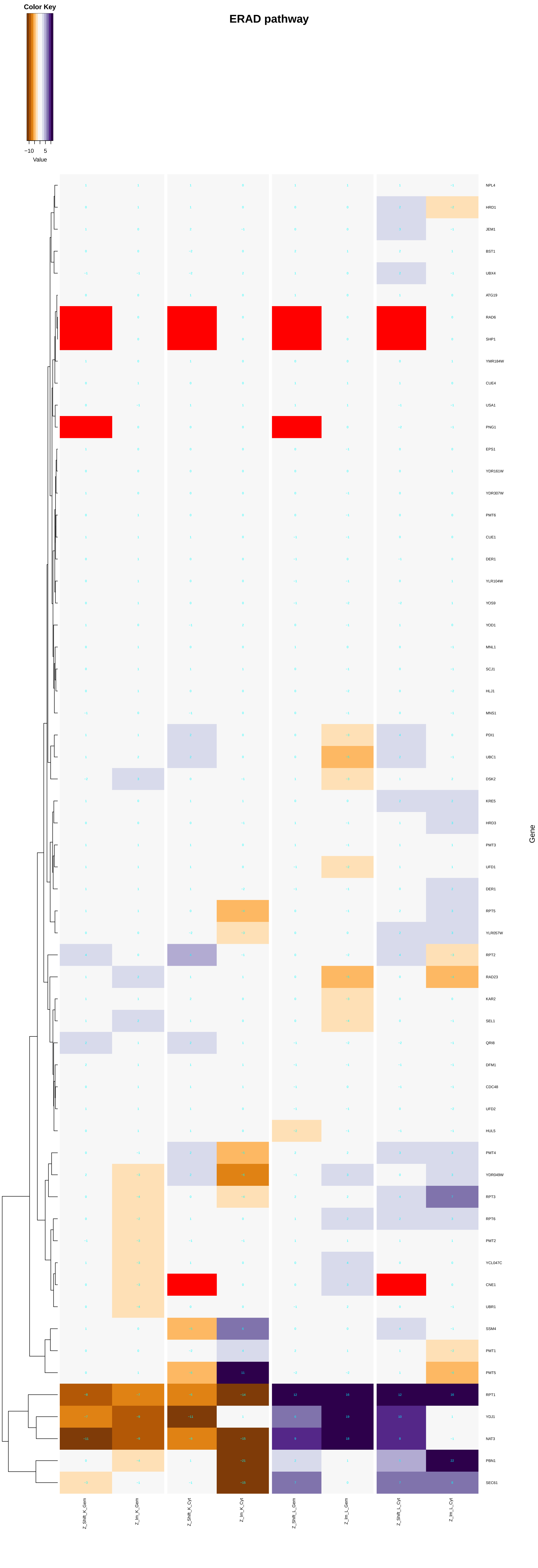

ERAD pathway
Color Key
−10
5
Value
Gene

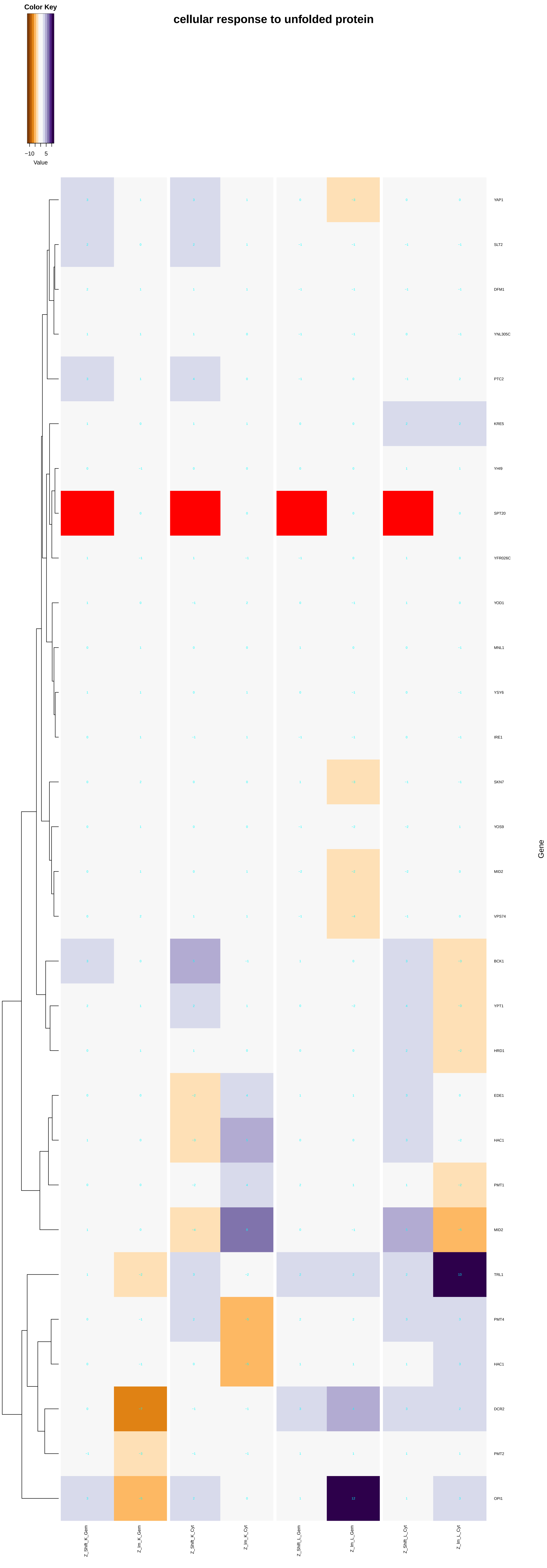

cellular response to unfolded protein
Color Key
−10 5
Value
YAP1
SLT2
DFM1
YNL305C
PTC2
KRE5
YHI9
SPT20
YFR026C
YOD1
MNL1
YSY6
IRE1
SKN7
YOS9
MID2
VPS74
BCK1
YPT1
HRD1
EDE1
HAC1
PMT1
MID2
TRL1
PMT4
HAC1
DCR2
PMT2
OPI1
Gene
Z_Shift_K_Gem
Z_lm_K_Gem
Z_Shift_K_Cyt
Z_lm_K_Cyt
Z_Shift_L_Gem
Z_lm_L_Gem
Z_Shift_L_Cyt
Z_lm_L_Cyt

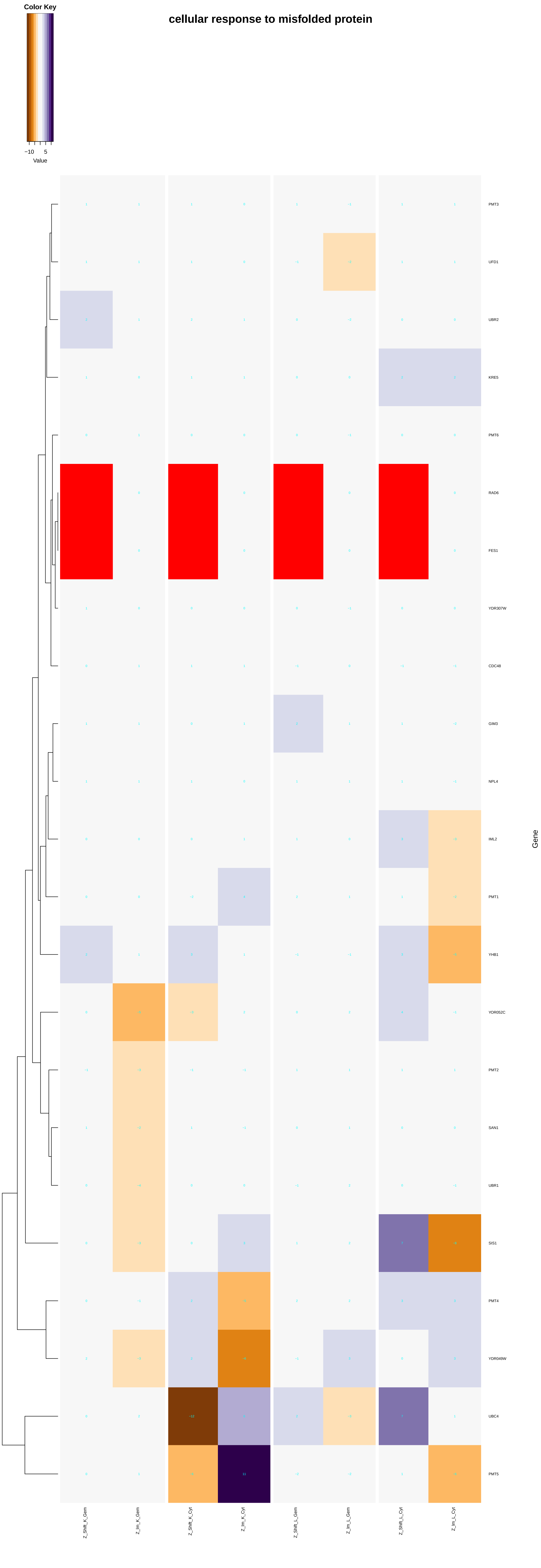

cellular response to misfolded protein
Color Key
−10
5
Value
PMT3
UFD1
UBR2
KRE5
PMT6
RAD6
FES1
YDR307W
CDC48
GIM3
NPL4
IML2
PMT1
YHB1
YDR052C
PMT2
SAN1
UBR1
SIS1
PMT4
YDR049W
UBC4
PMT5
Gene
Z_Shift_K_Gem
Z_Im_K_Gem
Z_Shift_K_Cyt
Z_Im_K_Cyt
Z_Shift_L_Gem
Z_Im_L_Gem
Z_Shift_L_Cyt
Z_Im_L_Cyt

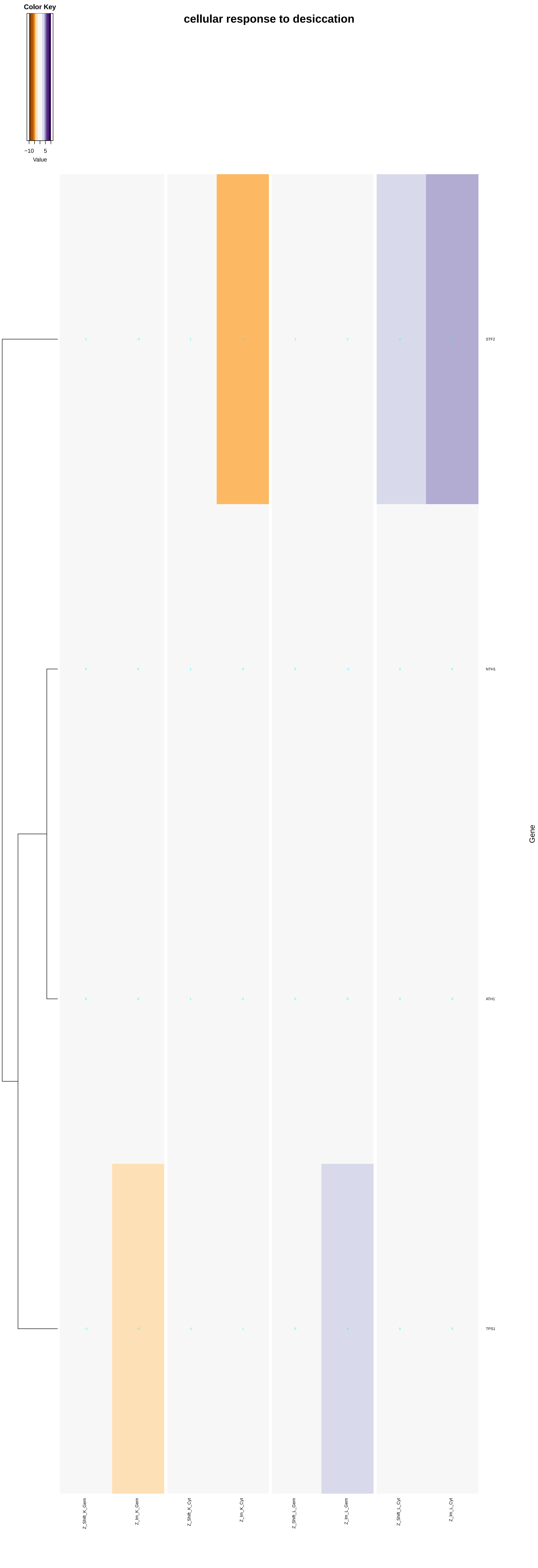
cellular response to desiccation
Color Key
−10
5
Value
STF2
NTH1
ATH1
TPS1
Gene
Z_Shift_K_Gem
Z_lm_K_Gem
Z_Shift_K_Cyt
Z_lm_K_Cyt
Z_Shift_L_Gem
Z_lm_L_Gem
Z_Shift_L_Cyt
Z_lm_L_Cyt

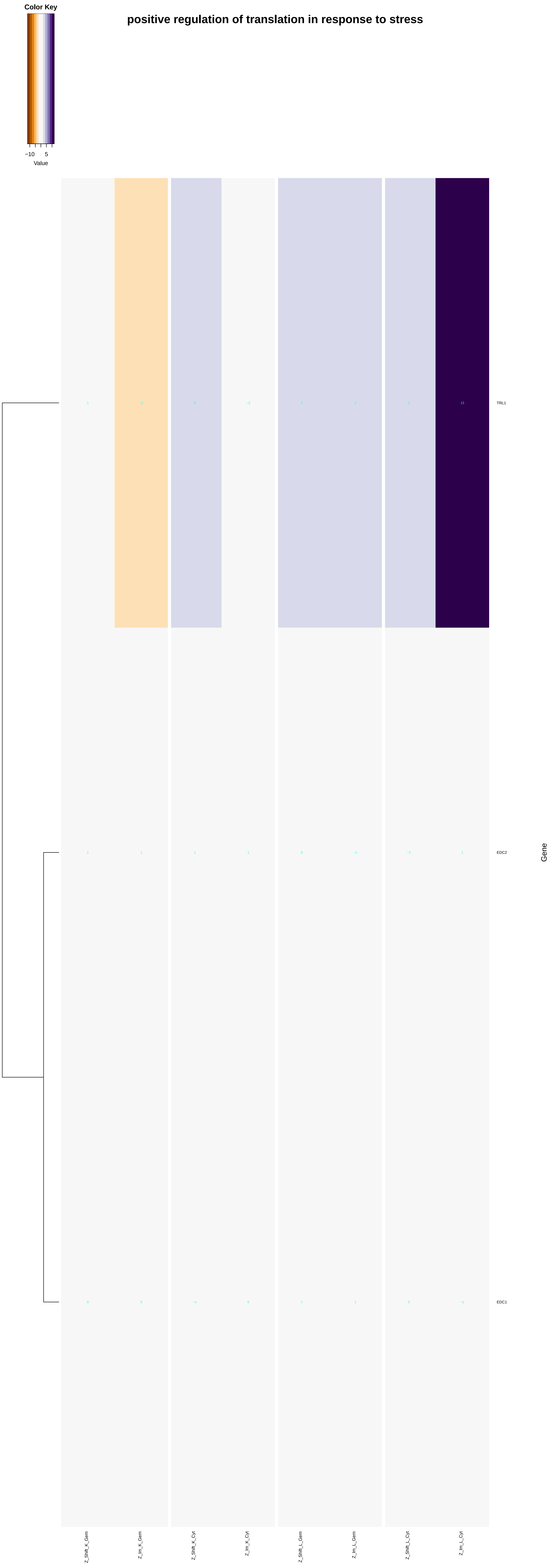
positive regulation of translation in response to stress
Color Key
−10
5
Value
TRL1
EDC2
EDC1
Gene
Z_Shift_K_Gem
Z_Im_K_Gem
Z_Shift_K_Cyt
Z_Im_K_Cyt
Z_Shift_L_Gem
Z_Im_L_Gem
Z_Shift_L_Cyt
Z_Im_L_Cyt

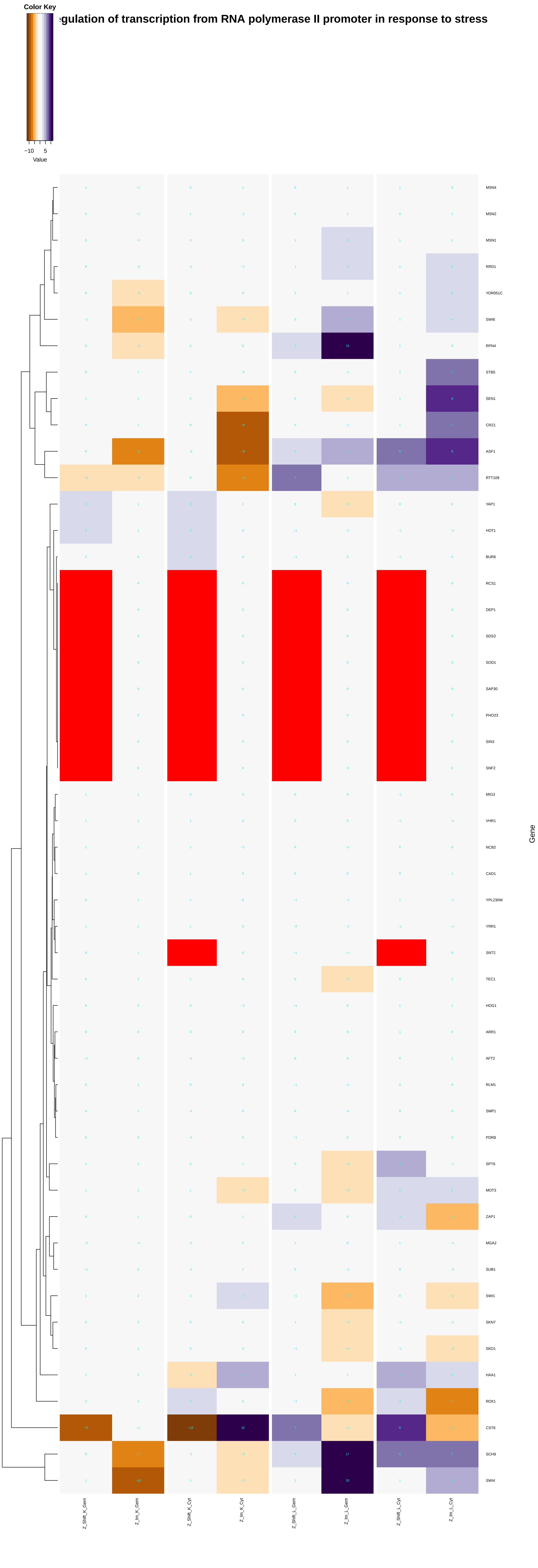

regulation of transcription from RNA polymerase II promoter in response to stress
Color Key
−10
5
Value
Gene
MSN4
MSN2
MSN1
RRD1
YOR051C
SWI6
RPN4
STB5
SEN1
CRZ1
ASF1
RTT109
YAP1
HOT1
BUR6
RCS1
DEP1
SDS3
SOD1
SAP30
PHO23
SIN3
SNF2
MIG3
VHR1
NCB2
CAD1
YPL230W
YRR1
SNT2
TEC1
HOG1
ARR1
AFT2
RLM1
SMP1
PDR8
SPT6
MOT3
ZAP1
MGA2
SUB1
SWI1
SKN7
SKO1
HAA1
ROX1
CST6
SCH9
SWI4
Z_Shift_K_Gem
Z_Im_K_Gem
Z_Shift_K_Cyt
Z_Im_K_Cyt
Z_Shift_L_Gem
Z_Im_L_Gem
Z_Shift_L_Cyt
Z_Im_L_Cyt

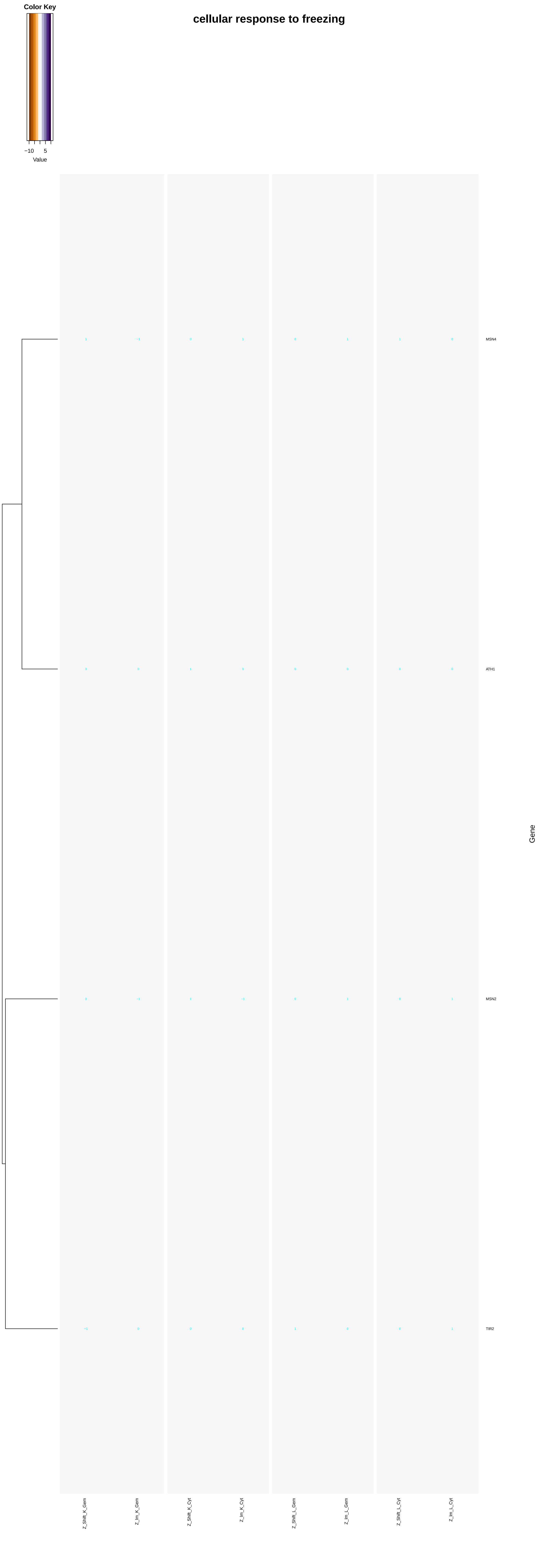

Color Key
−10
5
Value
cellular response to freezing
MSN4
ATH1
MSN2
TIR2
Gene
Z_Shift_K_Gem
Z_Im_K_Gem
Z_Shift_K_Cyt
Z_Im_K_Cyt
Z_Shift_L_Gem
Z_Im_L_Gem
Z_Shift_L_Cyt
Z_Im_L_Cyt

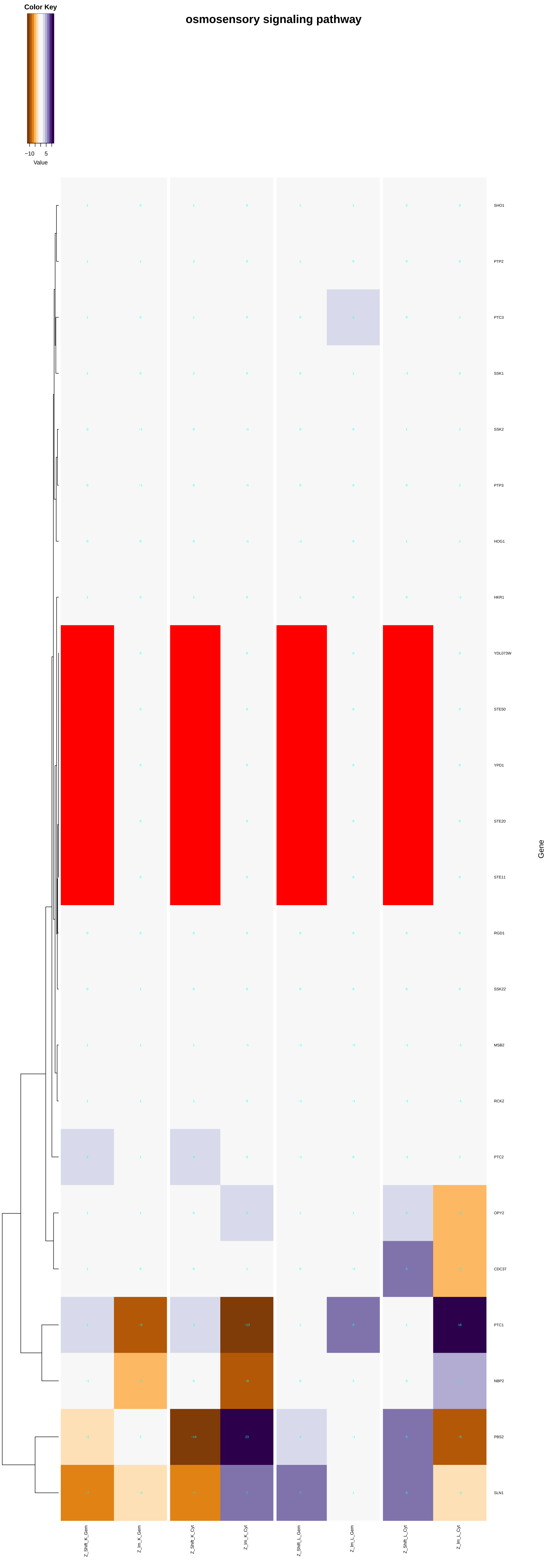

osmosensory signaling pathway
Color Key
−10
5
Value
SHO1
PTP2
PTC3
SSK1
SSK2
PTP3
HOG1
HKR1
YDL073W
STE50
YPD1
STE20
STE11
RGD1
SSK22
MSB2
RCK2
PTC2
OPY2
CDC37
PTC1
NBP2
PBS2
SLN1
Gene
Z_Shift_K_Gem
Z_Im_K_Gem
Z_Shift_K_Cyt
Z_Im_K_Cyt
Z_Shift_L_Gem
Z_Im_L_Gem
Z_Shift_L_Cyt
Z_Im_L_Cyt

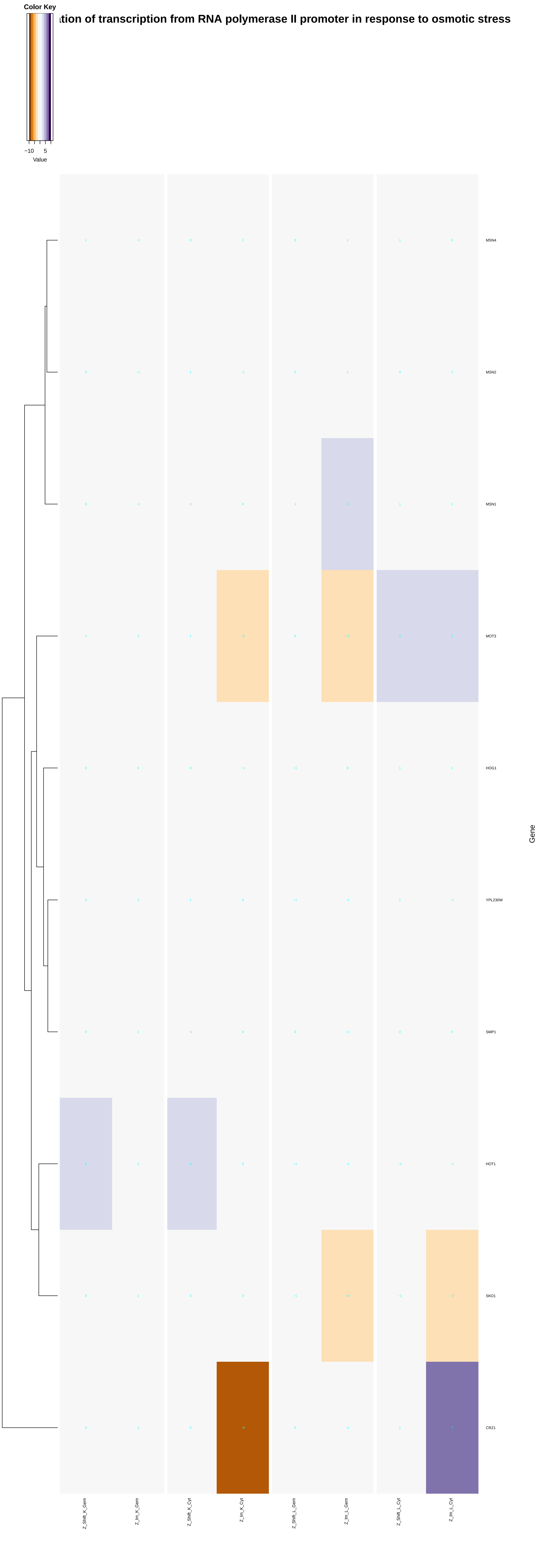
ıtion of transcription from RNA polymerase II promoter in response to osmotic stress
Color Key
−10
5
Value
MSN4
MSN2
MSN1
MOT3
HOG1
YPL230W
SMP1
HOT1
SKO1
CRZ1
Gene
Z_Shift_K_Gem
Z_Im_K_Gem
Z_Shift_K_Cyt
Z_Im_K_Cyt
Z_Shift_L_Gem
Z_Im_L_Gem
Z_Shift_L_Cyt
Z_Im_L_Cyt

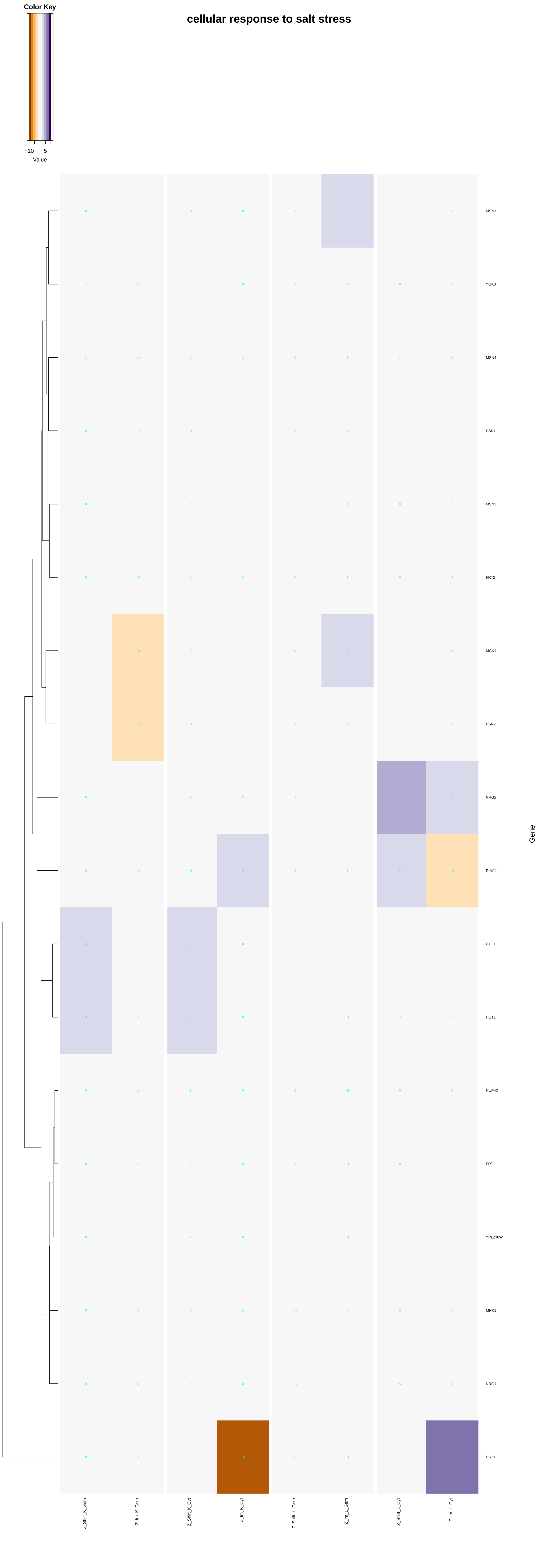
cellular response to salt stress
Color Key
−10
5
Value
MSN1
YGK3
MSN4
PSR1
MSN2
FRT2
MCK1
PSR2
NRG2
RIM11
CTT1
HOT1
NUP42
FRT1
YPL230W
MRK1
NRG1
CRZ1
Gene
Z_Shift_K_Gem
Z_Im_K_Gem
Z_Shift_K_Cyt
Z_Im_K_Cyt
Z_Shift_L_Gem
Z_Im_L_Gem
Z_Shift_L_Cyt
Z_Im_L_Cyt

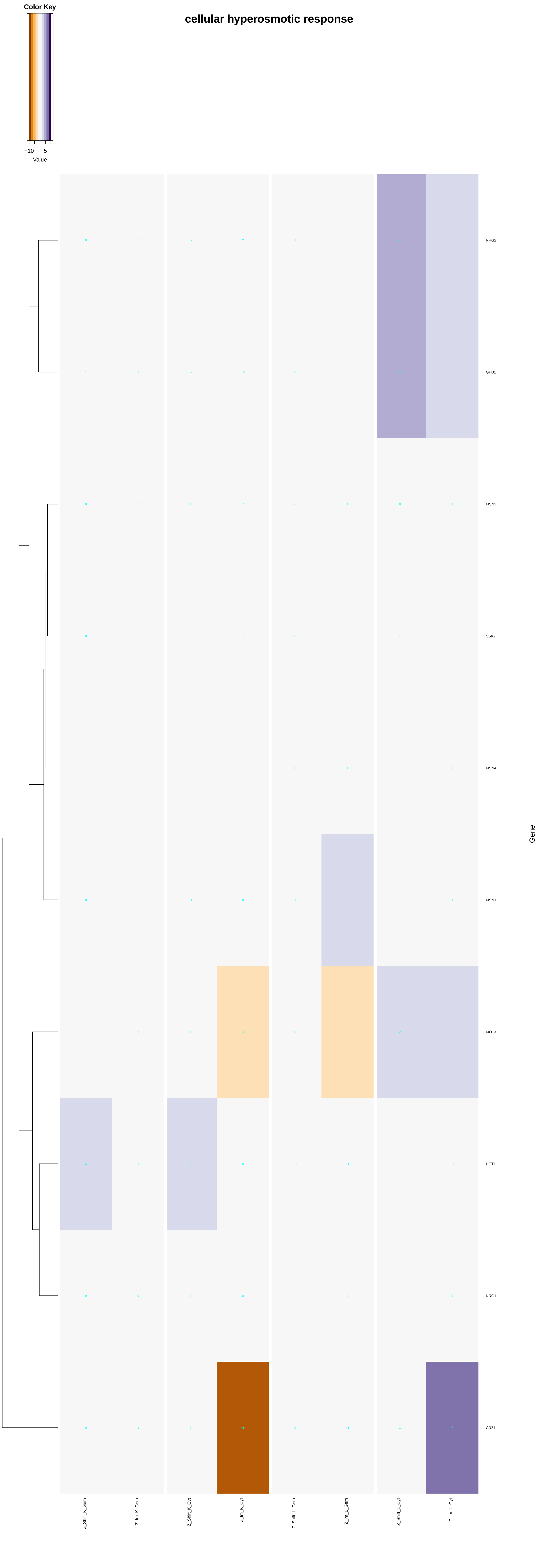

cellular hyperosmotic response
Color Key
−10
5
Value
NRG2
GPD1
MSN2
SSK2
MSN4
MSN1
MOT3
HOT1
NRG1
CRZ1
Gene
Z_Shift_K_Gem
Z_Im_K_Gem
Z_Shift_K_Cyt
Z_Im_K_Cyt
Z_Shift_L_Gem
Z_Im_L_Gem
Z_Shift_L_Cyt
Z_Im_L_Cyt

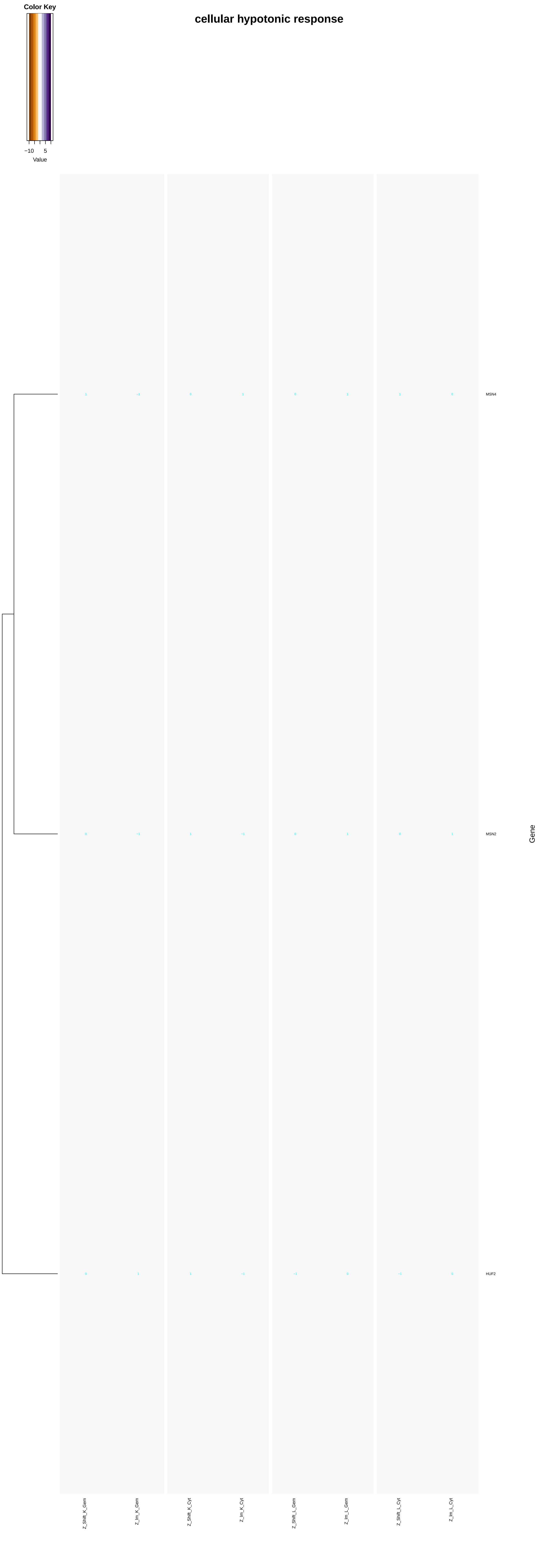
Color Key
−10
5
Value
cellular hypotonic response
MSN4
MSN2
HUF2
Gene
Z_Shift_K_Gem
Z_Im_K_Gem
Z_Shift_K_Cyt
Z_Im_K_Cyt
Z_Shift_L_Gem
Z_Im_L_Gem
Z_Shift_L_Cyt
Z_Im_L_Cyt

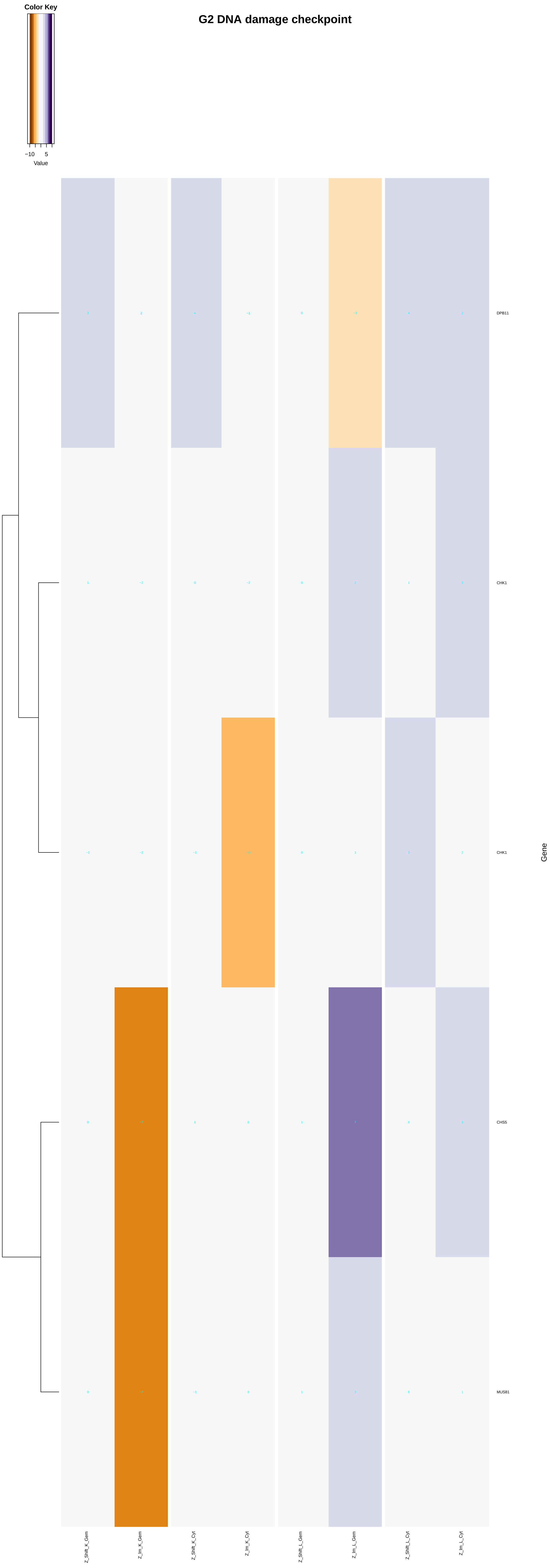
G2 DNA damage checkpoint
Color Key
−10
5
Value
DPB11
CHK1
CHK1
CHS5
MUS81
Gene
Z_Shift_K_Gem
Z_Im_K_Gem
Z_Shift_K_Cyt
Z_Im_K_Cyt
Z_Shift_L_Gem
Z_Im_L_Gem
Z_Shift_L_Cyt
Z_Im_L_Cyt

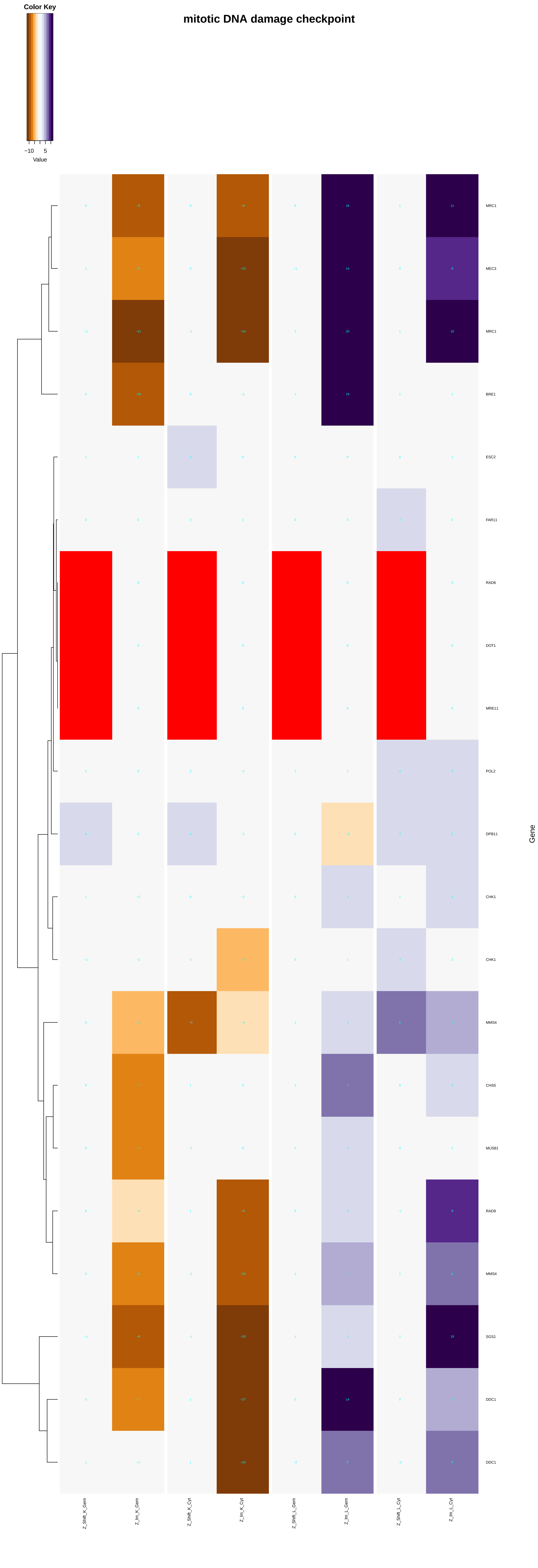

mitotic DNA damage checkpoint
Color Key
−10 5
Value
MRC1
MEC3
MRC1
BRE1
ESC2
FAR11
RAD6
DOT1
MRE11
POL2
DPB11
CHK1
CHK1
MMS4
CHS5
MUS81
RAD9
MMS4
SGS1
DDC1
DDC1
Gene
Z_Shift_K_Gem
Z_Im_K_Gem
Z_Shift_K_Cyt
Z_Im_K_Cyt
Z_Shift_L_Gem
Z_Im_L_Gem
Z_Shift_L_Cyt
Z_Im_L_Cyt

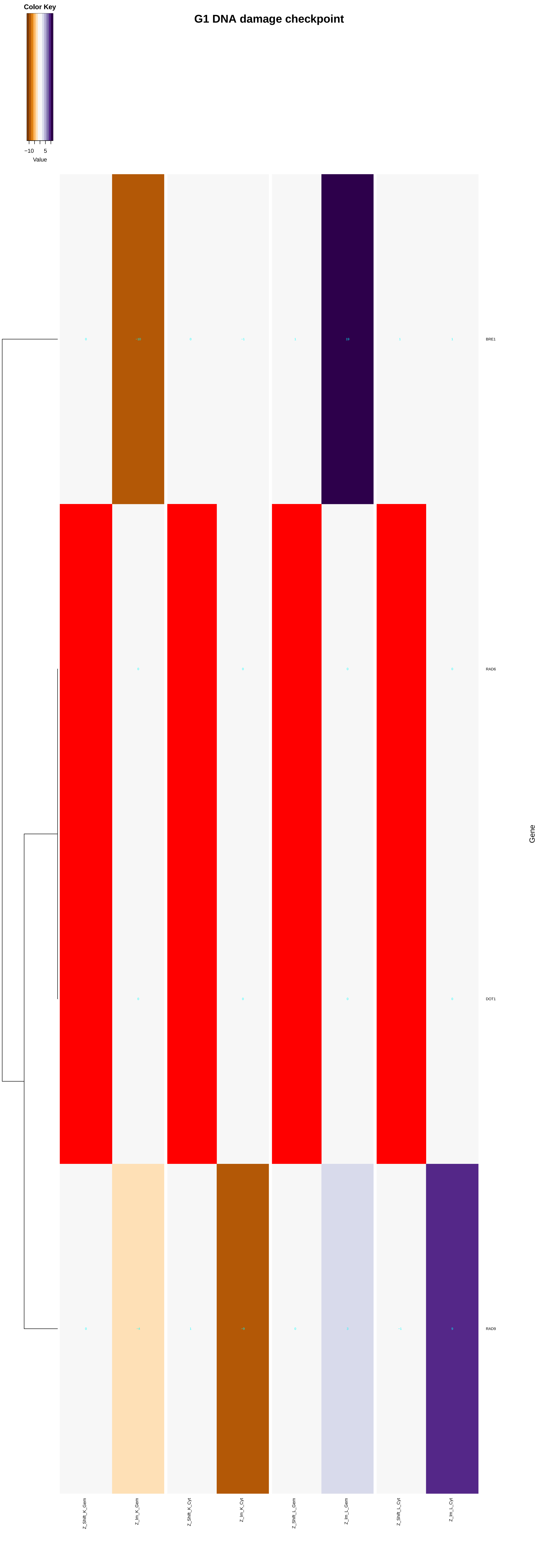

Color Key
−10
5
Value
G1 DNA damage checkpoint
BRE1
RAD6
DOT1
RAD9
Gene
Z_Shift_K_Gem
Z_Im_K_Gem
Z_Shift_K_Cyt
Z_Im_K_Cyt
Z_Shift_L_Gem
Z_Im_L_Gem
Z_Shift_L_Cyt
Z_Im_L_Cyt

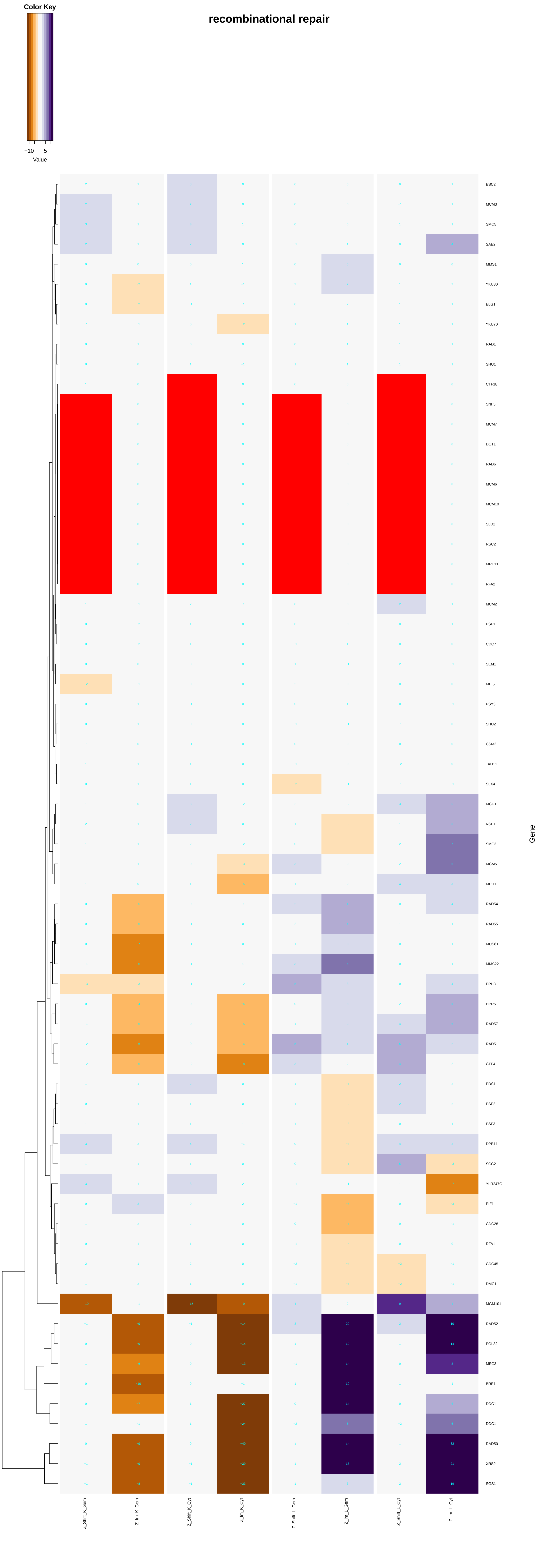
Color Key
−10
5
Value
recombinational repair
ESC2
MCM3
SMC5
SAE2
MMS1
YKU80
ELG1
YKU70
RAD1
SHU1
CTF18
SNF5
MCM7
DOT1
RAD6
MCM6
MCM10
SLD2
RSC2
MRE11
RFA2
MCM2
PSF1
CDC7
SEM1
MEI5
PSY3
SHU2
CSM2
TAH11
SLX4
MCD1
NSE1
SMC3
MCM5
MPH1
RAD54
RAD55
MUS81
MMS22
PPH3
HPR5
RAD57
RAD51
CTF4
PDS1
PSF2
PSF3
DPB11
SCC2
YLR247C
PIF1
CDC28
RFA1
CDC45
DMC1
MGM101
RAD52
POL32
MEC3
BRE1
DDC1
DDC1
RAD50
XRS2
SGS1
Gene
Z_Shift_K_Gem
Z_Im_K_Gem
Z_Shift_K_Cyt
Z_Im_K_Cyt
Z_Shift_L_Gem
Z_Im_L_Gem
Z_Shift_L_Cyt
Z_Im_L_Cyt

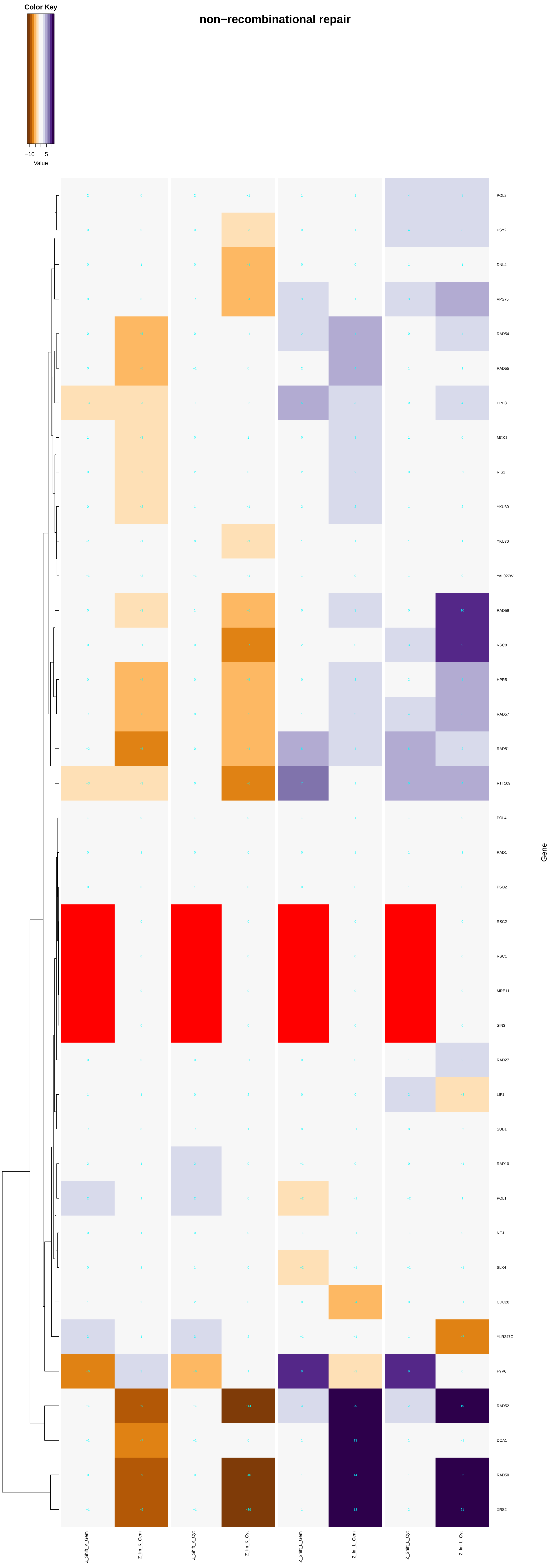

Color Key
−10
5
Value
non−recombinational repair
POL2
PSY2
DNL4
VPS75
RAD54
RAD55
PPH3
MCK1
RIS1
YKU80
YKU70
YAL027W
RAD59
RSC8
HPR5
RAD57
RAD51
RTT109
POL4
RAD1
PSO2
RSC2
RSC1
MRE11
SIN3
RAD27
LIF1
SUB1
RAD10
POL1
NEJ1
SLX4
CDC28
YLR247C
FYV6
RAD52
DOA1
RAD50
XRS2
Gene
Z_Shift_K_Gem
Z_Im_K_Gem
Z_Shift_K_Cyt
Z_Im_K_Cyt
Z_Shift_L_Gem
Z_Im_L_Gem
Z_Shift_L_Cyt
Z_Im_L_Cyt

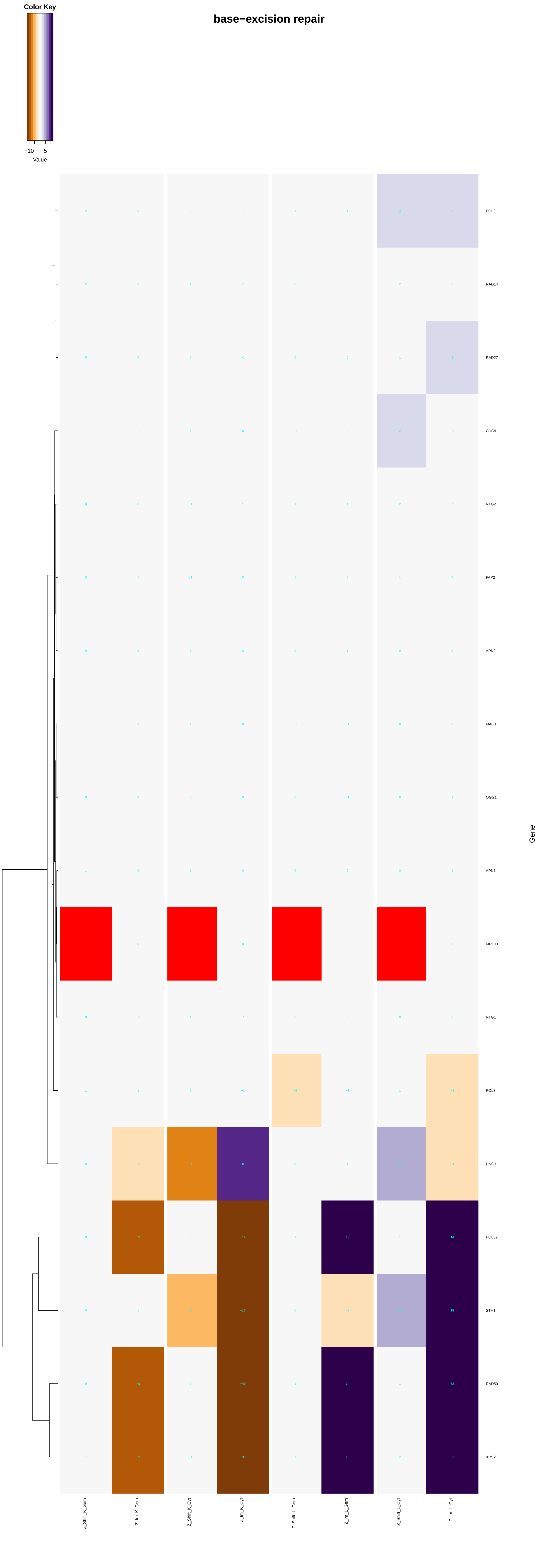
base−excision repair
Color Key
−10 5
Value
POL2
RAD14
RAD27
CDC9
NTG2
PAP2
APN2
MAG1
OGG1
APN1
MRE11
NTG1
POL3
UNG1
POL32
STH1
RAD50
XRS2
Gene
Z_Shift_K_Gem
Z_Im_K_Gem
Z_Shift_K_Cyt
Z_Im_K_Cyt
Z_Shift_L_Gem
Z_Im_L_Gem
Z_Shift_L_Cyt
Z_Im_L_Cyt

### nucleotide−excision repair

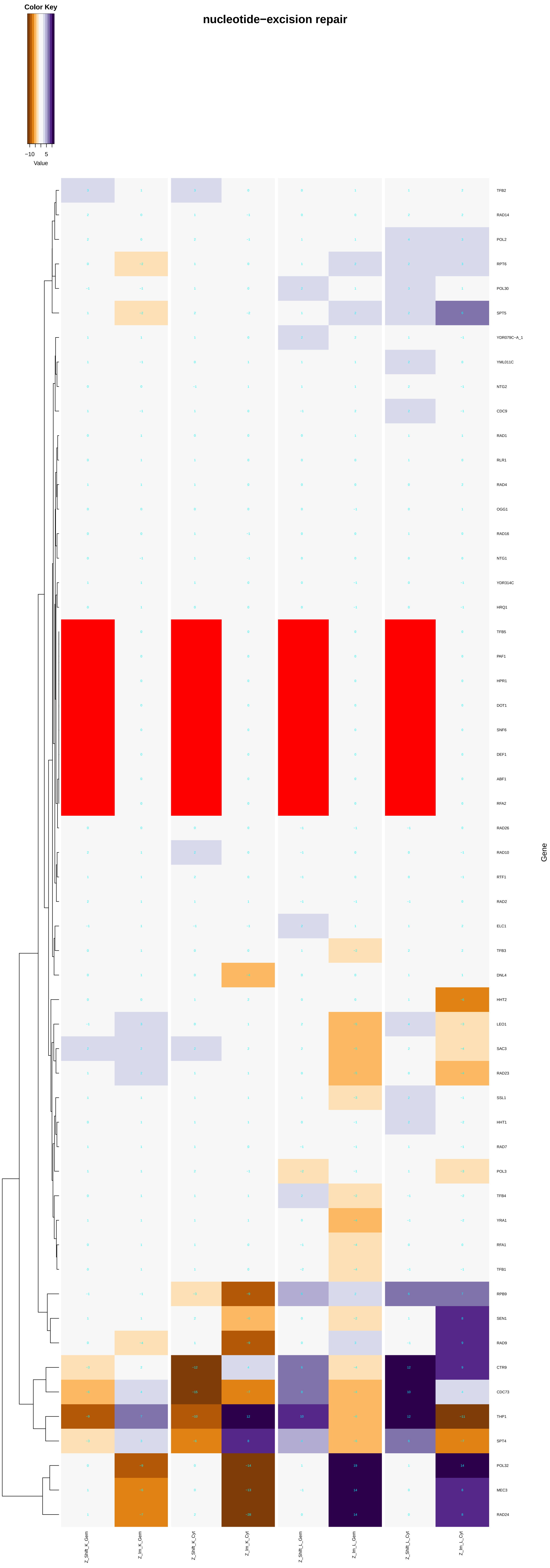

### mismatch repair

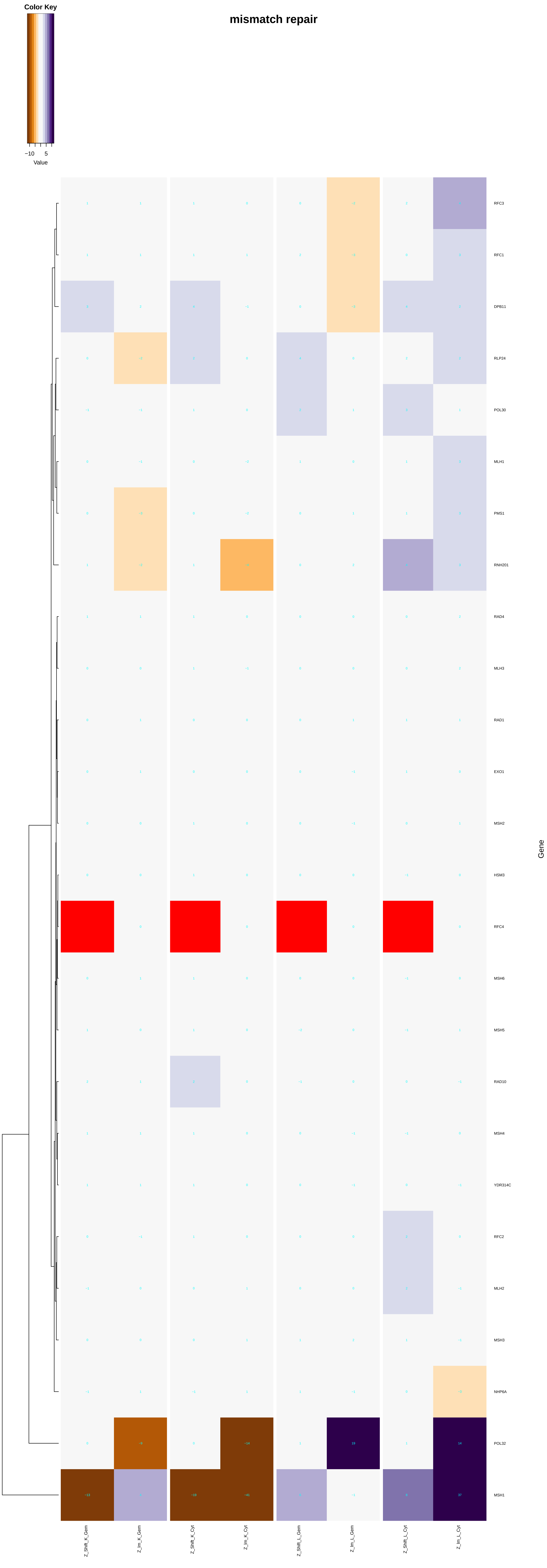

### postreplication repair

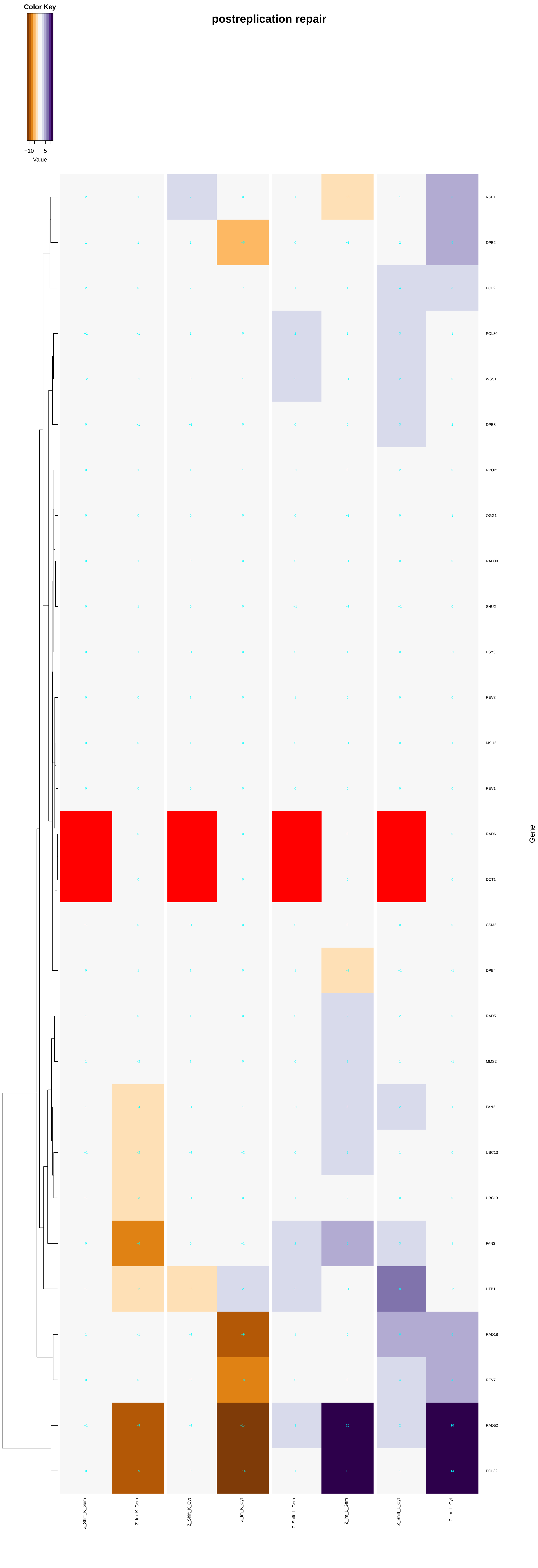

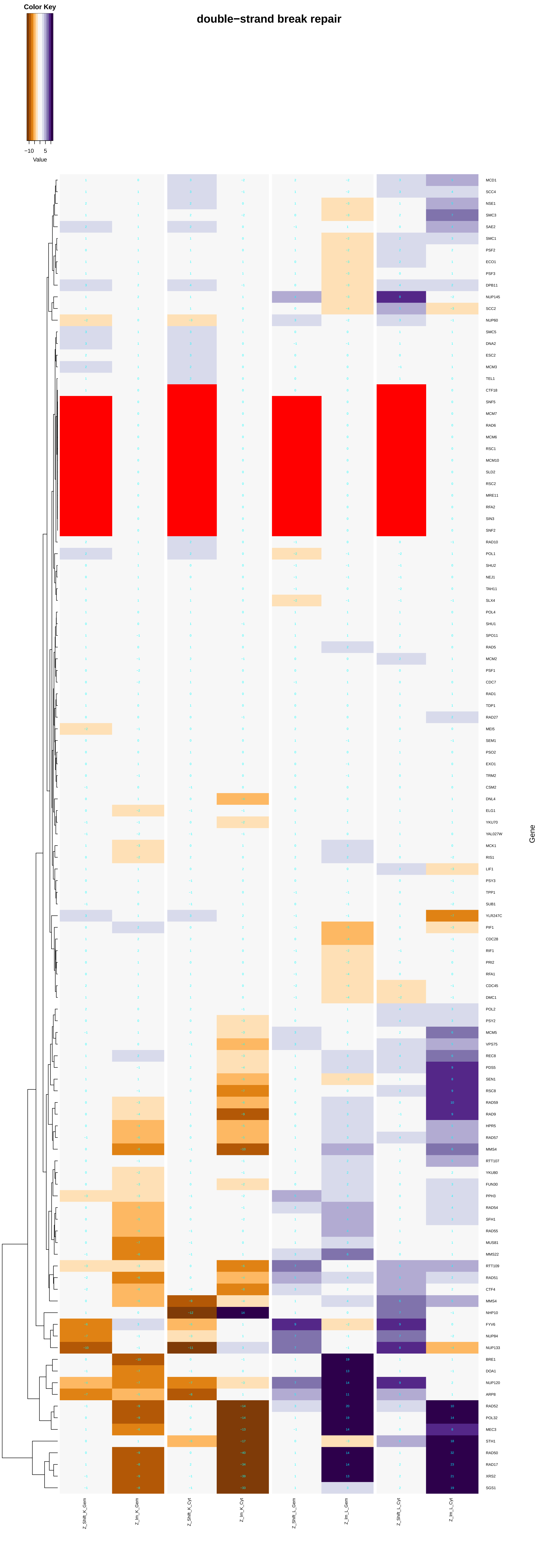
double−strand break repair
Color Key
−10
5
Value
Gene
MCD1
SCC4
NSE1
SMC3
SAE2
SMC1
PSF2
ECO1
PSF3
DPB11
NUP145
SCC2
NUP60
SMC5
DNA2
ESC2
MCM3
TEL1
CTF18
SNF5
MCM7
RAD6
MCM6
RSC1
MCM10
SLD2
RSC2
MRE11
RFA2
SIN3
SNF2
RAD10
POL1
SHU2
NEJ1
TAH11
SLX4
POL4
SHU1
SPO11
RAD5
MCM2
PSF1
CDC7
RAD1
TDP1
RAD27
MEI5
SEM1
PSO2
EXO1
TRM2
CSM2
DNL4
ELG1
YKU70
YAL027W
MCK1
RIS1
LIF1
PSY3
TPP1
SUB1
YLR247C
PIF1
CDC28
RIF1
PRI2
RFA1
CDC45
DMC1
POL2
PSY2
MCM5
VPS75
REC8
PDS5
SEN1
RSC8
RAD59
RAD9
HPR5
RAD57
MMS4
RTT107
YKU80
FUN30
PPH3
RAD54
SFH1
RAD55
MUS81
MMS22
RTT109
RAD51
CTF4
MMS4
NHP10
FYV6
NUP84
NUP133
BRE1
DOA1
NUP120
ARP8
RAD52
POL32
MEC3
STH1
RAD50
RAD17
XRS2
SGS1
Z_Shift_K_Gem
Z_Im_K_Gem
Z_Shift_K_Cyt
Z_Im_K_Cyt
Z_Shift_L_Gem
Z_Im_L_Gem
Z_Shift_L_Cyt
Z_Im_L_Cyt

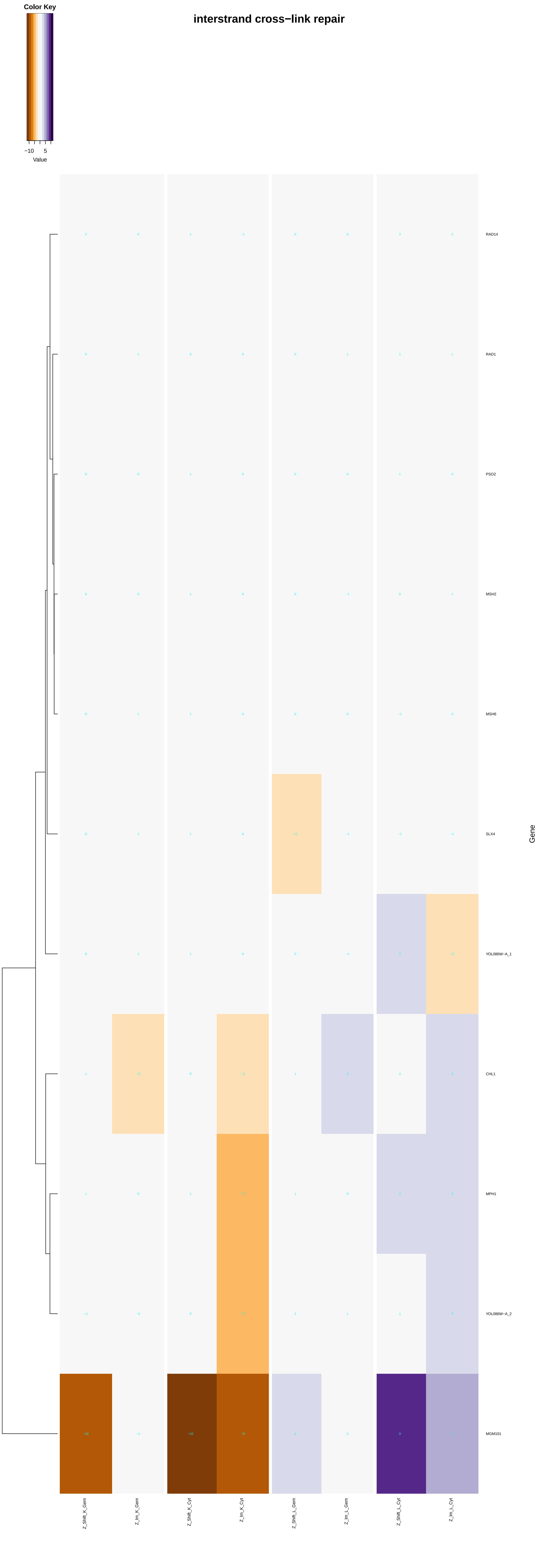
interstrand cross−link repair
Color Key
−10
5
Value
RAD14
RAD1
PSO2
MSH2
MSH6
SLX4
YOL086W−A_1
CHL1
MPH1
YOL086W−A_2
MGM101
Gene
Z_Shift_K_Gem
Z_Im_K_Gem
Z_Shift_K_Cyt
Z_Im_K_Cyt
Z_Shift_L_Gem
Z_Im_L_Gem
Z_Shift_L_Cyt
Z_Im_L_Cyt

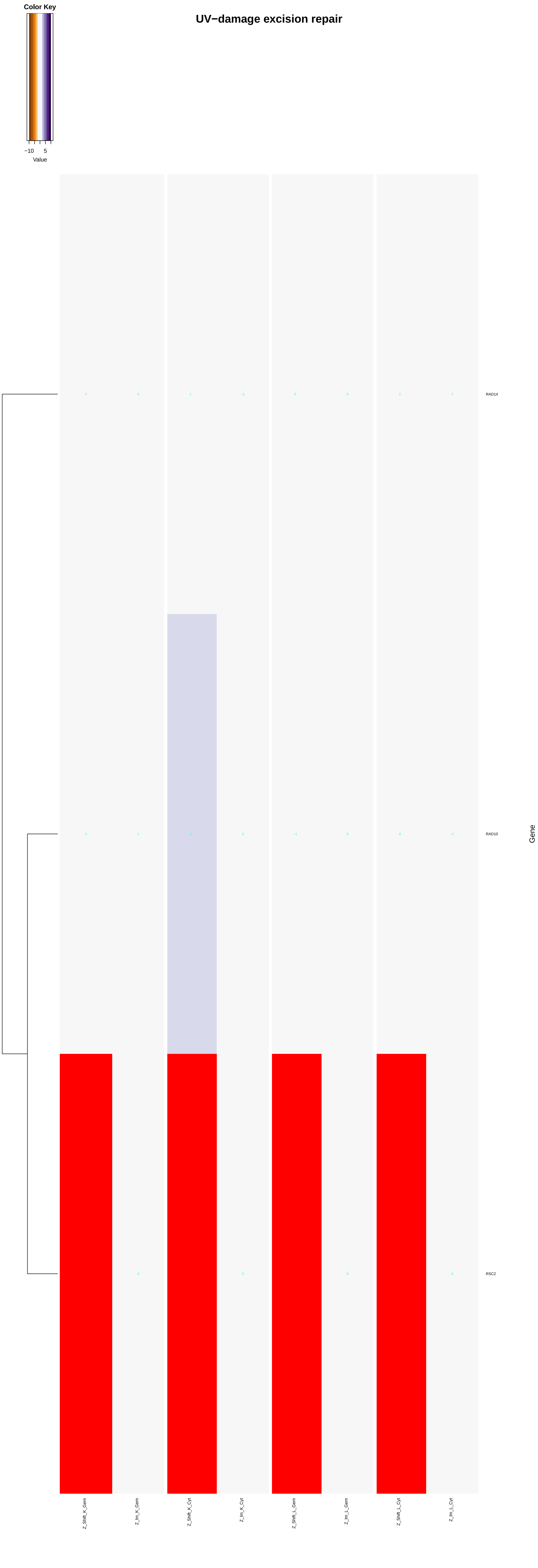
UV−damage excision repair
Color Key
−10
5
Value
RAD14
RAD10
RSC2
Gene
Z_Shift_K_Gem
Z_Im_K_Gem
Z_Shift_K_Cyt
Z_Im_K_Cyt
Z_Shift_L_Gem
Z_Im_L_Gem
Z_Shift_L_Cyt
Z_Im_L_Cyt

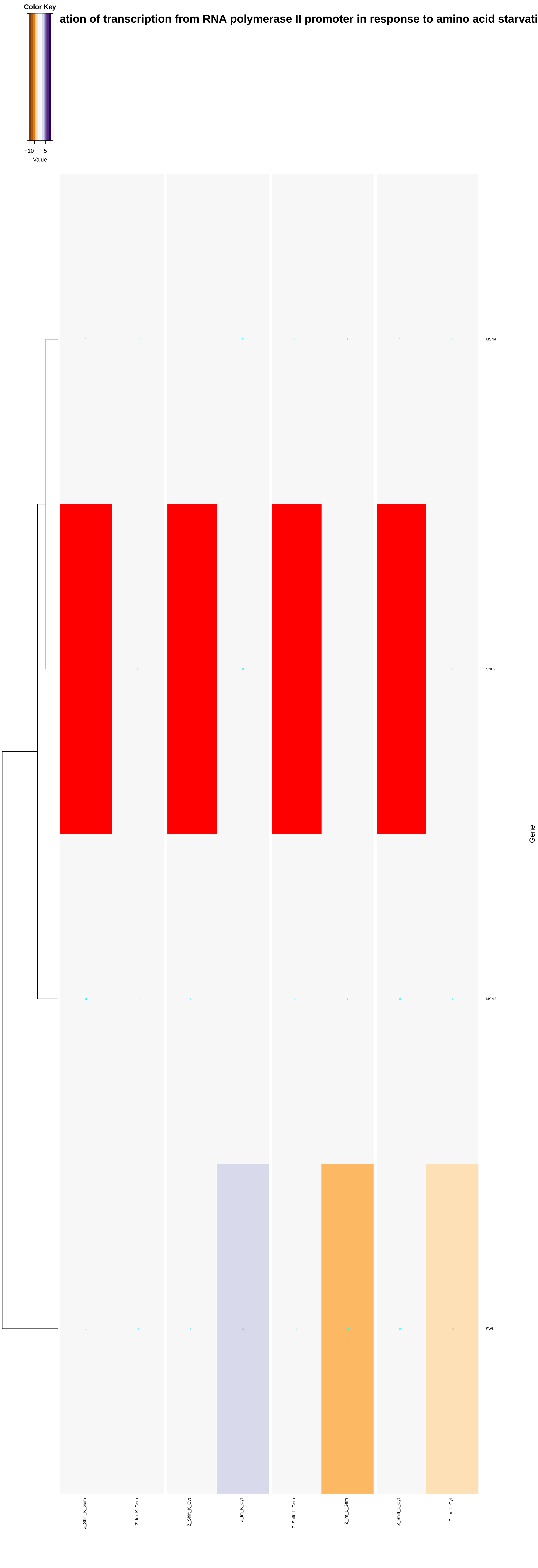
Color Key
−10
5
Value
ation of transcription from RNA polymerase II promoter in response to amino acid starvati
MSN4
SNF2
MSN2
SWI1
Gene
Z_Shift_K_Gem
Z_Im_K_Gem
Z_Shift_K_Cyt
Z_Im_K_Cyt
Z_Shift_L_Gem
Z_Im_L_Gem
Z_Shift_L_Cyt
Z_Im_L_Cyt

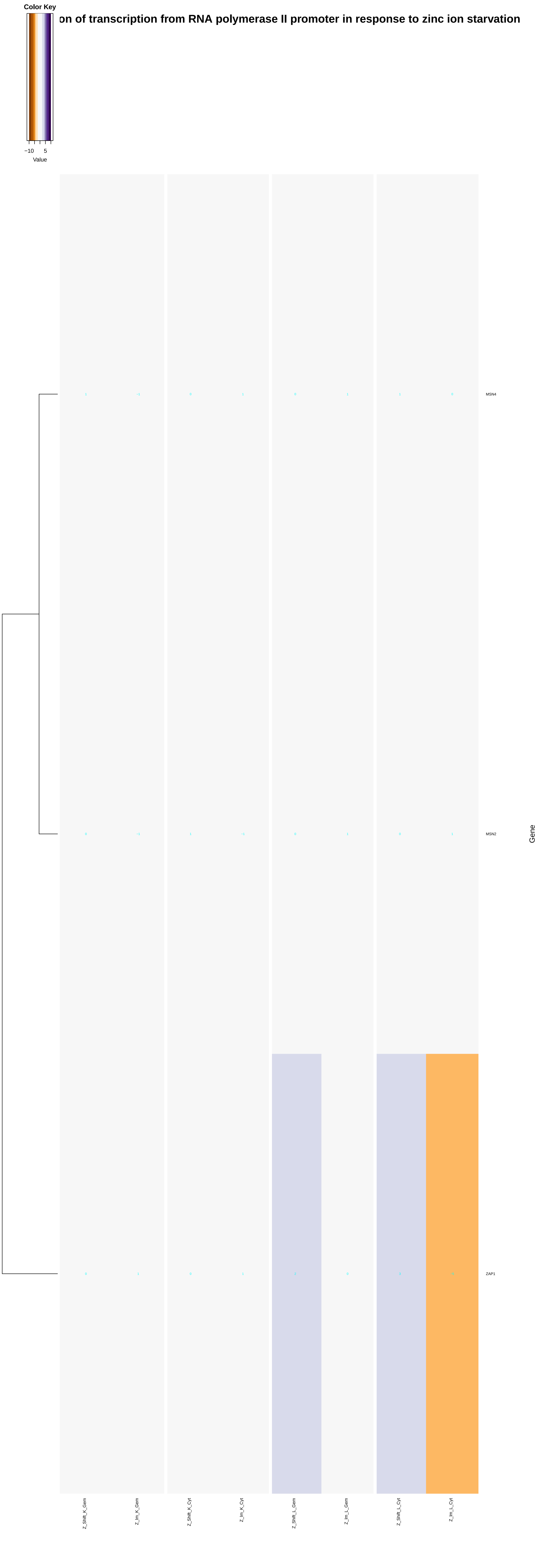
on of transcription from RNA polymerase II promoter in response to zinc ion starvation
Color Key
−10
5
Value
MSN4
MSN2
ZAP1
Gene
Z_Shift_K_Gem
Z_Im_K_Gem
Z_Shift_K_Cyt
Z_Im_K_Cyt
Z_Shift_L_Gem
Z_Im_L_Gem
Z_Shift_L_Cyt
Z_Im_L_Cyt

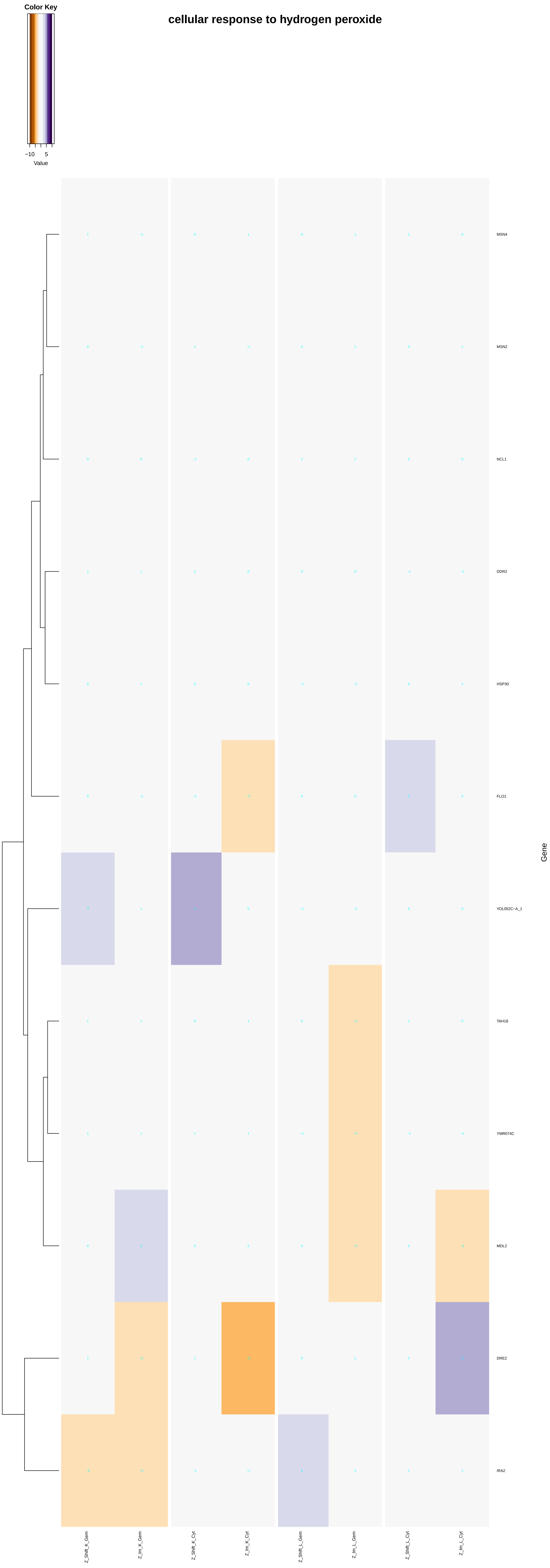
cellular response to hydrogen peroxide
Color Key
−10
5
Value
MSN4
MSN2
NCL1
DDR2
HSP30
FLO1
YOL052C−A_1
TAH18
YMR074C
MDL2
DRE2
IRA2
Gene
Z_Shift_K_Gem
Z_Im_K_Gem
Z_Shift_K_Cyt
Z_Im_K_Cyt
Z_Shift_L_Gem
Z_Im_L_Gem
Z_Shift_L_Cyt
Z_Im_L_Cyt

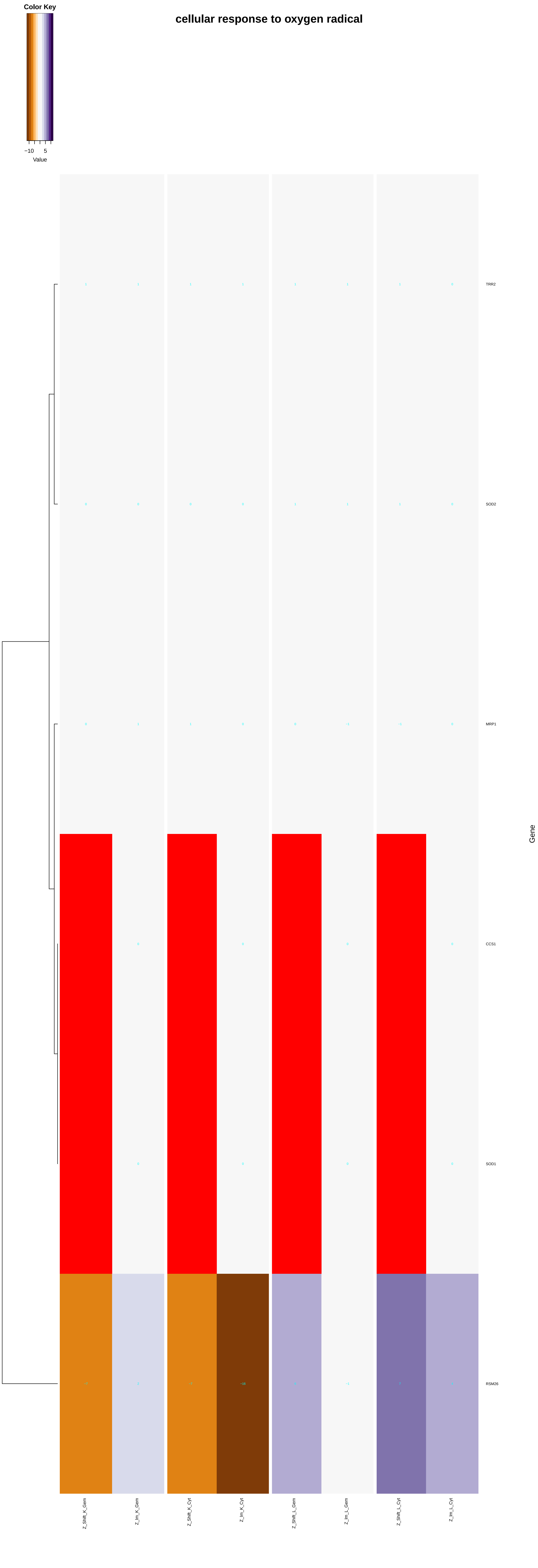
cellular response to oxygen radical
Color Key
−10
5
Value
TRR2
SOD2
MRP1
CCS1
SOD1
RSM26
Gene
Z_Shift_K_Gem
Z_Im_K_Gem
Z_Shift_K_Cyt
Z_Im_K_Cyt
Z_Shift_L_Gem
Z_Im_L_Gem
Z_Shift_L_Cyt
Z_Im_L_Cyt

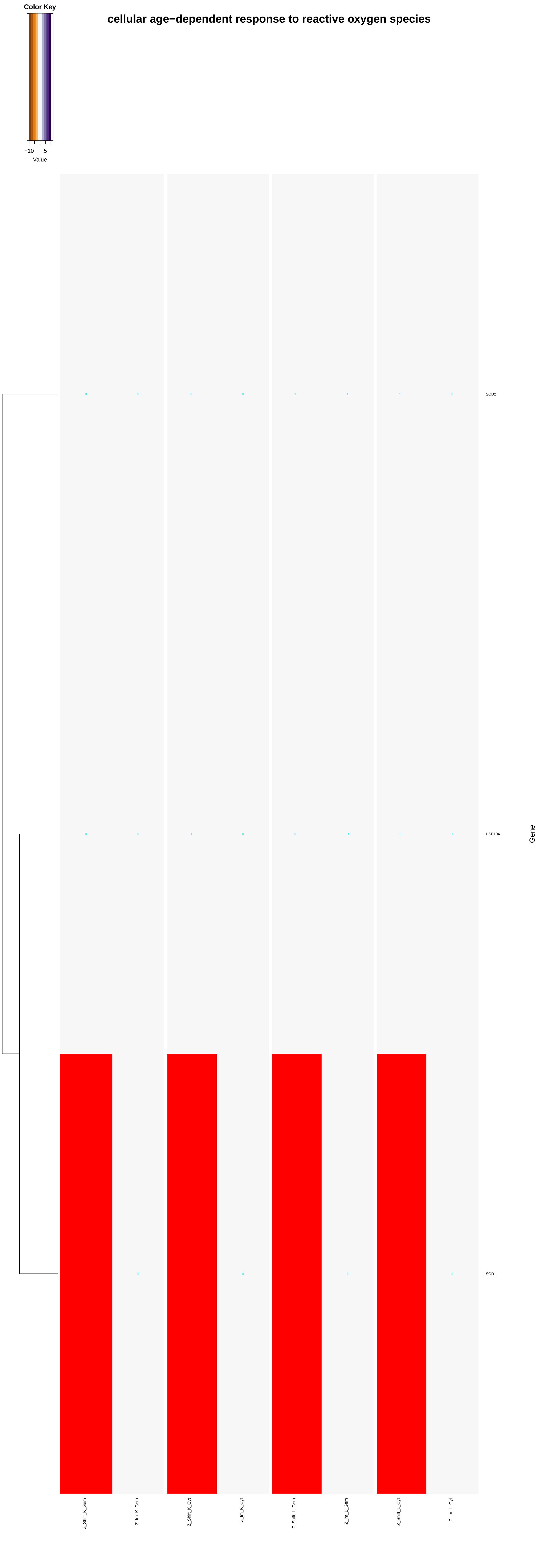
cellular age−dependent response to reactive oxygen species
Color Key
−10
5
Value
SOD2
HSP104
SOD1
Gene
Z_Shift_K_Gem
Z_Im_K_Gem
Z_Shift_K_Cyt
Z_Im_K_Cyt
Z_Shift_L_Gem
Z_Im_L_Gem
Z_Shift_L_Cyt
Z_Im_L_Cyt

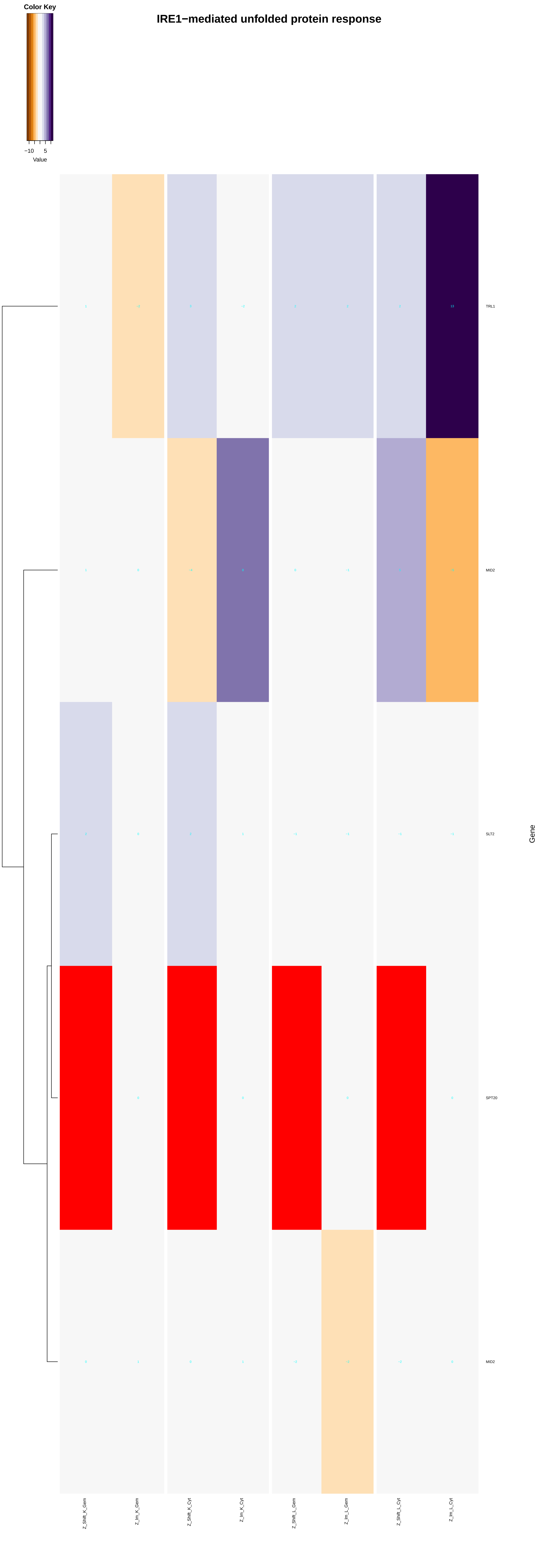
IRE1−mediated unfolded protein response
Color Key
−10
5
Value
TRL1
MID2
SLT2
SPT20
MID2
Gene
Z_Shift_K_Gem
Z_Im_K_Gem
Z_Shift_K_Cyt
Z_Im_K_Cyt
Z_Shift_L_Gem
Z_Im_L_Gem
Z_Shift_L_Cyt
Z_Im_L_Cyt

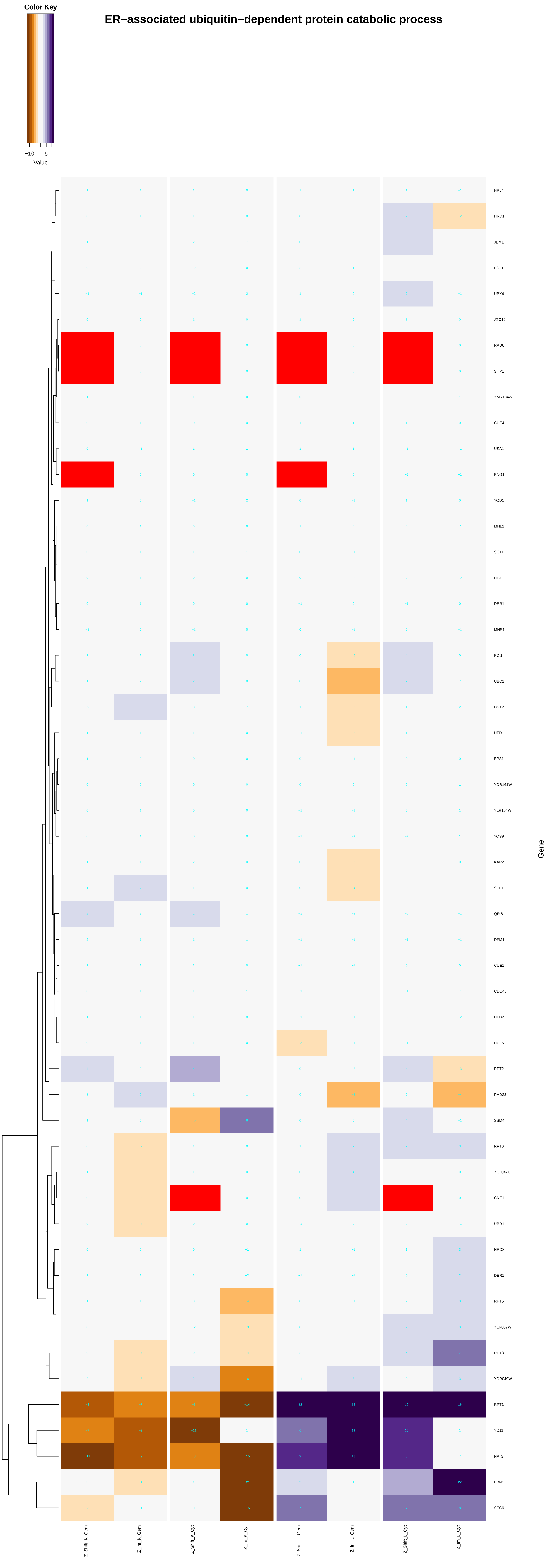
Color Key
−10
5
Value
ER−associated ubiquitin−dependent protein catabolic process
NPL4
HRD1
JEM1
BST1
UBX4
ATG19
RAD6
SHP1
YMR184W
CUE4
USA1
PNG1
YOD1
MNL1
SCJ1
HLJ1
DER1
MNS1
PDI1
UBC1
DSK2
UFD1
EPS1
YDR161W
YLR104W
YOS9
KAR2
SEL1
QRI8
DFM1
CUE1
CDC48
UFD2
HUL5
RPT2
RAD23
SSM4
RPT6
YCL047C
CNE1
UBR1
HRD3
DER1
RPT5
YLR057W
RPT3
YDR049W
RPT1
YDJ1
NAT3
PBN1
SEC61
Gene
Z_Shift_K_Gem
Z_Im_K_Gem
Z_Shift_K_Cyt
Z_Im_K_Cyt
Z_Shift_L_Gem
Z_Im_L_Gem
Z_Shift_L_Cyt
Z_Im_L_Cyt

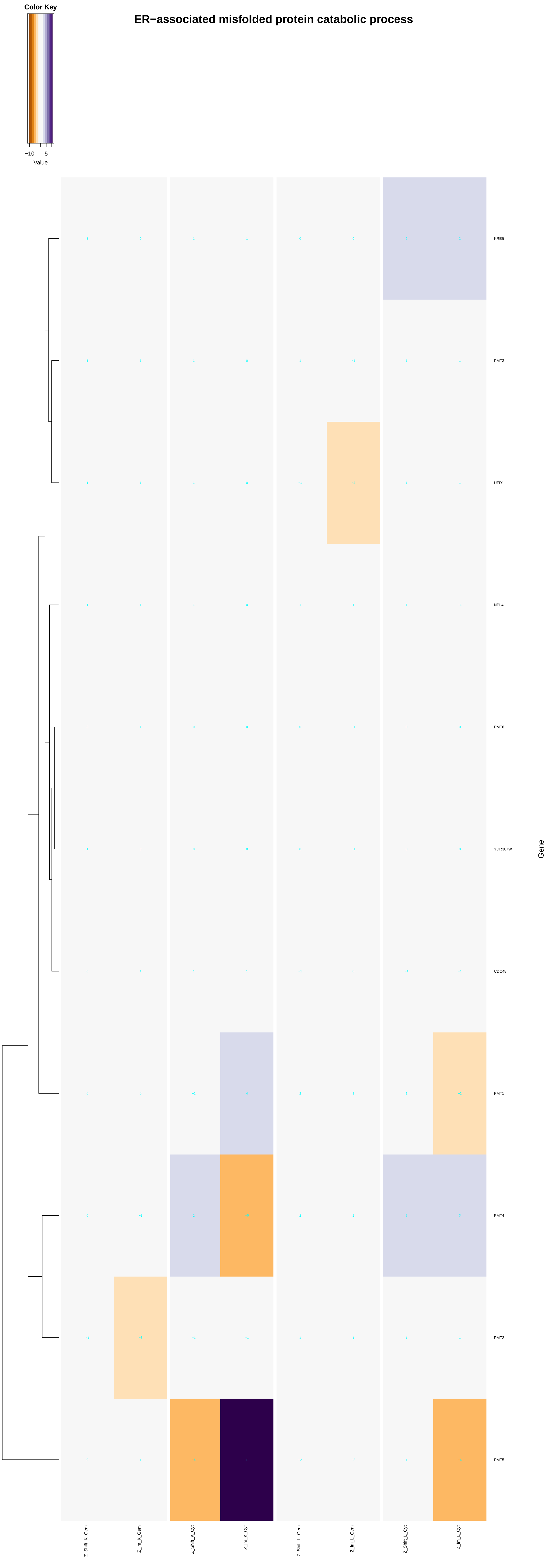
ER−associated misfolded protein catabolic process
Color Key
−10 5
Value
KRE5
PMT3
UFD1
NPL4
PMT6
YDR307W
CDC48
PMT1
PMT4
PMT2
PMT5
Gene
Z_Shift_K_Gem
Z_Im_K_Gem
Z_Shift_K_Cyt
Z_Im_K_Cyt
Z_Shift_L_Gem
Z_Im_L_Gem
Z_Shift_L_Cyt
Z_Im_L_Cyt

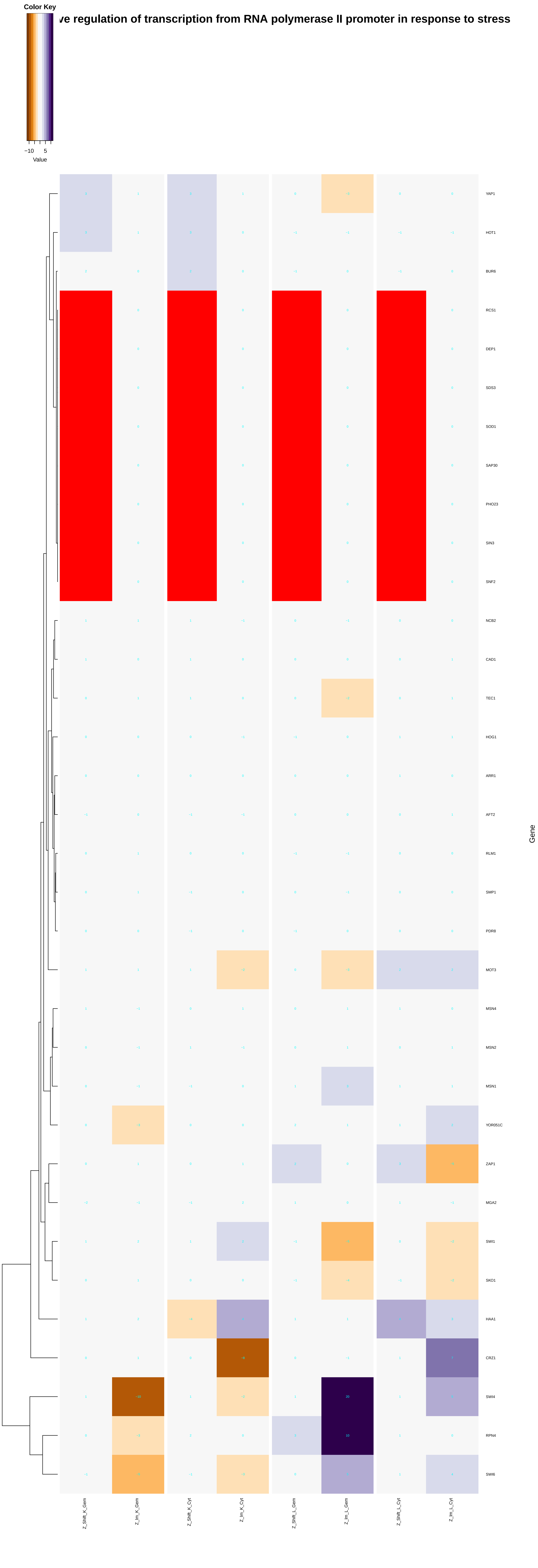

ve regulation of transcription from RNA polymerase II promoter in response to stress
Color Key
−10
5
Value
YAP1
HOT1
BUR6
RCS1
DEP1
SDS3
SOD1
SAP30
PHO23
SIN3
SNF2
NCB2
CAD1
TEC1
HOG1
ARR1
AFT2
RLM1
SMP1
PDR8
MOT3
MSN4
MSN2
MSN1
YOR051C
ZAP1
MGA2
SWI1
SKO1
HAA1
CRZ1
SWI4
RPN4
SWI6
Gene
Z_Shift_K_Gem
Z_Im_K_Gem
Z_Shift_K_Cyt
Z_Im_K_Cyt
Z_Shift_L_Gem
Z_Im_L_Gem
Z_Shift_L_Cyt
Z_Im_L_Cyt

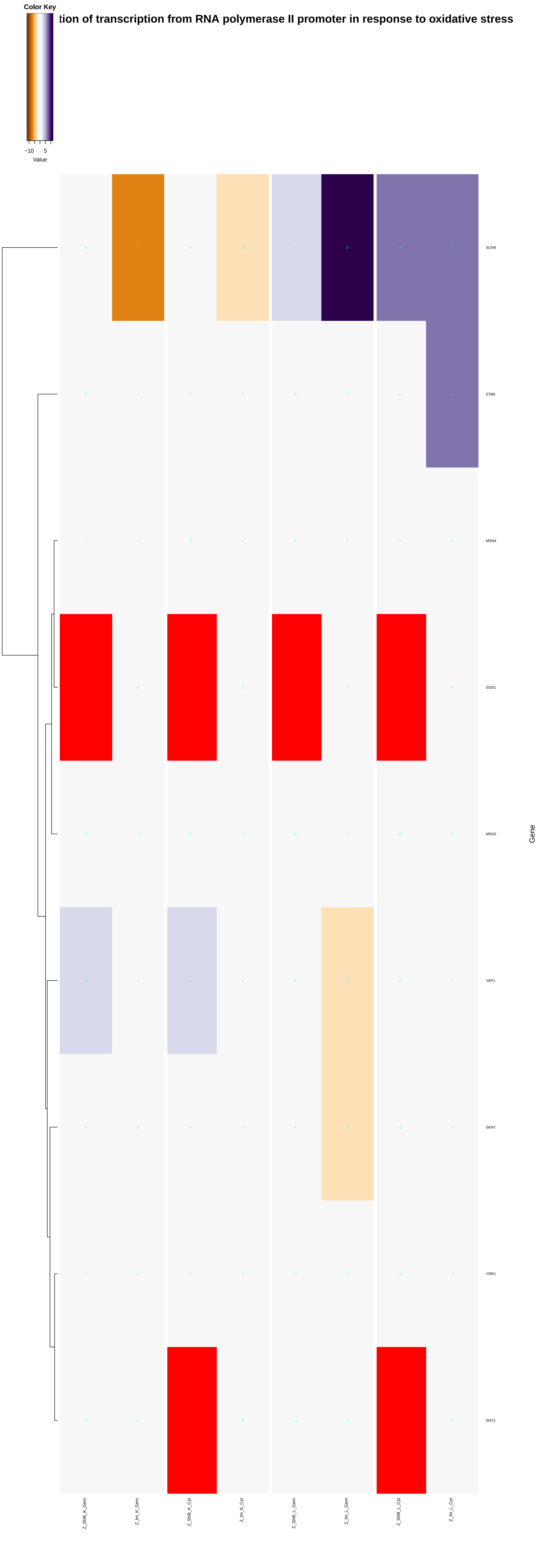
tion of transcription from RNA polymerase II promoter in response to oxidative stress
Color Key
−10
5
Value
SCH9
STB5
MSN4
SOD1
MSN2
YAP1
SKN7
YRR1
SNT2
Gene
Z_Shift_K_Gem
Z_Im_K_Gem
Z_Shift_K_Cyt
Z_Im_K_Cyt
Z_Shift_L_Gem
Z_Im_L_Gem
Z_Shift_L_Cyt
Z_Im_L_Cyt

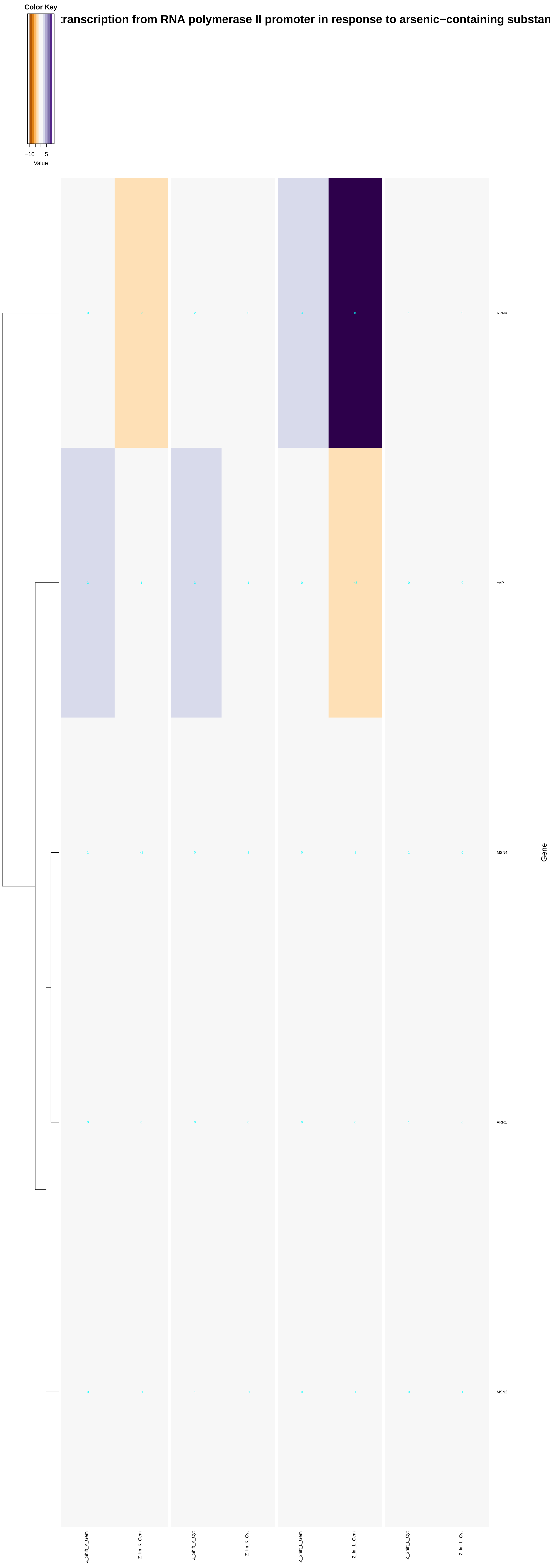

transcription from RNA polymerase II promoter in response to arsenic−containing substan
Color Key
−10
5
Value
0
−3
2
0
3
10
1
0
RPN4
3
1
3
1
0
−3
0
0
YAP1
1
−1
0
1
0
1
1
0
MSN4
0
0
0
0
0
0
1
0
ARR1
0
−1
1
−1
0
1
0
1
MSN2
Gene
Z_Shift_K_Gem
Z_lm_K_Gem
Z_Shift_K_Cyt
Z_lm_K_Cyt
Z_Shift_L_Gem
Z_lm_L_Gem
Z_Shift_L_Cyt
Z_lm_L_Cyt

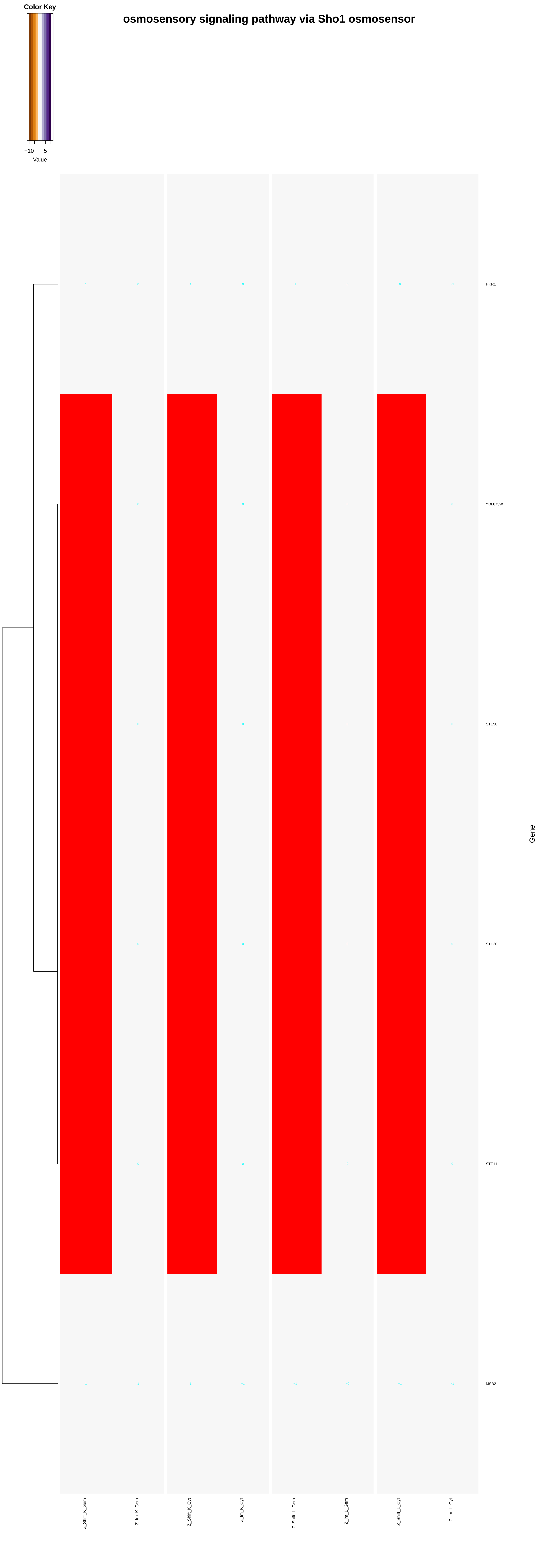
osmosensory signaling pathway via Sho1 osmosensor
Color Key
−10
5
Value
HKR1
YDL073W
STE50
STE20
STE11
MSB2
Gene
Z_Shift_K_Gem
Z_Im_K_Gem
Z_Shift_K_Cyt
Z_Im_K_Cyt
Z_Shift_L_Gem
Z_Im_L_Gem
Z_Shift_L_Cyt
Z_Im_L_Cyt

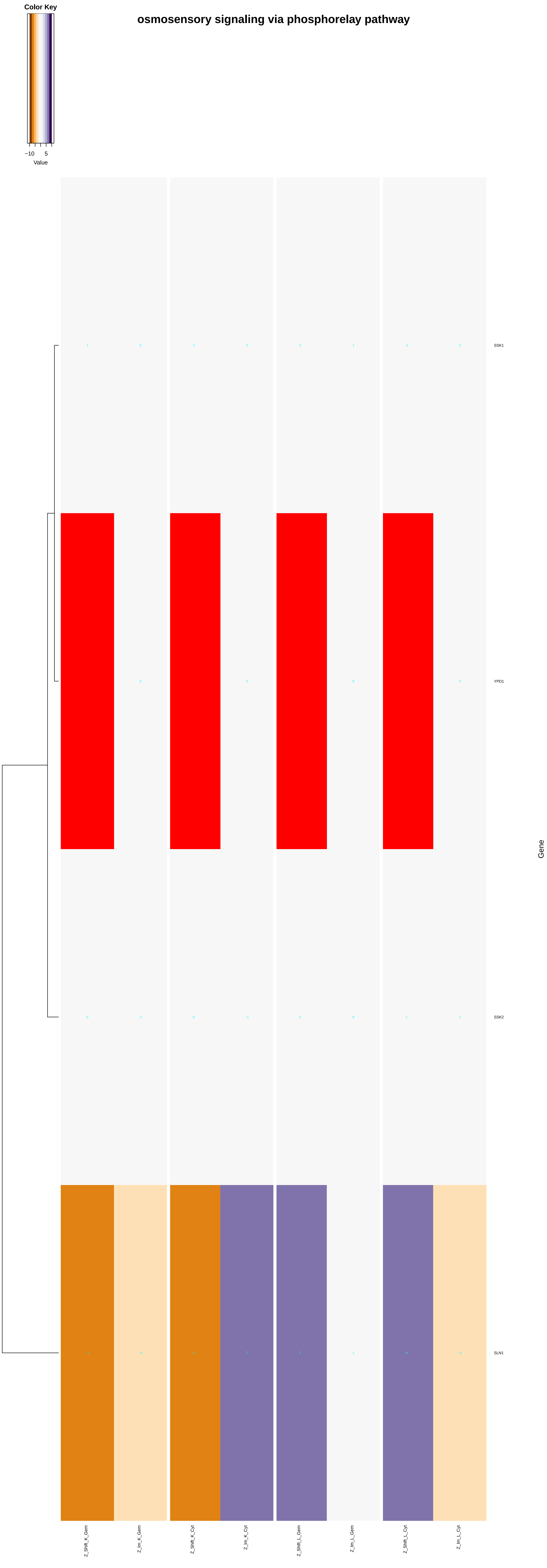

Color Key
−10
5
Value
osmosensory signaling via phosphorelay pathway
SSK1
YPD1
SSK2
SLN1
Gene
Z_Shift_K_Gem
Z_Im_K_Gem
Z_Shift_K_Cyt
Z_Im_K_Cyt
Z_Shift_L_Gem
Z_Im_L_Gem
Z_Shift_L_Cyt
Z_Im_L_Cyt

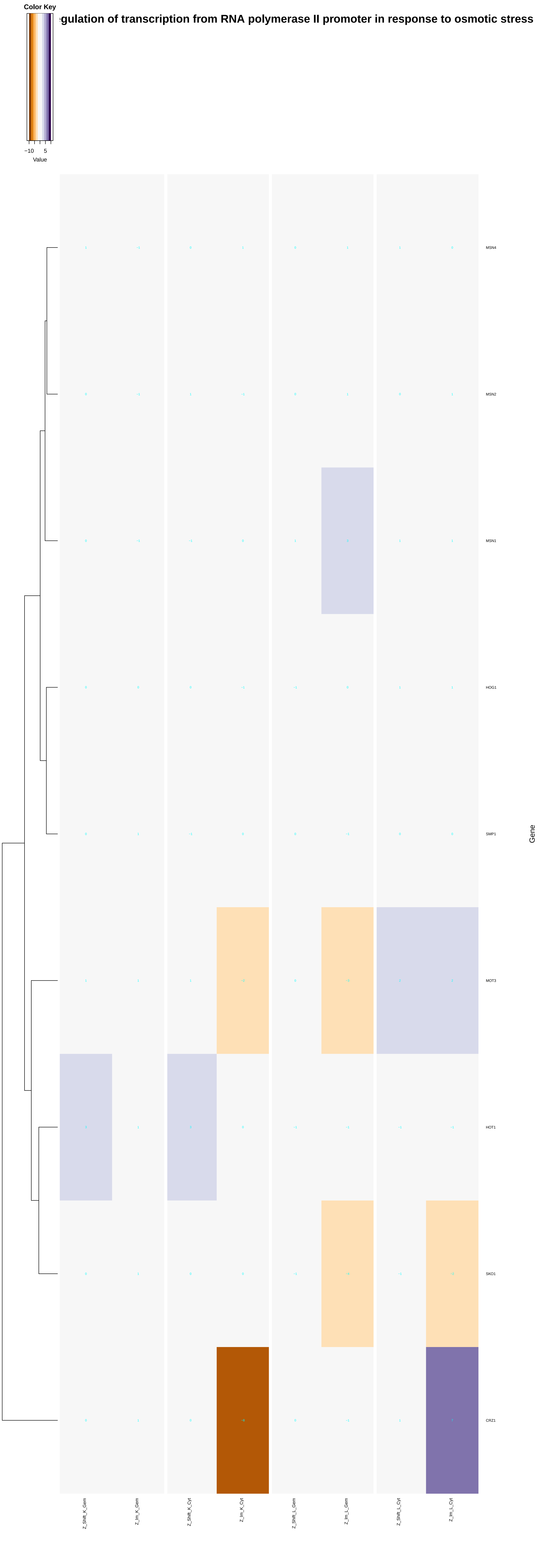
gulation of transcription from RNA polymerase II promoter in response to osmotic stress
Color Key
−10
5
Value
MSN4
MSN2
MSN1
HOG1
SMP1
MOT3
HOT1
SKO1
CRZ1
Gene
Z_Shift_K_Gem
Z_Im_K_Gem
Z_Shift_K_Cyt
Z_Im_K_Cyt
Z_Shift_L_Gem
Z_Im_L_Gem
Z_Shift_L_Cyt
Z_Im_L_Cyt

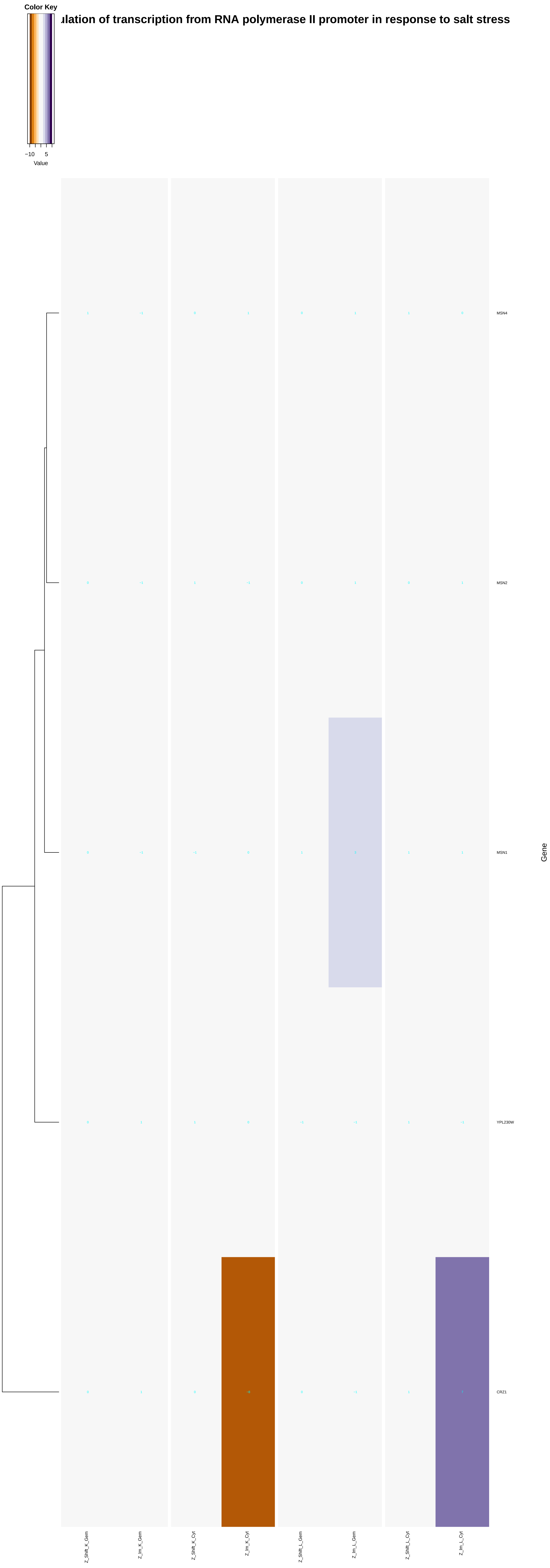

Color Key
−10
5
Value
ulation of transcription from RNA polymerase II promoter in response to salt stress
MSN4
MSN2
MSN1
YPL230W
CRZ1
Gene
Z_Shift_K_Gem
Z_Im_K_Gem
Z_Shift_K_Cyt
Z_Im_K_Cyt
Z_Shift_L_Gem
Z_Im_L_Gem
Z_Shift_L_Cyt
Z_Im_L_Cyt

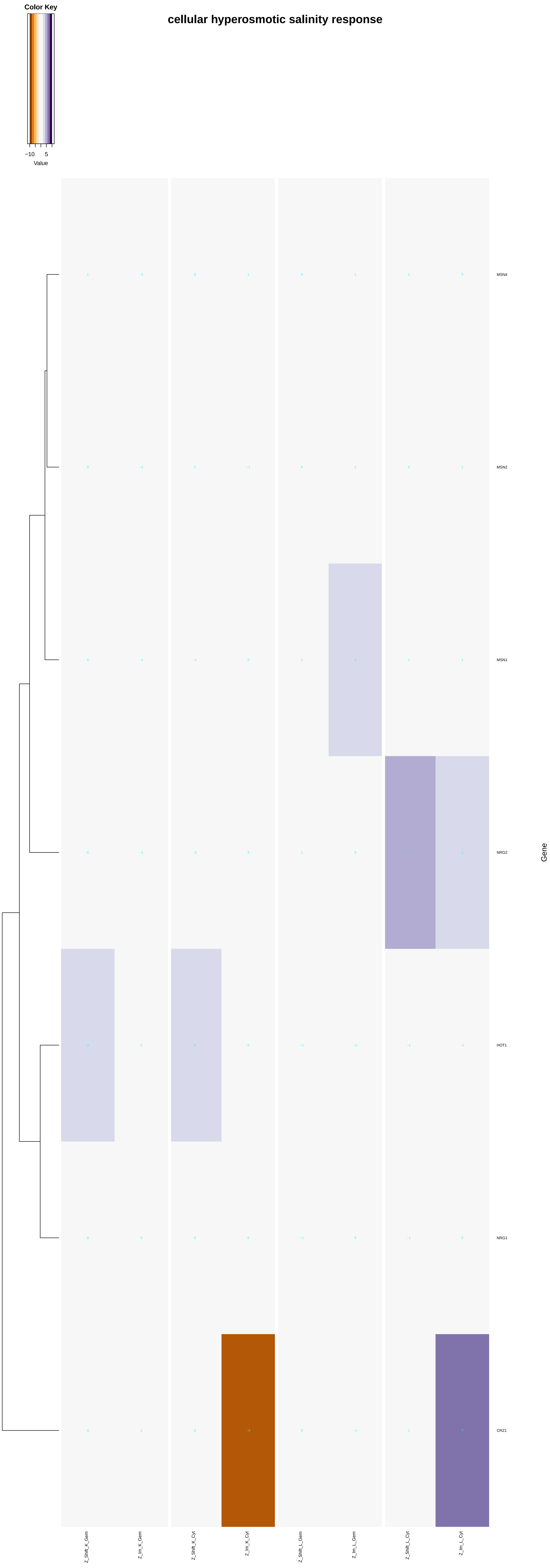
cellular hyperosmotic salinity response
Color Key
−10
5
Value
MSN4
MSN2
MSN1
NRG2
HOT1
NRG1
CRZ1
Gene
Z_Shift_K_Gem
Z_Im_K_Gem
Z_Shift_K_Cyt
Z_Im_K_Cyt
Z_Shift_L_Gem
Z_Im_L_Gem
Z_Shift_L_Cyt
Z_Im_L_Cyt

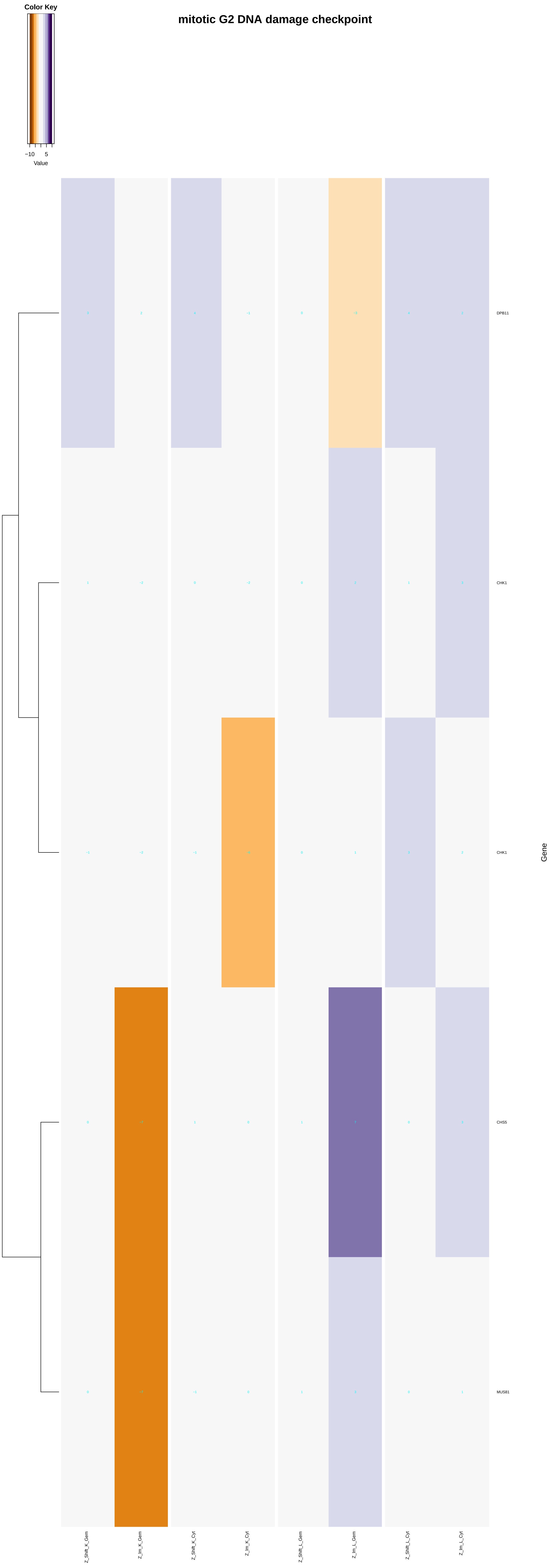

Color Key
−10
5
Value
mitotic G2 DNA damage checkpoint
DPB11
CHK1
CHK1
CHS5
MUS81
Gene
Z_Shift_K_Gem
Z_Im_K_Gem
Z_Shift_K_Cyt
Z_Im_K_Cyt
Z_Shift_L_Gem
Z_Im_L_Gem
Z_Shift_L_Cyt
Z_Im_L_Cyt

### mitotic G1 DNA damage checkpoint

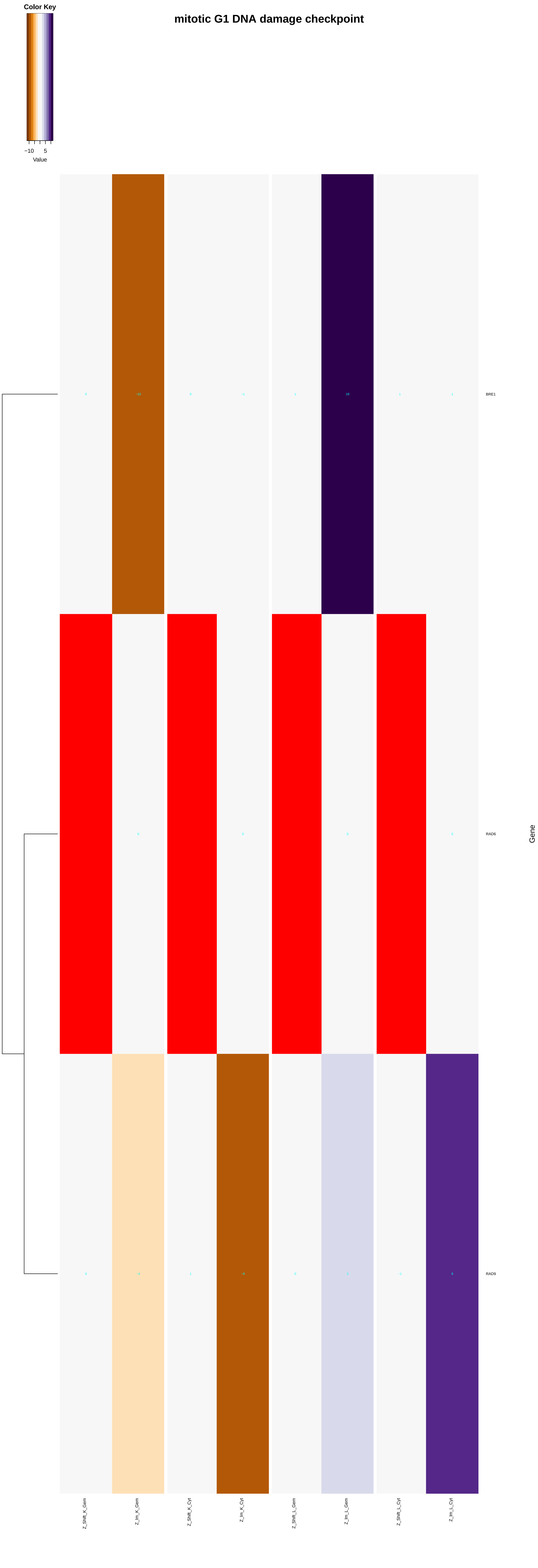

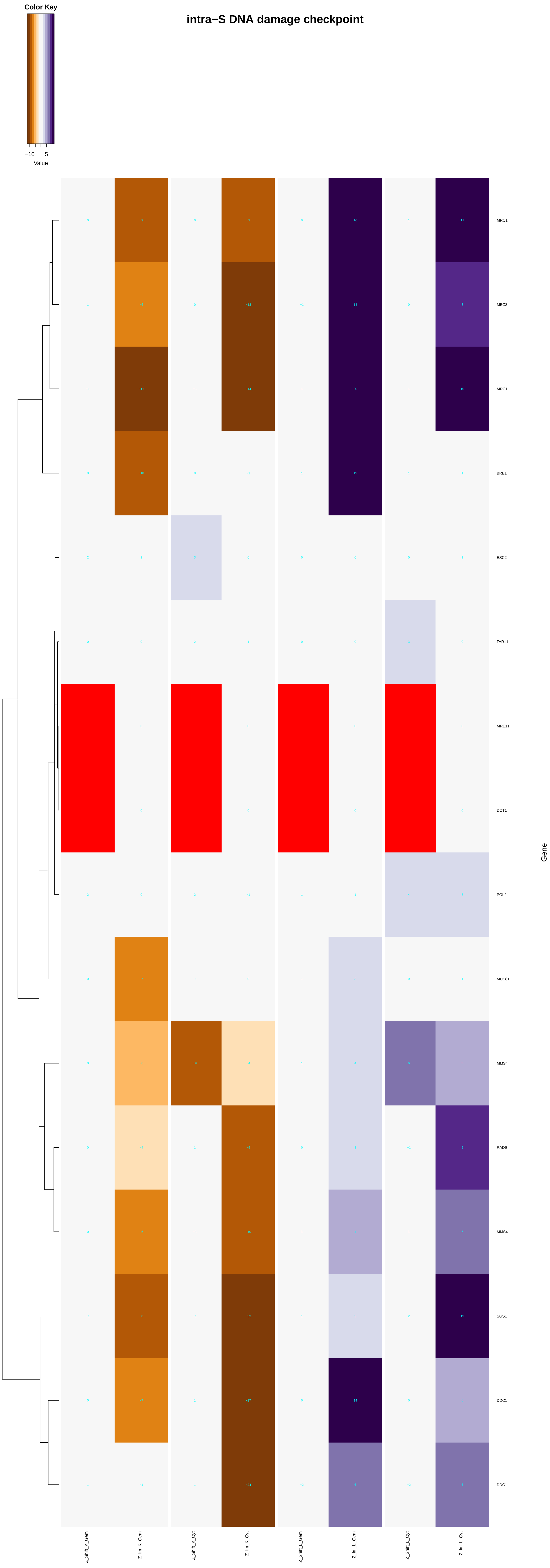
Color Key
−10 5
Value
intra−S DNA damage checkpoint
MRC1
MEC3
MRC1
BRE1
ESC2
FAR11
MRE11
DOT1
POL2
MUS81
MMS4
RAD9
MMS4
SGS1
DDC1
DDC1
Gene
Z_Shift_K_Gem
Z_Im_K_Gem
Z_Shift_K_Cyt
Z_Im_K_Cyt
Z_Shift_L_Gem
Z_Im_L_Gem
Z_Shift_L_Cyt
Z_Im_L_Cyt

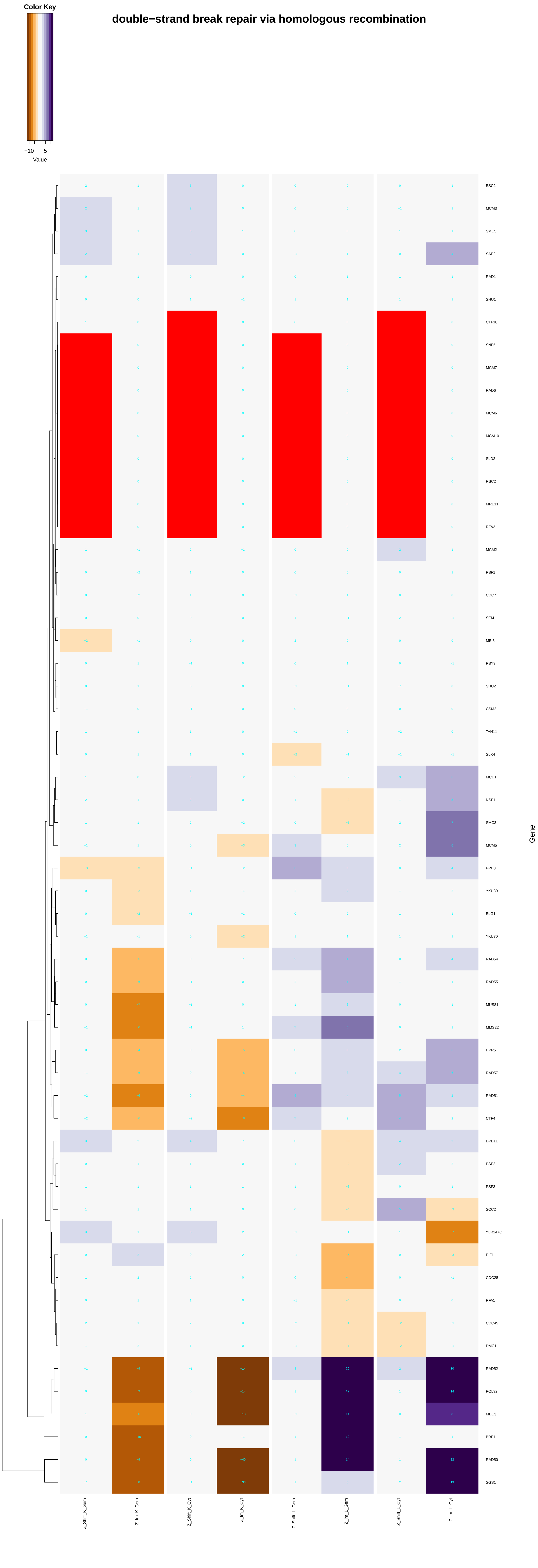
double−strand break repair via homologous recombination
Color Key
−10
5
Value
ESC2
MCM3
SMC5
SAE2
RAD1
SHU1
CTF18
SNF5
MCM7
RAD6
MCM6
MCM10
SLD2
RSC2
MRE11
RFA2
MCM2
PSF1
CDC7
SEM1
MEI5
PSY3
SHU2
CSM2
TAH11
SLX4
MCD1
NSE1
SMC3
MCM5
PPH3
YKU80
ELG1
YKU70
RAD54
RAD55
MUS81
MMS22
HPR5
RAD57
RAD51
CTF4
DPB11
PSF2
PSF3
SCC2
YLR247C
PIF1
CDC28
RFA1
CDC45
DMC1
RAD52
POL32
MEC3
BRE1
RAD50
SGS1
Gene
Z_Shift_K_Gem
Z_lm_K_Gem
Z_Shift_K_Cyt
Z_lm_K_Cyt
Z_Shift_L_Gem
Z_lm_L_Gem
Z_Shift_L_Cyt
Z_lm_L_Cyt

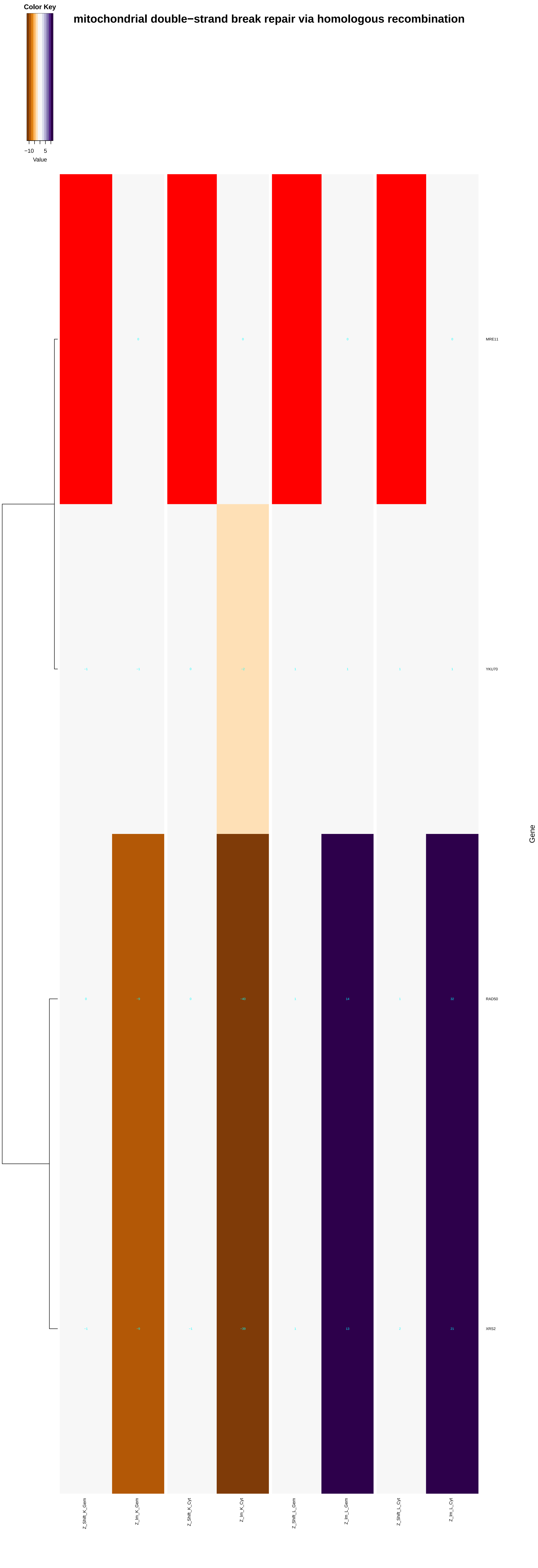

Color Key
−10
5
Value
mitochondrial double−strand break repair via homologous recombination
0
0
0
0
MRE11
−1
−1
0
−2
1
1
1
1
YKU70
0
−9
0
−40
1
14
1
32
RAD50
−1
−9
−1
−39
1
13
2
21
XRS2
Gene
Z_Shift_K_Gem
Z_Im_K_Gem
Z_Shift_K_Cyt
Z_Im_K_Cyt
Z_Shift_L_Gem
Z_Im_L_Gem
Z_Shift_L_Cyt
Z_Im_L_Cyt

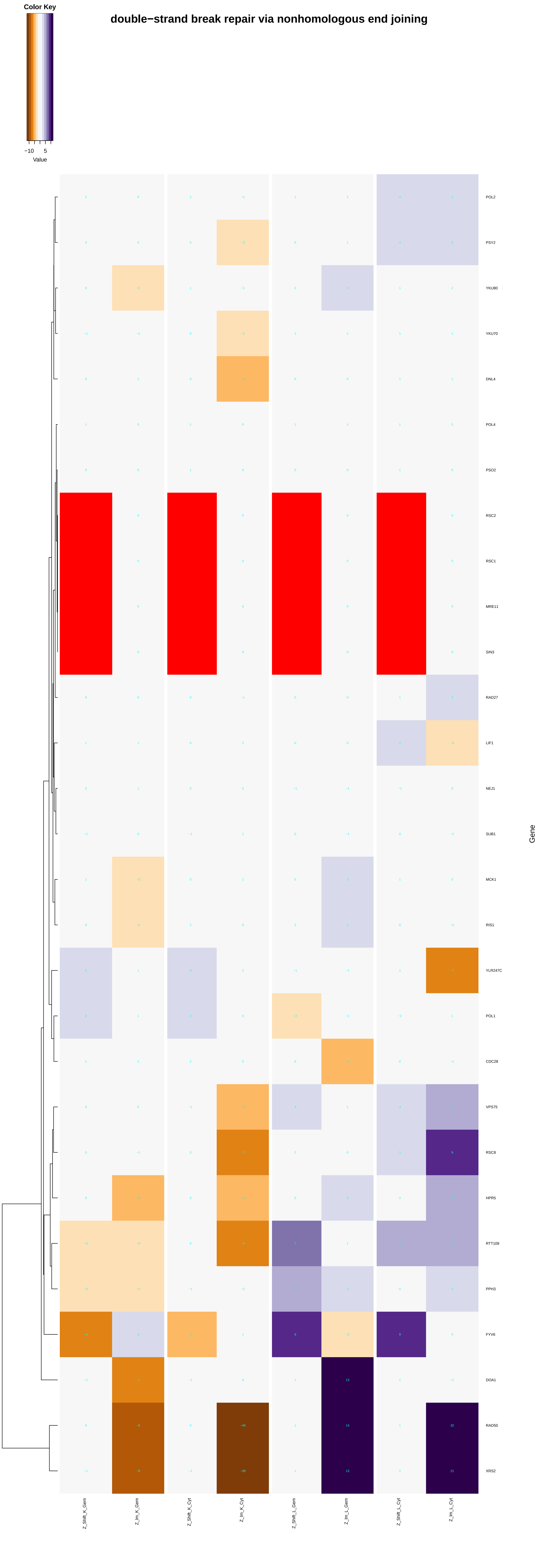
double−strand break repair via nonhomologous end joining
Color Key
−10
5
Value
POL2
PSY2
YKU80
YKU70
DNL4
POL4
PSO2
RSC2
RSC1
MRE11
SIN3
RAD27
LIF1
NEJ1
SUB1
MCK1
RIS1
YLR247C
POL1
CDC28
VPS75
RSC8
HPR5
RTT109
PPH3
FYV6
DOA1
RAD50
XRS2
Gene
Z_Shift_K_Gem
Z_lm_K_Gem
Z_Shift_K_Cyt
Z_lm_K_Cyt
Z_Shift_L_Gem
Z_lm_L_Gem
Z_Shift_L_Cyt
Z_lm_L_Cyt

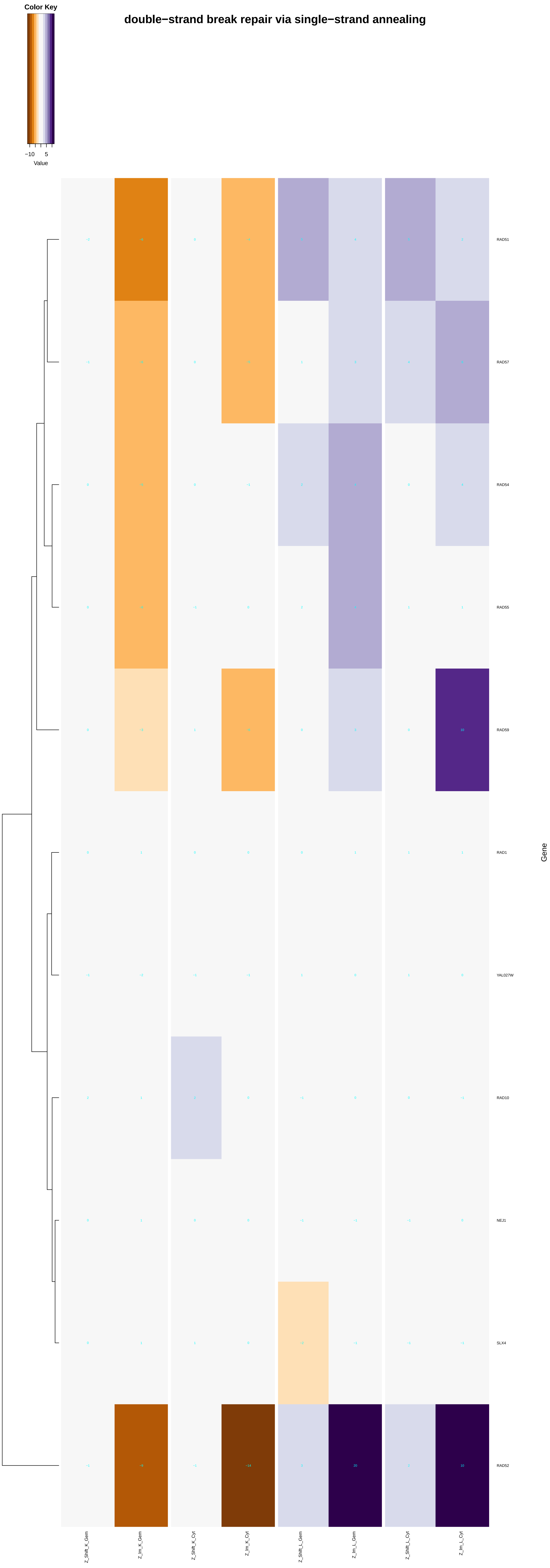

Color Key
−10
5
Value
double−strand break repair via single−strand annealing
RAD51
RAD57
RAD54
RAD55
RAD59
RAD1
YAL027W
RAD10
NEJ1
SLX4
RAD52
Gene
Z_Shift_K_Gem
Z_lm_K_Gem
Z_Shift_K_Cyt
Z_lm_K_Cyt
Z_Shift_L_Gem
Z_lm_L_Gem
Z_Shift_L_Cyt
Z_lm_L_Cyt

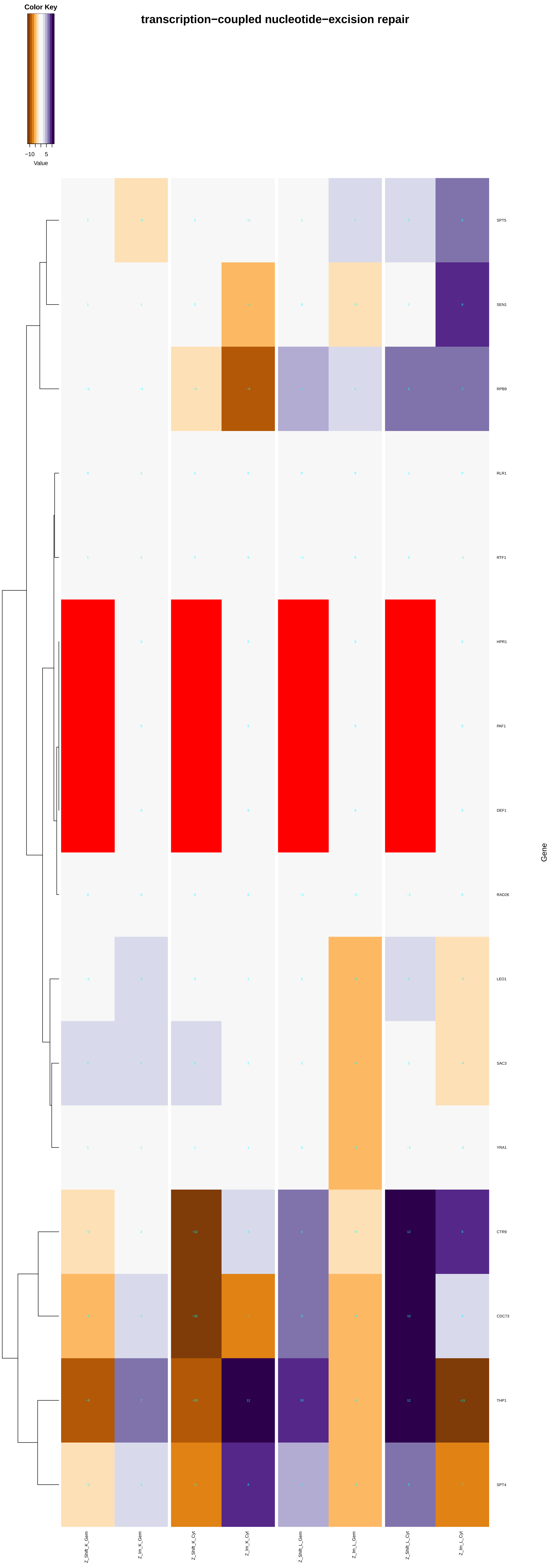
Color Key
−10
5
Value
transcription−coupled nucleotide−excision repair
SPT5
SEN1
RPB9
RLR1
RTF1
HPR1
PAF1
DEF1
RAD26
LEO1
SAC3
YRA1
CTR9
CDC73
THP1
SPT4
Gene
Z_Shift_K_Gem
Z_Im_K_Gem
Z_Shift_K_Cyt
Z_Im_K_Cyt
Z_Shift_L_Gem
Z_Im_L_Gem
Z_Shift_L_Cyt
Z_Im_L_Cyt

### global genome nucleotide−excision repair

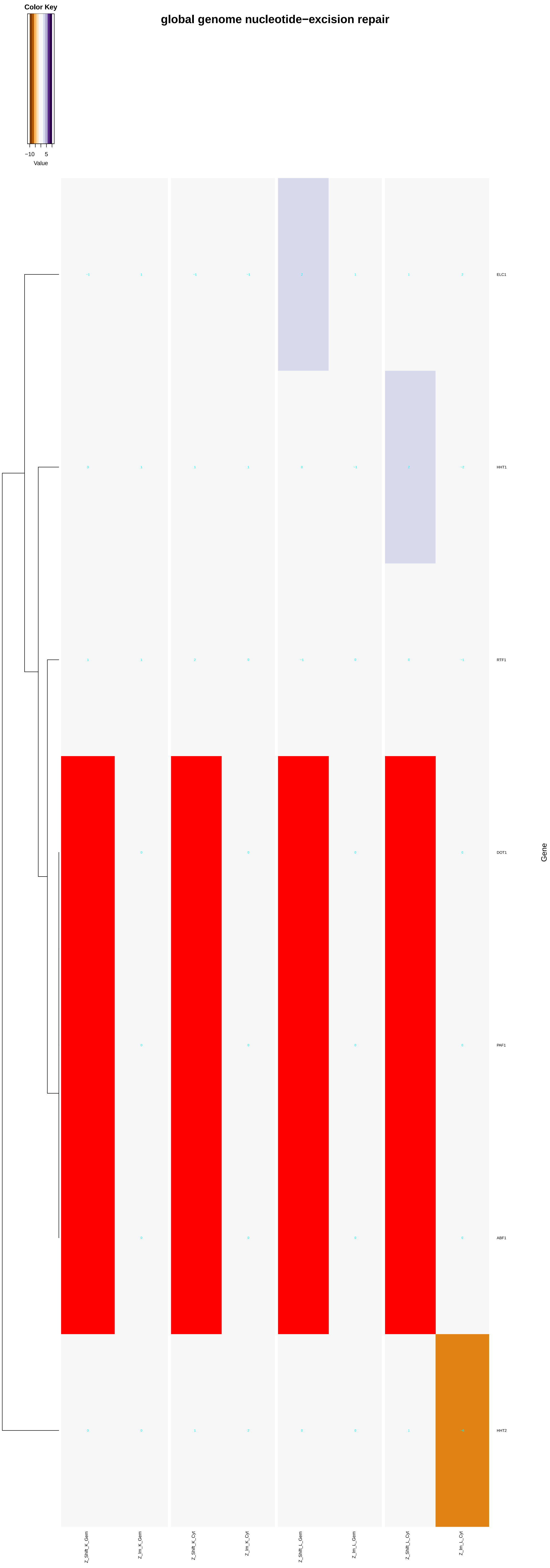

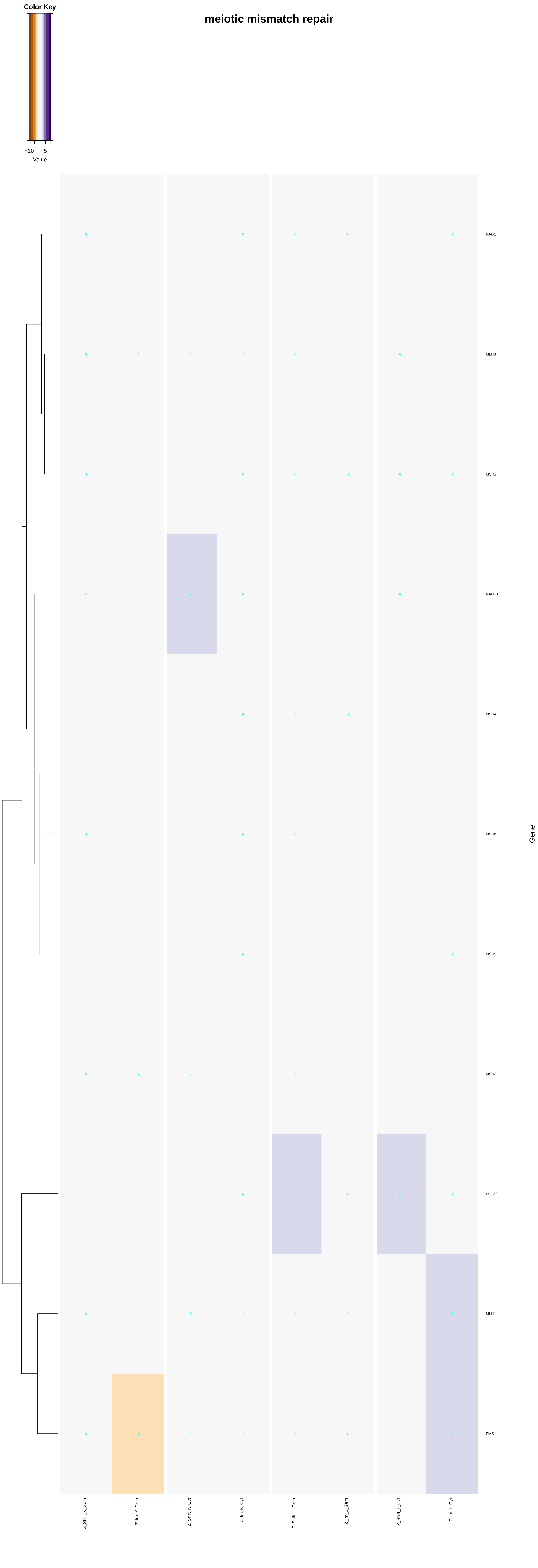

meiotic mismatch repair
Color Key
−10
5
Value
RAD1
MLH3
MSH2
RAD10
MSH4
MSH6
MSH5
MSH3
POL30
MLH1
PMS1
Gene
Z_Shift_K_Gem
Z_Im_K_Gem
Z_Shift_K_Cyt
Z_Im_K_Cyt
Z_Shift_L_Gem
Z_Im_L_Gem
Z_Shift_L_Cyt
Z_Im_L_Cyt

### translesion synthesis

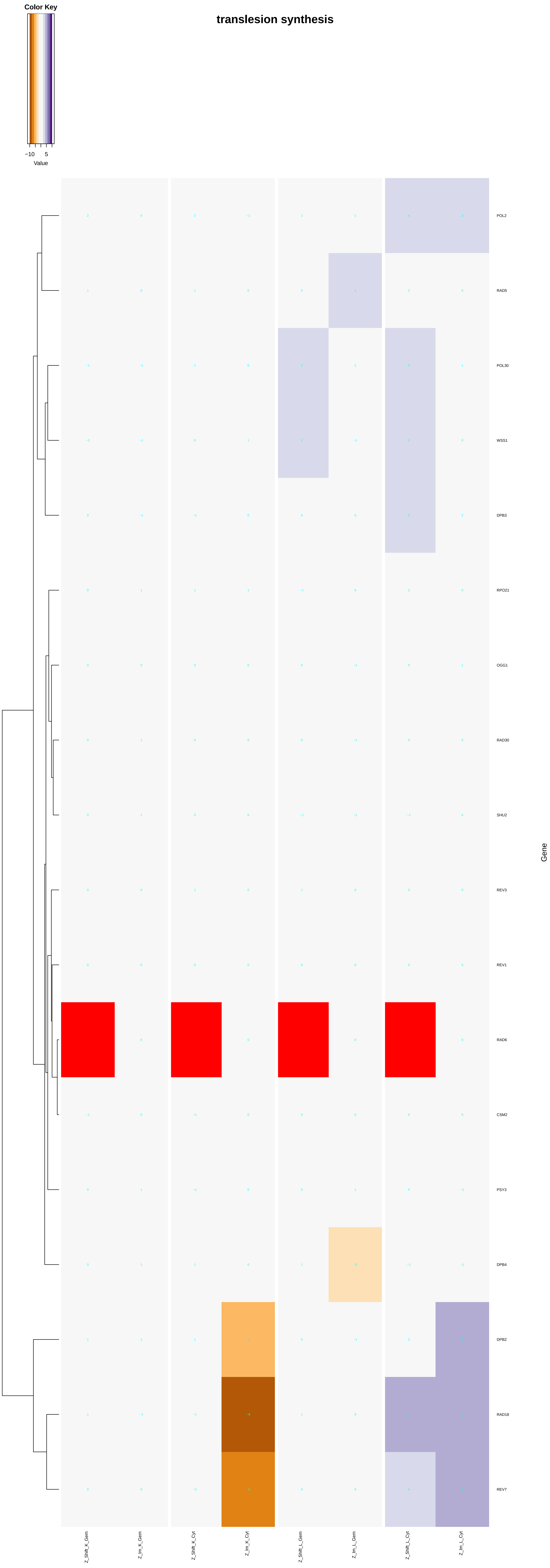

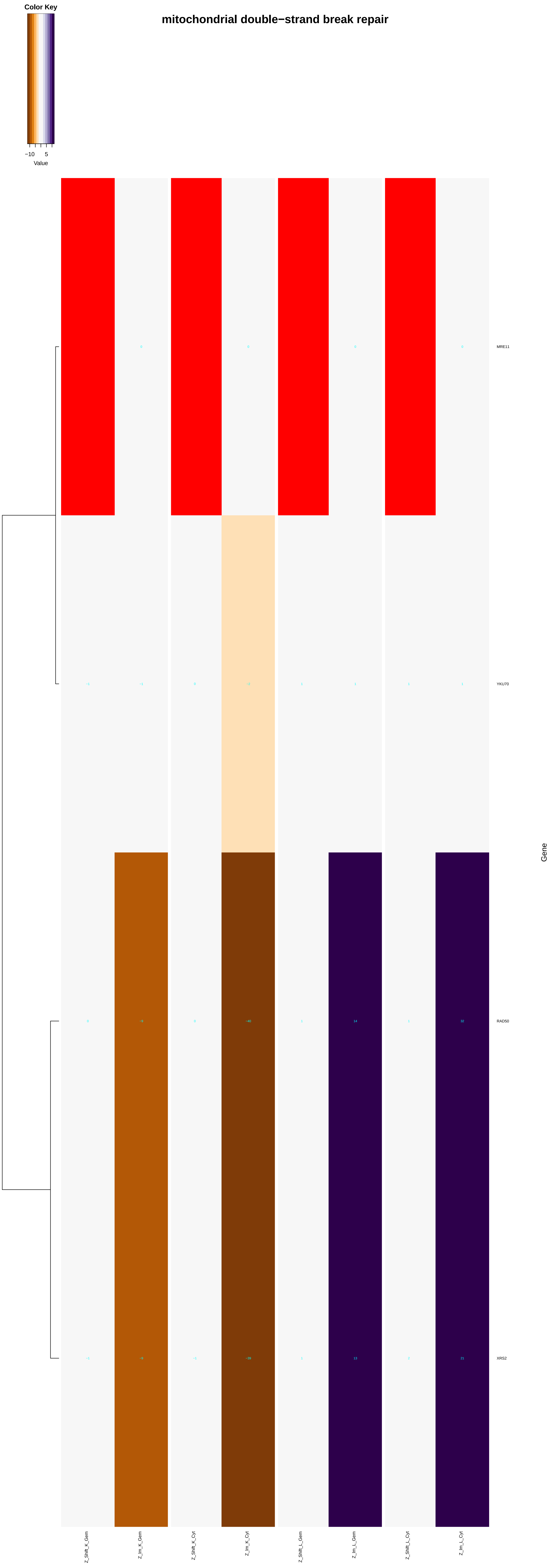

mitochondrial double−strand break repair
Color Key
−10
5
Value
MRE11
YKU70
RAD50
XRS2
Gene
Z_Shift_K_Gem
Z_Im_K_Gem
Z_Shift_K_Cyt
Z_Im_K_Cyt
Z_Shift_L_Gem
Z_Im_L_Gem
Z_Shift_L_Cyt
Z_Im_L_Cyt

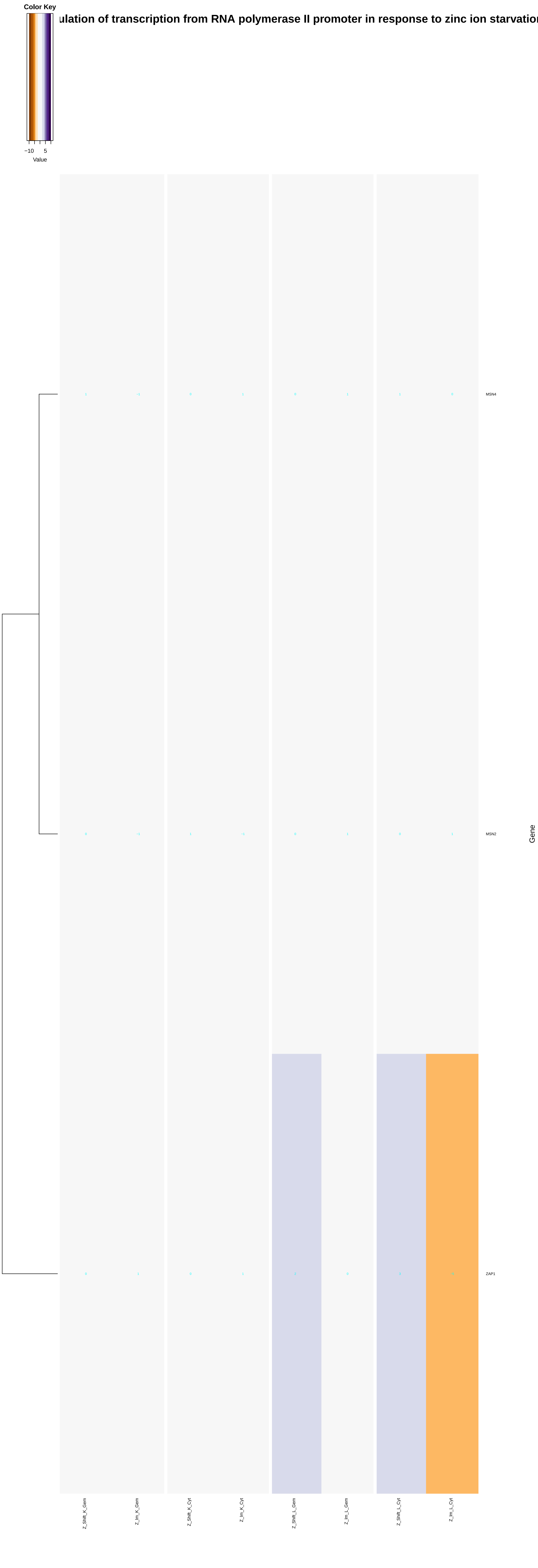

ulation of transcription from RNA polymerase II promoter in response to zinc ion starvation
Color Key
−10
5
Value
MSN4
MSN2
ZAP1
Gene
Z_Shift_K_Gem
Z_Im_K_Gem
Z_Shift_K_Cyt
Z_Im_K_Cyt
Z_Shift_L_Gem
Z_Im_L_Gem
Z_Shift_L_Cyt
Z_Im_L_Cyt

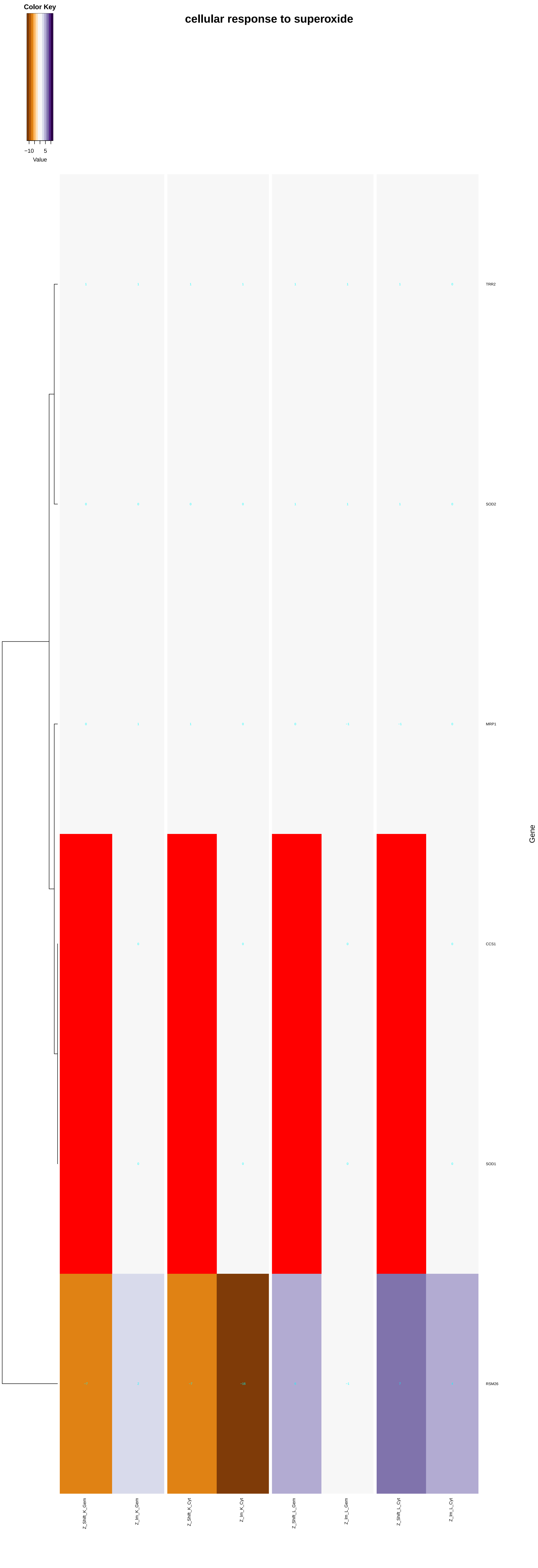

cellular response to superoxide
Color Key
−10
5
Value
TRR2
SOD2
MRP1
CCS1
SOD1
RSM26
Gene
Z_Shift_K_Gem
Z_Im_K_Gem
Z_Shift_K_Cyt
Z_Im_K_Cyt
Z_Shift_L_Gem
Z_Im_L_Gem
Z_Shift_L_Cyt
Z_Im_L_Cyt

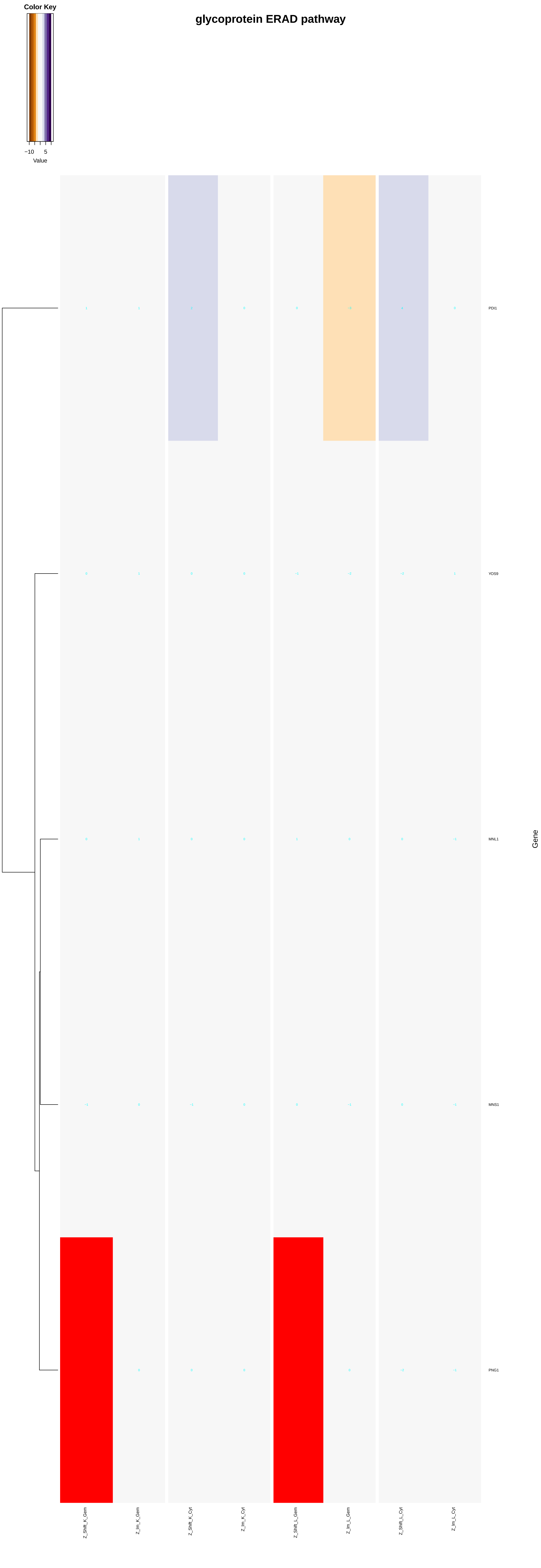
glycoprotein ERAD pathway
Color Key
−10
5
Value
PDI1
YOS9
MNL1
MNS1
PNG1
Gene
Z_Shift_K_Gem
Z_Im_K_Gem
Z_Shift_K_Cyt
Z_Im_K_Cyt
Z_Shift_L_Gem
Z_Im_L_Gem
Z_Shift_L_Cyt
Z_Im_L_Cyt

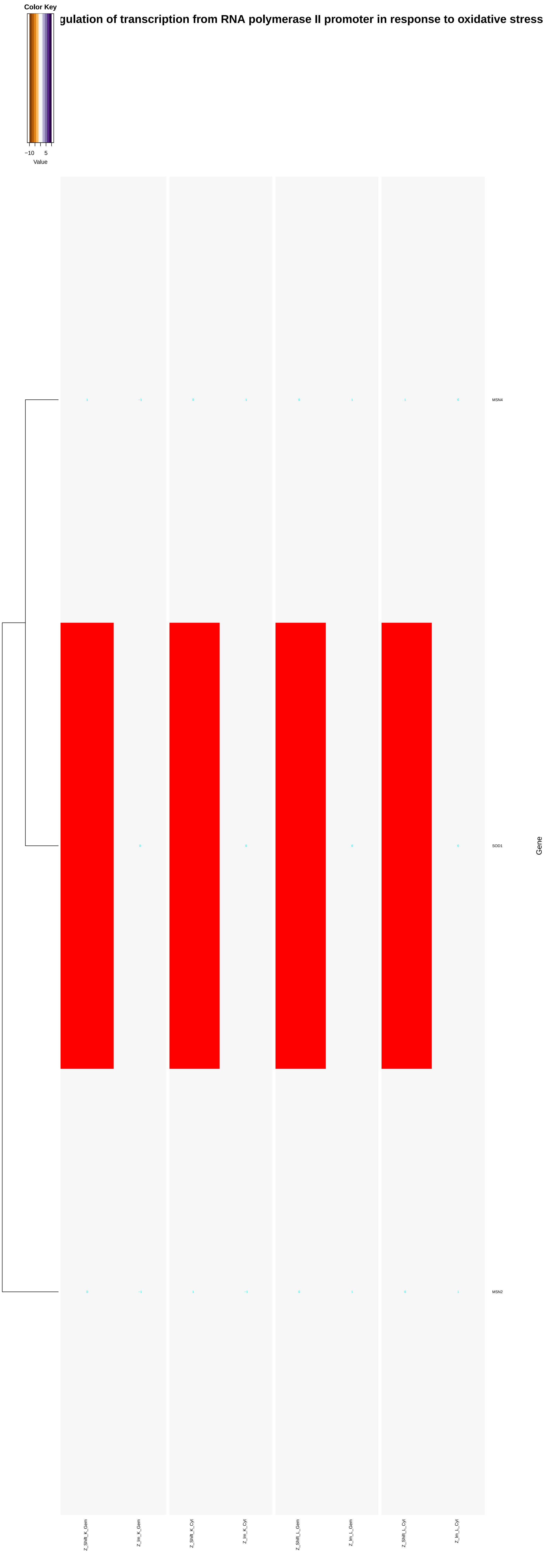

Color Key
−10
5
Value
gulation of transcription from RNA polymerase II promoter in response to oxidative stress
MSN4
SOD1
MSN2
Gene
Z_Shift_K_Gem
Z_Im_K_Gem
Z_Shift_K_Cyt
Z_Im_K_Cyt
Z_Shift_L_Gem
Z_Im_L_Gem
Z_Shift_L_Cyt
Z_Im_L_Cyt

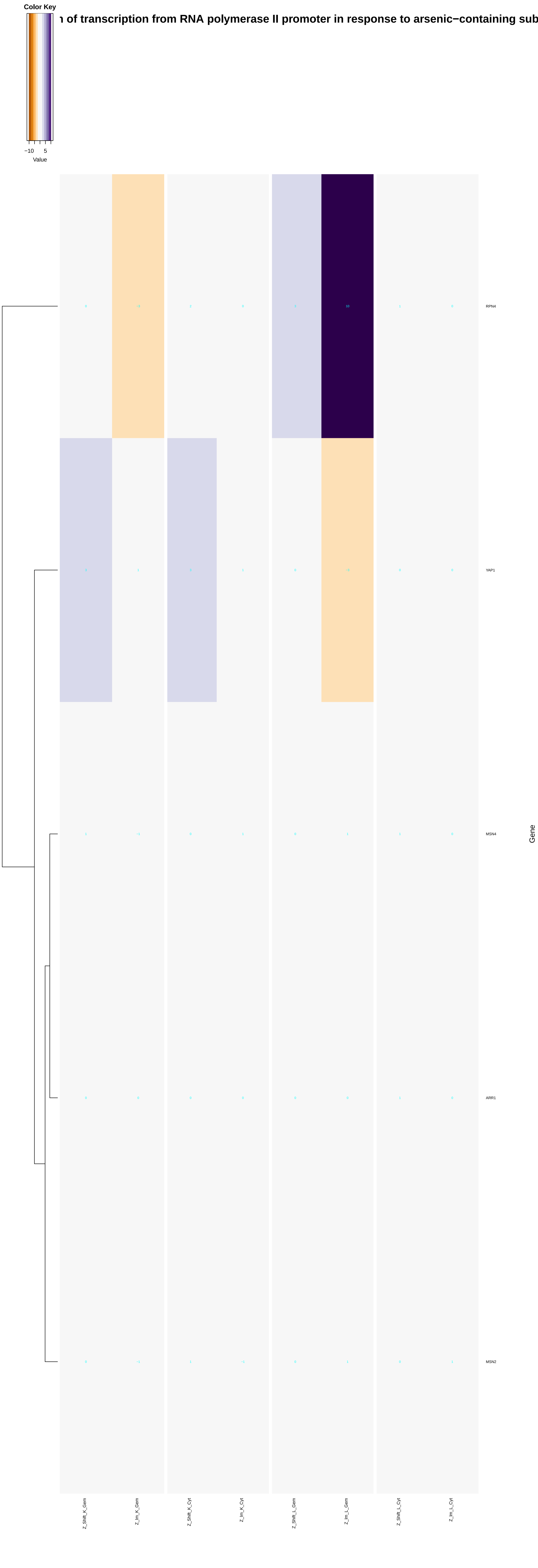
n of transcription from RNA polymerase II promoter in response to arsenic−containing sub
Color Key
−10
5
Value
RPN4
YAP1
MSN4
ARR1
MSN2
Gene
Z_Shift_K_Gem
Z_Im_K_Gem
Z_Shift_K_Cyt
Z_Im_K_Cyt
Z_Shift_L_Gem
Z_Im_L_Gem
Z_Shift_L_Cyt
Z_Im_L_Cyt

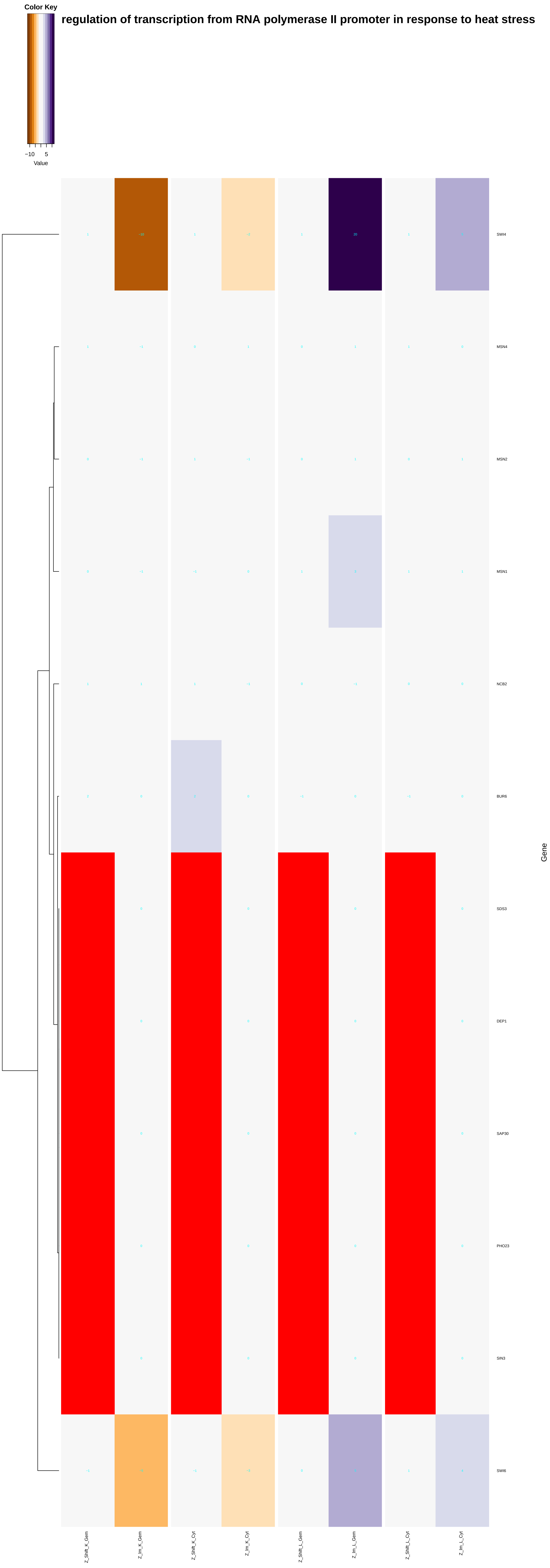
regulation of transcription from RNA polymerase II promoter in response to heat stress
Color Key
−10
5
Value
SWI4
MSN4
MSN2
MSN1
NCB2
BUR6
SDS3
DEP1
SAP30
PHO23
SIN3
SWI6
Gene
Z_Shift_K_Gem
Z_Im_K_Gem
Z_Shift_K_Cyt
Z_Im_K_Cyt
Z_Shift_L_Gem
Z_Im_L_Gem
Z_Shift_L_Cyt
Z_Im_L_Cyt

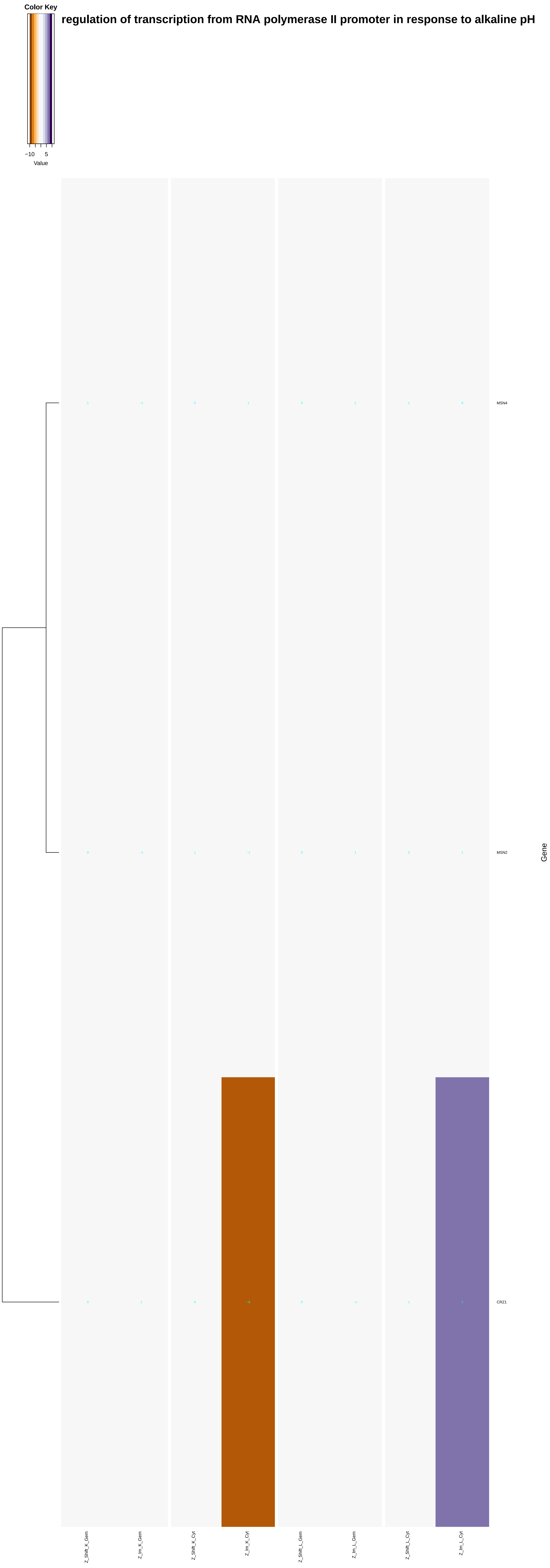
Color Key
−10
5
Value
regulation of transcription from RNA polymerase II promoter in response to alkaline pH
MSN4
MSN2
CRZ1
Gene
Z_Shift_K_Gem
Z_Im_K_Gem
Z_Shift_K_Cyt
Z_Im_K_Cyt
Z_Shift_L_Gem
Z_Im_L_Gem
Z_Shift_L_Cyt
Z_Im_L_Cyt

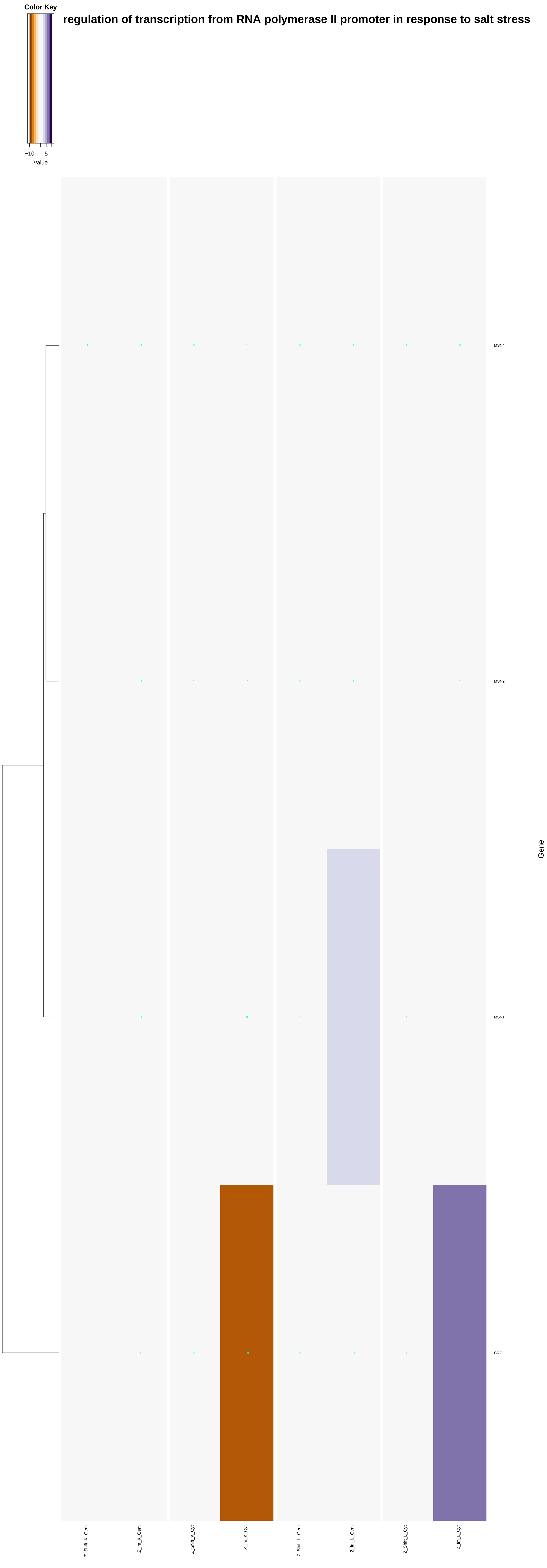
Color Key
−10
5
Value
regulation of transcription from RNA polymerase II promoter in response to salt stress
MSN4
MSN2
MSN1
CRZ1
Gene
Z_Shift_K_Gem
Z_Im_K_Gem
Z_Shift_K_Cyt
Z_Im_K_Cyt
Z_Shift_L_Gem
Z_Im_L_Gem
Z_Shift_L_Cyt
Z_Im_L_Cyt

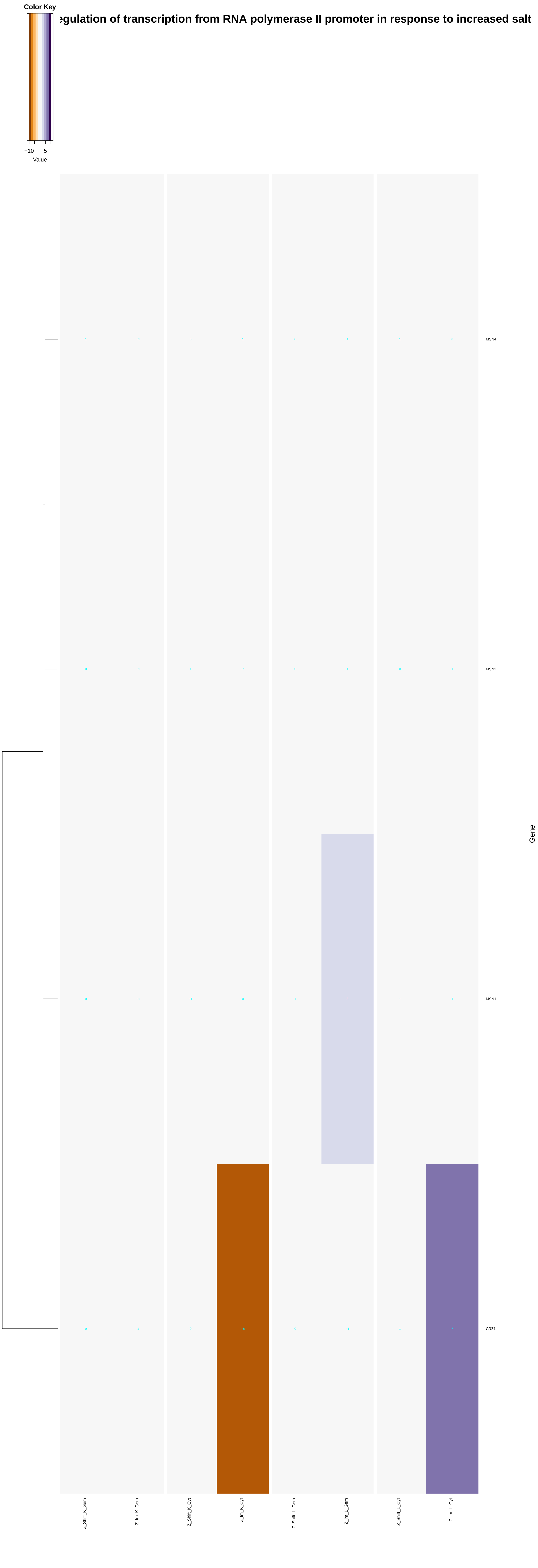
egulation of transcription from RNA polymerase II promoter in response to increased salt
Color Key
−10
5
Value
MSN4
MSN2
MSN1
CRZ1
Gene
Z_Shift_K_Gem
Z_Im_K_Gem
Z_Shift_K_Cyt
Z_Im_K_Cyt
Z_Shift_L_Gem
Z_Im_L_Gem
Z_Shift_L_Cyt
Z_Im_L_Cyt

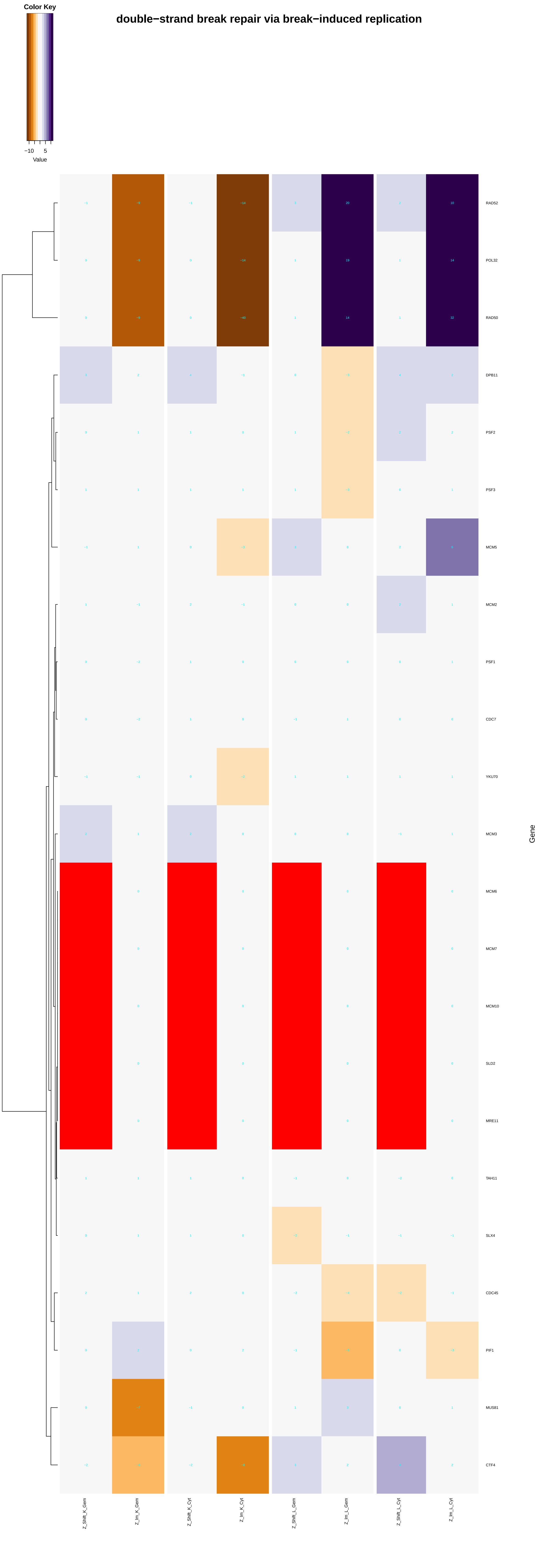

double−strand break repair via break−induced replication
Color Key
−10
5
Value
Gene
RAD52
POL32
RAD50
DPB11
PSF2
PSF3
MCM5
MCM2
PSF1
CDC7
YKU70
MCM3
MCM6
MCM7
MCM10
SLD2
MRE11
TAH11
SLX4
CDC45
PIF1
MUS81
CTF4
Z_Shift_K_Gem
Z_Im_K_Gem
Z_Shift_K_Cyt
Z_Im_K_Cyt
Z_Shift_L_Gem
Z_Im_L_Gem
Z_Shift_L_Cyt
Z_Im_L_Cyt

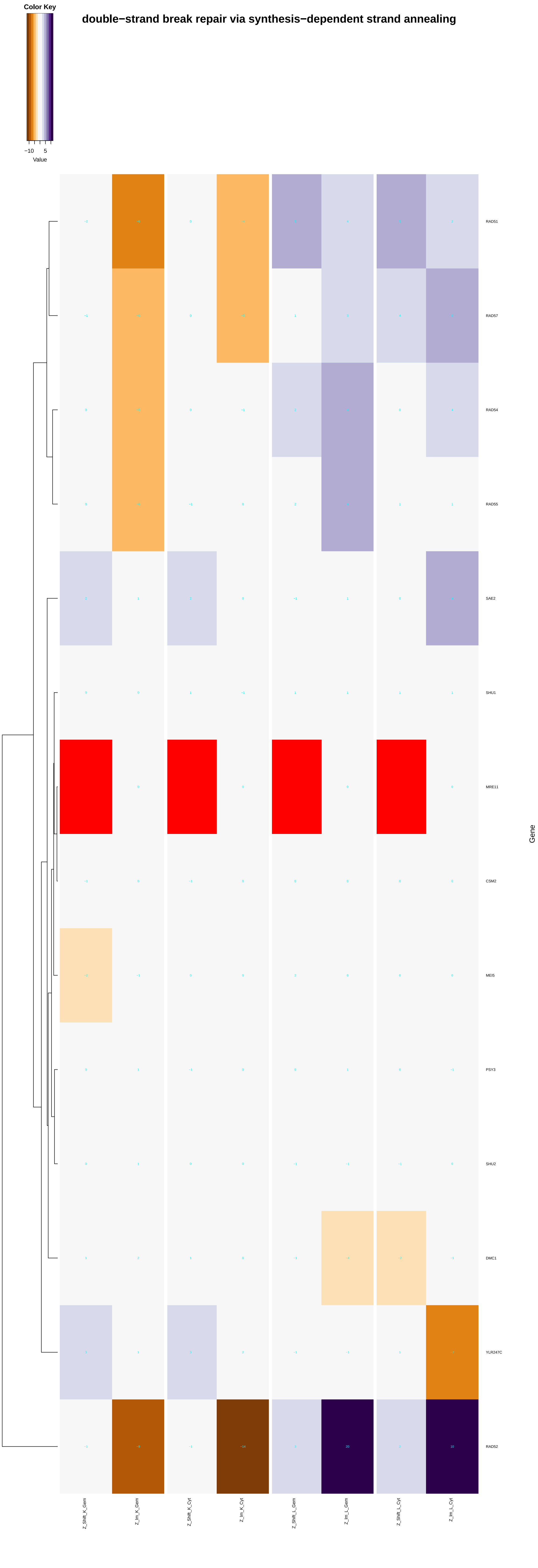
double−strand break repair via synthesis−dependent strand annealing
Color Key
−10
5
Value
RAD51
RAD57
RAD54
RAD55
SAE2
SHU1
MRE11
CSM2
MEI5
PSY3
SHU2
DMC1
YLR247C
RAD52
Gene
Z_Shift_K_Gem
Z_Im_K_Gem
Z_Shift_K_Cyt
Z_Im_K_Cyt
Z_Shift_L_Gem
Z_Im_L_Gem
Z_Shift_L_Cyt
Z_Im_L_Cyt

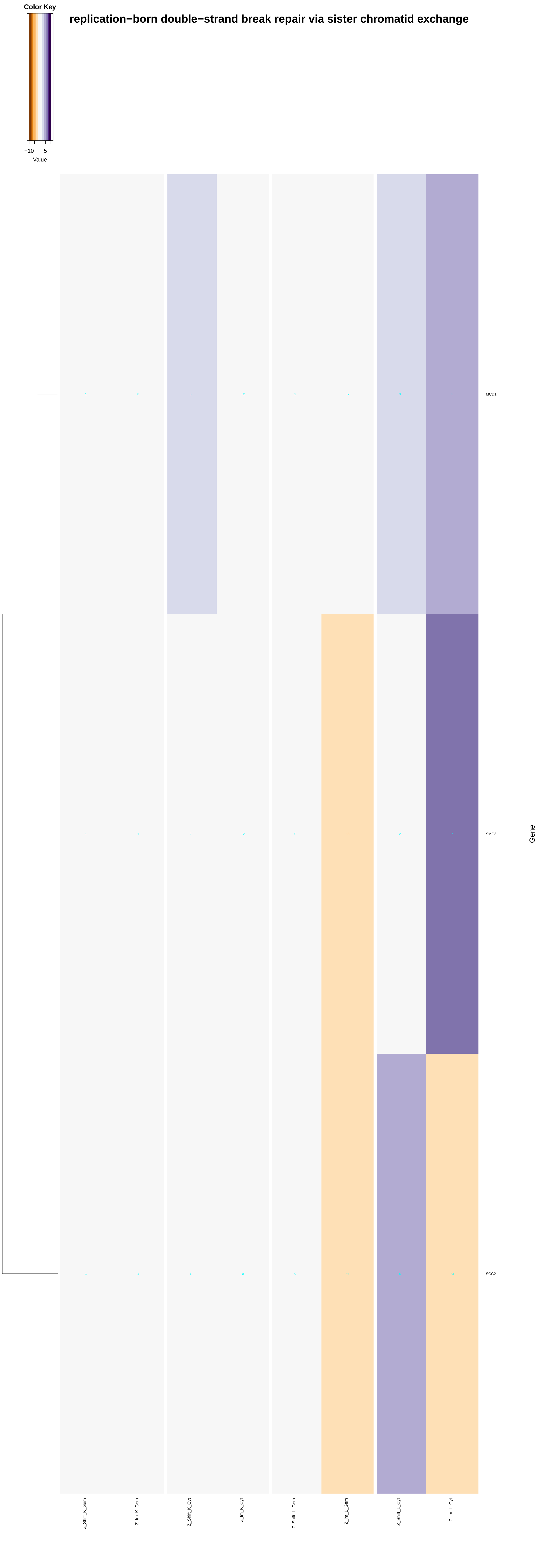
replication−born double−strand break repair via sister chromatid exchange
Color Key
−10
5
Value
MCD1
SMC3
SCC2
Gene
Z_Shift_K_Gem
Z_Im_K_Gem
Z_Shift_K_Cyt
Z_Im_K_Cyt
Z_Shift_L_Gem
Z_Im_L_Gem
Z_Shift_L_Cyt
Z_Im_L_Cyt

#### error−prone translesion synthesis

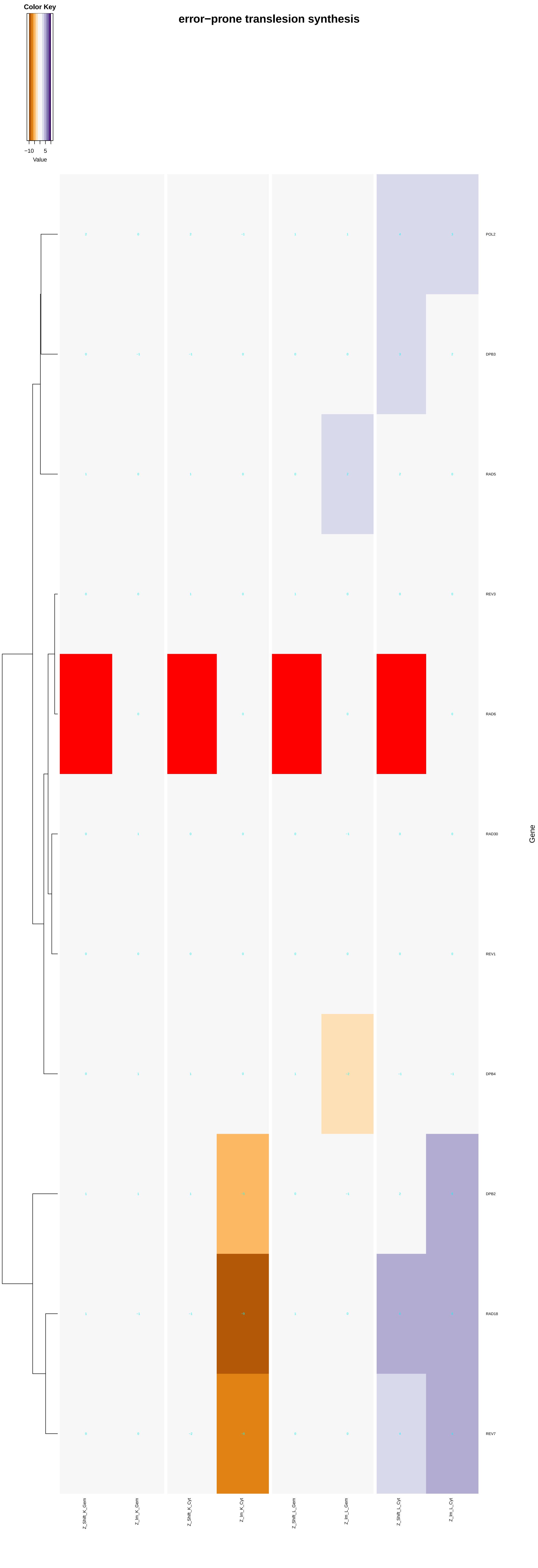

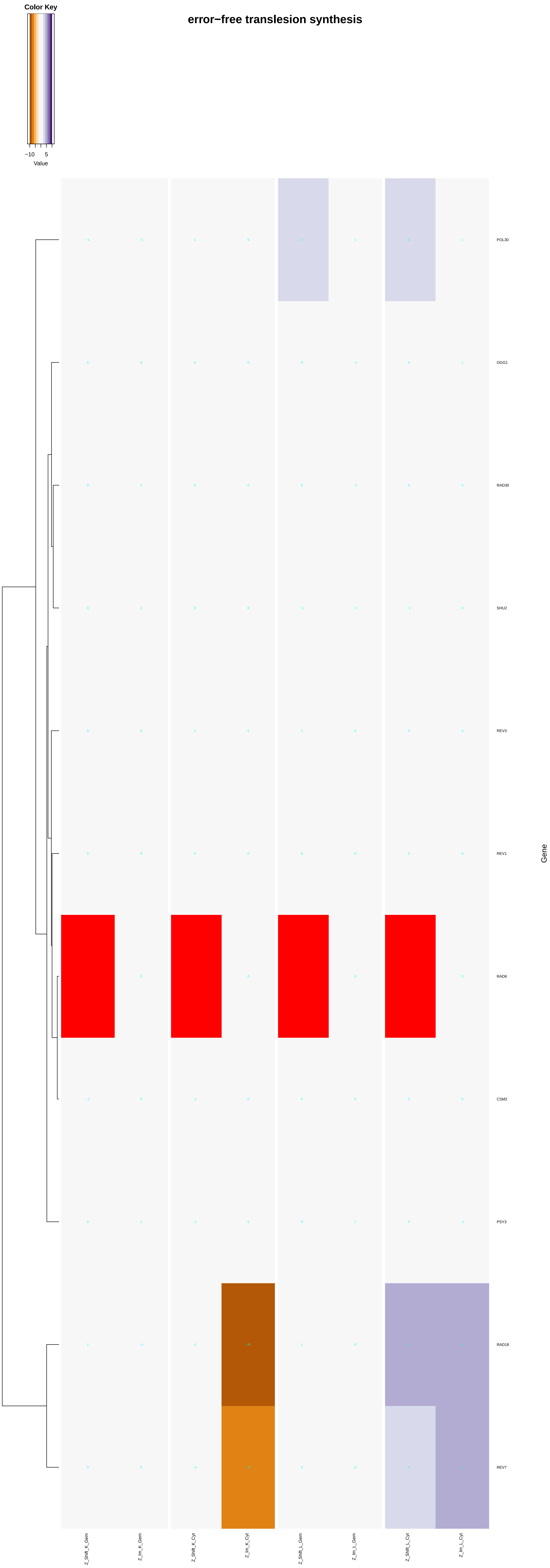

Color Key
−10
5
Value
error−free translesion synthesis
POL30
OGG1
RAD30
SHU2
REV3
REV1
RAD6
CSM2
PSY3
RAD18
REV7
Gene
Z_Shift_K_Gem
Z_Im_K_Gem
Z_Shift_K_Cyt
Z_Im_K_Cyt
Z_Shift_L_Gem
Z_Im_L_Gem
Z_Shift_L_Cyt
Z_Im_L_Cyt
