## Additional File 7 for "A humanized yeast phenomic model of deoxycytidine kinase to predict genetic buffering of nucleoside analog cytotoxicity": chromatin.pdf

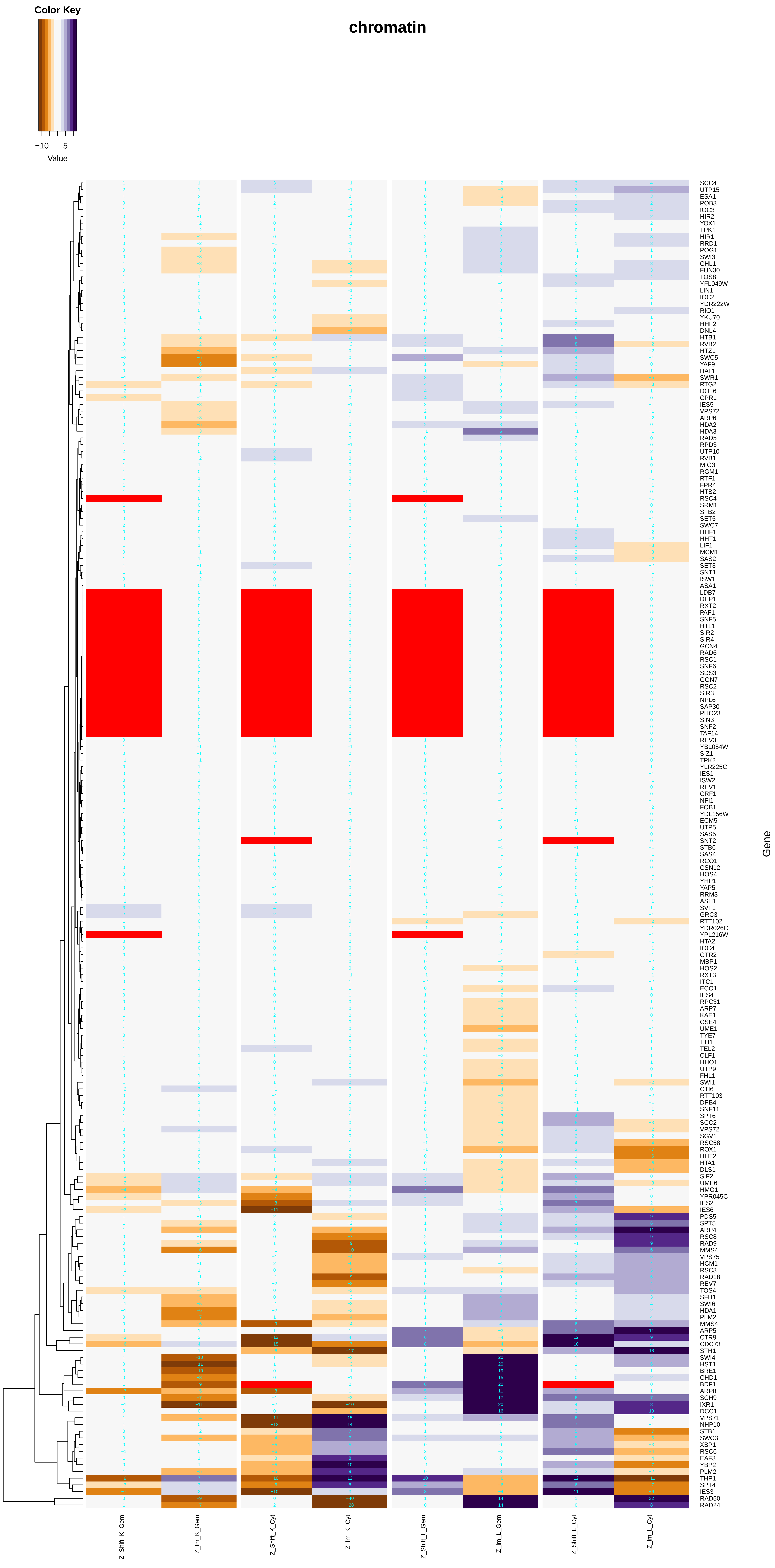
chromatin
Color Key
−10
5
Value
SCC4
UTP15
ESA1
POB3
IOC3
HIR2
YOX1
TPK1
HIR1
RRD1
POG1
SWI3
CHL1
FUN30
TOS8
YFL049W
LIN1
IOC2
YDR222W
RIO1
YKU70
HHF2
DNL4
HTB1
RVB2
HTZ1
SWC5
YAF9
HAT1
SWR1
RTG2
DOT6
CPR1
IES5
VPS72
ARP6
HDA2
HDA3
RAD5
RPD3
UTP10
RVB1
MIG3
RGM1
RTF1
FPR4
HTB2
RSC4
SRM1
STB2
SET5
SWC7
HHF1
HHT1
LIF1
MCM1
SAS2
SET3
SNT1
ISW1
ASA1
LDB7
DEP1
RXT2
PAF1
SNF5
HTL1
SIR2
SIR4
GCN4
RAD6
RSC1
SNF6
SDS3
GON7
RSC2
SIR3
NPL6
SAP30
PHO23
SIN3
SNF2
TAF14
REV3
YBL054W
SIZ1
TPK2
YLR225C
IES1
ISW2
REV1
CRF1
NFI1
FOB1
YDL156W
ECM5
UTP5
SAS5
SNT2
STB6
SAS4
RCO1
CSN12
HOS4
YHP1
YAP5
RRM3
ASH1
SVF1
GRC3
RTT102
YDR026C
YPL216W
HTA2
IOC4
GTR2
MBP1
HOS2
RXT3
ITC1
ECO1
IES4
RPC31
ARP7
KAE1
CSE4
UME1
TYE7
TTI1
TEL2
CLF1
HHO1
UTP9
FHL1
SWI1
CTI6
RTT103
DPB4
SNF11
SPT6
SCC2
VPS72
SGV1
RSC58
ROX1
HHT2
HTA1
DLS1
SIF2
UME6
HMO1
YPR045C
IES2
IES6
PDS5
SPT5
ARP4
RSC8
RAD9
MMS4
VPS75
HCM1
RSC3
RAD18
REV7
TOS4
SFH1
SWI6
HDA1
PLM2
MMS4
ARP5
CTR9
CDC73
STH1
SWI4
HST1
BRE1
CHD1
BDF1
ARP8
SCH9
IXR1
DCC1
VPS71
NHP10
STB1
SWC3
XBP1
RSC6
EAF3
YBP2
PLM2
THP1
SPT4
IES3
RAD50
RAD24
Gene
Z_Shift_K_Gem
Z_Im_K_Gem
Z_Shift_K_Cyt
Z_Im_K_Cyt
Z_Shift_L_Gem
Z_Im_L_Gem
Z_Shift_L_Cyt
Z_Im_L_Cyt

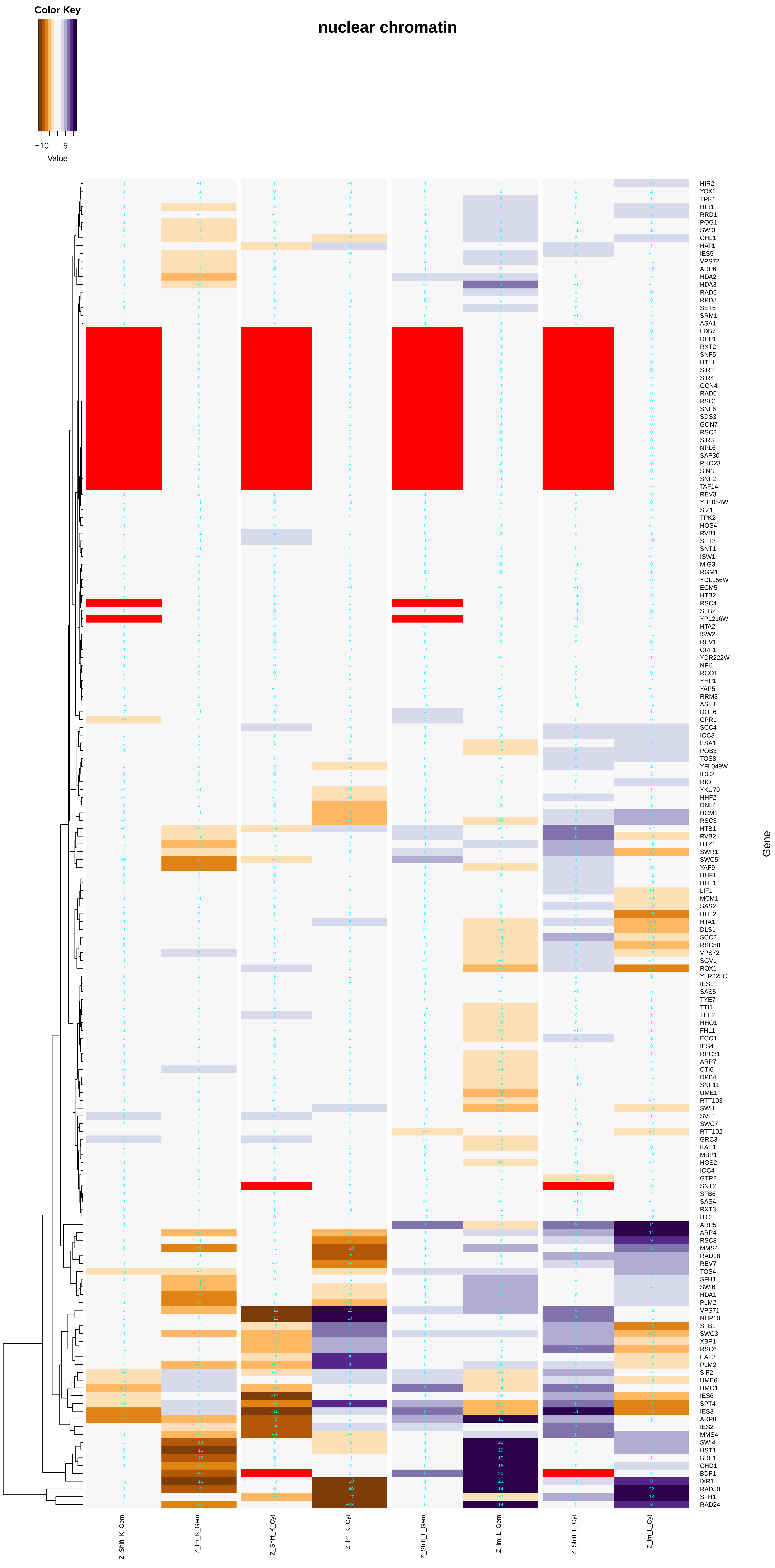
Color Key
−10
5
Value
nuclear chromatin
HIR2
YOX1
TPK1
HIR1
RRD1
POG1
SWI3
CHL1
HAT1
IES5
VPS72
ARP6
HDA2
HDA3
RAD5
RPD3
SET5
SRM1
ASA1
LDB7
DEP1
RXT2
SNF5
HTL1
SIR2
SIR4
GCN4
RAD6
RSC1
SNF6
SDS3
GON7
RSC2
SIR3
NPL6
SAP30
PHO23
SIN3
SNF2
TAF14
REV3
YBL054W
SIZ1
TPK2
HOS4
RVB1
SET3
SNT1
ISW1
MIG3
RGM1
YDL156W
ECM5
HTB2
RSC4
STB2
YPL216W
HTA2
ISW2
REV1
CRF1
YDR222W
NFI1
RCO1
YHP1
YAP5
RRM3
ASH1
DOT6
CPR1
SCC4
IOC3
ESA1
POB3
TOS8
YFL049W
IOC2
RIO1
YKU70
HHF2
DNL4
HCM1
RSC3
HTB1
RVB2
HTZ1
SWR1
SWC5
YAF9
HHF1
HHT1
LIF1
MCM1
SAS2
HHT2
HTA1
DLS1
SCC2
RSC58
VPS72
SGV1
ROX1
YLR225C
IES1
SAS5
TYE7
TTI1
TEL2
HHO1
FHL1
ECO1
IES4
RPC31
ARP7
CTI6
DPB4
SNF11
UME1
RTT103
SWI1
SVF1
SWC7
RTT102
GRC3
KAE1
MBP1
HOS2
IOC4
GTR2
SNT2
STB6
SAS4
RXT3
ITC1
ARP5
ARP4
RSC8
MMS4
RAD18
REV7
TOS4
SFH1
SWI6
HDA1
PLM2
VPS71
NHP10
STB1
SWC3
XBP1
RSC6
EAF3
PLM2
SIF2
UME6
HMO1
IES6
SPT4
IES3
ARP8
IES2
MMS4
SWI4
HST1
BRE1
CHD1
BDF1
IXR1
RAD50
STH1
RAD24
Gene
Z_Shift_K_Gem
Z_lm_K_Gem
Z_Shift_K_Cyt
Z_lm_K_Cyt
Z_Shift_L_Gem
Z_lm_L_Gem
Z_Shift_L_Cyt
Z_lm_L_Cyt

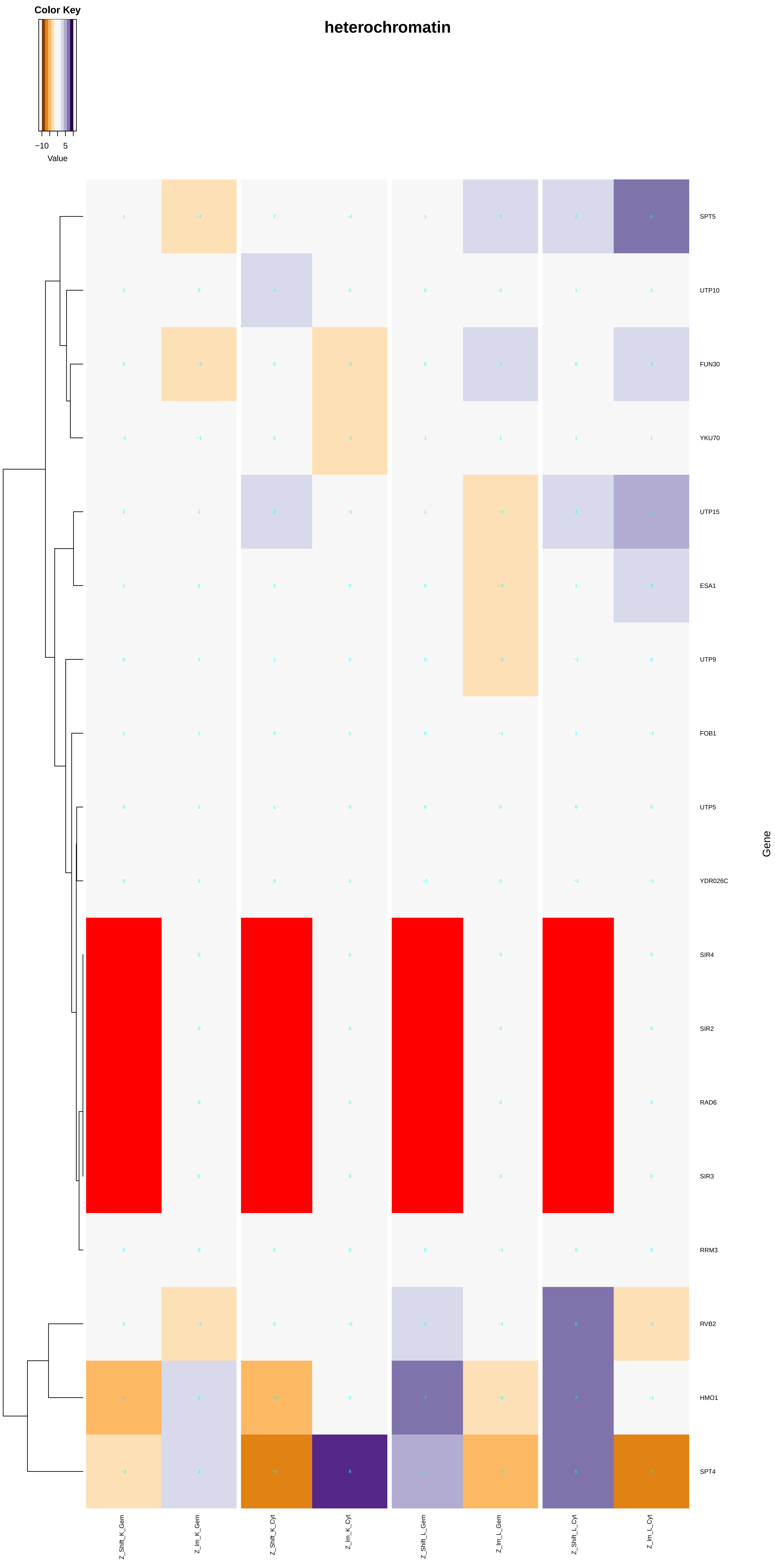
heterochromatin
Color Key
−10
5
Value
SPT5
UTP10
FUN30
YKU70
UTP15
ESA1
UTP9
FOB1
UTP5
YDR026C
SIR4
SIR2
RAD6
SIR3
RRM3
RVB2
HMO1
SPT4
Gene
Z_Shift_K_Gem
Z_Im_K_Gem
Z_Shift_K_Cyt
Z_Im_K_Cyt
Z_Shift_L_Gem
Z_Im_L_Gem
Z_Shift_L_Cyt
Z_Im_L_Cyt

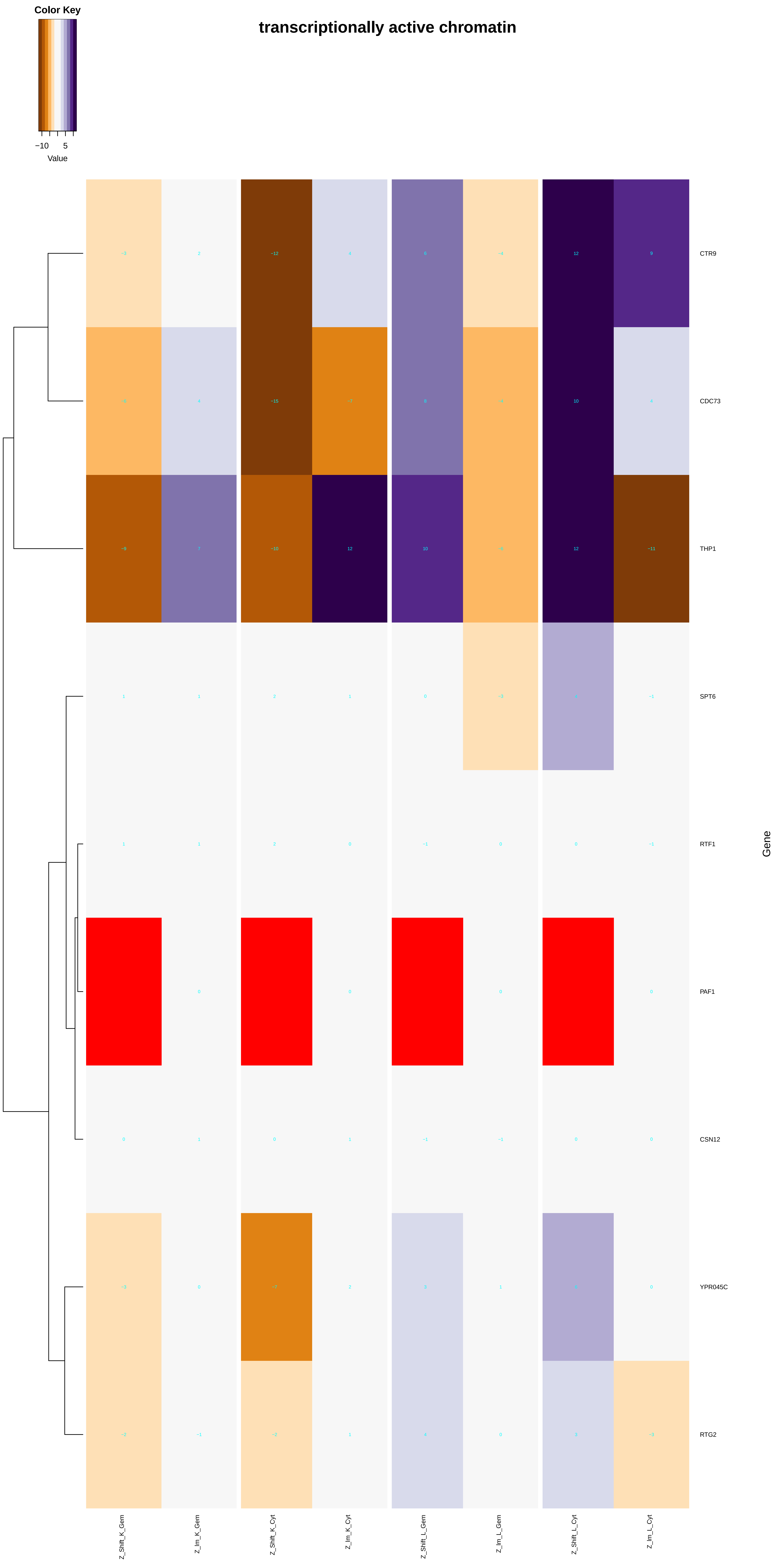
Color Key
−10
5
Value
transcriptionally active chromatin
CTR9
CDC73
THP1
SPT6
RTF1
PAF1
CSN12
YPR045C
RTG2
Gene
Z_Shift_K_Gem
Z_Im_K_Gem
Z_Shift_K_Cyt
Z_Im_K_Cyt
Z_Shift_L_Gem
Z_Im_L_Gem
Z_Shift_L_Cyt
Z_Im_L_Cyt

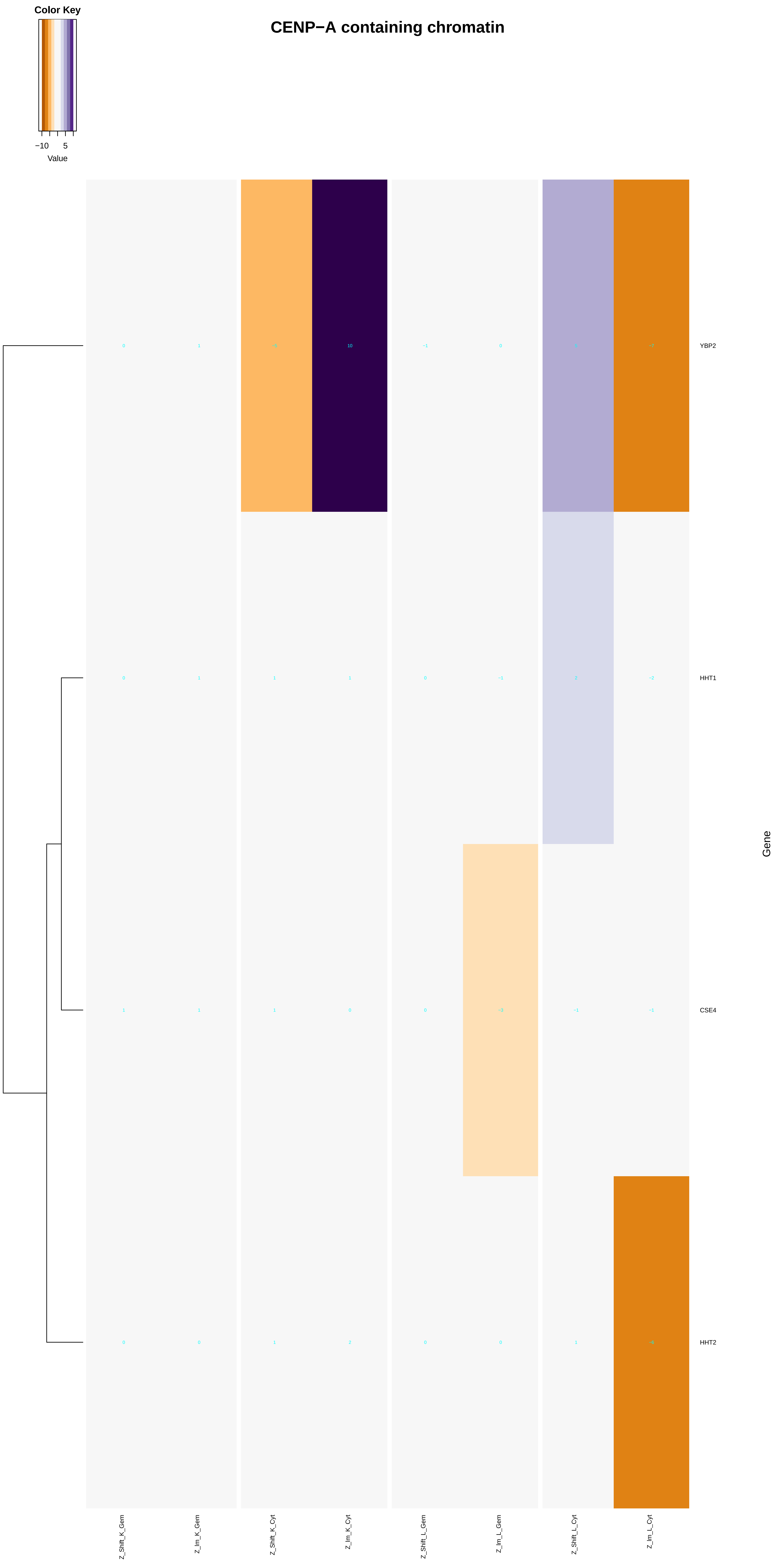
Color Key
−10
5
Value
CENP−A containing chromatin
YBP2
HHT1
CSE4
HHT2
Gene
Z_Shift_K_Gem
Z_Im_K_Gem
Z_Shift_K_Cyt
Z_Im_K_Cyt
Z_Shift_L_Gem
Z_Im_L_Gem
Z_Shift_L_Cyt
Z_Im_L_Cyt

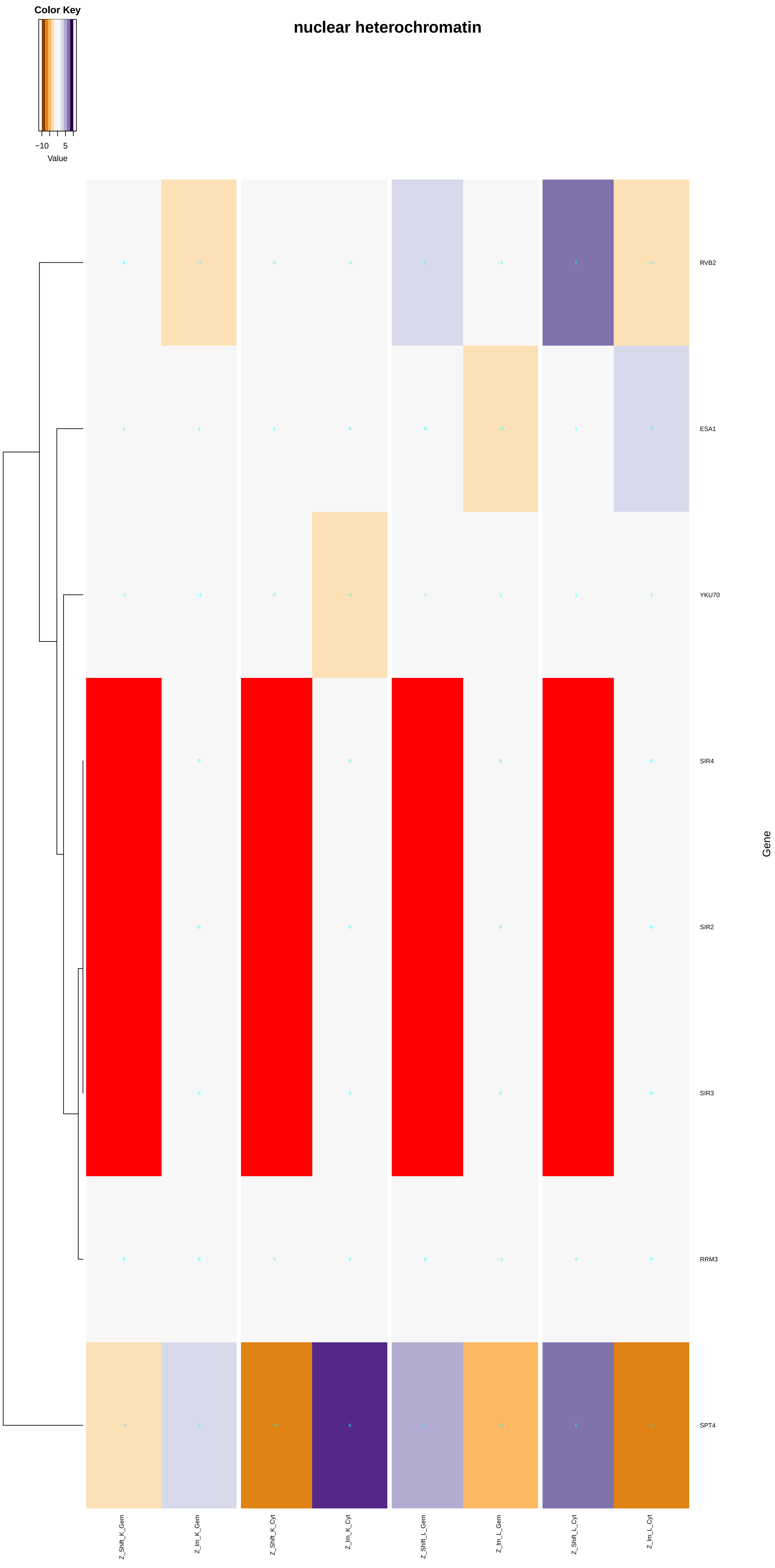

nuclear heterochromatin
Color Key
−10
5
Value
RVB2
ESA1
YKU70
SIR4
SIR2
SIR3
RRM3
SPT4
Gene
Z_Shift_K_Gem
Z_Im_K_Gem
Z_Shift_K_Cyt
Z_Im_K_Cyt
Z_Shift_L_Gem
Z_Im_L_Gem
Z_Shift_L_Cyt
Z_Im_L_Cyt

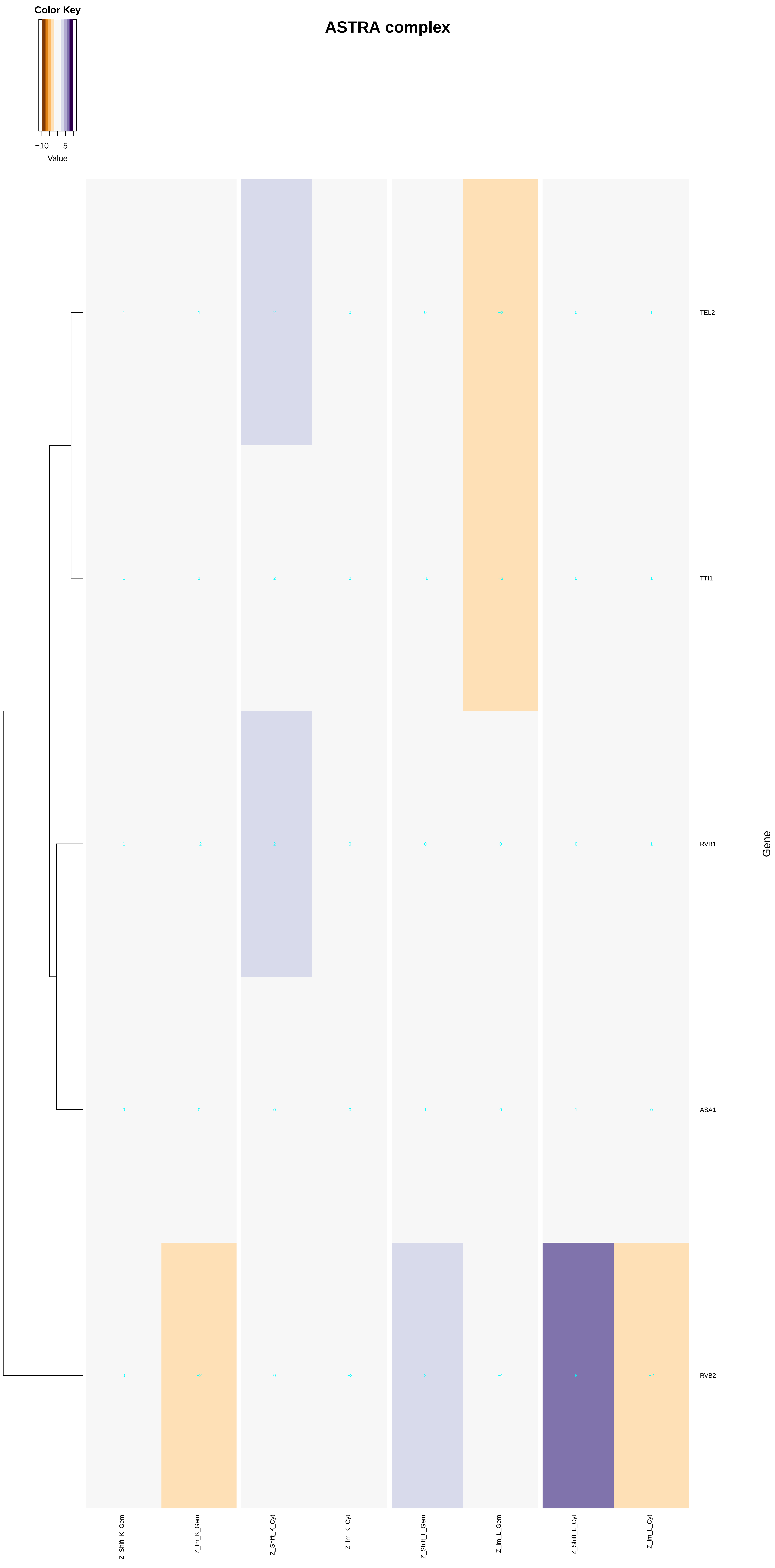
ASTRA complex
Color Key
−10
5
Value
TEL2
TTI1
RVB1
ASA1
RVB2
Gene
Z_Shift_K_Gem
Z_Im_K_Gem
Z_Shift_K_Cyt
Z_Im_K_Cyt
Z_Shift_L_Gem
Z_Im_L_Gem
Z_Shift_L_Cyt
Z_Im_L_Cyt

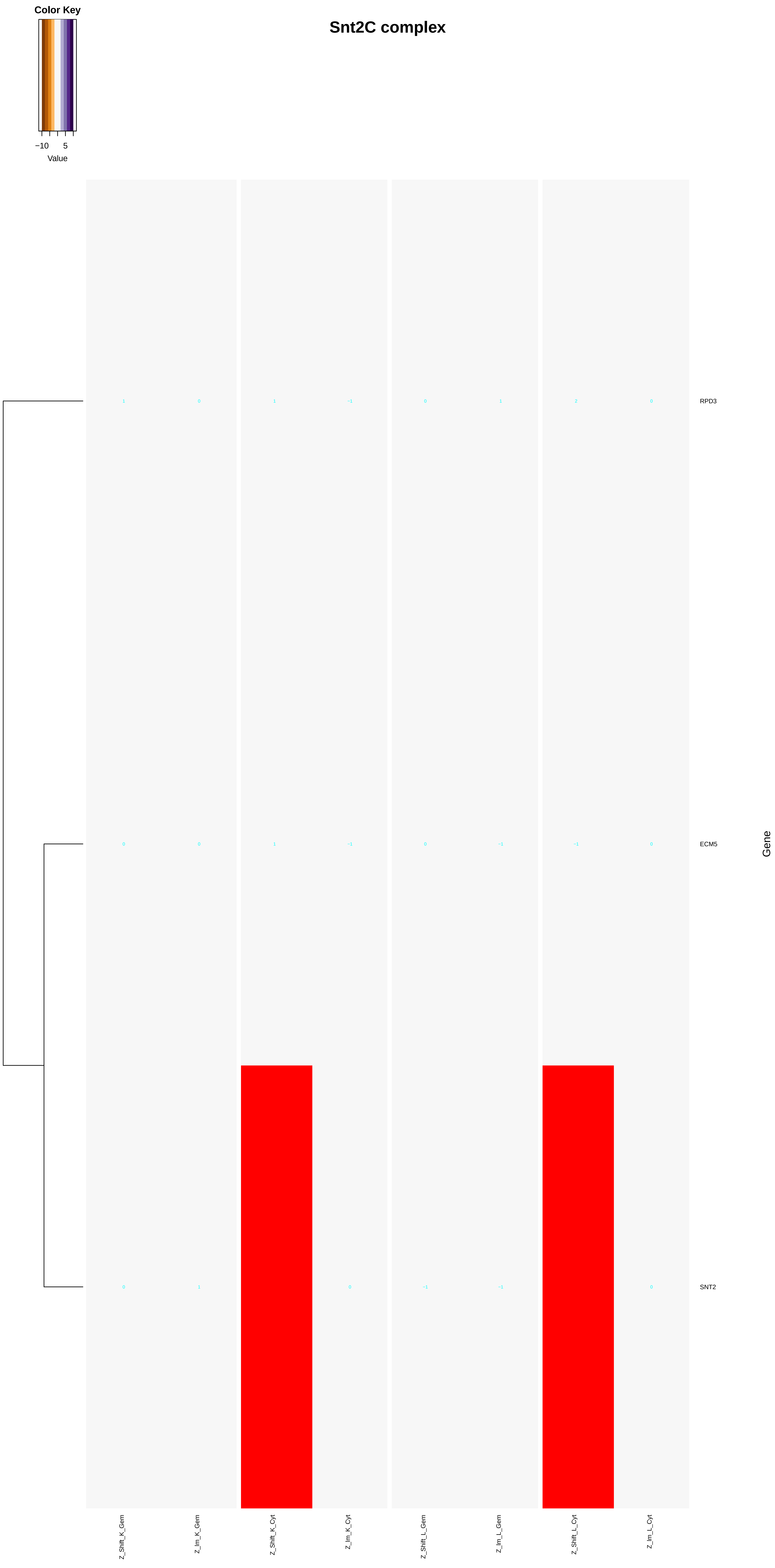
Color Key
−10
5
Value
Snt2C complex
RPD3
ECM5
SNT2
Gene
Z_Shift_K_Gem
Z_Im_K_Gem
Z_Shift_K_Cyt
Z_Im_K_Cyt
Z_Shift_L_Gem
Z_Im_L_Gem
Z_Shift_L_Cyt
Z_Im_L_Cyt

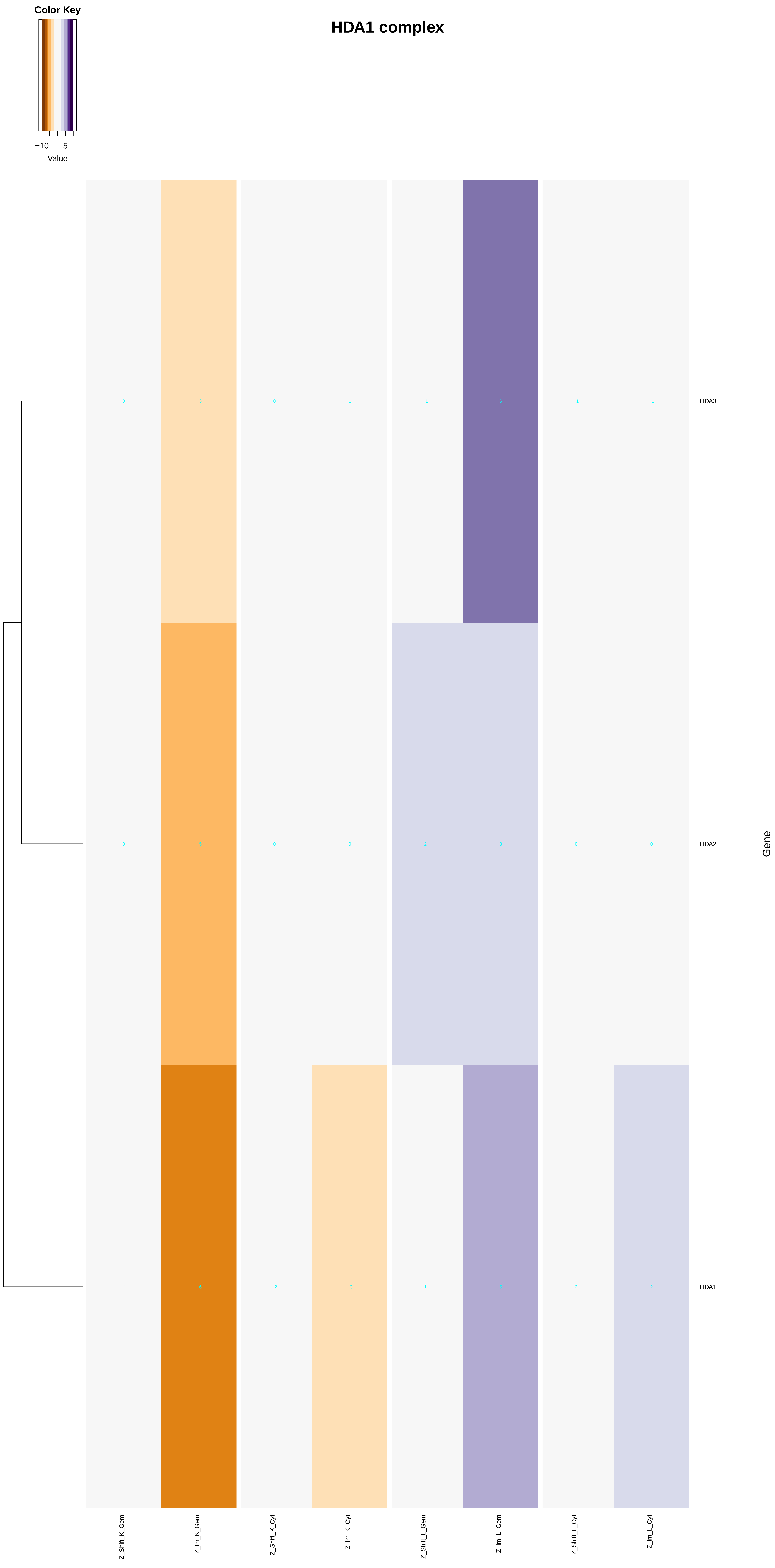
HDA1 complex
Color Key
−10
5
Value
HDA3
HDA2
HDA1
Gene
Z_Shift_K_Gem
Z_Im_K_Gem
Z_Shift_K_Cyt
Z_Im_K_Cyt
Z_Shift_L_Gem
Z_Im_L_Gem
Z_Shift_L_Cyt
Z_Im_L_Cyt

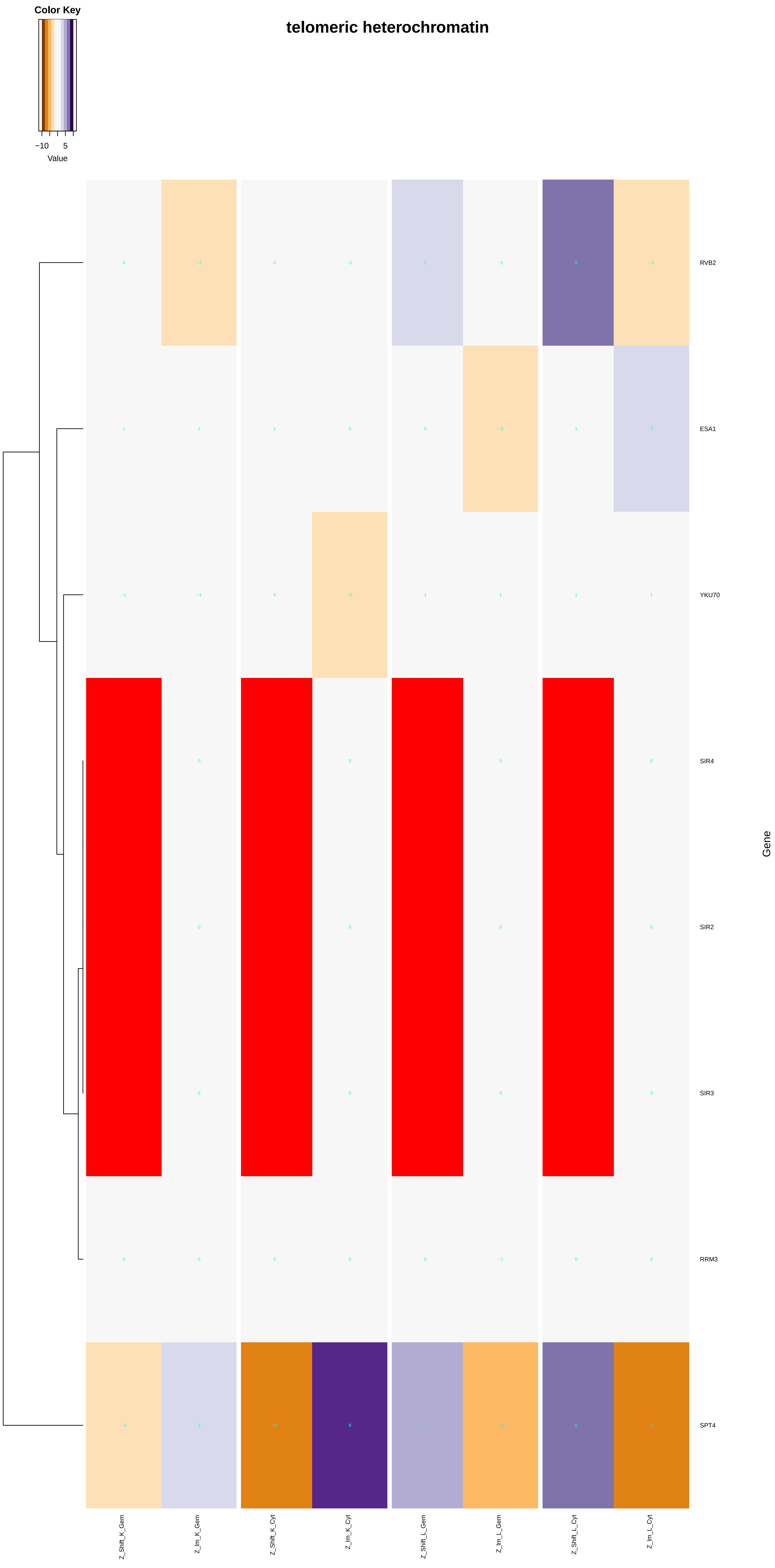

telomeric heterochromatin
Color Key
−10
5
Value
RVB2
ESA1
YKU70
SIR4
SIR2
SIR3
RRM3
SPT4
Gene
Z_Shift_K_Gem
Z_Im_K_Gem
Z_Shift_K_Cyt
Z_Im_K_Cyt
Z_Shift_L_Gem
Z_Im_L_Gem
Z_Shift_L_Cyt
Z_Im_L_Cyt

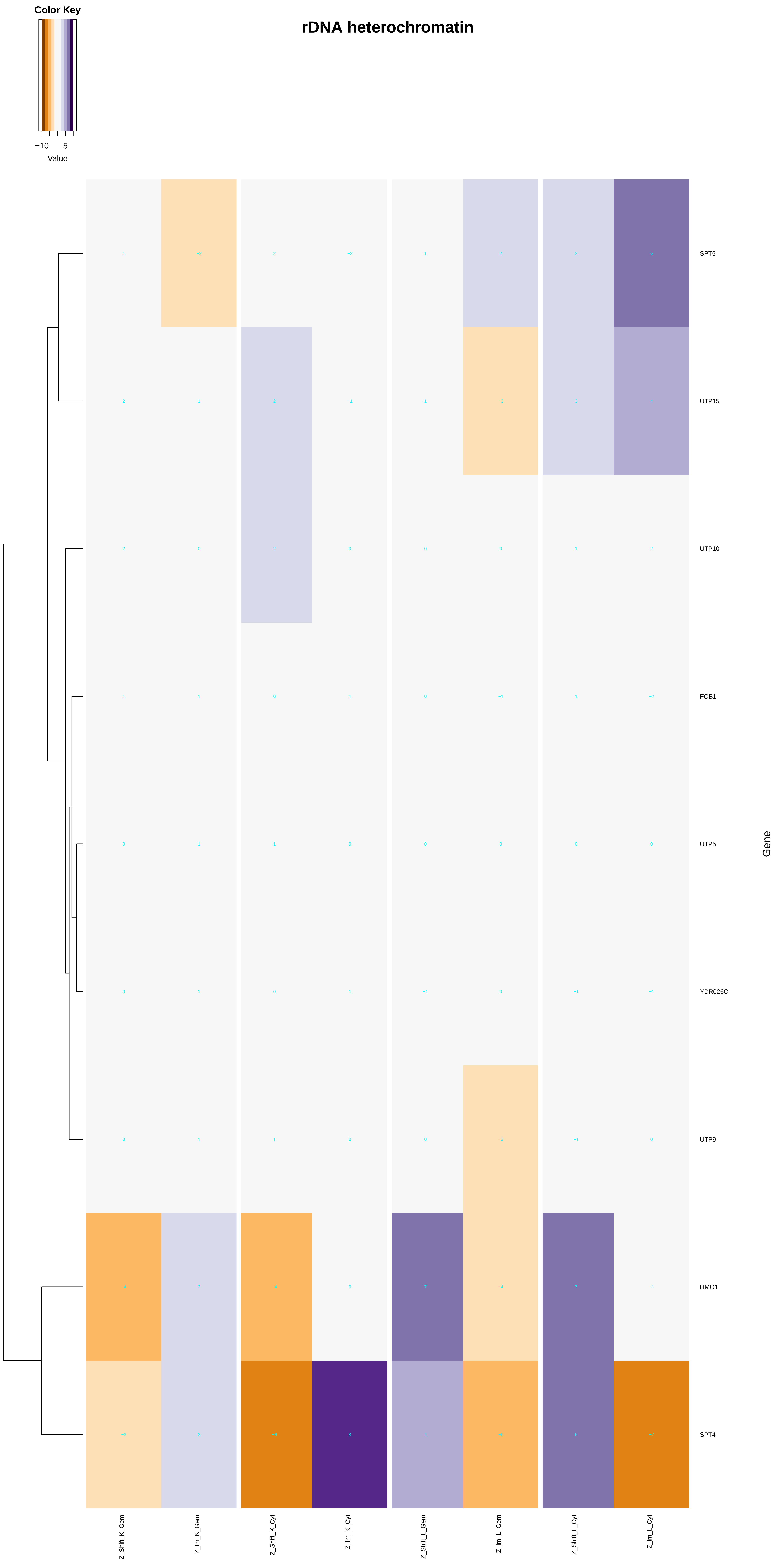

Color Key
−10
5
Value
rDNA heterochromatin
SPT5
UTP15
UTP10
FOB1
UTP5
YDR026C
UTP9
HMO1
SPT4
Gene
Z_Shift_K_Gem
Z_Im_K_Gem
Z_Shift_K_Cyt
Z_Im_K_Cyt
Z_Shift_L_Gem
Z_Im_L_Gem
Z_Shift_L_Cyt
Z_Im_L_Cyt

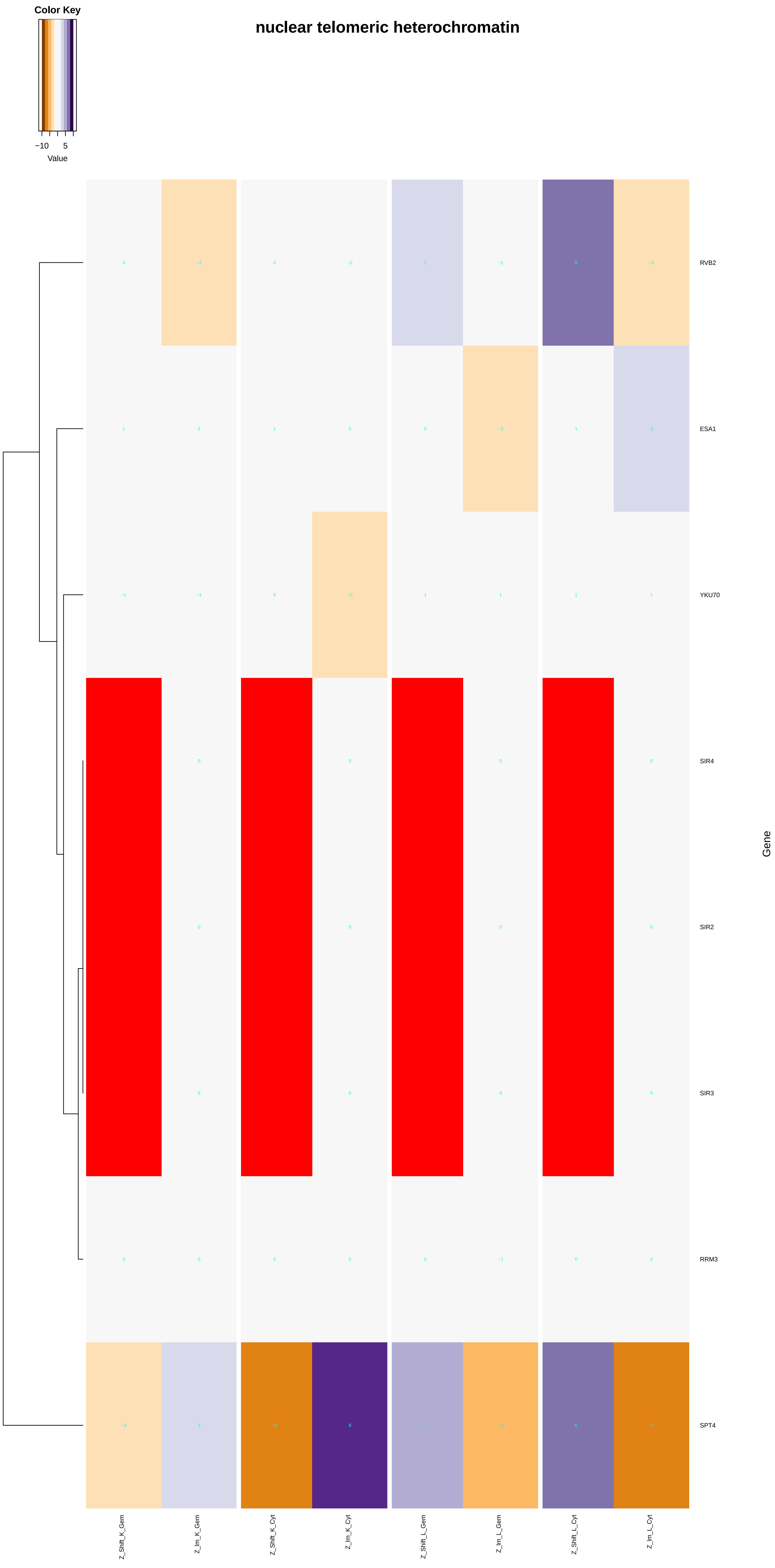

Color Key
−10
5
Value
nuclear telomeric heterochromatin
RVB2
ESA1
YKU70
SIR4
SIR2
SIR3
RRM3
SPT4
Gene
Z_Shift_K_Gem
Z_Im_K_Gem
Z_Shift_K_Cyt
Z_Im_K_Cyt
Z_Shift_L_Gem
Z_Im_L_Gem
Z_Shift_L_Cyt
Z_Im_L_Cyt
