## Additional File 7 for "A humanized yeast phenomic model of deoxycytidine kinase to predict genetic buffering of nucleoside analog cytotoxicity": chromatin_organization.pdf

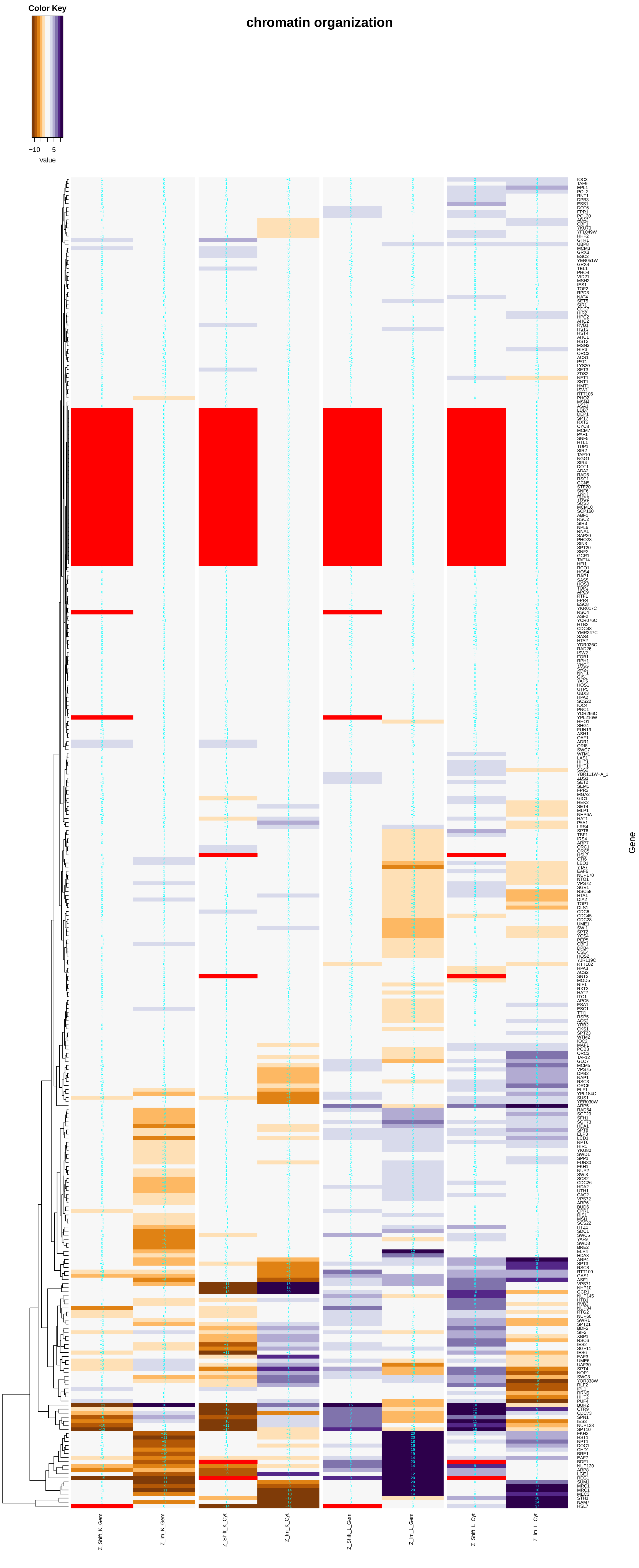
chromatin organization
Color Key
−10
5
Value
Gene
Z_Shift_K_Gem
Z_Im_K_Gem
Z_Shift_K_Cyt
Z_Im_K_Cyt
Z_Shift_L_Gem
Z_Im_L_Gem
Z_Shift_L_Cyt
Z_Im_L_Cyt

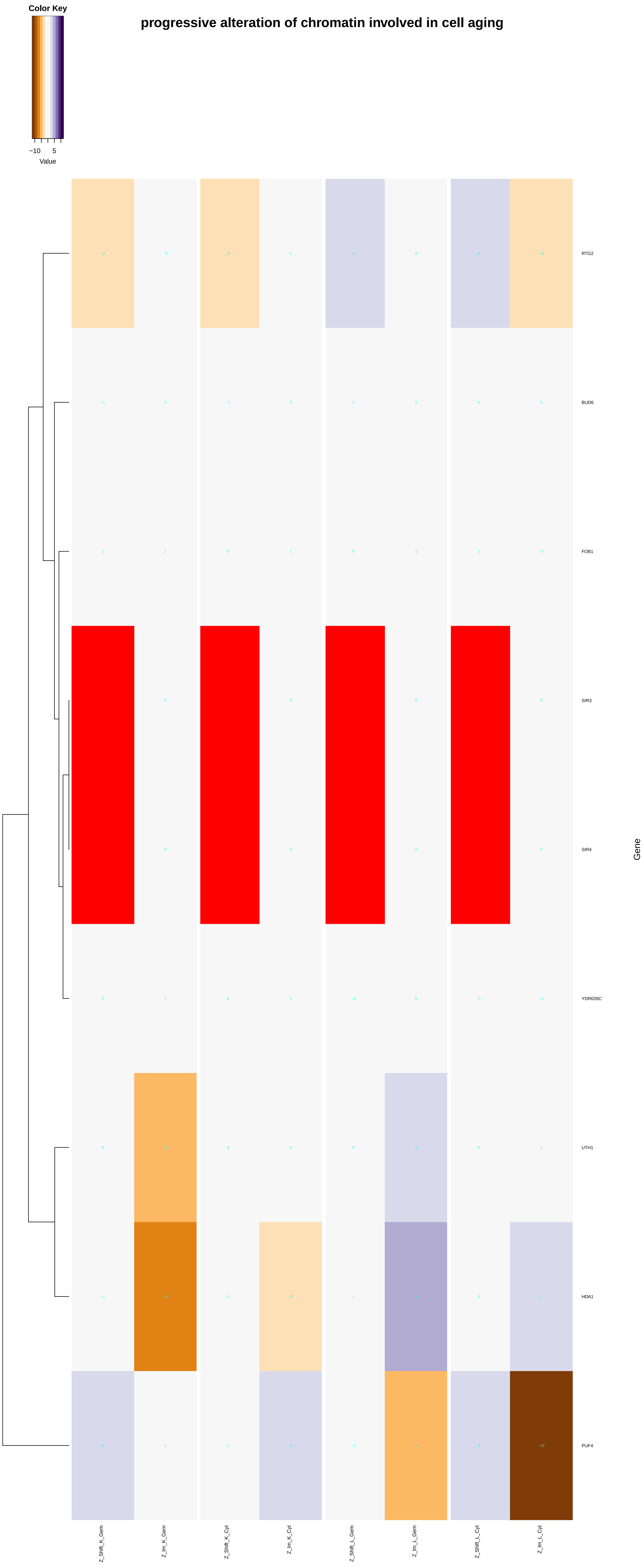

Color Key
−10
5
Value
progressive alteration of chromatin involved in cell aging
RTG2
BUD6
FOB1
SIR3
SIR4
YDR026C
UTH1
HDA1
PUF4
Gene
Z_Shift_K_Gem
Z_Im_K_Gem
Z_Shift_K_Cyt
Z_Im_K_Cyt
Z_Shift_L_Gem
Z_Im_L_Gem
Z_Shift_L_Cyt
Z_Im_L_Cyt

#### chromatin assembly or disassembly

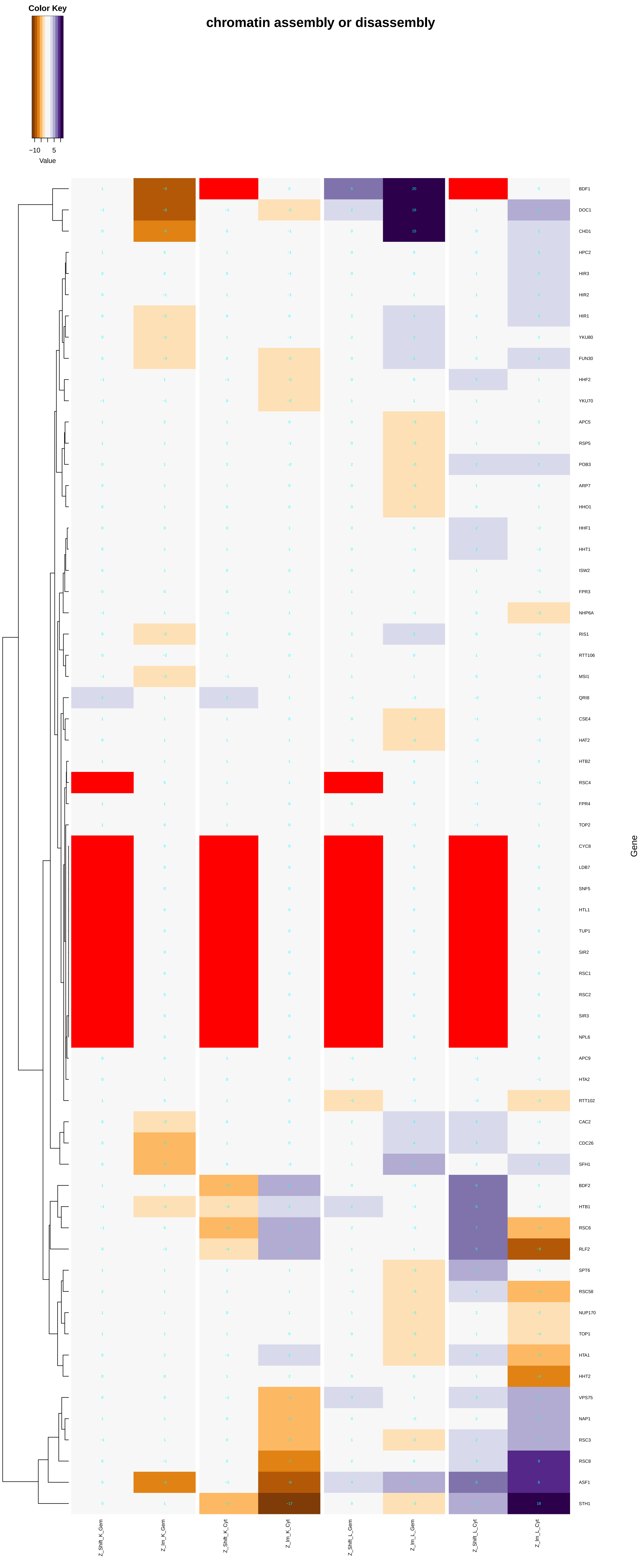

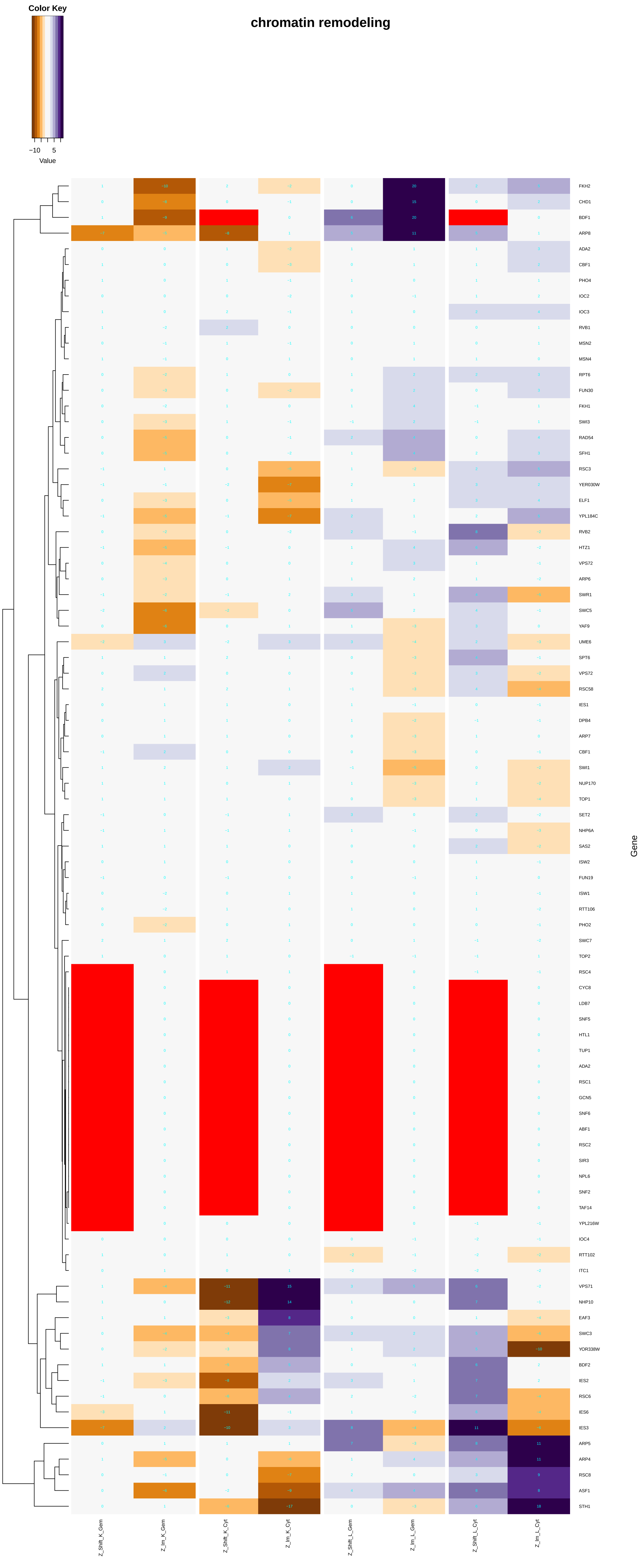
chromatin remodeling
Color Key
−10
5
Value
FKH2
CHD1
BDF1
ARP8
ADA2
CBF1
PHO4
IOC2
IOC3
RVB1
MSN2
MSN4
RPT6
FUN30
FKH1
SWI3
RAD54
SFH1
RSC3
YER030W
ELF1
YPL184C
RVB2
HTZ1
VPS72
ARP6
SWR1
SWC5
YAF9
UME6
SPT6
VPS72
RSC58
IES1
DPB4
ARP7
CBF1
SWI1
NUP170
TOP1
SET2
NHP6A
SAS2
ISW2
FUN19
ISW1
RTT106
PHO2
SWC7
TOP2
RSC4
CYC8
LDB7
SNF5
HTL1
TUP1
ADA2
RSC1
GCN5
SNF6
ABF1
RSC2
SIR3
NPL6
SNF2
TAF14
YPL216W
IOC4
RTT102
ITC1
VPS71
NHP10
EAF3
SWC3
YOR338W
BDF2
IES2
RSC6
IES6
IES3
ARP5
ARP4
RSC8
ASF1
STH1
Z_Shift_K_Gem
Z_Im_K_Gem
Z_Shift_K_Cyt
Z_Im_K_Cyt
Z_Shift_L_Gem
Z_Im_L_Gem
Z_Shift_L_Cyt
Z_Im_L_Cyt
Gene

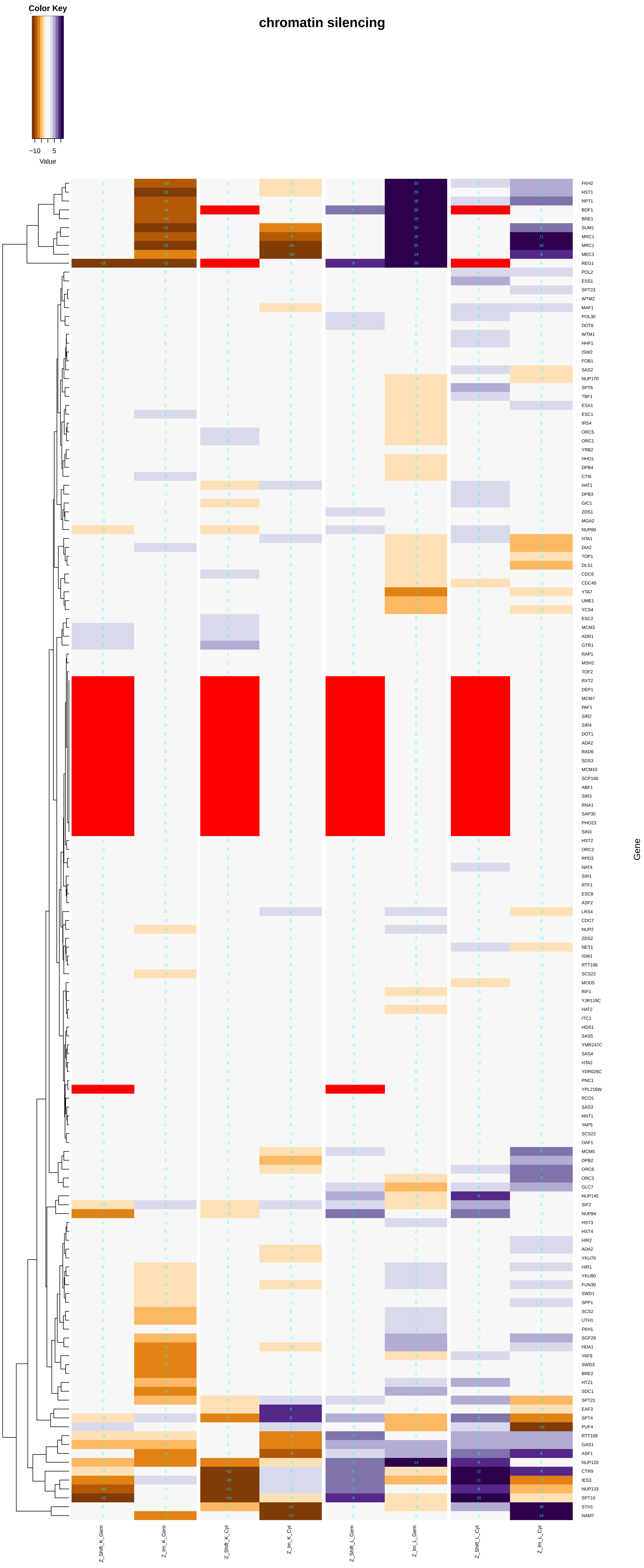
chromatin silencing
Color Key
−10
5
Value
FKH2
HST1
NPT1
BDF1
BRE1
SUM1
MRC1
MRC1
MEC3
REG1
POL2
ESS1
SPT23
WTM2
MAF1
POL30
DOT6
WTM1
HHF1
ISW2
FOB1
SAS2
NUP170
SPT6
TBF1
ESA1
ESC1
IRS4
ORC5
ORC1
YRB2
HHO1
DPB4
CTI6
HAT1
DPB3
GIC1
ZDS1
MGA2
NUP60
HTA1
DIA2
TOP1
DLS1
CDC6
CDC45
YTA7
UME1
YCS4
ESC2
MCM3
ADR1
GTR1
RAP1
MSH2
TOF2
RXT2
DEP1
MCM7
PAF1
SIR2
SIR4
DOT1
ADA2
RAD6
SDS3
MCM10
SCP160
ABF1
SIR3
RNA1
SAP30
PHO23
SIN3
HST2
ORC2
RPD3
NAT4
SIR1
RTF1
ESC8
ASF2
LRS4
CDC7
NUP2
ZDS2
NET1
ISW1
RTT106
SCS22
MOD5
RIF1
YJR119C
HAT2
ITC1
HOS1
SAS5
YMR247C
SAS4
HTA2
YDR026C
PNC1
YPL216W
RCO1
SAS3
NNT1
YAP5
SCS22
OAF1
MCM5
DPB2
ORC6
ORC3
GLC7
NUP145
SIF2
NUP84
HST3
HST4
HIR2
ADA2
YKU70
HIR1
YKU80
FUN30
SWD1
SPP1
SCS2
UTH1
FKH1
SGF29
HDA1
YAF9
SWD3
BRE2
HTZ1
SDC1
SPT21
EAF3
SPT4
PUF4
RTT109
GAS1
ASF1
NUP120
CTR9
IES3
NUP133
SPT10
STH1
NAM7
Gene
Z_Shift_K_Gem
Z_Im_K_Gem
Z_Shift_K_Cyt
Z_Im_K_Cyt
Z_Shift_L_Gem
Z_Im_L_Gem
Z_Shift_L_Cyt
Z_Im_L_Cyt

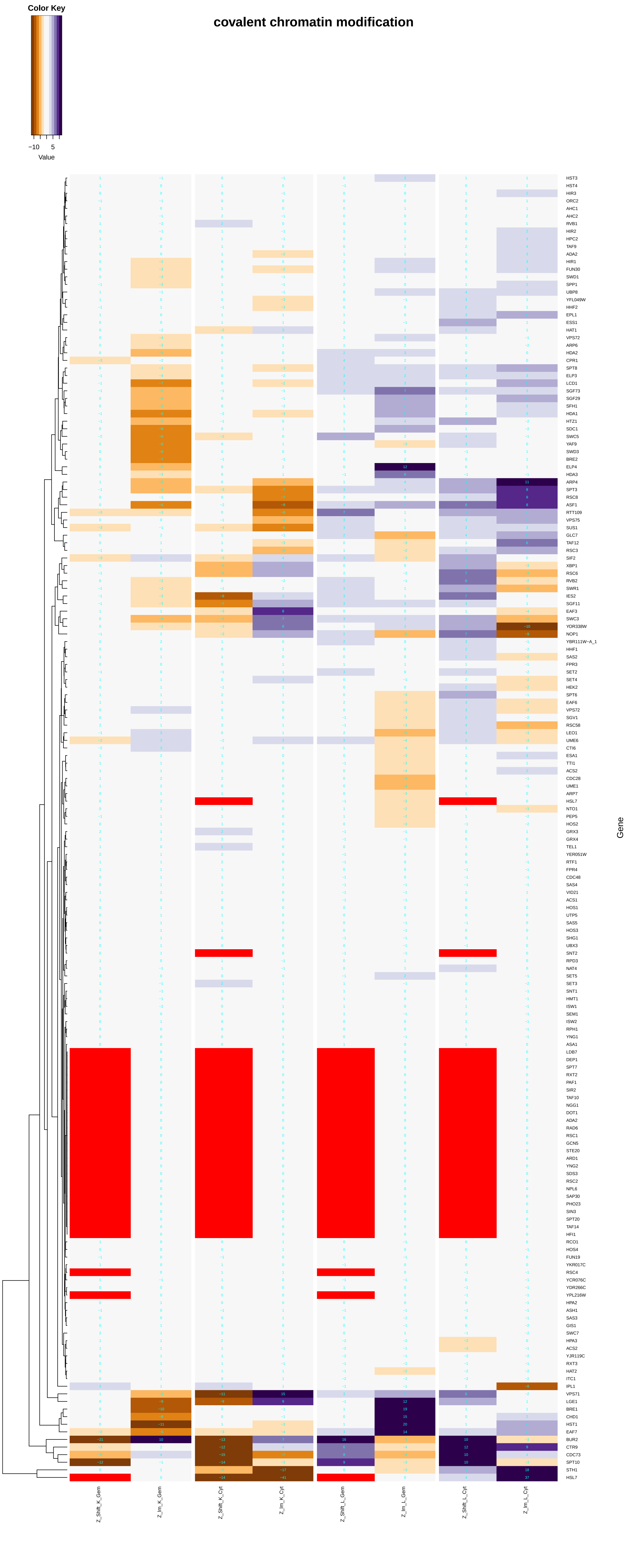

covalent chromatin modification
Color Key
−10
5
Value
Gene
HST3
HST4
HIR3
ORC2
AHC1
AHC2
RVB1
HIR2
HPC2
TAF9
ADA2
HIR1
FUN30
SWD1
SPP1
UBP8
YFL049W
HHF2
EPL1
ESS1
HAT1
VPS72
ARP6
HDA2
CPR1
SPT8
ELP3
LCD1
SGF73
SGF29
SFH1
HDA1
HTZ1
SDC1
SWC5
YAF9
SWD3
BRE2
ELP4
HDA3
ARP4
SPT3
RSC8
ASF1
RTT109
VPS75
SUS1
GLC7
TAF12
RSC3
SIF2
XBP1
RSC6
RVB2
SWR1
IES2
SGF11
EAF3
SWC3
YOR338W
NOP1
YBR111W−A_1
HHF1
SAS2
FPR3
SET2
SET4
HEK2
SPT6
EAF6
VPS72
SGV1
RSC58
LEO1
UME6
CTI6
ESA1
TTI1
ACS2
CDC28
UME1
ARP7
HSL7
NTO1
PEP5
HOS2
GRX3
GRX4
TEL1
YER051W
RTF1
FPR4
CDC48
SAS4
VID21
ACS1
HOS1
UTP5
SAS5
HOS3
SHG1
UBX3
SNT2
RPD3
NAT4
SET5
SET3
SNT1
HMT1
ISW1
SEM1
ISW2
RPH1
YNG1
ASA1
LDB7
DEP1
SPT7
RXT2
PAF1
SIR2
TAF10
NGG1
DOT1
ADA2
RAD6
RSC1
GCN5
STE20
ARD1
YNG2
SDS3
RSC2
NPL6
SAP30
PHO23
SIN3
SPT20
TAF14
HFI1
RCO1
HOS4
FUN19
YKR017C
RSC4
YCR076C
YDR266C
YPL216W
HPA2
ASH1
SAS3
GIS1
SWC7
HPA3
ACS2
YJR119C
RXT3
HAT2
ITC1
IPL1
VPS71
LGE1
BRE1
CHD1
HST1
EAF7
BUR2
CTR9
CDC73
SPT10
STH1
HSL7
Z_Shift_K_Gem
Z_Im_K_Gem
Z_Shift_K_Cyt
Z_Im_K_Cyt
Z_Shift_L_Gem
Z_Im_L_Gem
Z_Shift_L_Cyt
Z_Im_L_Cyt

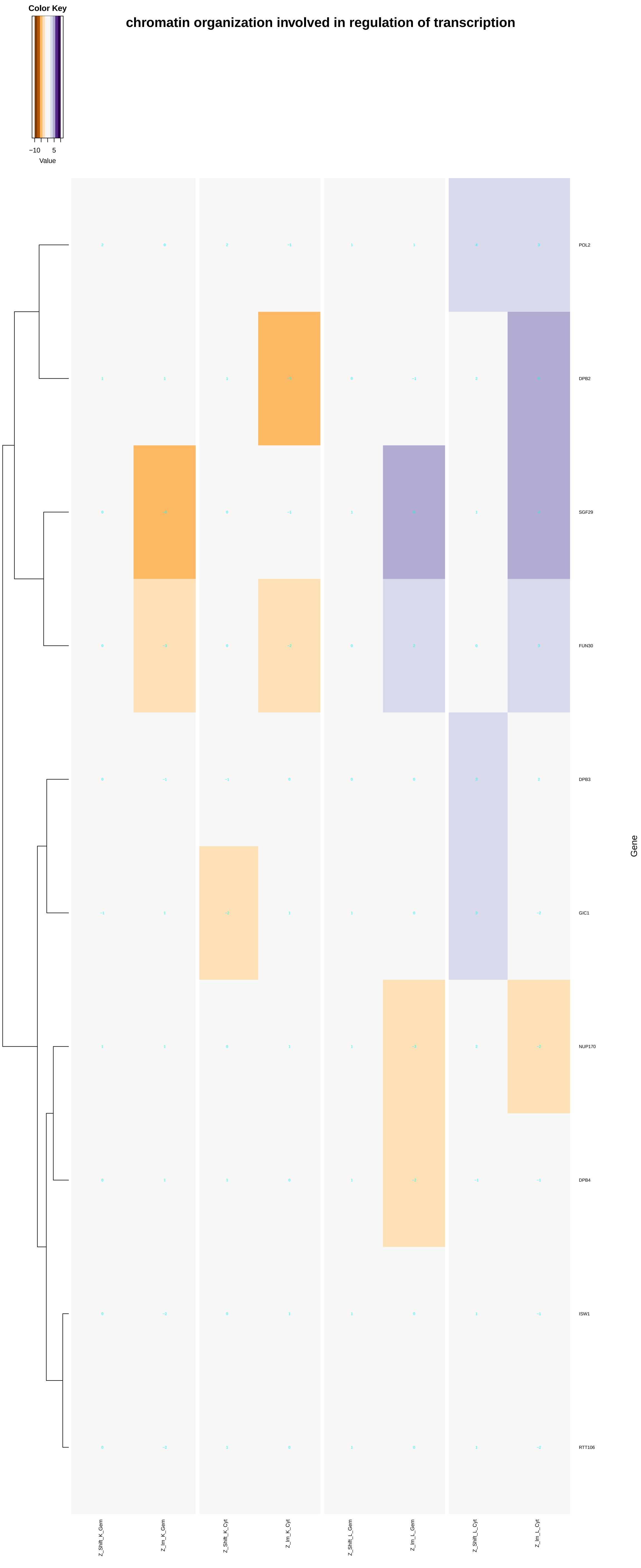
chromatin organization involved in regulation of transcription
Color Key
−10
5
Value
POL2
DPB2
SGF29
FUN30
DPB3
GIC1
NUP170
DPB4
ISW1
RTT106
Gene
Z_Shift_K_Gem
Z_Im_K_Gem
Z_Shift_K_Cyt
Z_Im_K_Cyt
Z_Shift_L_Gem
Z_Im_L_Gem
Z_Shift_L_Cyt
Z_Im_L_Cyt

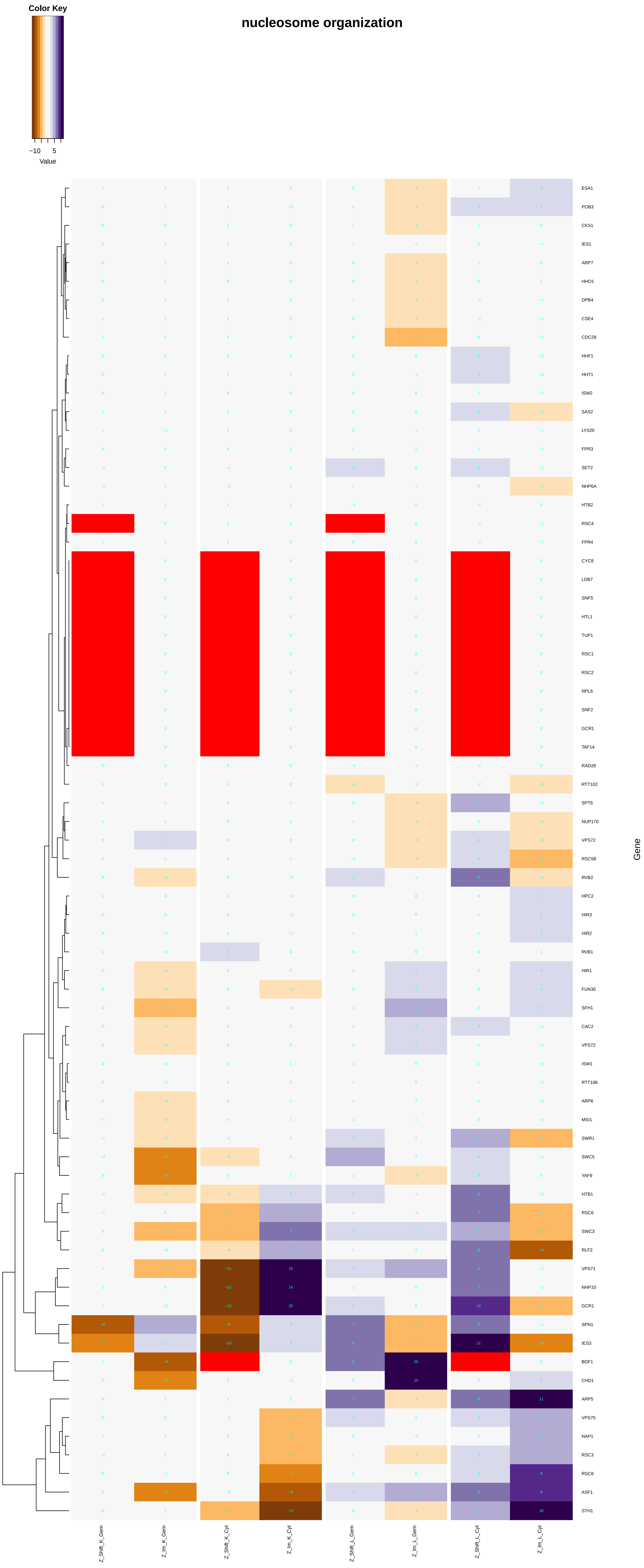

Color Key
−10
5
Value
nucleosome organization
ESA1
POB3
CKS1
IES1
ARP7
HHO1
DPB4
CSE4
CDC28
HHF1
HHT1
ISW2
SAS2
LYS20
FPR3
SET2
NHP6A
HTB2
RSC4
FPR4
CYC8
LDB7
SNF5
HTL1
TUP1
RSC1
RSC2
NPL6
SNF2
GCR1
TAF14
RAD26
RTT102
SPT6
NUP170
VPS72
RSC58
RVB2
HPC2
HIR3
HIR2
RVB1
HIR1
FUN30
SFH1
CAC2
VPS72
ISW1
RTT106
ARP6
MSI1
SWR1
SWC5
YAF9
HTB1
RSC6
SWC3
RLF2
VPS71
NHP10
GCR1
SPN1
IES3
BDF1
CHD1
ARP5
VPS75
NAP1
RSC3
RSC8
ASF1
STH1
Gene
Z_Shift_K_Gem
Z_Im_K_Gem
Z_Shift_K_Cyt
Z_Im_K_Cyt
Z_Shift_L_Gem
Z_Im_L_Gem
Z_Shift_L_Cyt
Z_Im_L_Cyt

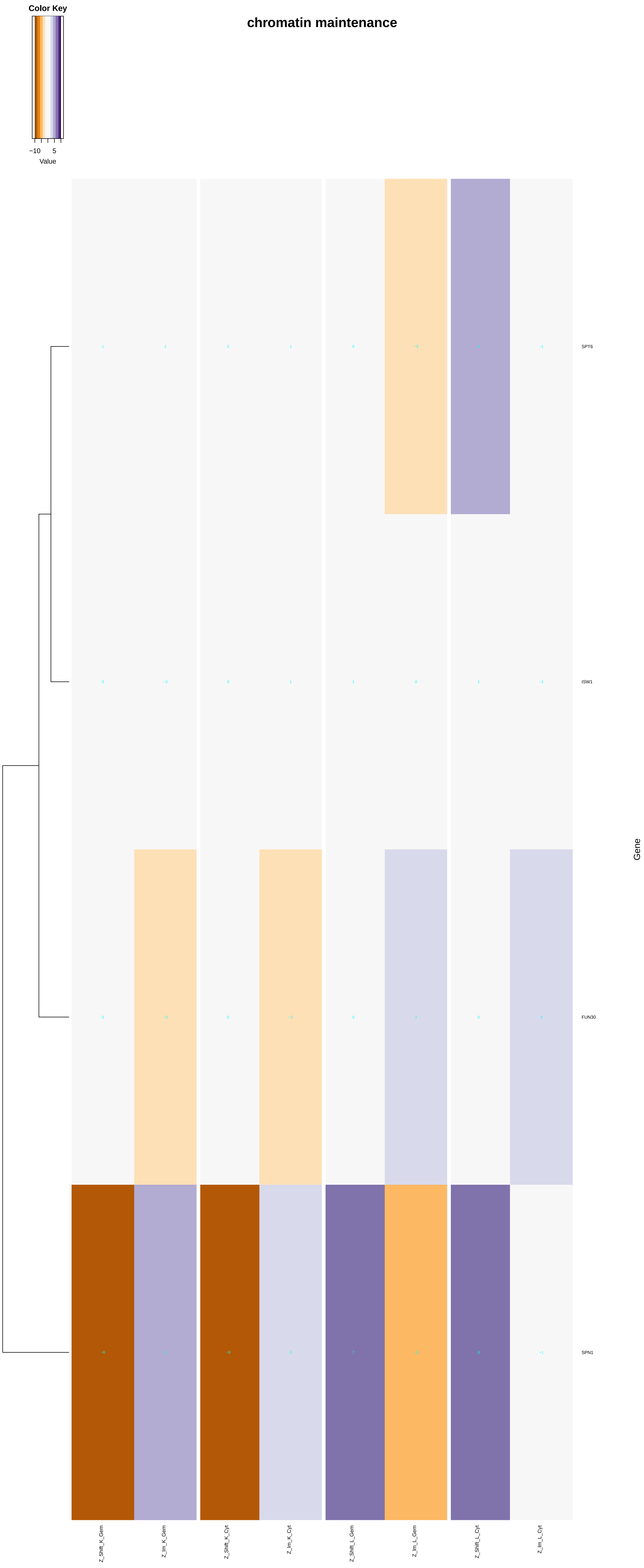
Color Key
−10
5
Value
chromatin maintenance
SPT6
ISW1
FUN30
SPN1
Gene
Z_Shift_K_Gem
Z_Im_K_Gem
Z_Shift_K_Cyt
Z_Im_K_Cyt
Z_Shift_L_Gem
Z_Im_L_Gem
Z_Shift_L_Cyt
Z_Im_L_Cyt

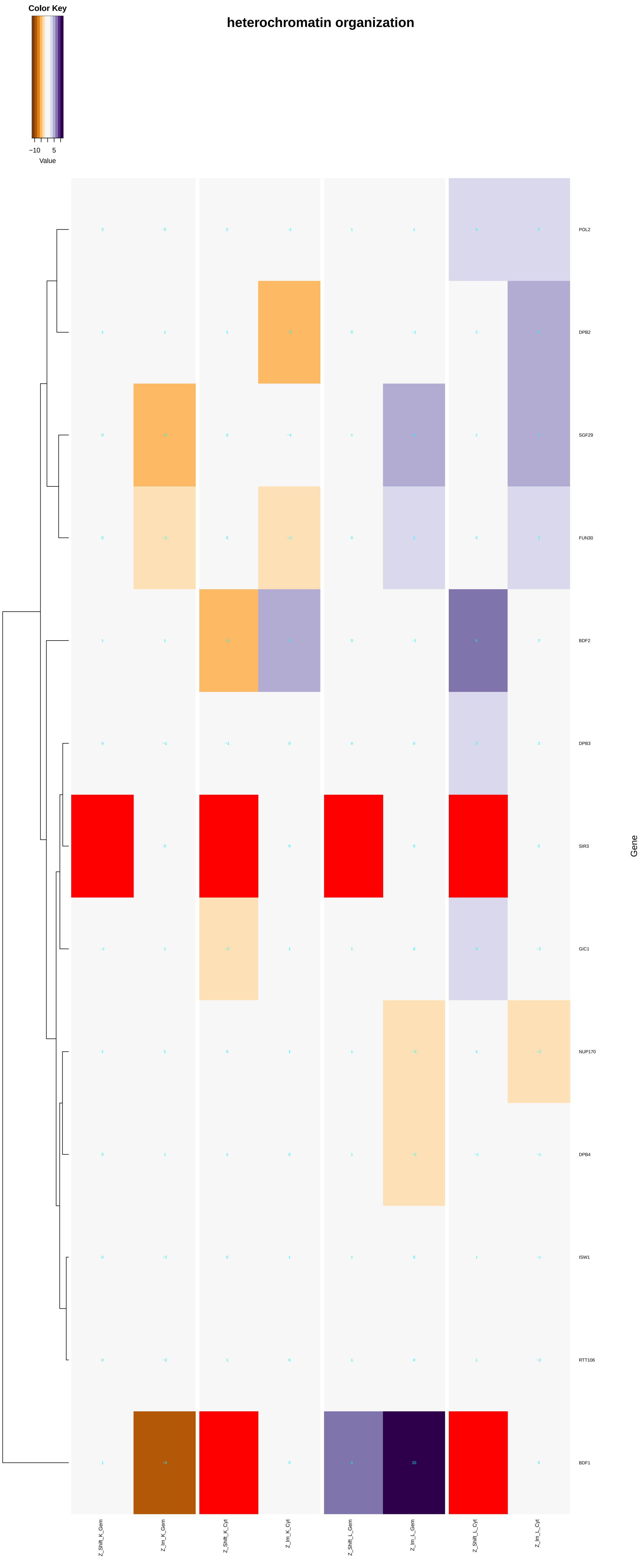
heterochromatin organization
Color Key
−10
5
Value
POL2
DPB2
SGF29
FUN30
BDF2
DPB3
SIR3
GIC1
NUP170
DPB4
ISW1
RTT106
BDF1
Gene
Z_Shift_K_Gem
Z_Im_K_Gem
Z_Shift_K_Cyt
Z_Im_K_Cyt
Z_Shift_L_Gem
Z_Im_L_Gem
Z_Shift_L_Cyt
Z_Im_L_Cyt

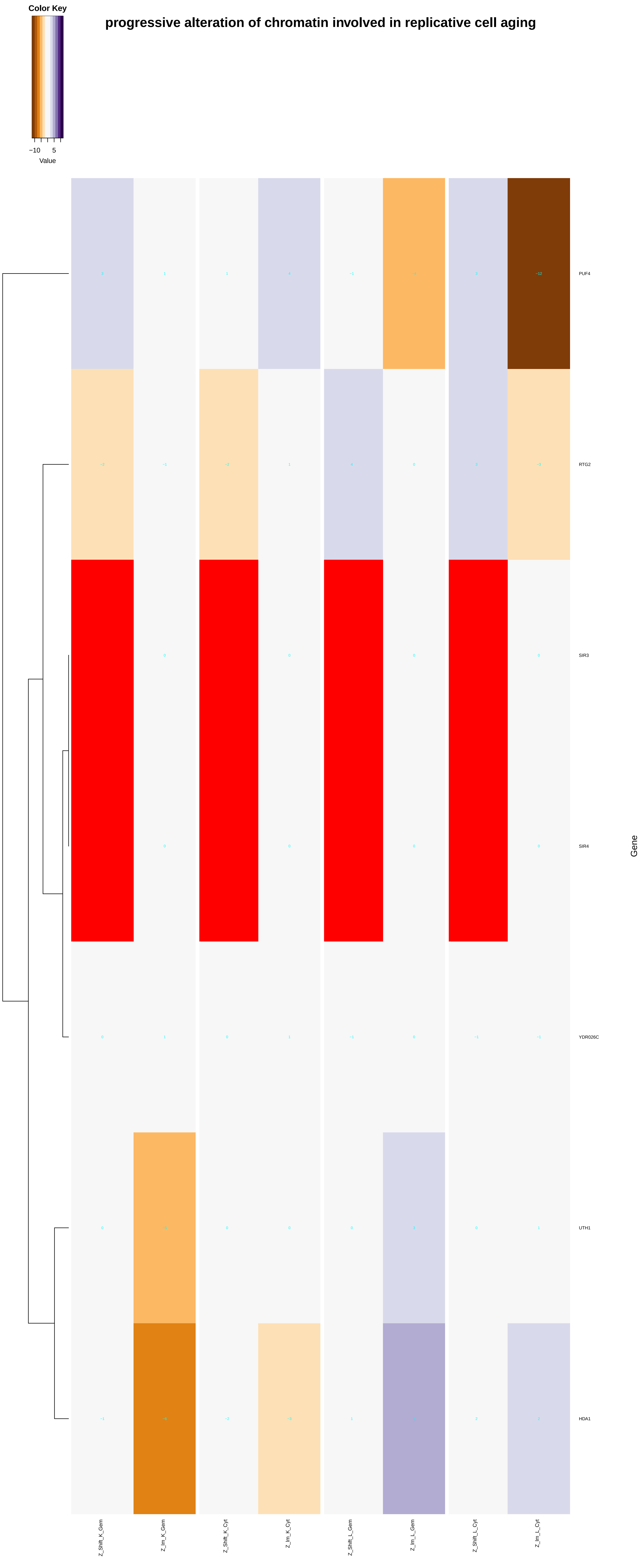

progressive alteration of chromatin involved in replicative cell aging
Color Key
−10
5
Value
PUF4
RTG2
SIR3
SIR4
YDR026C
UTH1
HDA1
Gene
Z_Shift_K_Gem
Z_Im_K_Gem
Z_Shift_K_Cyt
Z_Im_K_Cyt
Z_Shift_L_Gem
Z_Im_L_Gem
Z_Shift_L_Cyt
Z_Im_L_Cyt

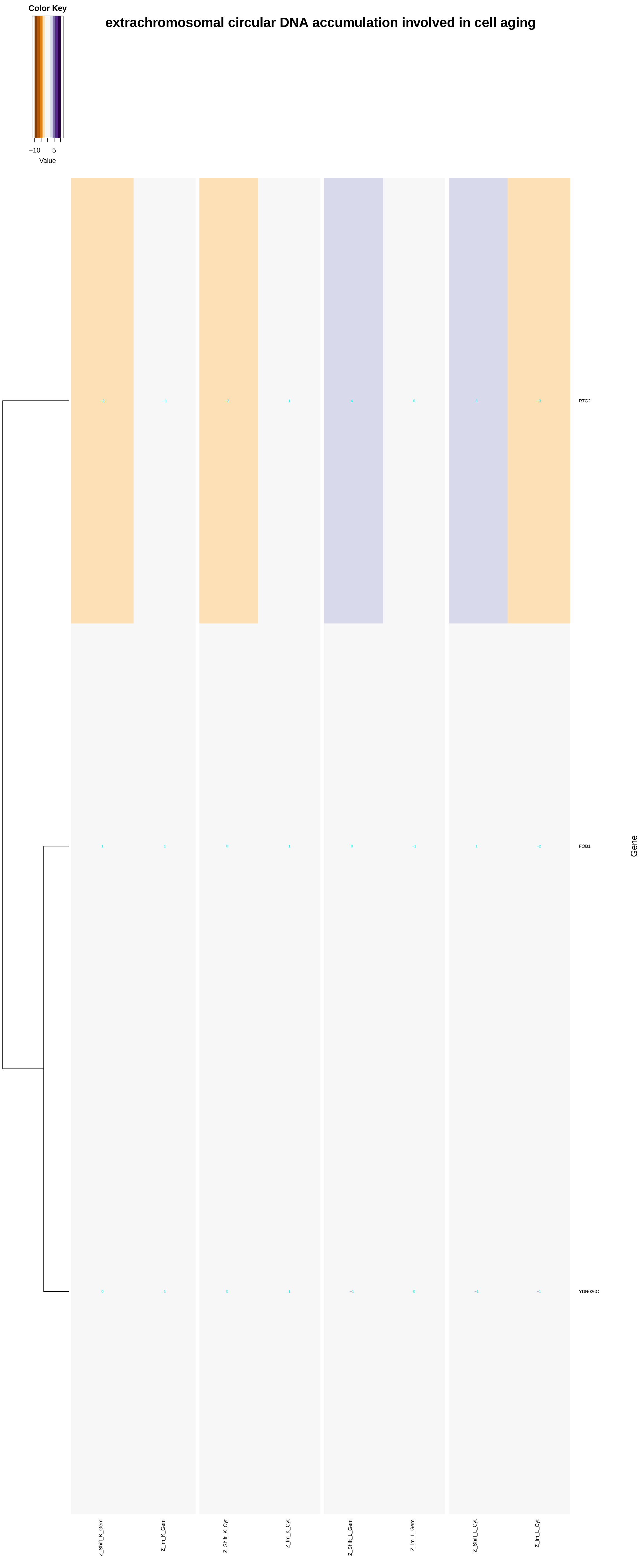

Color Key
−10
5
Value
extrachromosomal circular DNA accumulation involved in cell aging
RTG2
FOB1
YDR026C
Gene
Z_Shift_K_Gem
Z_Im_K_Gem
Z_Shift_K_Cyt
Z_Im_K_Cyt
Z_Shift_L_Gem
Z_Im_L_Gem
Z_Shift_L_Cyt
Z_Im_L_Cyt

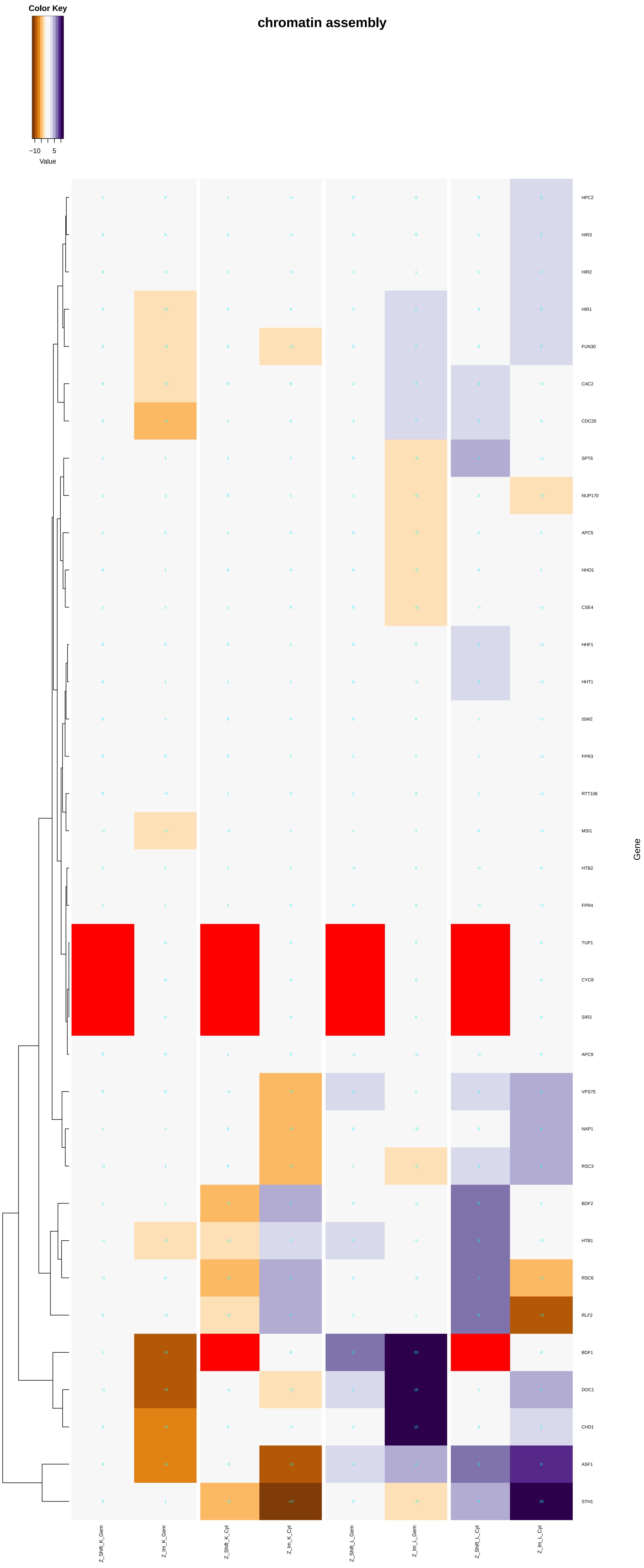

Color Key
−10
5
Value
chromatin assembly
HPC2
HIR3
HIR2
HIR1
FUN30
CAC2
CDC26
SPT6
NUP170
APC5
HHO1
CSE4
HHF1
HHT1
ISW2
FPR3
RTT106
MSI1
HTB2
FPR4
TUP1
CYC8
SIR3
APC9
VPS75
NAP1
RSC3
BDF2
HTB1
RSC6
RLF2
BDF1
DOC1
CHD1
ASF1
STH1
Gene
Z_Shift_K_Gem
Z_Im_K_Gem
Z_Shift_K_Cyt
Z_Im_K_Cyt
Z_Shift_L_Gem
Z_Im_L_Gem
Z_Shift_L_Cyt
Z_Im_L_Cyt

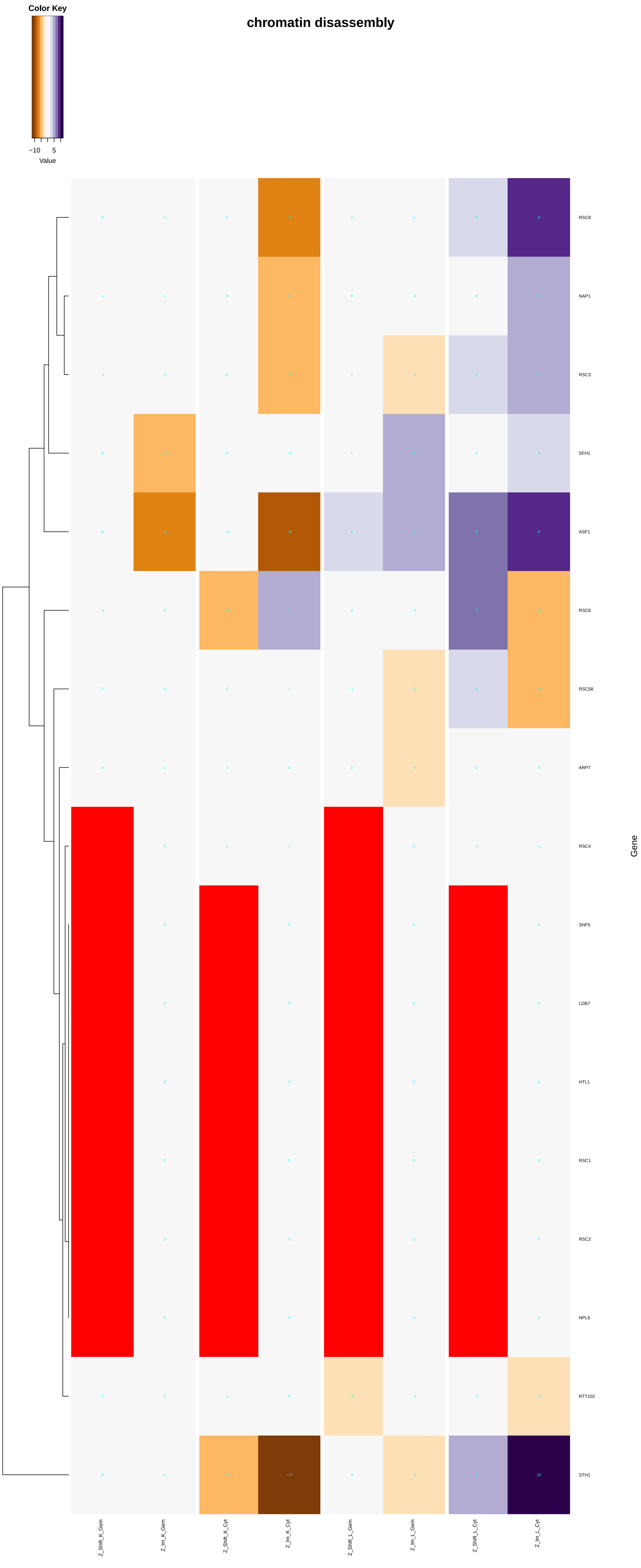

Color Key
−10
5
Value
chromatin disassembly
RSC8
NAP1
RSC3
SFH1
ASF1
RSC6
RSC58
ARP7
RSC4
SNF5
LDB7
HTL1
RSC1
RSC2
NPL6
RTT102
STH1
Gene
Z_Shift_K_Gem
Z_Im_K_Gem
Z_Shift_K_Cyt
Z_Im_K_Cyt
Z_Shift_L_Gem
Z_Im_L_Gem
Z_Shift_L_Cyt
Z_Im_L_Cyt

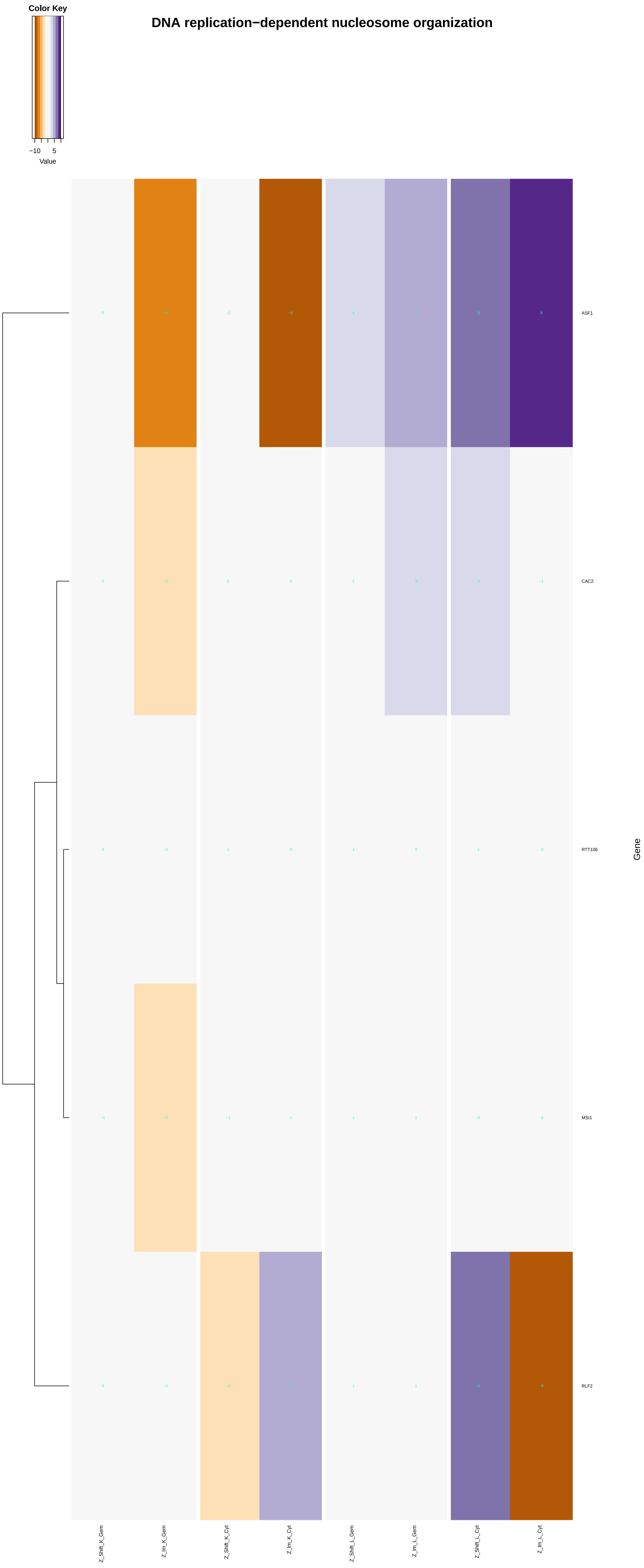
DNA replication−dependent nucleosome organization
Color Key
−10
5
Value
ASF1
CAC2
RTT106
MSI1
RLF2
Gene
Z_Shift_K_Gem
Z_Im_K_Gem
Z_Shift_K_Cyt
Z_Im_K_Cyt
Z_Shift_L_Gem
Z_Im_L_Gem
Z_Shift_L_Cyt
Z_Im_L_Cyt

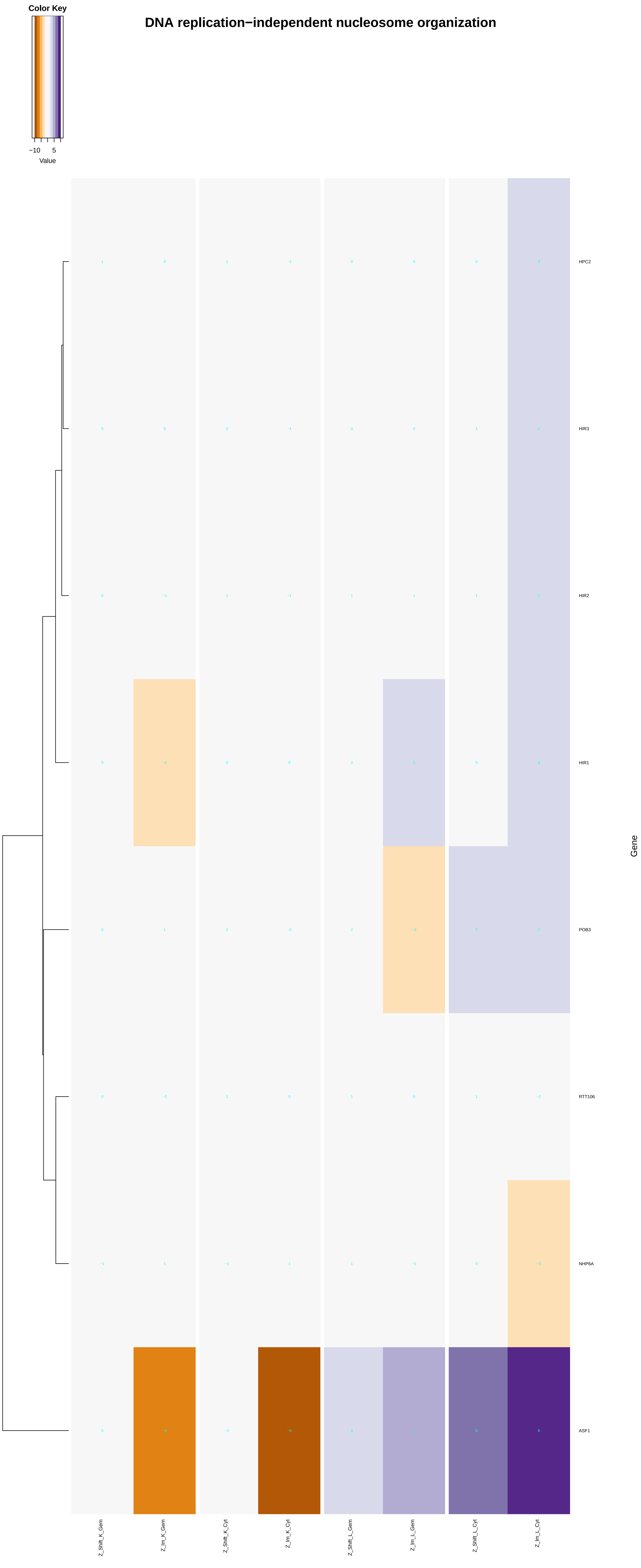

Color Key
−10
5
Value
DNA replication−independent nucleosome organization
HPC2
HIR3
HIR2
HIR1
POB3
RTT106
NHP6A
ASF1
Gene
Z_Shift_K_Gem
Z_lm_K_Gem
Z_Shift_K_Cyt
Z_lm_K_Cyt
Z_Shift_L_Gem
Z_lm_L_Gem
Z_Shift_L_Cyt
Z_lm_L_Cyt

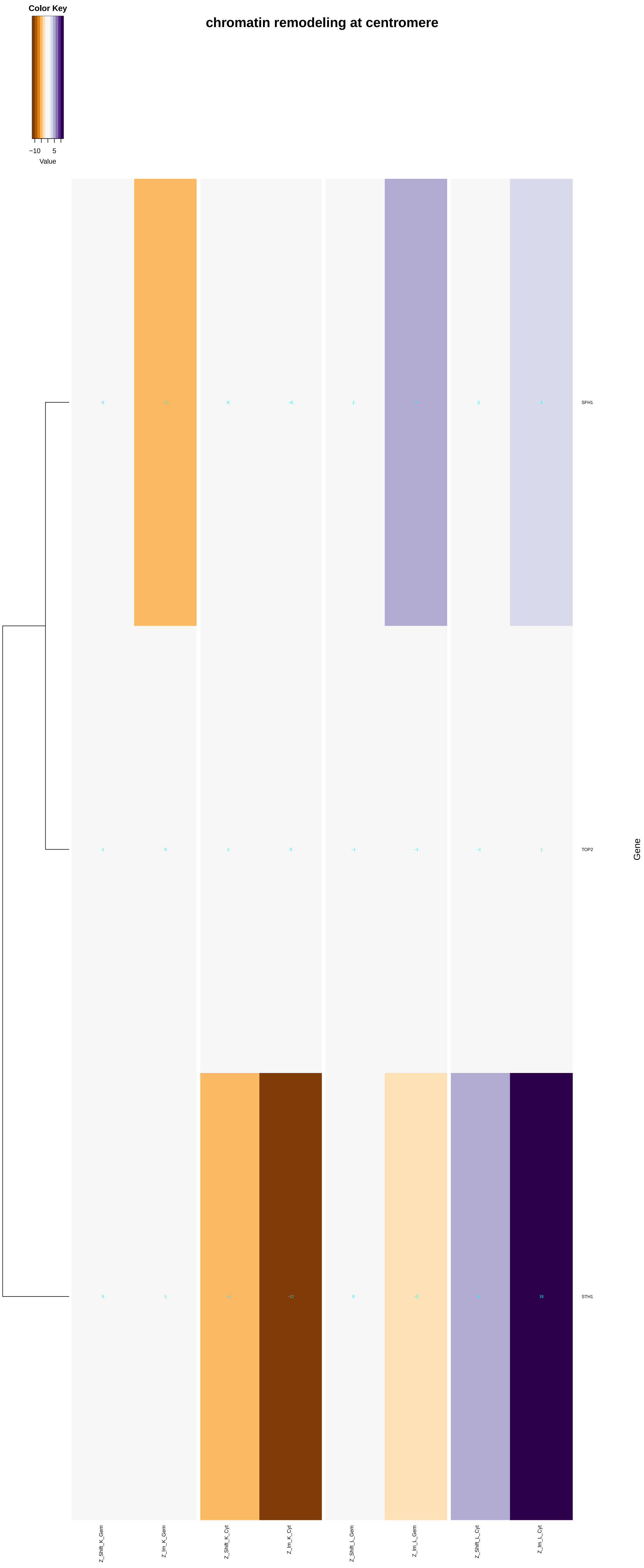
chromatin remodeling at centromere
Color Key
−10
5
Value
SFH1
TOP2
STH1
Gene
Z_Shift_K_Gem
Z_Im_K_Gem
Z_Shift_K_Cyt
Z_Im_K_Cyt
Z_Shift_L_Gem
Z_Im_L_Gem
Z_Shift_L_Cyt
Z_Im_L_Cyt

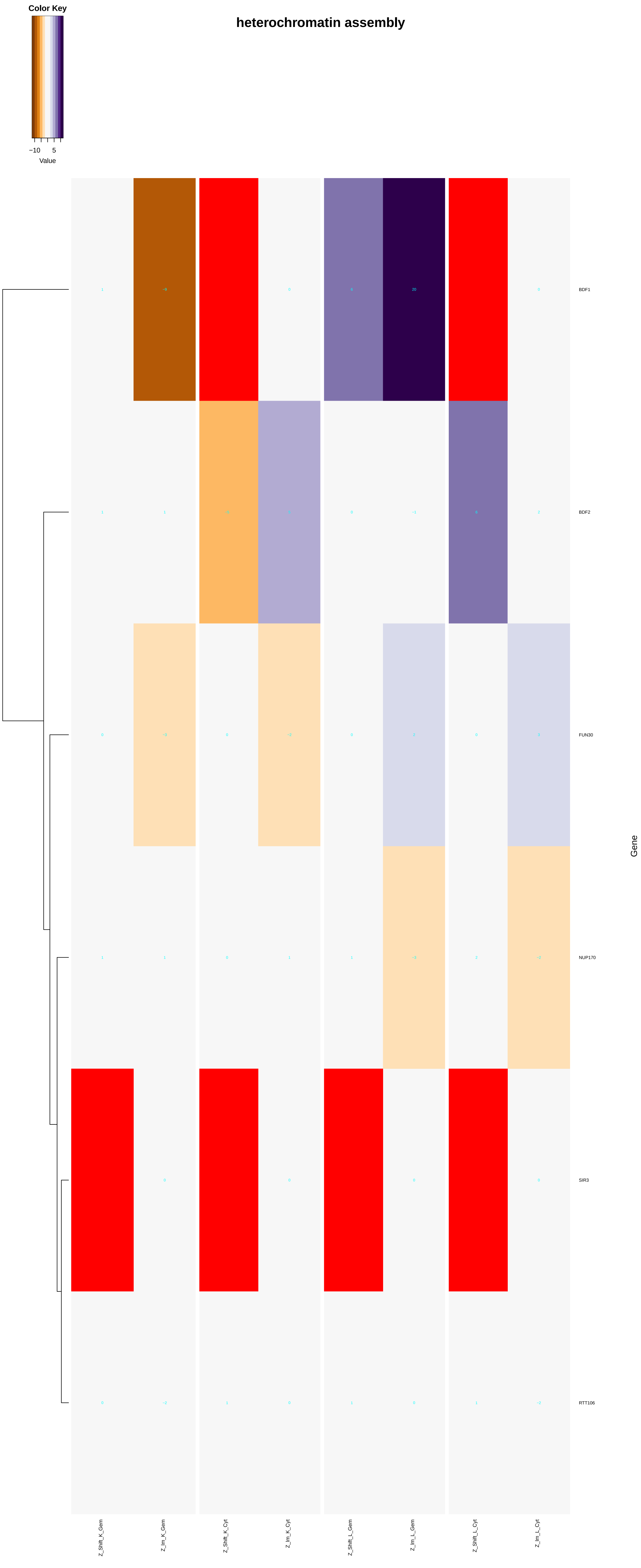
Color Key
−10 5
Value
heterochromatin assembly
BDF1
BDF2
FUN30
NUP170
SIR3
RTT106
Gene
Z_Shift_K_Gem
Z_Im_K_Gem
Z_Shift_K_Cyt
Z_Im_K_Cyt
Z_Shift_L_Gem
Z_Im_L_Gem
Z_Shift_L_Cyt
Z_Im_L_Cyt

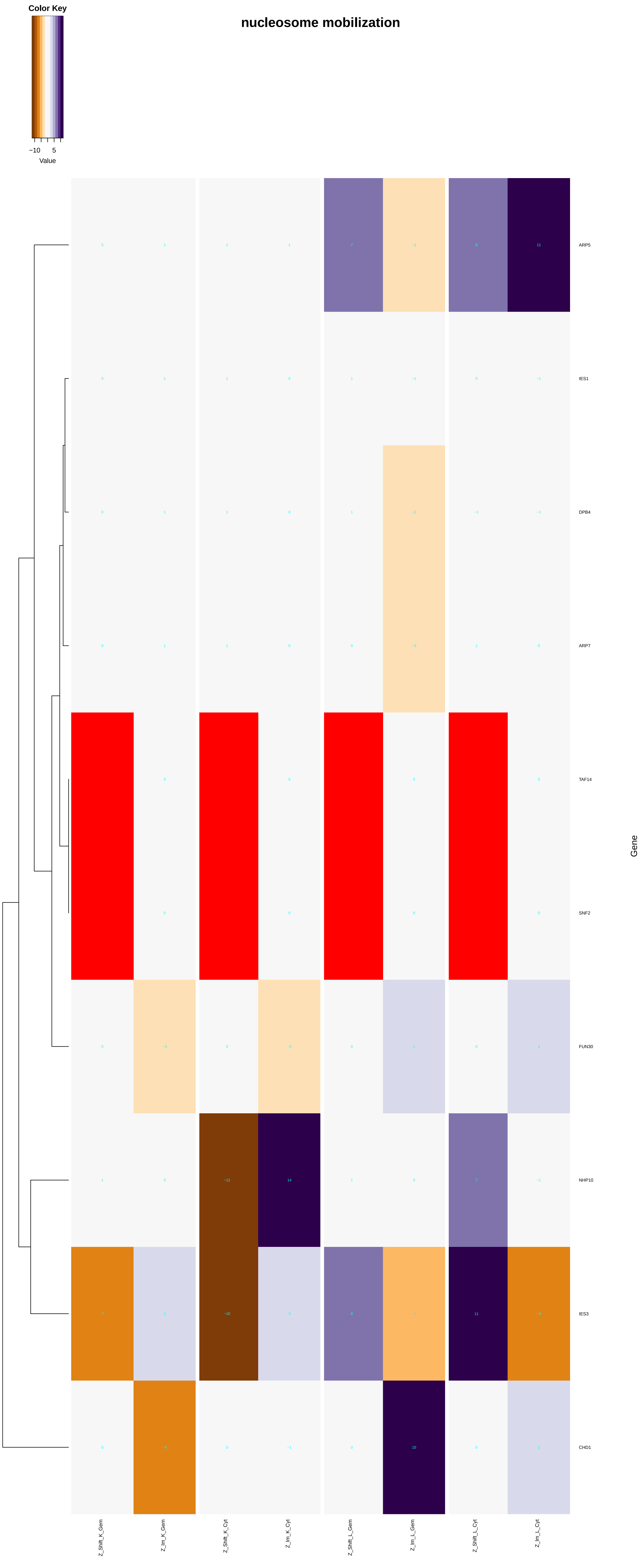

Color Key
−10
5
Value
nucleosome mobilization
ARP5
IES1
DPB4
ARP7
TAF14
SNF2
FUN30
NHP10
IES3
CHD1
Gene
Z_Shift_K_Gem
Z_Im_K_Gem
Z_Shift_K_Cyt
Z_Im_K_Cyt
Z_Shift_L_Gem
Z_Im_L_Gem
Z_Shift_L_Cyt
Z_Im_L_Cyt

#### ATP−dependent chromatin remodeling

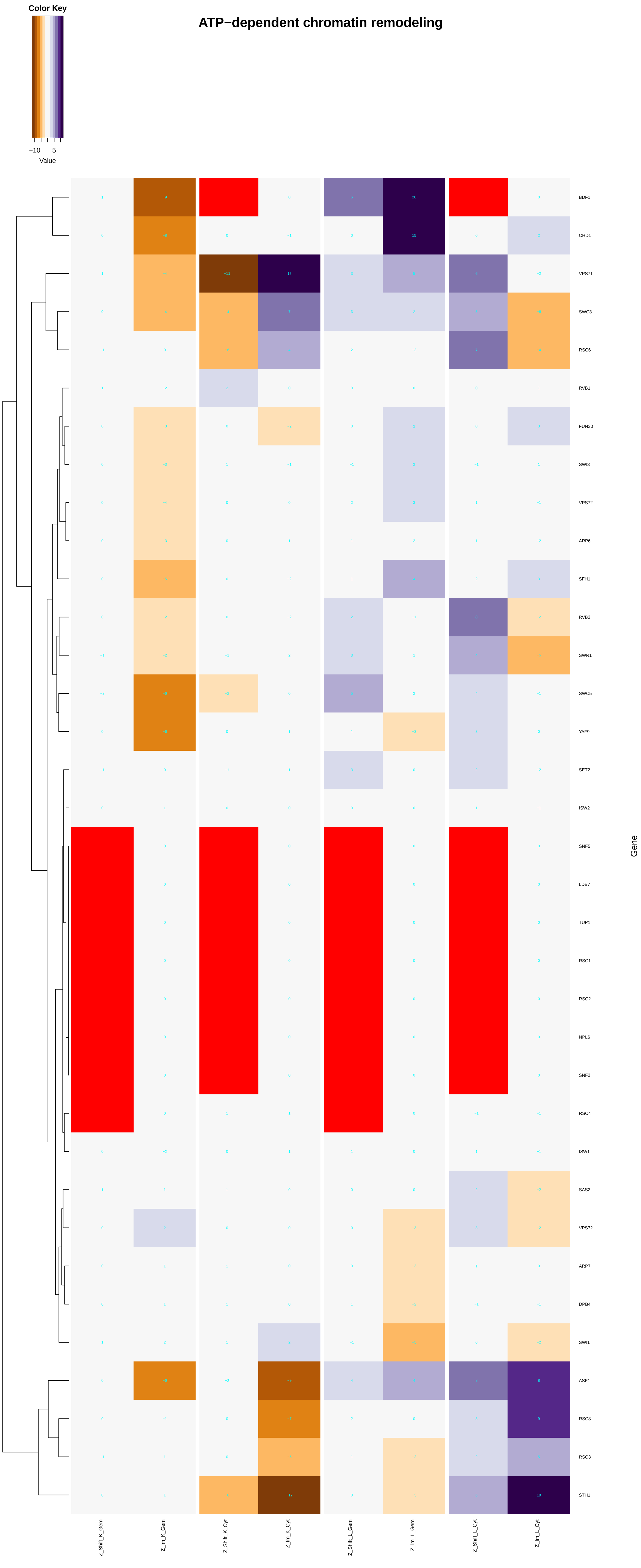

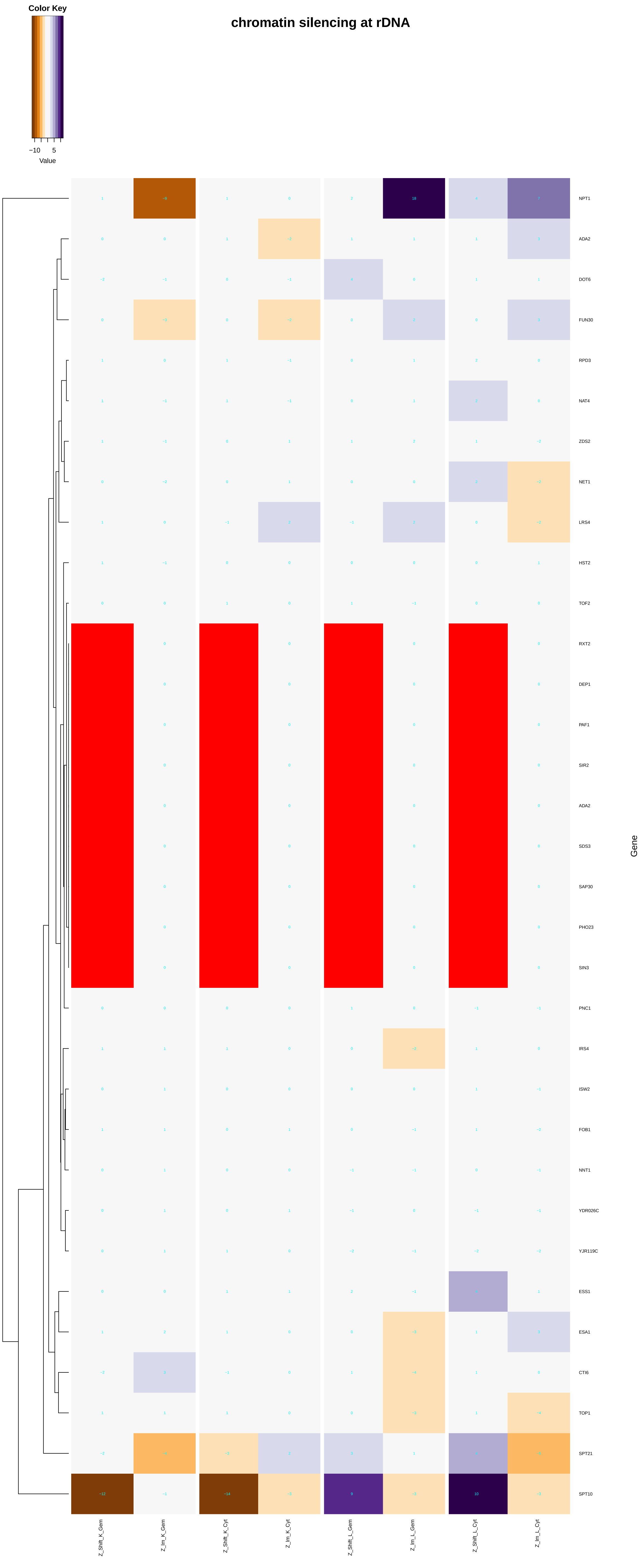

chromatin silencing at rDNA
Color Key
−10
5
Value
NPT1
ADA2
DOT6
FUN30
RPD3
NAT4
ZDS2
NET1
LRS4
HST2
TOF2
RXT2
DEP1
PAF1
SIR2
ADA2
SDS3
SAP30
PHO23
SIN3
PNC1
IRS4
ISW2
FOB1
NNT1
YDR026C
YJR119C
ESS1
ESA1
CTI6
TOP1
SPT21
SPT10
Gene
Z_Shift_K_Gem
Z_lm_K_Gem
Z_Shift_K_Cyt
Z_lm_K_Cyt
Z_Shift_L_Gem
Z_lm_L_Gem
Z_Shift_L_Cyt
Z_lm_L_Cyt

### chromatin silencing at telomere

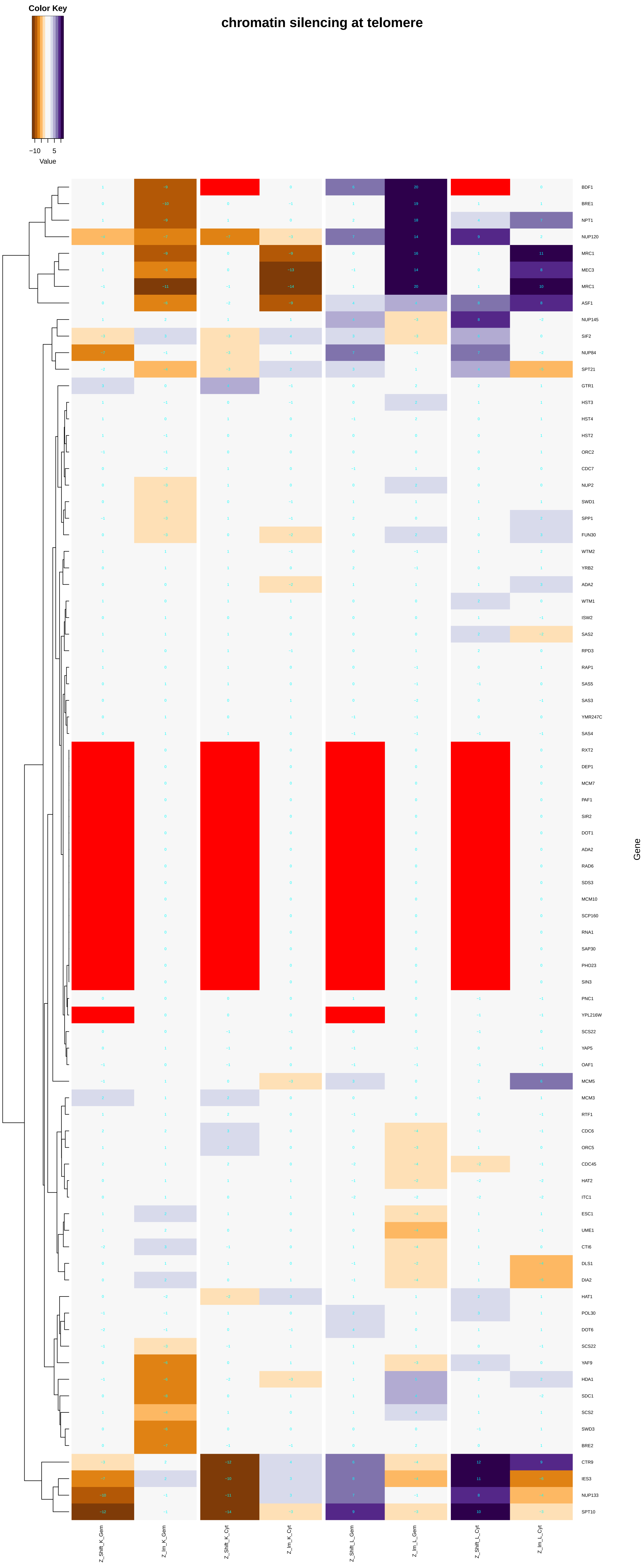

#### chromatin silencing at silent mating−type cassette

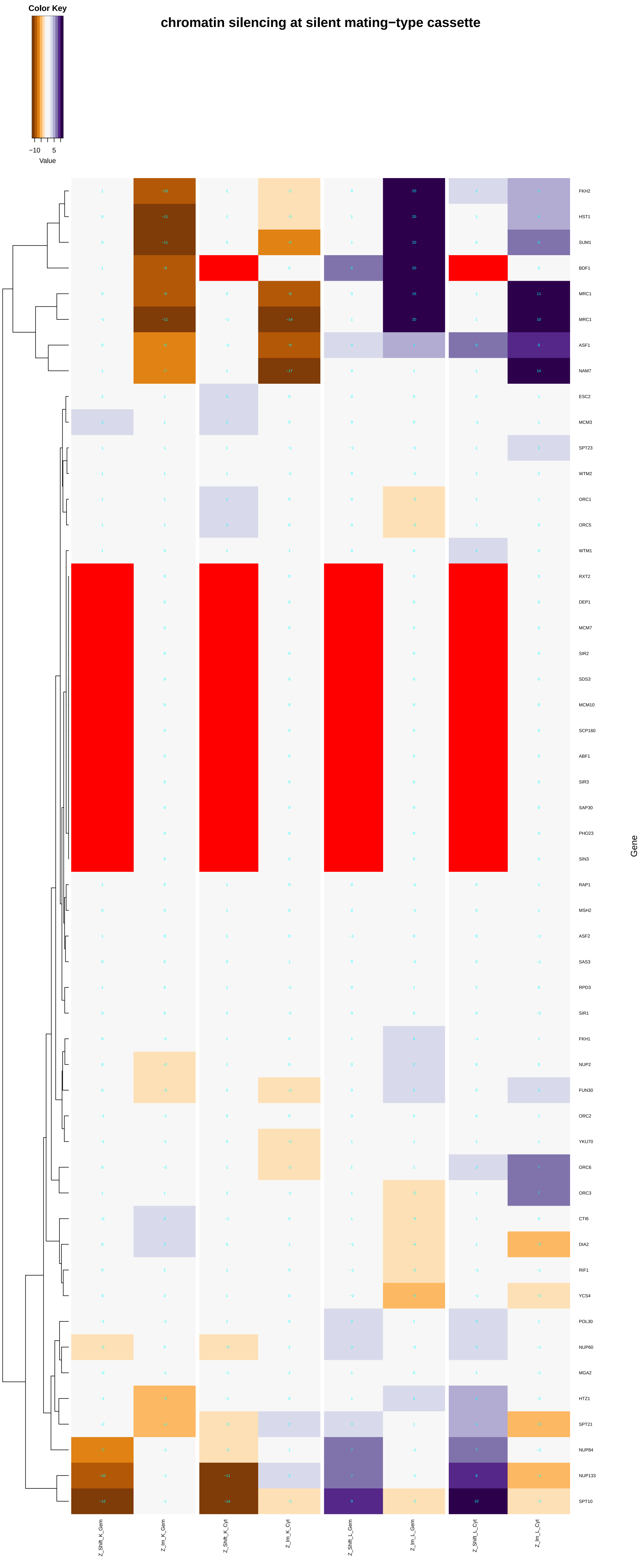

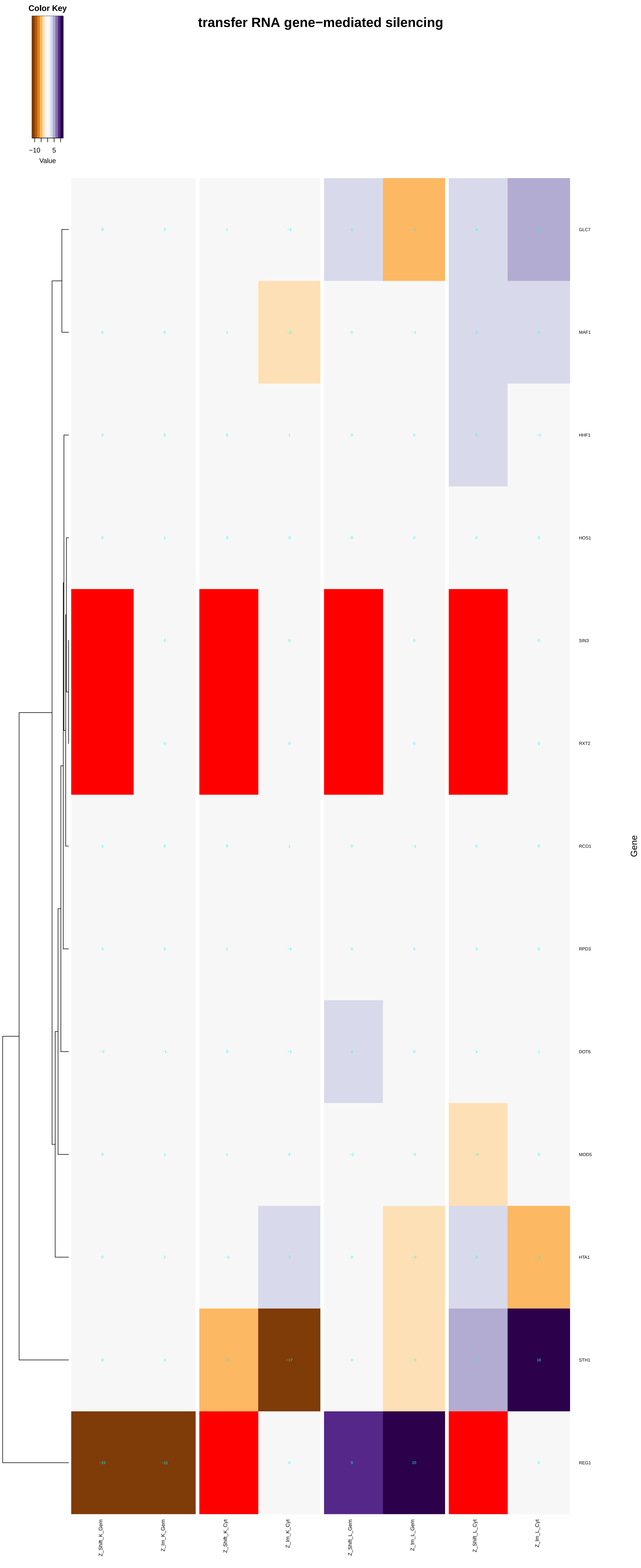

Color Key
−10
5
Value
transfer RNA gene−mediated silencing
GLC7
MAF1
HHF1
HOS1
SIN3
RXT2
RCO1
RPD3
DOT6
MOD5
HTA1
STH1
REG1
Gene
Z_Shift_K_Gem
Z_Im_K_Gem
Z_Shift_K_Cyt
Z_Im_K_Cyt
Z_Shift_L_Gem
Z_Im_L_Gem
Z_Shift_L_Cyt
Z_Im_L_Cyt

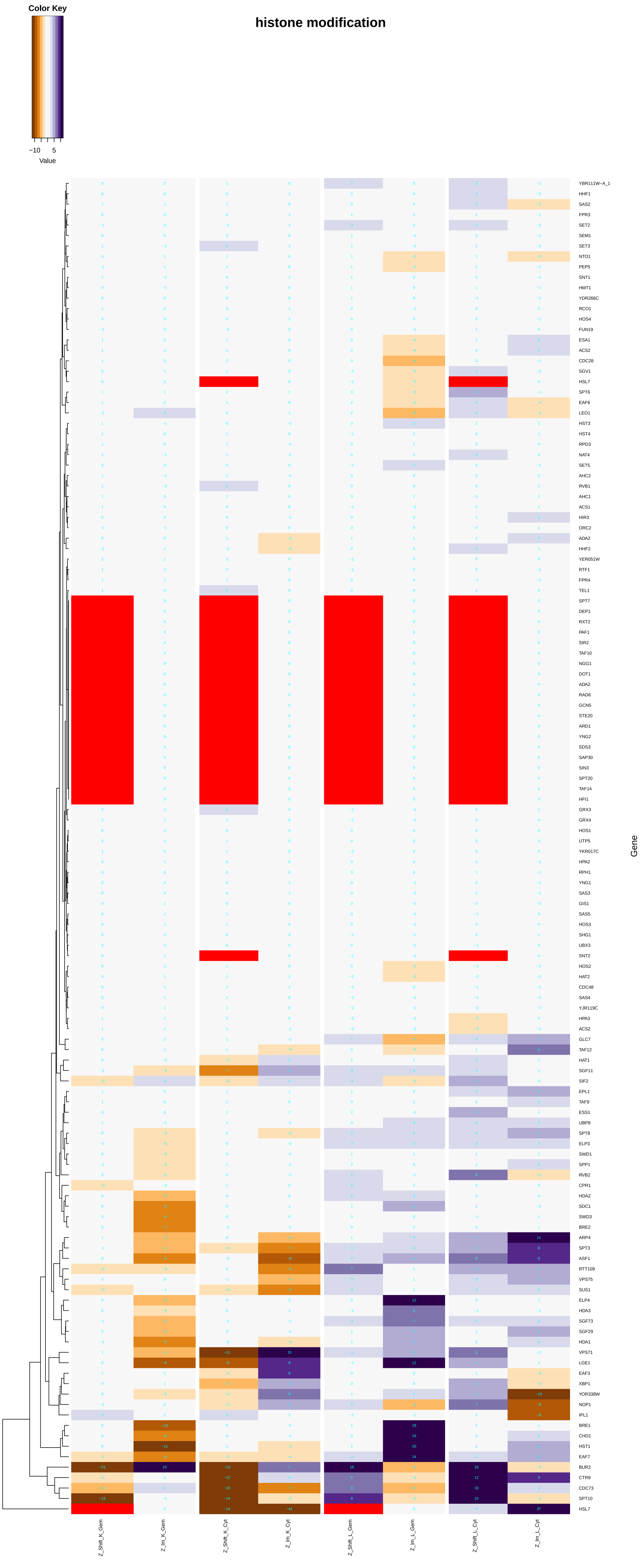
histone modification
Color Key
−10 5
Value
YBR111W−A_1
HHF1
SAS2
FPR3
SET2
SEM1
SET3
NTO1
PEP5
SNT1
HMT1
YDR266C
RCO1
HOS4
FUN19
ESA1
ACS2
CDC28
SGV1
HSL7
SPT6
EAF6
LEO1
HST3
HST4
RPD3
NAT4
SET5
AHC2
RVB1
AHC1
ACS1
HIR3
ORC2
ADA2
HHF2
YER051W
RTF1
FPR4
TEL1
SPT7
DEP1
RXT2
PAF1
SIR2
TAF10
NGG1
DOT1
ADA2
RAD6
GCN5
STE20
ARD1
YNG2
SDS3
SAP30
SIN3
SPT20
TAF14
HFI1
GRX3
GRX4
HOS1
UTP5
YKR017C
HPA2
RPH1
YNG1
SAS3
GIS1
SAS5
HOS3
SHG1
UBX3
SNT2
HOS2
HAT2
CDC48
SAS4
YJR119C
HPA3
ACS2
GLC7
TAF12
HAT1
SGF11
SIF2
EPL1
TAF9
ESS1
UBP8
SPT8
ELP3
SWD1
SPP1
RVB2
CPR1
HDA2
SDC1
SWD3
BRE1
ARP4
SPT3
ASF1
RTT109
VPS75
SUS1
ELP4
HDA3
SGF73
SGF29
HDA1
VPS71
LGE1
EAF3
XBP1
YOR338W
NOP1
IPL1
BRE1
CHD1
HST1
EAF7
BUR2
CTR9
CDC73
SPT10
HSL7
Gene
Z_Shift_K_Gem
Z_Im_K_Gem
Z_Shift_K_Cyt
Z_Im_K_Cyt
Z_Shift_L_Gem
Z_Im_L_Gem
Z_Shift_L_Cyt
Z_Im_L_Cyt

### chromatin organization involved in negative regulation of transcription

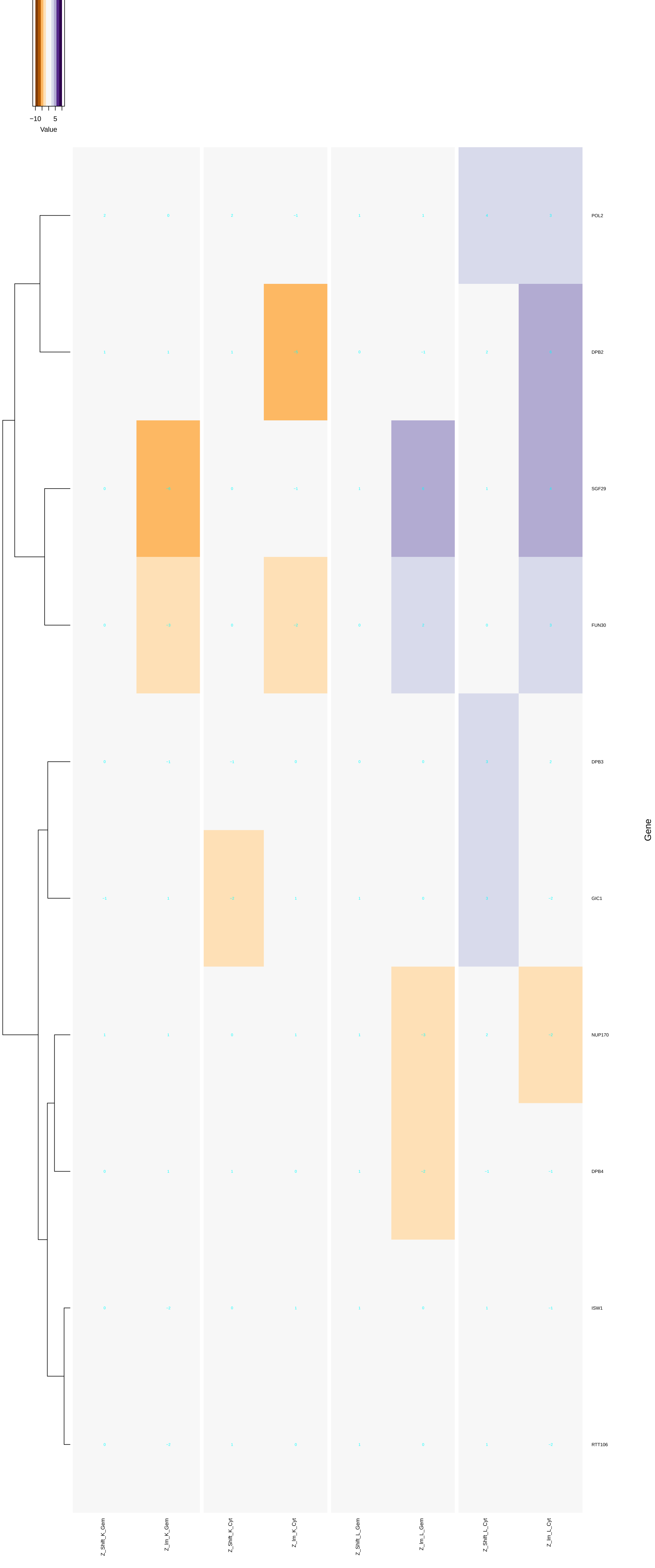

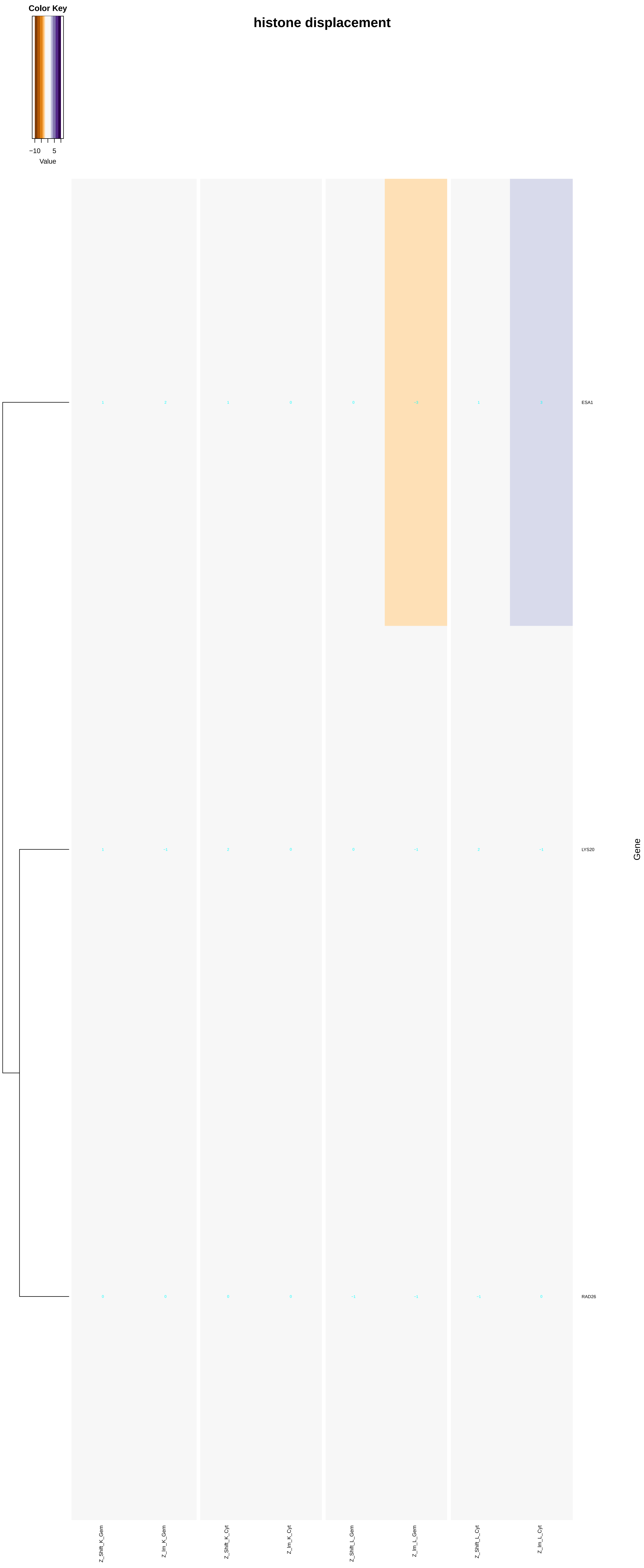
Color Key
−10
5
Value
histone displacement
1
2
1
0
0
−3
1
3
ESA1
1
−1
2
0
0
−1
2
−1
LYS20
0
0
0
0
−1
−1
−1
0
RAD26
Gene
Z_Shift_K_Gem
Z_Im_K_Gem
Z_Shift_K_Cyt
Z_Im_K_Cyt
Z_Shift_L_Gem
Z_Im_L_Gem
Z_Shift_L_Cyt
Z_Im_L_Cyt

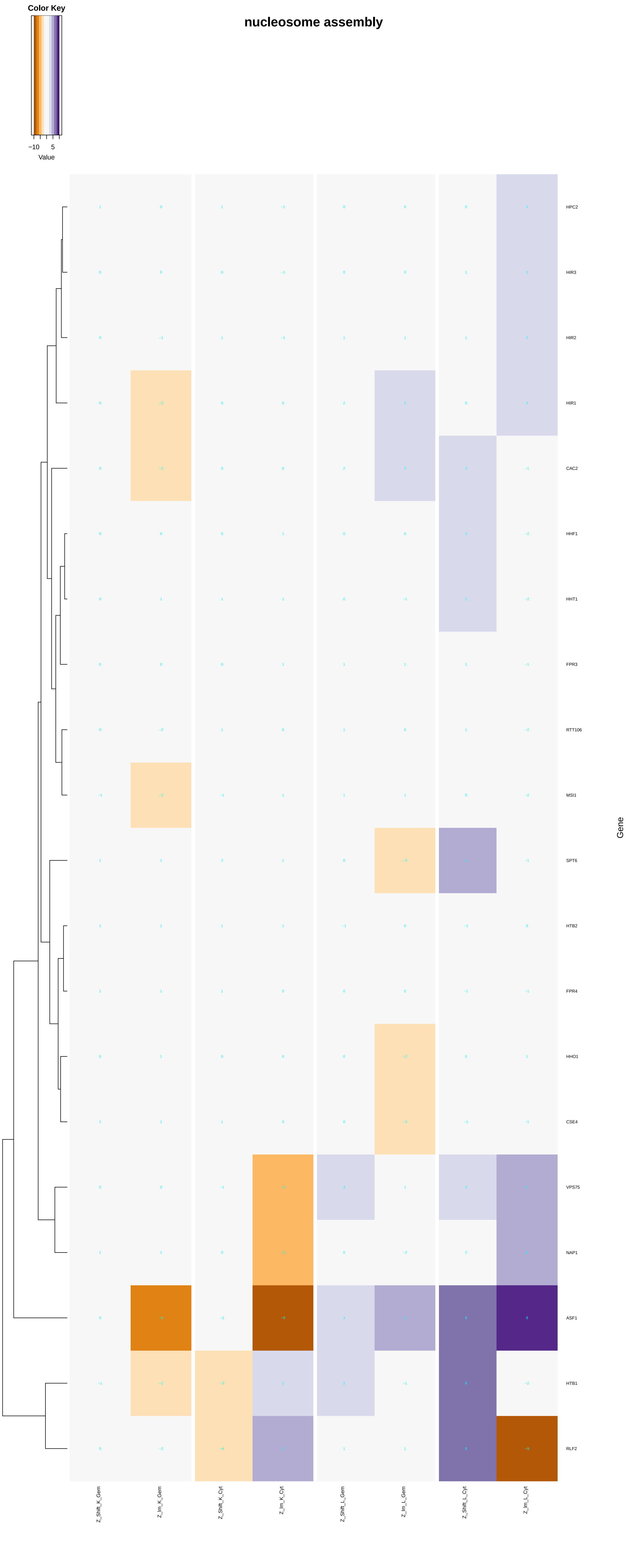
nucleosome assembly
Color Key
−10
5
Value
HPC2
HIR3
HIR2
HIR1
CAC2
HHF1
HHT1
FPR3
RTT106
MSI1
SPT6
HTB2
FPR4
HHO1
CSE4
VPS75
NAP1
ASF1
HTB1
RLF2
Gene
Z_Shift_K_Gem
Z_lm_K_Gem
Z_Shift_K_Cyt
Z_lm_K_Cyt
Z_Shift_L_Gem
Z_lm_L_Gem
Z_Shift_L_Cyt
Z_lm_L_Cyt

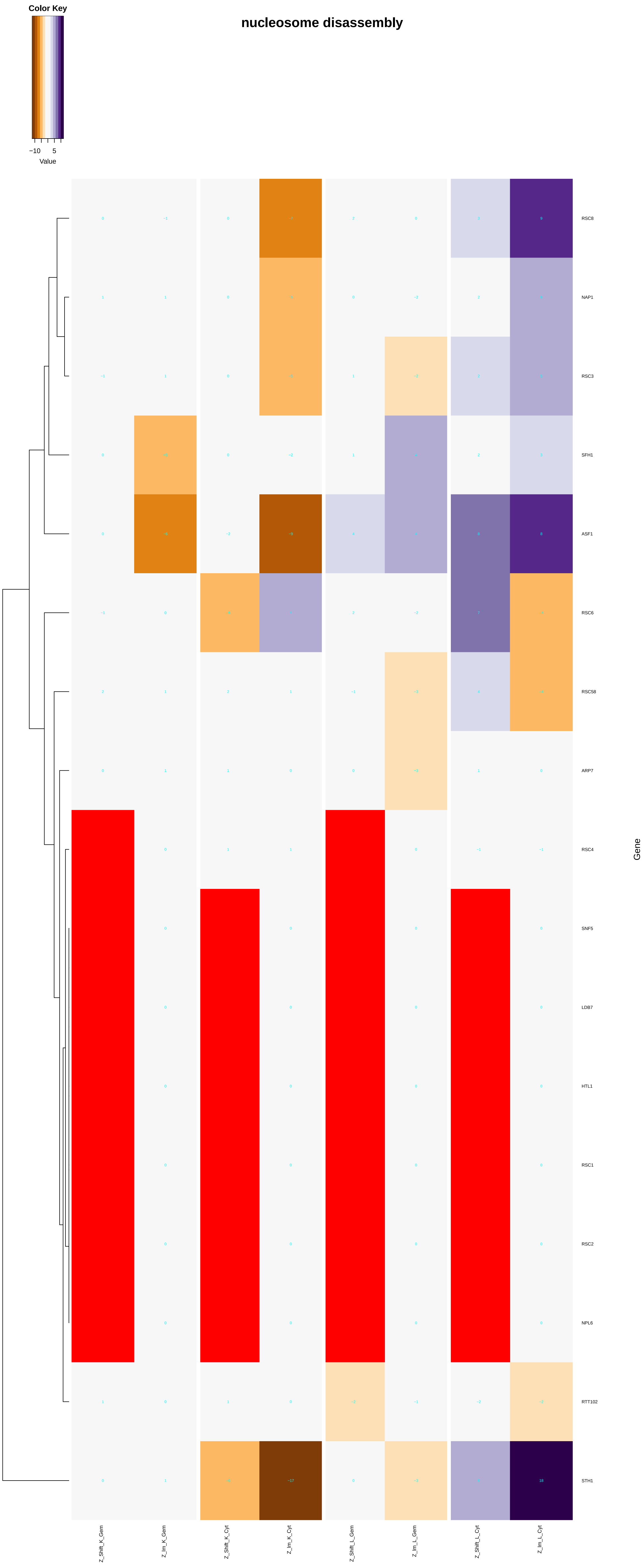

nucleosome disassembly
Color Key
−10
5
Value
RSC8
NAP1
RSC3
SFH1
ASF1
RSC6
RSC58
ARP7
RSC4
SNF5
LDB7
HTL1
RSC1
RSC2
NPL6
RTT102
STH1
Gene
Z_Shift_K_Gem
Z_Im_K_Gem
Z_Shift_K_Cyt
Z_Im_K_Cyt
Z_Shift_L_Gem
Z_Im_L_Gem
Z_Shift_L_Cyt
Z_Im_L_Cyt

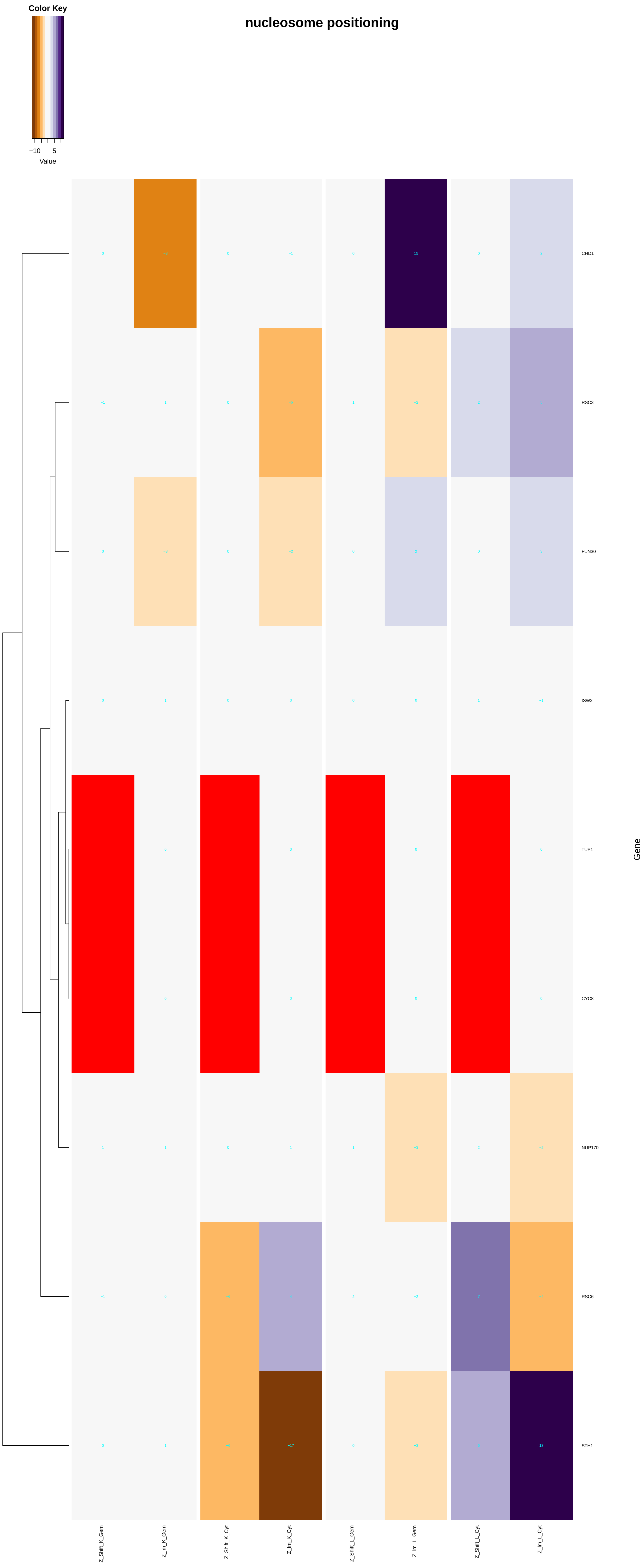

Color Key
−10
5
Value
nucleosome positioning
CHD1
RSC3
FUN30
ISW2
TUP1
CYC8
NUP170
RSC6
STH1
Gene
Z_Shift_K_Gem
Z_Im_K_Gem
Z_Shift_K_Cyt
Z_Im_K_Cyt
Z_Shift_L_Gem
Z_Im_L_Gem
Z_Shift_L_Cyt
Z_Im_L_Cyt

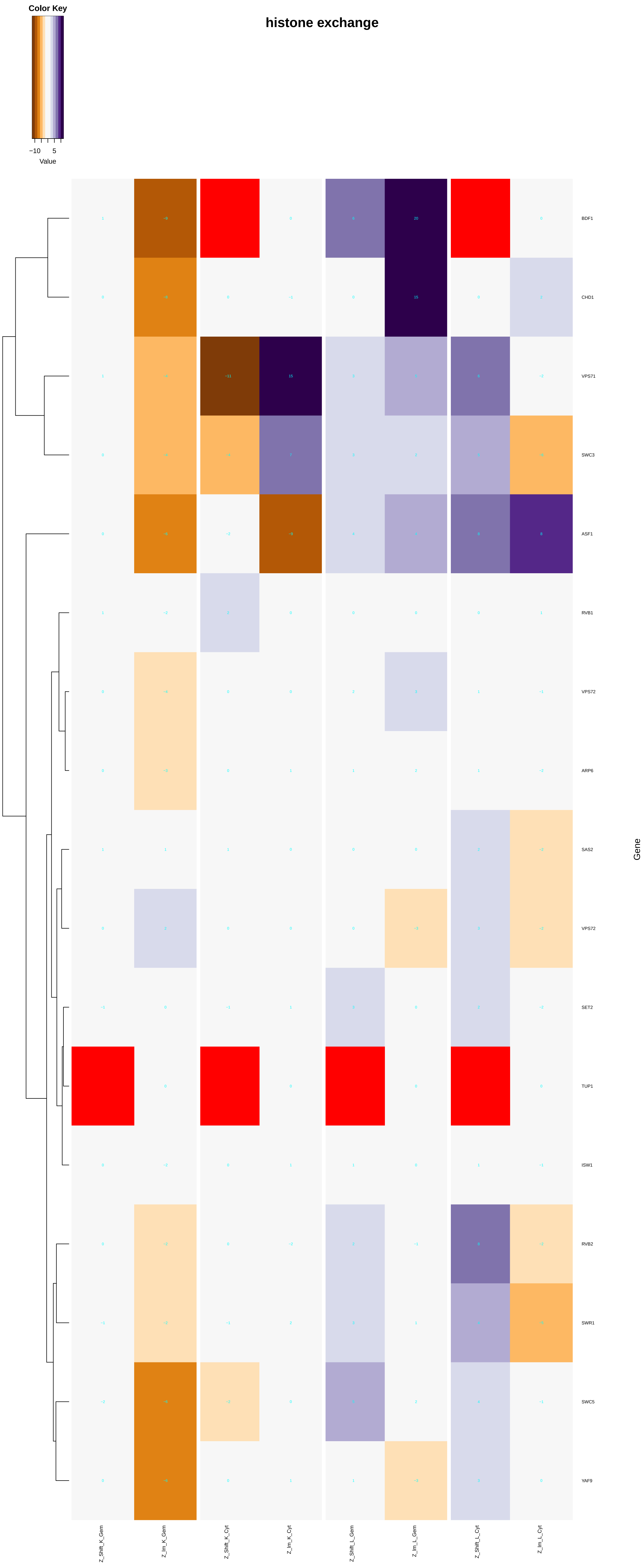
histone exchange
Color Key
−10
5
Value
BDF1
CHD1
VPS71
SWC3
ASF1
RVB1
VPS72
ARP6
SAS2
VPS72
SET2
TUP1
ISW1
RVB2
SWR1
SWC5
YAF9
Gene
Z_Shift_K_Gem
Z_Im_K_Gem
Z_Shift_K_Cyt
Z_Im_K_Cyt
Z_Shift_L_Gem
Z_Im_L_Gem
Z_Shift_L_Cyt
Z_Im_L_Cyt

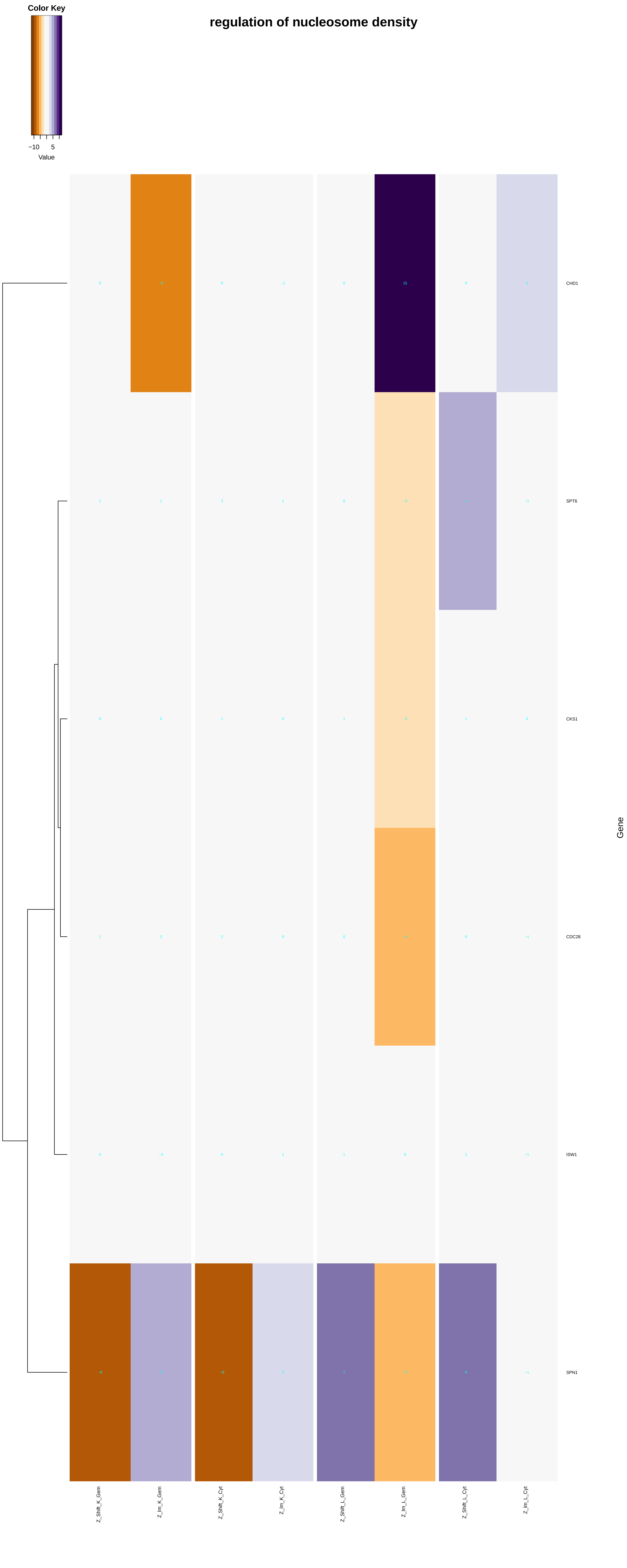
regulation of nucleosome density
Color Key
−10
5
Value
CHD1
SPT6
CKS1
CDC28
ISW1
SPN1
Gene
Z_Shift_K_Gem
Z_Im_K_Gem
Z_Shift_K_Cyt
Z_Im_K_Cyt
Z_Shift_L_Gem
Z_Im_L_Gem
Z_Shift_L_Cyt
Z_Im_L_Cyt

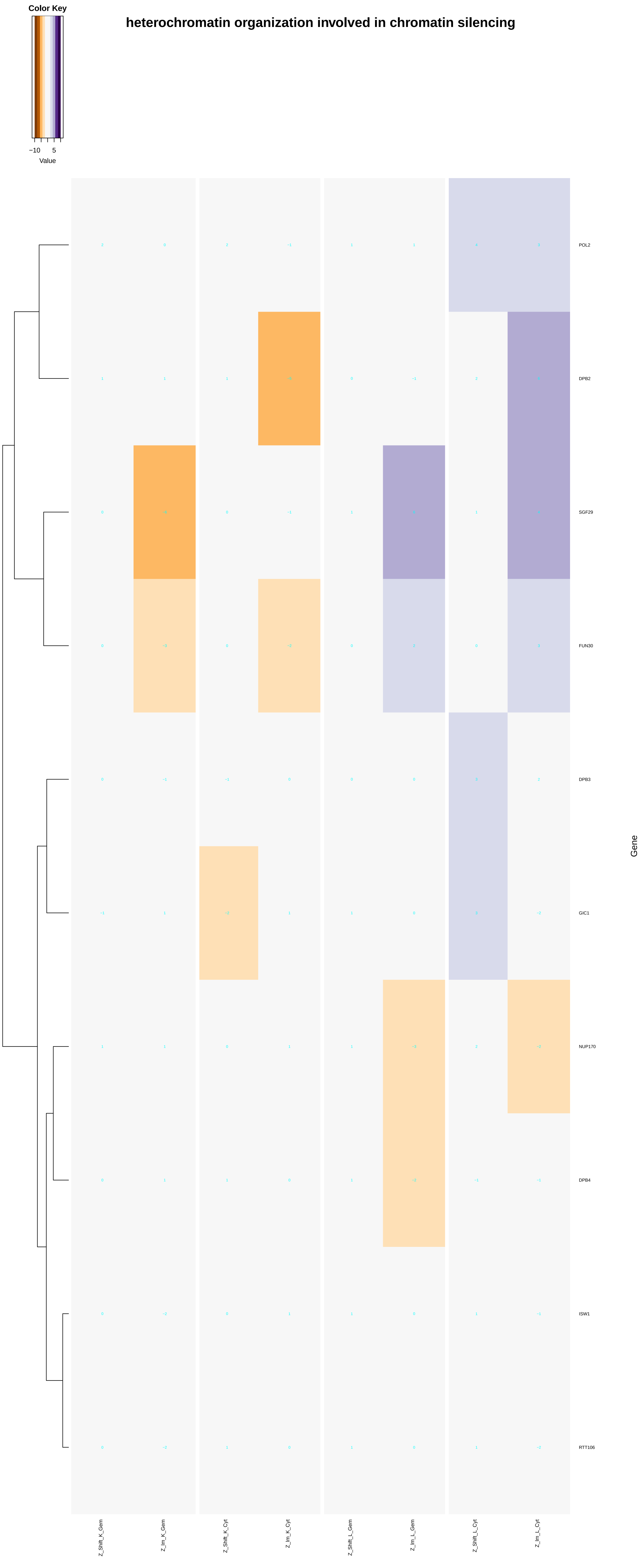
Color Key
−10
5
Value
heterochromatin organization involved in chromatin silencing
POL2
DPB2
SGF29
FUN30
DPB3
GIC1
NUP170
DPB4
ISW1
RTT106
Gene
Z_Shift_K_Gem
Z_lm_K_Gem
Z_Shift_K_Cyt
Z_lm_K_Cyt
Z_Shift_L_Gem
Z_lm_L_Gem
Z_Shift_L_Cyt
Z_lm_L_Cyt

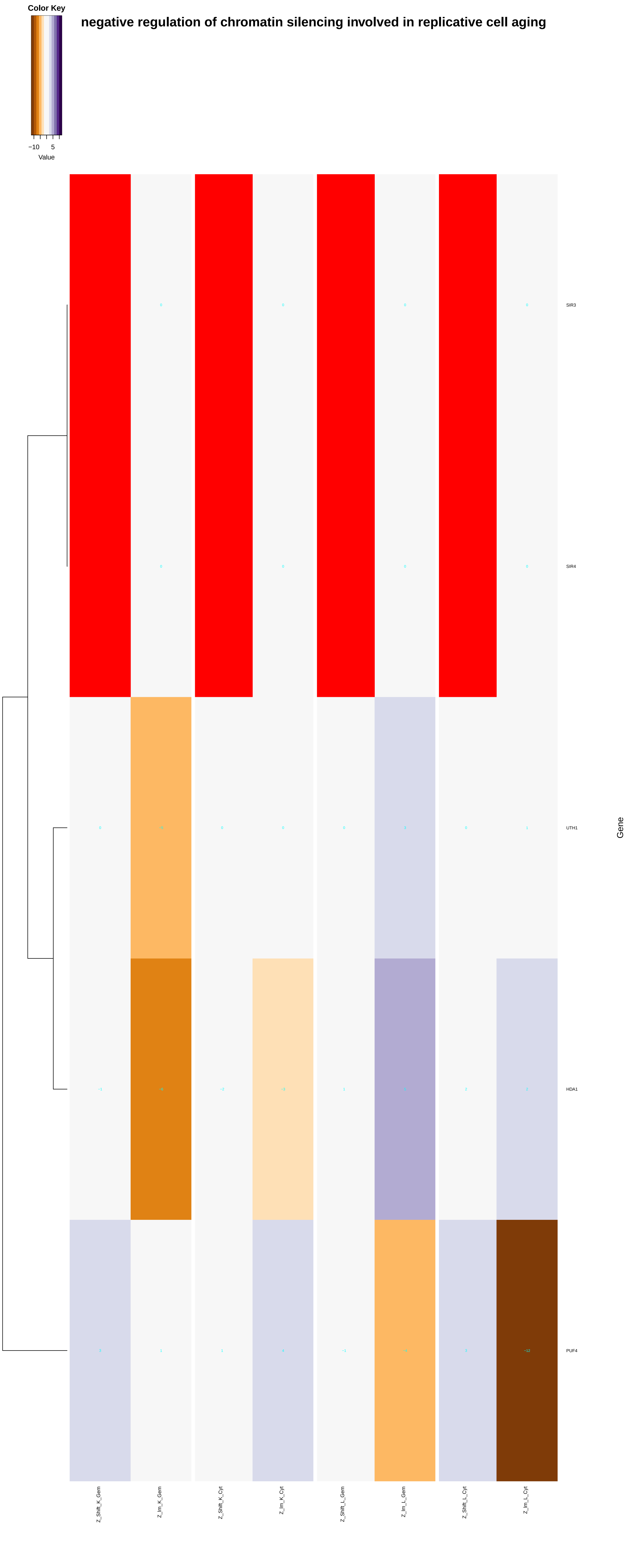

negative regulation of chromatin silencing involved in replicative cell aging
Color Key
−10
5
Value
SIR3
SIR4
UTH1
HDA1
PUF4
Gene
Z_Shift_K_Gem
Z_lm_K_Gem
Z_Shift_K_Cyt
Z_lm_K_Cyt
Z_Shift_L_Gem
Z_lm_L_Gem
Z_Shift_L_Cyt
Z_lm_L_Cyt

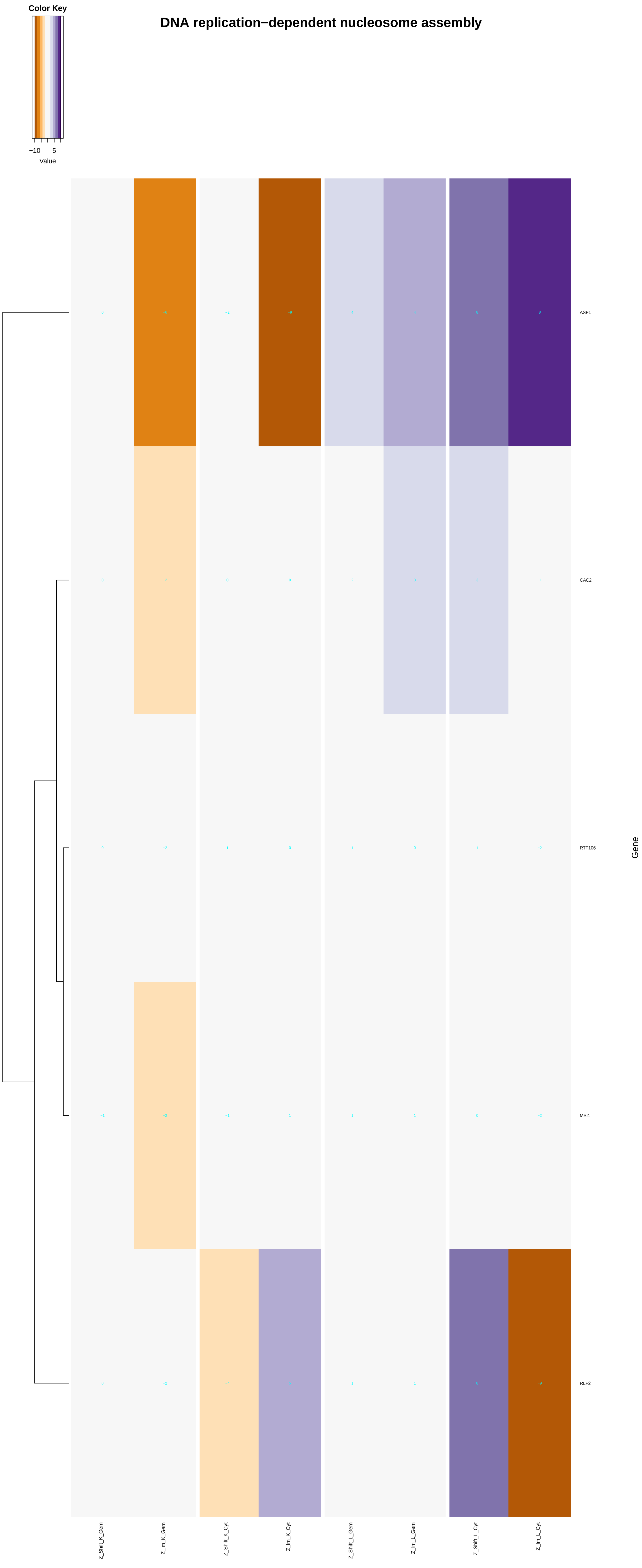

Color Key
−10
5
Value
DNA replication−dependent nucleosome assembly
ASF1
CAC2
RTT106
MSI1
RLF2
Gene
Z_Shift_K_Gem
Z_Im_K_Gem
Z_Shift_K_Cyt
Z_Im_K_Cyt
Z_Shift_L_Gem
Z_Im_L_Gem
Z_Shift_L_Cyt
Z_Im_L_Cyt

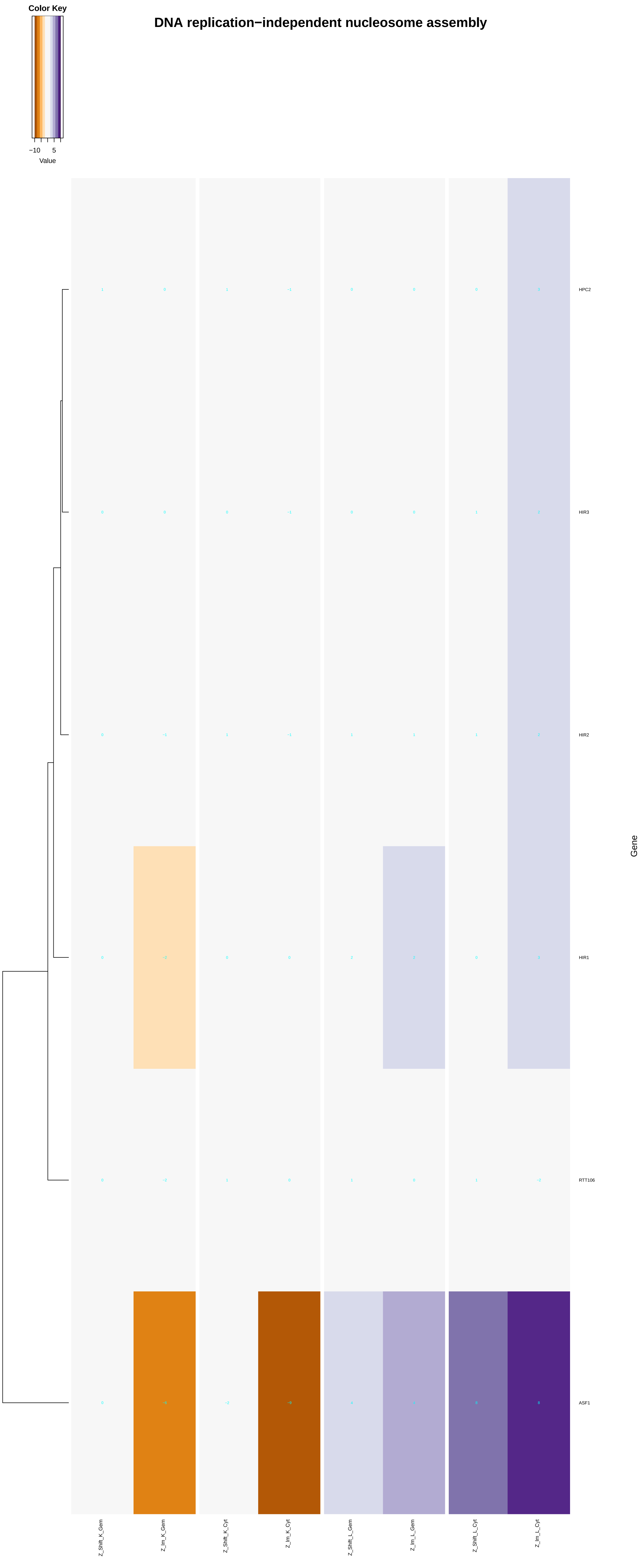

Color Key
−10
5
Value
DNA replication−independent nucleosome assembly
HPC2
HIR3
HIR2
HIR1
RTT106
ASF1
Gene
Z_Shift_K_Gem
Z_lm_K_Gem
Z_Shift_K_Cyt
Z_lm_K_Cyt
Z_Shift_L_Gem
Z_lm_L_Gem
Z_Shift_L_Cyt
Z_lm_L_Cyt

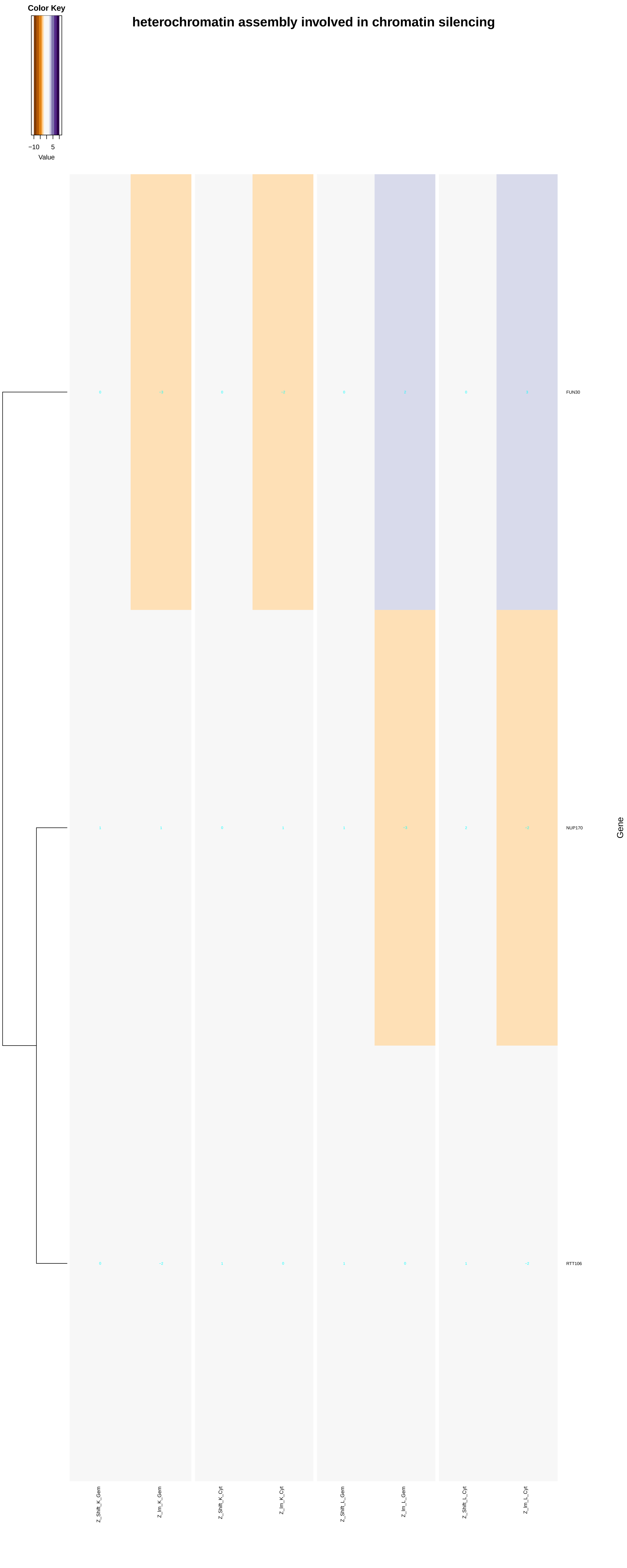
heterochromatin assembly involved in chromatin silencing
Color Key
−10
5
Value
FUN30
NUP170
RTT106
Gene
Z_Shift_K_Gem
Z_lm_K_Gem
Z_Shift_K_Cyt
Z_lm_K_Cyt
Z_Shift_L_Gem
Z_lm_L_Gem
Z_Shift_L_Cyt
Z_lm_L_Cyt

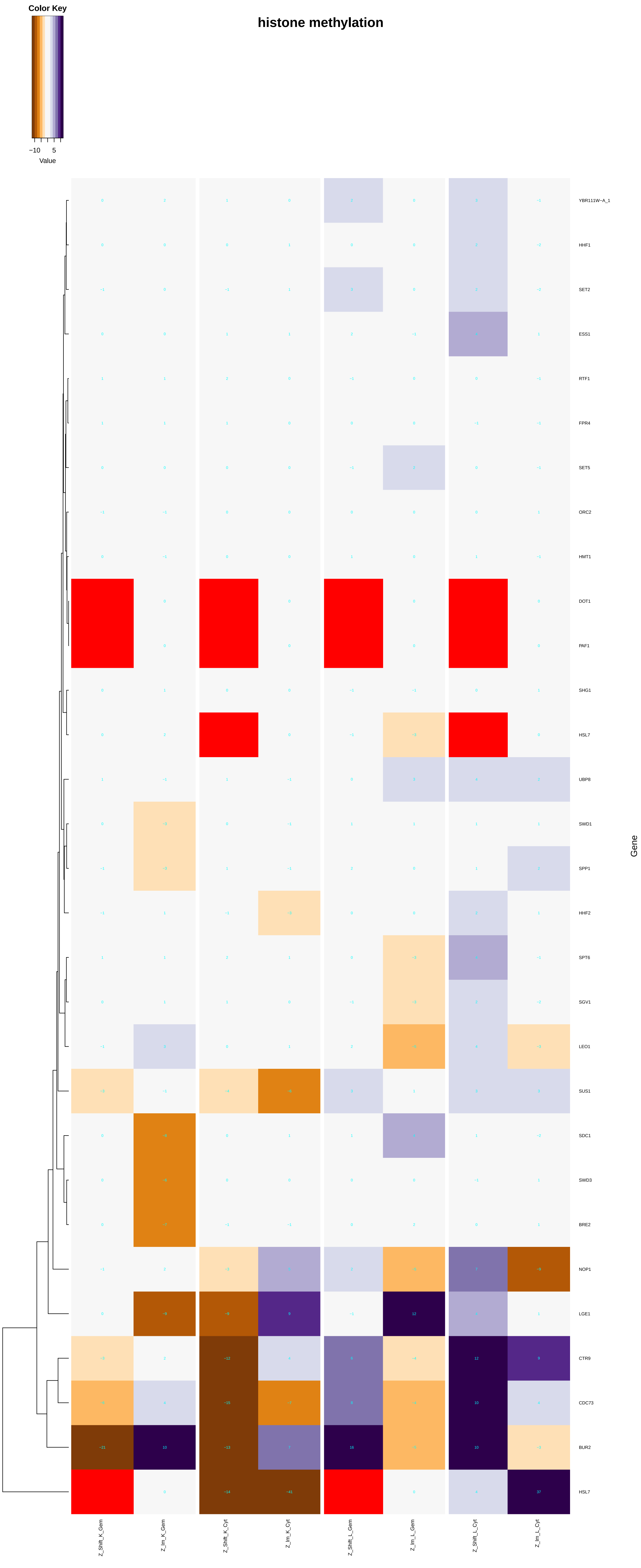

Color Key
−10
5
Value
histone methylation
YBR111W−A_1
HHF1
SET2
ESS1
RTF1
FPR4
SET5
ORC2
HMT1
DOT1
PAF1
SHG1
HSL7
UBP8
SWD1
SPP1
HHF2
SPT6
SGV1
LEO1
SUS1
SDC1
SWD3
BRE2
NOP1
LGE1
CTR9
CDC73
BUR2
HSL7
Gene
Z_Shift_K_Gem
Z_Im_K_Gem
Z_Shift_K_Cyt
Z_Im_K_Cyt
Z_Shift_L_Gem
Z_Im_L_Gem
Z_Shift_L_Cyt
Z_Im_L_Cyt

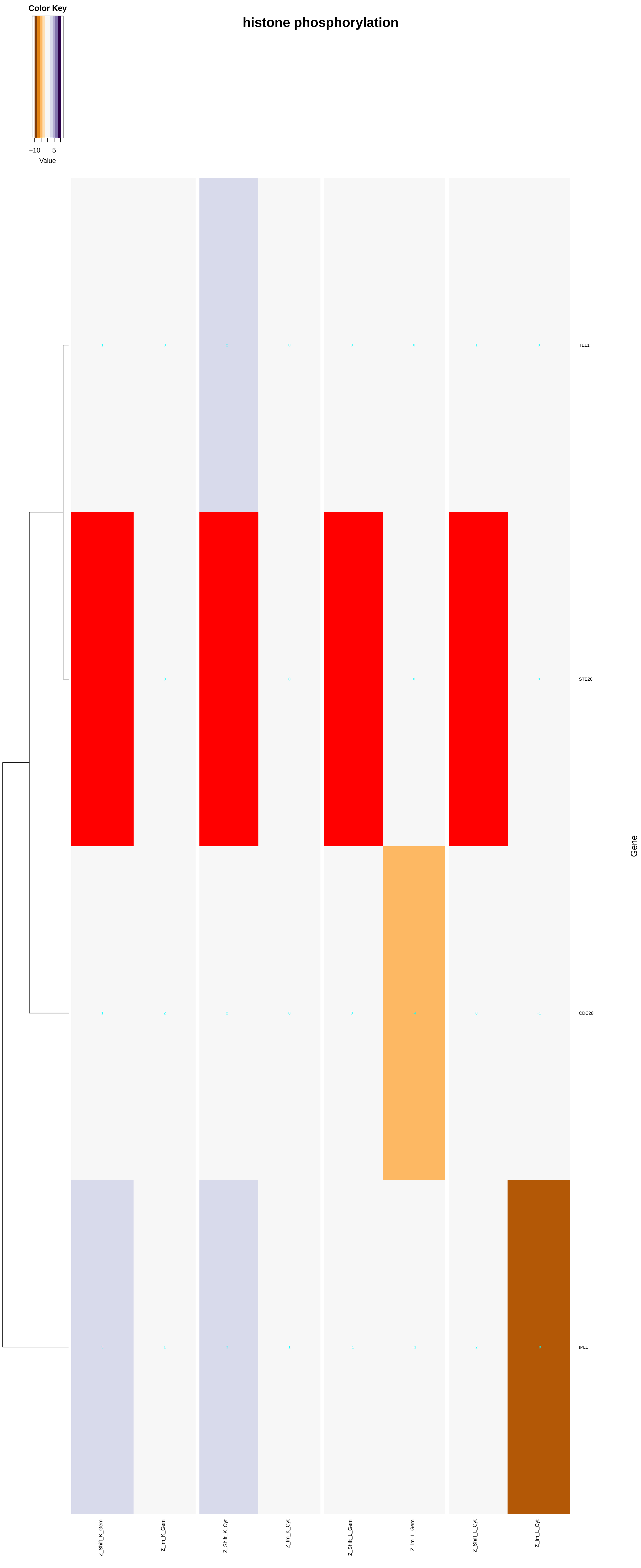
histone phosphorylation
Color Key
−10
5
Value
TEL1
STE20
CDC28
IPL1
Gene
Z_Shift_K_Gem
Z_Im_K_Gem
Z_Shift_K_Cyt
Z_Im_K_Cyt
Z_Shift_L_Gem
Z_Im_L_Gem
Z_Shift_L_Cyt
Z_Im_L_Cyt

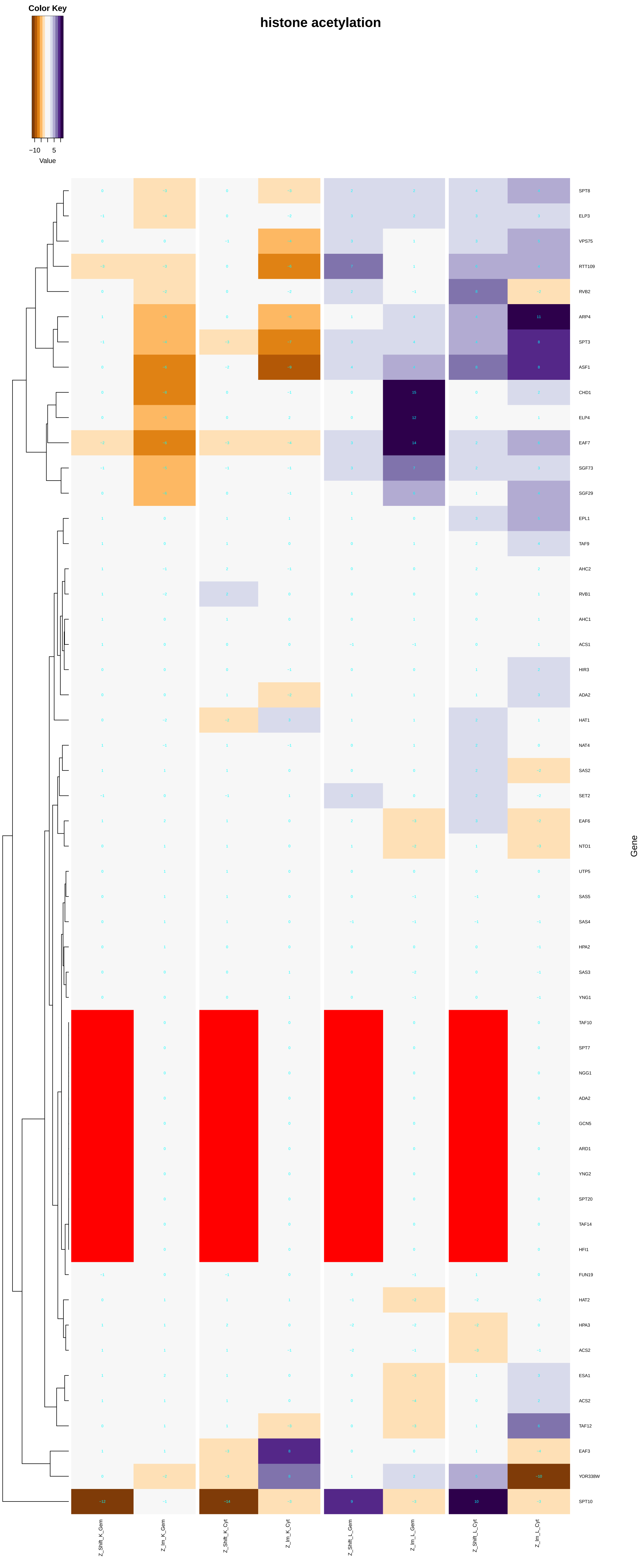

histone acetylation
Color Key
−10
5
Value
SPT8
ELP3
VPS75
RTT109
RVB2
ARP4
SPT3
ASF1
CHD1
ELP4
EAF7
SGF73
SGF29
EPL1
TAF9
AHC2
RVB1
AHC1
ACS1
HIR3
ADA2
HAT1
NAT4
SAS2
SET2
EAF6
NTO1
UTP5
SAS5
SAS4
HPA2
SAS3
YNG1
TAF10
SPT7
NGG1
ADA2
GCN5
ARD1
YNG2
SPT20
TAF14
HFI1
FUN19
HAT2
HPA3
ACS2
ESA1
ACS2
TAF12
EAF3
YOR338W
SPT10
Gene
Z_Shift_K_Gem
Z_Im_K_Gem
Z_Shift_K_Cyt
Z_Im_K_Cyt
Z_Shift_L_Gem
Z_Im_L_Gem
Z_Shift_L_Cyt
Z_Im_L_Cyt

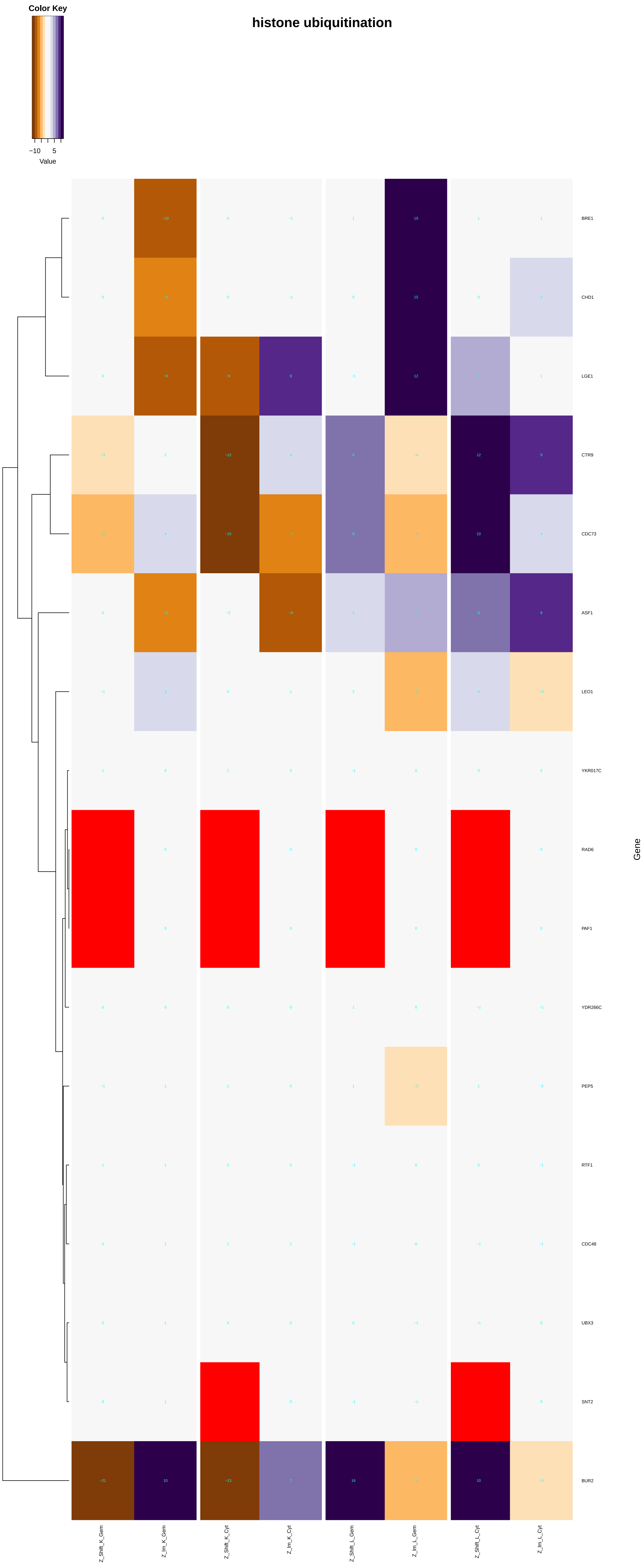

Color Key
−10 5
Value
histone ubiquitination
BRE1
CHD1
LGE1
CTR9
CDC73
ASF1
LEO1
YKR017C
RAD6
PAF1
YDR266C
PEP5
RTF1
CDC48
UBX3
SNT2
BUR2
Gene
Z_Shift_K_Gem
Z_Im_K_Gem
Z_Shift_K_Cyt
Z_Im_K_Cyt
Z_Shift_L_Gem
Z_Im_L_Gem
Z_Shift_L_Cyt
Z_Im_L_Cyt

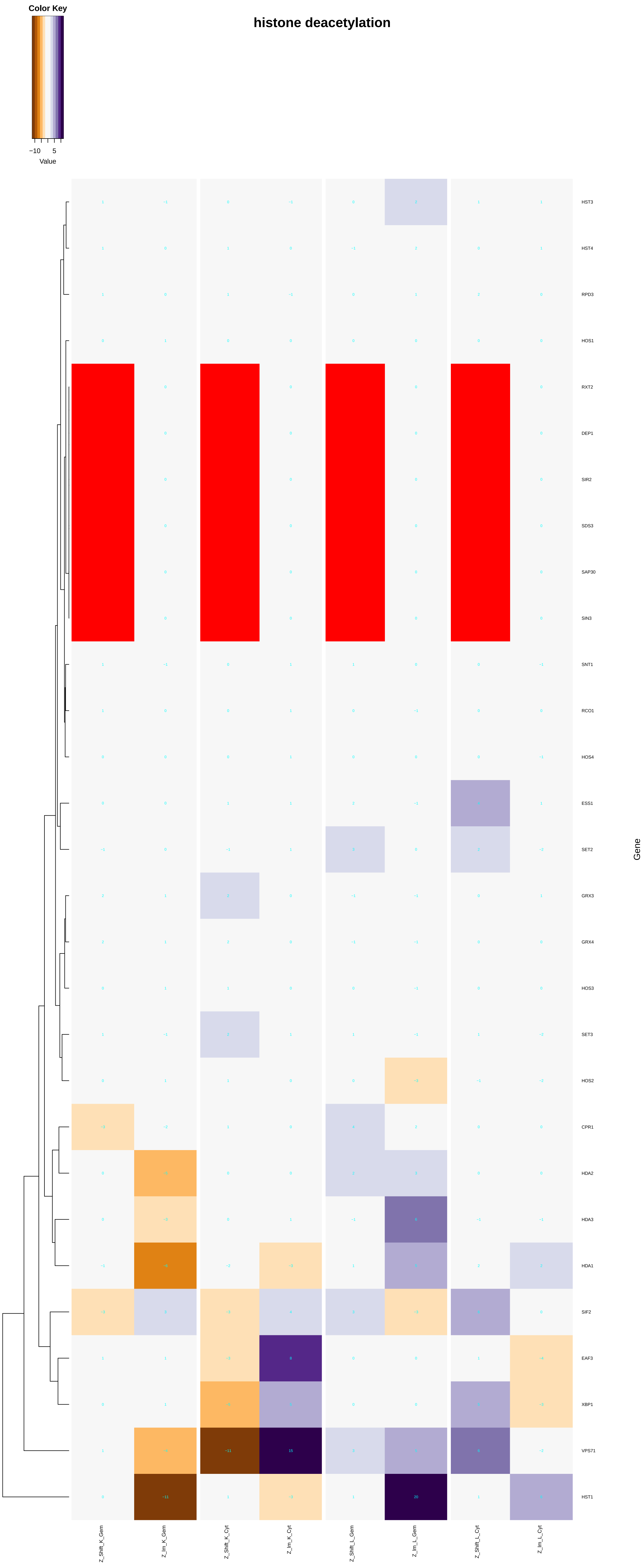
Color Key
−10 5
Value
histone deacetylation
HST3
HST4
RPD3
HOS1
RXT2
DEP1
SIR2
SDS3
SAP30
SIN3
SNT1
RCO1
HOS4
ESS1
SET2
GRX3
GRX4
HOS3
SET3
HOS2
CPR1
HDA2
HDA3
HDA1
SIF2
EAF3
XBP1
VPS71
HST1
Gene
Z_Shift_K_Gem
Z_Im_K_Gem
Z_Shift_K_Cyt
Z_Im_K_Cyt
Z_Shift_L_Gem
Z_Im_L_Gem
Z_Shift_L_Cyt
Z_Im_L_Cyt

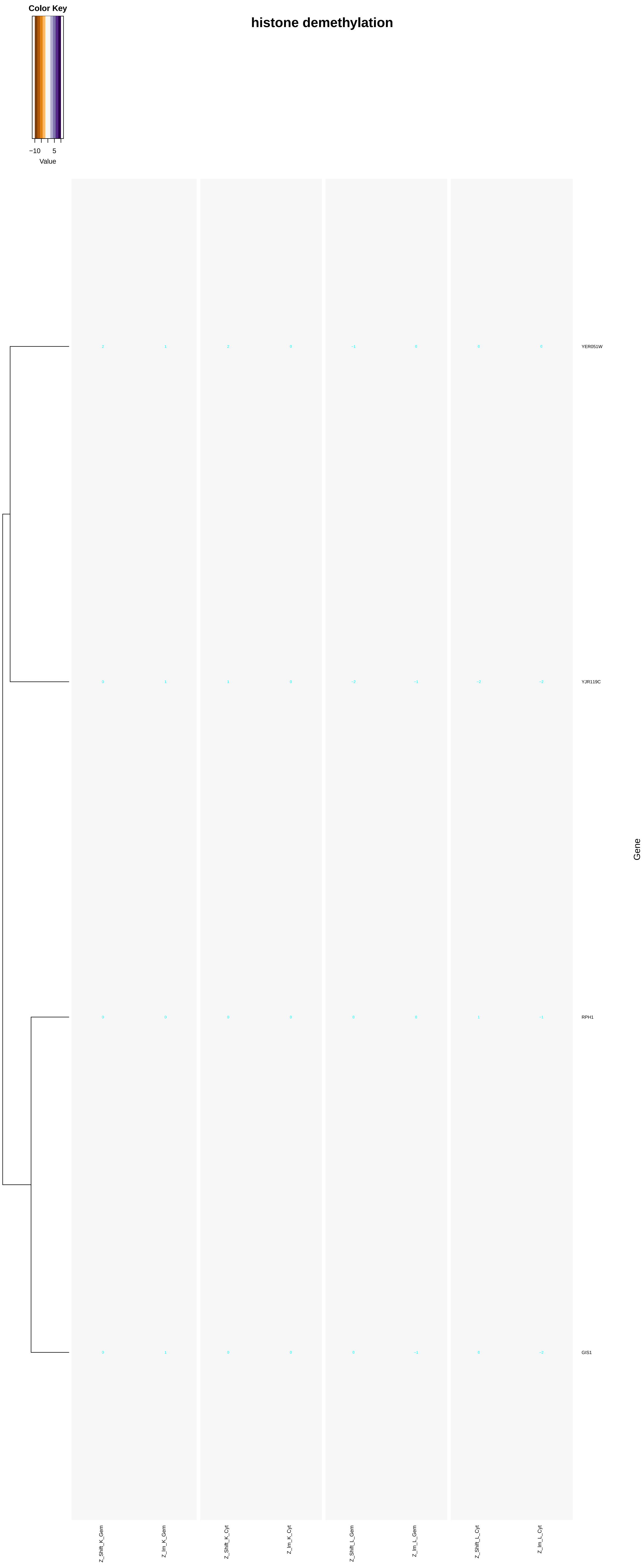

histone demethylation
Color Key
−10
5
Value
YER051W
YJR119C
RPH1
GIS1
Gene
Z_Shift_K_Gem
Z_lm_K_Gem
Z_Shift_K_Cyt
Z_lm_K_Cyt
Z_Shift_L_Gem
Z_lm_L_Gem
Z_Shift_L_Cyt
Z_lm_L_Cyt

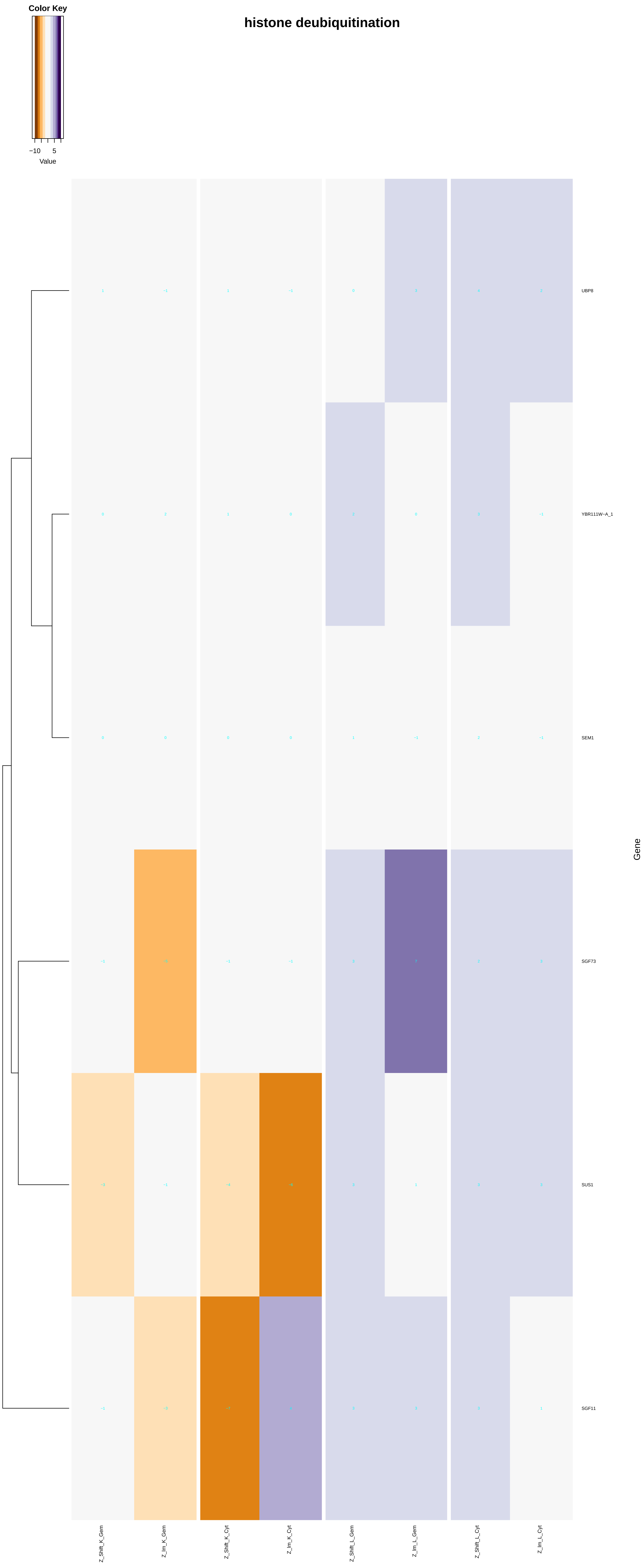

Color Key
−10
5
Value
histone deubiquitination
UBP8
YBR111W−A_1
SEM1
SGF73
SUS1
SGF11
Gene
Z_Shift_K_Gem
Z_Im_K_Gem
Z_Shift_K_Cyt
Z_Im_K_Cyt
Z_Shift_L_Gem
Z_Im_L_Gem
Z_Shift_L_Cyt
Z_Im_L_Cyt

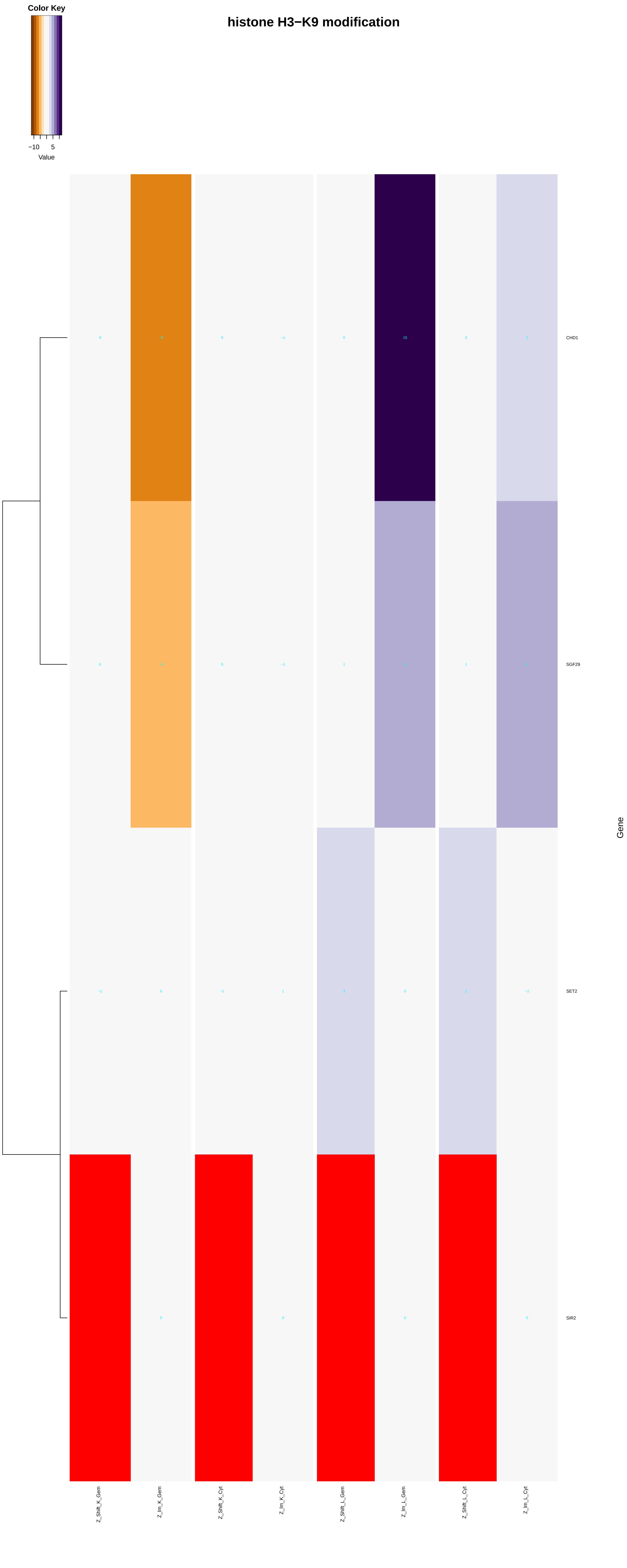

Color Key
−10
5
Value
histone H3−K9 modification
CHD1
SGF29
SET2
SIR2
Gene
Z_Shift_K_Gem
Z_Im_K_Gem
Z_Shift_K_Cyt
Z_Im_K_Cyt
Z_Shift_L_Gem
Z_Im_L_Gem
Z_Shift_L_Cyt
Z_Im_L_Cyt

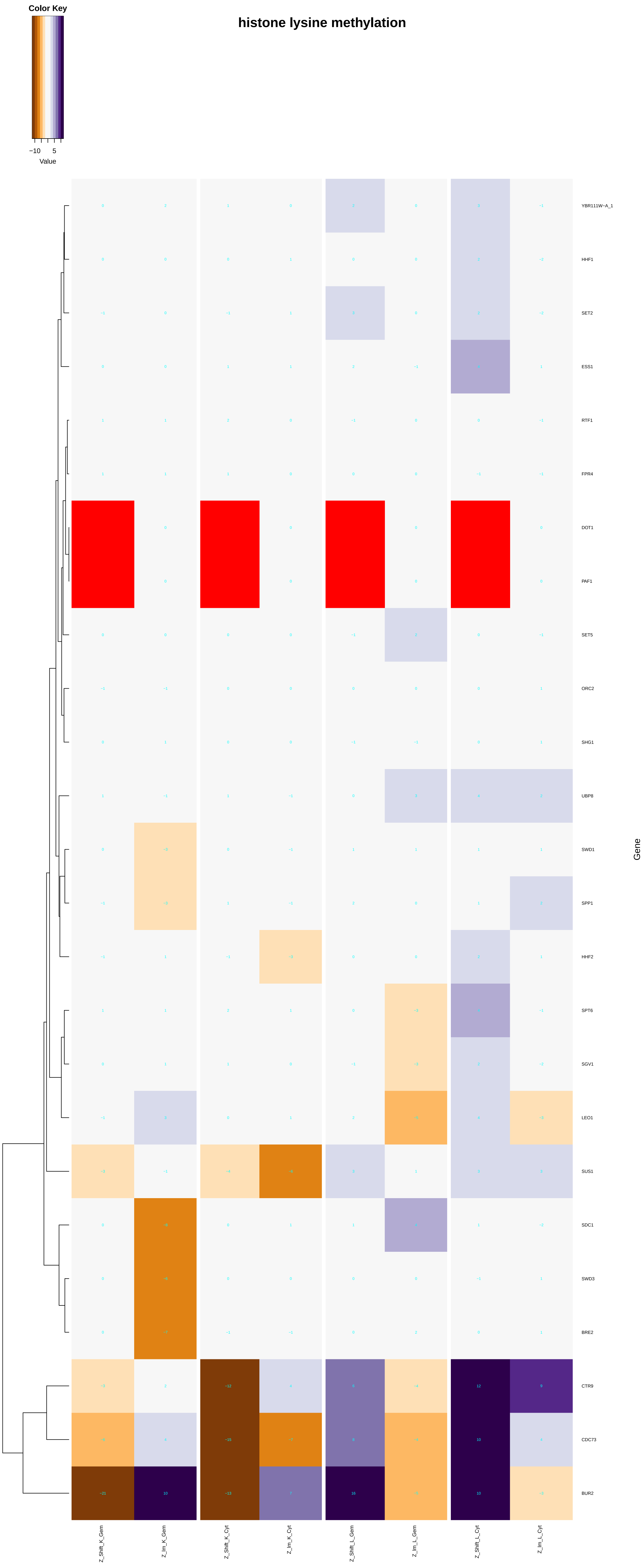
histone lysine methylation
Color Key
−10
5
Value
YBR111W−A_1
HHF1
SET2
ESS1
RTF1
FPR4
DOT1
PAF1
SET5
ORC2
SHG1
UBP8
SWD1
SPP1
HHF2
SPT6
SGV1
LEO1
SUS1
SDC1
SWD3
BRE2
CTR9
CDC73
BUR2
Gene
Z_Shift_K_Gem
Z_Im_K_Gem
Z_Shift_K_Cyt
Z_Im_K_Cyt
Z_Shift_L_Gem
Z_Im_L_Gem
Z_Shift_L_Cyt
Z_Im_L_Cyt

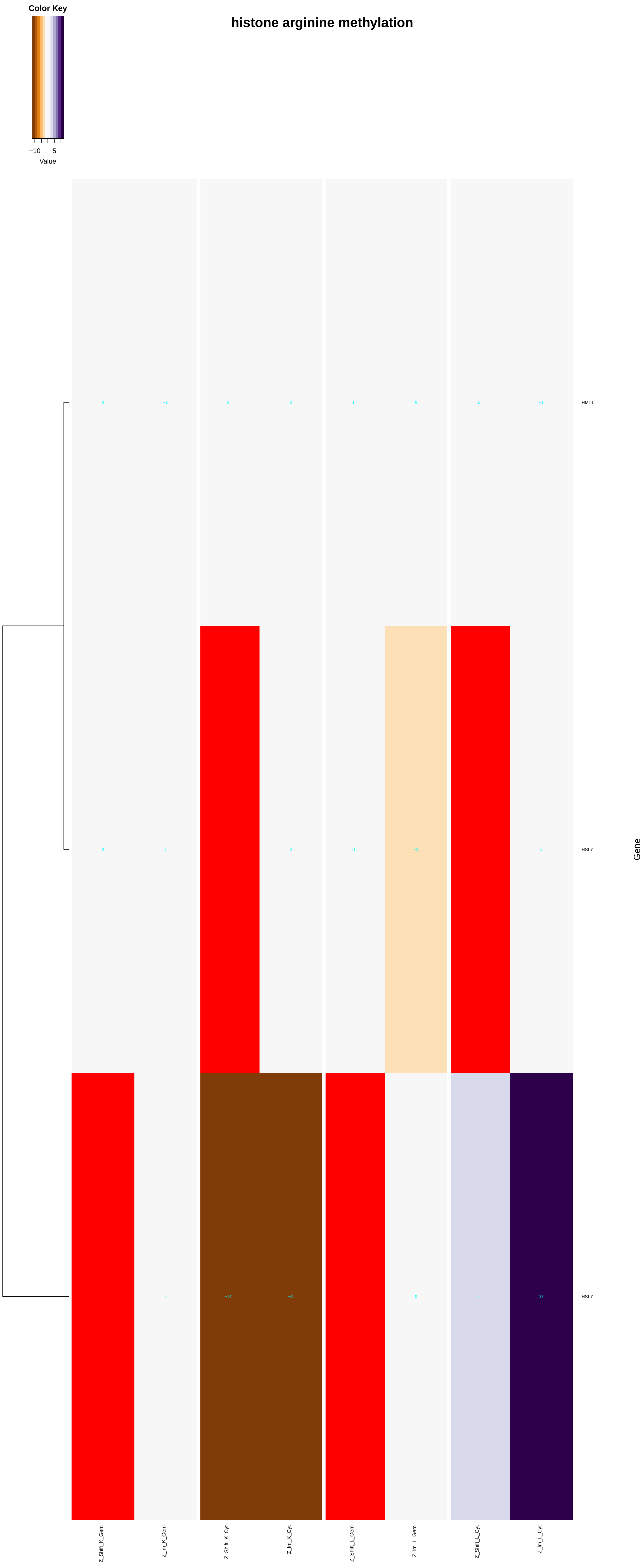
histone arginine methylation
Color Key
−10
5
Value
HMT1
HSL7
HSL7
Gene
Z_Shift_K_Gem
Z_Im_K_Gem
Z_Shift_K_Cyt
Z_Im_K_Cyt
Z_Shift_L_Gem
Z_Im_L_Gem
Z_Shift_L_Cyt
Z_Im_L_Cyt

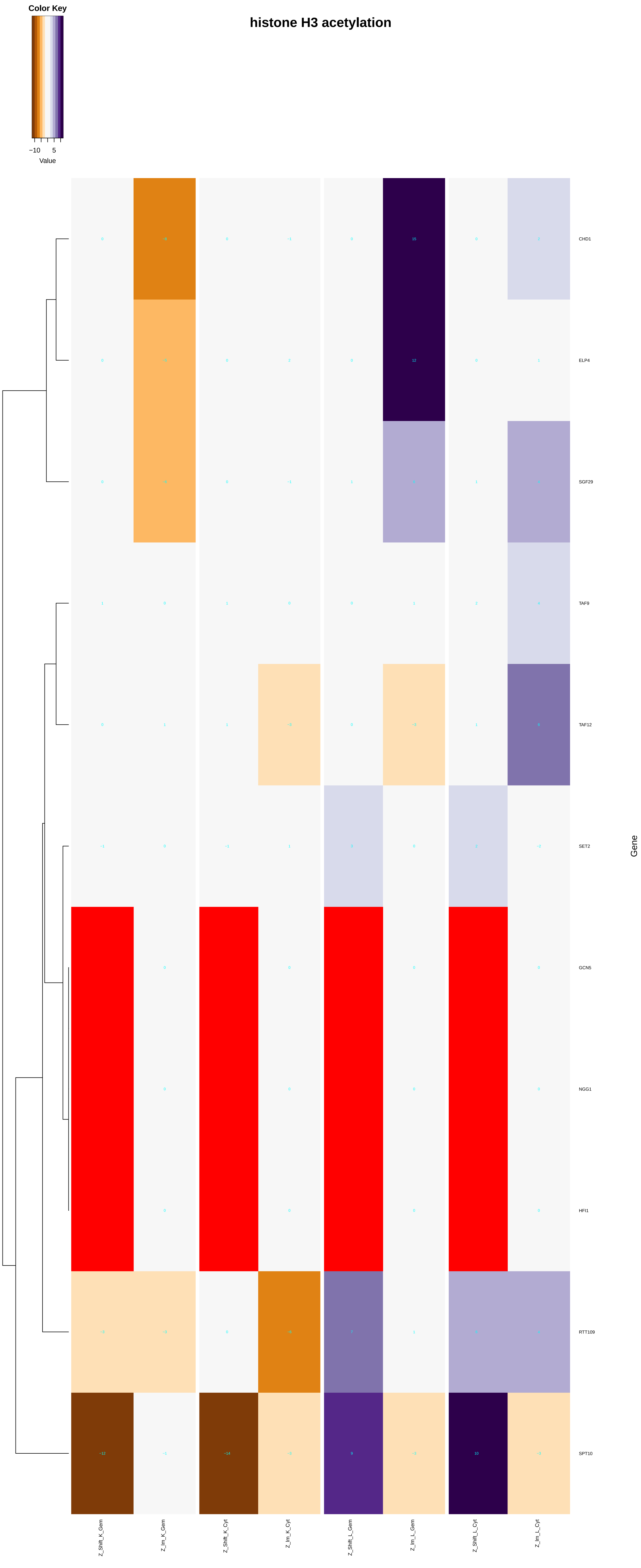

histone H3 acetylation
Color Key
−10
5
Value
CHD1
ELP4
SGF29
TAF9
TAF12
SET2
GCN5
NGG1
HFI1
RTT109
SPT10
Gene
Z_Shift_K_Gem
Z_Im_K_Gem
Z_Shift_K_Cyt
Z_Im_K_Cyt
Z_Shift_L_Gem
Z_Im_L_Gem
Z_Shift_L_Cyt
Z_Im_L_Cyt

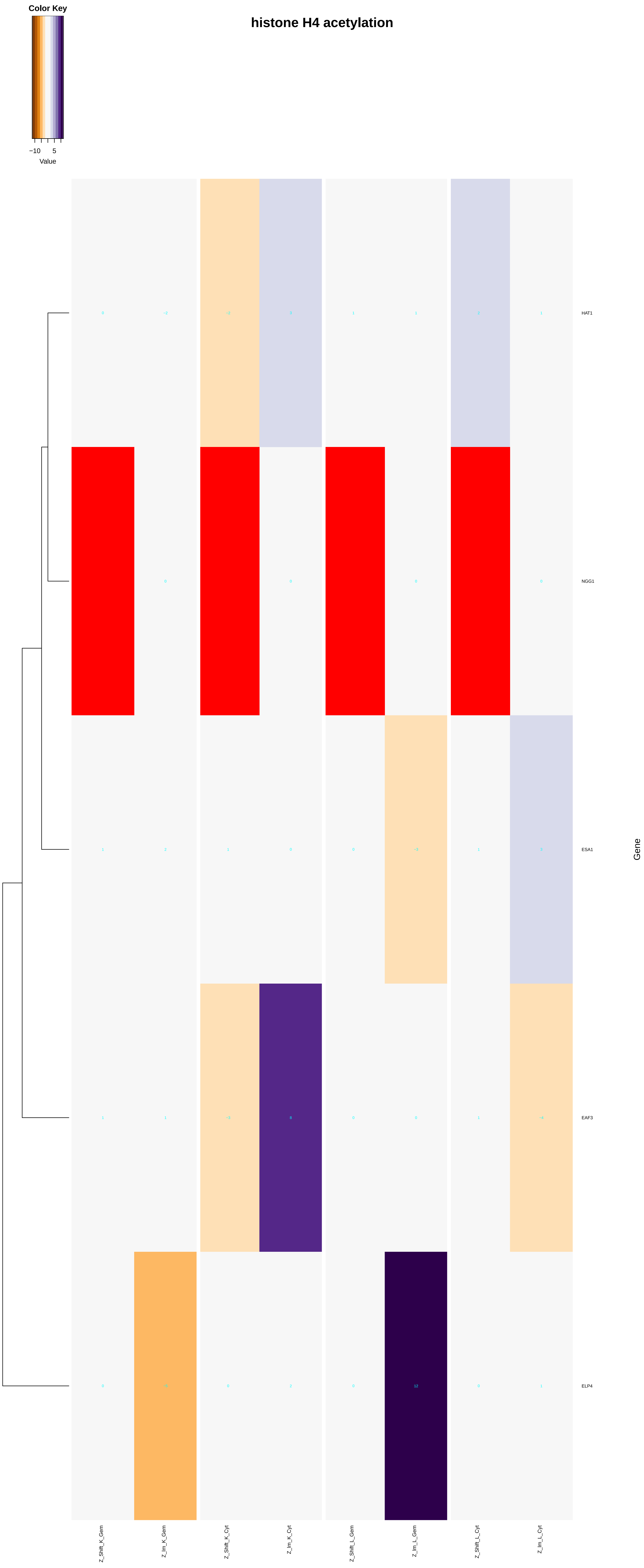
histone H4 acetylation
Color Key
−10
5
Value
HAT1
NGG1
ESA1
EAF3
ELP4
Gene
Z_Shift_K_Gem
Z_Im_K_Gem
Z_Shift_K_Cyt
Z_Im_K_Cyt
Z_Shift_L_Gem
Z_Im_L_Gem
Z_Shift_L_Cyt
Z_Im_L_Cyt

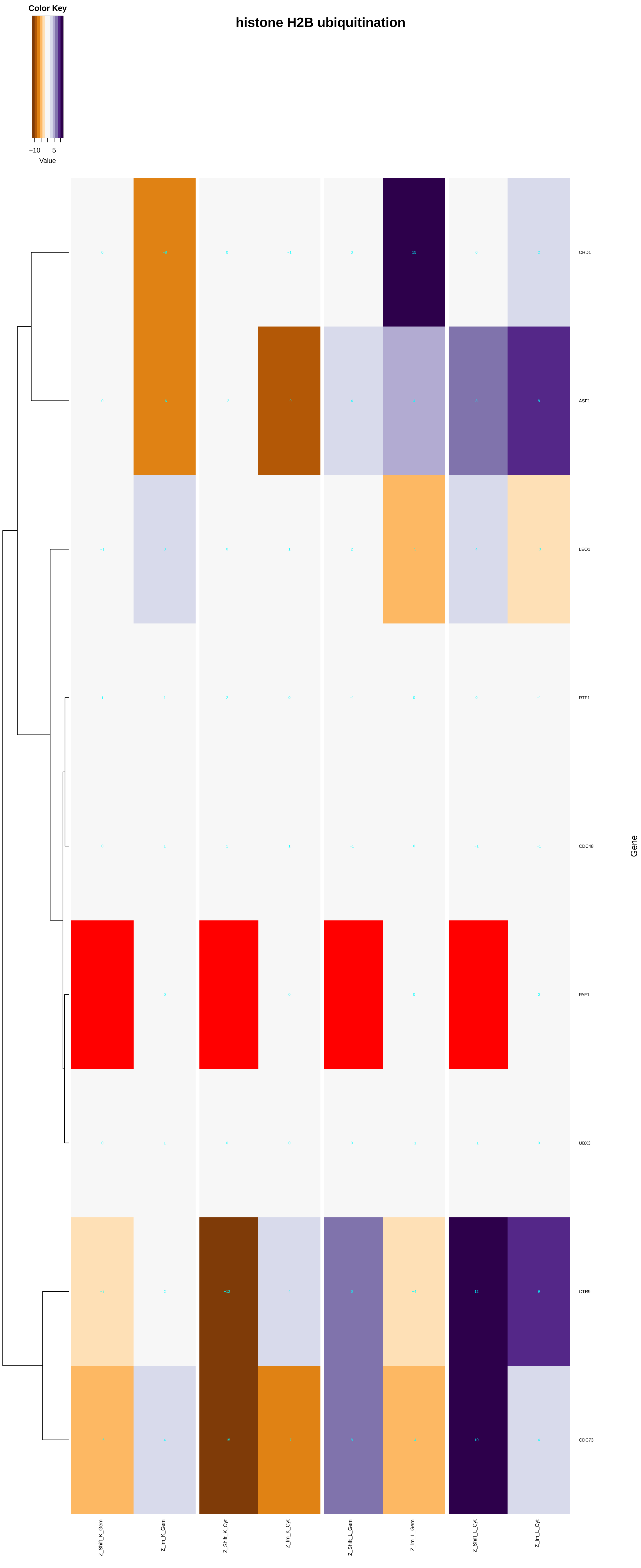
Color Key
−10 5
Value
histone H2B ubiquitination
CHD1
ASF1
LEO1
RTF1
CDC48
PAF1
UBX3
CTR9
CDC73
Gene
Z_Shift_K_Gem
Z_Im_K_Gem
Z_Shift_K_Cyt
Z_Im_K_Cyt
Z_Shift_L_Gem
Z_Im_L_Gem
Z_Shift_L_Cyt
Z_Im_L_Cyt

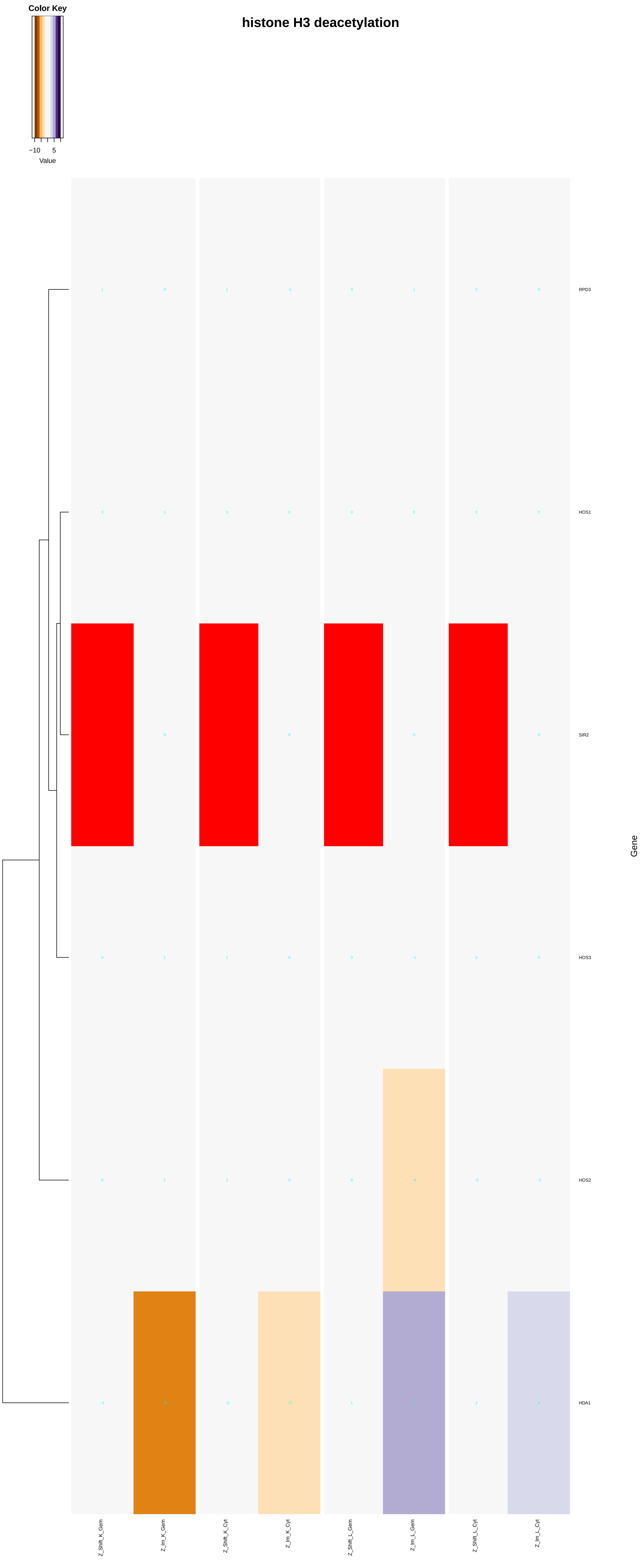
histone H3 deacetylation
Color Key
−10
5
Value
RPD3
HOS1
SIR2
HOS3
HOS2
HDA1
Gene
Z_Shift_K_Gem
Z_Im_K_Gem
Z_Shift_K_Cyt
Z_Im_K_Cyt
Z_Shift_L_Gem
Z_Im_L_Gem
Z_Shift_L_Cyt
Z_Im_L_Cyt

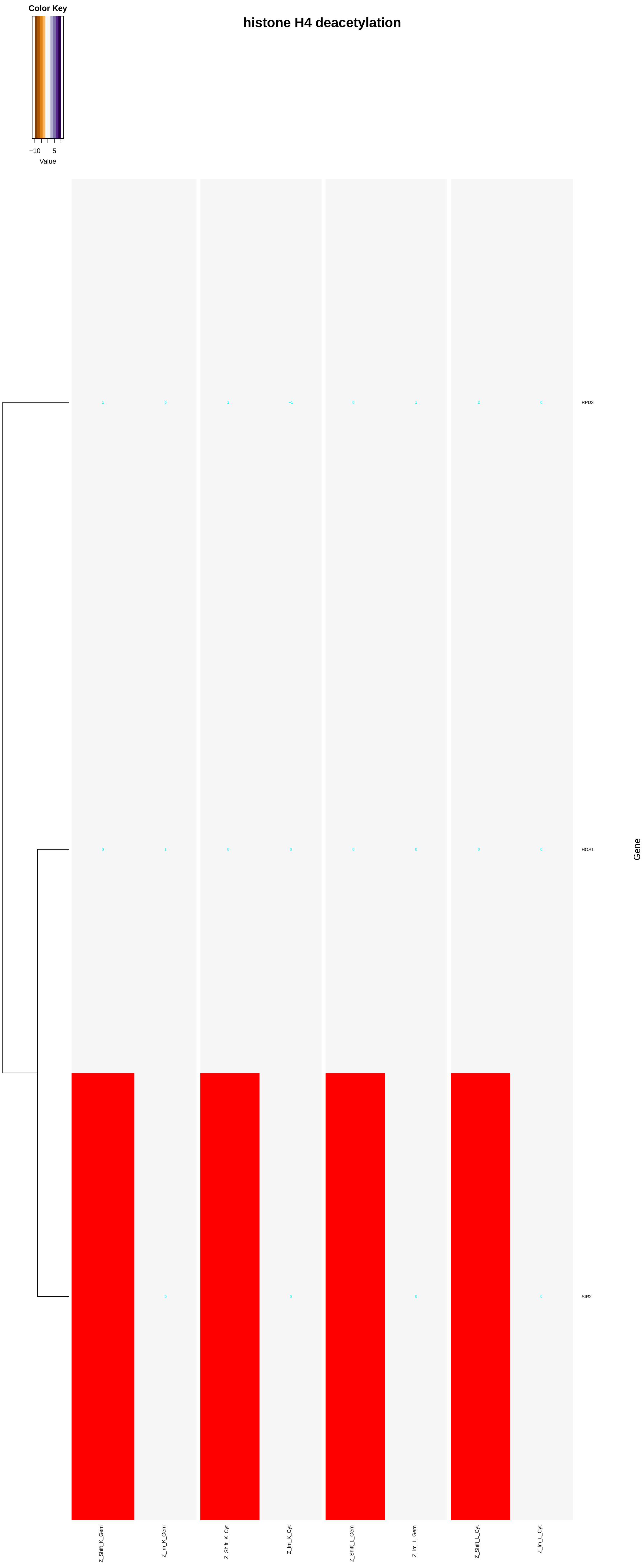

Color Key
−10
5
Value
histone H4 deacetylation
RPD3
HOS1
SIR2
Gene
Z_Shift_K_Gem
Z_Im_K_Gem
Z_Shift_K_Cyt
Z_Im_K_Cyt
Z_Shift_L_Gem
Z_Im_L_Gem
Z_Shift_L_Cyt
Z_Im_L_Cyt

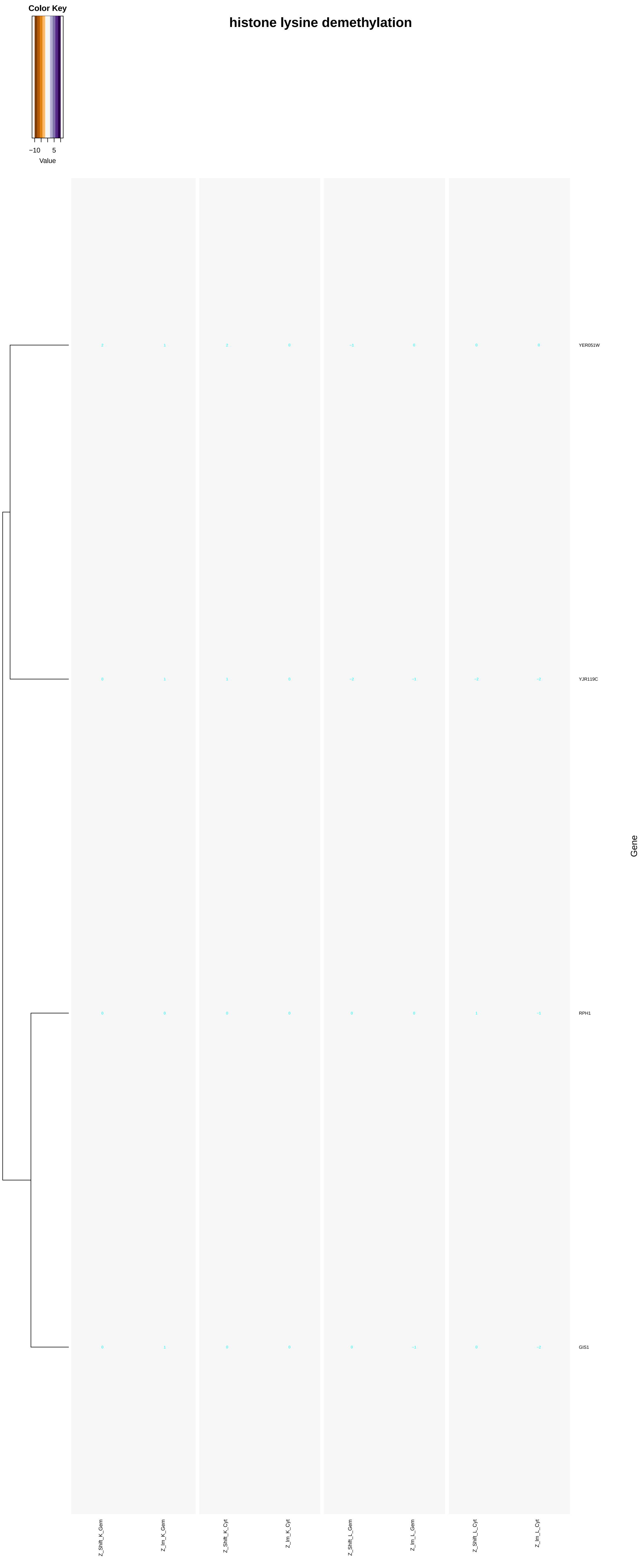
histone lysine demethylation
Color Key
−10
5
Value
YER051W
YJR119C
RPH1
GIS1
Gene
Z_Shift_K_Gem
Z_lm_K_Gem
Z_Shift_K_Cyt
Z_lm_K_Cyt
Z_Shift_L_Gem
Z_lm_L_Gem
Z_Shift_L_Cyt
Z_lm_L_Cyt

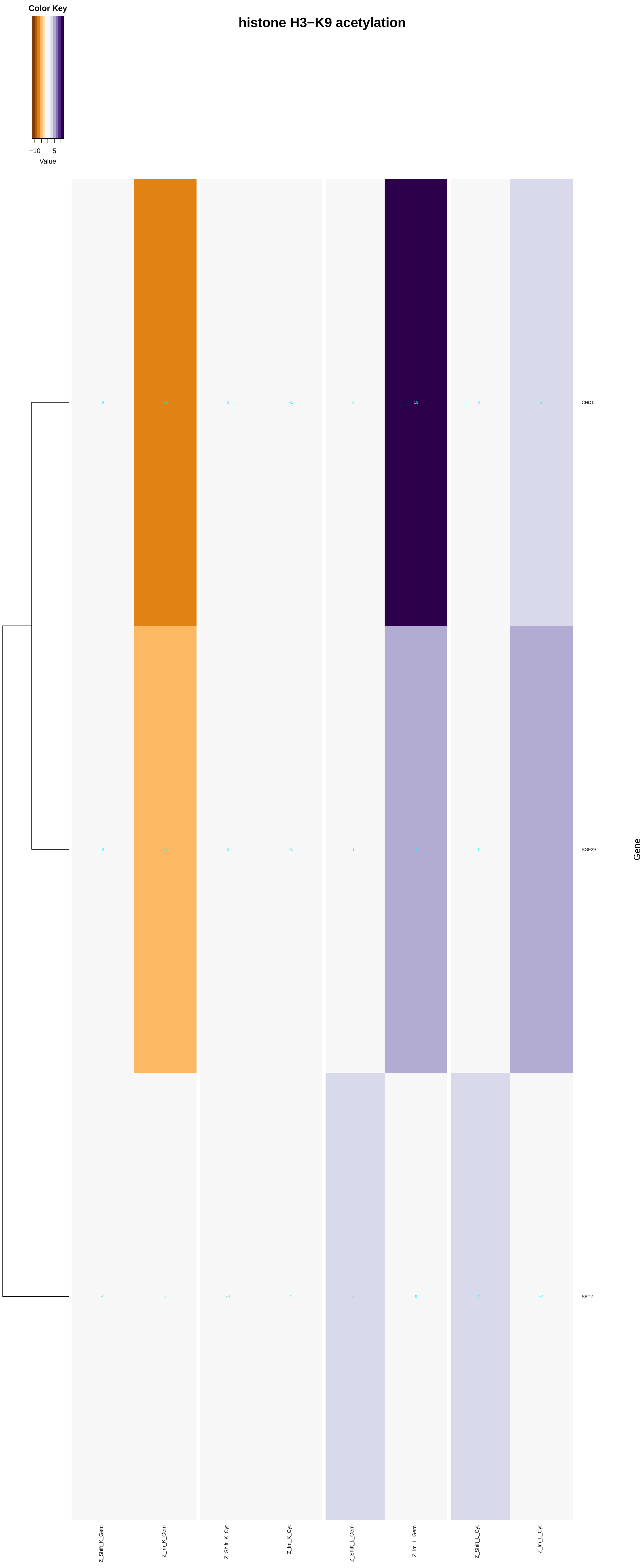

Color Key
−10
5
Value
histone H3−K9 acetylation
0
−8
0
−1
0
15
0
2
CHD1
0
−6
0
−1
1
6
1
4
SGF29
−1
0
−1
1
3
0
2
−2
SET2
Gene
Z_Shift_K_Gem
Z_lm_K_Gem
Z_Shift_K_Cyt
Z_lm_K_Cyt
Z_Shift_L_Gem
Z_lm_L_Gem
Z_Shift_L_Cyt
Z_lm_L_Cyt

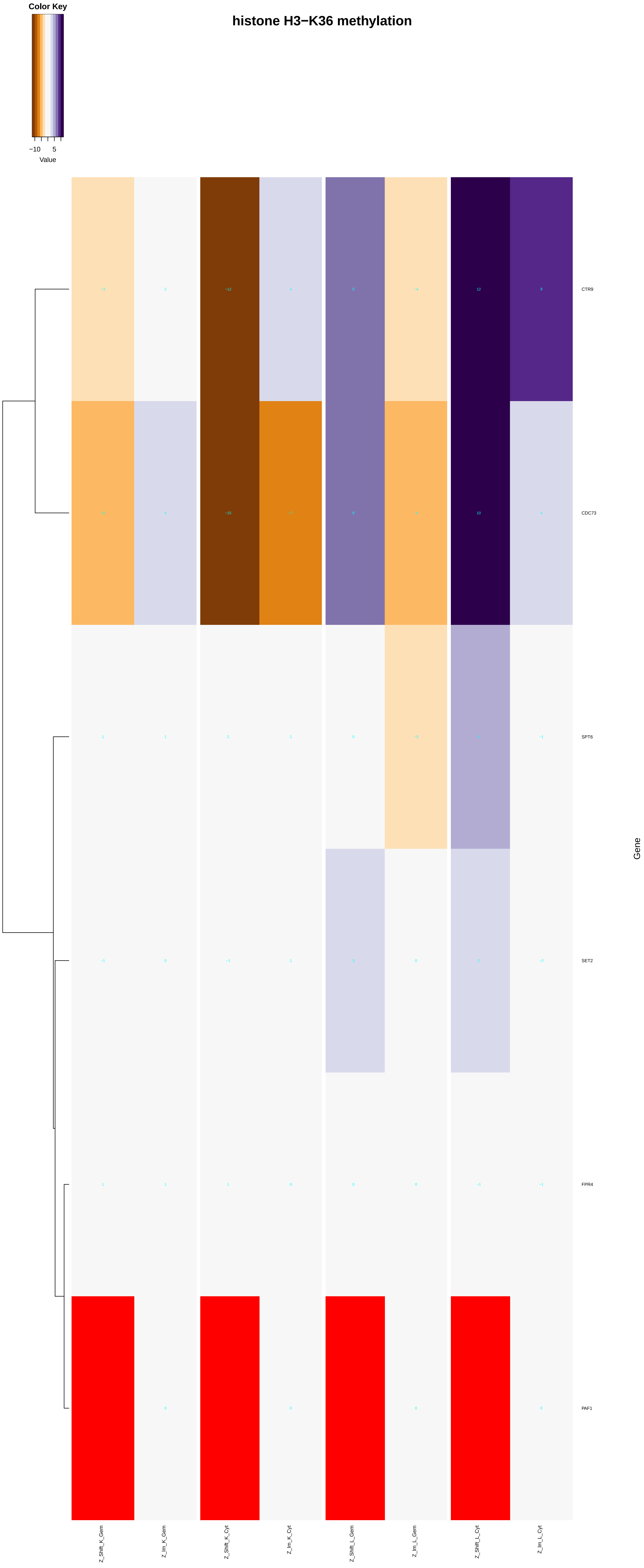
histone H3−K36 methylation
Color Key
−10
5
Value
CTR9
CDC73
SPT6
SET2
FPR4
PAF1
Gene
Z_Shift_K_Gem
Z_Im_K_Gem
Z_Shift_K_Cyt
Z_Im_K_Cyt
Z_Shift_L_Gem
Z_Im_L_Gem
Z_Shift_L_Cyt
Z_Im_L_Cyt

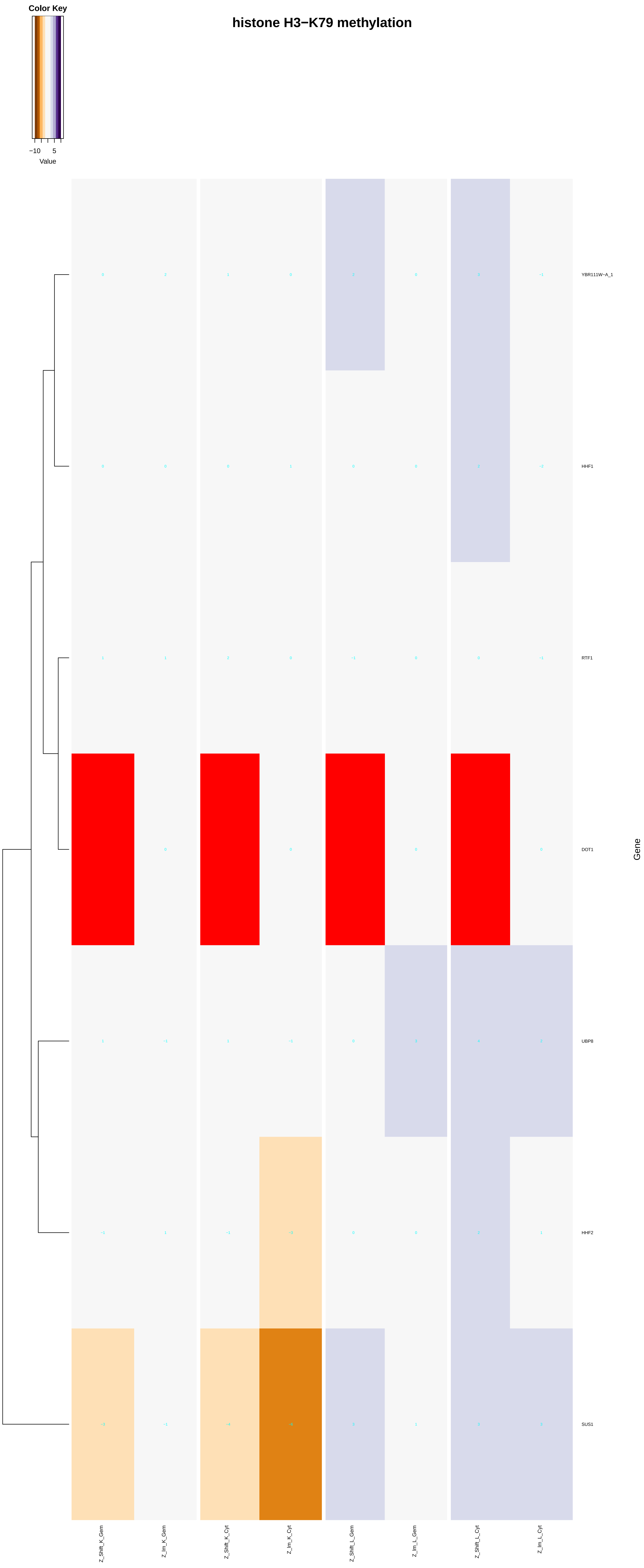

Color Key
−10
5
Value
histone H3−K79 methylation
YBR111W−A_1
HHF1
RTF1
DOT1
UBP8
HHF2
SUS1
Gene
Z_Shift_K_Gem
Z_Im_K_Gem
Z_Shift_K_Cyt
Z_Im_K_Cyt
Z_Shift_L_Gem
Z_Im_L_Gem
Z_Shift_L_Cyt
Z_Im_L_Cyt

#### histone H3−K4 methylation

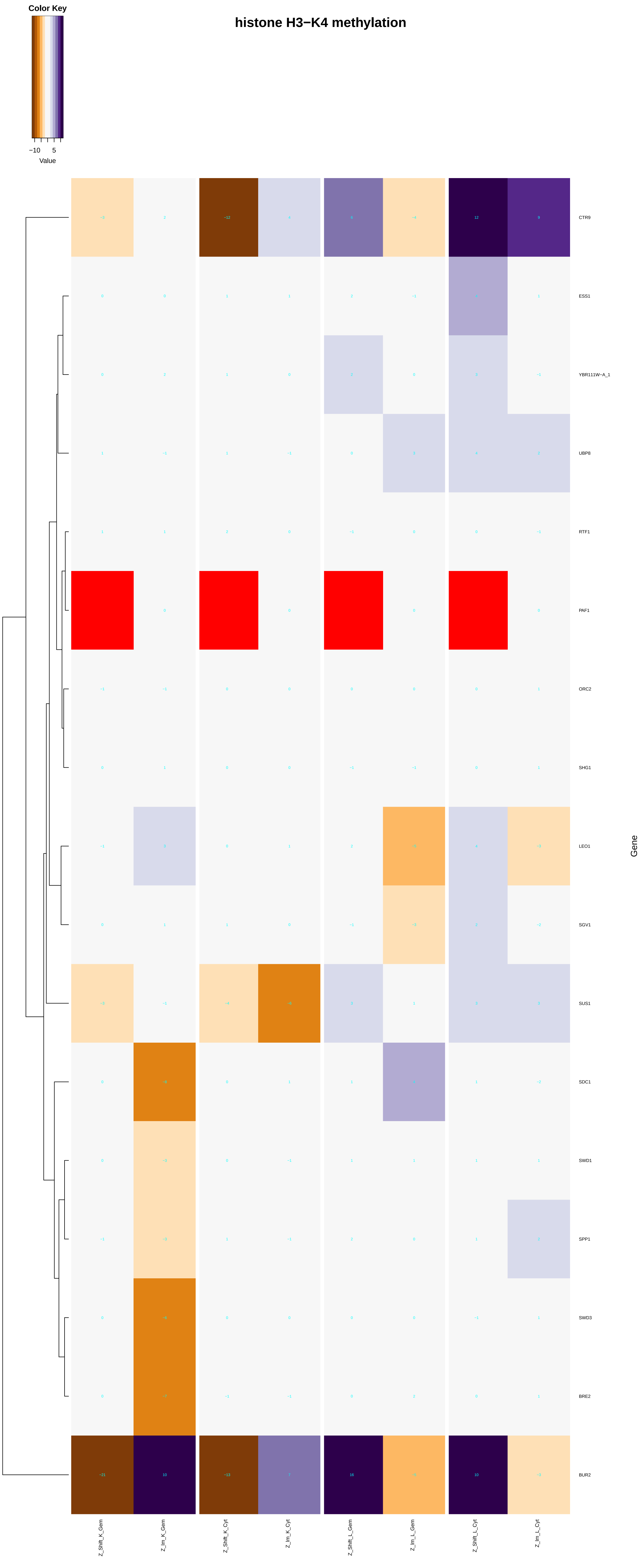

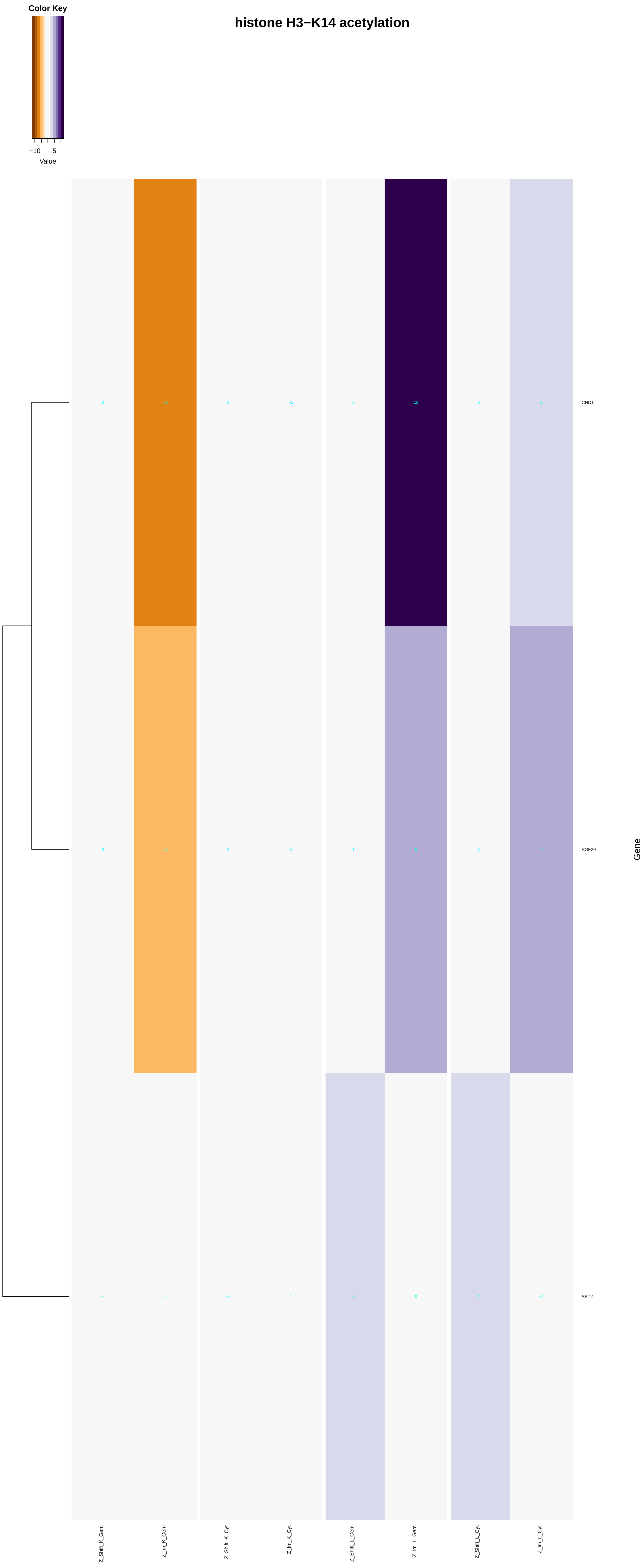
Color Key
−10
5
Value
histone H3−K14 acetylation
0
−8
0
−1
0
15
0
2
CHD1
0
−6
0
−1
1
6
1
4
SGF29
−1
0
−1
1
3
0
2
−2
SET2
Gene
Z_Shift_K_Gem
Z_Im_K_Gem
Z_Shift_K_Cyt
Z_Im_K_Cyt
Z_Shift_L_Gem
Z_Im_L_Gem
Z_Shift_L_Cyt
Z_Im_L_Cyt

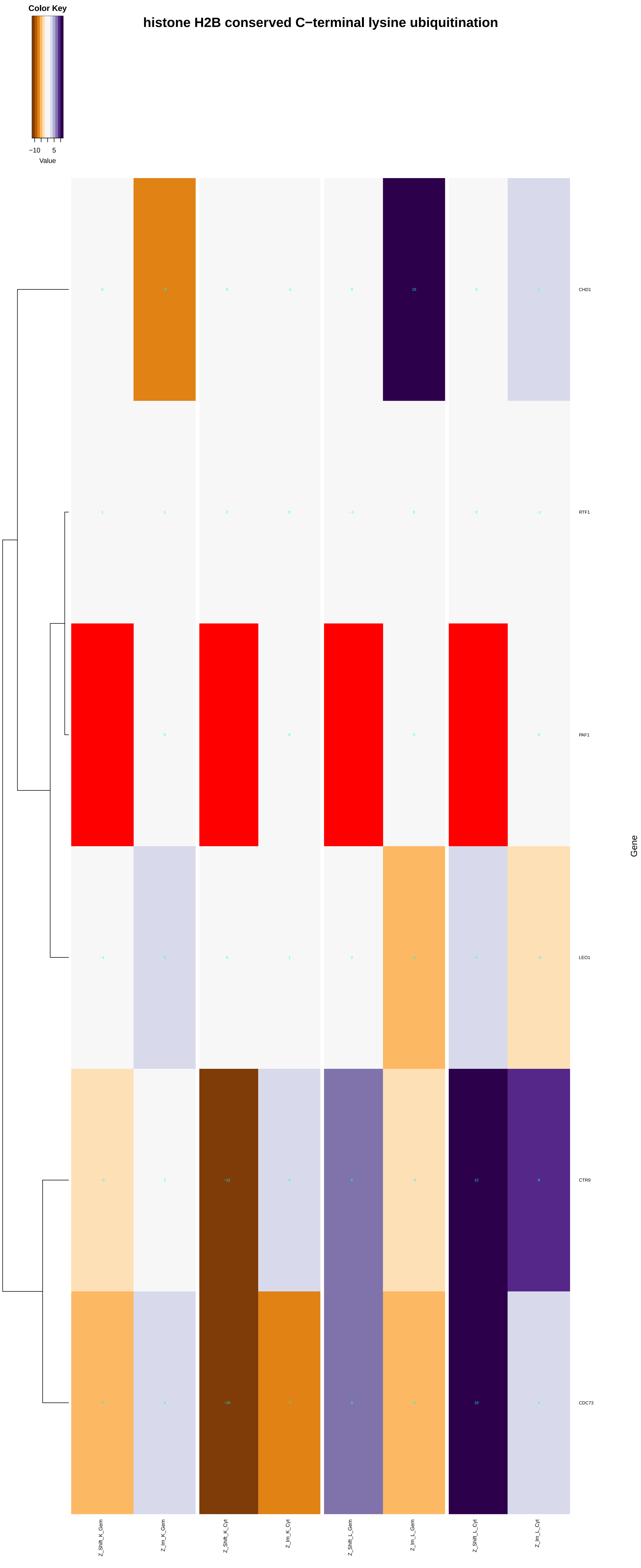

Color Key
−10
5
Value
histone H2B conserved C−terminal lysine ubiquitination
CHD1
RTF1
PAF1
LEO1
CTR9
CDC73
Gene
Z_Shift_K_Gem
Z_Im_K_Gem
Z_Shift_K_Cyt
Z_Im_K_Cyt
Z_Shift_L_Gem
Z_Im_L_Gem
Z_Shift_L_Cyt
Z_Im_L_Cyt

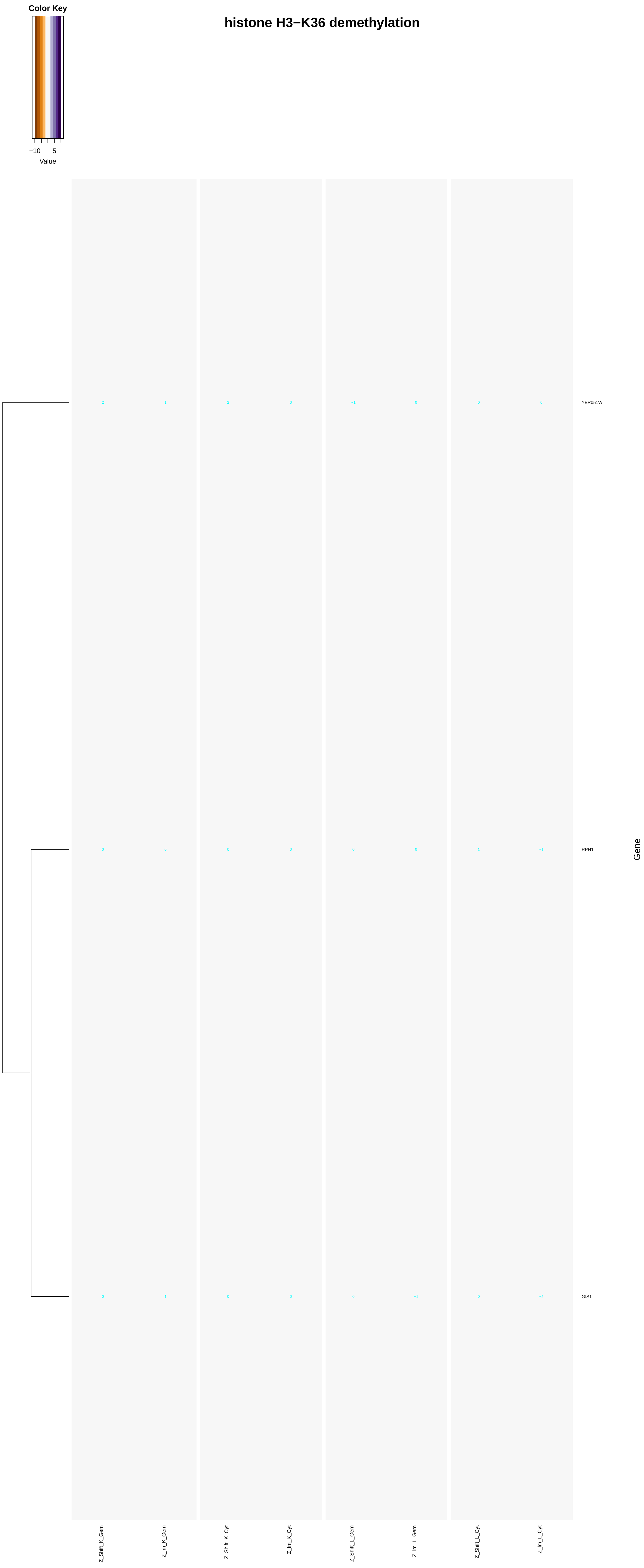
Color Key
−10
5
Value
histone H3−K36 demethylation
2 1 2 0 −1 0 0 0
YER051W
0 0 0 0 0 0 1 −1
RPH1
0 1 0 0 0 −1 0 −2
GIS1
Gene
Z_Shift_K_Gem
Z_lm_K_Gem
Z_Shift_K_Cyt
Z_lm_K_Cyt
Z_Shift_L_Gem
Z_lm_L_Gem
Z_Shift_L_Cyt
Z_lm_L_Cyt

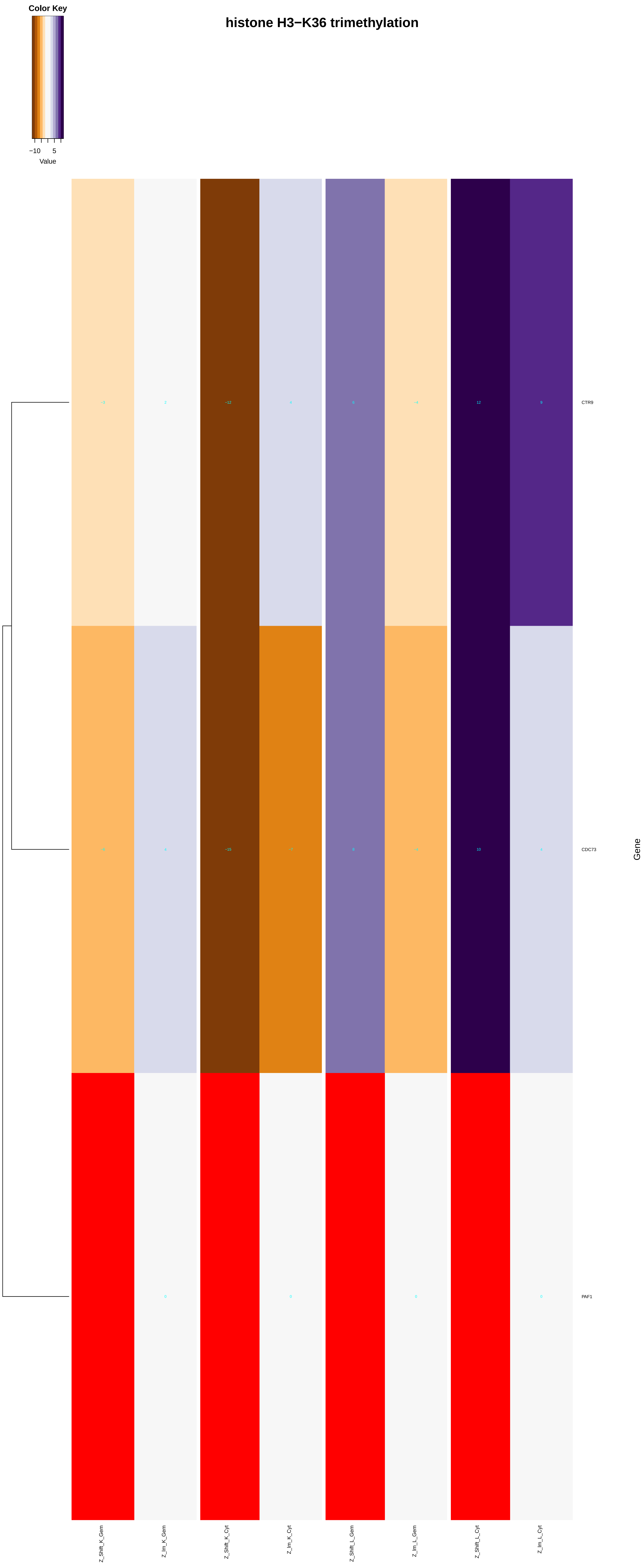
Color Key
−10
5
Value
histone H3−K36 trimethylation
−3
2
−12
4
6
−4
12
9
CTR9
−6
4
−15
−7
8
−4
10
4
CDC73
0
0
0
0
PAF1
Gene
Z_Shift_K_Gem
Z_Im_K_Gem
Z_Shift_K_Cyt
Z_Im_K_Cyt
Z_Shift_L_Gem
Z_Im_L_Gem
Z_Shift_L_Cyt
Z_Im_L_Cyt

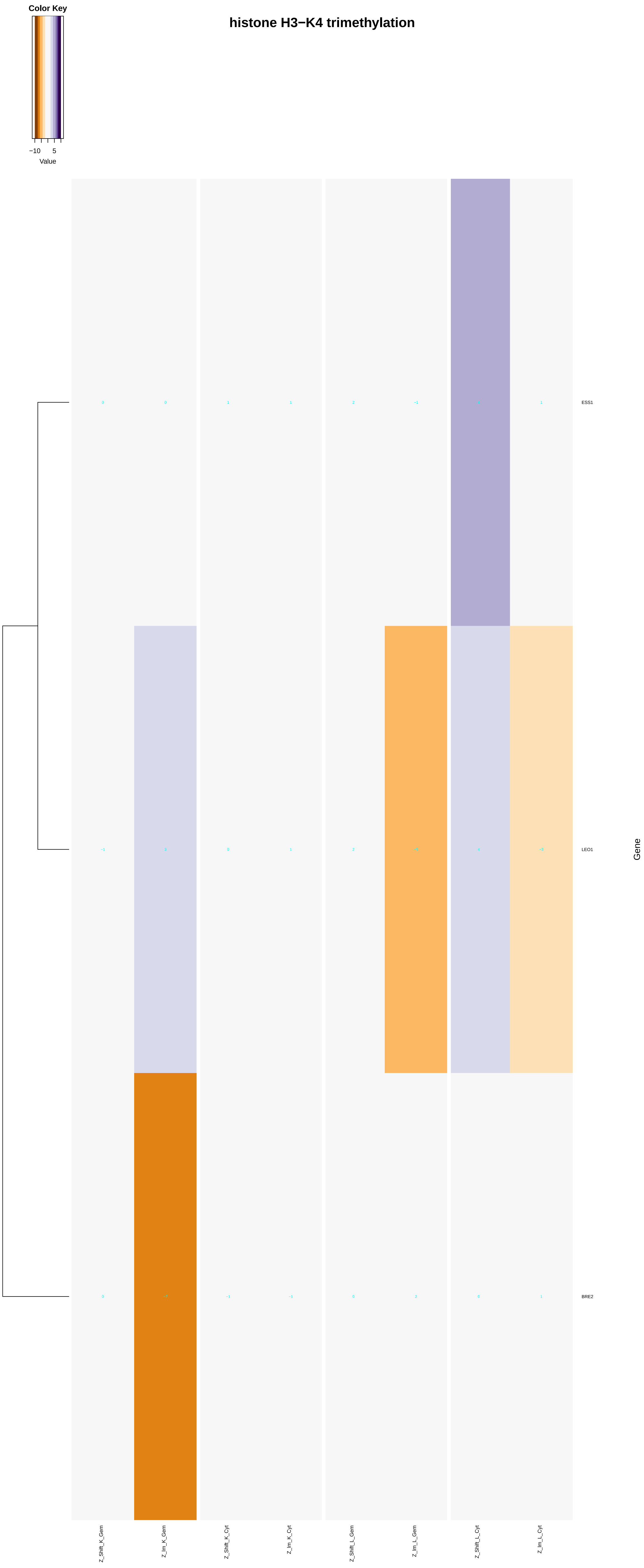
histone H3−K4 trimethylation
Color Key
−10
5
Value
ESS1
LEO1
BRE2
Gene
Z_Shift_K_Gem
Z_Im_K_Gem
Z_Shift_K_Cyt
Z_Im_K_Cyt
Z_Shift_L_Gem
Z_Im_L_Gem
Z_Shift_L_Cyt
Z_Im_L_Cyt
