## Additional File 7 for "A humanized yeast phenomic model of deoxycytidine kinase to predict genetic buffering of nucleoside analog cytotoxicity": chromosome_organization.pdf

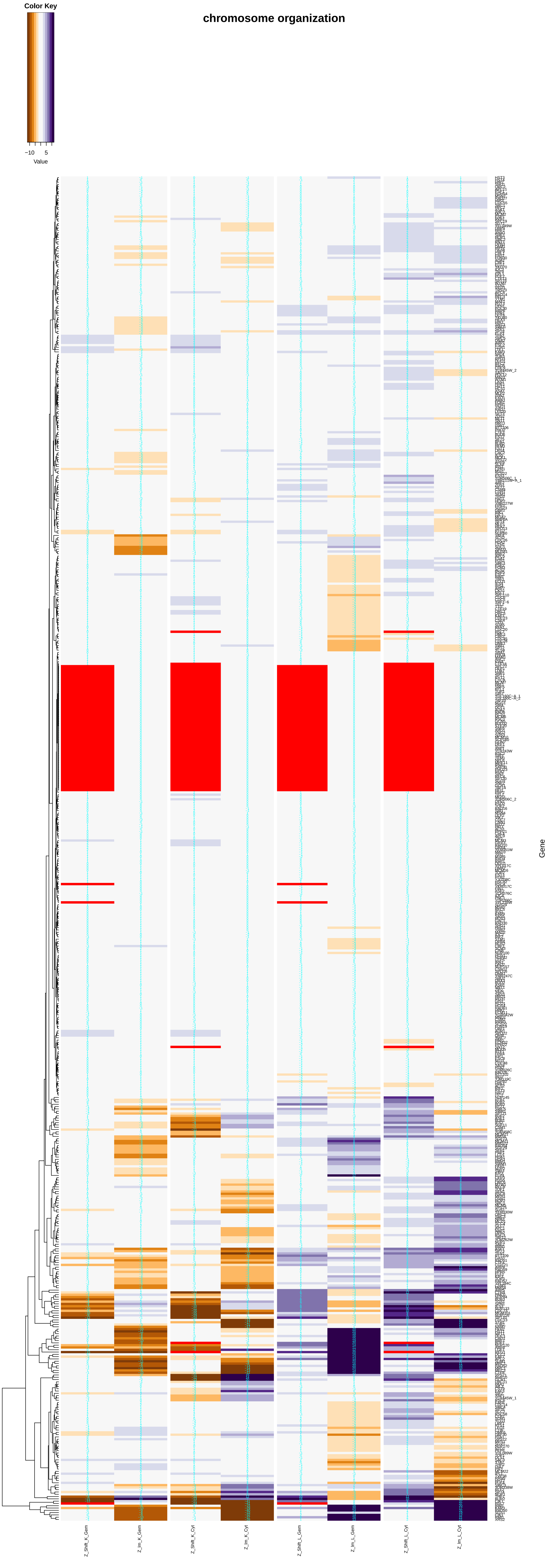

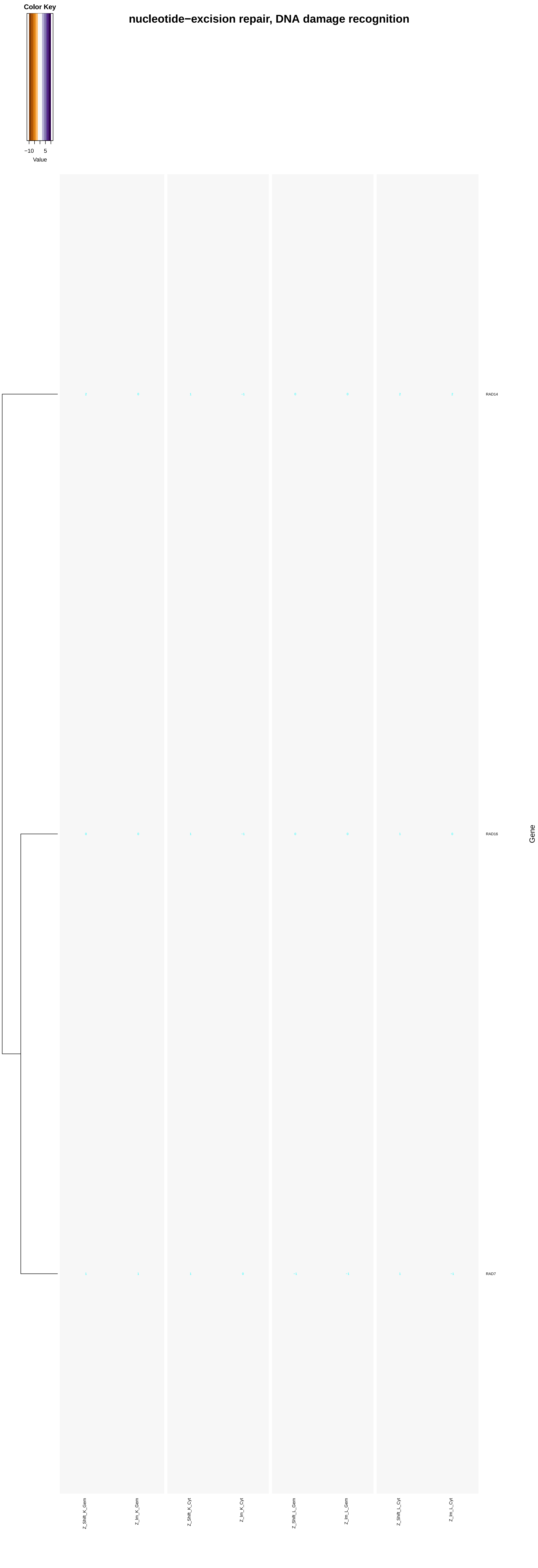
nucleotide−excision repair, DNA damage recognition
Color Key
−10
5
Value
2 0 1 −1 0 0 2 2
RAD14
0 0 1 −1 0 0 1 0
RAD16
1 1 1 0 −1 −1 1 −1
RAD7
Gene
Z_Shift_K_Gem
Z_Int_K_Gem
Z_Shift_K_Cyt
Z_Int_K_Cyt
Z_Shift_L_Gem
Z_Int_L_Gem
Z_Shift_L_Cyt
Z_Int_L_Cyt

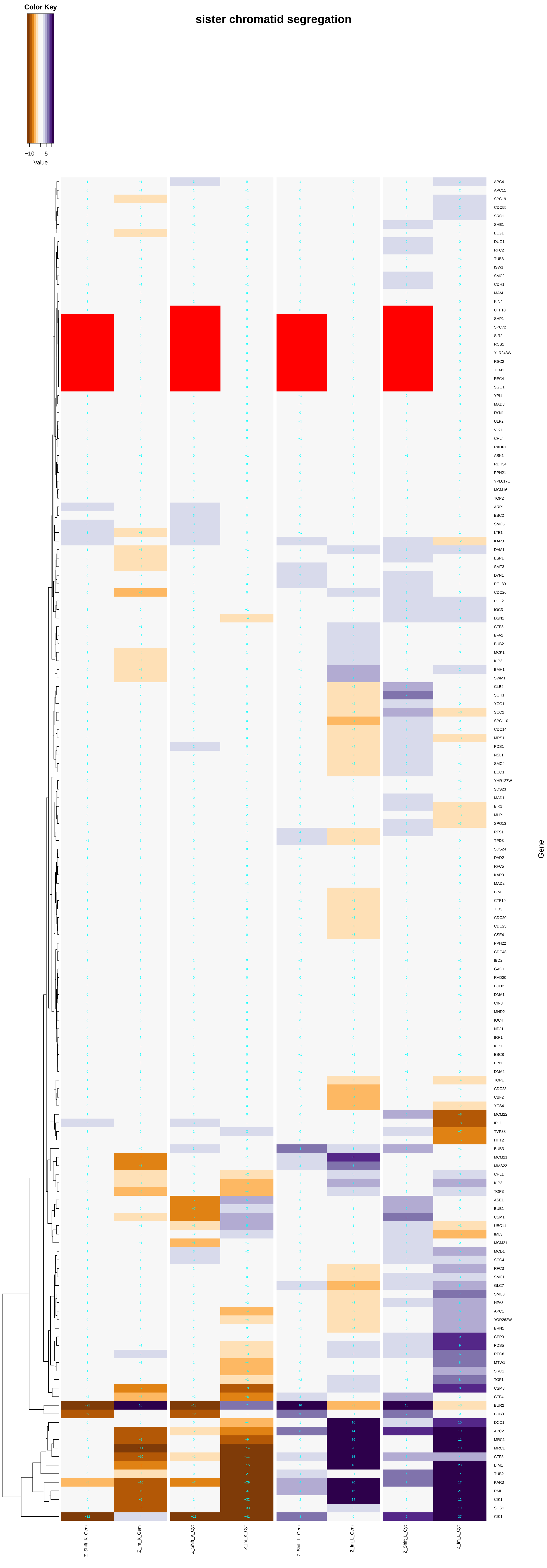
Color Key
−10
5
Value
sister chromatid segregation
APC4
APC11
SPC19
CDC55
SRC1
SHE1
ELG1
DUO1
RFC2
TUB3
ISW1
SMC2
CDH1
MAM1
KIN4
CTF18
SHP1
SPC72
SIR2
RCS1
YLR243W
RSC2
TEM1
RFC4
SGO1
YPI1
MAD3
DYN1
ULP2
VIK1
CHL4
RAD61
ASK1
RDH54
PPH21
YPL017C
MCM16
TOP2
ARP1
ESC2
SMC5
LTE1
KAR3
DAM1
ESP1
SMT3
DYN1
POL30
CDC26
POL2
IOC3
DSN1
CTF3
BFA1
BUB2
MCK1
KIP3
BMH1
SWM1
CLB2
SOH1
YCG1
SCC2
SPC110
CDC14
MPS1
PDS1
NSL1
SMC4
ECO1
YHR127W
SDS23
MAD1
BIK1
MLP1
SPO13
RTS1
TPD3
SDS24
DAD2
RFC5
KAR9
MAD2
BIM1
CTF19
TID3
CDC20
CDC23
CSE4
PPH22
CDC48
IBD2
GAC1
RAD30
BUD2
DMA1
CIN8
MND2
IOC4
NDJ1
IRR1
KIP1
ESC8
FIN1
DMA2
TOP1
CDC28
CBF2
YCS4
MCM22
IPL1
TVP38
HHT2
BUB3
MCM21
MMS22
CHL1
KIP3
TOP3
ASE1
BUB1
CSM1
UBC11
IML3
MCM21
MCD1
SCC4
RFC3
SMC1
GLC7
SMC3
NPA3
APC1
YOR262W
BRN1
CEP3
PDS5
REC8
MTW1
SRC1
TOF1
CSM3
CTF4
BUR2
BUB3
DCC1
APC2
MRC1
MRC1
CTF8
BIM1
TUB2
KAR3
RMI1
CIK1
SGS1
CIK1
Gene
Z_Shift_K_Gem
Z_Im_K_Gem
Z_Shift_K_Cyt
Z_Im_K_Cyt
Z_Shift_L_Gem
Z_Im_L_Gem
Z_Shift_L_Cyt
Z_Im_L_Cyt

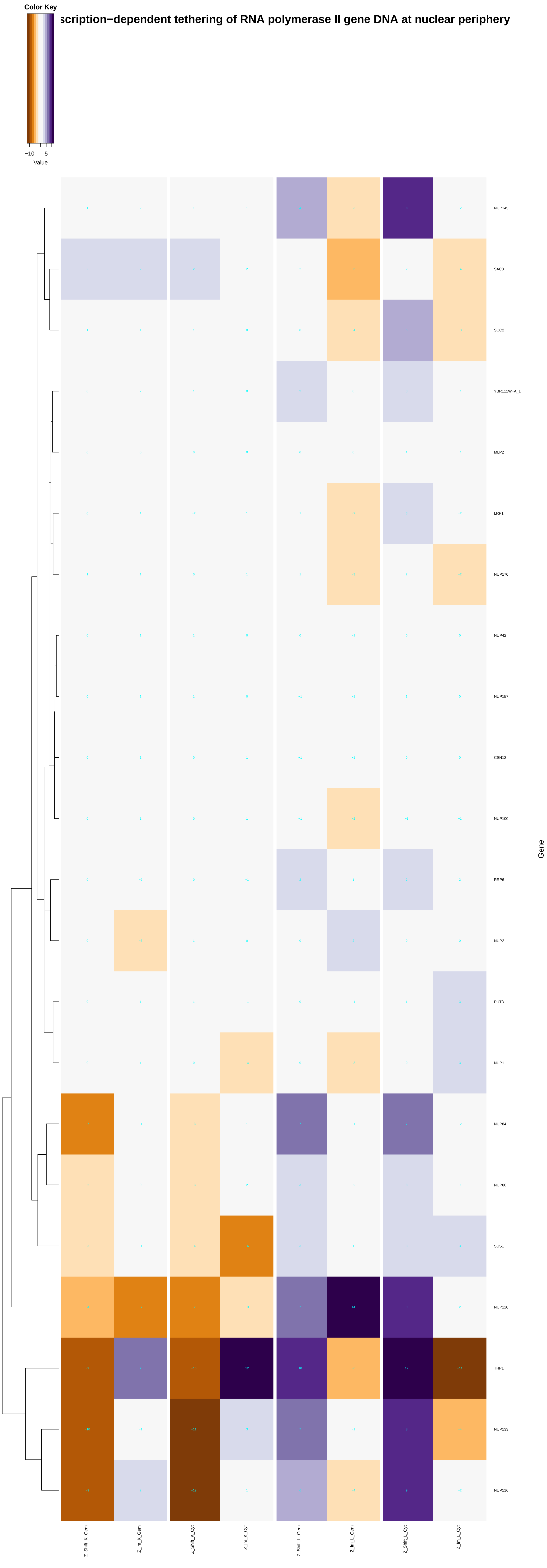

scription−dependent tethering of RNA polymerase II gene DNA at nuclear periphery
Color Key
−10
5
Value
NUP145
SAC3
SCC2
YBR111W−A_1
MLP2
LRP1
NUP170
NUP42
NUP157
CSN12
NUP100
RRP6
NUP2
PUT3
NUP1
NUP84
NUP60
SUS1
NUP120
THP1
NUP133
NUP116
Gene
Z_Shift_K_Gem
Z_Im_K_Gem
Z_Shift_K_Cyt
Z_Im_K_Cyt
Z_Shift_L_Gem
Z_Im_L_Gem
Z_Shift_L_Cyt
Z_Im_L_Cyt

### establishment of chromatin silencing

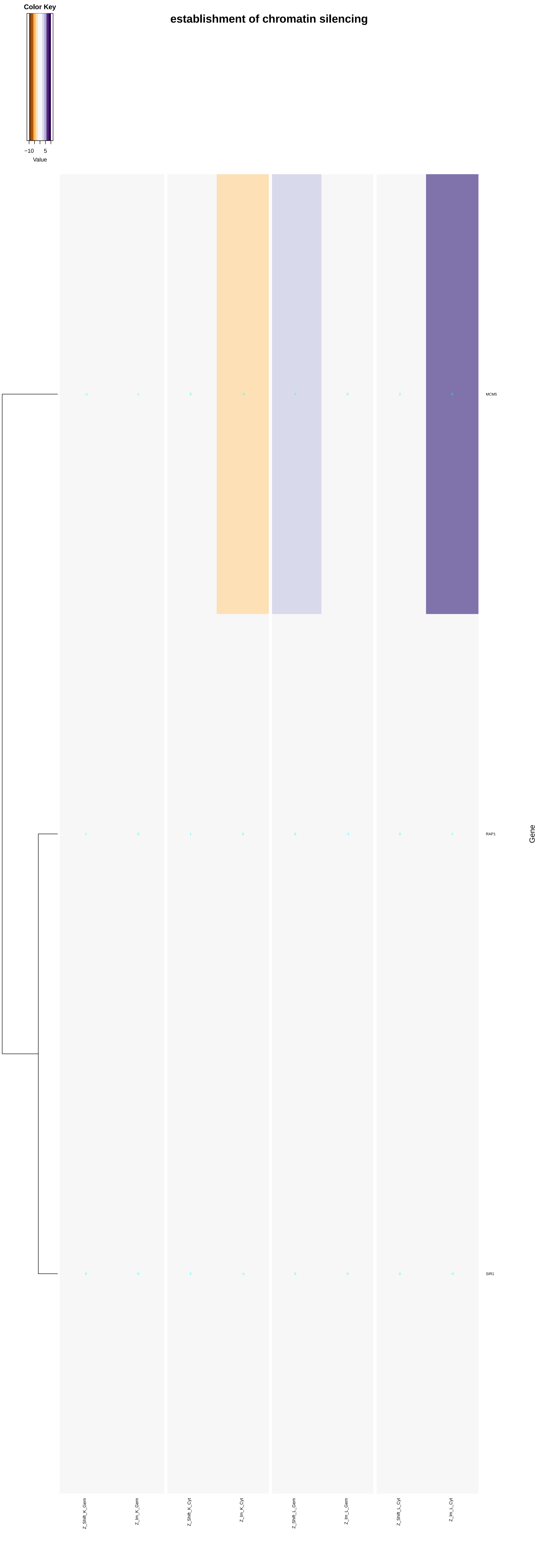

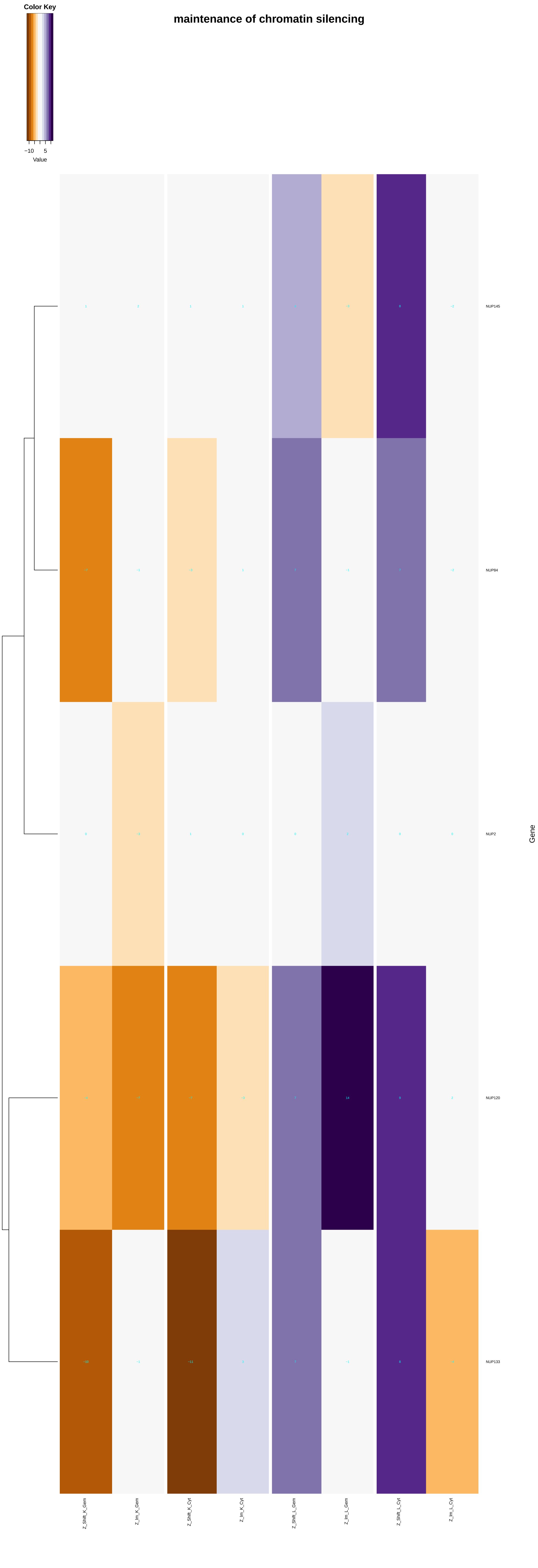
maintenance of chromatin silencing
Color Key
−10
5
Value
NUP145
NUP84
NUP2
NUP120
NUP133
Gene
Z_Shift_K_Gem
Z_Im_K_Gem
Z_Shift_K_Cyt
Z_Im_K_Cyt
Z_Shift_L_Gem
Z_Im_L_Gem
Z_Shift_L_Cyt
Z_Im_L_Cyt

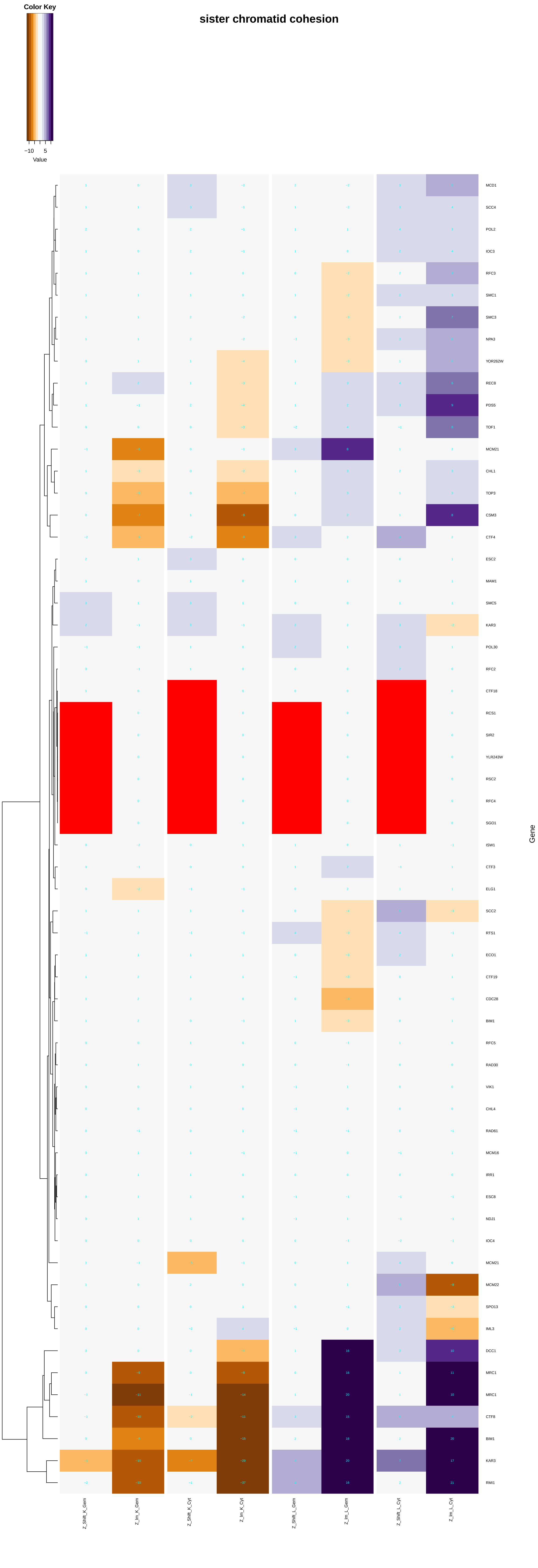

sister chromatid cohesion
Color Key
−10 5
Value
MCD1
SCC4
POL2
IOC3
RFC3
SMC1
SMC3
NPA3
YOR262W
REC8
PDS5
TOF1
MCM21
CHL1
TOP3
CSM3
CTF4
ESC2
MAM1
SMC5
KAR3
POL30
RFC2
CTF18
RCS1
SIR2
YLR243W
RSC2
RFC4
SGO1
ISW1
CTF3
ELG1
SCC2
RTS1
ECO1
CTF19
CDC28
BIM1
RFC5
RAD30
VIK1
CHL4
RAD61
MCM16
IRR1
ESC8
NDJ1
IOC4
MCM21
MCM22
SPO13
IML3
DCC1
MRC1
MRC1
CTF8
BIM1
KAR3
RMI1
Gene
Z_Shift_K_Gem
Z_lm_K_Gem
Z_Shift_K_Cyt
Z_lm_K_Cyt
Z_Shift_L_Gem
Z_lm_L_Gem
Z_Shift_L_Cyt
Z_lm_L_Cyt

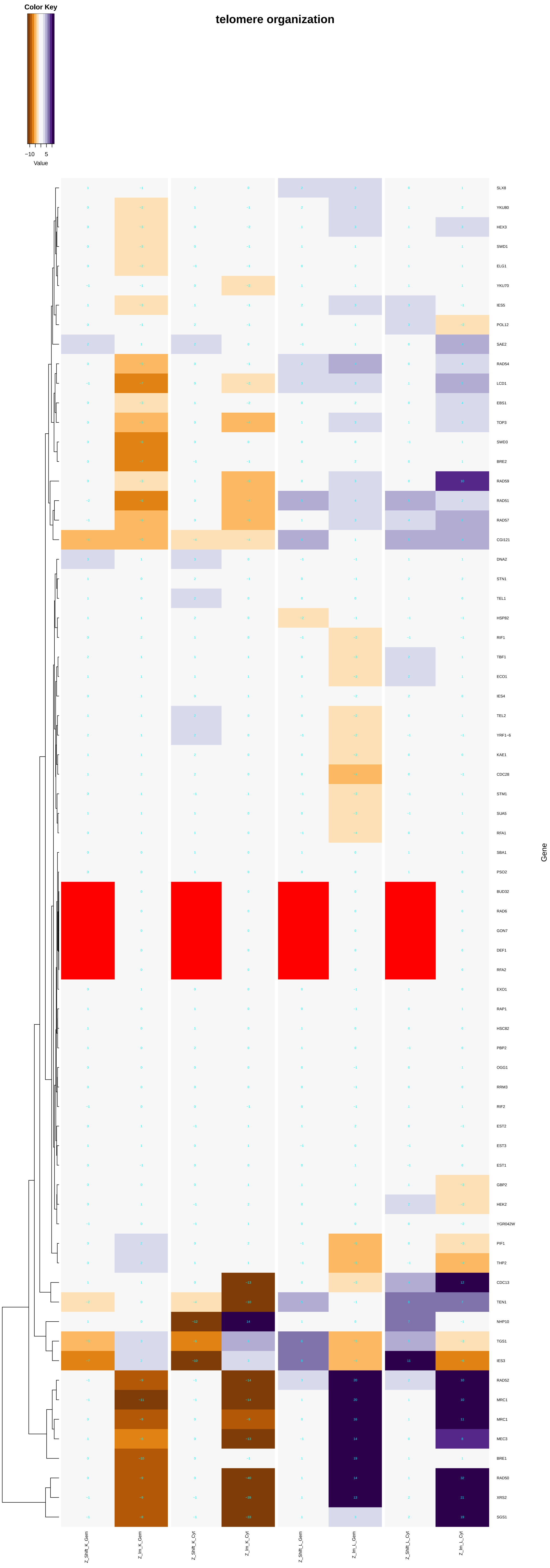
telomere organization
Color Key
−10
5
Value
SLX8
YKU80
HEX3
SWD1
ELG1
YKU70
IES5
POL12
SAE2
RAD54
LCD1
EBS1
TOP3
SWD3
BRE2
RAD59
RAD51
RAD57
CGI121
DNA2
STN1
TEL1
HSP82
RIF1
TBF1
ECO1
IES4
TEL2
YRF1−6
KAE1
CDC28
STM1
SUA5
RFA1
SBA1
PSO2
BUD32
RAD6
GON7
DEF1
RFA2
EXO1
RAP1
HSC82
PBP2
OGG1
RRM3
RIF2
EST2
EST3
EST1
GBP2
HEK2
YGR042W
PIF1
THP2
CDC13
TEN1
NHP10
TGS1
IES3
RAD52
MRC1
MRC1
MEC3
BRE1
RAD50
XRS2
SGS1
Gene
Z_Shift_K_Gem
Z_Im_K_Gem
Z_Shift_K_Cyt
Z_Im_K_Cyt
Z_Shift_L_Gem
Z_Im_L_Gem
Z_Shift_L_Cyt
Z_Im_L_Cyt

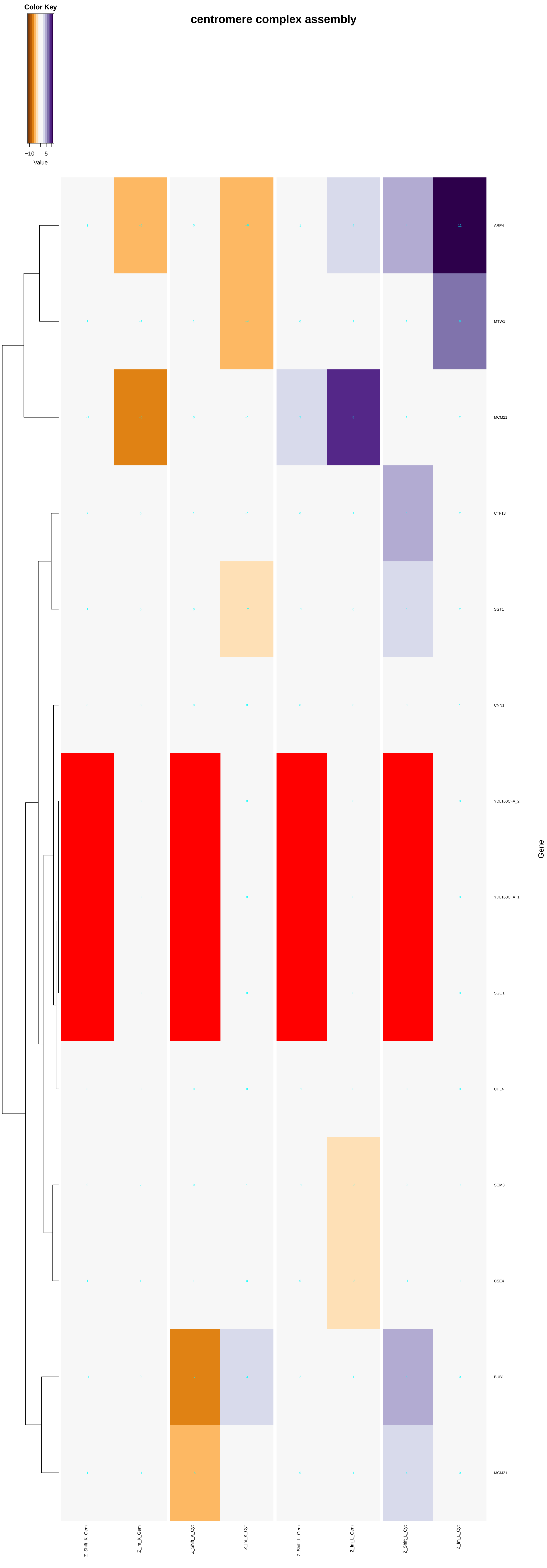
centromere complex assembly
Color Key
−10
5
Value
ARP4
MTW1
MCM21
CTF13
SGT1
CNN1
YDL160C−A_2
YDL160C−A_1
SGO1
CHL4
SCM3
CSE4
BUB1
MCM21
Gene
Z_Shift_K_Gem
Z_Im_K_Gem
Z_Shift_K_Cyt
Z_Im_K_Cyt
Z_Shift_L_Gem
Z_Im_L_Gem
Z_Shift_L_Cyt
Z_Im_L_Cyt

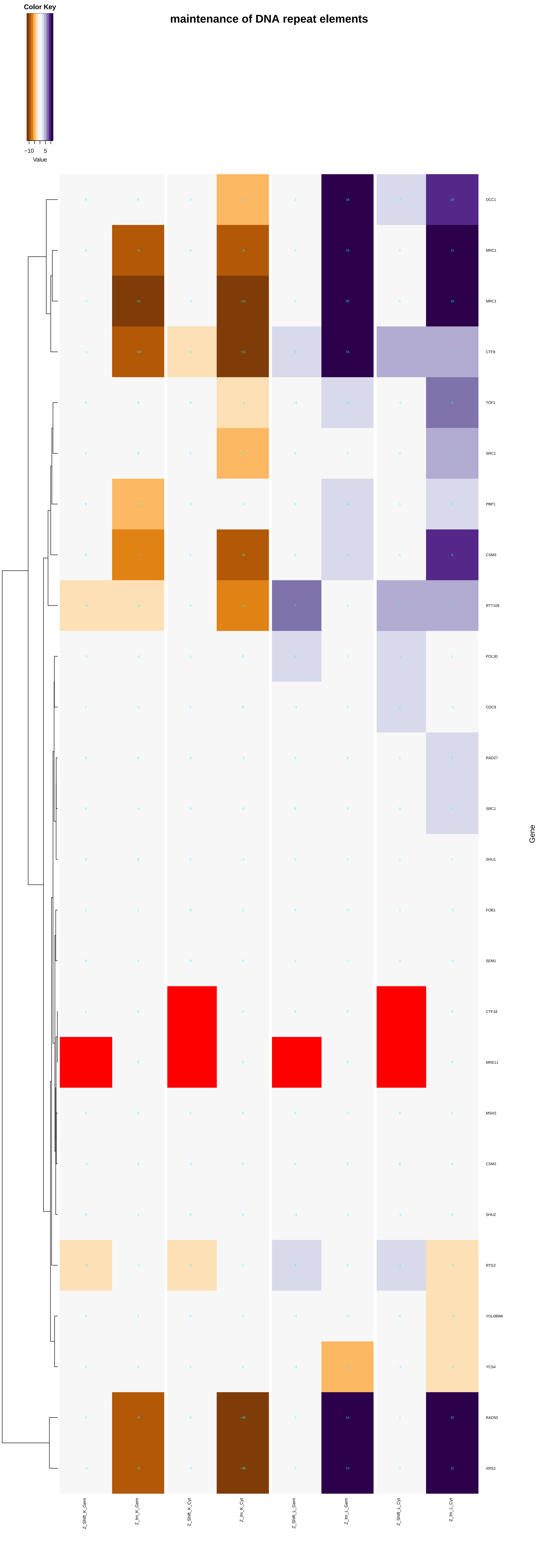
maintenance of DNA repeat elements
Color Key
−10
5
Value
DCC1
MRC1
MRC1
CTF8
TOF1
SRC1
PBP1
CSM3
RTT109
POL30
CDC9
RAD27
SRC1
SHU1
FOB1
SEM1
CTF18
MRE11
MSH2
CSM2
SHU2
RTG2
YDL089W
YCS4
RAD50
XRS2
Gene
Z_Shift_K_Gem
Z_lm_K_Gem
Z_Shift_K_Cyt
Z_lm_K_Cyt
Z_Shift_L_Gem
Z_lm_L_Gem
Z_Shift_L_Cyt
Z_lm_L_Cyt

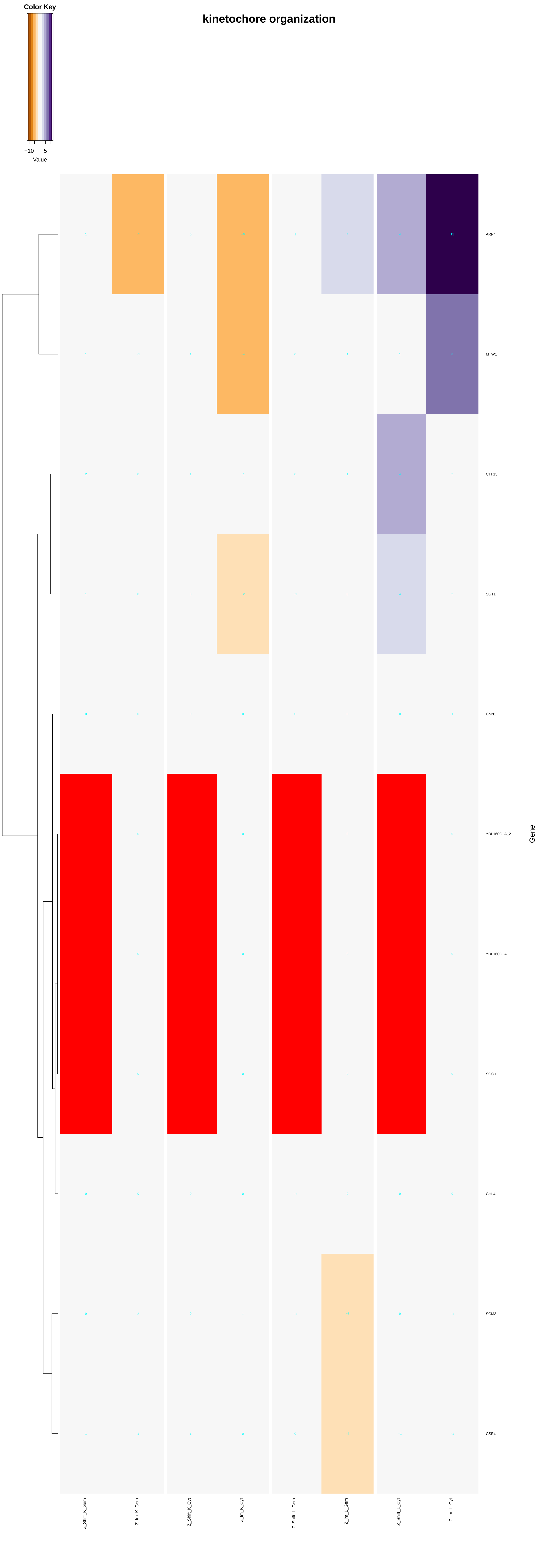

kinetochore organization
Color Key
−10
5
Value
ARP4
MTW1
CTF13
SGT1
CNN1
YDL160C−A_2
YDL160C−A_1
SGO1
CHL4
SCM3
CSE4
Gene
Z_Shift_K_Gem
Z_Im_K_Gem
Z_Shift_K_Cyt
Z_Im_K_Cyt
Z_Shift_L_Gem
Z_Im_L_Gem
Z_Shift_L_Cyt
Z_Im_L_Cyt

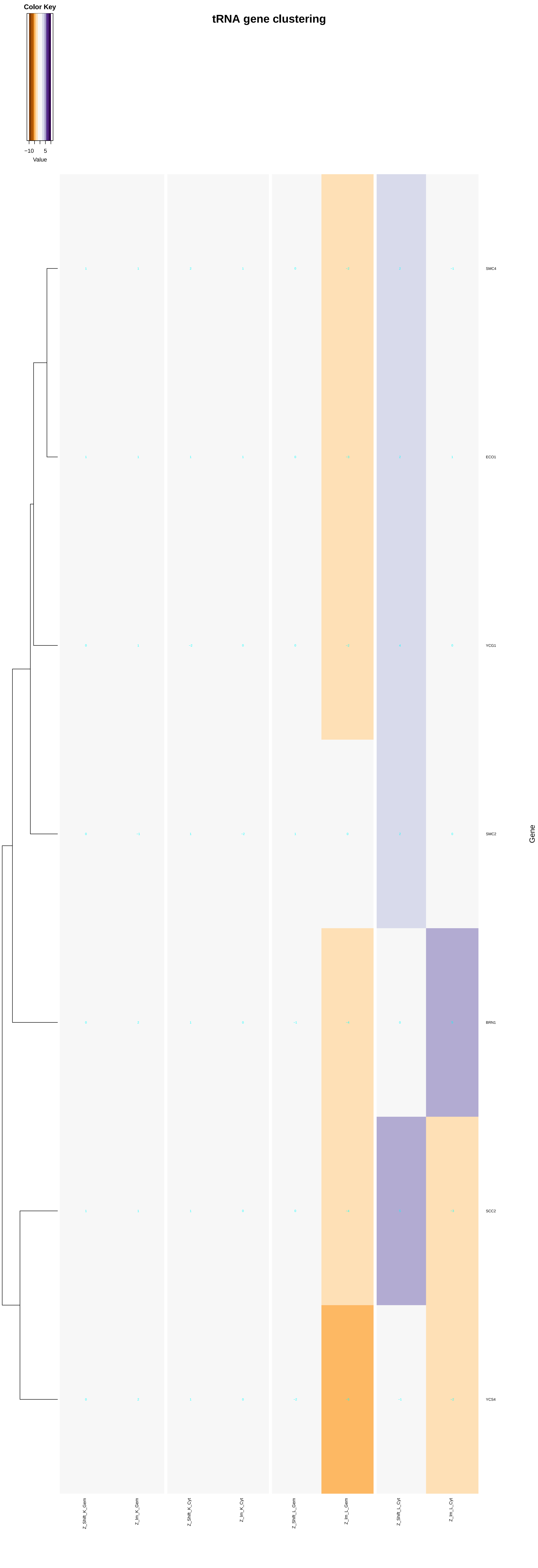

tRNA gene clustering
Color Key
−10
5
Value
SMC4
ECO1
YCG1
SMC2
BRN1
SCC2
YCS4
Gene
Z_Shift_K_Gem
Z_Im_K_Gem
Z_Shift_K_Cyt
Z_Im_K_Cyt
Z_Shift_L_Gem
Z_Im_L_Gem
Z_Shift_L_Cyt
Z_Im_L_Cyt

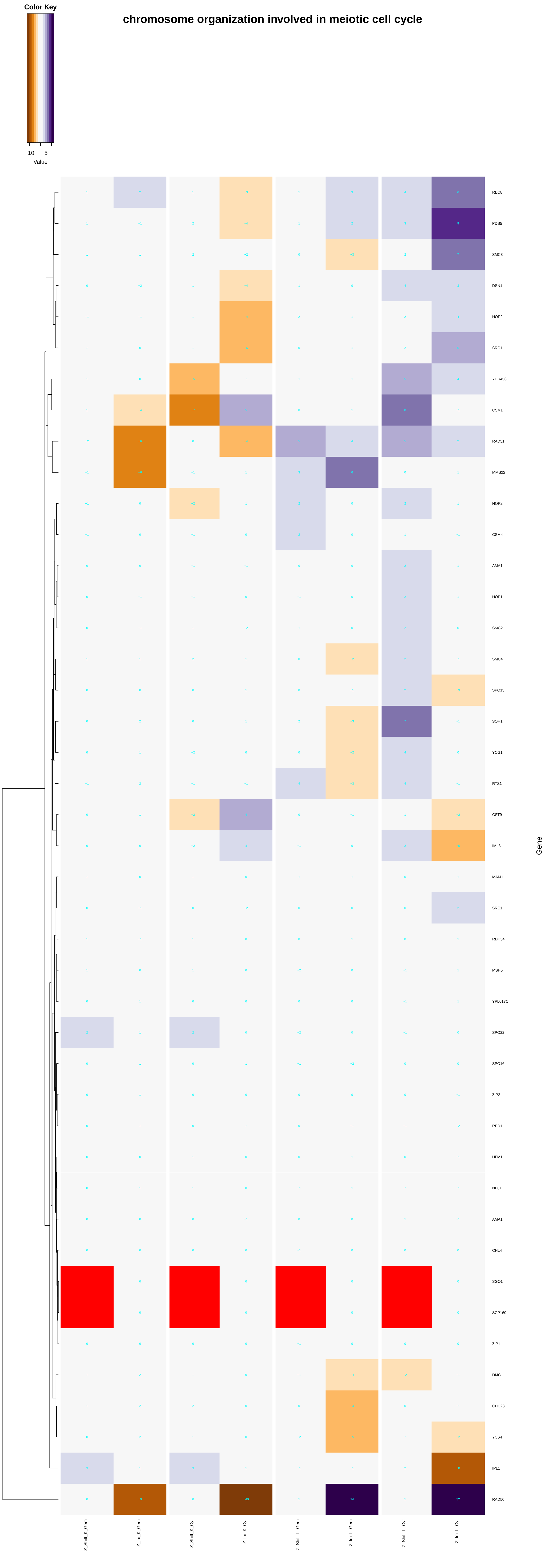
chromosome organization involved in meiotic cell cycle
Color Key
−10
5
Value
REC8
PDS5
SMC3
DSN1
HOP2
SRC1
YDR458C
CSM1
RAD51
MMS22
HOP2
CSM4
AMA1
HOP1
SMC2
SMC4
SPO13
SOH1
YCG1
RTS1
CST9
IML3
MAM1
SRC1
RDH54
MSH5
YPL017C
SPO22
SPO16
ZIP2
RED1
HFM1
NDJ1
AMA1
CHL4
SGO1
SCP160
ZIP1
DMC1
CDC28
YCS4
IPL1
RAD50
Gene
Z_Shift_K_Gem
Z_Im_K_Gem
Z_Shift_K_Cyt
Z_Im_K_Cyt
Z_Shift_L_Gem
Z_Im_L_Gem
Z_Shift_L_Cyt
Z_Im_L_Cyt

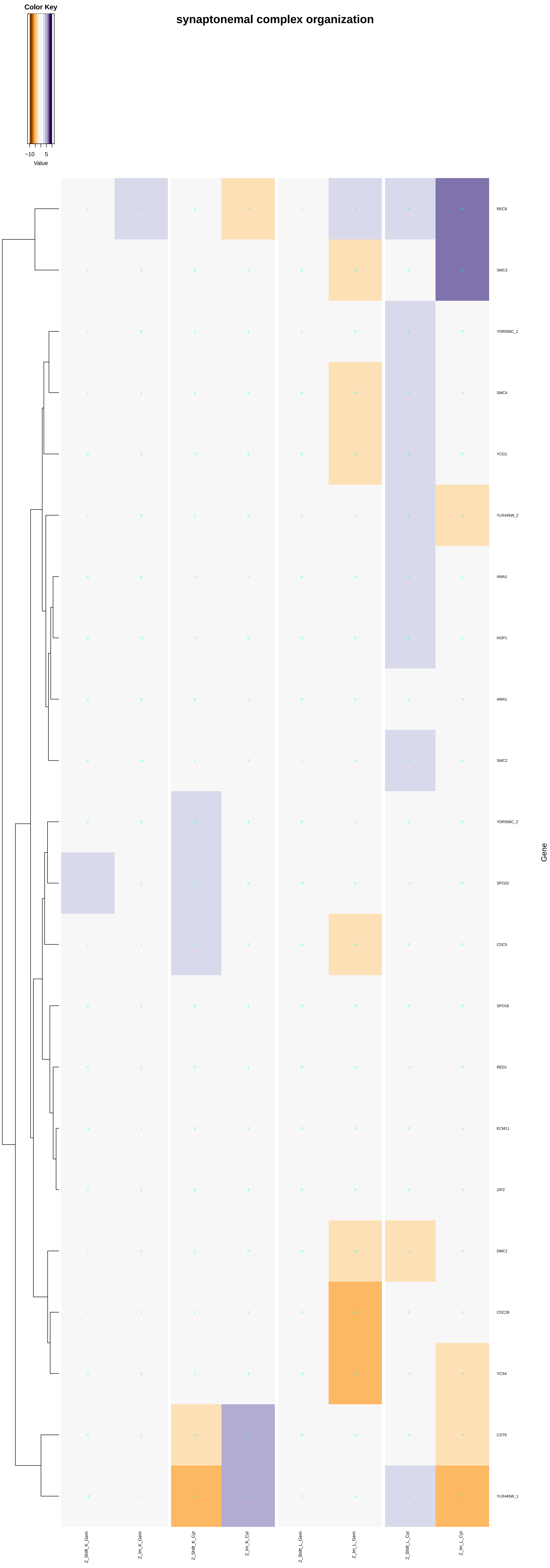
synaptonemal complex organization
Color Key
Value
Gene
REC8
SMC3
YDR506C_1
SMC4
YCG1
YLR445W_2
AMA1
HOP1
AMA1
SMC2
YDR506C_2
SPO22
CDC5
SPO16
RED1
ECM11
ZIP2
DMC1
CDC28
YCS4
CST9
YLR445W_1
Z_Shift_K_Gem
Z_Im_K_Gem
Z_Shift_K_Cyt
Z_Im_K_Cyt
Z_Shift_L_Gem
Z_Im_L_Gem
Z_Shift_L_Cyt
Z_Im_L_Cyt

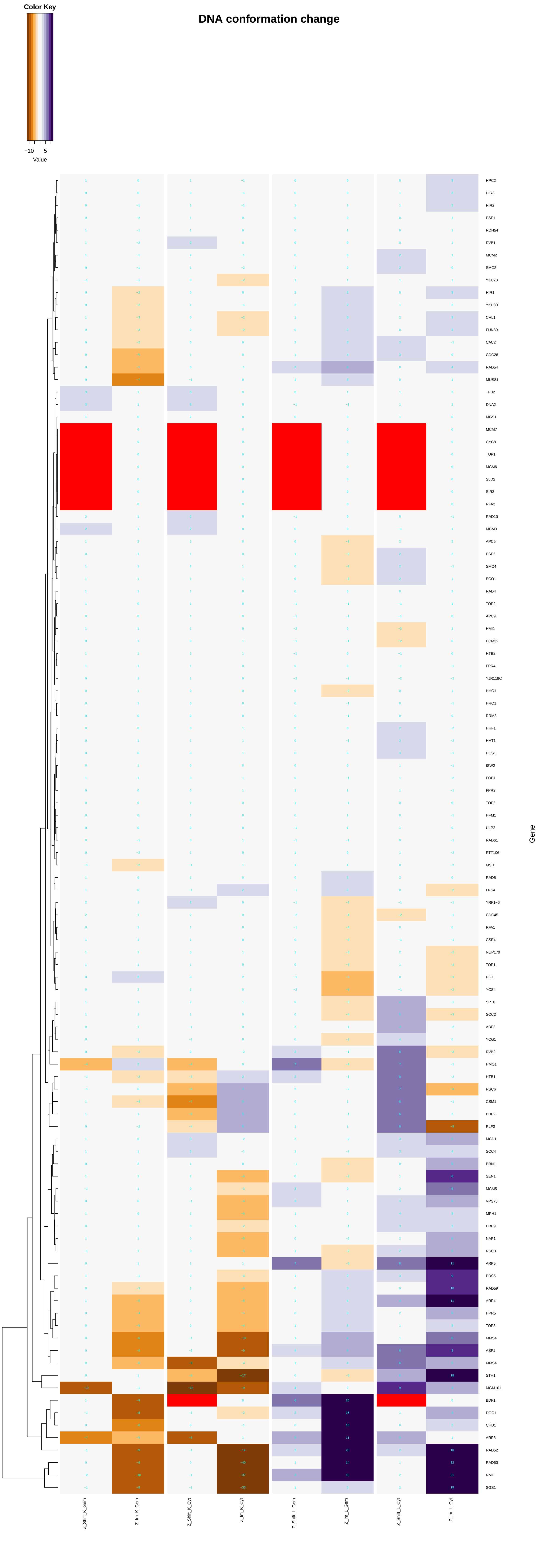
DNA conformation change
Color Key
−10
5
Value
HPC2
HIR3
HIR2
PSF1
RDH54
RVB1
MCM2
SMC2
YKU70
HIR1
YKU80
CHL1
FUN30
CAC2
CDC26
RAD54
MUS81
TFB2
DNA2
MGS1
MCM7
CYC8
TUP1
MCM6
SLD2
SIR3
RFA2
RAD10
MCM3
APC5
PSF2
SMC4
ECO1
RAD4
TOP2
APC9
HMI1
ECM32
HTB2
FPR4
YJR119C
HHO1
HRQ1
RRM3
HHF1
HHT1
HCS1
ISW2
FOB1
FPR3
TOF2
HFM1
ULP2
RAD61
RTT106
MSI1
RAD5
LRS4
YRF1−6
CDC45
RFA1
CSE4
NUP170
TOP1
PIF1
YCS4
SPT6
SCC2
ABF2
YCG1
RVB2
HMO1
HTB1
RSC6
CSM1
BDF2
RLF2
MCD1
SCC4
BRN1
SEN1
MCM5
VPS75
MPH1
DBP9
NAP1
RSC3
ARP5
PDS5
RAD59
ARP4
HPR5
TOP3
MMS4
ASF1
MMS4
STH1
MGM101
BDF1
DOC1
CHD1
ARP8
RAD52
RAD50
RMI1
SGS1
Gene
Z_Shift_K_Gem
Z_Im_K_Gem
Z_Shift_K_Cyt
Z_Im_K_Cyt
Z_Shift_L_Gem
Z_Im_L_Gem
Z_Shift_L_Cyt
Z_Im_L_Cyt

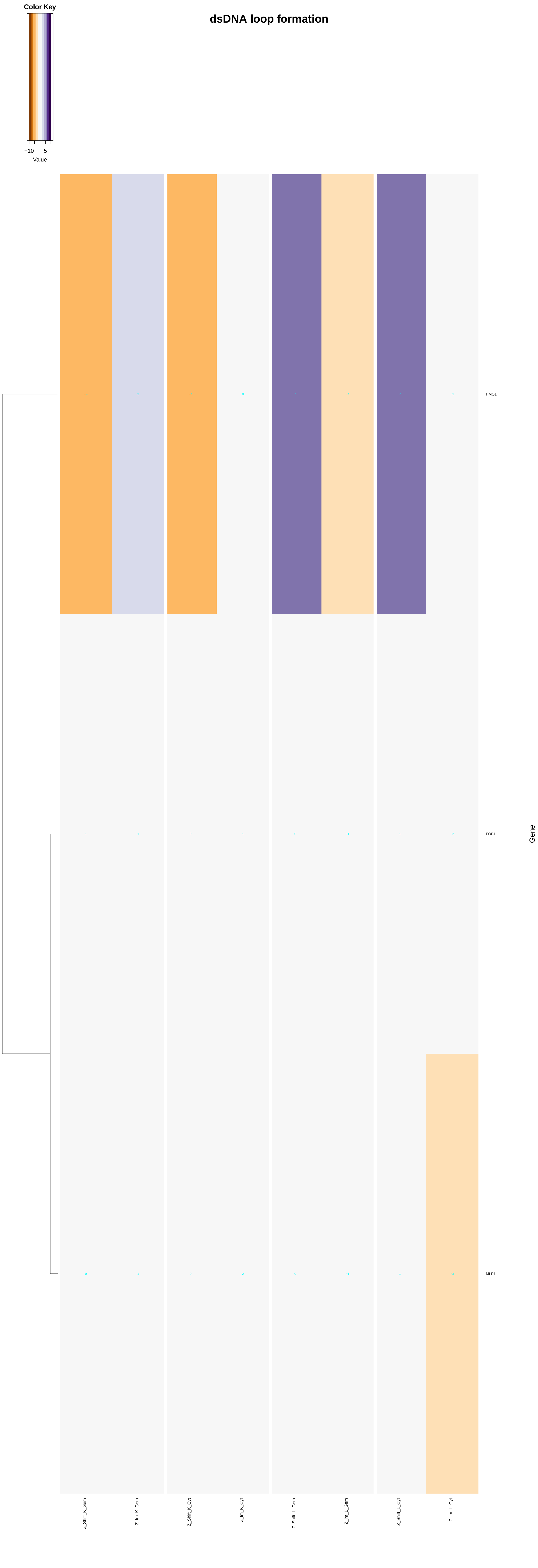

Color Key
−10
5
Value
dsDNA loop formation
HMO1
FOB1
MLP1
Gene
Z_Shift_K_Gem
Z_Im_K_Gem
Z_Shift_K_Cyt
Z_Im_K_Cyt
Z_Shift_L_Gem
Z_Im_L_Gem
Z_Shift_L_Cyt
Z_Im_L_Cyt

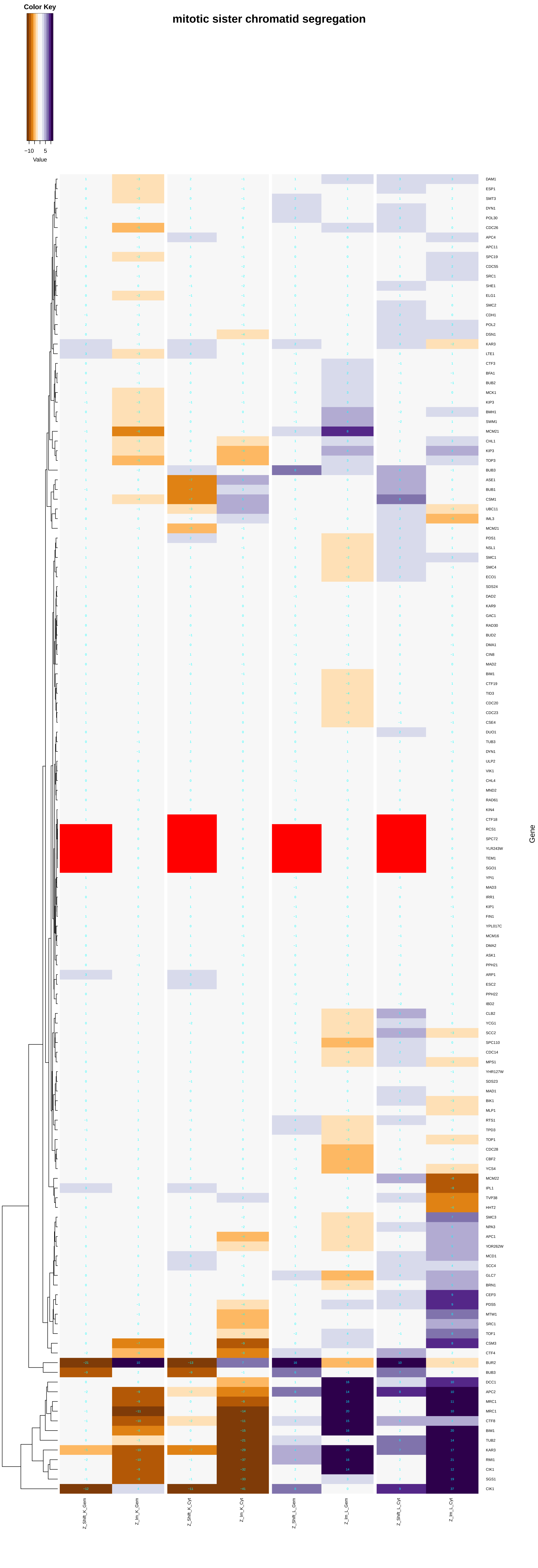
mitotic sister chromatid segregation
Color Key
−10
5
Value
Gene
DAM1
ESP1
SMT3
DYN1
POL30
CDC26
APC4
APC11
SPC19
CDC55
SRC1
SHE1
ELG1
SMC2
CDH1
POL2
DSN1
KAR3
LTE1
CTF3
BFA1
BUB2
MCK1
KIP3
BMH1
SWM1
MCM21
CHL1
KIP3
TOP3
BUB3
ASE1
BUB1
CSM1
UBC11
IML3
MCM21
PDS1
NSL1
SMC1
SMC4
ECO1
SDS24
DAD2
KAR9
GAC1
RAD30
BUD2
DMA1
CIN8
MAD2
BIM1
CTF19
TID3
CDC20
CDC23
CSE4
DUO1
TUB3
DYN1
ULP2
VIK1
CHL4
MND2
RAD61
KIN4
CTF18
RCS1
SPC72
YLR243W
TEM1
SGO1
YPI1
MAD3
IRR1
KIP1
FIN1
YPL017C
MCM16
DMA2
ASK1
PPH21
ARP1
ESC2
PPH22
IBD2
CLB2
YCG1
SCC2
SPC110
CDC14
MPS1
YHR127W
SDS23
MAD1
BIK1
MLP1
RTS1
TPD3
TOP1
CDC28
CBF2
YCS4
MCM22
IPL1
TVP38
HHT2
SMC3
NPA3
APC1
YOR262W
MCD1
SCC4
GLC7
BRN1
CEP3
PDS5
MTW1
SRC1
TOF1
CSM3
CTF4
BUR2
BUB3
DCC1
APC2
MRC1
MRC1
CTF8
BIM1
TUB2
KAR3
RMI1
CIK1
SGS1
CIK1
Z_Shift_K_Gem
Z_Im_K_Gem
Z_Shift_K_Cyt
Z_Im_K_Cyt
Z_Shift_L_Gem
Z_Im_L_Gem
Z_Shift_L_Cyt
Z_Im_L_Cyt

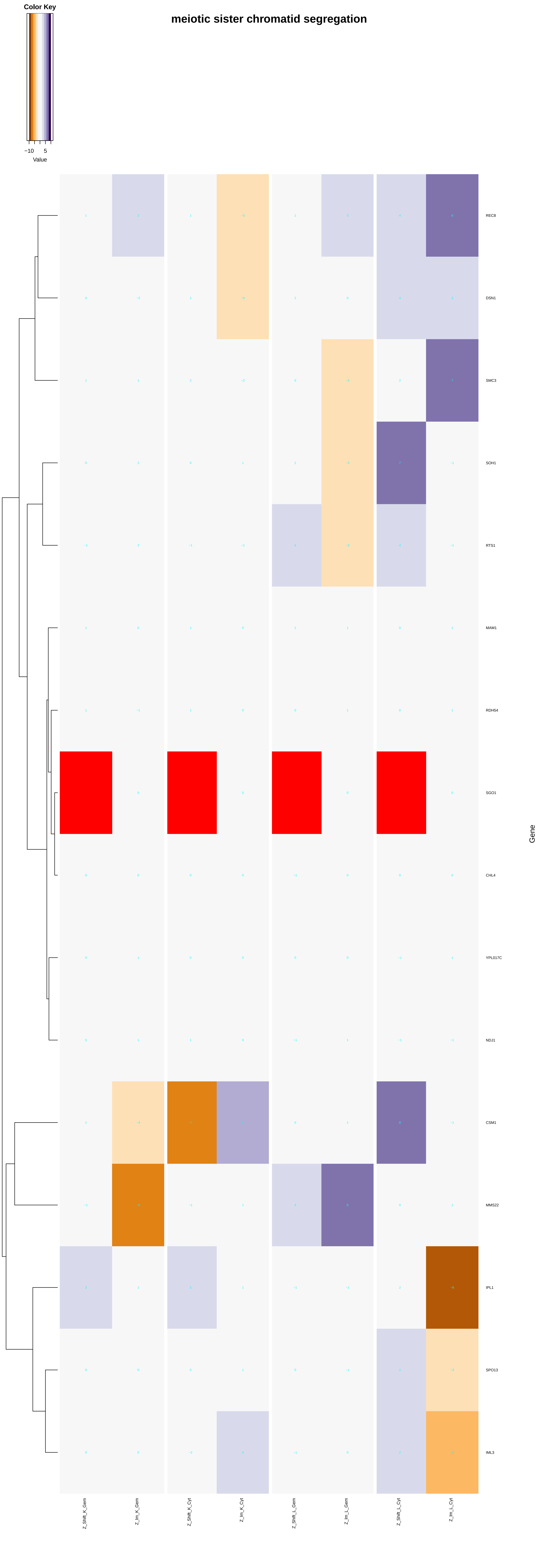

meiotic sister chromatid segregation
Color Key
−10
5
Value
REC8
DSN1
SMC3
SOH1
RTS1
MAM1
RDH54
SGO1
CHL4
YPL017C
NDJ1
CSM1
MMS22
IPL1
SPO13
IML3
Gene
Z_Shift_K_Gem
Z_Im_K_Gem
Z_Shift_K_Cyt
Z_Im_K_Cyt
Z_Shift_L_Gem
Z_Im_L_Gem
Z_Shift_L_Cyt
Z_Im_L_Cyt

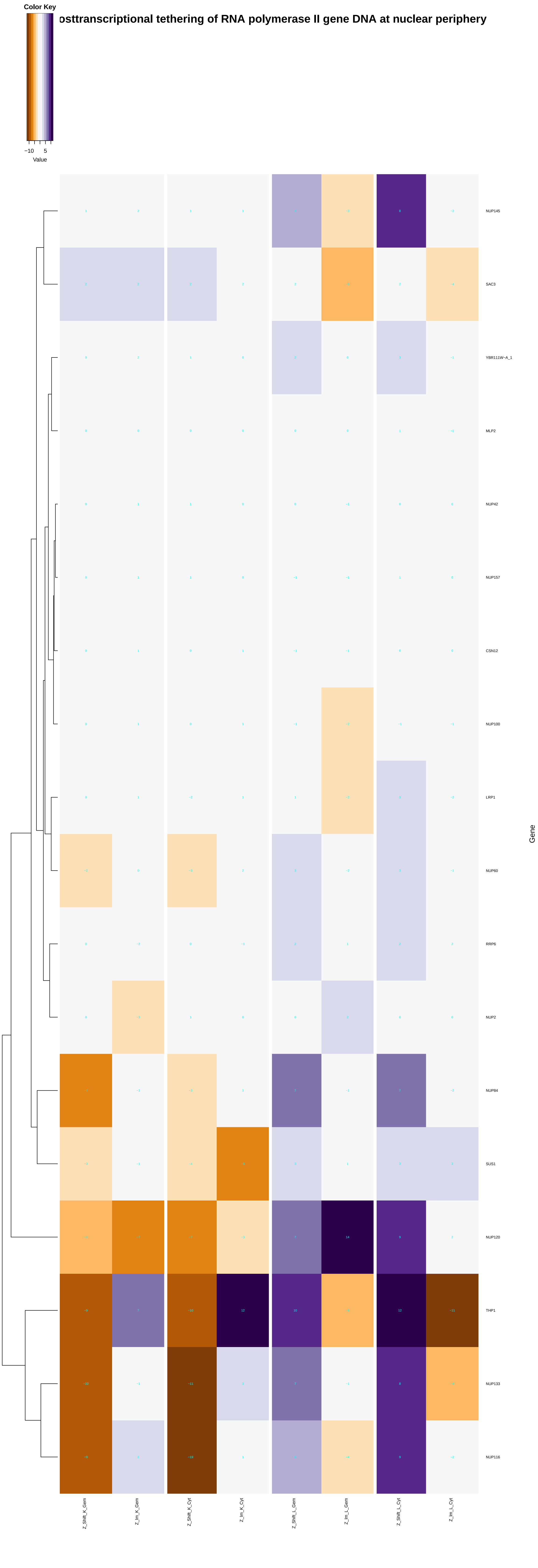

Color Key
−10
5
Value
osttranscriptional tethering of RNA polymerase II gene DNA at nuclear periphery
NUP145
SAC3
YBR111W−A_1
MLP2
NUP42
NUP157
CSN12
NUP100
LRP1
NUP60
RRP6
NUP2
NUP84
SUS1
NUP120
THP1
NUP133
NUP116
Gene
Z_Shift_K_Gem
Z_Im_K_Gem
Z_Shift_K_Cyt
Z_Im_K_Cyt
Z_Shift_L_Gem
Z_Im_L_Gem
Z_Shift_L_Cyt
Z_Im_L_Cyt

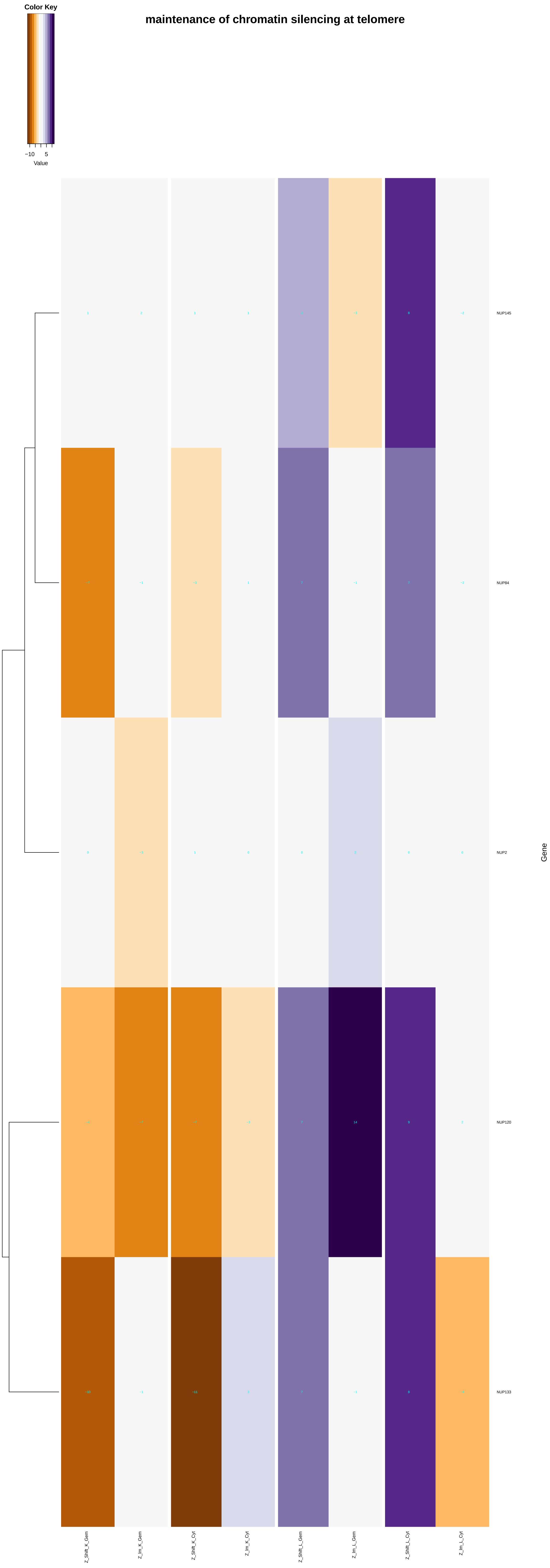
maintenance of chromatin silencing at telomere
Color Key
−10
5
Value
NUP145
NUP84
NUP2
NUP120
NUP133
Gene
Z_Shift_K_Gem
Z_Im_K_Gem
Z_Shift_K_Cyt
Z_Im_K_Cyt
Z_Shift_L_Gem
Z_Im_L_Gem
Z_Shift_L_Cyt
Z_Im_L_Cyt

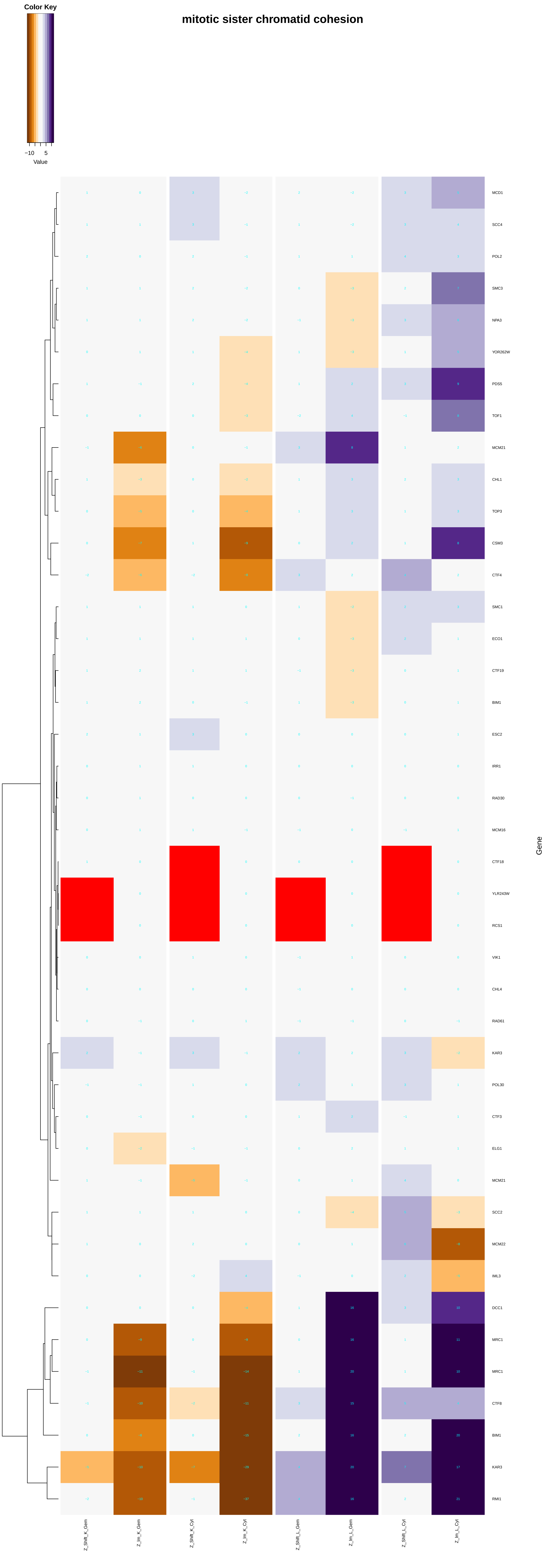
mitotic sister chromatid cohesion
Color Key
−10
5
Value
MCD1
SCC4
POL2
SMC3
NPA3
YOR262W
PDS5
TOF1
MCM21
CHL1
TOP3
CSM3
CTF4
SMC1
ECO1
CTF19
BIM1
ESC2
IRR1
RAD30
MCM16
CTF18
YLR243W
RCS1
VIK1
CHL4
RAD61
KAR3
POL30
CTF3
ELG1
MCM21
SCC2
MCM22
IML3
DCC1
MRC1
MRC1
CTF8
BIM1
KAR3
RMI1
Gene
Z_Shift_K_Gem
Z_Im_K_Gem
Z_Shift_K_Cyt
Z_Im_K_Cyt
Z_Shift_L_Gem
Z_Im_L_Gem
Z_Shift_L_Cyt
Z_Im_L_Cyt

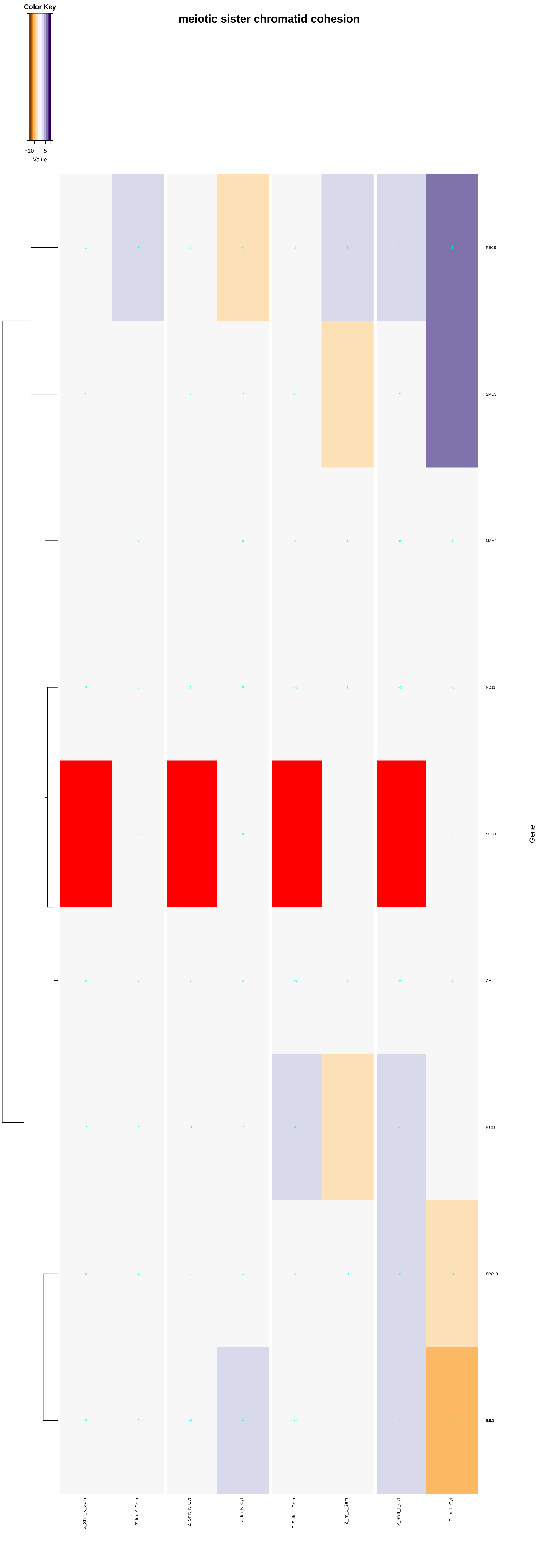
meiotic sister chromatid cohesion
Color Key
−10
5
Value
REC8
SMC3
MAM1
NDJ1
SGO1
CHL4
RTS1
SPO13
IML3
Gene
Z_Shift_K_Gem
Z_Im_K_Gem
Z_Shift_K_Cyt
Z_Im_K_Cyt
Z_Shift_L_Gem
Z_Im_L_Gem
Z_Shift_L_Cyt
Z_Im_L_Cyt

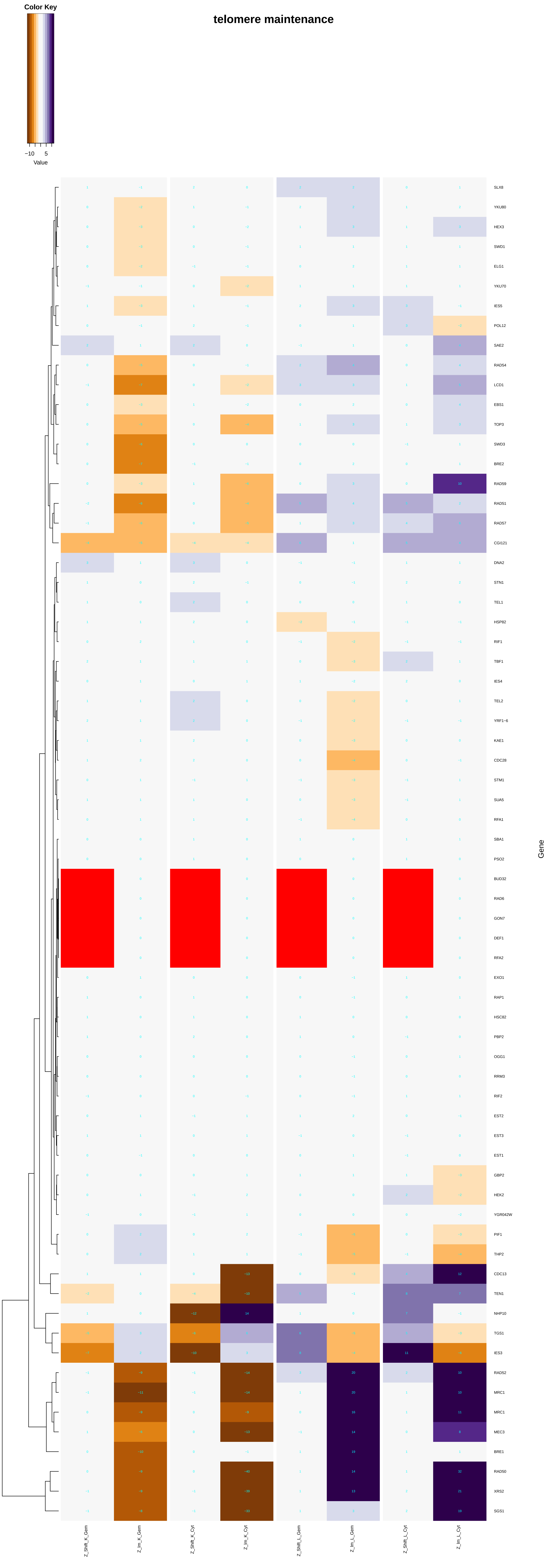

telomere maintenance
Color Key
−10
5
Value
Gene
SLX8
YKU80
HEX3
SWD1
ELG1
YKU70
IES5
POL12
SAE2
RAD54
LCD1
EBS1
TOP3
SWD3
BRE2
RAD59
RAD51
RAD57
CGI121
DNA2
STN1
TEL1
HSP82
RIF1
TBF1
IES4
TEL2
YRF1−6
KAE1
CDC28
STM1
SUA5
RFA1
SBA1
PSO2
BUD32
RAD6
GON7
DEF1
RFA2
EXO1
RAP1
HSC82
PBP2
OGG1
RRM3
RIF2
EST2
EST3
EST1
GBP2
HEK2
YGR042W
PIF1
THP2
CDC13
TEN1
NHP10
TGS1
IES3
RAD52
MRC1
MRC1
MEC3
BRE1
RAD50
XRS2
SGS1
Z_Shift_K_Gem
Z_lm_K_Gem
Z_Shift_K_Cyt
Z_lm_K_Cyt
Z_Shift_L_Gem
Z_lm_L_Gem
Z_Shift_L_Cyt
Z_lm_L_Cyt

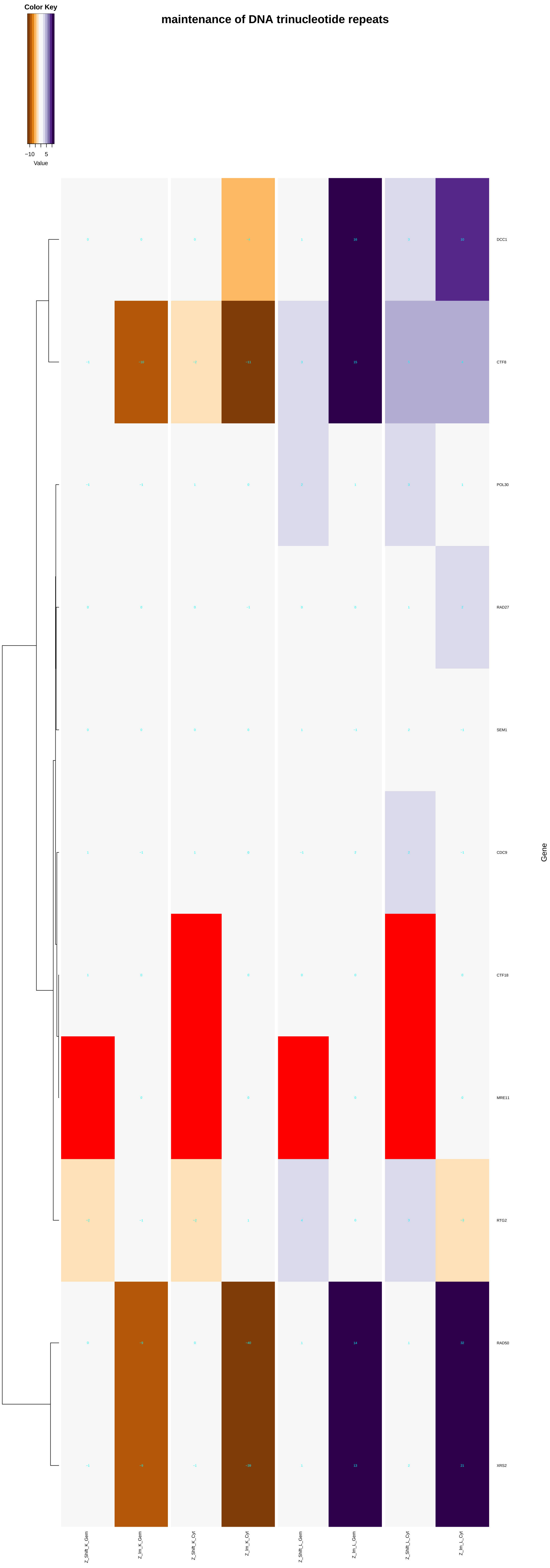
maintenance of DNA trinucleotide repeats
Color Key
−10
5
Value
DCC1
CTF8
POL30
RAD27
SEM1
CDC9
CTF18
MRE11
RTG2
RAD50
XRS2
Gene
Z_Shift_K_Gem
Z_Im_K_Gem
Z_Shift_K_Cyt
Z_Im_K_Cyt
Z_Shift_L_Gem
Z_Im_L_Gem
Z_Shift_L_Cyt
Z_Im_L_Cyt

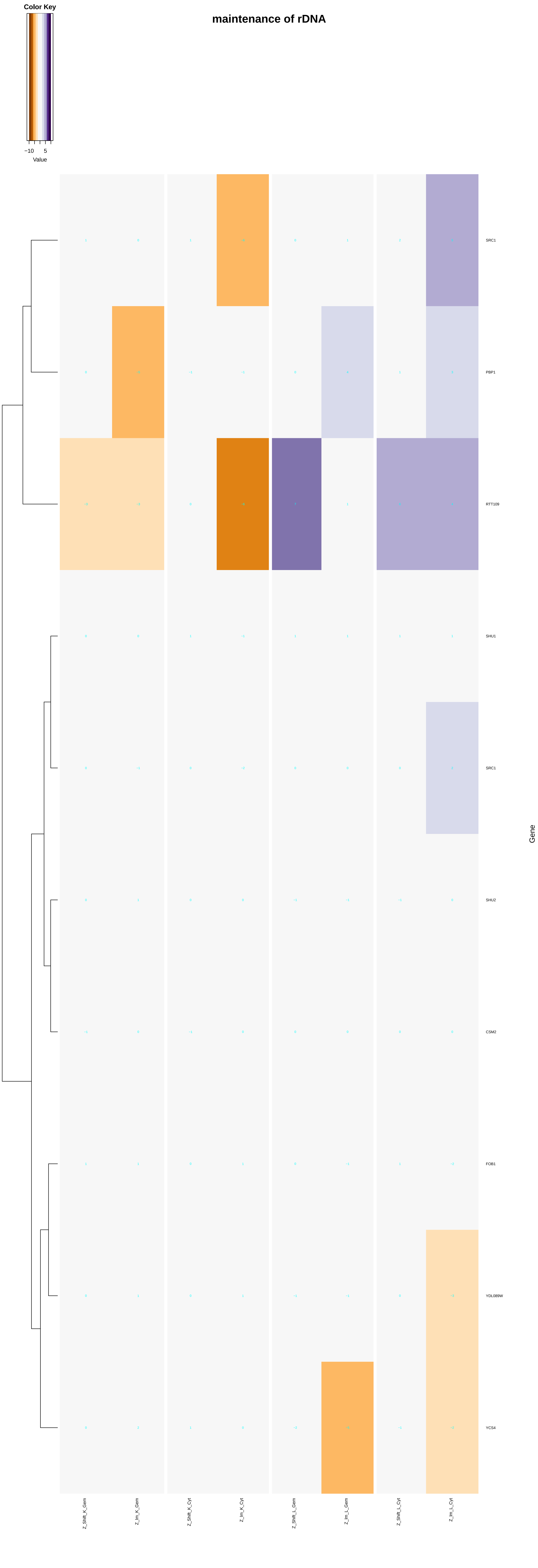

Color Key
−10
5
Value
maintenance of rDNA
SRC1
PBP1
RTT109
SHU1
SRC1
SHU2
CSM2
FOB1
YDL089W
YCS4
Gene
Z_Shift_K_Gem
Z_Im_K_Gem
Z_Shift_K_Cyt
Z_Im_K_Cyt
Z_Shift_L_Gem
Z_Im_L_Gem
Z_Shift_L_Cyt
Z_Im_L_Cyt

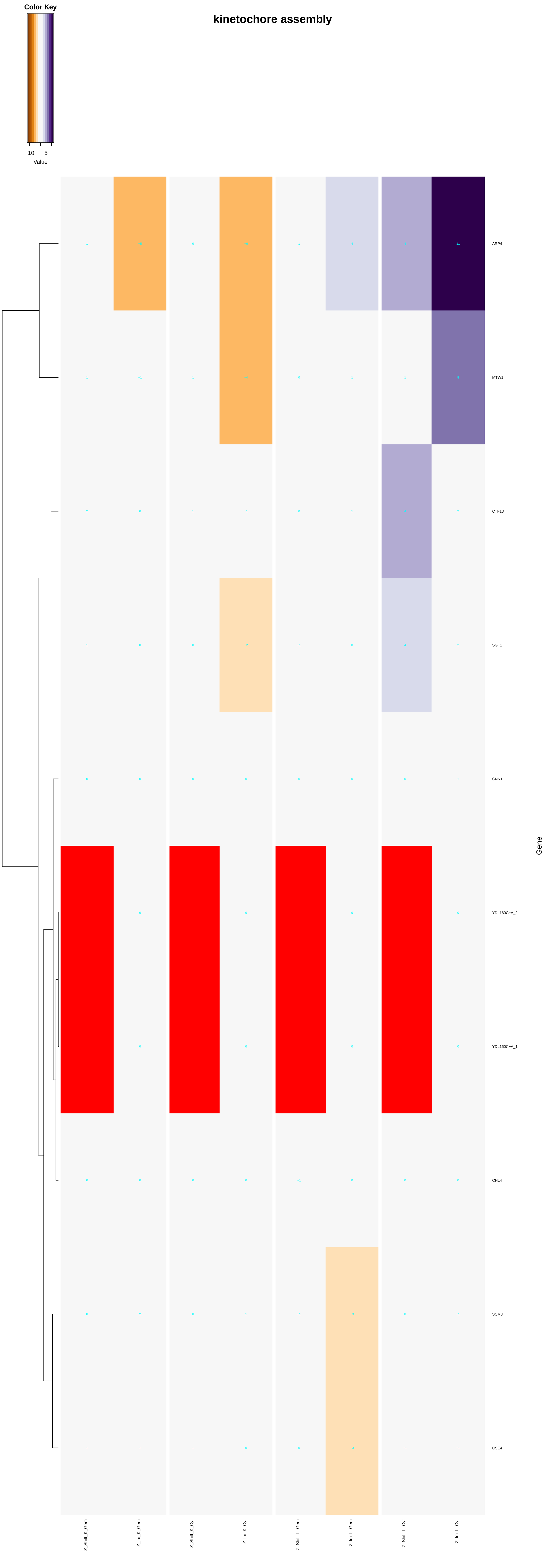
Color Key
−10
5
Value
kinetochore assembly
ARP4
MTW1
CTF13
SGT1
CNN1
YDL160C−A_2
YDL160C−A_1
CHL4
SCM3
CSE4
Gene
Z_Shift_K_Gem
Z_Im_K_Gem
Z_Shift_K_Cyt
Z_Im_K_Cyt
Z_Shift_L_Gem
Z_Im_L_Gem
Z_Shift_L_Cyt
Z_Im_L_Cyt

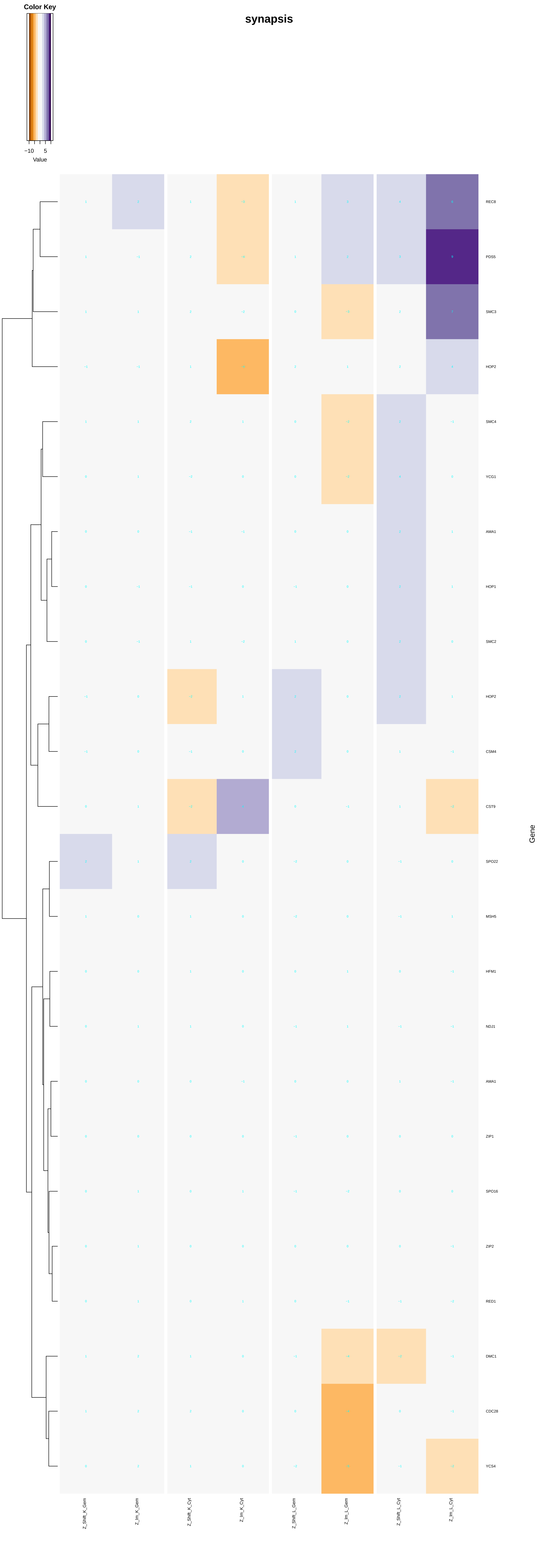
Color Key
−10
5
Value
synapsis
REC8
PDS5
SMC3
HOP2
SMC4
YCG1
AMA1
HOP1
SMC2
HOP2
CSM4
CST9
SPO22
MSH5
HFM1
NDJ1
AMA1
ZIP1
SPO16
ZIP2
RED1
DMC1
CDC28
YCS4
Gene
Z_Shift_K_Gem
Z_Im_K_Gem
Z_Shift_K_Cyt
Z_Im_K_Cyt
Z_Shift_L_Gem
Z_Im_L_Gem
Z_Shift_L_Cyt
Z_Im_L_Cyt

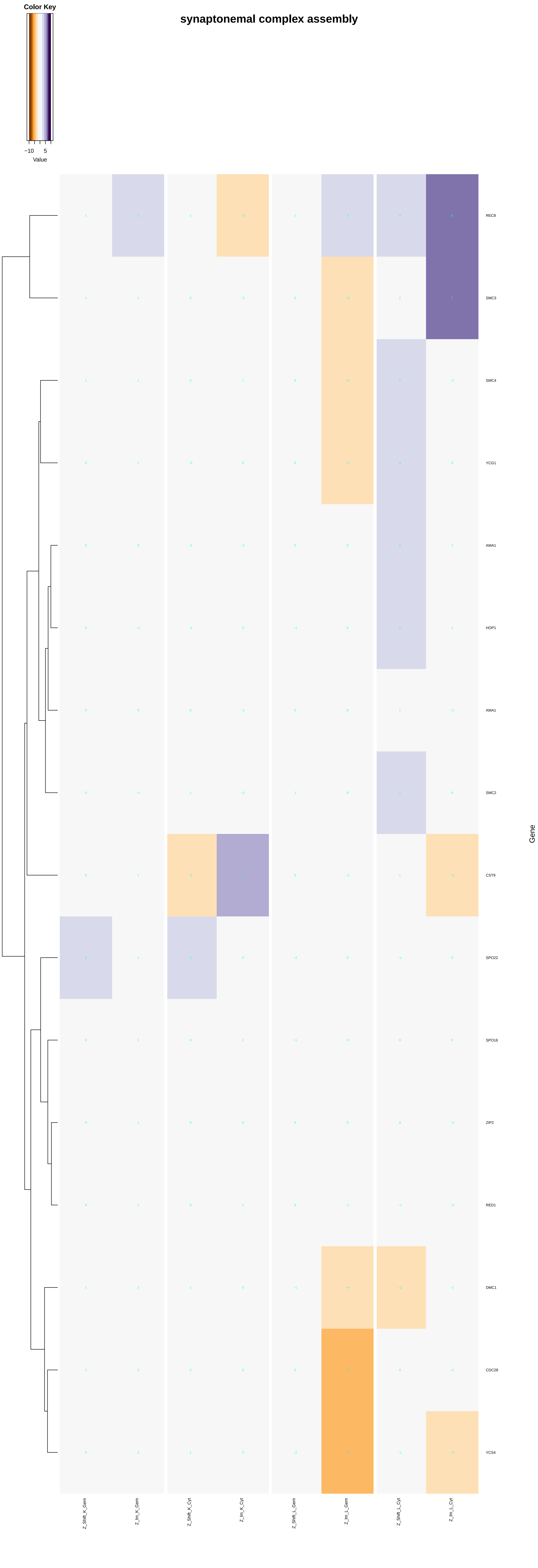
synaptonemal complex assembly
Color Key
−10
5
Value
REC8
SMC3
SMC4
YCG1
AMA1
HOP1
AMA1
SMC2
CST9
SPO22
SPO16
ZIP2
RED1
DMC1
CDC28
YCS4
Gene
Z_Shift_K_Gem
Z_Im_K_Gem
Z_Shift_K_Cyt
Z_Im_K_Cyt
Z_Shift_L_Gem
Z_Im_L_Gem
Z_Shift_L_Cyt
Z_Im_L_Cyt

### meiotic chromosome condensation

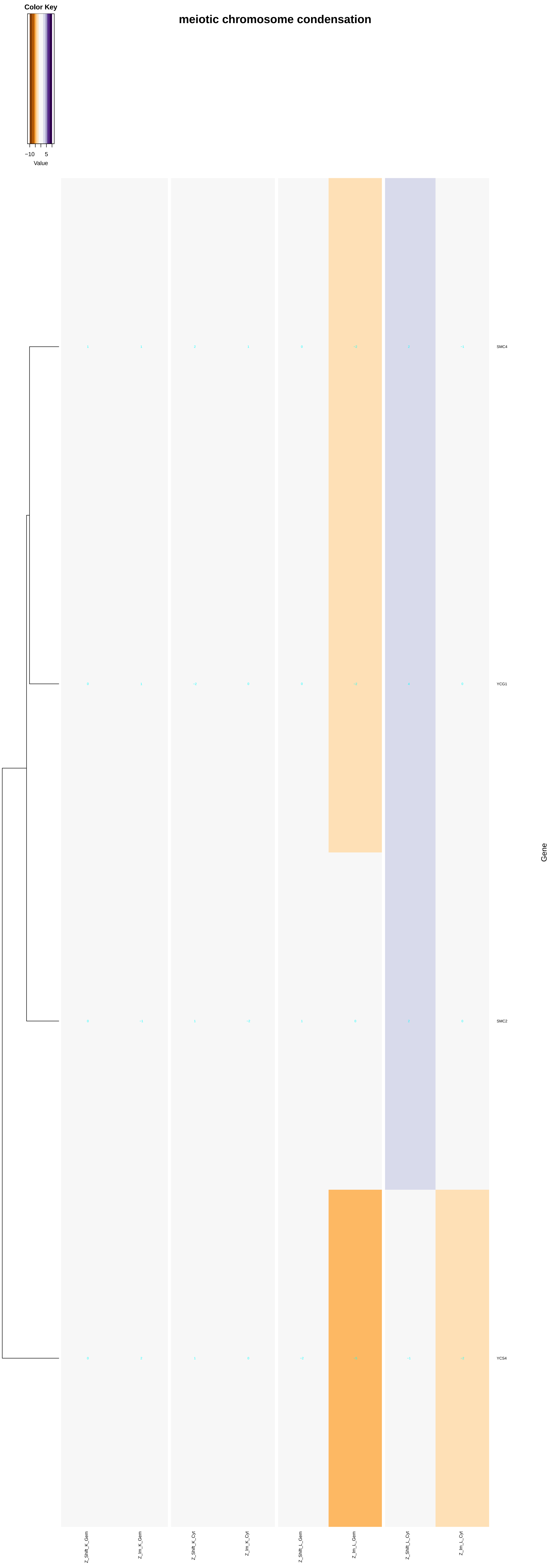

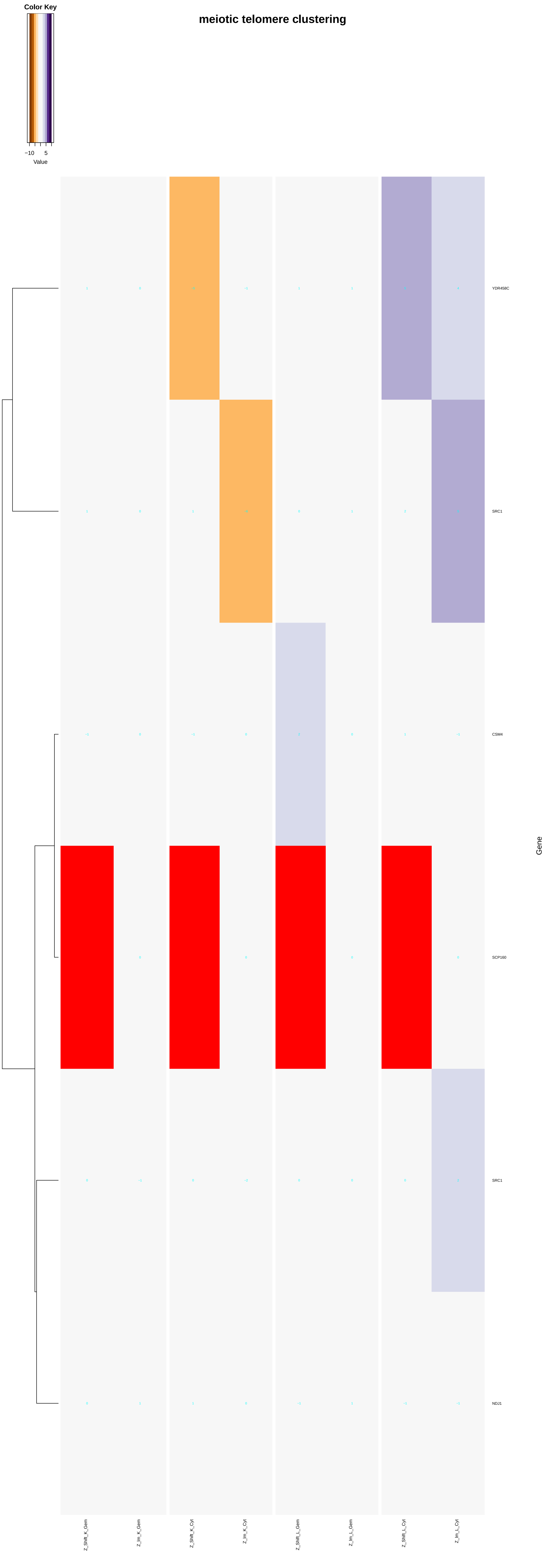
meiotic telomere clustering
Color Key
−10
5
Value
YDR458C
SRC1
CSM4
SCP160
SRC1
NDJ1
Gene
Z_Shift_K_Gem
Z_Im_K_Gem
Z_Shift_K_Cyt
Z_Im_K_Cyt
Z_Shift_L_Gem
Z_Im_L_Gem
Z_Shift_L_Cyt
Z_Im_L_Cyt

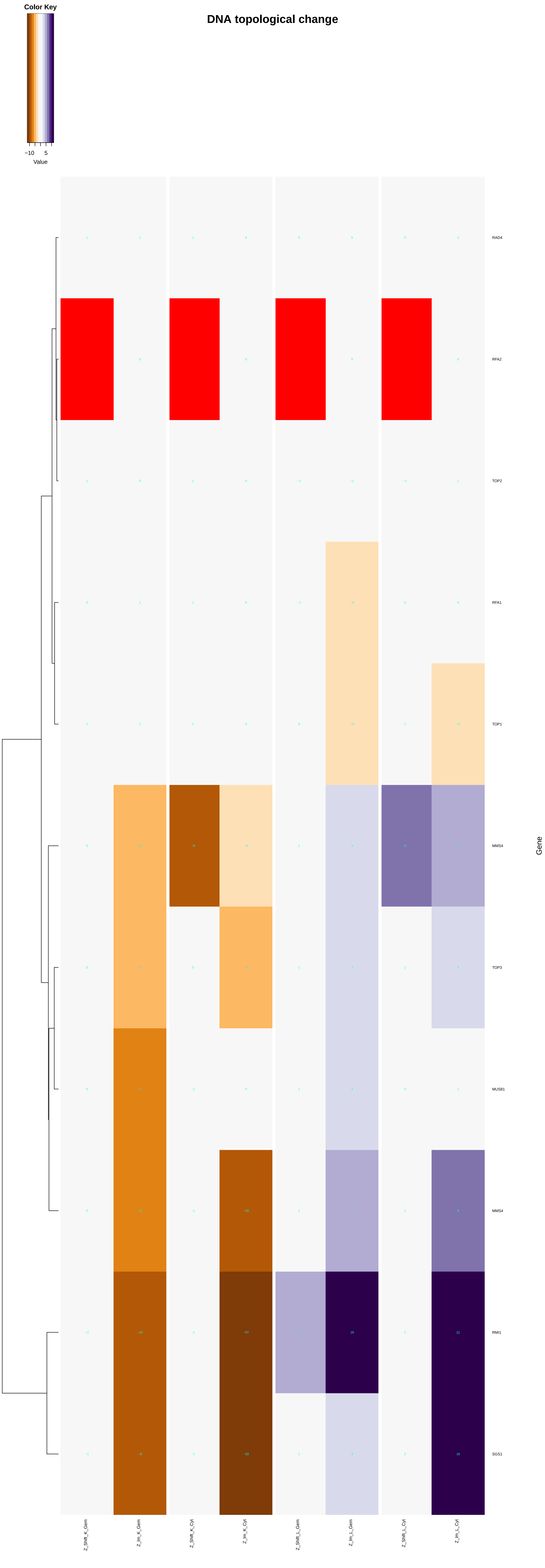
DNA topological change
Color Key
−10
5
Value
RAD4
RFA2
TOP2
RFA1
TOP1
MMS4
TOP3
MUS81
MMS4
RMI1
SGS1
Gene
Z_Shift_K_Gem
Z_Im_K_Gem
Z_Shift_K_Cyt
Z_Im_K_Cyt
Z_Shift_L_Gem
Z_Im_L_Gem
Z_Shift_L_Cyt
Z_Im_L_Cyt

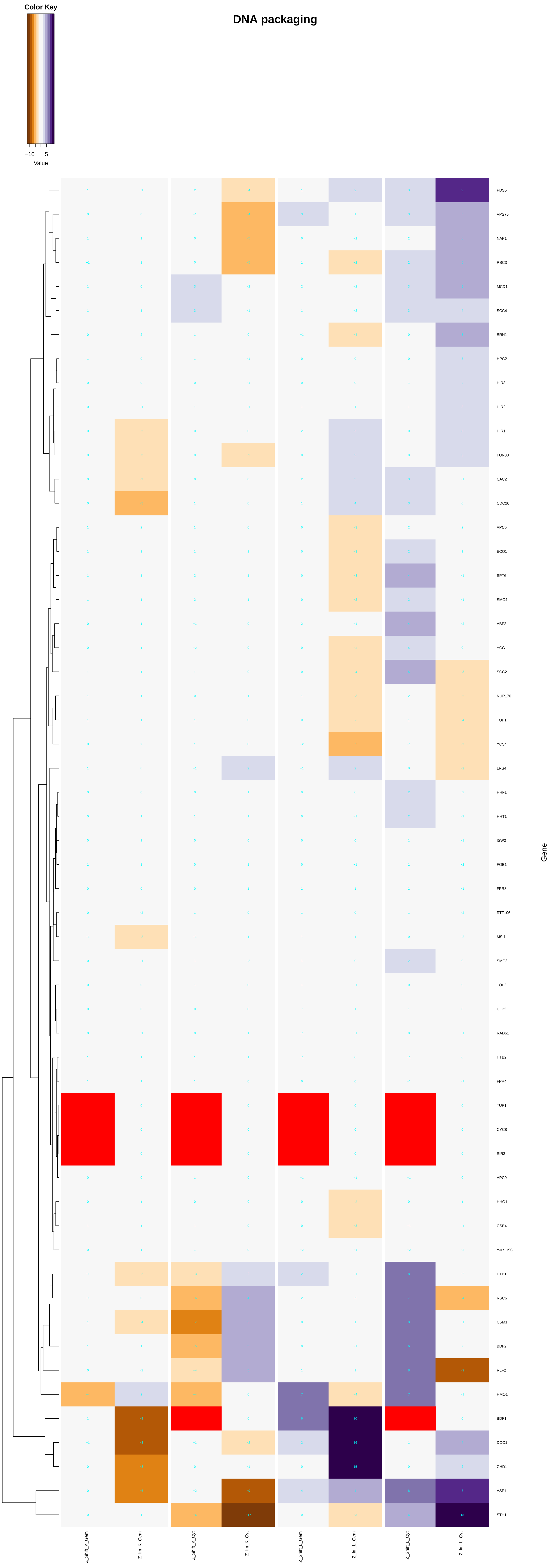

DNA packaging
Color Key
−10
5
Value
PDS5
VPS75
NAP1
RSC3
MCD1
SCC4
BRN1
HPC2
HIR3
HIR2
HIR1
FUN30
CAC2
CDC26
APC5
ECO1
SPT6
SMC4
ABF2
YCG1
SCC2
NUP170
TOP1
YCS4
LRS4
HHF1
HHT1
ISW2
FOB1
FPR3
RTT106
MSI1
SMC2
TOF2
ULP2
RAD61
HTB2
FPR4
TUP1
CYC8
SIR3
APC9
HHO1
CSE4
YJR119C
HTB1
RSC6
CSM1
BDF2
RLF2
HMO1
BDF1
DOC1
CHD1
ASF1
STH1
Gene
Z_Shift_K_Gem
Z_Im_K_Gem
Z_Shift_K_Cyt
Z_Im_K_Cyt
Z_Shift_L_Gem
Z_Im_L_Gem
Z_Shift_L_Cyt
Z_Im_L_Cyt

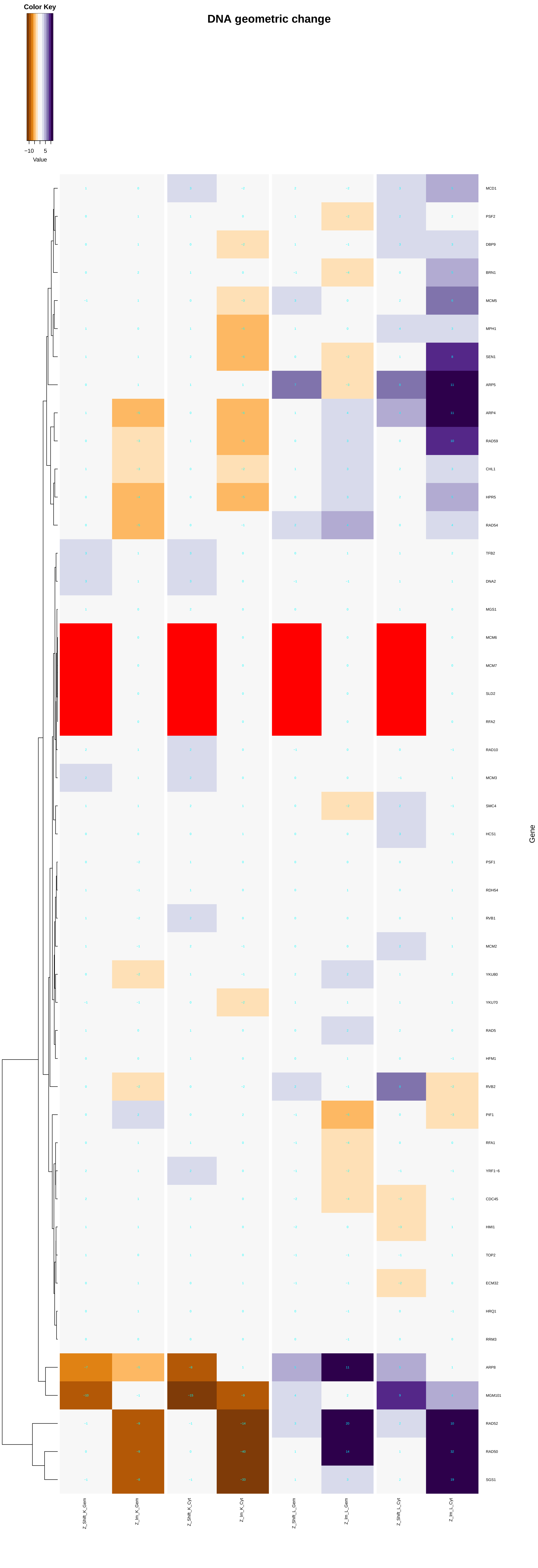

DNA geometric change
Color Key
−10
5
Value
Gene
MCD1
PSF2
DBP9
BRN1
MCM5
MPH1
SEN1
ARP5
ARP4
RAD59
CHL1
HPR5
RAD54
TFB2
DNA2
MGS1
MCM6
MCM7
SLD2
RFA2
RAD10
MCM3
SMC4
HCS1
PSF1
RDH54
RVB1
MCM2
YKU80
YKU70
RAD5
HFM1
RVB2
PIF1
RFA1
YRF1−6
CDC45
HMI1
TOP2
ECM32
HRQ1
RRM3
ARP8
MGM101
RAD52
RAD50
SGS1
Z_Shift_K_Gem
Z_Im_K_Gem
Z_Shift_K_Cyt
Z_Im_K_Cyt
Z_Shift_L_Gem
Z_Im_L_Gem
Z_Shift_L_Cyt
Z_Im_L_Cyt

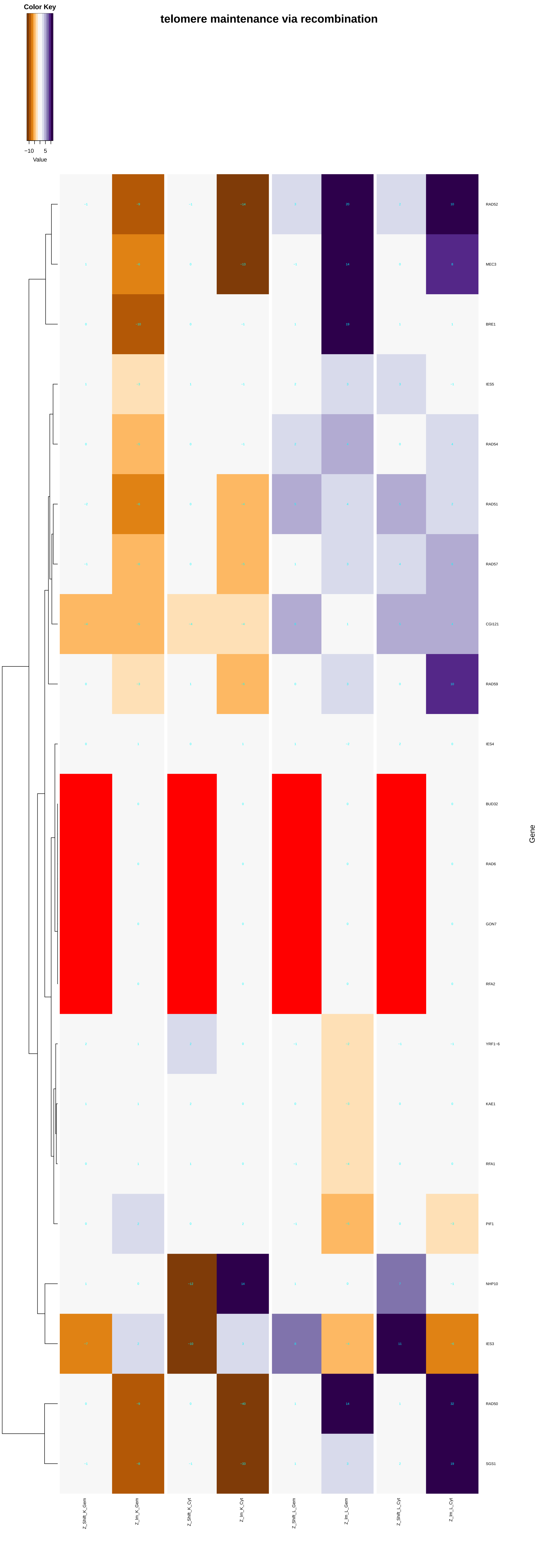

Color Key
−10
5
Value
telomere maintenance via recombination
RAD52
MEC3
BRE1
IES5
RAD54
RAD51
RAD57
CGI121
RAD59
IES4
BUD32
RAD6
GON7
RFA2
YRF1−6
KAE1
RFA1
PIF1
NHP10
IES3
RAD50
SGS1
Gene
Z_Shift_K_Gem
Z_Im_K_Gem
Z_Shift_K_Cyt
Z_Im_K_Cyt
Z_Shift_L_Gem
Z_Im_L_Gem
Z_Shift_L_Cyt
Z_Im_L_Cyt

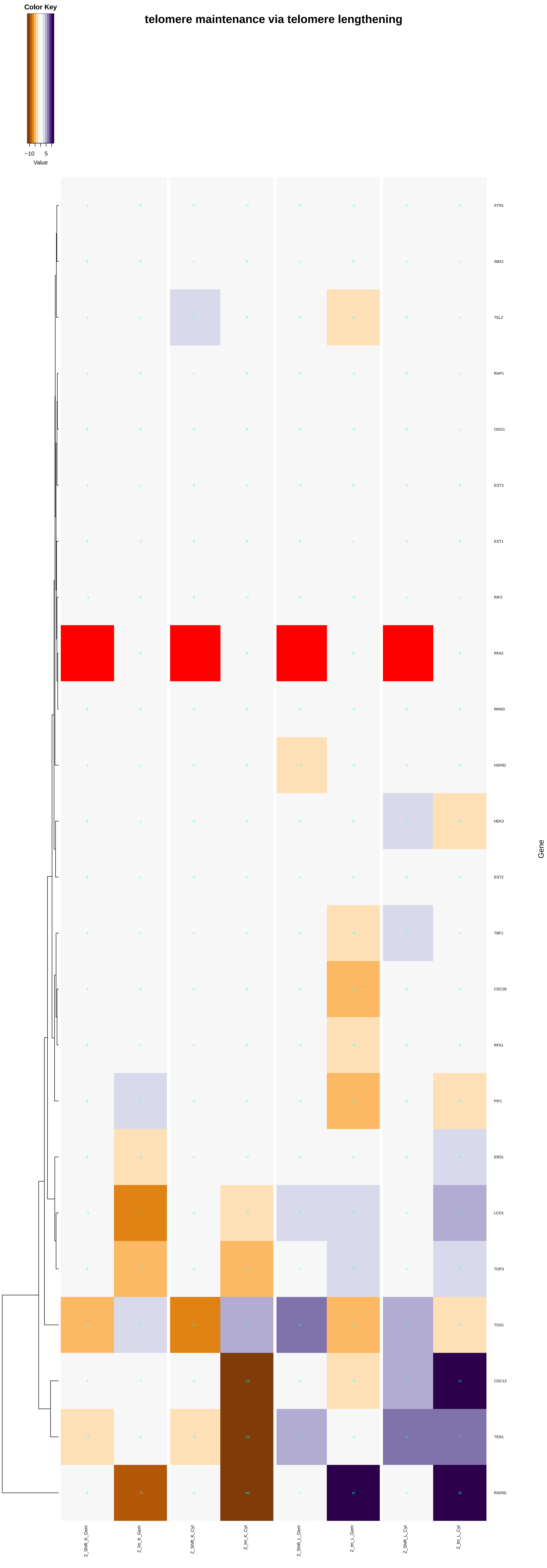
telomere maintenance via telomere lengthening
Color Key
−10
5
Value
STN1
SBA1
TEL2
RAP1
OGG1
EST3
EST1
RIF2
RFA2
RRM3
HSP82
HEK2
EST2
TBF1
CDC28
RFA1
PIF1
EBS1
LCD1
TOP3
TGS1
CDC13
TEN1
RAD50
Gene
Z_Shift_K_Gem
Z_lm_K_Gem
Z_Shift_K_Cyt
Z_lm_K_Cyt
Z_Shift_L_Gem
Z_lm_L_Gem
Z_Shift_L_Cyt
Z_lm_L_Cyt

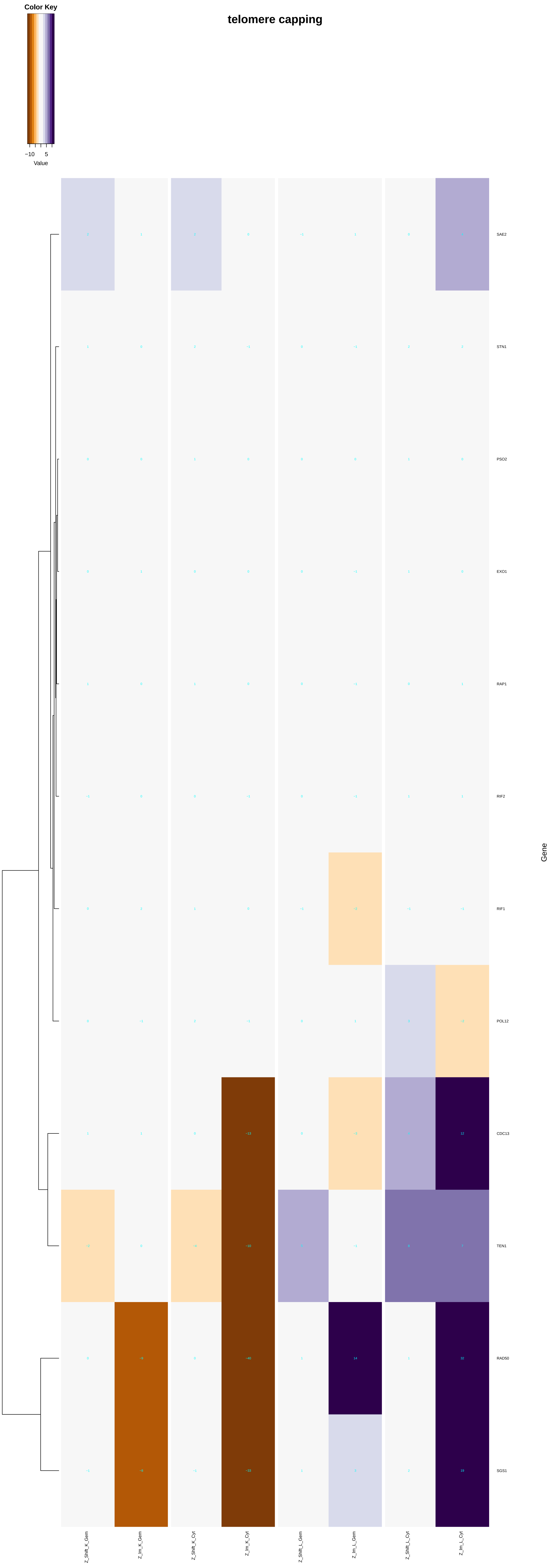

telomere capping
Color Key
−10
5
Value
SAE2
STN1
PSO2
EXO1
RAP1
RIF2
RIF1
POL12
CDC13
TEN1
RAD50
SGS1
Gene
Z_Shift_K_Gem
Z_Im_K_Gem
Z_Shift_K_Cyt
Z_Im_K_Cyt
Z_Shift_L_Gem
Z_Im_L_Gem
Z_Shift_L_Cyt
Z_Im_L_Cyt

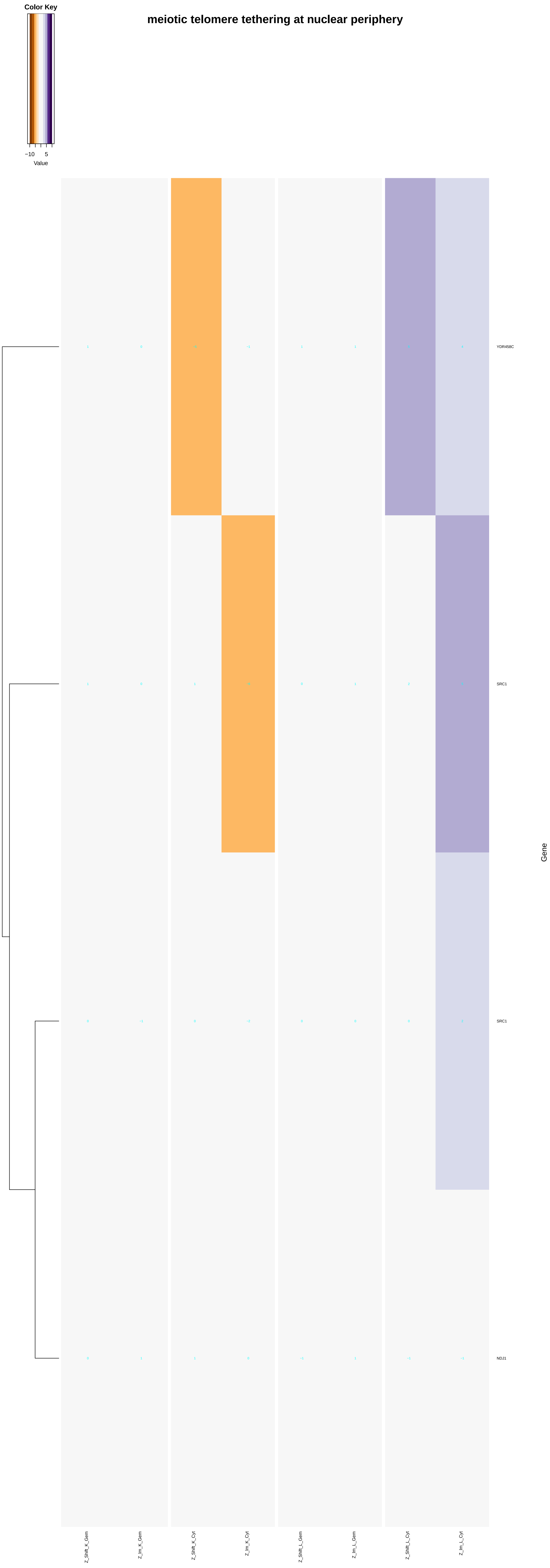
meiotic telomere tethering at nuclear periphery
Color Key
−10
5
Value
YDR458C
SRC1
SRC1
NDJ1
Gene
Z_Shift_K_Gem
Z_Im_K_Gem
Z_Shift_K_Cyt
Z_Im_K_Cyt
Z_Shift_L_Gem
Z_Im_L_Gem
Z_Shift_L_Cyt
Z_Im_L_Cyt

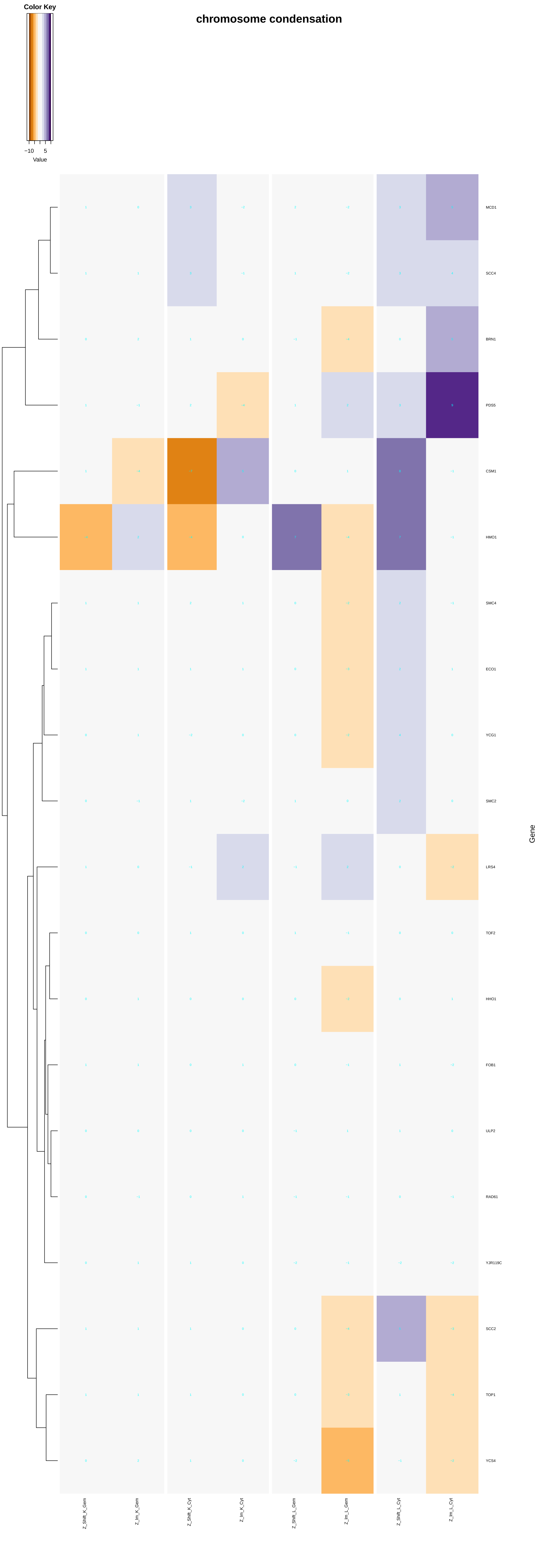
Color Key
−10
5
Value
chromosome condensation
MCD1
SCC4
BRN1
PDS5
CSM1
HMO1
SMC4
ECO1
YCG1
SMC2
LRS4
TOF2
HHO1
FOB1
ULP2
RAD61
YJR119C
SCC2
TOP1
YCS4
Gene
Z_Shift_K_Gem
Z_lm_K_Gem
Z_Shift_K_Cyt
Z_lm_K_Cyt
Z_Shift_L_Gem
Z_lm_L_Gem
Z_Shift_L_Cyt
Z_lm_L_Cyt

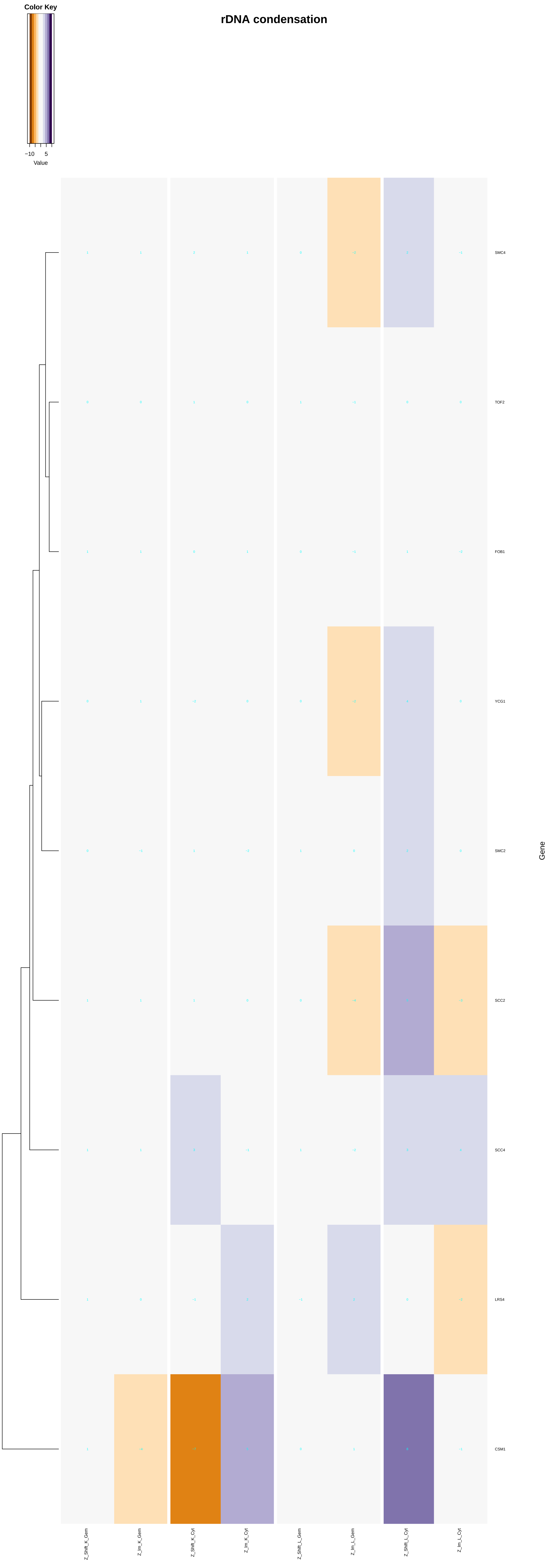
rDNA condensation
Color Key
−10
5
Value
SMC4
TOF2
FOB1
YCG1
SMC2
SCC2
SCC4
LRS4
CSM1
Gene
Z_Shift_K_Gem
Z_Im_K_Gem
Z_Shift_K_Cyt
Z_Im_K_Cyt
Z_Shift_L_Gem
Z_Im_L_Gem
Z_Shift_L_Cyt
Z_Im_L_Cyt

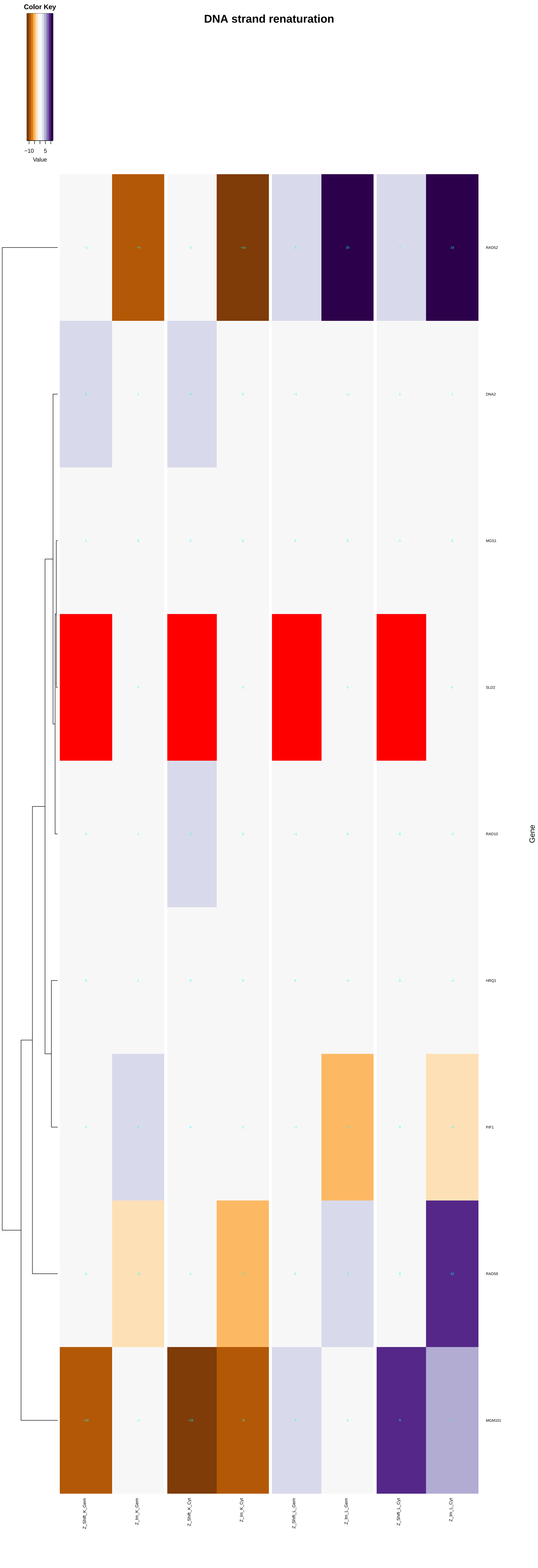
DNA strand renaturation
Color Key
−10
5
Value
RAD52
DNA2
MGS1
SLD2
RAD10
HRQ1
PIF1
RAD59
MGM101
Gene
Z_Shift_K_Gem
Z_Im_K_Gem
Z_Shift_K_Cyt
Z_Im_K_Cyt
Z_Shift_L_Gem
Z_Im_L_Gem
Z_Shift_L_Cyt
Z_Im_L_Cyt

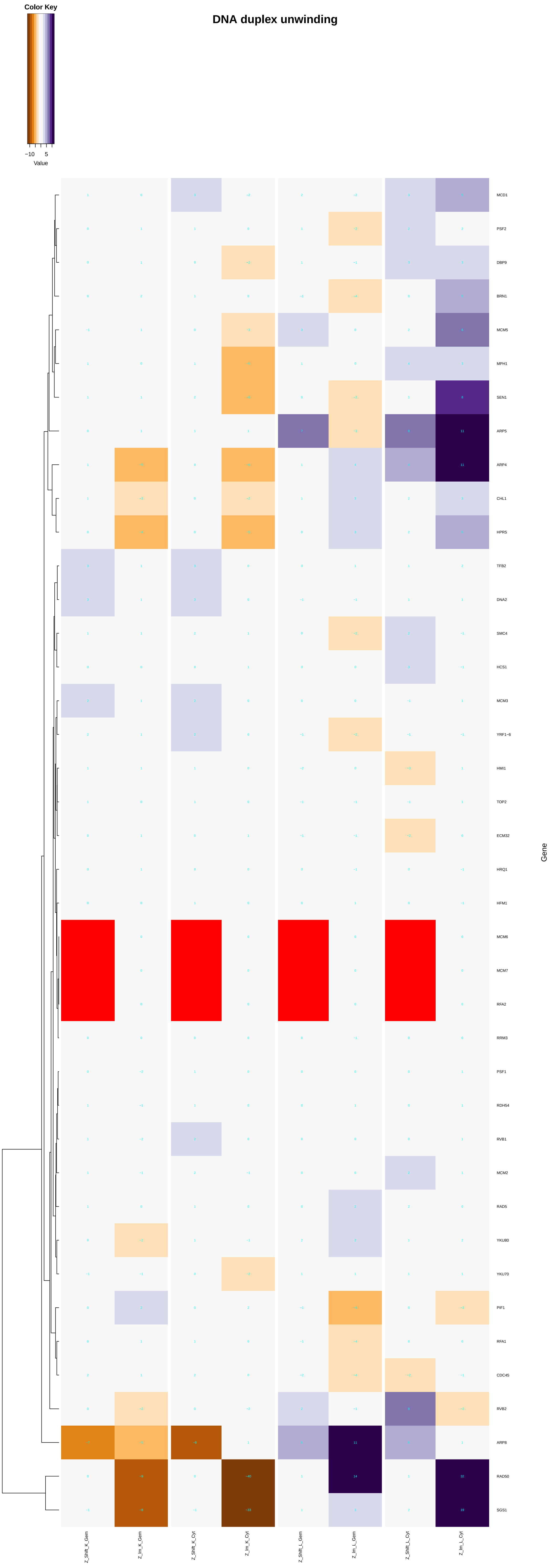

Color Key
−10
5
Value
DNA duplex unwinding
MCD1
PSF2
DBP9
BRN1
MCM5
MPH1
SEN1
ARP5
ARP4
CHL1
HPR5
TFB2
DNA2
SMC4
HCS1
MCM3
YRF1−6
HMI1
TOP2
ECM32
HRQ1
HFM1
MCM6
MCM7
RFA2
RRM3
PSF1
RDH54
RVB1
MCM2
RAD5
YKU80
YKU70
PIF1
RFA1
CDC45
RVB2
ARP8
RAD50
SGS1
Gene
Z_Shift_K_Gem
Z_Im_K_Gem
Z_Shift_K_Cyt
Z_Im_K_Cyt
Z_Shift_L_Gem
Z_Im_L_Gem
Z_Shift_L_Cyt
Z_Im_L_Cyt

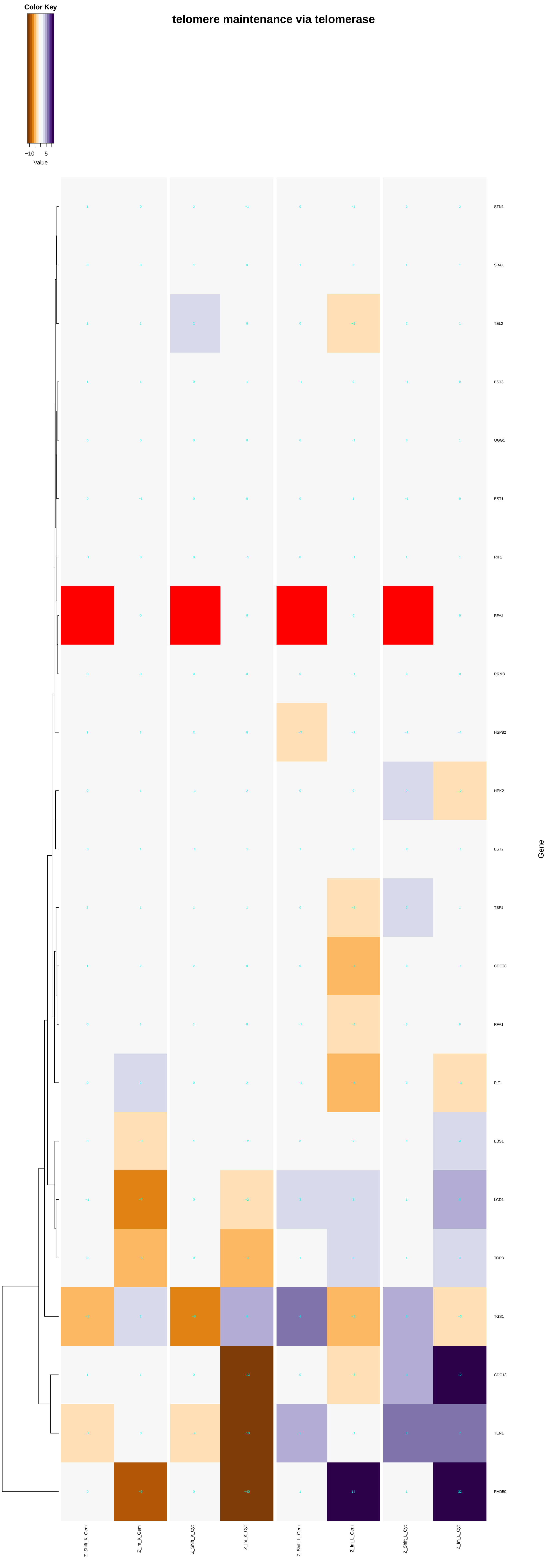
telomere maintenance via telomerase
Color Key
−10
5
Value
STN1
SBA1
TEL2
EST3
OGG1
EST1
RIF2
RFA2
RRM3
HSP82
HEK2
EST2
TBF1
CDC28
RFA1
PIF1
EBS1
LCD1
TOP3
TGS1
CDC13
TEN1
RAD50
Gene
Z_Shift_K_Gem
Z_Im_K_Gem
Z_Shift_K_Cyt
Z_Im_K_Cyt
Z_Shift_L_Gem
Z_Im_L_Gem
Z_Shift_L_Cyt
Z_Im_L_Cyt

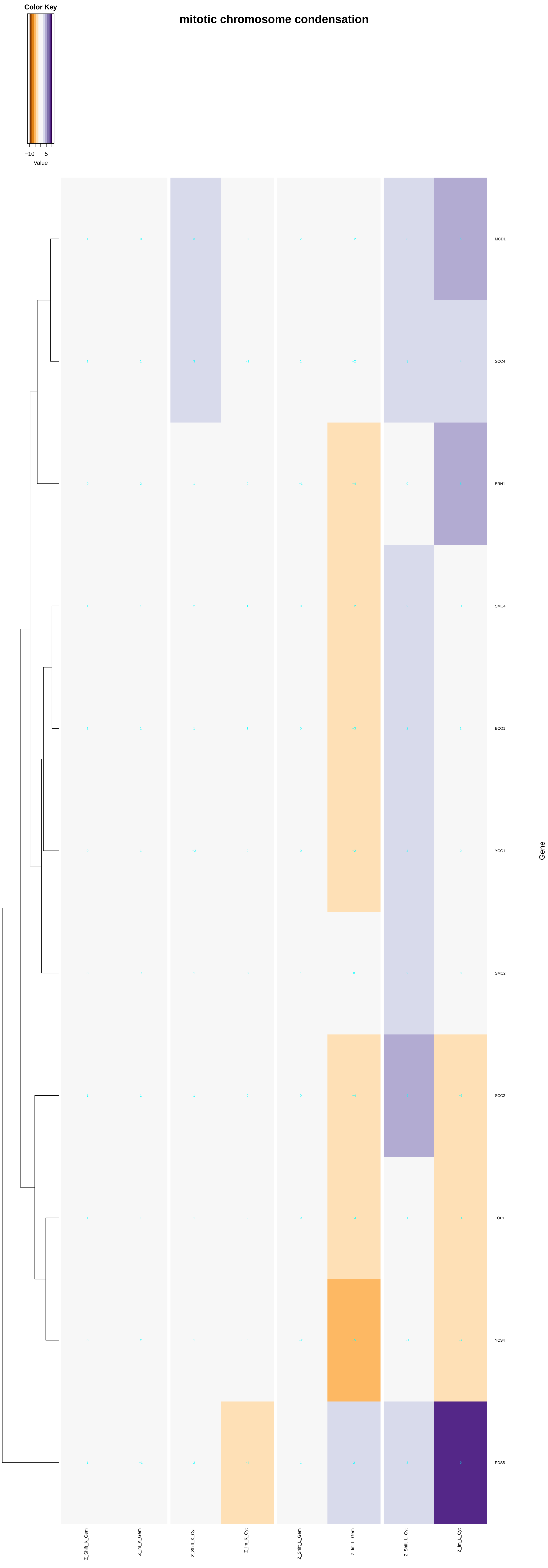
mitotic chromosome condensation
Color Key
−10
5
Value
MCD1
SCC4
BRN1
SMC4
ECO1
YCG1
SMC2
SCC2
TOP1
YCS4
PDS5
Gene
Z_Shift_K_Gem
Z_Im_K_Gem
Z_Shift_K_Cyt
Z_Im_K_Cyt
Z_Shift_L_Gem
Z_Im_L_Gem
Z_Shift_L_Cyt
Z_Im_L_Cyt

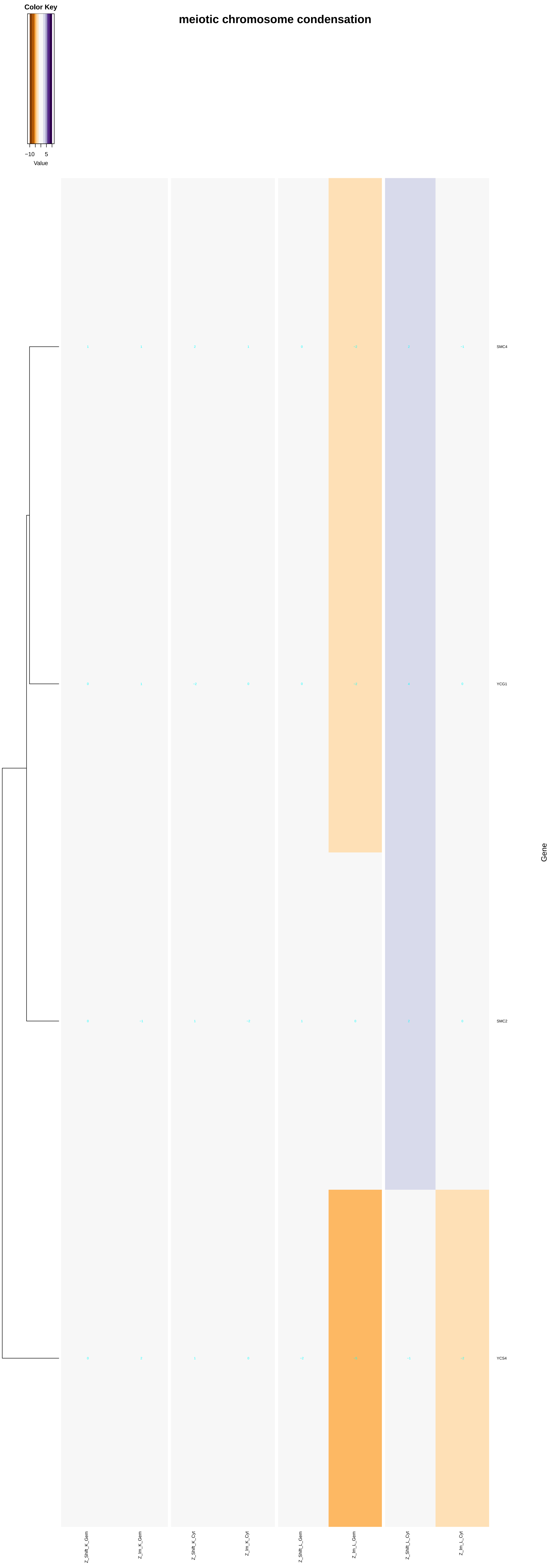
meiotic chromosome condensation
Color Key
−10
5
Value
SMC4
YCG1
SMC2
YCS4
Gene
Z_Shift_K_Gem
Z_Im_K_Gem
Z_Shift_K_Cyt
Z_Im_K_Cyt
Z_Shift_L_Gem
Z_Im_L_Gem
Z_Shift_L_Cyt
Z_Im_L_Cyt

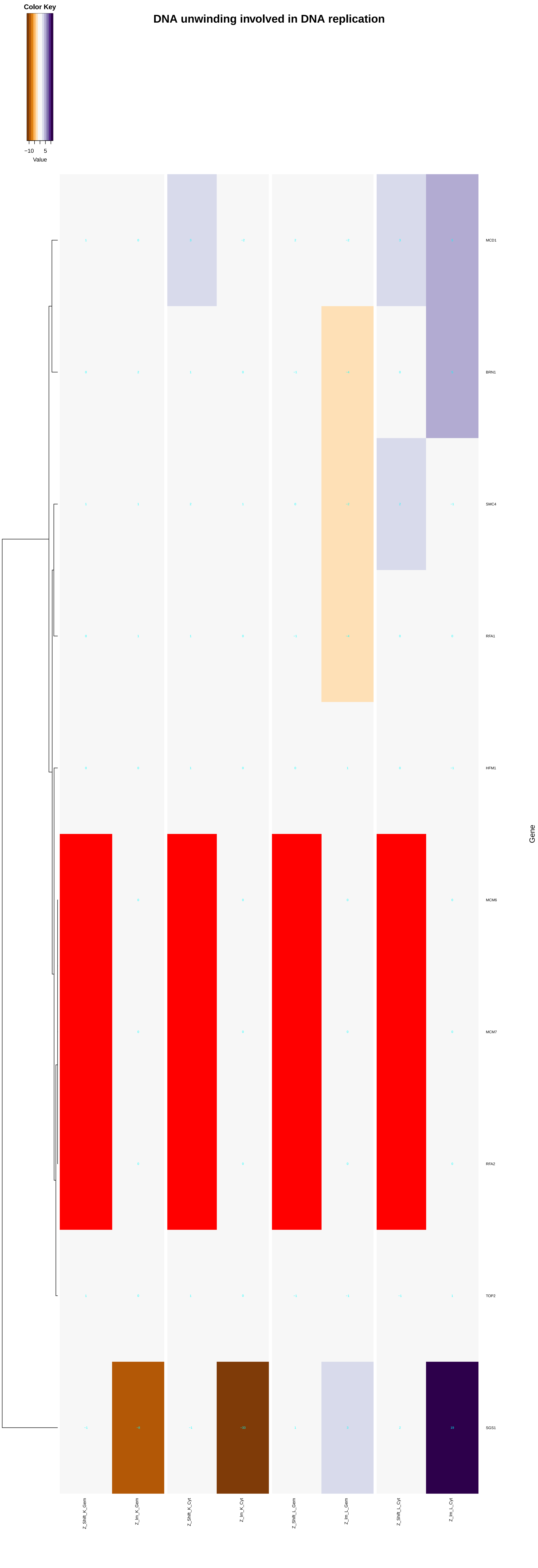
DNA unwinding involved in DNA replication
Color Key
−10
5
Value
MCD1
BRN1
SMC4
RFA1
HFM1
MCM6
MCM7
RFA2
TOP2
SGS1
Gene
Z_Shift_K_Gem
Z_Im_K_Gem
Z_Shift_K_Cyt
Z_Im_K_Cyt
Z_Shift_L_Gem
Z_Im_L_Gem
Z_Shift_L_Cyt
Z_Im_L_Cyt
