## Additional File 7 for "A humanized yeast phenomic model of deoxycytidine kinase to predict genetic buffering of nucleoside analog cytotoxicity": covalent_chromatin_modification.pdf

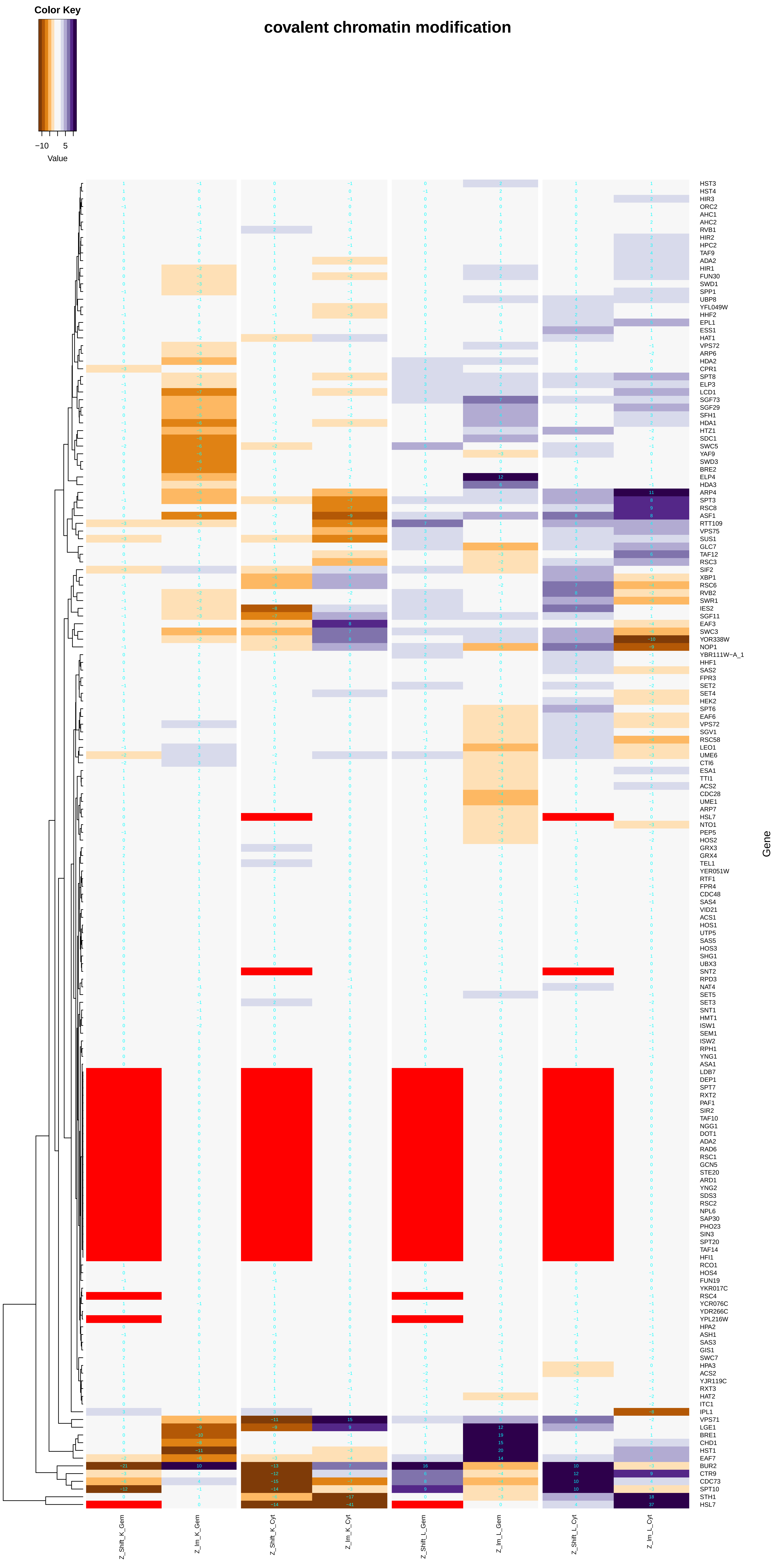

covalent chromatin modification
Color Key
−10 5
Value
HST3
HST4
HIR3
ORC2
AHC1
AHC2
RVB1
HIR2
HPC2
TAF9
ADA2
HIR1
FUN30
SWD1
SPP1
UBP8
YFL049W
HHF2
EPL1
ESS1
HAT1
VPS72
ARP6
HDA2
CPR1
SPT8
ELP3
LCD1
SGF73
SGF29
SFH1
HDA1
HTZ1
SDC1
SWC5
YAF9
SWD3
BRE2
ELP4
HDA3
ARP4
SPT3
RSC8
ASF1
RTT109
VPS75
SUS1
GLC7
TAF12
RSC3
SIF2
XBP1
RSC6
RVB2
SWR1
IES2
SGF11
EAF3
SWC3
YOR338W
NOP1
YBR111W−A_1
HHF1
SAS2
FPR3
SET2
SET4
HEK2
SPT6
EAF6
VPS72
SGV1
RSC58
LEO1
UME6
CTI6
ESA1
TTI1
ACS2
CDC28
UME1
ARP7
HSL7
NTO1
PEP5
HOS2
GRX3
GRX4
TEL1
YER051W
RTF1
FPR4
CDC48
SAS4
VID21
ACS1
HOS1
UTP5
SAS5
HOS3
SHG1
UBX3
SNT2
RPD3
NAT4
SET5
SET3
SNT1
HMT1
ISW1
SEM1
ISW2
RPH1
YNG1
ASA1
LDB7
DEP1
SPT7
RXT2
PAF1
SIR2
TAF10
NGG1
DOT1
ADA2
RAD6
RSC1
GCN5
STE20
ARD1
YNG2
SDS3
RSC2
NPL6
SAP30
PHO23
SIN3
SPT20
TAF14
HFI1
RCO1
HOS4
FUN19
YKR017C
RSC4
YCR076C
YDR266C
YPL216W
HPA2
ASH1
SAS3
GIS1
SWC7
HPA3
ACS2
YJR119C
RXT3
HAT2
ITC1
IPL1
VPS71
LGE1
BRE1
CHD1
HST1
EAF7
BUR2
CTR9
CDC73
SPT10
STH1
HSL7
Gene
Z_Shift_K_Gem
Z_Im_K_Gem
Z_Shift_K_Cyt
Z_Im_K_Cyt
Z_Shift_L_Gem
Z_Im_L_Gem
Z_Shift_L_Cyt
Z_Im_L_Cyt

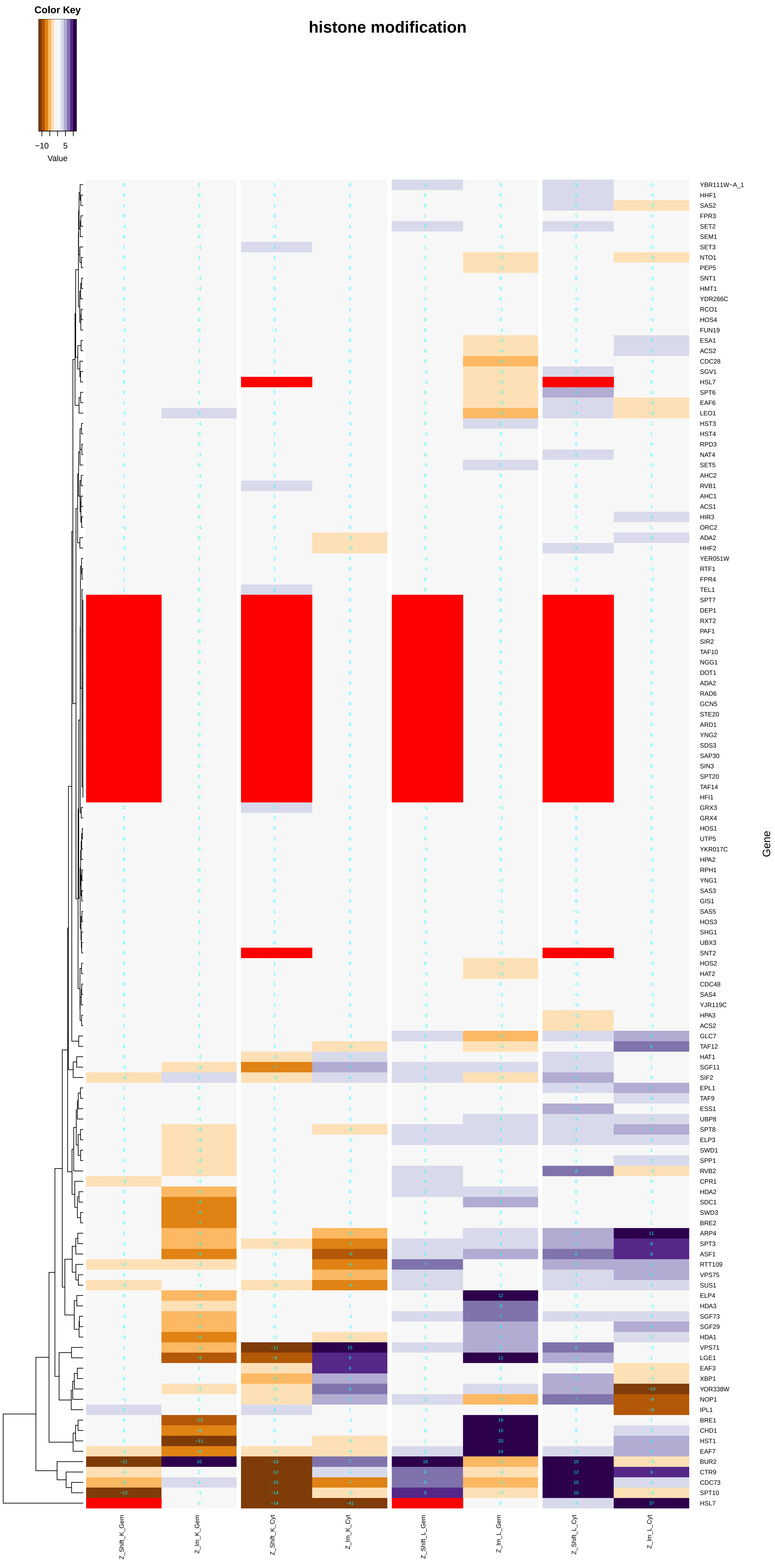
Color Key
−10 5
Value
histone modification
Gene
Z_Shift_K_Gem
Z_Im_K_Gem
Z_Shift_K_Cyt
Z_Im_K_Cyt
Z_Shift_L_Gem
Z_Im_L_Gem
Z_Shift_L_Cyt
Z_Im_L_Cyt
YBR111W−A_1
HHF1
SAS2
FPR3
SET2
SEM1
SET3
NTO1
PEP5
SNT1
HMT1
YDR266C
RCO1
HOS4
FUN19
ESA1
ACS2
CDC28
SGV1
HSL7
SPT6
EAF6
LEO1
HST3
HST4
RPD3
NAT4
SET5
AHC2
RVB1
AHC1
ACS1
HIR3
ORC2
ADA2
HHF2
YER051W
RTF1
FPR4
TEL1
SPT7
DEP1
RXT2
PAF1
SIR2
TAF10
NGG1
DOT1
ADA2
RAD6
GCN5
STE20
ARD1
YNG2
SDS3
SAP30
SIN3
SPT20
TAF14
HFI1
GRX3
GRX4
HOS1
UTP5
YKR017C
HPA2
RPH1
YNG1
SAS3
GIS1
SAS5
HOS3
SHG1
UBX3
SNT2
HOS2
HAT2
CDC48
SAS4
YJR119C
HPA3
ACS2
GLC7
TAF12
HAT1
SGF11
SIF2
EPL1
TAF9
ESS1
UBP8
SPT8
ELP3
SWD1
SPP1
RVB2
CPR1
HDA2
SDC1
SWD3
BRE2
ARP4
SPT3
ASF1
RTT109
VPS75
SUS1
ELP4
HDA3
SGF73
SGF29
HDA1
VPS71
LGE1
EAF3
XBP1
YOR338W
NOP1
IPL1
BRE1
CHD1
HST1
EAF7
BUR2
CTR9
CDC73
SPT10
HSL7

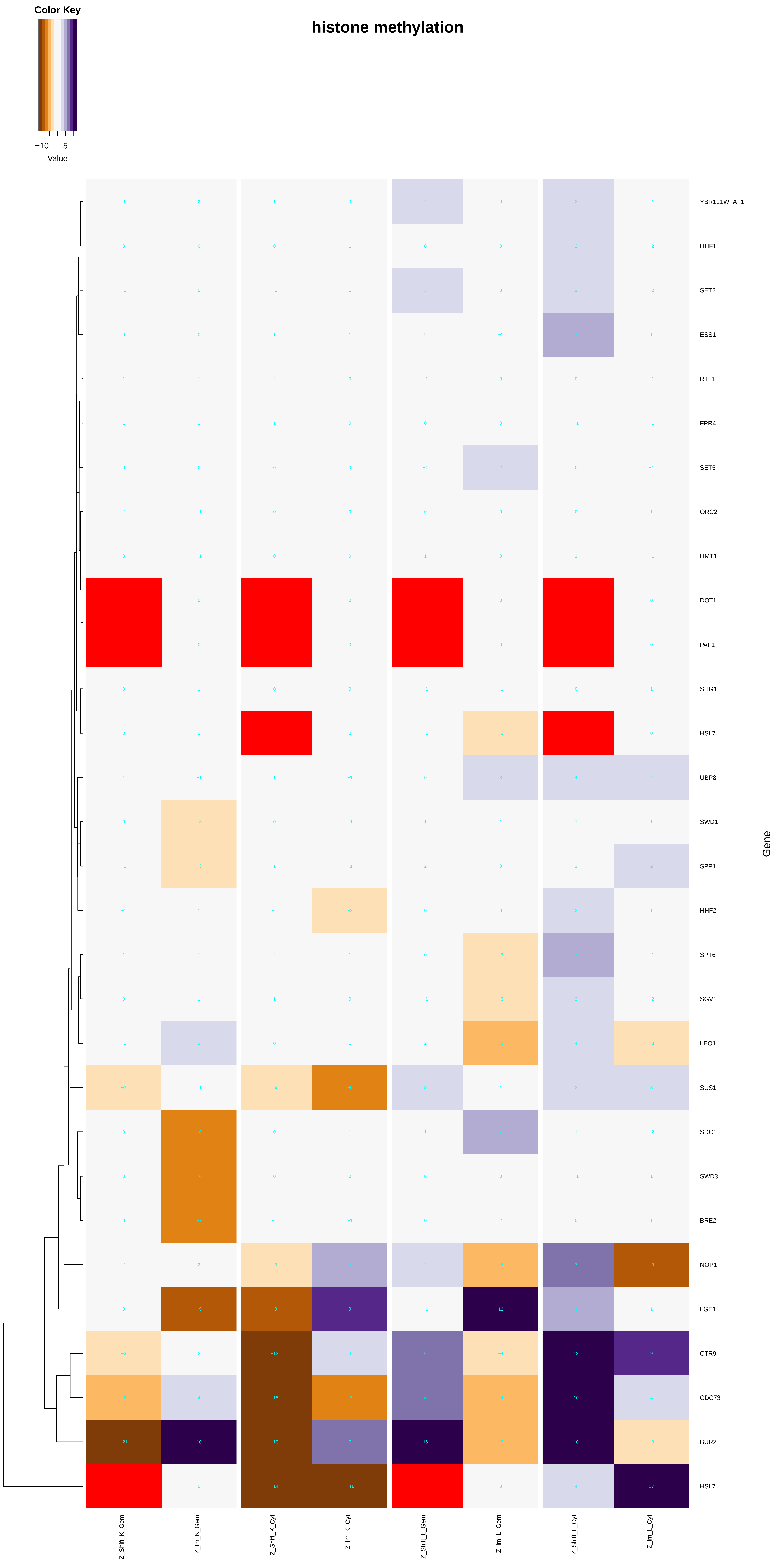

Color Key
−10
5
Value
histone methylation
YBR111W−A_1
HHF1
SET2
ESS1
RTF1
FPR4
SET5
ORC2
HMT1
DOT1
PAF1
SHG1
HSL7
UBP8
SWD1
SPP1
HHF2
SPT6
SGV1
LEO1
SUS1
SDC1
SWD3
BRE2
NOP1
LGE1
CTR9
CDC73
BUR2
HSL7
Gene
Z_Shift_K_Gem
Z_Im_K_Gem
Z_Shift_K_Cyt
Z_Im_K_Cyt
Z_Shift_L_Gem
Z_Im_L_Gem
Z_Shift_L_Cyt
Z_Im_L_Cyt

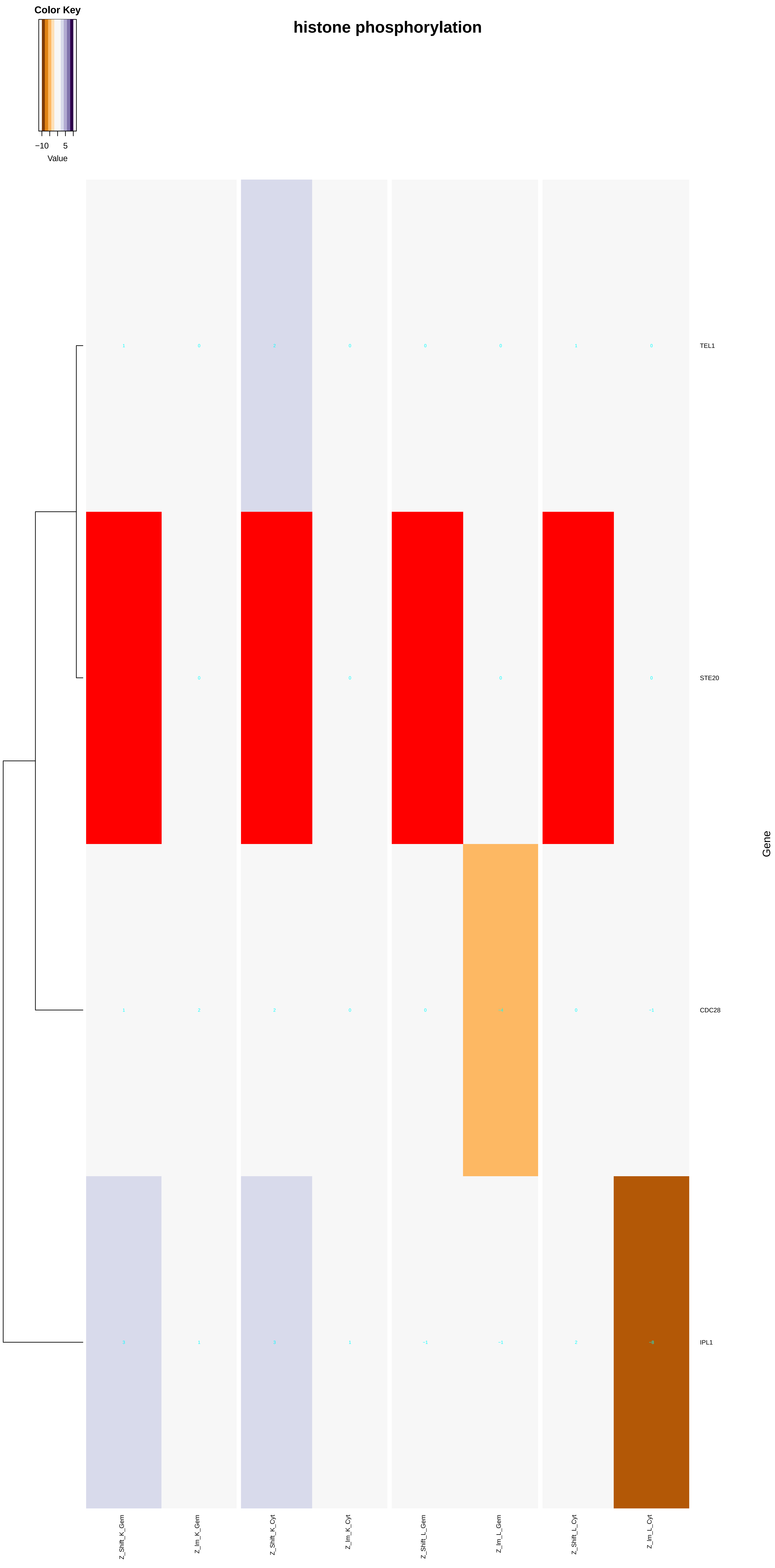

Color Key
−10
5
Value
histone phosphorylation
TEL1
STE20
CDC28
IPL1
Gene
Z_Shift_K_Gem
Z_Im_K_Gem
Z_Shift_K_Cyt
Z_Im_K_Cyt
Z_Shift_L_Gem
Z_Im_L_Gem
Z_Shift_L_Cyt
Z_Im_L_Cyt

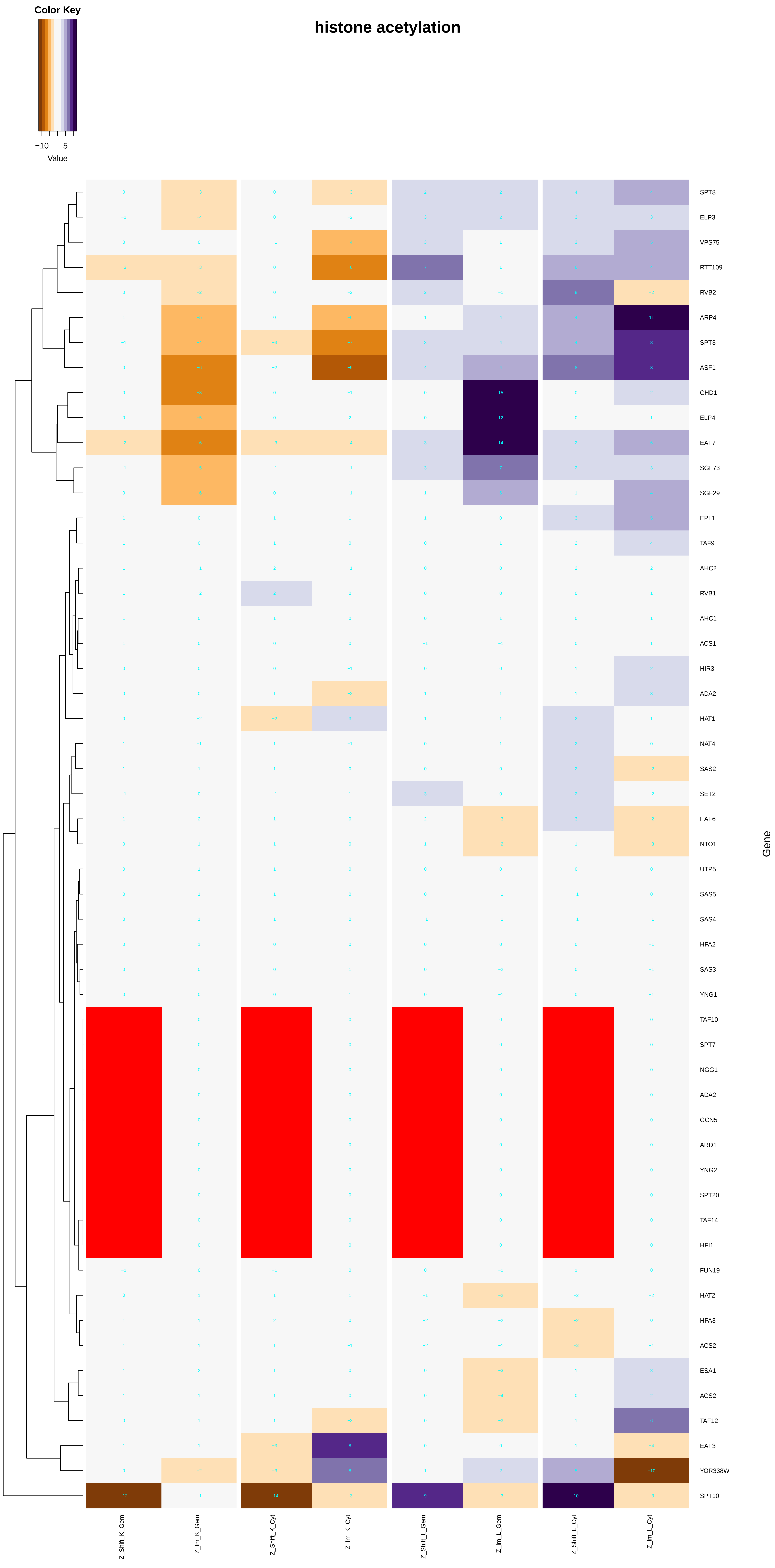
histone acetylation
Color Key
−10
5
Value
SPT8
ELP3
VPS75
RTT109
RVB2
ARP4
SPT3
ASF1
CHD1
ELP4
EAF7
SGF73
SGF29
EPL1
TAF9
AHC2
RVB1
AHC1
ACS1
HIR3
ADA2
HAT1
NAT4
SAS2
SET2
EAF6
NTO1
UTP5
SAS5
SAS4
HPA2
SAS3
YNG1
TAF10
SPT7
NGG1
ADA2
GCN5
ARD1
YNG2
SPT20
TAF14
HFI1
FUN19
HAT2
HPA3
ACS2
ESA1
ACS2
TAF12
EAF3
YOR338W
SPT10
Gene
Z_Shift_K_Gem
Z_Im_K_Gem
Z_Shift_K_Cyt
Z_Im_K_Cyt
Z_Shift_L_Gem
Z_Im_L_Gem
Z_Shift_L_Cyt
Z_Im_L_Cyt

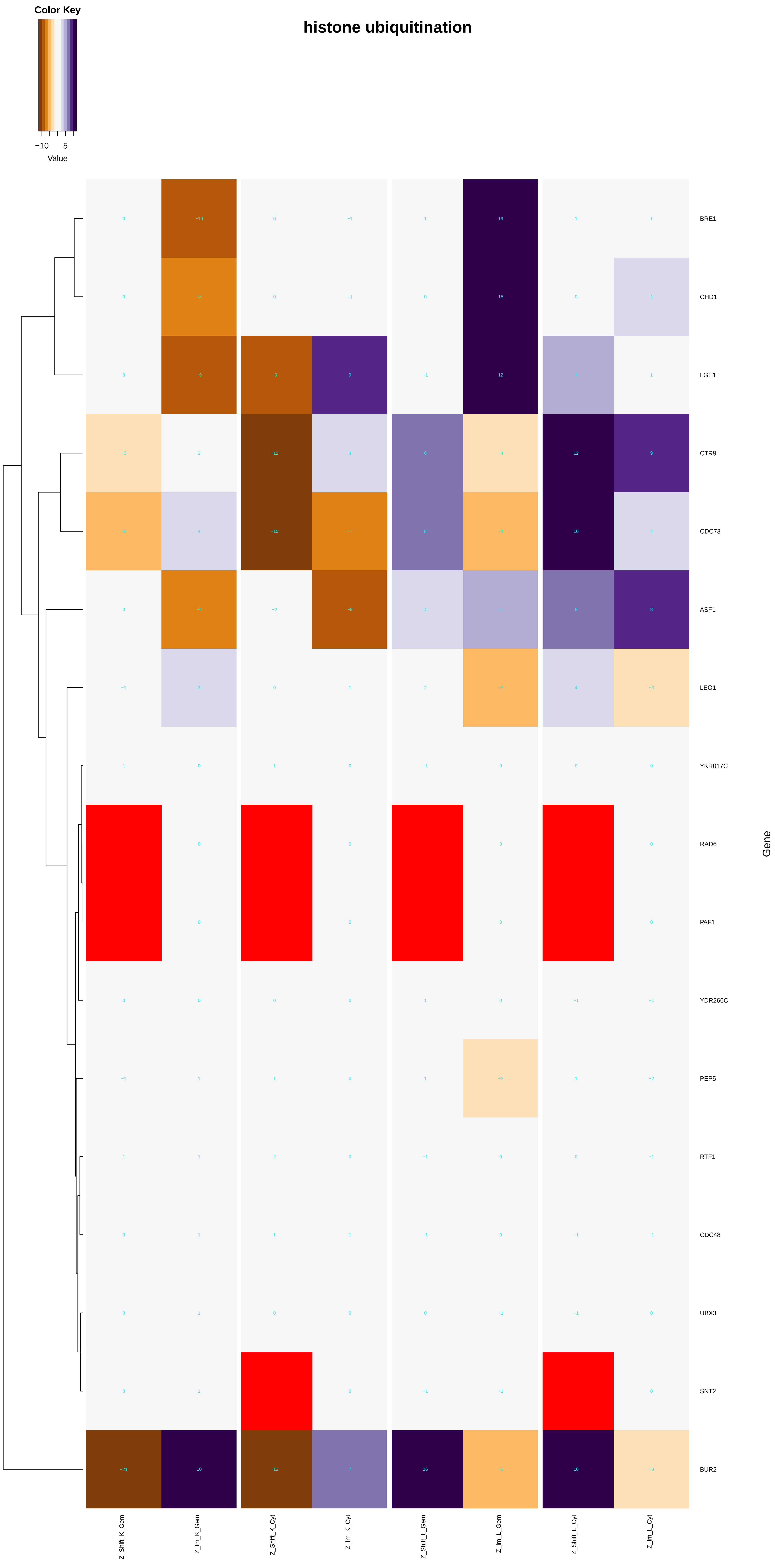

Color Key
−10
5
Value
histone ubiquitination
BRE1
CHD1
LGE1
CTR9
CDC73
ASF1
LEO1
YKR017C
RAD6
PAF1
YDR266C
PEP5
RTF1
CDC48
UBX3
SNT2
BUR2
Gene
Z_Shift_K_Gem
Z_Im_K_Gem
Z_Shift_K_Cyt
Z_Im_K_Cyt
Z_Shift_L_Gem
Z_Im_L_Gem
Z_Shift_L_Cyt
Z_Im_L_Cyt

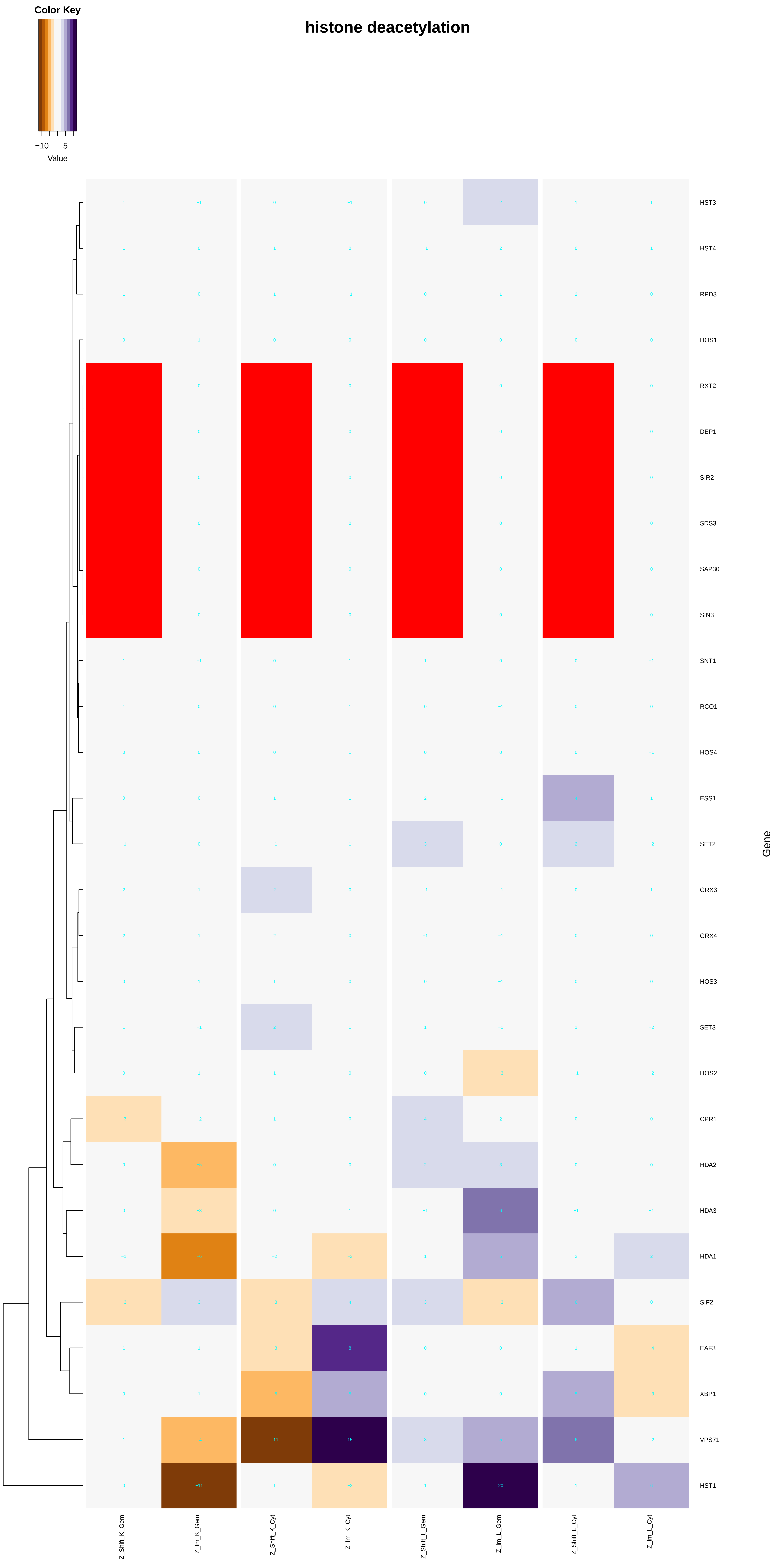

Color Key
−10
5
Value
histone deacetylation
HST3
HST4
RPD3
HOS1
RXT2
DEP1
SIR2
SDS3
SAP30
SIN3
SNT1
RCO1
HOS4
ESS1
SET2
GRX3
GRX4
HOS3
SET3
HOS2
CPR1
HDA2
HDA3
HDA1
SIF2
EAF3
XBP1
VPS71
HST1
Gene
Z_Shift_K_Gem
Z_Im_K_Gem
Z_Shift_K_Cyt
Z_Im_K_Cyt
Z_Shift_L_Gem
Z_Im_L_Gem
Z_Shift_L_Cyt
Z_Im_L_Cyt

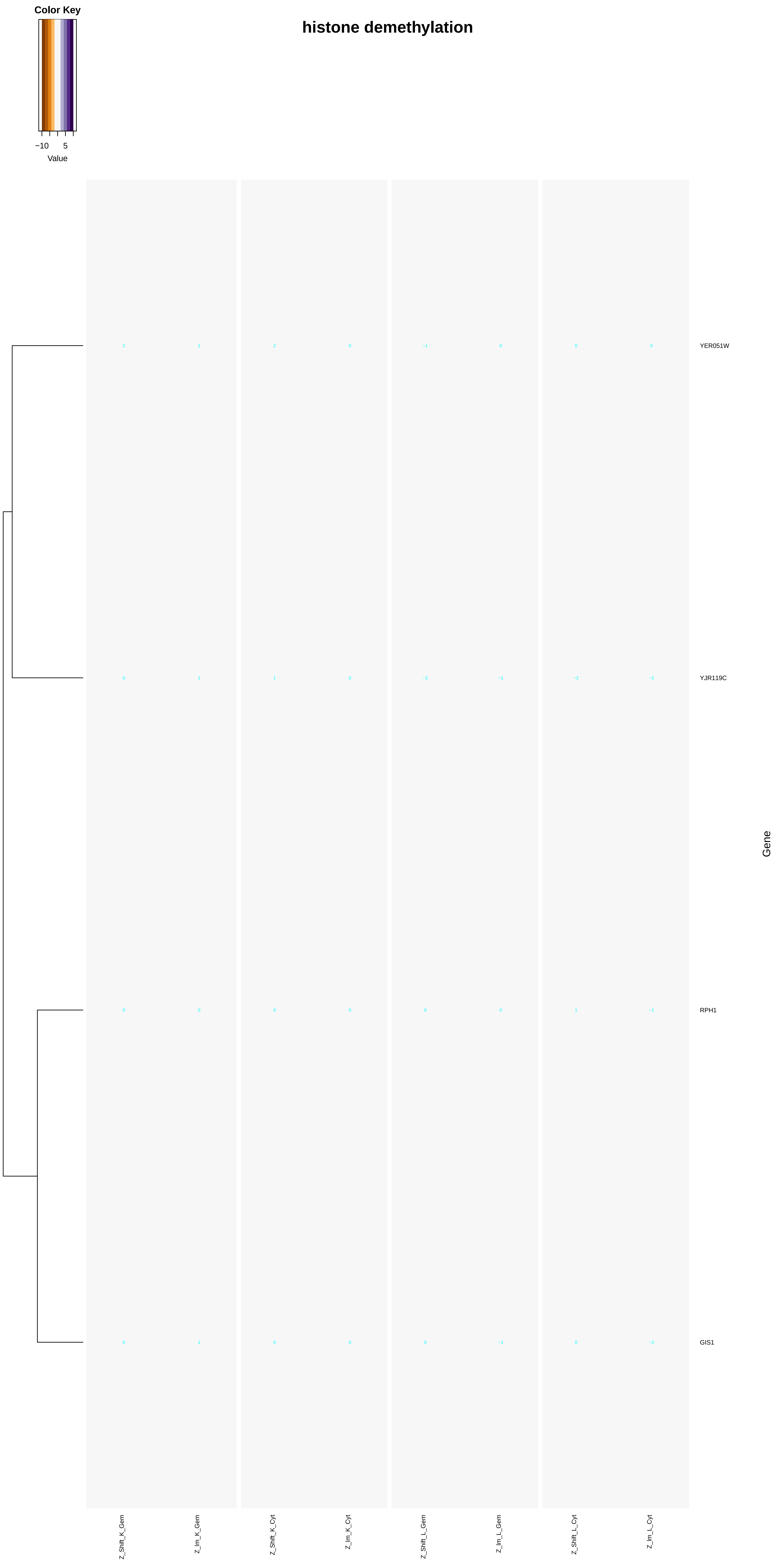
Color Key
−10
5
Value
histone demethylation
YER051W
YJR119C
RPH1
GIS1
Gene
Z_Shift_K_Gem
Z_Im_K_Gem
Z_Shift_K_Cyt
Z_Im_K_Cyt
Z_Shift_L_Gem
Z_Im_L_Gem
Z_Shift_L_Cyt
Z_Im_L_Cyt

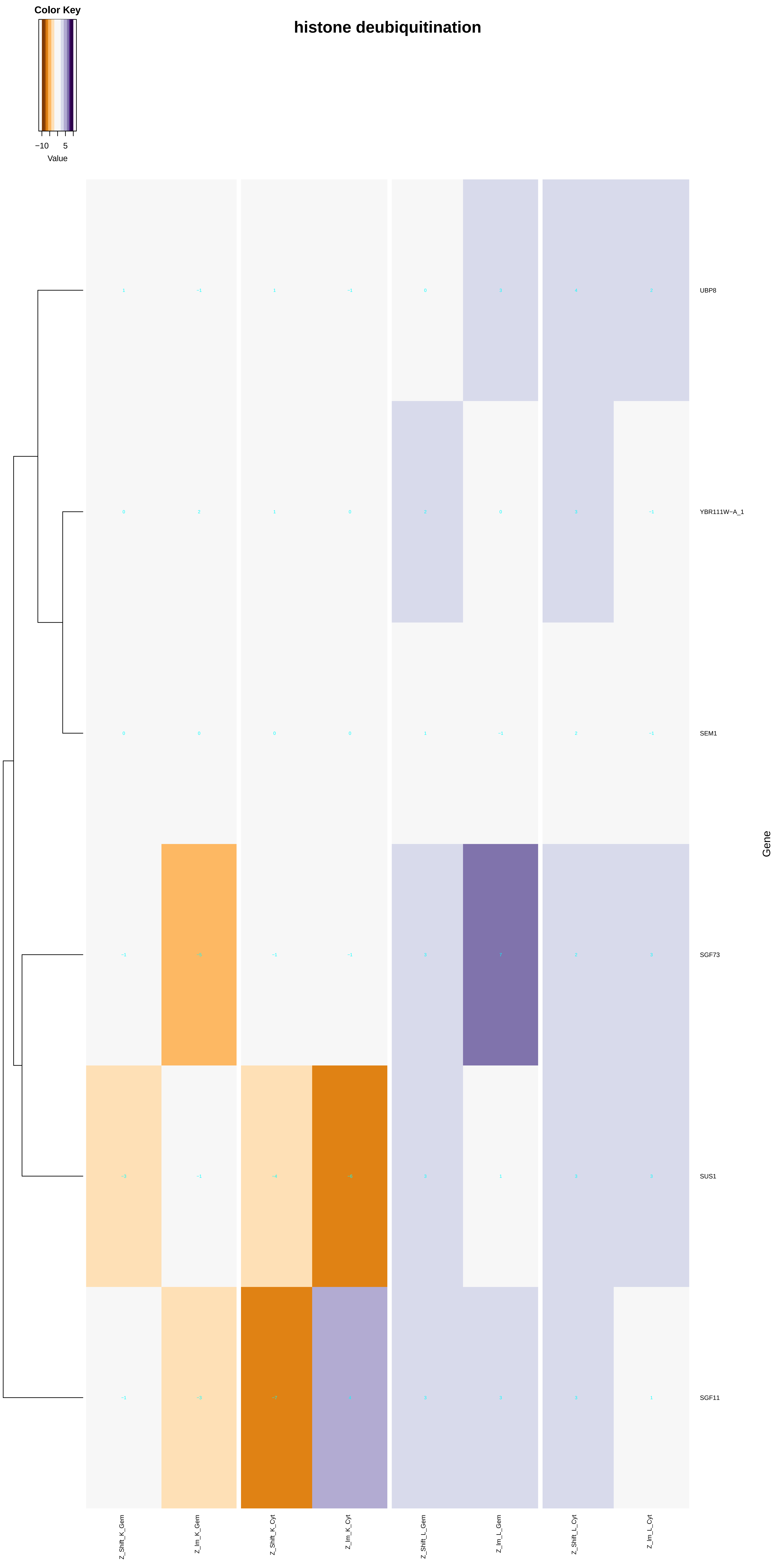

Color Key
−10
5
Value
histone deubiquitination
UBP8
YBR111W−A_1
SEM1
SGF73
SUS1
SGF11
Gene
Z_Shift_K_Gem
Z_Im_K_Gem
Z_Shift_K_Cyt
Z_Im_K_Cyt
Z_Shift_L_Gem
Z_Im_L_Gem
Z_Shift_L_Cyt
Z_Im_L_Cyt

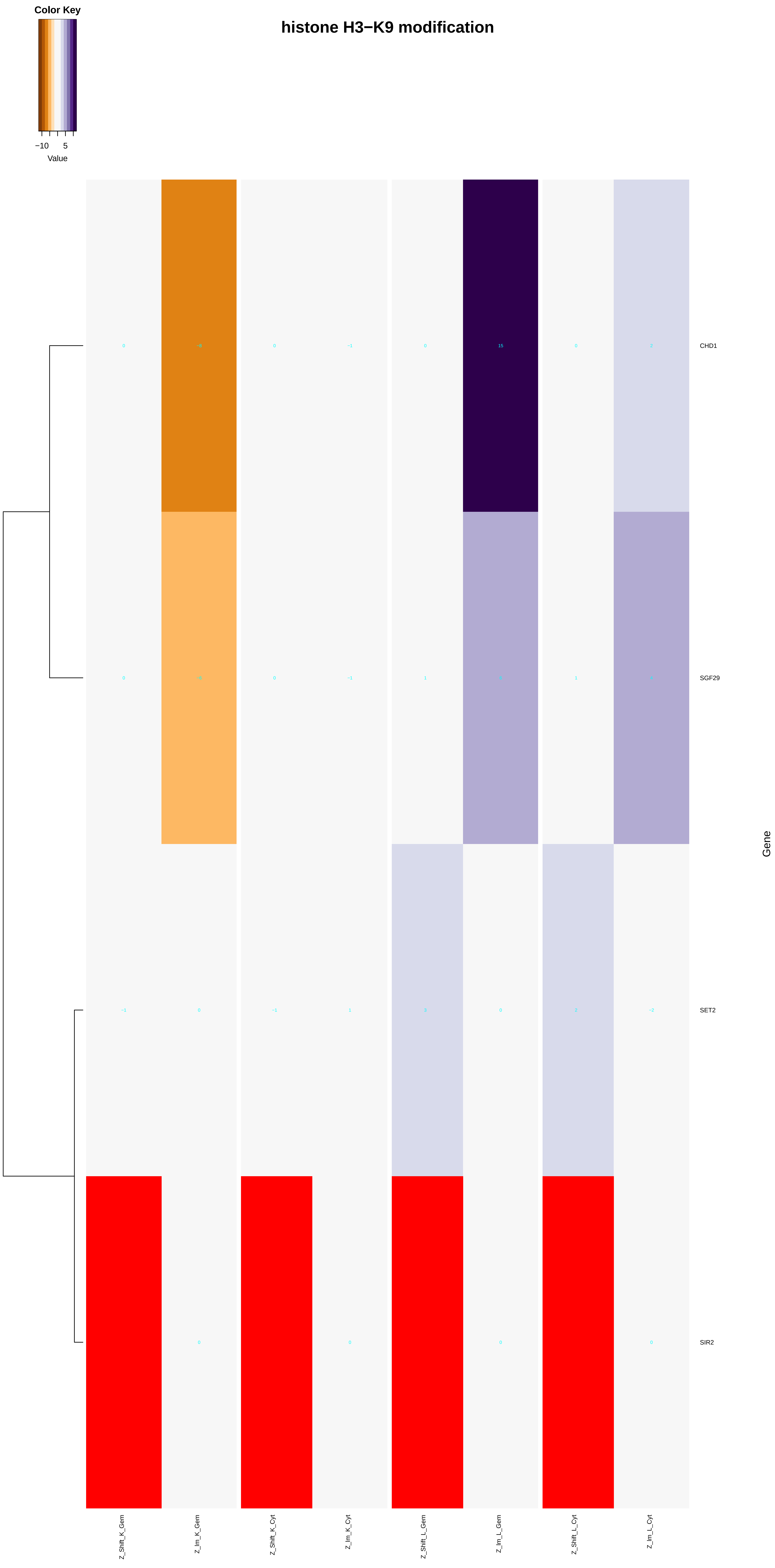
histone H3−K9 modification
Color Key
−10
5
Value
CHD1
SGF29
SET2
SIR2
Gene
Z_Shift_K_Gem
Z_Im_K_Gem
Z_Shift_K_Cyt
Z_Im_K_Cyt
Z_Shift_L_Gem
Z_Im_L_Gem
Z_Shift_L_Cyt
Z_Im_L_Cyt

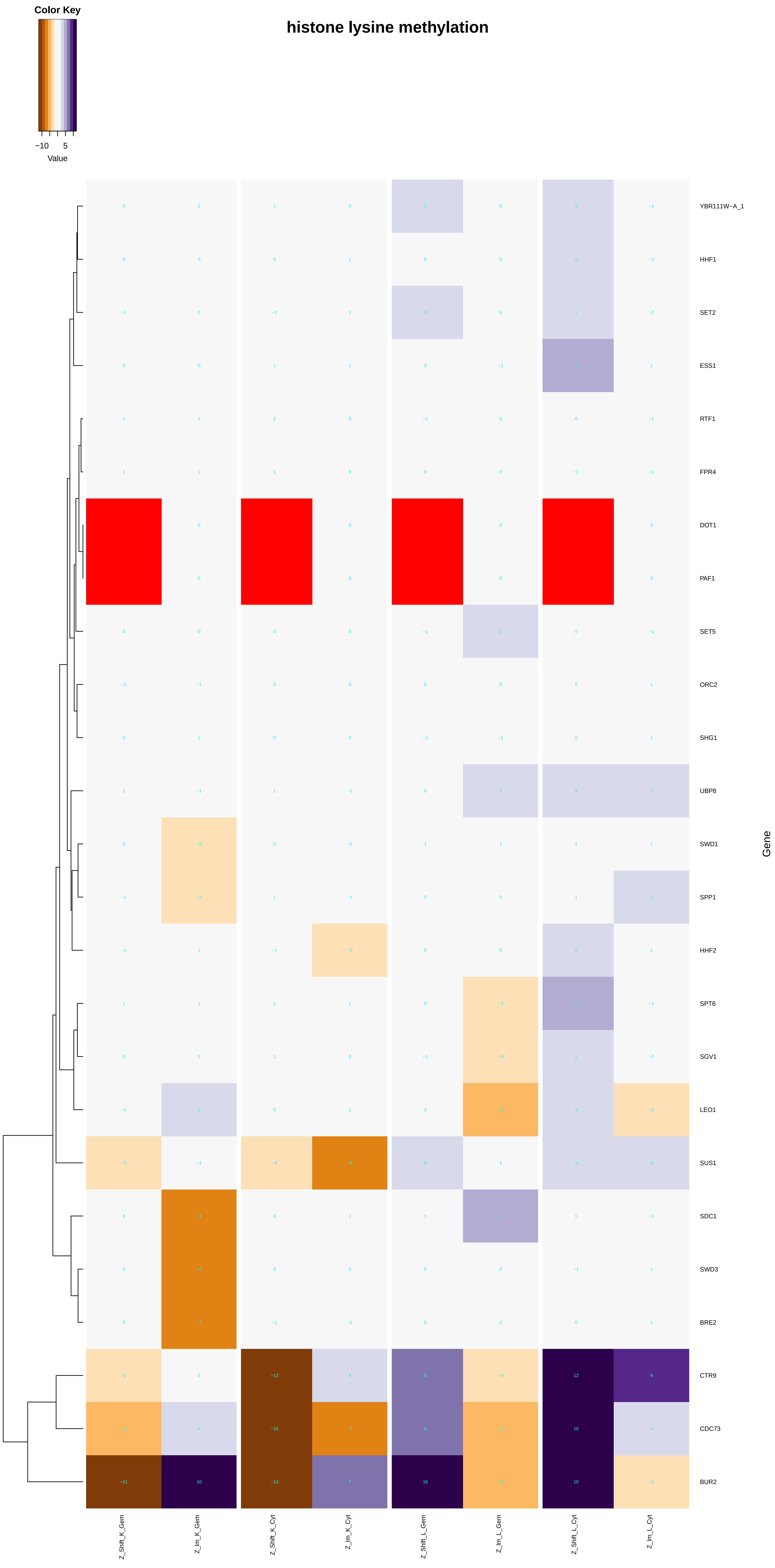
Color Key
−10 5
Value
histone lysine methylation
YBR111W−A_1
HHF1
SET2
ESS1
RTF1
FPR4
DOT1
PAF1
SET5
ORC2
SHG1
UBP8
SWD1
SPP1
HHF2
SPT6
SGV1
LEO1
SUS1
SDC1
SWD3
BRE2
CTR9
CDC73
BUR2
Gene
Z_Shift_K_Gem
Z_Im_K_Gem
Z_Shift_K_Cyt
Z_Im_K_Cyt
Z_Shift_L_Gem
Z_Im_L_Gem
Z_Shift_L_Cyt
Z_Im_L_Cyt

### histone arginine methylation

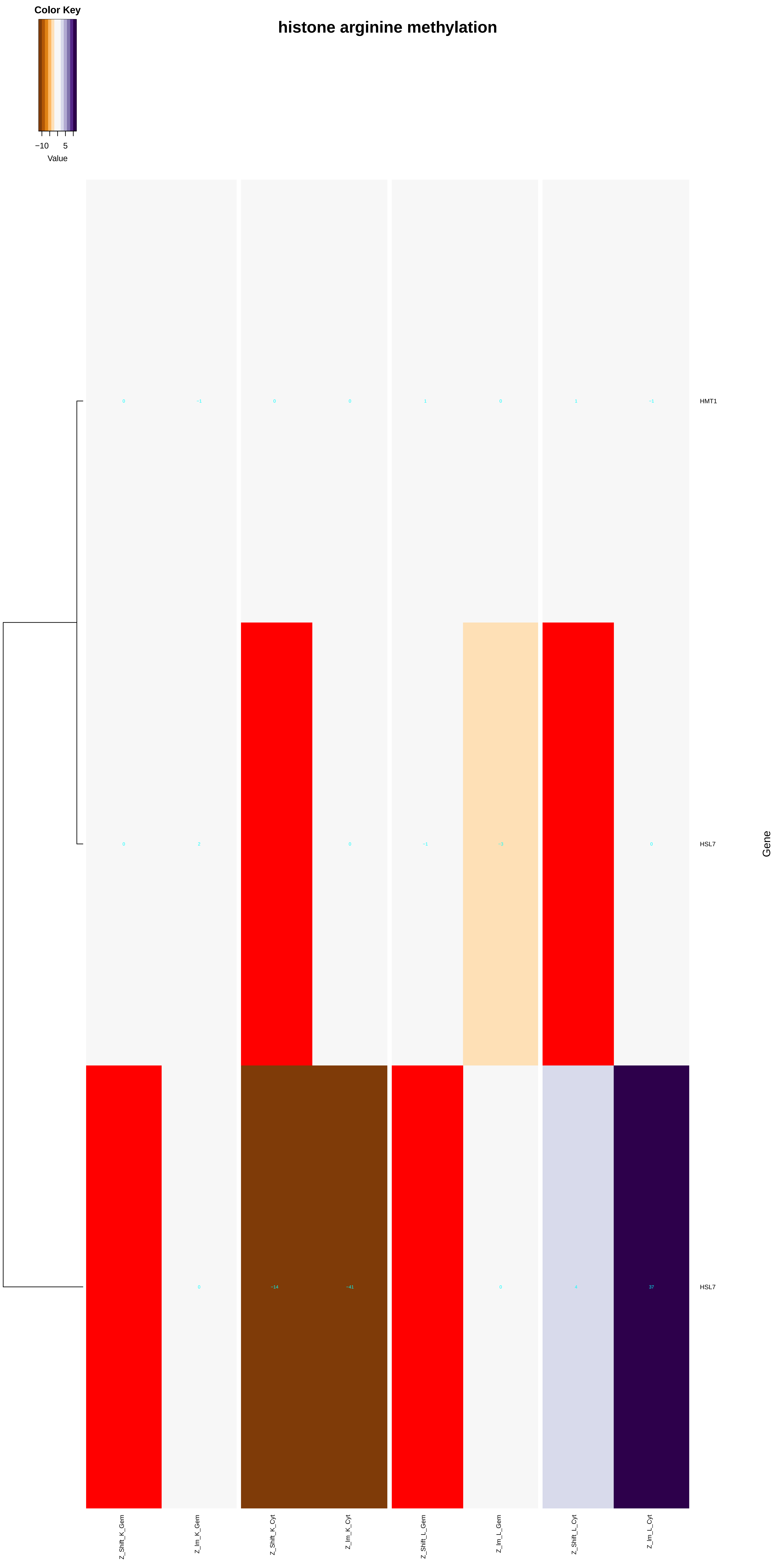

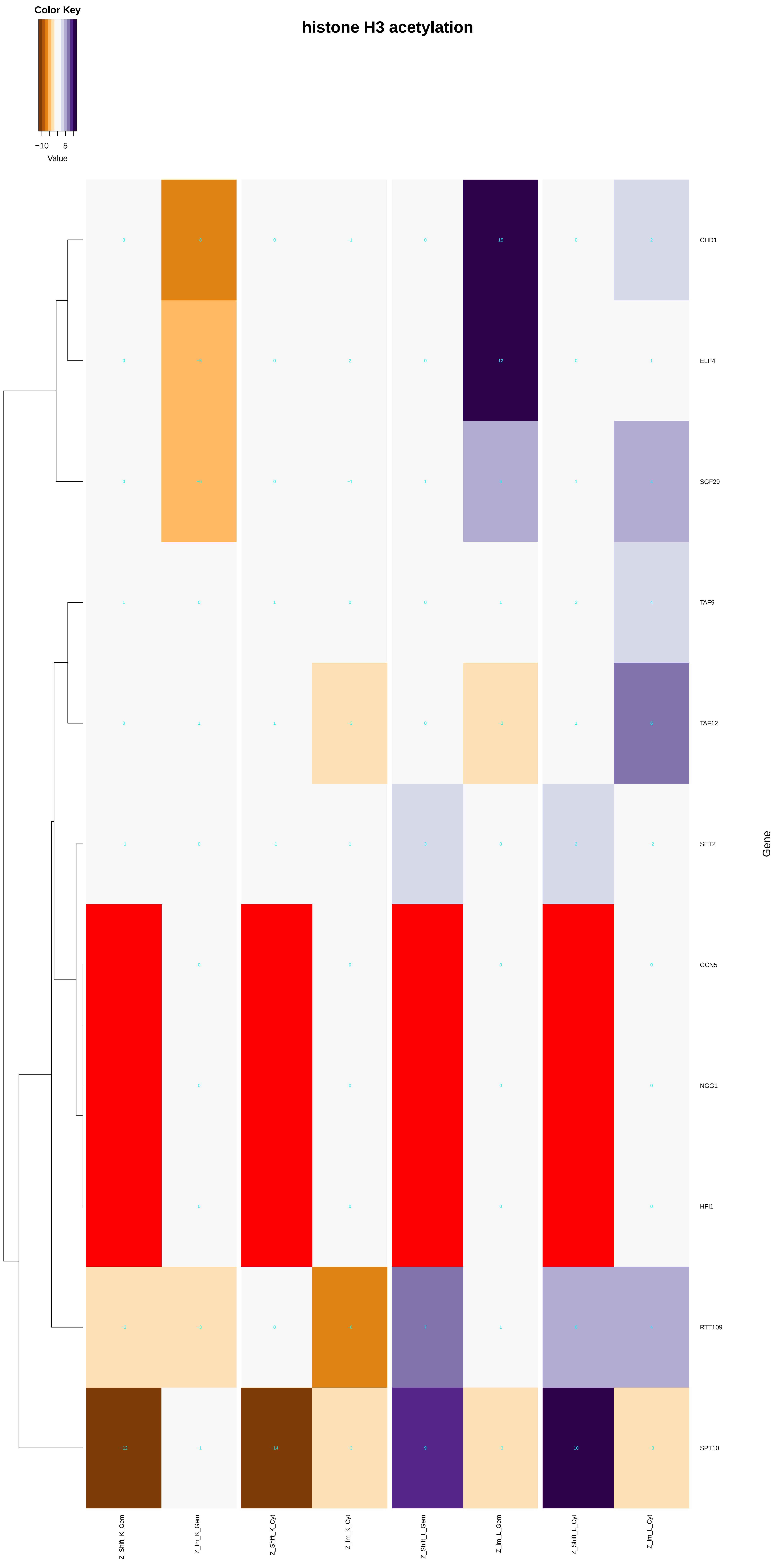

histone H3 acetylation
Color Key
−10
5
Value
CHD1
ELP4
SGF29
TAF9
TAF12
SET2
GCN5
NGG1
HFI1
RTT109
SPT10
Gene
Z_Shift_K_Gem
Z_Im_K_Gem
Z_Shift_K_Cyt
Z_Im_K_Cyt
Z_Shift_L_Gem
Z_Im_L_Gem
Z_Shift_L_Cyt
Z_Im_L_Cyt

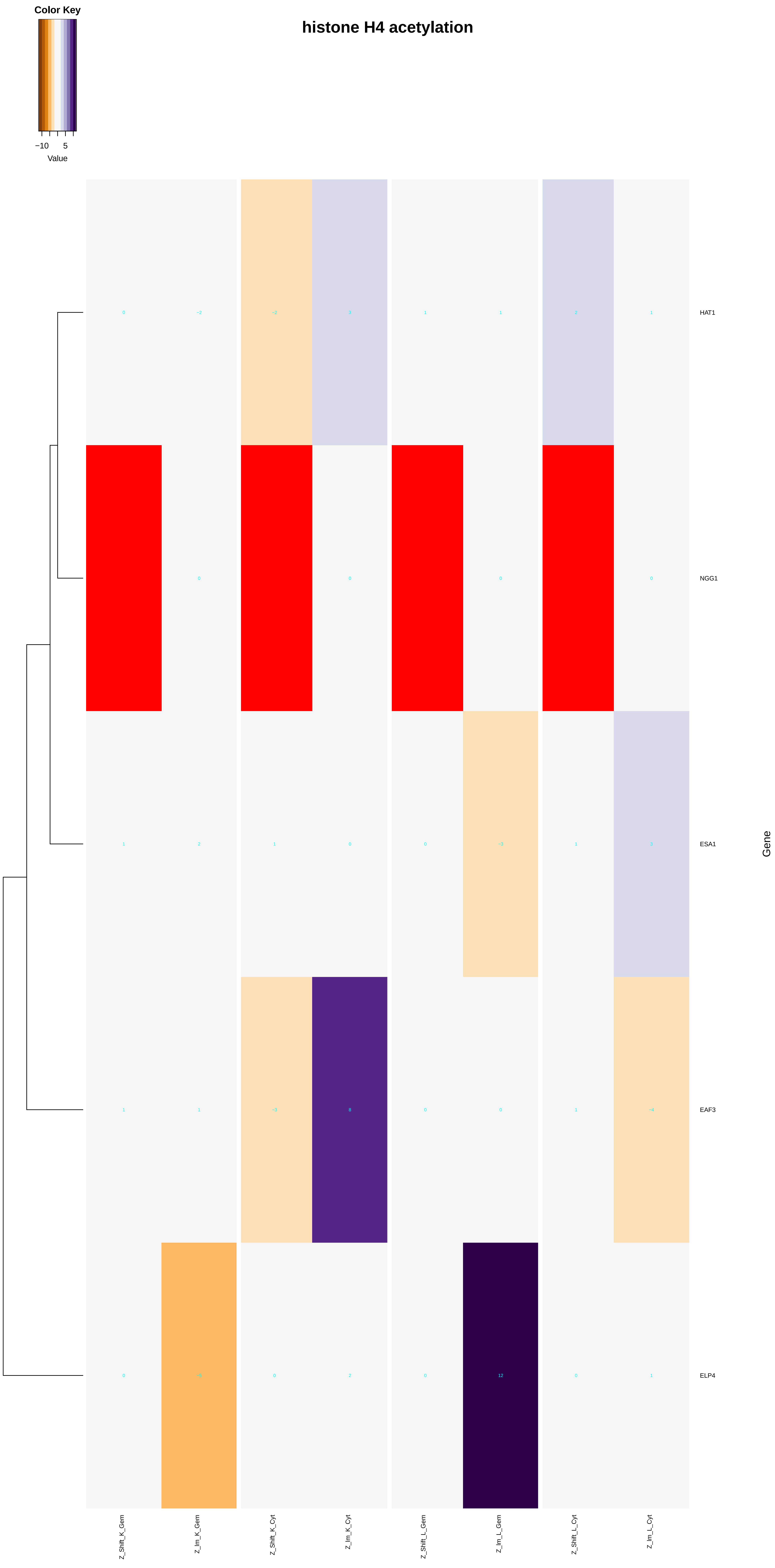

Color Key
−10
5
Value
histone H4 acetylation
HAT1
NGG1
ESA1
EAF3
ELP4
Gene
Z_Shift_K_Gem
Z_Im_K_Gem
Z_Shift_K_Cyt
Z_Im_K_Cyt
Z_Shift_L_Gem
Z_Im_L_Gem
Z_Shift_L_Cyt
Z_Im_L_Cyt

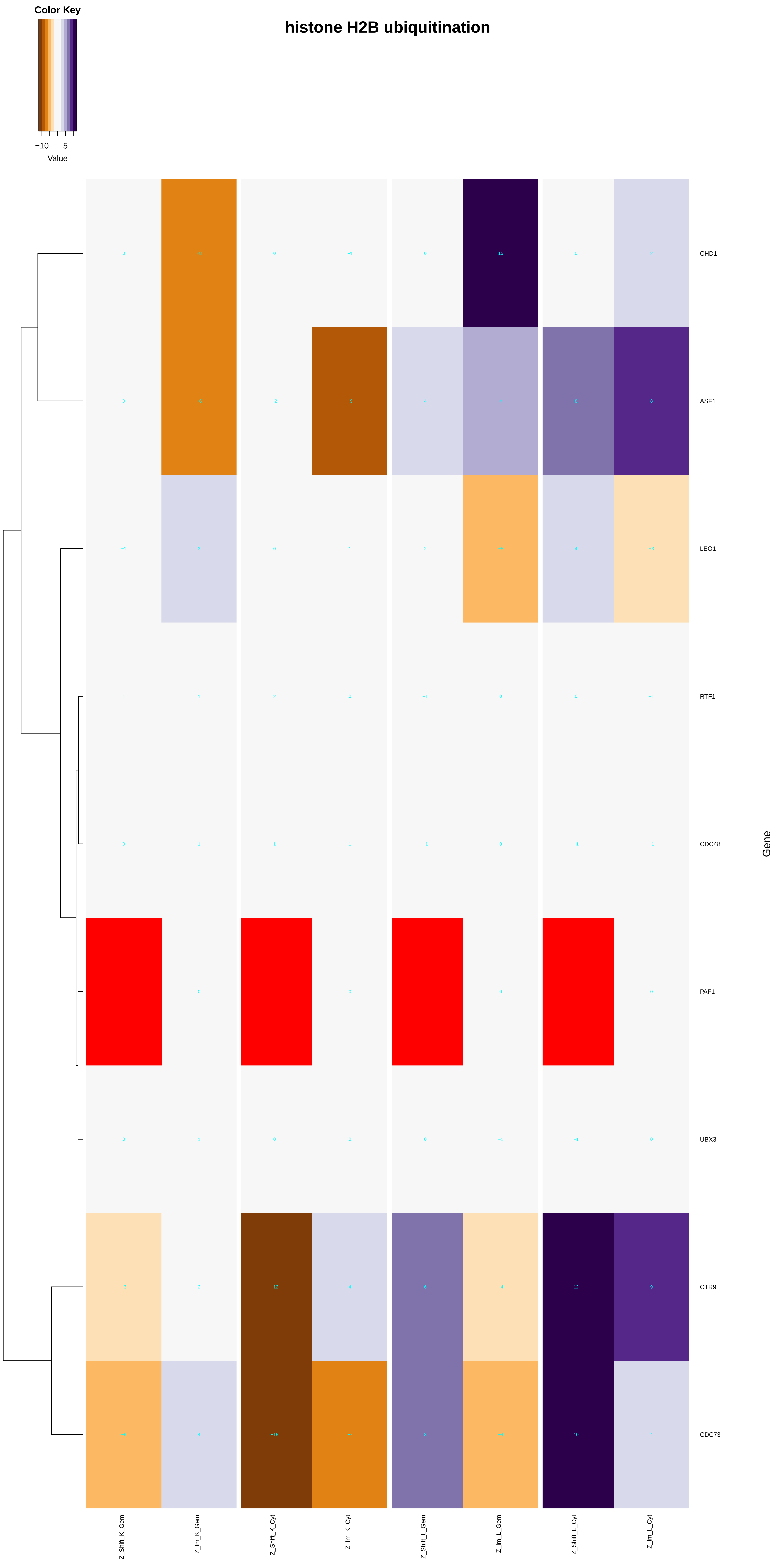

Color Key
−10 5
Value
histone H2B ubiquitination
CHD1
ASF1
LEO1
RTF1
CDC48
PAF1
UBX3
CTR9
CDC73
Gene
Z_Shift_K_Gem
Z_Im_K_Gem
Z_Shift_K_Cyt
Z_Im_K_Cyt
Z_Shift_L_Gem
Z_Im_L_Gem
Z_Shift_L_Cyt
Z_Im_L_Cyt

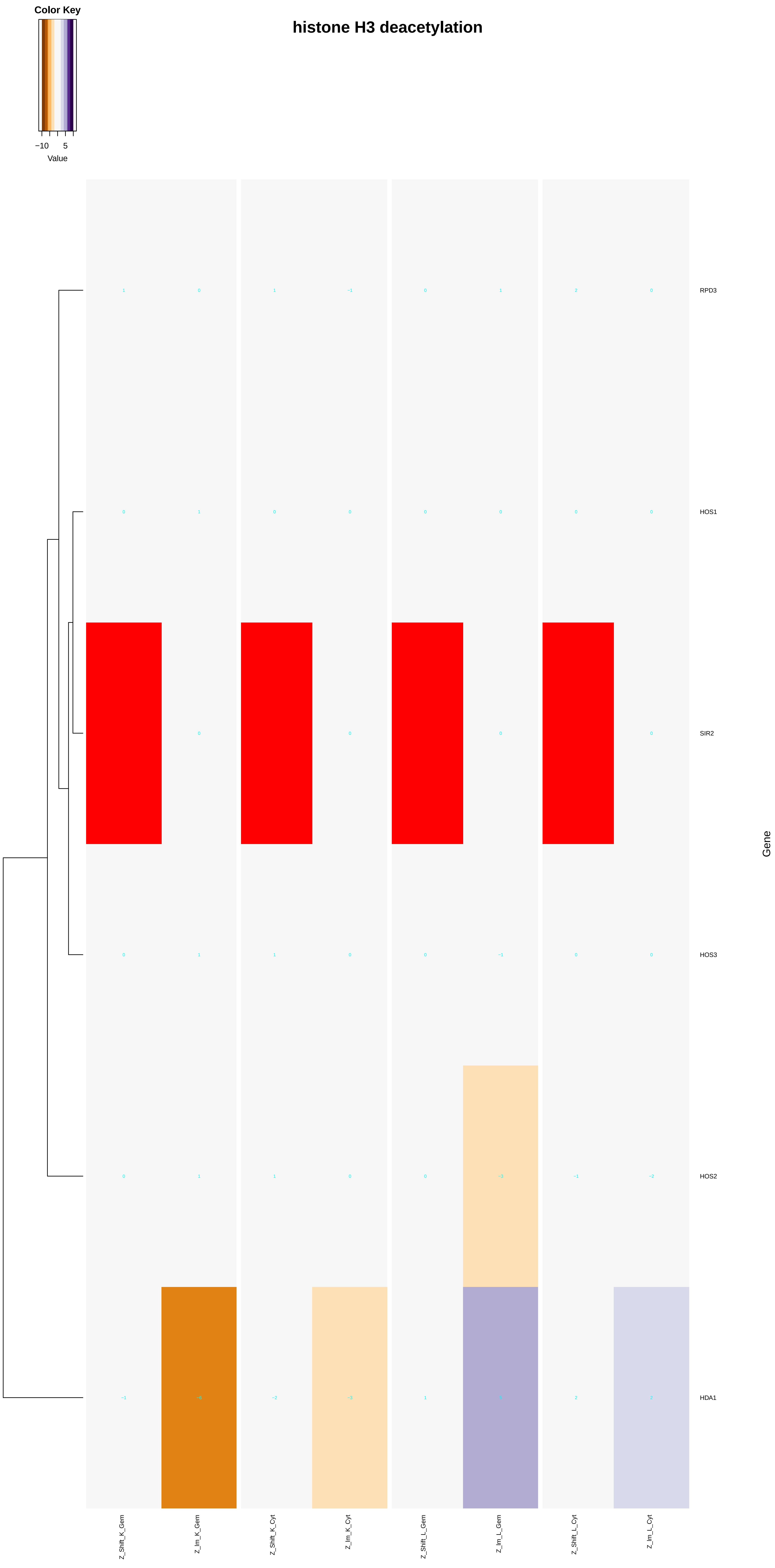
Color Key
−10
5
Value
histone H3 deacetylation
RPD3
HOS1
SIR2
HOS3
HOS2
HDA1
Gene
Z_Shift_K_Gem
Z_Im_K_Gem
Z_Shift_K_Cyt
Z_Im_K_Cyt
Z_Shift_L_Gem
Z_Im_L_Gem
Z_Shift_L_Cyt
Z_Im_L_Cyt

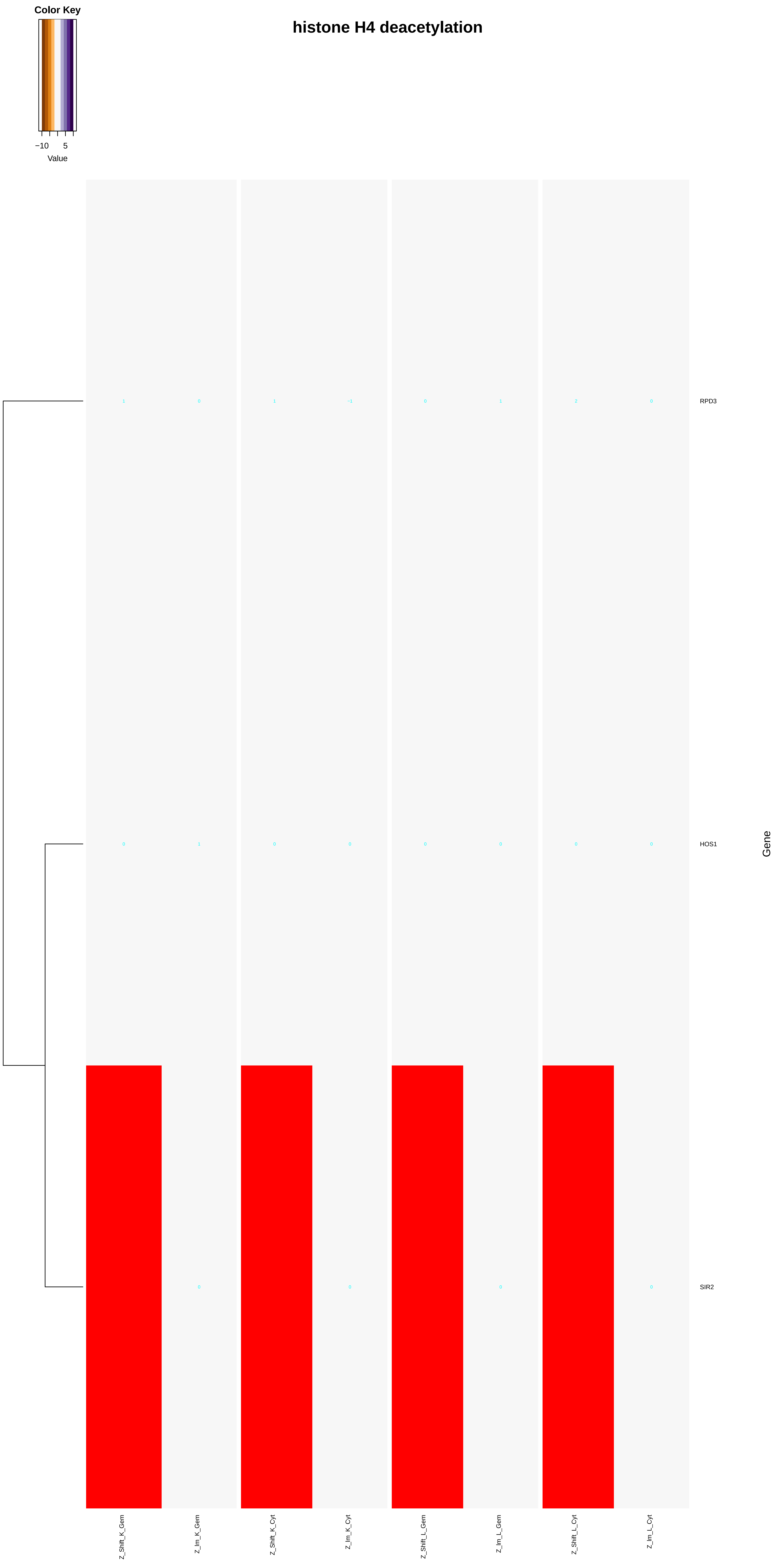

Color Key
−10
5
Value
histone H4 deacetylation
RPD3
HOS1
SIR2
Gene
Z_Shift_K_Gem
Z_Im_K_Gem
Z_Shift_K_Cyt
Z_Im_K_Cyt
Z_Shift_L_Gem
Z_Im_L_Gem
Z_Shift_L_Cyt
Z_Im_L_Cyt

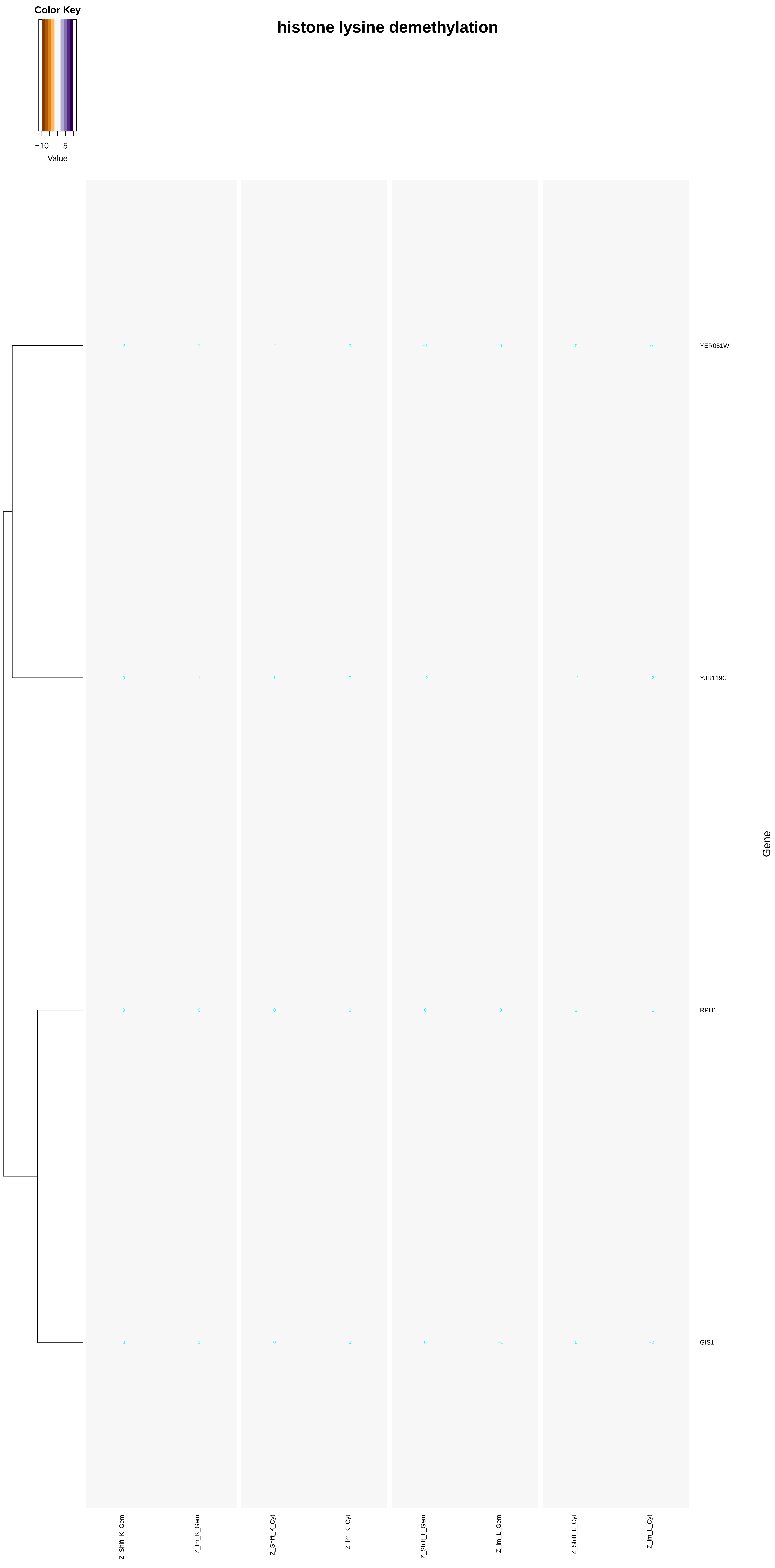

Color Key
−10
5
Value
histone lysine demethylation
YER051W
YJR119C
RPH1
GIS1
Gene
Z_Shift_K_Gem
Z_Im_K_Gem
Z_Shift_K_Cyt
Z_Im_K_Cyt
Z_Shift_L_Gem
Z_Im_L_Gem
Z_Shift_L_Cyt
Z_Im_L_Cyt

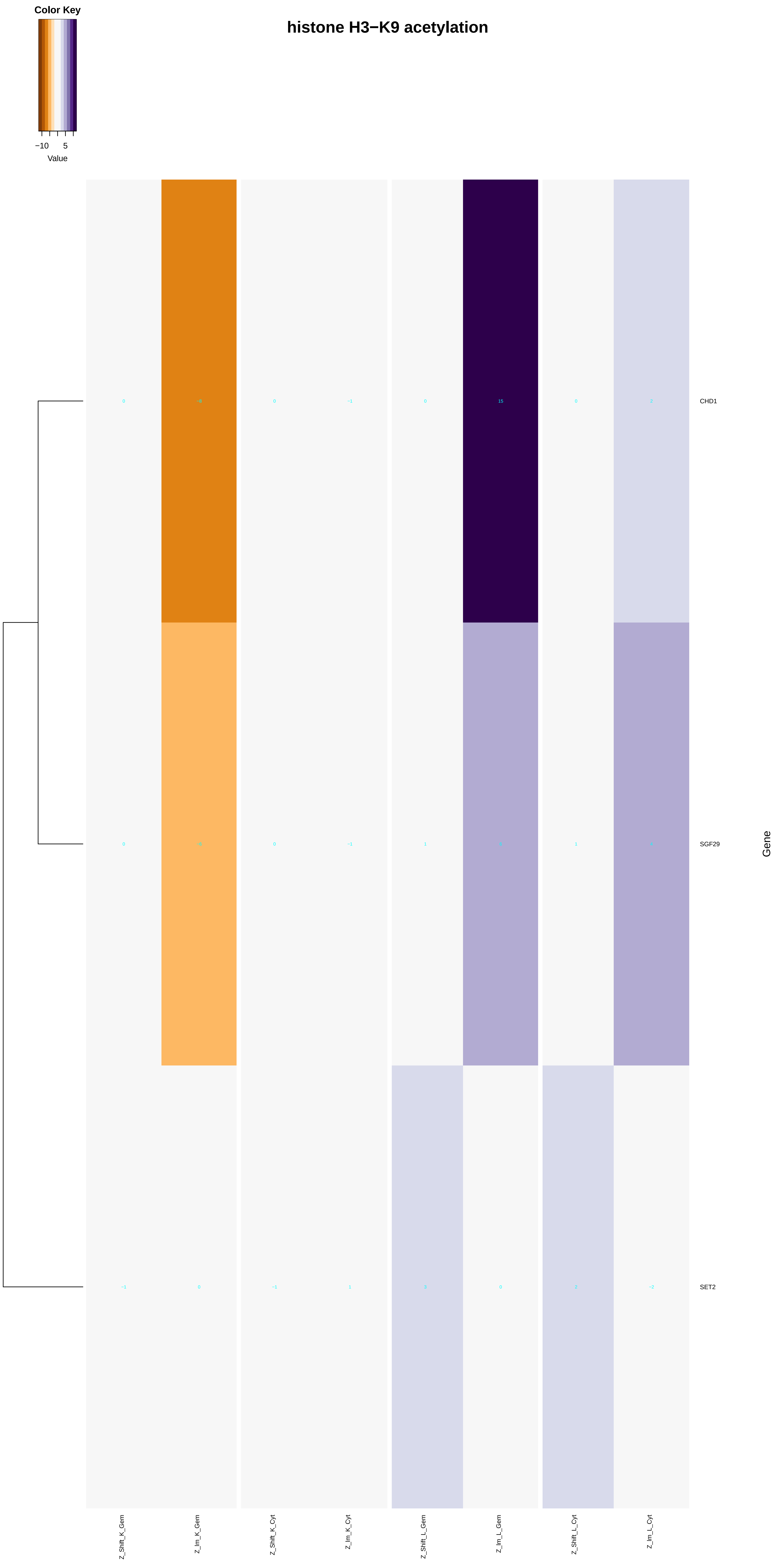

Color Key
−10
5
Value
histone H3−K9 acetylation
0
−8
0
−1
0
15
0
2
CHD1
0
−6
0
−1
1
6
1
4
SGF29
−1
0
−1
1
3
0
2
−2
SET2
Gene
Z_Shift_K_Gem
Z_Im_K_Gem
Z_Shift_K_Cyt
Z_Im_K_Cyt
Z_Shift_L_Gem
Z_Im_L_Gem
Z_Shift_L_Cyt
Z_Im_L_Cyt

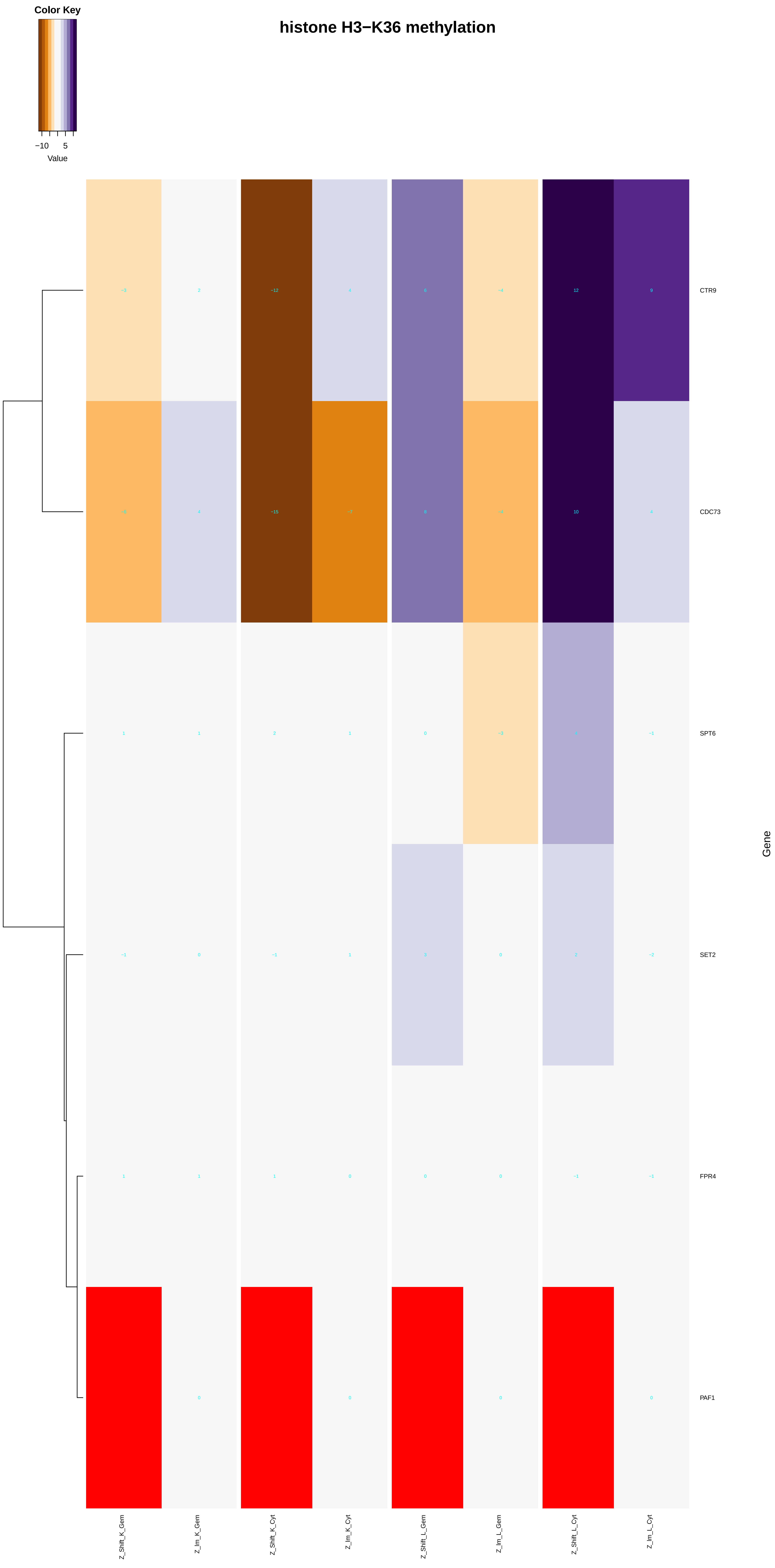
histone H3−K36 methylation
Color Key
−10
5
Value
CTR9
CDC73
SPT6
SET2
FPR4
PAF1
Gene
Z_Shift_K_Gem
Z_Im_K_Gem
Z_Shift_K_Cyt
Z_Im_K_Cyt
Z_Shift_L_Gem
Z_Im_L_Gem
Z_Shift_L_Cyt
Z_Im_L_Cyt

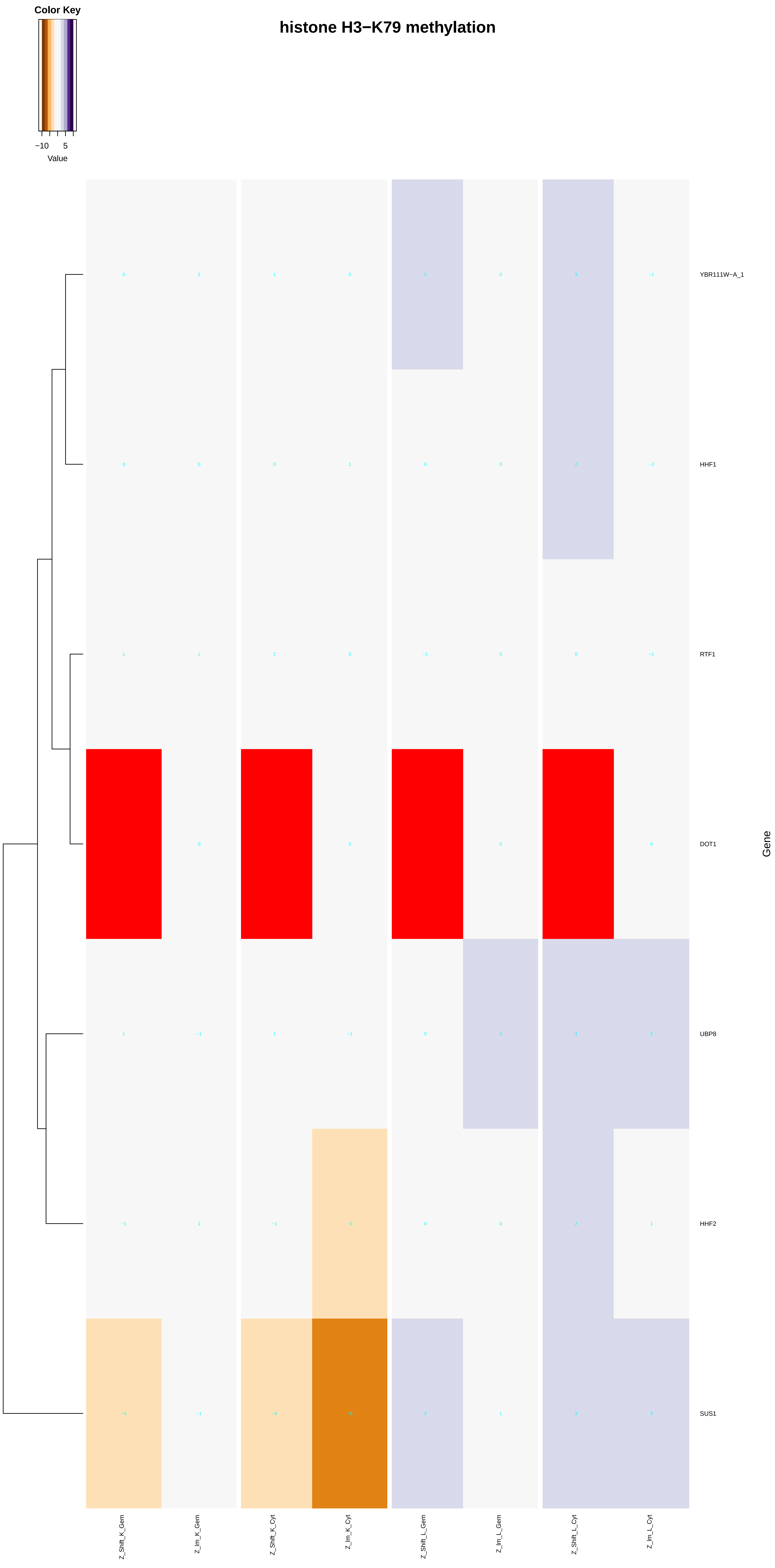

Color Key
−10
5
Value
histone H3−K79 methylation
YBR111W−A_1
HHF1
RTF1
DOT1
UBP8
HHF2
SUS1
Gene
Z_Shift_K_Gem
Z_Im_K_Gem
Z_Shift_K_Cyt
Z_Im_K_Cyt
Z_Shift_L_Gem
Z_Im_L_Gem
Z_Shift_L_Cyt
Z_Im_L_Cyt

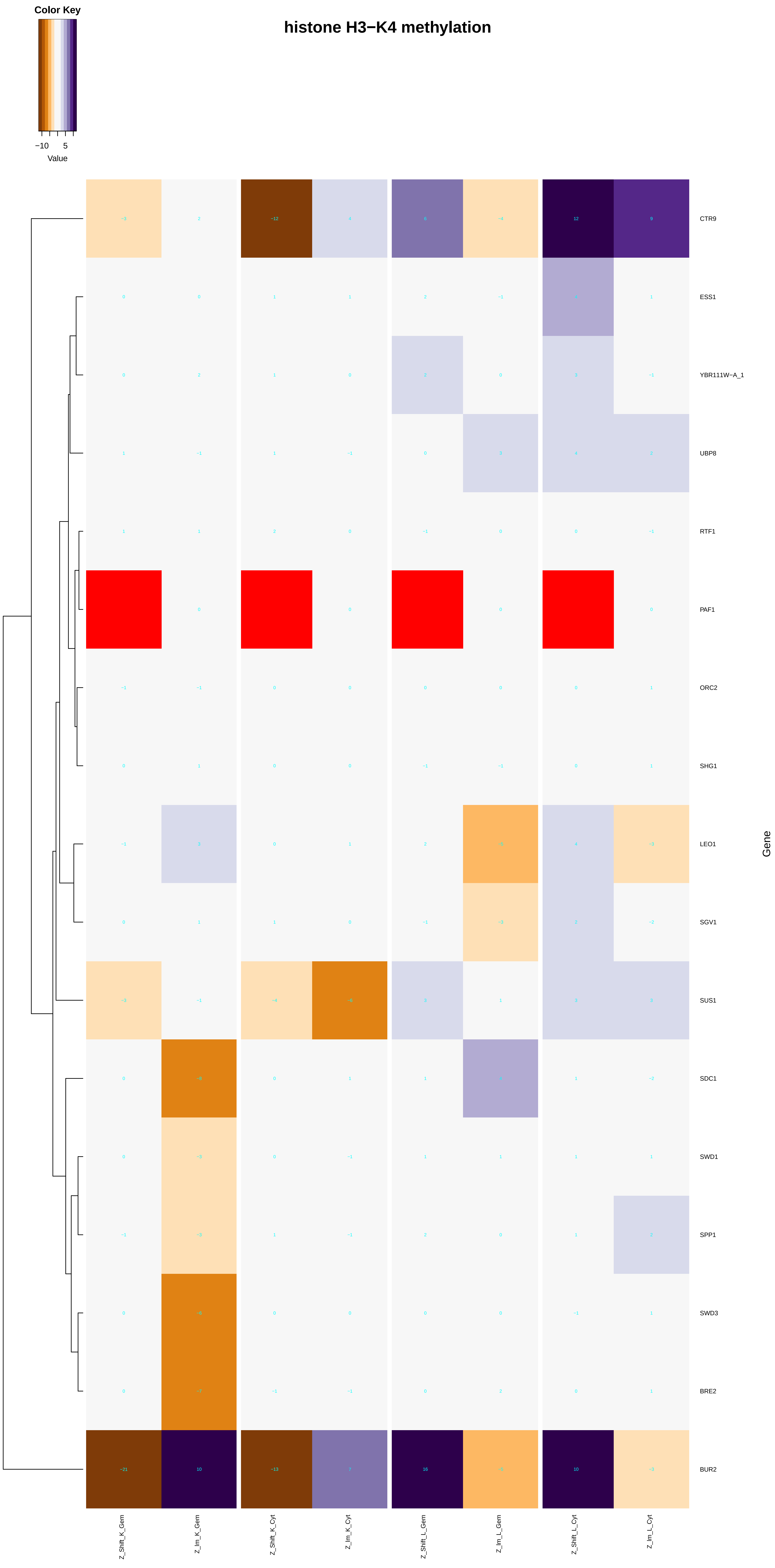

Color Key
−10
5
Value
histone H3−K4 methylation
CTR9
ESS1
YBR111W−A_1
UBP8
RTF1
PAF1
ORC2
SHG1
LEO1
SGV1
SUS1
SDC1
SWD1
SPP1
SWD3
BRE2
BUR2
Gene
Z_Shift_K_Gem
Z_Im_K_Gem
Z_Shift_K_Cyt
Z_Im_K_Cyt
Z_Shift_L_Gem
Z_Im_L_Gem
Z_Shift_L_Cyt
Z_Im_L_Cyt

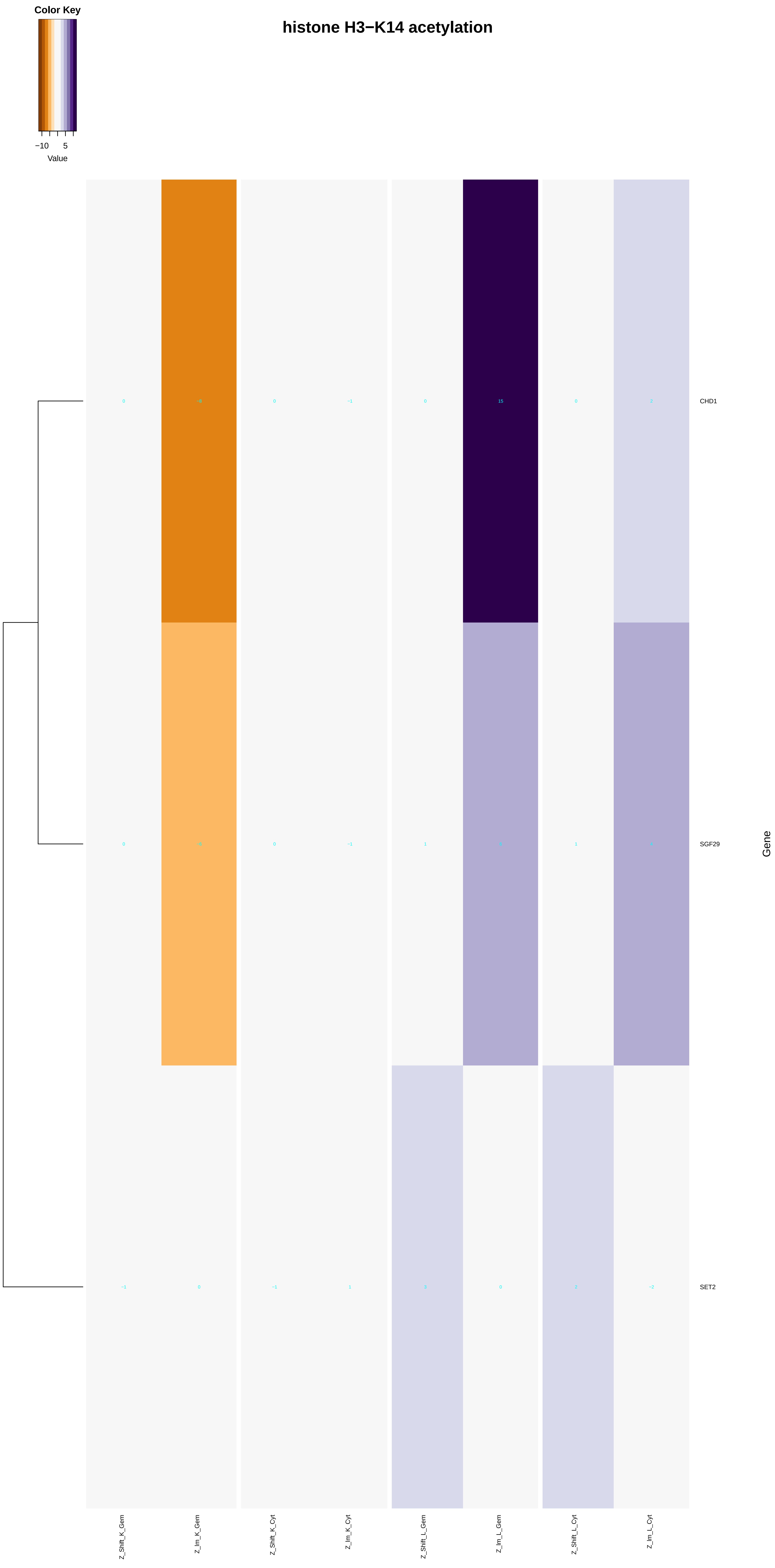

Color Key
−10
5
Value
histone H3−K14 acetylation
0
−8
0
−1
0
15
0
2
CHD1
0
−6
0
−1
1
6
1
4
SGF29
−1
0
−1
1
3
0
2
−2
SET2
Gene
Z_Shift_K_Gem
Z_Im_K_Gem
Z_Shift_K_Cyt
Z_Im_K_Cyt
Z_Shift_L_Gem
Z_Im_L_Gem
Z_Shift_L_Cyt
Z_Im_L_Cyt

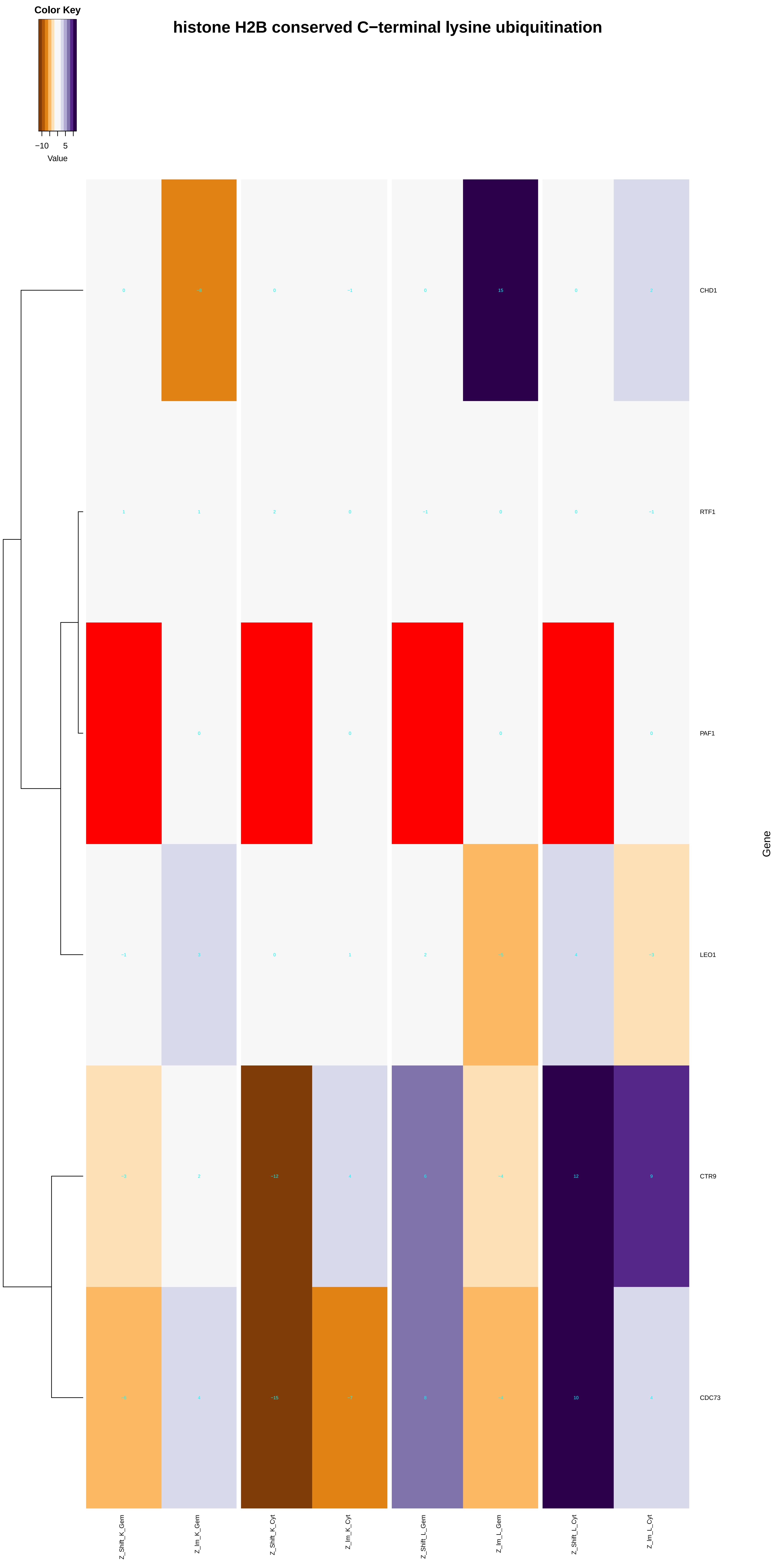

histone H2B conserved C−terminal lysine ubiquitination
Color Key
−10
5
Value
CHD1
RTF1
PAF1
LEO1
CTR9
CDC73
Gene
Z_Shift_K_Gem
Z_Im_K_Gem
Z_Shift_K_Cyt
Z_Im_K_Cyt
Z_Shift_L_Gem
Z_Im_L_Gem
Z_Shift_L_Cyt
Z_Im_L_Cyt

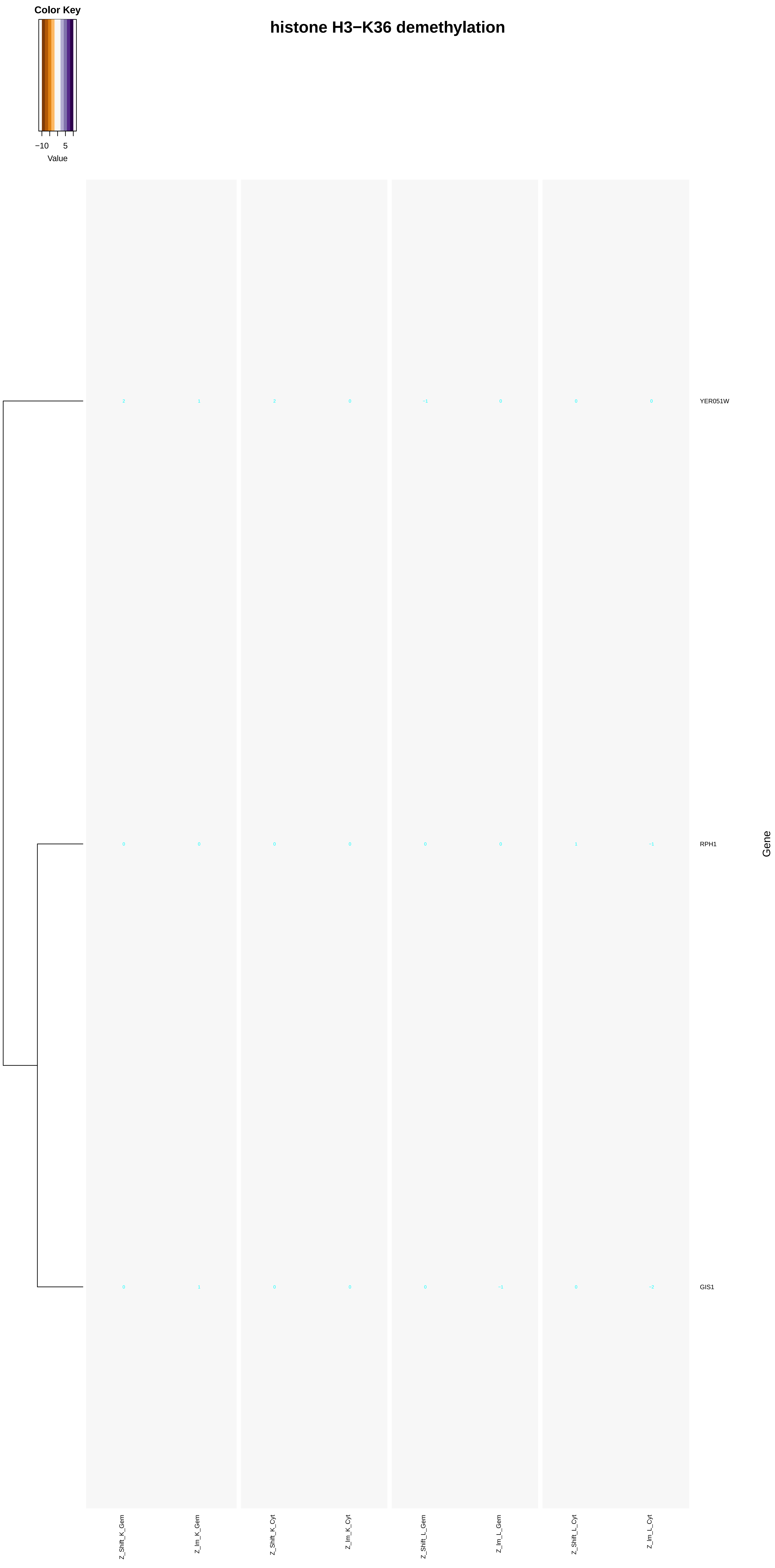
histone H3−K36 demethylation
Color Key
−10
5
Value
YER051W
RPH1
GIS1
Gene
2 1 2 0 −1 0 0 0
0 0 0 0 0 0 1 −1
0 1 0 0 0 −1 0 −2
Z_Shift_K_Gem
Z_Im_K_Gem
Z_Shift_K_Cyt
Z_Im_K_Cyt
Z_Shift_L_Gem
Z_Im_L_Gem
Z_Shift_L_Cyt
Z_Im_L_Cyt

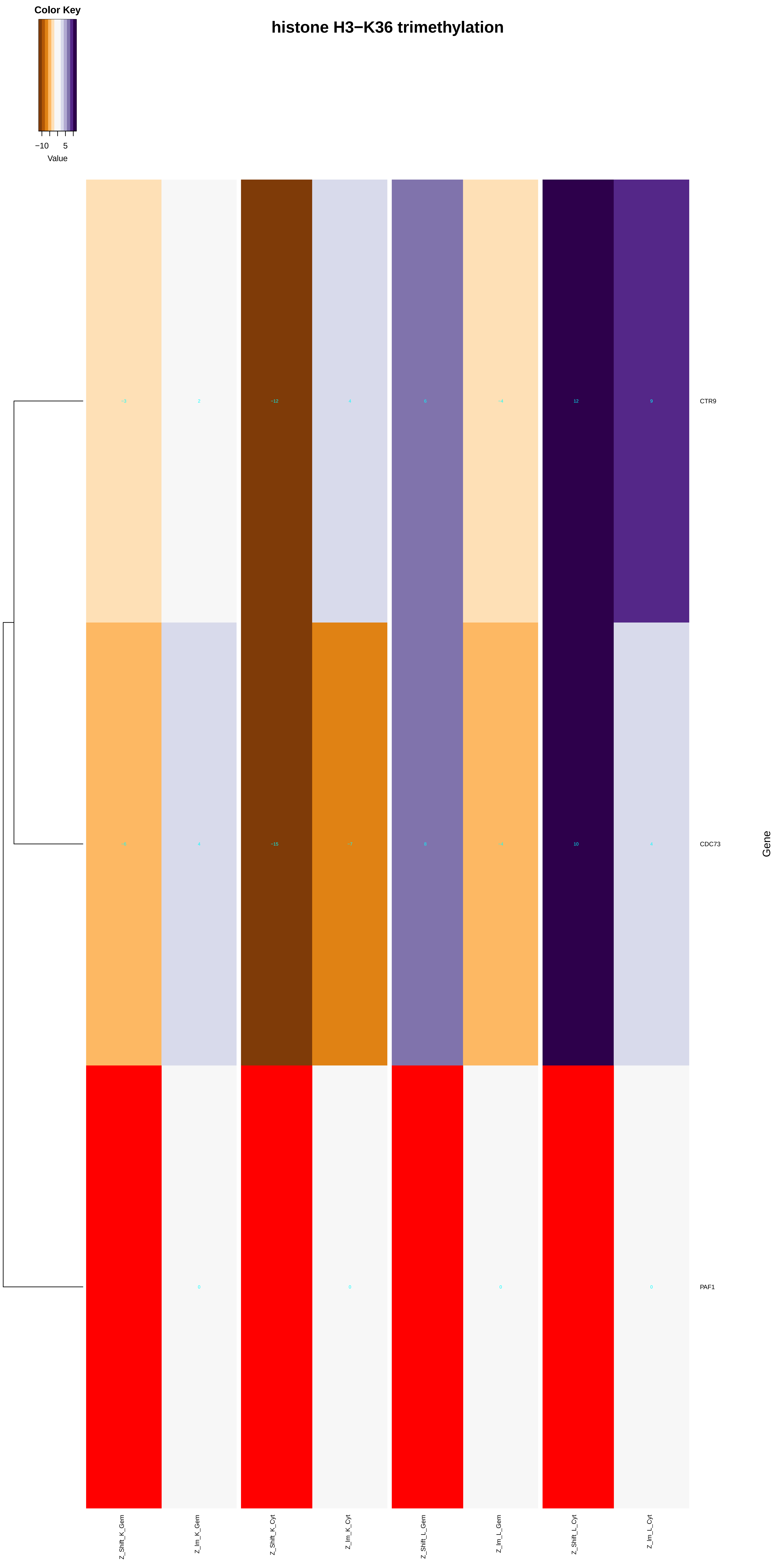

histone H3−K36 trimethylation
Color Key
−10
5
Value
−3
2
−12
4
6
−4
12
9
CTR9
−6
4
−15
−7
8
−4
10
4
CDC73
0
0
0
0
PAF1
Gene
Z_Shift_K_Gem
Z_Im_K_Gem
Z_Shift_K_Cyt
Z_Im_K_Cyt
Z_Shift_L_Gem
Z_Im_L_Gem
Z_Shift_L_Cyt
Z_Im_L_Cyt

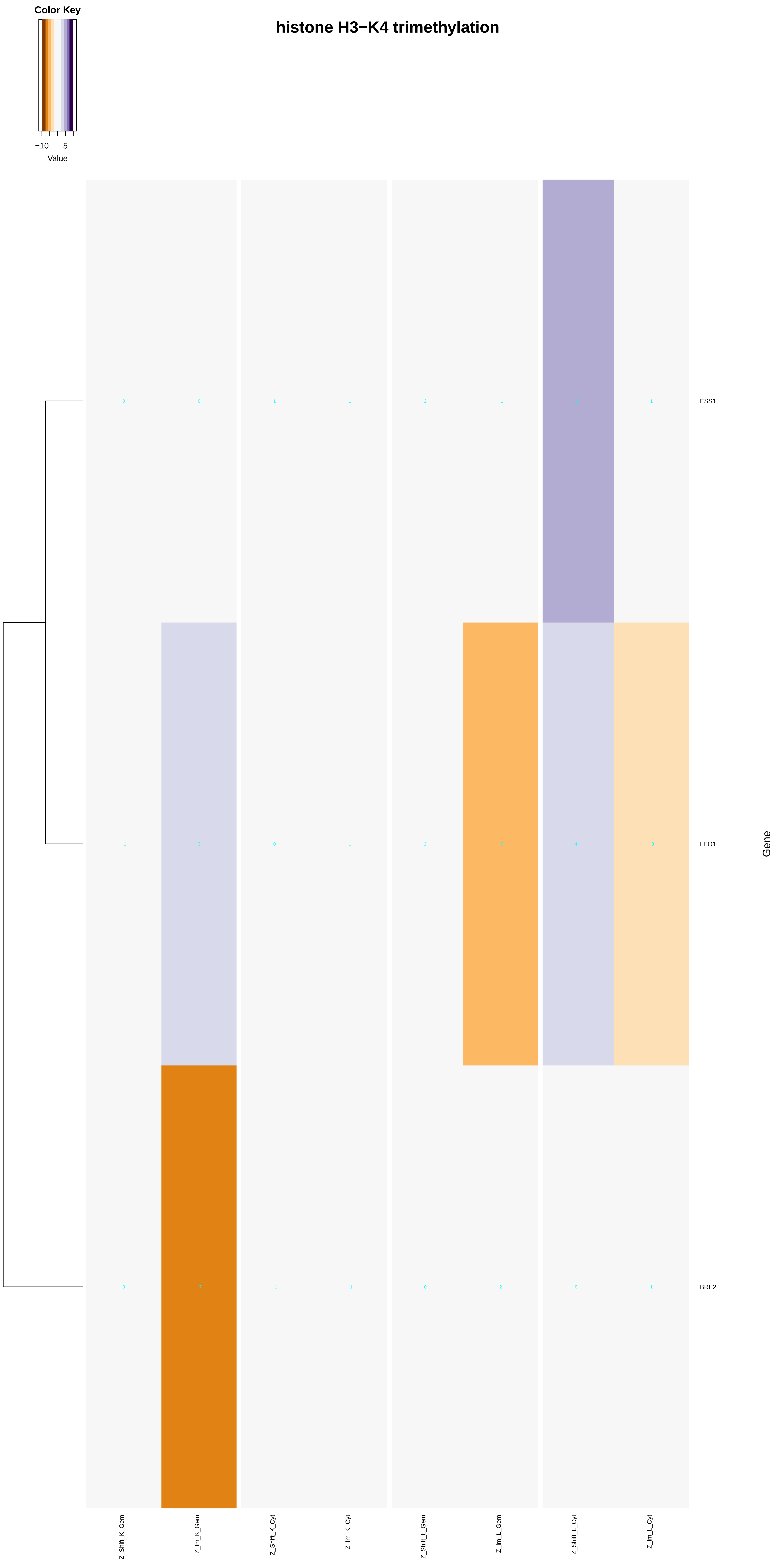

Color Key
−10
5
Value
histone H3−K4 trimethylation
ESS1
LEO1
BRE2
Gene
Z_Shift_K_Gem
Z_Im_K_Gem
Z_Shift_K_Cyt
Z_Im_K_Cyt
Z_Shift_L_Gem
Z_Im_L_Gem
Z_Shift_L_Cyt
Z_Im_L_Cyt
