## Additional File 7 for "A humanized yeast phenomic model of deoxycytidine kinase to predict genetic buffering of nucleoside analog cytotoxicity": cytoplasmic_vesicle.pdf

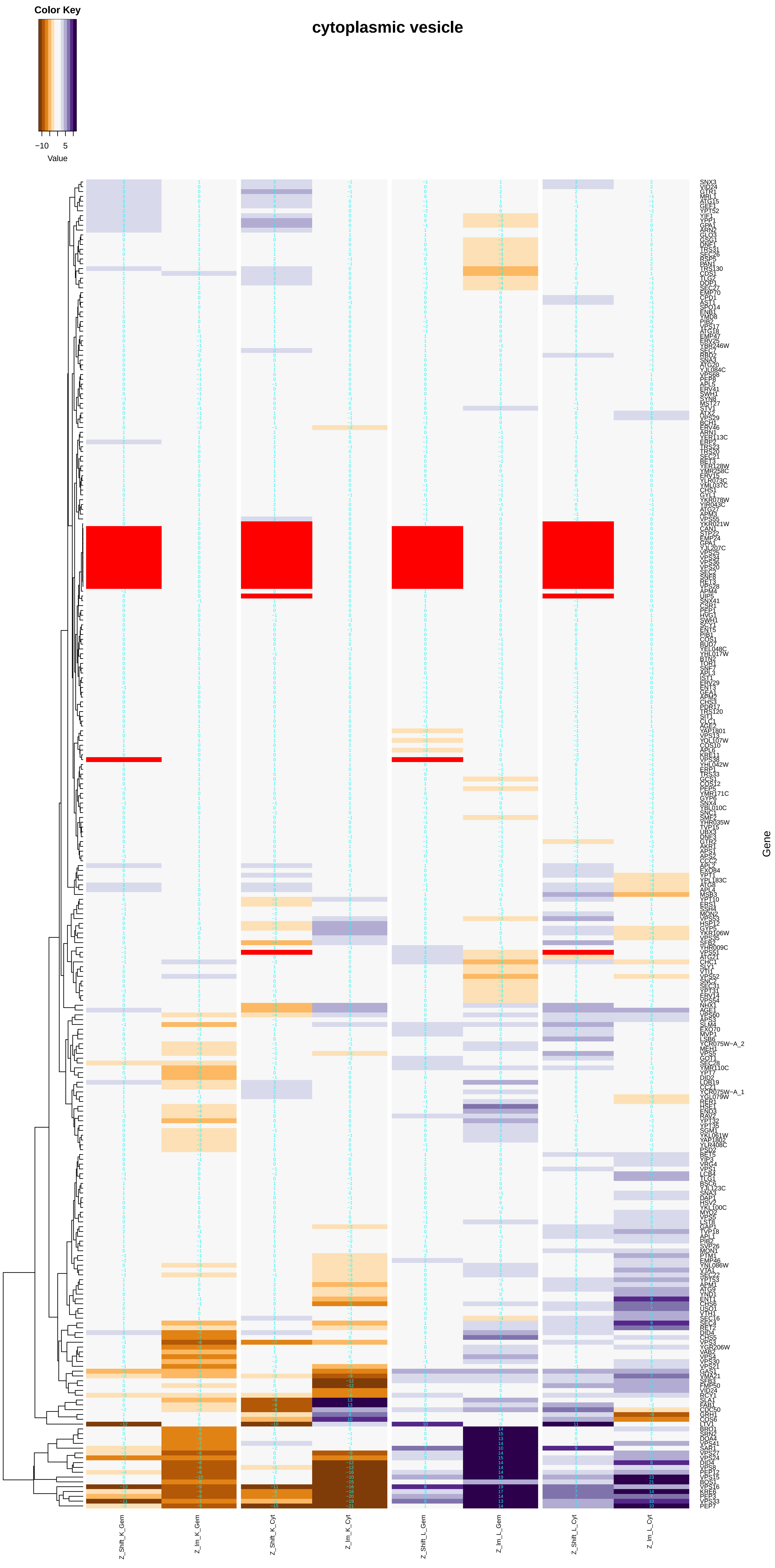
cytoplasmic vesicle
Color Key
−10
5
Value
Z_Shift_K_Gem
Z_Im_K_Gem
Z_Shift_K_Cyt
Z_Im_K_Cyt
Z_Shift_L_Gem
Z_Im_L_Gem
Z_Shift_L_Cyt
Z_Im_L_Cyt
Gene

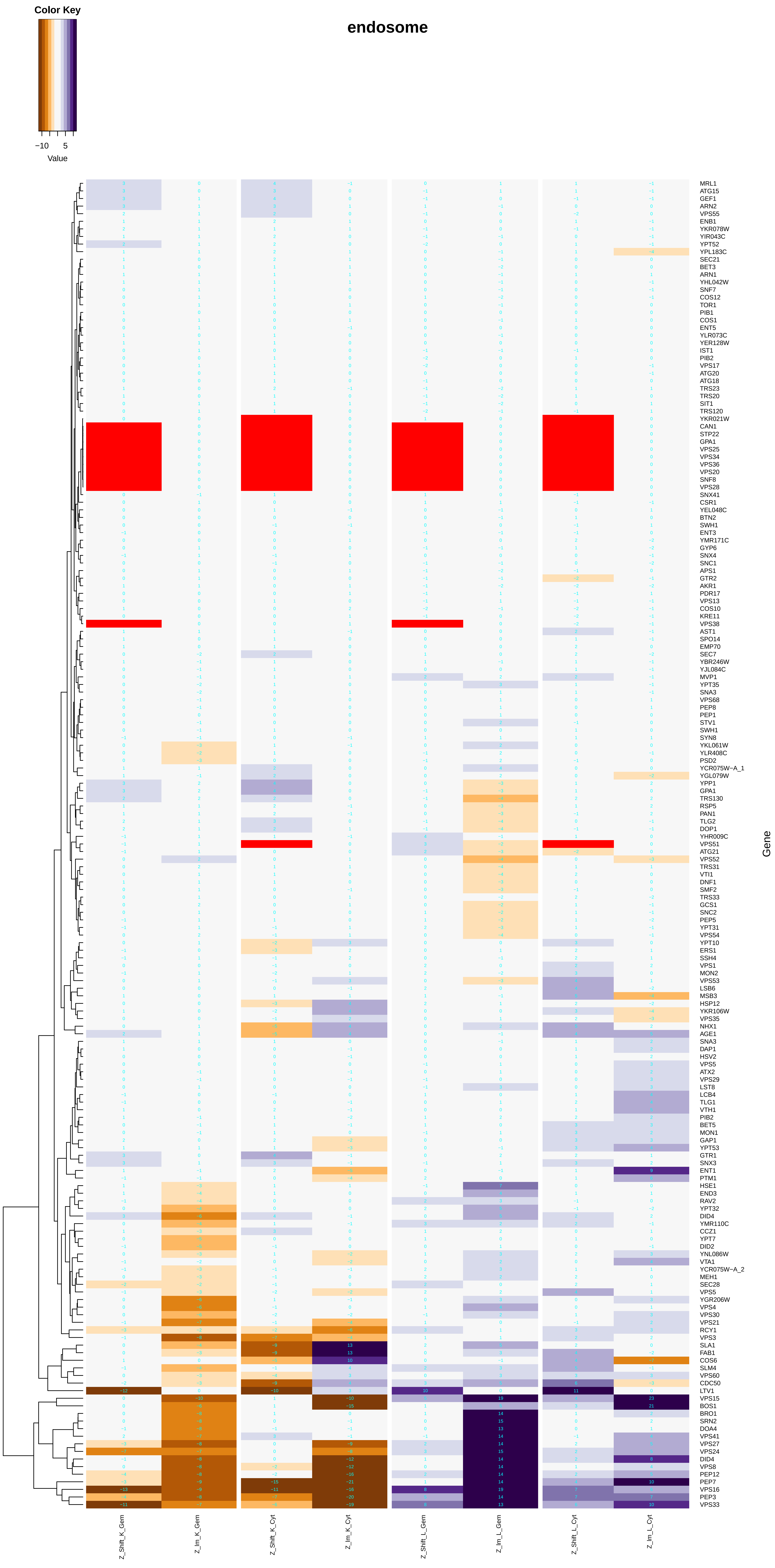
Color Key
−10
5
Value
endosome
Gene
MRL1
ATG15
GEF1
ARN2
VPS55
ENB1
YKR078W
YIR043C
YPT52
YPL183C
SEC21
BET3
ARN1
YHL042W
SNF7
COS12
TOR1
PIB1
COS1
ENT5
YLR073C
YER128W
IST1
PIB2
VPS17
ATG20
ATG18
TRS23
TRS20
SIT1
TRS120
YKR021W
CAN1
STP22
GPA1
VPS25
VPS34
VPS36
VPS20
SNF8
VPS28
SNX41
CSR1
YEL048C
BTN2
SWH1
ENT3
YMR171C
GYP6
SNX4
SNC1
APS1
GTR2
AKR1
PDR17
VPS13
COS10
KRE11
VPS38
AST1
SPO14
EMP70
SEC7
YBR246W
YJL084C
MVP1
YPT35
SNA3
VPS68
PEP8
PEP1
STV1
SWH1
SYN8
YKL061W
YLR408C
PSD2
YCR075W−A_1
YGL079W
YPP1
GPA1
TRS130
RSP5
PAN1
TLG2
DOP1
YHR009C
VPS51
ATG21
VPS52
TRS31
VTI1
DNF1
SMF2
TRS33
GCS1
SNC2
PEP5
YPT31
VPS54
YPT10
ERS1
SSH4
VPS1
MON2
VPS53
LSB6
MSB3
HSP12
YKR106W
VPS35
NHX1
AGE1
SNA3
DAP1
HSV2
VPS5
ATX2
VPS29
LST8
LCB4
TLG1
VTH1
PIB2
BET5
MON1
GAP1
YPT53
GTR1
SNX3
ENT1
PTM1
HSE1
END3
RAV2
YPT32
DID4
YMR110C
CCZ1
YPT7
DID2
YNL086W
VTA1
YCR075W−A_2
MEH1
SEC28
VPS5
YGR206W
VPS4
VPS30
VPS21
RCY1
VPS3
SLA1
FAB1
COS6
SLM4
VPS60
CDC50
LTV1
VPS15
BOS1
BRO1
SRN2
DOA4
VPS41
VPS27
VPS24
DID4
VPS8
PEP12
PEP7
VPS16
PEP3
VPS33
Z_Shift_K_Gem
Z_Im_K_Gem
Z_Shift_K_Cyt
Z_Im_K_Cyt
Z_Shift_L_Gem
Z_Im_L_Gem
Z_Shift_L_Cyt
Z_Im_L_Cyt

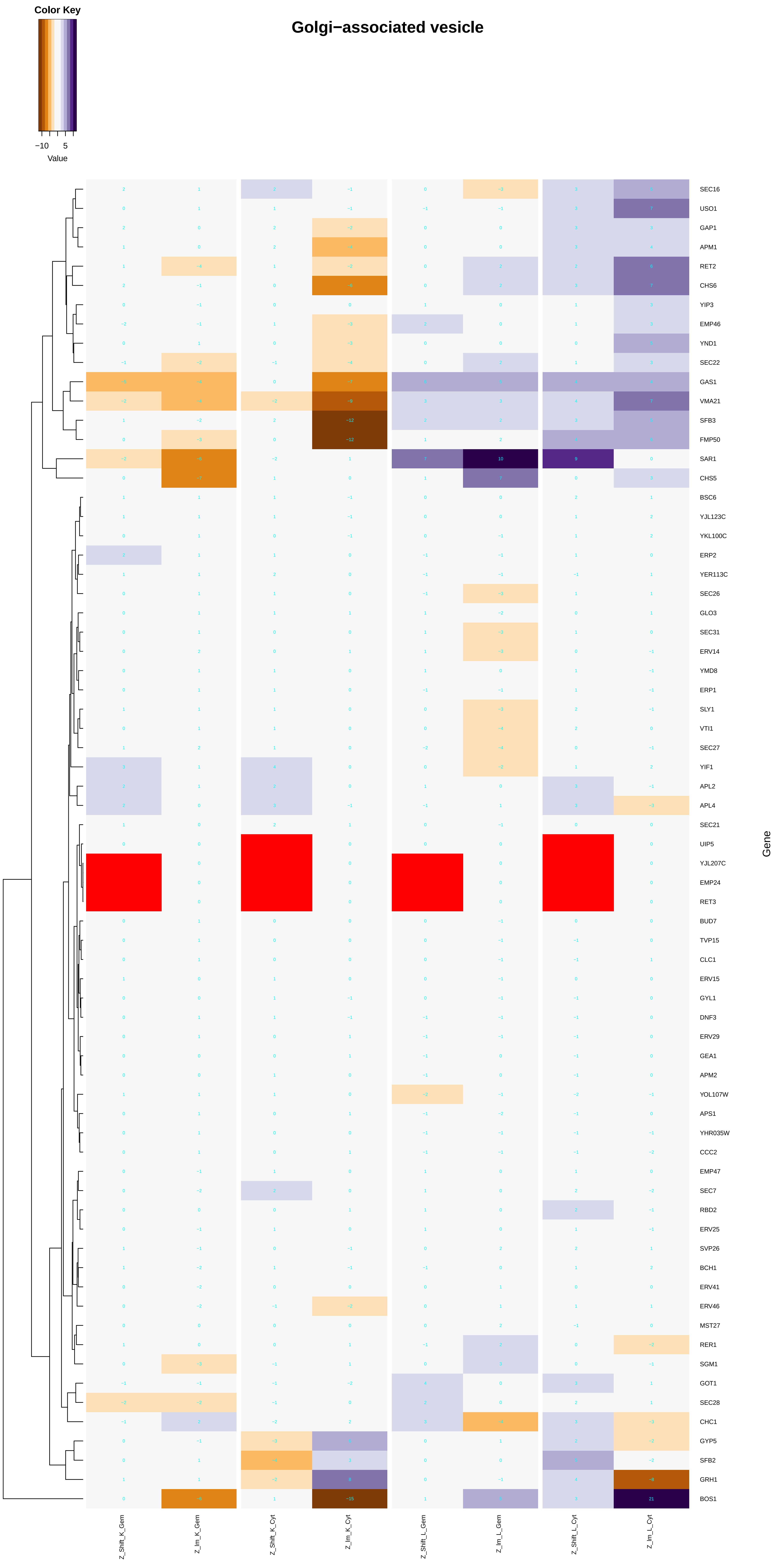
Golgi−associated vesicle
Color Key
−10
5
Value
Gene
SEC16
USO1
GAP1
APM1
RET2
CHS6
YIP3
EMP46
YND1
SEC22
GAS1
VMA21
SFB3
FMP50
SAR1
CHS5
BSC6
YJL123C
YKL100C
ERP2
YER113C
SEC26
GLO3
SEC31
ERV14
YMD8
ERP1
SLY1
VTI1
SEC27
YIF1
APL2
APL4
SEC21
UIP5
YJL207C
EMP24
RET3
BUD7
TVP15
CLC1
ERV15
GYL1
DNF3
ERV29
GEA1
APM2
YOL107W
APS1
YHR035W
CCC2
EMP47
SEC7
RBD2
ERV25
SVP26
BCH1
ERV41
ERV46
MST27
RER1
SGM1
GOT1
SEC28
CHC1
GYP5
SFB2
GRH1
BOS1
Z_Shift_K_Gem
Z_Im_K_Gem
Z_Shift_K_Cyt
Z_Im_K_Cyt
Z_Shift_L_Gem
Z_Im_L_Gem
Z_Shift_L_Cyt
Z_Im_L_Cyt

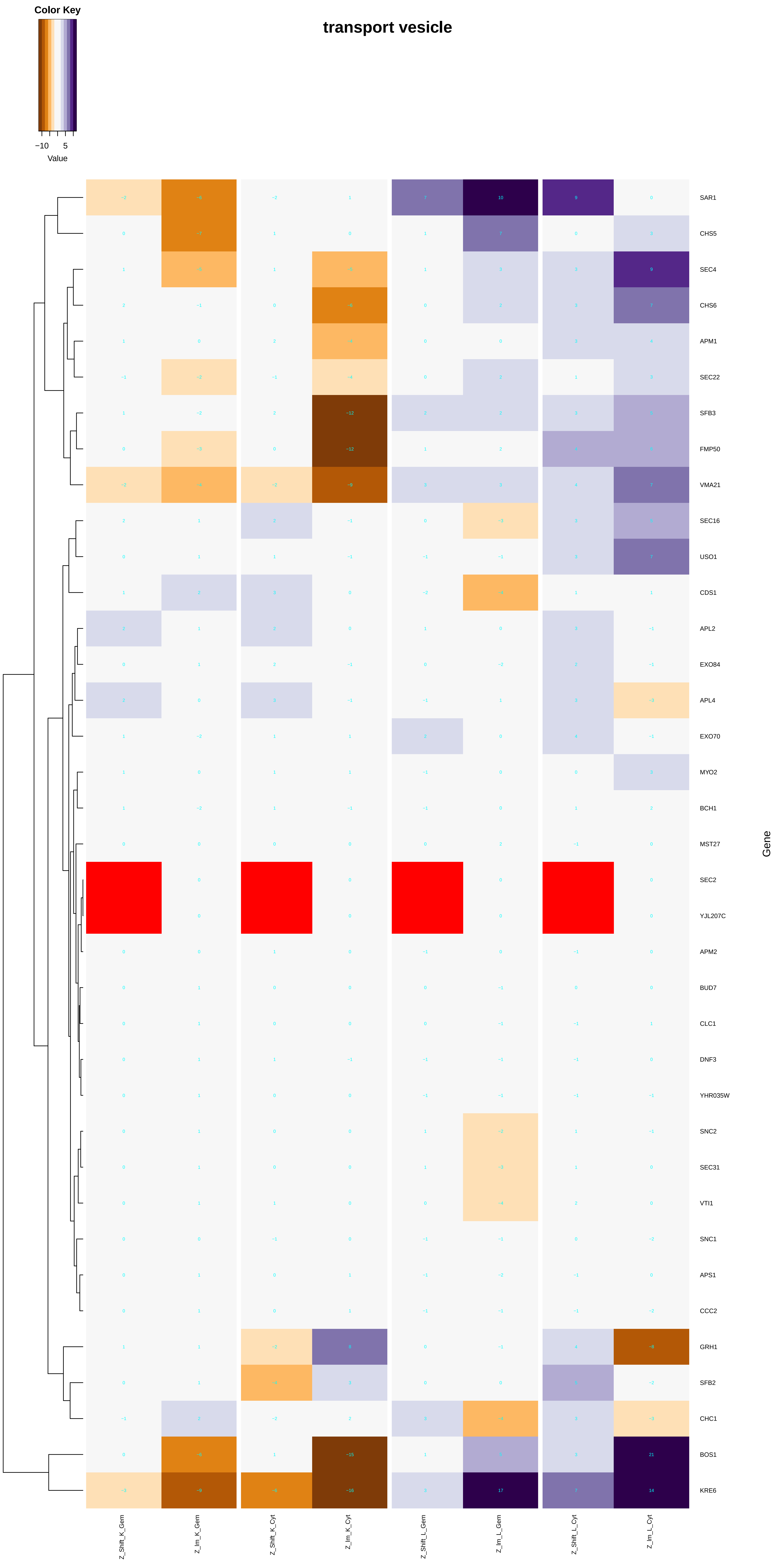

transport vesicle
Color Key
−10
5
Value
SAR1
CHS5
SEC4
CHS6
APM1
SEC22
SFB3
FMP50
VMA21
SEC16
USO1
CDS1
APL2
EXO84
APL4
EXO70
MYO2
BCH1
MST27
SEC2
YJL207C
APM2
BUD7
CLC1
DNF3
YHR035W
SNC2
SEC31
VTI1
SNC1
APS1
CCC2
GRH1
SFB2
CHC1
BOS1
KRE6
Gene
Z_Shift_K_Gem
Z_lm_K_Gem
Z_Shift_K_Cyt
Z_lm_K_Cyt
Z_Shift_L_Gem
Z_lm_L_Gem
Z_Shift_L_Cyt
Z_lm_L_Cyt

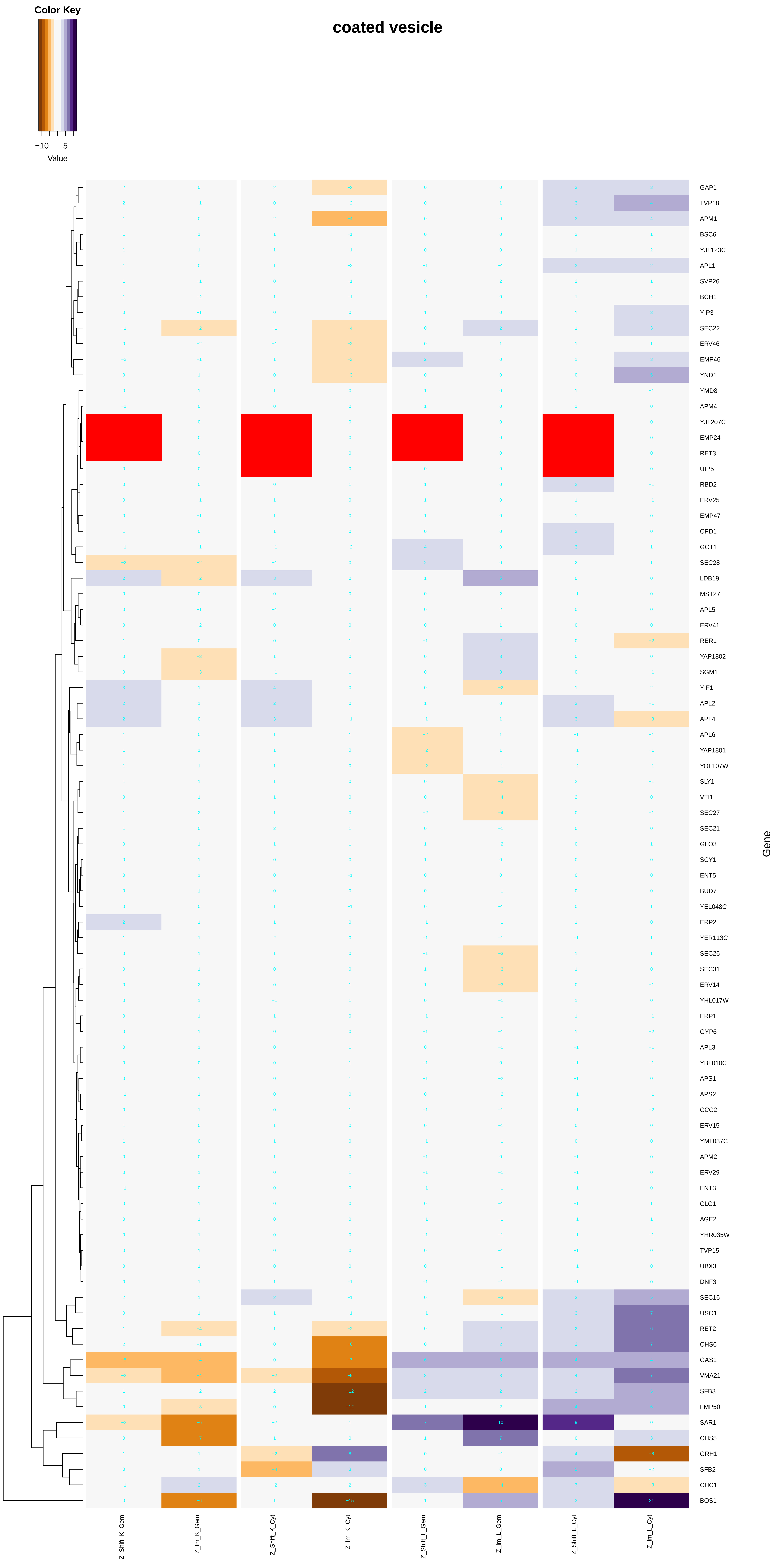
Color Key
−10 5
Value
coated vesicle
Gene
Z_Shift_K_Gem
Z_Im_K_Gem
Z_Shift_K_Cyt
Z_Im_K_Cyt
Z_Shift_L_Gem
Z_Im_L_Gem
Z_Shift_L_Cyt
Z_Im_L_Cyt
GAP1
TVP18
APM1
BSC6
YJL123C
APL1
SVP26
BCH1
YIP3
SEC22
ERV46
EMP46
YND1
YMD8
APM4
YJL207C
EMP24
RET3
UIP5
RBD2
ERV25
EMP47
CPD1
GOT1
SEC28
LDB19
MST27
APL5
ERV41
RER1
YAP1802
SGM1
YIF1
APL2
APL4
APL6
YAP1801
YOL107W
SLY1
VTI1
SEC27
SEC21
GLO3
SCY1
ENT5
BUD7
YEL048C
ERP2
YER113C
SEC26
SEC31
ERV14
YHL017W
ERP1
GYP6
APL3
YBL010C
APS1
APS2
CCC2
ERV15
YML037C
APM2
ERV29
ENT3
CLC1
AGE2
YHR035W
TVP15
UBX3
DNF3
SEC16
USO1
RET2
CHS6
GAS1
VMA21
SFB3
FMP50
SAR1
CHS5
GRH1
SFB2
CHC1
BOS1

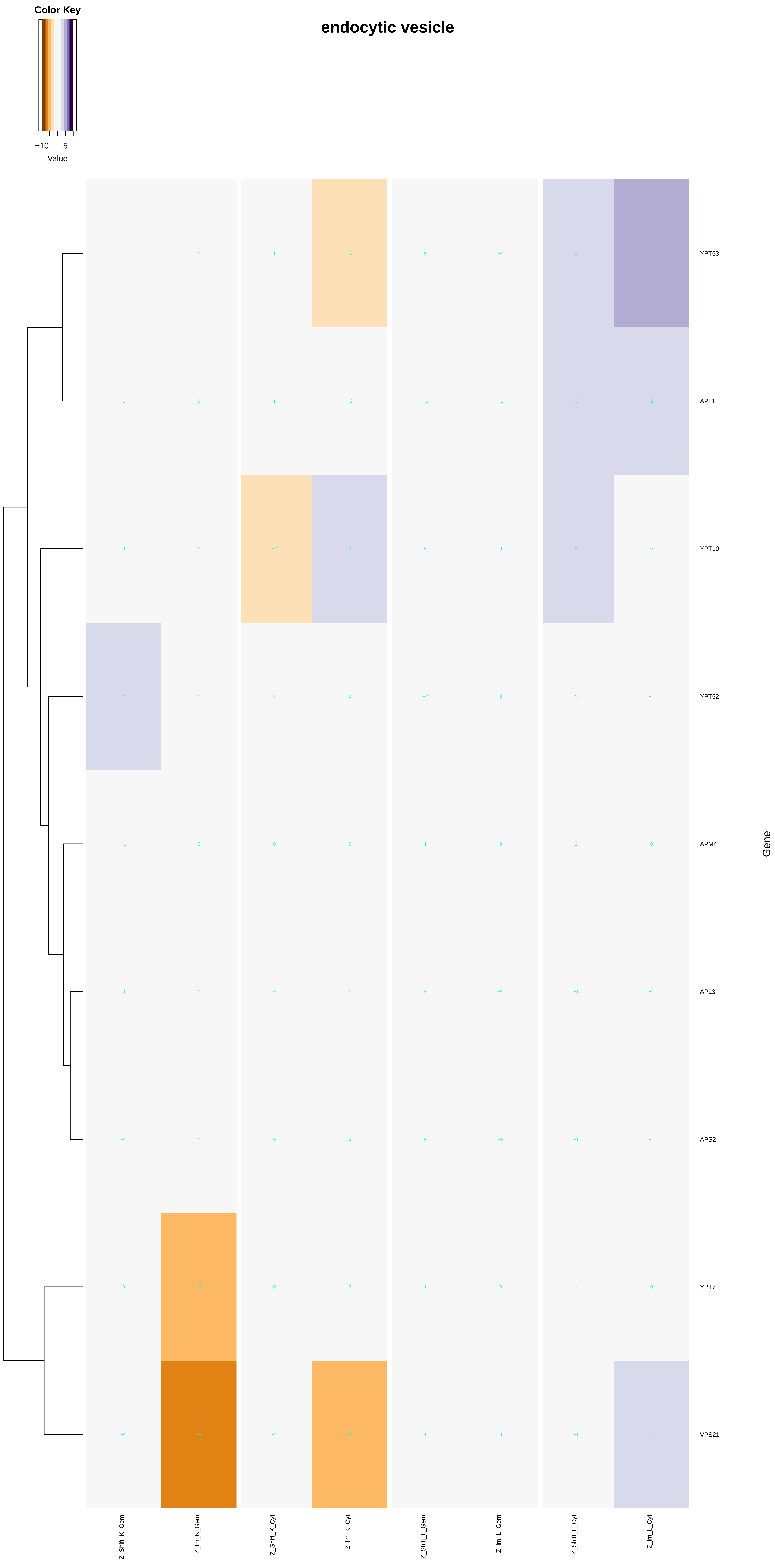

Color Key
−10
5
Value
endocytic vesicle
YPT53
APL1
YPT10
YPT52
APM4
APL3
APS2
YPT7
VPS21
Gene
Z_Shift_K_Gem
Z_Im_K_Gem
Z_Shift_K_Cyt
Z_Im_K_Cyt
Z_Shift_L_Gem
Z_Im_L_Gem
Z_Shift_L_Cyt
Z_Im_L_Cyt

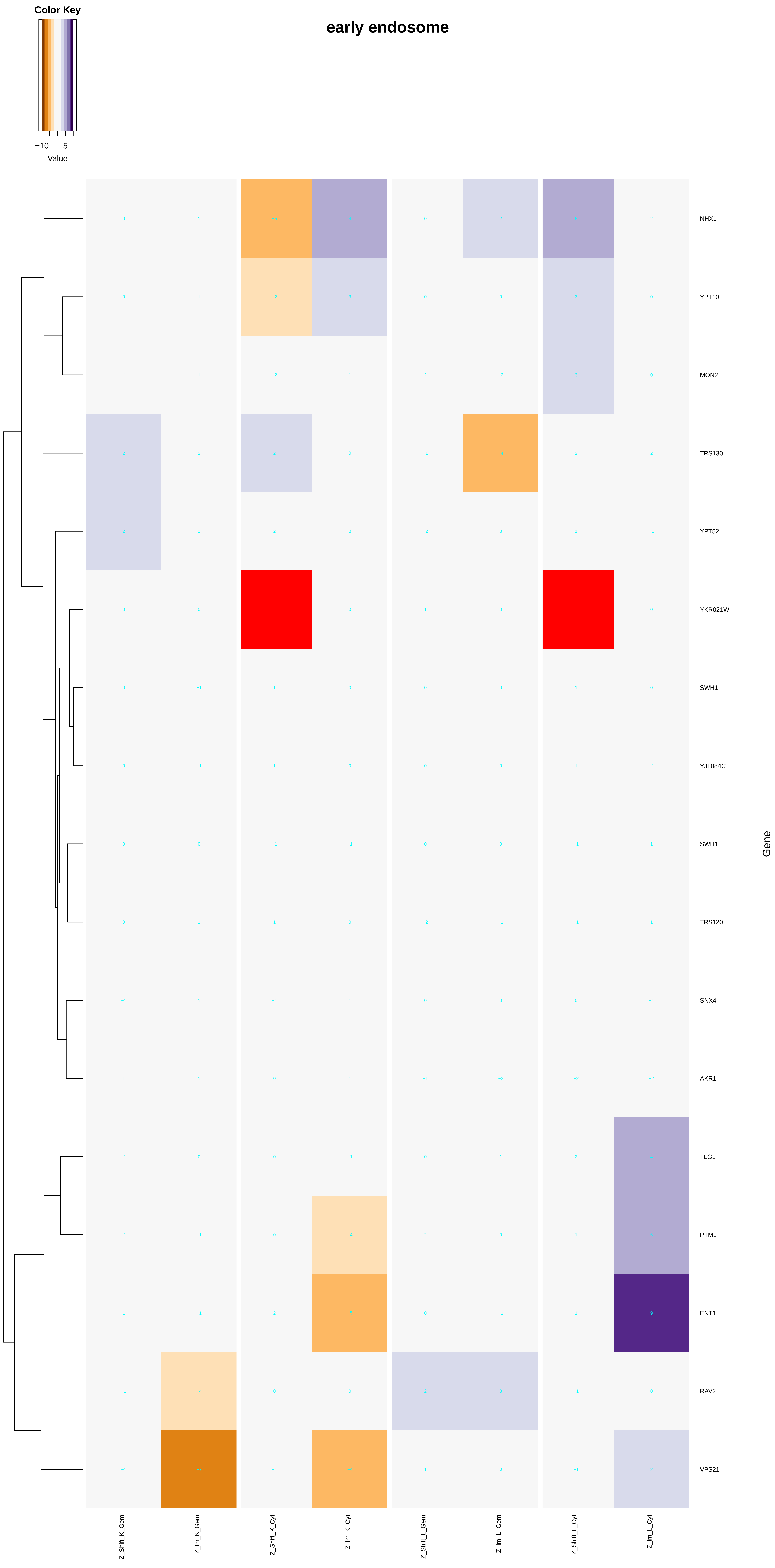

Color Key
−10
5
Value
early endosome
NHX1
YPT10
MON2
TRS130
YPT52
YKR021W
SWH1
YJL084C
SWH1
TRS120
SNX4
AKR1
TLG1
PTM1
ENT1
RAV2
VPS21
Gene
Z_Shift_K_Gem
Z_Im_K_Gem
Z_Shift_K_Cyt
Z_Im_K_Cyt
Z_Shift_L_Gem
Z_Im_L_Gem
Z_Shift_L_Cyt
Z_Im_L_Cyt

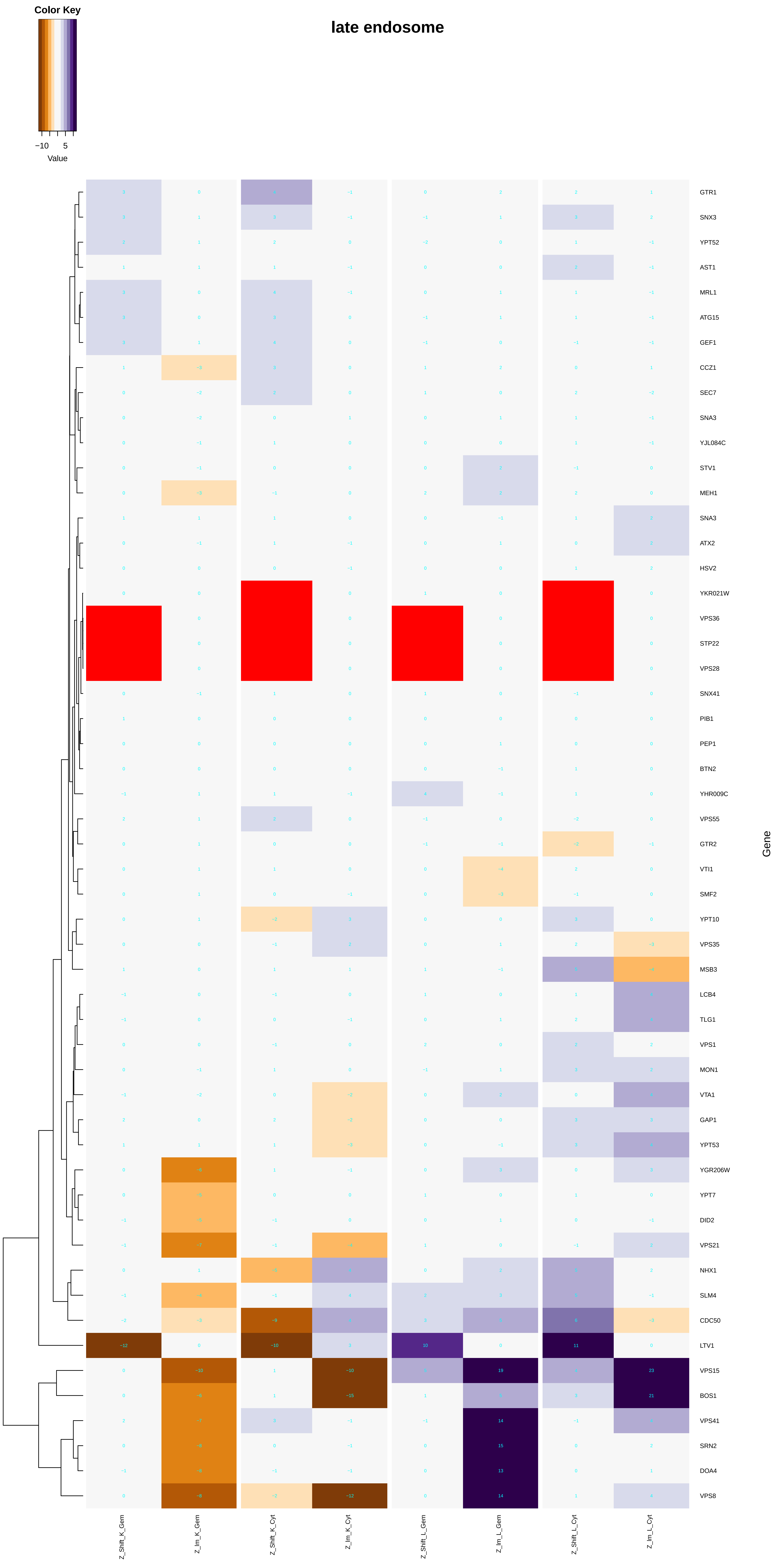
late endosome
Color Key
−10
5
Value
GTR1
SNX3
YPT52
AST1
MRL1
ATG15
GEF1
CCZ1
SEC7
SNA3
YJL084C
STV1
MEH1
SNA3
ATX2
HSV2
YKR021W
VPS36
STP22
VPS28
SNX41
PIB1
PEP1
BTN2
YHR009C
VPS55
GTR2
VTI1
SMF2
YPT10
VPS35
MSB3
LCB4
TLG1
VPS1
MON1
VTA1
GAP1
YPT53
YGR206W
YPT7
DID2
VPS21
NHX1
SLM4
CDC50
LTV1
VPS15
BOS1
VPS41
SRN2
DOA4
VPS8
Gene
Z_Shift_K_Gem
Z_lm_K_Gem
Z_Shift_K_Cyt
Z_lm_K_Cyt
Z_Shift_L_Gem
Z_lm_L_Gem
Z_Shift_L_Cyt
Z_lm_L_Cyt

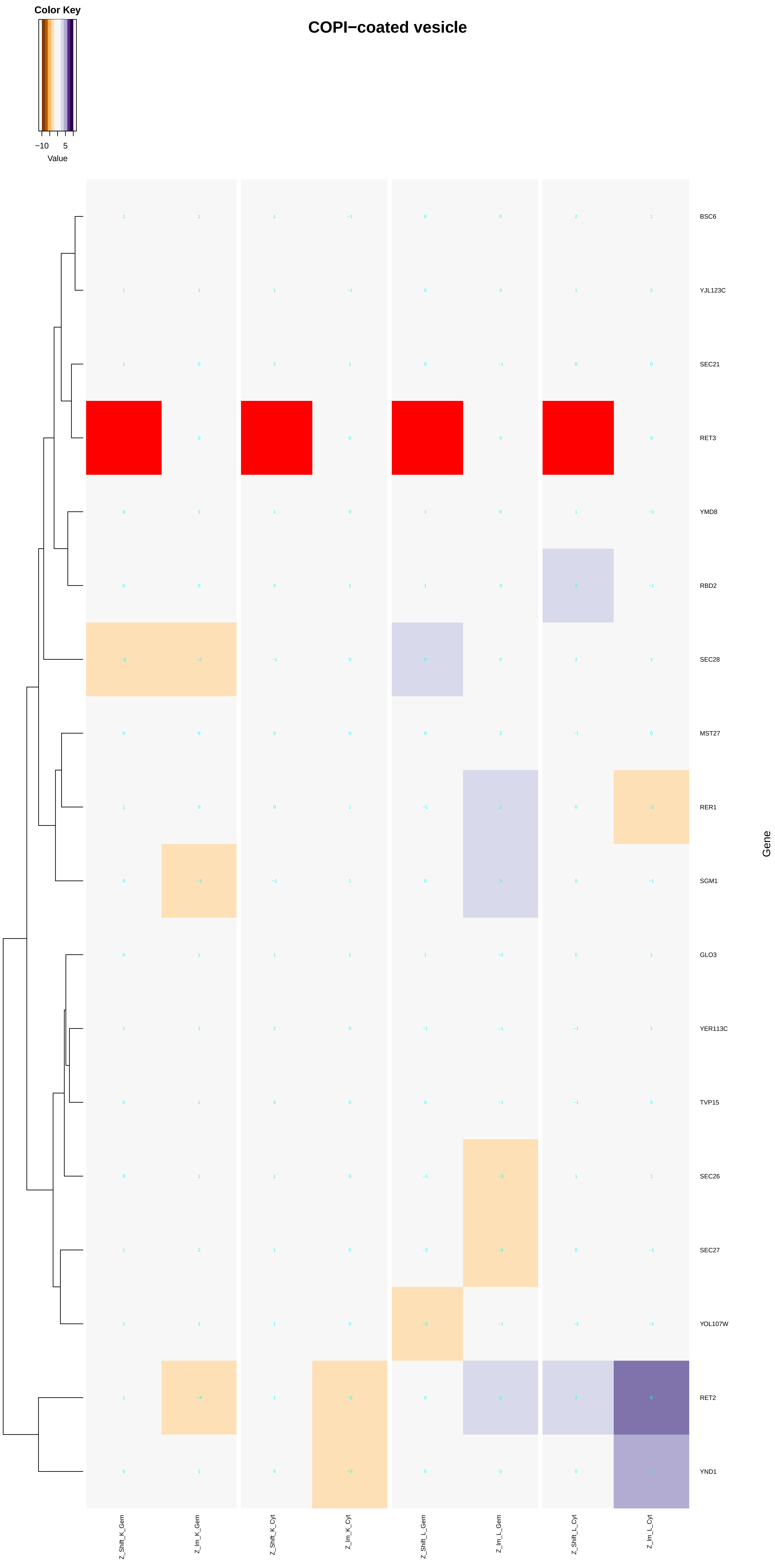
Color Key
−10
5
Value
COPI−coated vesicle
BSC6
YJL123C
SEC21
RET3
YMD8
RBD2
SEC28
MST27
RER1
SGM1
GLO3
YER113C
TVP15
SEC26
SEC27
YOL107W
RET2
YND1
Gene
Z_Shift_K_Gem
Z_Im_K_Gem
Z_Shift_K_Cyt
Z_Im_K_Cyt
Z_Shift_L_Gem
Z_Im_L_Gem
Z_Shift_L_Cyt
Z_Im_L_Cyt

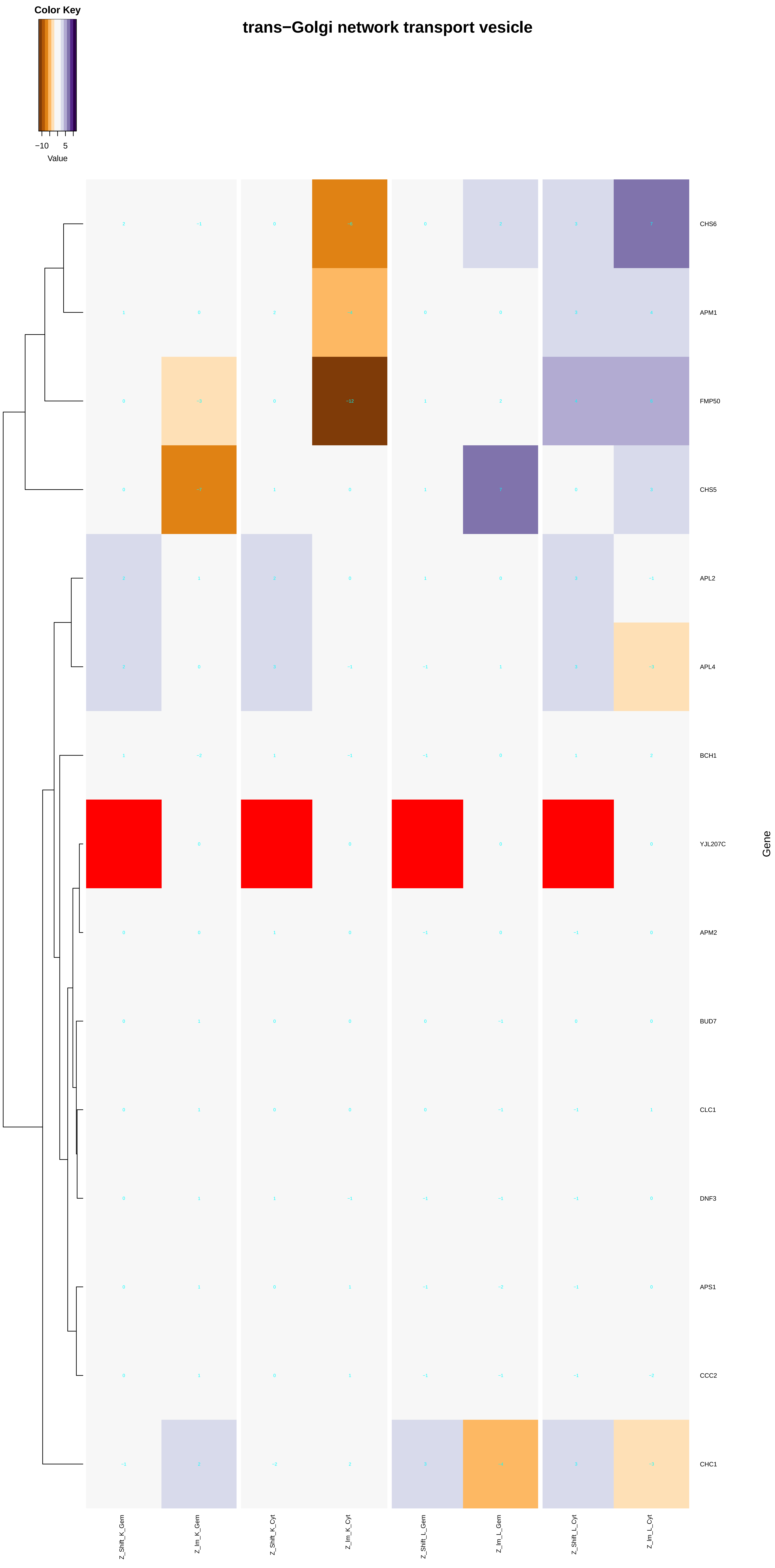

Color Key
−10
5
Value
trans−Golgi network transport vesicle
CHS6
APM1
FMP50
CHS5
APL2
APL4
BCH1
YJL207C
APM2
BUD7
CLC1
DNF3
APS1
CCC2
CHC1
Gene
Z_Shift_K_Gem
Z_Im_K_Gem
Z_Shift_K_Cyt
Z_Im_K_Cyt
Z_Shift_L_Gem
Z_Im_L_Gem
Z_Shift_L_Cyt
Z_Im_L_Cyt

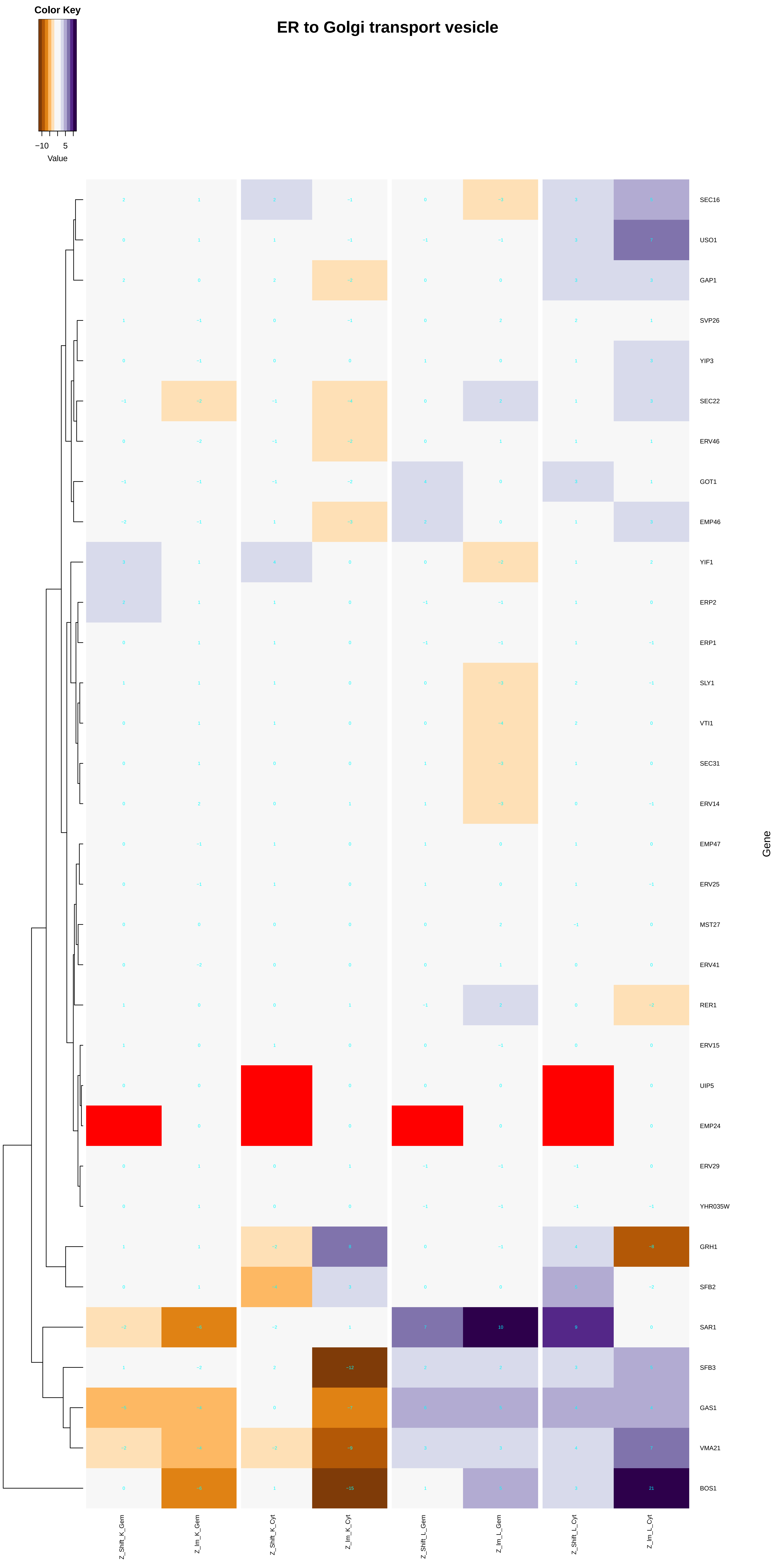

Color Key
−10
5
Value
ER to Golgi transport vesicle
SEC16
USO1
GAP1
SVP26
YIP3
SEC22
ERV46
GOT1
EMP46
YIF1
ERP2
ERP1
SLY1
VTI1
SEC31
ERV14
EMP47
ERV25
MST27
ERV41
RER1
ERV15
UIP5
EMP24
ERV29
YHR035W
GRH1
SFB2
SAR1
SFB3
GAS1
VMA21
BOS1
Gene
Z_Shift_K_Gem
Z_Im_K_Gem
Z_Shift_K_Cyt
Z_Im_K_Cyt
Z_Shift_L_Gem
Z_Im_L_Gem
Z_Shift_L_Cyt
Z_Im_L_Cyt

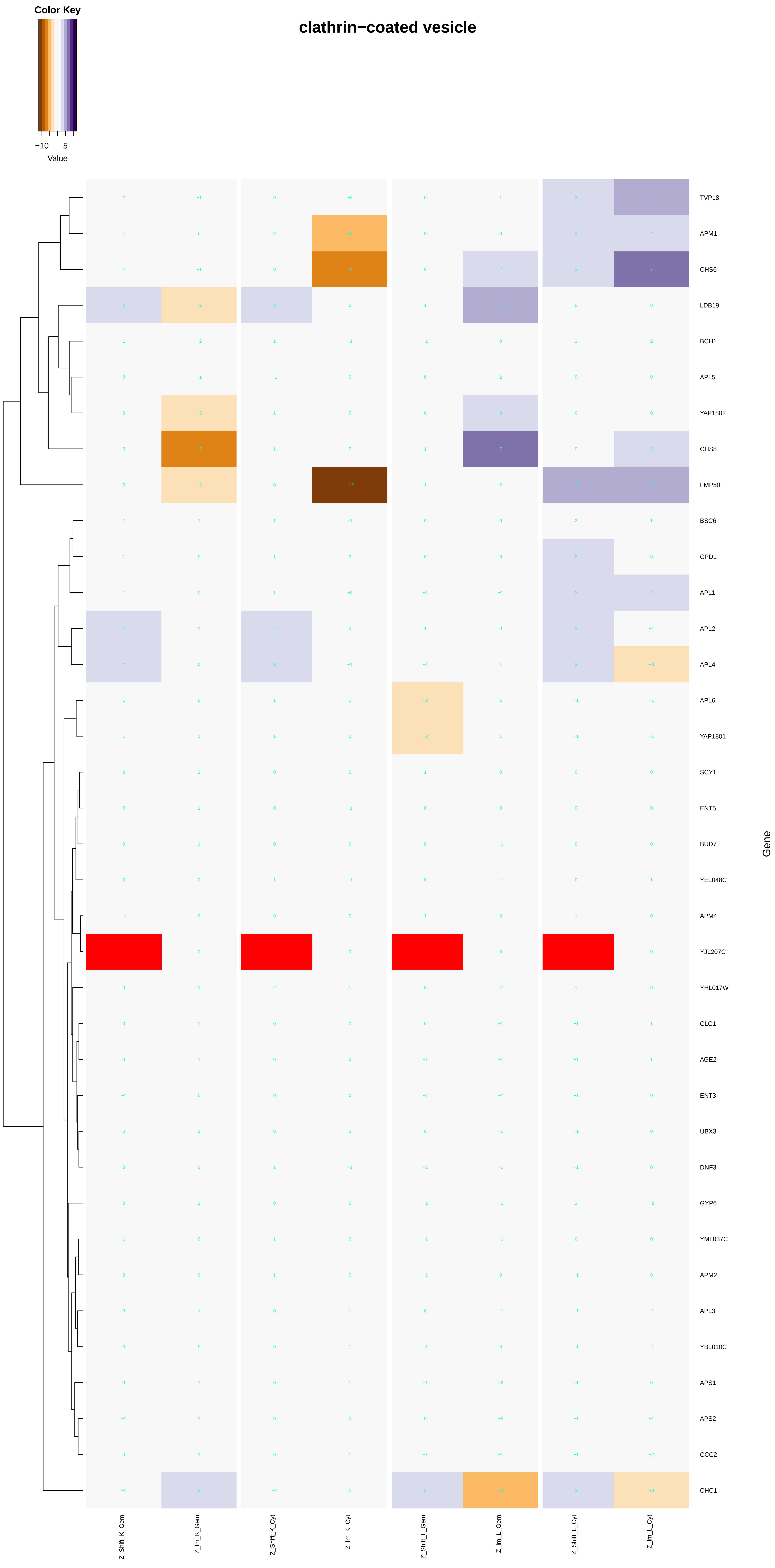

Color Key
−10
5
Value
clathrin−coated vesicle
TVP18
APM1
CHS6
LDB19
BCH1
APL5
YAP1802
CHS5
FMP50
BSC6
CPD1
APL1
APL2
APL4
APL6
YAP1801
SCY1
ENT5
BUD7
YEL048C
APM4
YJL207C
YHL017W
CLC1
AGE2
ENT3
UBX3
DNF3
GYP6
YML037C
APM2
APL3
YBL010C
APS1
APS2
CCC2
CHC1
Gene
Z_Shift_K_Gem
Z_Im_K_Gem
Z_Shift_K_Cyt
Z_Im_K_Cyt
Z_Shift_L_Gem
Z_Im_L_Gem
Z_Shift_L_Cyt
Z_Im_L_Cyt

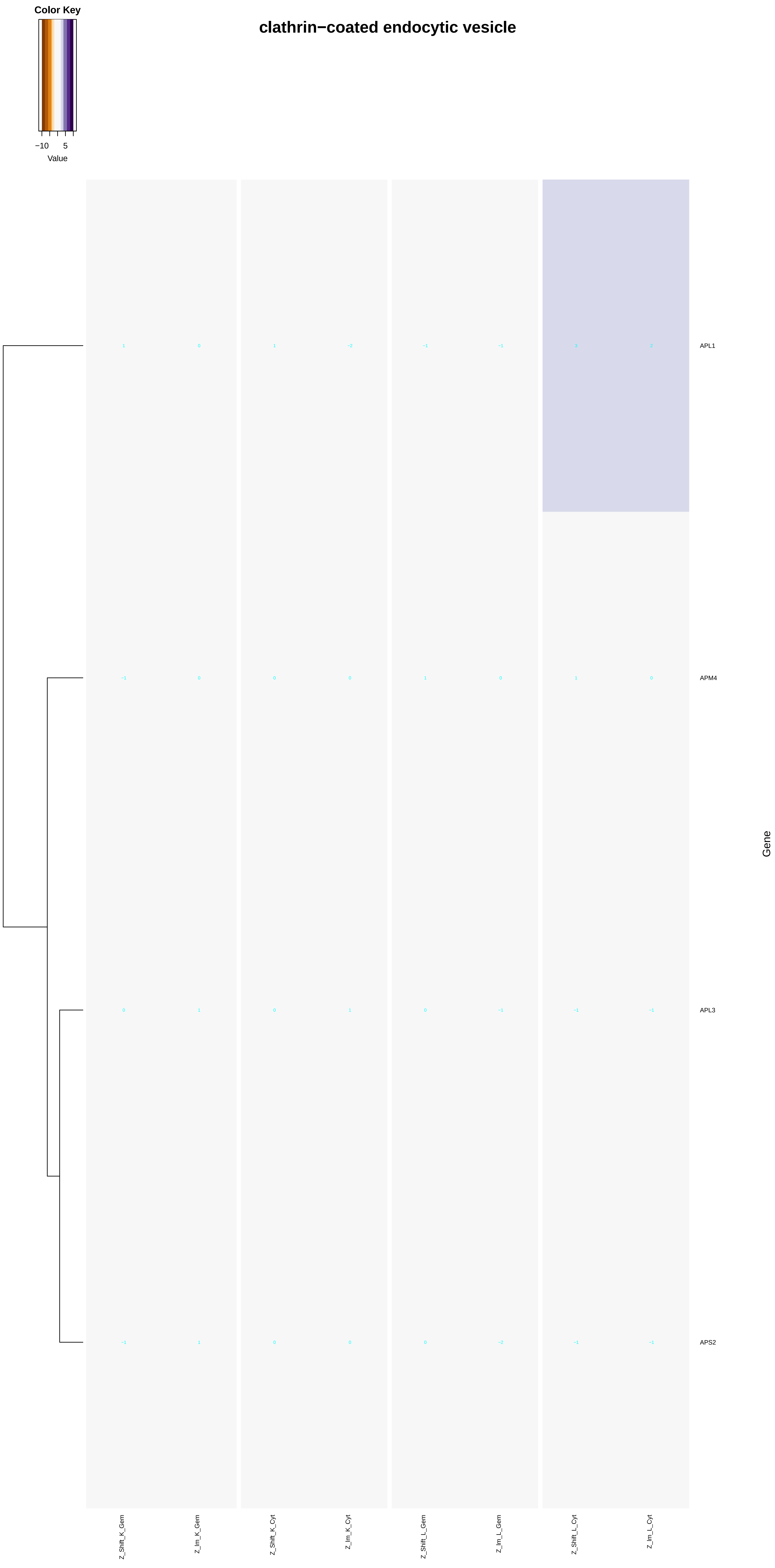
clathrin−coated endocytic vesicle
Color Key
−10
5
Value
APL1
APM4
APL3
APS2
Gene
Z_Shift_K_Gem
Z_Im_K_Gem
Z_Shift_K_Cyt
Z_Im_K_Cyt
Z_Shift_L_Gem
Z_Im_L_Gem
Z_Shift_L_Cyt
Z_Im_L_Cyt

### multivesicular body

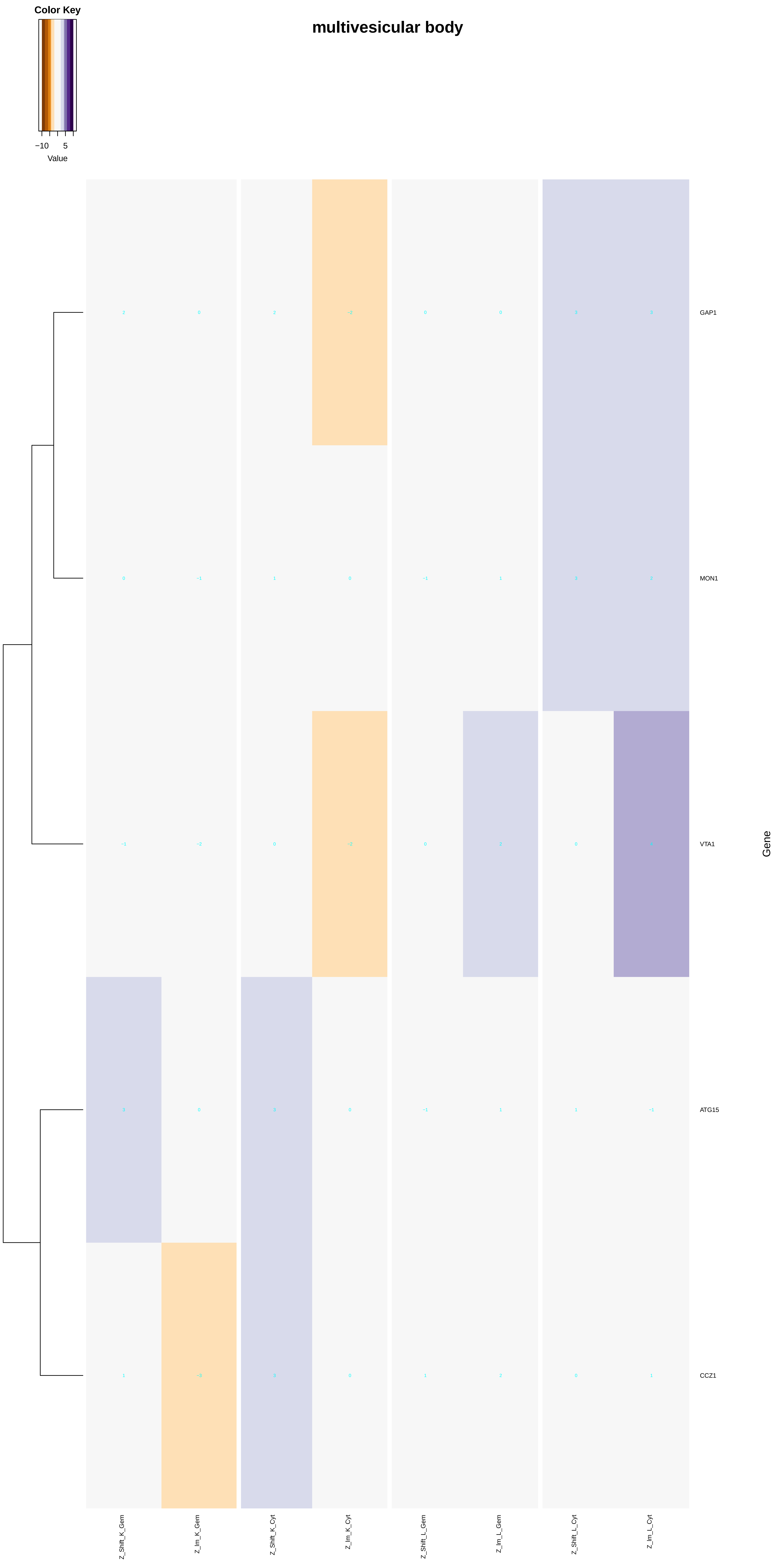
