## Additional File 7 for "A humanized yeast phenomic model of deoxycytidine kinase to predict genetic buffering of nucleoside analog cytotoxicity": DNA_metabolic_process.pdf

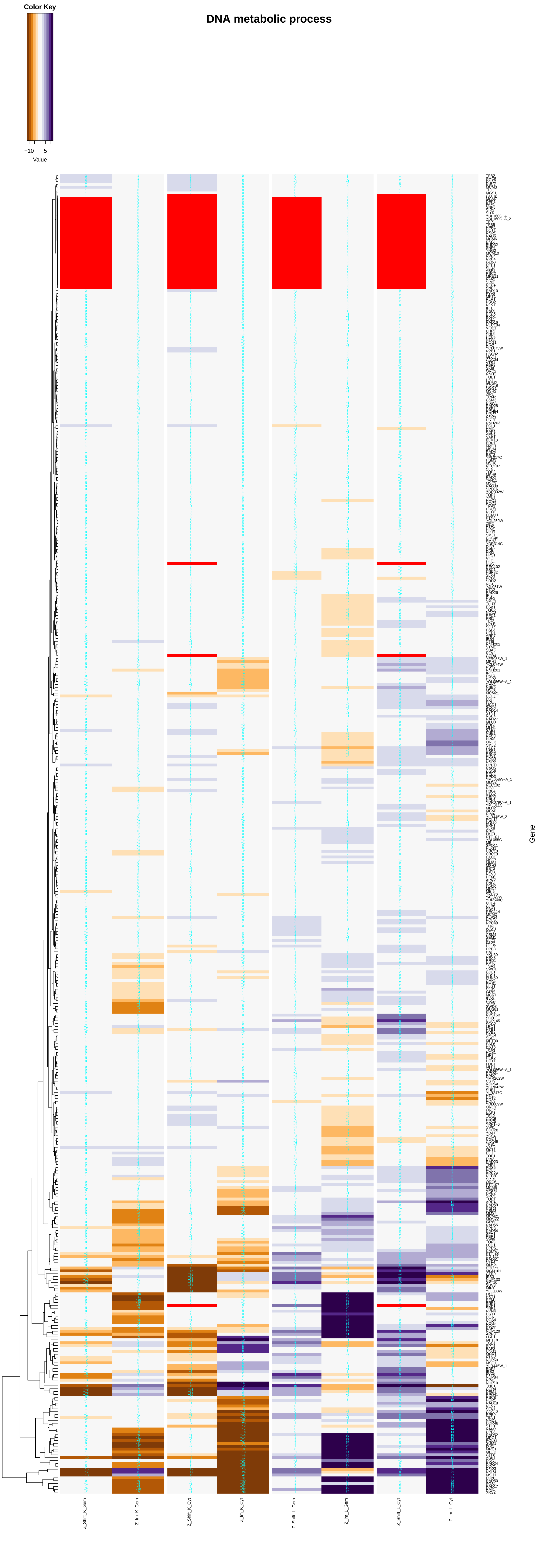
DNA metabolic process
Color Key
−10
5
Value
Gene
Z_Shift_K_Gem
Z_Im_K_Gem
Z_Shift_K_Cyt
Z_Im_K_Cyt
Z_Shift_L_Gem
Z_Im_L_Gem
Z_Shift_L_Cyt
Z_Im_L_Cyt

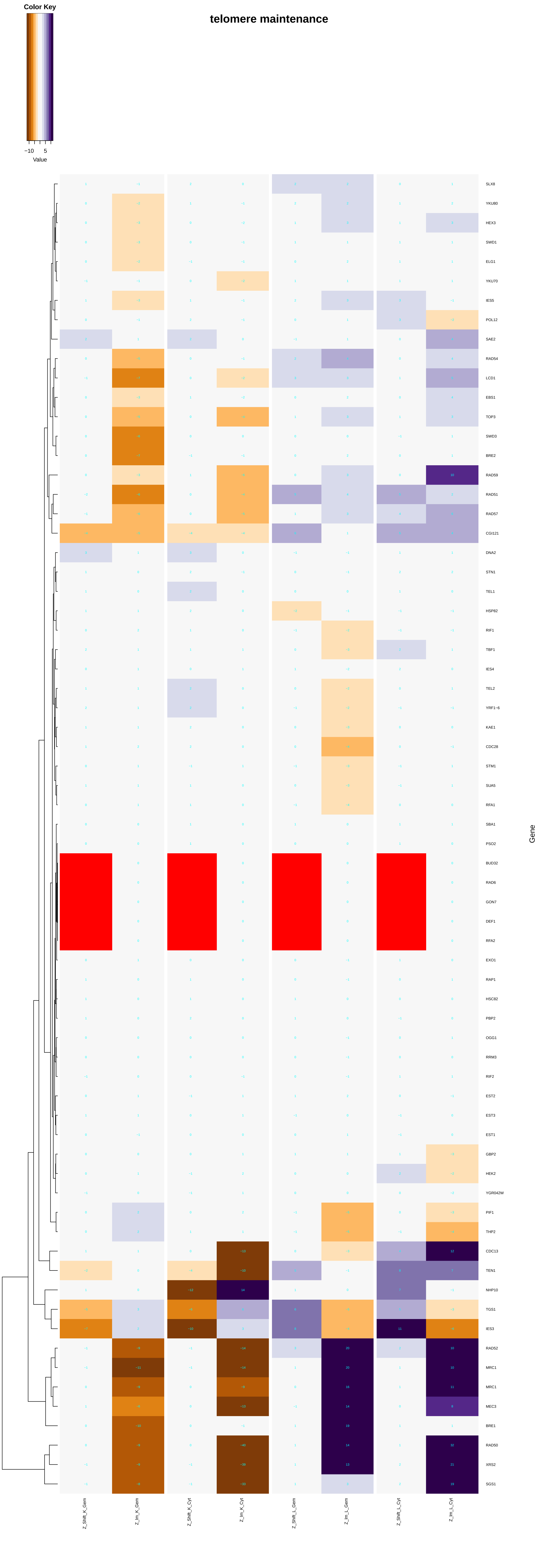
telomere maintenance
Color Key
−10 5
Value
SLX8
YKU80
HEX3
SWD1
ELG1
YKU70
IES5
POL12
SAE2
RAD54
LCD1
EBS1
TOP3
SWD3
BRE2
RAD59
RAD51
RAD57
CGI121
DNA2
STN1
TEL1
HSP82
RIF1
TBF1
IES4
TEL2
YRF1−6
KAE1
CDC28
STM1
SUA5
RFA1
SBA1
PSO2
BUD32
RAD6
GON7
DEF1
RFA2
EXO1
RAP1
HSC82
PBP2
OGG1
RRM3
RIF2
EST2
EST3
EST1
GBP2
HEK2
YGR042W
PIF1
THP2
CDC13
TEN1
NHP10
TGS1
IES3
RAD52
MRC1
MRC1
MEC3
BRE1
RAD50
XRS2
SGS1
Gene
Z_Shift_K_Gem
Z_Im_K_Gem
Z_Shift_K_Cyt
Z_Im_K_Cyt
Z_Shift_L_Gem
Z_Im_L_Gem
Z_Shift_L_Cyt
Z_Im_L_Cyt

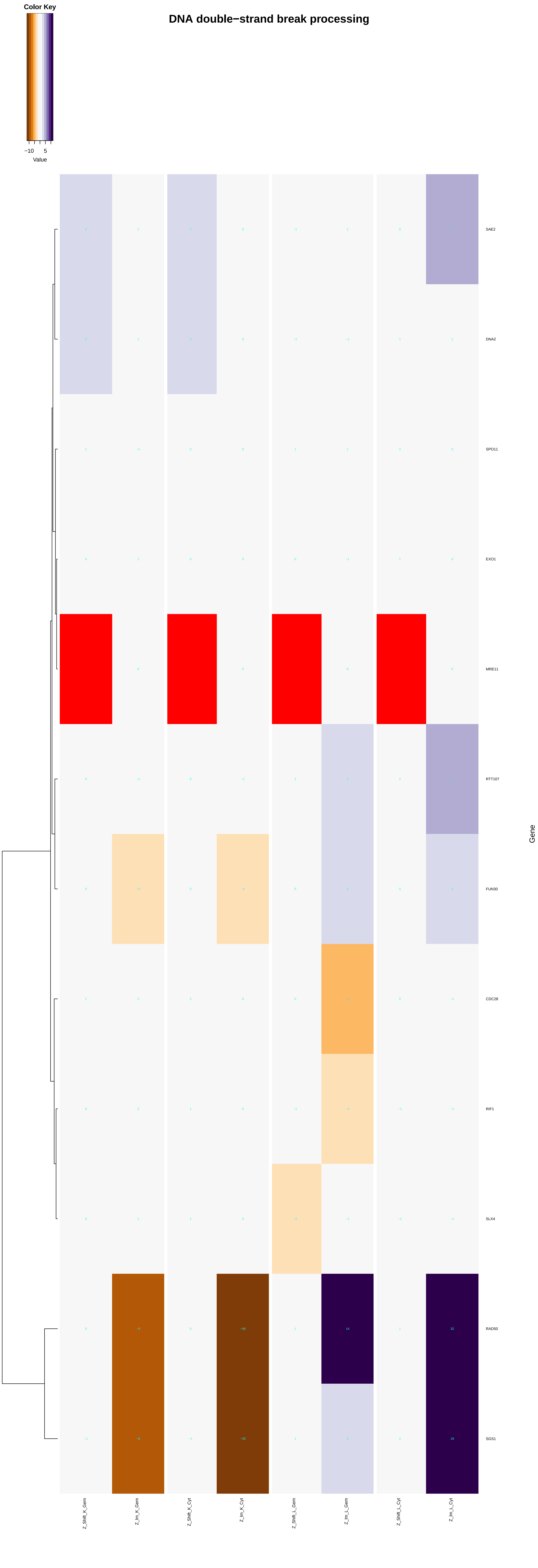
DNA double−strand break processing
Color Key
−10
5
Value
SAE2
DNA2
SPO11
EXO1
MRE11
RTT107
FUN30
CDC28
RIF1
SLX4
RAD50
SGS1
Gene
Z_Shift_K_Gem
Z_Im_K_Gem
Z_Shift_K_Cyt
Z_Im_K_Cyt
Z_Shift_L_Gem
Z_Im_L_Gem
Z_Shift_L_Cyt
Z_Im_L_Cyt

### DNA strand renaturation

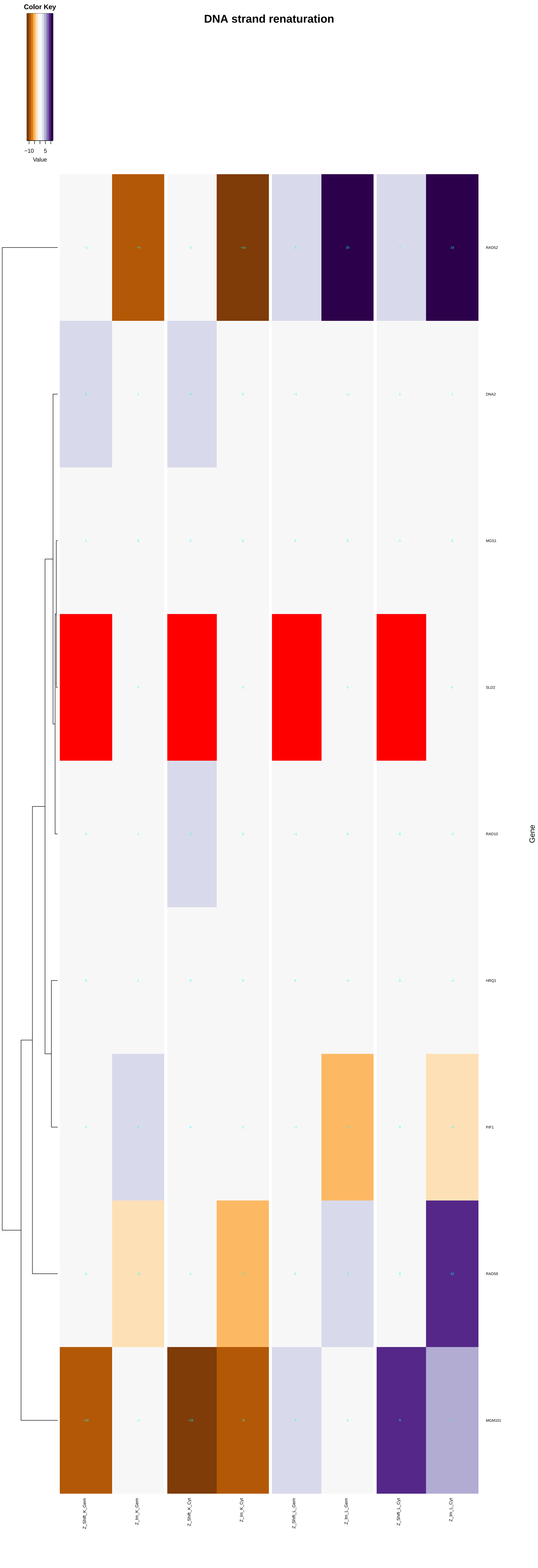

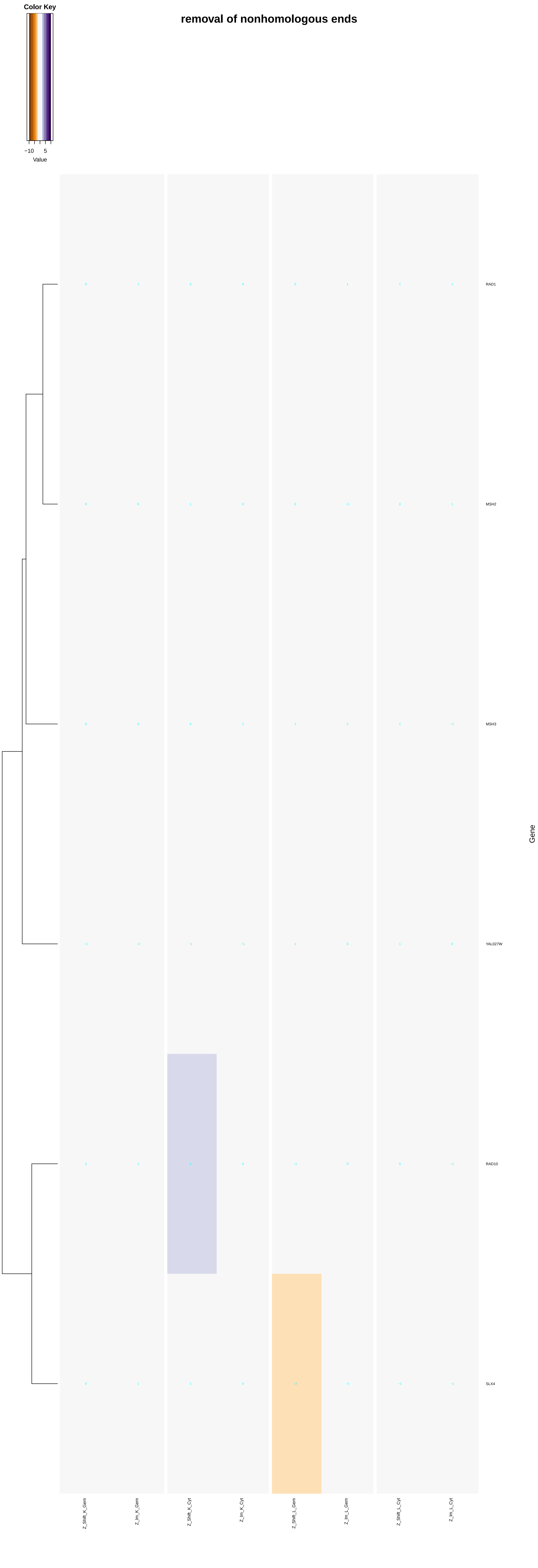

removal of nonhomologous ends
Color Key
−10
5
Value
RAD1
MSH2
MSH3
YAL027W
RAD10
SLX4
Gene
Z_Shift_K_Gem
Z_Im_K_Gem
Z_Shift_K_Cyt
Z_Im_K_Cyt
Z_Shift_L_Gem
Z_Im_L_Gem
Z_Shift_L_Cyt
Z_Im_L_Cyt

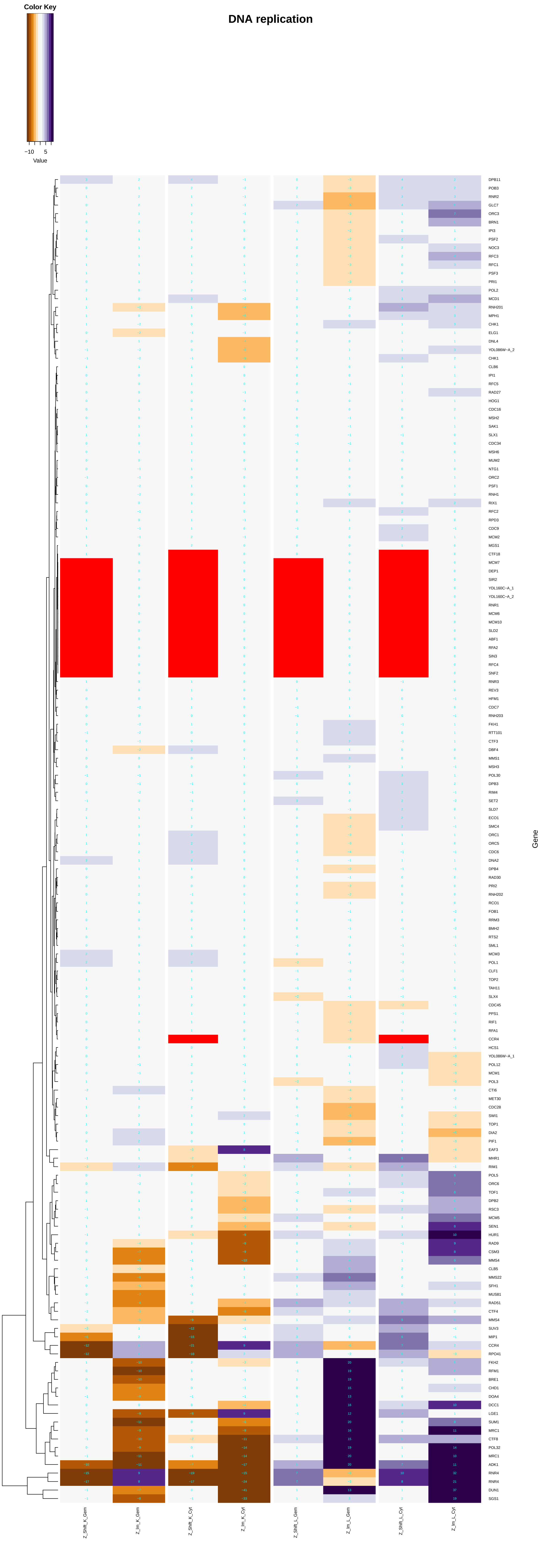
Color Key
−10
5
Value
DNA replication
DPB11
POB3
RNR2
GLC7
ORC3
BRN1
IPI3
PSF2
NOC3
RFC3
RFC1
PSF3
PRI1
POL2
MCD1
RNH201
MPH1
CHK1
ELG1
DNL4
YOL086W−A_2
CHK1
CLB6
IPI1
RFC5
RAD27
HOG1
CDC16
MSH2
SAK1
SLX1
CDC34
MSH6
MUM2
NTG1
ORC2
PSF1
RNH1
RIX1
RFC2
RPD3
CDC9
MCM2
MGS1
CTF18
MCM7
DEP1
SIR2
YDL160C−A_1
YDL160C−A_2
RNR1
MCM6
MCM10
SLD2
ABF1
RFA2
SIN3
RFC4
SNF2
RNR3
REV3
HFM1
CDC7
RNH203
FKH1
RTT101
CTF3
DBF4
MMS1
MSH3
POL30
DPB3
RIM4
SET2
SLD7
ECO1
SMC4
ORC1
ORC5
CDC6
DNA2
DPB4
RAD30
PRI2
RNH202
RCO1
FOB1
RRM3
BMH2
RTS2
SML1
MCM3
POL1
CLF1
TOP2
TAH11
SLX4
CDC45
PPS1
RIF1
RFA1
CCR4
HCS1
YOL086W−A_1
POL12
MCM1
POL3
CTI6
MET30
CDC28
SWI1
TOP1
DIA2
PIF1
EAF3
MHR1
RIM1
POL5
ORC6
TOF1
DPB2
RSC3
MCM5
SEN1
HUR1
RAD9
CSM3
MMS4
CLB5
MMS22
SFH1
MUS81
CTF4
RAD51
MMS4
SUV3
MIP1
CCR4
RPO41
FKH2
RFM1
BRE1
CHD1
DOA4
DCC1
LGE1
SUM1
MRC1
CTF8
POL32
MRC1
ADK1
RNR4
RNR4
DUN1
SGS1
Gene
Z_Shift_K_Gem
Z_Im_K_Gem
Z_Shift_K_Cyt
Z_Im_K_Cyt
Z_Shift_L_Gem
Z_Im_L_Gem
Z_Shift_L_Cyt
Z_Im_L_Cyt

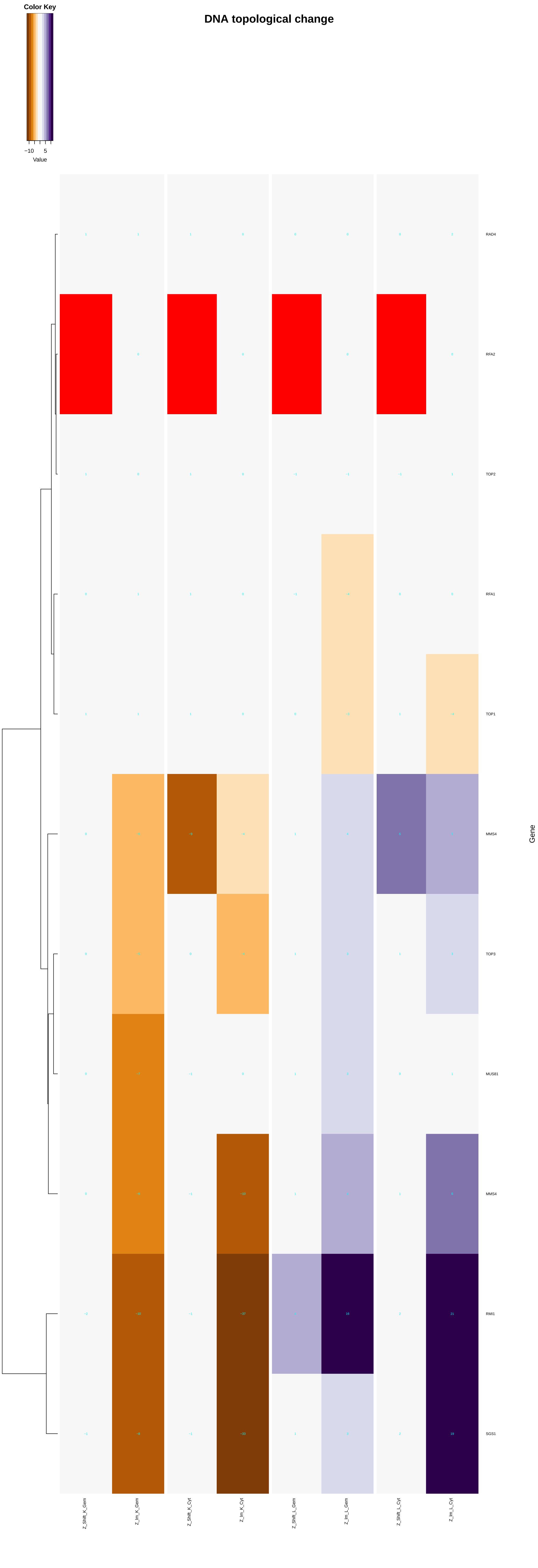
DNA topological change
Color Key
−10
5
Value
RAD4
RFA2
TOP2
RFA1
TOP1
MMS4
TOP3
MUS81
MMS4
RMI1
SGS1
Gene
Z_Shift_K_Gem
Z_Im_K_Gem
Z_Shift_K_Cyt
Z_Im_K_Cyt
Z_Shift_L_Gem
Z_Im_L_Gem
Z_Shift_L_Cyt
Z_Im_L_Cyt

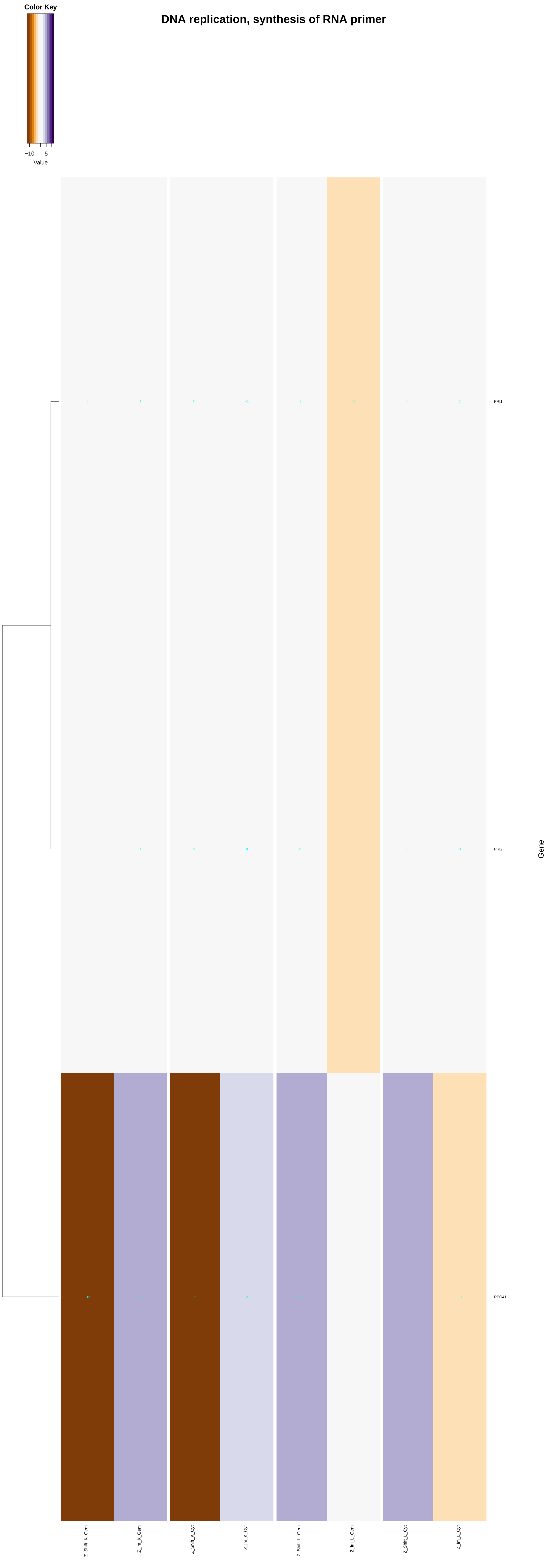
DNA replication, synthesis of RNA primer
Color Key
−10
5
Value
PRI1
PRI2
RPO41
Gene
Z_Shift_K_Gem
Z_Im_K_Gem
Z_Shift_K_Cyt
Z_Im_K_Cyt
Z_Shift_L_Gem
Z_Im_L_Gem
Z_Shift_L_Cyt
Z_Im_L_Cyt

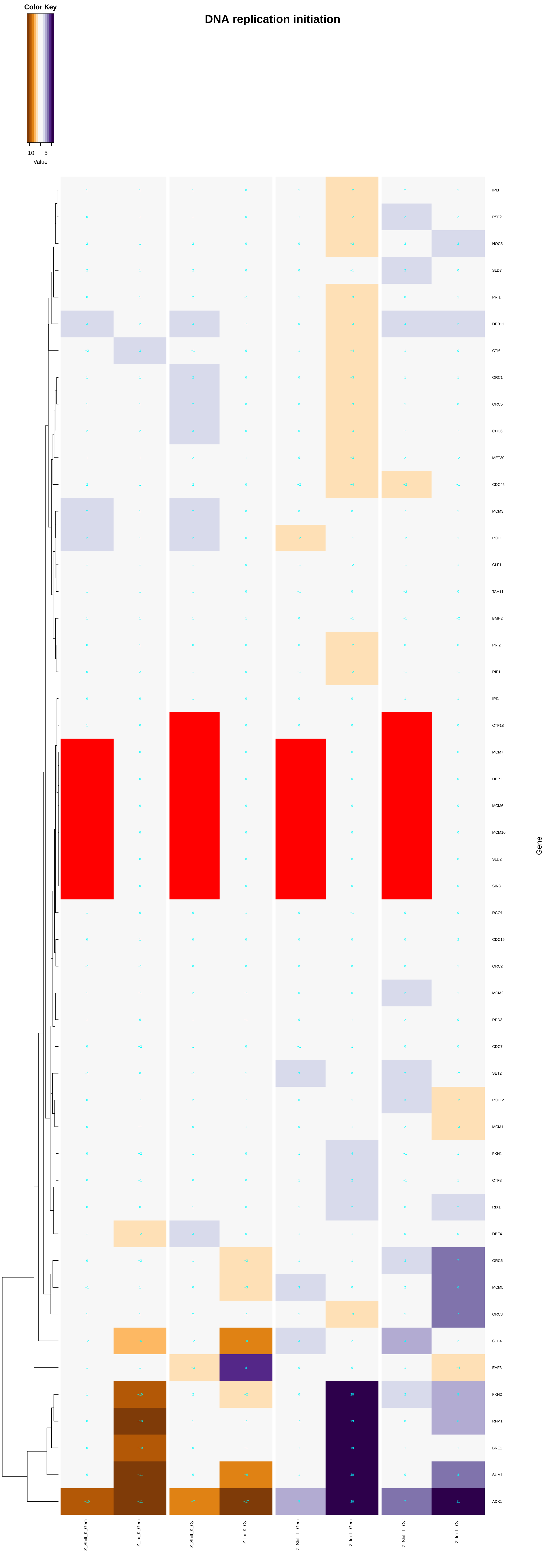
DNA replication initiation
Color Key
−10
5
Value
IPI3
PSF2
NOC3
SLD7
PRI1
DPB11
CTI6
ORC1
ORC5
CDC6
MET30
CDC45
MCM3
POL1
CLF1
TAH11
BMH2
PRI2
RIF1
IPI1
CTF18
MCM7
DEP1
MCM6
MCM10
SLD2
SIN3
RCO1
CDC16
ORC2
MCM2
RPD3
CDC7
SET2
POL12
MCM1
FKH1
CTF3
RIX1
DBF4
ORC6
MCM5
ORC3
CTF4
EAF3
FKH2
RFM1
BRE1
SUM1
ADK1
Gene
Z_Shift_K_Gem
Z_Im_K_Gem
Z_Shift_K_Cyt
Z_Im_K_Cyt
Z_Shift_L_Gem
Z_Im_L_Gem
Z_Shift_L_Cyt
Z_Im_L_Cyt

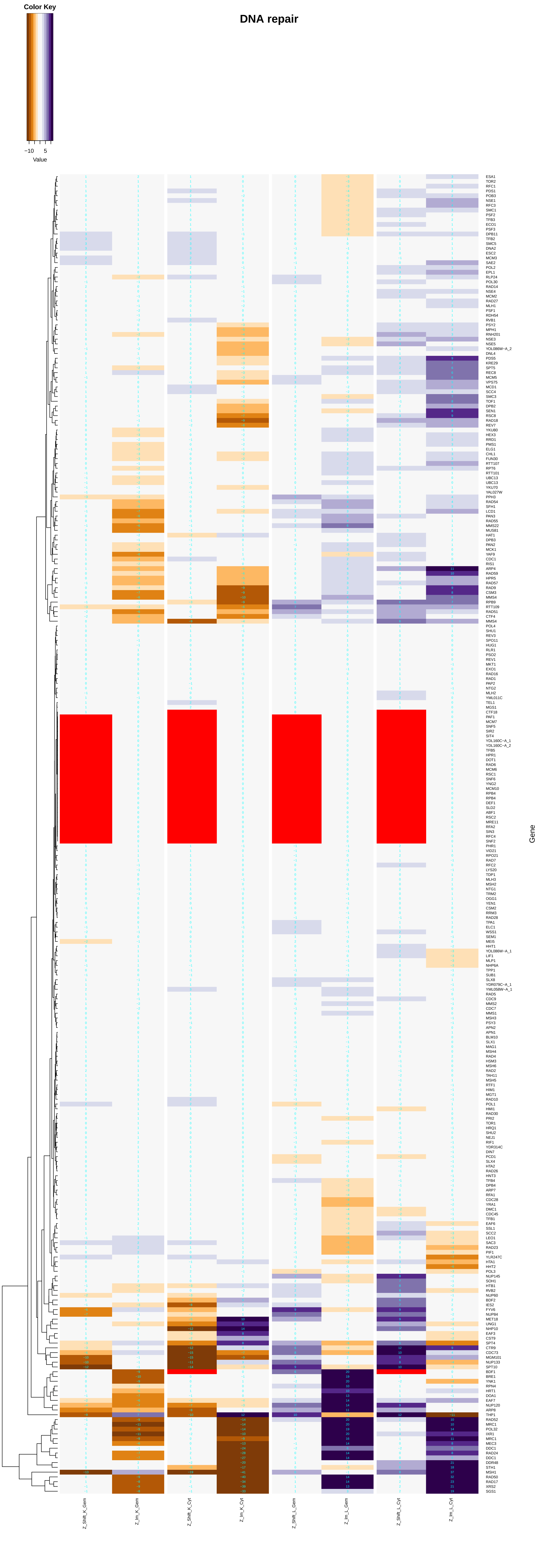

Color Key
−10
5
Value
DNA repair
Gene
Z_Shift_K_Gem
Z_Im_K_Gem
Z_Shift_K_Cyt
Z_Im_K_Cyt
Z_Shift_L_Gem
Z_Im_L_Gem
Z_Shift_L_Cyt
Z_Im_L_Cyt

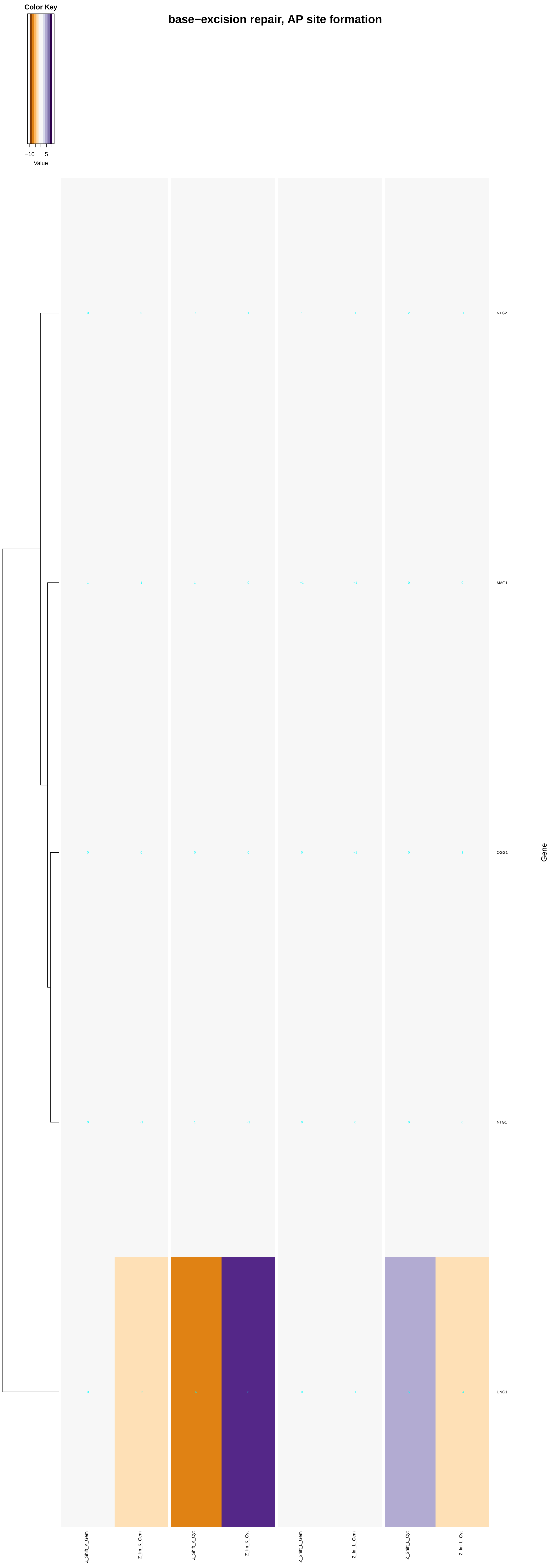

Color Key
−10
5
Value
base−excision repair, AP site formation
NTG2
MAG1
OGG1
NTG1
UNG1
Gene
Z_Shift_K_Gem
Z_Im_K_Gem
Z_Shift_K_Cyt
Z_Im_K_Cyt
Z_Shift_L_Gem
Z_Im_L_Gem
Z_Shift_L_Cyt
Z_Im_L_Cyt

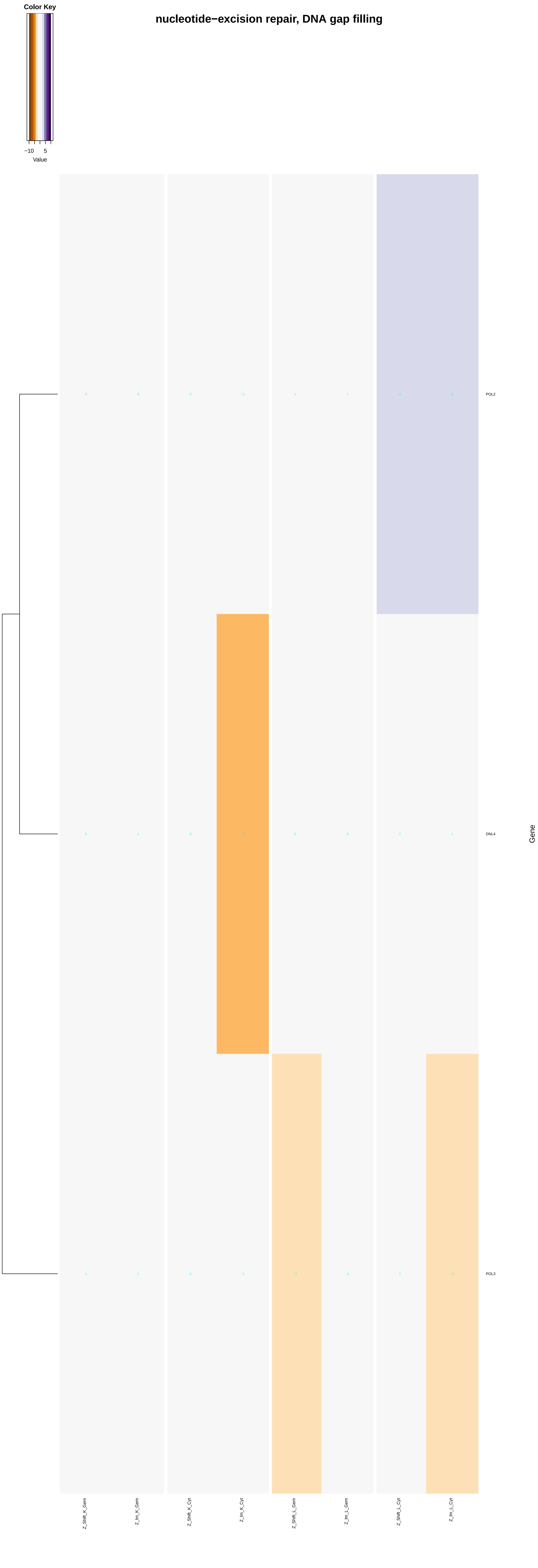

nucleotide−excision repair, DNA gap filling
Color Key
−10
5
Value
POL2
DNL4
POL3
Gene
Z_Shift_K_Gem
Z_Im_K_Gem
Z_Shift_K_Cyt
Z_Im_K_Cyt
Z_Shift_L_Gem
Z_Im_L_Gem
Z_Shift_L_Cyt
Z_Im_L_Cyt

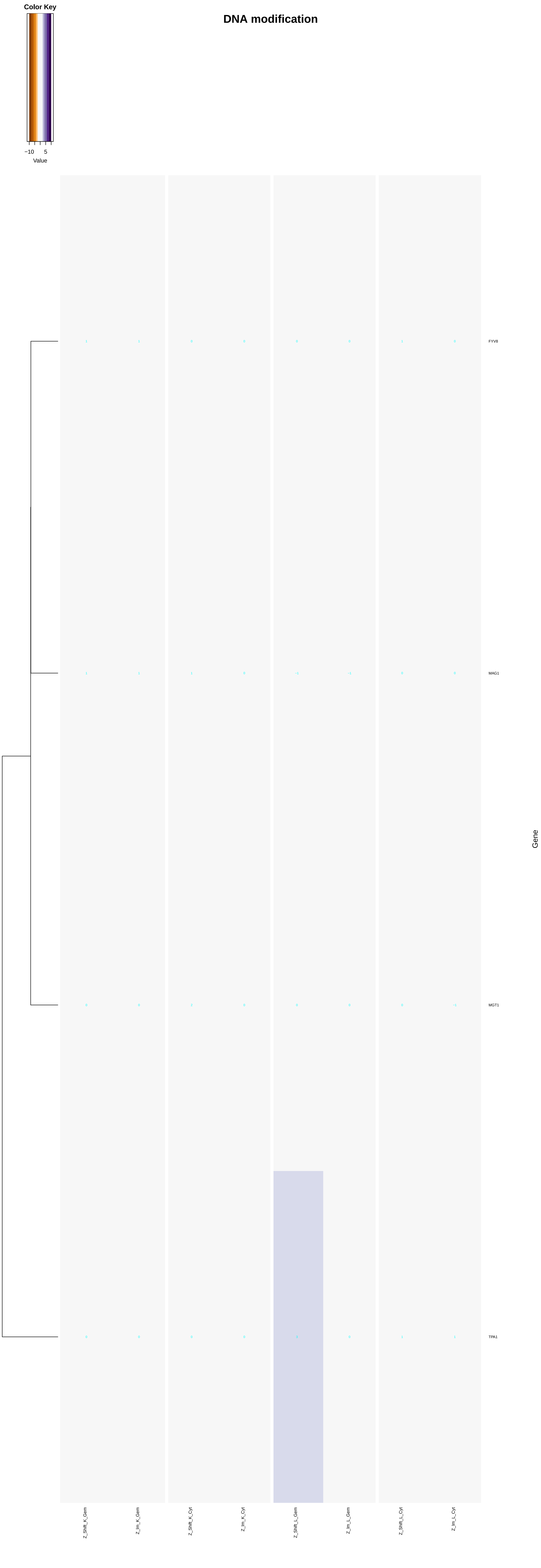
Color Key
−10
5
Value
DNA modification
FYV8
MAG1
MGT1
TPA1
Gene
Z_Shift_K_Gem
Z_Im_K_Gem
Z_Shift_K_Cyt
Z_Im_K_Cyt
Z_Shift_L_Gem
Z_Im_L_Gem
Z_Shift_L_Cyt
Z_Im_L_Cyt

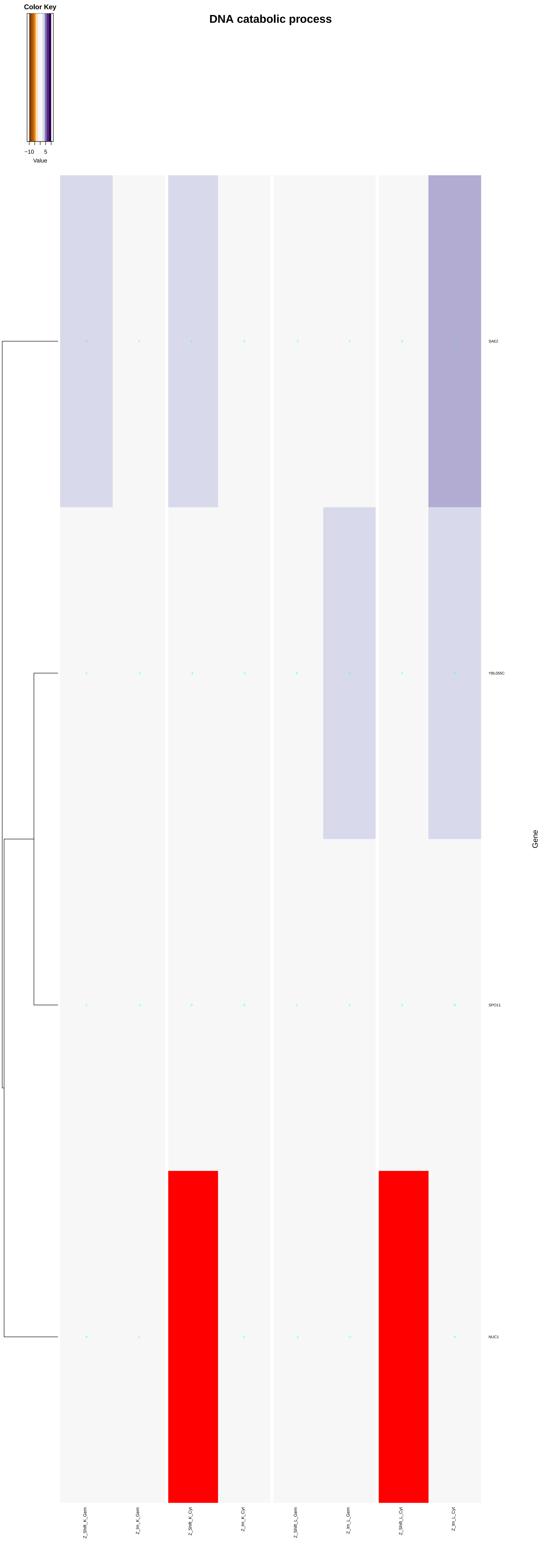
DNA catabolic process
Color Key
−10
5
Value
SAE2
YBL055C
SPO11
NUC1
Gene
Z_Shift_K_Gem
Z_Im_K_Gem
Z_Shift_K_Cyt
Z_Im_K_Cyt
Z_Shift_L_Gem
Z_Im_L_Gem
Z_Shift_L_Cyt
Z_Im_L_Cyt

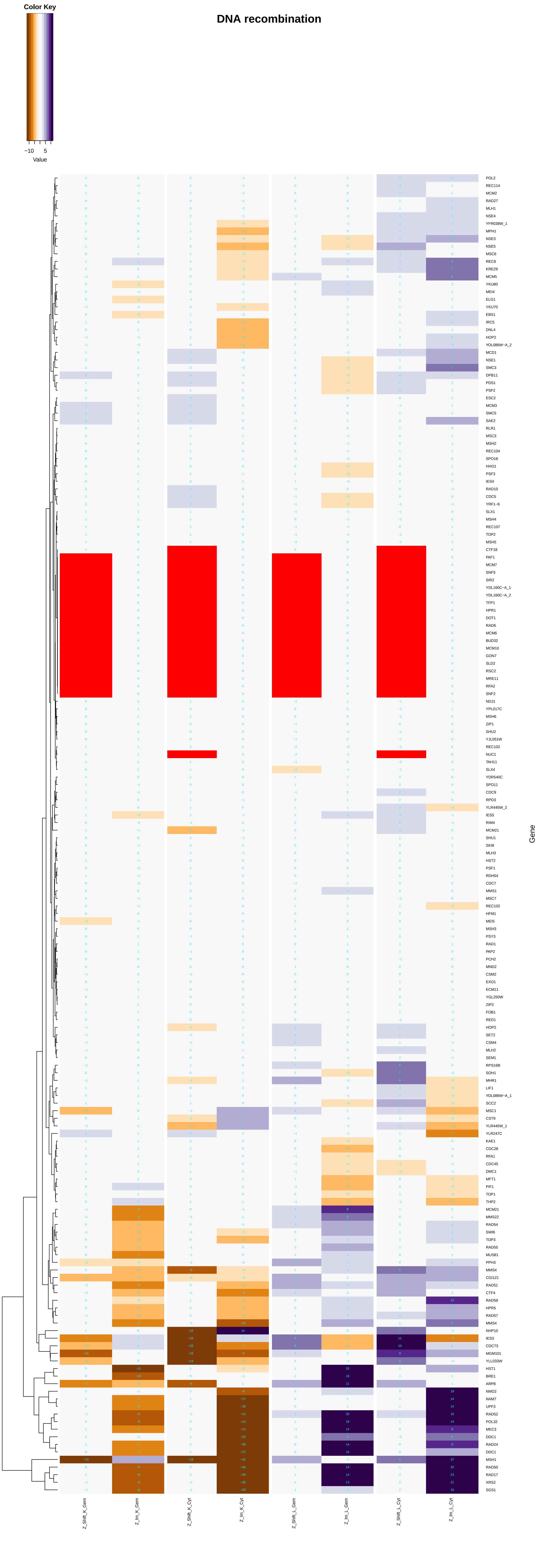

DNA recombination
Color Key
−10
5
Value
Gene
POL2
REC114
MCM2
RAD27
MLH1
NSE4
YFR038W_1
MPH1
NSE3
NSE5
MSC6
REC8
KRE29
MCM5
YKU80
MEI4
ELG1
YKU70
EBS1
IRC5
DNL4
HOP2
YOL086W−A_2
MCD1
NSE1
SMC3
DPB11
PDS1
PSF2
ESC2
MCM3
SMC5
SAE2
RLR1
MSC3
MSH2
REC104
SPO16
HHO1
PSF3
IES4
RAD10
CDC5
YRF1−6
SLX1
MSH4
REC107
TOP2
MSH5
CTF18
PAF1
MCM7
SNF5
SIR2
YDL160C−A_1
YDL160C−A_2
TFP1
HPR1
DOT1
RAD6
MCM6
BUD32
MCM10
GON7
SLD2
RSC2
MRE11
RFA2
SNF2
NDJ1
YPL017C
MSH6
ZIP1
SHU2
YJL051W
REC102
NUC1
TAH11
SLX4
YDR540C
SPO11
CDC9
RPD3
YLR445W_2
IES5
RIM4
MCM21
SHU1
SKI8
MLH3
HST2
PSF1
RDH54
CDC7
MMS1
MSC7
REC102
HFM1
MEI5
MSH3
PSY3
RAD1
PAP2
PCH2
MND2
CSM2
EXO1
ECM11
YGL250W
ZIP2
FOB1
RED1
HOP2
SET2
CSM4
MLH2
SEM1
RPS16B
SOH1
MHR1
LIF1
YOL086W−A_1
SCC2
MSC1
CST9
YLR445W_1
YLR247C
KAE1
CDC28
RFA1
CDC45
DMC1
MFT1
PIF1
TOP1
THP2
MCM21
MMS22
RAD54
SWI6
TOP3
RAD55
MUS81
PPH3
MMS4
CG121
RAD51
CTF4
RAD59
HPR5
RAD57
MMS4
NHP10
IES3
CDC73
MGM101
YLL033W
HST1
BRE1
ARP8
NMD2
NAM7
UPF3
RAD52
POL32
MEC3
DDC1
RAD24
DDC1
MSH1
RAD50
RAD17
XRS2
SGS1
Z_Shift_K_Gem
Z_Im_K_Gem
Z_Shift_K_Cyt
Z_Im_K_Cyt
Z_Shift_L_Gem
Z_Im_L_Gem
Z_Shift_L_Cyt
Z_Im_L_Cyt

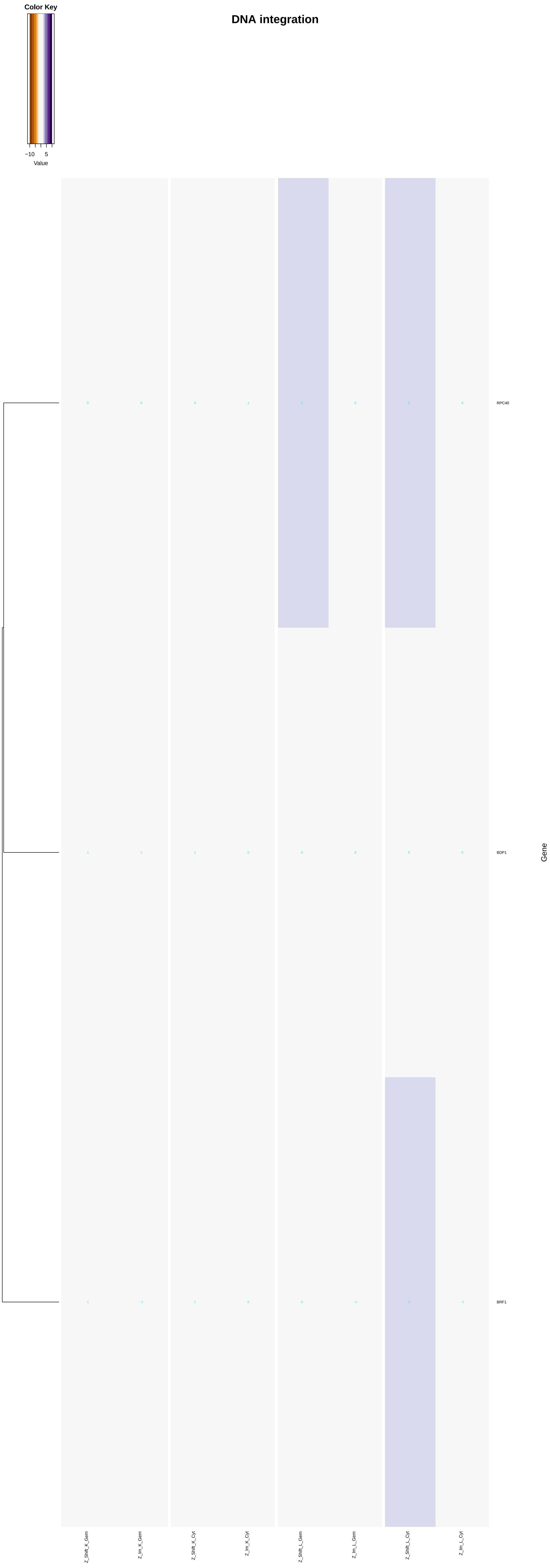
DNA integration
Color Key
−10
5
Value
RPC40
BDP1
BRF1
Gene
Z_Shift_K_Gem
Z_Im_K_Gem
Z_Shift_K_Cyt
Z_Im_K_Cyt
Z_Shift_L_Gem
Z_Im_L_Gem
Z_Shift_L_Cyt
Z_Im_L_Cyt

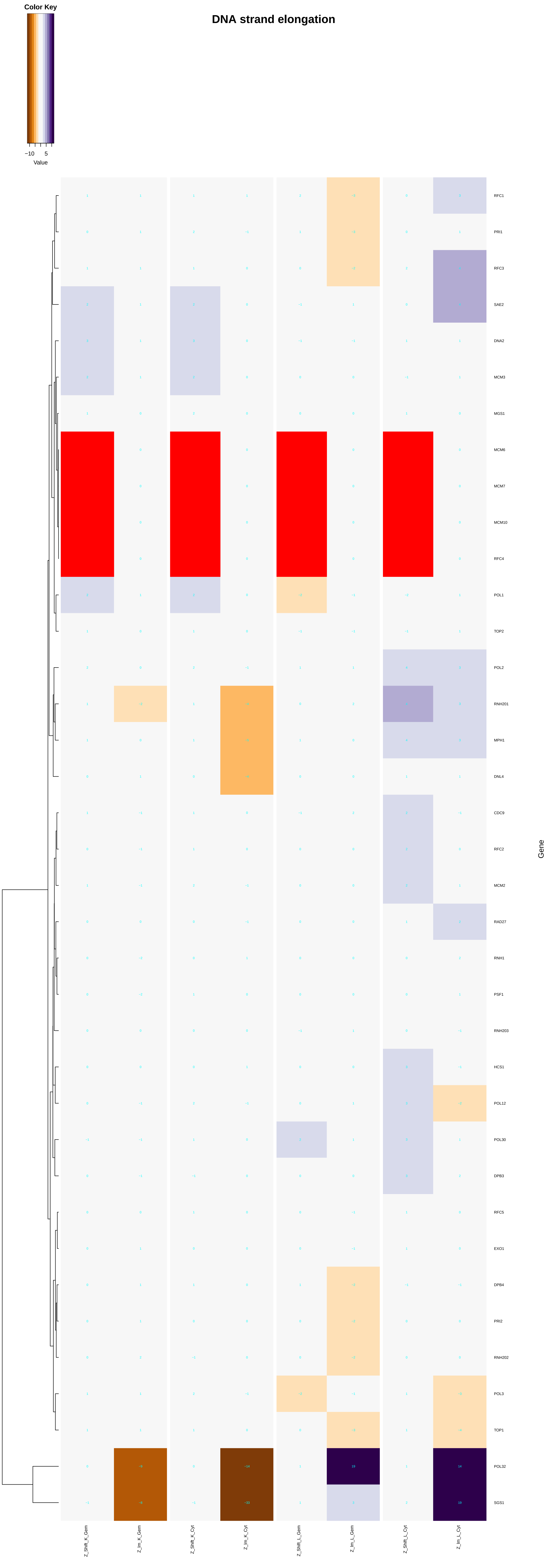
Color Key
−10 5
Value
DNA strand elongation
RFC1
PRI1
RFC3
SAE2
DNA2
MCM3
MGS1
MCM6
MCM7
MCM10
RFC4
POL1
TOP2
POL2
RNH201
MPH1
DNL4
CDC9
RFC2
MCM2
RAD27
RNH1
PSF1
RNH203
HCS1
POL12
POL30
DPB3
RFC5
EXO1
DPB4
PRI2
RNH202
POL3
TOP1
POL32
SGS1
Gene
Z_Shift_K_Gem
Z_Im_K_Gem
Z_Shift_K_Cyt
Z_Im_K_Cyt
Z_Shift_L_Gem
Z_Im_L_Gem
Z_Shift_L_Cyt
Z_Im_L_Cyt

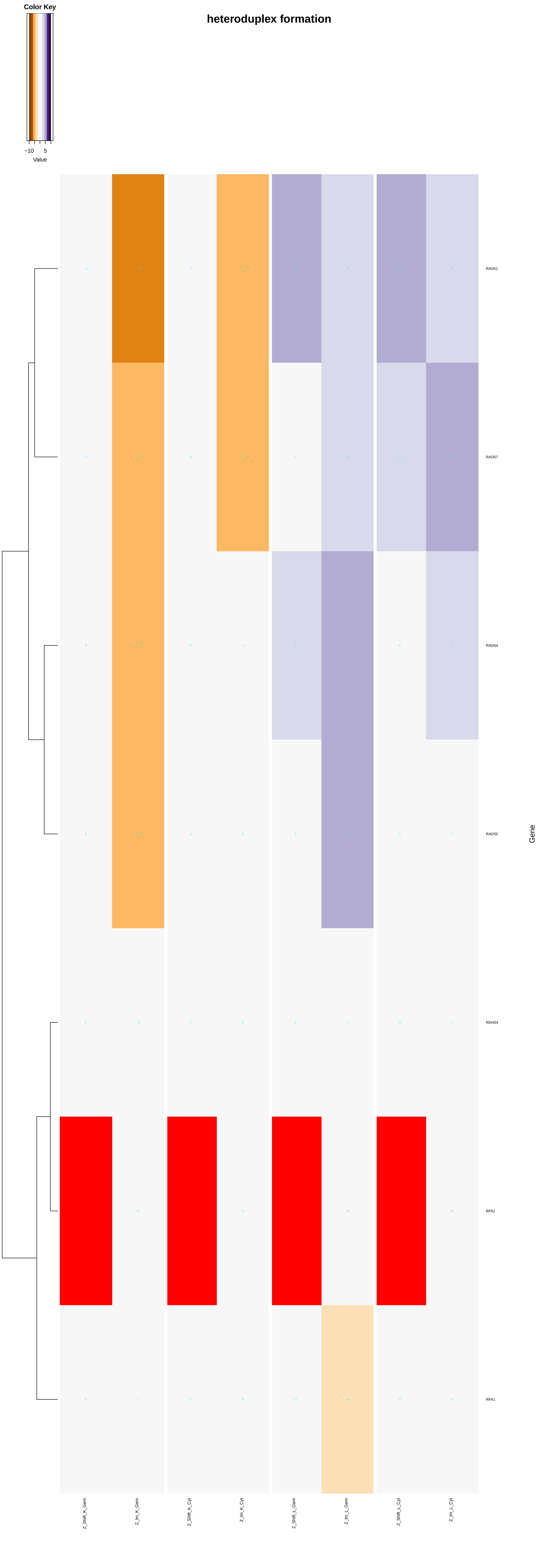
heteroduplex formation
Color Key
−10
5
Value
RAD51
RAD57
RAD54
RAD55
RDH54
RFA2
RFA1
Gene
Z_Shift_K_Gem
Z_Im_K_Gem
Z_Shift_K_Cyt
Z_Im_K_Cyt
Z_Shift_L_Gem
Z_Im_L_Gem
Z_Shift_L_Cyt
Z_Im_L_Cyt

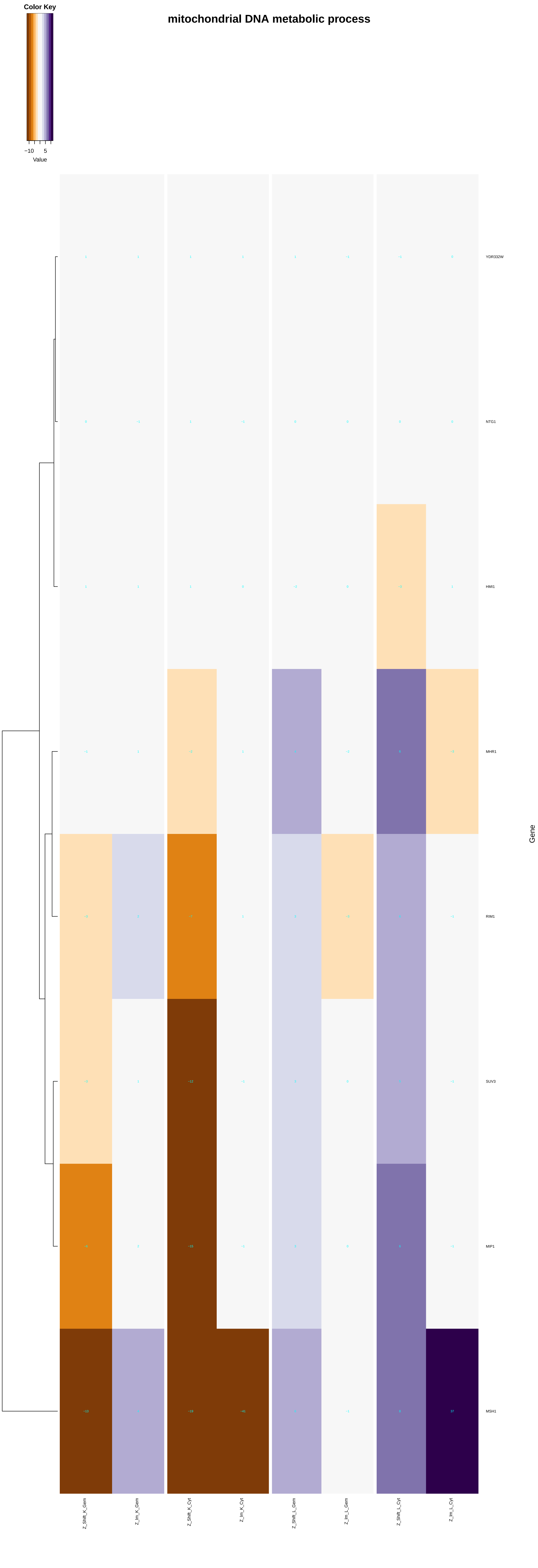
mitochondrial DNA metabolic process
Color Key
−10
5
Value
YDR332W
NTG1
HMI1
MHR1
RIM1
SUV3
MIP1
MSH1
Gene
Z_Shift_K_Gem
Z_Im_K_Gem
Z_Shift_K_Cyt
Z_Im_K_Cyt
Z_Shift_L_Gem
Z_Im_L_Gem
Z_Shift_L_Cyt
Z_Im_L_Cyt

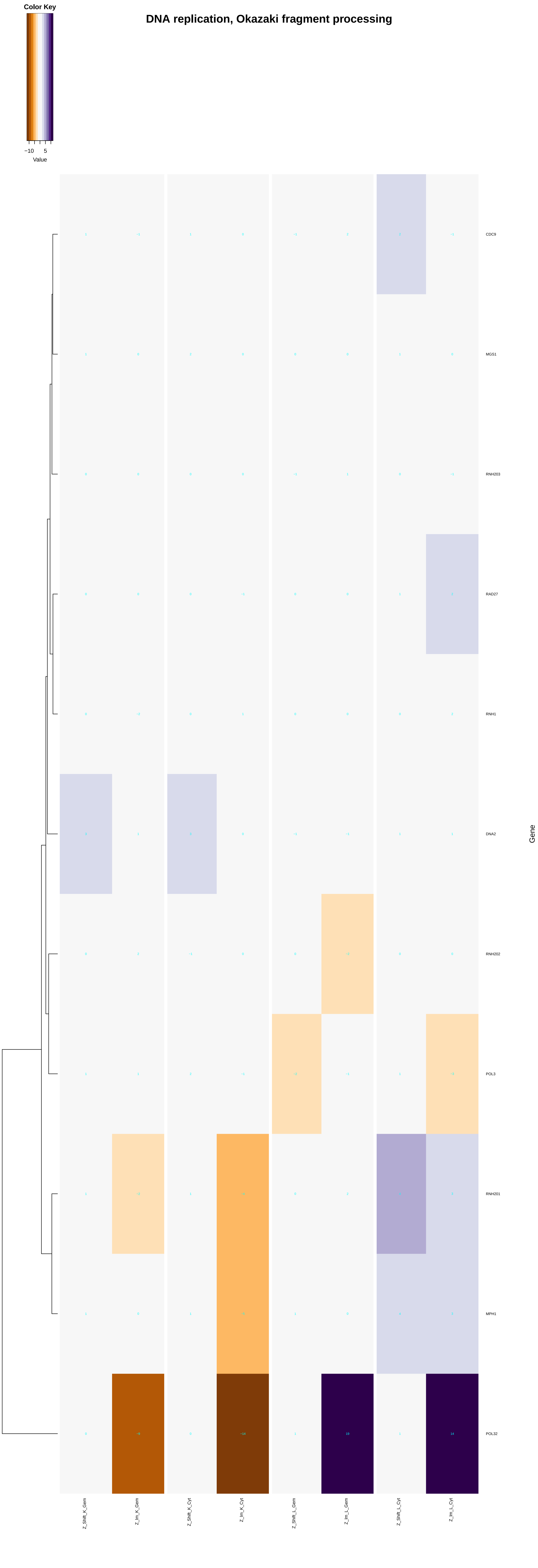
DNA replication, Okazaki fragment processing
Color Key
−10
5
Value
CDC9
MGS1
RNH203
RAD27
RNH1
DNA2
RNH202
POL3
RNH201
MPH1
POL32
Gene
Z_Shift_K_Gem
Z_Im_K_Gem
Z_Shift_K_Cyt
Z_Im_K_Cyt
Z_Shift_L_Gem
Z_Im_L_Gem
Z_Shift_L_Cyt
Z_Im_L_Cyt

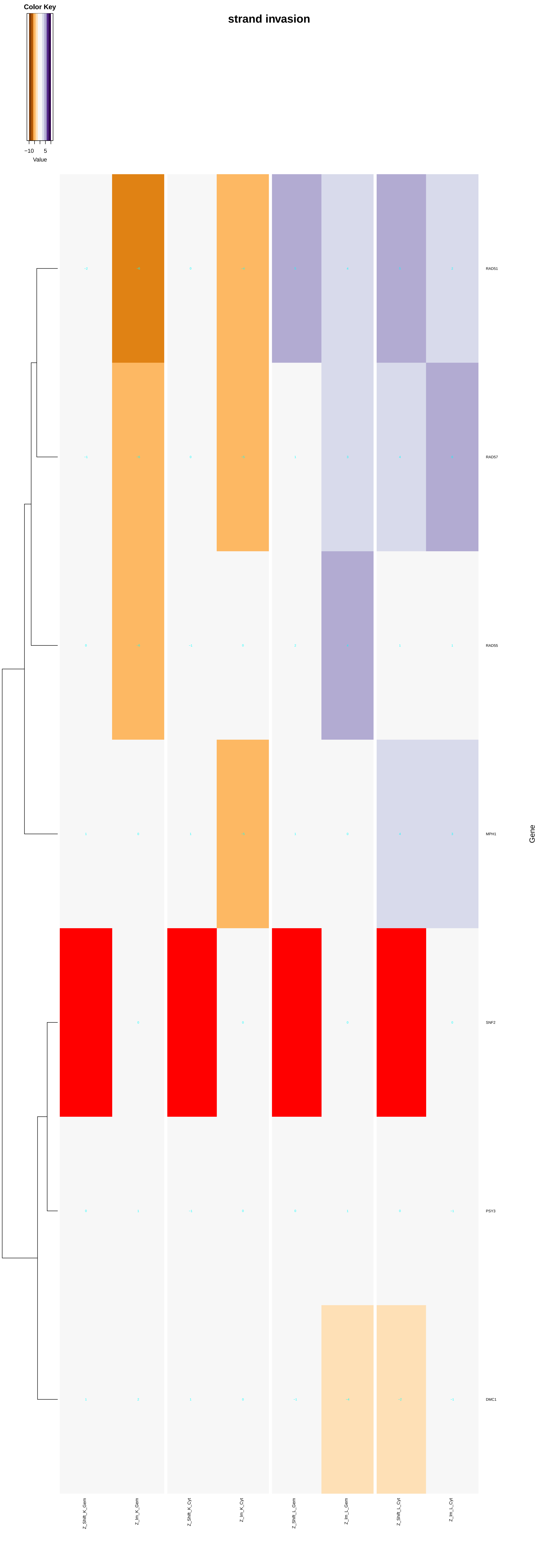
strand invasion
Color Key
−10
5
Value
RAD51
RAD57
RAD55
MPH1
SNF2
PSY3
DMC1
Gene
Z_Shift_K_Gem
Z_Im_K_Gem
Z_Shift_K_Cyt
Z_Im_K_Cyt
Z_Shift_L_Gem
Z_Im_L_Gem
Z_Shift_L_Cyt
Z_Im_L_Cyt

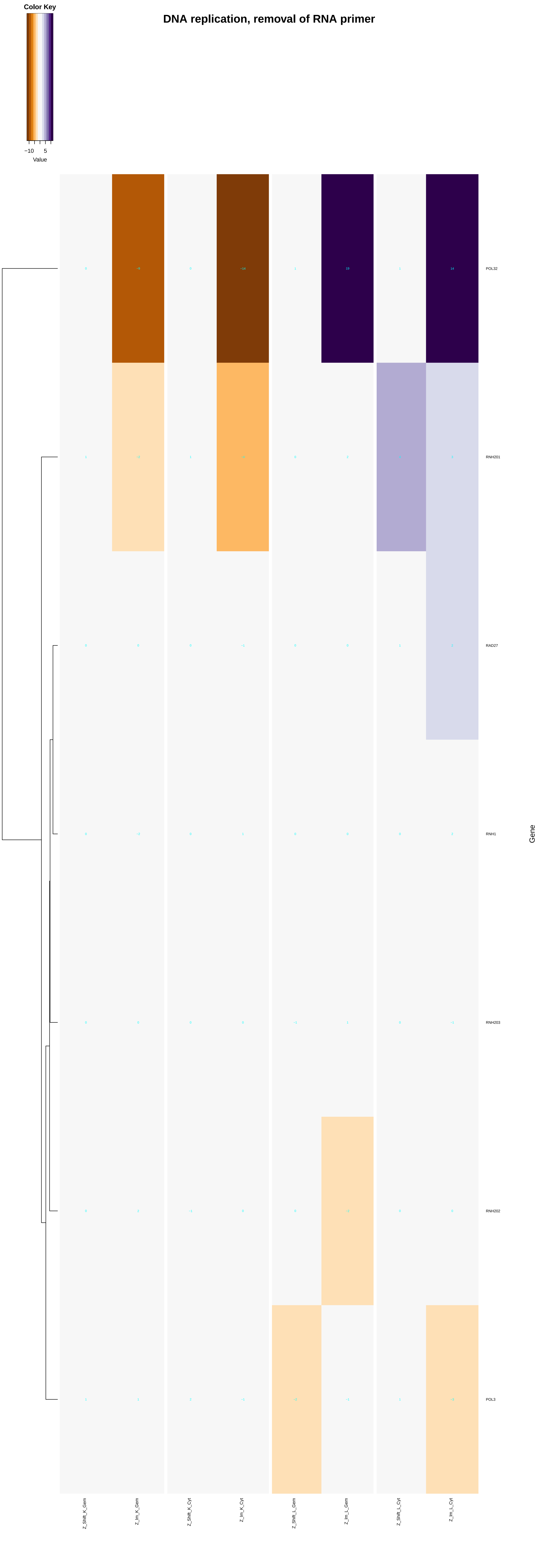
DNA replication, removal of RNA primer
Color Key
−10
5
Value
POL32
RNH201
RAD27
RNH1
RNH203
RNH202
POL3
Gene
Z_Shift_K_Gem
Z_Im_K_Gem
Z_Shift_K_Cyt
Z_Im_K_Cyt
Z_Shift_L_Gem
Z_Im_L_Gem
Z_Shift_L_Cyt
Z_Im_L_Cyt

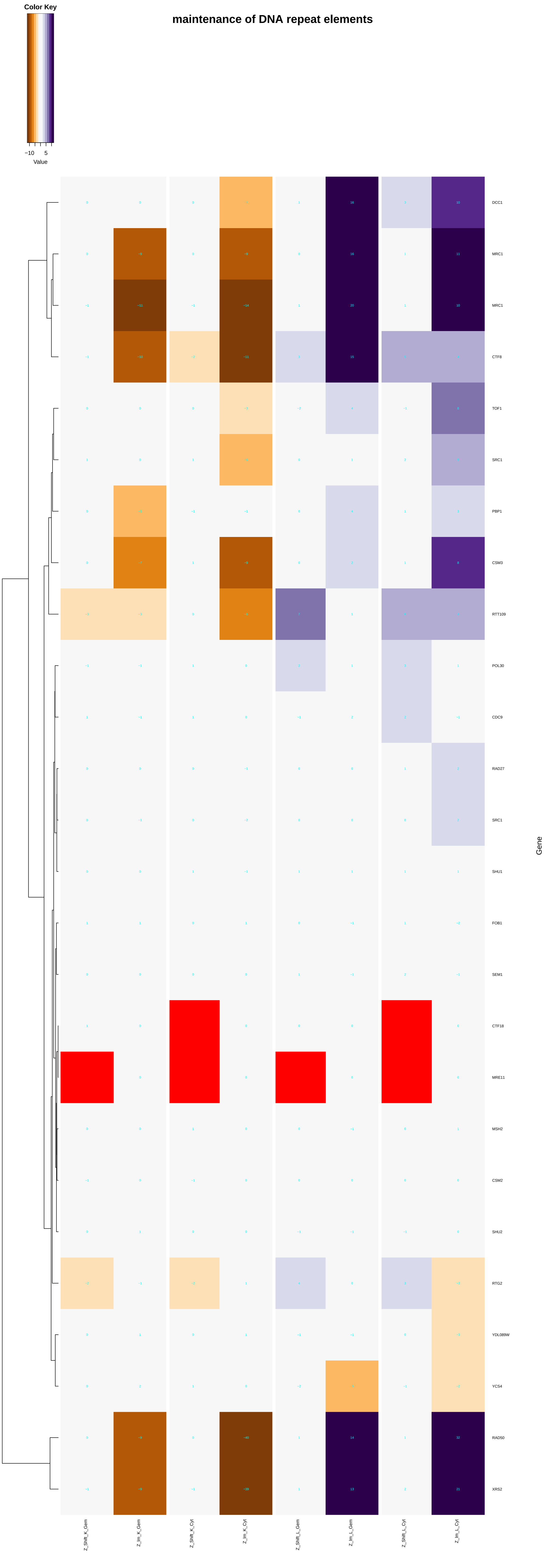
maintenance of DNA repeat elements
Color Key
−10
5
Value
DCC1
MRC1
MRC1
CTF8
TOF1
SRC1
PBP1
CSM3
RTT109
POL30
CDC9
RAD27
SRC1
SHU1
FOB1
SEM1
CTF18
MRE11
MSH2
CSM2
SHU2
RTG2
YDL089W
YCS4
RAD50
XRS2
Gene
Z_Shift_K_Gem
Z_Im_K_Gem
Z_Shift_K_Cyt
Z_Im_K_Cyt
Z_Shift_L_Gem
Z_Im_L_Gem
Z_Shift_L_Cyt
Z_Im_L_Cyt

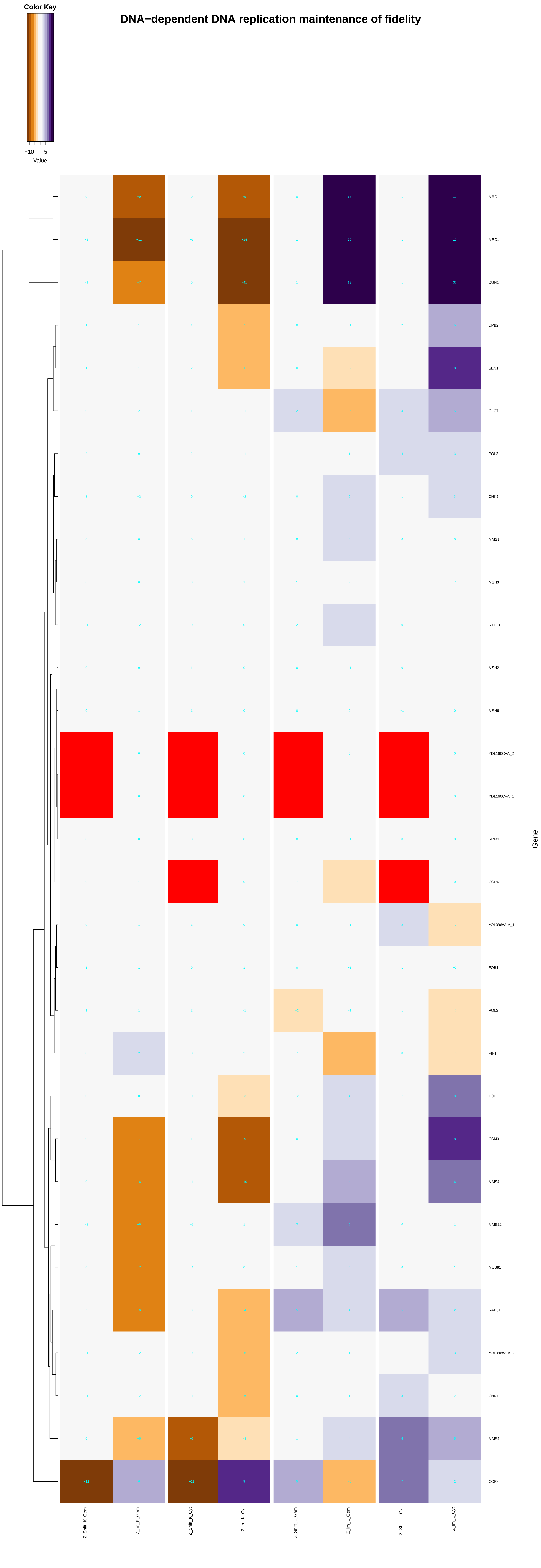

DNA−dependent DNA replication maintenance of fidelity
Color Key
−10
5
Value
MRC1
MRC1
DUN1
DPB2
SEN1
GLC7
POL2
CHK1
MMS1
MSH3
RTT101
MSH2
MSH6
YDL160C−A_2
YDL160C−A_1
RRM3
CCR4
YOL086W−A_1
FOB1
POL3
PIF1
TOF1
CSM3
MMS4
MMS22
MUS81
RAD51
YOL086W−A_2
CHK1
MMS4
CCR4
Gene
Z_Shift_K_Gem
Z_Im_K_Gem
Z_Shift_K_Cyt
Z_Im_K_Cyt
Z_Shift_L_Gem
Z_Im_L_Gem
Z_Shift_L_Cyt
Z_Im_L_Cyt

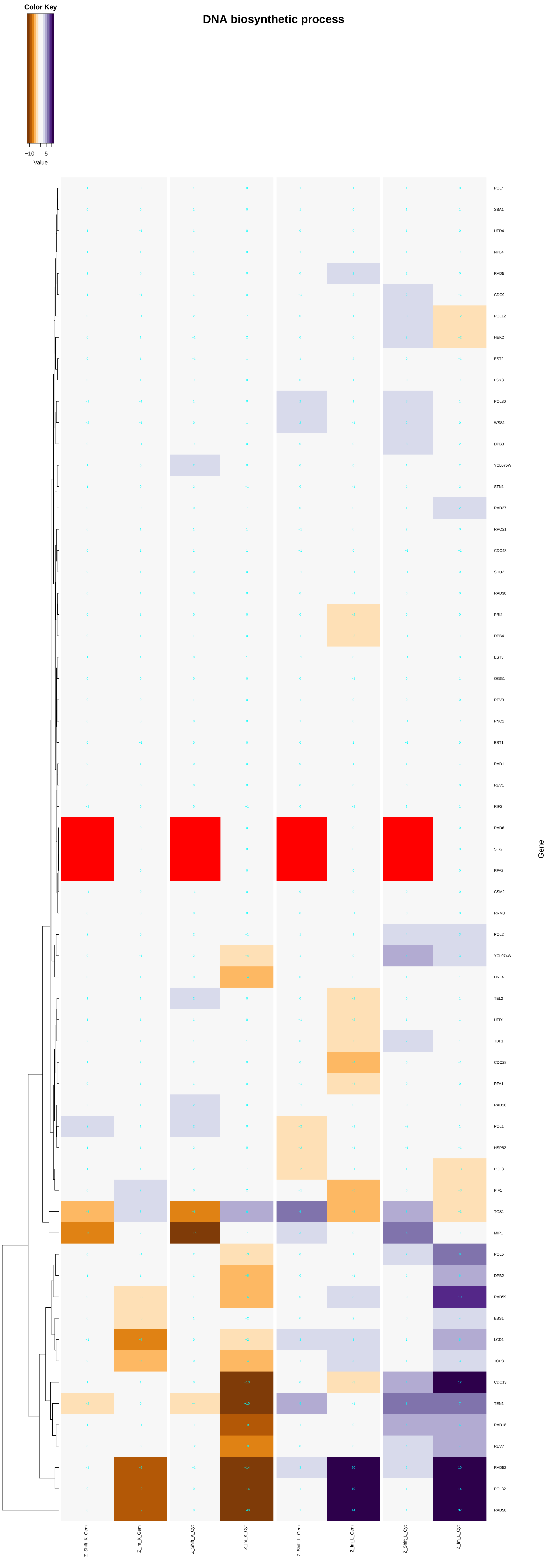
DNA biosynthetic process
Color Key
−10
5
Value
POL4
SBA1
UFD4
NPL4
RAD5
CDC9
POL12
HEK2
EST2
PSY3
POL30
WSS1
DPB3
YCL075W
STN1
RAD27
RPO21
CDC48
SHU2
RAD30
PRI2
DPB4
EST3
OGG1
REV3
PNC1
EST1
RAD1
REV1
RIF2
RAD6
SIR2
RFA2
CSM2
RRM3
POL2
YCL074W
DNL4
TEL2
UFD1
TBF1
CDC28
RFA1
RAD10
POL1
HSP82
POL3
PIF1
TGS1
MIP1
POL5
DPB2
RAD59
EBS1
LCD1
TOP3
CDC13
TEN1
RAD18
REV7
RAD52
POL32
RAD50
Gene
Z_Shift_K_Gem
Z_Im_K_Gem
Z_Shift_K_Cyt
Z_Im_K_Cyt
Z_Shift_L_Gem
Z_Im_L_Gem
Z_Shift_L_Cyt
Z_Im_L_Cyt

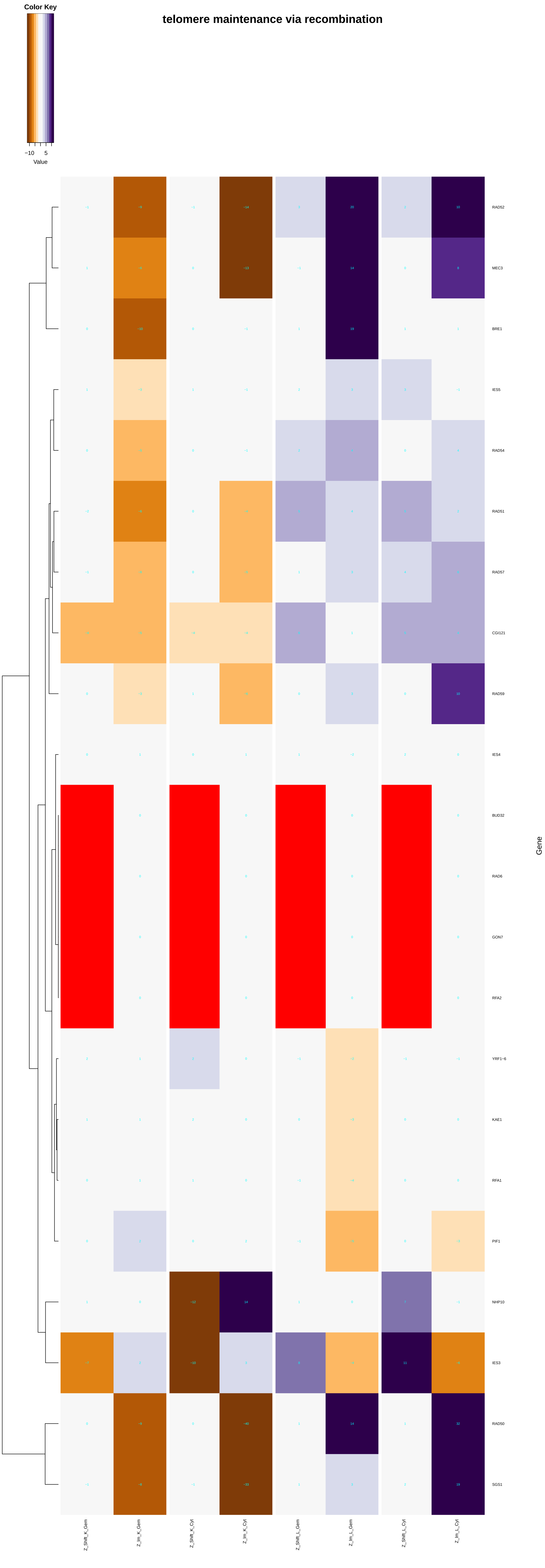

telomere maintenance via recombination
Color Key
−10
5
Value
RAD52
MEC3
BRE1
IES5
RAD54
RAD51
RAD57
CGI121
RAD59
IES4
BUD32
RAD6
GON7
RFA2
YRF1−6
KAE1
RFA1
PIF1
NHP10
IES3
RAD50
SGS1
Gene
Z_Shift_K_Gem
Z_Im_K_Gem
Z_Shift_K_Cyt
Z_Im_K_Cyt
Z_Shift_L_Gem
Z_Im_L_Gem
Z_Shift_L_Cyt
Z_Im_L_Cyt

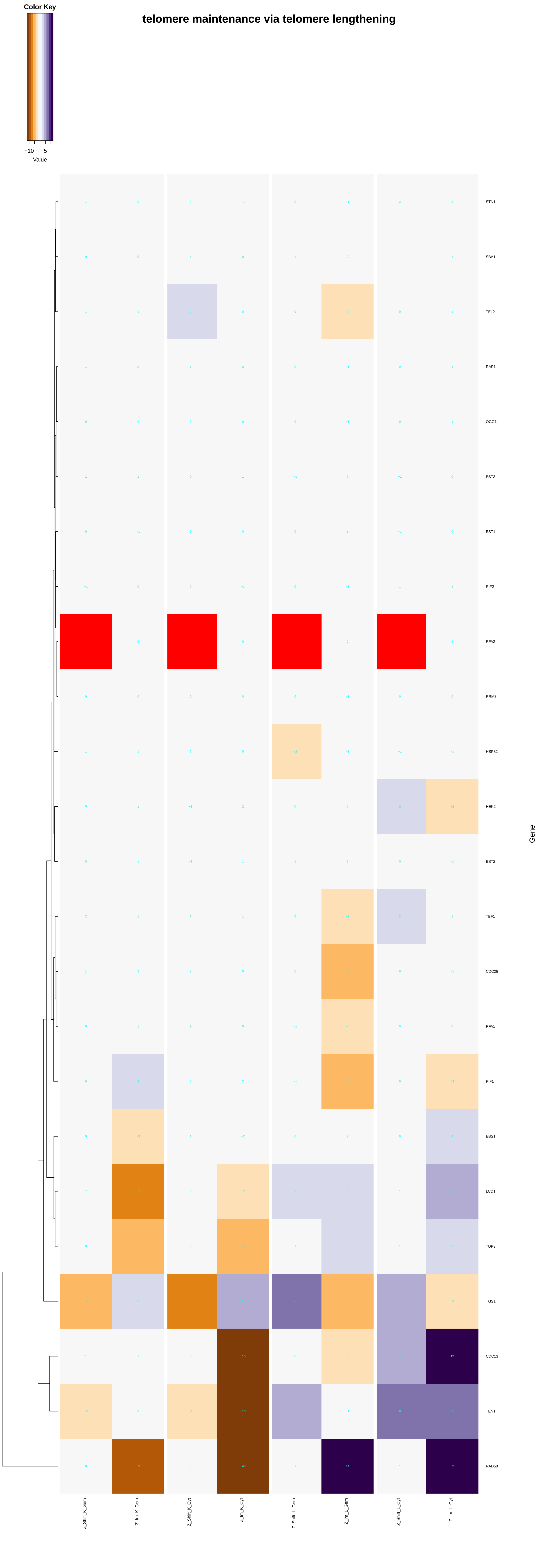

Color Key
−10
5
Value
telomere maintenance via telomere lengthening
STN1
SBA1
TEL2
RAP1
OGG1
EST3
EST1
RIF2
RFA2
RRM3
HSP82
HEK2
EST2
TBF1
CDC28
RFA1
PIF1
EBS1
LCD1
TOP3
TGS1
CDC13
TEN1
RAD50
Gene
Z_Shift_K_Gem
Z_Im_K_Gem
Z_Shift_K_Cyt
Z_Im_K_Cyt
Z_Shift_L_Gem
Z_Im_L_Gem
Z_Shift_L_Cyt
Z_Im_L_Cyt

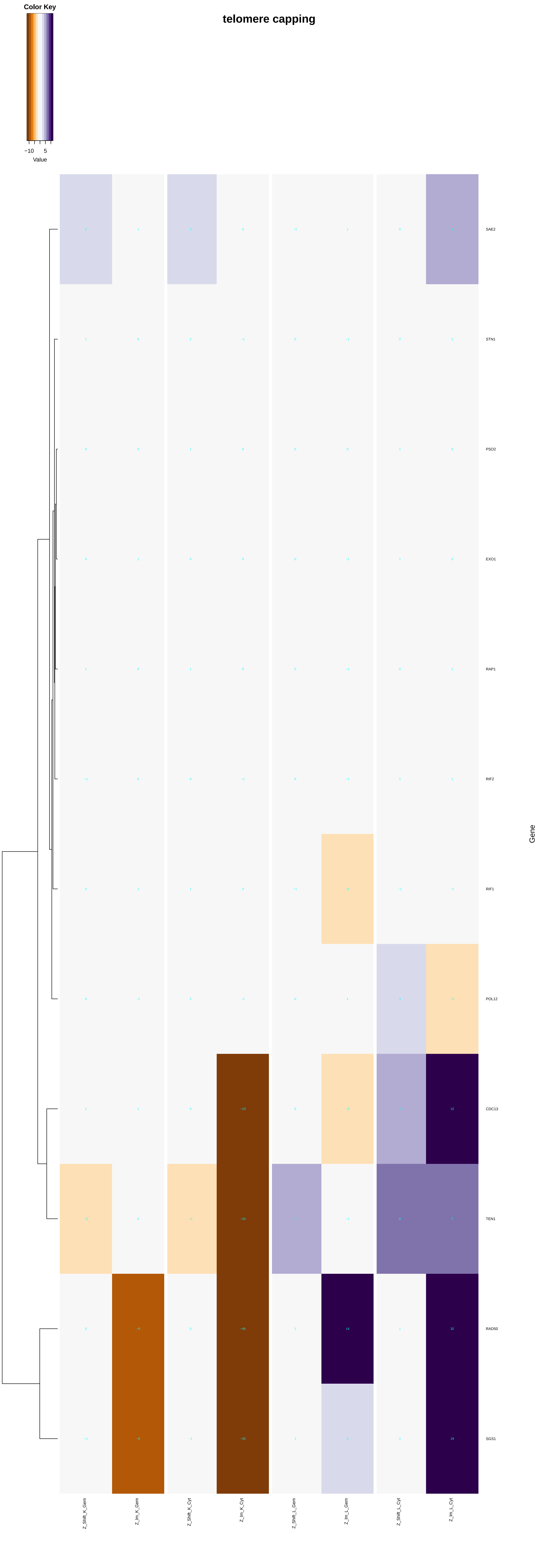
telomere capping
Color Key
−10
5
Value
SAE2
STN1
PSO2
EXO1
RAP1
RIF2
RIF1
POL12
CDC13
TEN1
RAD50
SGS1
Gene
Z_Shift_K_Gem
Z_lm_K_Gem
Z_Shift_K_Cyt
Z_lm_K_Cyt
Z_Shift_L_Gem
Z_lm_L_Gem
Z_Shift_L_Cyt
Z_lm_L_Cyt

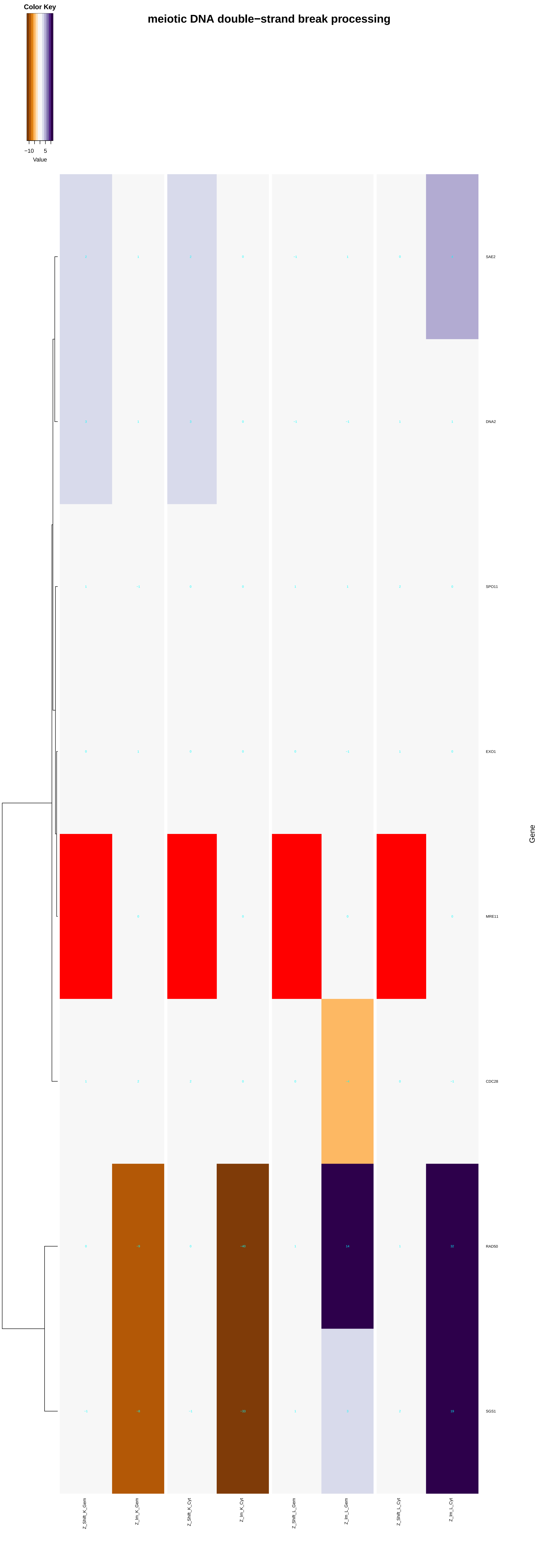

Color Key
−10
5
Value
meiotic DNA double−strand break processing
SAE2
DNA2
SPO11
EXO1
MRE11
CDC28
RAD50
SGS1
Gene
Z_Shift_K_Gem
Z_Im_K_Gem
Z_Shift_K_Cyt
Z_Im_K_Cyt
Z_Shift_L_Gem
Z_Im_L_Gem
Z_Shift_L_Cyt
Z_Im_L_Cyt

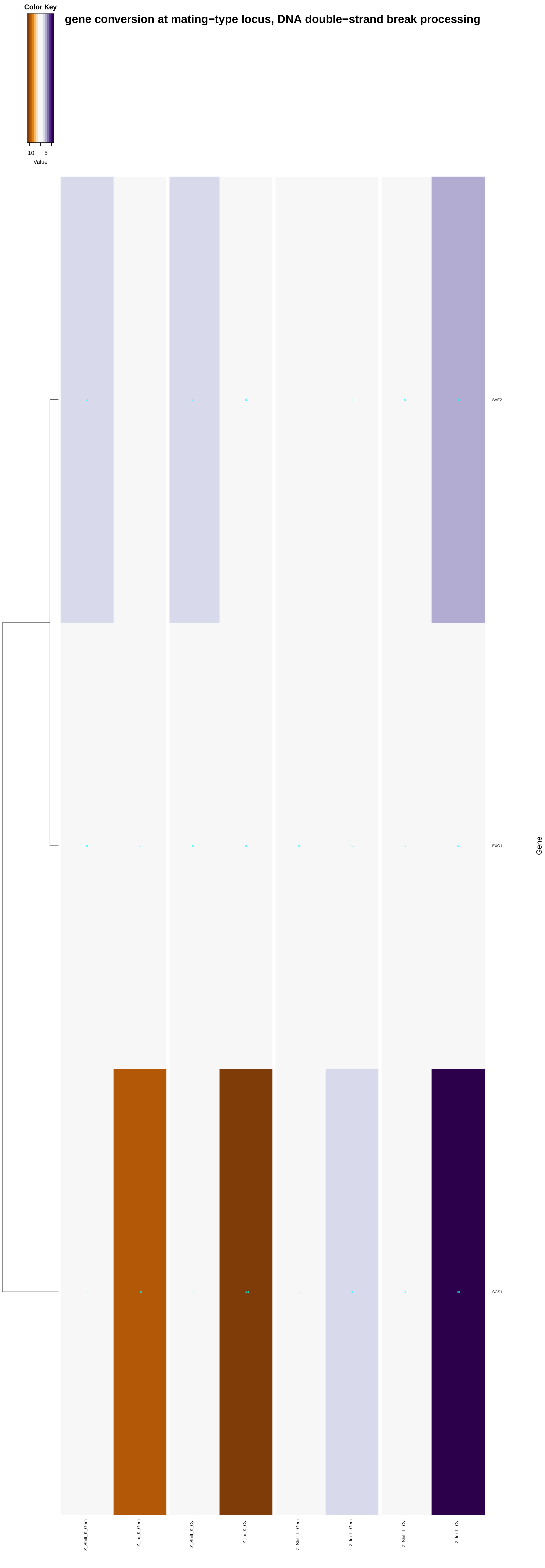
Color Key
−10
5
Value
gene conversion at mating−type locus, DNA double−strand break processing
2
1
2
0
−1
1
0
4
SAE2
0
1
0
0
0
−1
1
0
EXO1
−1
−8
−1
−33
1
3
2
19
SGS1
Gene
Z_Shift_K_Gem
Z_Im_K_Gem
Z_Shift_K_Cyt
Z_Im_K_Cyt
Z_Shift_L_Gem
Z_Im_L_Gem
Z_Shift_L_Cyt
Z_Im_L_Cyt

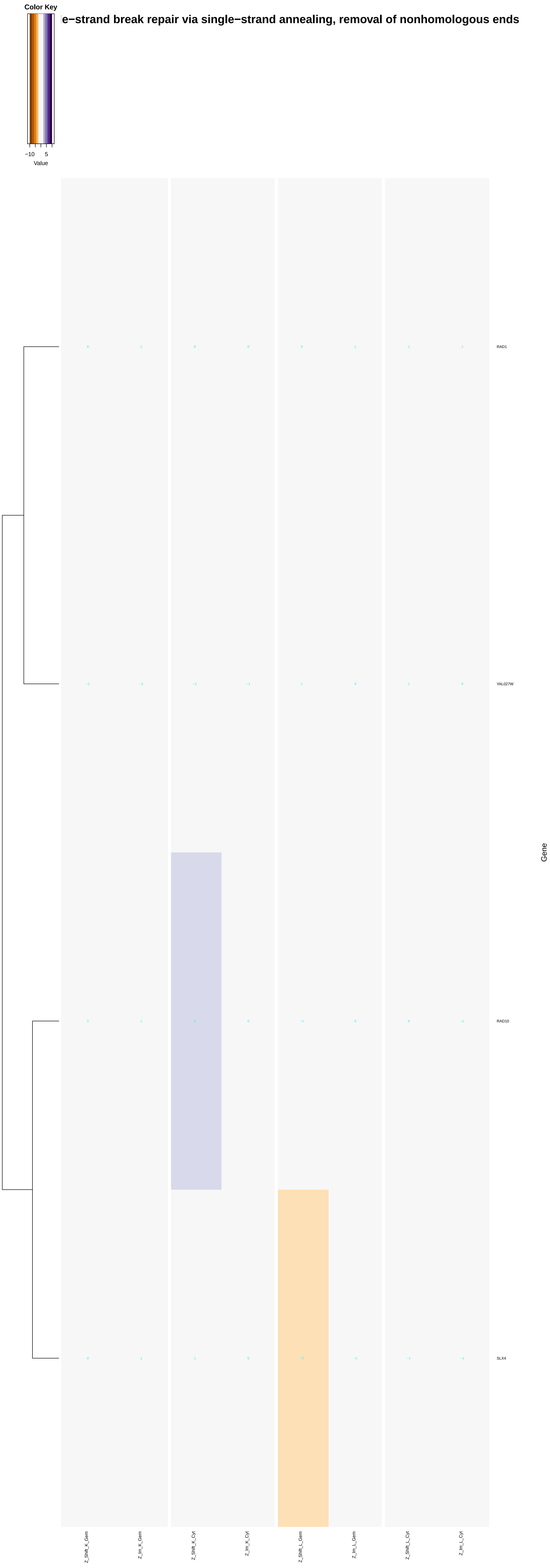

e−strand break repair via single−strand annealing, removal of nonhomologous ends
Color Key
−10
5
Value
RAD1
YAL027W
RAD10
SLX4
Gene
Z_Shift_K_Gem
Z_lm_K_Gem
Z_Shift_K_Cyt
Z_lm_K_Cyt
Z_Shift_L_Gem
Z_lm_L_Gem
Z_Shift_L_Cyt
Z_lm_L_Cyt

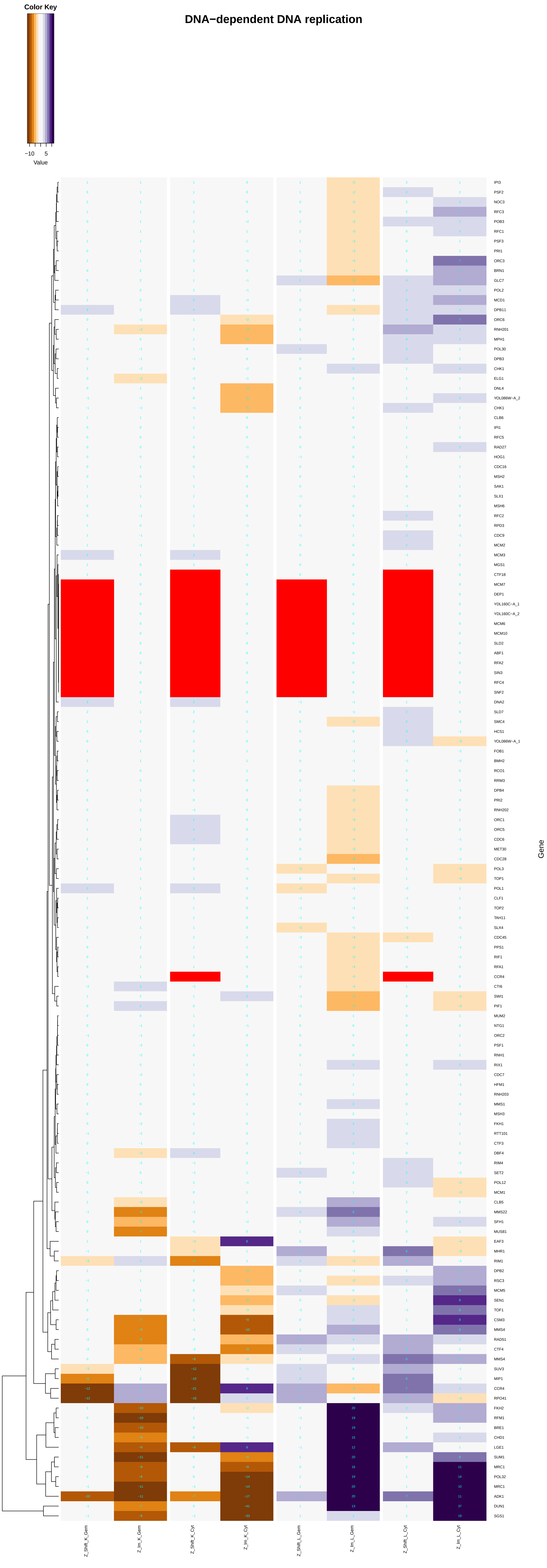
DNA−dependent DNA replication
Color Key
−10
5
Value
IPI3
PSF2
NOC3
RFC3
POB3
RFC1
PSF3
PRI1
ORC3
BRN1
GLC7
POL2
MCD1
DPB11
ORC6
RNH201
MPH1
POL30
DPB3
CHK1
ELG1
DNL4
YOL086W−A_2
CHK1
CLB6
IPI1
RFC5
RAD27
HOG1
CDC16
MSH2
SAK1
SLX1
MSH6
RFC2
RPD3
CDC9
MCM2
MCM3
MGS1
CTF18
MCM7
DEP1
YDL160C−A_1
YDL160C−A_2
MCM6
MCM10
SLD2
ABF1
RFA2
SIN3
RFC4
SNF2
DNA2
SLD7
SMC4
HCS1
YOL086W−A_1
FOB1
BMH2
RCO1
RRM3
DPB4
PRI2
RNH202
ORC1
ORC5
CDC6
MET30
CDC28
POL3
TOP1
POL1
CLF1
TOP2
TAH11
SLX4
CDC45
PPS1
RIF1
RFA1
CCR4
CTI6
SWI1
PIF1
MUM2
NTG1
ORC2
PSF1
RNH1
RIX1
CDC7
HFM1
RNH203
MMS1
MSH3
FKH1
RTT101
CTF3
DBF4
RIM4
SET2
POL12
MCM1
CLB5
MMS22
SFH1
MUS81
EAF3
MHR1
RIM1
DPB2
RSC3
MCM5
SEN1
TOF1
CSM3
MMS4
RAD51
CTF4
MMS4
SUV3
MIP1
CCR4
RPO41
FKH2
RFM1
BRE1
CHD1
LGE1
SUM1
MRC1
POL32
MRC1
ADK1
DUN1
SGS1
Gene
Z_Shift_K_Gem
Z_Im_K_Gem
Z_Shift_K_Cyt
Z_Im_K_Cyt
Z_Shift_L_Gem
Z_Im_L_Gem
Z_Shift_L_Cyt
Z_Im_L_Cyt

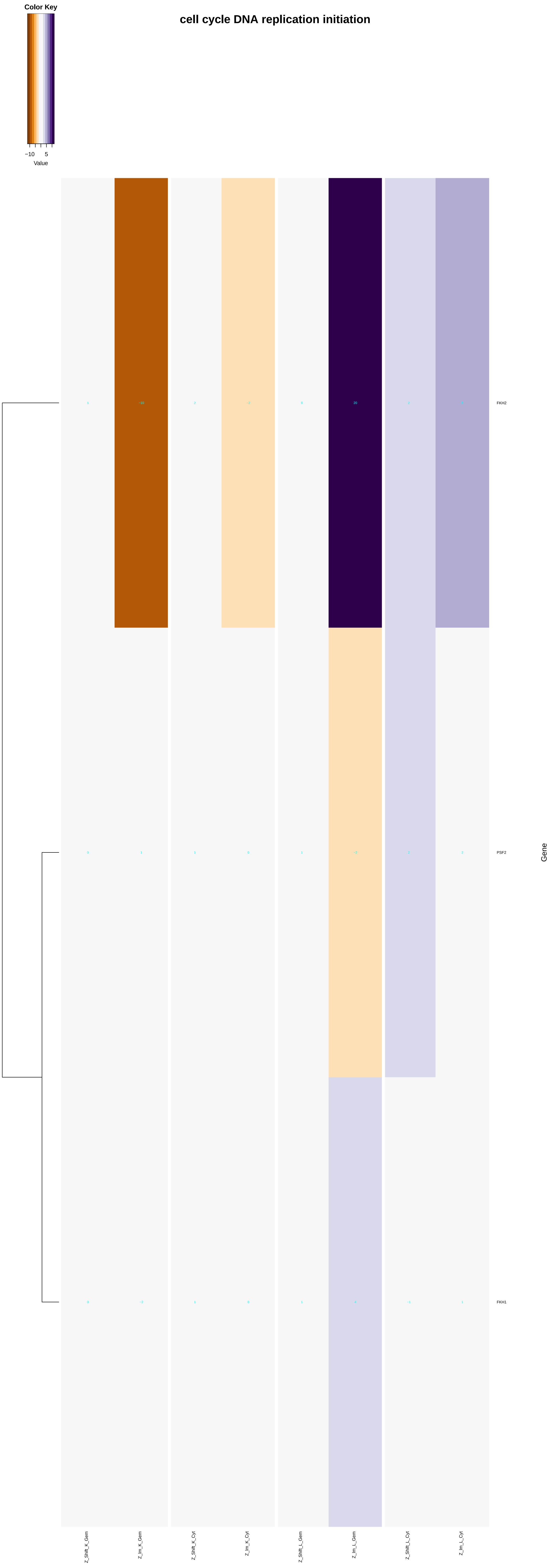
cell cycle DNA replication initiation
Color Key
−10
5
Value
FKH2
PSF2
FKH1
Gene
Z_Shift_K_Gem
Z_Im_K_Gem
Z_Shift_K_Cyt
Z_Im_K_Cyt
Z_Shift_L_Gem
Z_Im_L_Gem
Z_Shift_L_Cyt
Z_Im_L_Cyt

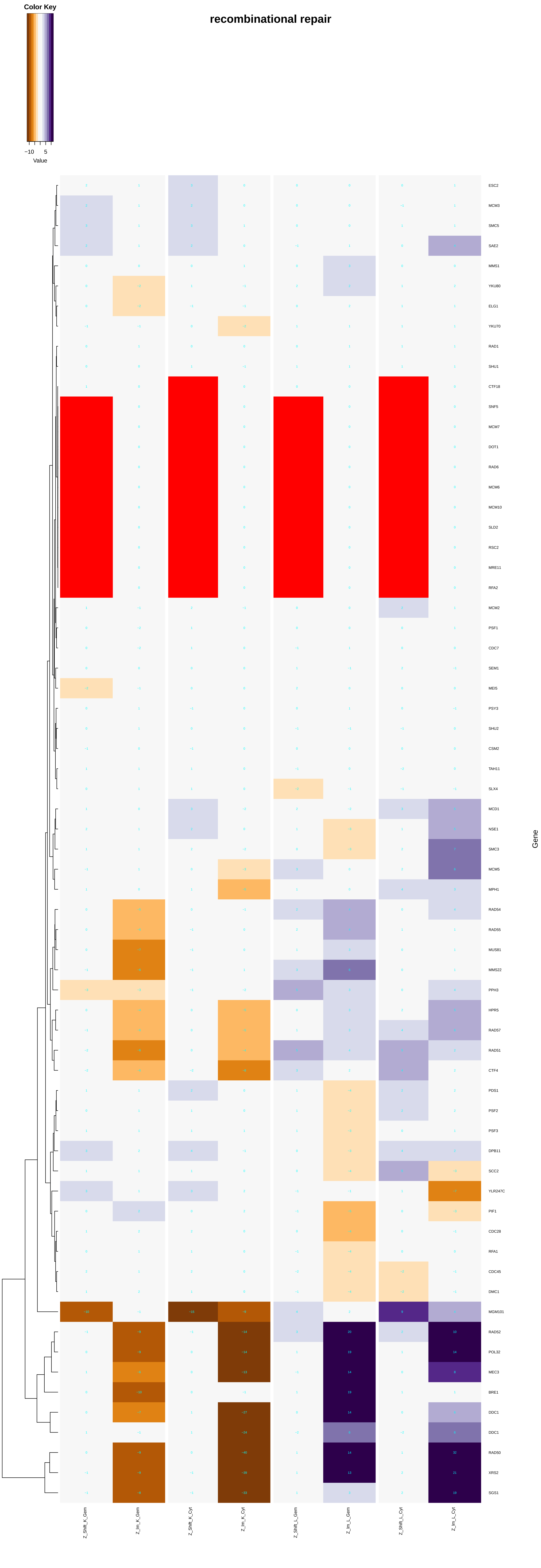

Color Key
−10
5
Value
recombinational repair
Gene

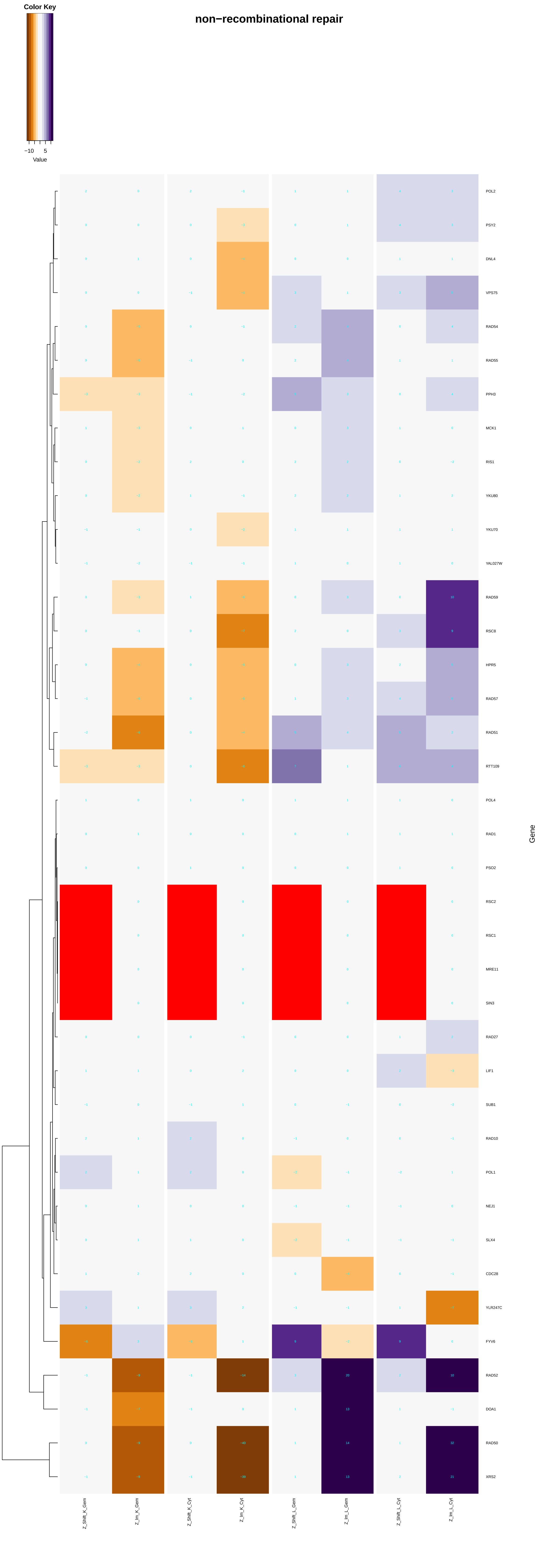

non−recombinational repair
Color Key
−10
5
Value
POL2
PSY2
DNL4
VPS75
RAD54
RAD55
PPH3
MCK1
RIS1
YKU80
YKU70
YAL027W
RAD59
RSC8
HPR5
RAD57
RAD51
RTT109
POL4
RAD1
PSO2
RSC2
RSC1
MRE11
SIN3
RAD27
LIF1
SUB1
RAD10
POL1
NEJ1
SLX4
CDC28
YLR247C
FYV6
RAD52
DOA1
RAD50
XRS2
Gene
Z_Shift_K_Gem
Z_Im_K_Gem
Z_Shift_K_Cyt
Z_Im_K_Cyt
Z_Shift_L_Gem
Z_Im_L_Gem
Z_Shift_L_Cyt
Z_Im_L_Cyt

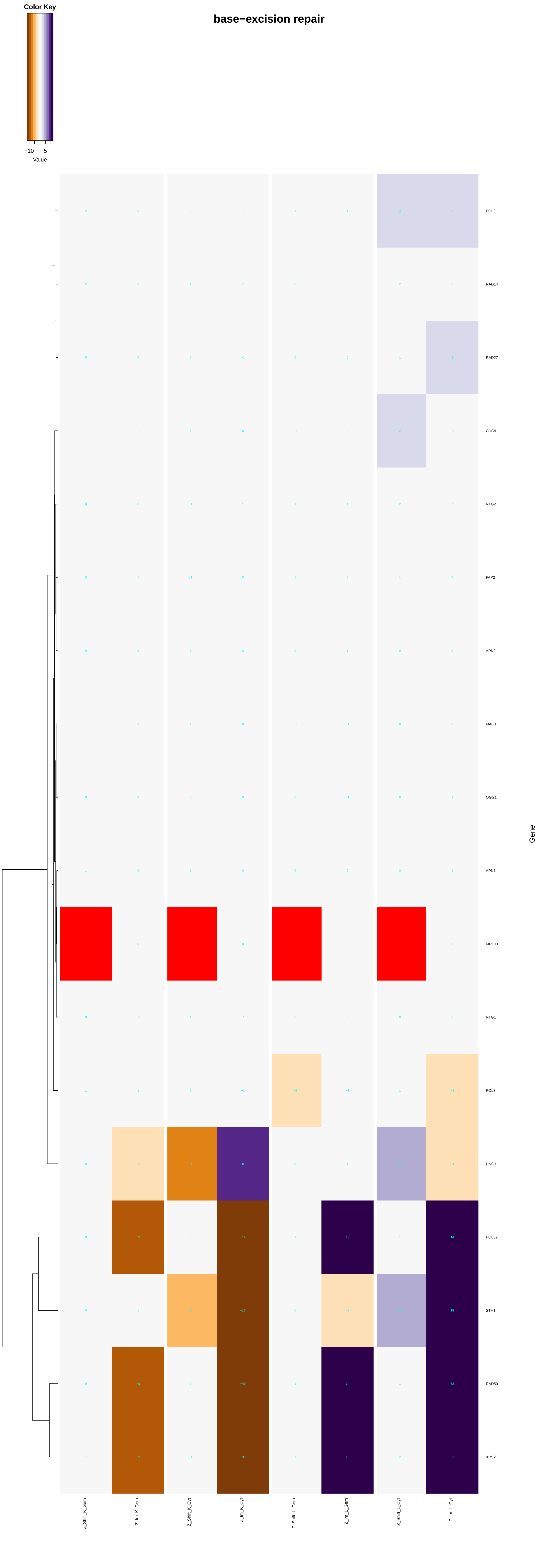
base−excision repair
Color Key
−10
5
Value
POL2
RAD14
RAD27
CDC9
NTG2
PAP2
APN2
MAG1
OGG1
APN1
MRE11
NTG1
POL3
UNG1
POL32
STH1
RAD50
XRS2
Gene
Z_Shift_K_Gem
Z_Im_K_Gem
Z_Shift_K_Cyt
Z_Im_K_Cyt
Z_Shift_L_Gem
Z_Im_L_Gem
Z_Shift_L_Cyt
Z_Im_L_Cyt

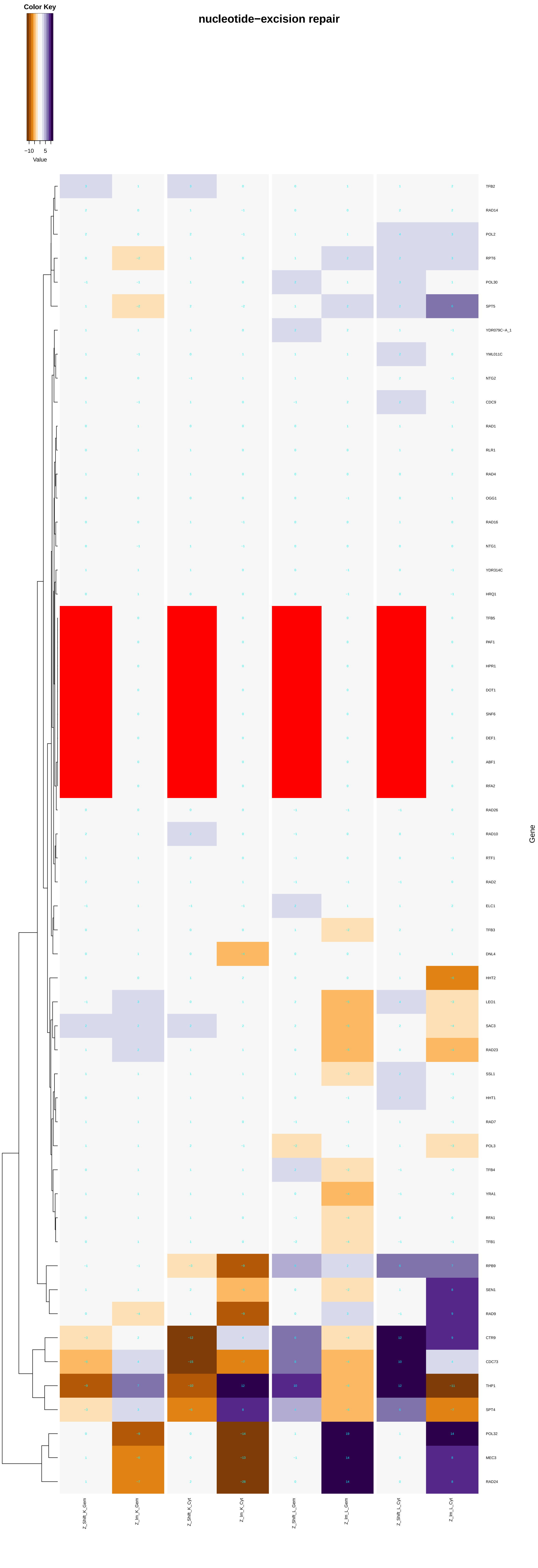
Color Key
−10
5
Value
nucleotide−excision repair
TFB2
RAD14
POL2
RPT6
POL30
SPT5
YDR079C−A_1
YML011C
NTG2
CDC9
RAD1
RLR1
RAD4
OGG1
RAD16
NTG1
YDR314C
HRQ1
TFB5
PAF1
HPR1
DOT1
SNF6
DEF1
ABF1
RFA2
RAD26
RAD10
RTF1
RAD2
ELC1
TFB3
DNL4
HHT2
LEO1
SAC3
RAD23
SSL1
HHT1
RAD7
POL3
TFB4
YRA1
RFA1
TFB1
RPB9
SEN1
RAD9
CTR9
CDC73
THP1
SPT4
POL32
MEC3
RAD24
Gene
Z_Shift_K_Gem
Z_Im_K_Gem
Z_Shift_K_Cyt
Z_Im_K_Cyt
Z_Shift_L_Gem
Z_Im_L_Gem
Z_Shift_L_Cyt
Z_Im_L_Cyt

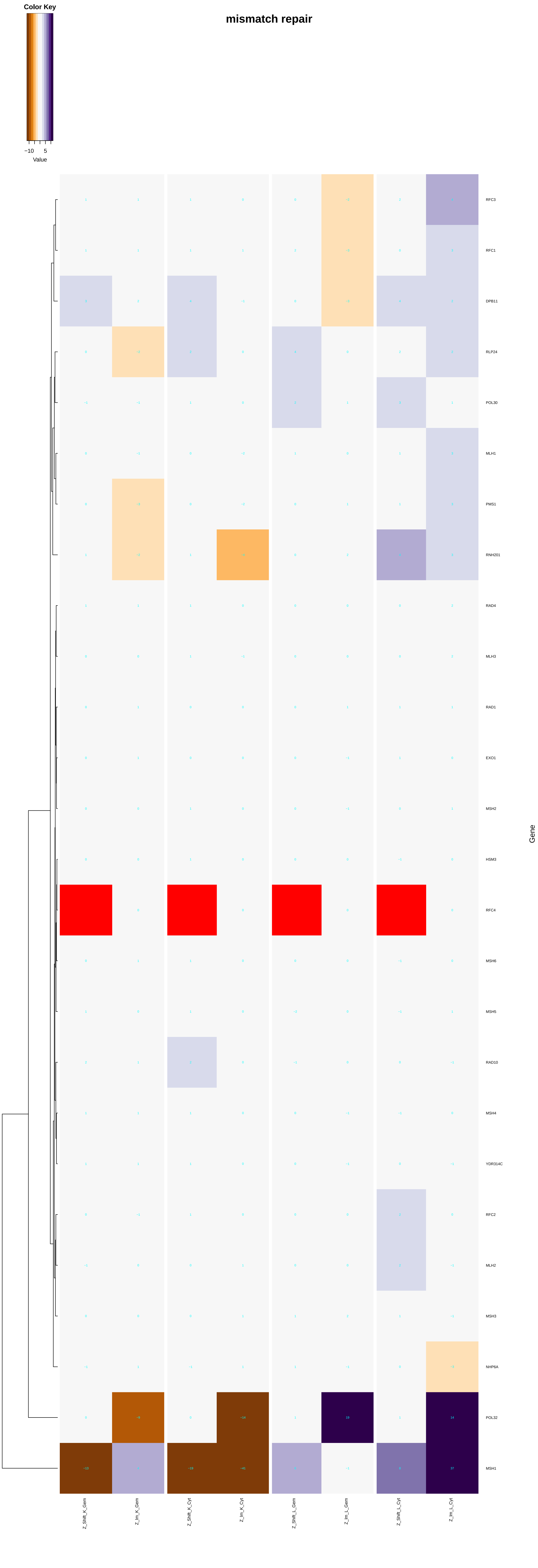
mismatch repair
Color Key
−10
5
Value
RFC3
RFC1
DPB11
RLP24
POL30
MLH1
PMS1
RNH201
RAD4
MLH3
RAD1
EXO1
MSH2
HSM3
RFC4
MSH6
MSH5
RAD10
MSH4
YDR314C
RFC2
MLH2
MSH3
NHP6A
POL32
MSH1
Gene
Z_Shift_K_Gem
Z_lm_K_Gem
Z_Shift_K_Cyt
Z_lm_K_Cyt
Z_Shift_L_Gem
Z_lm_L_Gem
Z_Shift_L_Cyt
Z_lm_L_Cyt

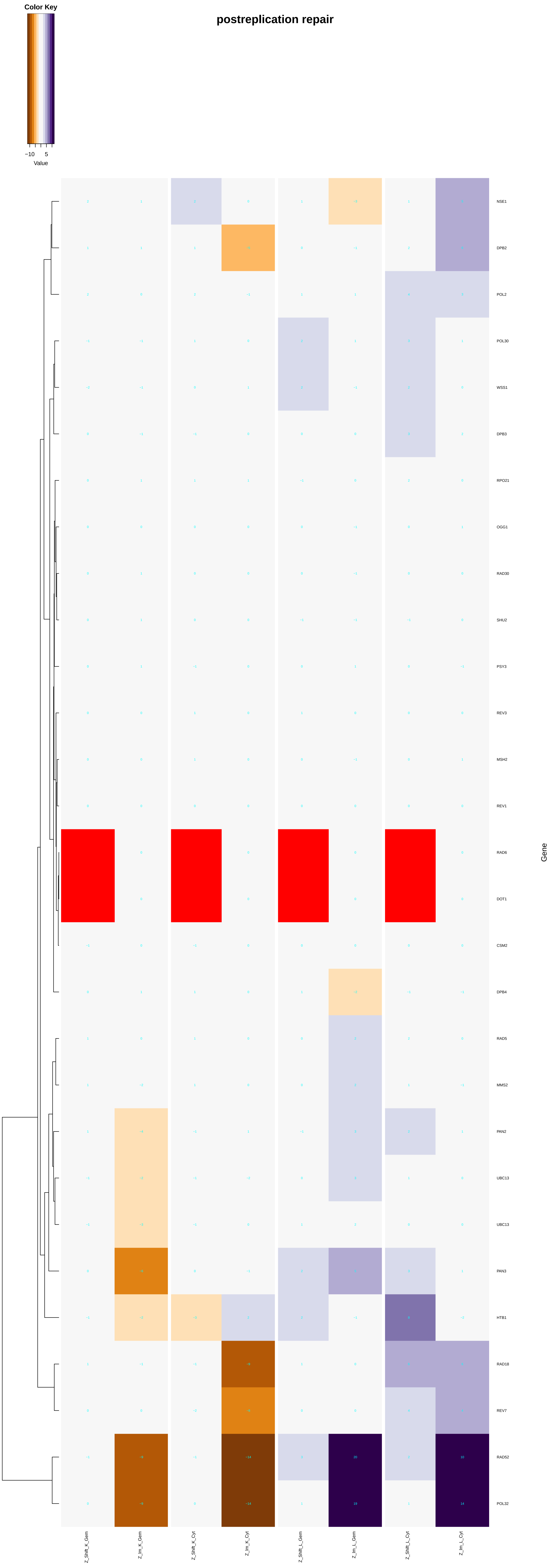
postreplication repair
Color Key
−10
5
Value
NSE1
DPB2
POL2
POL30
WSS1
DPB3
RPO21
OGG1
RAD30
SHU2
PSY3
REV3
MSH2
REV1
RAD6
DOT1
CSM2
DPB4
RAD5
MMS2
PAN2
UBC13
UBC13
PAN3
HTB1
RAD18
REV7
RAD52
POL32
Gene
Z_Shift_K_Gem
Z_Im_K_Gem
Z_Shift_K_Cyt
Z_Im_K_Cyt
Z_Shift_L_Gem
Z_Im_L_Gem
Z_Shift_L_Cyt
Z_Im_L_Cyt

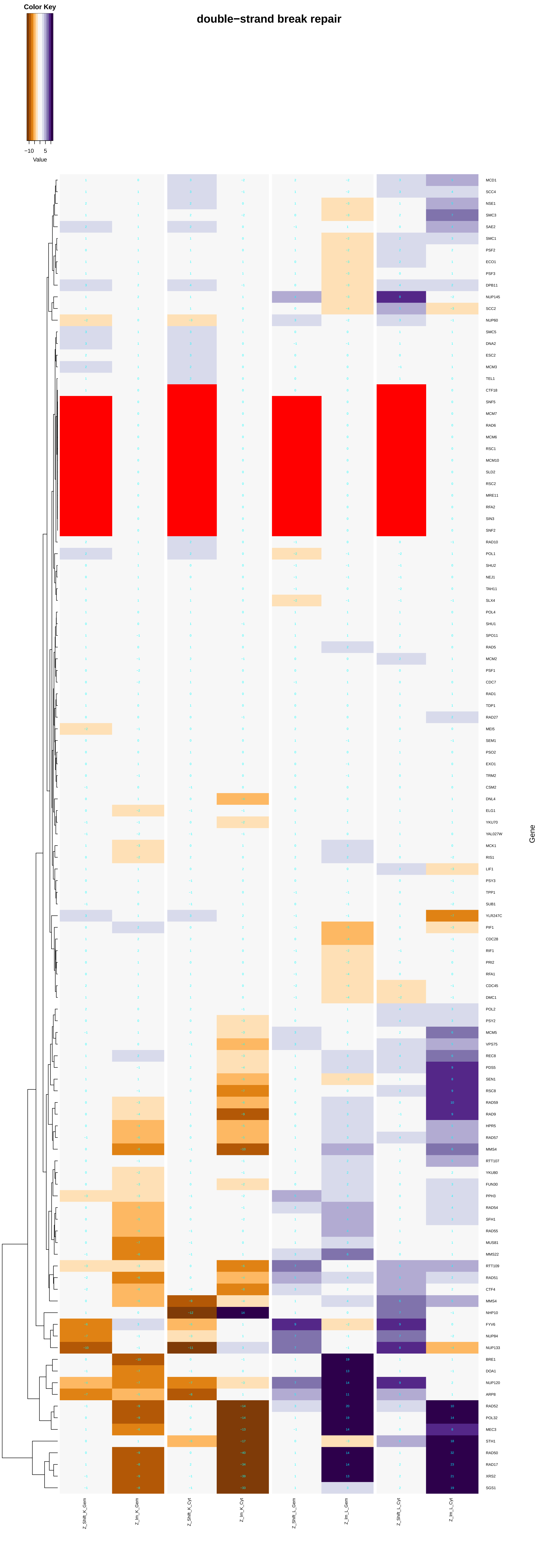

double−strand break repair
Color Key
−10
5
Value
MCD1
SCC4
NSE1
SMC3
SAE2
SMC1
PSF2
ECO1
PSF3
DPB11
NUP145
SCC2
NUP60
SMC5
DNA2
ESC2
MCM3
TEL1
CTF18
SNF5
MCM7
RAD6
MCM6
RSC1
MCM10
SLD2
RSC2
MRE11
RFA2
SIN3
SNF2
RAD10
POL1
SHU2
NEJ1
TAH11
SLX4
POL4
SHU1
SPO11
RAD5
MCM2
PSF1
CDC7
RAD1
TDP1
RAD27
MEI5
SEM1
PSO2
EXO1
TRM2
CSM2
DNL4
ELG1
YKU70
YAL027W
MCK1
RIS1
LIF1
PSY3
TPP1
SUB1
YLR247C
PIF1
CDC28
RIF1
PRI2
RFA1
CDC45
DMC1
POL2
PSY2
MCM5
VPS75
REC8
PDS5
SEN1
RSC8
RAD59
RAD9
HPR5
RAD57
MMS4
RTT107
YKU80
FUN30
PPH3
RAD54
SFH1
RAD55
MUS81
MMS22
RTT109
RAD51
CTF4
MMS4
NHP10
FYV6
NUP84
NUP133
BRE1
DOA1
NUP120
ARP8
RAD52
POL32
MEC3
STH1
RAD50
RAD17
XRS2
SGS1
Gene
Z_Shift_K_Gem
Z_Im_K_Gem
Z_Shift_K_Cyt
Z_Im_K_Cyt
Z_Shift_L_Gem
Z_Im_L_Gem
Z_Shift_L_Cyt
Z_Im_L_Cyt

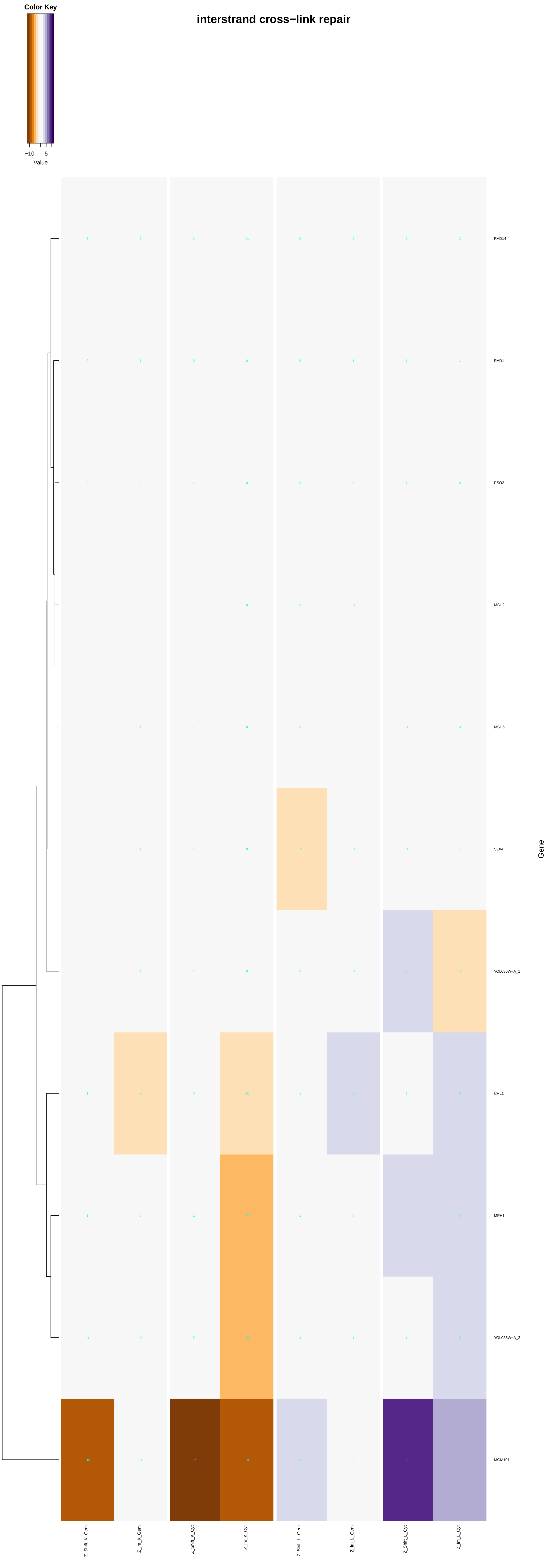

interstrand cross−link repair
Color Key
−10
5
Value
RAD14
RAD1
PSO2
MSH2
MSH6
SLX4
YOL086W−A_1
CHL1
MPH1
YOL086W−A_2
MGM101
Gene
Z_Shift_K_Gem
Z_Im_K_Gem
Z_Shift_K_Cyt
Z_Im_K_Cyt
Z_Shift_L_Gem
Z_Im_L_Gem
Z_Shift_L_Cyt
Z_Im_L_Cyt

#### UV−damage excision repair

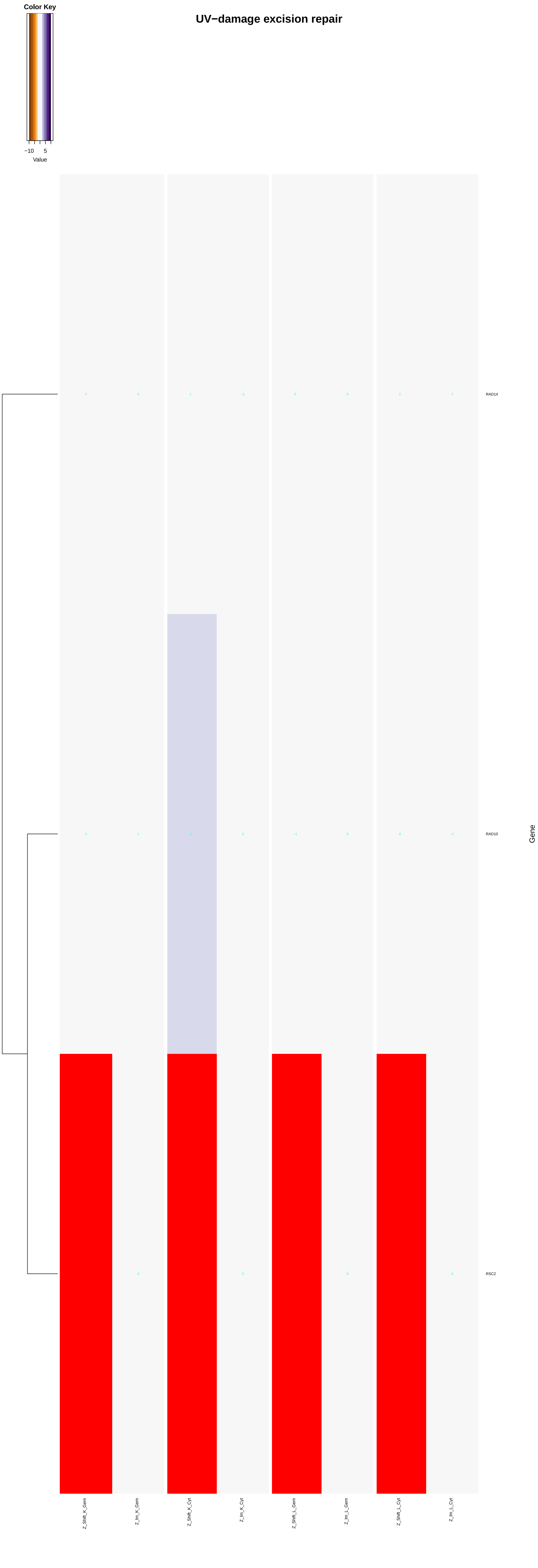

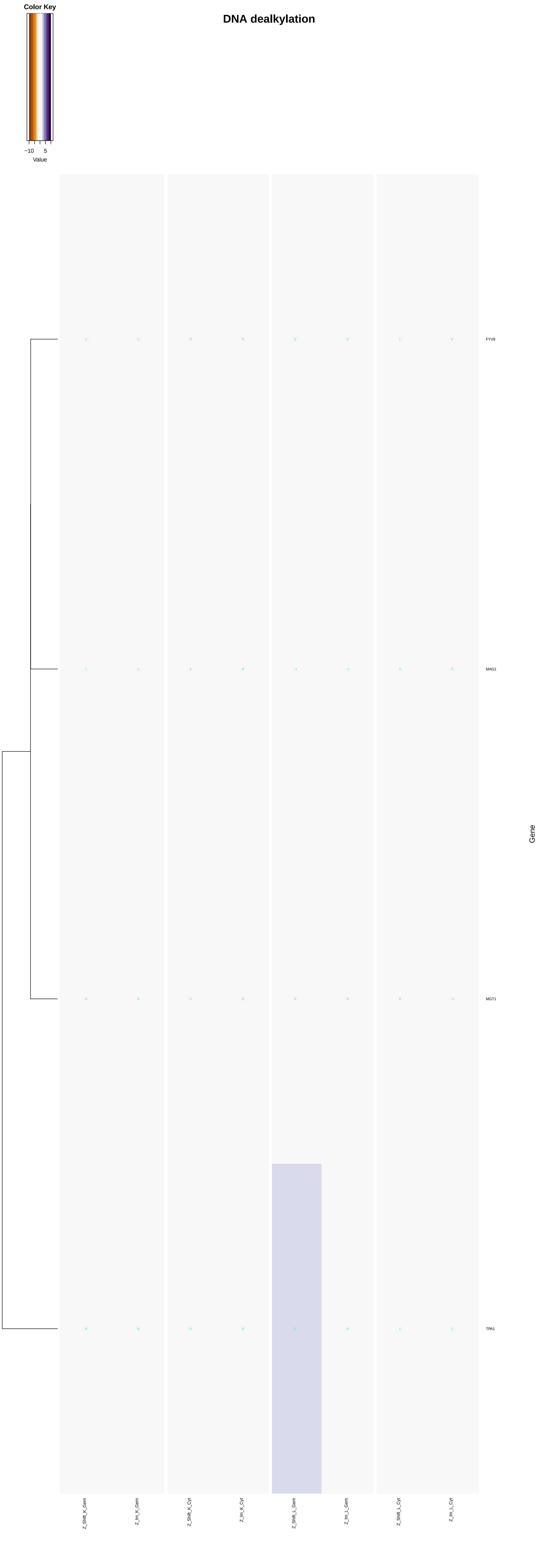
DNA dealkylation
Color Key
−10
5
Value
FYV8
MAG1
MGT1
TPA1
Gene
Z_Shift_K_Gem
Z_Im_K_Gem
Z_Shift_K_Cyt
Z_Im_K_Cyt
Z_Shift_L_Gem
Z_Im_L_Gem
Z_Shift_L_Cyt
Z_Im_L_Cyt

### DNA catabolic process, endonucleolytic

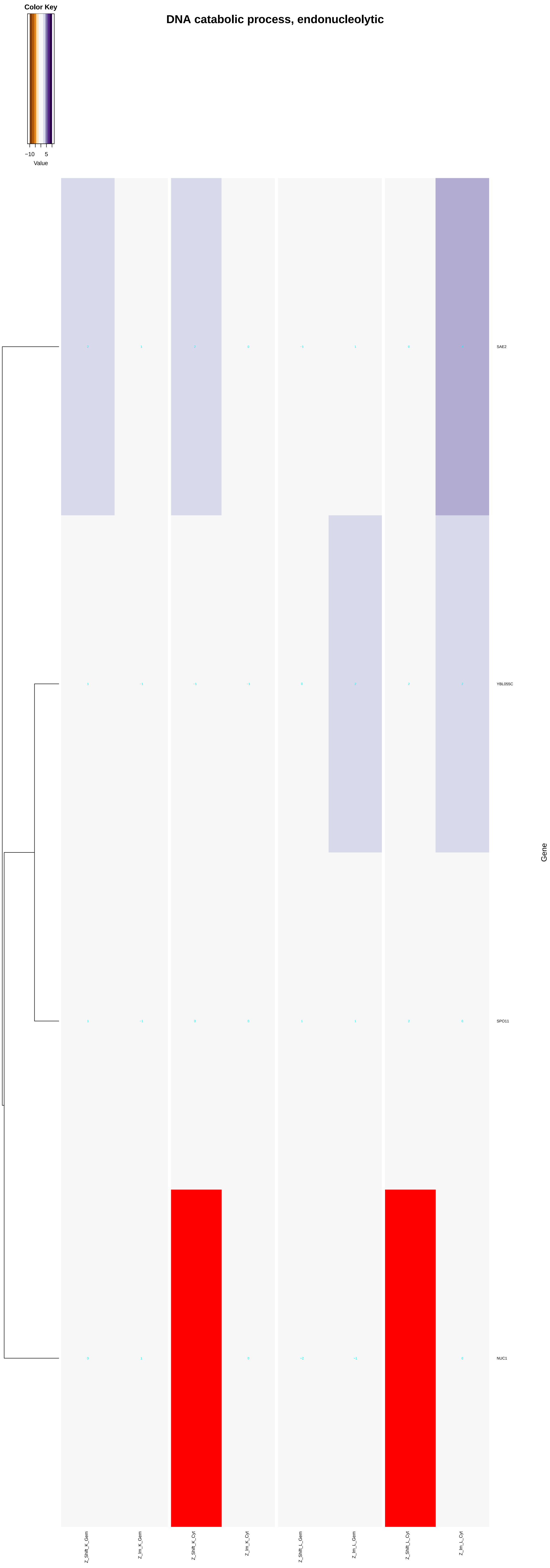

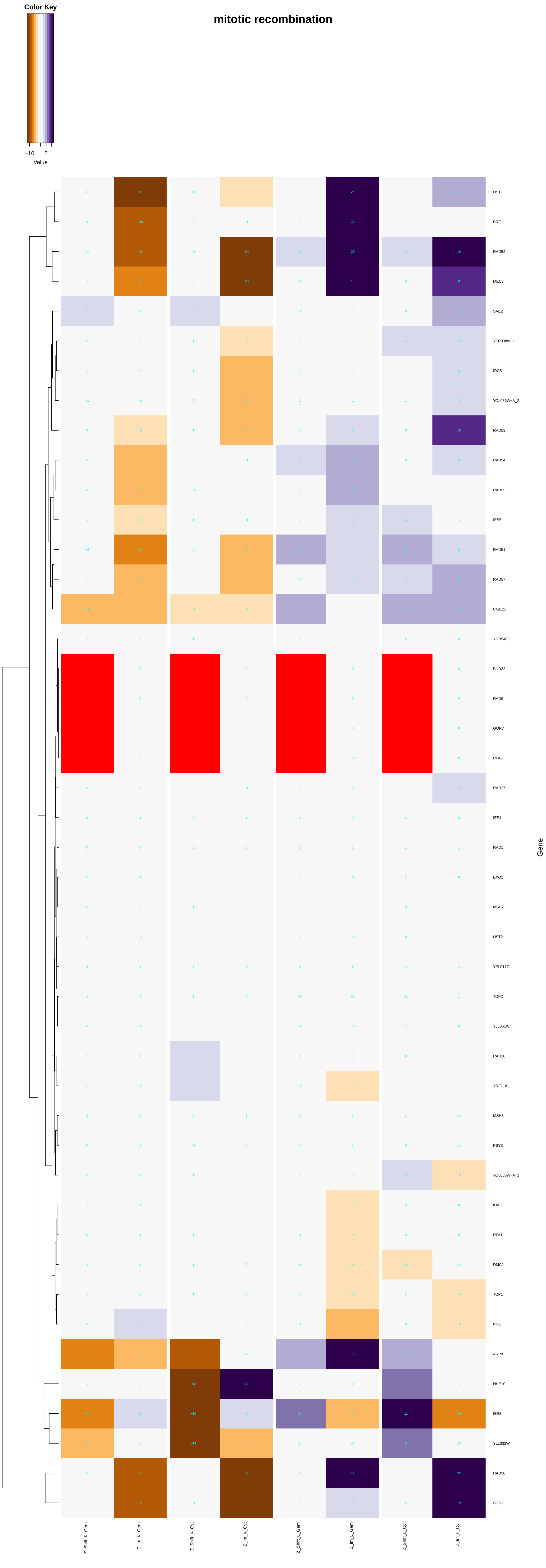
mitotic recombination
Color Key
−10
5
Value
HST1
BRE1
RAD52
MEC3
SAE2
YFR038W_1
IRC5
YOL086W−A_2
RAD59
RAD54
RAD55
IES5
RAD51
RAD57
CGI121
YDR540C
BUD32
RAD6
GON7
RFA2
RAD27
IES4
RAD1
EXO1
MSH2
HST2
YPL017C
TOP2
YJL051W
RAD10
YRF1−6
MSH3
PSY3
YOL086W−A_1
KAE1
RFA1
DMC1
TOP1
PIF1
ARP8
NHP10
IES3
YLL033W
RAD50
SGS1
Gene
Z_Shift_K_Gem
Z_Im_K_Gem
Z_Shift_K_Cyt
Z_Im_K_Cyt
Z_Shift_L_Gem
Z_Im_L_Gem
Z_Shift_L_Cyt
Z_Im_L_Cyt

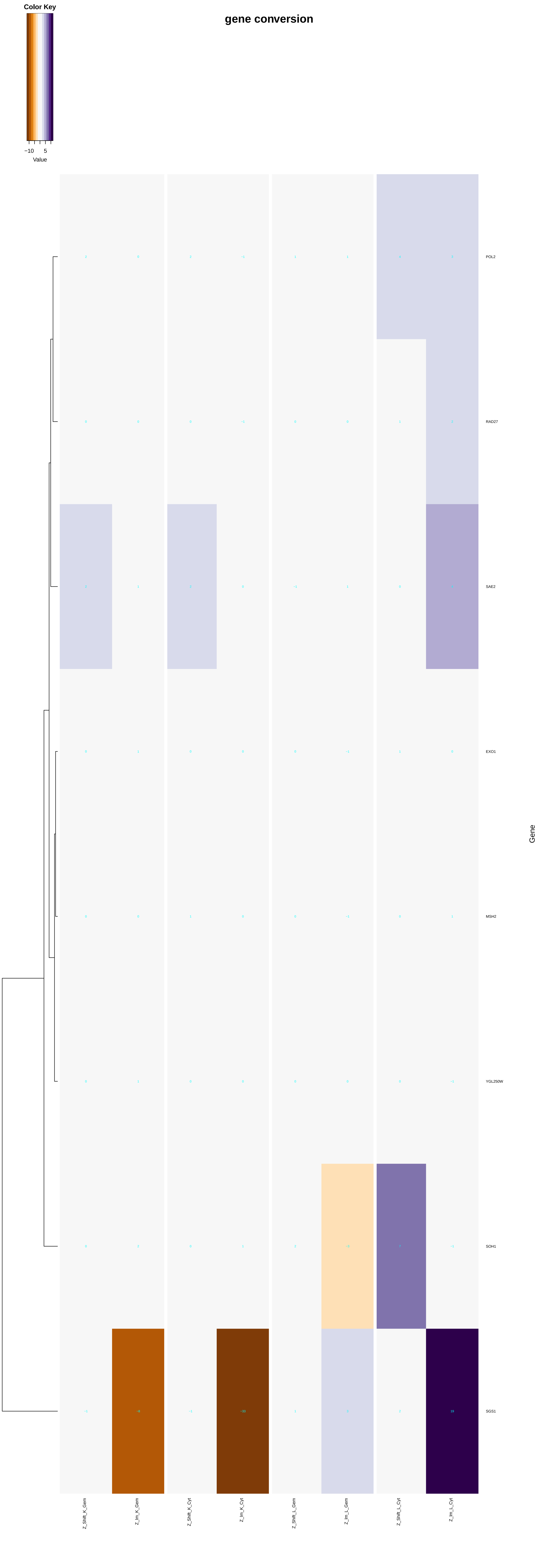
Color Key
−10
5
Value
gene conversion
POL2
RAD27
SAE2
EXO1
MSH2
YGL250W
SOH1
SGS1
Gene
Z_Shift_K_Gem
Z_Im_K_Gem
Z_Shift_K_Cyt
Z_Im_K_Cyt
Z_Shift_L_Gem
Z_Im_L_Gem
Z_Shift_L_Cyt
Z_Im_L_Cyt

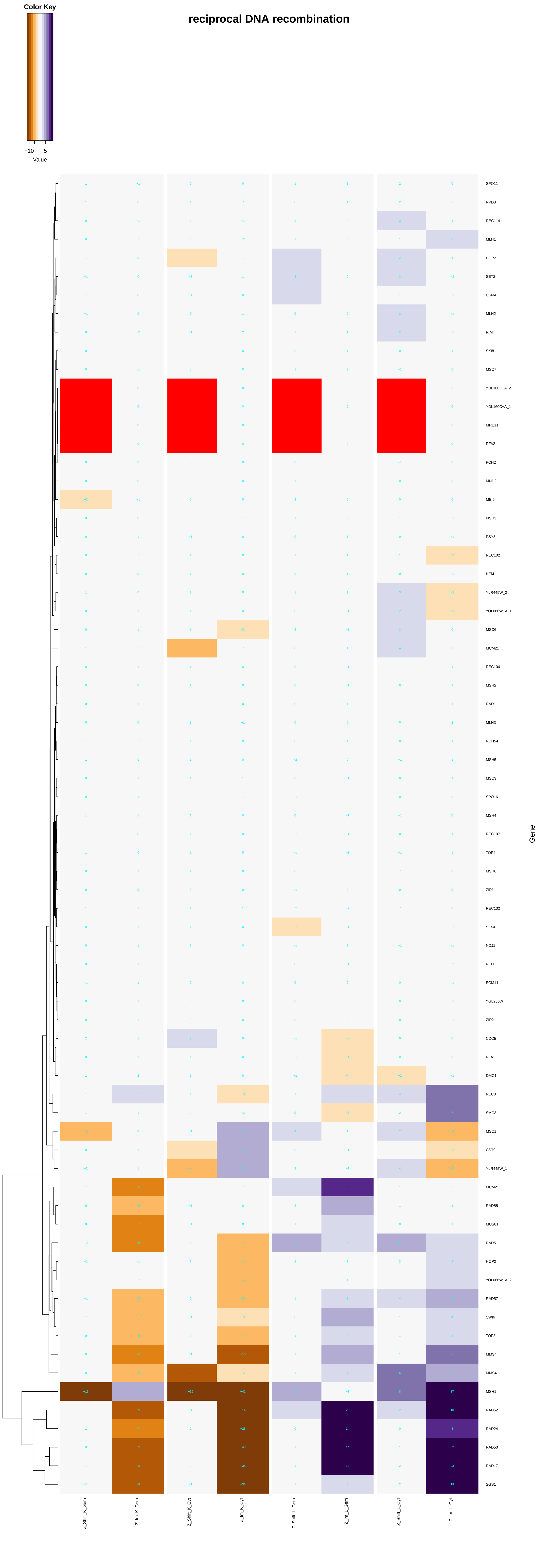
reciprocal DNA recombination
Color Key
−10
5
Value
SPO11
RPD3
REC114
MLH1
HOP2
SET2
CSM4
MLH2
RIM4
SKI8
MSC7
YDL160C−A_2
YDL160C−A_1
MRE11
RFA2
PCH2
MND2
MEI5
MSH3
PSY3
REC102
HFM1
YLR445W_2
YOL086W−A_1
MSC6
MCM21
REC104
MSH2
RAD1
MLH3
RDH54
MSH5
MSC3
SPO16
MSH4
REC107
TOP2
MSH6
ZIP1
REC102
SLX4
NDJ1
RED1
ECM11
YGL250W
ZIP2
CDC5
RFA1
DMC1
REC8
SMC3
MSC1
CST9
YLR445W_1
MCM21
RAD55
MUS81
RAD51
HOP2
YOL086W−A_2
RAD57
SWI6
TOP3
MMS4
MMS4
MSH1
RAD52
RAD24
RAD50
RAD17
SGS1
Gene
Z_Shift_K_Gem
Z_Im_K_Gem
Z_Shift_K_Cyt
Z_Im_K_Cyt
Z_Shift_L_Gem
Z_Im_L_Gem
Z_Shift_L_Cyt
Z_Im_L_Cyt

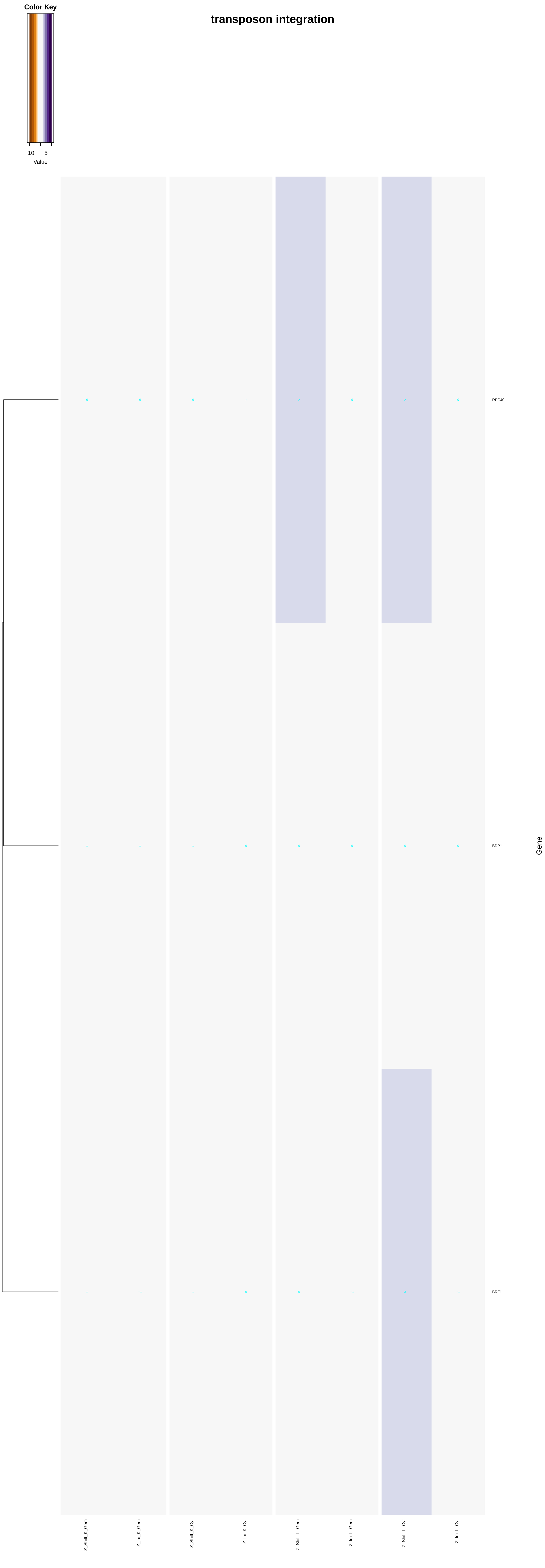
transposon integration
Color Key
−10
5
Value
RPC40
BDP1
BRF1
Gene
Z_Shift_K_Gem
Z_Im_K_Gem
Z_Shift_K_Cyt
Z_Im_K_Cyt
Z_Shift_L_Gem
Z_Im_L_Gem
Z_Shift_L_Cyt
Z_Im_L_Cyt

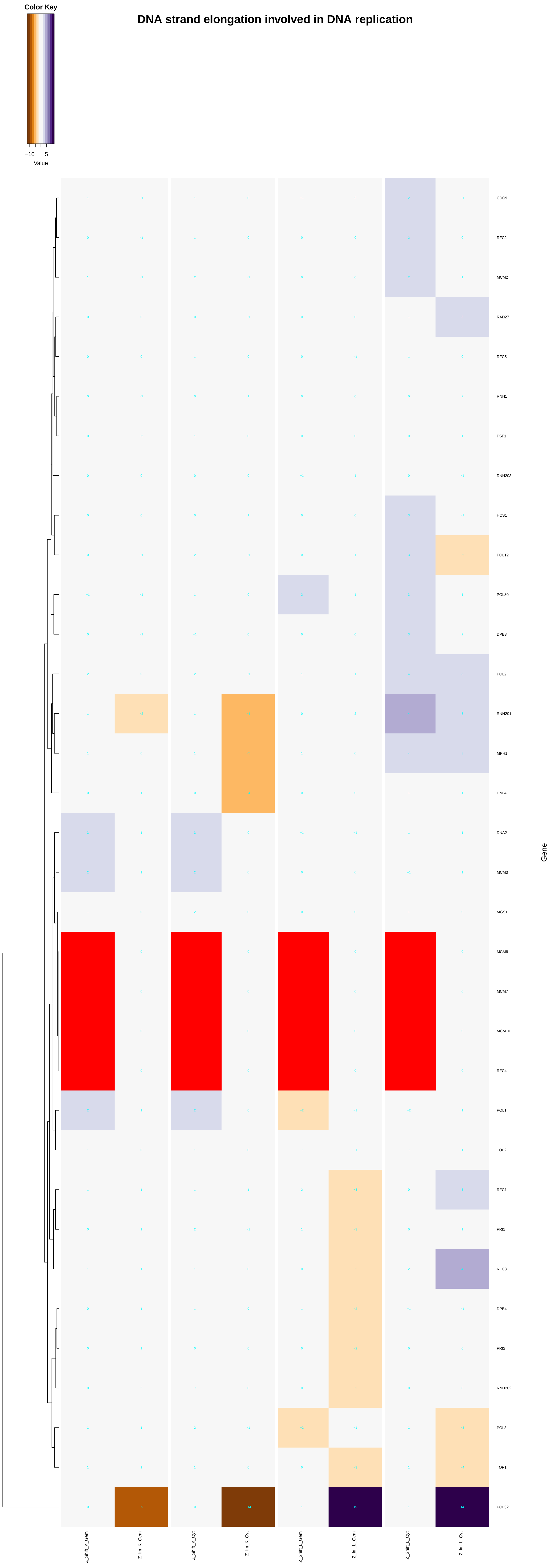
Color Key
−10
5
Value
DNA strand elongation involved in DNA replication
CDC9
RFC2
MCM2
RAD27
RFC5
RNH1
PSF1
RNH203
HCS1
POL12
POL30
DPB3
POL2
RNH201
MPH1
DNL4
DNA2
MCM3
MGS1
MCM6
MCM7
MCM10
RFC4
POL1
TOP2
RFC1
PRI1
RFC3
DPB4
PRI2
RNH202
POL3
TOP1
POL32
Gene
Z_Shift_K_Gem
Z_Im_K_Gem
Z_Shift_K_Cyt
Z_Im_K_Cyt
Z_Shift_L_Gem
Z_Im_L_Gem
Z_Shift_L_Cyt
Z_Im_L_Cyt

### telomeric 3' overhang formation

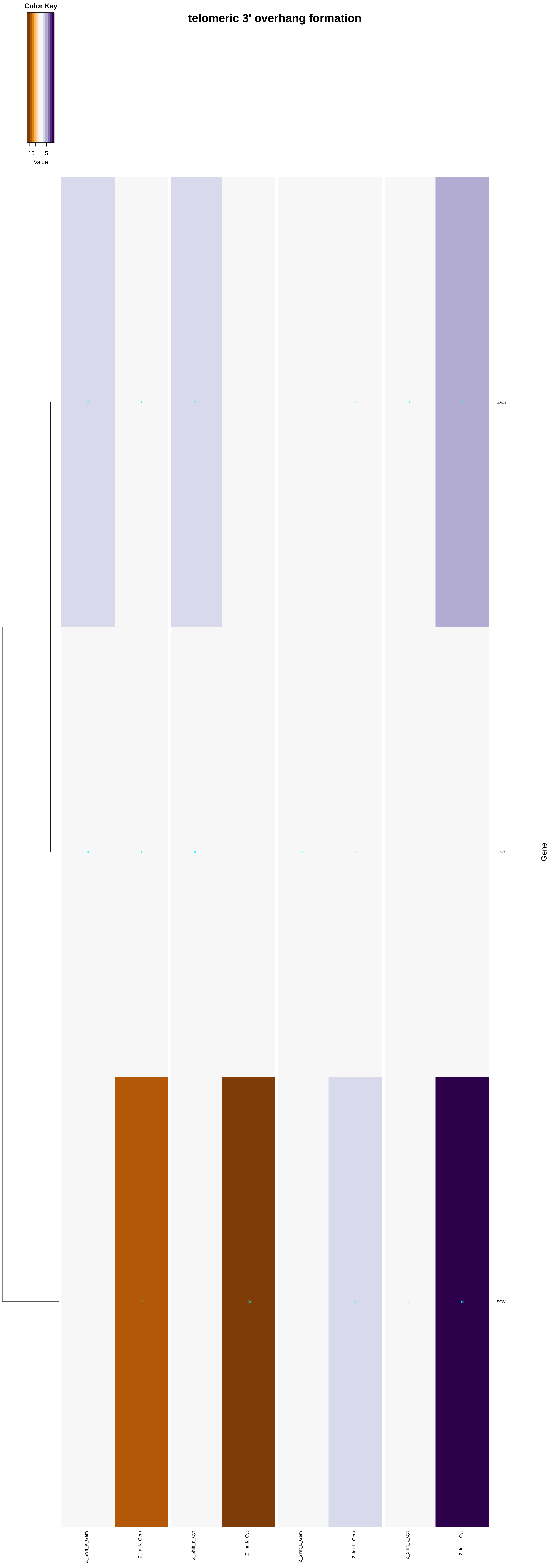

### mitochondrial DNA replication

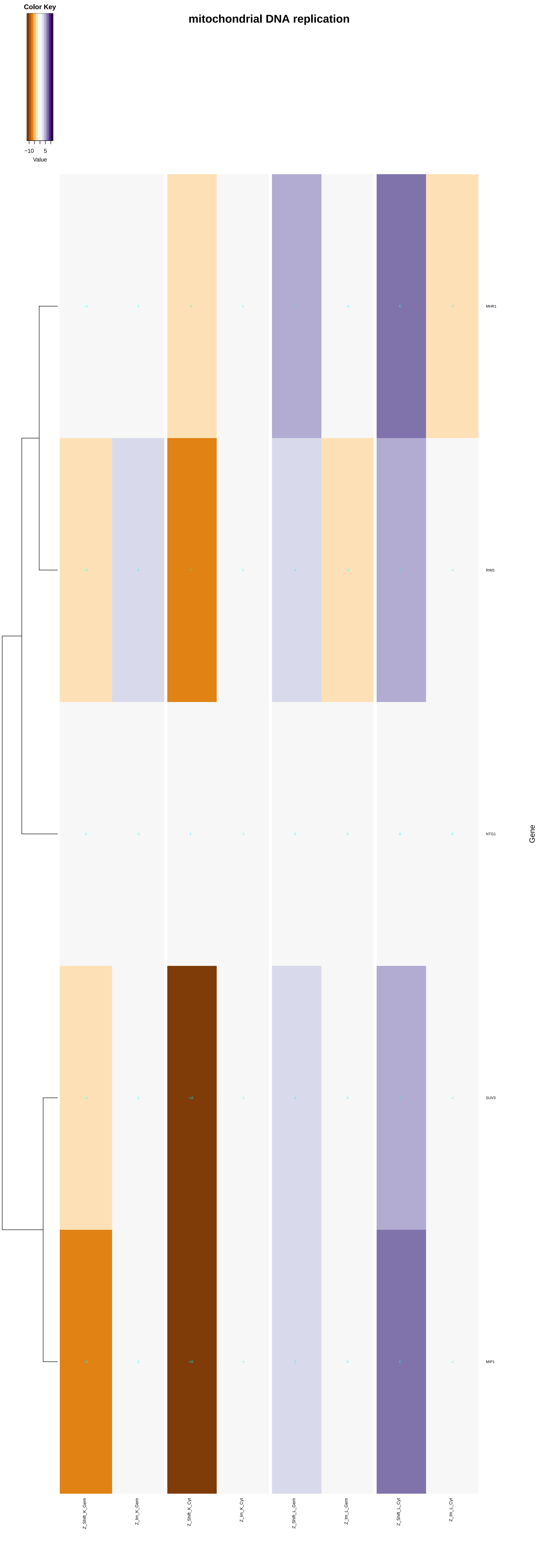

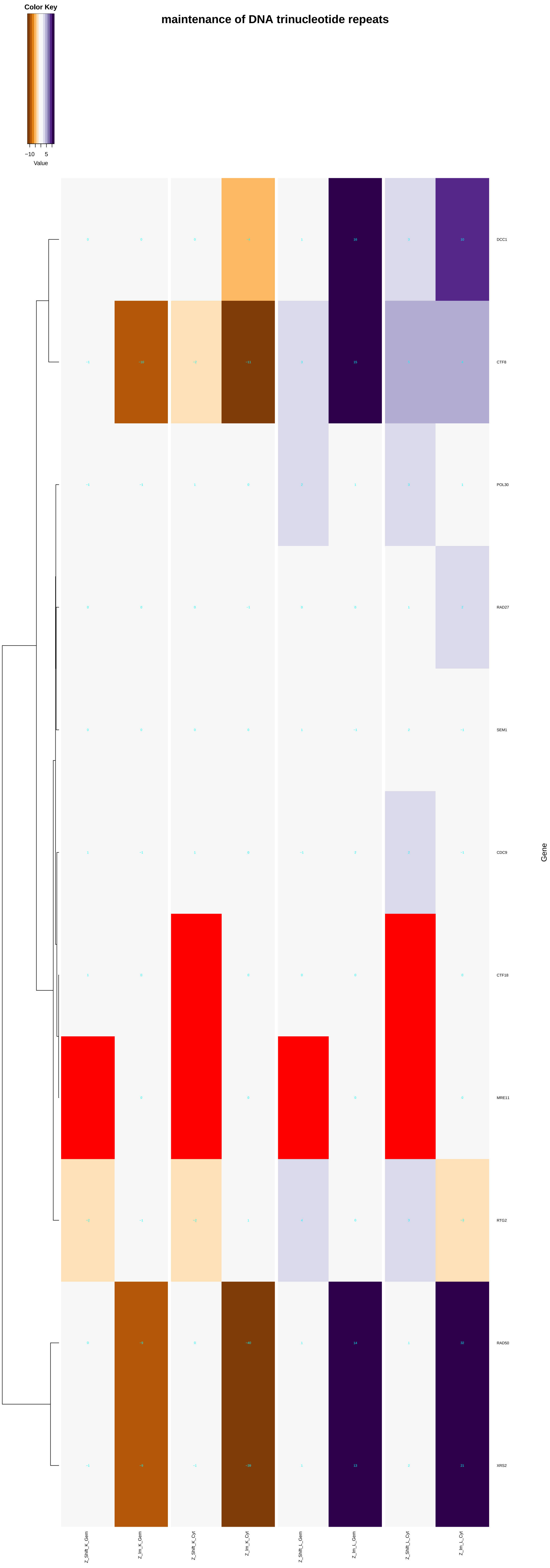

maintenance of DNA trinucleotide repeats
Color Key
−10
5
Value
DCC1
CTF8
POL30
RAD27
SEM1
CDC9
CTF18
MRE11
RTG2
RAD50
XRS2
Gene
Z_Shift_K_Gem
Z_Im_K_Gem
Z_Shift_K_Cyt
Z_Im_K_Cyt
Z_Shift_L_Gem
Z_Im_L_Gem
Z_Shift_L_Cyt
Z_Im_L_Cyt

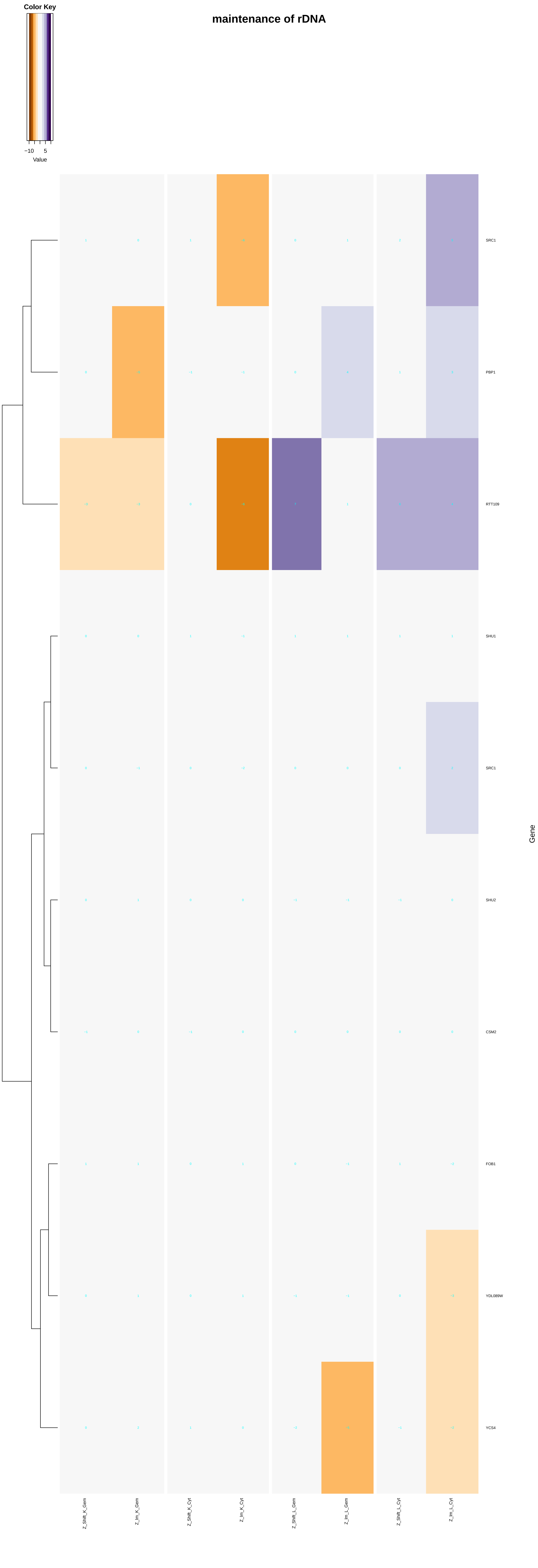

maintenance of rDNA
Color Key
−10
5
Value
SRC1
PBP1
RTT109
SHU1
SRC1
SHU2
CSM2
FOB1
YDL089W
YCS4
Gene
Z_Shift_K_Gem
Z_Im_K_Gem
Z_Shift_K_Cyt
Z_Im_K_Cyt
Z_Shift_L_Gem
Z_Im_L_Gem
Z_Shift_L_Cyt
Z_Im_L_Cyt

### replication fork processing

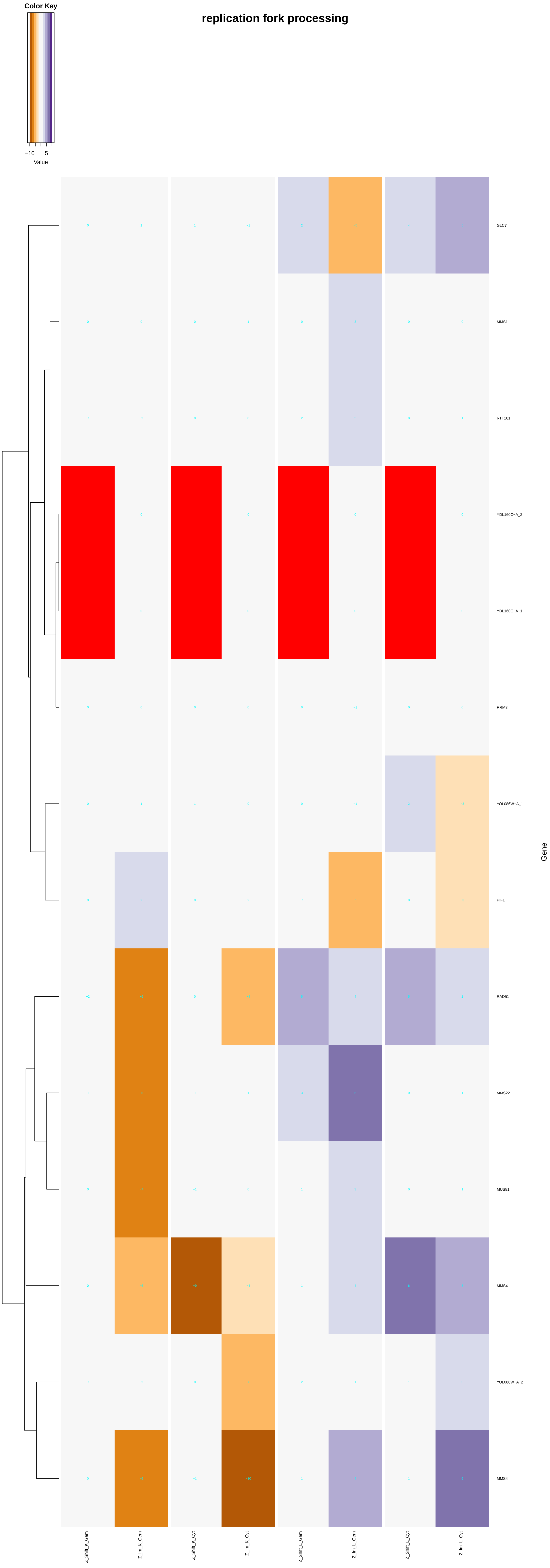

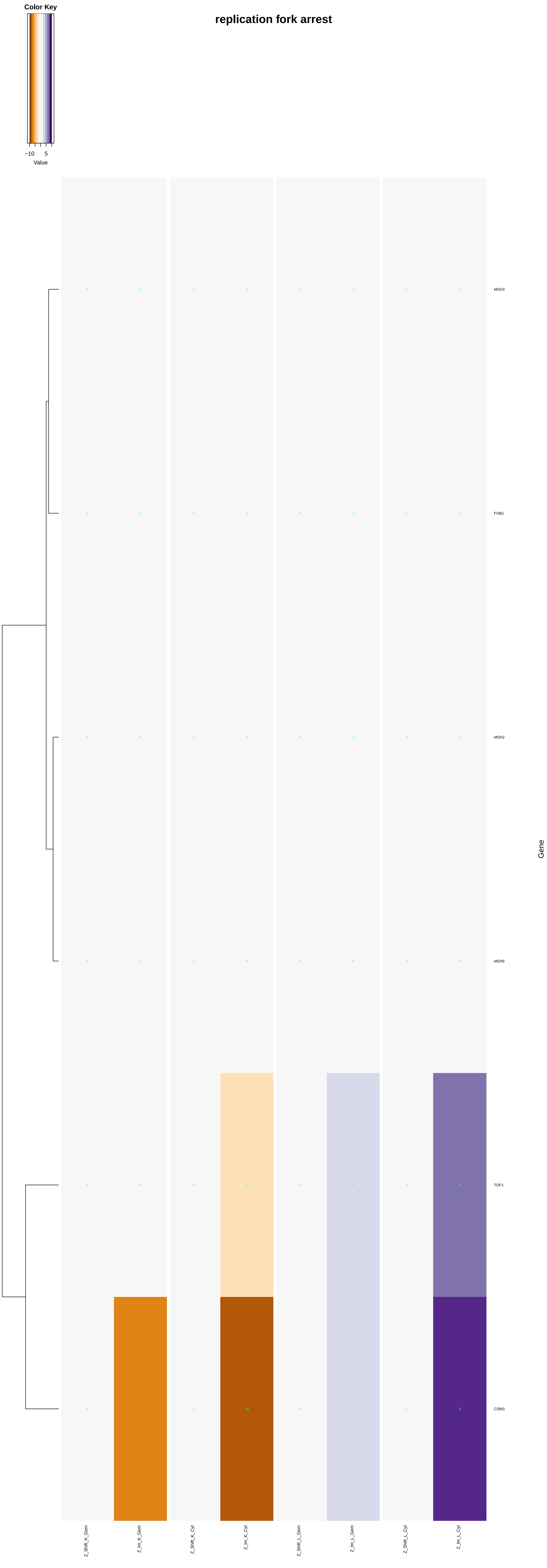
Color Key
−10
5
Value
replication fork arrest
MSH3
FOB1
MSH2
MSH6
TOF1
CSM3
Gene
Z_Shift_K_Gem
Z_Im_K_Gem
Z_Shift_K_Cyt
Z_Im_K_Cyt
Z_Shift_L_Gem
Z_Im_L_Gem
Z_Shift_L_Cyt
Z_Im_L_Cyt

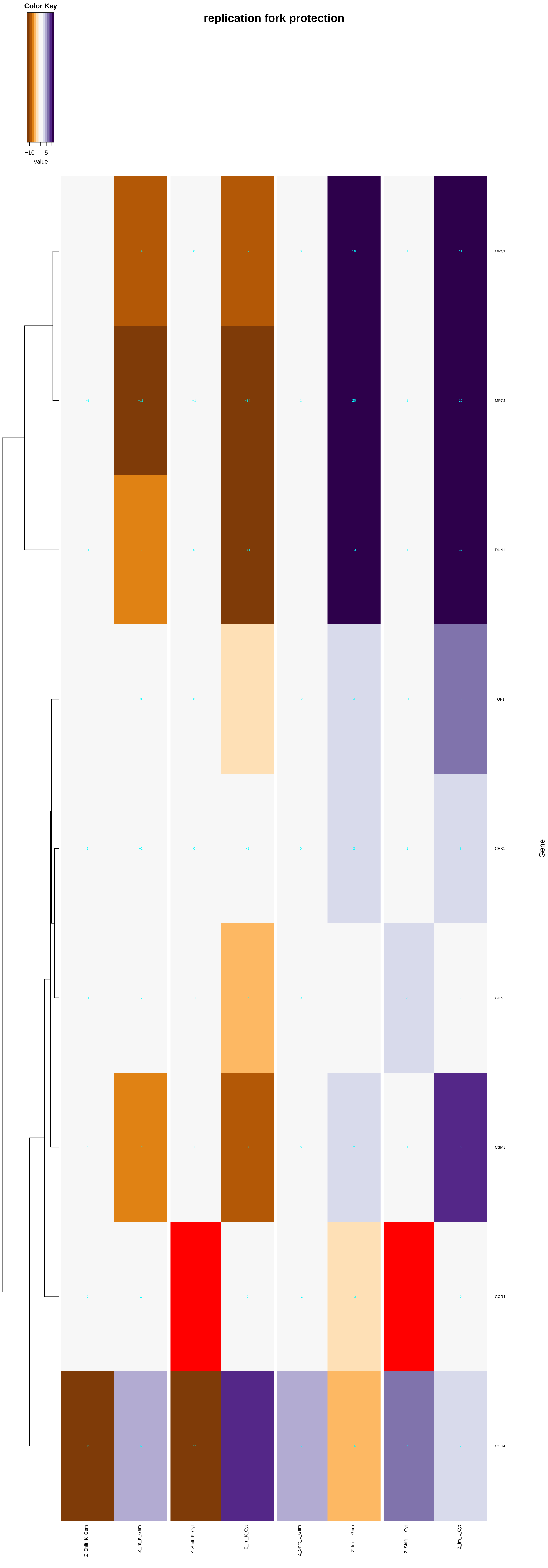

replication fork protection
Color Key
−10
5
Value
MRC1
MRC1
DUN1
TOF1
CHK1
CHK1
CSM3
CCR4
CCR4
Gene
Z_Shift_K_Gem
Z_Im_K_Gem
Z_Shift_K_Cyt
Z_Im_K_Cyt
Z_Shift_L_Gem
Z_Im_L_Gem
Z_Shift_L_Cyt
Z_Im_L_Cyt

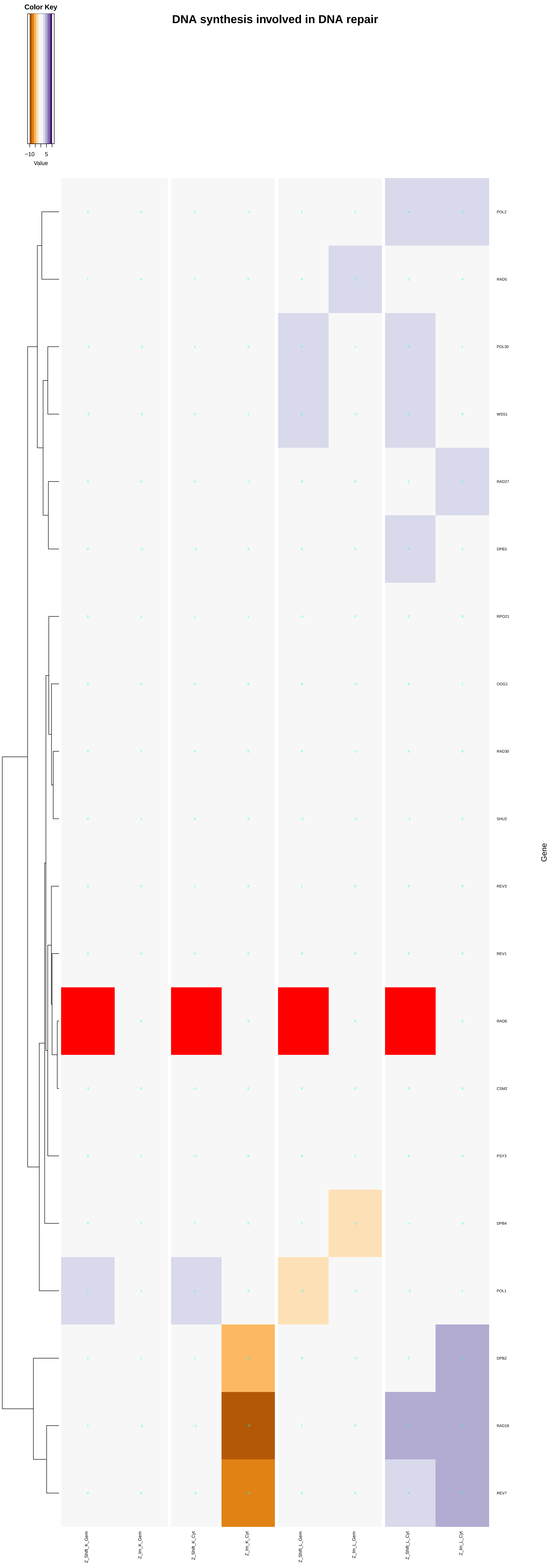

DNA synthesis involved in DNA repair
Color Key
−10 5
Value
POL2
RAD5
POL30
WSS1
RAD27
DPB3
RPO21
OGG1
RAD30
SHU2
REV3
REV1
RAD6
CSM2
PSY3
DPB4
POL1
DPB2
RAD18
REV7
Gene
Z_Shift_K_Gem
Z_Im_K_Gem
Z_Shift_K_Cyt
Z_Im_K_Cyt
Z_Shift_L_Gem
Z_Im_L_Gem
Z_Shift_L_Cyt
Z_Im_L_Cyt

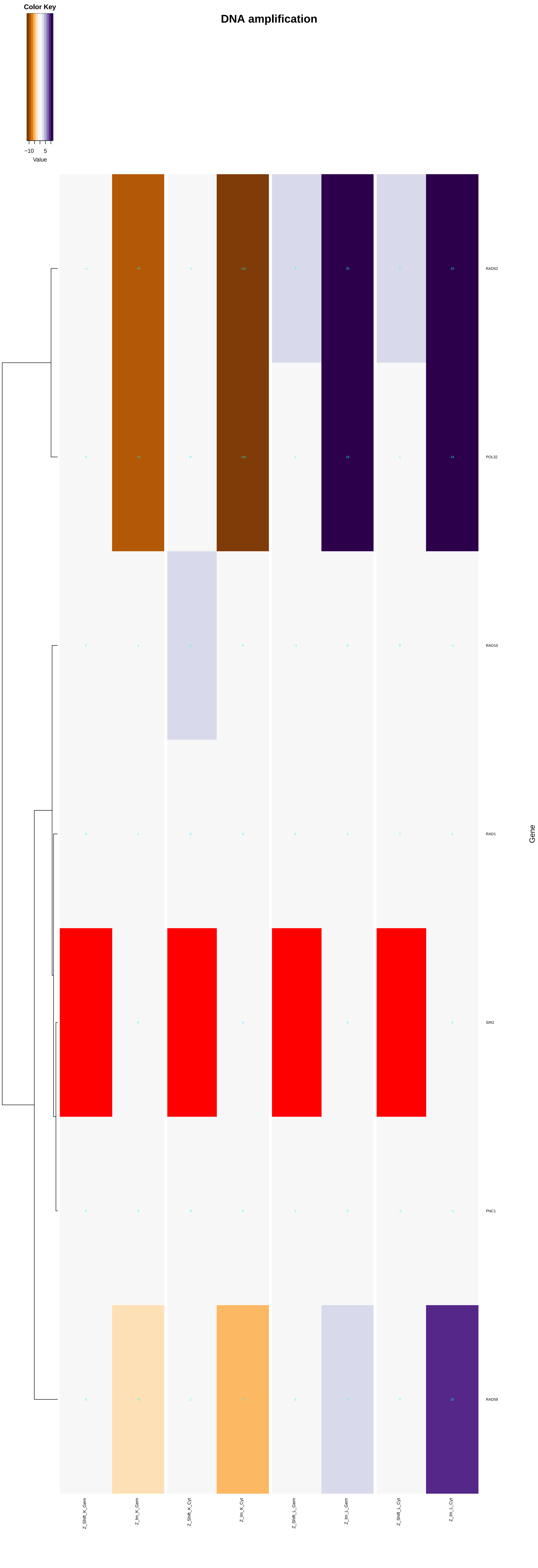
DNA amplification
Color Key
−10
5
Value
RAD52
POL32
RAD10
RAD1
SIR2
PNC1
RAD59
Gene
Z_Shift_K_Gem
Z_Im_K_Gem
Z_Shift_K_Cyt
Z_Im_K_Cyt
Z_Shift_L_Gem
Z_Im_L_Gem
Z_Shift_L_Cyt
Z_Im_L_Cyt

RNA−dependent DNA biosynthetic process

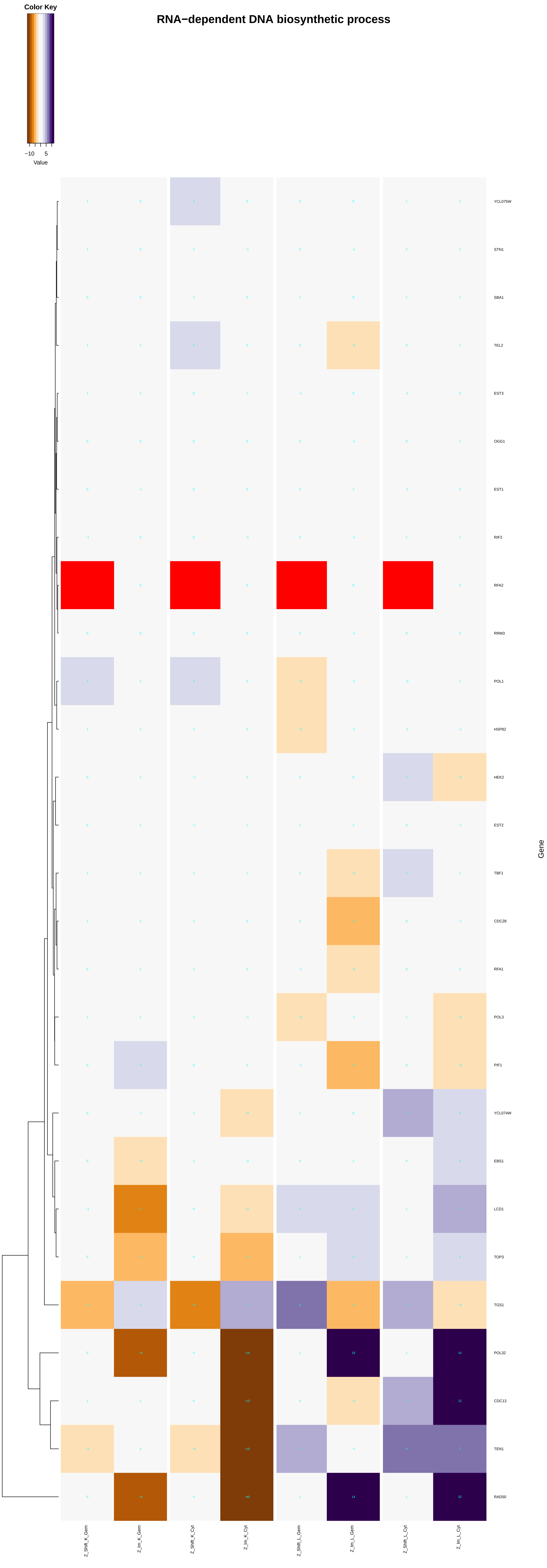

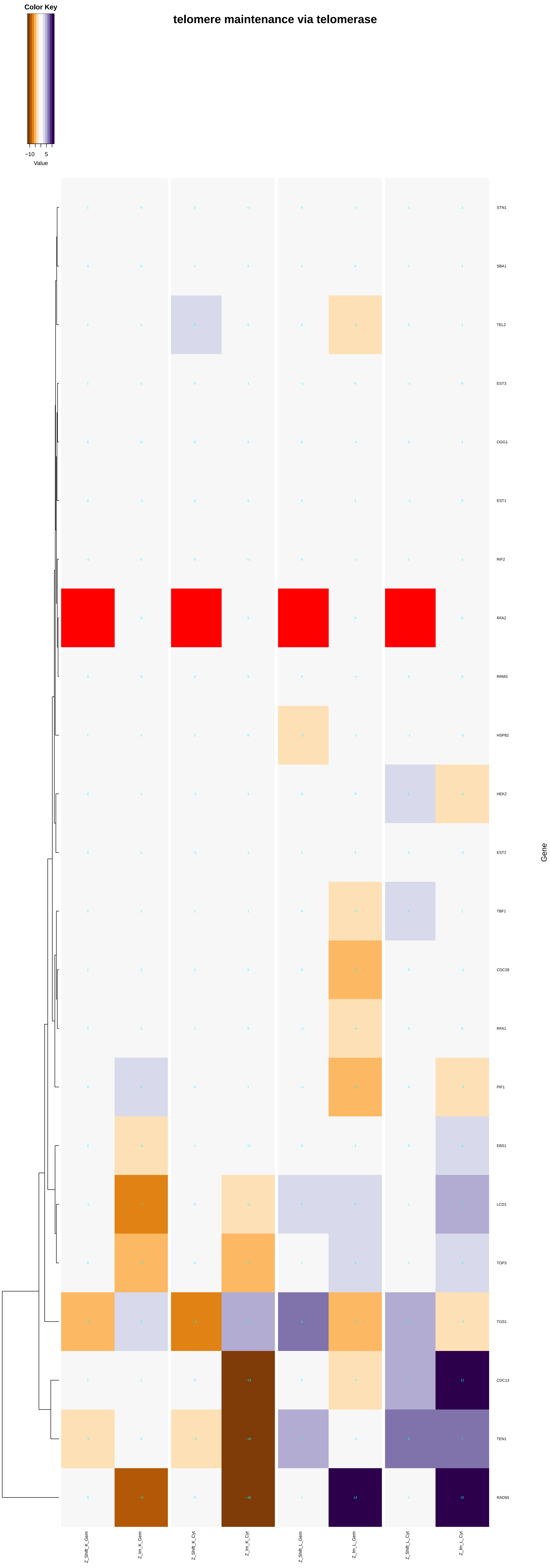

Color Key
−10
5
Value
telomere maintenance via telomerase
STN1
SBA1
TEL2
EST3
OGG1
EST1
RIF2
RFA2
RRM3
HSP82
HEK2
EST2
TBF1
CDC28
RFA1
PIF1
EBS1
LCD1
TOP3
TGS1
CDC13
TEN1
RAD50
Gene
Z_Shift_K_Gem
Z_lm_K_Gem
Z_Shift_K_Cyt
Z_lm_K_Cyt
Z_Shift_L_Gem
Z_lm_L_Gem
Z_Shift_L_Cyt
Z_lm_L_Cyt

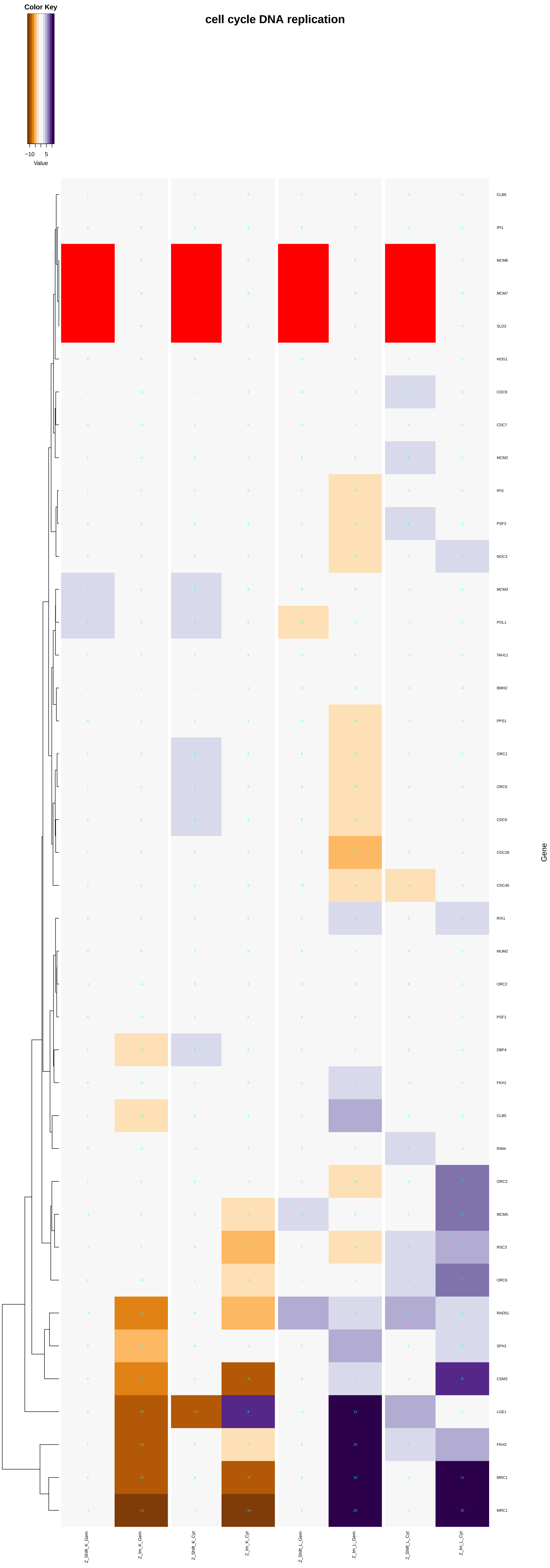

cell cycle DNA replication
Color Key
−10
5
Value
CLB6
IPI1
MCM6
MCM7
SLD2
HOG1
CDC9
CDC7
MCM2
IPI3
PSF2
NOC3
MCM3
POL1
TAH11
BMH2
PPS1
ORC1
ORC5
CDC6
CDC28
CDC45
RIX1
MUM2
ORC2
PSF1
DBF4
FKH1
CLB5
RIM4
ORC3
MCM5
RSC3
ORC6
RAD51
SFH1
CSM3
LGE1
FKH2
MRC1
MRC1
Gene
Z_Shift_K_Gem
Z_Im_K_Gem
Z_Shift_K_Cyt
Z_Im_K_Cyt
Z_Shift_L_Gem
Z_Im_L_Gem
Z_Shift_L_Cyt
Z_Im_L_Cyt

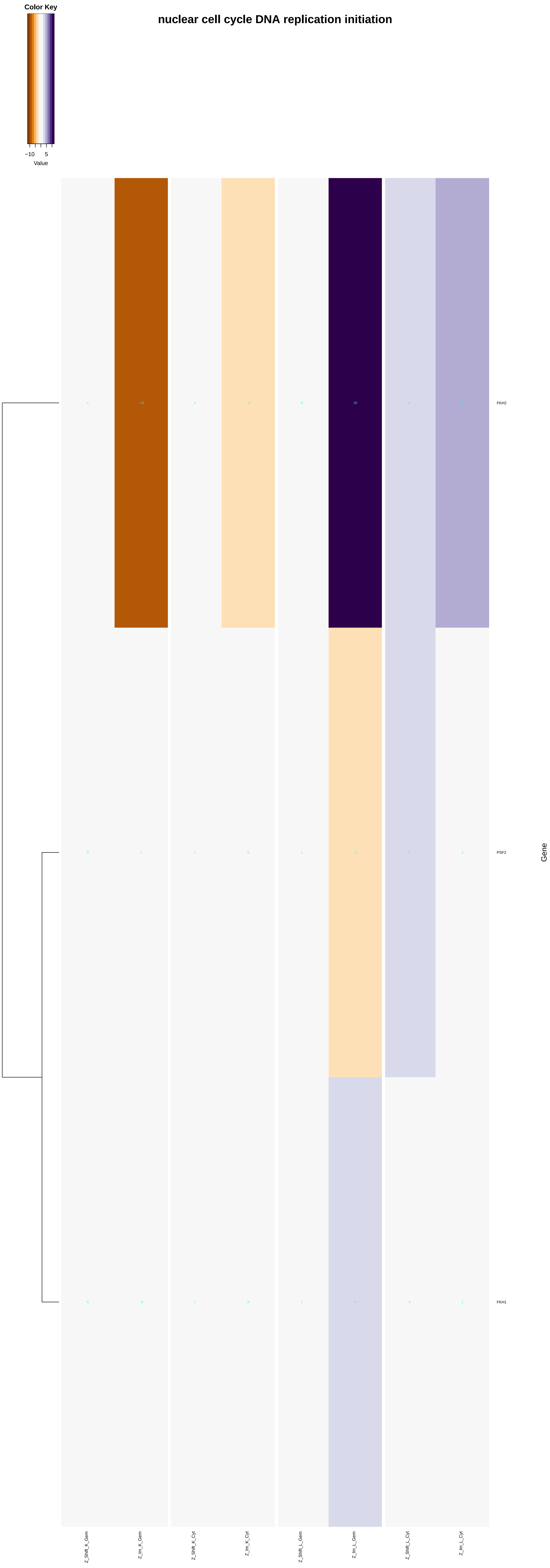
nuclear cell cycle DNA replication initiation
Color Key
−10
5
Value
FKH2
PSF2
FKH1
Gene
Z_Shift_K_Gem
Z_Im_K_Gem
Z_Shift_K_Cyt
Z_Im_K_Cyt
Z_Shift_L_Gem
Z_Im_L_Gem
Z_Shift_L_Cyt
Z_Im_L_Cyt

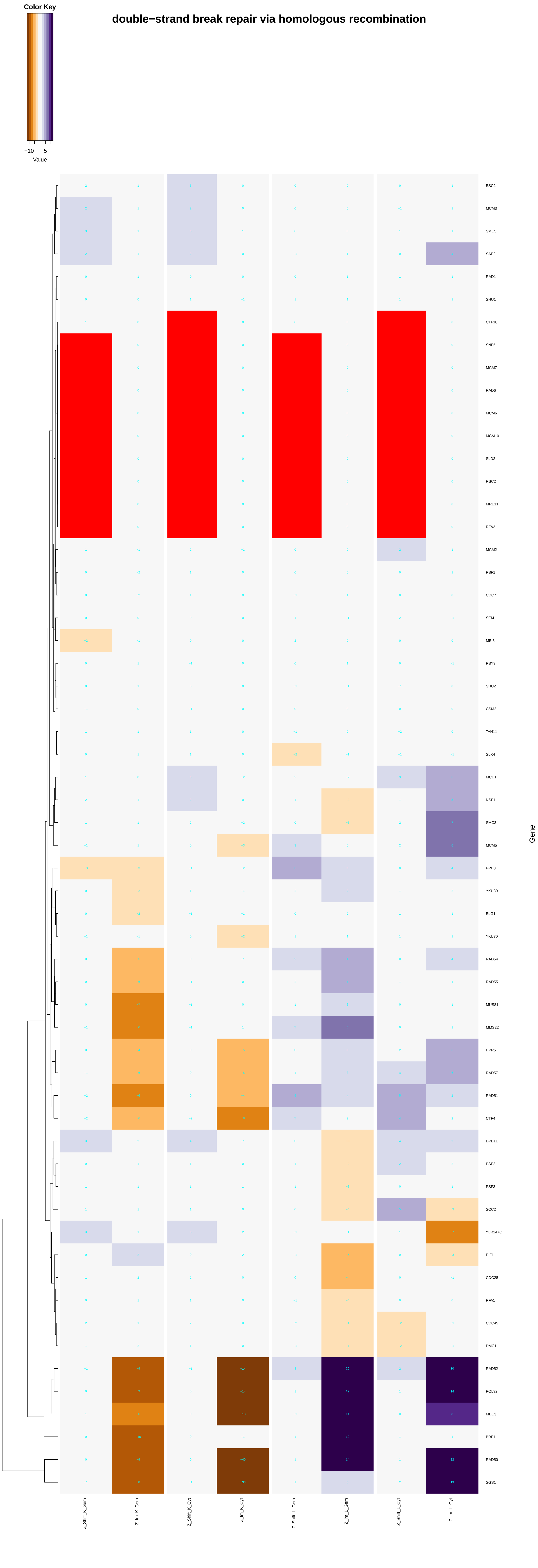
double−strand break repair via homologous recombination
Color Key
−10
5
Value
ESC2
MCM3
SMC5
SAE2
RAD1
SHU1
CTF18
SNF5
MCM7
RAD6
MCM6
MCM10
SLD2
RSC2
MRE11
RFA2
MCM2
PSF1
CDC7
SEM1
MEI5
PSY3
SHU2
CSM2
TAH11
SLX4
MCD1
NSE1
SMC3
MCM5
PPH3
YKU80
ELG1
YKU70
RAD54
RAD55
MUS81
MMS22
HPR5
RAD57
RAD51
CTF4
DPB11
PSF2
PSF3
SCC2
YLR247C
PIF1
CDC28
RFA1
CDC45
DMC1
RAD52
POL32
MEC3
BRE1
RAD50
SGS1
Gene
Z_Shift_K_Gem
Z_Im_K_Gem
Z_Shift_K_Cyt
Z_Im_K_Cyt
Z_Shift_L_Gem
Z_Im_L_Gem
Z_Shift_L_Cyt
Z_Im_L_Cyt

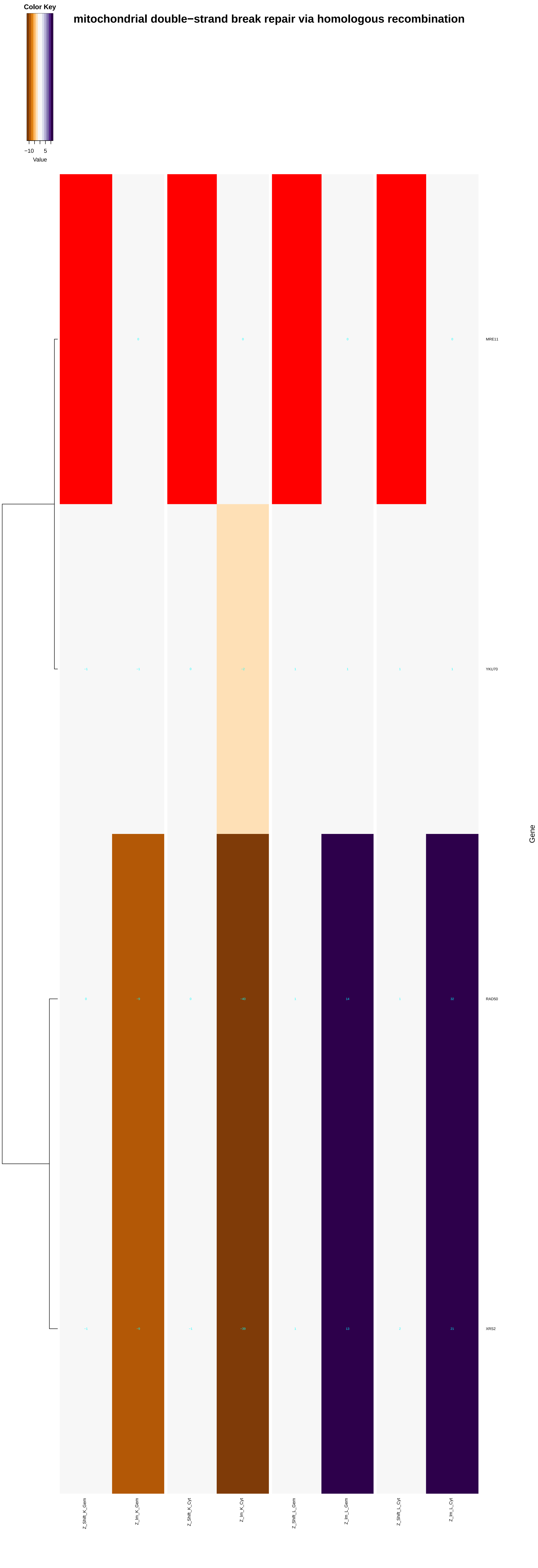
mitochondrial double−strand break repair via homologous recombination
Color Key
−10
5
Value
MRE11
YKU70
RAD50
XRS2
Gene
Z_Shift_K_Gem
Z_Im_K_Gem
Z_Shift_K_Cyt
Z_Im_K_Cyt
Z_Shift_L_Gem
Z_Im_L_Gem
Z_Shift_L_Cyt
Z_Im_L_Cyt

### double−strand break repair via nonhomologous end joining

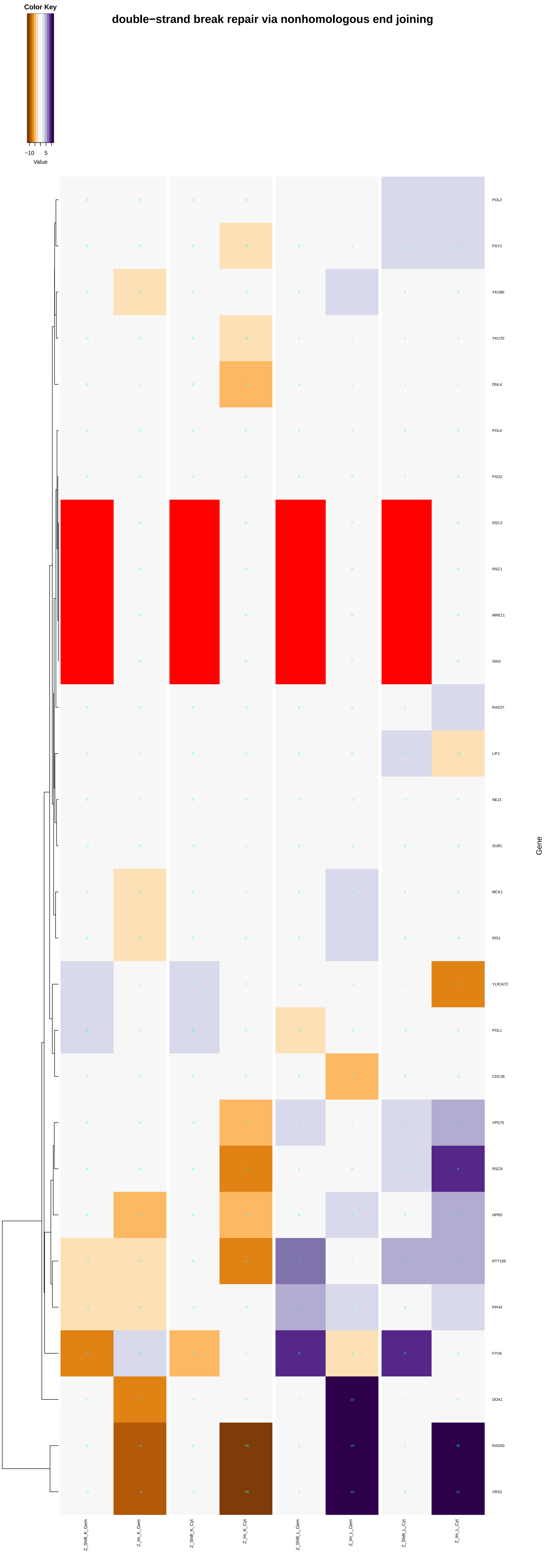

#### double−strand break repair via single−strand annealing

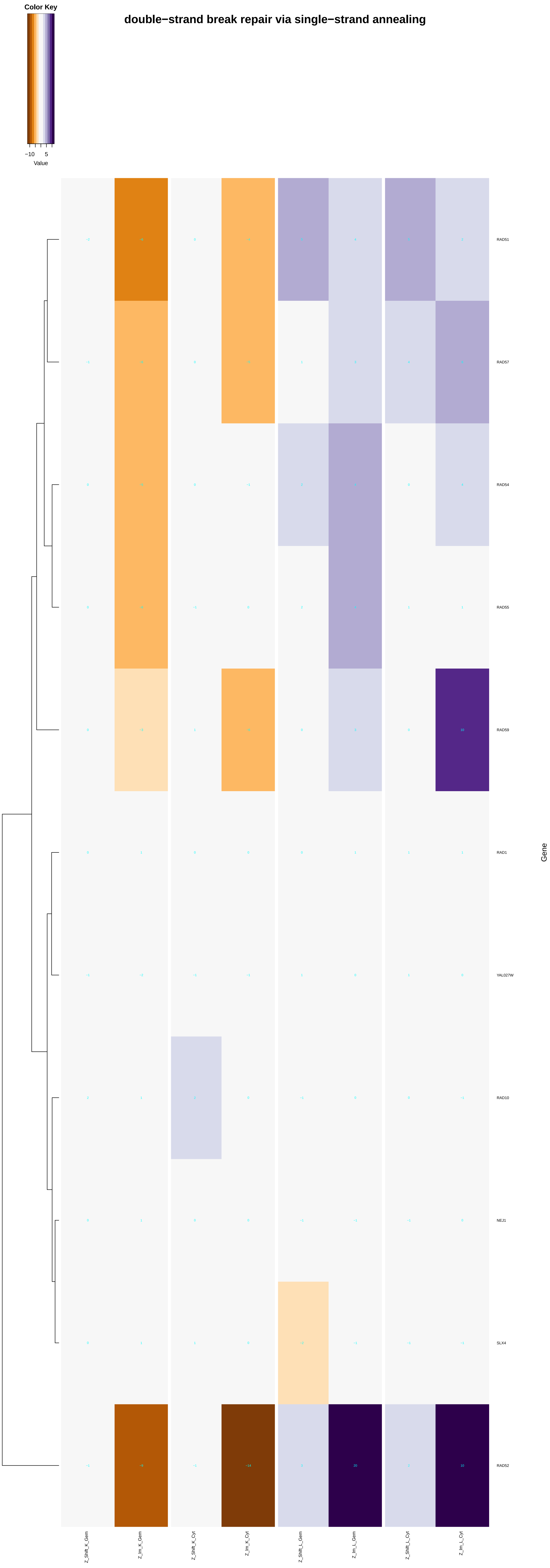

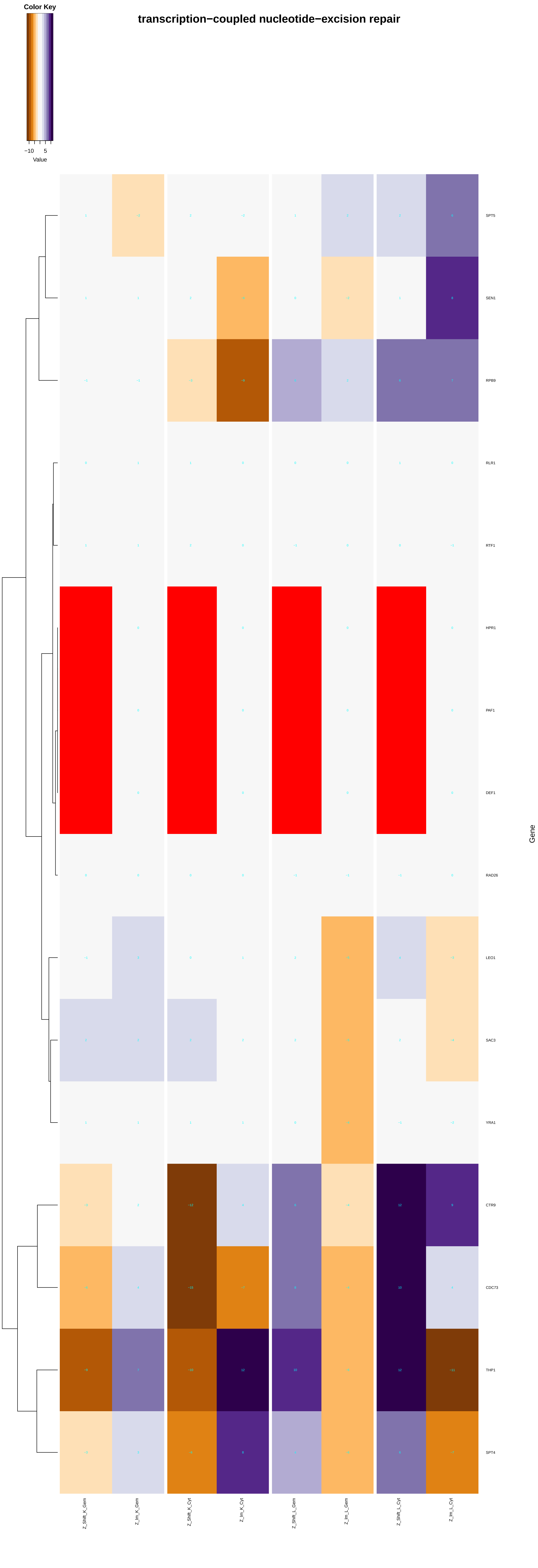
transcription−coupled nucleotide−excision repair
Color Key
−10
5
Value
SPT5
SEN1
RPB9
RLR1
RTF1
HPR1
PAF1
DEF1
RAD26
LEO1
SAC3
YRA1
CTR9
CDC73
THP1
SPT4
Gene
Z_Shift_K_Gem
Z_Im_K_Gem
Z_Shift_K_Cyt
Z_Im_K_Cyt
Z_Shift_L_Gem
Z_Im_L_Gem
Z_Shift_L_Cyt
Z_Im_L_Cyt

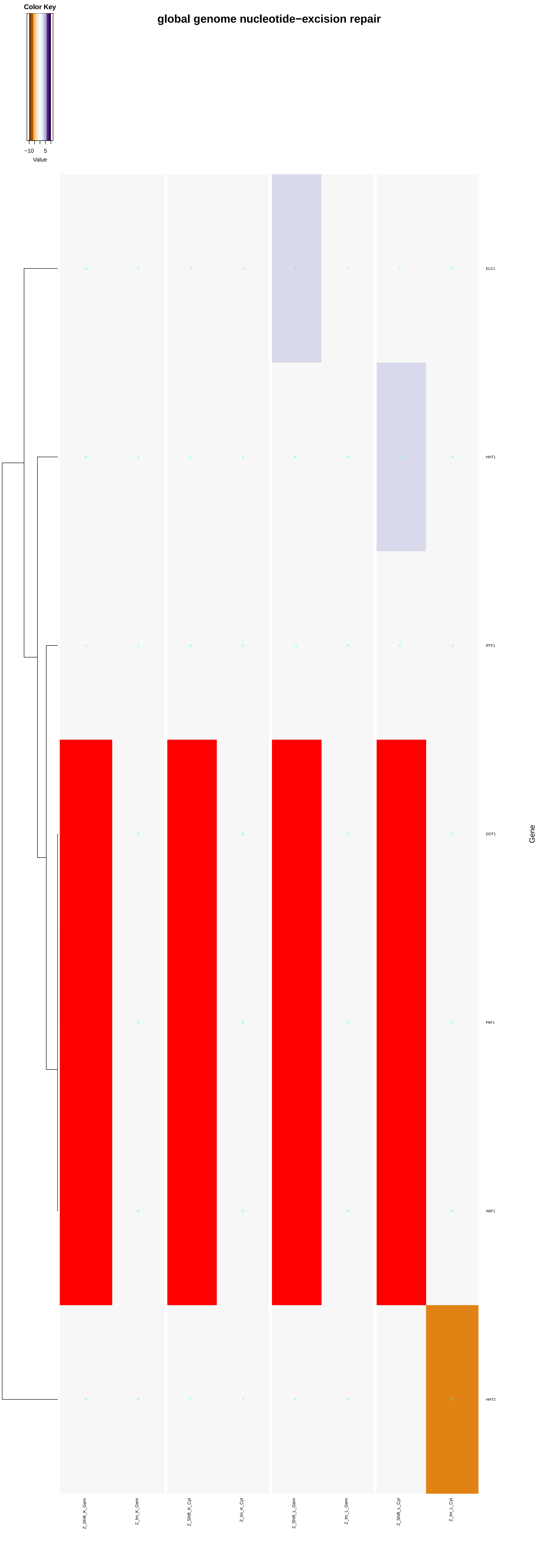
Color Key
−10
5
Value
global genome nucleotide−excision repair
ELC1
HHT1
RTF1
DOT1
PAF1
ABF1
HHT2
Gene
Z_Shift_K_Gem
Z_Im_K_Gem
Z_Shift_K_Cyt
Z_Im_K_Cyt
Z_Shift_L_Gem
Z_Im_L_Gem
Z_Shift_L_Cyt
Z_Im_L_Cyt

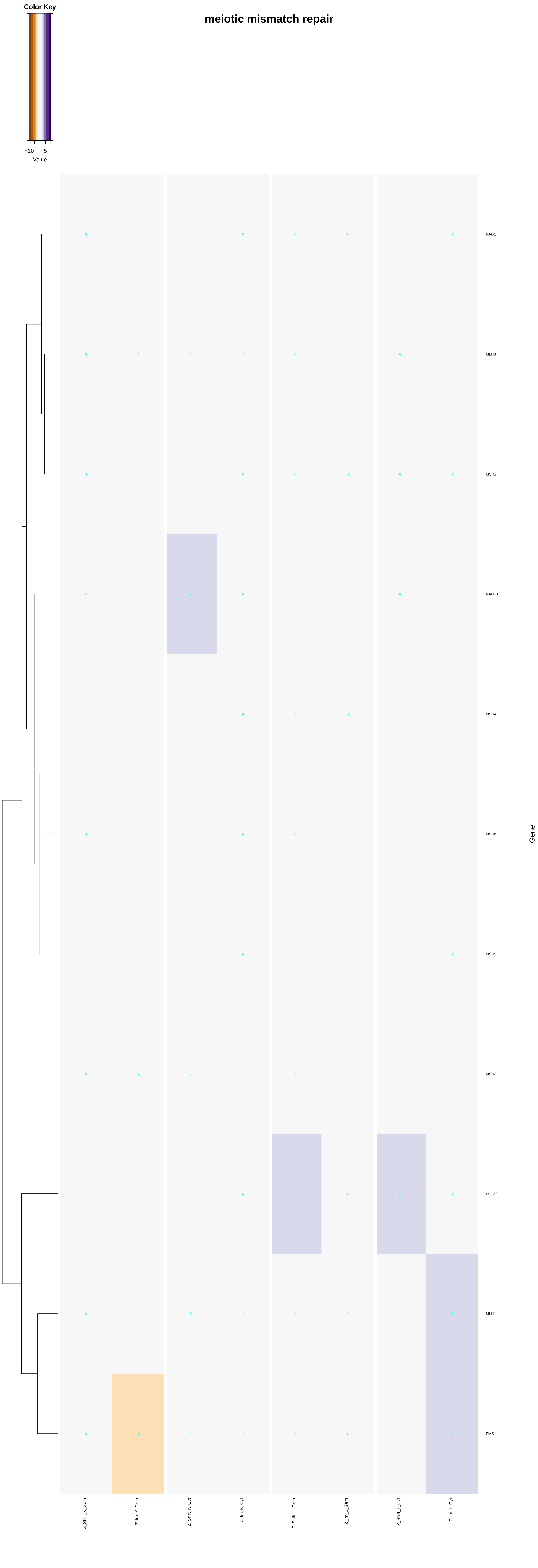
meiotic mismatch repair
Color Key
−10
5
Value
RAD1
MLH3
MSH2
RAD10
MSH4
MSH6
MSH5
MSH3
POL30
MLH1
PMS1
Gene
Z_Shift_K_Gem
Z_Im_K_Gem
Z_Shift_K_Cyt
Z_Im_K_Cyt
Z_Shift_L_Gem
Z_Im_L_Gem
Z_Shift_L_Cyt
Z_Im_L_Cyt

### translesion synthesis

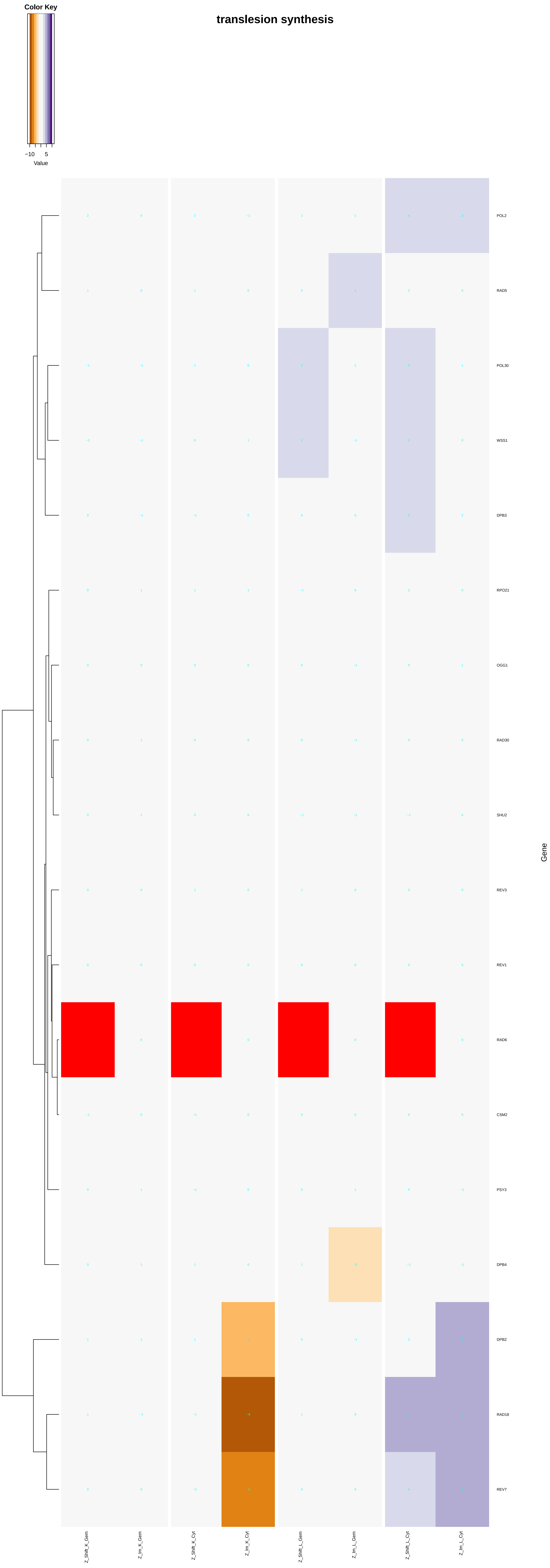

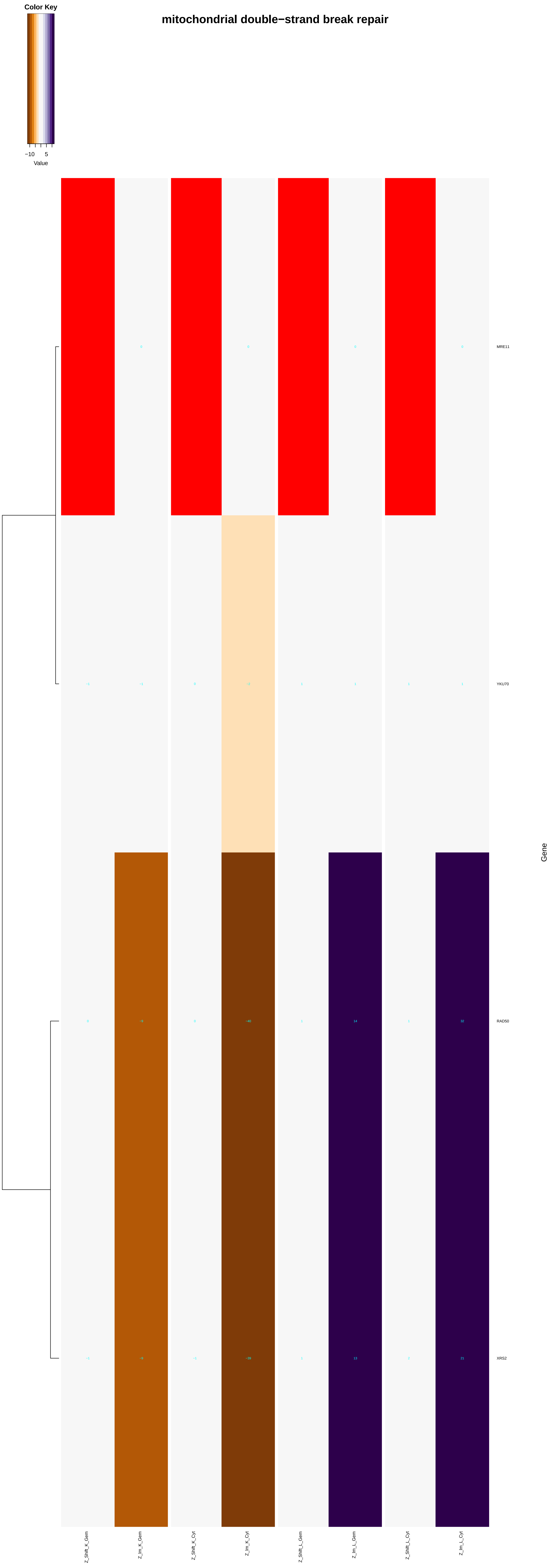
mitochondrial double−strand break repair
Color Key
−10
5
Value
MRE11
YKU70
RAD50
XRS2
Gene
Z_Shift_K_Gem
Z_Im_K_Gem
Z_Shift_K_Cyt
Z_Im_K_Cyt
Z_Shift_L_Gem
Z_Im_L_Gem
Z_Shift_L_Cyt
Z_Im_L_Cyt

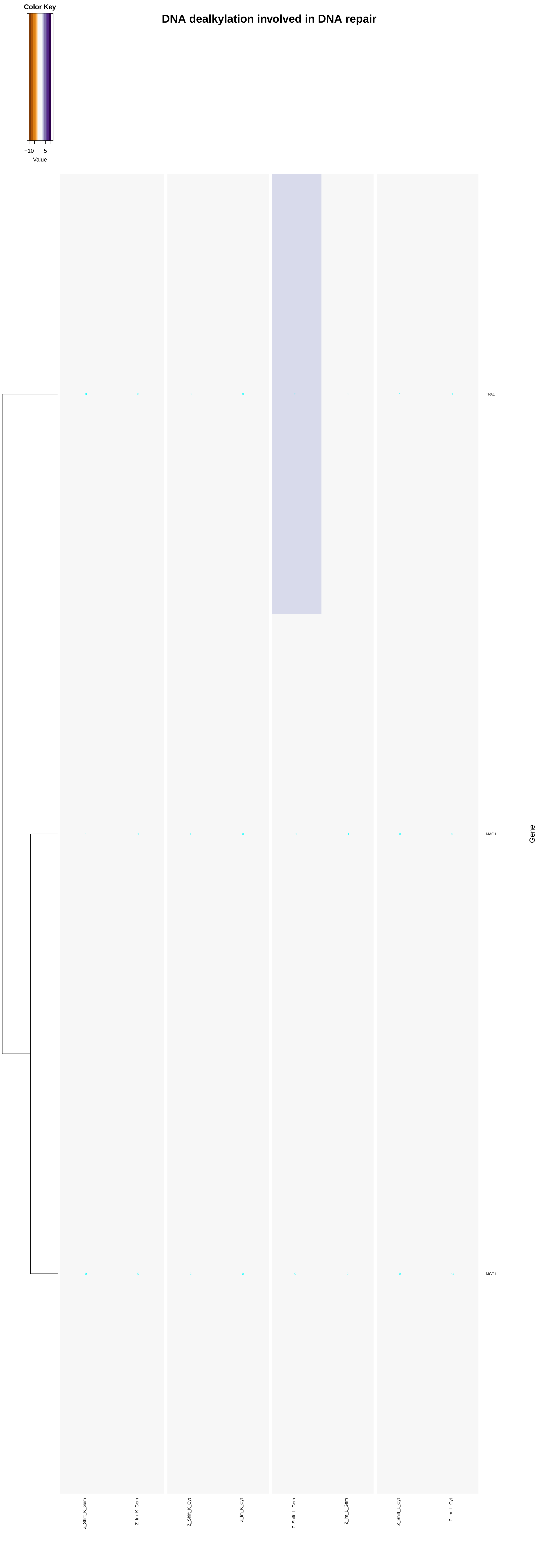
DNA dealkylation involved in DNA repair
Color Key
−10
5
Value
TPA1
MAG1
MGT1
Gene
Z_Shift_K_Gem
Z_Im_K_Gem
Z_Shift_K_Cyt
Z_Im_K_Cyt
Z_Shift_L_Gem
Z_Im_L_Gem
Z_Shift_L_Cyt
Z_Im_L_Cyt

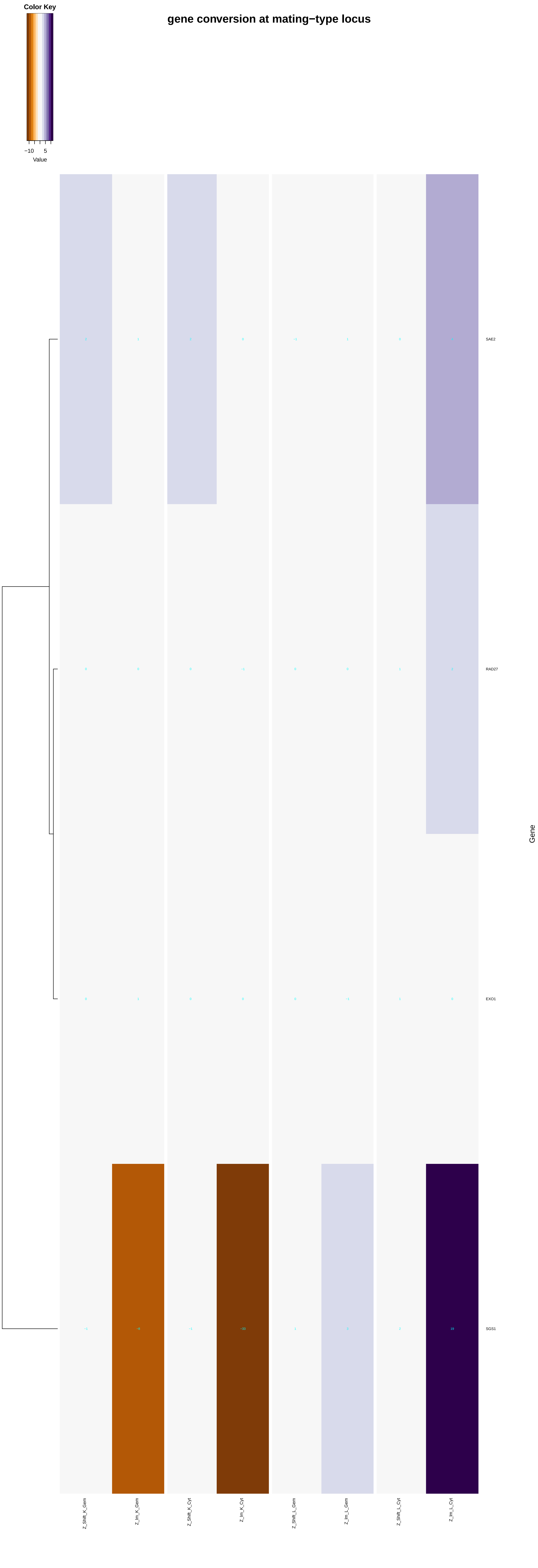
gene conversion at mating−type locus
Color Key
−10
5
Value
SAE2
RAD27
EXO1
SGS1
Gene
Z_Shift_K_Gem
Z_Im_K_Gem
Z_Shift_K_Cyt
Z_Im_K_Cyt
Z_Shift_L_Gem
Z_Im_L_Gem
Z_Shift_L_Cyt
Z_Im_L_Cyt

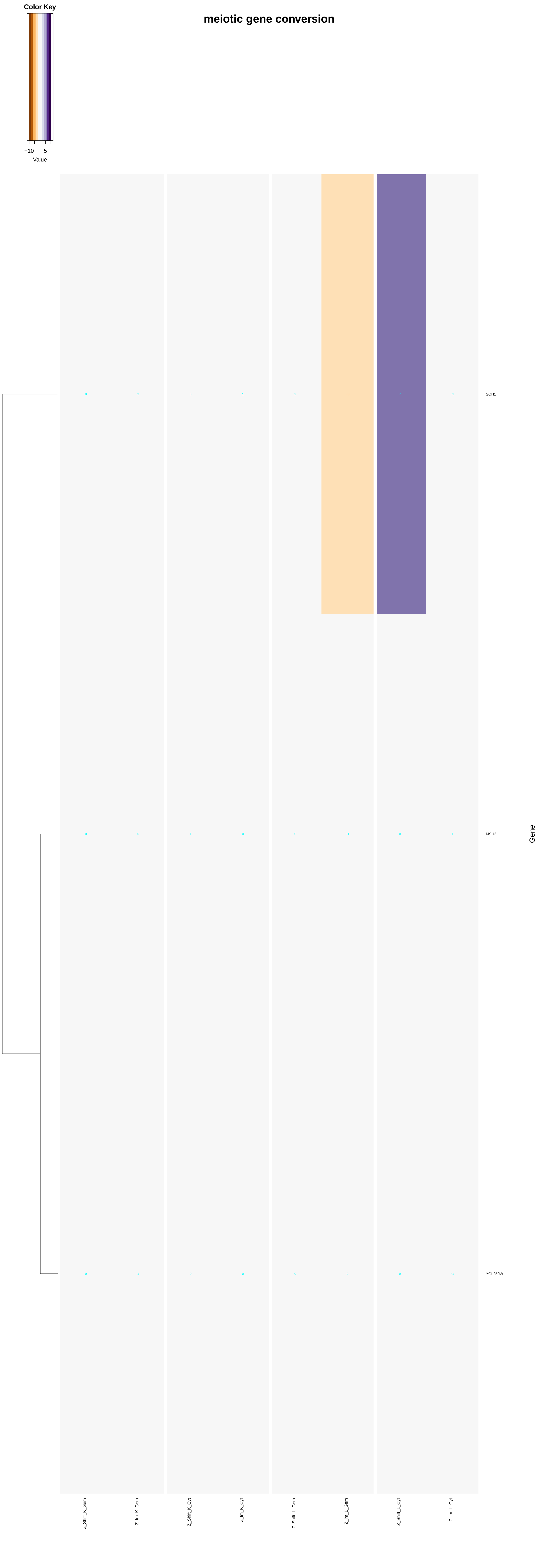
meiotic gene conversion
Color Key
−10
5
Value
SOH1
MSH2
YGL250W
Gene
Z_Shift_K_Gem
Z_Im_K_Gem
Z_Shift_K_Cyt
Z_Im_K_Cyt
Z_Shift_L_Gem
Z_Im_L_Gem
Z_Shift_L_Cyt
Z_Im_L_Cyt

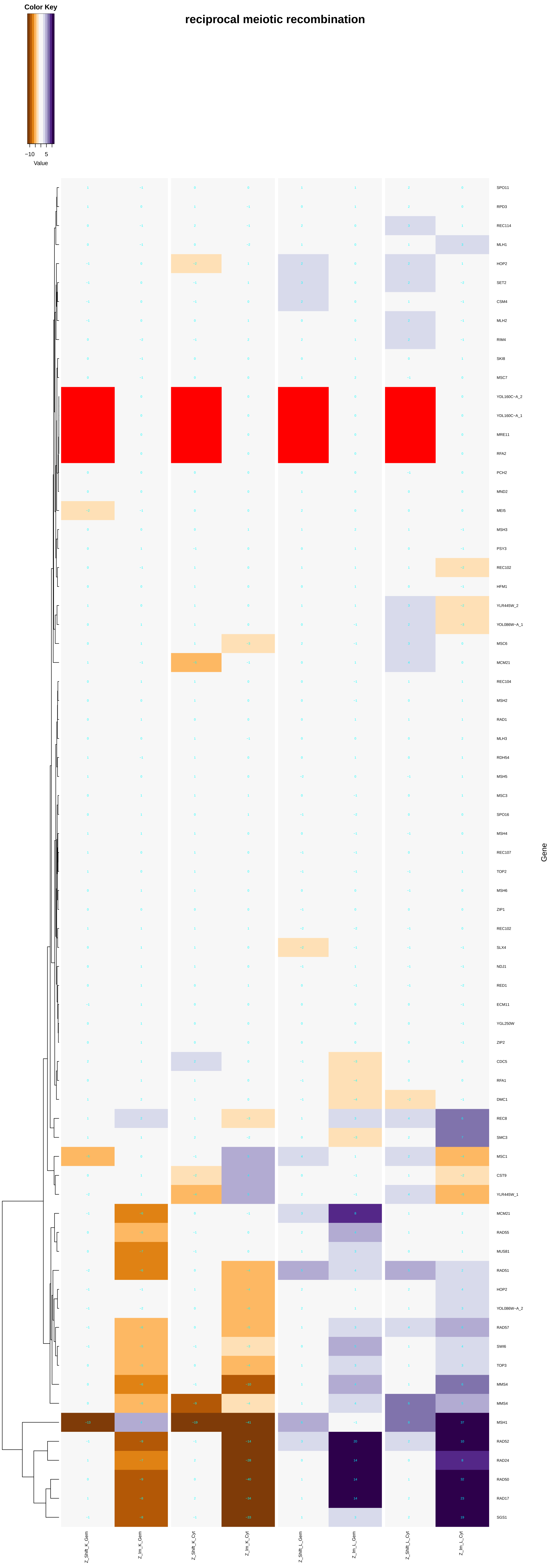
reciprocal meiotic recombination
Color Key
−10
5
Value
SPO11
RPD3
REC114
MLH1
HOP2
SET2
CSM4
MLH2
RIM4
SKI8
MSC7
YDL160C−A_2
YDL160C−A_1
MRE11
RFA2
PCH2
MND2
MEI5
MSH3
PSY3
REC102
HFM1
YLR445W_2
YOL086W−A_1
MSC6
MCM21
REC104
MSH2
RAD1
MLH3
RDH54
MSH5
MSC3
SPO16
MSH4
REC107
TOP2
MSH6
ZIP1
REC102
SLX4
NDJ1
RED1
ECM11
YGL250W
ZIP2
CDC5
RFA1
DMC1
REC8
SMC3
MSC1
CST9
YLR445W_1
MCM21
RAD55
MUS81
RAD51
HOP2
YOL086W−A_2
RAD57
SWI6
TOP3
MMS4
MMS4
MSH1
RAD52
RAD24
RAD50
RAD17
SGS1
Gene
Z_Shift_K_Gem
Z_Im_K_Gem
Z_Shift_K_Cyt
Z_Im_K_Cyt
Z_Shift_L_Gem
Z_Im_L_Gem
Z_Shift_L_Cyt
Z_Im_L_Cyt

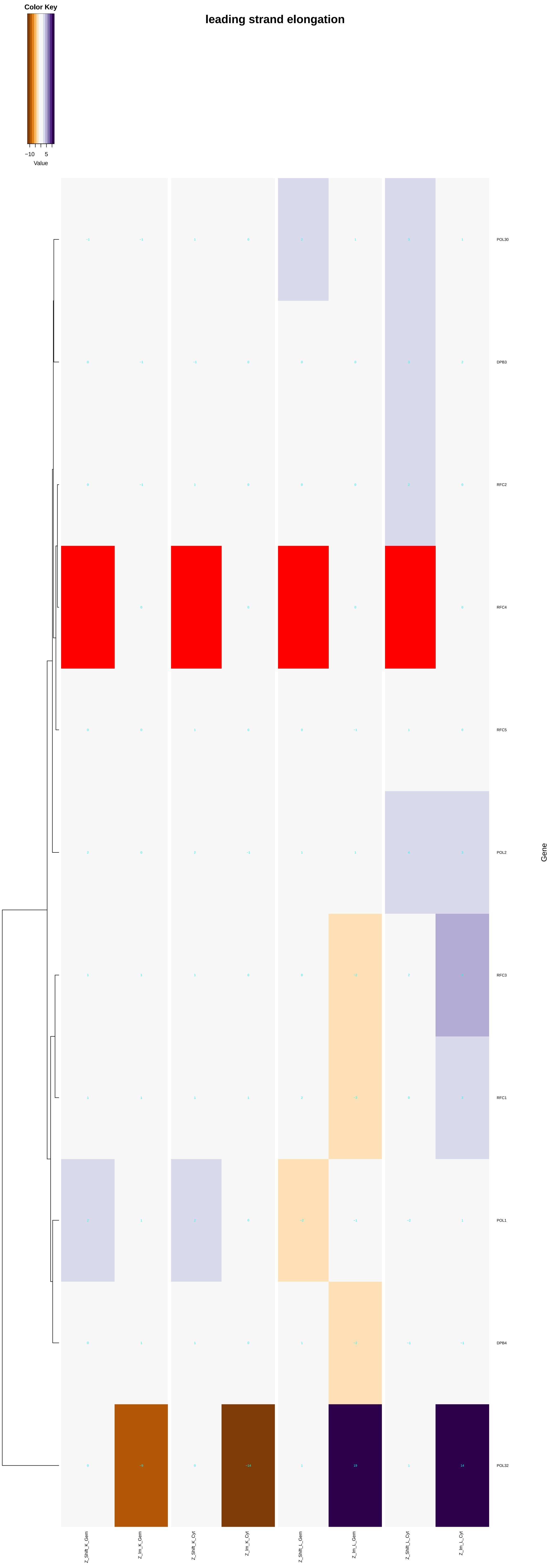
leading strand elongation
Color Key
−10
5
Value
POL30
DPB3
RFC2
RFC4
RFC5
POL2
RFC3
RFC1
POL1
DPB4
POL32
Gene
Z_Shift_K_Gem
Z_Im_K_Gem
Z_Shift_K_Cyt
Z_Im_K_Cyt
Z_Shift_L_Gem
Z_Im_L_Gem
Z_Shift_L_Cyt
Z_Im_L_Cyt

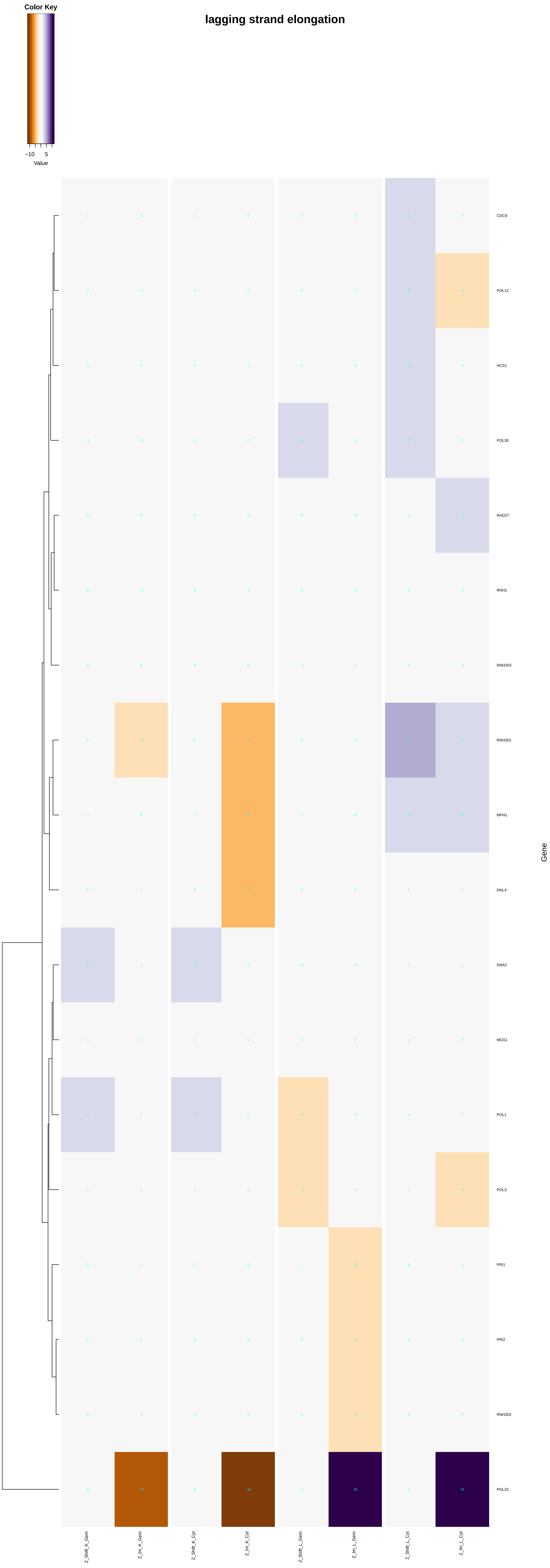

lagging strand elongation
Color Key
−10
5
Value
CDC9
POL12
HCS1
POL30
RAD27
RNH1
RNH203
RNH201
MPH1
DNL4
DNA2
MGS1
POL1
POL3
PRI1
PRI2
RNH202
POL32
Gene
Z_Shift_K_Gem
Z_Im_K_Gem
Z_Shift_K_Cyt
Z_Im_K_Cyt
Z_Shift_L_Gem
Z_Im_L_Gem
Z_Shift_L_Cyt
Z_Im_L_Cyt

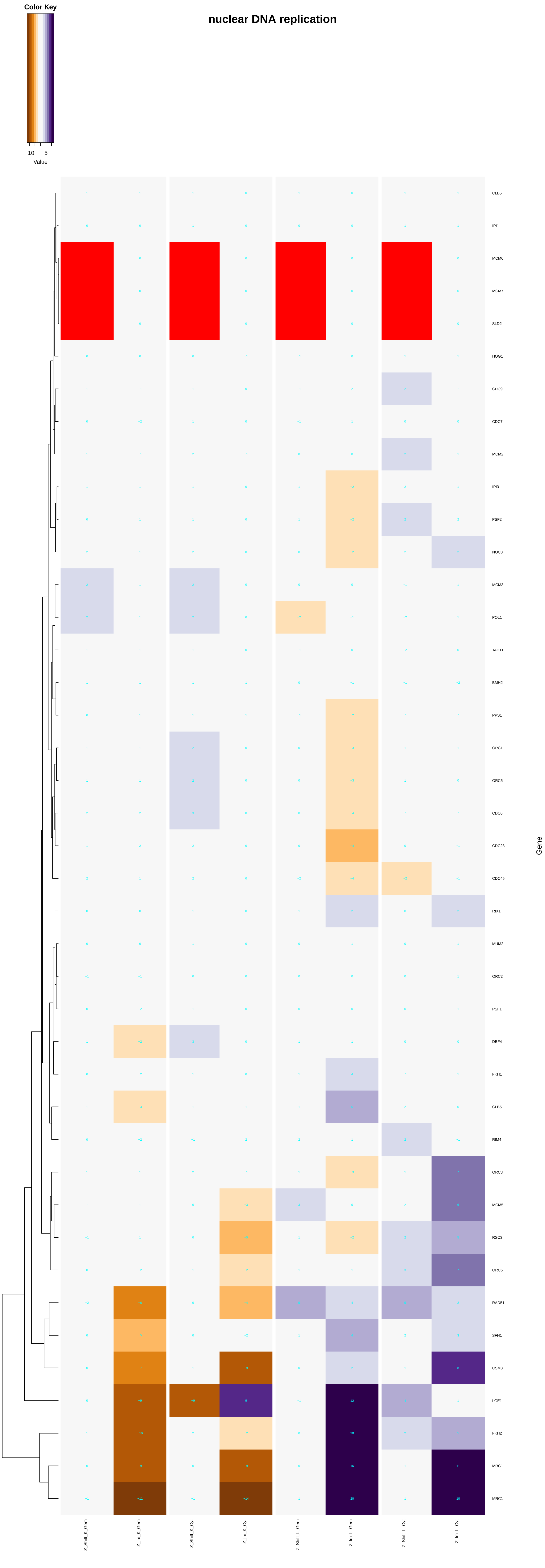

nuclear DNA replication
Color Key
−10
5
Value
Gene
CLB6
IPI1
MCM6
MCM7
SLD2
HOG1
CDC9
CDC7
MCM2
IPI3
PSF2
NOC3
MCM3
POL1
TAH11
BMH2
PPS1
ORC1
ORC5
CDC6
CDC28
CDC45
RIX1
MUM2
ORC2
PSF1
DBF4
FKH1
CLB5
RIM4
ORC3
MCM5
RSC3
ORC6
RAD51
SFH1
CSM3
LGE1
FKH2
MRC1
MRC1
Z_Shift_K_Gem
Z_Im_K_Gem
Z_Shift_K_Cyt
Z_Im_K_Cyt
Z_Shift_L_Gem
Z_Im_L_Gem
Z_Shift_L_Cyt
Z_Im_L_Cyt

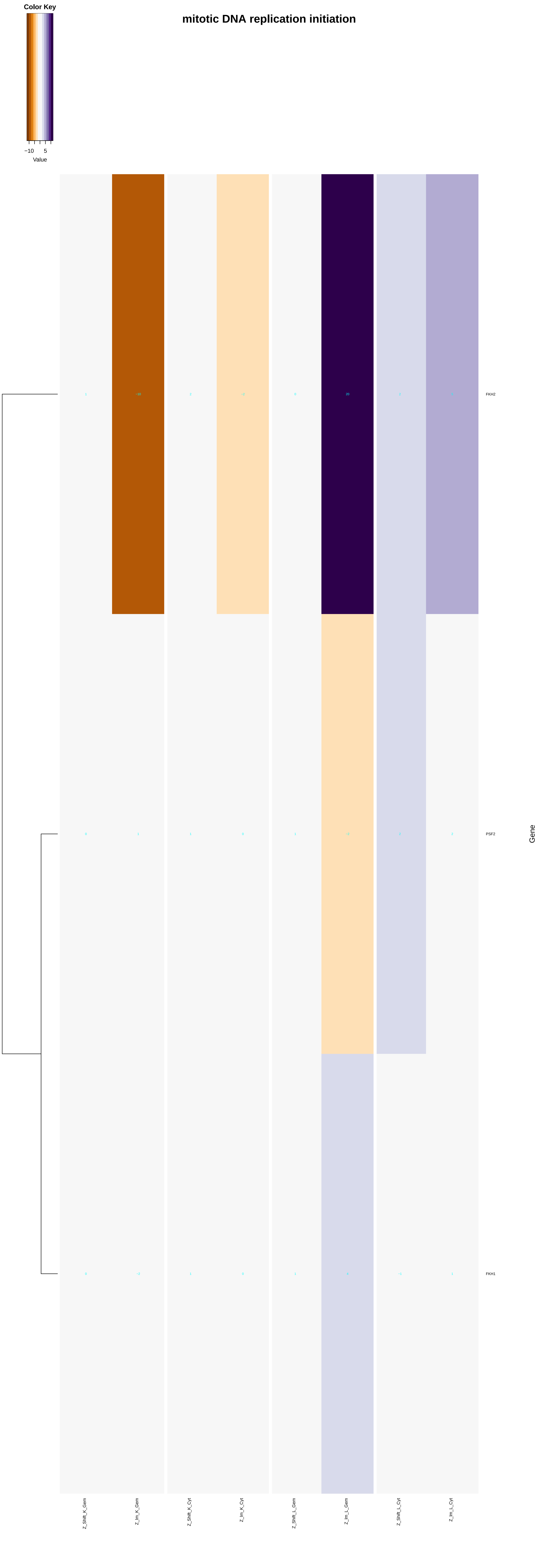
Color Key
−10
5
Value
mitotic DNA replication initiation
1
−10
2
−2
0
20
2
5
FKH2
0
1
1
0
1
−2
2
2
PSF2
0
−2
1
0
1
4
−1
1
FKH1
Gene
Z_Shift_K_Gem
Z_Int_K_Gem
Z_Shift_K_Cyt
Z_Int_K_Cyt
Z_Shift_L_Gem
Z_Int_L_Gem
Z_Shift_L_Cyt
Z_Int_L_Cyt

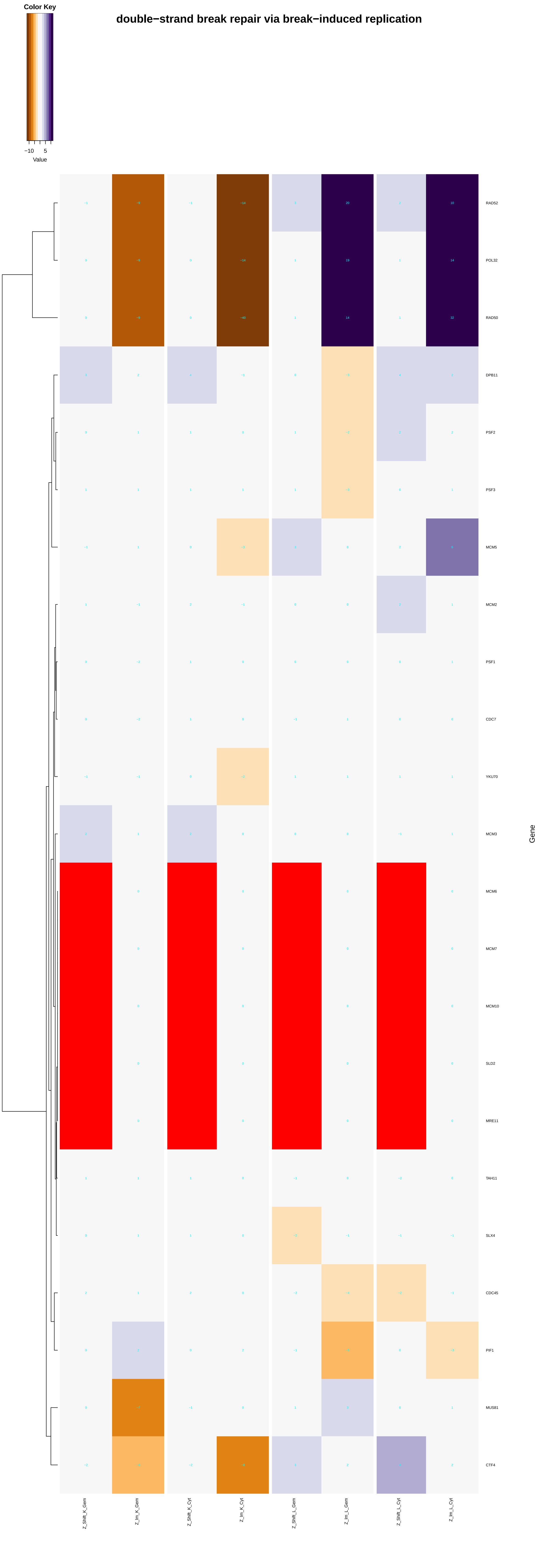

Color Key
−10
5
Value
double−strand break repair via break−induced replication
RAD52
POL32
RAD50
DPB11
PSF2
PSF3
MCM5
MCM2
PSF1
CDC7
YKU70
MCM3
MCM6
MCM7
MCM10
SLD2
MRE11
TAH11
SLX4
CDC45
PIF1
MUS81
CTF4
Gene
Z_Shift_K_Gem
Z_Im_K_Gem
Z_Shift_K_Cyt
Z_Im_K_Cyt
Z_Shift_L_Gem
Z_Im_L_Gem
Z_Shift_L_Cyt
Z_Im_L_Cyt

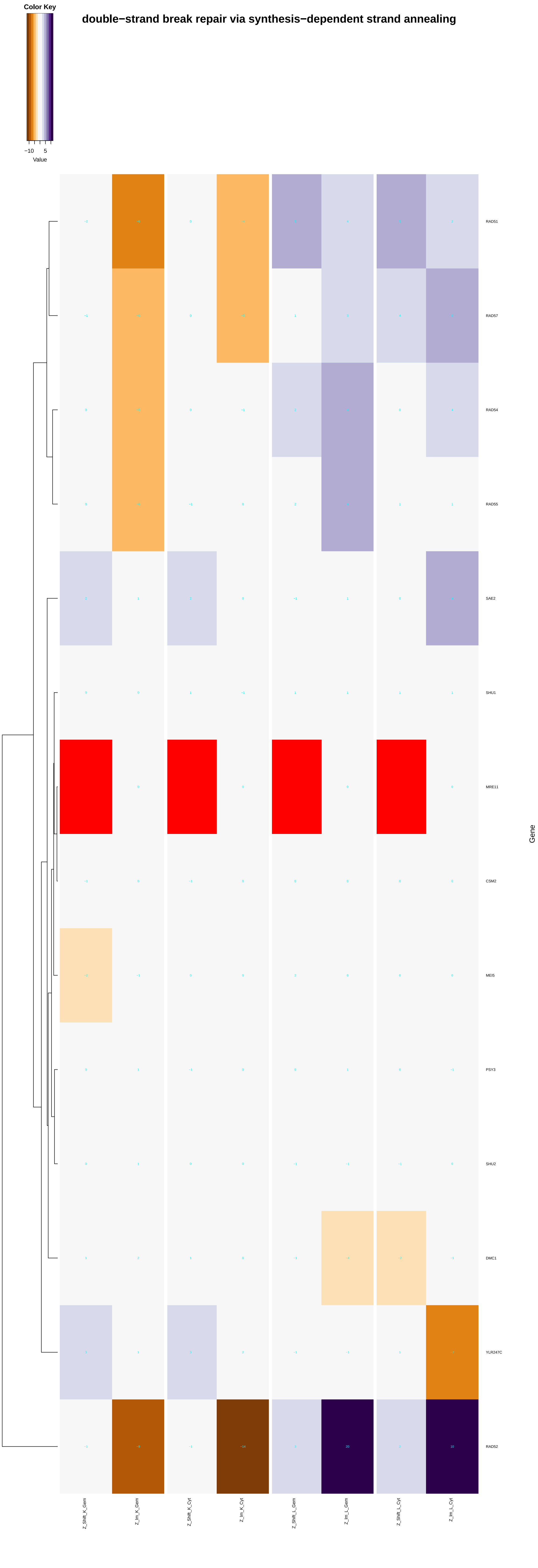

double−strand break repair via synthesis−dependent strand annealing
Color Key
−10
5
Value
RAD51
RAD57
RAD54
RAD55
SAE2
SHU1
MRE11
CSM2
MEI5
PSY3
SHU2
DMC1
YLR247C
RAD52
Gene
Z_Shift_K_Gem
Z_Im_K_Gem
Z_Shift_K_Cyt
Z_Im_K_Cyt
Z_Shift_L_Gem
Z_Im_L_Gem
Z_Shift_L_Cyt
Z_Im_L_Cyt

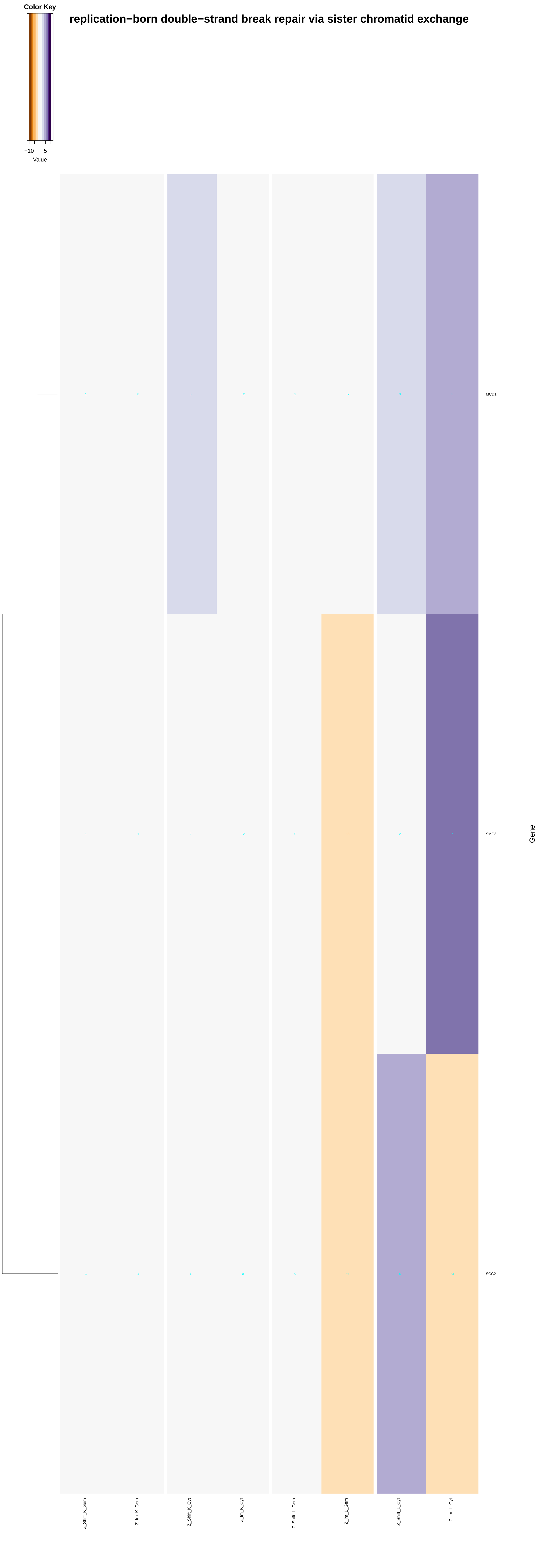

Color Key
−10
5
Value
replication−born double−strand break repair via sister chromatid exchange
MCD1
SMC3
SCC2
Gene
Z_Shift_K_Gem
Z_Im_K_Gem
Z_Shift_K_Cyt
Z_Im_K_Cyt
Z_Shift_L_Gem
Z_Im_L_Gem
Z_Shift_L_Cyt
Z_Im_L_Cyt

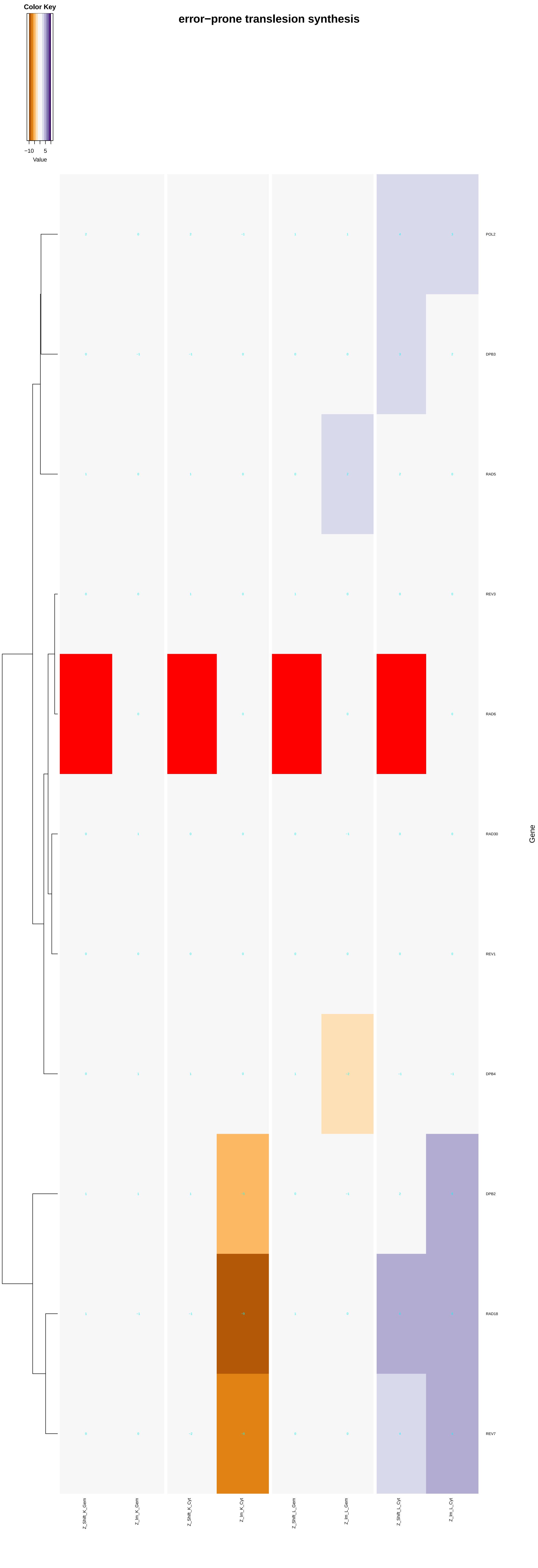

Color Key
−10
5
Value
error−prone translesion synthesis
POL2
DPB3
RAD5
REV3
RAD6
RAD30
REV1
DPB4
DPB2
RAD18
REV7
Gene
Z_Shift_K_Gem
Z_Im_K_Gem
Z_Shift_K_Cyt
Z_Im_K_Cyt
Z_Shift_L_Gem
Z_Im_L_Gem
Z_Shift_L_Cyt
Z_Im_L_Cyt

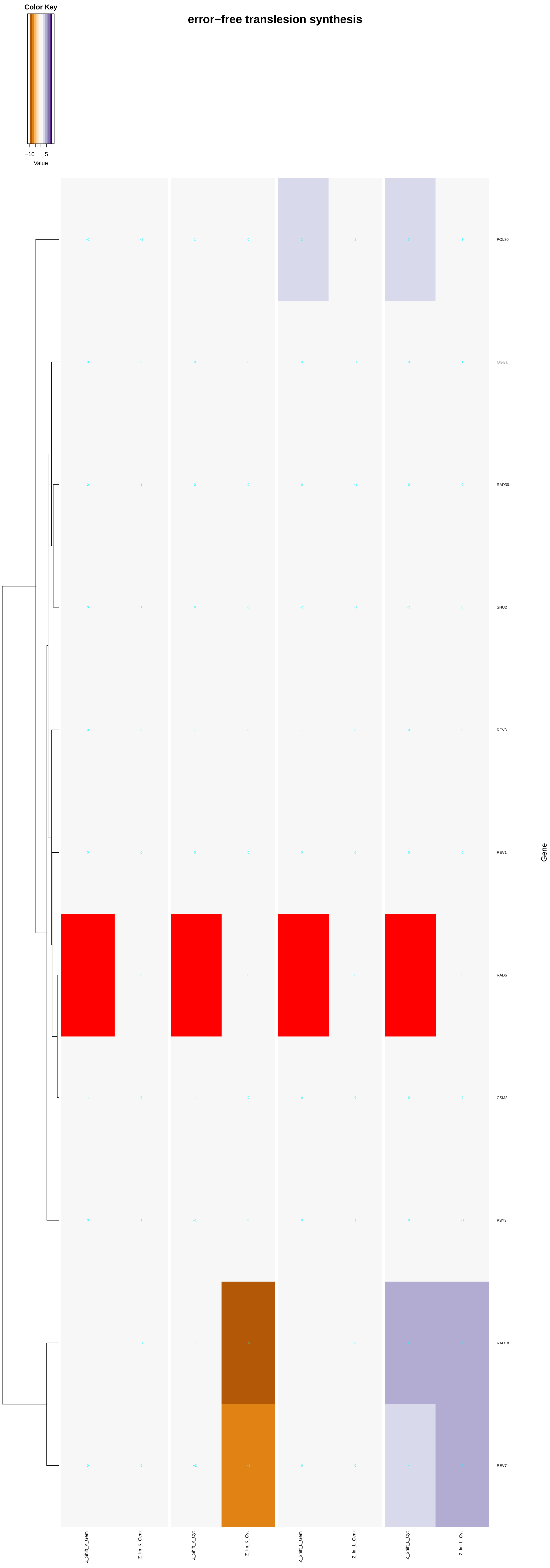
error−free translesion synthesis
Color Key
−10
5
Value
POL30
OGG1
RAD30
SHU2
REV3
REV1
RAD6
CSM2
PSY3
RAD18
REV7
Gene
Z_Shift_K_Gem
Z_Im_K_Gem
Z_Shift_K_Cyt
Z_Im_K_Cyt
Z_Shift_L_Gem
Z_Im_L_Gem
Z_Shift_L_Cyt
Z_Im_L_Cyt

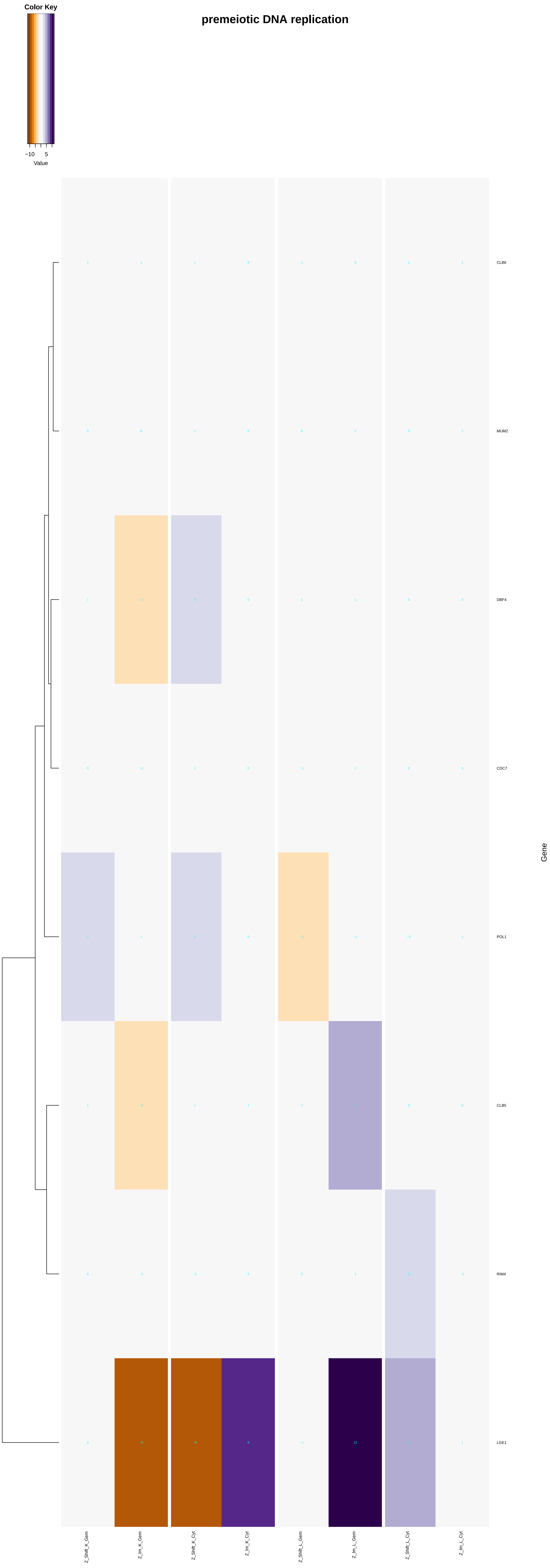
Color Key
−10
5
Value
premeiotic DNA replication
CLB6
MUM2
DBF4
CDC7
POL1
CLB5
RIM4
LGE1
Gene
Z_Shift_K_Gem
Z_Im_K_Gem
Z_Shift_K_Cyt
Z_Im_K_Cyt
Z_Shift_L_Gem
Z_Im_L_Gem
Z_Shift_L_Cyt
Z_Im_L_Cyt

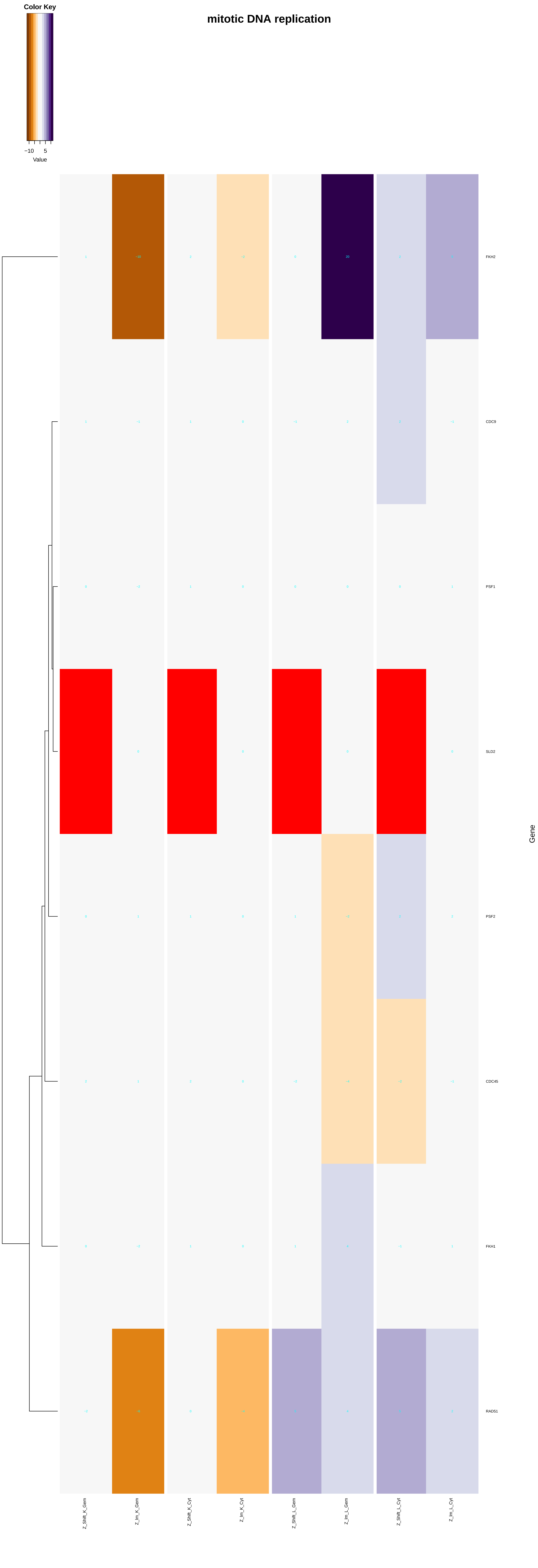

Color Key
−10
5
Value
mitotic DNA replication
FKH2
CDC9
PSF1
SLD2
PSF2
CDC45
FKH1
RAD51
Gene
Z_Shift_K_Gem
Z_Im_K_Gem
Z_Shift_K_Cyt
Z_Im_K_Cyt
Z_Shift_L_Gem
Z_Im_L_Gem
Z_Shift_L_Cyt
Z_Im_L_Cyt
