## Additional File 7 for "A humanized yeast phenomic model of deoxycytidine kinase to predict genetic buffering of nucleoside analog cytotoxicity": DNA_recombination.pdf

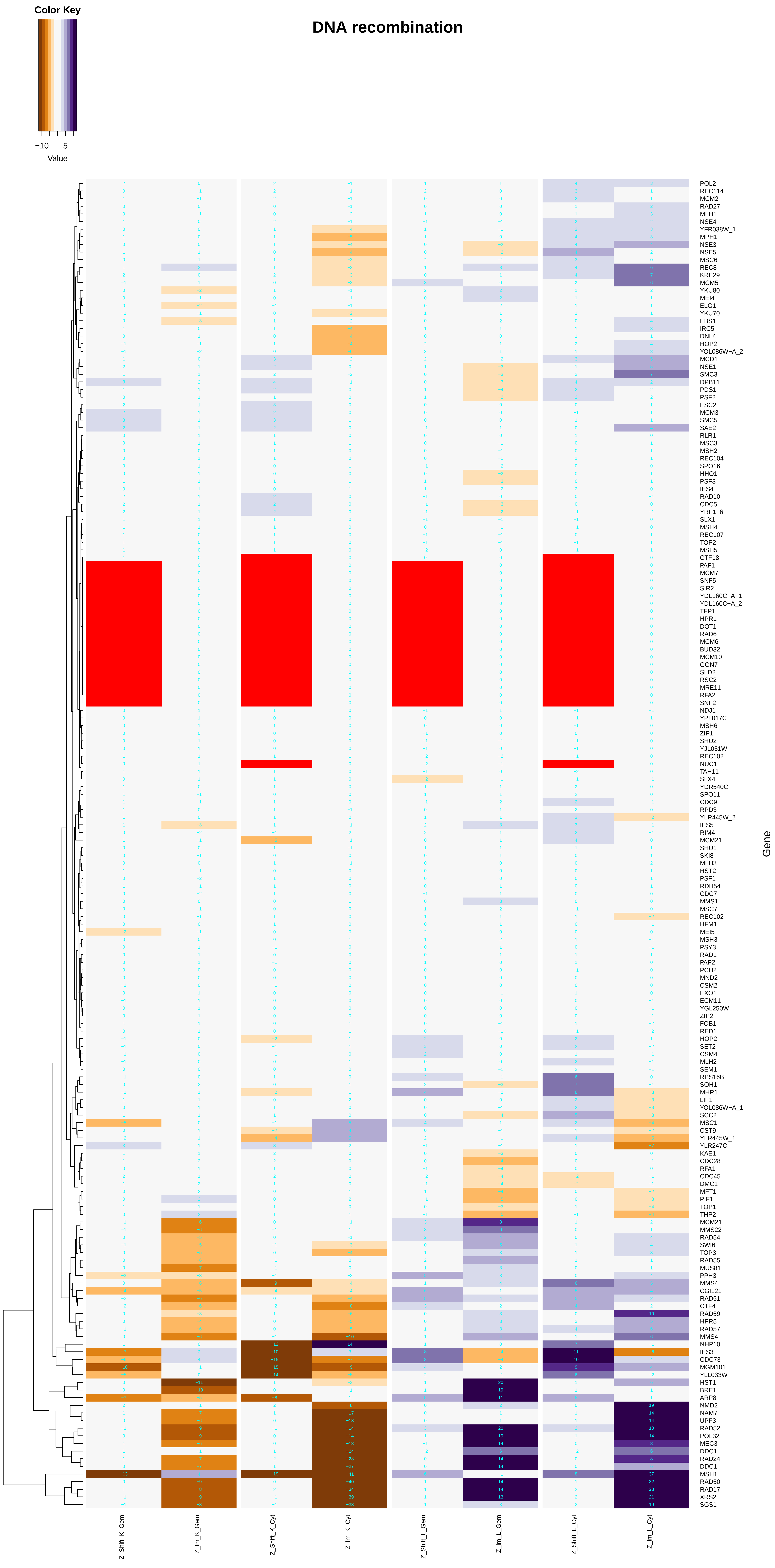
Color Key
−10
5
Value
DNA recombination
POL2
REC114
MCM2
RAD27
MLH1
NSE4
YFR038W_1
MPH1
NSE3
NSE5
MSC6
REC8
KRE29
MCM5
YKU80
MEI4
ELG1
YKU70
EBS1
IRC5
DNL4
HOP2
YOL086W−A_2
MCD1
NSE1
SMC3
DPB11
PDS1
PSF2
ESC2
MCM3
SMC5
SAE2
RLR1
MSC3
MSH2
REC104
SPO16
HHO1
PSF3
IES4
RAD10
CDC5
YRF1−6
SLX1
MSH4
REC107
TOP2
MSH5
CTF18
PAF1
MCM7
SNF5
SIR2
YDL160C−A_1
YDL160C−A_2
TFP1
HPR1
DOT1
RAD6
MCM6
BUD32
MCM10
GON7
SLD2
RSC2
MRE11
RFA2
SNF2
NDJ1
YPL017C
MSH6
ZIP1
SHU2
YJL051W
REC102
NUC1
TAH11
SLX4
YDR540C
SPO11
CDC9
RPD3
YLR445W_2
IES5
RIM4
MCM21
SHU1
SKI8
MLH3
HST2
PSF1
RDH54
CDC7
MMS1
MSC7
REC102
HFM1
MEI5
MSH3
PSY3
RAD1
PAP2
PCH2
MND2
CSM2
EXO1
ECM11
YGL250W
ZIP2
FOB1
RED1
HOP2
SET2
CSM4
MLH2
SEM1
RPS16B
SOH1
MHR1
LIF1
YOL086W−A_1
SCC2
MSC1
CST9
YLR445W_1
YLR247C
KAE1
CDC28
RFA1
CDC45
DMC1
MFT1
PIF1
TOP1
THP2
MCM21
MMS22
RAD54
SWI6
TOP3
RAD55
MUS81
PPH3
MMS4
CGI121
RAD51
CTF4
RAD59
HPR5
RAD57
MMS4
NHP10
IES3
CDC73
MGM101
YLL033W
HST1
BRE1
ARP8
NMD2
NAM7
UPF3
RAD52
POL32
MEC3
DDC1
RAD24
DDC1
MSH1
RAD50
RAD17
XRS2
SGS1
Gene
Z_Shift_K_Gem
Z_Im_K_Gem
Z_Shift_K_Cyt
Z_Im_K_Cyt
Z_Shift_L_Gem
Z_Im_L_Gem
Z_Shift_L_Cyt
Z_Im_L_Cyt

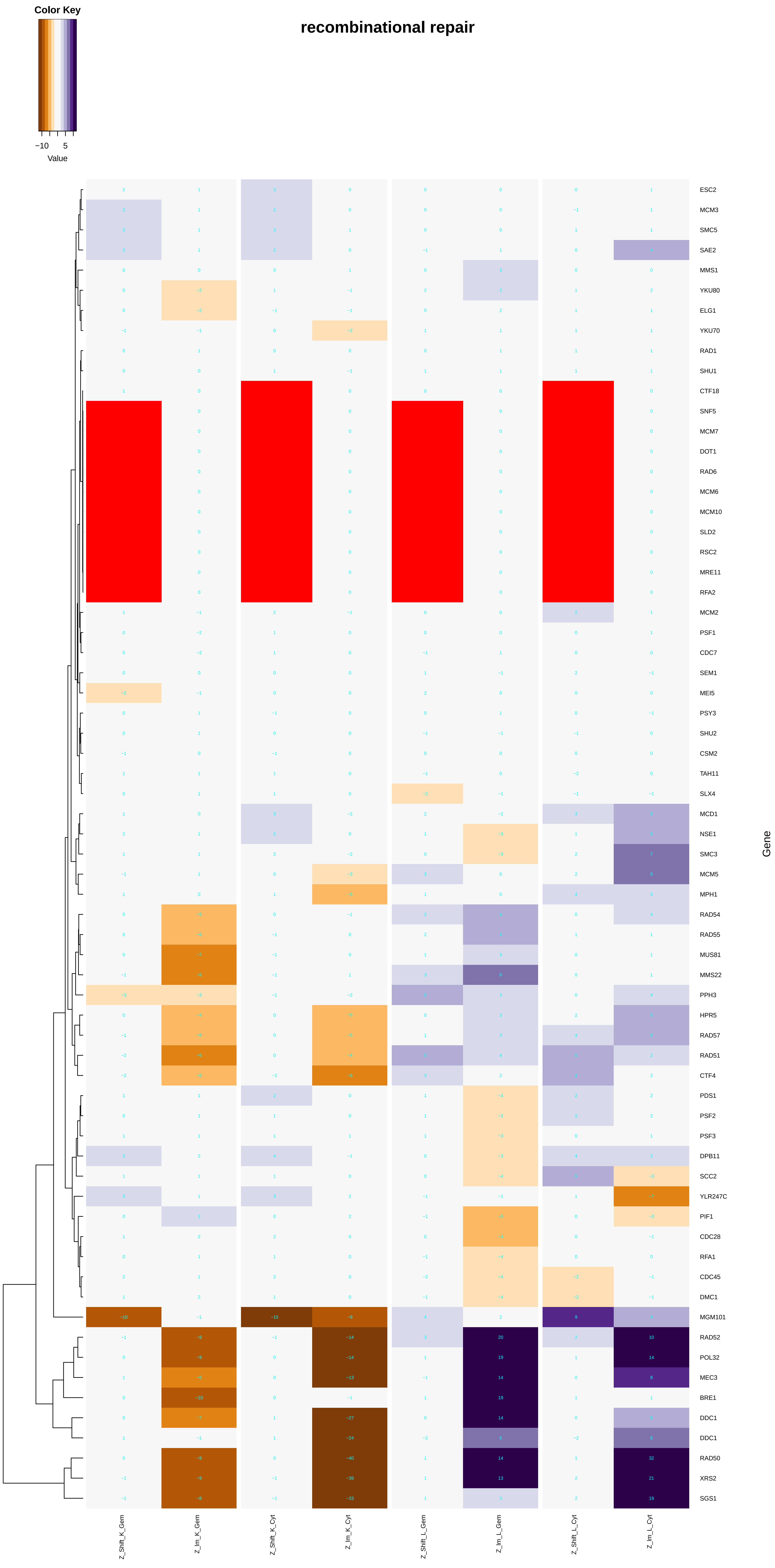

Color Key
−10
5
Value
recombinational repair
ESC2
MCM3
SMC5
SAE2
MMS1
YKU80
ELG1
YKU70
RAD1
SHU1
CTF18
SNF5
MCM7
DOT1
RAD6
MCM6
MCM10
SLD2
RSC2
MRE11
RFA2
MCM2
PSF1
CDC7
SEM1
MEI5
PSY3
SHU2
CSM2
TAH11
SLX4
MCD1
NSE1
SMC3
MCM5
MPH1
RAD54
RAD55
MUS81
MMS22
PPH3
HPR5
RAD57
RAD51
CTF4
PDS1
PSF2
PSF3
DPB11
SCC2
YLR247C
PIF1
CDC28
RFA1
CDC45
DMC1
MGM101
RAD52
POL32
MEC3
BRE1
DDC1
DDC1
RAD50
XRS2
SGS1
Gene
Z_Shift_K_Gem
Z_Im_K_Gem
Z_Shift_K_Cyt
Z_Im_K_Cyt
Z_Shift_L_Gem
Z_Im_L_Gem
Z_Shift_L_Cyt
Z_Im_L_Cyt

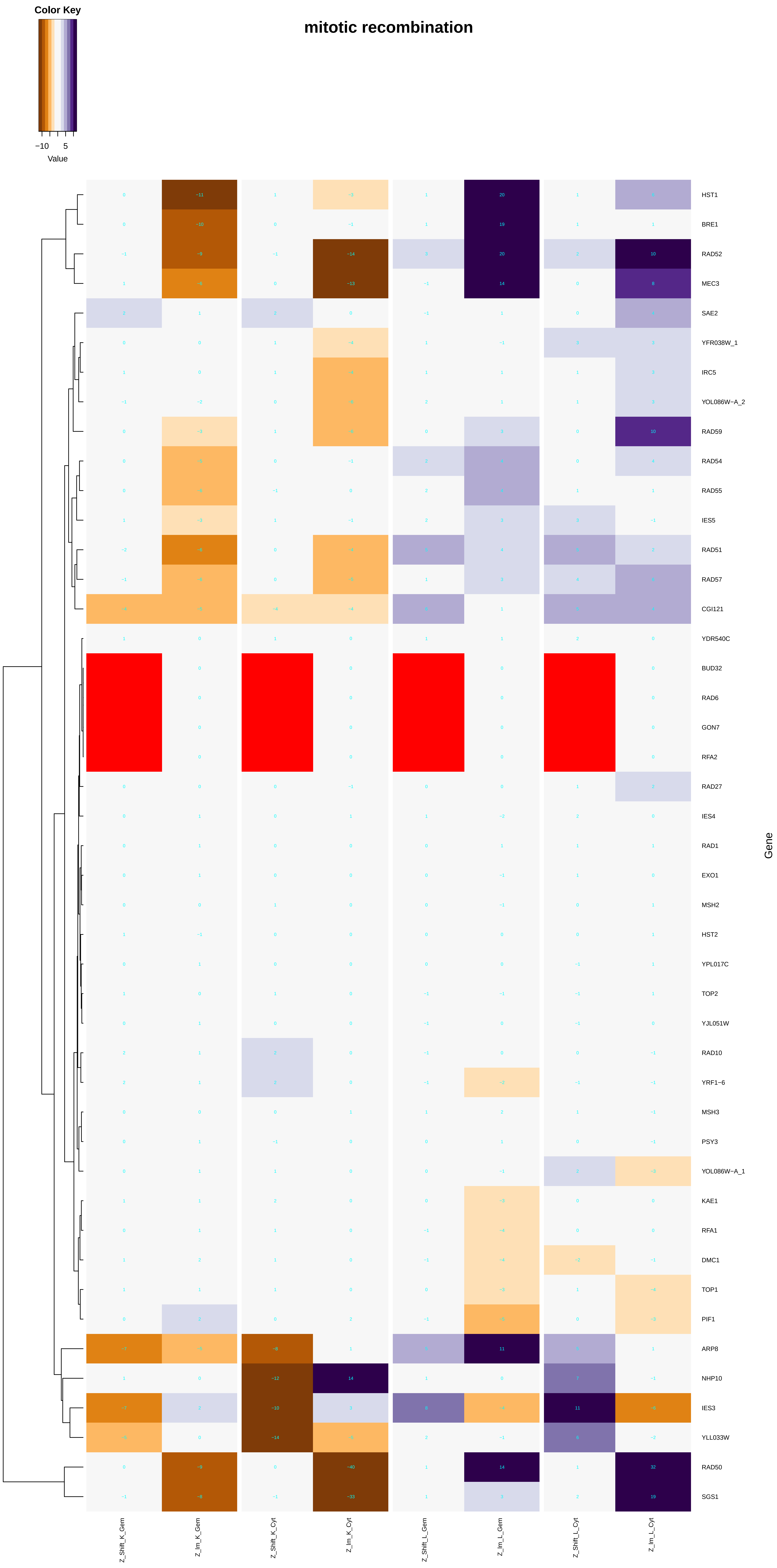
mitotic recombination
Color Key
−10
5
Value
HST1
BRE1
RAD52
MEC3
SAE2
YFR038W_1
IRC5
YOL086W−A_2
RAD59
RAD54
RAD55
IES5
RAD51
RAD57
CGI121
YDR540C
BUD32
RAD6
GON7
RFA2
RAD27
IES4
RAD1
EXO1
MSH2
HST2
YPL017C
TOP2
YJL051W
RAD10
YRF1−6
MSH3
PSY3
YOL086W−A_1
KAE1
RFA1
DMC1
TOP1
PIF1
ARP8
NHP10
IES3
YLL033W
RAD50
SGS1
Gene
Z_Shift_K_Gem
Z_Im_K_Gem
Z_Shift_K_Cyt
Z_Im_K_Cyt
Z_Shift_L_Gem
Z_Im_L_Gem
Z_Shift_L_Cyt
Z_Im_L_Cyt

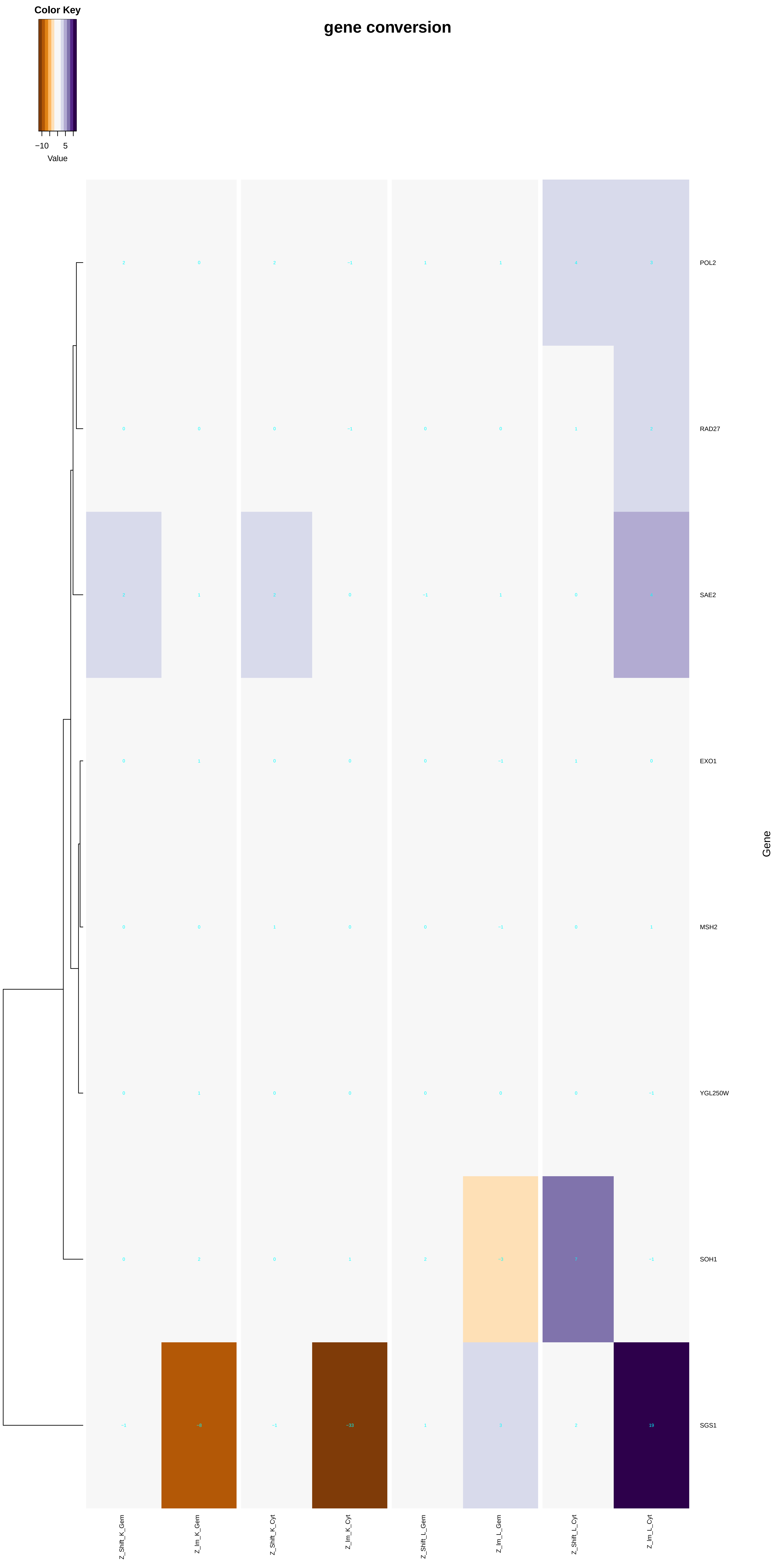
Color Key
−10
5
Value
gene conversion
POL2
RAD27
SAE2
EXO1
MSH2
YGL250W
SOH1
SGS1
Gene
Z_Shift_K_Gem
Z_Im_K_Gem
Z_Shift_K_Cyt
Z_Im_K_Cyt
Z_Shift_L_Gem
Z_Im_L_Gem
Z_Shift_L_Cyt
Z_Im_L_Cyt

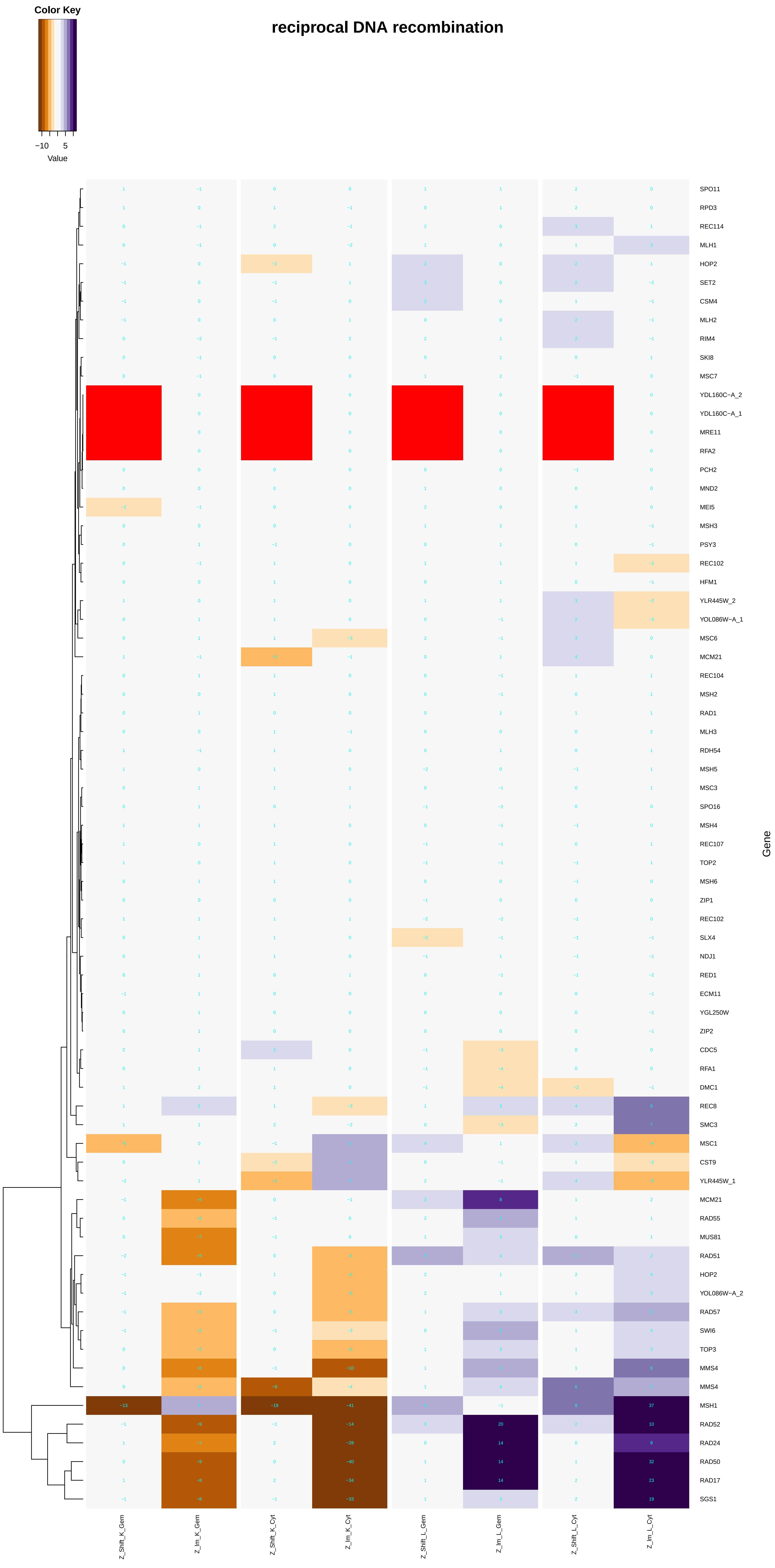
reciprocal DNA recombination
Color Key
−10
5
Value
SPO11
RPD3
REC114
MLH1
HOP2
SET2
CSM4
MLH2
RIM4
SKI8
MSC7
YDL160C−A_2
YDL160C−A_1
MRE11
RFA2
PCH2
MND2
MEI5
MSH3
PSY3
REC102
HFM1
YLR445W_2
YOL086W−A_1
MSC6
MCM21
REC104
MSH2
RAD1
MLH3
RDH54
MSH5
MSC3
SPO16
MSH4
REC107
TOP2
MSH6
ZIP1
REC102
SLX4
NDJ1
RED1
ECM11
YGL250W
ZIP2
CDC5
RFA1
DMC1
REC8
SMC3
MSC1
CST9
YLR445W_1
MCM21
RAD55
MUS81
RAD51
HOP2
YOL086W−A_2
RAD57
SWI6
TOP3
MMS4
MMS4
MSH1
RAD52
RAD24
RAD50
RAD17
SGS1
Gene
Z_Shift_K_Gem
Z_Im_K_Gem
Z_Shift_K_Cyt
Z_Im_K_Cyt
Z_Shift_L_Gem
Z_Im_L_Gem
Z_Shift_L_Cyt
Z_Im_L_Cyt

### double−strand break repair via homologous recombination

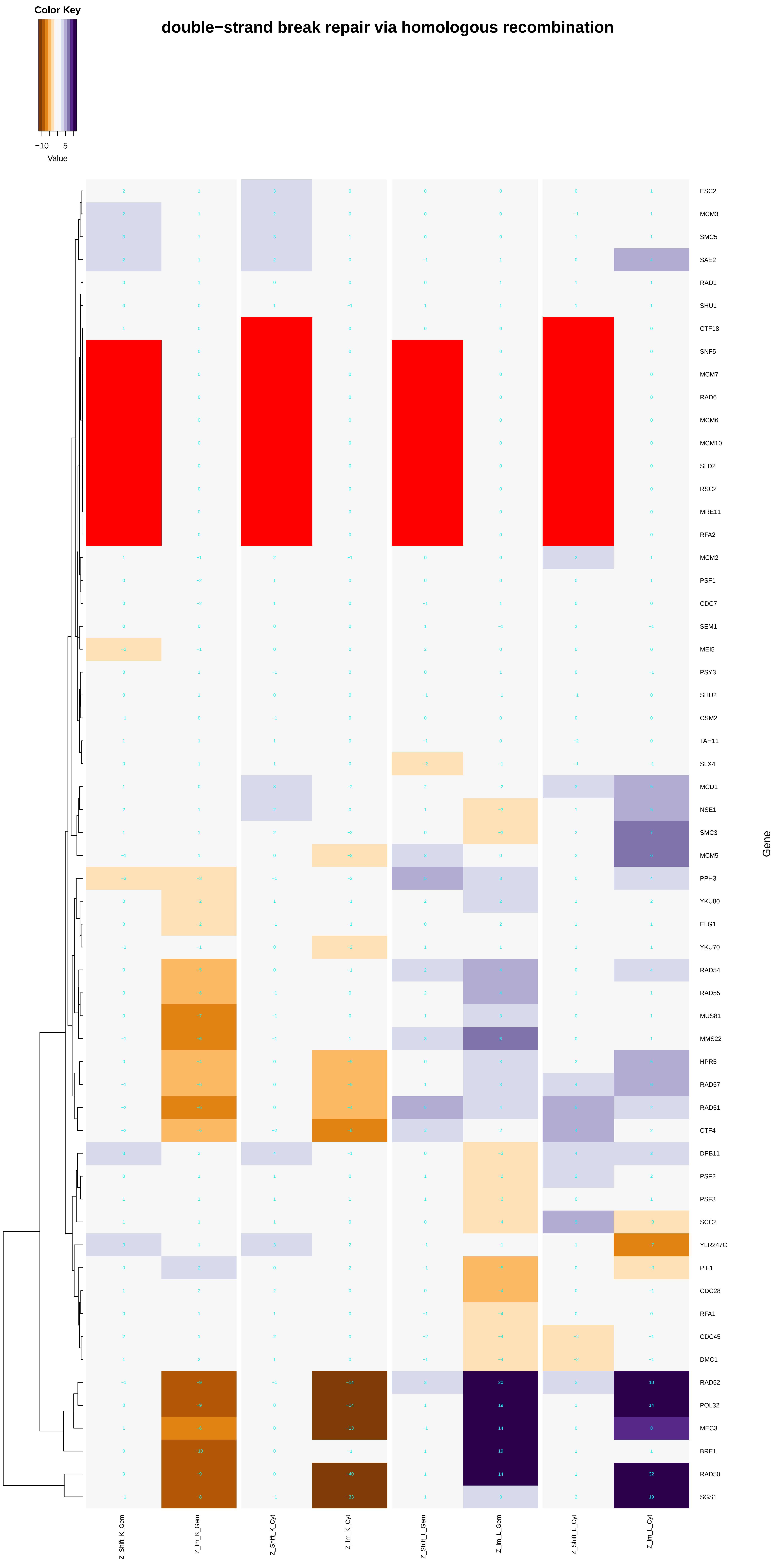

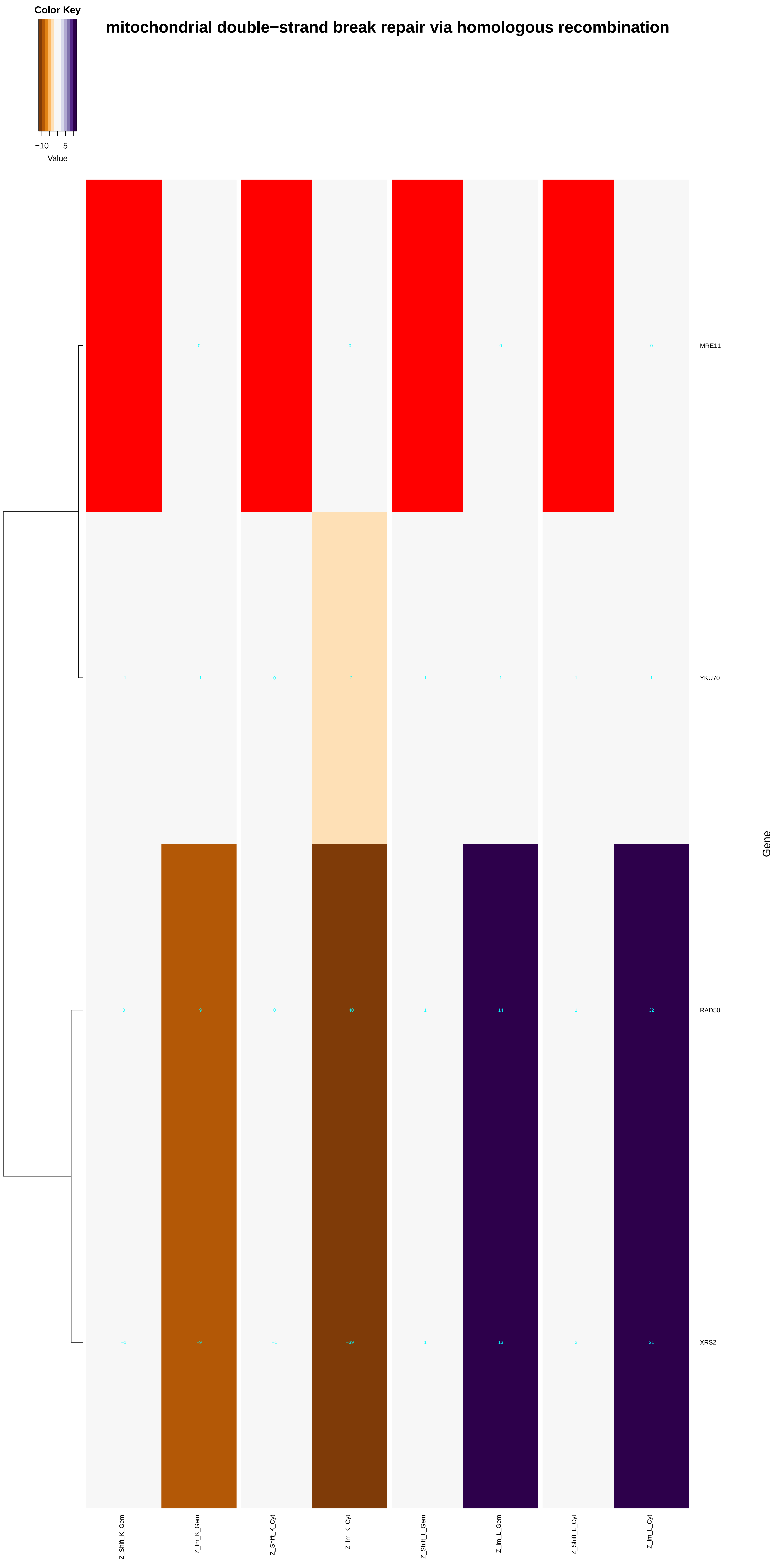
Color Key
−10
5
Value
mitochondrial double−strand break repair via homologous recombination
0
0
0
0
MRE11
−1
−1
0
−2
1
1
1
1
YKU70
0
−9
0
−40
1
14
1
32
RAD50
−1
−9
−1
−39
1
13
2
21
XRS2
Gene
Z_Shift_K_Gem
Z_Im_K_Gem
Z_Shift_K_Cyt
Z_Im_K_Cyt
Z_Shift_L_Gem
Z_Im_L_Gem
Z_Shift_L_Cyt
Z_Im_L_Cyt

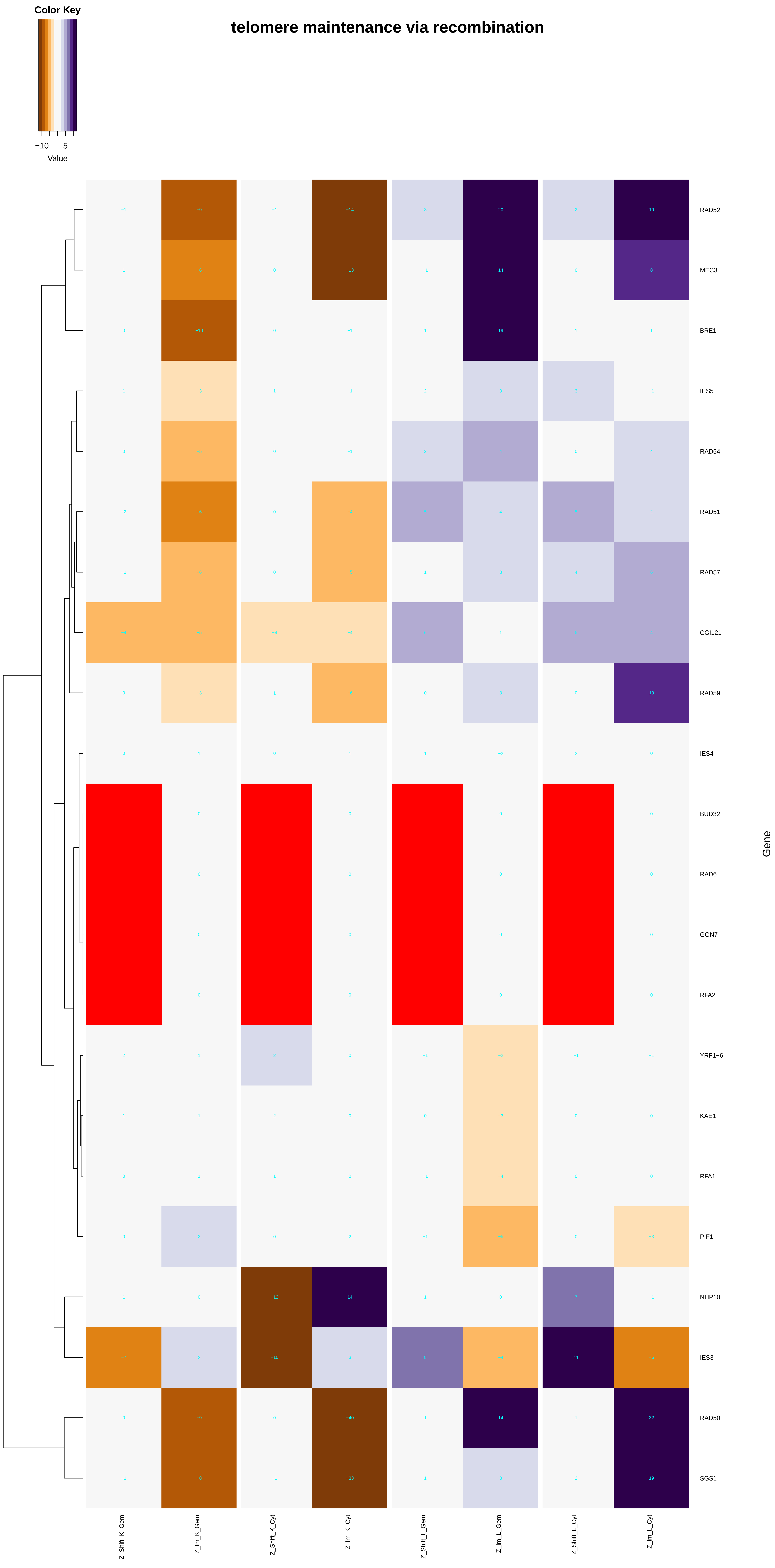

Color Key
−10
5
Value
telomere maintenance via recombination
RAD52
MEC3
BRE1
IES5
RAD54
RAD51
RAD57
CGI121
RAD59
IES4
BUD32
RAD6
GON7
RFA2
YRF1−6
KAE1
RFA1
PIF1
NHP10
IES3
RAD50
SGS1
Gene
Z_Shift_K_Gem
Z_Im_K_Gem
Z_Shift_K_Cyt
Z_Im_K_Cyt
Z_Shift_L_Gem
Z_Im_L_Gem
Z_Shift_L_Cyt
Z_Im_L_Cyt

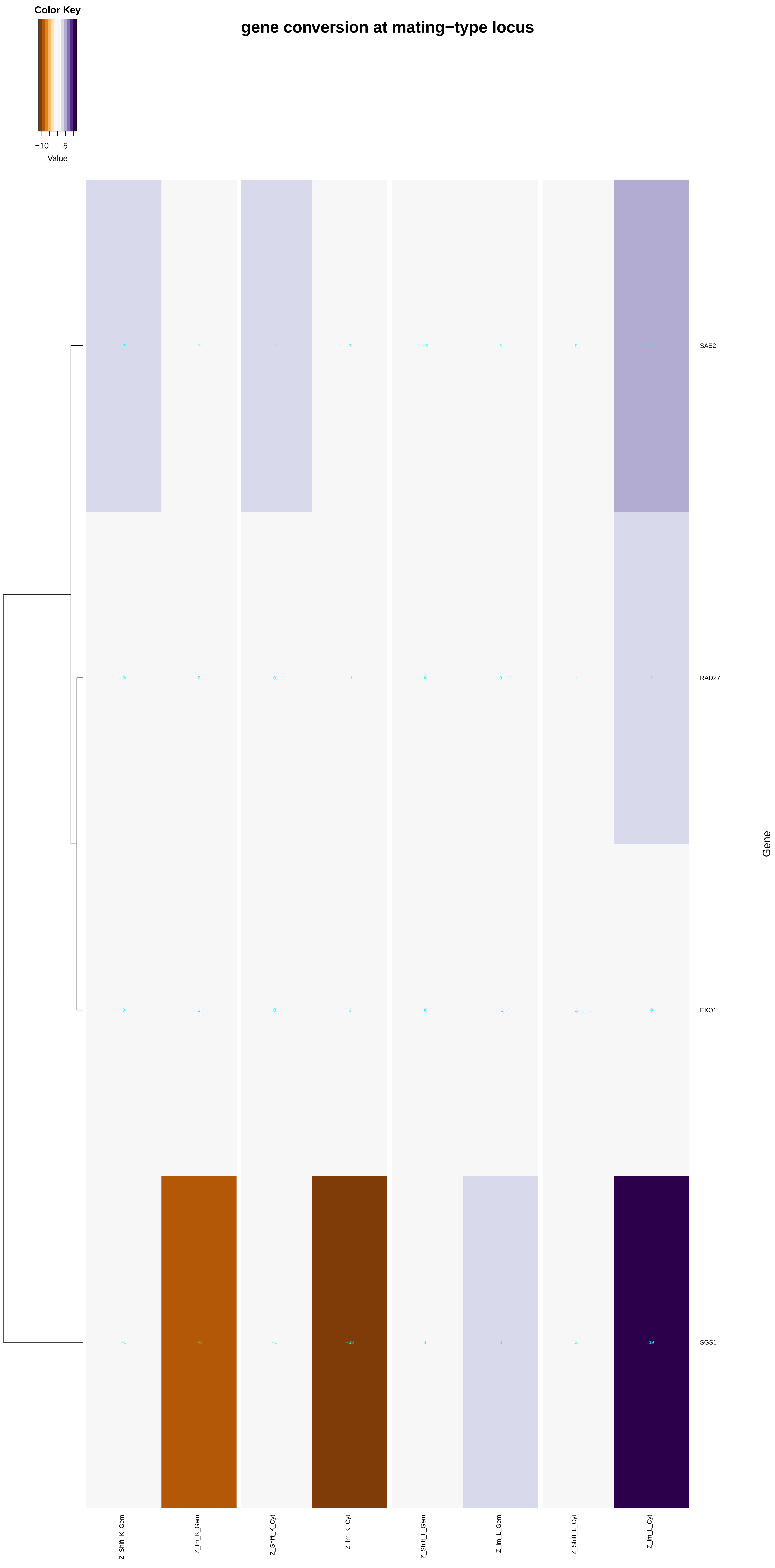

gene conversion at mating−type locus
Color Key
−10
5
Value
SAE2
RAD27
EXO1
SGS1
Gene
Z_Shift_K_Gem
Z_Im_K_Gem
Z_Shift_K_Cyt
Z_Im_K_Cyt
Z_Shift_L_Gem
Z_Im_L_Gem
Z_Shift_L_Cyt
Z_Im_L_Cyt

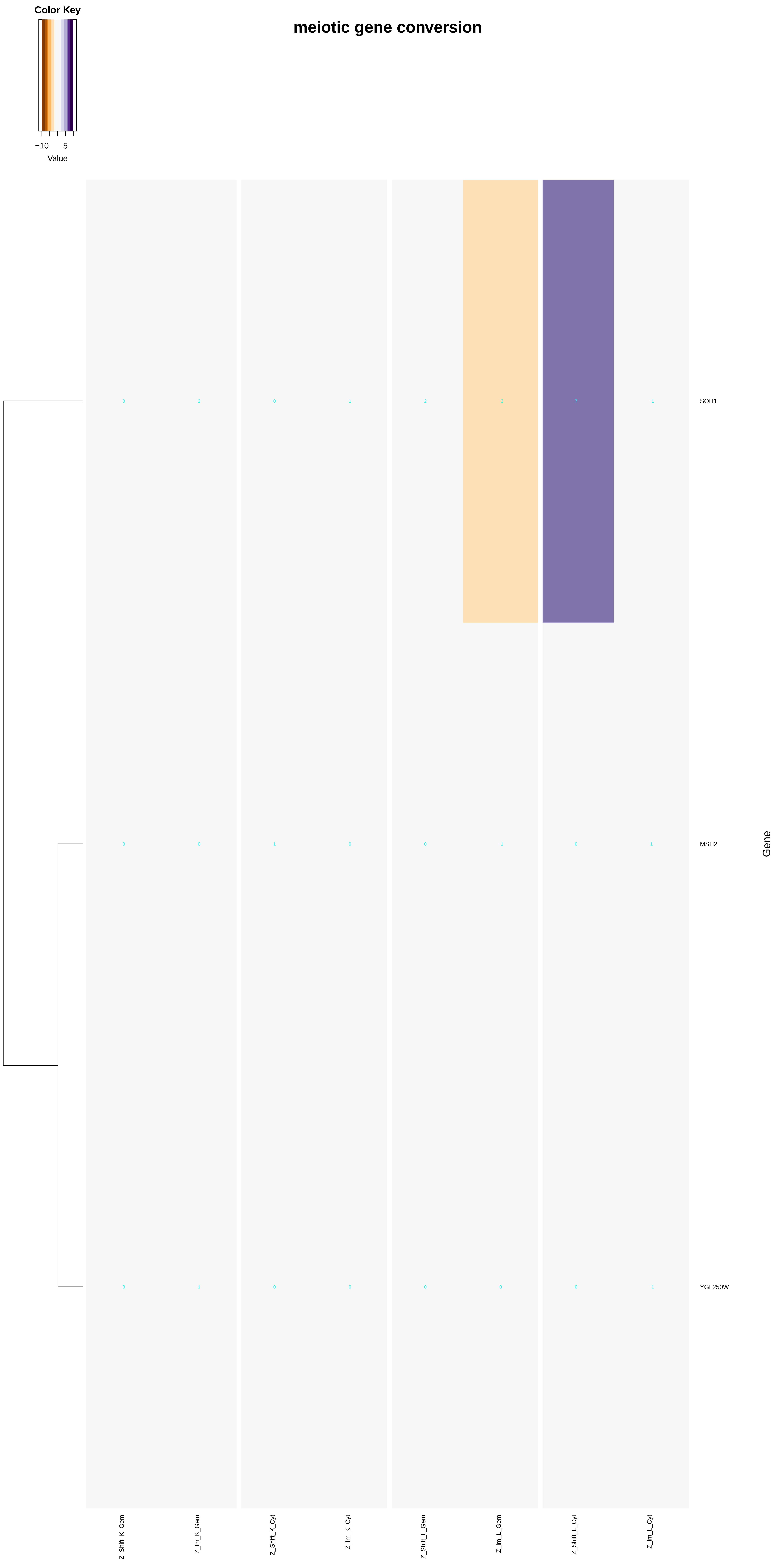
meiotic gene conversion
Color Key
−10
5
Value
SOH1
MSH2
YGL250W
Gene
Z_Shift_K_Gem
Z_Im_K_Gem
Z_Shift_K_Cyt
Z_Im_K_Cyt
Z_Shift_L_Gem
Z_Im_L_Gem
Z_Shift_L_Cyt
Z_Im_L_Cyt

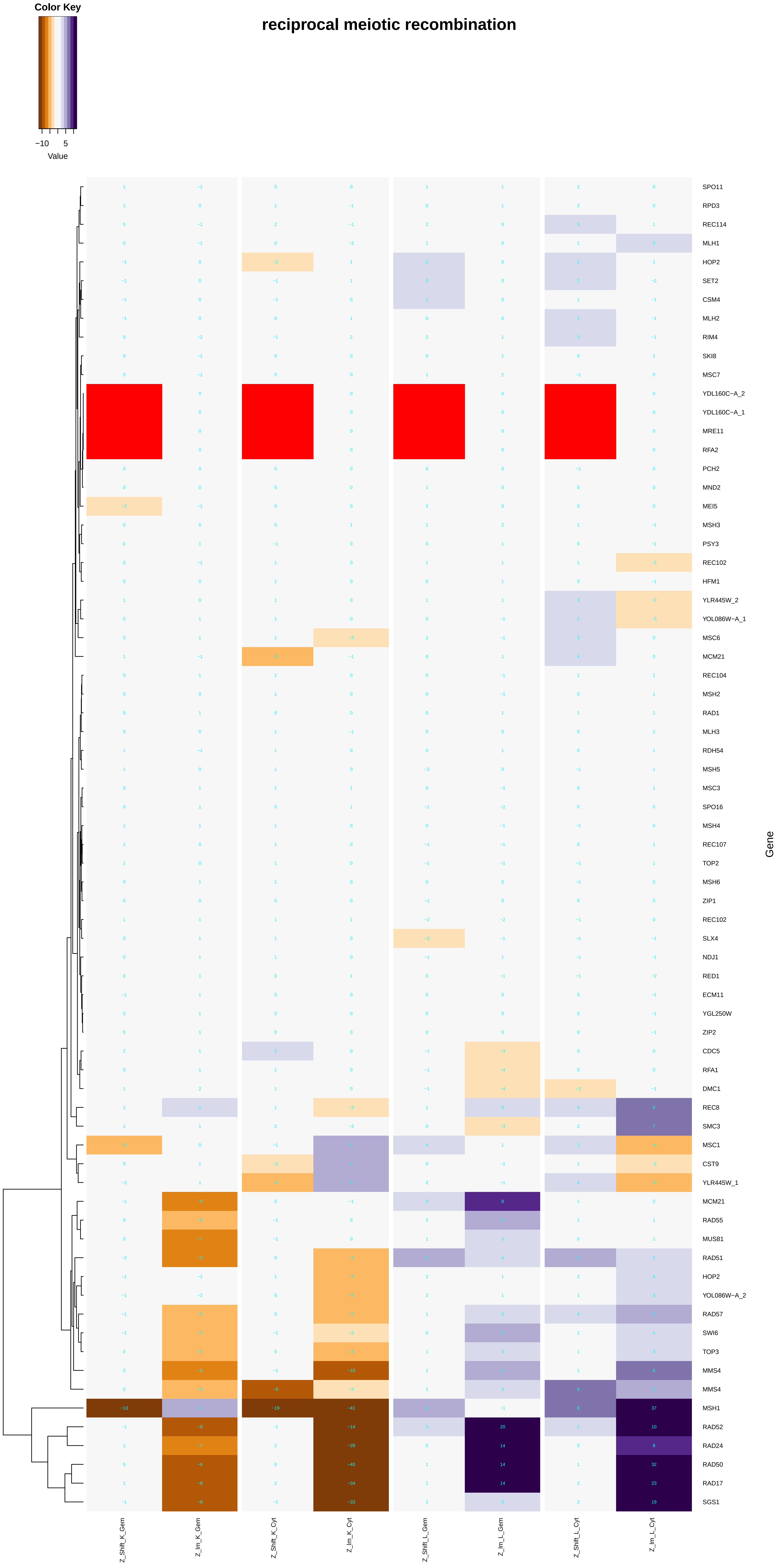

Color Key
−10 5
Value
reciprocal meiotic recombination
SPO11
RPD3
REC114
MLH1
HOP2
SET2
CSM4
MLH2
RIM4
SKI8
MSC7
YDL160C−A_2
YDL160C−A_1
MRE11
RFA2
PCH2
MND2
MEI5
MSH3
PSY3
REC102
HFM1
YLR445W_2
YOL086W−A_1
MSC6
MCM21
REC104
MSH2
RAD1
MLH3
RDH54
MSH5
MSC3
SPO16
MSH4
REC107
TOP2
MSH6
ZIP1
REC102
SLX4
NDJ1
RED1
ECM11
YGL250W
ZIP2
CDC5
RFA1
DMC1
REC8
SMC3
MSC1
CST9
YLR445W_1
MCM21
RAD55
MUS81
RAD51
HOP2
YOL086W−A_2
RAD57
SWI6
TOP3
MMS4
MMS4
MSH1
RAD52
RAD24
RAD50
RAD17
SGS1
Gene
Z_Shift_K_Gem
Z_Im_K_Gem
Z_Shift_K_Cyt
Z_Im_K_Cyt
Z_Shift_L_Gem
Z_Im_L_Gem
Z_Shift_L_Cyt
Z_Im_L_Cyt

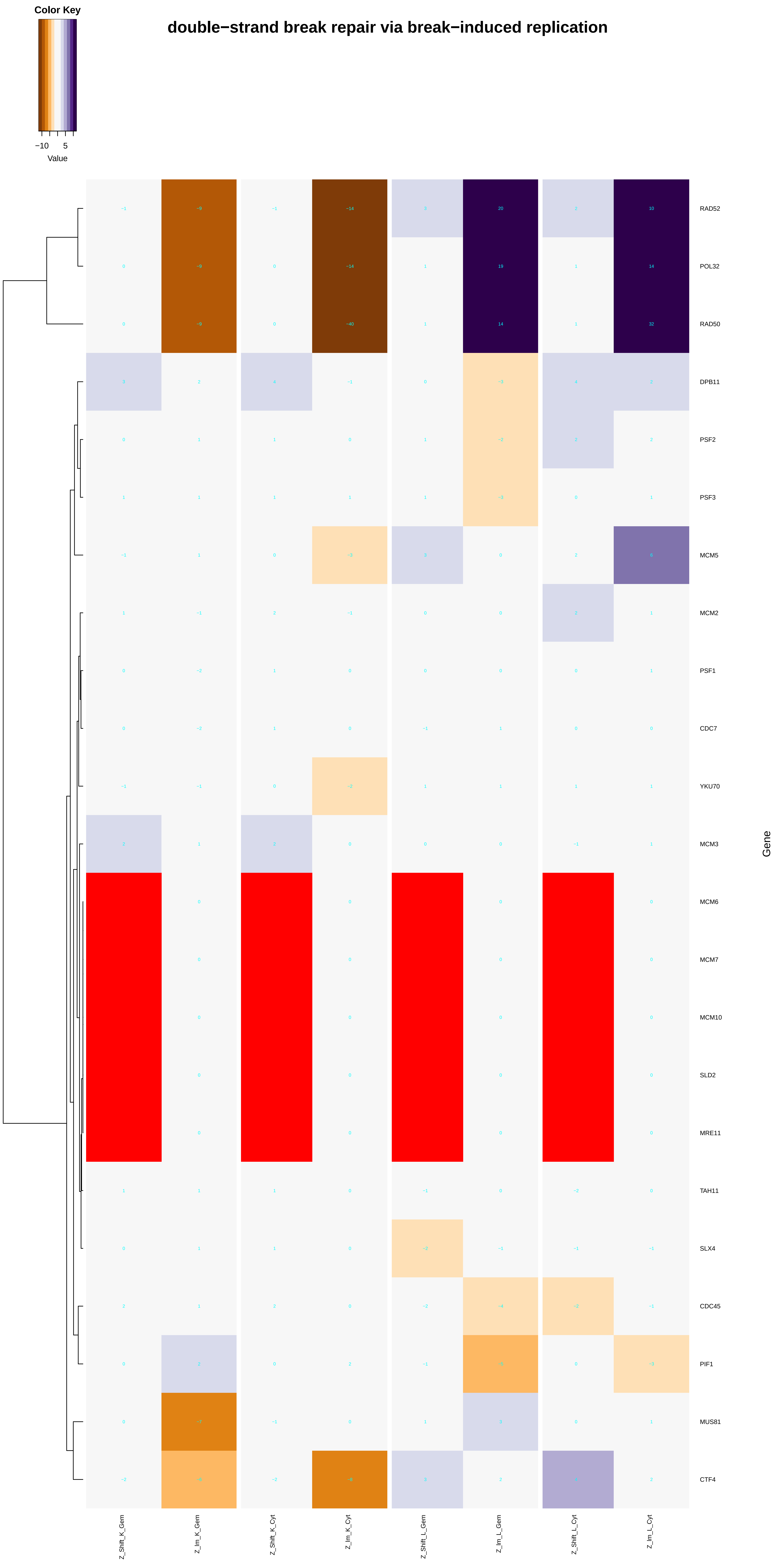

double−strand break repair via break−induced replication
Color Key
−10
5
Value
RAD52
POL32
RAD50
DPB11
PSF2
PSF3
MCM5
MCM2
PSF1
CDC7
YKU70
MCM3
MCM6
MCM7
MCM10
SLD2
MRE11
TAH11
SLX4
CDC45
PIF1
MUS81
CTF4
Gene
Z_Shift_K_Gem
Z_Im_K_Gem
Z_Shift_K_Cyt
Z_Im_K_Cyt
Z_Shift_L_Gem
Z_Im_L_Gem
Z_Shift_L_Cyt
Z_Im_L_Cyt

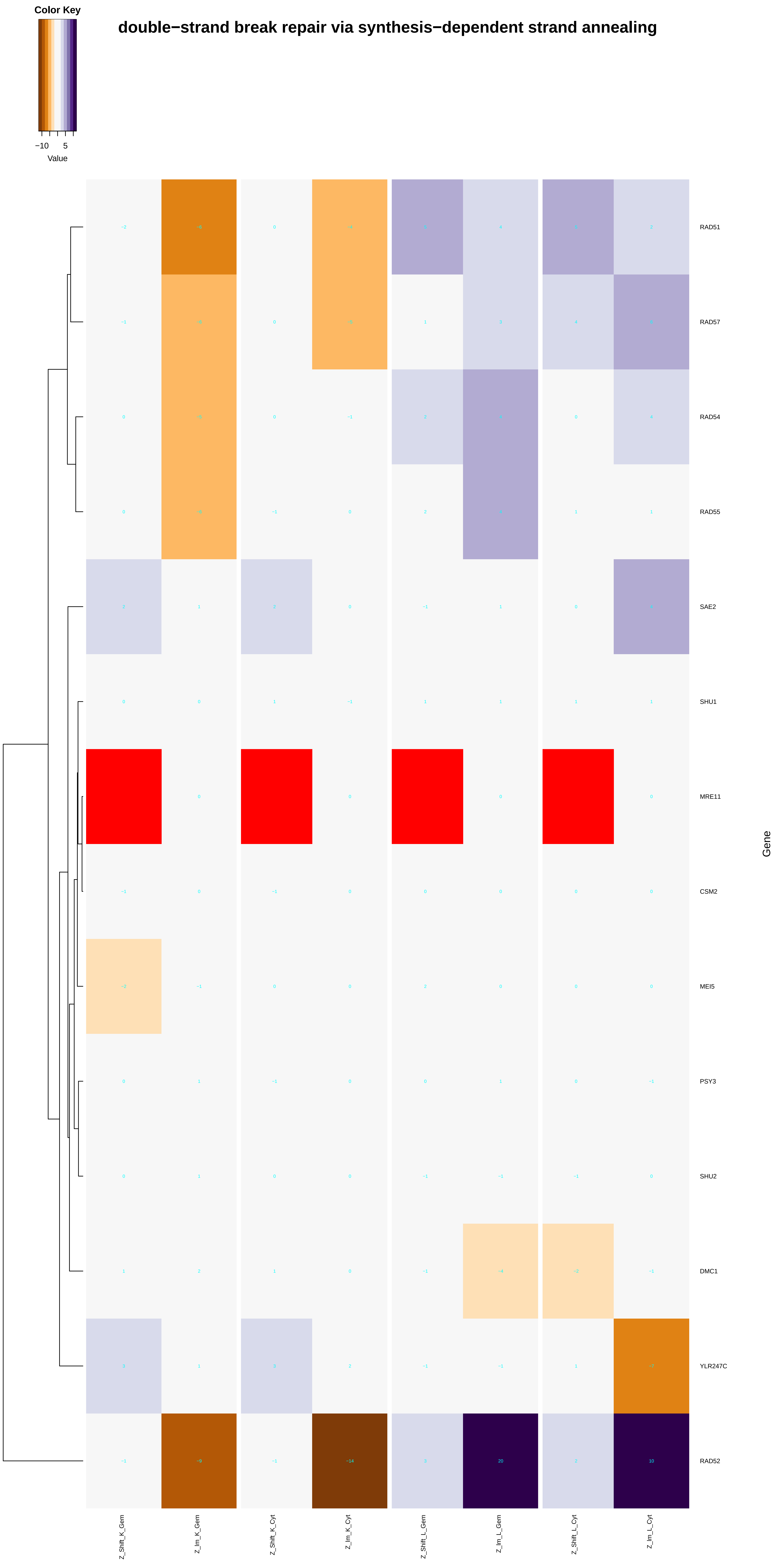

Color Key
−10
5
Value
double−strand break repair via synthesis−dependent strand annealing
RAD51
RAD57
RAD54
RAD55
SAE2
SHU1
MRE11
CSM2
MEI5
PSY3
SHU2
DMC1
YLR247C
RAD52
Gene
Z_Shift_K_Gem
Z_Im_K_Gem
Z_Shift_K_Cyt
Z_Im_K_Cyt
Z_Shift_L_Gem
Z_Im_L_Gem
Z_Shift_L_Cyt
Z_Im_L_Cyt

#### replication−born double−strand break repair via sister chromatid exchange

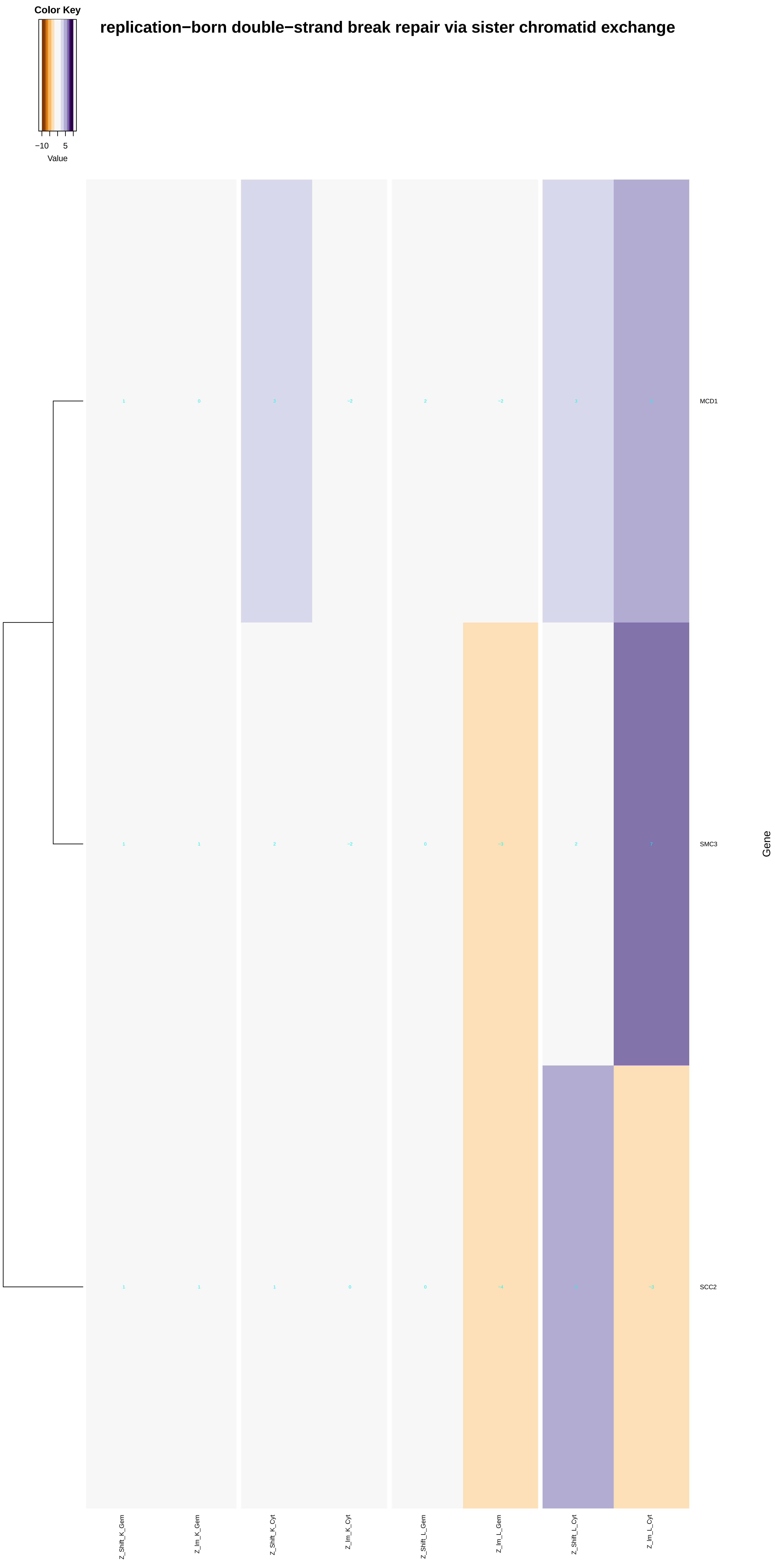
