## Additional File 7 for "A humanized yeast phenomic model of deoxycytidine kinase to predict genetic buffering of nucleoside analog cytotoxicity": DNA_repair.pdf

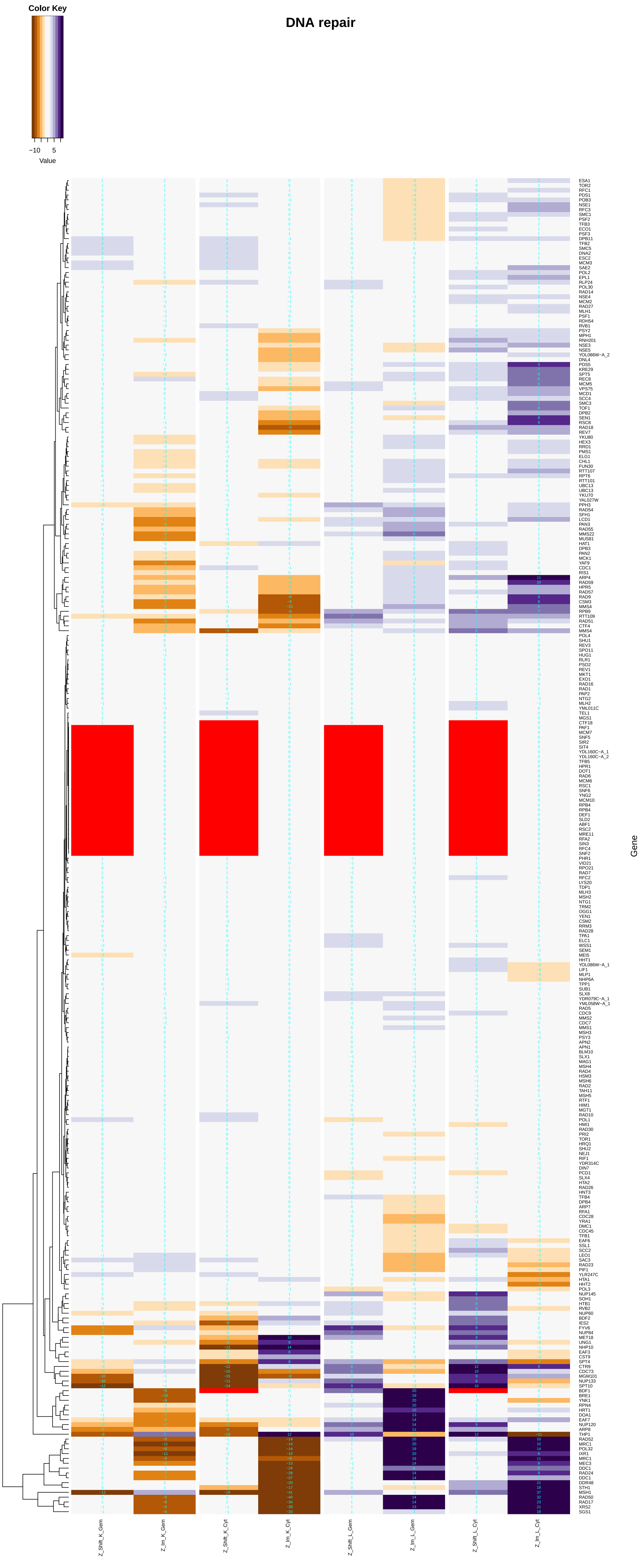

Color Key
−10 5
Value
Z_Shift_K_Gem
Z_Im_K_Gem
Z_Shift_K_Cyt
Z_Im_K_Cyt
Z_Shift_L_Gem
Z_Im_L_Gem
Z_Shift_L_Cyt
Z_Im_L_Cyt
Gene

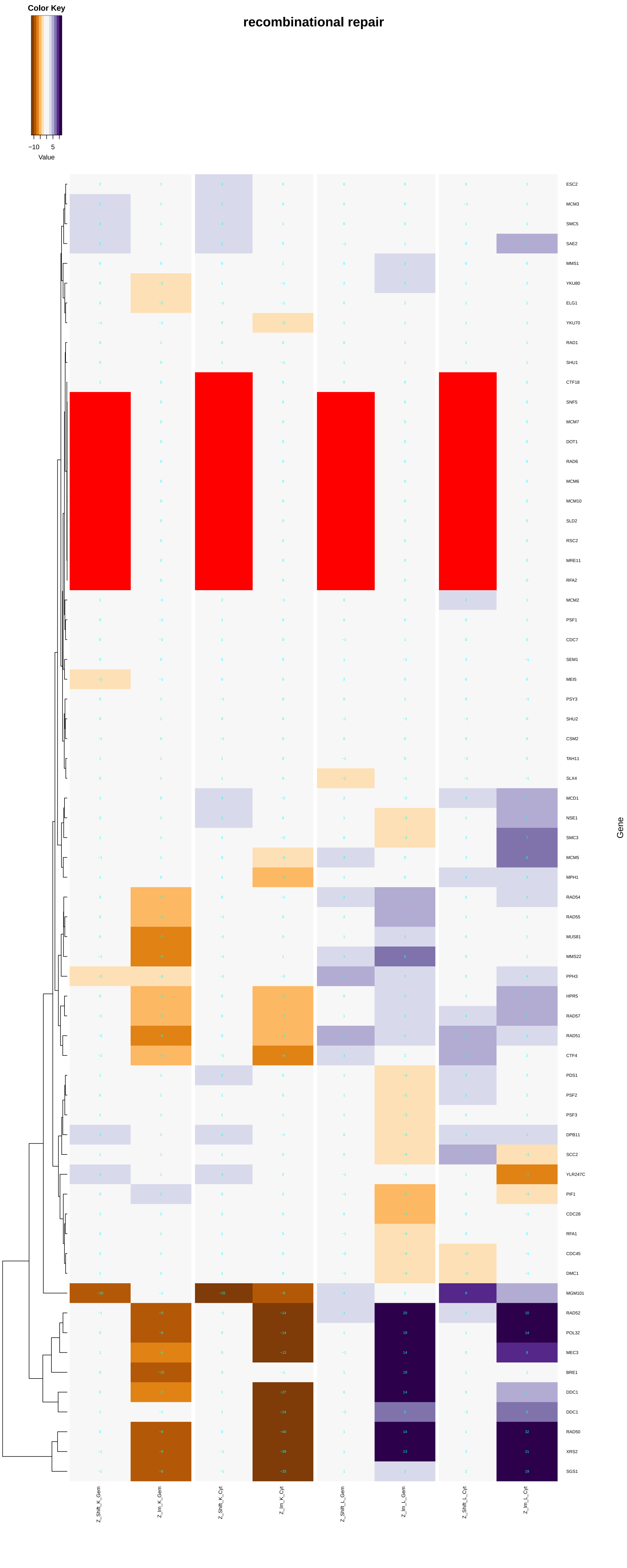

Color Key
−10
5
Value
recombinational repair
ESC2
MCM3
SMC5
SAE2
MMS1
YKU80
ELG1
YKU70
RAD1
SHU1
CTF18
SNF5
MCM7
DOT1
RAD6
MCM6
MCM10
SLD2
RSC2
MRE11
RFA2
MCM2
PSF1
CDC7
SEM1
MEI5
PSY3
SHU2
CSM2
TAH11
SLX4
MCD1
NSE1
SMC3
MCM5
MPH1
RAD54
RAD55
MUS81
MMS22
PPH3
HPR5
RAD57
RAD51
CTF4
PDS1
PSF2
PSF3
DPB11
SCC2
YLR247C
PIF1
CDC28
RFA1
CDC45
DMC1
MGM101
RAD52
POL32
MEC3
BRE1
DDC1
DDC1
RAD50
XRS2
SGS1
Gene
Z_Shift_K_Gem
Z_Im_K_Gem
Z_Shift_K_Cyt
Z_Im_K_Cyt
Z_Shift_L_Gem
Z_Im_L_Gem
Z_Shift_L_Cyt
Z_Im_L_Cyt

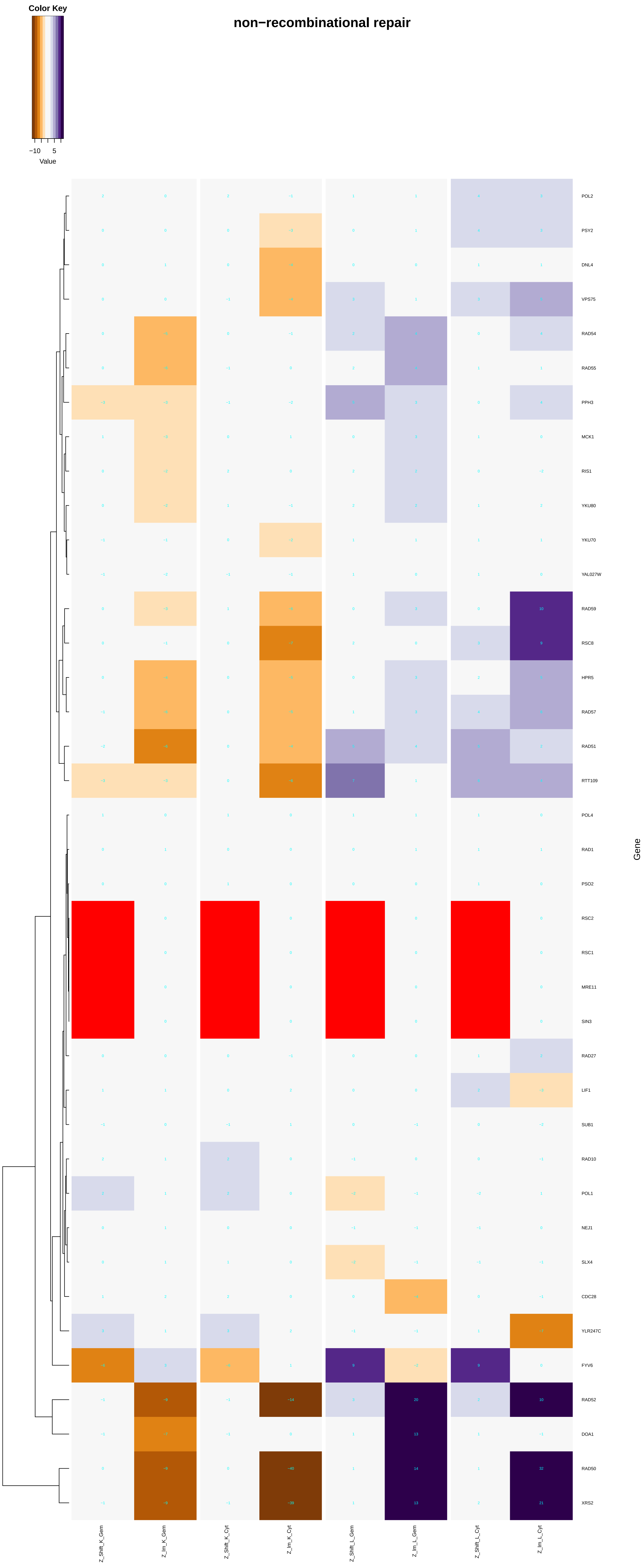
non−recombinational repair
Color Key
−10
5
Value
POL2
PSY2
DNL4
VPS75
RAD54
RAD55
PPH3
MCK1
RIS1
YKU80
YKU70
YAL027W
RAD59
RSC8
HPR5
RAD57
RAD51
RTT109
POL4
RAD1
PSO2
RSC2
RSC1
MRE11
SIN3
RAD27
LIF1
SUB1
RAD10
POL1
NEJ1
SLX4
CDC28
YLR247C
FYV6
RAD52
DOA1
RAD50
XRS2
Gene
Z_Shift_K_Gem
Z_Im_K_Gem
Z_Shift_K_Cyt
Z_Im_K_Cyt
Z_Shift_L_Gem
Z_Im_L_Gem
Z_Shift_L_Cyt
Z_Im_L_Cyt

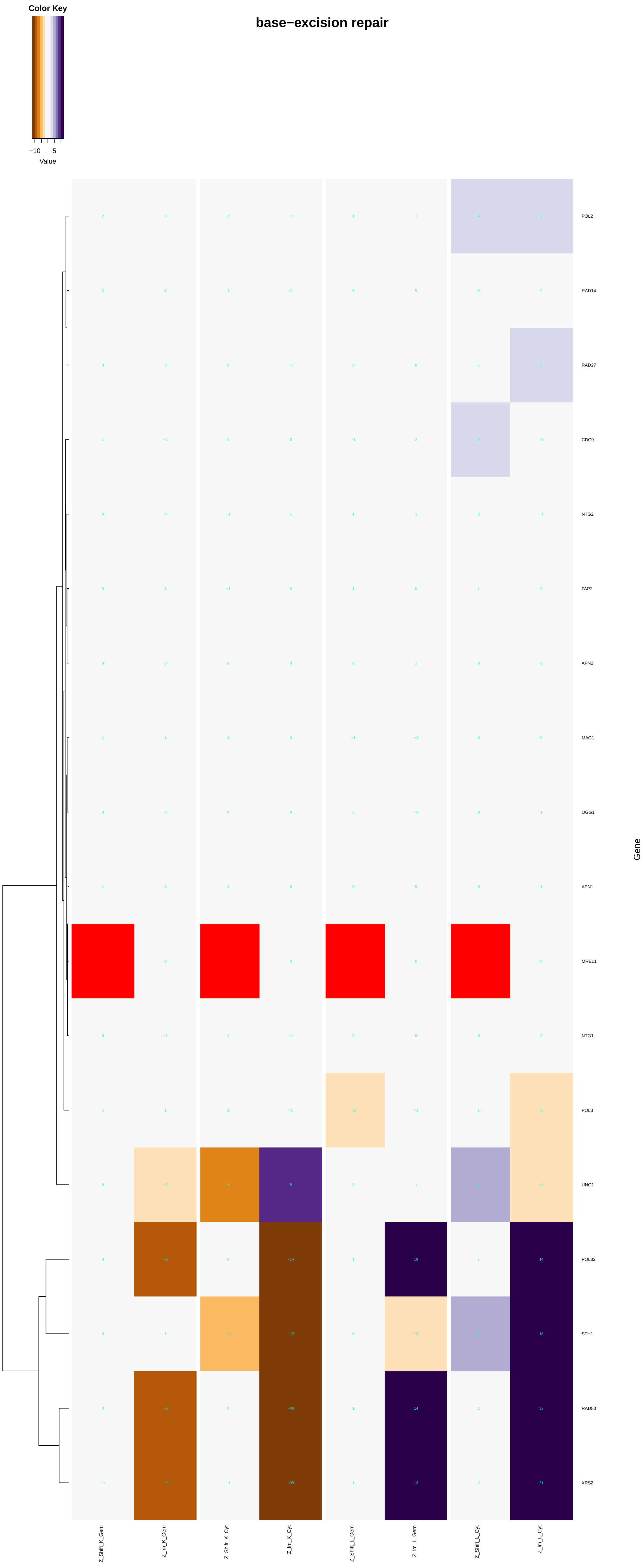

base−excision repair
Color Key
−10
5
Value
POL2
RAD14
RAD27
CDC9
NTG2
PAP2
APN2
MAG1
OGG1
APN1
MRE11
NTG1
POL3
UNG1
POL32
STH1
RAD50
XRS2
Gene
Z_Shift_K_Gem
Z_Im_K_Gem
Z_Shift_K_Cyt
Z_Im_K_Cyt
Z_Shift_L_Gem
Z_Im_L_Gem
Z_Shift_L_Cyt
Z_Im_L_Cyt

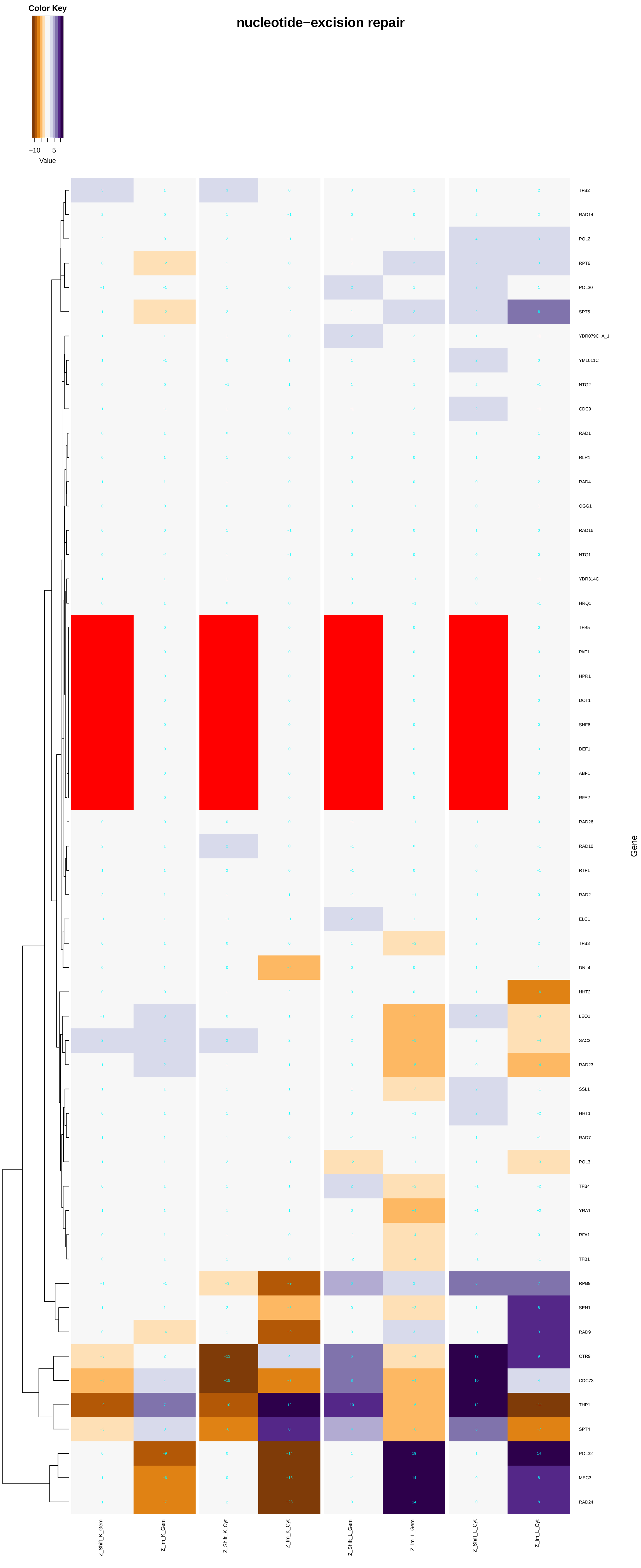

nucleotide−excision repair
Color Key
−10
5
Value
TFB2
RAD14
POL2
RPT6
POL30
SPT5
YDR079C−A_1
YML011C
NTG2
CDC9
RAD1
RLR1
RAD4
OGG1
RAD16
NTG1
YDR314C
HRQ1
TFB5
PAF1
HPR1
DOT1
SNF6
DEF1
ABF1
RFA2
RAD26
RAD10
RTF1
RAD2
ELC1
TFB3
DNL4
HHT2
LEO1
SAC3
RAD23
SSL1
HHT1
RAD7
POL3
TFB4
YRA1
RFA1
TFB1
RPB9
SEN1
RAD9
CTR9
CDC73
THP1
SPT4
POL32
MEC3
RAD24
Gene
Z_Shift_K_Gem
Z_Im_K_Gem
Z_Shift_K_Cyt
Z_Im_K_Cyt
Z_Shift_L_Gem
Z_Im_L_Gem
Z_Shift_L_Cyt
Z_Im_L_Cyt

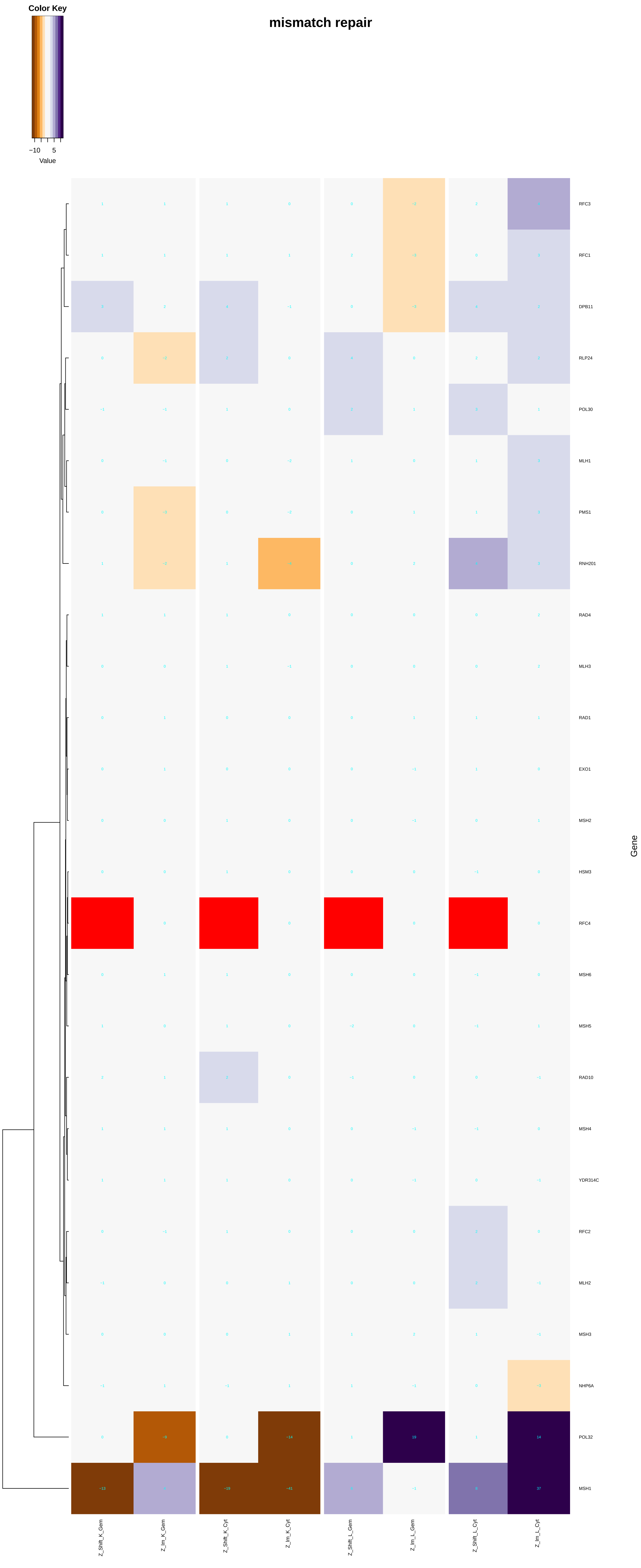

Color Key
−10
5
Value
mismatch repair
RFC3
RFC1
DPB11
RLP24
POL30
MLH1
PMS1
RNH201
RAD4
MLH3
RAD1
EXO1
MSH2
HSM3
RFC4
MSH6
MSH5
RAD10
MSH4
YDR314C
RFC2
MLH2
MSH3
NHP6A
POL32
MSH1
Gene
Z_Shift_K_Gem
Z_Im_K_Gem
Z_Shift_K_Cyt
Z_Im_K_Cyt
Z_Shift_L_Gem
Z_Im_L_Gem
Z_Shift_L_Cyt
Z_Im_L_Cyt

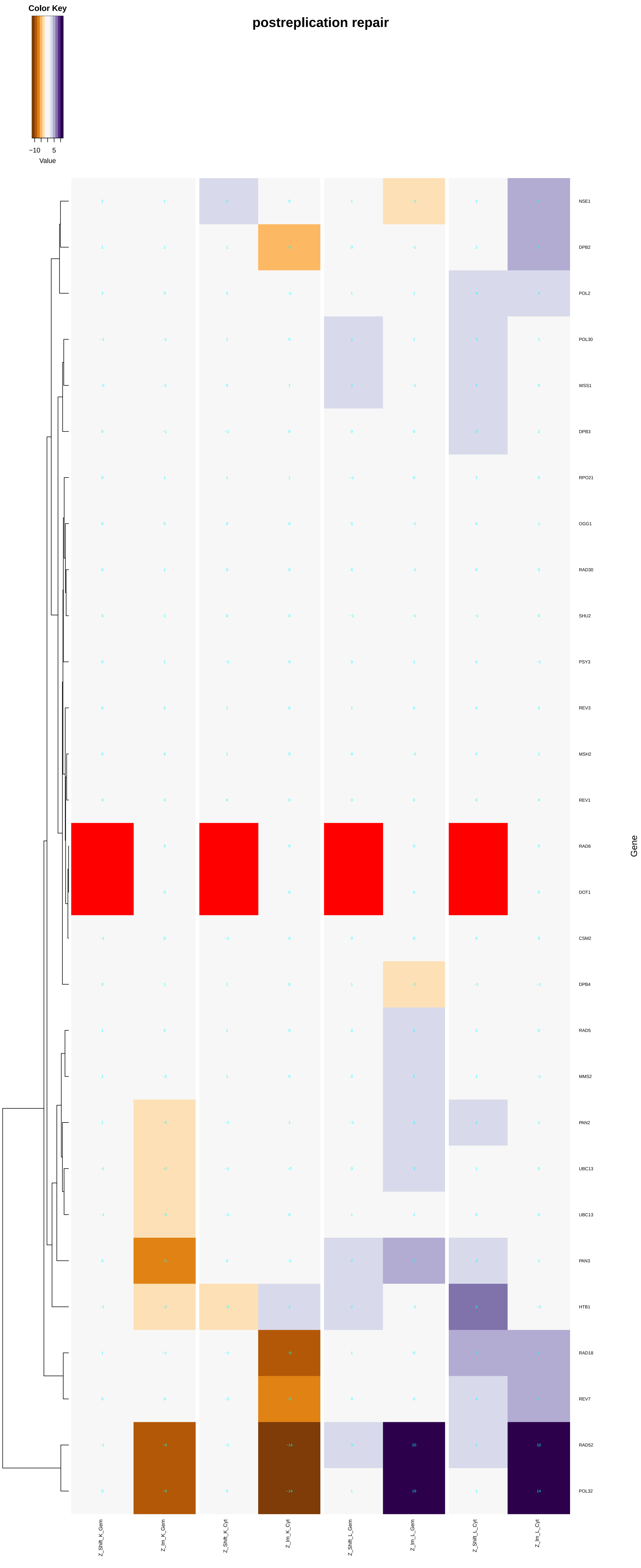
postreplication repair
Color Key
−10
5
Value
NSE1
DPB2
POL2
POL30
WSS1
DPB3
RPO21
OGG1
RAD30
SHU2
PSY3
REV3
MSH2
REV1
RAD6
DOT1
CSM2
DPB4
RAD5
MMS2
PAN2
UBC13
UBC13
PAN3
HTB1
RAD18
REV7
RAD52
POL32
Gene
Z_Shift_K_Gem
Z_Im_K_Gem
Z_Shift_K_Cyt
Z_Im_K_Cyt
Z_Shift_L_Gem
Z_Im_L_Gem
Z_Shift_L_Cyt
Z_Im_L_Cyt

double−strand break repair

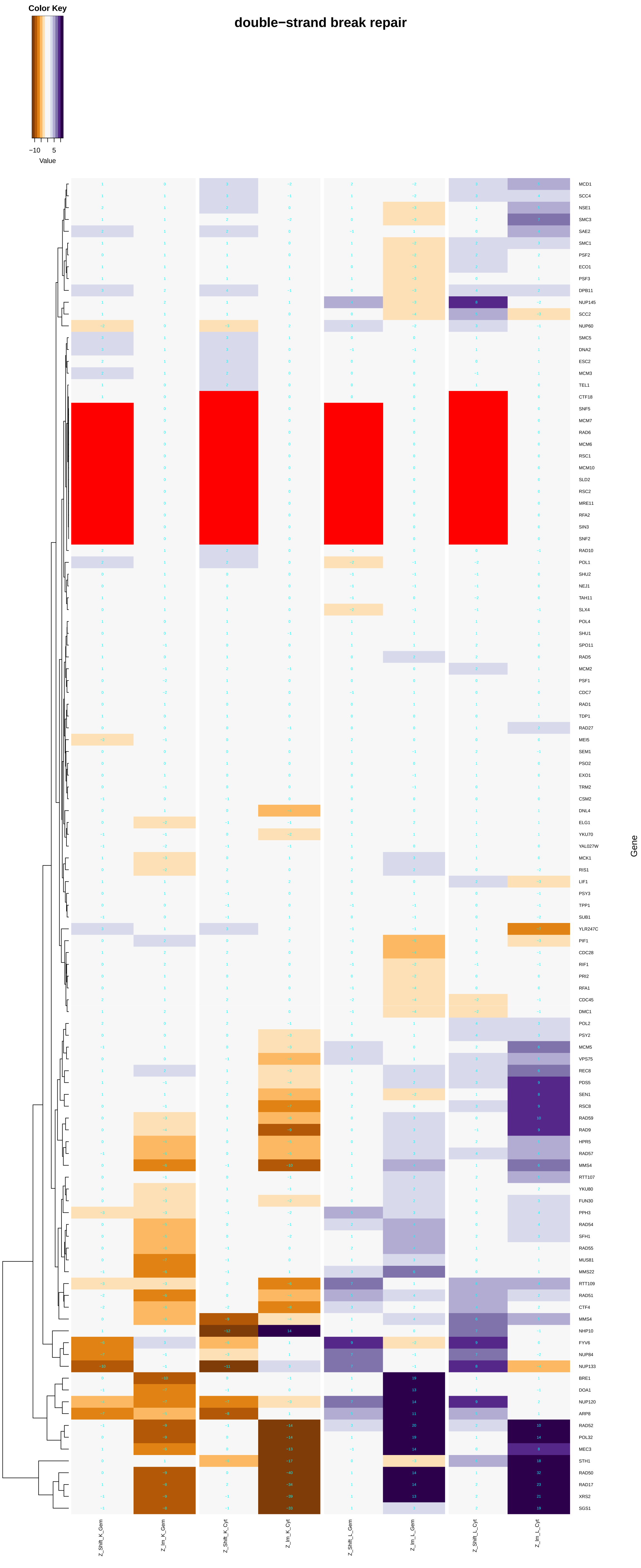

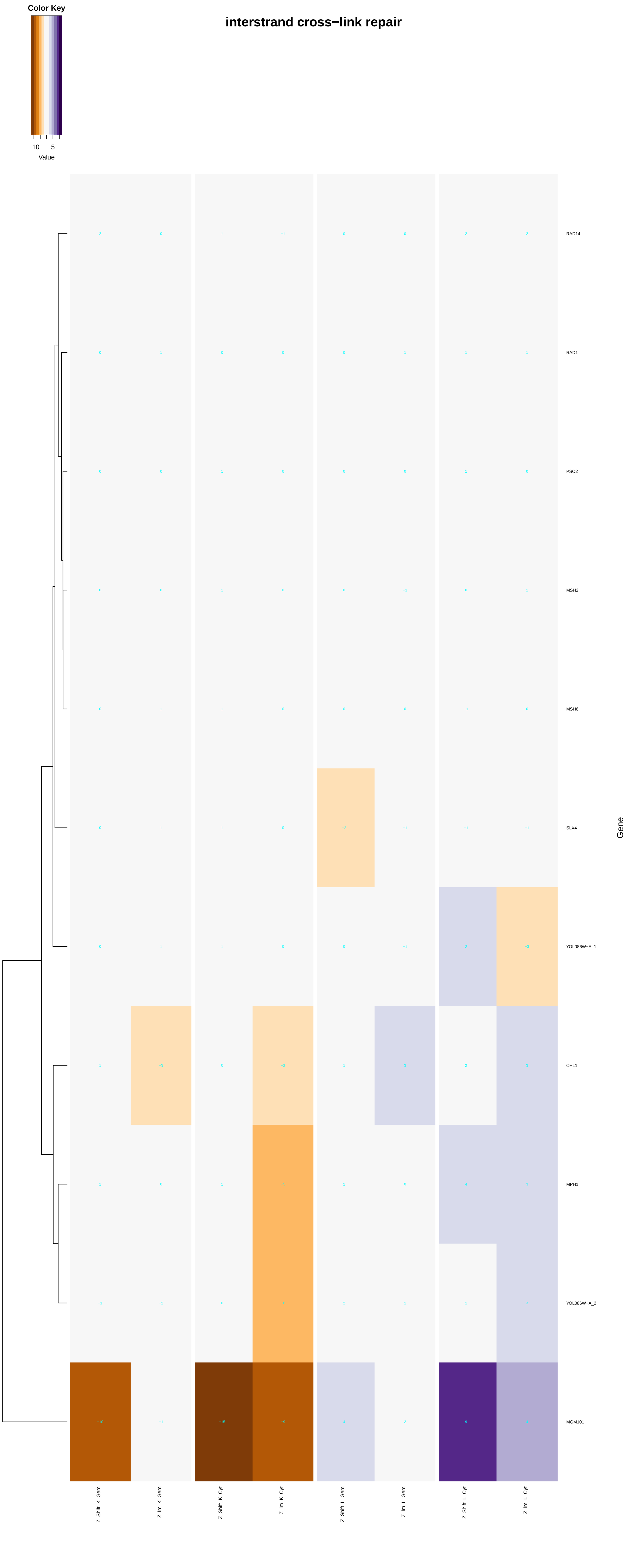
Color Key
−10
5
Value
interstrand cross−link repair
RAD14
RAD1
PSO2
MSH2
MSH6
SLX4
YOL086W−A_1
CHL1
MPH1
YOL086W−A_2
MGM101
Gene
Z_Shift_K_Gem
Z_Im_K_Gem
Z_Shift_K_Cyt
Z_Im_K_Cyt
Z_Shift_L_Gem
Z_Im_L_Gem
Z_Shift_L_Cyt
Z_Im_L_Cyt

### UV−damage excision repair

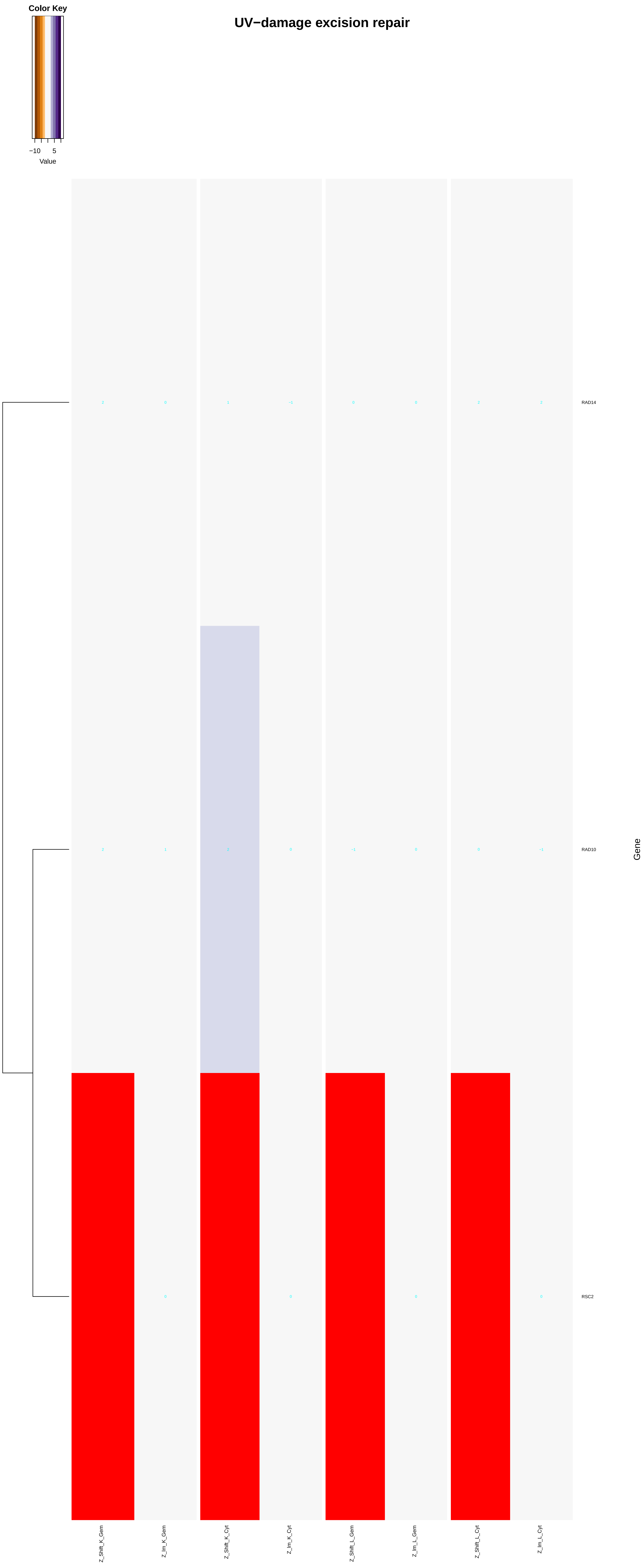

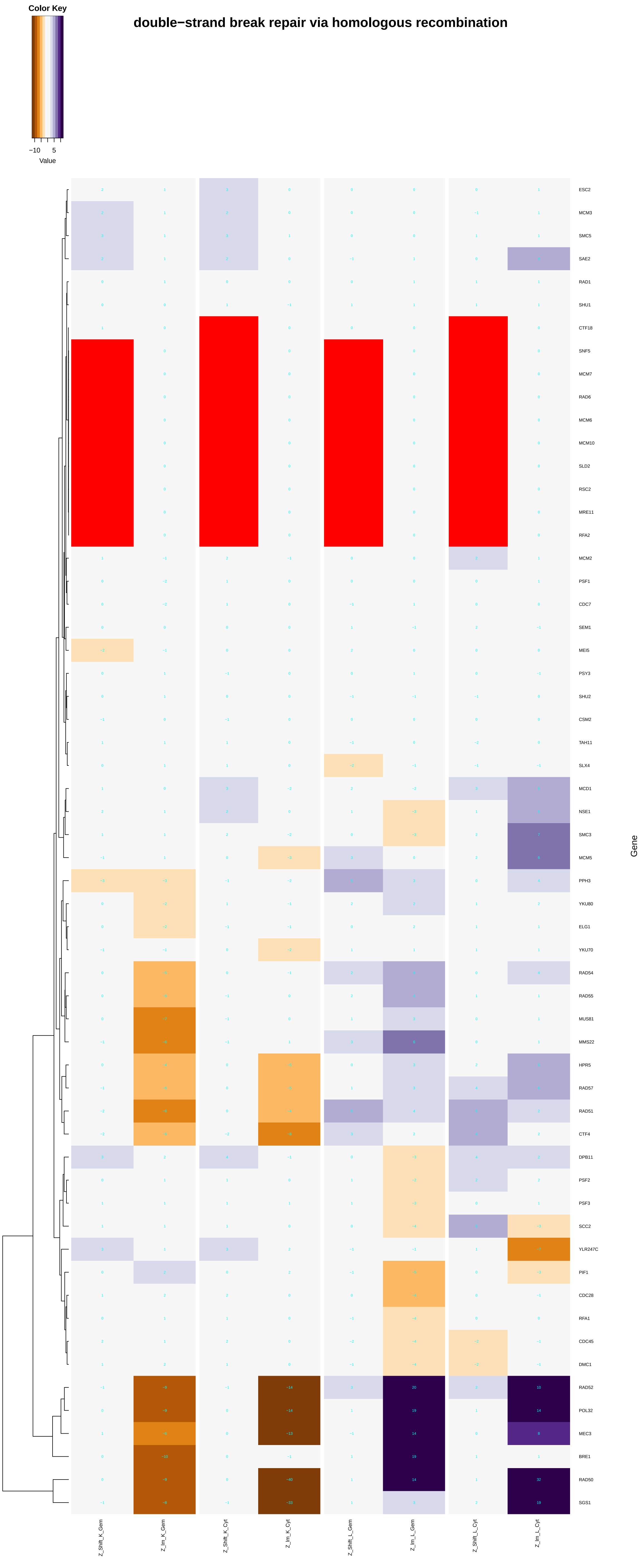
double−strand break repair via homologous recombination
Color Key
−10
5
Value
Gene
ESC2
MCM3
SMC5
SAE2
RAD1
SHU1
CTF18
SNF5
MCM7
RAD6
MCM6
MCM10
SLD2
RSC2
MRE11
RFA2
MCM2
PSF1
CDC7
SEM1
MEI5
PSY3
SHU2
CSM2
TAH11
SLX4
MCD1
NSE1
SMC3
MCM5
PPH3
YKU80
ELG1
YKU70
RAD54
RAD55
MUS81
MMS22
HPR5
RAD57
RAD51
CTF4
DPB11
PSF2
PSF3
SCC2
YLR247C
PIF1
CDC28
RFA1
CDC45
DMC1
RAD52
POL32
MEC3
BRE1
RAD50
SGS1
Z_Shift_K_Gem
Z_Im_K_Gem
Z_Shift_K_Cyt
Z_Im_K_Cyt
Z_Shift_L_Gem
Z_Im_L_Gem
Z_Shift_L_Cyt
Z_Im_L_Cyt

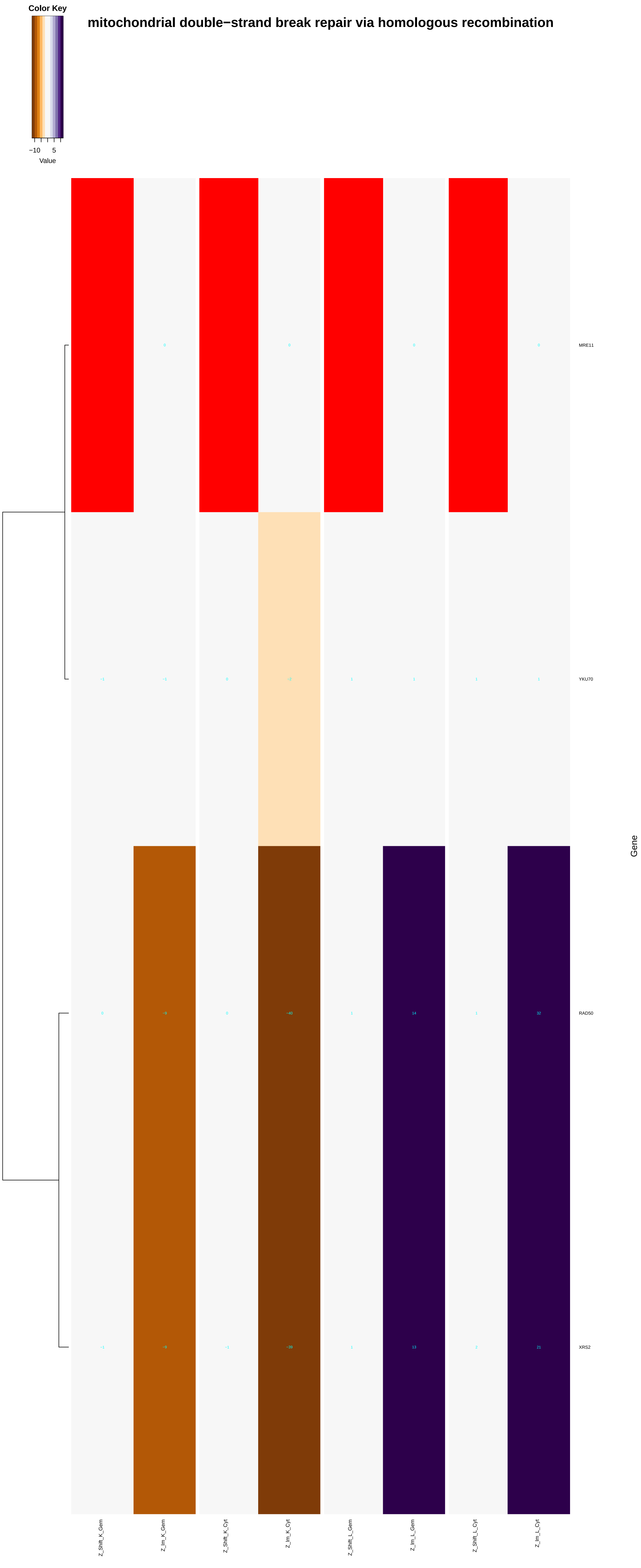
mitochondrial double−strand break repair via homologous recombination
Color Key
−10
5
Value
MRE11
YKU70
RAD50
XRS2
Gene
Z_Shift_K_Gem
Z_Im_K_Gem
Z_Shift_K_Cyt
Z_Im_K_Cyt
Z_Shift_L_Gem
Z_Im_L_Gem
Z_Shift_L_Cyt
Z_Im_L_Cyt

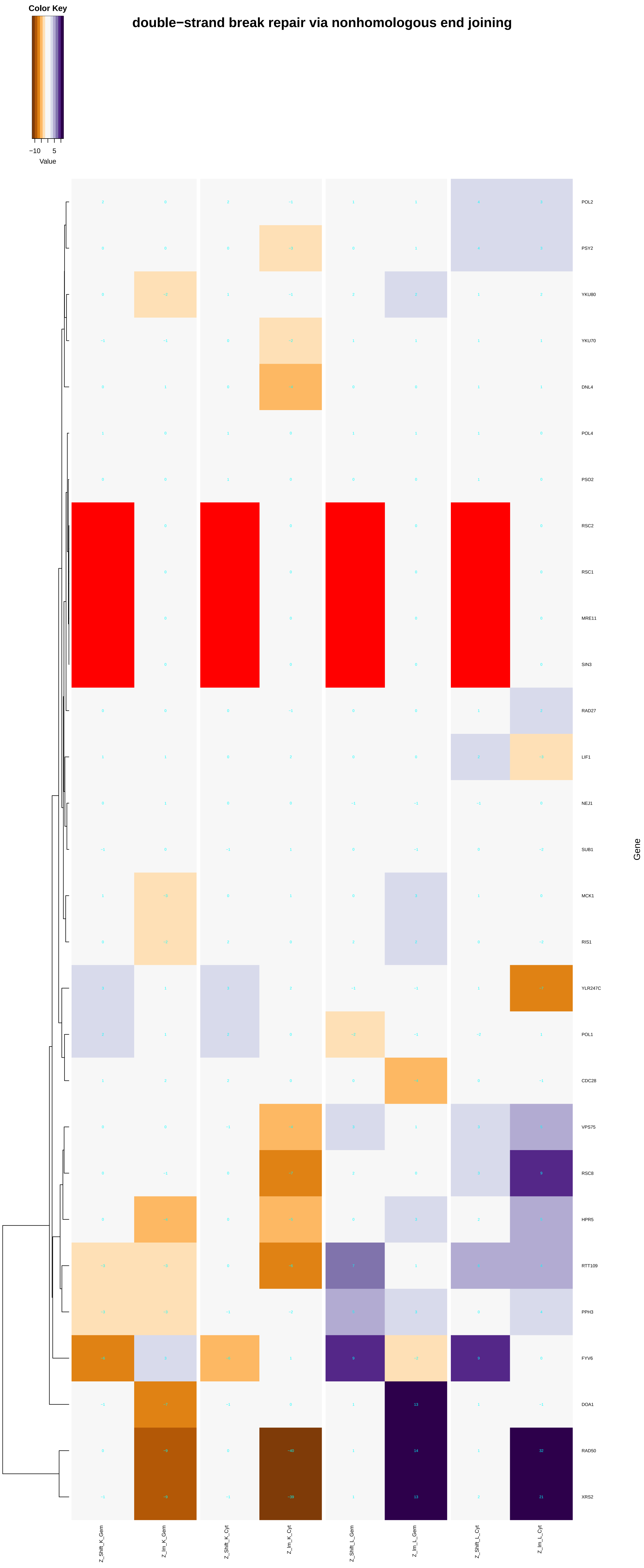

Color Key
−10
5
Value
double−strand break repair via nonhomologous end joining
POL2
PSY2
YKU80
YKU70
DNL4
POL4
PSO2
RSC2
RSC1
MRE11
SIN3
RAD27
LIF1
NEJ1
SUB1
MCK1
RIS1
YLR247C
POL1
CDC28
VPS75
RSC8
HPR5
RTT109
PPH3
FYV6
DOA1
RAD50
XRS2
Gene
Z_Shift_K_Gem
Z_Im_K_Gem
Z_Shift_K_Cyt
Z_Im_K_Cyt
Z_Shift_L_Gem
Z_Im_L_Gem
Z_Shift_L_Cyt
Z_Im_L_Cyt

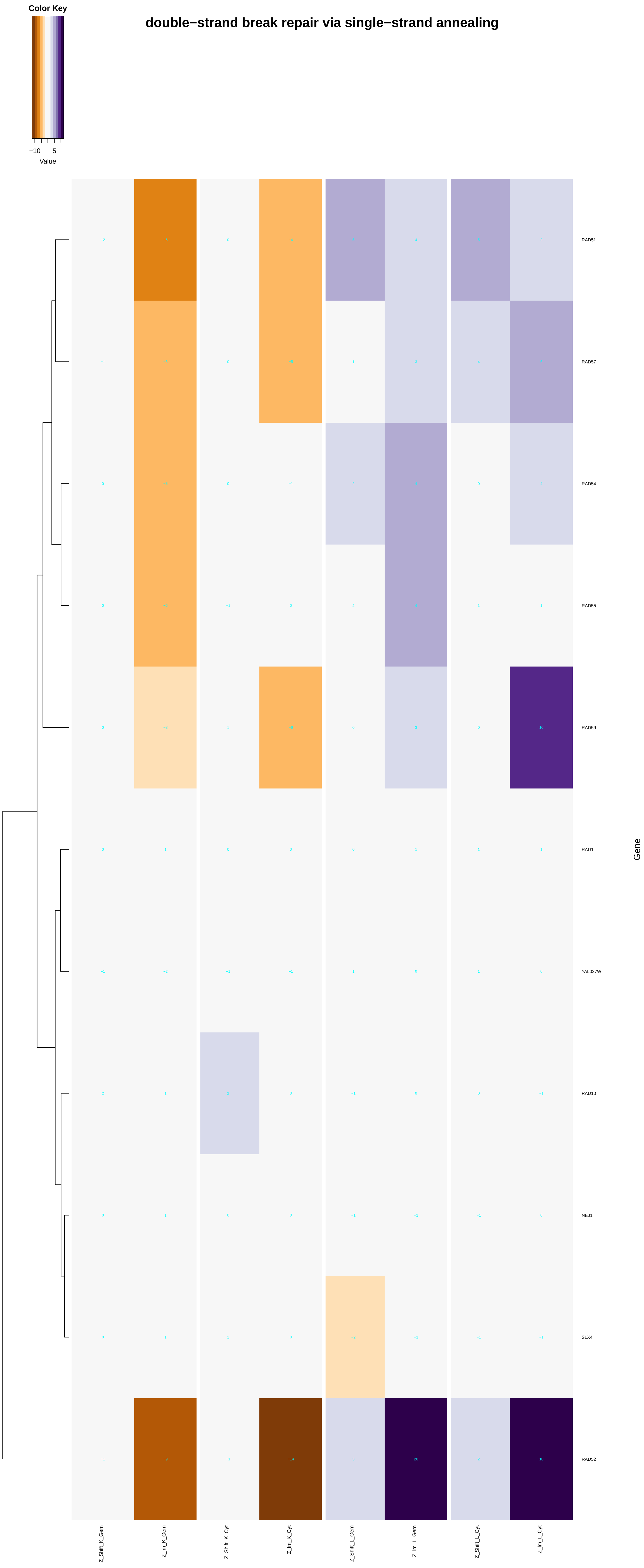
double−strand break repair via single−strand annealing
Color Key
−10
5
Value
RAD51
RAD57
RAD54
RAD55
RAD59
RAD1
YAL027W
RAD10
NEJ1
SLX4
RAD52
Gene
Z_Shift_K_Gem
Z_Im_K_Gem
Z_Shift_K_Cyt
Z_Im_K_Cyt
Z_Shift_L_Gem
Z_Im_L_Gem
Z_Shift_L_Cyt
Z_Im_L_Cyt

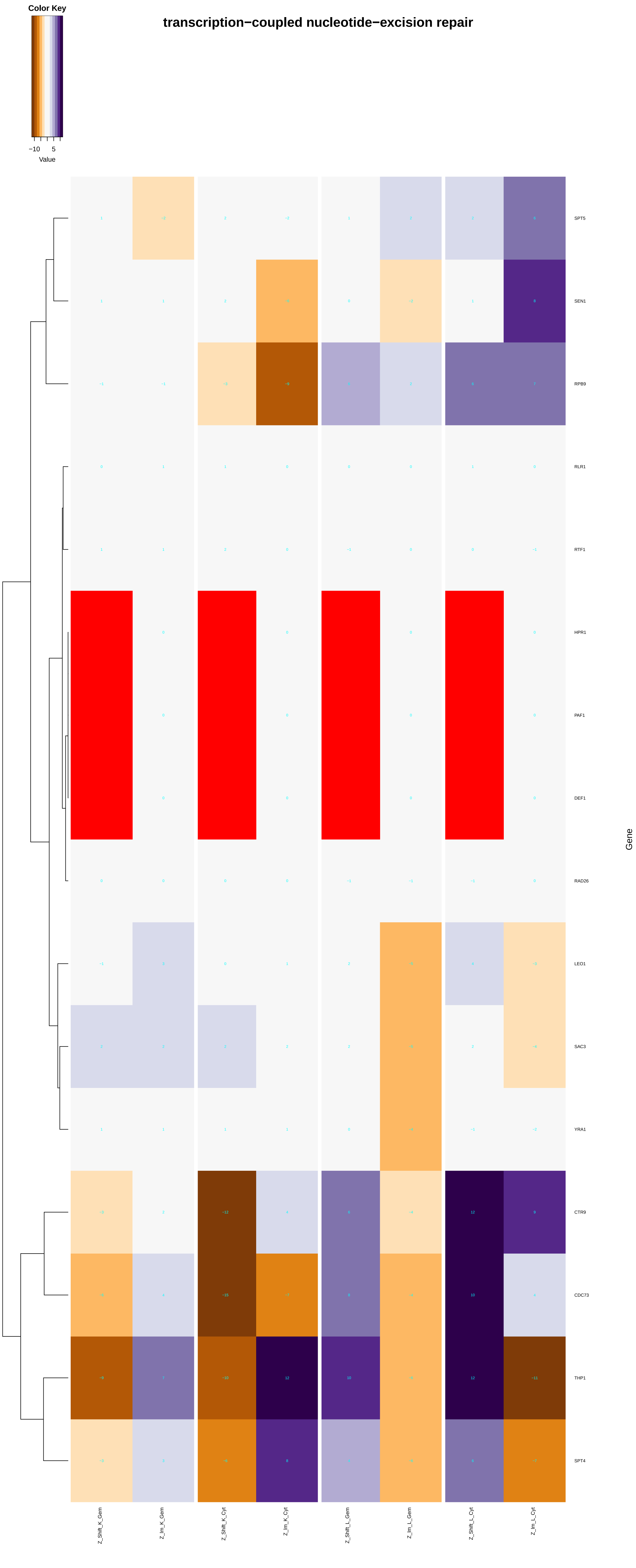
transcription−coupled nucleotide−excision repair
Color Key
−10
5
Value
SPT5
SEN1
RPB9
RLR1
RTF1
HPR1
PAF1
DEF1
RAD26
LEO1
SAC3
YRA1
CTR9
CDC73
THP1
SPT4
Gene
Z_Shift_K_Gem
Z_Im_K_Gem
Z_Shift_K_Cyt
Z_Im_K_Cyt
Z_Shift_L_Gem
Z_Im_L_Gem
Z_Shift_L_Cyt
Z_Im_L_Cyt

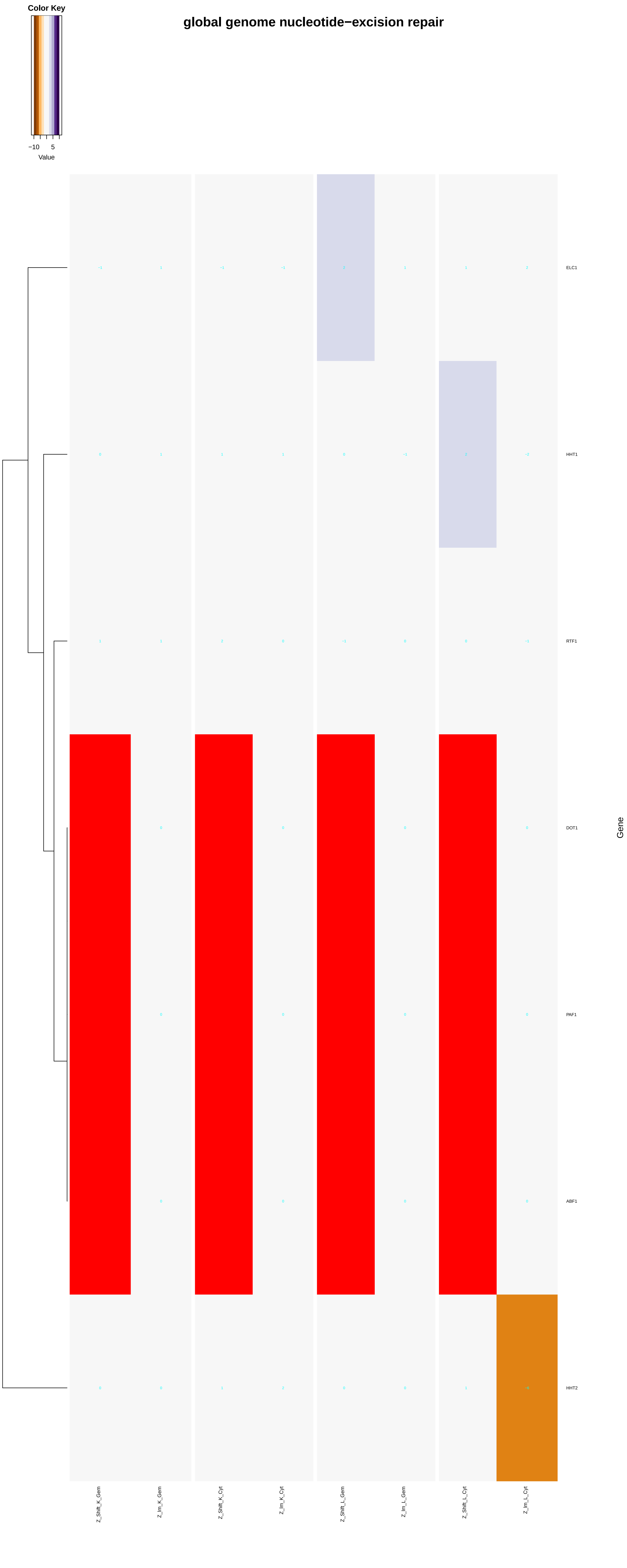

global genome nucleotide−excision repair
Color Key
−10
5
Value
ELC1
HHT1
RTF1
DOT1
PAF1
ABF1
HHT2
Gene
Z_Shift_K_Gem
Z_Im_K_Gem
Z_Shift_K_Cyt
Z_Im_K_Cyt
Z_Shift_L_Gem
Z_Im_L_Gem
Z_Shift_L_Cyt
Z_Im_L_Cyt

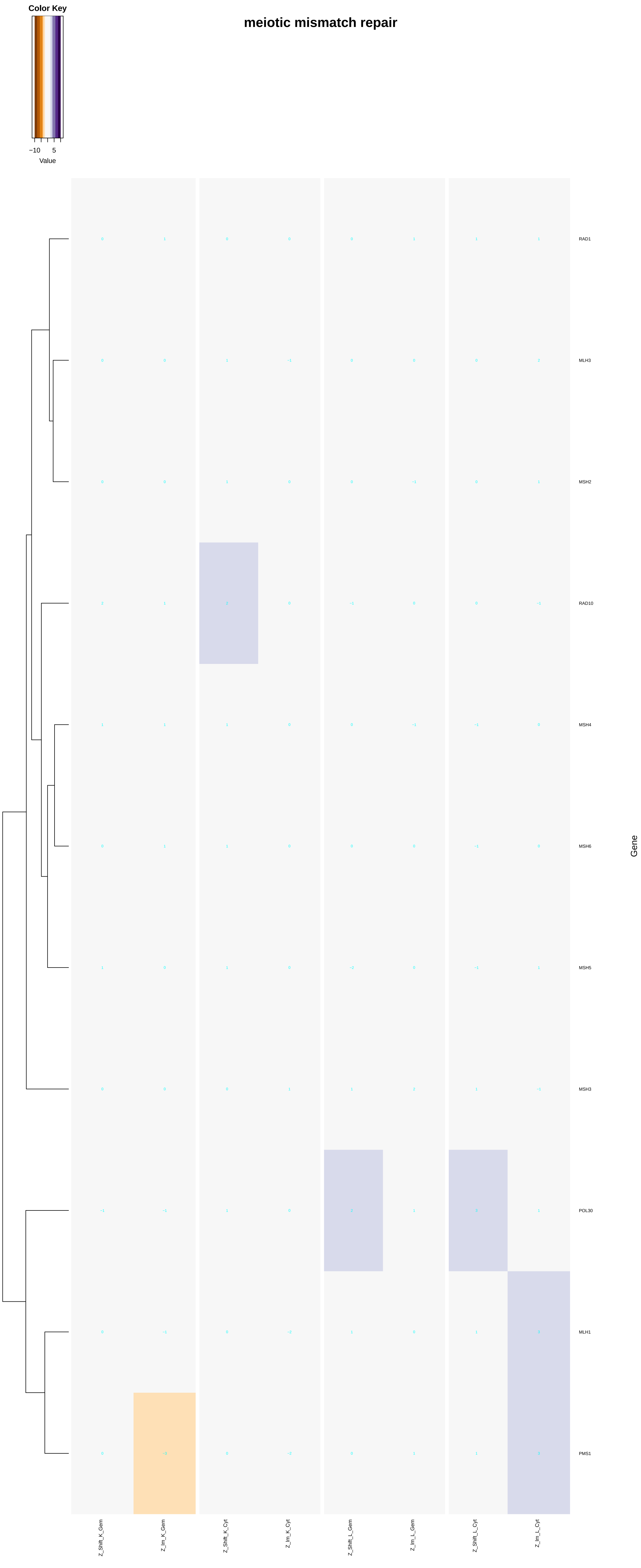
meiotic mismatch repair
Color Key
−10
5
Value
RAD1
MLH3
MSH2
RAD10
MSH4
MSH6
MSH5
MSH3
POL30
MLH1
PMS1
Gene
Z_Shift_K_Gem
Z_Im_K_Gem
Z_Shift_K_Cyt
Z_Im_K_Cyt
Z_Shift_L_Gem
Z_Im_L_Gem
Z_Shift_L_Cyt
Z_Im_L_Cyt

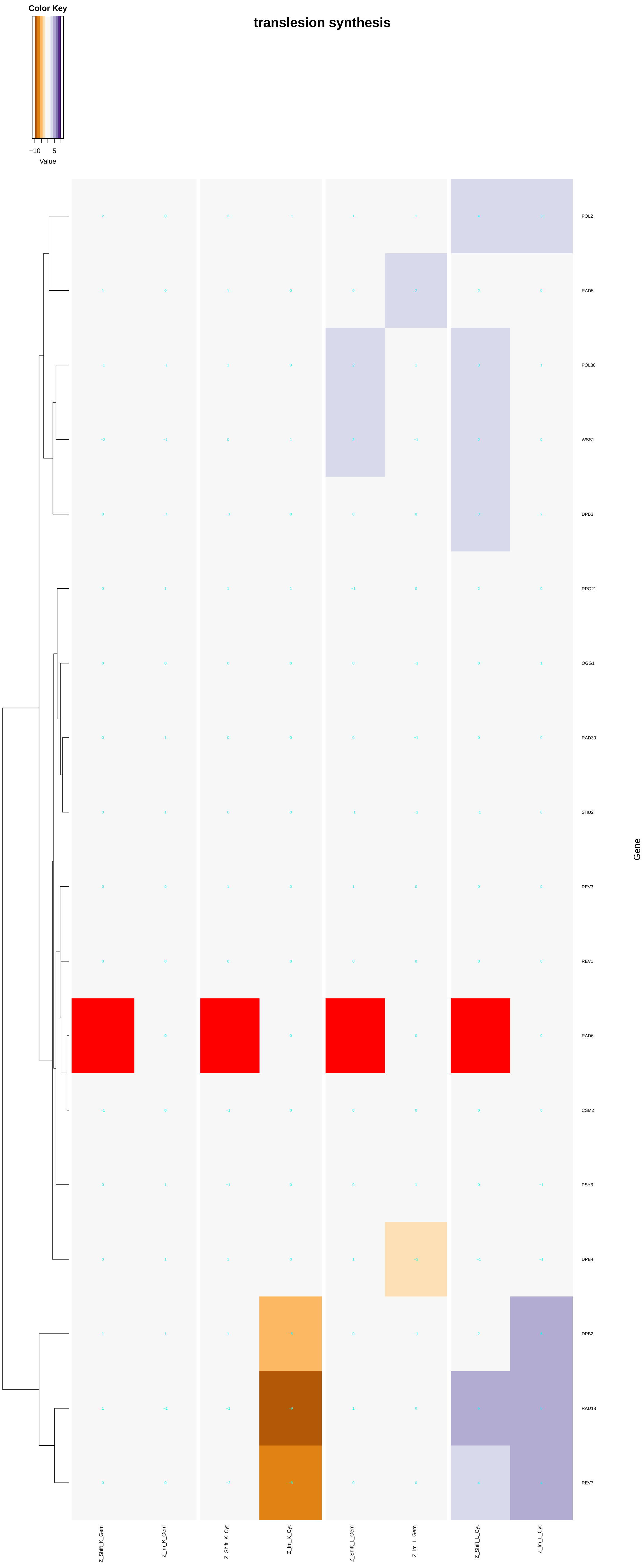

translesion synthesis
Color Key
−10
5
Value
POL2
RAD5
POL30
WSS1
DPB3
RPO21
OGG1
RAD30
SHU2
REV3
REV1
RAD6
CSM2
PSY3
DPB4
DPB2
RAD18
REV7
Gene
Z_Shift_K_Gem
Z_Im_K_Gem
Z_Shift_K_Cyt
Z_Im_K_Cyt
Z_Shift_L_Gem
Z_Im_L_Gem
Z_Shift_L_Cyt
Z_Im_L_Cyt

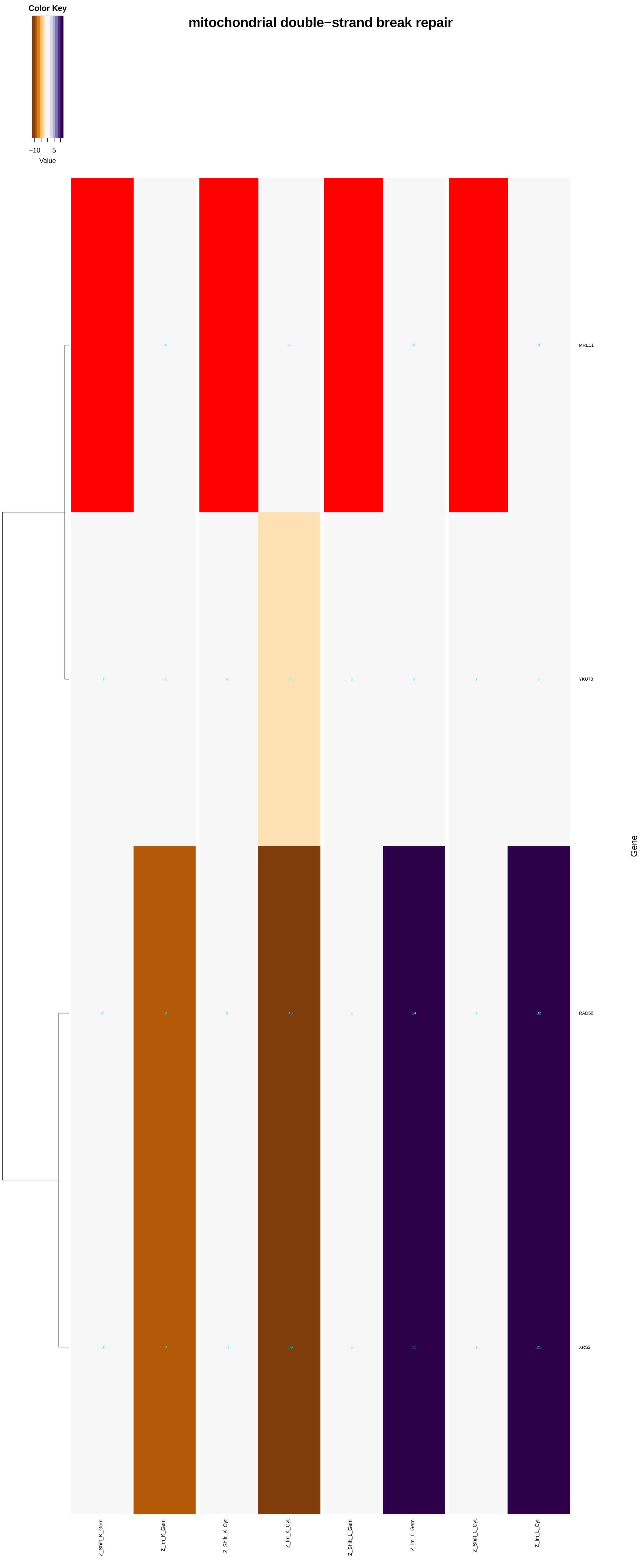

Color Key
−10
5
Value
mitochondrial double−strand break repair
MRE11
YKU70
RAD50
XRS2
Gene
Z_Shift_K_Gem
Z_Im_K_Gem
Z_Shift_K_Cyt
Z_Im_K_Cyt
Z_Shift_L_Gem
Z_Im_L_Gem
Z_Shift_L_Cyt
Z_Im_L_Cyt

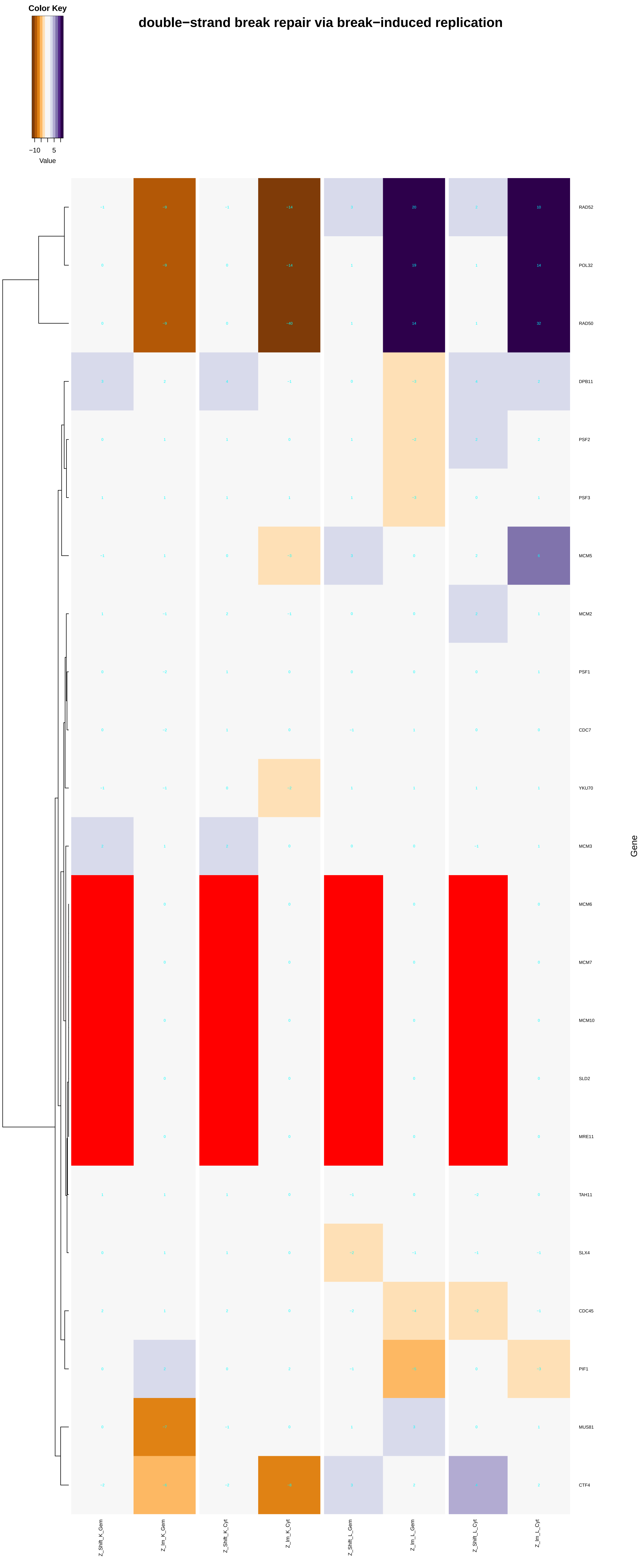
double−strand break repair via break−induced replication
Color Key
−10
5
Value
RAD52
POL32
RAD50
DPB11
PSF2
PSF3
MCM5
MCM2
PSF1
CDC7
YKU70
MCM3
MCM6
MCM7
MCM10
SLD2
MRE11
TAH11
SLX4
CDC45
PIF1
MUS81
CTF4
Gene
Z_Shift_K_Gem
Z_Im_K_Gem
Z_Shift_K_Cyt
Z_Im_K_Cyt
Z_Shift_L_Gem
Z_Im_L_Gem
Z_Shift_L_Cyt
Z_Im_L_Cyt

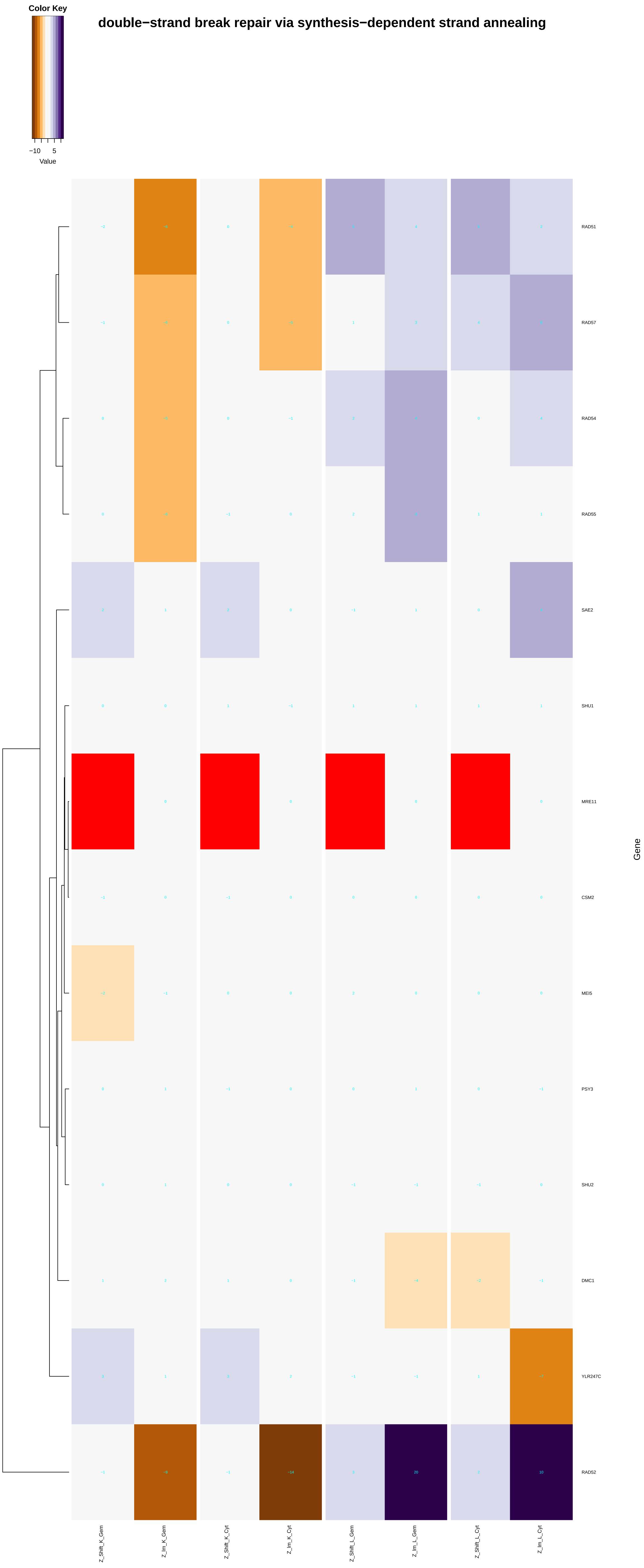
double−strand break repair via synthesis−dependent strand annealing
Color Key
−10
5
Value
RAD51
RAD57
RAD54
RAD55
SAE2
SHU1
MRE11
CSM2
MEI5
PSY3
SHU2
DMC1
YLR247C
RAD52
Gene
Z_Shift_K_Gem
Z_lm_K_Gem
Z_Shift_K_Cyt
Z_lm_K_Cyt
Z_Shift_L_Gem
Z_lm_L_Gem
Z_Shift_L_Cyt
Z_lm_L_Cyt

### replication−born double−strand break repair via sister chromatid exchange

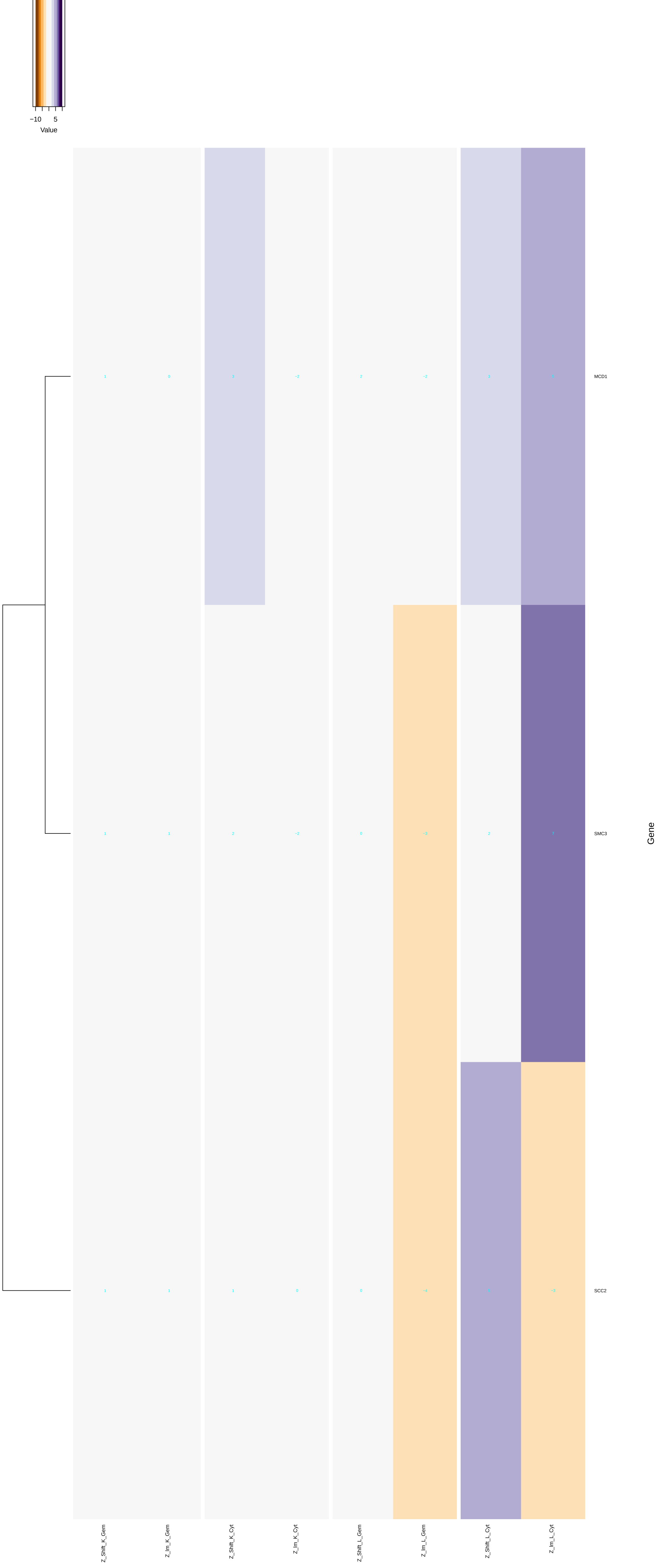

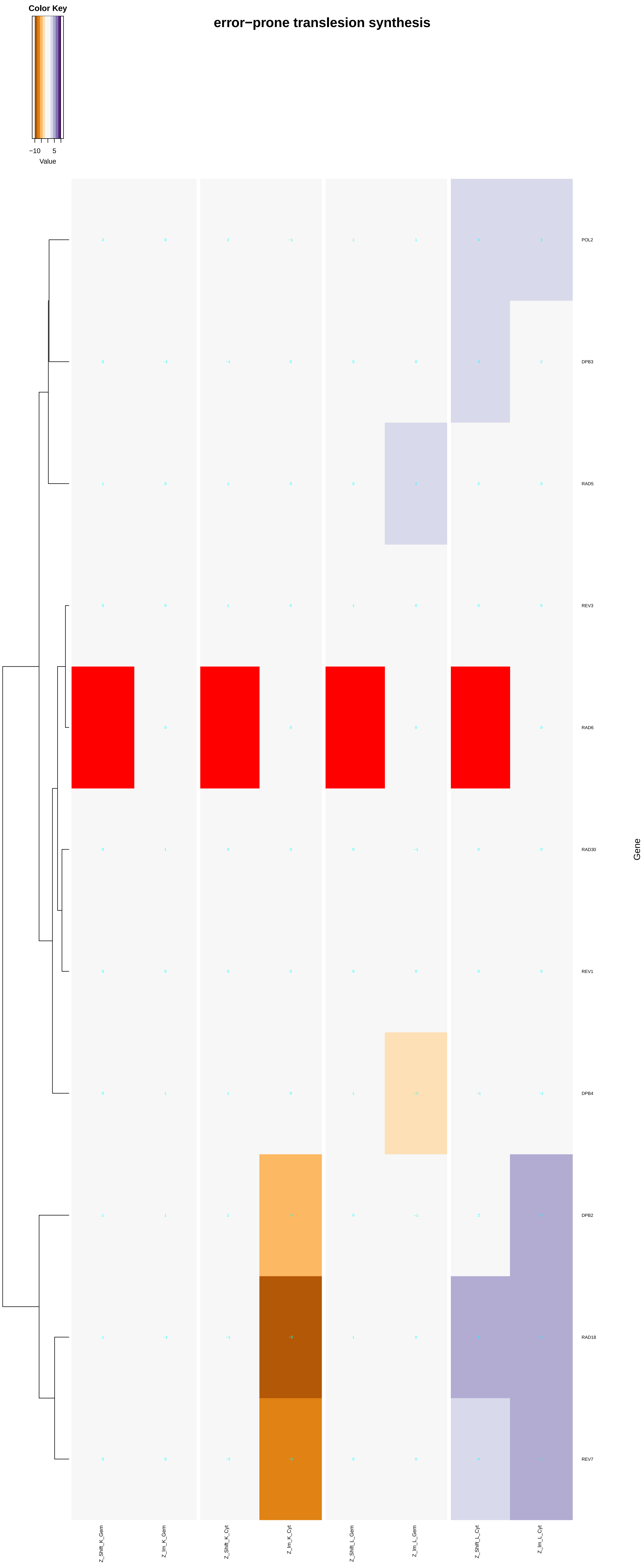
error−prone translesion synthesis
Color Key
−10
5
Value
POL2
DPB3
RAD5
REV3
RAD6
RAD30
REV1
DPB4
DPB2
RAD18
REV7
Gene
Z_Shift_K_Gem
Z_Im_K_Gem
Z_Shift_K_Cyt
Z_Im_K_Cyt
Z_Shift_L_Gem
Z_Im_L_Gem
Z_Shift_L_Cyt
Z_Im_L_Cyt

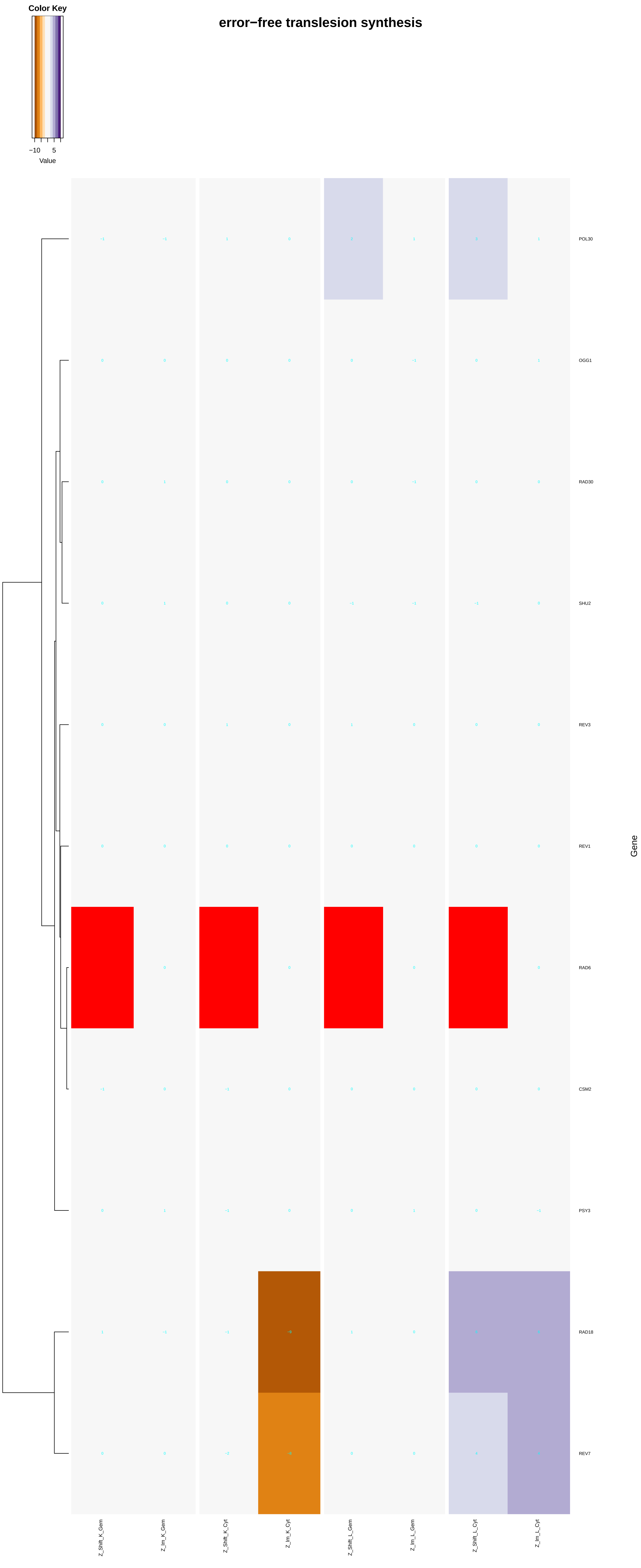
error−free translesion synthesis
Color Key
−10
5
Value
POL30
OGG1
RAD30
SHU2
REV3
REV1
RAD6
CSM2
PSY3
RAD18
REV7
Gene
Z_Shift_K_Gem
Z_Im_K_Gem
Z_Shift_K_Cyt
Z_Im_K_Cyt
Z_Shift_L_Gem
Z_Im_L_Gem
Z_Shift_L_Cyt
Z_Im_L_Cyt
