## Additional File 7 for "A humanized yeast phenomic model of deoxycytidine kinase to predict genetic buffering of nucleoside analog cytotoxicity": double-stranded_DNA_binding.pdf

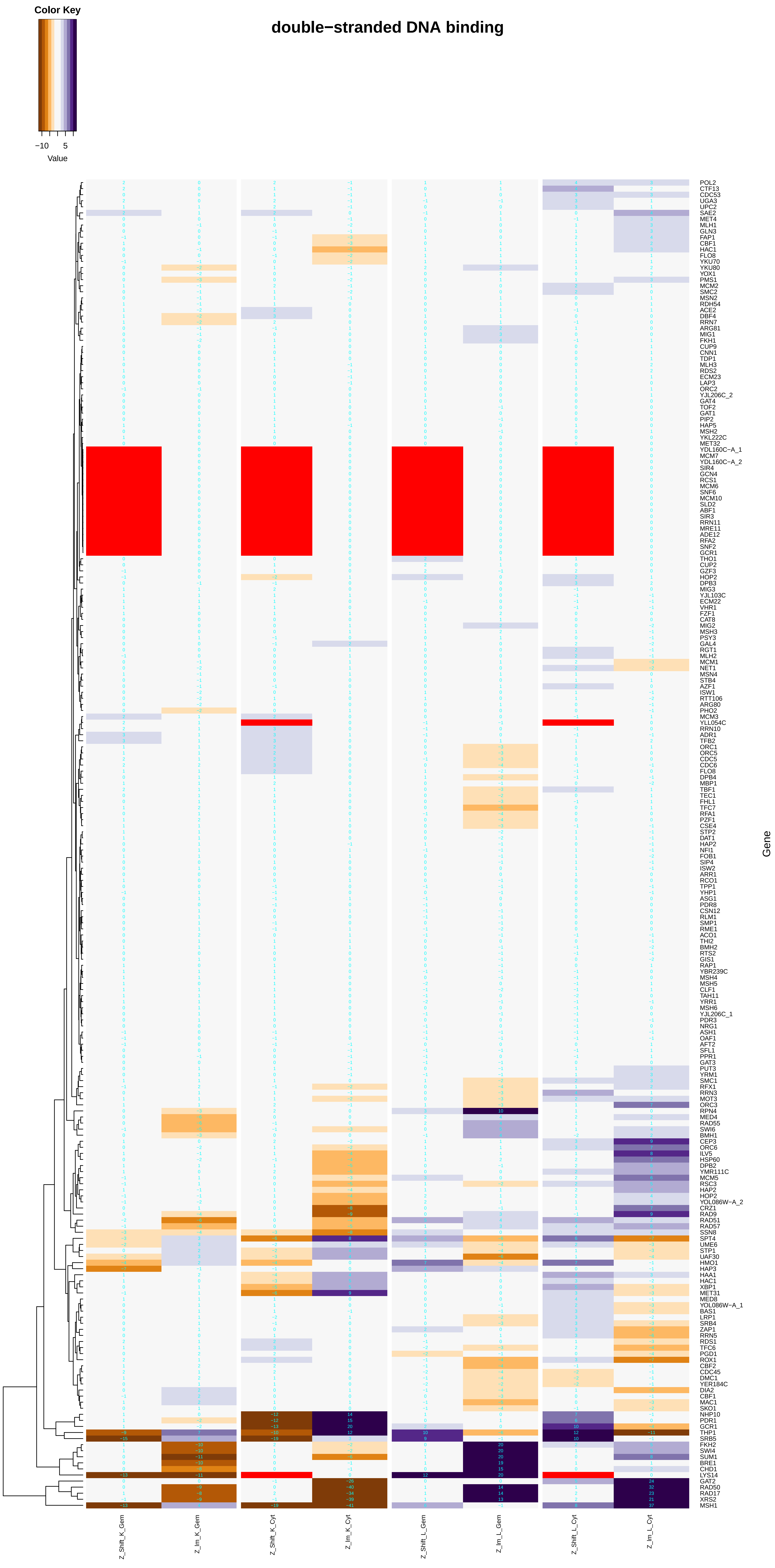

double−stranded DNA binding
Color Key
−10
5
Value
Gene
POL2
CTF13
CDC53
UGA3
UPC2
SAE2
MET4
MLH1
GLN3
FAP1
CBF1
HAC1
FLO8
YKU70
YKU80
YOX1
PMS1
MCM2
SMC2
MSN2
RDH54
ACE2
DBF4
RRN7
ARG81
MIG1
FKH1
CUP9
CNN1
TDP1
MLH3
RDS2
ECM23
LAP3
ORC2
YJL206C_2
GAT4
TOF2
GAT1
PIP2
HAP5
MSH2
YKL222C
MET32
YDL160C−A_1
MCM7
YDL160C−A_2
SIR4
GCN4
RCS1
MCM6
SNF6
MCM10
SLD2
ABF1
SIR3
RRN11
MRE11
ADE12
RFA2
SNF2
GCR1
THO1
CUP2
GZF3
HOP2
DPB3
MIG3
YJL103C
ECM22
VHR1
FZF1
CAT8
MIG2
MSH3
PSY3
GAL4
RGT1
MLH2
MCM1
NET1
MSN4
STB4
AZF1
ISW1
RTT106
ARG80
PHO2
MCM3
YLL054C
RRN10
ADR1
TFB2
ORC1
ORC5
CDC5
CDC6
FLO8
DPB4
MBP1
TBF1
TEC1
FHL1
TFC7
RFA1
PZF1
CSE4
STP2
DAT1
HAP2
NFI1
FOB1
SIP4
ISW2
ARR1
RCO1
TPP1
YHP1
ASG1
PDR8
CSN12
RLM1
SMP1
RME1
ACO1
THI2
BMH2
RTS2
GIS1
RAP1
YBR239C
MSH4
MSH5
CLF1
TAH11
YRR1
MSH6
YJL206C_1
PDR3
NRG1
ASH1
OAF1
AFT2
SFL1
PPR1
GAT3
PUT3
YRM1
SMC1
RFX1
RRN3
MOT3
ORC3
RPN4
MED4
RAD55
SWI6
BMH1
CEP3
ORC6
ILV5
HSP60
DPB2
YMR111C
MCM5
RSC3
HAP2
HOP2
YOL086W−A_2
CRZ1
RAD9
RAD51
RAD57
SSN8
SPT4
UME6
STP1
UAF30
HMO1
HAP3
HAA1
HAC1
XBP1
MET31
MED8
YOL086W−A_1
BAS1
LRP1
SRB4
ZAP1
RRN5
RDS1
TFC6
PGD1
ROX1
CBF2
CDC45
DMC1
YER184C
DIA2
CBF1
MAC1
SKO1
NHP10
PDR1
GCR1
THP1
SRB5
FKH2
SWI4
SUM1
BRE1
CHD1
LYS14
GAT2
RAD50
RAD17
XRS2
MSH1
Z_Shift_K_Gem
Z_Im_K_Gem
Z_Shift_K_Cyt
Z_Im_K_Cyt
Z_Shift_L_Gem
Z_Im_L_Gem
Z_Shift_L_Cyt
Z_Im_L_Cyt

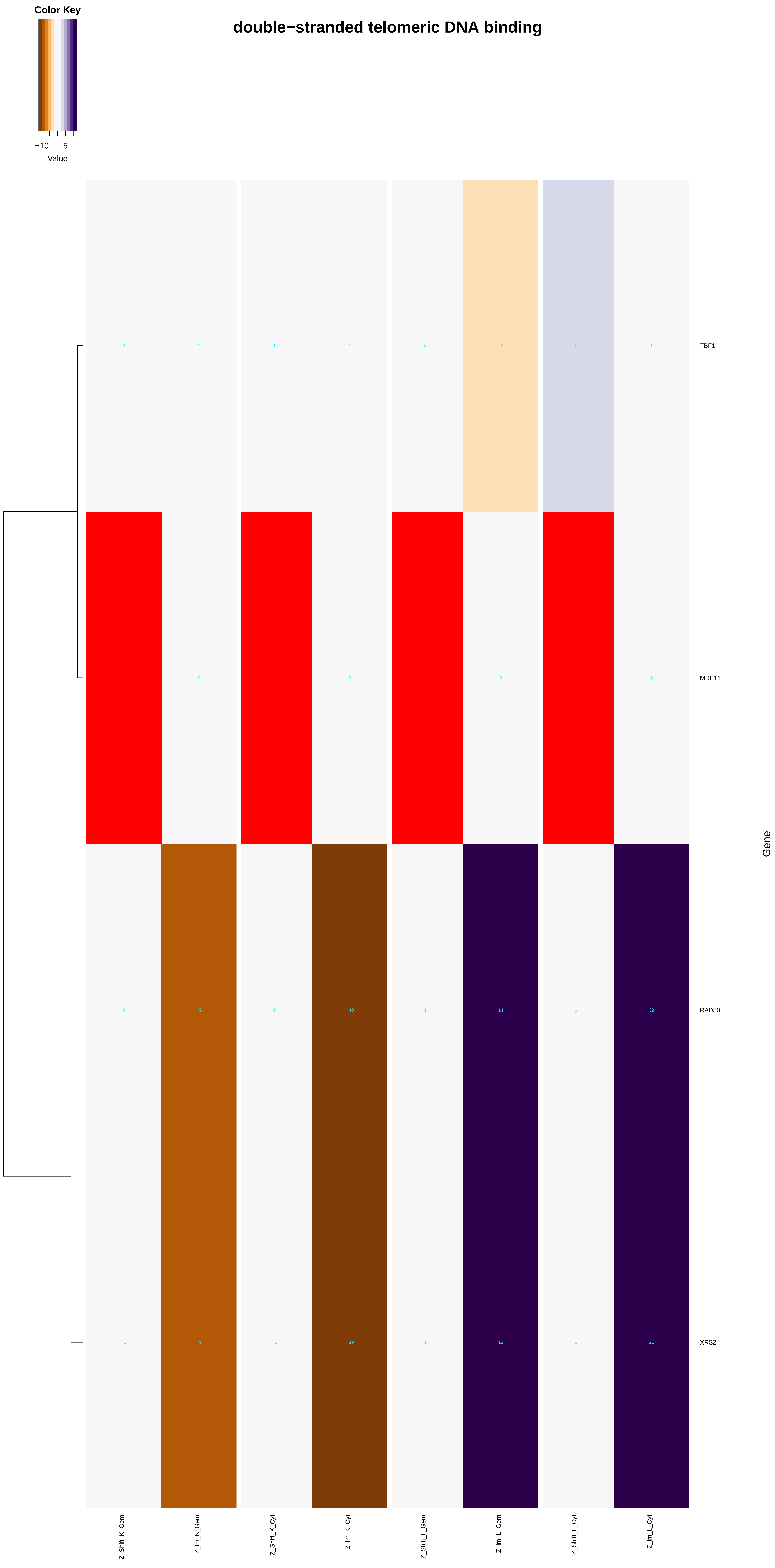

double−stranded telomeric DNA binding
Color Key
−10
5
Value
TBF1
MRE11
RAD50
XRS2
Gene
Z_Shift_K_Gem
Z_Im_K_Gem
Z_Shift_K_Cyt
Z_Im_K_Cyt
Z_Shift_L_Gem
Z_Im_L_Gem
Z_Shift_L_Cyt
Z_Im_L_Cyt

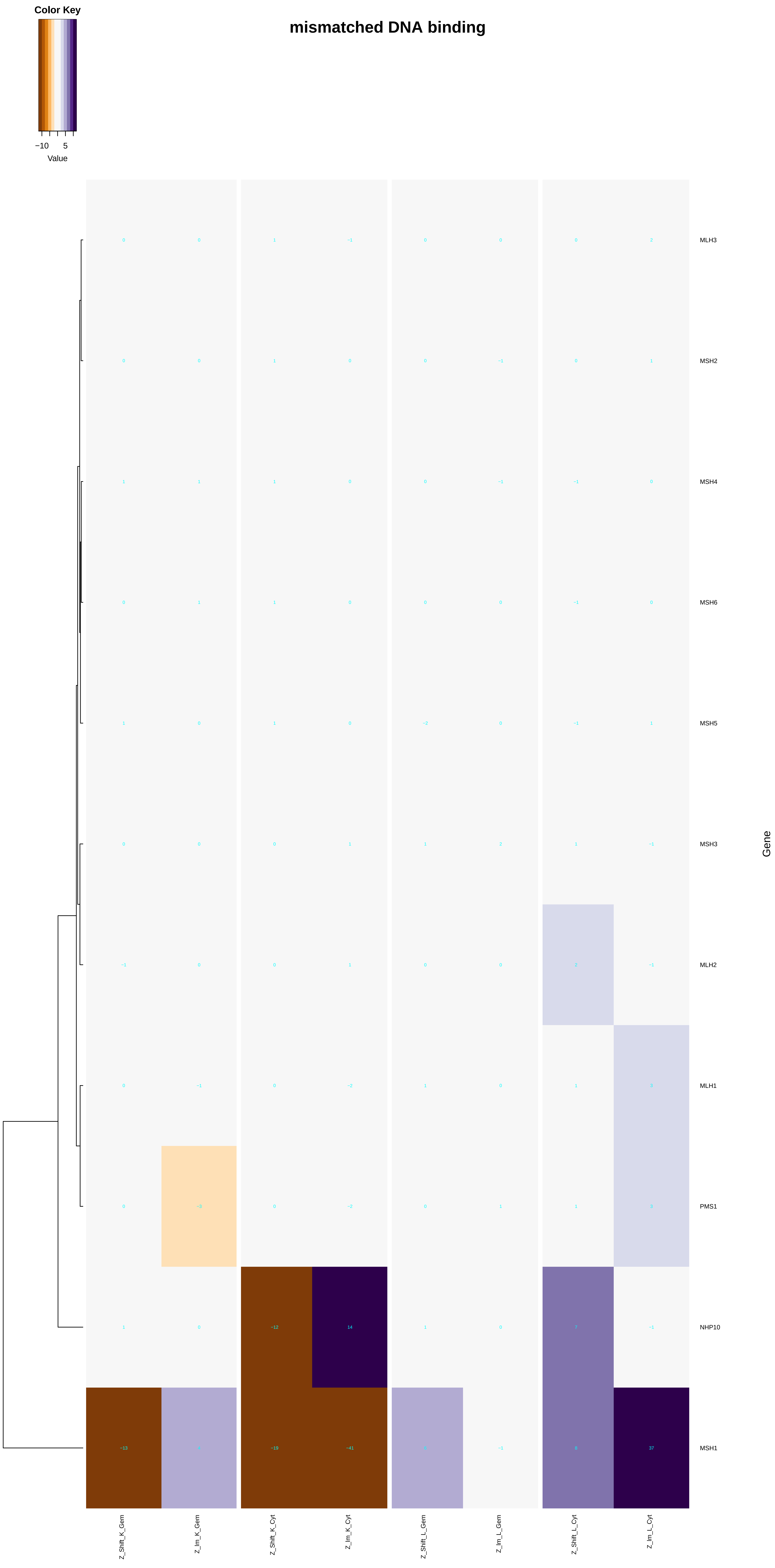

mismatched DNA binding
Color Key
−10
5
Value
Gene
MLH3
MSH2
MSH4
MSH6
MSH5
MSH3
MLH2
MLH1
PMS1
NHP10
MSH1
Z_Shift_K_Gem
Z_Im_K_Gem
Z_Shift_K_Cyt
Z_Im_K_Cyt
Z_Shift_L_Gem
Z_Im_L_Gem
Z_Shift_L_Cyt
Z_Im_L_Cyt

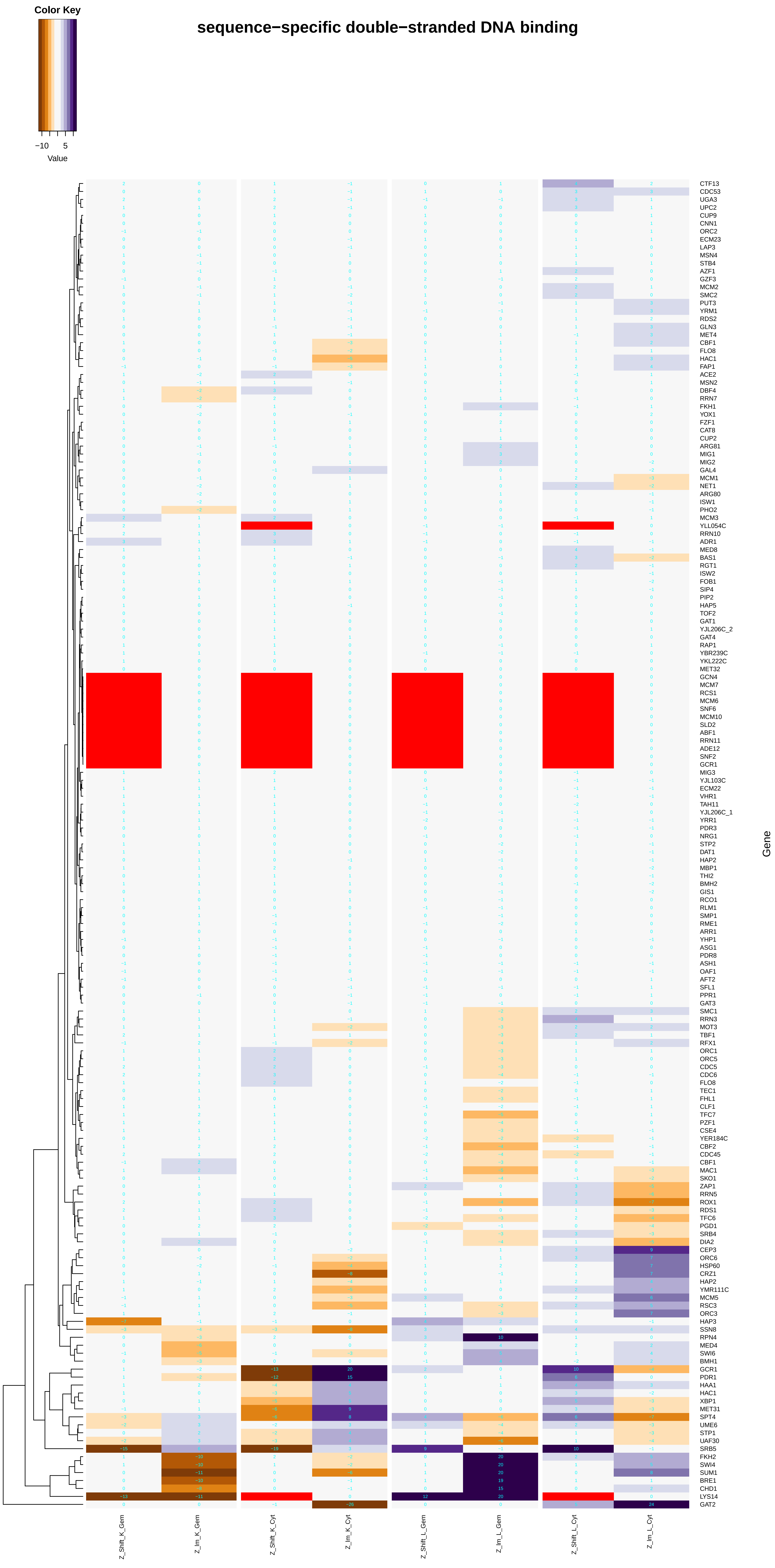

Color Key
−10
5
Value
sequence−specific double−stranded DNA binding
Z_Shift_K_Gem
Z_Im_K_Gem
Z_Shift_K_Cyt
Z_Im_K_Cyt
Z_Shift_L_Gem
Z_Im_L_Gem
Z_Shift_L_Cyt
Z_Im_L_Cyt
Gene
CTF13
CDC53
UGA3
UPC2
CUP9
CNN1
ORC2
ECM23
LAP3
MSN4
STB4
AZF1
GZF3
MCM2
SMC2
PUT3
YRM1
RDS2
GLN3
MET4
CBF1
FLO8
HAC1
FAP1
ACE2
MSN2
DBF4
RRN7
FKH1
YOX1
FZF1
CAT8
CUP2
ARG81
MIG1
MIG2
GAL4
MCM1
NET1
ARG80
ISW1
PHO2
MCM3
YLL054C
RRN10
ADR1
MED8
BAS1
RGT1
ISW2
FOB1
SIP4
PIP2
HAP5
TOF2
GAT1
YJL206C_2
GAT4
RAP1
YBR239C
YKL222C
MET32
GCN4
MCM7
RCS1
MCM6
SNF6
MCM10
SLD2
ABF1
RRN11
ADE12
SNF2
GCR1
MIG3
YJL103C
ECM22
VHR1
TAH11
YJL206C_1
YRR1
PDR3
NRG1
STP2
DAT1
HAP2
MBP1
THI2
BMH2
GIS1
RCO1
RLM1
SMP1
RME1
ARR1
YHP1
ASG1
PDR8
ASH1
OAF1
AFT2
SFL1
PPR1
GAT3
SMC1
RRN3
MOT3
TBF1
RFX1
ORC1
ORC5
CDC5
CDC6
FLO8
TEC1
FHL1
CLF1
TFC7
PZF1
CSE4
YER184C
CBF2
CDC45
CBF1
MAC1
SKO1
ZAP1
RRN5
ROX1
RDS1
TFC6
PGD1
SRB4
DIA2
CEP3
ORC6
HSP60
CRZ1
HAP2
YMR111C
MCM5
RSC3
ORC3
HAP3
SSN8
RPN4
MED4
SWI6
BMH1
GCR1
PDR1
HAA1
HAC1
XBP1
MET31
SPT4
UME6
STP1
UAF30
SRB5
FKH2
SWI4
SUM1
BRE1
CHD1
LYS14
GAT2

### DNA insertion or deletion binding

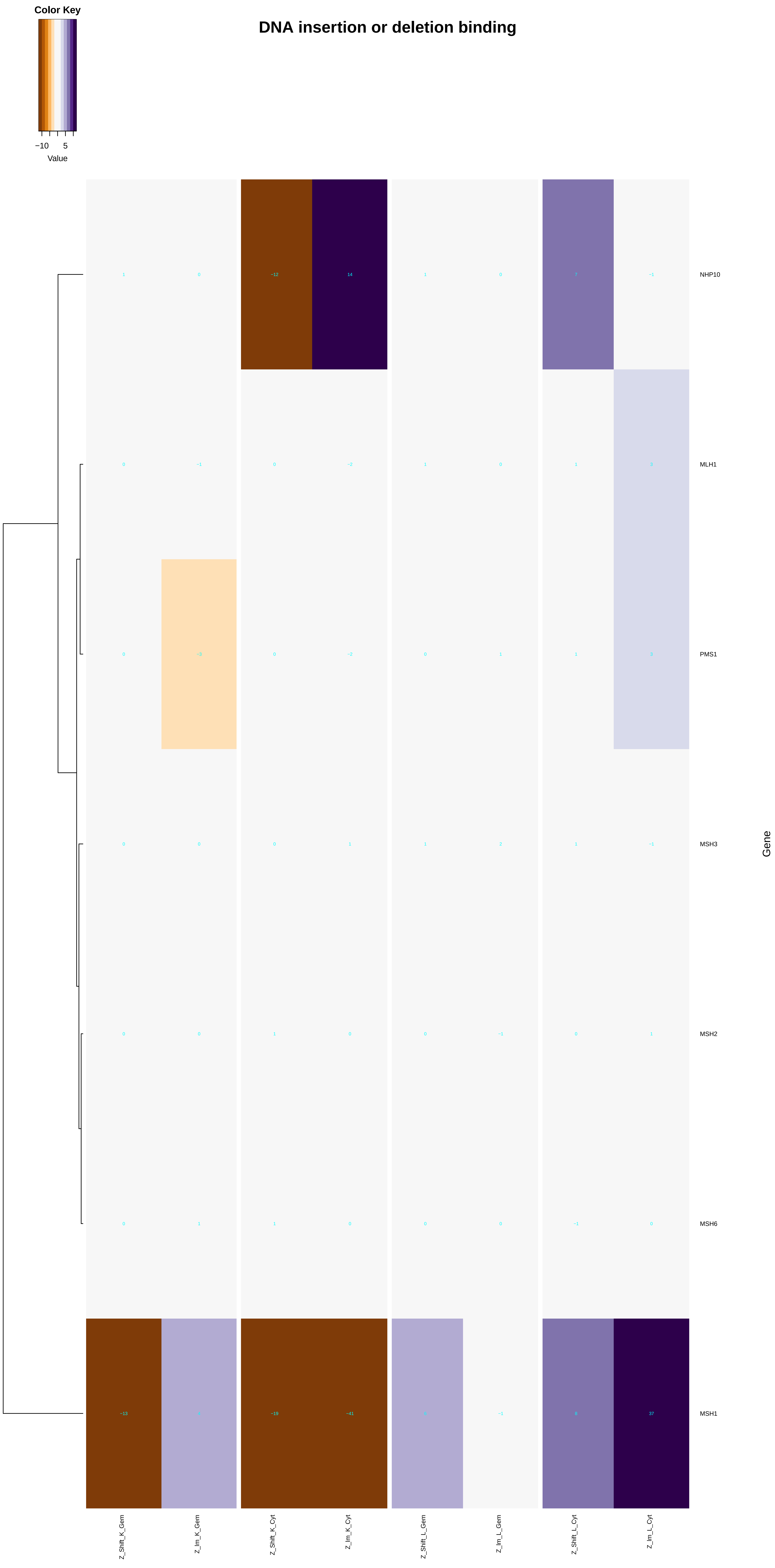

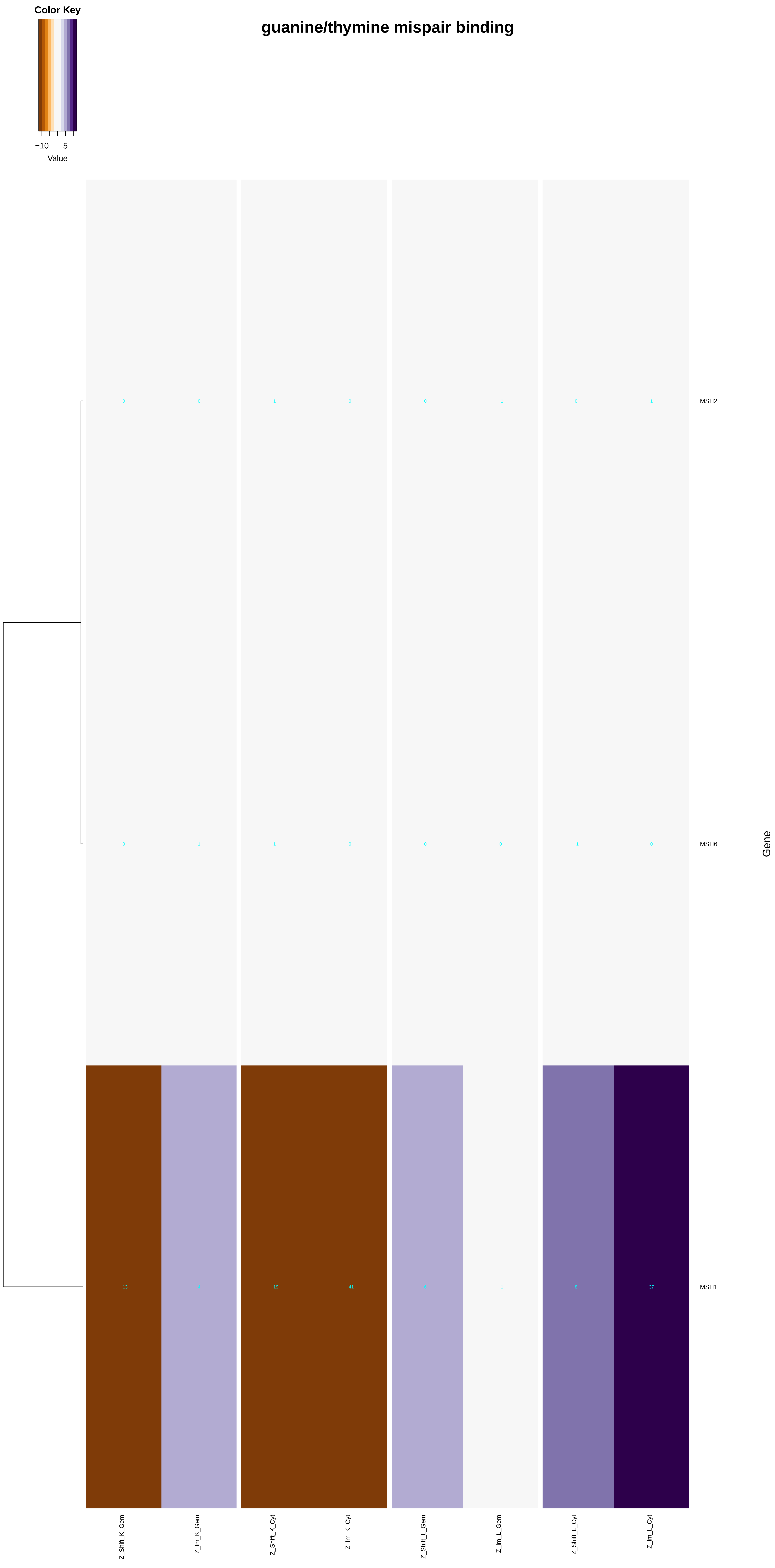
guanine/thymine mispair binding
Color Key
−10
5
Value
MSH2
MSH6
MSH1
Gene
Z_Shift_K_Gem
Z_Im_K_Gem
Z_Shift_K_Cyt
Z_Im_K_Cyt
Z_Shift_L_Gem
Z_Im_L_Gem
Z_Shift_L_Cyt
Z_Im_L_Cyt

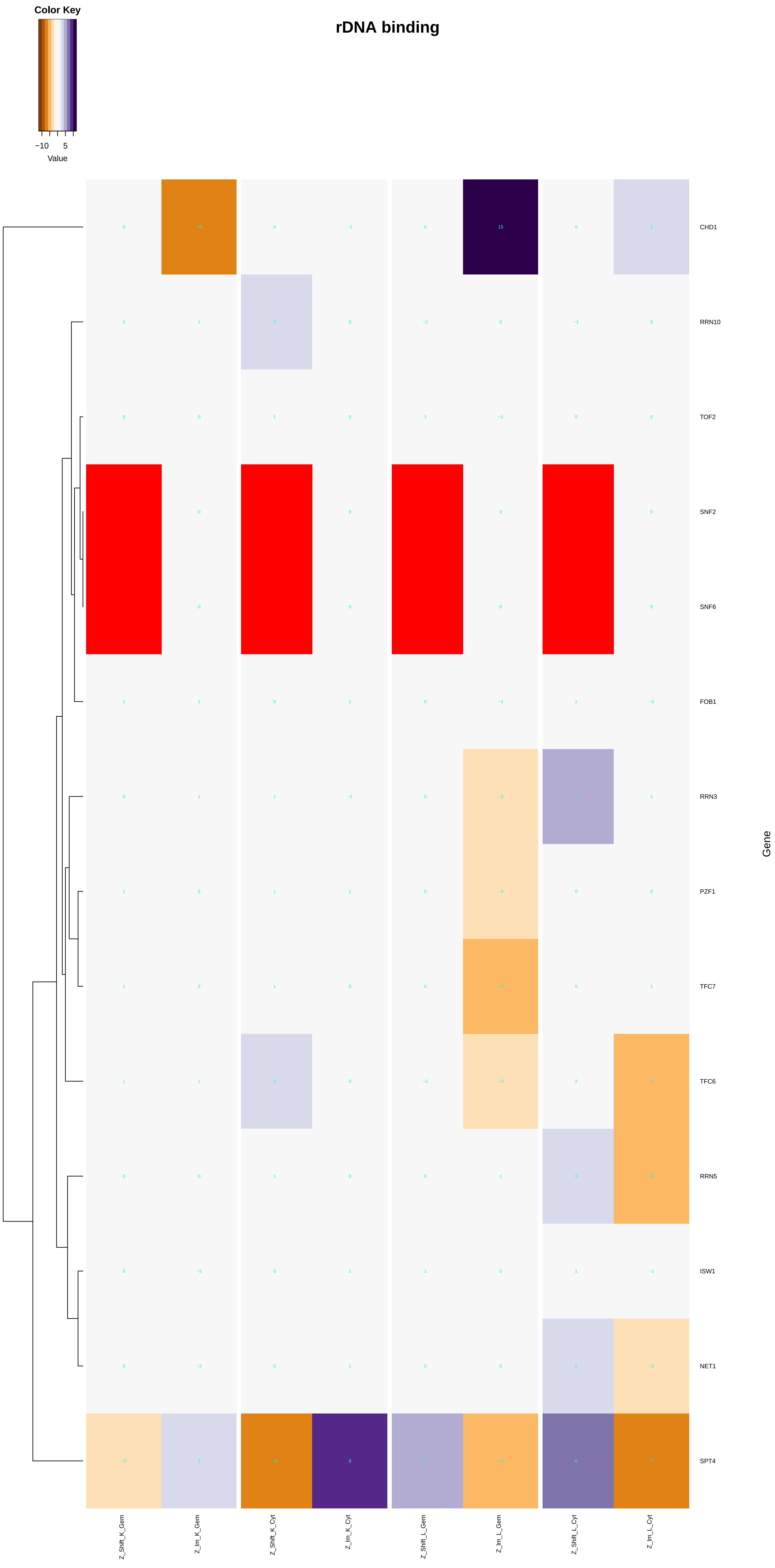
Color Key
−10
5
Value
rDNA binding
CHD1
RRN10
TOF2
SNF2
SNF6
FOB1
RRN3
PZF1
TFC7
TFC6
RRN5
ISW1
NET1
SPT4
Gene
Z_Shift_K_Gem
Z_Im_K_Gem
Z_Shift_K_Cyt
Z_Im_K_Cyt
Z_Shift_L_Gem
Z_Im_L_Gem
Z_Shift_L_Cyt
Z_Im_L_Cyt

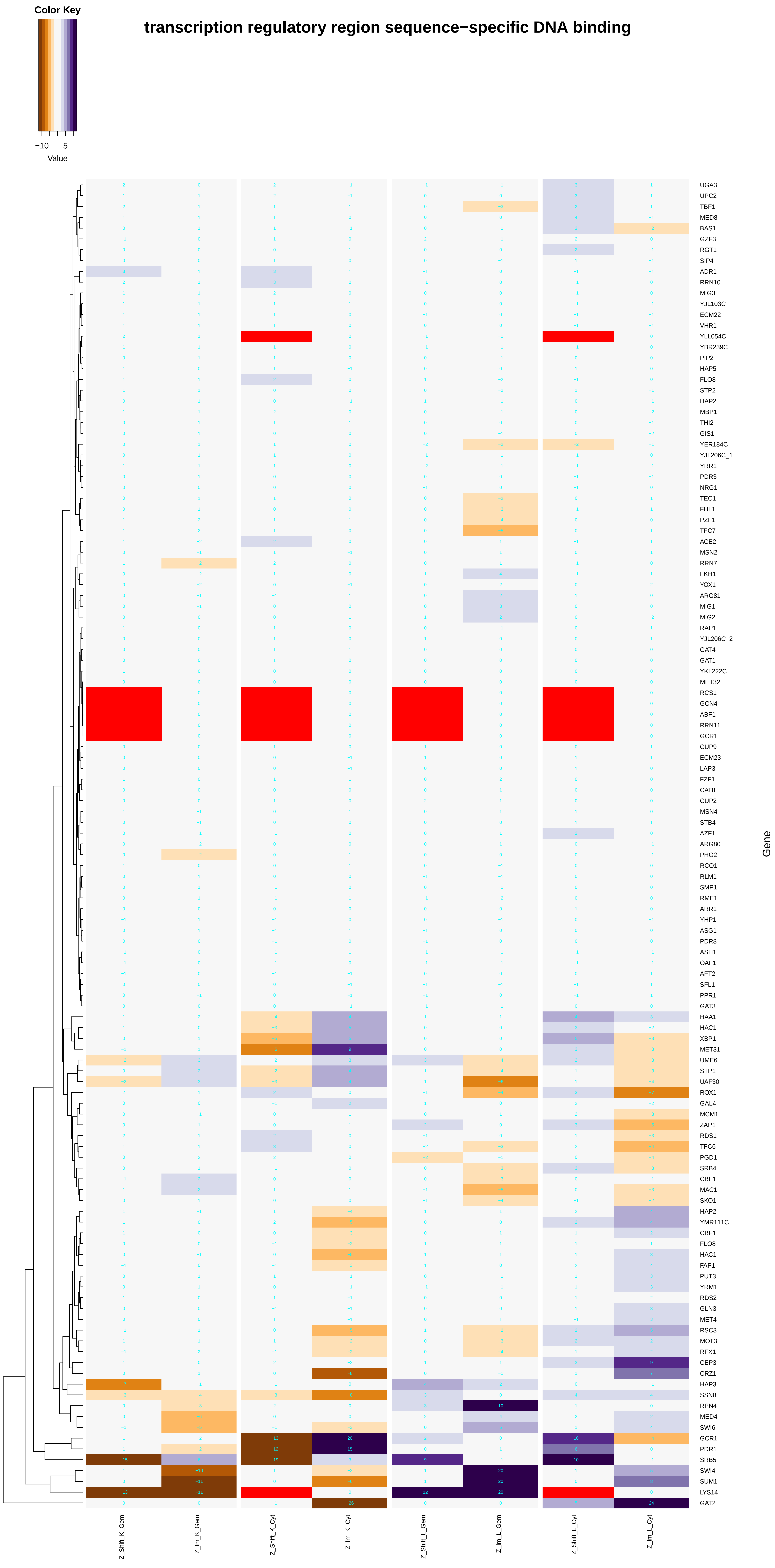
Color Key
−10
5
Value
transcription regulatory region sequence−specific DNA binding
UGA3
UPC2
TBF1
MED8
BAS1
GZF3
RGT1
SIP4
ADR1
RRN10
MIG3
YJL103C
ECM22
VHR1
YLL054C
YBR239C
PIP2
HAP5
FLO8
STP2
HAP2
MBP1
THI2
GIS1
YER184C
YJL206C_1
YRR1
PDR3
NRG1
TEC1
FHL1
PZF1
TFC7
ACE2
MSN2
RRN7
FKH1
YOX1
ARG81
MIG1
MIG2
RAP1
YJL206C_2
GAT4
GAT1
YKL222C
MET32
RCS1
GCN4
ABF1
RRN11
GCR1
CUP9
ECM23
LAP3
FZF1
CAT8
CUP2
MSN4
STB4
AZF1
ARG80
PHO2
RCO1
RLM1
SMP1
RME1
ARR1
YHP1
ASG1
PDR8
ASH1
OAF1
AFT2
SFL1
PPR1
GAT3
HAA1
HAC1
XBP1
MET31
UME6
STP1
UAF30
ROX1
GAL4
MCM1
ZAP1
RDS1
TFC6
PGD1
SRB4
CBF1
MAC1
SKO1
HAP2
YMR111C
CBF1
FLO8
HAC1
FAP1
PUT3
YRM1
RDS2
GLN3
MET4
RSC3
MOT3
RFX1
CEP3
CRZ1
HAP3
SSN8
RPN4
MED4
SWI6
GCR1
PDR1
SRB5
SWI4
SUM1
LYS14
GAT2
Gene
Z_Shift_K_Gem
Z_Im_K_Gem
Z_Shift_K_Cyt
Z_Im_K_Cyt
Z_Shift_L_Gem
Z_Im_L_Gem
Z_Shift_L_Cyt
Z_Im_L_Cyt

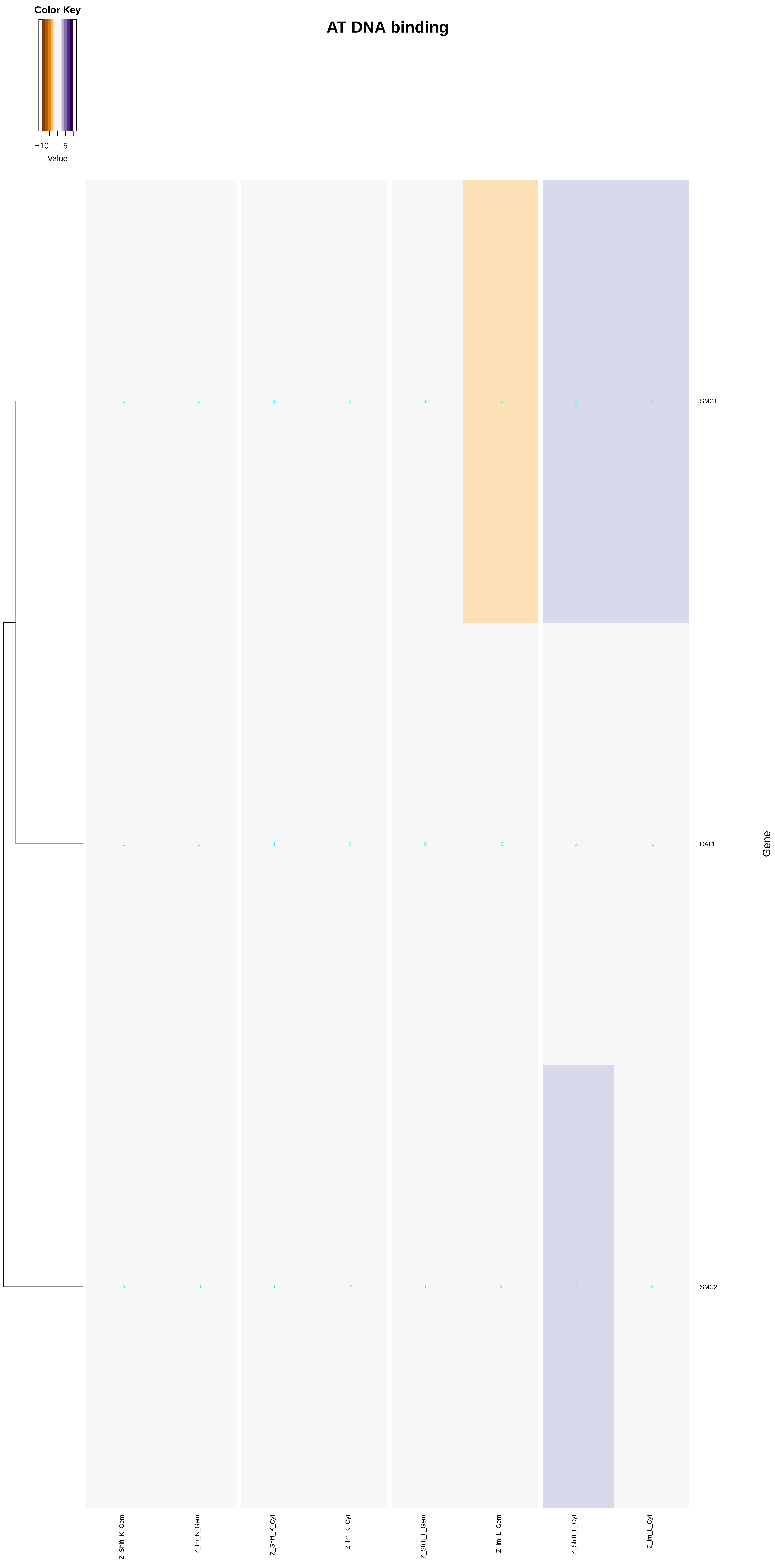

Color Key
−10
5
Value
AT DNA binding
SMC1
DAT1
SMC2
Gene
Z_Shift_K_Gem
Z_Im_K_Gem
Z_Shift_K_Cyt
Z_Im_K_Cyt
Z_Shift_L_Gem
Z_Im_L_Gem
Z_Shift_L_Cyt
Z_Im_L_Cyt

### DNA replication origin binding

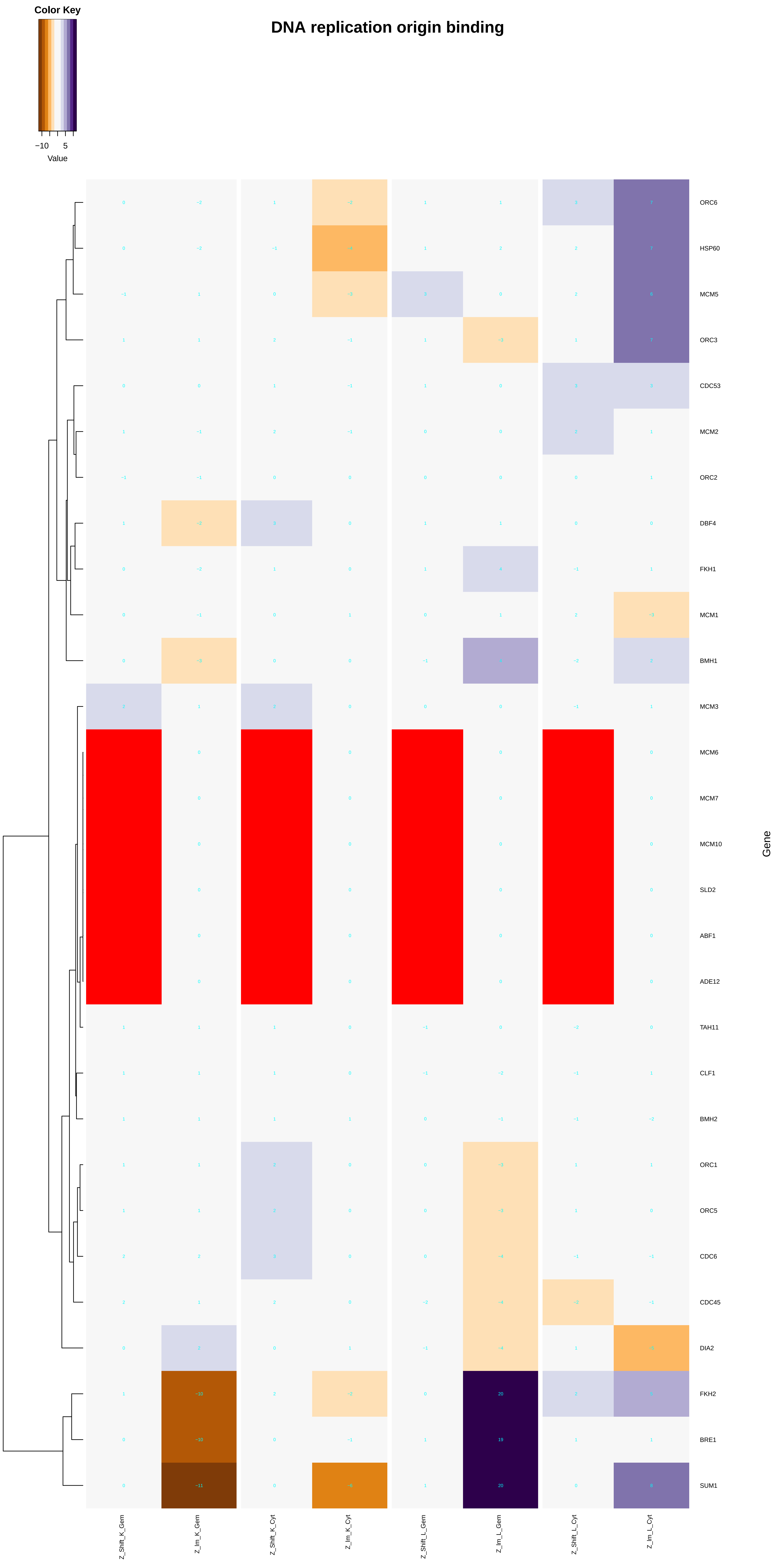

### centromeric DNA binding

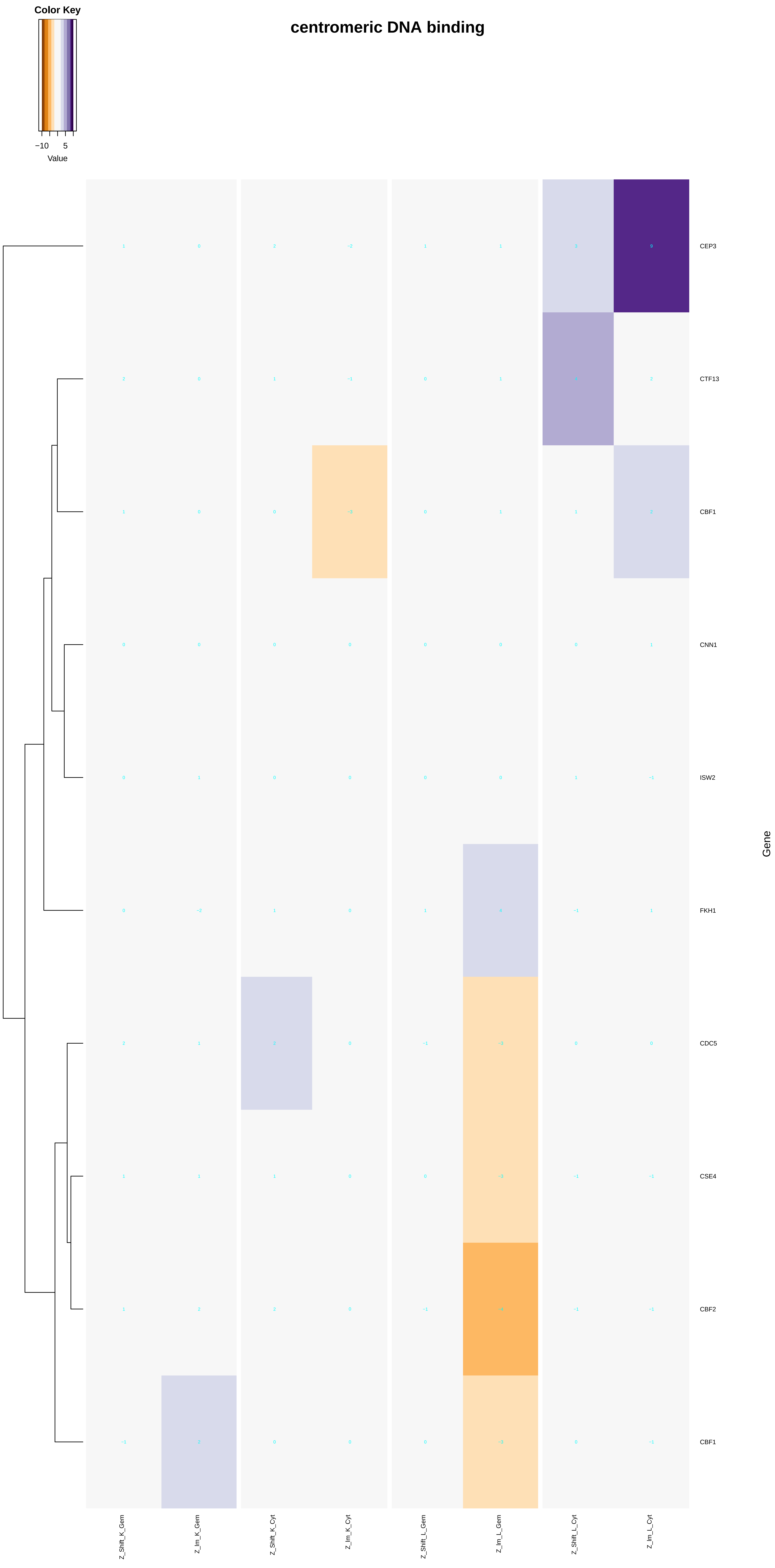

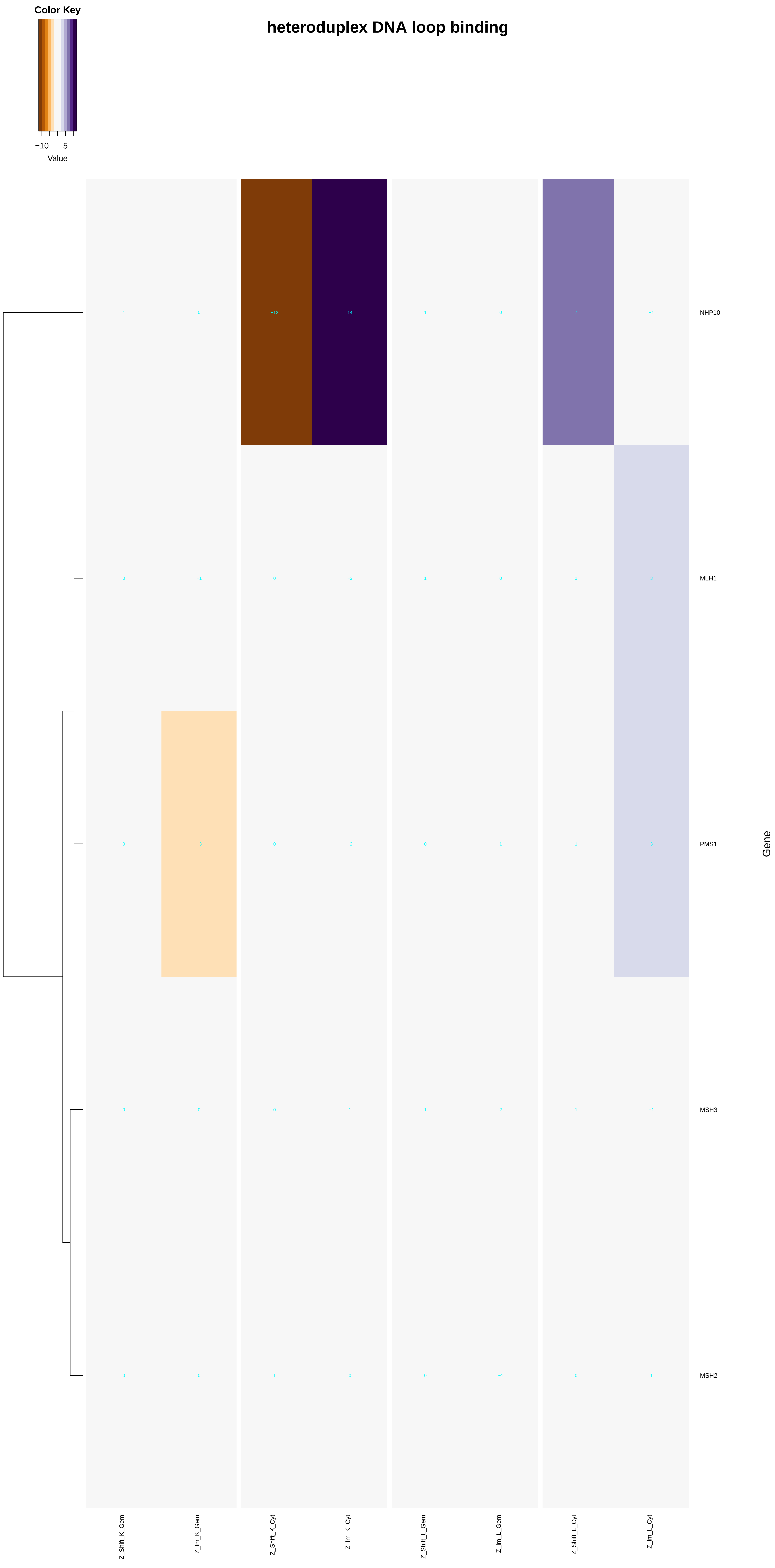
heteroduplex DNA loop binding
Color Key
−10
5
Value
NHP10
MLH1
PMS1
MSH3
MSH2
Gene
Z_Shift_K_Gem
Z_Im_K_Gem
Z_Shift_K_Cyt
Z_Im_K_Cyt
Z_Shift_L_Gem
Z_Im_L_Gem
Z_Shift_L_Cyt
Z_Im_L_Cyt

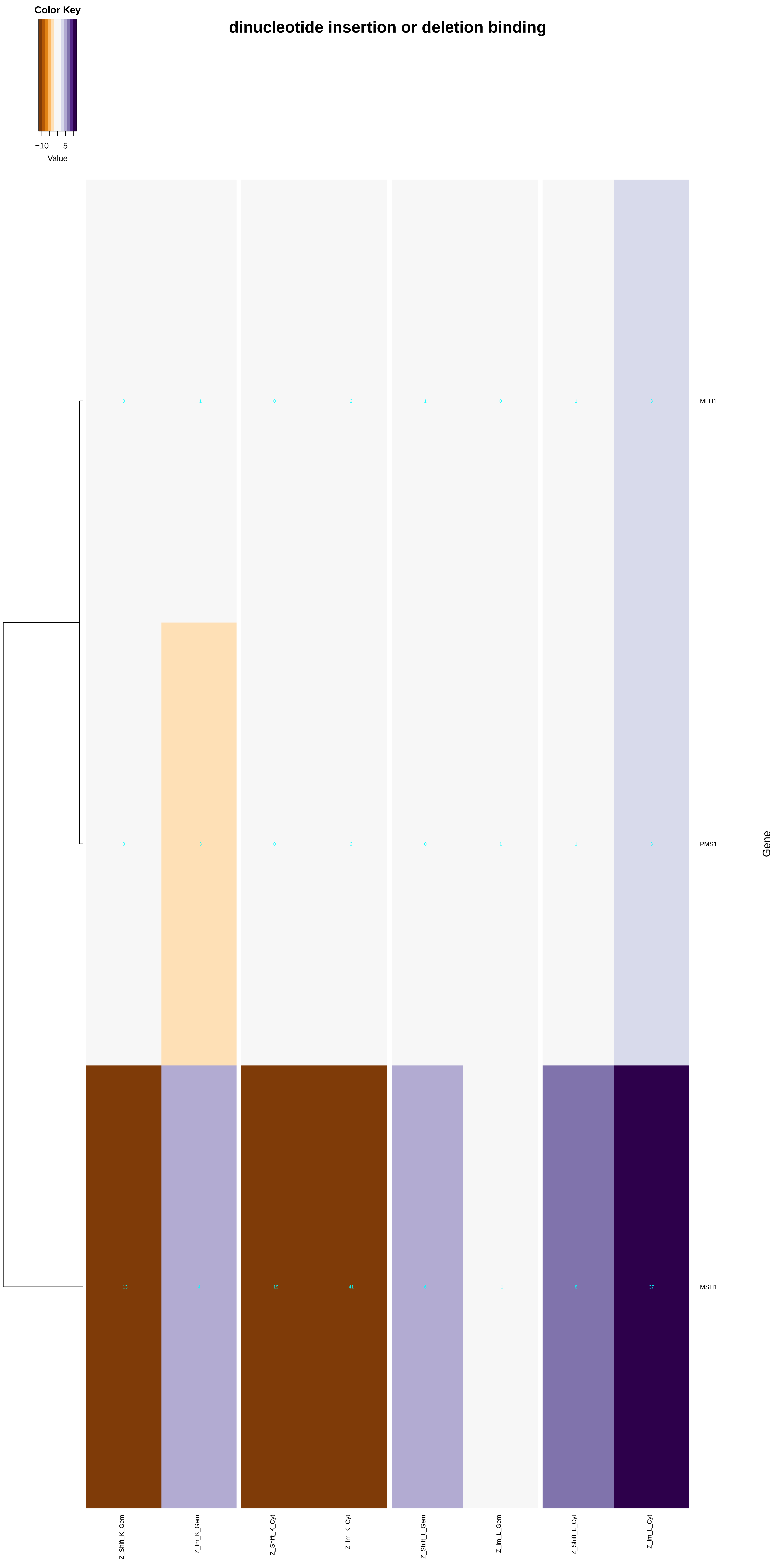
Color Key
−10
5
Value
dinucleotide insertion or deletion binding
MLH1
PMS1
MSH1
Gene
Z_Shift_K_Gem
Z_Im_K_Gem
Z_Shift_K_Cyt
Z_Im_K_Cyt
Z_Shift_L_Gem
Z_Im_L_Gem
Z_Shift_L_Cyt
Z_Im_L_Cyt

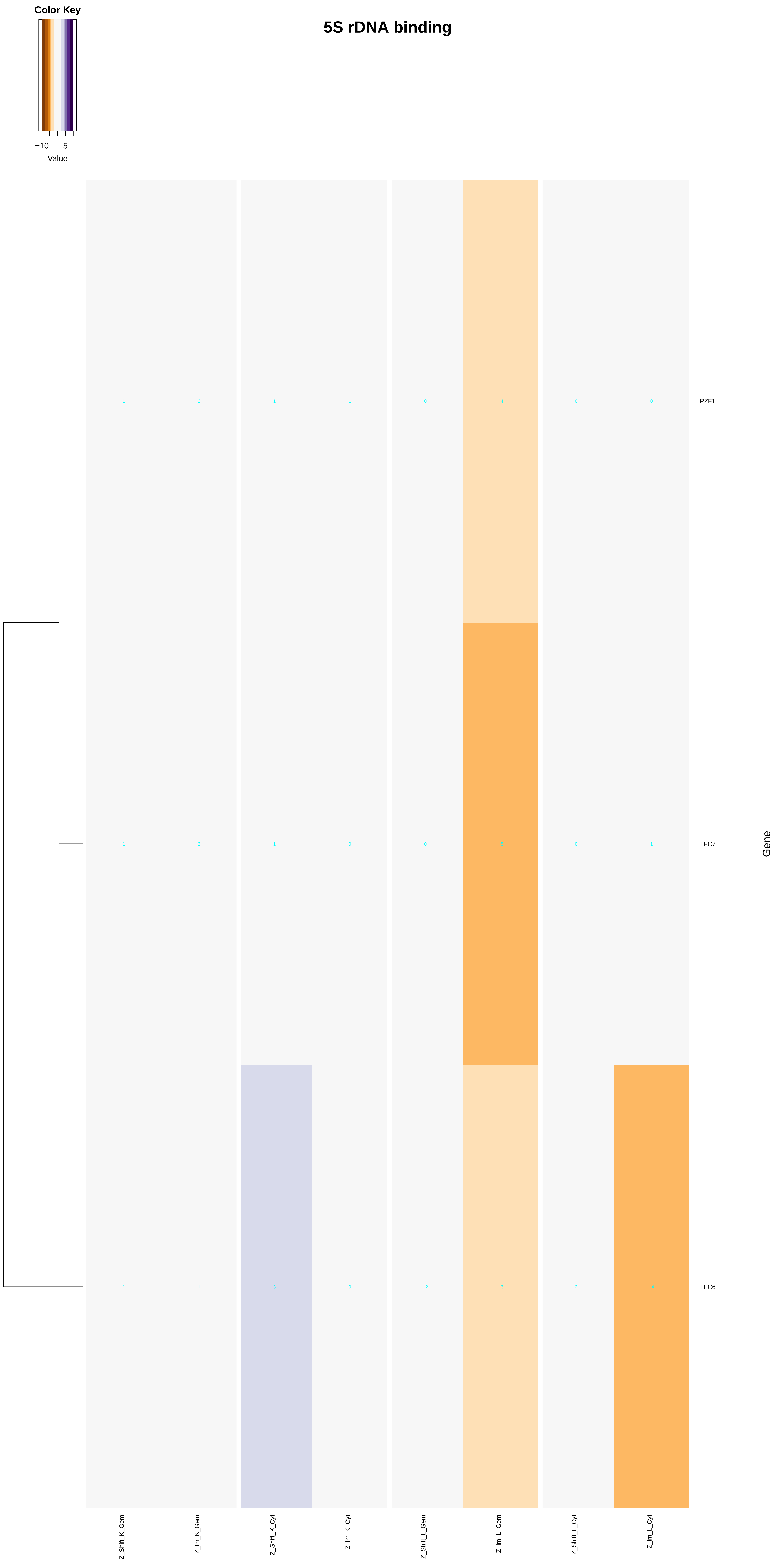

5S rDNA binding
Color Key
−10
5
Value
PZF1
TFC7
TFC6
Gene
Z_Shift_K_Gem
Z_Im_K_Gem
Z_Shift_K_Cyt
Z_Im_K_Cyt
Z_Shift_L_Gem
Z_Im_L_Gem
Z_Shift_L_Cyt
Z_Im_L_Cyt

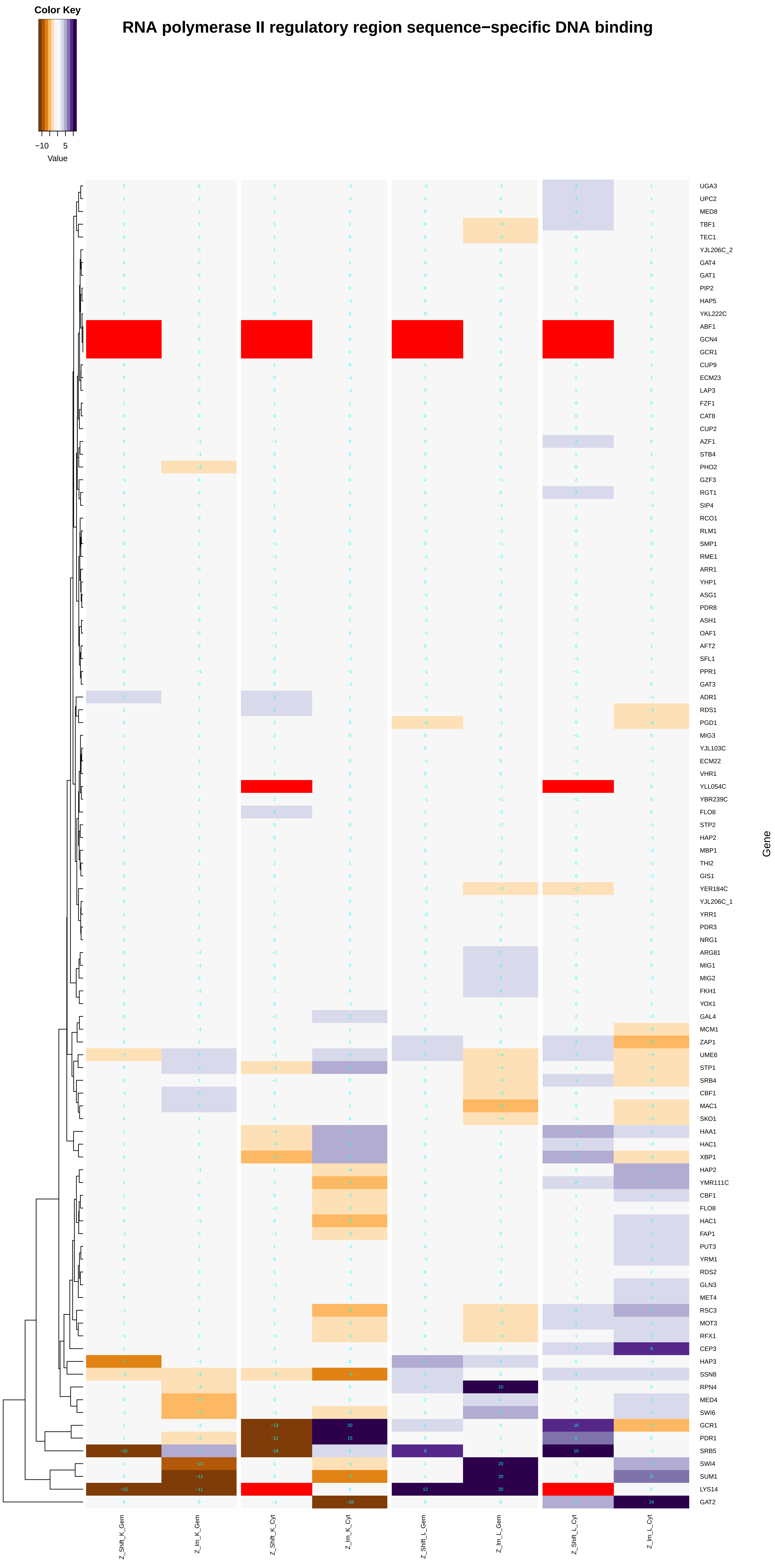

Color Key
−10
5
Value
RNA polymerase II regulatory region sequence−specific DNA binding
Gene
UGA3
UPC2
MED8
TBF1
TEC1
YJL206C_2
GAT4
GAT1
PIP2
HAP5
YKL222C
ABF1
GCN4
GCR1
CUP9
ECM23
LAP3
FZF1
CAT8
CUP2
AZF1
STB4
PHO2
GZF3
RGT1
SIP4
RCO1
RLM1
SMP1
RME1
ARR1
YHP1
ASG1
PDR8
ASH1
OAF1
AFT2
SFL1
PPR1
GAT3
ADR1
RDS1
PGD1
MIG3
YJL103C
ECM22
VHR1
YLL054C
YBR239C
FLO8
STP2
HAP2
MBP1
THI2
GIS1
YER184C
YJL206C_1
YRR1
PDR3
NRG1
ARG81
MIG1
MIG2
FKH1
YOX1
GAL4
MCM1
ZAP1
UME6
STP1
SRB4
CBF1
MAC1
SKO1
HAA1
HAC1
XBP1
HAP2
YMR111C
CBF1
FLO8
HAC1
FAP1
PUT3
YRM1
RDS2
GLN3
MET4
RSC3
MOT3
RFX1
CEP3
HAP3
SSN8
RPN4
MED4
SWI6
GCR1
PDR1
SRB5
SWI4
SUM1
LYS14
GAT2
Z_Shift_K_Gem
Z_Im_K_Gem
Z_Shift_K_Cyt
Z_Im_K_Cyt
Z_Shift_L_Gem
Z_Im_L_Gem
Z_Shift_L_Cyt
Z_Im_L_Cyt

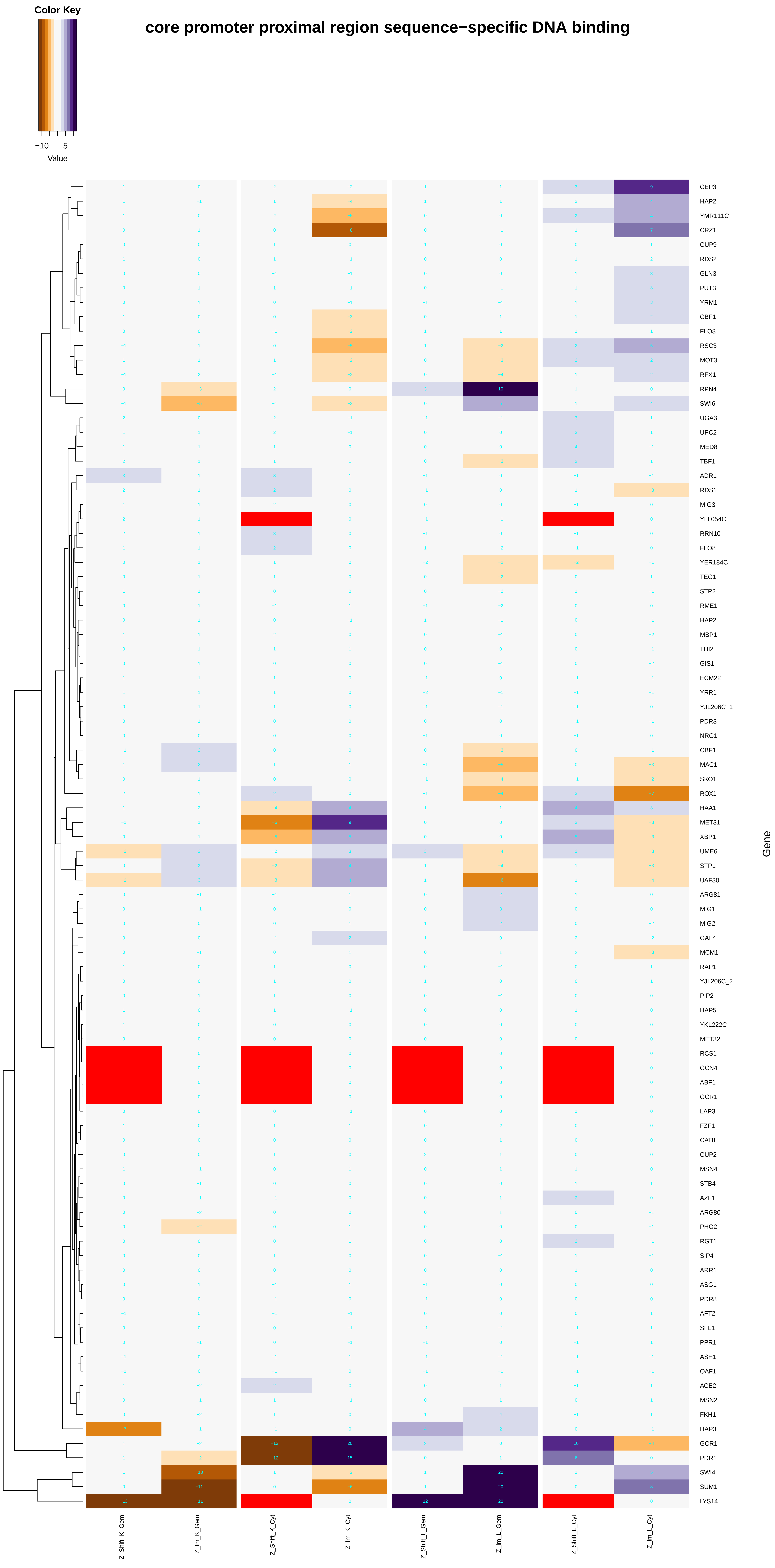
core promoter proximal region sequence−specific DNA binding
Color Key
−10
5
Value
CEP3
HAP2
YMR111C
CRZ1
CUP9
RDS2
GLN3
PUT3
YRM1
CBF1
FLO8
RSC3
MOT3
RFX1
RPN4
SWI6
UGA3
UPC2
MED8
TBF1
ADR1
RDS1
MIG3
YLL054C
RRN10
FLO8
YER184C
TEC1
STP2
RME1
HAP2
MBP1
THI2
GIS1
ECM22
YRR1
YJL206C_1
PDR3
NRG1
CBF1
MAC1
SKO1
ROX1
HAA1
MET31
XBP1
UME6
STP1
UAF30
ARG81
MIG1
MIG2
GAL4
MCM1
RAP1
YJL206C_2
PIP2
HAP5
YKL222C
MET32
RCS1
GCN4
ABF1
GCR1
LAP3
FZF1
CAT8
CUP2
MSN4
STB4
AZF1
ARG80
PHO2
RGT1
SIP4
ARR1
ASG1
PDR8
AFT2
SFL1
PPR1
ASH1
OAF1
ACE2
MSN2
FKH1
HAP3
GCR1
PDR1
SWI4
SUM1
LYS14
Gene
Z_Shift_K_Gem
Z_Im_K_Gem
Z_Shift_K_Cyt
Z_Im_K_Cyt
Z_Shift_L_Gem
Z_Im_L_Gem
Z_Shift_L_Cyt
Z_Im_L_Cyt

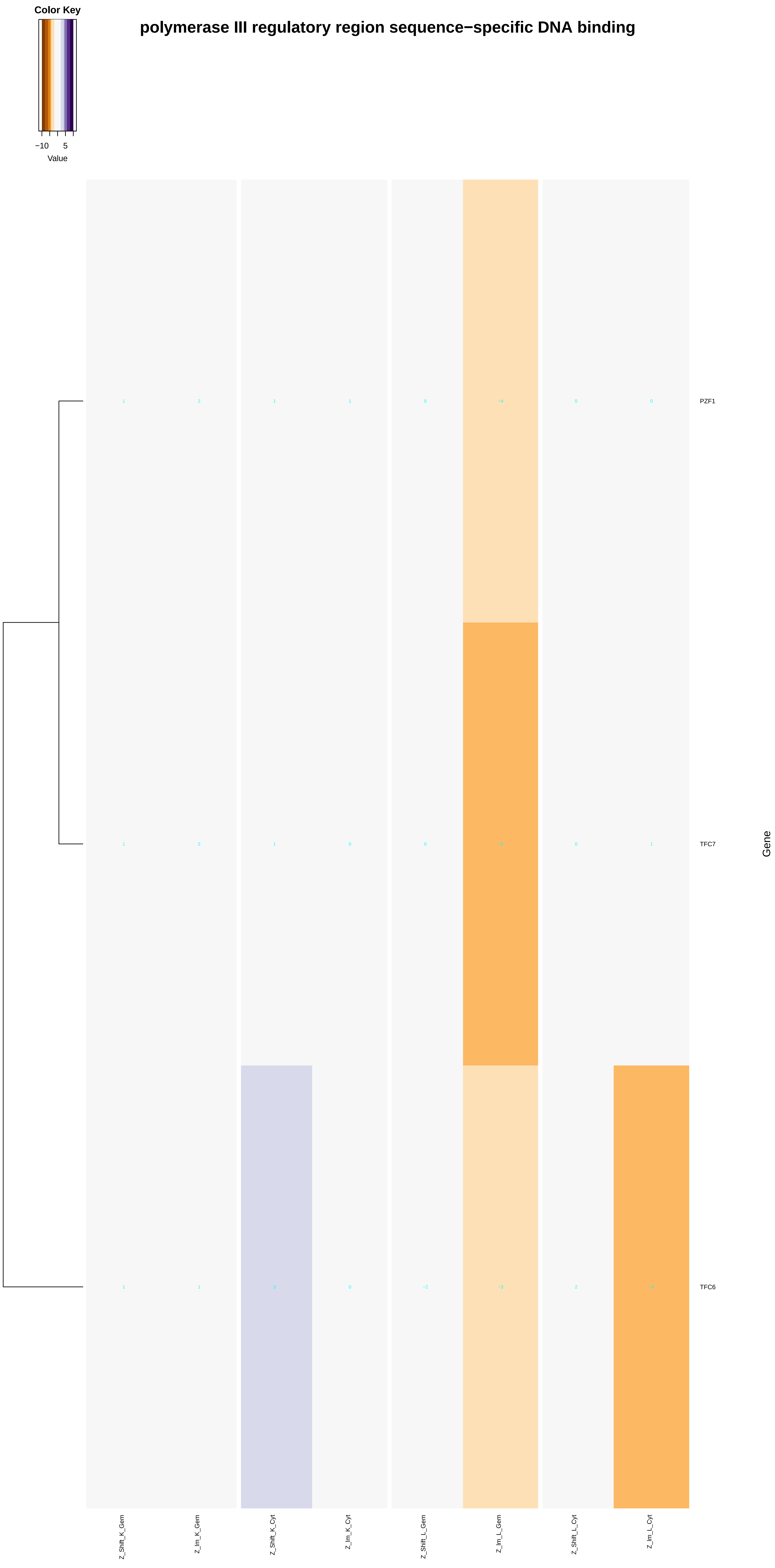
polymerase III regulatory region sequence−specific DNA binding
Color Key
−10
5
Value
1
2
1
1
0
−4
0
0
PZF1
1
2
1
0
0
−5
0
1
TFC7
1
1
3
0
−2
−3
2
−4
TFC6
Gene
Z_Shift_K_Gem
Z_lm_K_Gem
Z_Shift_K_Cyt
Z_lm_K_Cyt
Z_Shift_L_Gem
Z_lm_L_Gem
Z_Shift_L_Cyt
Z_lm_L_Cyt

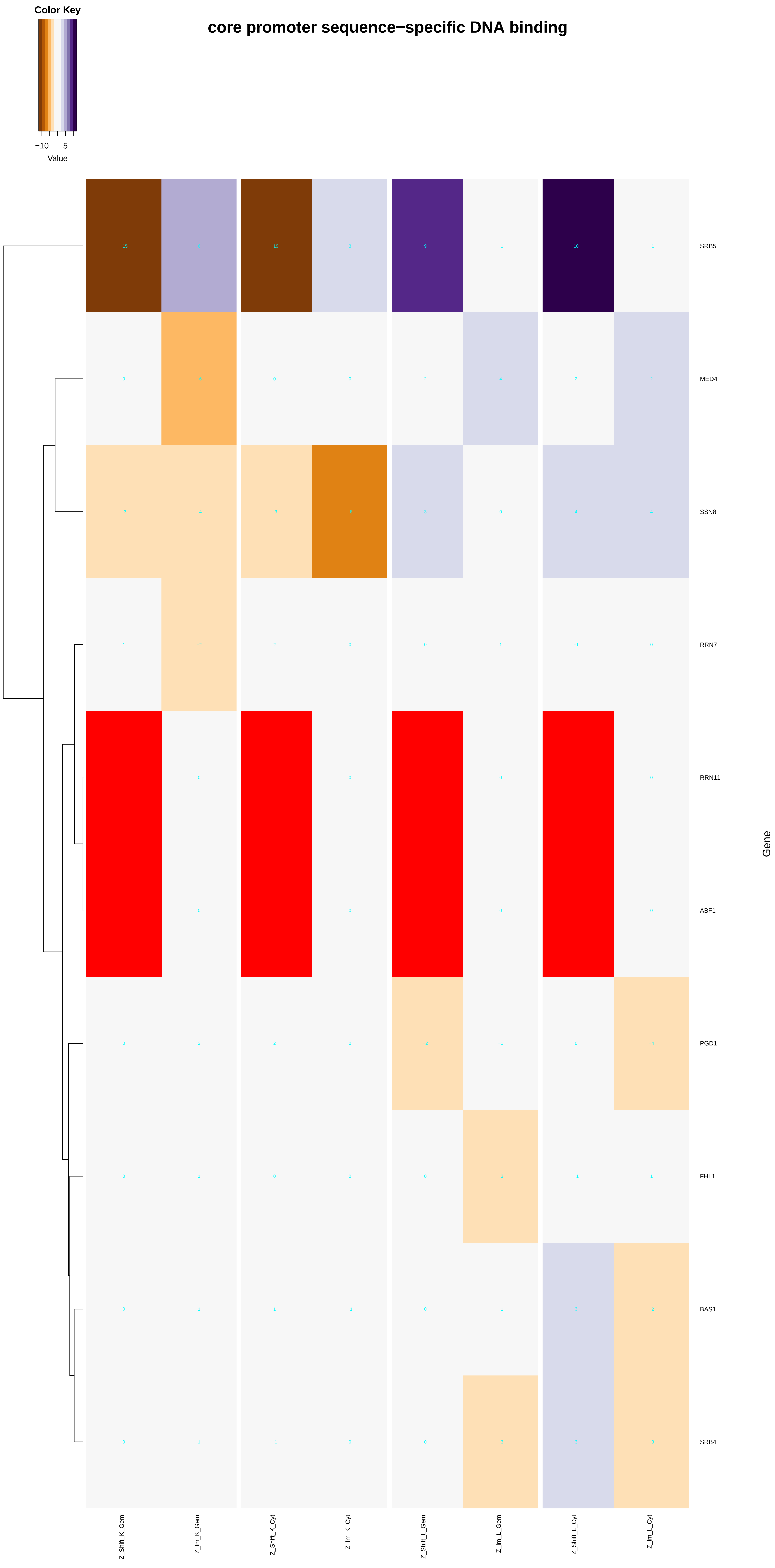
Color Key
−10
5
Value
core promoter sequence−specific DNA binding
SRB5
MED4
SSN8
RRN7
RRN11
ABF1
PGD1
FHL1
BAS1
SRB4
Gene
Z_Shift_K_Gem
Z_Im_K_Gem
Z_Shift_K_Cyt
Z_Im_K_Cyt
Z_Shift_L_Gem
Z_Im_L_Gem
Z_Shift_L_Cyt
Z_Im_L_Cyt

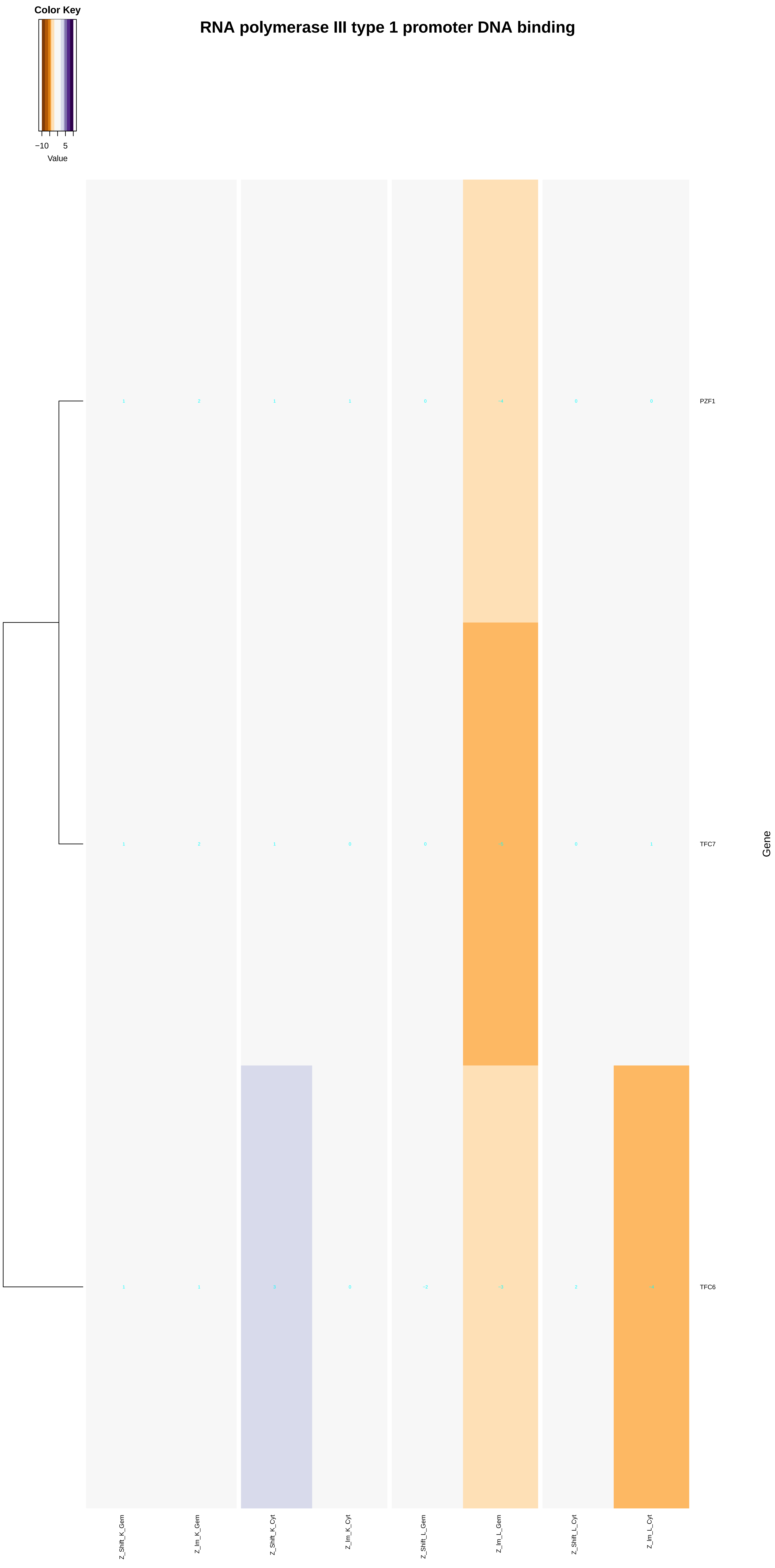

RNA polymerase III type 1 promoter DNA binding
Color Key
−10
5
Value
PZF1
TFC7
TFC6
Gene
Z_Shift_K_Gem
Z_Im_K_Gem
Z_Shift_K_Cyt
Z_Im_K_Cyt
Z_Shift_L_Gem
Z_Im_L_Gem
Z_Shift_L_Cyt
Z_Im_L_Cyt

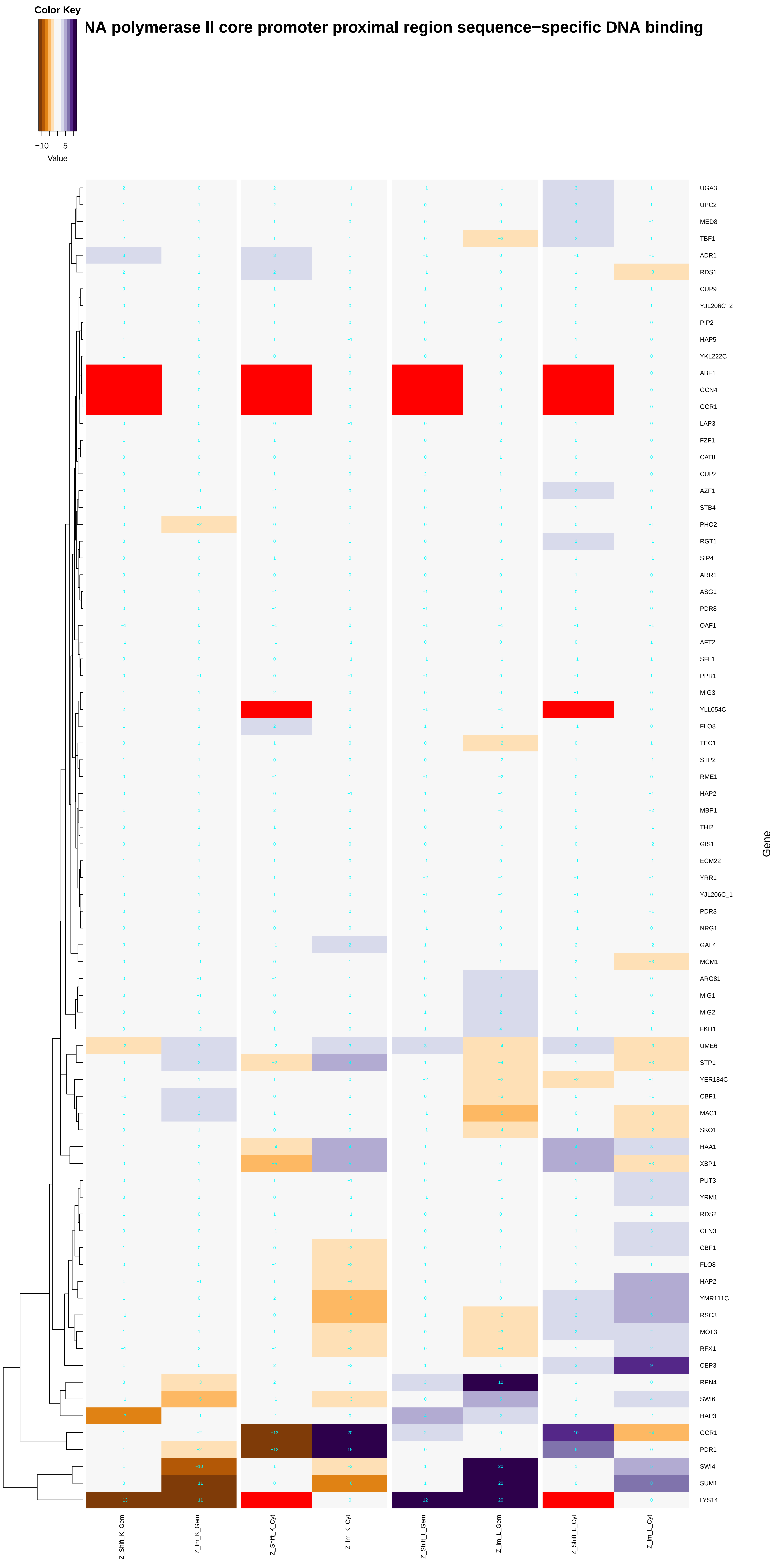

Color Key
−10 5
Value
NA polymerase II core promoter proximal region sequence−specific DNA binding
UGA3
UPC2
MED8
TBF1
ADR1
RDS1
CUP9
YJL206C_2
PIP2
HAP5
YKL222C
ABF1
GCN4
GCR1
LAP3
FZF1
CAT8
CUP2
AZF1
STB4
PHO2
RGT1
SIP4
ARR1
ASG1
PDR8
OAF1
AFT2
SFL1
PPR1
MIG3
YLL054C
FLO8
TEC1
STP2
RME1
HAP2
MBP1
THI2
GIS1
ECM22
YRR1
YJL206C_1
PDR3
NRG1
GAL4
MCM1
ARG81
MIG1
MIG2
FKH1
UME6
STP1
YER184C
CBF1
MAC1
SKO1
HAA1
XBP1
PUT3
YRM1
RDS2
GLN3
CBF1
FLO8
HAP2
YMR111C
RSC3
MOT3
RFX1
CEP3
RPN4
SWI6
HAP3
GCR1
PDR1
SWI4
SUM1
LYS14
Gene
Z_Shift_K_Gem
Z_Im_K_Gem
Z_Shift_K_Cyt
Z_Im_K_Cyt
Z_Shift_L_Gem
Z_Im_L_Gem
Z_Shift_L_Cyt
Z_Im_L_Cyt

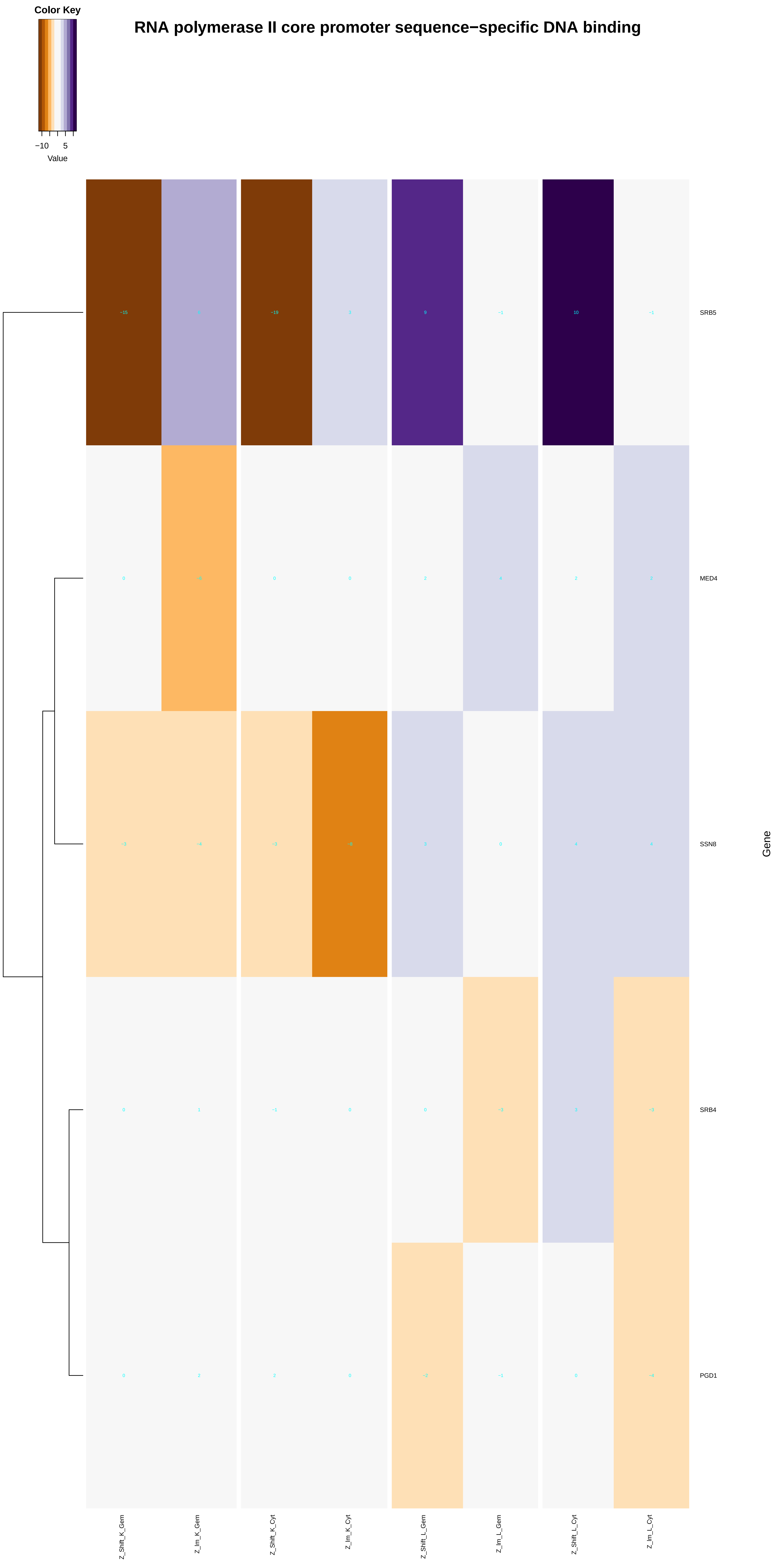
Color Key
−10 5
Value
RNA polymerase II core promoter sequence−specific DNA binding
−15
6
−19
3
9
−1
10
−1
SRB5
0
−6
0
0
2
4
2
2
MED4
−3
−4
−3
−8
3
0
4
4
SSN8
0
1
−1
0
0
−3
3
−3
SRB4
0
2
2
0
−2
−1
0
−4
PGD1
Gene
Z_Shift_K_Gem
Z_Im_K_Gem
Z_Shift_K_Cyt
Z_Im_K_Cyt
Z_Shift_L_Gem
Z_Im_L_Gem
Z_Shift_L_Cyt
Z_Im_L_Cyt

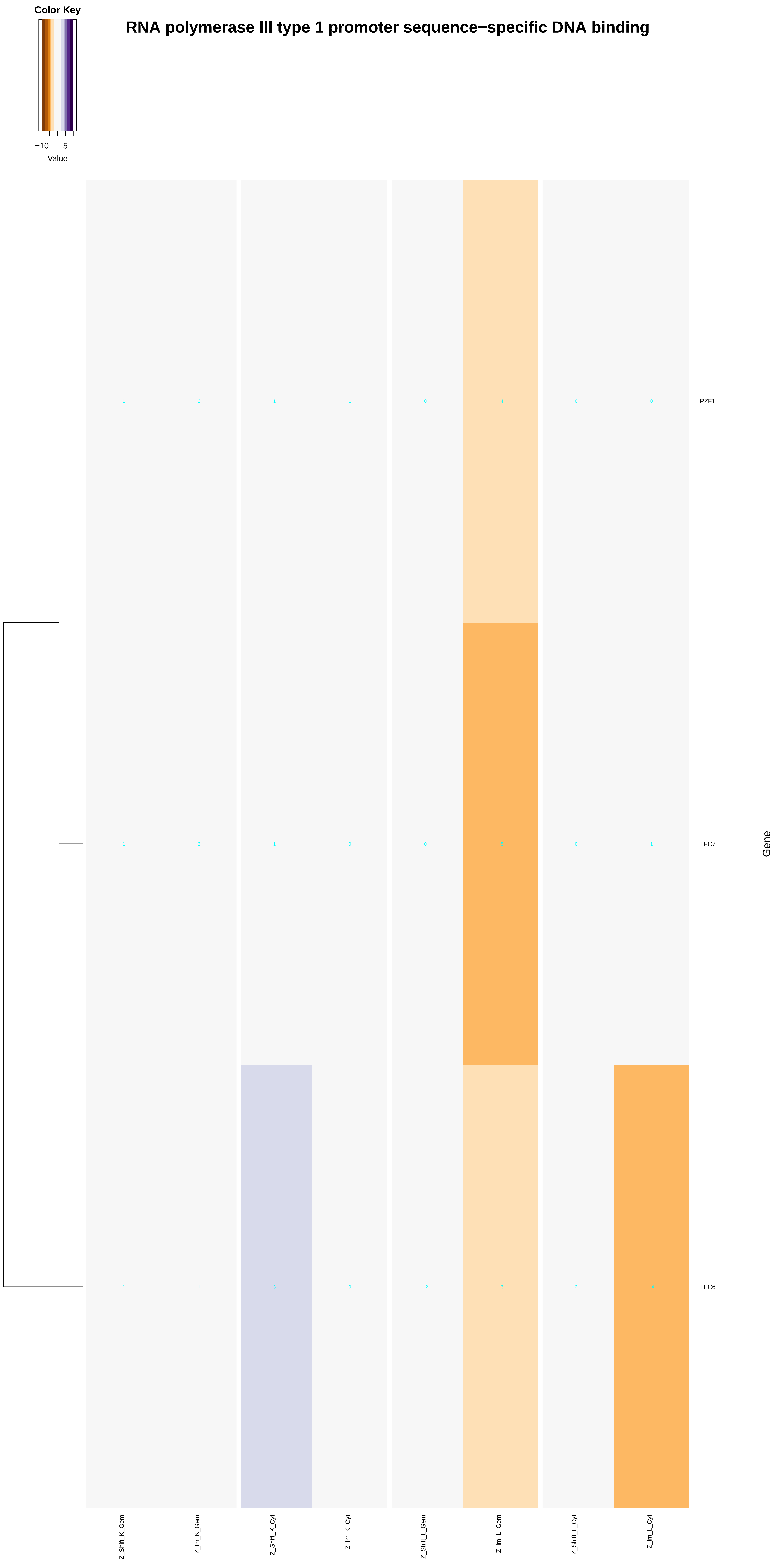

Color Key
−10
5
Value
RNA polymerase III type 1 promoter sequence−specific DNA binding
1
2
1
1
0
−4
0
0
PZF1
1
2
1
0
0
−5
0
1
TFC7
1
1
3
0
−2
−3
2
−4
TFC6
Gene
Z_Shift_K_Gem
Z_Im_K_Gem
Z_Shift_K_Cyt
Z_Im_K_Cyt
Z_Shift_L_Gem
Z_Im_L_Gem
Z_Shift_L_Cyt
Z_Im_L_Cyt
