## Additional File 7 for "A humanized yeast phenomic model of deoxycytidine kinase to predict genetic buffering of nucleoside analog cytotoxicity": endonuclease_activity.pdf

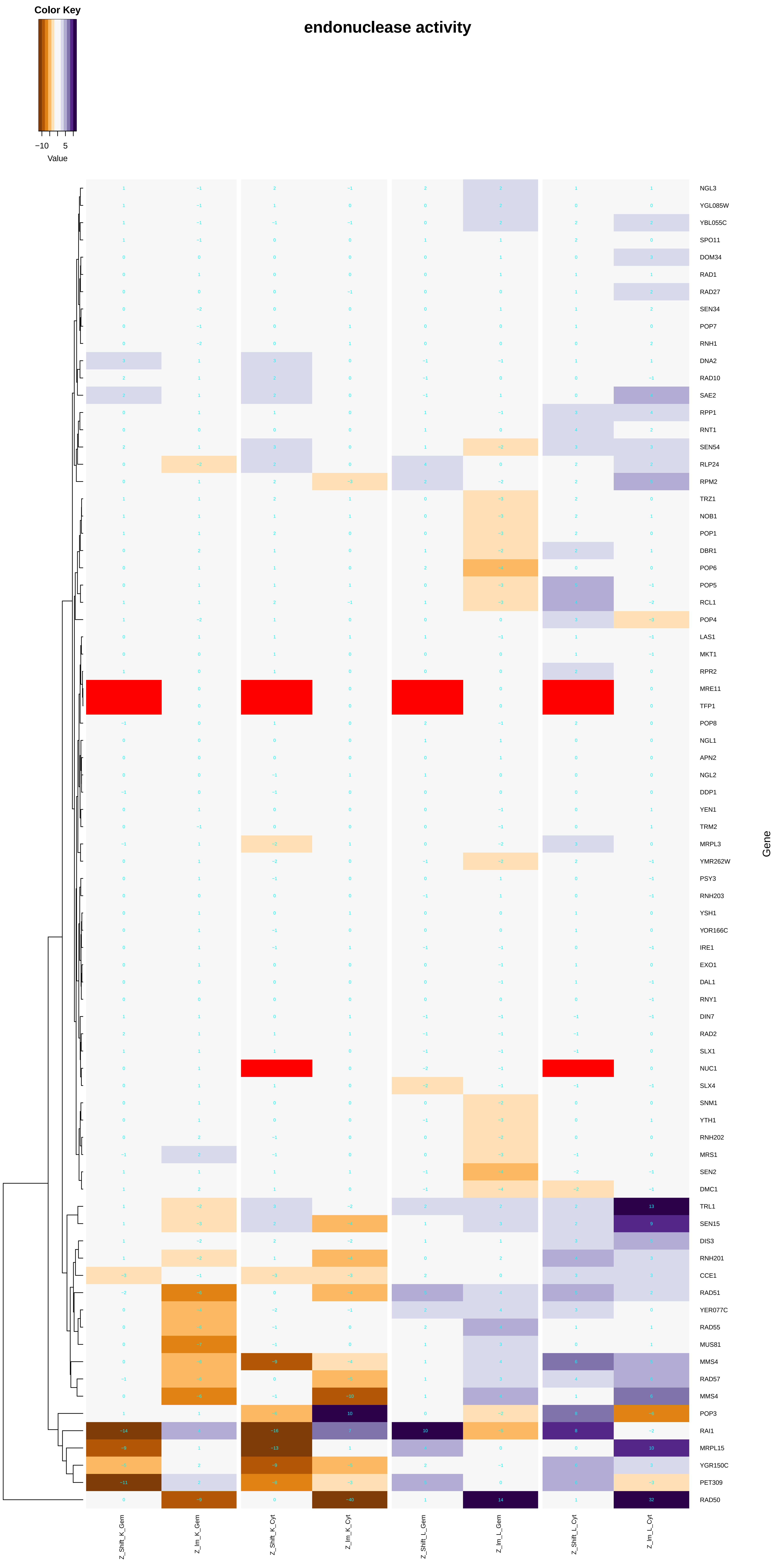

Color Key
−10
5
Value
endonuclease activity
Gene
NGL3
YGL085W
YBL055C
SPO11
DOM34
RAD1
RAD27
SEN34
POP7
RNH1
DNA2
RAD10
SAE2
RPP1
RNT1
SEN54
RLP24
RPM2
TRZ1
NOB1
POP1
DBR1
POP6
POP5
RCL1
POP4
LAS1
MKT1
RPR2
MRE11
TFP1
POP8
NGL1
APN2
NGL2
DDP1
YEN1
TRM2
MRPL3
YMR262W
PSY3
RNH203
YSH1
YOR166C
IRE1
EXO1
DAL1
RNY1
DIN7
RAD2
SLX1
NUC1
SLX4
SNM1
YTH1
RNH202
MRS1
SEN2
DMC1
TRL1
SEN15
DIS3
RNH201
CCE1
RAD51
YER077C
RAD55
MUS81
MMS4
RAD57
MMS4
POP3
RAI1
MRPL15
YGR150C
PET309
RAD50
Z_Shift_K_Gem
Z_Im_K_Gem
Z_Shift_K_Cyt
Z_Im_K_Cyt
Z_Shift_L_Gem
Z_Im_L_Gem
Z_Shift_L_Cyt
Z_Im_L_Cyt

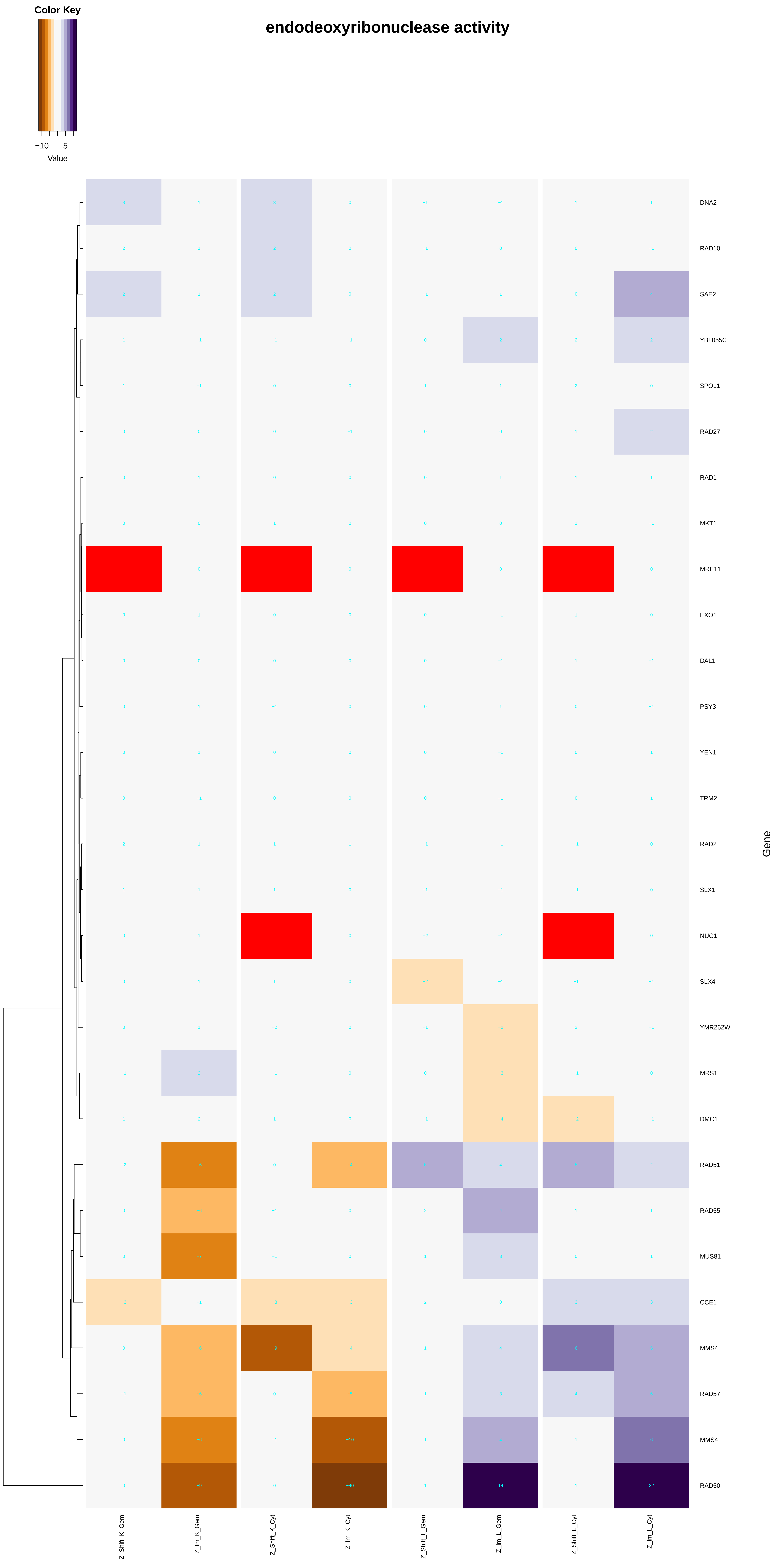
Color Key
−10
5
Value
endodeoxyribonuclease activity
DNA2
RAD10
SAE2
YBL055C
SPO11
RAD27
RAD1
MKT1
MRE11
EXO1
DAL1
PSY3
YEN1
TRM2
RAD2
SLX1
NUC1
SLX4
YMR262W
MRS1
DMC1
RAD51
RAD55
MUS81
CCE1
MMS4
RAD57
MMS4
RAD50
Gene
Z_Shift_K_Gem
Z_Im_K_Gem
Z_Shift_K_Cyt
Z_Im_K_Cyt
Z_Shift_L_Gem
Z_Im_L_Gem
Z_Shift_L_Cyt
Z_Im_L_Cyt

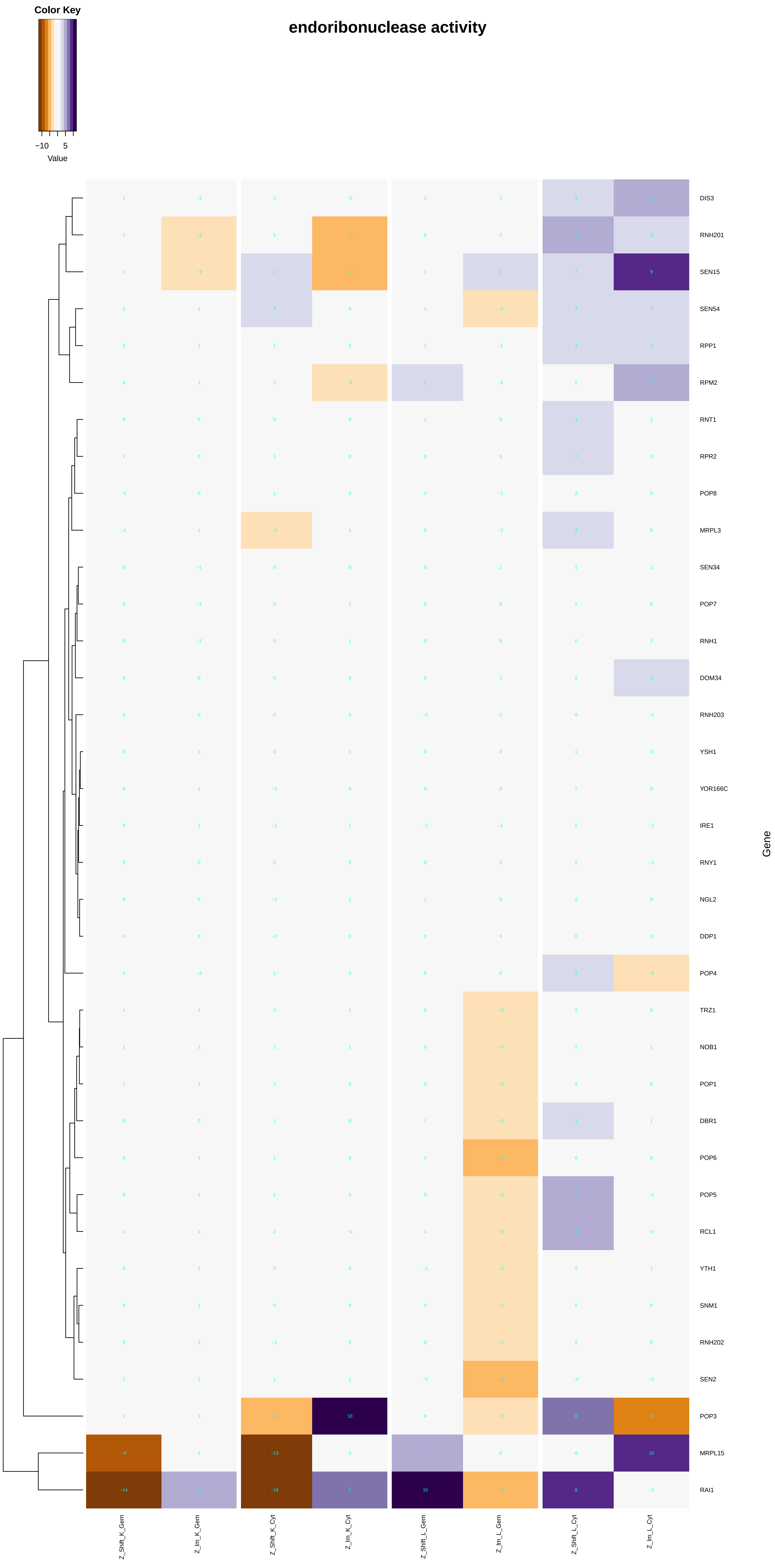

Color Key
−10
5
Value
endoribonuclease activity
DIS3
RNH201
SEN15
SEN54
RPP1
RPM2
RNT1
RPR2
POP8
MRPL3
SEN34
POP7
RNH1
DOM34
RNH203
YSH1
YOR166C
IRE1
RNY1
NGL2
DDP1
POP4
TRZ1
NOB1
POP1
DBR1
POP6
POP5
RCL1
YTH1
SNM1
RNH202
SEN2
POP3
MRPL15
RAI1
Gene
Z_Shift_K_Gem
Z_Im_K_Gem
Z_Shift_K_Cyt
Z_Im_K_Cyt
Z_Shift_L_Gem
Z_Im_L_Gem
Z_Shift_L_Cyt
Z_Im_L_Cyt

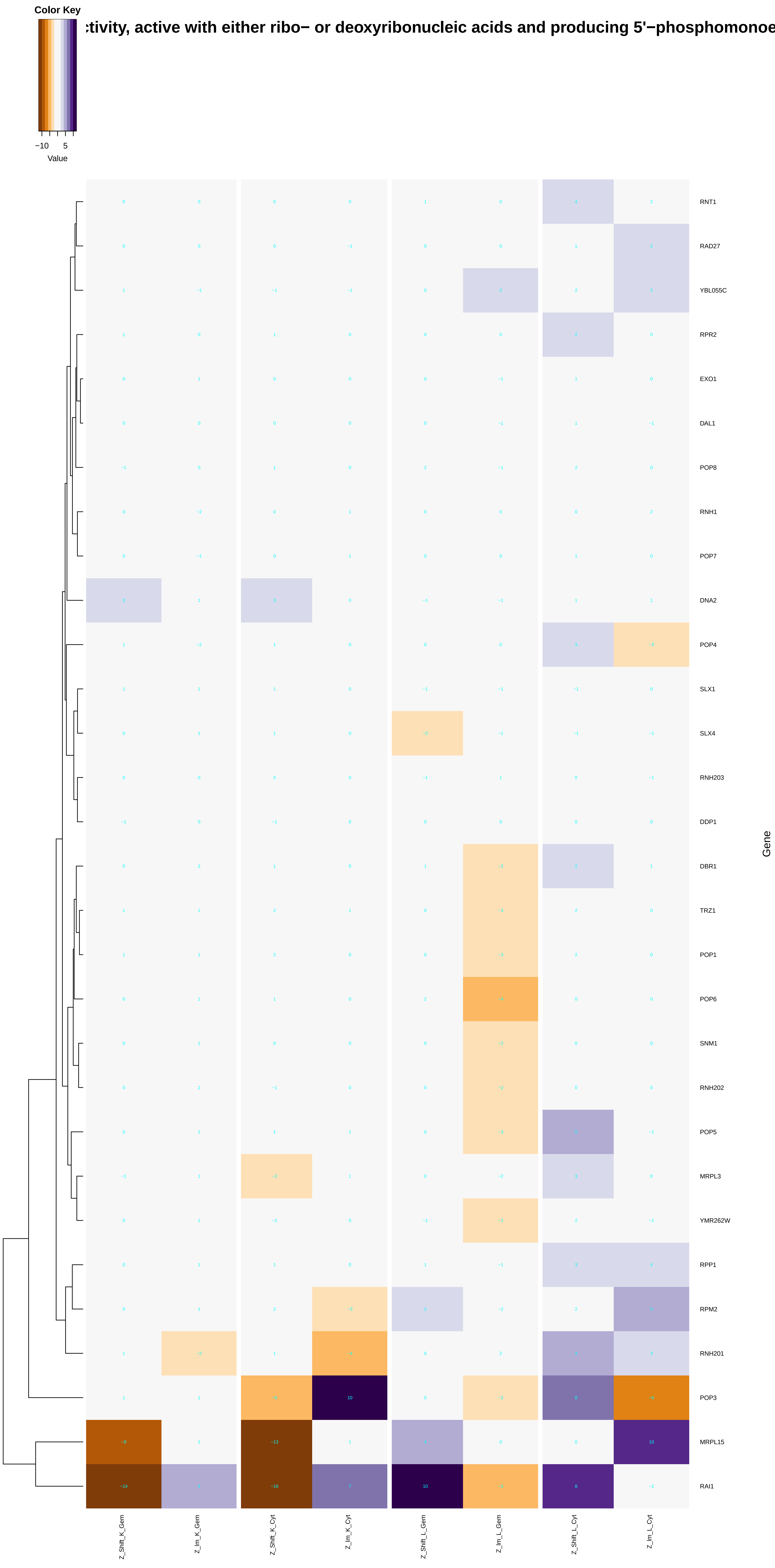
Color Key
−10 5
Value
:tivity, active with either ribo− or deoxyribonucleic acids and producing 5'−phosphomonoe
RNT1
RAD27
YBL055C
RPR2
EXO1
DAL1
POP8
RNH1
POP7
DNA2
POP4
SLX1
SLX4
RNH203
DDP1
DBR1
TRZ1
POP1
POP6
SNM1
RNH202
POP5
MRPL3
YMR262W
RPP1
RPM2
RNH201
POP3
MRPL15
RAI1
Gene
Z_Shift_K_Gem
Z_Im_K_Gem
Z_Shift_K_Cyt
Z_Im_K_Cyt
Z_Shift_L_Gem
Z_Im_L_Gem
Z_Shift_L_Cyt
Z_Im_L_Cyt

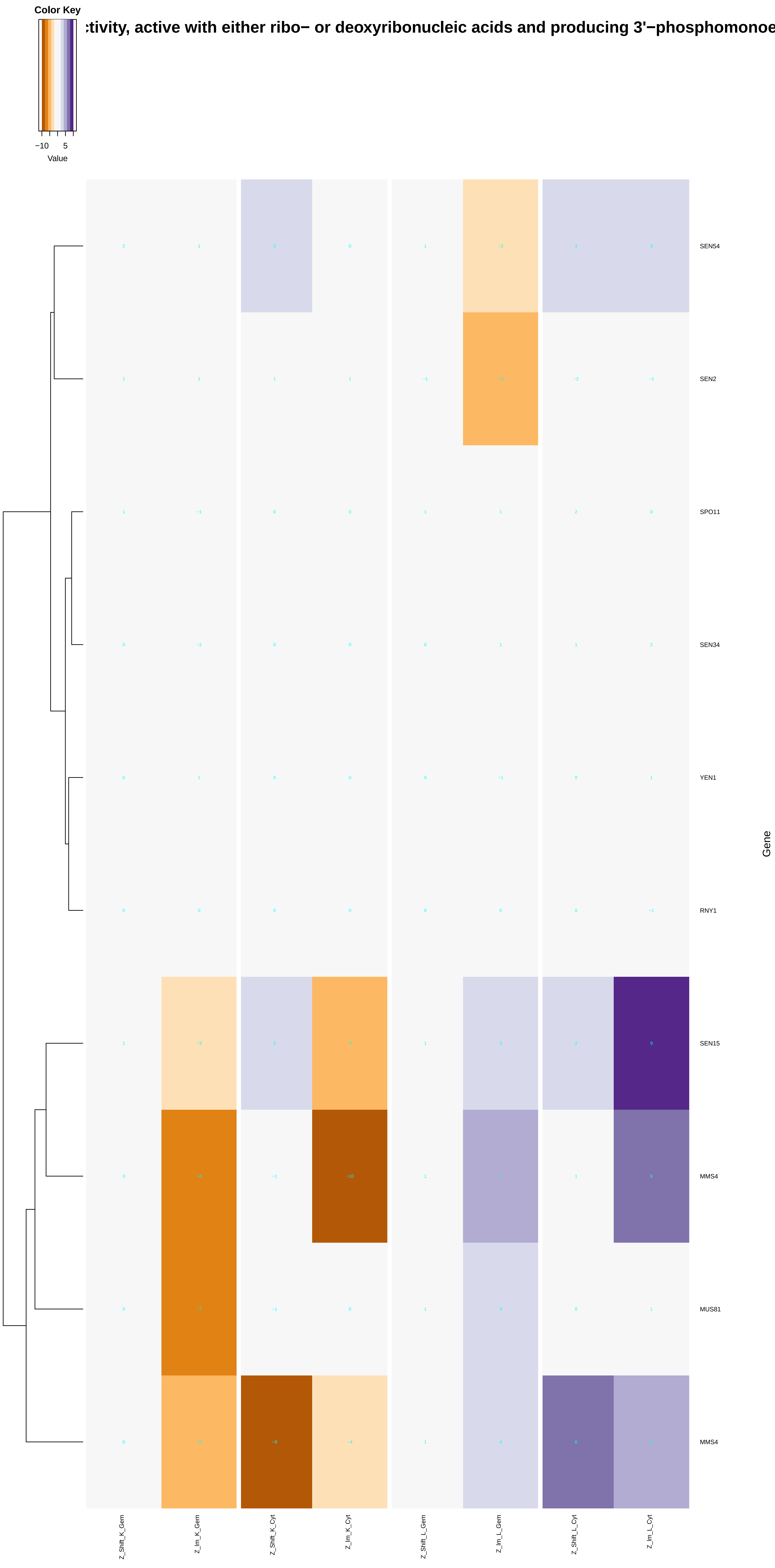

Color Key
−10
5
Value
:tivity, active with either ribo− or deoxyribonucleic acids and producing 3'−phosphomonoe
SEN54
SEN2
SPO11
SEN34
YEN1
RNY1
SEN15
MMS4
MUS81
MMS4
Gene
Z_Shift_K_Gem
Z_Im_K_Gem
Z_Shift_K_Cyt
Z_Im_K_Cyt
Z_Shift_L_Gem
Z_Im_L_Gem
Z_Shift_L_Cyt
Z_Im_L_Cyt

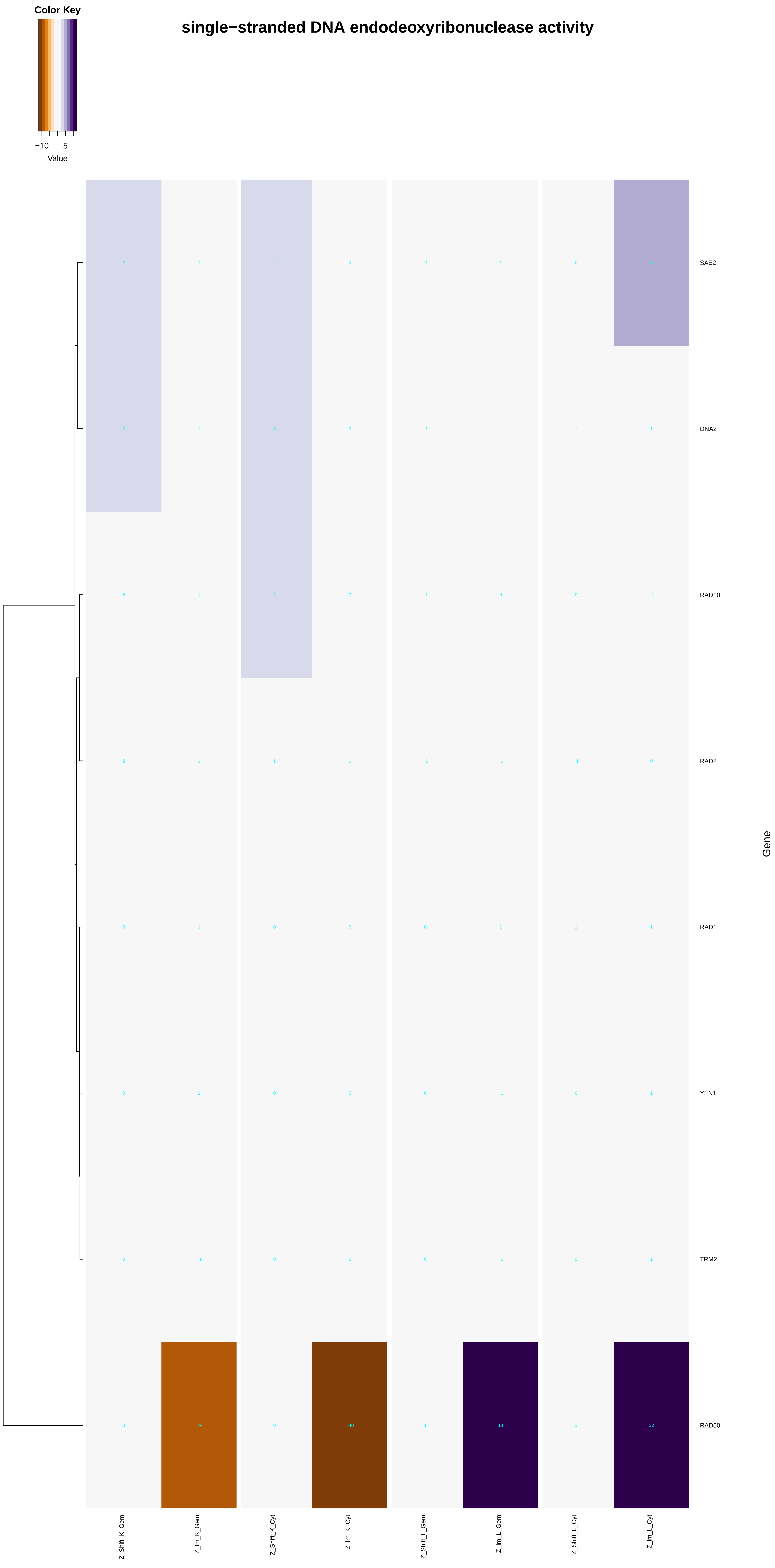
Color Key
−10
5
Value
single−stranded DNA endodeoxyribonuclease activity
SAE2
DNA2
RAD10
RAD2
RAD1
YEN1
TRM2
RAD50
Gene
Z_Shift_K_Gem
Z_Im_K_Gem
Z_Shift_K_Cyt
Z_Im_K_Cyt
Z_Shift_L_Gem
Z_Im_L_Gem
Z_Shift_L_Cyt
Z_Im_L_Cyt

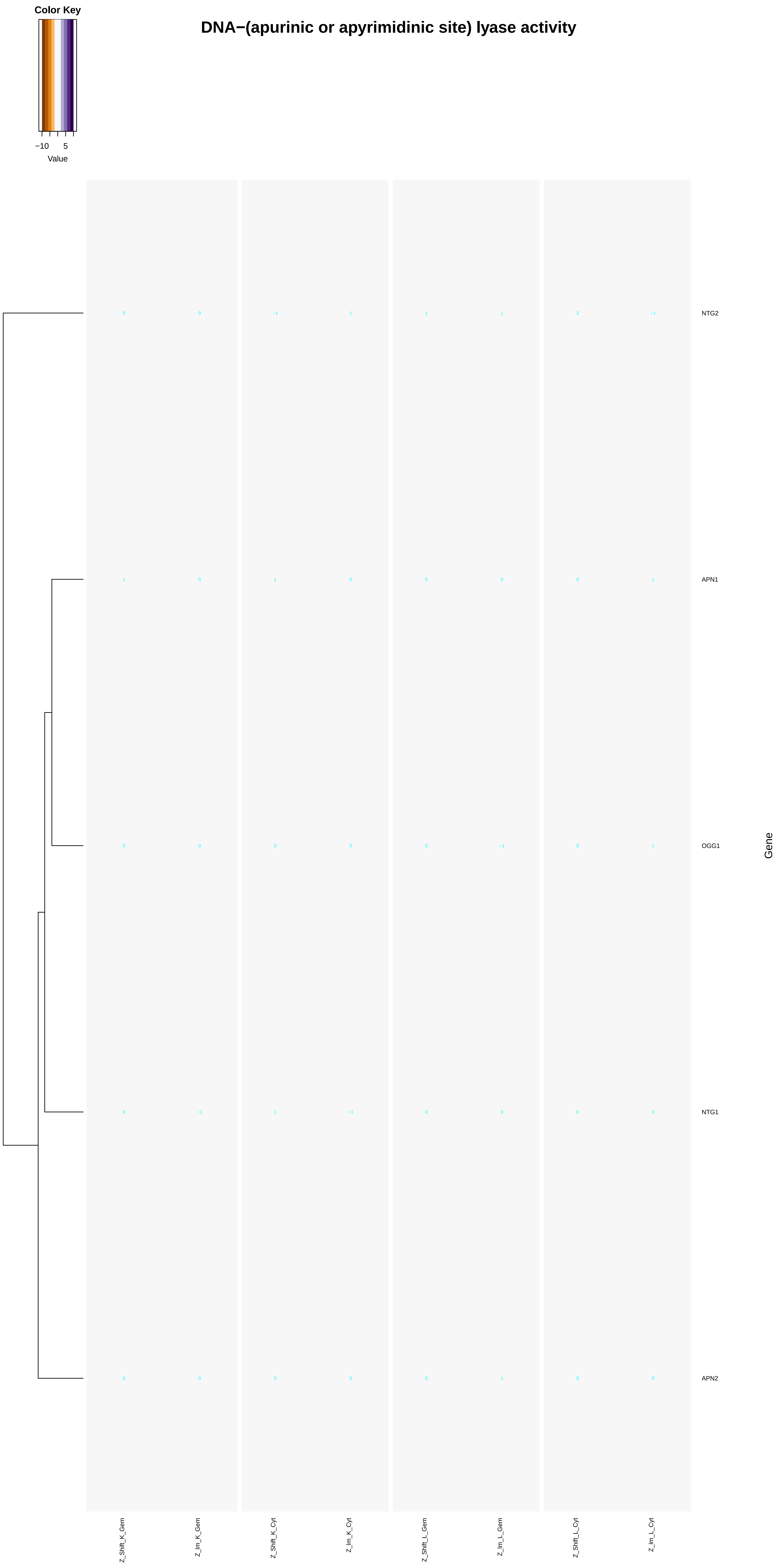

Color Key
−10
5
Value
DNA−(apurinic or apyrimidinic site) lyase activity
NTG2
APN1
OGG1
NTG1
APN2
Gene
Z_Shift_K_Gem
Z_Im_K_Gem
Z_Shift_K_Cyt
Z_Im_K_Cyt
Z_Shift_L_Gem
Z_Im_L_Gem
Z_Shift_L_Cyt
Z_Im_L_Cyt

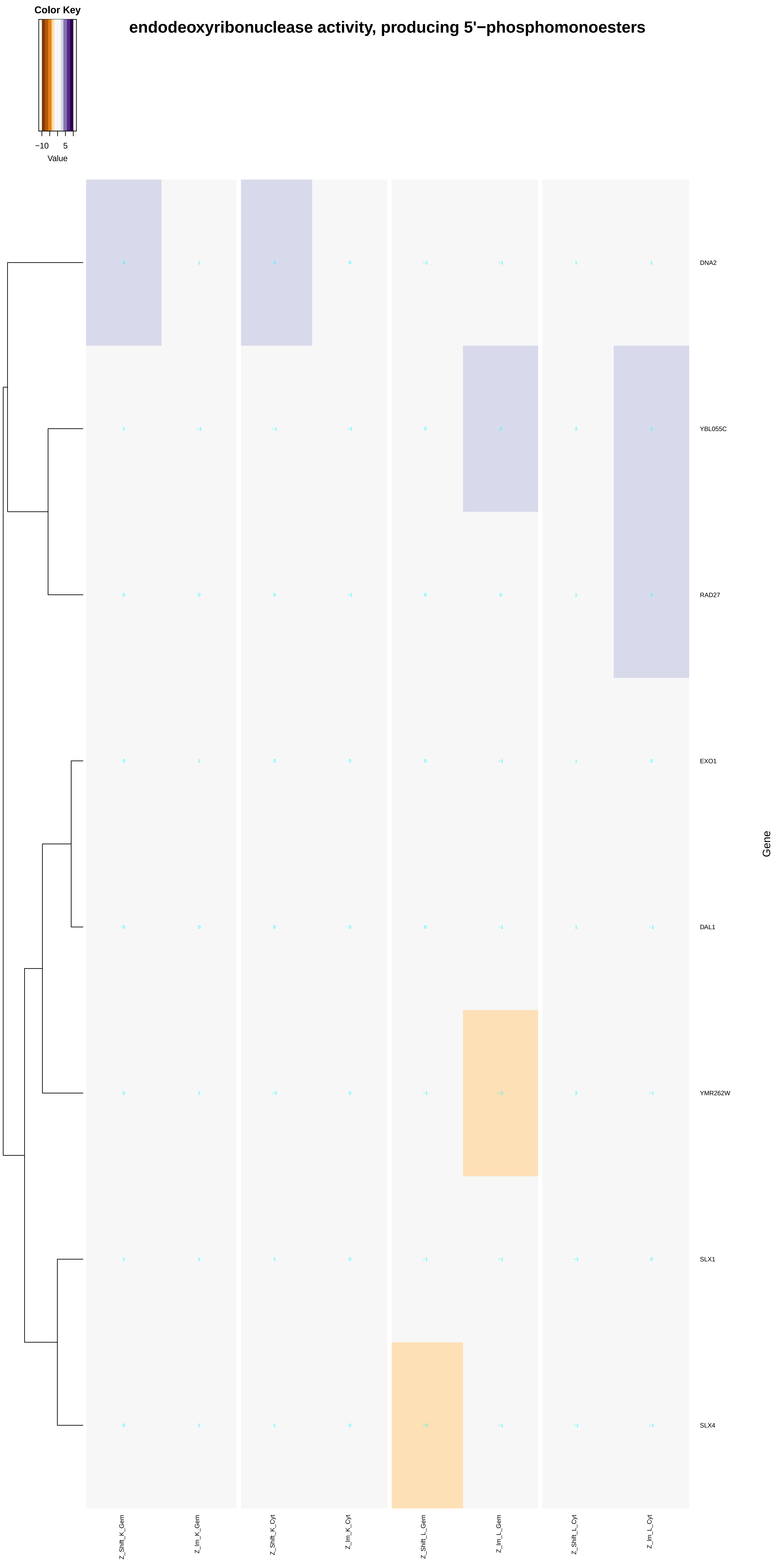

endodeoxyribonuclease activity, producing 5'−phosphomonoesters
Color Key
−10
5
Value
DNA2
YBL055C
RAD27
EXO1
DAL1
YMR262W
SLX1
SLX4
Gene
Z_Shift_K_Gem
Z_Im_K_Gem
Z_Shift_K_Cyt
Z_Im_K_Cyt
Z_Shift_L_Gem
Z_Im_L_Gem
Z_Shift_L_Cyt
Z_Im_L_Cyt

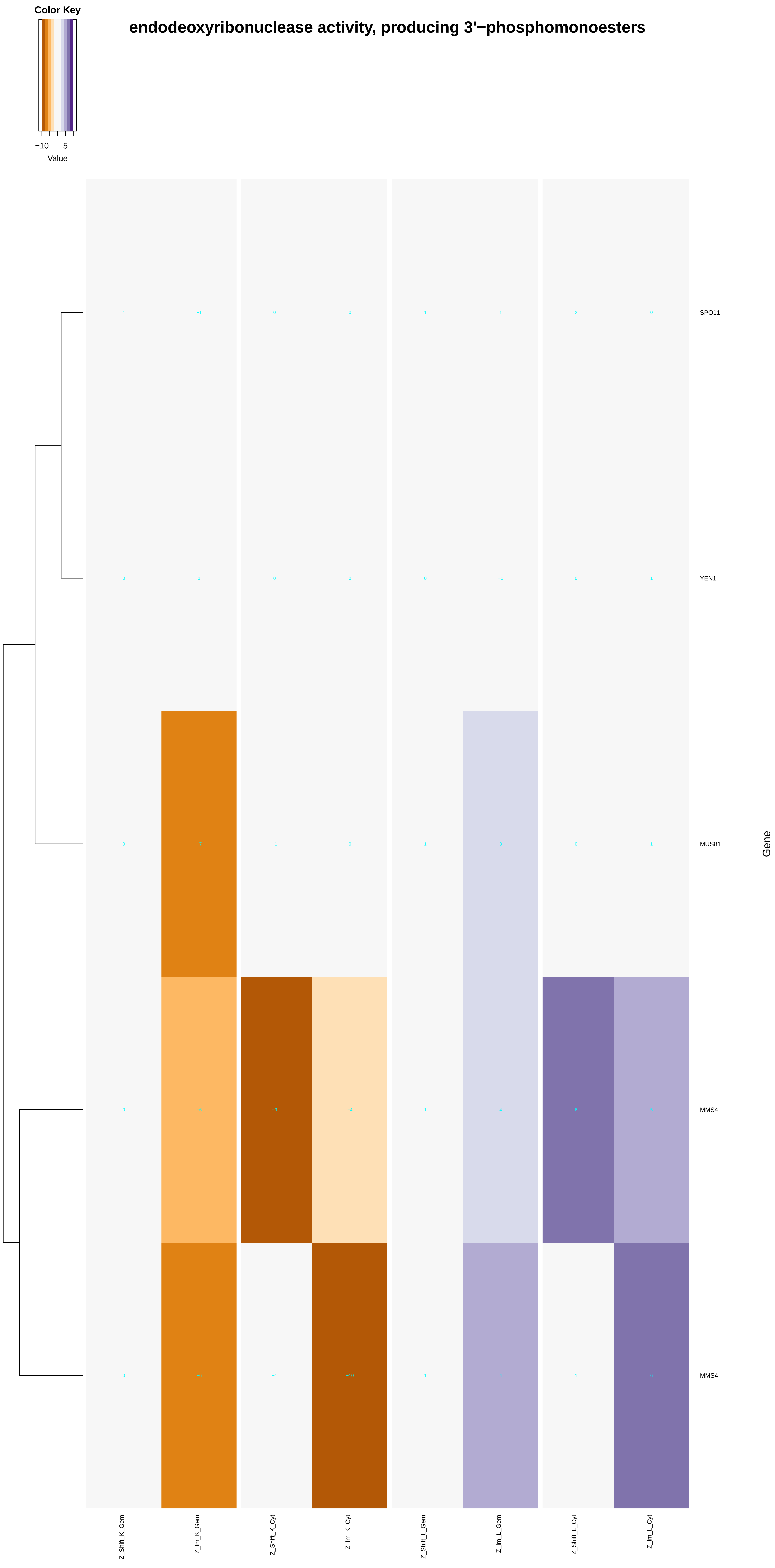
endodeoxyribonuclease activity, producing 3'−phosphomonoesters
Color Key
−10
5
Value
SPO11
YEN1
MUS81
MMS4
MMS4
Gene
Z_Shift_K_Gem
Z_Im_K_Gem
Z_Shift_K_Cyt
Z_Im_K_Cyt
Z_Shift_L_Gem
Z_Im_L_Gem
Z_Shift_L_Cyt
Z_Im_L_Cyt

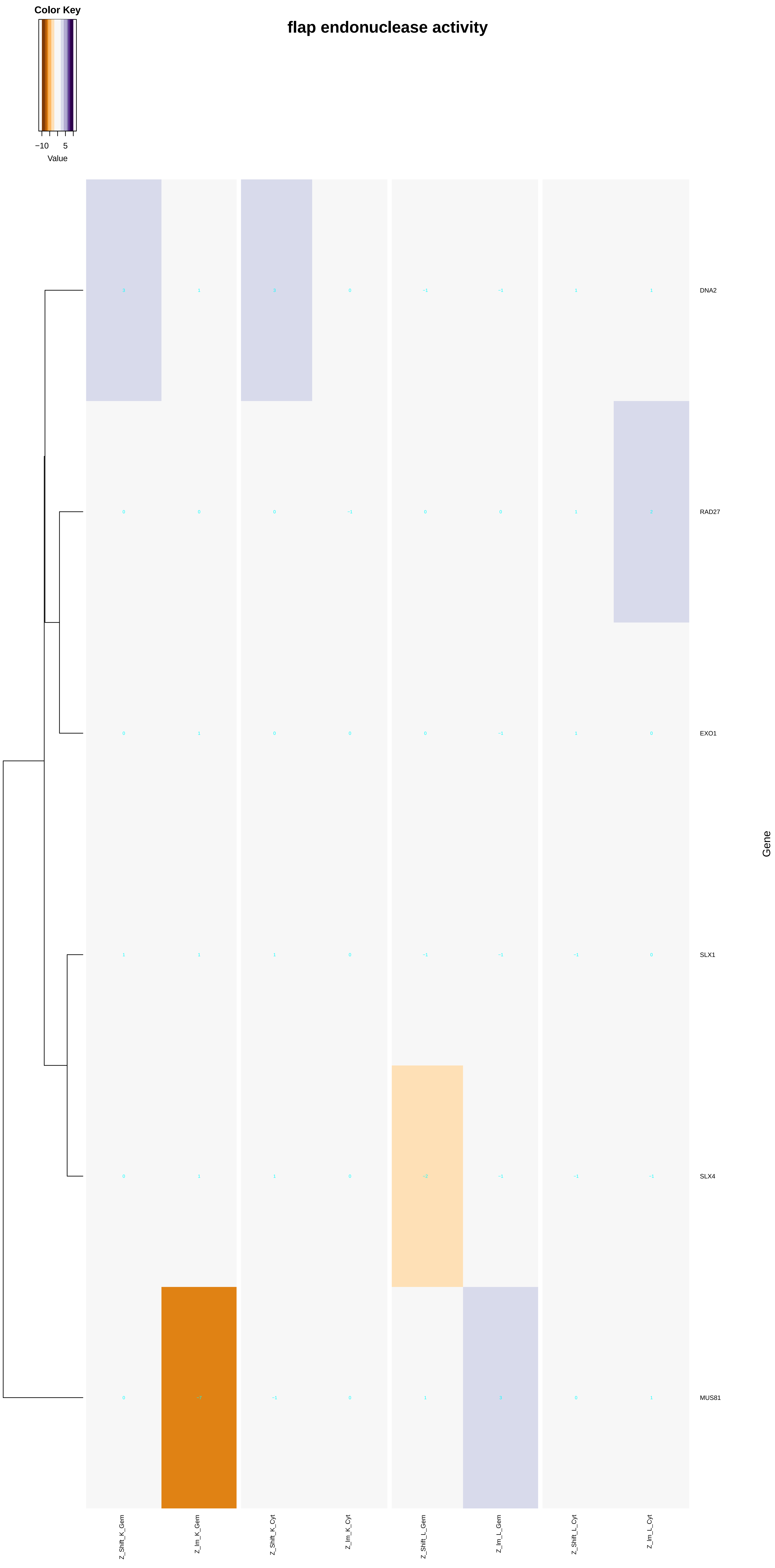

Color Key
−10
5
Value
flap endonuclease activity
DNA2
RAD27
EXO1
SLX1
SLX4
MUS81
Gene
Z_Shift_K_Gem
Z_Im_K_Gem
Z_Shift_K_Cyt
Z_Im_K_Cyt
Z_Shift_L_Gem
Z_Im_L_Gem
Z_Shift_L_Cyt
Z_Im_L_Cyt

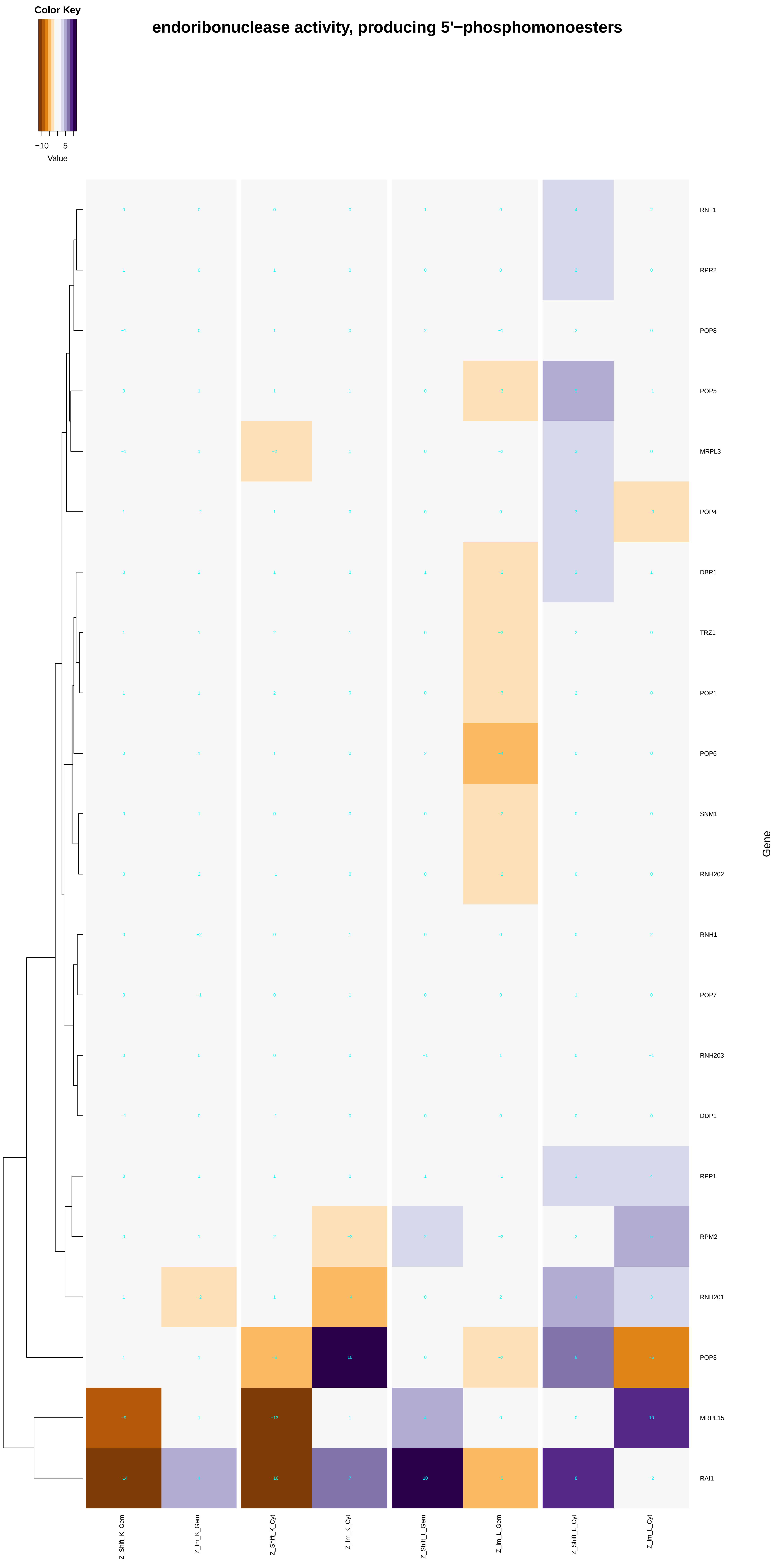
endoribonuclease activity, producing 5'−phosphomonoesters
Color Key
−10
5
Value
RNT1
RPR2
POP8
POP5
MRPL3
POP4
DBR1
TRZ1
POP1
POP6
SNM1
RNH202
RNH1
POP7
RNH203
DDP1
RPP1
RPM2
RNH201
POP3
MRPL15
RAI1
Gene
Z_Shift_K_Gem
Z_lm_K_Gem
Z_Shift_K_Cyt
Z_lm_K_Cyt
Z_Shift_L_Gem
Z_lm_L_Gem
Z_Shift_L_Cyt
Z_lm_L_Cyt

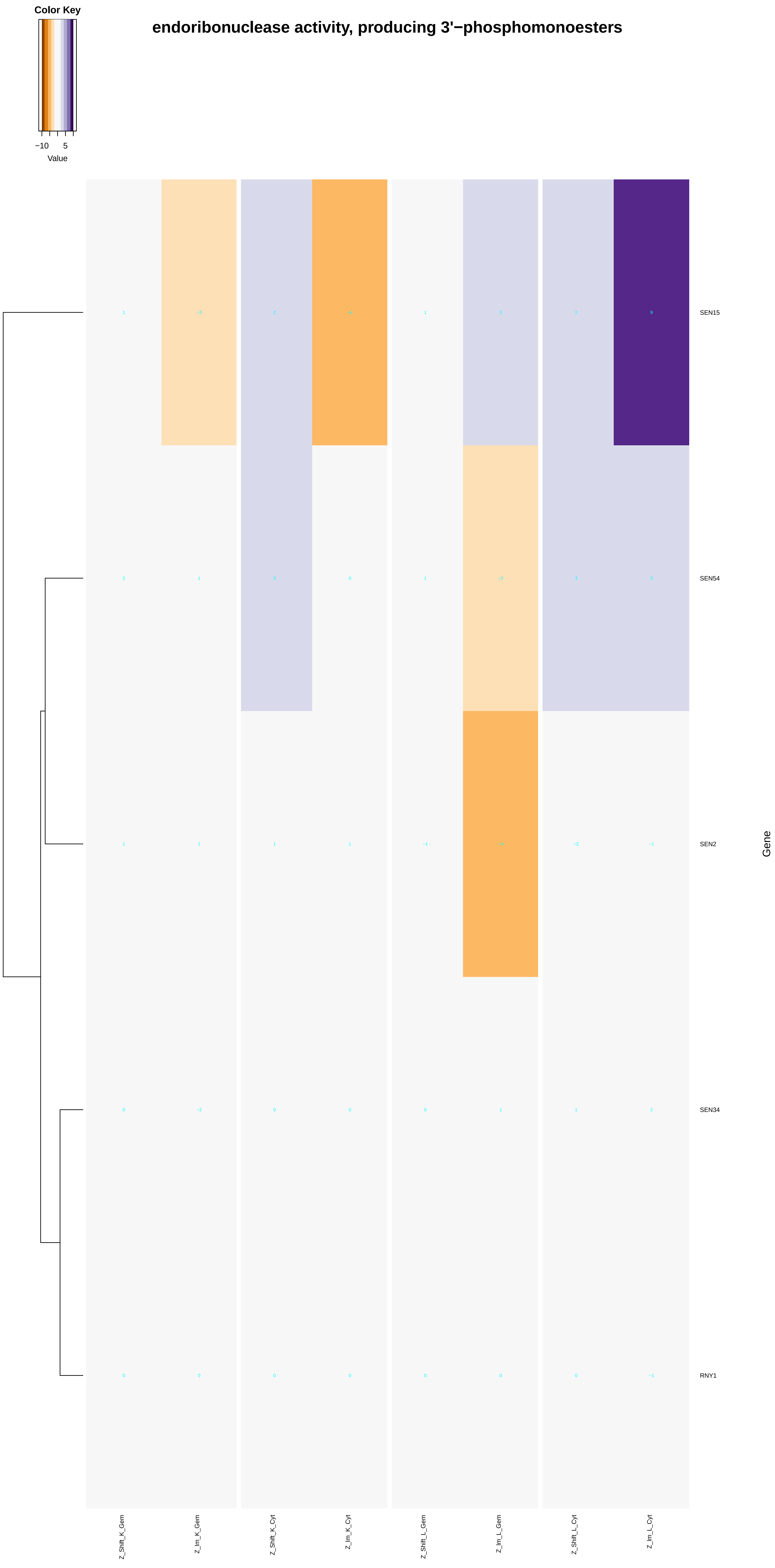
endoribonuclease activity, producing 3'−phosphomonoesters
Color Key
−10
5
Value
SEN15
SEN54
SEN2
SEN34
RNY1
Gene
Z_Shift_K_Gem
Z_Im_K_Gem
Z_Shift_K_Cyt
Z_Im_K_Cyt
Z_Shift_L_Gem
Z_Im_L_Gem
Z_Shift_L_Cyt
Z_Im_L_Cyt

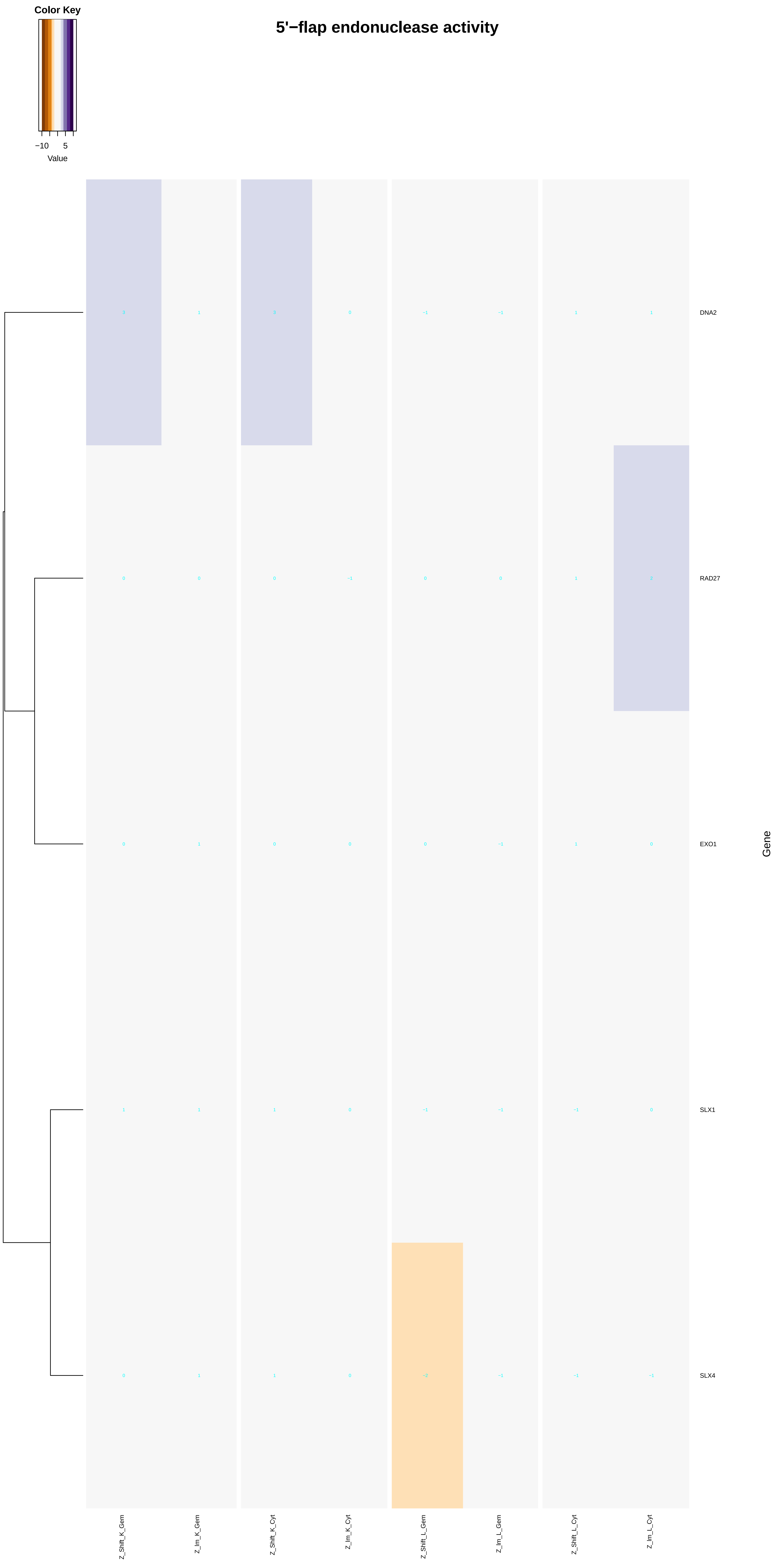
Color Key
−10
5
Value
5'−flap endonuclease activity
DNA2
RAD27
EXO1
SLX1
SLX4
Gene
Z_Shift_K_Gem
Z_Im_K_Gem
Z_Shift_K_Cyt
Z_Im_K_Cyt
Z_Shift_L_Gem
Z_Im_L_Gem
Z_Shift_L_Cyt
Z_Im_L_Cyt

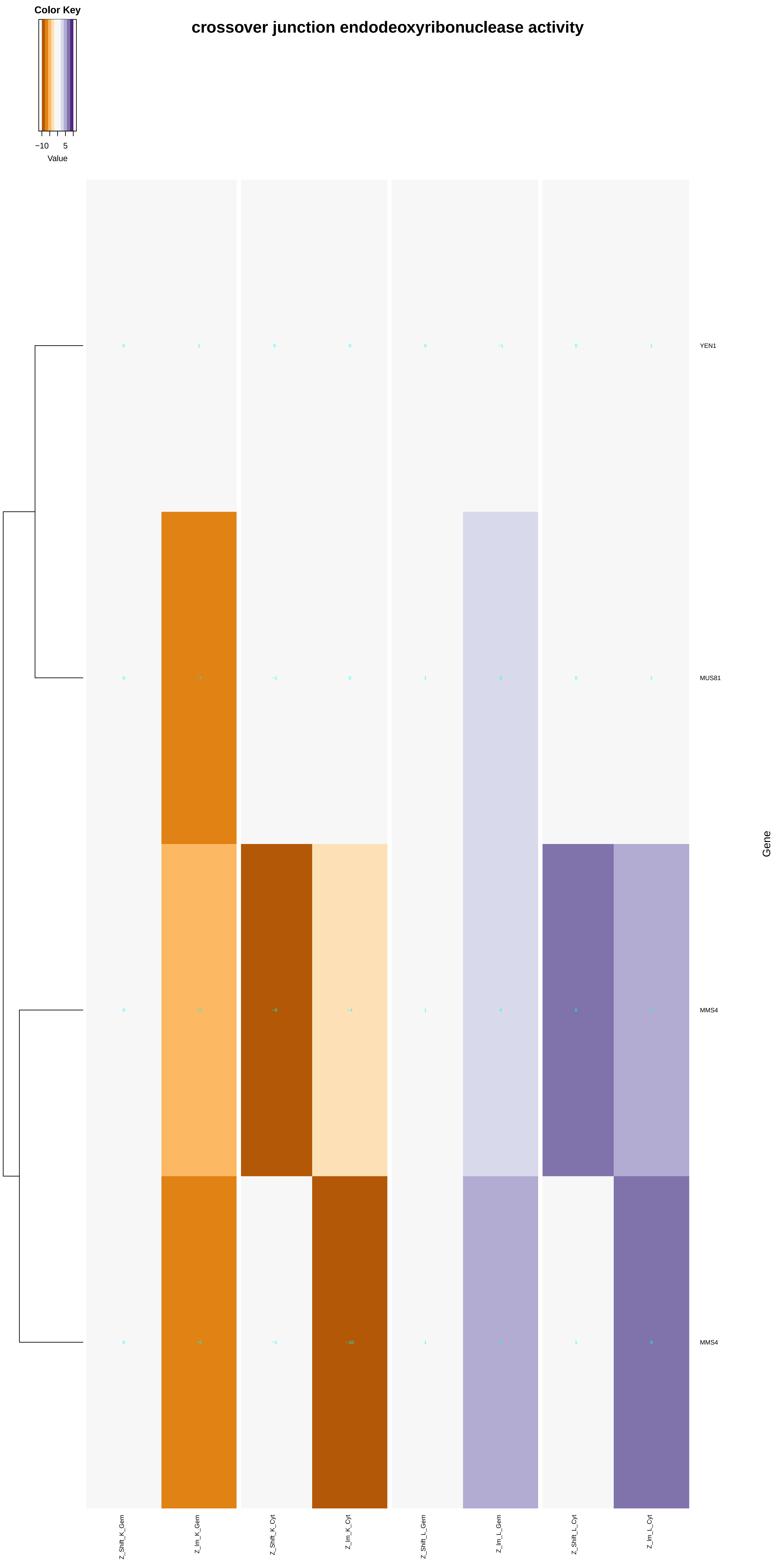
crossover junction endodeoxyribonuclease activity
Color Key
−10
5
Value
YEN1
MUS81
MMS4
MMS4
Gene
Z_Shift_K_Gem
Z_Im_K_Gem
Z_Shift_K_Cyt
Z_Im_K_Cyt
Z_Shift_L_Gem
Z_Im_L_Gem
Z_Shift_L_Cyt
Z_Im_L_Cyt

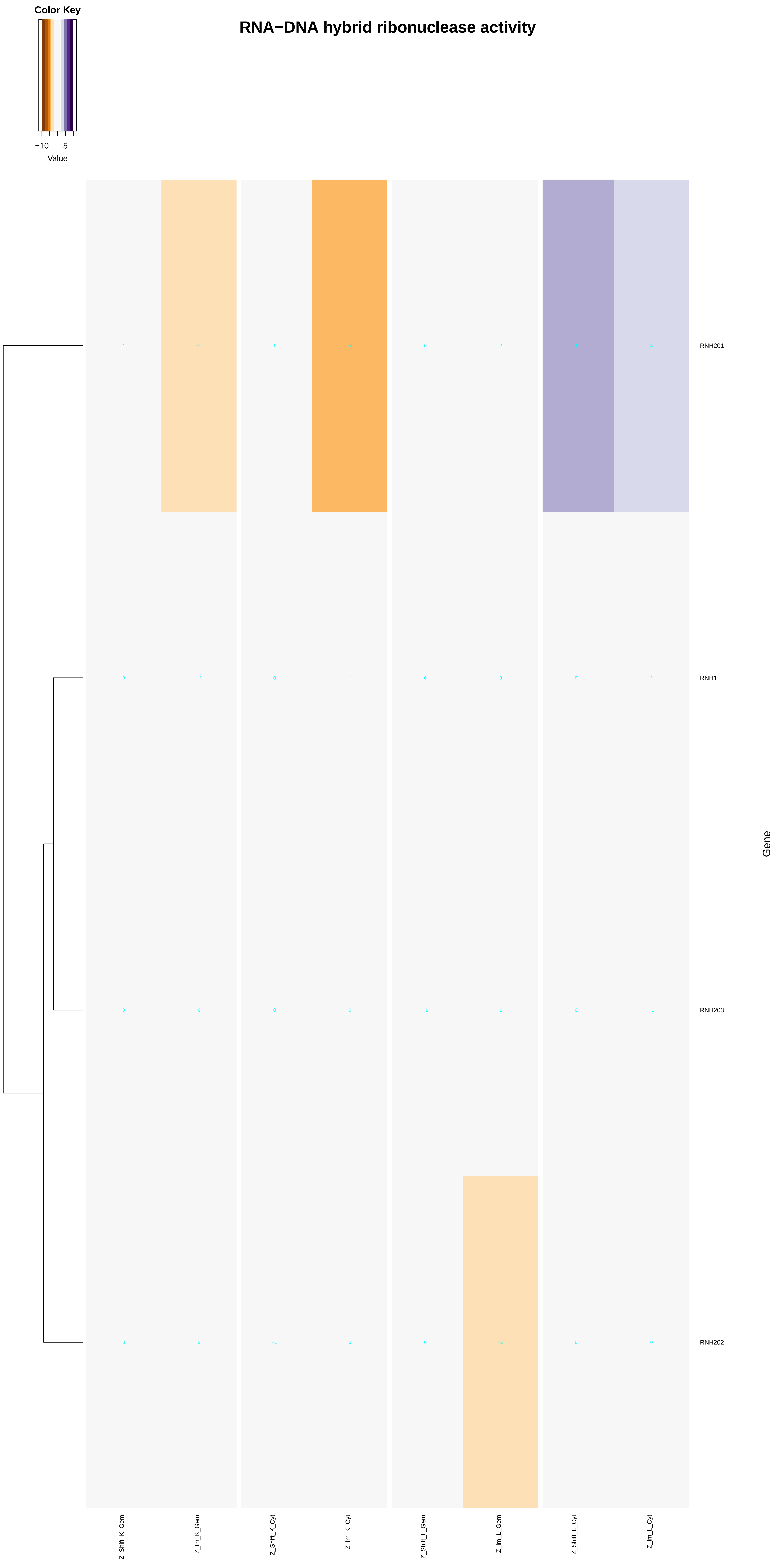
Color Key
−10
5
Value
RNA−DNA hybrid ribonuclease activity
RNH201
RNH1
RNH203
RNH202
Gene
Z_Shift_K_Gem
Z_Im_K_Gem
Z_Shift_K_Cyt
Z_Im_K_Cyt
Z_Shift_L_Gem
Z_Im_L_Gem
Z_Shift_L_Cyt
Z_Im_L_Cyt

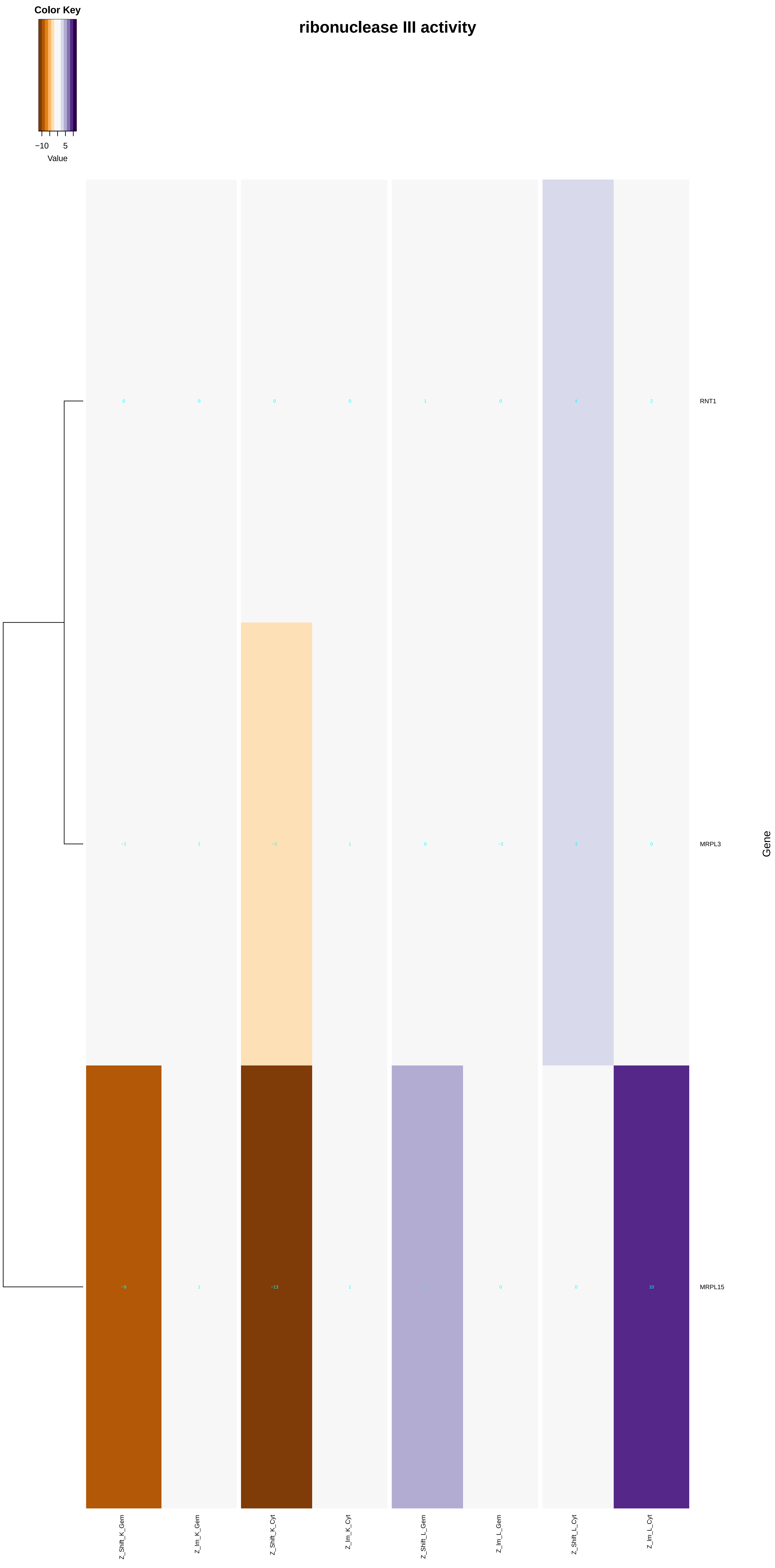

Color Key
−10
5
Value
ribonuclease III activity
RNT1
MRPL3
MRPL15
Gene
Z_Shift_K_Gem
Z_Im_K_Gem
Z_Shift_K_Cyt
Z_Im_K_Cyt
Z_Shift_L_Gem
Z_Im_L_Gem
Z_Shift_L_Cyt
Z_Im_L_Cyt

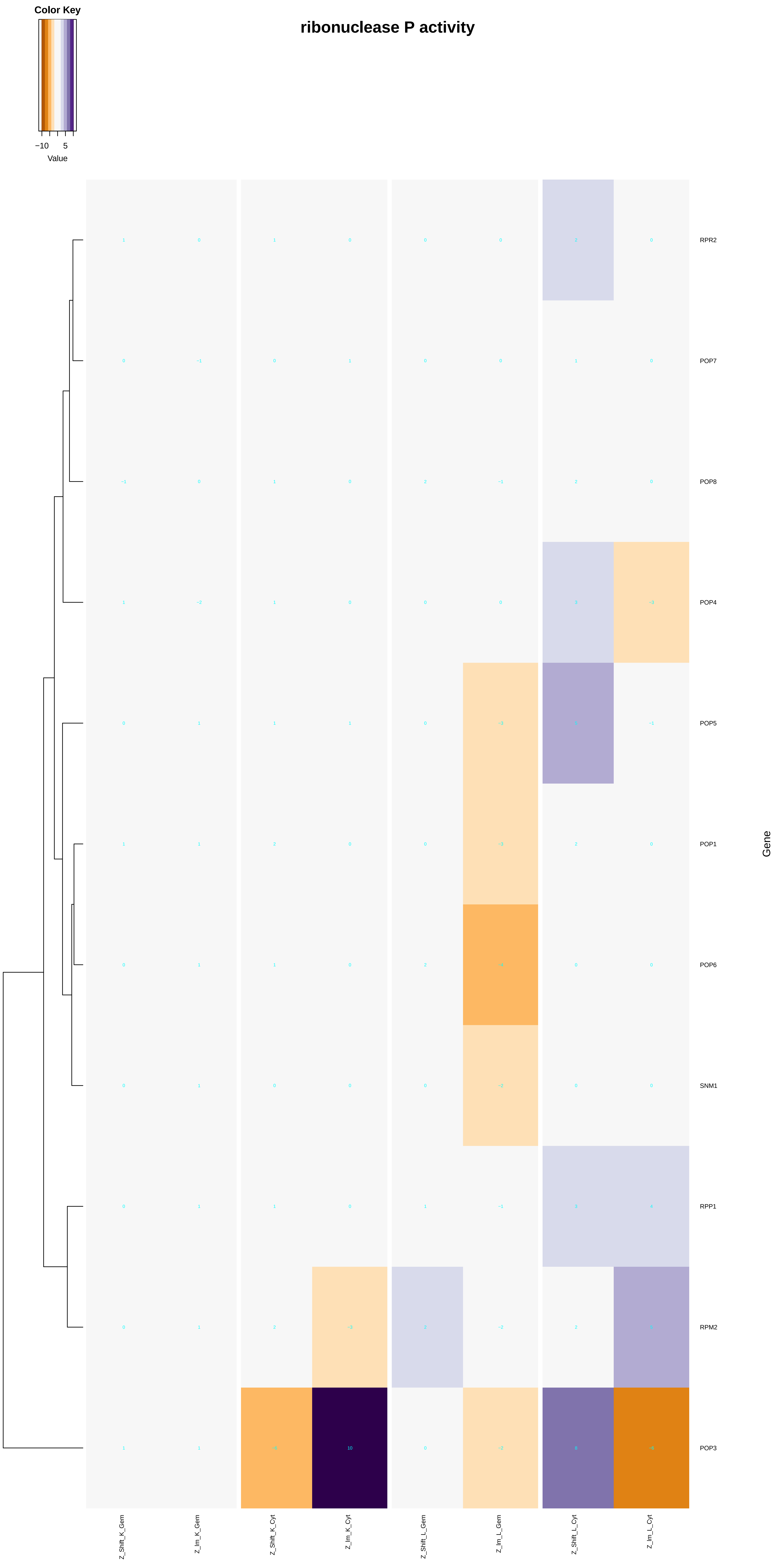
Color Key
−10 5
Value
ribonuclease P activity
RPR2
POP7
POP8
POP4
POP5
POP1
POP6
SNM1
RPP1
RPM2
POP3
Gene
Z_Shift_K_Gem
Z_Im_K_Gem
Z_Shift_K_Cyt
Z_Im_K_Cyt
Z_Shift_L_Gem
Z_Im_L_Gem
Z_Shift_L_Cyt
Z_Im_L_Cyt

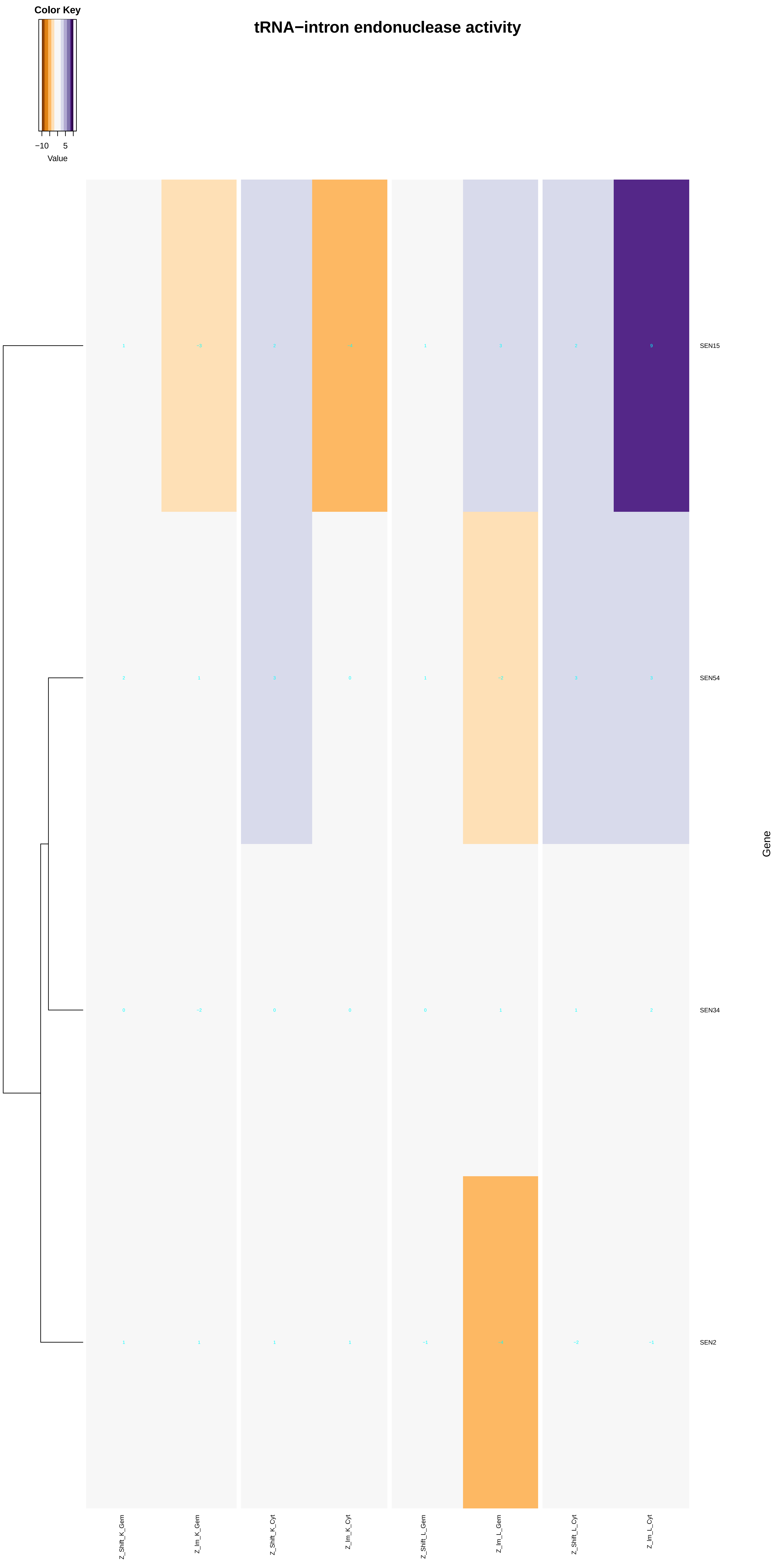

tRNA−intron endonuclease activity
Color Key
−10
5
Value
SEN15
SEN54
SEN34
SEN2
Gene
Z_Shift_K_Gem
Z_Im_K_Gem
Z_Shift_K_Cyt
Z_Im_K_Cyt
Z_Shift_L_Gem
Z_Im_L_Gem
Z_Shift_L_Cyt
Z_Im_L_Cyt
