## Additional File 7 for "A humanized yeast phenomic model of deoxycytidine kinase to predict genetic buffering of nucleoside analog cytotoxicity": enzyme_regulator_activity.pdf

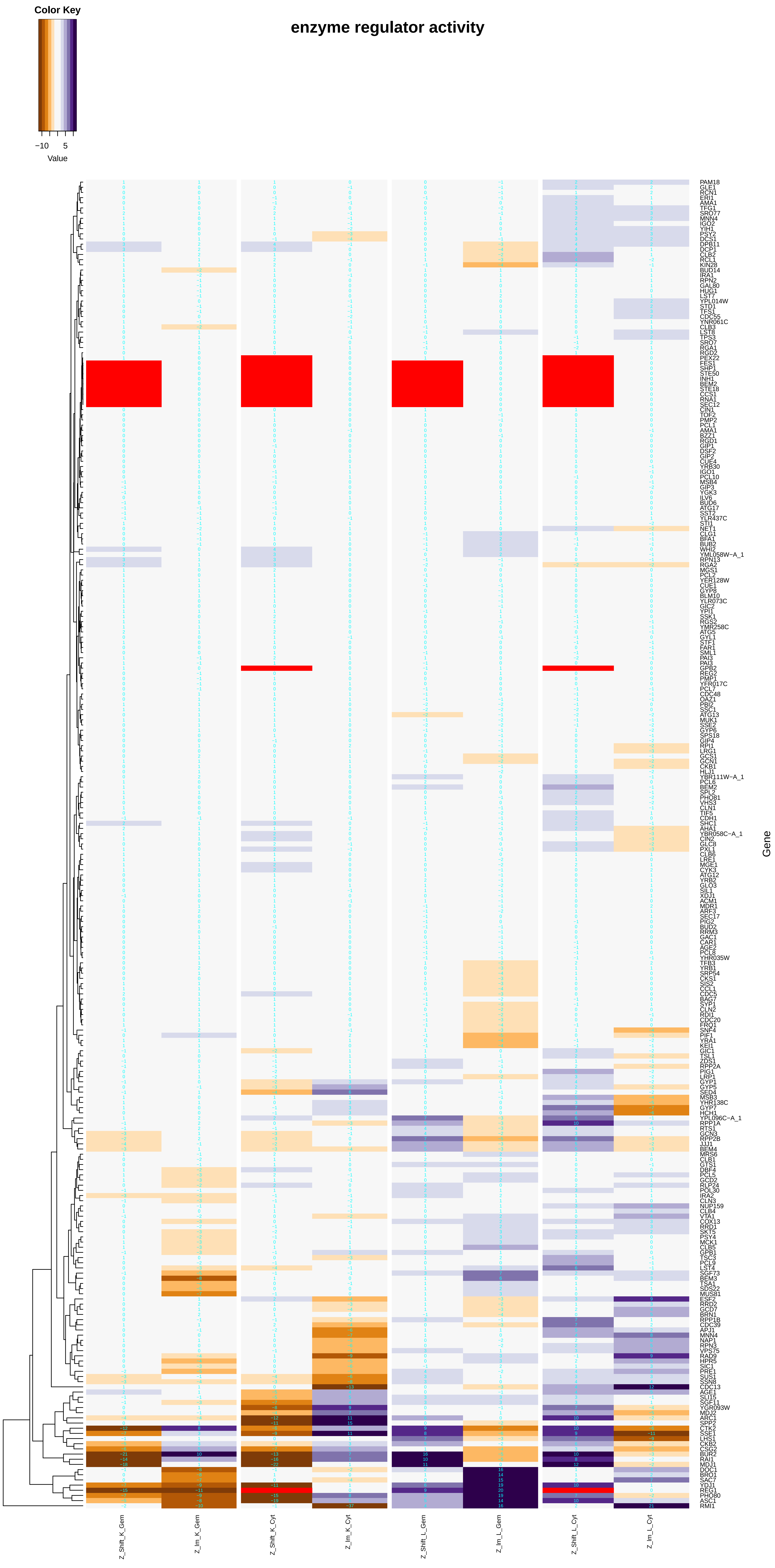
Color Key
−10
5
Value
enzyme regulator activity
Gene
Z_Shift_K_Gem
Z_Im_K_Gem
Z_Shift_K_Cyt
Z_Im_K_Cyt
Z_Shift_L_Gem
Z_Im_L_Gem
Z_Shift_L_Cyt
Z_Im_L_Cyt

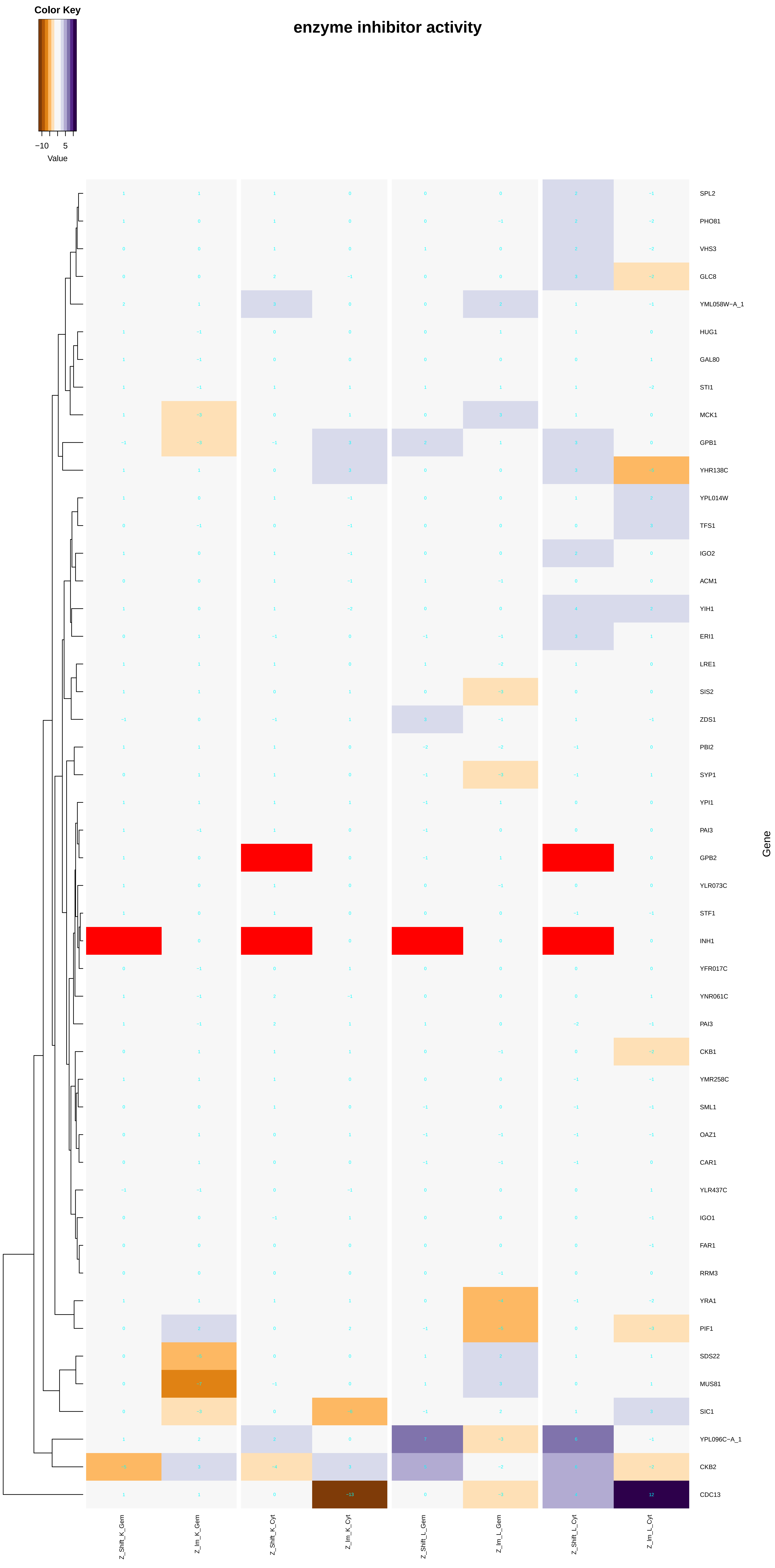
Color Key
−10
5
Value
enzyme inhibitor activity
SPL2
PHO81
VHS3
GLC8
YML058W−A_1
HUG1
GAL80
STI1
MCK1
GPB1
YHR138C
YPL014W
TFS1
IGO2
ACM1
YIH1
ERI1
LRE1
SIS2
ZDS1
PBI2
SYP1
YPI1
PAI3
GPB2
YLR073C
STF1
INH1
YFR017C
YNR061C
PAI3
CKB1
YMR258C
SML1
OAZ1
CAR1
YLR437C
IGO1
FAR1
RRM3
YRA1
PIF1
SDS22
MUS81
SIC1
YPL096C−A_1
CKB2
CDC13
Gene
Z_Shift_K_Gem
Z_Im_K_Gem
Z_Shift_K_Cyt
Z_Im_K_Cyt
Z_Shift_L_Gem
Z_Im_L_Gem
Z_Shift_L_Cyt
Z_Im_L_Cyt

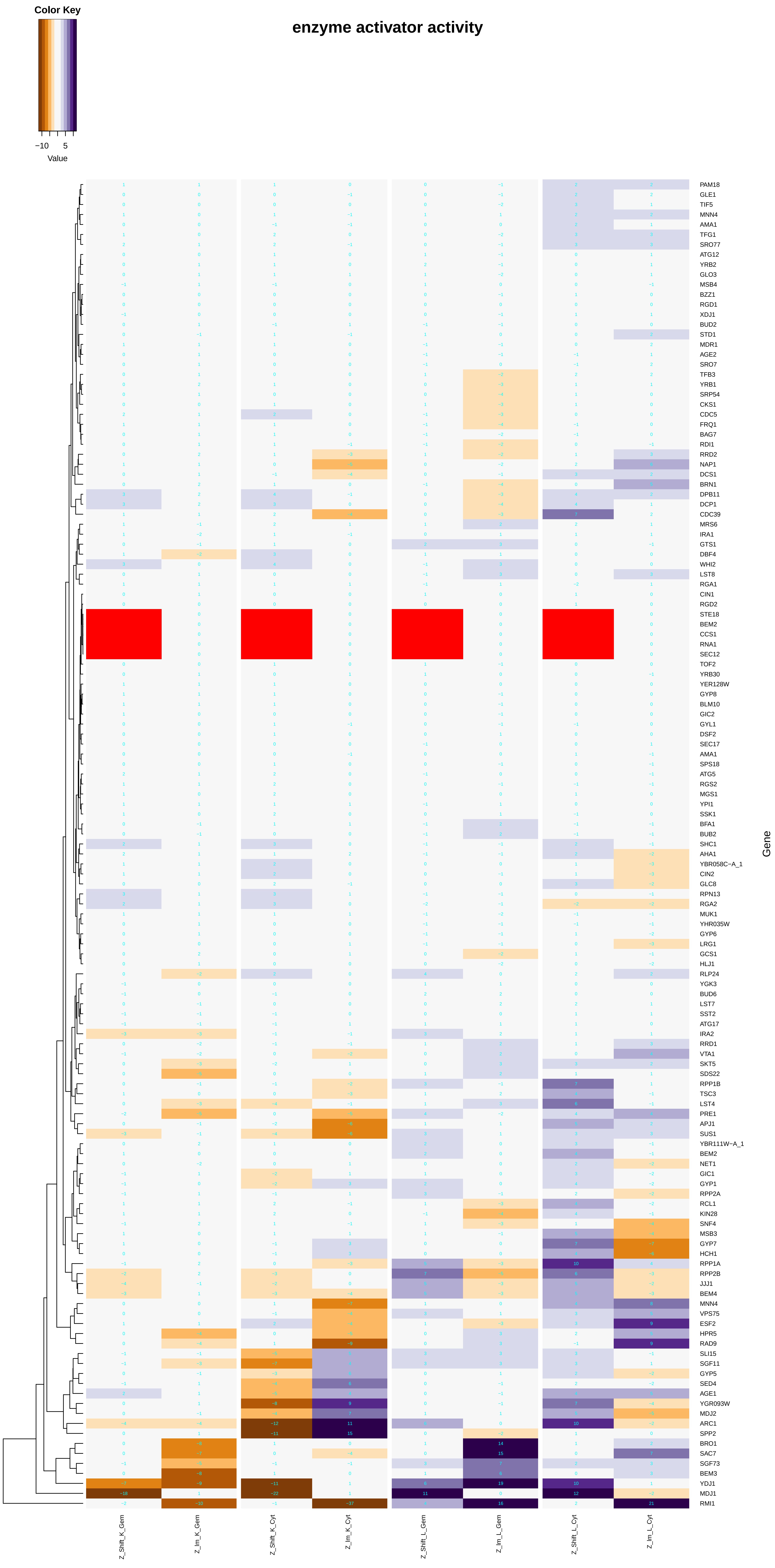
enzyme activator activity
Color Key
−10
5
Value
PAM18
GLE1
TIF5
MNN4
AMA1
TFG1
SRO77
ATG12
YRB2
GLO3
MSB4
BZZ1
RGD1
XDJ1
BUD2
STD1
MDR1
AGE2
SRO7
TFB3
YRB1
SRP54
CKS1
CDC5
FRQ1
BAG7
RDI1
RRD2
NAP1
DCS1
BRN1
DPB11
DCP1
CDC39
MRS6
IRA1
GTS1
DBF4
WHI2
LST8
RGA1
CIN1
RGD2
STE18
BEM2
CCS1
RNA1
SEC12
TOF2
YRB30
YER128W
GYP8
BLM10
GIC2
GYL1
DSF2
SEC17
AMA1
SPS18
ATG5
RGS2
MGS1
YPI1
SSK1
BFA1
BUB2
SHC1
AHA1
YBR058C−A_1
CIN2
GLC8
RPN13
RGA2
MUK1
YHR035W
GYP6
LRG1
GCS1
HLJ1
RLP24
YGK3
BUD6
LST7
SST2
ATG17
IRA2
RRD1
VTA1
SKT5
SDS22
RPP1B
TSC3
LST4
PRE1
APJ1
SUS1
YBR111W−A_1
BEM2
NET1
GIC1
GYP1
RPP2A
RCL1
KIN28
SNF4
MSB3
GYP7
HCH1
RPP1A
RPP2B
JJJ1
BEM4
MNN4
VPS75
ESF2
HPR5
RAD9
SLI15
SGF11
GYP5
SED4
AGE1
YGR093W
MDJ2
ARC1
SPP2
BRO1
SAC7
SGF73
BEM3
YDJ1
MDJ1
RMI1
Gene
Z_Shift_K_Gem
Z_Im_K_Gem
Z_Shift_K_Cyt
Z_Im_K_Cyt
Z_Shift_L_Gem
Z_Im_L_Gem
Z_Shift_L_Cyt
Z_Im_L_Cyt

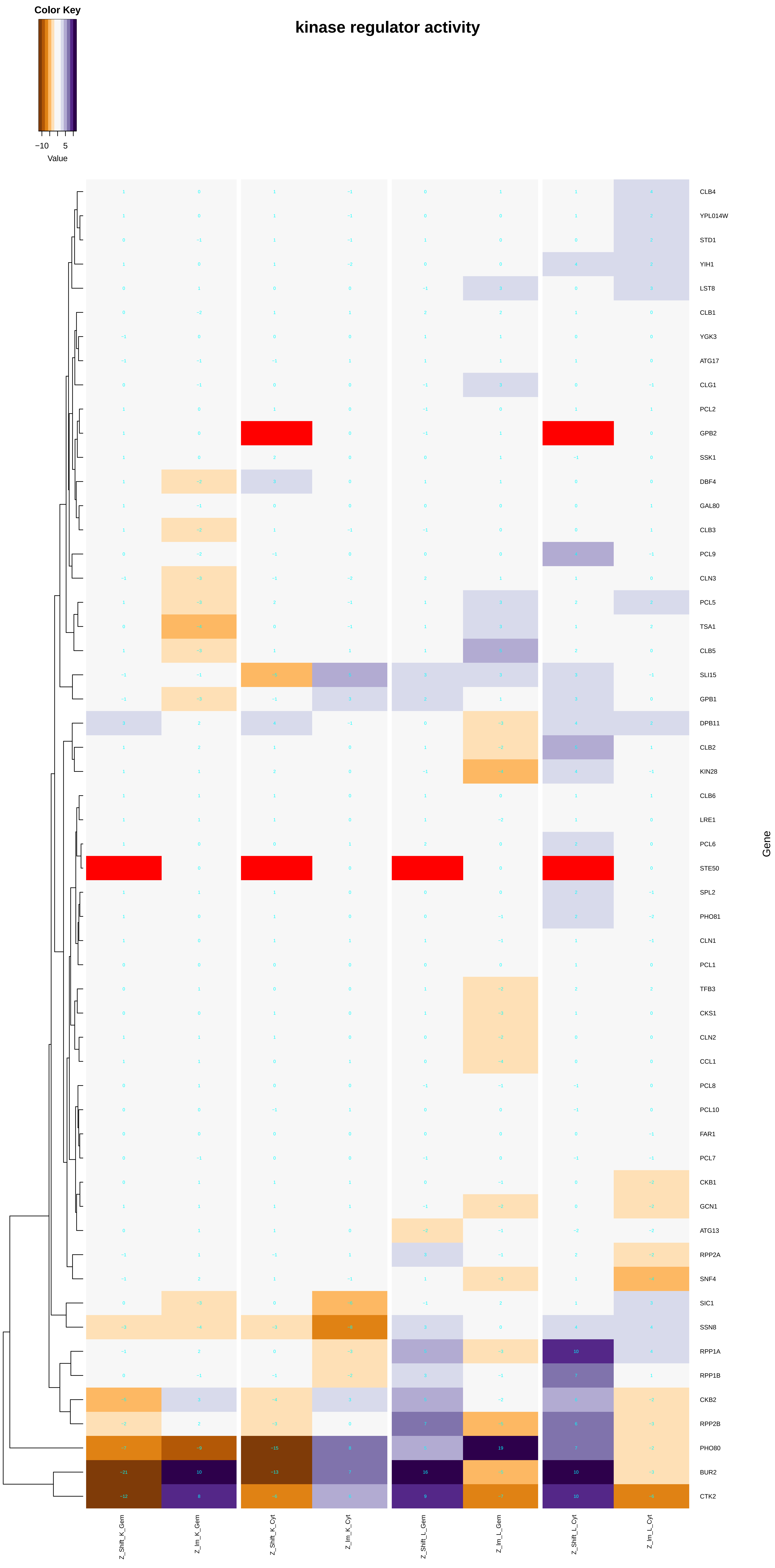

Color Key
−10
5
Value
kinase regulator activity
CLB4
YPL014W
STD1
YIH1
LST8
CLB1
YGK3
ATG17
CLG1
PCL2
GPB2
SSK1
DBF4
GAL80
CLB3
PCL9
CLN3
PCL5
TSA1
CLB5
SLI15
GPB1
DPB11
CLB2
KIN28
CLB6
LRE1
PCL6
STE50
SPL2
PHO81
CLN1
PCL1
TFB3
CKS1
CLN2
CCL1
PCL8
PCL10
FAR1
PCL7
CKB1
GCN1
ATG13
RPP2A
SNF4
SIC1
SSN8
RPP1A
RPP1B
CKB2
RPP2B
PHO80
BUR2
CTK2
Gene
Z_Shift_K_Gem
Z_Im_K_Gem
Z_Shift_K_Cyt
Z_Im_K_Cyt
Z_Shift_L_Gem
Z_Im_L_Gem
Z_Shift_L_Cyt
Z_Im_L_Cyt

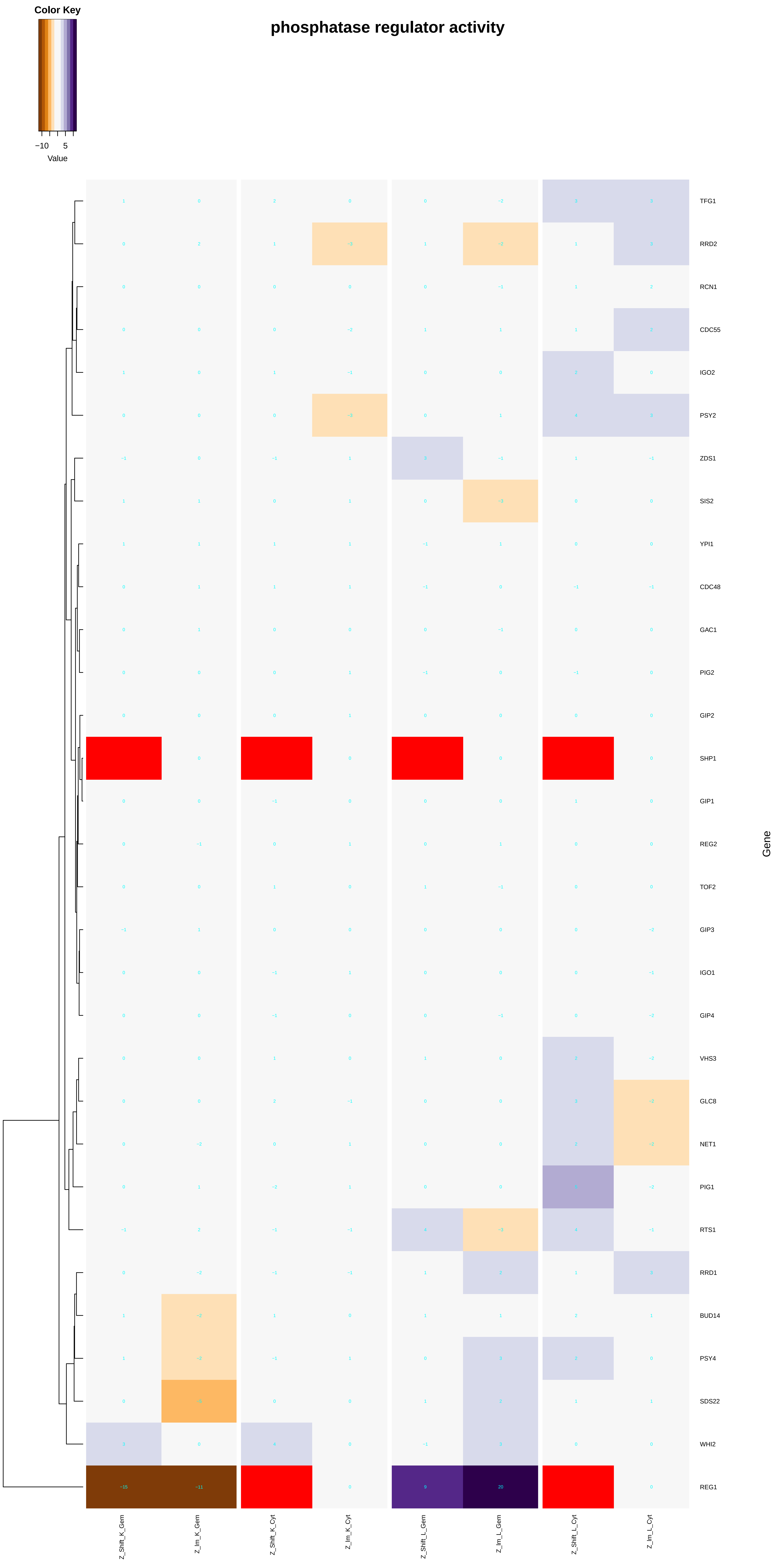
Color Key
phosphatase regulator activity
−10
5
Value
TFG1
RRD2
RCN1
CDC55
IGO2
PSY2
ZDS1
SIS2
YPI1
CDC48
GAC1
PIG2
GIP2
SHP1
GIP1
REG2
TOF2
GIP3
IGO1
GIP4
VHS3
GLC8
NET1
PIG1
RTS1
RRD1
BUD14
PSY4
SDS22
WHI2
REG1
Gene
Z_Shift_K_Gem
Z_Im_K_Gem
Z_Shift_K_Cyt
Z_Im_K_Cyt
Z_Shift_L_Gem
Z_Im_L_Gem
Z_Shift_L_Cyt
Z_Im_L_Cyt

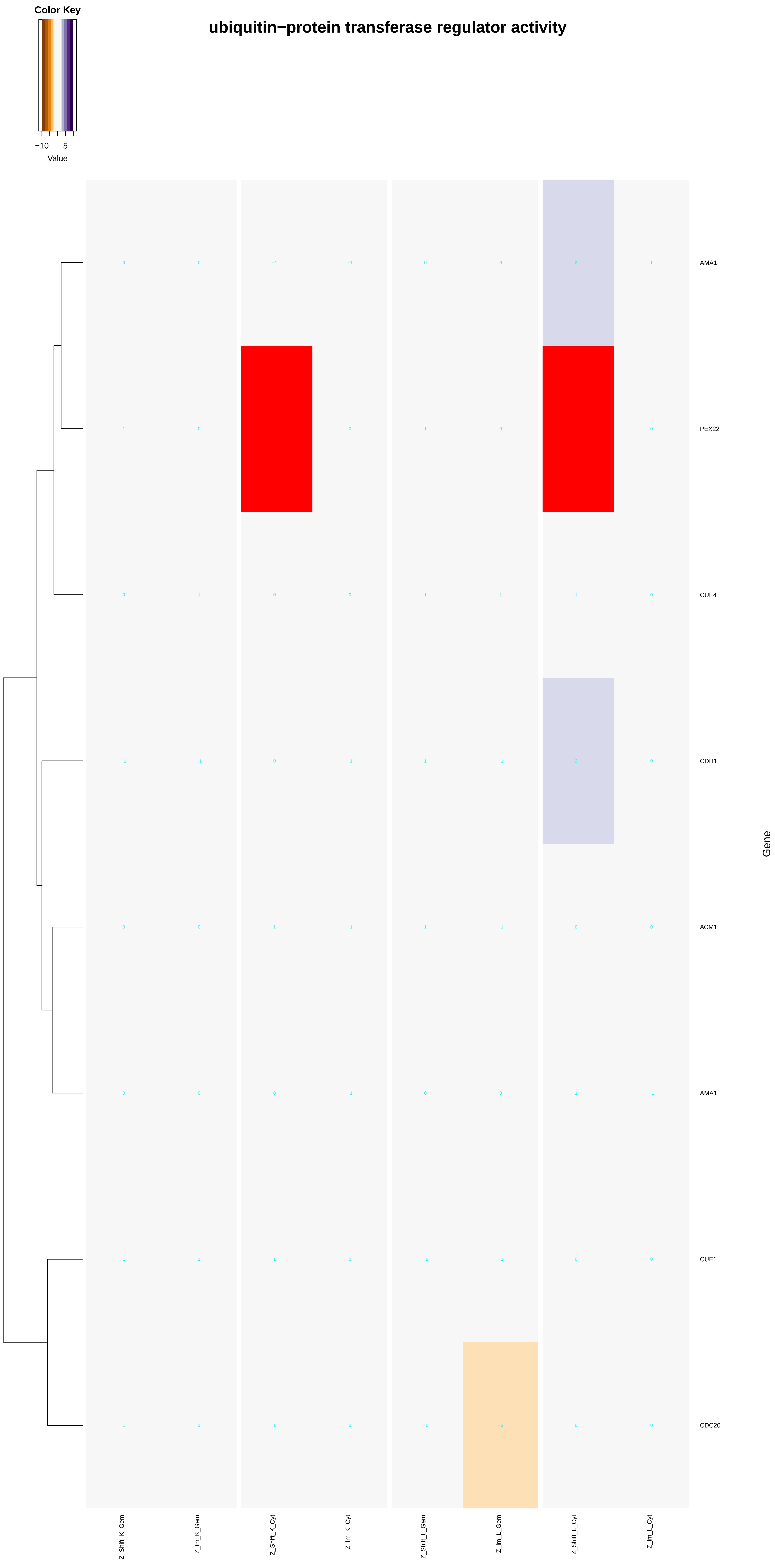
ubiquitin−protein transferase regulator activity
Color Key
−10
5
Value
AMA1
PEX22
CUE4
CDH1
ACM1
AMA1
CUE1
CDC20
Gene
Z_Shift_K_Gem
Z_Im_K_Gem
Z_Shift_K_Cyt
Z_Im_K_Cyt
Z_Shift_L_Gem
Z_Im_L_Gem
Z_Shift_L_Cyt
Z_Im_L_Cyt

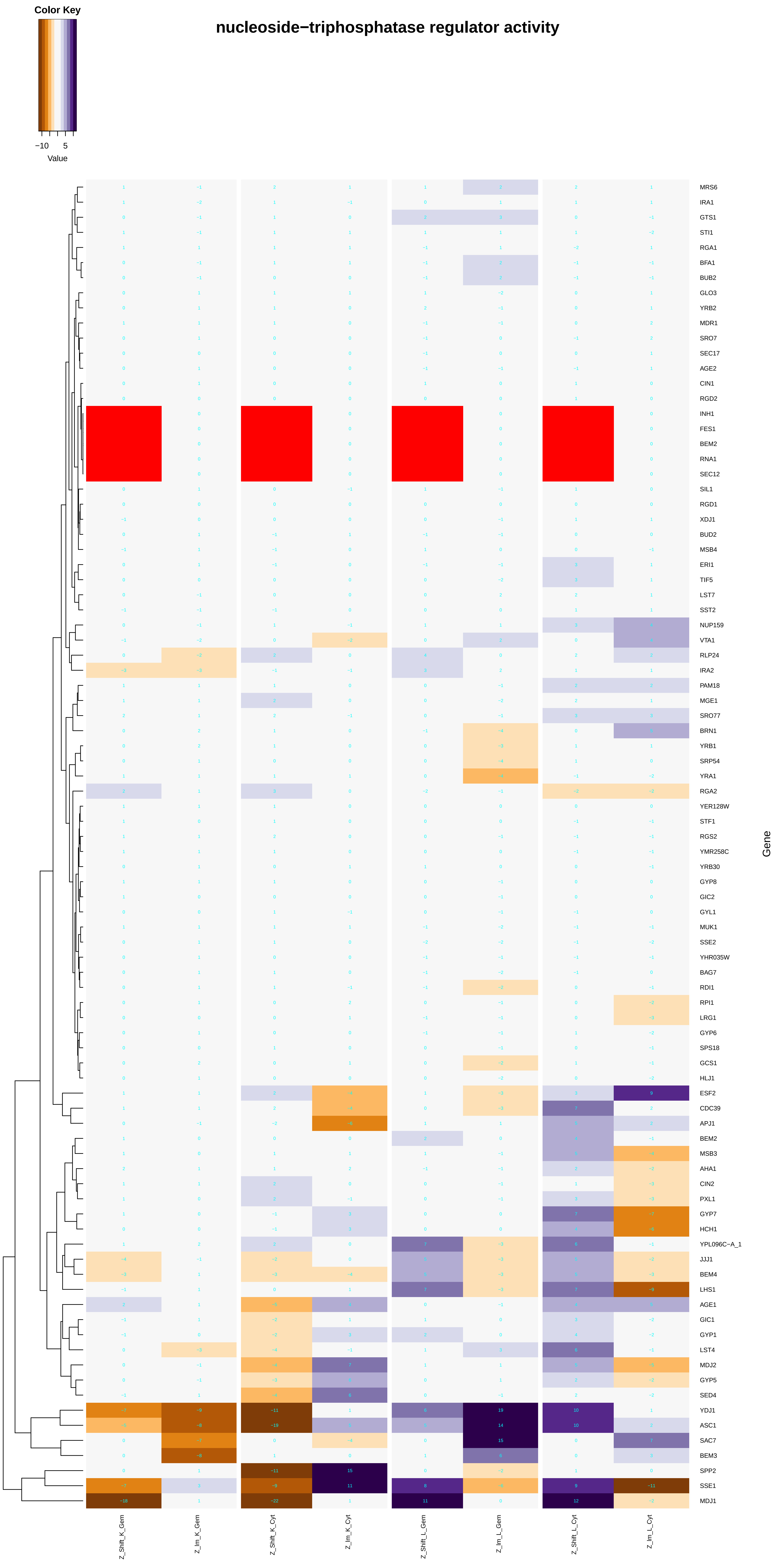
nucleoside−triphosphatase regulator activity
Color Key
−10
5
Value
MRS6
IRA1
GTS1
STI1
RGA1
BFA1
BUB2
GLO3
YRB2
MDR1
SRO7
SEC17
AGE2
CIN1
RGD2
INH1
FES1
BEM2
RNA1
SEC12
SIL1
RGD1
XDJ1
BUD2
MSB4
ERI1
TIF5
LST7
SST2
NUP159
VTA1
RLP24
IRA2
PAM18
MGE1
SRO77
BRN1
YRB1
SRP54
YRA1
RGA2
YER128W
STF1
RGS2
YMR258C
YRB30
GYP8
GIC2
GYL1
MUK1
SSE2
YHR035W
BAG7
RDI1
RPI1
LRG1
GYP6
SPS18
GCS1
HLJ1
ESF2
CDC39
APJ1
BEM2
MSB3
AHA1
CIN2
PXL1
GYP7
HCH1
YPL096C−A_1
JJJ1
BEM4
LHS1
AGE1
GIC1
GYP1
LST4
MDJ2
GYP5
SED4
YDJ1
ASC1
SAC7
BEM3
SPP2
SSE1
MDJ1
Gene
Z_Shift_K_Gem
Z_Im_K_Gem
Z_Shift_K_Cyt
Z_Im_K_Cyt
Z_Shift_L_Gem
Z_Im_L_Gem
Z_Shift_L_Cyt
Z_Im_L_Cyt

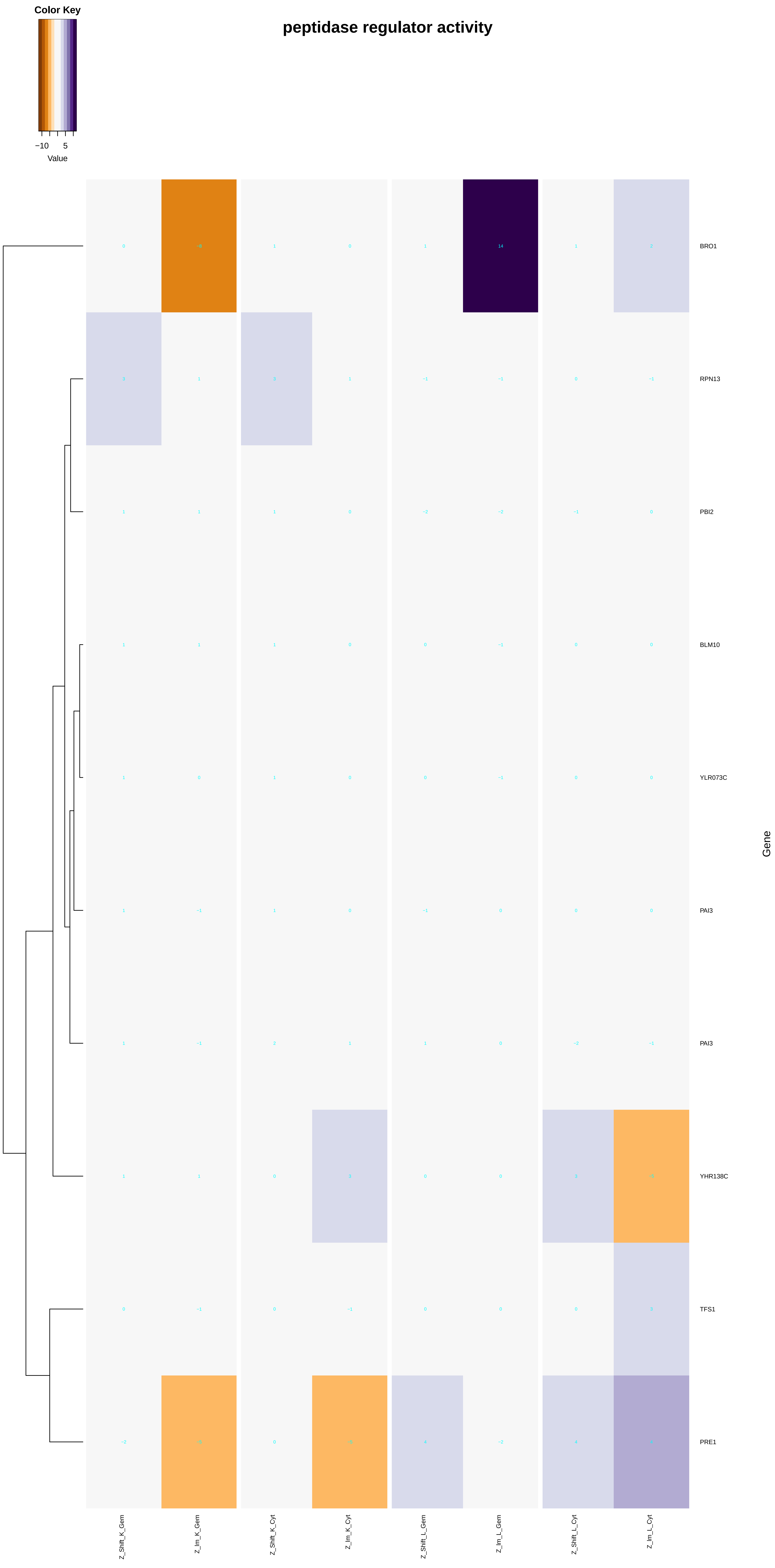
peptidase regulator activity
Color Key
−10
5
Value
BRO1
RPN13
PBI2
BLM10
YLR073C
PAI3
PAI3
YHR138C
TFS1
PRE1
Gene
Z_Shift_K_Gem
Z_Im_K_Gem
Z_Shift_K_Cyt
Z_Im_K_Cyt
Z_Shift_L_Gem
Z_Im_L_Gem
Z_Shift_L_Cyt
Z_Im_L_Cyt

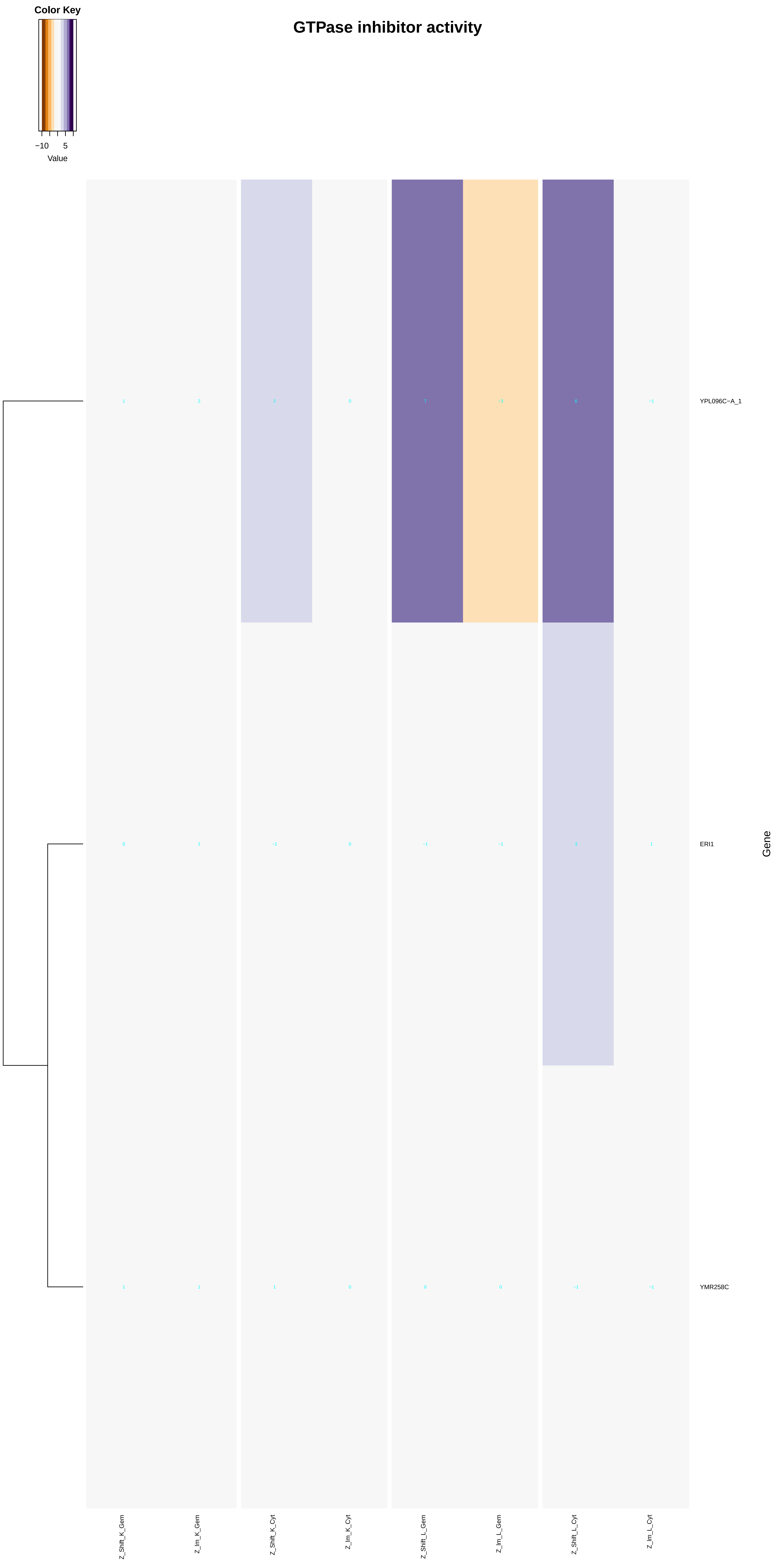
Color Key
−10
5
Value
GTPase inhibitor activity
YPL096C−A_1
ERI1
YMR258C
Gene
Z_Shift_K_Gem
Z_Im_K_Gem
Z_Shift_K_Cyt
Z_Im_K_Cyt
Z_Shift_L_Gem
Z_Im_L_Gem
Z_Shift_L_Cyt
Z_Im_L_Cyt

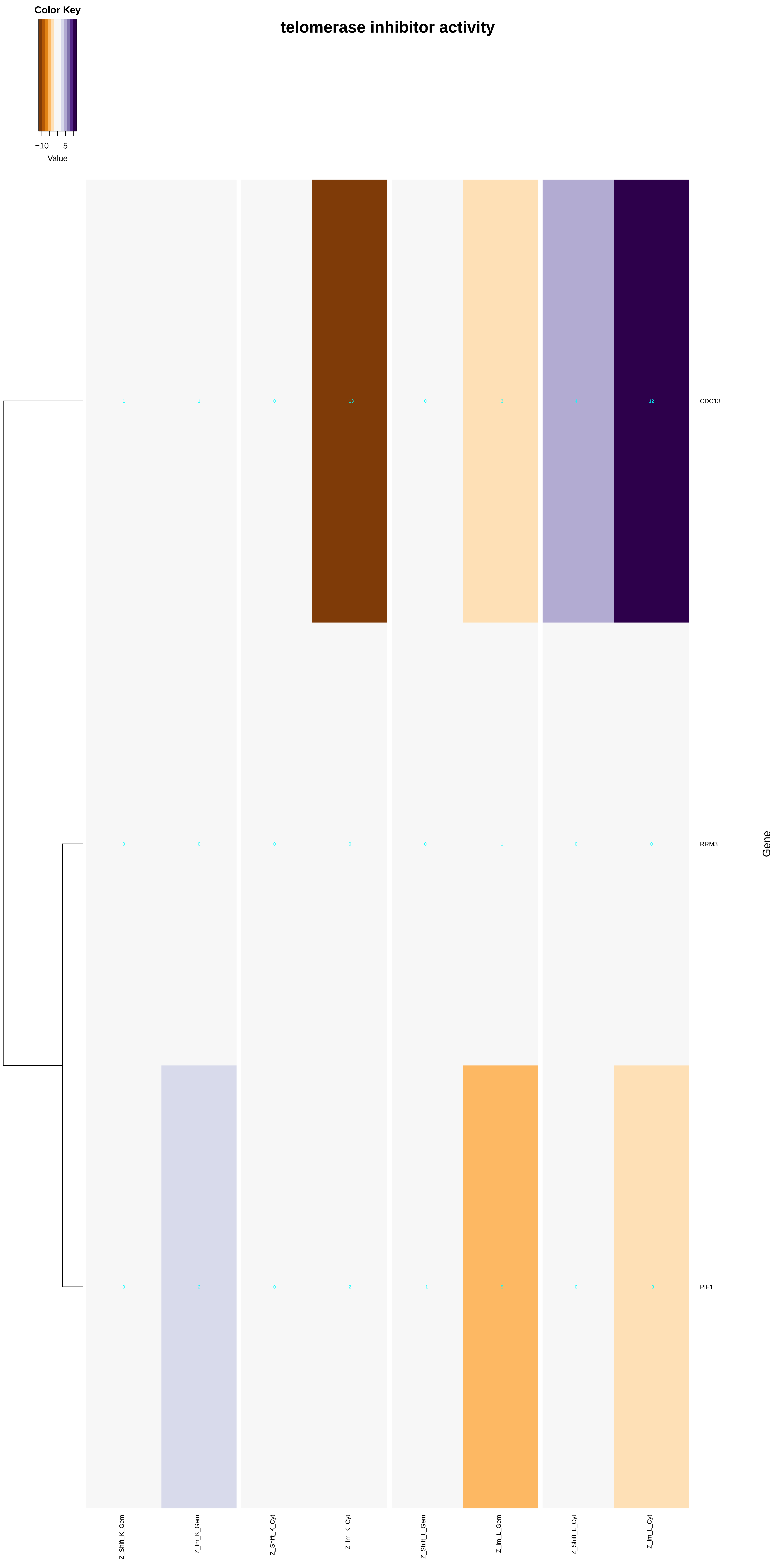

Color Key
−10
5
Value
telomerase inhibitor activity
CDC13
RRM3
PIF1
Gene
Z_Shift_K_Gem
Z_Im_K_Gem
Z_Shift_K_Cyt
Z_Im_K_Cyt
Z_Shift_L_Gem
Z_Im_L_Gem
Z_Shift_L_Cyt
Z_Im_L_Cyt

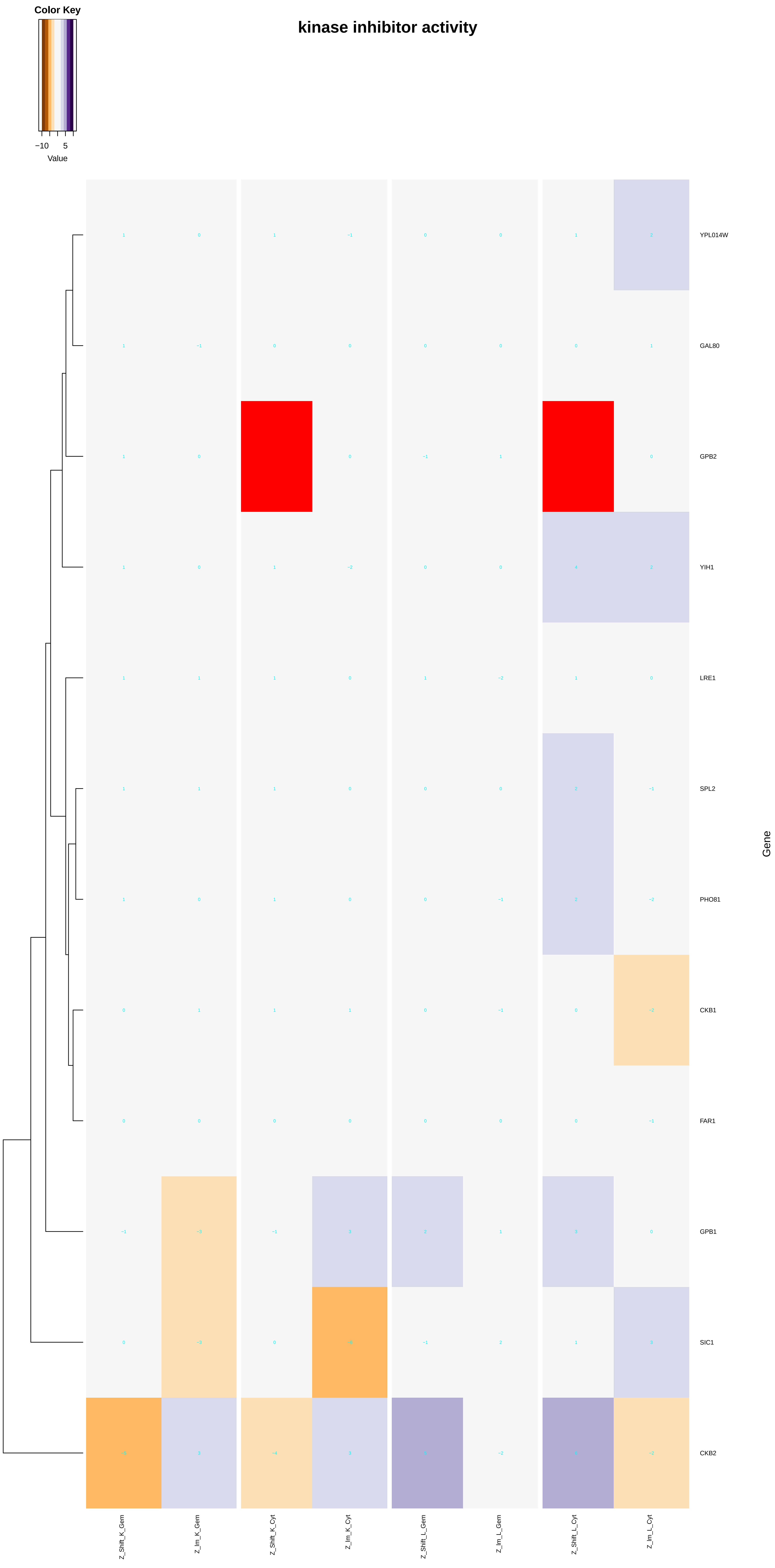
kinase inhibitor activity
Color Key
−10
5
Value
YPL014W
GAL80
GPB2
YIH1
LRE1
SPL2
PHO81
CKB1
FAR1
GPB1
SIC1
CKB2
Gene
Z_Shift_K_Gem
Z_lm_K_Gem
Z_Shift_K_Cyt
Z_lm_K_Cyt
Z_Shift_L_Gem
Z_lm_L_Gem
Z_Shift_L_Cyt
Z_lm_L_Cyt

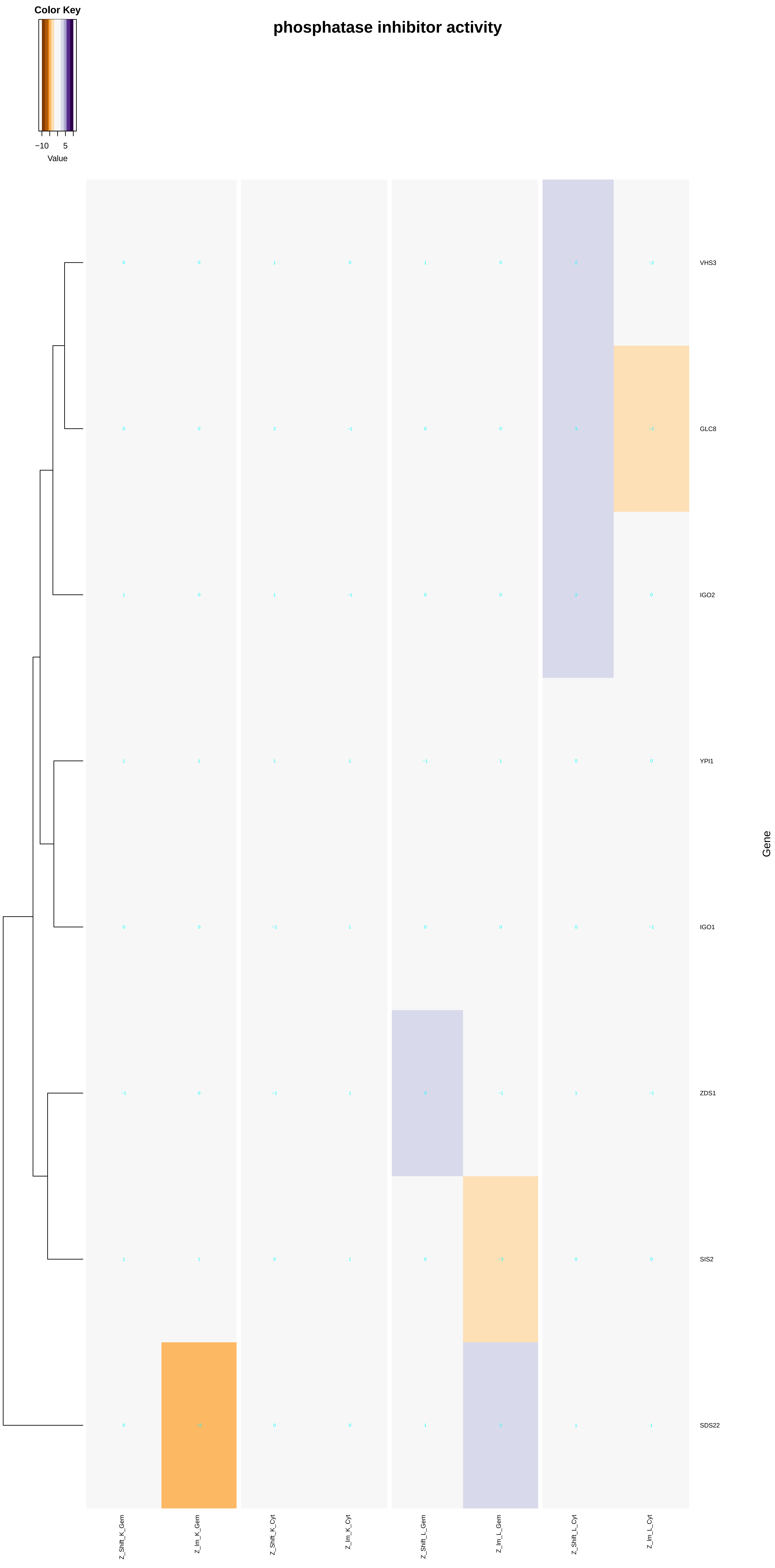

phosphatase inhibitor activity
Color Key
−10
5
Value
VHS3
GLC8
IGO2
YPI1
IGO1
ZDS1
SIS2
SDS22
Gene
Z_Shift_K_Gem
Z_Im_K_Gem
Z_Shift_K_Cyt
Z_Im_K_Cyt
Z_Shift_L_Gem
Z_Im_L_Gem
Z_Shift_L_Cyt
Z_Im_L_Cyt

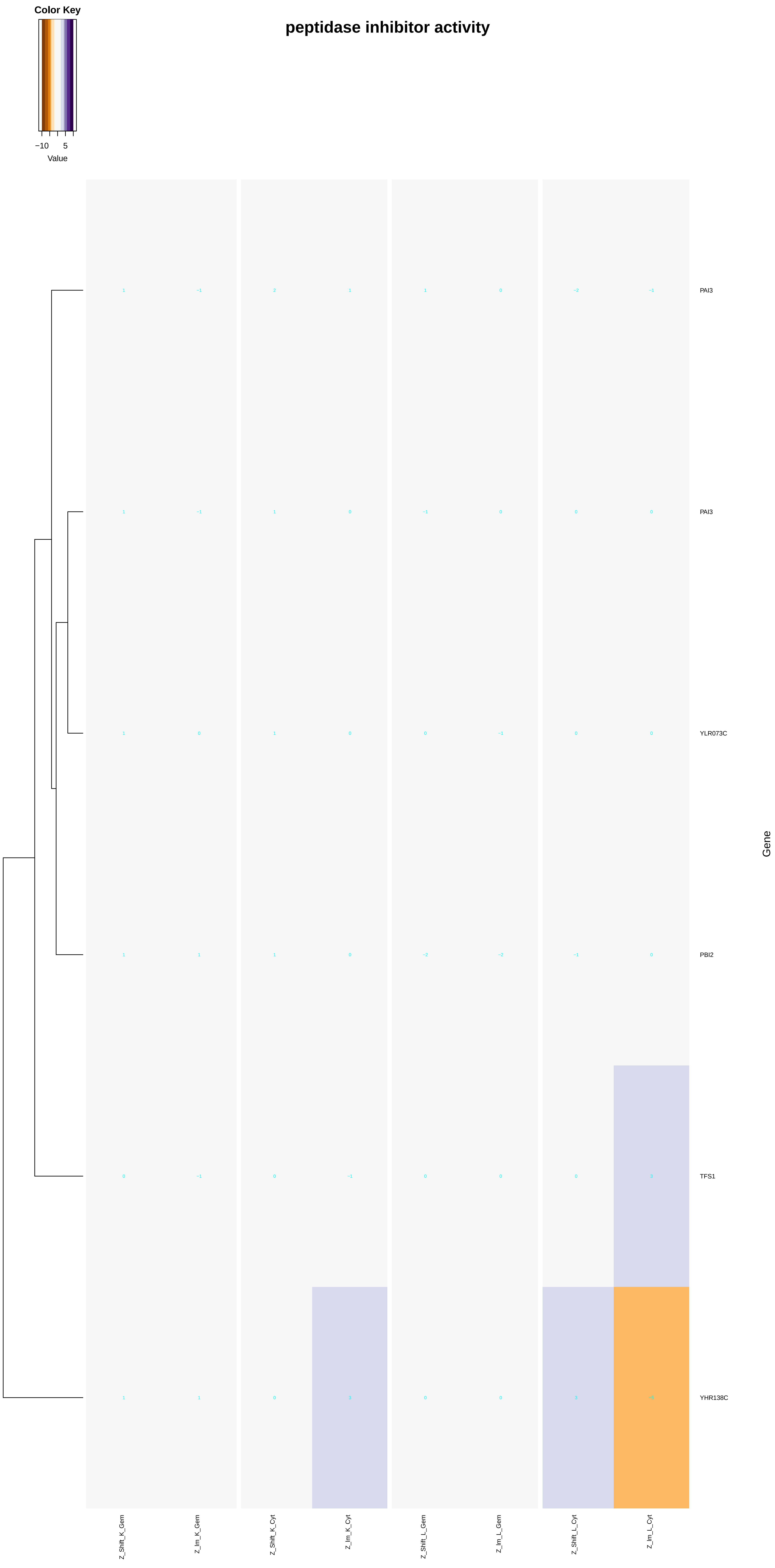

Color Key
−10
5
Value
peptidase inhibitor activity
PAI3
PAI3
YLR073C
PBI2
TFS1
YHR138C
Gene
Z_Shift_K_Gem
Z_Im_K_Gem
Z_Shift_K_Cyt
Z_Im_K_Cyt
Z_Shift_L_Gem
Z_Im_L_Gem
Z_Shift_L_Cyt
Z_Im_L_Cyt

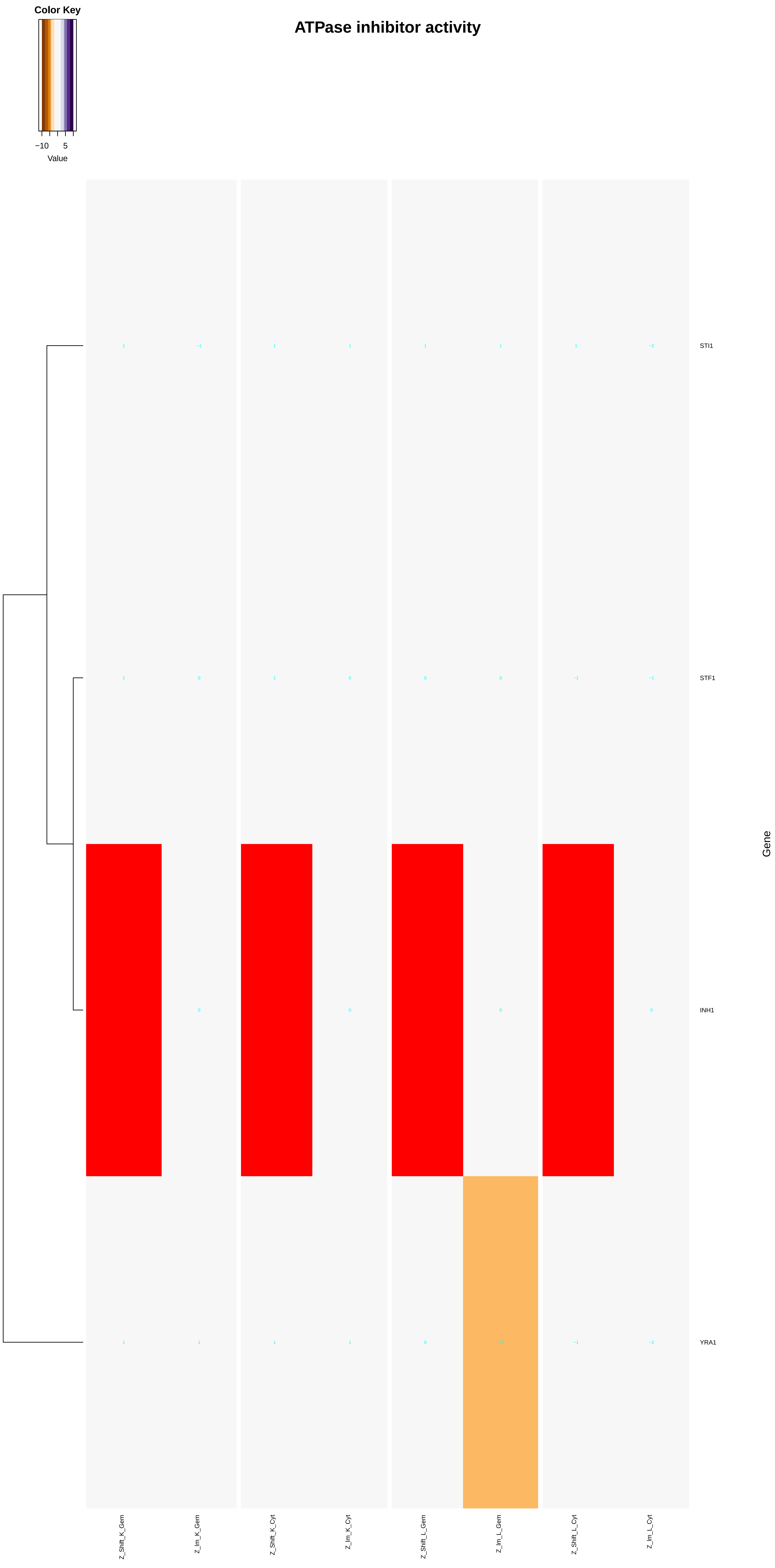
Color Key
−10
5
Value
ATPase inhibitor activity
STI1
STF1
INH1
YRA1
Gene
Z_Shift_K_Gem
Z_Im_K_Gem
Z_Shift_K_Cyt
Z_Im_K_Cyt
Z_Shift_L_Gem
Z_Im_L_Gem
Z_Shift_L_Cyt
Z_Im_L_Cyt

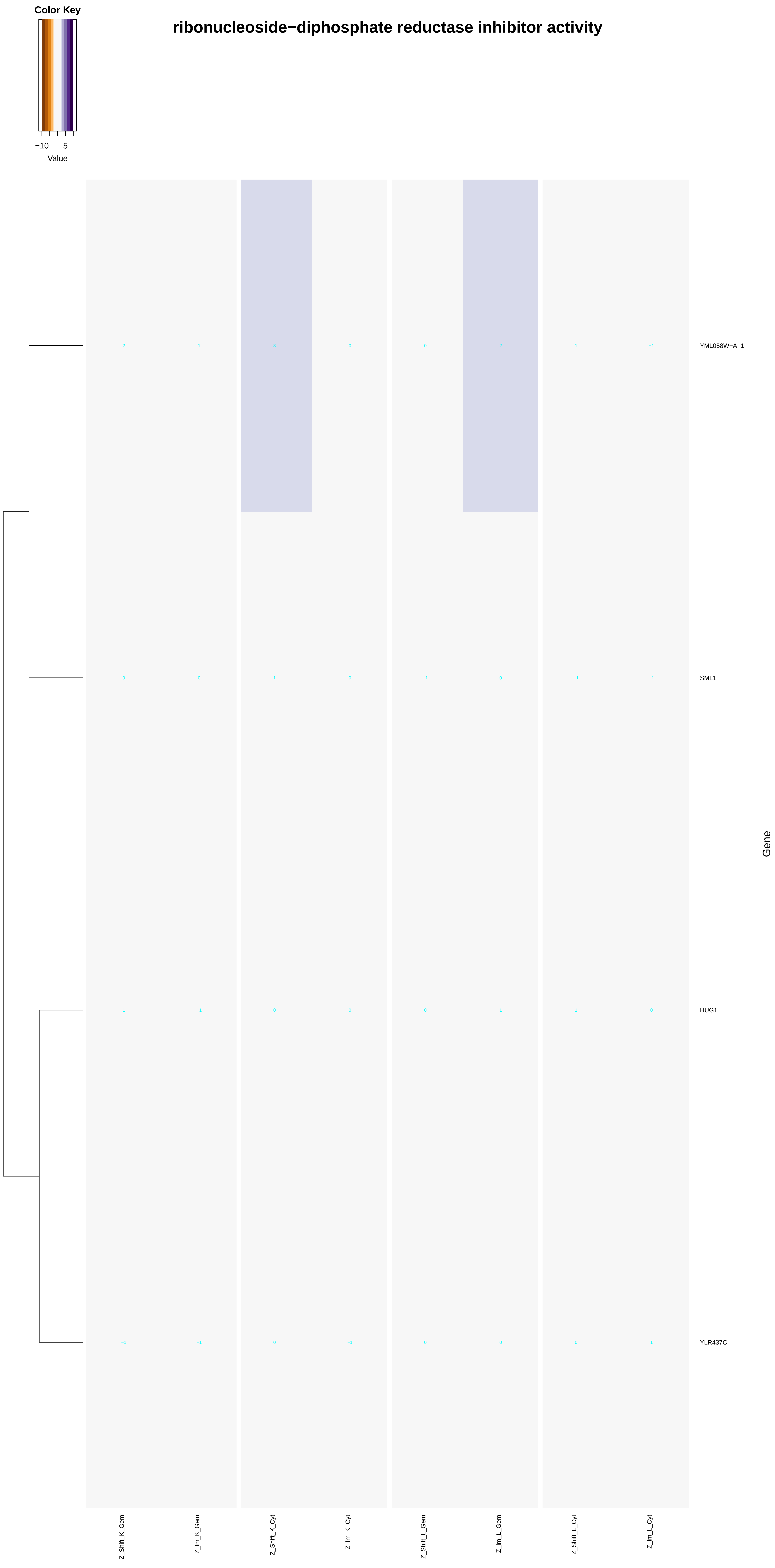
Color Key
−10
5
Value
ribonucleoside−diphosphate reductase inhibitor activity
YML058W−A_1
SML1
HUG1
YLR437C
Gene
Z_Shift_K_Gem
Z_Im_K_Gem
Z_Shift_K_Cyt
Z_Im_K_Cyt
Z_Shift_L_Gem
Z_Im_L_Gem
Z_Shift_L_Cyt
Z_Im_L_Cyt

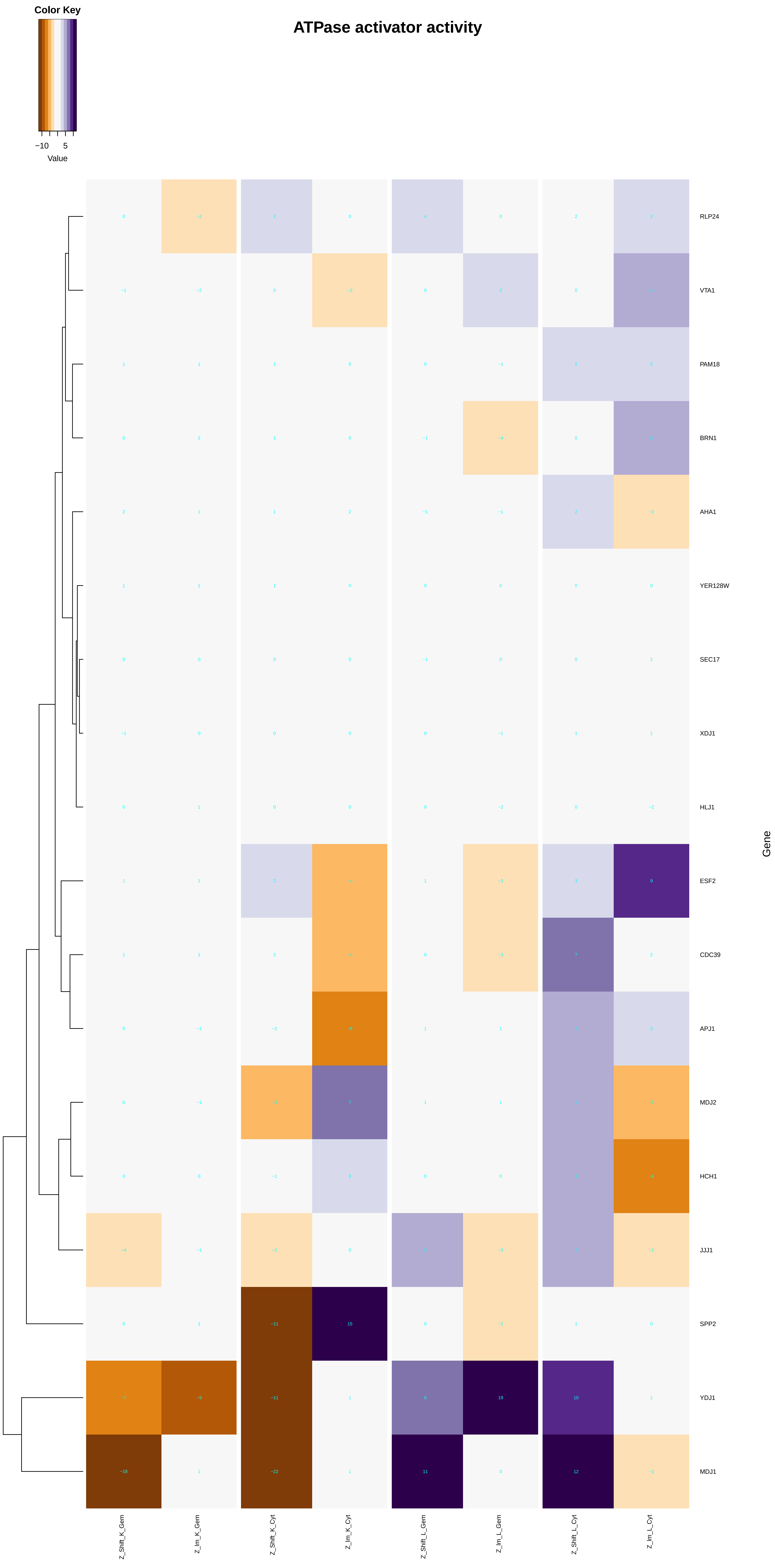
Color Key
−10
5
Value
ATPase activator activity
RLP24
VTA1
PAM18
BRN1
AHA1
YER128W
SEC17
XDJ1
HLJ1
ESF2
CDC39
APJ1
MDJ2
HCH1
JJJ1
SPP2
YDJ1
MDJ1
Gene
Z_Shift_K_Gem
Z_Im_K_Gem
Z_Shift_K_Cyt
Z_Im_K_Cyt
Z_Shift_L_Gem
Z_Im_L_Gem
Z_Shift_L_Cyt
Z_Im_L_Cyt

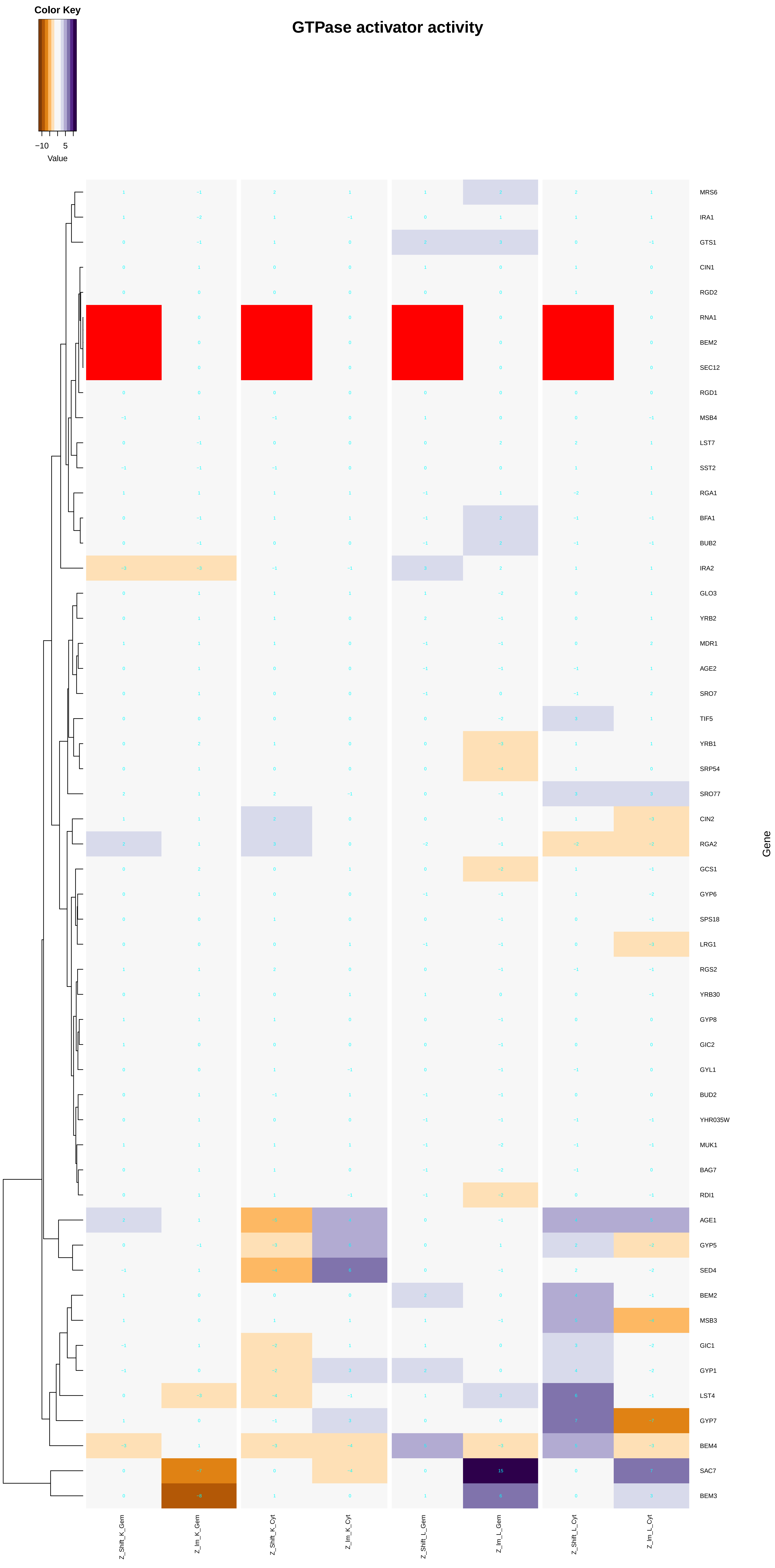
Color Key
−10
5
Value
GTPase activator activity
MRS6
IRA1
GTS1
CIN1
RGD2
RNA1
BEM2
SEC12
RGD1
MSB4
LST7
SST2
RGA1
BFA1
BUB2
IRA2
GLO3
YRB2
MDR1
AGE2
SRO7
TIF5
YRB1
SRP54
SRO77
CIN2
RGA2
GCS1
GYP6
SPS18
LRG1
RGS2
YRB30
GYP8
GIC2
GYL1
BUD2
YHR035W
MUK1
BAG7
RDI1
AGE1
GYP5
SED4
BEM2
MSB3
GIC1
GYP1
LST4
GYP7
BEM4
SAC7
BEM3
Gene
Z_Shift_K_Gem
Z_lm_K_Gem
Z_Shift_K_Cyt
Z_lm_K_Cyt
Z_Shift_L_Gem
Z_lm_L_Gem
Z_Shift_L_Cyt
Z_lm_L_Cyt

### peptidase activator activity

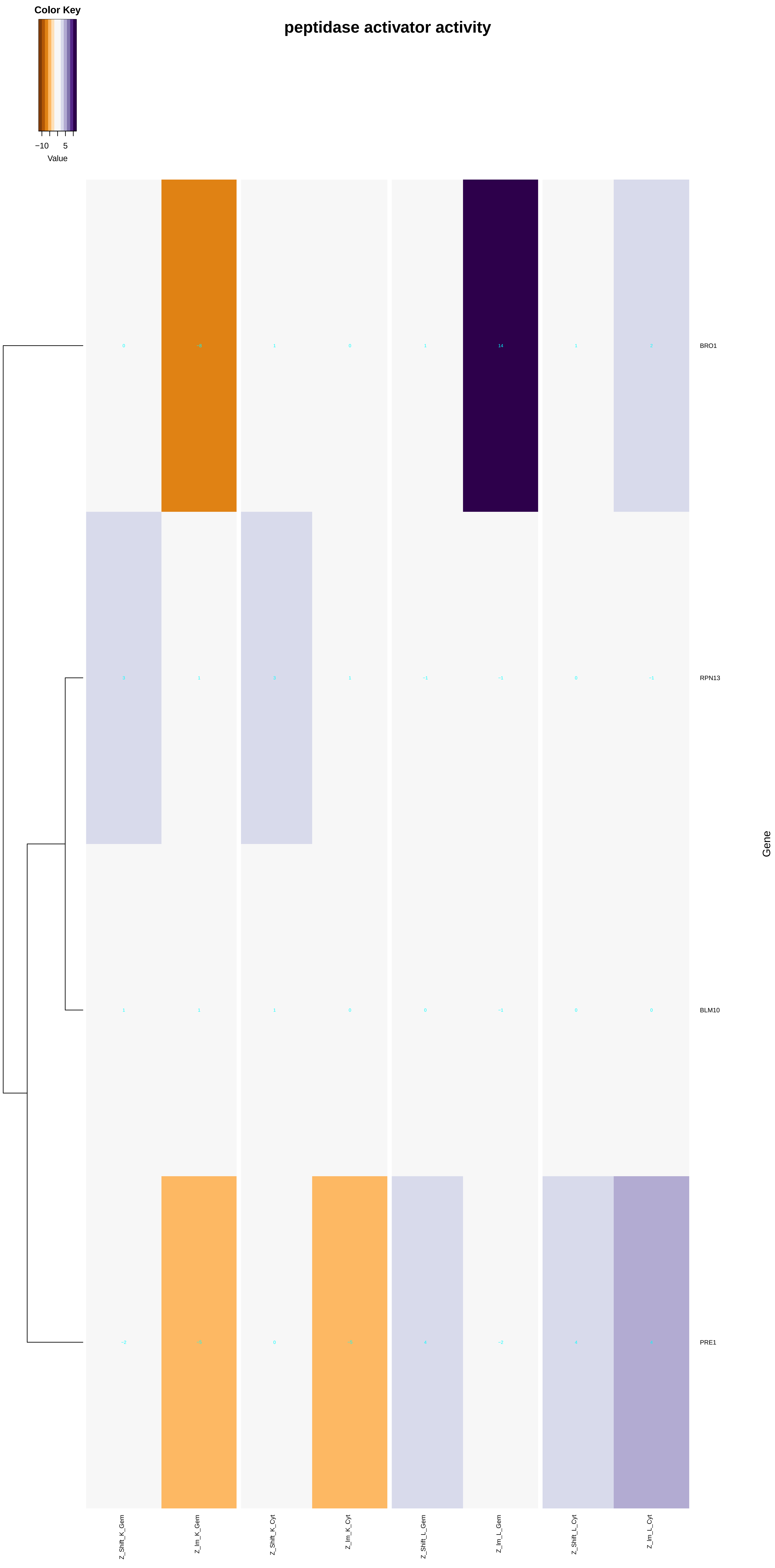

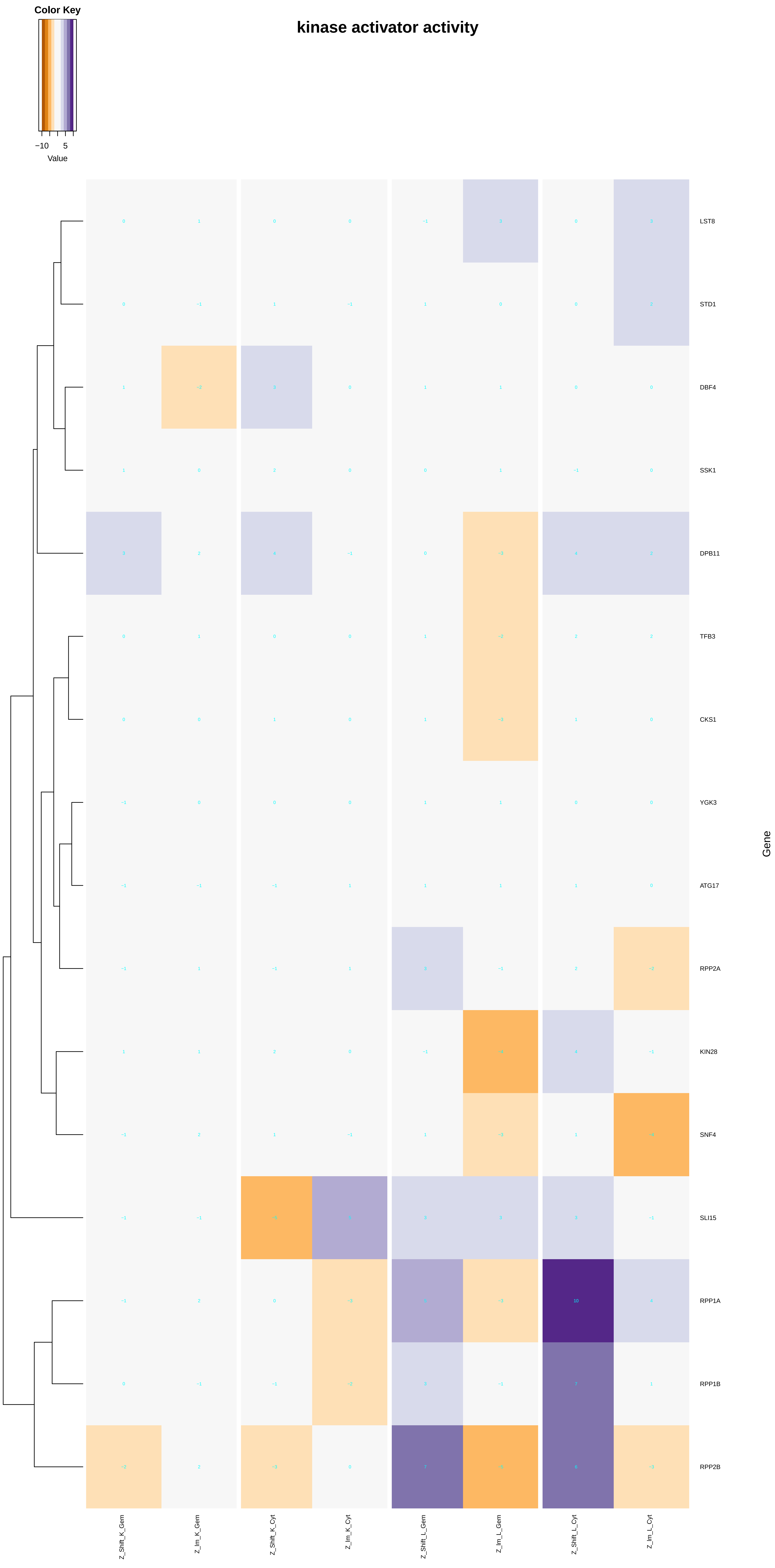
Color Key
−10
5
Value
kinase activator activity
LST8
STD1
DBF4
SSK1
DPB11
TFB3
CKS1
YGK3
ATG17
RPP2A
KIN28
SNF4
SLI15
RPP1A
RPP1B
RPP2B
Gene
Z_Shift_K_Gem
Z_Im_K_Gem
Z_Shift_K_Cyt
Z_Im_K_Cyt
Z_Shift_L_Gem
Z_Im_L_Gem
Z_Shift_L_Cyt
Z_Im_L_Cyt

#### phosphatase activator activity

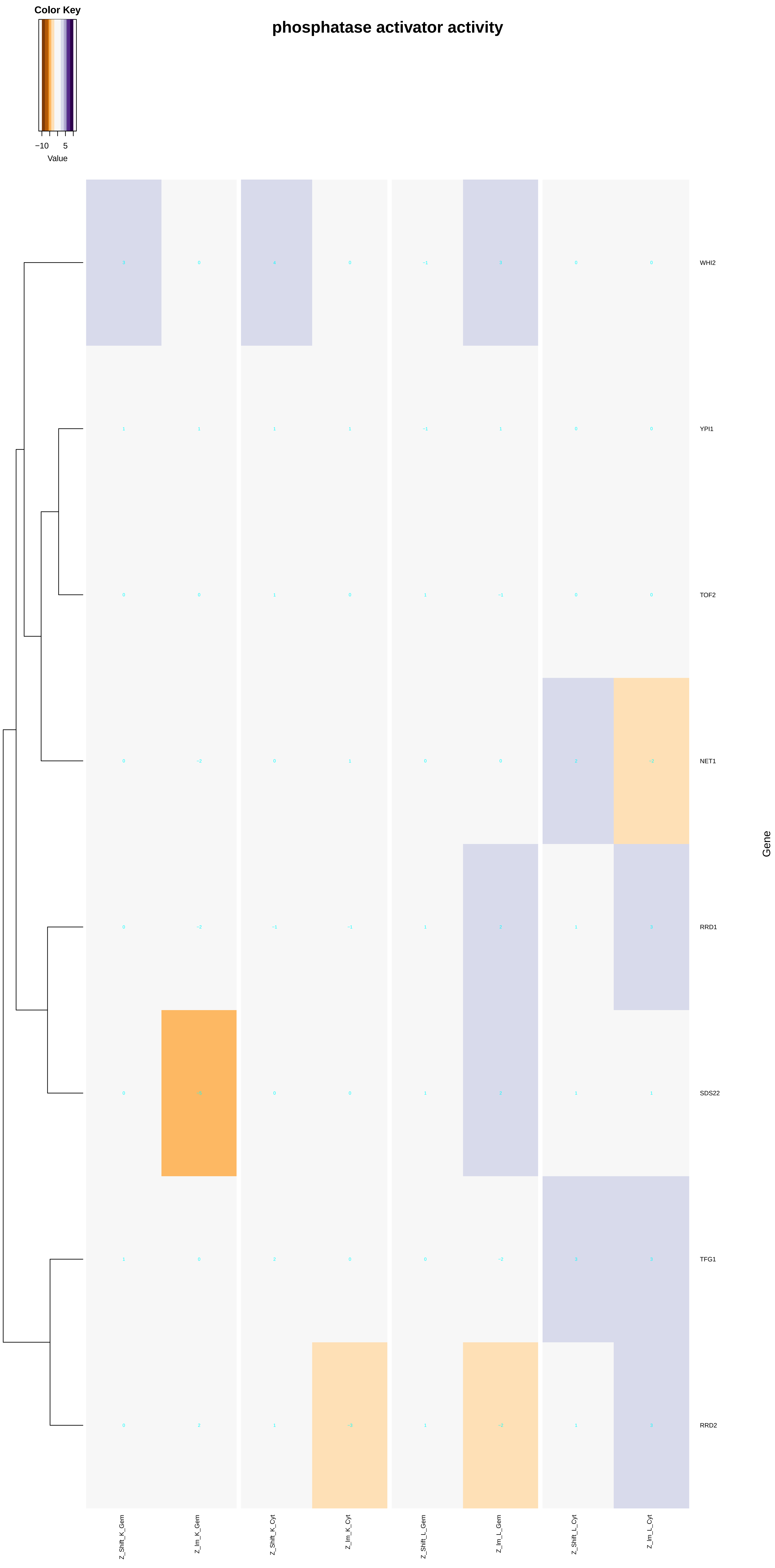

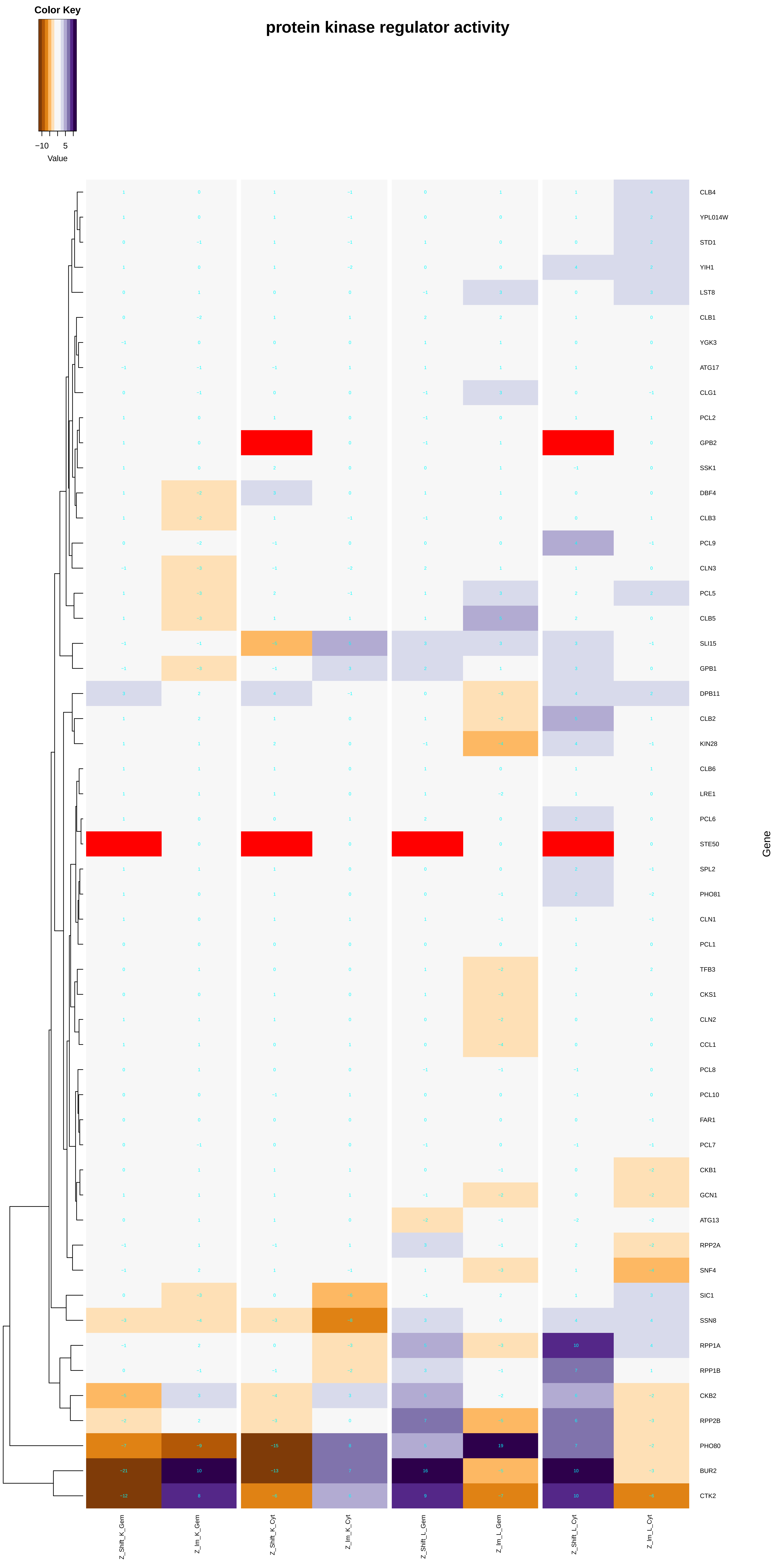

protein kinase regulator activity
Color Key
−10
5
Value
CLB4
YPL014W
STD1
YIH1
LST8
CLB1
YGK3
ATG17
CLG1
PCL2
GPB2
SSK1
DBF4
CLB3
PCL9
CLN3
PCL5
CLB5
SLI15
GPB1
DPB11
CLB2
KIN28
CLB6
LRE1
PCL6
STE50
SPL2
PHO81
CLN1
PCL1
TFB3
CKS1
CLN2
CCL1
PCL8
PCL10
FAR1
PCL7
CKB1
GCN1
ATG13
RPP2A
SNF4
SIC1
SSN8
RPP1A
RPP1B
CKB2
RPP2B
PHO80
BUR2
CTK2
Gene
Z_Shift_K_Gem
Z_Im_K_Gem
Z_Shift_K_Cyt
Z_Im_K_Cyt
Z_Shift_L_Gem
Z_Im_L_Gem
Z_Shift_L_Cyt
Z_Im_L_Cyt

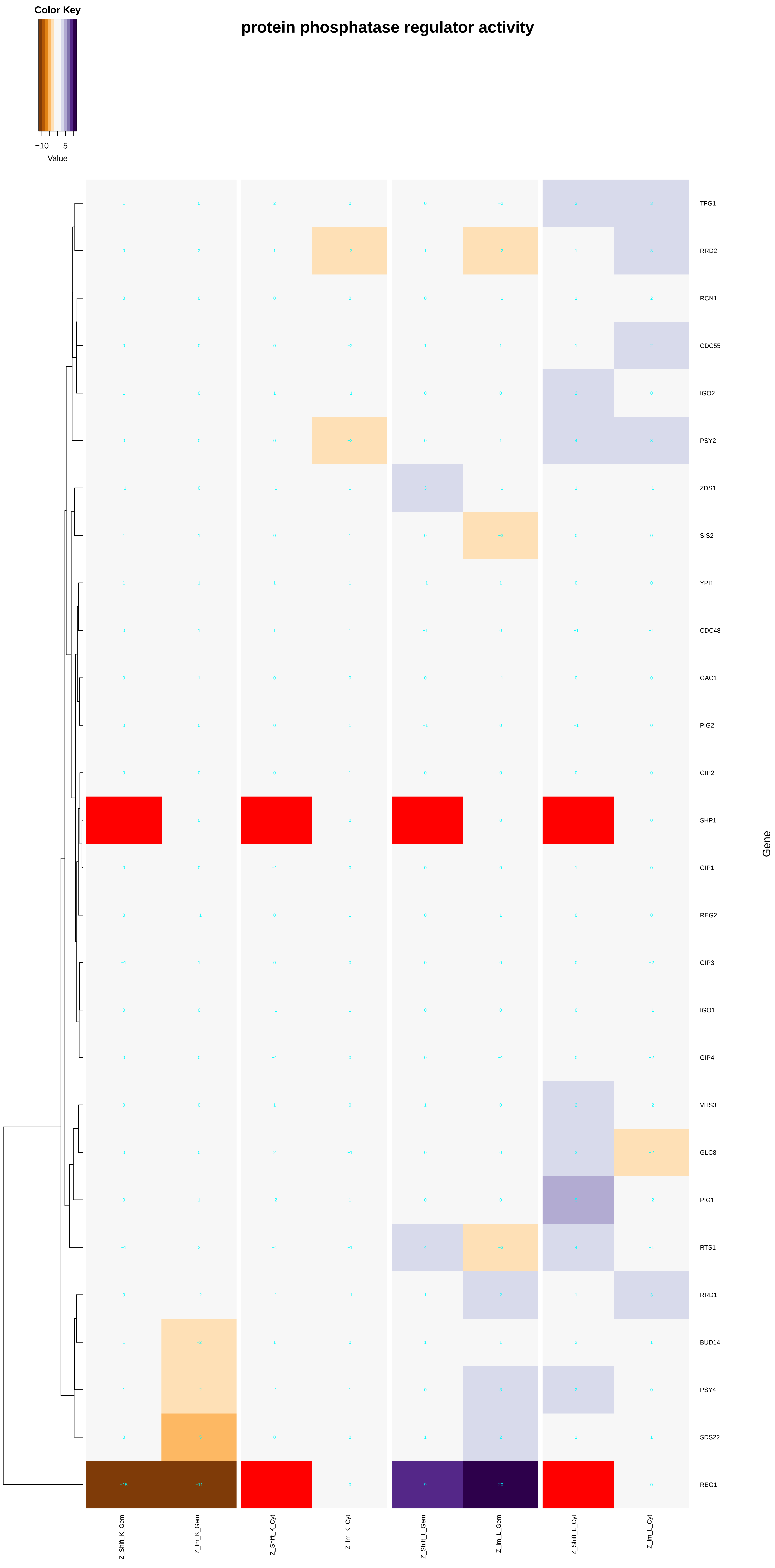

Color Key
−10
5
Value
protein phosphatase regulator activity
TFG1
RRD2
RCN1
CDC55
IGO2
PSY2
ZDS1
SIS2
YPI1
CDC48
GAC1
PIG2
GIP2
SHP1
GIP1
REG2
GIP3
IGO1
GIP4
VHS3
GLC8
PIG1
RTS1
RRD1
BUD14
PSY4
SDS22
REG1
Gene
Z_Shift_K_Gem
Z_Im_K_Gem
Z_Shift_K_Cyt
Z_Im_K_Cyt
Z_Shift_L_Gem
Z_Im_L_Gem
Z_Shift_L_Cyt
Z_Im_L_Cyt

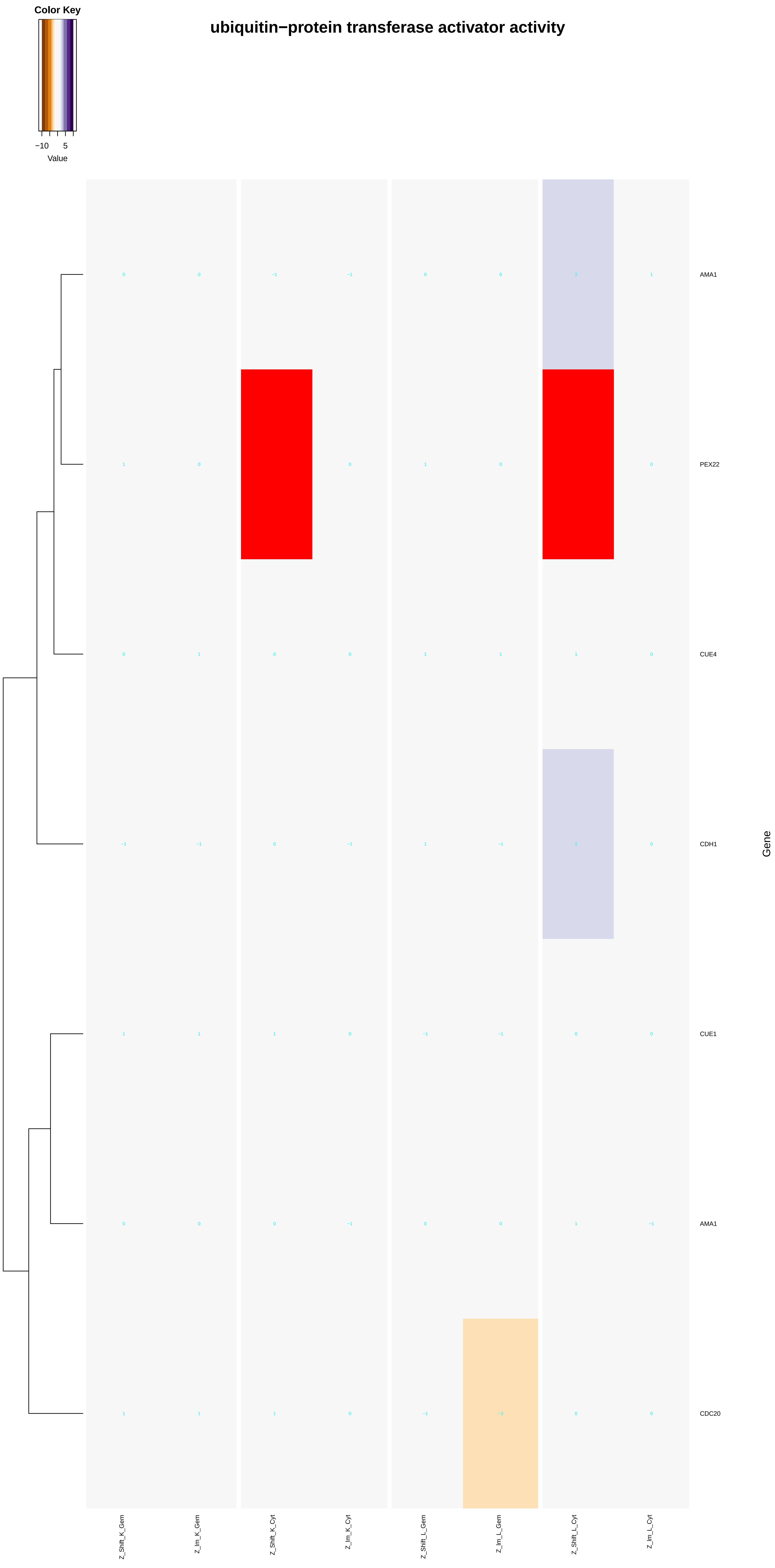

Color Key
−10
5
Value
ubiquitin−protein transferase activator activity
AMA1
PEX22
CUE4
CDH1
CUE1
AMA1
CDC20
Gene
Z_Shift_K_Gem
Z_lm_K_Gem
Z_Shift_K_Cyt
Z_lm_K_Cyt
Z_Shift_L_Gem
Z_lm_L_Gem
Z_Shift_L_Cyt
Z_lm_L_Cyt

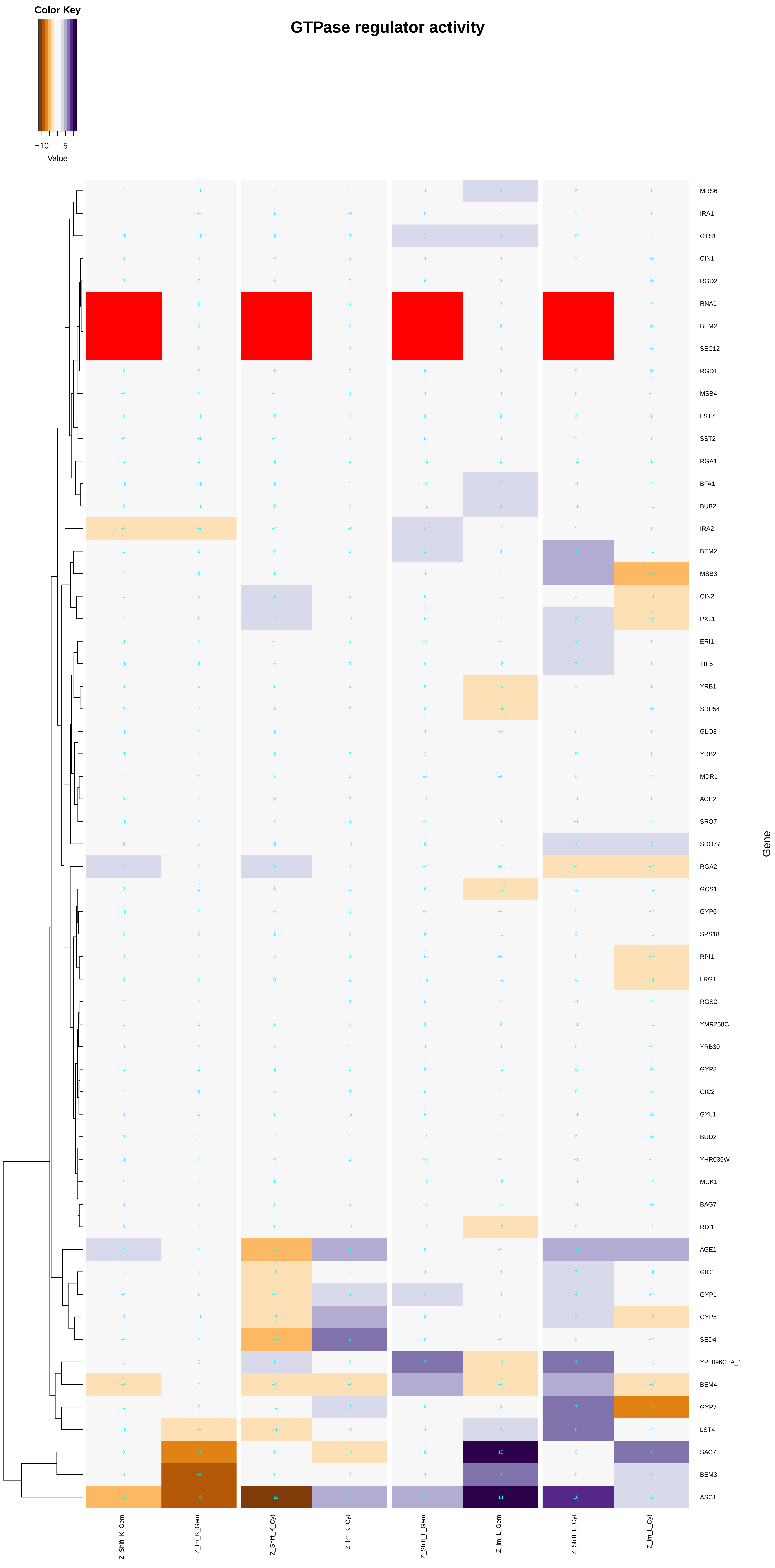

Color Key
−10
5
Value
GTPase regulator activity
MRS6
IRA1
GTS1
CIN1
RGD2
RNA1
BEM2
SEC12
RGD1
MSB4
LST7
SST2
RGA1
BFA1
BUB2
IRA2
BEM2
MSB3
CIN2
PXL1
ERI1
TIF5
YRB1
SRP54
GLO3
YRB2
MDR1
AGE2
SRO7
SRO77
RGA2
GCS1
GYP6
SPS18
RPI1
LRG1
RGS2
YMR258C
YRB30
GYP8
GIC2
GYL1
BUD2
YHR035W
MUK1
BAG7
RDI1
AGE1
GIC1
GYP1
GYP5
SED4
YPL096C−A_1
BEM4
GYP7
LST4
SAC7
BEM3
ASC1
Gene
Z_Shift_K_Gem
Z_Im_K_Gem
Z_Shift_K_Cyt
Z_Im_K_Cyt
Z_Shift_L_Gem
Z_Im_L_Gem
Z_Shift_L_Cyt
Z_Im_L_Cyt

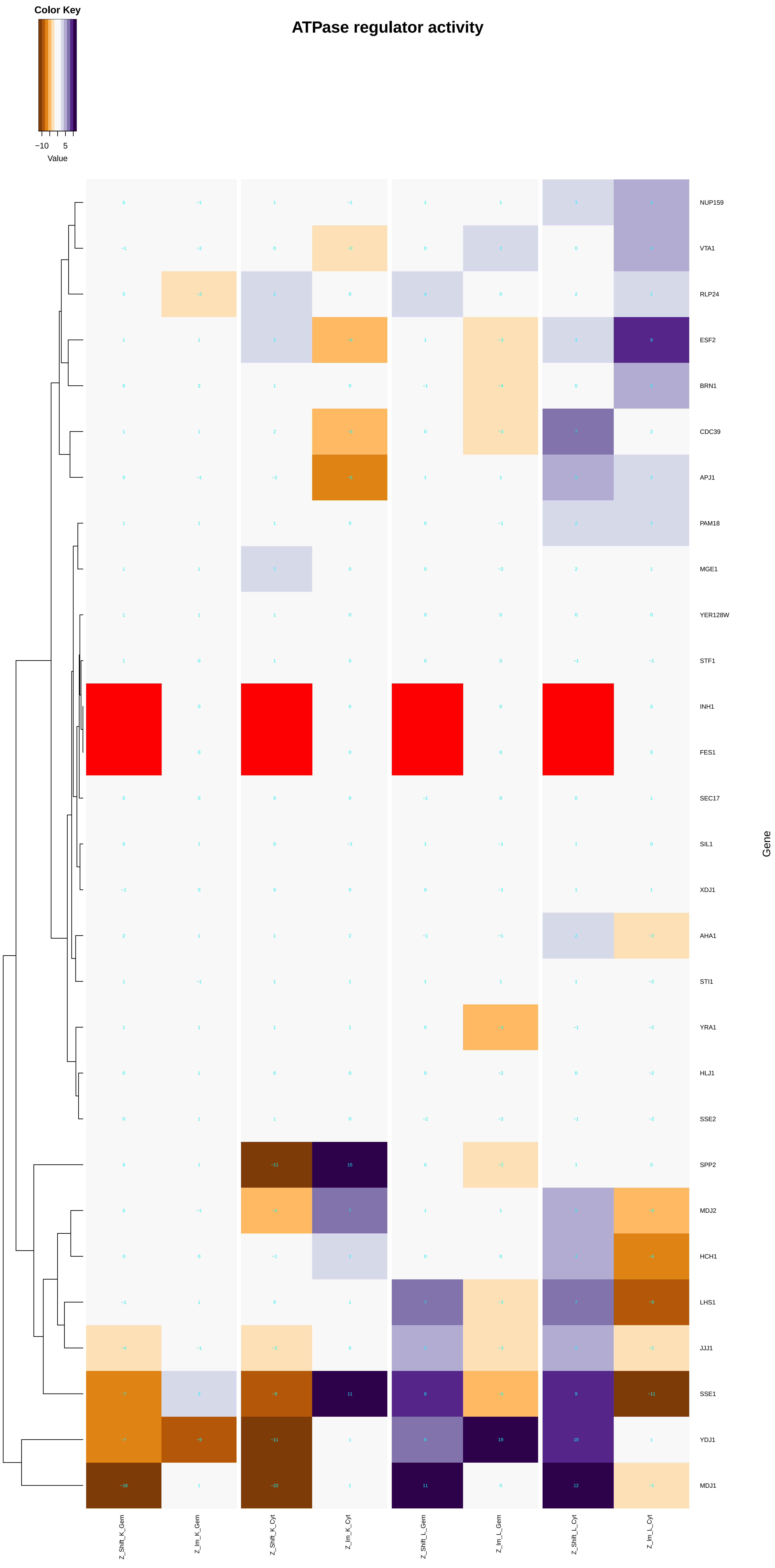
Color Key
−10
5
Value
ATPase regulator activity
NUP159
VTA1
RLP24
ESF2
BRN1
CDC39
APJ1
PAM18
MGE1
YER128W
STF1
INH1
FES1
SEC17
SIL1
XDJ1
AHA1
STI1
YRA1
HLJ1
SSE2
SPP2
MDJ2
HCH1
LHS1
JJJ1
SSE1
YDJ1
MDJ1
Gene
Z_Shift_K_Gem
Z_Im_K_Gem
Z_Shift_K_Cyt
Z_Im_K_Cyt
Z_Shift_L_Gem
Z_Im_L_Gem
Z_Shift_L_Cyt
Z_Im_L_Cyt

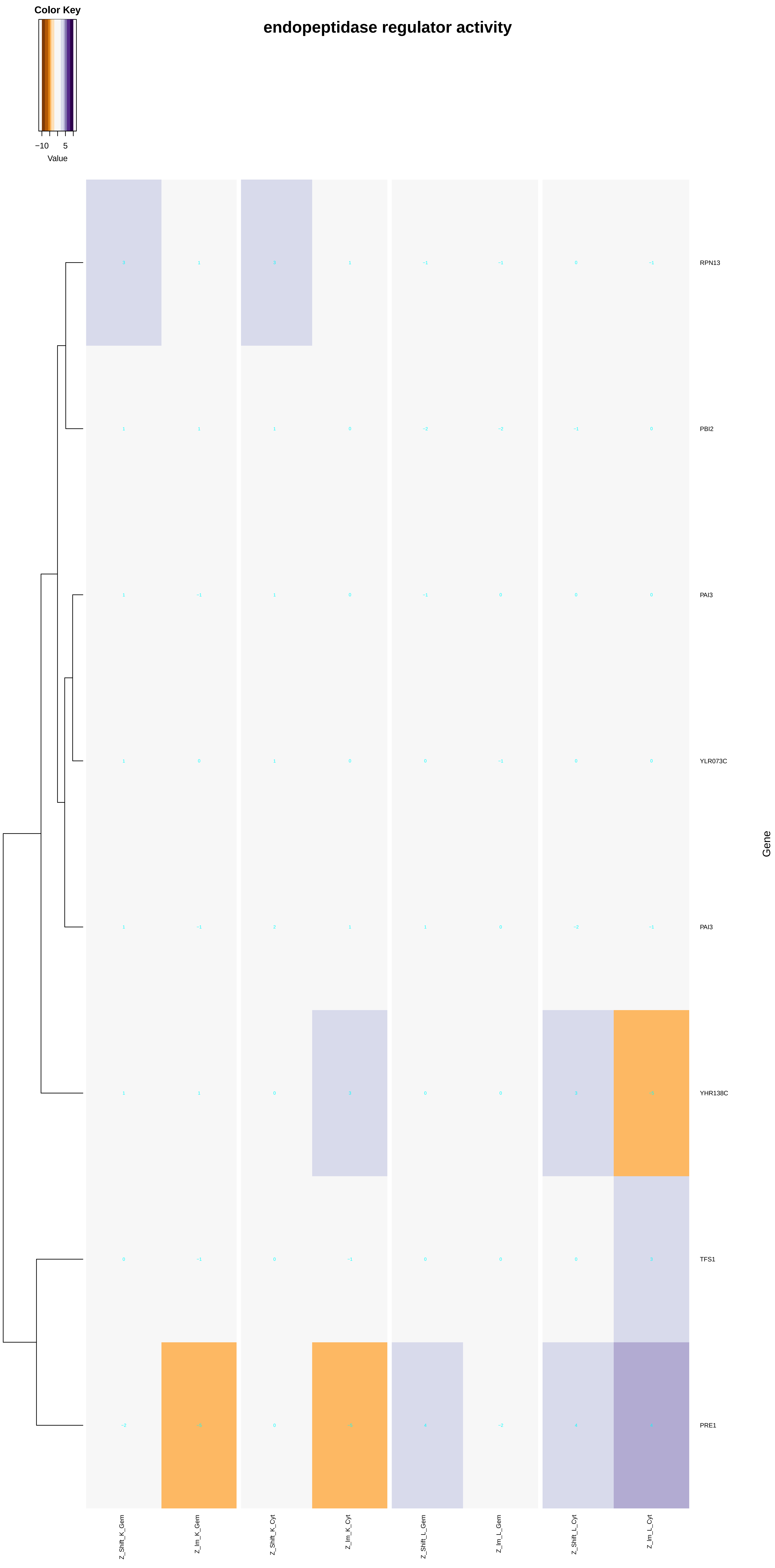
Color Key
−10
5
Value
endopeptidase regulator activity
RPN13
PBI2
PAI3
YLR073C
PAI3
YHR138C
TFS1
PRE1
Gene
Z_Shift_K_Gem
Z_Im_K_Gem
Z_Shift_K_Cyt
Z_Im_K_Cyt
Z_Shift_L_Gem
Z_Im_L_Gem
Z_Shift_L_Cyt
Z_Im_L_Cyt

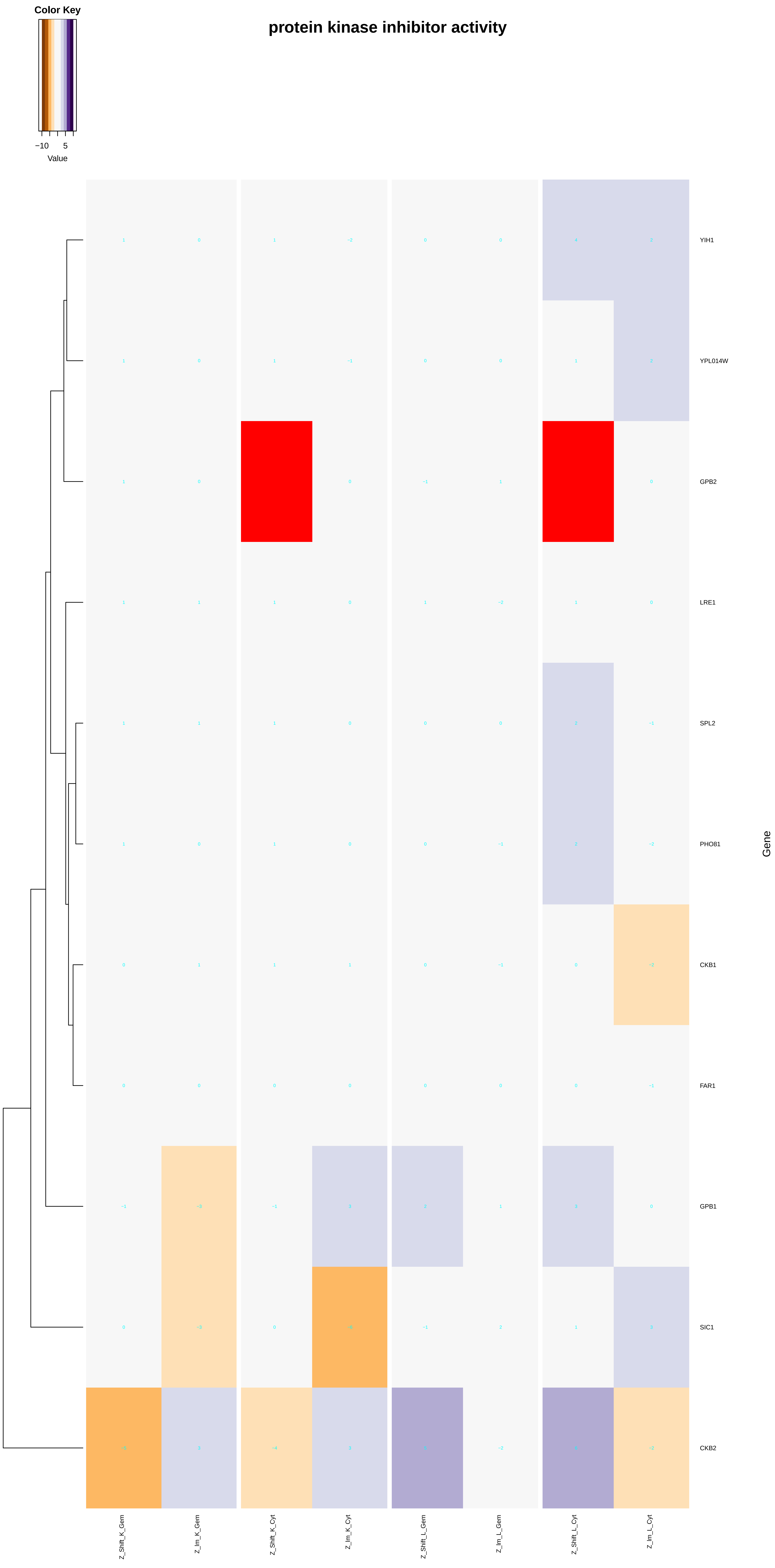
Color Key
−10
5
Value
protein kinase inhibitor activity
YIH1
YPL014W
GPB2
LRE1
SPL2
PHO81
CKB1
FAR1
GPB1
SIC1
CKB2
Gene
Z_Shift_K_Gem
Z_Im_K_Gem
Z_Shift_K_Cyt
Z_Im_K_Cyt
Z_Shift_L_Gem
Z_Im_L_Gem
Z_Shift_L_Cyt
Z_Im_L_Cyt

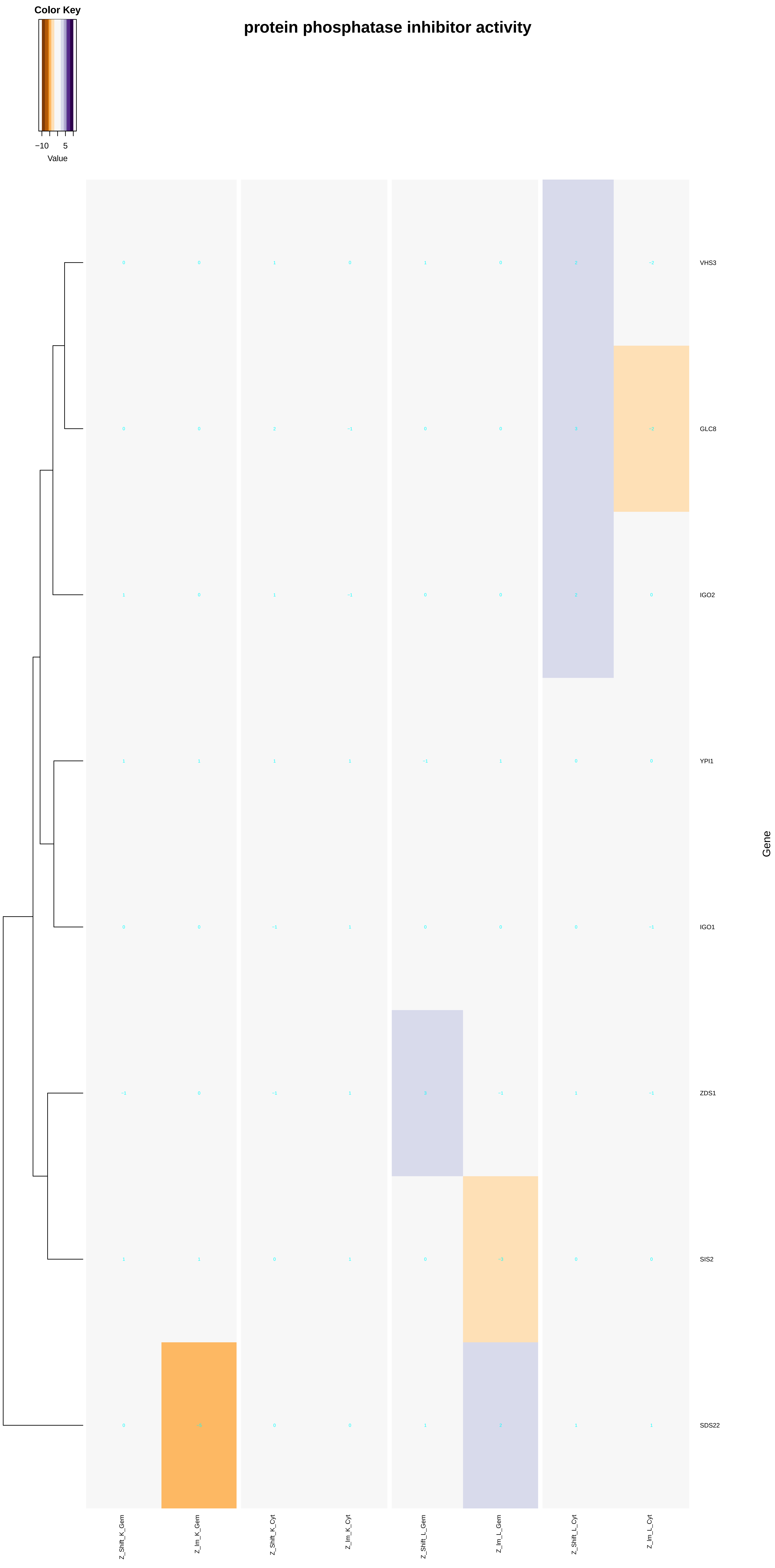
Color Key
−10
5
Value
protein phosphatase inhibitor activity
VHS3
GLC8
IGO2
YPI1
IGO1
ZDS1
SIS2
SDS22
Gene
Z_Shift_K_Gem
Z_Im_K_Gem
Z_Shift_K_Cyt
Z_Im_K_Cyt
Z_Shift_L_Gem
Z_Im_L_Gem
Z_Shift_L_Cyt
Z_Im_L_Cyt

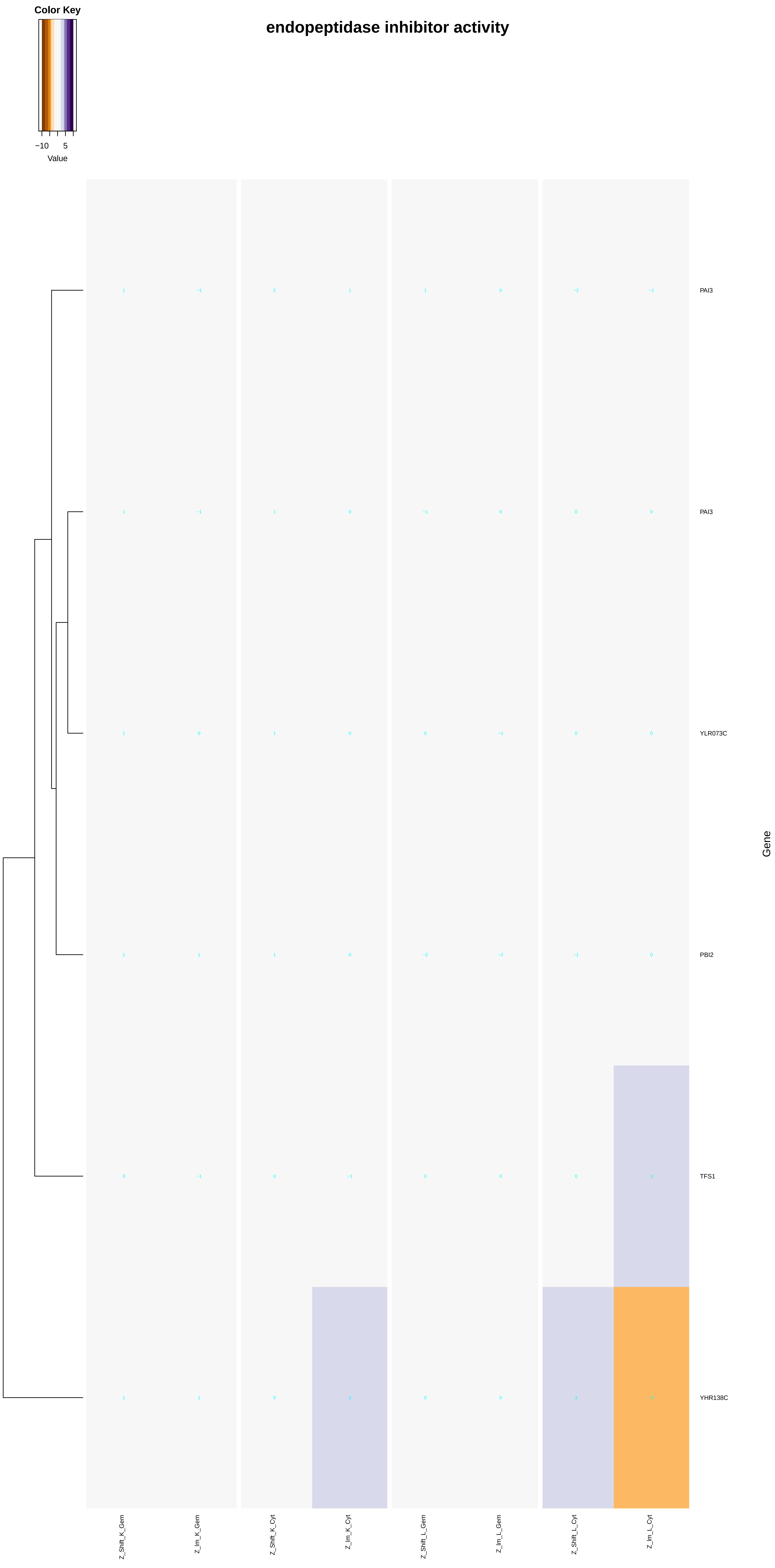
Color Key
−10
5
Value
endopeptidase inhibitor activity
PAI3
PAI3
YLR073C
PBI2
TFS1
YHR138C
Gene
Z_Shift_K_Gem
Z_Im_K_Gem
Z_Shift_K_Cyt
Z_Im_K_Cyt
Z_Shift_L_Gem
Z_Im_L_Gem
Z_Shift_L_Cyt
Z_Im_L_Cyt

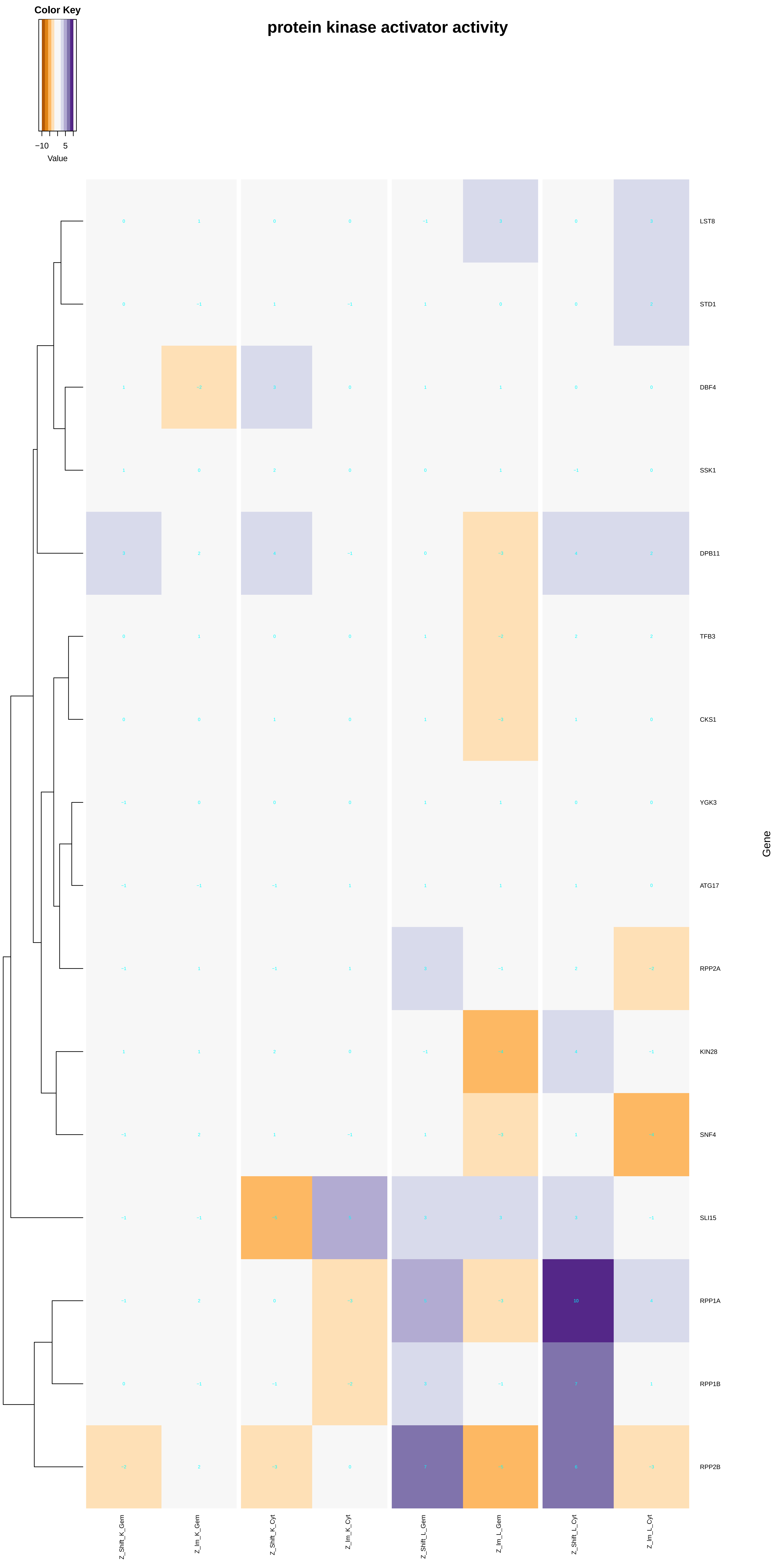

Color Key
−10
5
Value
protein kinase activator activity
LST8
STD1
DBF4
SSK1
DPB11
TFB3
CKS1
YGK3
ATG17
RPP2A
KIN28
SNF4
SLI15
RPP1A
RPP1B
RPP2B
Gene
Z_Shift_K_Gem
Z_Im_K_Gem
Z_Shift_K_Cyt
Z_Im_K_Cyt
Z_Shift_L_Gem
Z_Im_L_Gem
Z_Shift_L_Cyt
Z_Im_L_Cyt

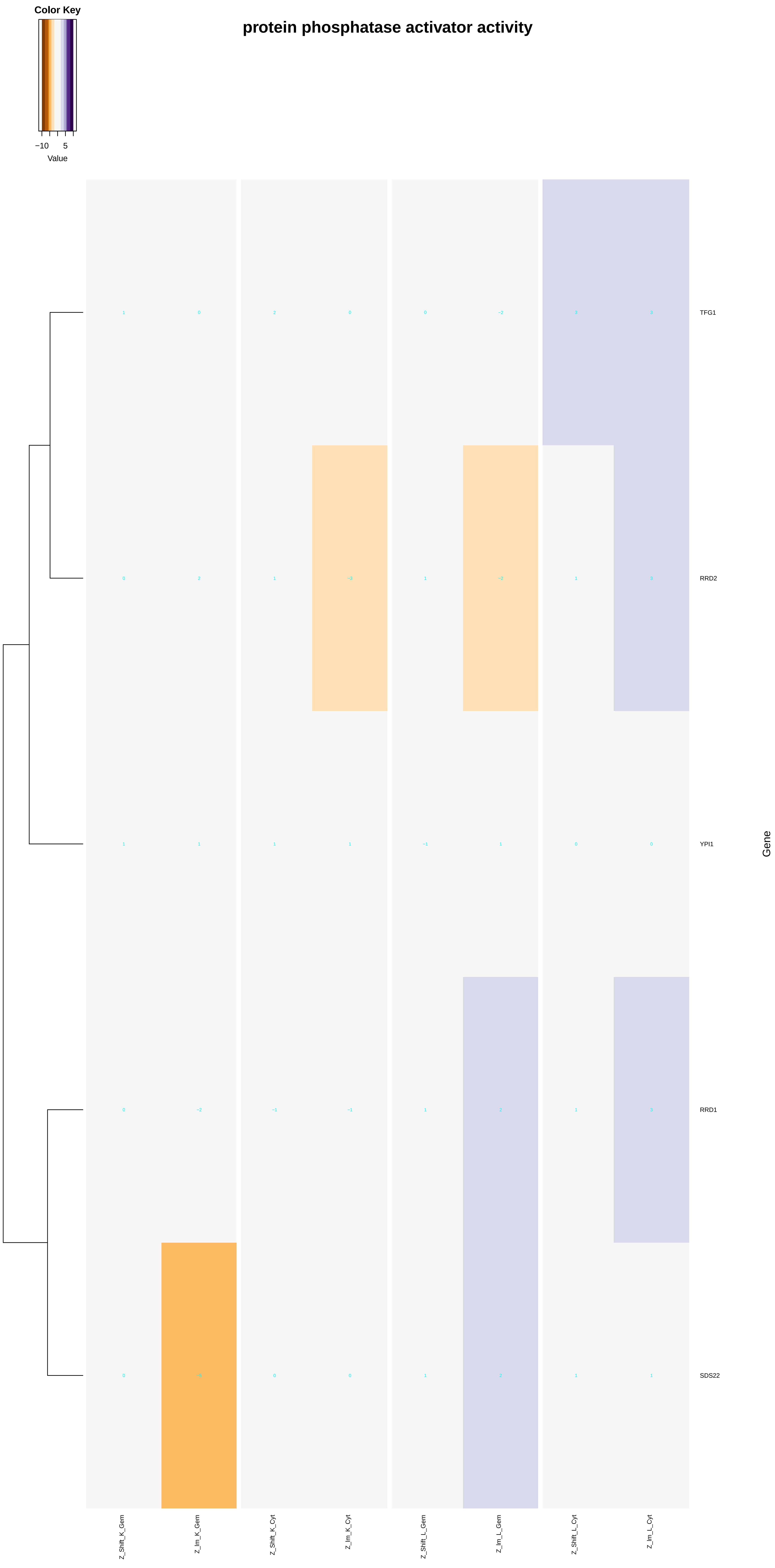

protein phosphatase activator activity
Color Key
−10
5
Value
TFG1
RRD2
YPI1
RRD1
SDS22
Gene
Z_Shift_K_Gem
Z_Im_K_Gem
Z_Shift_K_Cyt
Z_Im_K_Cyt
Z_Shift_L_Gem
Z_Im_L_Gem
Z_Shift_L_Cyt
Z_Im_L_Cyt

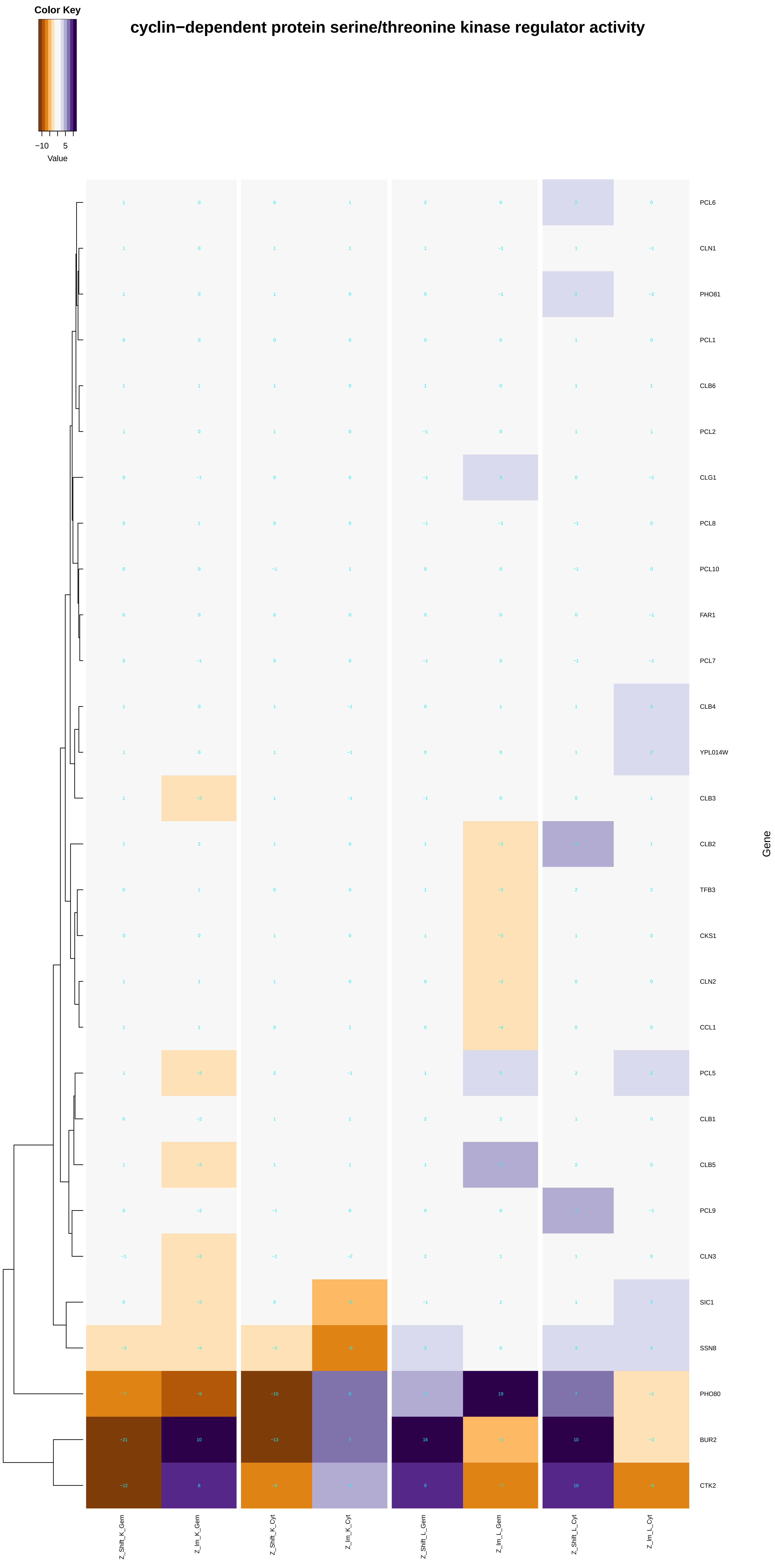

cyclin−dependent protein serine/threonine kinase regulator activity
Color Key
−10
5
Value
PCL6
CLN1
PHO81
PCL1
CLB6
PCL2
CLG1
PCL8
PCL10
FAR1
PCL7
CLB4
YPL014W
CLB3
CLB2
TFB3
CKS1
CLN2
CCL1
PCL5
CLB1
CLB5
PCL9
CLN3
SIC1
SSN8
PHO80
BUR2
CTK2
Gene
Z_Shift_K_Gem
Z_Im_K_Gem
Z_Shift_K_Cyt
Z_Im_K_Cyt
Z_Shift_L_Gem
Z_Im_L_Gem
Z_Shift_L_Cyt
Z_Im_L_Cyt

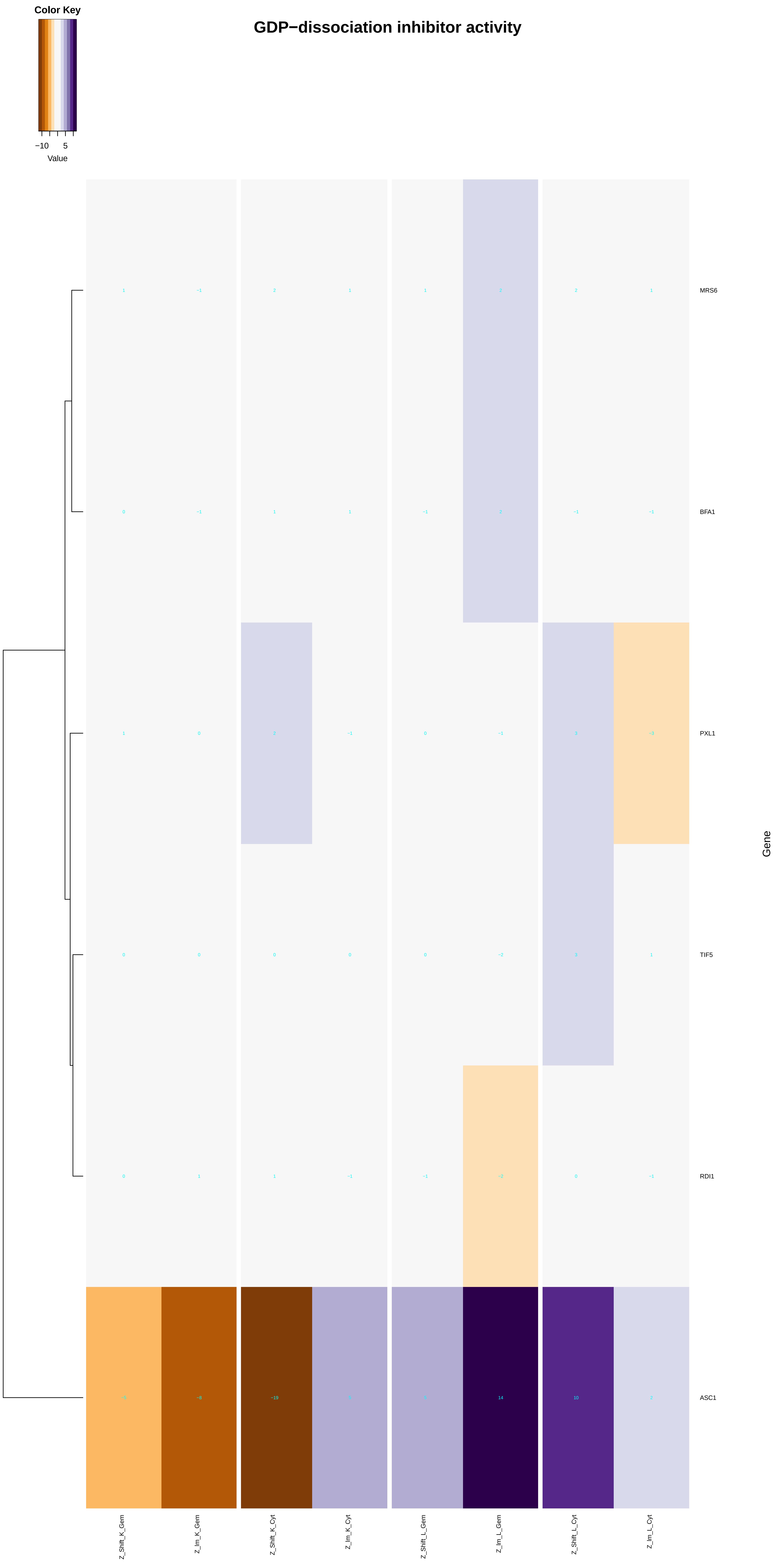

GDP−dissociation inhibitor activity
Color Key
−10
5
Value
MRS6
BFA1
PXL1
TIF5
RDI1
ASC1
Gene
Z_Shift_K_Gem
Z_Im_K_Gem
Z_Shift_K_Cyt
Z_Im_K_Cyt
Z_Shift_L_Gem
Z_Im_L_Gem
Z_Shift_L_Cyt
Z_Im_L_Cyt

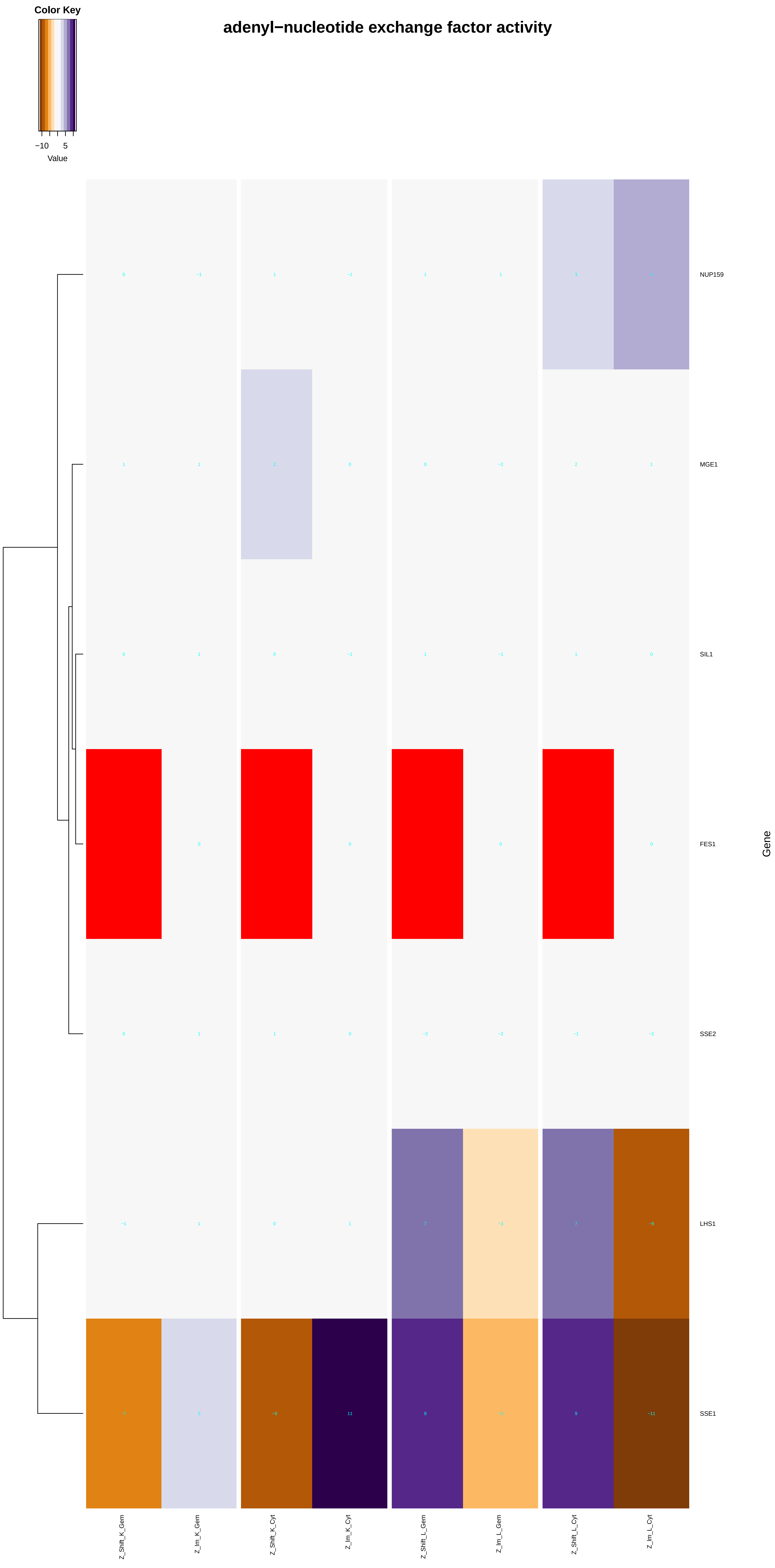

Color Key
−10
5
Value
adenyl−nucleotide exchange factor activity
NUP159
MGE1
SIL1
FES1
SSE2
LHS1
SSE1
Gene
Z_Shift_K_Gem
Z_Im_K_Gem
Z_Shift_K_Cyt
Z_Im_K_Cyt
Z_Shift_L_Gem
Z_Im_L_Gem
Z_Shift_L_Cyt
Z_Im_L_Cyt

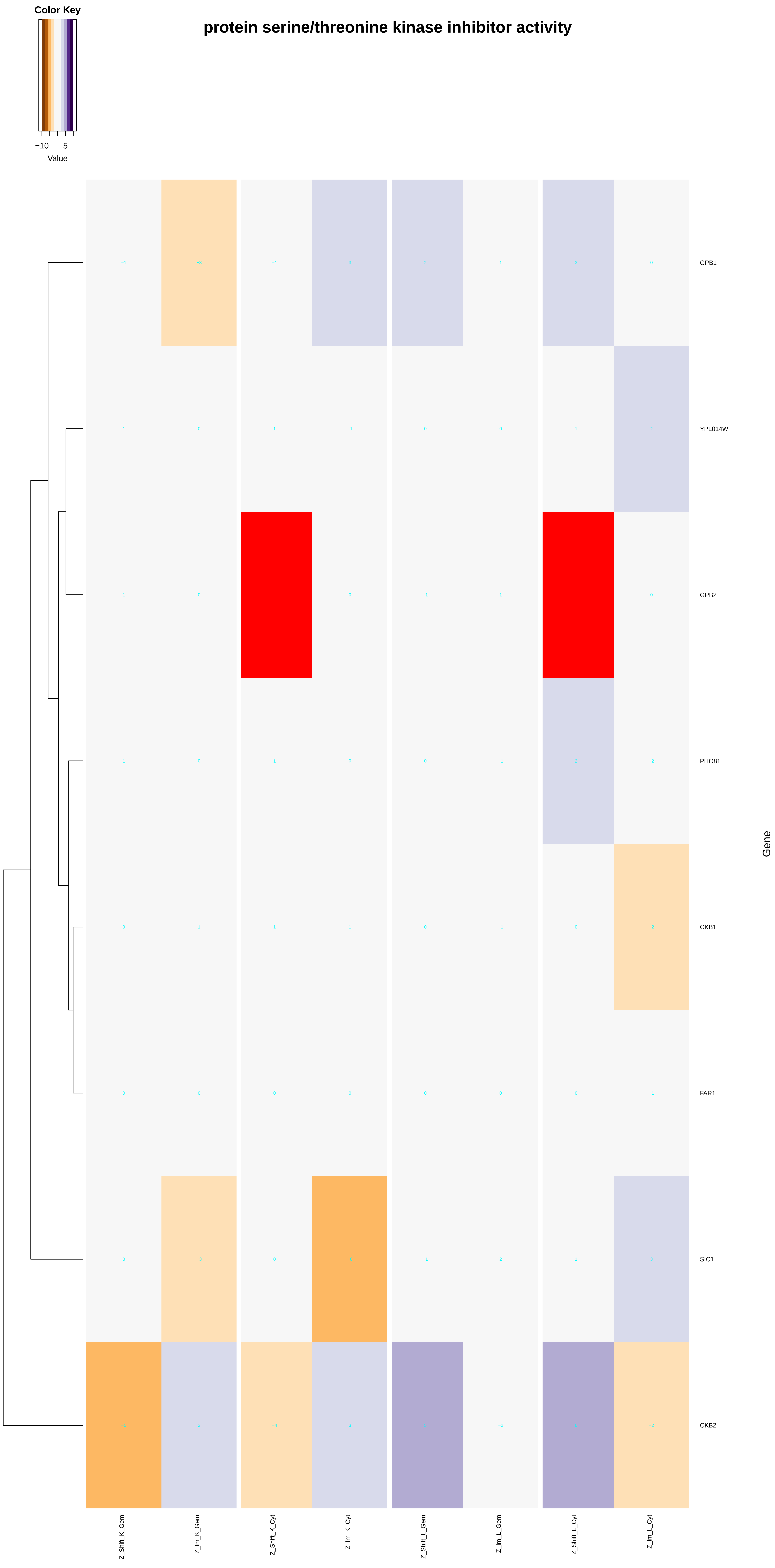

Color Key
−10
5
Value
protein serine/threonine kinase inhibitor activity
GPB1
YPL014W
GPB2
PHO81
CKB1
FAR1
SIC1
CKB2
Gene
Z_Shift_K_Gem
Z_Im_K_Gem
Z_Shift_K_Cyt
Z_Im_K_Cyt
Z_Shift_L_Gem
Z_Im_L_Gem
Z_Shift_L_Cyt
Z_Im_L_Cyt

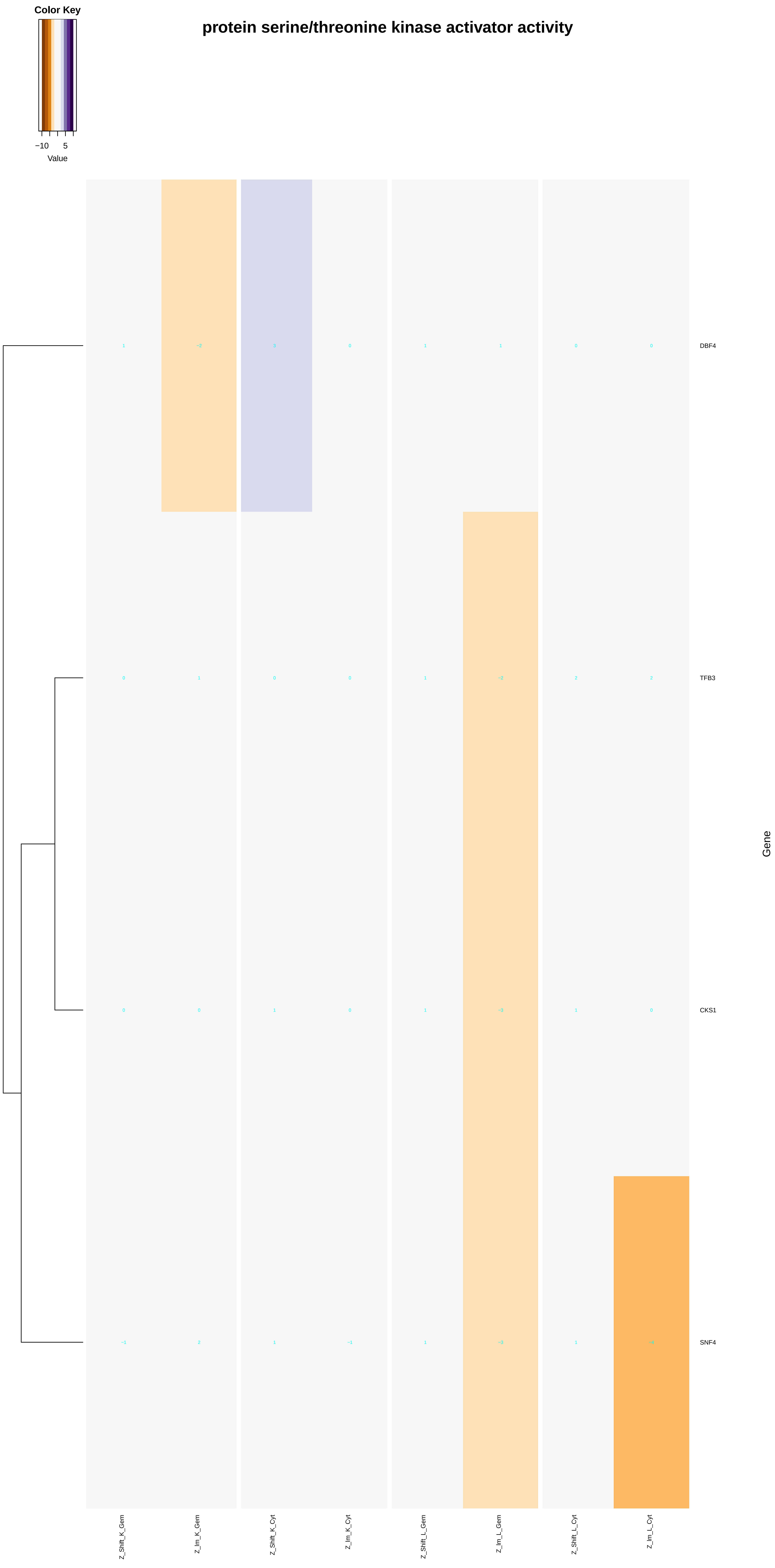

Color Key
−10
5
Value
protein serine/threonine kinase activator activity
DBF4
TFB3
CKS1
SNF4
Gene
Z_Shift_K_Gem
Z_Im_K_Gem
Z_Shift_K_Cyt
Z_Im_K_Cyt
Z_Shift_L_Gem
Z_Im_L_Gem
Z_Shift_L_Cyt
Z_Im_L_Cyt

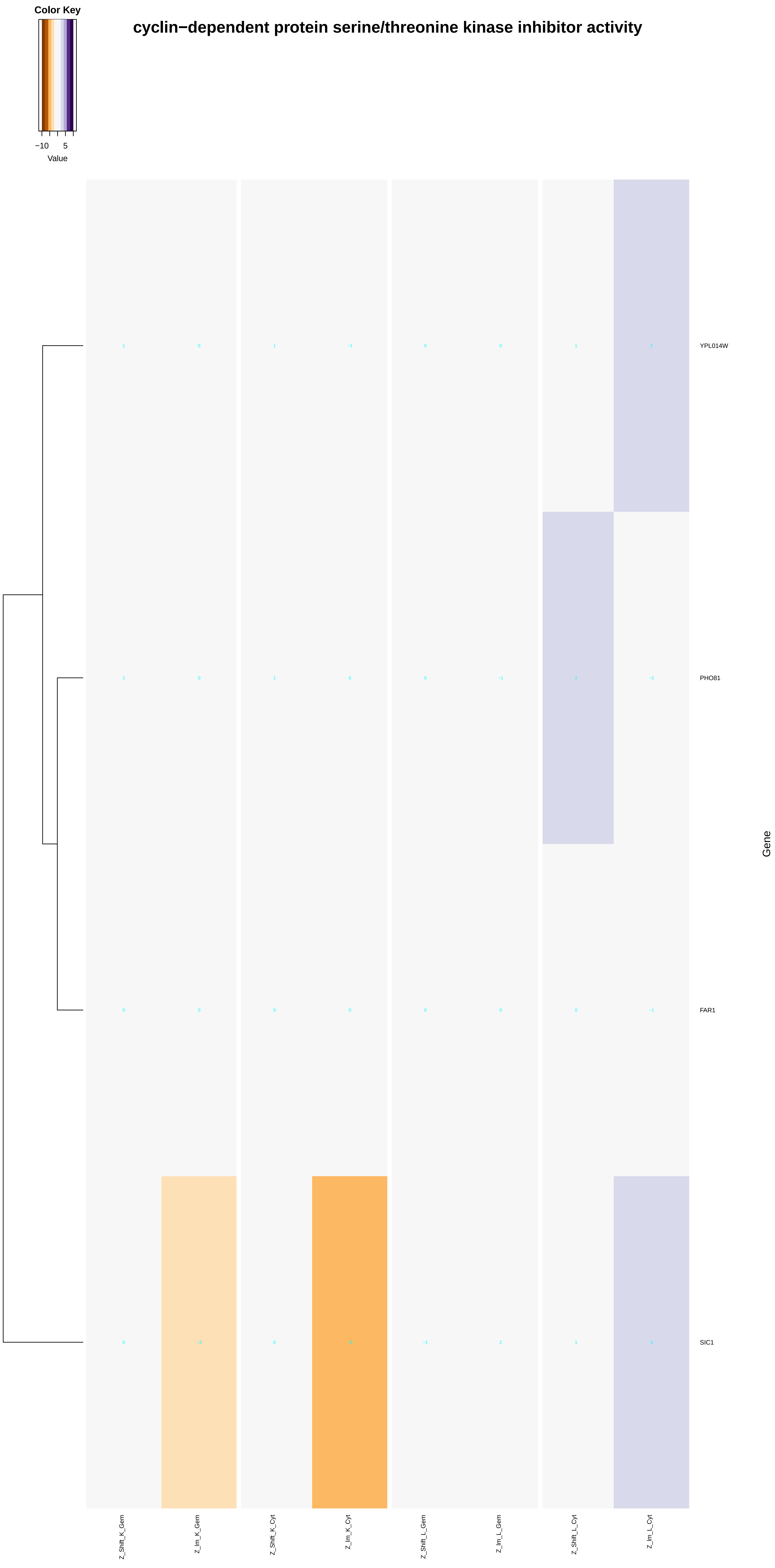
Color Key
−10
5
Value
cyclin−dependent protein serine/threonine kinase inhibitor activity
YPL014W
PHO81
FAR1
SIC1
Gene
Z_Shift_K_Gem
Z_Im_K_Gem
Z_Shift_K_Cyt
Z_Im_K_Cyt
Z_Shift_L_Gem
Z_Im_L_Gem
Z_Shift_L_Cyt
Z_Im_L_Cyt
