## Additional File 7 for "A humanized yeast phenomic model of deoxycytidine kinase to predict genetic buffering of nucleoside analog cytotoxicity": establishment_of_protein_localization.pdf

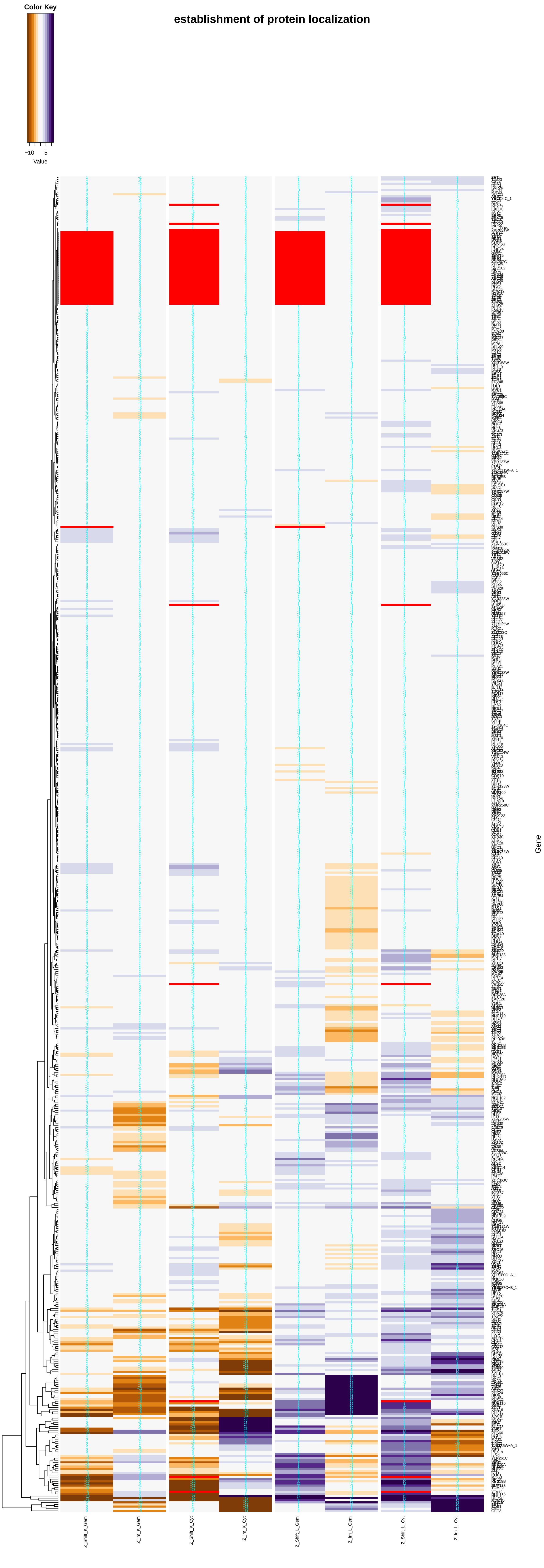
establishment of protein localization
Color Key
−10
5
Value
Gene
Z_Shift_K_Gem
Z_Im_K_Gem
Z_Shift_K_Cyt
Z_Im_K_Cyt
Z_Shift_L_Gem
Z_Im_L_Gem
Z_Shift_L_Cyt
Z_Im_L_Cyt

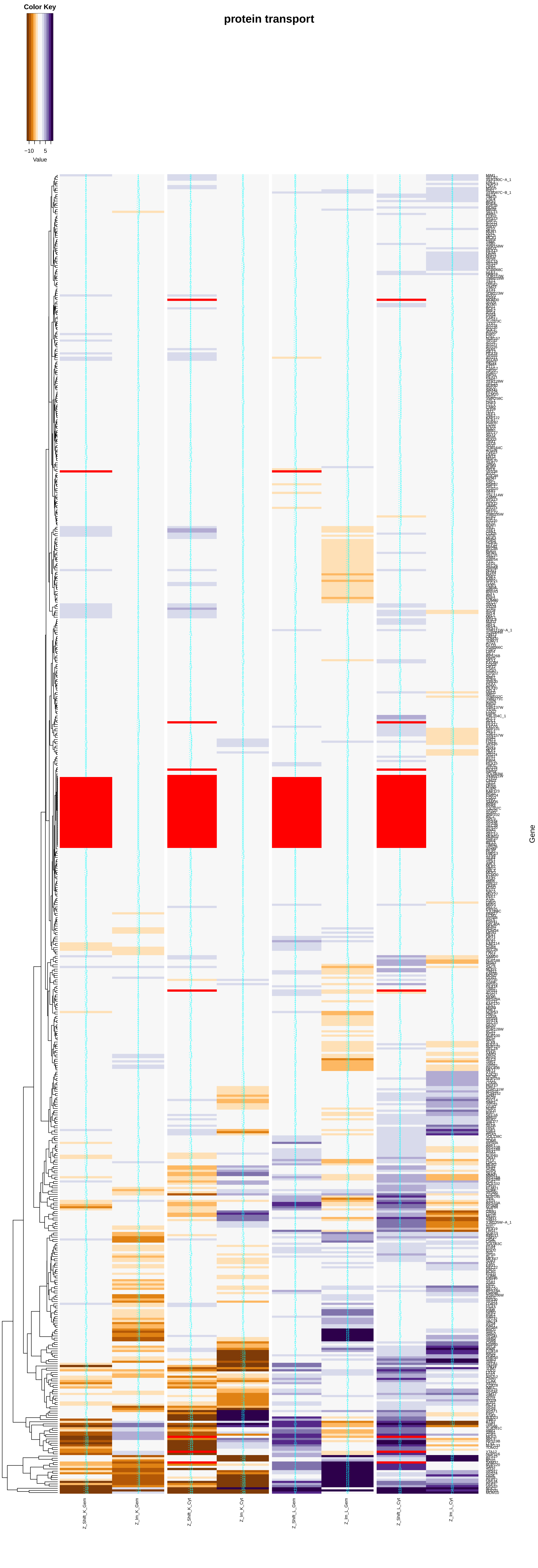

protein transport
Color Key
−10
5
Value
Gene
Z_Shift_K_Gem
Z_Im_K_Gem
Z_Shift_K_Cyt
Z_Im_K_Cyt
Z_Shift_L_Gem
Z_Im_L_Gem
Z_Shift_L_Cyt
Z_Im_L_Cyt

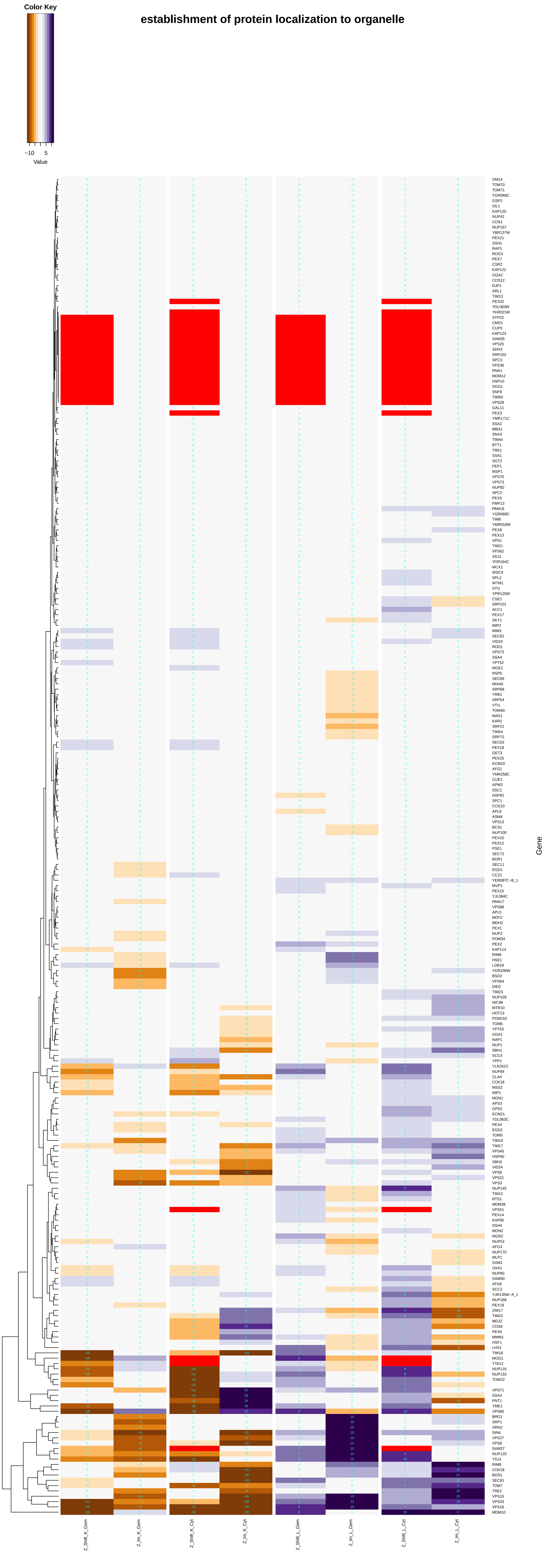
Color Key
−10
5
Value
establishment of protein localization to organelle
Z_Shift_K_Gem
Z_Im_K_Gem
Z_Shift_K_Cyt
Z_Im_K_Cyt
Z_Shift_L_Gem
Z_Im_L_Gem
Z_Shift_L_Cyt
Z_Im_L_Cyt
Gene

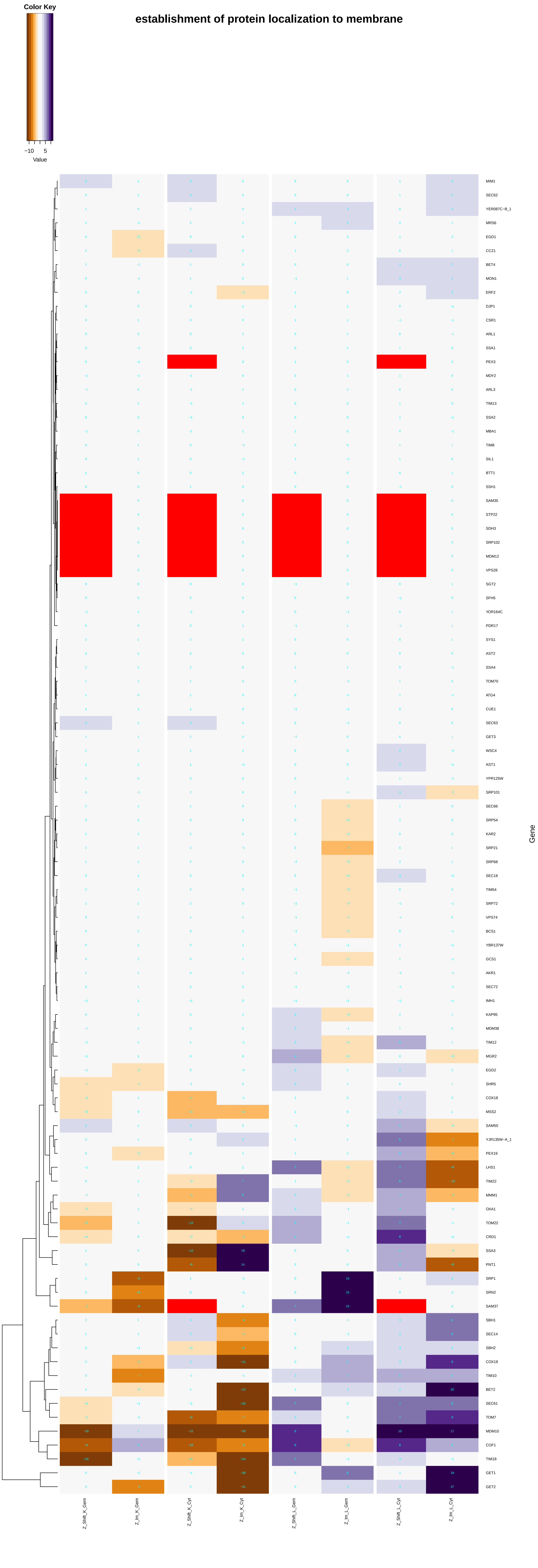
establishment of protein localization to membrane
Color Key
−10 5
Value
MIM1
SEC62
YER087C−B_1
MRS6
EGD1
CCZ1
BET4
MON1
ERF2
DJP1
CSR1
ARL1
SSA1
PEX3
MDY2
ARL3
TIM13
SSA2
MBA1
TIM8
SIL1
BTT1
SSH1
SAM35
STP22
SDH3
SRP102
MDM12
VPS28
SGT2
SFH5
YOR164C
PDR17
SYS1
AST2
SSA4
TOM70
ATG4
CUE1
SEC63
GET3
WSC4
AST1
YPR125W
SRP101
SEC66
SRP54
KAR2
SRP21
SRP68
SEC18
TIM54
SRP72
VPS74
BCS1
YBR137W
GCS1
AKR1
SEC72
IMH1
KAP95
MDM38
TIM12
MGR2
EGD2
SHR5
COX18
MSS2
SAM50
YJR135W−A_1
PEX19
LHS1
TIM22
MMM1
OXA1
TOM22
CRD1
SSA3
PNT1
SRP1
SRN2
SAM37
SBH1
SEC14
SBH2
COX18
TIM10
BET2
SEC61
TOM7
MDM10
COF1
TIM18
GET1
GET2
Gene
Z_Shift_K_Gem
Z_Im_K_Gem
Z_Shift_K_Cyt
Z_Im_K_Cyt
Z_Shift_L_Gem
Z_Im_L_Gem
Z_Shift_L_Cyt
Z_Im_L_Cyt

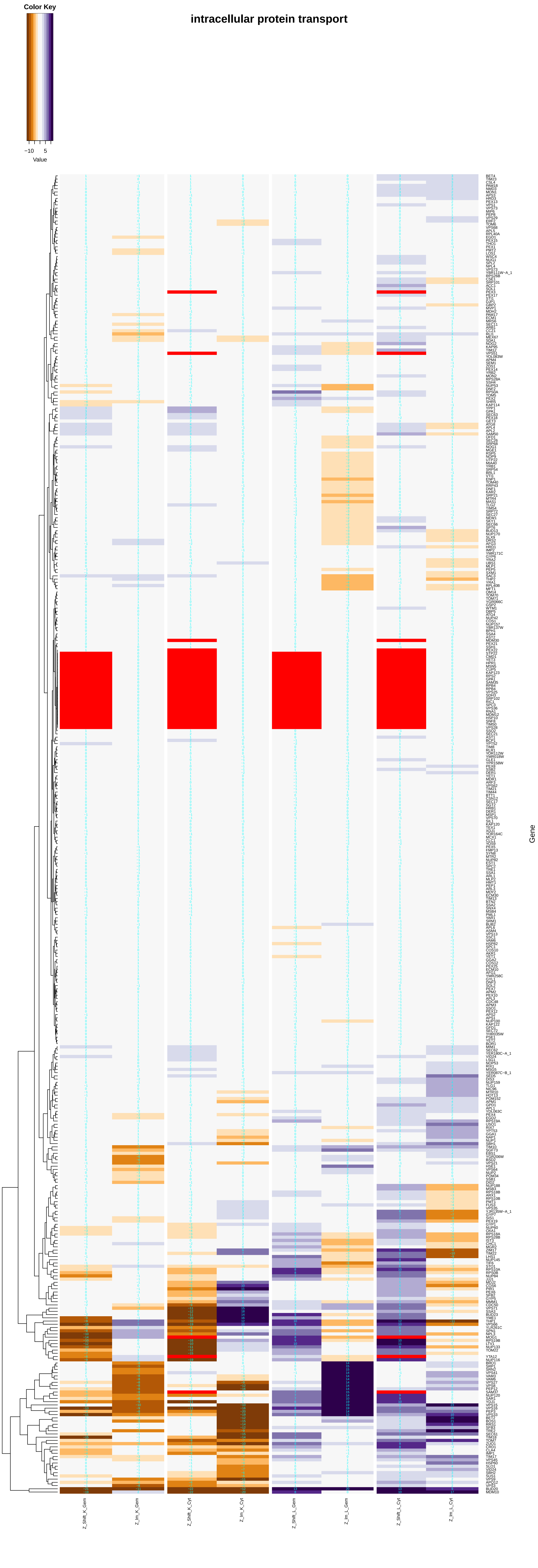
intracellular protein transport
Color Key
−10
5
Value
Gene
Z_Shift_K_Gem
Z_Im_K_Gem
Z_Shift_K_Cyt
Z_Im_K_Cyt
Z_Shift_L_Gem
Z_Im_L_Gem
Z_Shift_L_Cyt
Z_Im_L_Cyt

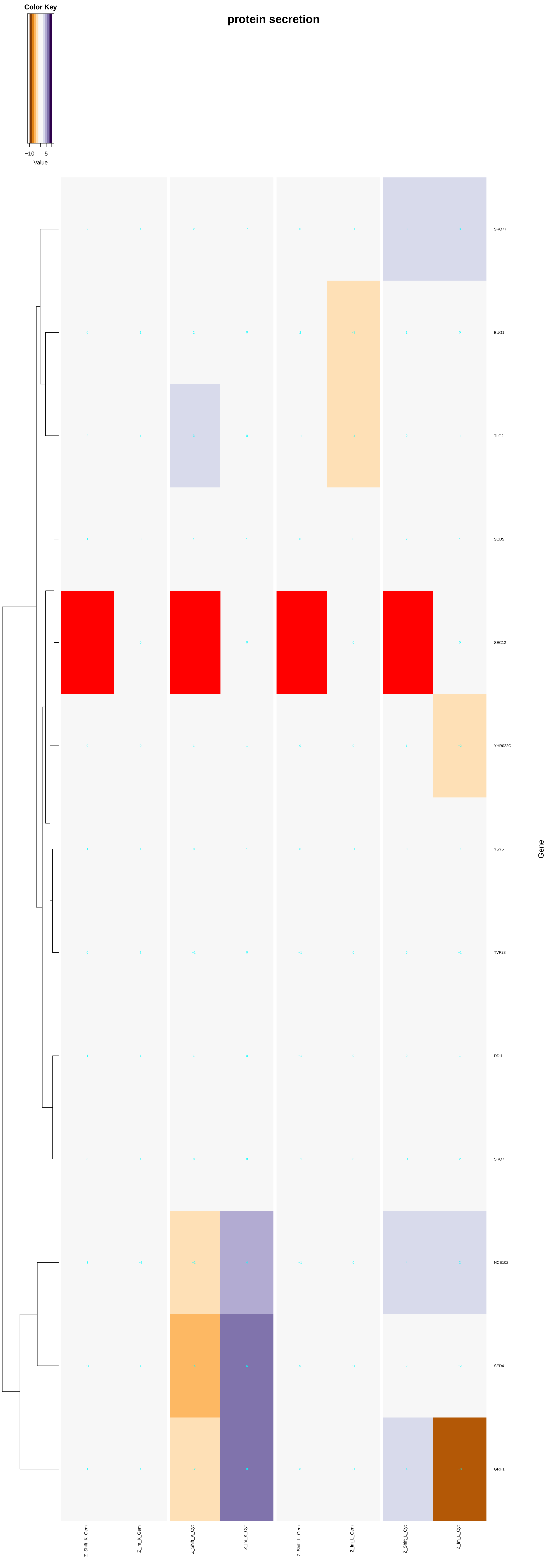
Color Key
−10
5
Value
protein secretion
SRO77
BUG1
TLG2
SCD5
SEC12
YHR022C
YSY6
TVP23
DDI1
SRO7
NCE102
SED4
GRH1
Gene
Z_Shift_K_Gem
Z_Im_K_Gem
Z_Shift_K_Cyt
Z_Im_K_Cyt
Z_Shift_L_Gem
Z_Im_L_Gem
Z_Shift_L_Cyt
Z_Im_L_Cyt

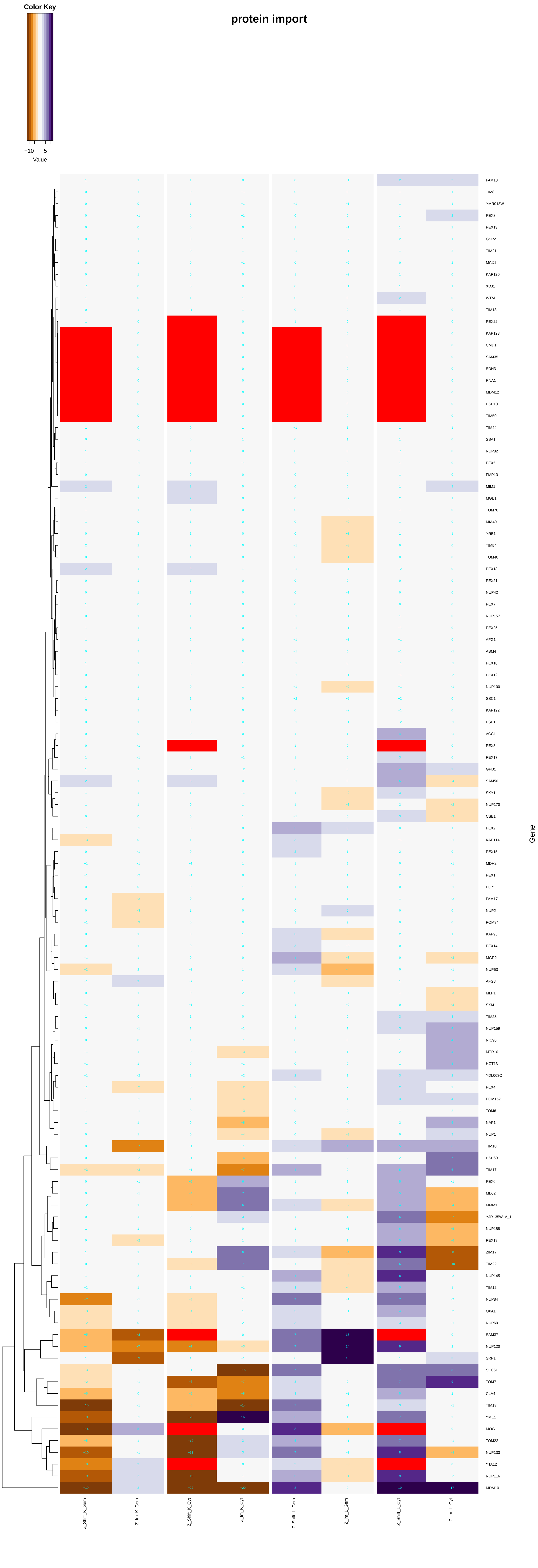
protein import
Color Key
−10
5
Value
PAM18
TIM8
YMR018W
PEX8
PEX13
GSP2
TIM21
MCX1
KAP120
XDJ1
WTM1
TIM13
PEX22
KAP123
CMD1
SAM35
SDH3
RNA1
MDM12
HSP10
TIM50
TIM44
SSA1
NUP82
PEX5
FMP13
MIM1
MGE1
TOM70
MIA40
YRB1
TIM54
TOM40
PEX18
PEX21
NUP42
PEX7
NUP157
PEX25
AFG1
ASM4
PEX10
PEX12
NUP100
SSC1
KAP122
PSE1
ACC1
PEX3
PEX17
GPD1
SAM50
SKY1
NUP170
CSE1
PEX2
KAP114
PEX15
MDH2
PEX1
DJP1
PAM17
NUP2
POM34
KAP95
PEX14
MGR2
NUP53
AFG3
MLP1
SXM1
TIM23
NUP159
NIC96
MTR10
HOT13
YDL063C
PEX4
POM152
TOM6
NAP1
NUP1
TIM10
HSP60
TIM17
PEX6
MDJ2
MMM1
YJR135W~A_1
NUP188
PEX19
ZIM17
TIM22
NUP145
TIM12
NUP84
OXA1
NUP60
SAM37
NUP120
SRP1
SEC61
TOM7
CLA4
TIM18
YME1
MOG1
TOM22
NUP133
YTA12
NUP116
MDM10
Gene
Z_Shift_K_Gem
Z_Im_K_Gem
Z_Shift_K_Cyt
Z_Im_K_Cyt
Z_Shift_L_Gem
Z_Im_L_Gem
Z_Shift_L_Cyt
Z_Im_L_Cyt

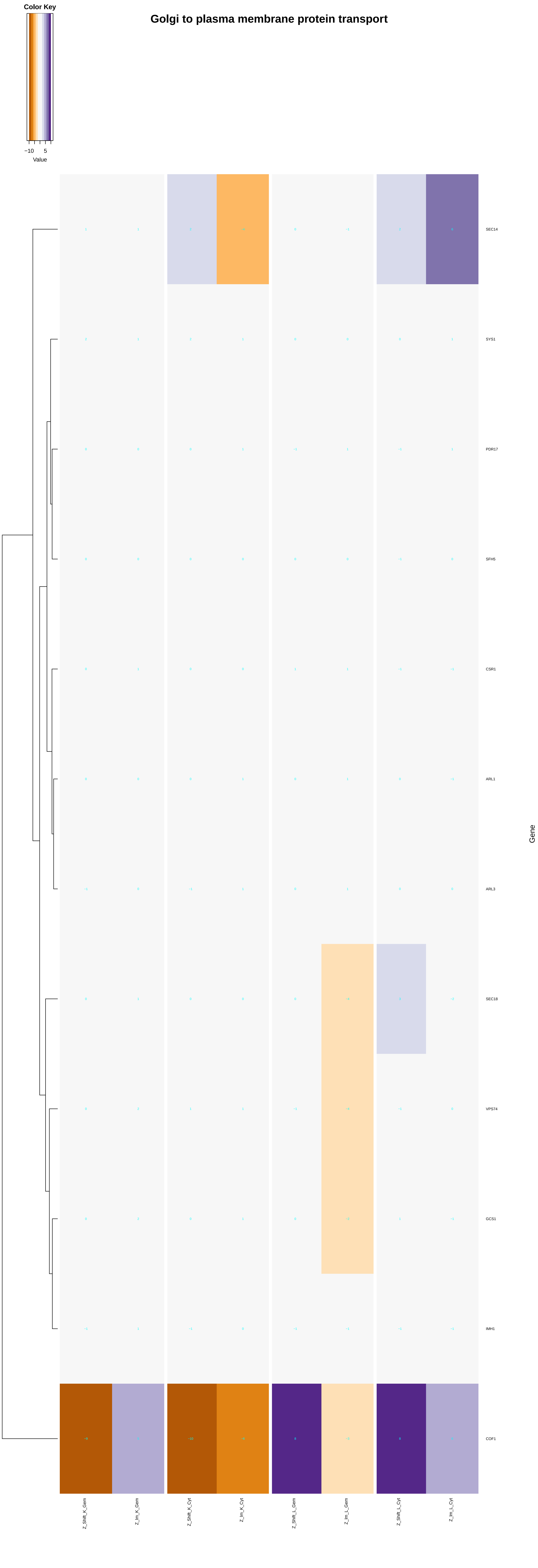
Golgi to plasma membrane protein transport
Color Key
−10
5
Value
SEC14
SYS1
PDR17
SFH5
CSR1
ARL1
ARL3
SEC18
VPS74
GCS1
IMH1
COF1
Gene
Z_Shift_K_Gem
Z_Im_K_Gem
Z_Shift_K_Cyt
Z_Im_K_Cyt
Z_Shift_L_Gem
Z_Im_L_Gem
Z_Shift_L_Cyt
Z_Im_L_Cyt

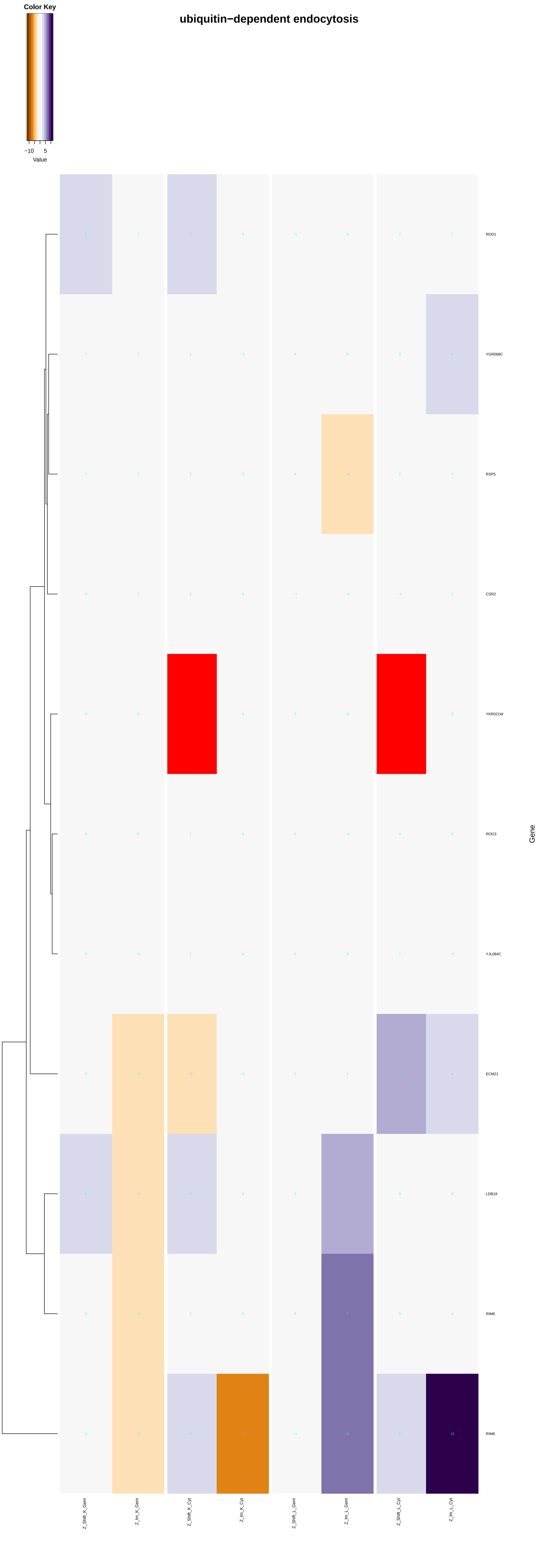

Color Key
−10
5
Value
ubiquitin−dependent endocytosis
ROD1
YGR068C
RSP5
CSR2
YKR021W
ROG3
YJL084C
ECM21
LDB19
RIM8
RIM8
Gene
Z_Shift_K_Gem
Z_Im_K_Gem
Z_Shift_K_Cyt
Z_Im_K_Cyt
Z_Shift_L_Gem
Z_Im_L_Gem
Z_Shift_L_Cyt
Z_Im_L_Cyt

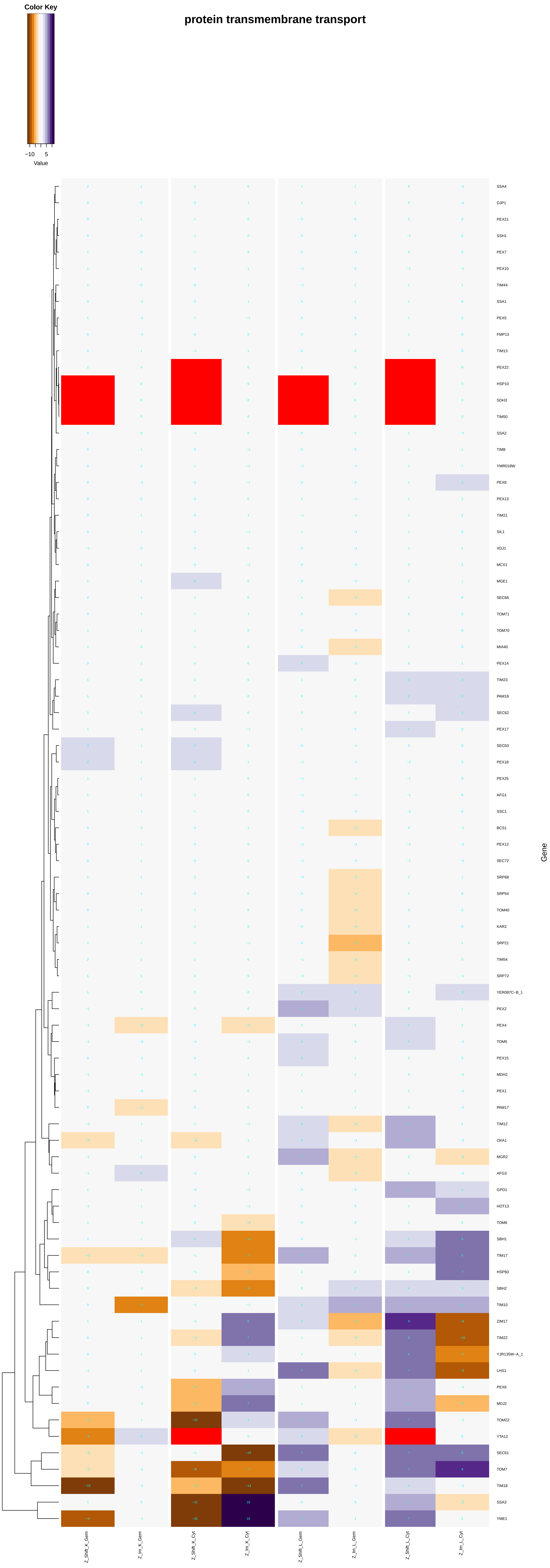

protein transmembrane transport
Color Key
−10
5
Value
SSA4
DJP1
PEX21
SSH1
PEX7
PEX10
TIM44
SSA1
PEX5
FMP13
TIM13
PEX22
HSP10
SDH3
TIM50
SSA2
TIM8
YMR018W
PEX8
PEX13
TIM21
SIL1
XDJ1
MCX1
MGE1
SEC66
TOM71
TOM70
MIA40
PEX14
TIM23
PAM18
SEC62
PEX17
SEC63
PEX18
PEX25
AFG1
SSC1
BCS1
PEX12
SEC72
SRP68
SRP54
TOM40
KAR2
SRP21
TIM54
SRP72
YER087C−B_1
PEX2
PEX4
TOM5
PEX15
MDH2
PEX1
PAM17
TIM12
OXA1
MGR2
AFG3
GPD1
HOT13
TOM6
SBH1
TIM17
HSP60
SBH2
TIM10
ZIM17
TIM22
YJR135W−A_1
LHS1
PEX6
MDJ2
TOM22
YTA12
SEC61
TOM7
TIM18
SSA3
YME1
Gene
Z_Shift_K_Gem
Z_Im_K_Gem
Z_Shift_K_Cyt
Z_Im_K_Cyt
Z_Shift_L_Gem
Z_Im_L_Gem
Z_Shift_L_Cyt
Z_Im_L_Cyt

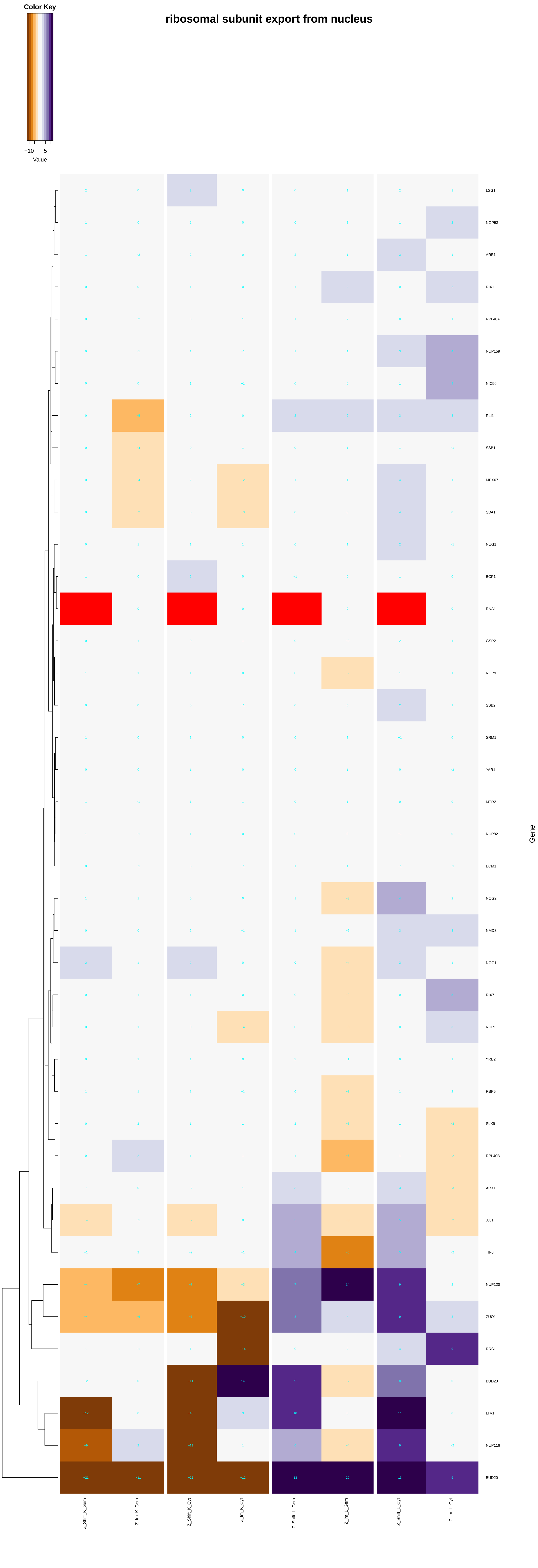

ribosomal subunit export from nucleus
Color Key
−10
5
Value
LSG1
NOP53
ARB1
RIX1
RPL40A
NUP159
NIC96
RLI1
SSB1
MEX67
SDA1
NUG1
BCP1
RNA1
GSP2
NOP9
SSB2
SRM1
YAR1
MTR2
NUP82
ECM1
NOG2
NMD3
NOG1
RIX7
NUP1
YRB2
RSP5
SLX9
RPL40B
ARX1
JJJ1
TIF6
NUP120
ZUO1
RRS1
BUD23
LTV1
NUP116
BUD20
Gene
Z_Shift_K_Gem
Z_Im_K_Gem
Z_Shift_K_Cyt
Z_Im_K_Cyt
Z_Shift_L_Gem
Z_Im_L_Gem
Z_Shift_L_Cyt
Z_Im_L_Cyt

#### protein import into nucleus

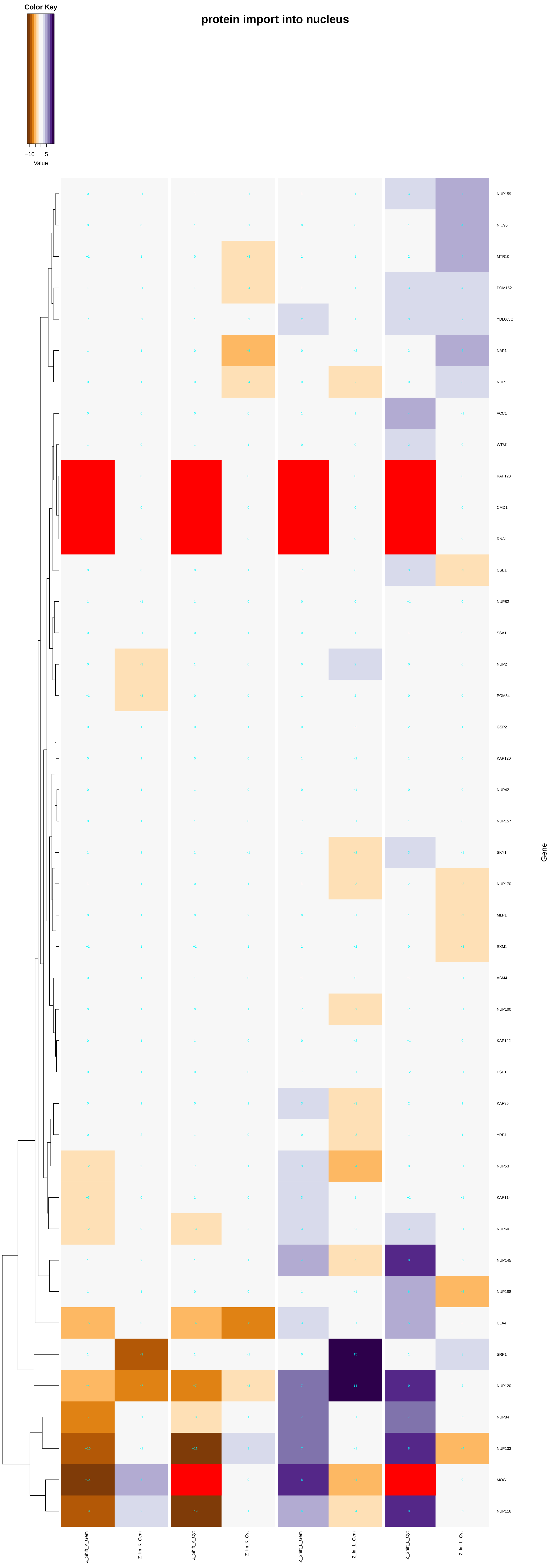

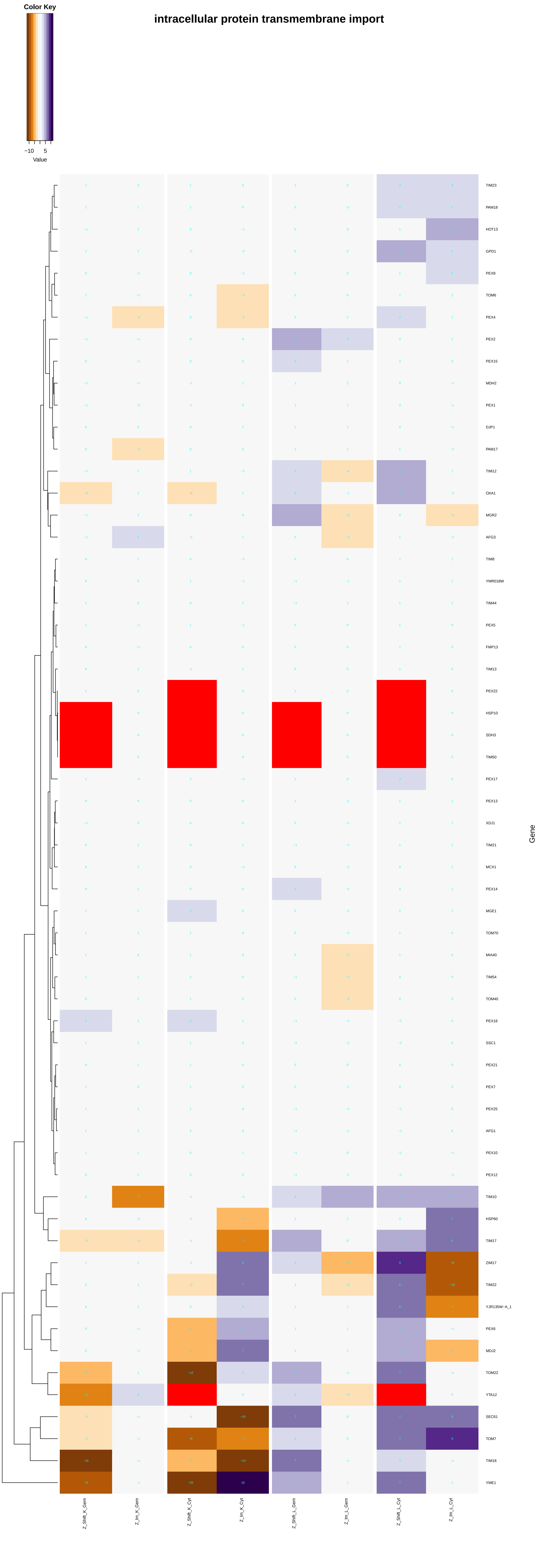

Color Key
−10
5
Value
intracellular protein transmembrane import
TIM23
PAM18
HOT13
GPD1
PEX8
TOM6
PEX4
PEX2
PEX15
MDH2
PEX1
DJP1
PAM17
TIM12
OXA1
MGR2
AFG3
TIM8
YMR018W
TIM44
PEX5
FMP13
TIM13
PEX22
HSP10
SDH3
TIM50
PEX17
PEX13
XDJ1
TIM21
MCX1
PEX14
MGE1
TOM70
MIA40
TIM54
TOM40
PEX18
SSC1
PEX21
PEX7
PEX25
AFG1
PEX10
PEX12
TIM10
HSP60
TIM17
ZIM17
TIM22
YJR135W−A_1
PEX6
MDJ2
TOM22
YTA12
SEC61
TOM7
TIM18
YME1
Gene
Z_Shift_K_Gem
Z_Im_K_Gem
Z_Shift_K_Cyt
Z_Im_K_Cyt
Z_Shift_L_Gem
Z_Im_L_Gem
Z_Shift_L_Cyt
Z_Im_L_Cyt

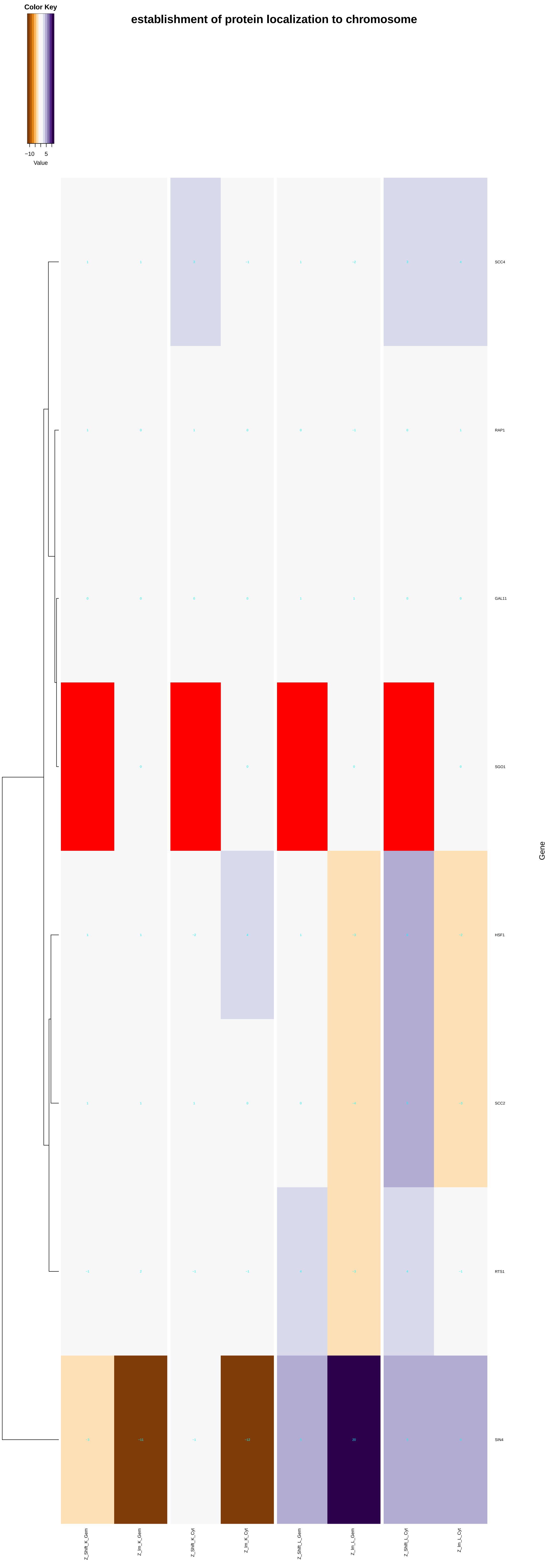
establishment of protein localization to chromosome
Color Key
−10
5
Value
SCC4
RAP1
GAL11
SGO1
HSF1
SCC2
RTS1
SIN4
Gene
Z_Shift_K_Gem
Z_lm_K_Gem
Z_Shift_K_Cyt
Z_lm_K_Cyt
Z_Shift_L_Gem
Z_lm_L_Gem
Z_Shift_L_Cyt
Z_lm_L_Cyt

### establishment of protein localization to endoplasmic reticulum

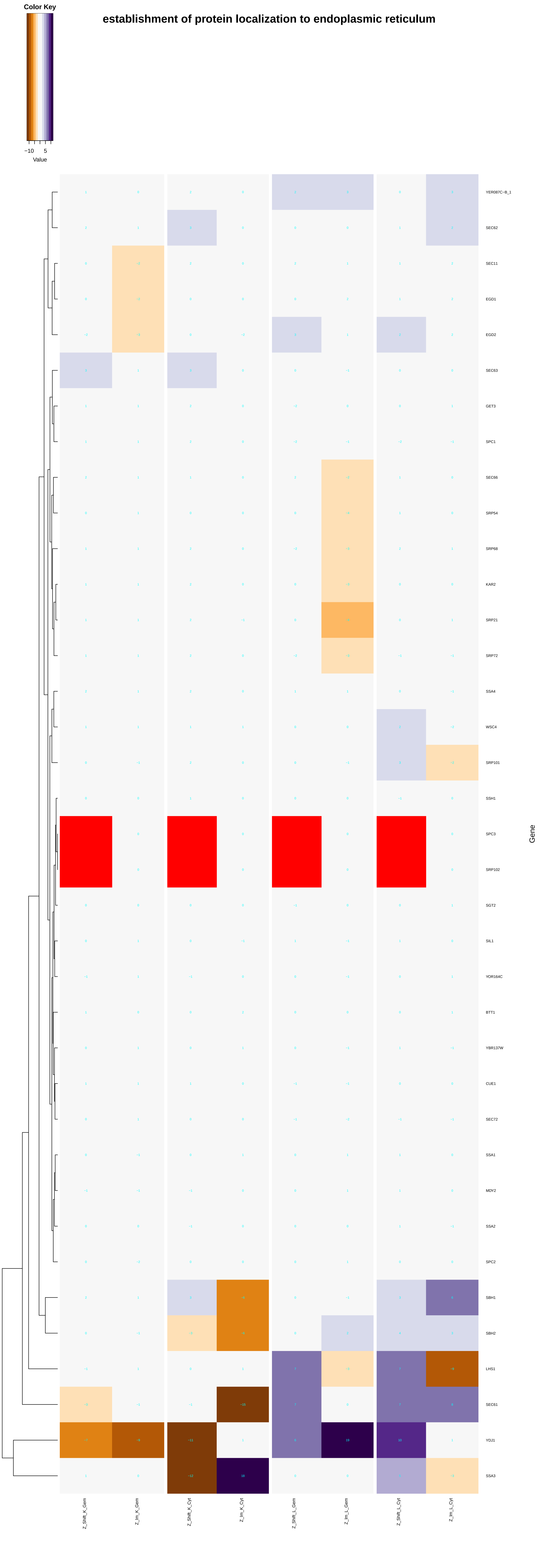

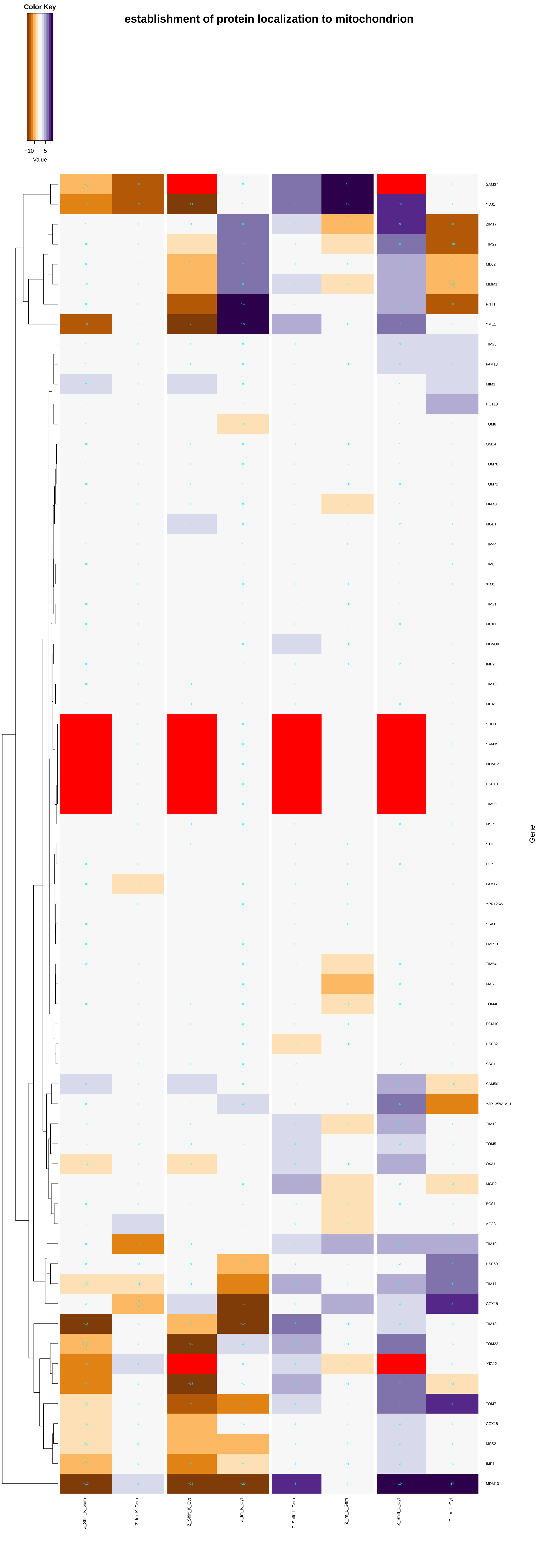

establishment of protein localization to mitochondrion
Color Key
−10
5
Value
SAM37
YDJ1
ZIM17
TIM22
MDJ2
MMM1
PNT1
YME1
TIM23
PAM18
MIM1
HOT13
TOM6
OM14
TOM70
TOM71
MIA40
MGE1
TIM44
TIM8
XDJ1
TIM21
MCX1
MDM38
IMP2
TIM13
MBA1
SDH3
SAM35
MDM12
HSP10
TIM50
MSP1
STI1
DJP1
PAM17
YPR125W
SSA1
FMP13
TIM54
MAS1
TOM40
ECM10
HSP82
SSC1
SAM50
YJR135W−A_1
TIM12
TOM5
OXA1
MGR2
BCS1
AFG3
TIM10
HSP60
TIM17
COX18
TIM18
TOM22
YTA12
TOM7
COX18
MSS2
IMP1
MDM10
Gene
Z_Shift_K_Gem
Z_Im_K_Gem
Z_Shift_K_Cyt
Z_Im_K_Cyt
Z_Shift_L_Gem
Z_Im_L_Gem
Z_Shift_L_Cyt
Z_Im_L_Cyt

#### establishment of protein localization to peroxisome

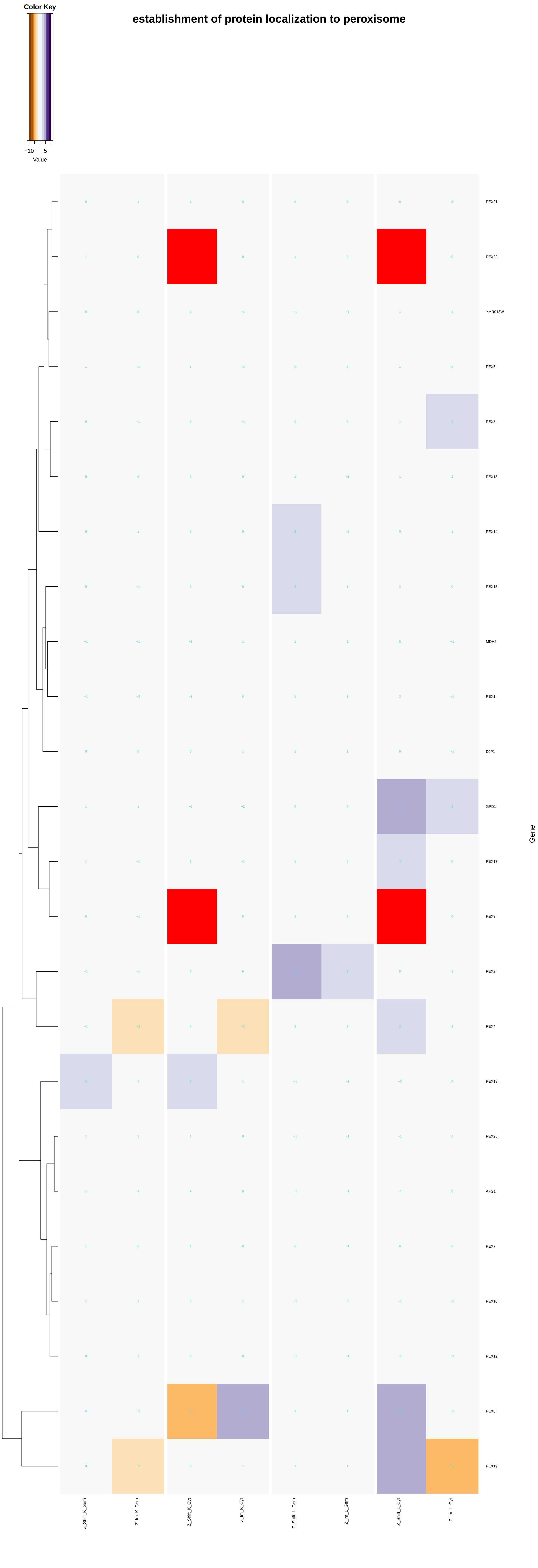

### establishment of protein localization to vacuole

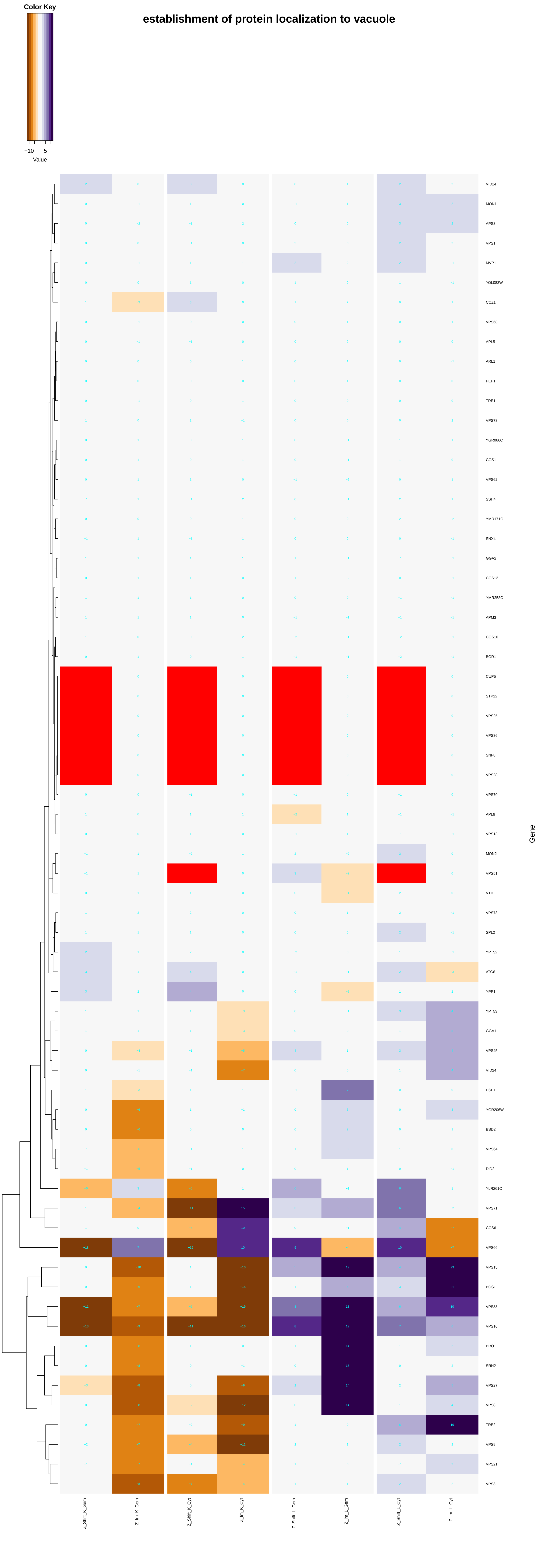

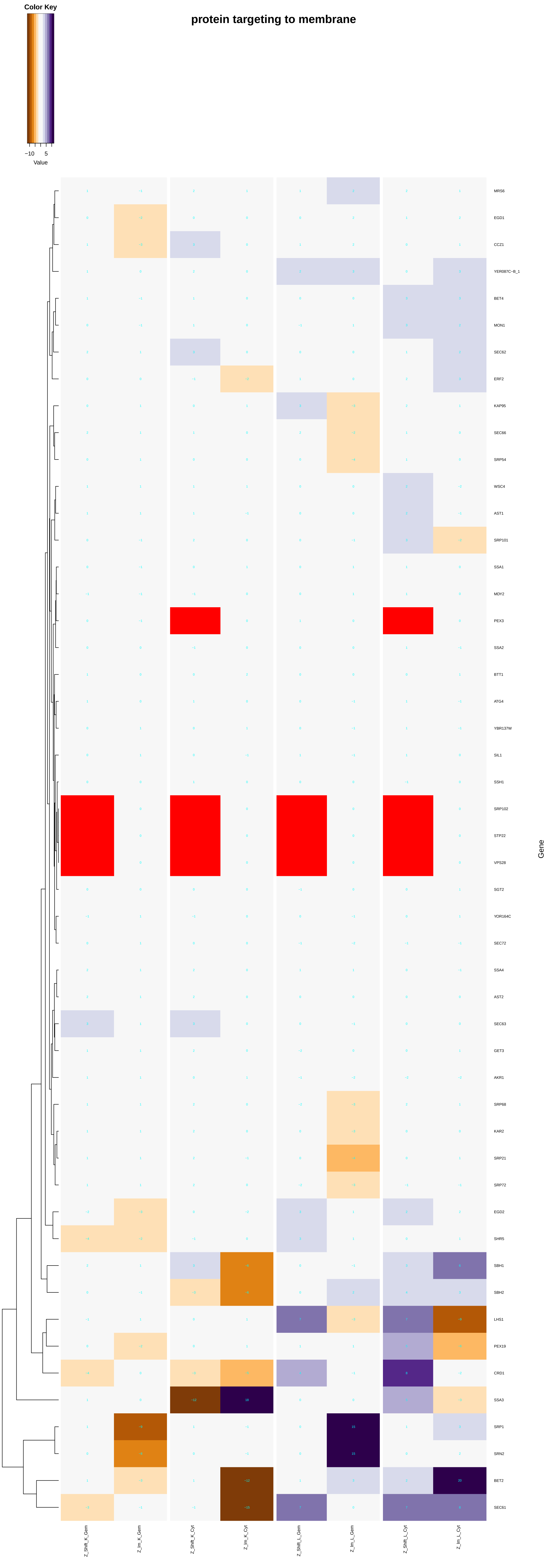

Color Key
−10
5
Value
protein targeting to membrane
MRS6
EGD1
CCZ1
YER087C−B_1
BET4
MON1
SEC62
ERF2
KAP95
SEC66
SRP54
WSC4
AST1
SRP101
SSA1
MDY2
PEX3
SSA2
BTT1
ATG4
YBR137W
SIL1
SSH1
SRP102
STP22
VPS28
SGT2
YOR164C
SEC72
SSA4
AST2
SEC63
GET3
AKR1
SRP68
KAR2
SRP21
SRP72
EGD2
SHR5
SBH1
SBH2
LHS1
PEX19
CRD1
SSA3
SRP1
SRN2
BET2
SEC61
Gene
Z_Shift_K_Gem
Z_Im_K_Gem
Z_Shift_K_Cyt
Z_Im_K_Cyt
Z_Shift_L_Gem
Z_Im_L_Gem
Z_Shift_L_Cyt
Z_Im_L_Cyt

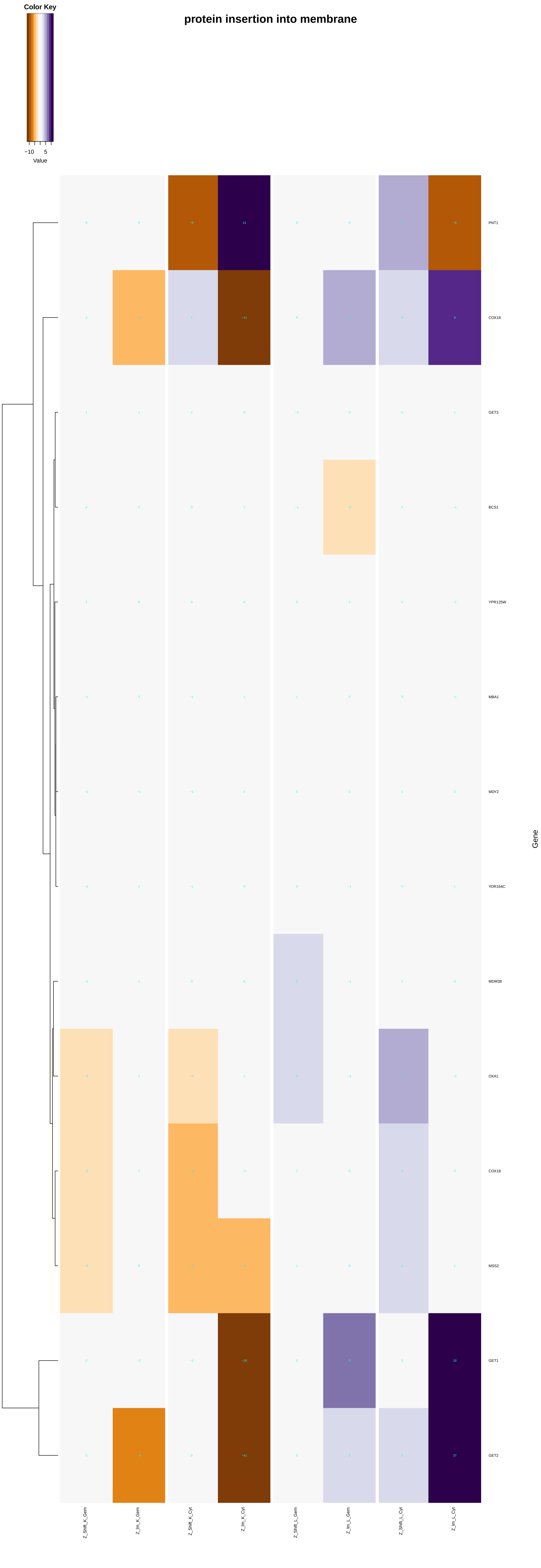
protein insertion into membrane
Color Key
−10
5
Value
PNT1
COX18
GET3
BCS1
YPR125W
MBA1
MDY2
YOR164C
MDM38
OXA1
COX18
MSS2
GET1
GET2
Gene
Z_Shift_K_Gem
Z_Im_K_Gem
Z_Shift_K_Cyt
Z_Im_K_Cyt
Z_Shift_L_Gem
Z_Im_L_Gem
Z_Shift_L_Cyt
Z_Im_L_Cyt

### establishment of protein localization to mitochondrial membrane

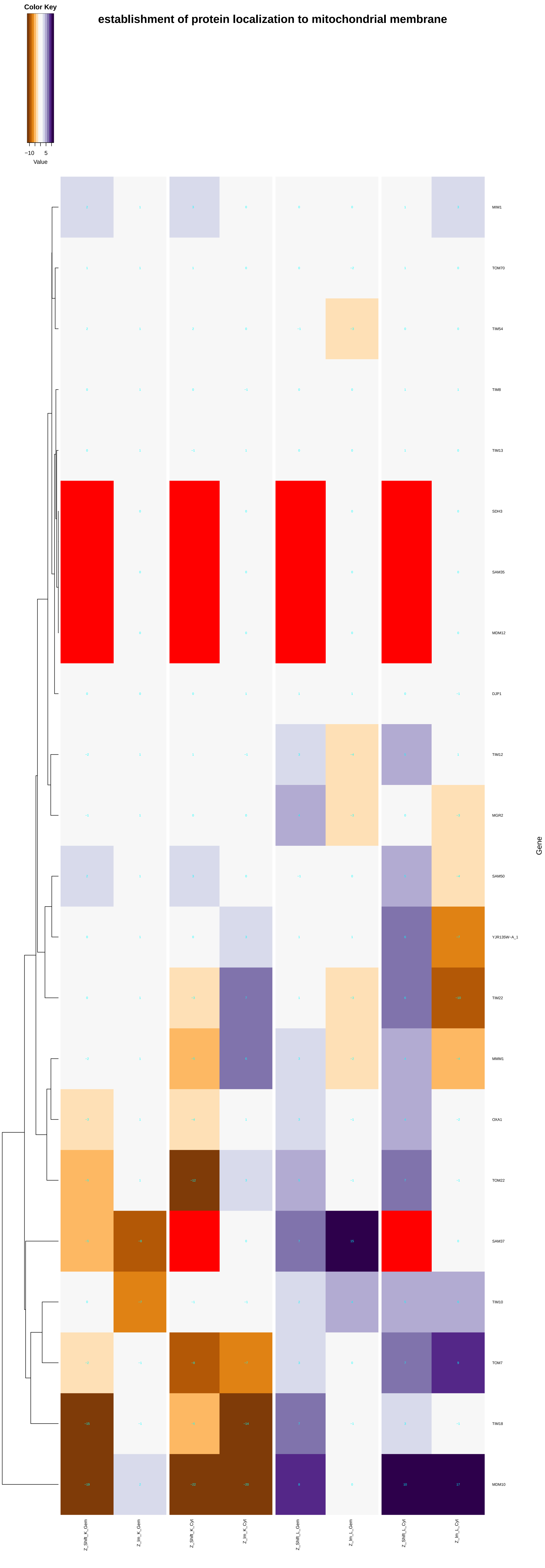

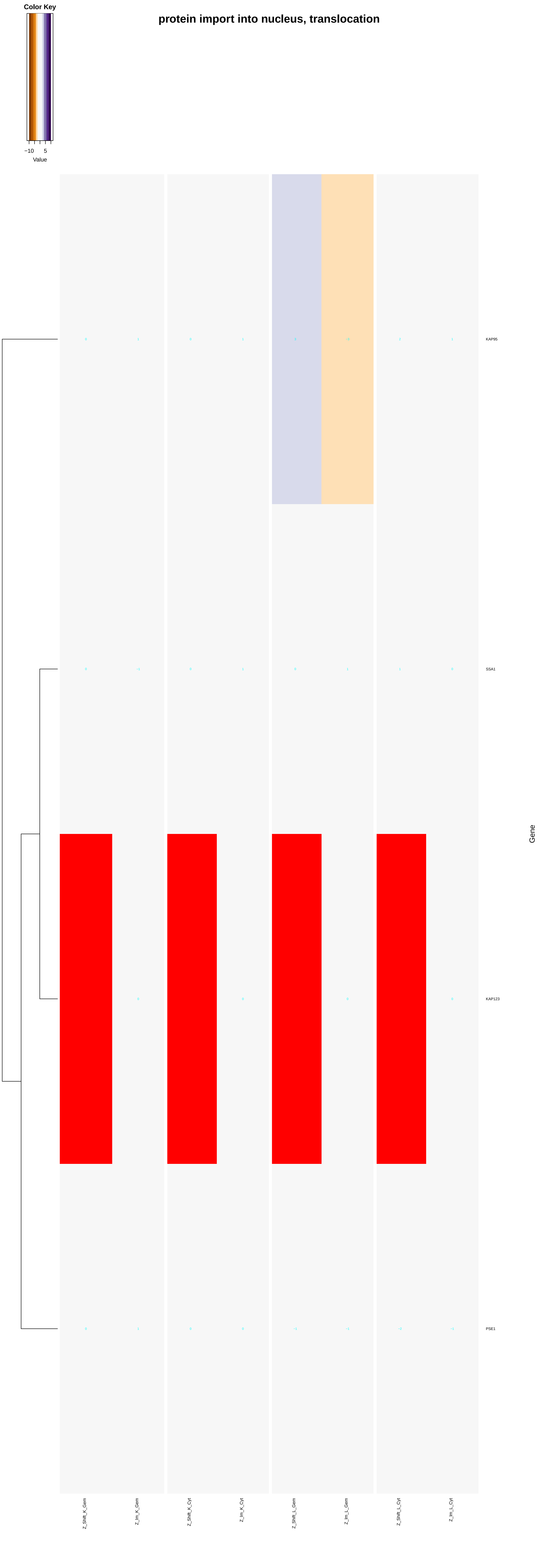
protein import into nucleus, translocation
Color Key
−10
5
Value
KAP95
SSA1
KAP123
PSE1
Gene
Z_Shift_K_Gem
Z_Im_K_Gem
Z_Shift_K_Cyt
Z_Im_K_Cyt
Z_Shift_L_Gem
Z_Im_L_Gem
Z_Shift_L_Cyt
Z_Im_L_Cyt

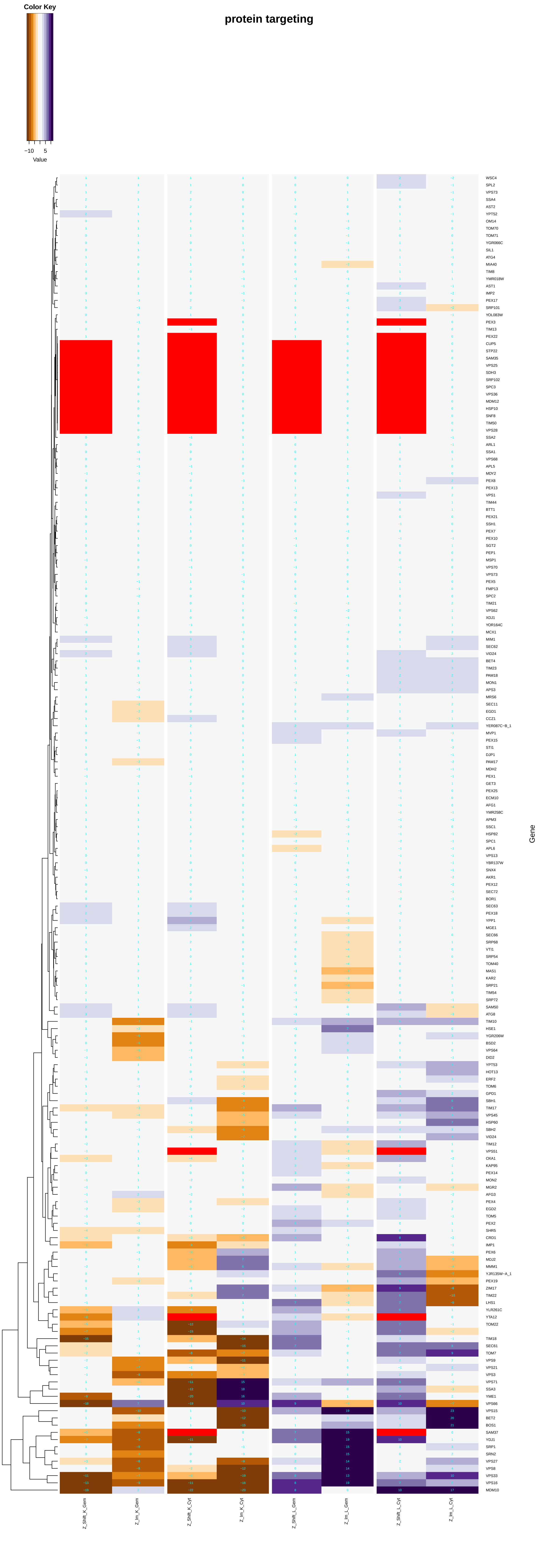
protein targeting
Color Key
−10
5
Value
Gene
WSC4
SPL2
VPS73
SSA4
AST2
YPT52
OM14
TOM70
TOM71
YGR066C
SIL1
ATG4
MIA40
TIM8
YMR018W
AST1
IMP2
PEX17
SRP101
YOL083W
PEX3
TIM13
PEX22
CUP5
STP22
SAM35
VPS25
SDH3
SRP102
SPC3
VPS36
MDM12
HSP10
SNF8
TIM50
VPS28
SSA2
ARL1
SSA1
VPS68
APL5
MDY2
PEX8
PEX13
VPS1
TIM44
BTT1
PEX21
SSH1
PEX7
PEX10
SGT2
PEP1
MSP1
VPS70
VPS73
PEX5
FMP13
SPC2
TIM21
VPS62
XDJ1
YOR164C
MCX1
MIM1
SEC62
VID24
BET4
TIM23
PAM18
MON1
APS3
MRS6
SEC11
EGD1
CCZ1
YER087C−B_1
MVP1
PEX15
STI1
DJP1
PAM17
MDH2
PEX1
GET3
PEX25
ECM10
AFG1
YMR258C
APM3
SSC1
HSP82
SPC1
APL6
VPS13
YBR137W
SNX4
AKR1
PEX12
SEC72
BOR1
SEC63
PEX18
YPP1
MGE1
SEC66
SRP68
VTI1
SRP54
TOM40
MAS1
KAR2
SRP21
TIM54
SRP72
SAM50
ATG8
TIM10
HSE1
YGR206W
BSD2
VPS64
DID2
YPT53
HOT13
ERF2
TOM6
GPD1
SBH1
TIM17
VPS45
HSP60
SBH2
VID24
TIM12
VPS51
OXA1
KAP95
PEX14
MON2
MGR2
AFG3
PEX4
EGD2
TOM5
PEX2
SHR5
CRD1
IMP1
PEX6
MDJ2
MMM1
YJR135W−A_1
PEX19
ZIM17
TIM22
LHS1
YLR261C
YTA12
TOM22
TIM18
SEC61
TOM7
VPS9
VPS21
VPS3
VPS71
YME1
SSA3
VPS66
VPS15
BET2
BOS1
SAM37
YDJ1
SRP1
SRN2
VPS27
VPS8
VPS33
VPS16
MDM10
Z_Shift_K_Gem
Z_Im_K_Gem
Z_Shift_K_Cyt
Z_Im_K_Cyt
Z_Shift_L_Gem
Z_Im_L_Gem
Z_Shift_L_Cyt
Z_Im_L_Cyt

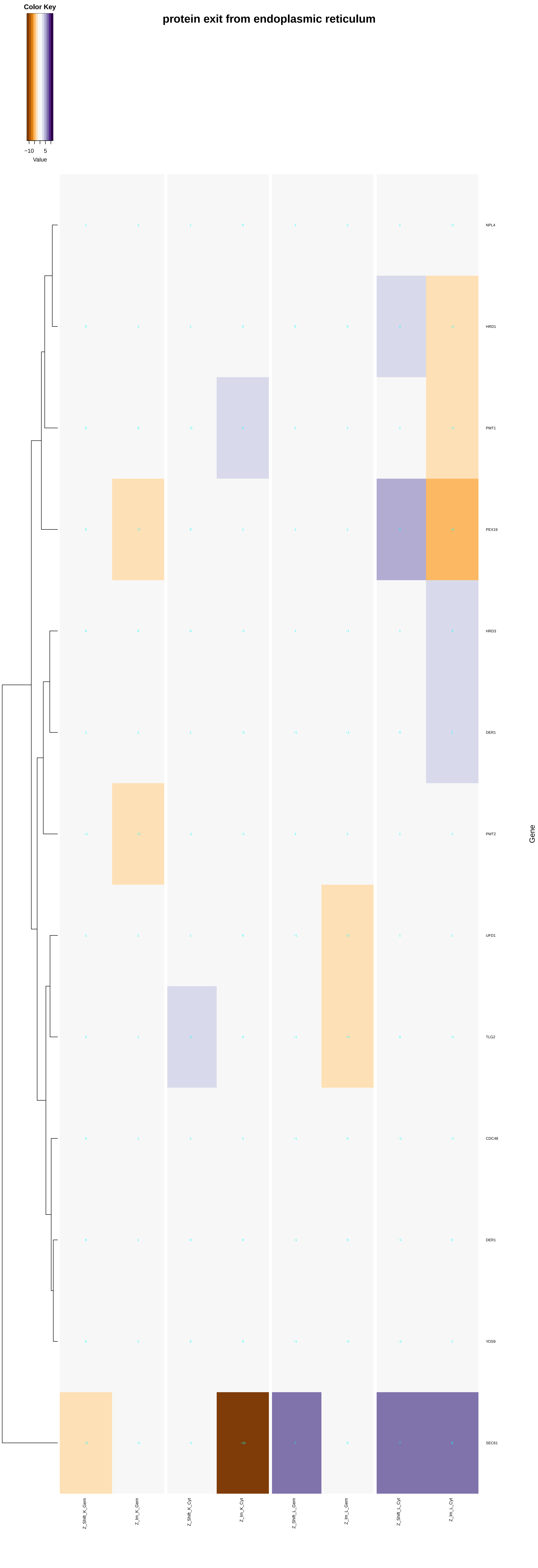
protein exit from endoplasmic reticulum
Color Key
−10
5
Value
NPL4
HRD1
PMT1
PEX19
HRD3
DER1
PMT2
UFD1
TLG2
CDC48
DER1
YOS9
SEC61
Gene
Z_Shift_K_Gem
Z_Im_K_Gem
Z_Shift_K_Cyt
Z_Im_K_Cyt
Z_Shift_L_Gem
Z_Im_L_Gem
Z_Shift_L_Cyt
Z_Im_L_Cyt

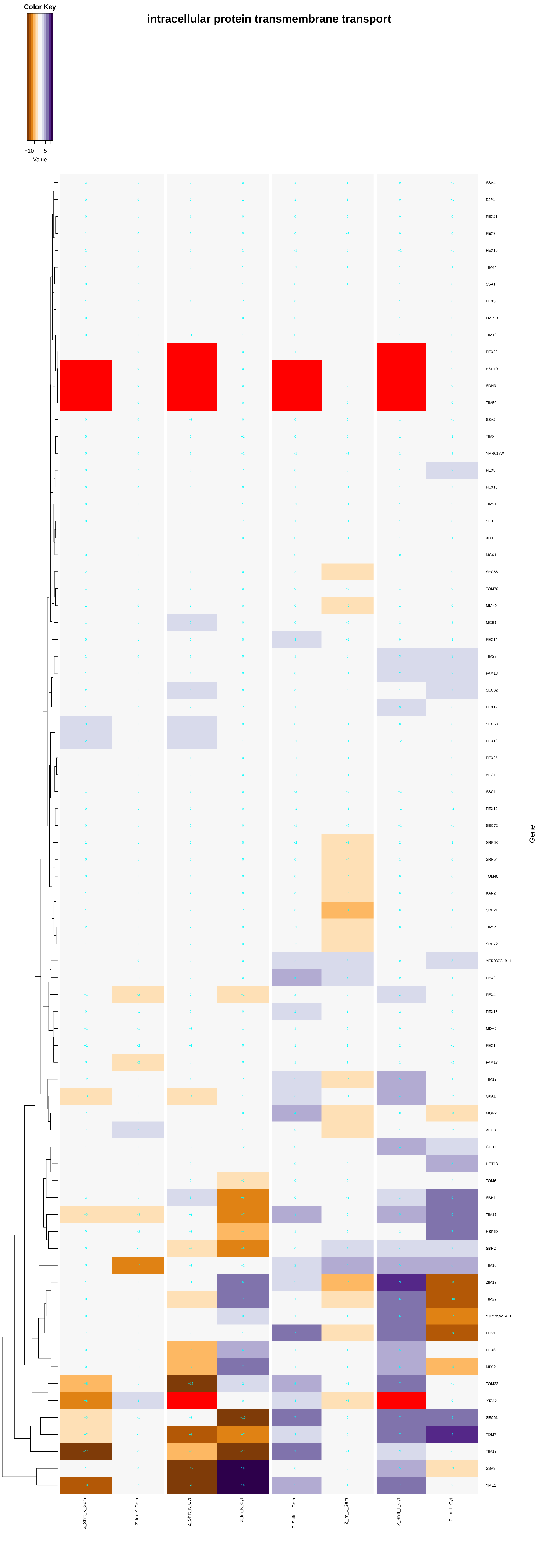

intracellular protein transmembrane transport
Color Key
−10
5
Value
Gene
SSA4
DJP1
PEX21
PEX7
PEX10
TIM44
SSA1
PEX5
FMP13
TIM13
PEX22
HSP10
SDH3
TIM50
SSA2
TIM8
YMR018W
PEX8
PEX13
TIM21
SIL1
XDJ1
MCX1
SEC66
TOM70
MIA40
MGE1
PEX14
TIM23
PAM18
SEC62
PEX17
SEC63
PEX18
PEX25
AFG1
SSC1
PEX12
SEC72
SRP68
SRP54
TOM40
KAR2
SRP21
TIM54
SRP72
YER087C−B_1
PEX2
PEX4
PEX15
MDH2
PEX1
PAM17
TIM12
OXA1
MGR2
AFG3
GPD1
HOT13
TOM6
SBH1
TIM17
HSP60
SBH2
TIM10
ZIM17
TIM22
YJR135W−A_1
LHS1
PEX6
MDJ2
TOM22
YTA12
SEC61
TOM7
TIM18
SSA3
YME1
Z_Shift_K_Gem
Z_Im_K_Gem
Z_Shift_K_Cyt
Z_Im_K_Cyt
Z_Shift_L_Gem
Z_Im_L_Gem
Z_Shift_L_Cyt
Z_Im_L_Cyt

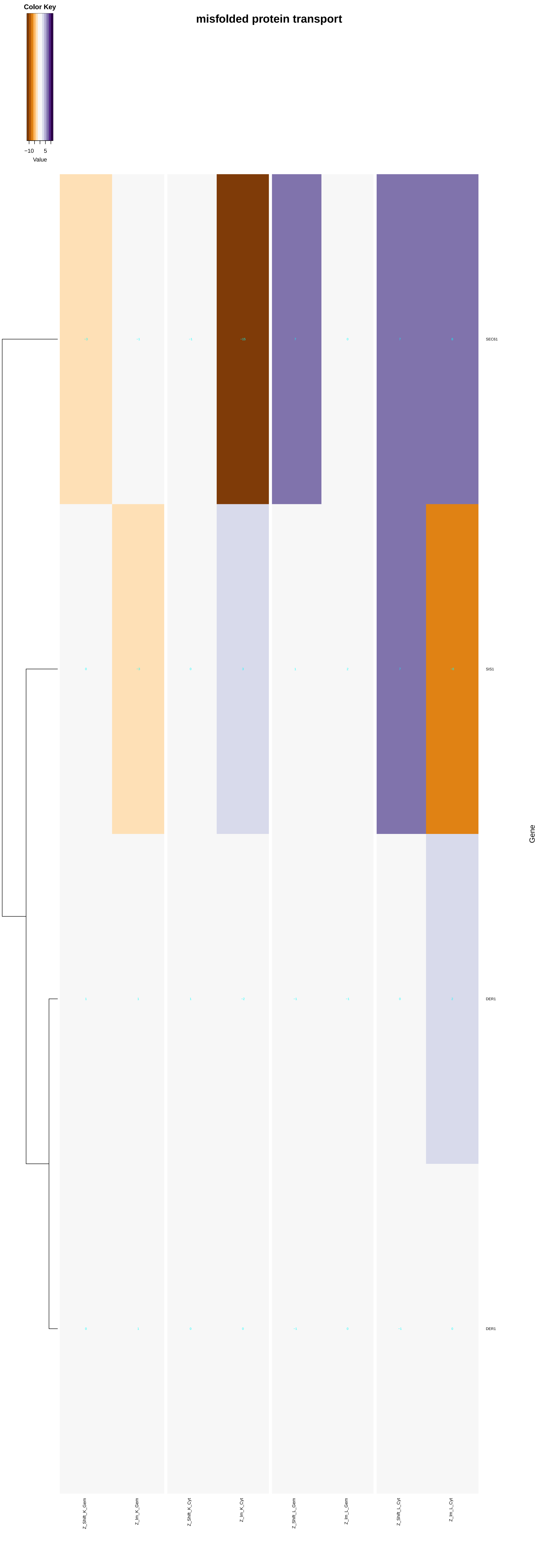

Color Key
−10
5
Value
misfolded protein transport
SEC61
SIS1
DER1
DER1
Gene
Z_Shift_K_Gem
Z_Im_K_Gem
Z_Shift_K_Cyt
Z_Im_K_Cyt
Z_Shift_L_Gem
Z_Im_L_Gem
Z_Shift_L_Cyt
Z_Im_L_Cyt

### ribosomal large subunit export from nucleus

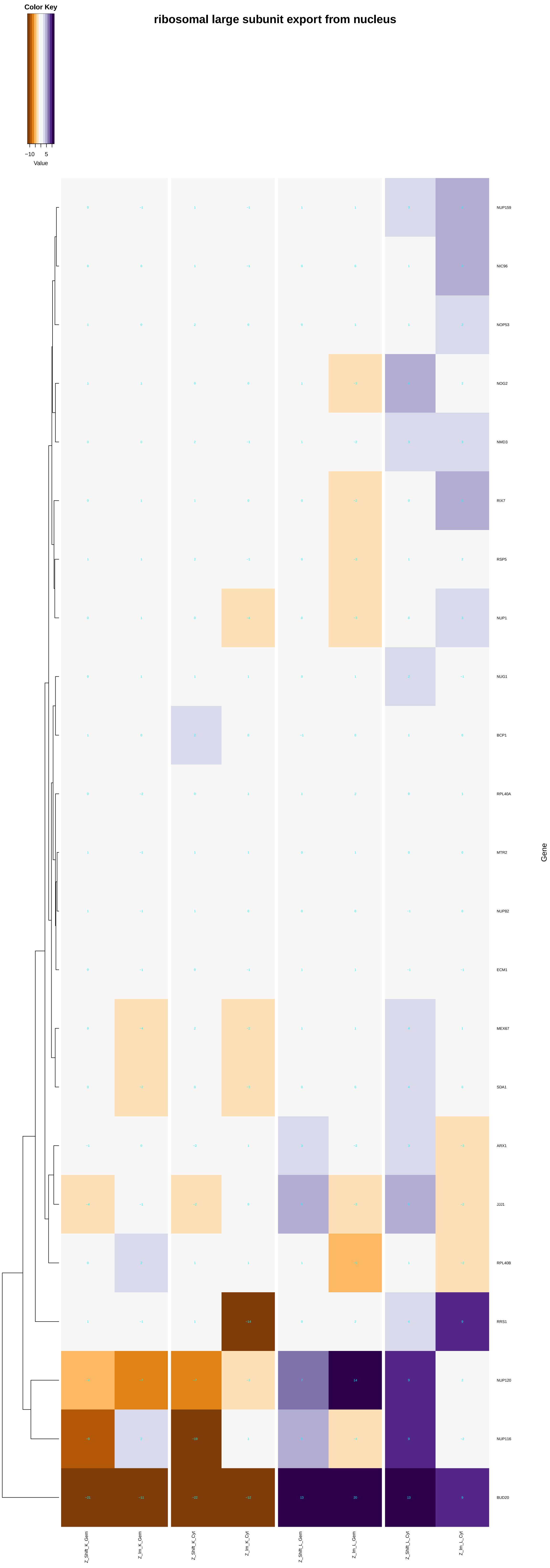

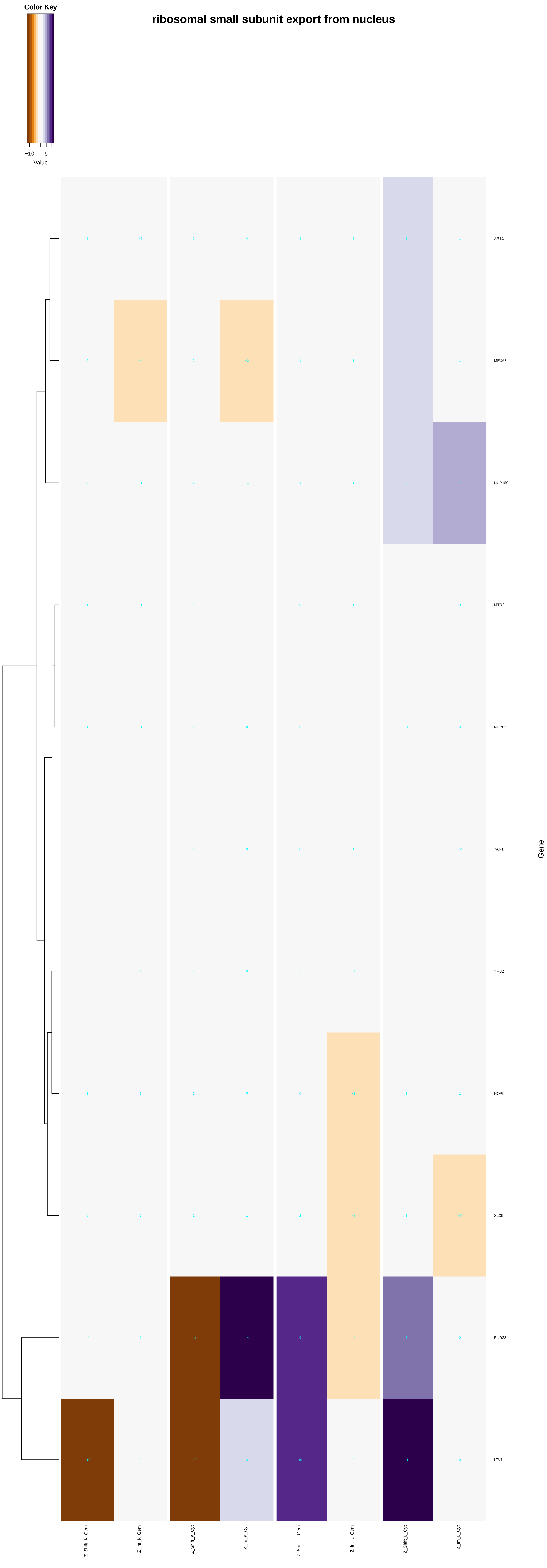

ribosomal small subunit export from nucleus
Color Key
Value
−10
5
ARB1
MEX67
NUP159
MTR2
NUP82
YAR1
YRB2
NOP9
SLX9
BUD23
LTV1
Gene
Z_Shift_K_Gem
Z_Im_K_Gem
Z_Shift_K_Cyt
Z_Im_K_Cyt
Z_Shift_L_Gem
Z_Im_L_Gem
Z_Shift_L_Cyt
Z_Im_L_Cyt

### NLS−bearing protein import into nucleus

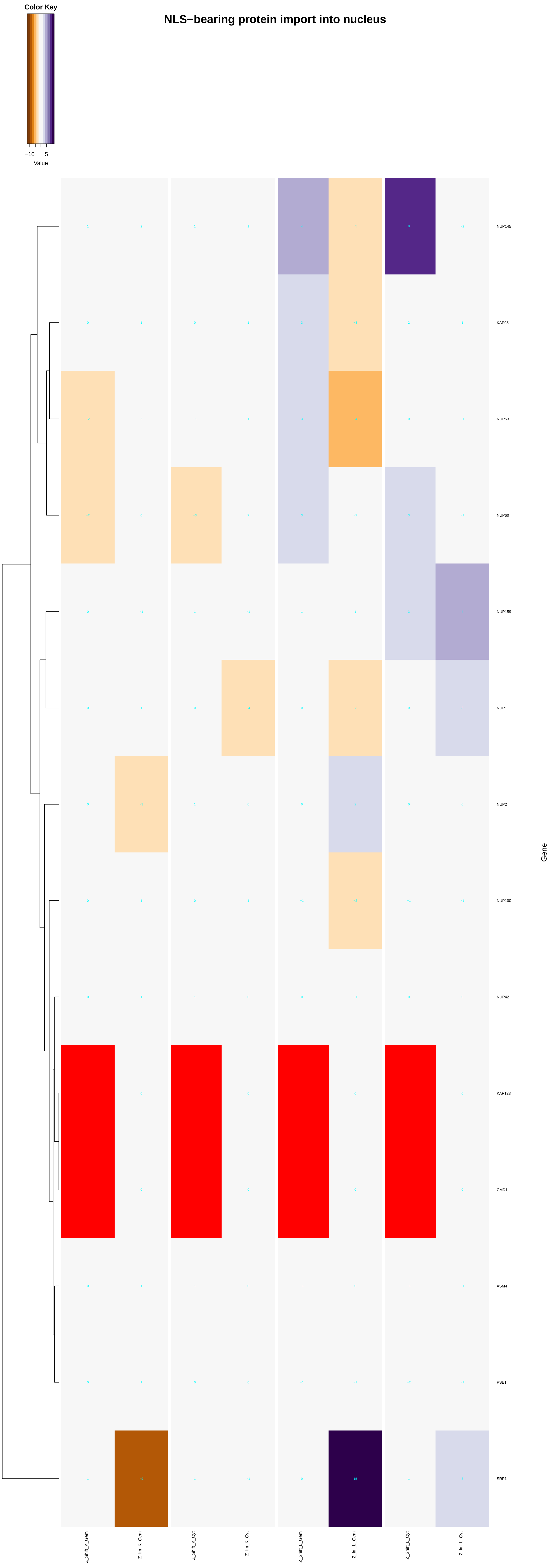

### ribosomal protein import into nucleus

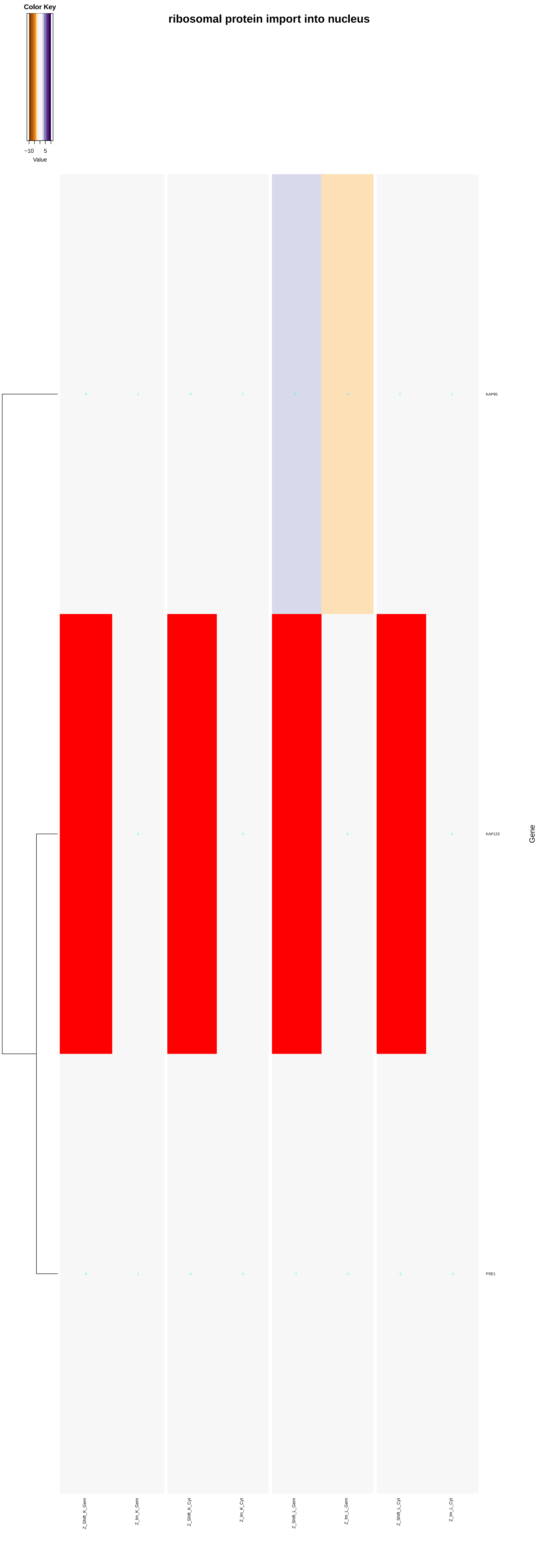

#### transcription factor import into nucleus

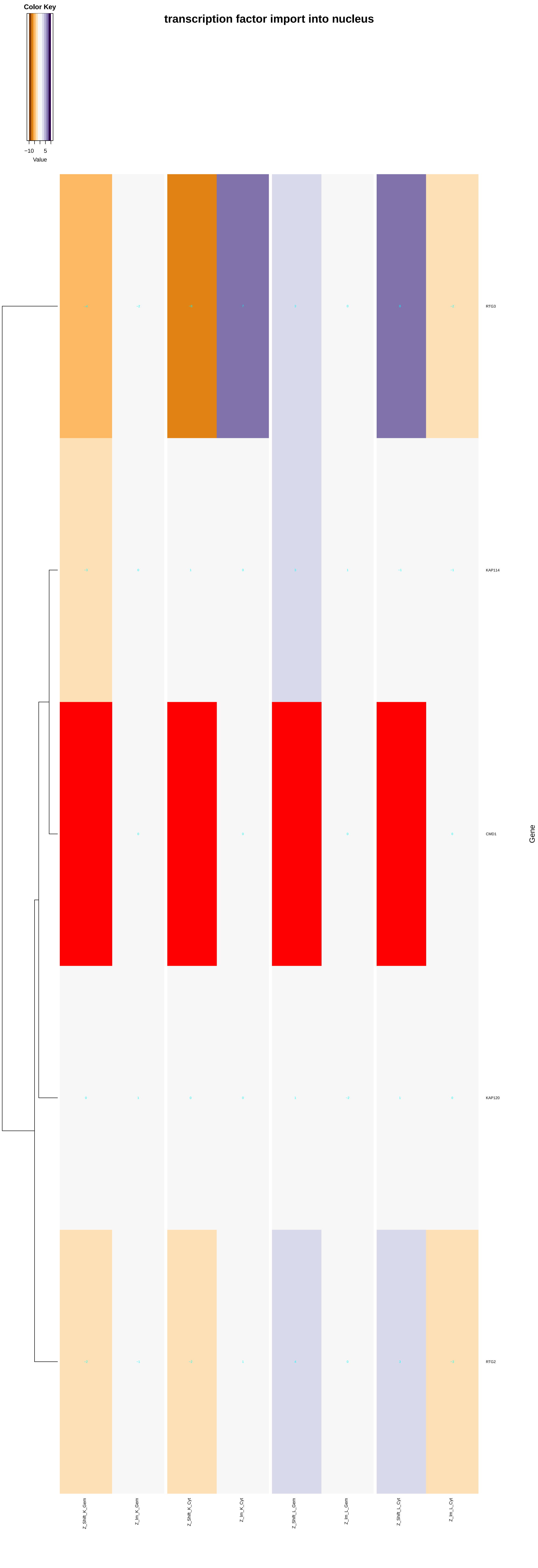

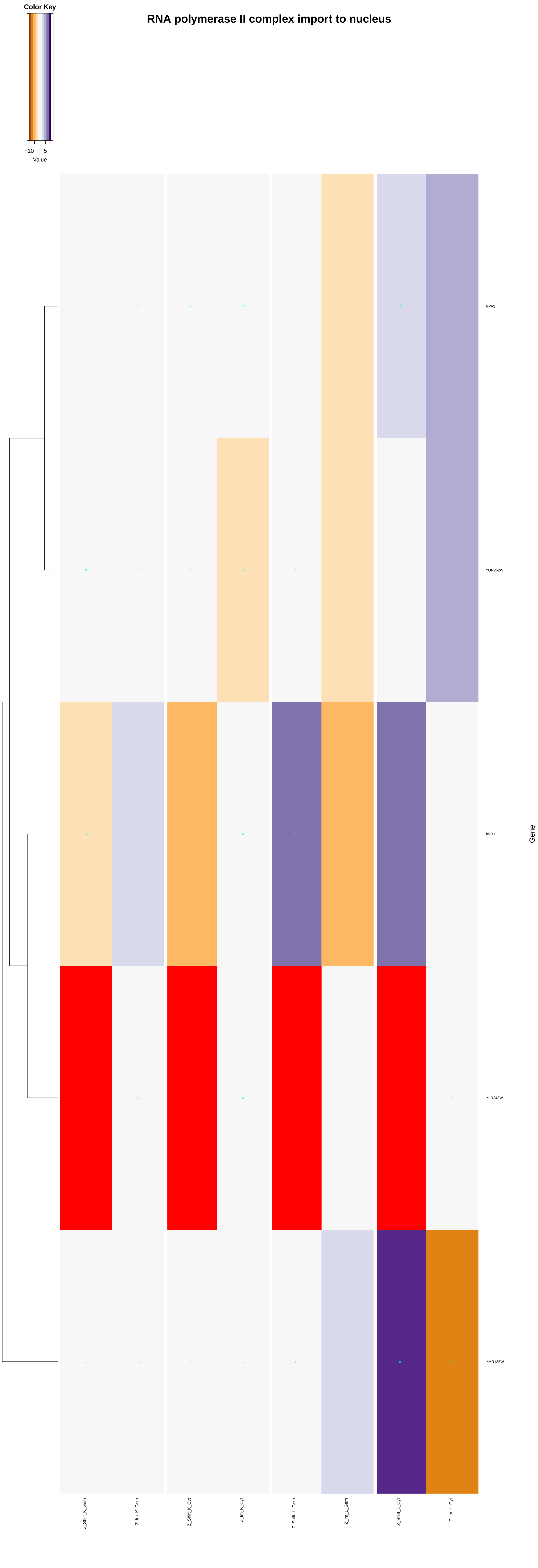
RNA polymerase II complex import to nucleus
Color Key
−10
5
Value
NPA3
YOR262W
IWR1
YLR243W
YMR185W
Gene
Z_Shift_K_Gem
Z_Im_K_Gem
Z_Shift_K_Cyt
Z_Im_K_Cyt
Z_Shift_L_Gem
Z_Im_L_Gem
Z_Shift_L_Cyt
Z_Im_L_Cyt

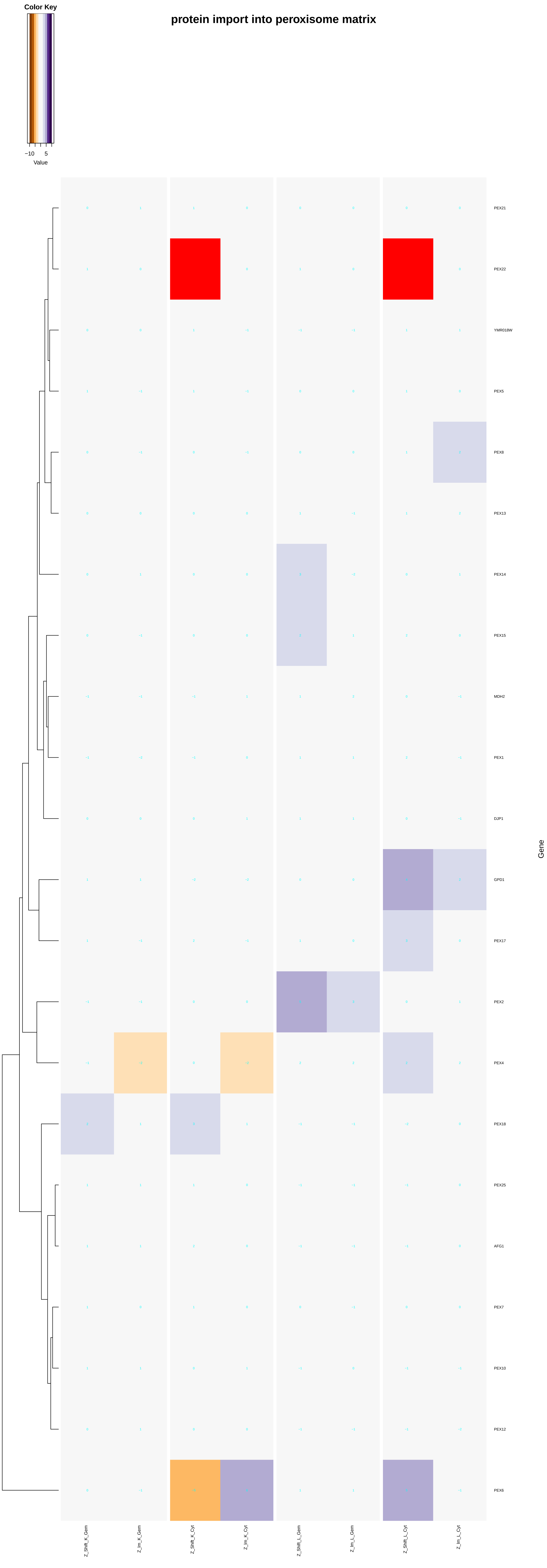

protein import into peroxisome matrix
Color Key
−10
5
Value
PEX21
PEX22
YMR018W
PEX5
PEX8
PEX13
PEX14
PEX15
MDH2
PEX1
DJP1
GPD1
PEX17
PEX2
PEX4
PEX18
PEX25
AFG1
PEX7
PEX10
PEX12
PEX6
Gene
Z_Shift_K_Gem
Z_Im_K_Gem
Z_Shift_K_Cyt
Z_Im_K_Cyt
Z_Shift_L_Gem
Z_Im_L_Gem
Z_Shift_L_Cyt
Z_Im_L_Cyt

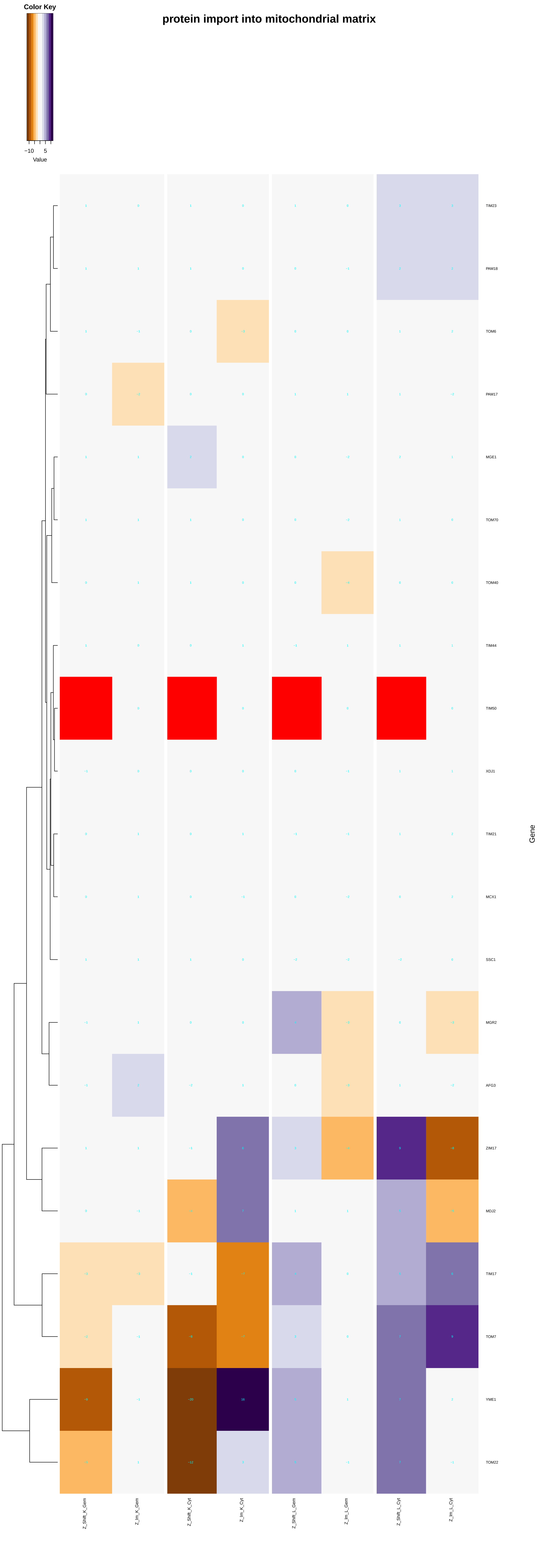
protein import into mitochondrial matrix
Color Key
−10
5
Value
TIM23
PAM18
TOM6
PAM17
MGE1
TOM70
TOM40
TIM44
TIM50
XDJ1
TIM21
MCX1
SSC1
MGR2
AFG3
ZIM17
MDJ2
TIM17
TOM7
YME1
TOM22
Gene
Z_Shift_K_Gem
Z_Im_K_Gem
Z_Shift_K_Cyt
Z_Im_K_Cyt
Z_Shift_L_Gem
Z_Im_L_Gem
Z_Shift_L_Cyt
Z_Im_L_Cyt

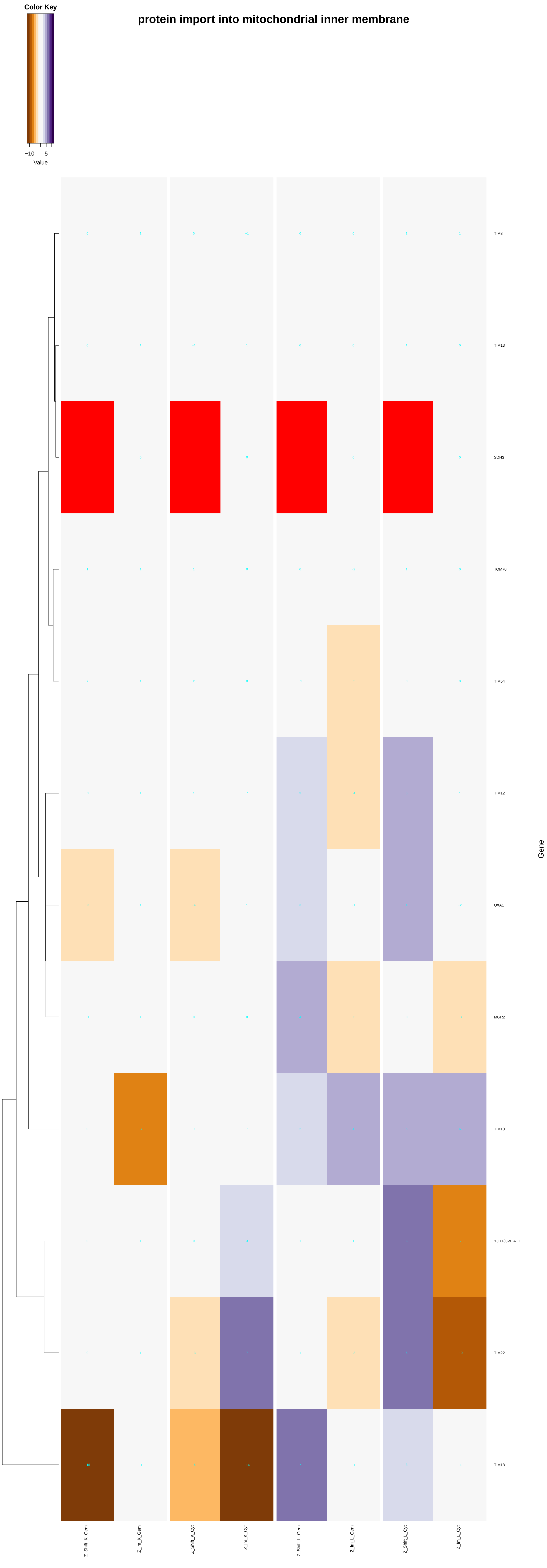
protein import into mitochondrial inner membrane
Color Key
−10
5
Value
TIM8
TIM13
SDH3
TOM70
TIM54
TIM12
OXA1
MGR2
TIM10
YJR135W−A_1
TIM22
TIM18
Gene
Z_Shift_K_Gem
Z_Im_K_Gem
Z_Shift_K_Cyt
Z_Im_K_Cyt
Z_Shift_L_Gem
Z_Im_L_Gem
Z_Shift_L_Cyt
Z_Im_L_Cyt

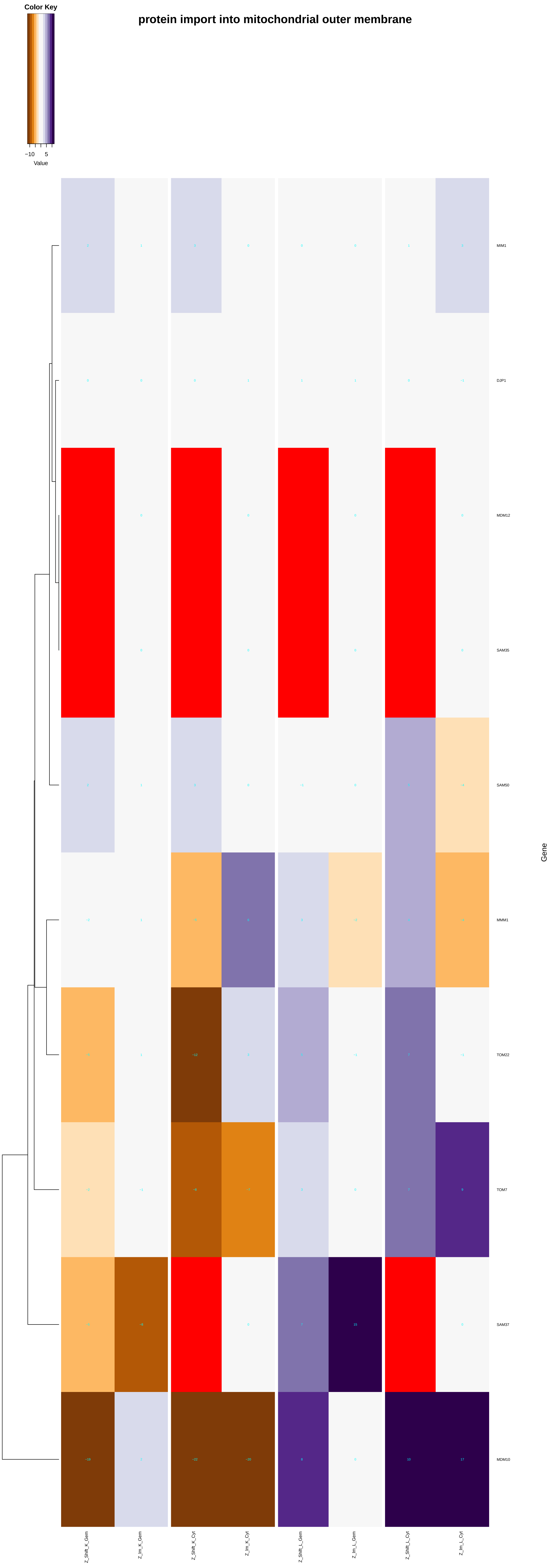
protein import into mitochondrial outer membrane
Color Key
−10
5
Value
MIM1
DJP1
MDM12
SAM35
SAM50
MMM1
TOM22
TOM7
SAM37
MDM10
Gene
Z_Shift_K_Gem
Z_Im_K_Gem
Z_Shift_K_Cyt
Z_Im_K_Cyt
Z_Shift_L_Gem
Z_Im_L_Gem
Z_Shift_L_Cyt
Z_Im_L_Cyt

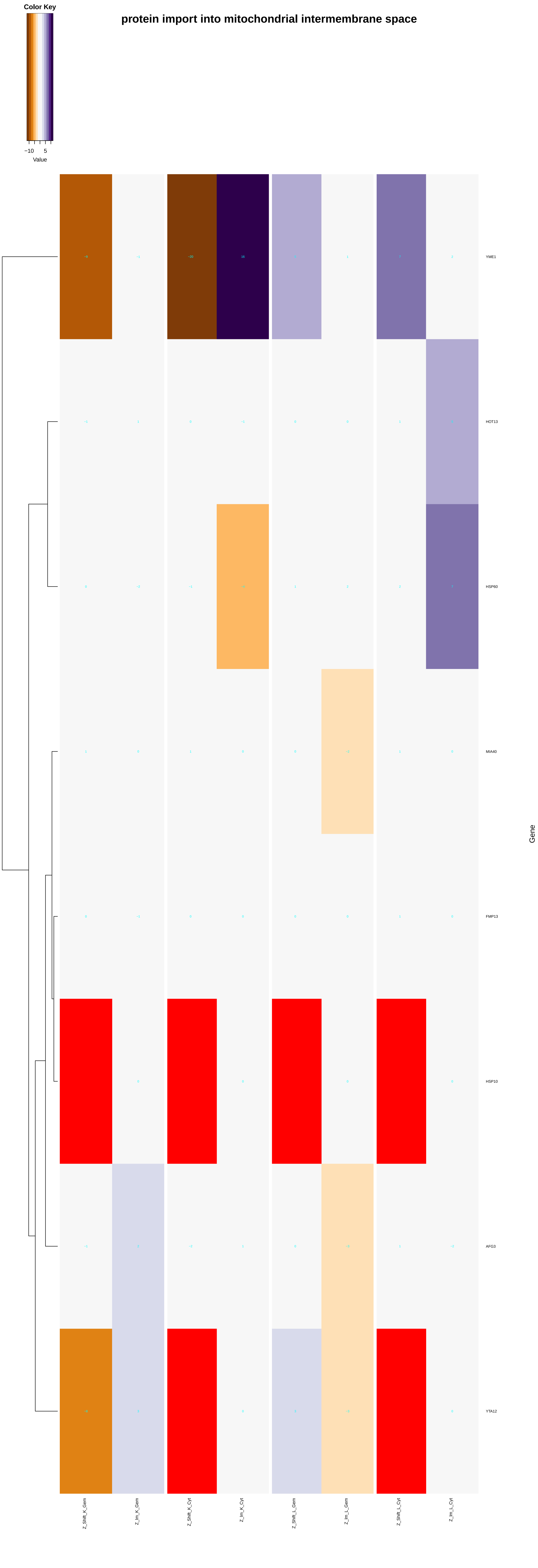
protein import into mitochondrial intermembrane space
Color Key
−10
5
Value
YME1
HOT13
HSP60
MIA40
FMP13
HSP10
AFG3
YTA12
Gene
Z_Shift_K_Gem
Z_Im_K_Gem
Z_Shift_K_Cyt
Z_Im_K_Cyt
Z_Shift_L_Gem
Z_Im_L_Gem
Z_Shift_L_Cyt
Z_Im_L_Cyt

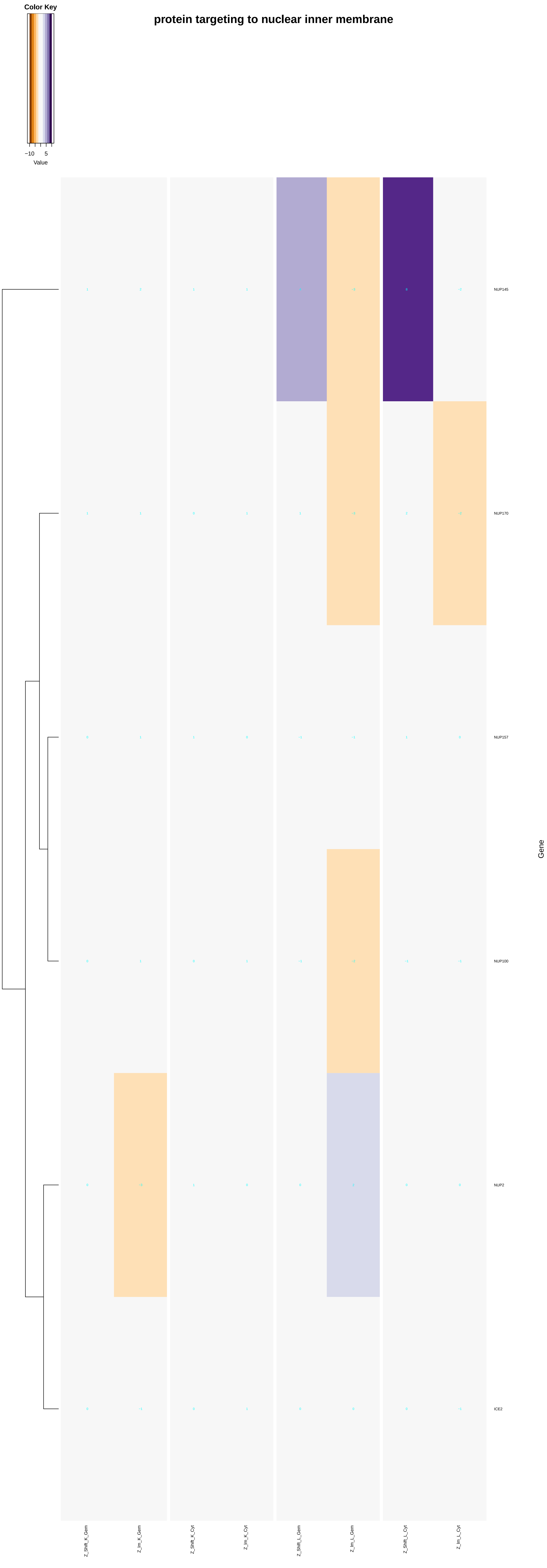
protein targeting to nuclear inner membrane
Color Key
−10
5
Value
NUP145
NUP170
NUP157
NUP100
NUP2
ICE2
Gene
Z_Shift_K_Gem
Z_Im_K_Gem
Z_Shift_K_Cyt
Z_Im_K_Cyt
Z_Shift_L_Gem
Z_Im_L_Gem
Z_Shift_L_Cyt
Z_Im_L_Cyt

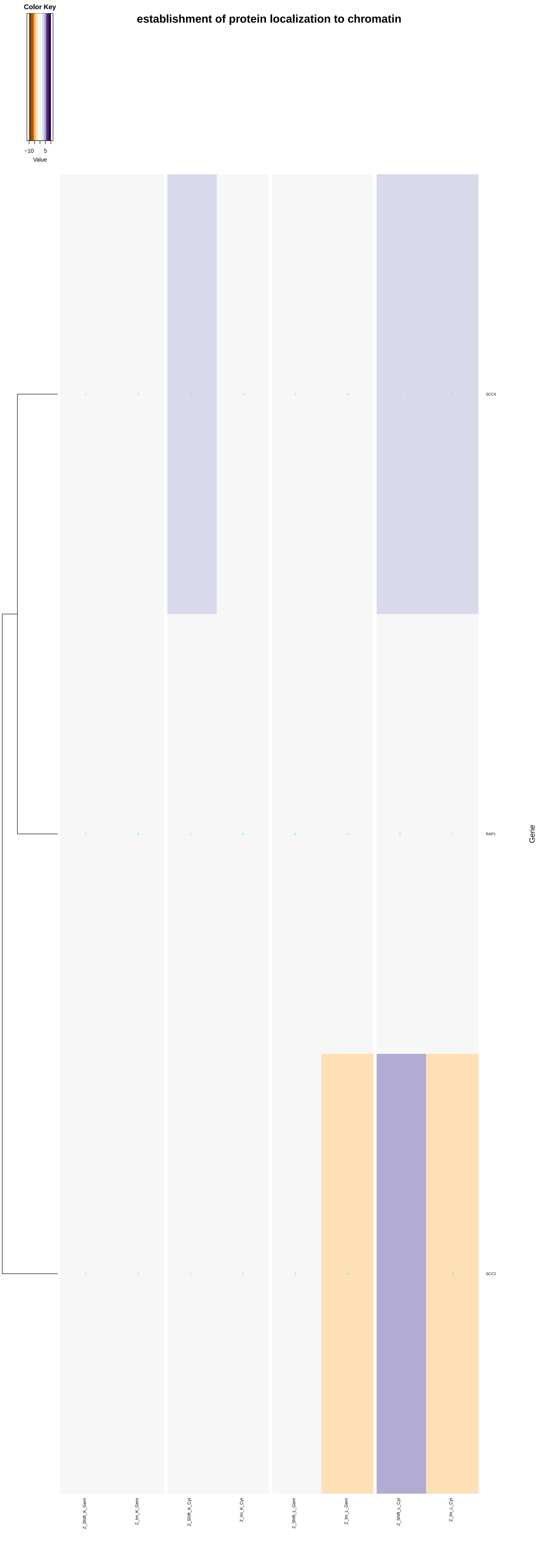
establishment of protein localization to chromatin
Color Key
−10
5
Value
SCC4
RAP1
SCC2
Gene
Z_Shift_K_Gem
Z_Im_K_Gem
Z_Shift_K_Cyt
Z_Im_K_Cyt
Z_Shift_L_Gem
Z_Im_L_Gem
Z_Shift_L_Cyt
Z_Im_L_Cyt

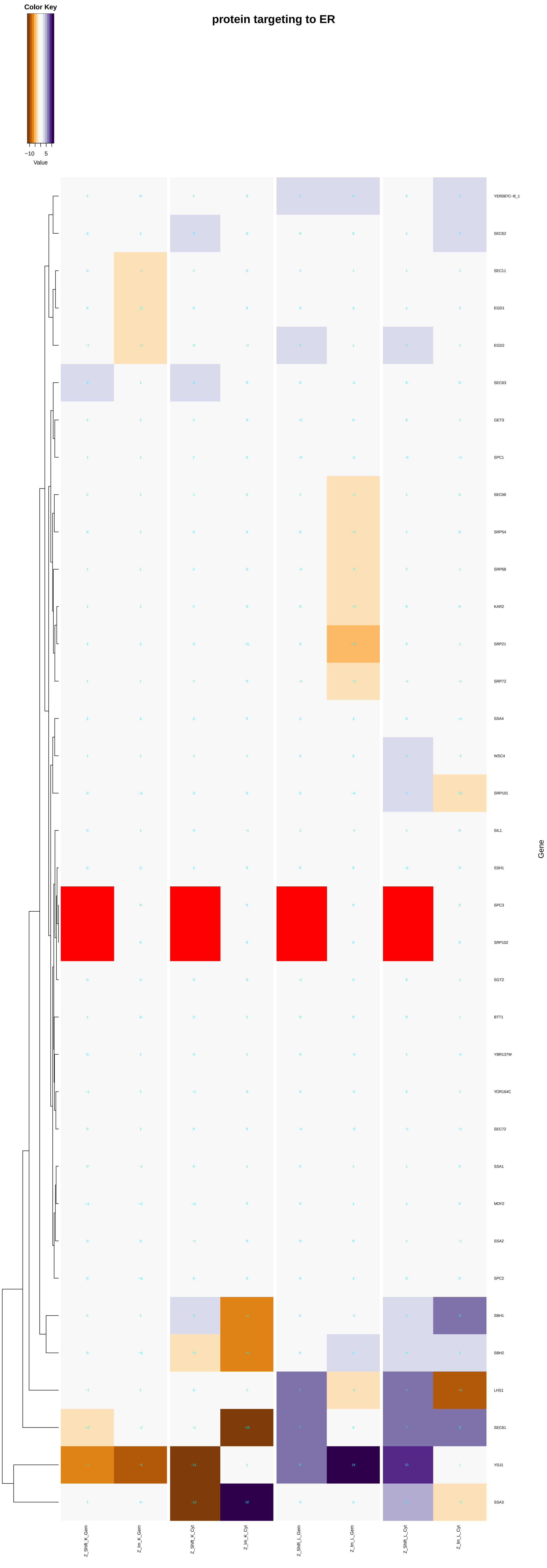
protein targeting to ER
Color Key
−10
5
Value
YER087C−B_1
SEC62
SEC11
EGD1
EGD2
SEC63
GET3
SPC1
SEC66
SRP54
SRP68
KAR2
SRP21
SRP72
SSA4
WSC4
SRP101
SIL1
SSH1
SPC3
SRP102
SGT2
BTT1
YBR137W
YOR164C
SEC72
SSA1
MDY2
SSA2
SPC2
SBH1
SBH2
LHS1
SEC61
YDJ1
SSA3
Gene
Z_Shift_K_Gem
Z_Im_K_Gem
Z_Shift_K_Cyt
Z_Im_K_Cyt
Z_Shift_L_Gem
Z_Im_L_Gem
Z_Shift_L_Cyt
Z_Im_L_Cyt

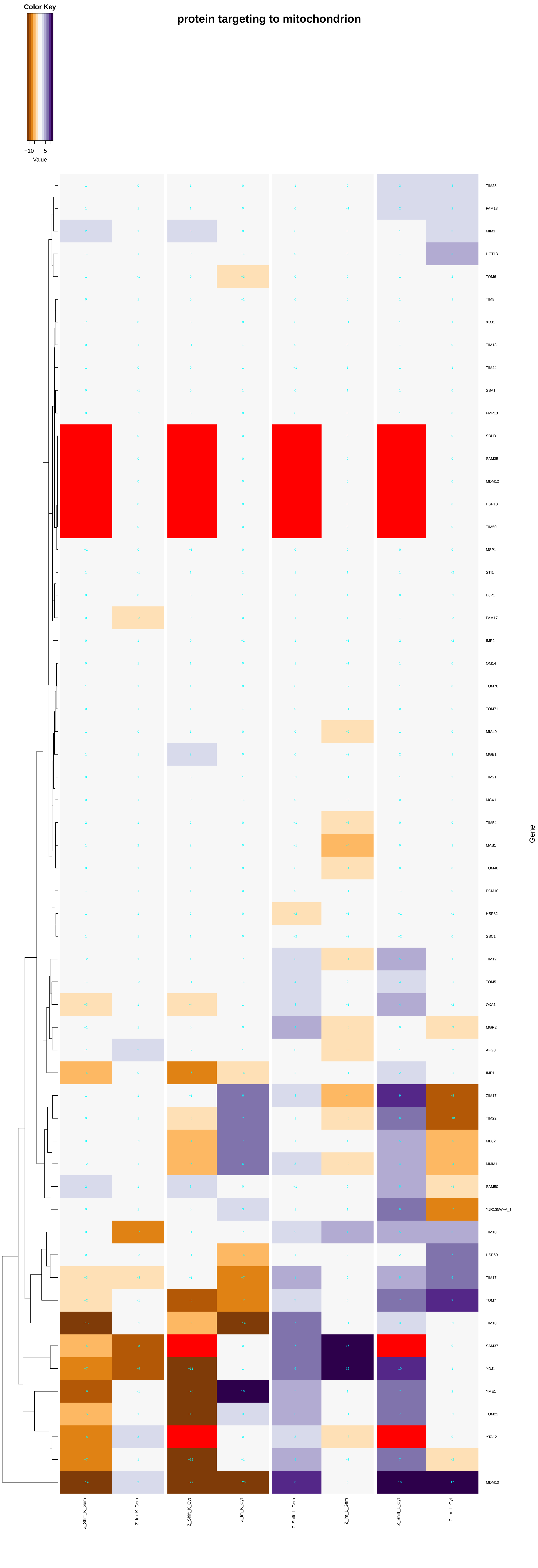
Color Key
−10
5
Value
protein targeting to mitochondrion
TIM23
PAM18
MIM1
HOT13
TOM6
TIM8
XDJ1
TIM13
TIM44
SSA1
FMP13
SDH3
SAM35
MDM12
HSP10
TIM50
MSP1
STI1
DJP1
PAM17
IMP2
OM14
TOM70
TOM71
MIA40
MGE1
TIM21
MCX1
TIM54
MAS1
TOM40
ECM10
HSP82
SSC1
TIM12
TOM5
OXA1
MGR2
AFG3
IMP1
ZIM17
TIM22
MDJ2
MMM1
SAM50
YJR135W−A_1
TIM10
HSP60
TIM17
TOM7
TIM18
SAM37
YDJ1
YME1
TOM22
YTA12
MDM10
Gene
Z_Shift_K_Gem
Z_Im_K_Gem
Z_Shift_K_Cyt
Z_Im_K_Cyt
Z_Shift_L_Gem
Z_Im_L_Gem
Z_Shift_L_Cyt
Z_Im_L_Cyt

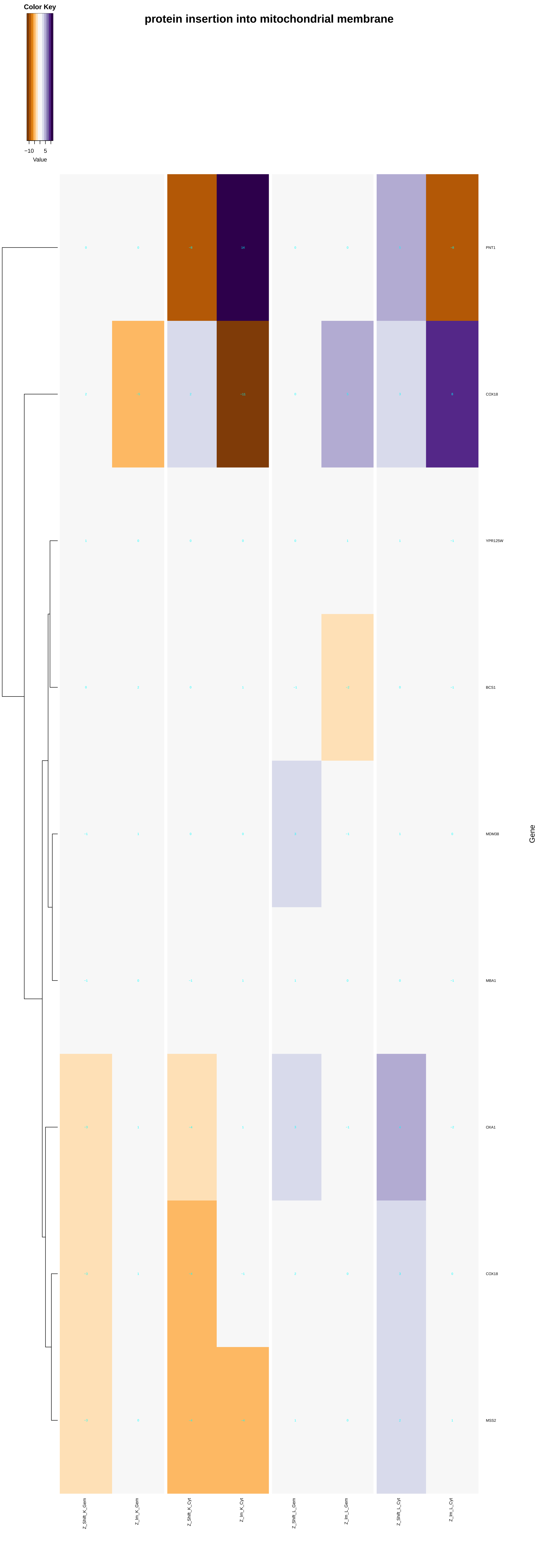
Color Key
−10
5
Value
protein insertion into mitochondrial membrane
PNT1
COX18
YPR125W
BCS1
MDM38
MBA1
OXA1
COX18
MSS2
Gene
Z_Shift_K_Gem
Z_Im_K_Gem
Z_Shift_K_Cyt
Z_Im_K_Cyt
Z_Shift_L_Gem
Z_Im_L_Gem
Z_Shift_L_Cyt
Z_Im_L_Cyt

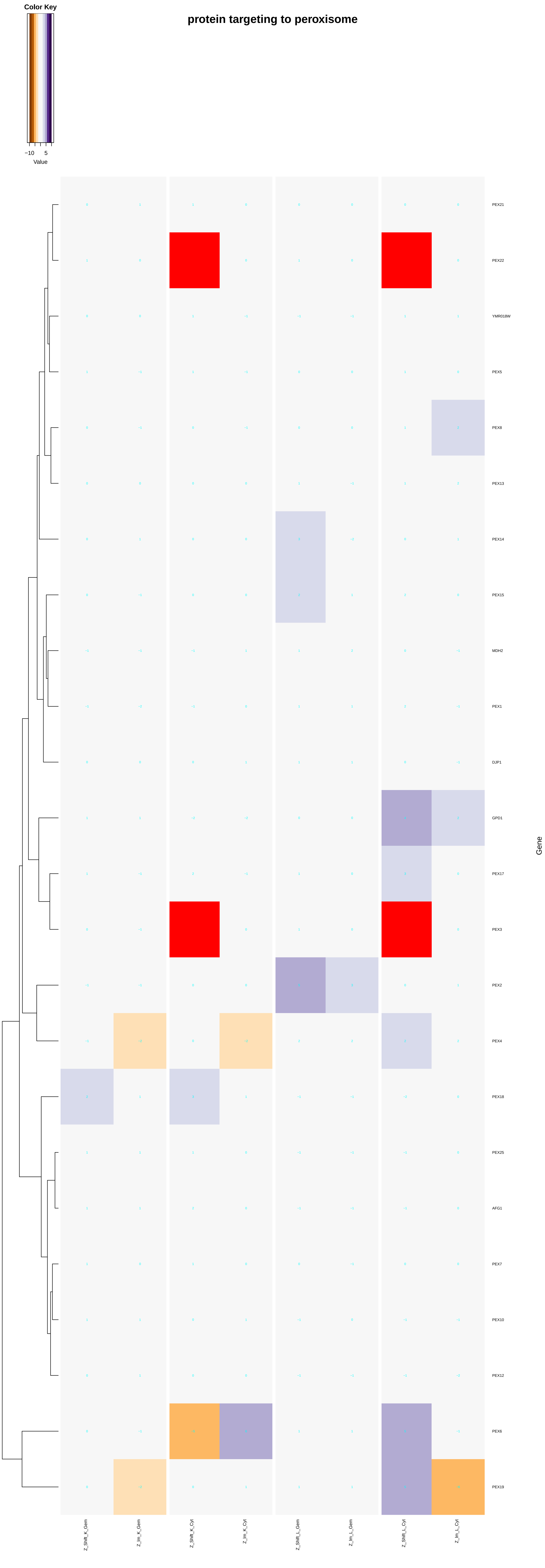

protein targeting to peroxisome
Color Key
−10
5
Value
PEX21
PEX22
YMR018W
PEX5
PEX8
PEX13
PEX14
PEX15
MDH2
PEX1
DJP1
GPD1
PEX17
PEX3
PEX2
PEX4
PEX18
PEX25
AFG1
PEX7
PEX10
PEX12
PEX6
PEX19
Gene
Z_Shift_K_Gem
Z_Im_K_Gem
Z_Shift_K_Cyt
Z_Im_K_Cyt
Z_Shift_L_Gem
Z_Im_L_Gem
Z_Shift_L_Cyt
Z_Im_L_Cyt

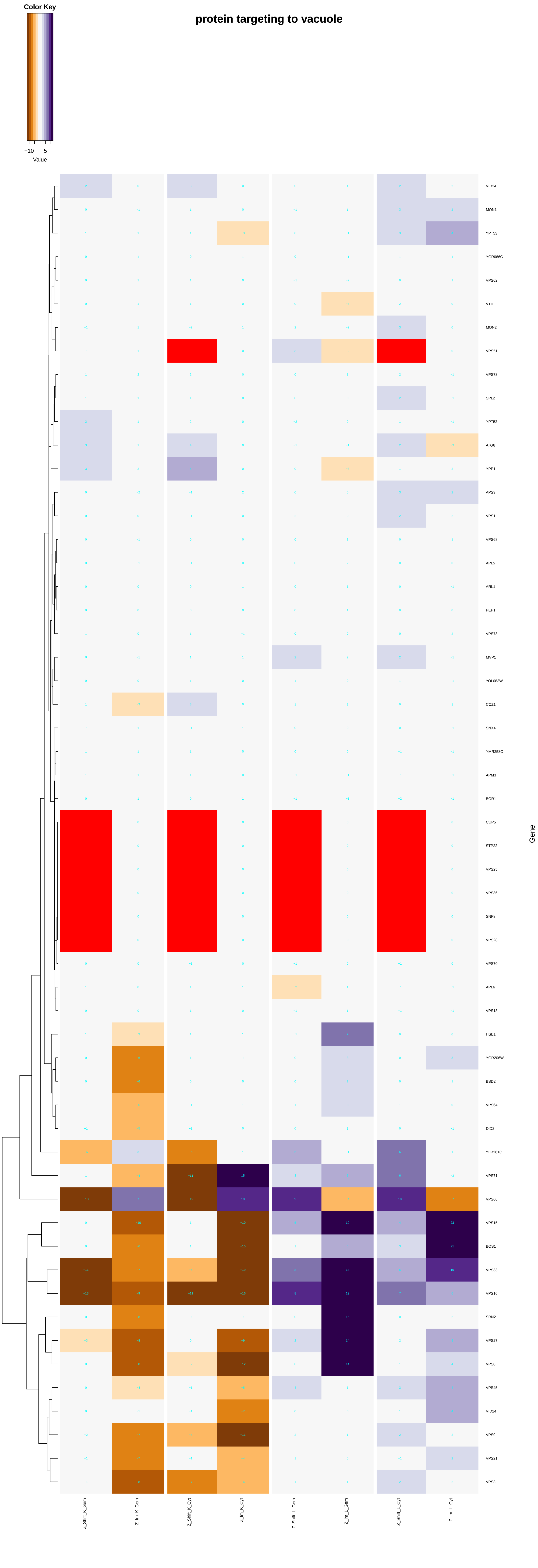

protein targeting to vacuole
Color Key
−10
5
Value
VID24
MON1
YPT53
YGR066C
VPS62
VTI1
MON2
VPS51
VPS73
SPL2
YPT52
ATG8
YPP1
APS3
VPS1
VPS68
APL5
ARL1
PEP1
VPS73
MVP1
YOL083W
CCZ1
SNX4
YMR258C
APM3
BOR1
CUP5
STP22
VPS25
VPS36
SNF8
VPS28
VPS70
APL6
VPS13
HSE1
YGR206W
BSD2
VPS64
DID2
YLR261C
VPS71
VPS66
VPS15
BOS1
VPS33
VPS16
SRN2
VPS27
VPS8
VPS45
VID24
VPS9
VPS21
VPS3
Gene
Z_Shift_K_Gem
Z_Im_K_Gem
Z_Shift_K_Cyt
Z_Im_K_Cyt
Z_Shift_L_Gem
Z_Im_L_Gem
Z_Shift_L_Cyt
Z_Im_L_Cyt

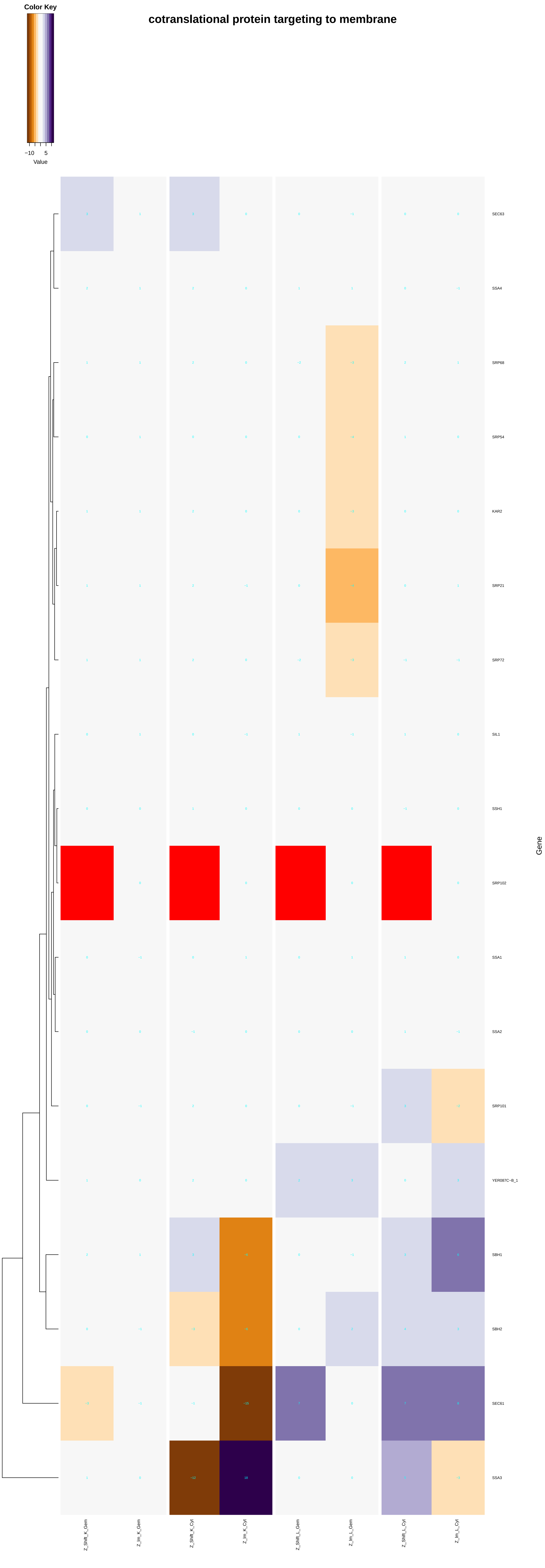
cotranslational protein targeting to membrane
Color Key
−10
5
Value
SEC63
SSA4
SRP68
SRP54
KAR2
SRP21
SRP72
SIL1
SSH1
SRP102
SSA1
SSA2
SRP101
YER087C−B_1
SBH1
SBH2
SEC61
SSA3
Gene
Z_Shift_K_Gem
Z_Im_K_Gem
Z_Shift_K_Cyt
Z_Im_K_Cyt
Z_Shift_L_Gem
Z_Im_L_Gem
Z_Shift_L_Cyt
Z_Im_L_Cyt

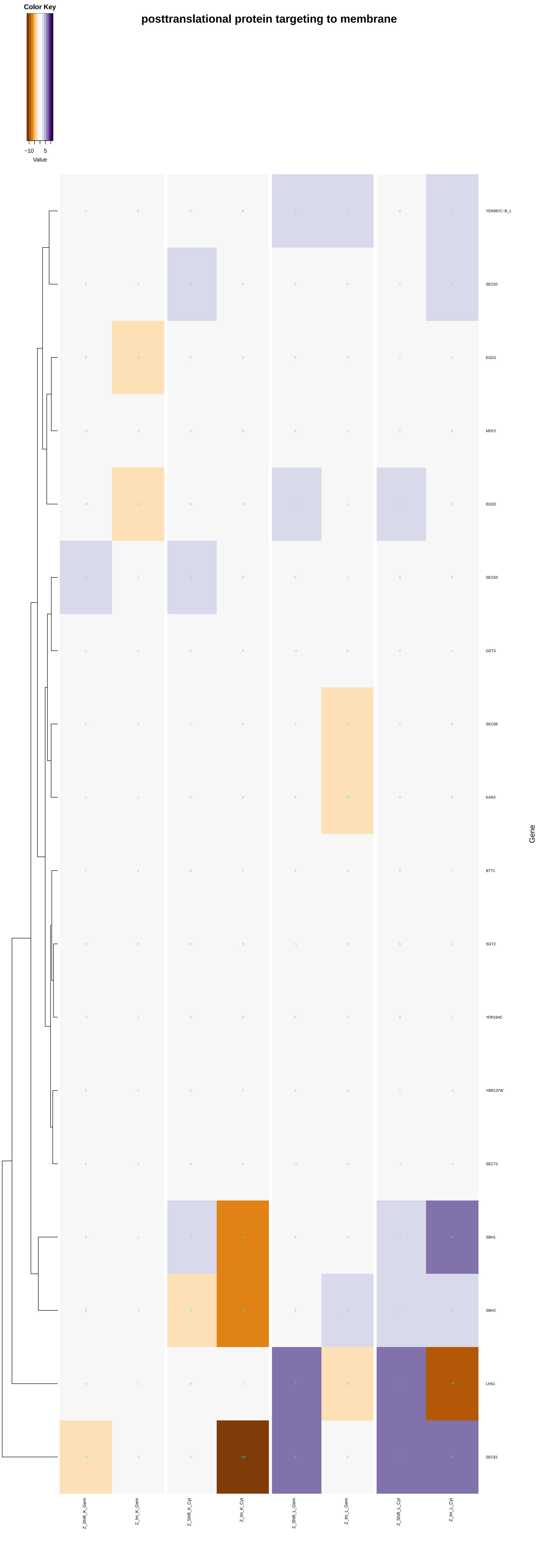

Color Key
−10
5
Value
posttranslational protein targeting to membrane
YER087C−B_1
SEC62
EGD1
MDY2
EGD2
SEC63
GET3
SEC66
KAR2
BTT1
SGT2
YOR164C
YBR137W
SEC72
SBH1
SBH2
LHS1
SEC61
Gene
Z_Shift_K_Gem
Z_Im_K_Gem
Z_Shift_K_Cyt
Z_Im_K_Cyt
Z_Shift_L_Gem
Z_Im_L_Gem
Z_Shift_L_Cyt
Z_Im_L_Cyt

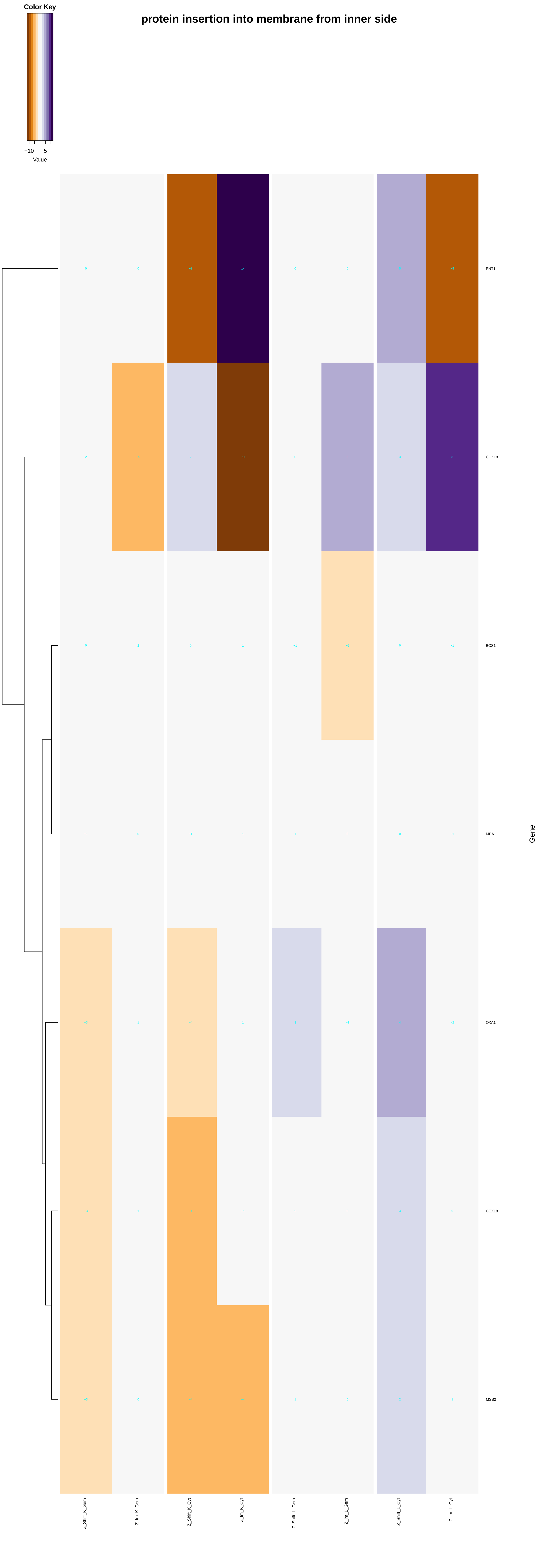

Color Key
−10
5
Value
protein insertion into membrane from inner side
PNT1
COX18
BCS1
MBA1
OXA1
COX18
MSS2
Gene
Z_Shift_K_Gem
Z_Im_K_Gem
Z_Shift_K_Cyt
Z_Im_K_Cyt
Z_Shift_L_Gem
Z_Im_L_Gem
Z_Shift_L_Cyt
Z_Im_L_Cyt

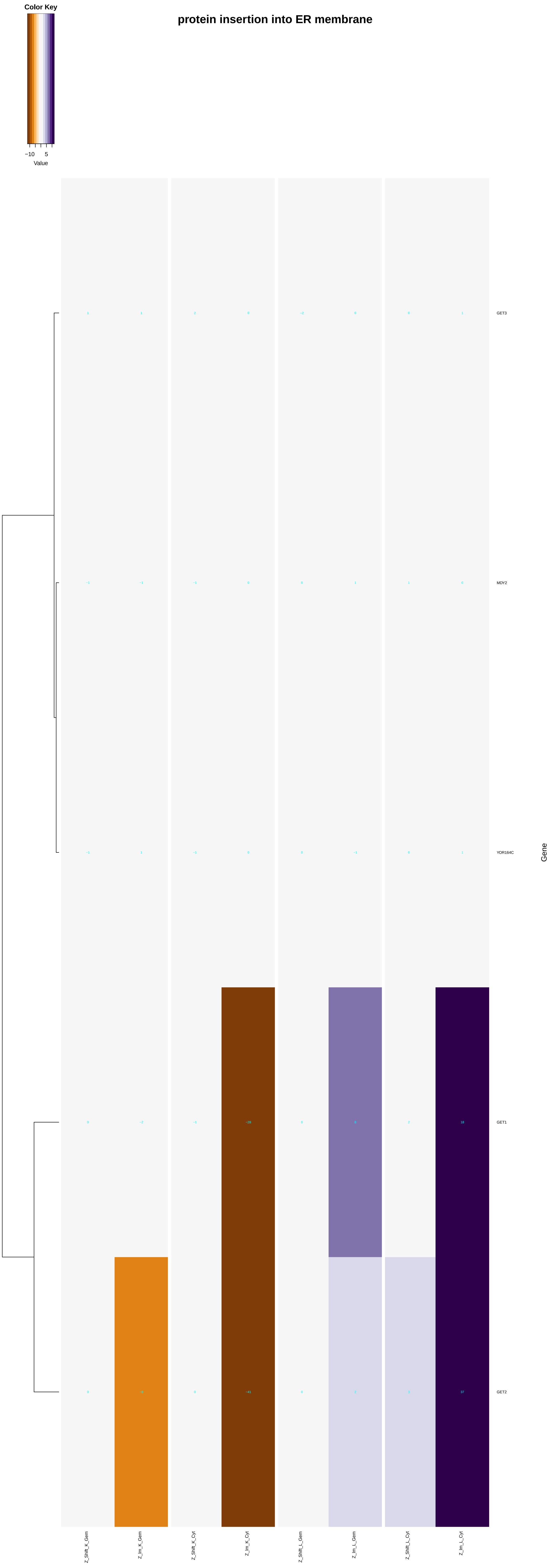

protein insertion into ER membrane
Color Key
−10
5
Value
GET3
MDY2
YOR164C
GET1
GET2
Gene
Z_Shift_K_Gem
Z_Im_K_Gem
Z_Shift_K_Cyt
Z_Im_K_Cyt
Z_Shift_L_Gem
Z_Im_L_Gem
Z_Shift_L_Cyt
Z_Im_L_Cyt

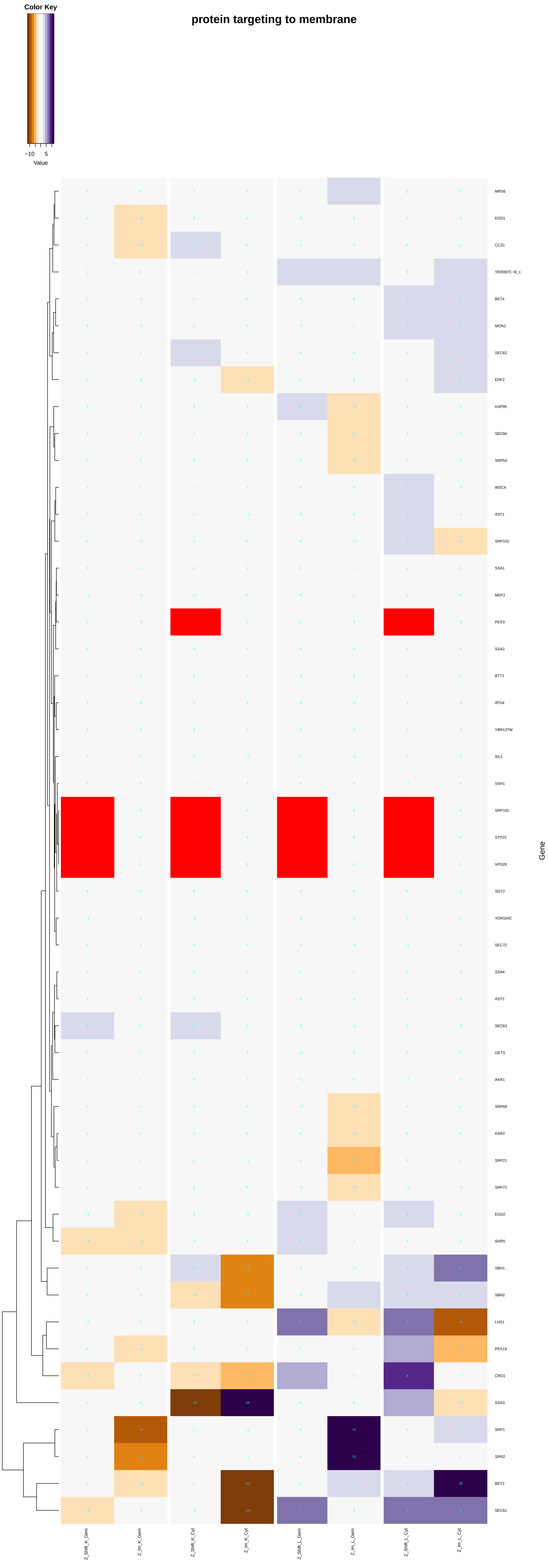
protein targeting to membrane
Color Key
−10
5
Value
MRS6
EGD1
CCZ1
YER087C−B_1
BET4
MON1
SEC62
ERF2
KAP95
SEC66
SRP54
WSC4
AST1
SRP101
SSA1
MDY2
PEX3
SSA2
BTT1
ATG4
YBR137W
SIL1
SSH1
SRP102
STP22
VPS28
SGT2
YOR164C
SEC72
SSA4
AST2
SEC63
GET3
AKR1
SRP68
KAR2
SRP21
SRP72
EGD2
SHR5
SBH1
SBH2
LHS1
PEX19
CRD1
SSA3
SRP1
SRN2
BET2
SEC61
Gene
Z_Shift_K_Gem
Z_Im_K_Gem
Z_Shift_K_Cyt
Z_Im_K_Cyt
Z_Shift_L_Gem
Z_Im_L_Gem
Z_Shift_L_Cyt
Z_Im_L_Cyt

#### retrograde protein transport, ER to cytosol

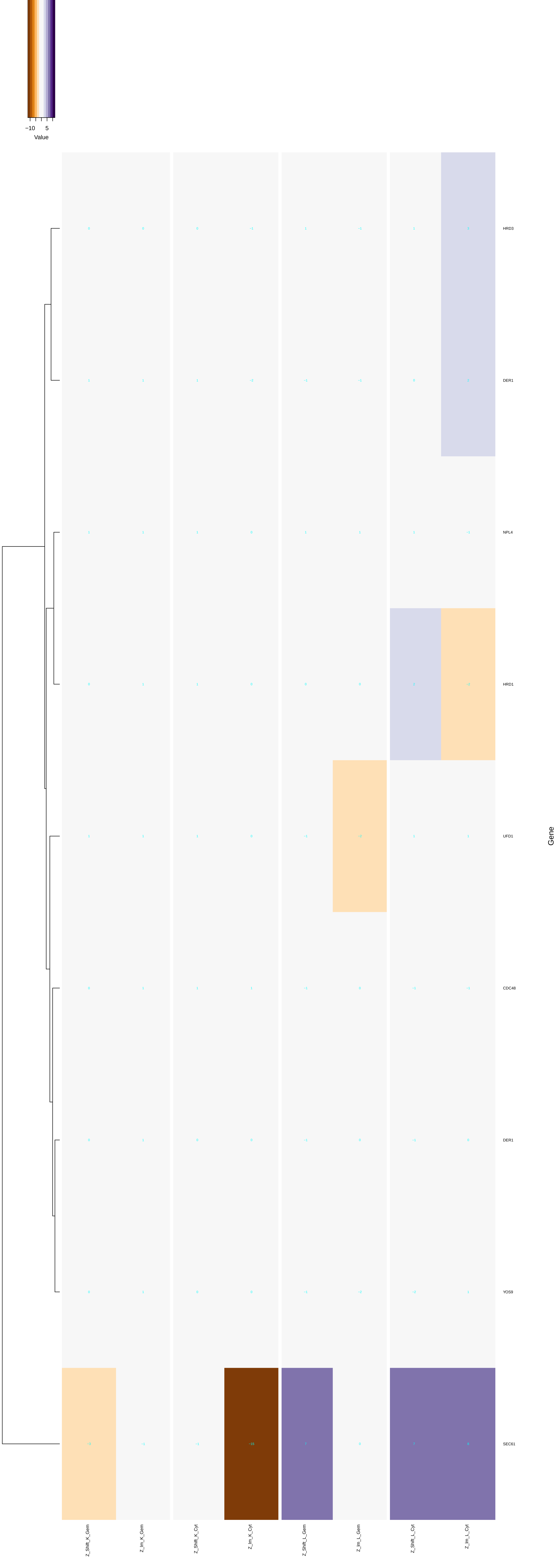

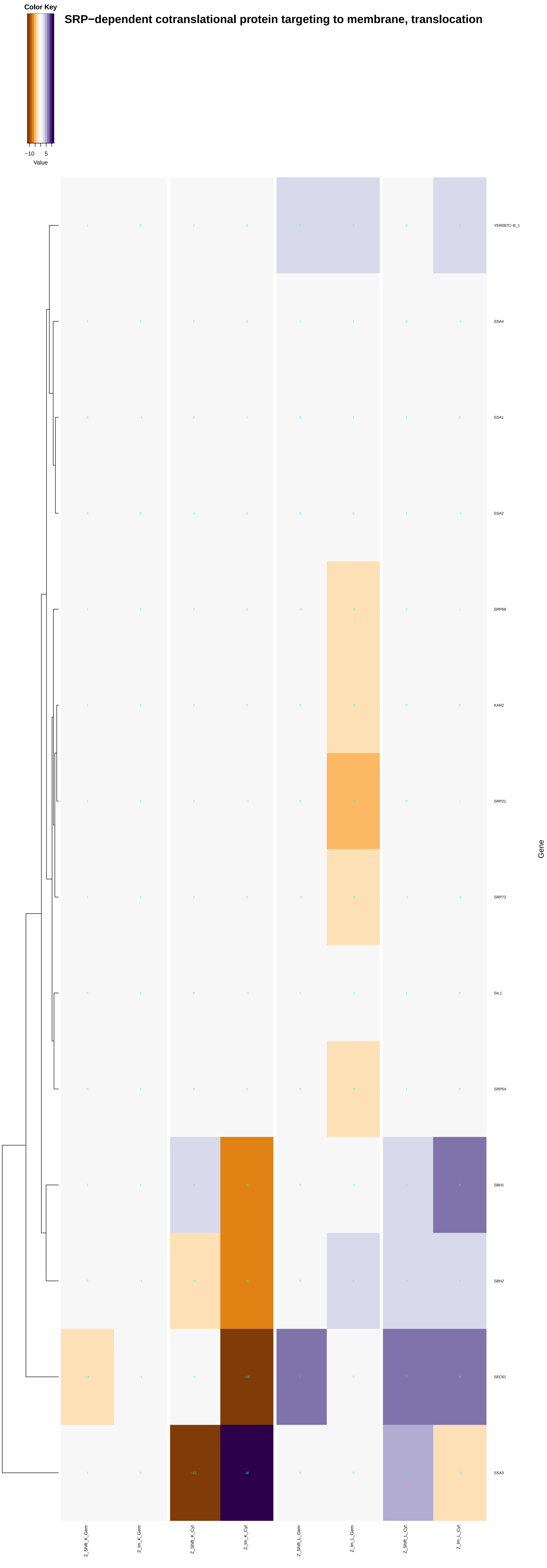
SRP−dependent cotranslational protein targeting to membrane, translocation
Color Key
−10
5
Value
YER087C−B_1
SSA4
SSA1
SSA2
SRP68
KAR2
SRP21
SRP72
SIL1
SRP54
SBH1
SBH2
SEC61
SSA3
Gene
Z_Shift_K_Gem
Z_Im_K_Gem
Z_Shift_K_Cyt
Z_Im_K_Cyt
Z_Shift_L_Gem
Z_Im_L_Gem
Z_Shift_L_Cyt
Z_Im_L_Cyt

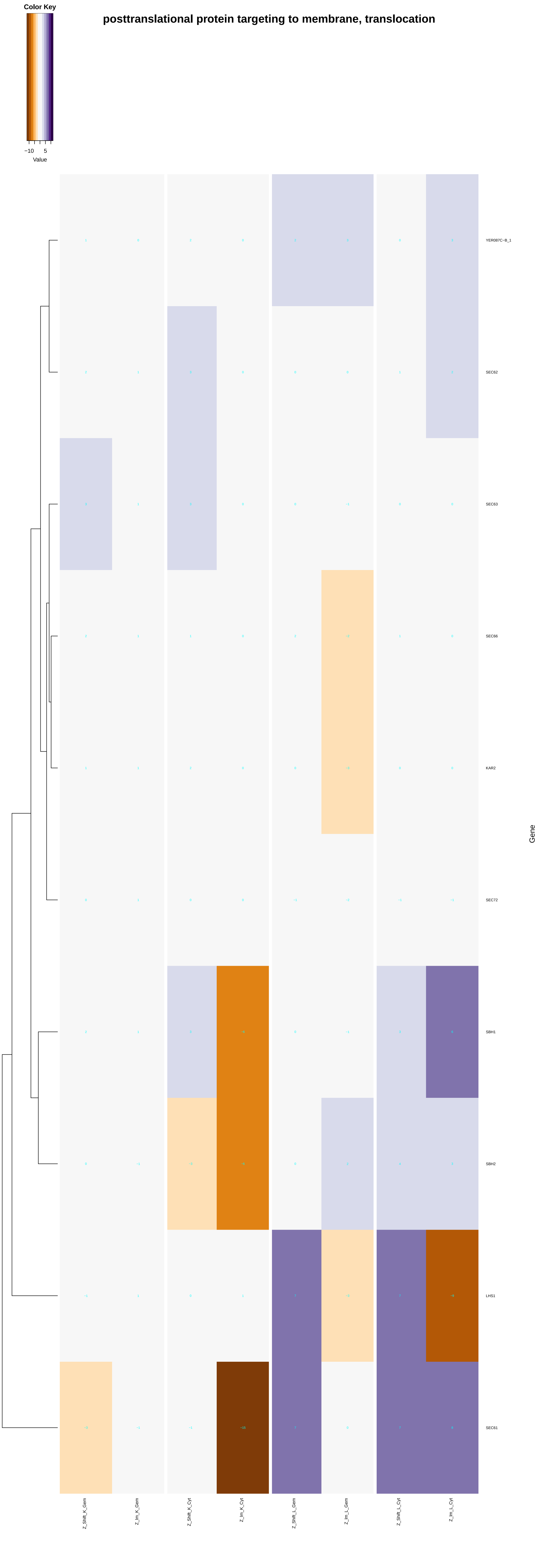
Color Key
−10
5
Value
posttranslational protein targeting to membrane, translocation
YER087C−B_1
SEC62
SEC63
SEC66
KAR2
SEC72
SBH1
SBH2
LHS1
SEC61
Gene
Z_Shift_K_Gem
Z_Im_K_Gem
Z_Shift_K_Cyt
Z_Im_K_Cyt
Z_Shift_L_Gem
Z_Im_L_Gem
Z_Shift_L_Cyt
Z_Im_L_Cyt

### intracellular protein transmembrane import

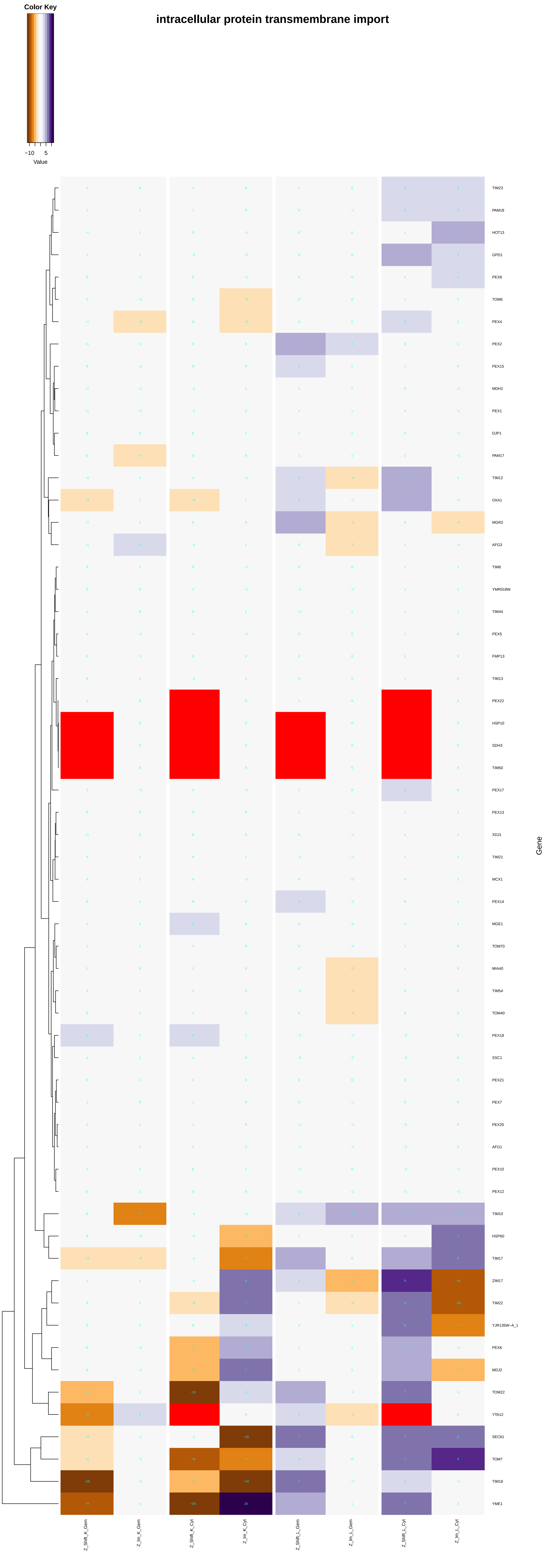

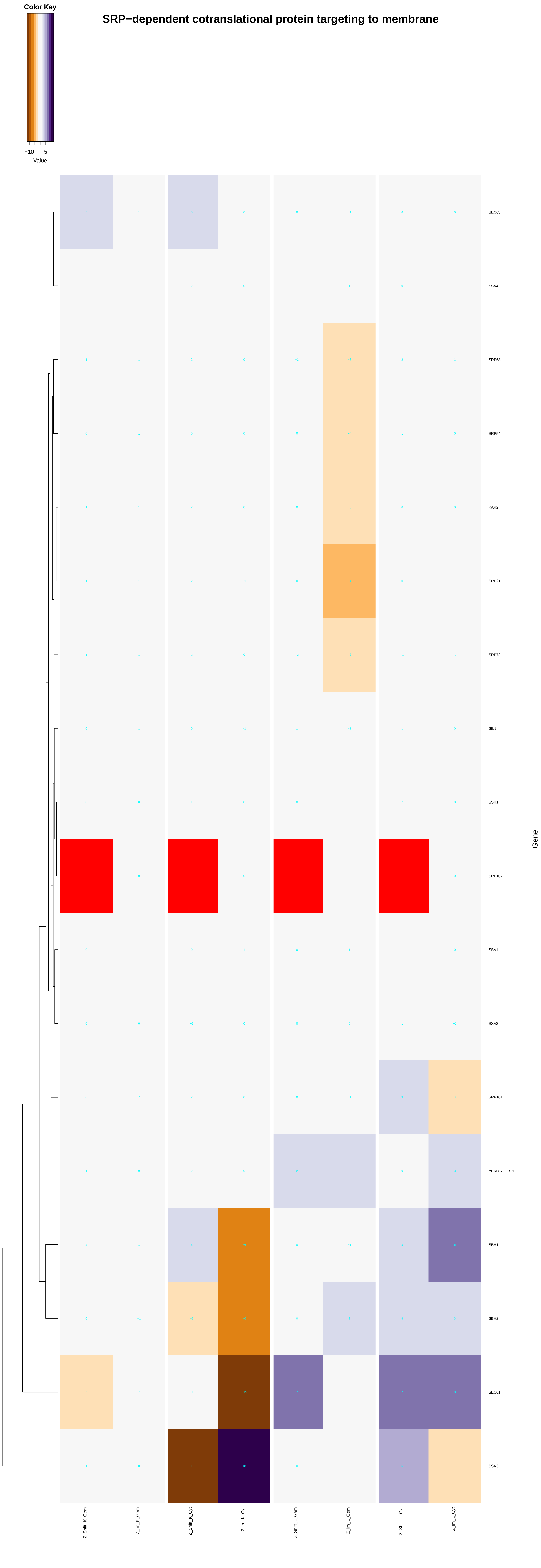
SRP−dependent cotranslational protein targeting to membrane
Color Key
−10
5
Value
SEC63
SSA4
SRP68
SRP54
KAR2
SRP21
SRP72
SIL1
SSH1
SRP102
SSA1
SSA2
SRP101
YER087C−B_1
SBH1
SBH2
SEC61
SSA3
Gene
Z_Shift_K_Gem
Z_Im_K_Gem
Z_Shift_K_Cyt
Z_Im_K_Cyt
Z_Shift_L_Gem
Z_Im_L_Gem
Z_Shift_L_Cyt
Z_Im_L_Cyt

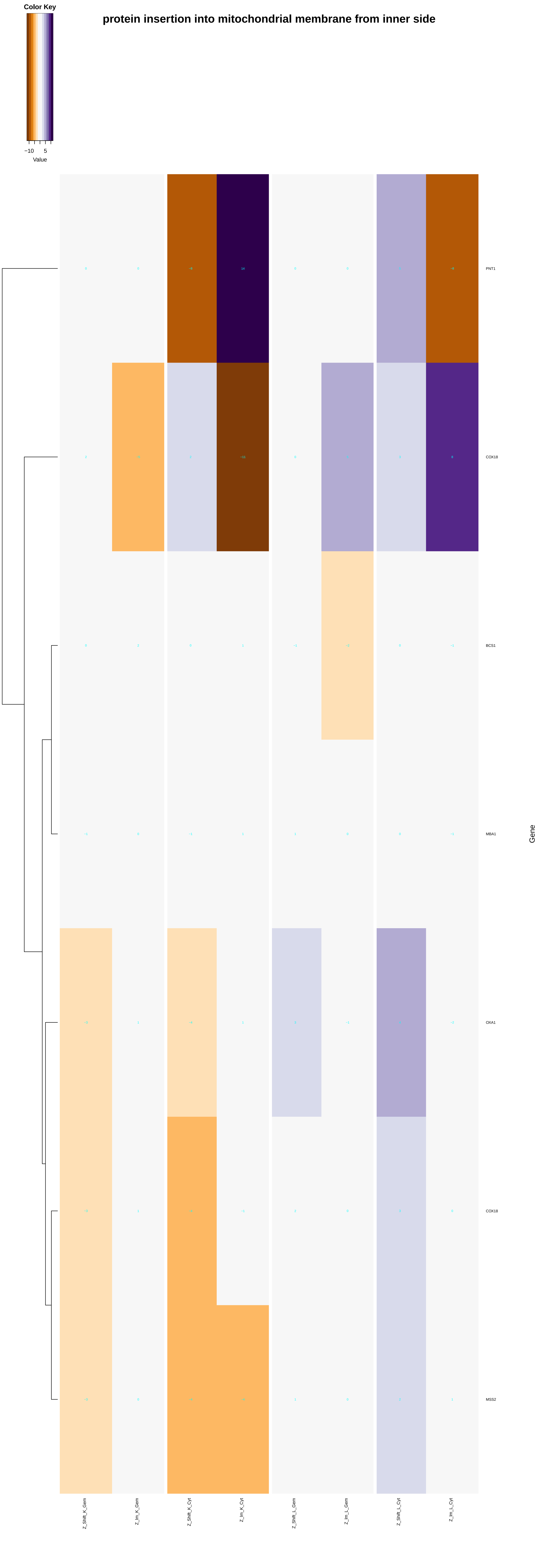

Color Key
−10
5
Value
protein insertion into mitochondrial membrane from inner side
PNT1
COX18
BCS1
MBA1
OXA1
COX18
MSS2
Gene
Z_Shift_K_Gem
Z_Im_K_Gem
Z_Shift_K_Cyt
Z_Im_K_Cyt
Z_Shift_L_Gem
Z_Im_L_Gem
Z_Shift_L_Cyt
Z_Im_L_Cyt

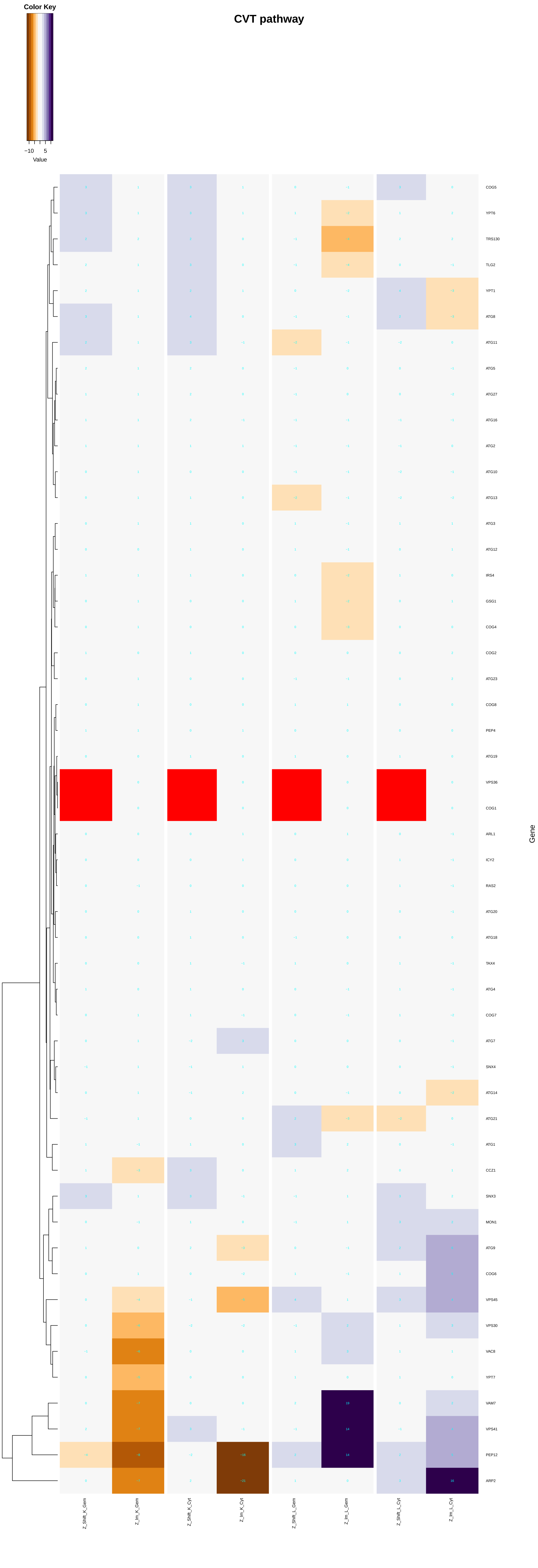
Color Key
−10
5
Value
CVT pathway
COG5
YPT6
TRS130
TLG2
YPT1
ATG8
ATG11
ATG5
ATG27
ATG16
ATG2
ATG10
ATG13
ATG3
ATG12
IRS4
GSG1
COG4
COG2
ATG23
COG8
PEP4
ATG19
VPS36
COG1
ARL1
ICY2
RAS2
ATG20
ATG18
TAX4
ATG4
COG7
ATG7
SNX4
ATG14
ATG21
ATG1
CCZ1
SNX3
MON1
ATG9
COG6
VPS45
VPS30
VAC8
YPT7
VAM7
VPS41
PEP12
ARP2
Gene
Z_Shift_K_Gem
Z_Im_K_Gem
Z_Shift_K_Cyt
Z_Im_K_Cyt
Z_Shift_L_Gem
Z_Im_L_Gem
Z_Shift_L_Cyt
Z_Im_L_Cyt

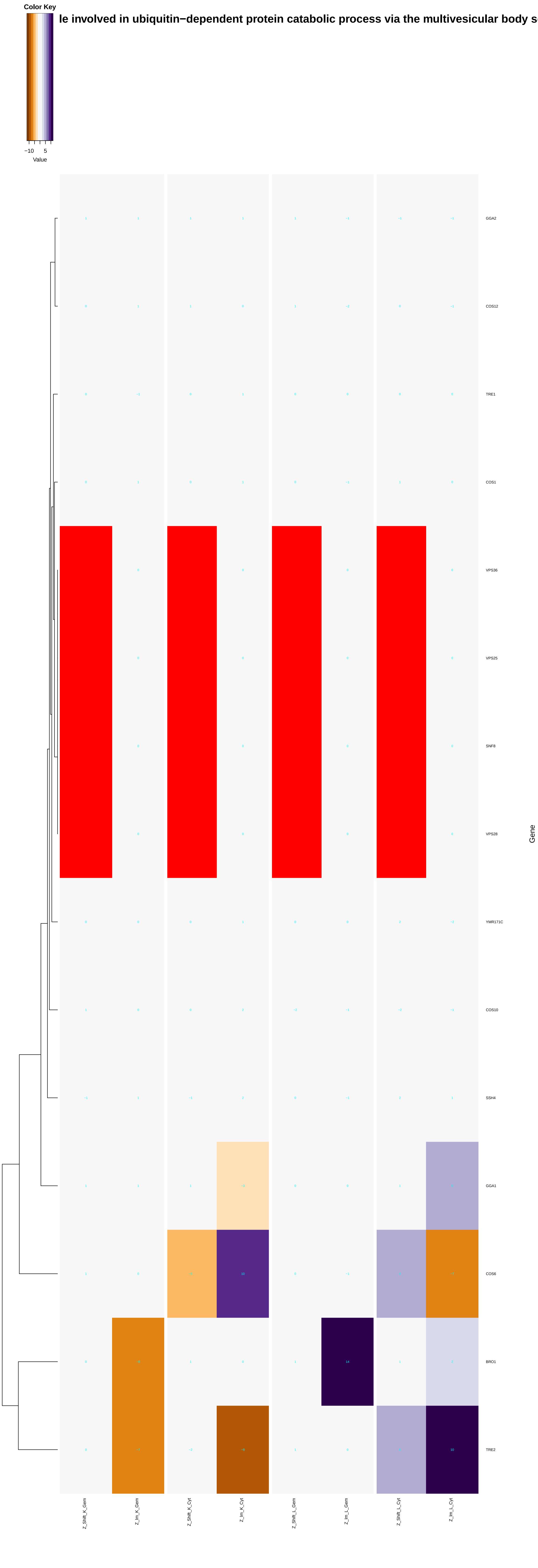

Color Key
−10
5
Value
le involved in ubiquitin−dependent protein catabolic process via the multivesicular body s
GGA2
COS12
TRE1
COS1
VPS36
VPS25
SNF8
VPS28
YMR171C
COS10
SSH4
GGA1
COS6
BRO1
TRE2
Gene
Z_Shift_K_Gem
Z_lm_K_Gem
Z_Shift_K_Cyt
Z_lm_K_Cyt
Z_Shift_L_Gem
Z_lm_L_Gem
Z_Shift_L_Cyt
Z_lm_L_Cyt

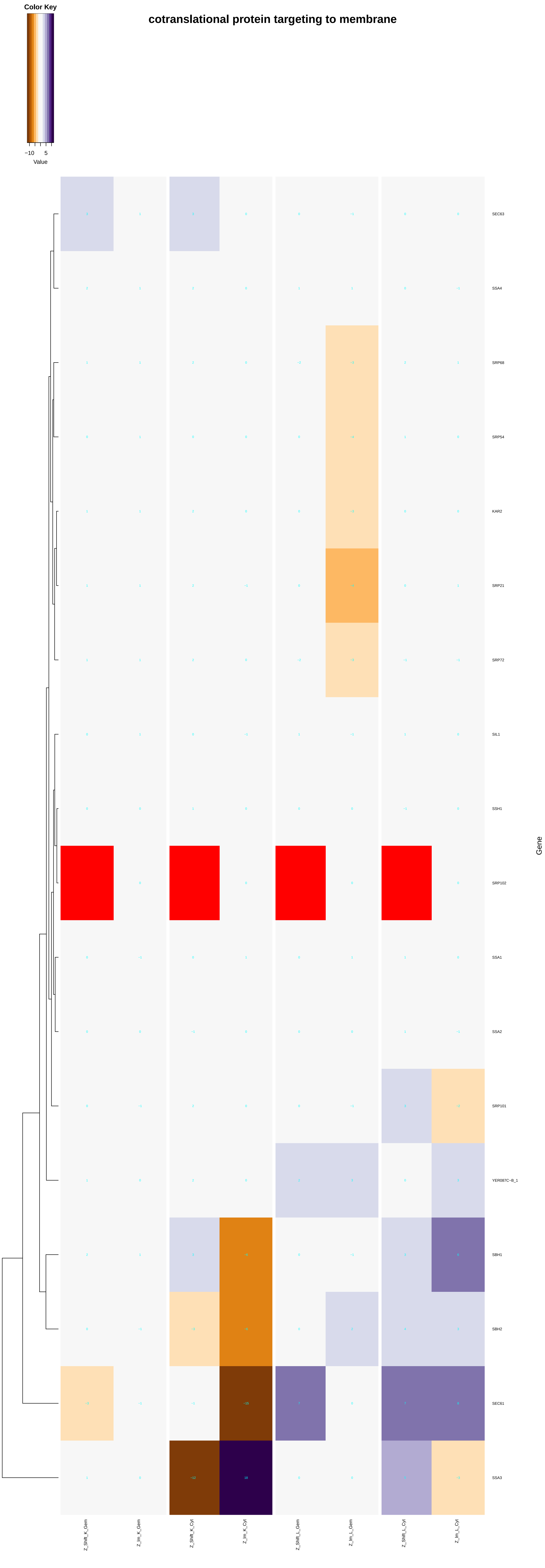
cotranslational protein targeting to membrane
Color Key
−10
5
Value
SEC63
SSA4
SRP68
SRP54
KAR2
SRP21
SRP72
SIL1
SSH1
SRP102
SSA1
SSA2
SRP101
YER087C−B_1
SBH1
SBH2
SEC61
SSA3
Gene
Z_Shift_K_Gem
Z_Int_K_Gem
Z_Shift_K_Cyt
Z_Int_K_Cyt
Z_Shift_L_Gem
Z_Int_L_Gem
Z_Shift_L_Cyt
Z_Int_L_Cyt

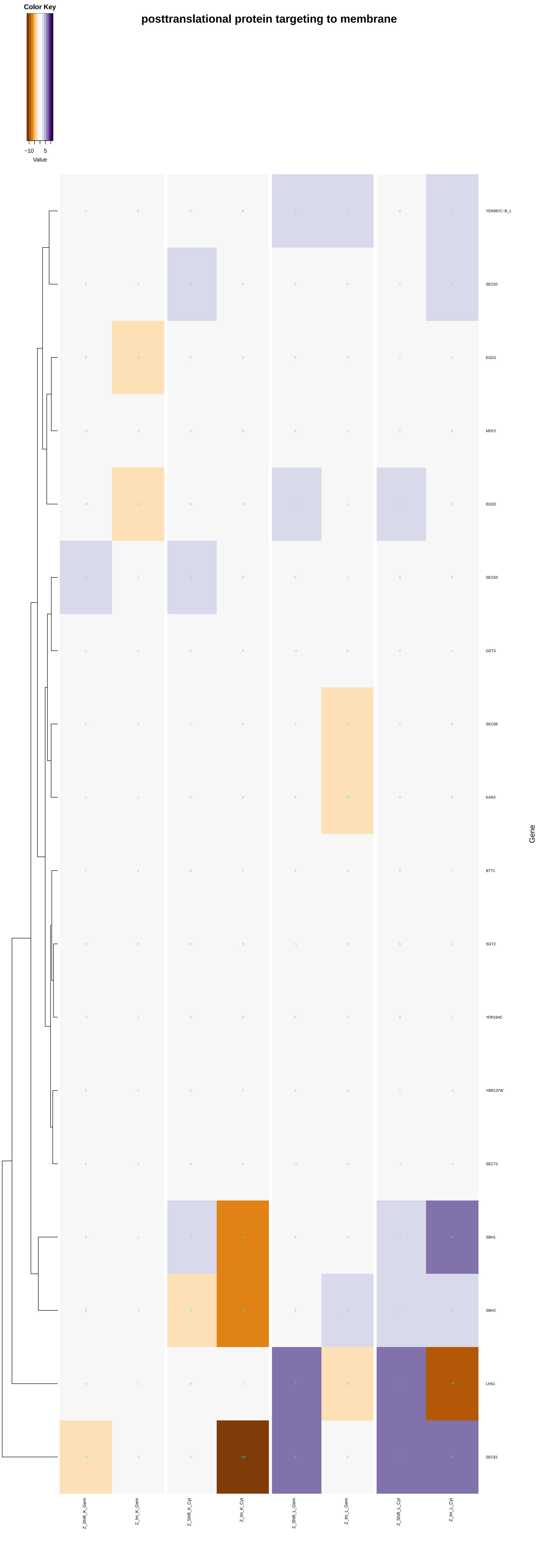

Color Key
−10
5
Value
posttranslational protein targeting to membrane
YER087C−B_1
SEC62
EGD1
MDY2
EGD2
SEC63
GET3
SEC66
KAR2
BTT1
SGT2
YOR164C
YBR137W
SEC72
SBH1
SBH2
LHS1
SEC61
Gene
Z_Shift_K_Gem
Z_Im_K_Gem
Z_Shift_K_Cyt
Z_Im_K_Cyt
Z_Shift_L_Gem
Z_Im_L_Gem
Z_Shift_L_Cyt
Z_Im_L_Cyt

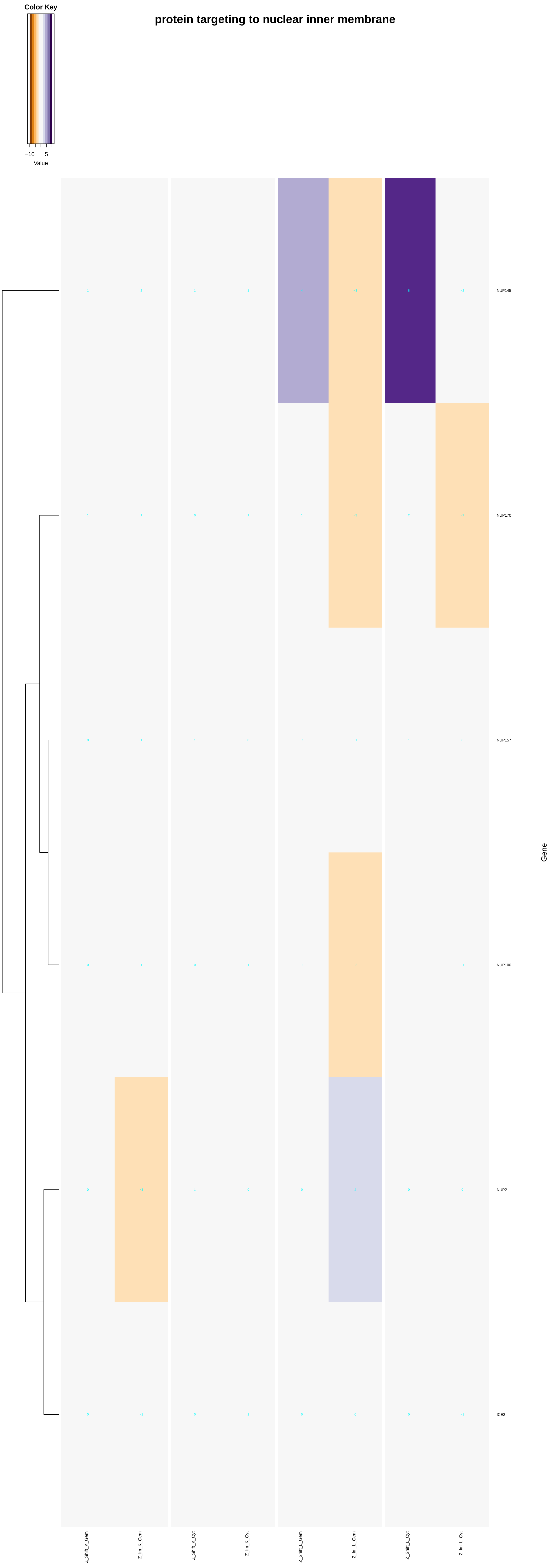

protein targeting to nuclear inner membrane
Color Key
−10
5
Value
NUP145
NUP170
NUP157
NUP100
NUP2
ICE2
Gene
Z_Shift_K_Gem
Z_lm_K_Gem
Z_Shift_K_Cyt
Z_lm_K_Cyt
Z_Shift_L_Gem
Z_lm_L_Gem
Z_Shift_L_Cyt
Z_lm_L_Cyt

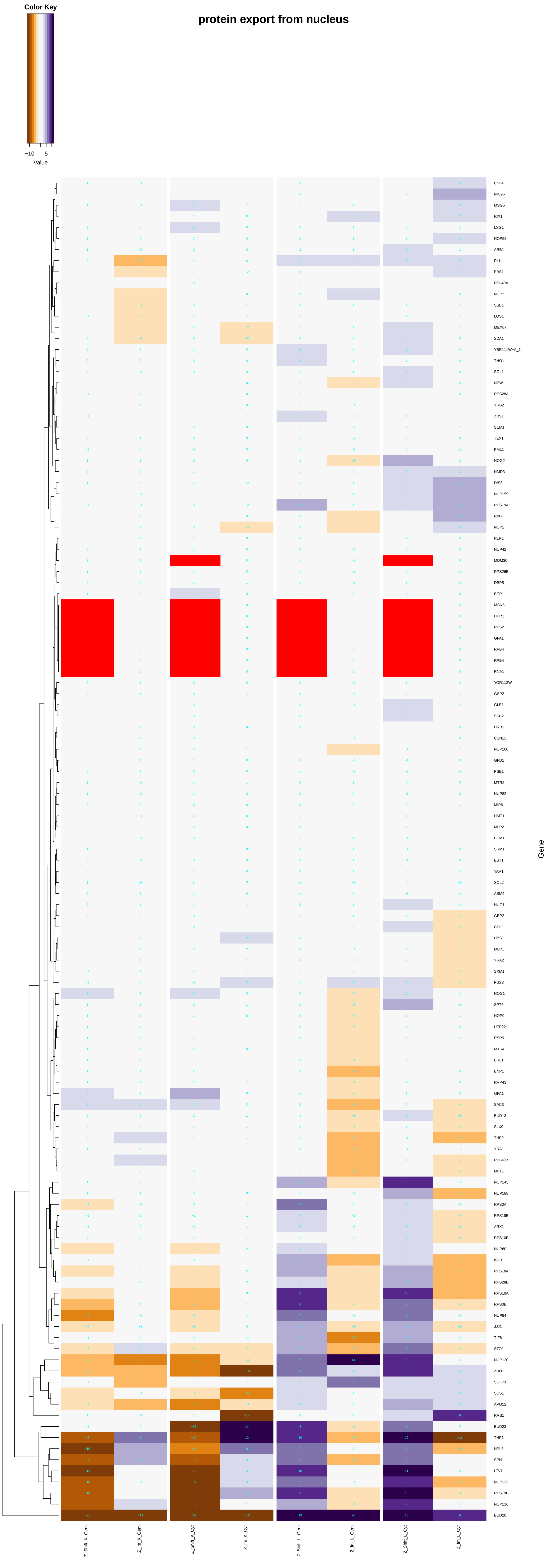

Color Key
−10
5
Value
protein export from nucleus
Gene
Z_Shift_K_Gem
Z_Im_K_Gem
Z_Shift_K_Cyt
Z_Im_K_Cyt
Z_Shift_L_Gem
Z_Im_L_Gem
Z_Shift_L_Cyt
Z_Im_L_Cyt
CSL4
NIC96
MSG5
RIX1
LSG1
NOP53
ARB1
RLI1
EBS1
RPL40A
NUP2
SSB1
LOS1
MEX67
SDA1
YBR111W-A_1
THO1
SOL1
NEW1
RPS28A
YRB2
ZDS1
SEM1
TEX1
PML1
NOG2
NMD3
DIS3
NUP159
RPS19A
RIX7
NUP1
RLR1
NUP42
MDM30
RPS26B
DBP5
BCP1
MSN5
HPR1
RPS2
GPA1
RPB4
RPB4
RNA1
YOR112W
GSP2
GLE1
SSB2
HRB1
CSN12
NUP100
GFD1
PSE1
MTR2
NUP82
MIP6
HMT1
MLP2
ECM1
SRM1
EST1
YAR1
SOL2
ASM4
NUG1
GBP2
CSE1
UBS1
MLP1
YRA2
SXM1
FUS3
NOG1
SPT6
NOP9
UTP22
RSP5
MTR4
BRL1
ENP1
RRP43
GPA1
SAC3
BUD13
SLX9
THP2
YRA1
RPL40B
MFT1
NUP145
NUP188
RPS0A
RPS18B
ARX1
RPS10B
NUP60
IST3
RPS18A
RPS28B
RPS10A
RPS0B
NUP84
JJJ1
TIF6
STO1
NUP120
ZUO1
SGF73
SUS1
APQ12
RRS1
BUD23
THP1
NPL3
SPN1
LTV1
NUP133
RPS19B
NUP116
BUD20

### RNA polymerase II complex import to nucleus

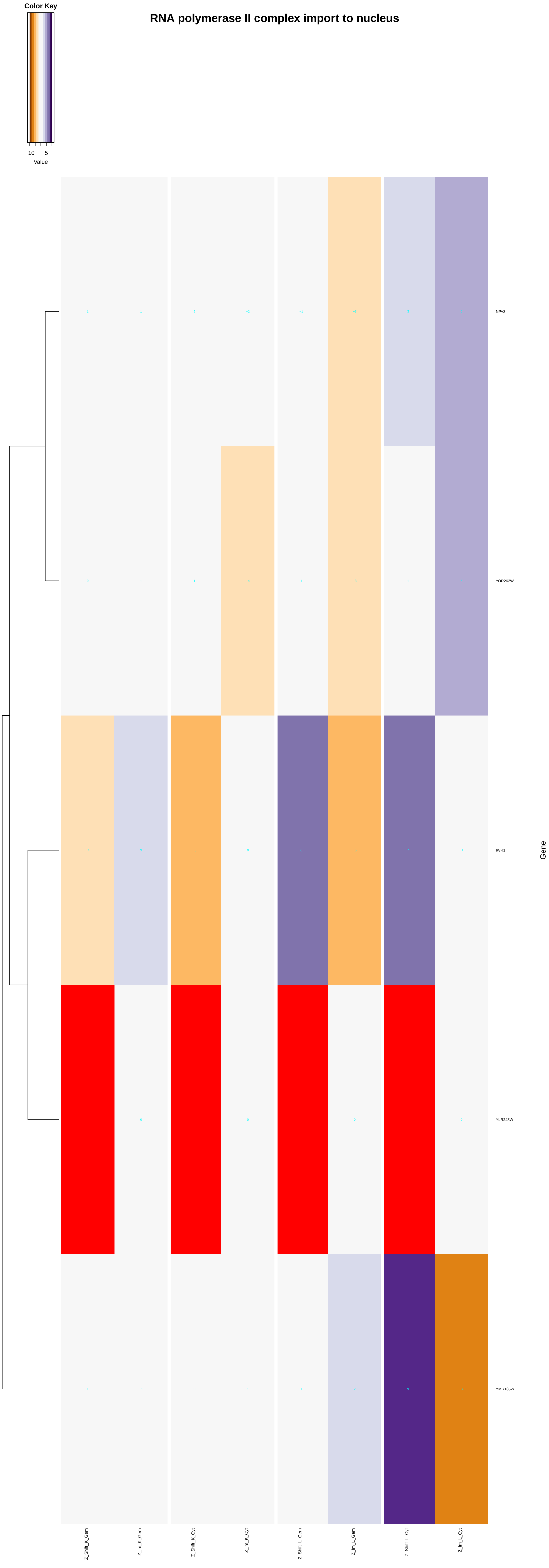

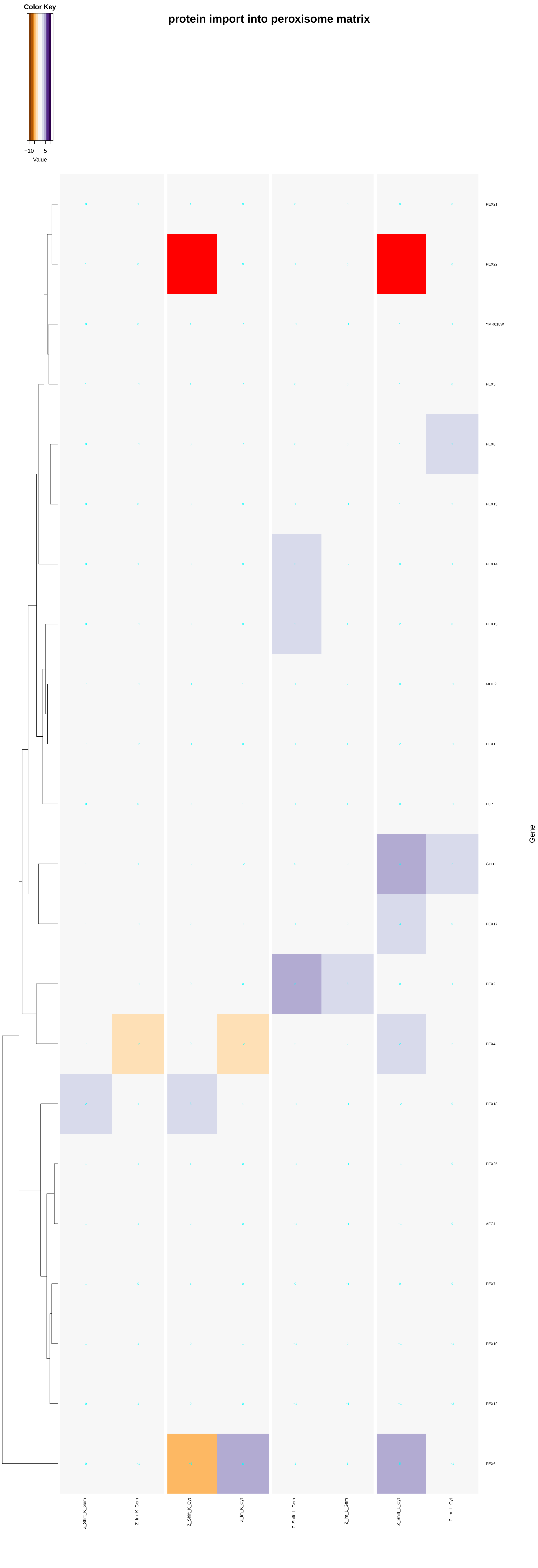

protein import into peroxisome matrix
Color Key
−10
5
Value
PEX21
PEX22
YMR018W
PEX5
PEX8
PEX13
PEX14
PEX15
MDH2
PEX1
DJP1
GPD1
PEX17
PEX2
PEX4
PEX18
PEX25
AFG1
PEX7
PEX10
PEX12
PEX6
Gene
Z_Shift_K_Gem
Z_lm_K_Gem
Z_Shift_K_Cyt
Z_lm_K_Cyt
Z_Shift_L_Gem
Z_lm_L_Gem
Z_Shift_L_Cyt
Z_lm_L_Cyt

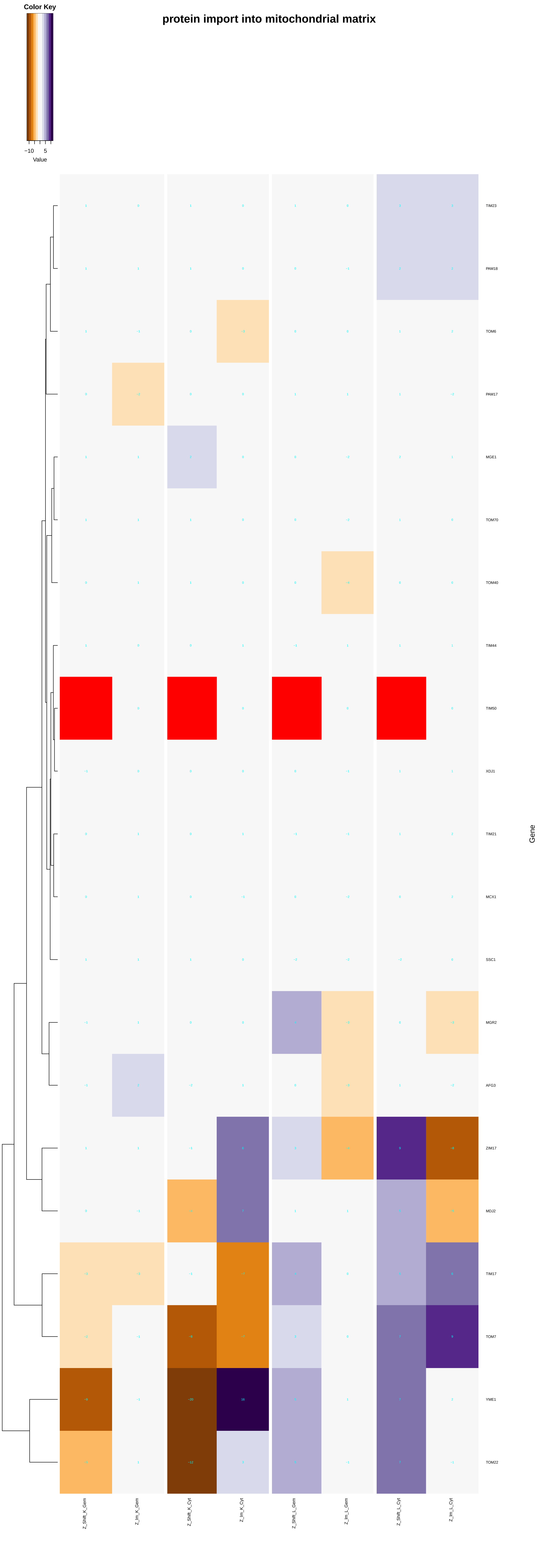

Color Key
−10
5
Value
protein import into mitochondrial matrix
TIM23
PAM18
TOM6
PAM17
MGE1
TOM70
TOM40
TIM44
TIM50
XDJ1
TIM21
MCX1
SSC1
MGR2
AFG3
ZIM17
MDJ2
TIM17
TOM7
YME1
TOM22
Gene
Z_Shift_K_Gem
Z_Im_K_Gem
Z_Shift_K_Cyt
Z_Im_K_Cyt
Z_Shift_L_Gem
Z_Im_L_Gem
Z_Shift_L_Cyt
Z_Im_L_Cyt

### protein import into mitochondrial inner membrane

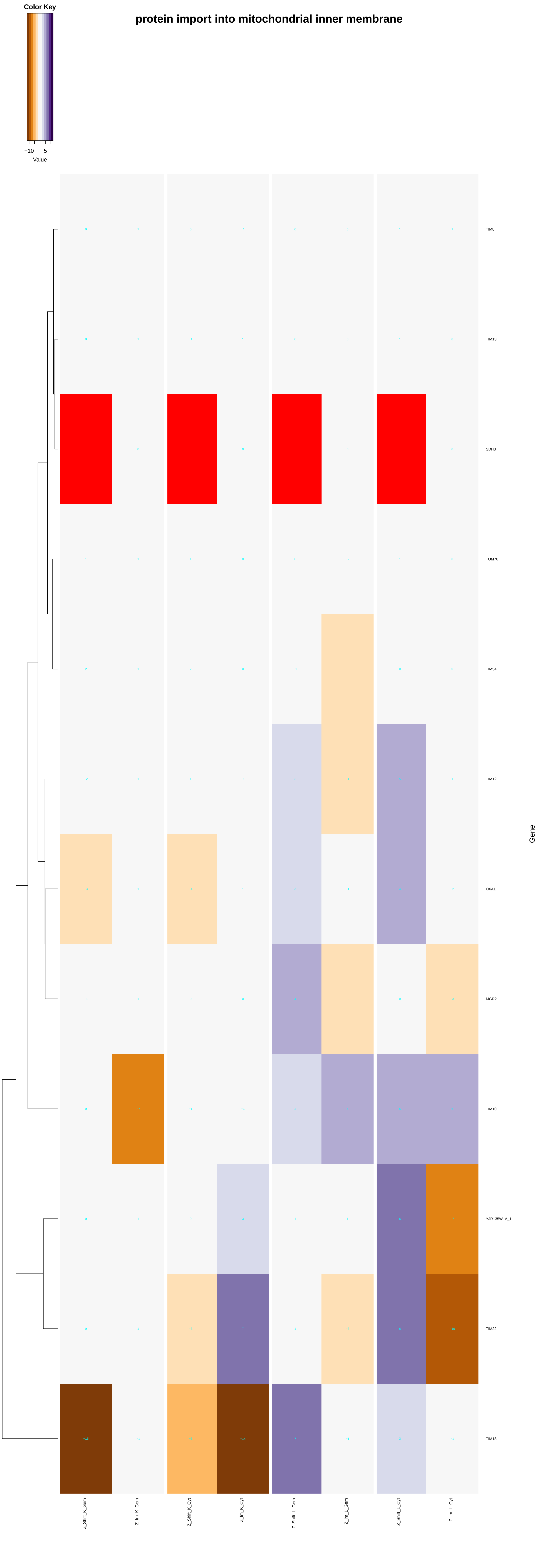

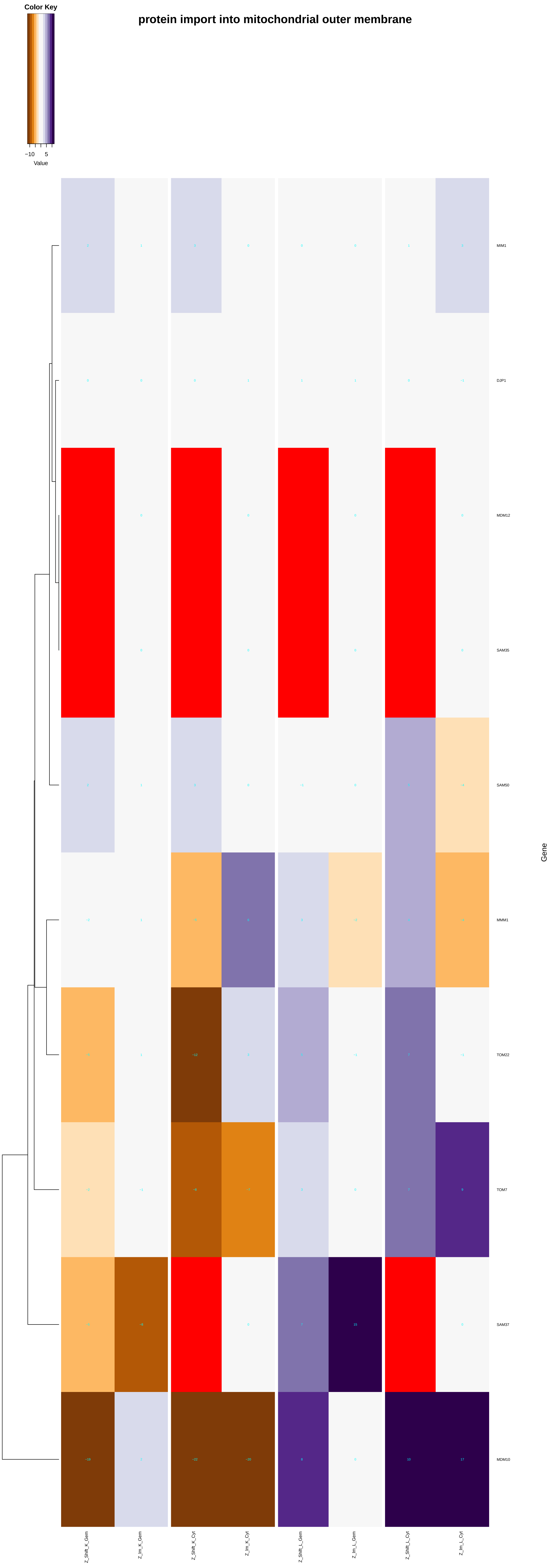
protein import into mitochondrial outer membrane
Color Key
−10
5
Value
MIM1
DJP1
MDM12
SAM35
SAM50
MMM1
TOM22
TOM7
SAM37
MDM10
Gene
Z_Shift_K_Gem
Z_Im_K_Gem
Z_Shift_K_Cyt
Z_Im_K_Cyt
Z_Shift_L_Gem
Z_Im_L_Gem
Z_Shift_L_Cyt
Z_Im_L_Cyt

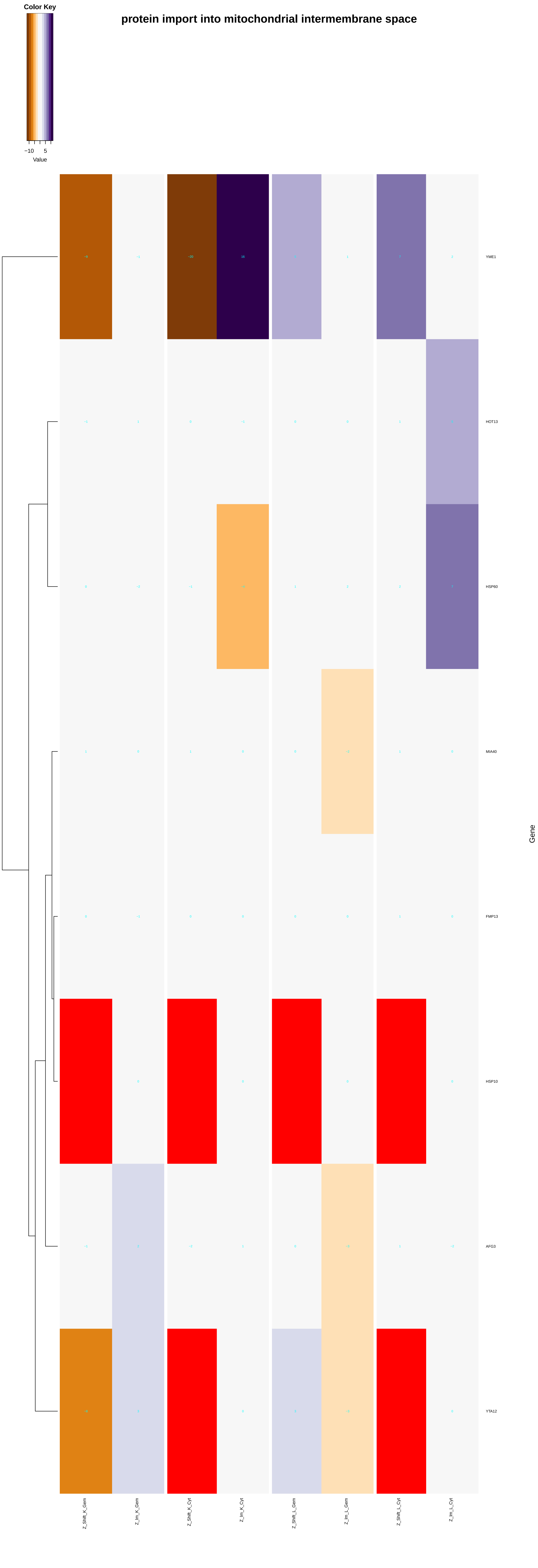
protein import into mitochondrial intermembrane space
Color Key
−10
5
Value
YME1
HOT13
HSP60
MIA40
FMP13
HSP10
AFG3
YTA12
Gene
Z_Shift_K_Gem
Z_Im_K_Gem
Z_Shift_K_Cyt
Z_Im_K_Cyt
Z_Shift_L_Gem
Z_Im_L_Gem
Z_Shift_L_Cyt
Z_Im_L_Cyt

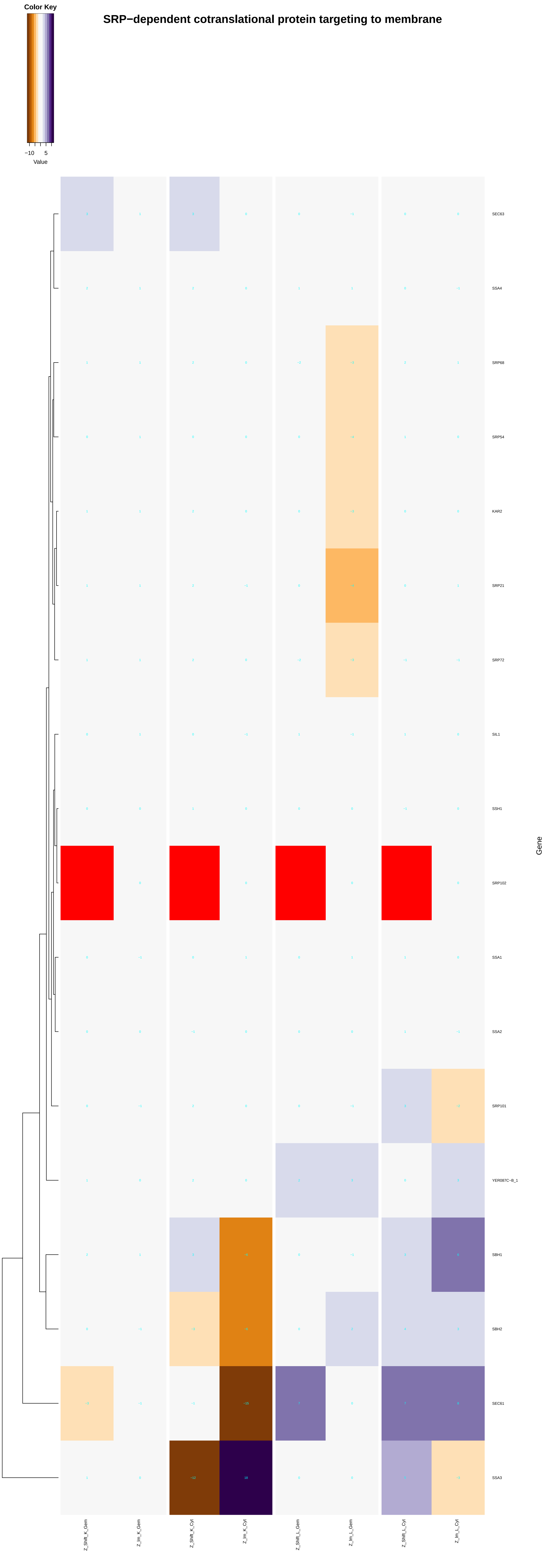

SRP−dependent cotranslational protein targeting to membrane
Color Key
−10
5
Value
SEC63
SSA4
SRP68
SRP54
KAR2
SRP21
SRP72
SIL1
SSH1
SRP102
SSA1
SSA2
SRP101
YER087C−B_1
SBH1
SBH2
SEC61
SSA3
Gene
Z_Shift_K_Gem
Z_Im_K_Gem
Z_Shift_K_Cyt
Z_Im_K_Cyt
Z_Shift_L_Gem
Z_Im_L_Gem
Z_Shift_L_Cyt
Z_Im_L_Cyt

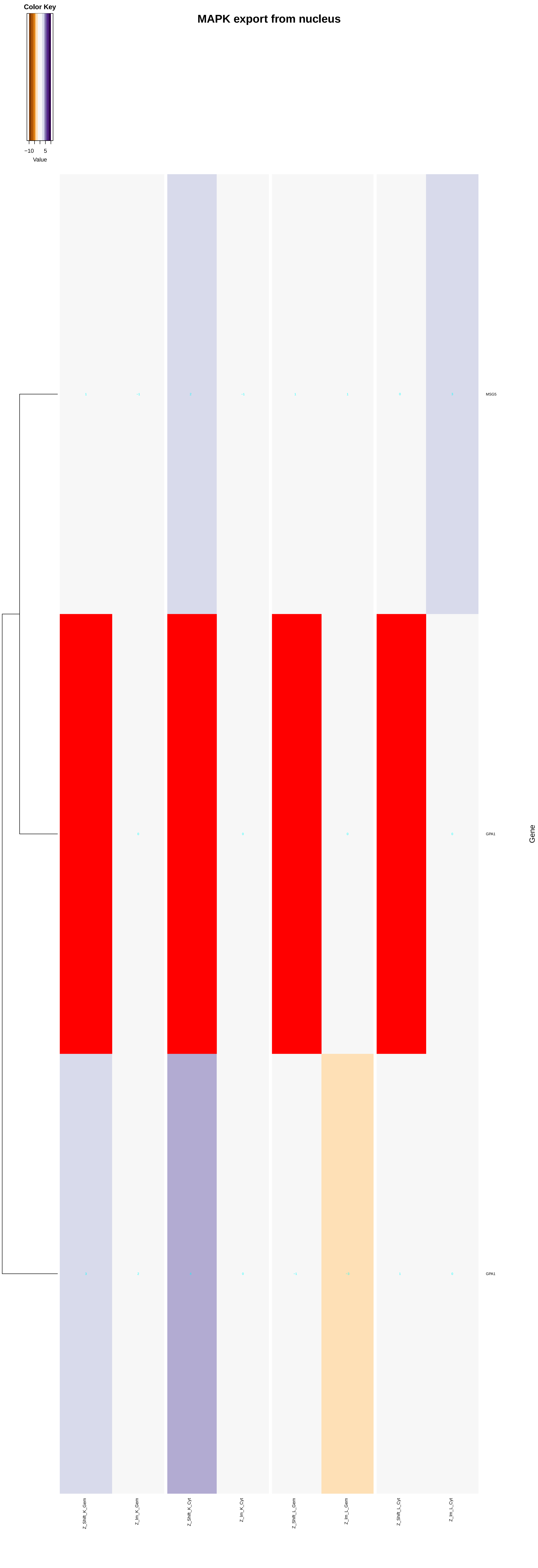

MAPK export from nucleus
Color Key
−10
5
Value
MSG5
GPA1
GPA1
Gene
Z_Shift_K_Gem
Z_Im_K_Gem
Z_Shift_K_Cyt
Z_Im_K_Cyt
Z_Shift_L_Gem
Z_Im_L_Gem
Z_Shift_L_Cyt
Z_Im_L_Cyt

### ribonucleoprotein complex export from nucleus

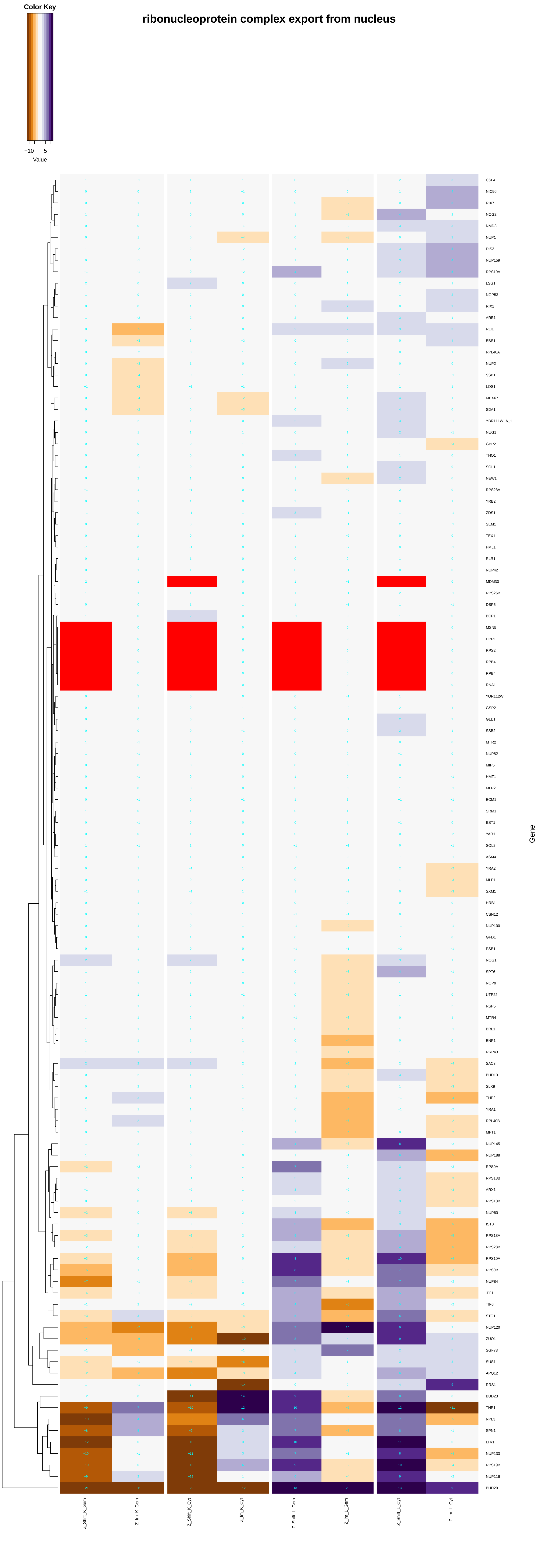

### mRNA−containing ribonucleoprotein complex export from nucleus

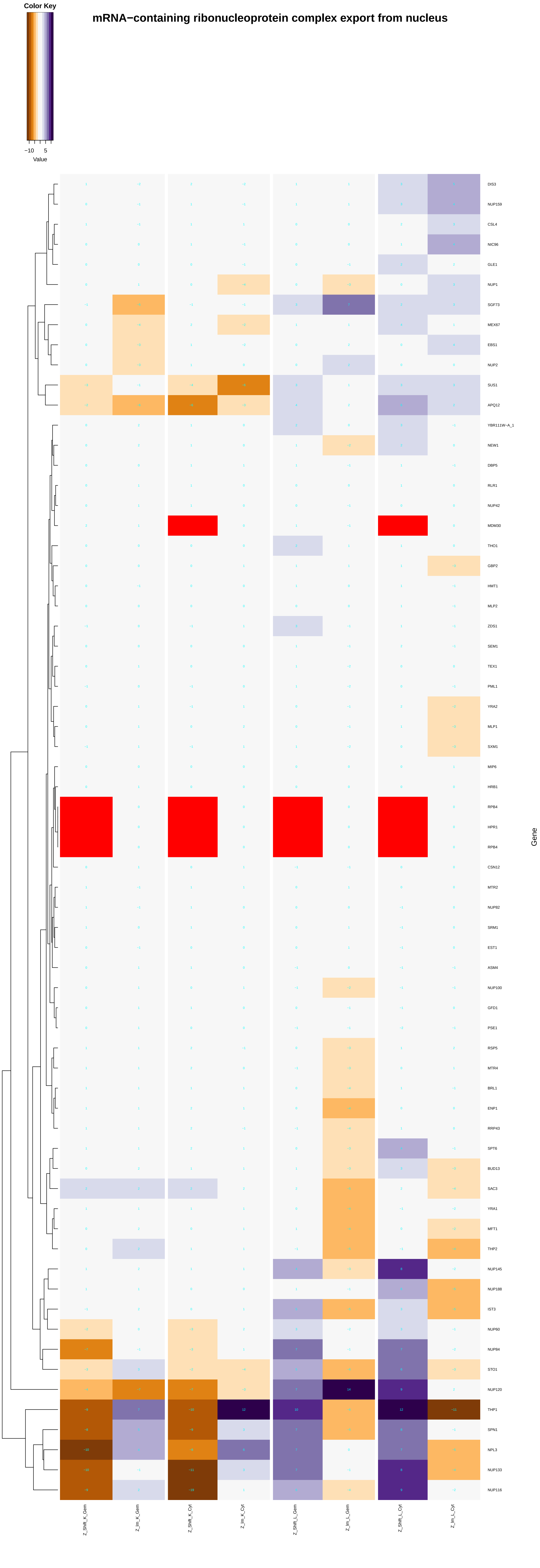

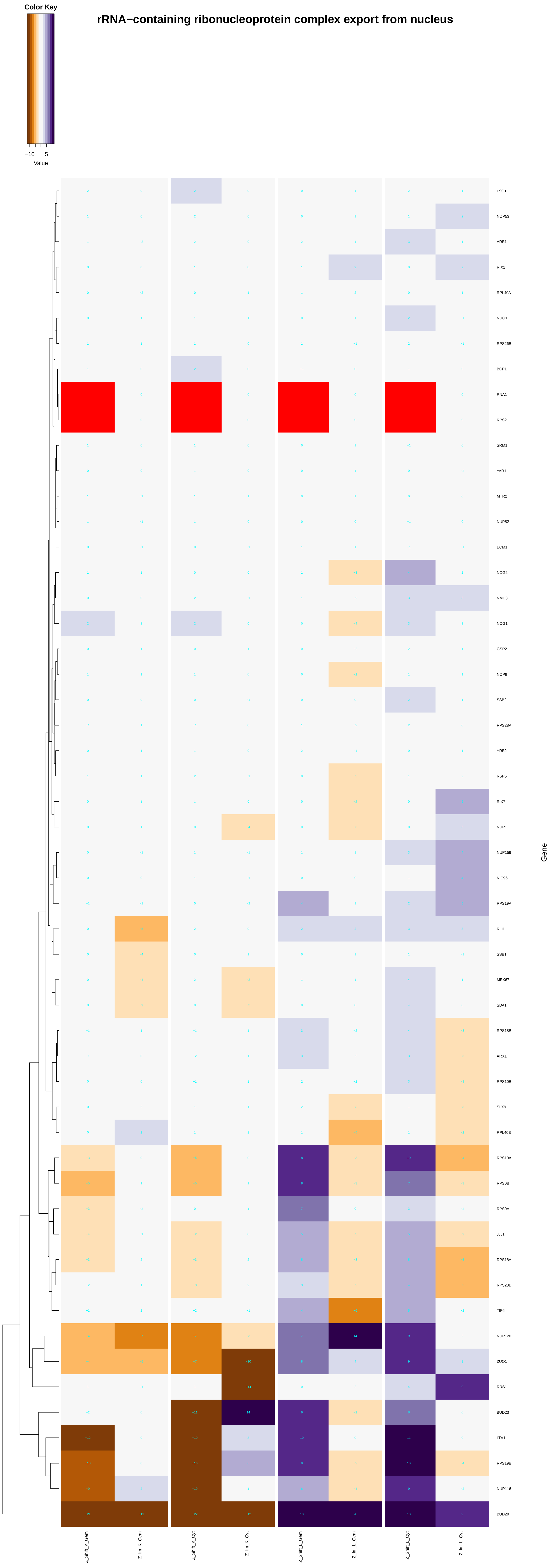

rRNA−containing ribonucleoprotein complex export from nucleus
Color Key
−10
5
Value
LSG1
NOP53
ARB1
RIX1
RPL40A
NUG1
RPS26B
BCP1
RNA1
RPS2
SRM1
YAR1
MTR2
NUP82
ECM1
NOG2
NMD3
NOG1
GSP2
NOP9
SSB2
RPS28A
YRB2
RSP5
RIX7
NUP1
NUP159
NIC96
RPS19A
RLI1
SSB1
MEX67
SDA1
RPS18B
ARX1
RPS10B
SLX9
RPL40B
RPS10A
RPS0B
RPS0A
JJJ1
RPS18A
RPS28B
TIF6
NUP120
ZUO1
RRS1
BUD23
LTV1
RPS19B
NUP116
BUD20
Gene
Z_Shift_K_Gem
Z_Im_K_Gem
Z_Shift_K_Cyt
Z_Im_K_Cyt
Z_Shift_L_Gem
Z_Im_L_Gem
Z_Shift_L_Cyt
Z_Im_L_Cyt

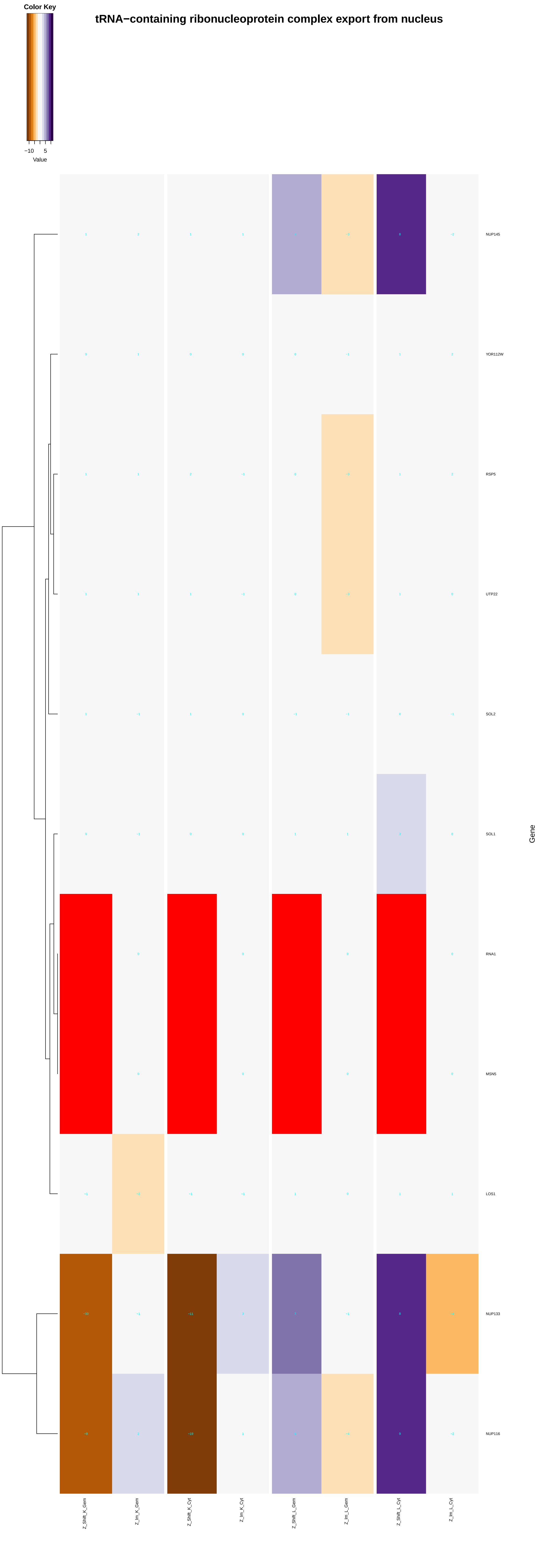
tRNA−containing ribonucleoprotein complex export from nucleus
Color Key
−10
5
Value
NUP145
YOR112W
RSP5
UTP22
SOL2
SOL1
RNA1
MSN5
LOS1
NUP133
NUP116
Gene
Z_Shift_K_Gem
Z_Im_K_Gem
Z_Shift_K_Cyt
Z_Im_K_Cyt
Z_Shift_L_Gem
Z_Im_L_Gem
Z_Shift_L_Cyt
Z_Im_L_Cyt

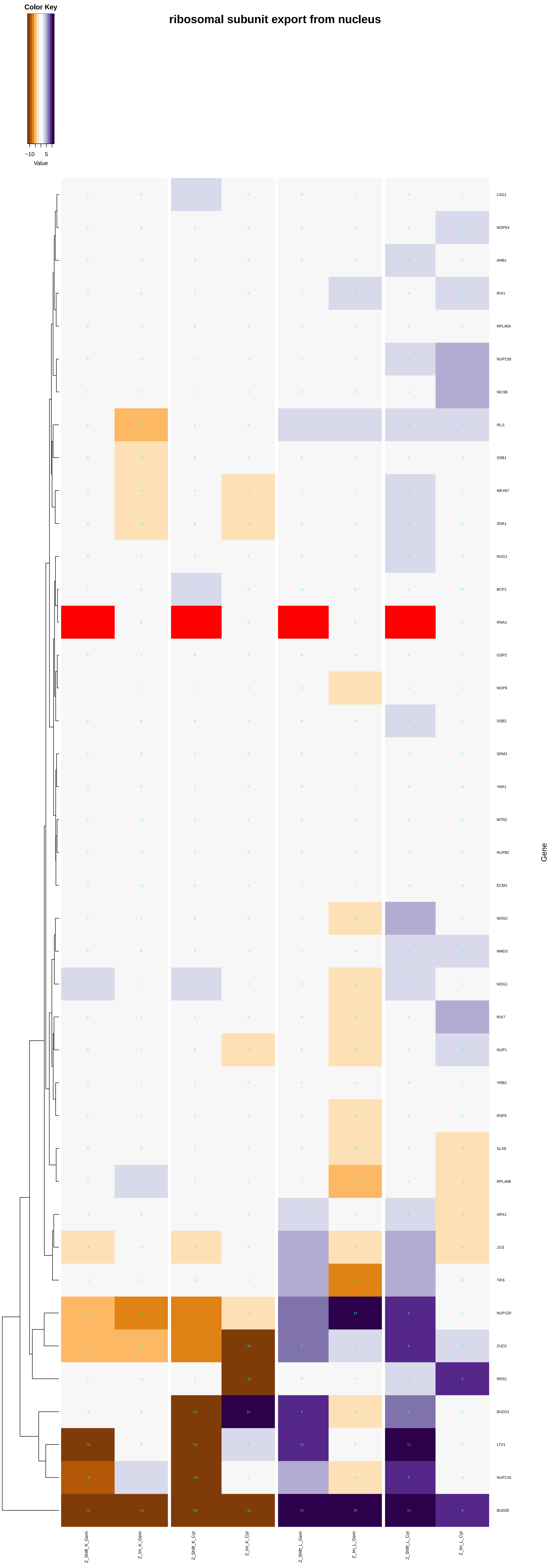
Color Key
−10
5
Value
ribosomal subunit export from nucleus
LSG1
NOP53
ARB1
RIX1
RPL40A
NUP159
NIC96
RLI1
SSB1
MEX67
SDA1
NUG1
BCP1
RNA1
GSP2
NOP9
SSB2
SRM1
YAR1
MTR2
NUP82
ECM1
NOG2
NMD3
NOG1
RIX7
NUP1
YRB2
RSP5
SLX9
RPL40B
ARX1
JJJ1
TIF6
NUP120
ZUO1
RRS1
BUD23
LTV1
NUP116
BUD20
Gene
Z_Shift_K_Gem
Z_Im_K_Gem
Z_Shift_K_Cyt
Z_Im_K_Cyt
Z_Shift_L_Gem
Z_Im_L_Gem
Z_Shift_L_Cyt
Z_Im_L_Cyt

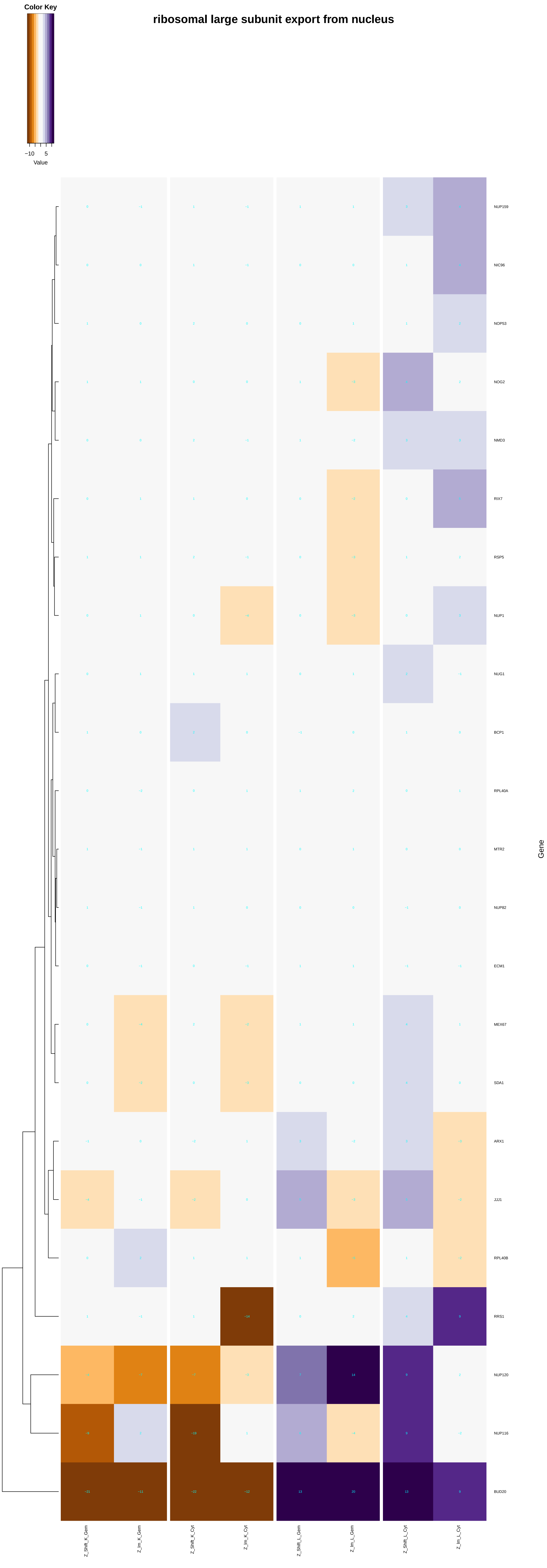
Color Key
−10
5
Value
ribosomal large subunit export from nucleus
NUP159
NIC96
NOP53
NOG2
NMD3
RIX7
RSP5
NUP1
NUG1
BCP1
RPL40A
MTR2
NUP82
ECM1
MEX67
SDA1
ARX1
JJJ1
RPL40B
RRS1
NUP120
NUP116
BUD20
Gene
Z_Shift_K_Gem
Z_Im_K_Gem
Z_Shift_K_Cyt
Z_Im_K_Cyt
Z_Shift_L_Gem
Z_Im_L_Gem
Z_Shift_L_Cyt
Z_Im_L_Cyt

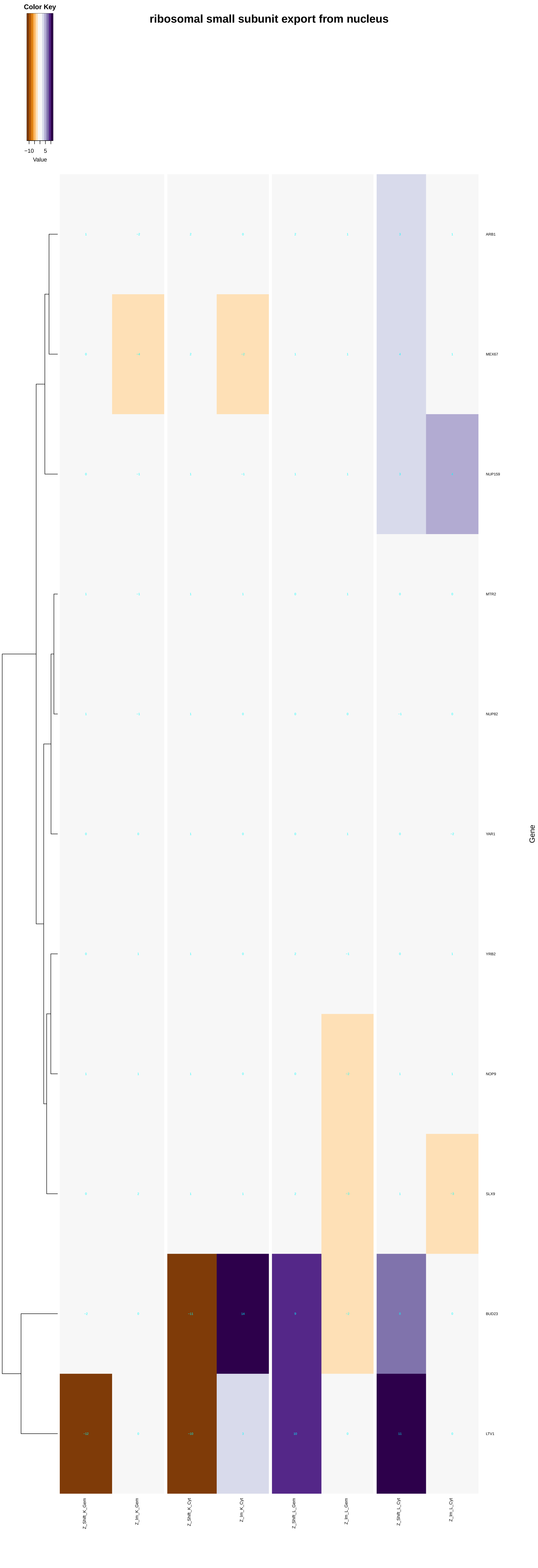
ribosomal small subunit export from nucleus
Color Key
−10
5
Value
ARB1
MEX67
NUP159
MTR2
NUP82
YAR1
YRB2
NOP9
SLX9
BUD23
LTV1
Gene
Z_Shift_K_Gem
Z_Im_K_Gem
Z_Shift_K_Cyt
Z_Im_K_Cyt
Z_Shift_L_Gem
Z_Im_L_Gem
Z_Shift_L_Cyt
Z_Im_L_Cyt
