## Additional File 7 for "A humanized yeast phenomic model of deoxycytidine kinase to predict genetic buffering of nucleoside analog cytotoxicity": gene_silencing.pdf

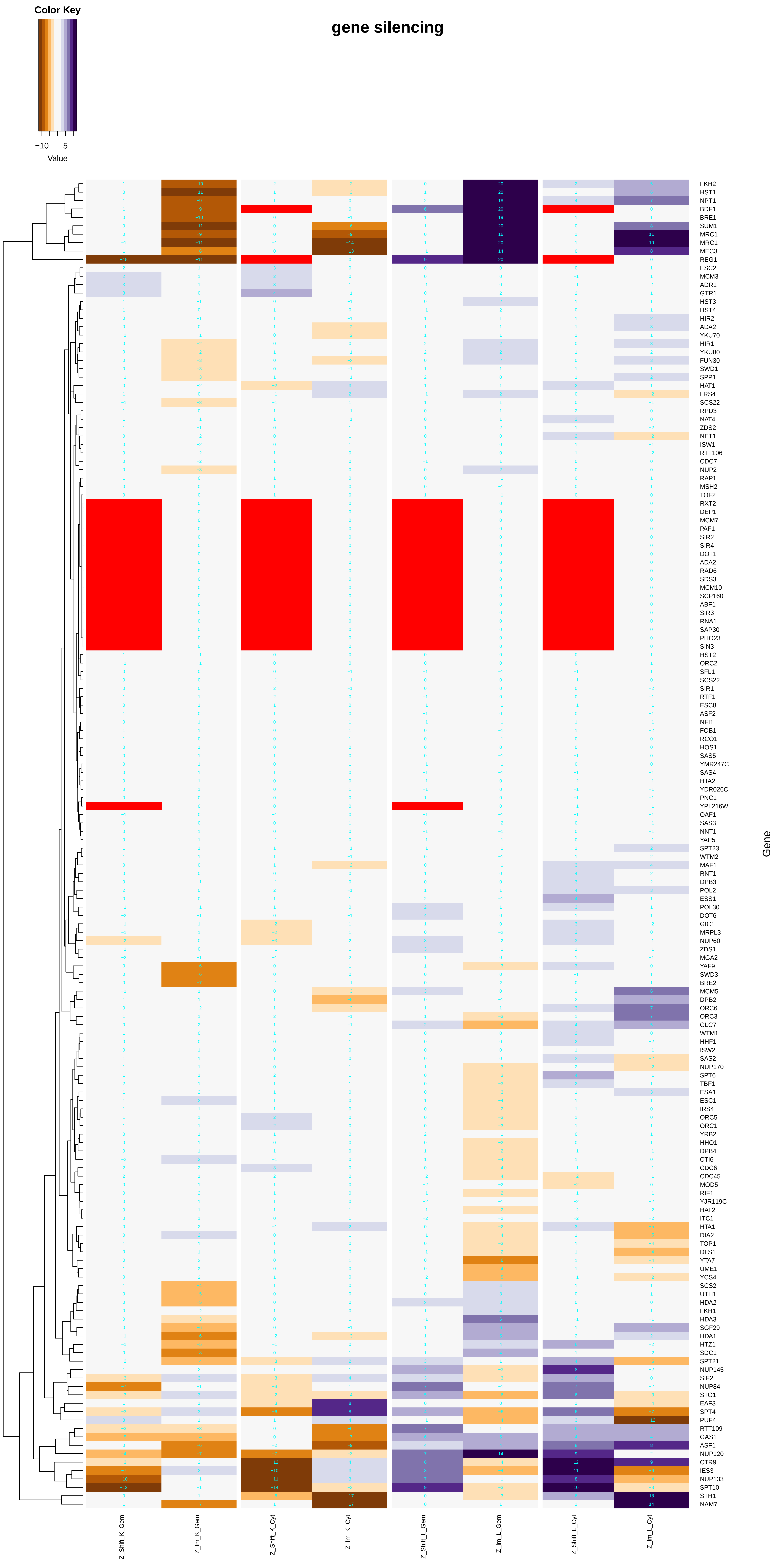

gene silencing
Color Key
−10
5
Value
FKH2
HST1
NPT1
BDF1
BRE1
SUM1
MRC1
MRC1
MEC3
REG1
ESC2
MCM3
ADR1
GTR1
HST3
HST4
HIR2
ADA2
YKU70
HIR1
YKU80
FUN30
SWD1
SPP1
HAT1
LRS4
SCS22
RPD3
NAT4
ZDS2
NET1
ISW1
RTT106
CDC7
NUP2
RAP1
MSH2
TOF2
RXT2
DEP1
MCM7
PAF1
SIR2
SIR4
DOT1
ADA2
RAD6
SDS3
MCM10
SCP160
ABF1
SIR3
RNA1
SAP30
PHO23
SIN3
HST2
ORC2
SFL1
SCS22
SIR1
RTF1
ESC8
ASF2
NFI1
FOB1
RCO1
HOS1
SAS5
YMR247C
SAS4
HTA2
YDR026C
PNC1
YPL216W
OAF1
SAS3
NNT1
YAP5
SPT23
WTM2
MAF1
RNT1
DPB3
POL2
ESS1
POL30
DOT6
GIC1
MRPL3
NUP60
ZDS1
MGA2
YAF9
SWD3
BRE2
MCM5
DPB2
ORC6
ORC3
GLC7
WTM1
HHF1
ISW2
SAS2
NUP170
SPT6
TBF1
ESA1
ESC1
IRS4
ORC5
ORC1
YRB2
HHO1
DPB4
CTI6
CDC6
CDC45
MOD5
RIF1
YJR119C
HAT2
ITC1
HTA1
DIA2
TOP1
DLS1
YTA7
UME1
YCS4
SCS2
UTH1
HDA2
FKH1
HDA3
SGF29
HDA1
HTZ1
SDC1
SPT21
NUP145
SIF2
NUP84
STO1
EAF3
SPT4
PUF4
RTT109
GAS1
ASF1
NUP120
CTR9
IES3
NUP133
SPT10
STH1
NAM7
Gene
Z_Shift_K_Gem
Z_Im_K_Gem
Z_Shift_K_Cyt
Z_Im_K_Cyt
Z_Shift_L_Gem
Z_Im_L_Gem
Z_Shift_L_Cyt
Z_Im_L_Cyt

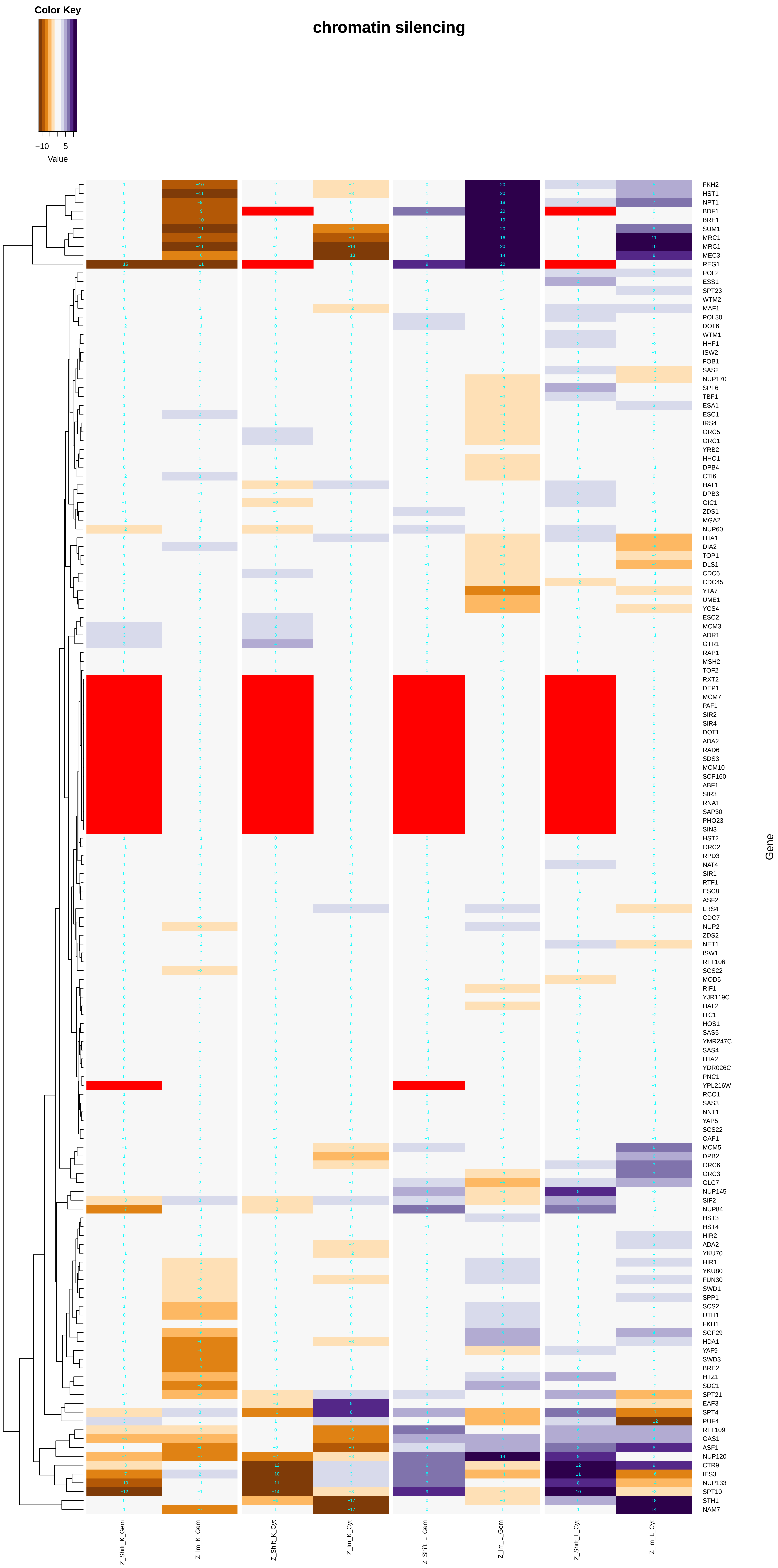

chromatin silencing
Color Key
−10
5
Value
FKH2
HST1
NPT1
BDF1
BRE1
SUM1
MRC1
MRC1
MEC3
REG1
POL2
ESS1
SPT23
WTM2
MAF1
POL30
DOT6
WTM1
HHF1
ISW2
FOB1
SAS2
NUP170
SPT6
TBF1
ESA1
ESC1
IRS4
ORC5
ORC1
YRB2
HHO1
DPB4
CTI6
HAT1
DPB3
GIC1
ZDS1
MGA2
NUP60
HTA1
DIA2
TOP1
DLS1
CDC6
CDC45
YTA7
UME1
YCS4
ESC2
MCM3
ADR1
GTR1
RAP1
MSH2
TOF2
RXT2
DEP1
MCM7
PAF1
SIR2
SIR4
DOT1
ADA2
RAD6
SDS3
MCM10
SCP160
ABF1
SIR3
RNA1
SAP30
PHO23
SIN3
HST2
ORC2
RPD3
NAT4
SIR1
RTF1
ESC8
ASF2
LRS4
CDC7
NUP2
ZDS2
NET1
ISW1
RTT106
SCS22
MOD5
RIF1
YJR119C
HAT2
ITC1
HOS1
SAS5
YMR247C
SAS4
HTA2
YDR026C
PNC1
YPL216W
RCO1
SAS3
NNT1
YAP5
SCS22
OAF1
MCM5
DPB2
ORC6
ORC3
GLC7
NUP145
SIF2
NUP84
HST3
HST4
HIR2
ADA2
YKU70
HIR1
YKU80
FUN30
SWD1
SPP1
SCS2
UTH1
FKH1
SGF29
HDA1
YAF9
SWD3
BRE2
HTZ1
SDC1
SPT21
EAF3
SPT4
PUF4
RTT109
GAS1
ASF1
NUP120
CTR9
IES3
NUP133
SPT10
STH1
NAM7
Gene
Z_Shift_K_Gem
Z_Im_K_Gem
Z_Shift_K_Cyt
Z_Im_K_Cyt
Z_Shift_L_Gem
Z_Im_L_Gem
Z_Shift_L_Cyt
Z_Im_L_Cyt

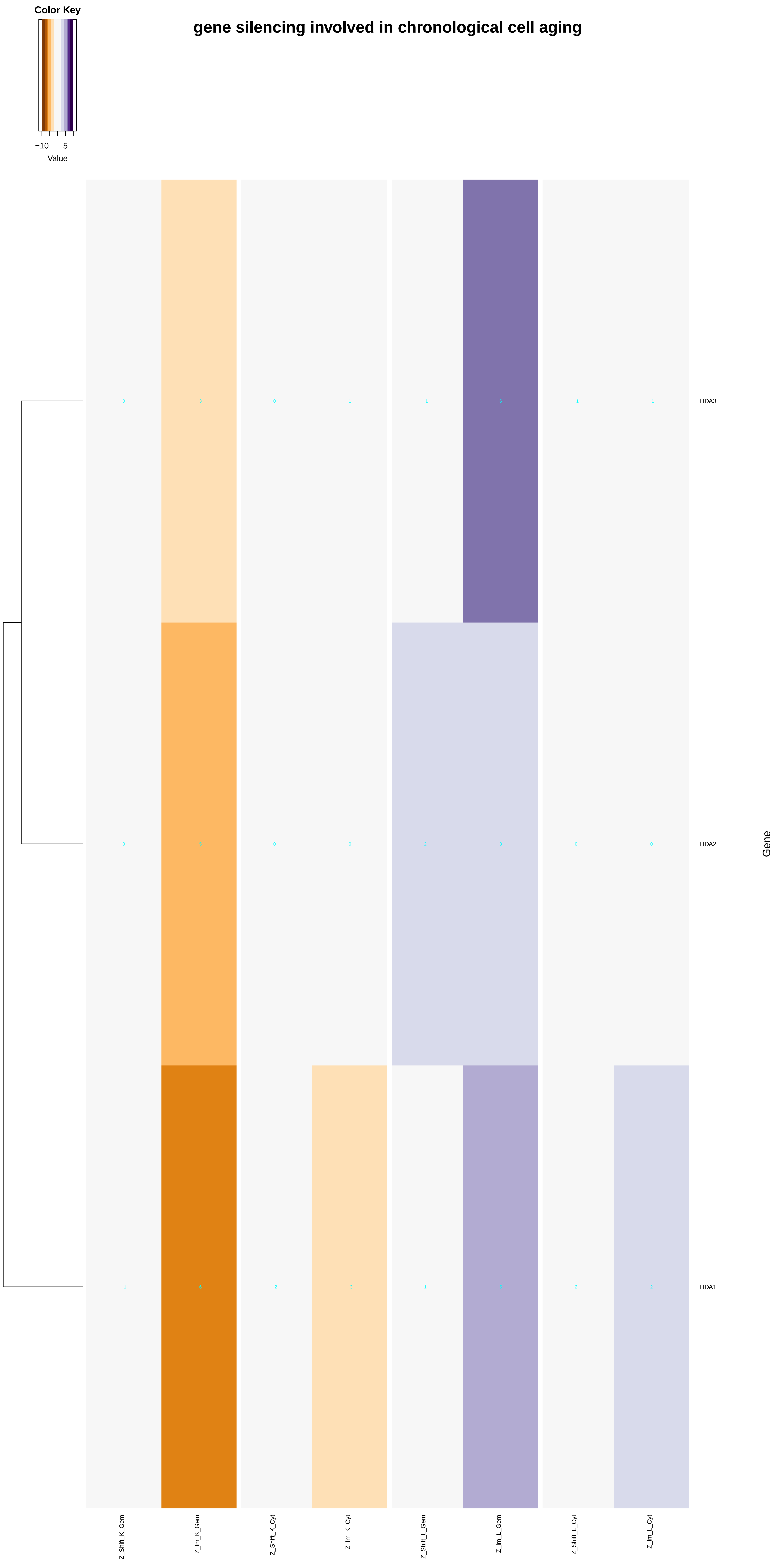
Color Key
−10
5
Value
gene silencing involved in chronological cell aging
HDA3
HDA2
HDA1
Gene
Z_Shift_K_Gem
Z_Im_K_Gem
Z_Shift_K_Cyt
Z_Im_K_Cyt
Z_Shift_L_Gem
Z_Im_L_Gem
Z_Shift_L_Cyt
Z_Im_L_Cyt

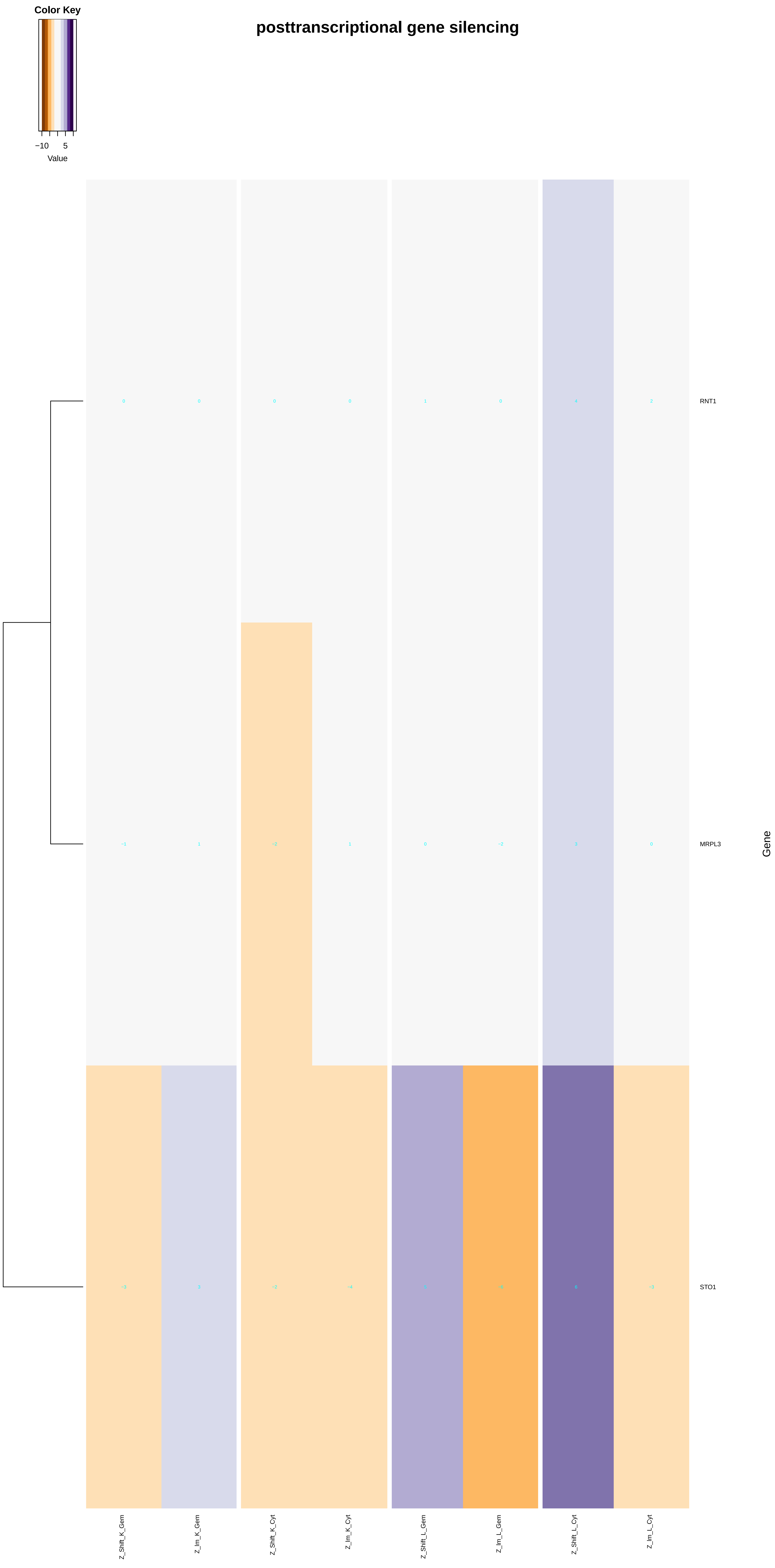
posttranscriptional gene silencing
Color Key
−10
5
Value
RNT1
MRPL3
STO1
Gene
Z_Shift_K_Gem
Z_Im_K_Gem
Z_Shift_K_Cyt
Z_Im_K_Cyt
Z_Shift_L_Gem
Z_Im_L_Gem
Z_Shift_L_Cyt
Z_Im_L_Cyt

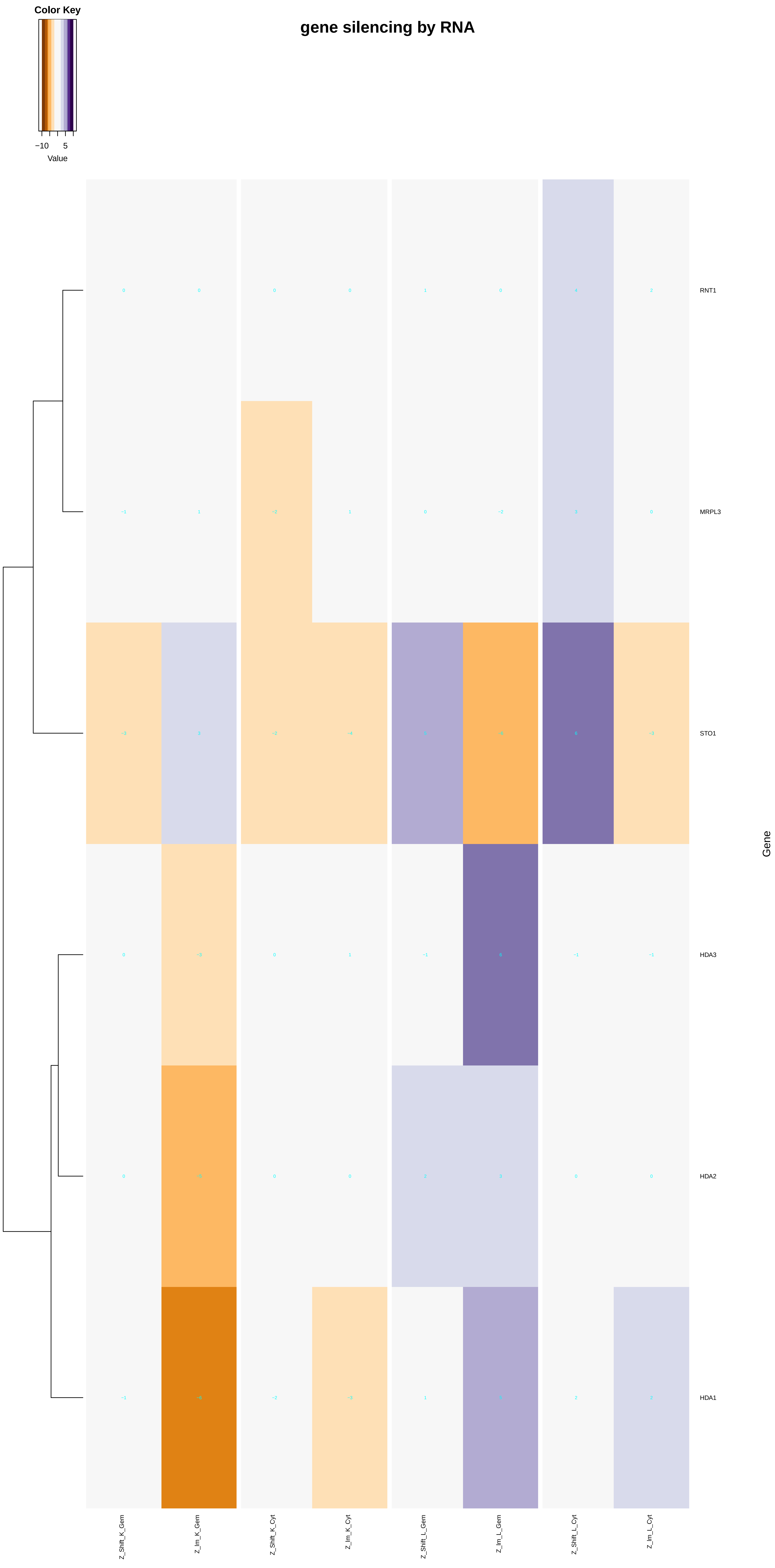
Color Key
−10
5
Value
gene silencing by RNA
RNT1
MRPL3
STO1
HDA3
HDA2
HDA1
Gene
Z_Shift_K_Gem
Z_Im_K_Gem
Z_Shift_K_Cyt
Z_Im_K_Cyt
Z_Shift_L_Gem
Z_Im_L_Gem
Z_Shift_L_Cyt
Z_Im_L_Cyt

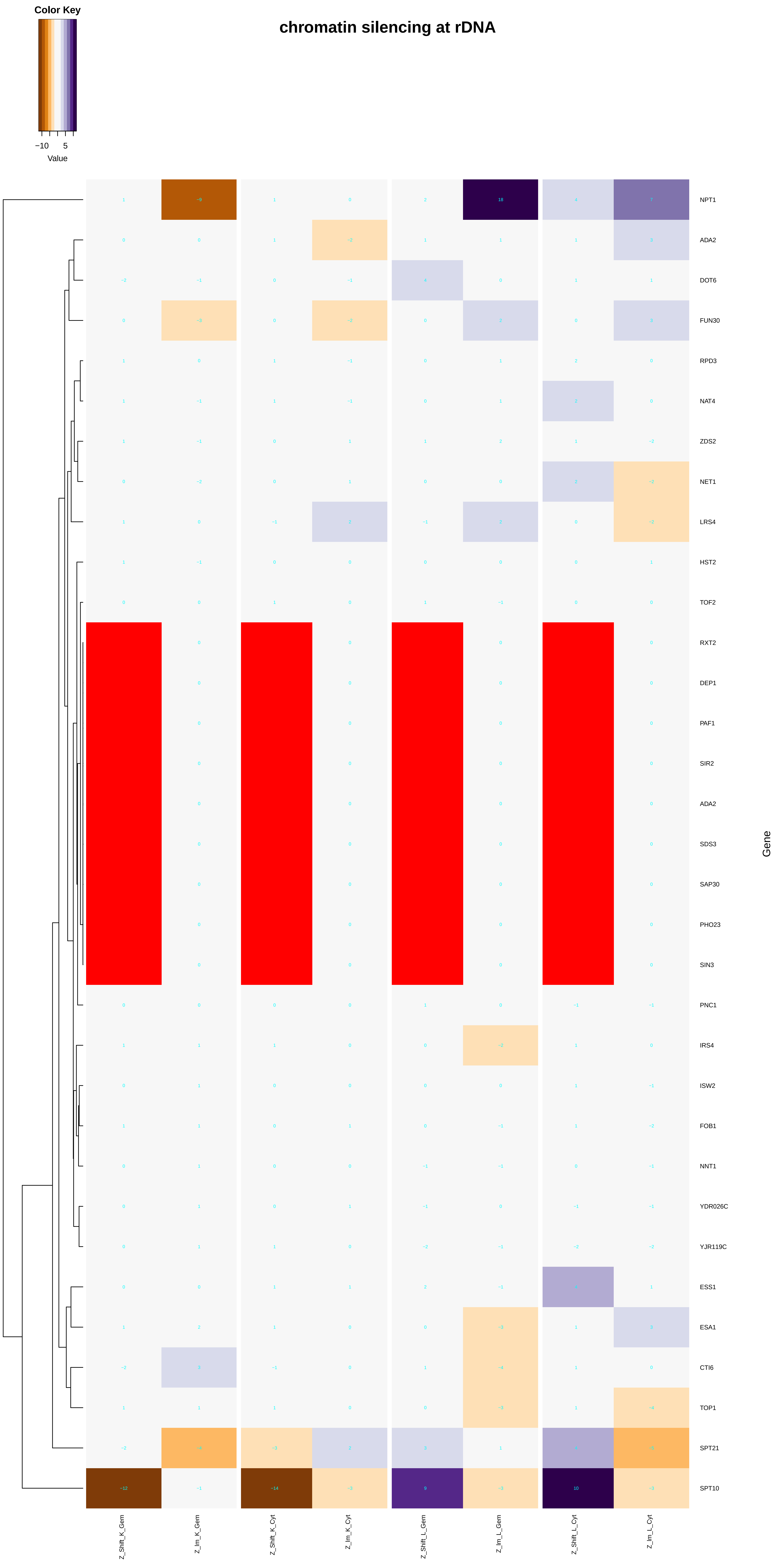

Color Key
−10
5
Value
chromatin silencing at rDNA
NPT1
ADA2
DOT6
FUN30
RPD3
NAT4
ZDS2
NET1
LRS4
HST2
TOF2
RXT2
DEP1
PAF1
SIR2
ADA2
SDS3
SAP30
PHO23
SIN3
PNC1
IRS4
ISW2
FOB1
NNT1
YDR026C
YJR119C
ESS1
ESA1
CTI6
TOP1
SPT21
SPT10
Gene
Z_Shift_K_Gem
Z_Im_K_Gem
Z_Shift_K_Cyt
Z_Im_K_Cyt
Z_Shift_L_Gem
Z_Im_L_Gem
Z_Shift_L_Cyt
Z_Im_L_Cyt

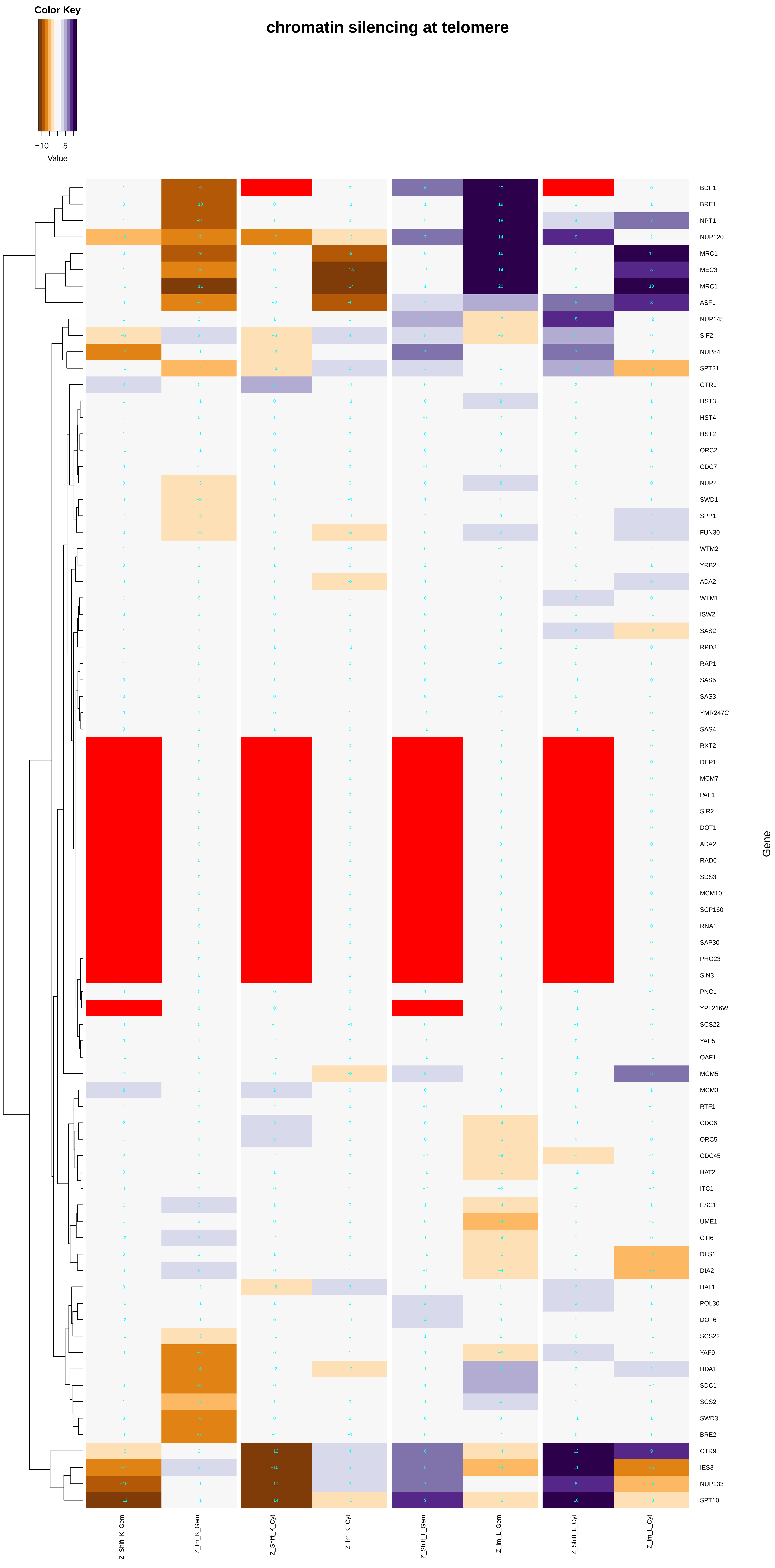

chromatin silencing at telomere
Color Key
−10
5
Value
Gene
BDF1
BRE1
NPT1
NUP120
MRC1
MEC3
MRC1
ASF1
NUP145
SIF2
NUP84
SPT21
GTR1
HST3
HST4
HST2
ORC2
CDC7
NUP2
SWD1
SPP1
FUN30
WTM2
YRB2
ADA2
WTM1
ISW2
SAS2
RPD3
RAP1
SAS5
SAS3
YMR247C
SAS4
RXT2
DEP1
MCM7
PAF1
SIR2
DOT1
ADA2
RAD6
SDS3
MCM10
SCP160
RNA1
SAP30
PHO23
SIN3
PNC1
YPL216W
SCS22
YAP5
OAF1
MCM5
MCM3
RTF1
CDC6
ORC5
CDC45
HAT2
ITC1
ESC1
UME1
CTI6
DLS1
DIA2
HAT1
POL30
DOT6
SCS22
YAF9
HDA1
SDC1
SCS2
SWD3
BRE2
CTR9
IES3
NUP133
SPT10
Z_Shift_K_Gem
Z_Im_K_Gem
Z_Shift_K_Cyt
Z_Im_K_Cyt
Z_Shift_L_Gem
Z_Im_L_Gem
Z_Shift_L_Cyt
Z_Im_L_Cyt

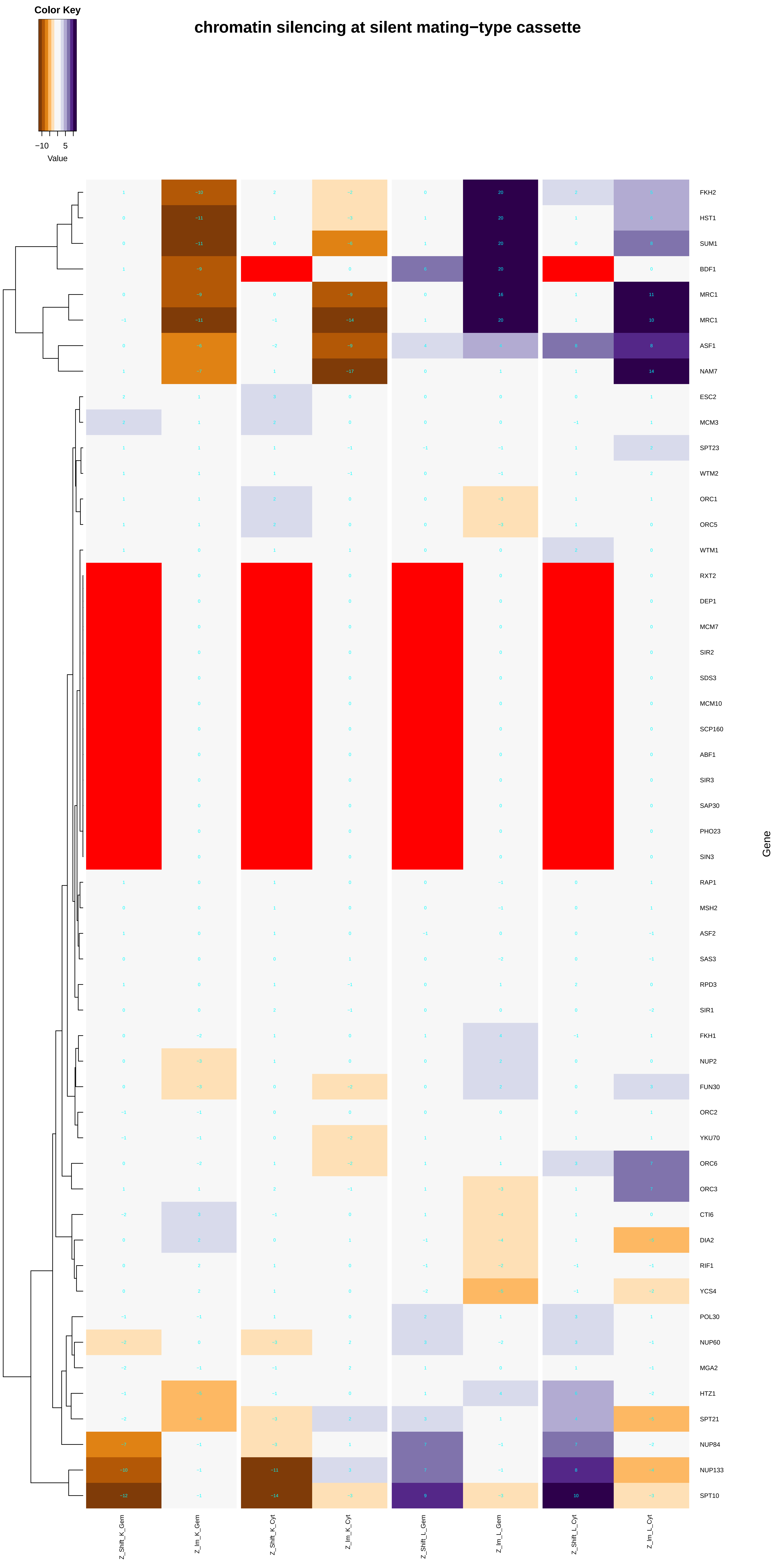
chromatin silencing at silent mating−type cassette
Color Key
−10
5
Value
FKH2
HST1
SUM1
BDF1
MRC1
MRC1
ASF1
NAM7
ESC2
MCM3
SPT23
WTM2
ORC1
ORC5
WTM1
RXT2
DEP1
MCM7
SIR2
SDS3
MCM10
SCP160
ABF1
SIR3
SAP30
PHO23
SIN3
RAP1
MSH2
ASF2
SAS3
RPD3
SIR1
FKH1
NUP2
FUN30
ORC2
YKU70
ORC6
ORC3
CTI6
DIA2
RIF1
YCS4
POL30
NUP60
MGA2
HTZ1
SPT21
NUP84
NUP133
SPT10
Gene
Z_Shift_K_Gem
Z_Im_K_Gem
Z_Shift_K_Cyt
Z_Im_K_Cyt
Z_Shift_L_Gem
Z_Im_L_Gem
Z_Shift_L_Cyt
Z_Im_L_Cyt

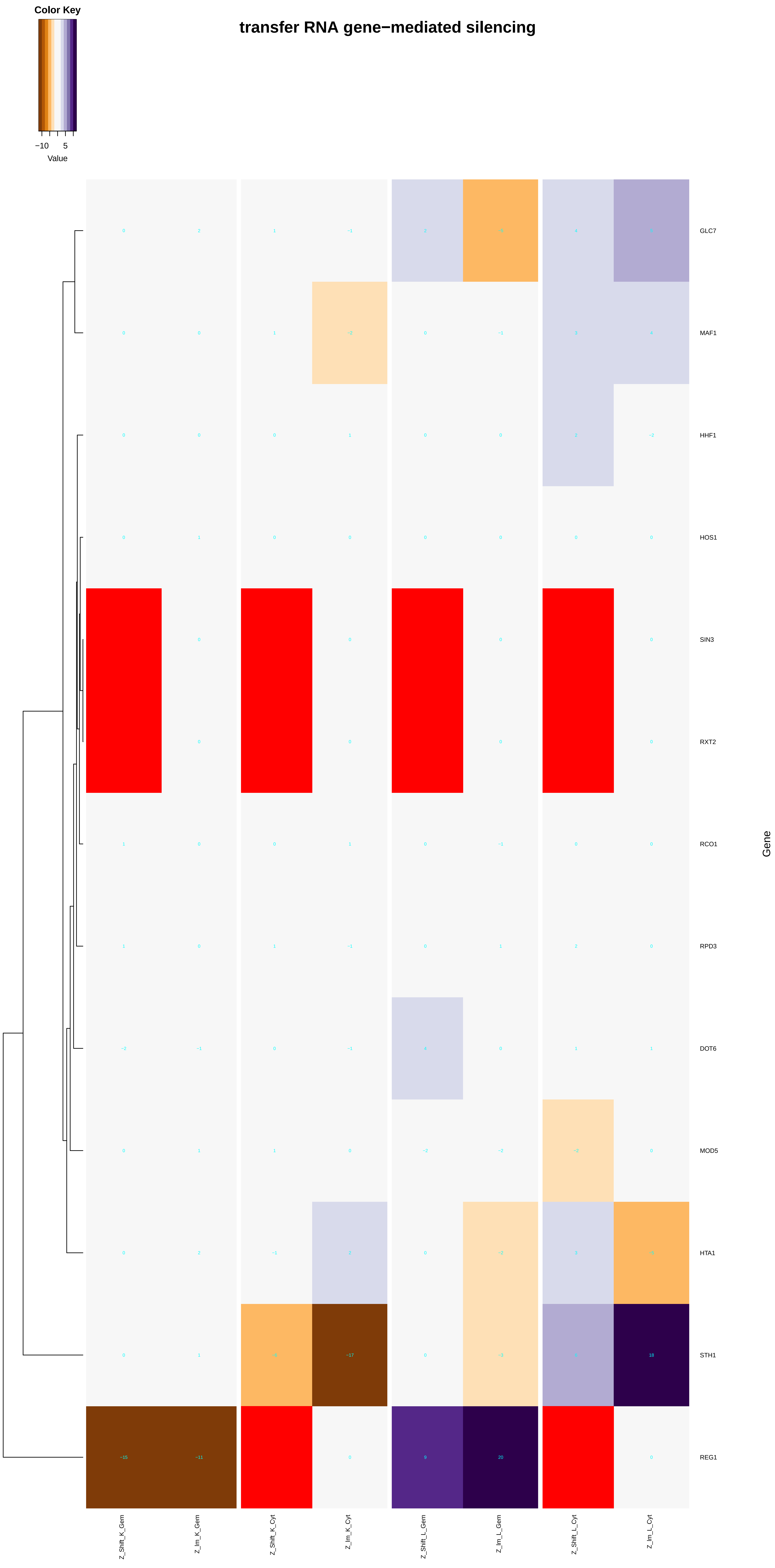
Color Key
−10 5
Value
transfer RNA gene−mediated silencing
GLC7
MAF1
HHF1
HOS1
SIN3
RXT2
RCO1
RPD3
DOT6
MOD5
HTA1
STH1
REG1
Gene
Z_Shift_K_Gem
Z_lm_K_Gem
Z_Shift_K_Cyt
Z_lm_K_Cyt
Z_Shift_L_Gem
Z_lm_L_Gem
Z_Shift_L_Cyt
Z_lm_L_Cyt

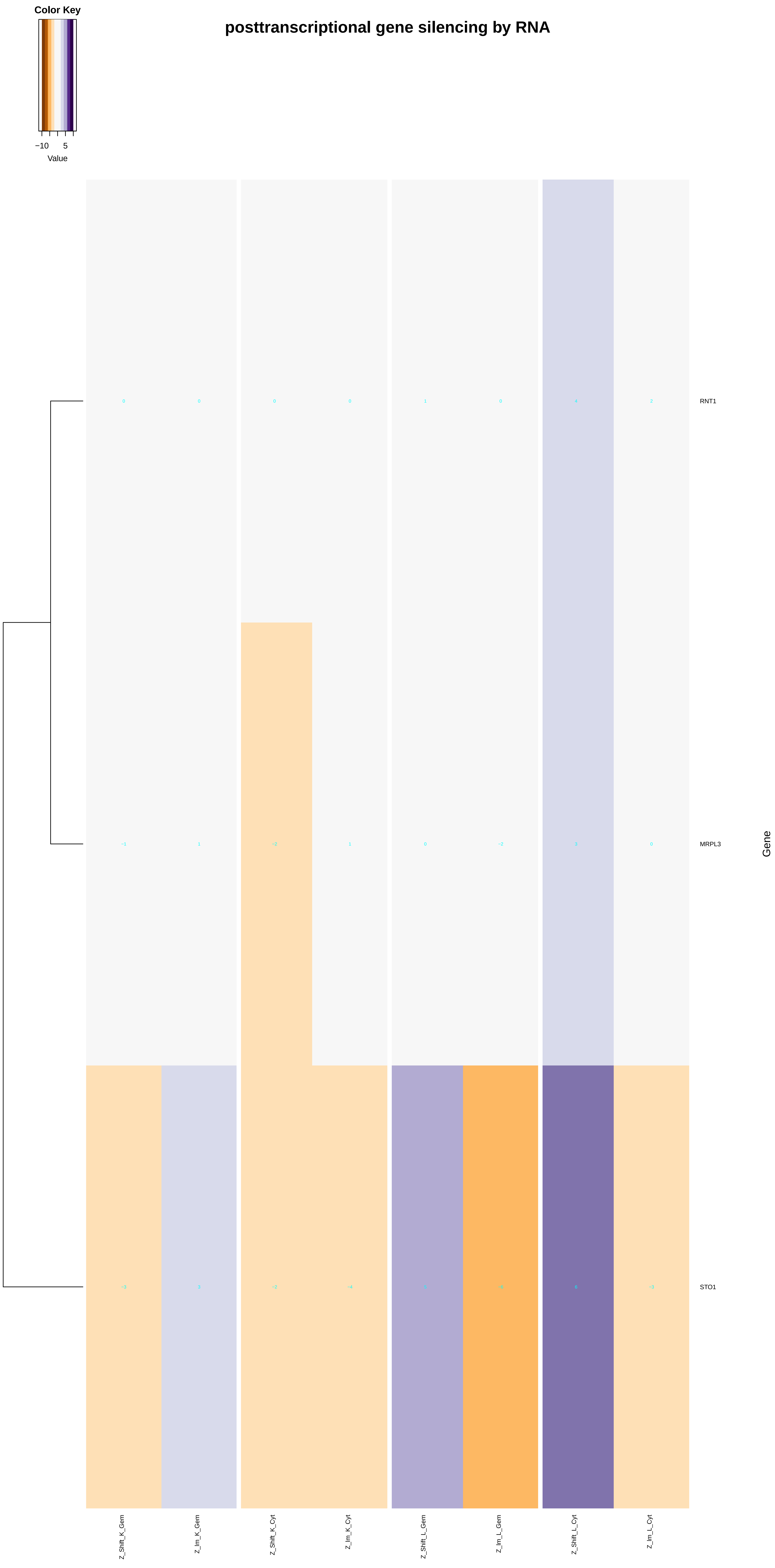

Color Key
−10
5
Value
posttranscriptional gene silencing by RNA
RNT1
MRPL3
STO1
Gene
Z_Shift_K_Gem
Z_Im_K_Gem
Z_Shift_K_Cyt
Z_Im_K_Cyt
Z_Shift_L_Gem
Z_Im_L_Gem
Z_Shift_L_Cyt
Z_Im_L_Cyt

### gene silencing by miRNA

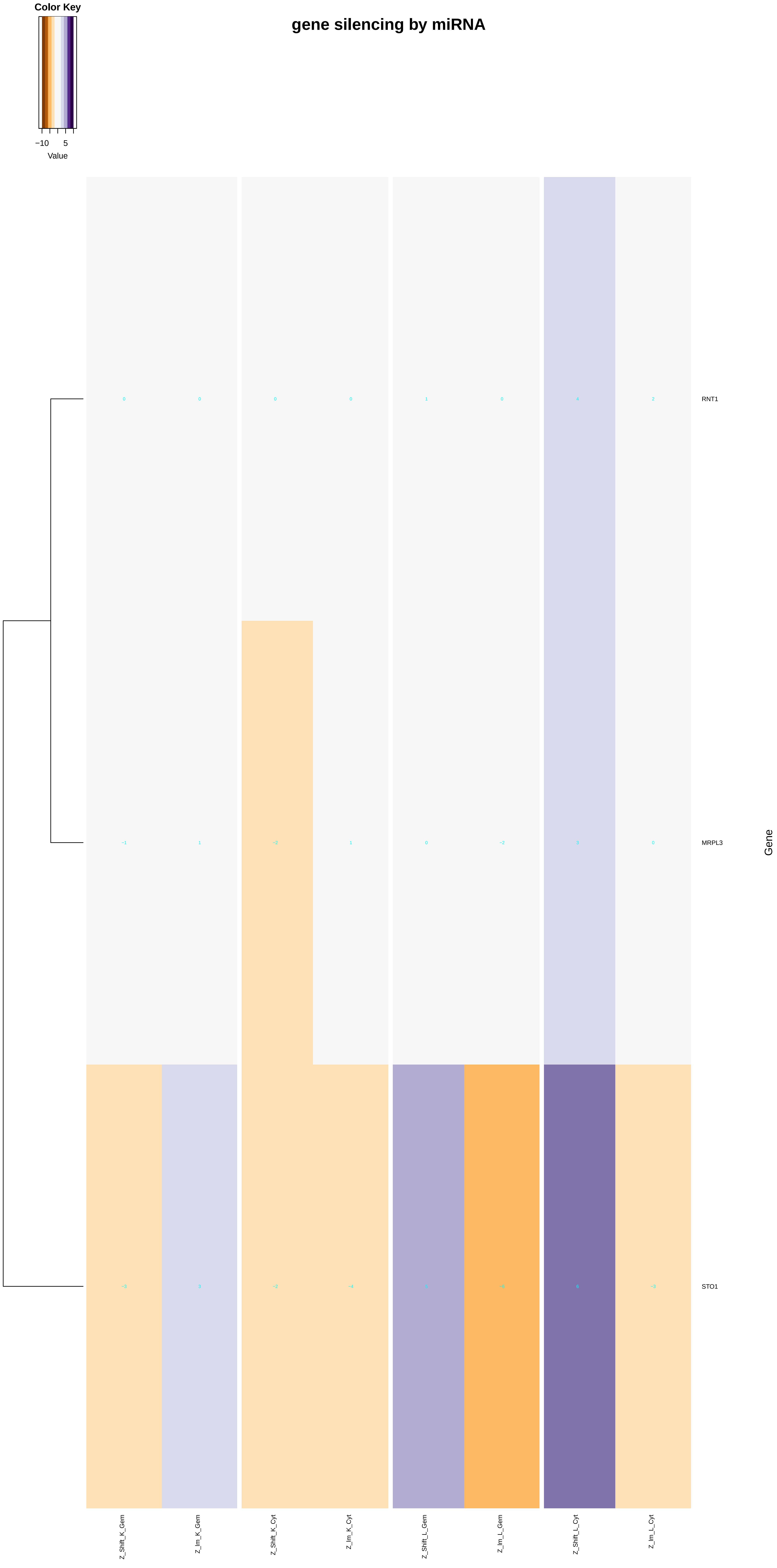
