## Additional File 7 for "A humanized yeast phenomic model of deoxycytidine kinase to predict genetic buffering of nucleoside analog cytotoxicity": helicase_activity.pdf

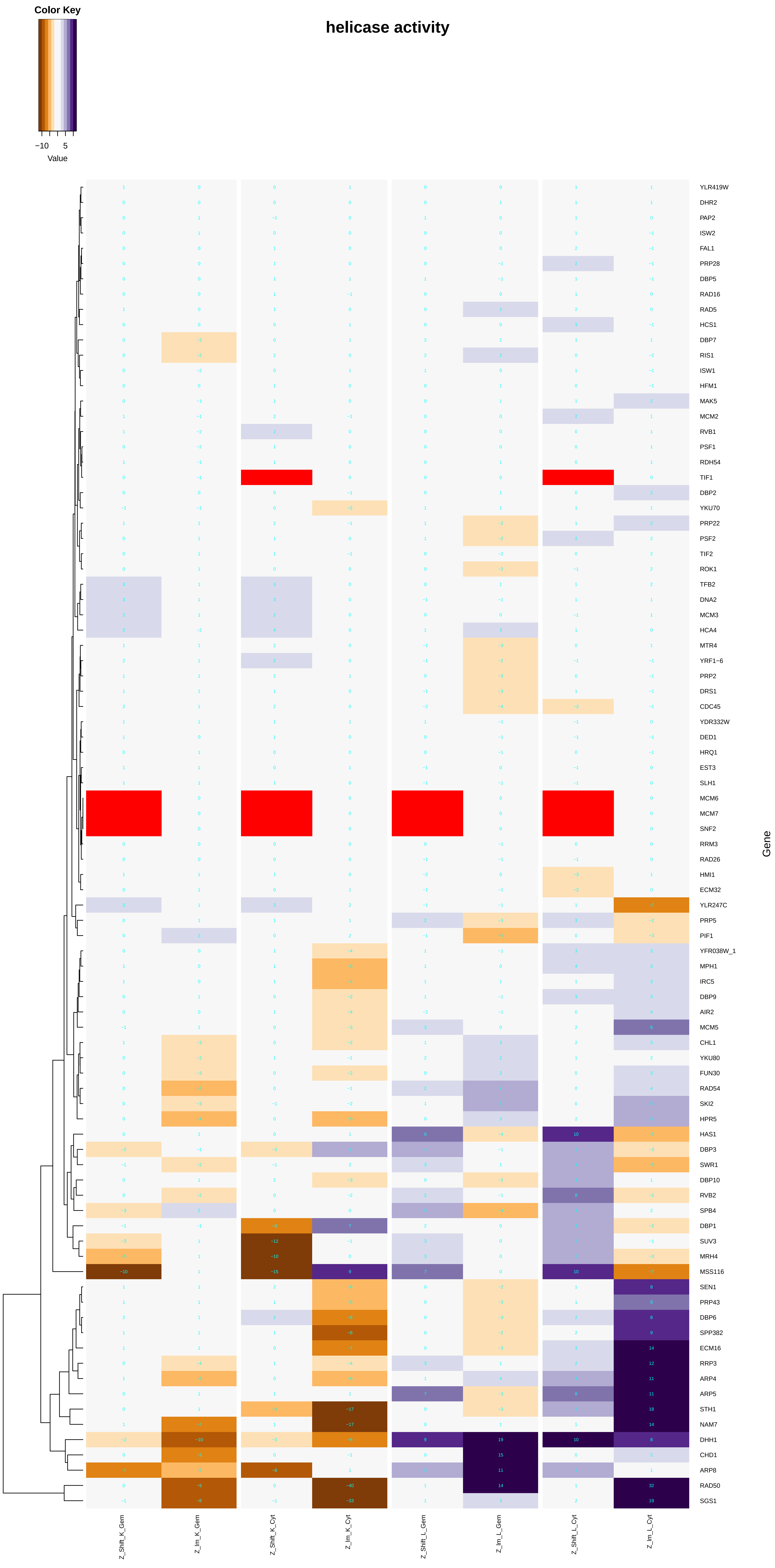
helicase activity
Color Key
−10 5
Value
YLR419W
DHR2
PAP2
ISW2
FAL1
PRP28
DBP5
RAD16
RAD5
HCS1
DBP7
RIS1
ISW1
HFM1
MAK5
MCM2
RVB1
PSF1
RDH54
TIF1
DBP2
YKU70
PRP22
PSF2
TIF2
ROK1
TFB2
DNA2
MCM3
HCA4
MTR4
YRF1−6
PRP2
DRS1
CDC45
YDR332W
DED1
HRQ1
EST3
SLH1
MCM6
MCM7
SNF2
RRM3
RAD26
HMI1
ECM32
YLR247C
PRP5
PIF1
YFR038W_1
MPH1
IRC5
DBP9
AIR2
MCM5
CHL1
YKU80
FUN30
RAD54
SKI2
HPR5
HAS1
DBP3
SWR1
DBP10
RVB2
SPB4
DBP1
SUV3
MRH4
MSS116
SEN1
PRP43
DBP6
SPP382
ECM16
RRP3
ARP4
ARP5
STH1
NAM7
DHH1
CHD1
ARP8
RAD50
SGS1
Gene
Z_Shift_K_Gem
Z_Im_K_Gem
Z_Shift_K_Cyt
Z_Im_K_Cyt
Z_Shift_L_Gem
Z_Im_L_Gem
Z_Shift_L_Cyt
Z_Im_L_Cyt

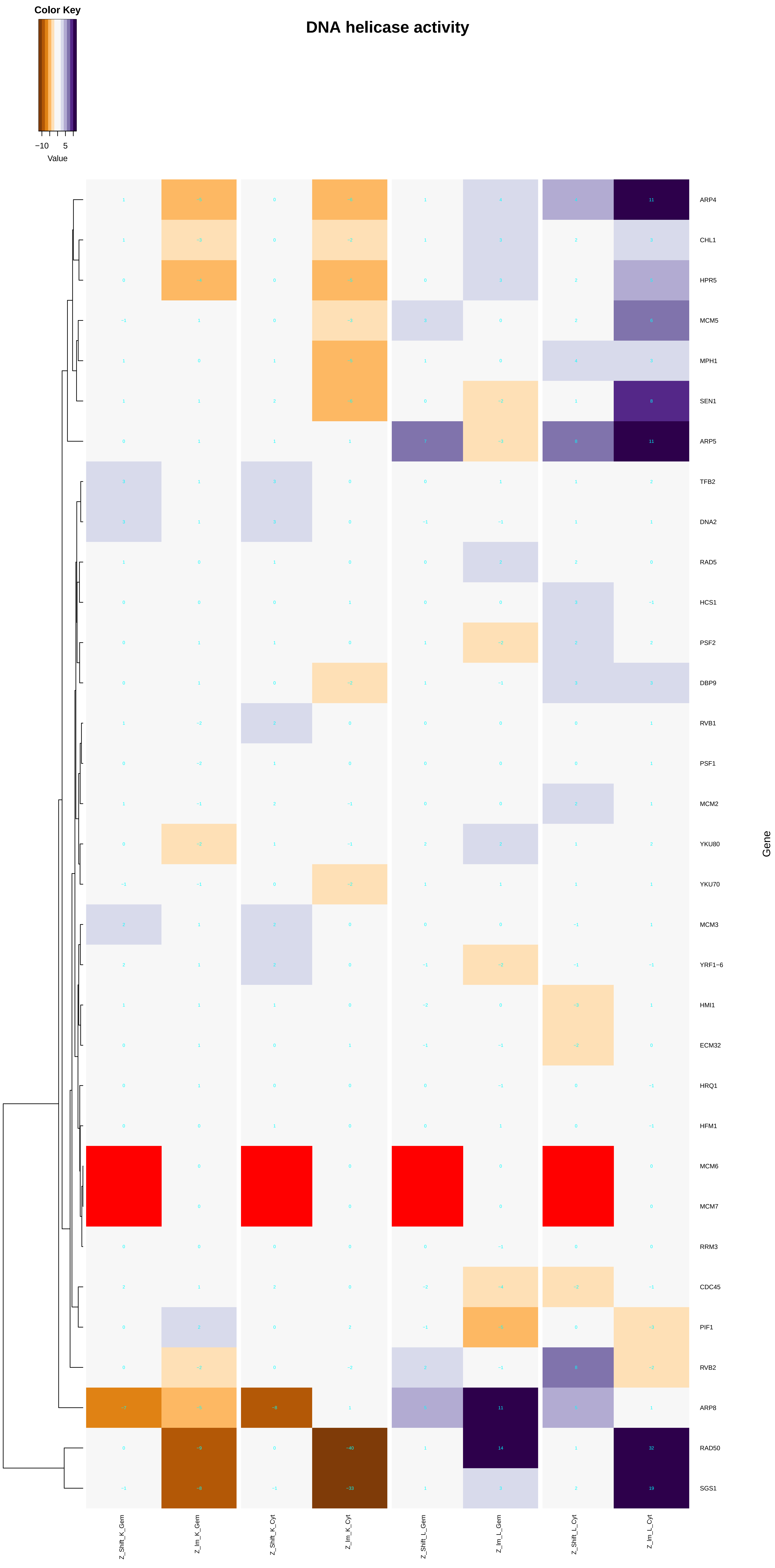

Color Key
−10
5
Value
DNA helicase activity
ARP4
CHL1
HPR5
MCM5
MPH1
SEN1
ARP5
TFB2
DNA2
RAD5
HCS1
PSF2
DBP9
RVB1
PSF1
MCM2
YKU80
YKU70
MCM3
YRF1−6
HMI1
ECM32
HRQ1
HFM1
MCM6
MCM7
RRM3
CDC45
PIF1
RVB2
ARP8
RAD50
SGS1
Gene
Z_Shift_K_Gem
Z_Im_K_Gem
Z_Shift_K_Cyt
Z_Im_K_Cyt
Z_Shift_L_Gem
Z_Im_L_Gem
Z_Shift_L_Cyt
Z_Im_L_Cyt

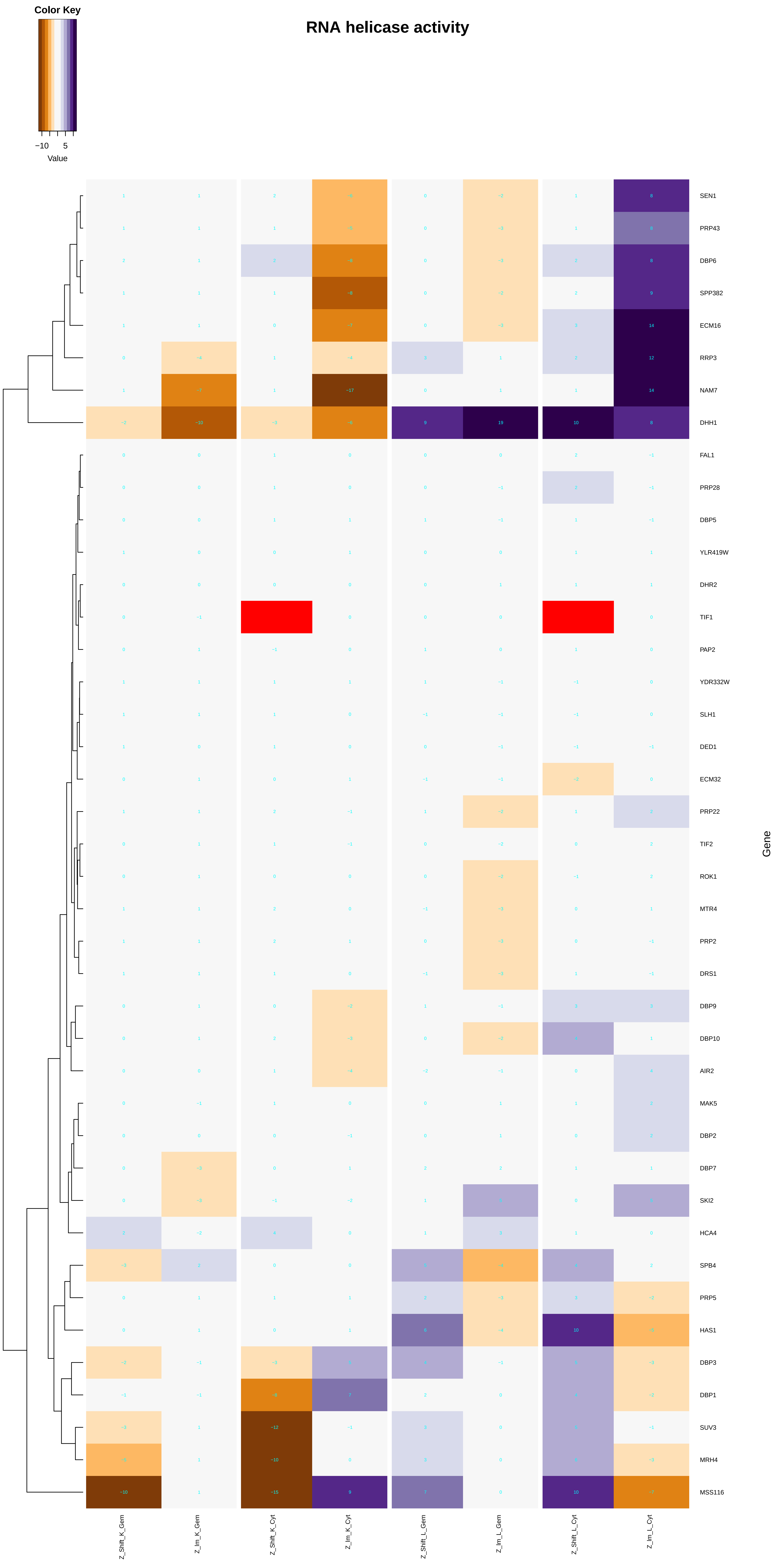

Color Key
−10
5
Value
RNA helicase activity
SEN1
PRP43
DBP6
SPP382
ECM16
RRP3
NAM7
DHH1
FAL1
PRP28
DBP5
YLR419W
DHR2
TIF1
PAP2
YDR332W
SLH1
DED1
ECM32
PRP22
TIF2
ROK1
MTR4
PRP2
DRS1
DBP9
DBP10
AIR2
MAK5
DBP2
DBP7
SKI2
HCA4
SPB4
PRP5
HAS1
DBP3
DBP1
SUV3
MRH4
MSS116
Gene
Z_Shift_K_Gem
Z_Im_K_Gem
Z_Shift_K_Cyt
Z_Im_K_Cyt
Z_Shift_L_Gem
Z_Im_L_Gem
Z_Shift_L_Cyt
Z_Im_L_Cyt

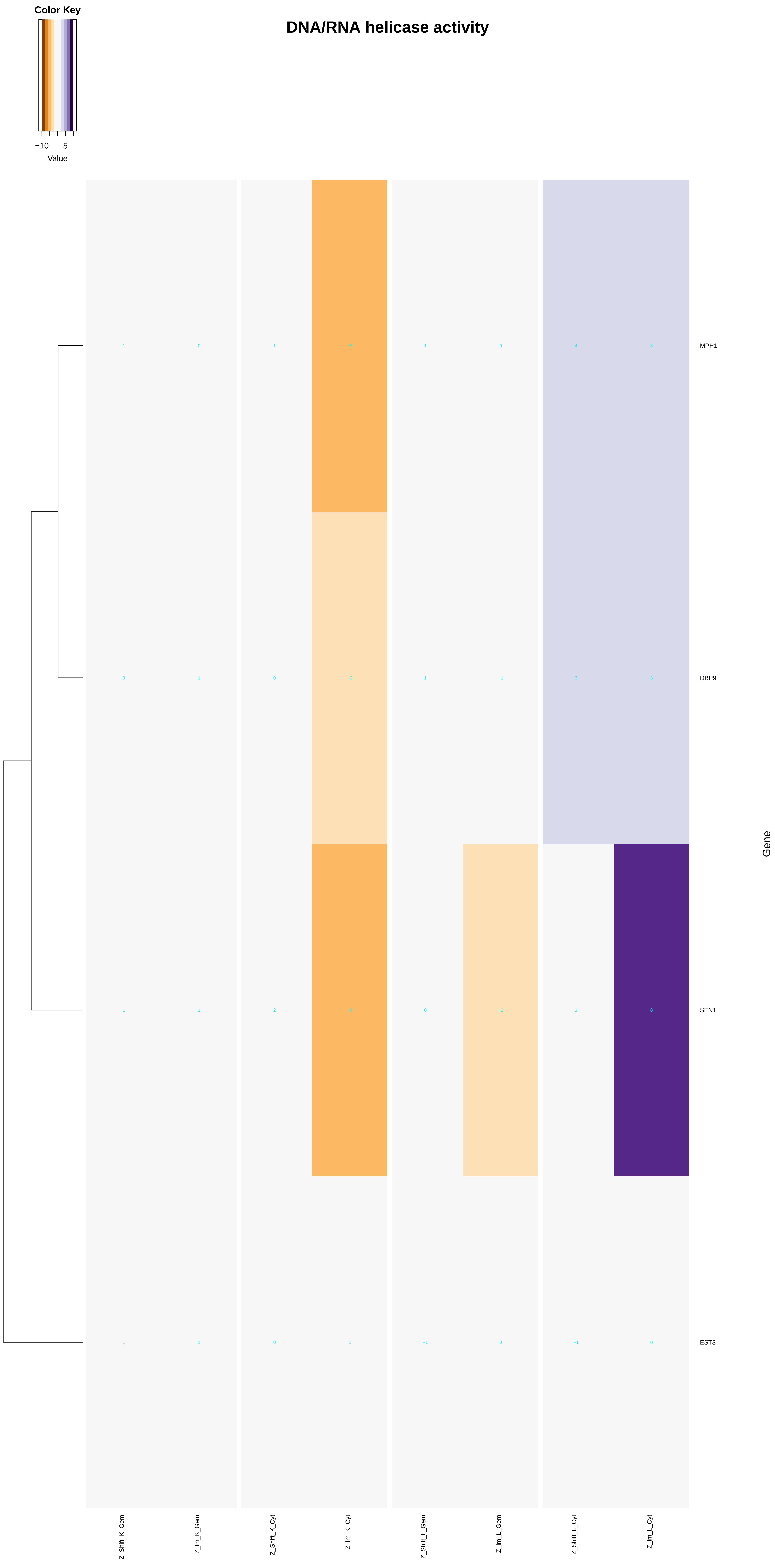
DNA/RNA helicase activity
Color Key
−10
5
Value
MPH1
DBP9
SEN1
EST3
Gene
Z_Shift_K_Gem
Z_Im_K_Gem
Z_Shift_K_Cyt
Z_Im_K_Cyt
Z_Shift_L_Gem
Z_Im_L_Gem
Z_Shift_L_Cyt
Z_Im_L_Cyt

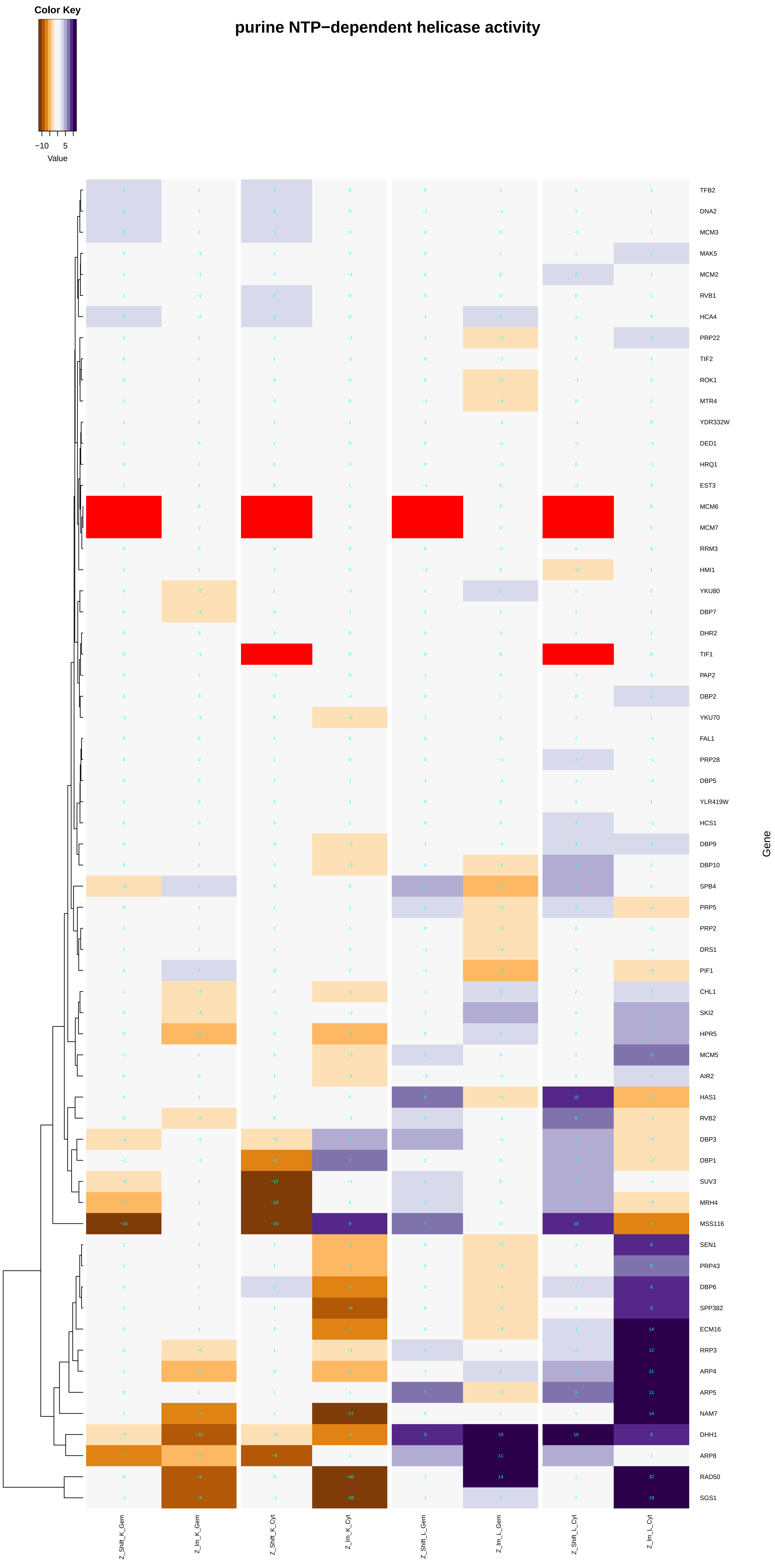
purine NTP−dependent helicase activity
Color Key
−10
5
Value
TFB2
DNA2
MCM3
MAK5
MCM2
RVB1
HCA4
PRP22
TIF2
ROK1
MTR4
YDR332W
DED1
HRQ1
EST3
MCM6
MCM7
RRM3
HMI1
YKU80
DBP7
DHR2
TIF1
PAP2
DBP2
YKU70
FAL1
PRP28
DBP5
YLR419W
HCS1
DBP9
DBP10
SPB4
PRP5
PRP2
DRS1
PIF1
CHL1
SKI2
HPR5
MCM5
AIR2
HAS1
RVB2
DBP3
DBP1
SUV3
MRH4
MSS116
SEN1
PRP43
DBP6
SPP382
ECM16
RRP3
ARP4
ARP5
NAM7
DHH1
ARP8
RAD50
SGS1
Gene
Z_Shift_K_Gem
Z_lm_K_Gem
Z_Shift_K_Cyt
Z_lm_K_Cyt
Z_Shift_L_Gem
Z_lm_L_Gem
Z_Shift_L_Cyt
Z_lm_L_Cyt

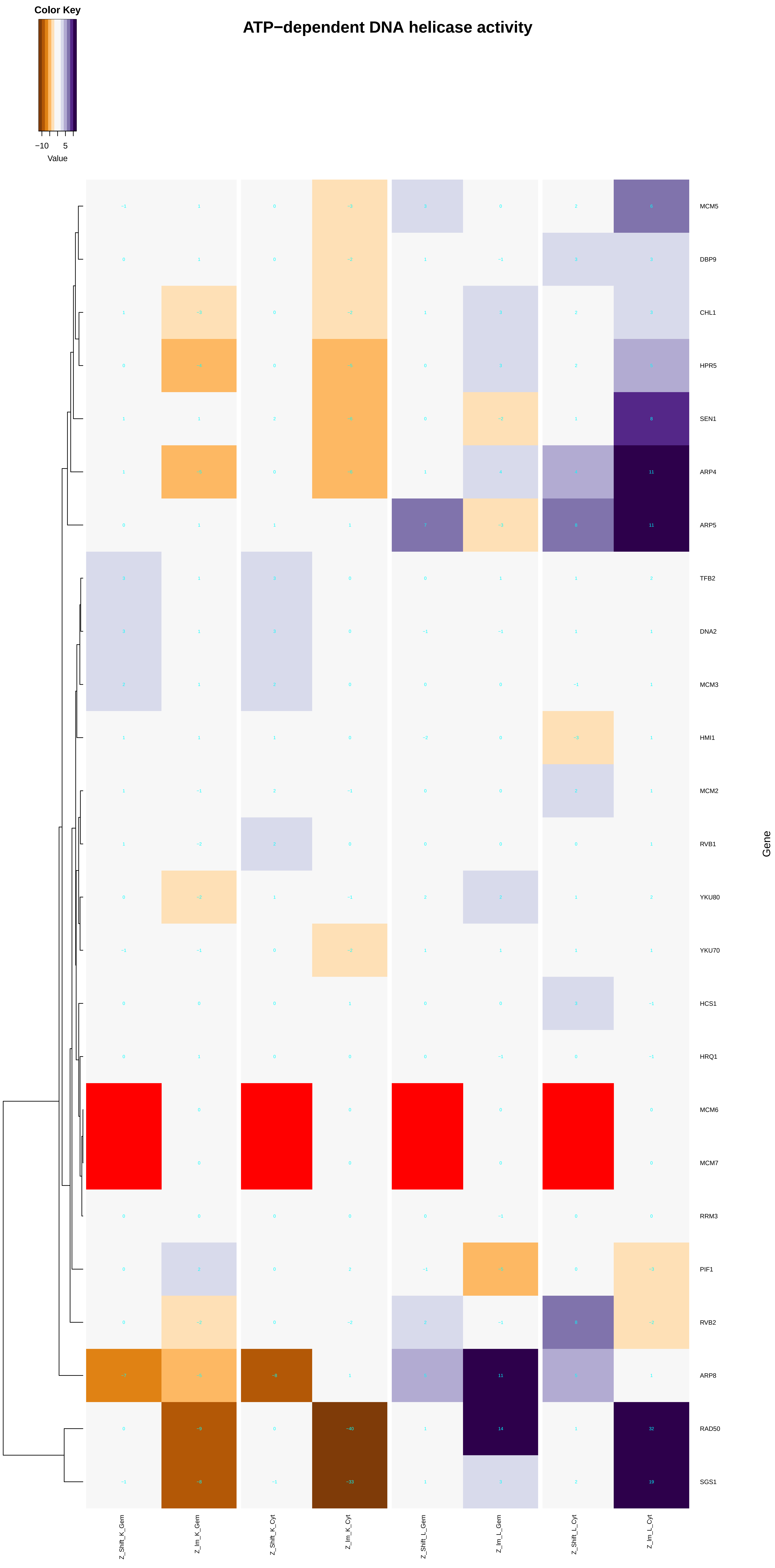

Color Key
−10
5
Value
ATP−dependent DNA helicase activity
MCM5
DBP9
CHL1
HPR5
SEN1
ARP4
ARP5
TFB2
DNA2
MCM3
HMI1
MCM2
RVB1
YKU80
YKU70
HCS1
HRQ1
MCM6
MCM7
RRM3
PIF1
RVB2
ARP8
RAD50
SGS1
Gene
Z_Shift_K_Gem
Z_Im_K_Gem
Z_Shift_K_Cyt
Z_Im_K_Cyt
Z_Shift_L_Gem
Z_Im_L_Gem
Z_Shift_L_Cyt
Z_Im_L_Cyt

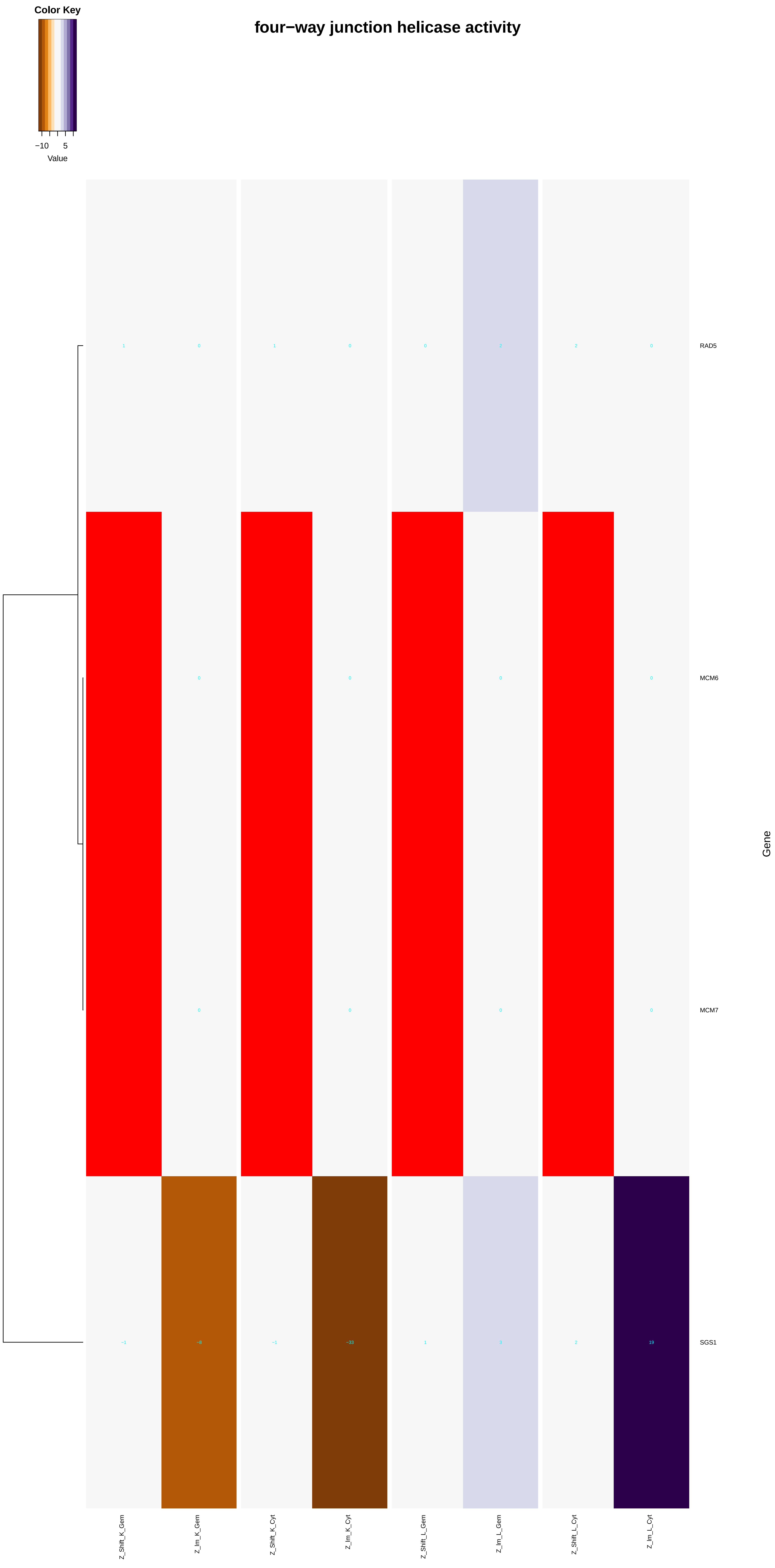
Color Key
−10
5
Value
four−way junction helicase activity
1
0
1
0
0
2
2
0
RAD5
0
0
0
0
MCM6
0
0
0
0
MCM7
−1
−8
−1
−33
1
3
2
19
SGS1
Gene
Z_Shift_K_Gem
Z_Im_K_Gem
Z_Shift_K_Cyt
Z_Im_K_Cyt
Z_Shift_L_Gem
Z_Im_L_Gem
Z_Shift_L_Cyt
Z_Im_L_Cyt

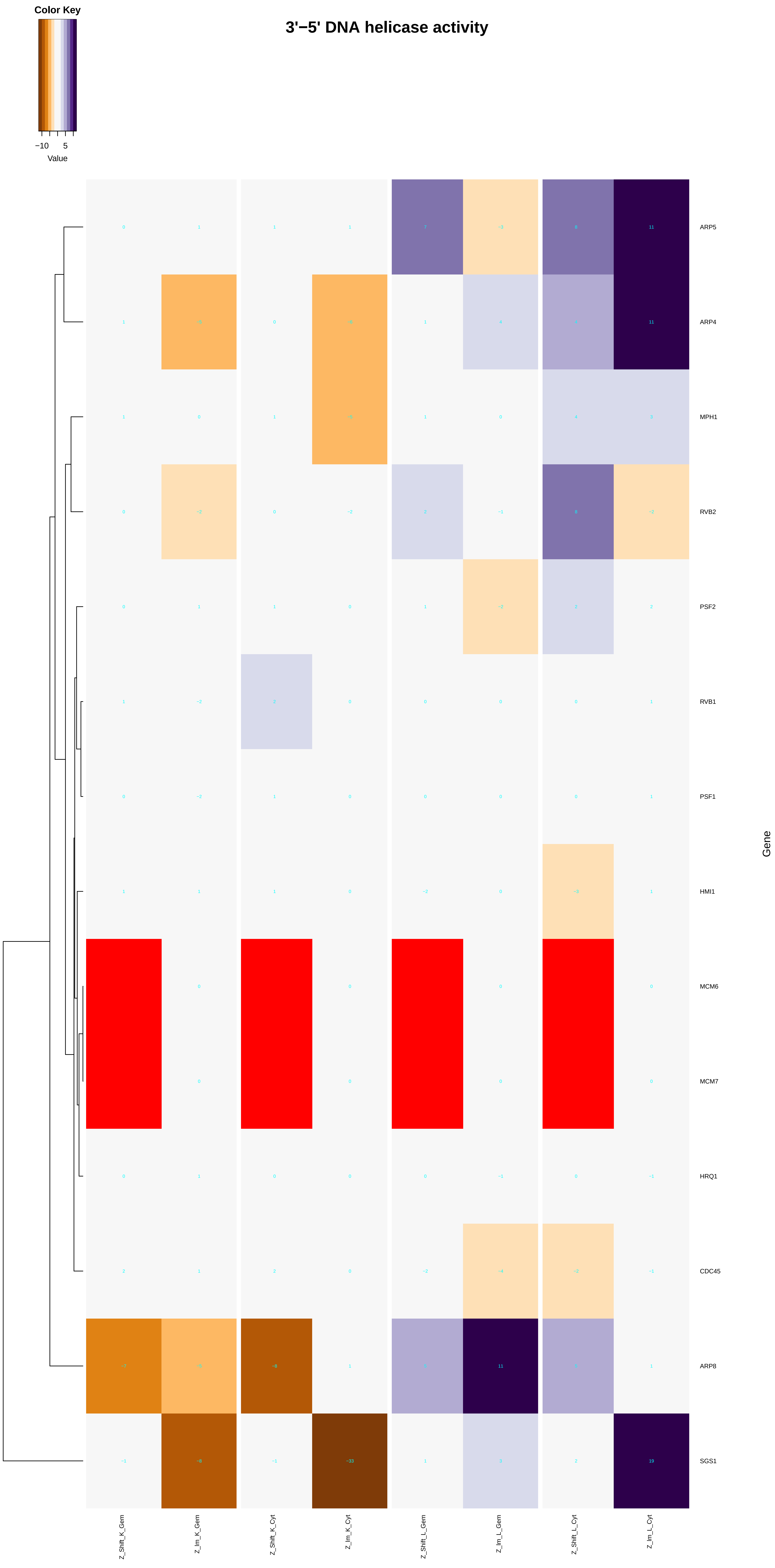
Color Key
−10
5
Value
3'−5' DNA helicase activity
ARP5
ARP4
MPH1
RVB2
PSF2
RVB1
PSF1
HMI1
MCM6
MCM7
HRQ1
CDC45
ARP8
SGS1
Gene
Z_Shift_K_Gem
Z_Im_K_Gem
Z_Shift_K_Cyt
Z_Im_K_Cyt
Z_Shift_L_Gem
Z_Im_L_Gem
Z_Shift_L_Cyt
Z_Im_L_Cyt

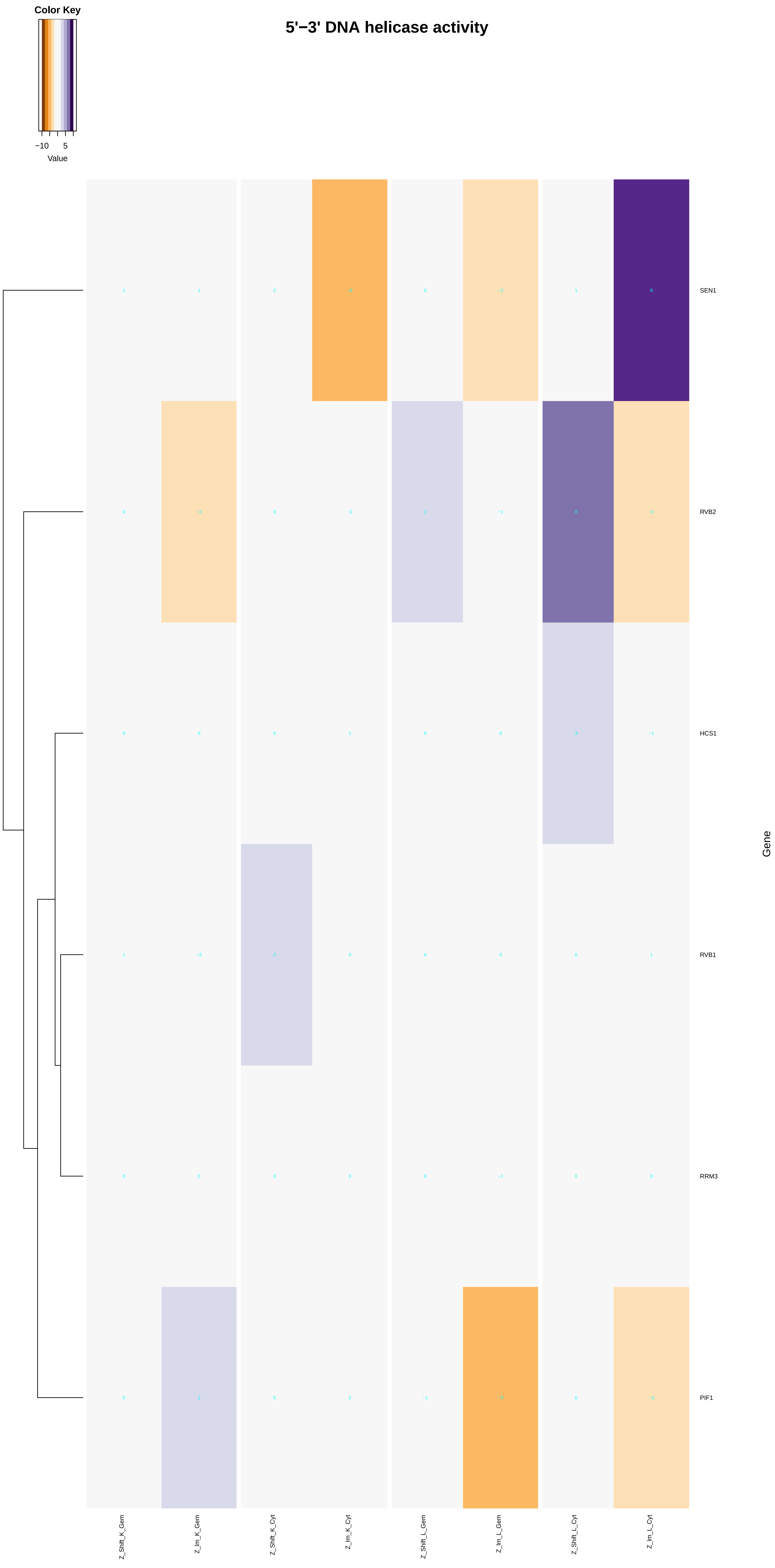
5'−3' DNA helicase activity
Color Key
−10
5
Value
SEN1
RVB2
HCS1
RVB1
RRM3
PIF1
Gene
Z_Shift_K_Gem
Z_Im_K_Gem
Z_Shift_K_Cyt
Z_Im_K_Cyt
Z_Shift_L_Gem
Z_Im_L_Gem
Z_Shift_L_Cyt
Z_Im_L_Cyt

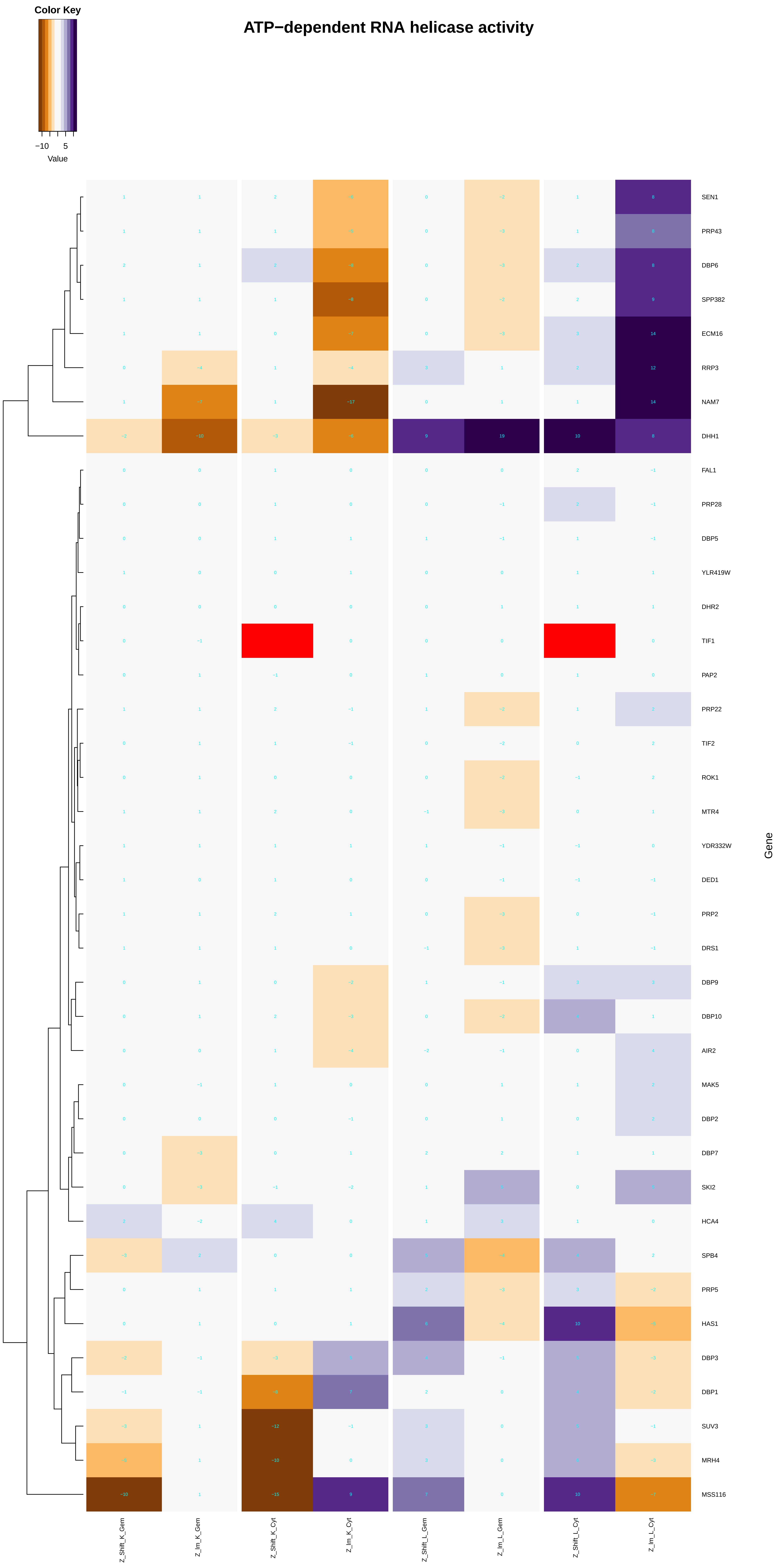
ATP−dependent RNA helicase activity
Color Key
−10
5
Value
SEN1
PRP43
DBP6
SPP382
ECM16
RRP3
NAM7
DHH1
FAL1
PRP28
DBP5
YLR419W
DHR2
TIF1
PAP2
PRP22
TIF2
ROK1
MTR4
YDR332W
DED1
PRP2
DRS1
DBP9
DBP10
AIR2
MAK5
DBP2
DBP7
SKI2
HCA4
SPB4
PRP5
HAS1
DBP3
DBP1
SUV3
MRH4
MSS116
Gene
Z_Shift_K_Gem
Z_Im_K_Gem
Z_Shift_K_Cyt
Z_Im_K_Cyt
Z_Shift_L_Gem
Z_Im_L_Gem
Z_Shift_L_Cyt
Z_Im_L_Cyt

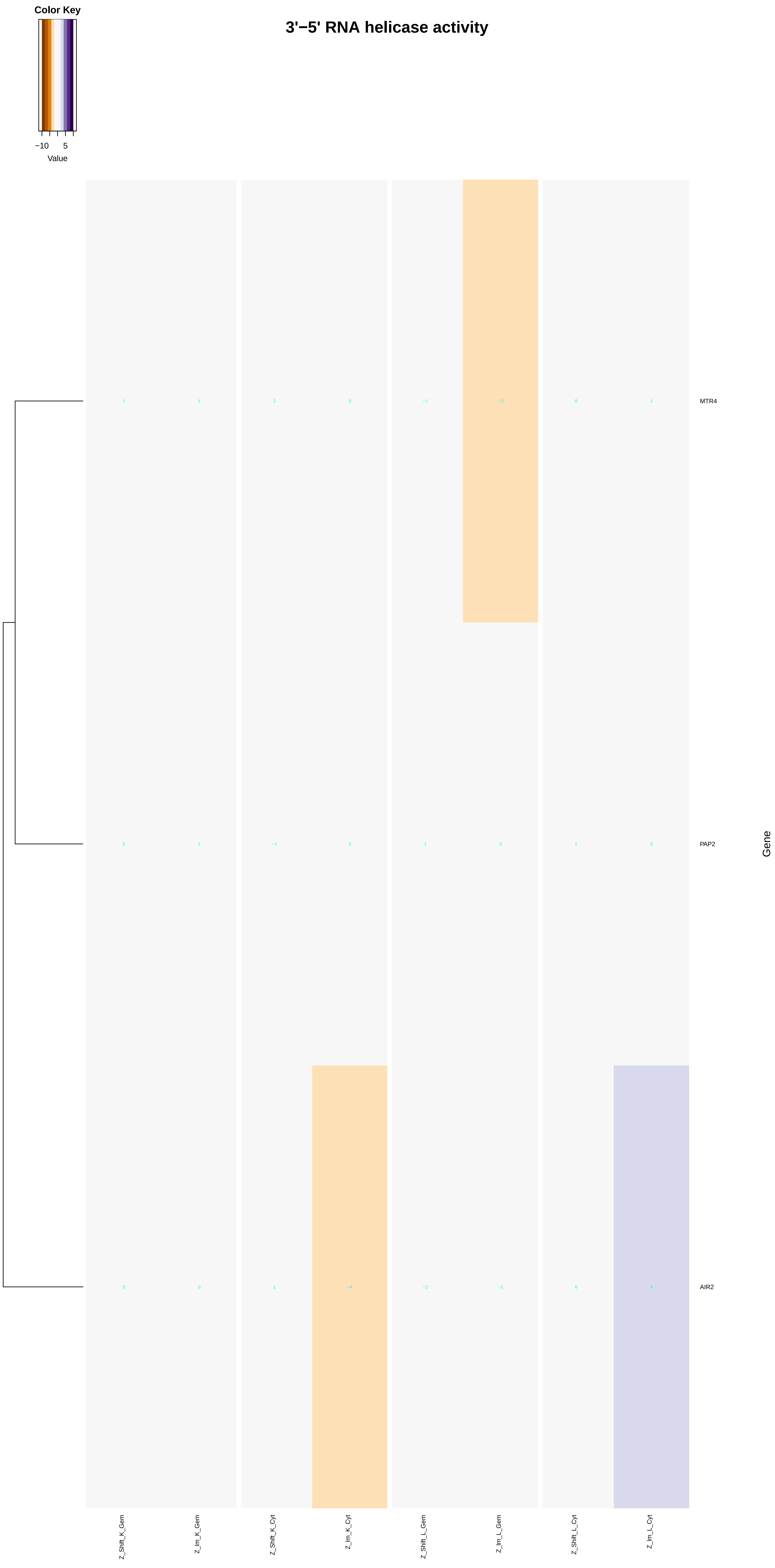
Color Key
−10
5
Value
3'−5' RNA helicase activity
1
1
2
0
−1
−3
0
1
MTR4
0
1
−1
0
1
0
1
0
PAP2
0
0
1
−4
−2
−1
0
4
AIR2
Gene
Z_Shift_K_Gem
Z_Im_K_Gem
Z_Shift_K_Cyt
Z_Im_K_Cyt
Z_Shift_L_Gem
Z_Im_L_Gem
Z_Shift_L_Cyt
Z_Im_L_Cyt

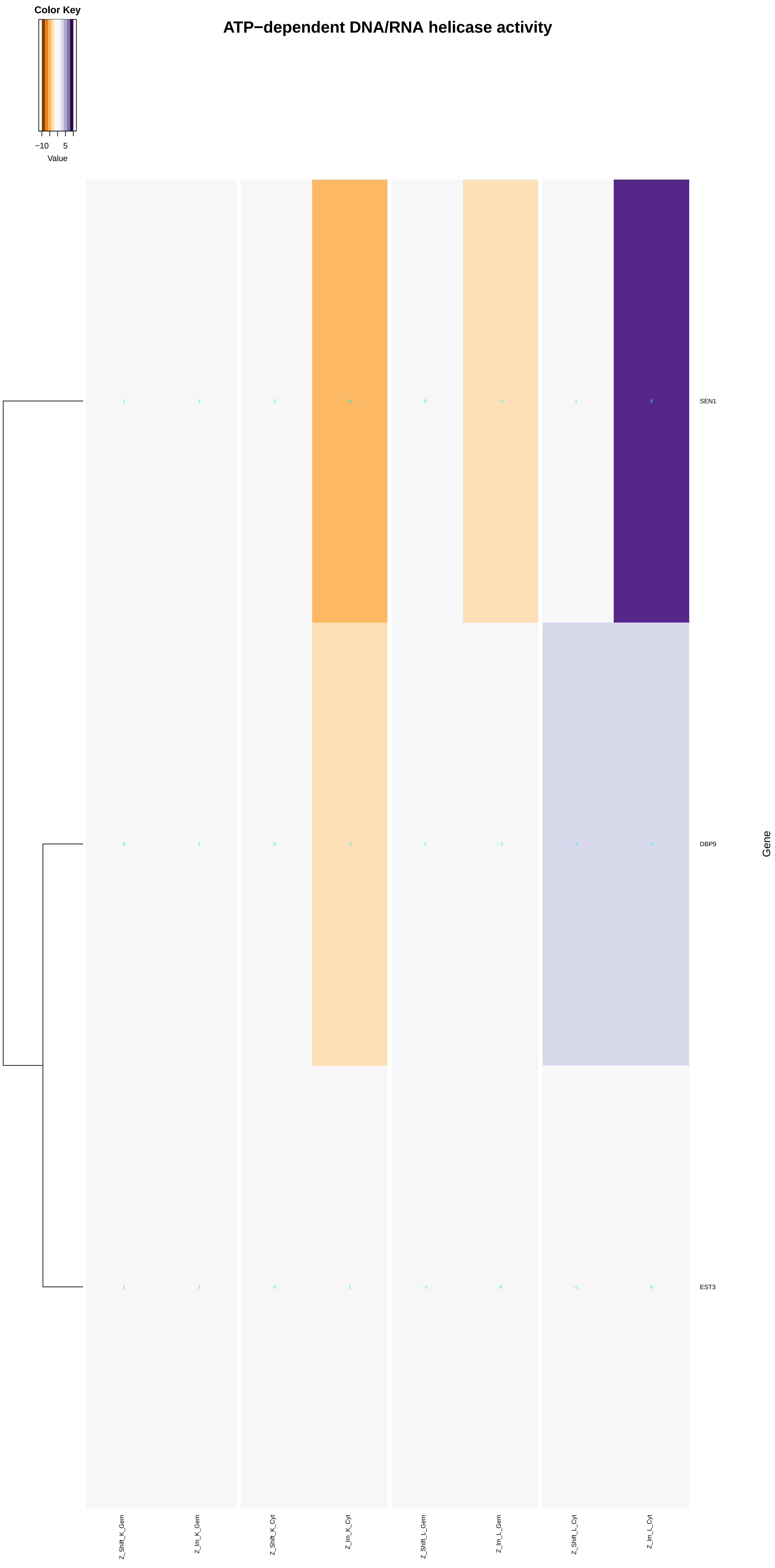
Color Key
−10
5
Value
ATP−dependent DNA/RNA helicase activity
SEN1
DBP9
EST3
Gene
Z_Shift_K_Gem
Z_Im_K_Gem
Z_Shift_K_Cyt
Z_Im_K_Cyt
Z_Shift_L_Gem
Z_Im_L_Gem
Z_Shift_L_Cyt
Z_Im_L_Cyt

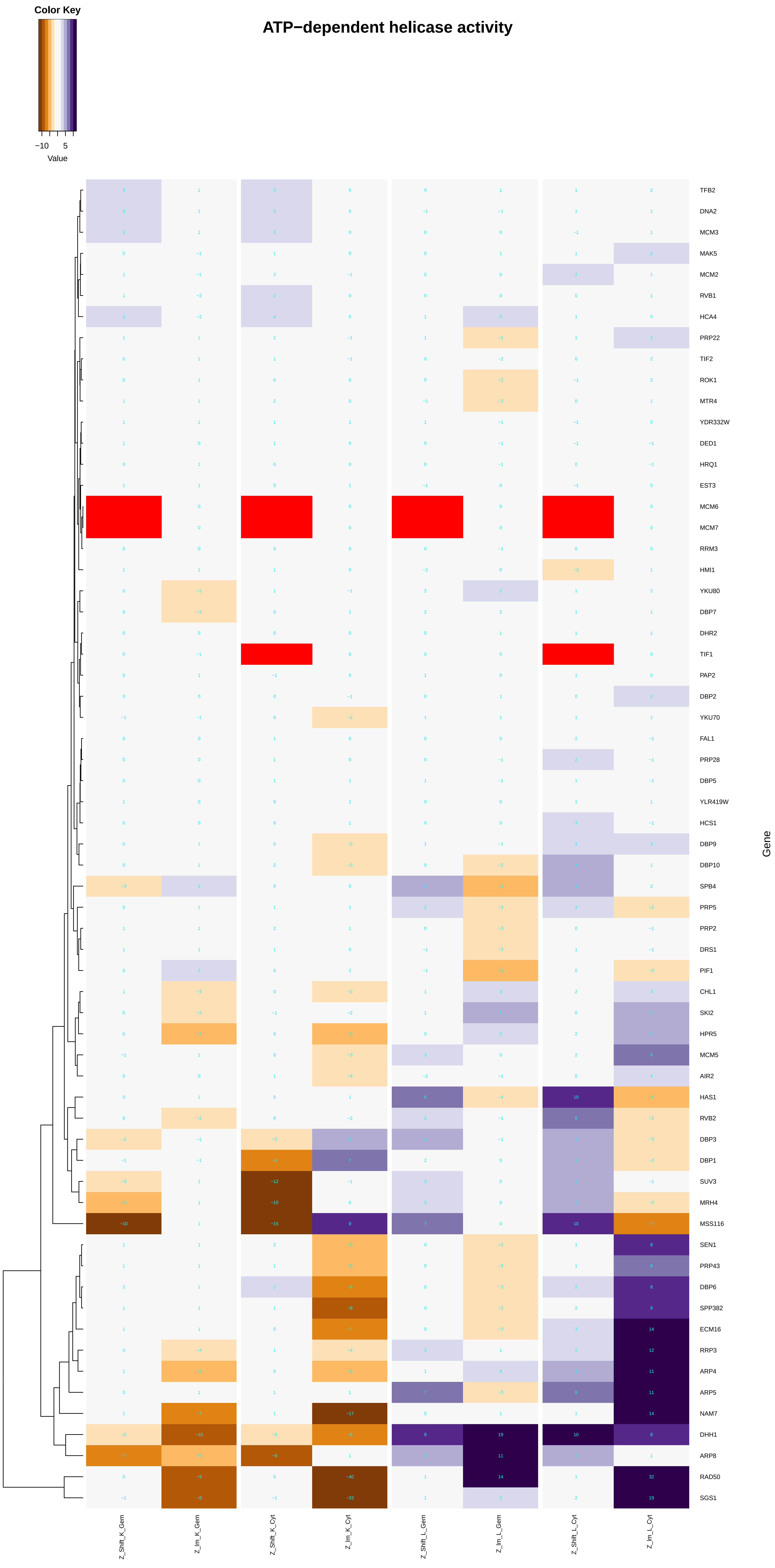
ATP−dependent helicase activity
Color Key
−10
5
Value
Gene
TFB2
DNA2
MCM3
MAK5
MCM2
RVB1
HCA4
PRP22
TIF2
ROK1
MTR4
YDR332W
DED1
HRQ1
EST3
MCM6
MCM7
RRM3
HMI1
YKU80
DBP7
DHR2
TIF1
PAP2
DBP2
YKU70
FAL1
PRP28
DBP5
YLR419W
HCS1
DBP9
DBP10
SPB4
PRP5
PRP2
DRS1
PIF1
CHL1
SKI2
HPR5
MCM5
AIR2
HAS1
RVB2
DBP3
DBP1
SUV3
MRH4
MSS116
SEN1
PRP43
DBP6
SPP382
ECM16
RRP3
ARP4
ARP5
NAM7
DHH1
ARP8
RAD50
SGS1
Z_Shift_K_Gem
Z_Im_K_Gem
Z_Shift_K_Cyt
Z_Im_K_Cyt
Z_Shift_L_Gem
Z_Im_L_Gem
Z_Shift_L_Cyt
Z_Im_L_Cyt

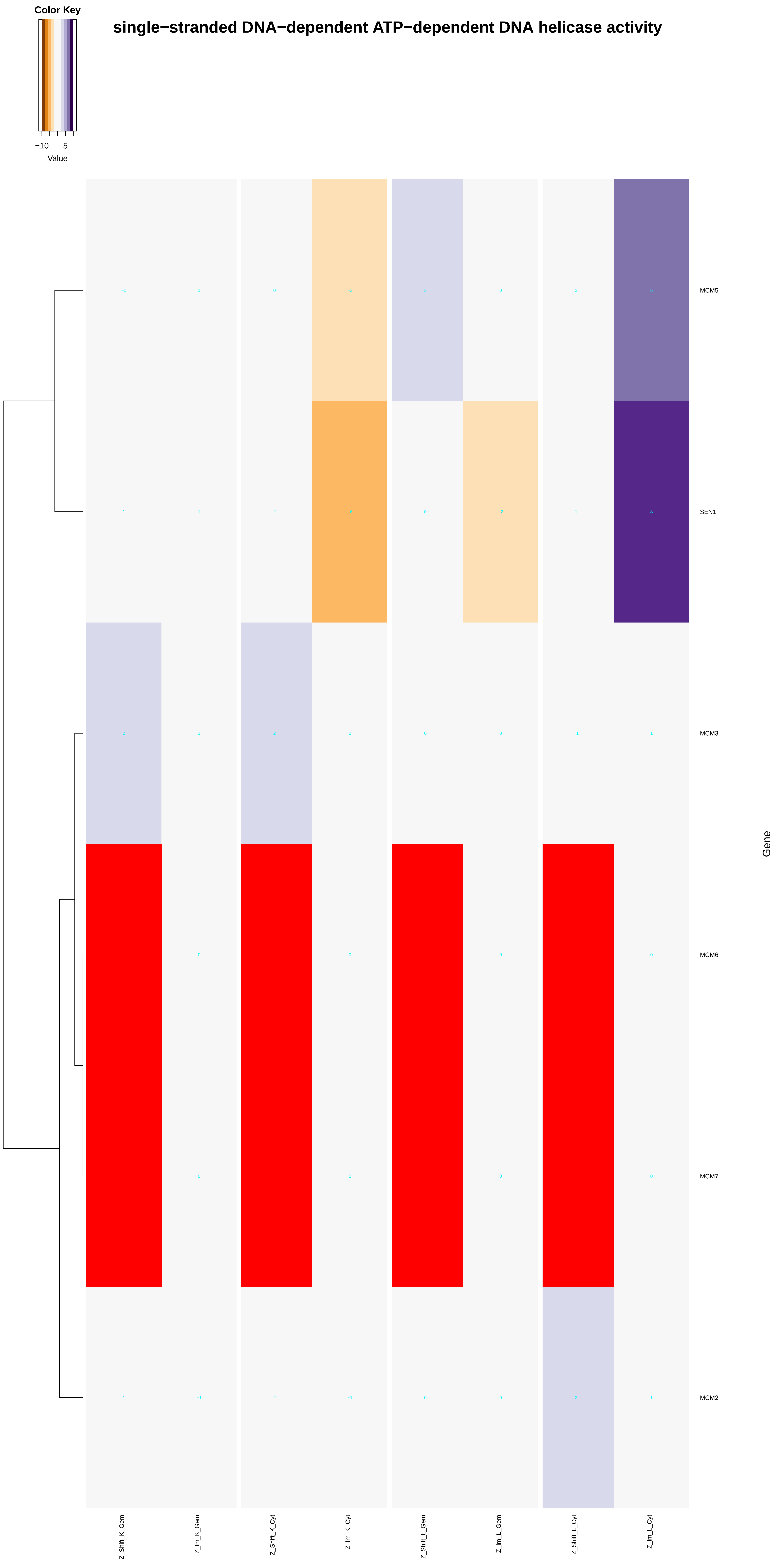

Color Key
−10
5
Value
single−stranded DNA−dependent ATP−dependent DNA helicase activity
MCM5
SEN1
MCM3
MCM6
MCM7
MCM2
Gene
Z_Shift_K_Gem
Z_Im_K_Gem
Z_Shift_K_Cyt
Z_Im_K_Cyt
Z_Shift_L_Gem
Z_Im_L_Gem
Z_Shift_L_Cyt
Z_Im_L_Cyt

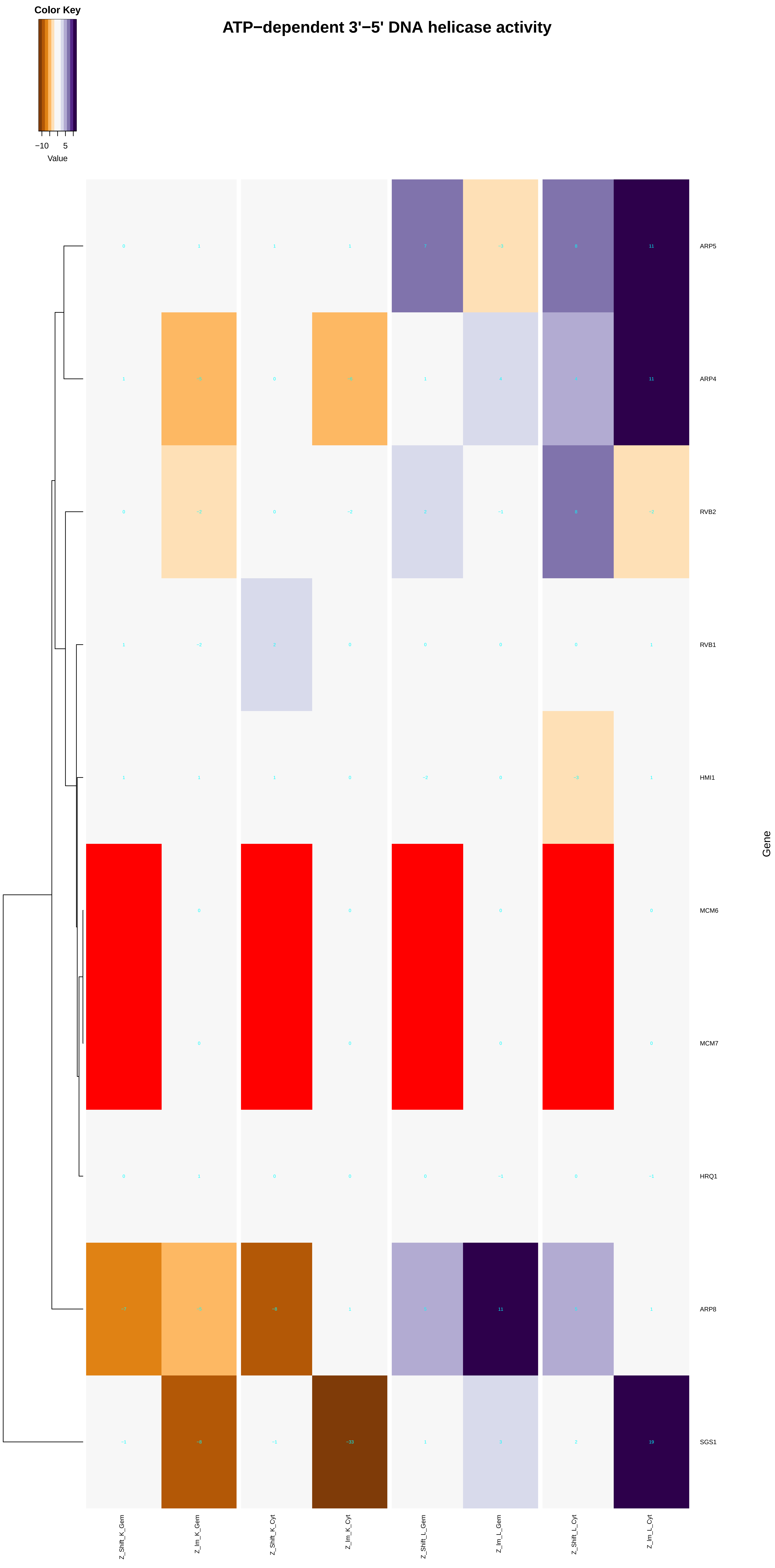
Color Key
−10
5
Value
ATP−dependent 3'−5' DNA helicase activity
ARP5
ARP4
RVB2
RVB1
HMI1
MCM6
MCM7
HRQ1
ARP8
SGS1
Gene
Z_Shift_K_Gem
Z_Im_K_Gem
Z_Shift_K_Cyt
Z_Im_K_Cyt
Z_Shift_L_Gem
Z_Im_L_Gem
Z_Shift_L_Cyt
Z_Im_L_Cyt

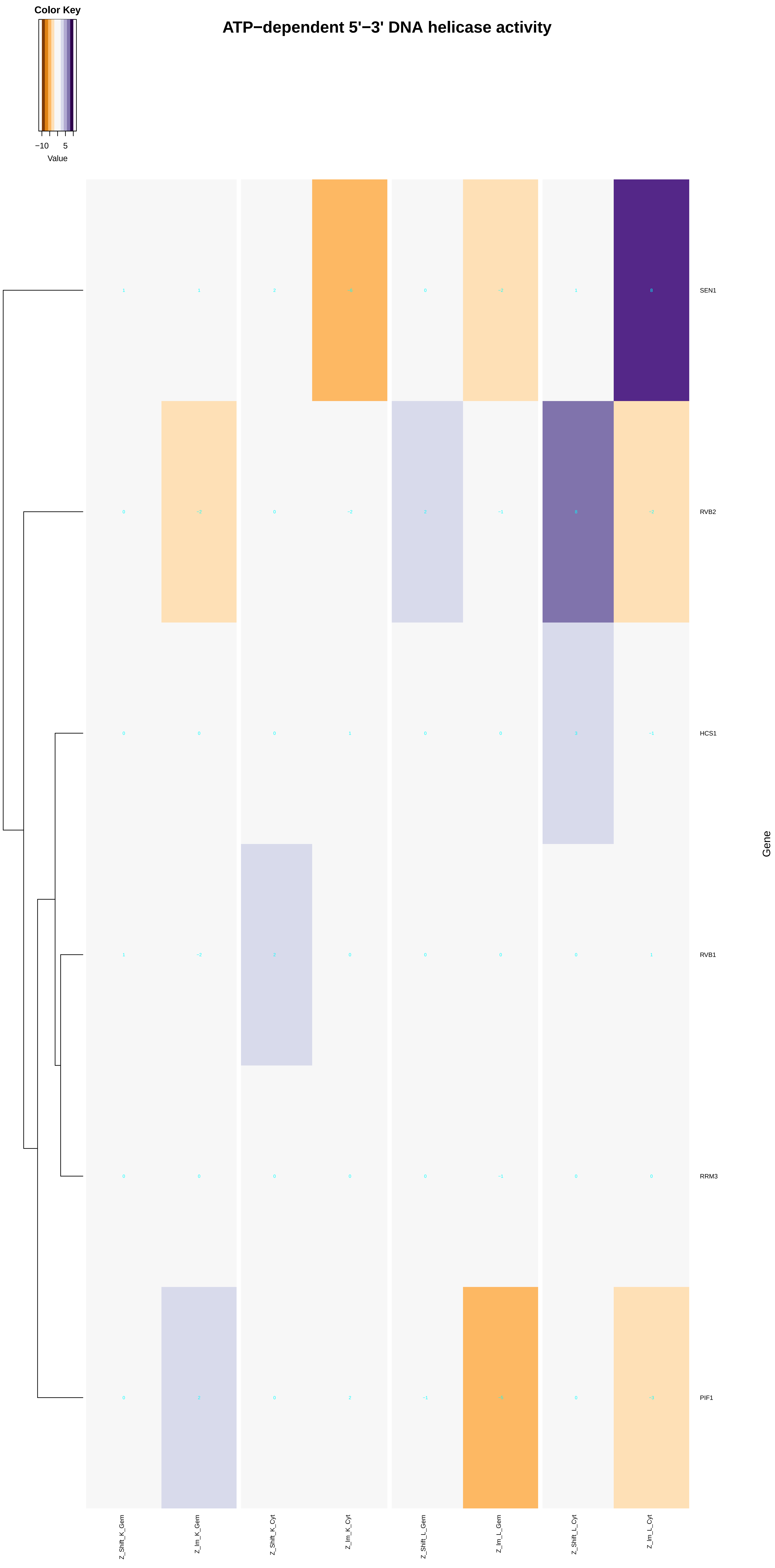
ATP−dependent 5'−3' DNA helicase activity
Color Key
−10
5
Value
SEN1
RVB2
HCS1
RVB1
RRM3
PIF1
Gene
Z_Shift_K_Gem
Z_Im_K_Gem
Z_Shift_K_Cyt
Z_Im_K_Cyt
Z_Shift_L_Gem
Z_Im_L_Gem
Z_Shift_L_Cyt
Z_Im_L_Cyt

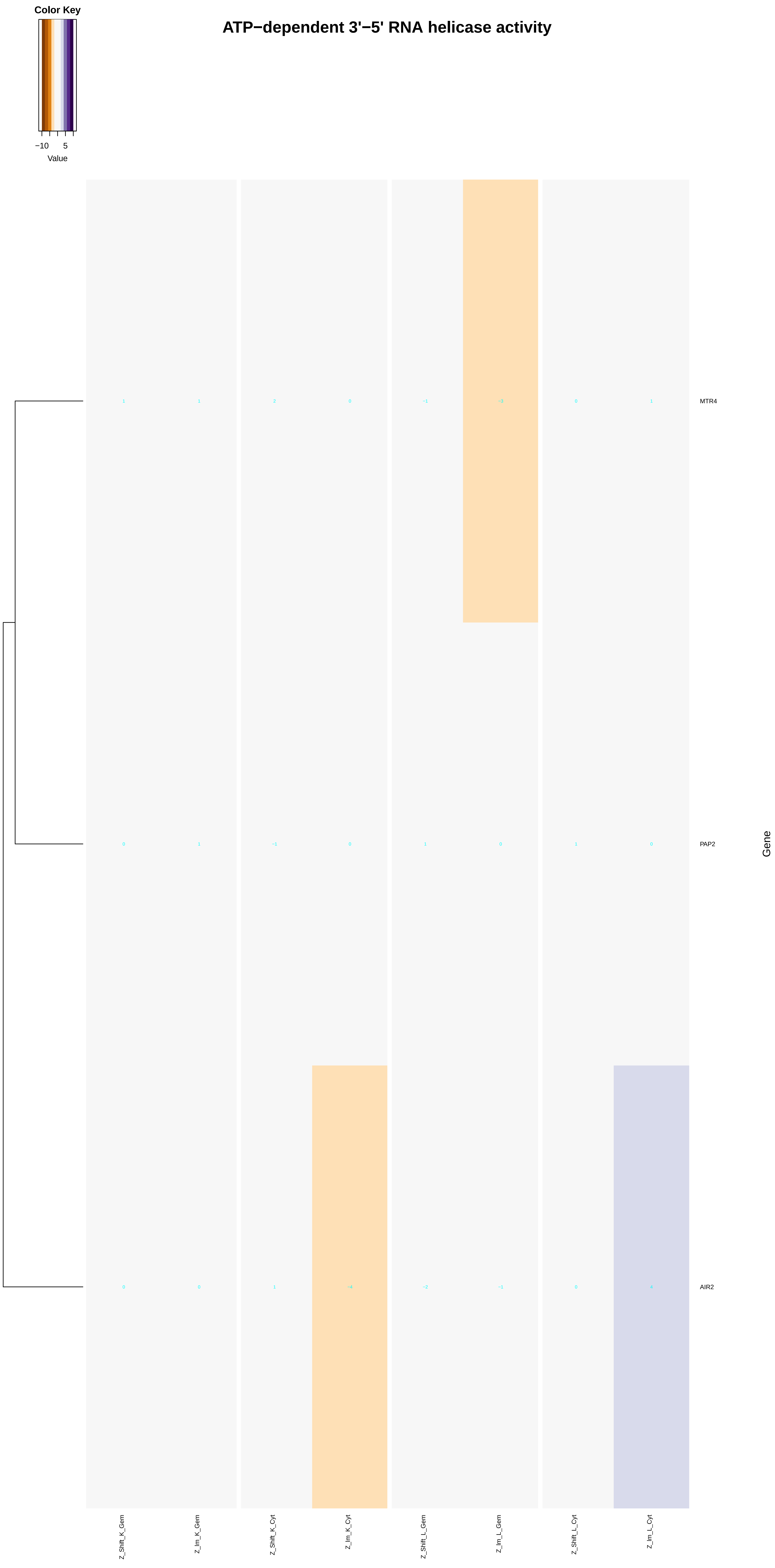
ATP−dependent 3'−5' RNA helicase activity
Color Key
−10
5
Value
MTR4
PAP2
AIR2
Gene
Z_Shift_K_Gem
Z_Im_K_Gem
Z_Shift_K_Cyt
Z_Im_K_Cyt
Z_Shift_L_Gem
Z_Im_L_Gem
Z_Shift_L_Cyt
Z_Im_L_Cyt
