## Additional File 7 for "A humanized yeast phenomic model of deoxycytidine kinase to predict genetic buffering of nucleoside analog cytotoxicity": hydrolase_activity,_acting_on_acid_anhydrides,_in_phosphorus-containing_anhydrides.pdf

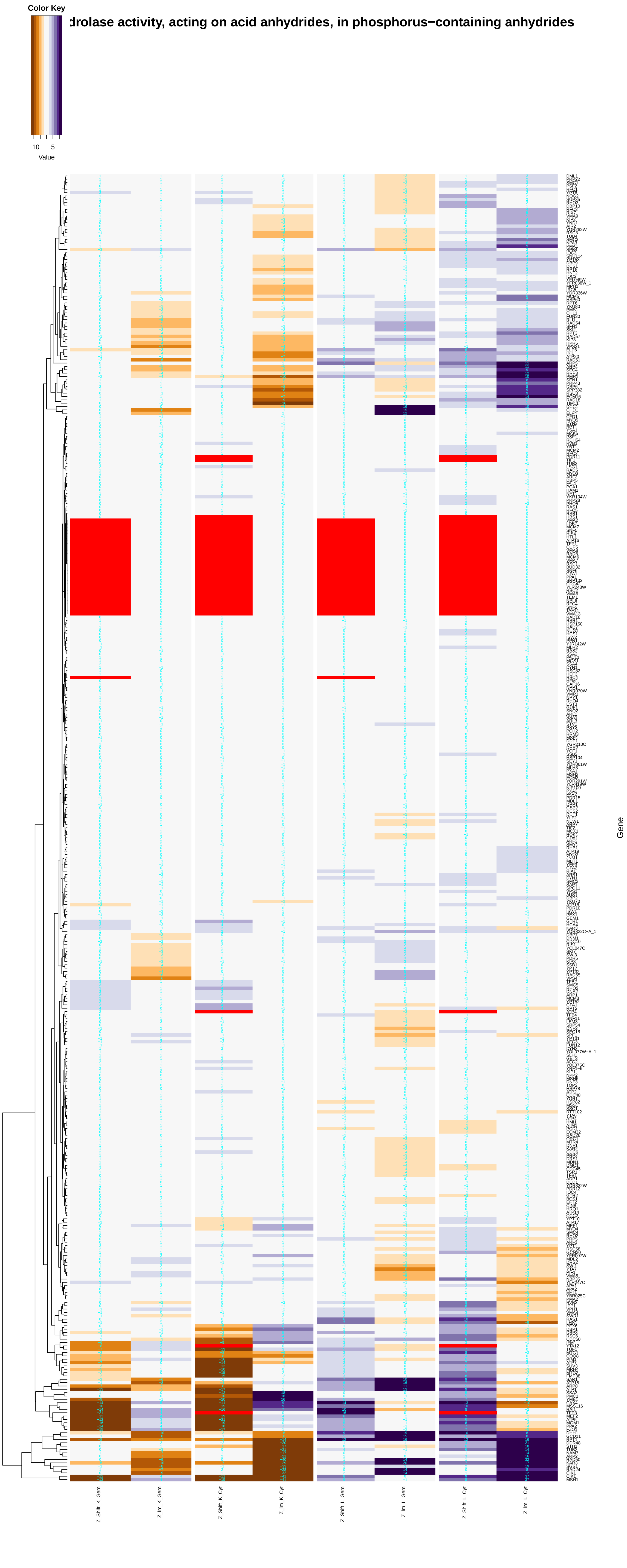
Color Key
−10
5
Value
drolase activity, acting on acid anhydrides, in phosphorus−containing anhydrides
Z_Shift_K_Gem
Z_Im_K_Gem
Z_Shift_K_Cyt
Z_Im_K_Cyt
Z_Shift_L_Gem
Z_Im_L_Gem
Z_Shift_L_Cyt
Z_Im_L_Cyt
Gene

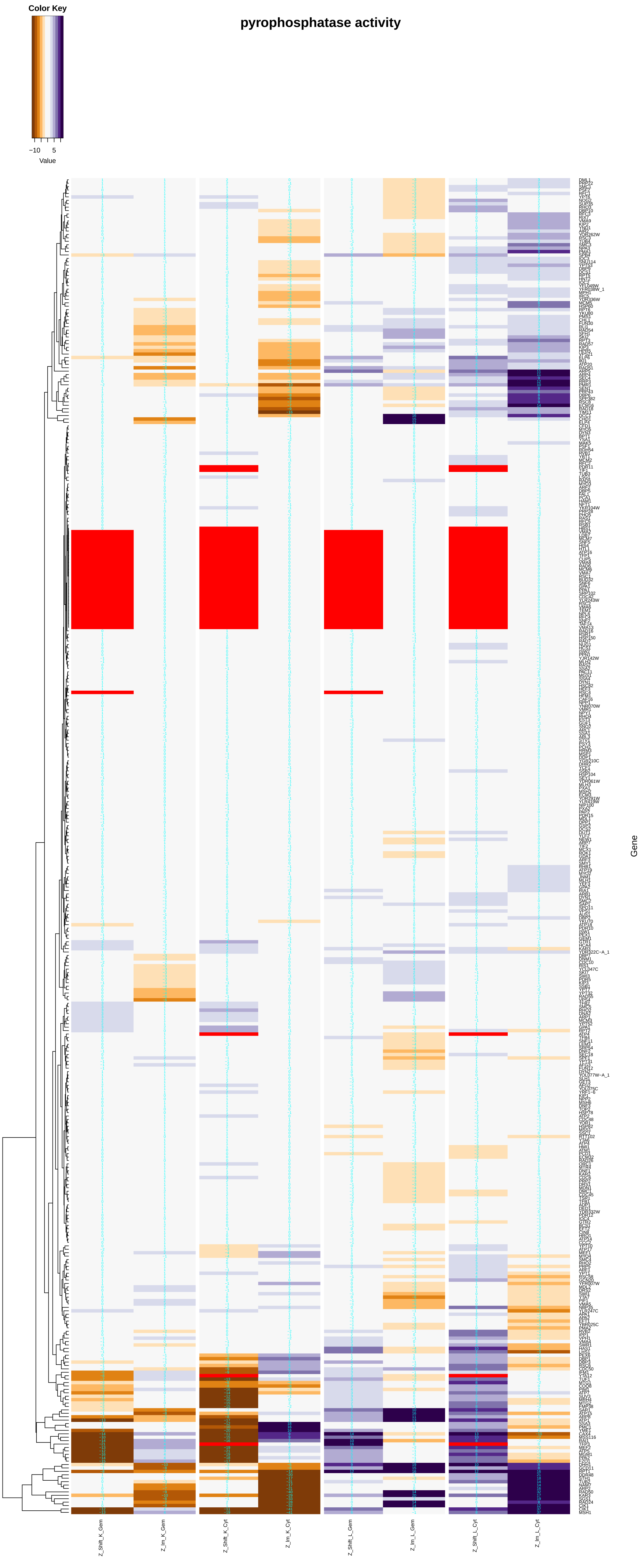

pyrophosphatase activity
Color Key
−10
5
Value
Gene
Z_Shift_K_Gem
Z_Im_K_Gem
Z_Shift_K_Cyt
Z_Im_K_Cyt
Z_Shift_L_Gem
Z_Im_L_Gem
Z_Shift_L_Cyt
Z_Im_L_Cyt

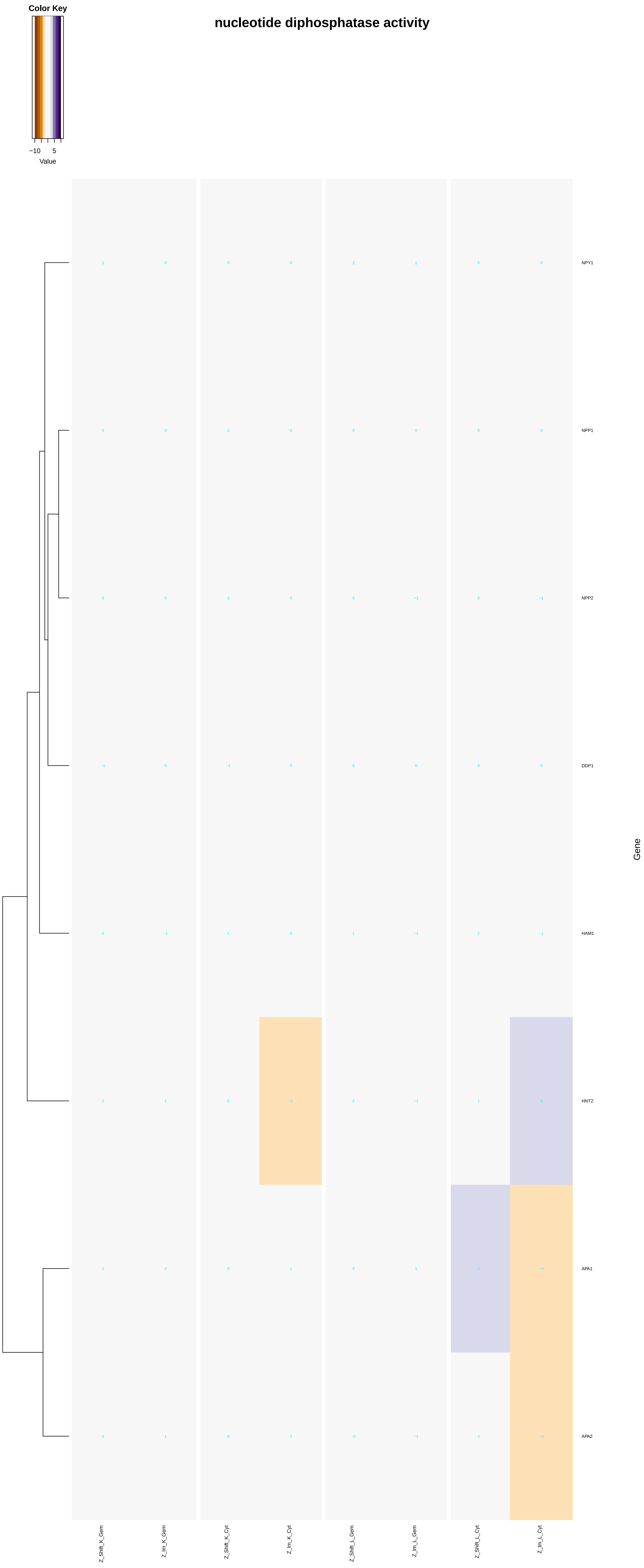
Color Key
−10
5
Value
nucleotide diphosphatase activity
NPY1
NPP1
NPP2
DDP1
HAM1
HNT2
APA1
APA2
Gene
Z_Shift_K_Gem
Z_lm_K_Gem
Z_Shift_K_Cyt
Z_lm_K_Cyt
Z_Shift_L_Gem
Z_lm_L_Gem
Z_Shift_L_Cyt
Z_lm_L_Cyt

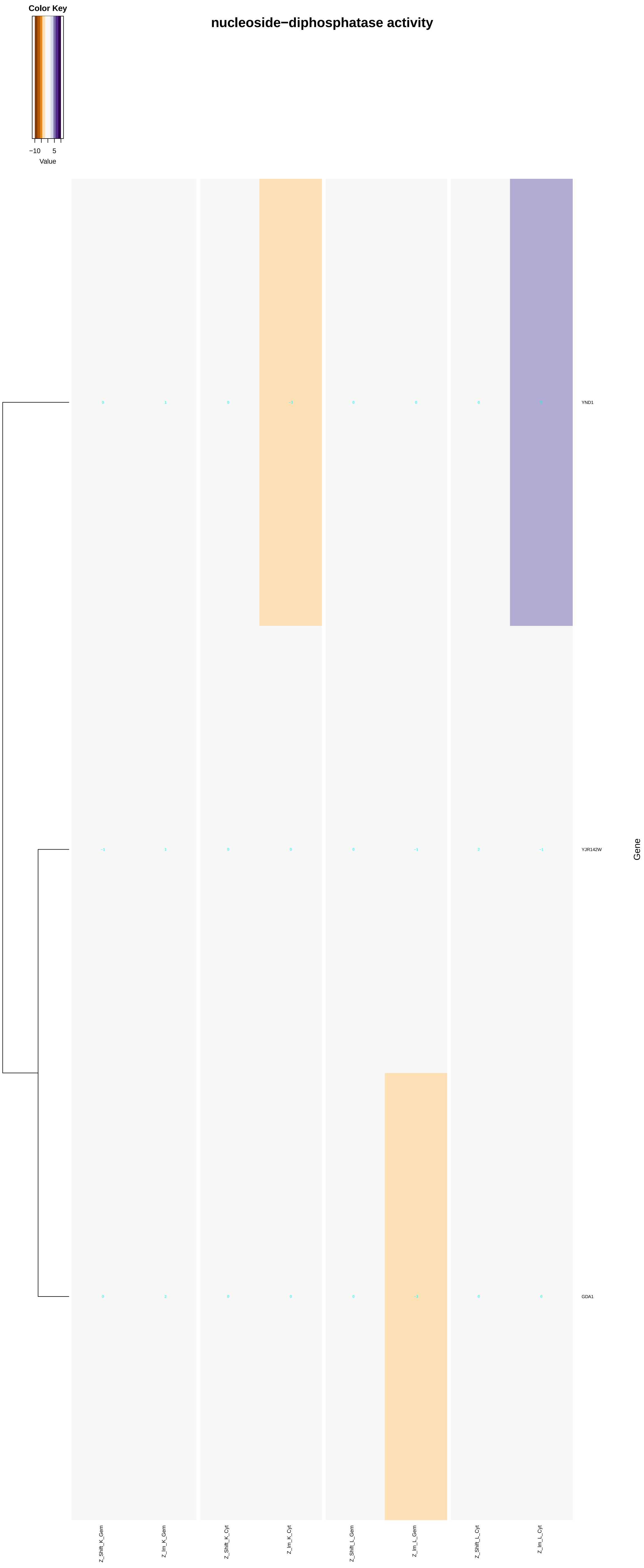
Color Key
−10
5
Value
nucleoside−diphosphatase activity
YND1
YJR142W
GDA1
Gene
Z_Shift_K_Gem
Z_Im_K_Gem
Z_Shift_K_Cyt
Z_Im_K_Cyt
Z_Shift_L_Gem
Z_Im_L_Gem
Z_Shift_L_Cyt
Z_Im_L_Cyt

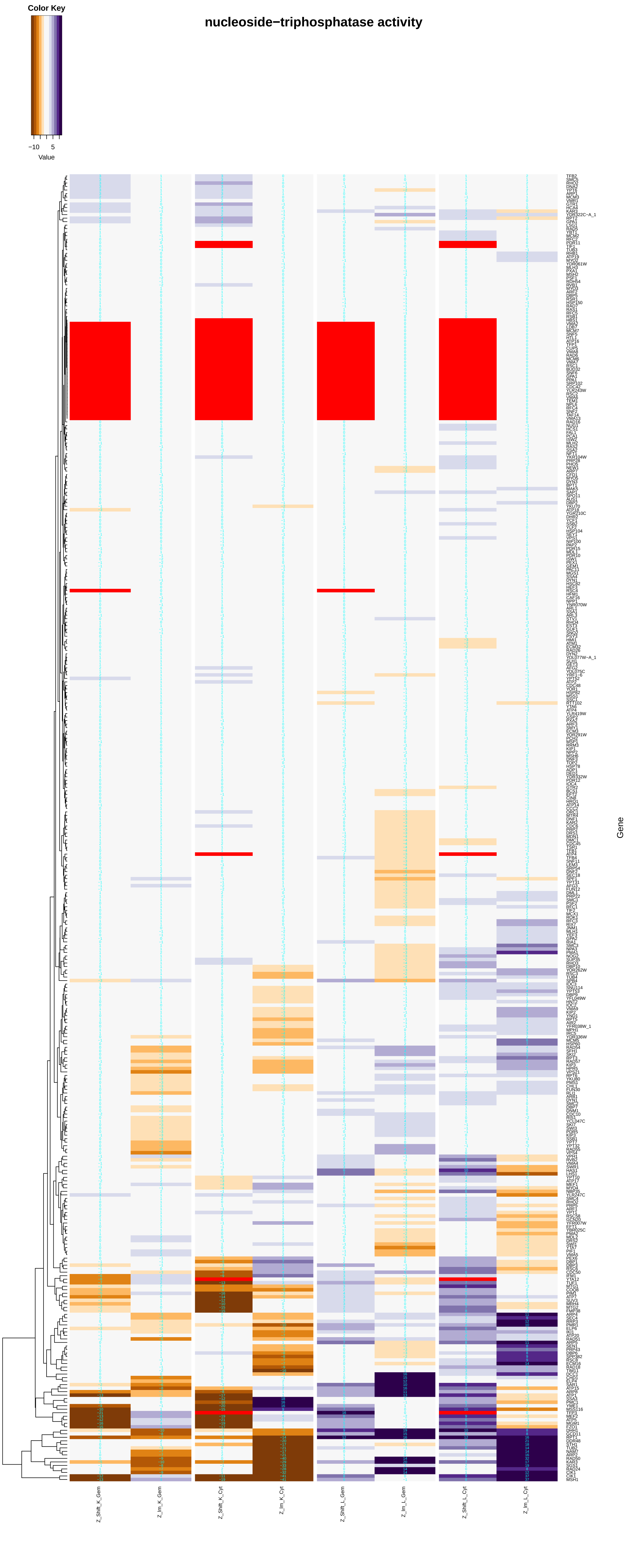
nucleoside−triphosphatase activity
Color Key
−10
5
Value
Z_Shift_K_Gem
Z_Im_K_Gem
Z_Shift_K_Cyt
Z_Im_K_Cyt
Z_Shift_L_Gem
Z_Im_L_Gem
Z_Shift_L_Cyt
Z_Im_L_Cyt
Gene

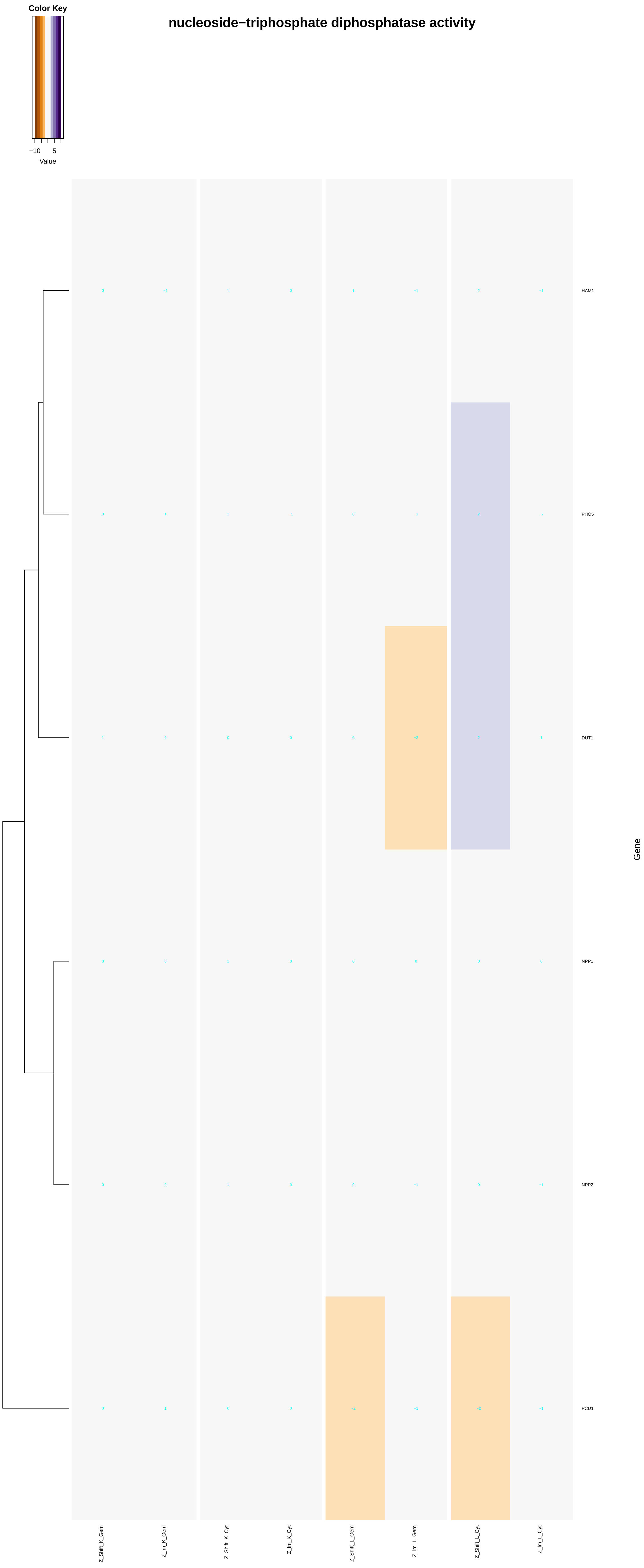
nucleoside−triphosphate diphosphatase activity
Color Key
−10
5
Value
HAM1
PHO5
DUT1
NPP1
NPP2
PCD1
Gene
Z_Shift_K_Gem
Z_Im_K_Gem
Z_Shift_K_Cyt
Z_Im_K_Cyt
Z_Shift_L_Gem
Z_Im_L_Gem
Z_Shift_L_Cyt
Z_Im_L_Cyt

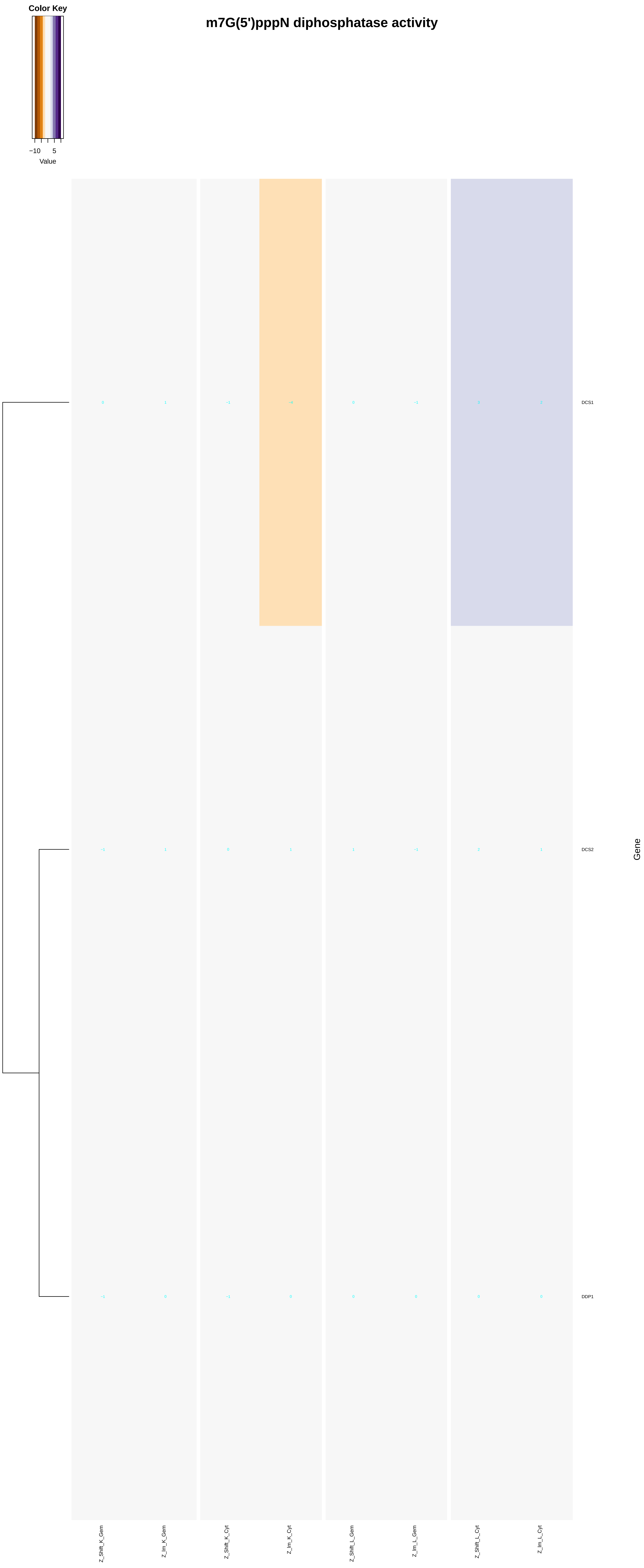

Color Key
−10
5
Value
m7G(5')pppN diphosphatase activity
DCS1
DCS2
DDP1
Gene
Z_Shift_K_Gem
Z_Im_K_Gem
Z_Shift_K_Cyt
Z_Im_K_Cyt
Z_Shift_L_Gem
Z_Im_L_Gem
Z_Shift_L_Cyt
Z_Im_L_Cyt

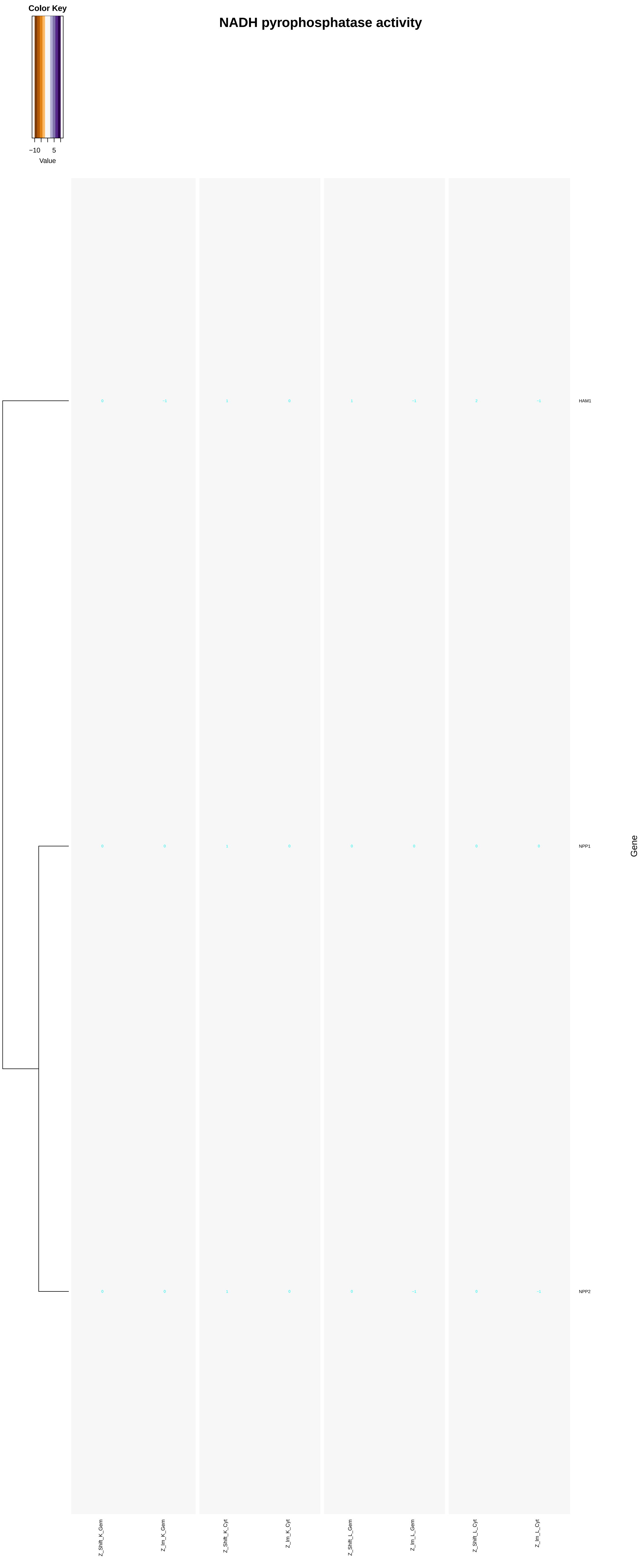
NADH pyrophosphatase activity
Color Key
−10
5
Value
HAM1
NPP1
NPP2
Gene
Z_Shift_K_Gem
Z_Im_K_Gem
Z_Shift_K_Cyt
Z_Im_K_Cyt
Z_Shift_L_Gem
Z_Im_L_Gem
Z_Shift_L_Cyt
Z_Im_L_Cyt

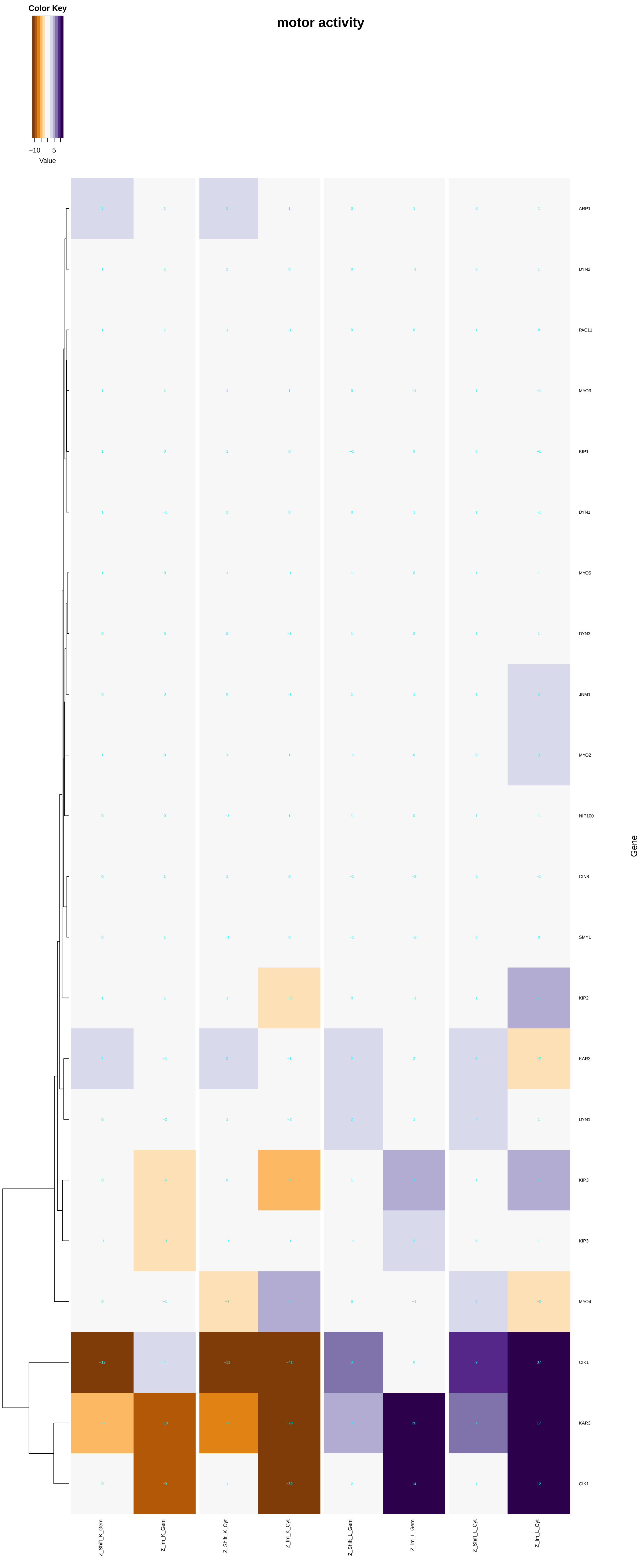
motor activity
Color Key
−10
5
Value
ARP1
DYN2
PAC11
MYO3
KIP1
DYN1
MYO5
DYN3
JNM1
MYO2
NIP100
CIN8
SMY1
KIP2
KAR3
DYN1
KIP3
KIP3
MYO4
CIK1
KAR3
CIK1
Gene
Z_Shift_K_Gem
Z_Im_K_Gem
Z_Shift_K_Cyt
Z_Im_K_Cyt
Z_Shift_L_Gem
Z_Im_L_Gem
Z_Shift_L_Cyt
Z_Im_L_Cyt

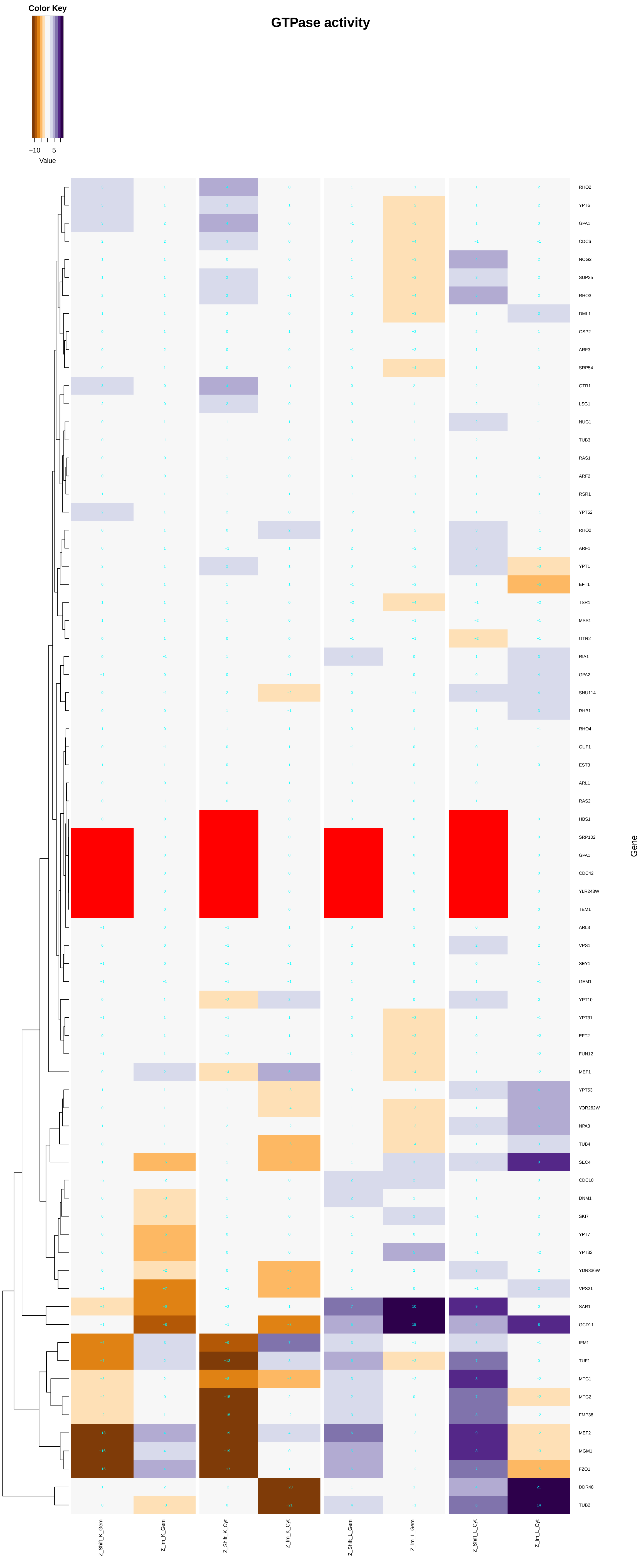
Color Key
−10
5
Value
GTPase activity
Gene
Z_Shift_K_Gem
Z_Im_K_Gem
Z_Shift_K_Cyt
Z_Im_K_Cyt
Z_Shift_L_Gem
Z_Im_L_Gem
Z_Shift_L_Cyt
Z_Im_L_Cyt
RHO2
YPT6
GPA1
CDC6
NOG2
SUP35
RHO3
DML1
GSP2
ARF3
SRP54
GTR1
LSG1
NUG1
TUB3
RAS1
ARF2
RSR1
YPT52
RHO2
ARF1
YPT1
EFT1
TSR1
MSS1
GTR2
RIA1
GPA2
SNU114
RHB1
RHO4
GUF1
EST3
ARL1
RAS2
HBS1
SRP102
GPA1
CDC42
YLR243W
TEM1
ARL3
VPS1
SEY1
GEM1
YPT10
YPT31
EFT2
FUN12
MEF1
YPT53
YOR262W
NPA3
TUB4
SEC4
CDC10
DNM1
SKI7
YPT7
YPT32
YDR336W
VPS21
SAR1
GCD11
IFM1
TUF1
MTG1
MTG2
FMP38
MEF2
MGM1
FZO1
DDR48
TUB2

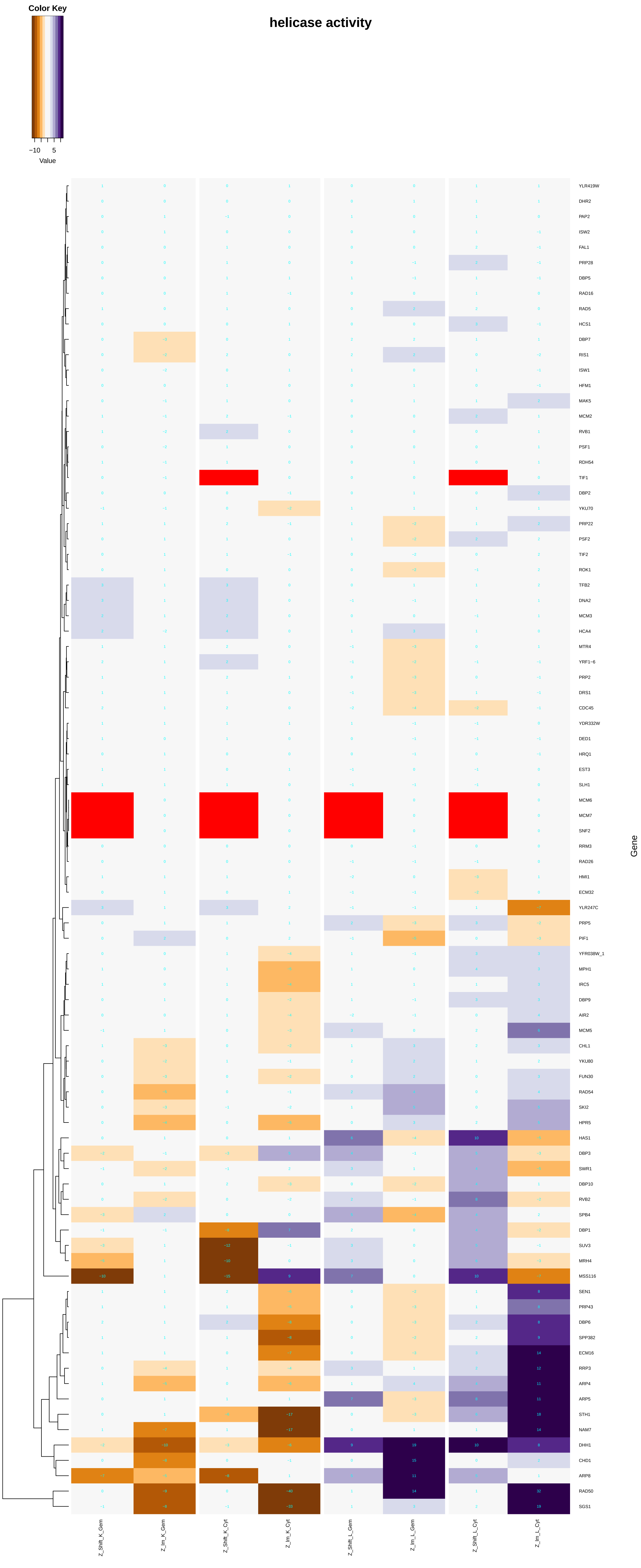
helicase activity
Color Key
−10
5
Value
YLR419W
DHR2
PAP2
ISW2
FAL1
PRP28
DBP5
RAD16
RAD5
HCS1
DBP7
RIS1
ISW1
HFM1
MAK5
MCM2
RVB1
PSF1
RDH54
TIF1
DBP2
YKU70
PRP22
PSF2
TIF2
ROK1
TFB2
DNA2
MCM3
HCA4
MTR4
YRF1−6
PRP2
DRS1
CDC45
YDR332W
DED1
HRQ1
EST3
SLH1
MCM6
MCM7
SNF2
RRM3
RAD26
HMI1
ECM32
YLR247C
PRP5
PIF1
YFR038W_1
MPH1
IRC5
DBP9
AIR2
MCM5
CHL1
YKU80
FUN30
RAD54
SKI2
HPR5
HAS1
DBP3
SWR1
DBP10
RVB2
SPB4
DBP1
SUV3
MRH4
MSS116
SEN1
PRP43
DBP6
SPP382
ECM16
RRP3
ARP4
ARP5
STH1
NAM7
DHH1
CHD1
ARP8
RAD50
SGS1
Gene
Z_Shift_K_Gem
Z_Im_K_Gem
Z_Shift_K_Cyt
Z_Im_K_Cyt
Z_Shift_L_Gem
Z_Im_L_Gem
Z_Shift_L_Cyt
Z_Im_L_Cyt

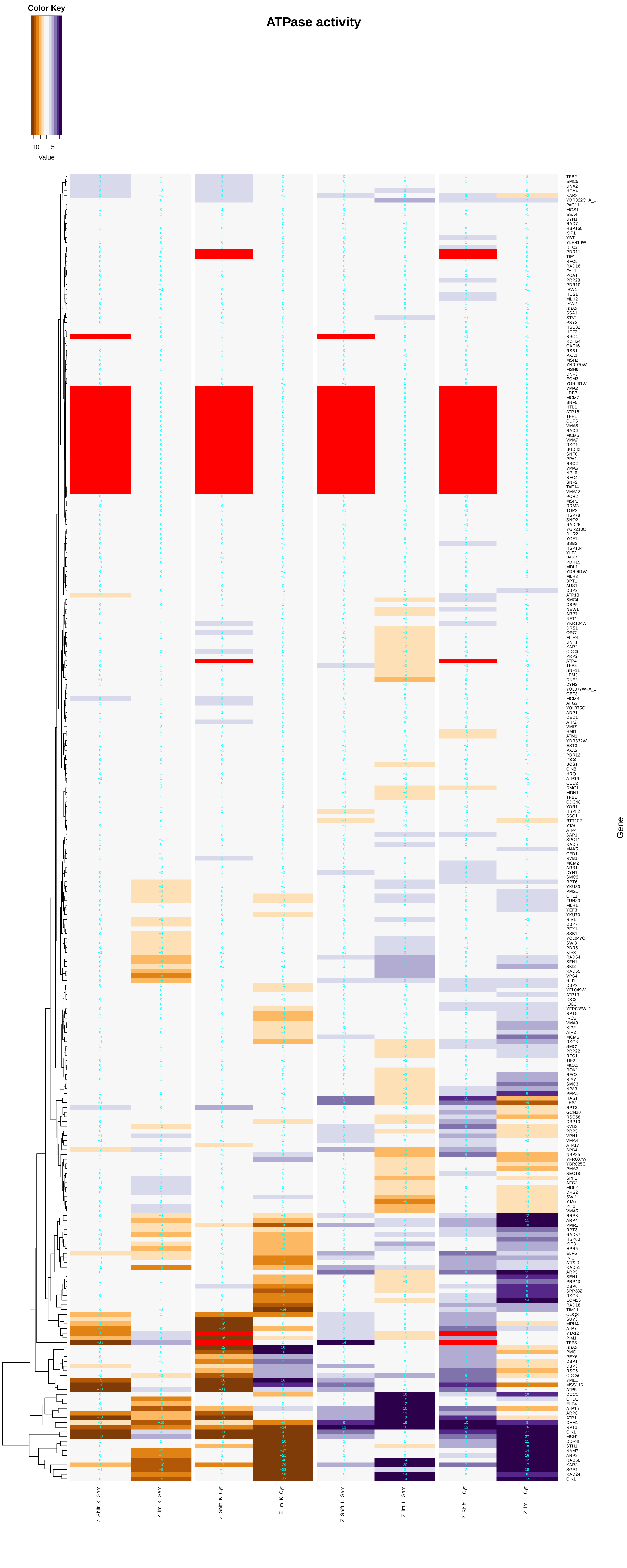
ATPase activity
Color Key
−10 5
Value
Gene
Z_Shift_K_Gem
Z_Im_K_Gem
Z_Shift_K_Cyt
Z_Im_K_Cyt
Z_Shift_L_Gem
Z_Im_L_Gem
Z_Shift_L_Cyt
Z_Im_L_Cyt

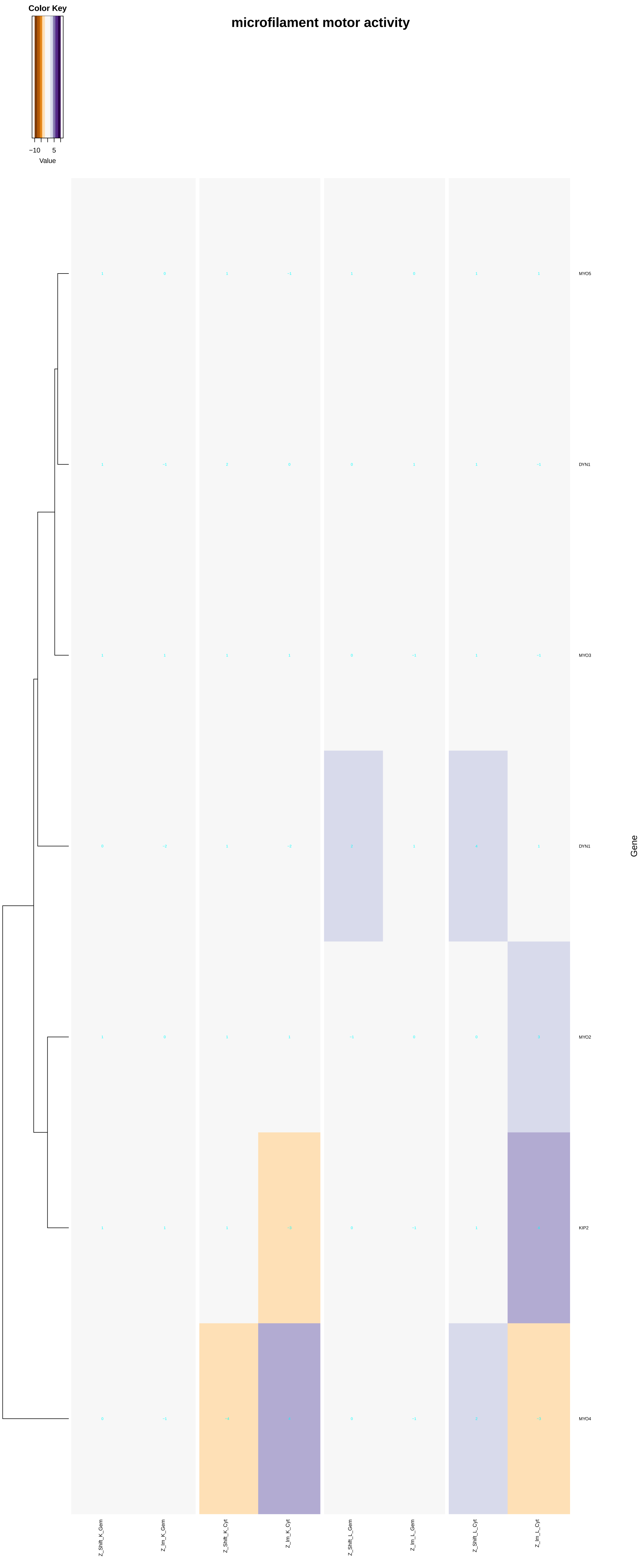
microfilament motor activity
Color Key
−10
5
Value
MYO5
DYN1
MYO3
DYN1
MYO2
KIP2
MYO4
Gene
Z_Shift_K_Gem
Z_Im_K_Gem
Z_Shift_K_Cyt
Z_Im_K_Cyt
Z_Shift_L_Gem
Z_Im_L_Gem
Z_Shift_L_Cyt
Z_Im_L_Cyt

microtubule motor activity

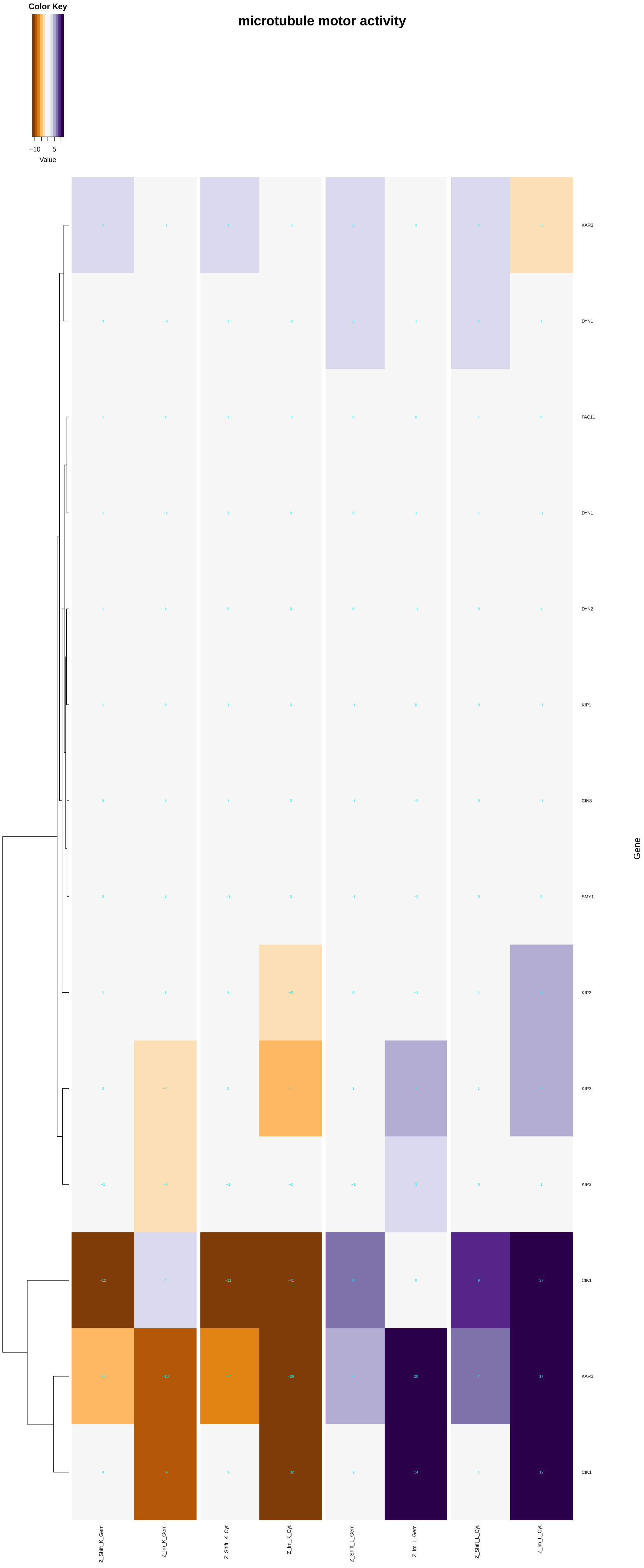

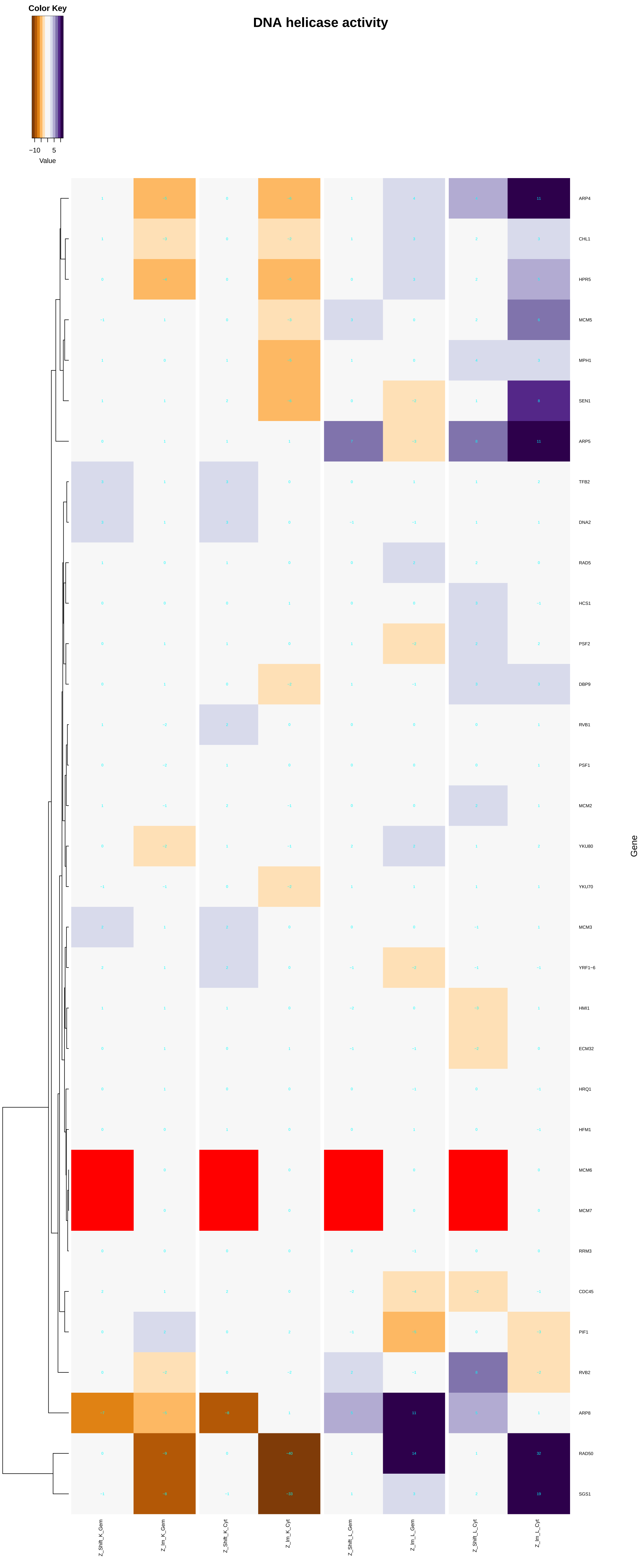
DNA helicase activity
Color Key
−10
5
Value
ARP4
CHL1
HPR5
MCM5
MPH1
SEN1
ARP5
TFB2
DNA2
RAD5
HCS1
PSF2
DBP9
RVB1
PSF1
MCM2
YKU80
YKU70
MCM3
YRF1−6
HMI1
ECM32
HRQ1
HFM1
MCM6
MCM7
RRM3
CDC45
PIF1
RVB2
ARP8
RAD50
SGS1
Gene
Z_Shift_K_Gem
Z_Im_K_Gem
Z_Shift_K_Cyt
Z_Im_K_Cyt
Z_Shift_L_Gem
Z_Im_L_Gem
Z_Shift_L_Cyt
Z_Im_L_Cyt

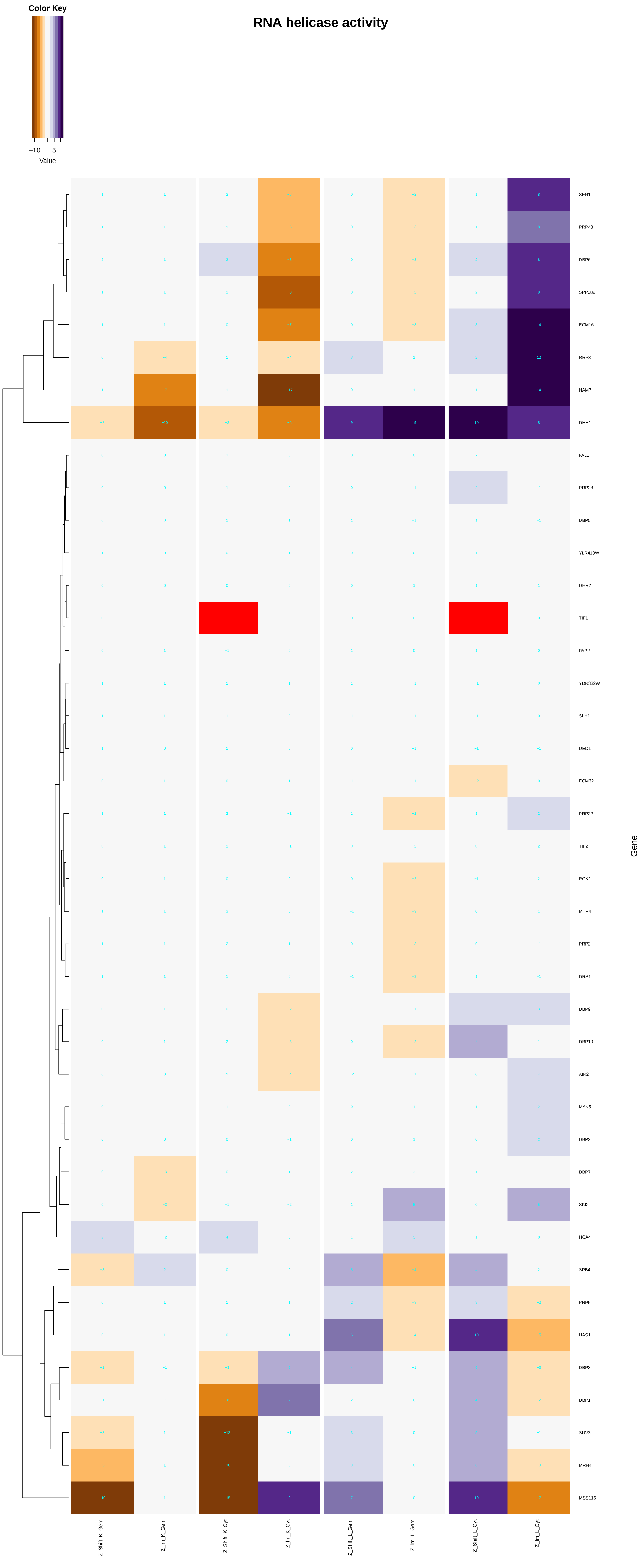
RNA helicase activity
Color Key
−10
5
Value
SEN1
PRP43
DBP6
SPP382
ECM16
RRP3
NAM7
DHH1
FAL1
PRP28
DBP5
YLR419W
DHR2
TIF1
PAP2
YDR332W
SLH1
DED1
ECM32
PRP22
TIF2
ROK1
MTR4
PRP2
DRS1
DBP9
DBP10
AIR2
MAK5
DBP2
DBP7
SKI2
HCA4
SPB4
PRP5
HAS1
DBP3
DBP1
SUV3
MRH4
MSS116
Gene
Z_Shift_K_Gem
Z_Im_K_Gem
Z_Shift_K_Cyt
Z_Im_K_Cyt
Z_Shift_L_Gem
Z_Im_L_Gem
Z_Shift_L_Cyt
Z_Im_L_Cyt

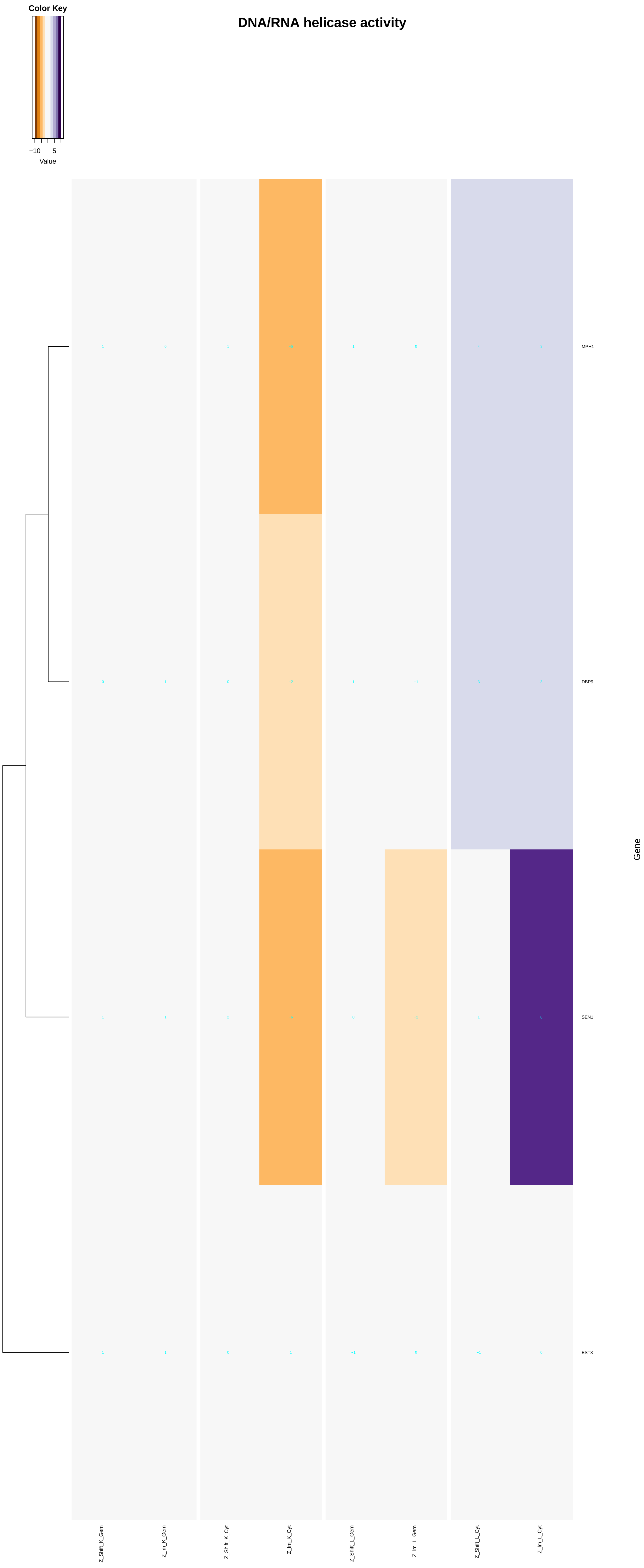
DNA/RNA helicase activity
Color Key
−10
5
Value
MPH1
DBP9
SEN1
EST3
Gene
Z_Shift_K_Gem
Z_Im_K_Gem
Z_Shift_K_Cyt
Z_Im_K_Cyt
Z_Shift_L_Gem
Z_Im_L_Gem
Z_Shift_L_Cyt
Z_Im_L_Cyt

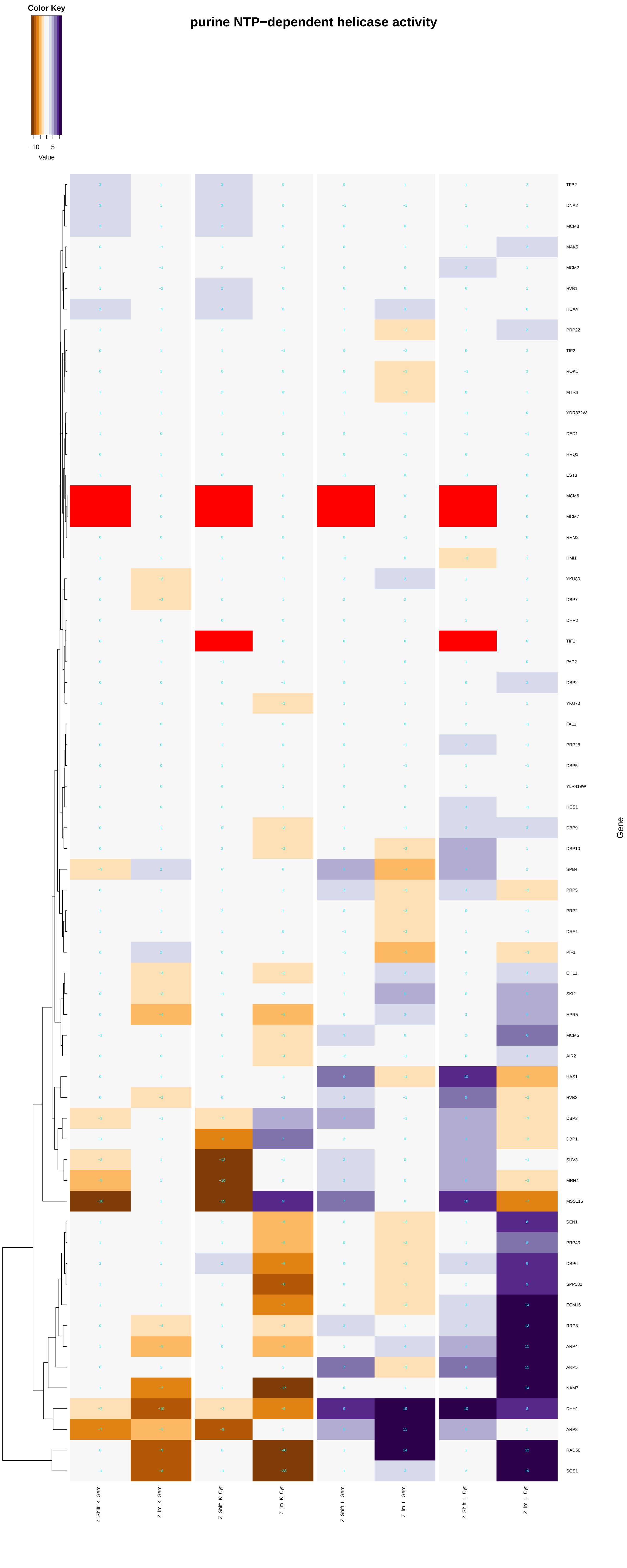
purine NTP−dependent helicase activity
Color Key
−10 5
Value
TFB2
DNA2
MCM3
MAK5
MCM2
RVB1
HCA4
PRP22
TIF2
ROK1
MTR4
YDR332W
DED1
HRQ1
EST3
MCM6
MCM7
RRM3
HMI1
YKU80
DBP7
DHR2
TIF1
PAP2
DBP2
YKU70
FAL1
PRP28
DBP5
YLR419W
HCS1
DBP9
DBP10
SPB4
PRP5
PRP2
DRS1
PIF1
CHL1
SKI2
HPR5
MCM5
AIR2
HAS1
RVB2
DBP3
DBP1
SUV3
MRH4
MSS116
SEN1
PRP43
DBP6
SPP382
ECM16
RRP3
ARP4
ARP5
NAM7
DHH1
ARP8
RAD50
SGS1
Gene
Z_Shift_K_Gem
Z_Im_K_Gem
Z_Shift_K_Cyt
Z_Im_K_Cyt
Z_Shift_L_Gem
Z_Im_L_Gem
Z_Shift_L_Cyt
Z_Im_L_Cyt

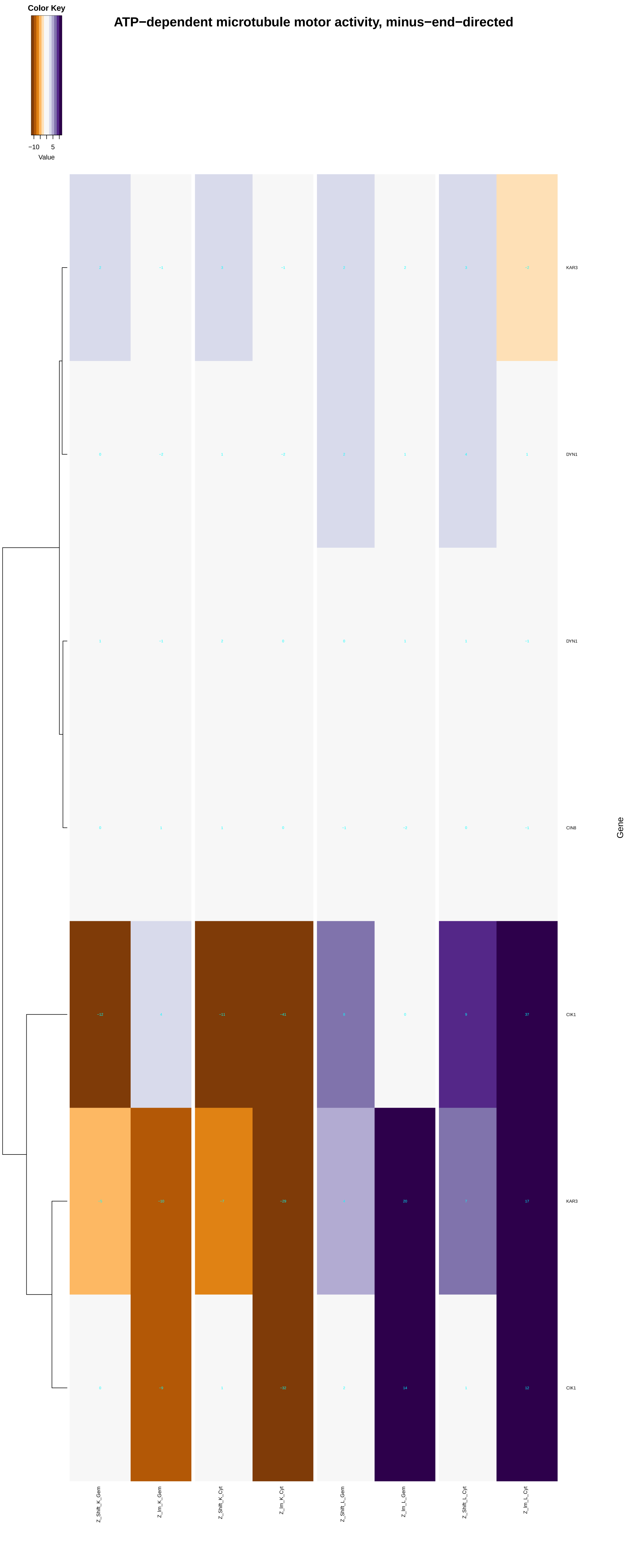

Color Key
−10
5
Value
ATP−dependent microtubule motor activity, minus−end−directed
KAR3
DYN1
DYN1
CIN8
CIK1
KAR3
CIK1
Gene
Z_Shift_K_Gem
Z_Im_K_Gem
Z_Shift_K_Cyt
Z_Im_K_Cyt
Z_Shift_L_Gem
Z_Im_L_Gem
Z_Shift_L_Cyt
Z_Im_L_Cyt

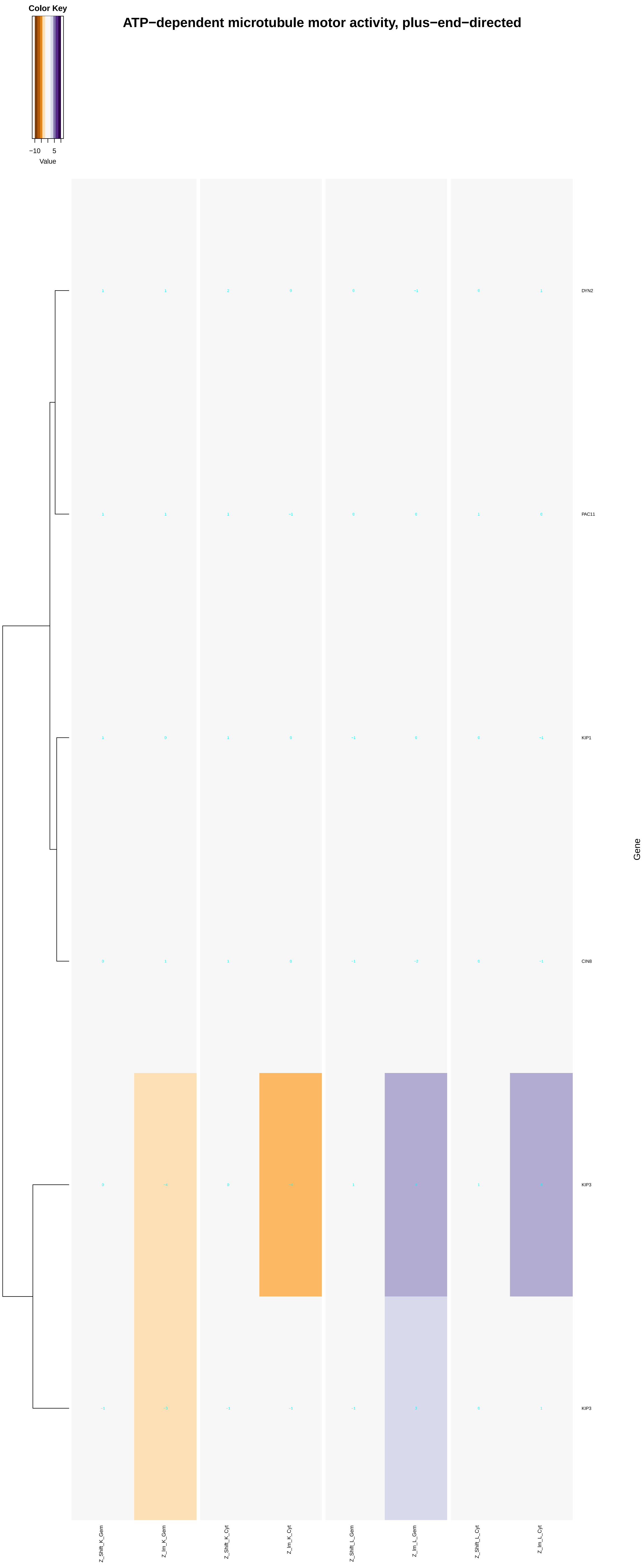
ATP−dependent microtubule motor activity, plus−end−directed
Color Key
−10
5
Value
DYN2
PAC11
KIP1
CIN8
KIP3
KIP3
Gene
Z_Shift_K_Gem
Z_Im_K_Gem
Z_Shift_K_Cyt
Z_Im_K_Cyt
Z_Shift_L_Gem
Z_Im_L_Gem
Z_Shift_L_Cyt
Z_Im_L_Cyt

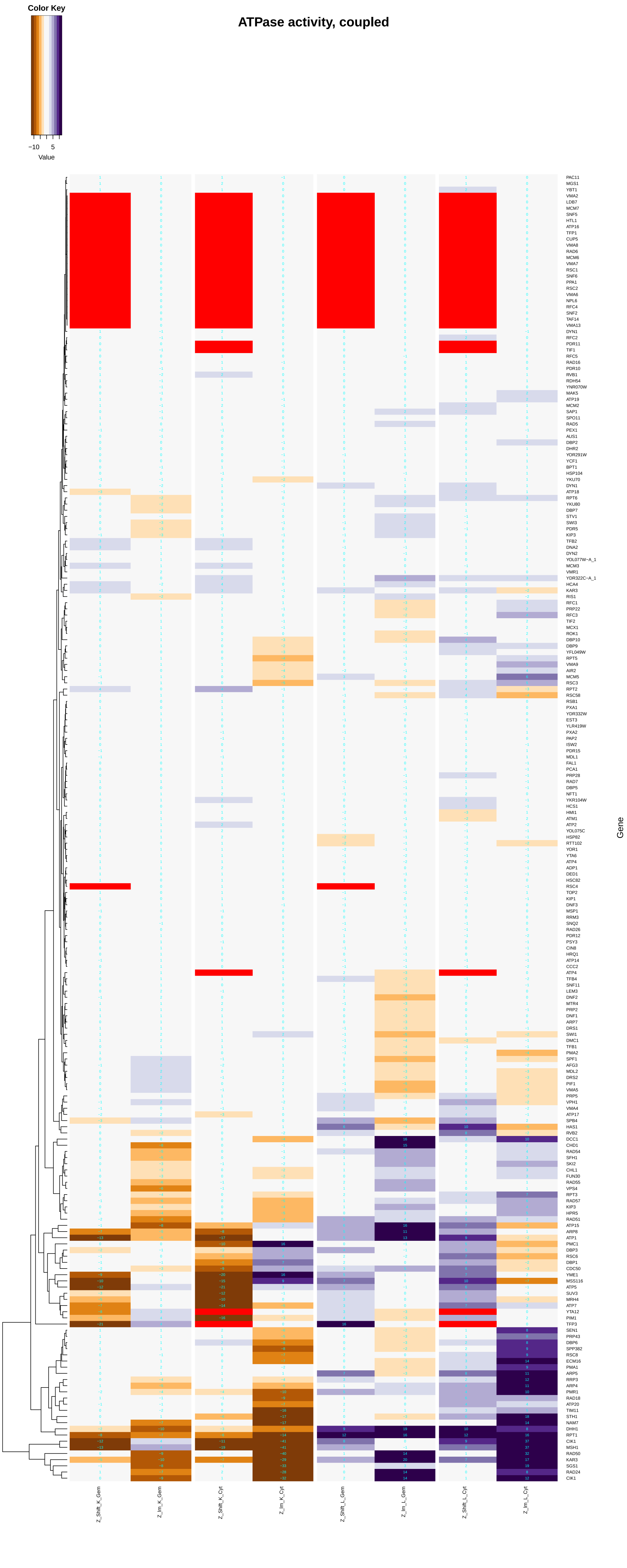

ATPase activity, coupled
Color Key
−10 5
Value
Gene
Z_Shift_K_Gem
Z_Im_K_Gem
Z_Shift_K_Cyt
Z_Im_K_Cyt
Z_Shift_L_Gem
Z_Im_L_Gem
Z_Shift_L_Cyt
Z_Im_L_Cyt

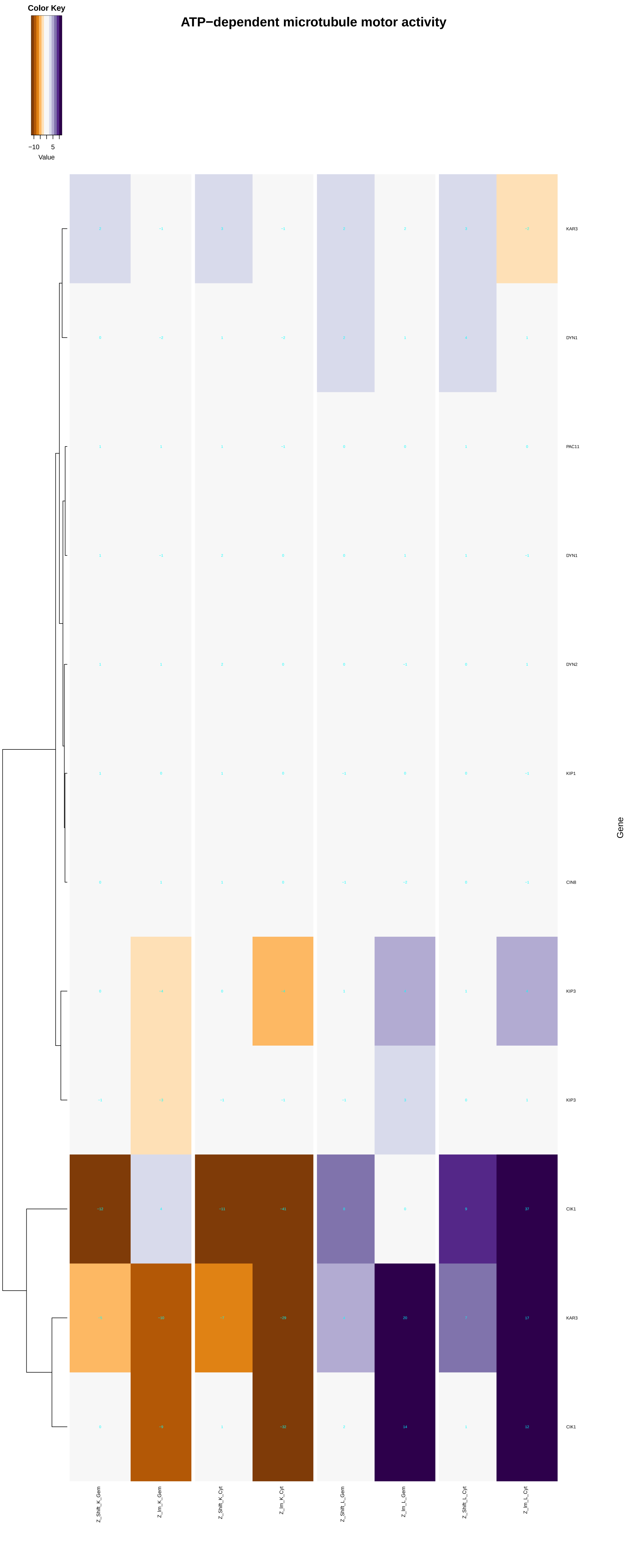
Color Key
−10
5
Value
ATP−dependent microtubule motor activity
KAR3
DYN1
PAC11
DYN1
DYN2
KIP1
CIN8
KIP3
KIP3
CIK1
KAR3
CIK1
Gene
Z_Shift_K_Gem
Z_Im_K_Gem
Z_Shift_K_Cyt
Z_Im_K_Cyt
Z_Shift_L_Gem
Z_Im_L_Gem
Z_Shift_L_Cyt
Z_Im_L_Cyt

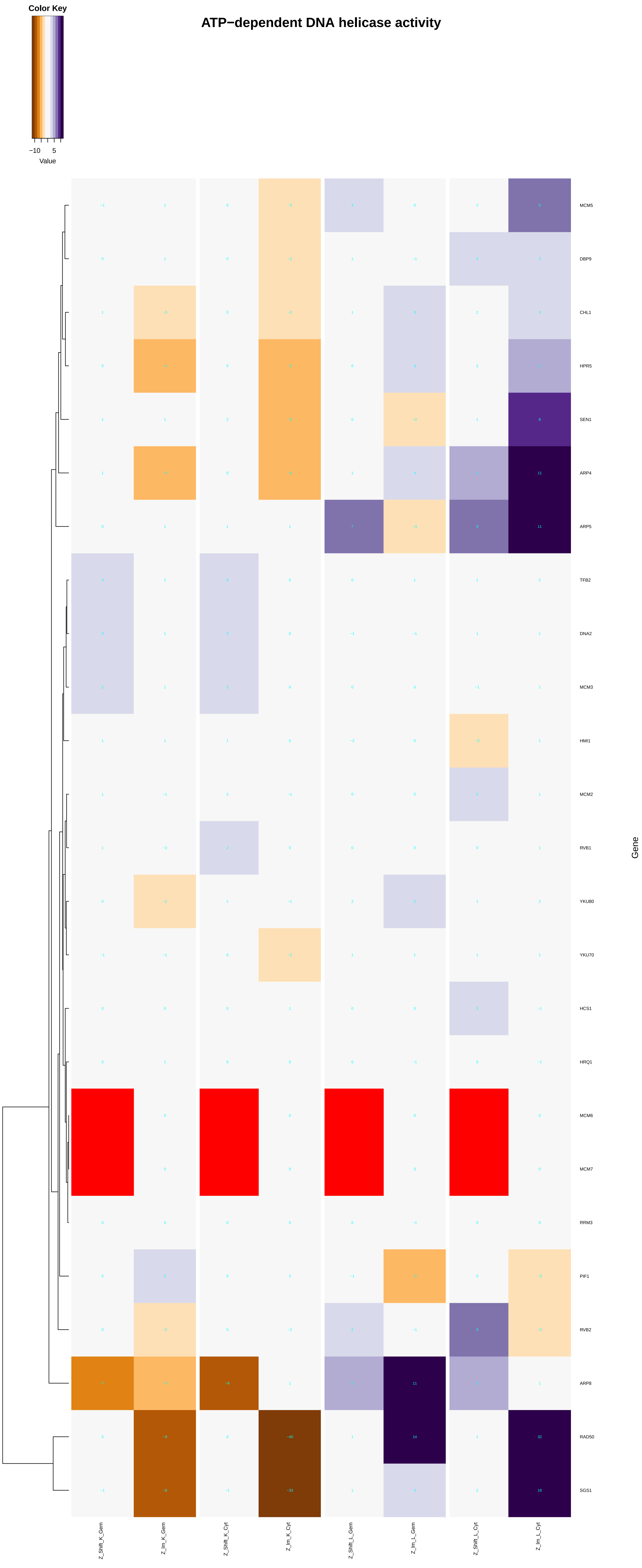

ATP−dependent DNA helicase activity
Color Key
−10
5
Value
MCM5
DBP9
CHL1
HPR5
SEN1
ARP4
ARP5
TFB2
DNA2
MCM3
HMI1
MCM2
RVB1
YKU80
YKU70
HCS1
HRQ1
MCM6
MCM7
RRM3
PIF1
RVB2
ARP8
RAD50
SGS1
Gene
Z_Shift_K_Gem
Z_Im_K_Gem
Z_Shift_K_Cyt
Z_Im_K_Cyt
Z_Shift_L_Gem
Z_Im_L_Gem
Z_Shift_L_Cyt
Z_Im_L_Cyt

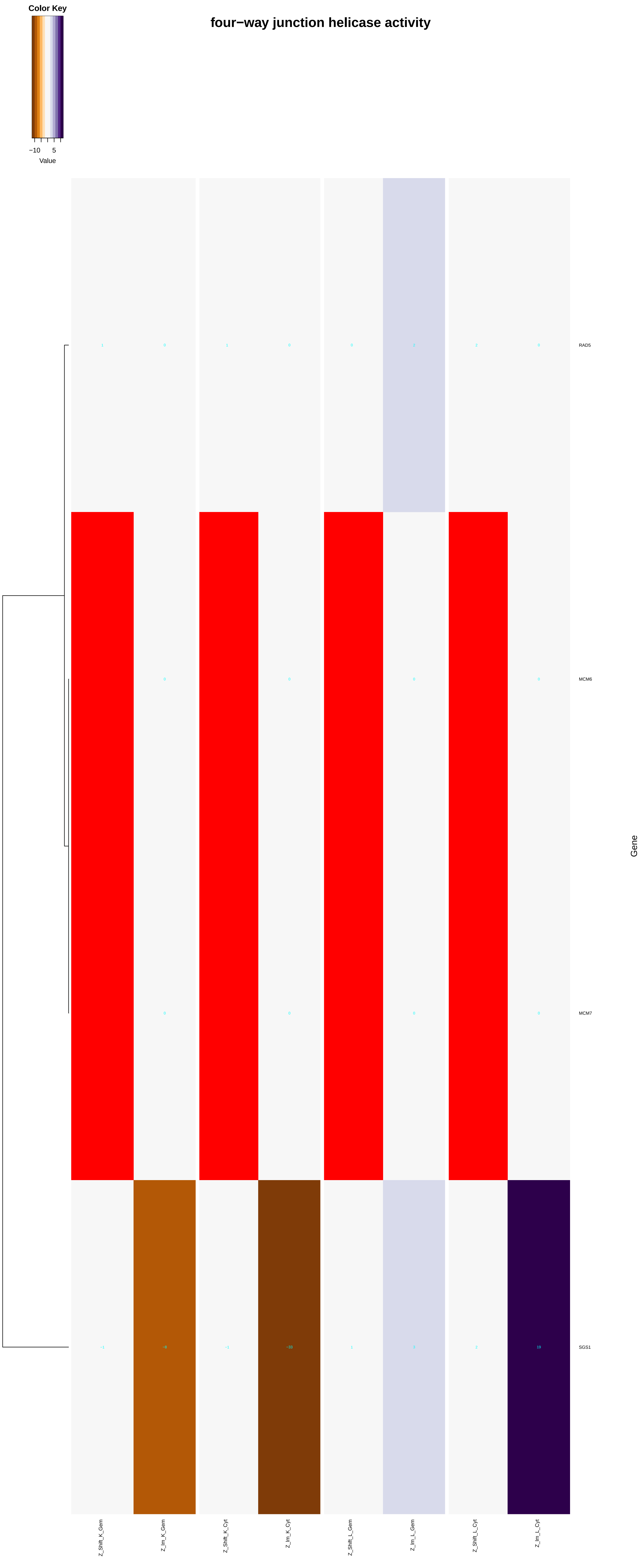
four−way junction helicase activity
Color Key
−10
5
Value
RAD5
MCM6
MCM7
SGS1
Gene
Z_Shift_K_Gem
Z_Im_K_Gem
Z_Shift_K_Cyt
Z_Im_K_Cyt
Z_Shift_L_Gem
Z_Im_L_Gem
Z_Shift_L_Cyt
Z_Im_L_Cyt

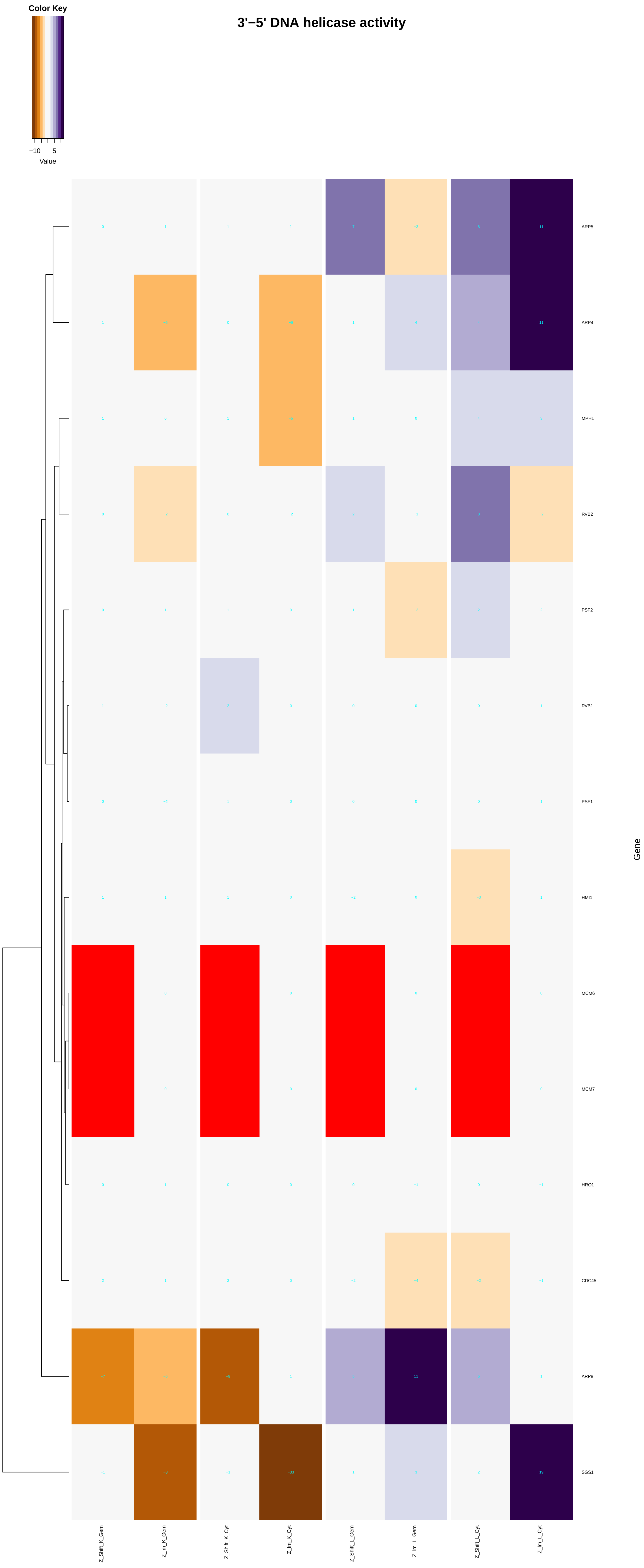
3'−5' DNA helicase activity
Color Key
−10
5
Value
ARP5
ARP4
MPH1
RVB2
PSF2
RVB1
PSF1
HMI1
MCM6
MCM7
HRQ1
CDC45
ARP8
SGS1
Gene
Z_Shift_K_Gem
Z_lm_K_Gem
Z_Shift_K_Cyt
Z_lm_K_Cyt
Z_Shift_L_Gem
Z_lm_L_Gem
Z_Shift_L_Cyt
Z_lm_L_Cyt

#### 5'−3' DNA helicase activity

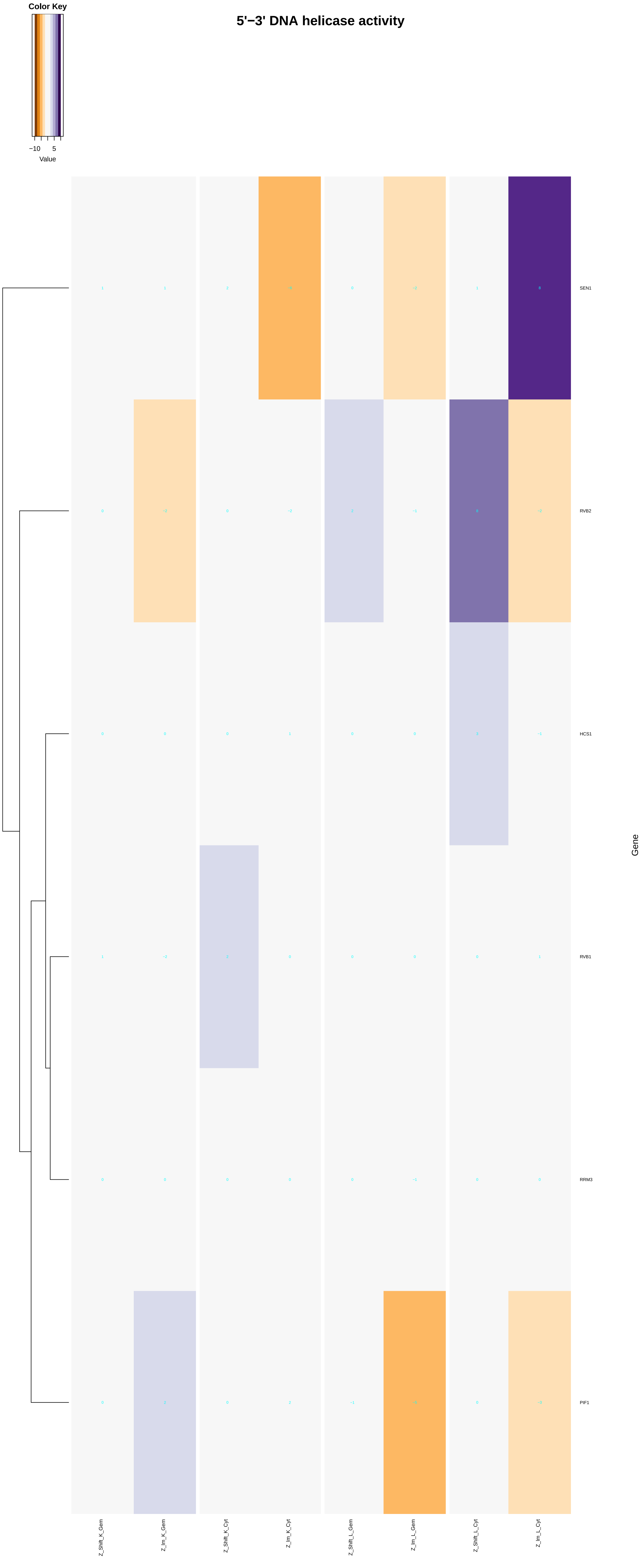

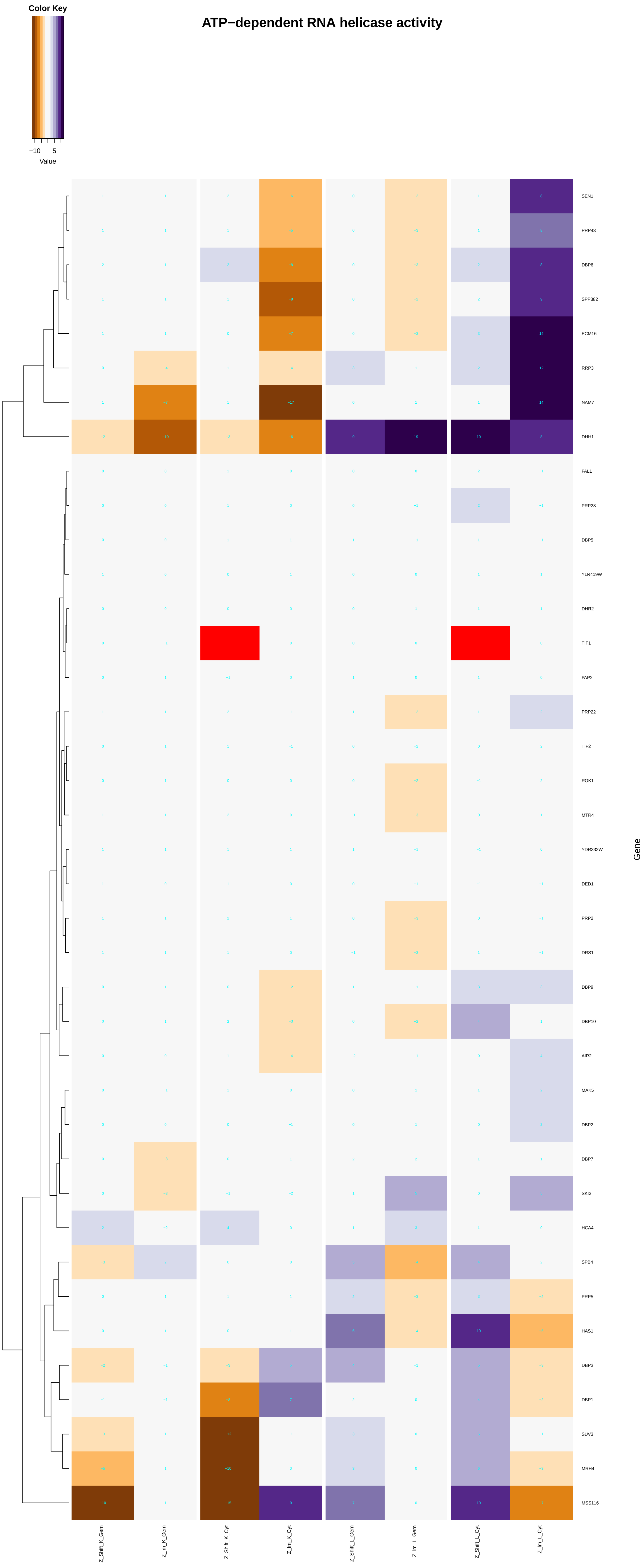
ATP−dependent RNA helicase activity
Color Key
−10
5
Value
SEN1
PRP43
DBP6
SPP382
ECM16
RRP3
NAM7
DHH1
FAL1
PRP28
DBP5
YLR419W
DHR2
TIF1
PAP2
PRP22
TIF2
ROK1
MTR4
YDR332W
DED1
PRP2
DRS1
DBP9
DBP10
AIR2
MAK5
DBP2
DBP7
SKI2
HCA4
SPB4
PRP5
HAS1
DBP3
DBP1
SUV3
MRH4
MSS116
Gene
Z_Shift_K_Gem
Z_Im_K_Gem
Z_Shift_K_Cyt
Z_Im_K_Cyt
Z_Shift_L_Gem
Z_Im_L_Gem
Z_Shift_L_Cyt
Z_Im_L_Cyt

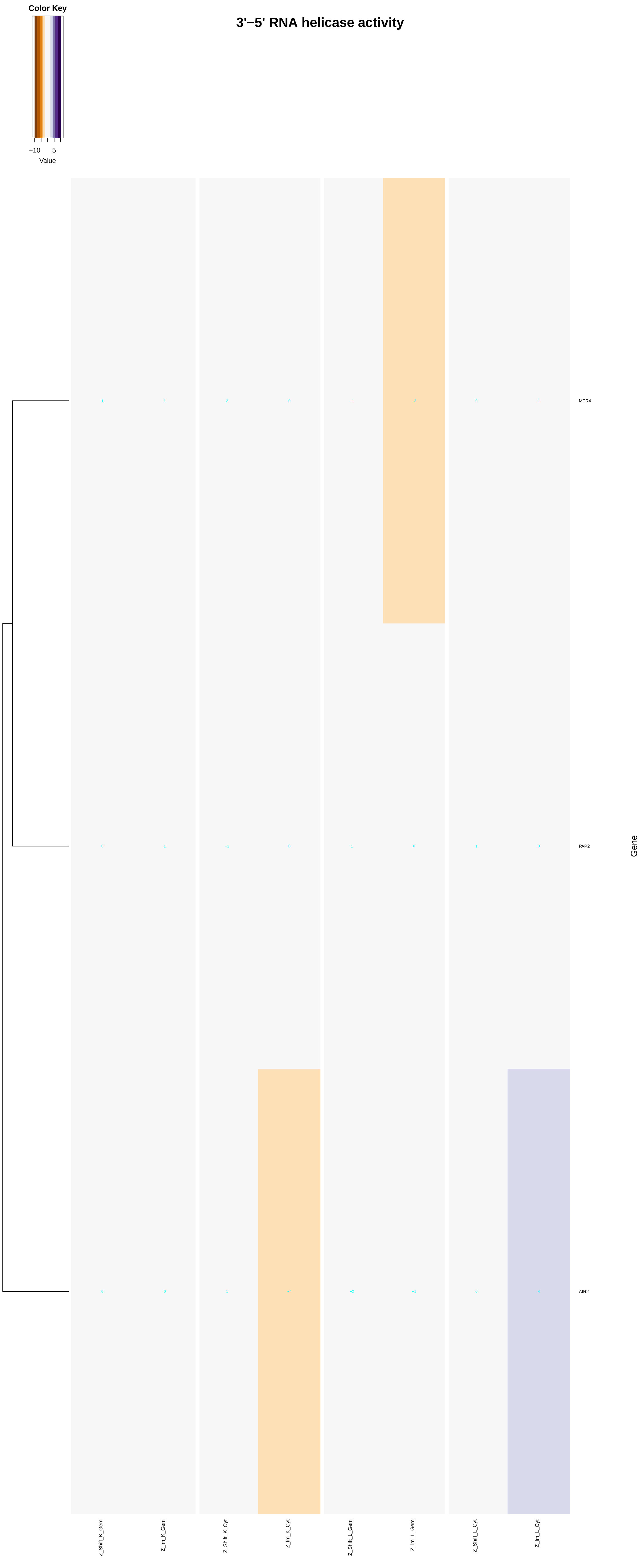
3'−5' RNA helicase activity
Color Key
−10
5
Value
MTR4
PAP2
AIR2
Gene
Z_Shift_K_Gem
Z_Im_K_Gem
Z_Shift_K_Cyt
Z_Im_K_Cyt
Z_Shift_L_Gem
Z_Im_L_Gem
Z_Shift_L_Cyt
Z_Im_L_Cyt

### ATP−dependent DNA/RNA helicase activity

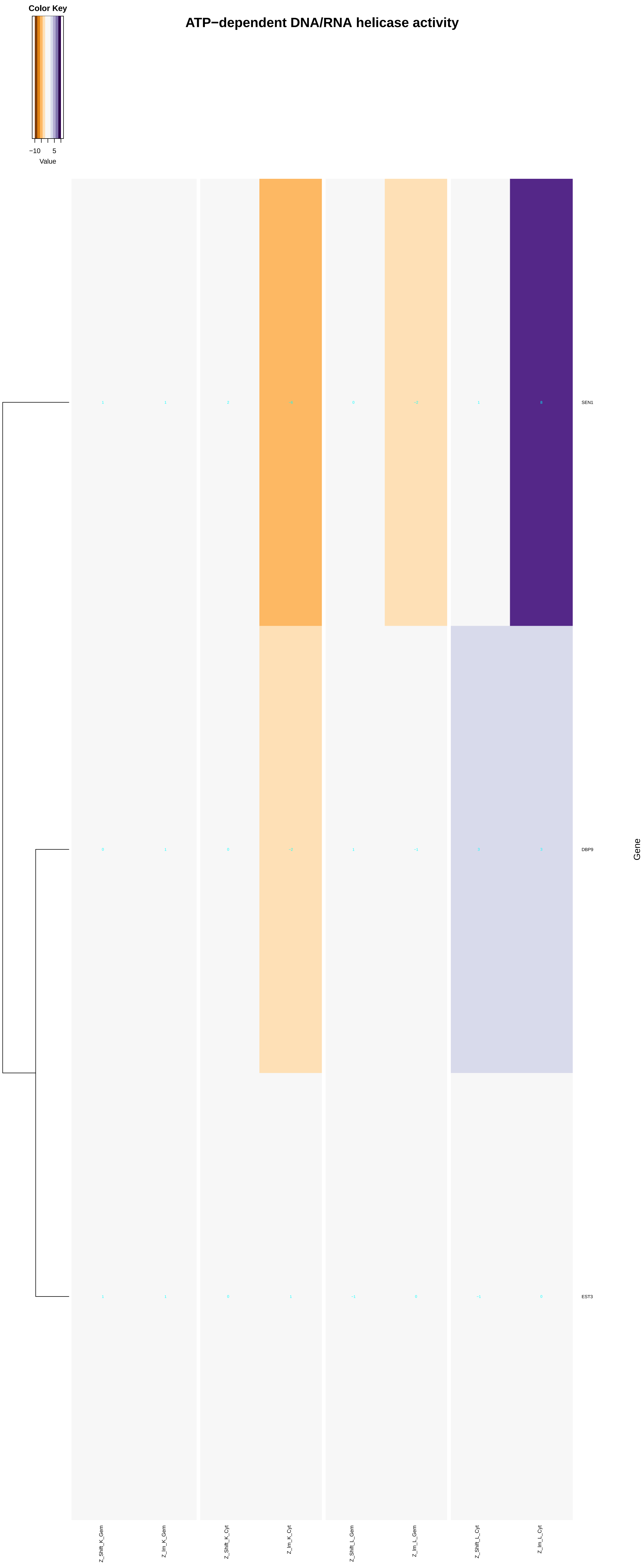

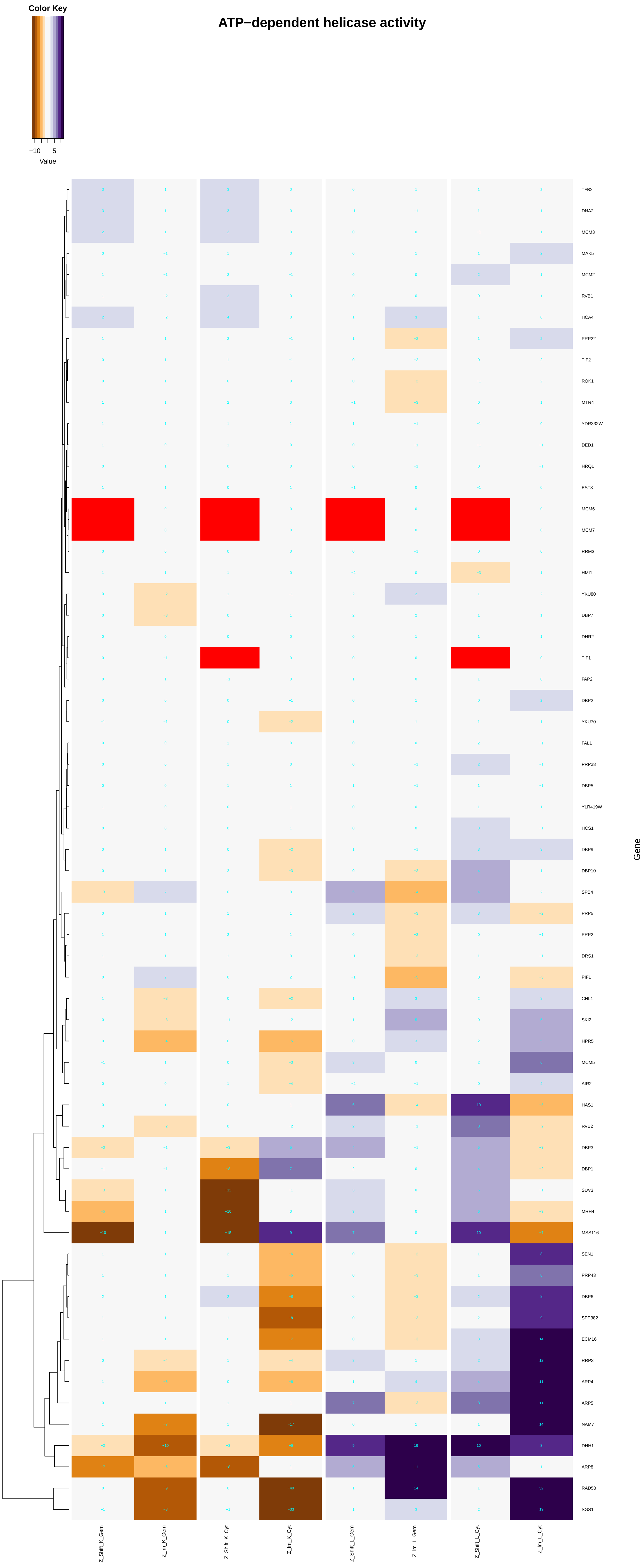
ATP−dependent helicase activity
Color Key
−10
5
Value
TFB2
DNA2
MCM3
MAK5
MCM2
RVB1
HCA4
PRP22
TIF2
ROK1
MTR4
YDR332W
DED1
HRQ1
EST3
MCM6
MCM7
RRM3
HMI1
YKU80
DBP7
DHR2
TIF1
PAP2
DBP2
YKU70
FAL1
PRP28
DBP5
YLR419W
HCS1
DBP9
DBP10
SPB4
PRP5
PRP2
DRS1
PIF1
CHL1
SKI2
HPR5
MCM5
AIR2
HAS1
RVB2
DBP3
DBP1
SUV3
MRH4
MSS116
SEN1
PRP43
DBP6
SPP382
ECM16
RRP3
ARP4
ARP5
NAM7
DHH1
ARP8
RAD50
SGS1
Gene
Z_Shift_K_Gem
Z_Im_K_Gem
Z_Shift_K_Cyt
Z_Im_K_Cyt
Z_Shift_L_Gem
Z_Im_L_Gem
Z_Shift_L_Cyt
Z_Im_L_Cyt

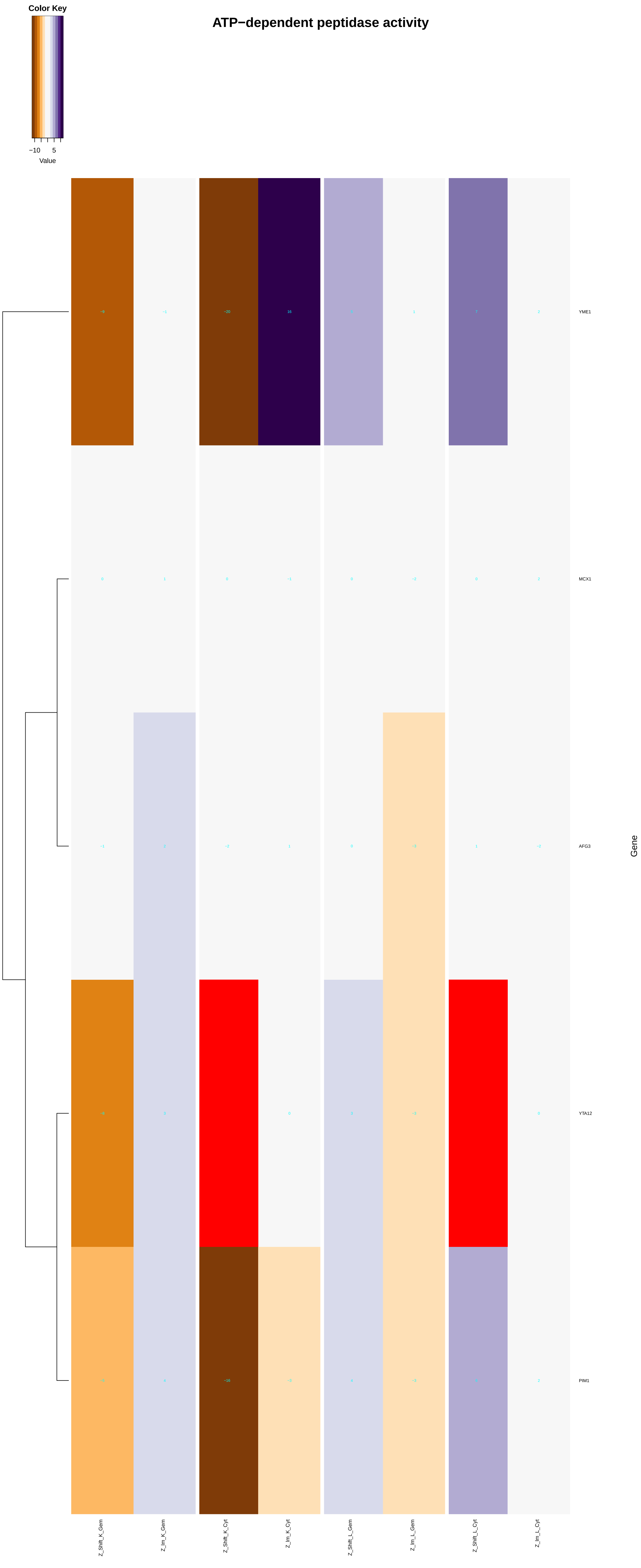

Color Key
−10 5
Value
ATP−dependent peptidase activity
YME1
MCX1
AFG3
YTA12
PIM1
Gene
Z_Shift_K_Gem
Z_Im_K_Gem
Z_Shift_K_Cyt
Z_Im_K_Cyt
Z_Shift_L_Gem
Z_Im_L_Gem
Z_Shift_L_Cyt
Z_Im_L_Cyt

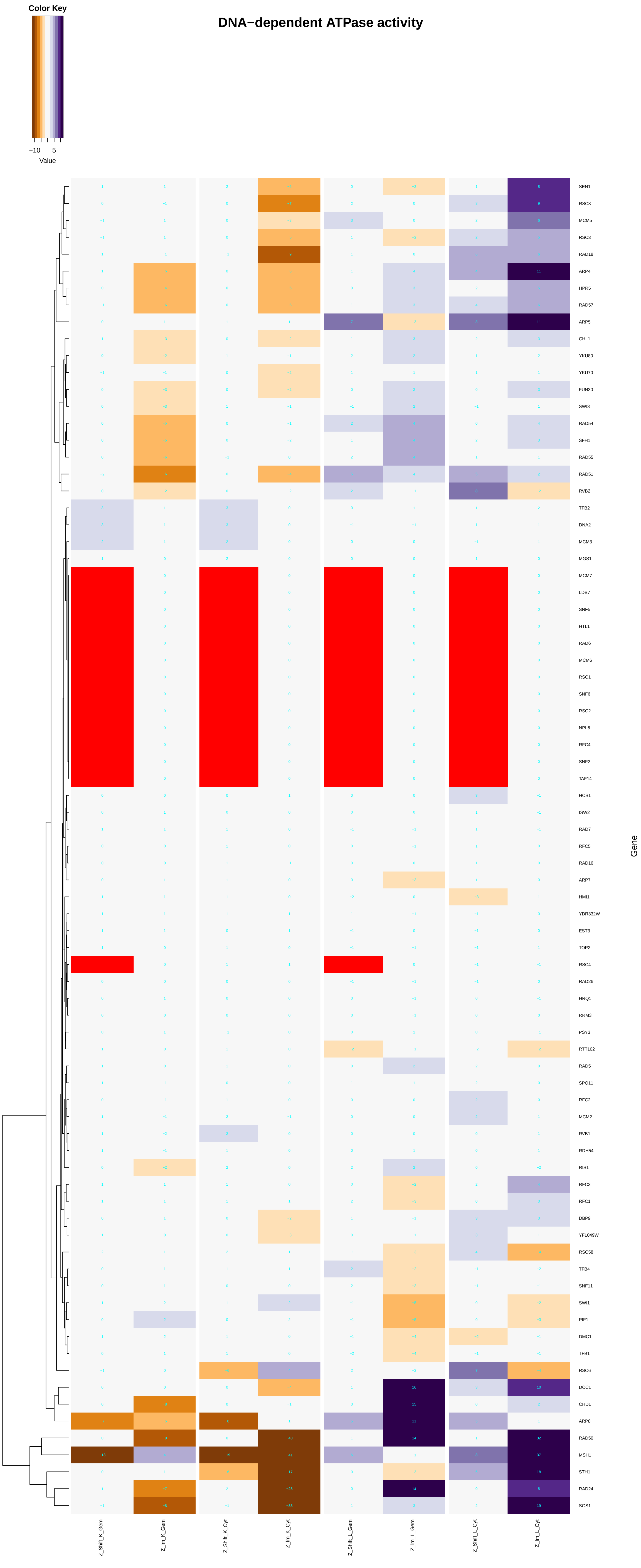
DNA−dependent ATPase activity
Color Key
−10
5
Value
Gene
Z_Shift_K_Gem
Z_Im_K_Gem
Z_Shift_K_Cyt
Z_Im_K_Cyt
Z_Shift_L_Gem
Z_Im_L_Gem
Z_Shift_L_Cyt
Z_Im_L_Cyt

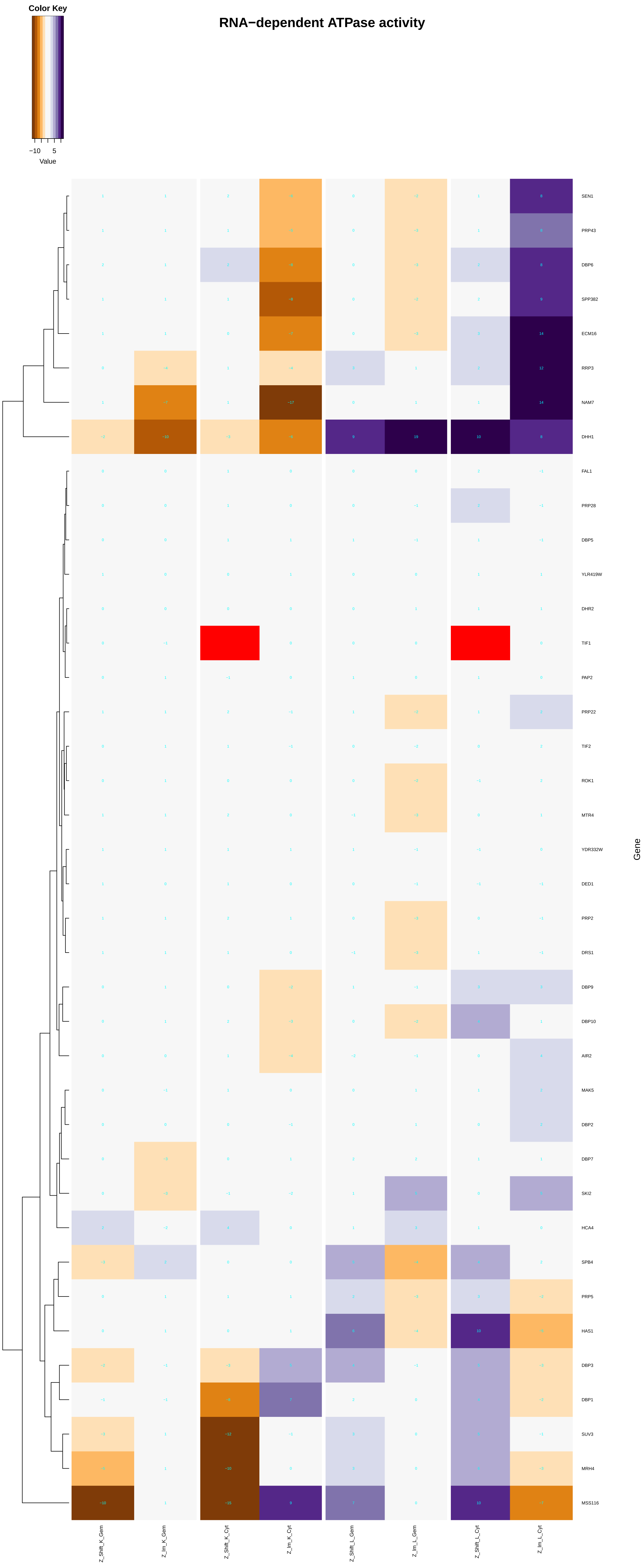

RNA−dependent ATPase activity
Color Key
−10 5
Value
SEN1
PRP43
DBP6
SPP382
ECM16
RRP3
NAM7
DHH1
FAL1
PRP28
DBP5
YLR419W
DHR2
TIF1
PAP2
PRP22
TIF2
ROK1
MTR4
YDR332W
DED1
PRP2
DRS1
DBP9
DBP10
AIR2
MAK5
DBP2
DBP7
SKI2
HCA4
SPB4
PRP5
HAS1
DBP3
DBP1
SUV3
MRH4
MSS116
Z_Shift_K_Gem
Z_Im_K_Gem
Z_Shift_K_Cyt
Z_Im_K_Cyt
Z_Shift_L_Gem
Z_Im_L_Gem
Z_Shift_L_Cyt
Z_Im_L_Cyt
Gene

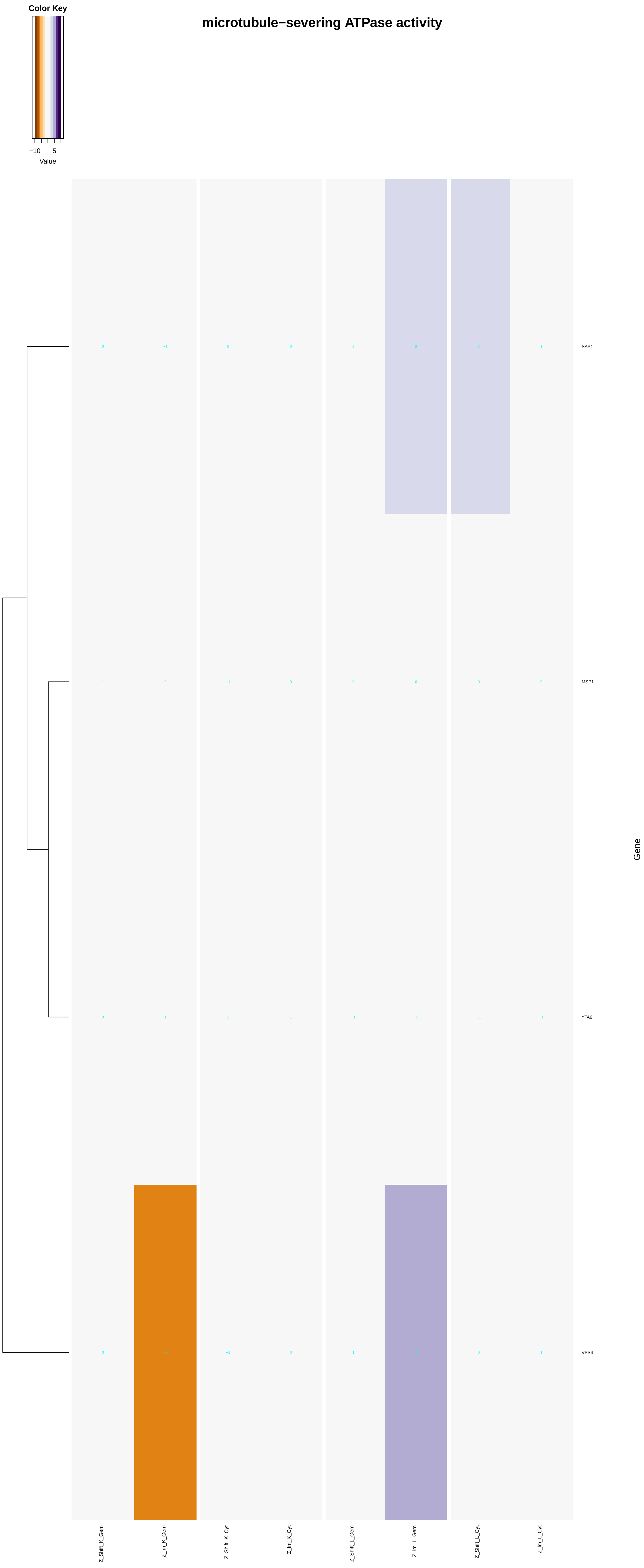
microtubule−severing ATPase activity
Color Key
−10
5
Value
SAP1
MSP1
YTA6
VPS4
Gene
Z_Shift_K_Gem
Z_Im_K_Gem
Z_Shift_K_Cyt
Z_Im_K_Cyt
Z_Shift_L_Gem
Z_Im_L_Gem
Z_Shift_L_Cyt
Z_Im_L_Cyt

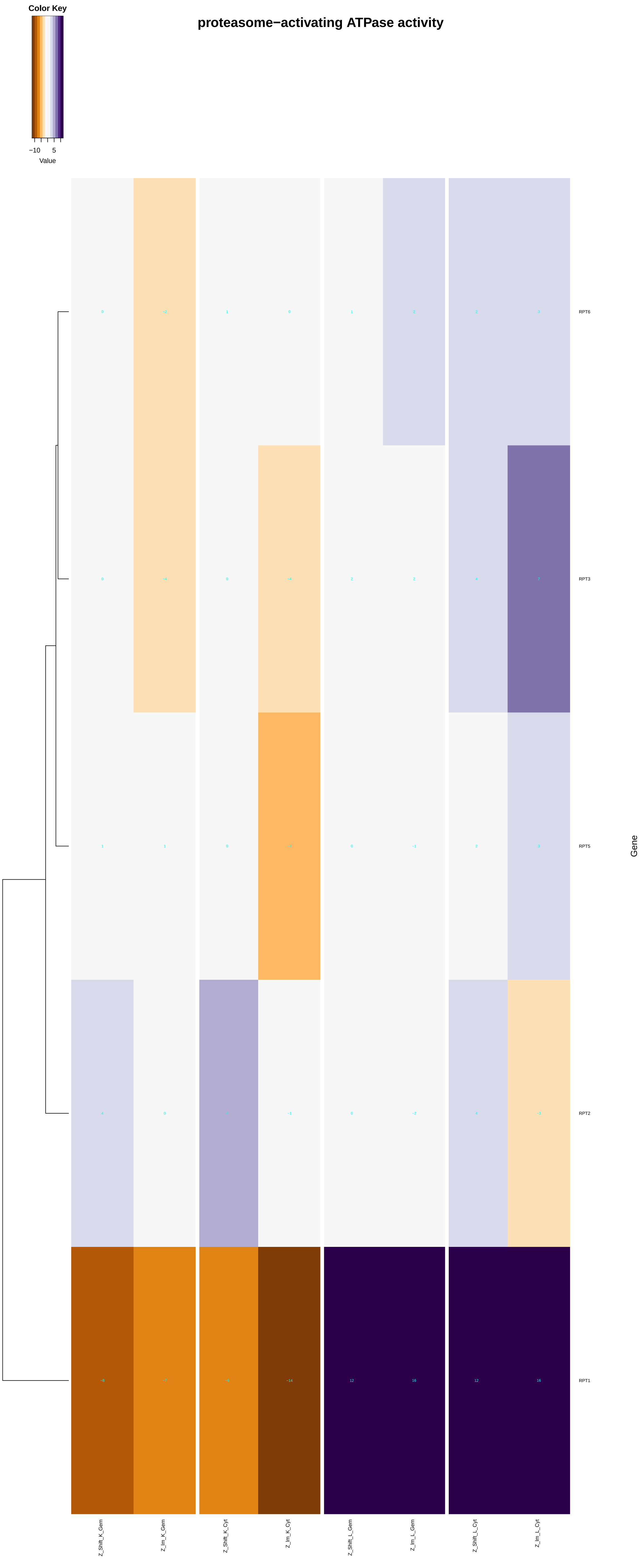
proteasome−activating ATPase activity
Color Key
−10
5
Value
RPT6
RPT3
RPT5
RPT2
RPT1
Gene
Z_Shift_K_Gem
Z_Im_K_Gem
Z_Shift_K_Cyt
Z_Im_K_Cyt
Z_Shift_L_Gem
Z_Im_L_Gem
Z_Shift_L_Cyt
Z_Im_L_Cyt

#### ATPase activity, coupled to movement of substances

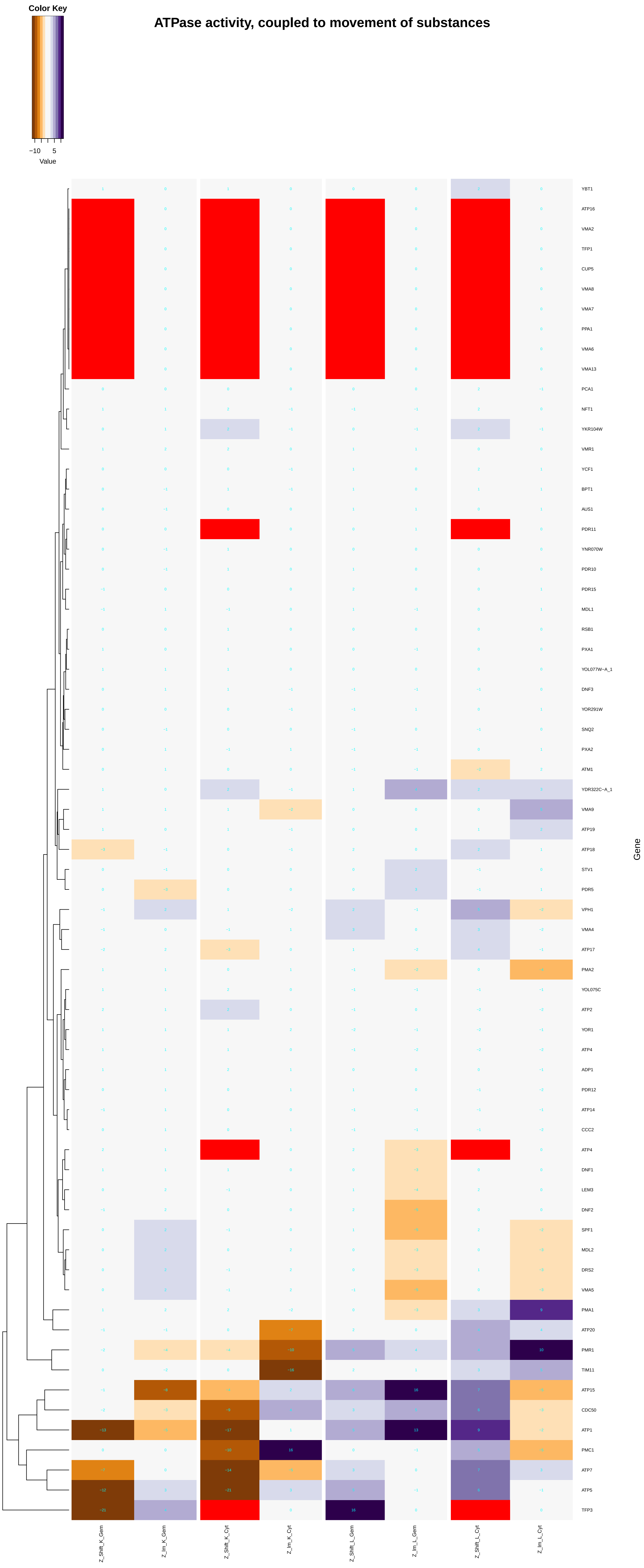

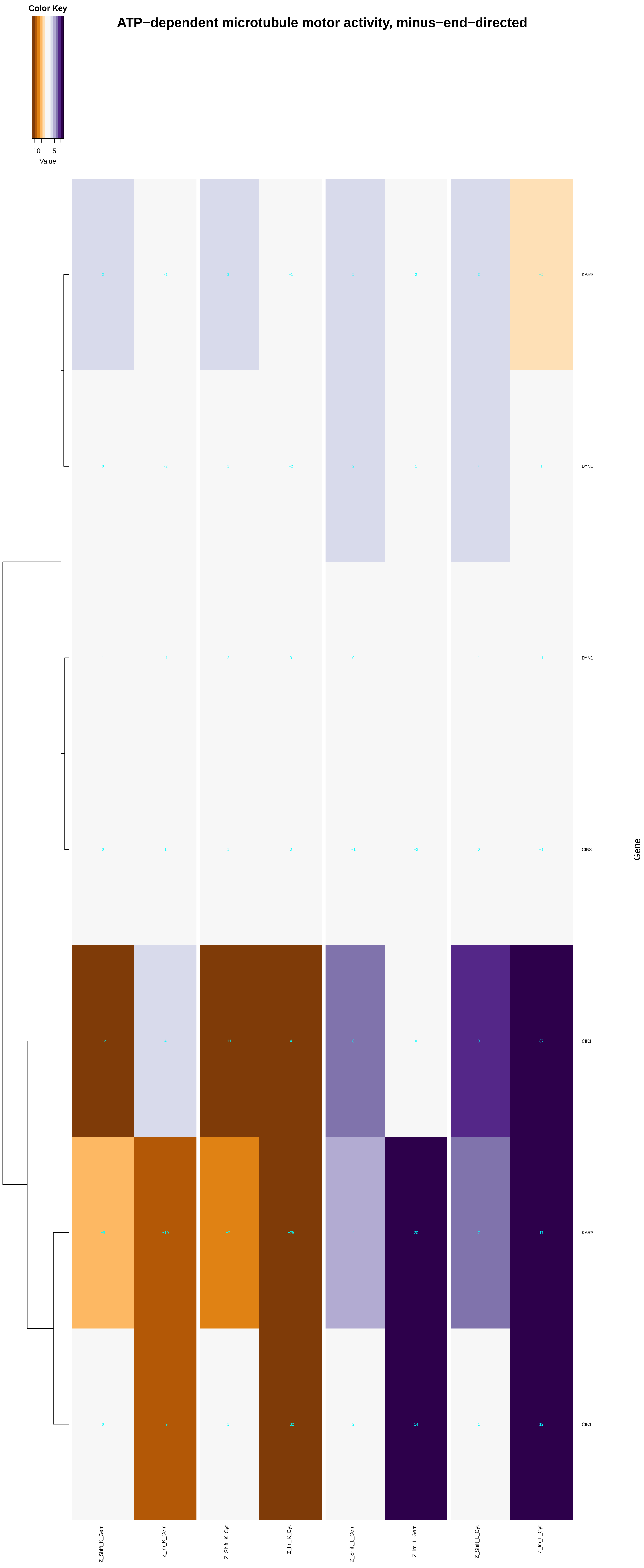
ATP−dependent microtubule motor activity, minus−end−directed
Color Key
−10
5
Value
KAR3
DYN1
DYN1
CIN8
CIK1
KAR3
CIK1
Gene
Z_Shift_K_Gem
Z_Im_K_Gem
Z_Shift_K_Cyt
Z_Im_K_Cyt
Z_Shift_L_Gem
Z_Im_L_Gem
Z_Shift_L_Cyt
Z_Im_L_Cyt

### ATP−dependent microtubule motor activity, plus−end−directed

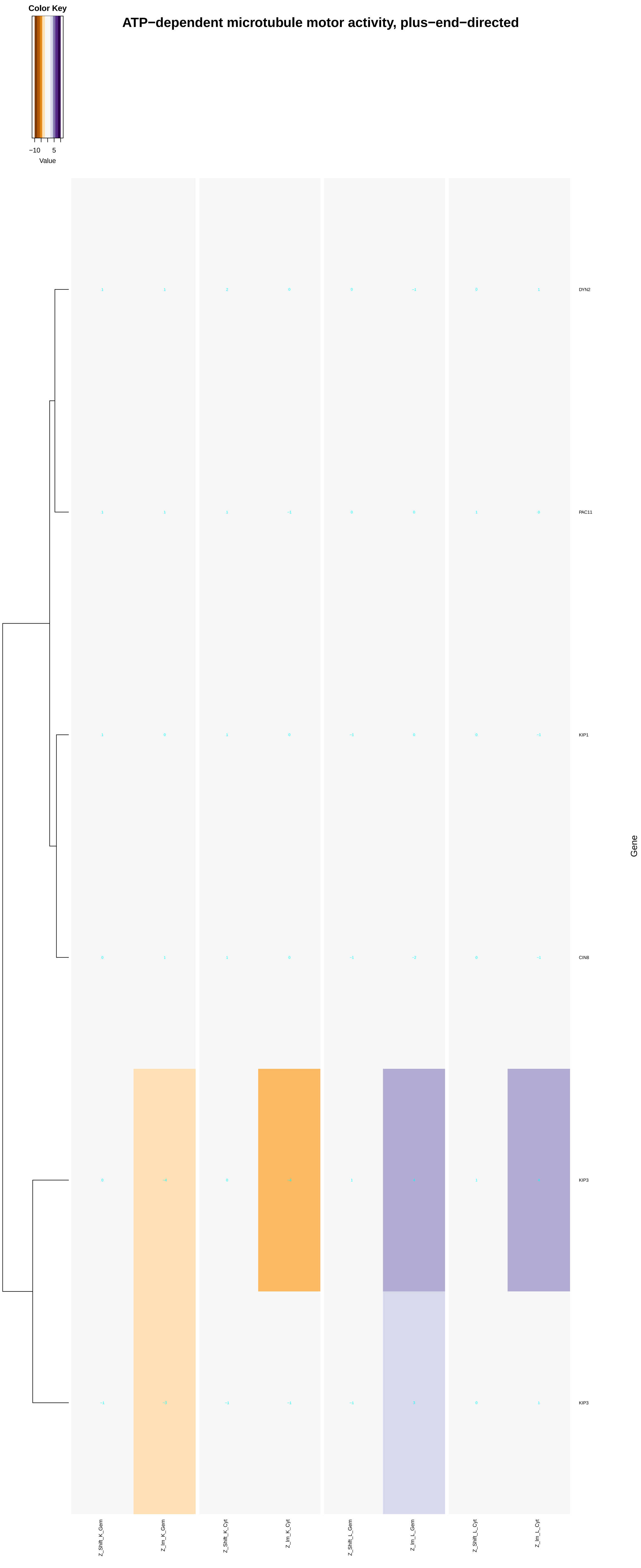

### single−stranded DNA−dependent ATP−dependent DNA helicase activity

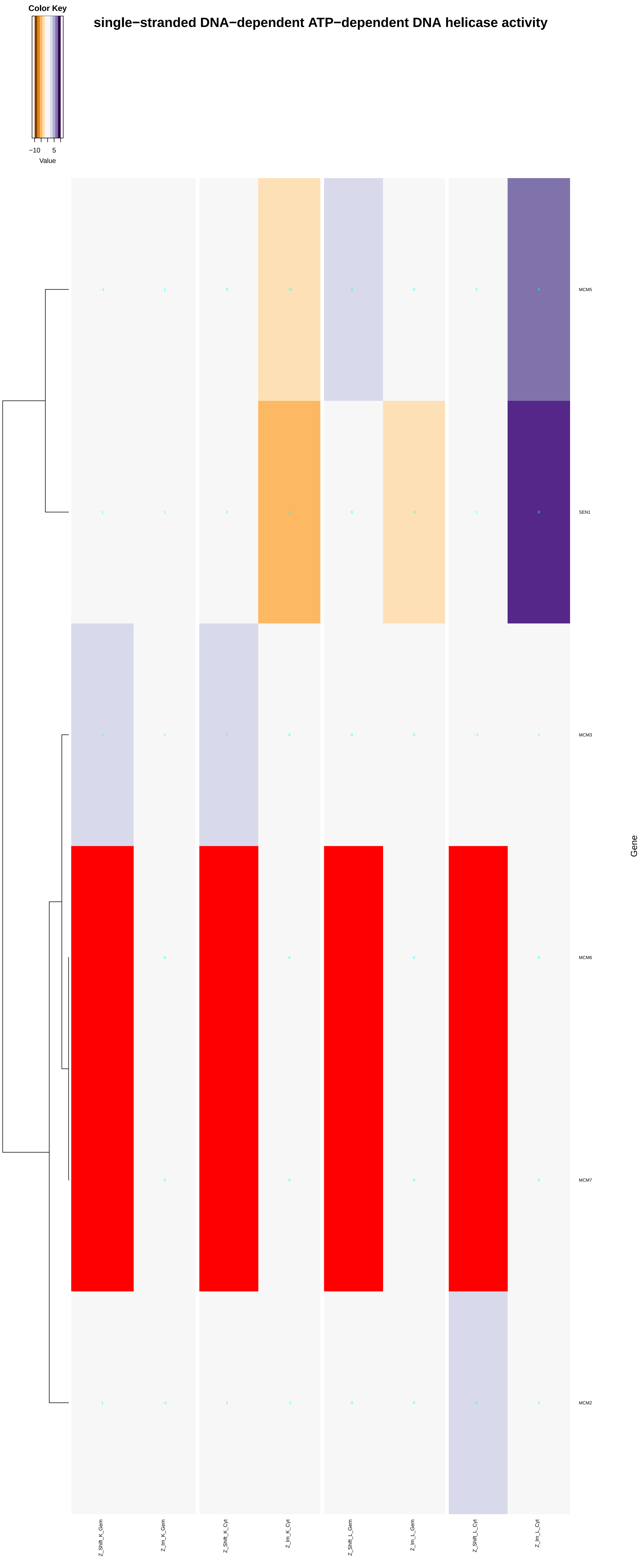

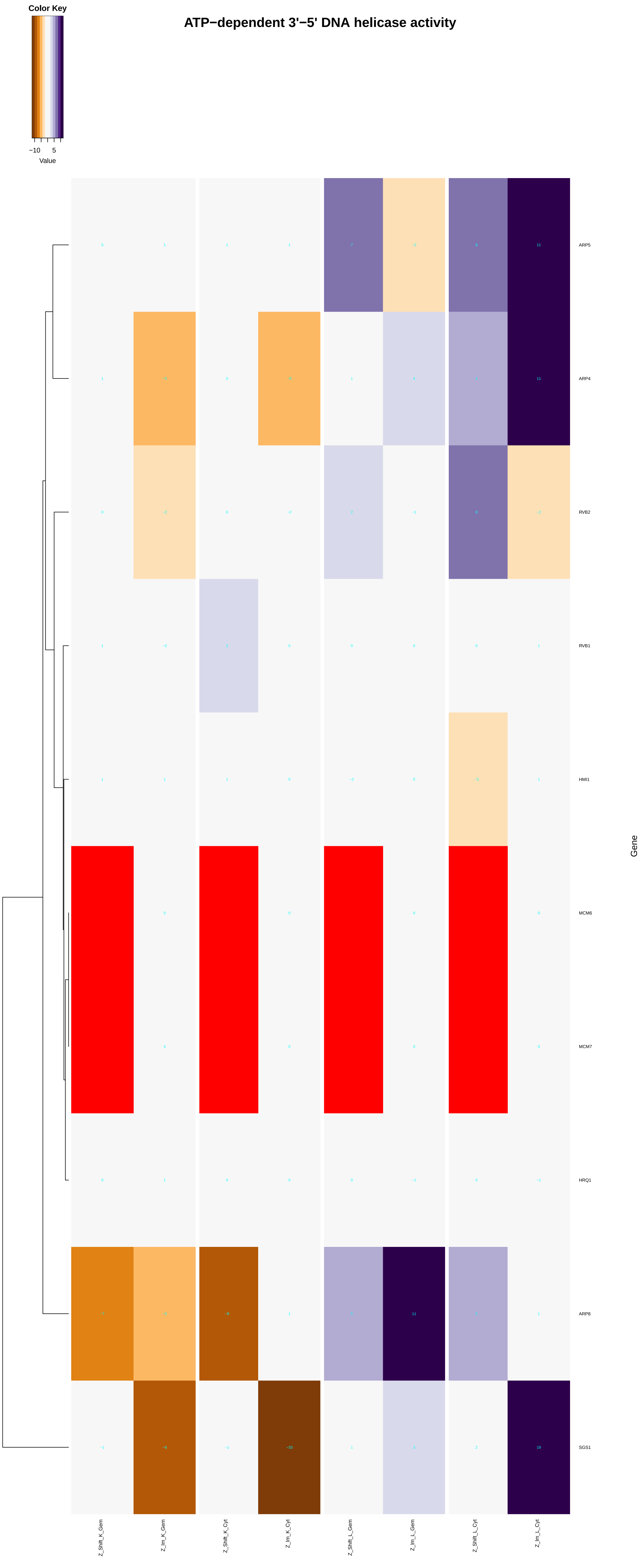

Color Key
−10
5
Value
ATP−dependent 3'−5' DNA helicase activity
ARP5
ARP4
RVB2
RVB1
HMI1
MCM6
MCM7
HRQ1
ARP8
SGS1
Gene
Z_Shift_K_Gem
Z_lm_K_Gem
Z_Shift_K_Cyt
Z_lm_K_Cyt
Z_Shift_L_Gem
Z_lm_L_Gem
Z_Shift_L_Cyt
Z_lm_L_Cyt

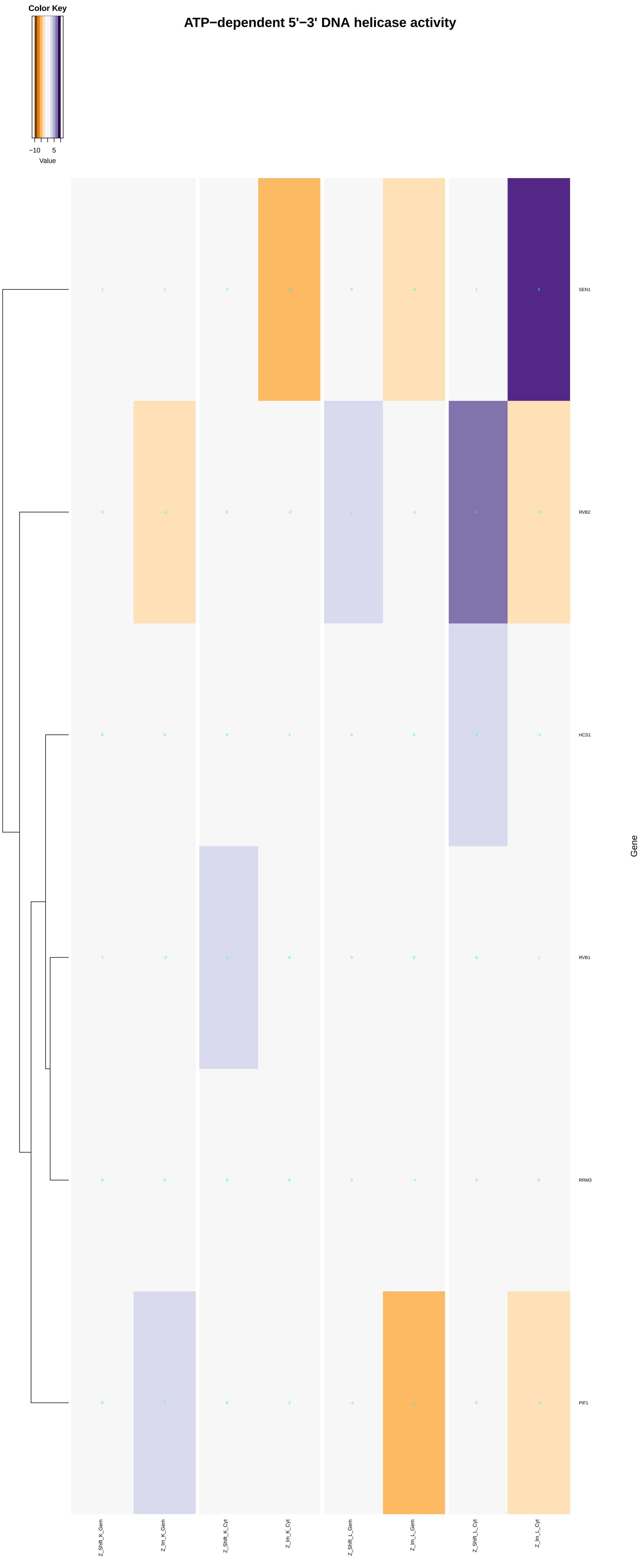

ATP−dependent 5'−3' DNA helicase activity
Color Key
−10
5
Value
SEN1
RVB2
HCS1
RVB1
RRM3
PIF1
Gene
Z_Shift_K_Gem
Z_lm_K_Gem
Z_Shift_K_Cyt
Z_lm_K_Cyt
Z_Shift_L_Gem
Z_lm_L_Gem
Z_Shift_L_Cyt
Z_lm_L_Cyt

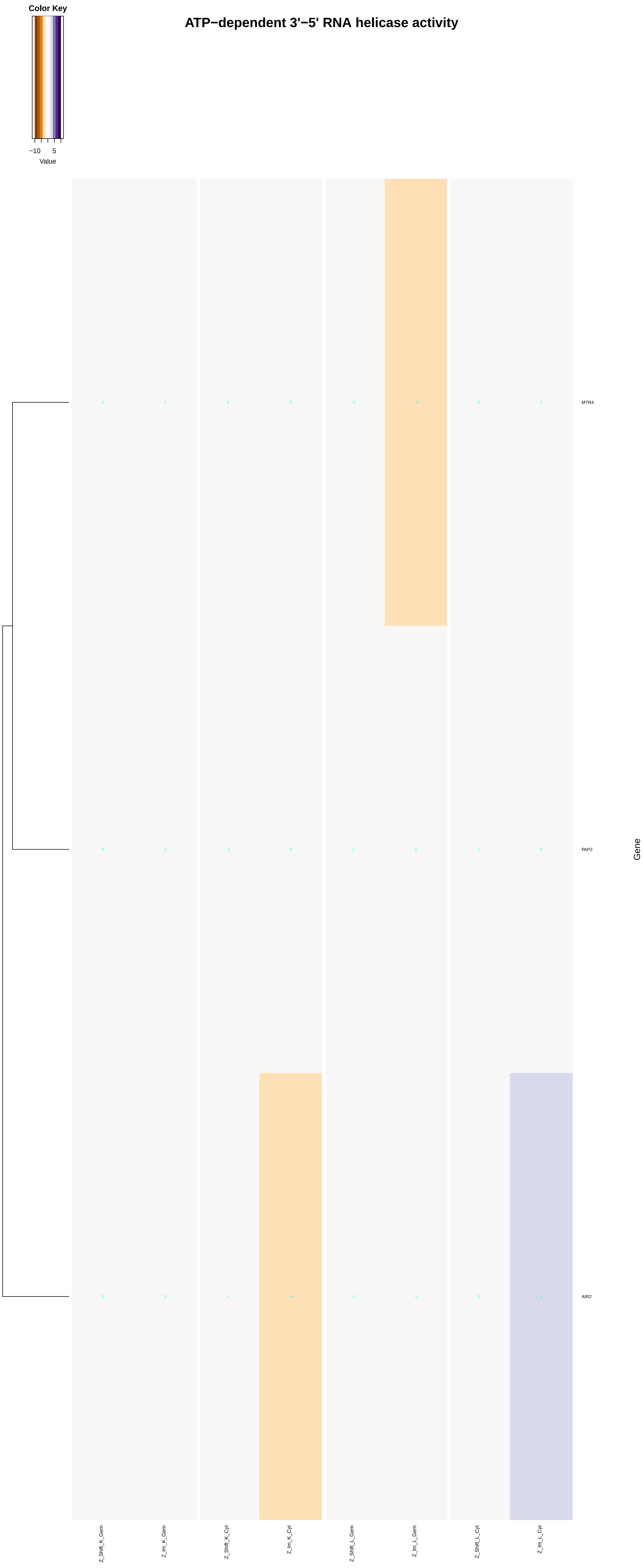

Color Key
−10
5
Value
ATP−dependent 3'−5' RNA helicase activity
MTR4
PAP2
AIR2
Gene
Z_Shift_K_Gem
Z_Im_K_Gem
Z_Shift_K_Cyt
Z_Im_K_Cyt
Z_Shift_L_Gem
Z_Im_L_Gem
Z_Shift_L_Cyt
Z_Im_L_Cyt

### DNA translocase activity

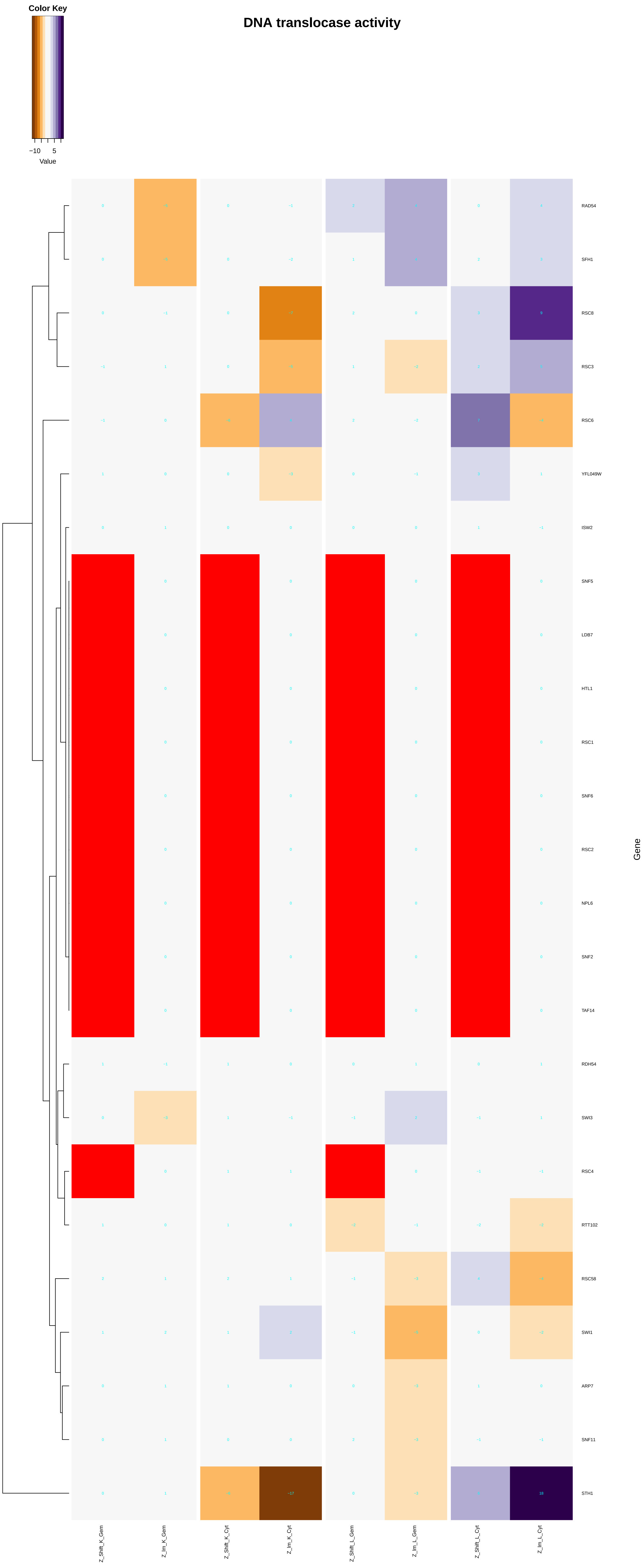

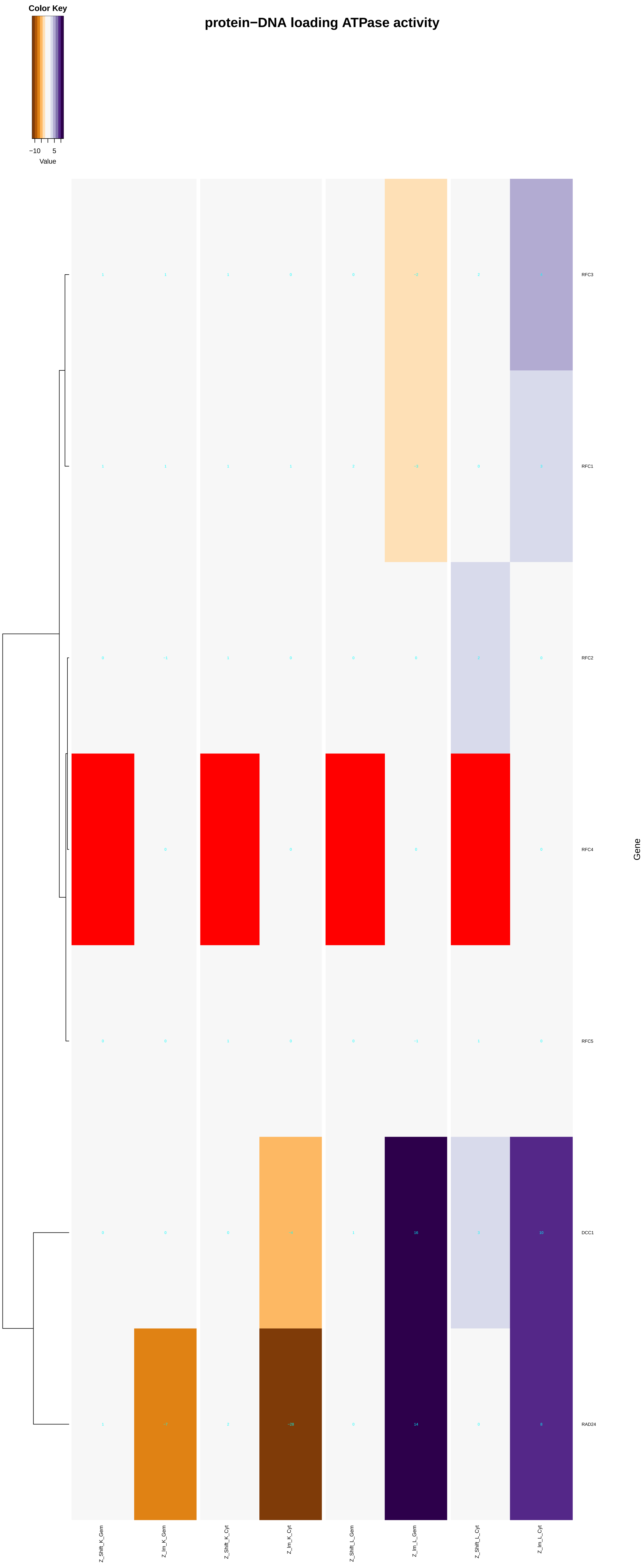

Color Key
−10
5
Value
protein−DNA loading ATPase activity
RFC3
RFC1
RFC2
RFC4
RFC5
DCC1
RAD24
Gene
Z_Shift_K_Gem
Z_Im_K_Gem
Z_Shift_K_Cyt
Z_Im_K_Cyt
Z_Shift_L_Gem
Z_Im_L_Gem
Z_Shift_L_Cyt
Z_Im_L_Cyt

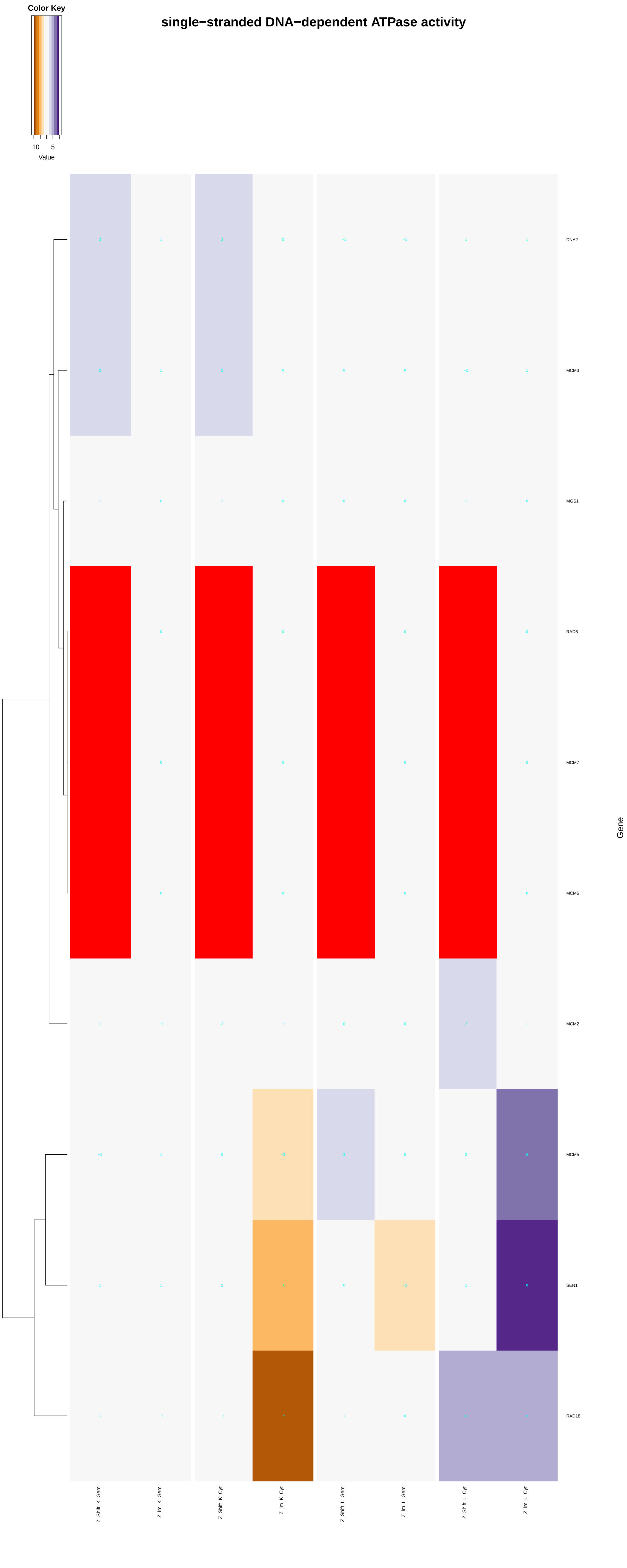

single−stranded DNA−dependent ATPase activity
Color Key
−10
5
Value
DNA2
MCM3
MGS1
RAD6
MCM7
MCM6
MCM2
MCM5
SEN1
RAD18
Gene
Z_Shift_K_Gem
Z_Im_K_Gem
Z_Shift_K_Cyt
Z_Im_K_Cyt
Z_Shift_L_Gem
Z_Im_L_Gem
Z_Shift_L_Cyt
Z_Im_L_Cyt

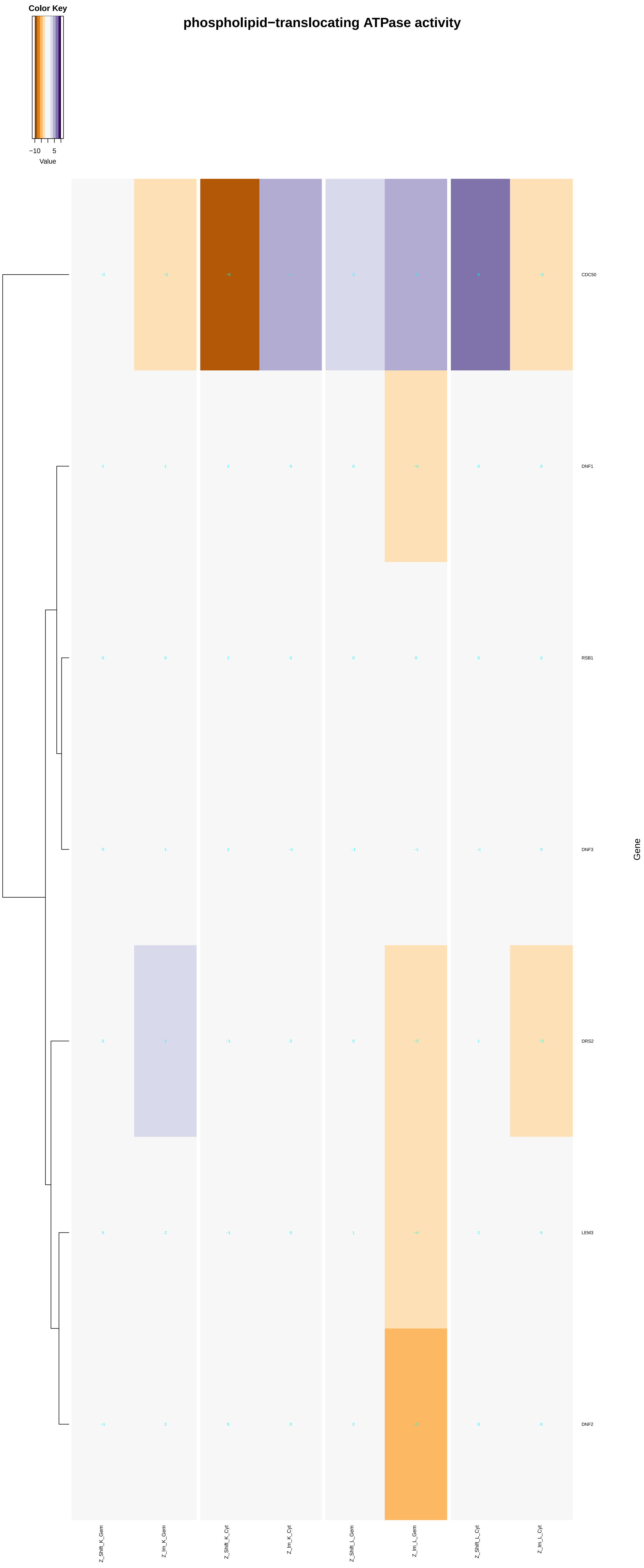
Color Key
−10
5
Value
phospholipid−translocating ATPase activity
CDC50
DNF1
RSB1
DNF3
DRS2
LEM3
DNF2
Gene
Z_Shift_K_Gem
Z_Im_K_Gem
Z_Shift_K_Cyt
Z_Im_K_Cyt
Z_Shift_L_Gem
Z_Im_L_Gem
Z_Shift_L_Cyt
Z_Im_L_Cyt

### ATPase activity, coupled to transmembrane movement of substances

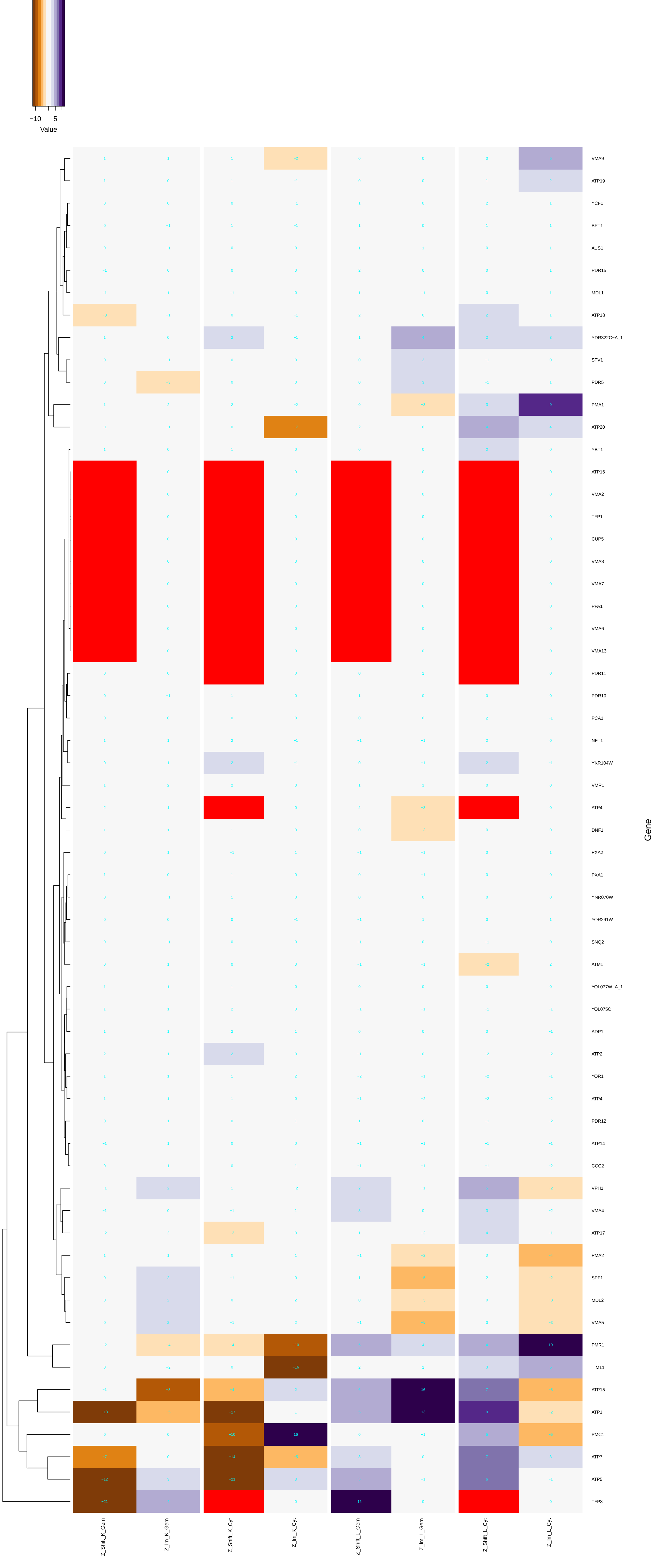

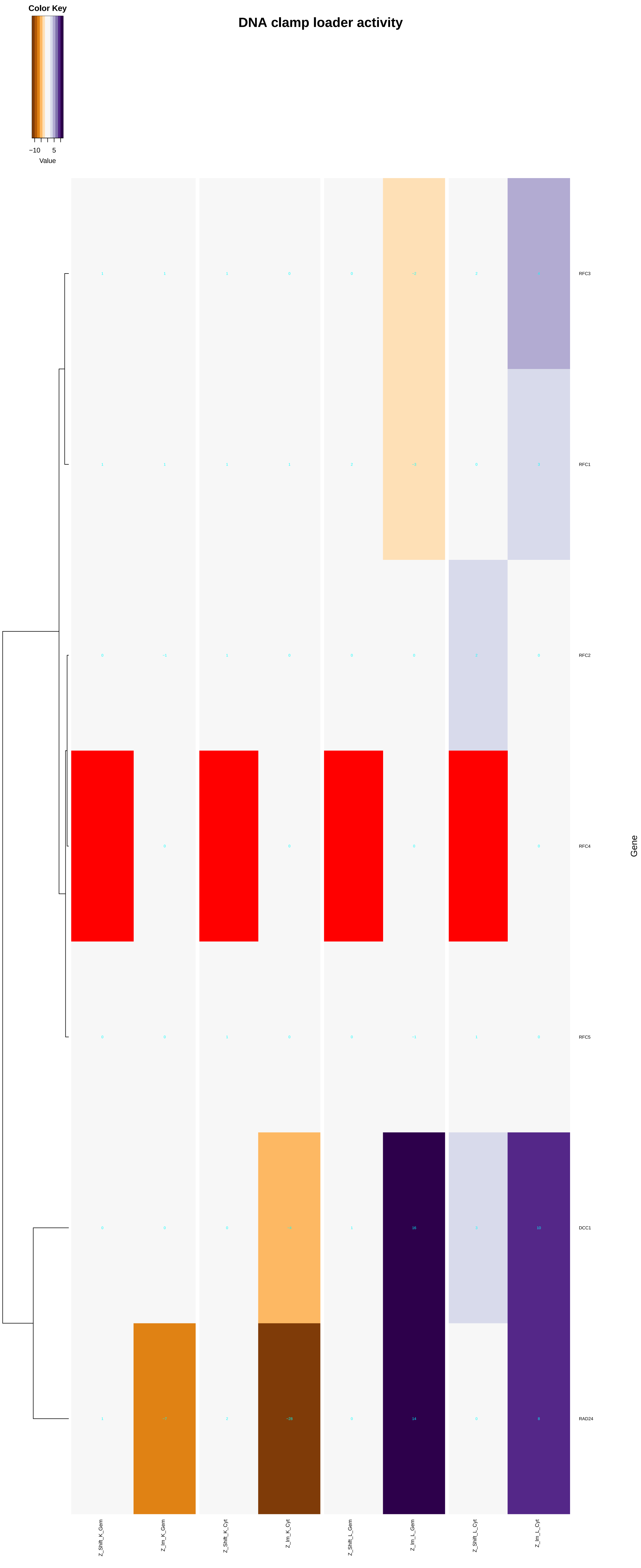
DNA clamp loader activity
Color Key
−10
5
Value
RFC3
RFC1
RFC2
RFC4
RFC5
DCC1
RAD24
Gene
Z_Shift_K_Gem
Z_Im_K_Gem
Z_Shift_K_Cyt
Z_Im_K_Cyt
Z_Shift_L_Gem
Z_Im_L_Gem
Z_Shift_L_Cyt
Z_Im_L_Cyt

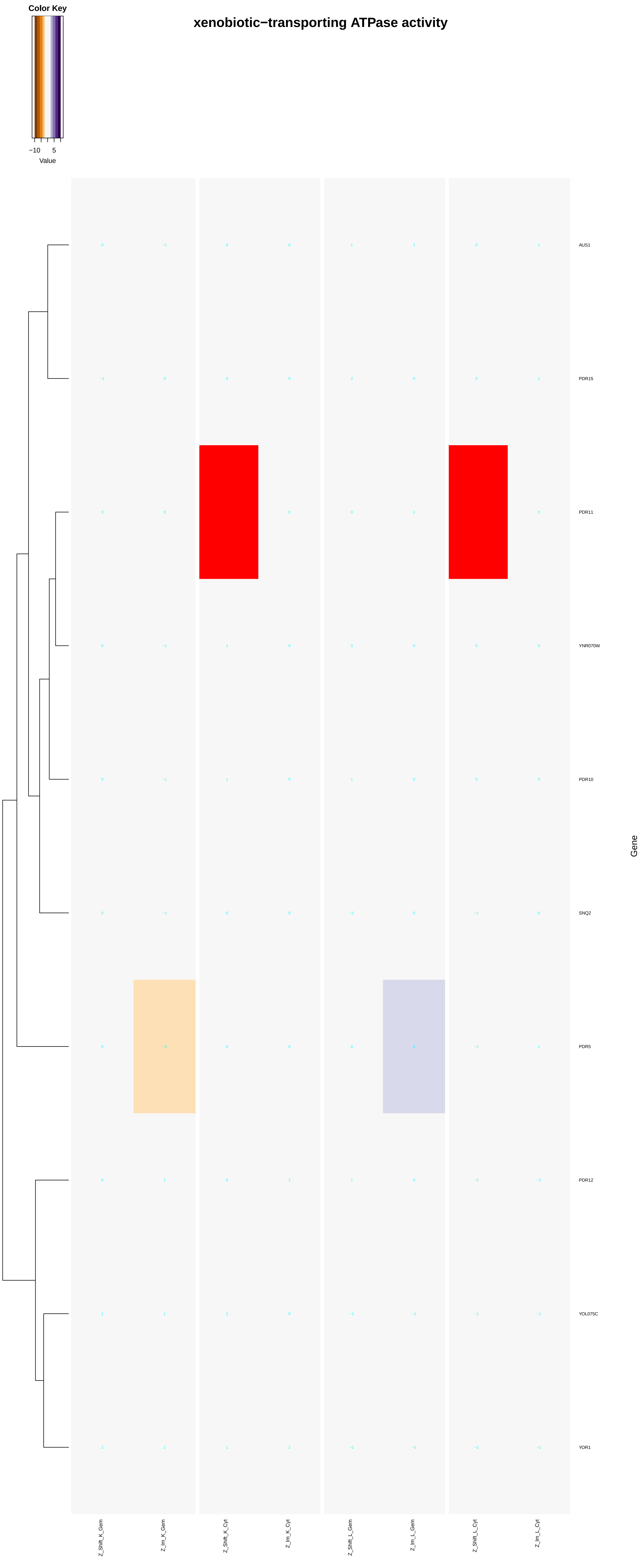
Color Key
−10
5
Value
xenobiotic−transporting ATPase activity
AUS1
PDR15
PDR11
YNR070W
PDR10
SNQ2
PDR5
PDR12
YOL075C
YOR1
Gene
Z_Shift_K_Gem
Z_Im_K_Gem
Z_Shift_K_Cyt
Z_Im_K_Cyt
Z_Shift_L_Gem
Z_Im_L_Gem
Z_Shift_L_Cyt
Z_Im_L_Cyt

### ATPase coupled ion transmembrane transporter activity

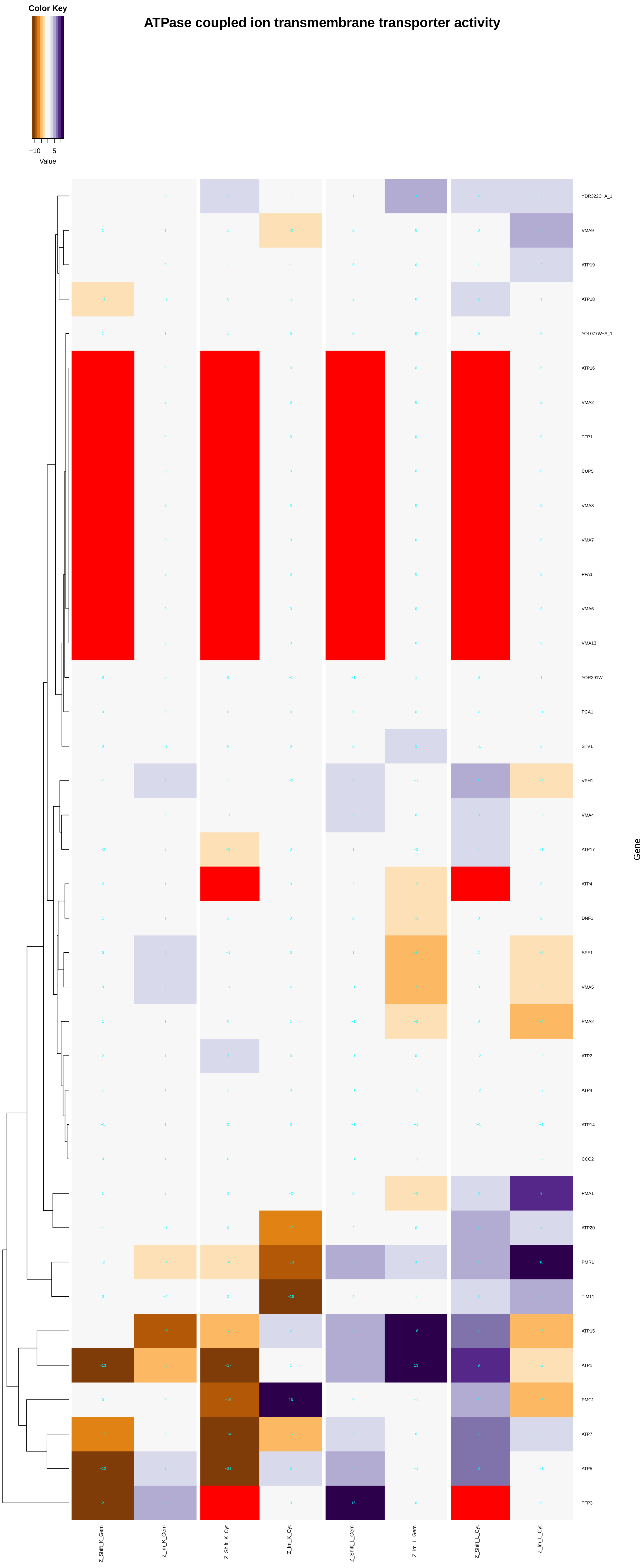

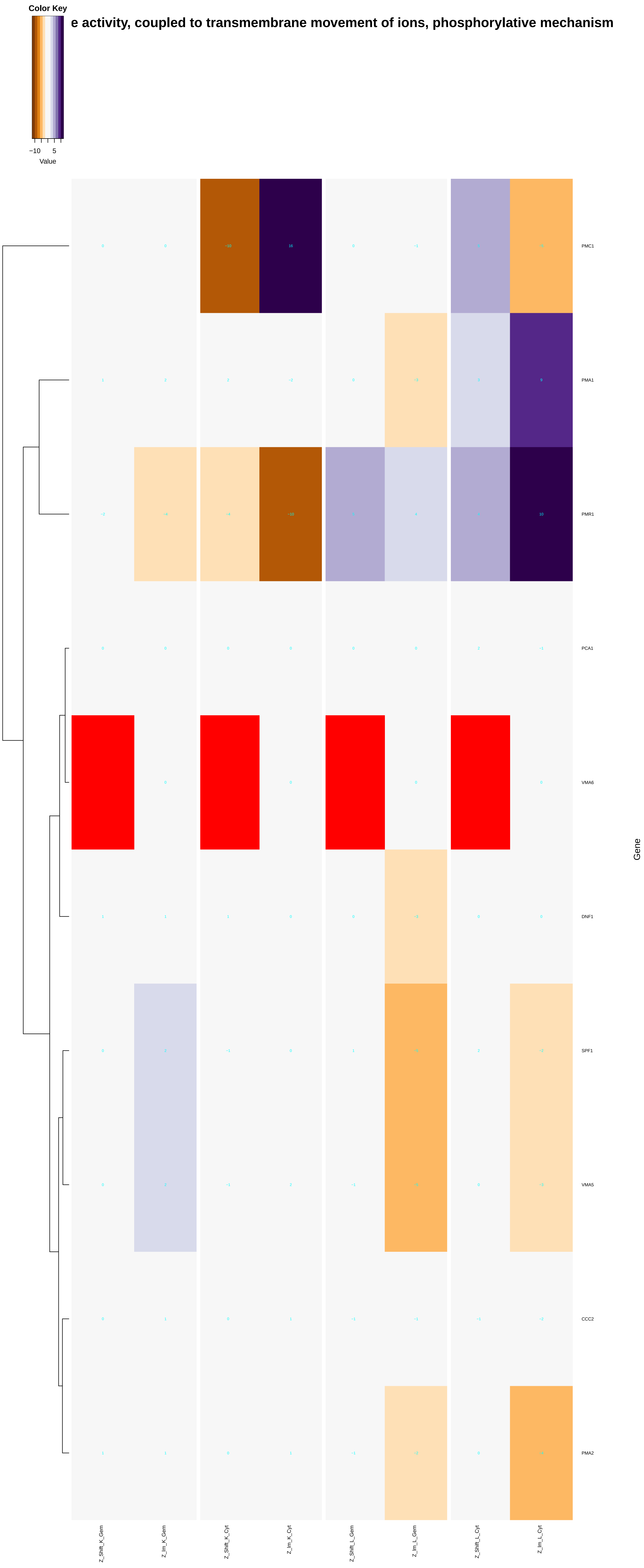
Color Key
−10
5
Value
e activity, coupled to transmembrane movement of ions, phosphorylative mechanism
PMC1
PMA1
PMR1
PCA1
VMA6
DNF1
SPF1
VMA5
CCC2
PMA2
Gene
Z_Shift_K_Gem
Z_Im_K_Gem
Z_Shift_K_Cyt
Z_Im_K_Cyt
Z_Shift_L_Gem
Z_Im_L_Gem
Z_Shift_L_Cyt
Z_Im_L_Cyt

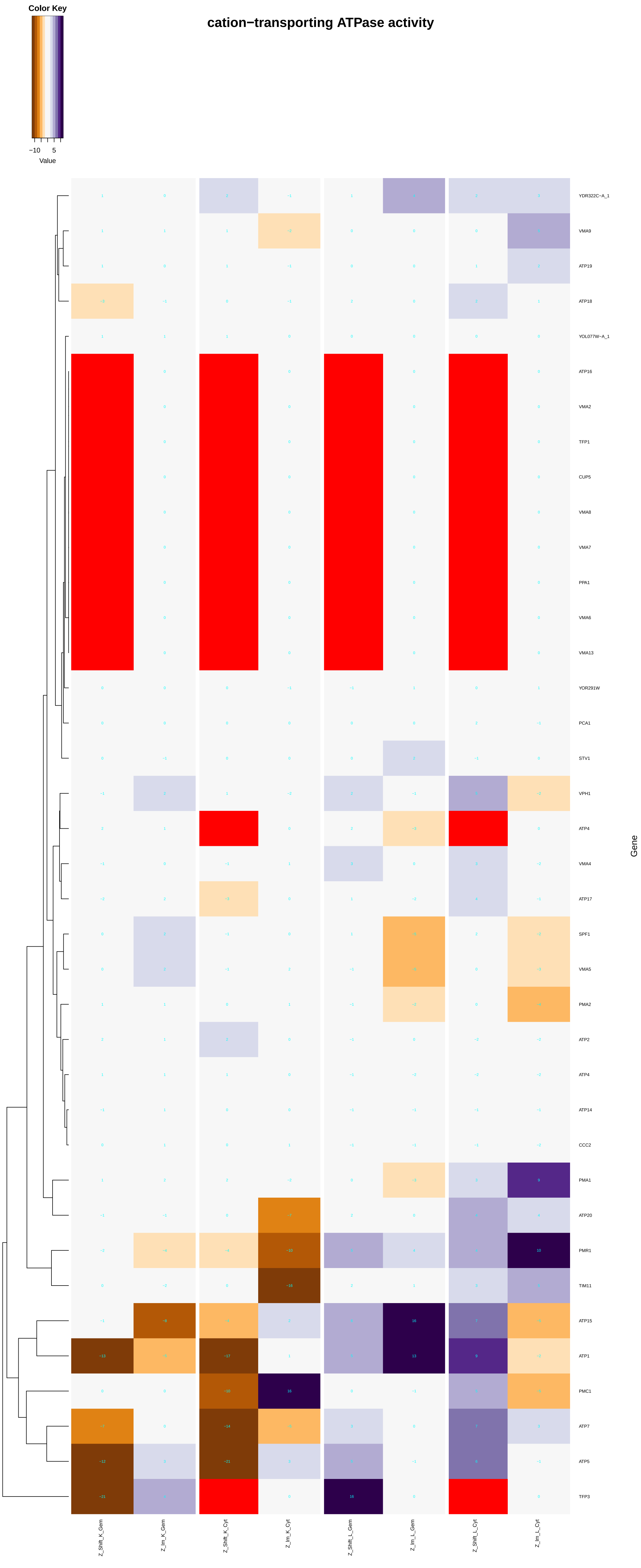

cation−transporting ATPase activity
Color Key
−10
5
Value
YDR322C−A_1
VMA9
ATP19
ATP18
YOL077W−A_1
ATP16
VMA2
TFP1
CUP5
VMA8
VMA7
PPA1
VMA6
VMA13
YOR291W
PCA1
STV1
VPH1
ATP4
VMA4
ATP17
SPF1
VMA5
PMA2
ATP2
ATP4
ATP14
CCC2
PMA1
ATP20
PMR1
TIM11
ATP15
ATP1
PMC1
ATP7
ATP5
TFP3
Gene
Z_Shift_K_Gem
Z_Im_K_Gem
Z_Shift_K_Cyt
Z_Im_K_Cyt
Z_Shift_L_Gem
Z_Im_L_Gem
Z_Shift_L_Cyt
Z_Im_L_Cyt

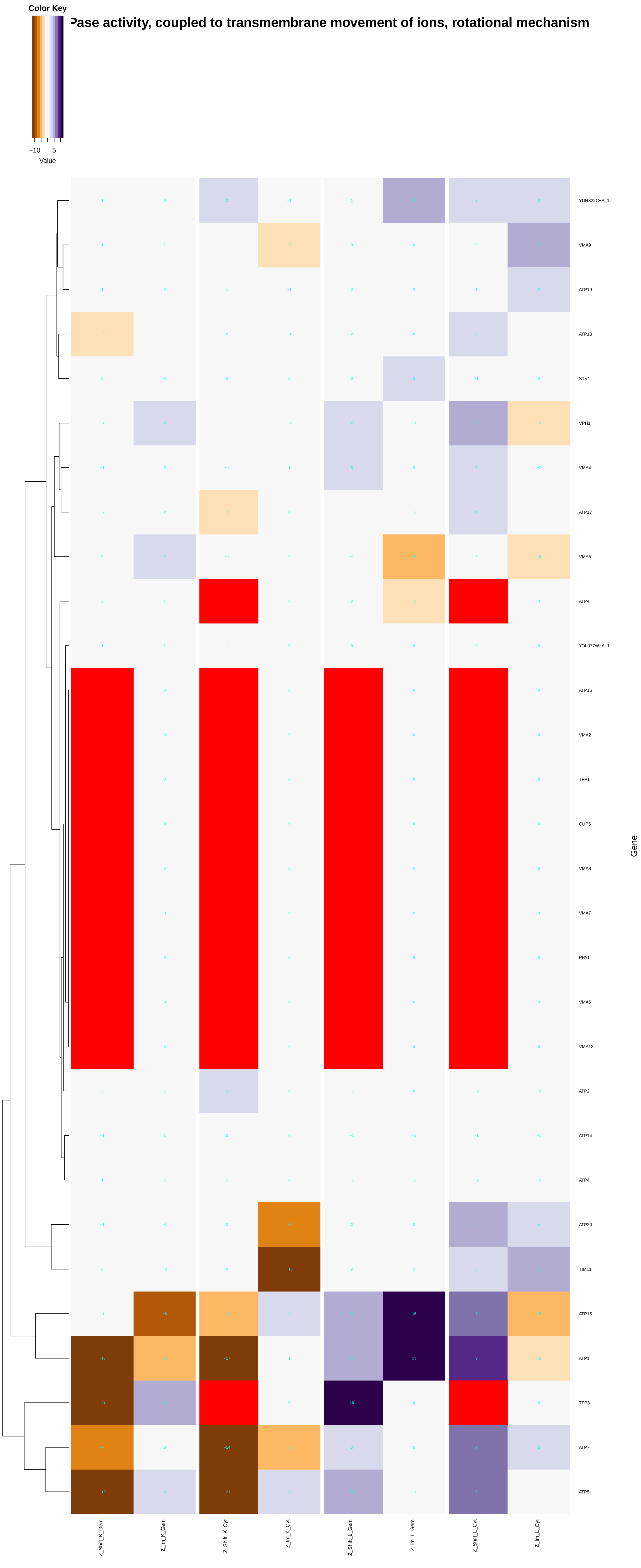
Color Key
−10
5
Value
Pase activity, coupled to transmembrane movement of ions, rotational mechanism
YDR322C−A_1
VMA9
ATP19
ATP18
STV1
VPH1
VMA4
ATP17
VMA5
ATP4
YOL077W−A_1
ATP16
VMA2
TFP1
CUP5
VMA8
VMA7
PPA1
VMA6
VMA13
ATP2
ATP14
ATP4
ATP20
TIM11
ATP15
ATP1
TFP3
ATP7
ATP5
Gene
Z_Shift_K_Gem
Z_Im_K_Gem
Z_Shift_K_Cyt
Z_Im_K_Cyt
Z_Shift_L_Gem
Z_Im_L_Gem
Z_Shift_L_Cyt
Z_Im_L_Cyt

### hydrogen−exporting ATPase activity, phosphorylative mechanism

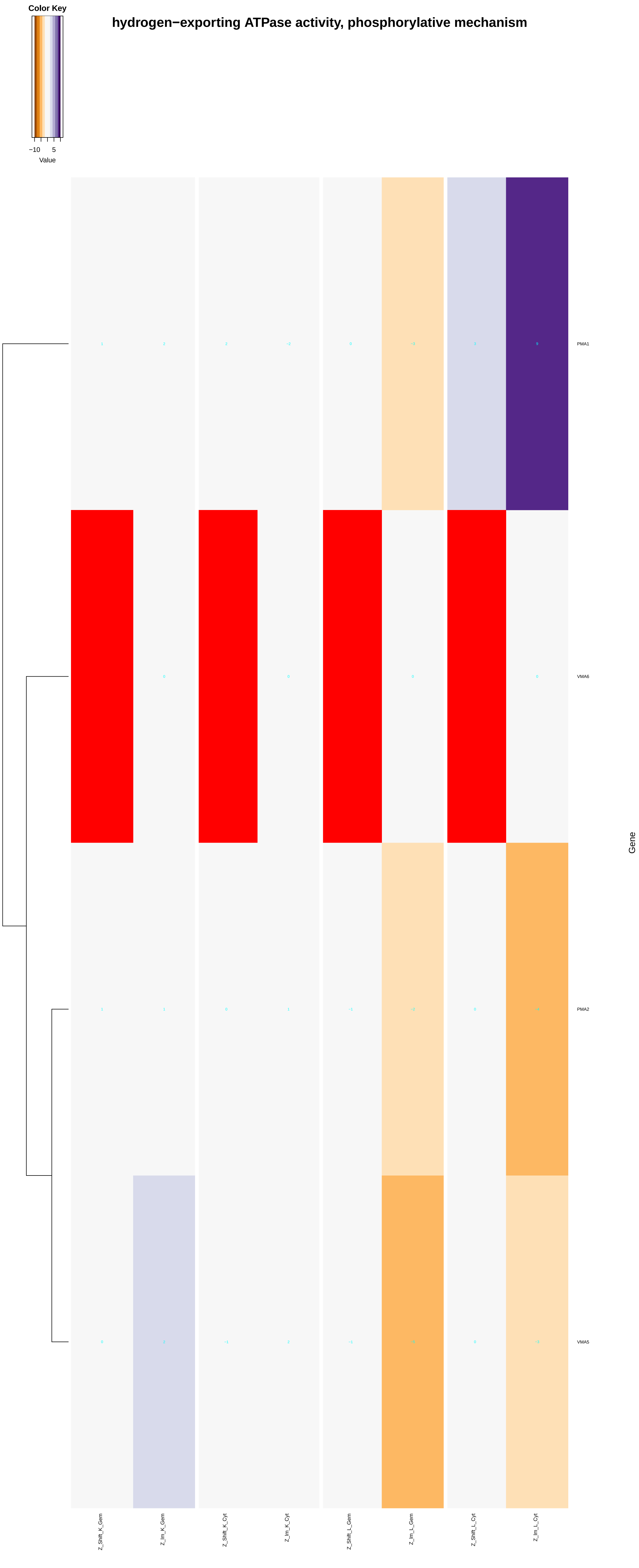

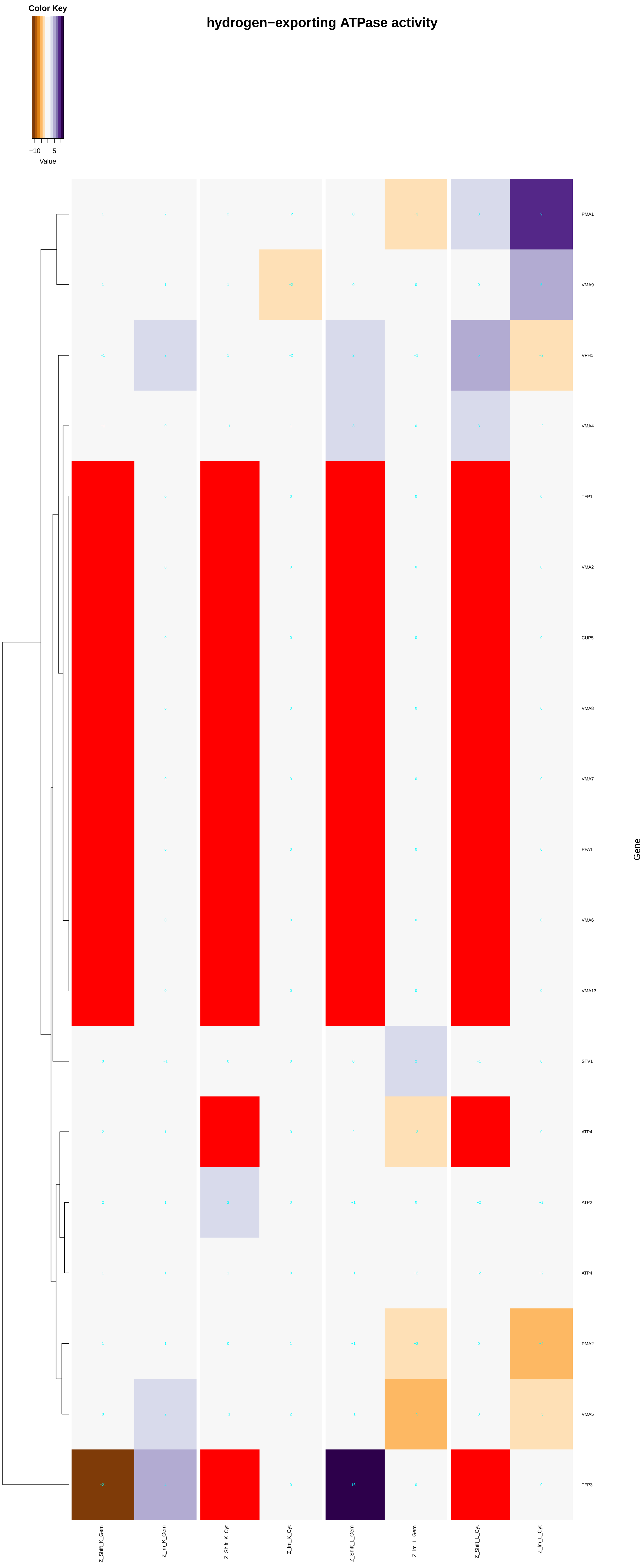
hydrogen−exporting ATPase activity
Color Key
−10 5
Value
Gene
PMA1
VMA9
VPH1
VMA4
TFP1
VMA2
CUP5
VMA8
VMA7
PPA1
VMA6
VMA13
STV1
ATP4
ATP2
ATP4
PMA2
VMA5
TFP3
Z_Shift_K_Gem
Z_Im_K_Gem
Z_Shift_K_Cyt
Z_Im_K_Cyt
Z_Shift_L_Gem
Z_Im_L_Gem
Z_Shift_L_Cyt
Z_Im_L_Cyt

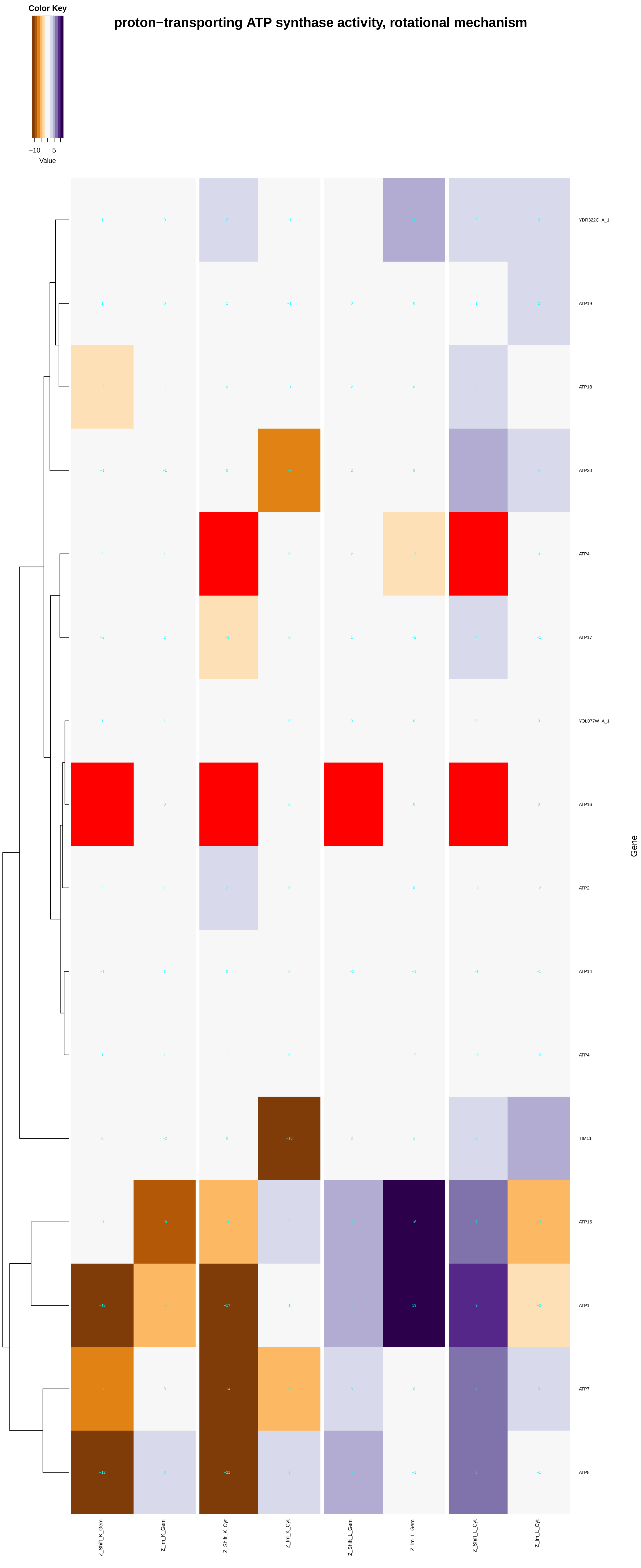

Color Key
−10
5
Value
proton−transporting ATP synthase activity, rotational mechanism
YDR322C−A_1
ATP19
ATP18
ATP20
ATP4
ATP17
YOL077W−A_1
ATP16
ATP2
ATP14
ATP4
TIM11
ATP15
ATP1
ATP7
ATP5
Gene
Z_Shift_K_Gem
Z_Im_K_Gem
Z_Shift_K_Cyt
Z_Im_K_Cyt
Z_Shift_L_Gem
Z_Im_L_Gem
Z_Shift_L_Cyt
Z_Im_L_Cyt

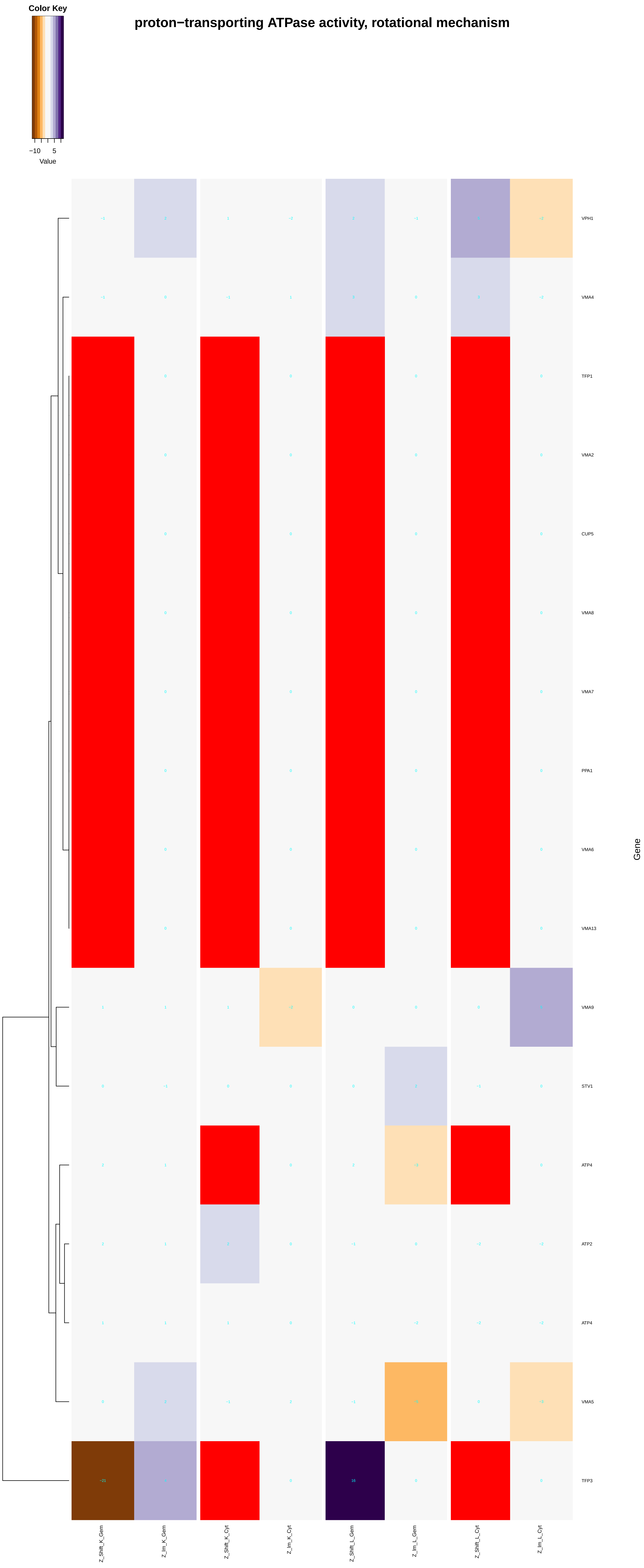

Color Key
−10
5
Value
proton−transporting ATPase activity, rotational mechanism
VPH1
VMA4
TFP1
VMA2
CUP5
VMA8
VMA7
PPA1
VMA6
VMA13
VMA9
STV1
ATP4
ATP2
ATP4
VMA5
TFP3
Gene
Z_Shift_K_Gem
Z_Im_K_Gem
Z_Shift_K_Cyt
Z_Im_K_Cyt
Z_Shift_L_Gem
Z_Im_L_Gem
Z_Shift_L_Cyt
Z_Im_L_Cyt
