## Additional File 7 for "A humanized yeast phenomic model of deoxycytidine kinase to predict genetic buffering of nucleoside analog cytotoxicity": hydrolase_activity,_acting_on_acid_anhydrides.pdf

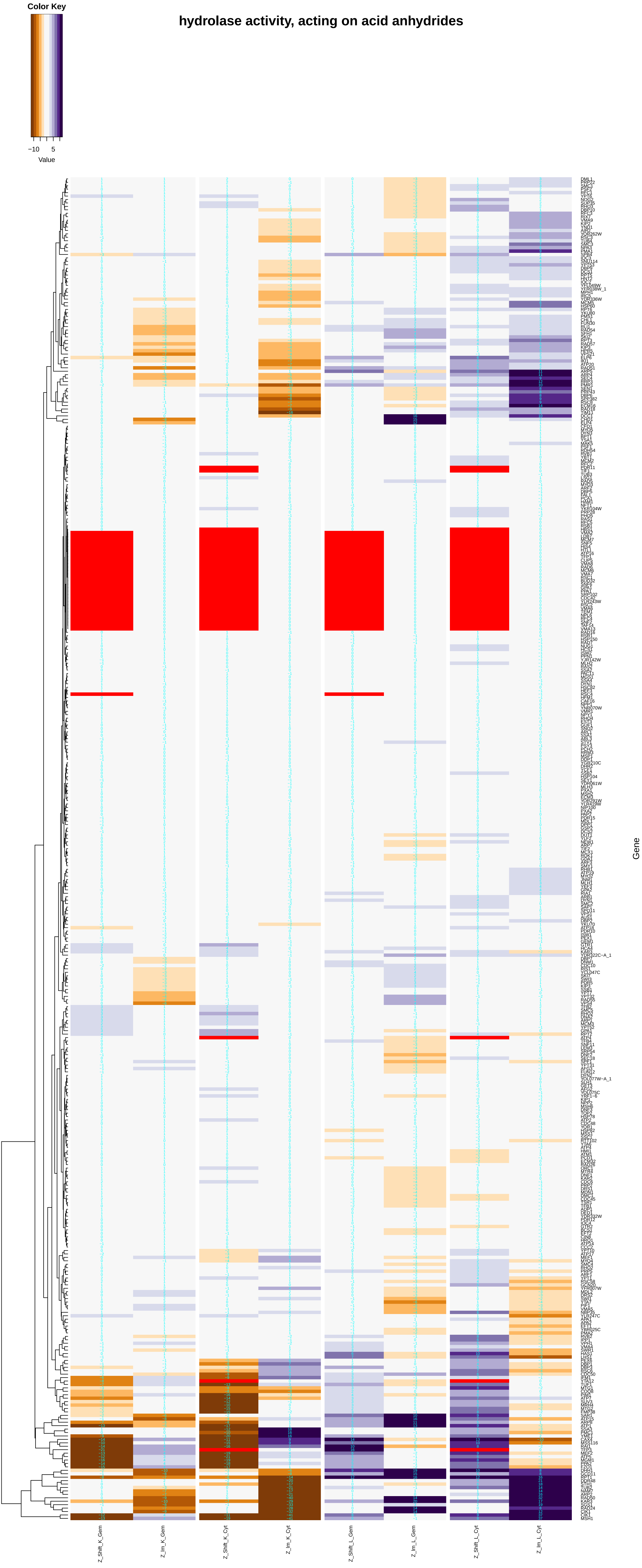
hydrolase activity, acting on acid anhydrides
Color Key
−10 5
Value
Gene
Z_Shift_K_Gem
Z_Im_K_Gem
Z_Shift_K_Cyt
Z_Im_K_Cyt
Z_Shift_L_Gem
Z_Im_L_Gem
Z_Shift_L_Cyt
Z_Im_L_Cyt

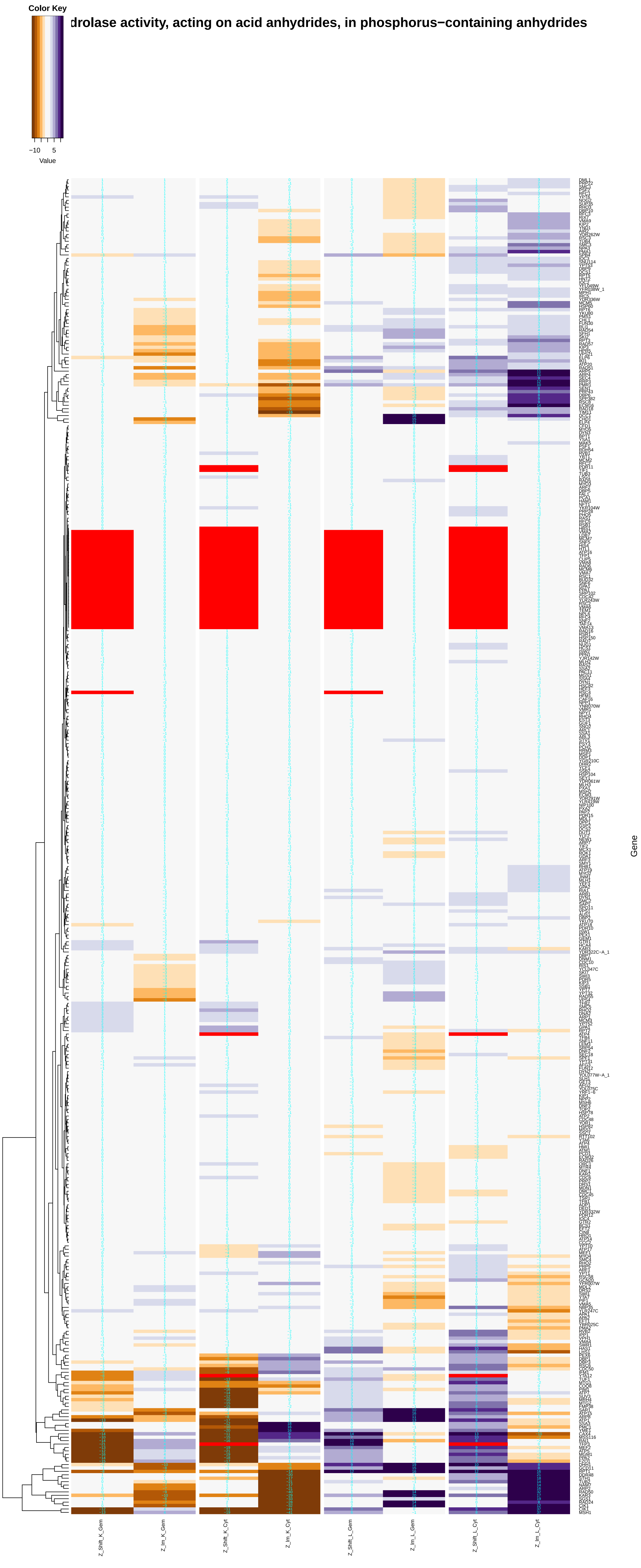
Color Key
−10
5
Value
drolase activity, acting on acid anhydrides, in phosphorus−containing anhydrides
Gene
Z_Shift_K_Gem
Z_Im_K_Gem
Z_Shift_K_Cyt
Z_Im_K_Cyt
Z_Shift_L_Gem
Z_Im_L_Gem
Z_Shift_L_Cyt
Z_Im_L_Cyt

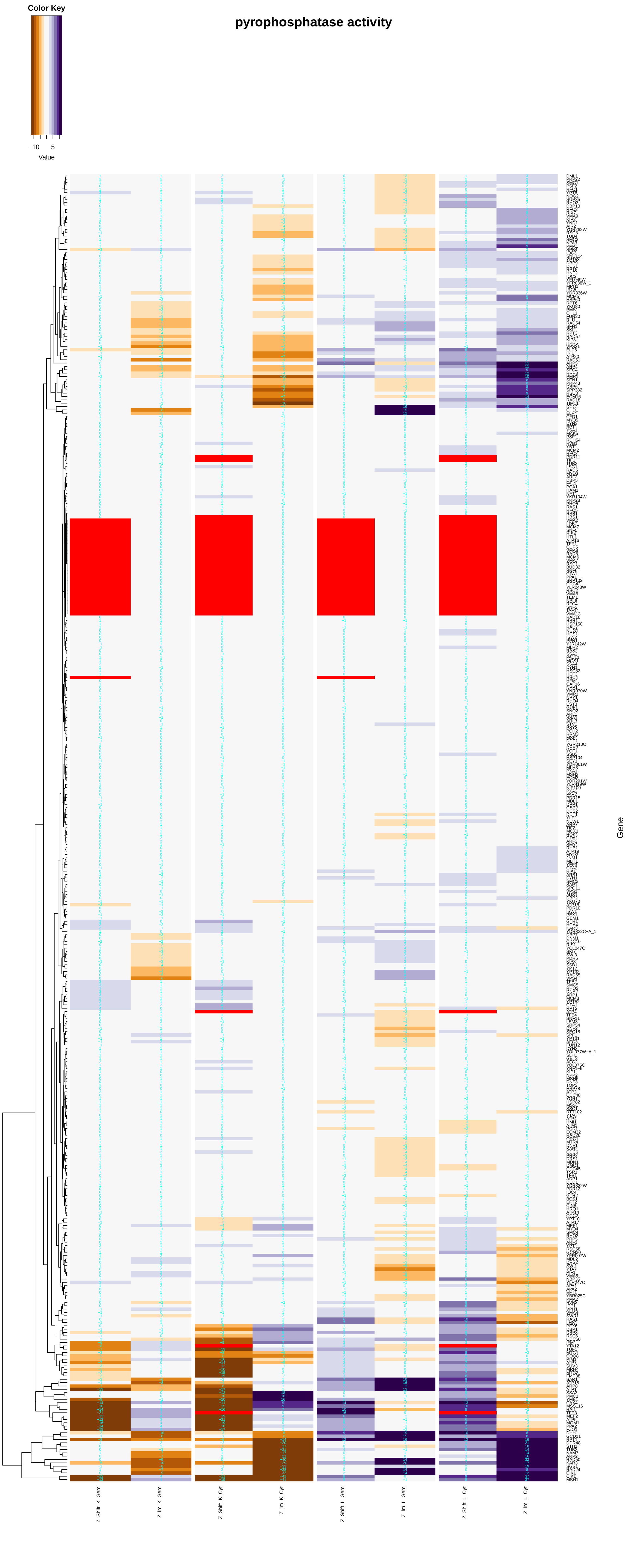

Color Key
−10
5
Value
pyrophosphatase activity
Z_Shift_K_Gem
Z_Im_K_Gem
Z_Shift_K_Cyt
Z_Im_K_Cyt
Z_Shift_L_Gem
Z_Im_L_Gem
Z_Shift_L_Cyt
Z_Im_L_Cyt
Gene

### ATPase activity, coupled to transmembrane movement of substances

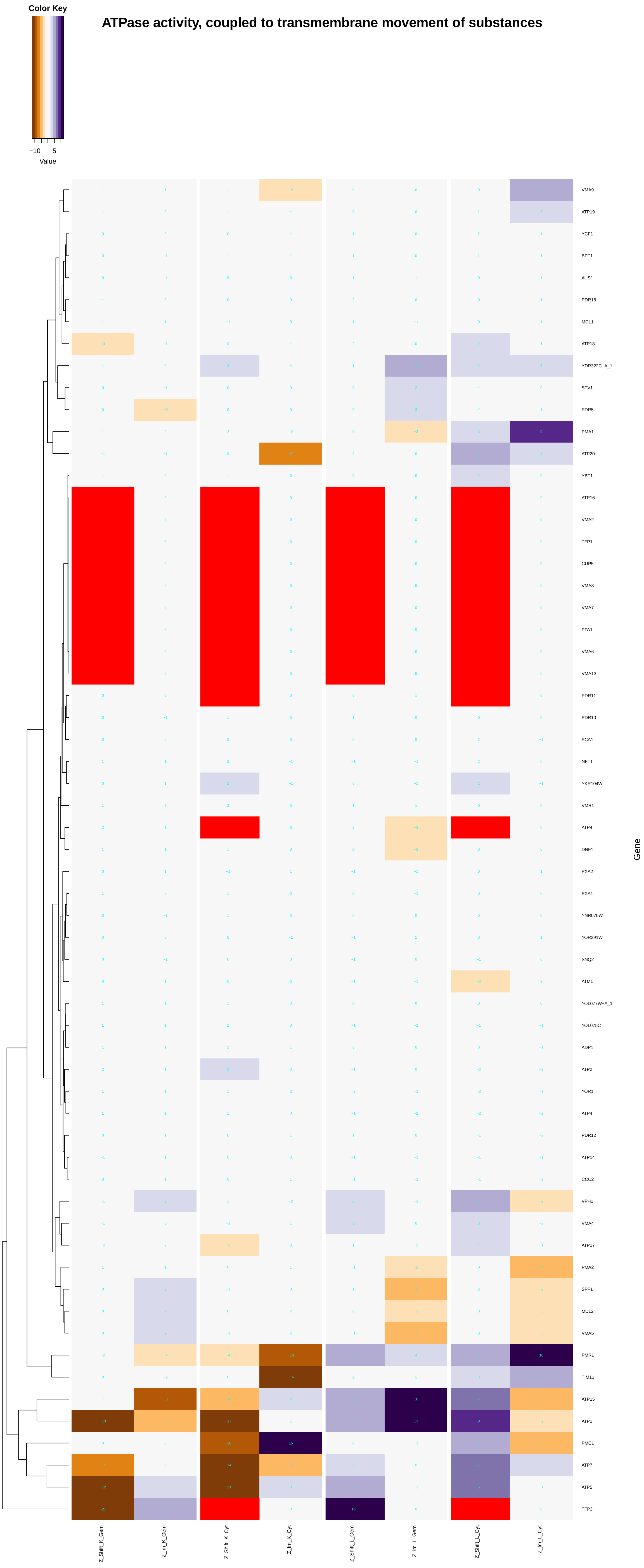

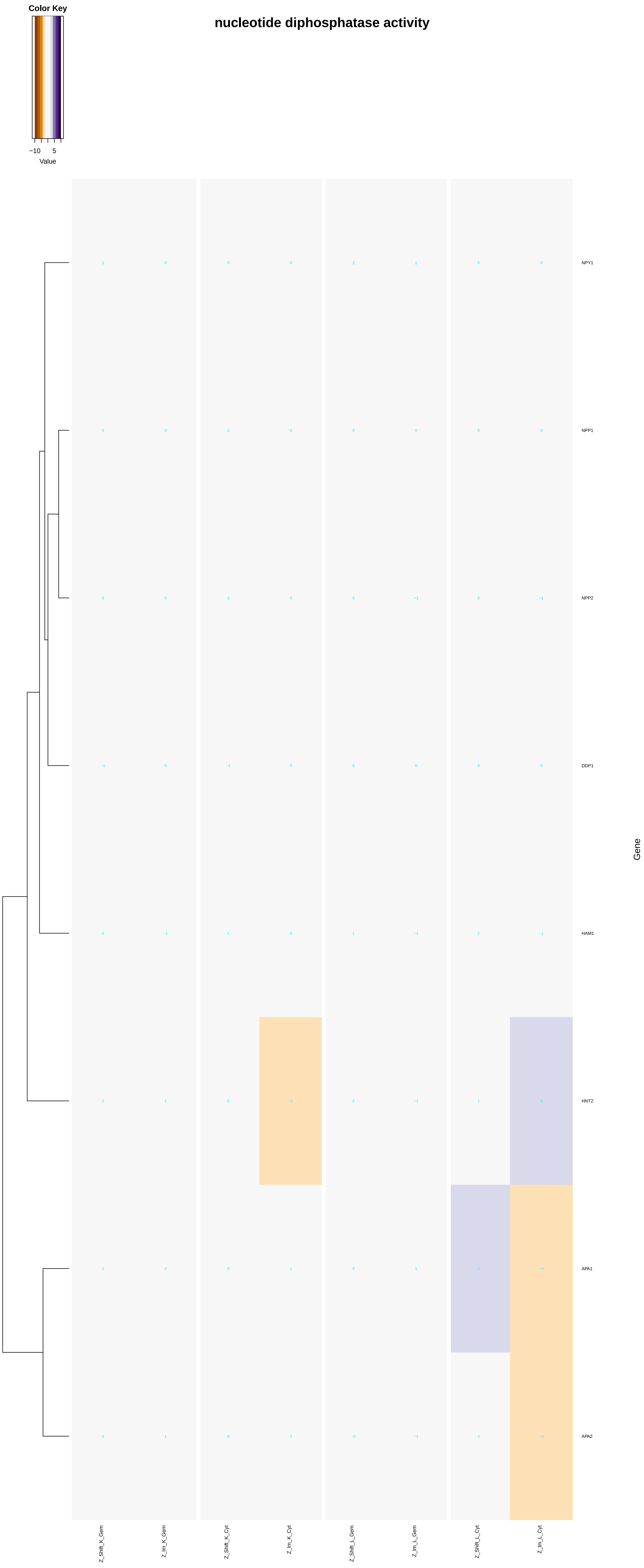
nucleotide diphosphatase activity
Color Key
−10
5
Value
NPY1
NPP1
NPP2
DDP1
HAM1
HNT2
APA1
APA2
Gene
Z_Shift_K_Gem
Z_Im_K_Gem
Z_Shift_K_Cyt
Z_Im_K_Cyt
Z_Shift_L_Gem
Z_Im_L_Gem
Z_Shift_L_Cyt
Z_Im_L_Cyt

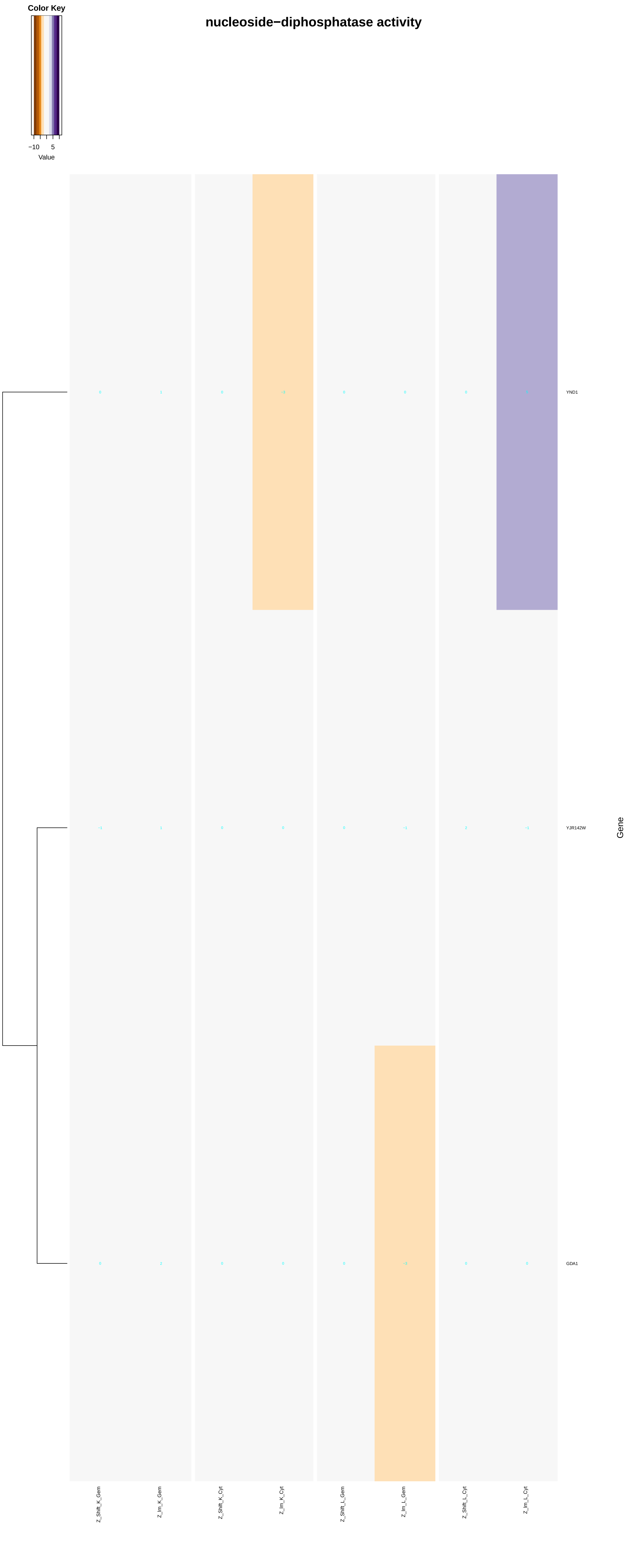
nucleoside−diphosphatase activity
Color Key
−10
5
Value
YND1
YJR142W
GDA1
Gene
Z_Shift_K_Gem
Z_Im_K_Gem
Z_Shift_K_Cyt
Z_Im_K_Cyt
Z_Shift_L_Gem
Z_Im_L_Gem
Z_Shift_L_Cyt
Z_Im_L_Cyt

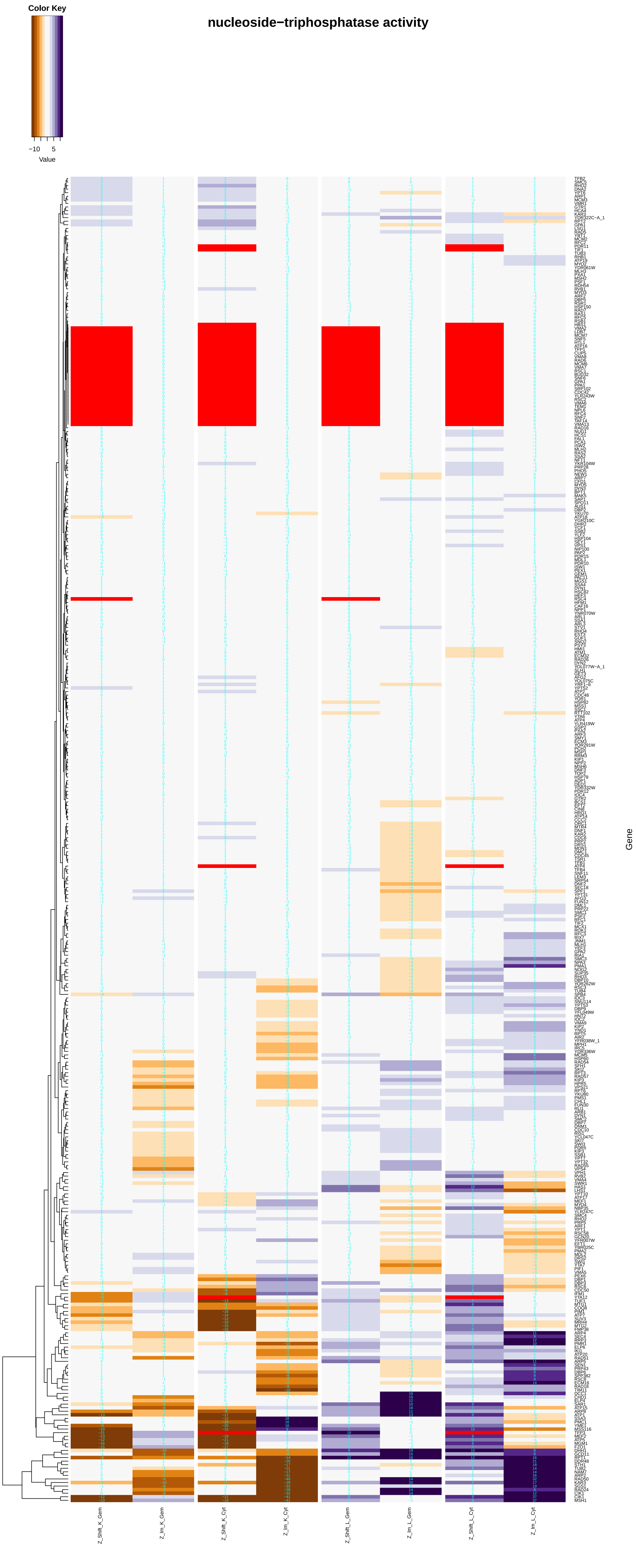
nucleoside−triphosphatase activity
Color Key
−10
5
Value
Z_Shift_K_Gem
Z_Im_K_Gem
Z_Shift_K_Cyt
Z_Im_K_Cyt
Z_Shift_L_Gem
Z_Im_L_Gem
Z_Shift_L_Cyt
Z_Im_L_Cyt
Gene

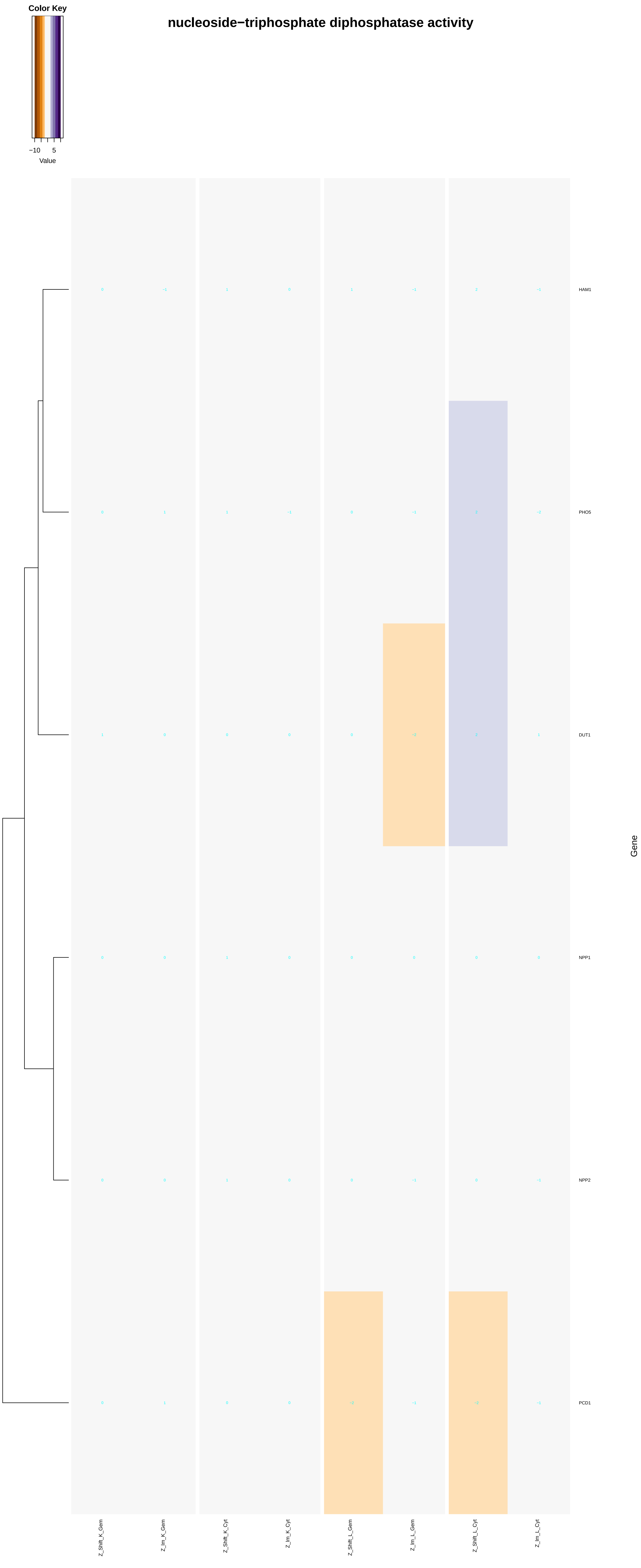
Color Key
−10
5
Value
nucleoside−triphosphate diphosphatase activity
HAM1
PHO5
DUT1
NPP1
NPP2
PCD1
Gene
Z_Shift_K_Gem
Z_Im_K_Gem
Z_Shift_K_Cyt
Z_Im_K_Cyt
Z_Shift_L_Gem
Z_Im_L_Gem
Z_Shift_L_Cyt
Z_Im_L_Cyt

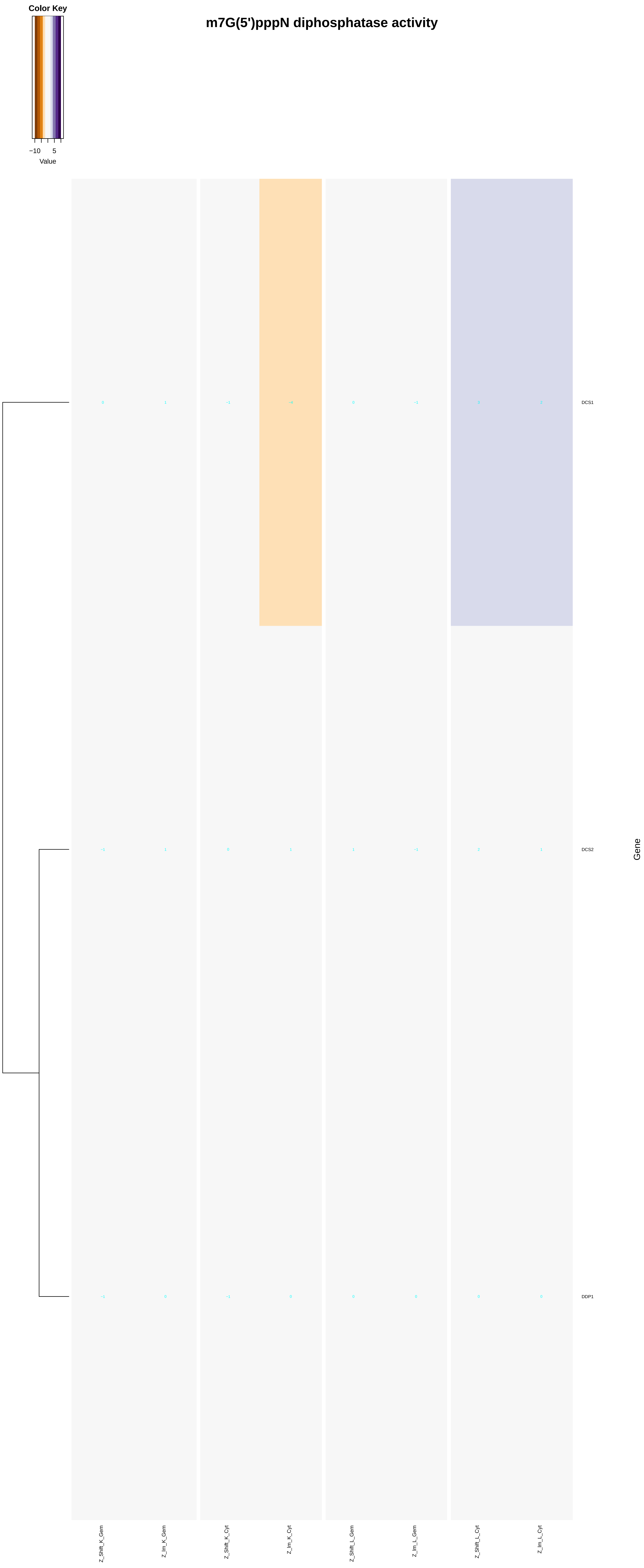

m7G(5')pppN diphosphatase activity
Color Key
−10
5
Value
DCS1
DCS2
DDP1
Gene
Z_Shift_K_Gem
Z_Im_K_Gem
Z_Shift_K_Cyt
Z_Im_K_Cyt
Z_Shift_L_Gem
Z_Im_L_Gem
Z_Shift_L_Cyt
Z_Im_L_Cyt

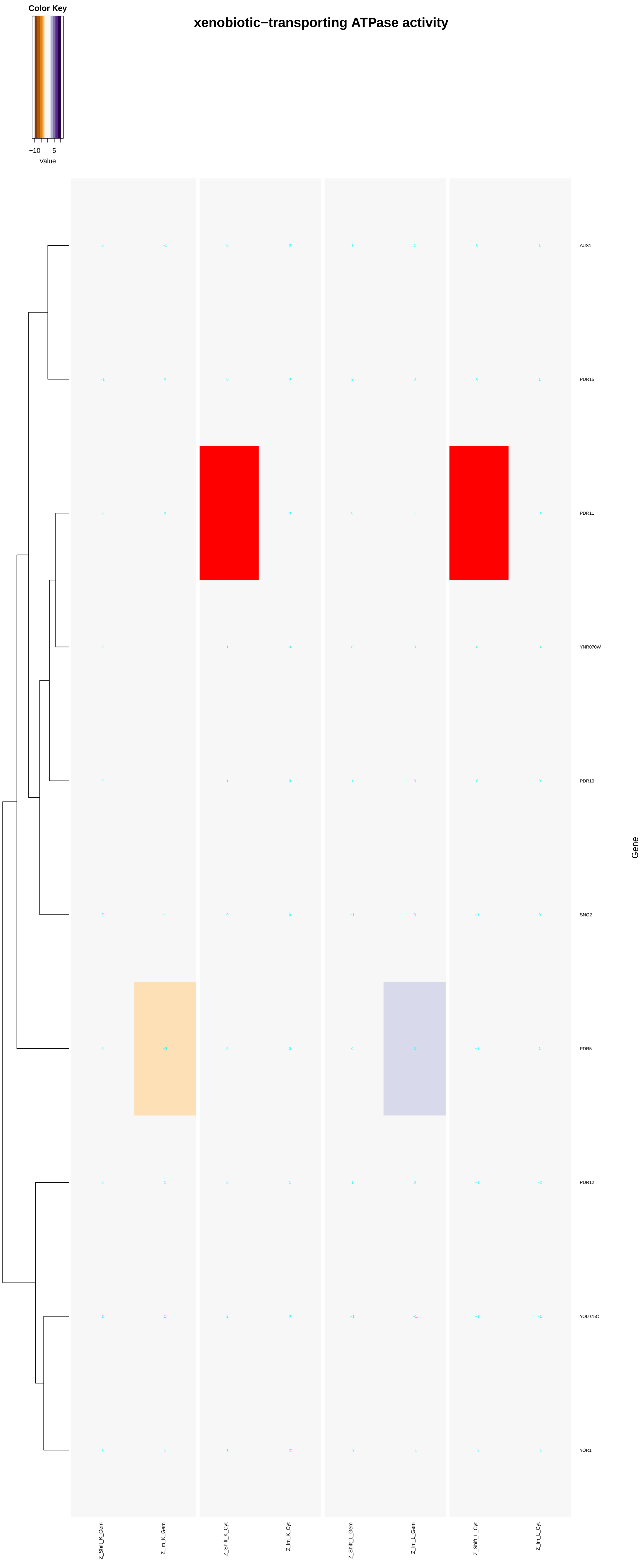

Color Key
−10 5
Value
xenobiotic−transporting ATPase activity
AUS1
PDR15
PDR11
YNR070W
PDR10
SNQ2
PDR5
PDR12
YOL075C
YOR1
Gene
Z_Shift_K_Gem
Z_Im_K_Gem
Z_Shift_K_Cyt
Z_Im_K_Cyt
Z_Shift_L_Gem
Z_Im_L_Gem
Z_Shift_L_Cyt
Z_Im_L_Cyt

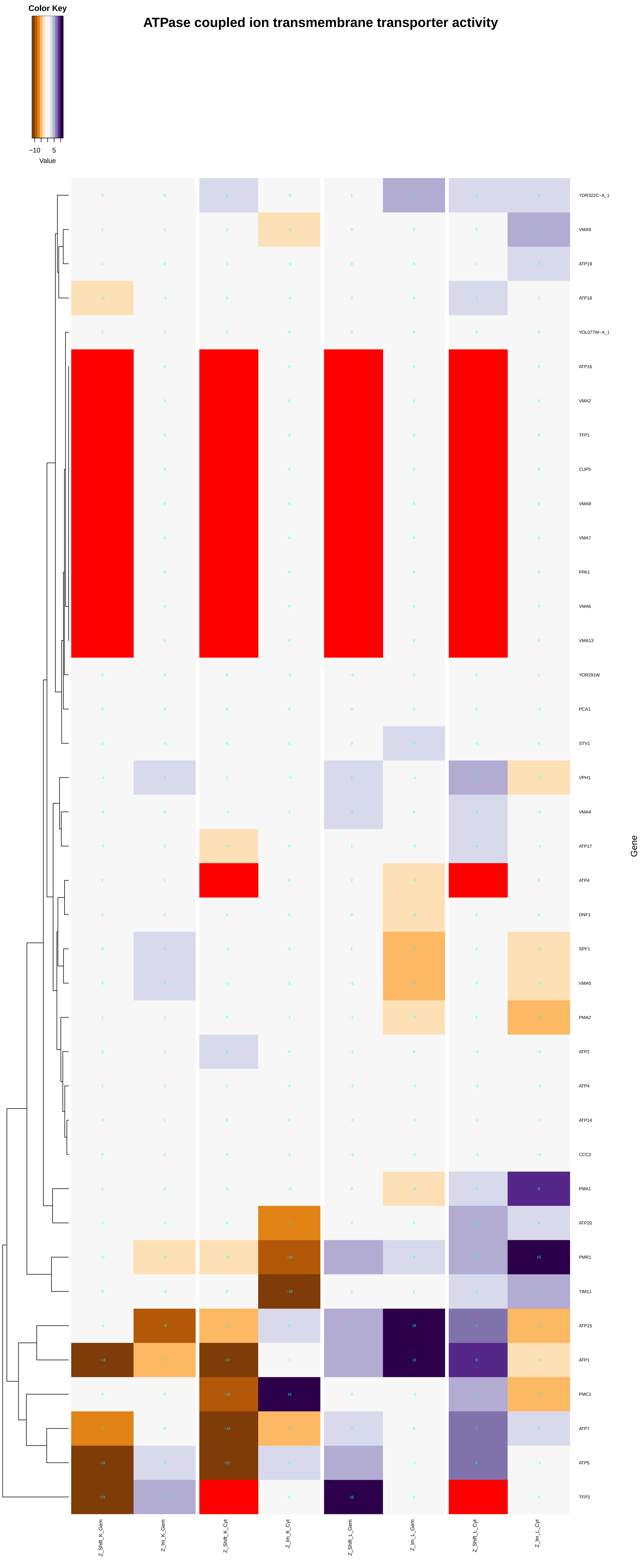
Color Key
−10
5
Value
ATPase coupled ion transmembrane transporter activity
YDR322C−A_1
VMA9
ATP19
ATP18
YOL077W−A_1
ATP16
VMA2
TFP1
CUP5
VMA8
VMA7
PPA1
VMA6
VMA13
YOR291W
PCA1
STV1
VPH1
VMA4
ATP17
ATP4
DNF1
SPF1
VMA5
PMA2
ATP2
ATP4
ATP14
CCC2
PMA1
ATP20
PMR1
TIM11
ATP15
ATP1
PMC1
ATP7
ATP5
TFP3
Gene
Z_Shift_K_Gem
Z_Im_K_Gem
Z_Shift_K_Cyt
Z_Im_K_Cyt
Z_Shift_L_Gem
Z_Im_L_Gem
Z_Shift_L_Cyt
Z_Im_L_Cyt

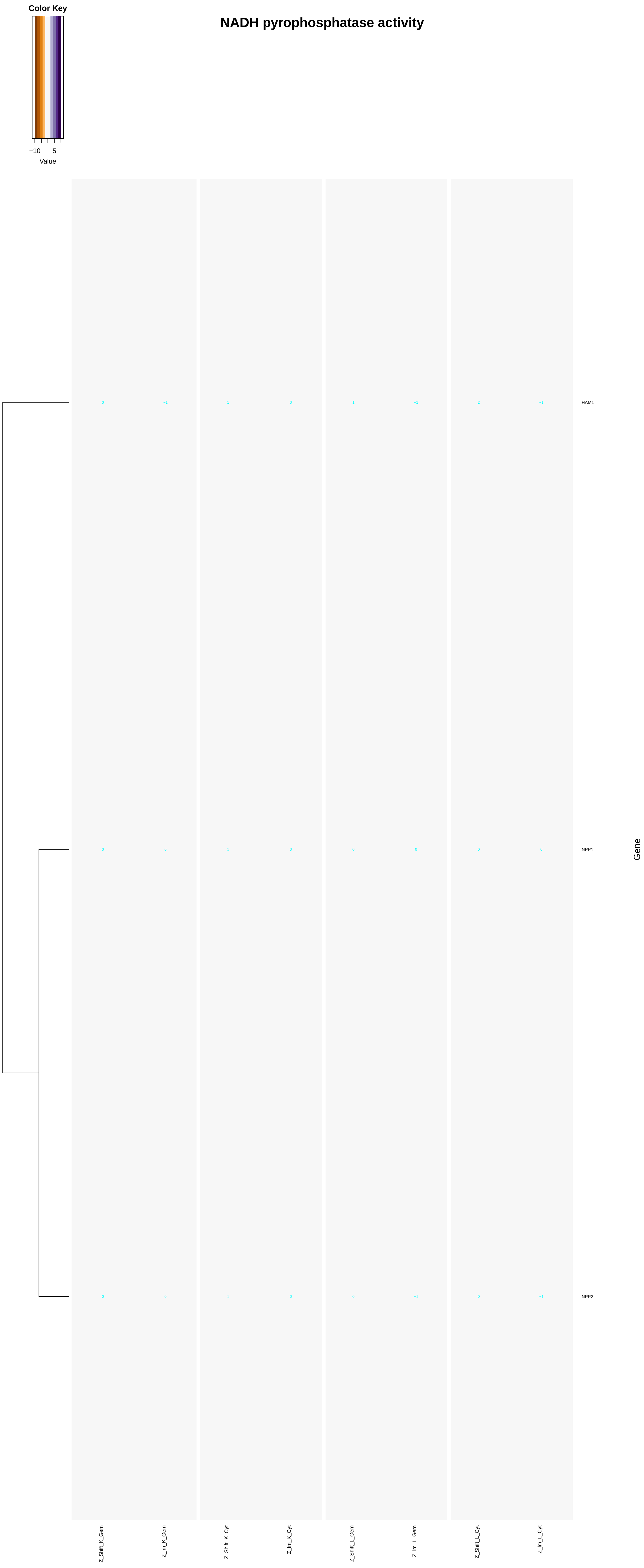

Color Key
−10
5
Value
NADH pyrophosphatase activity
HAM1
NPP1
NPP2
Gene
Z_Shift_K_Gem
Z_Im_K_Gem
Z_Shift_K_Cyt
Z_Im_K_Cyt
Z_Shift_L_Gem
Z_Im_L_Gem
Z_Shift_L_Cyt
Z_Im_L_Cyt

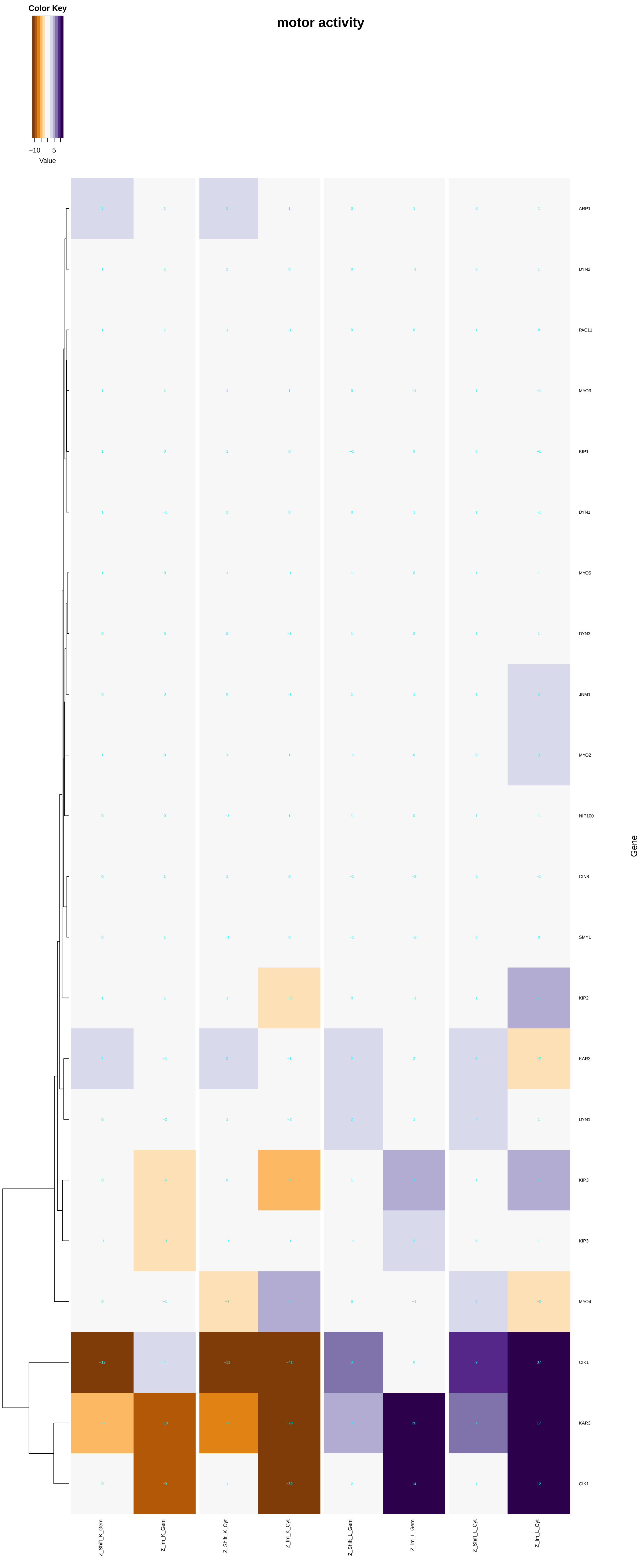
Color Key
−10
5
Value
motor activity
ARP1
DYN2
PAC11
MYO3
KIP1
DYN1
MYO5
DYN3
JNM1
MYO2
NIP100
CIN8
SMY1
KIP2
KAR3
DYN1
KIP3
KIP3
MYO4
CIK1
KAR3
CIK1
Gene
Z_Shift_K_Gem
Z_Im_K_Gem
Z_Shift_K_Cyt
Z_Im_K_Cyt
Z_Shift_L_Gem
Z_Im_L_Gem
Z_Shift_L_Cyt
Z_Im_L_Cyt

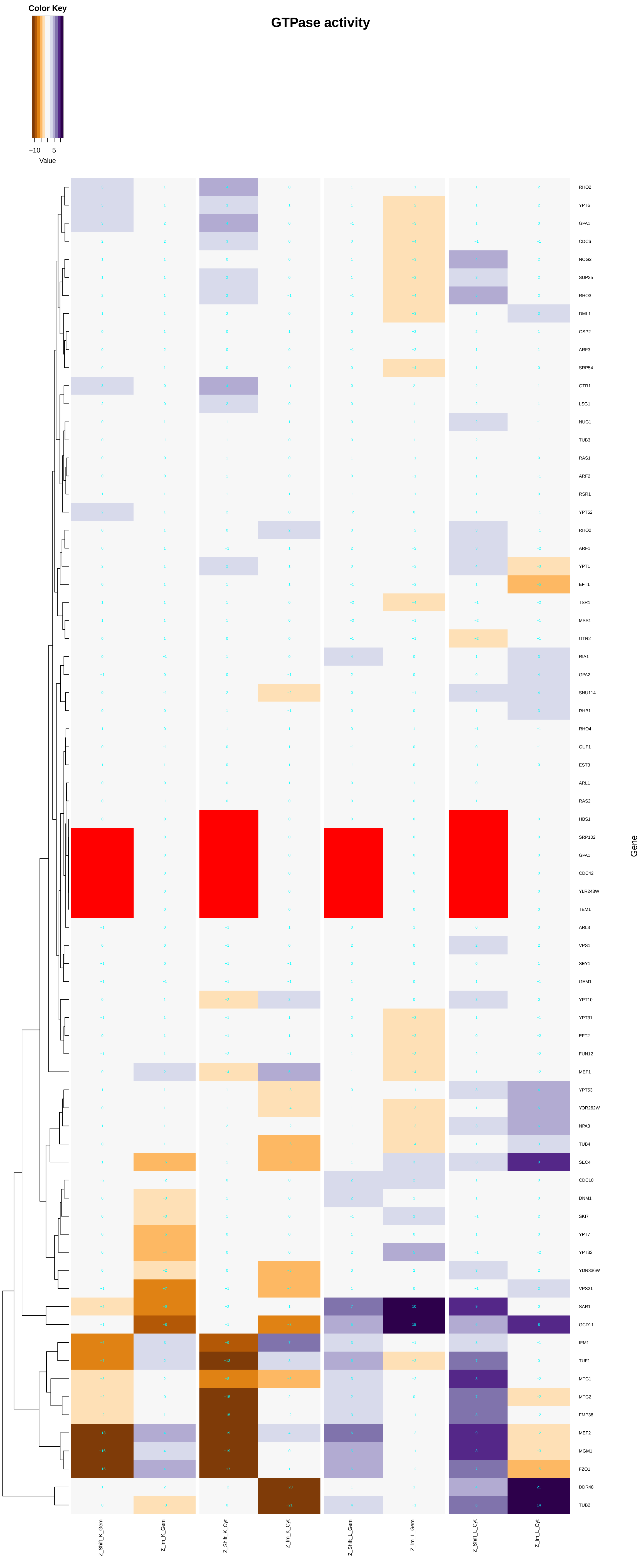

Color Key
−10
5
Value
GTPase activity
RHO2
YPT6
GPA1
CDC6
NOG2
SUP35
RHO3
DML1
GSP2
ARF3
SRP54
GTR1
LSG1
NUG1
TUB3
RAS1
ARF2
RSR1
YPT52
RHO2
ARF1
YPT1
EFT1
TSR1
MSS1
GTR2
RIA1
GPA2
SNU114
RHB1
RHO4
GUF1
EST3
ARL1
RAS2
HBS1
SRP102
GPA1
CDC42
YLR243W
TEM1
ARL3
VPS1
SEY1
GEM1
YPT10
YPT31
EFT2
FUN12
MEF1
YPT53
YOR262W
NPA3
TUB4
SEC4
CDC10
DNM1
SKI7
YPT7
YPT32
YDR336W
VPS21
SAR1
GCD11
IFM1
TUF1
MTG1
MTG2
FMP38
MEF2
MGM1
FZO1
DDR48
TUB2
Gene
Z_Shift_K_Gem
Z_Im_K_Gem
Z_Shift_K_Cyt
Z_Im_K_Cyt
Z_Shift_L_Gem
Z_Im_L_Gem
Z_Shift_L_Cyt
Z_Im_L_Cyt

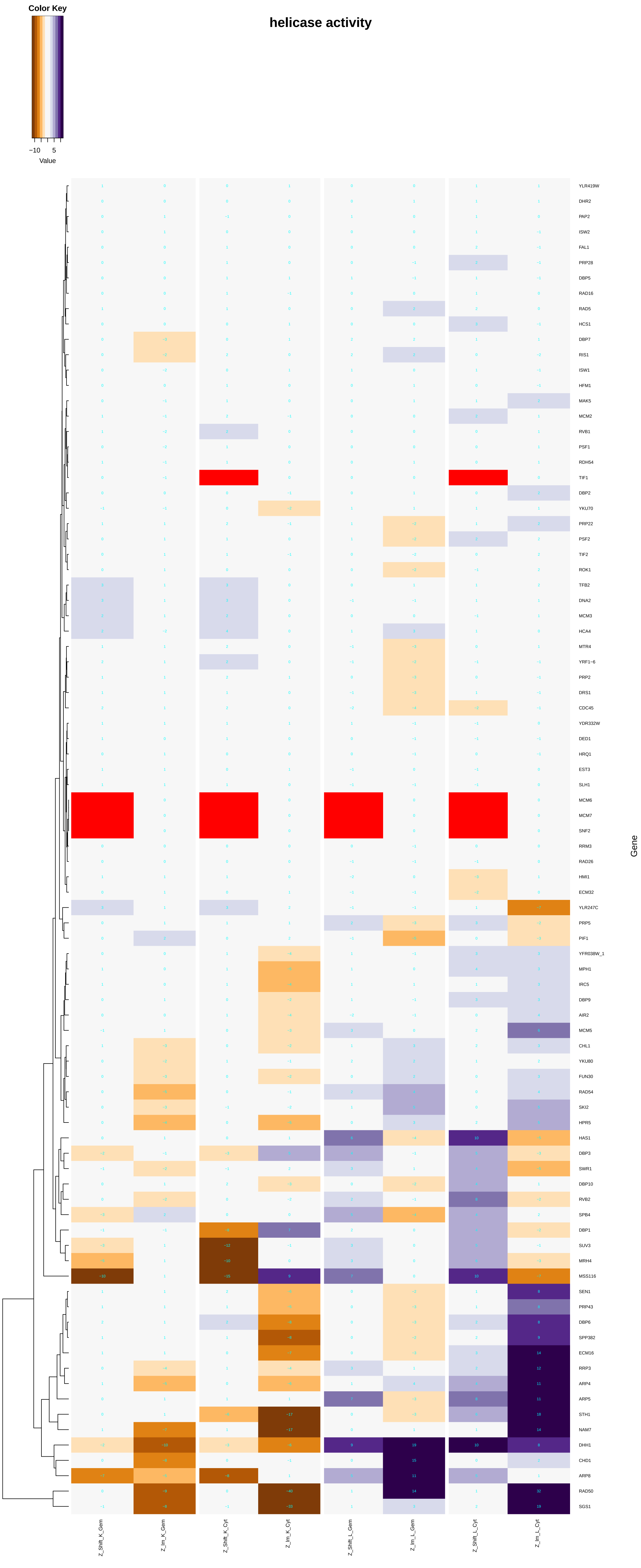

Color Key
−10 5
Value
helicase activity
YLR419W
DHR2
PAP2
ISW2
FAL1
PRP28
DBP5
RAD16
RAD5
HCS1
DBP7
RIS1
ISW1
HFM1
MAK5
MCM2
RVB1
PSF1
RDH54
TIF1
DBP2
YKU70
PRP22
PSF2
TIF2
ROK1
TFB2
DNA2
MCM3
HCA4
MTR4
YRF1−6
PRP2
DRS1
CDC45
YDR332W
DED1
HRQ1
EST3
SLH1
MCM6
MCM7
SNF2
RRM3
RAD26
HMI1
ECM32
YLR247C
PRP5
PIF1
YFR038W_1
MPH1
IRC5
DBP9
AIR2
MCM5
CHL1
YKU80
FUN30
RAD54
SKI2
HPR5
HAS1
DBP3
SWR1
DBP10
RVB2
SPB4
DBP1
SUV3
MRH4
MSS116
SEN1
PRP43
DBP6
SPP382
ECM16
RRP3
ARP4
ARP5
STH1
NAM7
DHH1
CHD1
ARP8
RAD50
SGS1
Gene
Z_Shift_K_Gem
Z_Im_K_Gem
Z_Shift_K_Cyt
Z_Im_K_Cyt
Z_Shift_L_Gem
Z_Im_L_Gem
Z_Shift_L_Cyt
Z_Im_L_Cyt

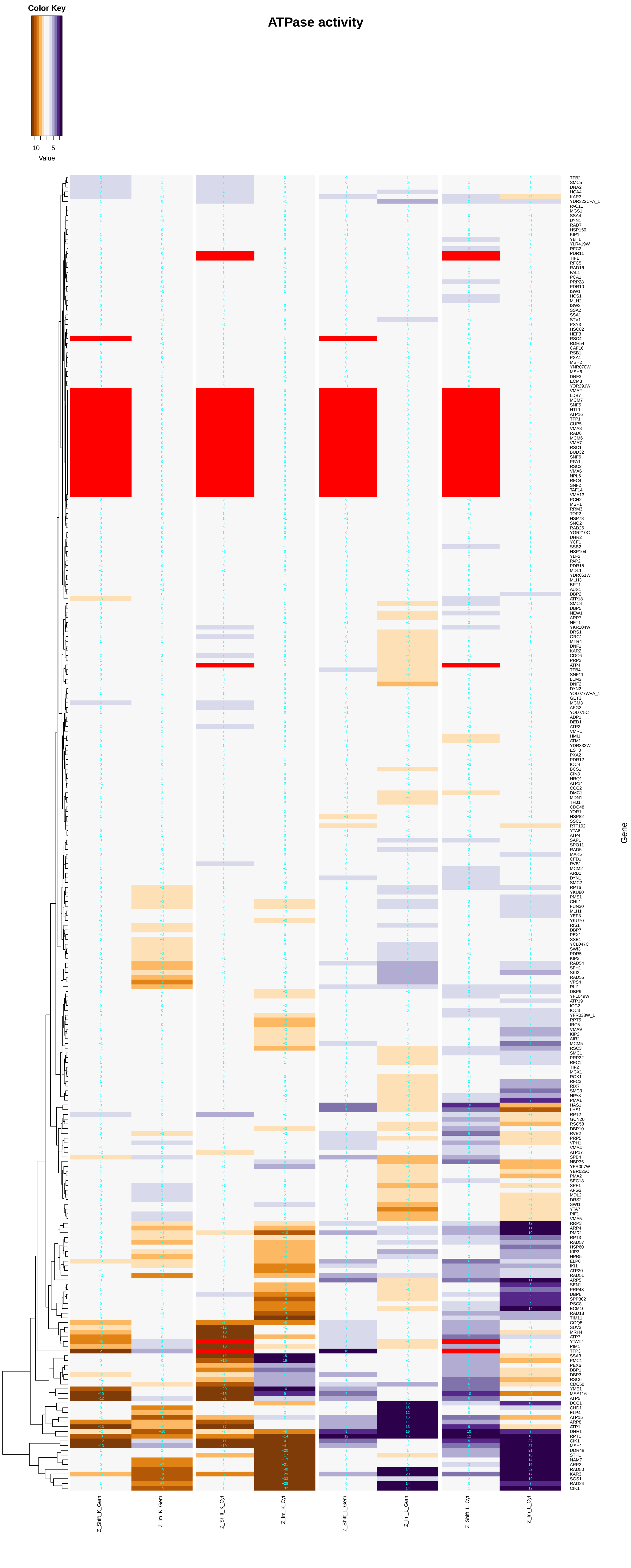

ATPase activity
Color Key
−10
5
Value
Z_Shift_K_Gem
Z_Im_K_Gem
Z_Shift_K_Cyt
Z_Im_K_Cyt
Z_Shift_L_Gem
Z_Im_L_Gem
Z_Shift_L_Cyt
Z_Im_L_Cyt
Gene

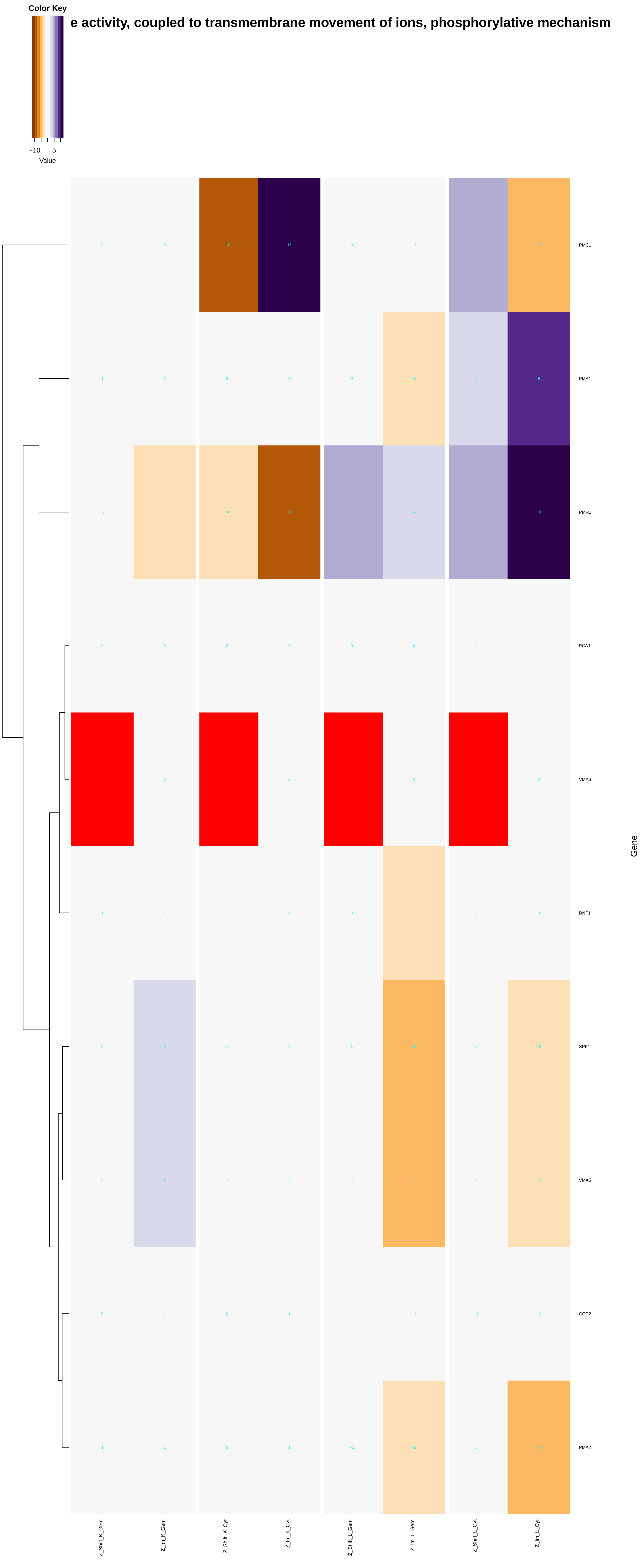

Color Key
−10
5
Value
e activity, coupled to transmembrane movement of ions, phosphorylative mechanism
PMC1
PMA1
PMR1
PCA1
VMA6
DNF1
SPF1
VMA5
CCC2
PMA2
Gene
Z_Shift_K_Gem
Z_Im_K_Gem
Z_Shift_K_Cyt
Z_Im_K_Cyt
Z_Shift_L_Gem
Z_Im_L_Gem
Z_Shift_L_Cyt
Z_Im_L_Cyt

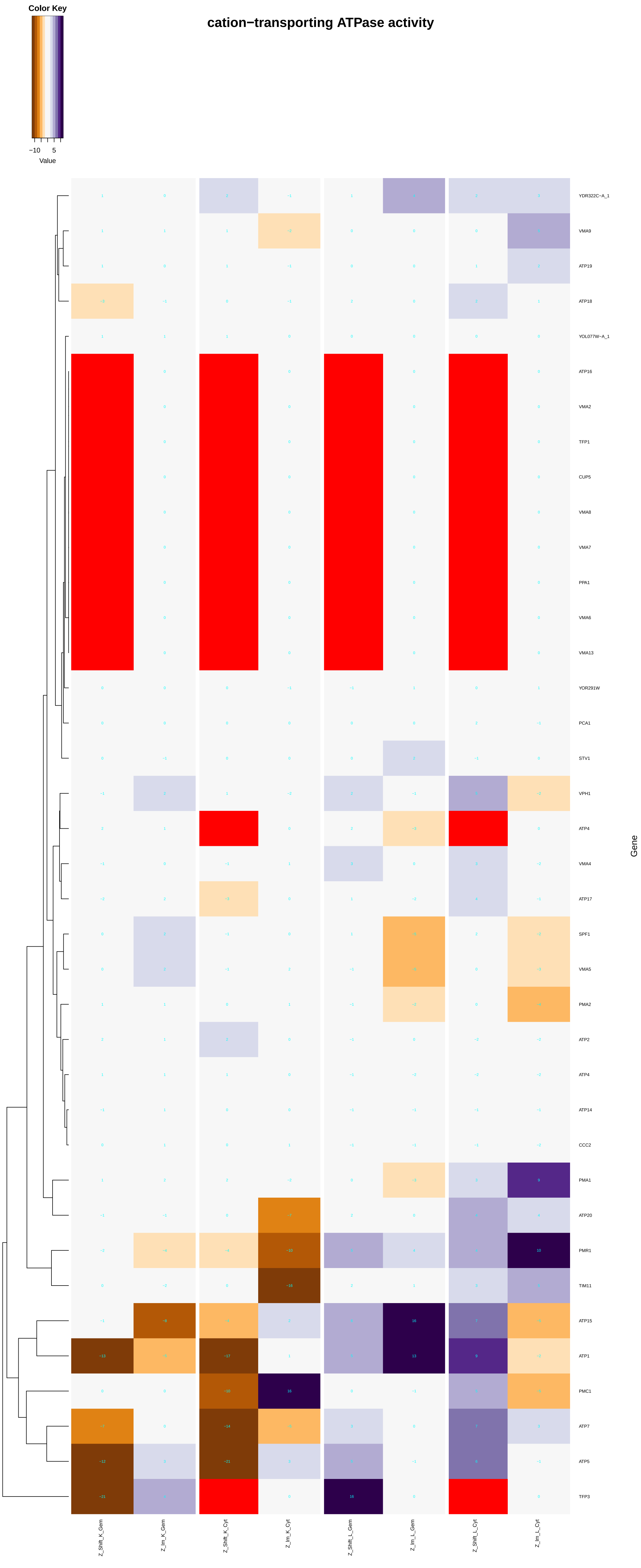
cation−transporting ATPase activity
Color Key
−10
5
Value
YDR322C−A_1
VMA9
ATP19
ATP18
YOL077W−A_1
ATP16
VMA2
TFP1
CUP5
VMA8
VMA7
PPA1
VMA6
VMA13
YOR291W
PCA1
STV1
VPH1
ATP4
VMA4
ATP17
SPF1
VMA5
PMA2
ATP2
ATP4
ATP14
CCC2
PMA1
ATP20
PMR1
TIM11
ATP15
ATP1
PMC1
ATP7
ATP5
TFP3
Gene
Z_Shift_K_Gem
Z_Im_K_Gem
Z_Shift_K_Cyt
Z_Im_K_Cyt
Z_Shift_L_Gem
Z_Im_L_Gem
Z_Shift_L_Cyt
Z_Im_L_Cyt

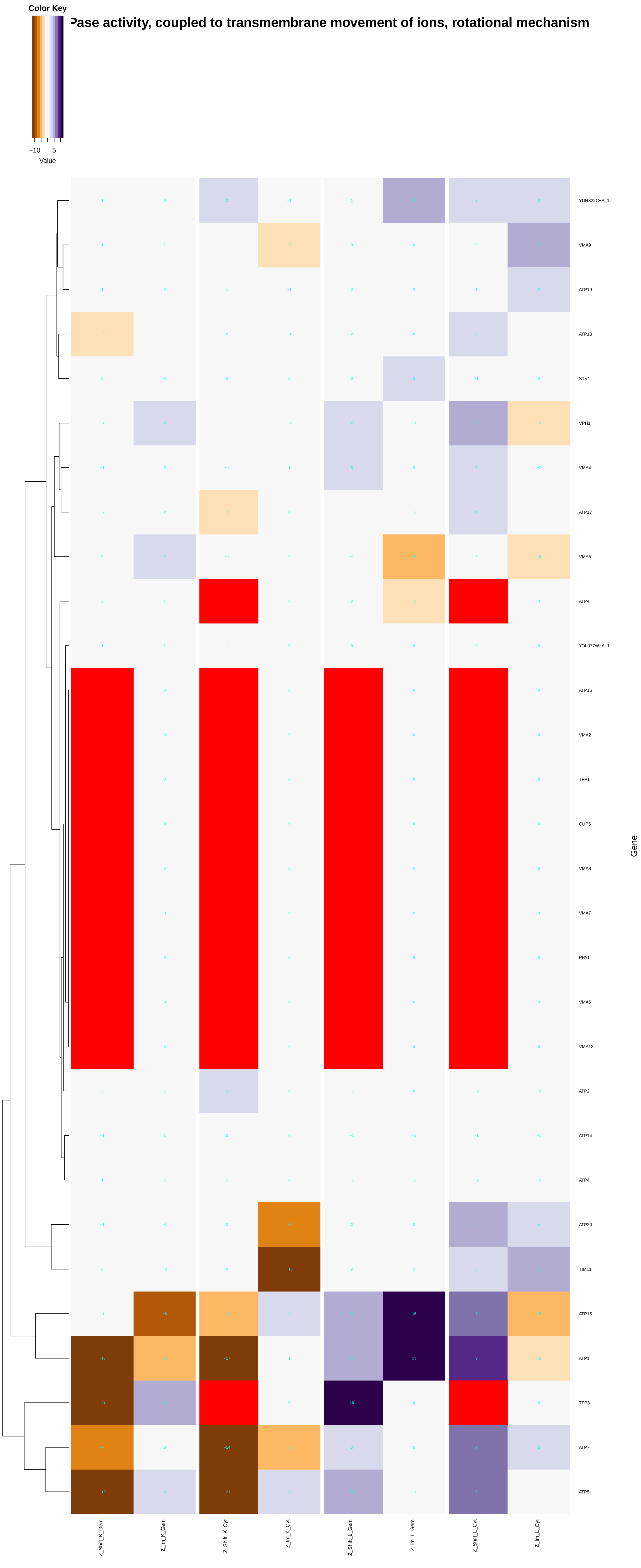
Pase activity, coupled to transmembrane movement of ions, rotational mechanism
Color Key
−10 5
Value
YDR322C−A_1
VMA9
ATP19
ATP18
STV1
VPH1
VMA4
ATP17
VMA5
ATP4
YOL077W−A_1
ATP16
VMA2
TFP1
CUP5
VMA8
VMA7
PPA1
VMA6
VMA13
ATP2
ATP14
ATP4
ATP20
TIM11
ATP15
ATP1
TFP3
ATP7
ATP5
Gene
Z_Shift_K_Gem
Z_Im_K_Gem
Z_Shift_K_Cyt
Z_Im_K_Cyt
Z_Shift_L_Gem
Z_Im_L_Gem
Z_Shift_L_Cyt
Z_Im_L_Cyt

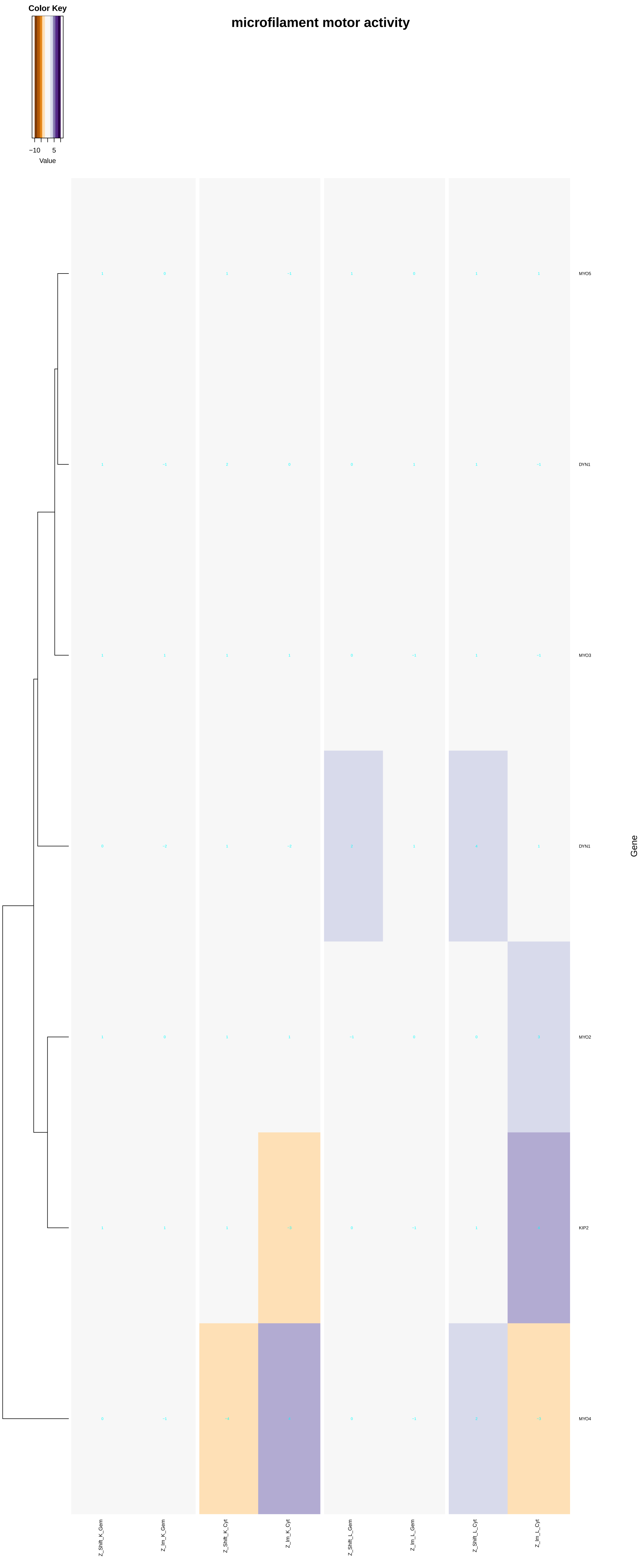
microfilament motor activity
Color Key
−10
5
Value
MYO5
DYN1
MYO3
DYN1
MYO2
KIP2
MYO4
Gene
Z_Shift_K_Gem
Z_Im_K_Gem
Z_Shift_K_Cyt
Z_Im_K_Cyt
Z_Shift_L_Gem
Z_Im_L_Gem
Z_Shift_L_Cyt
Z_Im_L_Cyt

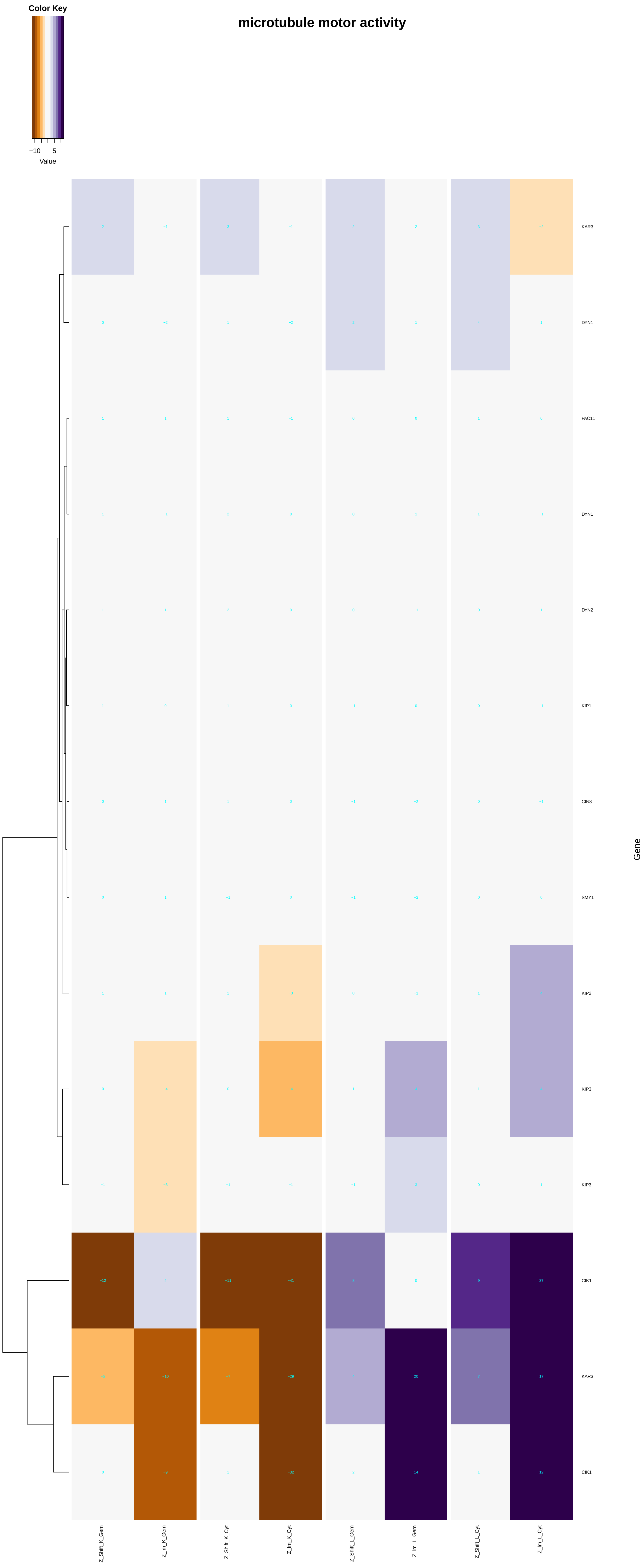

Color Key
−10
5
Value
microtubule motor activity
KAR3
DYN1
PAC11
DYN1
DYN2
KIP1
CIN8
SMY1
KIP2
KIP3
KIP3
CIK1
KAR3
CIK1
Gene
Z_Shift_K_Gem
Z_Im_K_Gem
Z_Shift_K_Cyt
Z_Im_K_Cyt
Z_Shift_L_Gem
Z_Im_L_Gem
Z_Shift_L_Cyt
Z_Im_L_Cyt

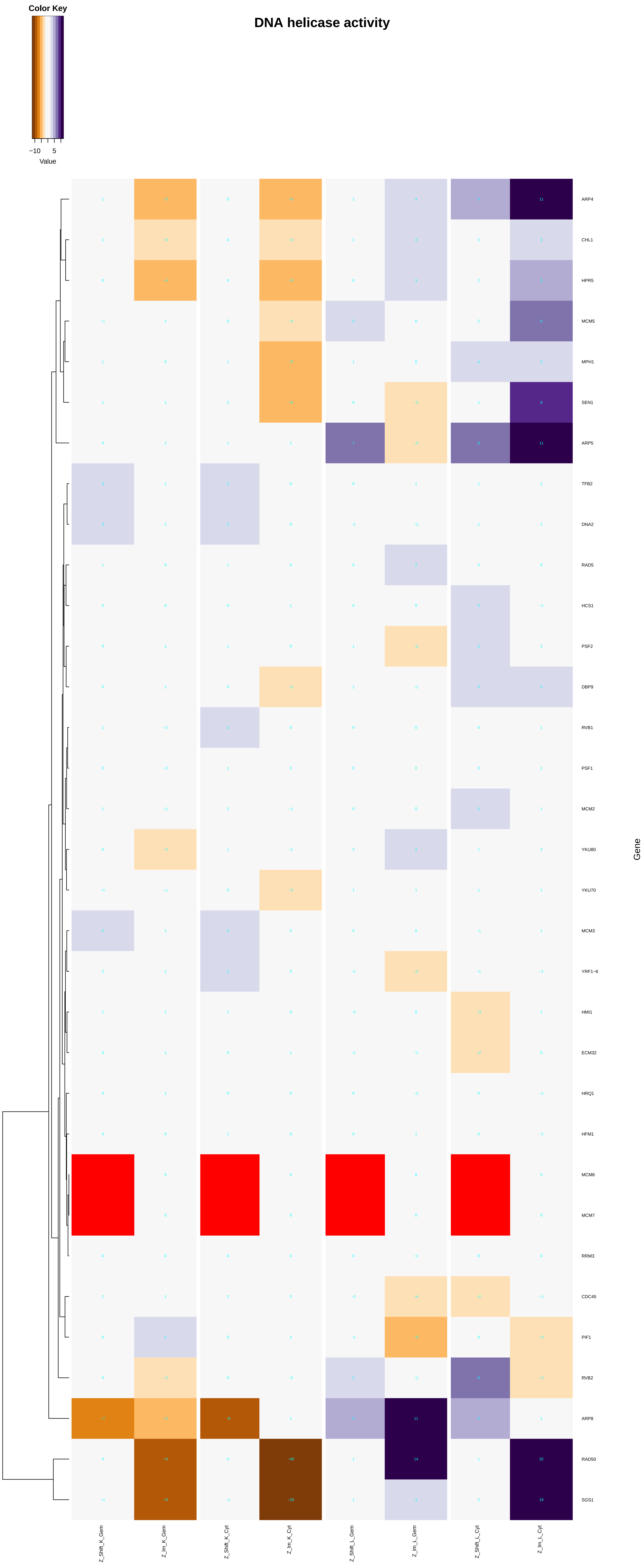
Color Key
−10
5
Value
DNA helicase activity
ARP4
CHL1
HPR5
MCM5
MPH1
SEN1
ARP5
TFB2
DNA2
RAD5
HCS1
PSF2
DBP9
RVB1
PSF1
MCM2
YKU80
YKU70
MCM3
YRF1−6
HMI1
ECM32
HRQ1
HFM1
MCM6
MCM7
RRM3
CDC45
PIF1
RVB2
ARP8
RAD50
SGS1
Gene
Z_Shift_K_Gem
Z_Im_K_Gem
Z_Shift_K_Cyt
Z_Im_K_Cyt
Z_Shift_L_Gem
Z_Im_L_Gem
Z_Shift_L_Cyt
Z_Im_L_Cyt

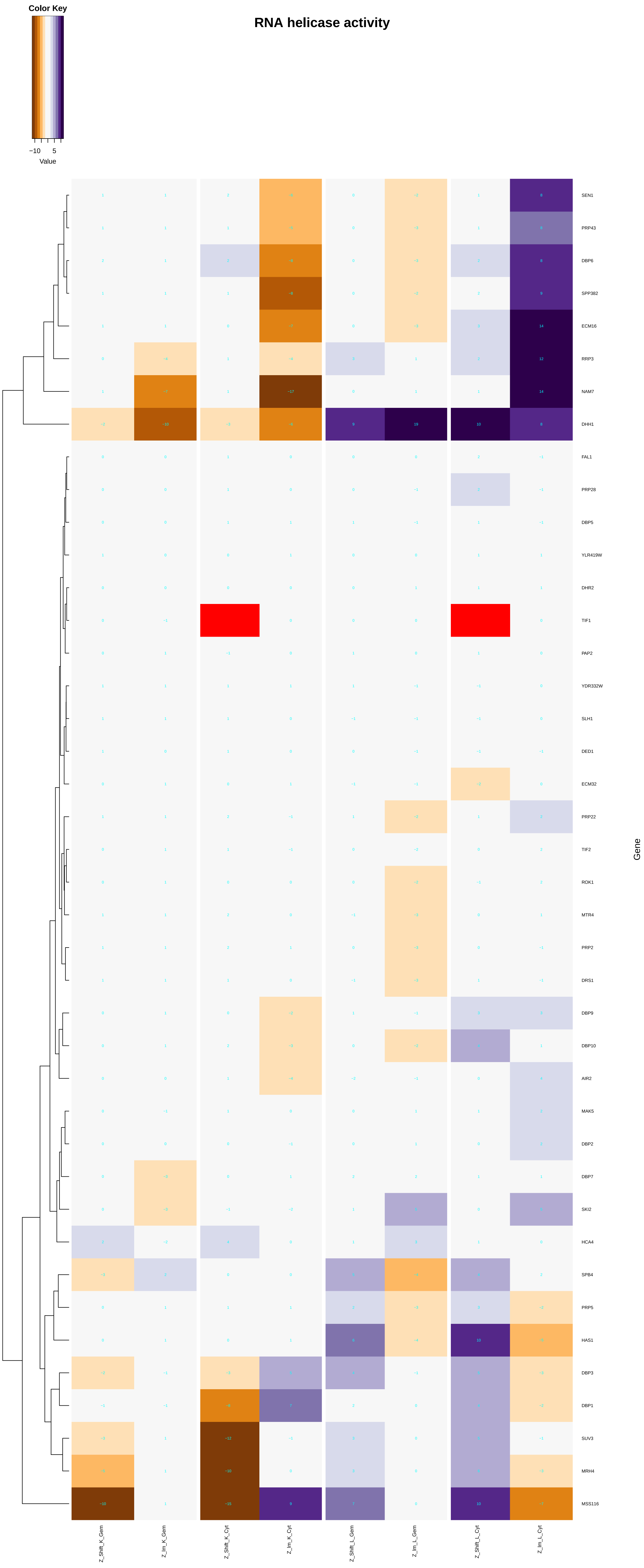
Color Key
−10
5
Value
RNA helicase activity
SEN1
PRP43
DBP6
SPP382
ECM16
RRP3
NAM7
DHH1
FAL1
PRP28
DBP5
YLR419W
DHR2
TIF1
PAP2
YDR332W
SLH1
DED1
ECM32
PRP22
TIF2
ROK1
MTR4
PRP2
DRS1
DBP9
DBP10
AIR2
MAK5
DBP2
DBP7
SKI2
HCA4
SPB4
PRP5
HAS1
DBP3
DBP1
SUV3
MRH4
MSS116
Gene
Z_Shift_K_Gem
Z_Im_K_Gem
Z_Shift_K_Cyt
Z_Im_K_Cyt
Z_Shift_L_Gem
Z_Im_L_Gem
Z_Shift_L_Cyt
Z_Im_L_Cyt

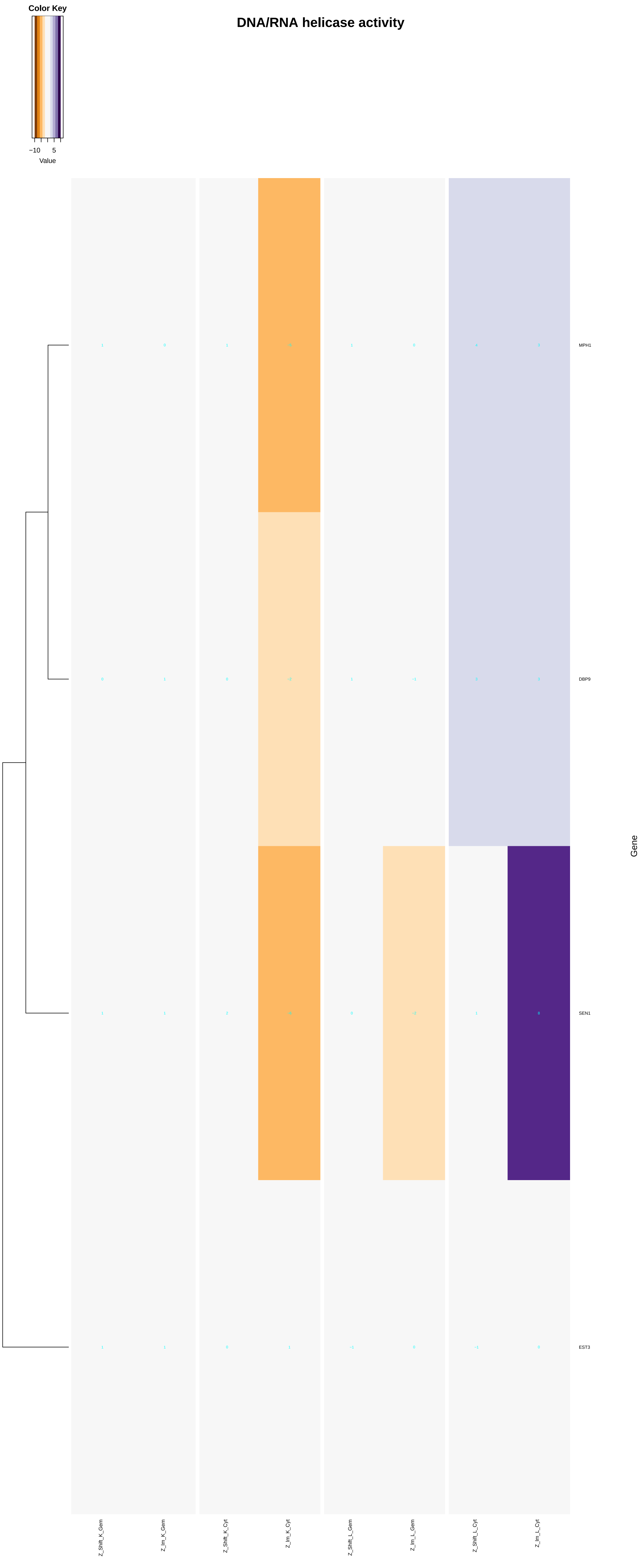
DNA/RNA helicase activity
Color Key
−10
5
Value
MPH1
DBP9
SEN1
EST3
Gene
Z_Shift_K_Gem
Z_Im_K_Gem
Z_Shift_K_Cyt
Z_Im_K_Cyt
Z_Shift_L_Gem
Z_Im_L_Gem
Z_Shift_L_Cyt
Z_Im_L_Cyt

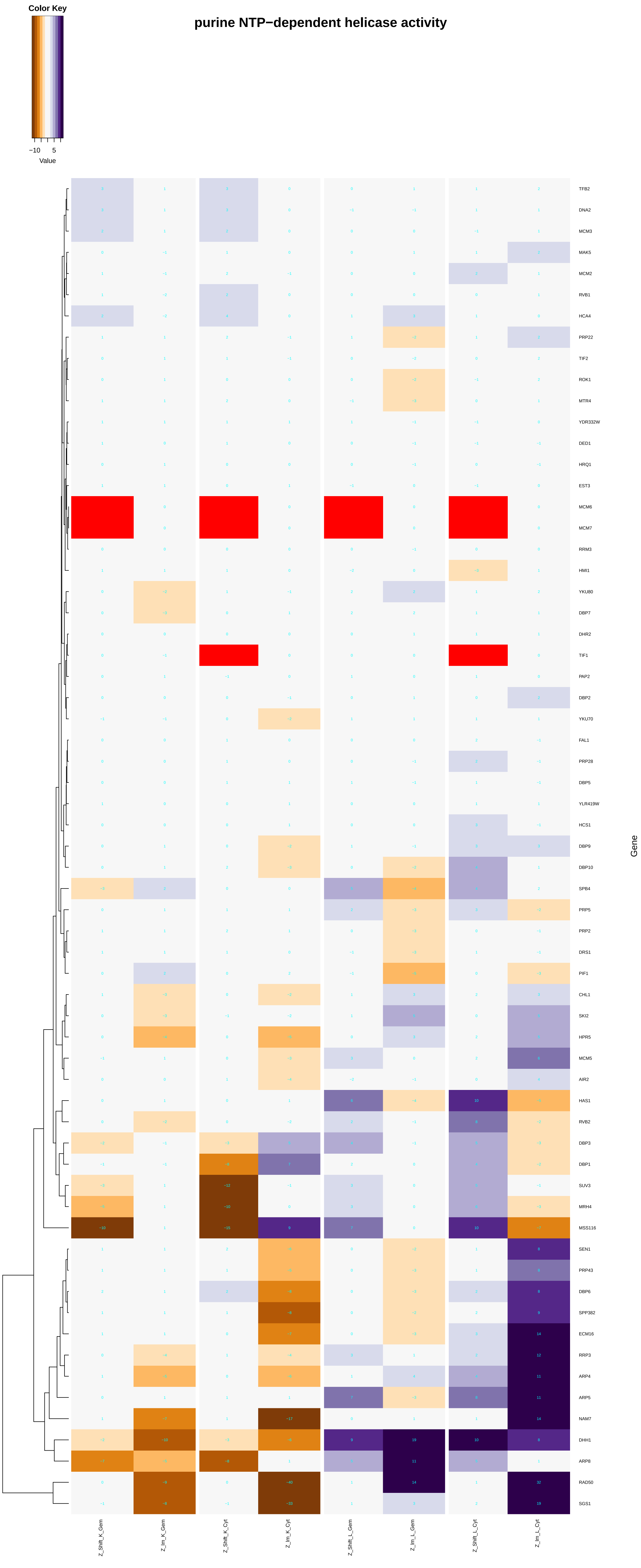

Color Key
−10
5
Value
purine NTP−dependent helicase activity
TFB2
DNA2
MCM3
MAK5
MCM2
RVB1
HCA4
PRP22
TIF2
ROK1
MTR4
YDR332W
DED1
HRQ1
EST3
MCM6
MCM7
RRM3
HMI1
YKU80
DBP7
DHR2
TIF1
PAP2
DBP2
YKU70
FAL1
PRP28
DBP5
YLR419W
HCS1
DBP9
DBP10
SPB4
PRP5
PRP2
DRS1
PIF1
CHL1
SKI2
HPR5
MCM5
AIR2
HAS1
RVB2
DBP3
DBP1
SUV3
MRH4
MSS116
SEN1
PRP43
DBP6
SPP382
ECM16
RRP3
ARP4
ARP5
NAM7
DHH1
ARP8
RAD50
SGS1
Gene
Z_Shift_K_Gem
Z_Im_K_Gem
Z_Shift_K_Cyt
Z_Im_K_Cyt
Z_Shift_L_Gem
Z_Im_L_Gem
Z_Shift_L_Cyt
Z_Im_L_Cyt

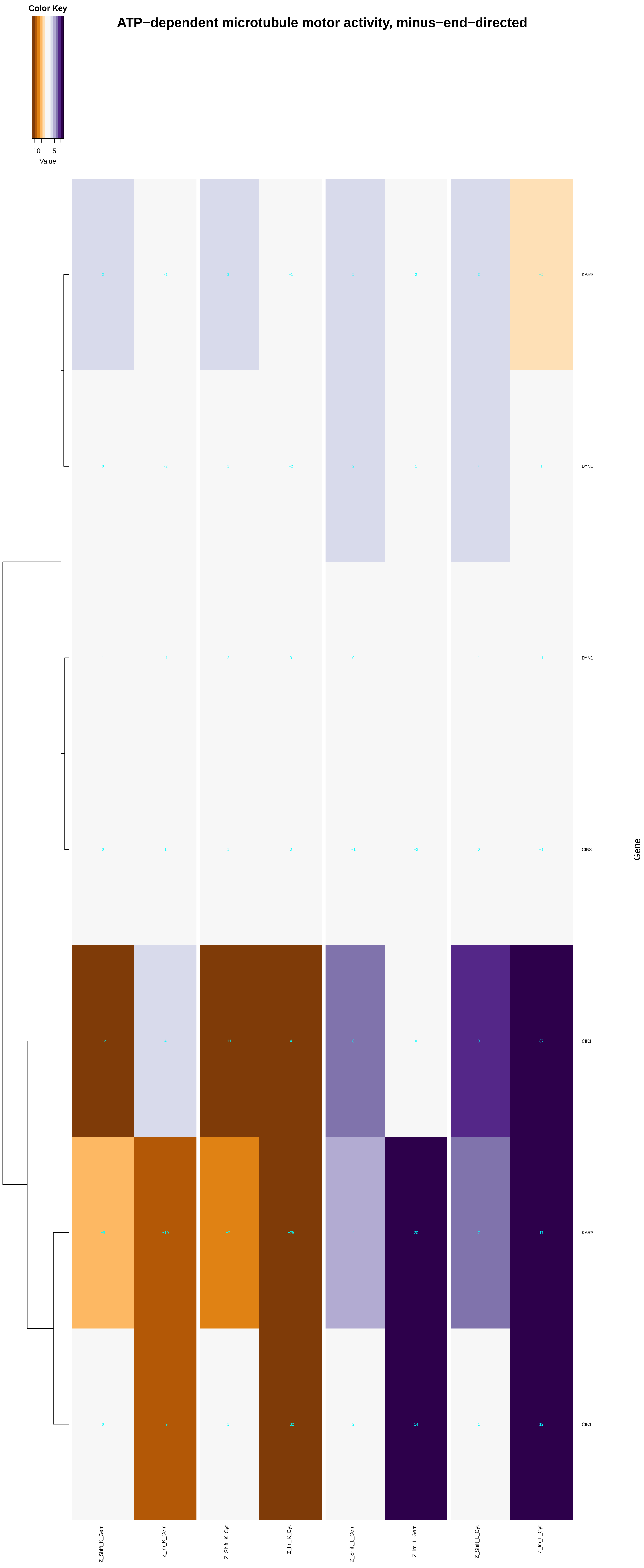
Color Key
−10
5
Value
ATP−dependent microtubule motor activity, minus−end−directed
KAR3
DYN1
DYN1
CIN8
CIK1
KAR3
CIK1
Gene
Z_Shift_K_Gem
Z_Im_K_Gem
Z_Shift_K_Cyt
Z_Im_K_Cyt
Z_Shift_L_Gem
Z_Im_L_Gem
Z_Shift_L_Cyt
Z_Im_L_Cyt

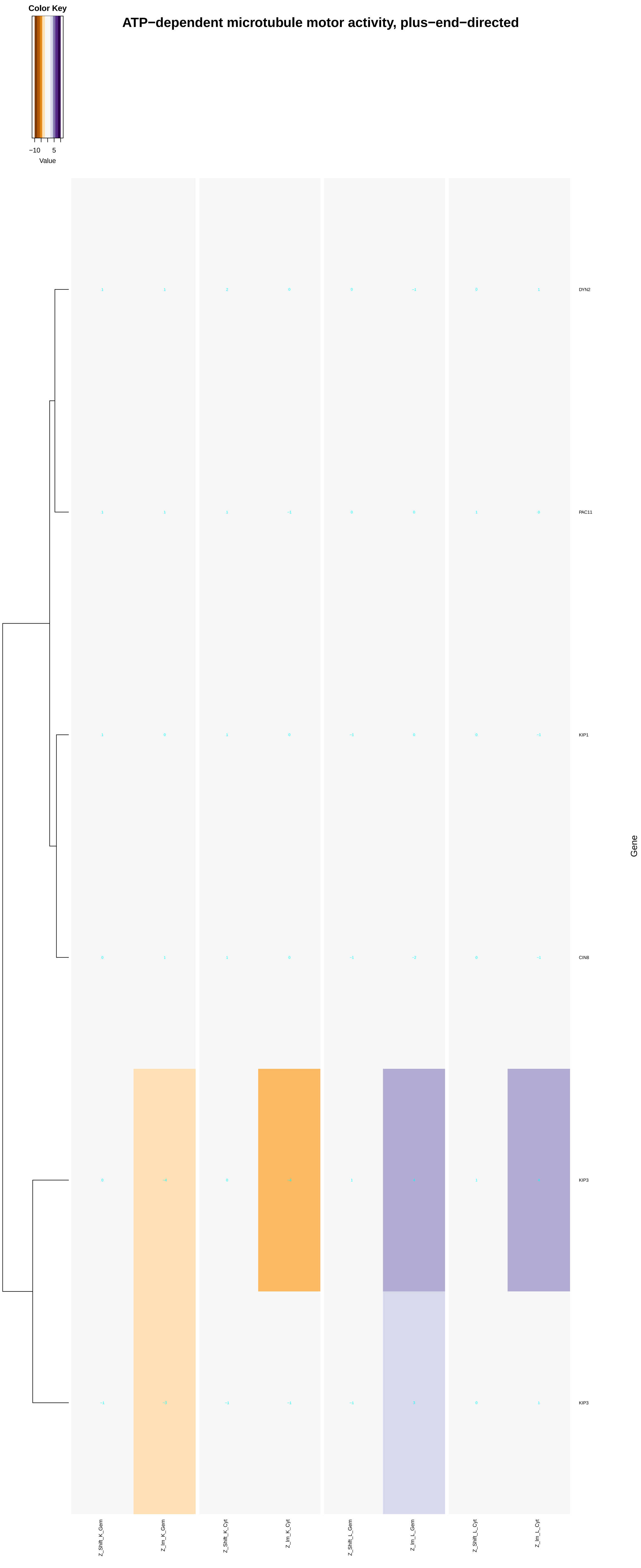
ATP−dependent microtubule motor activity, plus−end−directed
Color Key
−10
5
Value
DYN2
PAC11
KIP1
CIN8
KIP3
KIP3
Gene
Z_Shift_K_Gem
Z_Im_K_Gem
Z_Shift_K_Cyt
Z_Im_K_Cyt
Z_Shift_L_Gem
Z_Im_L_Gem
Z_Shift_L_Cyt
Z_Im_L_Cyt

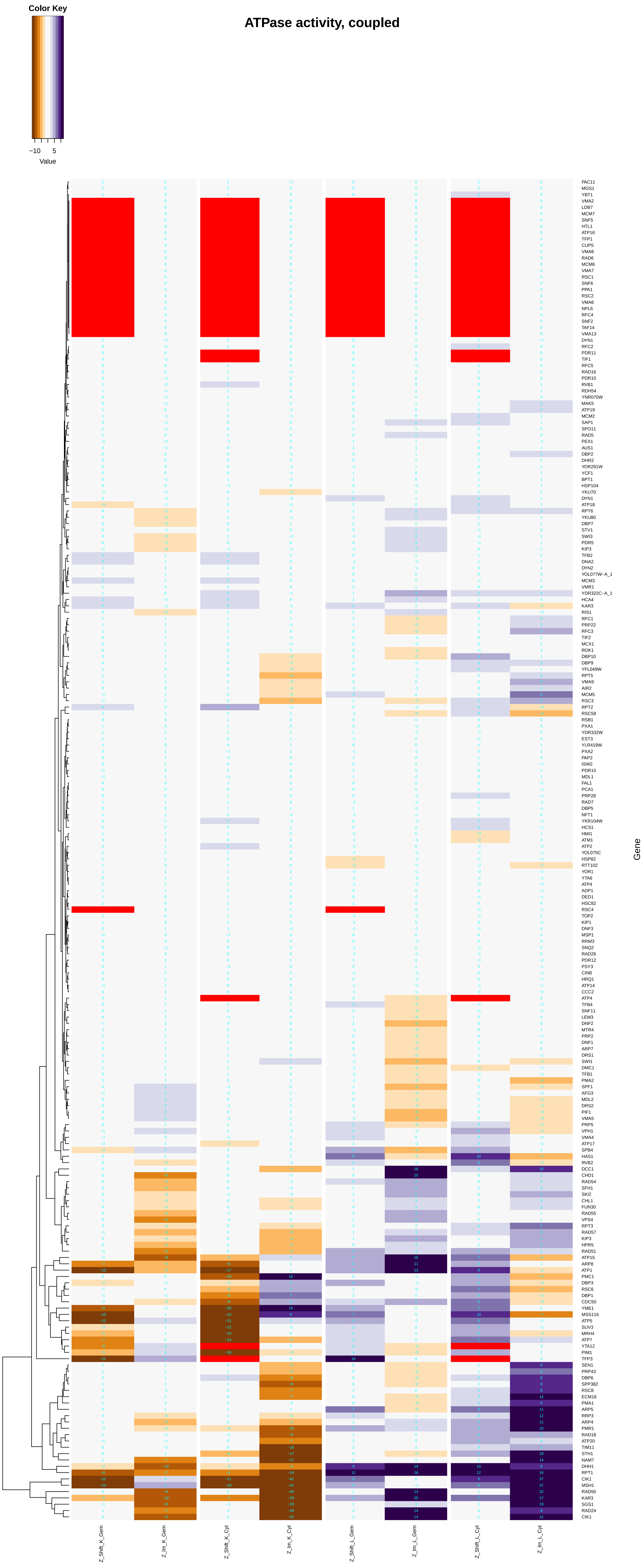

Color Key
−10
5
Value
ATPase activity, coupled
Gene
Z_Shift_K_Gem
Z_Im_K_Gem
Z_Shift_K_Cyt
Z_Im_K_Cyt
Z_Shift_L_Gem
Z_Im_L_Gem
Z_Shift_L_Cyt
Z_Im_L_Cyt
PAC11
MGS1
YBT1
VMA2
LDB7
MCM7
SNF5
HTL1
ATP16
TFP1
CUP5
VMA8
RAD6
MCM6
VMA7
RSC1
SNF6
PPA1
RSC2
VMA6
NPL6
RFC4
SNF2
TAF14
VMA13
DYN1
RFC2
PDR11
TIF1
RFC5
RAD16
PDR10
RVB1
RDH54
YNR070W
MAK5
ATP19
MCM2
SAP1
SPO11
RAD5
PEX1
AUS1
DBP2
DHR2
YOR291W
YCF1
BPT1
HSP104
YKU70
DYN1
ATP18
RPT6
YKU80
DBP7
STV1
SWI3
PDR5
KIP3
TFB2
DNA2
DYN2
YOL077W−A_1
MCM3
VMR1
YDR322C−A_1
HCA4
KAR3
RIS1
RFC1
PRP22
RFC3
TIF2
MCX1
ROK1
DBP10
DBP9
YFL049W
RPT5
VMA9
AIR2
MCM5
RSC3
RPT2
RSC58
RSB1
PXA1
YDR332W
EST3
YLR419W
PXA2
PAP2
ISW2
PDR15
MDL1
FAL1
PCA1
PRP28
RAD7
DBP5
NFT1
YKR104W
HCS1
HMI1
ATM1
ATP2
YOL075C
HSP82
RTT102
YOR1
YTA6
ATP4
ADP1
DED1
HSC82
RSC4
TOP2
KIP1
DNF3
MSP1
RRM3
SNQ2
RAD26
PDR12
PSY3
CIN8
HRQ1
ATP14
CCC2
ATP4
TFB4
SNF11
LEM3
DNF2
MTR4
PRP2
DNF1
ARP7
DRS1
SWI1
DMC1
TFB1
PMA2
SPF1
AFG3
MDL2
DRS2
PIF1
VMA5
PRP5
VPH1
VMA4
ATP17
SPB4
HAS1
RVB2
DCC1
CHD1
RAD54
SFH1
SKI2
CHL1
FUN30
RAD55
VPS4
RPT3
RAD57
KIP3
HPR5
RAD51
ATP15
ARP8
ATP1
PMC1
DBP3
RSC6
DBP1
CDC50
YME1
MSS116
ATP5
SUV3
MRH4
ATP7
YTA12
PIM1
TFP3
SEN1
PRP43
DBP6
SPP382
RSC8
ECM16
PMA1
ARP5
RRP3
ARP4
PMR1
RAD18
ATP20
TIM11
STH1
NAM7
DHH1
RPT1
CIK1
MSH1
RAD50
KAR3
SGS1
RAD24
CIK1

### hydrogen−exporting ATPase activity, phosphorylative mechanism

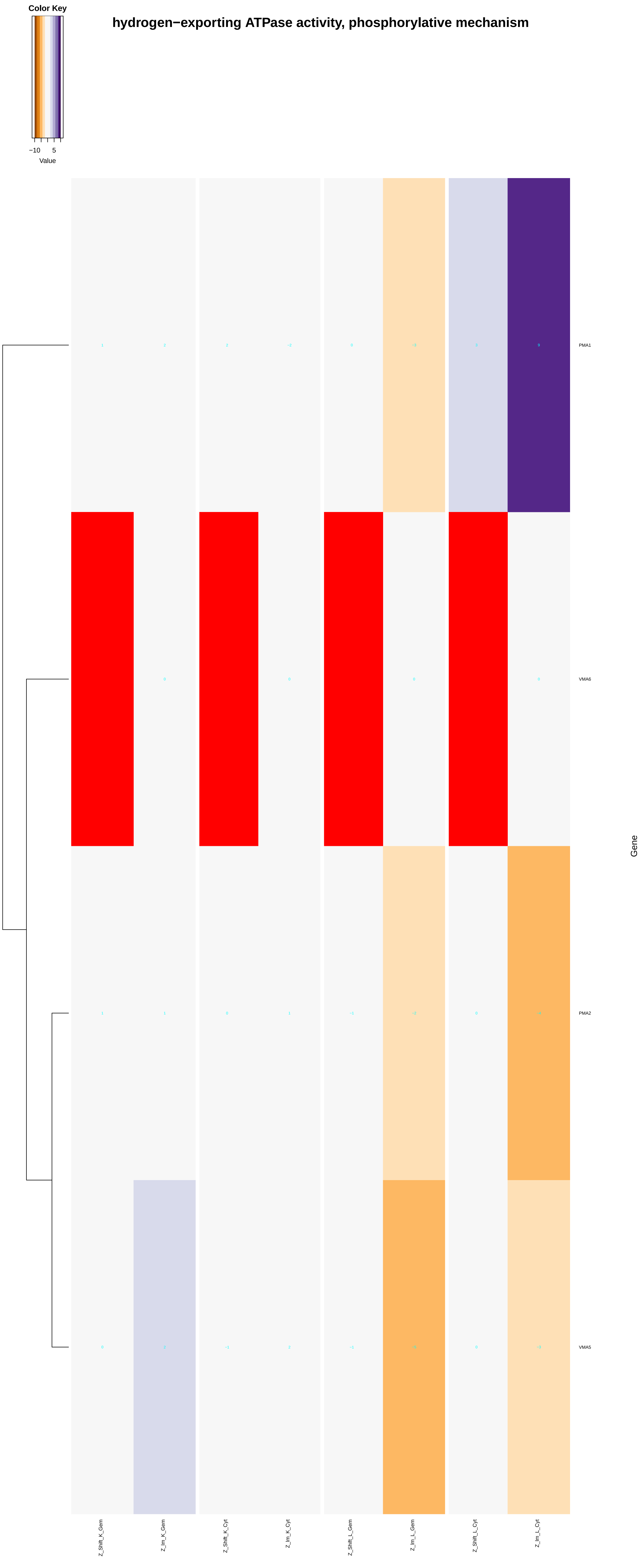

#### hydrogen−exporting ATPase activity

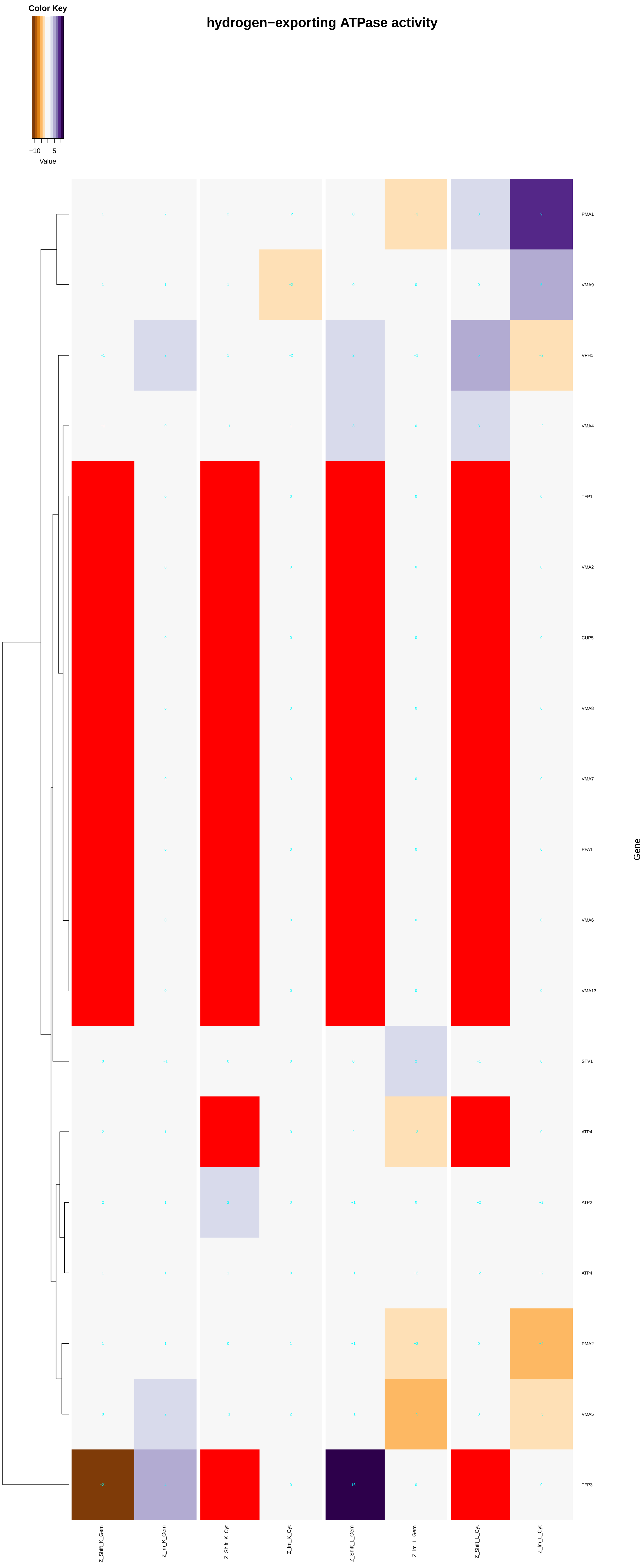

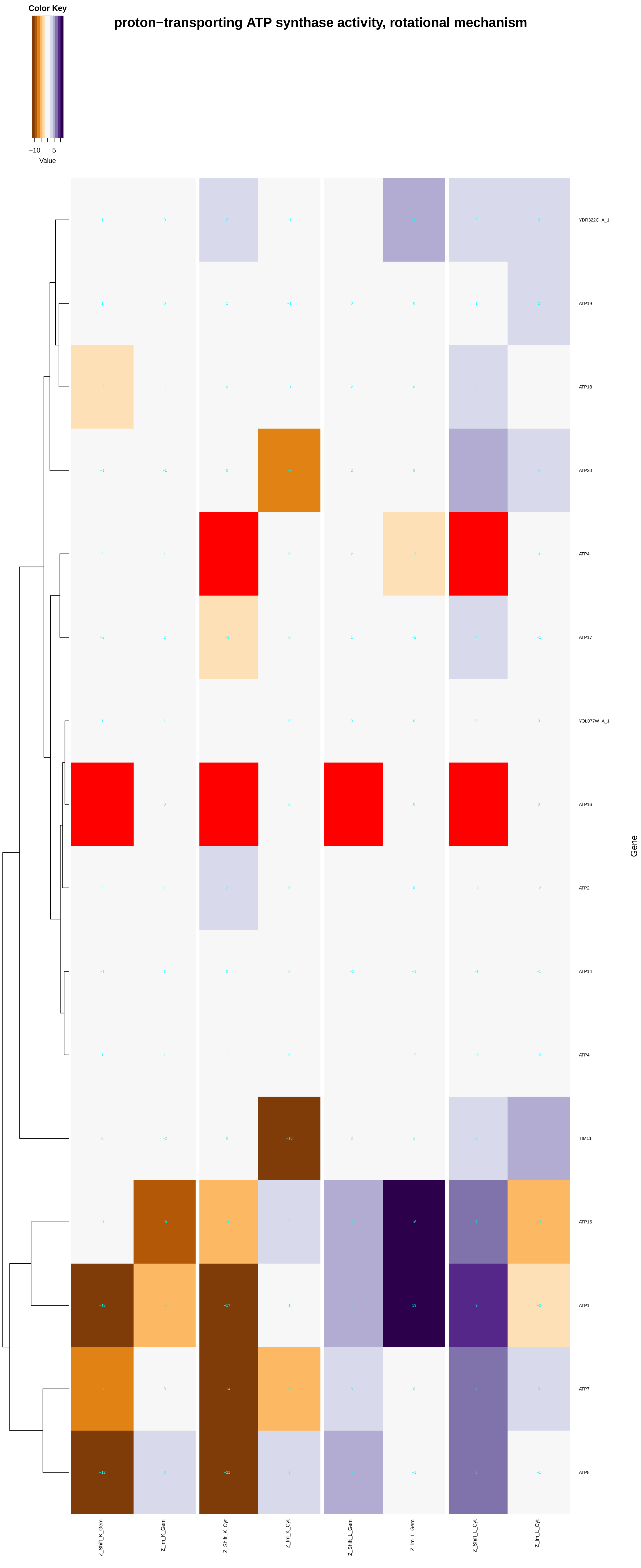
Color Key
−10
5
Value
proton−transporting ATP synthase activity, rotational mechanism
YDR322C−A_1
ATP19
ATP18
ATP20
ATP4
ATP17
YOL077W−A_1
ATP16
ATP2
ATP14
ATP4
TIM11
ATP15
ATP1
ATP7
ATP5
Gene
Z_Shift_K_Gem
Z_Im_K_Gem
Z_Shift_K_Cyt
Z_Im_K_Cyt
Z_Shift_L_Gem
Z_Im_L_Gem
Z_Shift_L_Cyt
Z_Im_L_Cyt

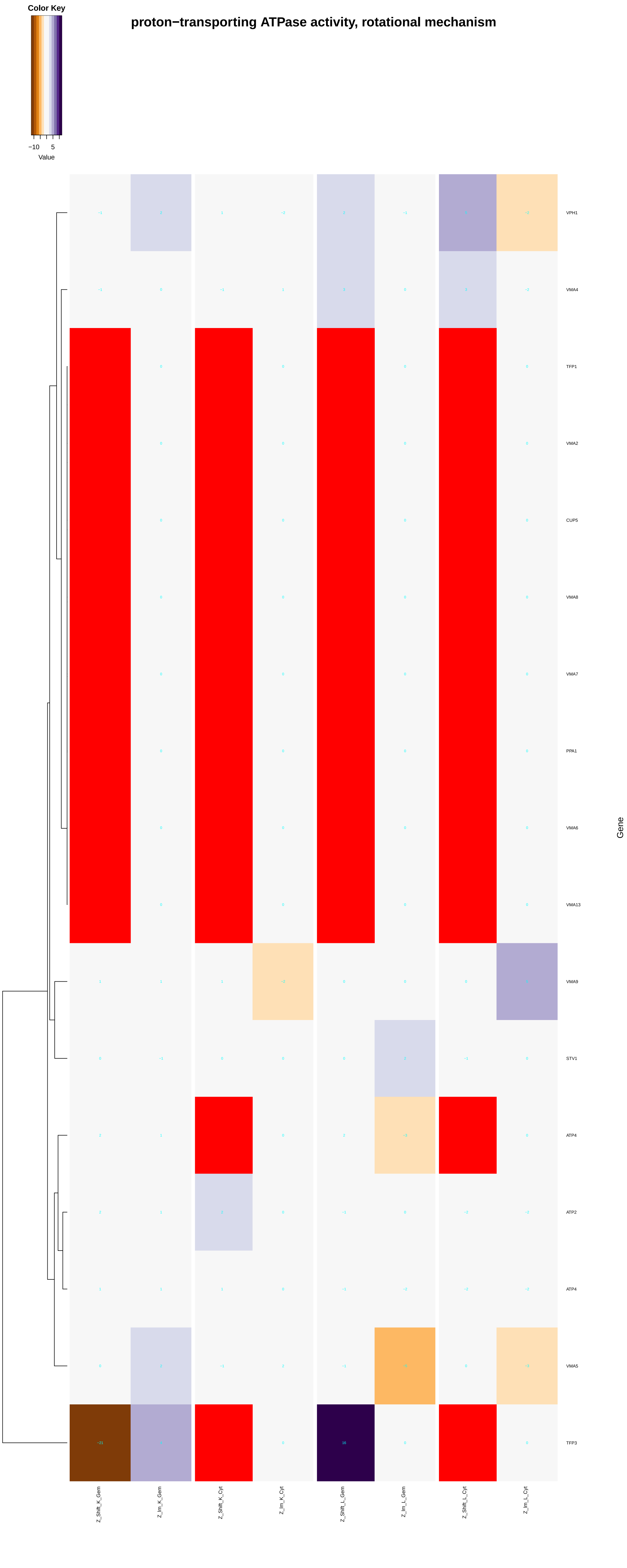
Color Key
−10
5
Value
proton−transporting ATPase activity, rotational mechanism
VPH1
VMA4
TFP1
VMA2
CUP5
VMA8
VMA7
PPA1
VMA6
VMA13
VMA9
STV1
ATP4
ATP2
ATP4
VMA5
TFP3
Gene
Z_Shift_K_Gem
Z_Im_K_Gem
Z_Shift_K_Cyt
Z_Im_K_Cyt
Z_Shift_L_Gem
Z_Im_L_Gem
Z_Shift_L_Cyt
Z_Im_L_Cyt

### ATP−dependent microtubule motor activity

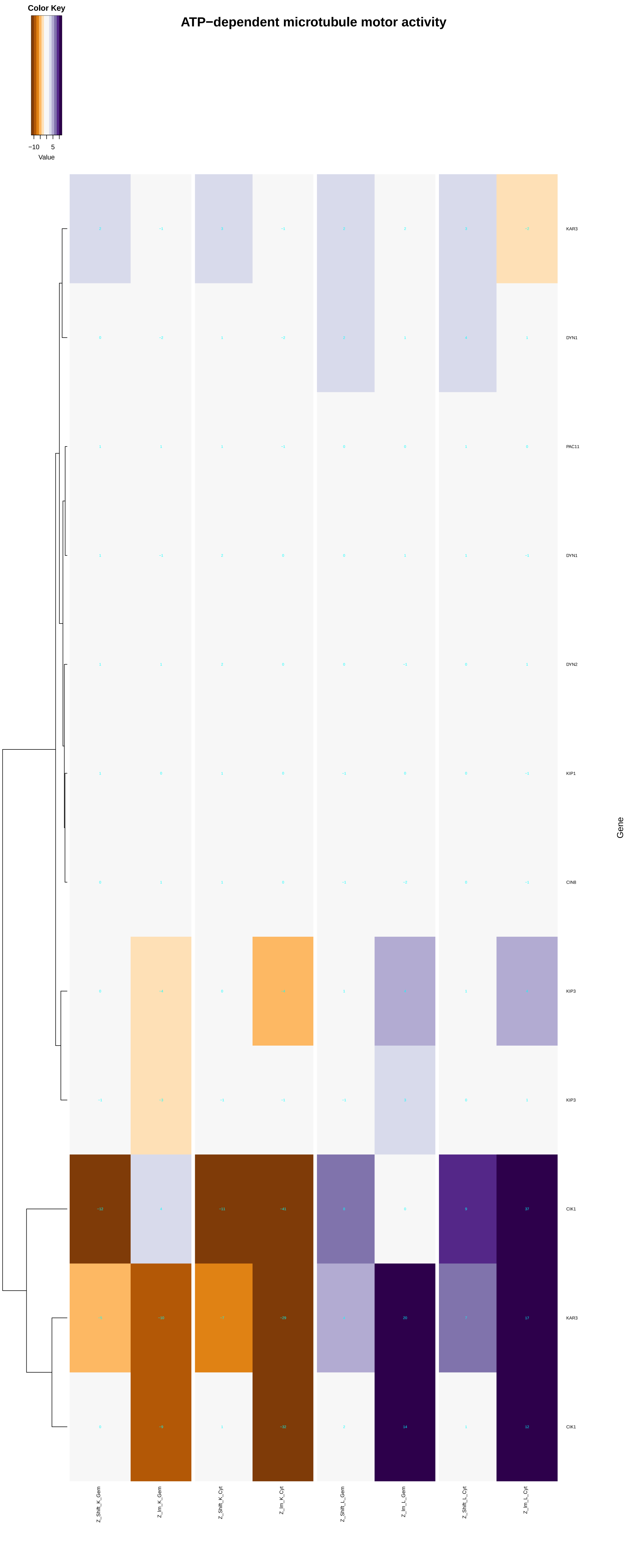

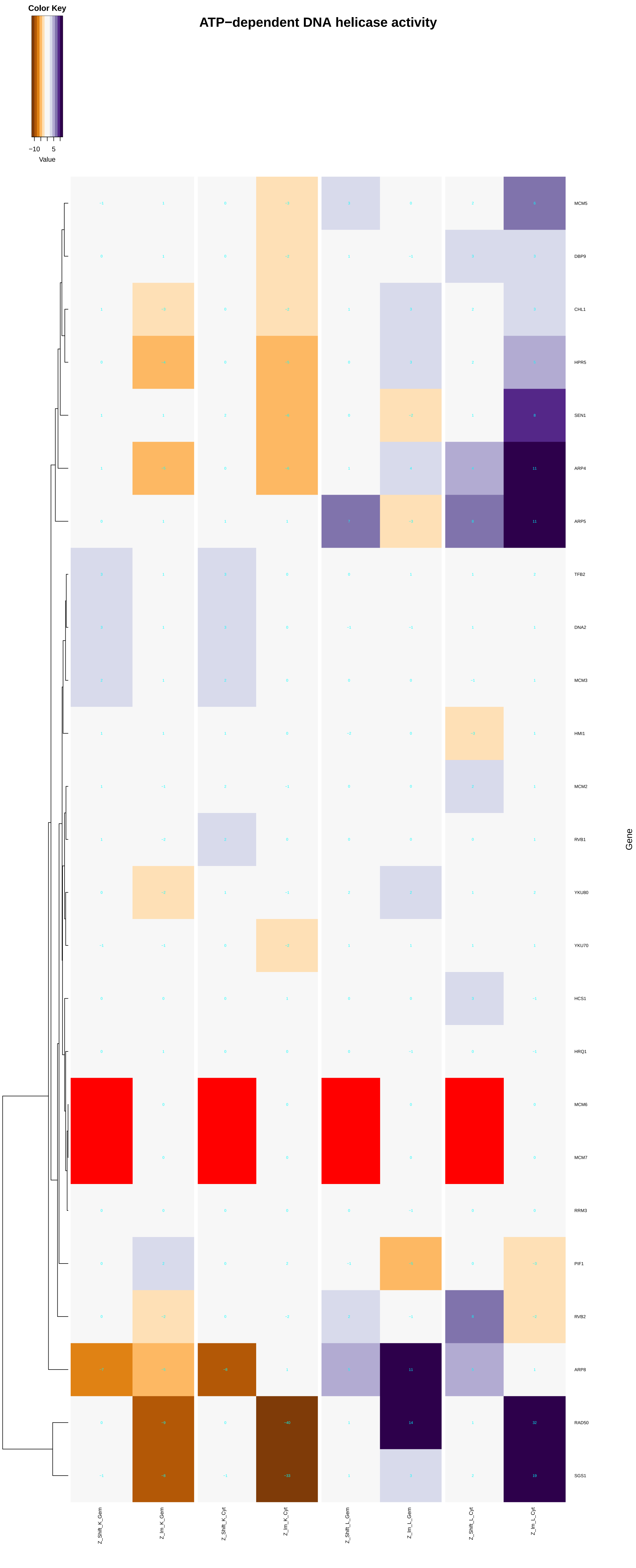

Color Key
−10
5
Value
ATP−dependent DNA helicase activity
MCM5
DBP9
CHL1
HPR5
SEN1
ARP4
ARP5
TFB2
DNA2
MCM3
HMI1
MCM2
RVB1
YKU80
YKU70
HCS1
HRQ1
MCM6
MCM7
RRM3
PIF1
RVB2
ARP8
RAD50
SGS1
Gene
Z_Shift_K_Gem
Z_Im_K_Gem
Z_Shift_K_Cyt
Z_Im_K_Cyt
Z_Shift_L_Gem
Z_Im_L_Gem
Z_Shift_L_Cyt
Z_Im_L_Cyt

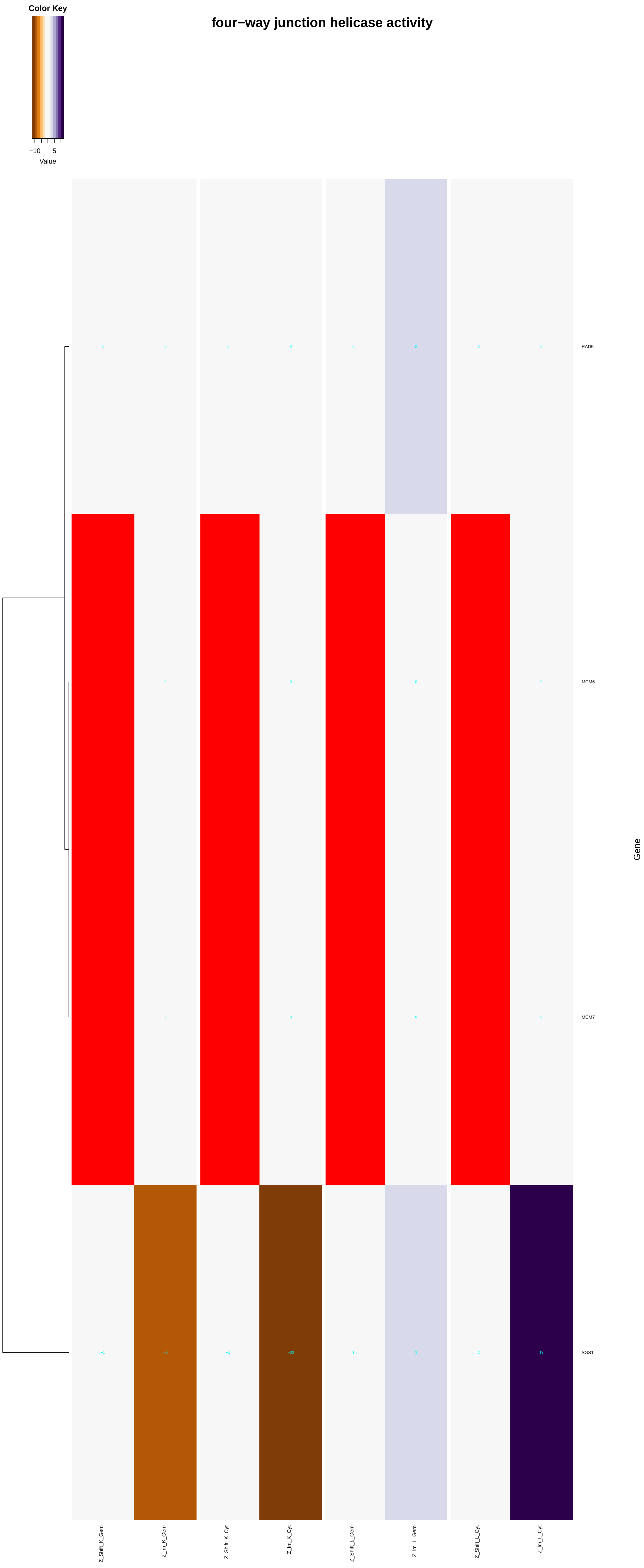
four−way junction helicase activity
Color Key
−10
5
Value
1
0
1
0
0
2
2
0
RAD5
0
0
0
0
MCM6
0
0
0
0
MCM7
−1
−8
−1
−33
1
3
2
19
SGS1
Z_Shift_K_Gem
Z_Im_K_Gem
Z_Shift_K_Cyt
Z_Im_K_Cyt
Z_Shift_L_Gem
Z_Im_L_Gem
Z_Shift_L_Cyt
Z_Im_L_Cyt
Gene

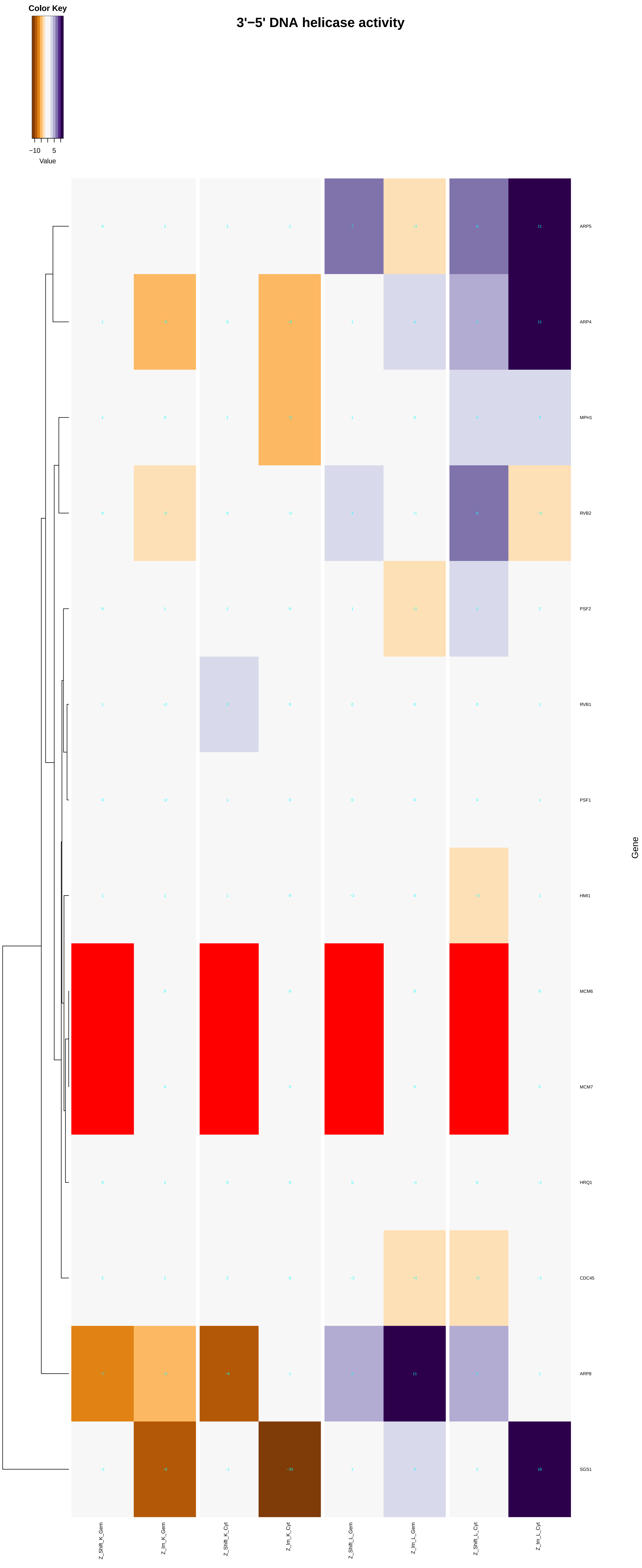

3'−5' DNA helicase activity
Color Key
−10
5
Value
ARP5
ARP4
MPH1
RVB2
PSF2
RVB1
PSF1
HMI1
MCM6
MCM7
HRQ1
CDC45
ARP8
SGS1
Gene
Z_Shift_K_Gem
Z_lm_K_Gem
Z_Shift_K_Cyt
Z_lm_K_Cyt
Z_Shift_L_Gem
Z_lm_L_Gem
Z_Shift_L_Cyt
Z_lm_L_Cyt

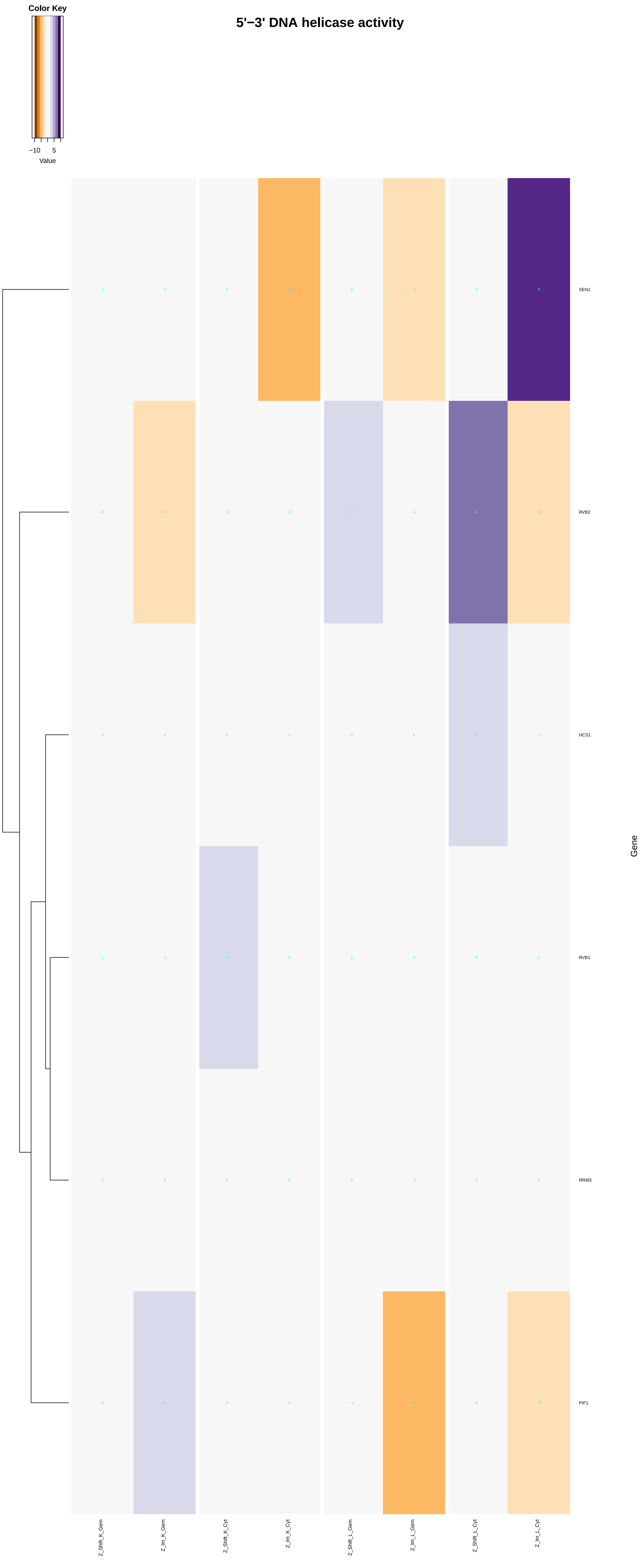

Color Key
−10
5
Value
5'−3' DNA helicase activity
SEN1
RVB2
HCS1
RVB1
RRM3
PIF1
Gene
Z_Shift_K_Gem
Z_Im_K_Gem
Z_Shift_K_Cyt
Z_Im_K_Cyt
Z_Shift_L_Gem
Z_Im_L_Gem
Z_Shift_L_Cyt
Z_Im_L_Cyt

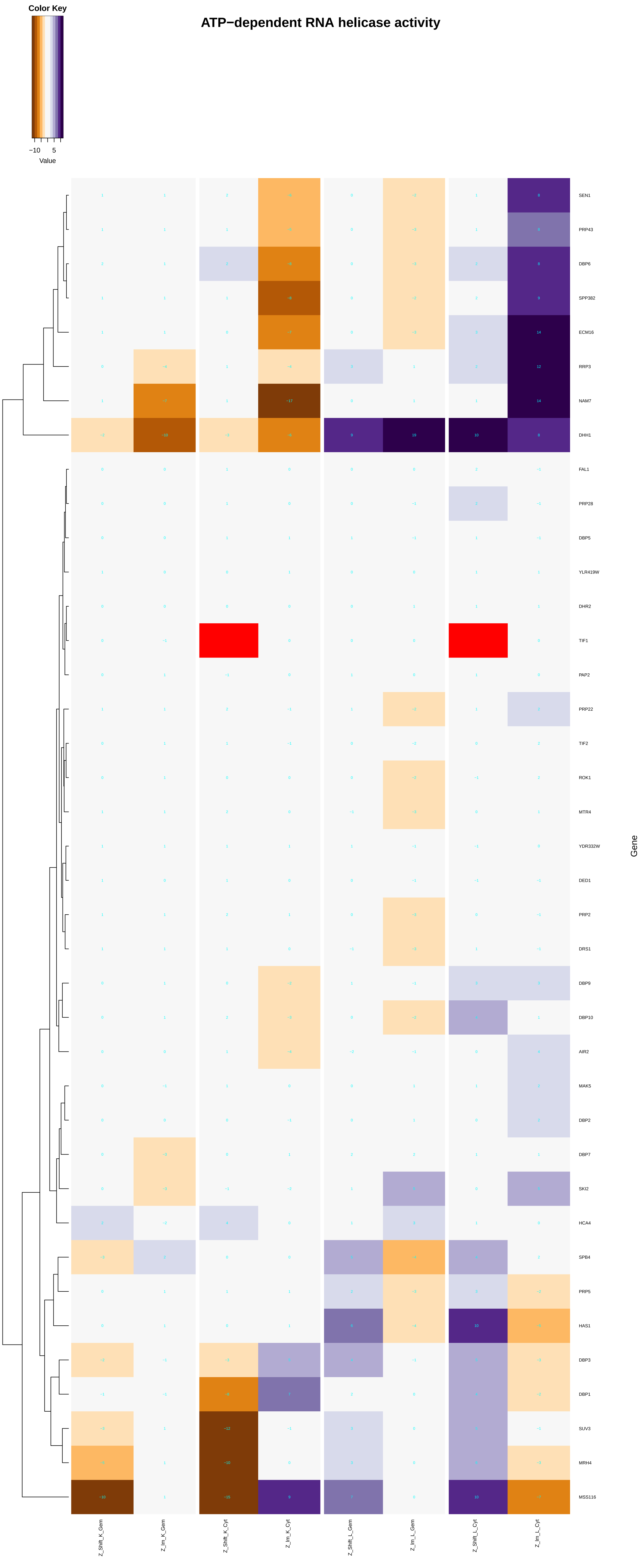
ATP−dependent RNA helicase activity
Color Key
−10
5
Value
SEN1
PRP43
DBP6
SPP382
ECM16
RRP3
NAM7
DHH1
FAL1
PRP28
DBP5
YLR419W
DHR2
TIF1
PAP2
PRP22
TIF2
ROK1
MTR4
YDR332W
DED1
PRP2
DRS1
DBP9
DBP10
AIR2
MAK5
DBP2
DBP7
SKI2
HCA4
SPB4
PRP5
HAS1
DBP3
DBP1
SUV3
MRH4
MSS116
Gene
Z_Shift_K_Gem
Z_Im_K_Gem
Z_Shift_K_Cyt
Z_Im_K_Cyt
Z_Shift_L_Gem
Z_Im_L_Gem
Z_Shift_L_Cyt
Z_Im_L_Cyt

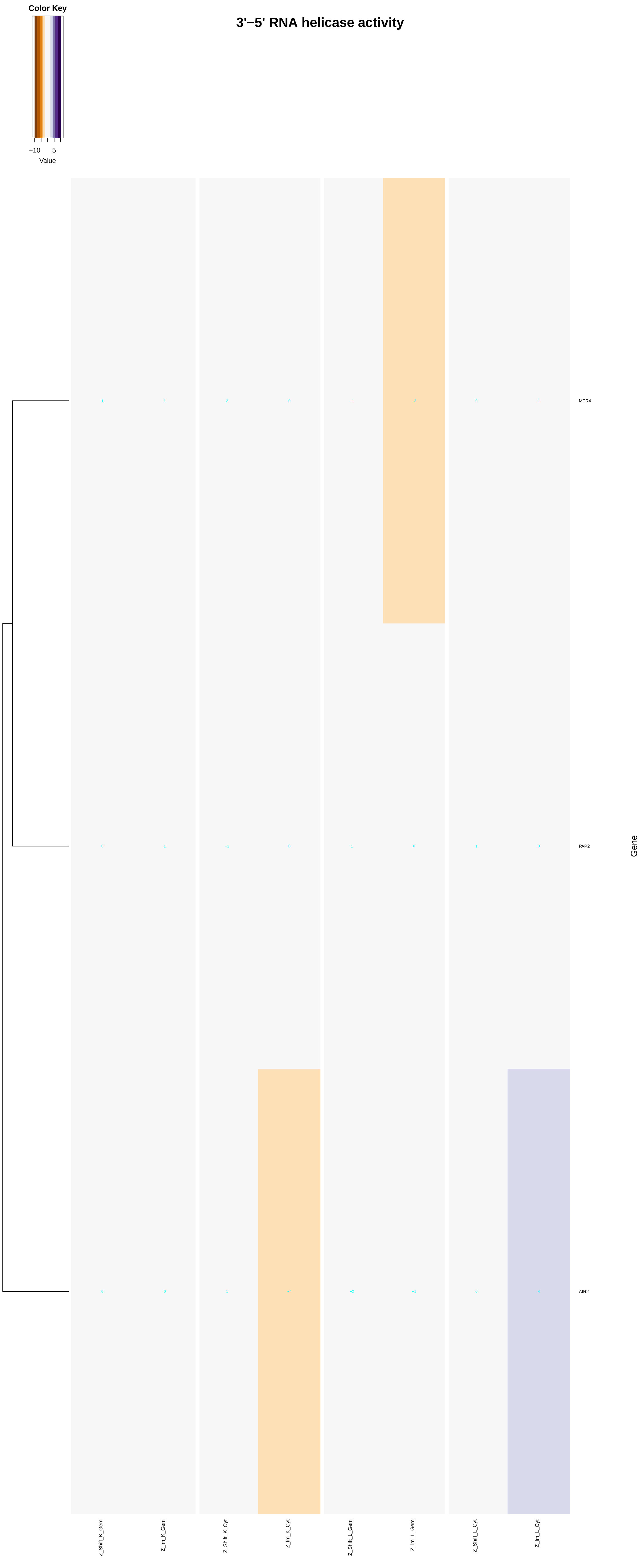
3'−5' RNA helicase activity
Color Key
−10
5
Value
MTR4
PAP2
AIR2
Gene
Z_Shift_K_Gem
Z_Im_K_Gem
Z_Shift_K_Cyt
Z_Im_K_Cyt
Z_Shift_L_Gem
Z_Im_L_Gem
Z_Shift_L_Cyt
Z_Im_L_Cyt

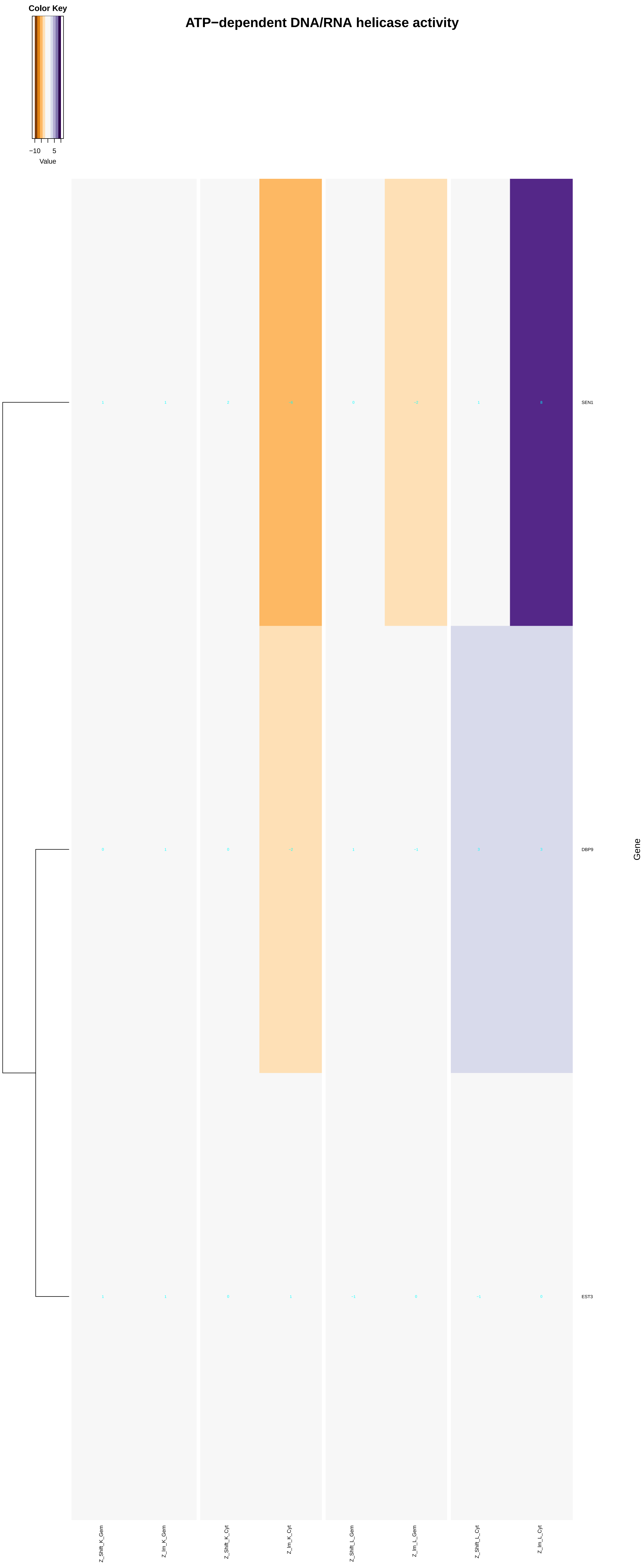
Color Key
−10 5
Value
ATP−dependent DNA/RNA helicase activity
SEN1
DBP9
EST3
Gene
Z_Shift_K_Gem
Z_Im_K_Gem
Z_Shift_K_Cyt
Z_Im_K_Cyt
Z_Shift_L_Gem
Z_Im_L_Gem
Z_Shift_L_Cyt
Z_Im_L_Cyt

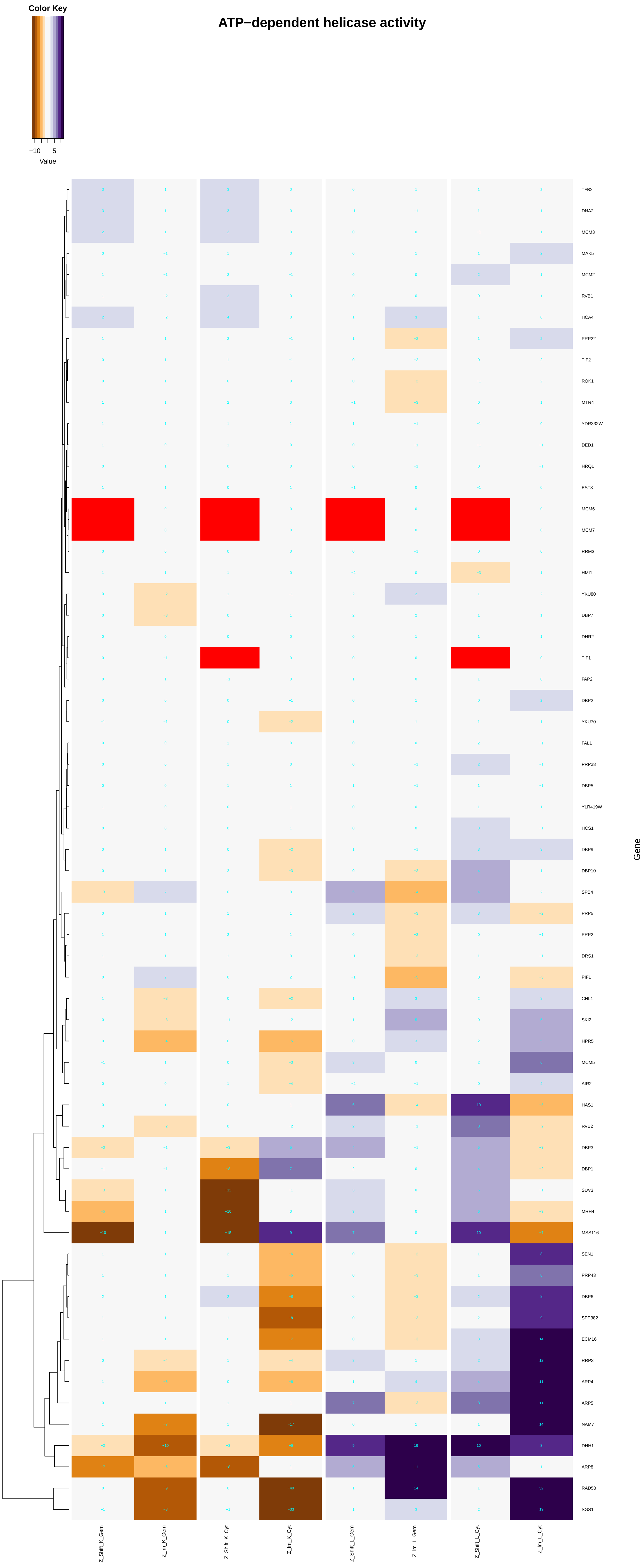

ATP−dependent helicase activity
Color Key
−10
5
Value
TFB2
DNA2
MCM3
MAK5
MCM2
RVB1
HCA4
PRP22
TIF2
ROK1
MTR4
YDR332W
DED1
HRQ1
EST3
MCM6
MCM7
RRM3
HMI1
YKU80
DBP7
DHR2
TIF1
PAP2
DBP2
YKU70
FAL1
PRP28
DBP5
YLR419W
HCS1
DBP9
DBP10
SPB4
PRP5
PRP2
DRS1
PIF1
CHL1
SKI2
HPR5
MCM5
AIR2
HAS1
RVB2
DBP3
DBP1
SUV3
MRH4
MSS116
SEN1
PRP43
DBP6
SPP382
ECM16
RRP3
ARP4
ARP5
NAM7
DHH1
ARP8
RAD50
SGS1
Gene
Z_Shift_K_Gem
Z_Im_K_Gem
Z_Shift_K_Cyt
Z_Im_K_Cyt
Z_Shift_L_Gem
Z_Im_L_Gem
Z_Shift_L_Cyt
Z_Im_L_Cyt

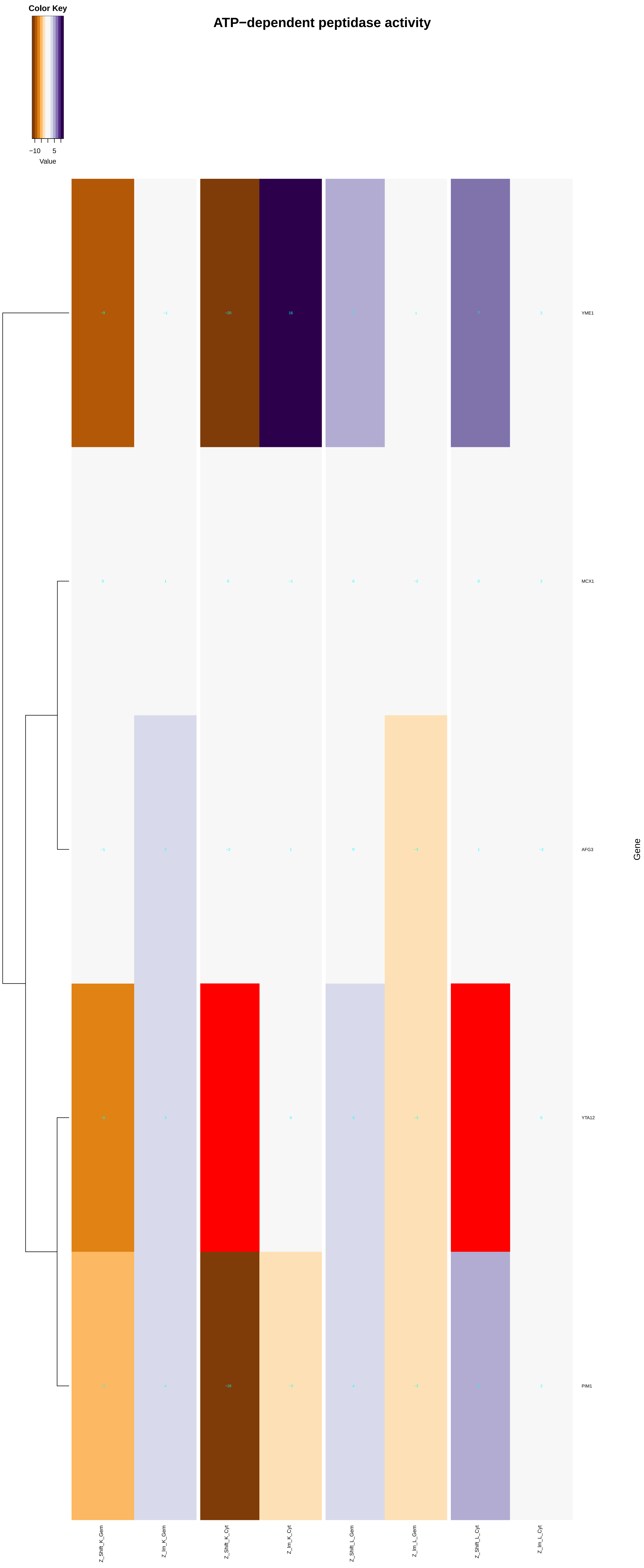

ATP−dependent peptidase activity
Color Key
−10
5
Value
YME1
MCX1
AFG3
YTA12
PIM1
Gene
Z_Shift_K_Gem
Z_Im_K_Gem
Z_Shift_K_Cyt
Z_Im_K_Cyt
Z_Shift_L_Gem
Z_Im_L_Gem
Z_Shift_L_Cyt
Z_Im_L_Cyt

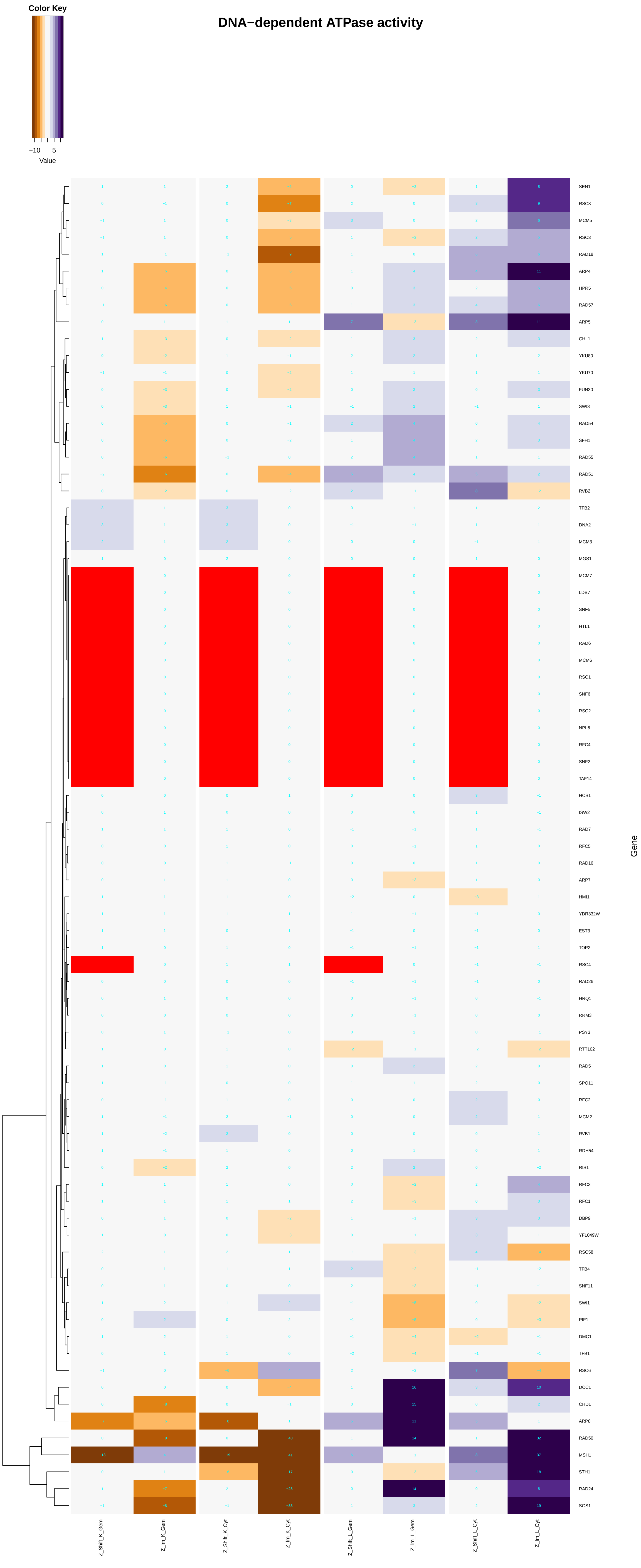

Color Key
−10 5
Value
DNA−dependent ATPase activity
SEN1
RSC8
MCM5
RSC3
RAD18
ARP4
HPR5
RAD57
ARP5
CHL1
YKU80
YKU70
FUN30
SWI3
RAD54
SFH1
RAD55
RAD51
RVB2
TFB2
DNA2
MCM3
MGS1
MCM7
LDB7
SNF5
HTL1
RAD6
MCM6
RSC1
SNF6
RSC2
NPL6
RFC4
SNF2
TAF14
HCS1
ISW2
RAD7
RFC5
RAD16
ARP7
HMI1
YDR332W
EST3
TOP2
RSC4
RAD26
HRQ1
RRM3
PSY3
RTT102
RAD5
SPO11
RFC2
MCM2
RVB1
RDH54
RIS1
RFC3
RFC1
DBP9
YFL049W
RSC58
TFB4
SNF11
SWI1
PIF1
DMC1
TFB1
RSC6
DCC1
CHD1
ARP8
RAD50
MSH1
STH1
RAD24
SGS1
Gene
Z_Shift_K_Gem
Z_Im_K_Gem
Z_Shift_K_Cyt
Z_Im_K_Cyt
Z_Shift_L_Gem
Z_Im_L_Gem
Z_Shift_L_Cyt
Z_Im_L_Cyt

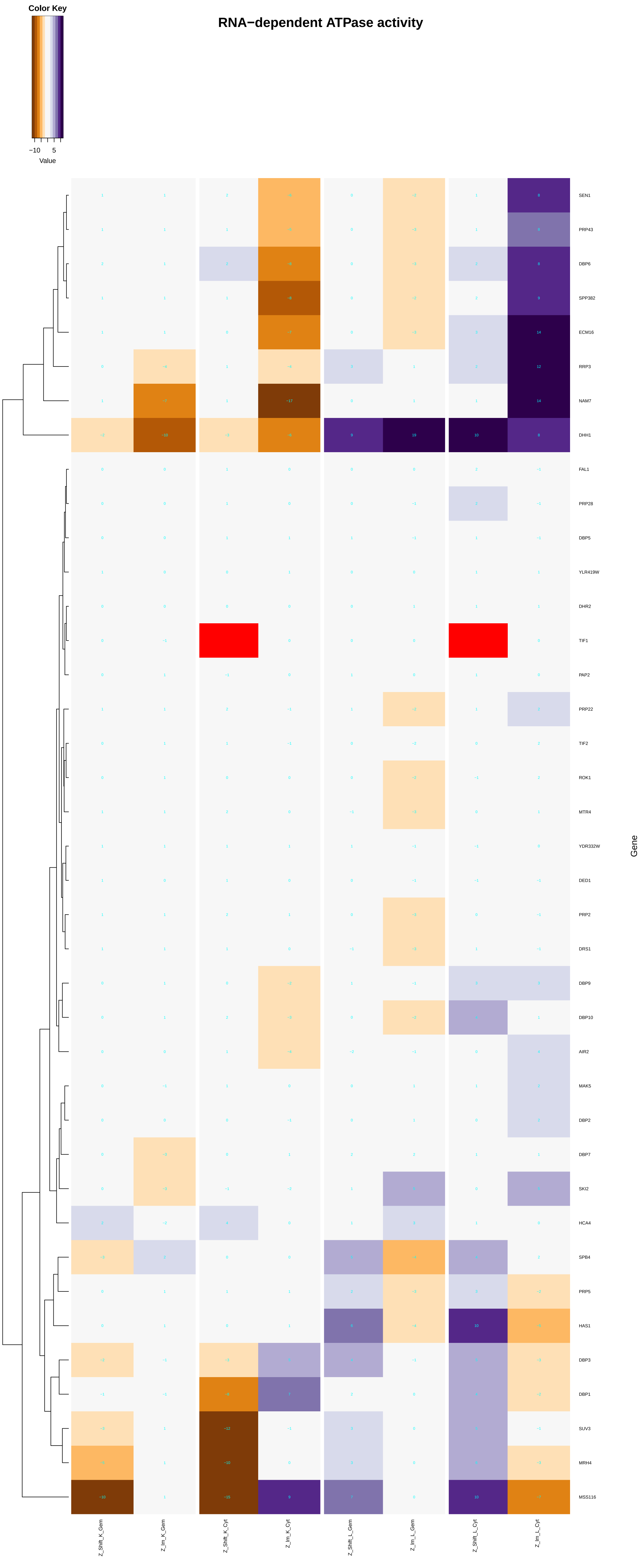
RNA−dependent ATPase activity
Color Key
−10
5
Value
SEN1
PRP43
DBP6
SPP382
ECM16
RRP3
NAM7
DHH1
FAL1
PRP28
DBP5
YLR419W
DHR2
TIF1
PAP2
PRP22
TIF2
ROK1
MTR4
YDR332W
DED1
PRP2
DRS1
DBP9
DBP10
AIR2
MAK5
DBP2
DBP7
SKI2
HCA4
SPB4
PRP5
HAS1
DBP3
DBP1
SUV3
MRH4
MSS116
Gene
Z_Shift_K_Gem
Z_Im_K_Gem
Z_Shift_K_Cyt
Z_Im_K_Cyt
Z_Shift_L_Gem
Z_Im_L_Gem
Z_Shift_L_Cyt
Z_Im_L_Cyt

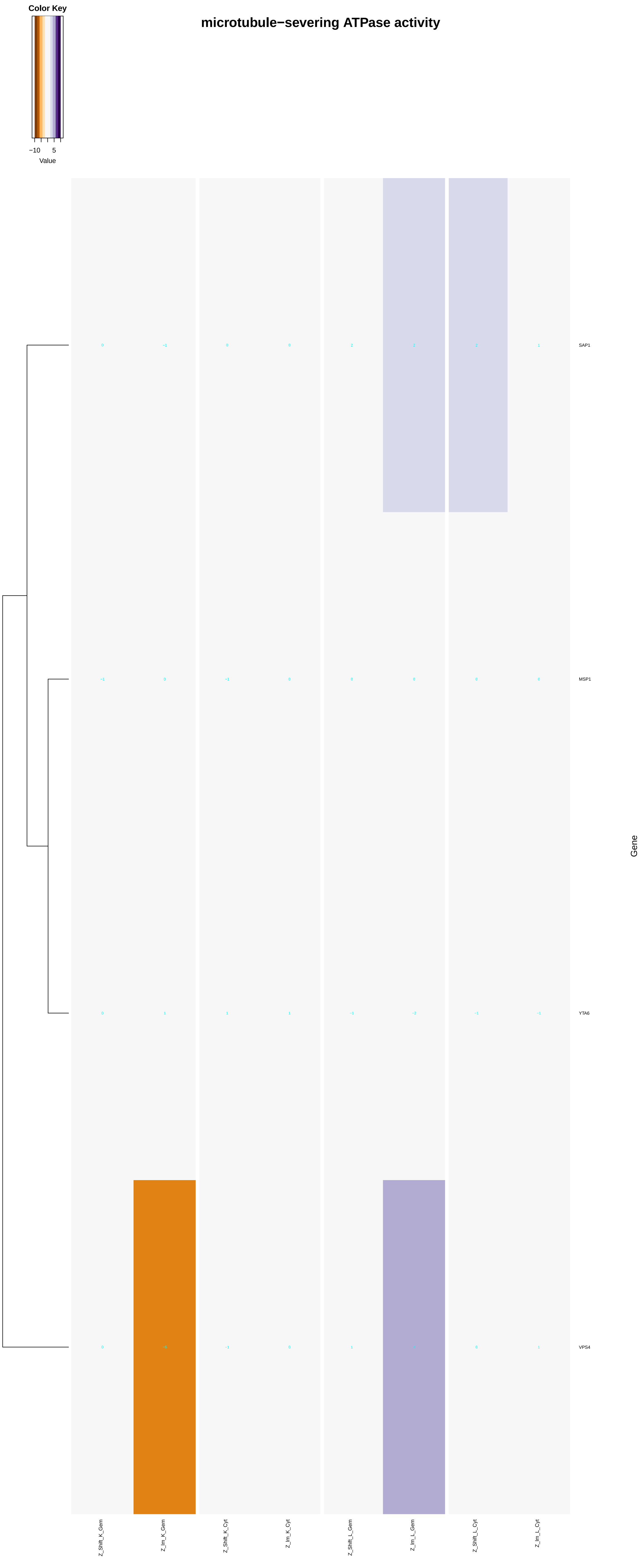

Color Key
−10
5
Value
microtubule−severing ATPase activity
SAP1
MSP1
YTA6
VPS4
Gene
Z_Shift_K_Gem
Z_Im_K_Gem
Z_Shift_K_Cyt
Z_Im_K_Cyt
Z_Shift_L_Gem
Z_Im_L_Gem
Z_Shift_L_Cyt
Z_Im_L_Cyt

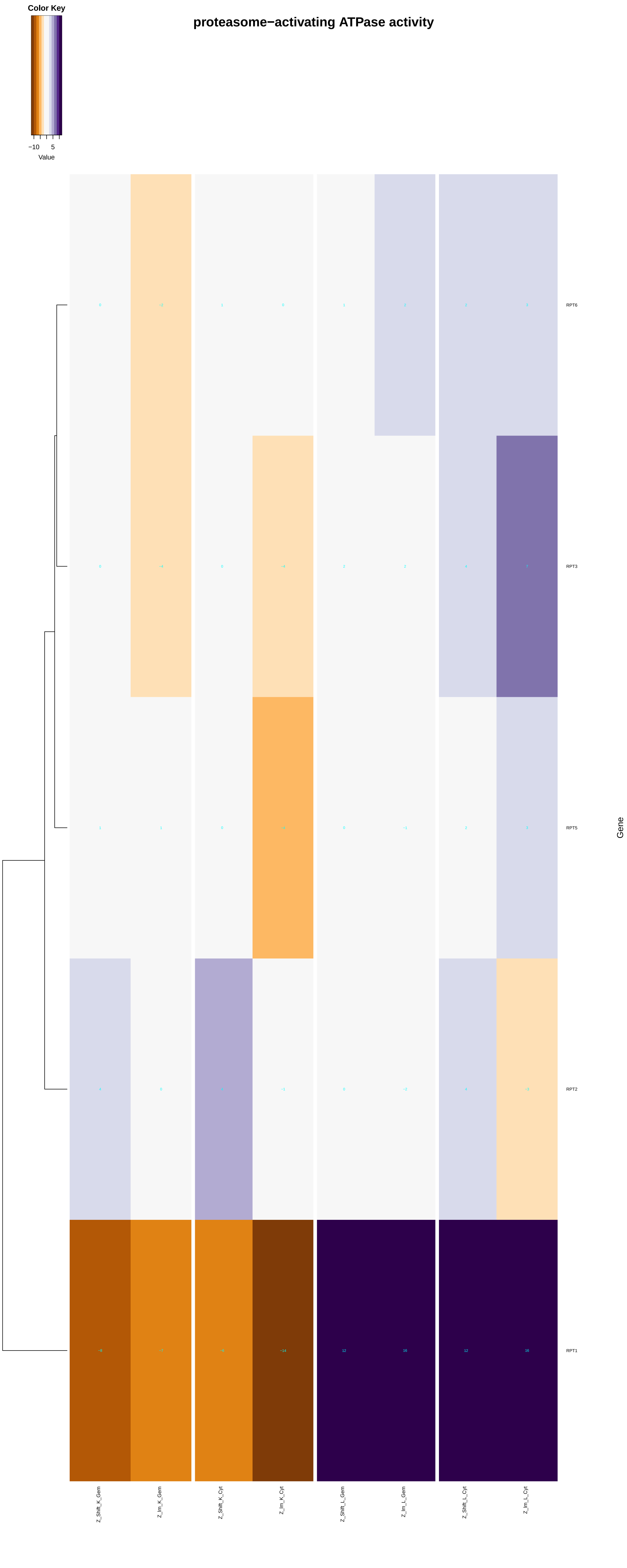

proteasome−activating ATPase activity
Color Key
−10
5
Value
RPT6
RPT3
RPT5
RPT2
RPT1
Gene
Z_Shift_K_Gem
Z_Im_K_Gem
Z_Shift_K_Cyt
Z_Im_K_Cyt
Z_Shift_L_Gem
Z_Im_L_Gem
Z_Shift_L_Cyt
Z_Im_L_Cyt

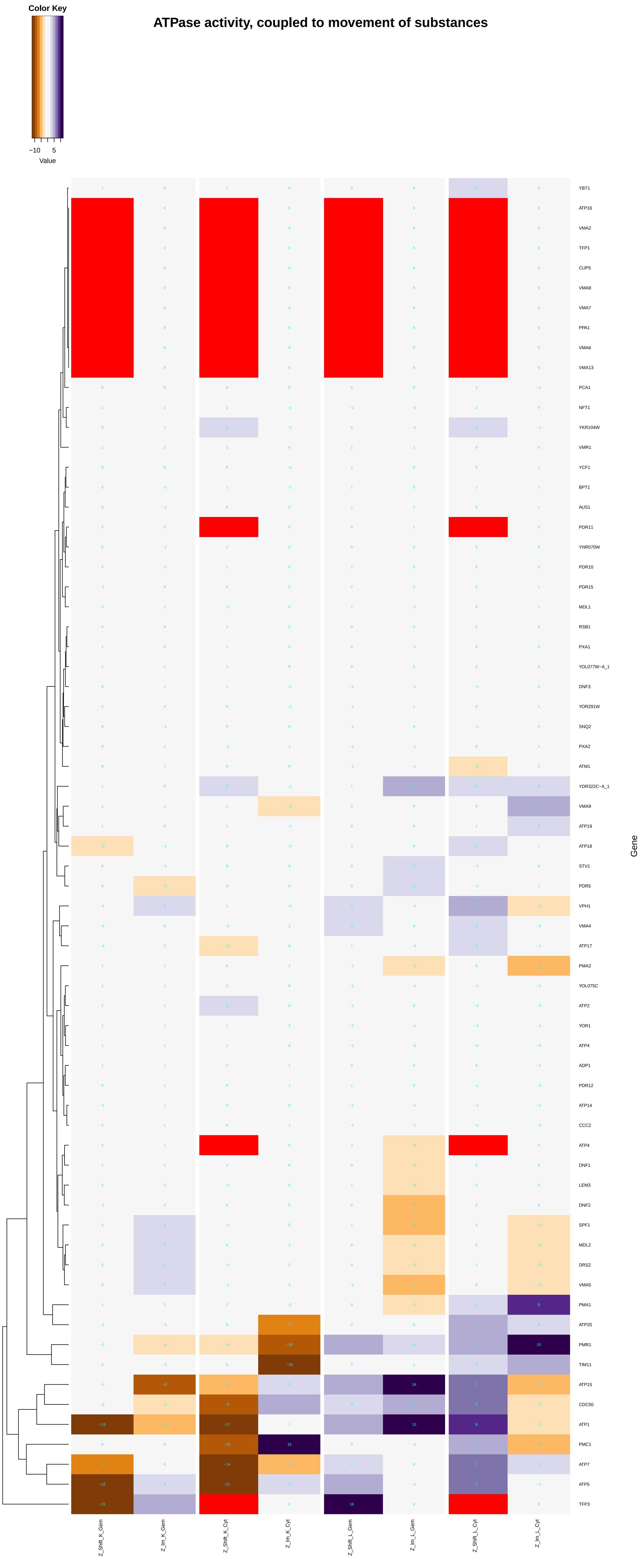

ATPase activity, coupled to movement of substances
Color Key
−10
5
Value
YBT1
ATP16
VMA2
TFP1
CUP5
VMA8
VMA7
PPA1
VMA6
VMA13
PCA1
NFT1
YKR104W
VMR1
YCF1
BPT1
AUS1
PDR11
YNR070W
PDR10
PDR15
MDL1
RSB1
PXA1
YOL077W−A_1
DNF3
YOR291W
SNQ2
PXA2
ATM1
YDR322C−A_1
VMA9
ATP19
ATP18
STV1
PDR5
VPH1
VMA4
ATP17
PMA2
YOL075C
ATP2
YOR1
ATP4
ADP1
PDR12
ATP14
CCC2
ATP4
DNF1
LEM3
DNF2
SPF1
MDL2
DRS2
VMA5
PMA1
ATP20
PMR1
TIM11
ATP15
CDC50
ATP1
PMC1
ATP7
ATP5
TFP3
Gene
Z_Shift_K_Gem
Z_Im_K_Gem
Z_Shift_K_Cyt
Z_Im_K_Cyt
Z_Shift_L_Gem
Z_Im_L_Gem
Z_Shift_L_Cyt
Z_Im_L_Cyt

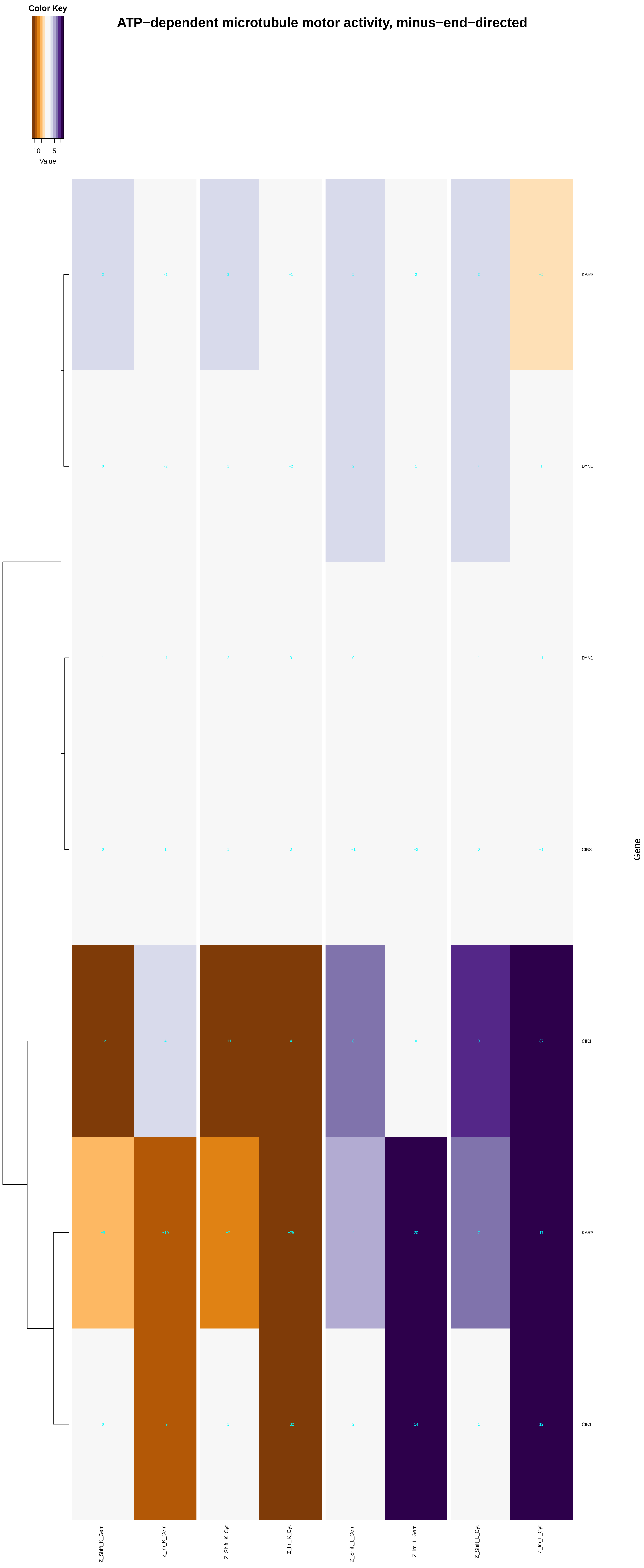
Color Key
−10
5
Value
ATP−dependent microtubule motor activity, minus−end−directed
KAR3
DYN1
DYN1
CIN8
CIK1
KAR3
CIK1
Gene
Z_Shift_K_Gem
Z_Im_K_Gem
Z_Shift_K_Cyt
Z_Im_K_Cyt
Z_Shift_L_Gem
Z_Im_L_Gem
Z_Shift_L_Cyt
Z_Im_L_Cyt

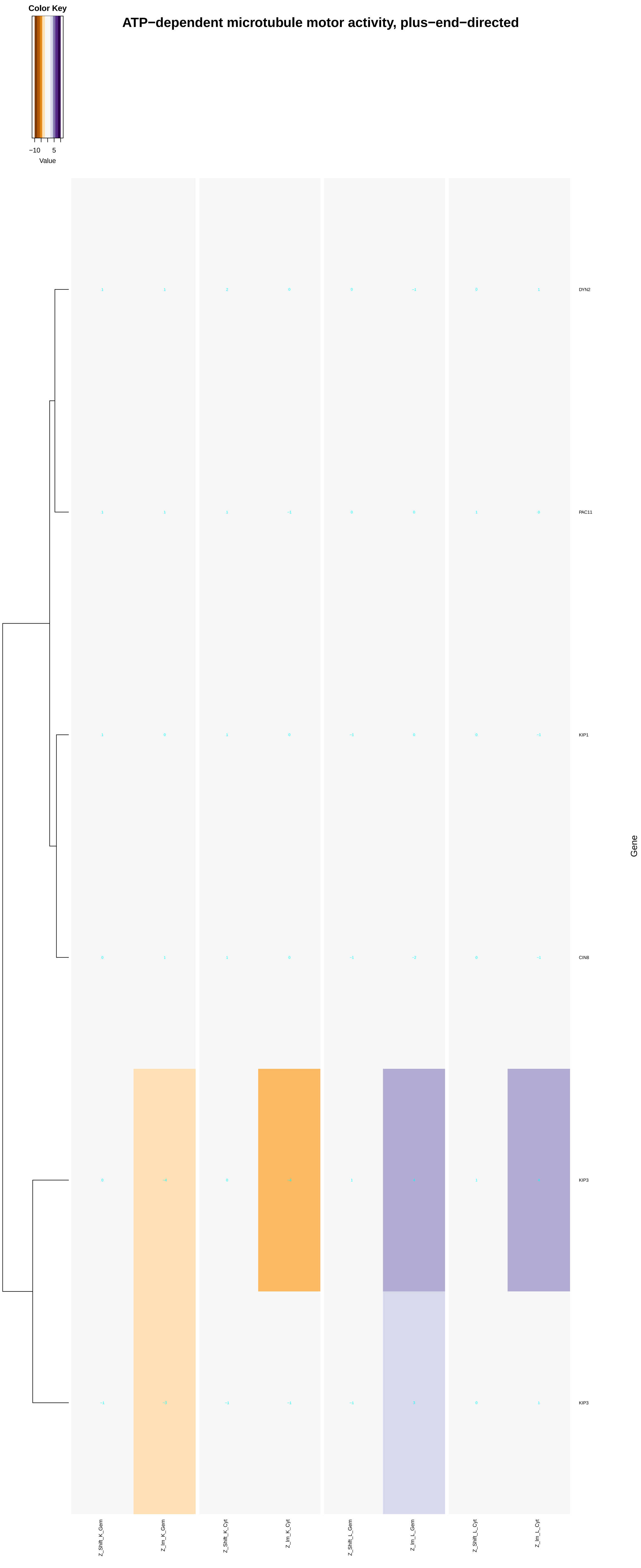
ATP−dependent microtubule motor activity, plus−end−directed
Color Key
−10
5
Value
DYN2
PAC11
KIP1
CIN8
KIP3
KIP3
Gene
Z_Shift_K_Gem
Z_Im_K_Gem
Z_Shift_K_Cyt
Z_Im_K_Cyt
Z_Shift_L_Gem
Z_Im_L_Gem
Z_Shift_L_Cyt
Z_Im_L_Cyt

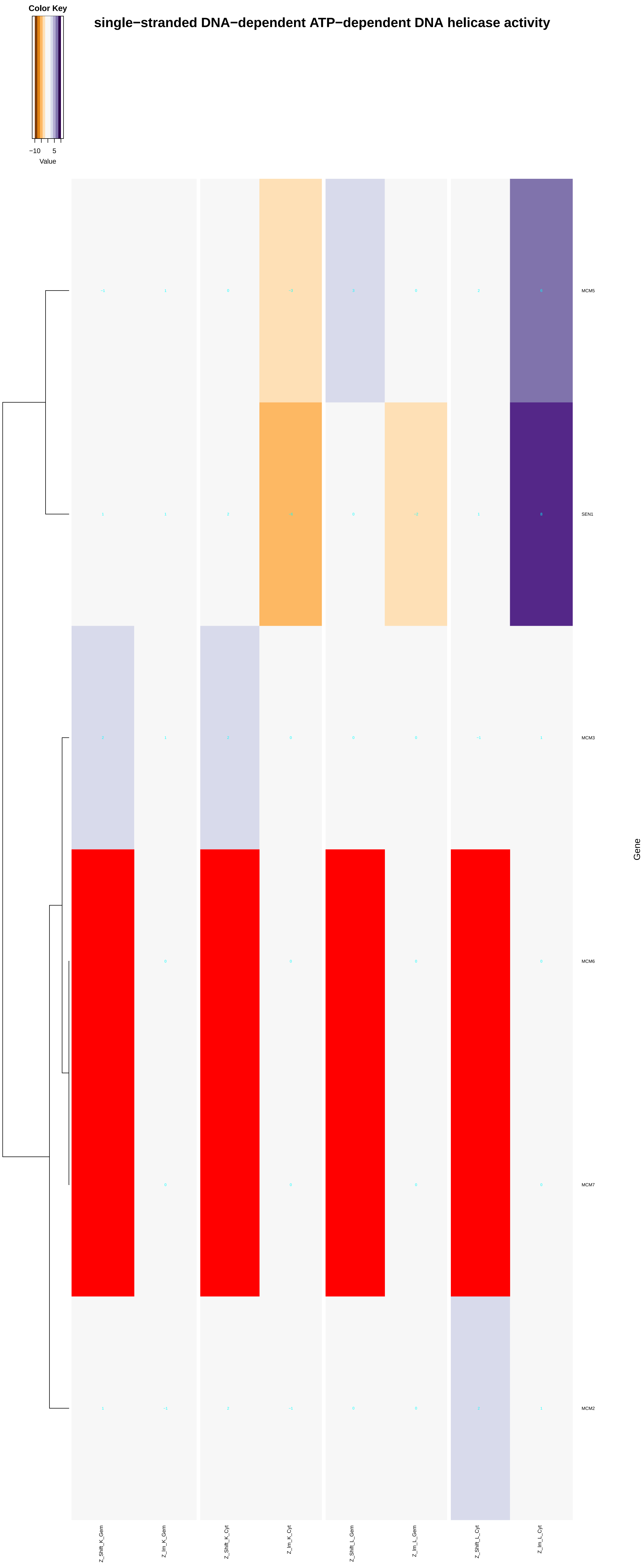

single−stranded DNA−dependent ATP−dependent DNA helicase activity
Color Key
−10
5
Value
MCM5
SEN1
MCM3
MCM6
MCM7
MCM2
Gene
Z_Shift_K_Gem
Z_Im_K_Gem
Z_Shift_K_Cyt
Z_Im_K_Cyt
Z_Shift_L_Gem
Z_Im_L_Gem
Z_Shift_L_Cyt
Z_Im_L_Cyt

### ATP−dependent 3'−5' DNA helicase activity

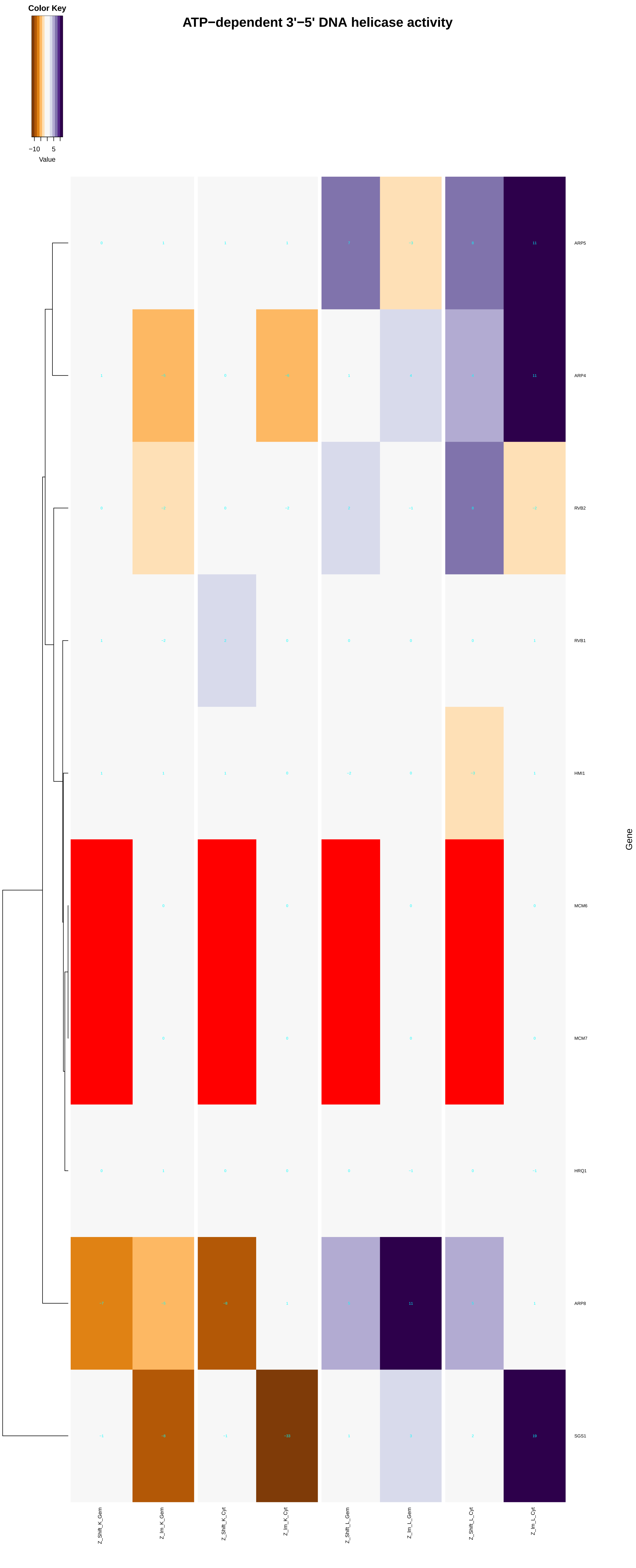

#### ATP−dependent 5'−3' DNA helicase activity

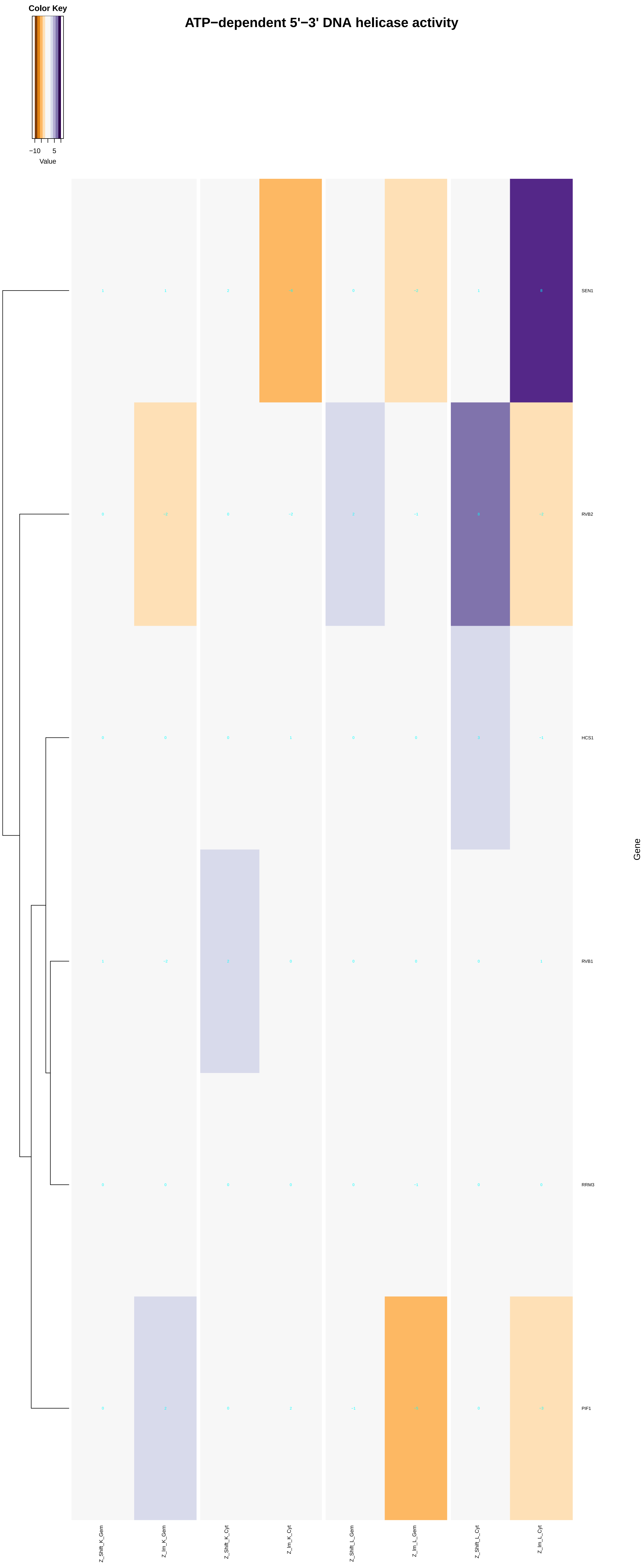

### ATP−dependent 3'−5' RNA helicase activity

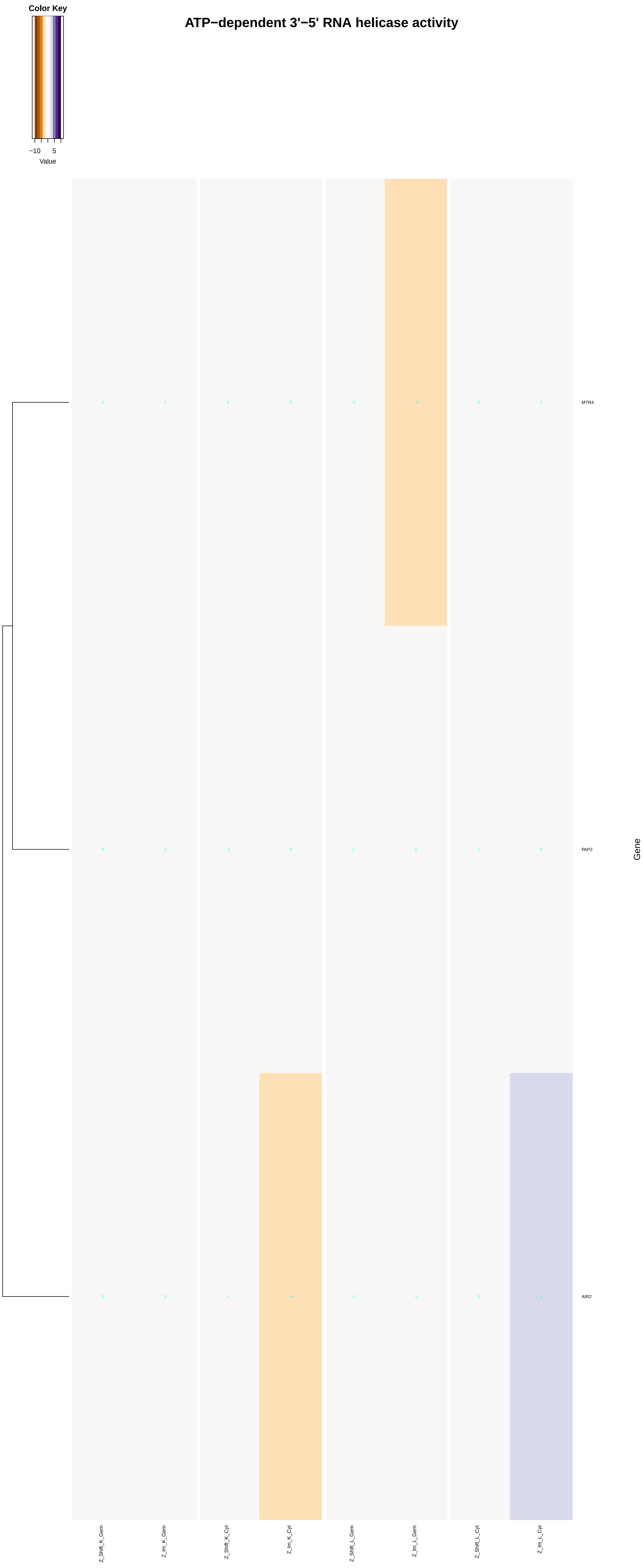

DNA translocase activity

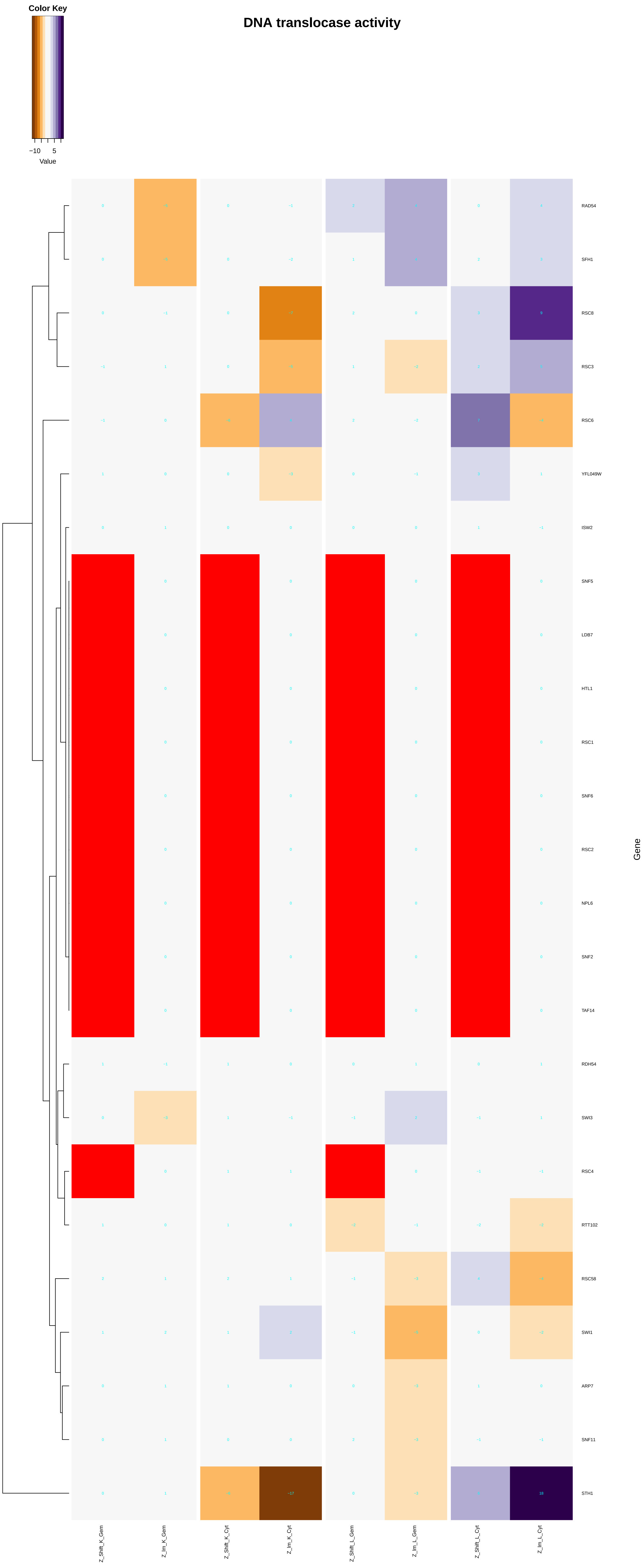

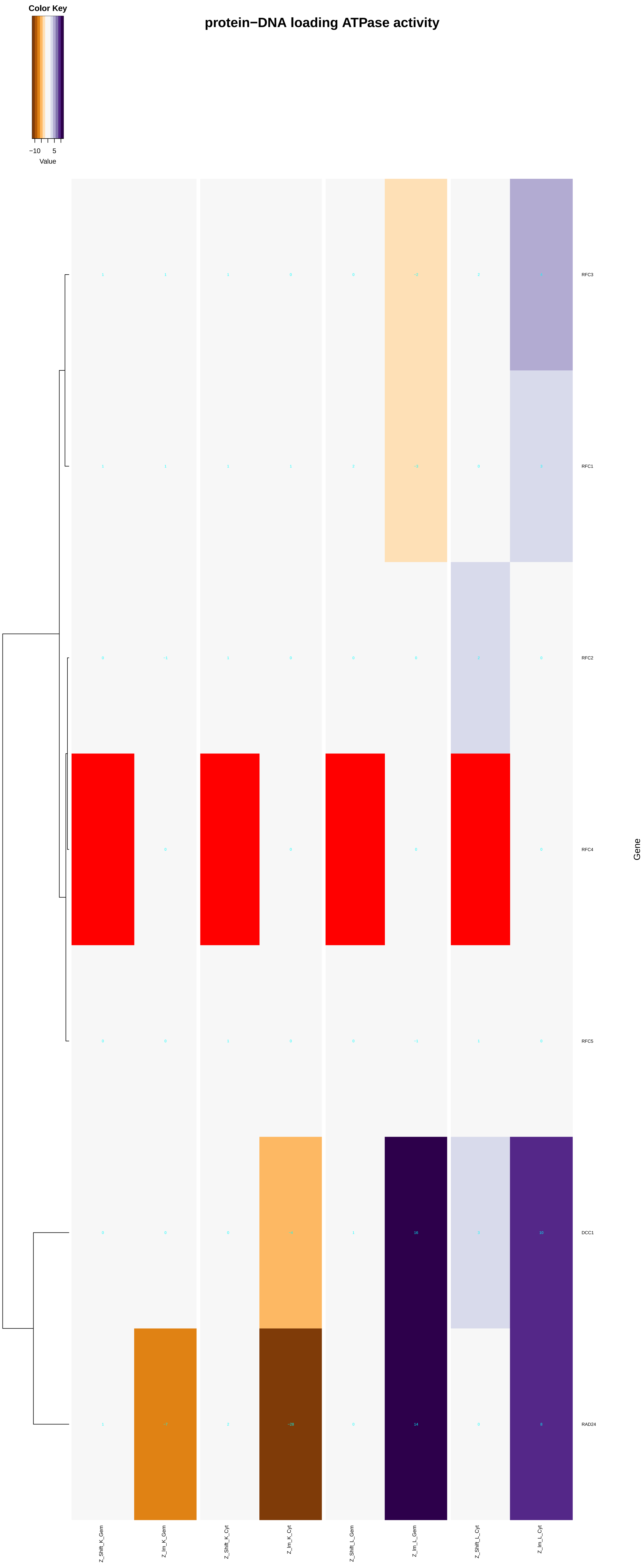

Color Key
−10
5
Value
protein−DNA loading ATPase activity
RFC3
RFC1
RFC2
RFC4
RFC5
DCC1
RAD24
Gene
Z_Shift_K_Gem
Z_Im_K_Gem
Z_Shift_K_Cyt
Z_Im_K_Cyt
Z_Shift_L_Gem
Z_Im_L_Gem
Z_Shift_L_Cyt
Z_Im_L_Cyt

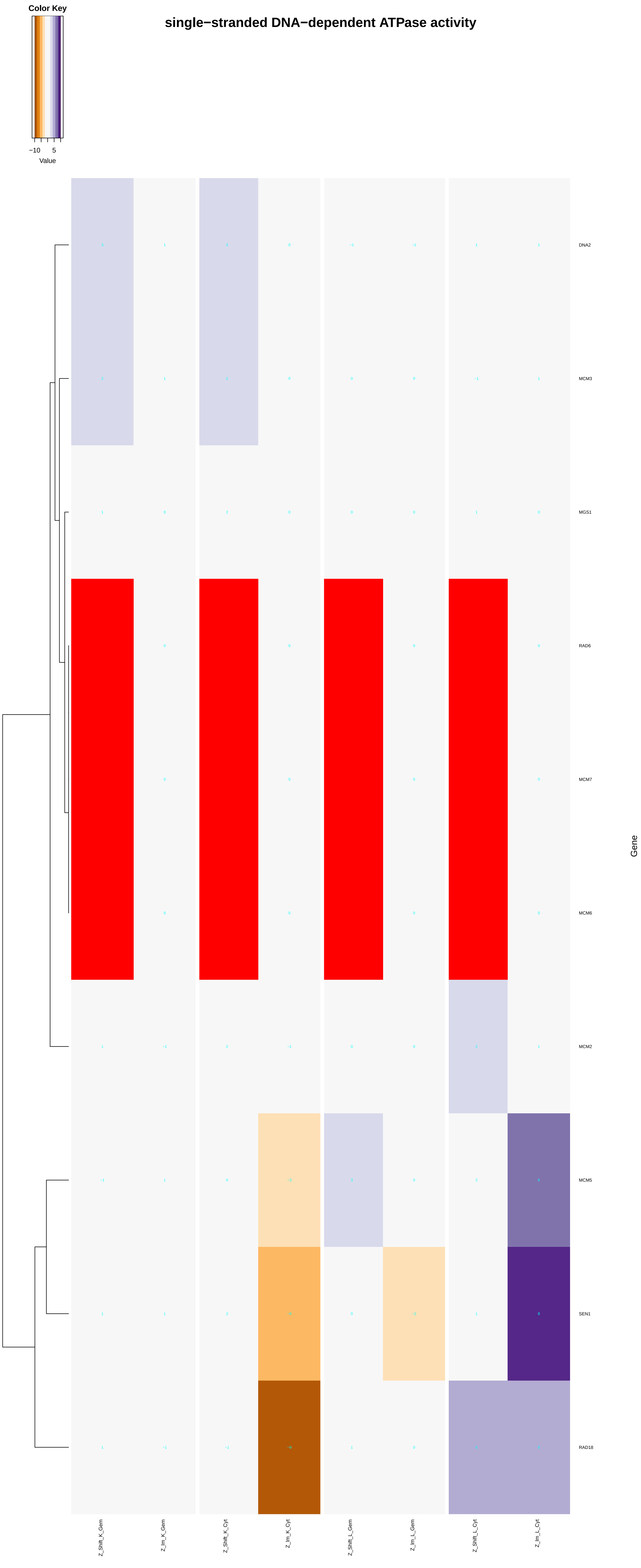

single−stranded DNA−dependent ATPase activity
Color Key
−10
5
Value
DNA2
MCM3
MGS1
RAD6
MCM7
MCM6
MCM2
MCM5
SEN1
RAD18
Gene
Z_Shift_K_Gem
Z_Im_K_Gem
Z_Shift_K_Cyt
Z_Im_K_Cyt
Z_Shift_L_Gem
Z_Im_L_Gem
Z_Shift_L_Cyt
Z_Im_L_Cyt

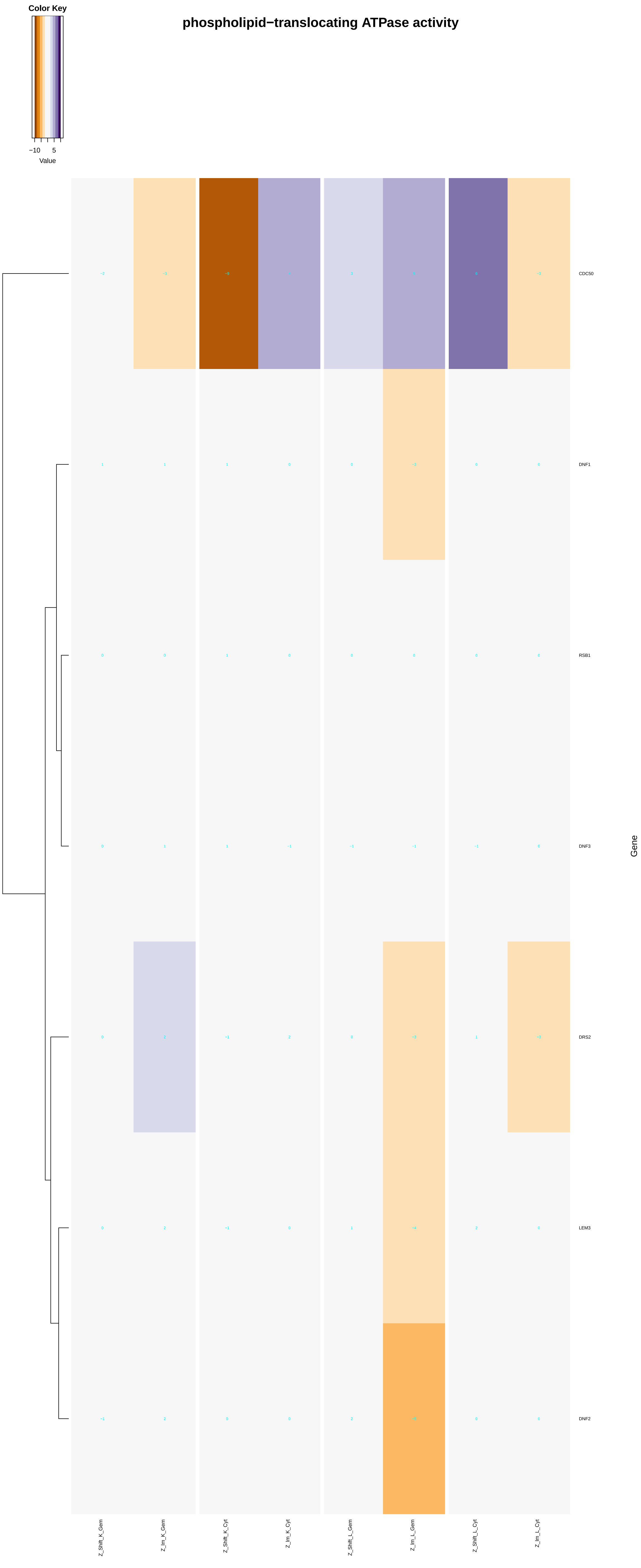
phospholipid−translocating ATPase activity
Color Key
−10
5
Value
CDC50
DNF1
RSB1
DNF3
DRS2
LEM3
DNF2
Gene
Z_Shift_K_Gem
Z_Im_K_Gem
Z_Shift_K_Cyt
Z_Im_K_Cyt
Z_Shift_L_Gem
Z_Im_L_Gem
Z_Shift_L_Cyt
Z_Im_L_Cyt

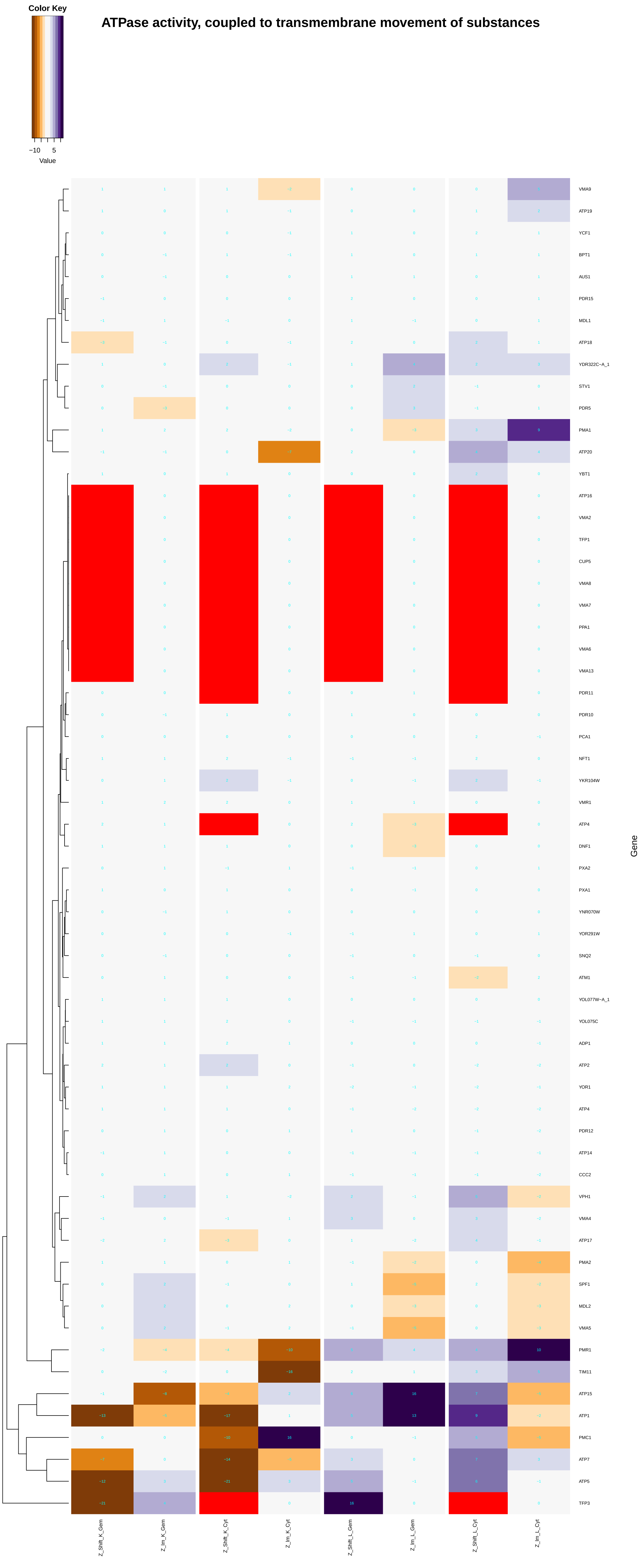
Color Key
−10
5
Value
ATPase activity, coupled to transmembrane movement of substances
VMA9
ATP19
YCF1
BPT1
AUS1
PDR15
MDL1
ATP18
YDR322C−A_1
STV1
PDR5
PMA1
ATP20
YBT1
ATP16
VMA2
TFP1
CUP5
VMA8
VMA7
PPA1
VMA6
VMA13
PDR11
PDR10
PCA1
NFT1
YKR104W
VMR1
ATP4
DNF1
PXA2
PXA1
YNR070W
YOR291W
SNQ2
ATM1
YOL077W−A_1
YOL075C
ADP1
ATP2
YOR1
ATP4
PDR12
ATP14
CCC2
VPH1
VMA4
ATP17
PMA2
SPF1
MDL2
VMA5
PMR1
TIM11
ATP15
ATP1
PMC1
ATP7
ATP5
TFP3
Gene
Z_Shift_K_Gem
Z_lm_K_Gem
Z_Shift_K_Cyt
Z_lm_K_Cyt
Z_Shift_L_Gem
Z_lm_L_Gem
Z_Shift_L_Cyt
Z_lm_L_Cyt

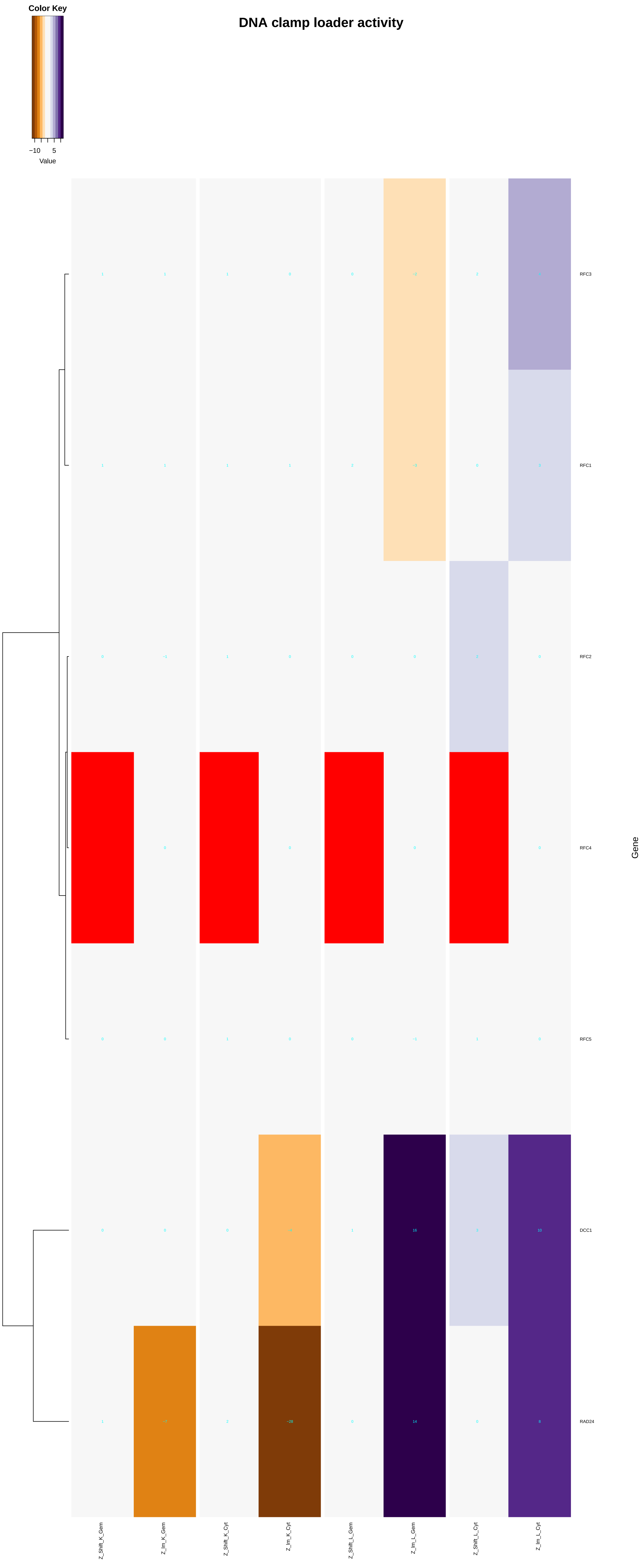

Color Key
−10
5
Value
DNA clamp loader activity
RFC3
RFC1
RFC2
RFC4
RFC5
DCC1
RAD24
Gene
Z_Shift_K_Gem
Z_Im_K_Gem
Z_Shift_K_Cyt
Z_Im_K_Cyt
Z_Shift_L_Gem
Z_Im_L_Gem
Z_Shift_L_Cyt
Z_Im_L_Cyt

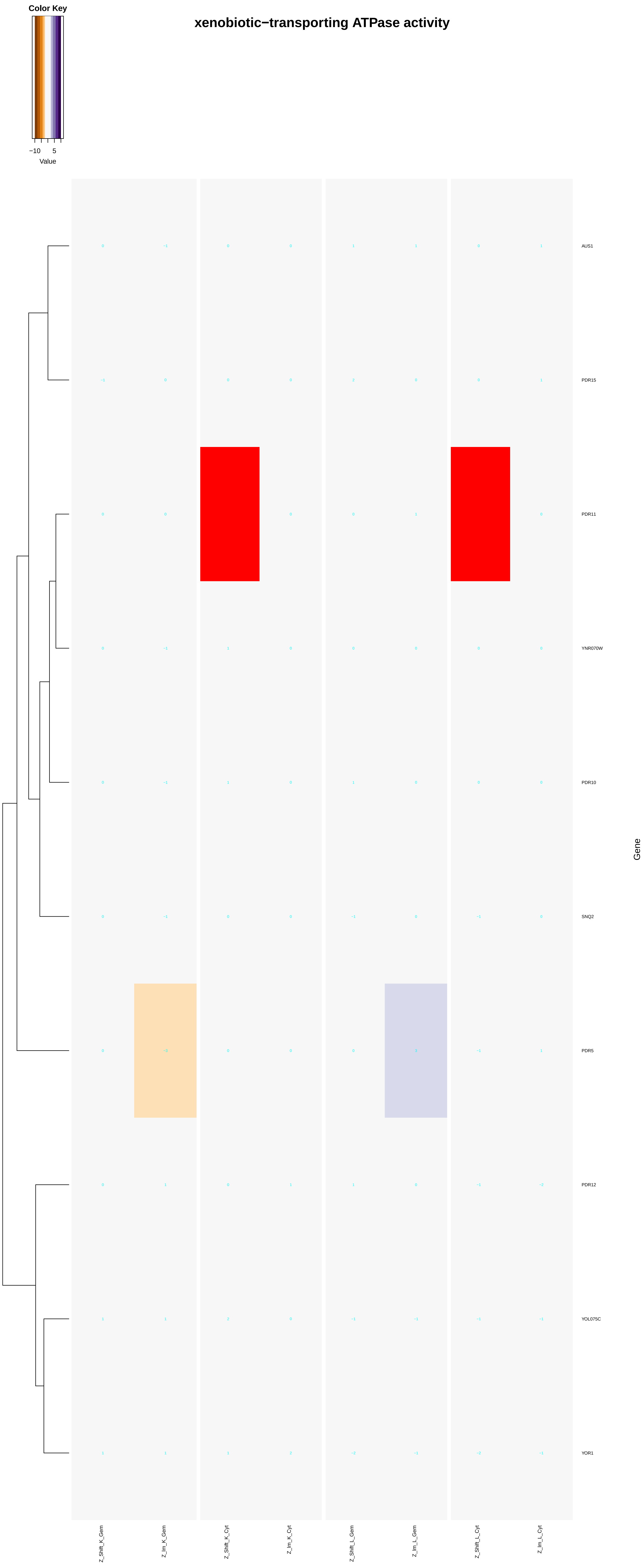
Color Key
−10
5
Value
xenobiotic−transporting ATPase activity
AUS1
PDR15
PDR11
YNR070W
PDR10
SNQ2
PDR5
PDR12
YOL075C
YOR1
Gene
Z_Shift_K_Gem
Z_Im_K_Gem
Z_Shift_K_Cyt
Z_Im_K_Cyt
Z_Shift_L_Gem
Z_Im_L_Gem
Z_Shift_L_Cyt
Z_Im_L_Cyt

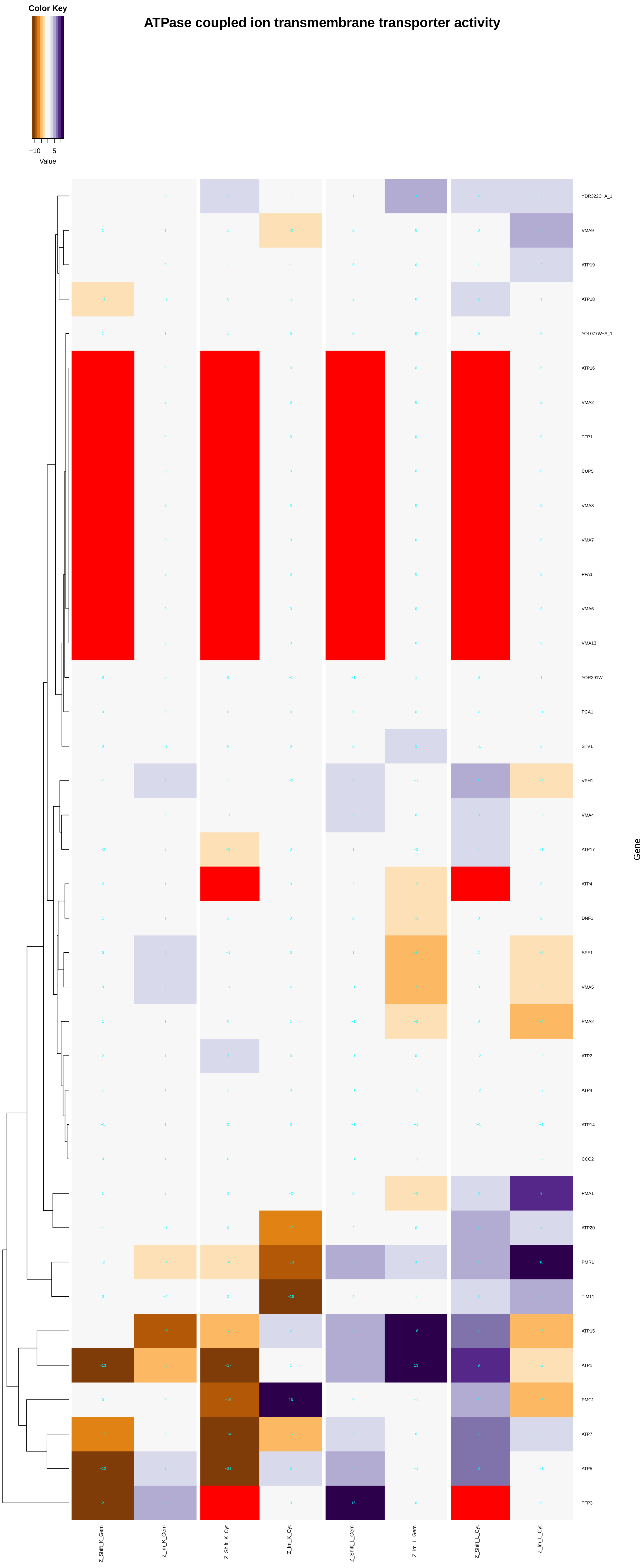
Color Key
−10
5
Value
ATPase coupled ion transmembrane transporter activity
YDR322C−A_1
VMA9
ATP19
ATP18
YOL077W−A_1
ATP16
VMA2
TFP1
CUP5
VMA8
VMA7
PPA1
VMA6
VMA13
YOR291W
PCA1
STV1
VPH1
VMA4
ATP17
ATP4
DNF1
SPF1
VMA5
PMA2
ATP2
ATP4
ATP14
CCC2
PMA1
ATP20
PMR1
TIM11
ATP15
ATP1
PMC1
ATP7
ATP5
TFP3
Gene
Z_Shift_K_Gem
Z_Im_K_Gem
Z_Shift_K_Cyt
Z_Im_K_Cyt
Z_Shift_L_Gem
Z_Im_L_Gem
Z_Shift_L_Cyt
Z_Im_L_Cyt

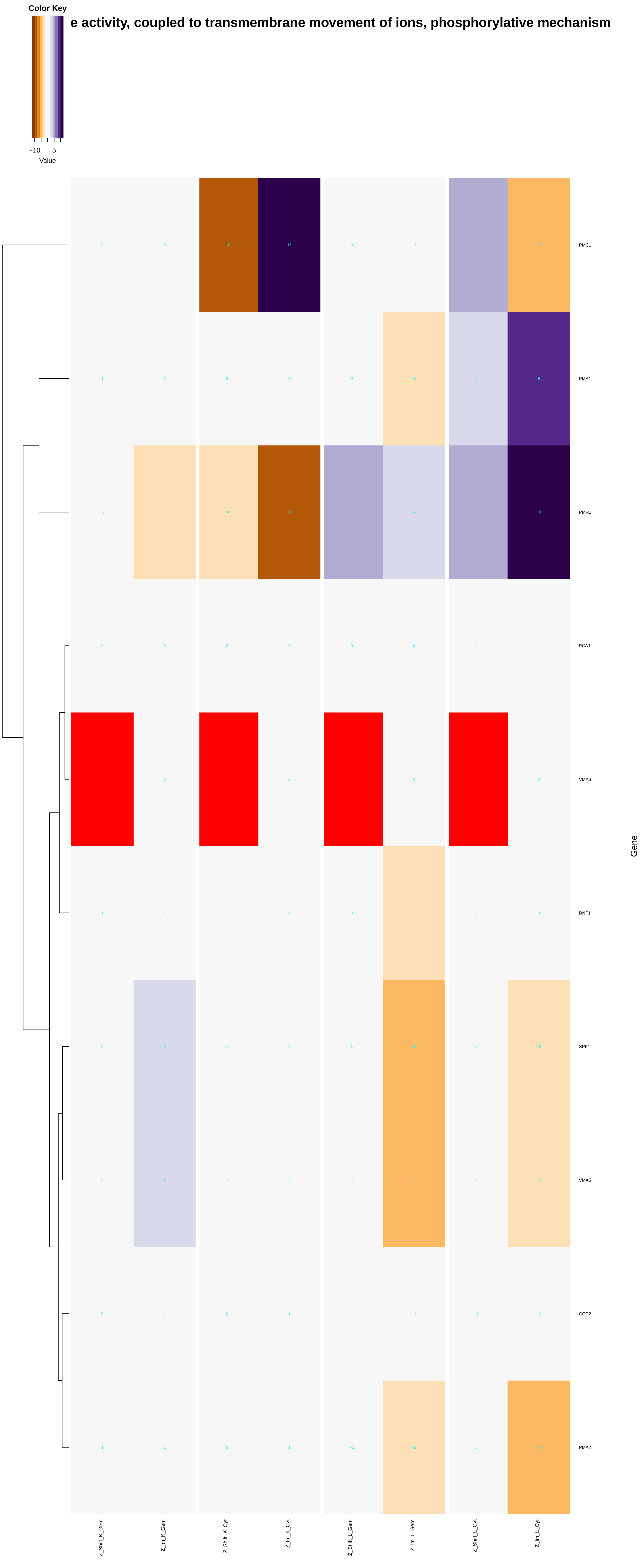
e activity, coupled to transmembrane movement of ions, phosphorylative mechanism
Color Key
−10
5
Value
PMC1
PMA1
PMR1
PCA1
VMA6
DNF1
SPF1
VMA5
CCC2
PMA2
Gene
Z_Shift_K_Gem
Z_Im_K_Gem
Z_Shift_K_Cyt
Z_Im_K_Cyt
Z_Shift_L_Gem
Z_Im_L_Gem
Z_Shift_L_Cyt
Z_Im_L_Cyt

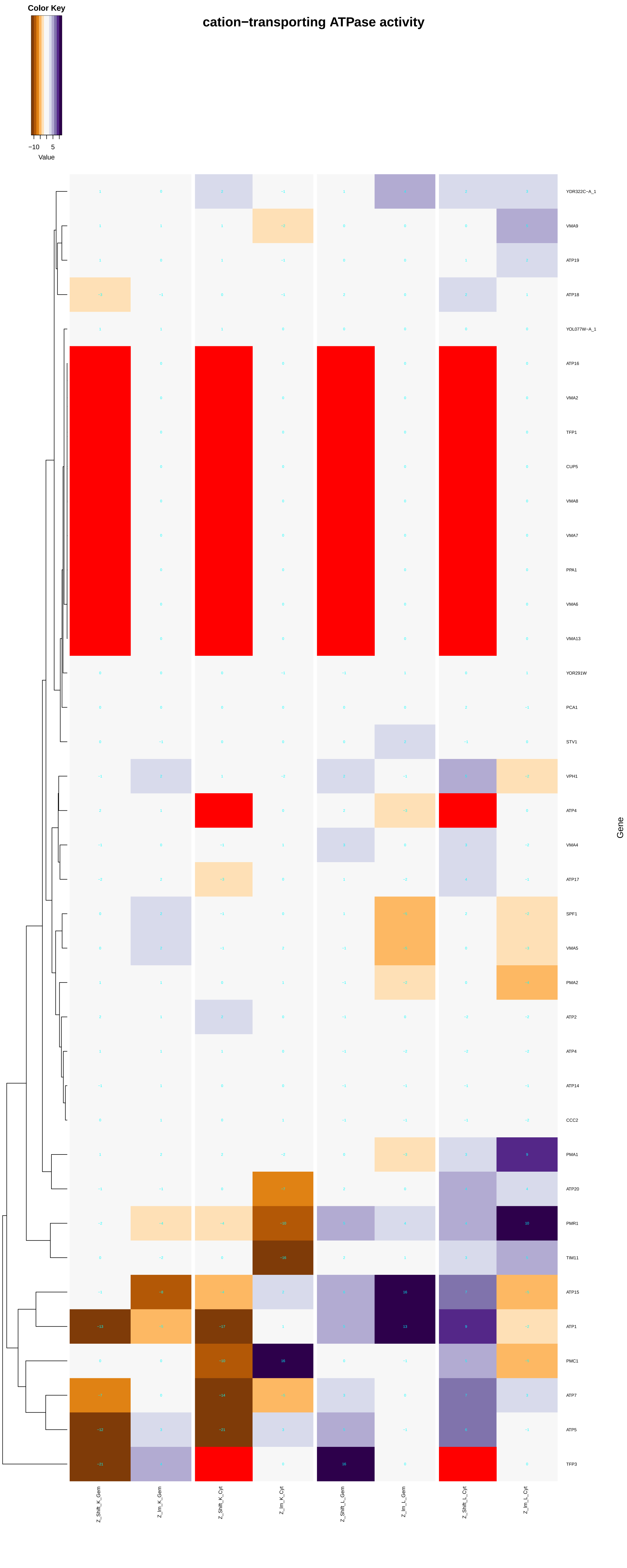

cation−transporting ATPase activity
Color Key
−10
5
Value
YDR322C−A_1
VMA9
ATP19
ATP18
YOL077W−A_1
ATP16
VMA2
TFP1
CUP5
VMA8
VMA7
PPA1
VMA6
VMA13
YOR291W
PCA1
STV1
VPH1
ATP4
VMA4
ATP17
SPF1
VMA5
PMA2
ATP2
ATP4
ATP14
CCC2
PMA1
ATP20
PMR1
TIM11
ATP15
ATP1
PMC1
ATP7
ATP5
TFP3
Gene
Z_Shift_K_Gem
Z_Im_K_Gem
Z_Shift_K_Cyt
Z_Im_K_Cyt
Z_Shift_L_Gem
Z_Im_L_Gem
Z_Shift_L_Cyt
Z_Im_L_Cyt

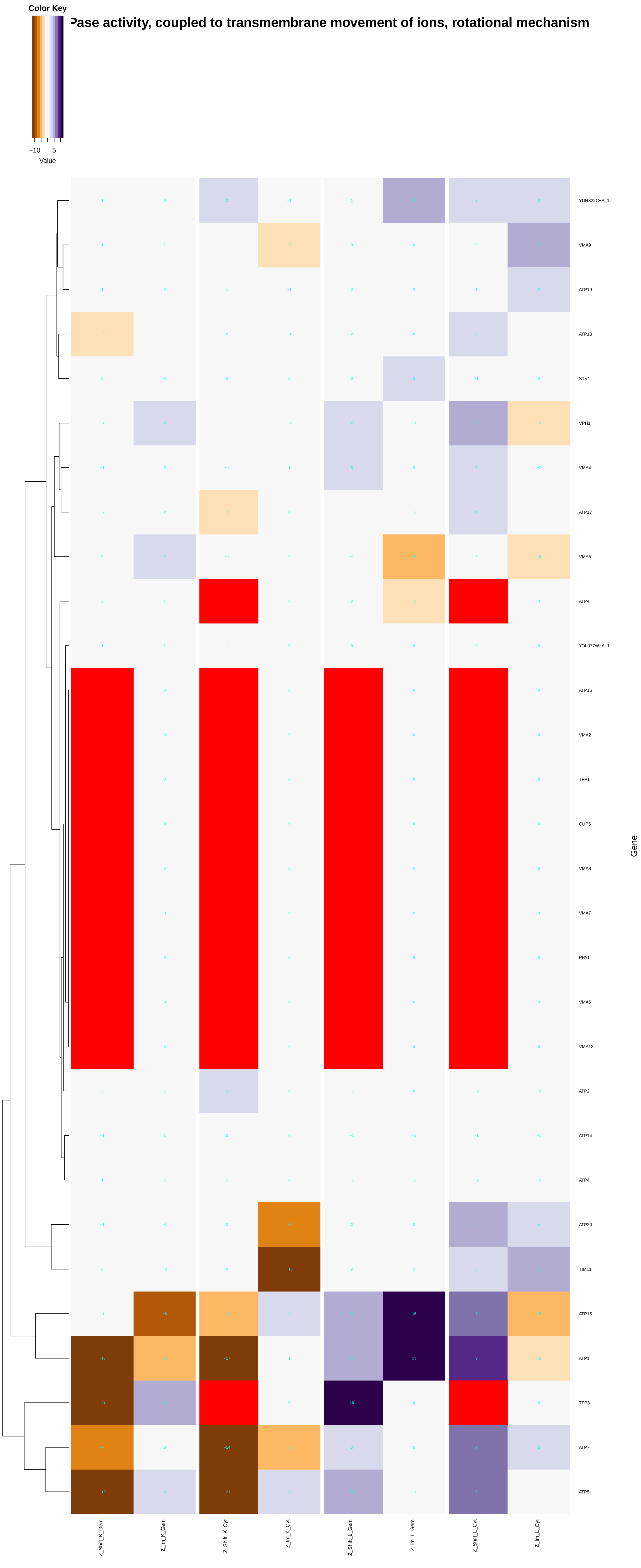

Color Key
−10 5
Value
Pase activity, coupled to transmembrane movement of ions, rotational mechanism
YDR322C−A_1
VMA9
ATP19
ATP18
STV1
VPH1
VMA4
ATP17
VMA5
ATP4
YOL077W−A_1
ATP16
VMA2
TFP1
CUP5
VMA8
VMA7
PPA1
VMA6
VMA13
ATP2
ATP14
ATP4
ATP20
TIM11
ATP15
ATP1
TFP3
ATP7
ATP5
Gene
Z_Shift_K_Gem
Z_Im_K_Gem
Z_Shift_K_Cyt
Z_Im_K_Cyt
Z_Shift_L_Gem
Z_Im_L_Gem
Z_Shift_L_Cyt
Z_Im_L_Cyt

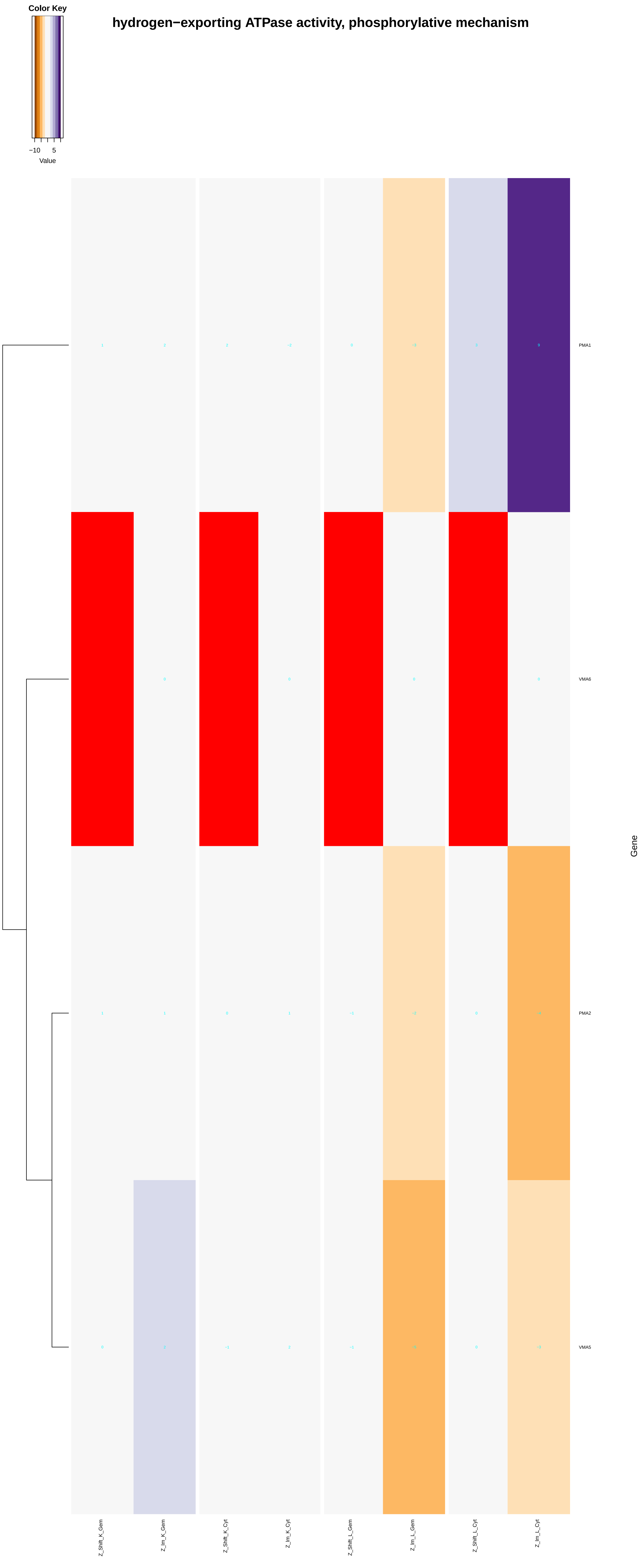

Color Key
−10
5
Value
hydrogen−exporting ATPase activity, phosphorylative mechanism
PMA1
VMA6
PMA2
VMA5
Gene
Z_Shift_K_Gem
Z_Im_K_Gem
Z_Shift_K_Cyt
Z_Im_K_Cyt
Z_Shift_L_Gem
Z_Im_L_Gem
Z_Shift_L_Cyt
Z_Im_L_Cyt

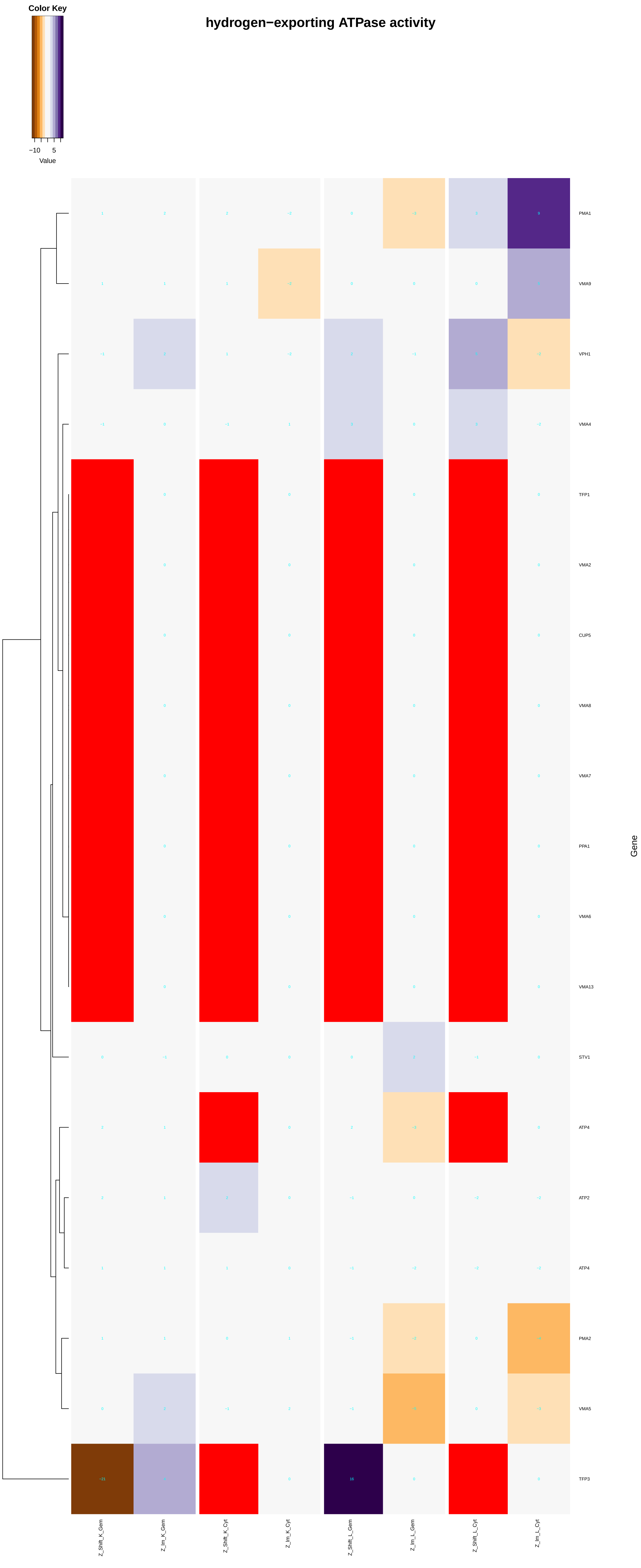
hydrogen−exporting ATPase activity
Color Key
−10
5
Value
PMA1
VMA9
VPH1
VMA4
TFP1
VMA2
CUP5
VMA8
VMA7
PPA1
VMA6
VMA13
STV1
ATP4
ATP2
ATP4
PMA2
VMA5
TFP3
Gene
Z_Shift_K_Gem
Z_Im_K_Gem
Z_Shift_K_Cyt
Z_Im_K_Cyt
Z_Shift_L_Gem
Z_Im_L_Gem
Z_Shift_L_Cyt
Z_Im_L_Cyt

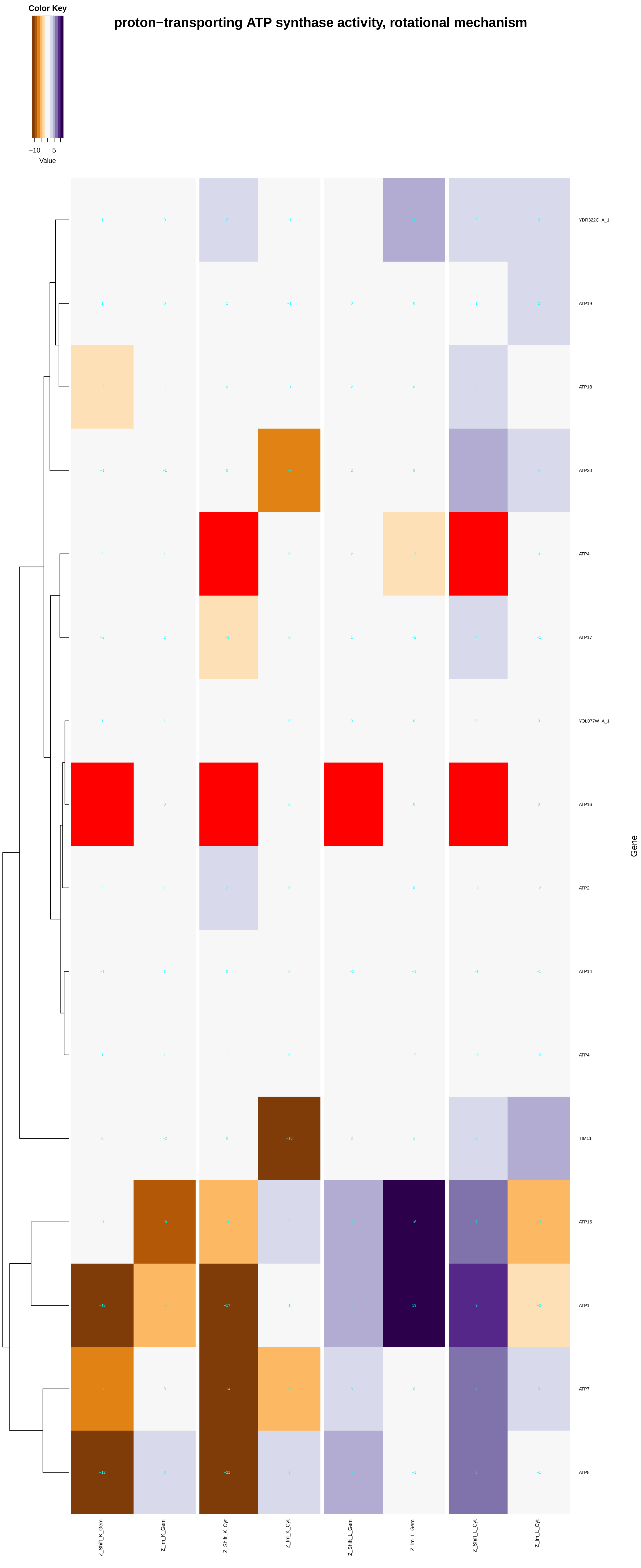
Color Key
−10
5
Value
proton−transporting ATP synthase activity, rotational mechanism
YDR322C−A_1
ATP19
ATP18
ATP20
ATP4
ATP17
YOL077W−A_1
ATP16
ATP2
ATP14
ATP4
TIM11
ATP15
ATP1
ATP7
ATP5
Gene
Z_Shift_K_Gem
Z_Im_K_Gem
Z_Shift_K_Cyt
Z_Im_K_Cyt
Z_Shift_L_Gem
Z_Im_L_Gem
Z_Shift_L_Cyt
Z_Im_L_Cyt

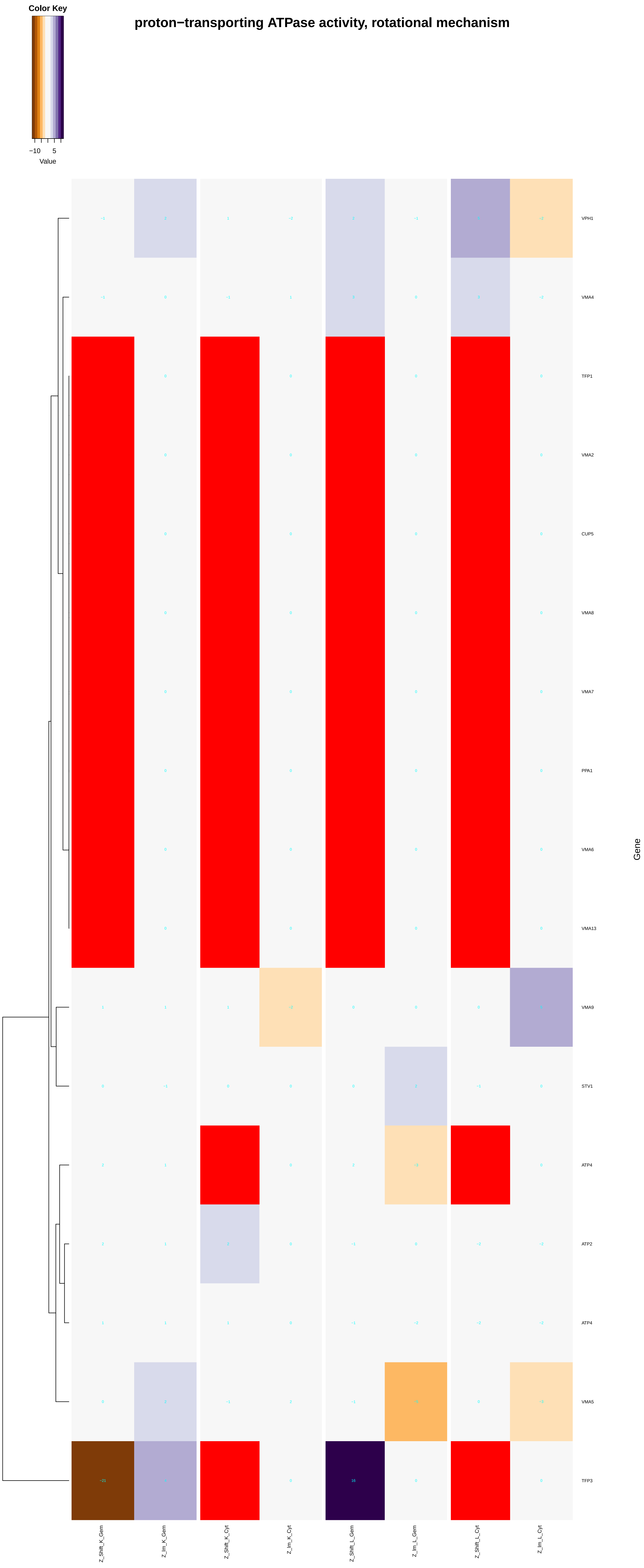

proton−transporting ATPase activity, rotational mechanism
Color Key
−10
5
Value
VPH1
VMA4
TFP1
VMA2
CUP5
VMA8
VMA7
PPA1
VMA6
VMA13
VMA9
STV1
ATP4
ATP2
ATP4
VMA5
TFP3
Gene
Z_Shift_K_Gem
Z_Im_K_Gem
Z_Shift_K_Cyt
Z_Im_K_Cyt
Z_Shift_L_Gem
Z_Im_L_Gem
Z_Shift_L_Cyt
Z_Im_L_Cyt
