## Additional File 7 for "A humanized yeast phenomic model of deoxycytidine kinase to predict genetic buffering of nucleoside analog cytotoxicity": intracellular_protein_transport.pdf

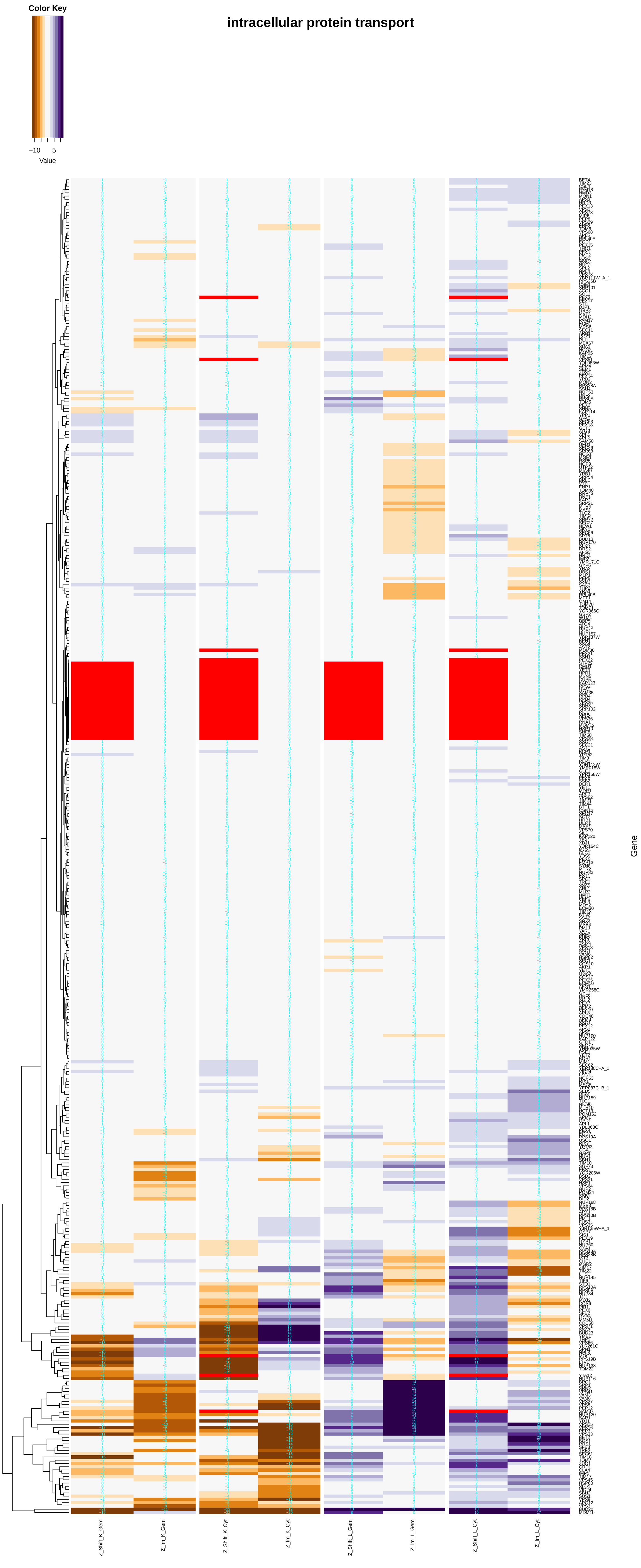

intracellular protein transport
Color Key
−10
5
Value
Gene
Z_Shift_K_Gem
Z_Im_K_Gem
Z_Shift_K_Cyt
Z_Im_K_Cyt
Z_Shift_L_Gem
Z_Im_L_Gem
Z_Shift_L_Cyt
Z_Im_L_Cyt

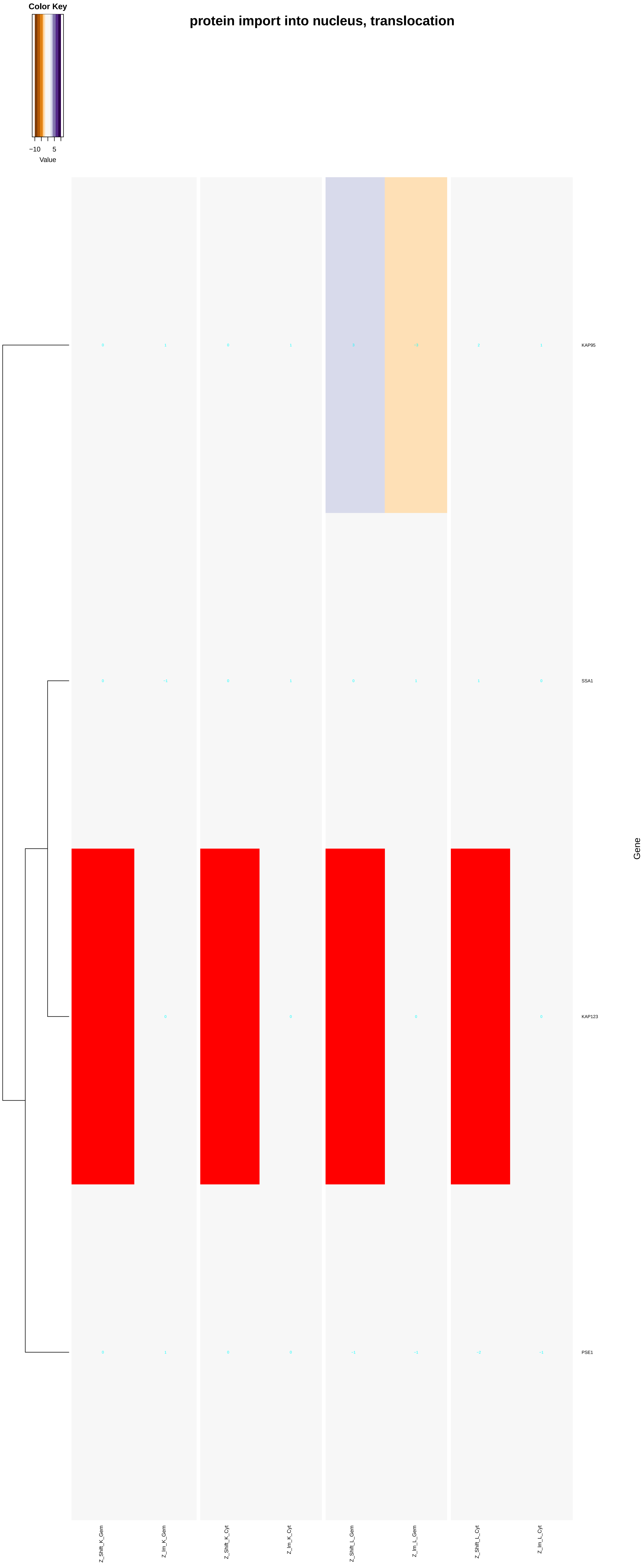
protein import into nucleus, translocation
Color Key
−10
5
Value
KAP95
SSA1
KAP123
PSE1
Gene
Z_Shift_K_Gem
Z_Im_K_Gem
Z_Shift_K_Cyt
Z_Im_K_Cyt
Z_Shift_L_Gem
Z_Im_L_Gem
Z_Shift_L_Cyt
Z_Im_L_Cyt

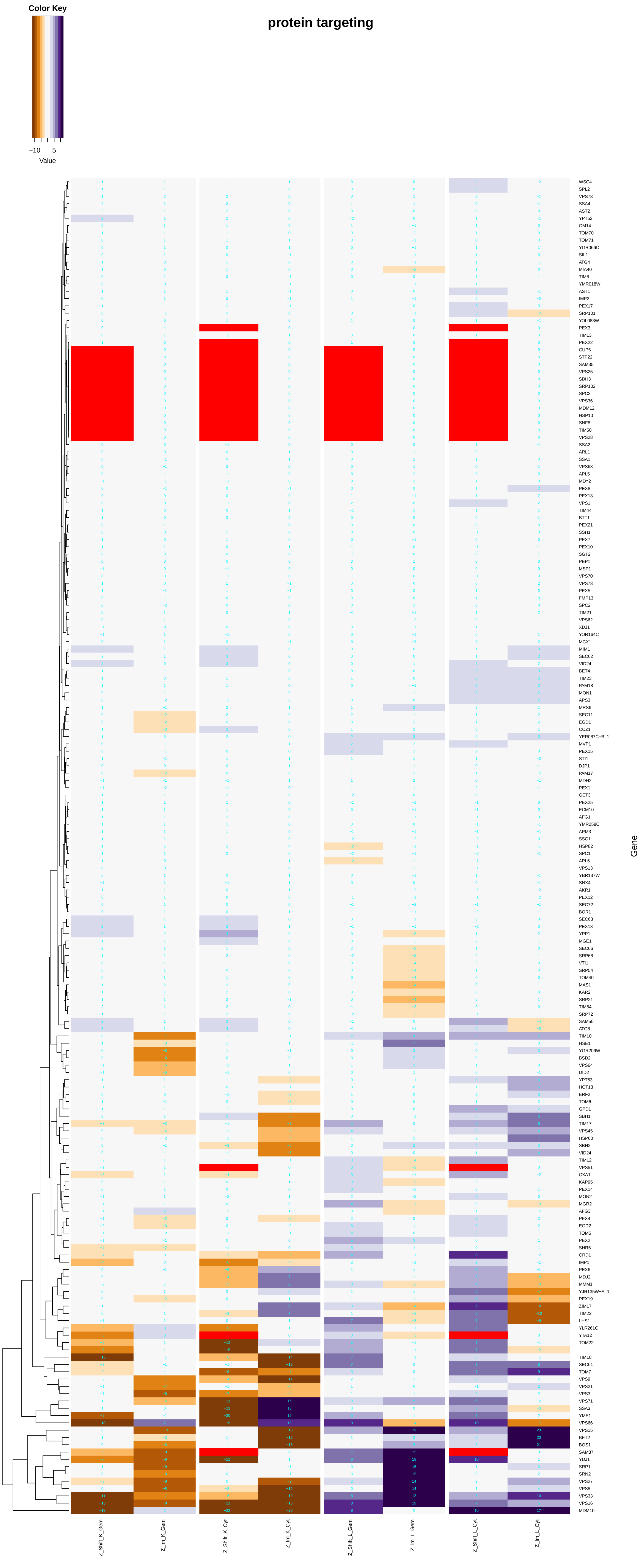
protein targeting
Color Key
−10
5
Value
Gene
WSC4
SPL2
VPS73
SSA4
AST2
YPT52
OM14
TOM70
TOM71
YGR066C
SIL1
ATG4
MIA40
TIM8
YMR018W
AST1
IMP2
PEX17
SRP101
YOL083W
PEX3
TIM13
PEX22
CUP5
STP22
SAM35
VPS25
SDH3
SRP102
SPC3
VPS36
MDM12
HSP10
SNF8
TIM50
VPS28
SSA2
ARL1
SSA1
VPS68
APL5
MDY2
PEX8
PEX13
VPS1
TIM44
BTT1
PEX21
SSH1
PEX7
PEX10
SGT2
PEP1
MSP1
VPS70
VPS73
PEX5
FMP13
SPC2
TIM21
VPS62
XDJ1
YOR164C
MCX1
MIM1
SEC62
VID24
BET4
TIM23
PAM18
MON1
APS3
MRS6
SEC11
EGD1
CCZ1
YER087C−B_1
MVP1
PEX15
STI1
DJP1
PAM17
MDH2
PEX1
GET3
PEX25
ECM10
AFG1
YMR258C
APM3
SSC1
HSP82
SPC1
APL6
VPS13
YBR137W
SNX4
AKR1
PEX12
SEC72
BOR1
SEC63
PEX18
YPP1
MGE1
SEC66
SRP68
VTI1
SRP54
TOM40
MAS1
KAR2
SRP21
TIM54
SRP72
SAM50
ATG8
TIM10
HSE1
YGR206W
BSD2
VPS64
DID2
YPT53
HOT13
ERF2
TOM6
GPD1
SBH1
TIM17
VPS45
HSP60
SBH2
VID24
TIM12
VPS51
OXA1
KAP95
PEX14
MON2
MGR2
AFG3
PEX4
EGD2
TOM5
PEX2
SHR5
CRD1
IMP1
PEX6
MDJ2
MMM1
YJR135W−A_1
PEX19
ZIM17
TIM22
LHS1
YLR261C
YTA12
TOM22
TIM18
SEC61
TOM7
VPS9
VPS21
VPS3
VPS71
SSA3
YME1
VPS66
VPS15
BET2
BOS1
SAM37
YDJ1
SRP1
SRN2
VPS27
VPS8
VPS33
VPS16
MDM10
Z_Shift_K_Gem
Z_Im_K_Gem
Z_Shift_K_Cyt
Z_Im_K_Cyt
Z_Shift_L_Gem
Z_Im_L_Gem
Z_Shift_L_Cyt
Z_Im_L_Cyt

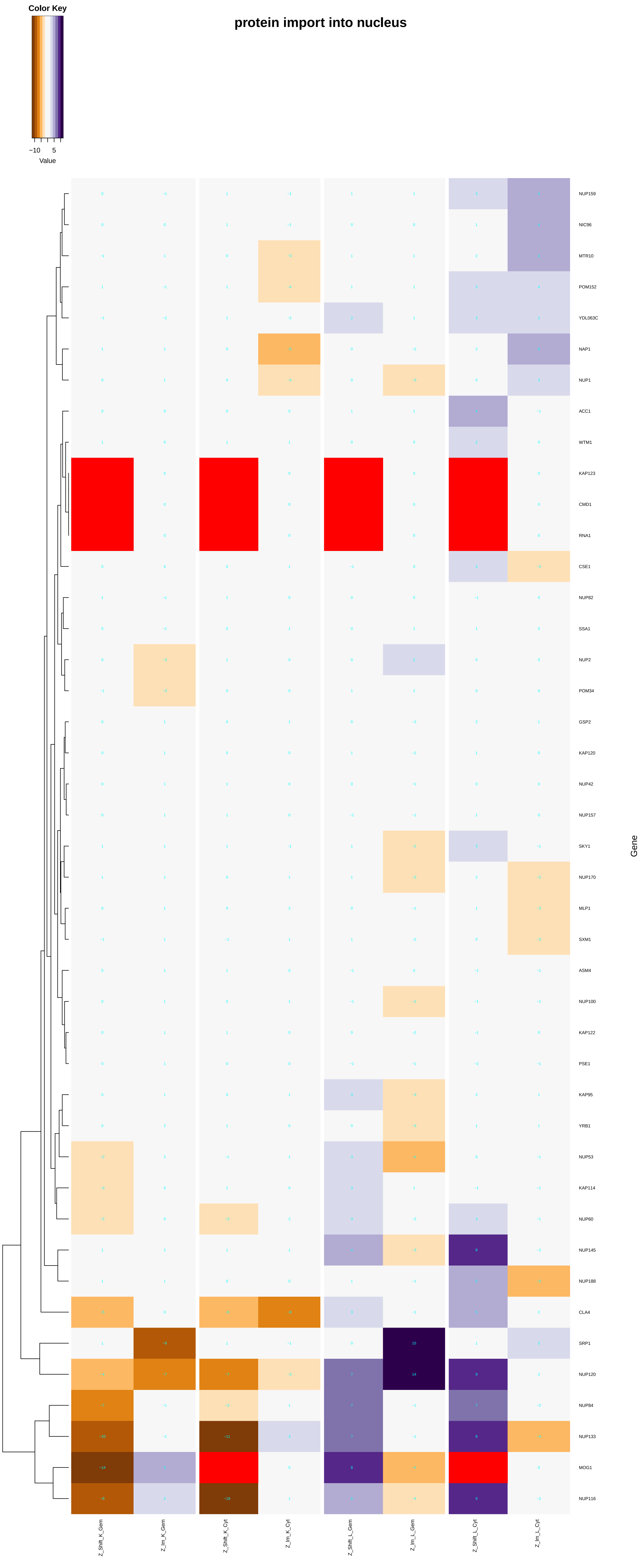
Color Key
−10 5
Value
protein import into nucleus
NUP159
NIC96
MTR10
POM152
YDL063C
NAP1
NUP1
ACC1
WTM1
KAP123
CMD1
RNA1
CSE1
NUP82
SSA1
NUP2
POM34
GSP2
KAP120
NUP42
NUP157
SKY1
NUP170
MLP1
SXM1
ASM4
NUP100
KAP122
PSE1
KAP95
YRB1
NUP53
KAP114
NUP60
NUP145
NUP188
CLA4
SRP1
NUP120
NUP84
NUP133
MOG1
NUP116
Gene
Z_Shift_K_Gem
Z_Im_K_Gem
Z_Shift_K_Cyt
Z_Im_K_Cyt
Z_Shift_L_Gem
Z_Im_L_Gem
Z_Shift_L_Cyt
Z_Im_L_Cyt

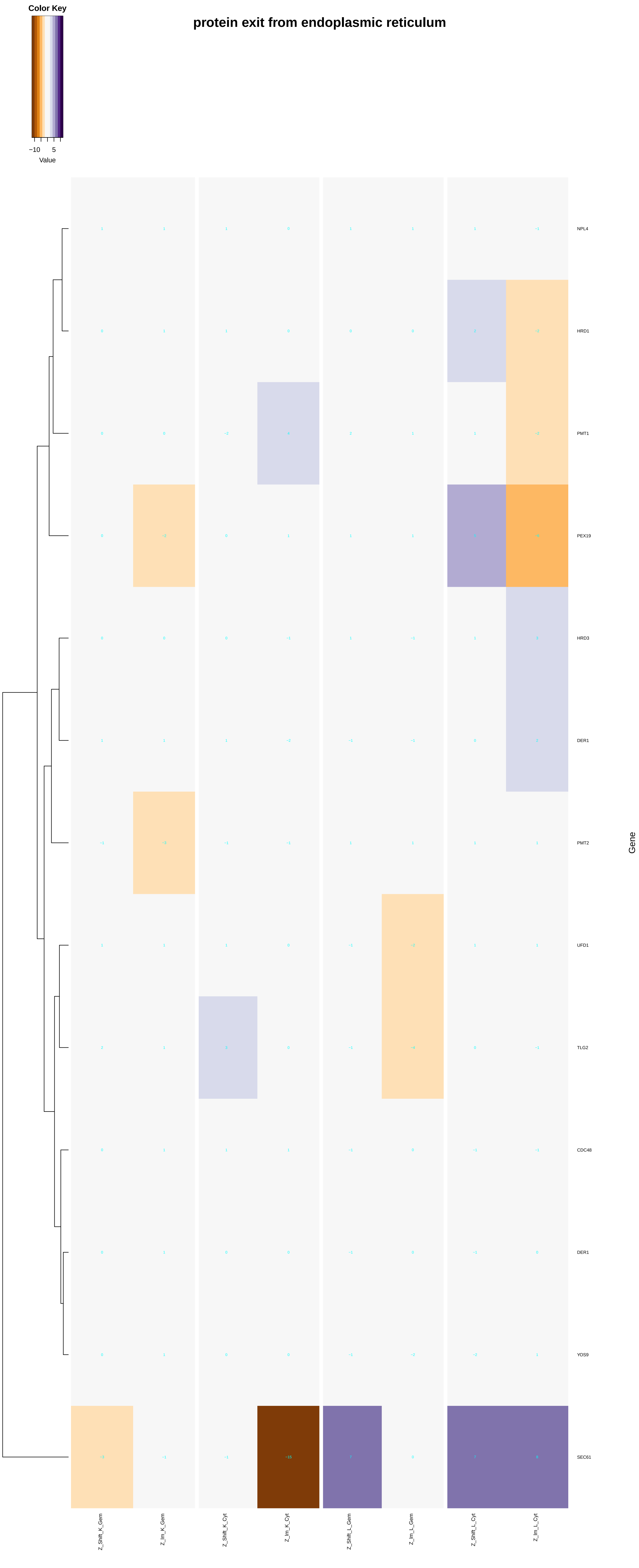
Color Key
−10
5
Value
protein exit from endoplasmic reticulum
NPL4
HRD1
PMT1
PEX19
HRD3
DER1
PMT2
UFD1
TLG2
CDC48
DER1
YOS9
SEC61
Gene
Z_Shift_K_Gem
Z_Im_K_Gem
Z_Shift_K_Cyt
Z_Im_K_Cyt
Z_Shift_L_Gem
Z_Im_L_Gem
Z_Shift_L_Cyt
Z_Im_L_Cyt

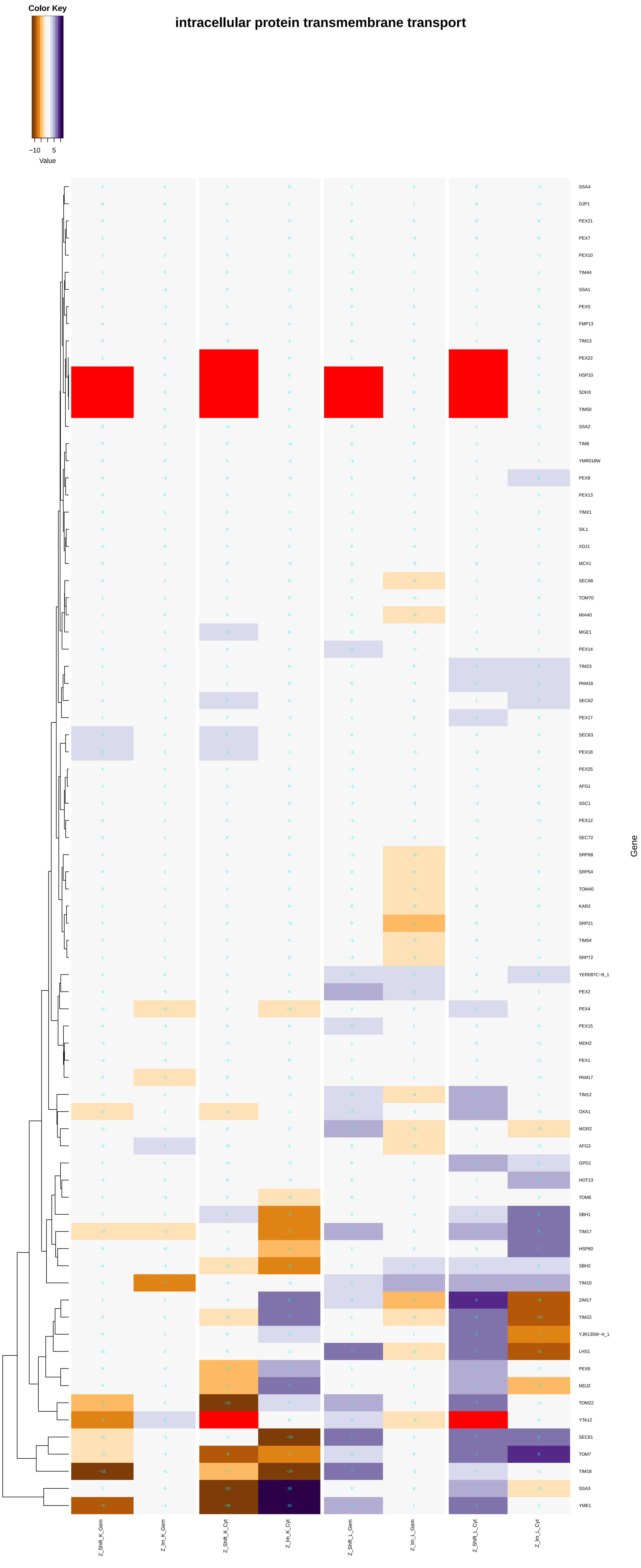

intracellular protein transmembrane transport
Color Key
−10
5
Value
Gene
SSA4
DJP1
PEX21
PEX7
PEX10
TIM44
SSA1
PEX5
FMP13
TIM13
PEX22
HSP10
SDH3
TIM50
SSA2
TIM8
YMR018W
PEX8
PEX13
TIM21
SIL1
XDJ1
MCX1
SEC66
TOM70
MIA40
MGE1
PEX14
TIM23
PAM18
SEC62
PEX17
SEC63
PEX18
PEX25
AFG1
SSC1
PEX12
SEC72
SRP68
SRP54
TOM40
KAR2
SRP21
TIM54
SRP72
YER087C−B_1
PEX2
PEX4
PEX15
MDH2
PEX1
PAM17
TIM12
OXA1
MGR2
AFG3
GPD1
HOT13
TOM6
SBH1
TIM17
HSP60
SBH2
TIM10
ZIM17
TIM22
YJR135W−A_1
LHS1
PEX6
MDJ2
TOM22
YTA12
SEC61
TOM7
TIM18
SSA3
YME1
Z_Shift_K_Gem
Z_Im_K_Gem
Z_Shift_K_Cyt
Z_Im_K_Cyt
Z_Shift_L_Gem
Z_Im_L_Gem
Z_Shift_L_Cyt
Z_Im_L_Cyt

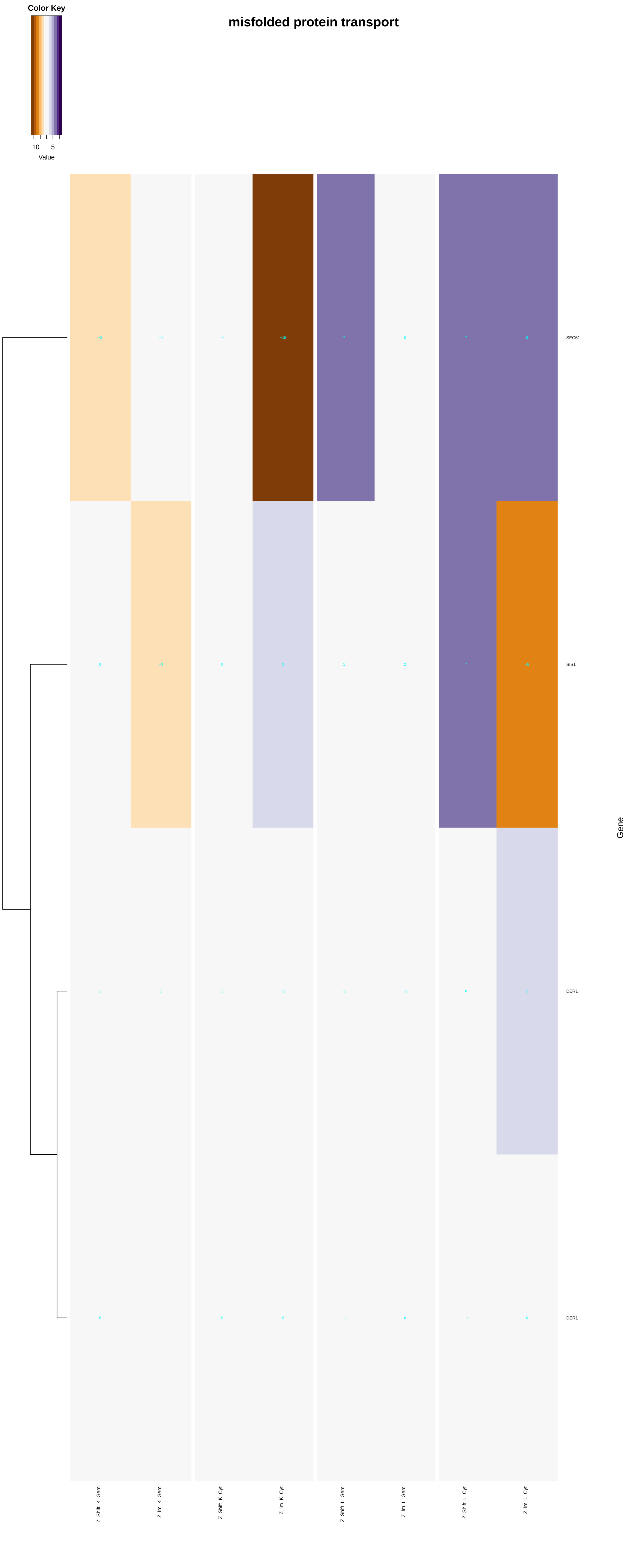
Color Key
−10
5
Value
misfolded protein transport
SEC61
SIS1
DER1
DER1
Gene
Z_Shift_K_Gem
Z_Im_K_Gem
Z_Shift_K_Cyt
Z_Im_K_Cyt
Z_Shift_L_Gem
Z_Im_L_Gem
Z_Shift_L_Cyt
Z_Im_L_Cyt

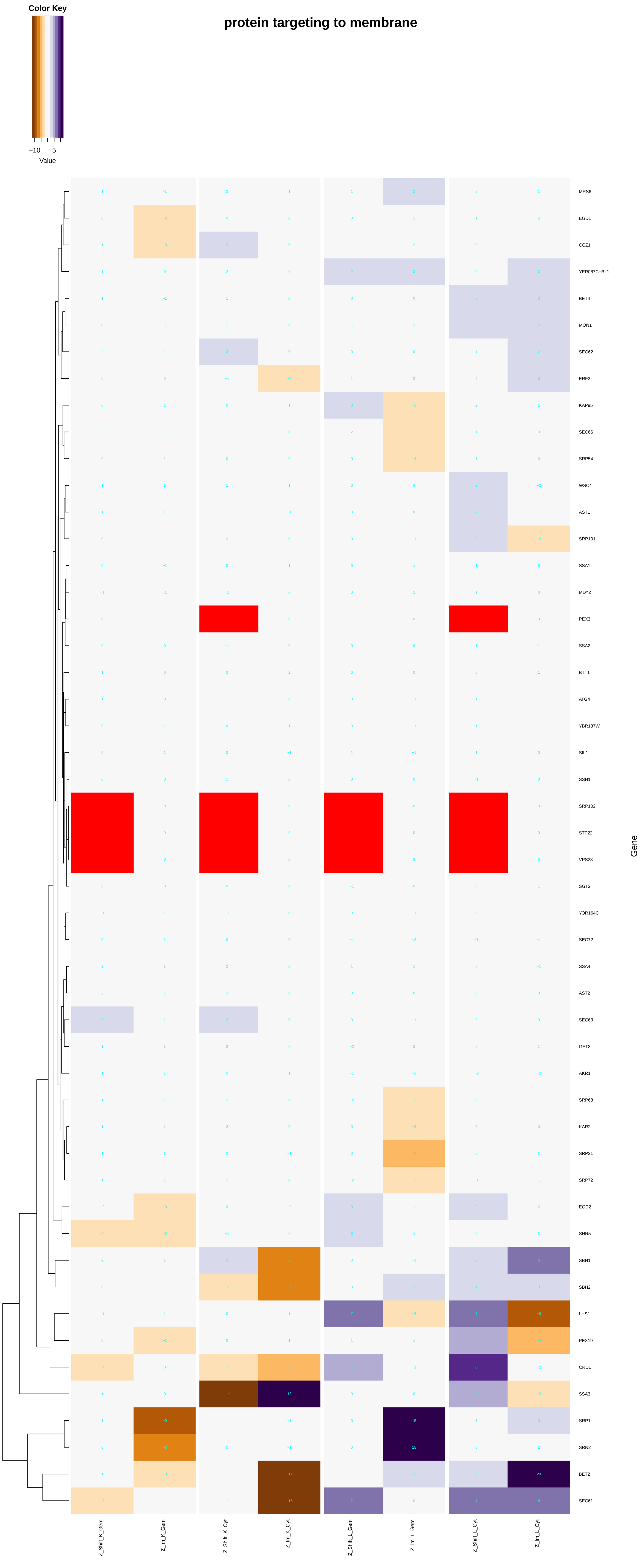

Color Key
−10
5
Value
protein targeting to membrane
MRS6
EGD1
CCZ1
YER087C−B_1
BET4
MON1
SEC62
ERF2
KAP95
SEC66
SRP54
WSC4
AST1
SRP101
SSA1
MDY2
PEX3
SSA2
BTT1
ATG4
YBR137W
SIL1
SSH1
SRP102
STP22
VPS28
SGT2
YOR164C
SEC72
SSA4
AST2
SEC63
GET3
AKR1
SRP68
KAR2
SRP21
SRP72
EGD2
SHR5
SBH1
SBH2
LHS1
PEX19
CRD1
SSA3
SRP1
SRN2
BET2
SEC61
Gene
Z_Shift_K_Gem
Z_Im_K_Gem
Z_Shift_K_Cyt
Z_Im_K_Cyt
Z_Shift_L_Gem
Z_Im_L_Gem
Z_Shift_L_Cyt
Z_Im_L_Cyt

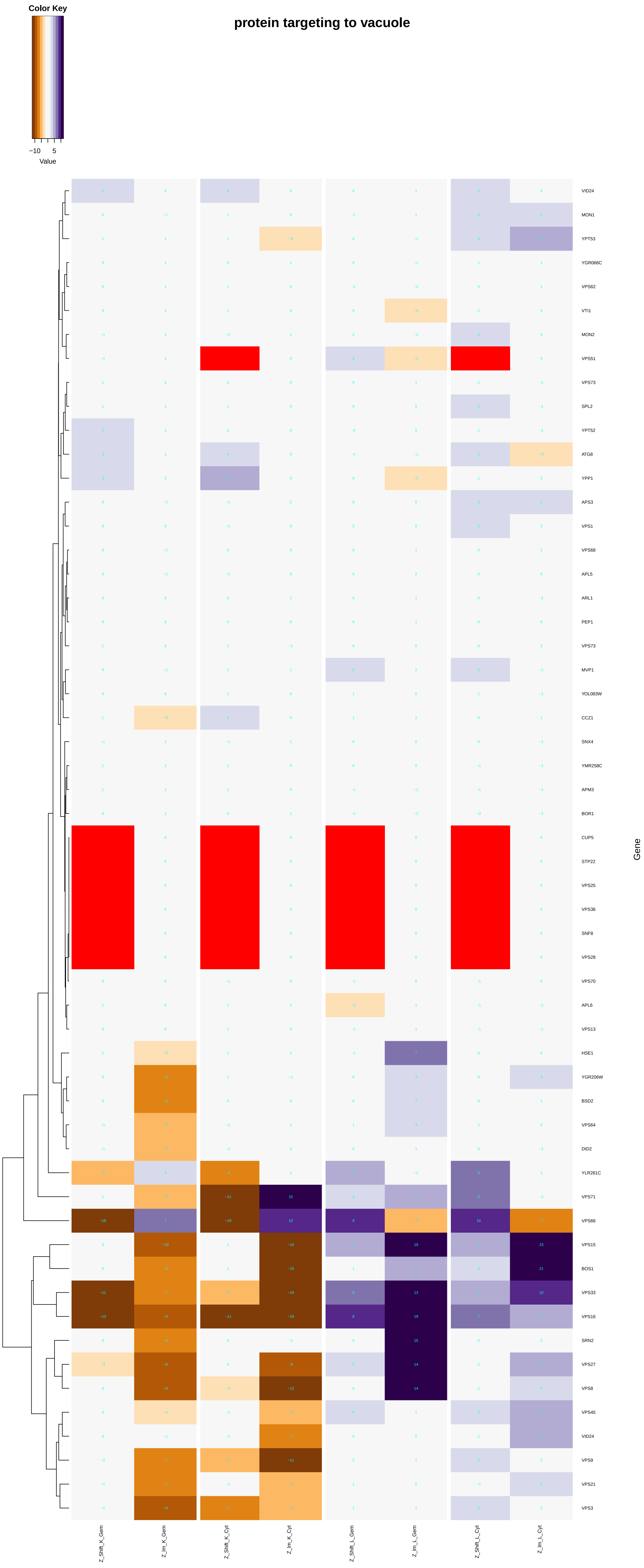

Color Key
−10
5
Value
protein targeting to vacuole
VID24
MON1
YPT53
YGR066C
VPS62
VTI1
MON2
VPS51
VPS73
SPL2
YPT52
ATG8
YPP1
APS3
VPS1
VPS68
APL5
ARL1
PEP1
VPS73
MVP1
YOL083W
CCZ1
SNX4
YMR258C
APM3
BOR1
CUP5
STP22
VPS25
VPS36
SNF8
VPS28
VPS70
APL6
VPS13
HSE1
YGR206W
BSD2
VPS64
DID2
YLR261C
VPS71
VPS66
VPS15
BOS1
VPS33
VPS16
SRN2
VPS27
VPS8
VPS45
VID24
VPS9
VPS21
VPS3
Gene
Z_Shift_K_Gem
Z_Im_K_Gem
Z_Shift_K_Cyt
Z_Im_K_Cyt
Z_Shift_L_Gem
Z_Im_L_Gem
Z_Shift_L_Cyt
Z_Im_L_Cyt

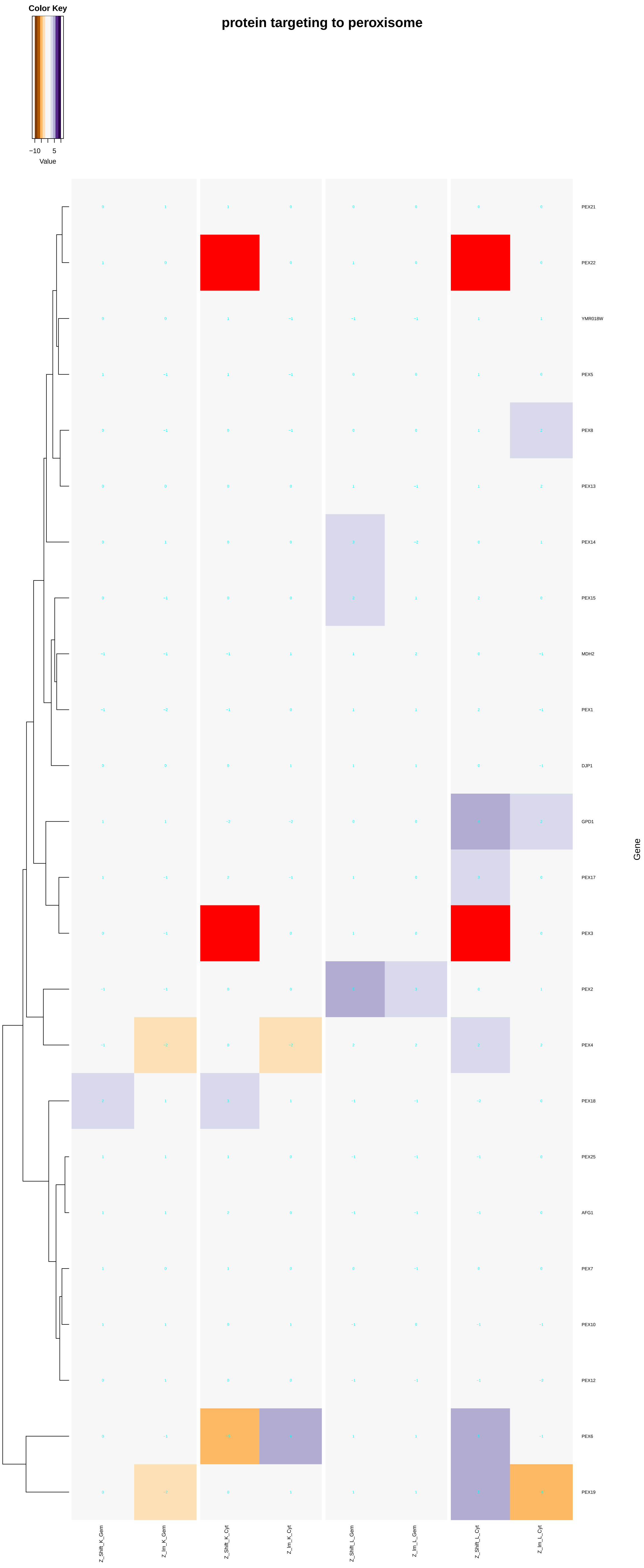

Color Key
−10
5
Value
protein targeting to peroxisome
PEX21
PEX22
YMR018W
PEX5
PEX8
PEX13
PEX14
PEX15
MDH2
PEX1
DJP1
GPD1
PEX17
PEX3
PEX2
PEX4
PEX18
PEX25
AFG1
PEX7
PEX10
PEX12
PEX6
PEX19
Gene
Z_Shift_K_Gem
Z_Im_K_Gem
Z_Shift_K_Cyt
Z_Im_K_Cyt
Z_Shift_L_Gem
Z_Im_L_Gem
Z_Shift_L_Cyt
Z_Im_L_Cyt

**protein targeting to mitochondrion**

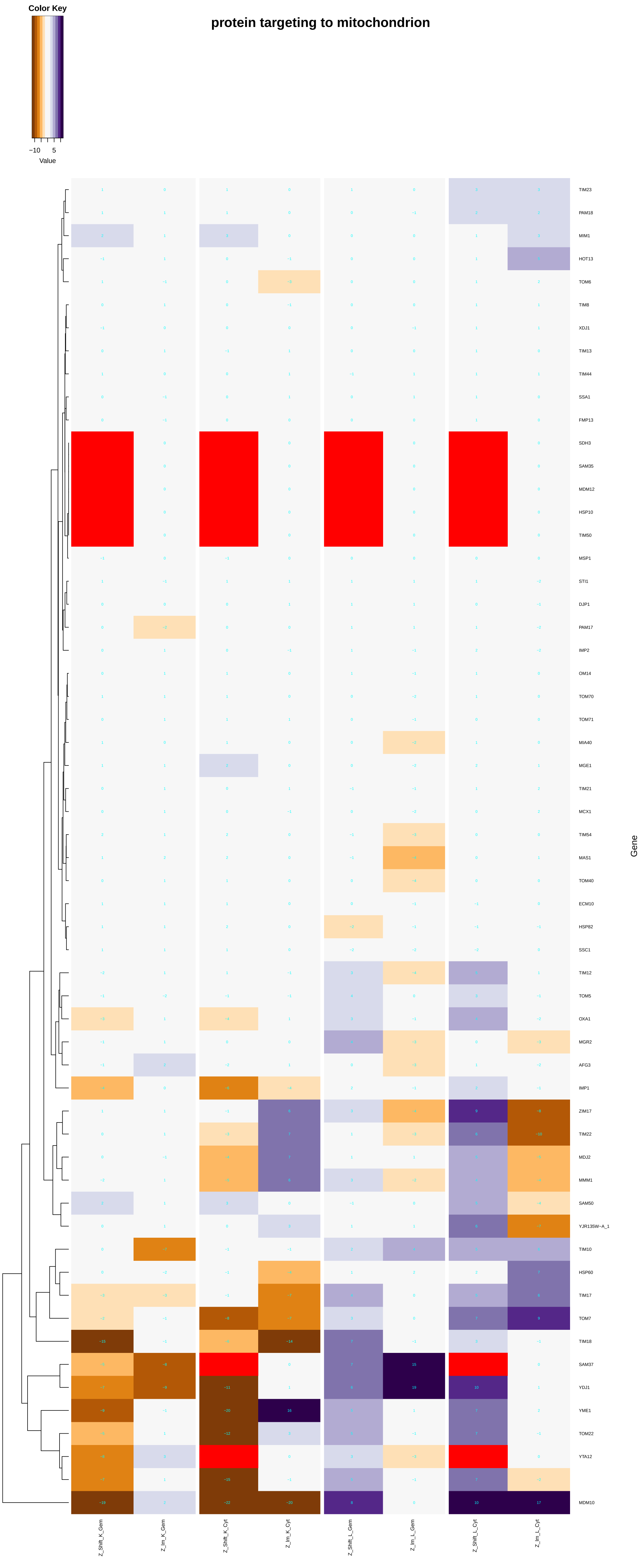

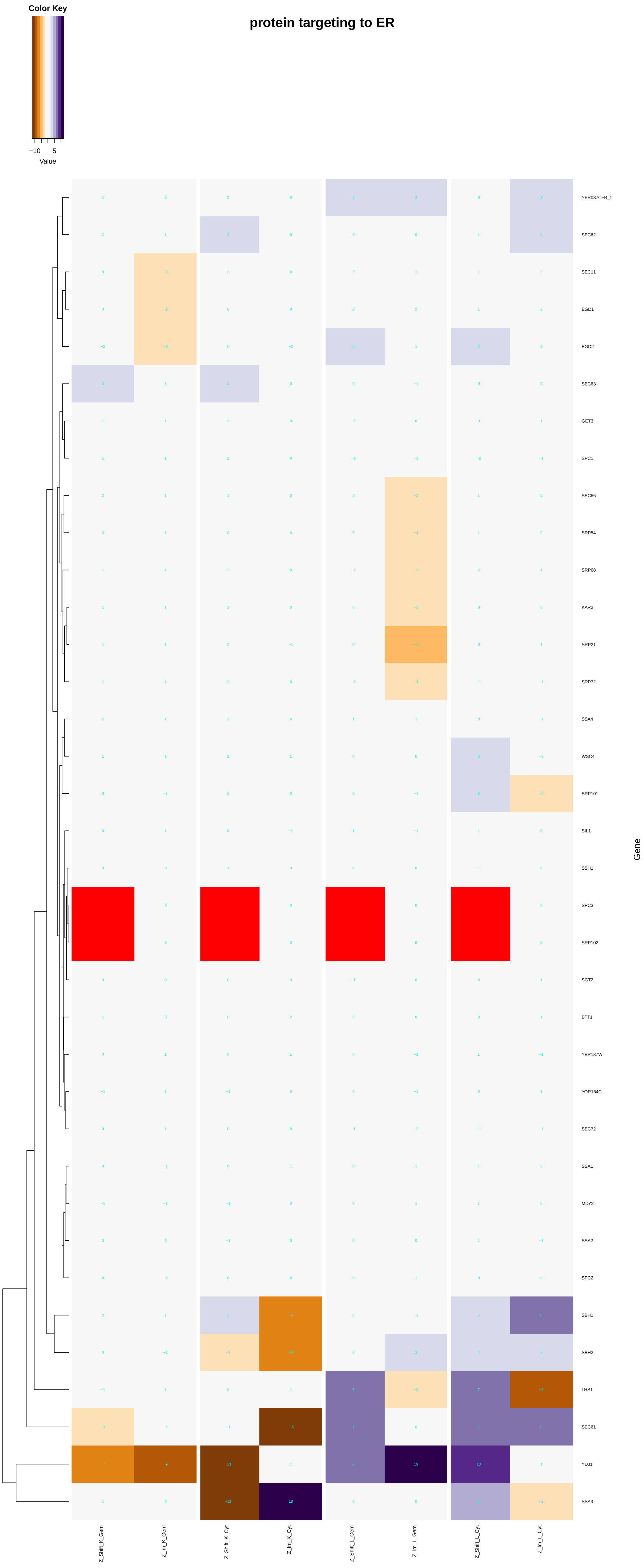

protein targeting to ER
Color Key
−10
5
Value
Gene
YER087C−B_1
SEC62
SEC11
EGD1
EGD2
SEC63
GET3
SPC1
SEC66
SRP54
SRP68
KAR2
SRP21
SRP72
SSA4
WSC4
SRP101
SIL1
SSH1
SPC3
SRP102
SGT2
BTT1
YBR137W
YOR164C
SEC72
SSA1
MDY2
SSA2
SPC2
SBH1
SBH2
LHS1
SEC61
YDJ1
SSA3
Z_Shift_K_Gem
Z_Im_K_Gem
Z_Shift_K_Cyt
Z_Im_K_Cyt
Z_Shift_L_Gem
Z_Im_L_Gem
Z_Shift_L_Cyt
Z_Im_L_Cyt

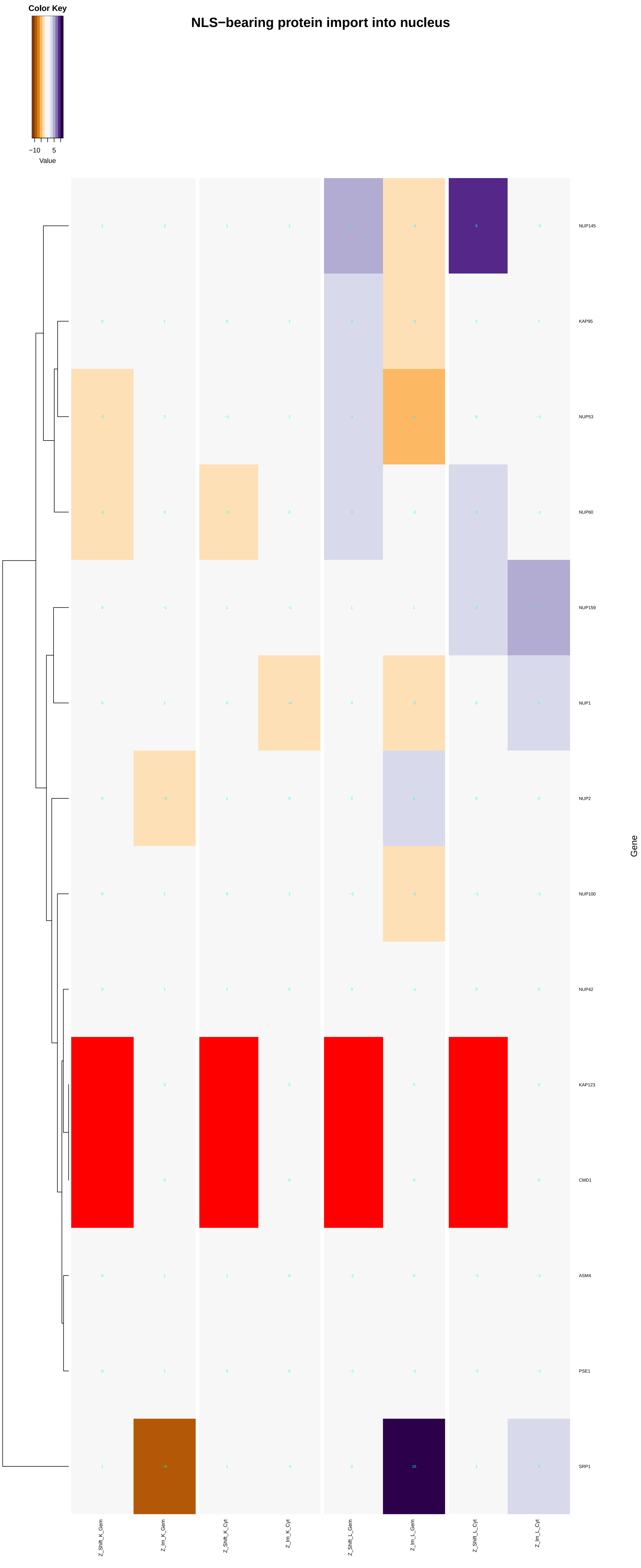

Color Key
−10
5
Value
NLS−bearing protein import into nucleus
NUP145
KAP95
NUP53
NUP60
NUP159
NUP1
NUP2
NUP100
NUP42
KAP123
CMD1
ASM4
PSE1
SRP1
Gene
Z_Shift_K_Gem
Z_Im_K_Gem
Z_Shift_K_Cyt
Z_Im_K_Cyt
Z_Shift_L_Gem
Z_Im_L_Gem
Z_Shift_L_Cyt
Z_Im_L_Cyt

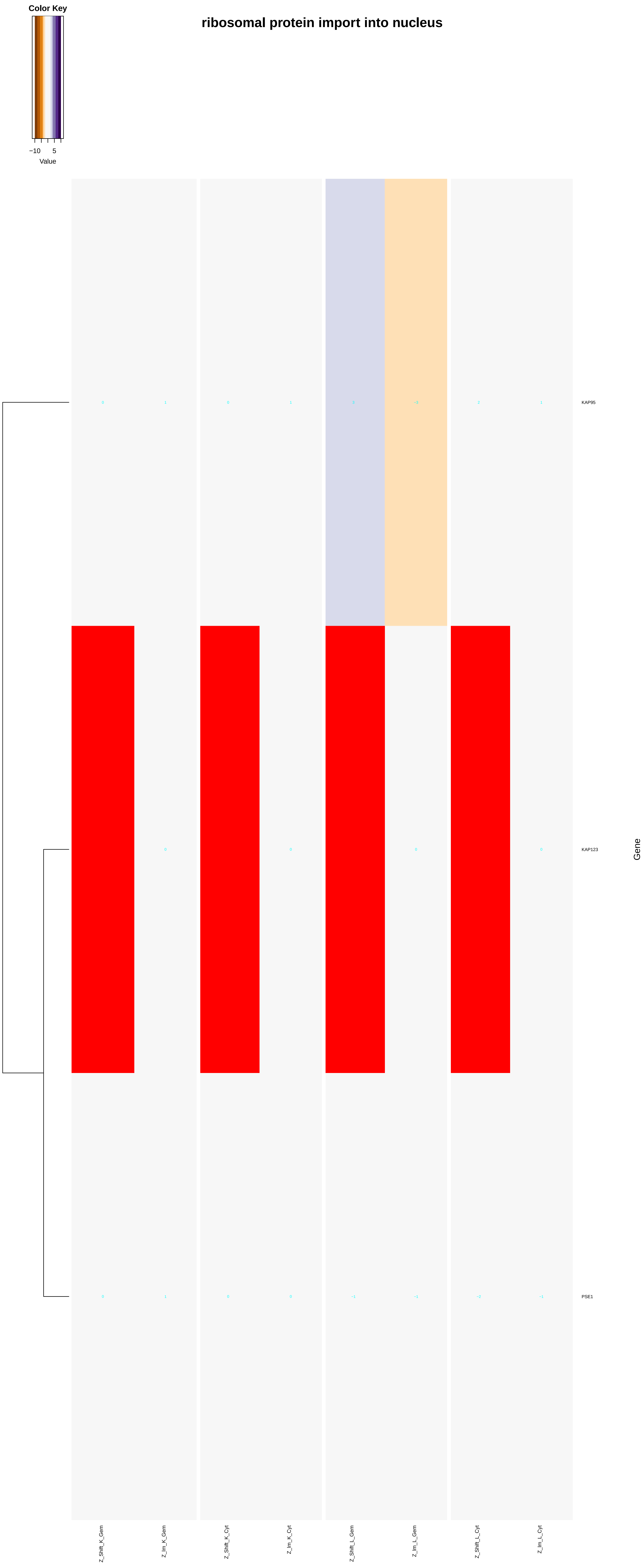
Color Key
−10
5
Value
ribosomal protein import into nucleus
KAP95
KAP123
PSE1
Gene
Z_Shift_K_Gem
Z_Im_K_Gem
Z_Shift_K_Cyt
Z_Im_K_Cyt
Z_Shift_L_Gem
Z_Im_L_Gem
Z_Shift_L_Cyt
Z_Im_L_Cyt

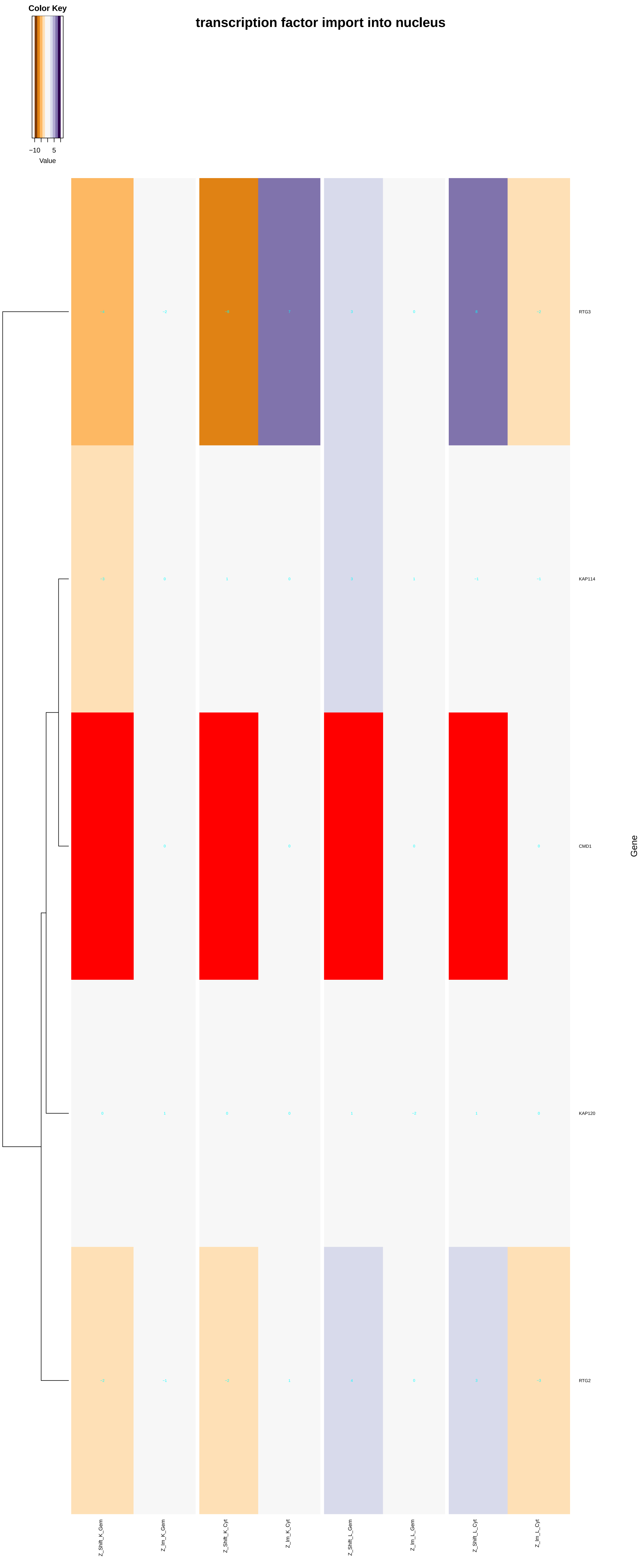
Color Key
−10
5
Value
transcription factor import into nucleus
RTG3
KAP114
CMD1
KAP120
RTG2
Gene
Z_Shift_K_Gem
Z_Im_K_Gem
Z_Shift_K_Cyt
Z_Im_K_Cyt
Z_Shift_L_Gem
Z_Im_L_Gem
Z_Shift_L_Cyt
Z_Im_L_Cyt

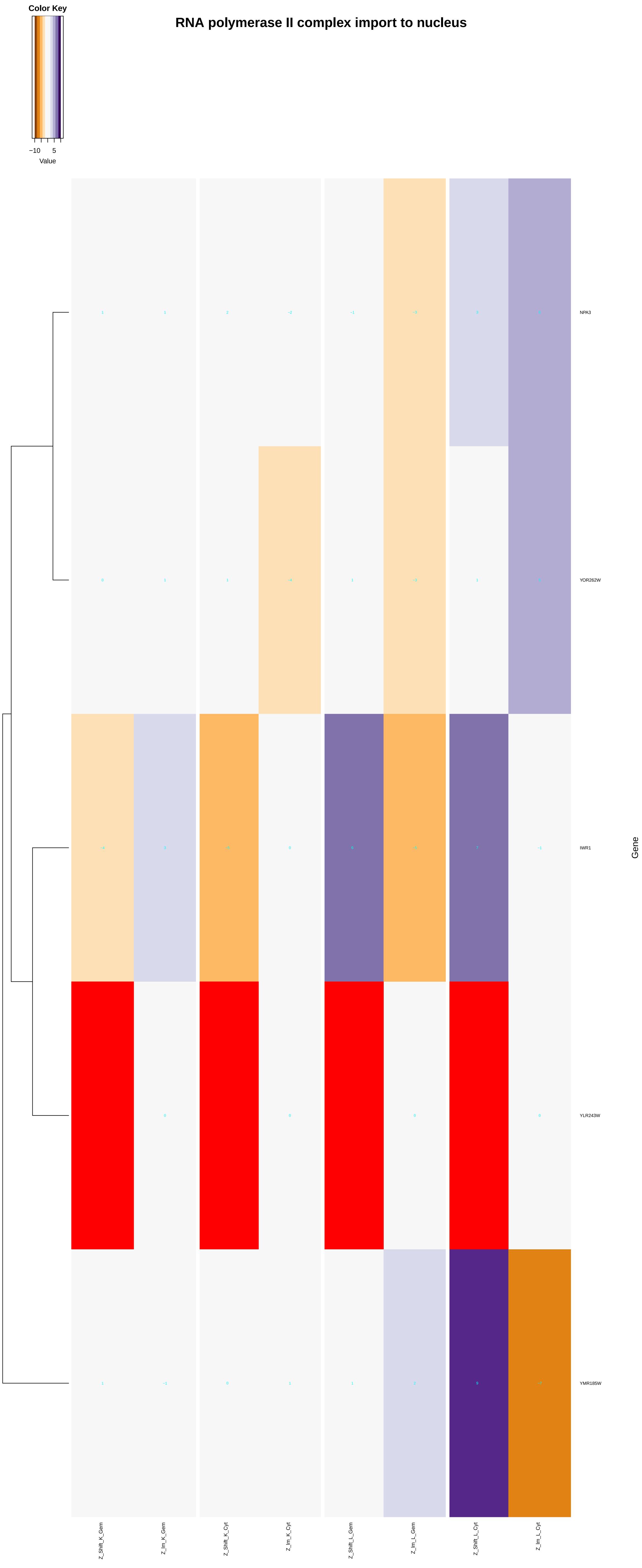
RNA polymerase II complex import to nucleus
Color Key
−10
5
Value
NPA3
YOR262W
IWR1
YLR243W
YMR185W
Gene
Z_Shift_K_Gem
Z_Im_K_Gem
Z_Shift_K_Cyt
Z_Im_K_Cyt
Z_Shift_L_Gem
Z_Im_L_Gem
Z_Shift_L_Cyt
Z_Im_L_Cyt

### retrograde protein transport, ER to cytosol

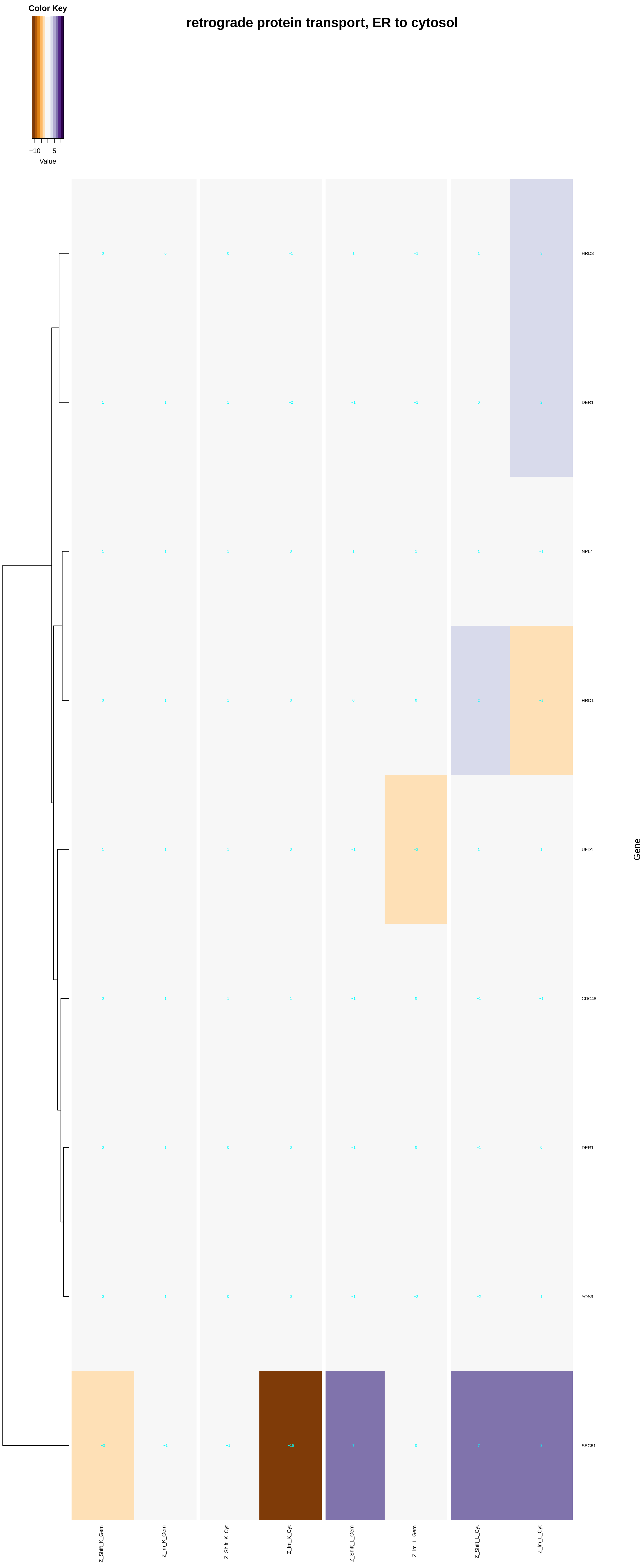

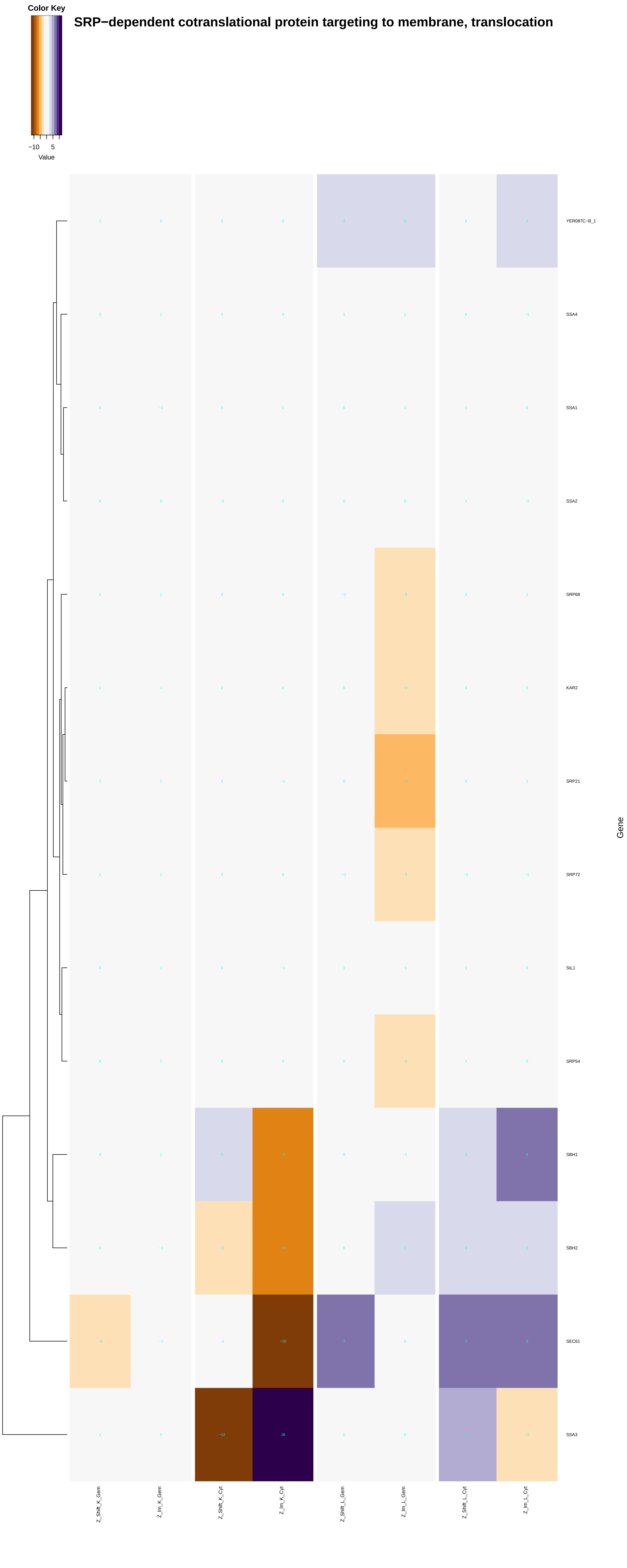
Color Key
−10
5
Value
SRP−dependent cotranslational protein targeting to membrane, translocation
YER087C−B_1
SSA4
SSA1
SSA2
SRP68
KAR2
SRP21
SRP72
SIL1
SRP54
SBH1
SBH2
SEC61
SSA3
Gene
Z_Shift_K_Gem
Z_Im_K_Gem
Z_Shift_K_Cyt
Z_Im_K_Cyt
Z_Shift_L_Gem
Z_Im_L_Gem
Z_Shift_L_Cyt
Z_Im_L_Cyt

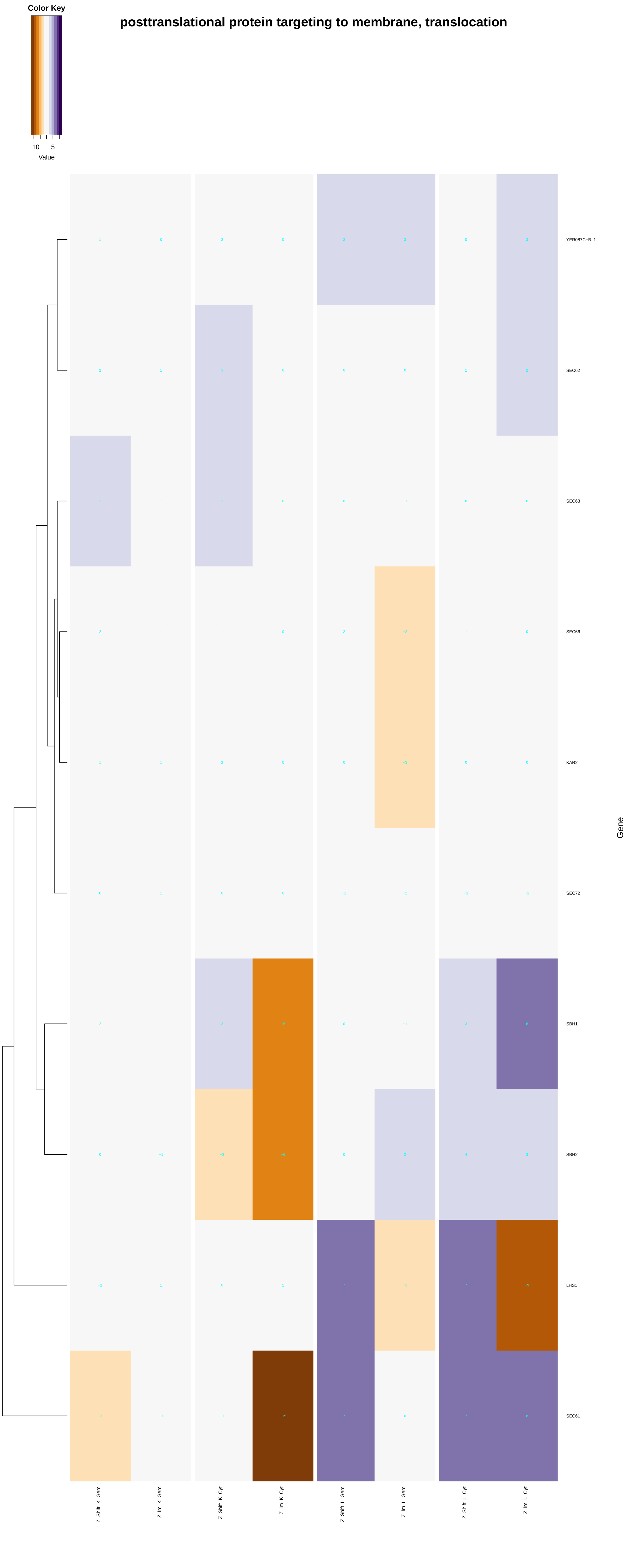
posttranslational protein targeting to membrane, translocation
Color Key
−10
5
Value
YER087C−B_1
SEC62
SEC63
SEC66
KAR2
SEC72
SBH1
SBH2
LHS1
SEC61
Gene
Z_Shift_K_Gem
Z_Im_K_Gem
Z_Shift_K_Cyt
Z_Im_K_Cyt
Z_Shift_L_Gem
Z_Im_L_Gem
Z_Shift_L_Cyt
Z_Im_L_Cyt

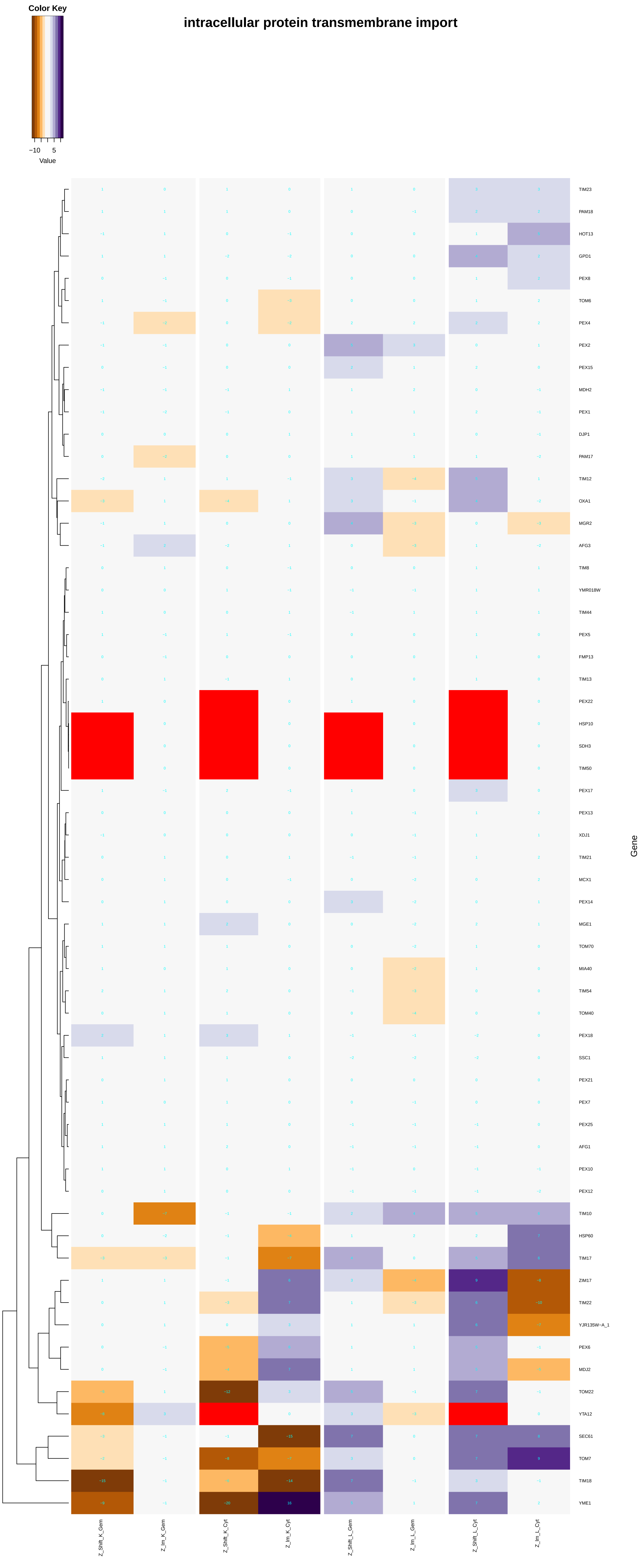

intracellular protein transmembrane import
Color Key
−10
5
Value
Gene
TIM23
PAM18
HOT13
GPD1
PEX8
TOM6
PEX4
PEX2
PEX15
MDH2
PEX1
DJP1
PAM17
TIM12
OXA1
MGR2
AFG3
TIM8
YMR018W
TIM44
PEX5
FMP13
TIM13
PEX22
HSP10
SDH3
TIM50
PEX17
PEX13
XDJ1
TIM21
MCX1
PEX14
MGE1
TOM70
MIA40
TIM54
TOM40
PEX18
SSC1
PEX21
PEX7
PEX25
AFG1
PEX10
PEX12
TIM10
HSP60
TIM17
ZIM17
TIM22
YJR135W−A_1
PEX6
MDJ2
TOM22
YTA12
SEC61
TOM7
TIM18
YME1
Z_Shift_K_Gem
Z_Im_K_Gem
Z_Shift_K_Cyt
Z_Im_K_Cyt
Z_Shift_L_Gem
Z_Im_L_Gem
Z_Shift_L_Cyt
Z_Im_L_Cyt

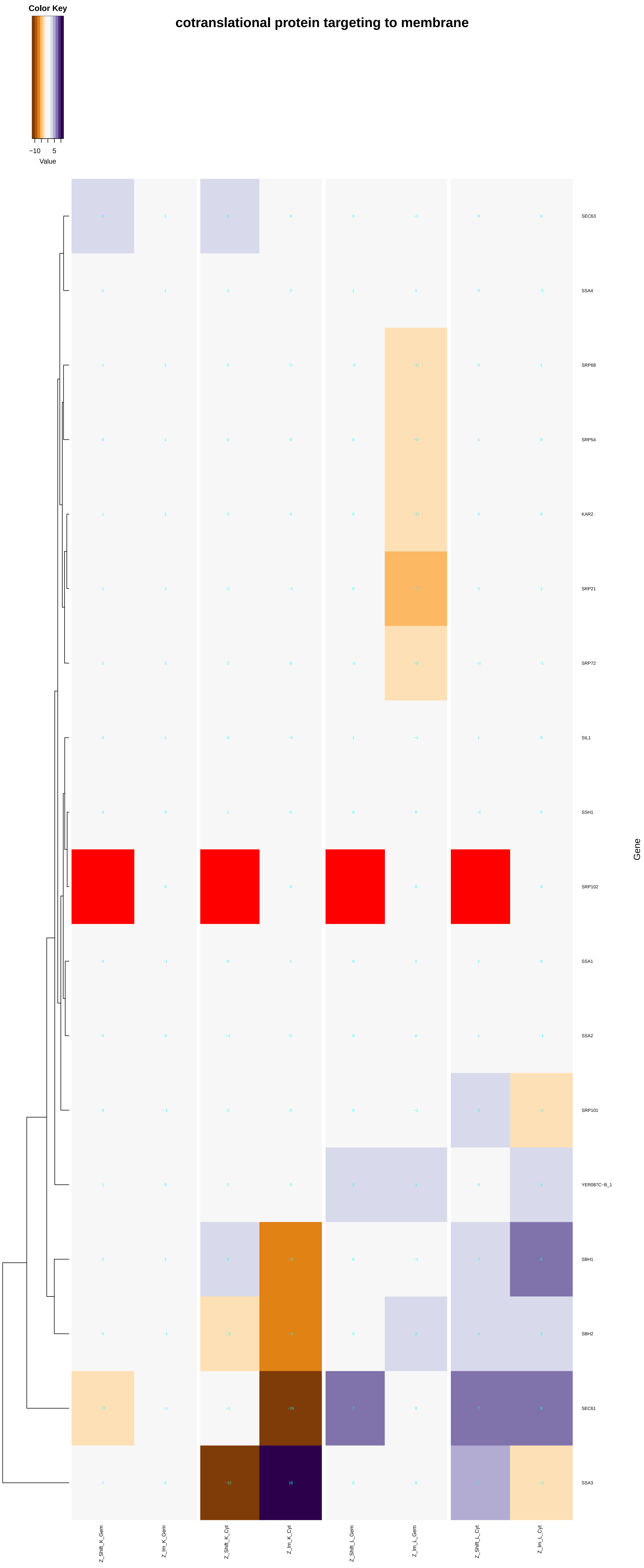
Color Key
−10
5
Value
cotranslational protein targeting to membrane
SEC63
SSA4
SRP68
SRP54
KAR2
SRP21
SRP72
SIL1
SSH1
SRP102
SSA1
SSA2
SRP101
YER087C−B_1
SBH1
SBH2
SEC61
SSA3
Gene
Z_Shift_K_Gem
Z_Im_K_Gem
Z_Shift_K_Cyt
Z_Im_K_Cyt
Z_Shift_L_Gem
Z_Im_L_Gem
Z_Shift_L_Cyt
Z_Im_L_Cyt

posttranslational protein targeting to membrane

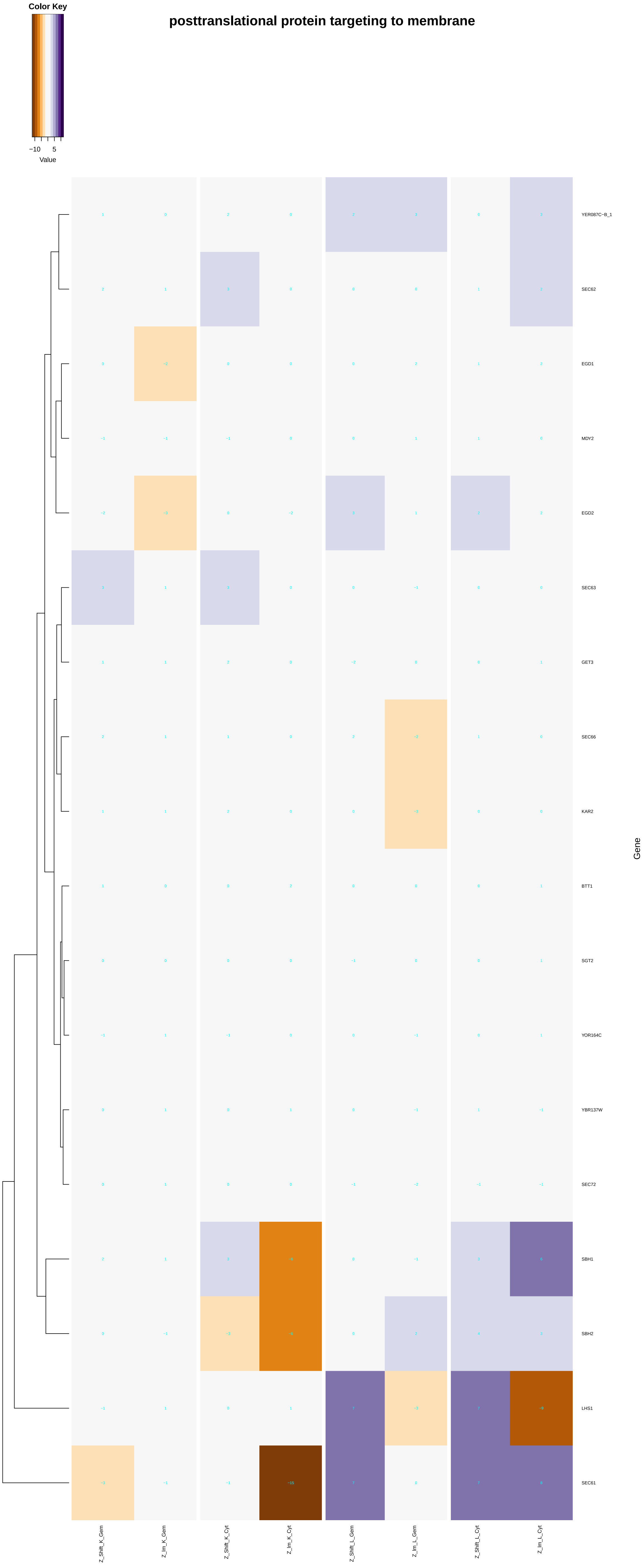

#### protein targeting to nuclear inner membrane

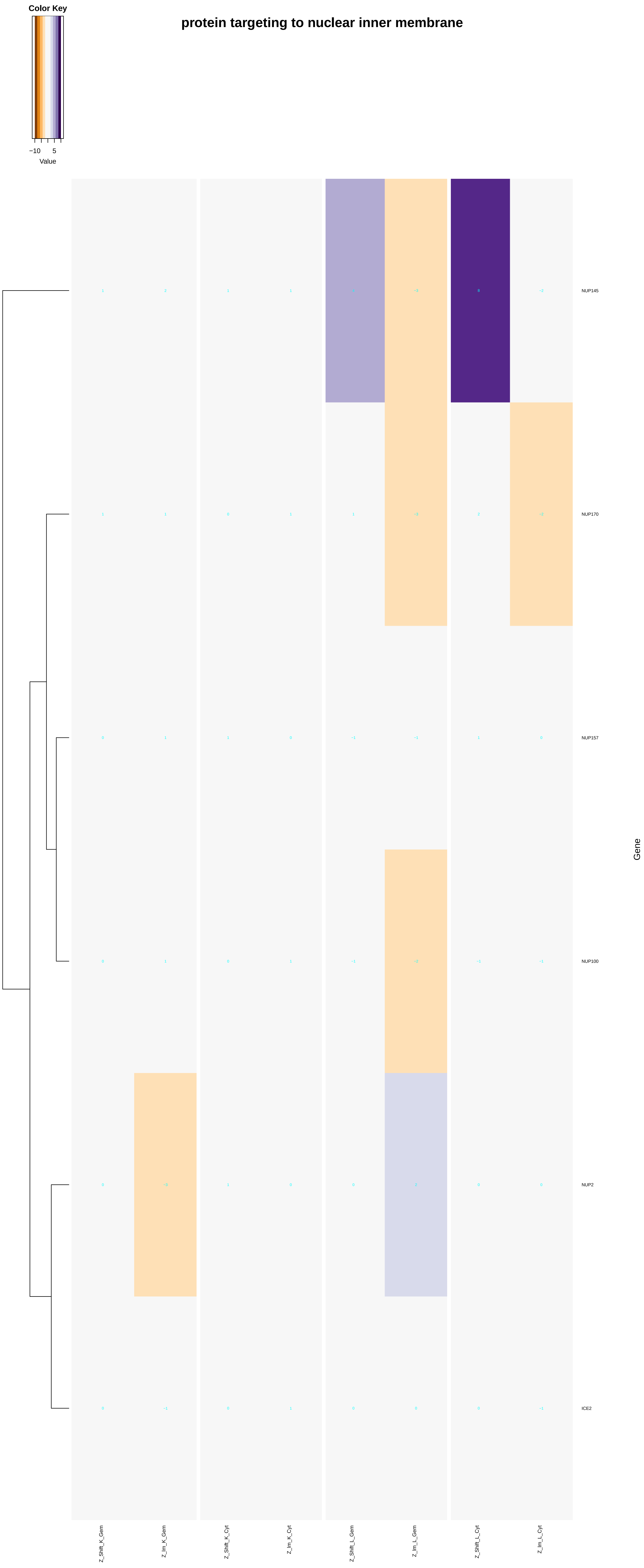

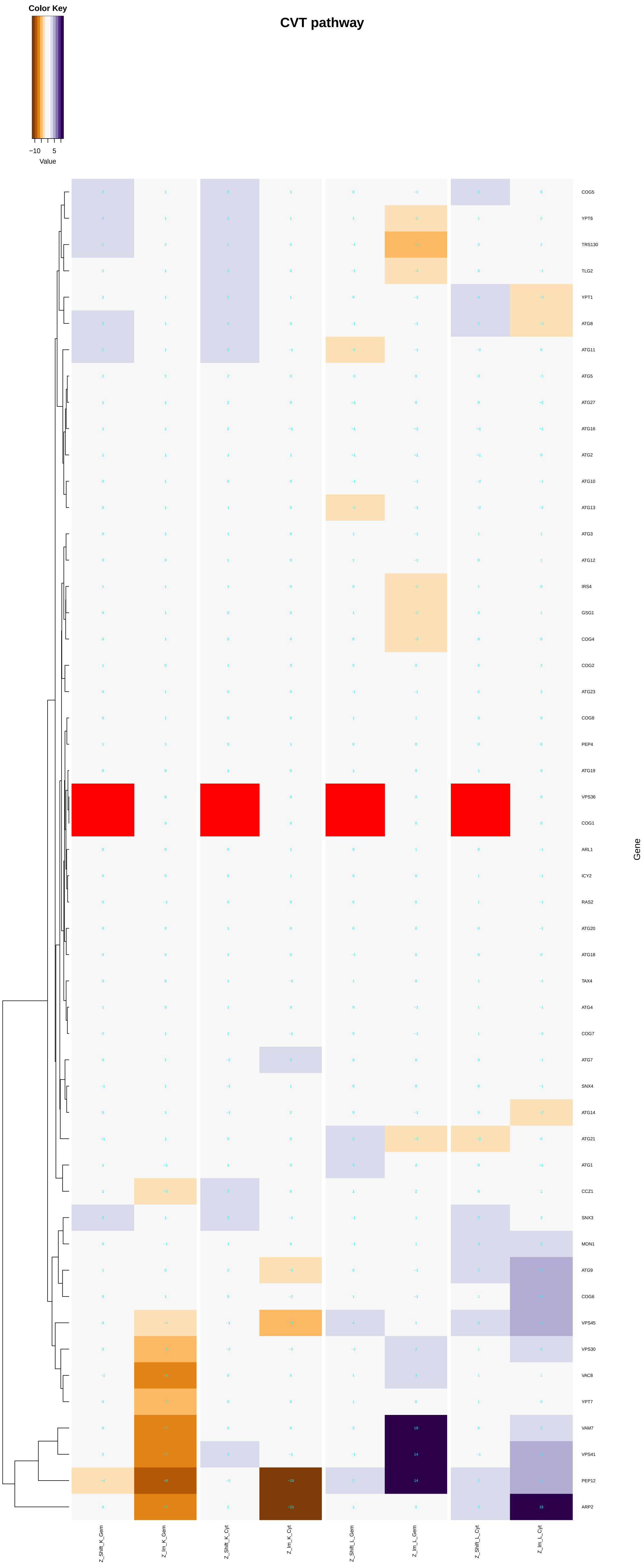

CVT pathway
Color Key
−10
5
Value
COG5
YPT6
TRS130
TLG2
YPT1
ATG8
ATG11
ATG5
ATG27
ATG16
ATG2
ATG10
ATG13
ATG3
ATG12
IRS4
GSG1
COG4
COG2
ATG23
COG8
PEP4
ATG19
VPS36
COG1
ARL1
ICY2
RAS2
ATG20
ATG18
TAX4
ATG4
COG7
ATG7
SNX4
ATG14
ATG21
ATG1
CCZ1
SNX3
MON1
ATG9
COG6
VPS45
VPS30
VAC8
YPT7
VAM7
VPS41
PEP12
ARP2
Gene
Z_Shift_K_Gem
Z_Im_K_Gem
Z_Shift_K_Cyt
Z_Im_K_Cyt
Z_Shift_L_Gem
Z_Im_L_Gem
Z_Shift_L_Cyt
Z_Im_L_Cyt

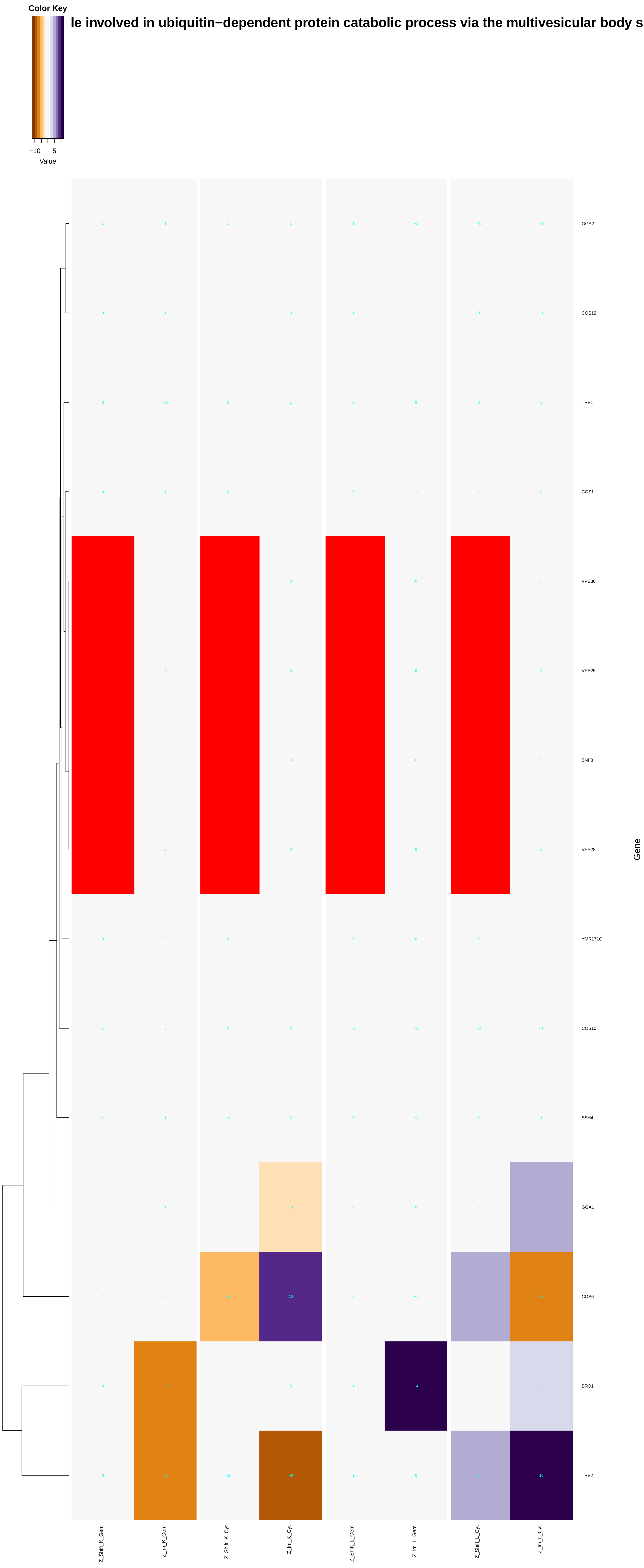
Color Key
−10
5
Value
le involved in ubiquitin−dependent protein catabolic process via the multivesicular body s
GGA2
COS12
TRE1
COS1
VPS36
VPS25
SNF8
VPS28
YMR171C
COS10
SSH4
GGA1
COS6
BRO1
TRE2
Gene
Z_Shift_K_Gem
Z_Im_K_Gem
Z_Shift_K_Cyt
Z_Im_K_Cyt
Z_Shift_L_Gem
Z_Im_L_Gem
Z_Shift_L_Cyt
Z_Im_L_Cyt

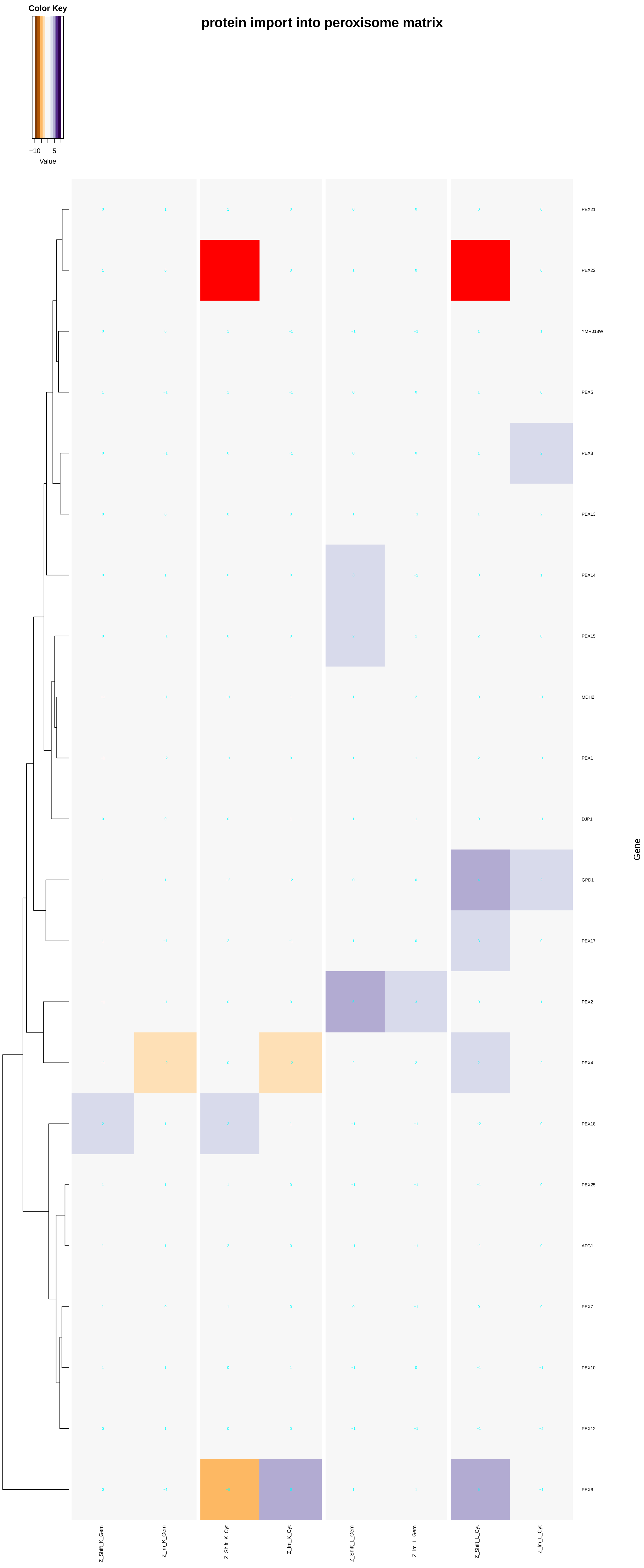

Color Key
−10
5
Value
protein import into peroxisome matrix
PEX21
PEX22
YMR018W
PEX5
PEX8
PEX13
PEX14
PEX15
MDH2
PEX1
DJP1
GPD1
PEX17
PEX2
PEX4
PEX18
PEX25
AFG1
PEX7
PEX10
PEX12
PEX6
Gene
Z_Shift_K_Gem
Z_Im_K_Gem
Z_Shift_K_Cyt
Z_Im_K_Cyt
Z_Shift_L_Gem
Z_Im_L_Gem
Z_Shift_L_Cyt
Z_Im_L_Cyt

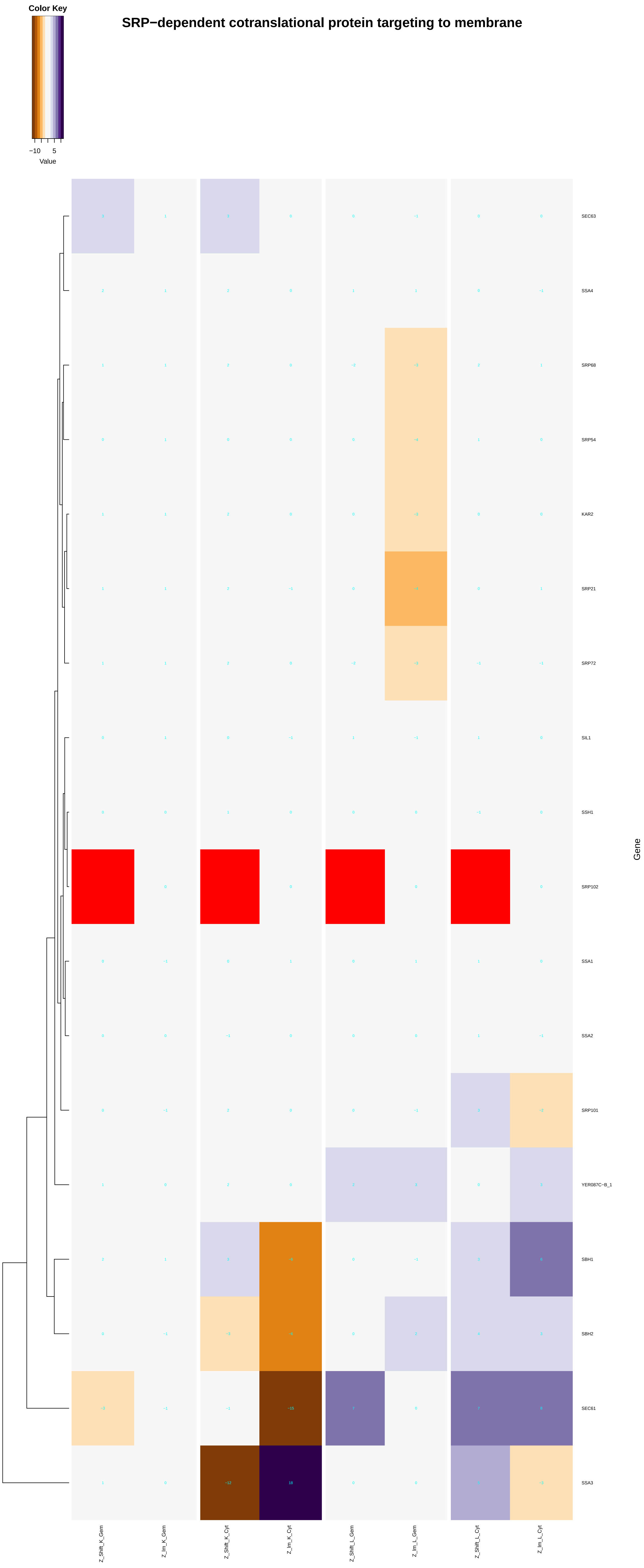
Color Key
−10
5
Value
SRP−dependent cotranslational protein targeting to membrane
SEC63
SSA4
SRP68
SRP54
KAR2
SRP21
SRP72
SIL1
SSH1
SRP102
SSA1
SSA2
SRP101
YER087C−B_1
SBH1
SBH2
SEC61
SSA3
Gene
Z_Shift_K_Gem
Z_Im_K_Gem
Z_Shift_K_Cyt
Z_Im_K_Cyt
Z_Shift_L_Gem
Z_Im_L_Gem
Z_Shift_L_Cyt
Z_Im_L_Cyt

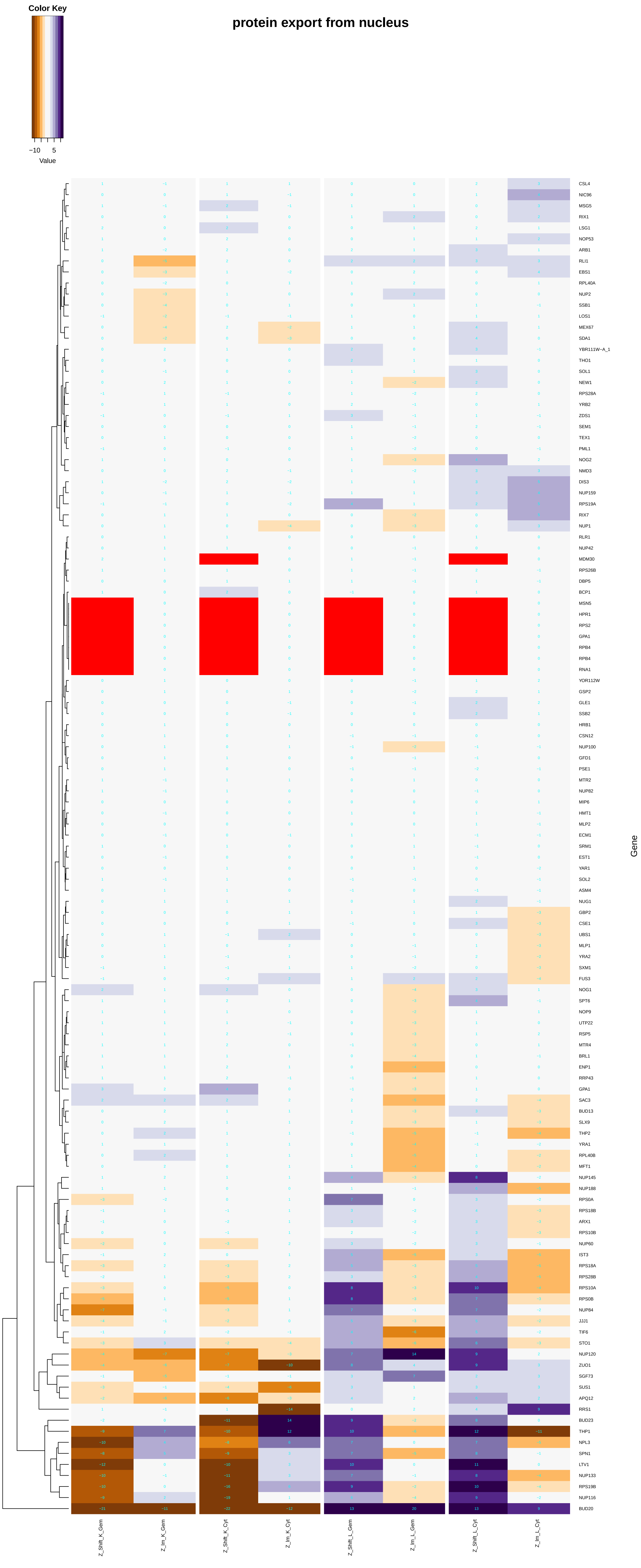
protein export from nucleus
Color Key
−10
5
Value
Gene
CSL4
NIC96
MSG5
RIX1
LSG1
NOP53
ARB1
RLI1
EBS1
RPL40A
NUP2
SSB1
LOS1
MEX67
SDA1
YBR111W−A_1
THO1
SOL1
NEW1
RPS28A
YRB2
ZDS1
SEM1
TEX1
PML1
NOG2
NMD3
DIS3
NUP159
RPS19A
RIX7
NUP1
RLR1
NUP42
MDM30
RPS26B
DBP5
BCP1
MSN5
HPR1
RPS2
GPA1
RPB4
RPB4
RNA1
YOR112W
GSP2
GLE1
SSB2
HRB1
CSN12
NUP100
GFD1
PSE1
MTR2
NUP82
MIP6
HMT1
MLP2
ECM1
SRM1
EST1
YAR1
SOL2
ASM4
NUG1
GBP2
CSE1
UBS1
MLP1
YRA2
SXM1
FUS3
NOG1
SPT6
NOP9
UTP22
RSP5
MTR4
BRL1
ENP1
RRP43
GPA1
SAC3
BUD13
SLX9
THP2
YRA1
RPL40B
MFT1
NUP145
NUP188
RPS0A
RPS18B
ARX1
RPS10B
NUP60
IST3
RPS18A
RPS28B
RPS10A
RPS0B
NUP84
JJJ1
TIF6
STO1
NUP120
ZUO1
SGF73
SUS1
APQ12
RRS1
BUD23
THP1
NPL3
SPN1
LTV1
NUP133
RPS19B
NUP116
BUD20
Z_Shift_K_Gem
Z_Im_K_Gem
Z_Shift_K_Cyt
Z_Im_K_Cyt
Z_Shift_L_Gem
Z_Im_L_Gem
Z_Shift_L_Cyt
Z_Im_L_Cyt

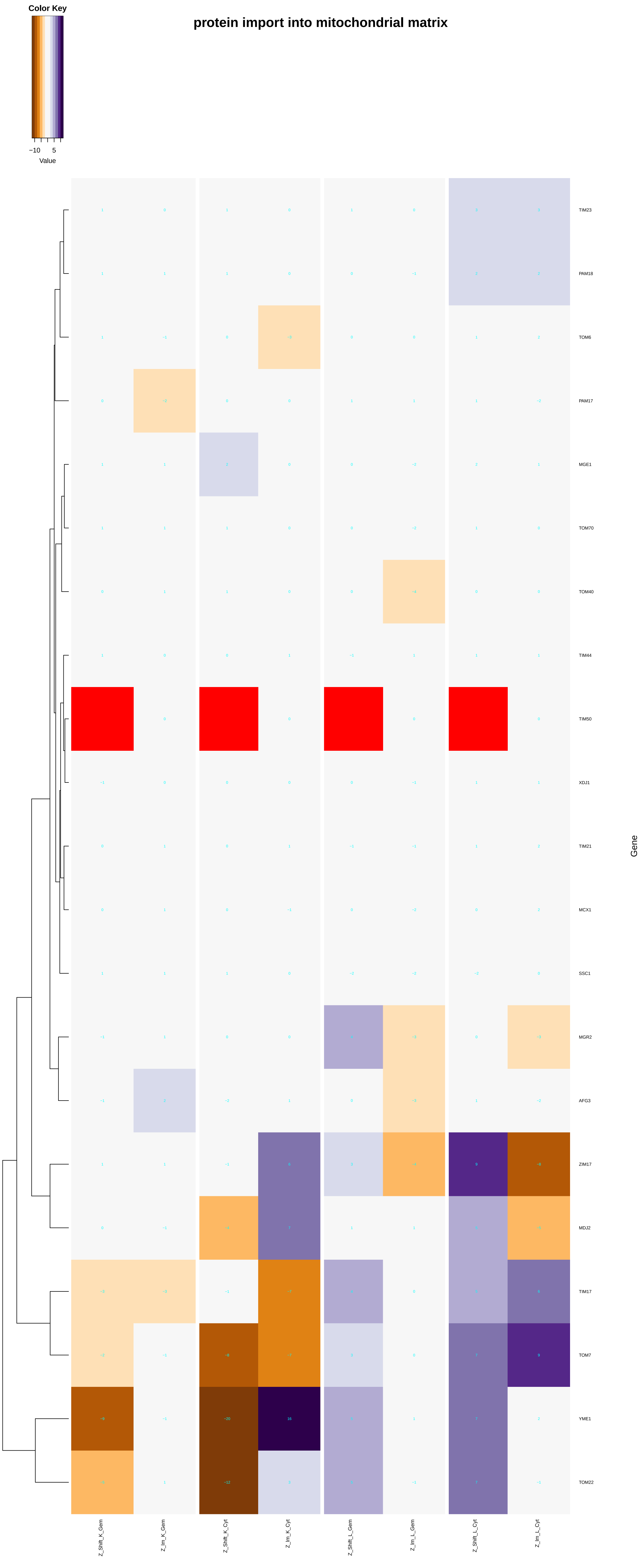

Color Key
−10
5
Value
protein import into mitochondrial matrix
TIM23
PAM18
TOM6
PAM17
MGE1
TOM70
TOM40
TIM44
TIM50
XDJ1
TIM21
MCX1
SSC1
MGR2
AFG3
ZIM17
MDJ2
TIM17
TOM7
YME1
TOM22
Gene
Z_Shift_K_Gem
Z_Im_K_Gem
Z_Shift_K_Cyt
Z_Im_K_Cyt
Z_Shift_L_Gem
Z_Im_L_Gem
Z_Shift_L_Cyt
Z_Im_L_Cyt

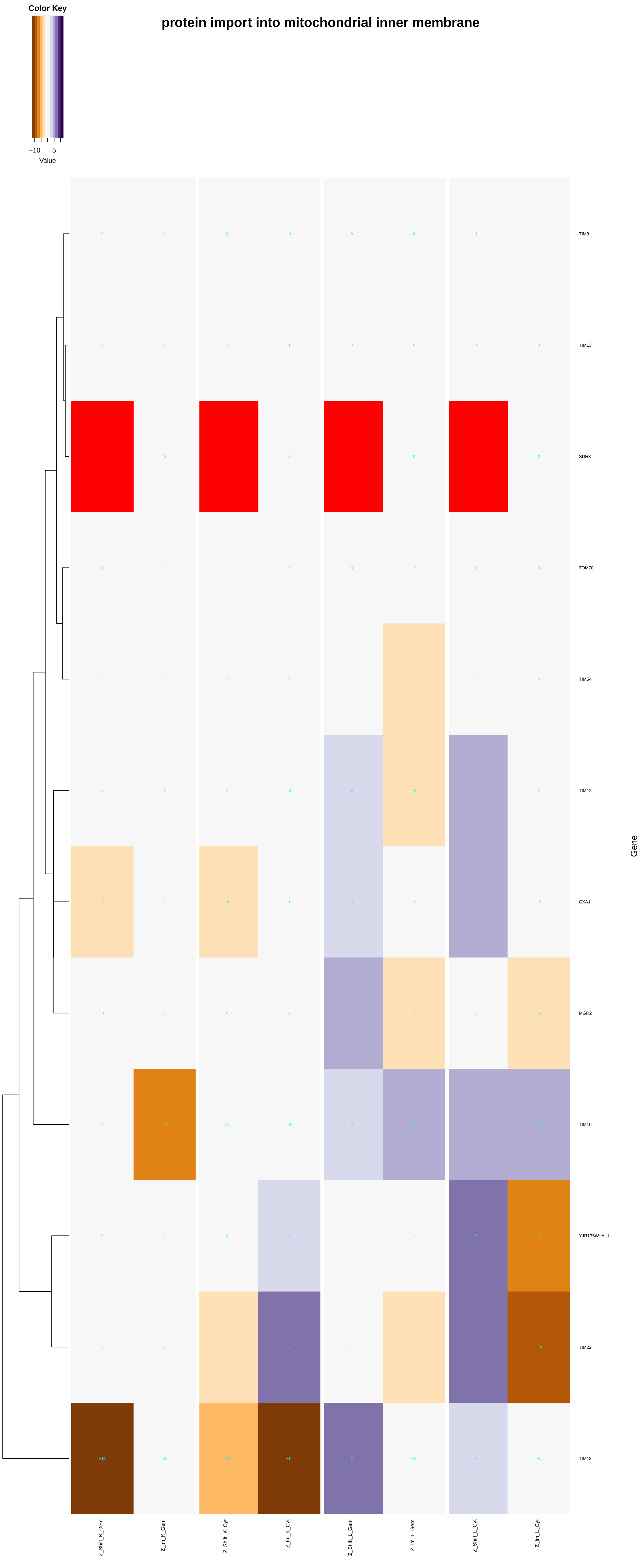
Color Key
−10
5
Value
protein import into mitochondrial inner membrane
TIM8
TIM13
SDH3
TOM70
TIM54
TIM12
OXA1
MGR2
TIM10
YJR135W−A_1
TIM22
TIM18
Gene
Z_Shift_K_Gem
Z_Im_K_Gem
Z_Shift_K_Cyt
Z_Im_K_Cyt
Z_Shift_L_Gem
Z_Im_L_Gem
Z_Shift_L_Cyt
Z_Im_L_Cyt

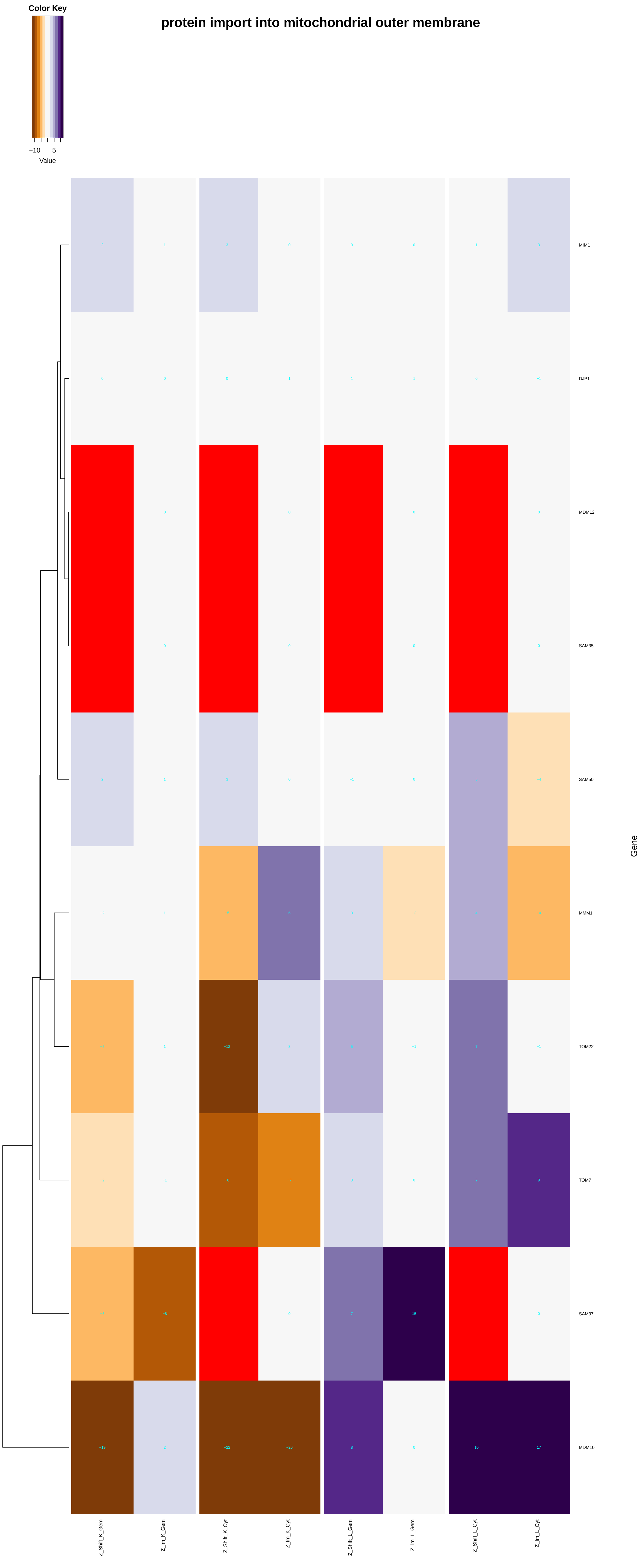

Color Key
−10
5
Value
protein import into mitochondrial outer membrane
MIM1
DJP1
MDM12
SAM35
SAM50
MMM1
TOM22
TOM7
SAM37
MDM10
Gene
Z_Shift_K_Gem
Z_Im_K_Gem
Z_Shift_K_Cyt
Z_Im_K_Cyt
Z_Shift_L_Gem
Z_Im_L_Gem
Z_Shift_L_Cyt
Z_Im_L_Cyt

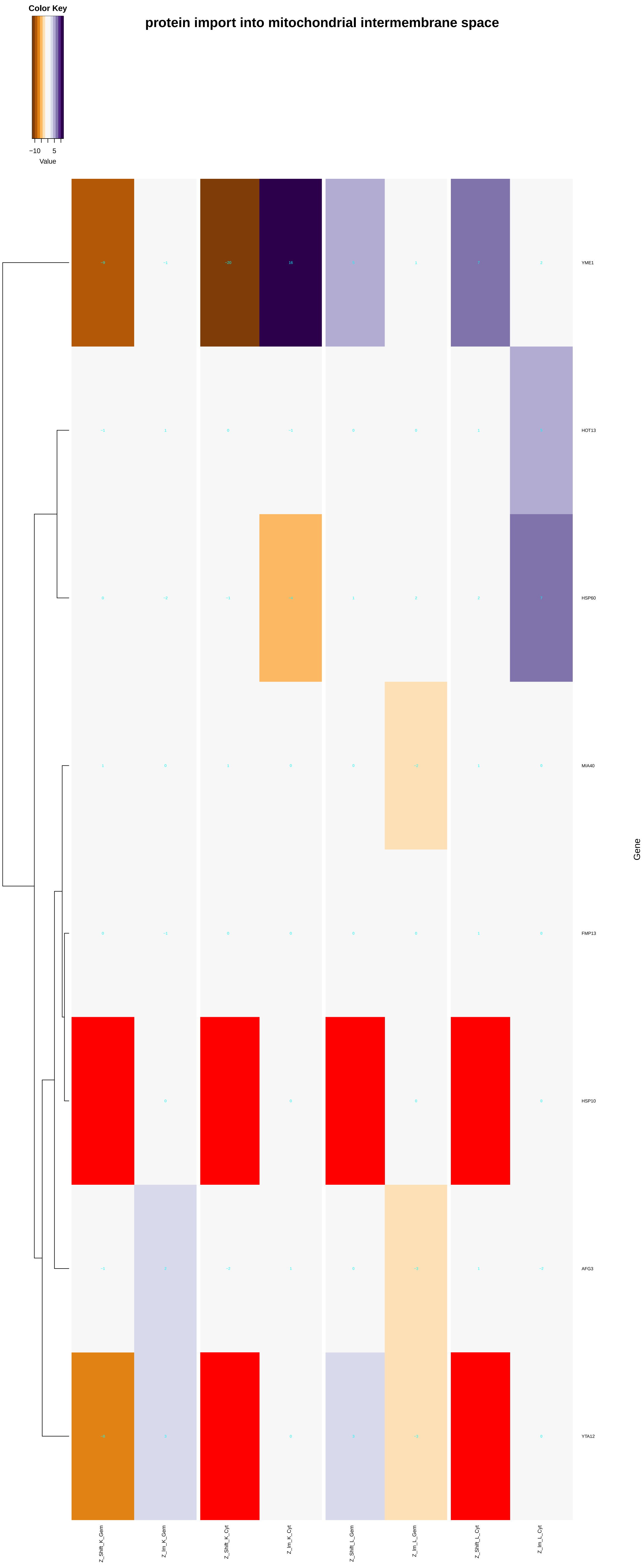

protein import into mitochondrial intermembrane space
Color Key
−10
5
Value
YME1
HOT13
HSP60
MIA40
FMP13
HSP10
AFG3
YTA12
Gene
Z_Shift_K_Gem
Z_Im_K_Gem
Z_Shift_K_Cyt
Z_Im_K_Cyt
Z_Shift_L_Gem
Z_Im_L_Gem
Z_Shift_L_Cyt
Z_Im_L_Cyt

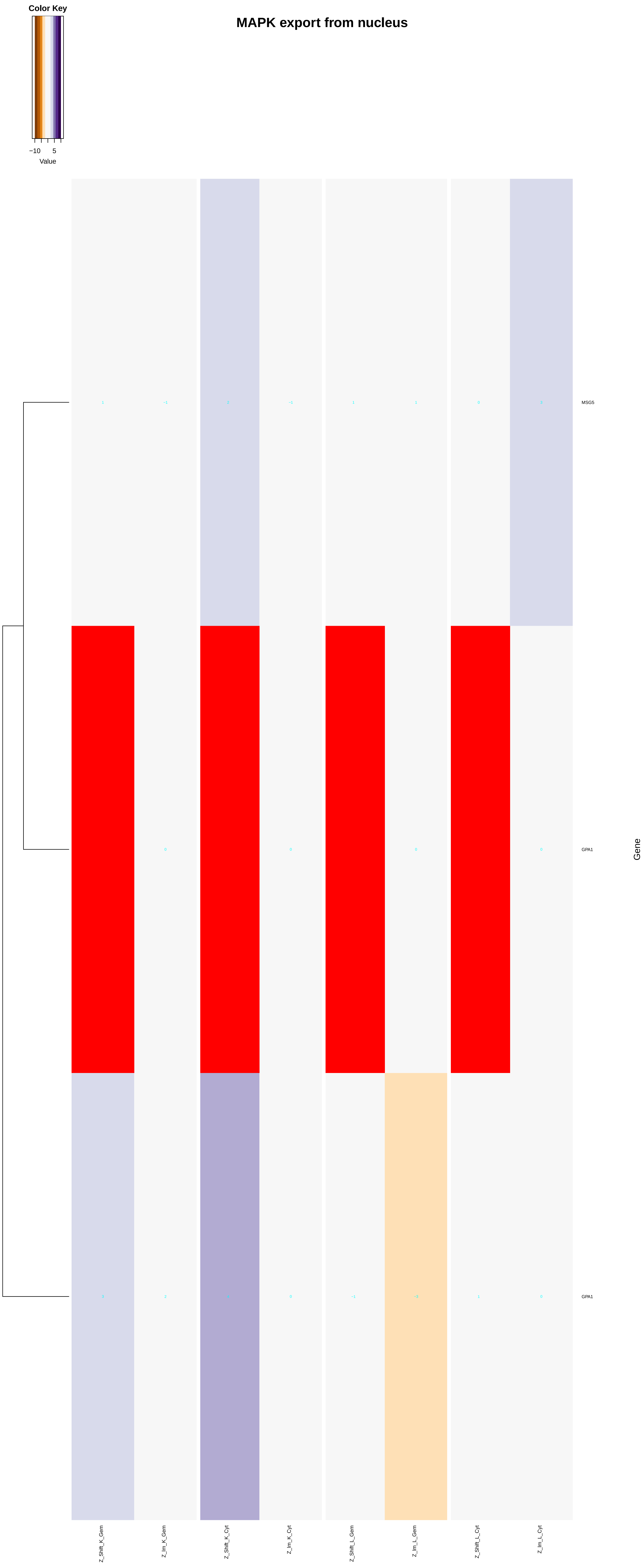

Color Key
−10
5
Value
MAPK export from nucleus
MSG5
GPA1
GPA1
Gene
Z_Shift_K_Gem
Z_Im_K_Gem
Z_Shift_K_Cyt
Z_Im_K_Cyt
Z_Shift_L_Gem
Z_Im_L_Gem
Z_Shift_L_Cyt
Z_Im_L_Cyt

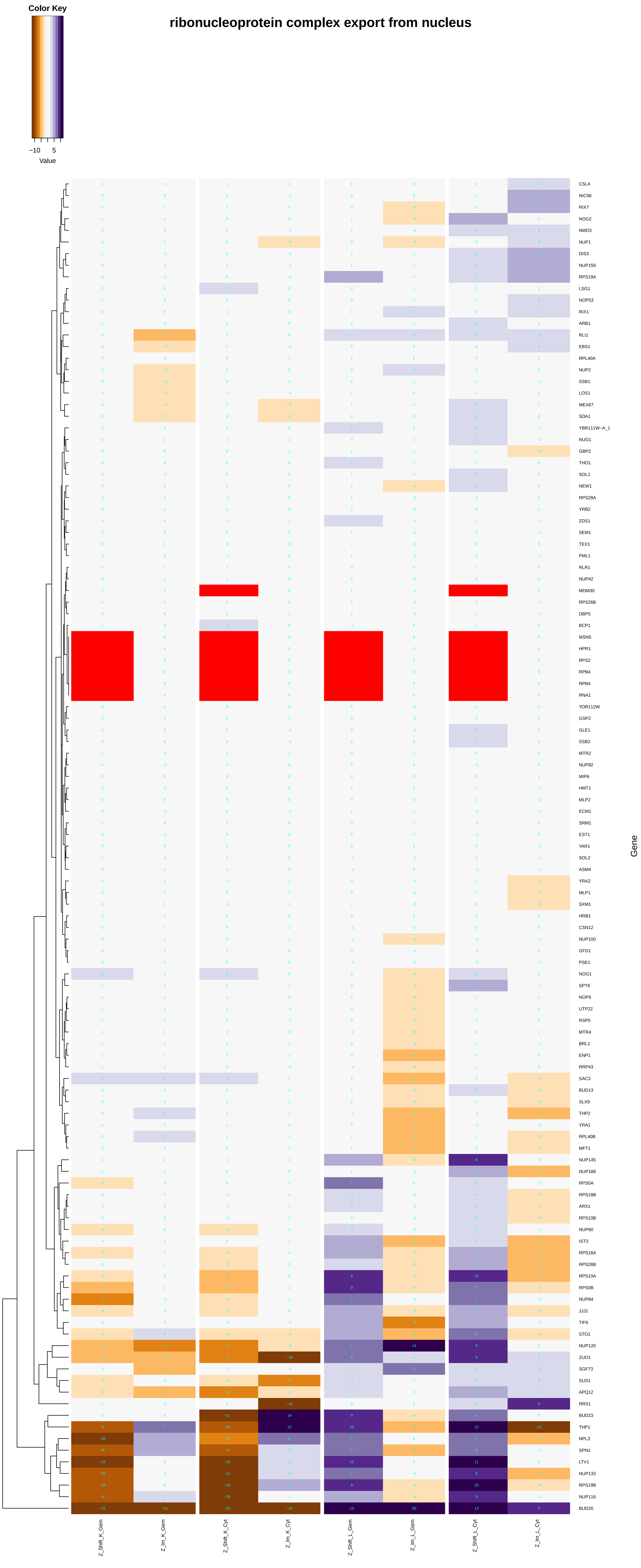

Color Key
−10
5
Value
ribonucleoprotein complex export from nucleus
CSL4
NIC96
RIX7
NOG2
NMD3
NUP1
DIS3
NUP159
RPS19A
LSG1
NOP53
RIX1
ARB1
RLI1
EBS1
RPL40A
NUP2
SSB1
LOS1
MEX67
SDA1
YBR111W−A_1
NUG1
GBP2
THO1
SOL1
NEW1
RPS28A
YRB2
ZDS1
SEM1
TEX1
PML1
RLR1
NUP42
MDM30
RPS26B
DBP5
BCP1
MSN5
HPR1
RPS2
RPB4
RPB4
RNA1
YOR112W
GSP2
GLE1
SSB2
MTR2
NUP82
MIP6
HMT1
MLP2
ECM1
SRM1
EST1
YAR1
SOL2
ASM4
YRA2
MLP1
SXM1
HRB1
CSN12
NUP100
GFD1
PSE1
NOG1
SPT6
NOP9
UTP22
RSP5
MTR4
BRL1
ENP1
RRP43
SAC3
BUD13
SLX9
THP2
YRA1
RPL40B
MFT1
NUP145
NUP188
RPS0A
RPS18B
ARX1
RPS10B
NUP60
IST3
RPS18A
RPS28B
RPS10A
RPS0B
NUP84
JJJ1
TIF6
STO1
NUP120
ZUO1
SGF73
SUS1
APQ12
RRS1
BUD23
THP1
NPL3
SPN1
LTV1
NUP133
RPS19B
NUP116
BUD20
Gene
Z_Shift_K_Gem
Z_Im_K_Gem
Z_Shift_K_Cyt
Z_Im_K_Cyt
Z_Shift_L_Gem
Z_Im_L_Gem
Z_Shift_L_Cyt
Z_Im_L_Cyt

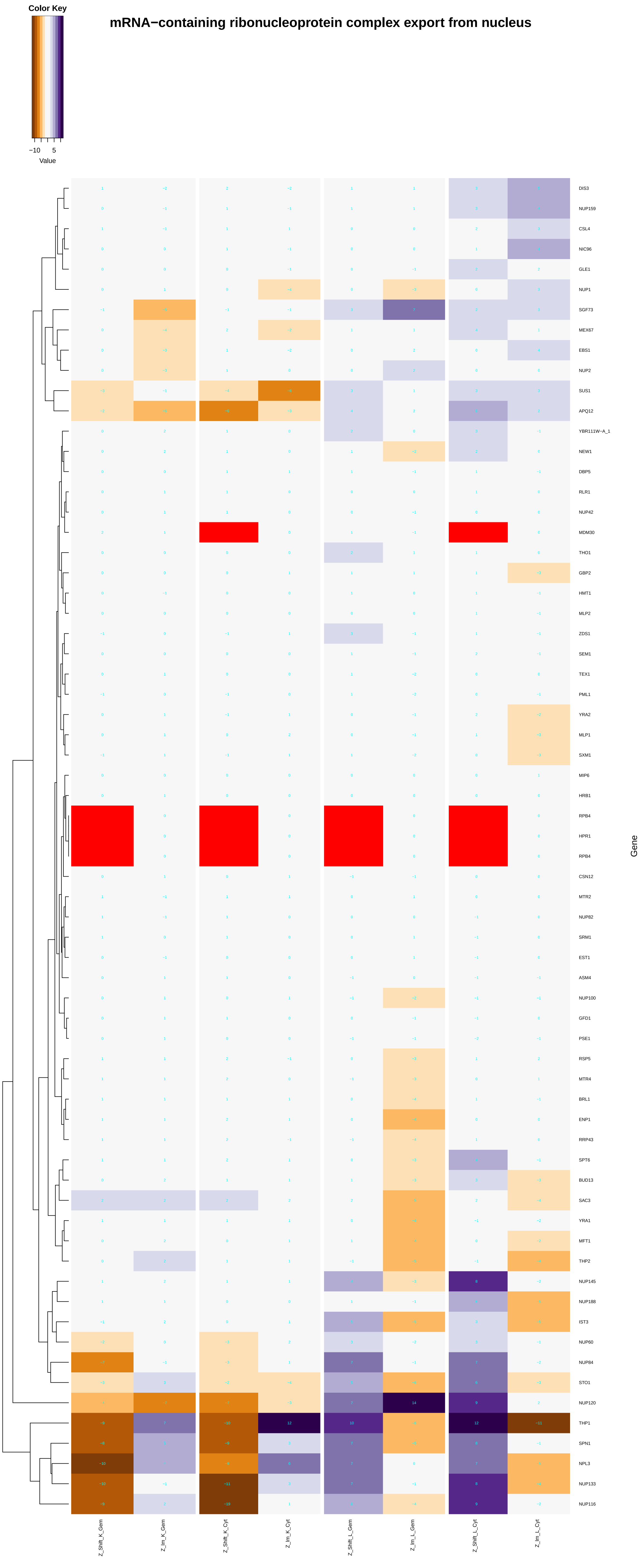
mRNA−containing ribonucleoprotein complex export from nucleus
Color Key
−10
5
Value
DIS3
NUP159
CSL4
NIC96
GLE1
NUP1
SGF73
MEX67
EBS1
NUP2
SUS1
APQ12
YBR111W−A_1
NEW1
DBP5
RLR1
NUP42
MDM30
THO1
GBP2
HMT1
MLP2
ZDS1
SEM1
TEX1
PML1
YRA2
MLP1
SXM1
MIP6
HRB1
RPB4
HPR1
RPB4
CSN12
MTR2
NUP82
SRM1
EST1
ASM4
NUP100
GFD1
PSE1
RSP5
MTR4
BRL1
ENP1
RRP43
SPT6
BUD13
SAC3
YRA1
MFT1
THP2
NUP145
NUP188
IST3
NUP60
NUP84
STO1
NUP120
THP1
SPN1
NPL3
NUP133
NUP116
Gene
Z_Shift_K_Gem
Z_Im_K_Gem
Z_Shift_K_Cyt
Z_Im_K_Cyt
Z_Shift_L_Gem
Z_Im_L_Gem
Z_Shift_L_Cyt
Z_Im_L_Cyt

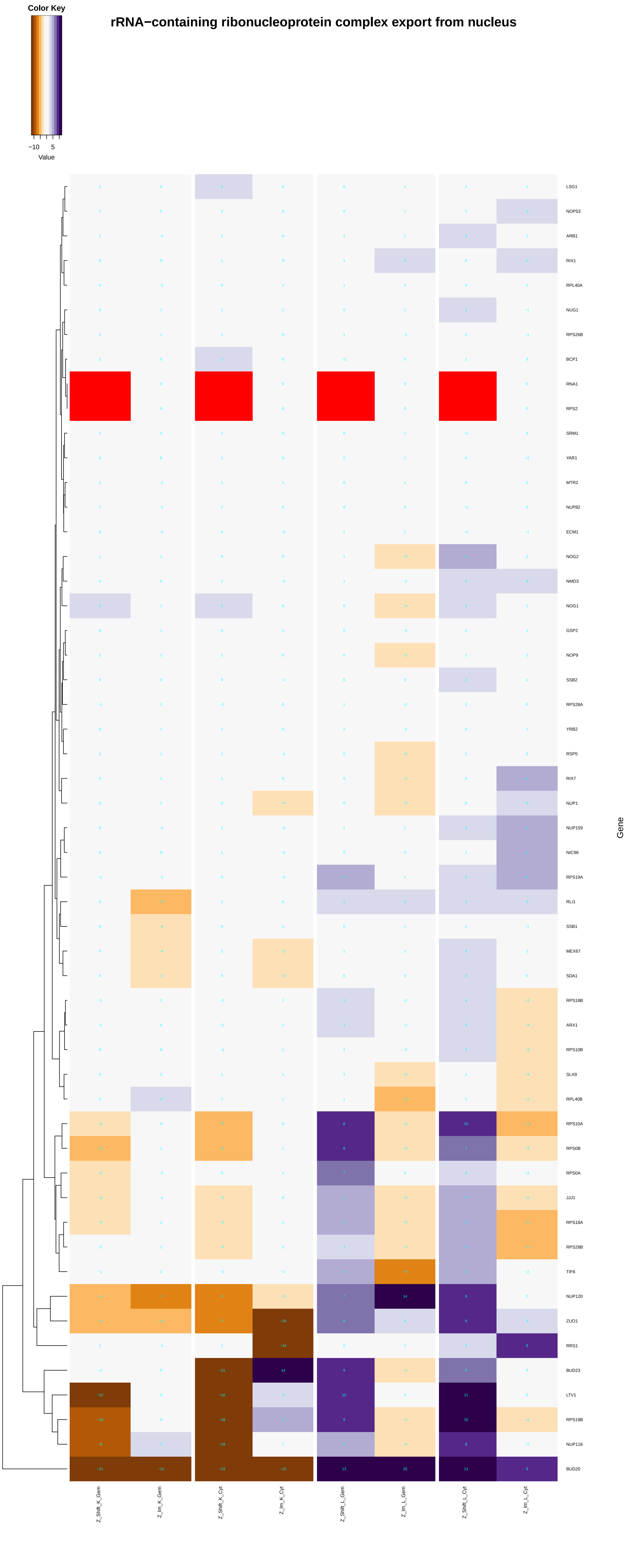
rRNA−containing ribonucleoprotein complex export from nucleus
Color Key
−10
5
Value
LSG1
NOP53
ARB1
RIX1
RPL40A
NUG1
RPS26B
BCP1
RNA1
RPS2
SRM1
YAR1
MTR2
NUP82
ECM1
NOG2
NMD3
NOG1
GSP2
NOP9
SSB2
RPS28A
YRB2
RSP5
RIX7
NUP1
NUP159
NIC96
RPS19A
RLI1
SSB1
MEX67
SDA1
RPS18B
ARX1
RPS10B
SLX9
RPL40B
RPS10A
RPS0B
RPS0A
JJJ1
RPS18A
RPS28B
TIF6
NUP120
ZUO1
RRS1
BUD23
LTV1
RPS19B
NUP116
BUD20
Gene
Z_Shift_K_Gem
Z_Im_K_Gem
Z_Shift_K_Cyt
Z_Im_K_Cyt
Z_Shift_L_Gem
Z_Im_L_Gem
Z_Shift_L_Cyt
Z_Im_L_Cyt

### tRNA−containing ribonucleoprotein complex export from nucleus

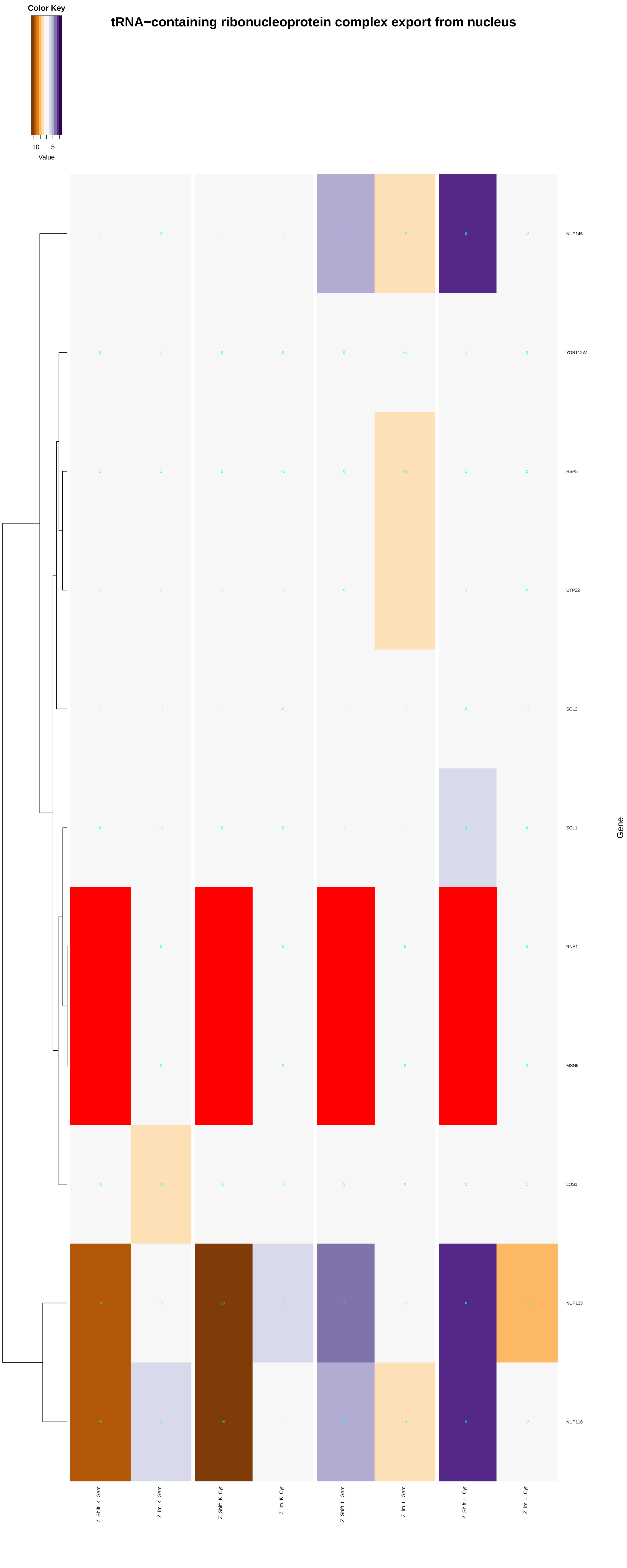

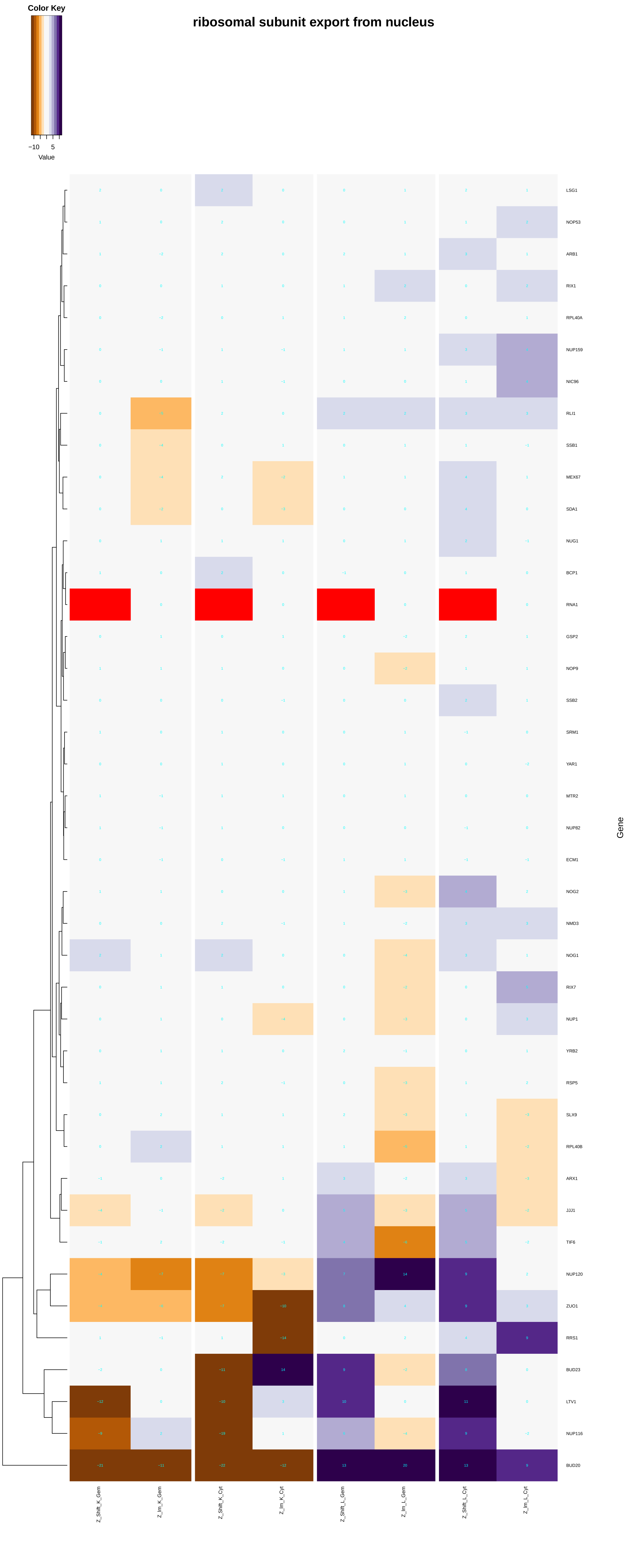

Color Key
−10
5
Value
ribosomal subunit export from nucleus
LSG1
NOP53
ARB1
RIX1
RPL40A
NUP159
NIC96
RLI1
SSB1
MEX67
SDA1
NUG1
BCP1
RNA1
GSP2
NOP9
SSB2
SRM1
YAR1
MTR2
NUP82
ECM1
NOG2
NMD3
NOG1
RIX7
NUP1
YRB2
RSP5
SLX9
RPL40B
ARX1
JJJ1
TIF6
NUP120
ZUO1
RRS1
BUD23
LTV1
NUP116
BUD20
Gene
Z_Shift_K_Gem
Z_Im_K_Gem
Z_Shift_K_Cyt
Z_Im_K_Cyt
Z_Shift_L_Gem
Z_Im_L_Gem
Z_Shift_L_Cyt
Z_Im_L_Cyt

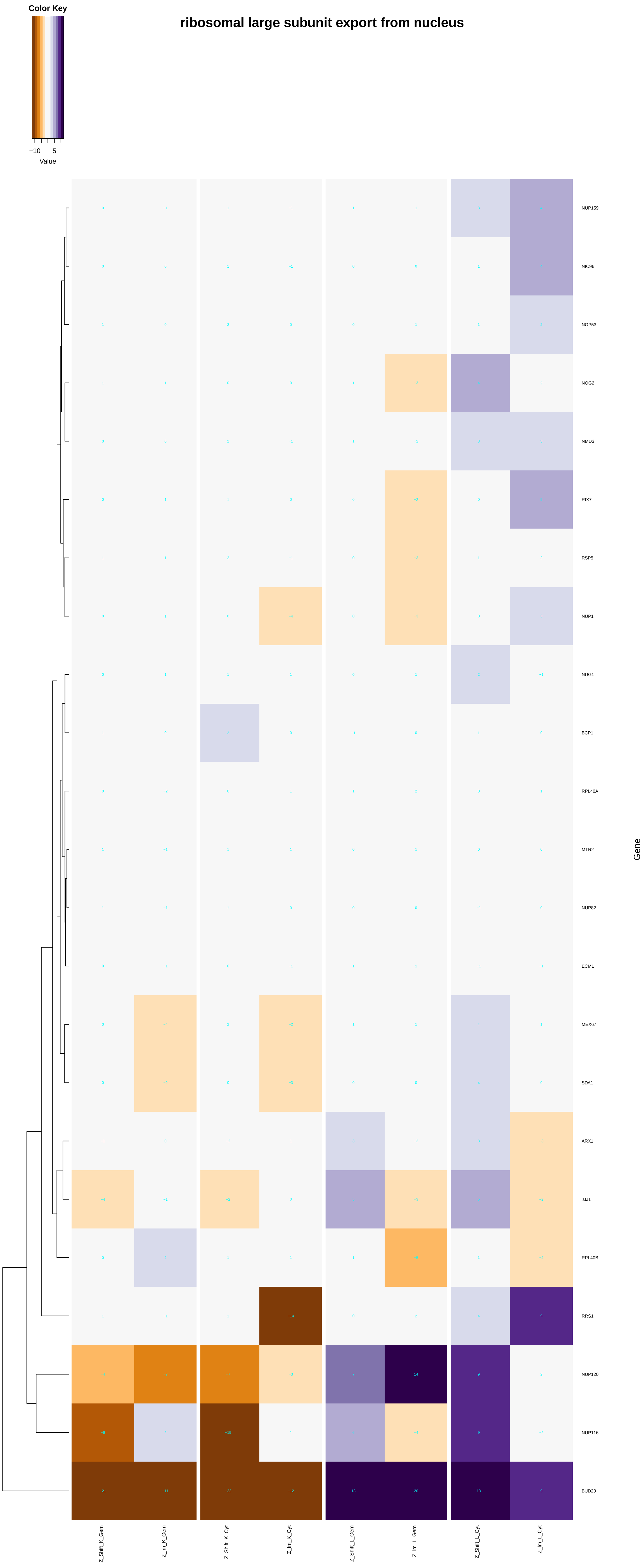
Color Key
−10
5
Value
ribosomal large subunit export from nucleus
NUP159
NIC96
NOP53
NOG2
NMD3
RIX7
RSP5
NUP1
NUG1
BCP1
RPL40A
MTR2
NUP82
ECM1
MEX67
SDA1
ARX1
JJJ1
RPL40B
RRS1
NUP120
NUP116
BUD20
Gene
Z_Shift_K_Gem
Z_Im_K_Gem
Z_Shift_K_Cyt
Z_Im_K_Cyt
Z_Shift_L_Gem
Z_Im_L_Gem
Z_Shift_L_Cyt
Z_Im_L_Cyt

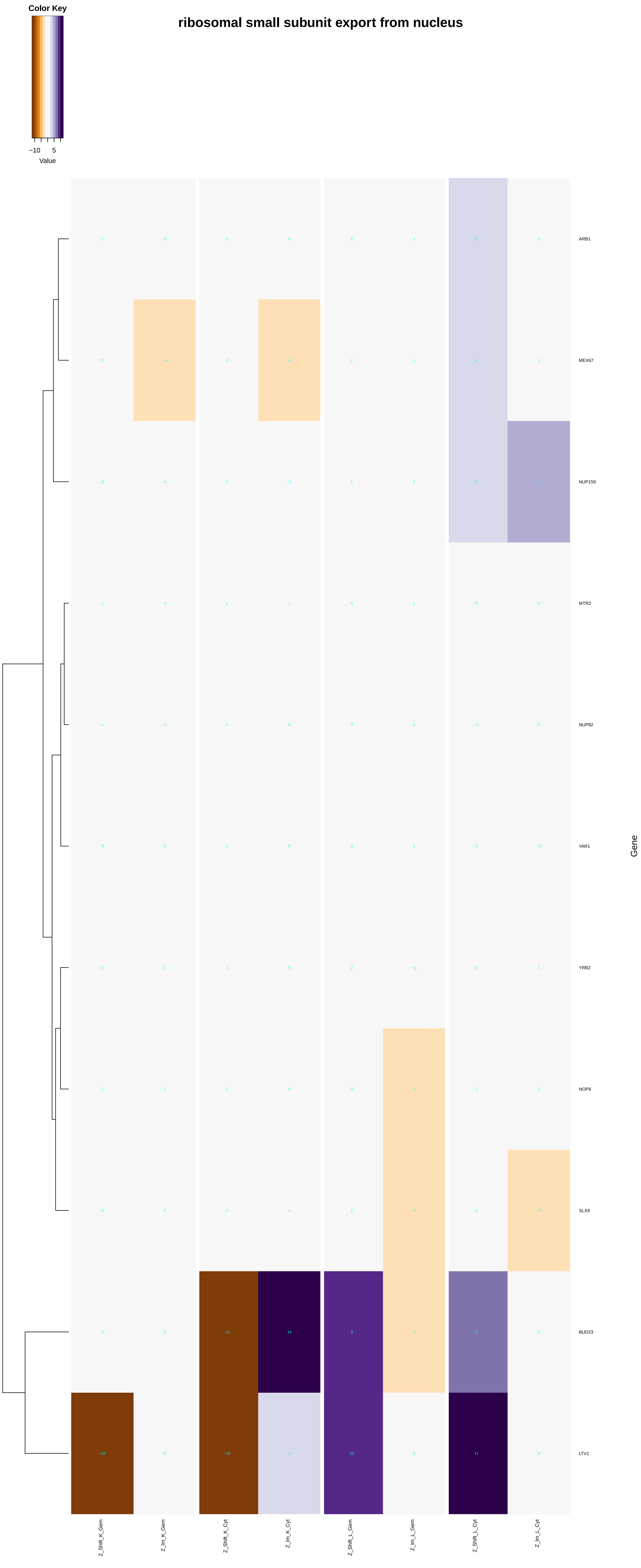
Color Key
−10 5
Value
ribosomal small subunit export from nucleus
ARB1
MEX67
NUP159
MTR2
NUP82
YAR1
YRB2
NOP9
SLX9
BUD23
LTV1
Gene
Z_Shift_K_Gem
Z_Im_K_Gem
Z_Shift_K_Cyt
Z_Im_K_Cyt
Z_Shift_L_Gem
Z_Im_L_Gem
Z_Shift_L_Cyt
Z_Im_L_Cyt
