## Additional File 7 for "A humanized yeast phenomic model of deoxycytidine kinase to predict genetic buffering of nucleoside analog cytotoxicity": intracellular_transport.pdf

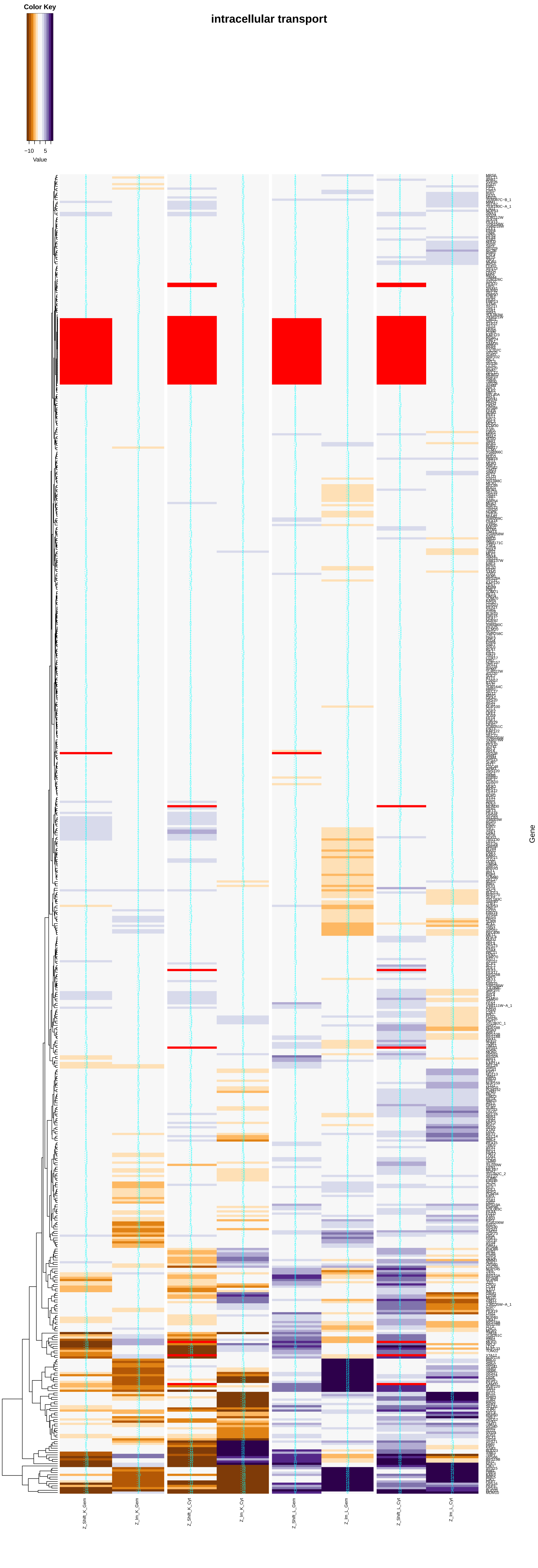
intracellular transport
Color Key
−10
5
Value
Gene
Z_Shift_K_Gem
Z_Im_K_Gem
Z_Shift_K_Cyt
Z_Im_K_Cyt
Z_Shift_L_Gem
Z_Im_L_Gem
Z_Shift_L_Cyt
Z_Im_L_Cyt

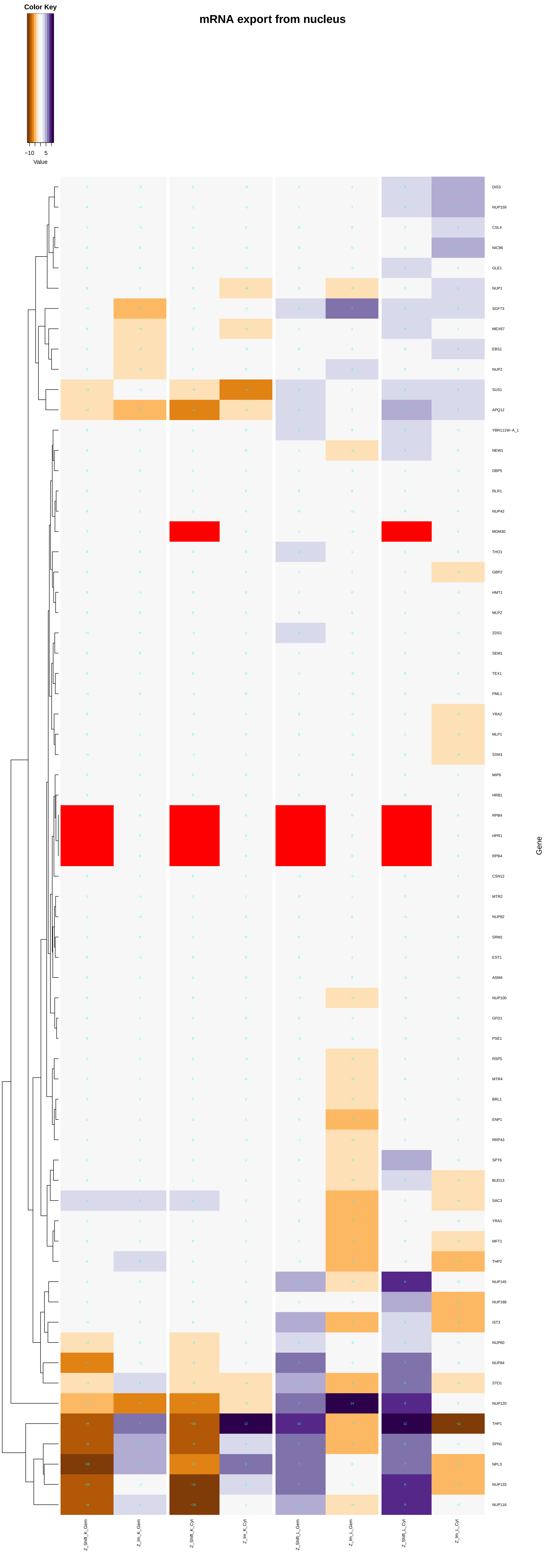
mRNA export from nucleus
Color Key
−10
5
Value
DIS3
NUP159
CSL4
NIC96
GLE1
NUP1
SGF73
MEX67
EBS1
NUP2
SUS1
APQ12
YBR111W−A_1
NEW1
DBP5
RLR1
NUP42
MDM30
THO1
GBP2
HMT1
MLP2
ZDS1
SEM1
TEX1
PML1
YRA2
MLP1
SXM1
MIP6
HRB1
RPB4
HPR1
RPB4
CSN12
MTR2
NUP82
SRM1
EST1
ASM4
NUP100
GFD1
PSE1
RSP5
MTR4
BRL1
ENP1
RRP43
SPT6
BUD13
SAC3
YRA1
MFT1
THP2
NUP145
NUP188
IST3
NUP60
NUP84
STO1
NUP120
THP1
SPN1
NPL3
NUP133
NUP116
Gene
Z_Shift_K_Gem
Z_Im_K_Gem
Z_Shift_K_Cyt
Z_Im_K_Cyt
Z_Shift_L_Gem
Z_Im_L_Gem
Z_Shift_L_Cyt
Z_Im_L_Cyt

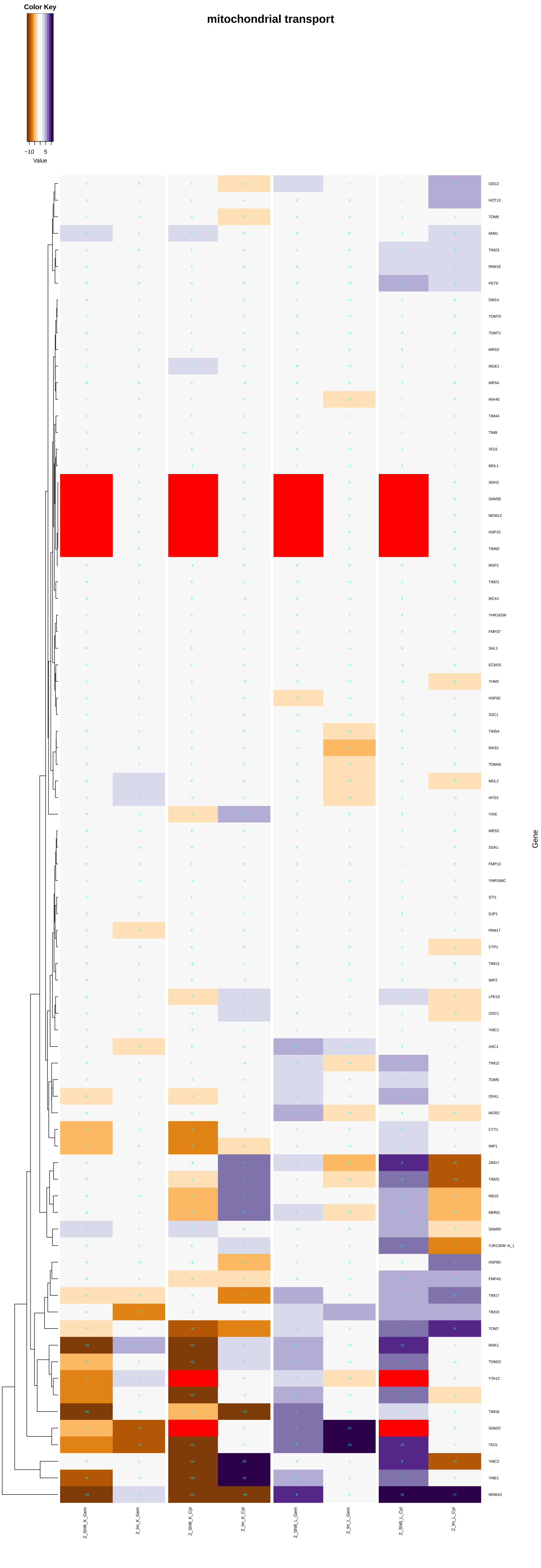
mitochondrial transport
Color Key
−10 5
Value
Gene
ODC2
HOT13
TOM6
MIM1
TIM23
PAM18
PET9
OM14
TOM70
TOM71
MRS3
MGE1
MRS4
MIA40
TIM44
TIM8
XDJ1
MDL1
SDH3
SAM35
MDM12
HSP10
TIM50
MSP1
TIM21
MCX1
YHR162W
FMP37
SAL1
ECM10
YHM2
HSP82
SSC1
TIM54
MAS1
TOM40
MDL2
AFG3
YIA6
MRS2
SSA1
FMP13
YMR166C
STI1
DJP1
PAM17
CTP1
TIM13
IMP2
LPE10
ODC1
YMC1
AAC1
TIM12
TOM5
OXA1
MGR2
CYT1
IMP1
ZIM17
TIM22
MDJ2
MMM1
SAM50
YJR135W−A_1
HSP60
FMP43
TIM17
TIM10
TOM7
MSK1
TOM22
YTA12
TIM18
SAM37
YDJ1
YMC2
YME1
MDM10
Z_Shift_K_Gem
Z_Im_K_Gem
Z_Shift_K_Cyt
Z_Im_K_Cyt
Z_Shift_L_Gem
Z_Im_L_Gem
Z_Shift_L_Cyt
Z_Im_L_Cyt

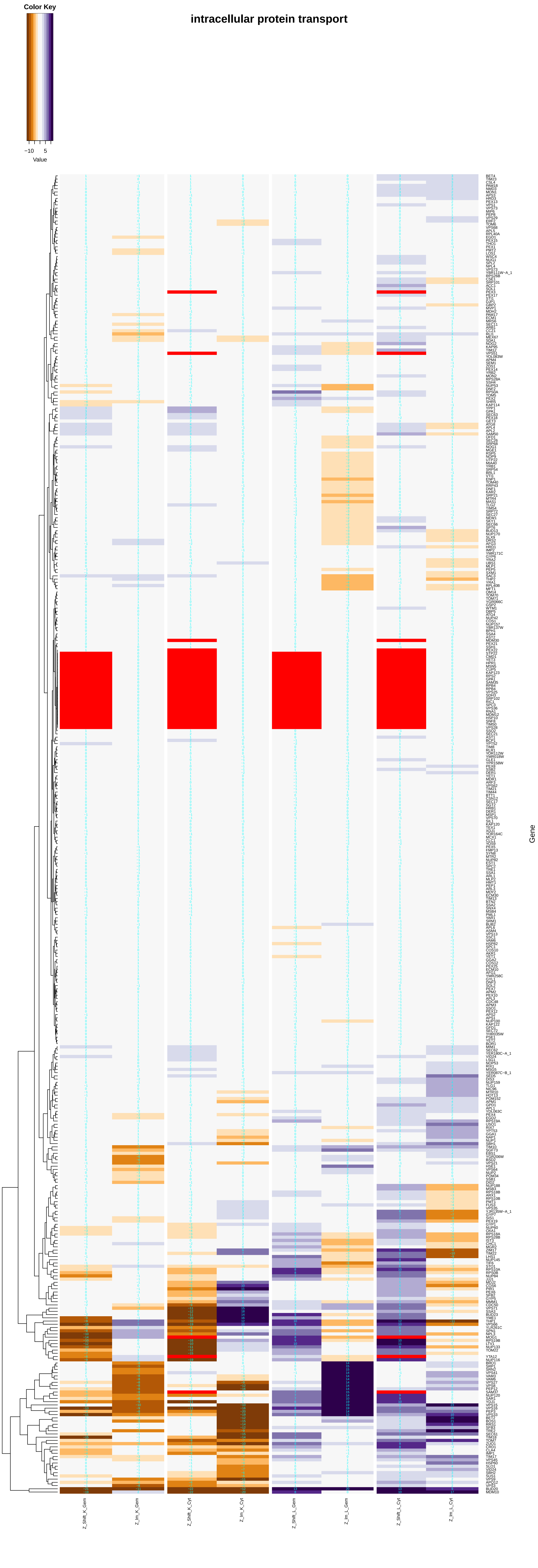

Color Key
−10
5
Value
intracellular protein transport
Gene
Z_Shift_K_Gem
Z_Im_K_Gem
Z_Shift_K_Cyt
Z_Im_K_Cyt
Z_Shift_L_Gem
Z_Im_L_Gem
Z_Shift_L_Cyt
Z_Im_L_Cyt

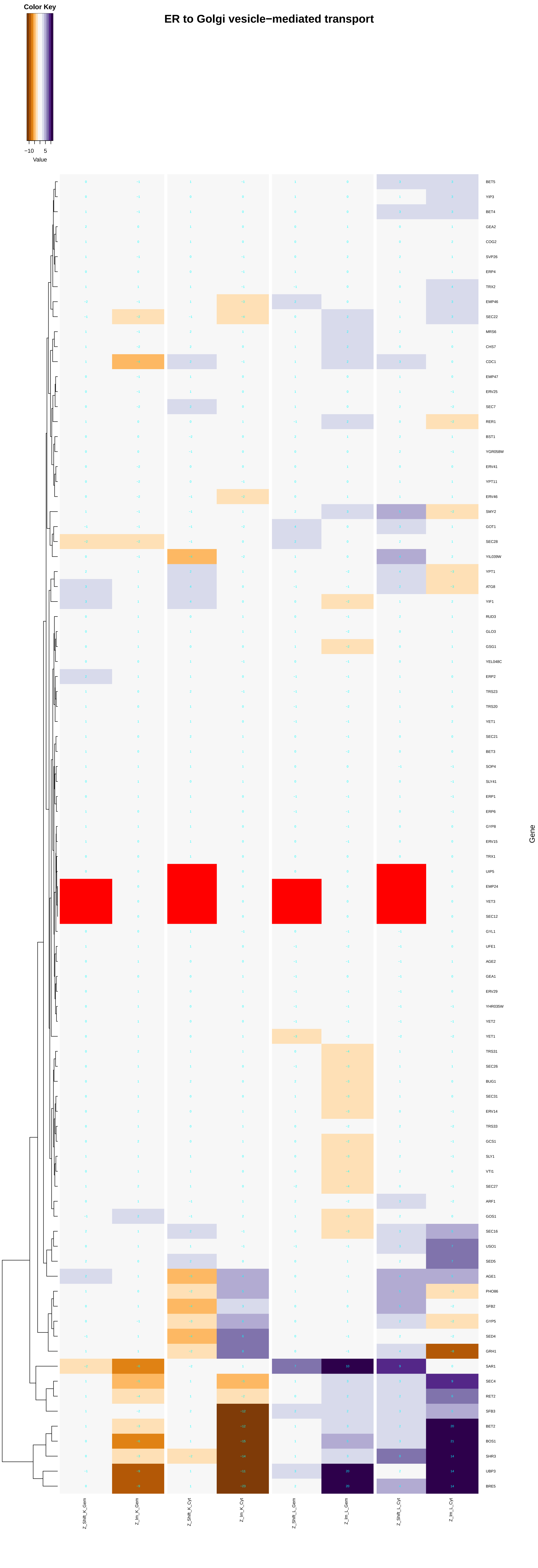

Color Key
−10
5
Value
ER to Golgi vesicle−mediated transport
BET5
YIP3
BET4
GEA2
COG2
SVP26
ERP4
TRX2
EMP46
SEC22
MRS6
CHS7
CDC1
EMP47
ERV25
SEC7
RER1
BST1
YGR058W
ERV41
YPT11
ERV46
SMY2
GOT1
SEC28
YIL039W
YPT1
ATG8
YIF1
RUD3
GLO3
GSG1
YEL048C
ERP2
TRS23
TRS20
YET1
SEC21
BET3
SOP4
SLY41
ERP1
ERP6
GYP8
ERV15
TRX1
UIP5
EMP24
YET3
SEC12
GYL1
UFE1
AGE2
GEA1
ERV29
YHR035W
YET2
YET1
TRS31
SEC26
BUG1
SEC31
ERV14
TRS33
GCS1
SLY1
VTI1
SEC27
ARF1
GOS1
SEC16
USO1
SED5
AGE1
PHO86
SFB2
GYP5
SED4
GRH1
SAR1
SEC4
RET2
SFB3
BET2
BOS1
SHR3
UBP3
BRE5
Gene
Z_Shift_K_Gem
Z_Im_K_Gem
Z_Shift_K_Cyt
Z_Im_K_Cyt
Z_Shift_L_Gem
Z_Im_L_Gem
Z_Shift_L_Cyt
Z_Im_L_Cyt

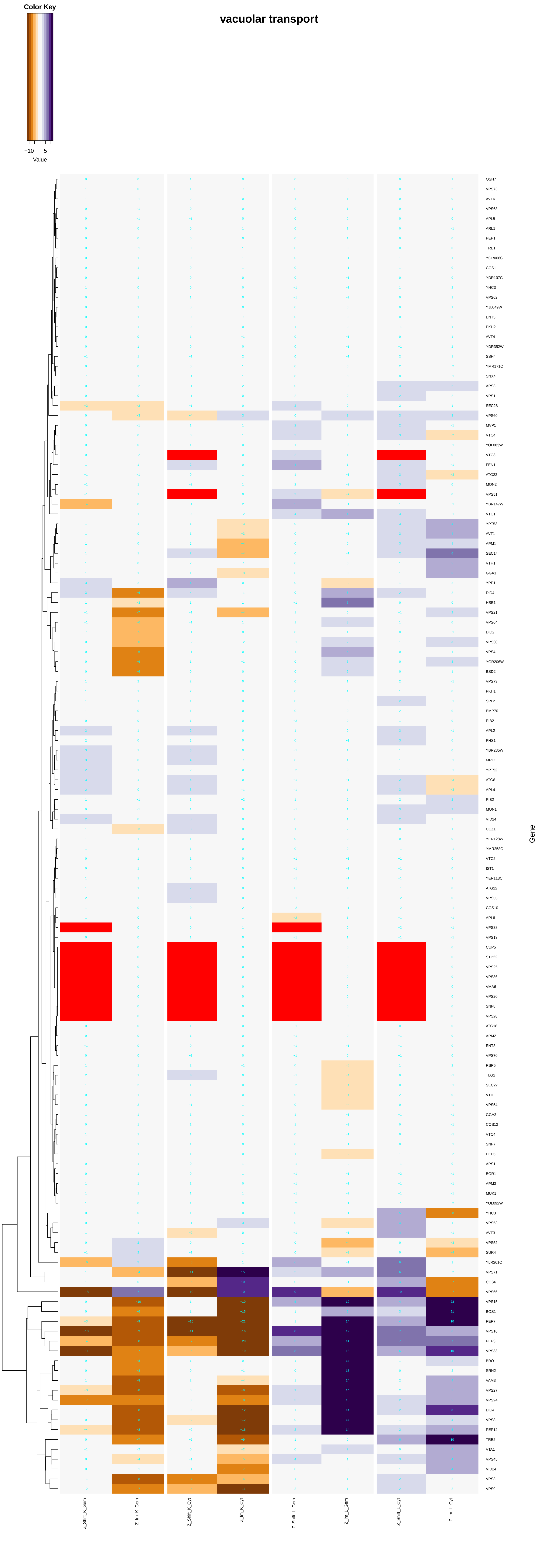

vacuolar transport
Color Key
−10
5
Value
Gene
OSH7
VPS73
AVT6
VPS68
APL5
ARL1
PEP1
TRE1
YGR066C
COS1
YDR107C
YHC3
VPS62
YJL049W
ENT5
PKH2
AVT4
YDR352W
SSH4
YMR171C
SNX4
APS3
VPS1
SEC28
VPS60
MVP1
VTC4
YOL083W
VTC3
FEN1
ATG22
MON2
VPS51
YBR147W
VTC1
YPT53
AVT1
APM1
SEC14
VTH1
GGA1
YPP1
DID4
HSE1
VPS21
VPS64
DID2
VPS30
VPS4
YGR206W
BSD2
VPS73
PKH1
SPL2
EMP70
PIB2
APL2
PHS1
YBR235W
MRL1
YPT52
ATG8
APL4
PIB2
MON1
VID24
CCZ1
YER128W
YMR258C
VTC2
IST1
YER113C
ATG22
VPS55
COS10
APL6
VPS38
VPS13
CUP5
STP22
VPS25
VPS36
VMA6
VPS20
SNF8
VPS28
ATG18
APM2
ENT3
VPS70
RSP5
TLG2
SEC27
VTI1
VPS54
GGA2
COS12
VTC4
SNF7
PEP5
APS1
BOR1
APM3
MUK1
YOL092W
YHC3
VPS53
AVT3
VPS52
SUR4
YLR261C
VPS71
COS6
VPS66
VPS15
BOS1
PEP7
VPS16
PEP3
VPS33
BRO1
SRN2
VAM3
VPS27
VPS24
DID4
VPS8
PEP12
TRE2
VTA1
VPS45
VID24
VPS3
VPS9
Z_Shift_K_Gem
Z_Im_K_Gem
Z_Shift_K_Cyt
Z_Im_K_Cyt
Z_Shift_L_Gem
Z_Im_L_Gem
Z_Shift_L_Cyt
Z_Im_L_Cyt

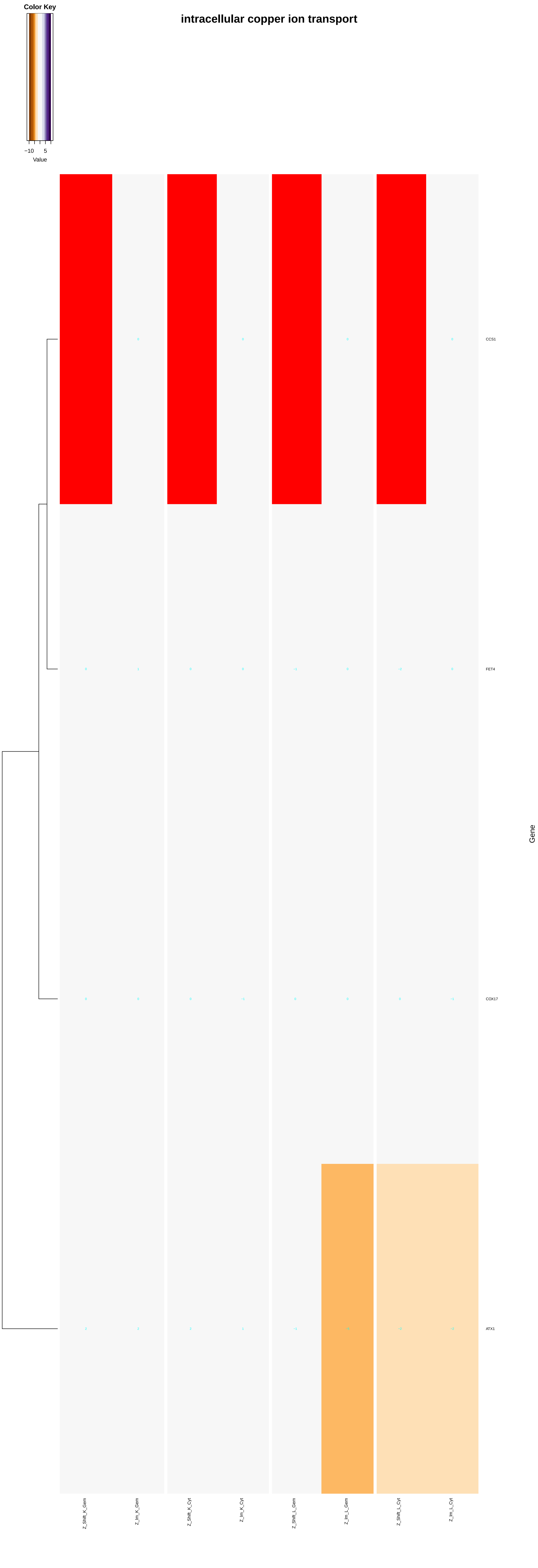
intracellular copper ion transport
Color Key
−10
5
Value
CCS1
FET4
COX17
ATX1
Gene
Z_Shift_K_Gem
Z_Im_K_Gem
Z_Shift_K_Cyt
Z_Im_K_Cyt
Z_Shift_L_Gem
Z_Im_L_Gem
Z_Shift_L_Cyt
Z_Im_L_Cyt

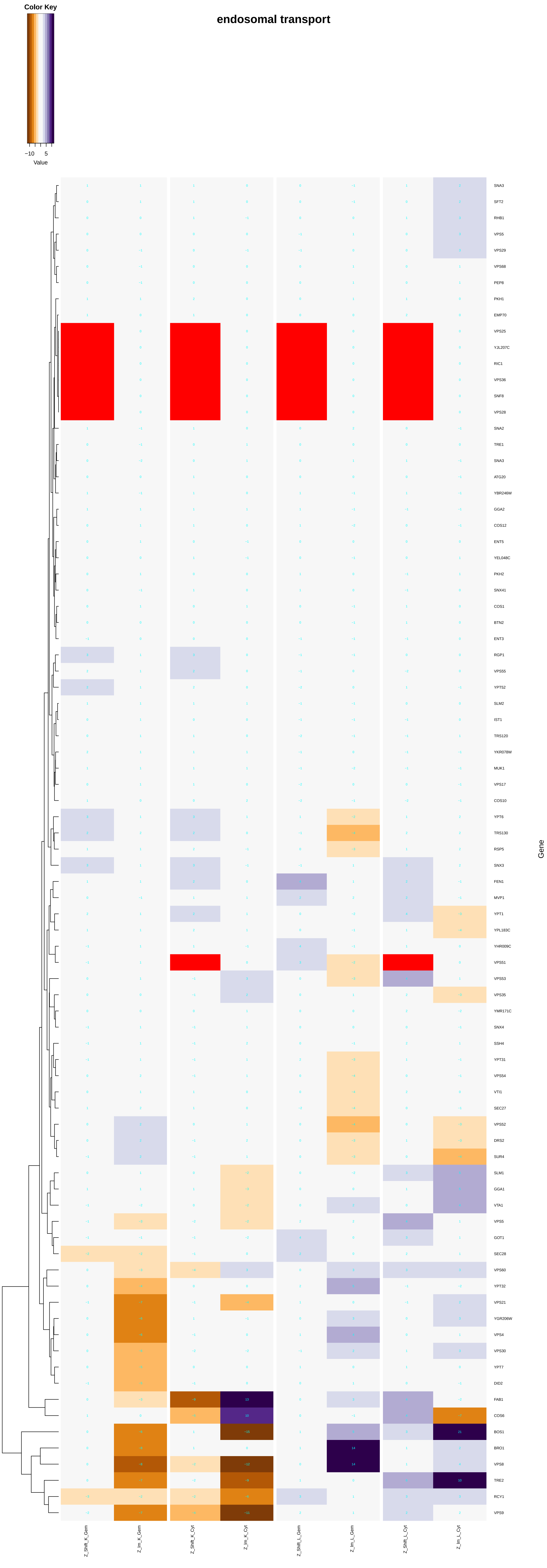
endosomal transport
Color Key
−10
5
Value
SNA3
SFT2
RHB1
VPS5
VPS29
VPS68
PEP8
PKH1
EMP70
VPS25
YJL207C
RIC1
VPS36
SNF8
VPS28
SNA2
TRE1
SNA3
ATG20
YBR246W
GGA2
COS12
ENT5
YEL048C
PKH2
SNX41
COS1
BTN2
ENT3
RGP1
VPS55
YPT52
SLM2
IST1
TRS120
YKR078W
MUK1
VPS17
COS10
YPT6
TRS130
RSP5
SNX3
FEN1
MVP1
YPT1
YPL183C
YHR009C
VPS51
VPS53
VPS35
YMR171C
SNX4
SSH4
YPT31
VPS54
VTI1
SEC27
VPS52
DRS2
SUR4
SLM1
GGA1
VTA1
VPS5
GOT1
SEC28
VPS60
YPT32
VPS21
YGR206W
VPS4
VPS30
YPT7
DID2
FAB1
COS6
BOS1
BRO1
VPS8
TRE2
RCY1
VPS9
Gene
Z_Shift_K_Gem
Z_Im_K_Gem
Z_Shift_K_Cyt
Z_Im_K_Cyt
Z_Shift_L_Gem
Z_Im_L_Gem
Z_Shift_L_Cyt
Z_Im_L_Cyt

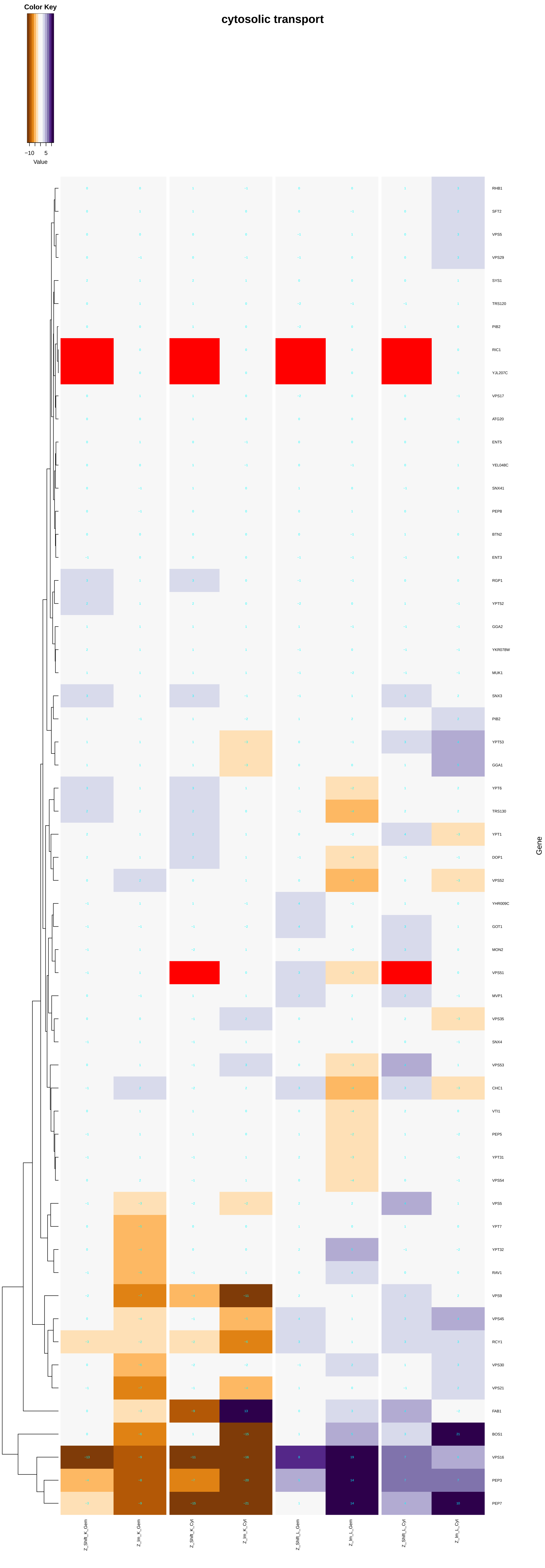
Color Key
−10
5
Value
cytosolic transport
RHB1
SFT2
VPS5
VPS29
SYS1
TRS120
PIB2
RIC1
YJL207C
VPS17
ATG20
ENT5
YEL048C
SNX41
PEP8
BTN2
ENT3
RGP1
YPT52
GGA2
YKR078W
MUK1
SNX3
PIB2
YPT53
GGA1
YPT6
TRS130
YPT1
DOP1
VPS52
YHR009C
GOT1
MON2
VPS51
MVP1
VPS35
SNX4
VPS53
CHC1
VTI1
PEP5
YPT31
VPS54
VPS5
YPT7
YPT32
RAV1
VPS9
VPS45
RCY1
VPS30
VPS21
FAB1
BOS1
VPS16
PEP3
PEP7
Gene
Z_Shift_K_Gem
Z_Im_K_Gem
Z_Shift_K_Cyt
Z_Im_K_Cyt
Z_Shift_L_Gem
Z_Im_L_Gem
Z_Shift_L_Cyt
Z_Im_L_Cyt

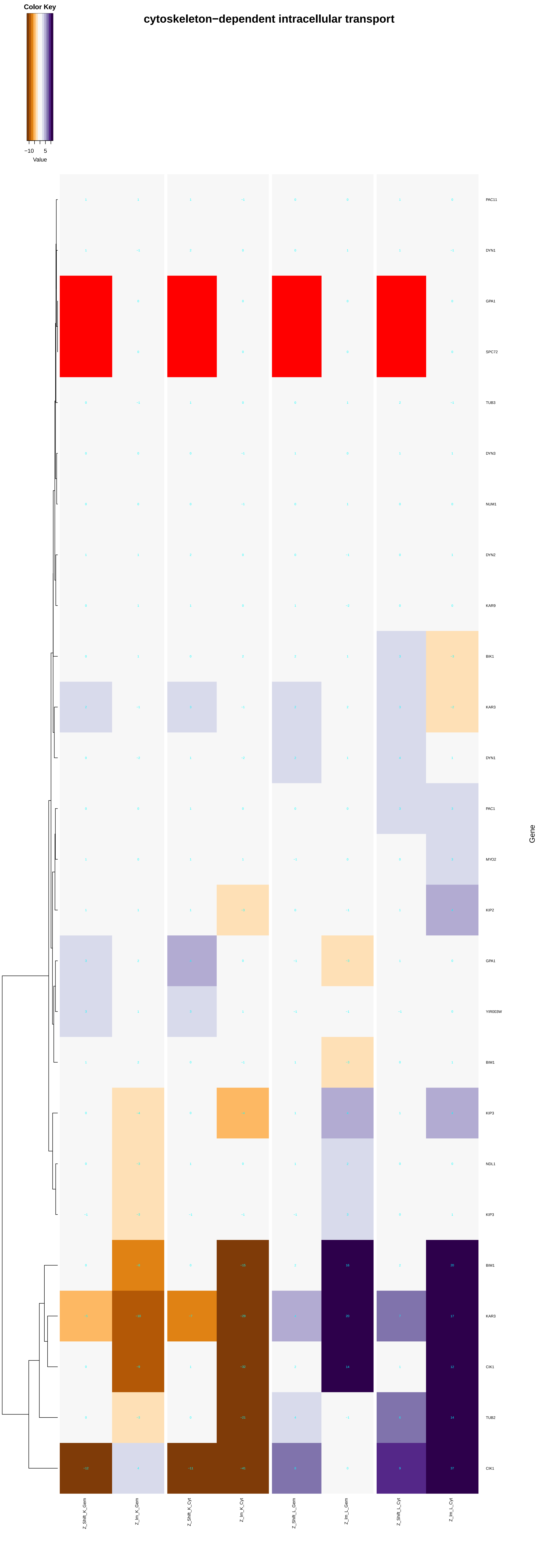

cytoskeleton−dependent intracellular transport
Color Key
Value
Gene
PAC11
DYN1
GPA1
SPC72
TUB3
DYN3
NUM1
DYN2
KAR9
BIK1
KAR3
DYN1
PAC1
MYO2
KIP2
GPA1
YIR003W
BIM1
KIP3
NDL1
KIP3
BIM1
KAR3
CIK1
TUB2
CIK1
Z_Shift_K_Gem
Z_Im_K_Gem
Z_Shift_K_Cyt
Z_Im_K_Cyt
Z_Shift_L_Gem
Z_Im_L_Gem
Z_Shift_L_Cyt
Z_Im_L_Cyt

### peroxisomal transport

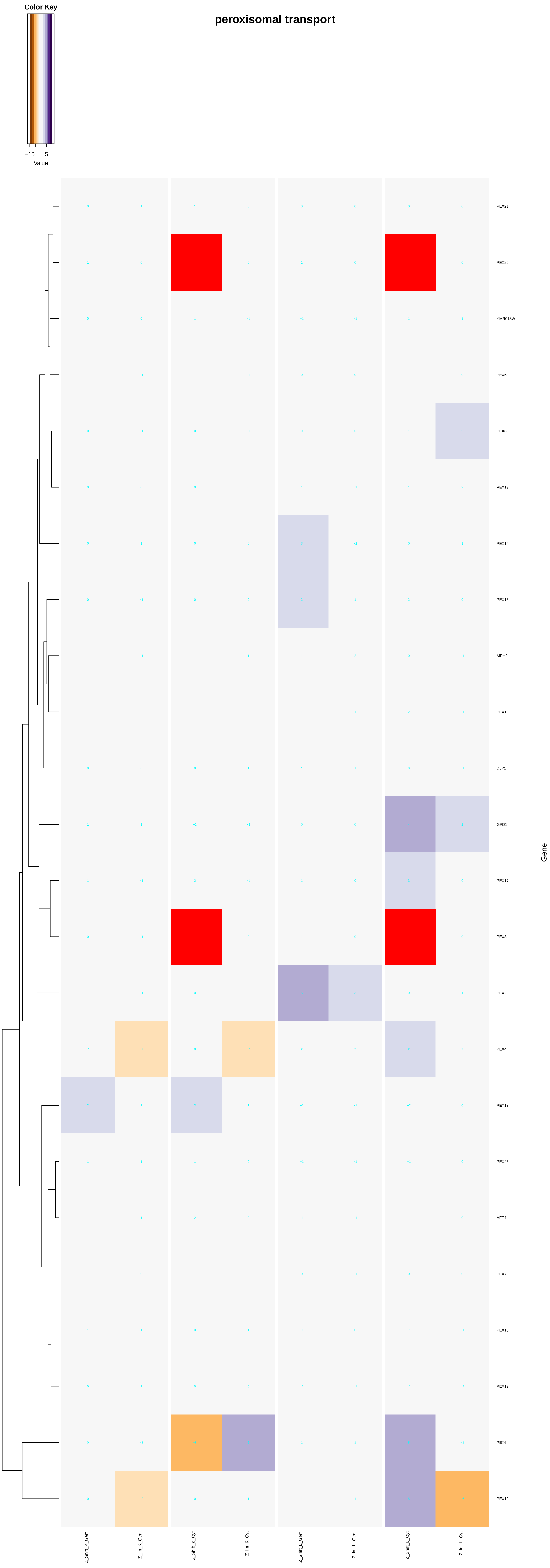

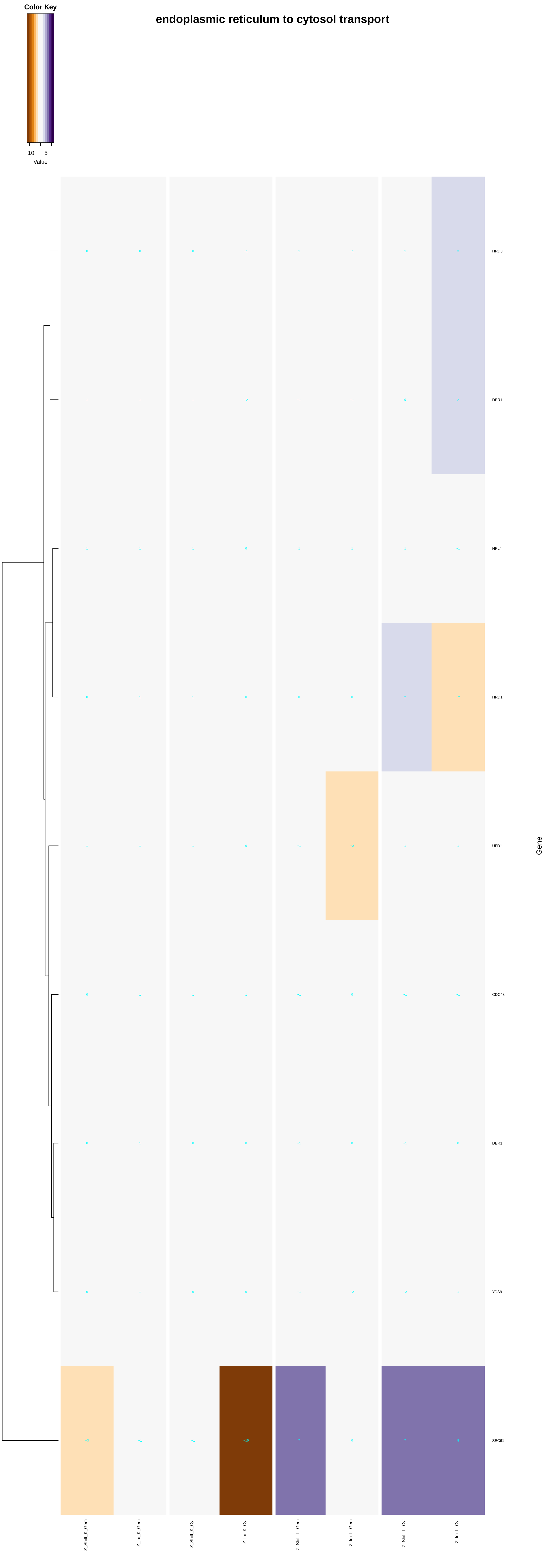
endoplasmic reticulum to cytosol transport
Color Key
−10
5
Value
HRD3
DER1
NPL4
HRD1
UFD1
CDC48
DER1
YOS9
SEC61
Gene
Z_Shift_K_Gem
Z_Im_K_Gem
Z_Shift_K_Cyt
Z_Im_K_Cyt
Z_Shift_L_Gem
Z_Im_L_Gem
Z_Shift_L_Cyt
Z_Im_L_Cyt

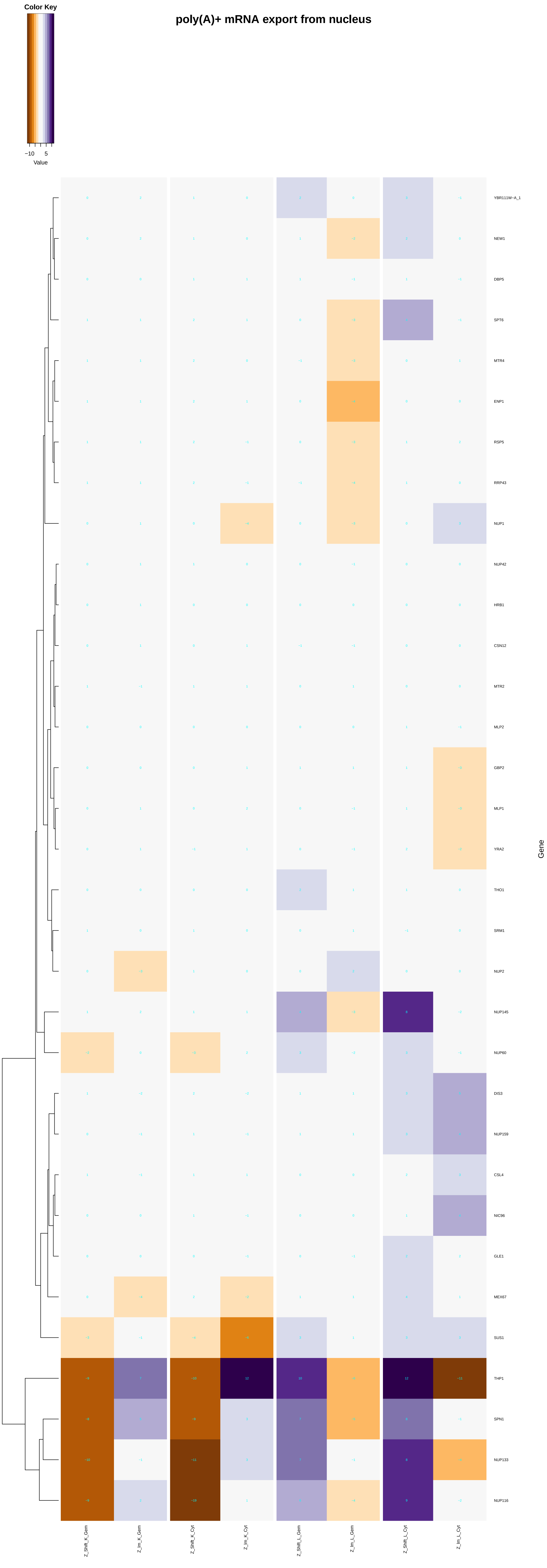
poly(A)+ mRNA export from nucleus
Color Key
−10
5
Value
YBR111W−A_1
NEW1
DBP5
SPT6
MTR4
ENP1
RSP5
RRP43
NUP1
NUP42
HRB1
CSN12
MTR2
MLP2
GBP2
MLP1
YRA2
THO1
SRM1
NUP2
NUP145
NUP60
DIS3
NUP159
CSL4
NIC96
GLE1
MEX67
SUS1
THP1
SPN1
NUP133
NUP116
Gene
Z_Shift_K_Gem
Z_Im_K_Gem
Z_Shift_K_Cyt
Z_Im_K_Cyt
Z_Shift_L_Gem
Z_Im_L_Gem
Z_Shift_L_Cyt
Z_Im_L_Cyt

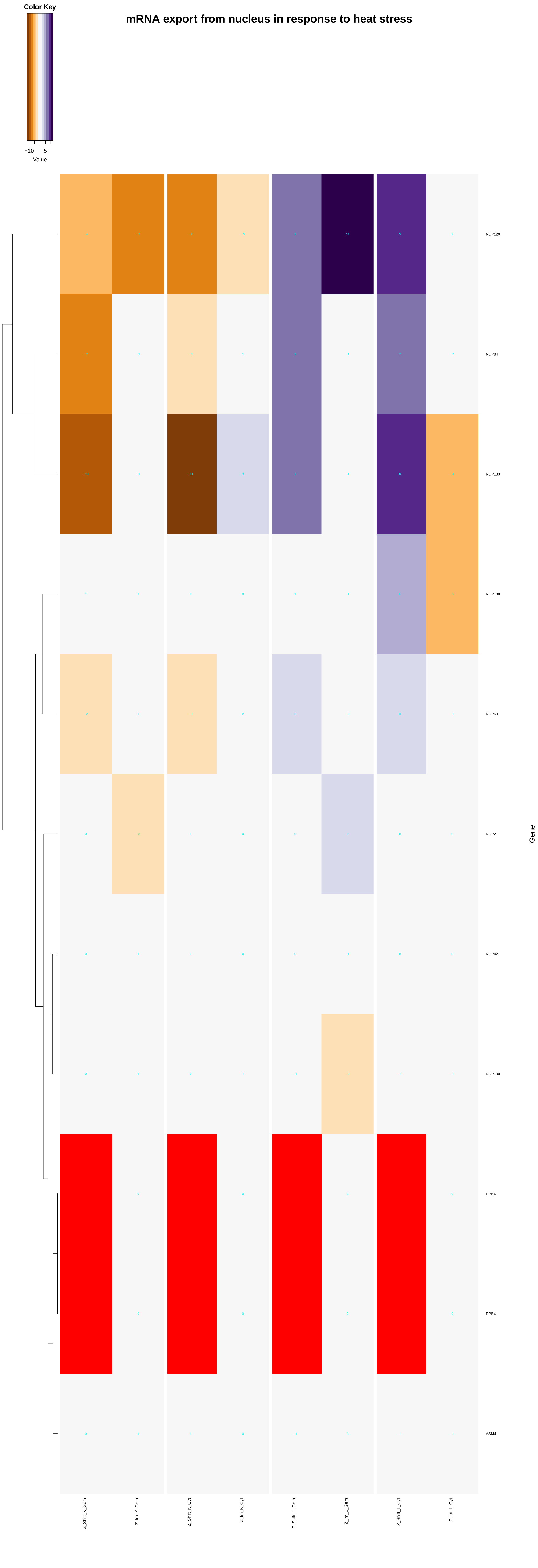
mRNA export from nucleus in response to heat stress
Color Key
−10
5
Value
NUP120
NUP84
NUP133
NUP188
NUP60
NUP2
NUP42
NUP100
RPB4
RPB4
ASM4
Gene
Z_Shift_K_Gem
Z_Im_K_Gem
Z_Shift_K_Cyt
Z_Im_K_Cyt
Z_Shift_L_Gem
Z_Im_L_Gem
Z_Shift_L_Cyt
Z_Im_L_Cyt

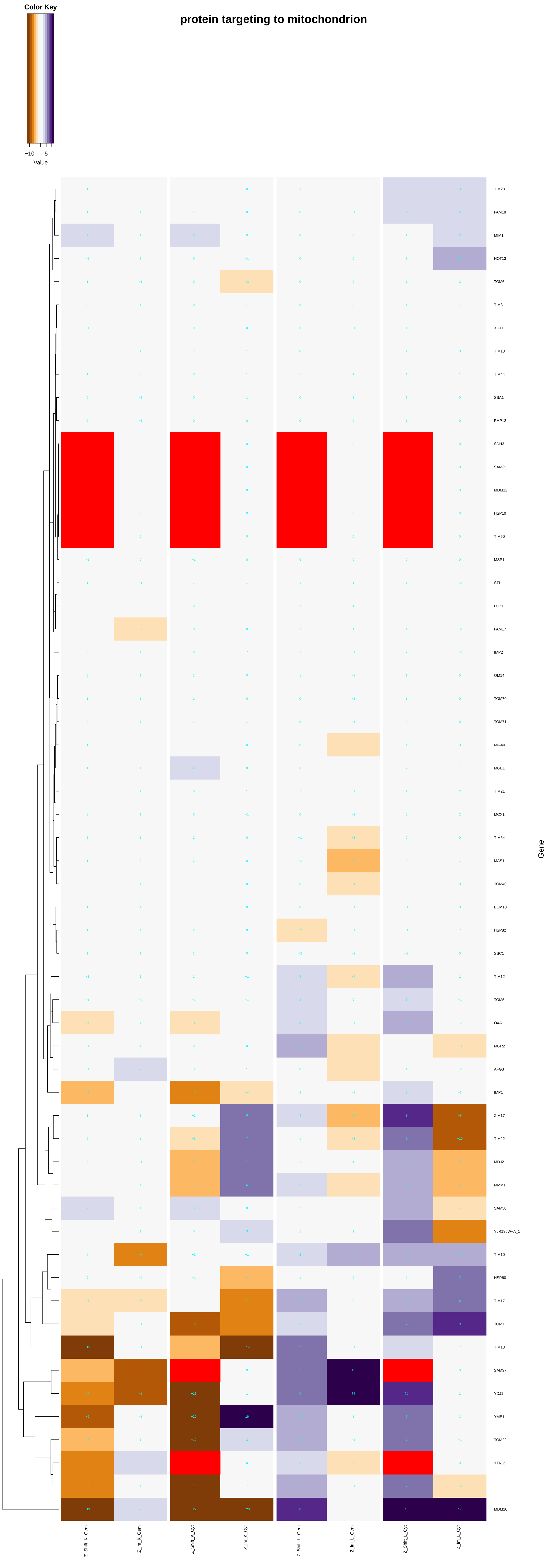
Color Key
−10
5
Value
protein targeting to mitochondrion
TIM23
PAM18
MIM1
HOT13
TOM6
TIM8
XDJ1
TIM13
TIM44
SSA1
FMP13
SDH3
SAM35
MDM12
HSP10
TIM50
MSP1
STI1
DJP1
PAM17
IMP2
OM14
TOM70
TOM71
MIA40
MGE1
TIM21
MCX1
TIM54
MAS1
TOM40
ECM10
HSP82
SSC1
TIM12
TOM5
OXA1
MGR2
AFG3
IMP1
ZIM17
TIM22
MDJ2
MMM1
SAM50
YJR135W−A_1
TIM10
HSP60
TIM17
TOM7
TIM18
SAM37
YDJ1
YME1
TOM22
YTA12
MDM10
Gene
Z_Shift_K_Gem
Z_Im_K_Gem
Z_Shift_K_Cyt
Z_Im_K_Cyt
Z_Shift_L_Gem
Z_Im_L_Gem
Z_Shift_L_Cyt
Z_Im_L_Cyt

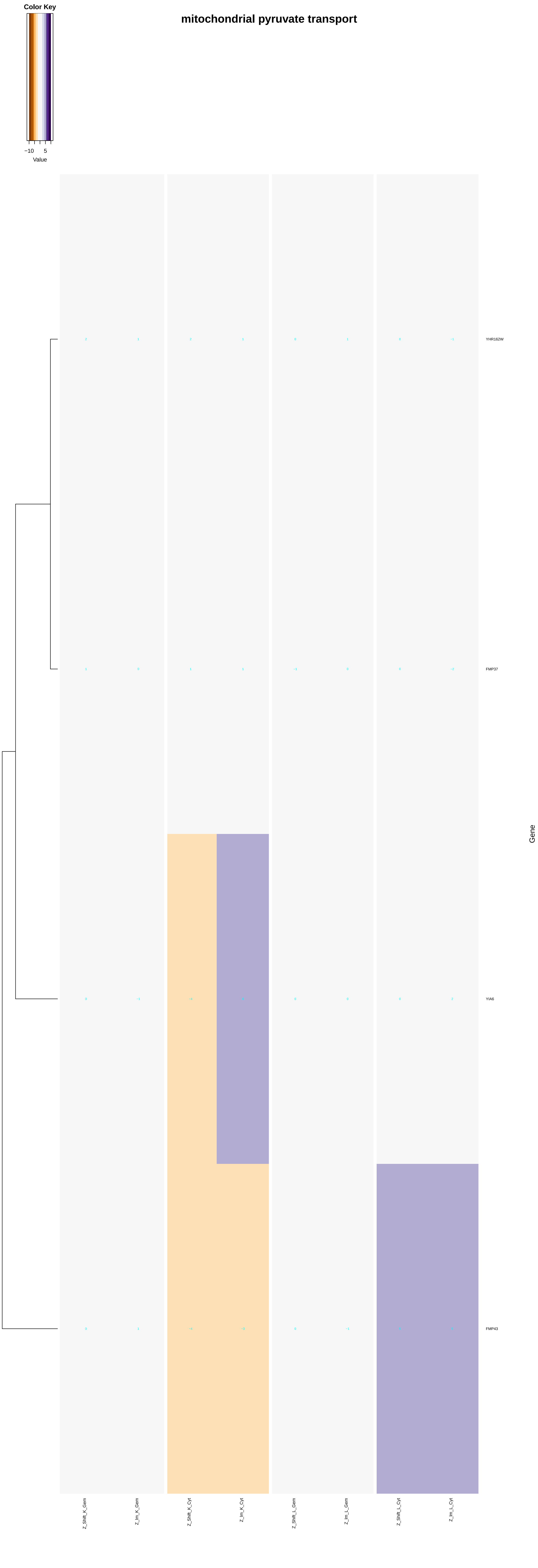

mitochondrial pyruvate transport
Color Key
−10
5
Value
YHR162W
FMP37
YIA6
FMP43
Gene
Z_Shift_K_Gem
Z_Im_K_Gem
Z_Shift_K_Cyt
Z_Im_K_Cyt
Z_Shift_L_Gem
Z_Im_L_Gem
Z_Shift_L_Cyt
Z_Im_L_Cyt

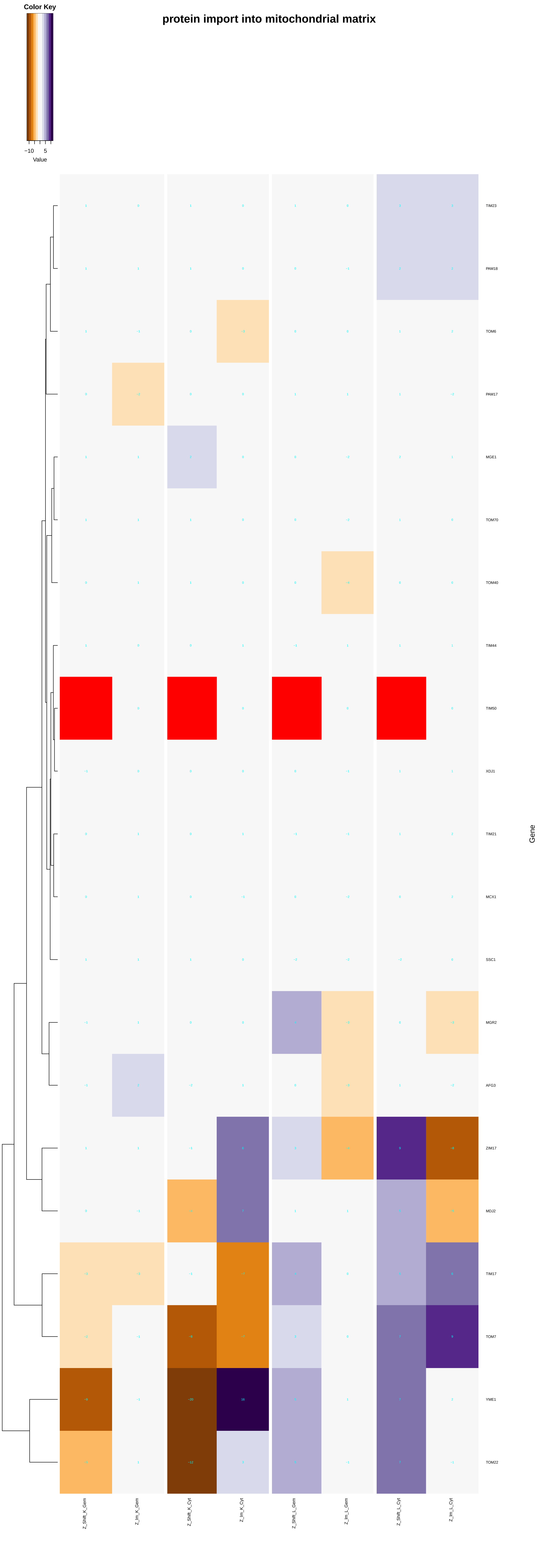
protein import into mitochondrial matrix
Color Key
−10
5
Value
TIM23
PAM18
TOM6
PAM17
MGE1
TOM70
TOM40
TIM44
TIM50
XDJ1
TIM21
MCX1
SSC1
MGR2
AFG3
ZIM17
MDJ2
TIM17
TOM7
YME1
TOM22
Gene
Z_Shift_K_Gem
Z_Im_K_Gem
Z_Shift_K_Cyt
Z_Im_K_Cyt
Z_Shift_L_Gem
Z_Im_L_Gem
Z_Shift_L_Cyt
Z_Im_L_Cyt

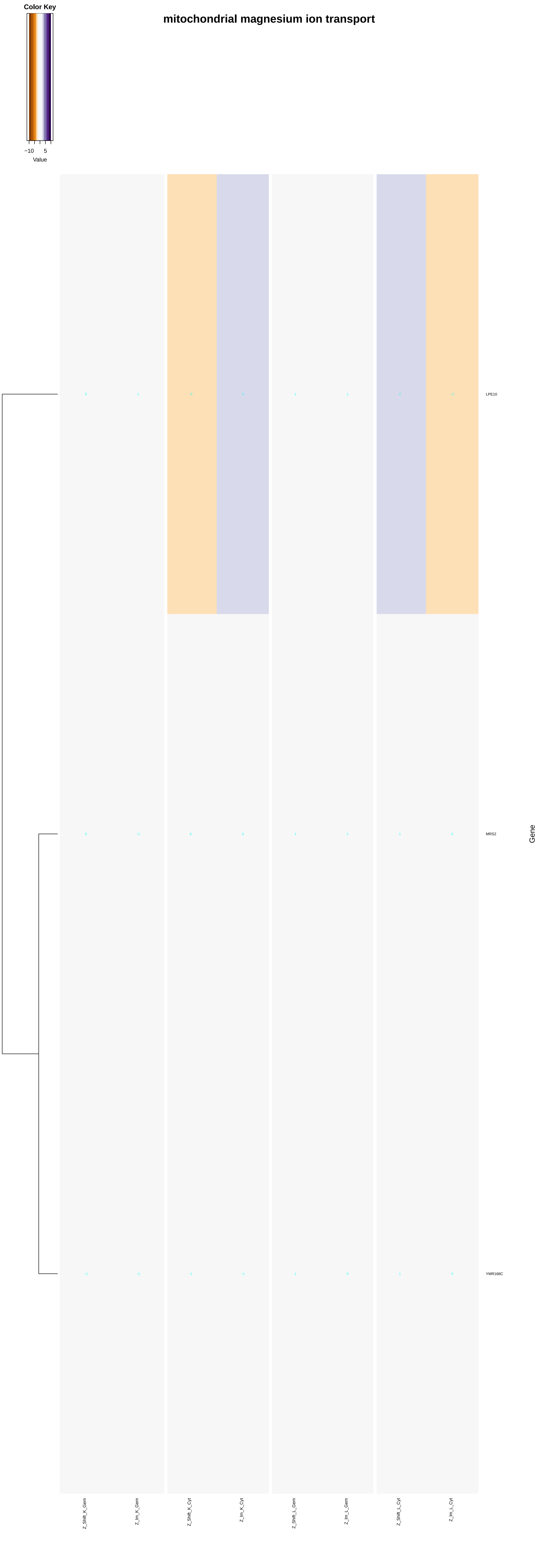
mitochondrial magnesium ion transport
Color Key
−10
5
Value
LPE10
MRS2
YMR166C
Gene
Z_Shift_K_Gem
Z_Im_K_Gem
Z_Shift_K_Cyt
Z_Im_K_Cyt
Z_Shift_L_Gem
Z_Im_L_Gem
Z_Shift_L_Cyt
Z_Im_L_Cyt

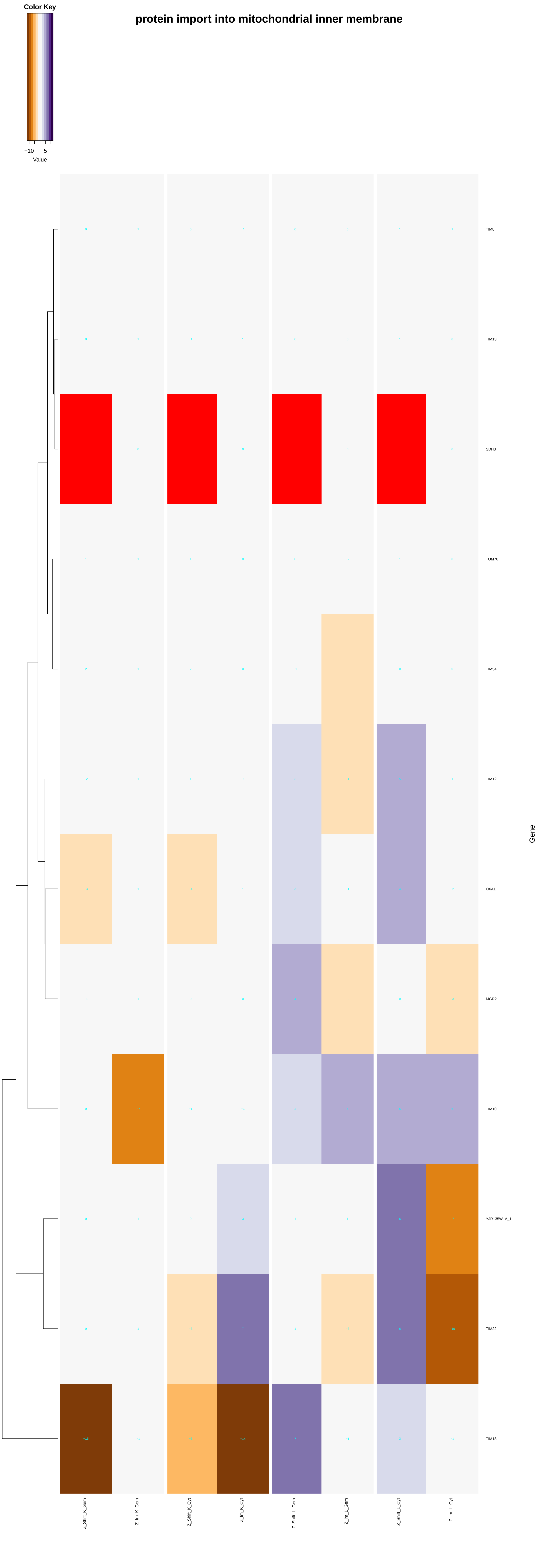
protein import into mitochondrial inner membrane
Color Key
−10
5
Value
TIM8
TIM13
SDH3
TOM70
TIM54
TIM12
OXA1
MGR2
TIM10
YJR135W−A_1
TIM22
TIM18
Gene
Z_Shift_K_Gem
Z_Im_K_Gem
Z_Shift_K_Cyt
Z_Im_K_Cyt
Z_Shift_L_Gem
Z_Im_L_Gem
Z_Shift_L_Cyt
Z_Im_L_Cyt

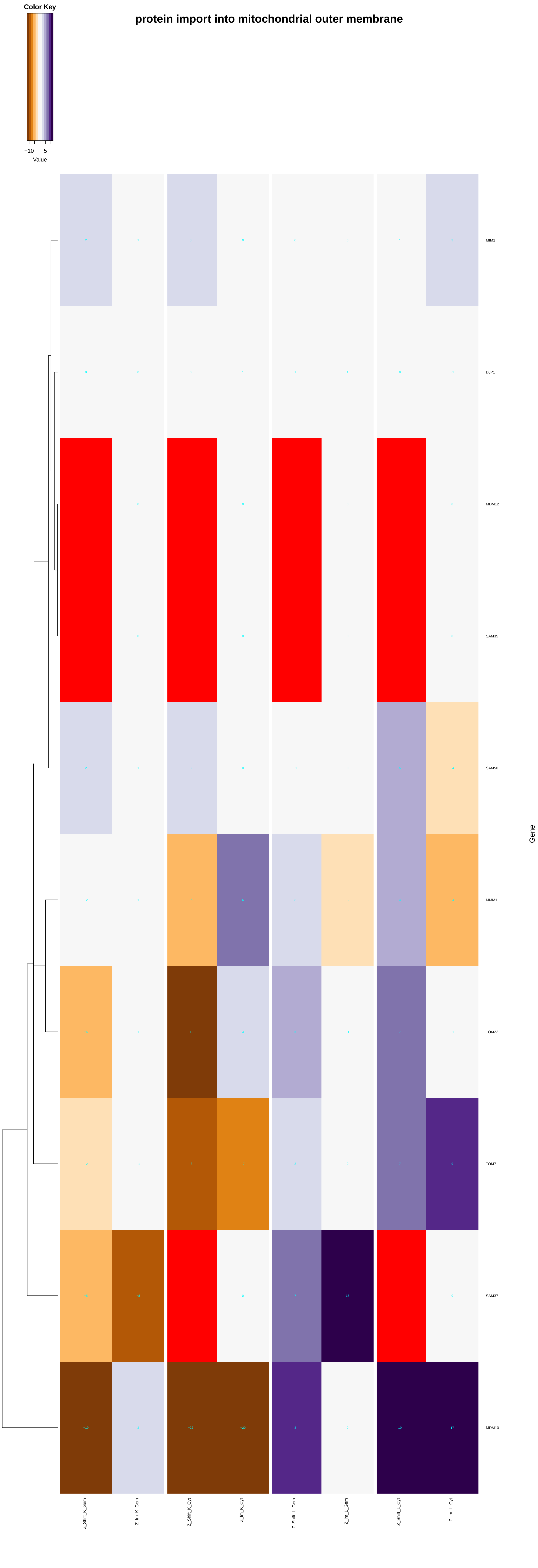

Color Key
−10
5
Value
protein import into mitochondrial outer membrane
MIM1
DJP1
MDM12
SAM35
SAM50
MMM1
TOM22
TOM7
SAM37
MDM10
Gene
Z_Shift_K_Gem
Z_Im_K_Gem
Z_Shift_K_Cyt
Z_Im_K_Cyt
Z_Shift_L_Gem
Z_Im_L_Gem
Z_Shift_L_Cyt
Z_Im_L_Cyt

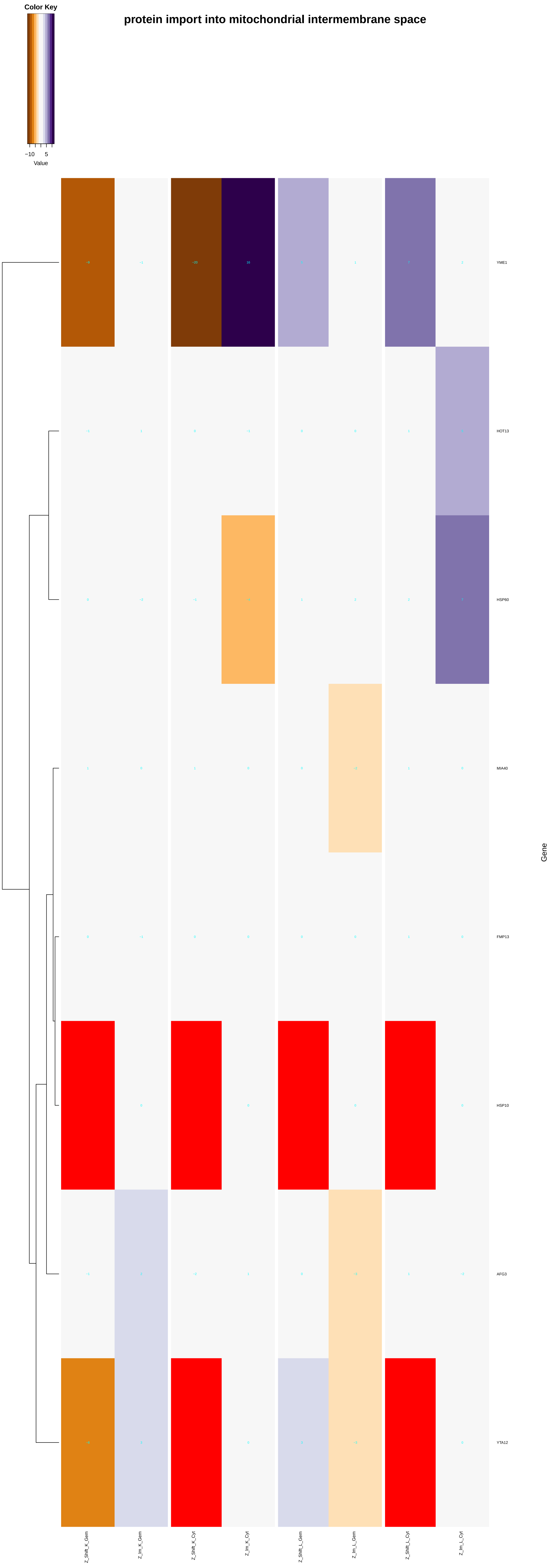
protein import into mitochondrial intermembrane space
Color Key
−10
5
Value
YME1
HOT13
HSP60
MIA40
FMP13
HSP10
AFG3
YTA12
Gene
Z_Shift_K_Gem
Z_Im_K_Gem
Z_Shift_K_Cyt
Z_Im_K_Cyt
Z_Shift_L_Gem
Z_Im_L_Gem
Z_Shift_L_Cyt
Z_Im_L_Cyt

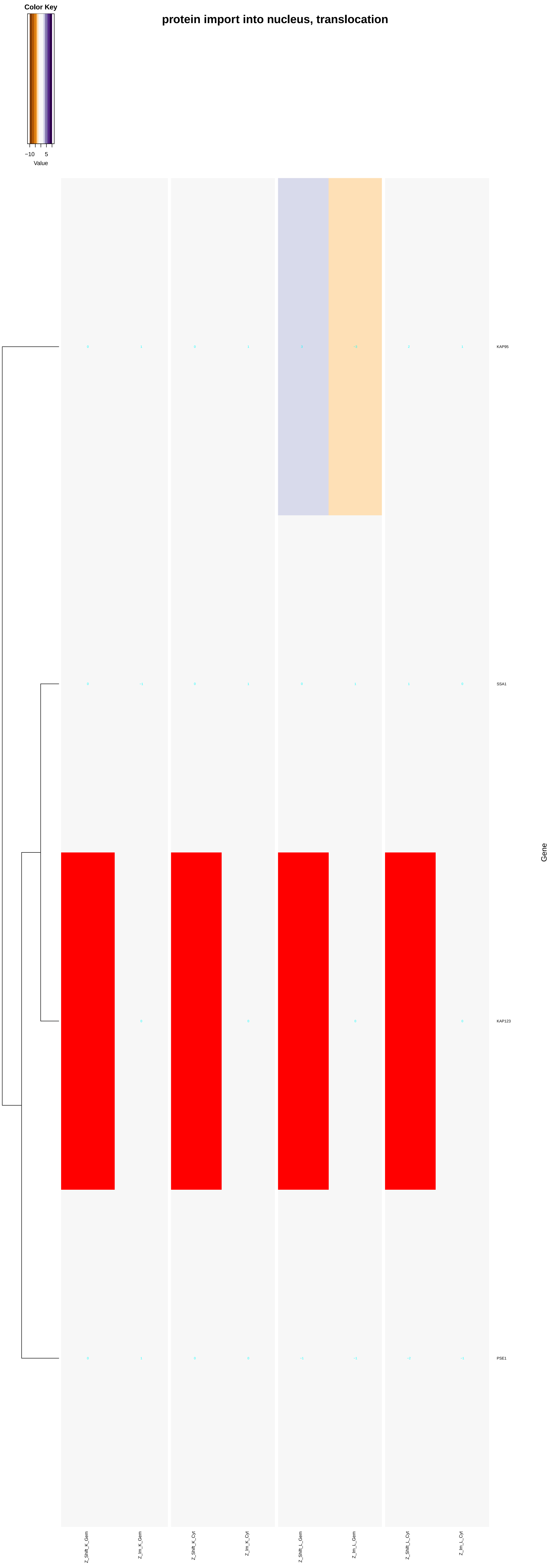

Color Key
−10
5
Value
protein import into nucleus, translocation
KAP95
SSA1
KAP123
PSE1
Gene
Z_Shift_K_Gem
Z_Im_K_Gem
Z_Shift_K_Cyt
Z_Im_K_Cyt
Z_Shift_L_Gem
Z_Im_L_Gem
Z_Shift_L_Cyt
Z_Im_L_Cyt

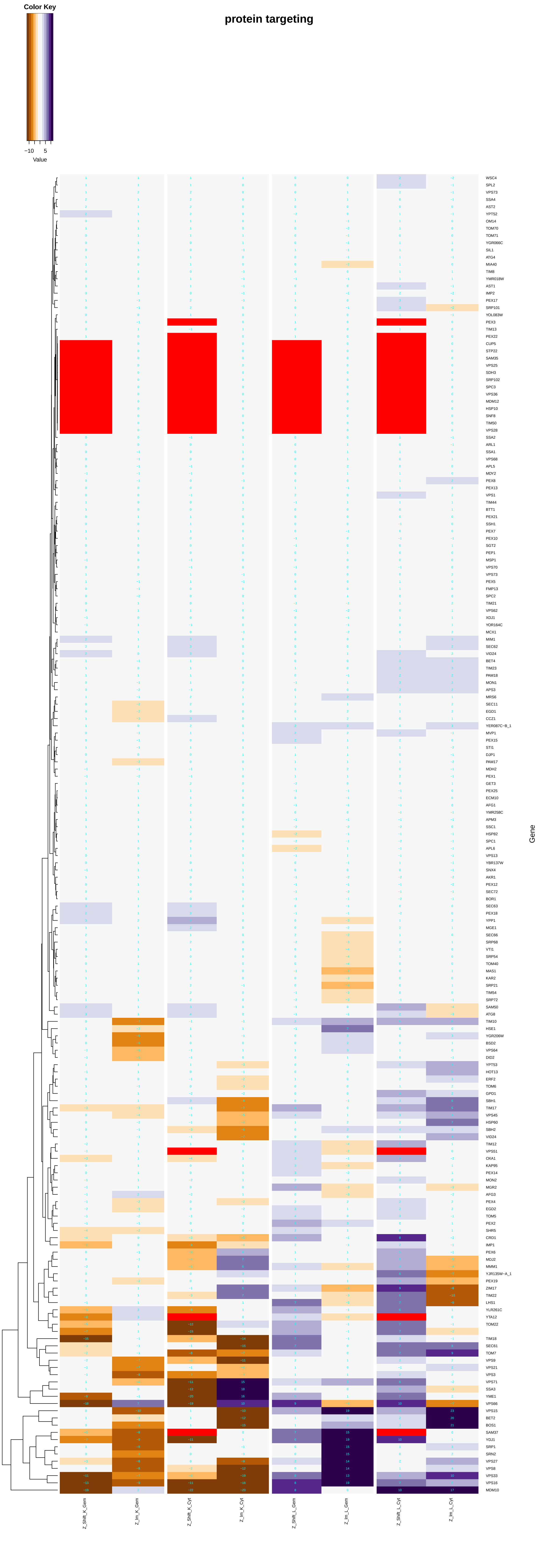
protein targeting
Color Key
−10
5
Value
Gene
Z_Shift_K_Gem
Z_Im_K_Gem
Z_Shift_K_Cyt
Z_Im_K_Cyt
Z_Shift_L_Gem
Z_Im_L_Gem
Z_Shift_L_Cyt
Z_Im_L_Cyt

### protein import into nucleus

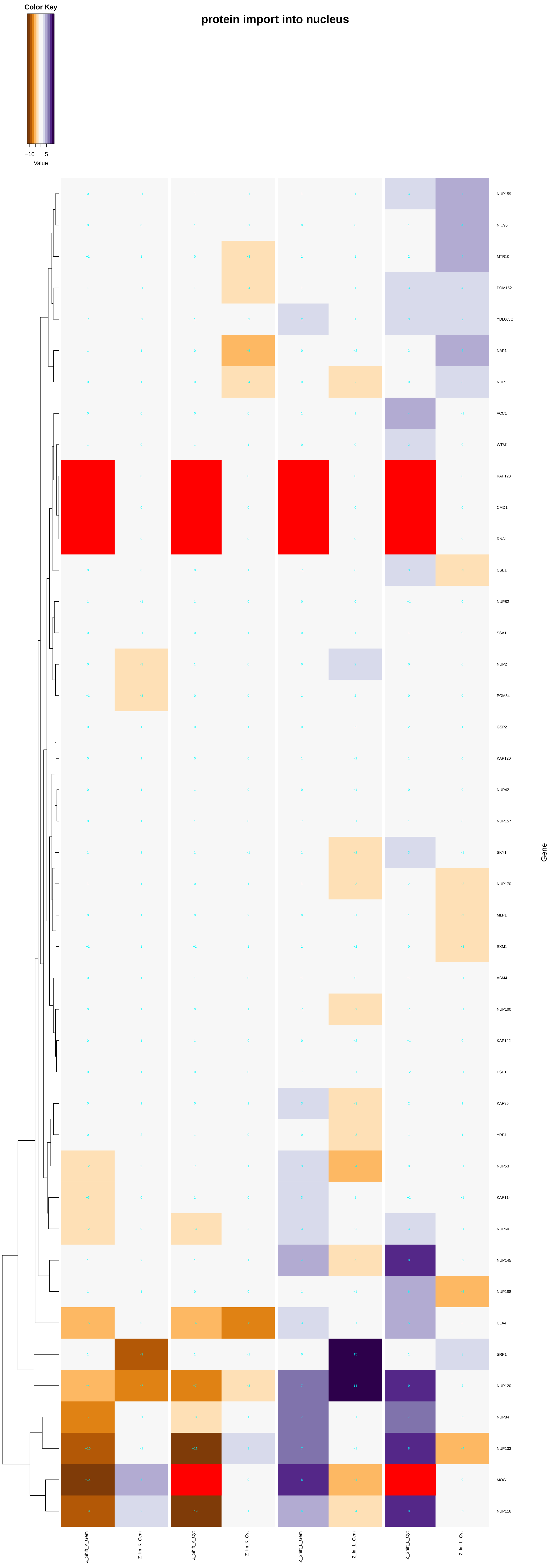

#### protein exit from endoplasmic reticulum

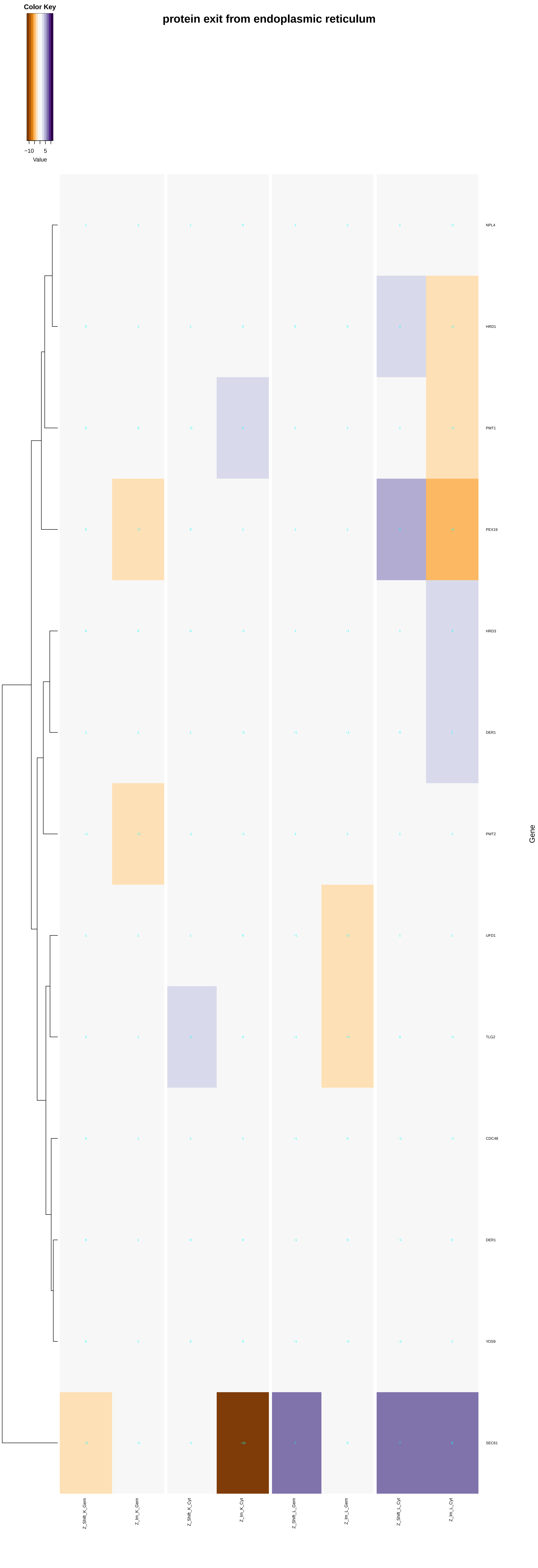

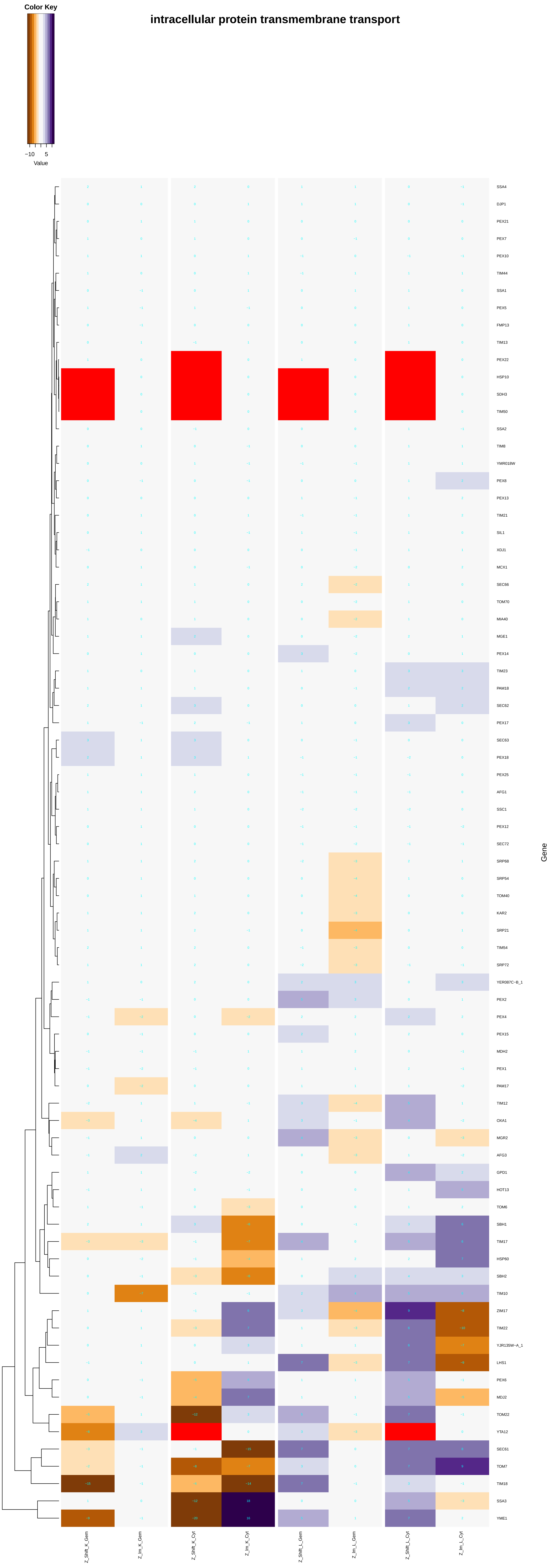
intracellular protein transmembrane transport
Color Key
−10
5
Value
SSA4
DJP1
PEX21
PEX7
PEX10
TIM44
SSA1
PEX5
FMP13
TIM13
PEX22
HSP10
SDH3
TIM50
SSA2
TIM8
YMR018W
PEX8
PEX13
TIM21
SIL1
XDJ1
MCX1
SEC66
TOM70
MIA40
MGE1
PEX14
TIM23
PAM18
SEC62
PEX17
SEC63
PEX18
PEX25
AFG1
SSC1
PEX12
SEC72
SRP68
SRP54
TOM40
KAR2
SRP21
TIM54
SRP72
YER087C−B_1
PEX2
PEX4
PEX15
MDH2
PEX1
PAM17
TIM12
OXA1
MGR2
AFG3
GPD1
HOT13
TOM6
SBH1
TIM17
HSP60
SBH2
TIM10
ZIM17
TIM22
YJR135W−A_1
LHS1
PEX6
MDJ2
TOM22
YTA12
SEC61
TOM7
TIM18
SSA3
YME1
Gene
Z_Shift_K_Gem
Z_Im_K_Gem
Z_Shift_K_Cyt
Z_Im_K_Cyt
Z_Shift_L_Gem
Z_Im_L_Gem
Z_Shift_L_Cyt
Z_Im_L_Cyt

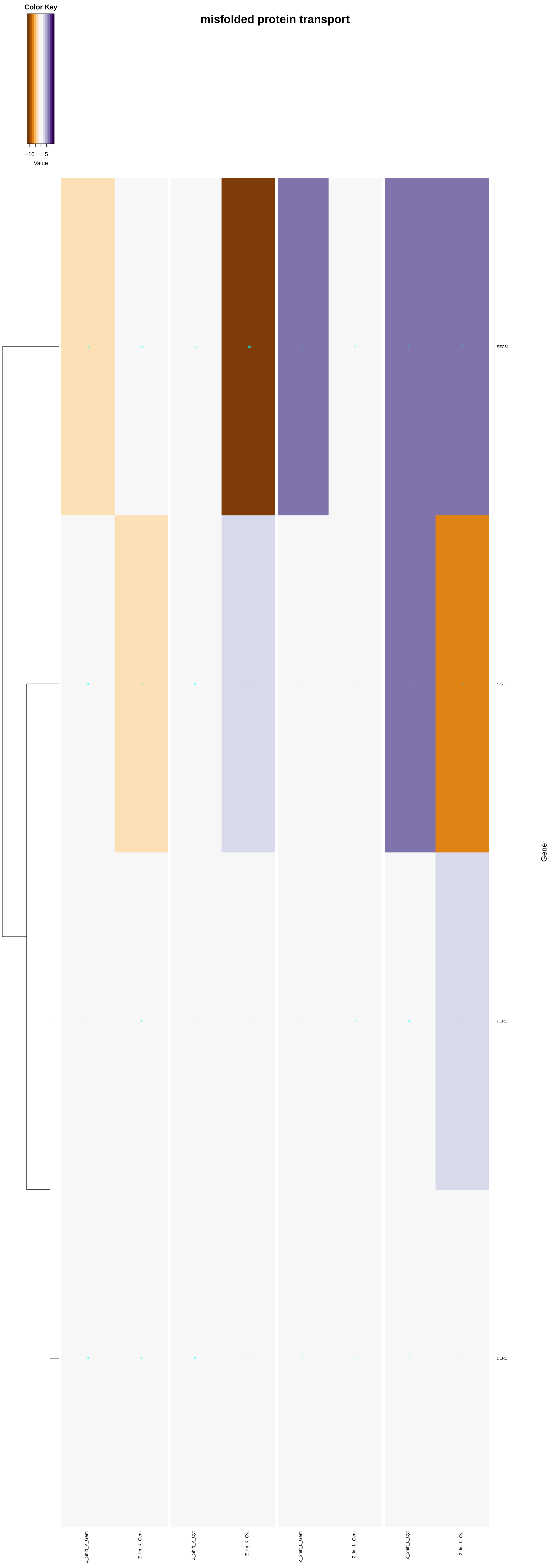
misfolded protein transport
Color Key
−10
5
Value
SEC61
SIS1
DER1
DER1
Gene
Z_Shift_K_Gem
Z_Im_K_Gem
Z_Shift_K_Cyt
Z_Im_K_Cyt
Z_Shift_L_Gem
Z_Im_L_Gem
Z_Shift_L_Cyt
Z_Im_L_Cyt

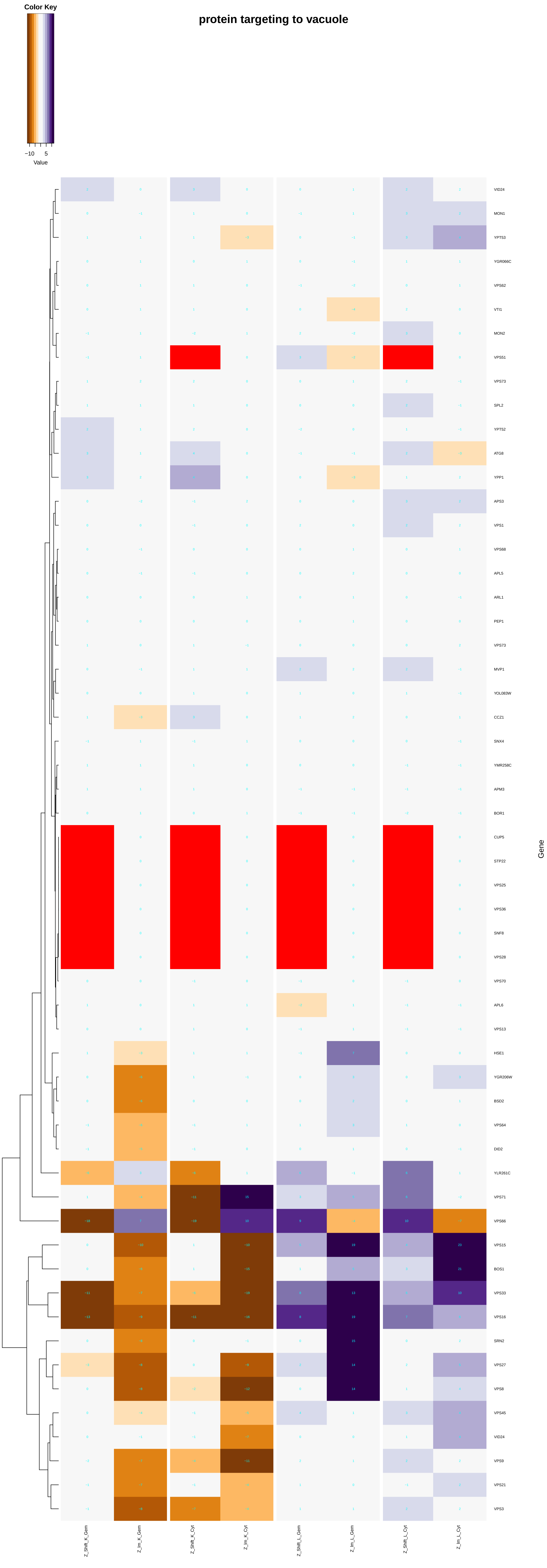
protein targeting to vacuole
Color Key
−10
5
Value
VID24
MON1
YPT53
YGR066C
VPS62
VTI1
MON2
VPS51
VPS73
SPL2
YPT52
ATG8
YPP1
APS3
VPS1
VPS68
APL5
ARL1
PEP1
VPS73
MVP1
YOL083W
CCZ1
SNX4
YMR258C
APM3
BOR1
CUP5
STP22
VPS25
VPS36
SNF8
VPS28
VPS70
APL6
VPS13
HSE1
YGR206W
BSD2
VPS64
DID2
YLR261C
VPS71
VPS66
VPS15
BOS1
VPS33
VPS16
SRN2
VPS27
VPS8
VPS45
VID24
VPS9
VPS21
VPS3
Gene
Z_Shift_K_Gem
Z_Im_K_Gem
Z_Shift_K_Cyt
Z_Im_K_Cyt
Z_Shift_L_Gem
Z_Im_L_Gem
Z_Shift_L_Cyt
Z_Im_L_Cyt

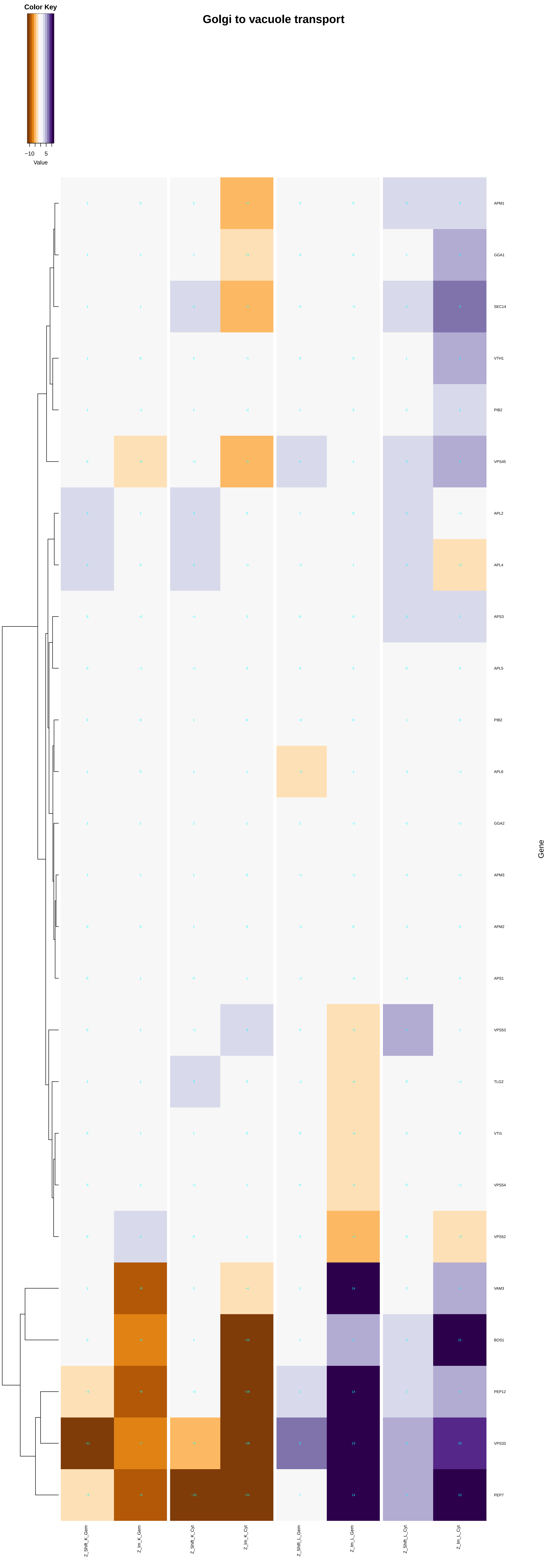
Golgi to vacuole transport
Color Key
−10
5
Value
APM1
GGA1
SEC14
VTH1
PIB2
VPS45
APL2
APL4
APS3
APL5
PIB2
APL6
GGA2
APM3
APM2
APS1
VPS53
TLG2
VTI1
VPS54
VPS52
VAM3
BOS1
PEP12
VPS33
PEP7
Gene
Z_Shift_K_Gem
Z_Im_K_Gem
Z_Shift_K_Cyt
Z_Im_K_Cyt
Z_Shift_L_Gem
Z_Im_L_Gem
Z_Shift_L_Cyt
Z_Im_L_Cyt

#### late endosome to vacuole transport

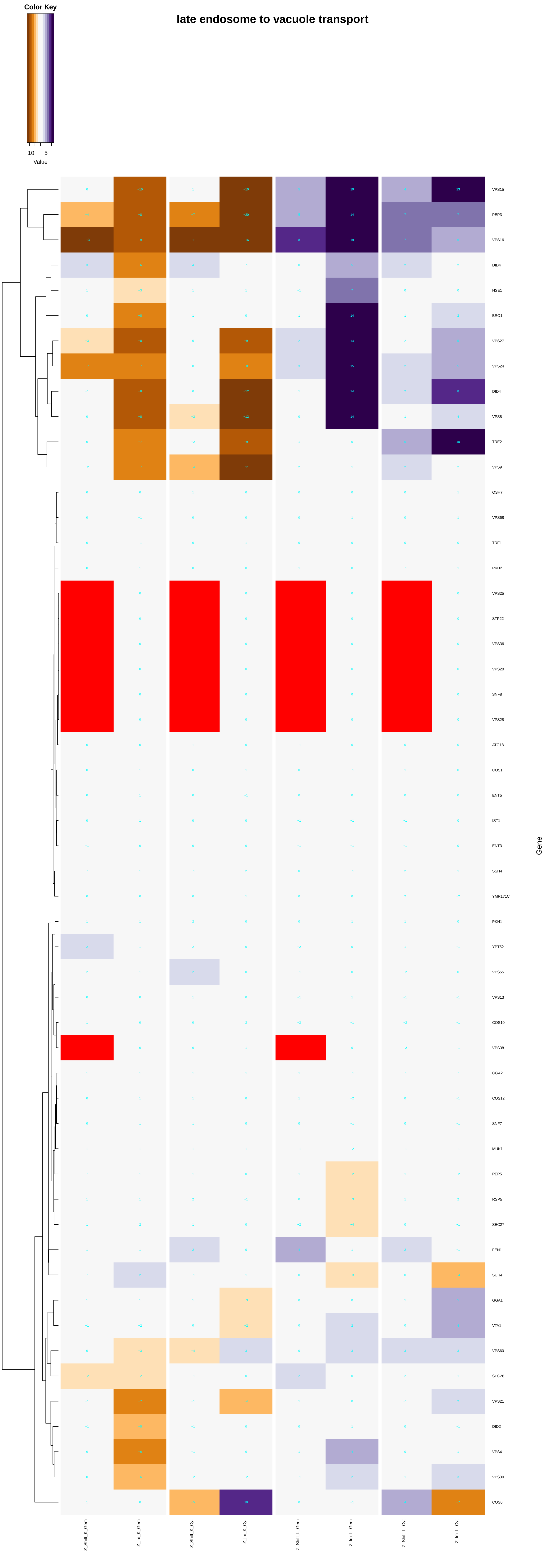

### Golgi to endosome transport

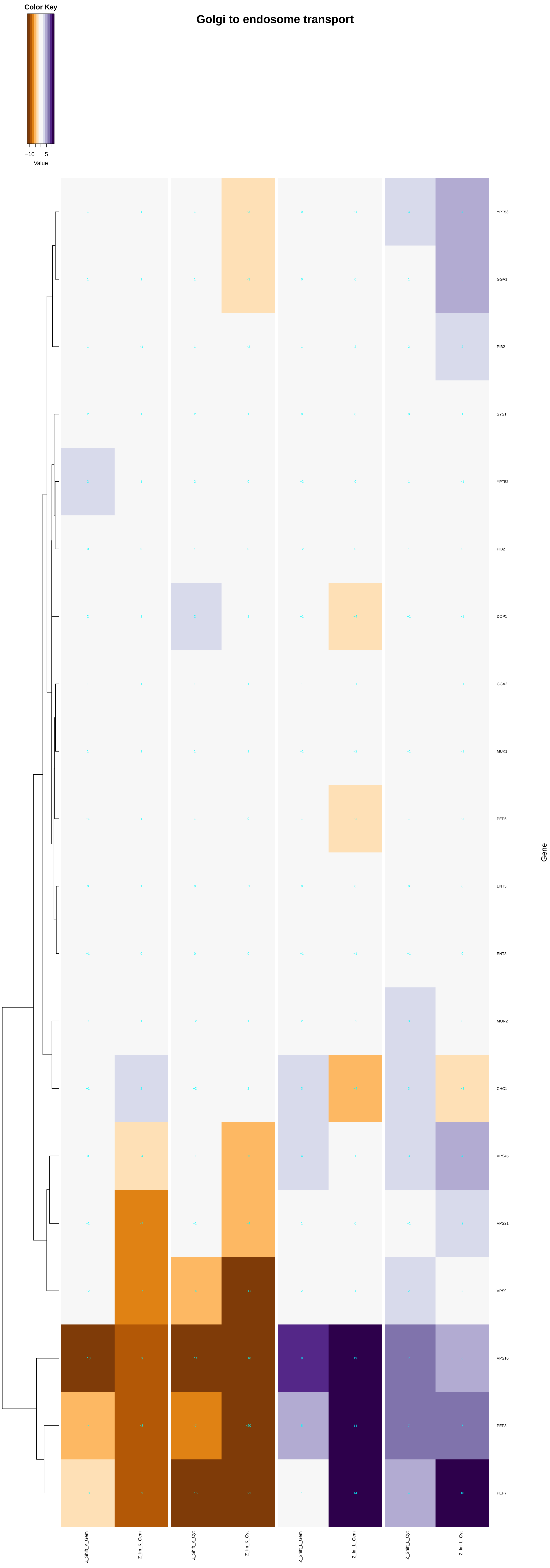

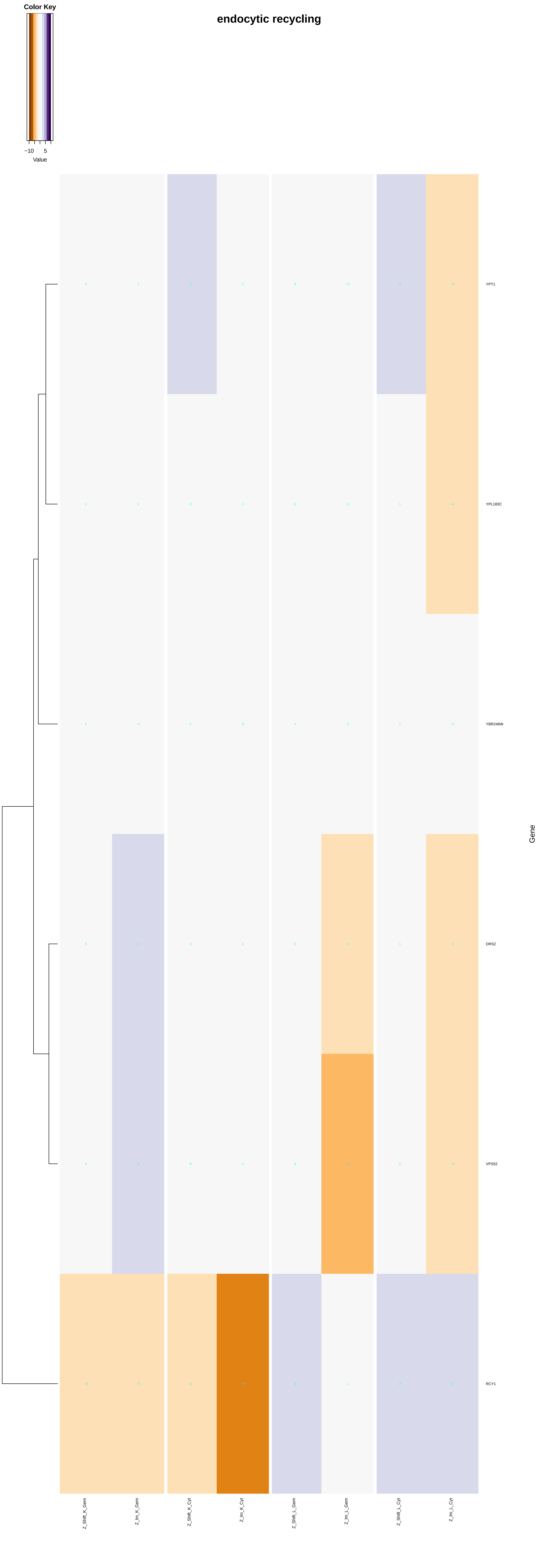

endocytic recycling
Color Key
−10
5
Value
YPT1
YPL183C
YBR246W
DRS2
VPS52
RCY1
Gene
Z_Shift_K_Gem
Z_Im_K_Gem
Z_Shift_K_Cyt
Z_Im_K_Cyt
Z_Shift_L_Gem
Z_Im_L_Gem
Z_Shift_L_Cyt
Z_Im_L_Cyt

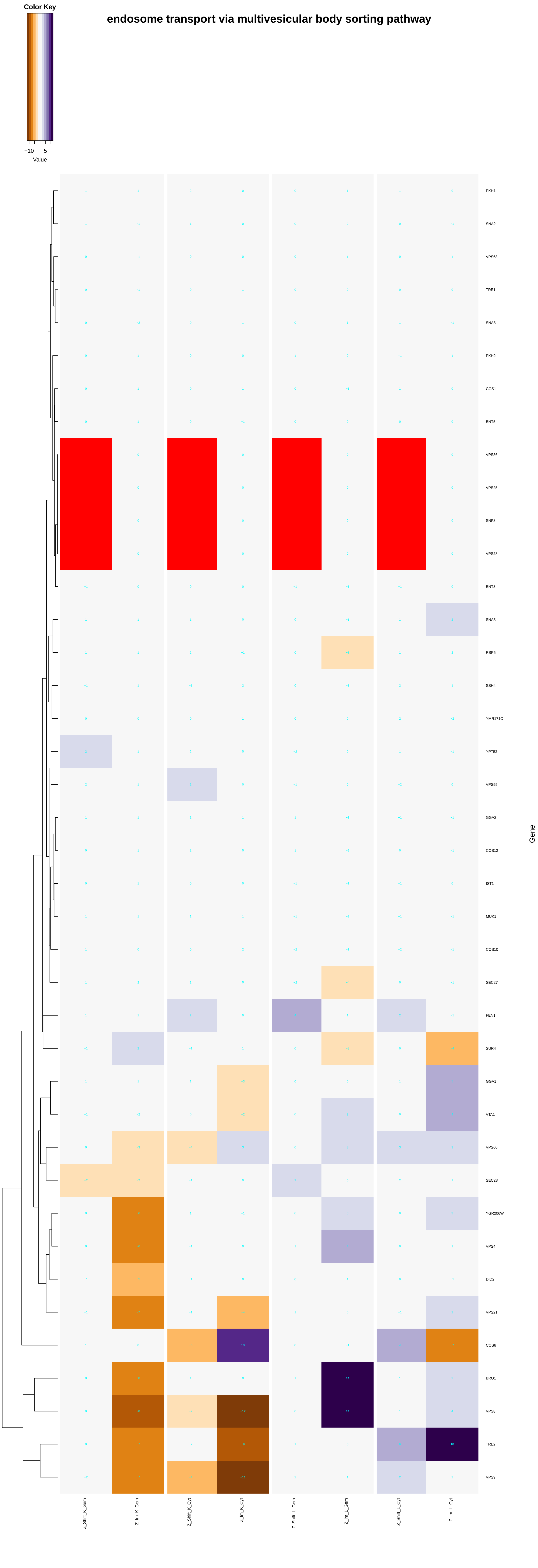
endosome transport via multivesicular body sorting pathway
Color Key
−10
5
Value
PKH1
SNA2
VPS68
TRE1
SNA3
PKH2
COS1
ENT5
VPS36
VPS25
SNF8
VPS28
ENT3
SNA3
RSP5
SSH4
YMR171C
YPT52
VPS55
GGA2
COS12
IST1
MUK1
COS10
SEC27
FEN1
SUR4
GGA1
VTA1
VPS60
SEC28
YGR206W
VPS4
DID2
VPS21
COS6
BRO1
VPS8
TRE2
VPS9
Gene
Z_Shift_K_Gem
Z_Im_K_Gem
Z_Shift_K_Cyt
Z_Im_K_Cyt
Z_Shift_L_Gem
Z_Im_L_Gem
Z_Shift_L_Cyt
Z_Im_L_Cyt

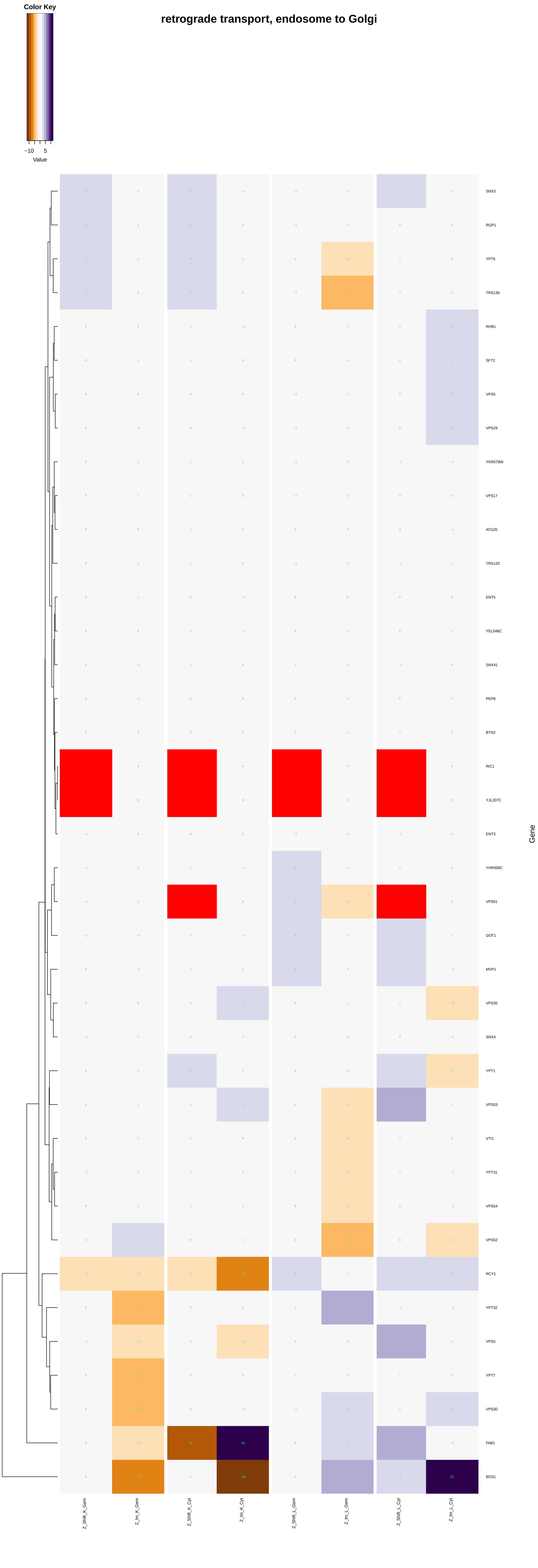

Color Key
−10
5
Value
retrograde transport, endosome to Golgi
SNX3
RGP1
YPT6
TRS130
RHB1
SFT2
VPS5
VPS29
YKR078W
VPS17
ATG20
TRS120
ENT5
YEL048C
SNX41
PEP8
BTN2
RIC1
YJL207C
ENT3
YHR009C
VPS51
GOT1
MVP1
VPS35
SNX4
YPT1
VPS53
VTI1
YPT31
VPS54
VPS52
RCY1
YPT32
VPS5
YPT7
VPS30
FAB1
BOS1
Gene
Z_Shift_K_Gem
Z_Im_K_Gem
Z_Shift_K_Cyt
Z_Im_K_Cyt
Z_Shift_L_Gem
Z_Im_L_Gem
Z_Shift_L_Cyt
Z_Im_L_Cyt

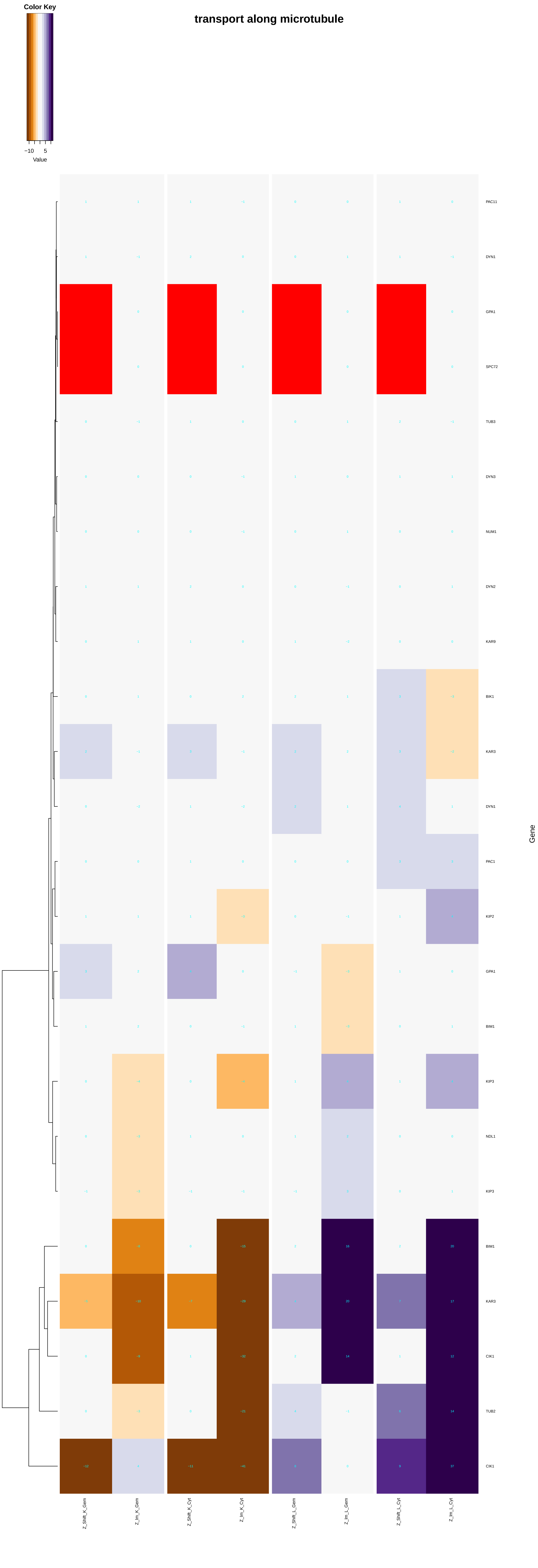
Color Key
−10
5
Value
transport along microtubule
PAC11
DYN1
GPA1
SPC72
TUB3
DYN3
NUM1
DYN2
KAR9
BIK1
KAR3
DYN1
PAC1
KIP2
GPA1
BIM1
KIP3
NDL1
KIP3
BIM1
KAR3
CIK1
TUB2
CIK1
Gene
Z_Shift_K_Gem
Z_Im_K_Gem
Z_Shift_K_Cyt
Z_Im_K_Cyt
Z_Shift_L_Gem
Z_Im_L_Gem
Z_Shift_L_Cyt
Z_Im_L_Cyt

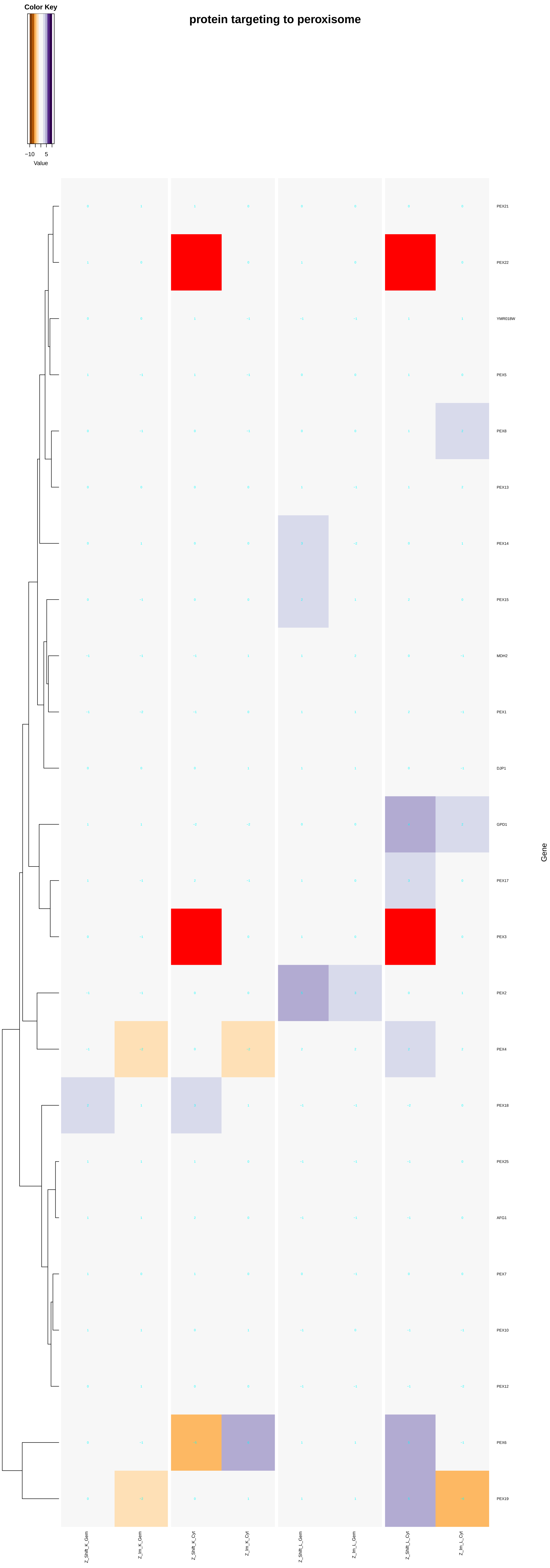

Color Key
−10
5
Value
protein targeting to peroxisome
PEX21
PEX22
YMR018W
PEX5
PEX8
PEX13
PEX14
PEX15
MDH2
PEX1
DJP1
GPD1
PEX17
PEX3
PEX2
PEX4
PEX18
PEX25
AFG1
PEX7
PEX10
PEX12
PEX6
PEX19
Gene
Z_Shift_K_Gem
Z_Int_K_Gem
Z_Shift_K_Cyt
Z_Int_K_Cyt
Z_Shift_L_Gem
Z_Int_L_Gem
Z_Shift_L_Cyt
Z_Int_L_Cyt

#### peroxisomal membrane transport

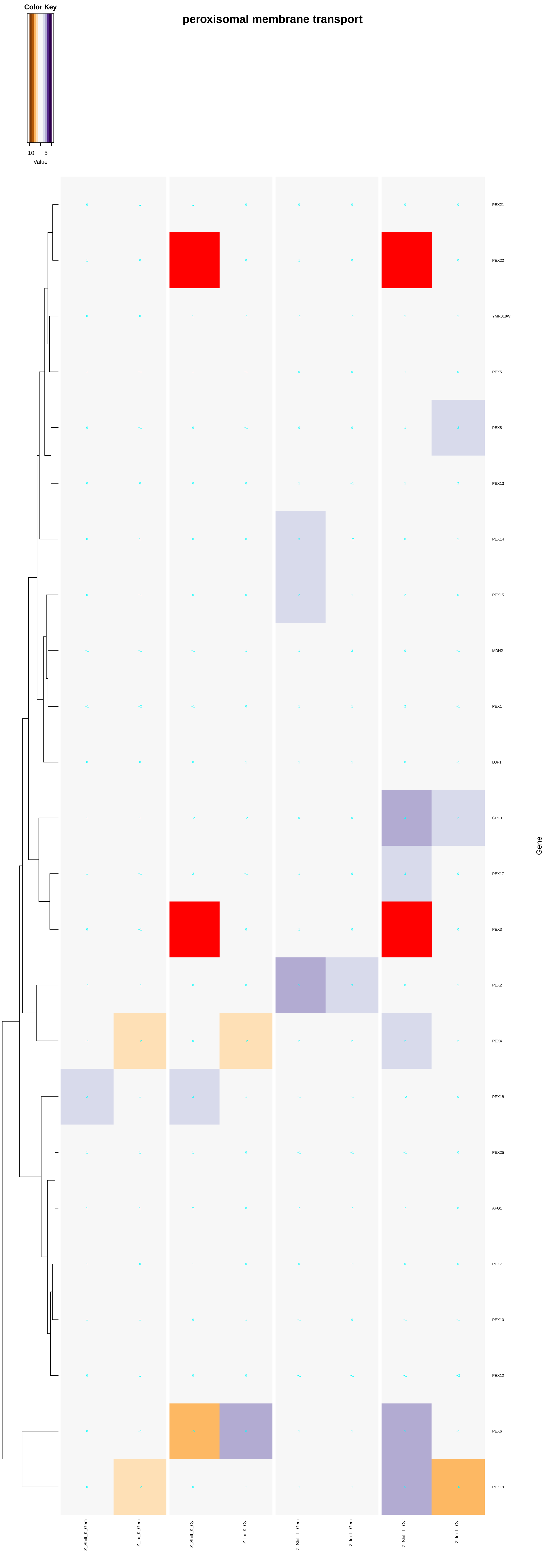

### rRNA export from nucleus

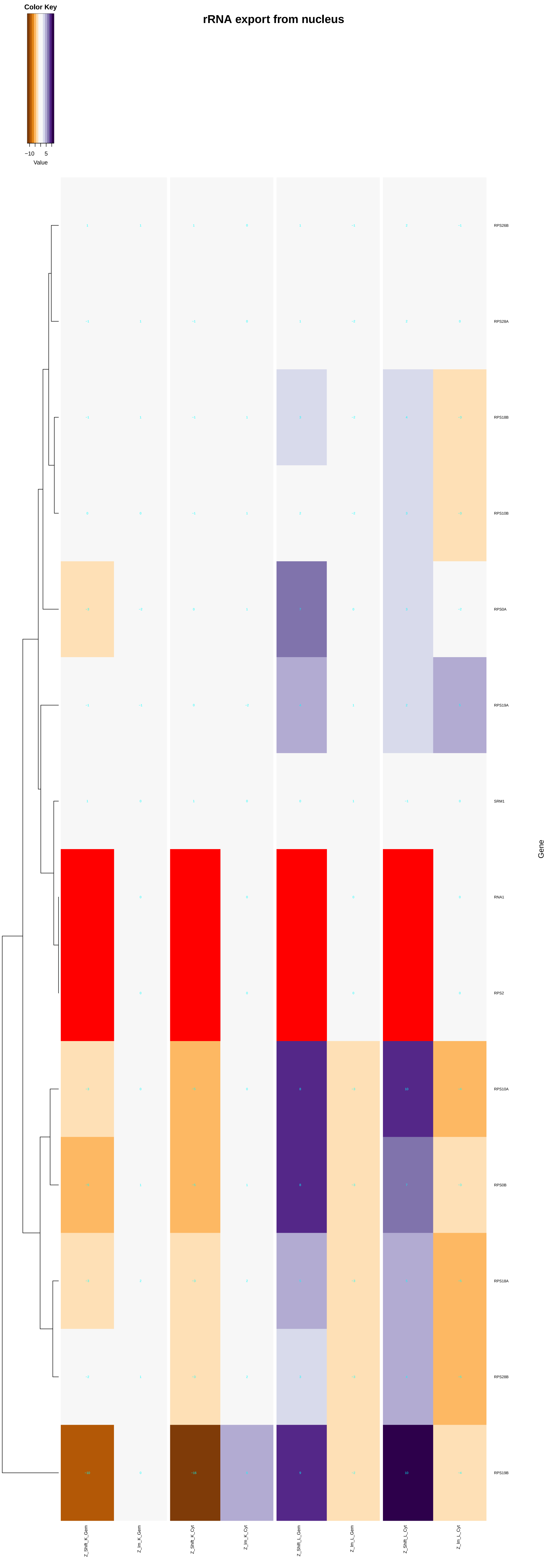

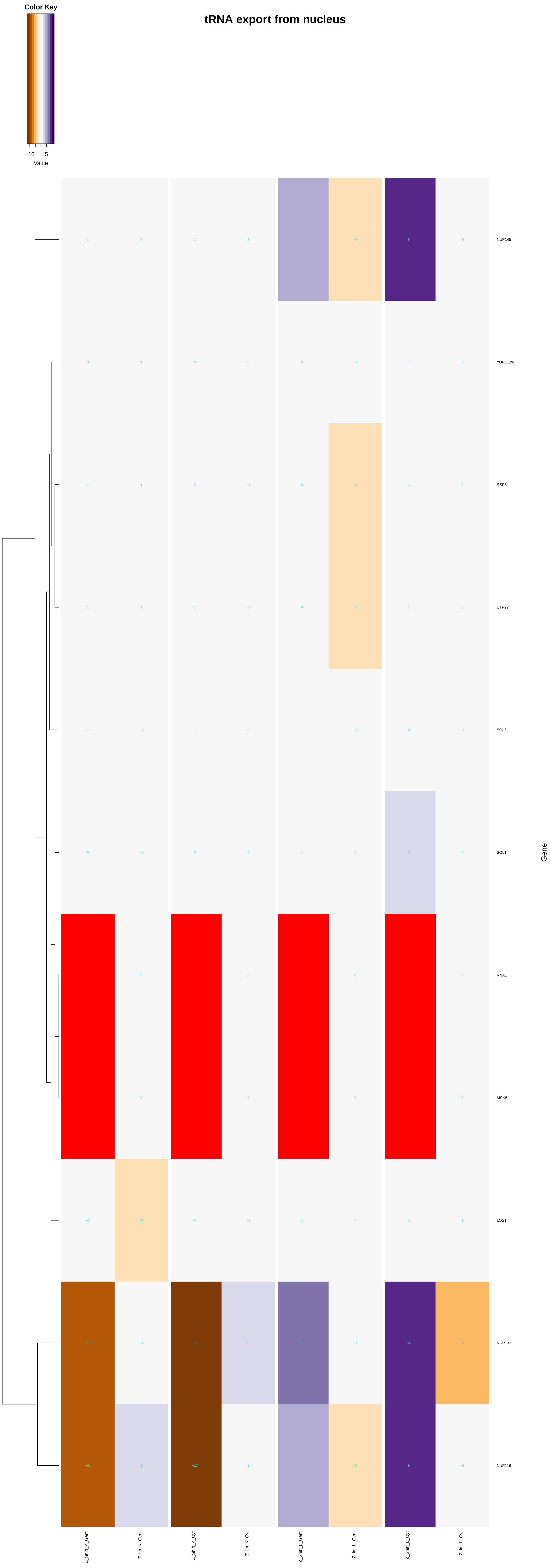

Color Key
−10
5
Value
tRNA export from nucleus
NUP145
YOR112W
RSP5
UTP22
SOL2
SOL1
RNA1
MSN5
LOS1
NUP133
NUP116
Gene
Z_Shift_K_Gem
Z_Im_K_Gem
Z_Shift_K_Cyt
Z_Im_K_Cyt
Z_Shift_L_Gem
Z_Im_L_Gem
Z_Shift_L_Cyt
Z_Im_L_Cyt

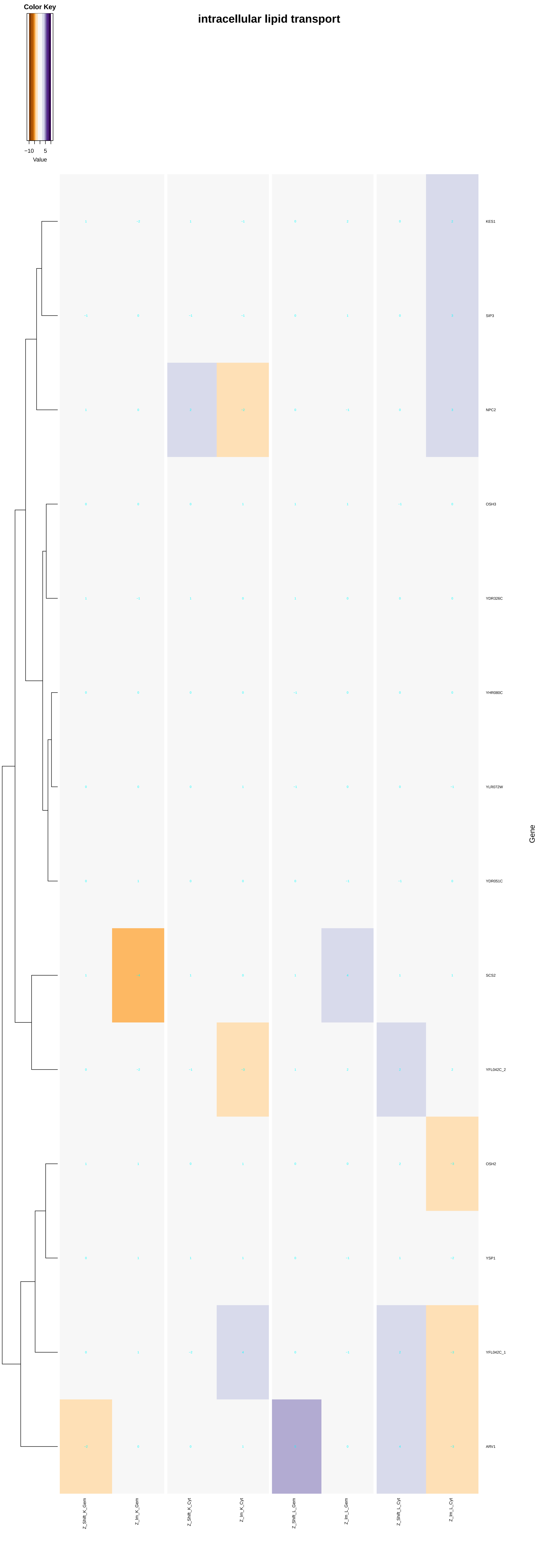

intracellular lipid transport
Color Key
−10
5
Value
KES1
SIP3
NPC2
OSH3
YDR326C
YHR080C
YLR072W
YDR051C
SCS2
YFL042C_2
OSH2
YSP1
YFL042C_1
ARV1
Gene
Z_Shift_K_Gem
Z_Im_K_Gem
Z_Shift_K_Cyt
Z_Im_K_Cyt
Z_Shift_L_Gem
Z_Im_L_Gem
Z_Shift_L_Cyt
Z_Im_L_Cyt

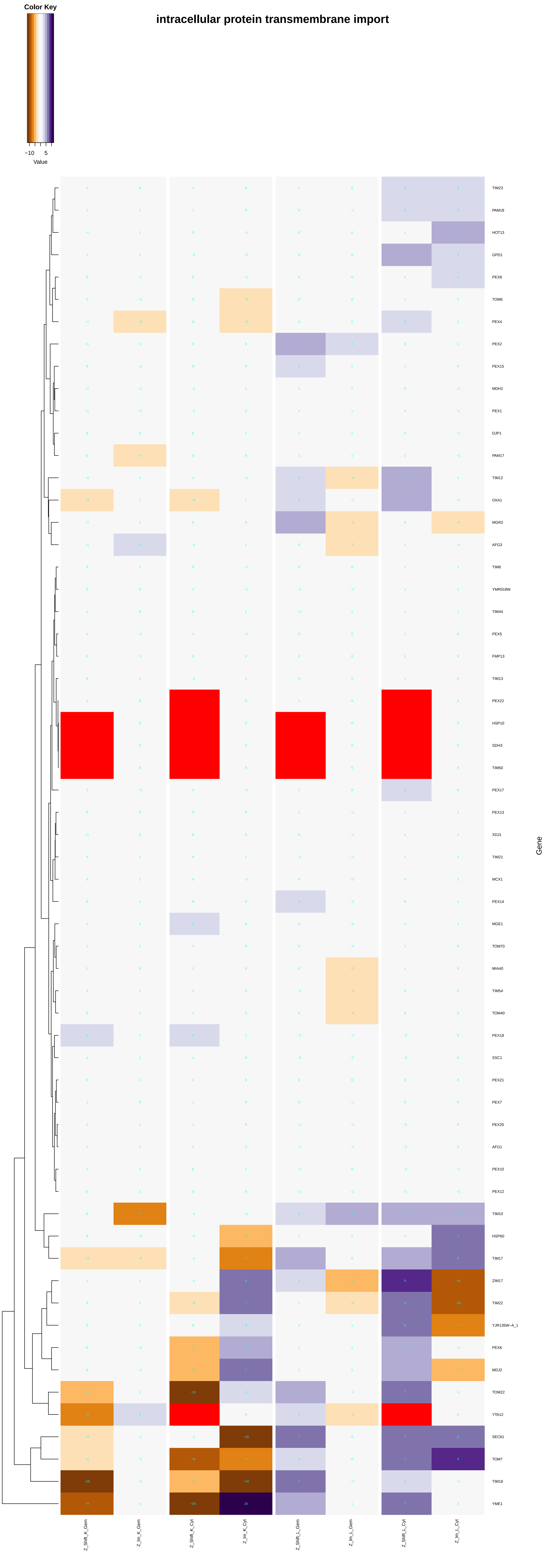
intracellular protein transmembrane import
Color Key
−10
5
Value
TIM23
PAM18
HOT13
GPD1
PEX8
TOM6
PEX4
PEX2
PEX15
MDH2
PEX1
DJP1
PAM17
TIM12
OXA1
MGR2
AFG3
TIM8
YMR018W
TIM44
PEX5
FMP13
TIM13
PEX22
HSP10
SDH3
TIM50
PEX17
PEX13
XDJ1
TIM21
MCX1
PEX14
MGE1
TOM70
MIA40
TIM54
TOM40
PEX18
SSC1
PEX21
PEX7
PEX25
AFG1
PEX10
PEX12
TIM10
HSP60
TIM17
ZIM17
TIM22
YJR135W-A_1
PEX6
MDJ2
TOM22
YTA12
SEC61
TOM7
TIM18
YME1
Gene
Z_Shift_K_Gem
Z_Im_K_Gem
Z_Shift_K_Cyt
Z_Im_K_Cyt
Z_Shift_L_Gem
Z_Im_L_Gem
Z_Shift_L_Cyt
Z_Im_L_Cyt

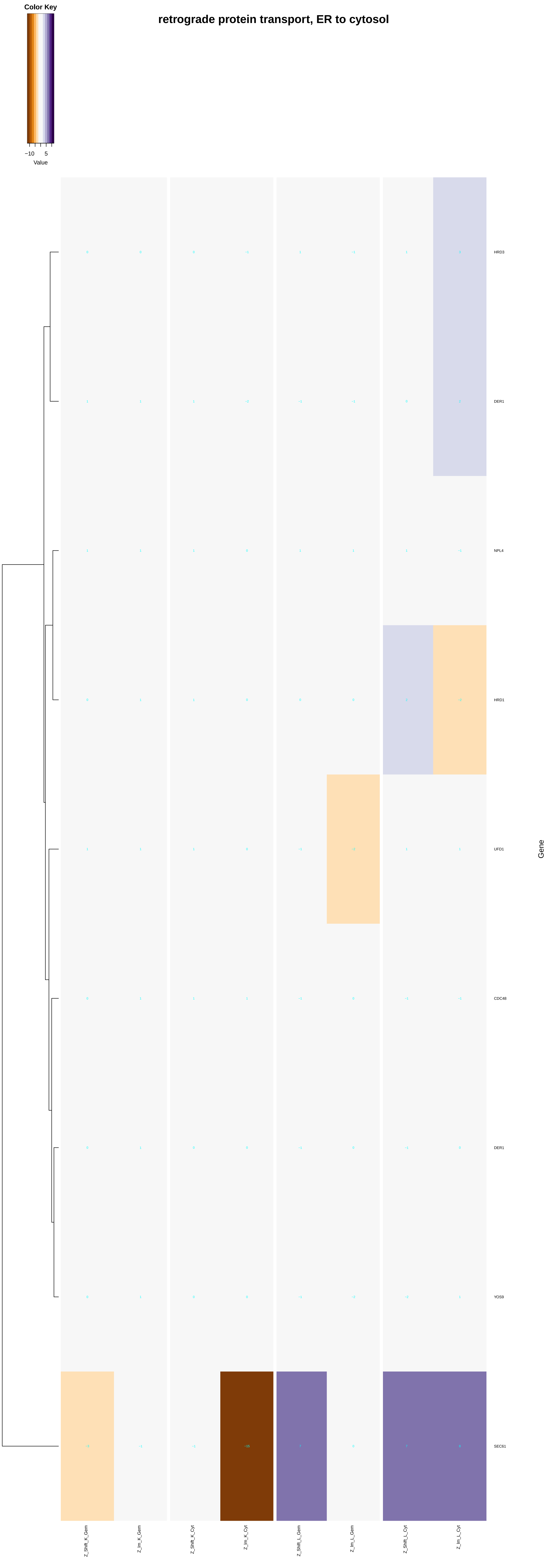
retrograde protein transport, ER to cytosol
Color Key
−10
5
Value
HRD3
DER1
NPL4
HRD1
UFD1
CDC48
DER1
YOS9
SEC61
Gene
Z_Shift_K_Gem
Z_Im_K_Gem
Z_Shift_K_Cyt
Z_Im_K_Cyt
Z_Shift_L_Gem
Z_Im_L_Gem
Z_Shift_L_Cyt
Z_Im_L_Cyt

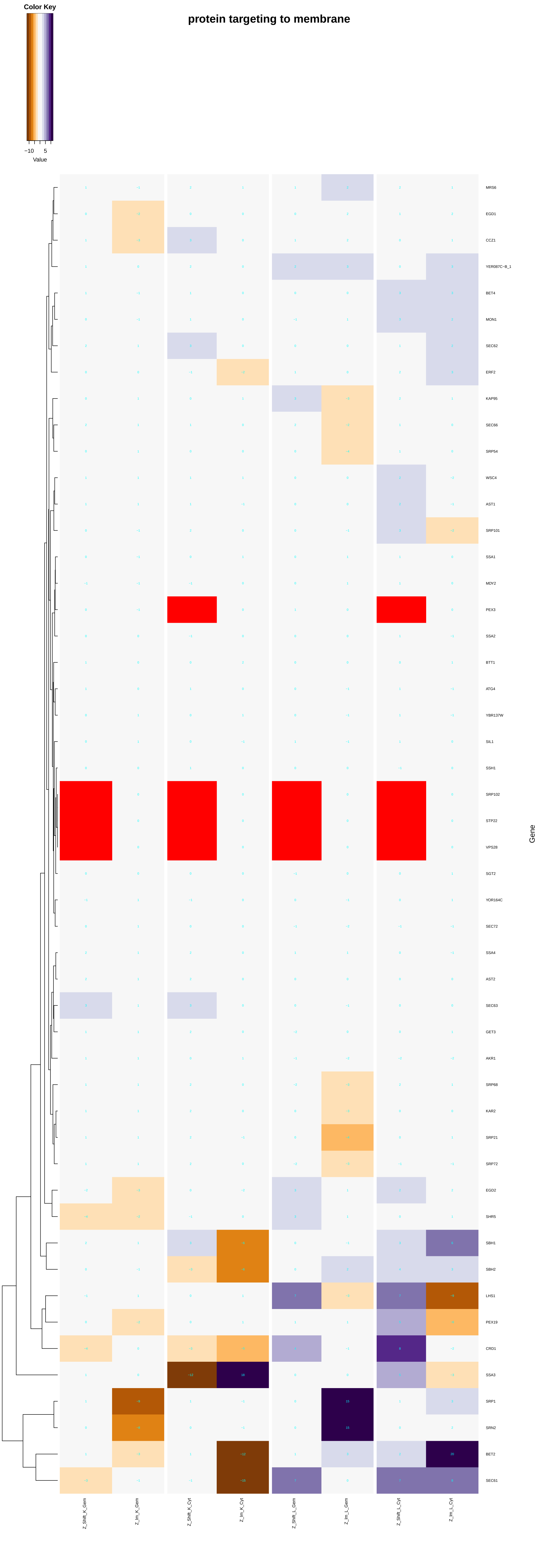
protein targeting to membrane
Color Key
−10
5
Value
MRS6
EGD1
CCZ1
YER087C−B_1
BET4
MON1
SEC62
ERF2
KAP95
SEC66
SRP54
WSC4
AST1
SRP101
SSA1
MDY2
PEX3
SSA2
BTT1
ATG4
YBR137W
SIL1
SSH1
SRP102
STP22
VPS28
SGT2
YOR164C
SEC72
SSA4
AST2
SEC63
GET3
AKR1
SRP68
KAR2
SRP21
SRP72
EGD2
SHR5
SBH1
SBH2
LHS1
PEX19
CRD1
SSA3
SRP1
SRN2
BET2
SEC61
Gene
Z_Shift_K_Gem
Z_Im_K_Gem
Z_Shift_K_Cyt
Z_Im_K_Cyt
Z_Shift_L_Gem
Z_Im_L_Gem
Z_Shift_L_Cyt
Z_Im_L_Cyt

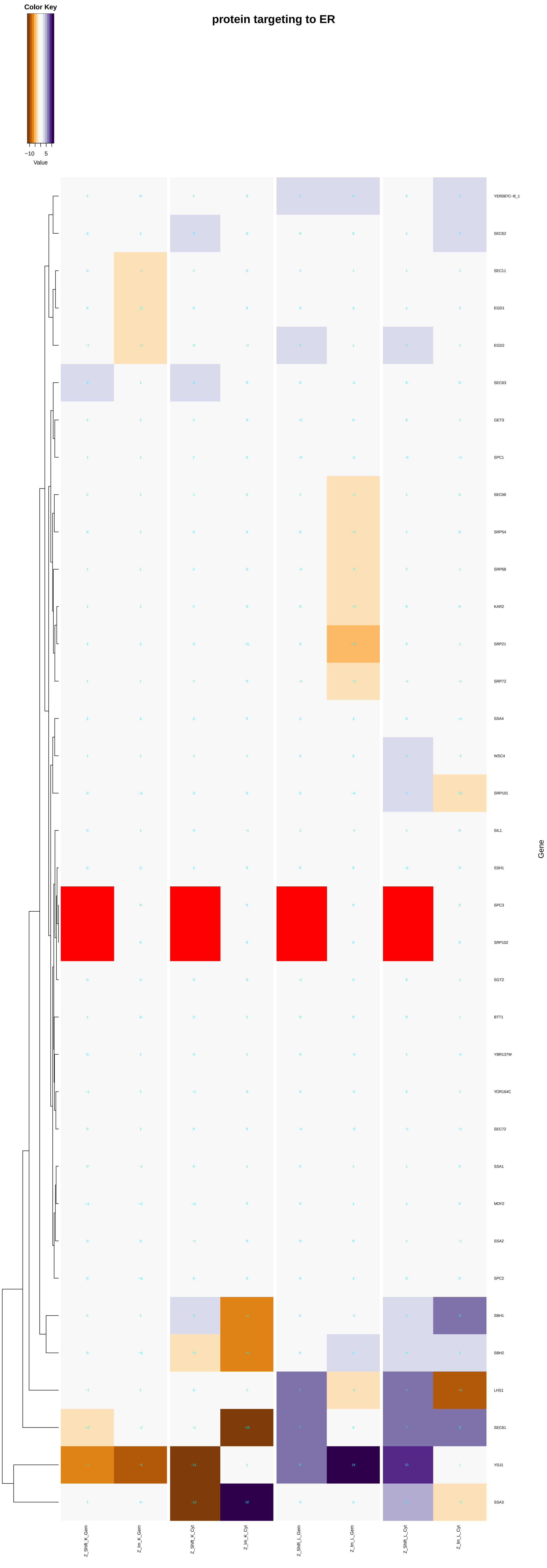

Color Key
−10
5
Value
protein targeting to ER
YER087C−B_1
SEC62
SEC11
EGD1
EGD2
SEC63
GET3
SPC1
SEC66
SRP54
SRP68
KAR2
SRP21
SRP72
SSA4
WSC4
SRP101
SIL1
SSH1
SPC3
SRP102
SGT2
BTT1
YBR137W
YOR164C
SEC72
SSA1
MDY2
SSA2
SPC2
SBH1
SBH2
LHS1
SEC61
YDJ1
SSA3
Gene
Z_Shift_K_Gem
Z_Im_K_Gem
Z_Shift_K_Cyt
Z_Im_K_Cyt
Z_Shift_L_Gem
Z_Im_L_Gem
Z_Shift_L_Cyt
Z_Im_L_Cyt

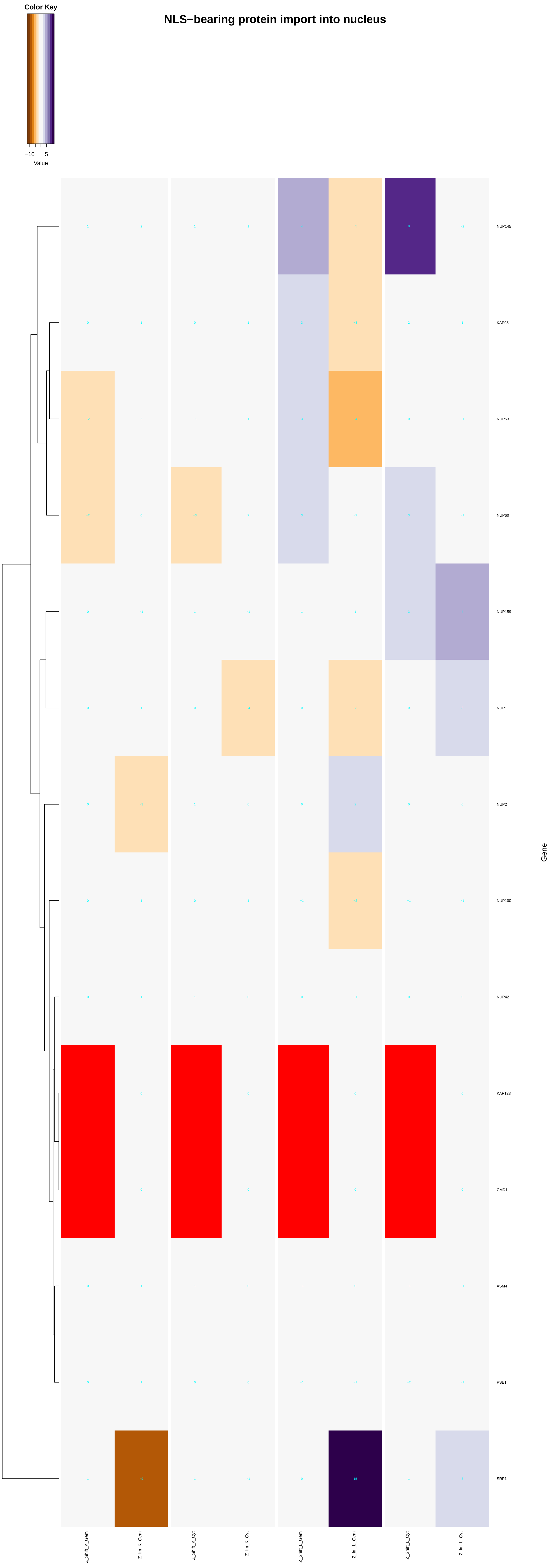
NLS−bearing protein import into nucleus
Color Key
−10
5
Value
NUP145
KAP95
NUP53
NUP60
NUP159
NUP1
NUP2
NUP100
NUP42
KAP123
CMD1
ASM4
PSE1
SRP1
Gene
Z_Shift_K_Gem
Z_Im_K_Gem
Z_Shift_K_Cyt
Z_Im_K_Cyt
Z_Shift_L_Gem
Z_Im_L_Gem
Z_Shift_L_Cyt
Z_Im_L_Cyt

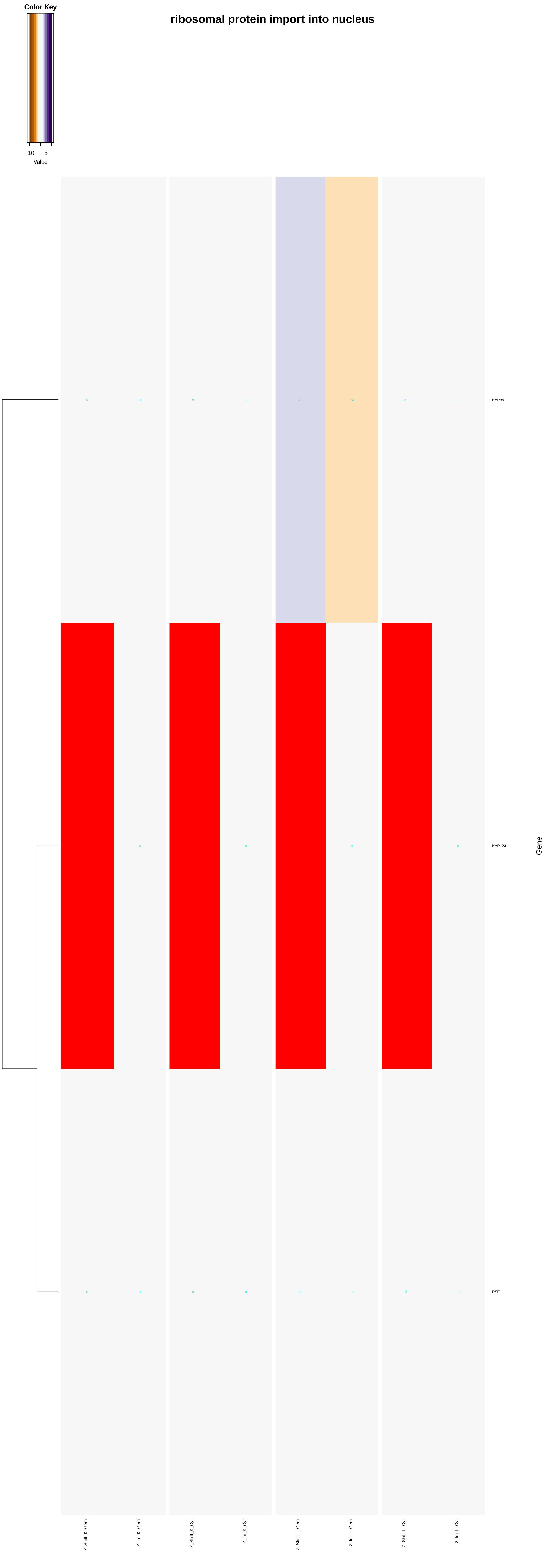
ribosomal protein import into nucleus
Color Key
−10
5
Value
KAP95
KAP123
PSE1
Gene
Z_Shift_K_Gem
Z_Im_K_Gem
Z_Shift_K_Cyt
Z_Im_K_Cyt
Z_Shift_L_Gem
Z_Im_L_Gem
Z_Shift_L_Cyt
Z_Im_L_Cyt

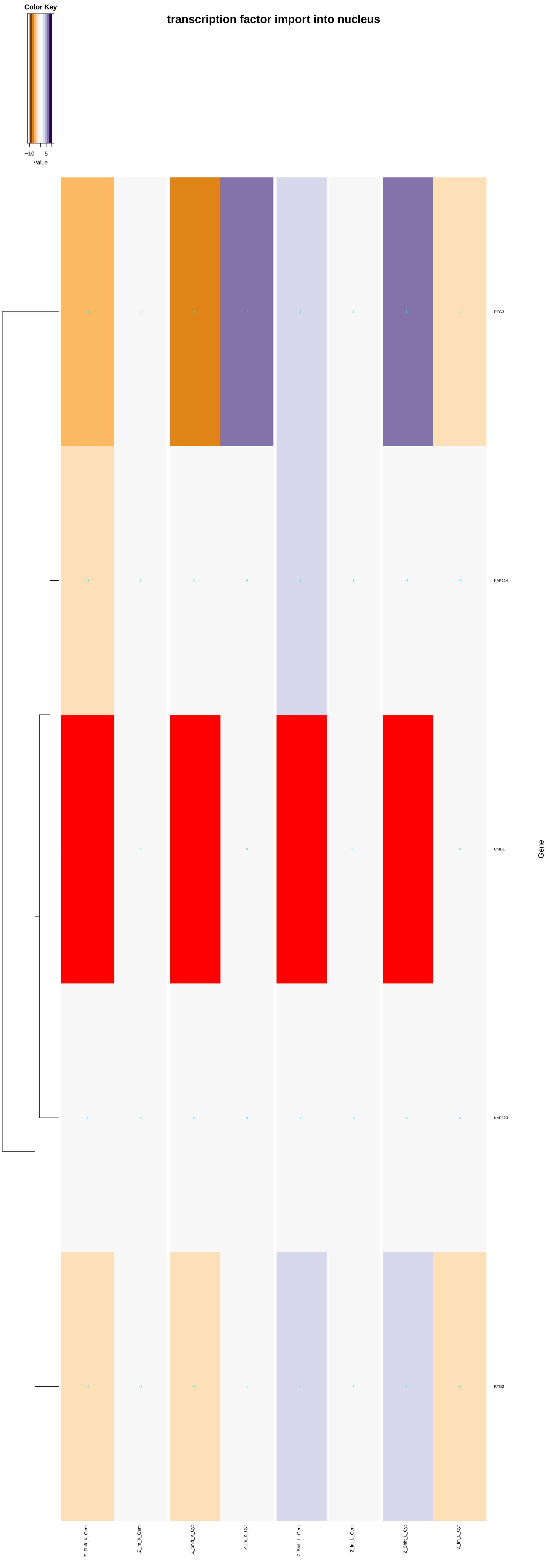
transcription factor import into nucleus
Color Key
−10
5
Value
RTG3
KAP114
CMD1
KAP120
RTG2
Gene
Z_Shift_K_Gem
Z_Im_K_Gem
Z_Shift_K_Cyt
Z_Im_K_Cyt
Z_Shift_L_Gem
Z_Im_L_Gem
Z_Shift_L_Cyt
Z_Im_L_Cyt

### RNA polymerase II complex import to nucleus

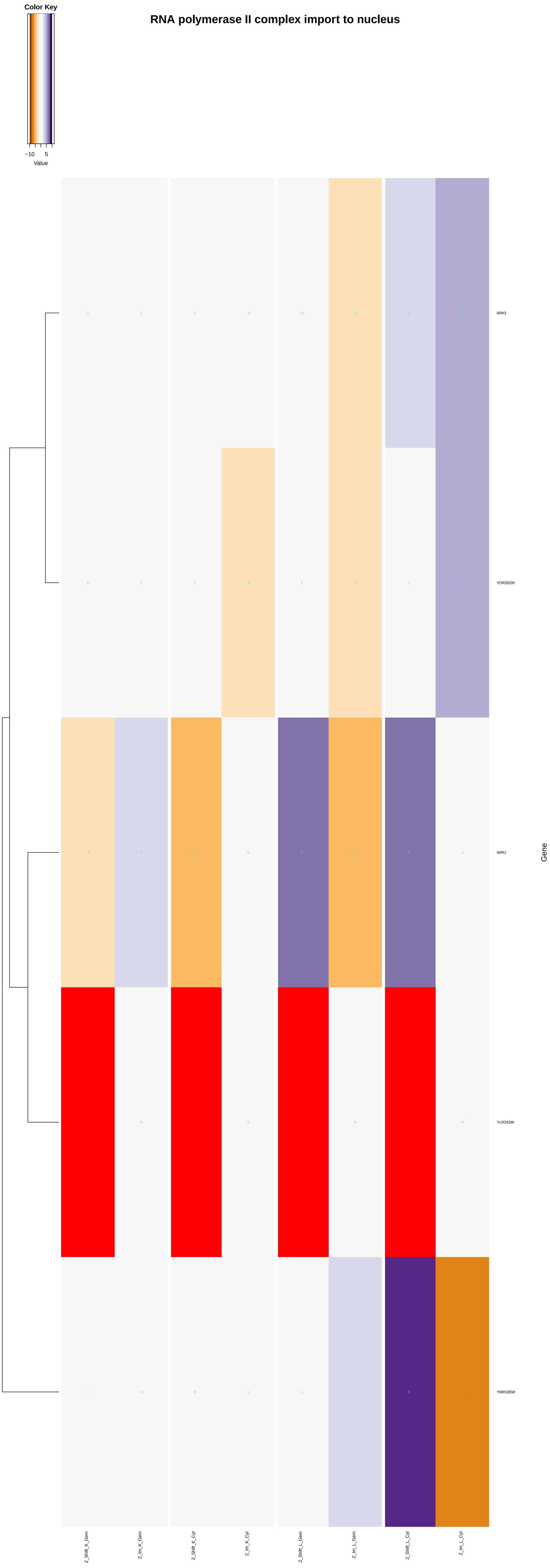

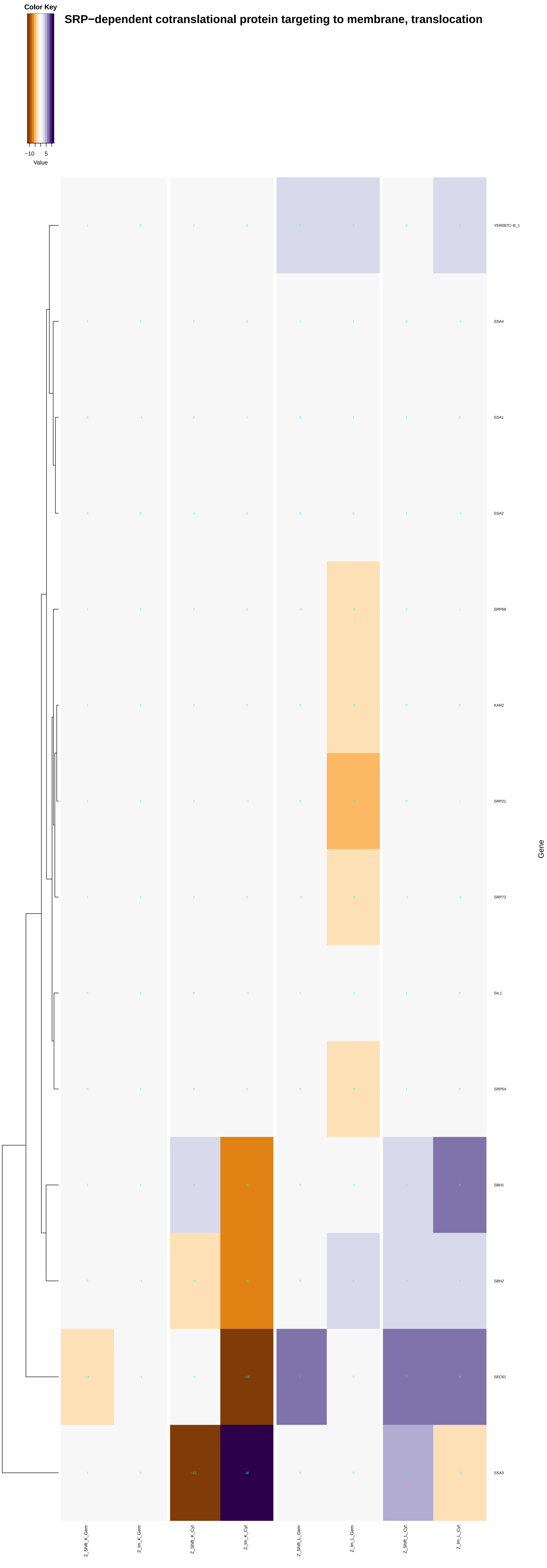

SRP−dependent cotranslational protein targeting to membrane, translocation
Color Key
−10
5
Value
YER087C−B_1
SSA4
SSA1
SSA2
SRP68
KAR2
SRP21
SRP72
SIL1
SRP54
SBH1
SBH2
SEC61
SSA3
Gene
Z_Shift_K_Gem
Z_Im_K_Gem
Z_Shift_K_Cyt
Z_Im_K_Cyt
Z_Shift_L_Gem
Z_Im_L_Gem
Z_Shift_L_Cyt
Z_Im_L_Cyt

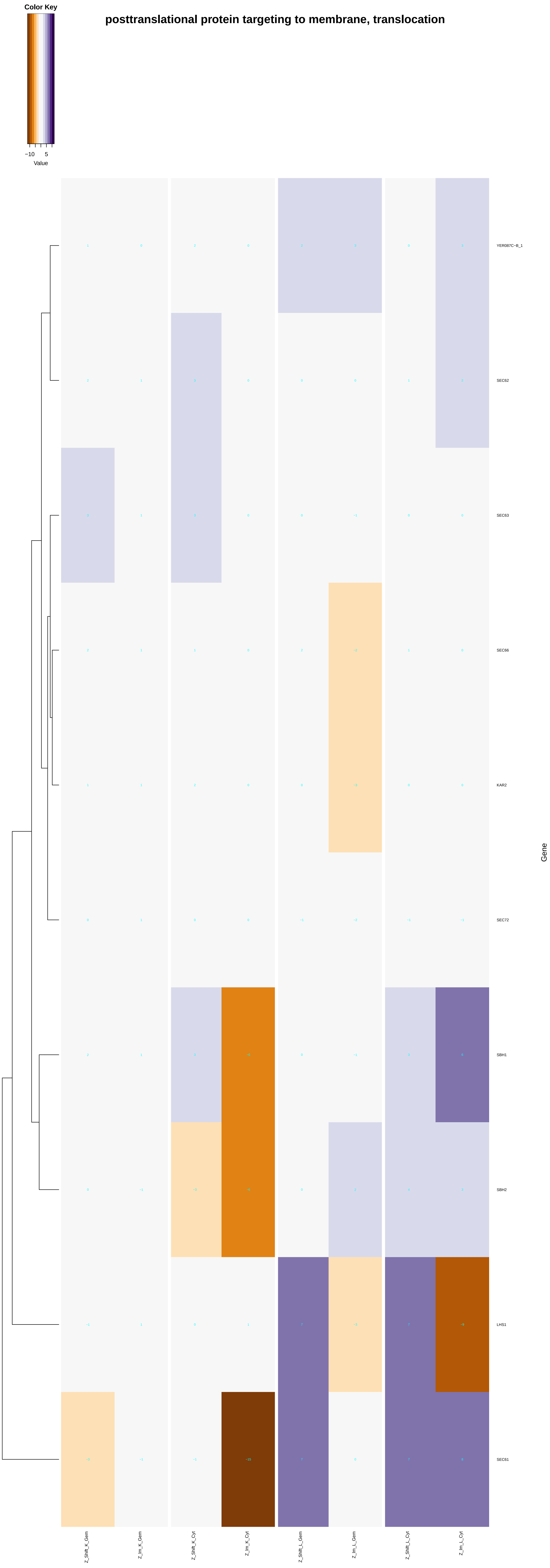

Color Key
−10
5
Value
posttranslational protein targeting to membrane, translocation
YER087C−B_1
SEC62
SEC63
SEC66
KAR2
SEC72
SBH1
SBH2
LHS1
SEC61
Gene
Z_Shift_K_Gem
Z_Im_K_Gem
Z_Shift_K_Cyt
Z_Im_K_Cyt
Z_Shift_L_Gem
Z_Im_L_Gem
Z_Shift_L_Cyt
Z_Im_L_Cyt

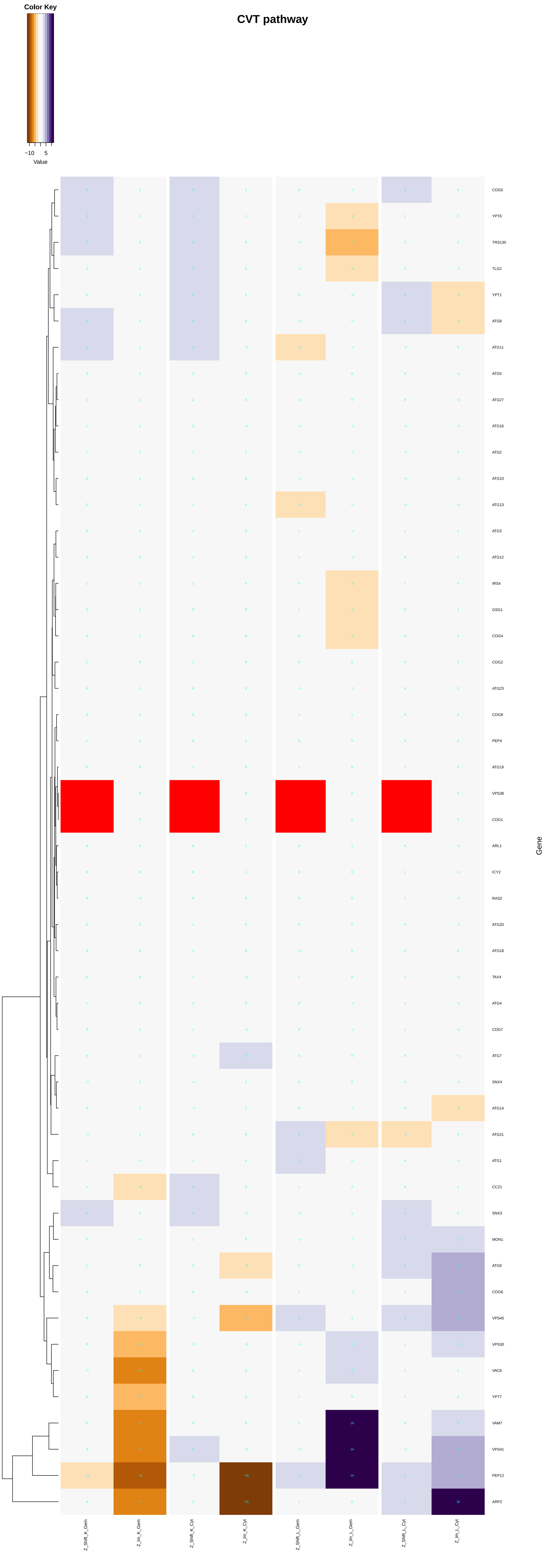
CVT pathway
Color Key
−10
5
Value
COG5
YPT6
TRS130
TLG2
YPT1
ATG8
ATG11
ATG5
ATG27
ATG16
ATG2
ATG10
ATG13
ATG3
ATG12
IRS4
GSG1
COG4
COG2
ATG23
COG8
PEP4
ATG19
VPS36
COG1
ARL1
ICY2
RAS2
ATG20
ATG18
TAX4
ATG4
COG7
ATG7
SNX4
ATG14
ATG21
ATG1
CCZ1
SNX3
MON1
ATG9
COG6
VPS45
VPS30
VAC8
YPT7
VAM7
VPS41
PEP12
ARP2
Gene
Z_Shift_K_Gem
Z_Im_K_Gem
Z_Shift_K_Cyt
Z_Im_K_Cyt
Z_Shift_L_Gem
Z_Im_L_Gem
Z_Shift_L_Cyt
Z_Im_L_Cyt

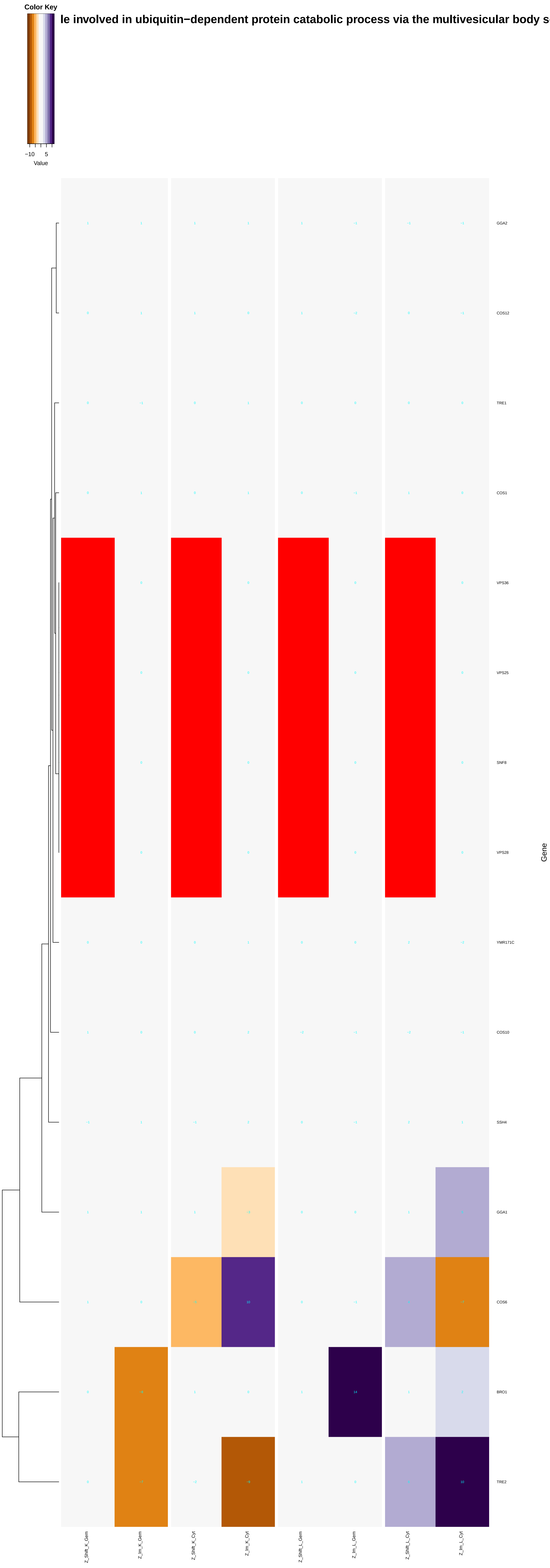

le involved in ubiquitin−dependent protein catabolic process via the multivesicular body s
Color Key
−10
5
Value
GGA2
COS12
TRE1
COS1
VPS36
VPS25
SNF8
VPS28
YMR171C
COS10
SSH4
GGA1
COS6
BRO1
TRE2
Gene
Z_Shift_K_Gem
Z_Im_K_Gem
Z_Shift_K_Cyt
Z_Im_K_Cyt
Z_Shift_L_Gem
Z_Im_L_Gem
Z_Shift_L_Cyt
Z_Im_L_Cyt

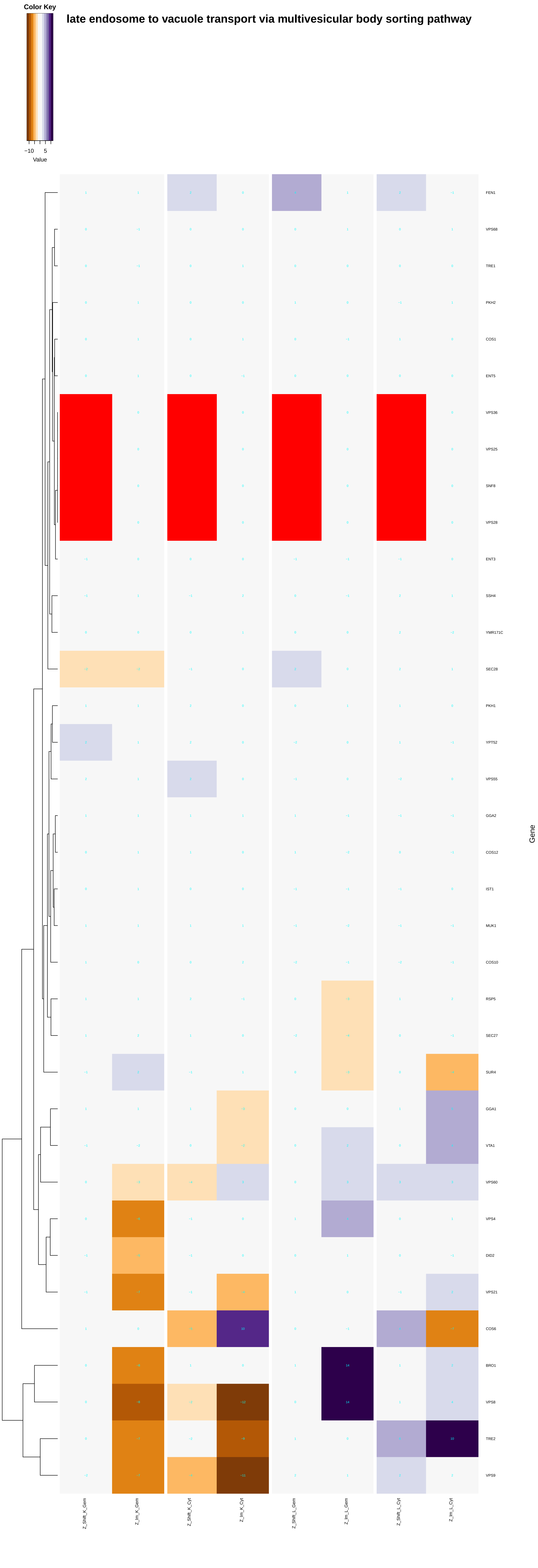

Color Key
−10
5
Value
late endosome to vacuole transport via multivesicular body sorting pathway
FEN1
VPS68
TRE1
PKH2
COS1
ENT5
VPS36
VPS25
SNF8
VPS28
ENT3
SSH4
YMR171C
SEC28
PKH1
YPT52
VPS55
GGA2
COS12
IST1
MUK1
COS10
RSP5
SEC27
SUR4
GGA1
VTA1
VPS60
VPS4
DID2
VPS21
COS6
BRO1
VPS8
TRE2
VPS9
Gene
Z_Shift_K_Gem
Z_lm_K_Gem
Z_Shift_K_Cyt
Z_lm_K_Cyt
Z_Shift_L_Gem
Z_lm_L_Gem
Z_Shift_L_Cyt
Z_lm_L_Cyt

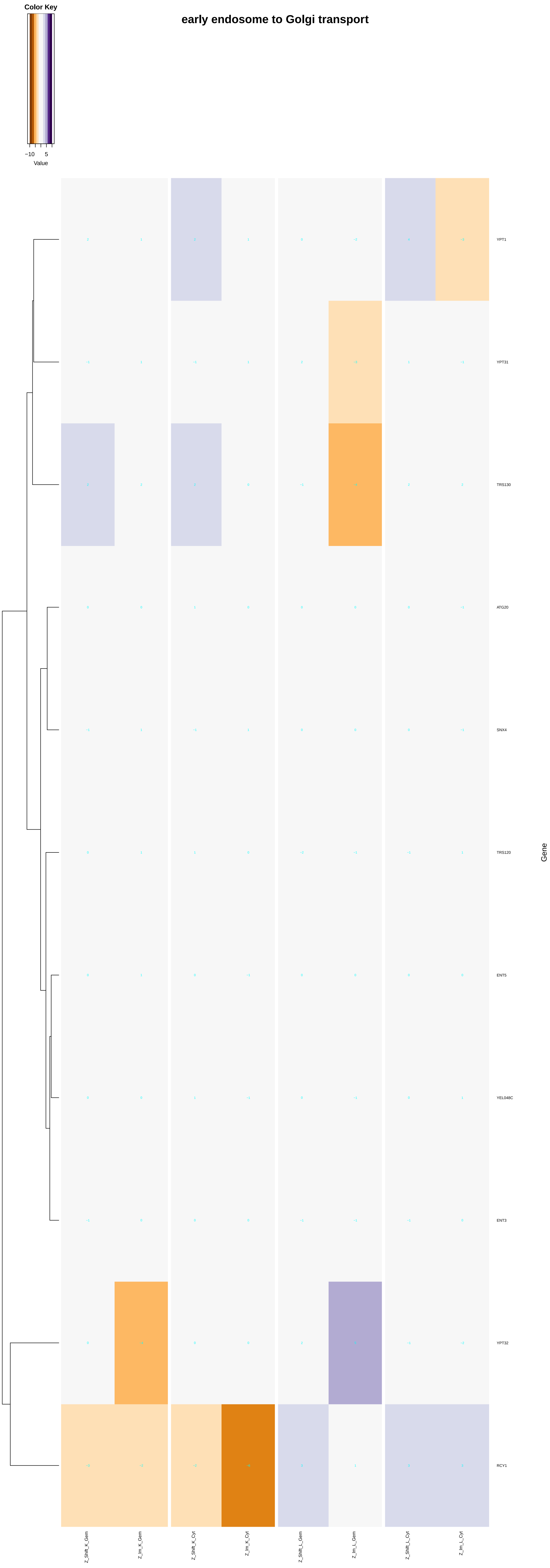
Color Key
−10
5
Value
early endosome to Golgi transport
YPT1
YPT31
TRS130
ATG20
SNX4
TRS120
ENT5
YEL048C
ENT3
YPT32
RCY1
Gene
Z_Shift_K_Gem
Z_Im_K_Gem
Z_Shift_K_Cyt
Z_Im_K_Cyt
Z_Shift_L_Gem
Z_Im_L_Gem
Z_Shift_L_Cyt
Z_Im_L_Cyt

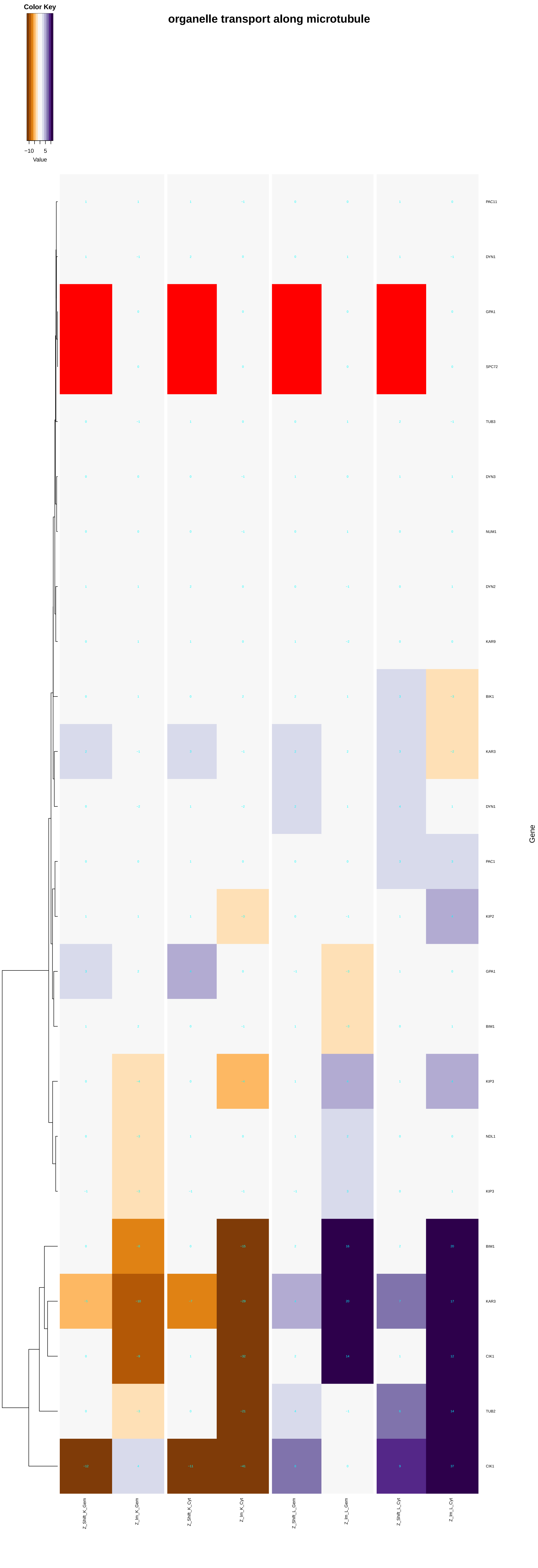

organelle transport along microtubule
Color Key
−10
5
Value
PAC11
DYN1
GPA1
SPC72
TUB3
DYN3
NUM1
DYN2
KAR9
BIK1
KAR3
DYN1
PAC1
KIP2
GPA1
BIM1
KIP3
NDL1
KIP3
BIM1
KAR3
CIK1
TUB2
CIK1
Gene
Z_Shift_K_Gem
Z_Im_K_Gem
Z_Shift_K_Cyt
Z_Im_K_Cyt
Z_Shift_L_Gem
Z_Im_L_Gem
Z_Shift_L_Cyt
Z_Im_L_Cyt

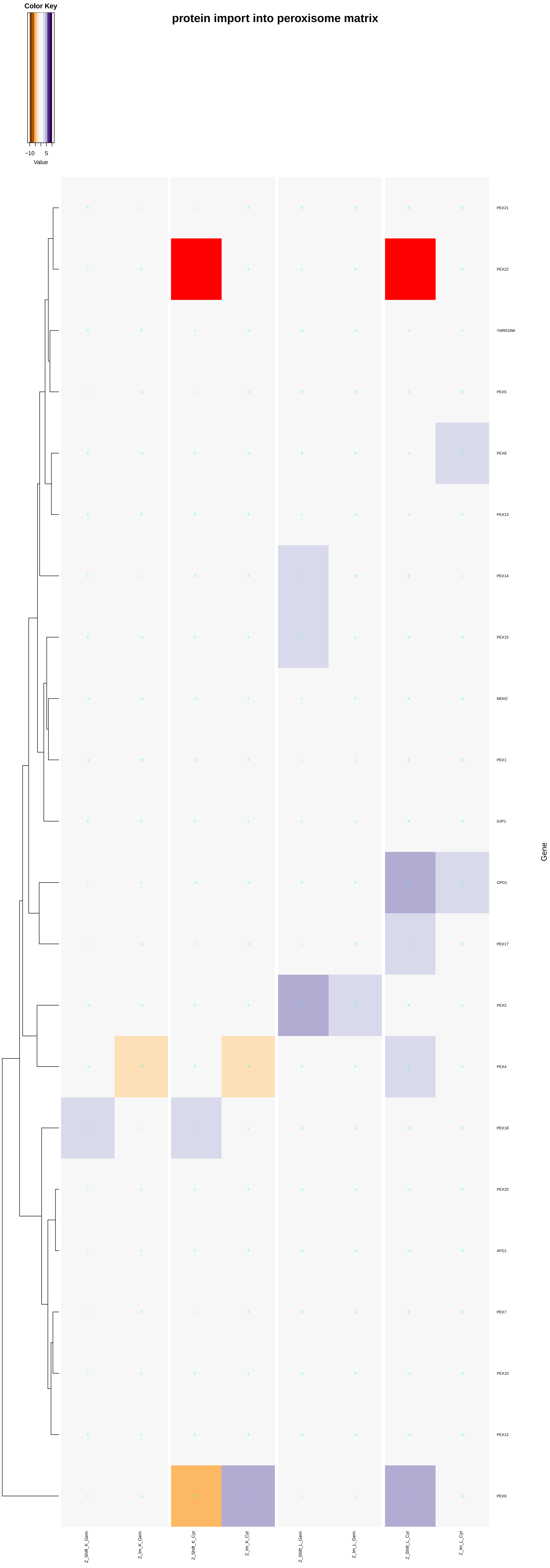
protein import into peroxisome matrix
Color Key
−10
5
Value
PEX21
PEX22
YMR018W
PEX5
PEX8
PEX13
PEX14
PEX15
MDH2
PEX1
DJP1
GPD1
PEX17
PEX2
PEX4
PEX18
PEX25
AFG1
PEX7
PEX10
PEX12
PEX6
Gene
Z_Shift_K_Gem
Z_Im_K_Gem
Z_Shift_K_Cyt
Z_Im_K_Cyt
Z_Shift_L_Gem
Z_Im_L_Gem
Z_Shift_L_Cyt
Z_Im_L_Cyt

### intracellular sterol transport

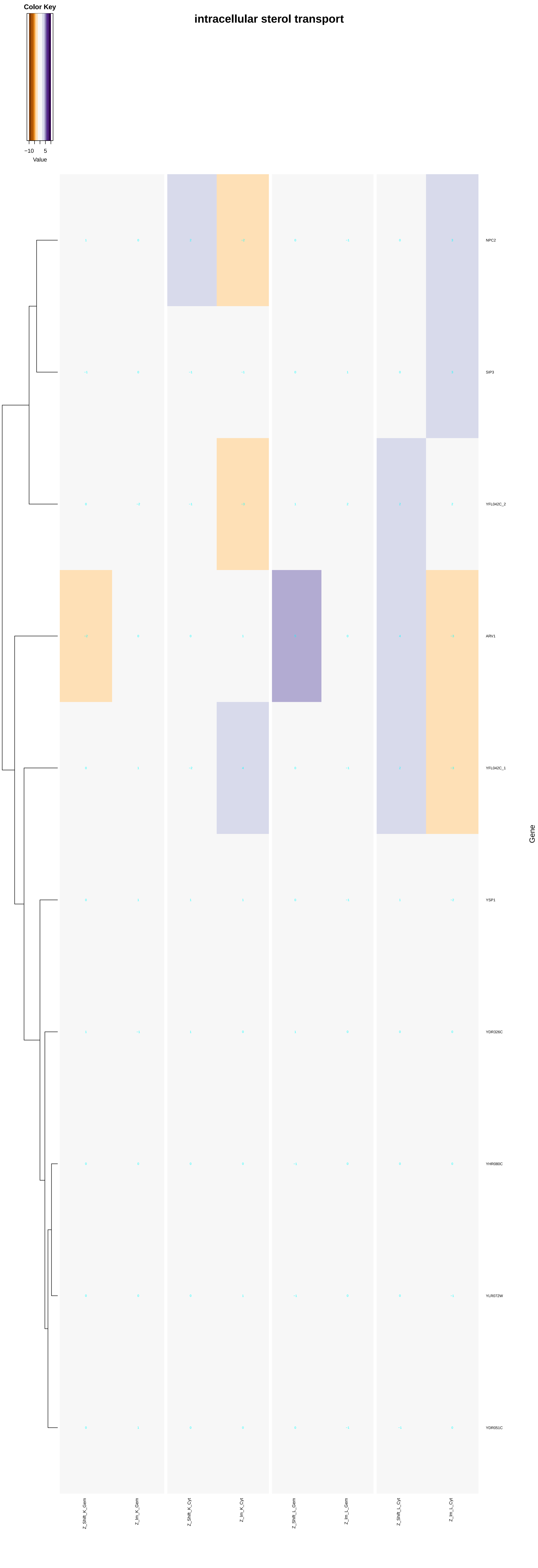

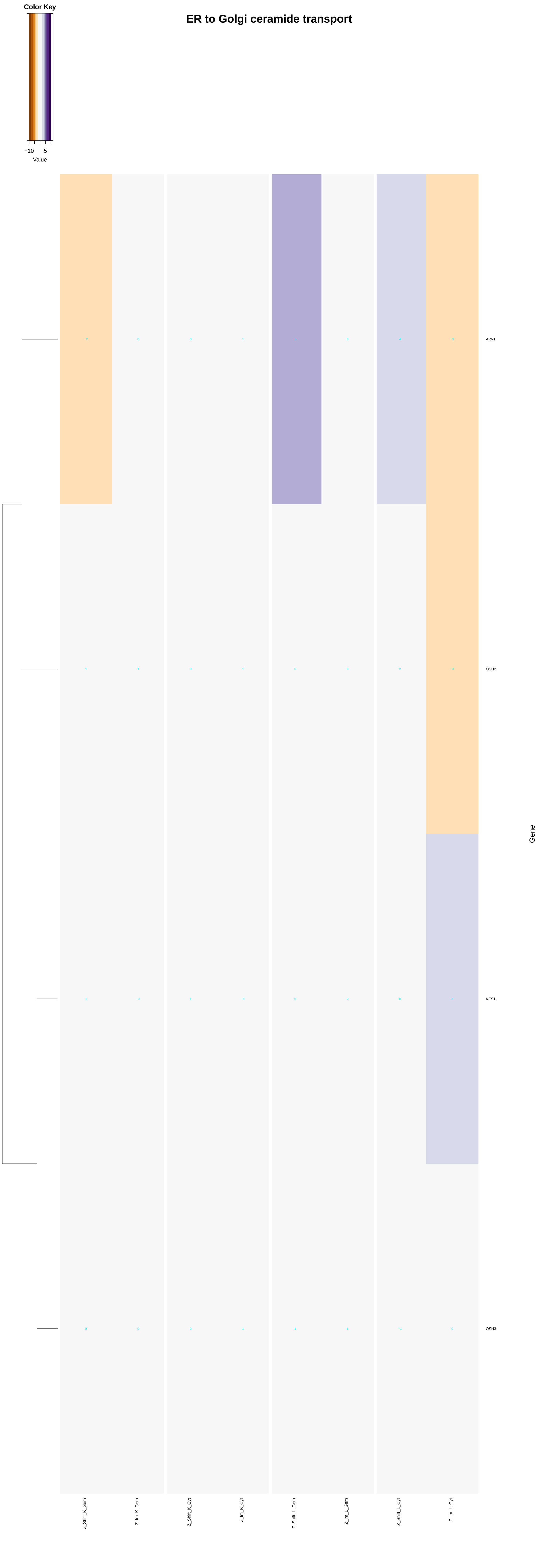

ER to Golgi ceramide transport
Color Key
−10
5
Value
ARV1
OSH2
KES1
OSH3
Gene
Z_Shift_K_Gem
Z_Im_K_Gem
Z_Shift_K_Cyt
Z_Im_K_Cyt
Z_Shift_L_Gem
Z_Im_L_Gem
Z_Shift_L_Cyt
Z_Im_L_Cyt

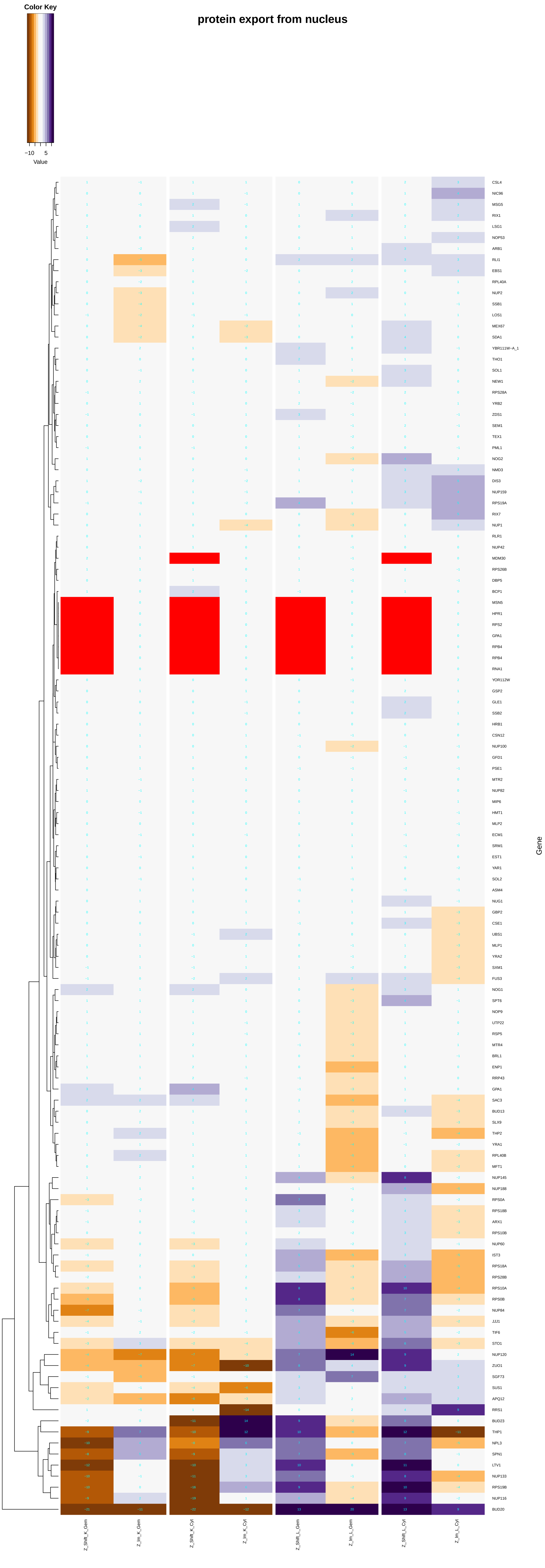
protein export from nucleus
Color Key
−10
5
Value
CSL4
NIC96
MSG5
RIX1
LSG1
NOP53
ARB1
RLI1
EBS1
RPL40A
NUP2
SSB1
LOS1
MEX67
SDA1
YBR111W−A_1
THO1
SOL1
NEW1
RPS28A
YRB2
ZDS1
SEM1
TEX1
PML1
NOG2
NMD3
DIS3
NUP159
RPS19A
RIX7
NUP1
RLR1
NUP42
MDM30
RPS26B
DBP5
BCP1
MSN5
HPR1
RPS2
GPA1
RPB4
RPB4
RNA1
YOR112W
GSP2
GLE1
SSB2
HRB1
CSN12
NUP100
GFD1
PSE1
MTR2
NUP82
MIP6
HMT1
MLP2
ECM1
SRM1
EST1
YAR1
SOL2
ASM4
NUG1
GBP2
CSE1
UBS1
MLP1
YRA2
SXM1
FUS3
NOG1
SPT6
NOP9
UTP22
RSP5
MTR4
BRL1
ENP1
RRP43
GPA1
SAC3
BUD13
SLX9
THP2
YRA1
RPL40B
MFT1
NUP145
NUP188
RPS0A
RPS18B
ARX1
RPS10B
NUP60
IST3
RPS18A
RPS28B
RPS10A
RPS0B
NUP84
JJJ1
TIF6
STO1
NUP120
ZUO1
SGF73
SUS1
APQ12
RRS1
BUD23
THP1
NPL3
SPN1
LTV1
NUP133
RPS19B
NUP116
BUD20
Gene
Z_Shift_K_Gem
Z_Im_K_Gem
Z_Shift_K_Cyt
Z_Im_K_Cyt
Z_Shift_L_Gem
Z_Im_L_Gem
Z_Shift_L_Cyt
Z_Im_L_Cyt

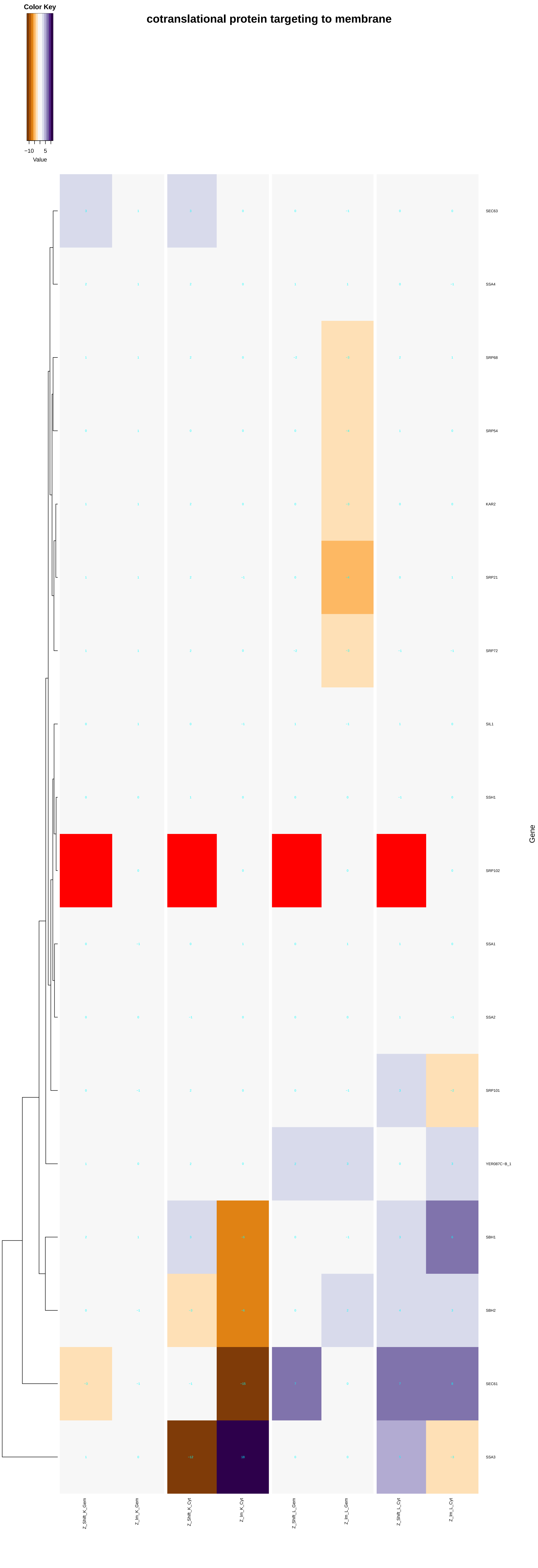
cotranslational protein targeting to membrane
Color Key
−10
5
Value
SEC63
SSA4
SRP68
SRP54
KAR2
SRP21
SRP72
SIL1
SSH1
SRP102
SSA1
SSA2
SRP101
YER087C−B_1
SBH1
SBH2
SEC61
SSA3
Gene
Z_Shift_K_Gem
Z_Im_K_Gem
Z_Shift_K_Cyt
Z_Im_K_Cyt
Z_Shift_L_Gem
Z_Im_L_Gem
Z_Shift_L_Cyt
Z_Im_L_Cyt

#### posttranslational protein targeting to membrane

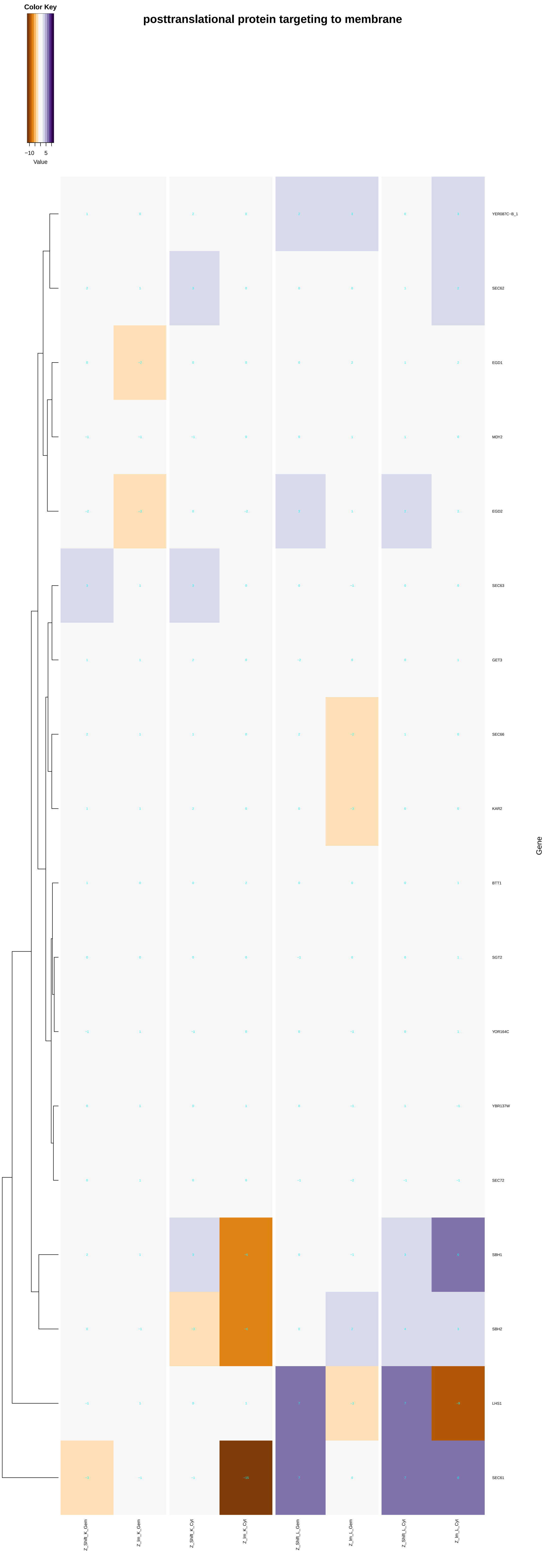

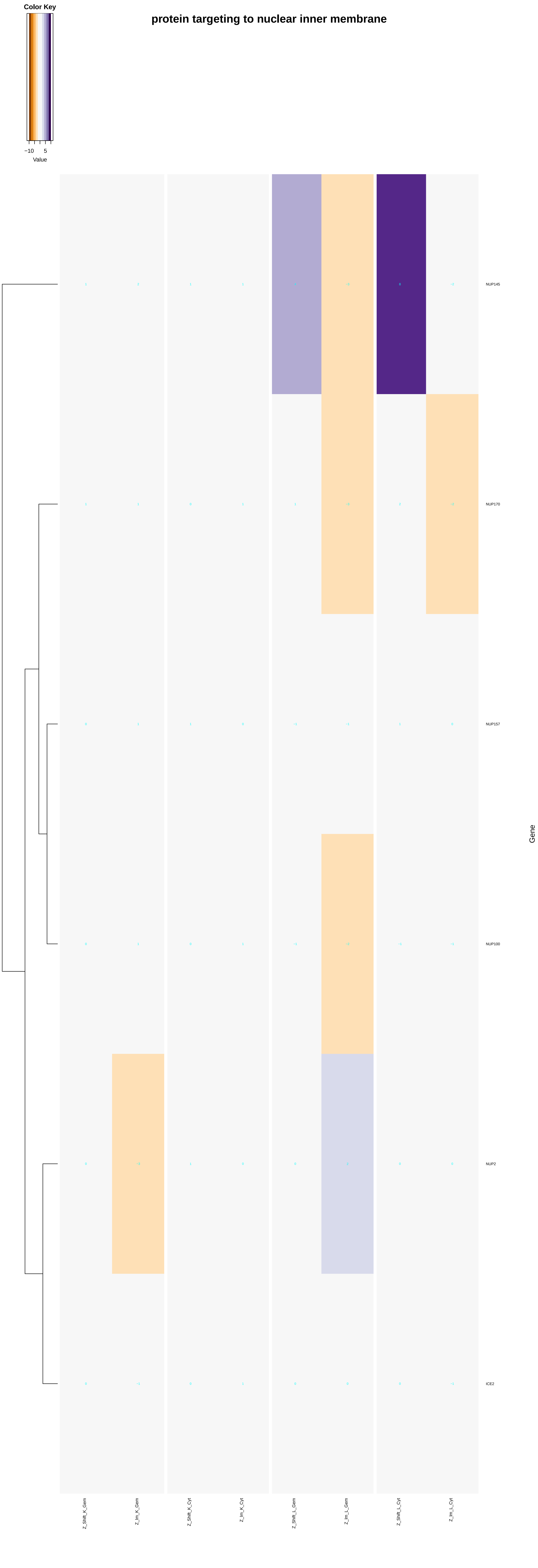
protein targeting to nuclear inner membrane
Color Key
−10
5
Value
NUP145
NUP170
NUP157
NUP100
NUP2
ICE2
Gene
Z_Shift_K_Gem
Z_lm_K_Gem
Z_Shift_K_Cyt
Z_lm_K_Cyt
Z_Shift_L_Gem
Z_lm_L_Gem
Z_Shift_L_Cyt
Z_lm_L_Cyt

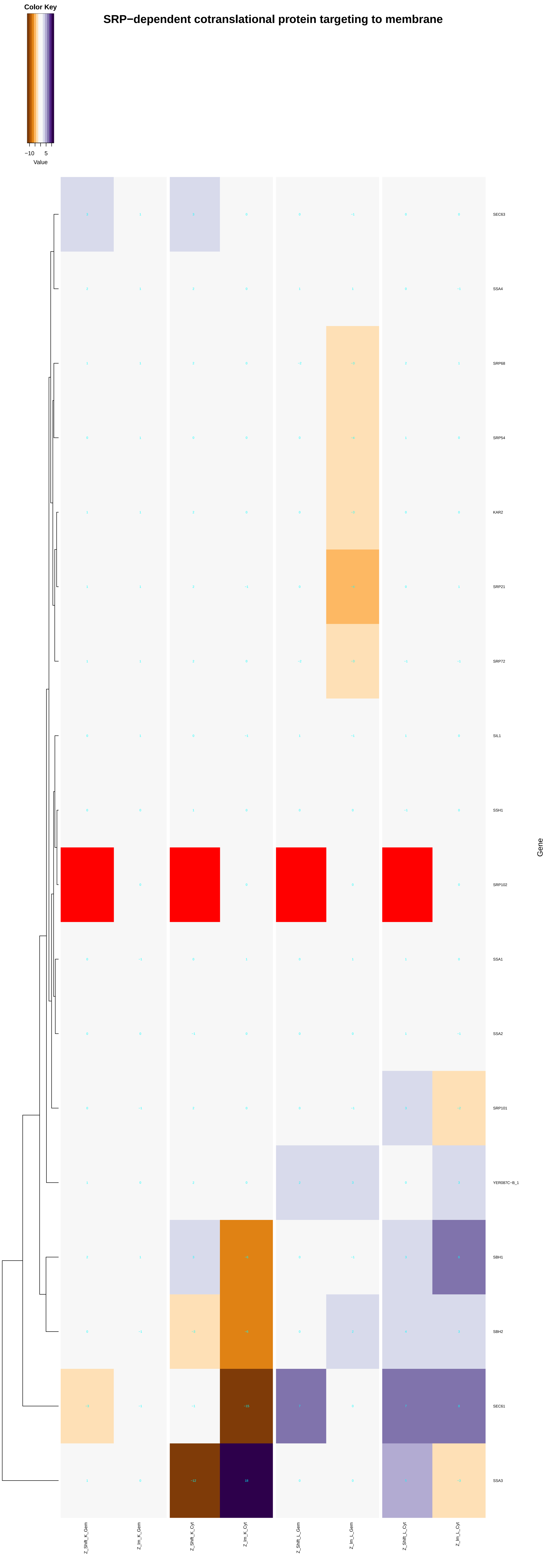
SRP−dependent cotranslational protein targeting to membrane
Color Key
−10
5
Value
SEC63
SSA4
SRP68
SRP54
KAR2
SRP21
SRP72
SIL1
SSH1
SRP102
SSA1
SSA2
SRP101
YER087C−B_1
SBH1
SBH2
SEC61
SSA3
Gene
Z_Shift_K_Gem
Z_Im_K_Gem
Z_Shift_K_Cyt
Z_Im_K_Cyt
Z_Shift_L_Gem
Z_Im_L_Gem
Z_Shift_L_Cyt
Z_Im_L_Cyt

### nuclear migration along microtubule

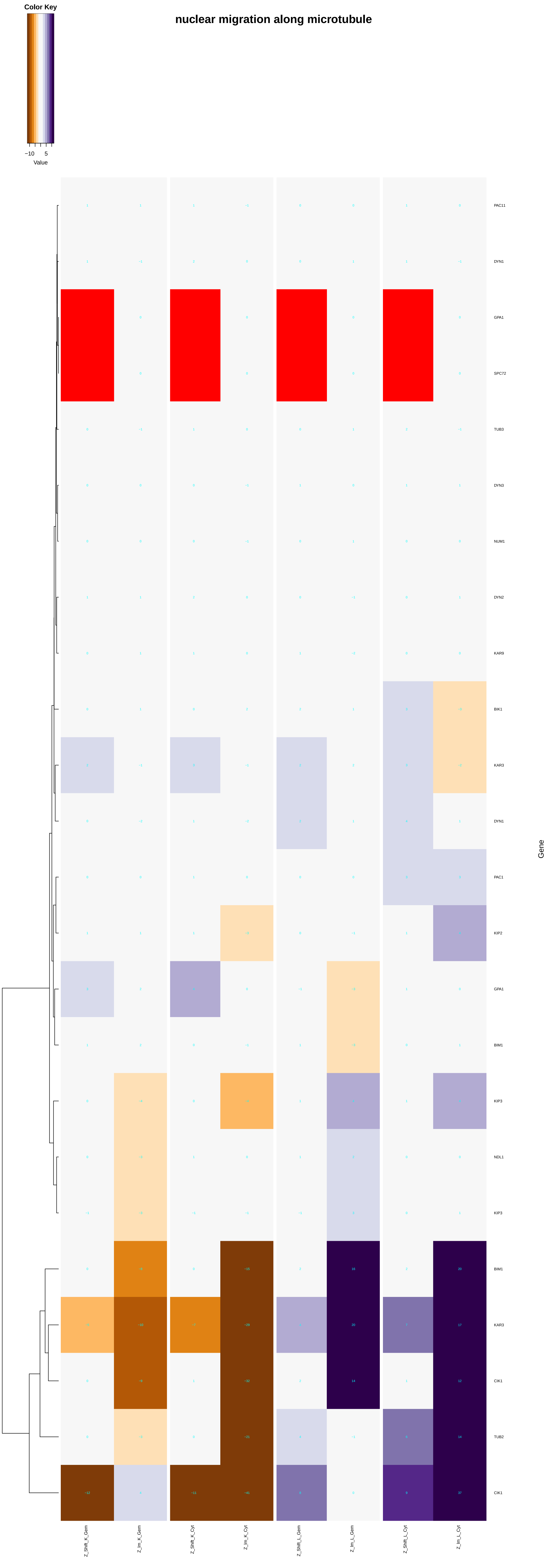

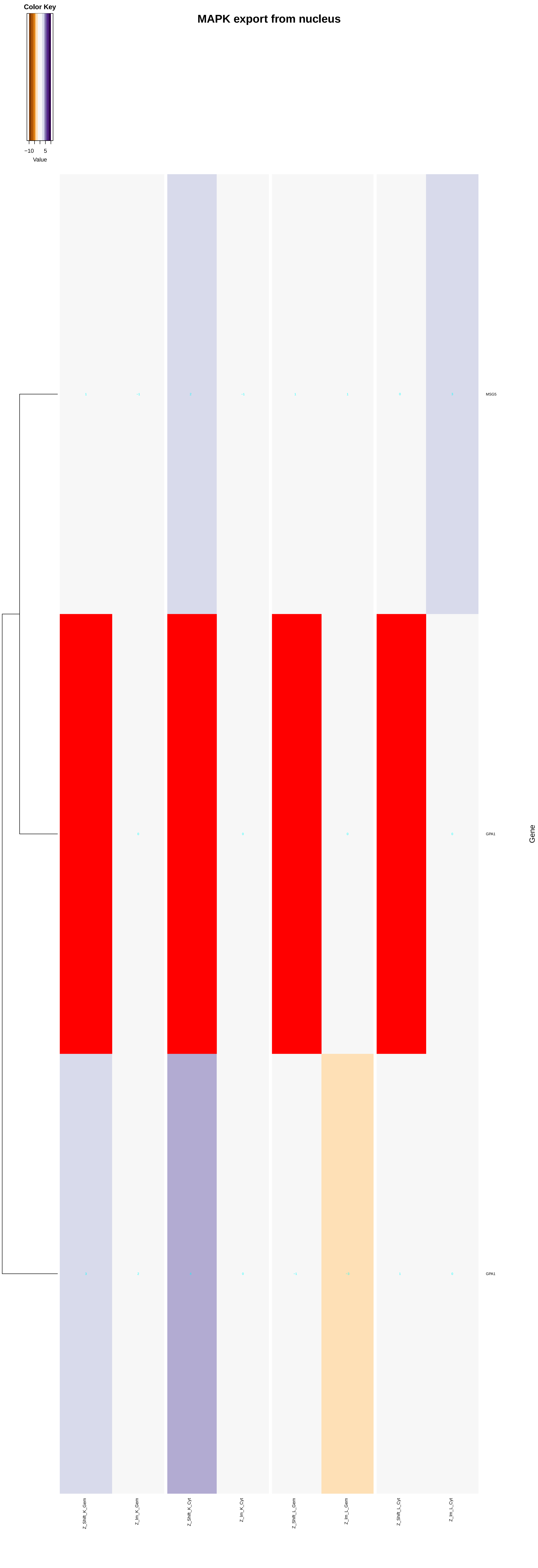
MAPK export from nucleus
Color Key
−10
5
Value
MSG5
GPA1
GPA1
Gene
Z_Shift_K_Gem
Z_Im_K_Gem
Z_Shift_K_Cyt
Z_Im_K_Cyt
Z_Shift_L_Gem
Z_Im_L_Gem
Z_Shift_L_Cyt
Z_Im_L_Cyt

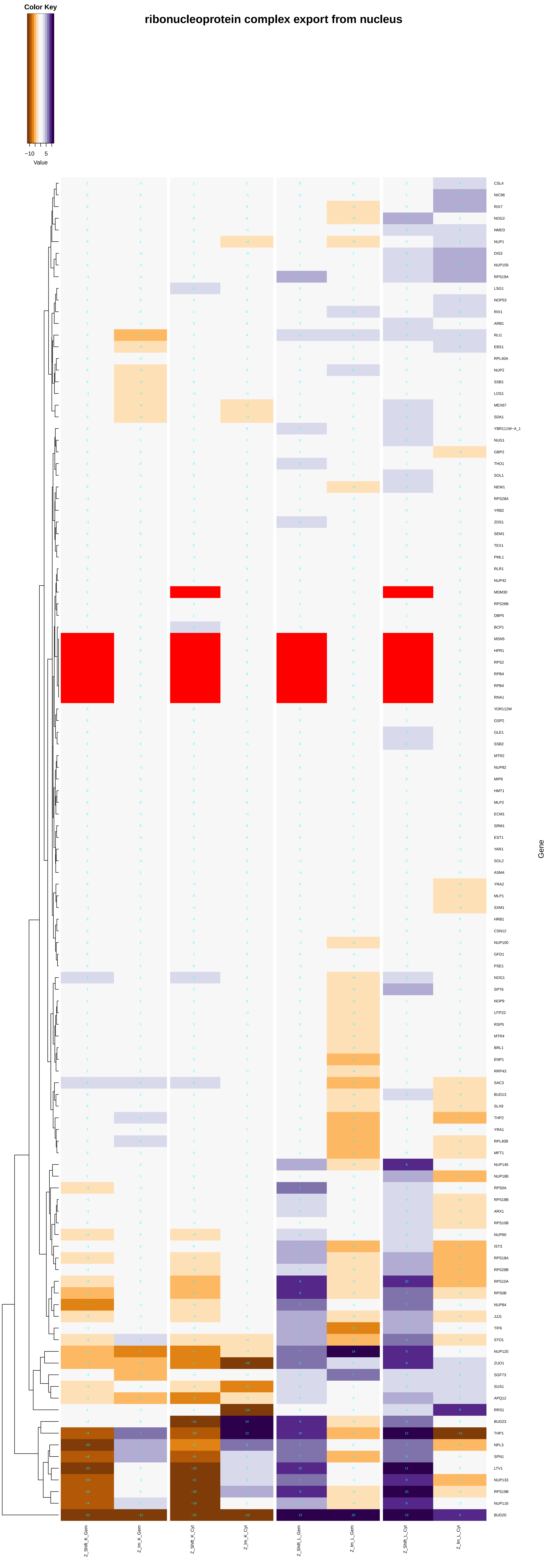

ribonucleoprotein complex export from nucleus
Color Key
−10 5
Value
Gene
CSL4
NIC96
RIX7
NOG2
NMD3
NUP1
DIS3
NUP159
RPS19A
LSG1
NOP53
RIX1
ARB1
RLI1
EBS1
RPL40A
NUP2
SSB1
LOS1
MEX67
SDA1
YBR111W−A_1
NUG1
GBP2
THO1
SOL1
NEW1
RPS28A
YRB2
ZDS1
SEM1
TEX1
PML1
RLR1
NUP42
MDM30
RPS26B
DBP5
BCP1
MSN5
HPR1
RPS2
RPB4
RPB4
RNA1
YOR112W
GSP2
GLE1
SSB2
MTR2
NUP82
MIP6
HMT1
MLP2
ECM1
SRM1
EST1
YAR1
SOL2
ASM4
YRA2
MLP1
SXM1
HRB1
CSN12
NUP100
GFD1
PSE1
NOG1
SPT6
NOP9
UTP22
RSP5
MTR4
BRL1
ENP1
RRP43
SAC3
BUD13
SLX9
THP2
YRA1
RPL40B
MFT1
NUP145
NUP188
RPS0A
RPS18B
ARX1
RPS10B
NUP60
IST3
RPS18A
RPS28B
RPS10A
RPS0B
NUP84
JJJ1
TIF6
STO1
NUP120
ZUO1
SGF73
SUS1
APQ12
RRS1
BUD23
THP1
NPL3
SPN1
LTV1
NUP133
RPS19B
NUP116
BUD20
Z_Shift_K_Gem
Z_Im_K_Gem
Z_Shift_K_Cyt
Z_Im_K_Cyt
Z_Shift_L_Gem
Z_Im_L_Gem
Z_Shift_L_Cyt
Z_Im_L_Cyt

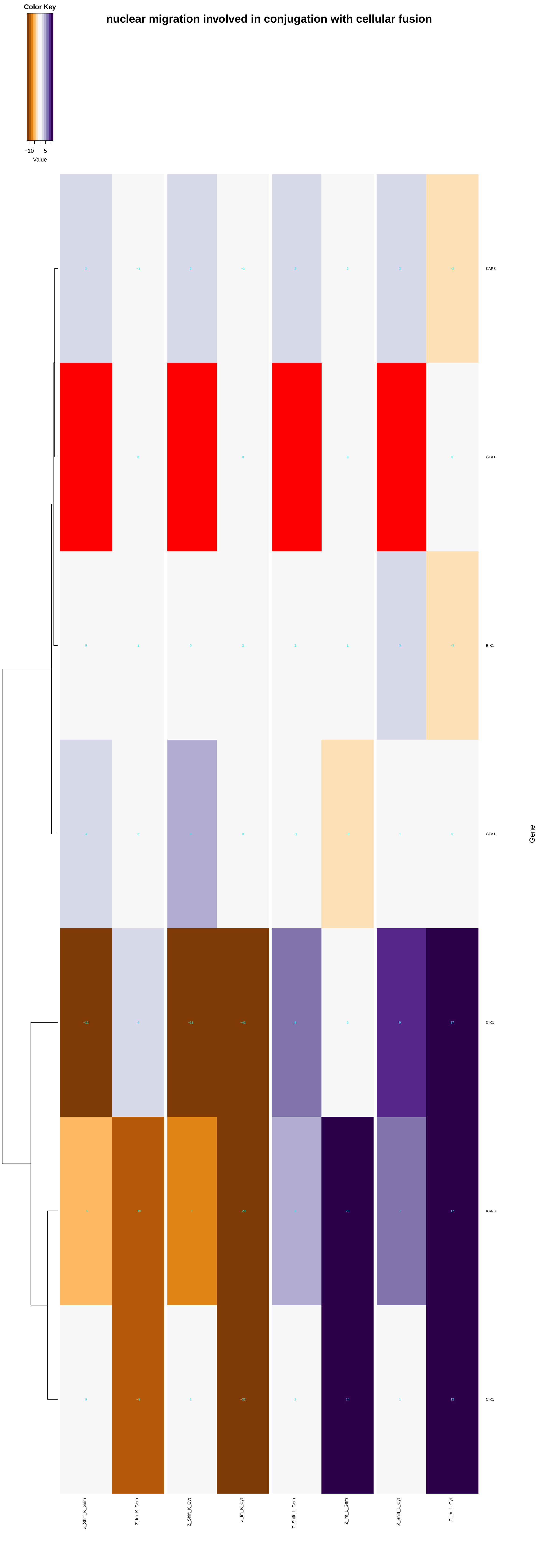
Color Key
−10
5
Value
nuclear migration involved in conjugation with cellular fusion
KAR3
GPA1
BIK1
GPA1
CIK1
KAR3
CIK1
Gene
Z_Shift_K_Gem
Z_Im_K_Gem
Z_Shift_K_Cyt
Z_Im_K_Cyt
Z_Shift_L_Gem
Z_Im_L_Gem
Z_Shift_L_Cyt
Z_Im_L_Cyt

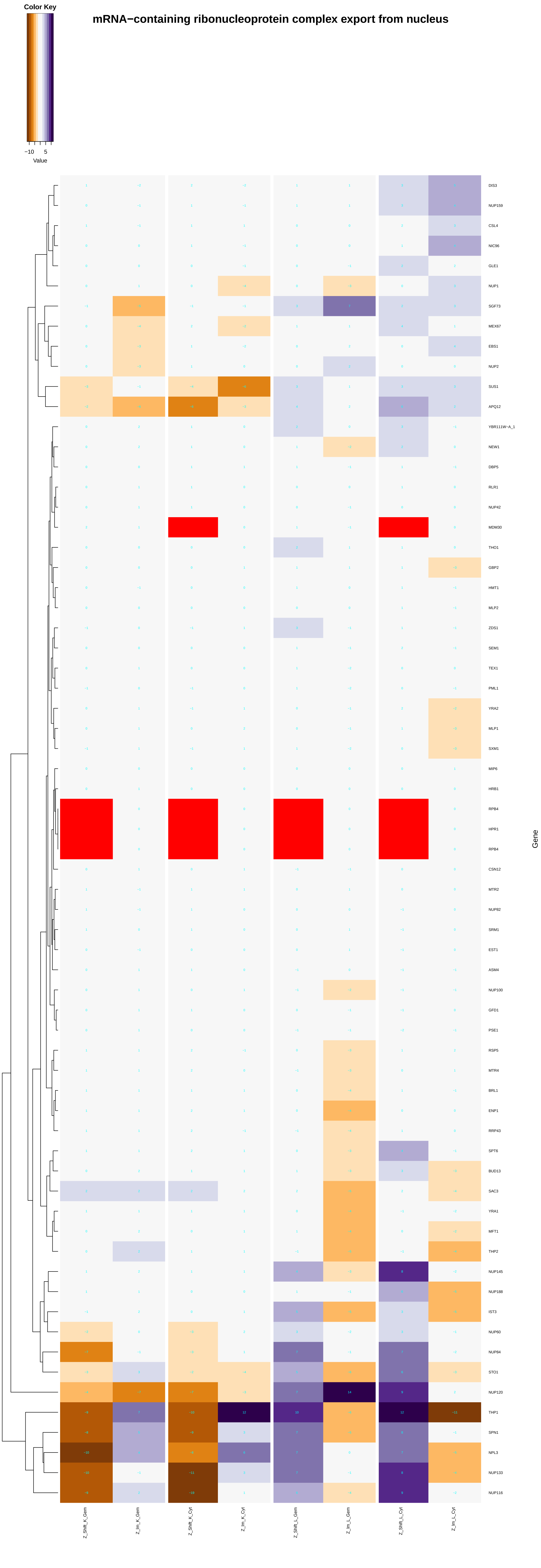
mRNA−containing ribonucleoprotein complex export from nucleus
Color Key
−10
5
Value
DIS3
NUP159
CSL4
NIC96
GLE1
NUP1
SGF73
MEX67
EBS1
NUP2
SUS1
APQ12
YBR111W−A_1
NEW1
DBP5
RLR1
NUP42
MDM30
THO1
GBP2
HMT1
MLP2
ZDS1
SEM1
TEX1
PML1
YRA2
MLP1
SXM1
MIP6
HRB1
RPB4
HPR1
RPB4
CSN12
MTR2
NUP82
SRM1
EST1
ASM4
NUP100
GFD1
PSE1
RSP5
MTR4
BRL1
ENP1
RRP43
SPT6
BUD13
SAC3
YRA1
MFT1
THP2
NUP145
NUP188
IST3
NUP60
NUP84
STO1
NUP120
THP1
SPN1
NPL3
NUP133
NUP116
Gene
Z_Shift_K_Gem
Z_Im_K_Gem
Z_Shift_K_Cyt
Z_Im_K_Cyt
Z_Shift_L_Gem
Z_Im_L_Gem
Z_Shift_L_Cyt
Z_Im_L_Cyt

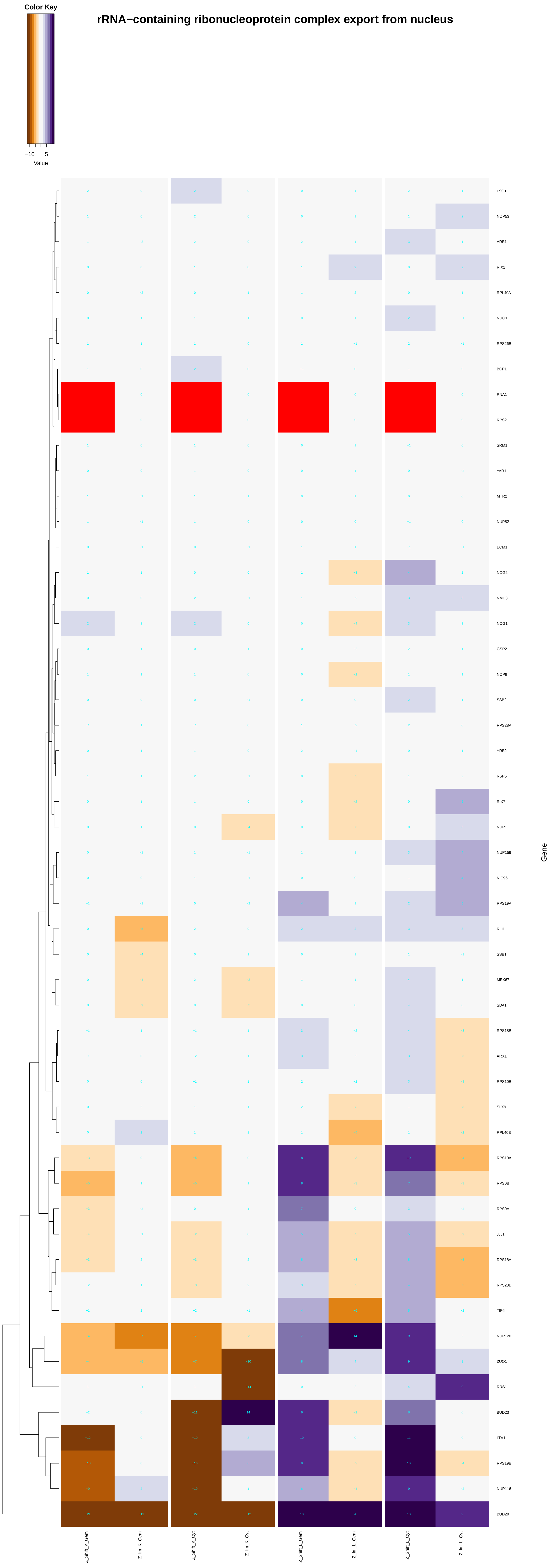
Color Key
−10
5
Value
rRNA−containing ribonucleoprotein complex export from nucleus
LSG1
NOP53
ARB1
RIX1
RPL40A
NUG1
RPS26B
BCP1
RNA1
RPS2
SRM1
YAR1
MTR2
NUP82
ECM1
NOG2
NMD3
NOG1
GSP2
NOP9
SSB2
RPS28A
YRB2
RSP5
RIX7
NUP1
NUP159
NIC96
RPS19A
RLI1
SSB1
MEX67
SDA1
RPS18B
ARX1
RPS10B
SLX9
RPL40B
RPS10A
RPS0B
RPS0A
JJJ1
RPS18A
RPS28B
TIF6
NUP120
ZUO1
RRS1
BUD23
LTV1
RPS19B
NUP116
BUD20
Gene
Z_Shift_K_Gem
Z_Im_K_Gem
Z_Shift_K_Cyt
Z_Im_K_Cyt
Z_Shift_L_Gem
Z_Im_L_Gem
Z_Shift_L_Cyt
Z_Im_L_Cyt

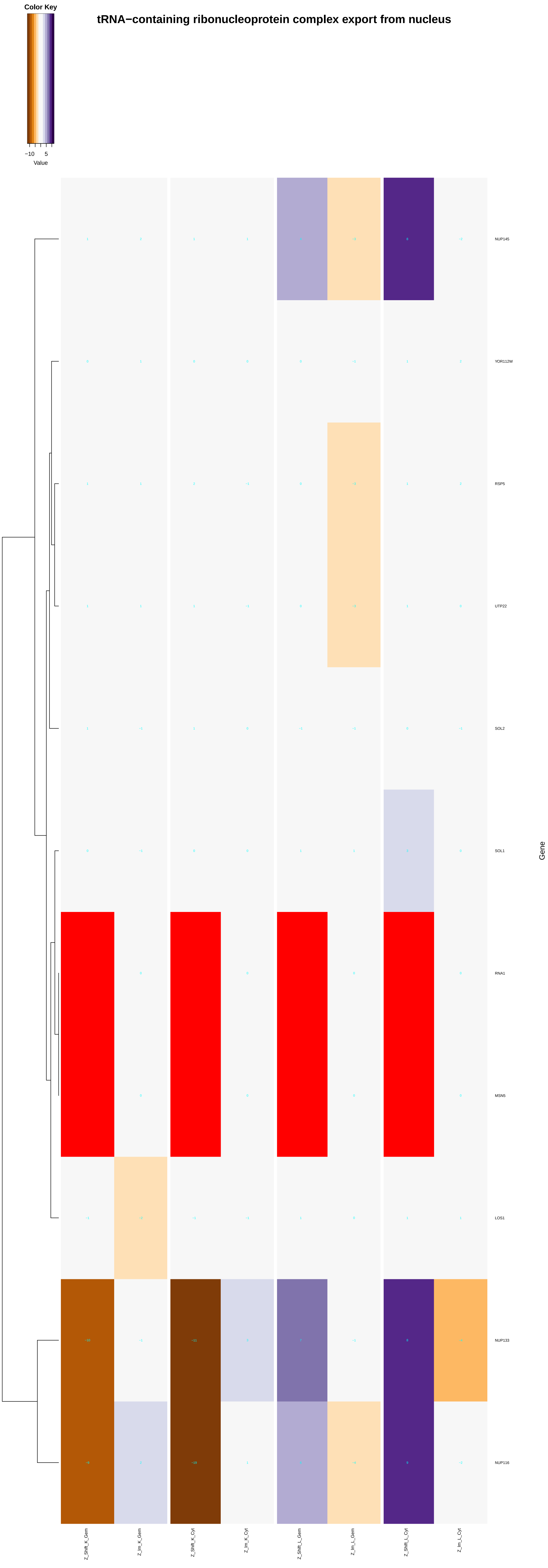
tRNA−containing ribonucleoprotein complex export from nucleus
Color Key
−10
5
Value
NUP145
YOR112W
RSP5
UTP22
SOL2
SOL1
RNA1
MSN5
LOS1
NUP133
NUP116
Gene
Z_Shift_K_Gem
Z_Im_K_Gem
Z_Shift_K_Cyt
Z_Im_K_Cyt
Z_Shift_L_Gem
Z_Im_L_Gem
Z_Shift_L_Cyt
Z_Im_L_Cyt

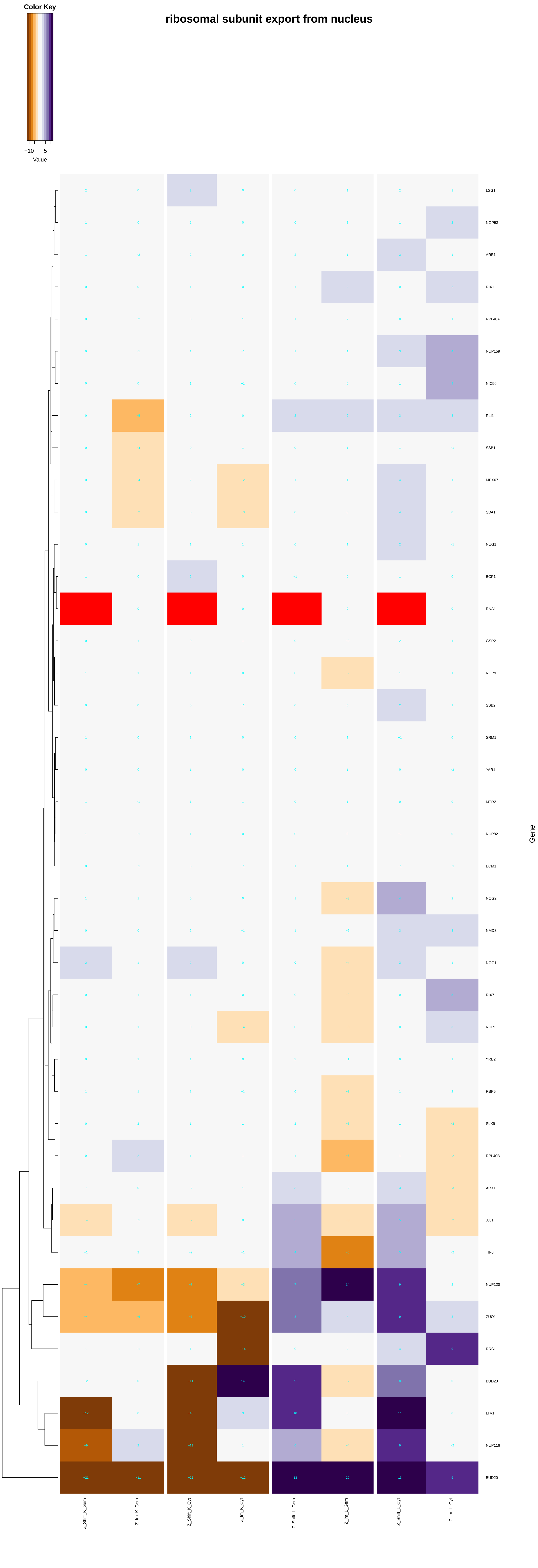
ribosomal subunit export from nucleus
Color Key
−10
5
Value
LSG1
NOP53
ARB1
RIX1
RPL40A
NUP159
NIC96
RLI1
SSB1
MEX67
SDA1
NUG1
BCP1
RNA1
GSP2
NOP9
SSB2
SRM1
YAR1
MTR2
NUP82
ECM1
NOG2
NMD3
NOG1
RIX7
NUP1
YRB2
RSP5
SLX9
RPL40B
ARX1
JJJ1
TIF6
NUP120
ZUO1
RRS1
BUD23
LTV1
NUP116
BUD20
Z_Shift_K_Gem
Z_Im_K_Gem
Z_Shift_K_Cyt
Z_Im_K_Cyt
Z_Shift_L_Gem
Z_Im_L_Gem
Z_Shift_L_Cyt
Z_Im_L_Cyt
Gene

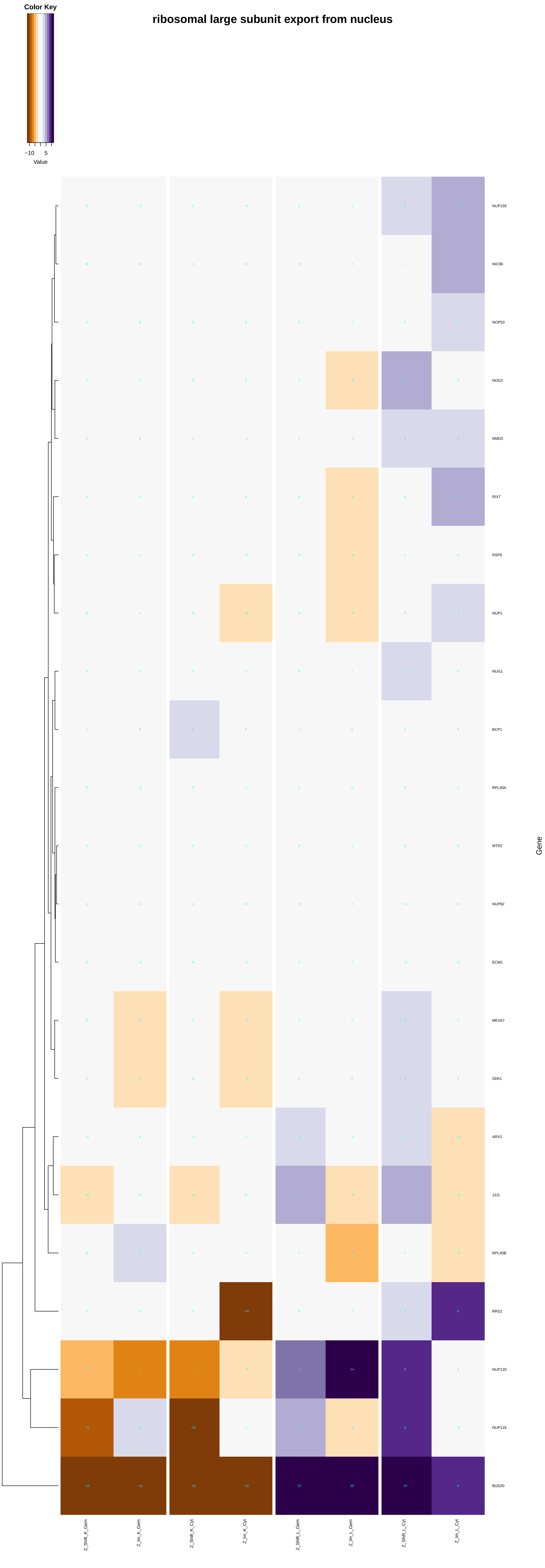

Color Key
−10
5
Value
ribosomal large subunit export from nucleus
NUP159
NIC96
NOP53
NOG2
NMD3
RIX7
RSP5
NUP1
NUG1
BCP1
RPL40A
MTR2
NUP82
ECM1
MEX67
SDA1
ARX1
JJJ1
RPL40B
RRS1
NUP120
NUP116
BUD20
Gene
Z_Shift_K_Gem
Z_Im_K_Gem
Z_Shift_K_Cyt
Z_Im_K_Cyt
Z_Shift_L_Gem
Z_Im_L_Gem
Z_Shift_L_Cyt
Z_Im_L_Cyt

### ribosomal small subunit export from nucleus

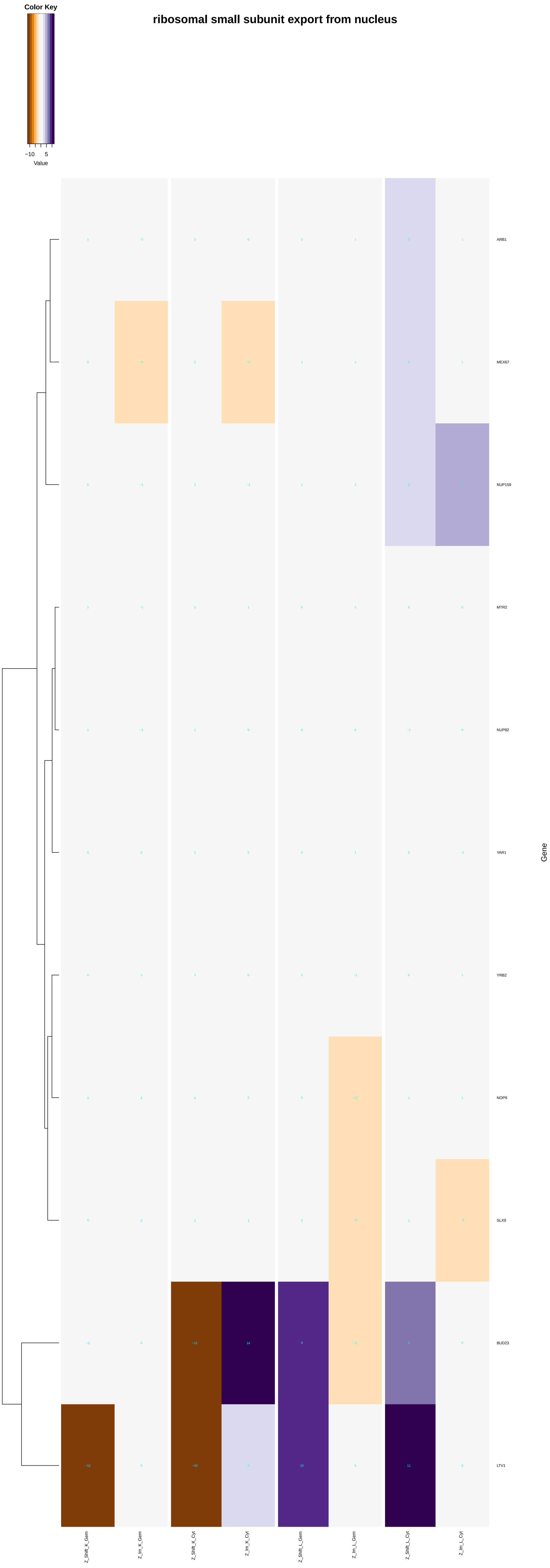
