## Additional File 7 for "A humanized yeast phenomic model of deoxycytidine kinase to predict genetic buffering of nucleoside analog cytotoxicity": lipid_binding.pdf

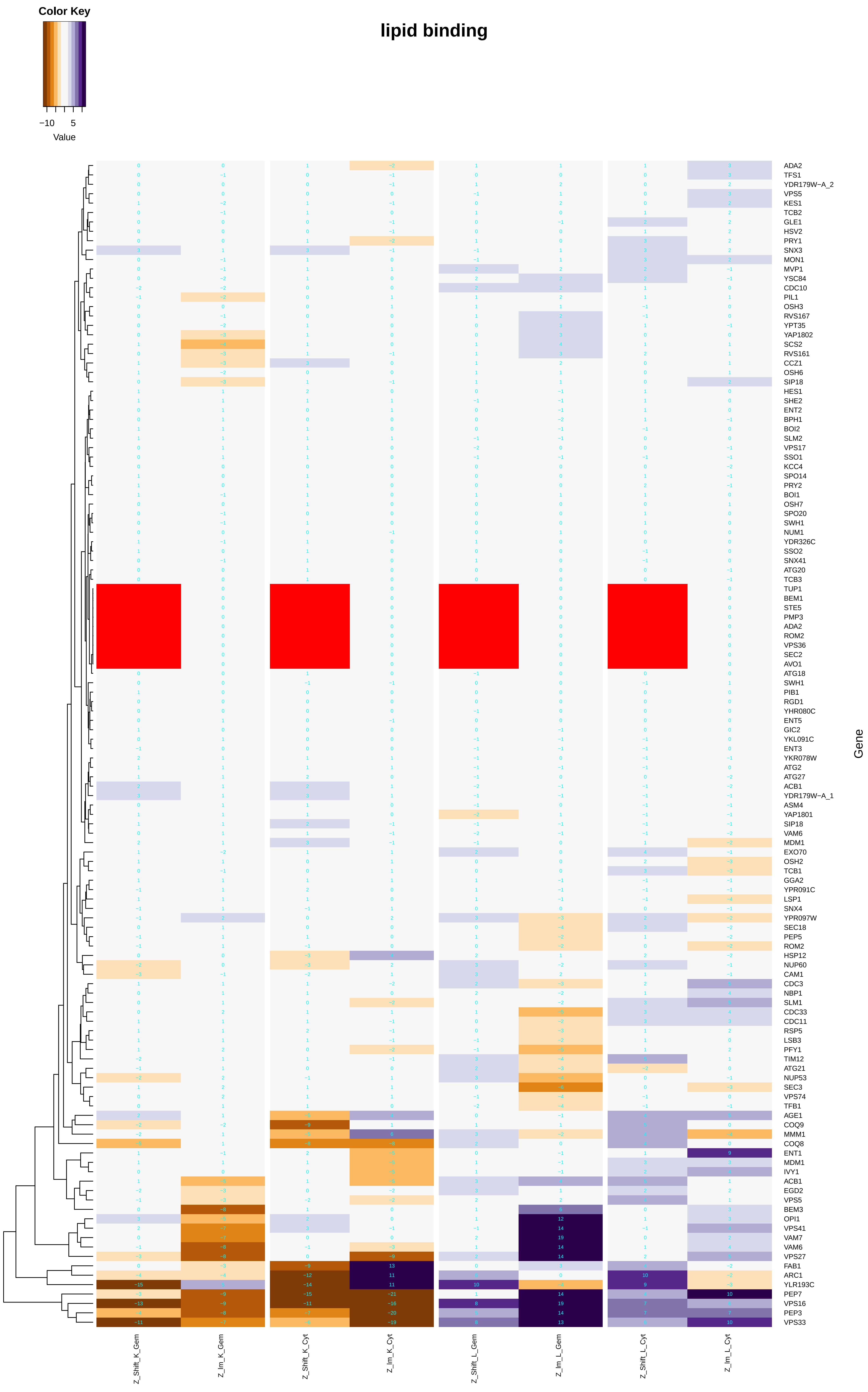
Color Key
−10
5
Value
lipid binding
ADA2
TFS1
YDR179W−A_2
VPS5
KES1
TCB2
GLE1
HSV2
PRY1
SNX3
MON1
MVP1
YSC84
CDC10
PIL1
OSH3
RVS167
YPT35
YAP1802
SCS2
RVS161
CCZ1
OSH6
SIP18
HES1
SHE2
ENT2
BPH1
BOI2
SLM2
VPS17
SSO1
KCC4
SPO14
PRY2
BOI1
OSH7
SPO20
SWH1
NUM1
YDR326C
SSO2
SNX41
ATG20
TCB3
TUP1
BEM1
STE5
PMP3
ADA2
ROM2
VPS36
SEC2
AVO1
ATG18
SWH1
PIB1
RGD1
YHR080C
ENT5
GIC2
YKL091C
ENT3
YKR078W
ATG2
ATG27
ACB1
YDR179W−A_1
ASM4
YAP1801
SIP18
VAM6
MDM1
EXO70
OSH2
TCB1
GGA2
YPR091C
LSP1
SNX4
YPR097W
SEC18
PEP5
ROM2
HSP12
NUP60
CAM1
CDC3
NBP1
SLM1
CDC33
CDC11
RSP5
LSB3
PFY1
TIM12
ATG21
NUP53
SEC3
VPS74
TFB1
AGE1
COQ9
MMM1
COQ8
ENT1
MDM1
IVY1
ACB1
EGD2
VPS5
BEM3
OPI1
VPS41
VAM7
VAM6
VPS27
FAB1
ARC1
YLR193C
PEP7
VPS16
PEP3
VPS33
Gene
Z_Shift_K_Gem
Z_Im_K_Gem
Z_Shift_K_Cyt
Z_Im_K_Cyt
Z_Shift_L_Gem
Z_Im_L_Gem
Z_Shift_L_Cyt
Z_Im_L_Cyt

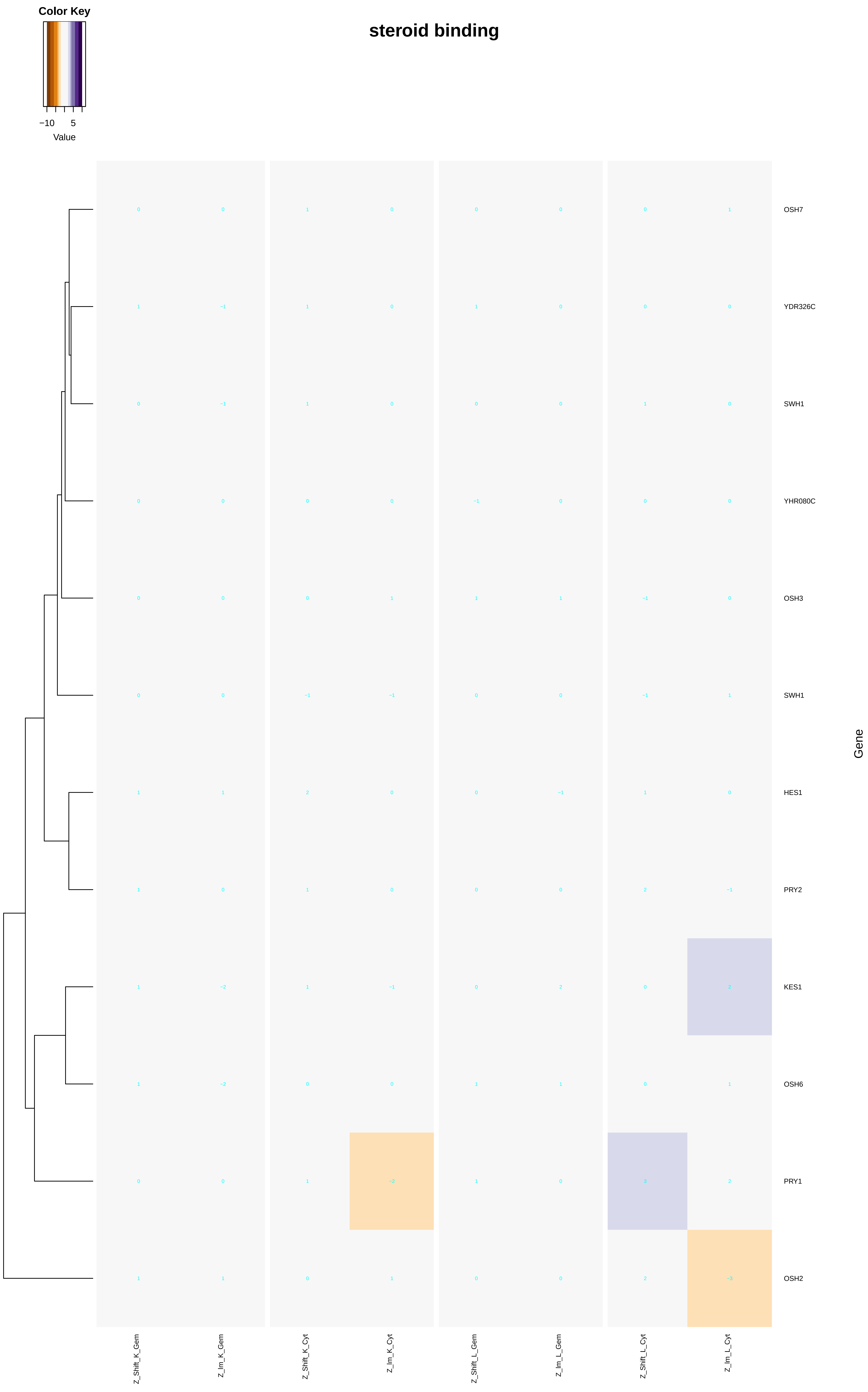

Color Key
−10
5
Value
steroid binding
OSH7
YDR326C
SWH1
YHR080C
OSH3
SWH1
HES1
PRY2
KES1
OSH6
PRY1
OSH2
Gene
Z_Shift_K_Gem
Z_Im_K_Gem
Z_Shift_K_Cyt
Z_Im_K_Cyt
Z_Shift_L_Gem
Z_Im_L_Gem
Z_Shift_L_Cyt
Z_Im_L_Cyt

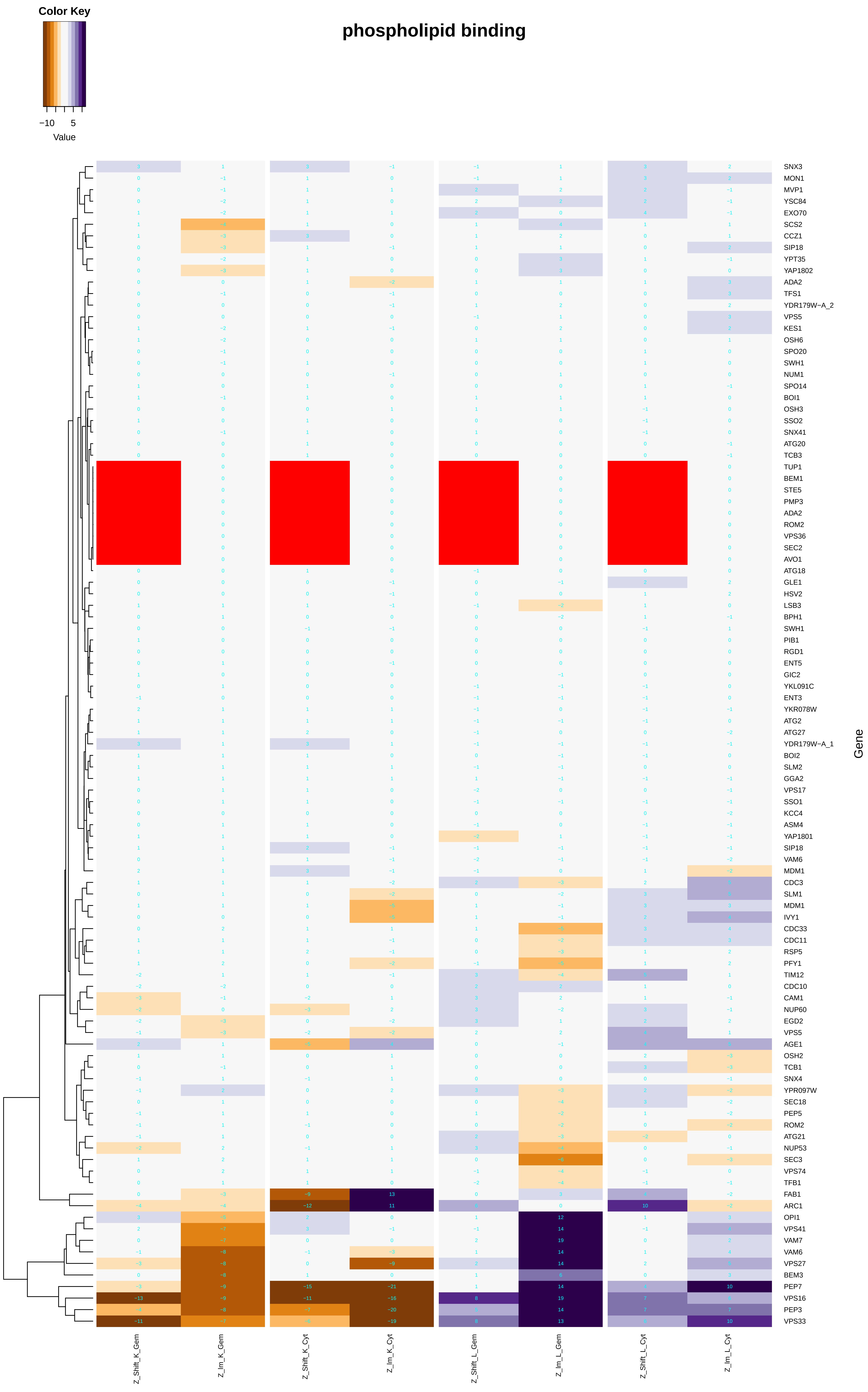
phospholipid binding
Color Key
−10
5
Value
Gene
SNX3
MON1
MVP1
YSC84
EXO70
SCS2
CCZ1
SIP18
YPT35
YAP1802
ADA2
TFS1
YDR179W−A_2
VPS5
KES1
OSH6
SPO20
SWH1
NUM1
SPO14
BOI1
OSH3
SSO2
SNX41
ATG20
TCB3
TUP1
BEM1
STE5
PMP3
ADA2
ROM2
VPS36
SEC2
AVO1
ATG18
GLE1
HSV2
LSB3
BPH1
SWH1
PIB1
RGD1
ENT5
GIC2
YKL091C
ENT3
YKR078W
ATG2
ATG27
YDR179W−A_1
BOI2
SLM2
GGA2
VPS17
SSO1
KCC4
ASM4
YAP1801
SIP18
VAM6
MDM1
CDC3
SLM1
MDM1
IVY1
CDC33
CDC11
RSP5
PFY1
TIM12
CDC10
CAM1
NUP60
EGD2
VPS5
AGE1
OSH2
TCB1
SNX4
YPR097W
SEC18
PEP5
ROM2
ATG21
NUP53
SEC3
VPS74
TFB1
FAB1
ARC1
OPI1
VPS41
VAM7
VAM6
VPS27
BEM3
PEP7
VPS16
PEP3
VPS33
Z_Shift_K_Gem
Z_Im_K_Gem
Z_Shift_K_Cyt
Z_Im_K_Cyt
Z_Shift_L_Gem
Z_Im_L_Gem
Z_Shift_L_Cyt
Z_Im_L_Cyt

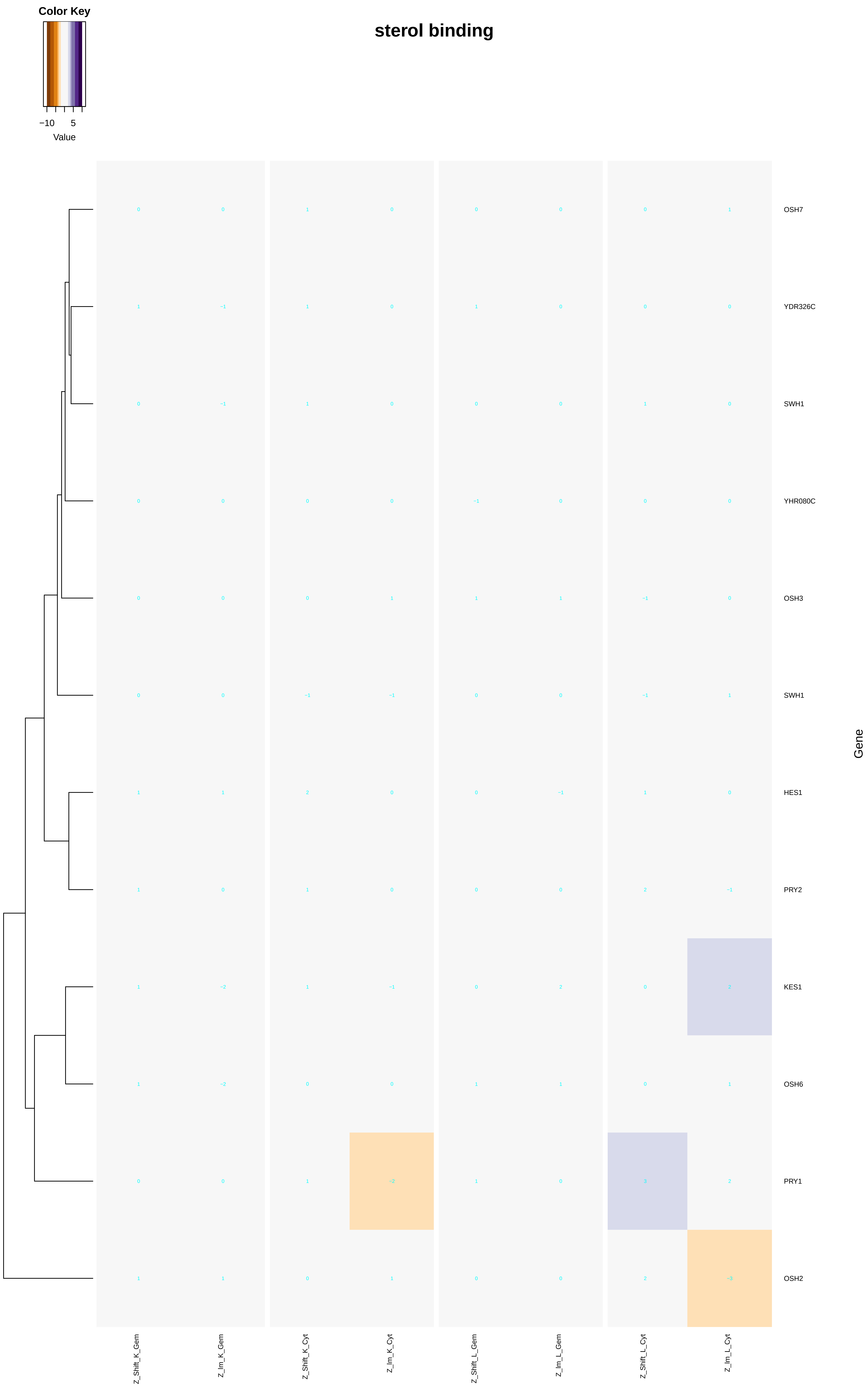
Color Key
−10 5
Value
sterol binding
OSH7
YDR326C
SWH1
YHR080C
OSH3
SWH1
HES1
PRY2
KES1
OSH6
PRY1
OSH2
Gene
Z_Shift_K_Gem
Z_Im_K_Gem
Z_Shift_K_Cyt
Z_Im_K_Cyt
Z_Shift_L_Gem
Z_Im_L_Gem
Z_Shift_L_Cyt
Z_Im_L_Cyt

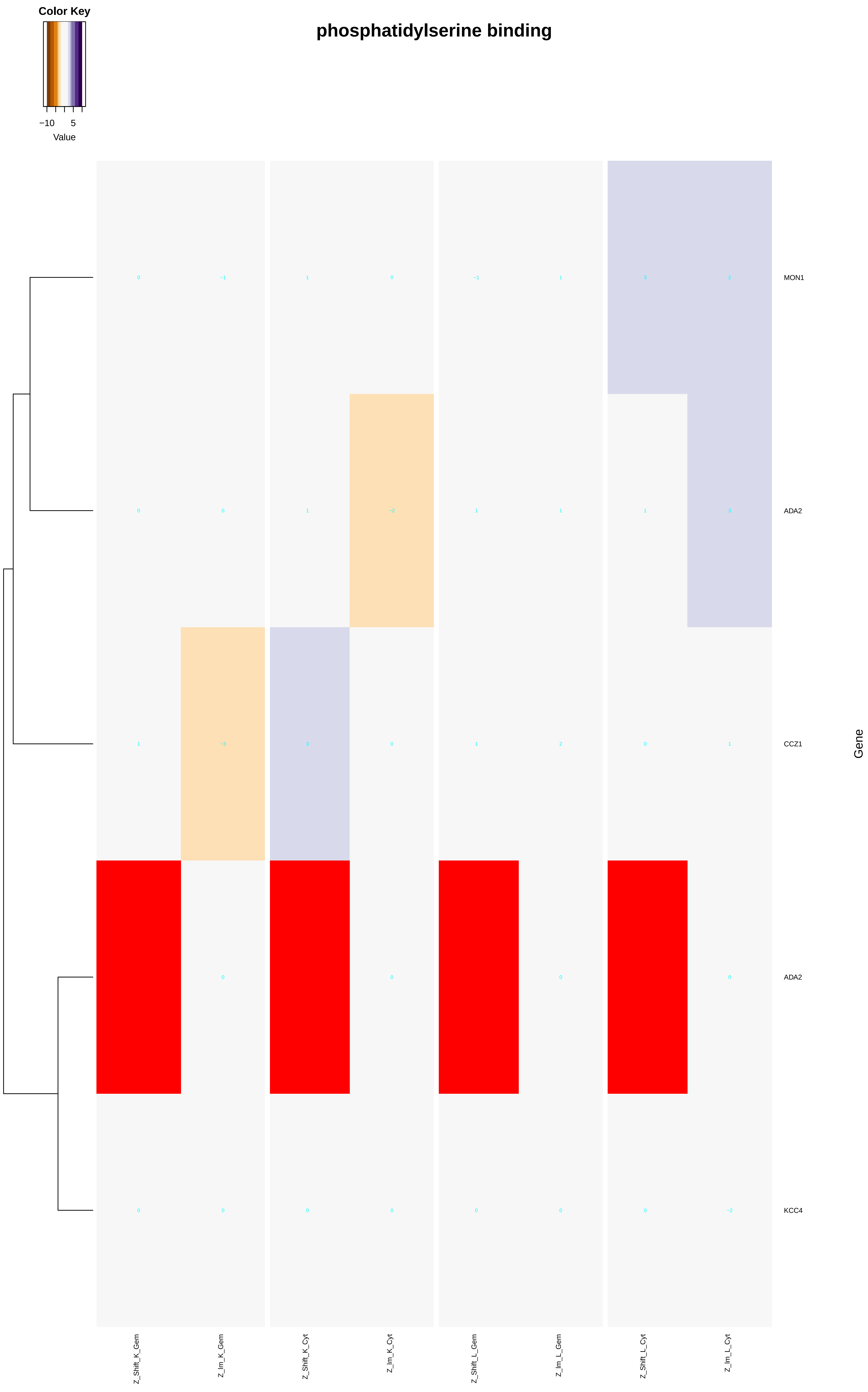

Color Key
−10
5
Value
phosphatidylserine binding
MON1
ADA2
CCZ1
ADA2
KCC4
Gene
Z_Shift_K_Gem
Z_Im_K_Gem
Z_Shift_K_Cyt
Z_Im_K_Cyt
Z_Shift_L_Gem
Z_Im_L_Gem
Z_Shift_L_Cyt
Z_Im_L_Cyt

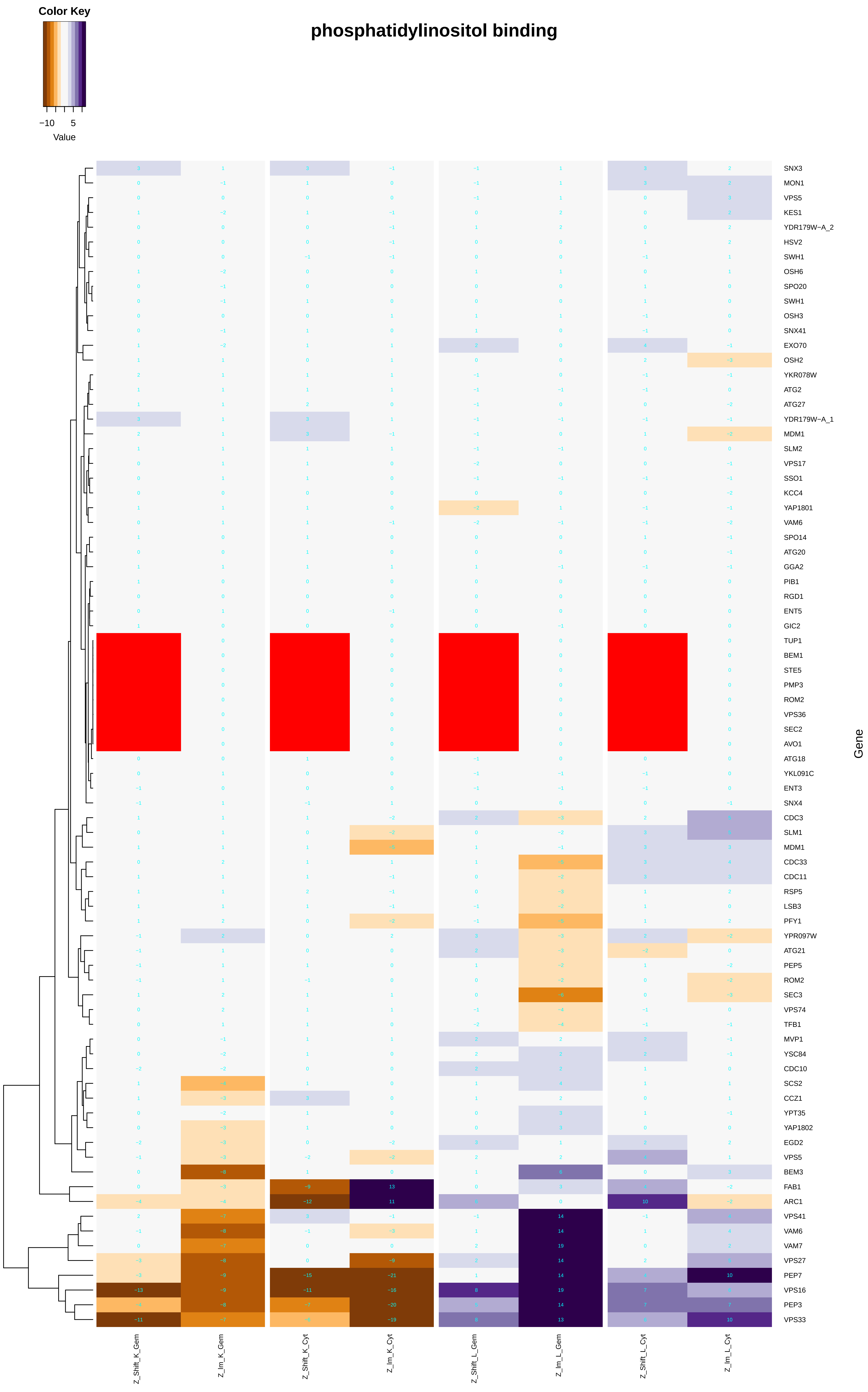
Color Key
−10
5
Value
phosphatidylinositol binding
SNX3
MON1
VPS5
KES1
YDR179W−A_2
HSV2
SWH1
OSH6
SPO20
SWH1
OSH3
SNX41
EXO70
OSH2
YKR078W
ATG2
ATG27
YDR179W−A_1
MDM1
SLM2
VPS17
SSO1
KCC4
YAP1801
VAM6
SPO14
ATG20
GGA2
PIB1
RGD1
ENT5
GIC2
TUP1
BEM1
STE5
PMP3
ROM2
VPS36
SEC2
AVO1
ATG18
YKL091C
ENT3
SNX4
CDC3
SLM1
MDM1
CDC33
CDC11
RSP5
LSB3
PFY1
YPR097W
ATG21
PEP5
ROM2
SEC3
VPS74
TFB1
MVP1
YSC84
CDC10
SCS2
CCZ1
YPT35
YAP1802
EGD2
VPS5
BEM3
FAB1
ARC1
VPS41
VAM6
VAM7
VPS27
PEP7
VPS16
PEP3
VPS33
Gene
Z_Shift_K_Gem
Z_Im_K_Gem
Z_Shift_K_Cyt
Z_Im_K_Cyt
Z_Shift_L_Gem
Z_Im_L_Gem
Z_Shift_L_Cyt
Z_Im_L_Cyt

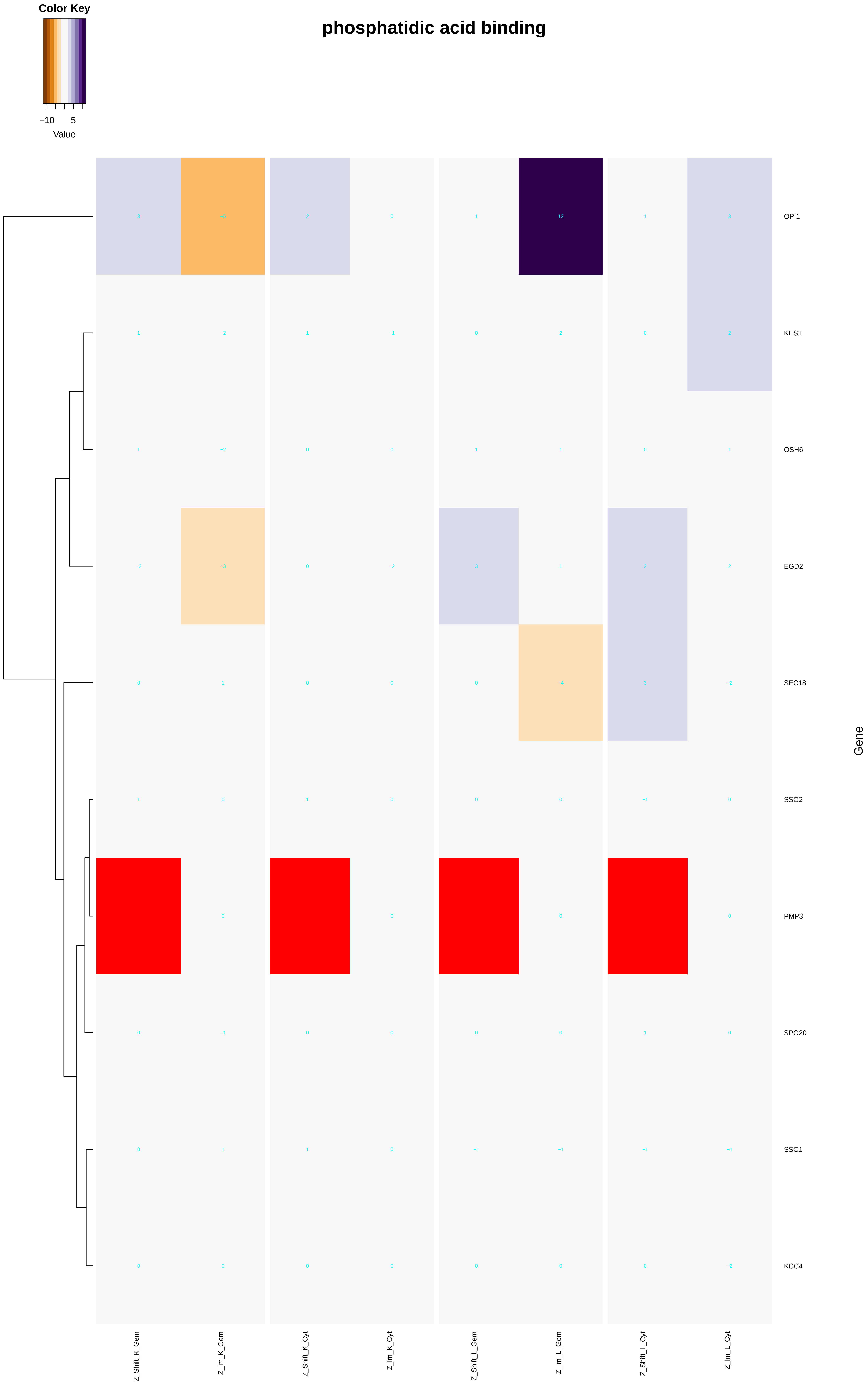

Color Key
−10
5
Value
phosphatidic acid binding
OPI1
KES1
OSH6
EGD2
SEC18
SSO2
PMP3
SPO20
SSO1
KCC4
Gene
Z_Shift_K_Gem
Z_Im_K_Gem
Z_Shift_K_Cyt
Z_Im_K_Cyt
Z_Shift_L_Gem
Z_Im_L_Gem
Z_Shift_L_Cyt
Z_Im_L_Cyt

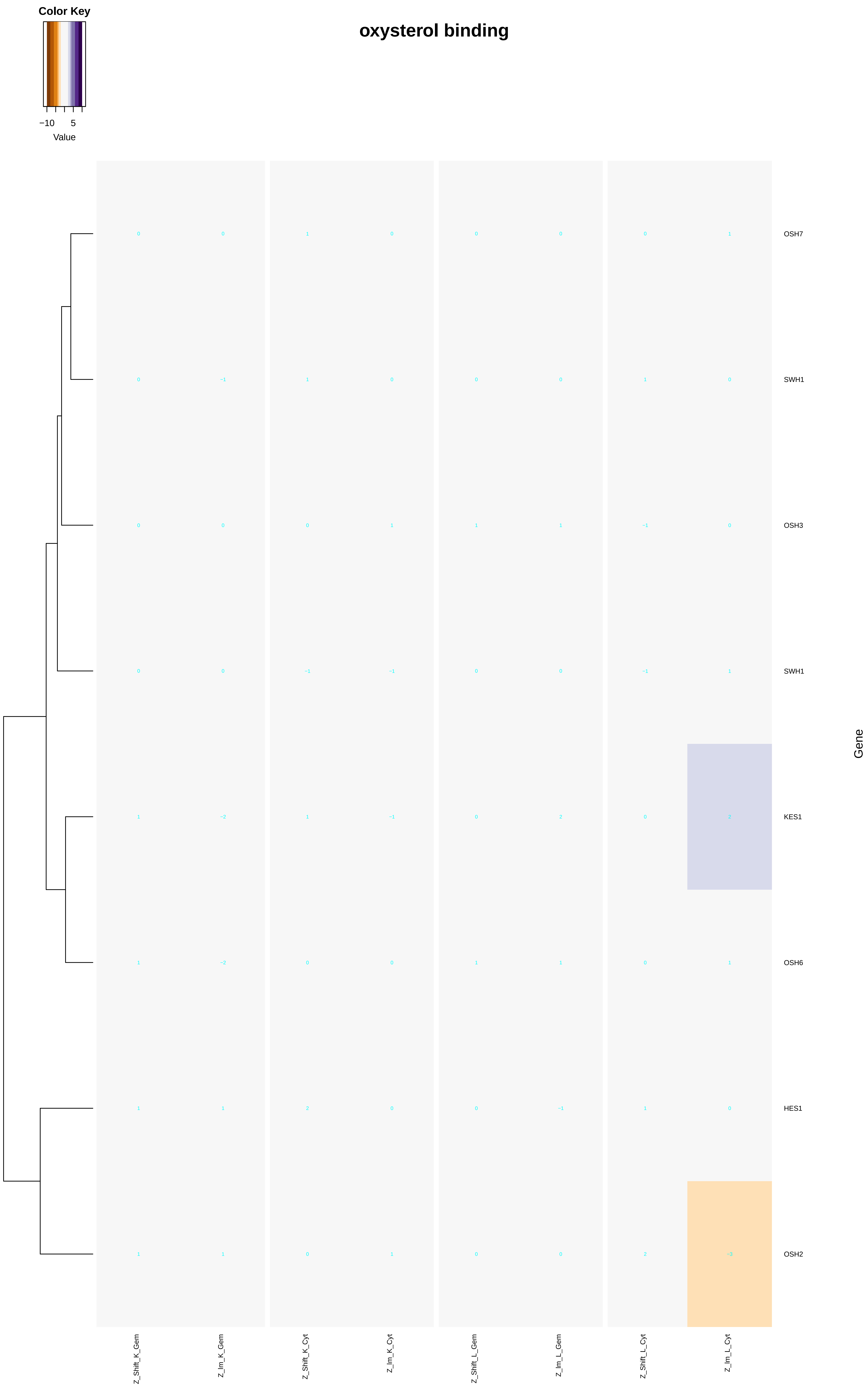

Color Key
−10
5
Value
oxysterol binding
OSH7
SWH1
OSH3
SWH1
KES1
OSH6
HES1
OSH2
Gene
Z_Shift_K_Gem
Z_Im_K_Gem
Z_Shift_K_Cyt
Z_Im_K_Cyt
Z_Shift_L_Gem
Z_Im_L_Gem
Z_Shift_L_Cyt
Z_Im_L_Cyt

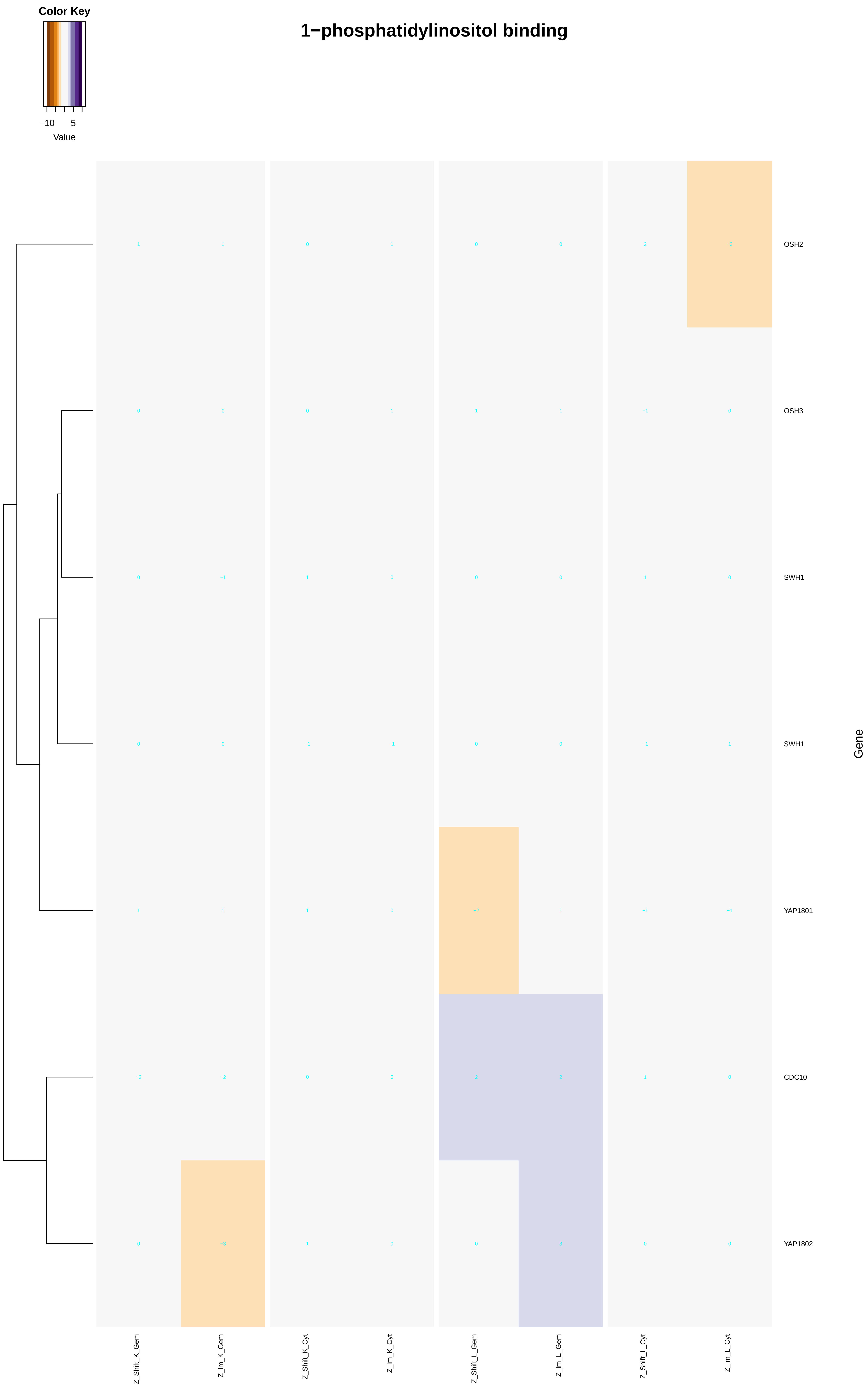

1−phosphatidylinositol binding
Color Key
−10
5
Value
OSH2
OSH3
SWH1
SWH1
YAP1801
CDC10
YAP1802
Gene
Z_Shift_K_Gem
Z_Im_K_Gem
Z_Shift_K_Cyt
Z_Im_K_Cyt
Z_Shift_L_Gem
Z_Im_L_Gem
Z_Shift_L_Cyt
Z_Im_L_Cyt

### phosphatidylinositol phosphate binding

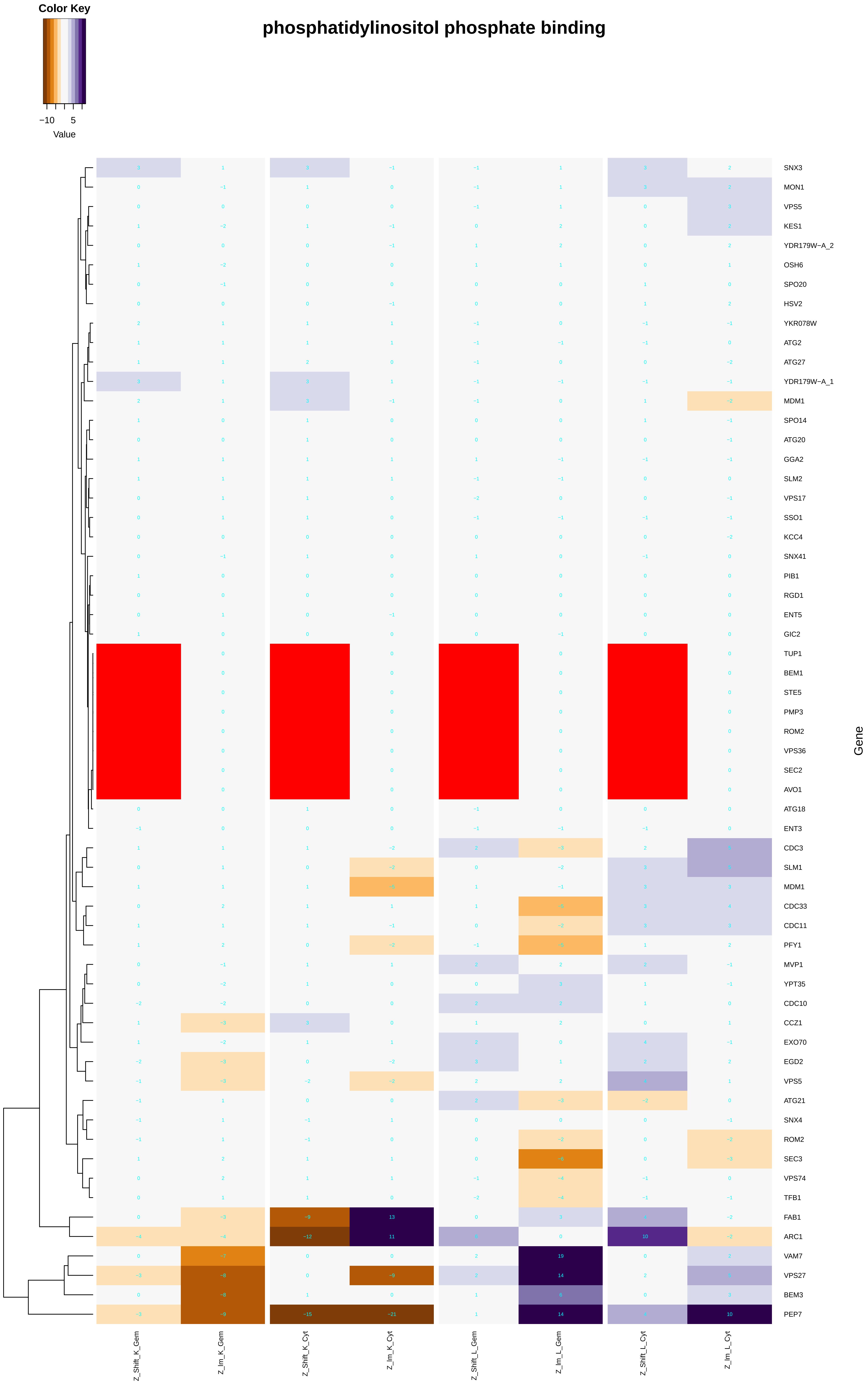

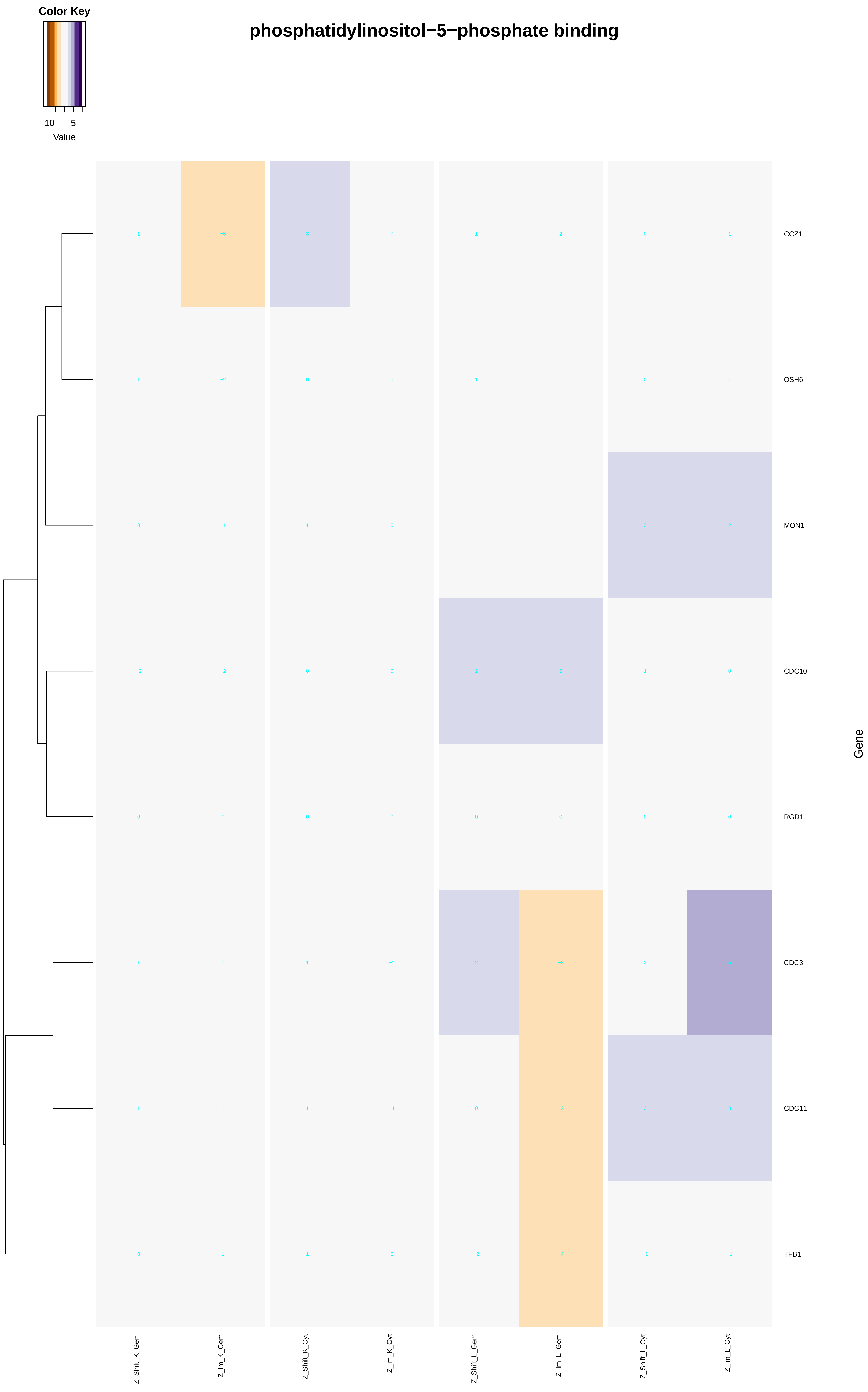

phosphatidylinositol−5−phosphate binding
Color Key
−10
5
Value
CCZ1
OSH6
MON1
CDC10
RGD1
CDC3
CDC11
TFB1
Gene
Z_Shift_K_Gem
Z_Im_K_Gem
Z_Shift_K_Cyt
Z_Im_K_Cyt
Z_Shift_L_Gem
Z_Im_L_Gem
Z_Shift_L_Cyt
Z_Im_L_Cyt

### phosphatidylinositol−3−phosphate binding

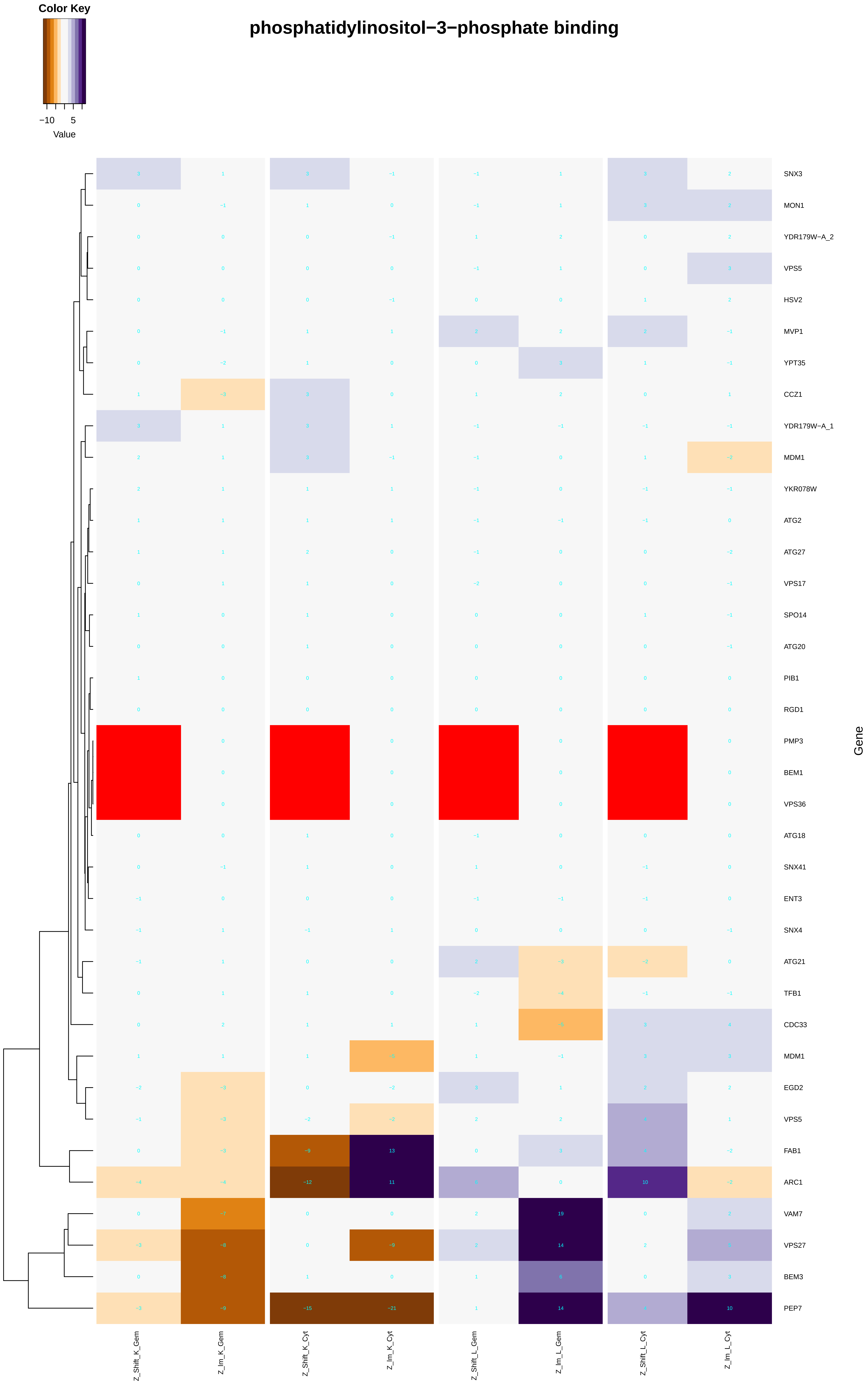

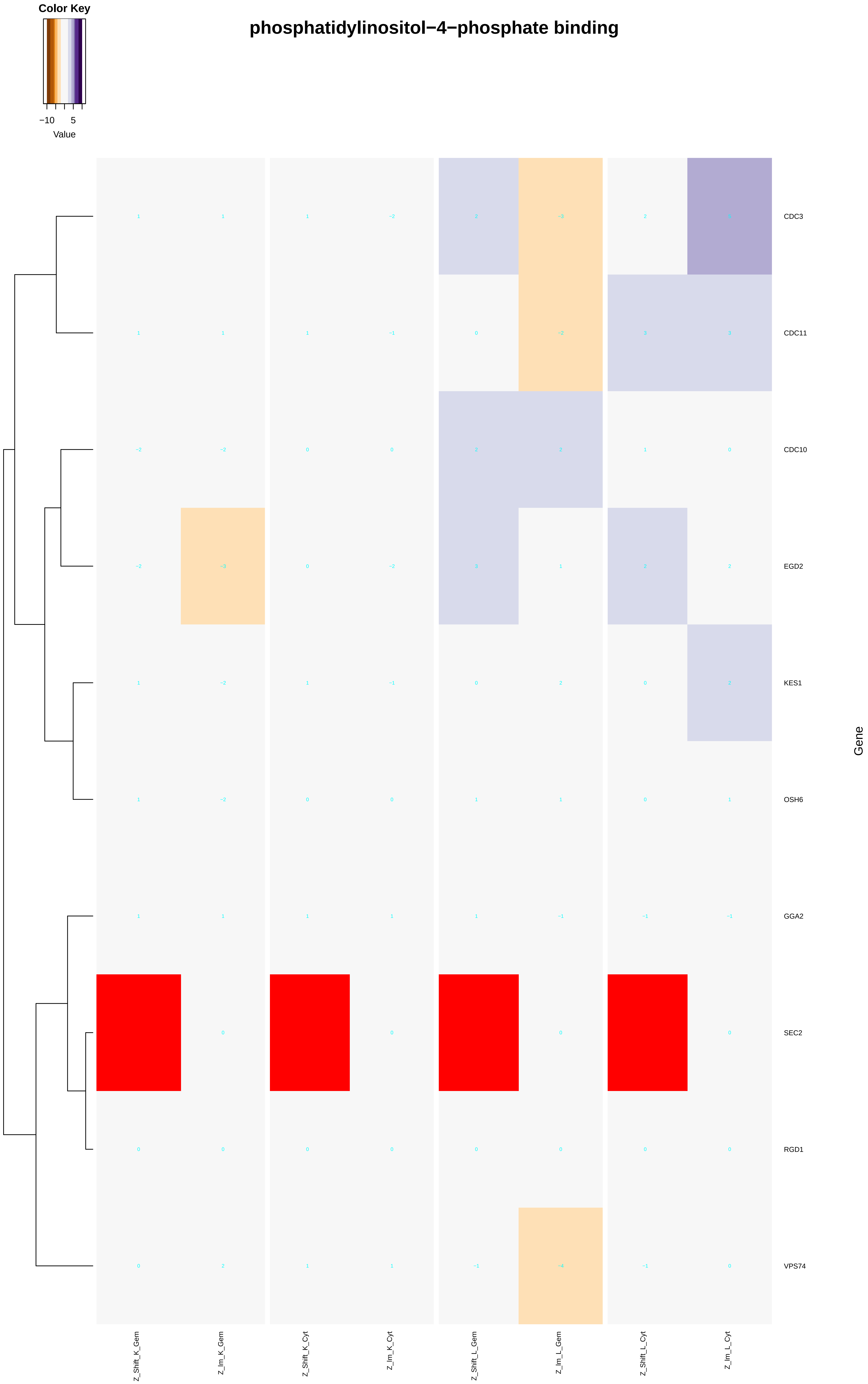
Color Key
−10
5
Value
phosphatidylinositol−4−phosphate binding
CDC3
CDC11
CDC10
EGD2
KES1
OSH6
GGA2
SEC2
RGD1
VPS74
Gene
Z_Shift_K_Gem
Z_Im_K_Gem
Z_Shift_K_Cyt
Z_Im_K_Cyt
Z_Shift_L_Gem
Z_Im_L_Gem
Z_Shift_L_Cyt
Z_Im_L_Cyt

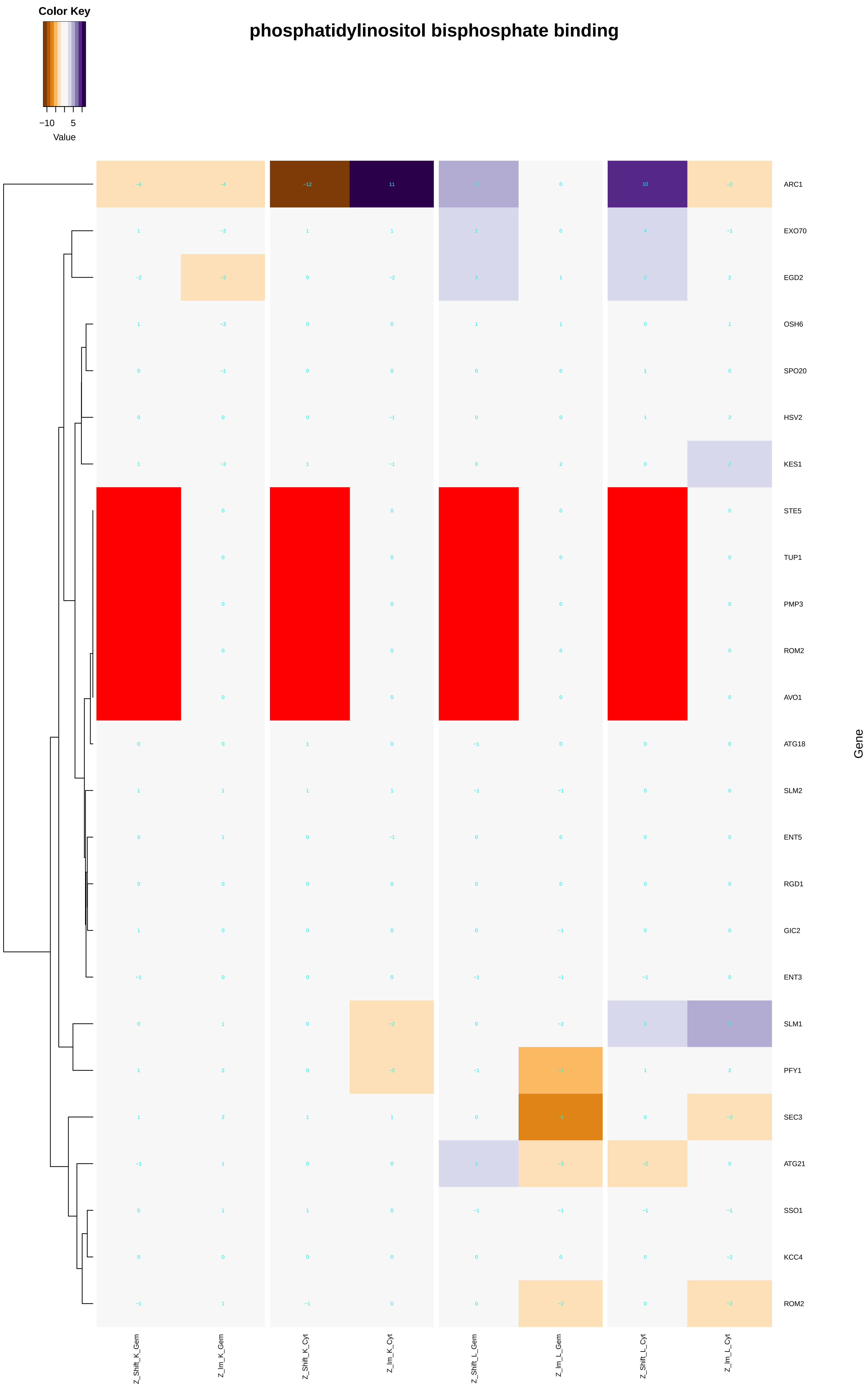

phosphatidylinositol bisphosphate binding
Color Key
−10
5
Value
ARC1
EXO70
EGD2
OSH6
SPO20
HSV2
KES1
STE5
TUP1
PMP3
ROM2
AVO1
ATG18
SLM2
ENT5
RGD1
GIC2
ENT3
SLM1
PFY1
SEC3
ATG21
SSO1
KCC4
ROM2
Gene
Z_Shift_K_Gem
Z_Im_K_Gem
Z_Shift_K_Cyt
Z_Im_K_Cyt
Z_Shift_L_Gem
Z_Im_L_Gem
Z_Shift_L_Cyt
Z_Im_L_Cyt

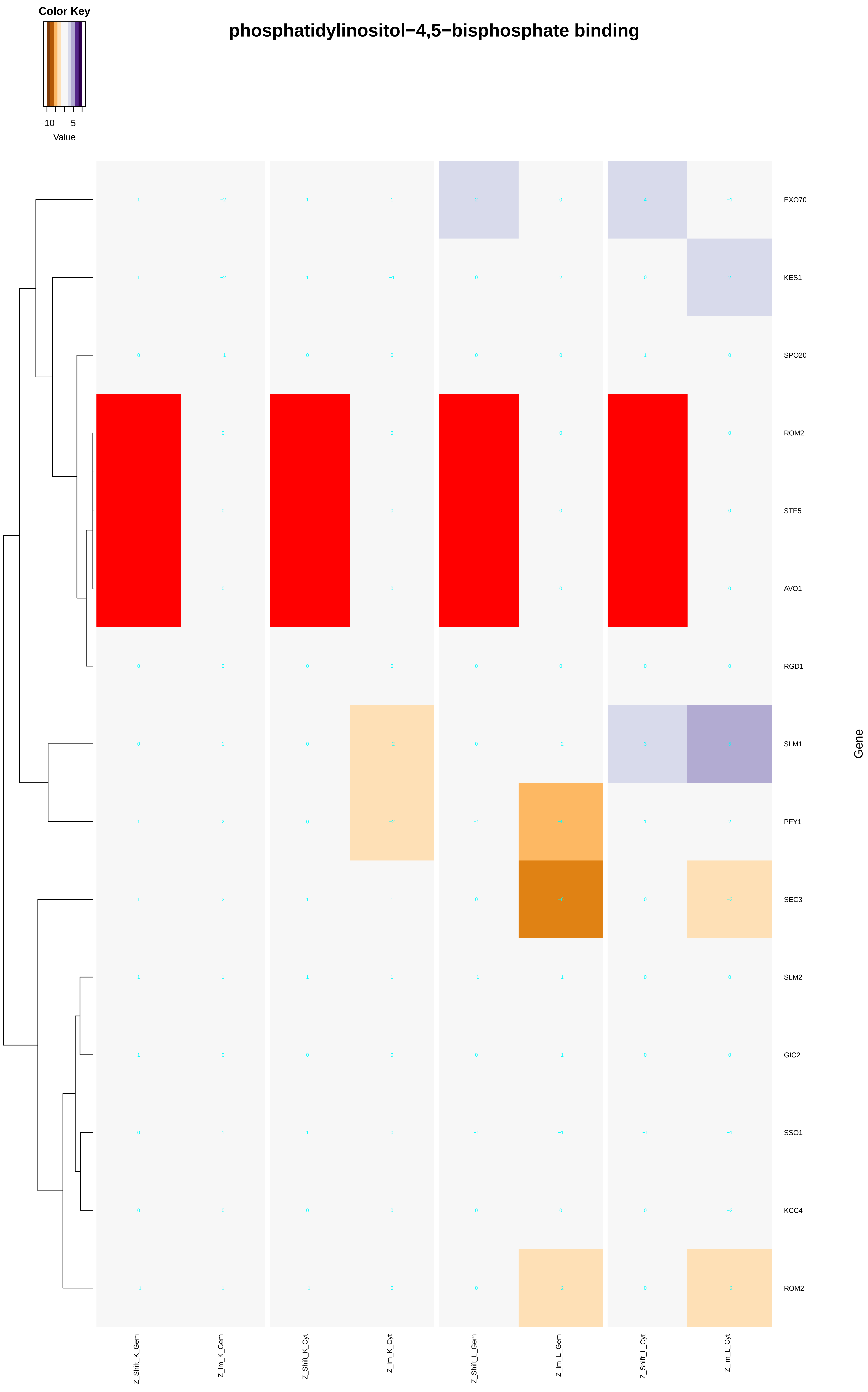

Color Key
−10
5
Value
phosphatidylinositol−4,5−bisphosphate binding
EXO70
KES1
SPO20
ROM2
STE5
AVO1
RGD1
SLM1
PFY1
SEC3
SLM2
GIC2
SSO1
KCC4
ROM2
Gene
Z_Shift_K_Gem
Z_Im_K_Gem
Z_Shift_K_Cyt
Z_Im_K_Cyt
Z_Shift_L_Gem
Z_Im_L_Gem
Z_Shift_L_Cyt
Z_Im_L_Cyt

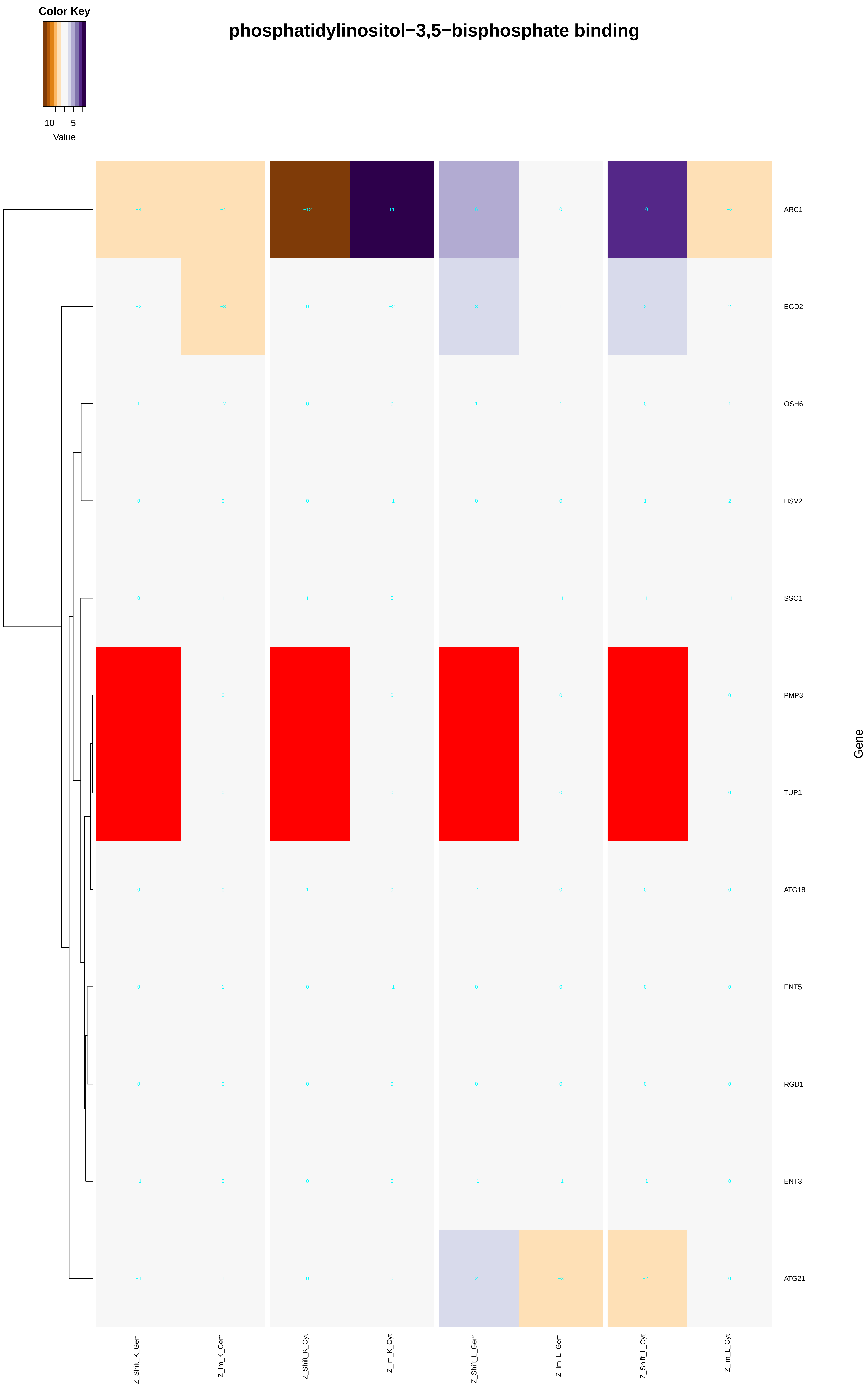
Color Key
−10
5
Value
phosphatidylinositol−3,5−bisphosphate binding
ARC1
EGD2
OSH6
HSV2
SSO1
PMP3
TUP1
ATG18
ENT5
RGD1
ENT3
ATG21
Gene
Z_Shift_K_Gem
Z_Im_K_Gem
Z_Shift_K_Cyt
Z_Im_K_Cyt
Z_Shift_L_Gem
Z_Im_L_Gem
Z_Shift_L_Cyt
Z_Im_L_Cyt
