## Additional File 7 for "A humanized yeast phenomic model of deoxycytidine kinase to predict genetic buffering of nucleoside analog cytotoxicity": membrane_organization.pdf

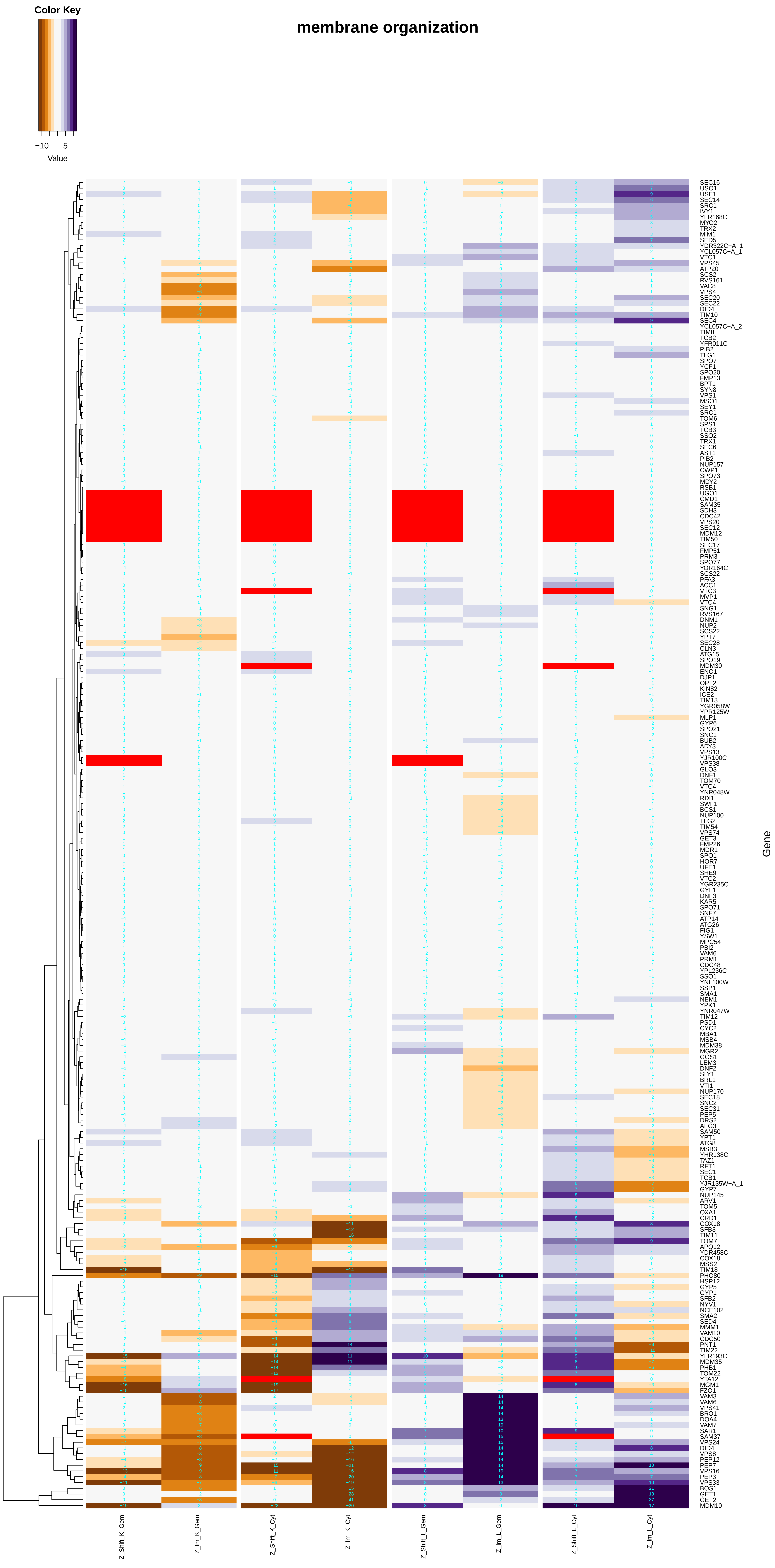
Color Key
−10 5
Value
membrane organization
SEC16
USO1
USE1
SEC14
SRC1
IVY1
YLR168C
MYO2
TRX2
MIM1
SED5
YDR322C−A_1
YCL057C−A_1
VTC1
VPS45
ATP20
SCS2
RVS161
VAC8
VPS4
SEC20
SEC22
DID4
TIM10
SEC4
YCL057C−A_2
TIM8
TCB2
YFR011C
PIB2
TLG1
SPO7
YCF1
SPO20
FMP13
BPT1
SYN8
VPS1
MSO1
SEY1
SRC1
TOM6
SPS1
TCB3
SSO2
TRX1
SEC6
AST1
PIB2
NUP157
CWP1
SPO73
MDY2
RSB1
UGO1
CMD1
SAM35
SDH3
CDC42
VPS20
SEC12
MDM12
TIM50
SEC17
FMP51
PRM3
SPO77
YOR164C
SCS22
PFA3
ACC1
VTC3
MVP1
VTC4
SNG1
RVS167
DNM1
NUP2
SCS22
YPT7
SEC28
CLN3
ATG15
SPO19
MDM30
ENO1
DJP1
OPT2
KIN82
ICE2
TIM13
YGR058W
YPR125W
MLP1
GYP6
SPO21
SNC1
BUB2
ADY3
VPS13
YJR100C
VPS38
GLO3
DNF1
TOM70
VTC4
YNR048W
RDI1
SWF1
BCS1
NUP100
TLG2
TIM54
VPS74
GET3
FMP26
MDR1
SPO1
HOR7
UFE1
SHE9
VTC2
YGR235C
GYL1
DNF3
KAR5
SPO71
SNF7
ATP14
ATG26
FIG1
YSW1
MPC54
PBI2
VAM6
PRM1
CDC48
YPL236C
SSO1
YNL100W
SSP1
SMA1
NEM1
YPK1
YNR047W
TIM12
PSD1
CYC2
MBA1
MSB4
MDM38
MGR2
GOS1
LEM3
DNF2
SLY1
BRL1
VTI1
NUP170
SEC18
SNC2
SEC31
PEP5
DRS2
AFG3
SAM50
YPT1
ATG8
MSB3
YHR138C
TAZ1
RFT1
SEC1
TCB1
YJR135W−A_1
GYP7
NUP145
ARV1
TOM5
OXA1
CRD1
COX18
SFB3
TIM11
TOM7
APQ12
YDR458C
COX18
MSS2
TIM18
PHO80
HSP12
GYP5
GYP1
SFB2
NYV1
NCE102
SMA2
SED4
MMM1
VAM10
CDC50
PNT1
TIM22
YLR193C
MDM35
PHB1
TOM22
YTA12
MGM1
FZO1
VAM3
VAM6
VPS41
BRO1
DOA4
VAM7
SAR1
SAM37
VPS24
DID4
VPS8
PEP12
PEP7
VPS16
PEP3
VPS33
BOS1
GET1
GET2
MDM10
Gene
Z_Shift_K_Gem
Z_Im_K_Gem
Z_Shift_K_Cyt
Z_Im_K_Cyt
Z_Shift_L_Gem
Z_Im_L_Gem
Z_Shift_L_Cyt
Z_Im_L_Cyt

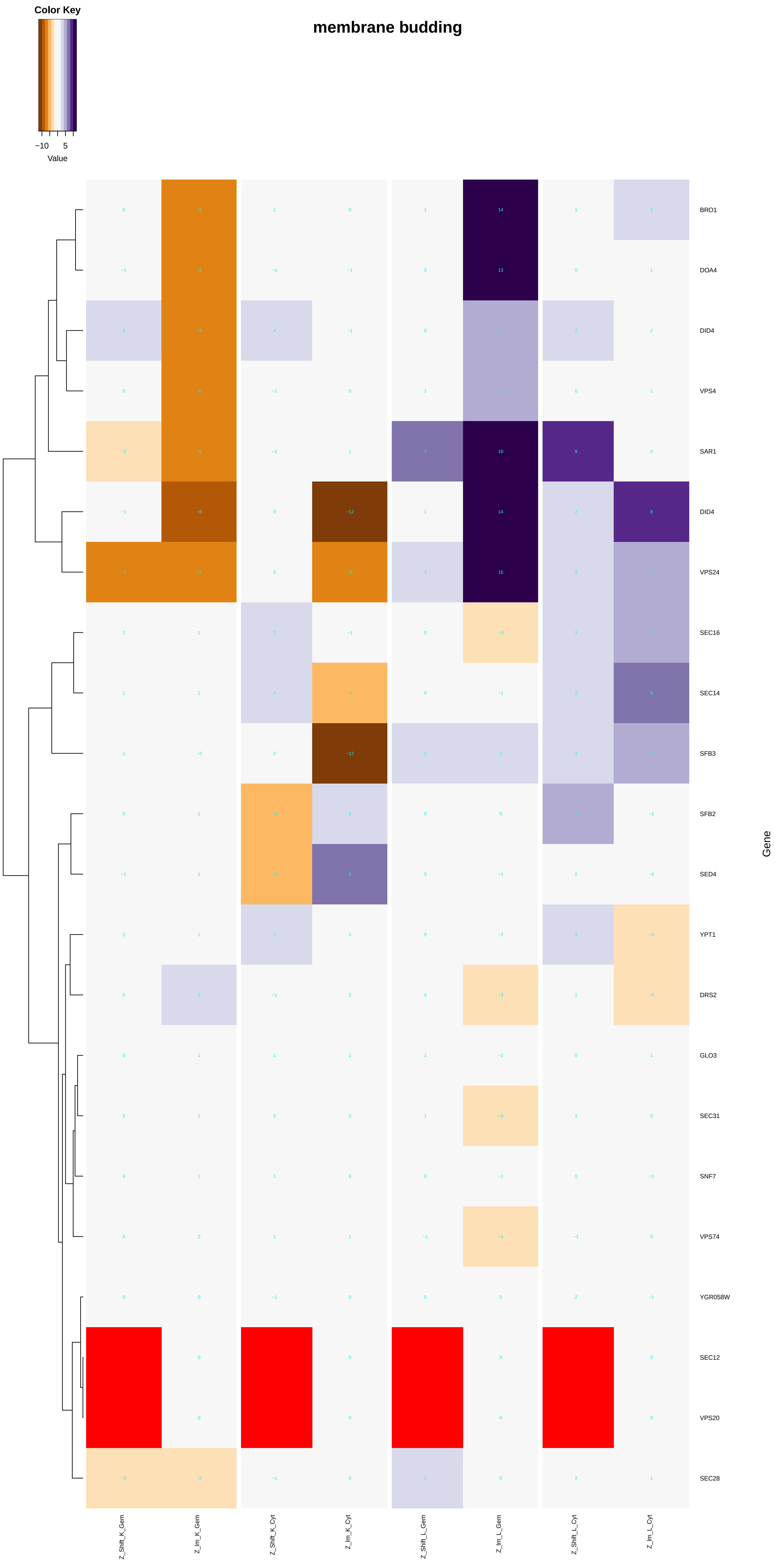
Color Key
−10
5
Value
membrane budding
BRO1
DOA4
DID4
VPS4
SAR1
DID4
VPS24
SEC16
SEC14
SFB3
SFB2
SED4
YPT1
DRS2
GLO3
SEC31
SNF7
VPS74
YGR058W
SEC12
VPS20
SEC28
Gene
Z_Shift_K_Gem
Z_Im_K_Gem
Z_Shift_K_Cyt
Z_Im_K_Cyt
Z_Shift_L_Gem
Z_Im_L_Gem
Z_Shift_L_Cyt
Z_Im_L_Cyt

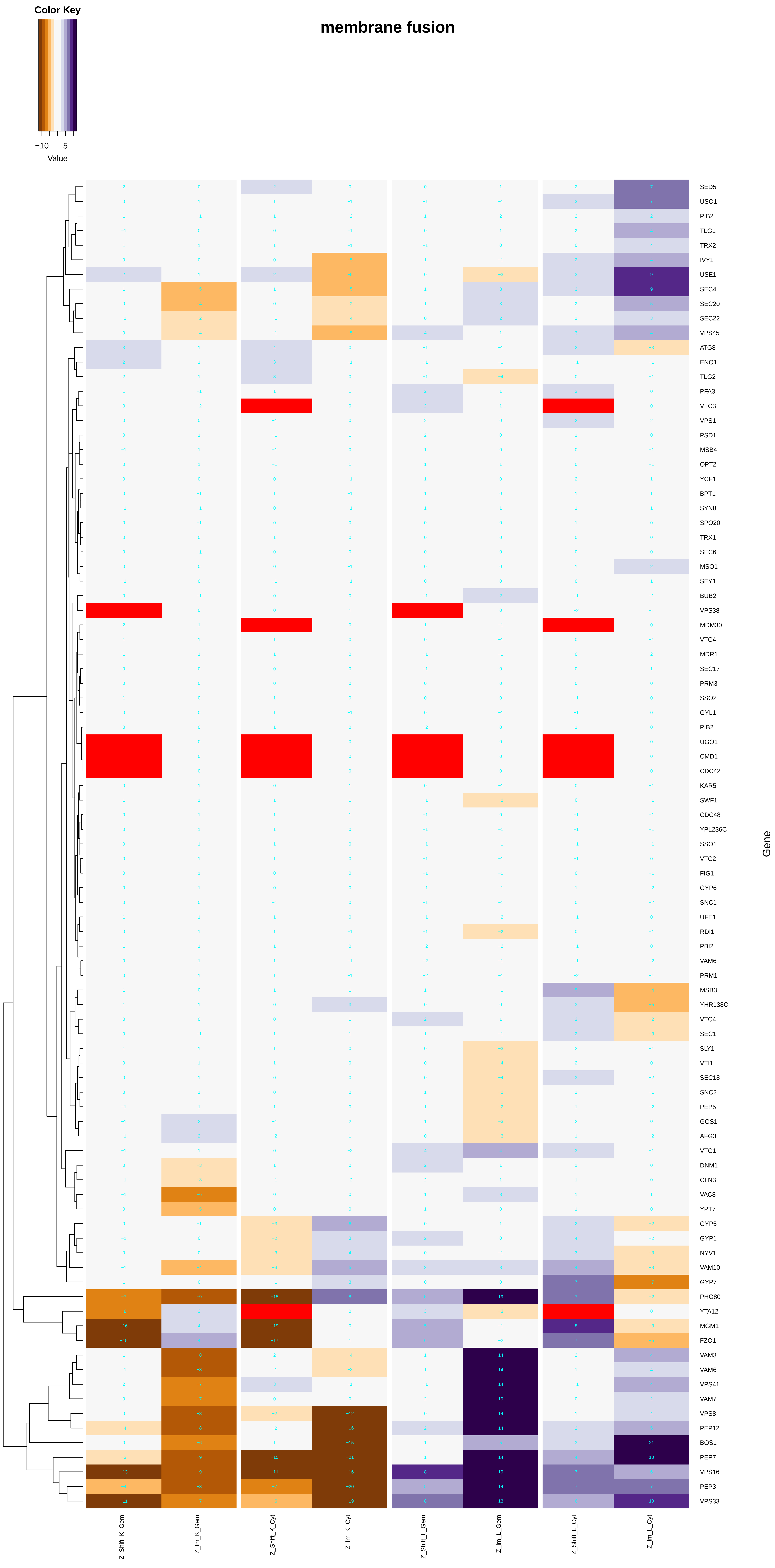
Color Key
−10
5
Value
membrane fusion
SED5
USO1
PIB2
TLG1
TRX2
IVY1
USE1
SEC4
SEC20
SEC22
VPS45
ATG8
ENO1
TLG2
PFA3
VTC3
VPS1
PSD1
MSB4
OPT2
YCF1
BPT1
SYN8
SPO20
TRX1
SEC6
MSO1
SEY1
BUB2
VPS38
MDM30
VTC4
MDR1
SEC17
PRM3
SSO2
GYL1
PIB2
UGO1
CMD1
CDC42
KAR5
SWF1
CDC48
YPL236C
SSO1
VTC2
FIG1
GYP6
SNC1
UFE1
RDI1
PBI2
VAM6
PRM1
MSB3
YHR138C
VTC4
SEC1
SLY1
VTI1
SEC18
SNC2
PEP5
GOS1
AFG3
VTC1
DNM1
CLN3
VAC8
YPT7
GYP5
GYP1
NYV1
VAM10
GYP7
PHO80
YTA12
MGM1
FZO1
VAM3
VAM6
VPS41
VAM7
VPS8
PEP12
BOS1
PEP7
VPS16
PEP3
VPS33
Gene
Z_Shift_K_Gem
Z_lm_K_Gem
Z_Shift_K_Cyt
Z_lm_K_Cyt
Z_Shift_L_Gem
Z_lm_L_Gem
Z_Shift_L_Cyt
Z_lm_L_Cyt

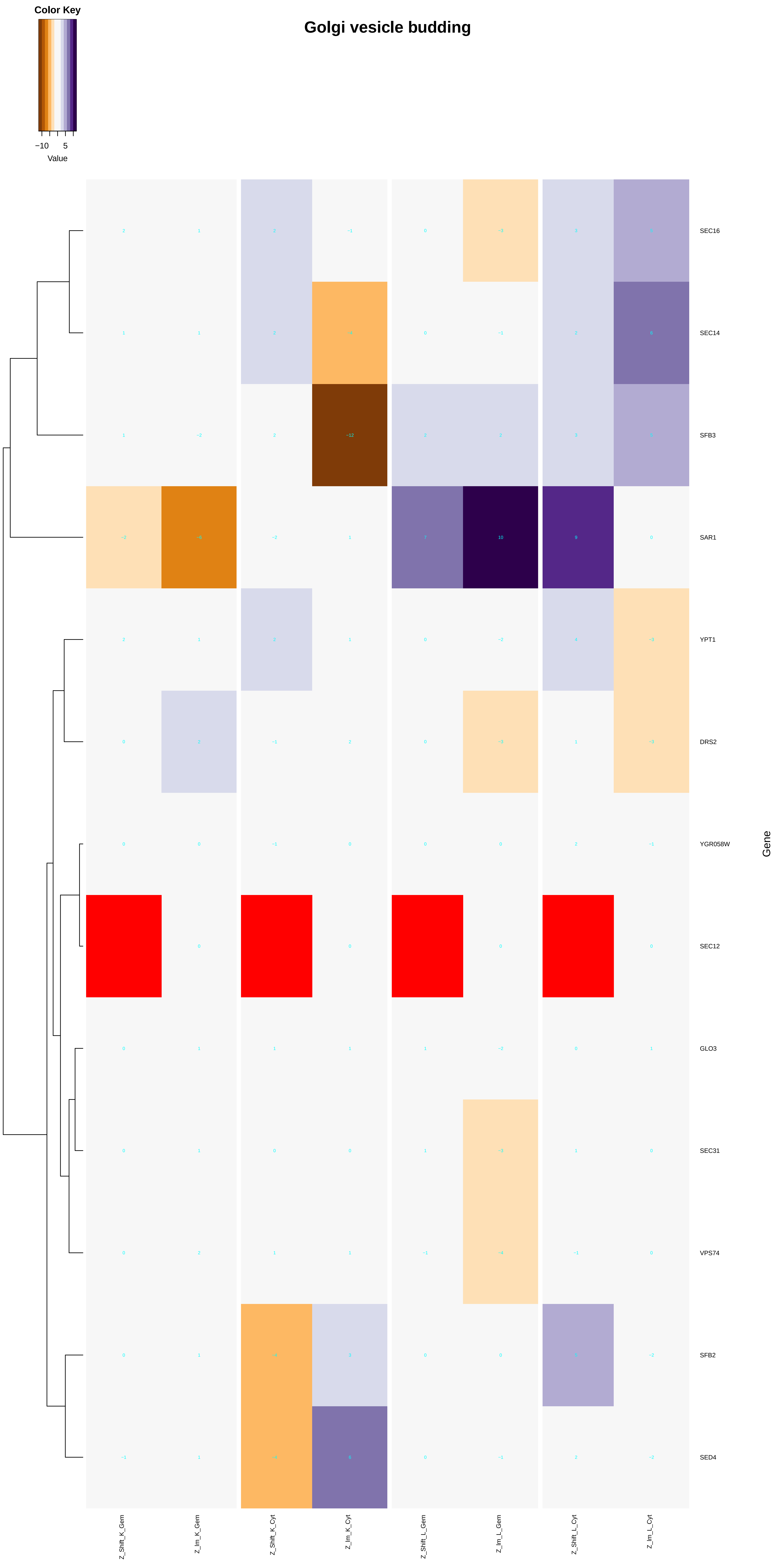

Color Key
−10
5
Value
Golgi vesicle budding
SEC16
SEC14
SFB3
SAR1
YPT1
DRS2
YGR058W
SEC12
GLO3
SEC31
VPS74
SFB2
SED4
Gene
Z_Shift_K_Gem
Z_Im_K_Gem
Z_Shift_K_Cyt
Z_Im_K_Cyt
Z_Shift_L_Gem
Z_Im_L_Gem
Z_Shift_L_Cyt
Z_Im_L_Cyt

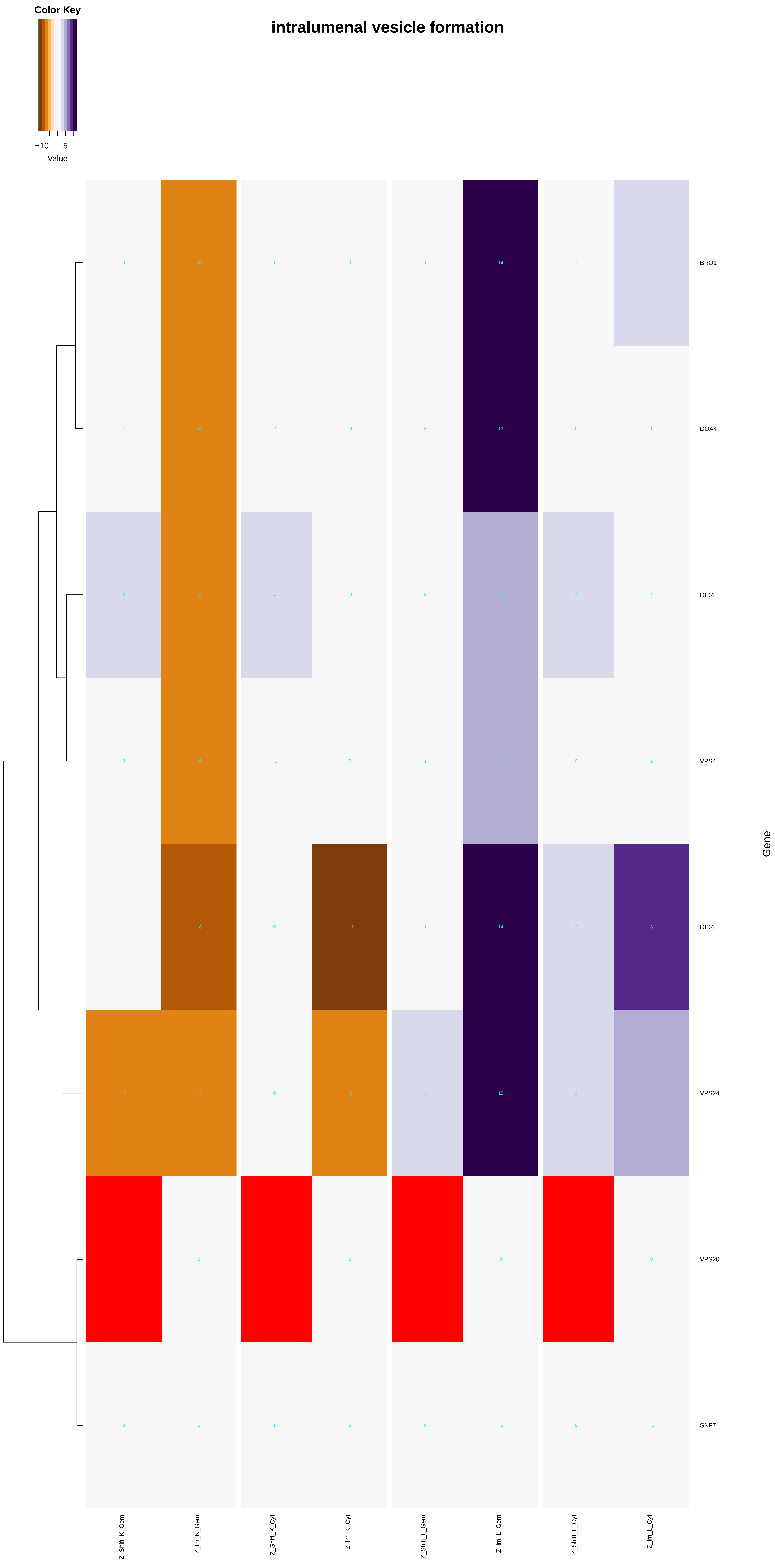

Color Key
−10
5
Value
intralumenal vesicle formation
BRO1
DOA4
DID4
VPS4
DID4
VPS24
VPS20
SNF7
Gene
Z_Shift_K_Gem
Z_Im_K_Gem
Z_Shift_K_Cyt
Z_Im_K_Cyt
Z_Shift_L_Gem
Z_Im_L_Gem
Z_Shift_L_Cyt
Z_Im_L_Cyt

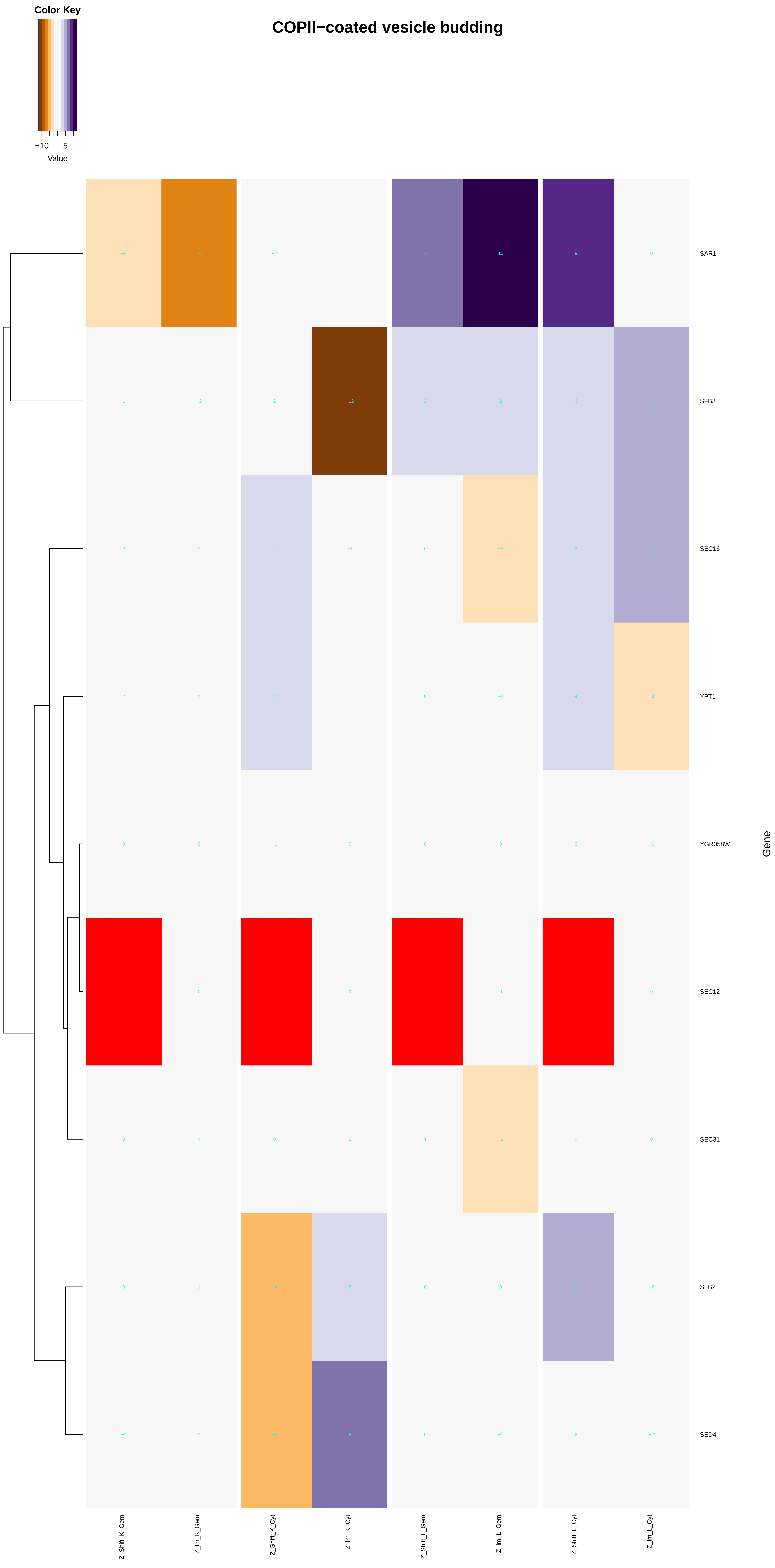
Color Key
−10
5
Value
COPII−coated vesicle budding
SAR1
SFB3
SEC16
YPT1
YGR058W
SEC12
SEC31
SFB2
SED4
Gene
Z_Shift_K_Gem
Z_Im_K_Gem
Z_Shift_K_Cyt
Z_Im_K_Cyt
Z_Shift_L_Gem
Z_Im_L_Gem
Z_Shift_L_Cyt
Z_Im_L_Cyt

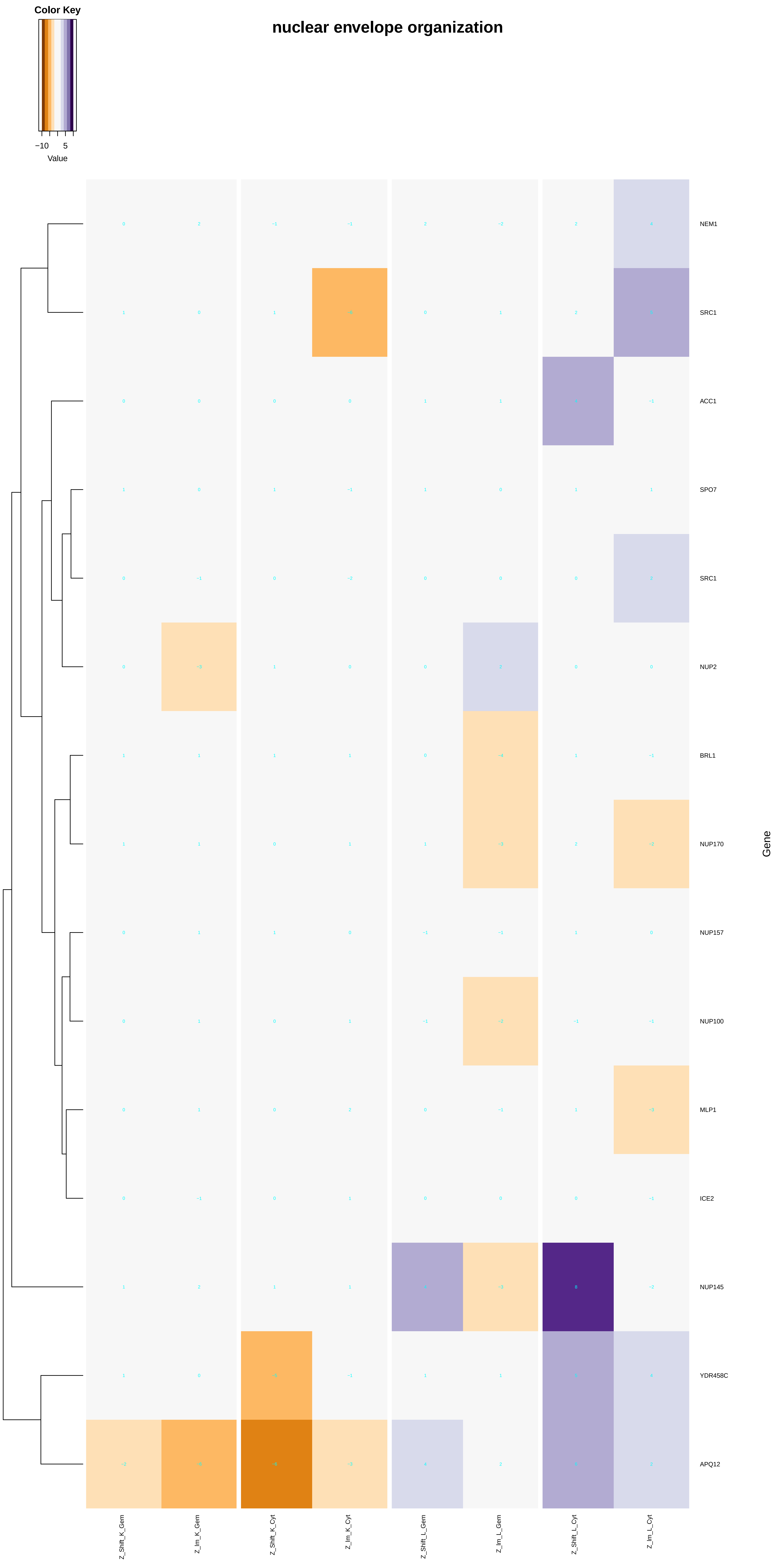

Color Key
−10
5
Value
nuclear envelope organization
NEM1
SRC1
ACC1
SPO7
SRC1
NUP2
BRL1
NUP170
NUP157
NUP100
MLP1
ICE2
NUP145
YDR458C
APQ12
Gene
Z_Shift_K_Gem
Z_Im_K_Gem
Z_Shift_K_Cyt
Z_Im_K_Cyt
Z_Shift_L_Gem
Z_Im_L_Gem
Z_Shift_L_Cyt
Z_Im_L_Cyt

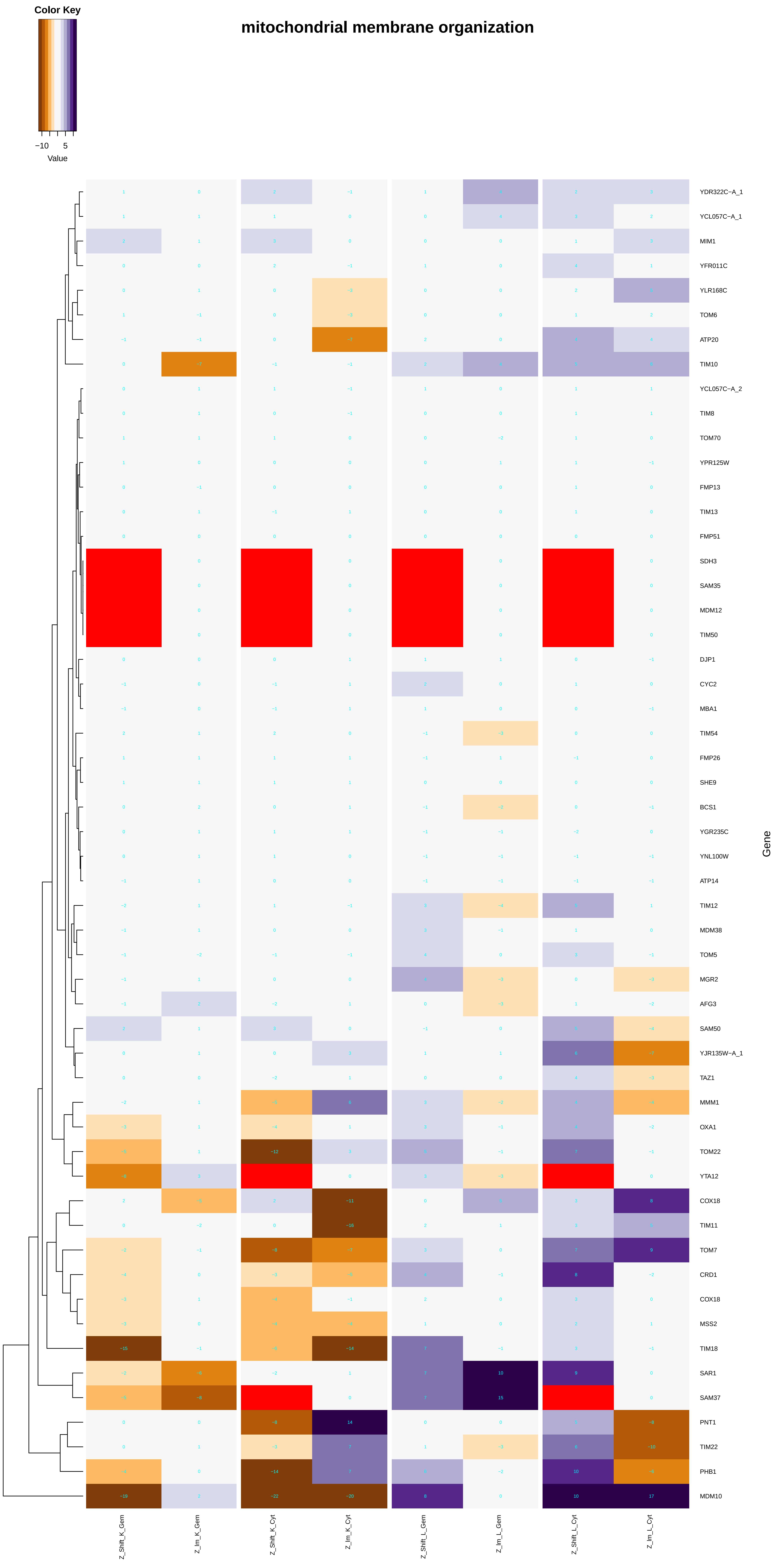

Color Key
−10
5
Value
mitochondrial membrane organization
YDR322C−A_1
YCL057C−A_1
MIM1
YFR011C
YLR168C
TOM6
ATP20
TIM10
YCL057C−A_2
TIM8
TOM70
YPR125W
FMP13
TIM13
FMP51
SDH3
SAM35
MDM12
TIM50
DJP1
CYC2
MBA1
TIM54
FMP26
SHE9
BCS1
YGR235C
YNL100W
ATP14
TIM12
MDM38
TOM5
MGR2
AFG3
SAM50
YJR135W−A_1
TAZ1
MMM1
OXA1
TOM22
YTA12
COX18
TIM11
TOM7
CRD1
COX18
MSS2
TIM18
SAR1
SAM37
PNT1
TIM22
PHB1
MDM10
Gene
Z_Shift_K_Gem
Z_Im_K_Gem
Z_Shift_K_Cyt
Z_Im_K_Cyt
Z_Shift_L_Gem
Z_Im_L_Gem
Z_Shift_L_Cyt
Z_Im_L_Cyt

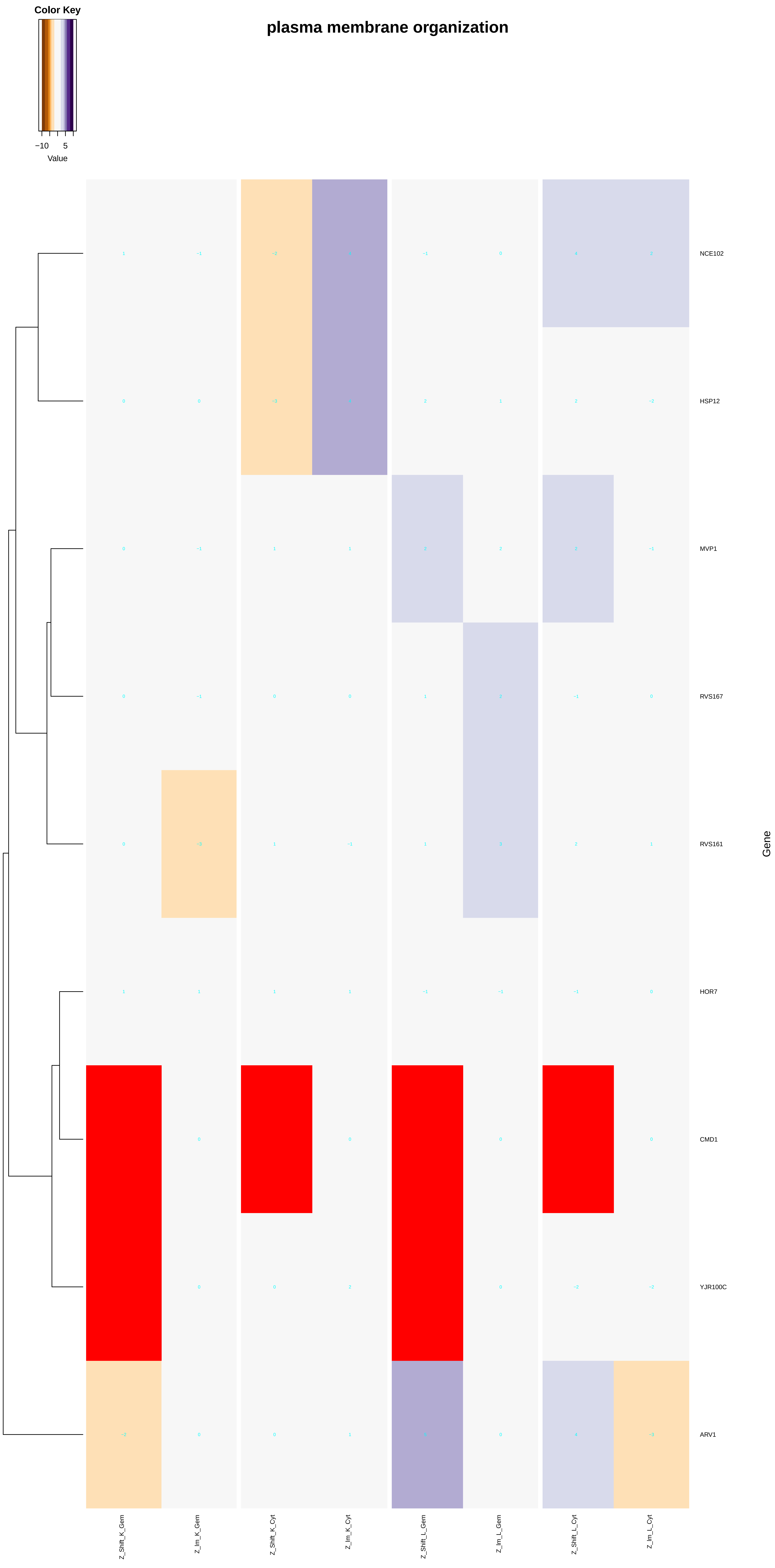

Color Key
−10
5
Value
plasma membrane organization
NCE102
HSP12
MVP1
RVS167
RVS161
HOR7
CMD1
YJR100C
ARV1
Gene
Z_Shift_K_Gem
Z_Im_K_Gem
Z_Shift_K_Cyt
Z_Im_K_Cyt
Z_Shift_L_Gem
Z_Im_L_Gem
Z_Shift_L_Cyt
Z_Im_L_Cyt

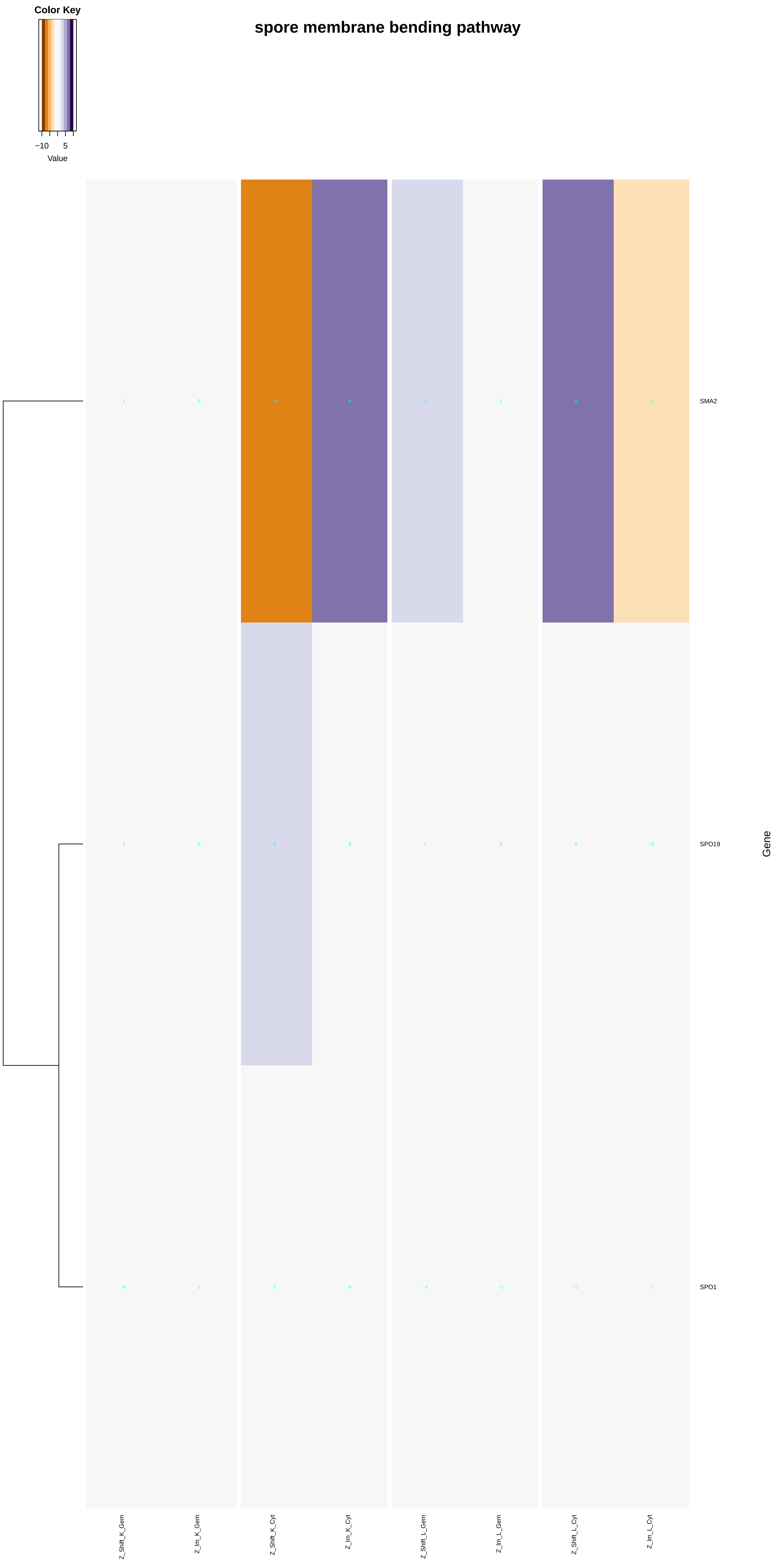
Color Key
−10
5
Value
spore membrane bending pathway
1
0
−8
8
2
1
8
−2
SMA2
1
0
2
0
1
0
0
−2
SPO19
0
1
1
0
−2
−1
−1
1
SPO1
Gene
Z_Shift_K_Gem
Z_Im_K_Gem
Z_Shift_K_Cyt
Z_Im_K_Cyt
Z_Shift_L_Gem
Z_Im_L_Gem
Z_Shift_L_Cyt
Z_Im_L_Cyt

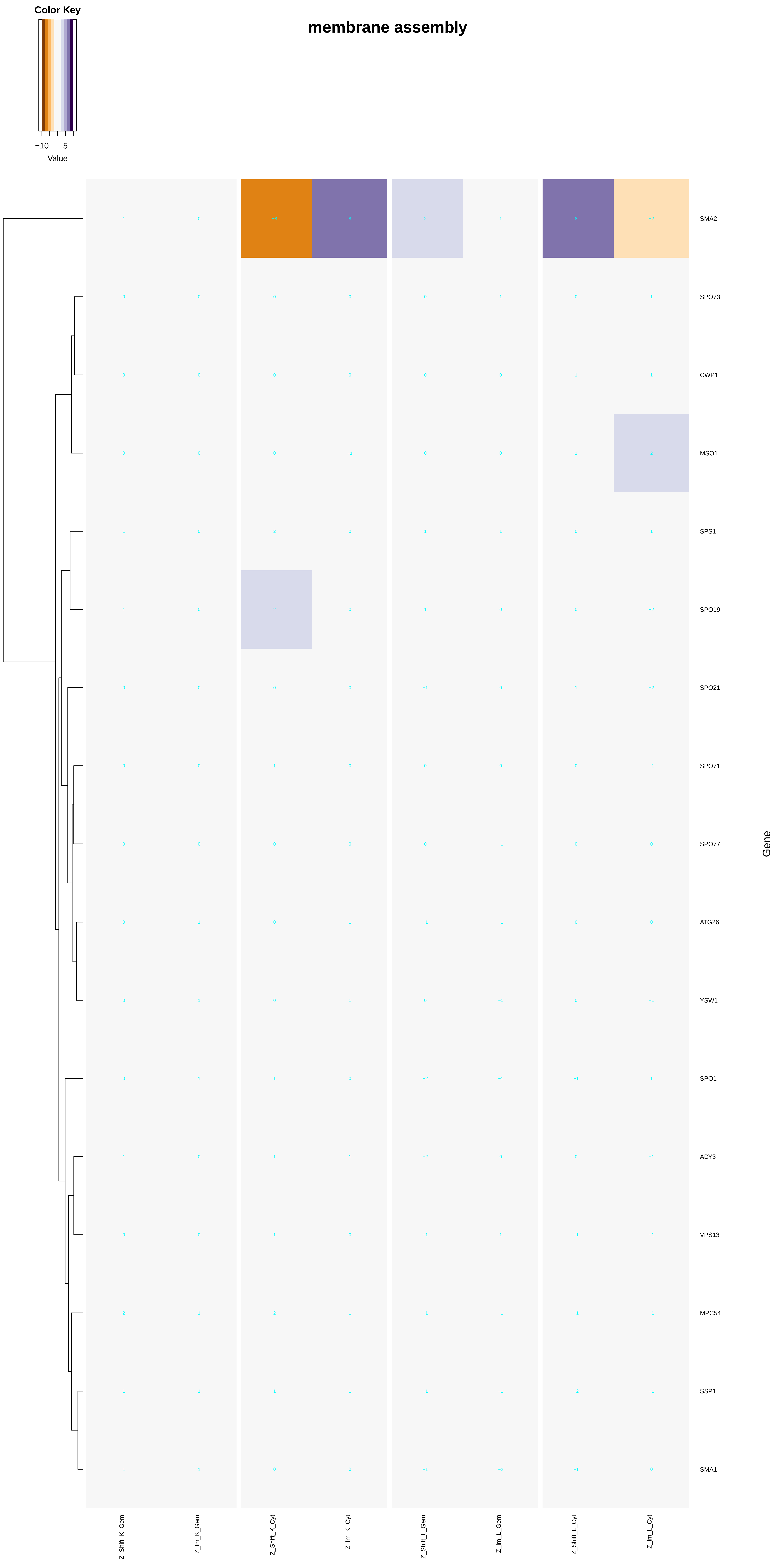
Color Key
−10
5
Value
membrane assembly
SMA2
SPO73
CWP1
MSO1
SPS1
SPO19
SPO21
SPO71
SPO77
ATG26
YSW1
SPO1
ADY3
VPS13
MPC54
SSP1
SMA1
Gene
Z_Shift_K_Gem
Z_lm_K_Gem
Z_Shift_K_Cyt
Z_lm_K_Cyt
Z_Shift_L_Gem
Z_lm_L_Gem
Z_Shift_L_Cyt
Z_lm_L_Cyt

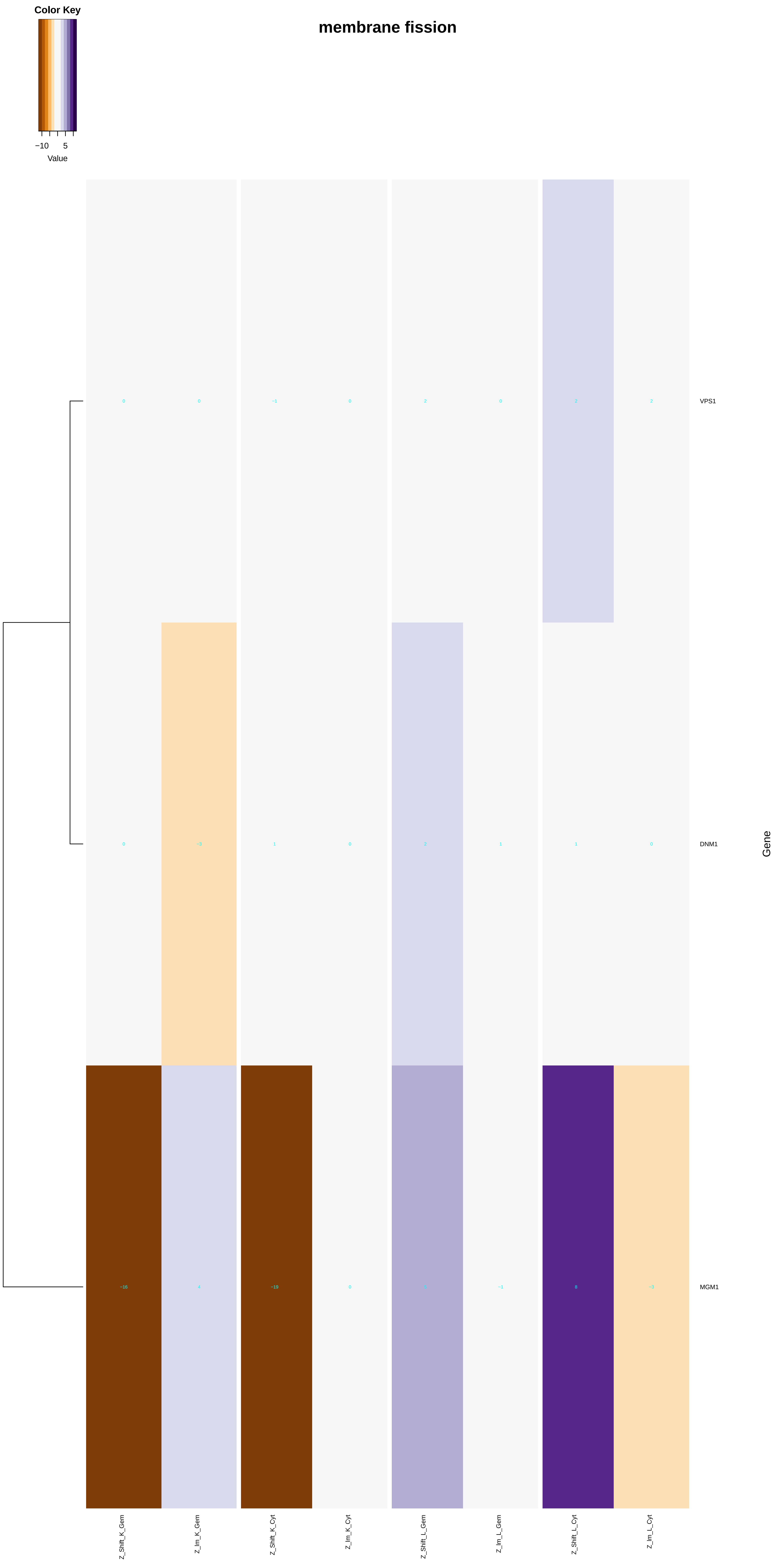
Color Key
−10
5
Value
membrane fission
0
0
−1
0
2
0
2
2
VPS1
0
−3
1
0
2
1
1
0
DNM1
−16
4
−19
0
5
−1
8
−3
MGM1
Gene
Z_Shift_K_Gem
Z_Im_K_Gem
Z_Shift_K_Cyt
Z_Im_K_Cyt
Z_Shift_L_Gem
Z_Im_L_Gem
Z_Shift_L_Cyt
Z_Im_L_Cyt

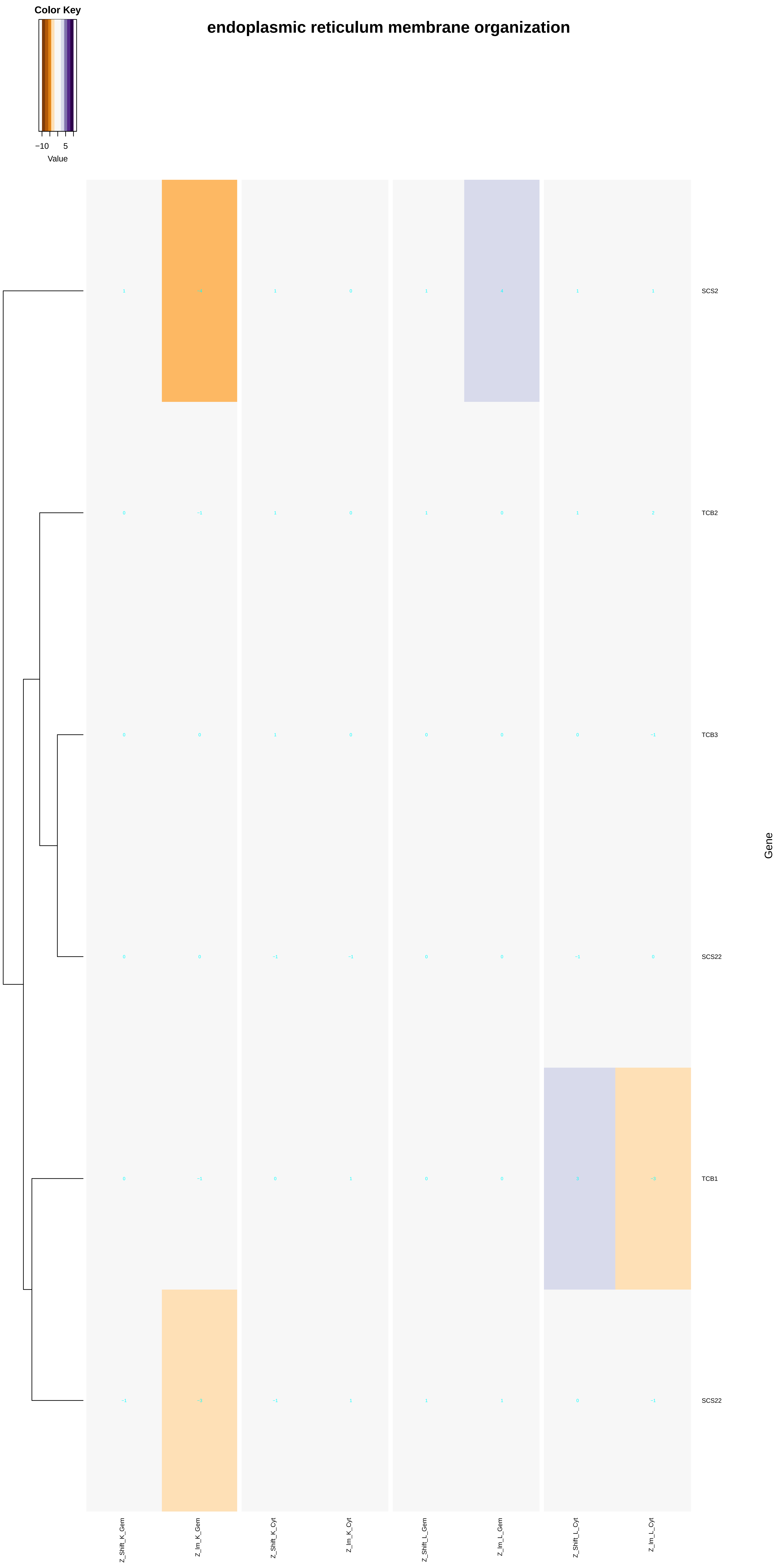
Color Key
−10
5
Value
endoplasmic reticulum membrane organization
SCS2
TCB2
TCB3
SCS22
TCB1
SCS22
Gene
Z_Shift_K_Gem
Z_Im_K_Gem
Z_Shift_K_Cyt
Z_Im_K_Cyt
Z_Shift_L_Gem
Z_Im_L_Gem
Z_Shift_L_Cyt
Z_Im_L_Cyt

### regulation of membrane lipid distribution

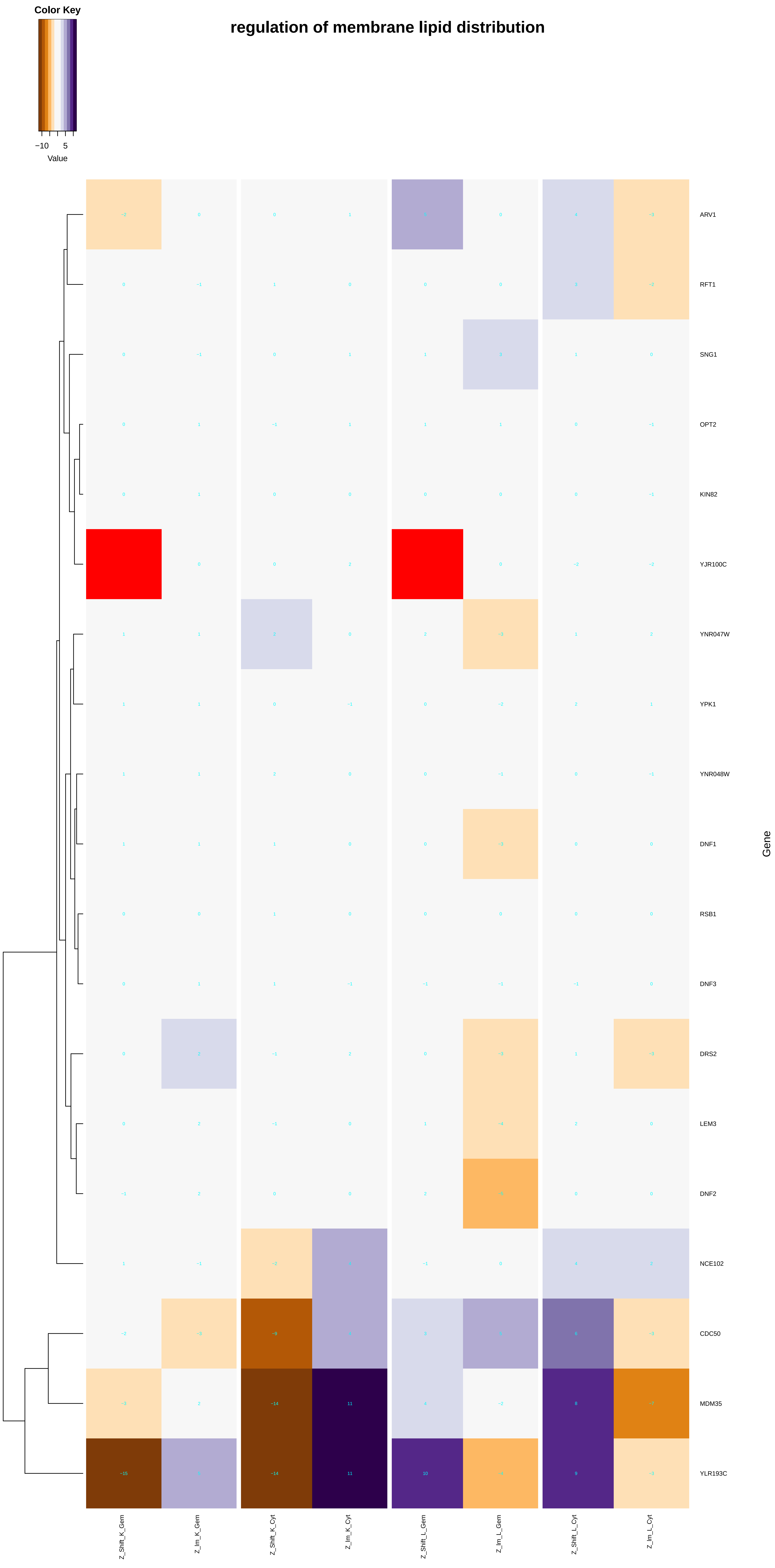

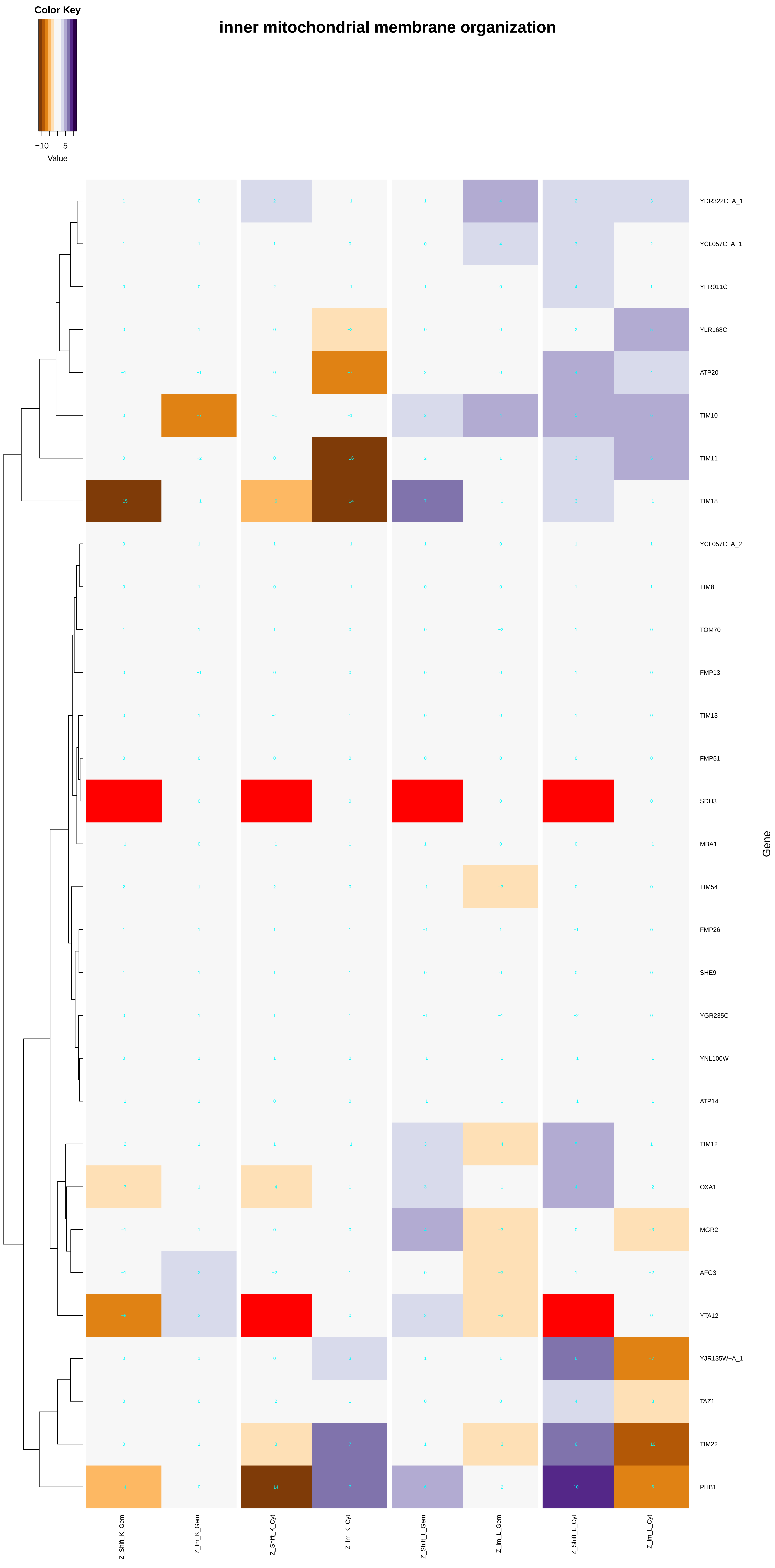
Color Key
−10 5
Value
inner mitochondrial membrane organization
YDR322C−A_1
YCL057C−A_1
YFR011C
YLR168C
ATP20
TIM10
TIM11
TIM18
YCL057C−A_2
TIM8
TOM70
FMP13
TIM13
FMP51
SDH3
MBA1
TIM54
FMP26
SHE9
YGR235C
YNL100W
ATP14
TIM12
OXA1
MGR2
AFG3
YTA12
YJR135W−A_1
TAZ1
TIM22
PHB1
Gene
Z_Shift_K_Gem
Z_Im_K_Gem
Z_Shift_K_Cyt
Z_Im_K_Cyt
Z_Shift_L_Gem
Z_Im_L_Gem
Z_Shift_L_Cyt
Z_Im_L_Cyt

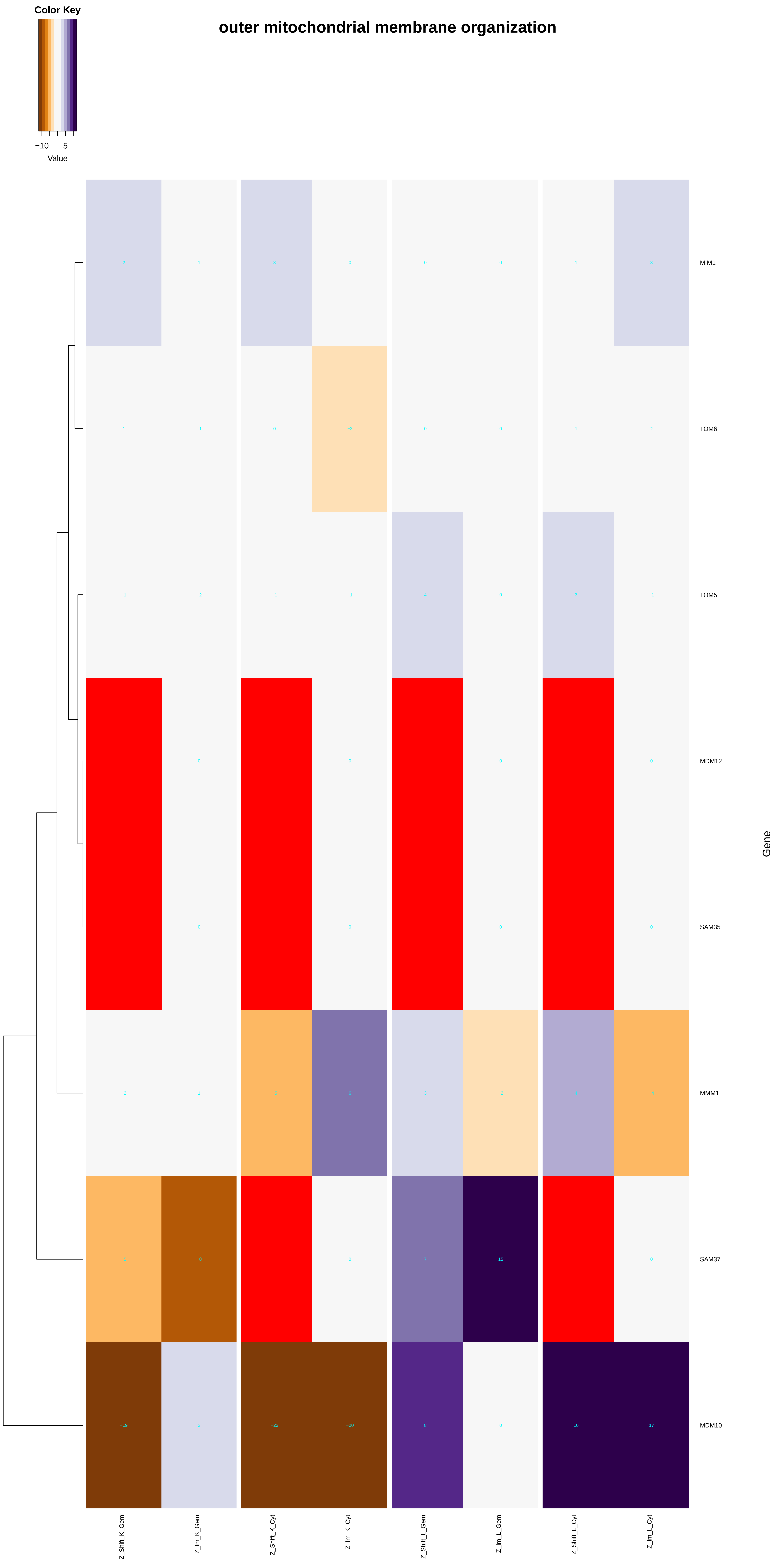
Color Key
−10
5
Value
outer mitochondrial membrane organization
MIM1
TOM6
TOM5
MDM12
SAM35
MMM1
SAM37
MDM10
Gene
Z_Shift_K_Gem
Z_Im_K_Gem
Z_Shift_K_Cyt
Z_Im_K_Cyt
Z_Shift_L_Gem
Z_Im_L_Gem
Z_Shift_L_Cyt
Z_Im_L_Cyt

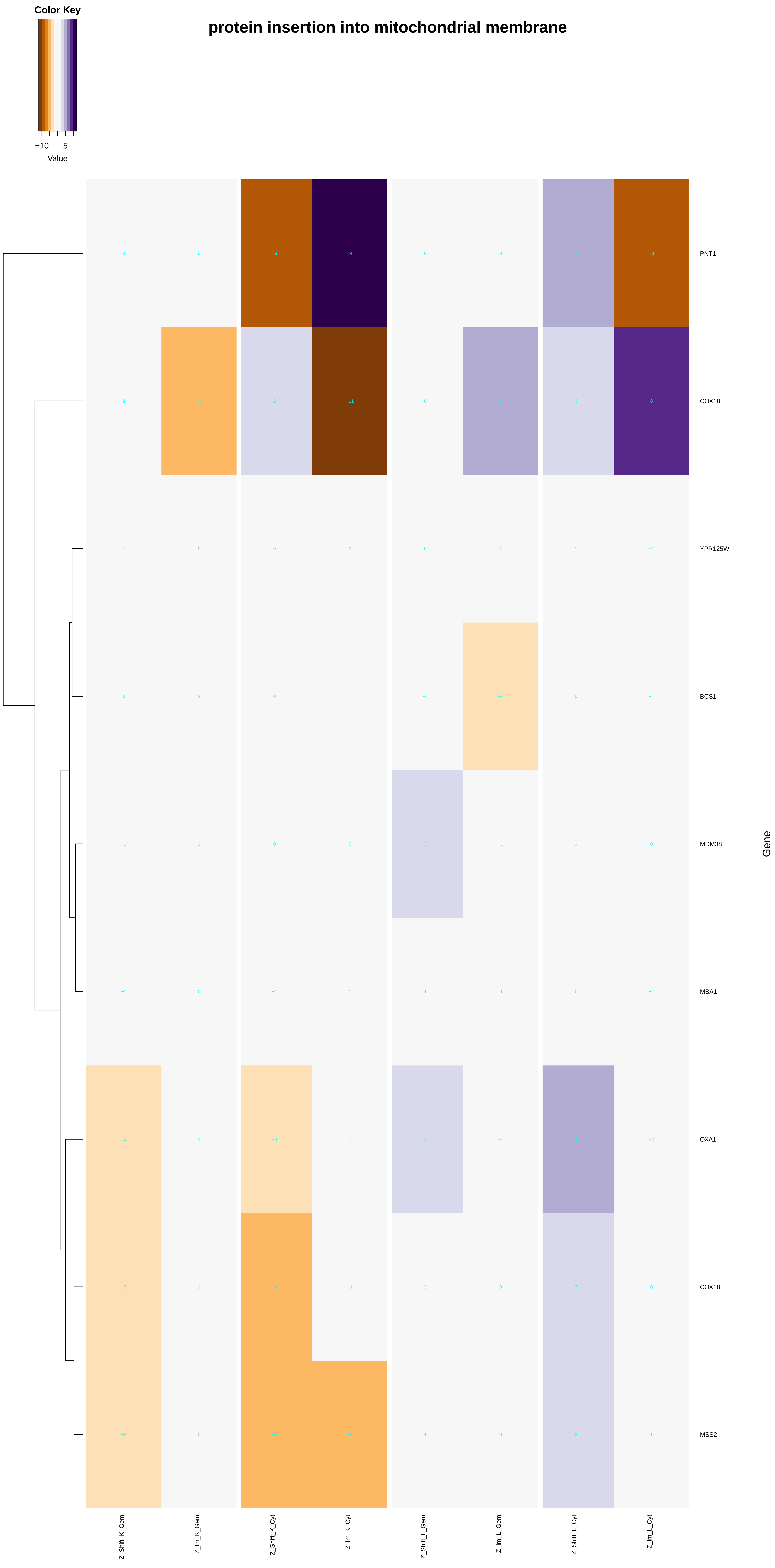

Color Key
−10 5
Value
protein insertion into mitochondrial membrane
PNT1
COX18
YPR125W
BCS1
MDM38
MBA1
OXA1
COX18
MSS2
Gene
Z_Shift_K_Gem
Z_Im_K_Gem
Z_Shift_K_Cyt
Z_Im_K_Cyt
Z_Shift_L_Gem
Z_Im_L_Gem
Z_Shift_L_Cyt
Z_Im_L_Cyt

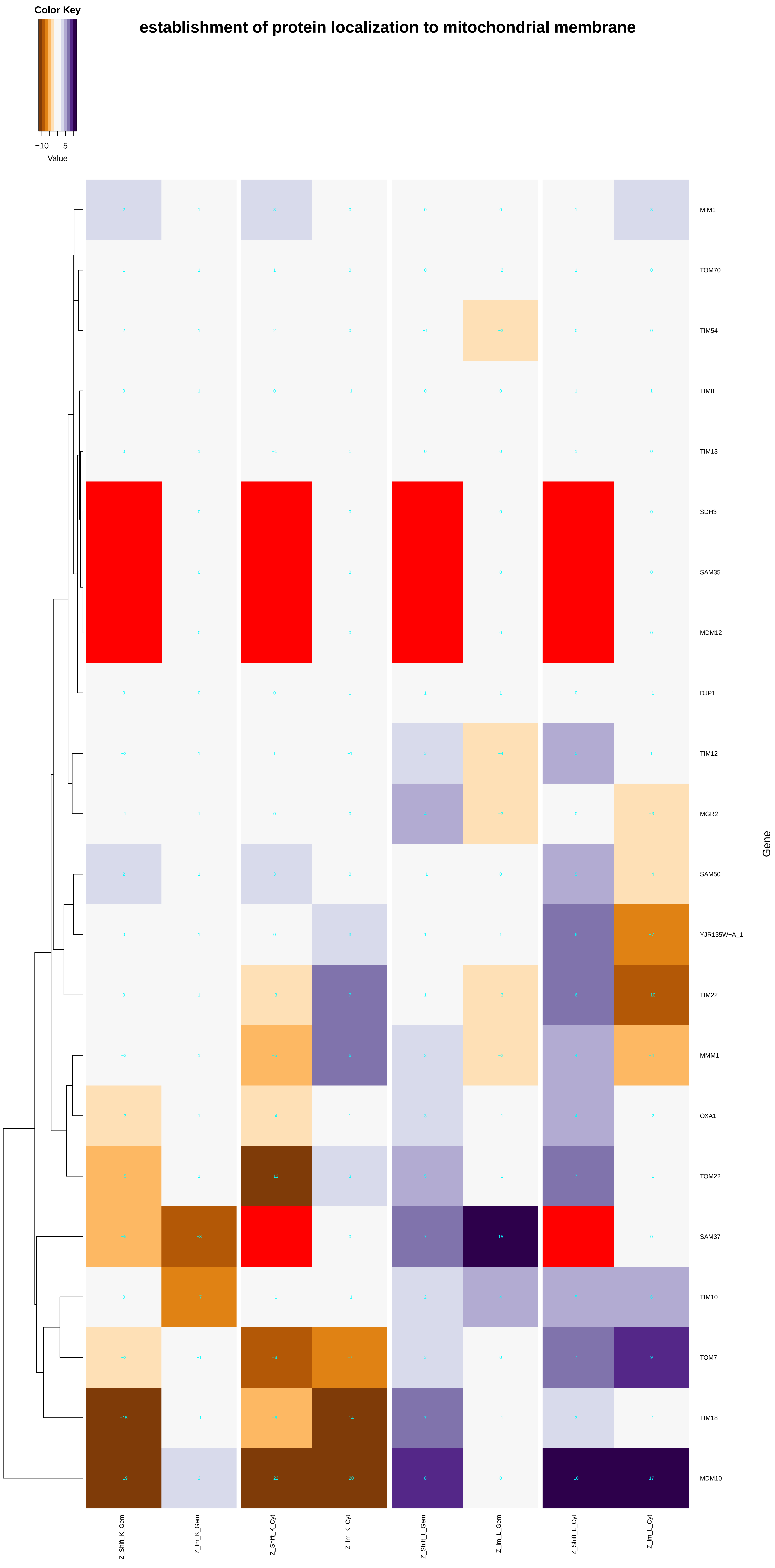

Color Key
−10 5
Value
establishment of protein localization to mitochondrial membrane
MIM1
TOM70
TIM54
TIM8
TIM13
SDH3
SAM35
MDM12
DJP1
TIM12
MGR2
SAM50
YJR135W−A_1
TIM22
MMM1
OXA1
TOM22
SAM37
TIM10
TOM7
TIM18
MDM10
Gene
Z_Shift_K_Gem
Z_Im_K_Gem
Z_Shift_K_Cyt
Z_Im_K_Cyt
Z_Shift_L_Gem
Z_Im_L_Gem
Z_Shift_L_Cyt
Z_Im_L_Cyt

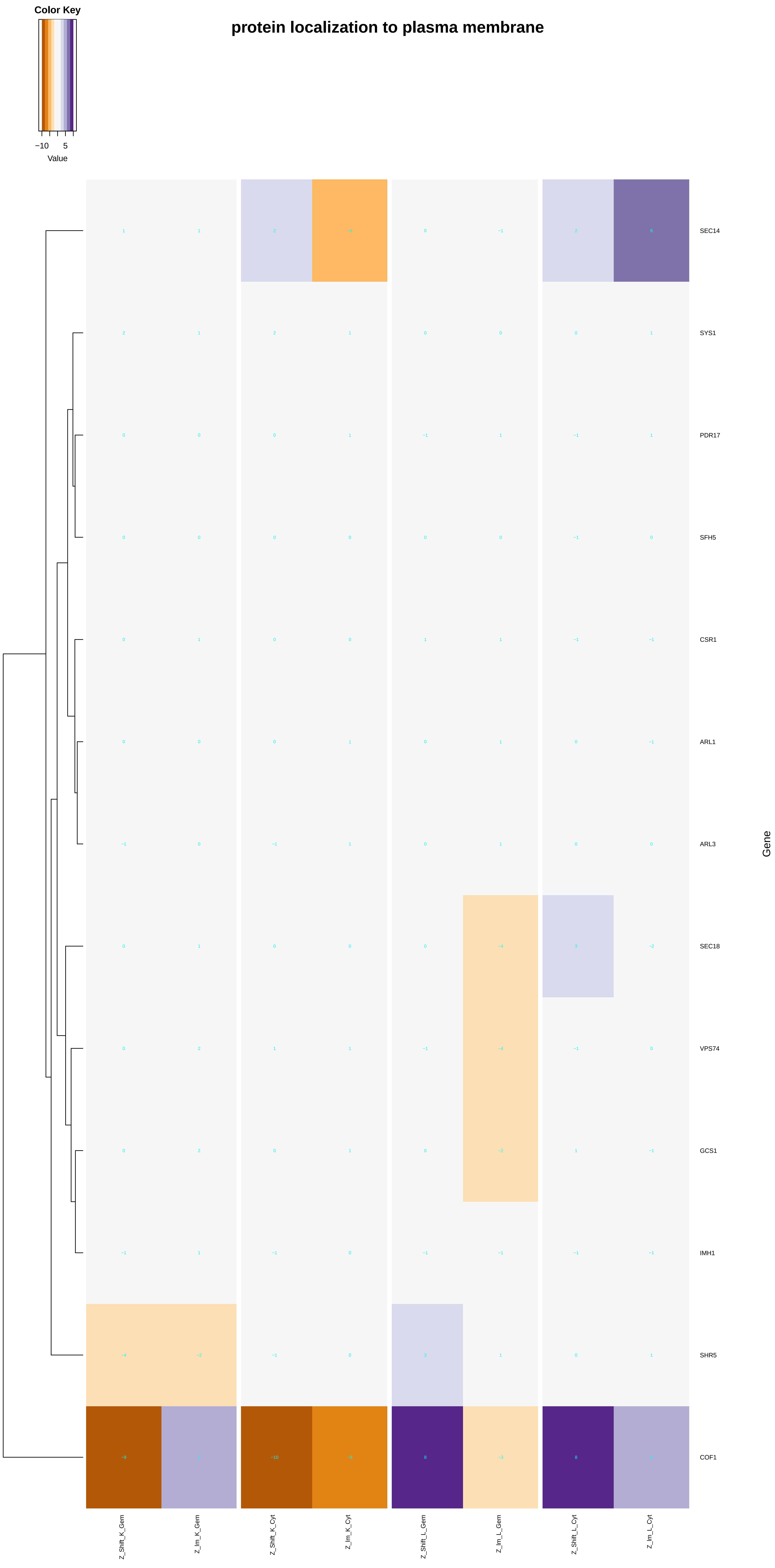
Color Key
−10
5
Value
protein localization to plasma membrane
SEC14
SYS1
PDR17
SFH5
CSR1
ARL1
ARL3
SEC18
VPS74
GCS1
IMH1
SHR5
COF1
Gene
Z_Shift_K_Gem
Z_Im_K_Gem
Z_Shift_K_Cyt
Z_Im_K_Cyt
Z_Shift_L_Gem
Z_Im_L_Gem
Z_Shift_L_Cyt
Z_Im_L_Cyt

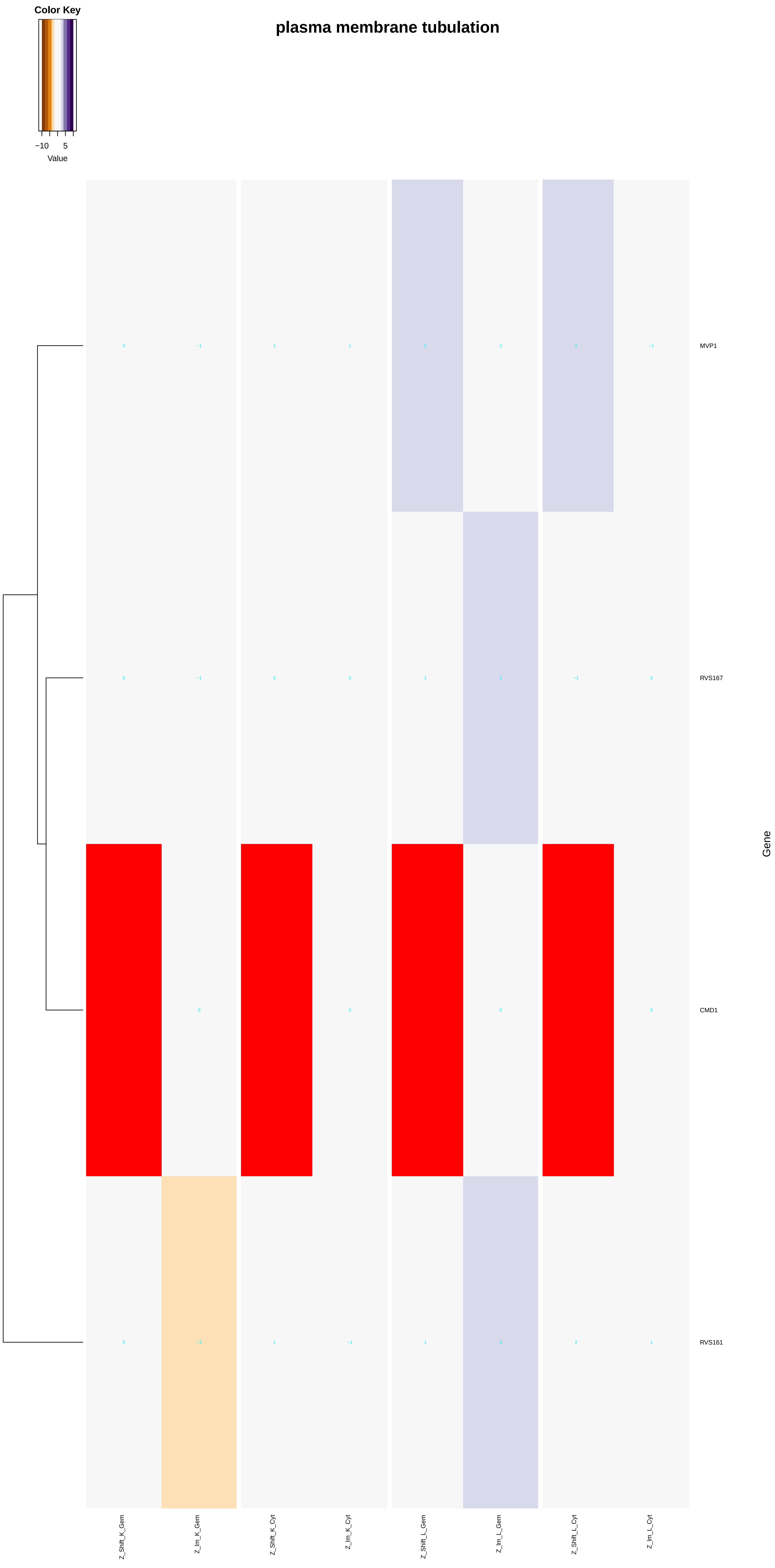
plasma membrane tubulation
Color Key
−10
5
Value
MVP1
RVS167
CMD1
RVS161
Gene
Z_Shift_K_Gem
Z_Im_K_Gem
Z_Shift_K_Cyt
Z_Im_K_Cyt
Z_Shift_L_Gem
Z_Im_L_Gem
Z_Shift_L_Cyt
Z_Im_L_Cyt

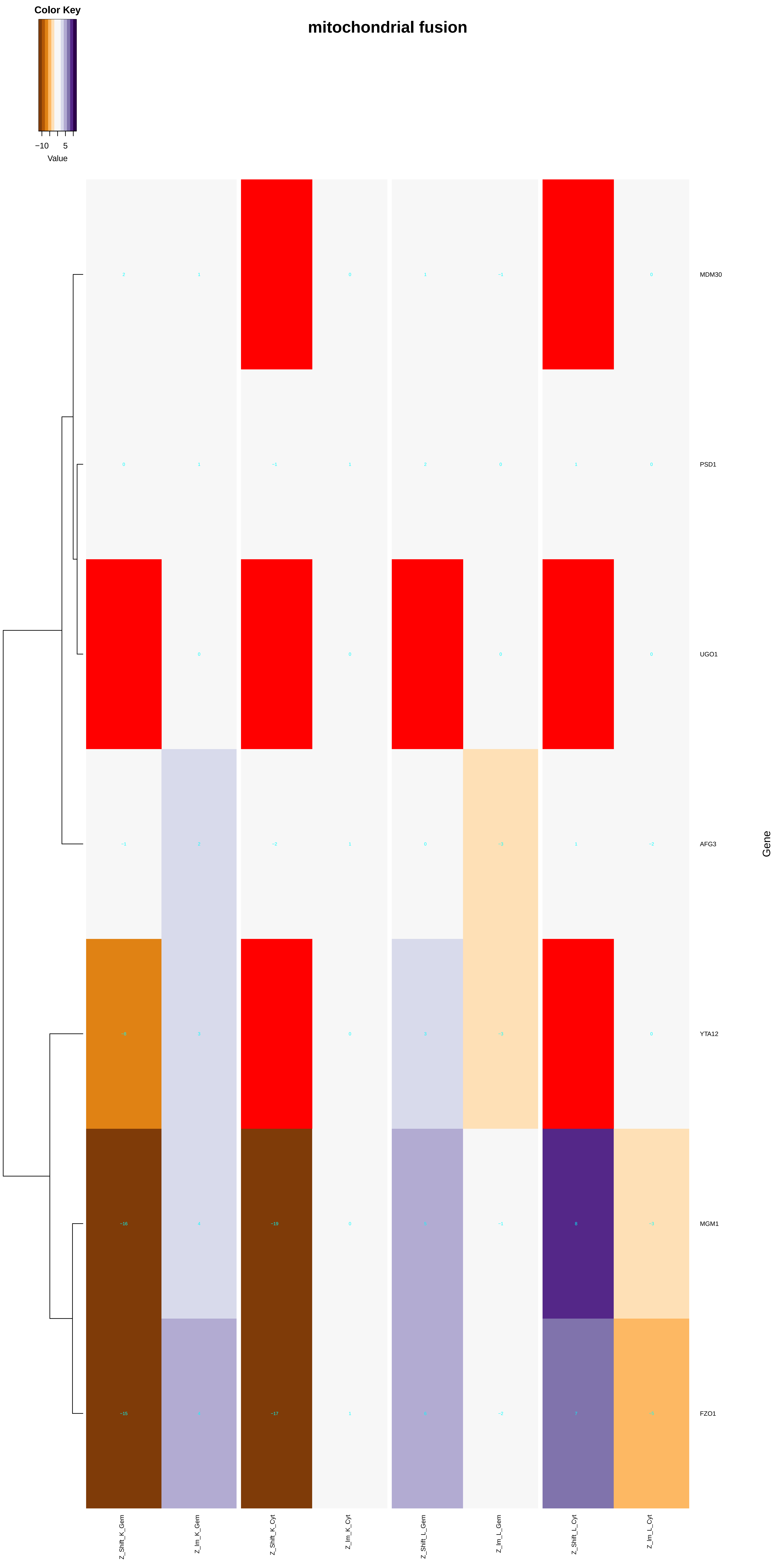
Color Key
−10
5
Value
mitochondrial fusion
MDM30
PSD1
UGO1
AFG3
YTA12
MGM1
FZO1
Gene
Z_Shift_K_Gem
Z_Im_K_Gem
Z_Shift_K_Cyt
Z_Im_K_Cyt
Z_Shift_L_Gem
Z_Im_L_Gem
Z_Shift_L_Cyt
Z_Im_L_Cyt

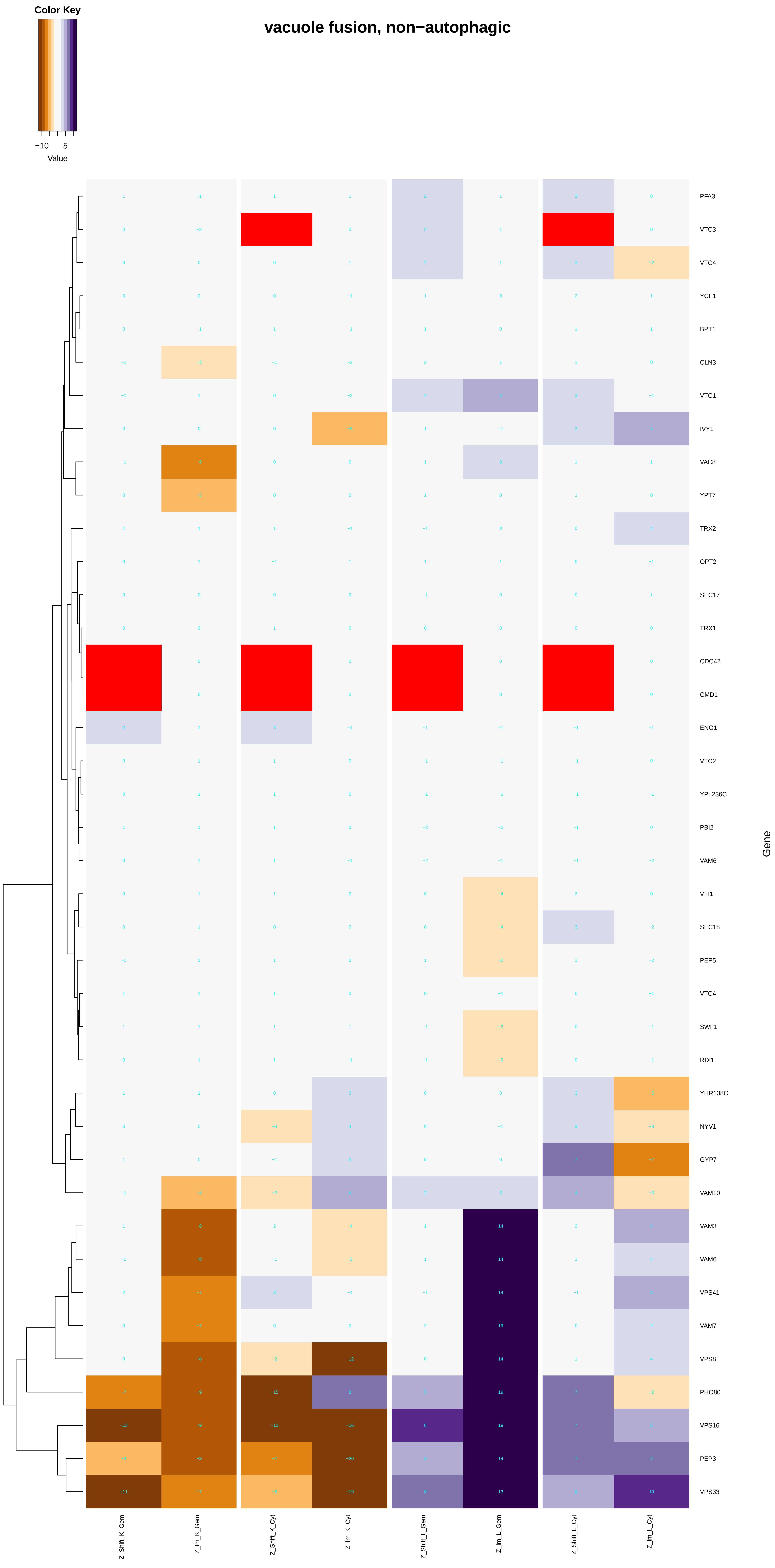

Color Key
−10 5
Value
vacuole fusion, non−autophagic
PFA3
VTC3
VTC4
YCF1
BPT1
CLN3
VTC1
IVY1
VAC8
YPT7
TRX2
OPT2
SEC17
TRX1
CDC42
CMD1
ENO1
VTC2
YPL236C
PBI2
VAM6
VTI1
SEC18
PEP5
VTC4
SWF1
RDI1
YHR138C
NYV1
GYP7
VAM10
VAM3
VAM6
VPS41
VAM7
VPS8
PHO80
VPS16
PEP3
VPS33
Gene
Z_Shift_K_Gem
Z_Im_K_Gem
Z_Shift_K_Cyt
Z_Im_K_Cyt
Z_Shift_L_Gem
Z_Im_L_Gem
Z_Shift_L_Cyt
Z_Im_L_Cyt

### organelle membrane fusion

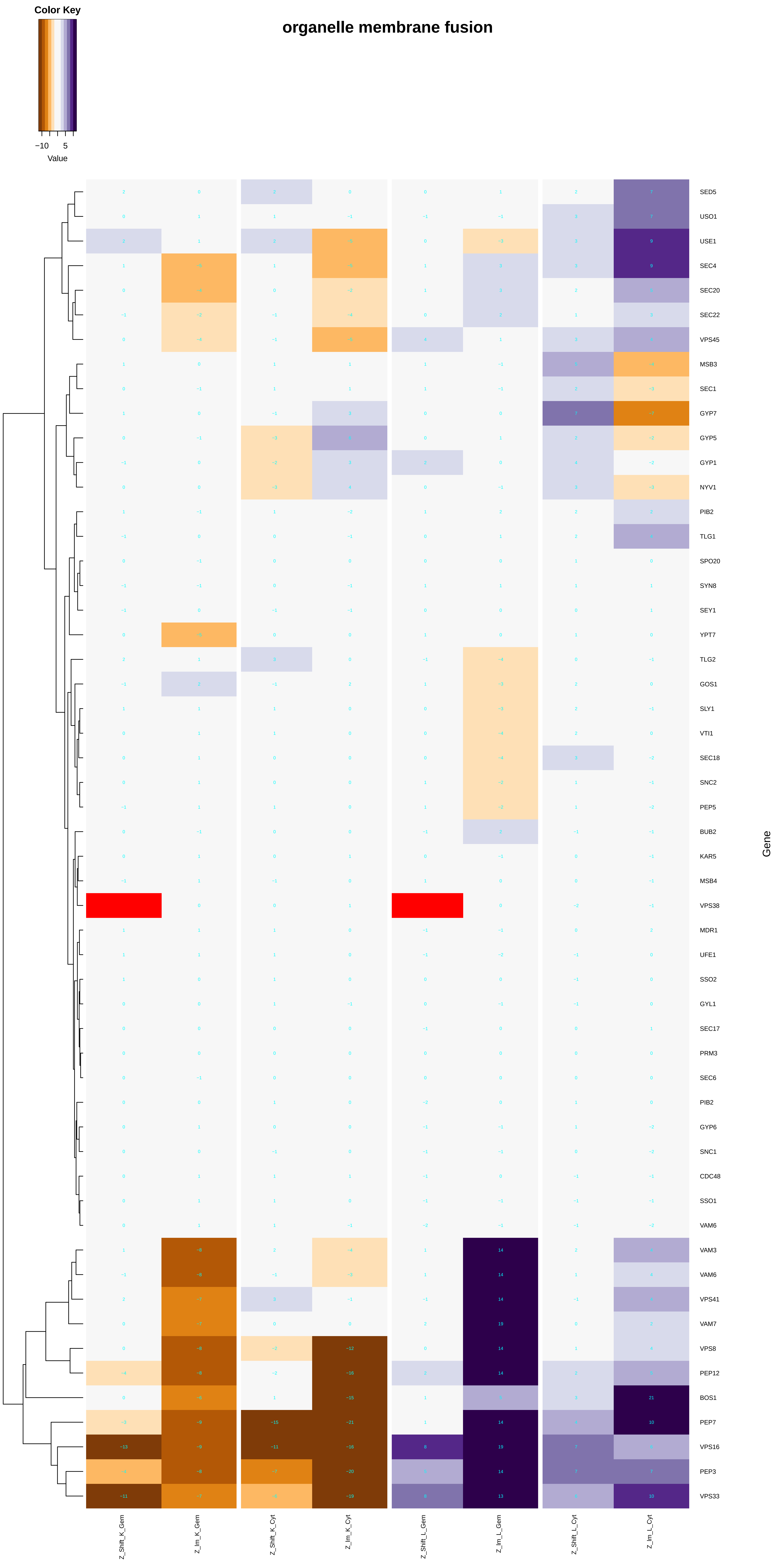

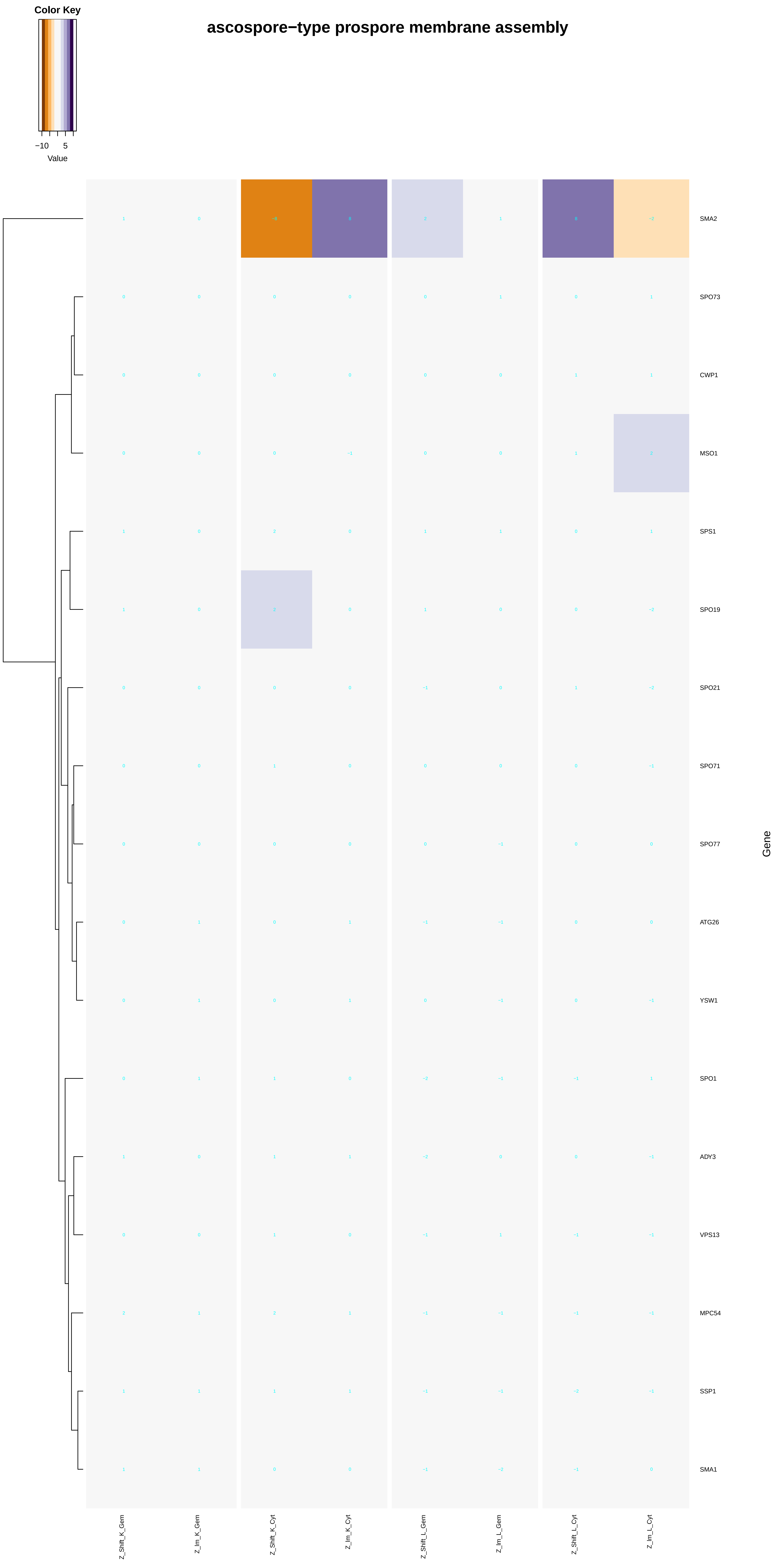
Color Key
−10
5
Value
ascospore−type prospore membrane assembly
SMA2
SPO73
CWP1
MSO1
SPS1
SPO19
SPO21
SPO71
SPO77
ATG26
YSW1
SPO1
ADY3
VPS13
MPC54
SSP1
SMA1
Gene
Z_Shift_K_Gem
Z_lm_K_Gem
Z_Shift_K_Cyt
Z_lm_K_Cyt
Z_Shift_L_Gem
Z_lm_L_Gem
Z_Shift_L_Cyt
Z_lm_L_Cyt

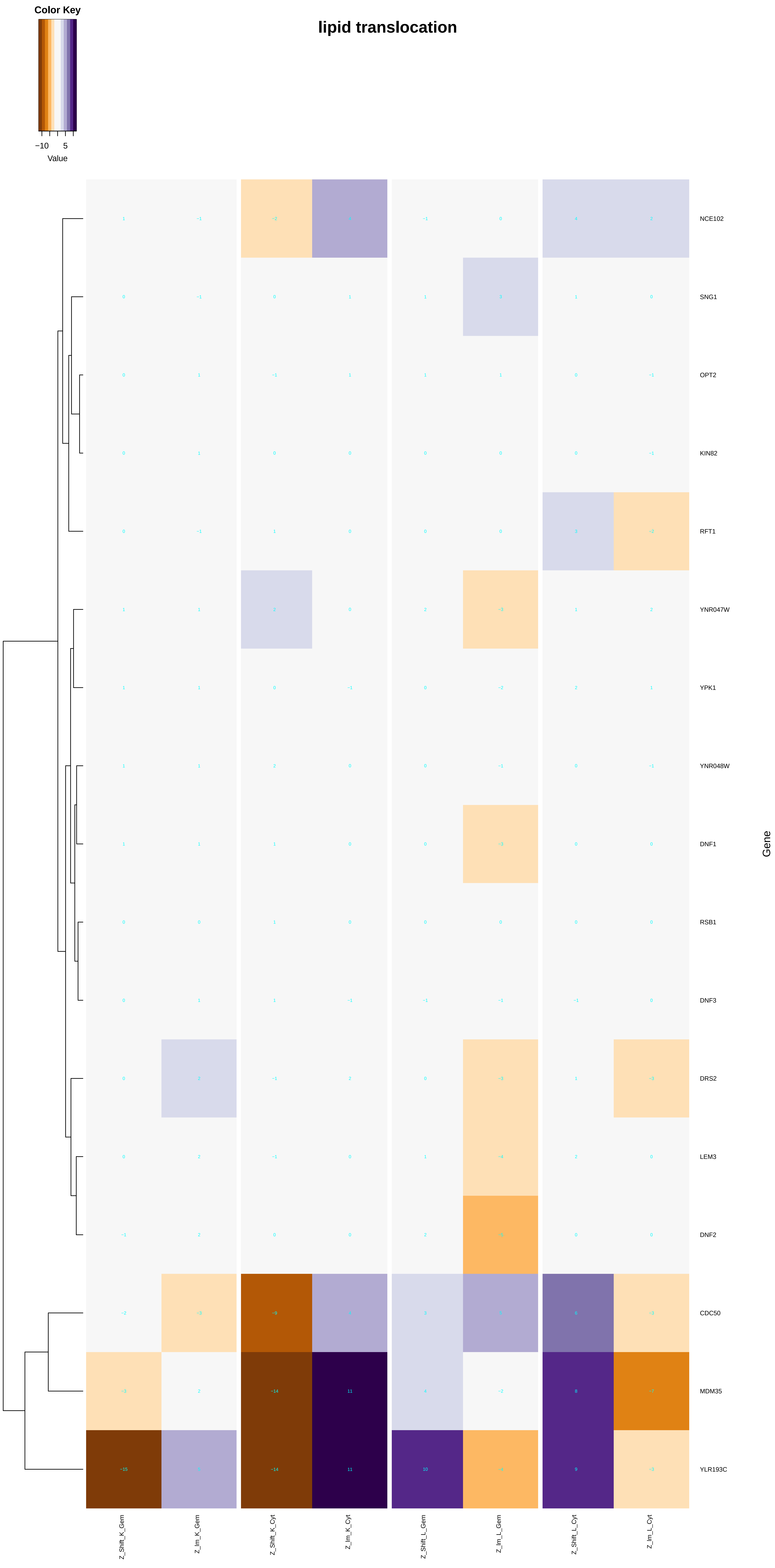
Color Key
−10
5
Value
lipid translocation
NCE102
SNG1
OPT2
KIN82
RFT1
YNR047W
YPK1
YNR048W
DNF1
RSB1
DNF3
DRS2
LEM3
DNF2
CDC50
MDM35
YLR193C
Gene
Z_Shift_K_Gem
Z_Im_K_Gem
Z_Shift_K_Cyt
Z_Im_K_Cyt
Z_Shift_L_Gem
Z_Im_L_Gem
Z_Shift_L_Cyt
Z_Im_L_Cyt

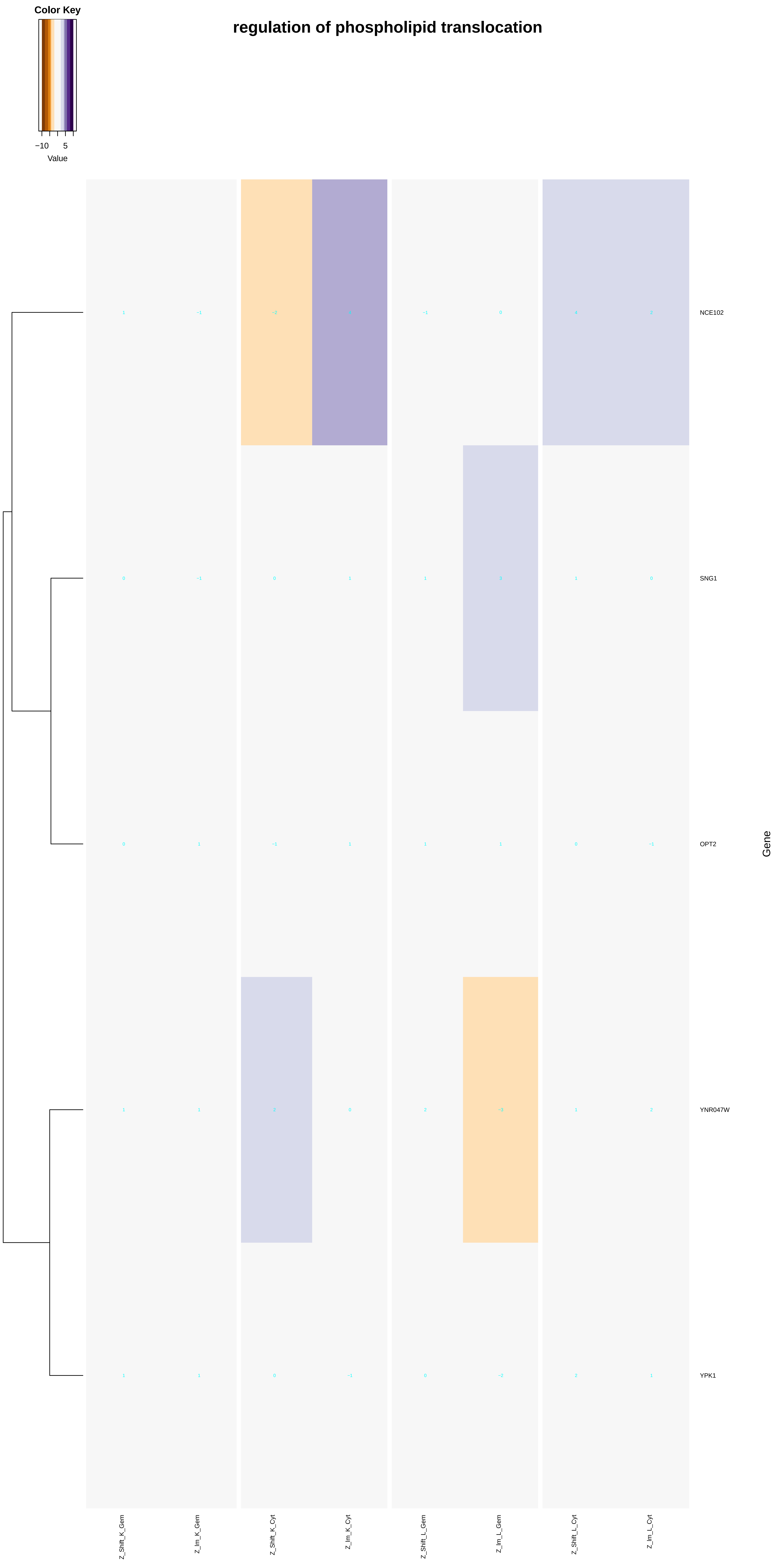

Color Key
−10
5
Value
regulation of phospholipid translocation
NCE102
SNG1
OPT2
YNR047W
YPK1
Gene
Z_Shift_K_Gem
Z_Im_K_Gem
Z_Shift_K_Cyt
Z_Im_K_Cyt
Z_Shift_L_Gem
Z_Im_L_Gem
Z_Shift_L_Cyt
Z_Im_L_Cyt

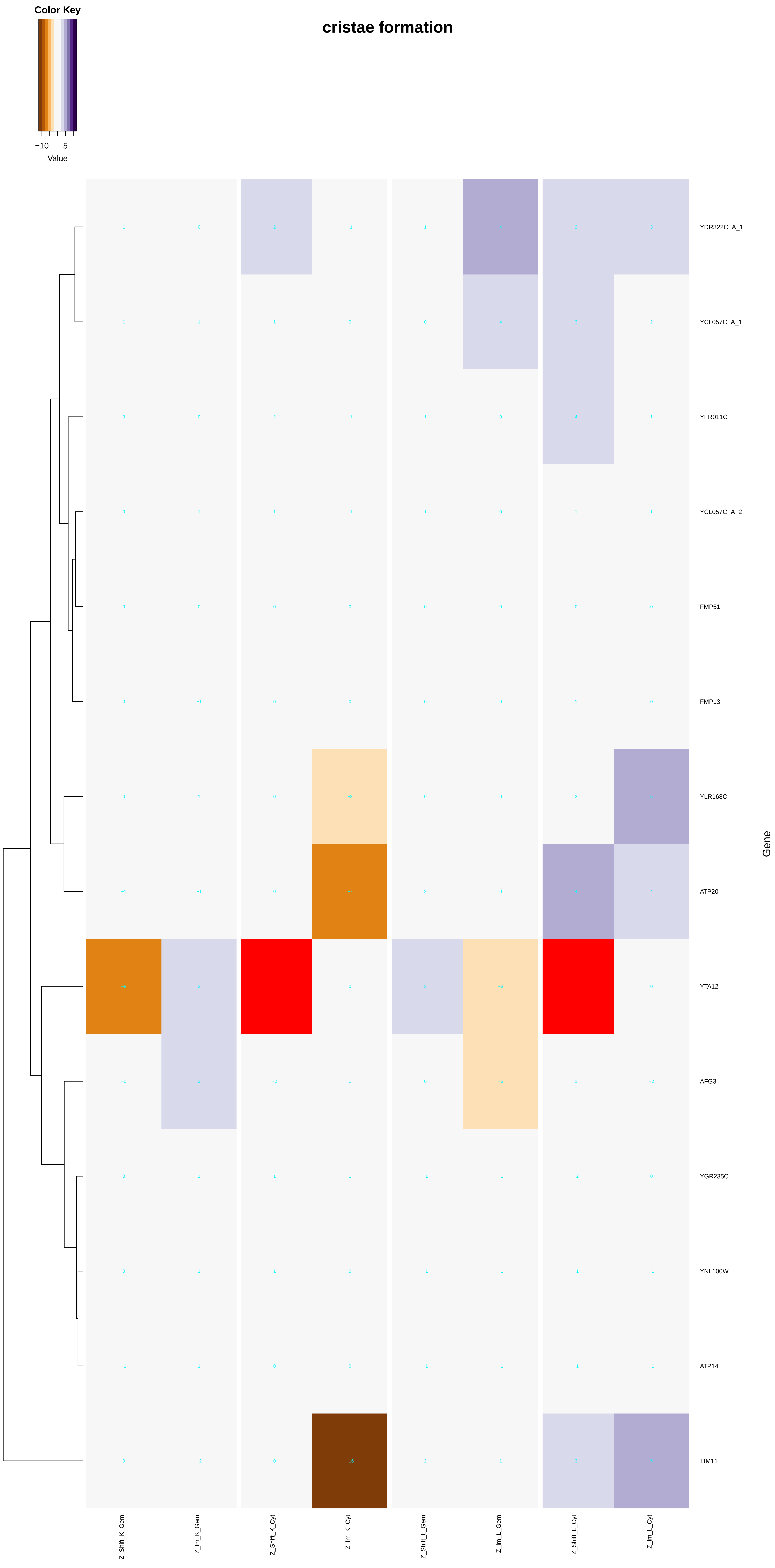
cristae formation
Color Key
−10
5
Value
YDR322C−A_1
YCL057C−A_1
YFR011C
YCL057C−A_2
FMP51
FMP13
YLR168C
ATP20
YTA12
AFG3
YGR235C
YNL100W
ATP14
TIM11
Gene
Z_Shift_K_Gem
Z_Im_K_Gem
Z_Shift_K_Cyt
Z_Im_K_Cyt
Z_Shift_L_Gem
Z_Im_L_Gem
Z_Shift_L_Cyt
Z_Im_L_Cyt

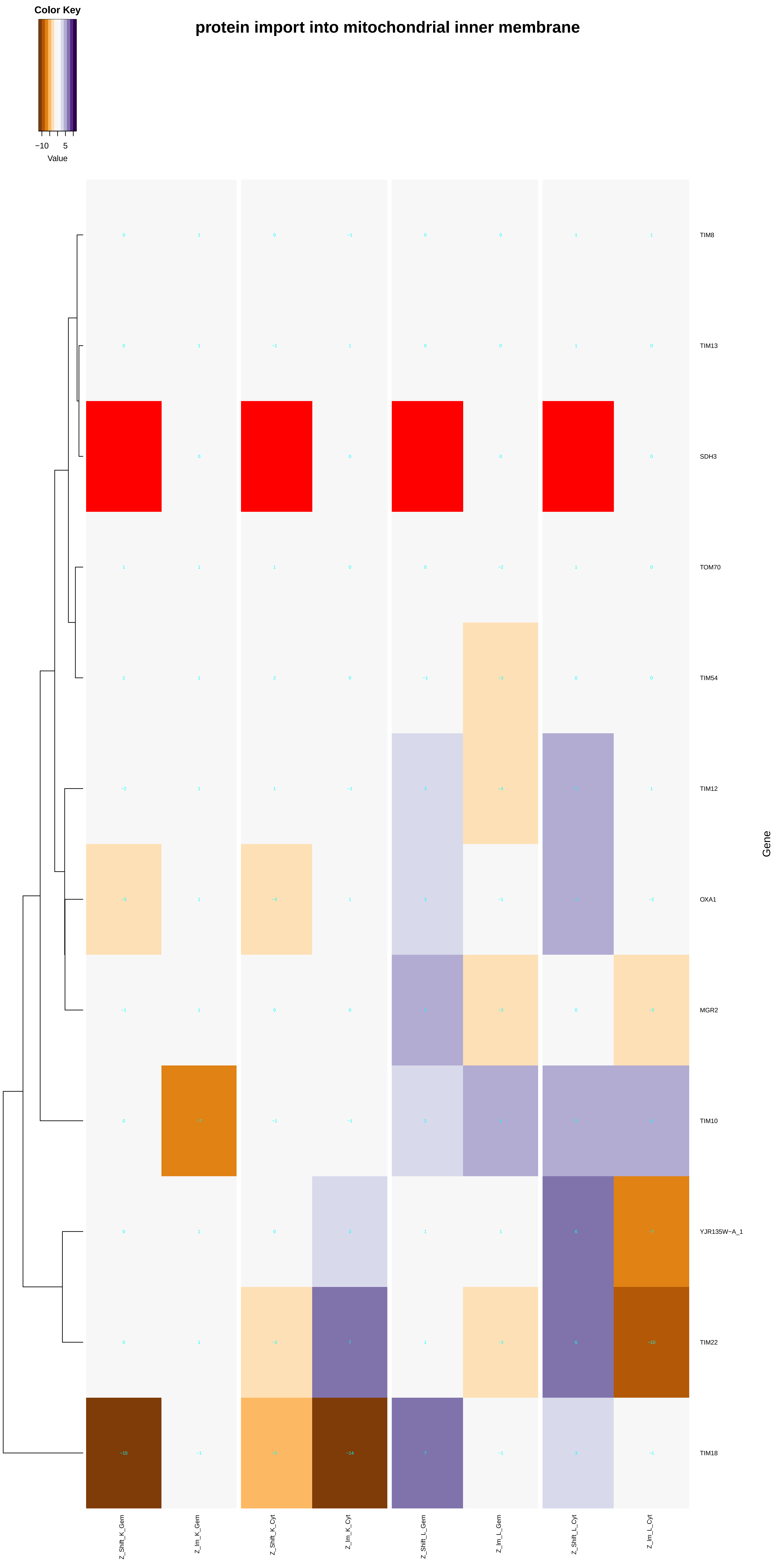
Color Key
−10
5
Value
protein import into mitochondrial inner membrane
TIM8
TIM13
SDH3
TOM70
TIM54
TIM12
OXA1
MGR2
TIM10
YJR135W−A_1
TIM22
TIM18
Gene
Z_Shift_K_Gem
Z_Im_K_Gem
Z_Shift_K_Cyt
Z_Im_K_Cyt
Z_Shift_L_Gem
Z_Im_L_Gem
Z_Shift_L_Cyt
Z_Im_L_Cyt

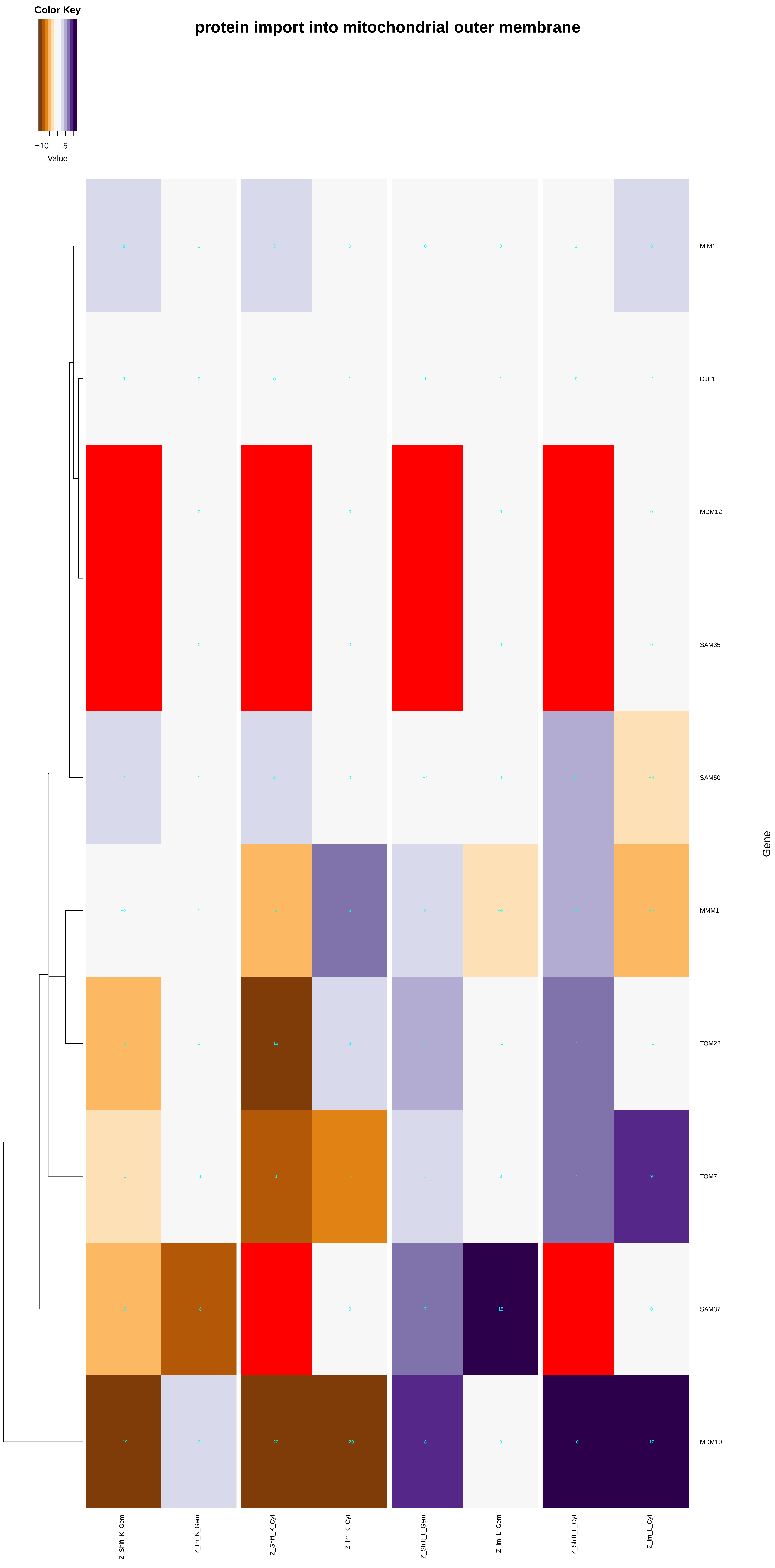

protein import into mitochondrial outer membrane
Color Key
−10
5
Value
MIM1
DJP1
MDM12
SAM35
SAM50
MMM1
TOM22
TOM7
SAM37
MDM10
Gene
Z_Shift_K_Gem
Z_Im_K_Gem
Z_Shift_K_Cyt
Z_Im_K_Cyt
Z_Shift_L_Gem
Z_Im_L_Gem
Z_Shift_L_Cyt
Z_Im_L_Cyt

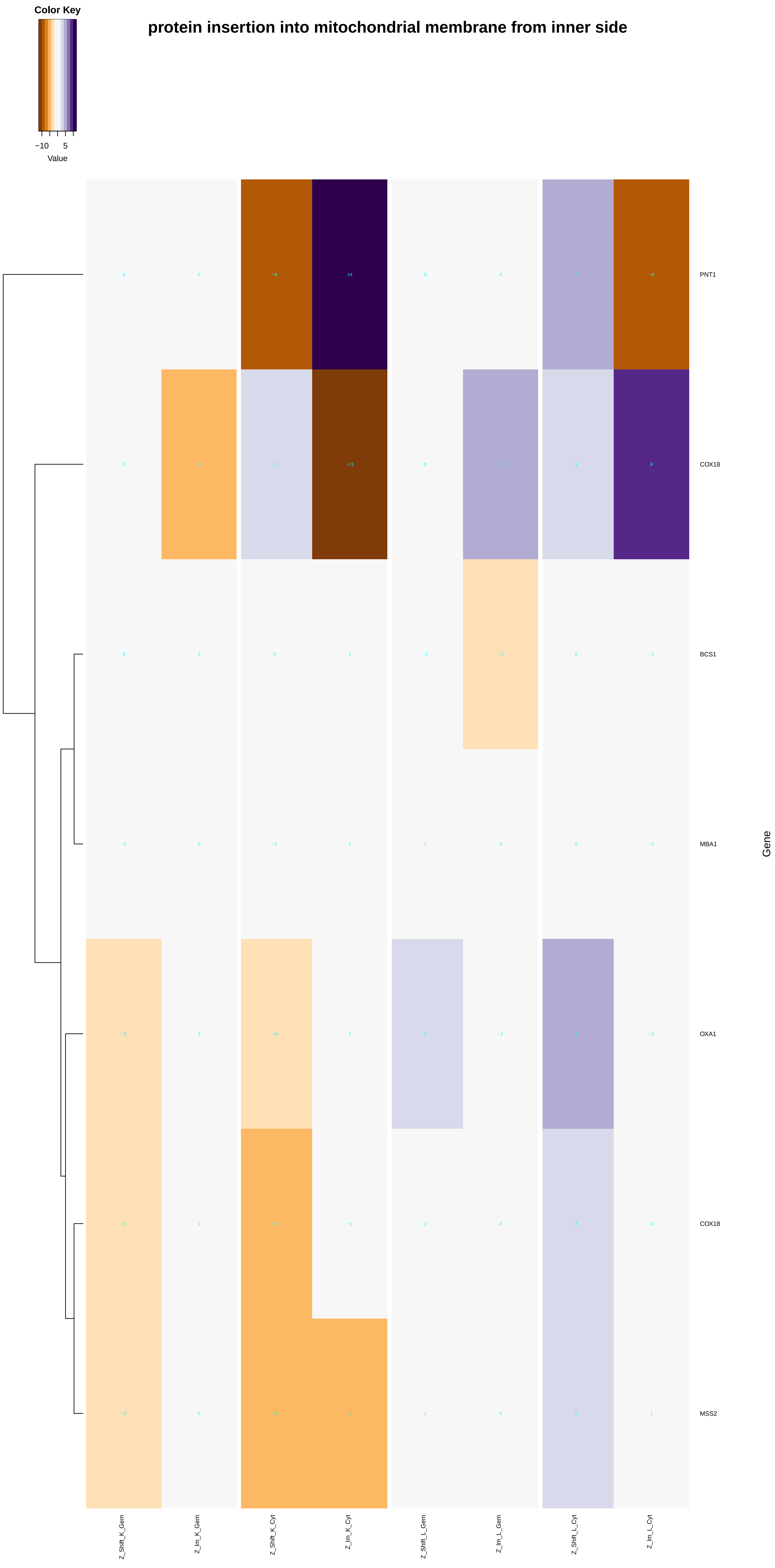

protein insertion into mitochondrial membrane from inner side
Color Key
−10
5
Value
PNT1
COX18
BCS1
MBA1
OXA1
COX18
MSS2
Gene
Z_Shift_K_Gem
Z_Im_K_Gem
Z_Shift_K_Cyt
Z_Im_K_Cyt
Z_Shift_L_Gem
Z_Im_L_Gem
Z_Shift_L_Cyt
Z_Im_L_Cyt

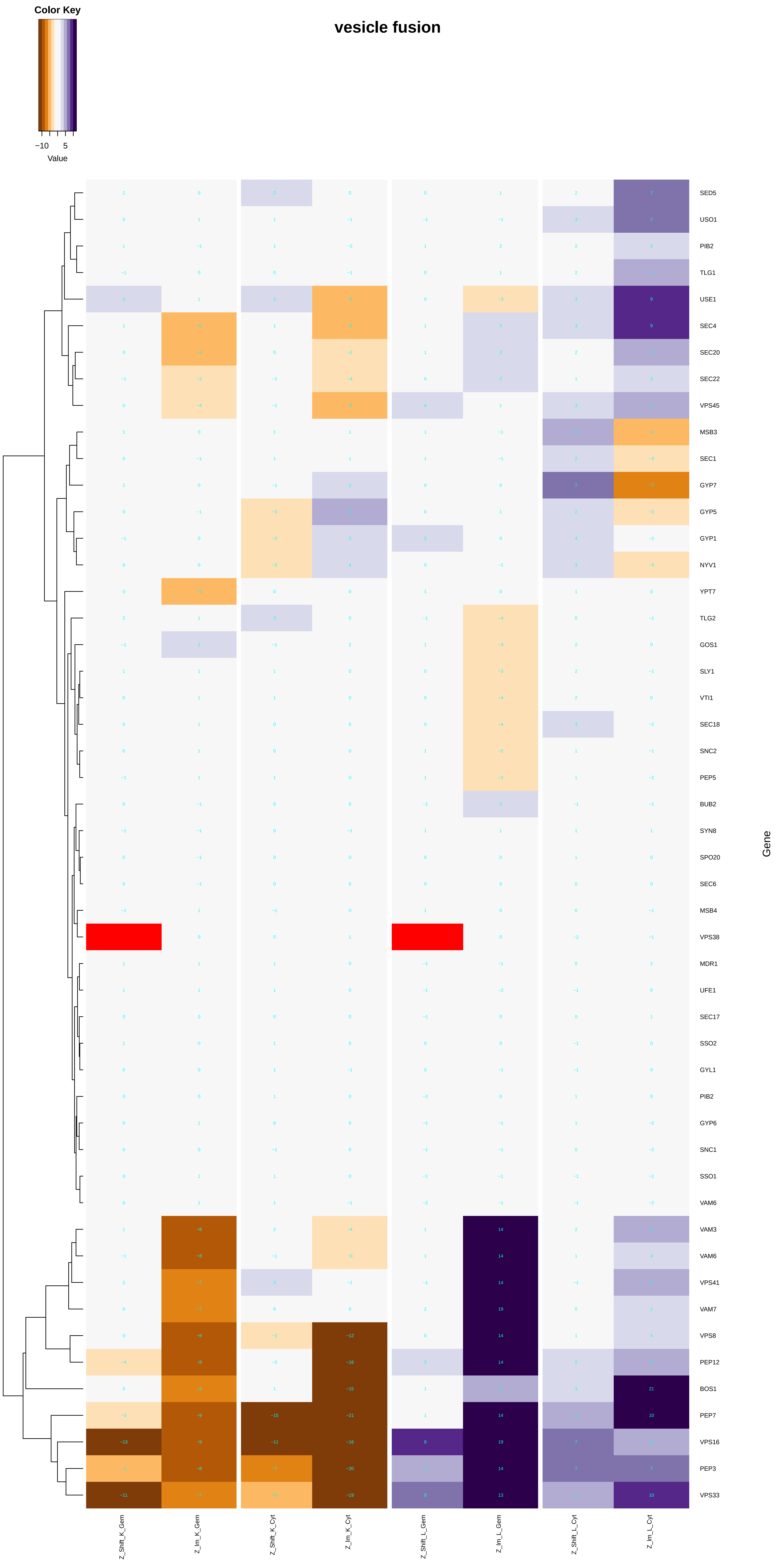

Color Key
−10
5
Value
vesicle fusion
SED5
USO1
PIB2
TLG1
USE1
SEC4
SEC20
SEC22
VPS45
MSB3
SEC1
GYP7
GYP5
GYP1
NYV1
YPT7
TLG2
GOS1
SLY1
VTI1
SEC18
SNC2
PEP5
BUB2
SYN8
SPO20
SEC6
MSB4
VPS38
MDR1
UFE1
SEC17
SSO2
GYL1
PIB2
GYP6
SNC1
SSO1
VAM6
VAM3
VAM6
VPS41
VAM7
VPS8
PEP12
BOS1
PEP7
VPS16
PEP3
VPS33
Gene
Z_Shift_K_Gem
Z_Im_K_Gem
Z_Shift_K_Cyt
Z_Im_K_Cyt
Z_Shift_L_Gem
Z_Im_L_Gem
Z_Shift_L_Cyt
Z_Im_L_Cyt

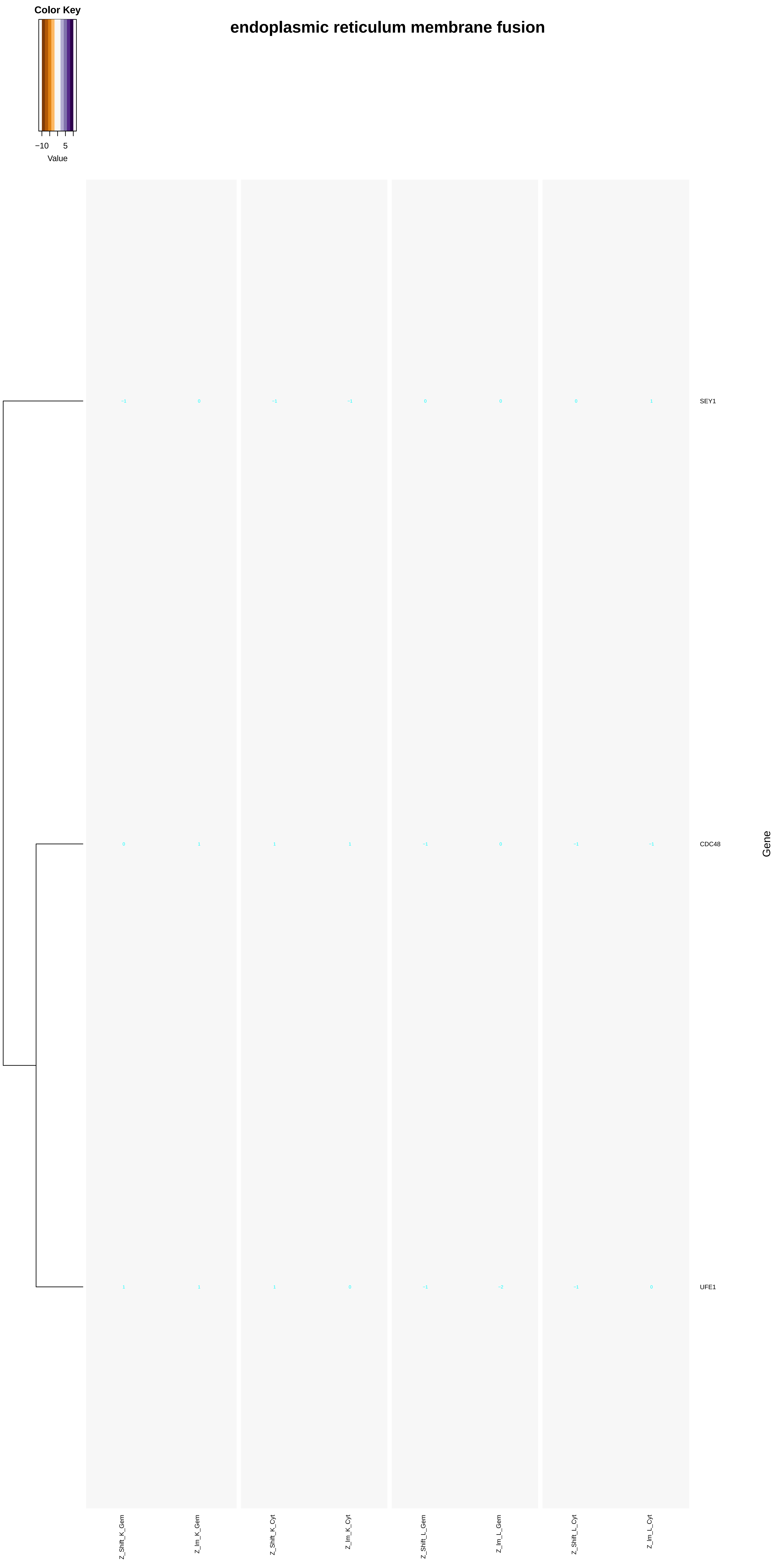
Color Key
−10
5
Value
endoplasmic reticulum membrane fusion
SEY1
CDC48
UFE1
Gene
Z_Shift_K_Gem
Z_Im_K_Gem
Z_Shift_K_Cyt
Z_Im_K_Cyt
Z_Shift_L_Gem
Z_Im_L_Gem
Z_Shift_L_Cyt
Z_Im_L_Cyt

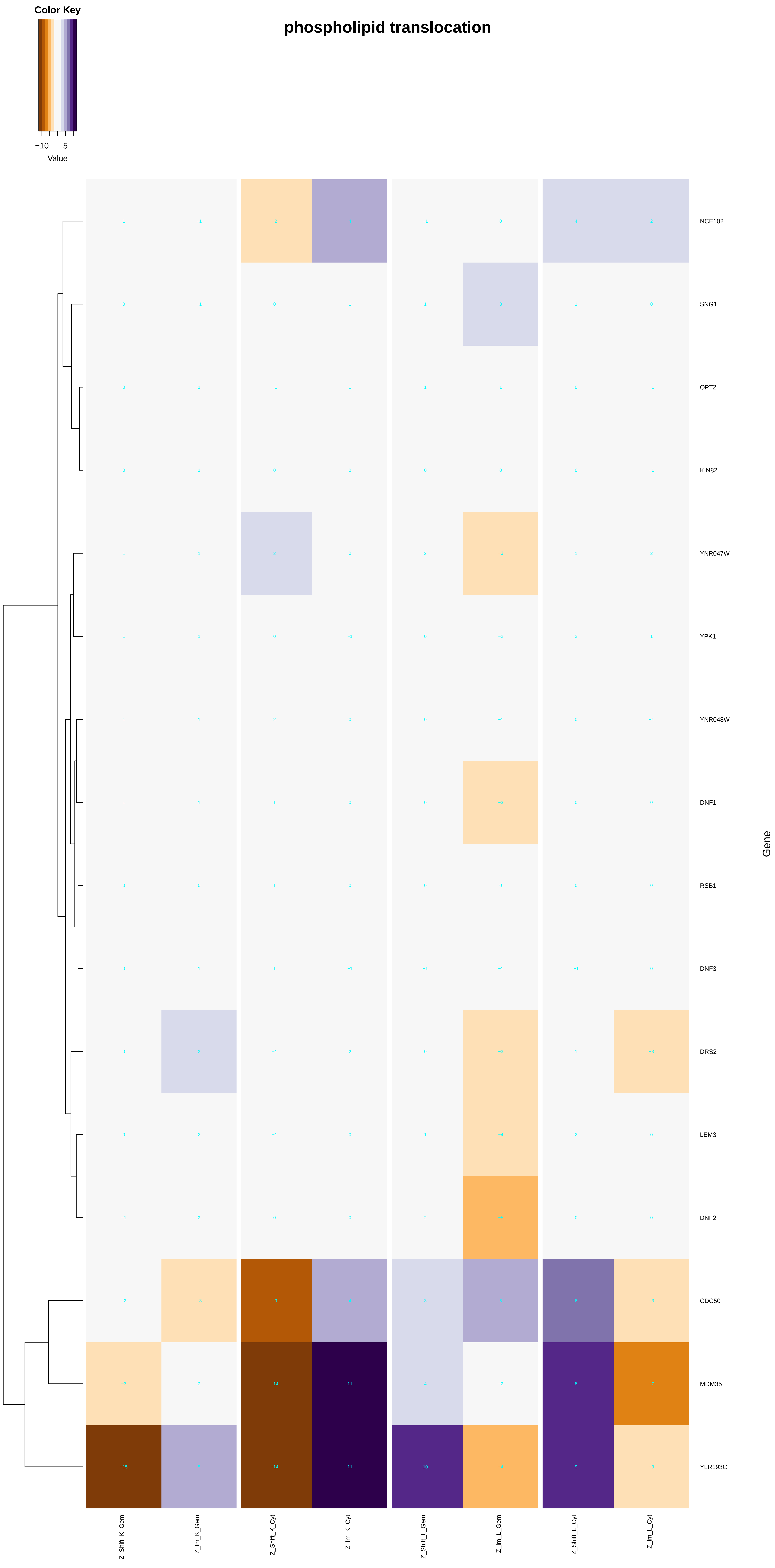
Color Key
−10
5
Value
phospholipid translocation
NCE102
SNG1
OPT2
KIN82
YNR047W
YPK1
YNR048W
DNF1
RSB1
DNF3
DRS2
LEM3
DNF2
CDC50
MDM35
YLR193C
Gene
Z_Shift_K_Gem
Z_Im_K_Gem
Z_Shift_K_Cyt
Z_Im_K_Cyt
Z_Shift_L_Gem
Z_Im_L_Gem
Z_Shift_L_Cyt
Z_Im_L_Cyt

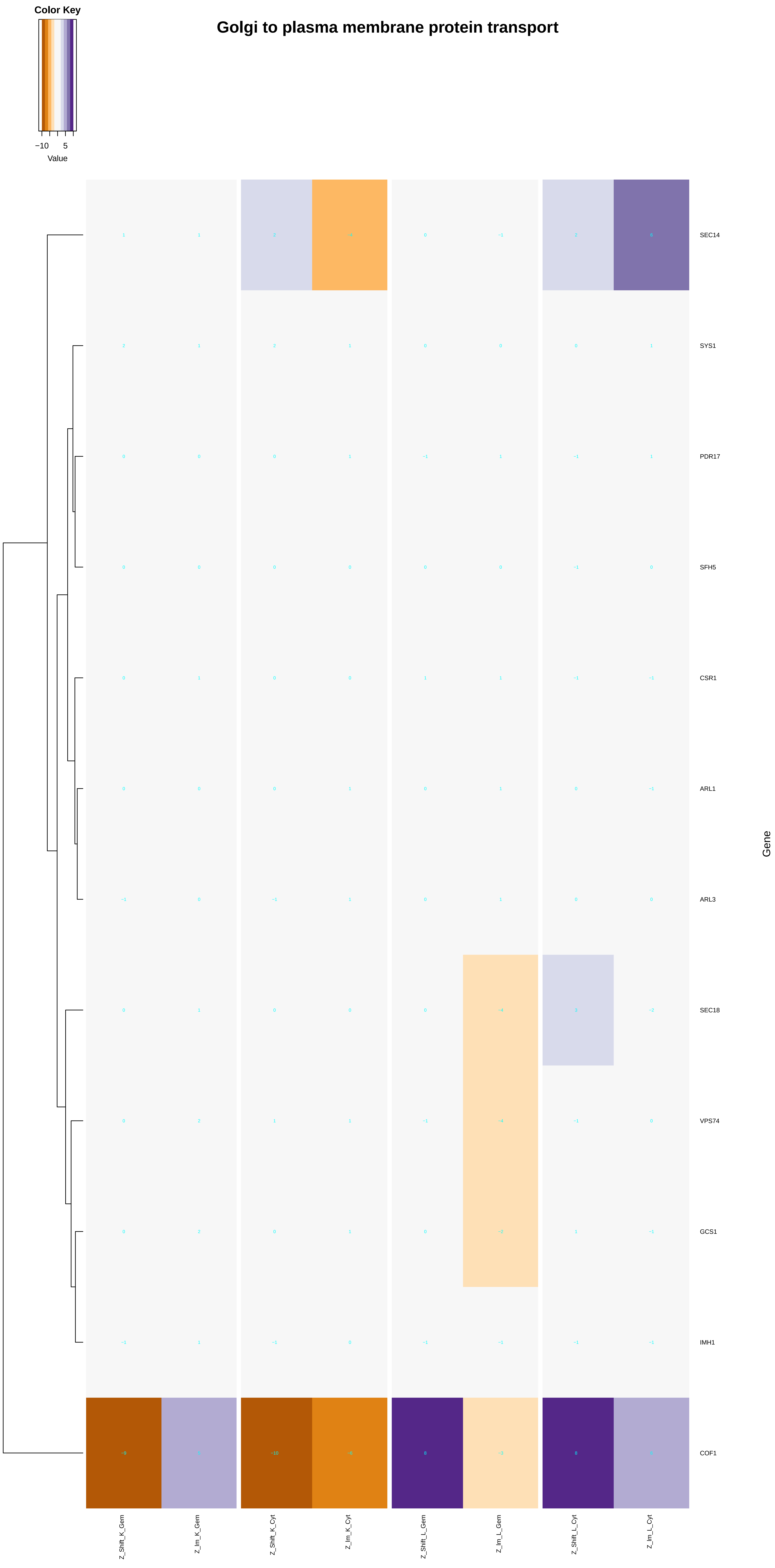

Golgi to plasma membrane protein transport
Color Key
−10
5
Value
SEC14
SYS1
PDR17
SFH5
CSR1
ARL1
ARL3
SEC18
VPS74
GCS1
IMH1
COF1
Gene
Z_Shift_K_Gem
Z_lm_K_Gem
Z_Shift_K_Cyt
Z_lm_K_Cyt
Z_Shift_L_Gem
Z_lm_L_Gem
Z_Shift_L_Cyt
Z_lm_L_Cyt

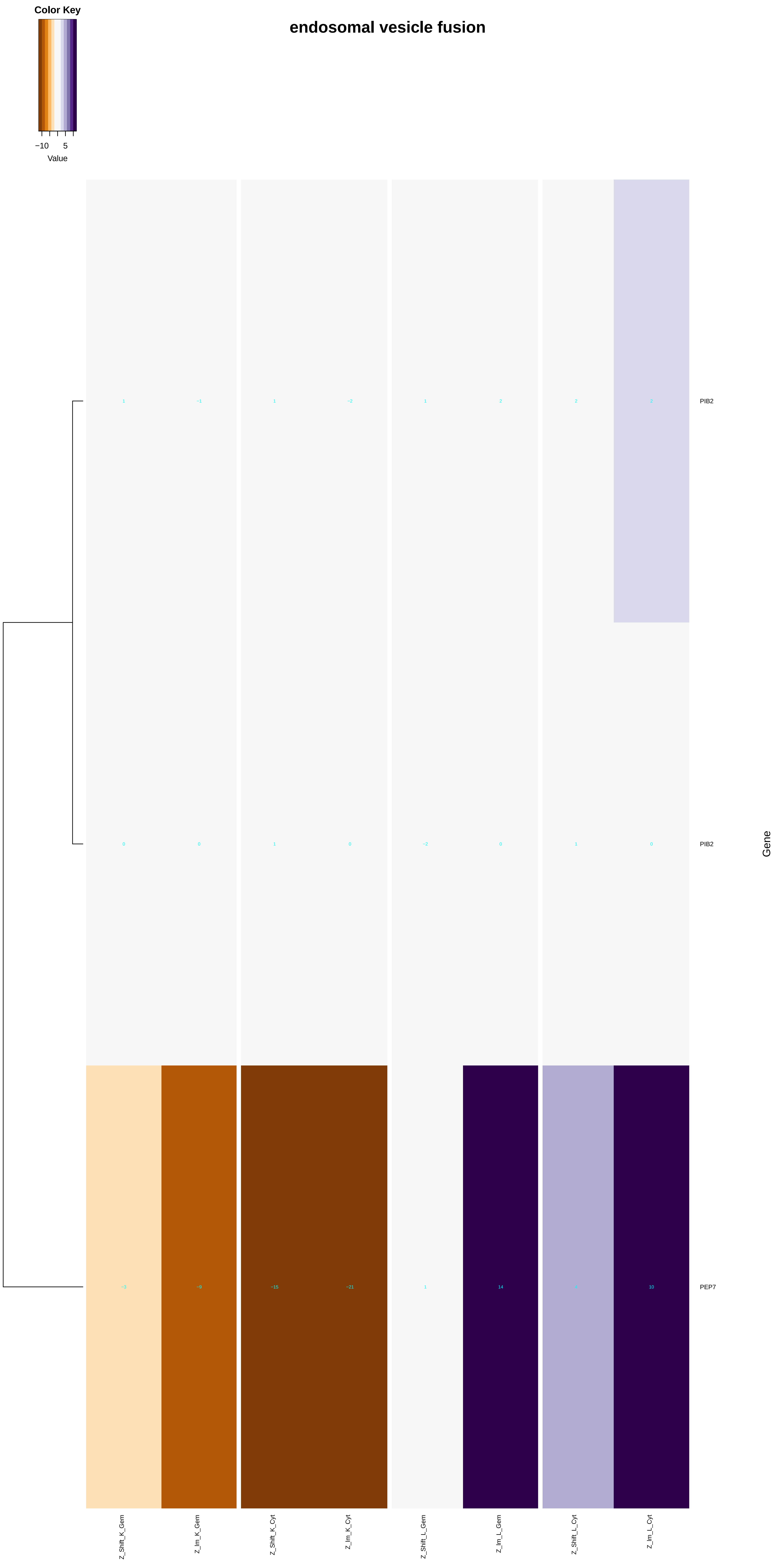

Color Key
−10
5
Value
endosomal vesicle fusion
PIB2
PIB2
PEP7
Gene
Z_Shift_K_Gem
Z_Im_K_Gem
Z_Shift_K_Cyt
Z_Im_K_Cyt
Z_Shift_L_Gem
Z_Im_L_Gem
Z_Shift_L_Cyt
Z_Im_L_Cyt

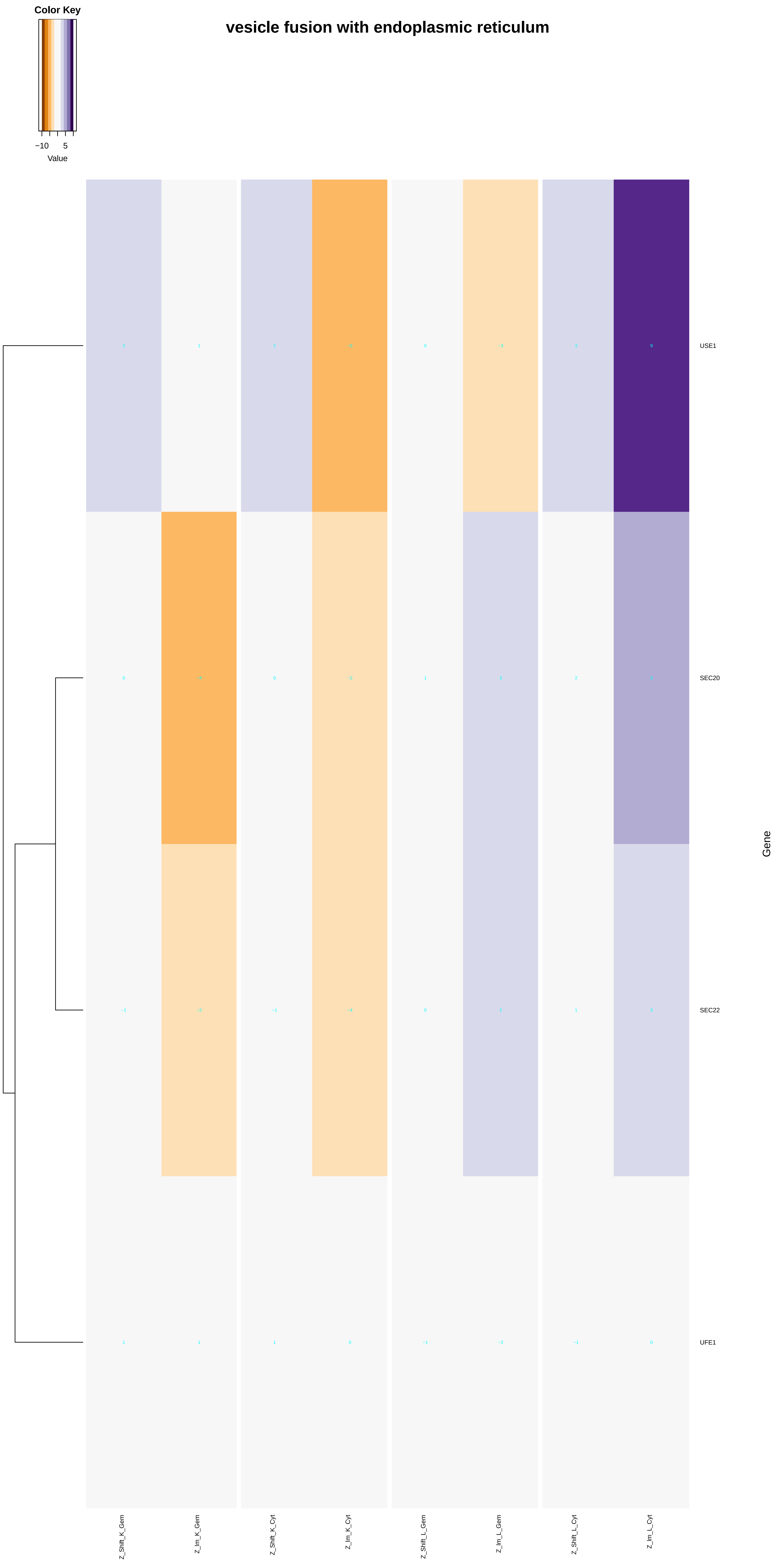

Color Key
−10
5
Value
vesicle fusion with endoplasmic reticulum
USE1
SEC20
SEC22
UFE1
Gene
Z_Shift_K_Gem
Z_Im_K_Gem
Z_Shift_K_Cyt
Z_Im_K_Cyt
Z_Shift_L_Gem
Z_Im_L_Gem
Z_Shift_L_Cyt
Z_Im_L_Cyt

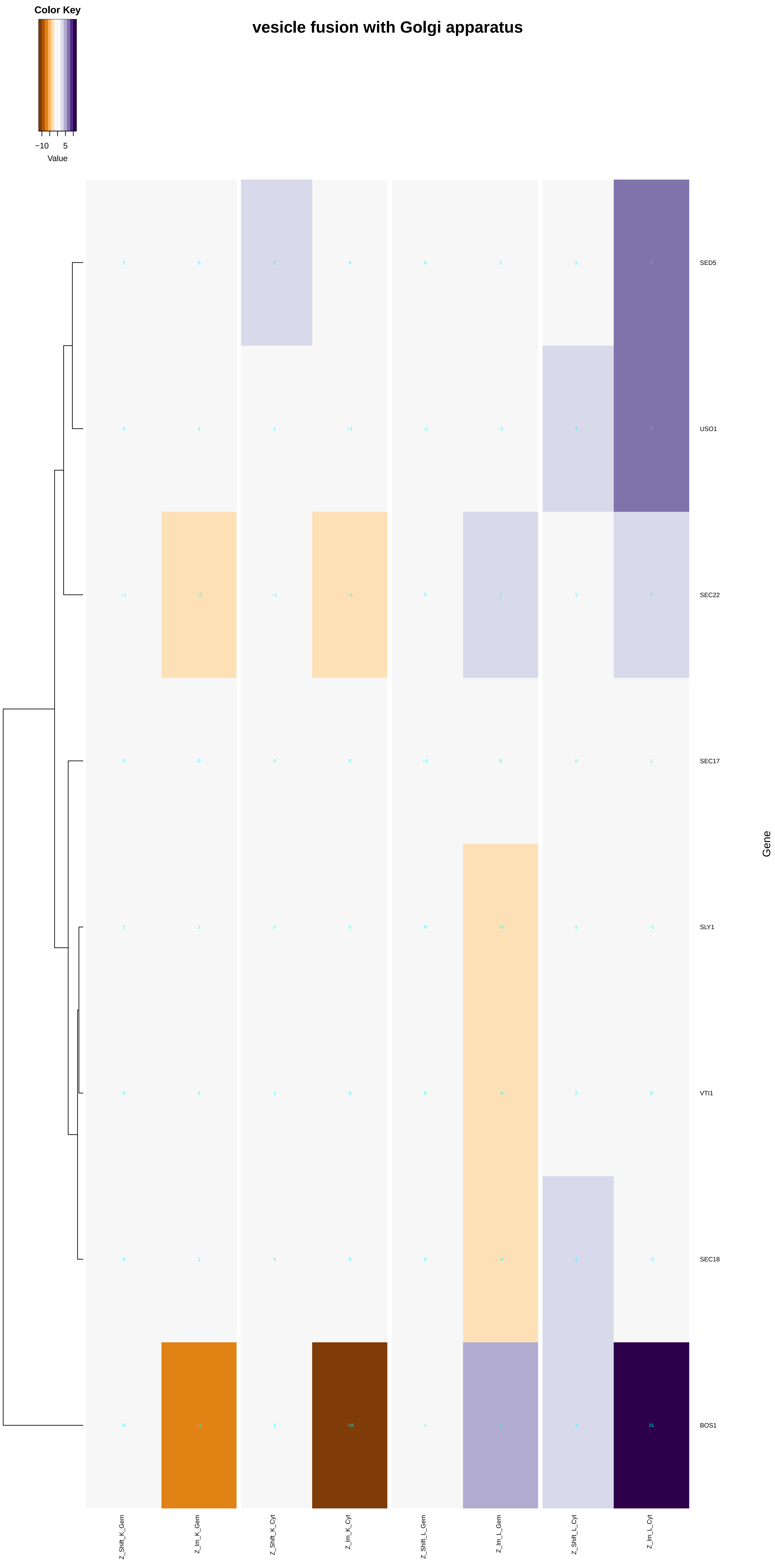

vesicle fusion with Golgi apparatus
Color Key
−10
5
Value
SED5
USO1
SEC22
SEC17
SLY1
VTI1
SEC18
BOS1
Gene
Z_Shift_K_Gem
Z_Im_K_Gem
Z_Shift_K_Cyt
Z_Im_K_Cyt
Z_Shift_L_Gem
Z_Im_L_Gem
Z_Shift_L_Cyt
Z_Im_L_Cyt
