## Additional File 7 for "A humanized yeast phenomic model of deoxycytidine kinase to predict genetic buffering of nucleoside analog cytotoxicity": mRNA_catabolic_process.pdf

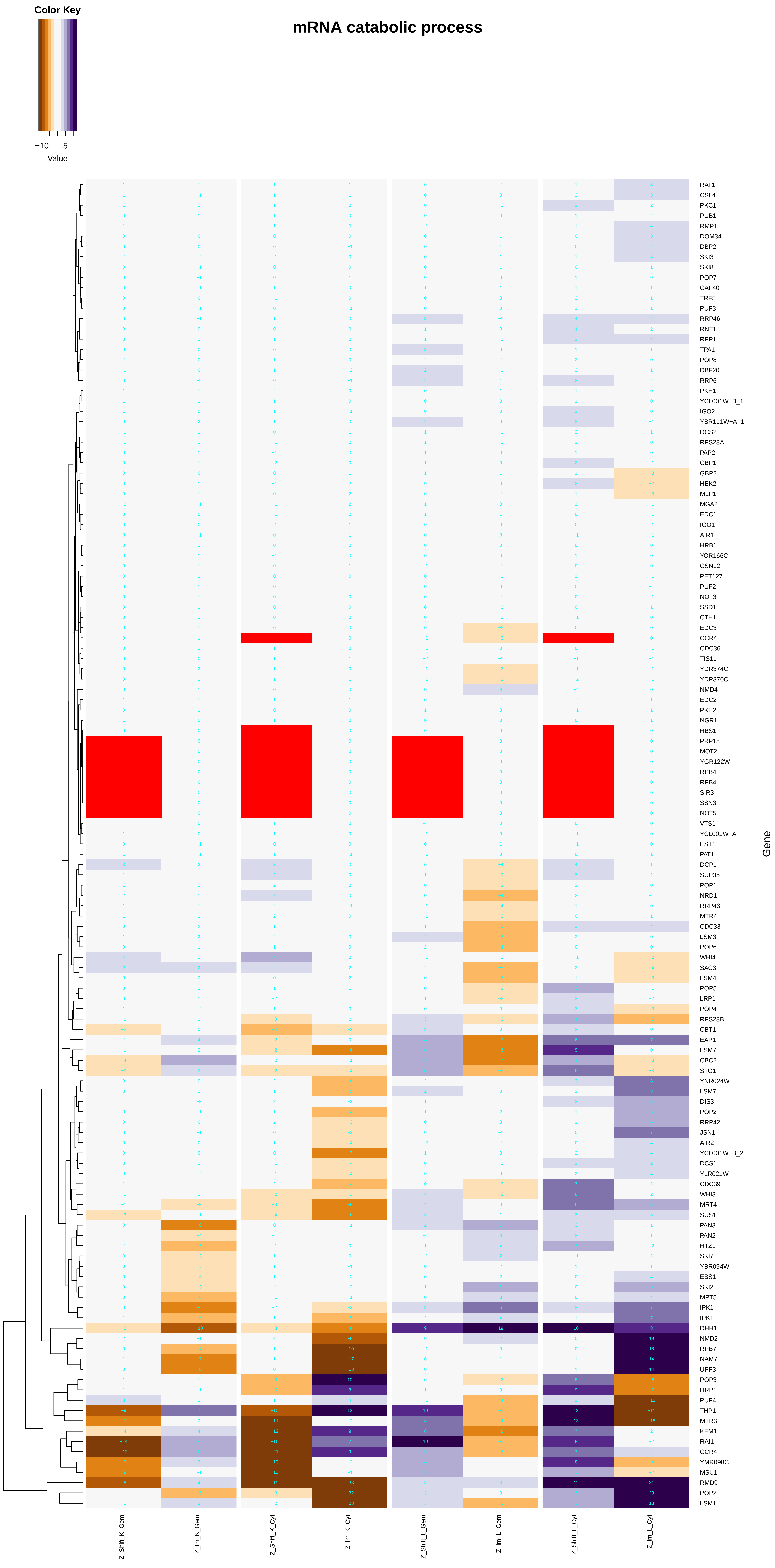

Color Key
−10
5
Value
mRNA catabolic process
Gene
RAT1
CSL4
PKC1
PUB1
RMP1
DOM34
DBP2
SKI3
SKI8
POP7
CAF40
TRF5
PUF3
RRP46
RNT1
RPP1
TPA1
POP8
DBF20
RRP6
PKH1
YCL001W−B_1
IGO2
YBR111W−A_1
DCS2
RPS28A
PAP2
CBP1
GBP2
HEK2
MLP1
MGA2
EDC1
IGO1
AIR1
HRB1
YOR166C
CSN12
PET127
PUF2
NOT3
SSD1
CTH1
EDC3
CCR4
CDC36
TIS11
YDR374C
YDR370C
NMD4
EDC2
PKH2
NGR1
HBS1
PRP18
MOT2
YGR122W
RPB4
RPB4
SIR3
SSN3
NOT5
VTS1
YCL001W−A
EST1
PAT1
DCP1
SUP35
POP1
NRD1
RRP43
MTR4
CDC33
LSM3
POP6
WHI4
SAC3
LSM4
POP5
LRP1
POP4
RPS28B
CBT1
EAP1
LSM7
CBC2
STO1
YNR024W
LSM7
DIS3
POP2
RRP42
JSN1
AIR2
YCL001W−B_2
DCS1
YLR021W
CDC39
WHI3
MRT4
SUS1
PAN3
PAN2
HTZ1
SKI7
YBR094W
EBS1
SKI2
MPT5
IPK1
IPK1
DHH1
NMD2
RPB7
NAM7
UPF3
POP3
HRP1
PUF4
THP1
MTR3
KEM1
RAI1
CCR4
YMR098C
MSU1
RMD9
POP2
LSM1
Z_Shift_K_Gem
Z_Im_K_Gem
Z_Shift_K_Cyt
Z_Im_K_Cyt
Z_Shift_L_Gem
Z_Im_L_Gem
Z_Shift_L_Cyt
Z_Im_L_Cyt

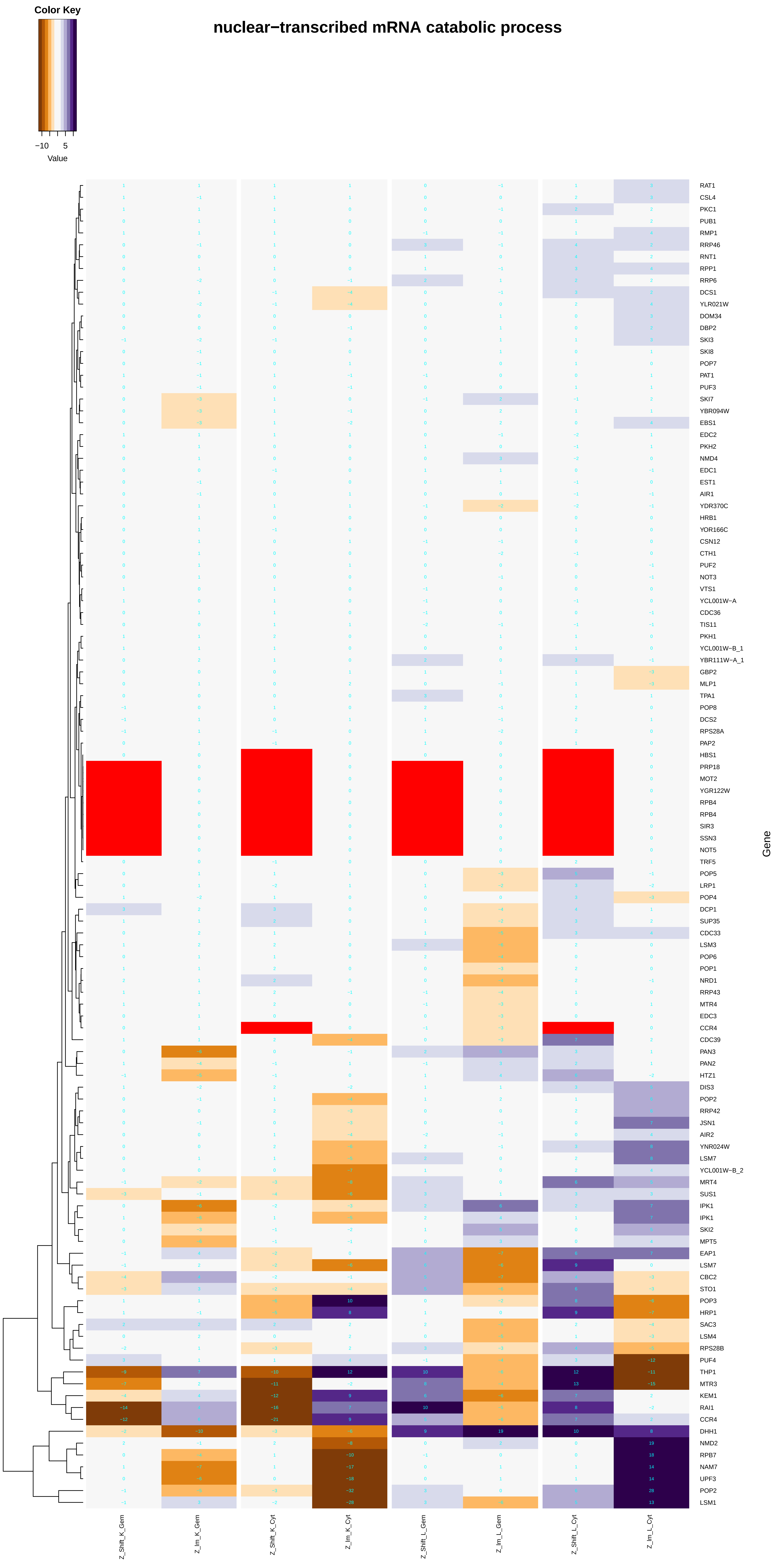
nuclear−transcribed mRNA catabolic process
Color Key
−10 5
Value
Gene
RAT1
CSL4
PKC1
PUB1
RMP1
RRP46
RNT1
RPP1
RRP6
DCS1
YLR021W
DOM34
DBP2
SKI3
SKI8
POP7
PAT1
PUF3
SKI7
YBR094W
EBS1
EDC2
PKH2
NMD4
EDC1
EST1
AIR1
YDR370C
HRB1
YOR166C
CSN12
CTH1
PUF2
NOT3
VTS1
YCL001W−A
CDC36
TIS11
PKH1
YCL001W−B_1
YBR111W−A_1
GBP2
MLP1
TPA1
POP8
DCS2
RPS28A
PAP2
HBS1
PRP18
MOT2
YGR122W
RPB4
RPB4
SIR3
SSN3
NOT5
TRF5
POP5
LRP1
POP4
DCP1
SUP35
CDC33
LSM3
POP6
POP1
NRD1
RRP43
MTR4
EDC3
CCR4
CDC39
PAN3
PAN2
HTZ1
DIS3
POP2
RRP42
JSN1
AIR2
YNR024W
LSM7
YCL001W−B_2
MRT4
SUS1
IPK1
IPK1
SKI2
MPT5
EAP1
LSM7
CBC2
STO1
POP3
HRP1
SAC3
LSM4
RPS28B
PUF4
THP1
MTR3
KEM1
RAI1
CCR4
DHH1
NMD2
RPB7
NAM7
UPF3
POP2
LSM1
Z_Shift_K_Gem
Z_Im_K_Gem
Z_Shift_K_Cyt
Z_Im_K_Cyt
Z_Shift_L_Gem
Z_Im_L_Gem
Z_Shift_L_Cyt
Z_Im_L_Cyt

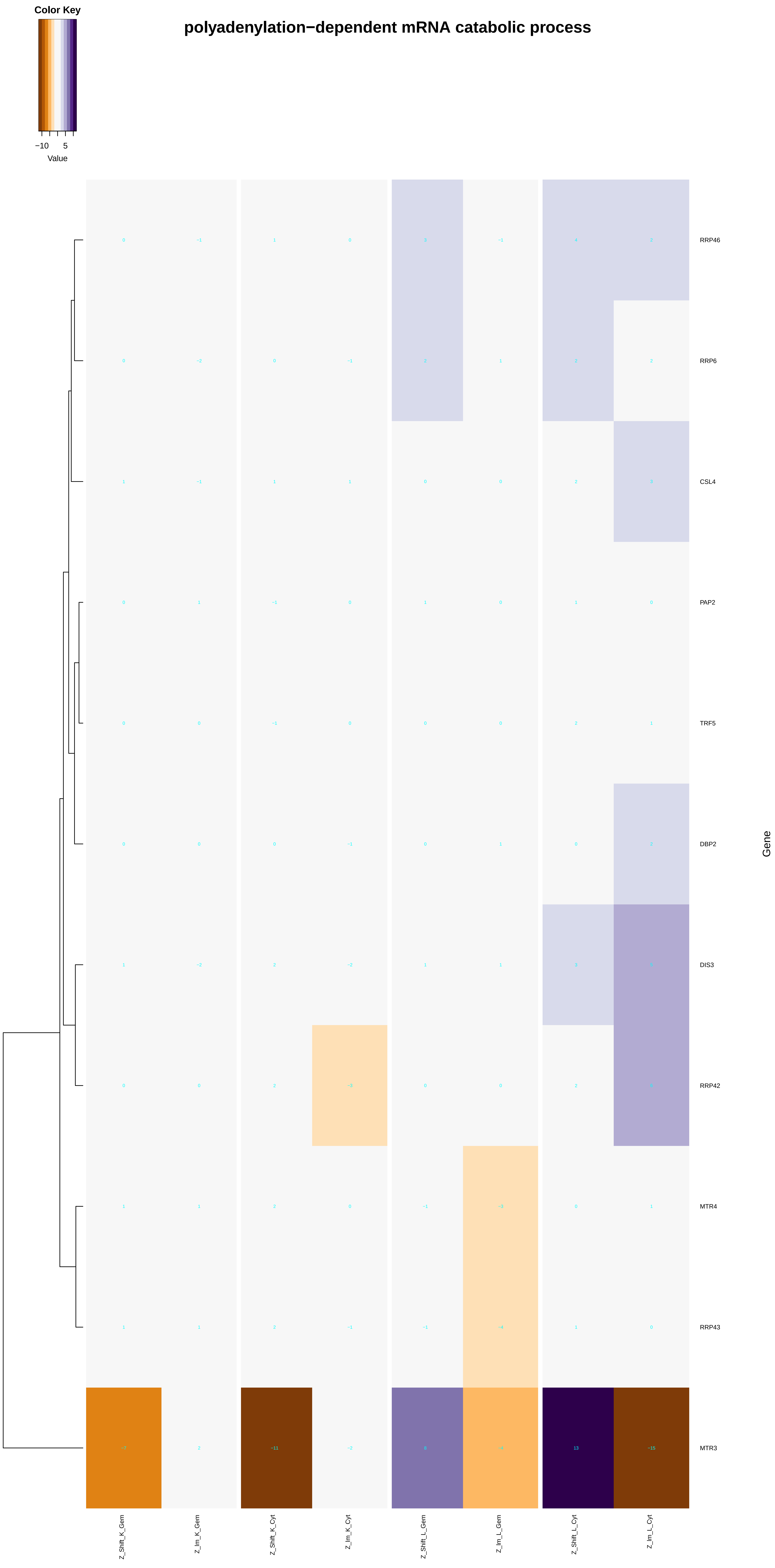
Color Key
−10
5
Value
polyadenylation−dependent mRNA catabolic process
RRP46
RRP6
CSL4
PAP2
TRF5
DBP2
DIS3
RRP42
MTR4
RRP43
MTR3
Gene
Z_Shift_K_Gem
Z_Im_K_Gem
Z_Shift_K_Cyt
Z_Im_K_Cyt
Z_Shift_L_Gem
Z_Im_L_Gem
Z_Shift_L_Cyt
Z_Im_L_Cyt

### nuclear−transcribed mRNA catabolic process, nonsense−mediated decay

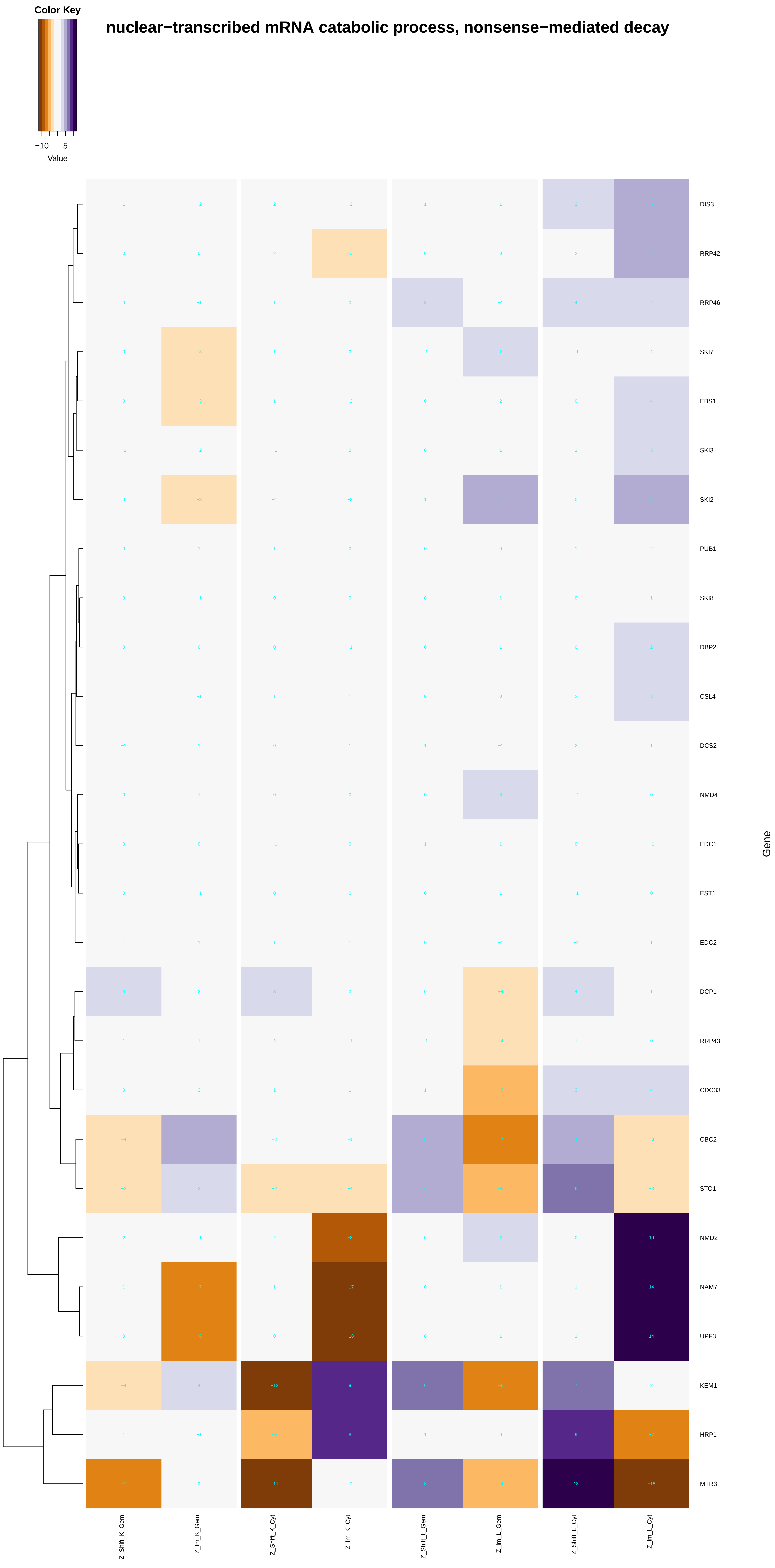

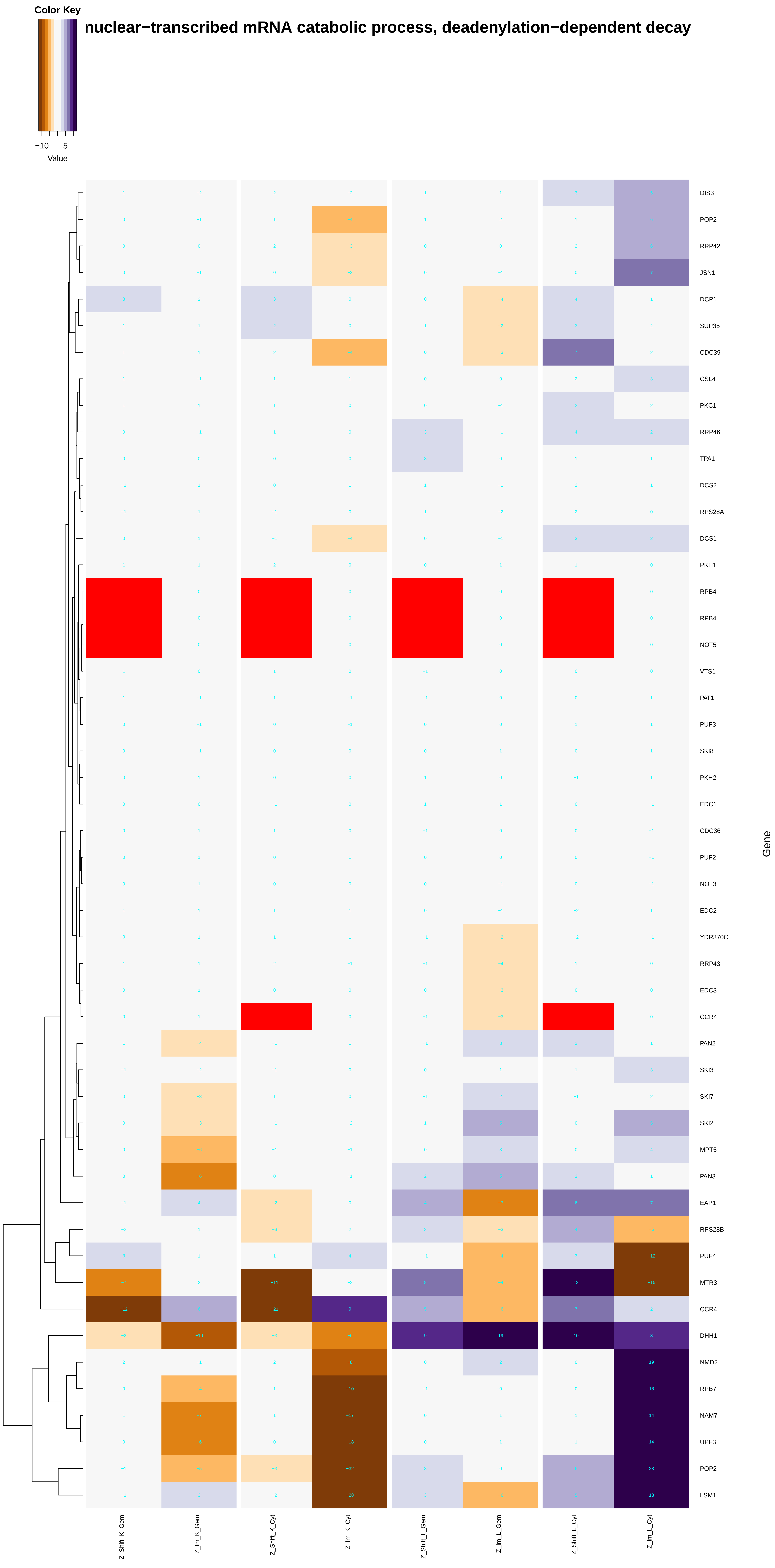
Color Key
−10
5
Value
nuclear−transcribed mRNA catabolic process, deadenylation−dependent decay
DIS3
POP2
RRP42
JSN1
DCP1
SUP35
CDC39
CSL4
PKC1
RRP46
TPA1
DCS2
RPS28A
DCS1
PKH1
RPB4
RPB4
NOT5
VTS1
PAT1
PUF3
SKI8
PKH2
EDC1
CDC36
PUF2
NOT3
EDC2
YDR370C
RRP43
EDC3
CCR4
PAN2
SKI3
SKI7
SKI2
MPT5
PAN3
EAP1
RPS28B
PUF4
MTR3
CCR4
DHH1
NMD2
RPB7
NAM7
UPF3
POP2
LSM1
Gene
Z_Shift_K_Gem
Z_Im_K_Gem
Z_Shift_K_Cyt
Z_Im_K_Cyt
Z_Shift_L_Gem
Z_Im_L_Gem
Z_Shift_L_Cyt
Z_Im_L_Cyt

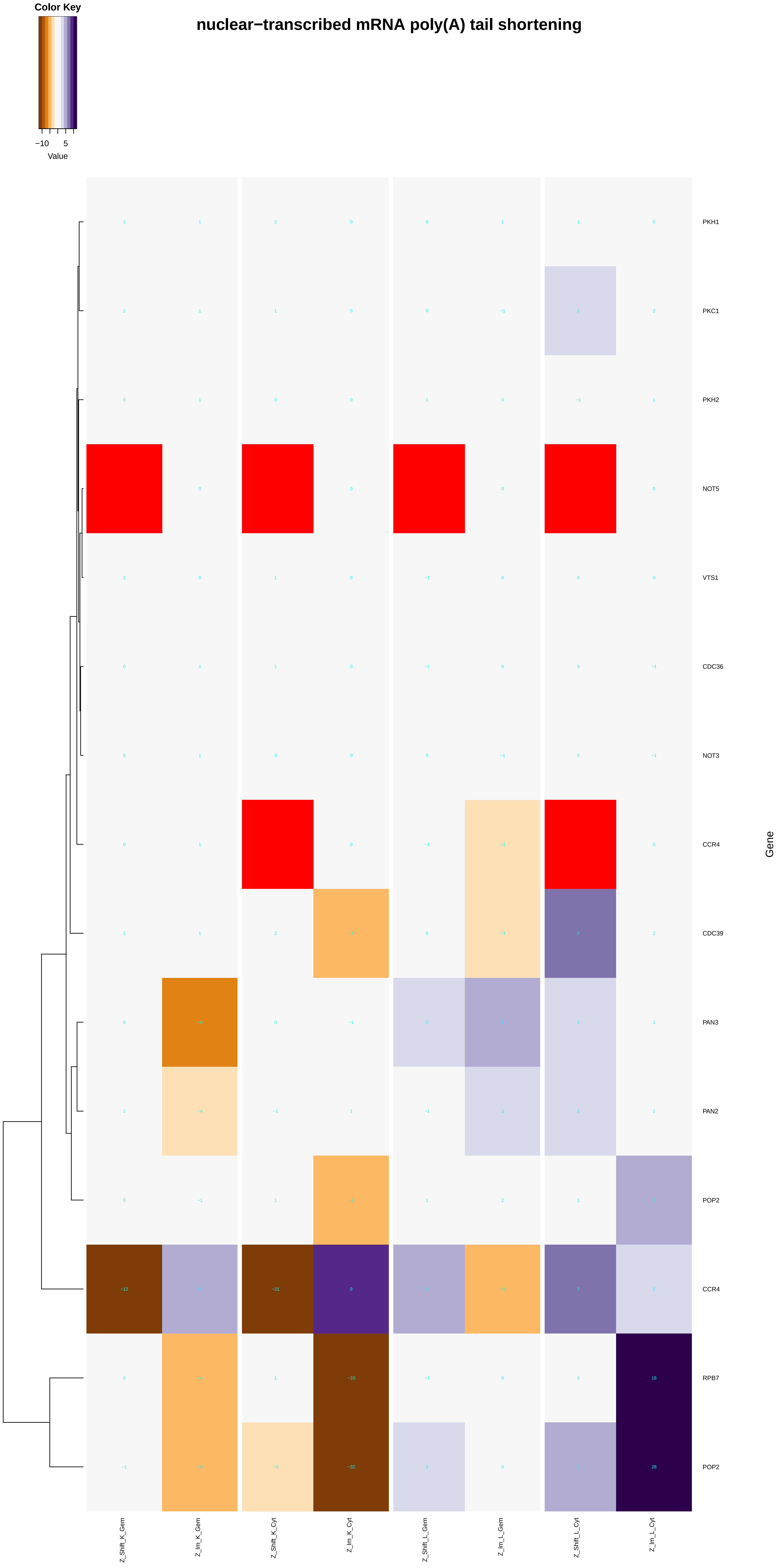
nuclear−transcribed mRNA poly(A) tail shortening
Color Key
−10
5
Value
PKH1
PKC1
PKH2
NOT5
VTS1
CDC36
NOT3
CCR4
CDC39
PAN3
PAN2
POP2
CCR4
RPB7
POP2
Gene
Z_Shift_K_Gem
Z_Im_K_Gem
Z_Shift_K_Cyt
Z_Im_K_Cyt
Z_Shift_L_Gem
Z_Im_L_Gem
Z_Shift_L_Cyt
Z_Im_L_Cyt

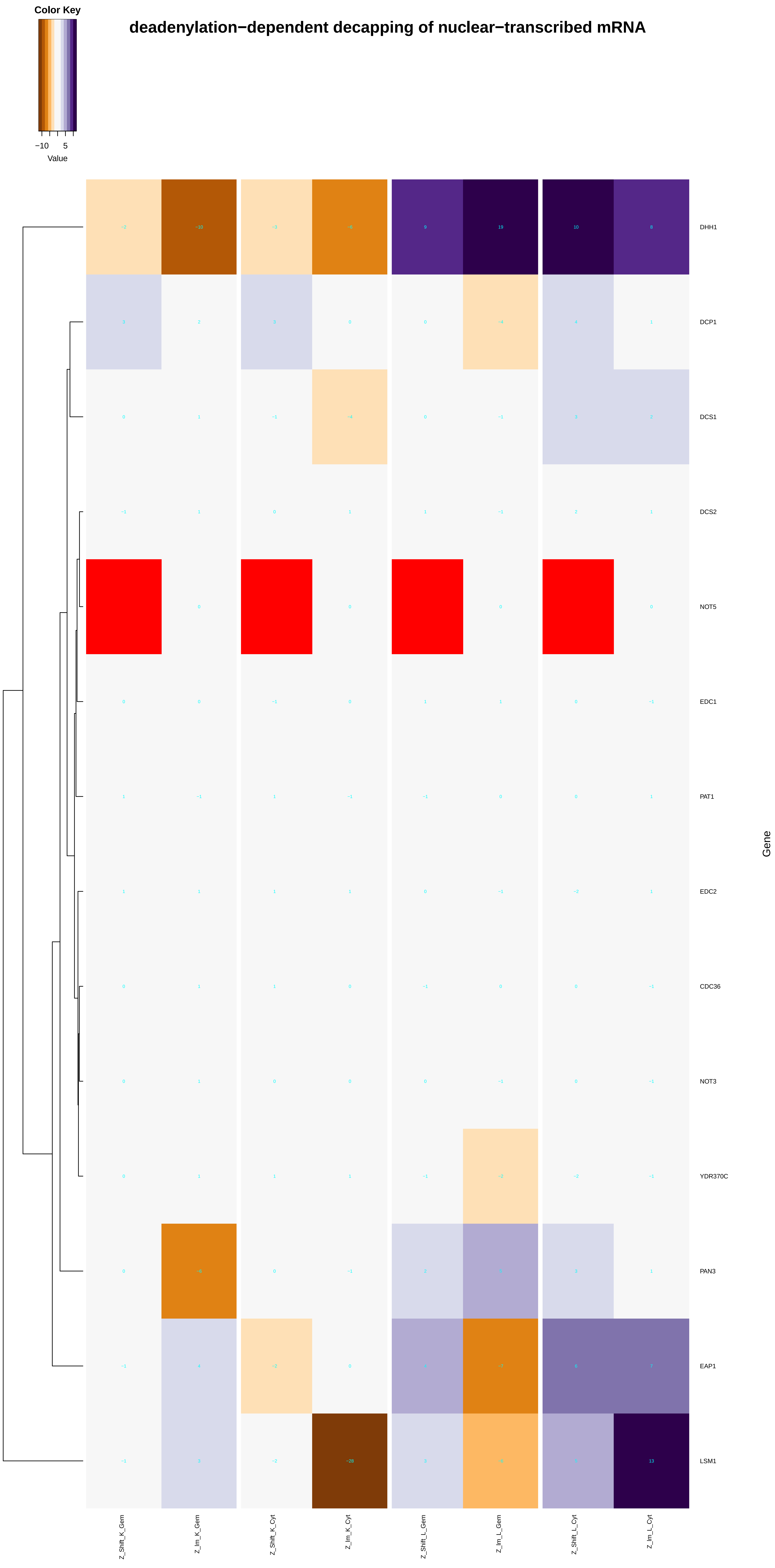

Color Key
−10 5
Value
deadenylation−dependent decapping of nuclear−transcribed mRNA
DHH1
DCP1
DCS1
DCS2
NOT5
EDC1
PAT1
EDC2
CDC36
NOT3
YDR370C
PAN3
EAP1
LSM1
Gene
Z_Shift_K_Gem
Z_Im_K_Gem
Z_Shift_K_Cyt
Z_Im_K_Cyt
Z_Shift_L_Gem
Z_Im_L_Gem
Z_Shift_L_Cyt
Z_Im_L_Cyt

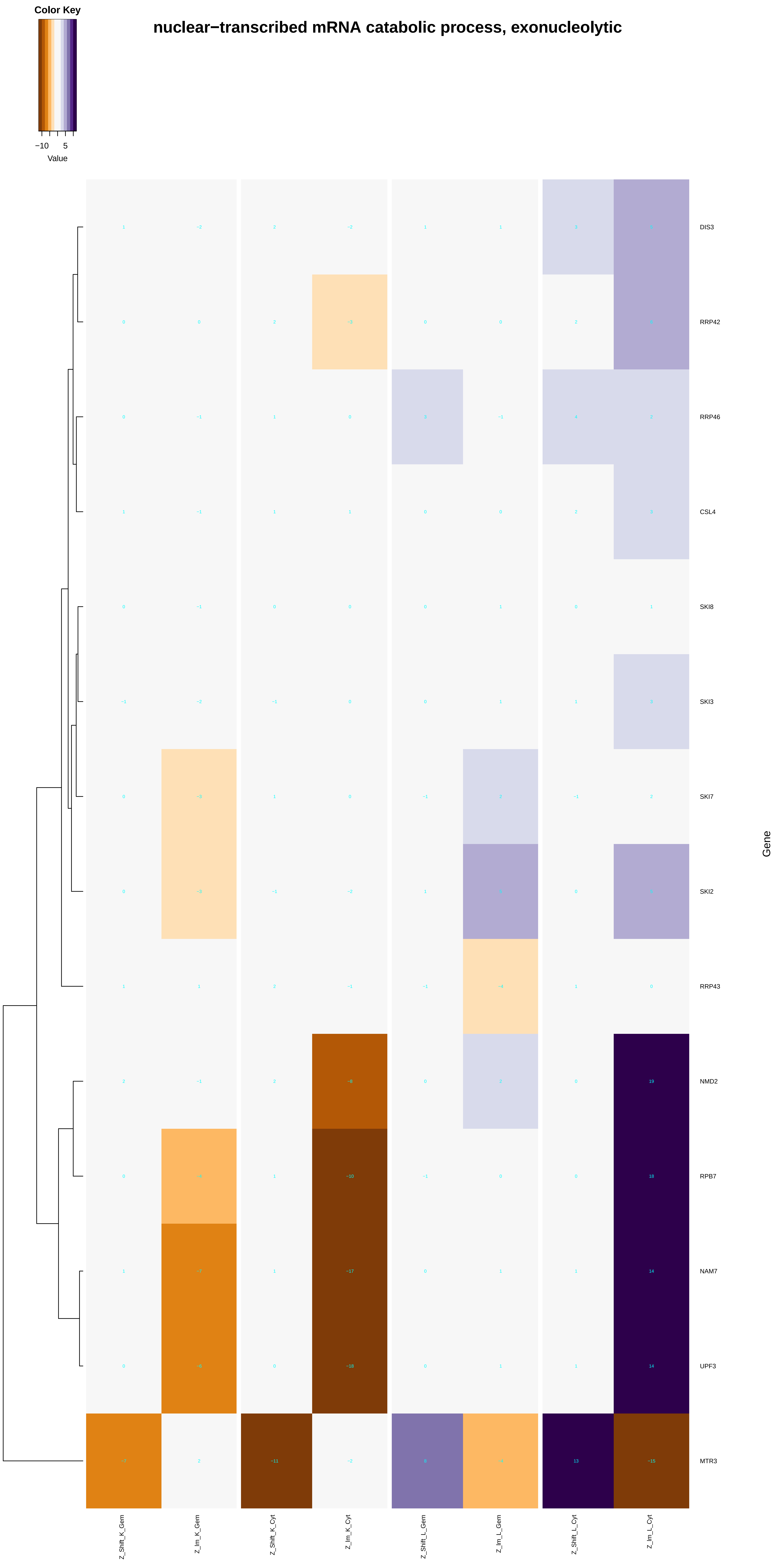

Color Key
−10
5
Value
nuclear−transcribed mRNA catabolic process, exonucleolytic
DIS3
RRP42
RRP46
CSL4
SKI8
SKI3
SKI7
SKI2
RRP43
NMD2
RPB7
NAM7
UPF3
MTR3
Gene
Z_Shift_K_Gem
Z_Im_K_Gem
Z_Shift_K_Cyt
Z_Im_K_Cyt
Z_Shift_L_Gem
Z_Im_L_Gem
Z_Shift_L_Cyt
Z_Im_L_Cyt

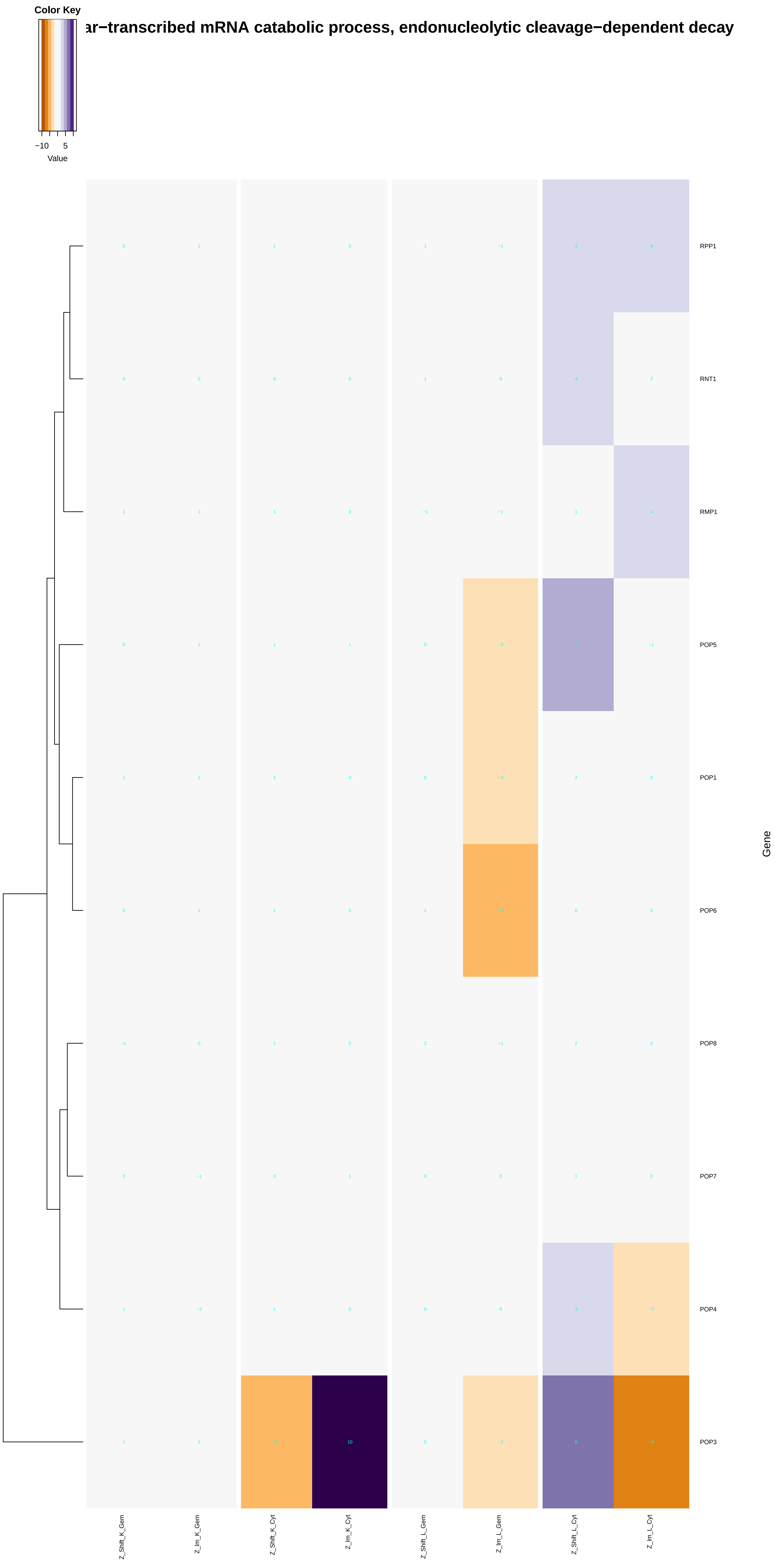
Color Key
−10
5
Value
ar−transcribed mRNA catabolic process, endonucleolytic cleavage−dependent decay
RPP1
RNT1
RMP1
POP5
POP1
POP6
POP8
POP7
POP4
POP3
Gene
Z_Shift_K_Gem
Z_Im_K_Gem
Z_Shift_K_Cyt
Z_Im_K_Cyt
Z_Shift_L_Gem
Z_Im_L_Gem
Z_Shift_L_Cyt
Z_Im_L_Cyt

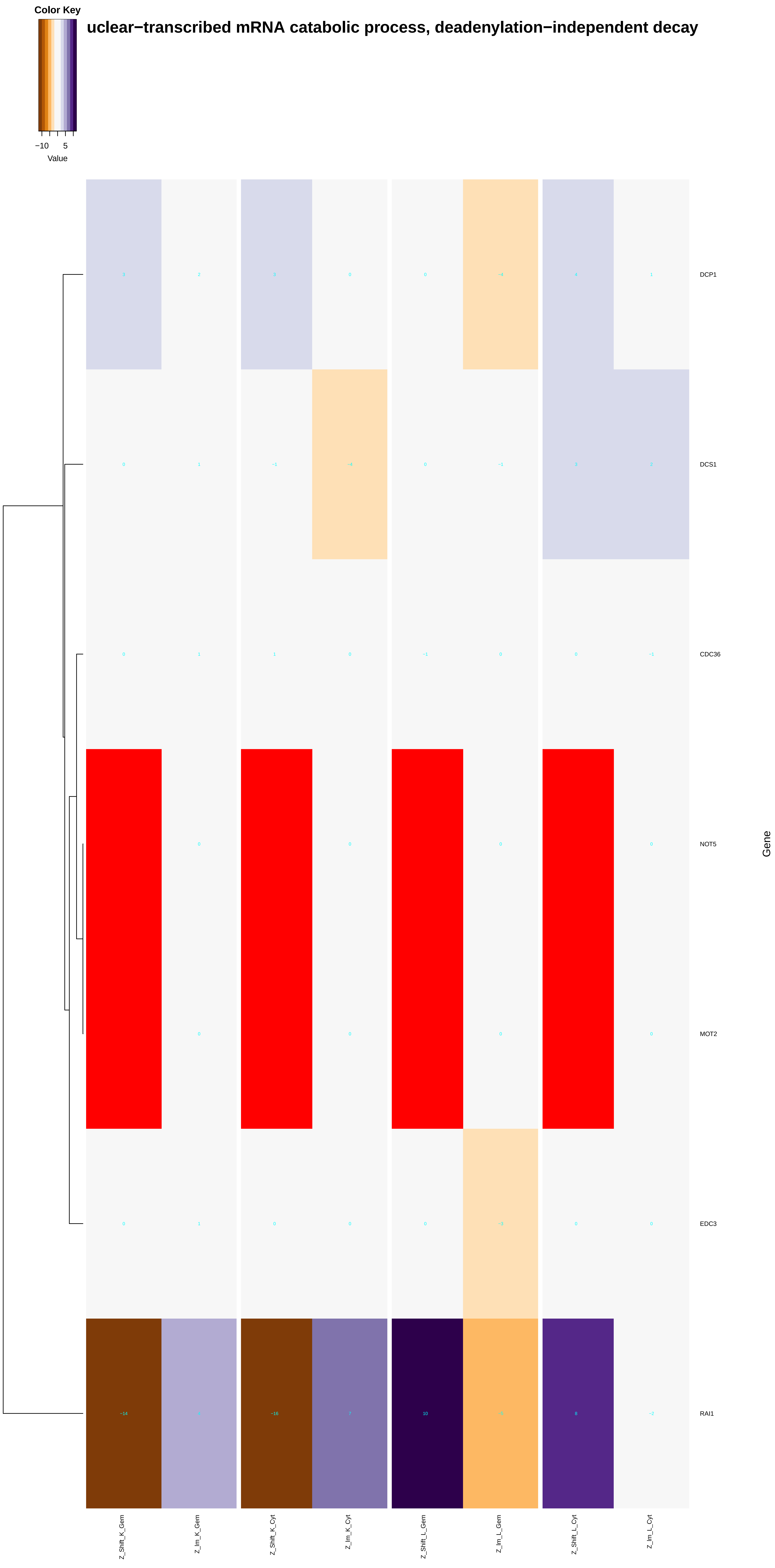
Color Key
−10
5
Value
uclear−transcribed mRNA catabolic process, deadenylation−independent decay
DCP1
DCS1
CDC36
NOT5
MOT2
EDC3
RAI1
Gene
Z_Shift_K_Gem
Z_Im_K_Gem
Z_Shift_K_Cyt
Z_Im_K_Cyt
Z_Shift_L_Gem
Z_Im_L_Gem
Z_Shift_L_Cyt
Z_Im_L_Cyt

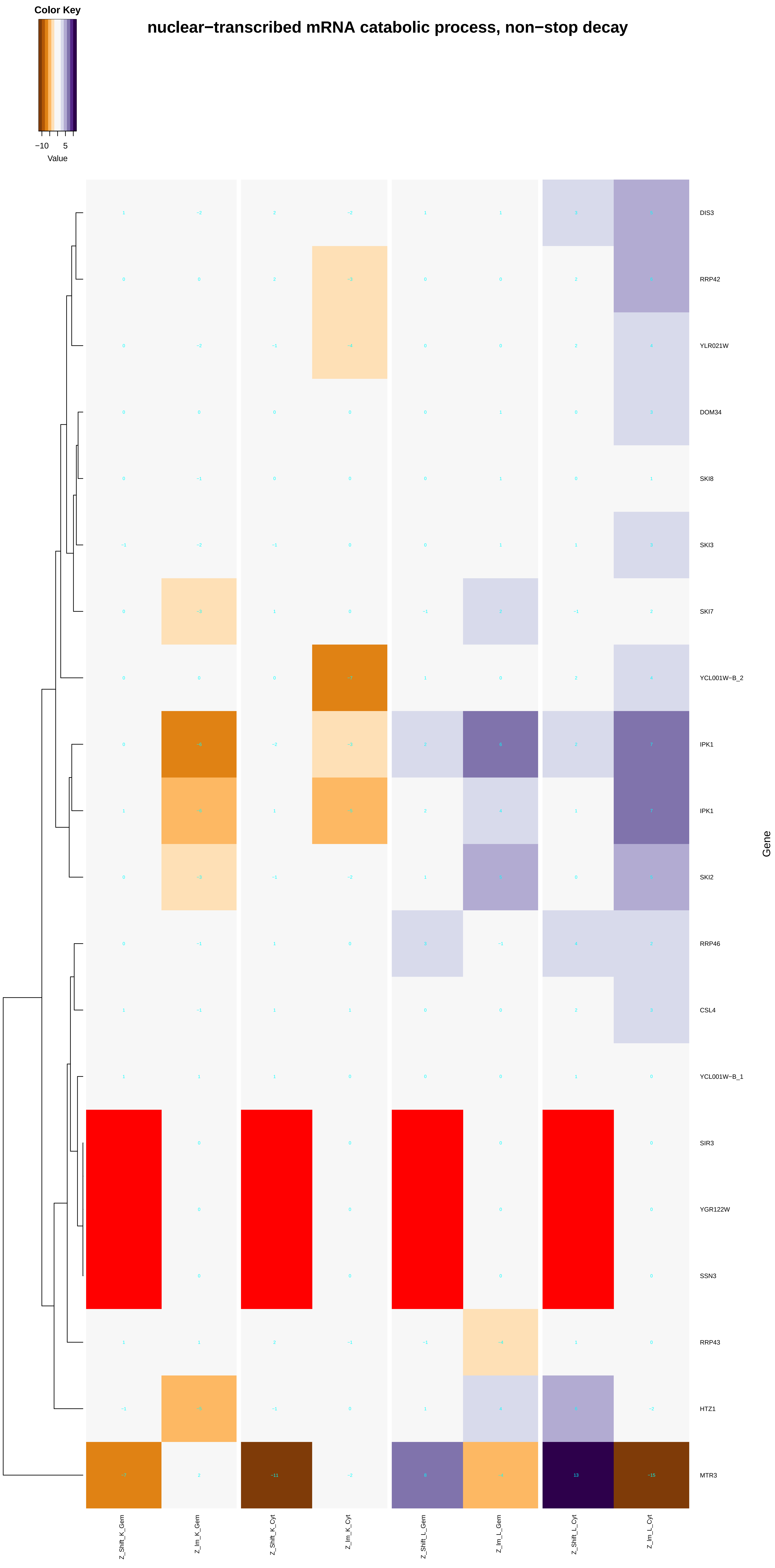
Color Key
−10
5
Value
nuclear−transcribed mRNA catabolic process, non−stop decay
DIS3
RRP42
YLR021W
DOM34
SKI8
SKI3
SKI7
YCL001W−B_2
IPK1
IPK1
SKI2
RRP46
CSL4
YCL001W−B_1
SIR3
YGR122W
SSN3
RRP43
HTZ1
MTR3
Gene
Z_Shift_K_Gem
Z_lm_K_Gem
Z_Shift_K_Cyt
Z_lm_K_Cyt
Z_Shift_L_Gem
Z_lm_L_Gem
Z_Shift_L_Cyt
Z_lm_L_Cyt

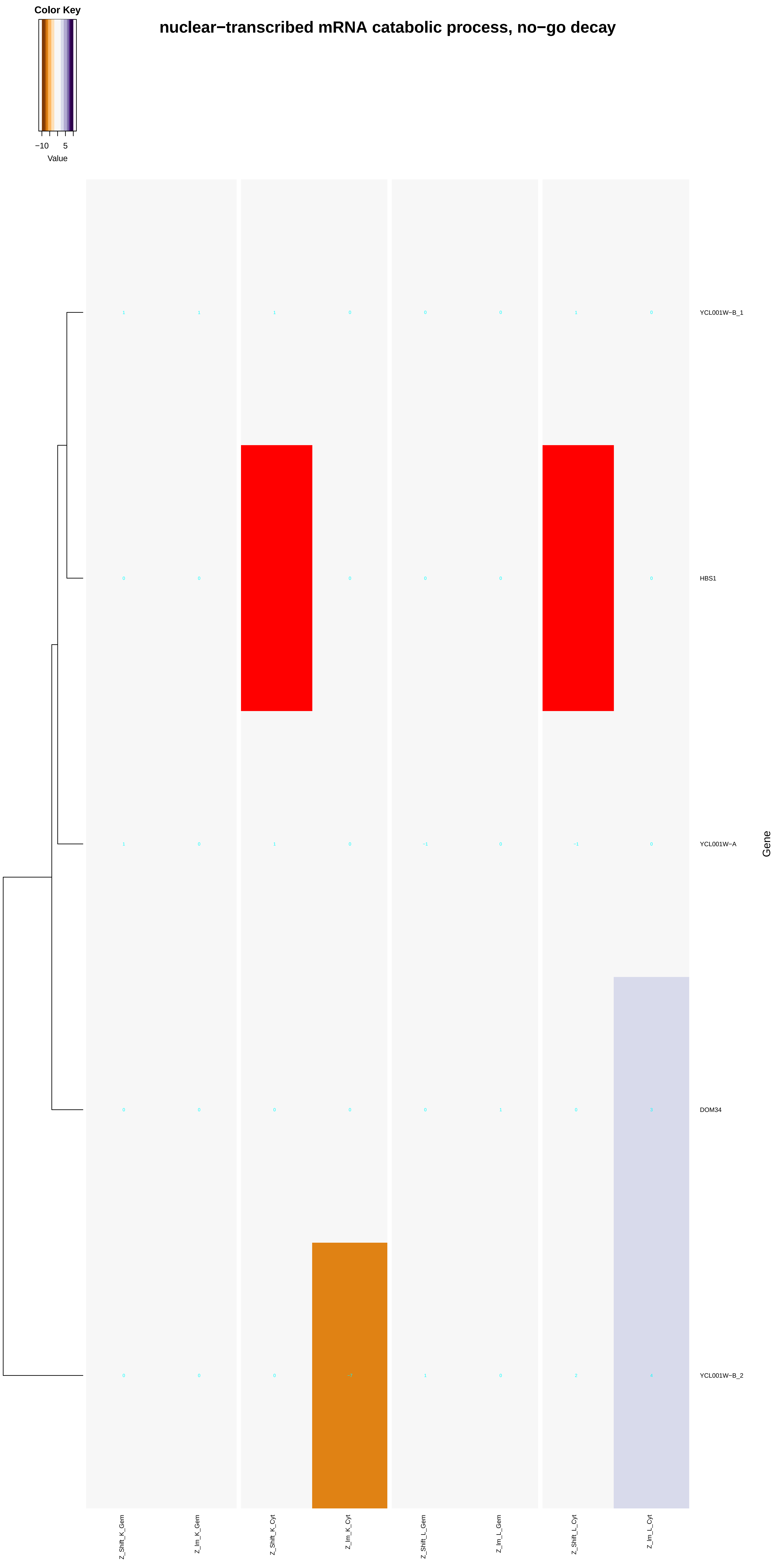

Color Key
−10
5
Value
nuclear−transcribed mRNA catabolic process, no−go decay
YCL001W−B_1
HBS1
YCL001W−A
DOM34
YCL001W−B_2
Gene
Z_Shift_K_Gem
Z_Im_K_Gem
Z_Shift_K_Cyt
Z_Im_K_Cyt
Z_Shift_L_Gem
Z_Im_L_Gem
Z_Shift_L_Cyt
Z_Im_L_Cyt

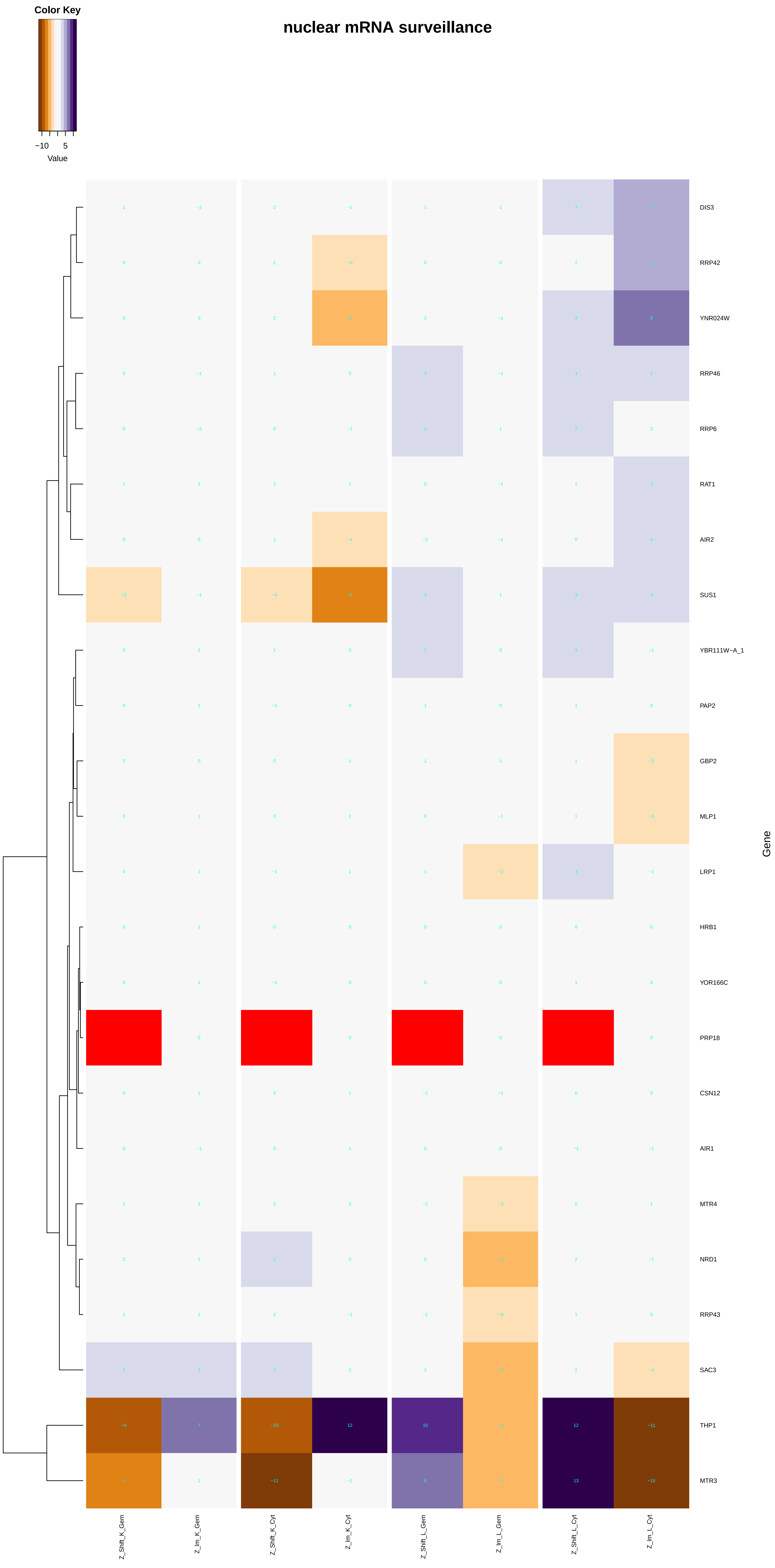

Color Key
−10
5
Value
nuclear mRNA surveillance
DIS3
RRP42
YNR024W
RRP46
RRP6
RAT1
AIR2
SUS1
YBR111W−A_1
PAP2
GBP2
MLP1
LRP1
HRB1
YOR166C
PRP18
CSN12
AIR1
MTR4
NRD1
RRP43
SAC3
THP1
MTR3
Gene
Z_Shift_K_Gem
Z_Im_K_Gem
Z_Shift_K_Cyt
Z_Im_K_Cyt
Z_Shift_L_Gem
Z_Im_L_Gem
Z_Shift_L_Cyt
Z_Im_L_Cyt

### histone mRNA catabolic process

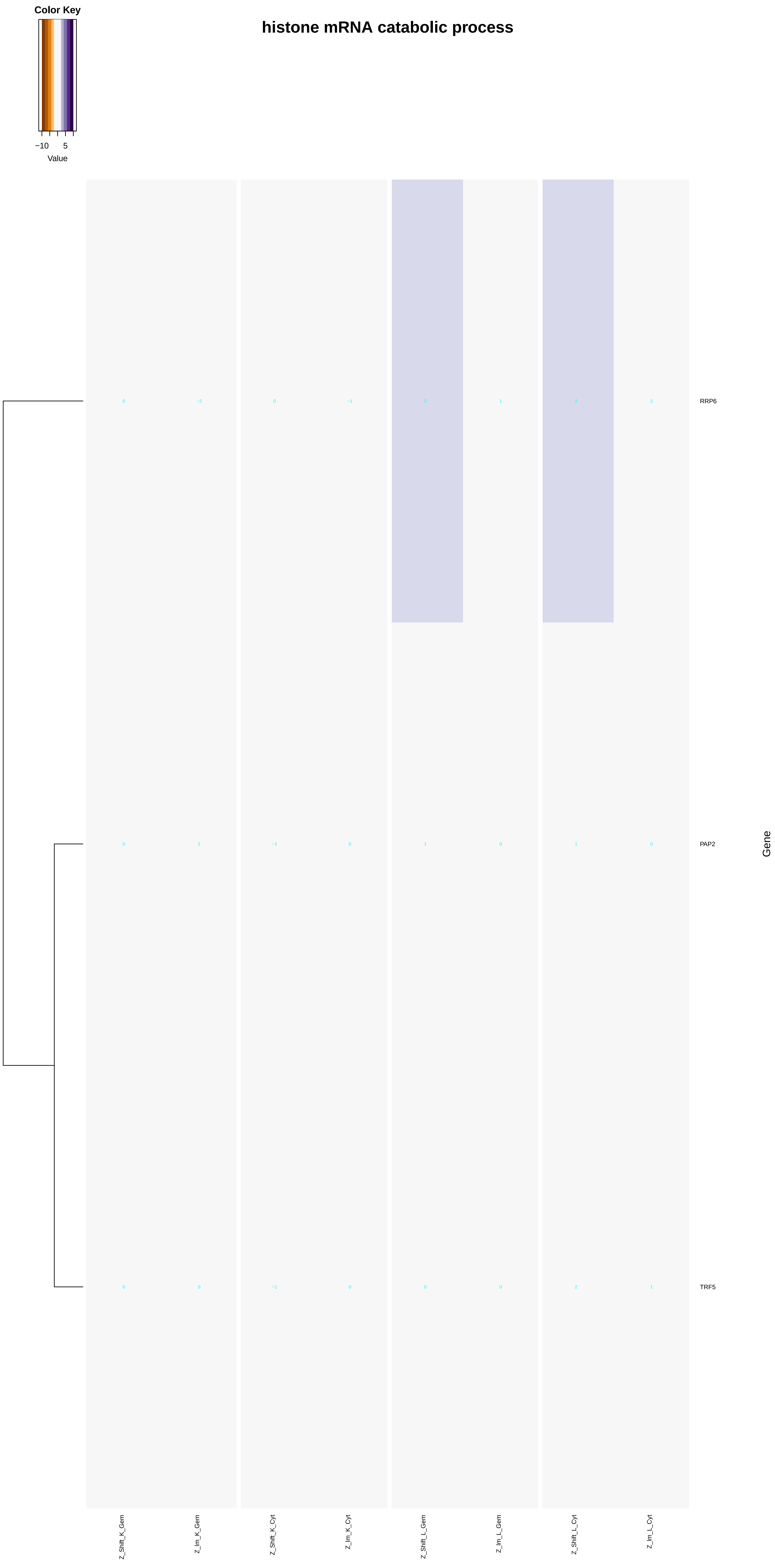

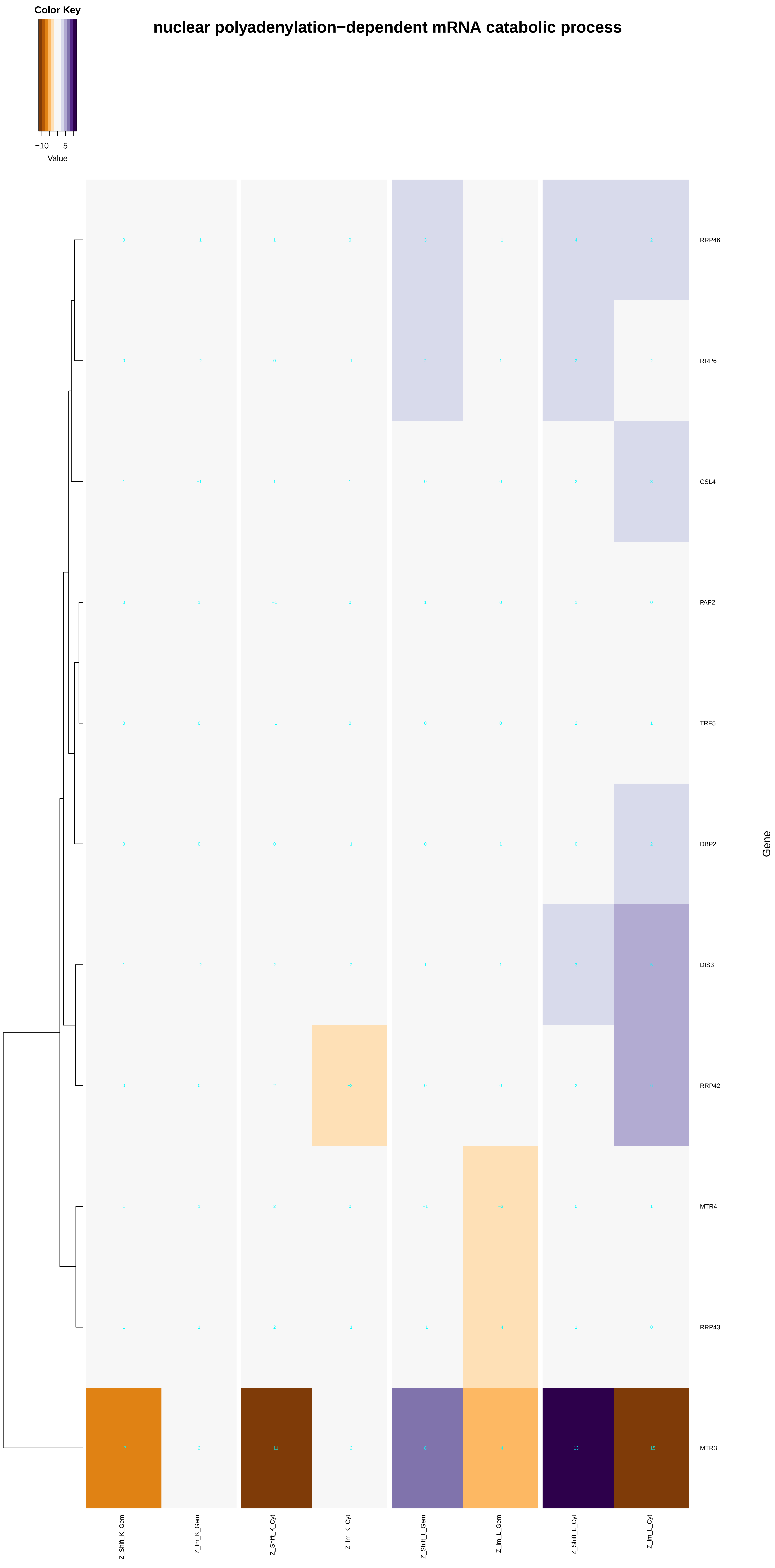

Color Key
−10
5
Value
nuclear polyadenylation−dependent mRNA catabolic process
RRP46
RRP6
CSL4
PAP2
TRF5
DBP2
DIS3
RRP42
MTR4
RRP43
MTR3
Gene
Z_Shift_K_Gem
Z_Im_K_Gem
Z_Shift_K_Cyt
Z_Im_K_Cyt
Z_Shift_L_Gem
Z_Im_L_Gem
Z_Shift_L_Cyt
Z_Im_L_Cyt

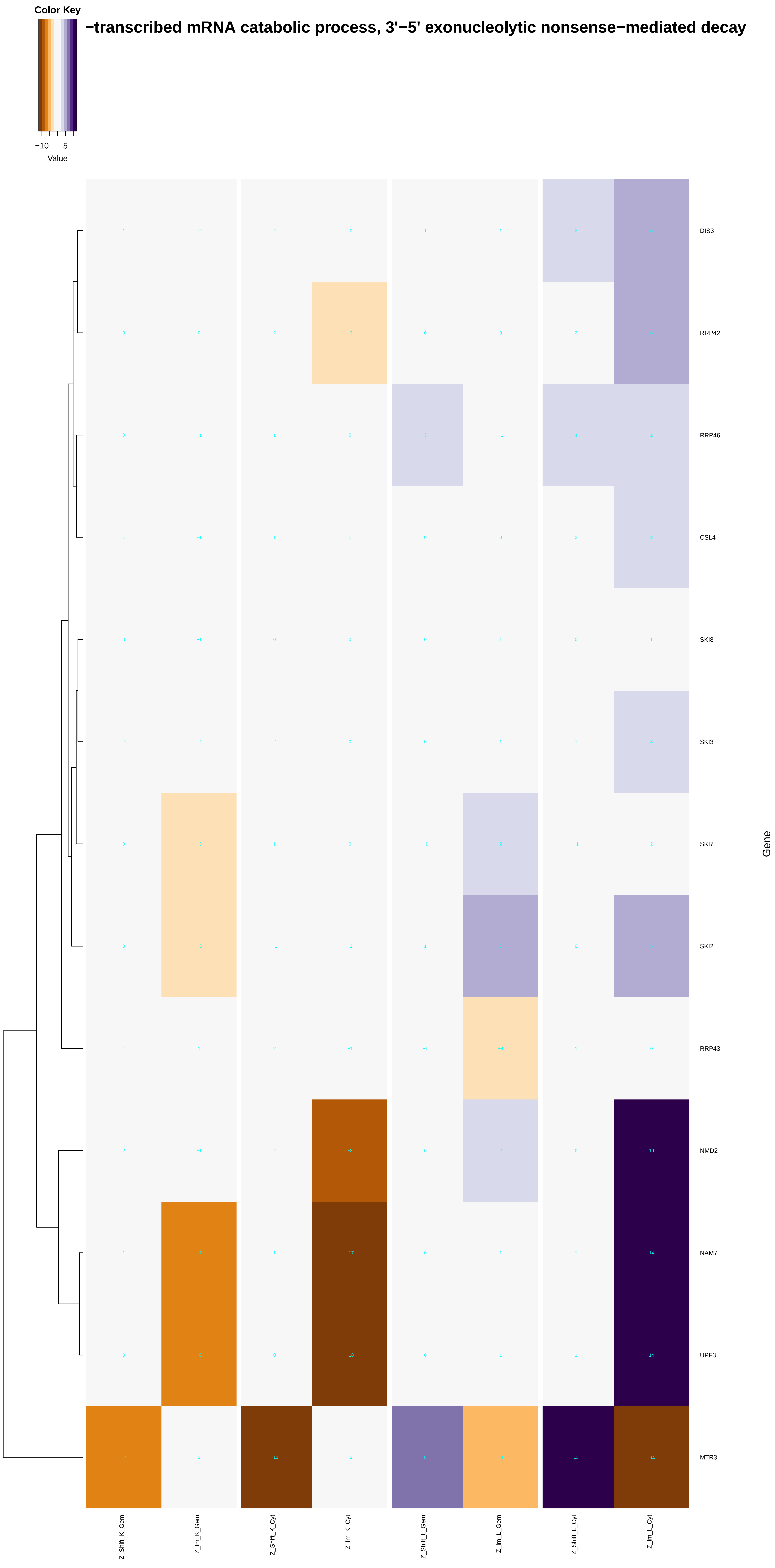
Color Key
−10
5
Value
−transcribed mRNA catabolic process, 3'−5' exonucleolytic nonsense−mediated decay
DIS3
RRP42
RRP46
CSL4
SKI8
SKI3
SKI7
SKI2
RRP43
NMD2
NAM7
UPF3
MTR3
Gene
Z_Shift_K_Gem
Z_Im_K_Gem
Z_Shift_K_Cyt
Z_Im_K_Cyt
Z_Shift_L_Gem
Z_Im_L_Gem
Z_Shift_L_Cyt
Z_Im_L_Cyt

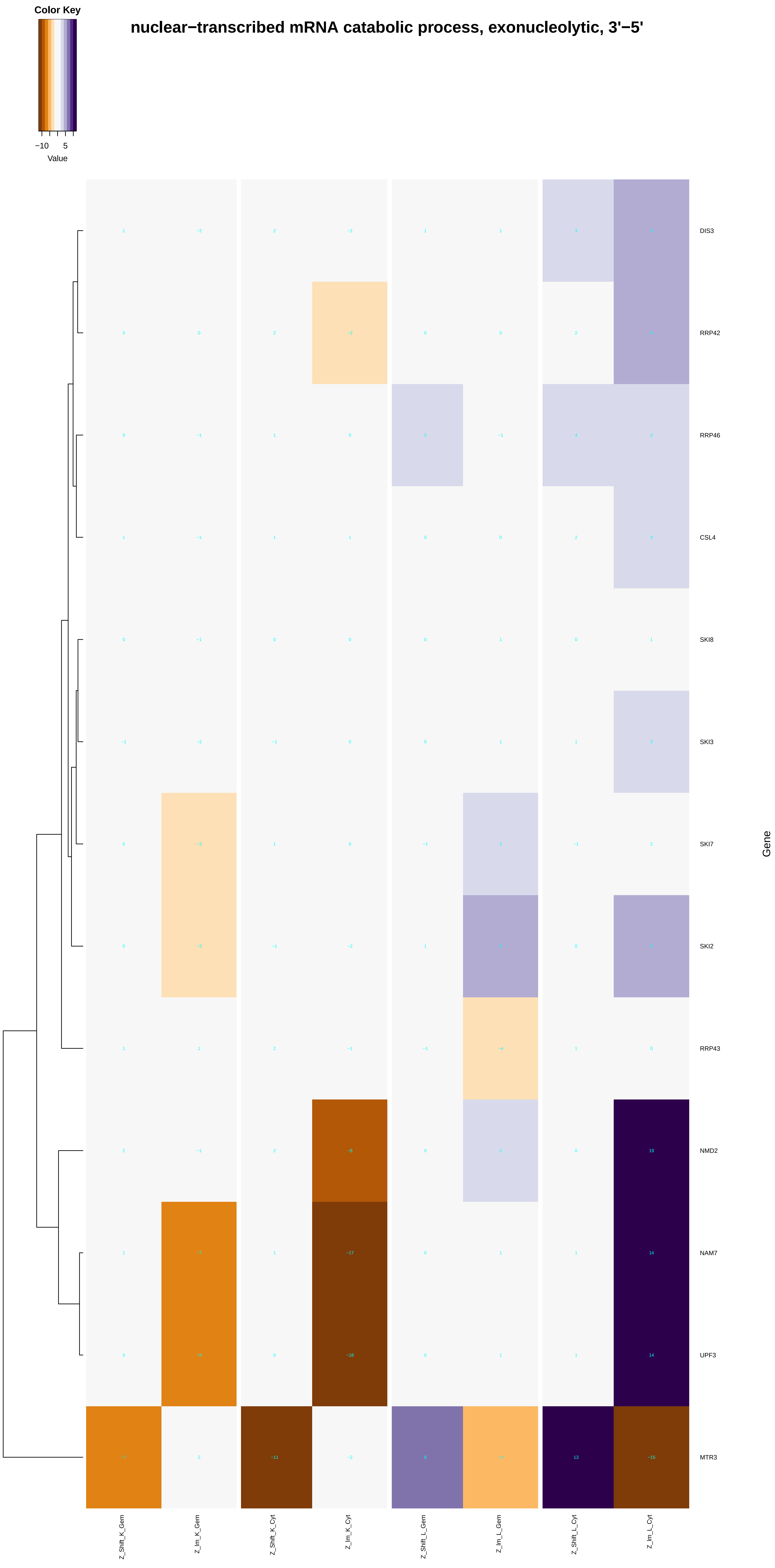

Color Key
−10
5
Value
nuclear−transcribed mRNA catabolic process, exonucleolytic, 3'−5'
DIS3
RRP42
RRP46
CSL4
SKI8
SKI3
SKI7
SKI2
RRP43
NMD2
NAM7
UPF3
MTR3
Gene
Z_Shift_K_Gem
Z_lm_K_Gem
Z_Shift_K_Cyt
Z_lm_K_Cyt
Z_Shift_L_Gem
Z_lm_L_Gem
Z_Shift_L_Cyt
Z_lm_L_Cyt

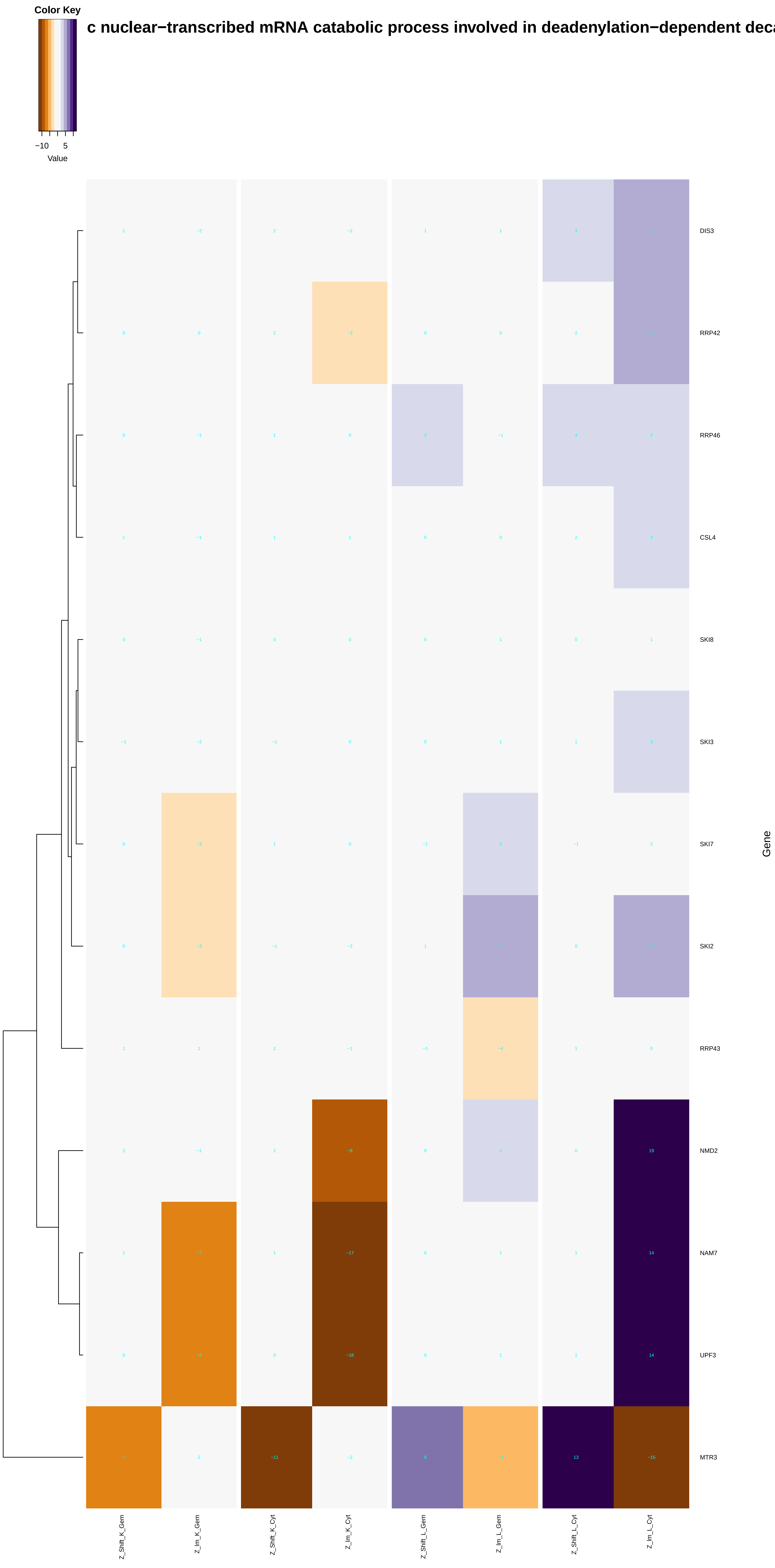

Color Key
−10
5
Value
c nuclear−transcribed mRNA catabolic process involved in deadenylation−dependent deca
DIS3
RRP42
RRP46
CSL4
SKI8
SKI3
SKI7
SKI2
RRP43
NMD2
NAM7
UPF3
MTR3
Gene
Z_Shift_K_Gem
Z_Im_K_Gem
Z_Shift_K_Cyt
Z_Im_K_Cyt
Z_Shift_L_Gem
Z_Im_L_Gem
Z_Shift_L_Cyt
Z_Im_L_Cyt

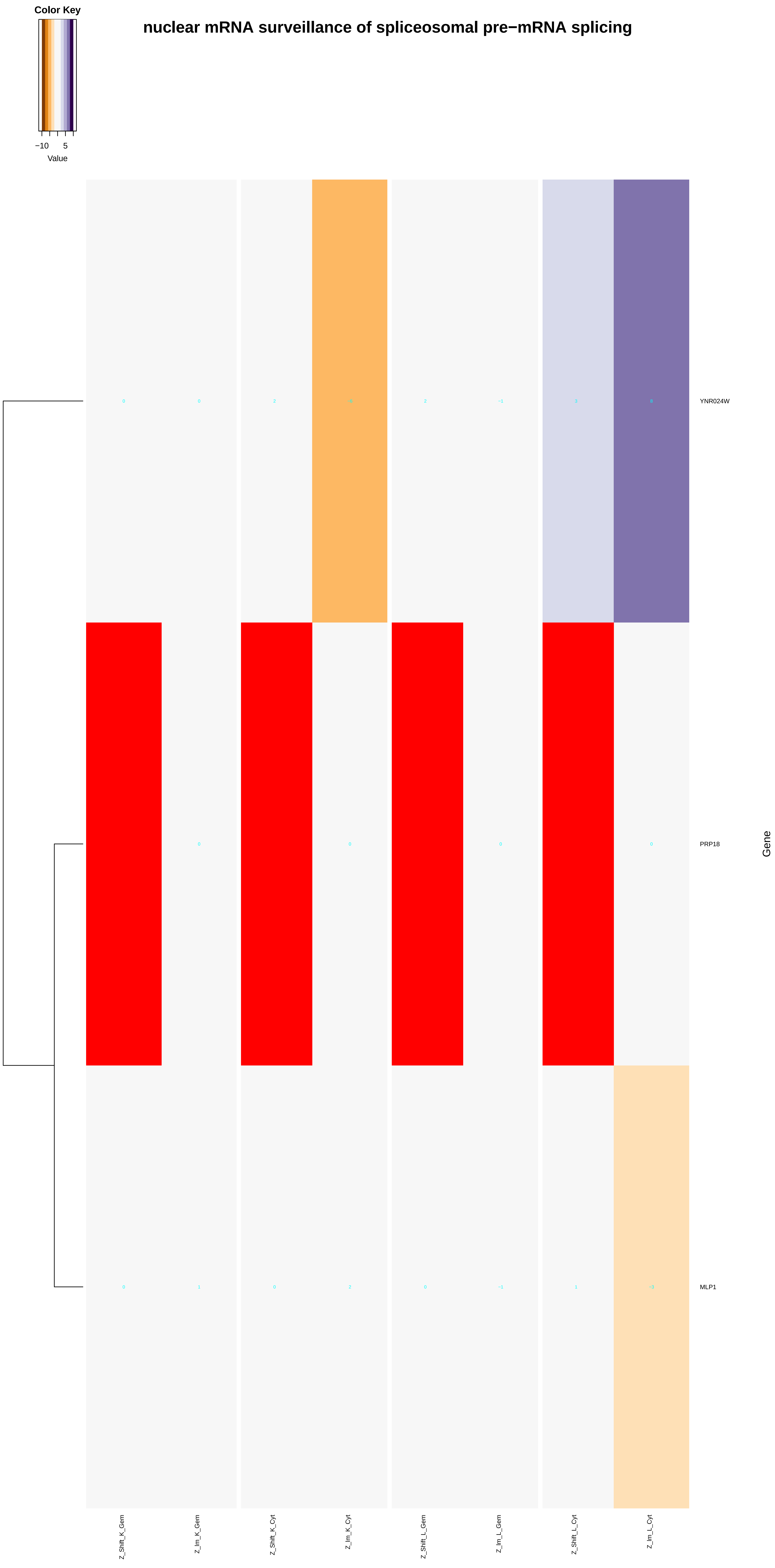
Color Key
−10
5
Value
nuclear mRNA surveillance of spliceosomal pre−mRNA splicing
YNR024W
PRP18
MLP1
Gene
Z_Shift_K_Gem
Z_Im_K_Gem
Z_Shift_K_Cyt
Z_Im_K_Cyt
Z_Shift_L_Gem
Z_Im_L_Gem
Z_Shift_L_Cyt
Z_Im_L_Cyt

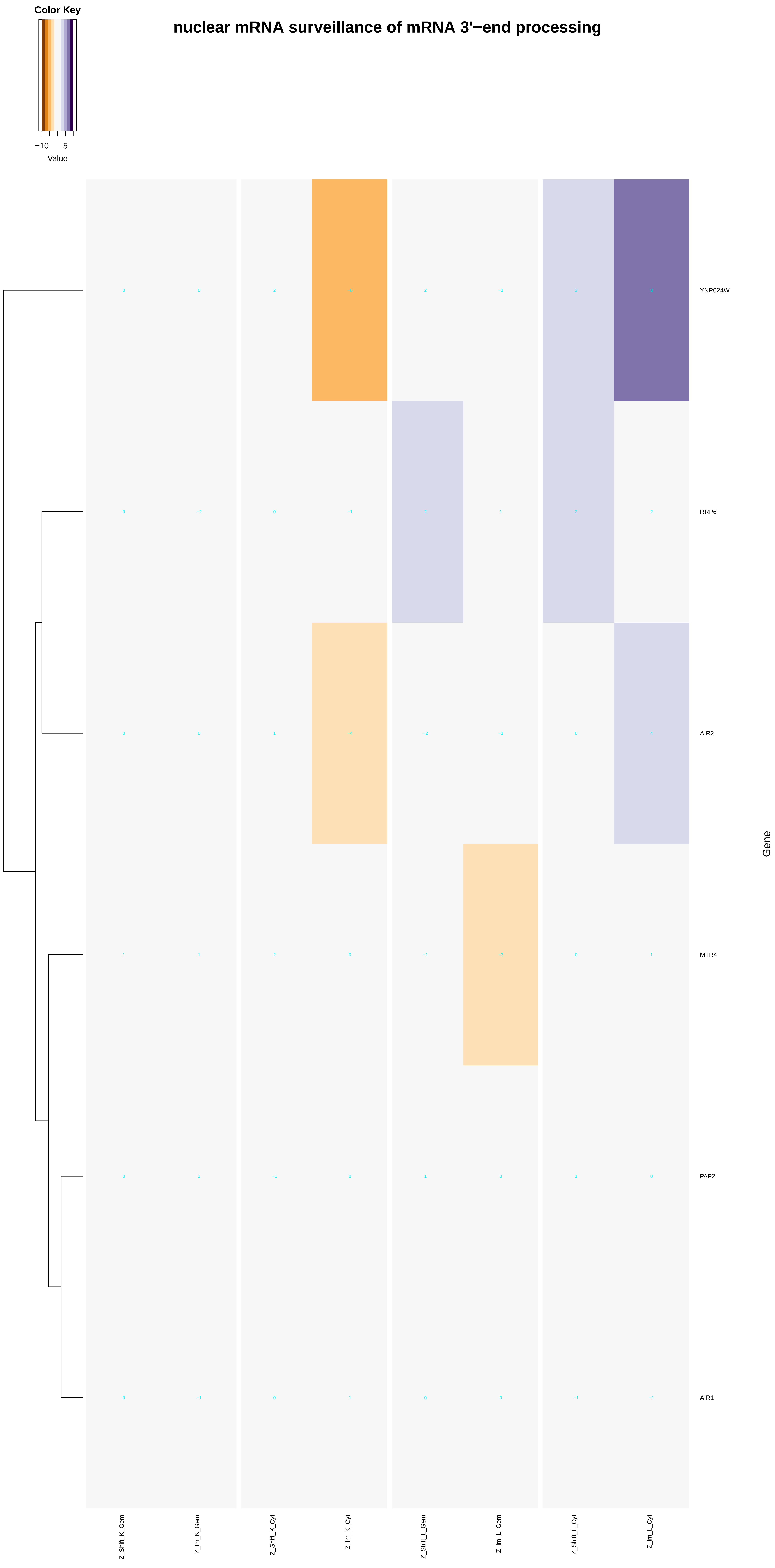

Color Key
−10
5
Value
nuclear mRNA surveillance of mRNA 3'−end processing
YNR024W
RRP6
AIR2
MTR4
PAP2
AIR1
Gene
Z_Shift_K_Gem
Z_Im_K_Gem
Z_Shift_K_Cyt
Z_Im_K_Cyt
Z_Shift_L_Gem
Z_Im_L_Gem
Z_Shift_L_Cyt
Z_Im_L_Cyt
