## Additional File 7 for "A humanized yeast phenomic model of deoxycytidine kinase to predict genetic buffering of nucleoside analog cytotoxicity": mRNA_metabolic_process.pdf

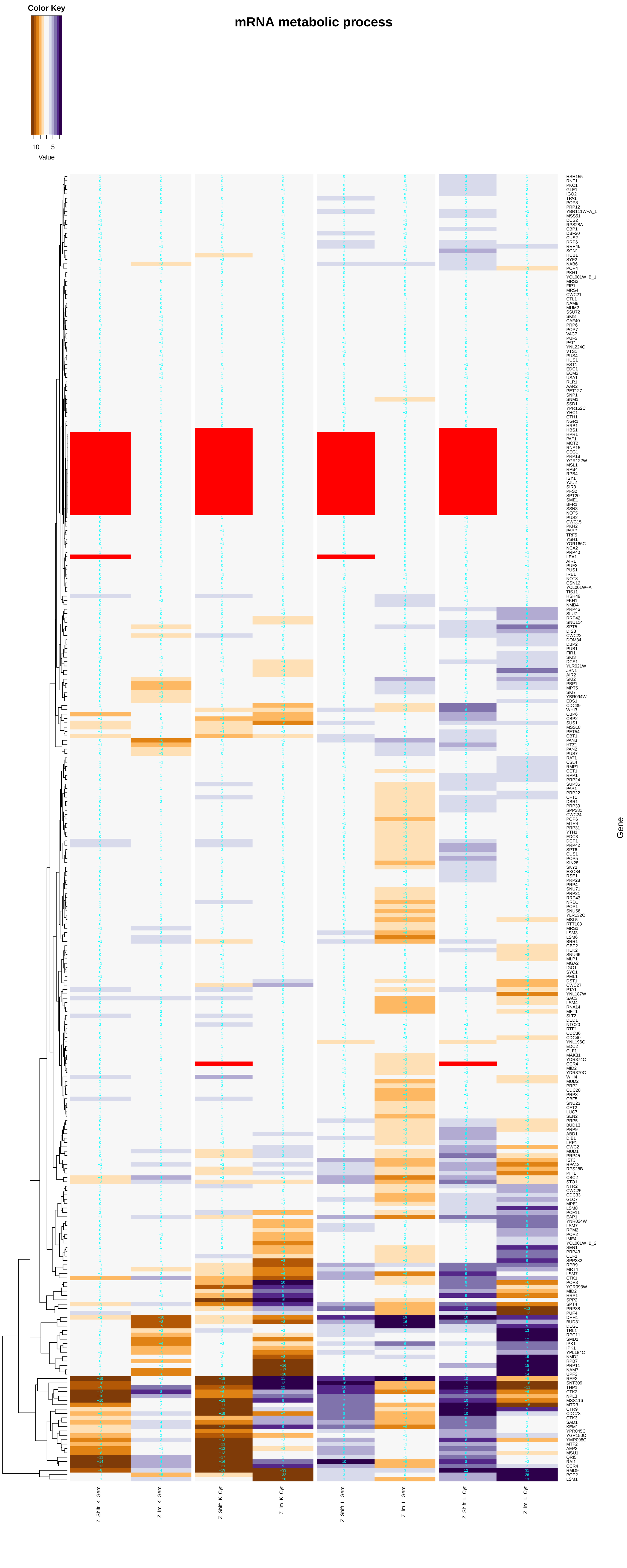
mRNA metabolic process
Color Key
−10 5
Value
Gene
Z_Shift_K_Gem
Z_Im_K_Gem
Z_Shift_K_Cyt
Z_Im_K_Cyt
Z_Shift_L_Gem
Z_Im_L_Gem
Z_Shift_L_Cyt
Z_Im_L_Cyt

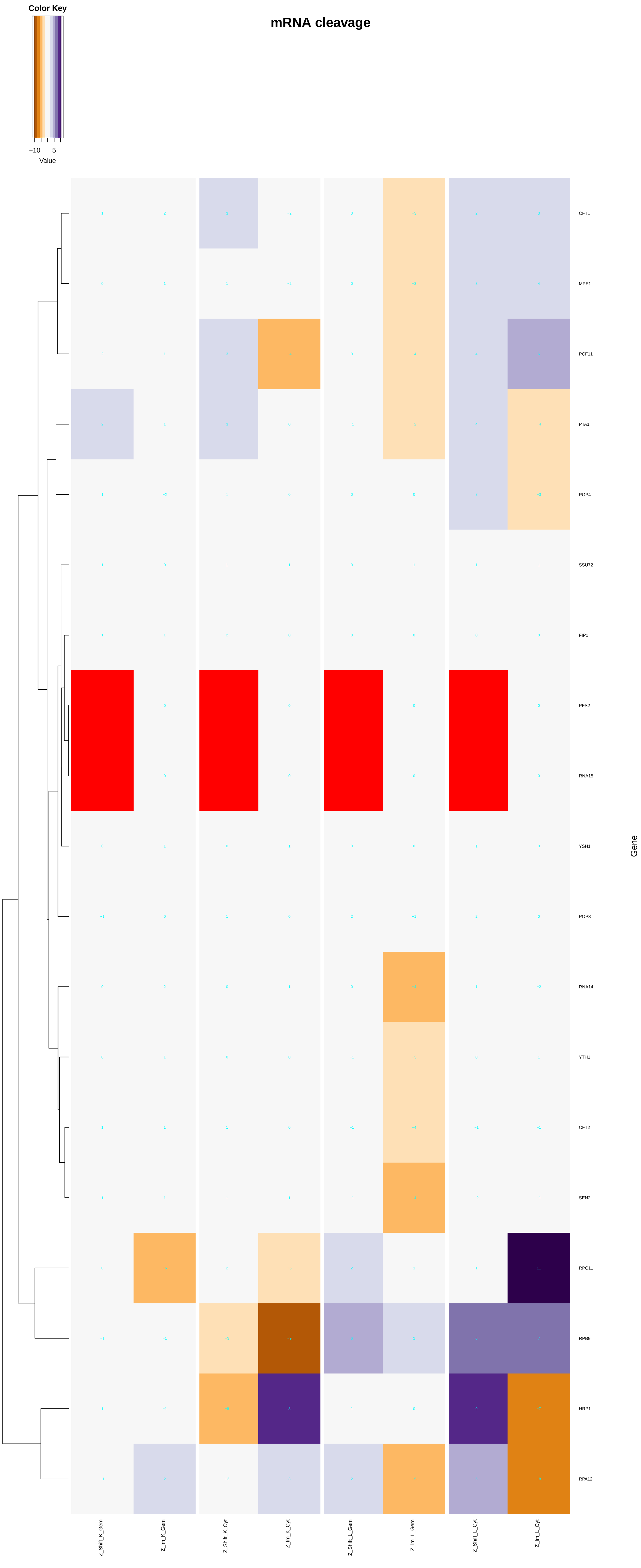
Color Key
−10
5
Value
mRNA cleavage
CFT1
MPE1
PCF11
PTA1
POP4
SSU72
FIP1
PFS2
RNA15
YSH1
POP8
RNA14
YTH1
CFT2
SEN2
RPC11
RPB9
HRP1
RPA12
Gene
Z_Shift_K_Gem
Z_Im_K_Gem
Z_Shift_K_Cyt
Z_Im_K_Cyt
Z_Shift_L_Gem
Z_Im_L_Gem
Z_Shift_L_Cyt
Z_Im_L_Cyt

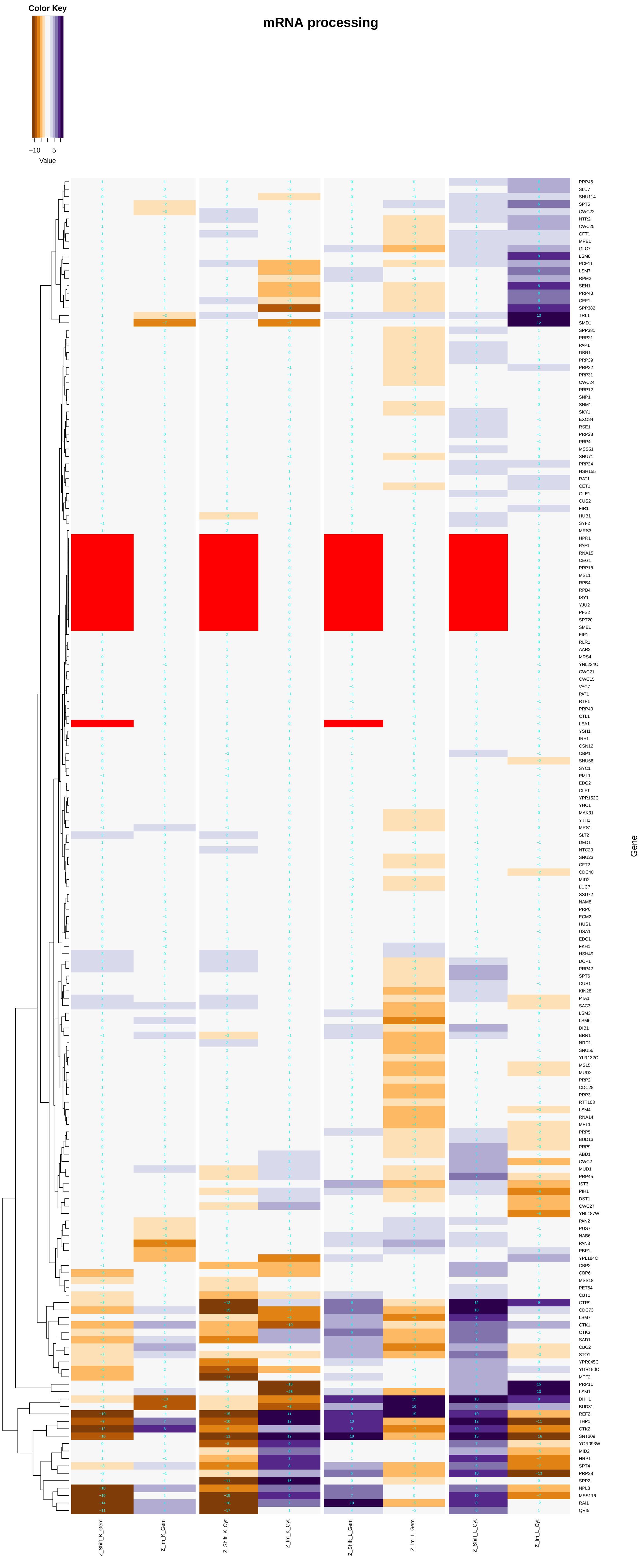

mRNA processing
Color Key
−10
5
Value
Gene
Z_Shift_K_Gem
Z_Im_K_Gem
Z_Shift_K_Cyt
Z_Im_K_Cyt
Z_Shift_L_Gem
Z_Im_L_Gem
Z_Shift_L_Cyt
Z_Im_L_Cyt
PRP46
SLU7
SNU114
SPT5
CWC22
NTR2
CWC25
CFT1
MPE1
GLC7
LSM8
PCF11
LSM7
RPM2
SEN1
PRP43
CEF1
SPP382
TRL1
SMD1
SPP381
PRP21
PAP1
DBR1
PRP39
PRP22
PRP31
CWC24
PRP12
SNP1
SNM1
SKY1
EXO84
RSE1
PRP28
PRP4
MSS51
SNU71
PRP24
HSH155
RAT1
CET1
GLE1
CUS2
FIR1
HUB1
SYF2
MRS3
HPR1
PAF1
RNA15
CEG1
PRP18
MSL1
RPB4
RPB4
ISY1
YJU2
PFS2
SPT20
SME1
FIP1
RLR1
AAR2
MRS4
YNL224C
CWC21
CWC15
VAC7
PAT1
RTF1
PRP40
CTL1
LEA1
YSH1
IRE1
CSN12
CBP1
SNU66
SYC1
PML1
EDC2
CLF1
YPR152C
YHC1
MAK31
YTH1
MRS1
SLT2
DED1
NTC20
SNU23
CFT2
CDC40
MID2
LUC7
SSU72
NAM8
PRP6
ECM2
HUS1
USA1
EDC1
FKH1
HSH49
DCP1
PRP42
SPT6
CUS1
KIN28
PTA1
SAC3
LSM3
LSM6
DIB1
BRR1
NRD1
SNU56
YLR132C
MSL5
MUD2
PRP2
CDC28
PRP3
RTT103
LSM4
RNA14
MFT1
PRP5
BUD13
PRP9
ABD1
CWC2
MUD1
PRP45
IST3
PIH1
DST1
CWC27
YNL187W
PAN2
PUS7
NAB6
PAN3
PBP1
YPL184C
CBP2
CBP6
MSS18
PET54
CBT1
CTR9
CDC73
LSM7
CTK1
CTK3
SAD1
CBC2
STO1
YPR045C
YGR150C
MTF2
PRP11
LSM1
DHH1
BUD31
REF2
THP1
CTK2
SNT309
YGR093W
MID2
HRP1
SPT4
PRP38
SPP2
NPL3
MSS116
RAI1
QRI5

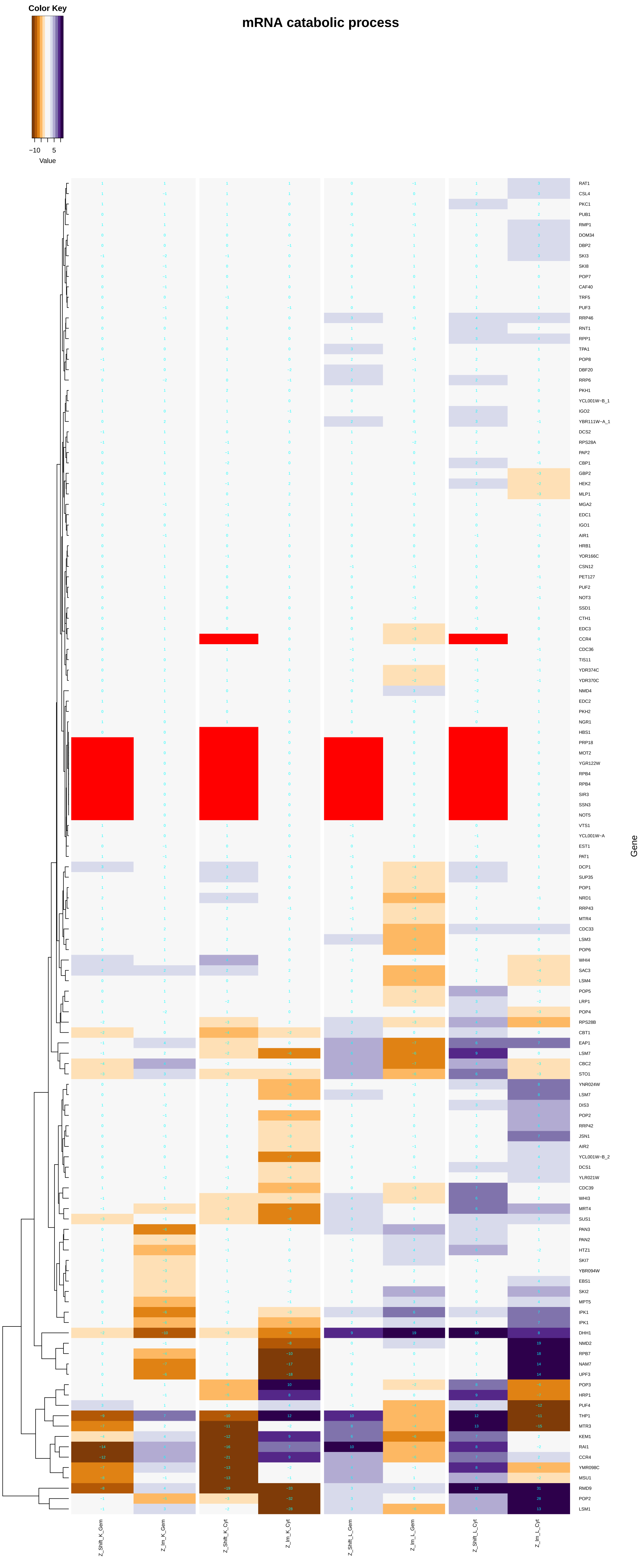
Color Key
mRNA catabolic process
−10
5
Value
Gene
RAT1
CSL4
PKC1
PUB1
RMP1
DOM34
DBP2
SKI3
SKI8
POP7
CAF40
TRF5
PUF3
RRP46
RNT1
RPP1
TPA1
POP8
DBF20
RRP6
PKH1
YCL001W−B_1
IGO2
YBR111W−A_1
DCS2
RPS28A
PAP2
CBP1
GBP2
HEK2
MLP1
MGA2
EDC1
IGO1
AIR1
HRB1
YOR166C
CSN12
PET127
PUF2
NOT3
SSD1
CTH1
EDC3
CCR4
CDC36
TIS11
YDR374C
YDR370C
NMD4
EDC2
PKH2
NGR1
HBS1
PRP18
MOT2
YGR122W
RPB4
RPB4
SIR3
SSN3
NOT5
VTS1
YCL001W−A
EST1
PAT1
DCP1
SUP35
POP1
NRD1
RRP43
MTR4
CDC33
LSM3
POP6
WHI4
SAC3
LSM4
POP5
LRP1
POP4
RPS28B
CBT1
EAP1
LSM7
CBC2
STO1
YNR024W
LSM5
DIS3
POP2
RRP42
JSN1
AIR2
YCL001W−B_2
DCS1
YLR021W
CDC39
WHI3
MRT4
SUS1
PAN3
PAN2
HTZ1
SKI7
YBR094W
EBS1
SKI2
MPT5
IPK1
IPK1
DHH1
NMD2
RPB7
NAM7
UPF3
POP3
HRP1
PUF4
THP1
MTR3
KEM1
RAI1
CCR4
YMR098C
MSU1
RMD9
POP2
LSM1
Z_Shift_K_Gem
Z_Im_K_Gem
Z_Shift_K_Cyt
Z_Im_K_Cyt
Z_Shift_L_Gem
Z_Im_L_Gem
Z_Shift_L_Cyt
Z_Im_L_Cyt

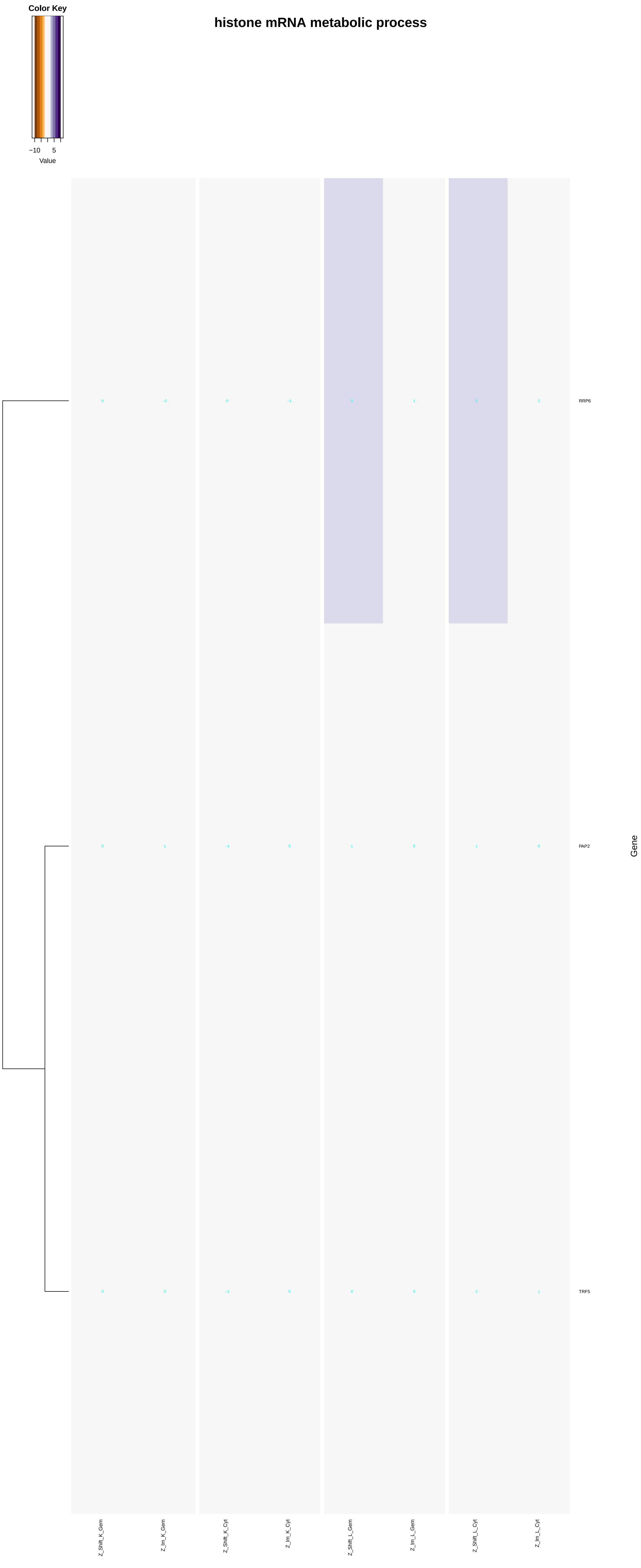

Color Key
−10
5
Value
histone mRNA metabolic process
0
−2
0
−1
2
1
2
2
RRP6
0
1
−1
0
1
0
1
0
PAP2
0
0
−1
0
0
0
2
1
TRF5
Gene
Z_Shift_K_Gem
Z_Im_K_Gem
Z_Shift_K_Cyt
Z_Im_K_Cyt
Z_Shift_L_Gem
Z_Im_L_Gem
Z_Shift_L_Cyt
Z_Im_L_Cyt

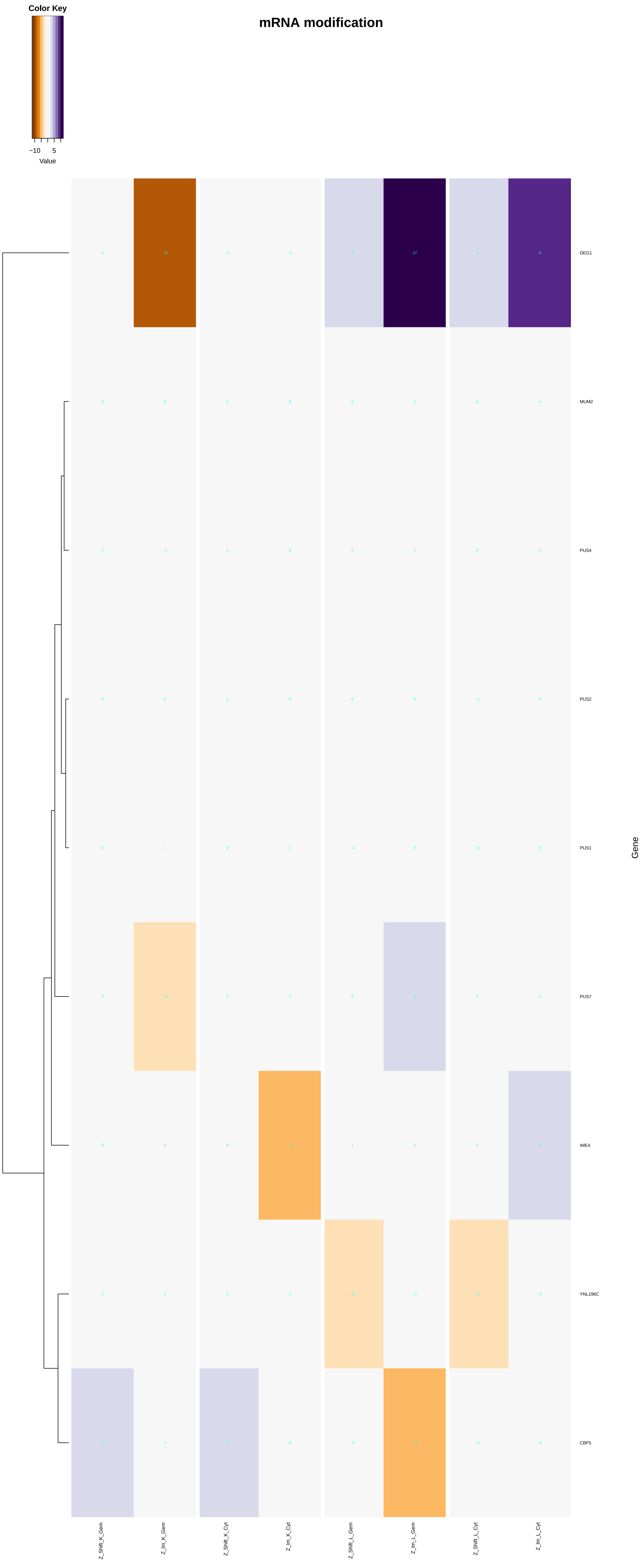
mRNA modification
Color Key
−10
5
Value
DEG1
MUM2
PUS4
PUS2
PUS1
PUS7
IME4
YNL196C
CBF5
Gene
Z_Shift_K_Gem
Z_lm_K_Gem
Z_Shift_K_Cyt
Z_lm_K_Cyt
Z_Shift_L_Gem
Z_lm_L_Gem
Z_Shift_L_Cyt
Z_lm_L_Cyt

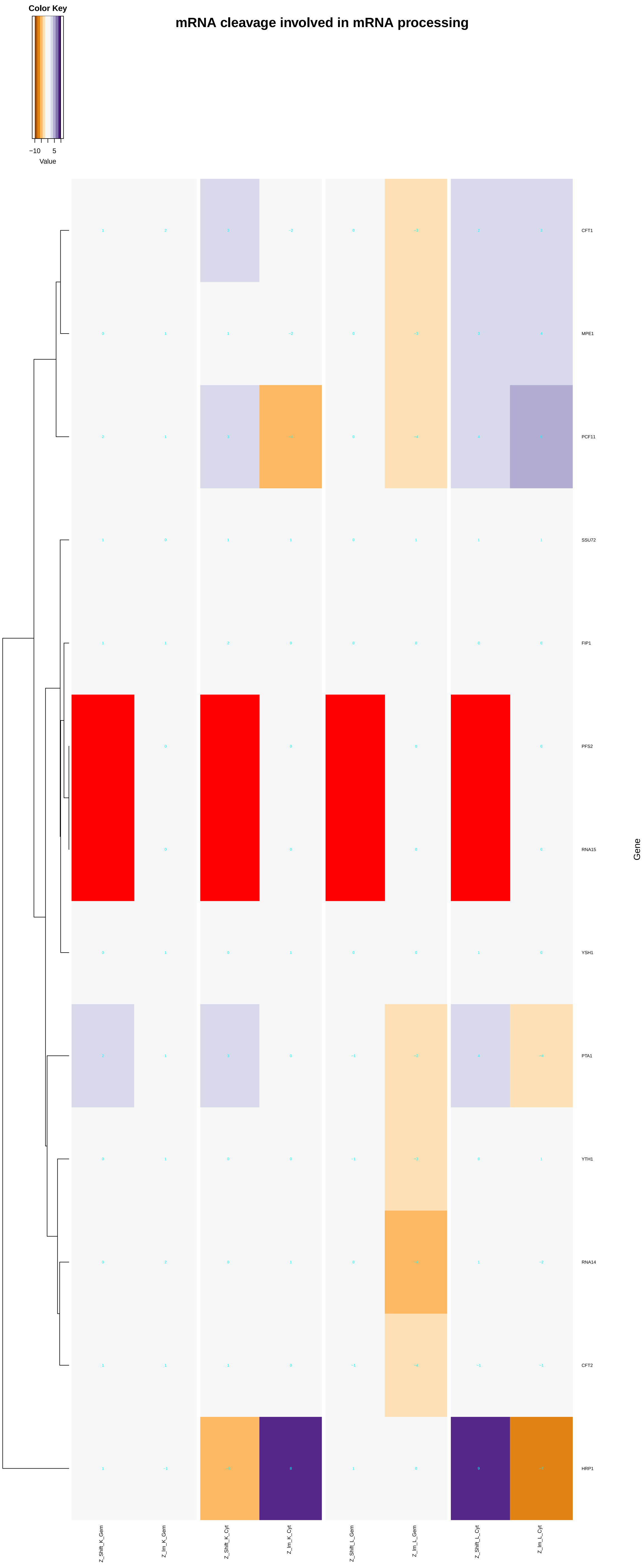
Color Key
−10
5
Value
mRNA cleavage involved in mRNA processing
CFT1
MPE1
PCF11
SSU72
FIP1
PFS2
RNA15
YSH1
PTA1
YTH1
RNA14
CFT2
HRP1
Gene
Z_Shift_K_Gem
Z_Im_K_Gem
Z_Shift_K_Cyt
Z_Im_K_Cyt
Z_Shift_L_Gem
Z_Im_L_Gem
Z_Shift_L_Cyt
Z_Im_L_Cyt

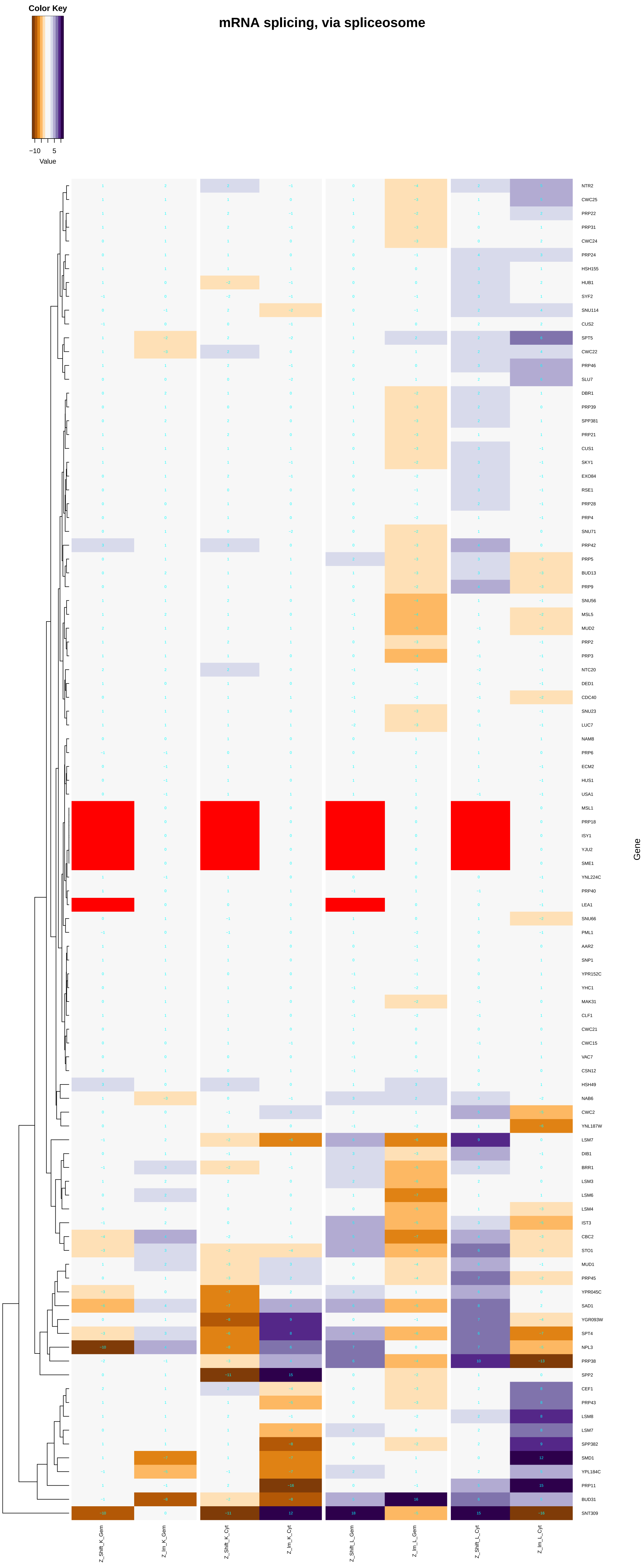

Color Key
−10 5
Value
mRNA splicing, via spliceosome
NTR2
CWC25
PRP22
PRP31
CWC24
PRP24
HSH155
HUB1
SYF2
SNU114
CUS2
SPT5
CWC22
PRP46
SLU7
DBR1
PRP39
SPP381
PRP21
CUS1
SKY1
EXO84
RSE1
PRP28
PRP4
SNU71
PRP42
PRP5
BUD13
PRP9
SNU56
MSL5
MUD2
PRP2
PRP3
NTC20
DED1
CDC40
SNU23
LUC7
NAM8
PRP6
ECM2
HUS1
USA1
MSL1
PRP18
ISY1
YJU2
SME1
YNL224C
PRP40
LEA1
SNU66
PML1
AAR2
SNP1
YPR152C
YHC1
MAK31
CLF1
CWC21
CWC15
VAC7
CSN12
HSH49
NAB6
CWC2
YNL187W
LSM7
DIB1
BRR1
LSM3
LSM6
LSM4
IST3
CBC2
STO1
MUD1
PRP45
YPR045C
SAD1
YGR093W
SPT4
NPL3
PRP38
SPP2
CEF1
PRP43
LSM8
LSM7
SPP382
SMD1
YPL184C
PRP11
BUD31
SNT309
Gene
Z_Shift_K_Gem
Z_Im_K_Gem
Z_Shift_K_Cyt
Z_Im_K_Cyt
Z_Shift_L_Gem
Z_Im_L_Gem
Z_Shift_L_Cyt
Z_Im_L_Cyt

### 7−methylguanosine mRNA capping

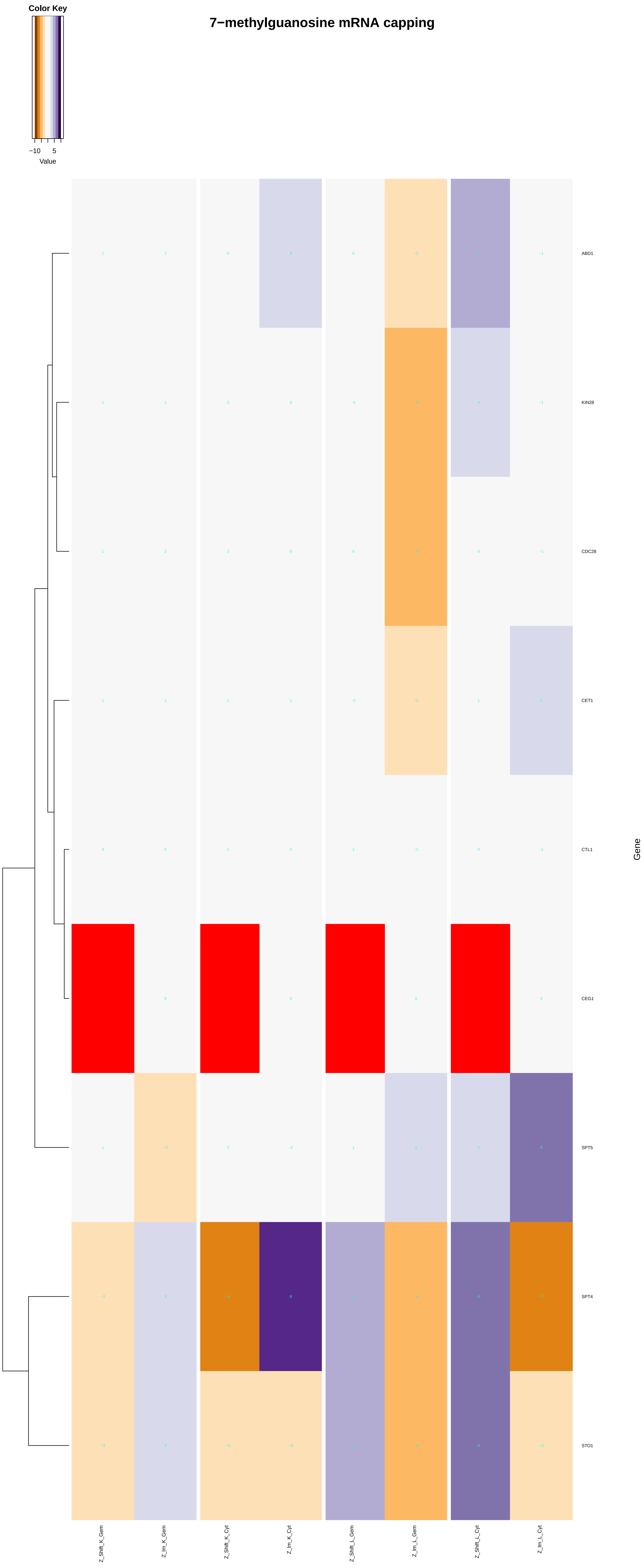

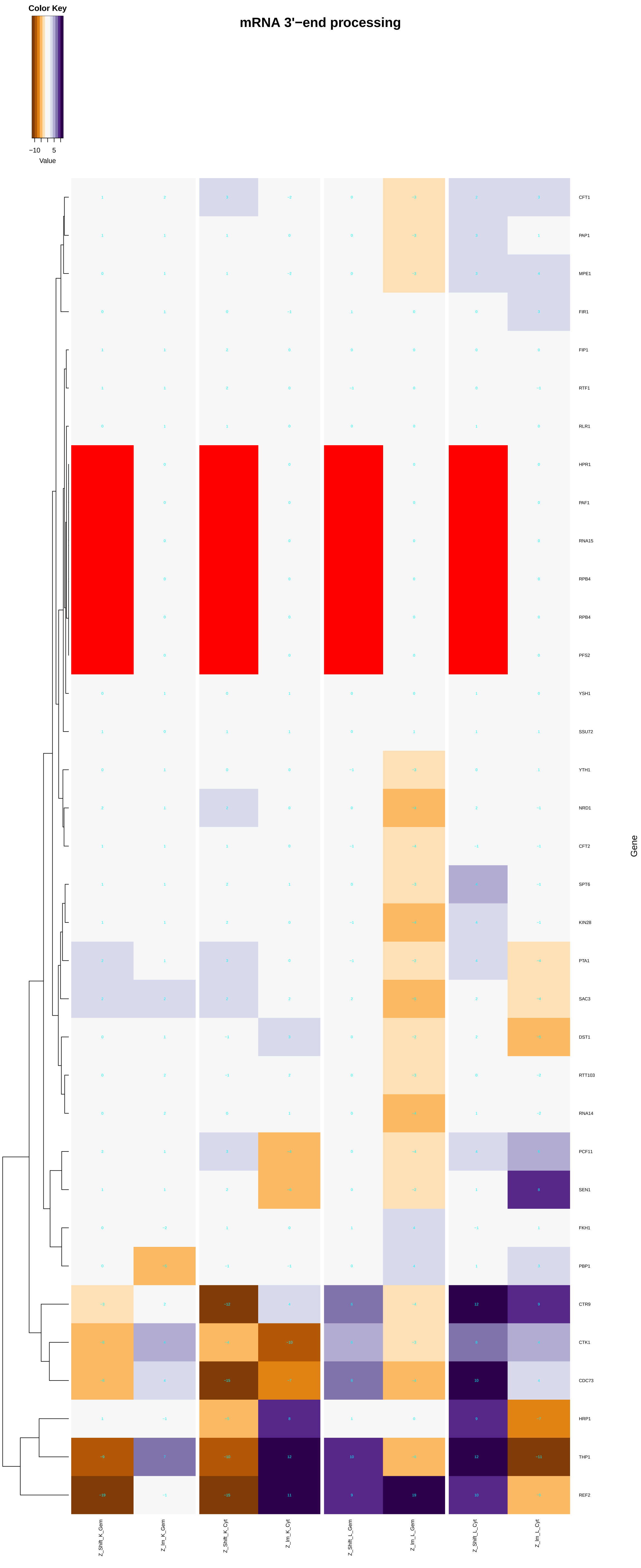

Color Key
−10
5
Value
mRNA 3'−end processing
CFT1
PAP1
MPE1
FIR1
FIP1
RTF1
RLR1
HPR1
PAF1
RNA15
RPB4
RPB4
PFS2
YSH1
SSU72
YTH1
NRD1
CFT2
SPT6
KIN28
PTA1
SAC3
DST1
RTT103
RNA14
PCF11
SEN1
FKH1
PBP1
CTR9
CTK1
CDC73
HRP1
THP1
REF2
Gene
Z_Shift_K_Gem
Z_Im_K_Gem
Z_Shift_K_Cyt
Z_Im_K_Cyt
Z_Shift_L_Gem
Z_Im_L_Gem
Z_Shift_L_Cyt
Z_Im_L_Cyt

### mRNA splicing, via endonucleolytic cleavage and ligation

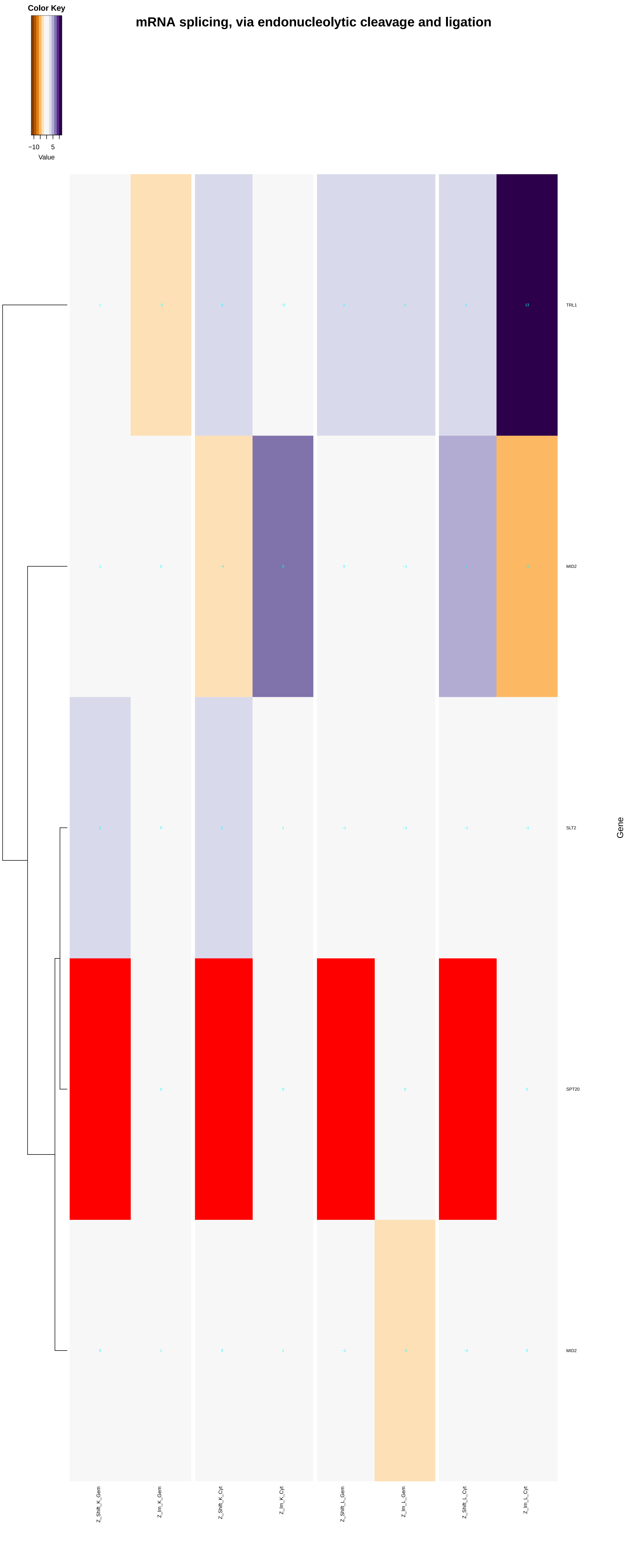

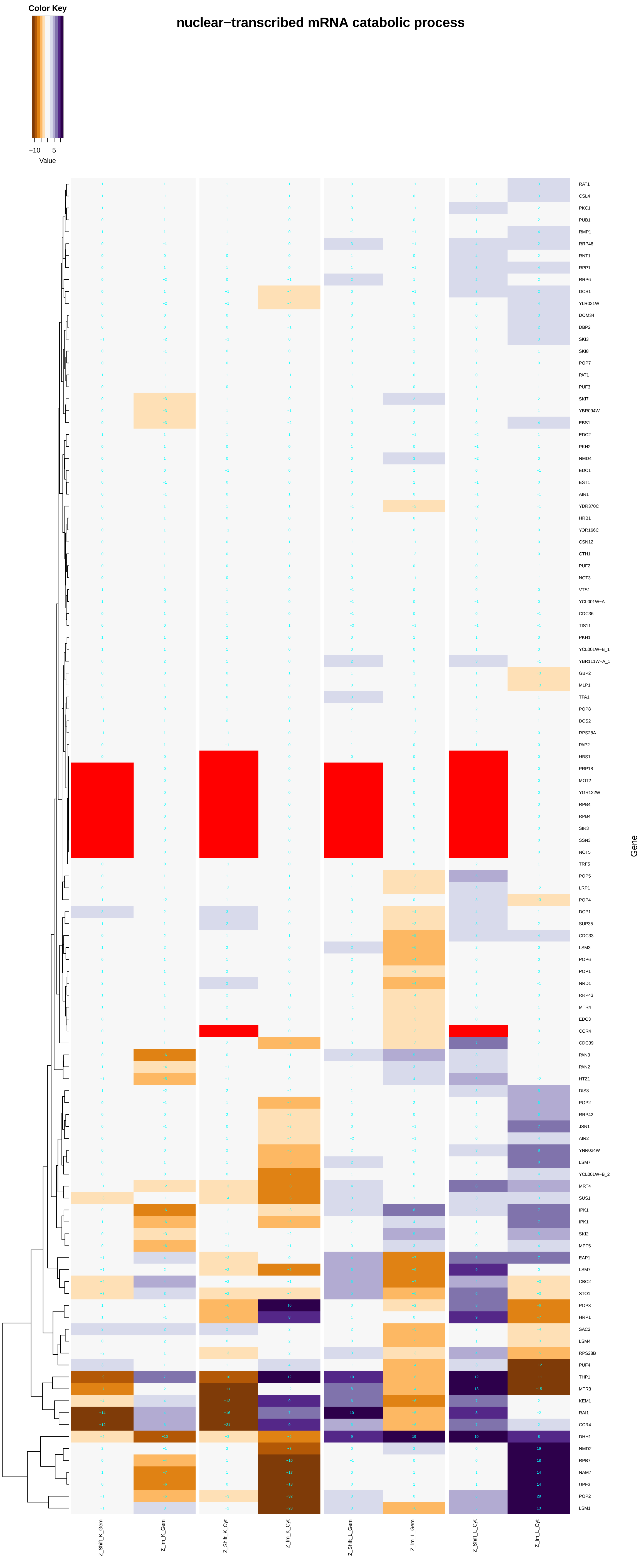
Color Key
−10 5
Value
nuclear−transcribed mRNA catabolic process
RAT1
CSL4
PKC1
PUB1
RMP1
RRP46
RNT1
RPP1
RRP6
DCS1
YLR021W
DOM34
DBP2
SKI3
SKI8
POP7
PAT1
PUF3
SKI7
YBR094W
EBS1
EDC2
PKH2
NMD4
EDC1
EST1
AIR1
YDR370C
HRB1
YOR166C
CSN12
CTH1
PUF2
NOT3
VTS1
YCL001W−A
CDC36
TIS11
PKH1
YCL001W−B_1
YBR111W−A_1
GBP2
MLP1
TPA1
POP8
DCS2
RPS28A
PAP2
HBS1
PRP18
MOT2
YGR122W
RPB4
RPB4
SIR3
SSN3
NOT5
TRF5
POP5
LRP1
POP4
DCP1
SUP35
CDC33
LSM3
POP6
POP1
NRD1
RRP43
MTR4
EDC3
CCR4
CDC39
PAN3
PAN2
HTZ1
DIS3
POP2
RRP42
JSN1
AIR2
YNR024W
LSM7
YCL001W−B_2
MRT4
SUS1
IPK1
IPK1
SKI2
MPT5
EAP1
LSM7
CBC2
STO1
POP3
HRP1
SAC3
LSM4
RPS28B
PUF4
THP1
MTR3
KEM1
RAI1
CCR4
DHH1
NMD2
RPB7
NAM7
UPF3
POP2
LSM1
Gene
Z_Shift_K_Gem
Z_Im_K_Gem
Z_Shift_K_Cyt
Z_Im_K_Cyt
Z_Shift_L_Gem
Z_Im_L_Gem
Z_Shift_L_Cyt
Z_Im_L_Cyt

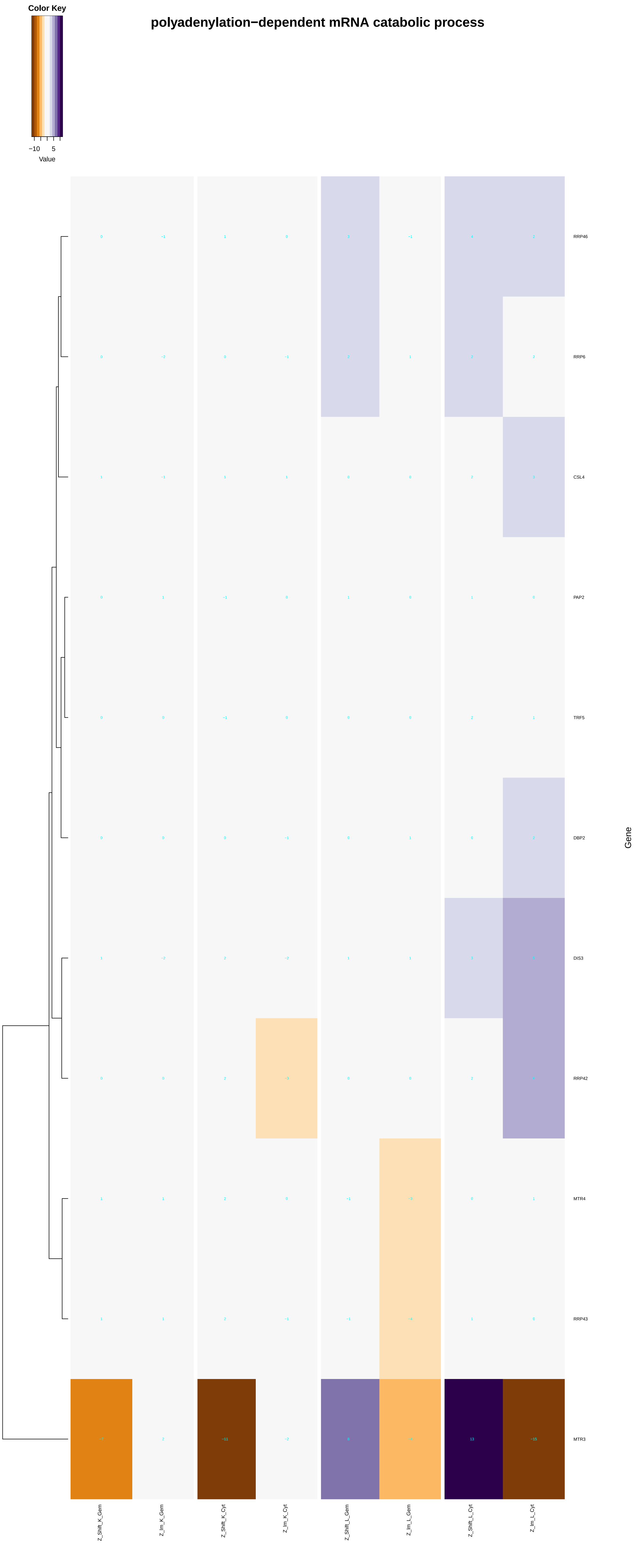
polyadenylation−dependent mRNA catabolic process
Color Key
−10
5
Value
RRP46
RRP6
CSL4
PAP2
TRF5
DBP2
DIS3
RRP42
MTR4
RRP43
MTR3
Gene
Z_Shift_K_Gem
Z_Im_K_Gem
Z_Shift_K_Cyt
Z_Im_K_Cyt
Z_Shift_L_Gem
Z_Im_L_Gem
Z_Shift_L_Cyt
Z_Im_L_Cyt

### histone mRNA catabolic process

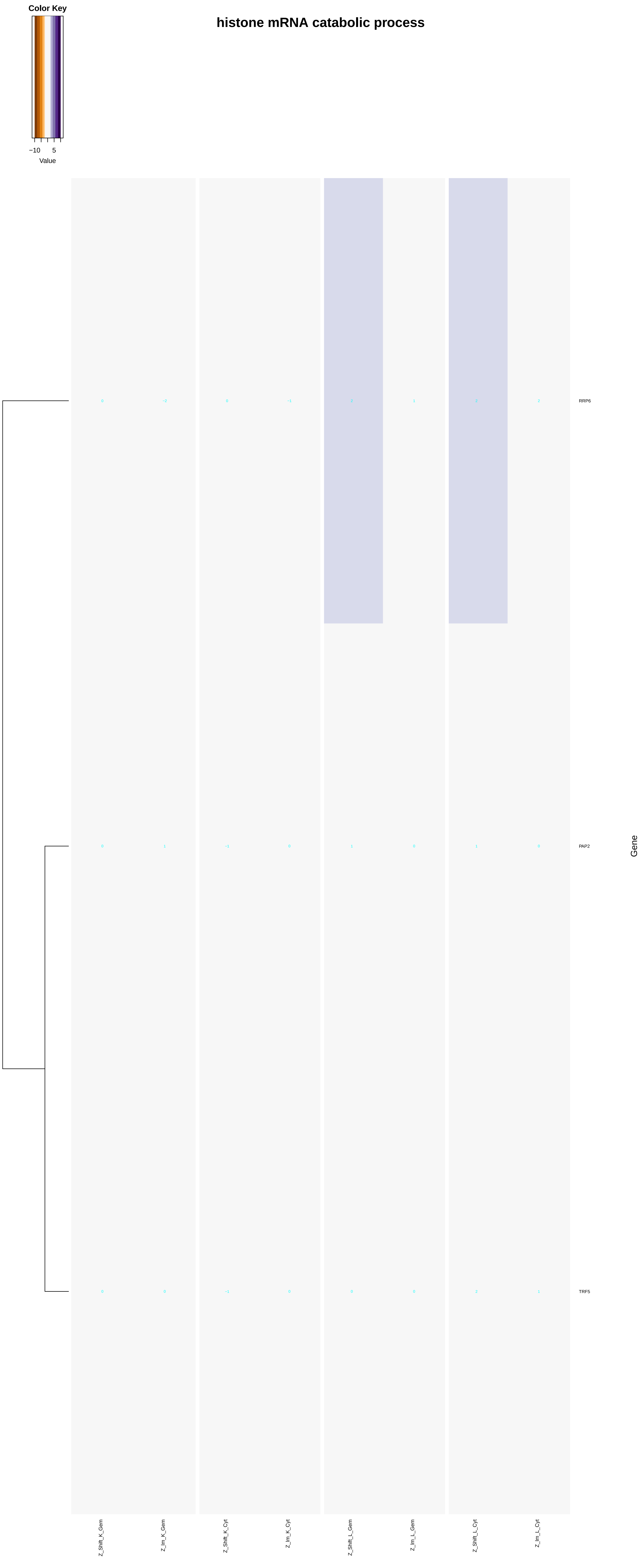

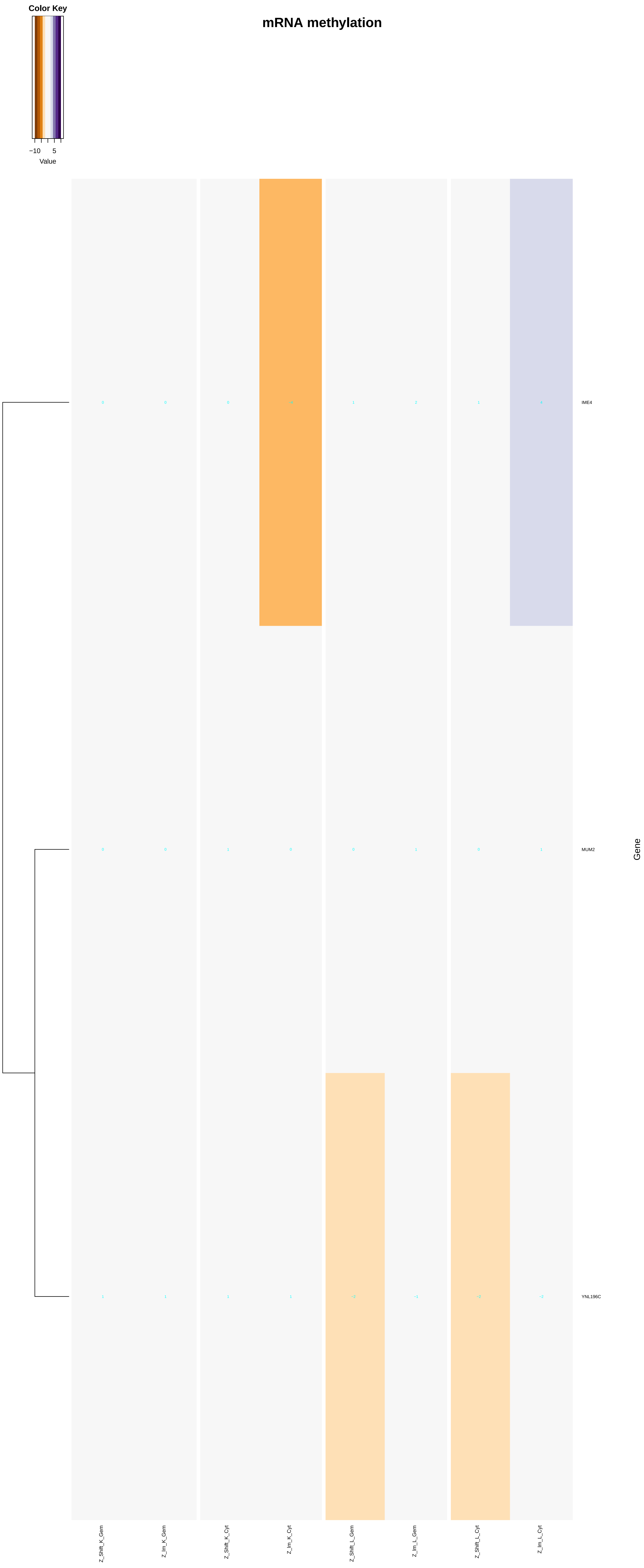

Color Key
−10
5
Value
mRNA methylation
IME4
MUM2
YNL196C
Gene
Z_Shift_K_Gem
Z_Im_K_Gem
Z_Shift_K_Cyt
Z_Im_K_Cyt
Z_Shift_L_Gem
Z_Im_L_Gem
Z_Shift_L_Cyt
Z_Im_L_Cyt

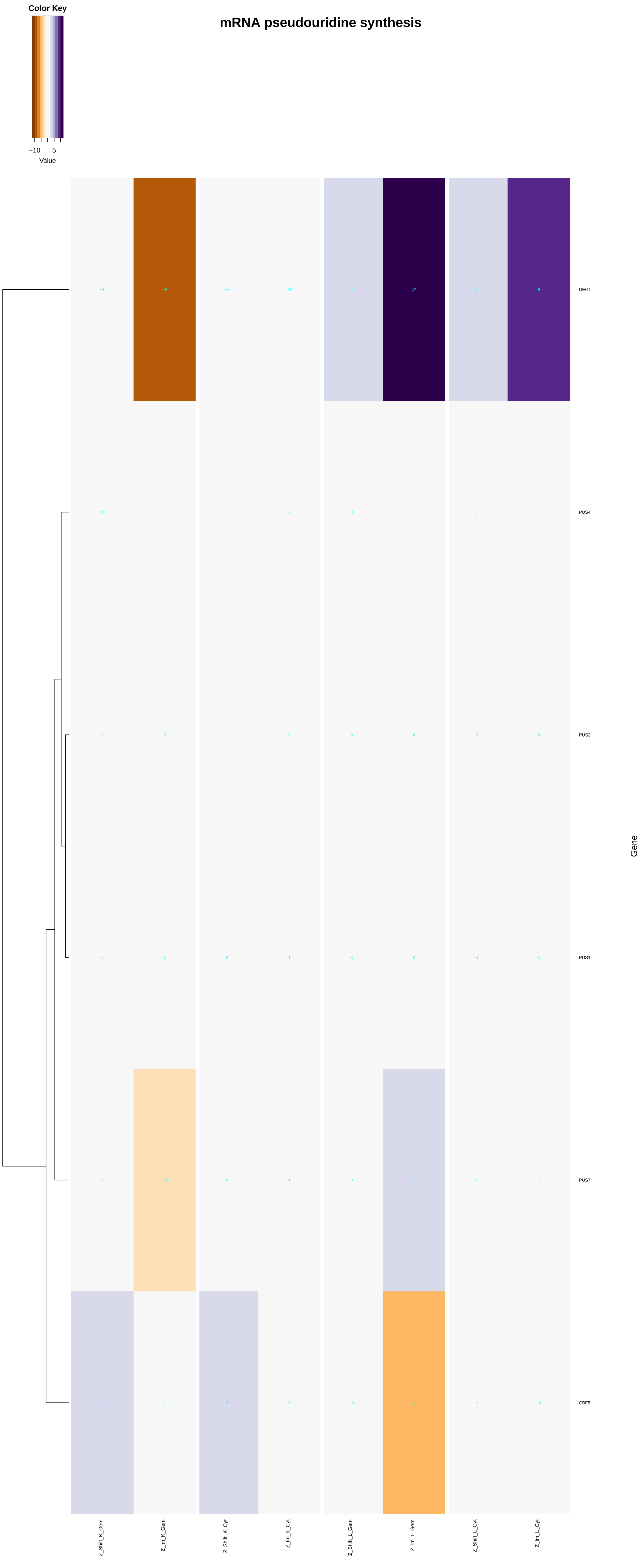

Color Key
−10
5
Value
mRNA pseudouridine synthesis
DEG1
PUS4
PUS2
PUS1
PUS7
CBF5
Gene
Z_Shift_K_Gem
Z_lm_K_Gem
Z_Shift_K_Cyt
Z_lm_K_Cyt
Z_Shift_L_Gem
Z_lm_L_Gem
Z_Shift_L_Cyt
Z_lm_L_Cyt

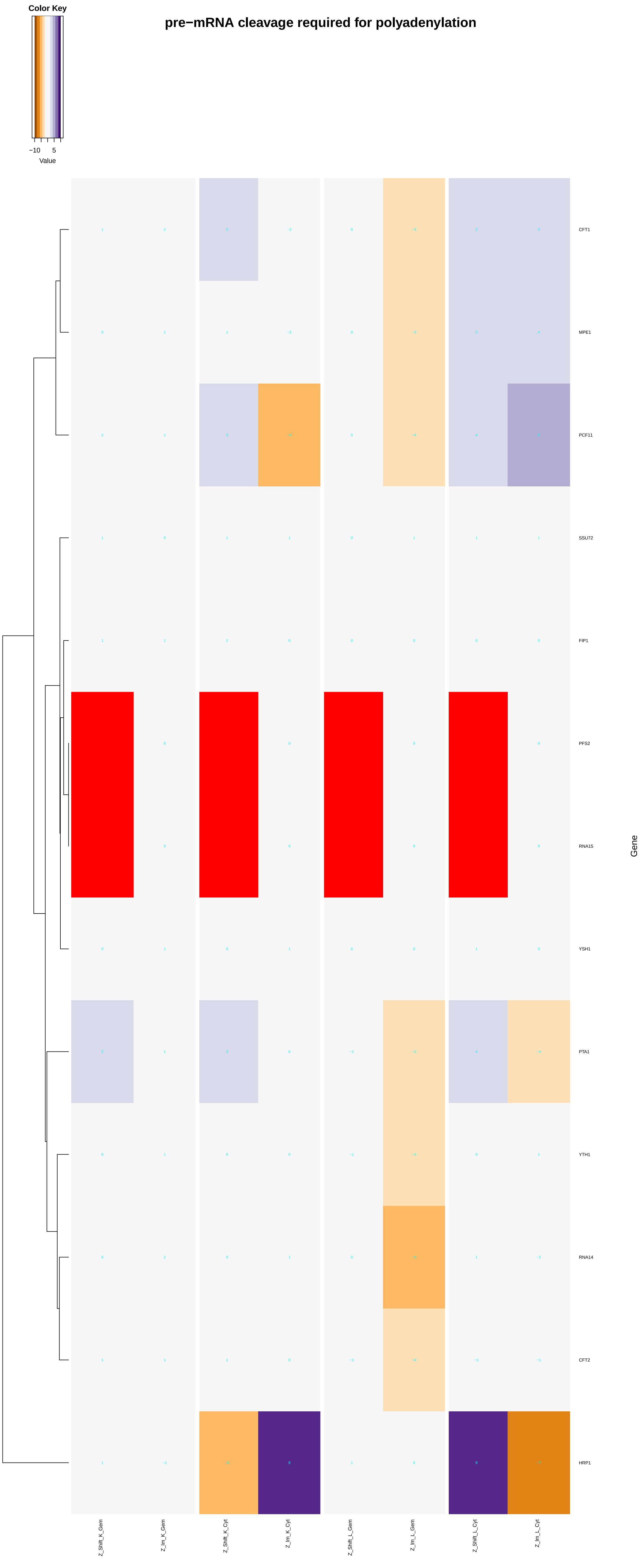
Color Key
−10 5
Value
pre−mRNA cleavage required for polyadenylation
CFT1
MPE1
PCF11
SSU72
FIP1
PFS2
RNA15
YSH1
PTA1
YTH1
RNA14
CFT2
HRP1
Gene
Z_Shift_K_Gem
Z_Im_K_Gem
Z_Shift_K_Cyt
Z_Im_K_Cyt
Z_Shift_L_Gem
Z_Im_L_Gem
Z_Shift_L_Cyt
Z_Im_L_Cyt

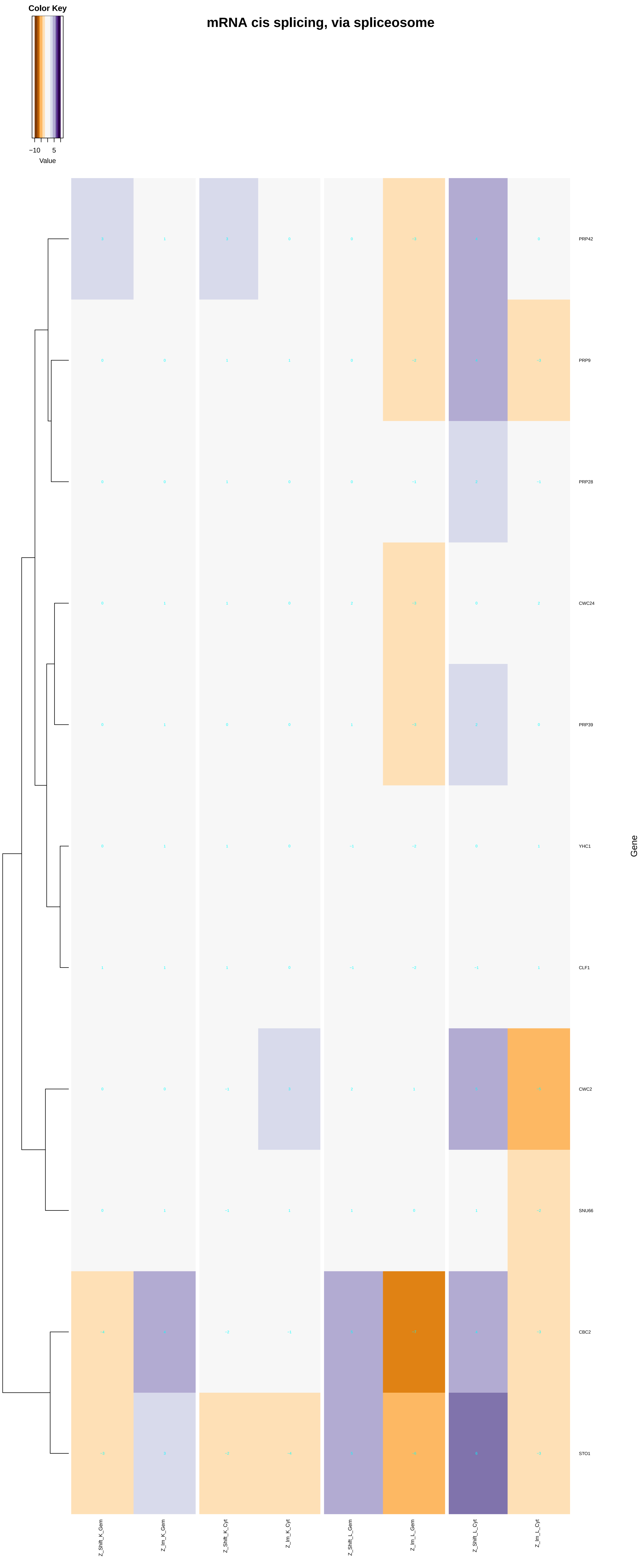
Color Key
−10
5
Value
mRNA cis splicing, via spliceosome
PRP42
PRP9
PRP28
CWC24
PRP39
YHC1
CLF1
CWC2
SNU66
CBC2
STO1
Gene
Z_Shift_K_Gem
Z_Im_K_Gem
Z_Shift_K_Cyt
Z_Im_K_Cyt
Z_Shift_L_Gem
Z_Im_L_Gem
Z_Shift_L_Cyt
Z_Im_L_Cyt

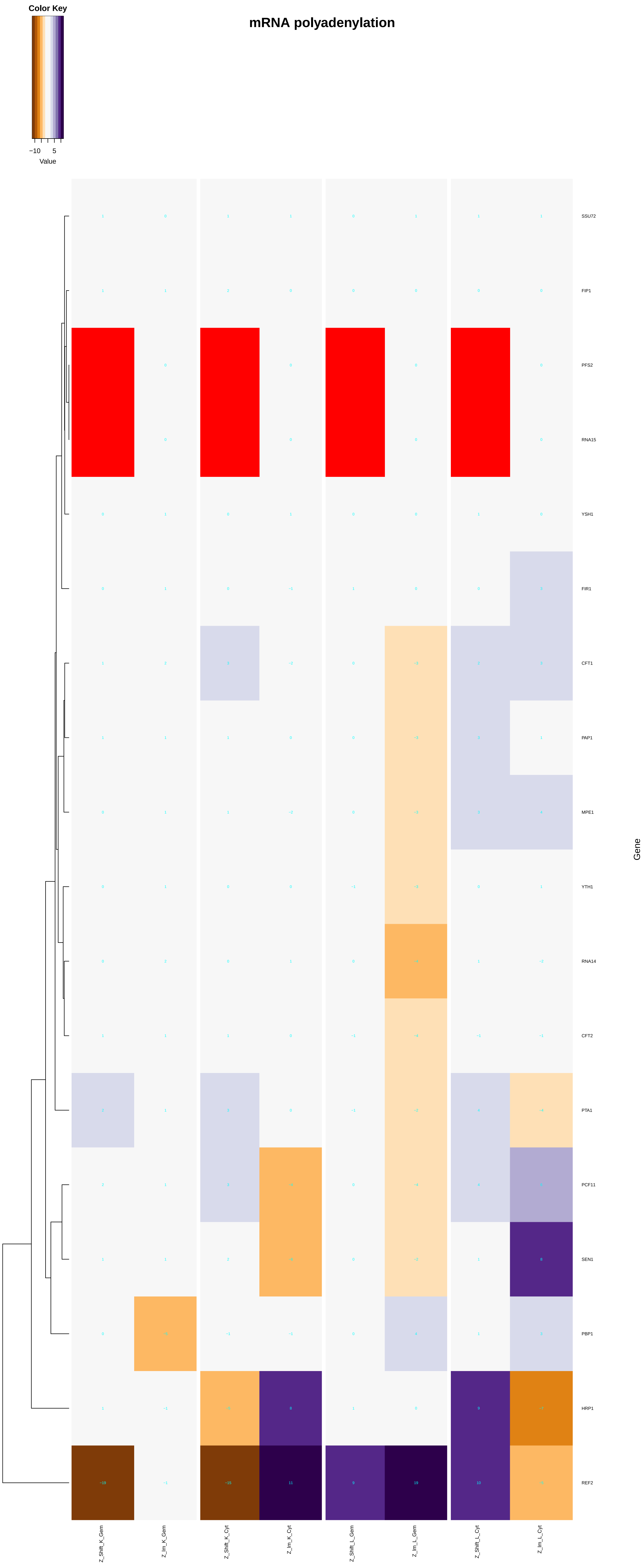

Color Key
−10
5
Value
mRNA polyadenylation
SSU72
FIP1
PFS2
RNA15
YSH1
FIR1
CFT1
PAP1
MPE1
YTH1
RNA14
CFT2
PTA1
PCF11
SEN1
PBP1
HRP1
REF2
Z_Shift_K_Gem
Z_Im_K_Gem
Z_Shift_K_Cyt
Z_Im_K_Cyt
Z_Shift_L_Gem
Z_Im_L_Gem
Z_Shift_L_Cyt
Z_Im_L_Cyt
Gene

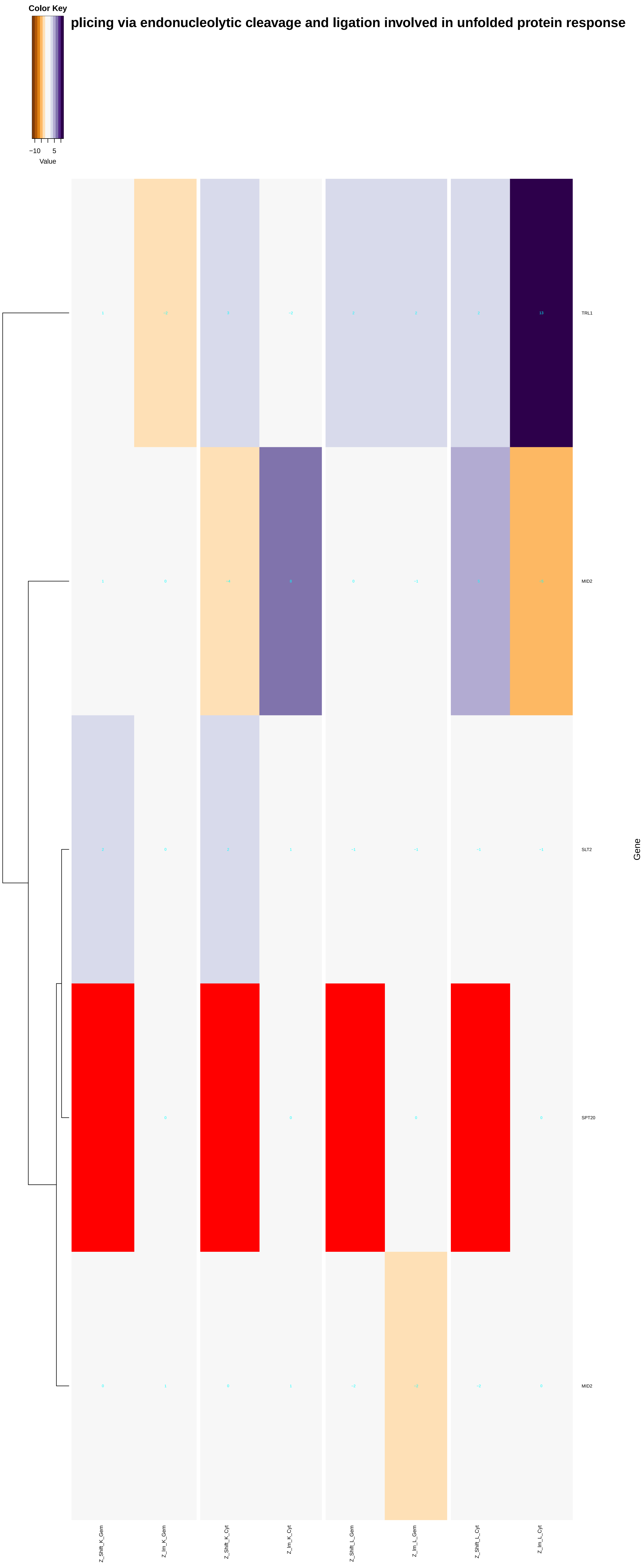
Color Key
−10
5
Value
plicing via endonucleolytic cleavage and ligation involved in unfolded protein response
TRL1
MID2
SLT2
SPT20
MID2
Gene
Z_Shift_K_Gem
Z_Im_K_Gem
Z_Shift_K_Cyt
Z_Im_K_Cyt
Z_Shift_L_Gem
Z_Im_L_Gem
Z_Shift_L_Cyt
Z_Im_L_Cyt

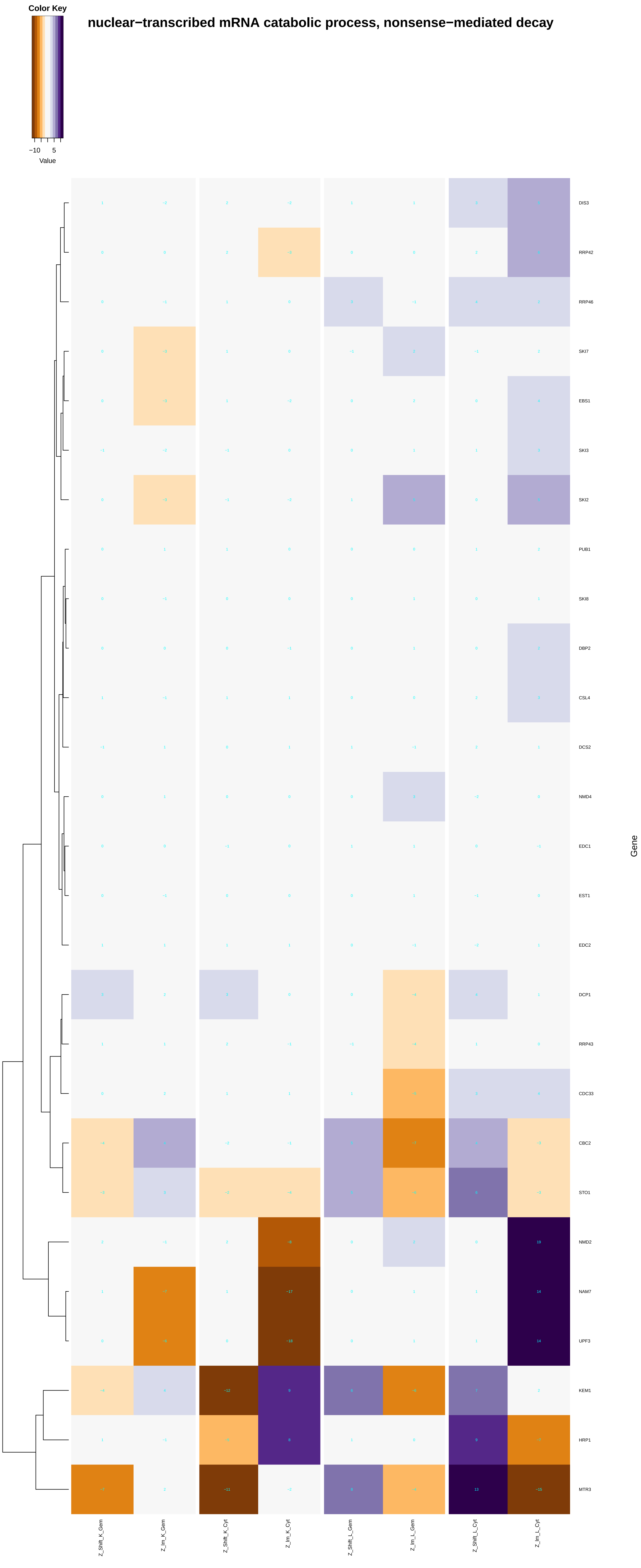
Color Key
−10
5
Value
nuclear−transcribed mRNA catabolic process, nonsense−mediated decay
DIS3
RRP42
RRP46
SKI7
EBS1
SKI3
SKI2
PUB1
SKI8
DBP2
CSL4
DCS2
NMD4
EDC1
EST1
EDC2
DCP1
RRP43
CDC33
CBC2
STO1
NMD2
NAM7
UPF3
KEM1
HRP1
MTR3
Gene
Z_Shift_K_Gem
Z_lm_K_Gem
Z_Shift_K_Cyt
Z_lm_K_Cyt
Z_Shift_L_Gem
Z_lm_L_Gem
Z_Shift_L_Cyt
Z_lm_L_Cyt

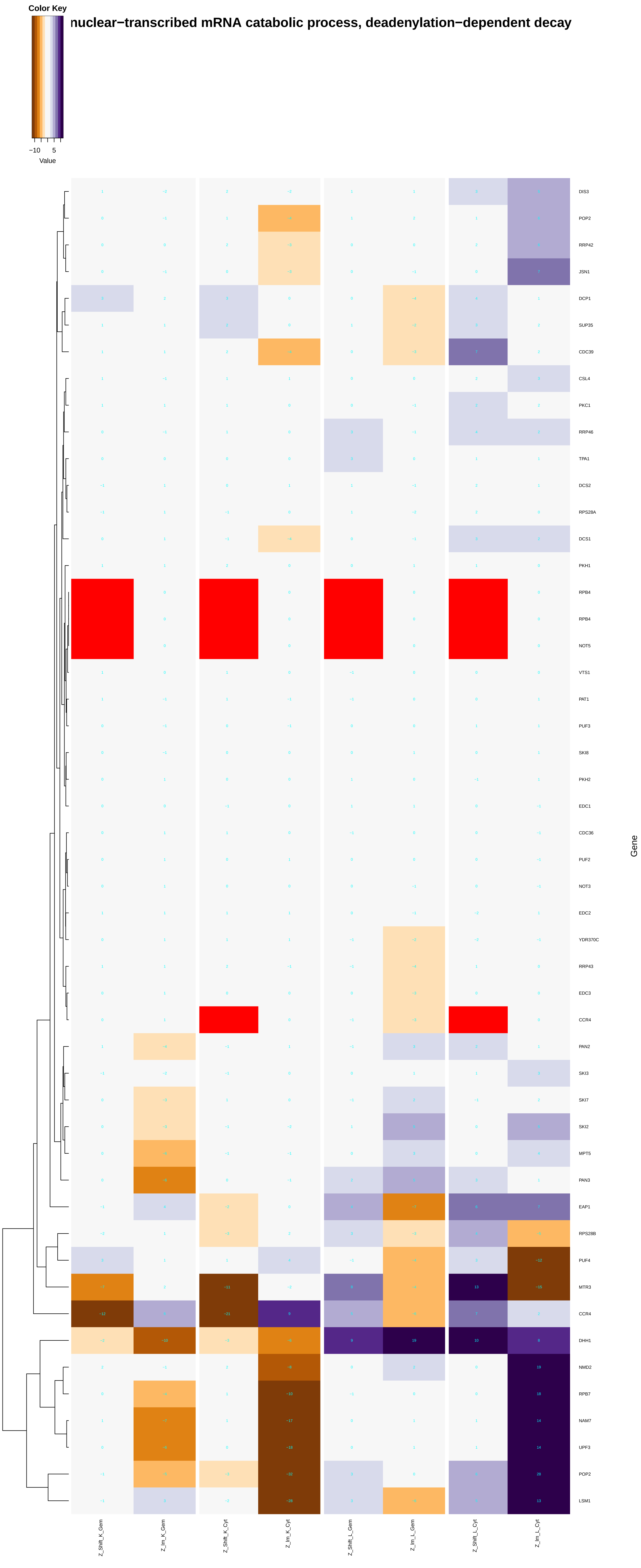

Color Key
−10
5
Value
nuclear−transcribed mRNA catabolic process, deadenylation−dependent decay
Gene
DIS3
POP2
RRP42
JSN1
DCP1
SUP35
CDC39
CSL4
PKC1
RRP46
TPA1
DCS2
RPS28A
DCS1
PKH1
RPB4
RPB4
NOT5
VTS1
PAT1
PUF3
SKI8
PKH2
EDC1
CDC36
PUF2
NOT3
EDC2
YDR370C
RRP43
EDC3
CCR4
PAN2
SKI3
SKI7
SKI2
MPT5
PAN3
EAP1
RPS28B
PUF4
MTR3
CCR4
DHH1
NMD2
RPB7
NAM7
UPF3
POP2
LSM1
Z_Shift_K_Gem
Z_Im_K_Gem
Z_Shift_K_Cyt
Z_Im_K_Cyt
Z_Shift_L_Gem
Z_Im_L_Gem
Z_Shift_L_Cyt
Z_Im_L_Cyt

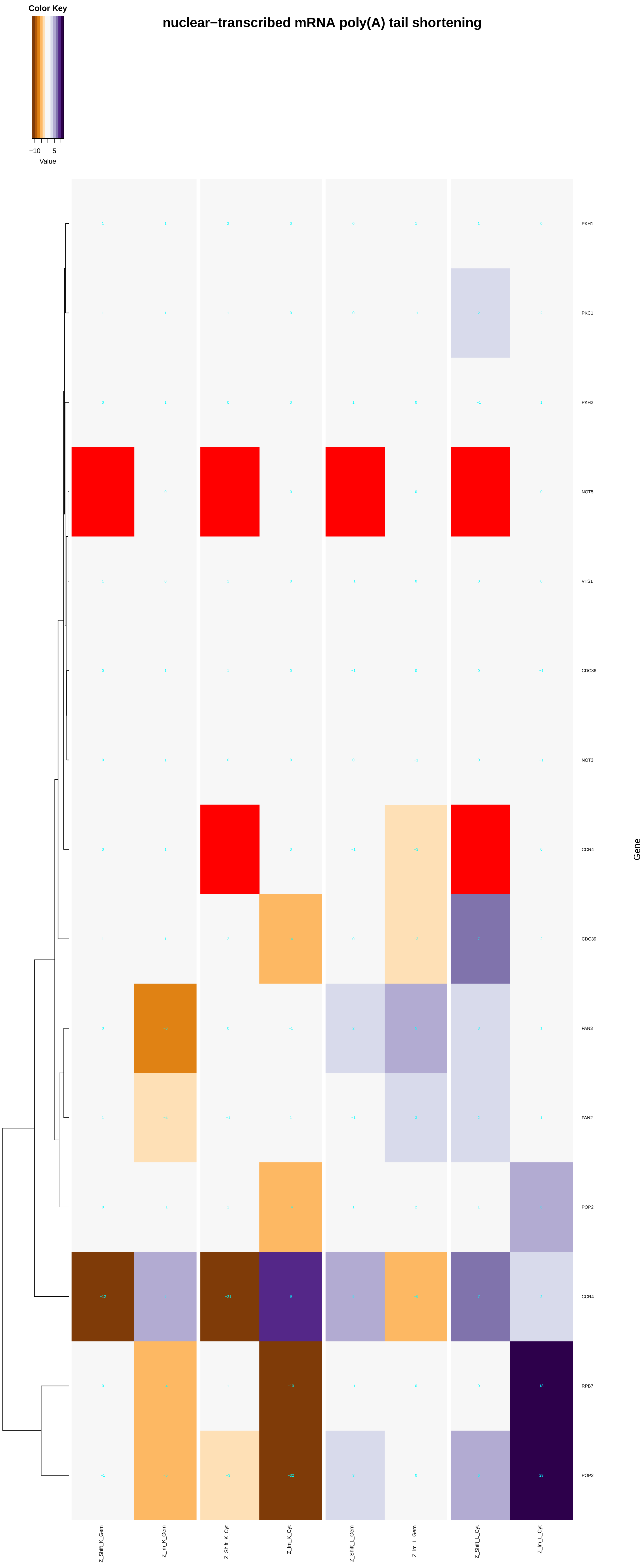

Color Key
−10 5
Value
nuclear−transcribed mRNA poly(A) tail shortening
PKH1
PKC1
PKH2
NOT5
VTS1
CDC36
NOT3
CCR4
CDC39
PAN3
PAN2
POP2
CCR4
RPB7
POP2
Gene
Z_Shift_K_Gem
Z_Im_K_Gem
Z_Shift_K_Cyt
Z_Im_K_Cyt
Z_Shift_L_Gem
Z_Im_L_Gem
Z_Shift_L_Cyt
Z_Im_L_Cyt

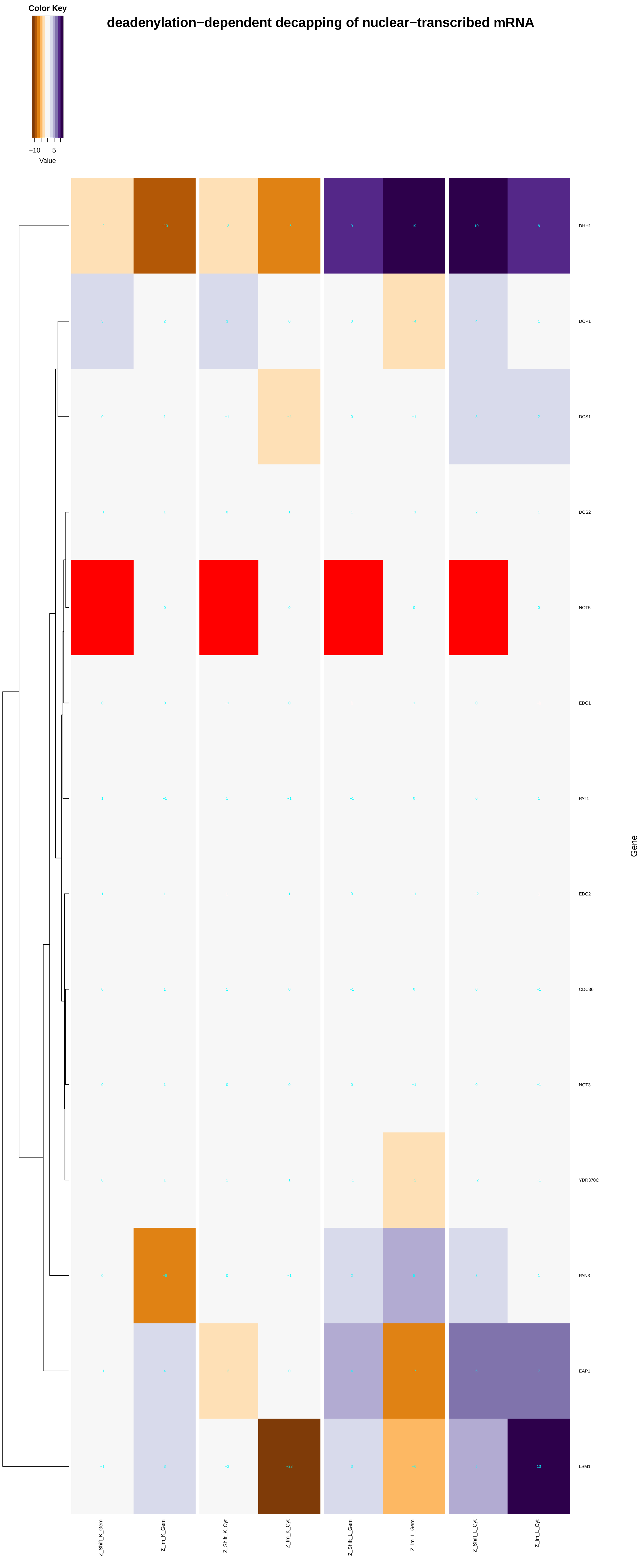
Color Key
−10
5
Value
deadenylation−dependent decapping of nuclear−transcribed mRNA
DHH1
DCP1
DCS1
DCS2
NOT5
EDC1
PAT1
EDC2
CDC36
NOT3
YDR370C
PAN3
EAP1
LSM1
Gene
Z_Shift_K_Gem
Z_Im_K_Gem
Z_Shift_K_Cyt
Z_Im_K_Cyt
Z_Shift_L_Gem
Z_Im_L_Gem
Z_Shift_L_Cyt
Z_Im_L_Cyt

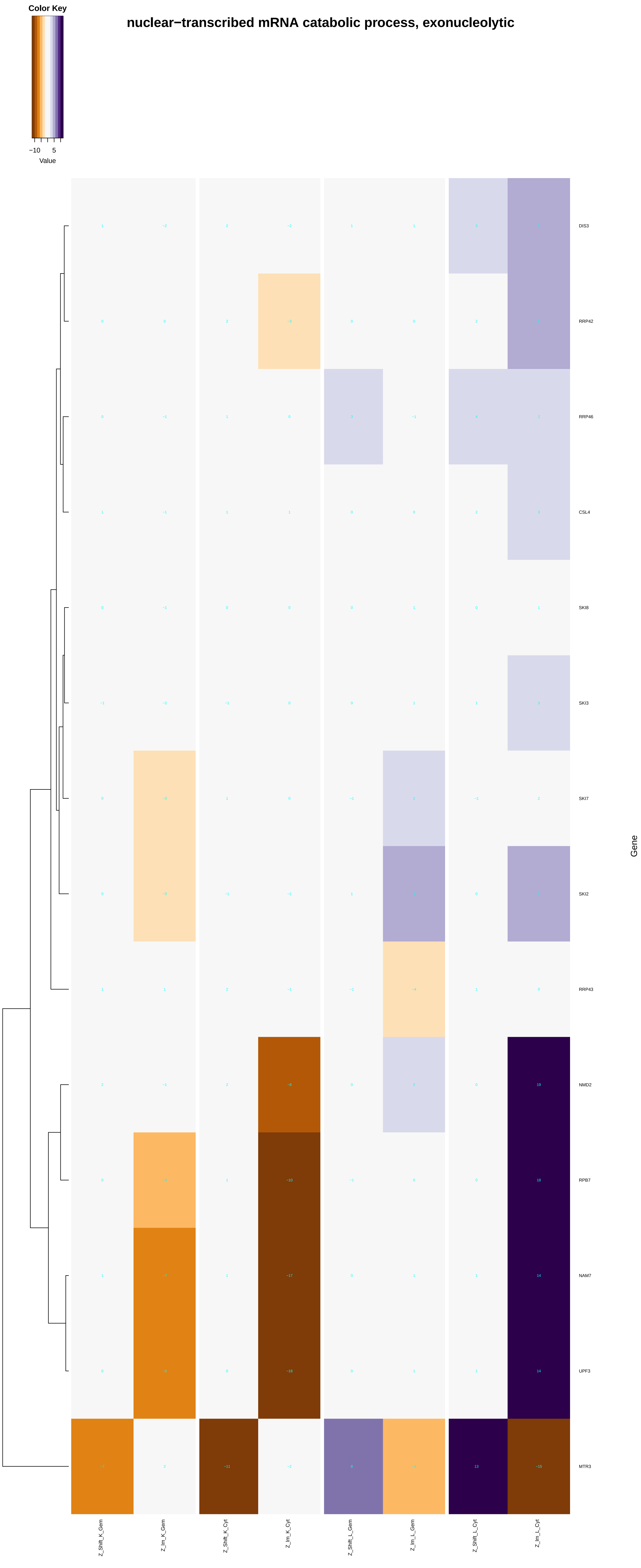
Color Key
−10
5
Value
nuclear−transcribed mRNA catabolic process, exonucleolytic
DIS3
RRP42
RRP46
CSL4
SKI8
SKI3
SKI7
SKI2
RRP43
NMD2
RPB7
NAM7
UPF3
MTR3
Gene
Z_Shift_K_Gem
Z_lm_K_Gem
Z_Shift_K_Cyt
Z_lm_K_Cyt
Z_Shift_L_Gem
Z_lm_L_Gem
Z_Shift_L_Cyt
Z_lm_L_Cyt

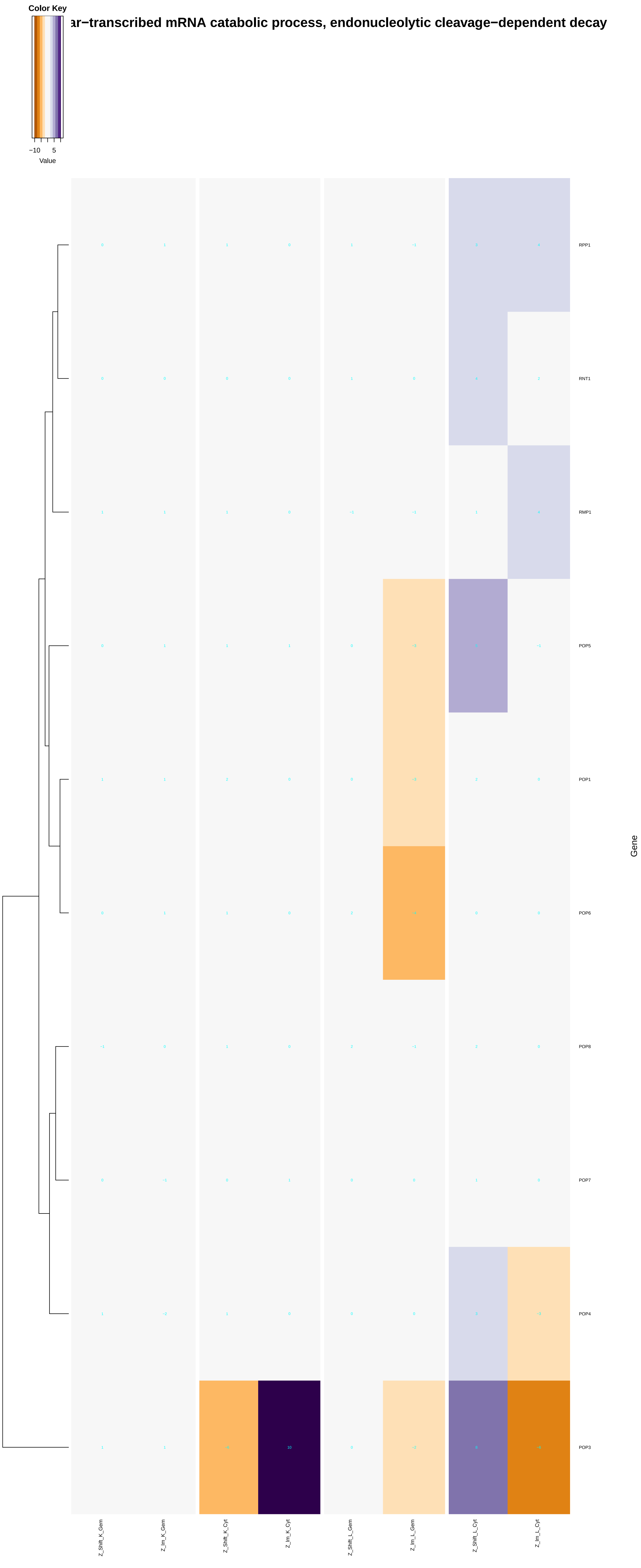
ar−transcribed mRNA catabolic process, endonucleolytic cleavage−dependent decay
Color Key
−10
5
Value
RPP1
RNT1
RMP1
POP5
POP1
POP6
POP8
POP7
POP4
POP3
Gene
Z_Shift_K_Gem
Z_Im_K_Gem
Z_Shift_K_Cyt
Z_Im_K_Cyt
Z_Shift_L_Gem
Z_Im_L_Gem
Z_Shift_L_Cyt
Z_Im_L_Cyt

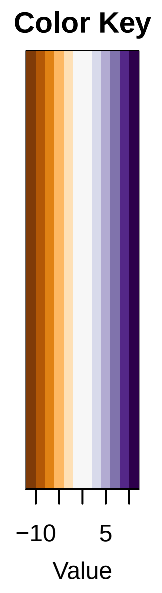


**uclear−transcribed mRNA catabolic process, deadenylation−independent decay**

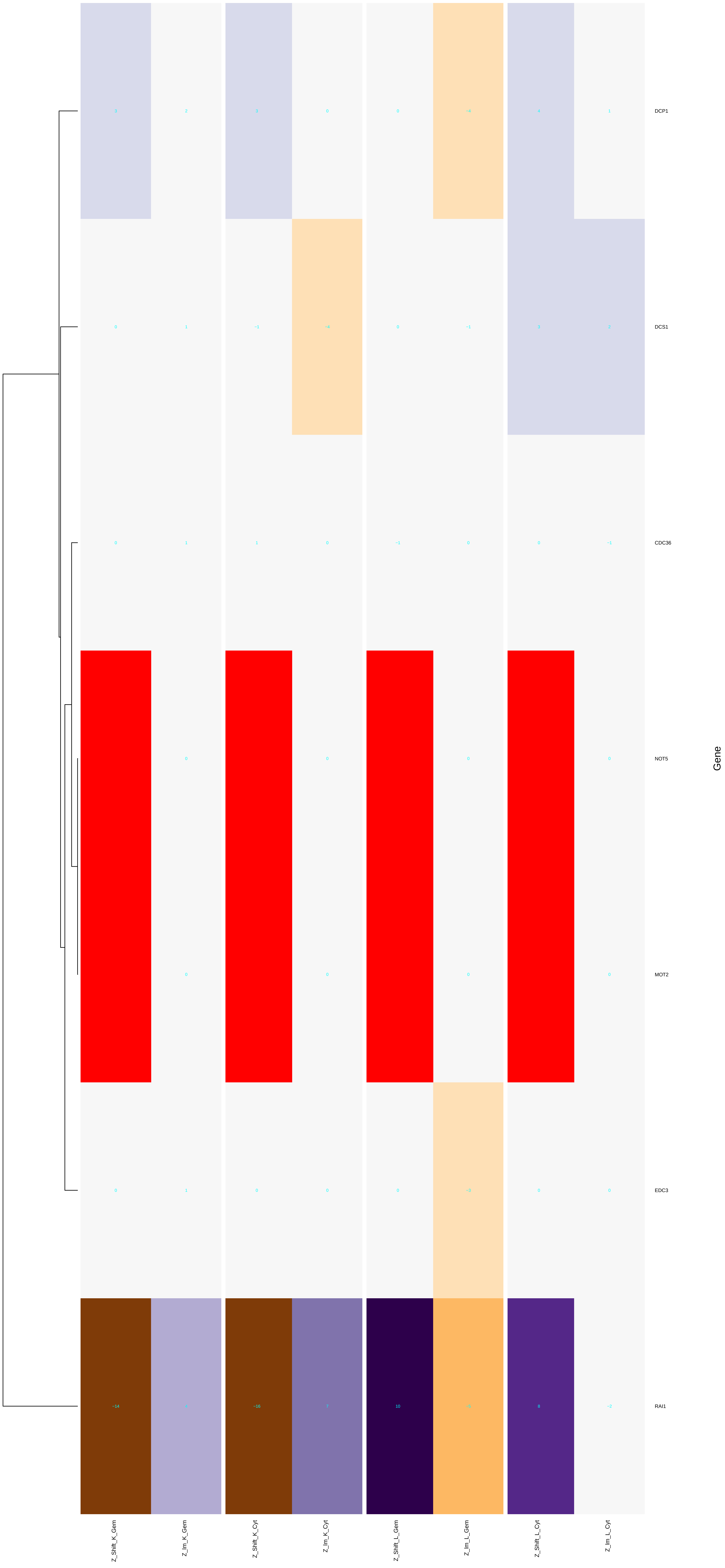

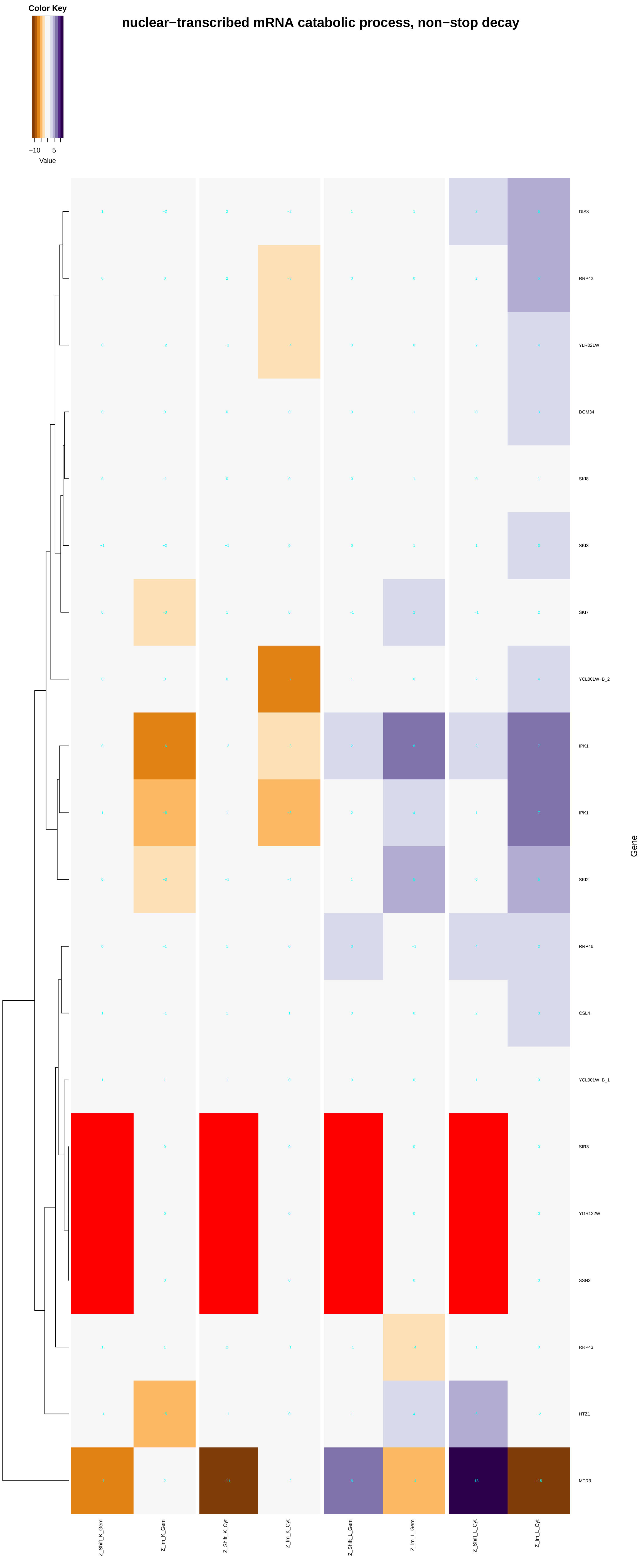

Color Key
−10 5
Value
nuclear−transcribed mRNA catabolic process, non−stop decay
DIS3
RRP42
YLR021W
DOM34
SKI8
SKI3
SKI7
YCL001W−B_2
IPK1
IPK1
SKI2
RRP46
CSL4
YCL001W−B_1
SIR3
YGR122W
SSN3
RRP43
HTZ1
MTR3
Gene
Z_Shift_K_Gem
Z_Im_K_Gem
Z_Shift_K_Cyt
Z_Im_K_Cyt
Z_Shift_L_Gem
Z_Im_L_Gem
Z_Shift_L_Cyt
Z_Im_L_Cyt

### nuclear−transcribed mRNA catabolic process, no−go decay

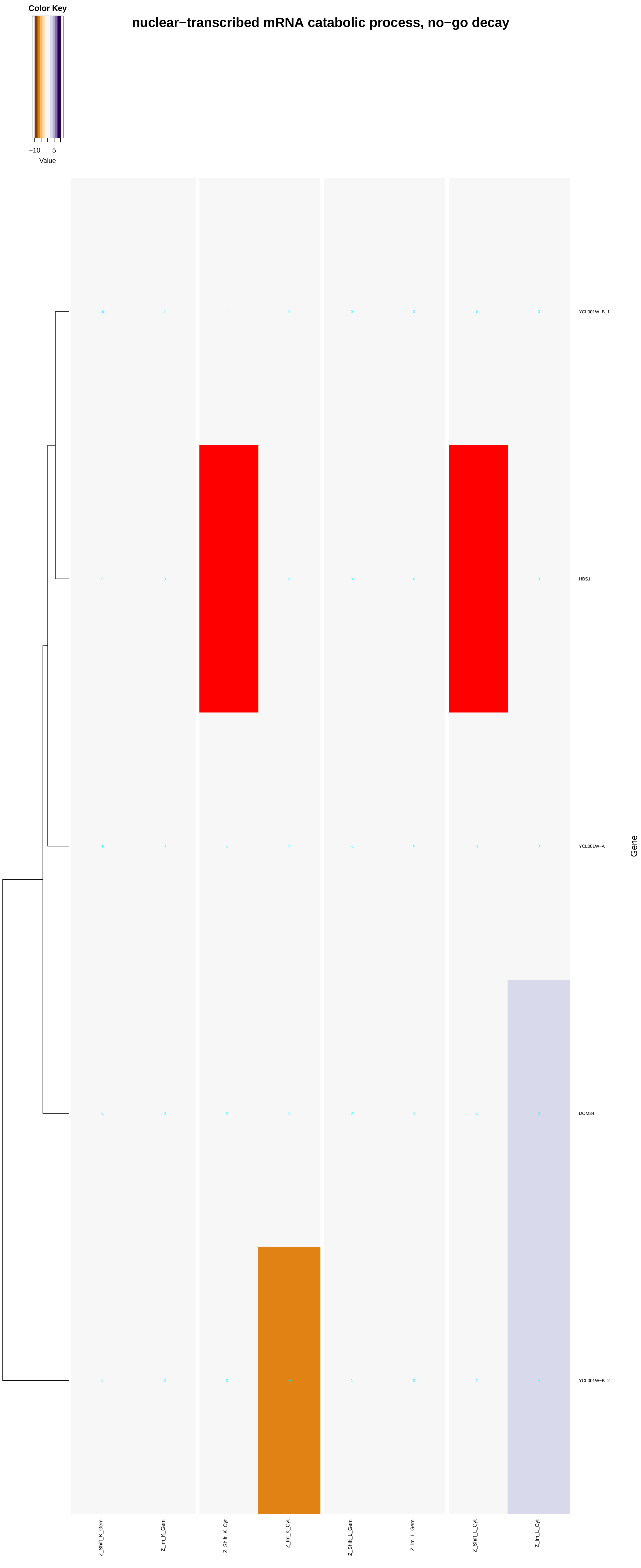

### nuclear mRNA surveillance

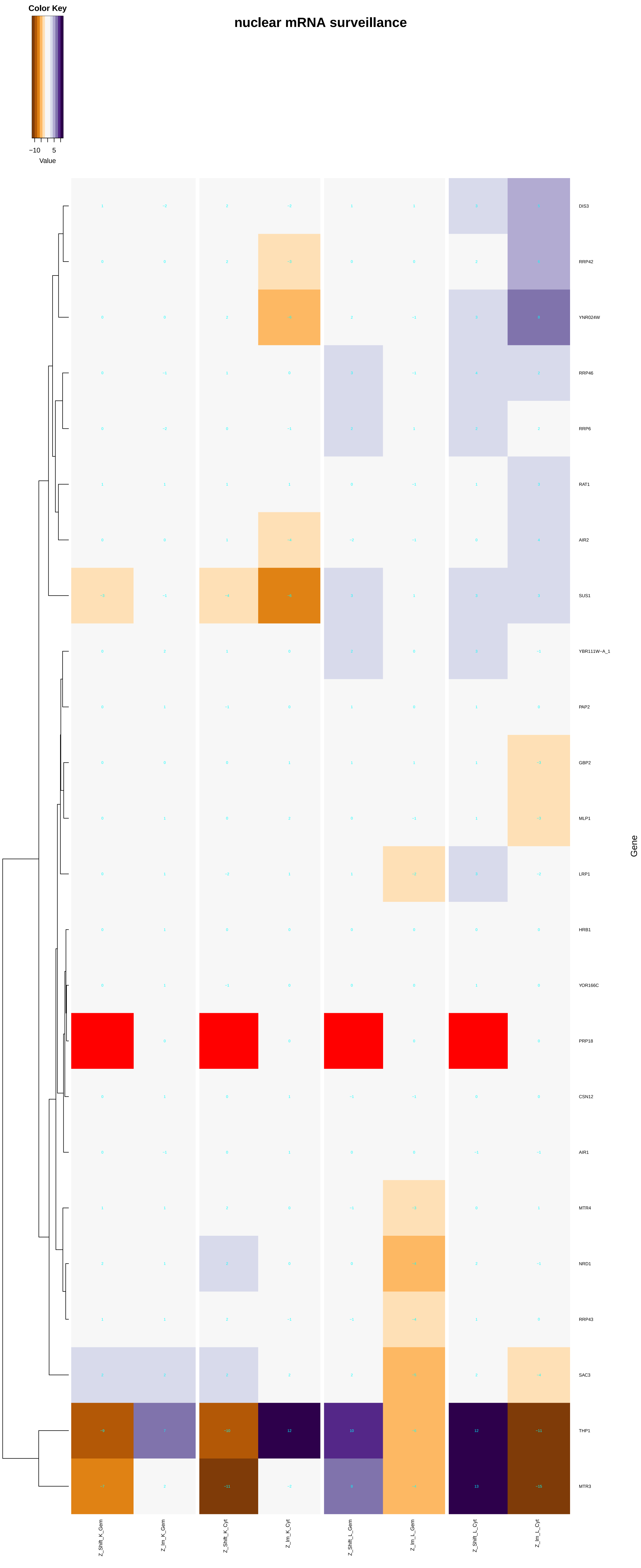

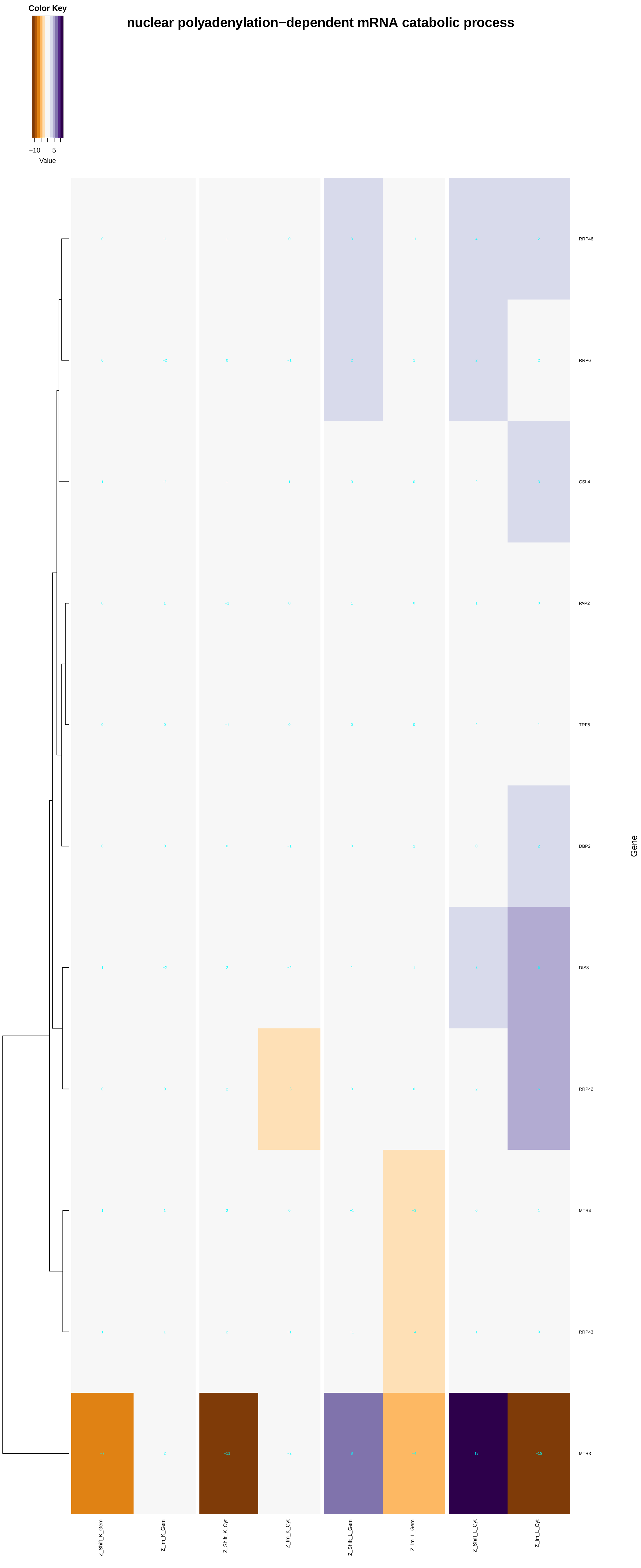
nuclear polyadenylation−dependent mRNA catabolic process
Color Key
−10
5
Value
RRP46
RRP6
CSL4
PAP2
TRF5
DBP2
DIS3
RRP42
MTR4
RRP43
MTR3
Gene
Z_Shift_K_Gem
Z_Im_K_Gem
Z_Shift_K_Cyt
Z_Im_K_Cyt
Z_Shift_L_Gem
Z_Im_L_Gem
Z_Shift_L_Cyt
Z_Im_L_Cyt

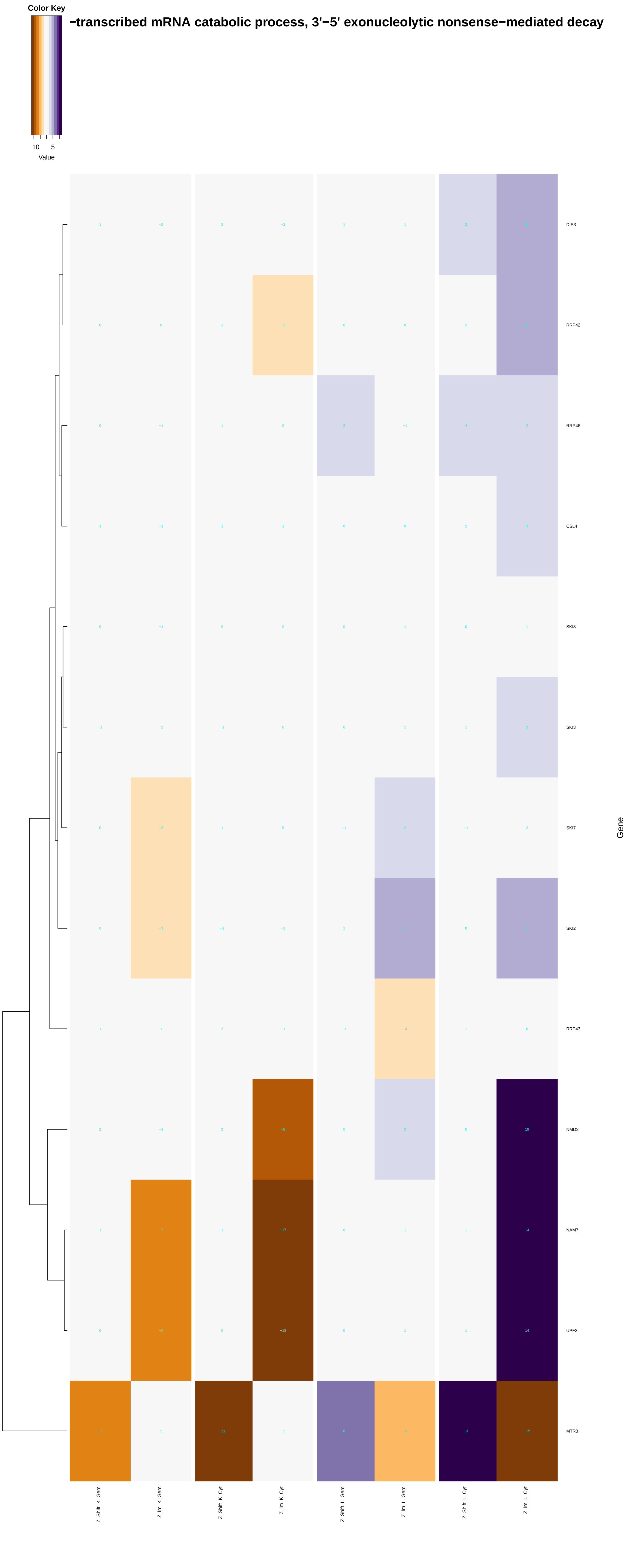

Color Key
−10 5
Value
−transcribed mRNA catabolic process, 3'−5' exonucleolytic nonsense−mediated decay
DIS3
RRP42
RRP46
CSL4
SKI8
SKI3
SKI7
SKI2
RRP43
NMD2
NAM7
UPF3
MTR3
Gene
Z_Shift_K_Gem
Z_Im_K_Gem
Z_Shift_K_Cyt
Z_Im_K_Cyt
Z_Shift_L_Gem
Z_Im_L_Gem
Z_Shift_L_Cyt
Z_Im_L_Cyt

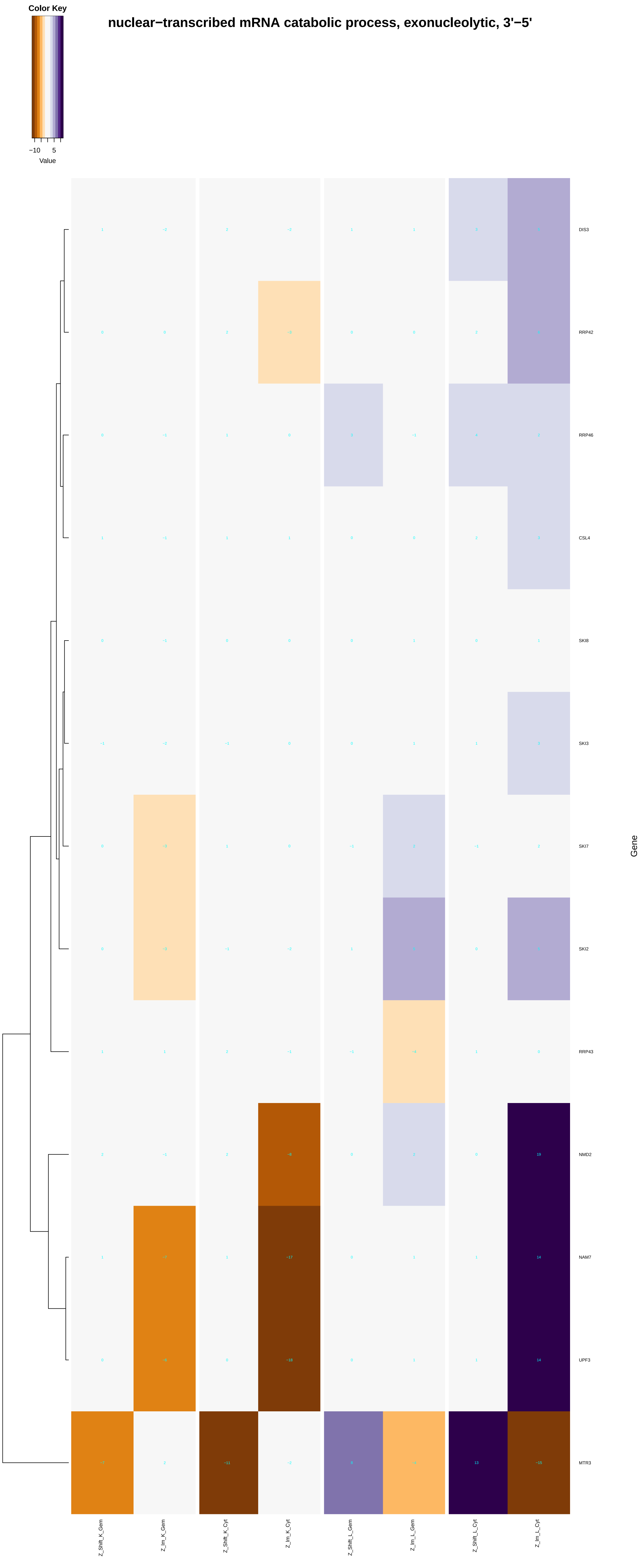

Color Key
−10
5
Value
nuclear−transcribed mRNA catabolic process, exonucleolytic, 3'−5'
DIS3
RRP42
RRP46
CSL4
SKI8
SKI3
SKI7
SKI2
RRP43
NMD2
NAM7
UPF3
MTR3
Gene
Z_Shift_K_Gem
Z_lm_K_Gem
Z_Shift_K_Cyt
Z_lm_K_Cyt
Z_Shift_L_Gem
Z_lm_L_Gem
Z_Shift_L_Cyt
Z_lm_L_Cyt

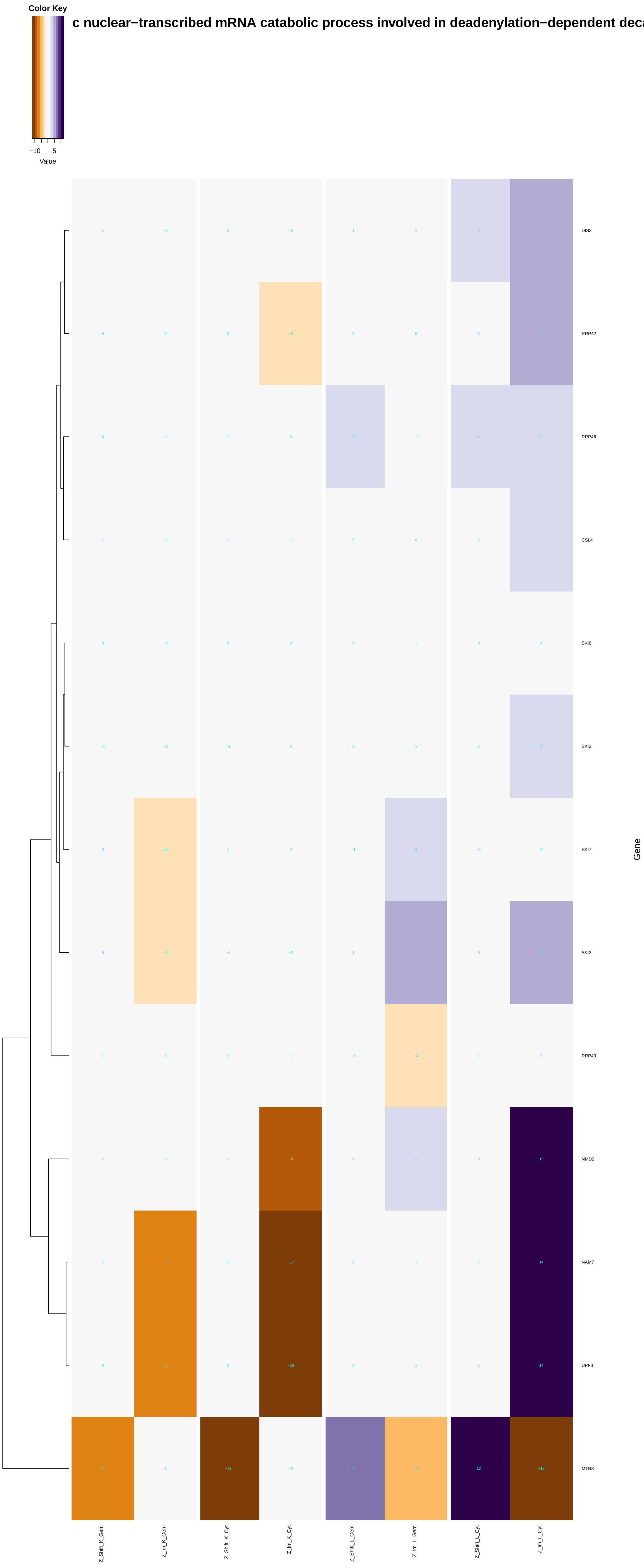
Color Key
−10
5
Value
c nuclear−transcribed mRNA catabolic process involved in deadenylation−dependent deca
DIS3
RRP42
RRP46
CSL4
SKI8
SKI3
SKI7
SKI2
RRP43
NMD2
NAM7
UPF3
MTR3
Gene
Z_Shift_K_Gem
Z_Im_K_Gem
Z_Shift_K_Cyt
Z_Im_K_Cyt
Z_Shift_L_Gem
Z_Im_L_Gem
Z_Shift_L_Cyt
Z_Im_L_Cyt

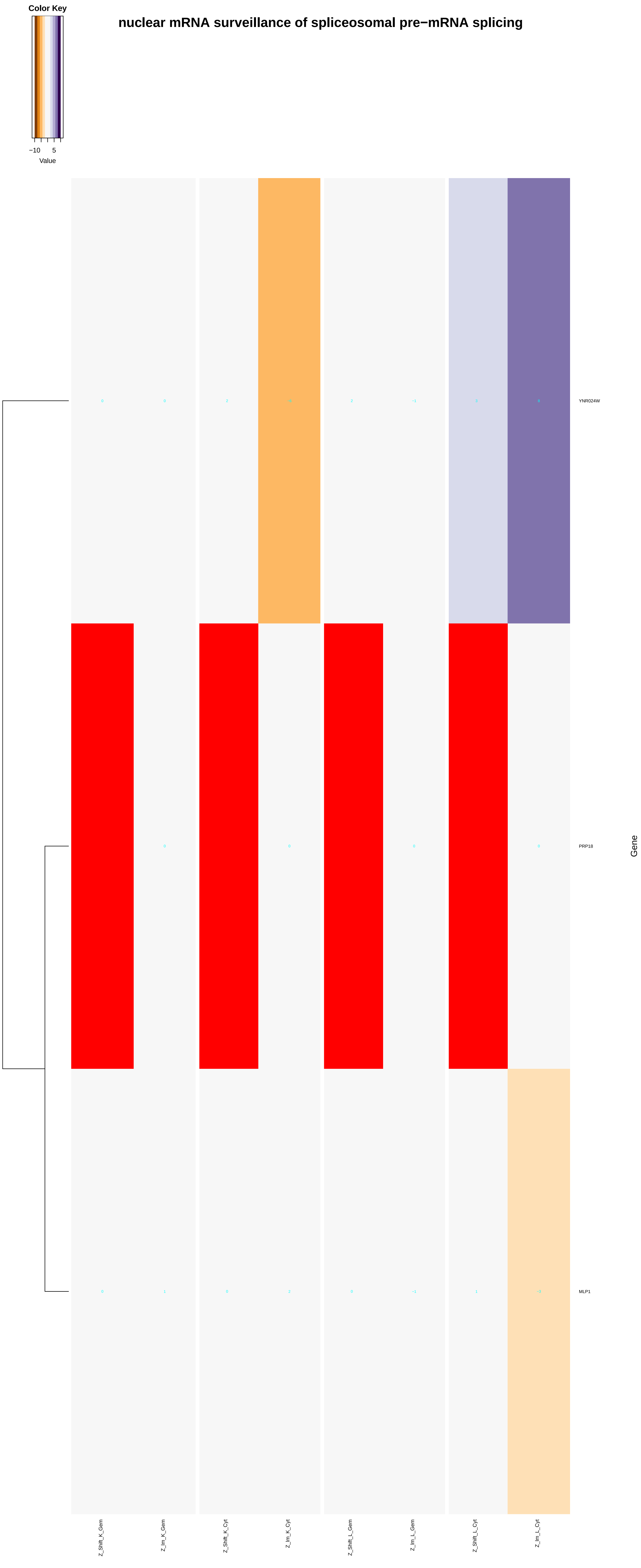
Color Key
−10
5
Value
nuclear mRNA surveillance of spliceosomal pre−mRNA splicing
YNR024W
PRP18
MLP1
Gene
Z_Shift_K_Gem
Z_Im_K_Gem
Z_Shift_K_Cyt
Z_Im_K_Cyt
Z_Shift_L_Gem
Z_Im_L_Gem
Z_Shift_L_Cyt
Z_Im_L_Cyt

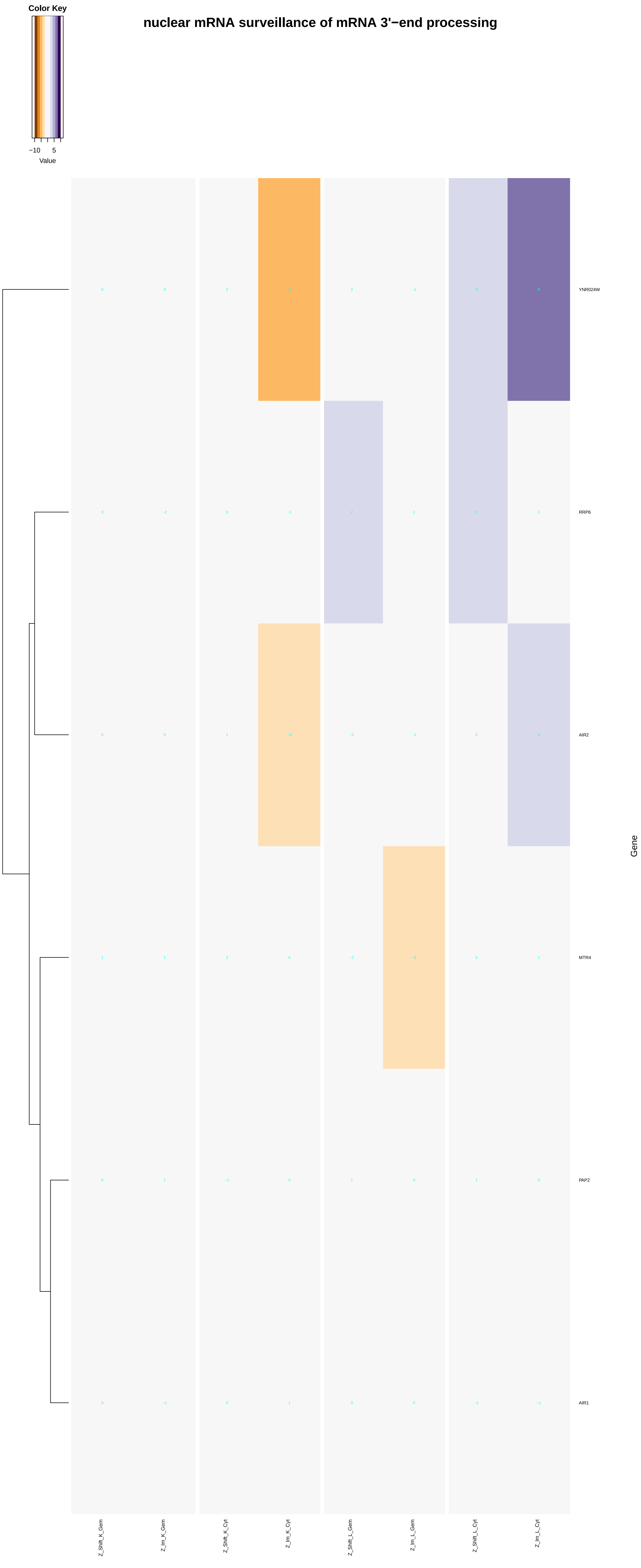
nuclear mRNA surveillance of mRNA 3'−end processing
Color Key
−10
5
Value
YNR024W
RRP6
AIR2
MTR4
PAP2
AIR1
Gene
Z_Shift_K_Gem
Z_Im_K_Gem
Z_Shift_K_Cyt
Z_Im_K_Cyt
Z_Shift_L_Gem
Z_Im_L_Gem
Z_Shift_L_Cyt
Z_Im_L_Cyt
