## Additional File 7 for "A humanized yeast phenomic model of deoxycytidine kinase to predict genetic buffering of nucleoside analog cytotoxicity": ncRNA_3'-end_processing.pdf

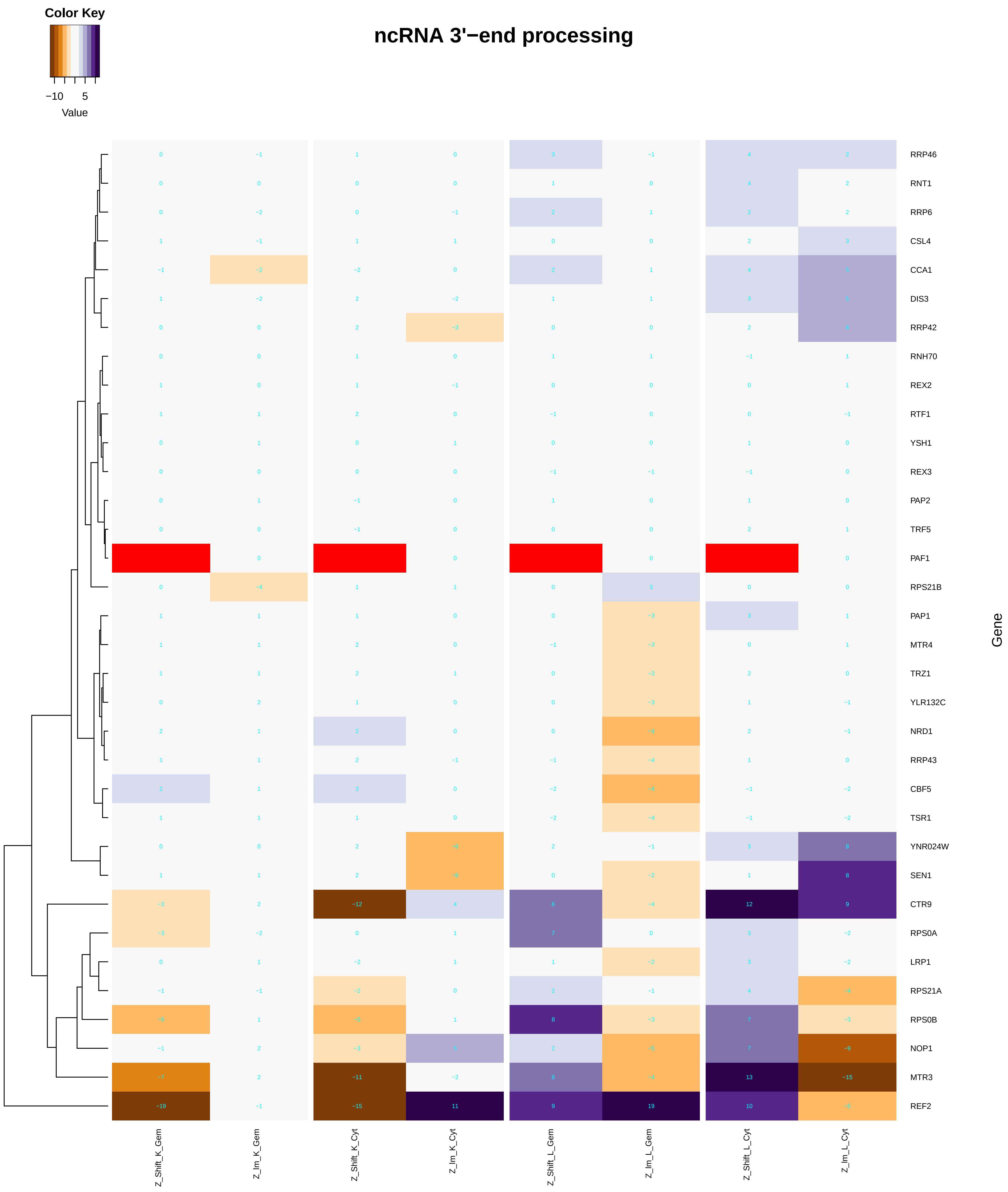

Color Key
−10
5
Value
ncRNA 3'−end processing
Gene
RRP46
RNT1
RRP6
CSL4
CCA1
DIS3
RRP42
RNH70
REX2
RTF1
YSH1
REX3
PAP2
TRF5
PAF1
RPS21B
PAP1
MTR4
TRZ1
YLR132C
NRD1
RRP43
CBF5
TSR1
YNR024W
SEN1
CTR9
RPS0A
LRP1
RPS21A
RPS0B
NOP1
MTR3
REF2
Z_Shift_K_Gem
Z_Im_K_Gem
Z_Shift_K_Cyt
Z_Im_K_Cyt
Z_Shift_L_Gem
Z_Im_L_Gem
Z_Shift_L_Cyt
Z_Im_L_Cyt

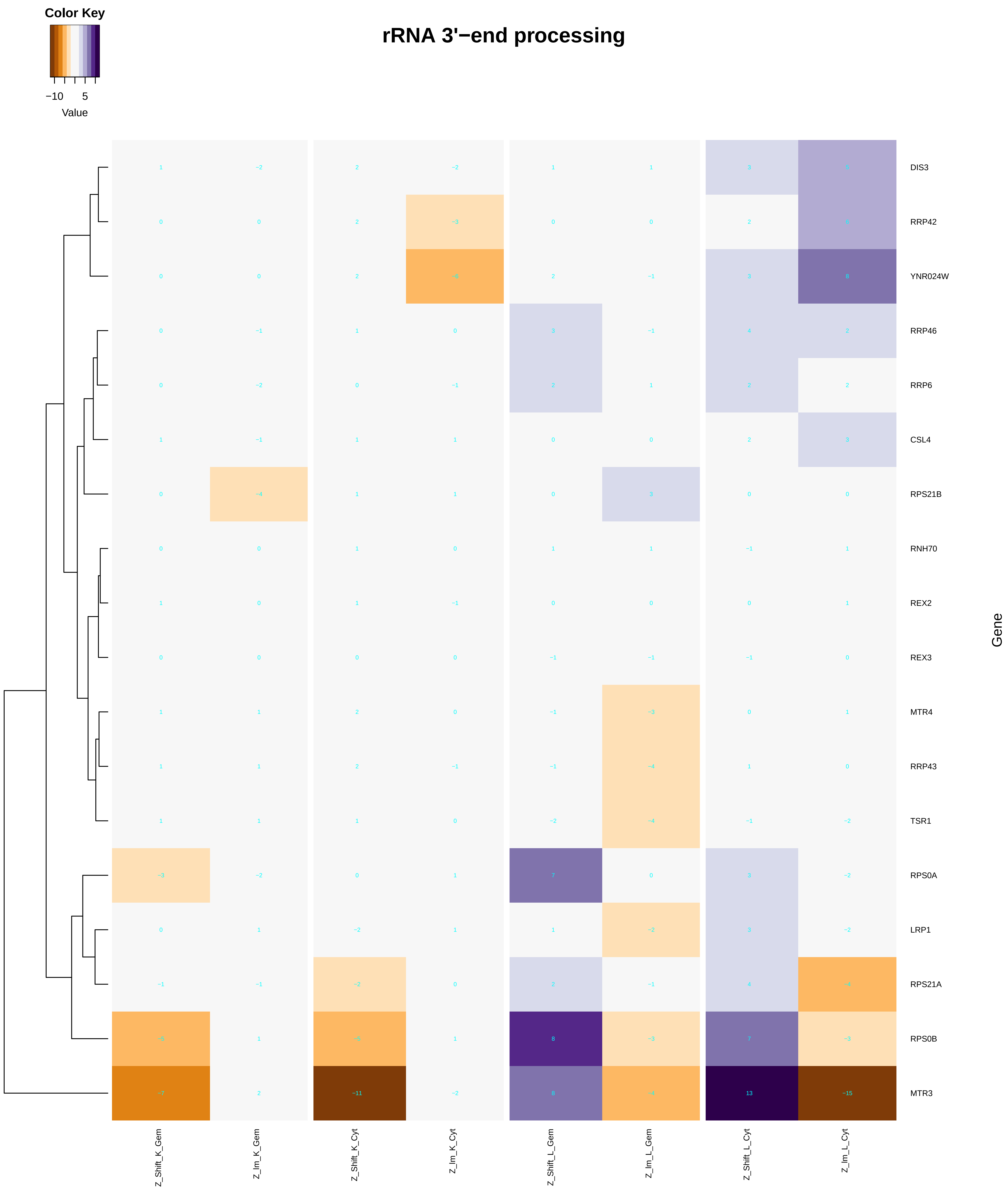
Color Key
−10
5
Value
rRNA 3'−end processing
DIS3
RRP42
YNR024W
RRP46
RRP6
CSL4
RPS21B
RNH70
REX2
REX3
MTR4
RRP43
TSR1
RPS0A
LRP1
RPS21A
RPS0B
MTR3
Gene
Z_Shift_K_Gem
Z_Im_K_Gem
Z_Shift_K_Cyt
Z_Im_K_Cyt
Z_Shift_L_Gem
Z_Im_L_Gem
Z_Shift_L_Cyt
Z_Im_L_Cyt

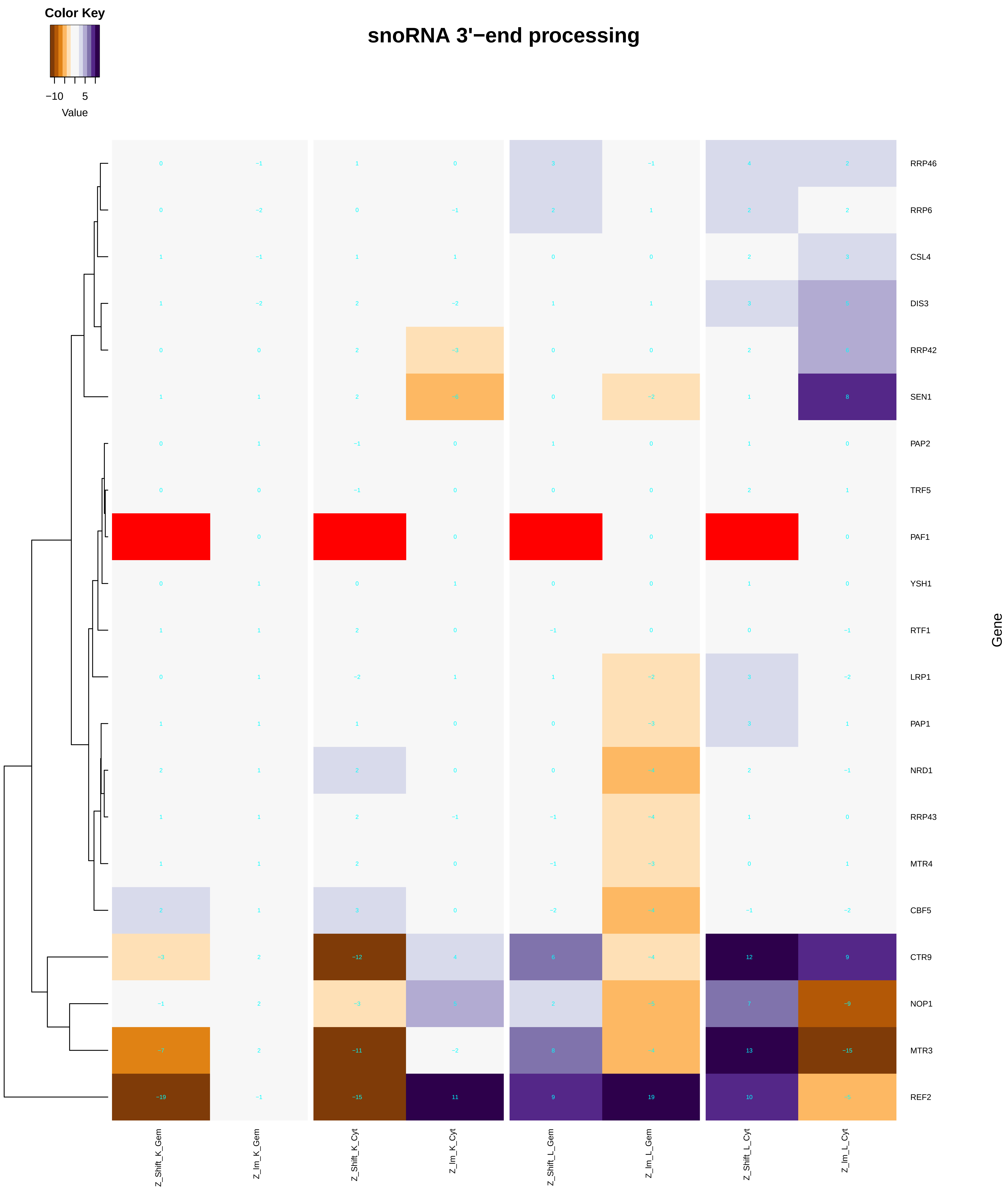

Color Key
−10 5
Value
snoRNA 3'−end processing
RRP46
RRP6
CSL4
DIS3
RRP42
SEN1
PAP2
TRF5
PAF1
YSH1
RTF1
LRP1
PAP1
NRD1
RRP43
MTR4
CBF5
CTR9
NOP1
MTR3
REF2
Gene
Z_Shift_K_Gem
Z_Im_K_Gem
Z_Shift_K_Cyt
Z_Im_K_Cyt
Z_Shift_L_Gem
Z_Im_L_Gem
Z_Shift_L_Cyt
Z_Im_L_Cyt

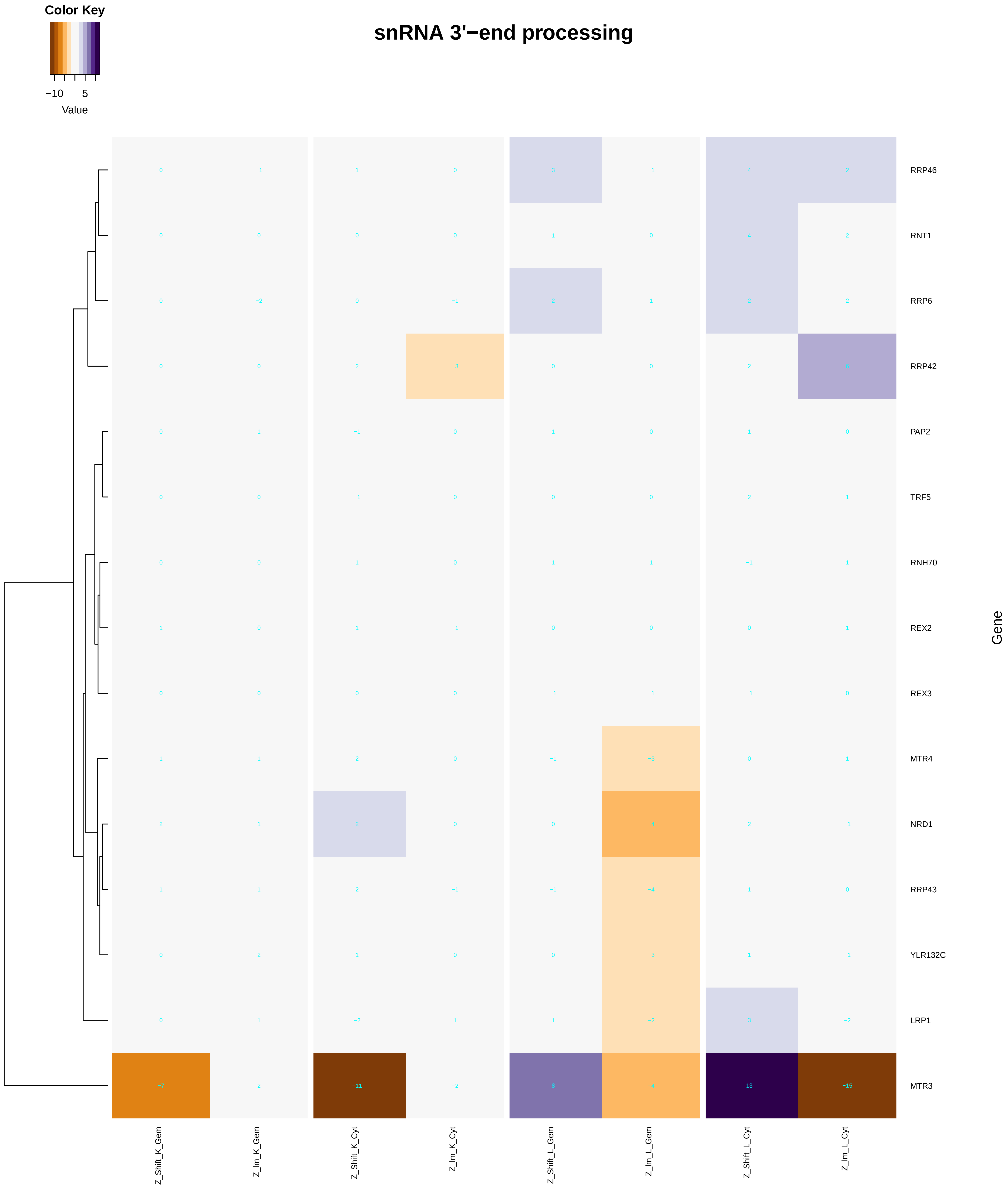
Color Key
−10
5
Value
snRNA 3'−end processing
RRP46
RNT1
RRP6
RRP42
PAP2
TRF5
RNH70
REX2
REX3
MTR4
NRD1
RRP43
YLR132C
LRP1
MTR3
Gene
Z_Shift_K_Gem
Z_Im_K_Gem
Z_Shift_K_Cyt
Z_Im_K_Cyt
Z_Shift_L_Gem
Z_Im_L_Gem
Z_Shift_L_Cyt
Z_Im_L_Cyt

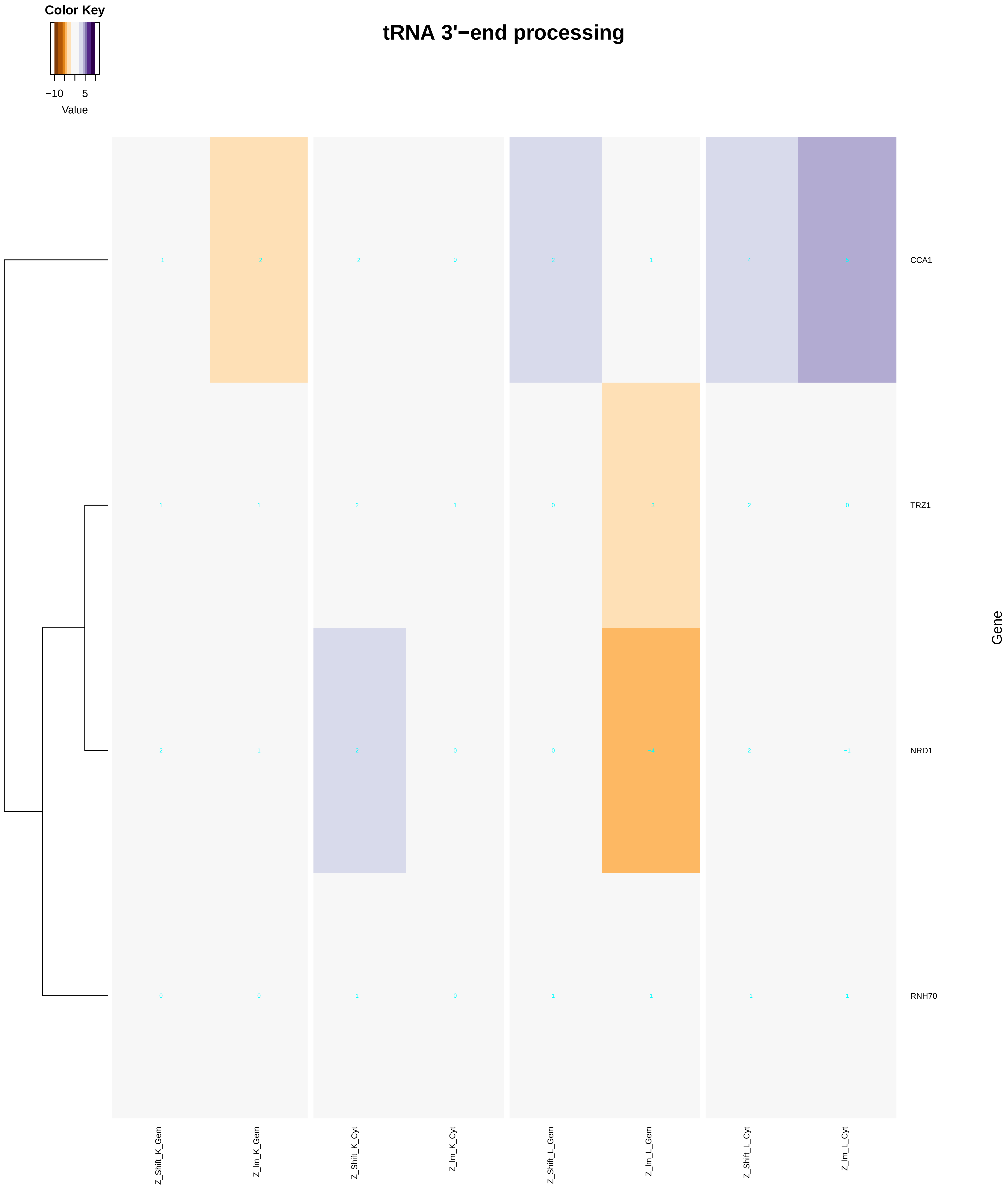

Color Key
−10
5
Value
tRNA 3'−end processing
−1 −2 −2 0 2 1 4 5
CCA1
1 1 2 1 0 −3 2 0
TRZ1
2 1 2 0 0 −4 2 −1
NRD1
0 0 1 0 1 1 −1 1
RNH70
Gene
Z_Shift_K_Gem
Z_Im_K_Gem
Z_Shift_K_Cyt
Z_Im_K_Cyt
Z_Shift_L_Gem
Z_Im_L_Gem
Z_Shift_L_Cyt
Z_Im_L_Cyt

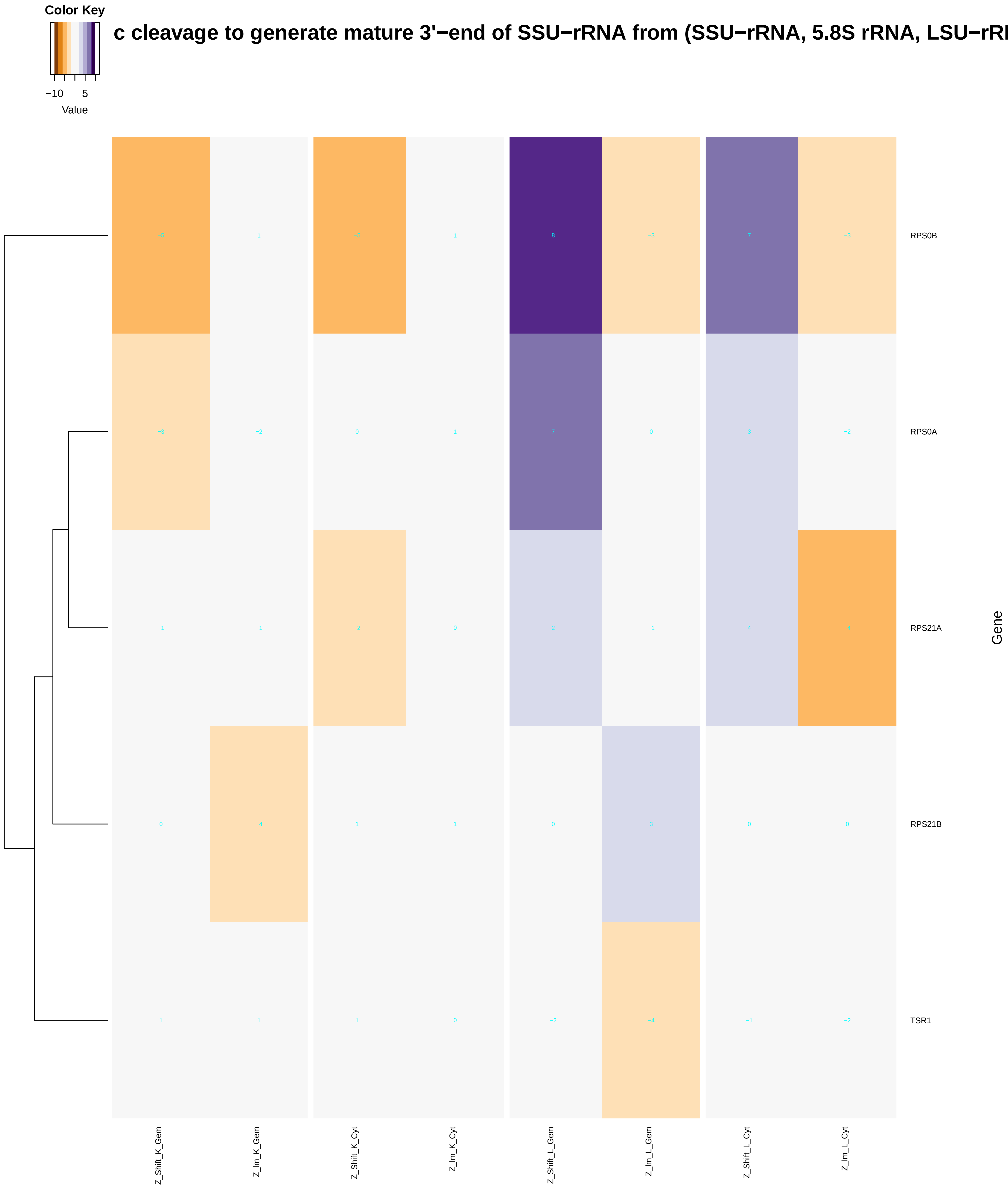

Color Key
−10
5
Value
c cleavage to generate mature 3'−end of SSU−rRNA from (SSU−rRNA, 5.8S rRNA, LSU−rR
RPS0B
RPS0A
RPS21A
RPS21B
TSR1
Gene
Z_Shift_K_Gem
Z_Im_K_Gem
Z_Shift_K_Cyt
Z_Im_K_Cyt
Z_Shift_L_Gem
Z_Im_L_Gem
Z_Shift_L_Cyt
Z_Im_L_Cyt

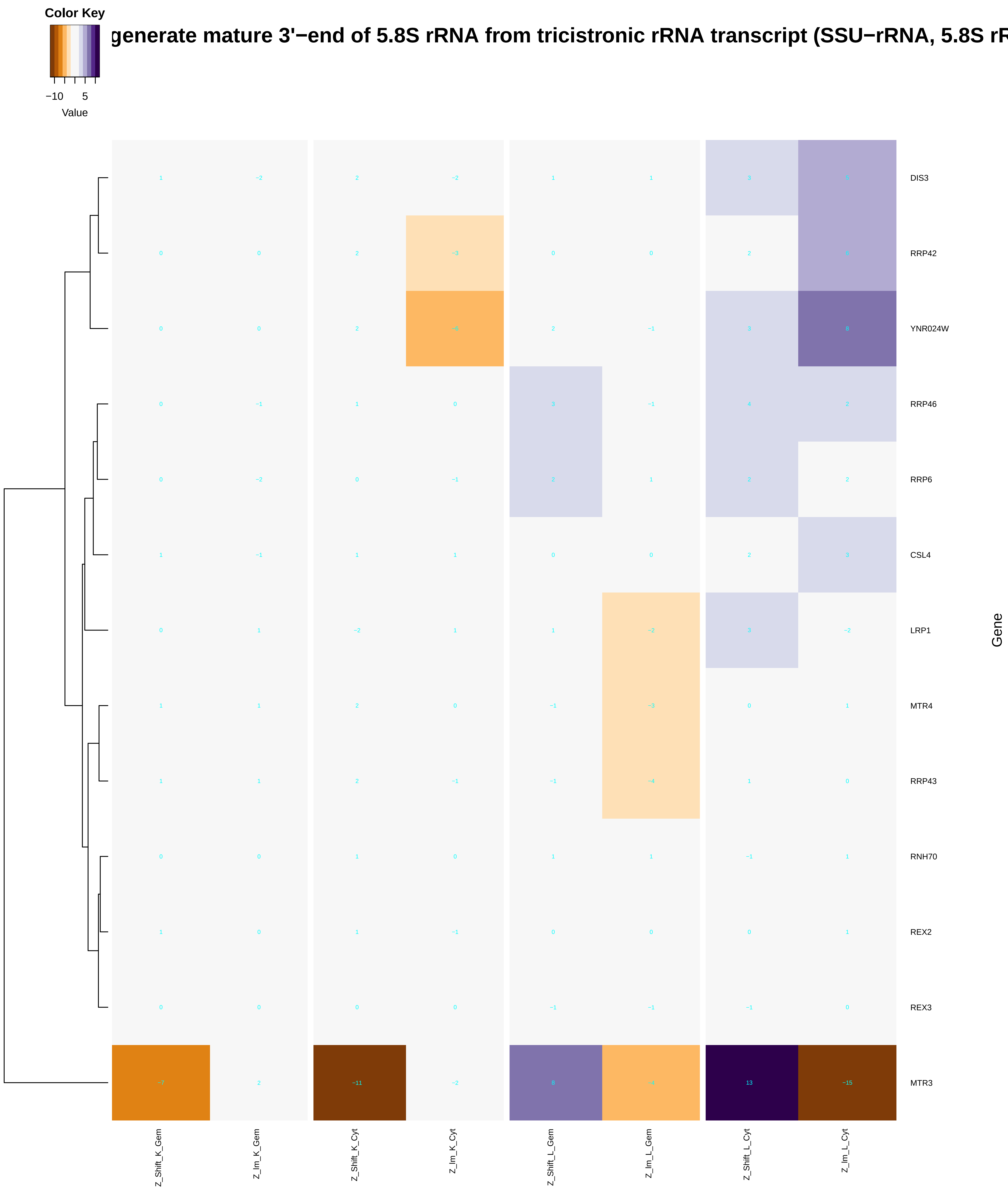
Color Key
−10
5
Value
generate mature 3'−end of 5.8S rRNA from tricistronic rRNA transcript (SSU−rRNA, 5.8S rR
DIS3
RRP42
YNR024W
RRP46
RRP6
CSL4
LRP1
MTR4
RRP43
RNH70
REX2
REX3
MTR3
Gene
Z_Shift_K_Gem
Z_Im_K_Gem
Z_Shift_K_Cyt
Z_Im_K_Cyt
Z_Shift_L_Gem
Z_Im_L_Gem
Z_Shift_L_Cyt
Z_Im_L_Cyt

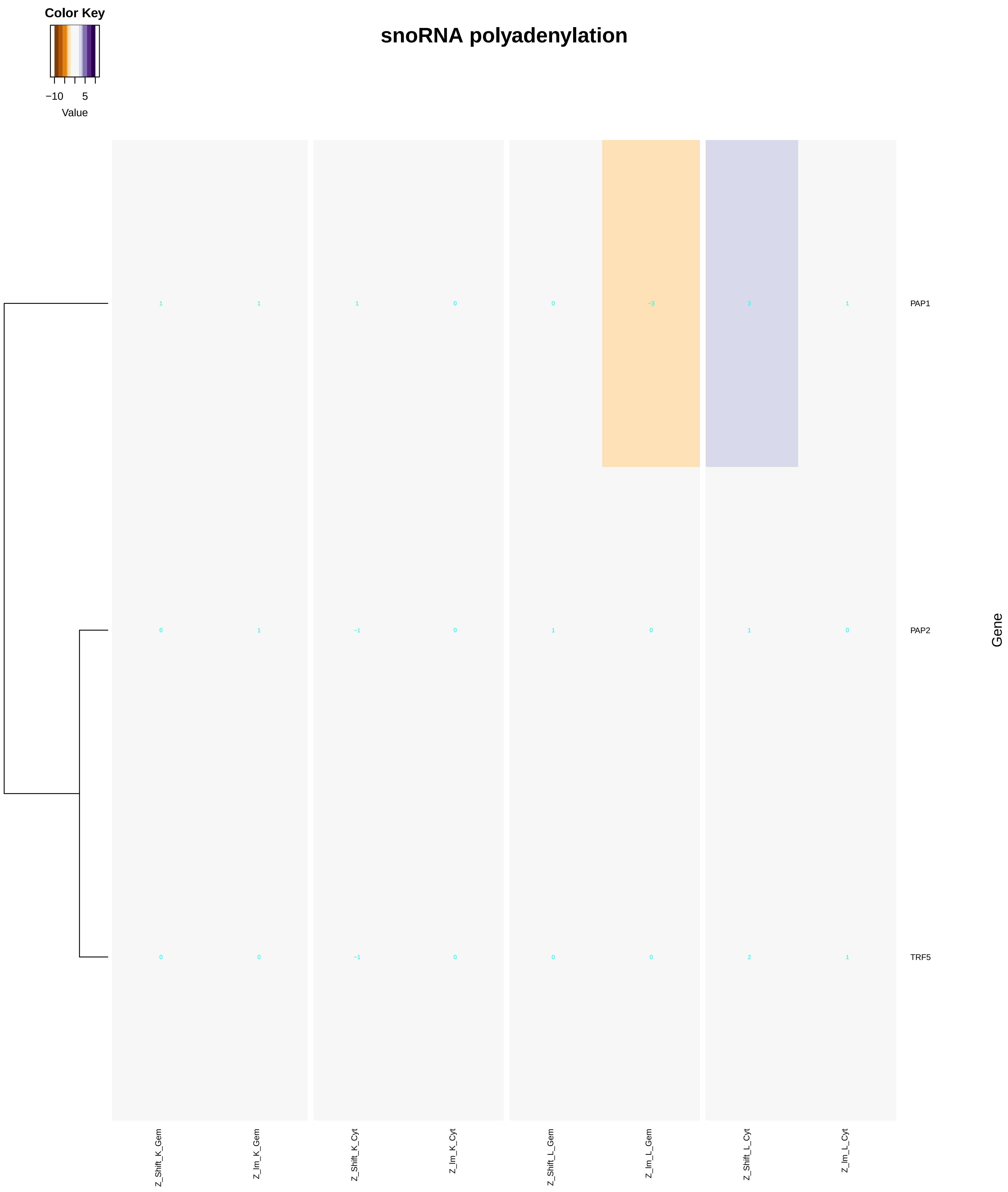
Color Key
−10
5
Value
snoRNA polyadenylation
PAP1
PAP2
TRF5
Gene
Z_Shift_K_Gem
Z_Im_K_Gem
Z_Shift_K_Cyt
Z_Im_K_Cyt
Z_Shift_L_Gem
Z_Im_L_Gem
Z_Shift_L_Cyt
Z_Im_L_Cyt

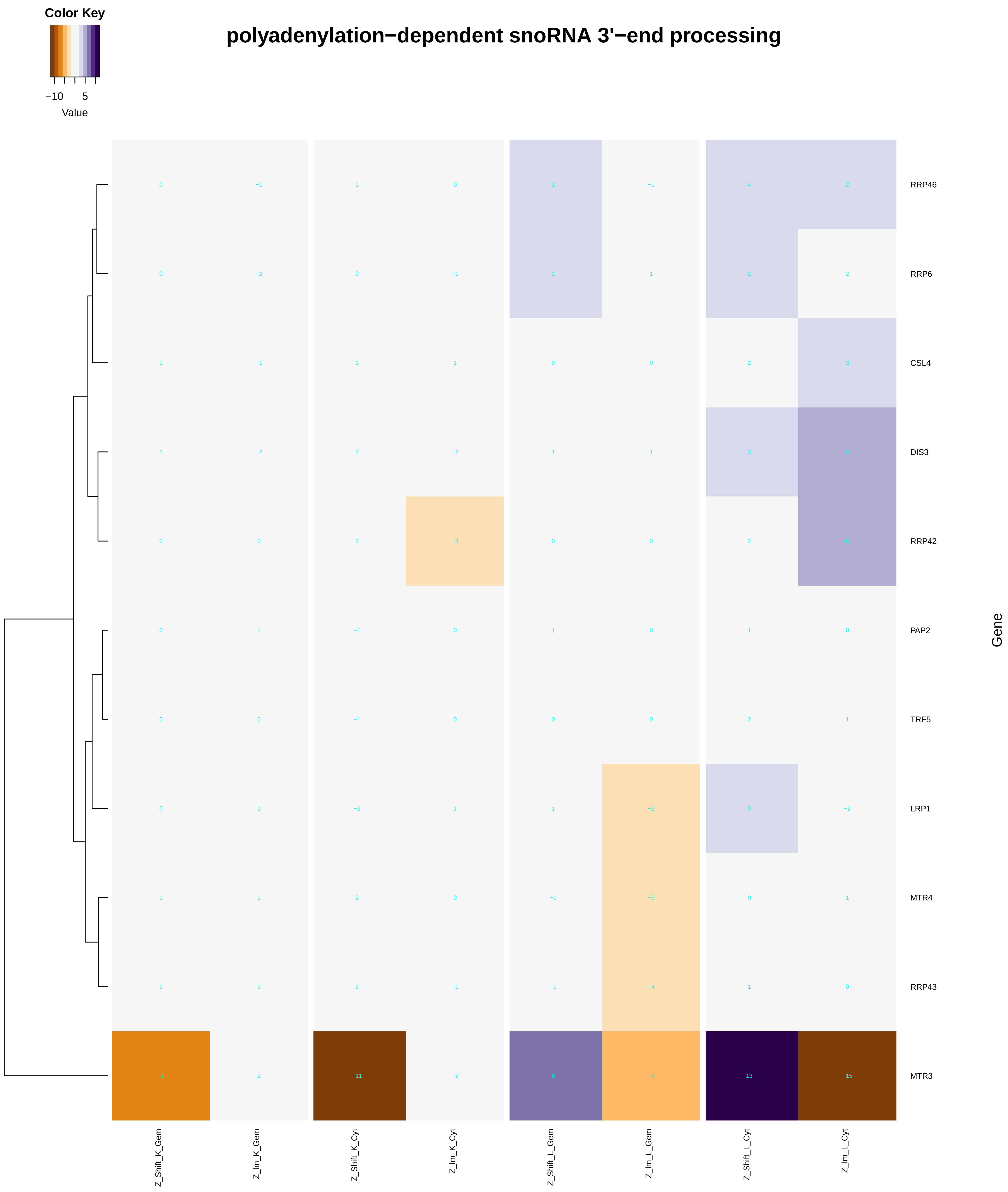

polyadenylation−dependent snoRNA 3'−end processing
Color Key
−10
5
Value
RRP46
RRP6
CSL4
DIS3
RRP42
PAP2
TRF5
LRP1
MTR4
RRP43
MTR3
Gene
Z_Shift_K_Gem
Z_Im_K_Gem
Z_Shift_K_Cyt
Z_Im_K_Cyt
Z_Shift_L_Gem
Z_Im_L_Gem
Z_Shift_L_Cyt
Z_Im_L_Cyt

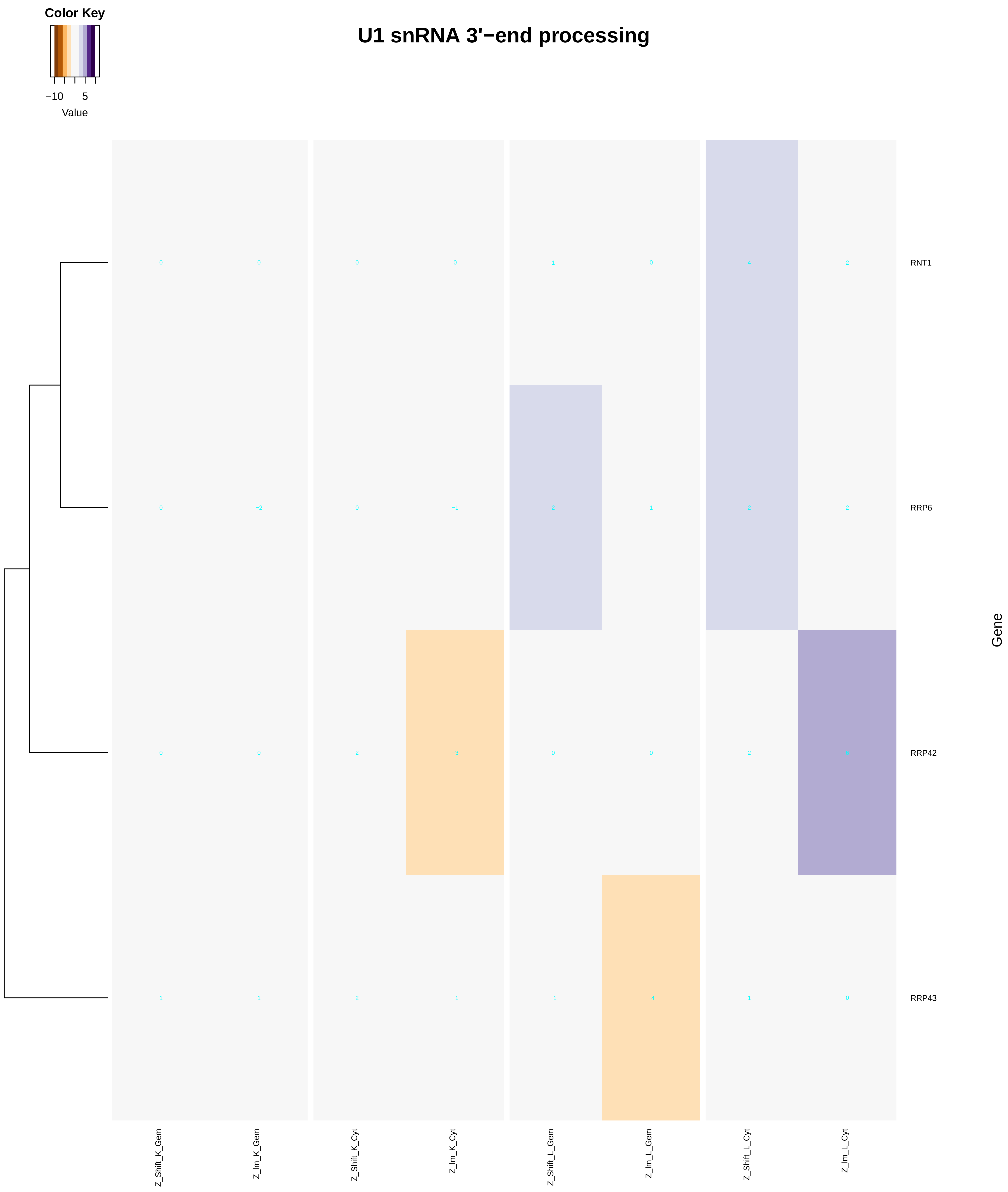
U1 snRNA 3'−end processing
Color Key
−10
5
Value
RNT1
RRP6
RRP42
RRP43
Gene
Z_Shift_K_Gem
Z_Im_K_Gem
Z_Shift_K_Cyt
Z_Im_K_Cyt
Z_Shift_L_Gem
Z_Im_L_Gem
Z_Shift_L_Cyt
Z_Im_L_Cyt

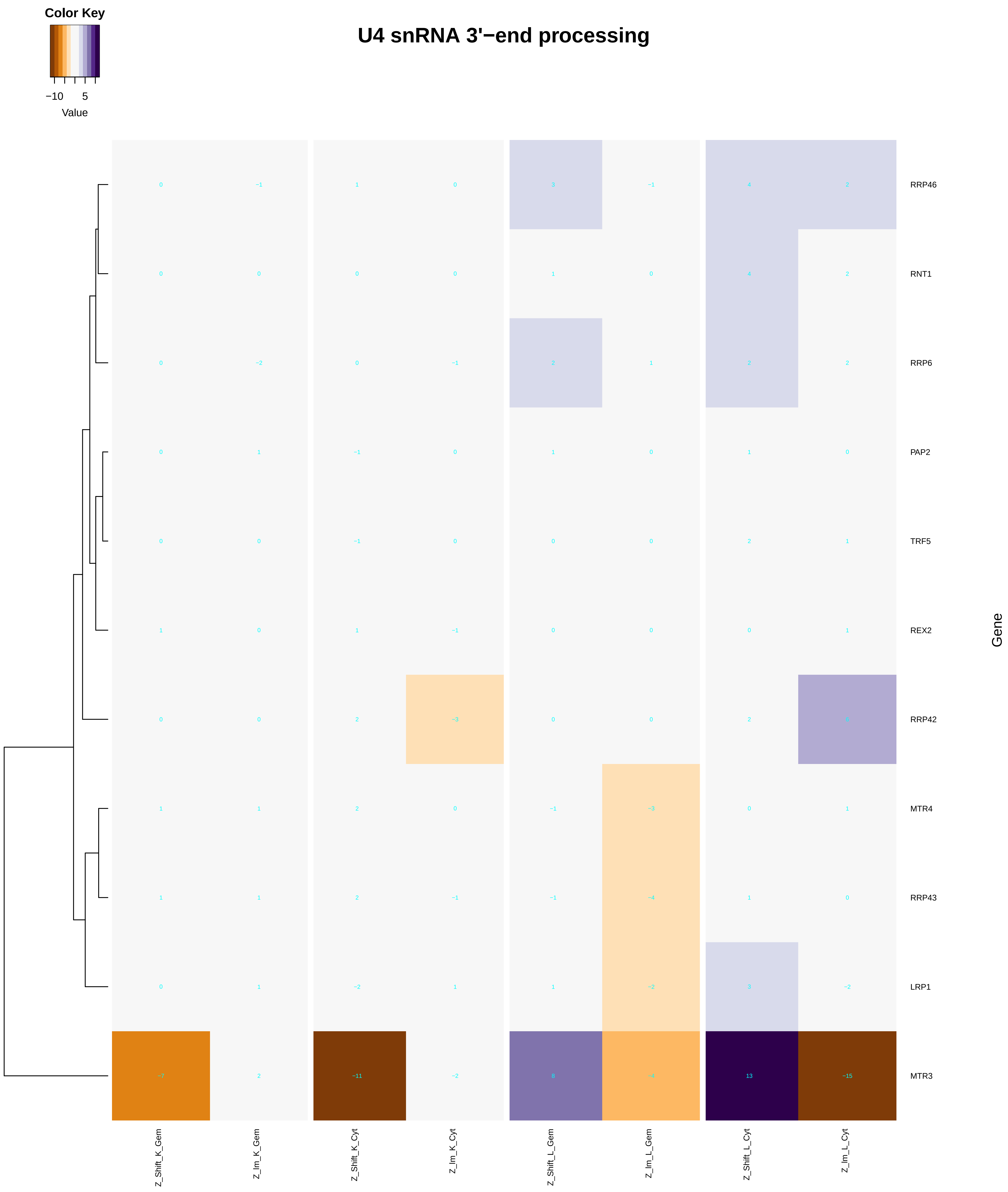
U4 snRNA 3'−end processing
Color Key
−10
5
Value
RRP46
RNT1
RRP6
PAP2
TRF5
REX2
RRP42
MTR4
RRP43
LRP1
MTR3
Gene
Z_Shift_K_Gem
Z_Im_K_Gem
Z_Shift_K_Cyt
Z_Im_K_Cyt
Z_Shift_L_Gem
Z_Im_L_Gem
Z_Shift_L_Cyt
Z_Im_L_Cyt

### U5 snRNA 3'−end processing

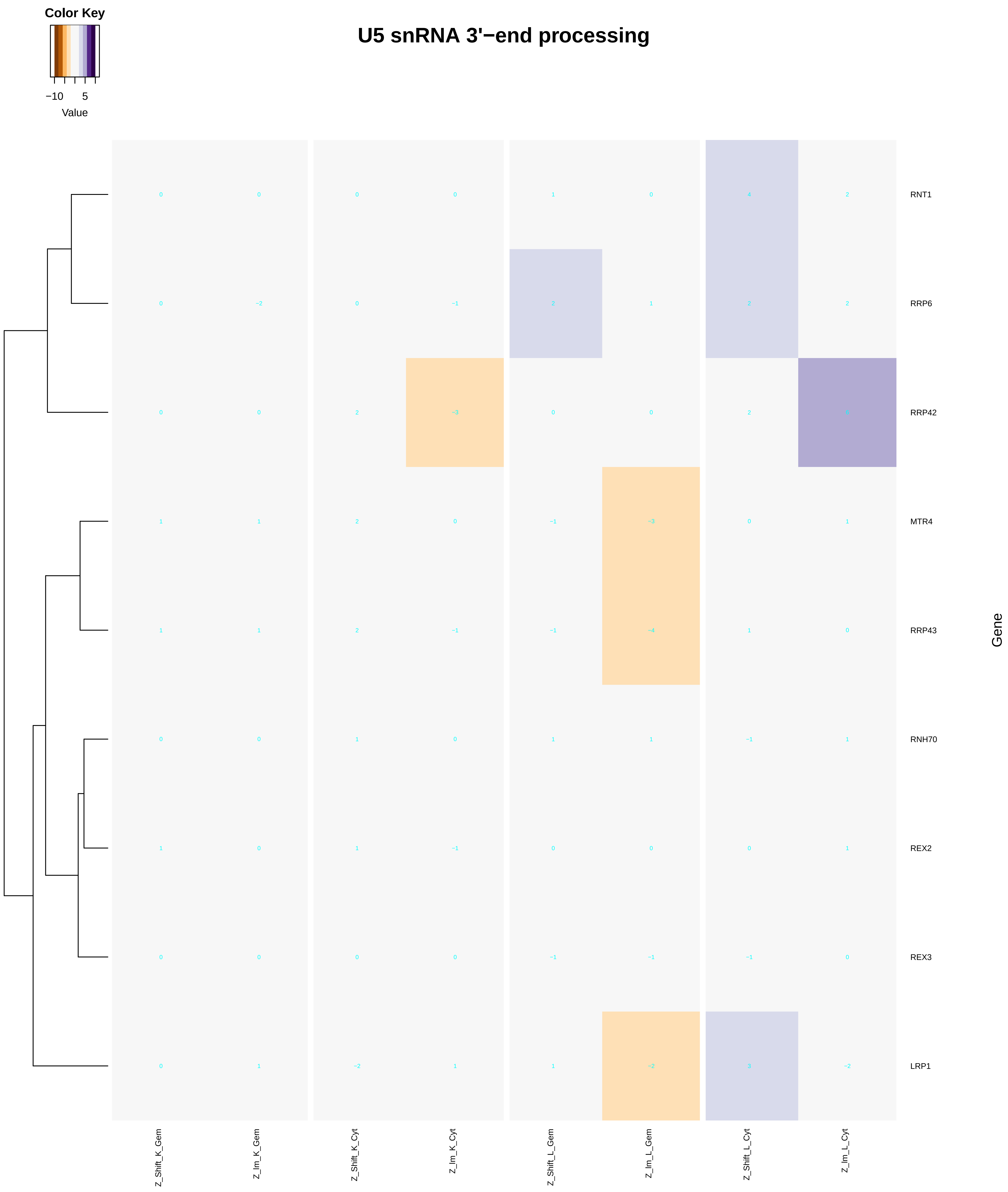
