## Additional File 7 for "A humanized yeast phenomic model of deoxycytidine kinase to predict genetic buffering of nucleoside analog cytotoxicity": ncRNA_processing.pdf

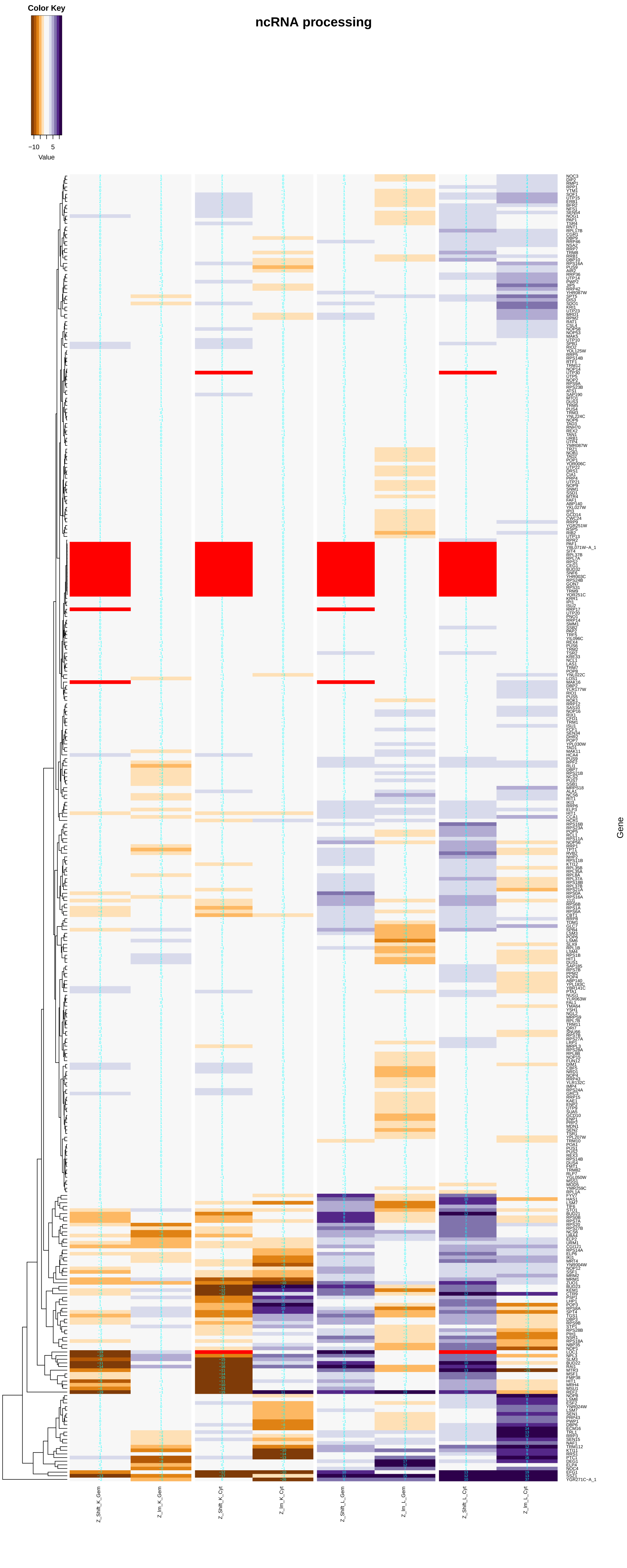

ncRNA processing
Color Key
−10
5
Value
Z_Shift_K_Gem
Z_Im_K_Gem
Z_Shift_K_Cyt
Z_Im_K_Cyt
Z_Shift_L_Gem
Z_Im_L_Gem
Z_Shift_L_Cyt
Z_Im_L_Cyt
Gene

### rRNA processing

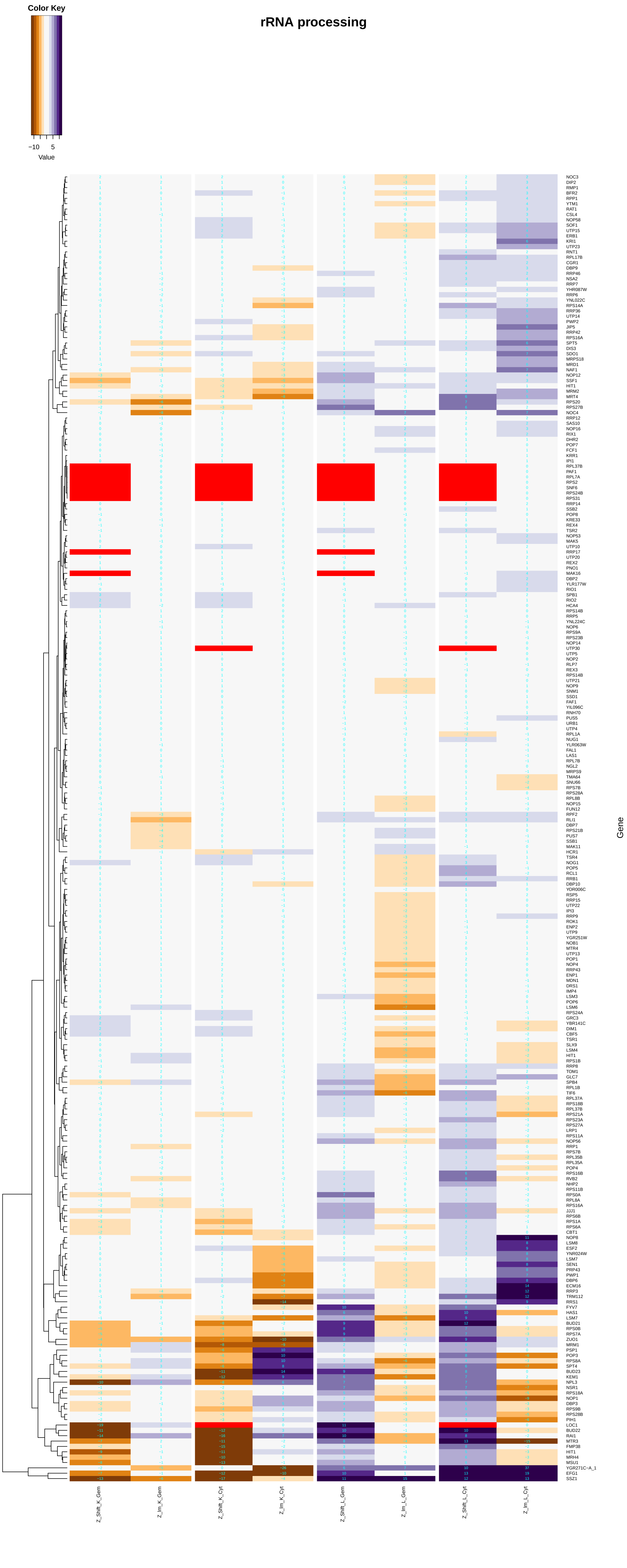

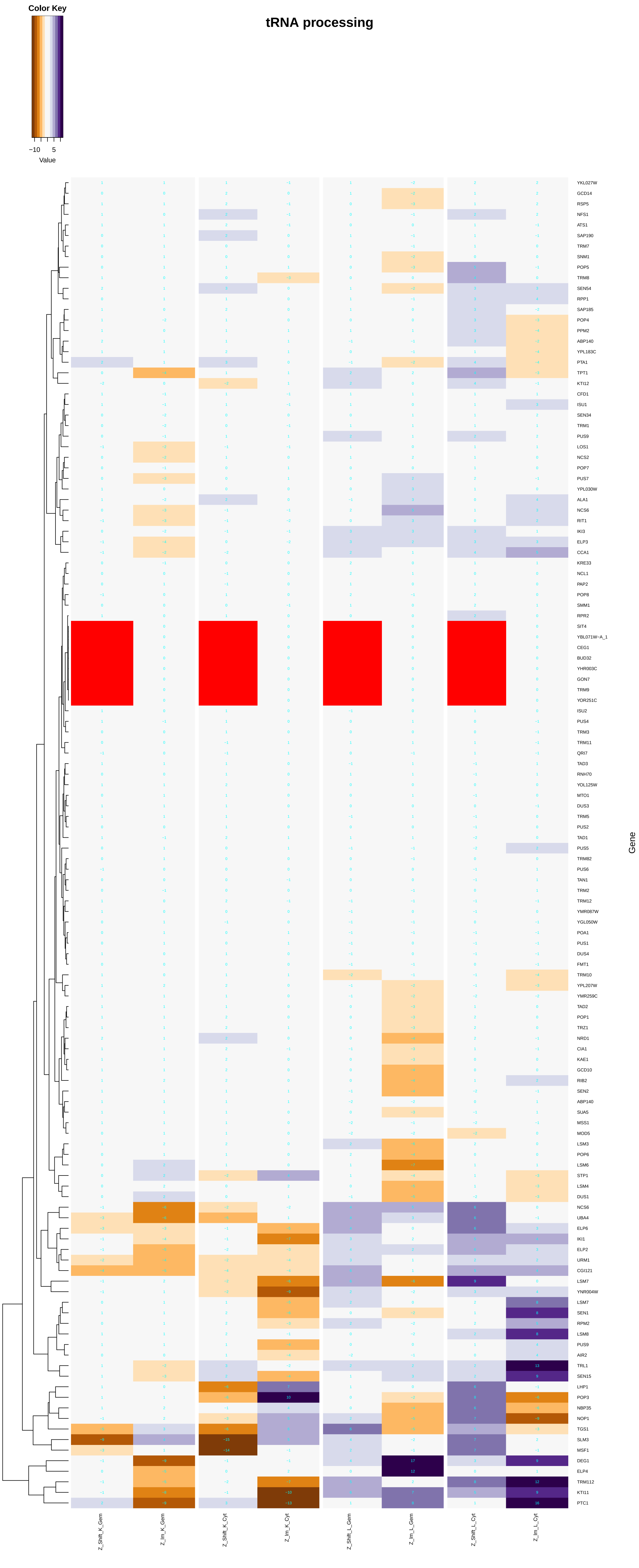

tRNA processing
Color Key
−10
5
Value
YKL027W
GCD14
RSP5
NFS1
ATS1
SAP190
TRM7
SNM1
POP5
TRM8
SEN54
RPP1
SAP185
POP4
PPM2
ABP140
YPL183C
PTA1
TPT1
KTI12
CFD1
ISU1
SEN34
TRM1
PUS9
LOS1
NCS2
POP7
PUS7
YPL030W
ALA1
NCS6
RIT1
IKI3
ELP3
CCA1
KRE33
NCL1
PAP2
POP8
SMM1
RPR2
SIT4
YBL071W−A_1
CEG1
BUD32
YHR003C
GON7
TRM9
YOR251C
ISU2
PUS4
TRM3
TRM11
QRI7
TAD3
RNH70
YOL125W
MTO1
DUS3
TRM5
PUS2
TAD1
PUS5
TRM82
PUS6
TAN1
TRM2
TRM12
YMR087W
YGL050W
POA1
PUS1
DUS4
FMT1
TRM10
YPL207W
YMR259C
TAD2
POP1
TRZ1
NRD1
CIA1
KAE1
GCD10
RIB2
SEN2
ABP140
SUA5
MSS1
MOD5
LSM3
POP6
LSM6
STP1
LSM4
DUS1
NCS6
UBA4
ELP6
IKI1
ELP2
URM1
CGI121
LSM7
YNR004W
LSM7
SEN1
RPM2
LSM8
PUS9
AIR2
TRL1
SEN15
LHP1
POP3
NBP35
NOP1
TGS1
SLM3
MSF1
DEG1
ELP4
TRM112
KTI11
PTC1
Gene
Z_Shift_K_Gem
Z_Im_K_Gem
Z_Shift_K_Cyt
Z_Im_K_Cyt
Z_Shift_L_Gem
Z_Im_L_Gem
Z_Shift_L_Cyt
Z_Im_L_Cyt

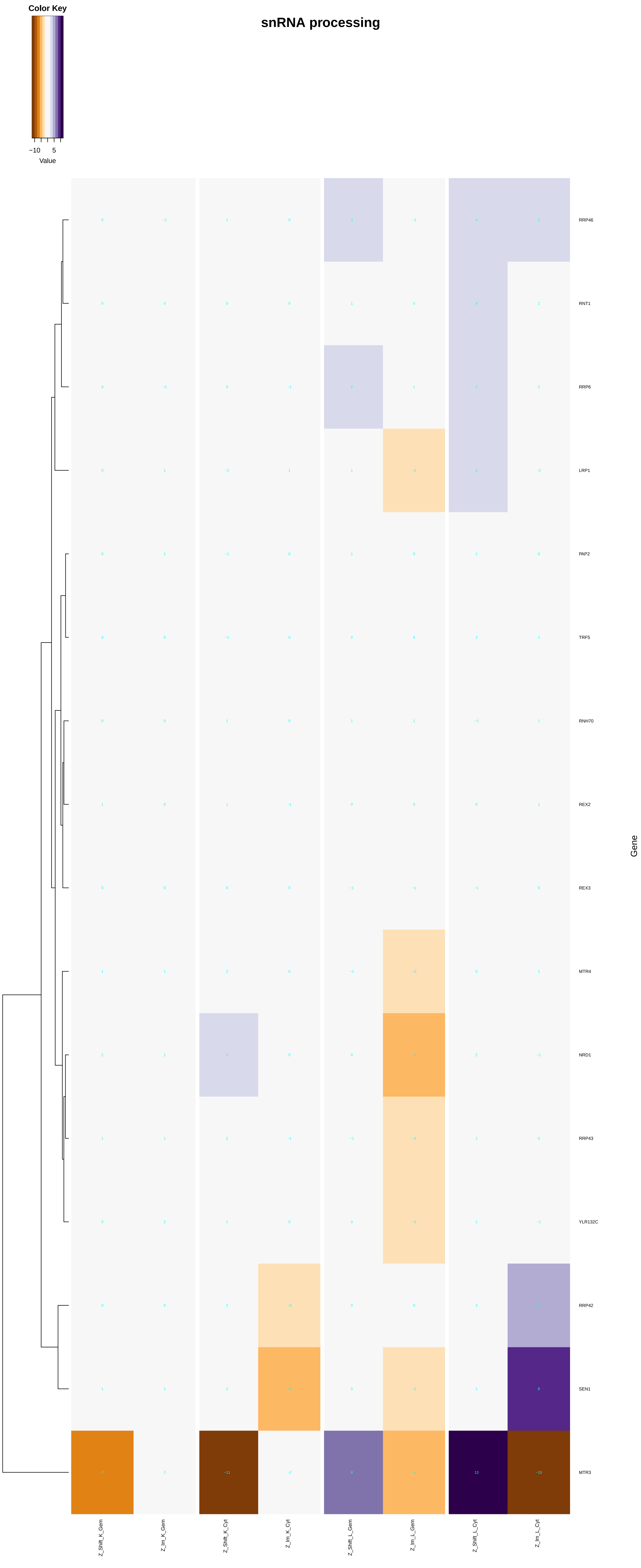
Color Key
−10
5
Value
snRNA processing
RRP46
RNT1
RRP6
LRP1
PAP2
TRF5
RNH70
REX2
REX3
MTR4
NRD1
RRP43
YLR132C
RRP42
SEN1
MTR3
Gene
Z_Shift_K_Gem
Z_Im_K_Gem
Z_Shift_K_Cyt
Z_Im_K_Cyt
Z_Shift_L_Gem
Z_Im_L_Gem
Z_Shift_L_Cyt
Z_Im_L_Cyt

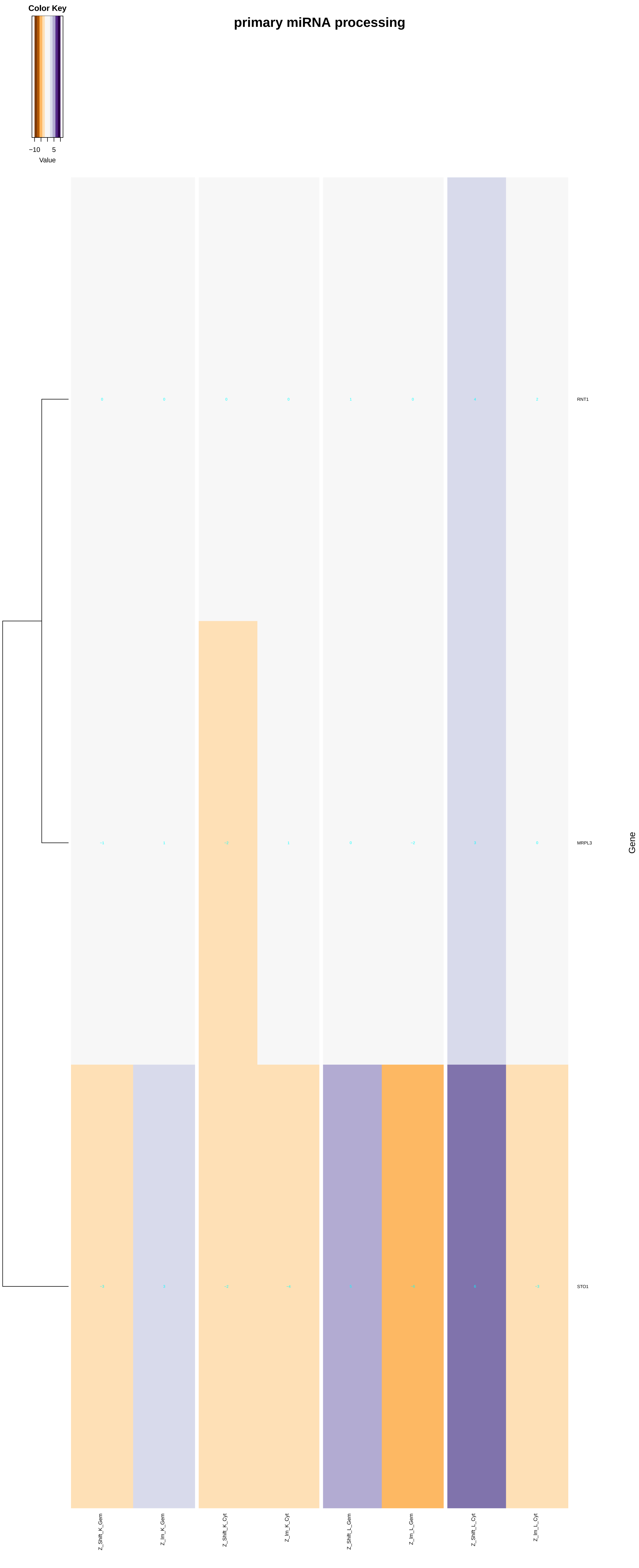
primary miRNA processing
Color Key
−10
5
Value
RNT1
MRPL3
STO1
Gene
Z_Shift_K_Gem
Z_Im_K_Gem
Z_Shift_K_Cyt
Z_Im_K_Cyt
Z_Shift_L_Gem
Z_Im_L_Gem
Z_Shift_L_Cyt
Z_Im_L_Cyt

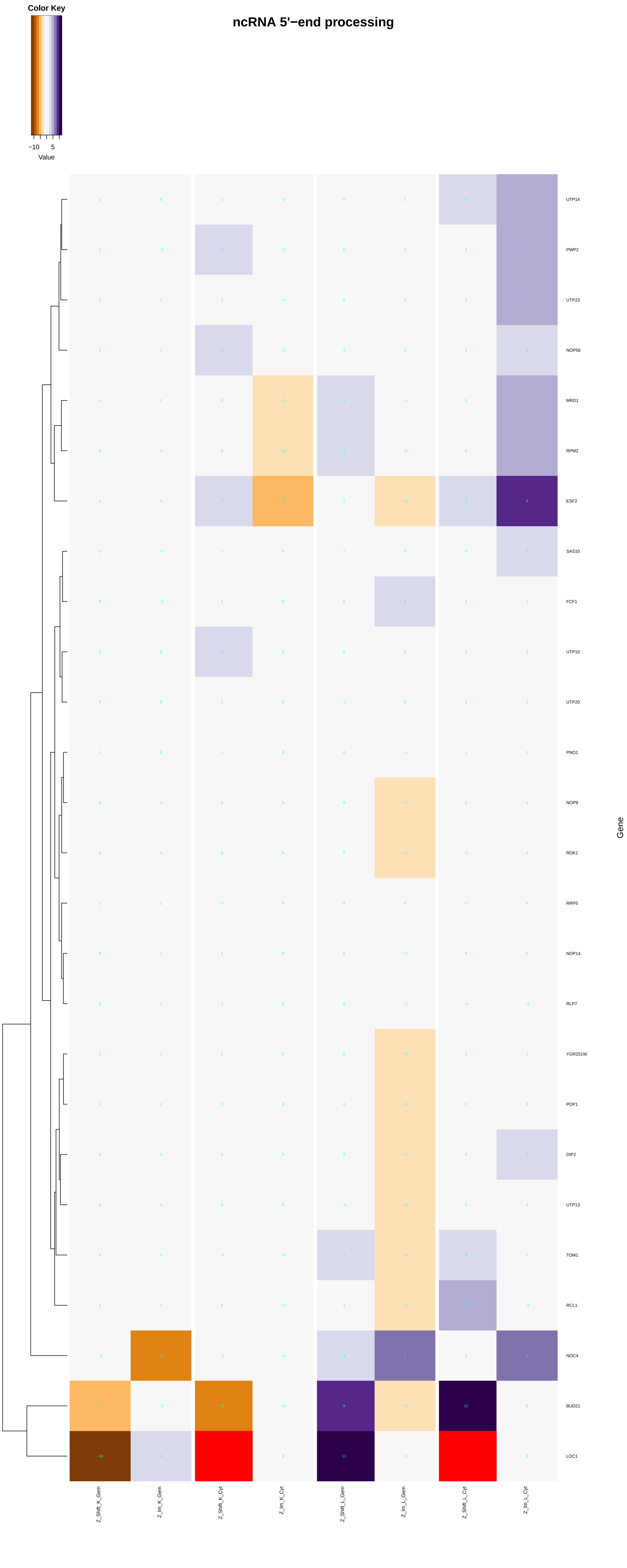
ncRNA 5'−end processing
Color Key
−10
5
Value
UTP14
PWP2
UTP23
NOP58
MRD1
RPM2
ESF2
SAS10
FCF1
UTP10
UTP20
PNO1
NOP9
ROK1
RRP5
NOP14
RLP7
YGR251W
POP1
DIP2
UTP13
TOM1
RCL1
NOC4
BUD21
LOC1
Gene
Z_Shift_K_Gem
Z_Im_K_Gem
Z_Shift_K_Cyt
Z_Im_K_Cyt
Z_Shift_L_Gem
Z_Im_L_Gem
Z_Shift_L_Cyt
Z_Im_L_Cyt

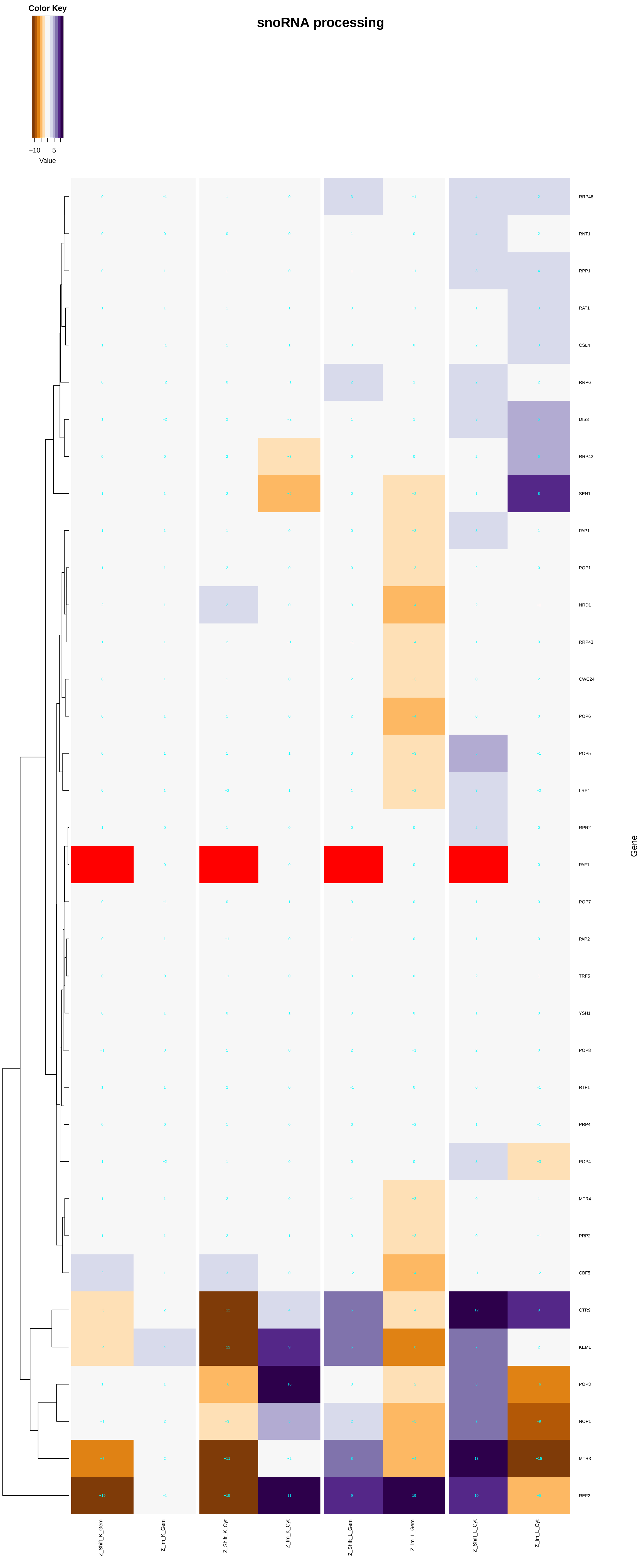

Color Key
−10
5
Value
snoRNA processing
RRP46
RNT1
RPP1
RAT1
CSL4
RRP6
DIS3
RRP42
SEN1
PAP1
POP1
NRD1
RRP43
CWC24
POP6
POP5
LRP1
RPR2
PAF1
POP7
PAP2
TRF5
YSH1
POP8
RTF1
PRP4
POP4
MTR4
PRP2
CBF5
CTR9
KEM1
POP3
NOP1
MTR3
REF2
Gene
Z_Shift_K_Gem
Z_Im_K_Gem
Z_Shift_K_Cyt
Z_Im_K_Cyt
Z_Shift_L_Gem
Z_Im_L_Gem
Z_Shift_L_Cyt
Z_Im_L_Cyt

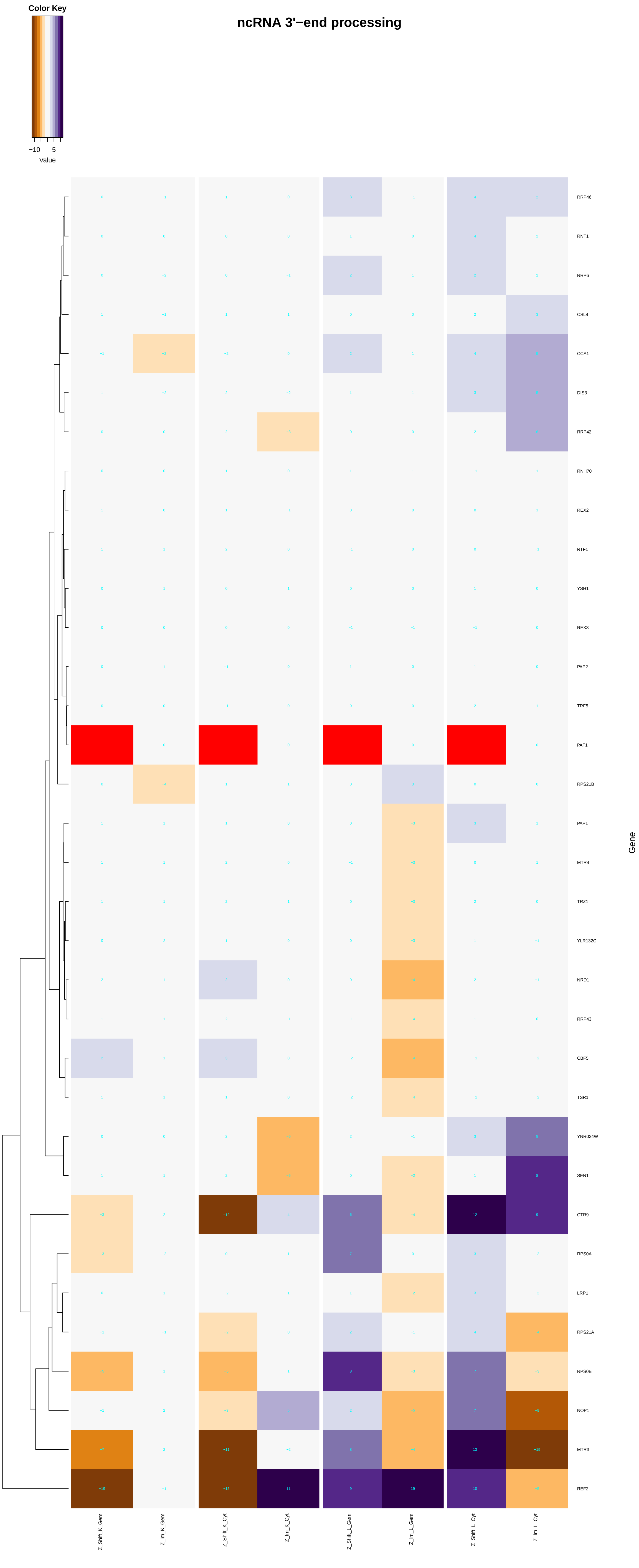

ncRNA 3'−end processing
Color Key
−10 5
Value
RRP46
RNT1
RRP6
CSL4
CCA1
DIS3
RRP42
RNH70
REX2
RTF1
YSH1
REX3
PAP2
TRF5
PAF1
RPS21B
PAP1
MTR4
TRZ1
YLR132C
NRD1
RRP43
CBF5
TSR1
YNR024W
SEN1
CTR9
RPS0A
LRP1
RPS21A
RPS0B
NOP1
MTR3
REF2
Gene
Z_Shift_K_Gem
Z_Im_K_Gem
Z_Shift_K_Cyt
Z_Im_K_Cyt
Z_Shift_L_Gem
Z_Im_L_Gem
Z_Shift_L_Cyt
Z_Im_L_Cyt

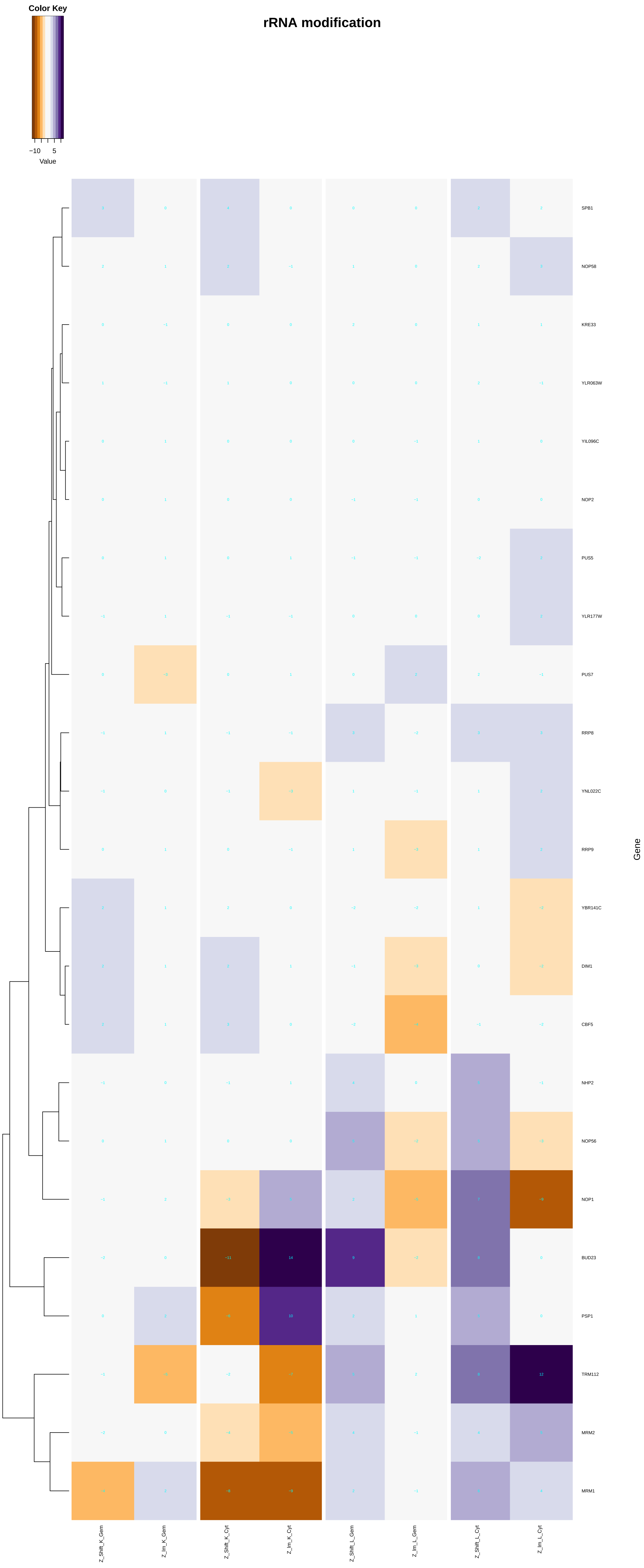

Color Key
−10
5
Value
rRNA modification
SPB1
NOP58
KRE33
YLR063W
YIL096C
NOP2
PUS5
YLR177W
PUS7
RRP8
YNL022C
RRP9
YBR141C
DIM1
CBF5
NHP2
NOP56
NOP1
BUD23
PSP1
TRM112
MRM2
MRM1
Gene
Z_Shift_K_Gem
Z_lm_K_Gem
Z_Shift_K_Cyt
Z_lm_K_Cyt
Z_Shift_L_Gem
Z_lm_L_Gem
Z_Shift_L_Cyt
Z_lm_L_Cyt

### maturation of 5.8S rRNA

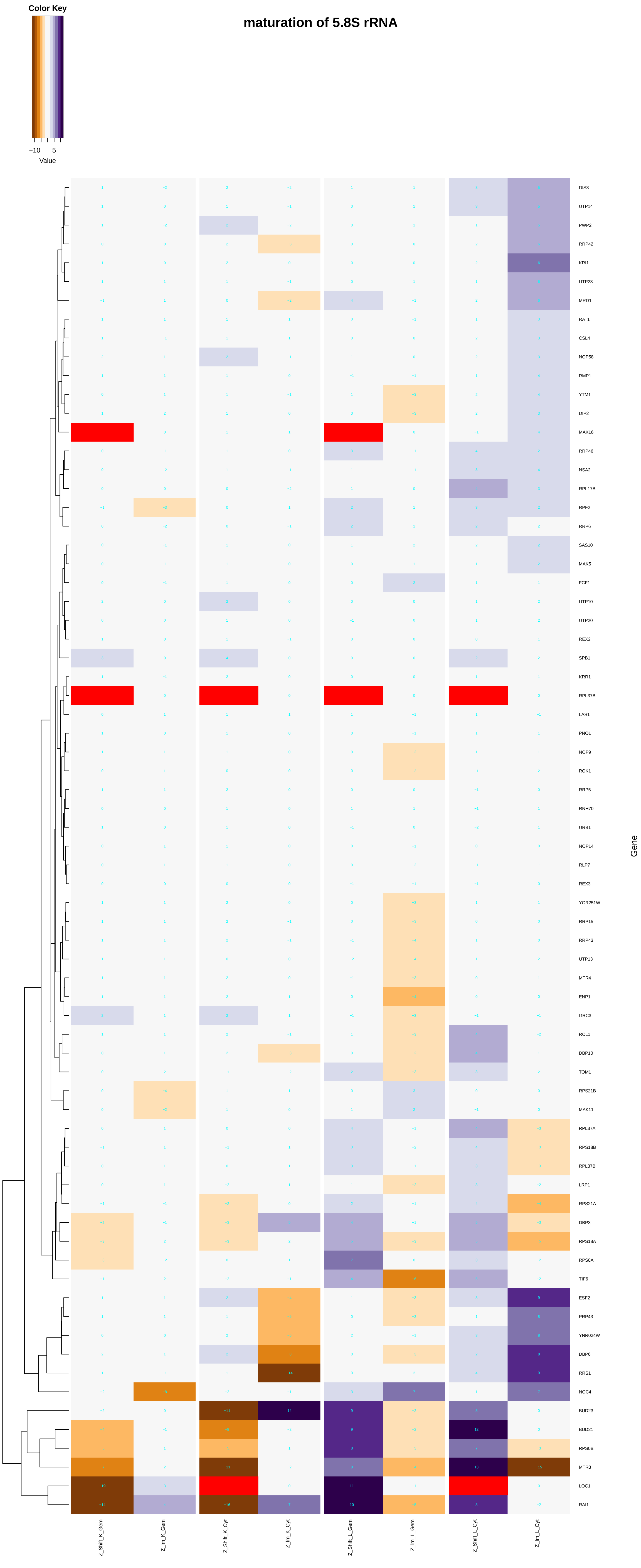

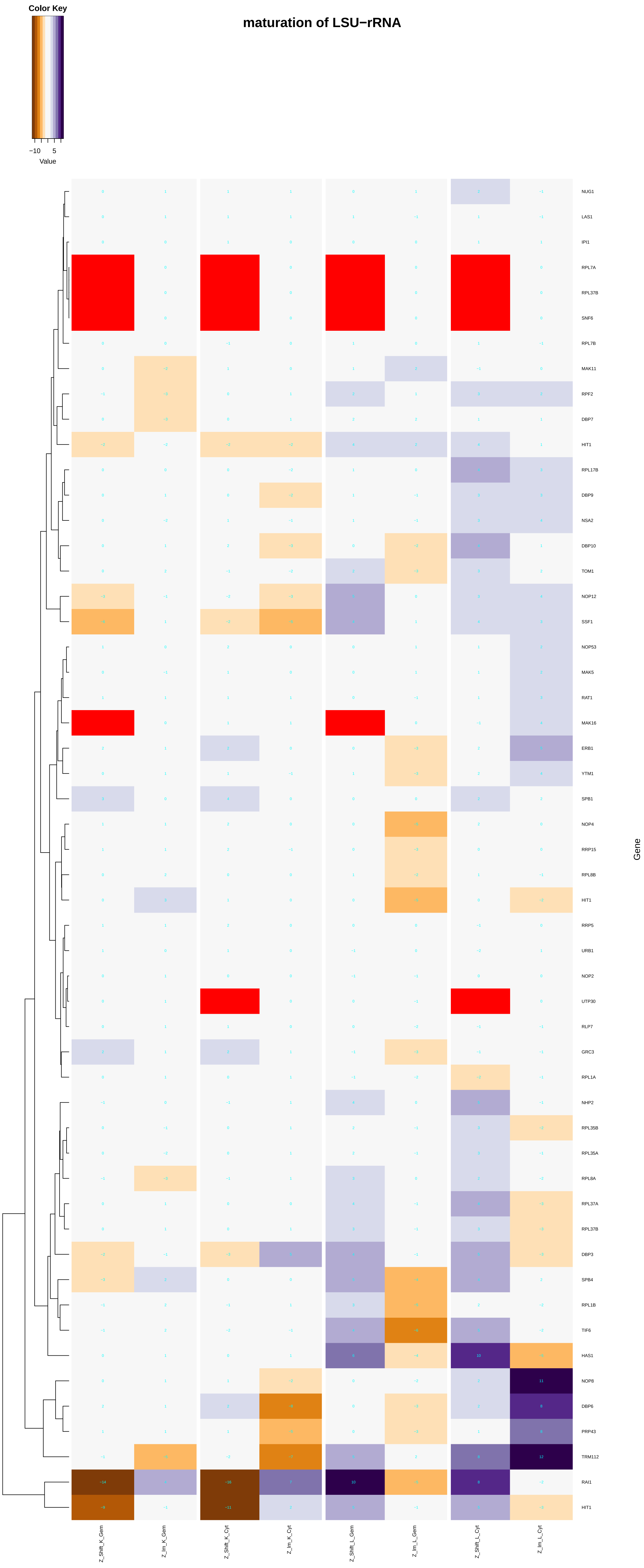

maturation of LSU−rRNA
Color Key
−10
5
Value
NUG1
LAS1
IPI1
RPL7A
RPL37B
SNF6
RPL7B
MAK11
RPF2
DBP7
HIT1
RPL17B
DBP9
NSA2
DBP10
TOM1
NOP12
SSF1
NOP53
MAK5
RAT1
MAK16
ERB1
YTM1
SPB1
NOP4
RRP15
RPL8B
HIT1
RRP5
URB1
NOP2
UTP30
RLP7
GRC3
RPL1A
NHP2
RPL35B
RPL35A
RPL8A
RPL37A
RPL37B
DBP3
SPB4
RPL1B
TIF6
HAS1
NOP8
DBP6
PRP43
TRM112
RAI1
HIT1
Gene
Z_Shift_K_Gem
Z_lm_K_Gem
Z_Shift_K_Cyt
Z_lm_K_Cyt
Z_Shift_L_Gem
Z_lm_L_Gem
Z_Shift_L_Cyt
Z_lm_L_Cyt

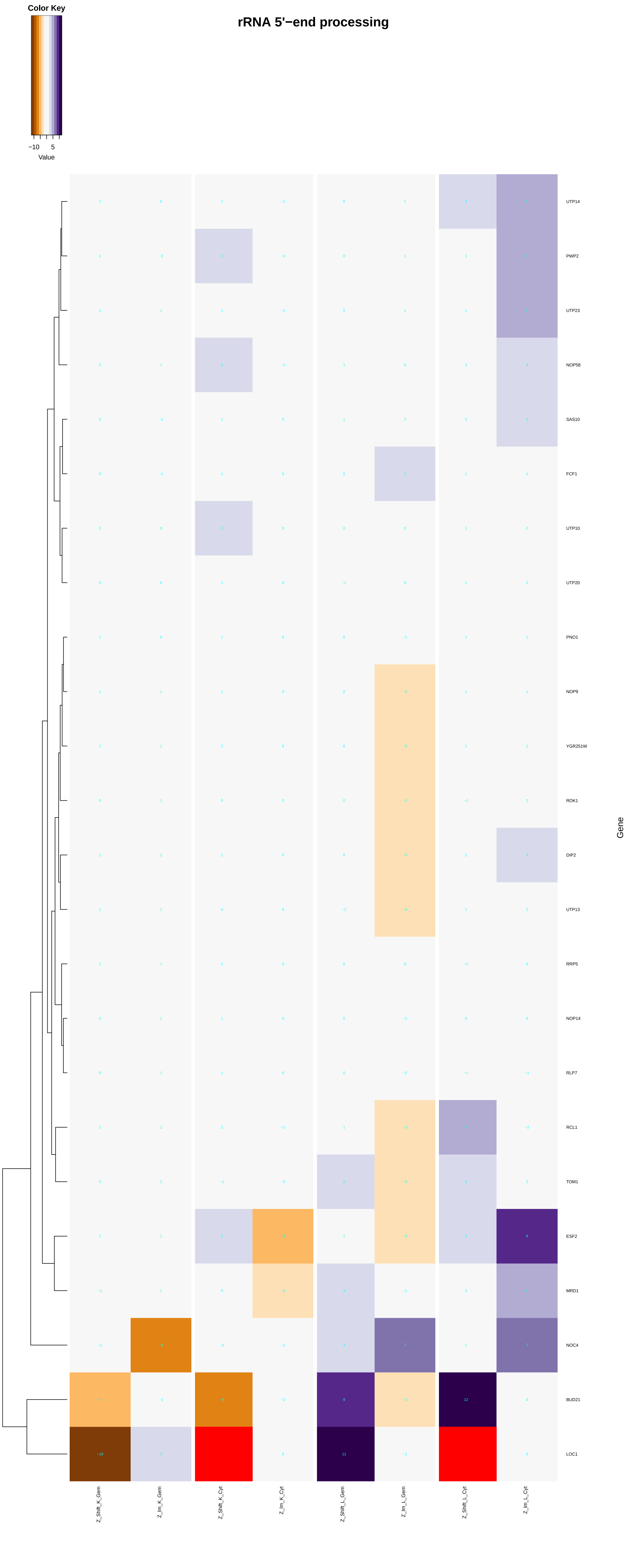

Color Key
−10
5
Value
rRNA 5'−end processing
UTP14
PWP2
UTP23
NOP58
SAS10
FCF1
UTP10
UTP20
PNO1
NOP9
YGR251W
ROK1
DIP2
UTP13
RRP5
NOP14
RLP7
RCL1
TOM1
ESF2
MRD1
NOC4
BUD21
LOC1
Gene
Z_Shift_K_Gem
Z_Im_K_Gem
Z_Shift_K_Cyt
Z_Im_K_Cyt
Z_Shift_L_Gem
Z_Im_L_Gem
Z_Shift_L_Cyt
Z_Im_L_Cyt

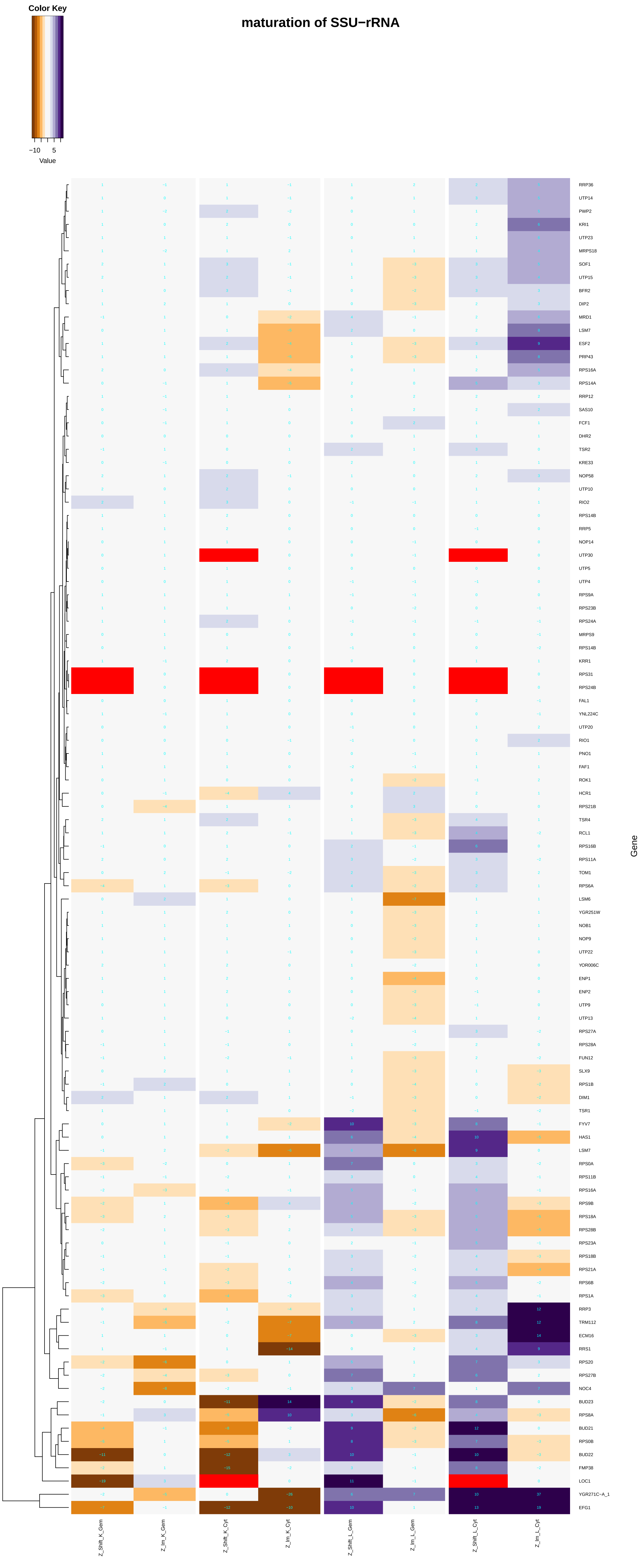

maturation of SSU−rRNA
Color Key
−10
5
Value
Gene
RRP36
UTP14
PWP2
KRI1
UTP23
MRPS18
SOF1
UTP15
BFR2
DIP2
MRD1
LSM7
ESF2
PRP43
RPS16A
RPS14A
RRP12
SAS10
FCF1
DHR2
TSR2
KRE33
NOP58
UTP10
RIO2
RPS14B
RRP5
NOP14
UTP30
UTP5
UTP4
RPS9A
RPS23B
RPS24A
MRPS9
RPS14B
KRR1
RPS31
RPS24B
FAL1
YNL224C
UTP20
RIO1
PNO1
FAF1
ROK1
HCR1
RPS21B
TSR4
RCL1
RPS16B
RPS11A
TOM1
RPS6A
LSM6
YGR251W
NOB1
NOP9
UTP22
YOR006C
ENP1
ENP2
UTP9
UTP13
RPS27A
RPS28A
FUN12
SLX9
RPS1B
DIM1
TSR1
FYV7
HAS1
LSM7
RPS0A
RPS11B
RPS16A
RPS9B
RPS18A
RPS28B
RPS23A
RPS18B
RPS21A
RPS6B
RPS1A
RRP3
TRM112
ECM16
RRS1
RPS20
RPS27B
NOC4
BUD23
RPS8A
BUD21
RPS0B
BUD22
FMP38
LOC1
YGR271C−A_1
EFG1
Z_Shift_K_Gem
Z_Im_K_Gem
Z_Shift_K_Cyt
Z_Im_K_Cyt
Z_Shift_L_Gem
Z_Im_L_Gem
Z_Shift_L_Cyt
Z_Im_L_Cyt

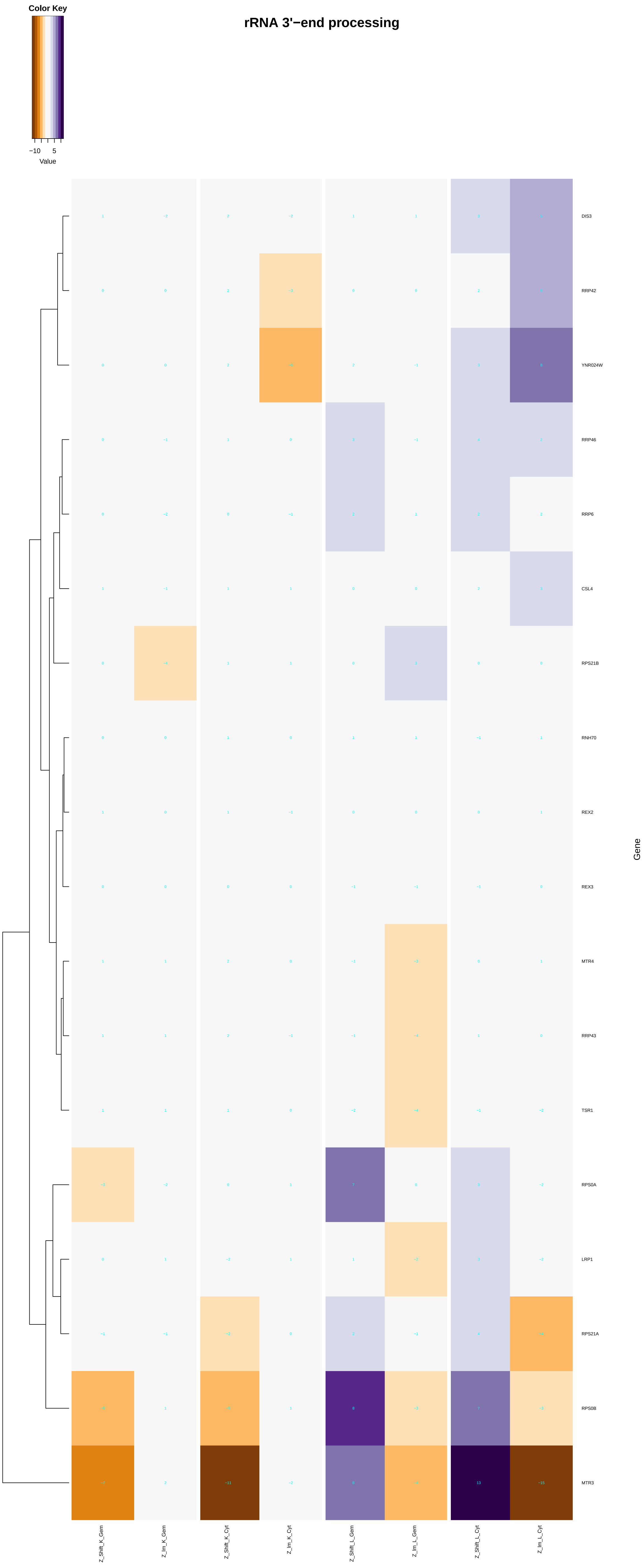
Color Key
−10
5
Value
rRNA 3'−end processing
DIS3
RRP42
YNR024W
RRP46
RRP6
CSL4
RPS21B
RNH70
REX2
REX3
MTR4
RRP43
TSR1
RPS0A
LRP1
RPS21A
RPS0B
MTR3
Gene
Z_Shift_K_Gem
Z_Im_K_Gem
Z_Shift_K_Cyt
Z_Im_K_Cyt
Z_Shift_L_Gem
Z_Im_L_Gem
Z_Shift_L_Cyt
Z_Im_L_Cyt

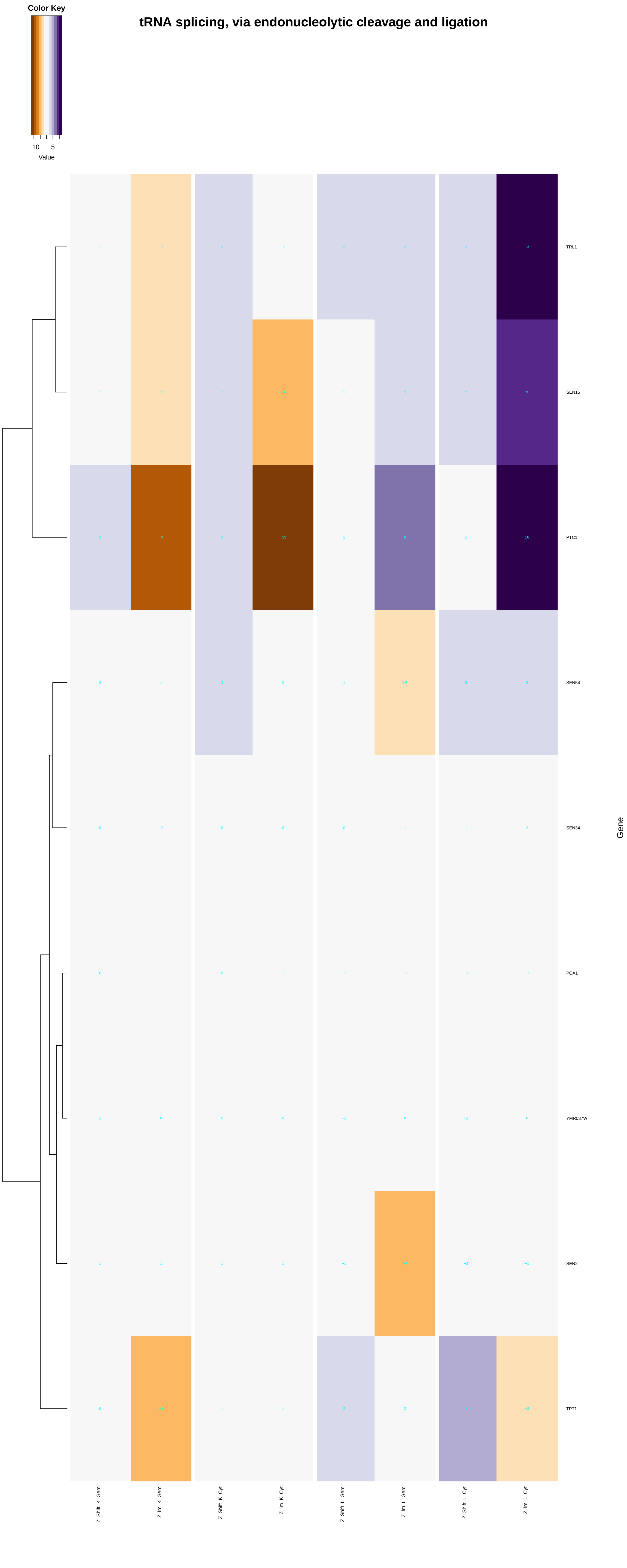
Color Key
−10
5
Value
tRNA splicing, via endonucleolytic cleavage and ligation
TRL1
SEN15
PTC1
SEN54
SEN34
POA1
YMR087W
SEN2
TPT1
Gene
Z_Shift_K_Gem
Z_Im_K_Gem
Z_Shift_K_Cyt
Z_Im_K_Cyt
Z_Shift_L_Gem
Z_Im_L_Gem
Z_Shift_L_Cyt
Z_Im_L_Cyt

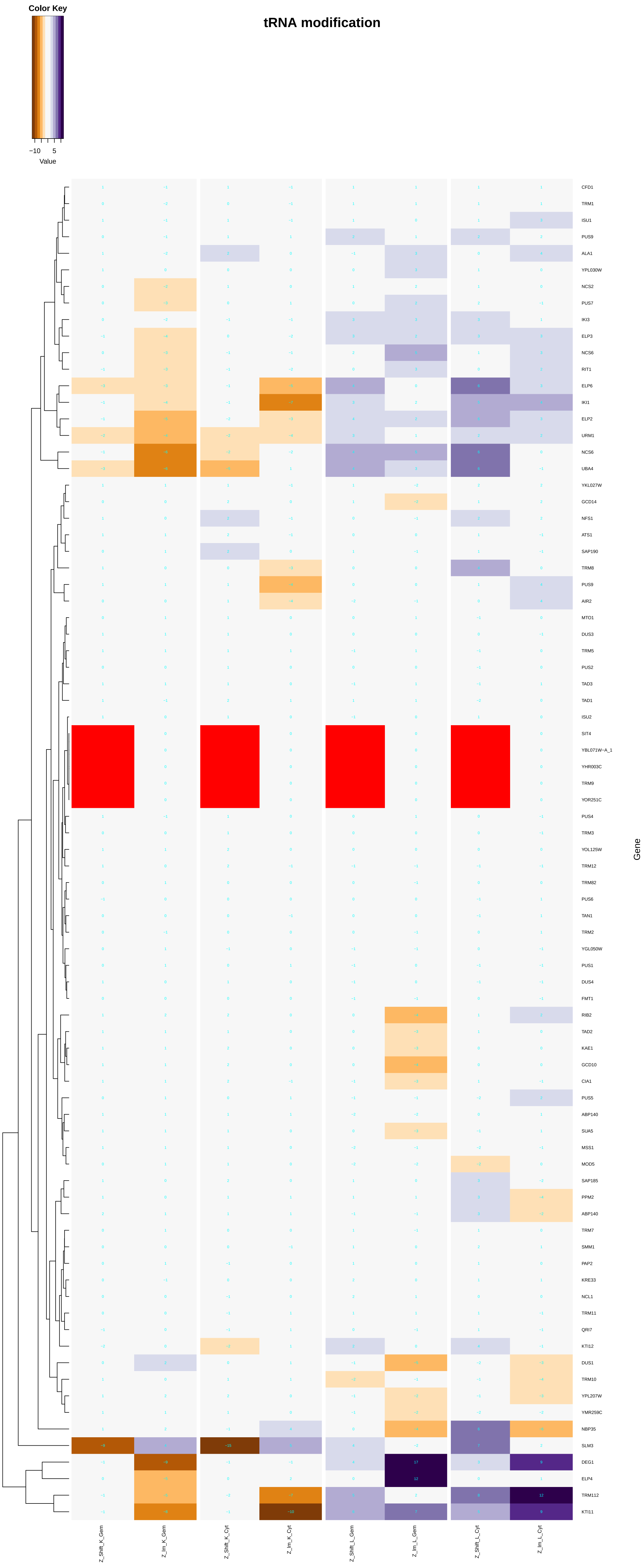
tRNA modification
Color Key
−10
5
Value
CFD1
TRM1
ISU1
PUS9
ALA1
YPL030W
NCS2
PUS7
IKI3
ELP3
NCS6
RIT1
ELP6
IKI1
ELP2
URM1
NCS6
UBA4
YKL027W
GCD14
NFS1
ATS1
SAP190
TRM8
PUS9
AIR2
MTO1
DUS3
TRM5
PUS2
TAD3
TAD1
ISU2
SIT4
YBL071W-A_1
YHR003C
TRM9
YOR251C
PUS4
TRM3
YOL125W
TRM12
TRM82
PUS6
TAN1
TRM2
YGL050W
PUS1
DUS4
FMT1
RIB2
TAD2
KAE1
GCD10
CIA1
PUS5
ABP140
SUA5
MSS1
MOD5
SAP185
PPM2
ABP140
TRM7
SMM1
PAP2
KRE33
NCL1
TRM11
QRI7
KTI12
DUS1
TRM10
YPL207W
YMR259C
NBP35
SLM3
DEG1
ELP4
TRM112
KTI11
Gene
Z_Shift_K_Gem
Z_lm_K_Gem
Z_Shift_K_Cyt
Z_lm_K_Cyt
Z_Shift_L_Gem
Z_lm_L_Gem
Z_Shift_L_Cyt
Z_lm_L_Cyt

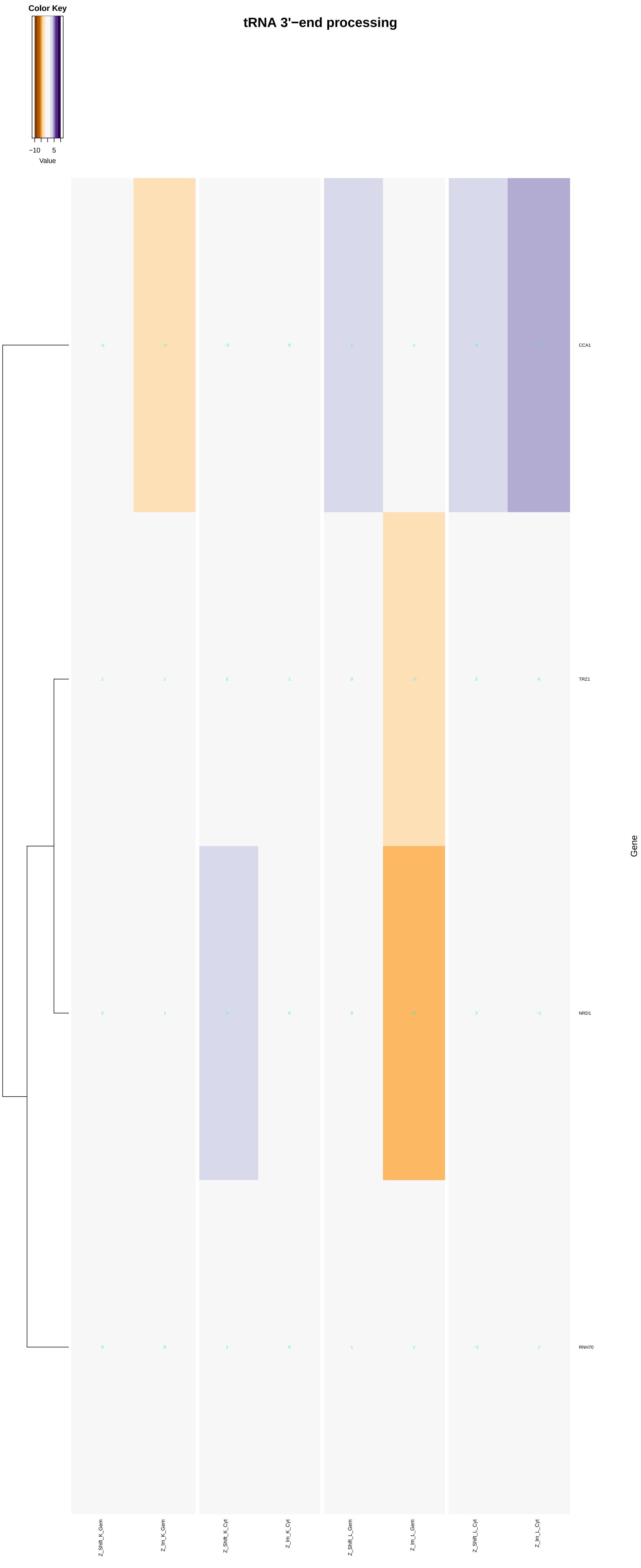
tRNA 3'−end processing
Color Key
−10
5
Value
CCA1
TRZ1
NRD1
RNH70
Gene
Z_Shift_K_Gem
Z_Im_K_Gem
Z_Shift_K_Cyt
Z_Im_K_Cyt
Z_Shift_L_Gem
Z_Im_L_Gem
Z_Shift_L_Cyt
Z_Im_L_Cyt

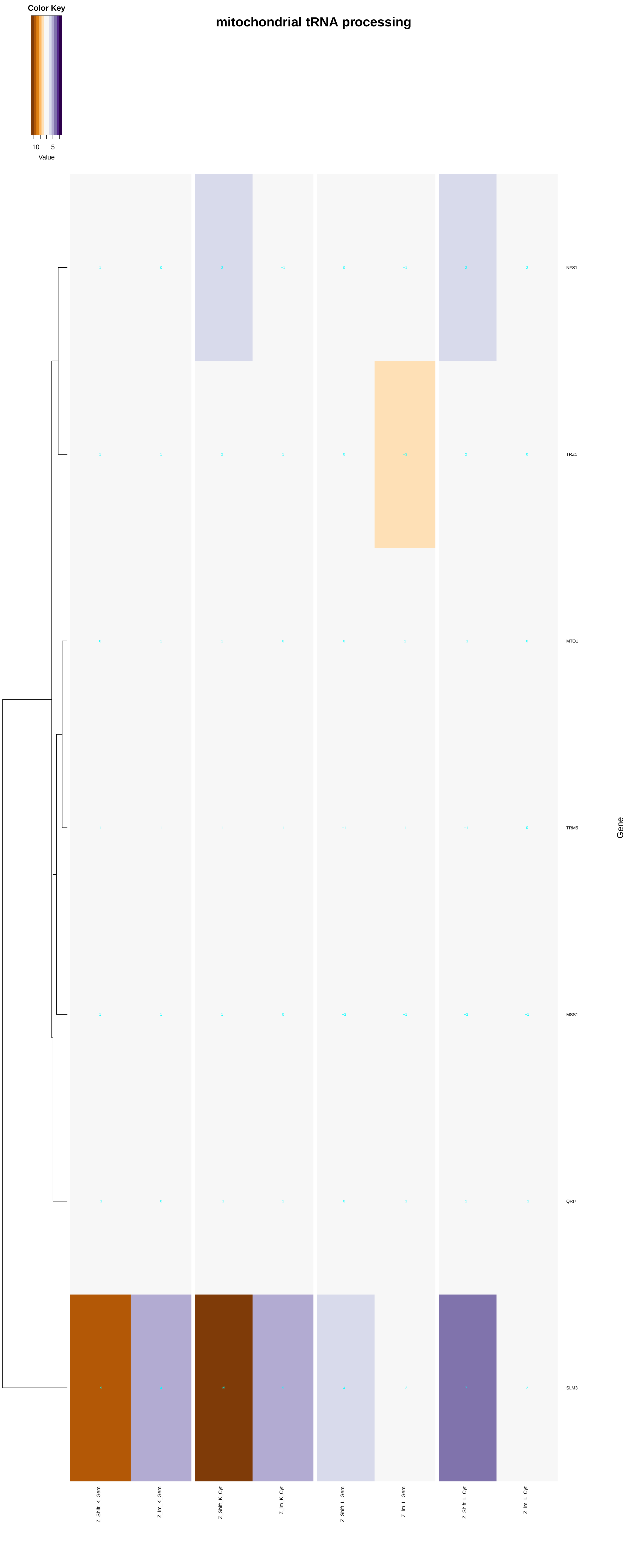

mitochondrial tRNA processing
Color Key
−10
5
Value
NFS1
TRZ1
MTO1
TRM5
MSS1
QRI7
SLM3
Gene
Z_Shift_K_Gem
Z_Im_K_Gem
Z_Shift_K_Cyt
Z_Im_K_Cyt
Z_Shift_L_Gem
Z_Im_L_Gem
Z_Shift_L_Cyt
Z_Im_L_Cyt

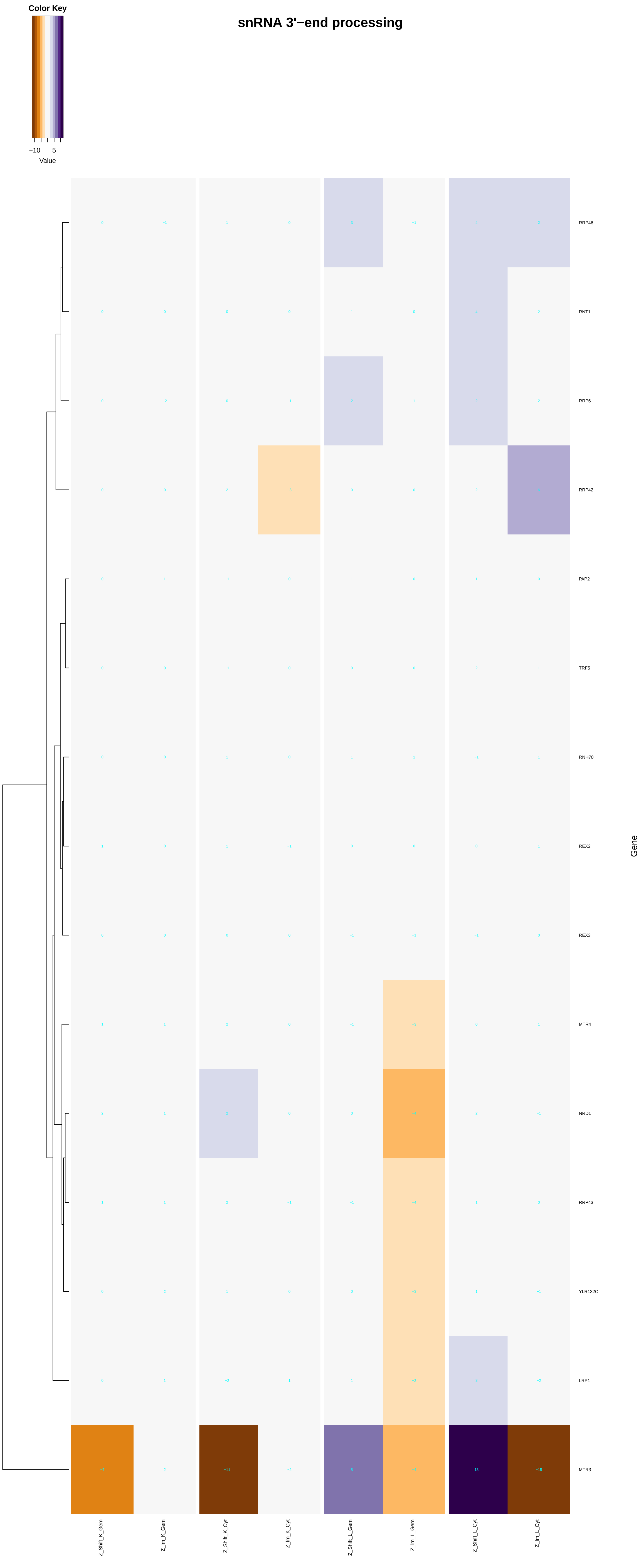
snRNA 3'−end processing
Color Key
−10
5
Value
RRP46
RNT1
RRP6
RRP42
PAP2
TRF5
RNH70
REX2
REX3
MTR4
NRD1
RRP43
YLR132C
LRP1
MTR3
Gene
Z_Shift_K_Gem
Z_Im_K_Gem
Z_Shift_K_Cyt
Z_Im_K_Cyt
Z_Shift_L_Gem
Z_Im_L_Gem
Z_Shift_L_Cyt
Z_Im_L_Cyt

### intronic snoRNA processing

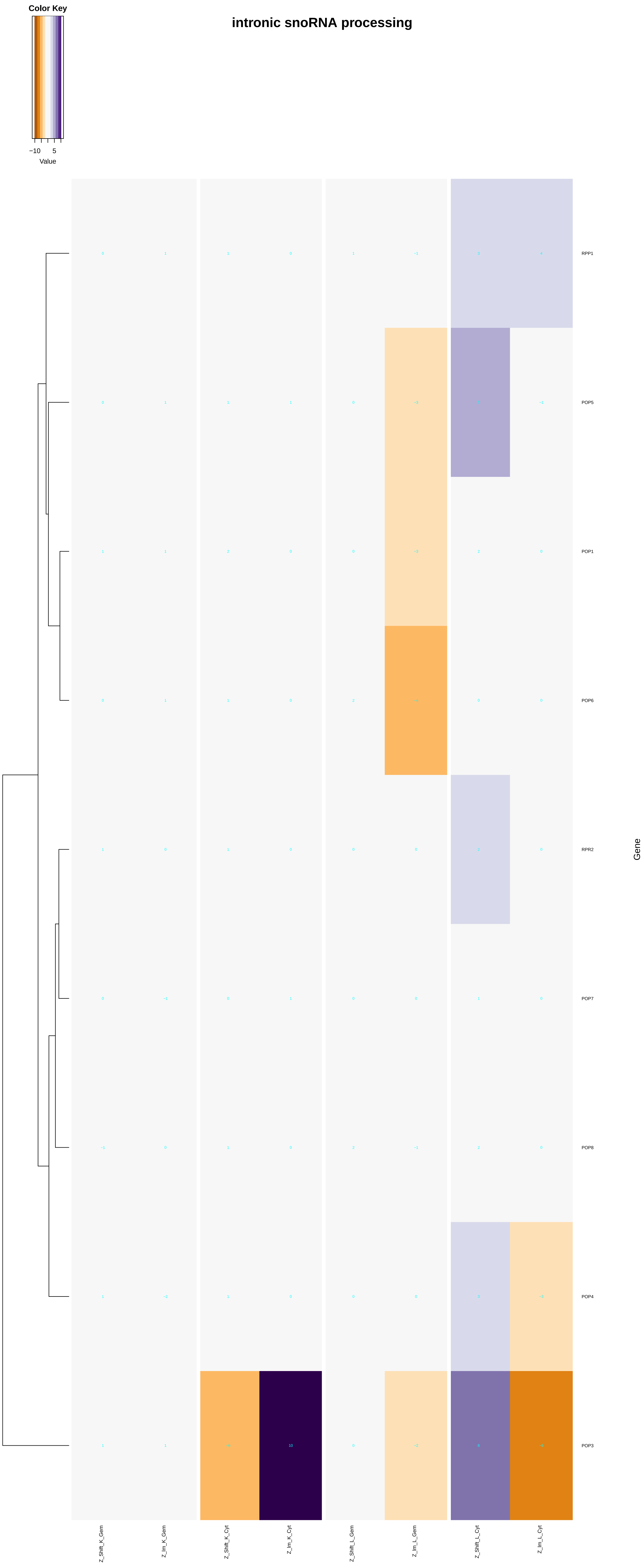

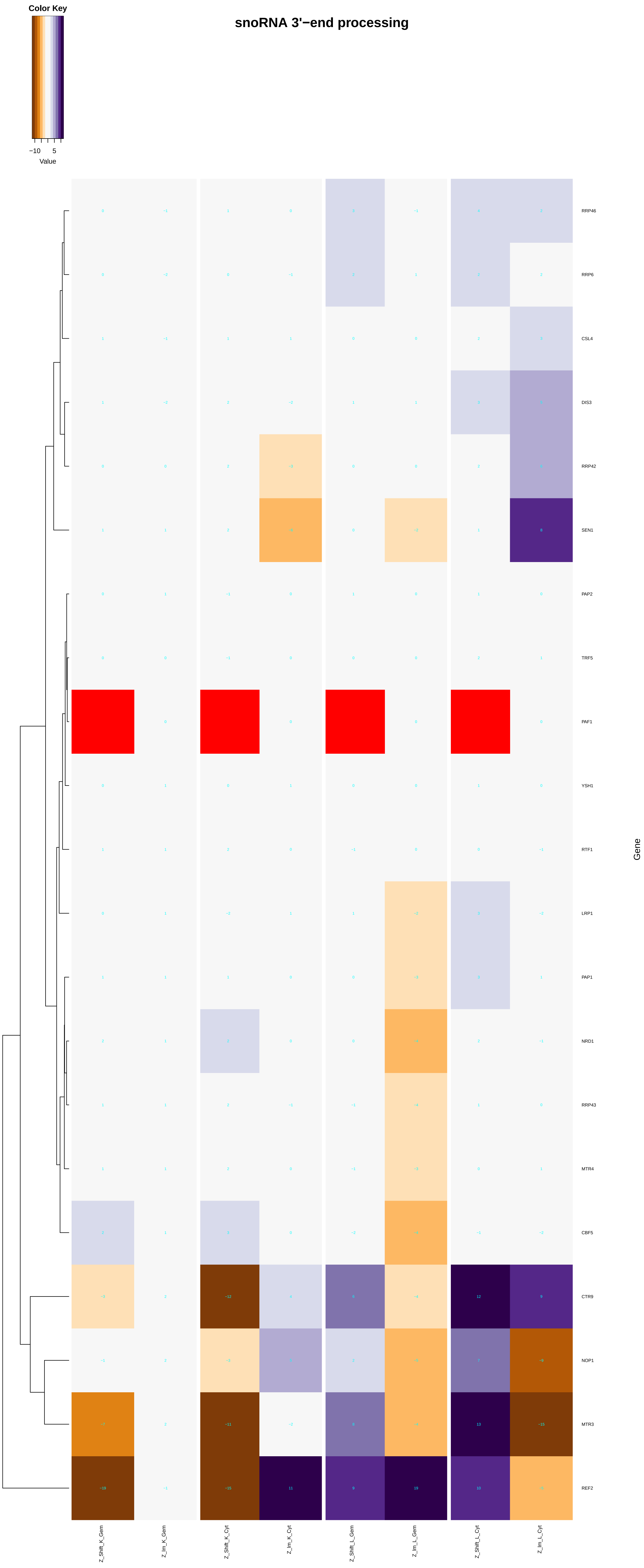

snoRNA 3'−end processing
Color Key
−10
5
Value
RRP46
RRP6
CSL4
DIS3
RRP42
SEN1
PAP2
TRF5
PAF1
YSH1
RTF1
LRP1
PAP1
NRD1
RRP43
MTR4
CBF5
CTR9
NOP1
MTR3
REF2
Gene
Z_Shift_K_Gem
Z_Im_K_Gem
Z_Shift_K_Cyt
Z_Im_K_Cyt
Z_Shift_L_Gem
Z_Im_L_Gem
Z_Shift_L_Cyt
Z_Im_L_Cyt

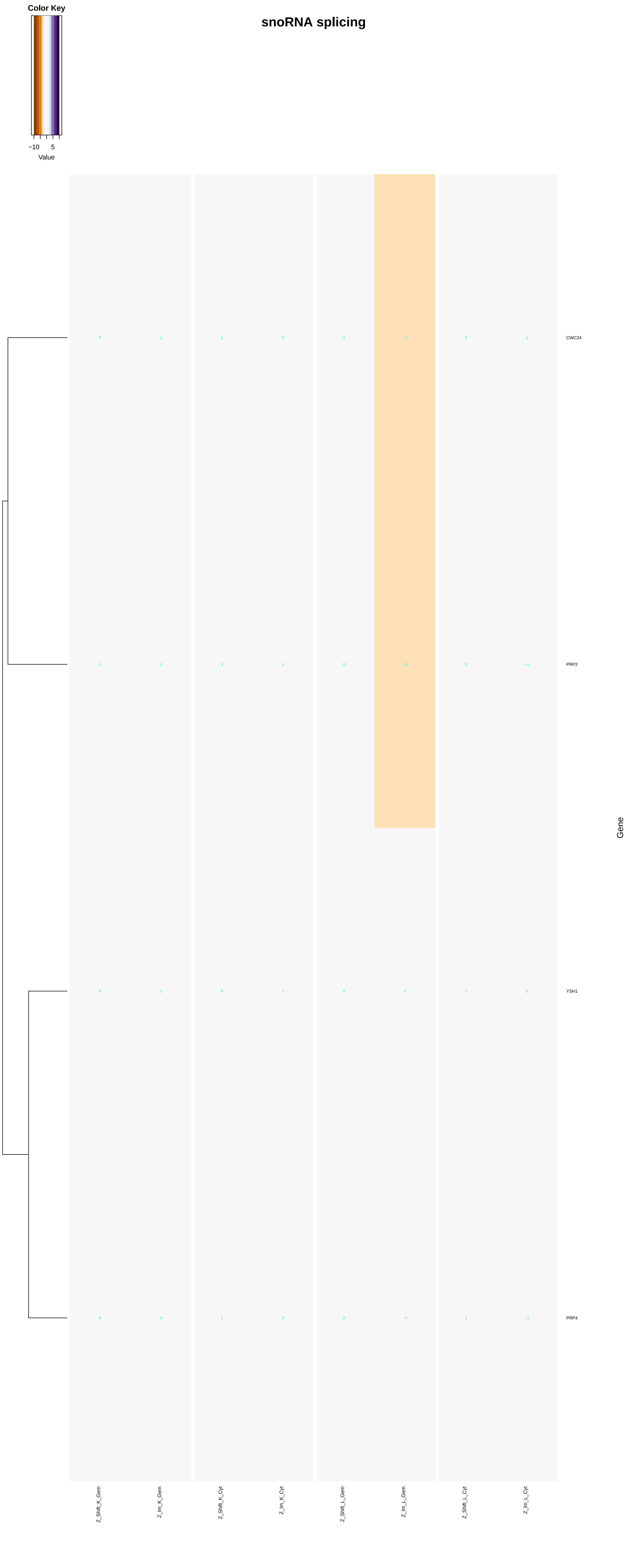
Color Key
−10
5
Value
snoRNA splicing
0 1 1 0 2 −3 0 2
CWC24
1 1 2 1 0 −3 0 −1
PRP2
0 1 0 1 0 0 1 0
YSH1
0 0 1 0 0 −2 1 −1
PRP4
Gene
Z_Shift_K_Gem
Z_Im_K_Gem
Z_Shift_K_Cyt
Z_Im_K_Cyt
Z_Shift_L_Gem
Z_Im_L_Gem
Z_Shift_L_Cyt
Z_Im_L_Cyt

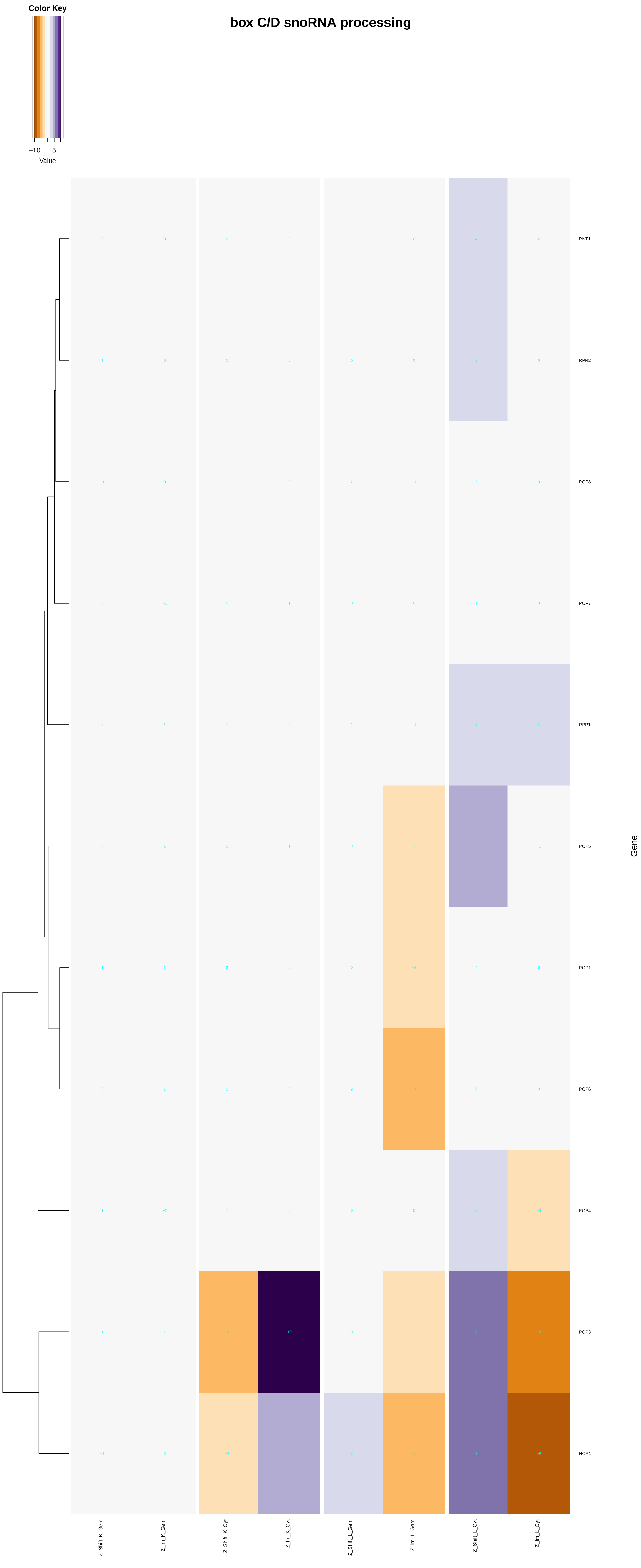
box C/D snoRNA processing
Color Key
−10
5
Value
RNT1
RPR2
POP8
POP7
RPP1
POP5
POP1
POP6
POP4
POP3
NOP1
Gene
Z_Shift_K_Gem
Z_Im_K_Gem
Z_Shift_K_Cyt
Z_Im_K_Cyt
Z_Shift_L_Gem
Z_Im_L_Gem
Z_Shift_L_Cyt
Z_Im_L_Cyt

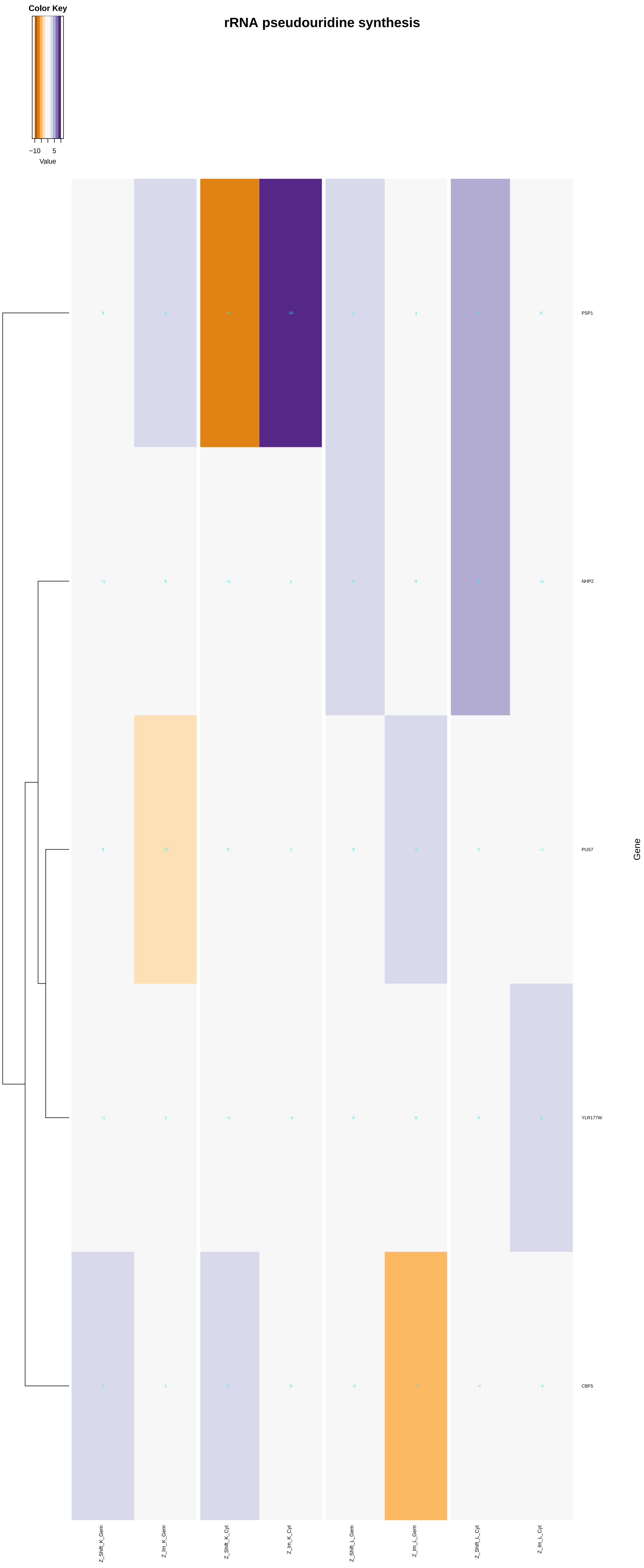
rRNA pseudouridine synthesis
Color Key
−10
5
Value
PSP1
NHP2
PUS7
YLR177W
CBF5
Gene
Z_Shift_K_Gem
Z_Im_K_Gem
Z_Shift_K_Cyt
Z_Im_K_Cyt
Z_Shift_L_Gem
Z_Im_L_Gem
Z_Shift_L_Cyt
Z_Im_L_Cyt

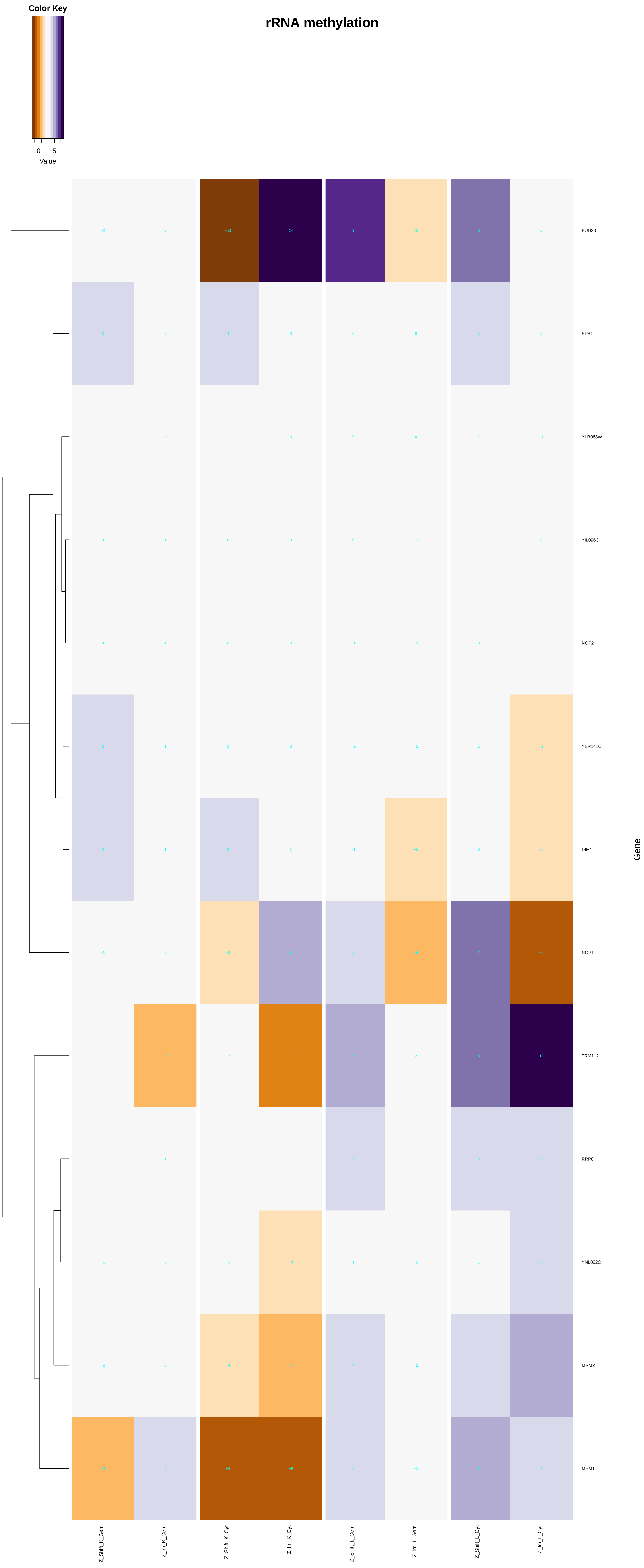

Color Key
−10
5
Value
rRNA methylation
BUD23
SPB1
YLR063W
YIL096C
NOP2
YBR141C
DIM1
NOP1
TRM112
RRP8
YNL022C
MRM2
MRM1
Gene
Z_Shift_K_Gem
Z_Im_K_Gem
Z_Shift_K_Cyt
Z_Im_K_Cyt
Z_Shift_L_Gem
Z_Im_L_Gem
Z_Shift_L_Cyt
Z_Im_L_Cyt

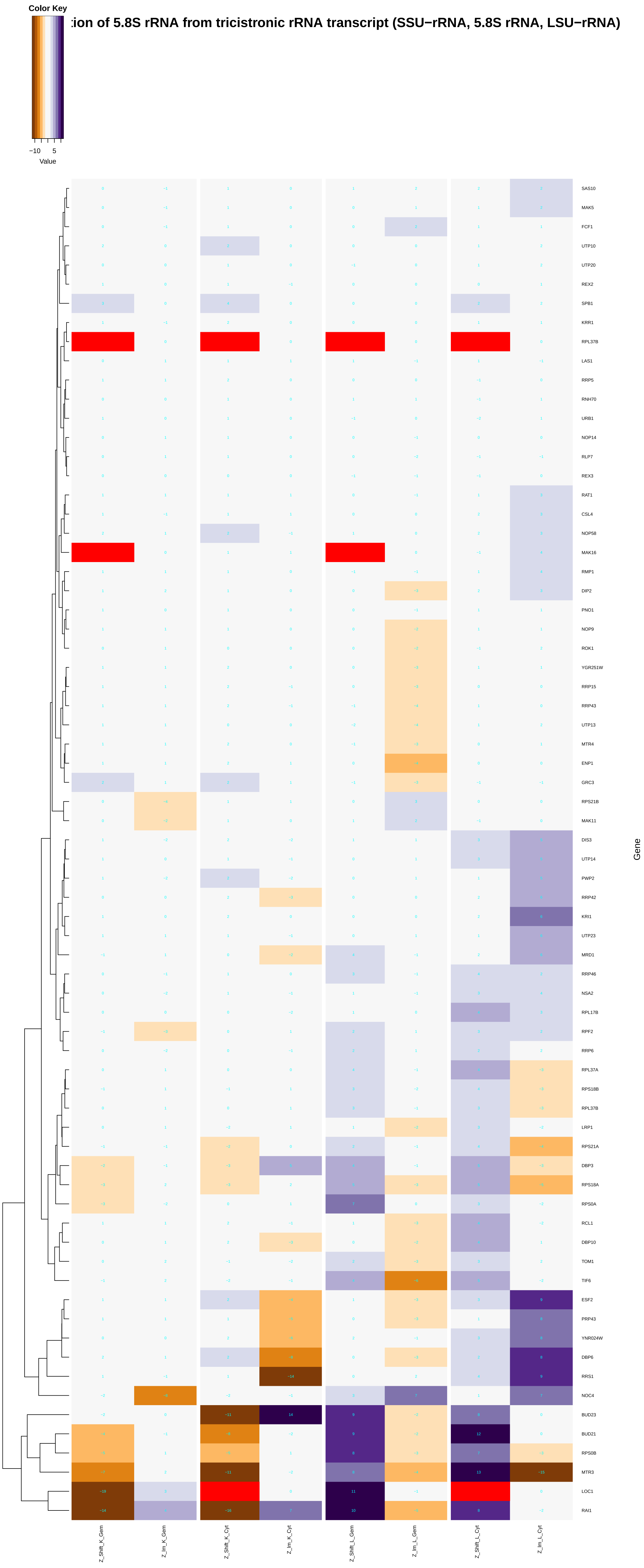

Color Key
−10 5
Value
tion of 5.8S rRNA from tricistronic rRNA transcript (SSU−rRNA, 5.8S rRNA, LSU−rRNA)
SAS10
MAK5
FCF1
UTP10
UTP20
REX2
SPB1
KRR1
RPL37B
LAS1
RRP5
RNH70
URB1
NOP14
RLP7
REX3
RAT1
CSL4
NOP58
MAK16
RMP1
DIP2
PNO1
NOP9
ROK1
YGR251W
RRP15
RRP43
UTP13
MTR4
ENP1
GRC3
RPS21B
MAK11
DIS3
UTP14
PWP2
RRP42
KRI1
UTP23
MRD1
RRP46
NSA2
RPL17B
RPF2
RRP6
RPL37A
RPS18B
RPL37B
LRP1
RPS21A
DBP3
RPS18A
RPS0A
RCL1
DBP10
TOM1
TIF6
ESF2
PRP43
YNR024W
DBP6
RRS1
NOC4
BUD23
BUD21
RPS0B
MTR3
LOC1
RAI1
Gene
Z_Shift_K_Gem
Z_Im_K_Gem
Z_Shift_K_Cyt
Z_Im_K_Cyt
Z_Shift_L_Gem
Z_Im_L_Gem
Z_Shift_L_Cyt
Z_Im_L_Cyt

ion of LSU−rRNA from tricistronic rRNA transcript (SSU−rRNA, 5.8S rRNA, LSU−rRNA)

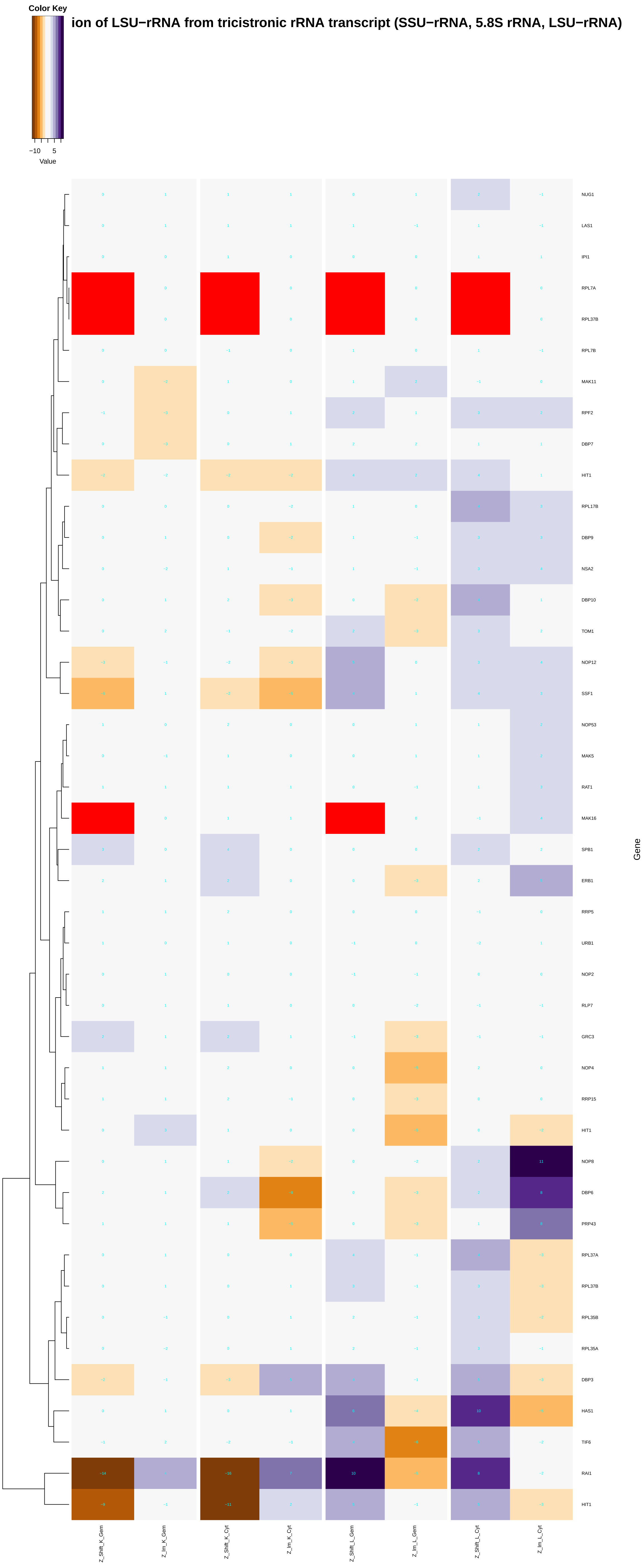

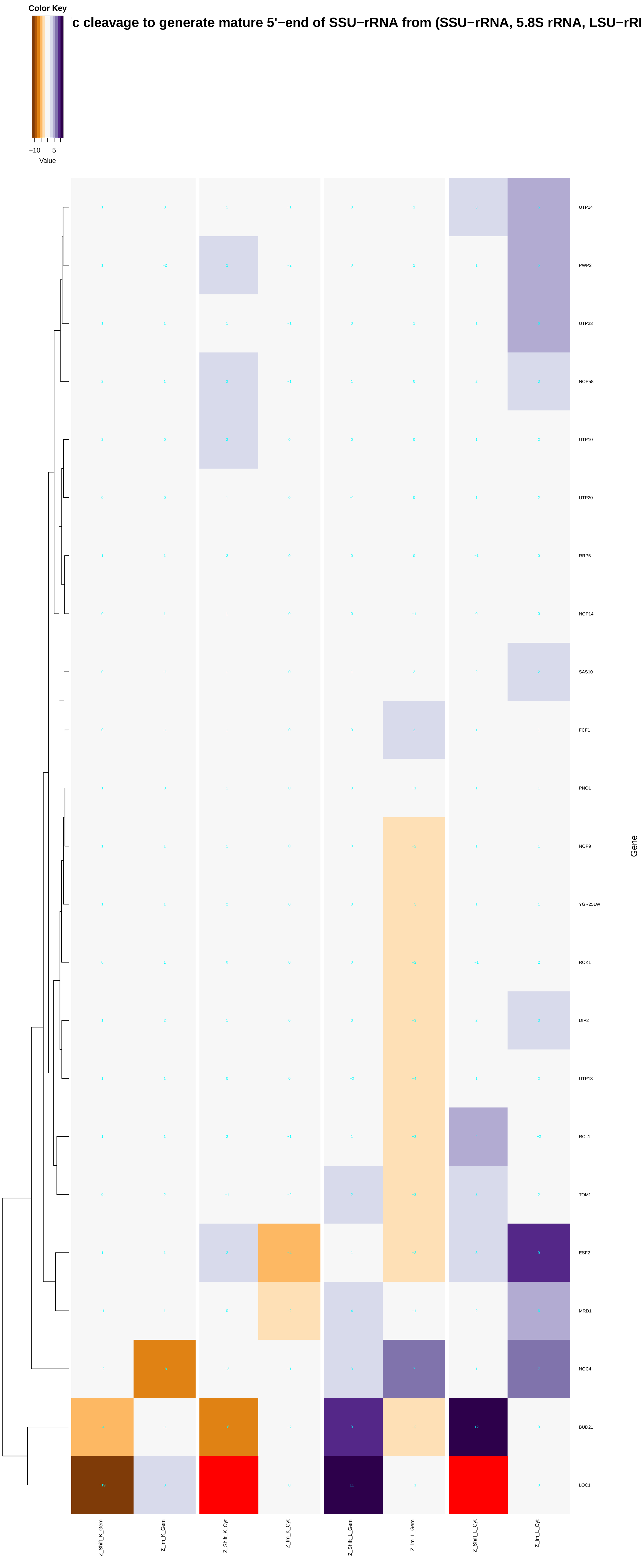

Color Key
−10
5
Value
c cleavage to generate mature 5'−end of SSU−rRNA from (SSU−rRNA, 5.8S rRNA, LSU−rR
UTP14
PWP2
UTP23
NOP58
UTP10
UTP20
RRP5
NOP14
SAS10
FCF1
PNO1
NOP9
YGR251W
ROK1
DIP2
UTP13
RCL1
TOM1
ESF2
MRD1
NOC4
BUD21
LOC1
Gene
Z_Shift_K_Gem
Z_Im_K_Gem
Z_Shift_K_Cyt
Z_Im_K_Cyt
Z_Shift_L_Gem
Z_Im_L_Gem
Z_Shift_L_Cyt
Z_Im_L_Cyt

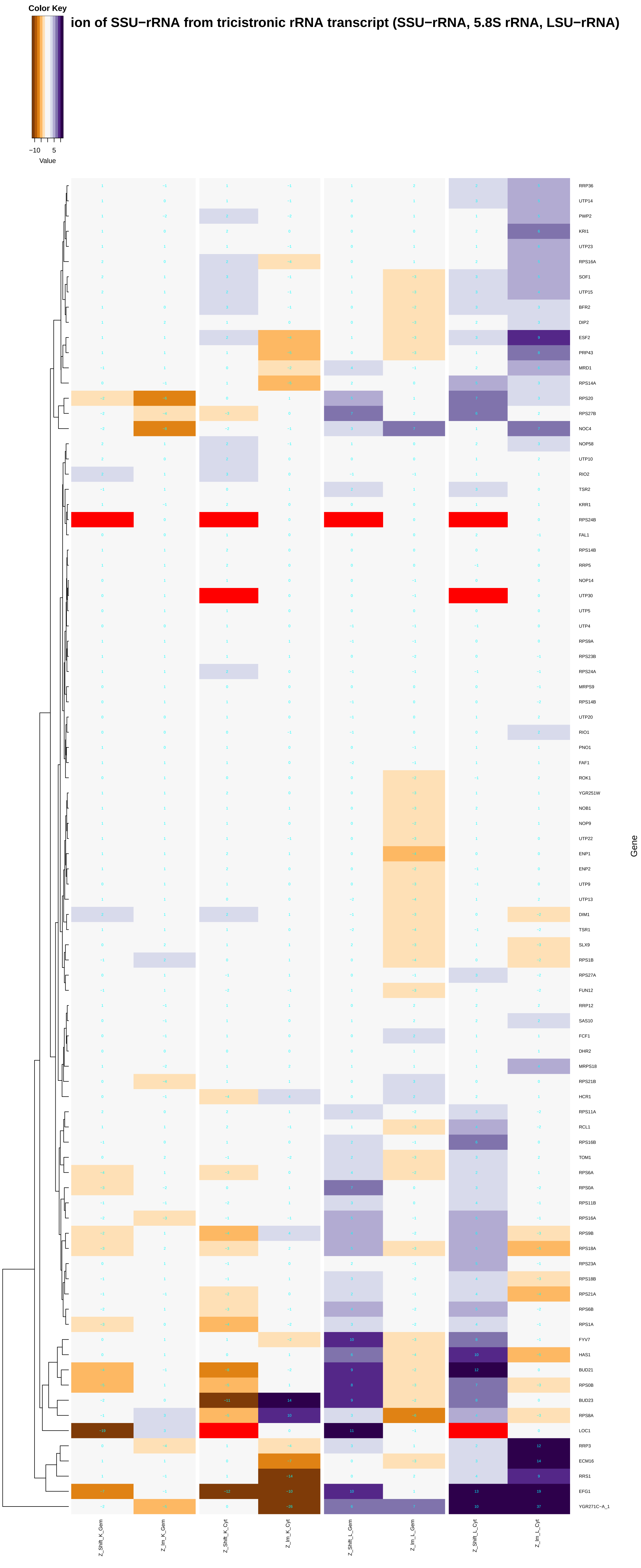
ion of SSU−rRNA from tricistronic rRNA transcript (SSU−rRNA, 5.8S rRNA, LSU−rRNA)
Color Key
−10
5
Value
Gene
RRP36
UTP14
PWP2
KRI1
UTP23
RPS16A
SOF1
UTP15
BFR2
DIP2
ESF2
PRP43
MRD1
RPS14A
RPS20
RPS27B
NOC4
NOP58
UTP10
RIO2
TSR2
KRR1
RPS24B
FAL1
RPS14B
RRP5
NOP14
UTP30
UTP5
UTP4
RPS9A
RPS23B
RPS24A
MRPS9
RPS14B
UTP20
RIO1
PNO1
FAF1
ROK1
YGR251W
NOB1
NOP9
UTP22
ENP1
ENP2
UTP9
UTP13
DIM1
TSR1
SLX9
RPS1B
RPS27A
FUN12
RRP12
SAS10
FCF1
DHR2
MRPS18
RPS21B
HCR1
RPS11A
RCL1
RPS16B
TOM1
RPS6A
RPS0A
RPS11B
RPS16A
RPS9B
RPS18A
RPS23A
RPS18B
RPS21A
RPS6B
RPS1A
FYV7
HAS1
BUD21
RPS0B
BUD23
RPS8A
LOC1
RRP3
ECM16
RRS1
EFG1
YGR271C−A_1
Z_Shift_K_Gem
Z_Im_K_Gem
Z_Shift_K_Cyt
Z_Im_K_Cyt
Z_Shift_L_Gem
Z_Im_L_Gem
Z_Shift_L_Cyt
Z_Im_L_Cyt

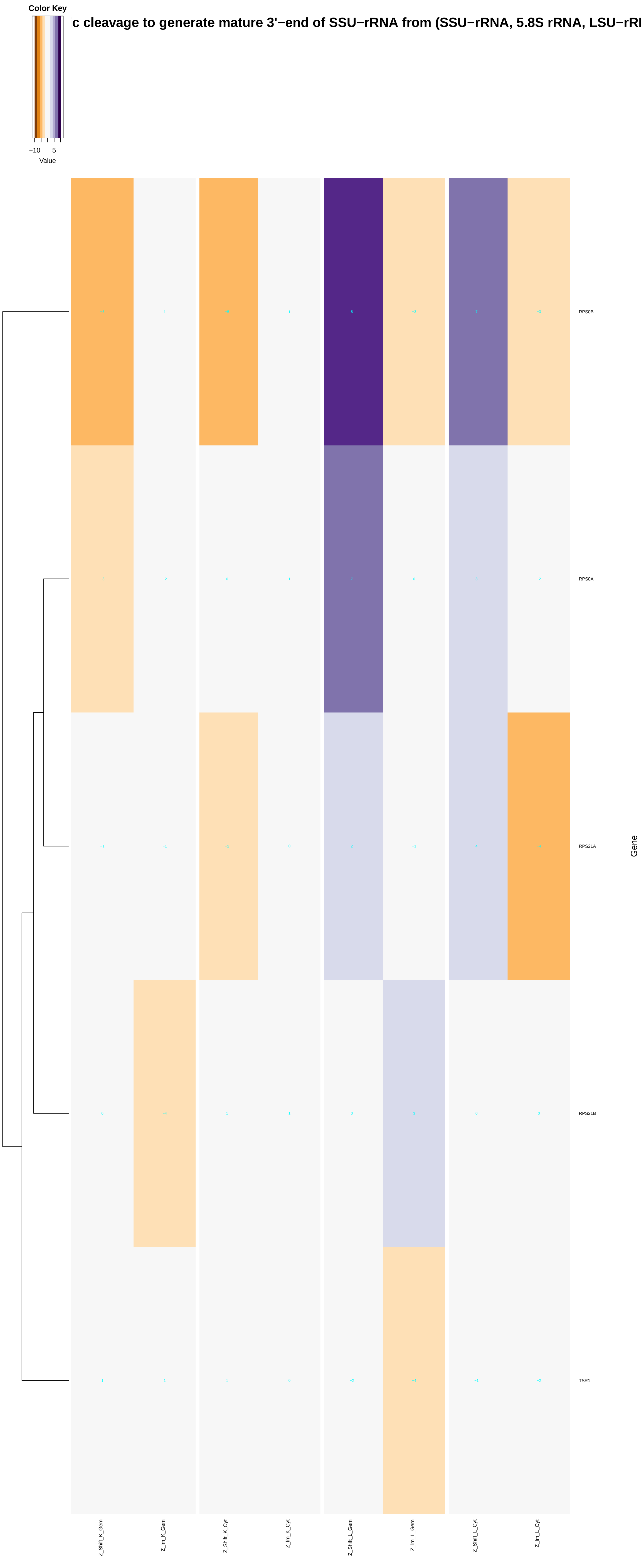
c cleavage to generate mature 3'−end of SSU−rRNA from (SSU−rRNA, 5.8S rRNA, LSU−rRI
Color Key
−10
5
Value
RPS0B
RPS0A
RPS21A
RPS21B
TSR1
Gene
Z_Shift_K_Gem
Z_Im_K_Gem
Z_Shift_K_Cyt
Z_Im_K_Cyt
Z_Shift_L_Gem
Z_Im_L_Gem
Z_Shift_L_Cyt
Z_Im_L_Cyt

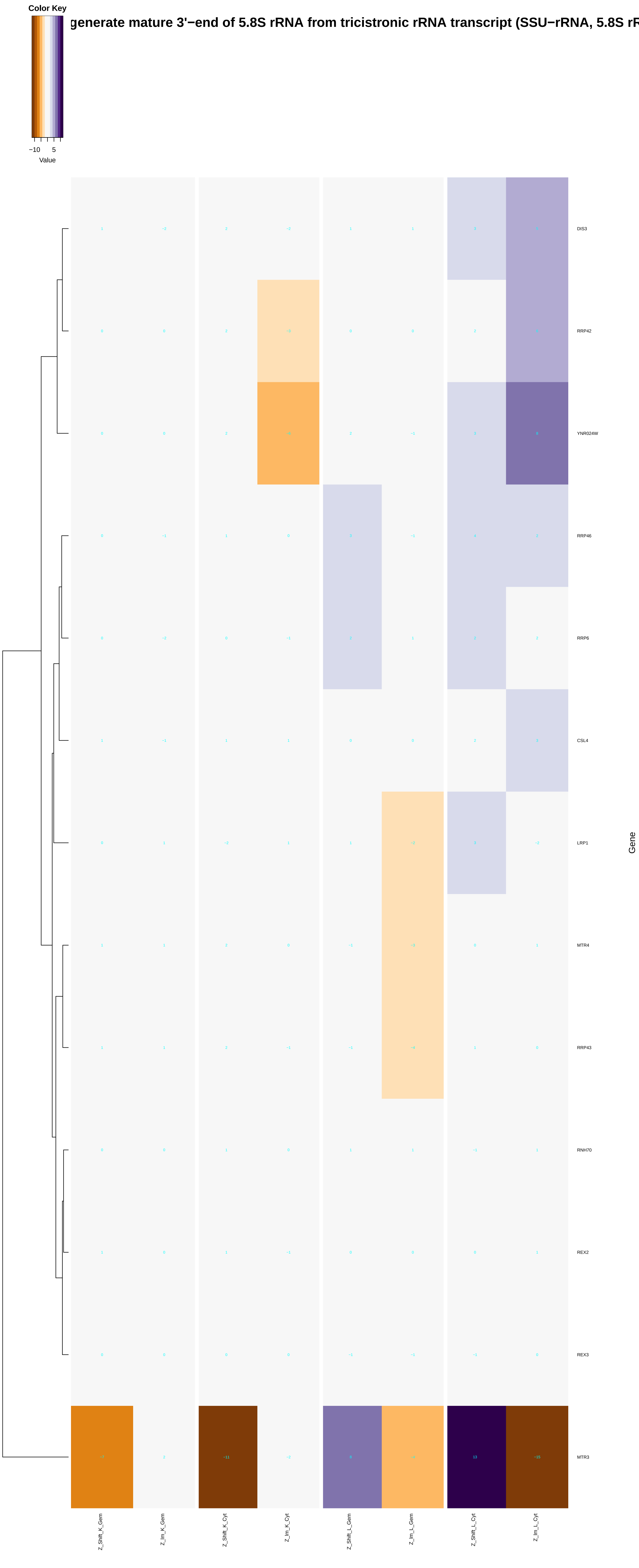

Color Key
−10
5
Value
generate mature 3'−end of 5.8S rRNA from tricistronic rRNA transcript (SSU−rRNA, 5.8S rR
DIS3
RRP42
YNR024W
RRP46
RRP6
CSL4
LRP1
MTR4
RRP43
RNH70
REX2
REX3
MTR3
Gene
Z_Shift_K_Gem
Z_Im_K_Gem
Z_Shift_K_Cyt
Z_Im_K_Cyt
Z_Shift_L_Gem
Z_Im_L_Gem
Z_Shift_L_Cyt
Z_Im_L_Cyt

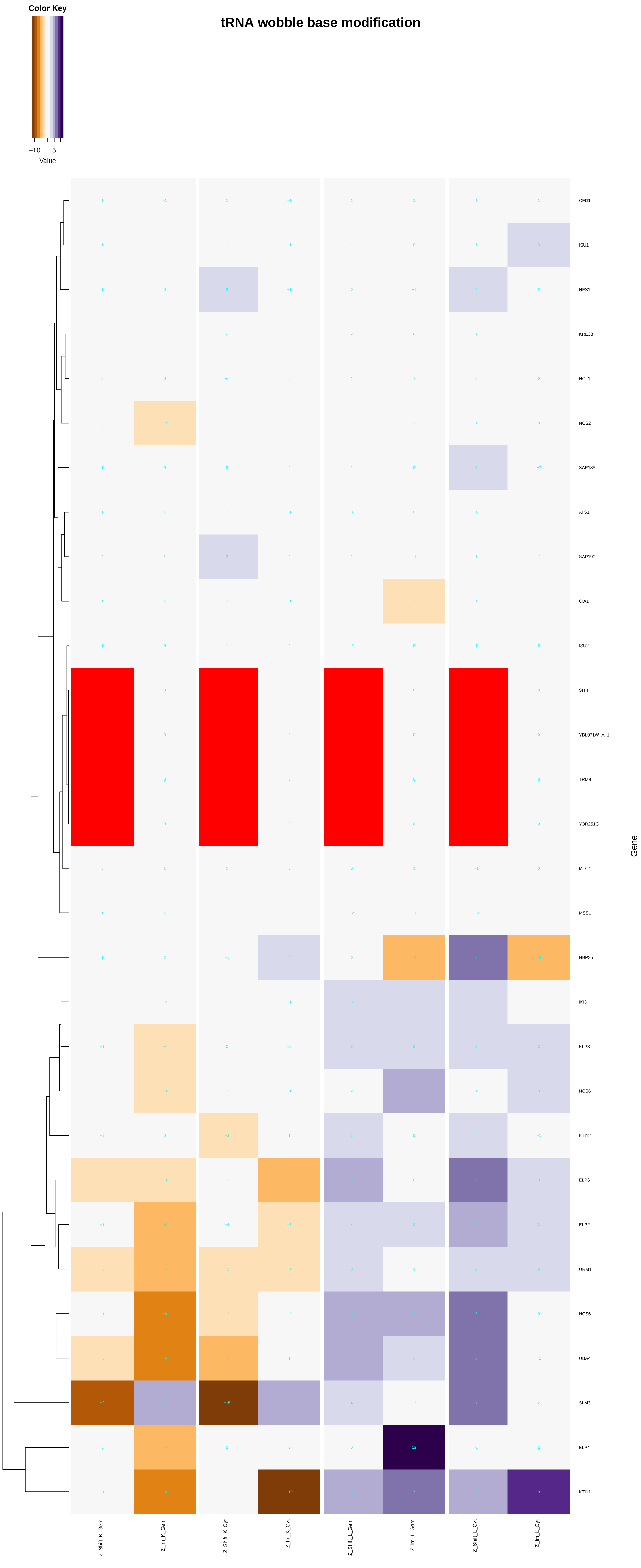

tRNA wobble base modification
Color Key
−10
5
Value
CFD1
ISU1
NFS1
KRE33
NCL1
NCS2
SAP185
ATS1
SAP190
CIA1
ISU2
SIT4
YBL071W−A_1
TRM9
YOR251C
MTO1
MSS1
NBP35
IKI3
ELP3
NCS6
KTI12
ELP6
ELP2
URM1
NCS6
UBA4
SLM3
ELP4
KTI11
Gene
Z_Shift_K_Gem
Z_Im_K_Gem
Z_Shift_K_Cyt
Z_Im_K_Cyt
Z_Shift_L_Gem
Z_Im_L_Gem
Z_Shift_L_Cyt
Z_Im_L_Cyt

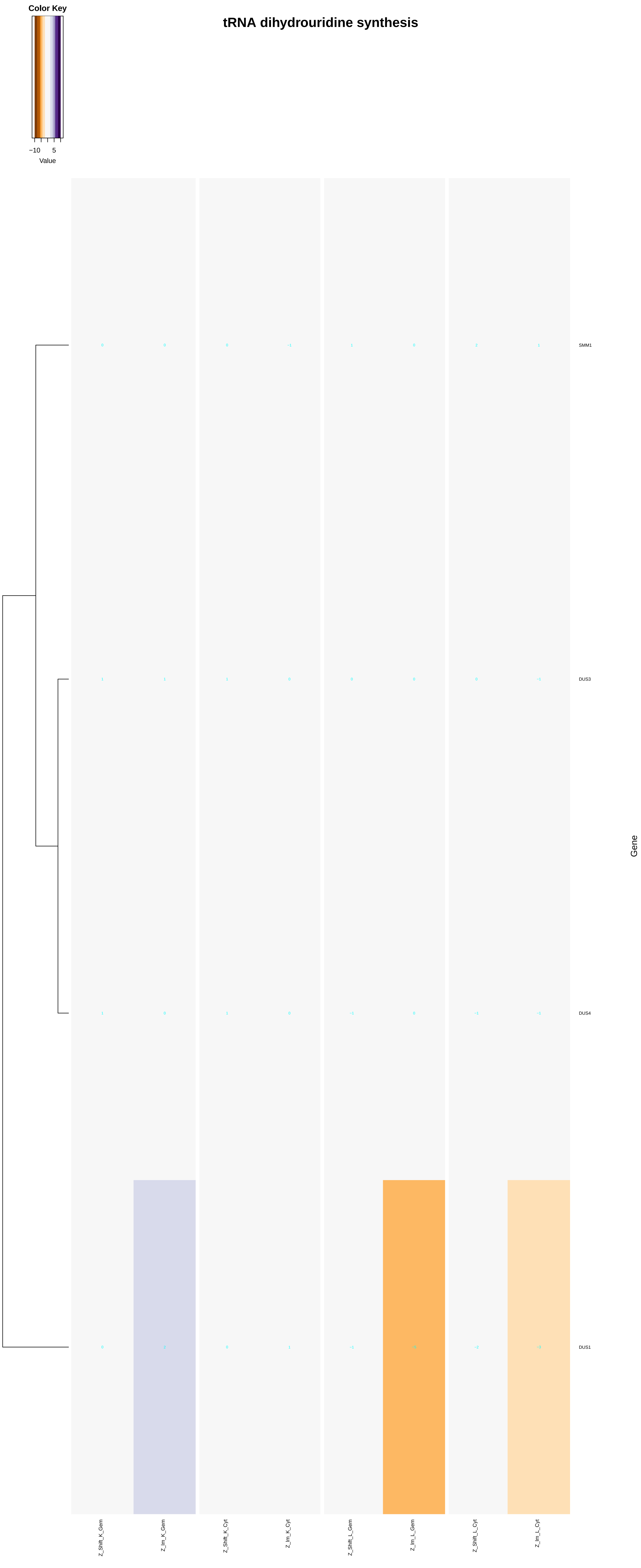
tRNA dihydrouridine synthesis
Color Key
−10
5
Value
SMM1
DUS3
DUS4
DUS1
Gene
Z_Shift_K_Gem
Z_Im_K_Gem
Z_Shift_K_Cyt
Z_Im_K_Cyt
Z_Shift_L_Gem
Z_Im_L_Gem
Z_Shift_L_Cyt
Z_Im_L_Cyt

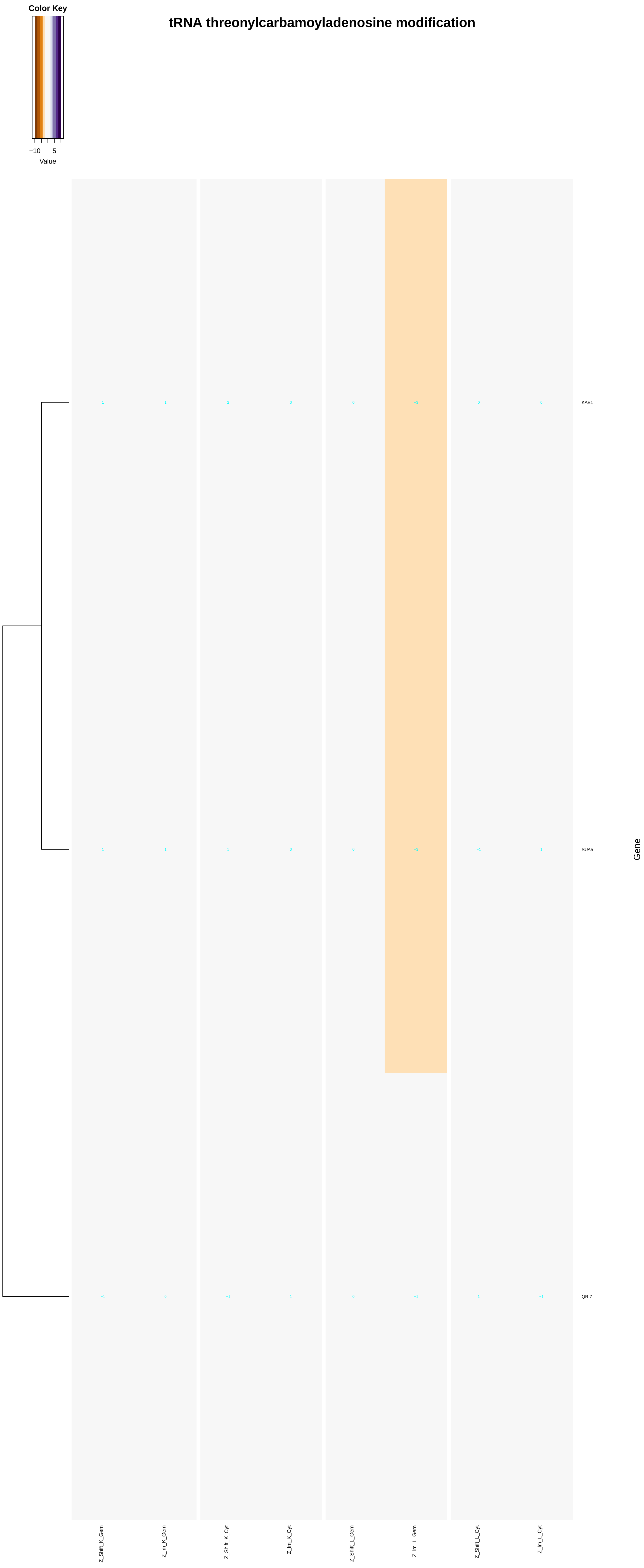

Color Key
−10
5
Value
tRNA threonylcarbamoyladenosine modification
1 1 2 0 0 −3 0 0 KAE1
1 1 1 0 0 −3 −1 1 SUA5
−1 0 −1 1 0 −1 1 −1 QRI7
Gene
Z_Shift_K_Gem
Z_Im_K_Gem
Z_Shift_K_Cyt
Z_Im_K_Cyt
Z_Shift_L_Gem
Z_Im_L_Gem
Z_Shift_L_Cyt
Z_Im_L_Cyt

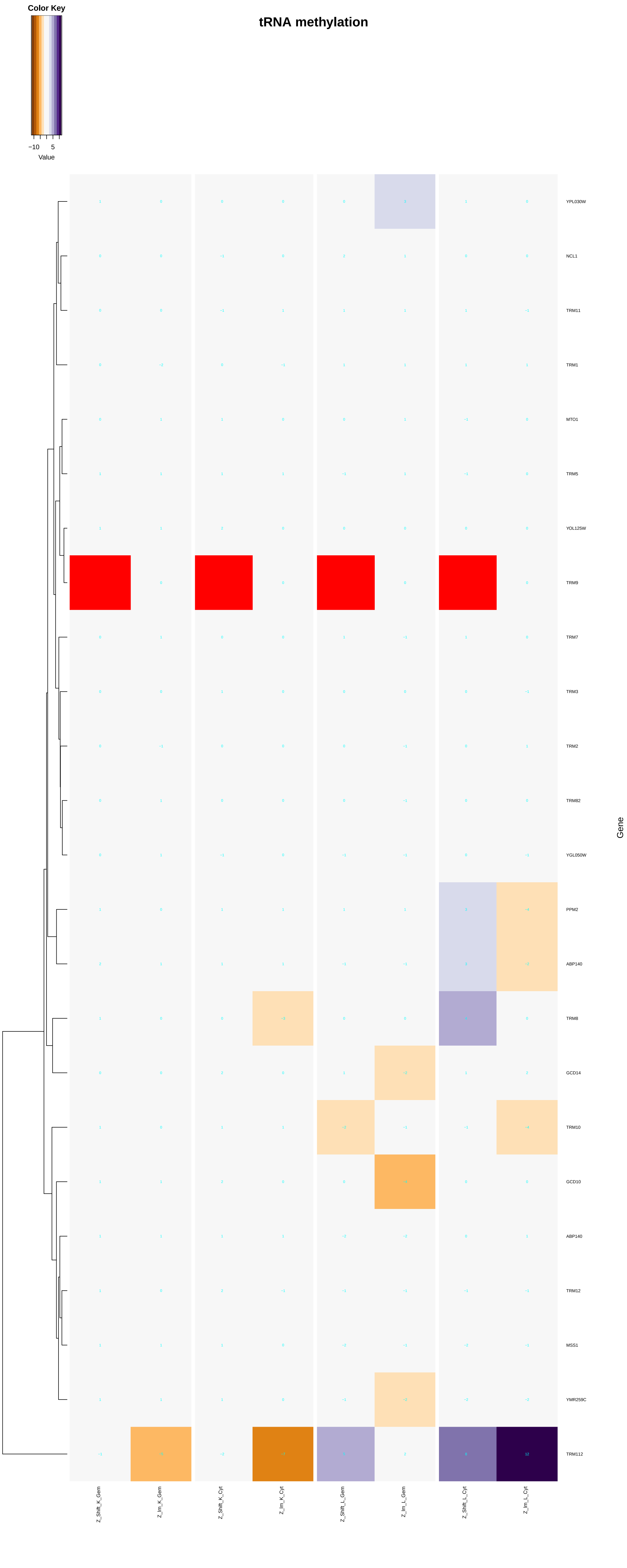
tRNA methylation
Color Key
−10
5
Value
YPL030W
NCL1
TRM11
TRM1
MTO1
TRM5
YOL125W
TRM9
TRM7
TRM3
TRM2
TRM82
YGL050W
PPM2
ABP140
TRM8
GCD14
TRM10
GCD10
ABP140
TRM12
MSS1
YMR259C
TRM112
Gene
Z_Shift_K_Gem
Z_Im_K_Gem
Z_Shift_K_Cyt
Z_Im_K_Cyt
Z_Shift_L_Gem
Z_Im_L_Gem
Z_Shift_L_Cyt
Z_Im_L_Cyt

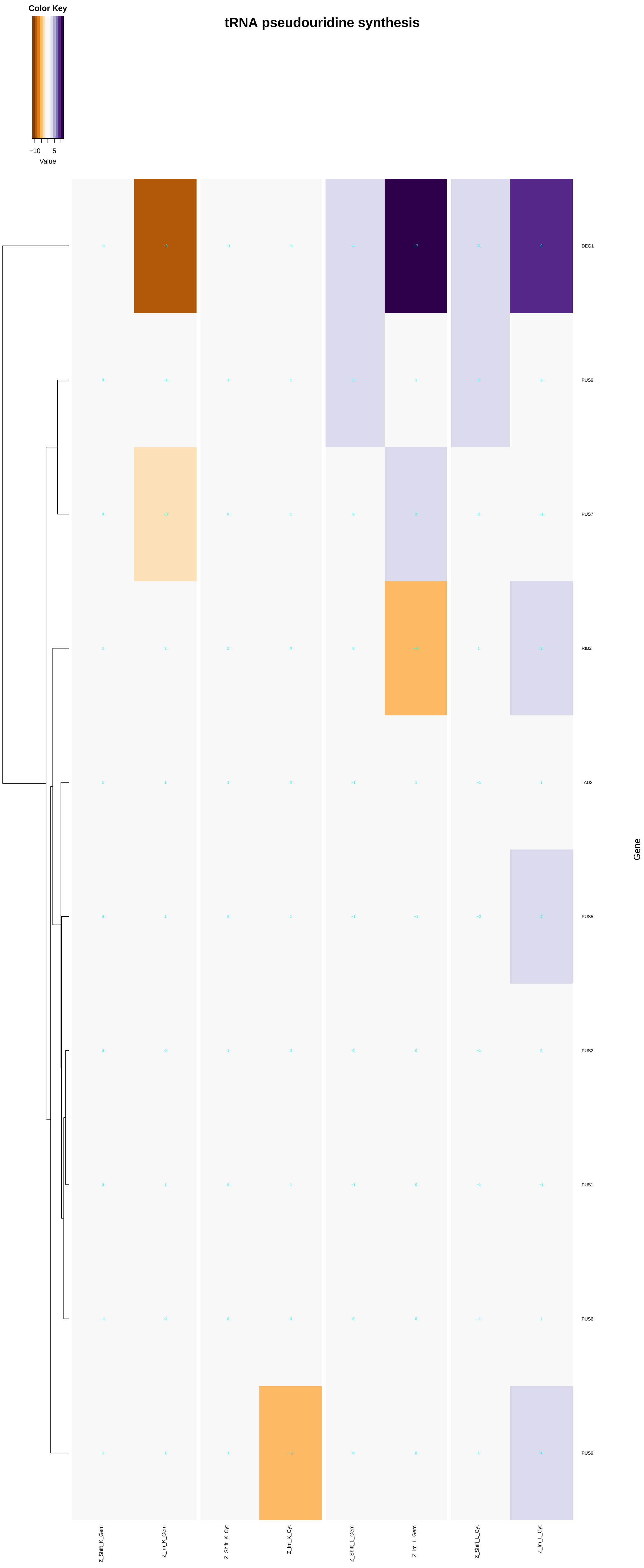
tRNA pseudouridine synthesis
Color Key
−10
5
Value
DEG1
PUS9
PUS7
RIB2
TAD3
PUS5
PUS2
PUS1
PUS6
PUS9
Gene
Z_Shift_K_Gem
Z_Im_K_Gem
Z_Shift_K_Cyt
Z_Im_K_Cyt
Z_Shift_L_Gem
Z_Im_L_Gem
Z_Shift_L_Cyt
Z_Im_L_Cyt

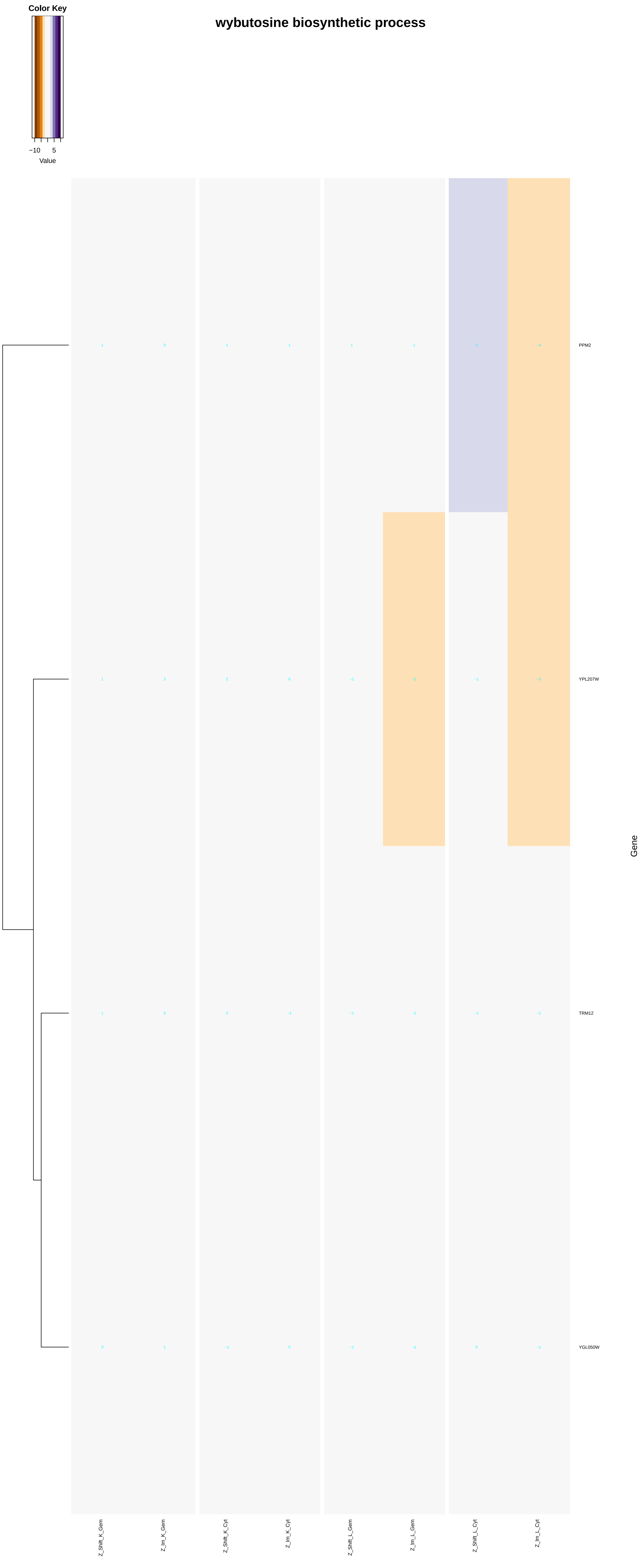
Color Key
−10
5
Value
wybutosine biosynthetic process
PPM2
YPL207W
TRM12
YGL050W
Gene
Z_Shift_K_Gem
Z_Im_K_Gem
Z_Shift_K_Cyt
Z_Im_K_Cyt
Z_Shift_L_Gem
Z_Im_L_Gem
Z_Shift_L_Cyt
Z_Im_L_Cyt

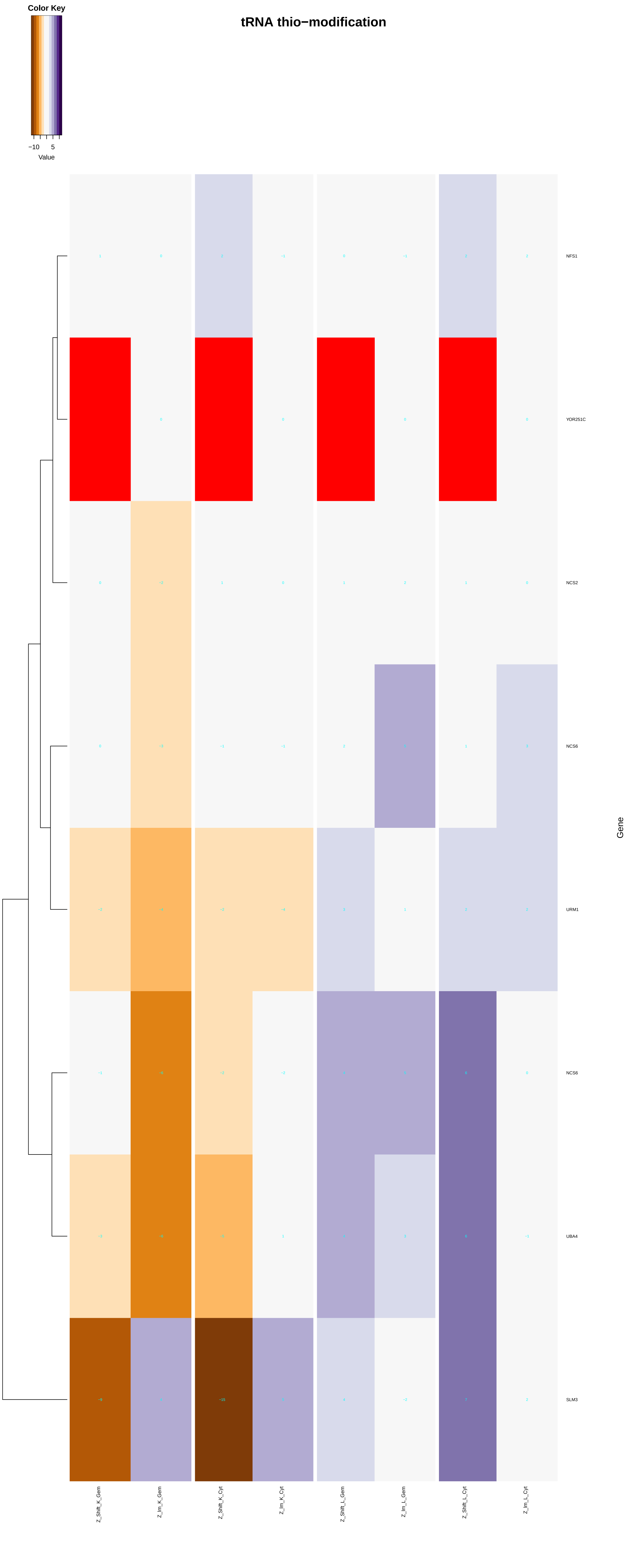
Color Key
−10
5
Value
tRNA thio−modification
NFS1
YOR251C
NCS2
NCS6
URM1
NCS6
UBA4
SLM3
Gene
Z_Shift_K_Gem
Z_Im_K_Gem
Z_Shift_K_Cyt
Z_Im_K_Cyt
Z_Shift_L_Gem
Z_Im_L_Gem
Z_Shift_L_Cyt
Z_Im_L_Cyt

### mitochondrial tRNA modification

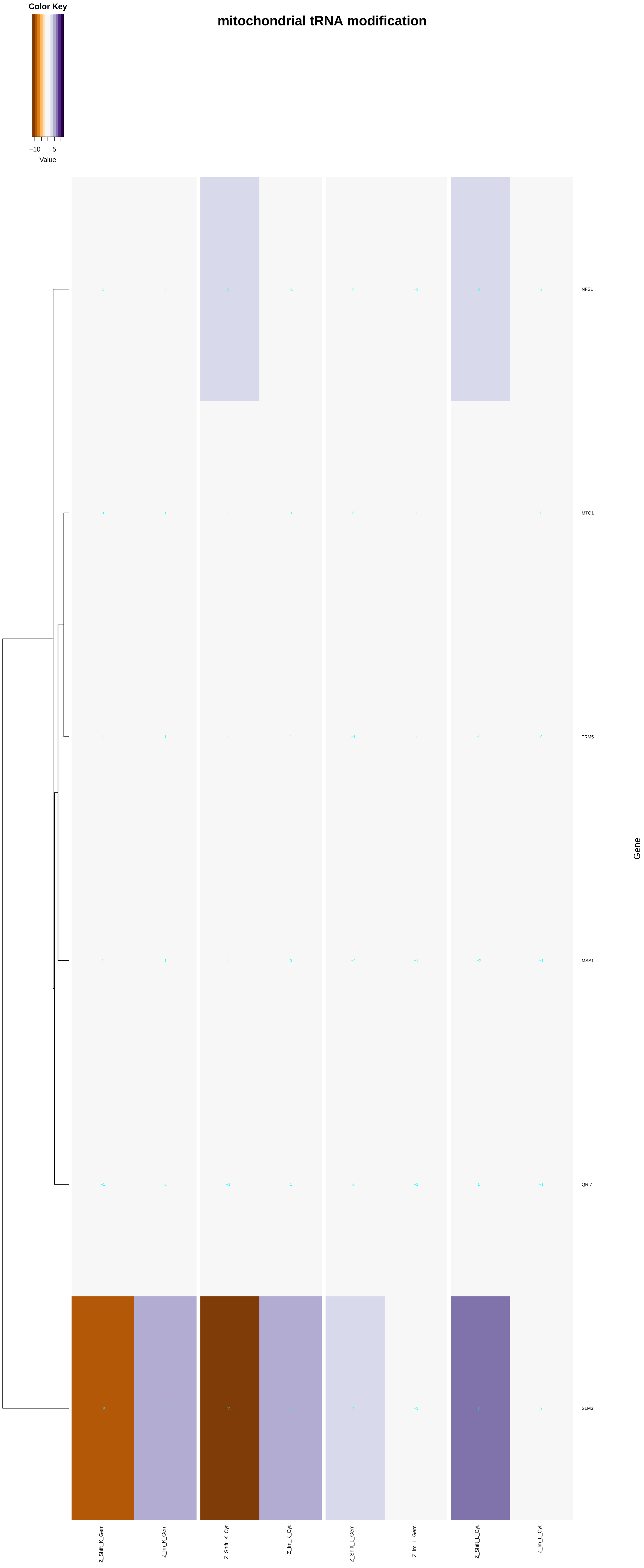

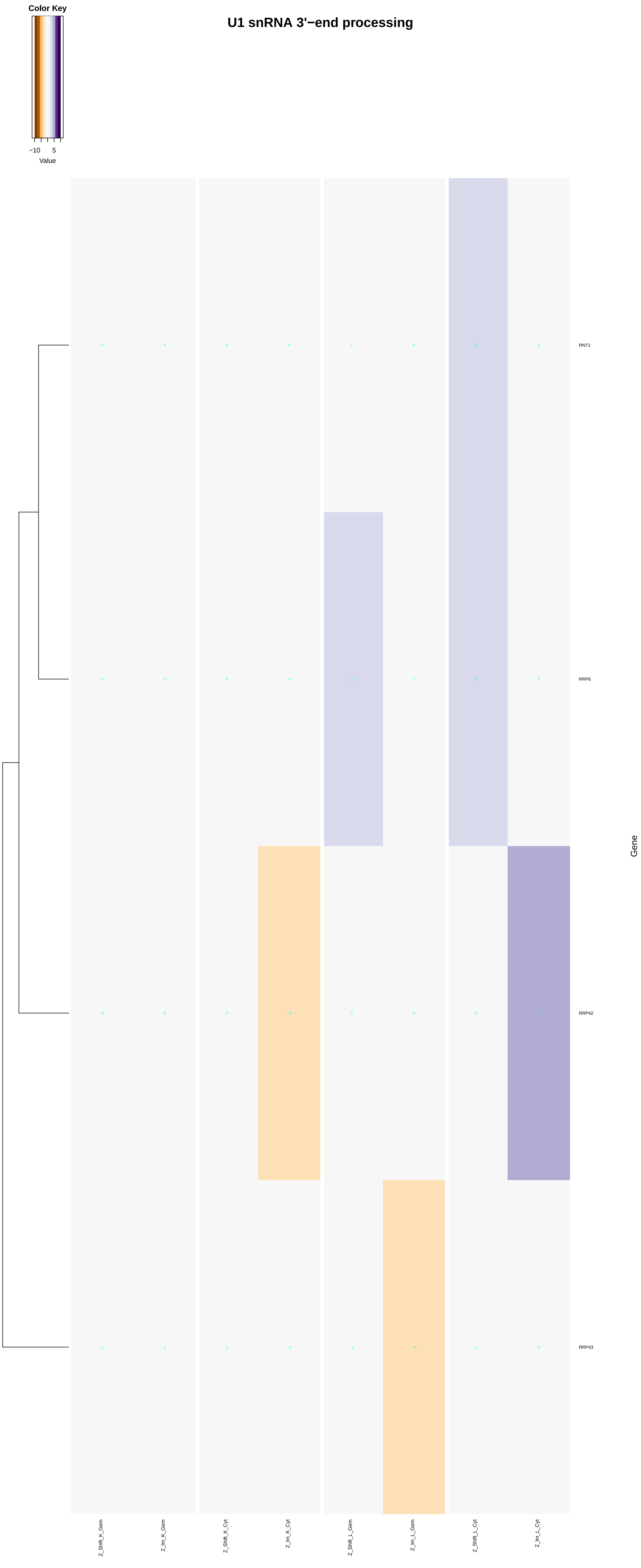

Color Key
−10
5
Value
U1 snRNA 3'−end processing
RNT1
RRP6
RRP42
RRP43
Gene
Z_Shift_K_Gem
Z_Im_K_Gem
Z_Shift_K_Cyt
Z_Im_K_Cyt
Z_Shift_L_Gem
Z_Im_L_Gem
Z_Shift_L_Cyt
Z_Im_L_Cyt

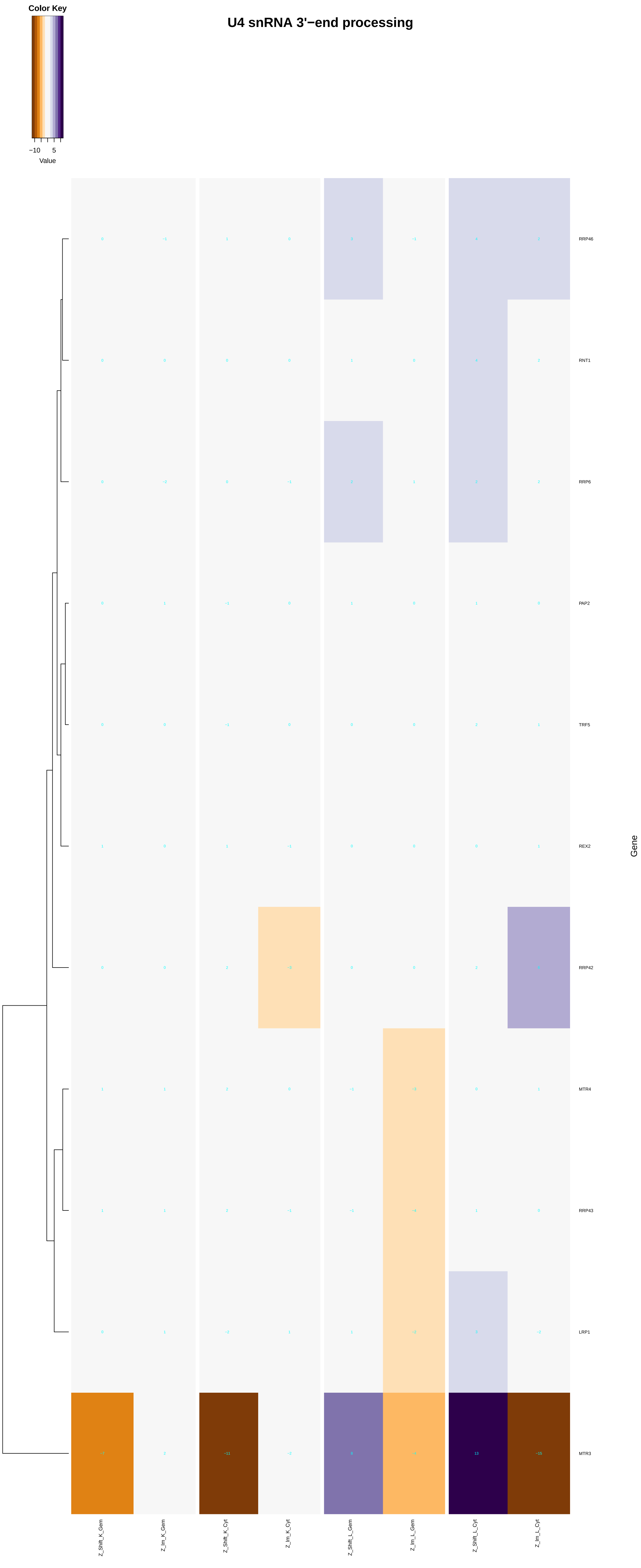
U4 snRNA 3'−end processing
Color Key
−10 5
Value
RRP46
RNT1
RRP6
PAP2
TRF5
REX2
RRP42
MTR4
RRP43
LRP1
MTR3
Gene
Z_Shift_K_Gem
Z_Im_K_Gem
Z_Shift_K_Cyt
Z_Im_K_Cyt
Z_Shift_L_Gem
Z_Im_L_Gem
Z_Shift_L_Cyt
Z_Im_L_Cyt

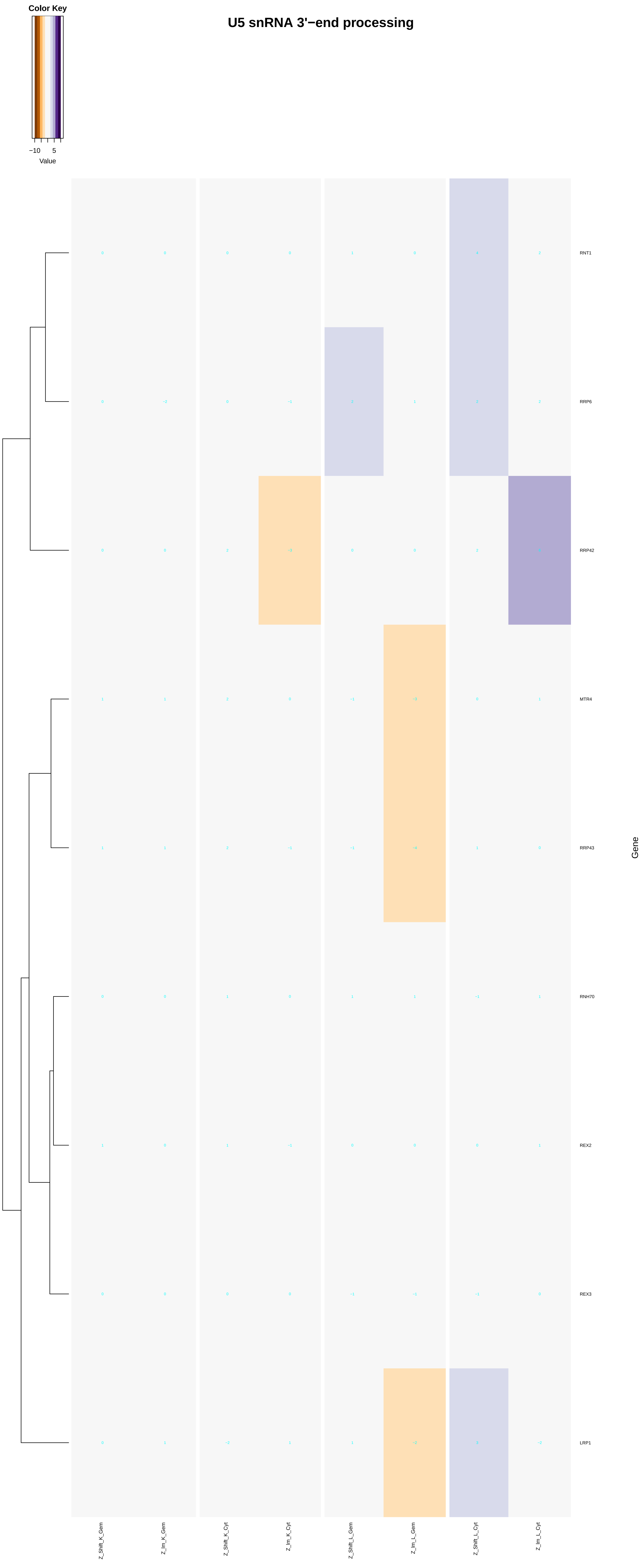

U5 snRNA 3'−end processing
Color Key
−10
5
Value
RNT1
RRP6
RRP42
MTR4
RRP43
RNH70
REX2
REX3
LRP1
Gene
Z_Shift_K_Gem
Z_Im_K_Gem
Z_Shift_K_Cyt
Z_Im_K_Cyt
Z_Shift_L_Gem
Z_Im_L_Gem
Z_Shift_L_Cyt
Z_Im_L_Cyt

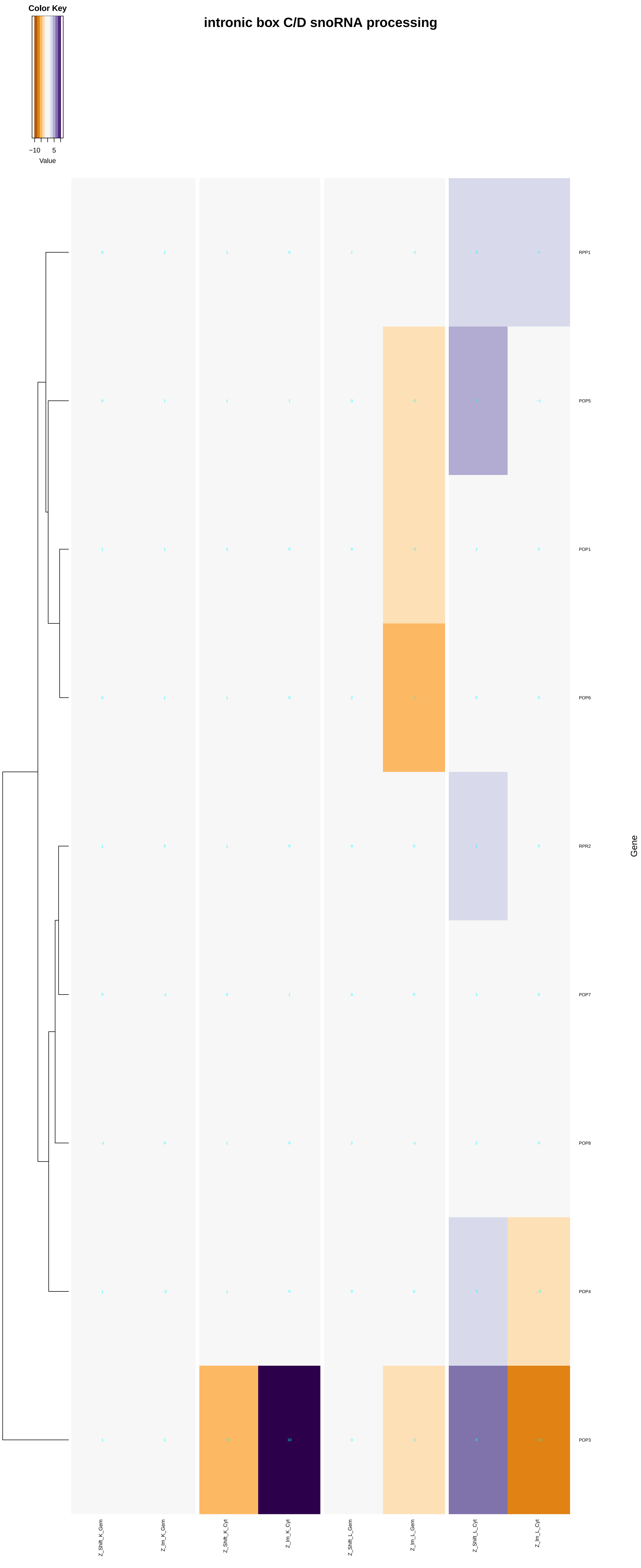
intronic box C/D snoRNA processing
Color Key
−10
5
Value
RPP1
POP5
POP1
POP6
RPR2
POP7
POP8
POP4
POP3
Gene
Z_Shift_K_Gem
Z_lm_K_Gem
Z_Shift_K_Cyt
Z_lm_K_Cyt
Z_Shift_L_Gem
Z_lm_L_Gem
Z_Shift_L_Cyt
Z_lm_L_Cyt

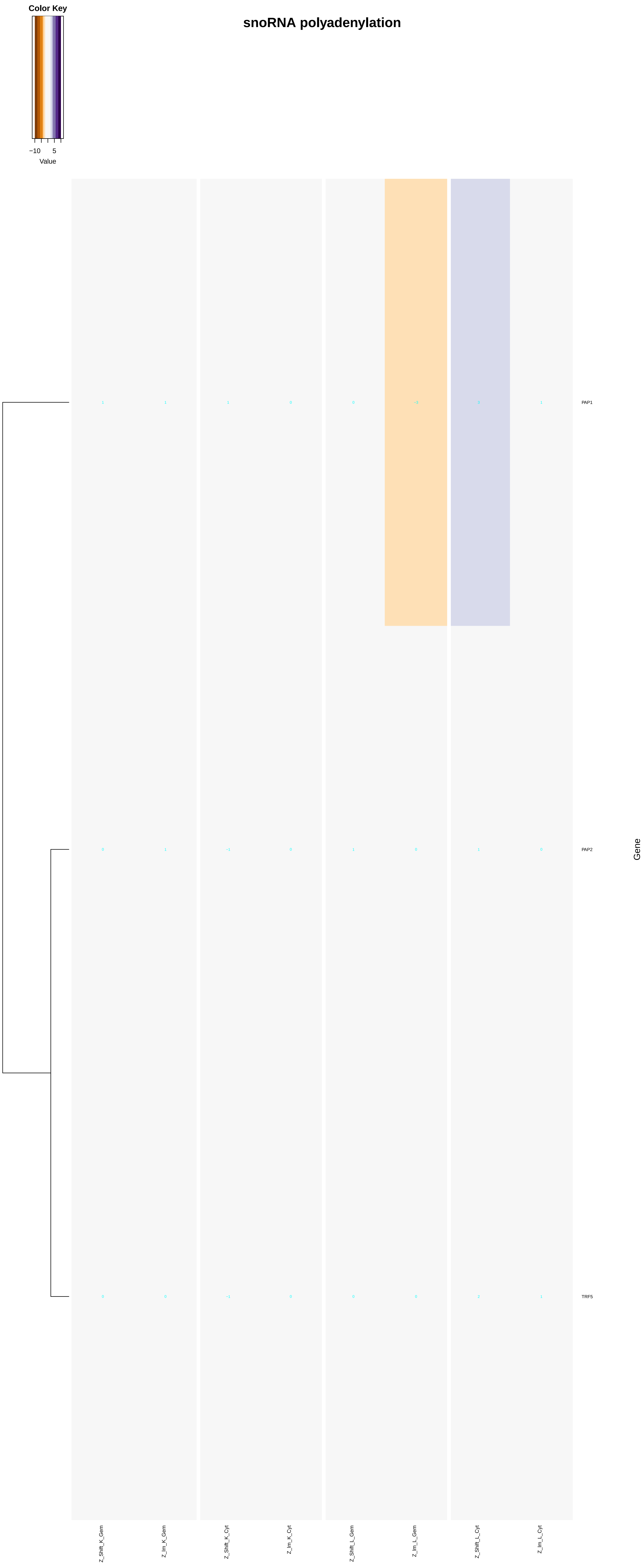
Color Key
−10
5
Value
snoRNA polyadenylation
PAP1
PAP2
TRF5
Gene
Z_Shift_K_Gem
Z_Im_K_Gem
Z_Shift_K_Cyt
Z_Im_K_Cyt
Z_Shift_L_Gem
Z_Im_L_Gem
Z_Shift_L_Cyt
Z_Im_L_Cyt

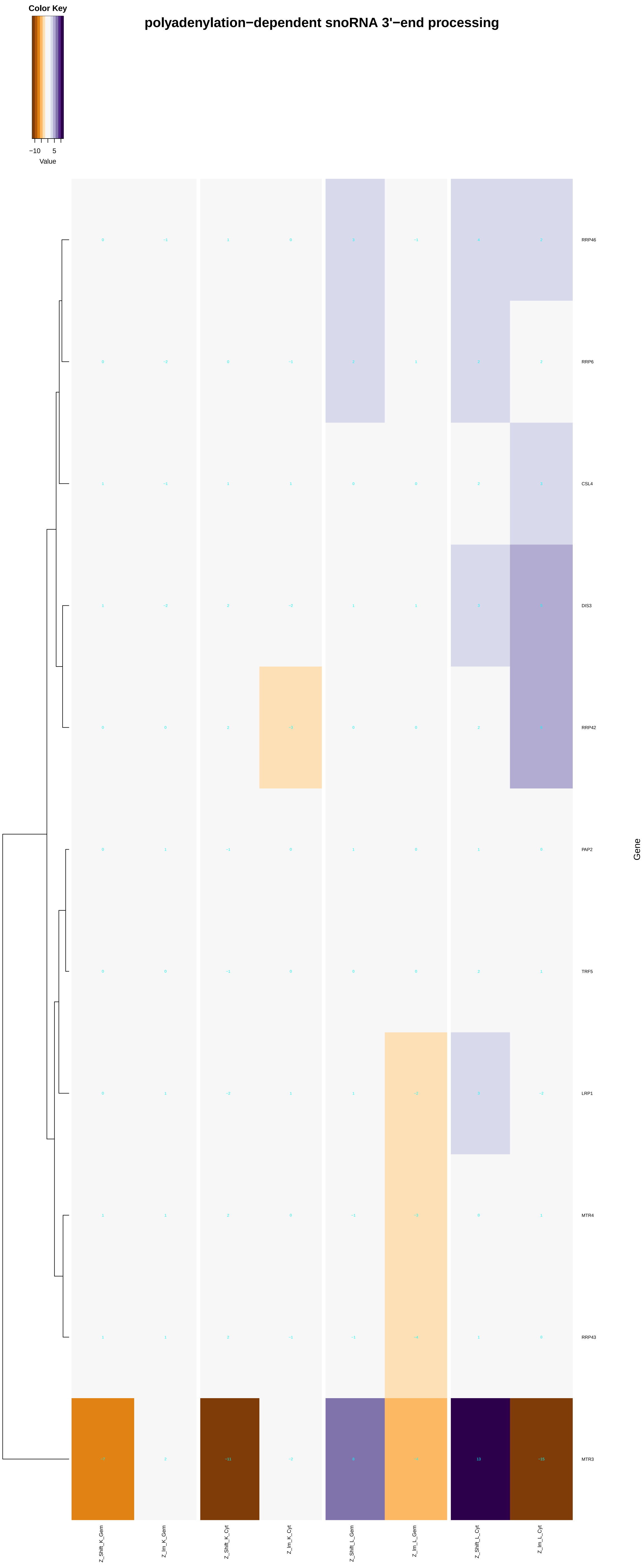
polyadenylation−dependent snoRNA 3'−end processing
Color Key
−10
5
Value
RRP46
RRP6
CSL4
DIS3
RRP42
PAP2
TRF5
LRP1
MTR4
RRP43
MTR3
Gene
Z_Shift_K_Gem
Z_Im_K_Gem
Z_Shift_K_Cyt
Z_Im_K_Cyt
Z_Shift_L_Gem
Z_Im_L_Gem
Z_Shift_L_Cyt
Z_Im_L_Cyt

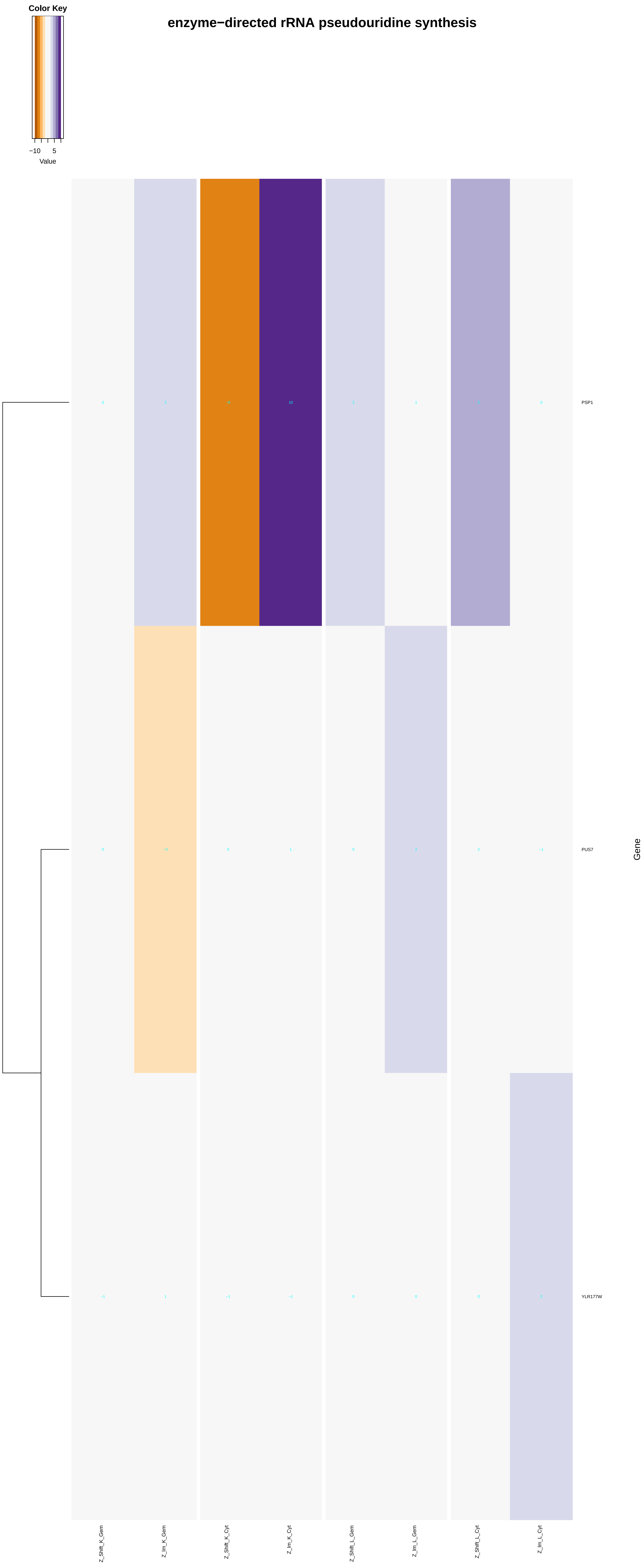

Color Key
−10
5
Value
enzyme−directed rRNA pseudouridine synthesis
0
2
−6
10
2
1
5
0
PSP1
0
−3
0
1
0
2
2
−1
PUS7
−1
1
−1
−1
0
0
0
2
YLR177W
Gene
Z_Shift_K_Gem
Z_Im_K_Gem
Z_Shift_K_Cyt
Z_Im_K_Cyt
Z_Shift_L_Gem
Z_Im_L_Gem
Z_Shift_L_Cyt
Z_Im_L_Cyt

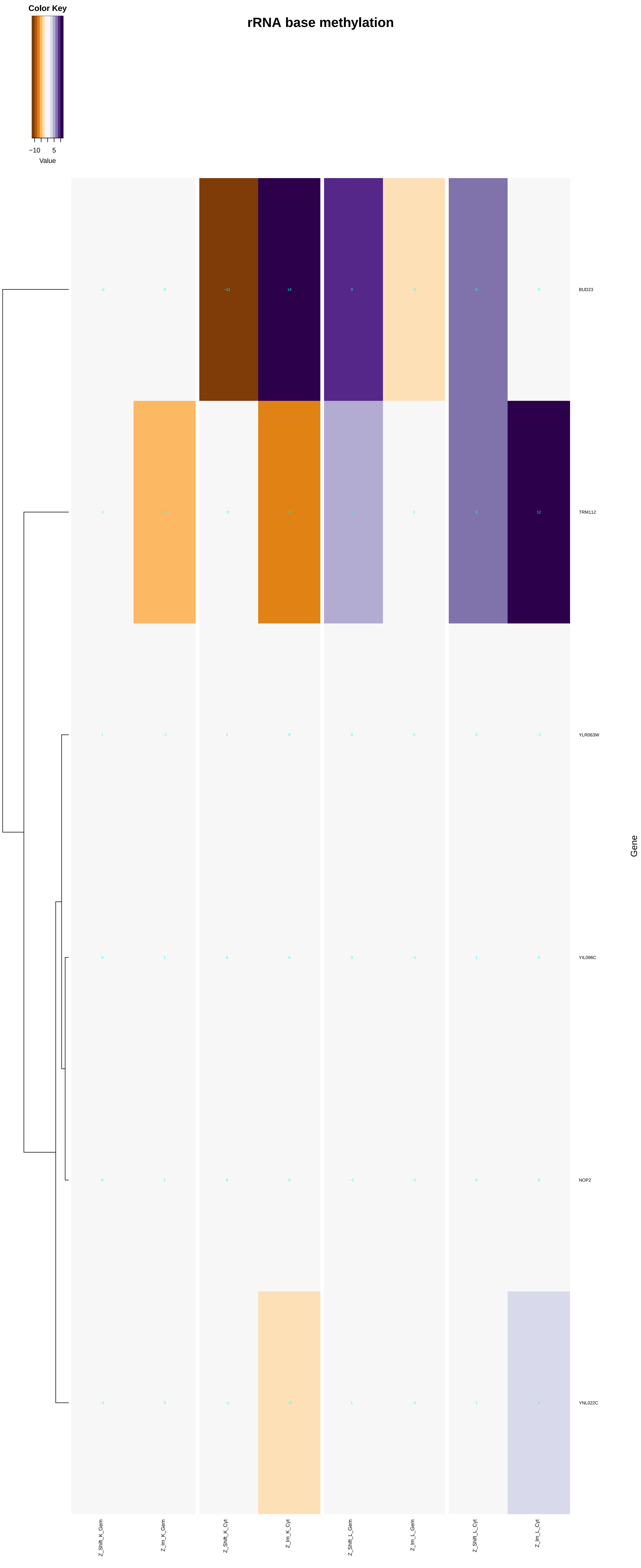
rRNA base methylation
Color Key
−10
5
Value
BUD23
TRM112
YLR063W
YIL096C
NOP2
YNL022C
Gene
Z_Shift_K_Gem
Z_Im_K_Gem
Z_Shift_K_Cyt
Z_Im_K_Cyt
Z_Shift_L_Gem
Z_Im_L_Gem
Z_Shift_L_Cyt
Z_Im_L_Cyt

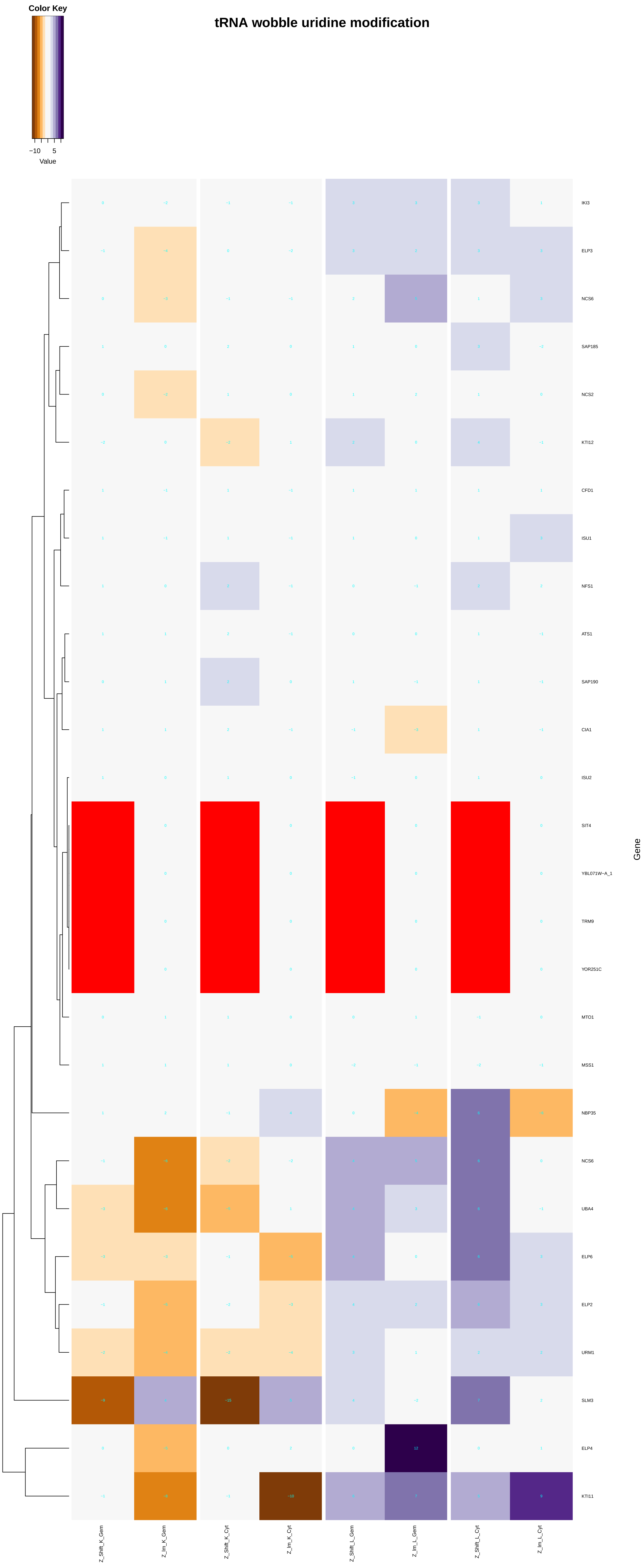

tRNA wobble uridine modification
Color Key
−10
5
Value
IKI3
ELP3
NCS6
SAP185
NCS2
KTI12
CFD1
ISU1
NFS1
ATS1
SAP190
CIA1
ISU2
SIT4
YBL071W−A_1
TRM9
YOR251C
MTO1
MSS1
NBP35
NCS6
UBA4
ELP6
ELP2
URM1
SLM3
ELP4
KTI11
Gene
Z_Shift_K_Gem
Z_Im_K_Gem
Z_Shift_K_Cyt
Z_Im_K_Cyt
Z_Shift_L_Gem
Z_Im_L_Gem
Z_Shift_L_Cyt
Z_Im_L_Cyt

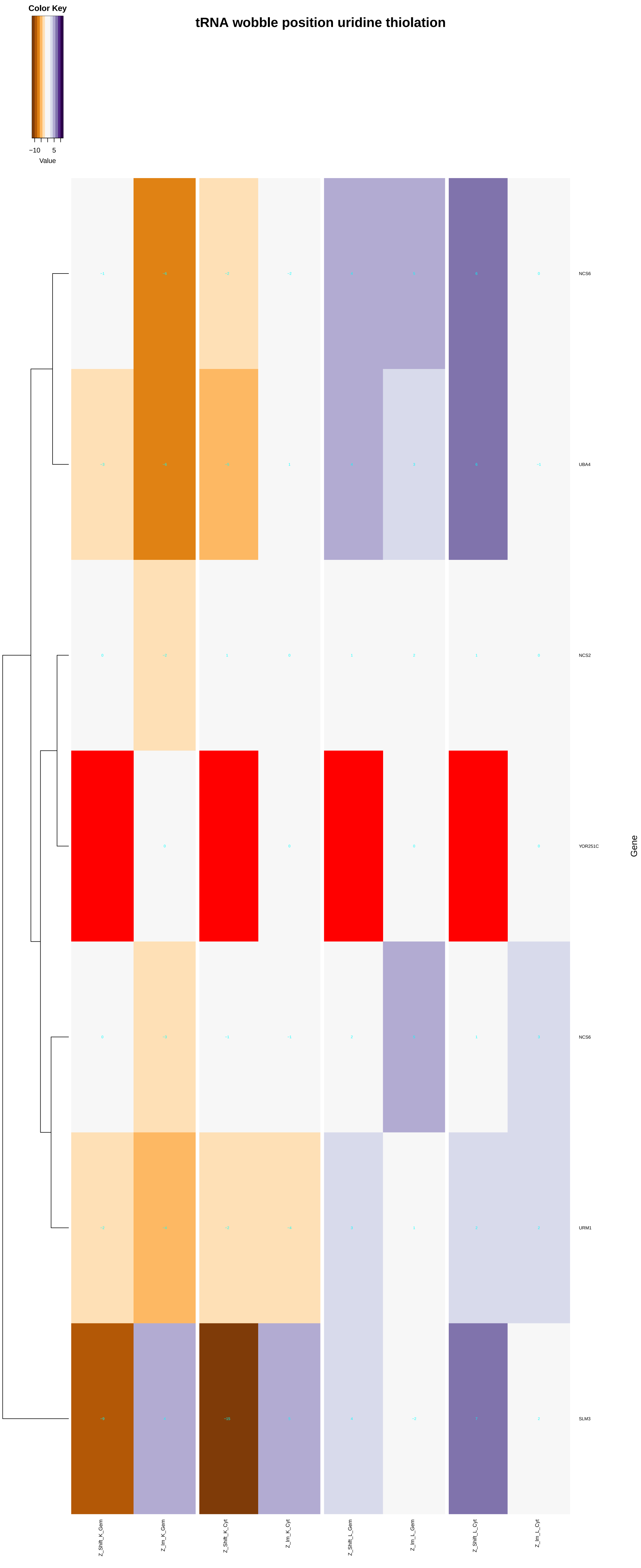

tRNA wobble position uridine thiolation
Color Key
−10
5
Value
NCS6
UBA4
NCS2
YOR251C
NCS6
URM1
SLM3
Gene
Z_Shift_K_Gem
Z_Im_K_Gem
Z_Shift_K_Cyt
Z_Im_K_Cyt
Z_Shift_L_Gem
Z_Im_L_Gem
Z_Shift_L_Cyt
Z_Im_L_Cyt

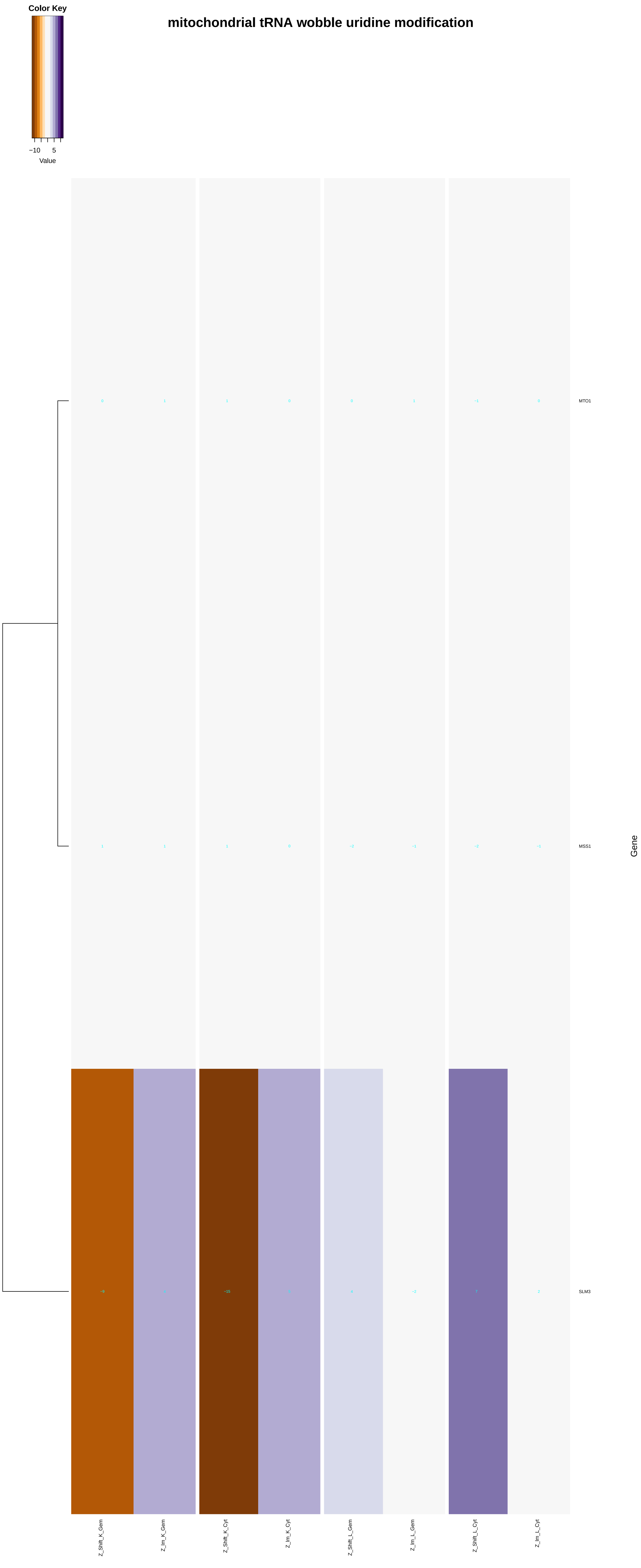
Color Key
−10 5
Value
mitochondrial tRNA wobble uridine modification
MTO1
MSS1
SLM3
Gene
Z_Shift_K_Gem
Z_Im_K_Gem
Z_Shift_K_Cyt
Z_Im_K_Cyt
Z_Shift_L_Gem
Z_Im_L_Gem
Z_Shift_L_Cyt
Z_Im_L_Cyt
