## Additional File 7 for "A humanized yeast phenomic model of deoxycytidine kinase to predict genetic buffering of nucleoside analog cytotoxicity": negative_regulation_of_gene_expression.pdf

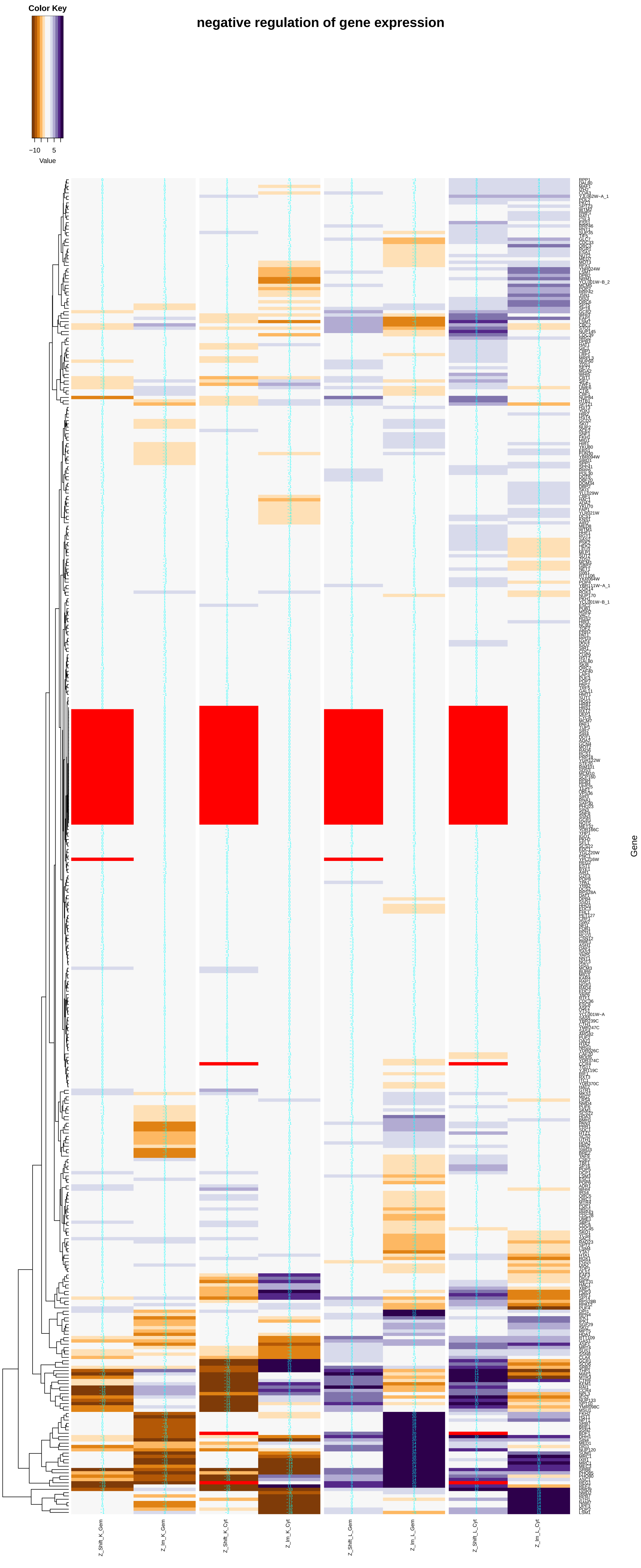

Color Key
−10 5
Value
negative regulation of gene expression
Gene
Z_Shift_K_Gem
Z_Im_K_Gem
Z_Shift_K_Cyt
Z_Im_K_Cyt
Z_Shift_L_Gem
Z_Im_L_Gem
Z_Shift_L_Cyt
Z_Im_L_Cyt

### mRNA catabolic process

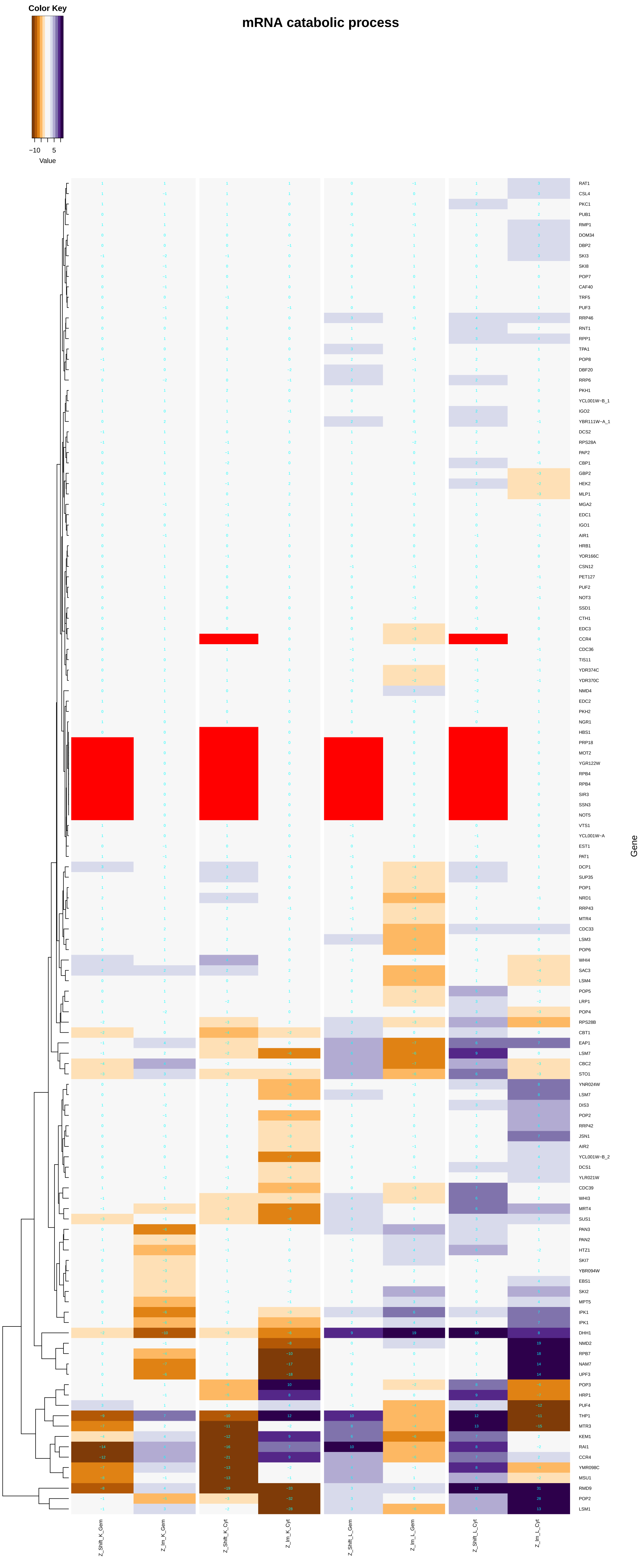

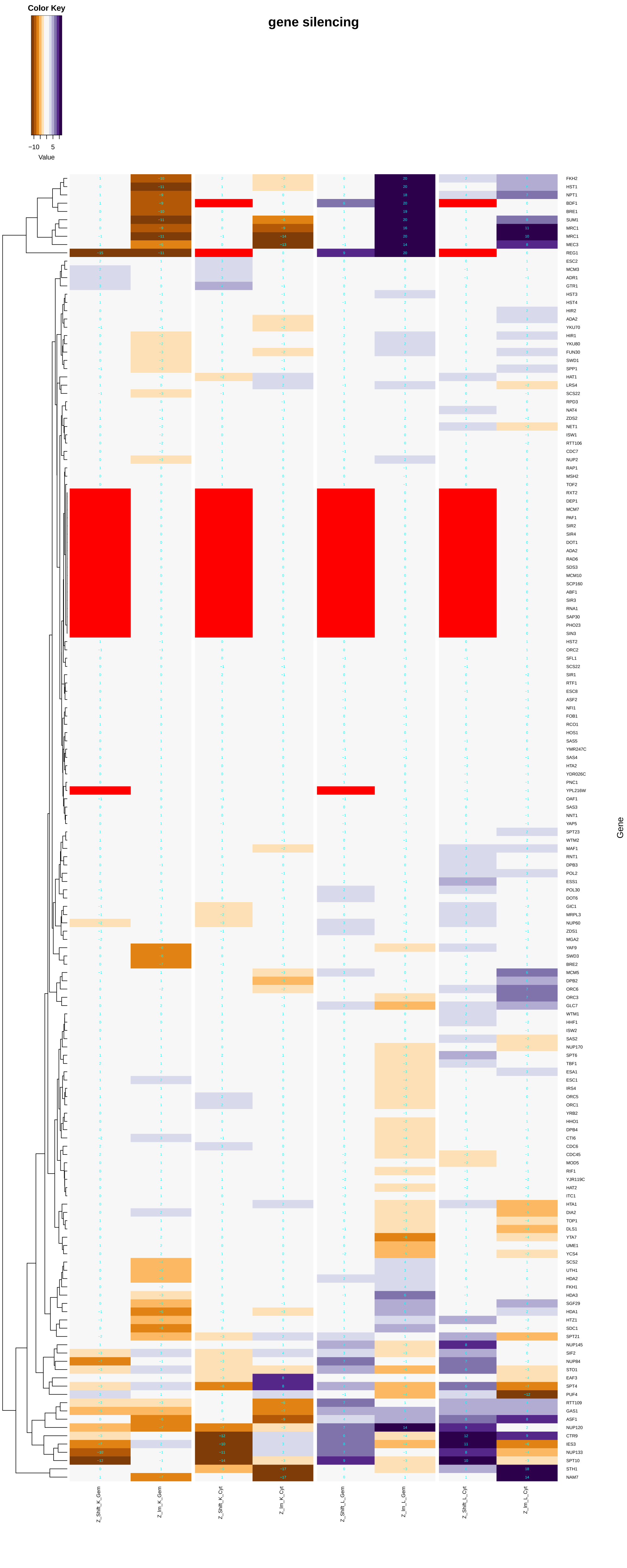

gene silencing
Color Key
−10 5
Value
FKH2
HST1
NPT1
BDF1
BRE1
SUM1
MRC1
MRC1
MEC3
REG1
ESC2
MCM3
ADR1
GTR1
HST3
HST4
HIR2
ADA2
YKU70
HIR1
YKU80
FUN30
SWD1
SPP1
HAT1
LRS4
SCS22
RPD3
NAT4
ZDS2
NET1
ISW1
RTT106
CDC7
NUP2
RAP1
MSH2
TOF2
RXT2
DEP1
MCM7
PAF1
SIR2
SIR4
DOT1
ADA2
RAD6
SDS3
MCM10
SCP160
ABF1
SIR3
RNA1
SAP30
PHO23
SIN3
HST2
ORC2
SFL1
SCS22
SIR1
RTF1
ESC8
ASF2
NFI1
FOB1
RCO1
HOS1
SAS5
YMR247C
SAS4
HTA2
YDR026C
PNC1
YPL216W
OAF1
SAS3
NNT1
YAP5
SPT23
WTM2
MAF1
RNT1
DPB3
POL2
ESS1
POL30
DOT6
GIC1
MRPL3
NUP60
ZDS1
MGA2
YAF9
SWD3
BRE2
MCM5
DPB2
ORC6
ORC3
GLC7
WTM1
HHF1
ISW2
SAS2
NUP170
SPT6
TBF1
ESA1
ESC1
IRS4
ORC5
ORC1
YRB2
HHO1
DPB4
CTI6
CDC6
CDC45
MOD5
RIF1
YJR119C
HAT2
ITC1
HTA1
DIA2
TOP1
DLS1
YTA7
UME1
YCS4
SCS2
UTH1
HDA2
FKH1
HDA3
SGF29
HDA1
HTZ1
SDC1
SPT21
NUP145
SIF2
NUP84
STO1
EAF3
SPT4
PUF4
RTT109
GAS1
ASF1
NUP120
CTR9
IES3
NUP133
SPT10
STH1
NAM7
Gene
Z_Shift_K_Gem
Z_Im_K_Gem
Z_Shift_K_Cyt
Z_Im_K_Cyt
Z_Shift_L_Gem
Z_Im_L_Gem
Z_Shift_L_Cyt
Z_Im_L_Cyt

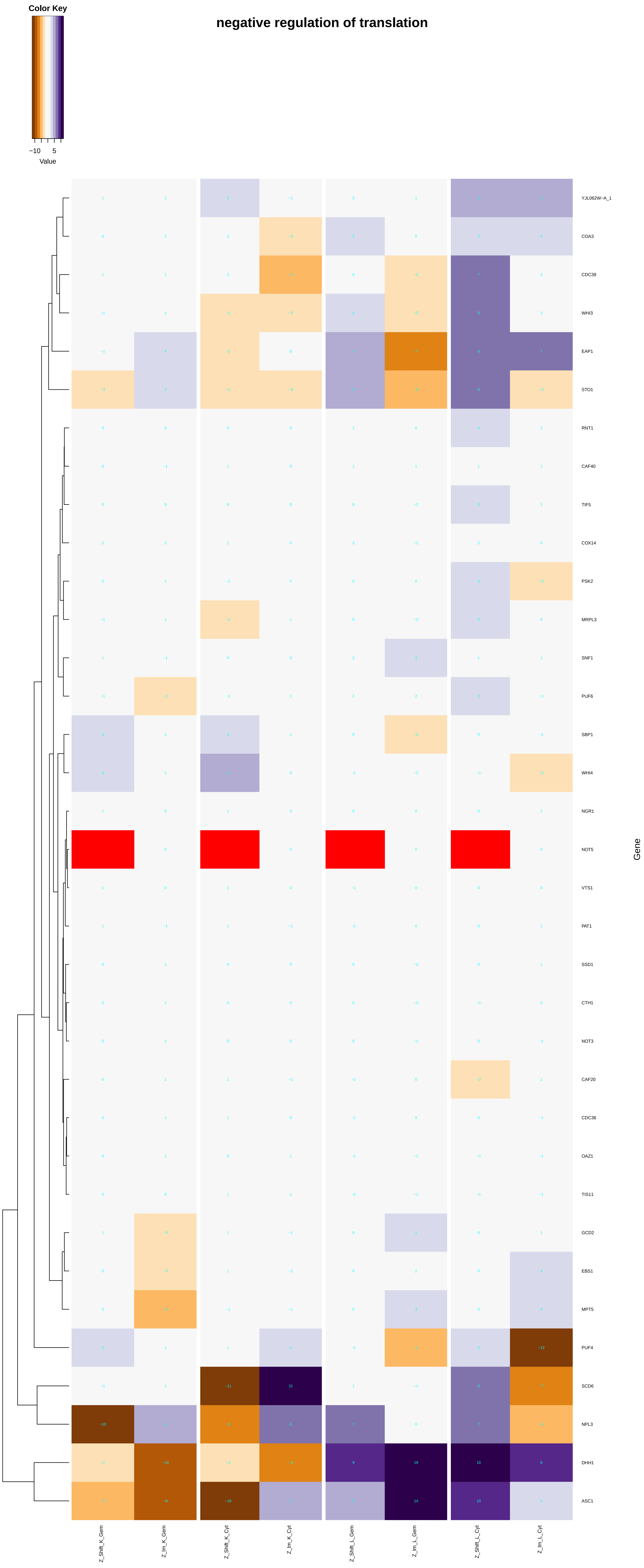
Color Key
−10
5
Value
negative regulation of translation
YJL062W−A_1
COA3
CDC39
WHI3
EAP1
STO1
RNT1
CAF40
TIF5
COX14
PSK2
MRPL3
SNF1
PUF6
SBP1
WHI4
NGR1
NOT5
VTS1
PAT1
SSD1
CTH1
NOT3
CAF20
CDC36
OAZ1
TIS11
GCD2
EBS1
MPT5
PUF4
SCD6
NPL3
DHH1
ASC1
Gene
Z_Shift_K_Gem
Z_Im_K_Gem
Z_Shift_K_Cyt
Z_Im_K_Cyt
Z_Shift_L_Gem
Z_Im_L_Gem
Z_Shift_L_Cyt
Z_Im_L_Cyt

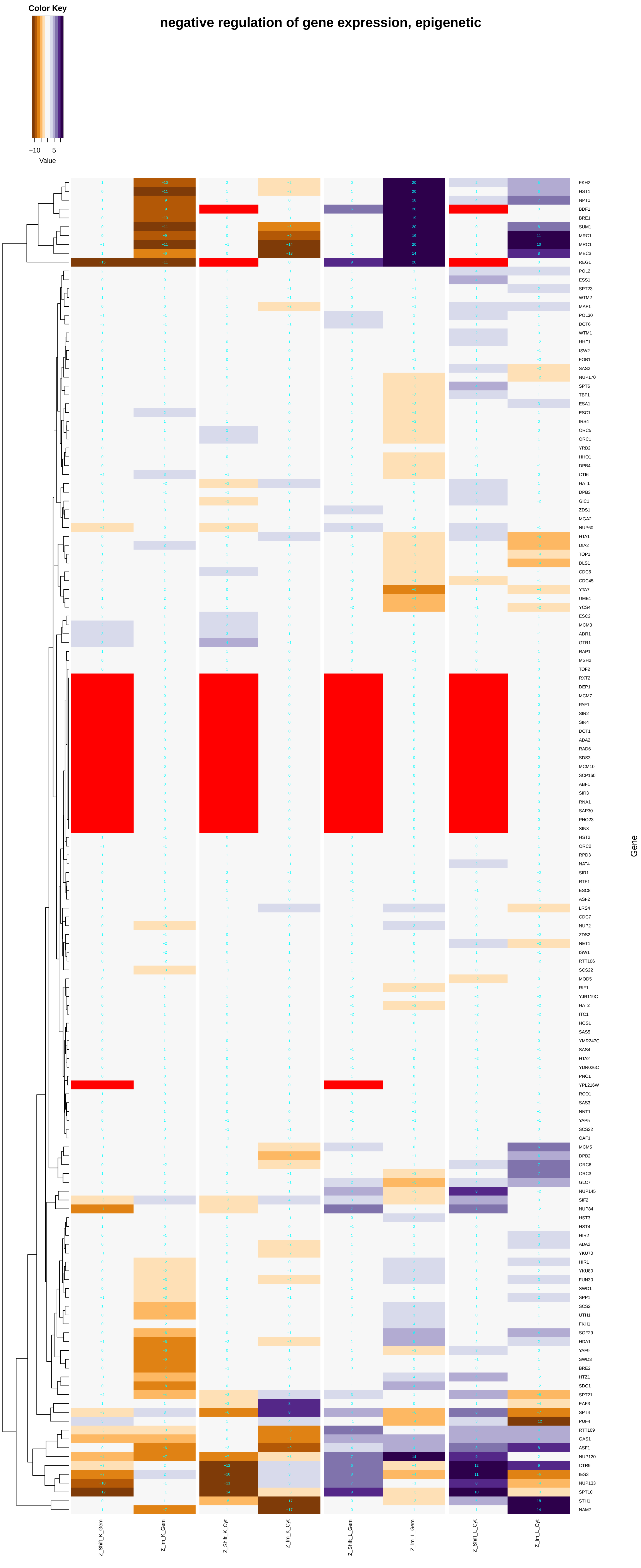

negative regulation of gene expression, epigenetic
Color Key
−10
5
Value
Gene
FKH2
HST1
NPT1
BDF1
BRE1
SUM1
MRC1
MRC1
MEC3
REG1
POL2
ESS1
SPT23
WTM2
MAF1
POL30
DOT6
WTM1
HHF1
ISW2
FOB1
SAS2
NUP170
SPT6
TBF1
ESA1
ESC1
IRS4
ORC5
ORC1
YRB2
HHO1
DPB4
CTI6
HAT1
DPB3
GIC1
ZDS1
MGA2
NUP60
HTA1
DIA2
TOP1
DLS1
CDC6
CDC45
YTA7
UME1
YCS4
ESC2
MCM3
ADR1
GTR1
RAP1
MSH2
TOF2
RXT2
DEP1
MCM7
PAF1
SIR2
SIR4
DOT1
ADA2
RAD6
SDS3
MCM10
SCP160
ABF1
SIR3
RNA1
SAP30
PHO23
SIN3
HST2
ORC2
RPD3
NAT4
SIR1
RTF1
ESC8
ASF2
LRS4
CDC7
NUP2
ZDS2
NET1
ISW1
RTT106
SCS22
MOD5
RIF1
YJR119C
HAT2
ITC1
HOS1
SAS5
YMR247C
SAS4
HTA2
YDR026C
PNC1
YPL216W
RCO1
SAS3
NNT1
YAP5
SCS22
OAF1
MCM5
DPB2
ORC6
ORC3
GLC7
NUP145
SIF2
NUP84
HST3
HST4
HIR2
ADA2
YKU70
HIR1
YKU80
FUN30
SWD1
SPP1
SCS2
UTH1
FKH1
SGF29
HDA1
YAF9
SWD3
BRE2
HTZ1
SDC1
SPT21
EAF3
SPT4
PUF4
RTT109
GAS1
ASF1
NUP120
CTR9
IES3
NUP133
SPT10
STH1
NAM7
Z_Shift_K_Gem
Z_Im_K_Gem
Z_Shift_K_Cyt
Z_Im_K_Cyt
Z_Shift_L_Gem
Z_Im_L_Gem
Z_Shift_L_Cyt
Z_Im_L_Cyt

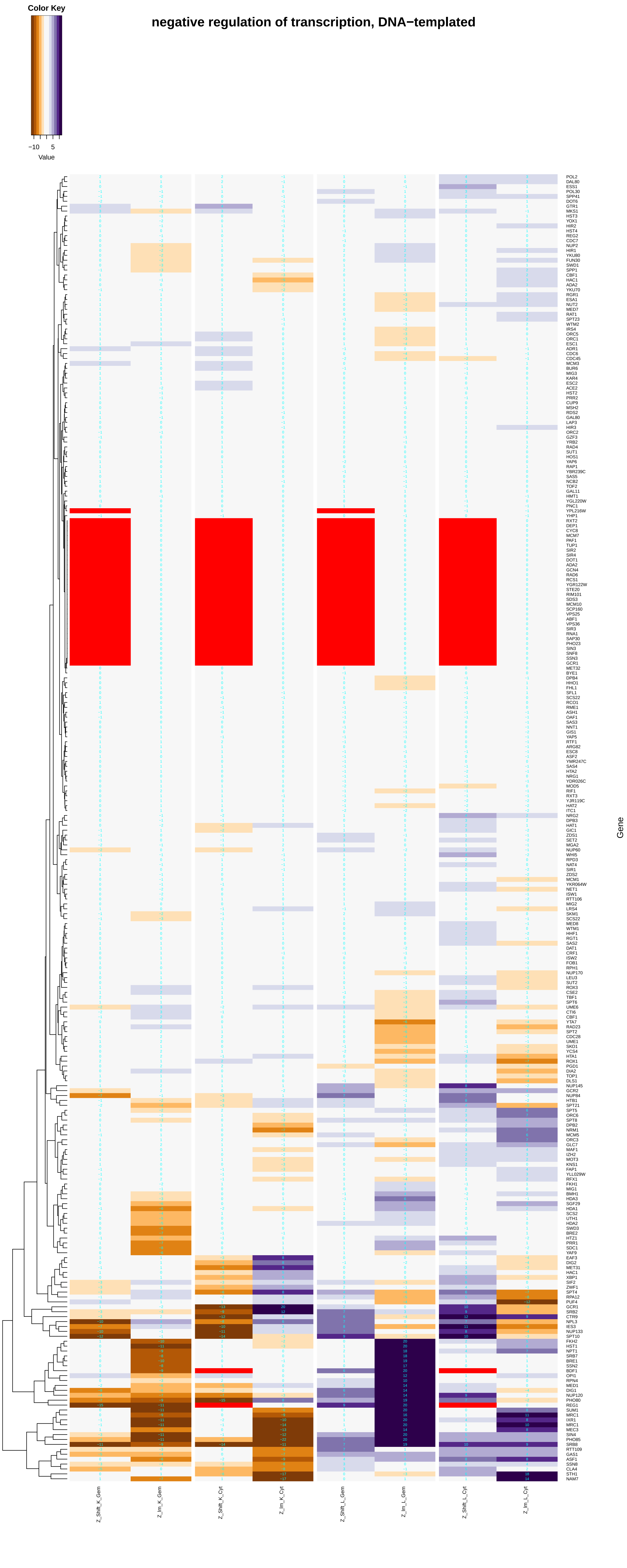
Color Key
−10
5
Value
negative regulation of transcription, DNA−templated
POL2
DAL80
ESS1
POL30
SPP41
DOT6
GTR1
MKS1
HST3
YOX1
HIR2
HST4
REG2
CDC7
NUP2
HIR1
YKU80
FUN30
SWD1
SPP1
CBF1
HAC1
ADA2
YKU70
RGR1
ESA1
NUT2
MED7
RAT1
SPT23
WTM2
IRS4
ORC5
ORC1
ESC1
ADR1
CDC6
CDC45
MCM3
BUR6
MIG3
KAR4
ESC2
ACE2
HST2
PRR2
CUP9
MSH2
RDS2
GAL80
LAP3
HIR3
ORC2
GZF3
YRB2
RAD4
SUT1
HOS1
YAP6
RAP1
YBR239C
SAS5
NCB2
TOF2
GAL11
HMT1
YGL220W
PNC1
YPL216W
YHP1
RXT2
DEP1
CYC8
MCM7
PAF1
TUP1
SIR2
SIR4
DOT1
ADA2
GCN4
RAD6
RCS1
YGR122W
STE20
RIM101
SDS3
MCM10
SCP160
VPS25
ABF1
VPS36
SIR3
RNA1
SAP30
PHO23
SIN3
SNF8
SSN3
GCR1
MET32
BYE1
DPB4
HHO1
FHL1
SFL1
SCS22
RCO1
RME1
ASH1
OAF1
SAS3
NNT1
GIS1
YAP5
RTF1
ARG82
ESC8
ASF2
YMR247C
SAS4
HTA2
NRG1
YDR026C
MOD5
RIF1
RXT3
YJR119C
HAT2
ITC1
NRG2
DPB3
HAT1
GIC1
ZDS1
SET2
MGA2
NUP60
WHI5
RPD3
NAT4
SIR1
ZDS2
MCM1
YKR064W
NET1
ISW1
RTT106
MIG2
LRS4
SKM1
SCS22
MED8
WTM1
HHF1
RGT1
SAS2
DAT1
CRF1
ISW2
FOB1
RPH1
NUP170
LEU3
SUT2
ROX3
CSE2
TBF1
SPT6
UME6
CTI6
CBF1
YTA7
RAD23
SPT2
CDC28
UME1
SKO1
YCS4
HTA1
ROX1
PGD1
DIA2
TOP1
DLS1
NUP145
GCR2
NUP84
HTB1
SPT21
SPT5
ORC6
SPT8
DPB2
NRM1
MCM5
ORC3
GLC7
MAF1
IZH2
MOT3
KNS1
FAP1
YLL029W
RFX1
FKH1
MIG1
BMH1
HDA3
SGF29
HDA1
SCS2
UTH1
HDA2
SWD3
BRE2
HTZ1
PRR1
SDC1
YAF9
EAF3
DIG2
MET31
HAC1
XBP1
SIF2
ZWF1
SPT4
RPA12
PUF4
GCR1
SRB2
CTR9
NPL3
IES3
NUP133
SPT10
FKH2
HST1
NPT1
SRB7
BRE1
SSN2
BDF1
OPI1
RPN4
MED1
DIG1
NUP120
PHO80
REG1
SUM1
MRC1
IXR1
MRC1
MEC3
SIN4
PHO85
SRB8
RTT109
GAL11
ASF1
SSN8
CLA4
STH1
NAM7
Gene
Z_Shift_K_Gem
Z_Im_K_Gem
Z_Shift_K_Cyt
Z_Im_K_Cyt
Z_Shift_L_Gem
Z_Im_L_Gem
Z_Shift_L_Cyt
Z_Im_L_Cyt

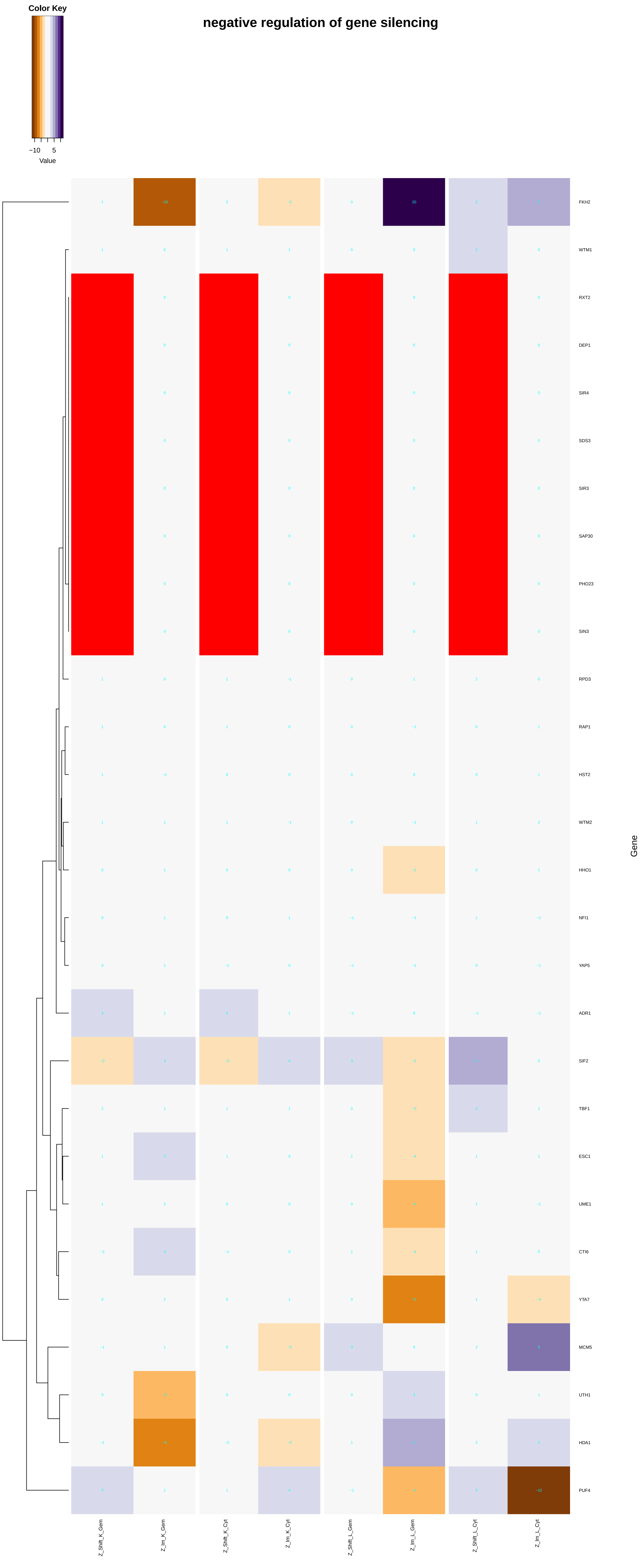

Color Key
−10
5
Value
negative regulation of gene silencing
FKH2
WTM1
RXT2
DEP1
SIR4
SDS3
SIR3
SAP30
PHO23
SIN3
RPD3
RAP1
HST2
WTM2
HHO1
NFI1
YAP5
ADR1
SIF2
TBF1
ESC1
UME1
CTI6
YTA7
MCM5
UTH1
HDA1
PUF4
Gene
Z_Shift_K_Gem
Z_Im_K_Gem
Z_Shift_K_Cyt
Z_Im_K_Cyt
Z_Shift_L_Gem
Z_Im_L_Gem
Z_Shift_L_Cyt
Z_Im_L_Cyt

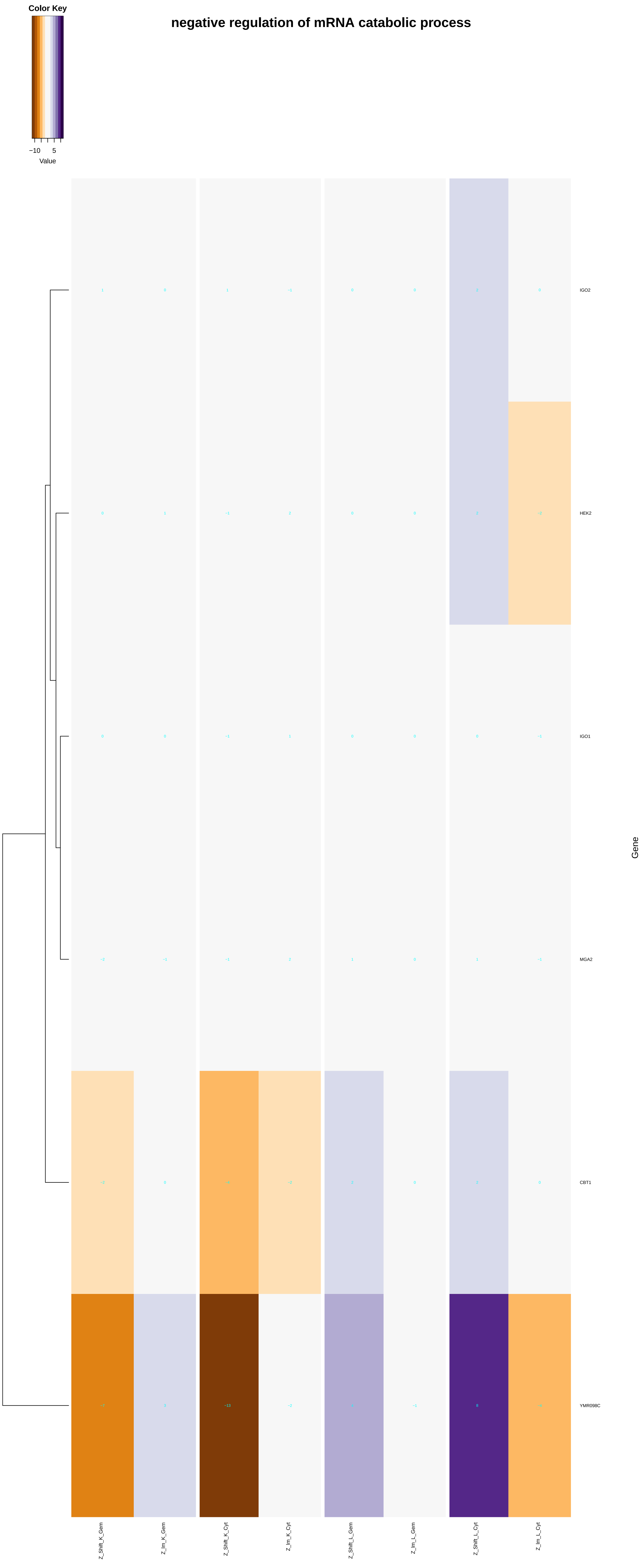
negative regulation of mRNA catabolic process
Color Key
−10
5
Value
IGO2
HEK2
IGO1
MGA2
CBT1
YMR098C
Gene
Z_Shift_K_Gem
Z_Im_K_Gem
Z_Shift_K_Cyt
Z_Im_K_Cyt
Z_Shift_L_Gem
Z_Im_L_Gem
Z_Shift_L_Cyt
Z_Im_L_Cyt

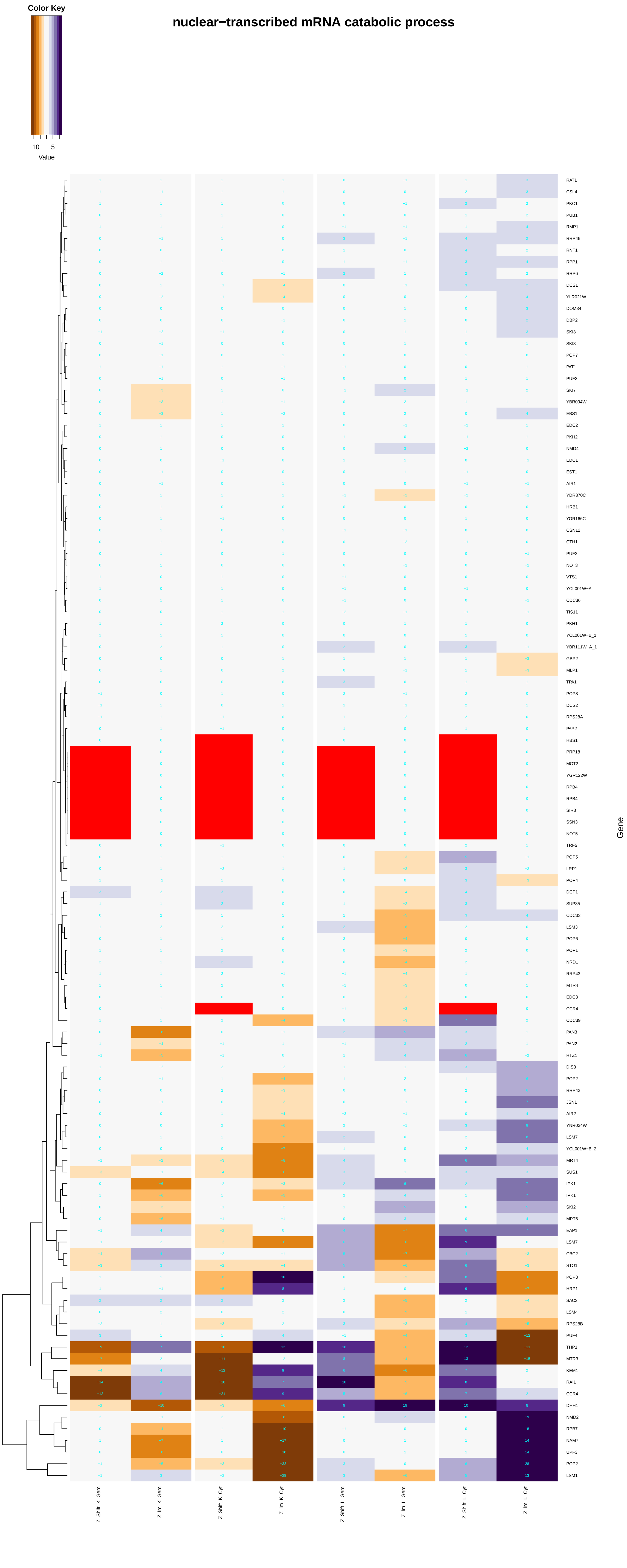
nuclear−transcribed mRNA catabolic process
Color Key
−10 5
Value
Gene
Z_Shift_K_Gem
Z_Im_K_Gem
Z_Shift_K_Cyt
Z_Im_K_Cyt
Z_Shift_L_Gem
Z_Im_L_Gem
Z_Shift_L_Cyt
Z_Im_L_Cyt
RAT1
CSL4
PKC1
PUB1
RMP1
RRP46
RNT1
RPP1
RRP6
DCS1
YLR021W
DOM34
DBP2
SKI3
SKI8
POP7
PAT1
PUF3
SKI7
YBR094W
EBS1
EDC2
PKH2
NMD4
EDC1
EST1
AIR1
YDR370C
HRB1
YOR166C
CSN12
CTH1
PUF2
NOT3
VTS1
YCL001W−A
CDC36
TIS11
PKH1
YCL001W−B_1
YBR111W−A_1
GBP2
MLP1
TPA1
POP8
DCS2
RPS28A
PAP2
HBS1
PRP18
MOT2
YGR122W
RPB4
RPB4
SIR3
SSN3
NOT5
TRF5
POP5
LRP1
POP4
DCP1
SUP35
CDC33
LSM3
POP6
POP1
NRD1
RRP43
MTR4
EDC3
CCR4
CDC39
PAN3
PAN2
HTZ1
DIS3
POP2
RRP42
JSN1
AIR2
YNR024W
LSM7
YCL001W−B_2
MRT4
SUS1
IPK1
IPK1
SKI2
MPT5
EAP1
LSM7
CBC2
STO1
POP3
HRP1
SAC3
LSM4
RPS28B
PUF4
THP1
MTR3
KEM1
RAI1
CCR4
DHH1
NMD2
RPB7
NAM7
UPF3
POP2
LSM1

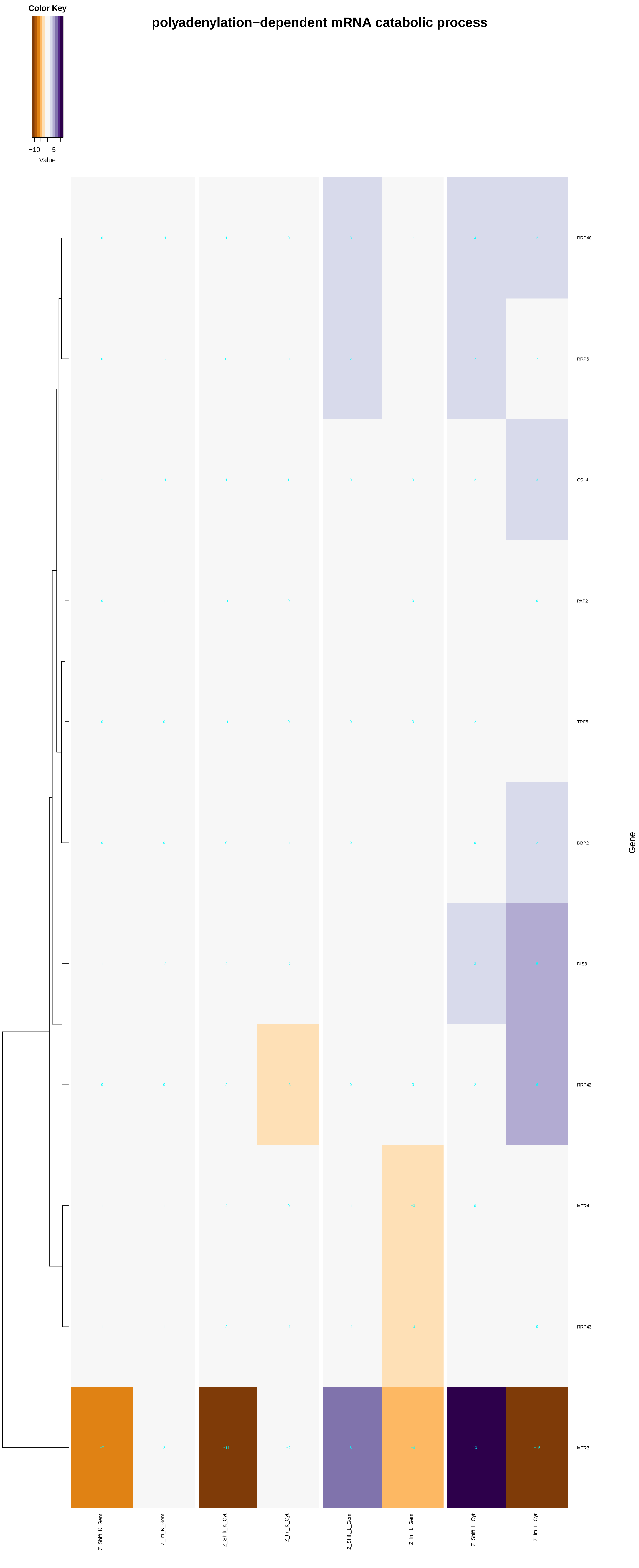

polyadenylation−dependent mRNA catabolic process
Color Key
−10
5
Value
RRP46
RRP6
CSL4
PAP2
TRF5
DBP2
DIS3
RRP42
MTR4
RRP43
MTR3
Gene
Z_Shift_K_Gem
Z_Im_K_Gem
Z_Shift_K_Cyt
Z_Im_K_Cyt
Z_Shift_L_Gem
Z_Im_L_Gem
Z_Shift_L_Cyt
Z_Im_L_Cyt

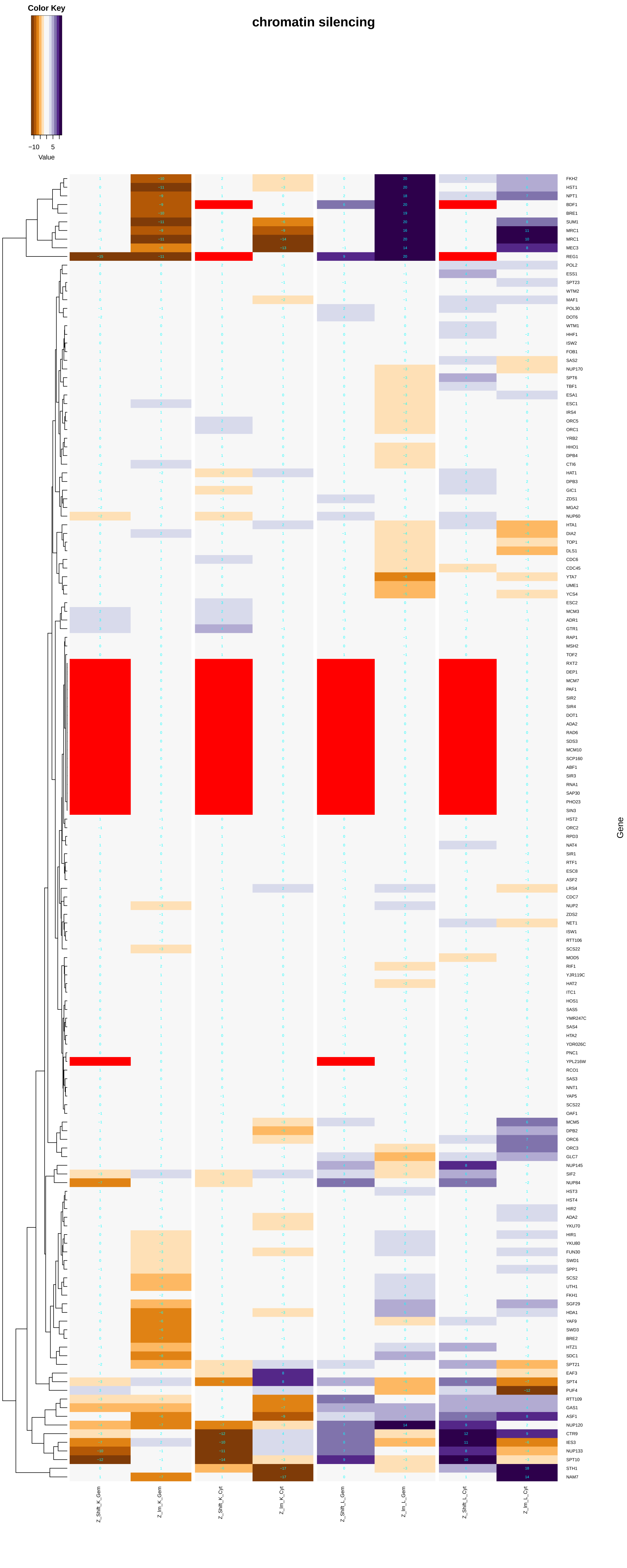

chromatin silencing
Color Key
−10
5
Value
FKH2
HST1
NPT1
BDF1
BRE1
SUM1
MRC1
MRC1
MEC3
REG1
POL2
ESS1
SPT23
WTM2
MAF1
POL30
DOT6
WTM1
HHF1
ISW2
FOB1
SAS2
NUP170
SPT6
TBF1
ESA1
ESC1
IRS4
ORC5
ORC1
YRB2
HHO1
DPB4
CTI6
HAT1
DPB3
GIC1
ZDS1
MGA2
NUP60
HTA1
DIA2
TOP1
DLS1
CDC6
CDC45
YTA7
UME1
YCS4
ESC2
MCM3
ADR1
GTR1
RAP1
MSH2
TOF2
RXT2
DEP1
MCM7
PAF1
SIR2
SIR4
DOT1
ADA2
RAD6
SDS3
MCM10
SCP160
ABF1
SIR3
RNA1
SAP30
PHO23
SIN3
HST2
ORC2
RPD3
NAT4
SIR1
RTF1
ESC8
ASF2
LRS4
CDC7
NUP2
ZDS2
NET1
ISW1
RTT106
SCS22
MOD5
RIF1
YJR119C
HAT2
ITC1
HOS1
SAS5
YMR247C
SAS4
HTA2
YDR026C
PNC1
YPL216W
RCO1
SAS3
NNT1
YAP5
SCS22
OAF1
MCM5
DPB2
ORC6
ORC3
GLC7
NUP145
SIF2
NUP84
HST3
HST4
HIR2
ADA2
YKU70
HIR1
YKU80
FUN30
SWD1
SPP1
SCS2
UTH1
FKH1
SGF29
HDA1
YAF9
SWD3
BRE2
HTZ1
SDC1
SPT21
EAF3
SPT4
PUF4
RTT109
GAS1
ASF1
NUP120
CTR9
IES3
NUP133
SPT10
STH1
NAM7
Gene
Z_Shift_K_Gem
Z_Im_K_Gem
Z_Shift_K_Cyt
Z_Im_K_Cyt
Z_Shift_L_Gem
Z_Im_L_Gem
Z_Shift_L_Cyt
Z_Im_L_Cyt

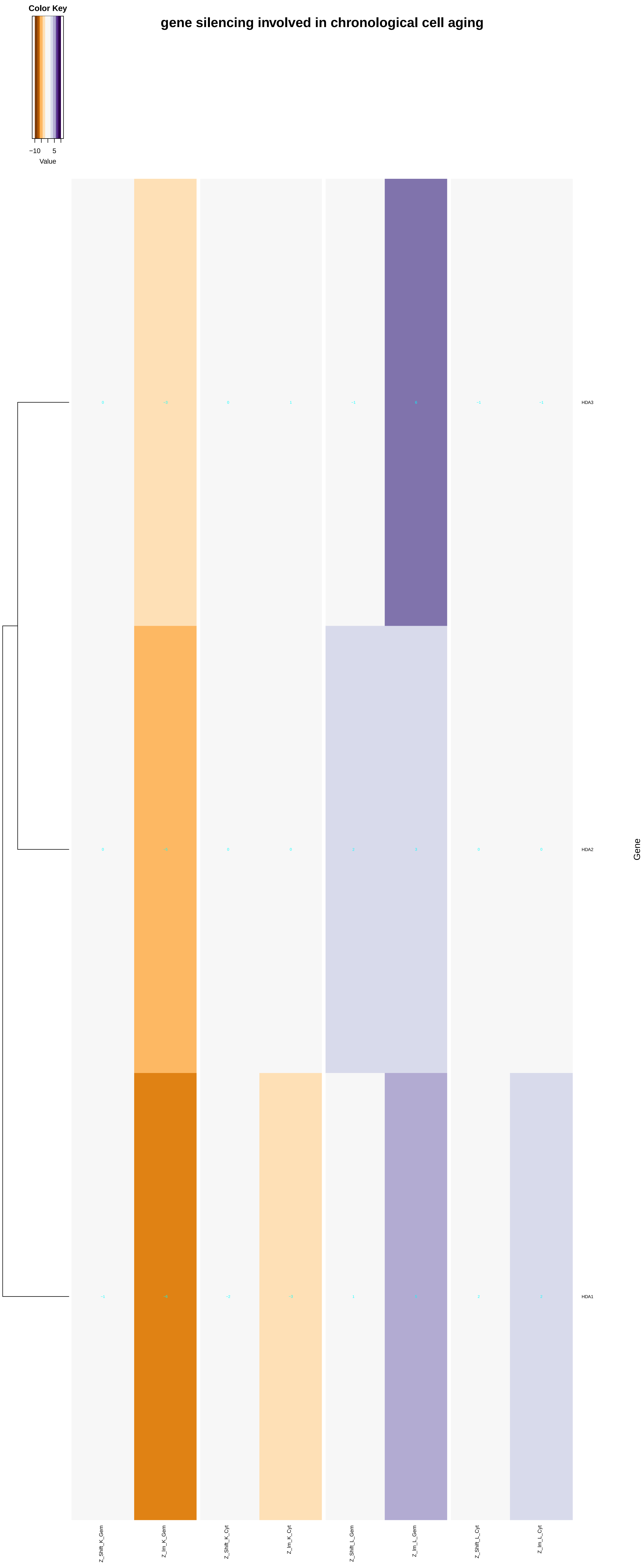

gene silencing involved in chronological cell aging
Color Key
−10
5
Value
HDA3
HDA2
HDA1
Gene
Z_Shift_K_Gem
Z_lm_K_Gem
Z_Shift_K_Cyt
Z_lm_K_Cyt
Z_Shift_L_Gem
Z_lm_L_Gem
Z_Shift_L_Cyt
Z_lm_L_Cyt

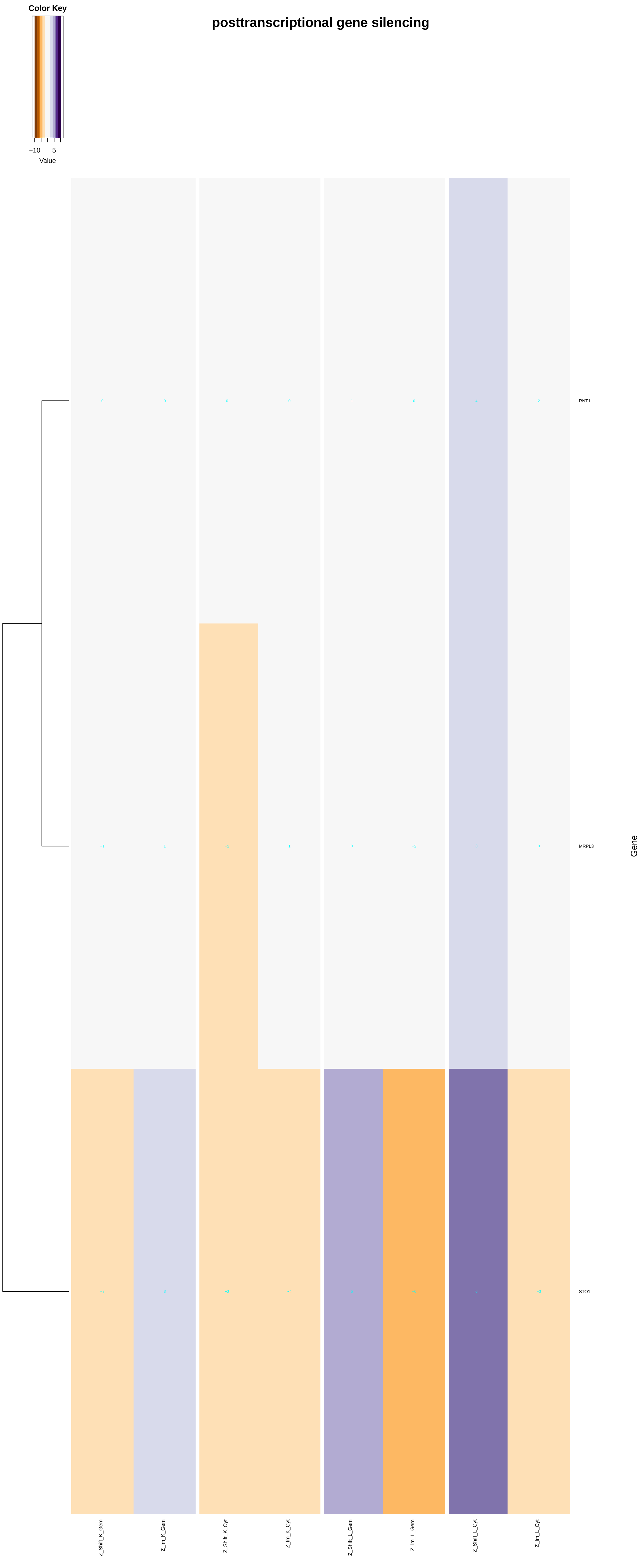
posttranscriptional gene silencing
Color Key
−10
5
Value
RNT1
MRPL3
STO1
Gene
Z_Shift_K_Gem
Z_Im_K_Gem
Z_Shift_K_Cyt
Z_Im_K_Cyt
Z_Shift_L_Gem
Z_Im_L_Gem
Z_Shift_L_Cyt
Z_Im_L_Cyt

### gene silencing by RNA

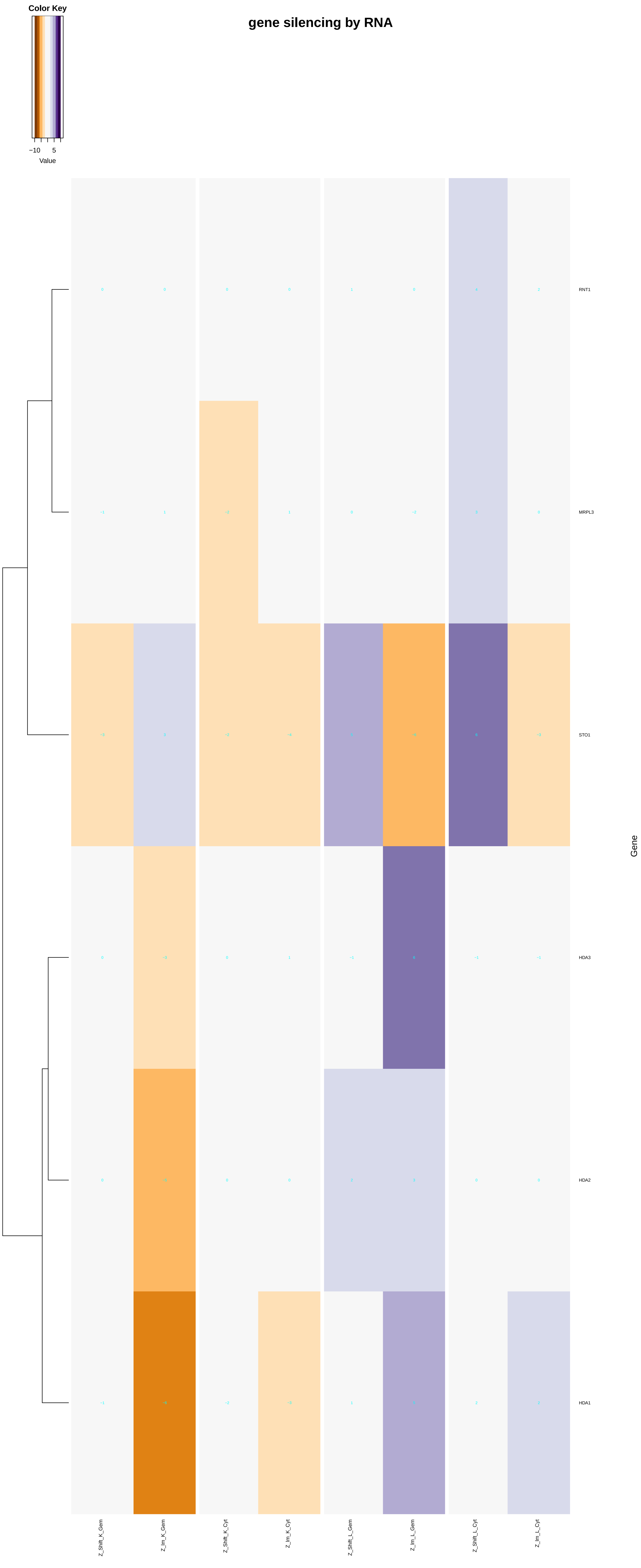

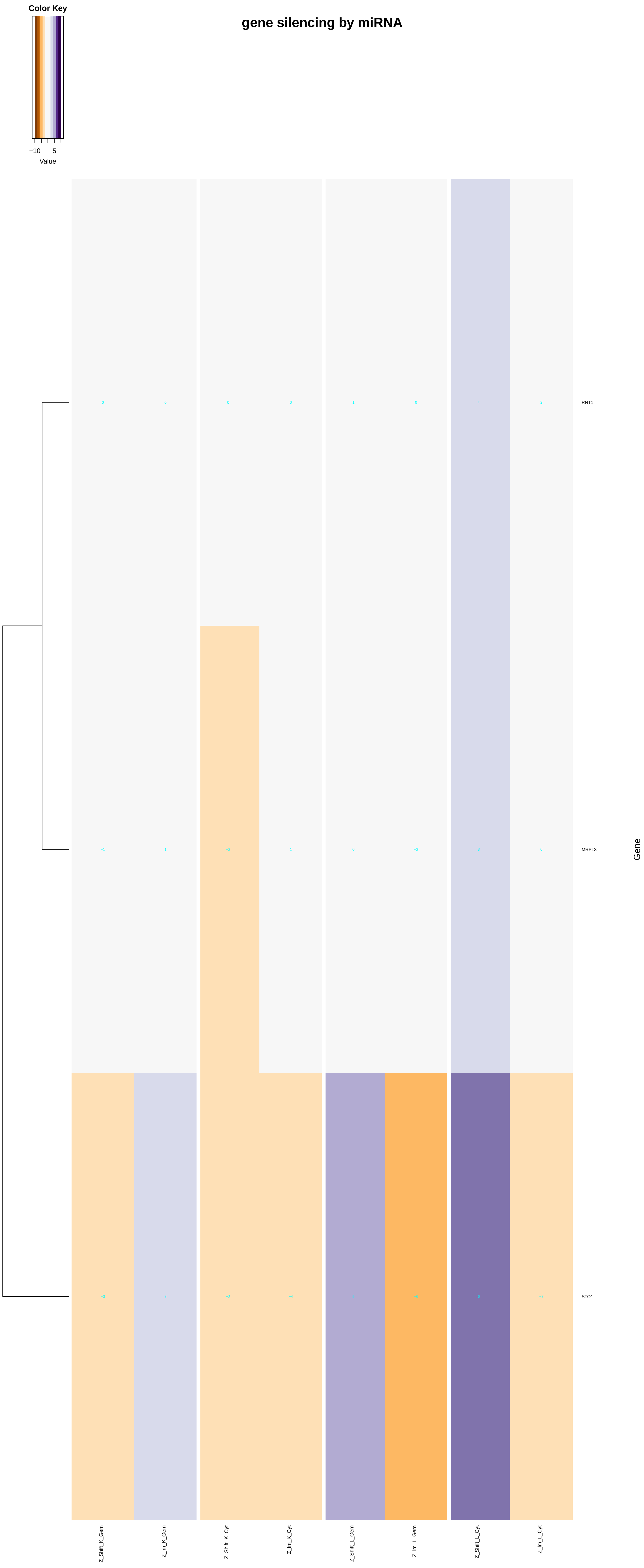
gene silencing by miRNA
Color Key
−10
5
Value
RNT1
MRPL3
STO1
Gene
Z_Shift_K_Gem
Z_Im_K_Gem
Z_Shift_K_Cyt
Z_Im_K_Cyt
Z_Shift_L_Gem
Z_Im_L_Gem
Z_Shift_L_Cyt
Z_Im_L_Cyt

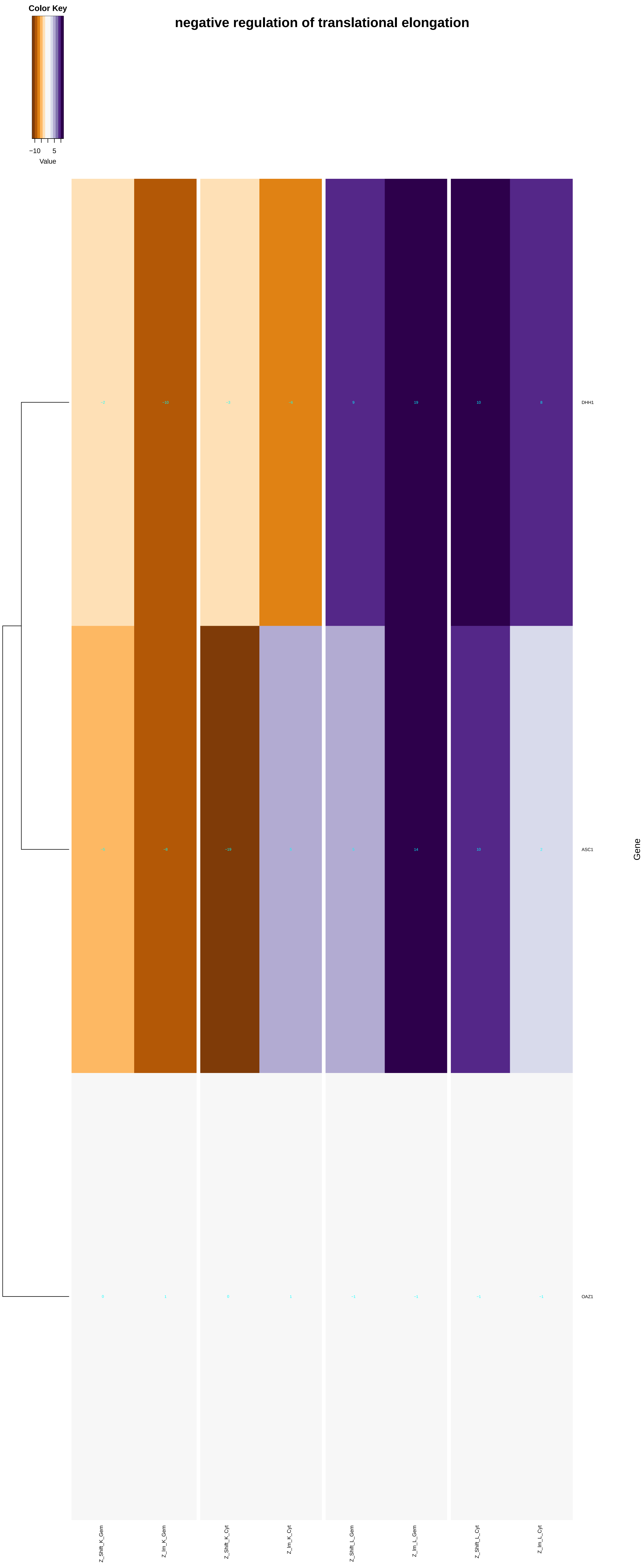

negative regulation of translational elongation
Color Key
−10
5
Value
−2
−10
−3
−6
9
19
10
8
DHH1
−5
−8
−19
5
5
14
10
2
ASC1
0
1
0
1
−1
−1
−1
−1
OAZ1
Gene
Z_Shift_K_Gem
Z_Im_K_Gem
Z_Shift_K_Cyt
Z_Im_K_Cyt
Z_Shift_L_Gem
Z_Im_L_Gem
Z_Shift_L_Cyt
Z_Im_L_Cyt

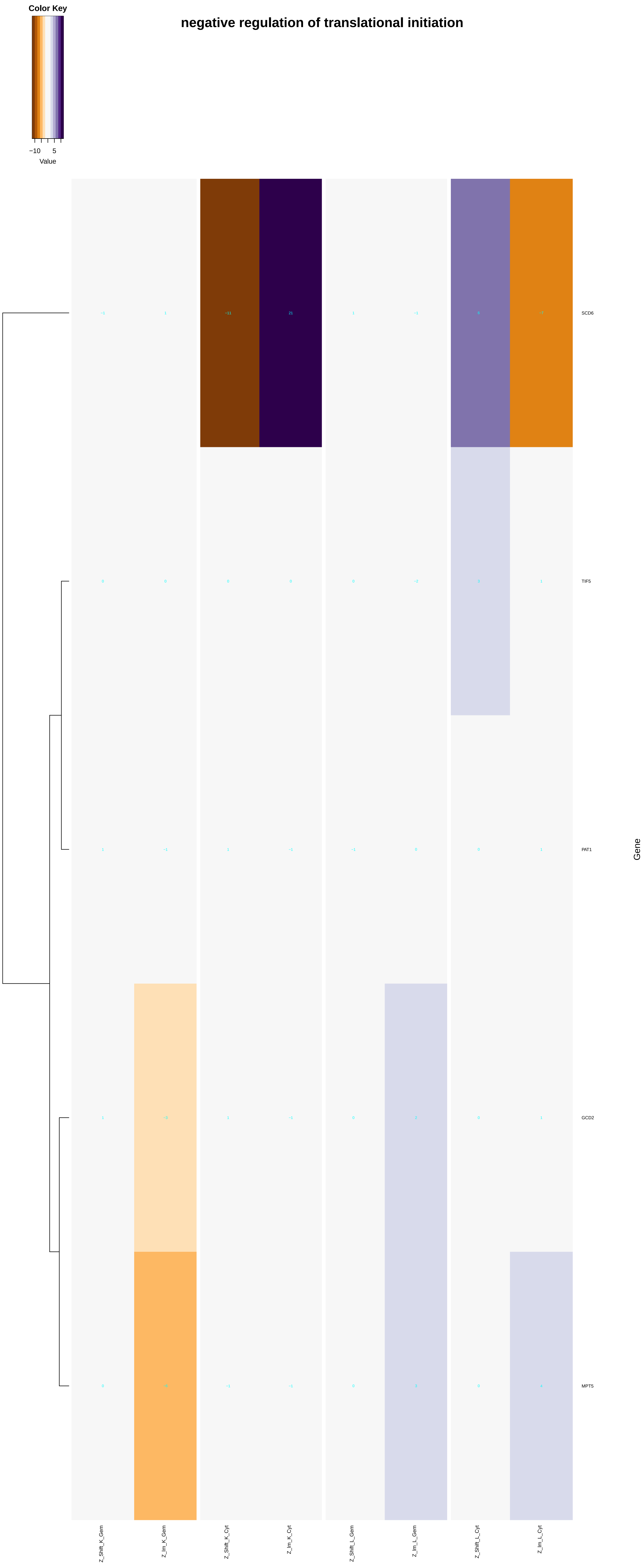

Color Key
−10
5
Value
negative regulation of translational initiation
SCD6
TIF5
PAT1
GCD2
MPT5
Gene
Z_Shift_K_Gem
Z_Im_K_Gem
Z_Shift_K_Cyt
Z_Im_K_Cyt
Z_Shift_L_Gem
Z_Im_L_Gem
Z_Shift_L_Cyt
Z_Im_L_Cyt

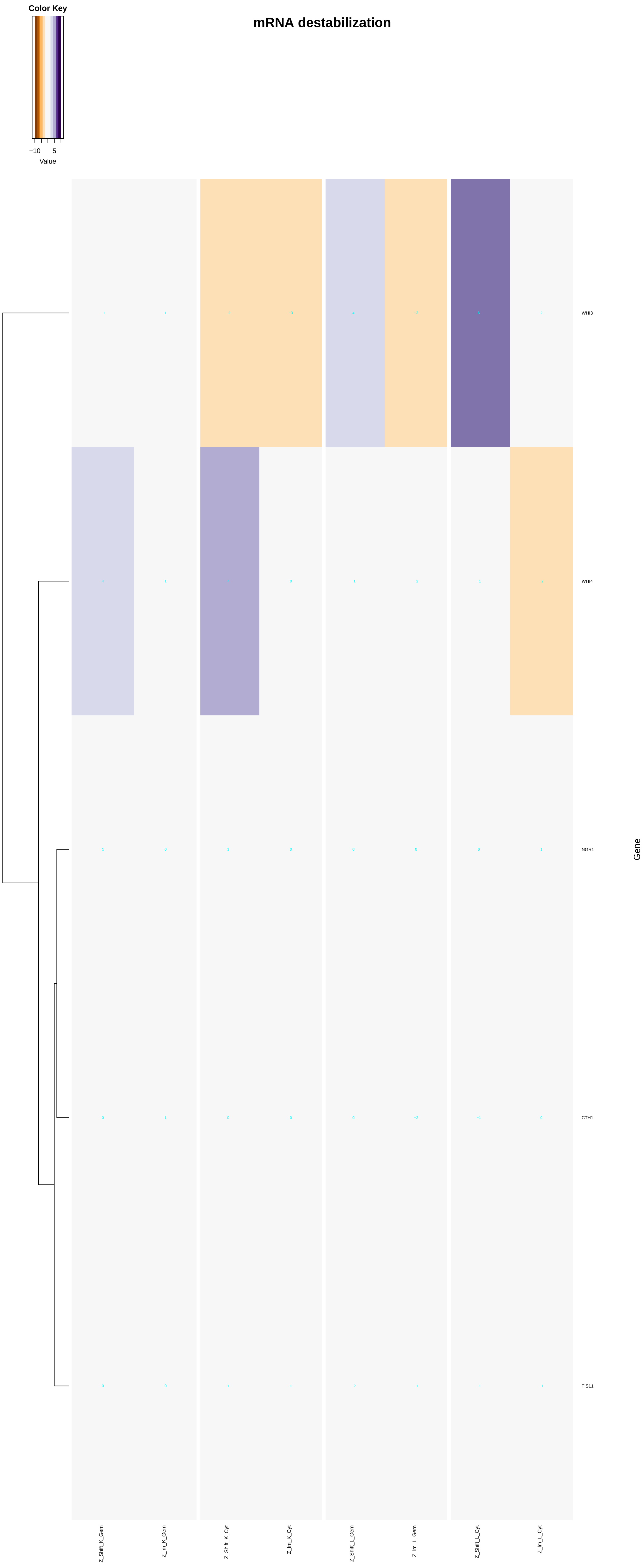
mRNA destabilization
Color Key
−10
5
Value
WHI3
WHI4
NGR1
CTH1
TIS11
Gene
Z_Shift_K_Gem
Z_Im_K_Gem
Z_Shift_K_Cyt
Z_Im_K_Cyt
Z_Shift_L_Gem
Z_Im_L_Gem
Z_Shift_L_Cyt
Z_Im_L_Cyt

### negative regulation of mitochondrial translation

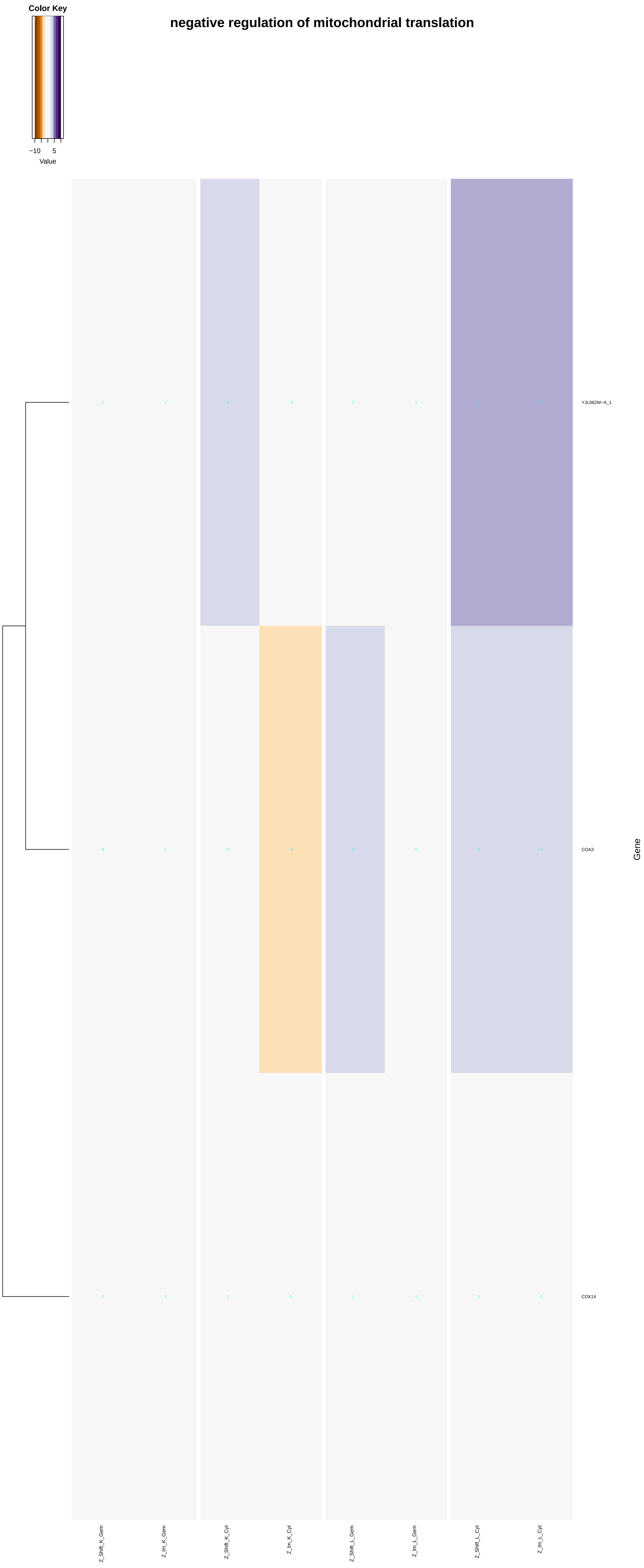

### negative regulation of transcription from RNA polymerase II promoter

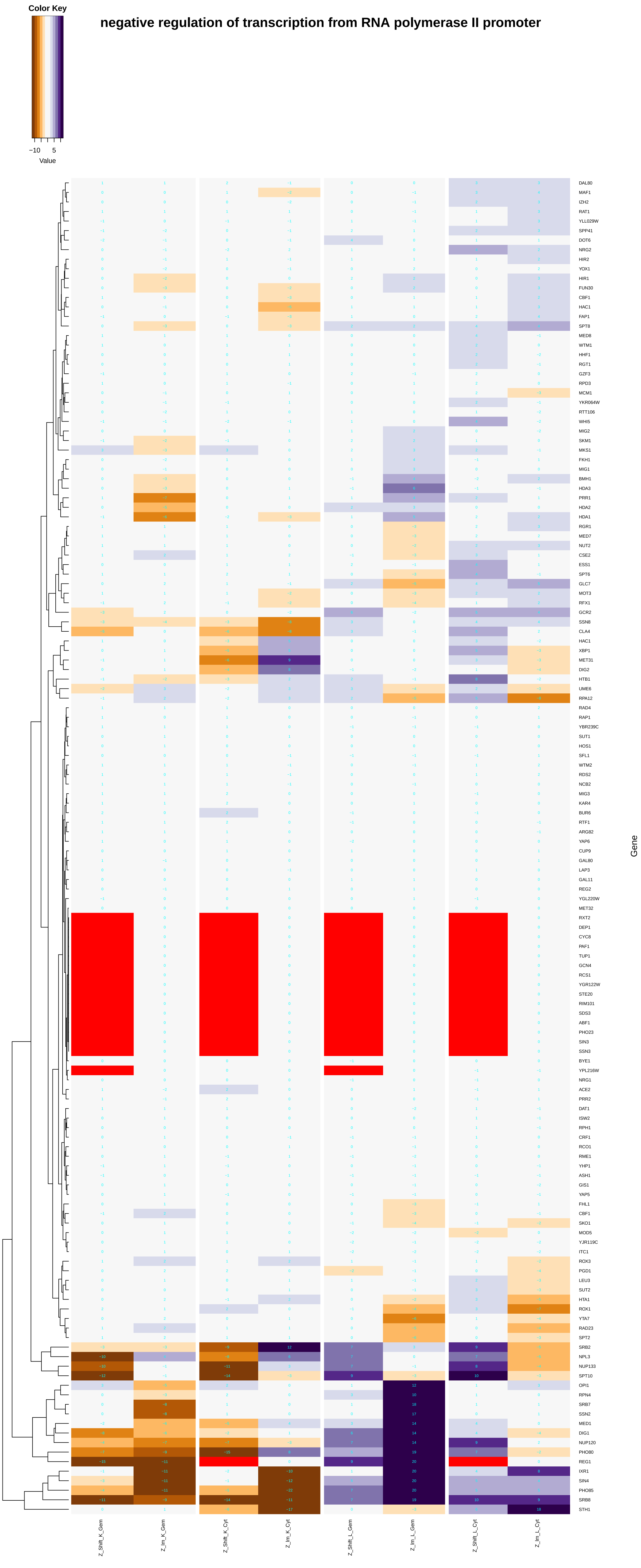

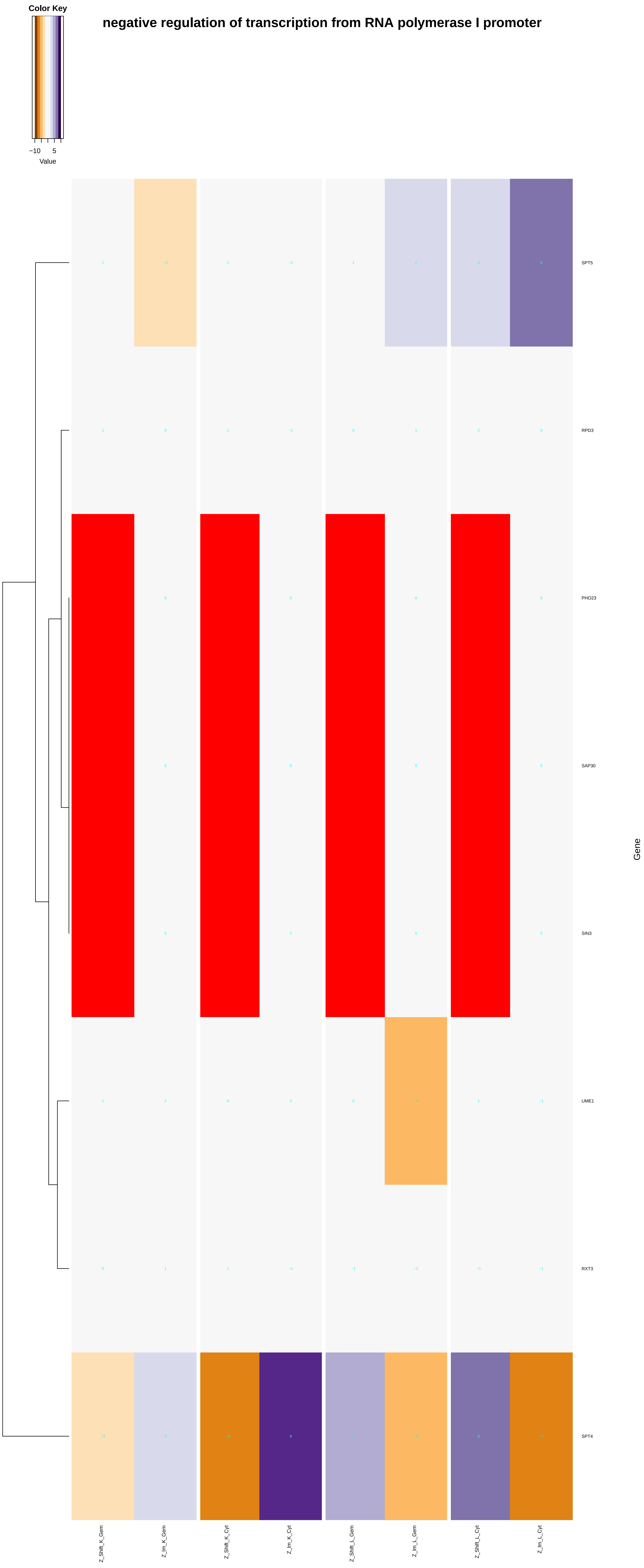

Color Key
−10
5
Value
negative regulation of transcription from RNA polymerase I promoter
SPT5
RPD3
PHO23
SAP30
SIN3
UME1
RXT3
SPT4
Gene
Z_Shift_K_Gem
Z_Im_K_Gem
Z_Shift_K_Cyt
Z_Im_K_Cyt
Z_Shift_L_Gem
Z_Im_L_Gem
Z_Shift_L_Cyt
Z_Im_L_Cyt

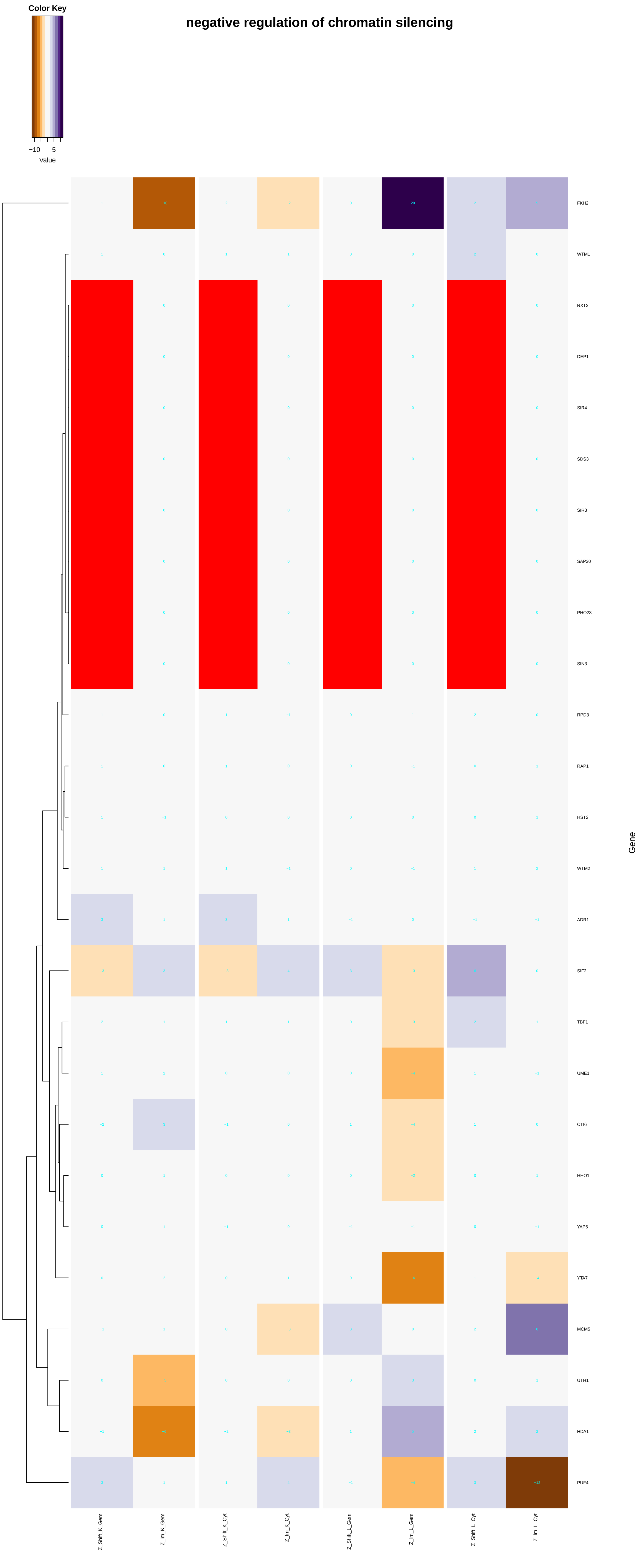
Color Key
−10
5
Value
negative regulation of chromatin silencing
FKH2
WTM1
RXT2
DEP1
SIR4
SDS3
SIR3
SAP30
PHO23
SIN3
RPD3
RAP1
HST2
WTM2
ADR1
SIF2
TBF1
UME1
CTI6
HHO1
YAP5
YTA7
MCM5
UTH1
HDA1
PUF4
Gene
Z_Shift_K_Gem
Z_lm_K_Gem
Z_Shift_K_Cyt
Z_lm_K_Cyt
Z_Shift_L_Gem
Z_lm_L_Gem
Z_Shift_L_Cyt
Z_lm_L_Cyt

### negative regulation of DNA−templated transcription, elongation

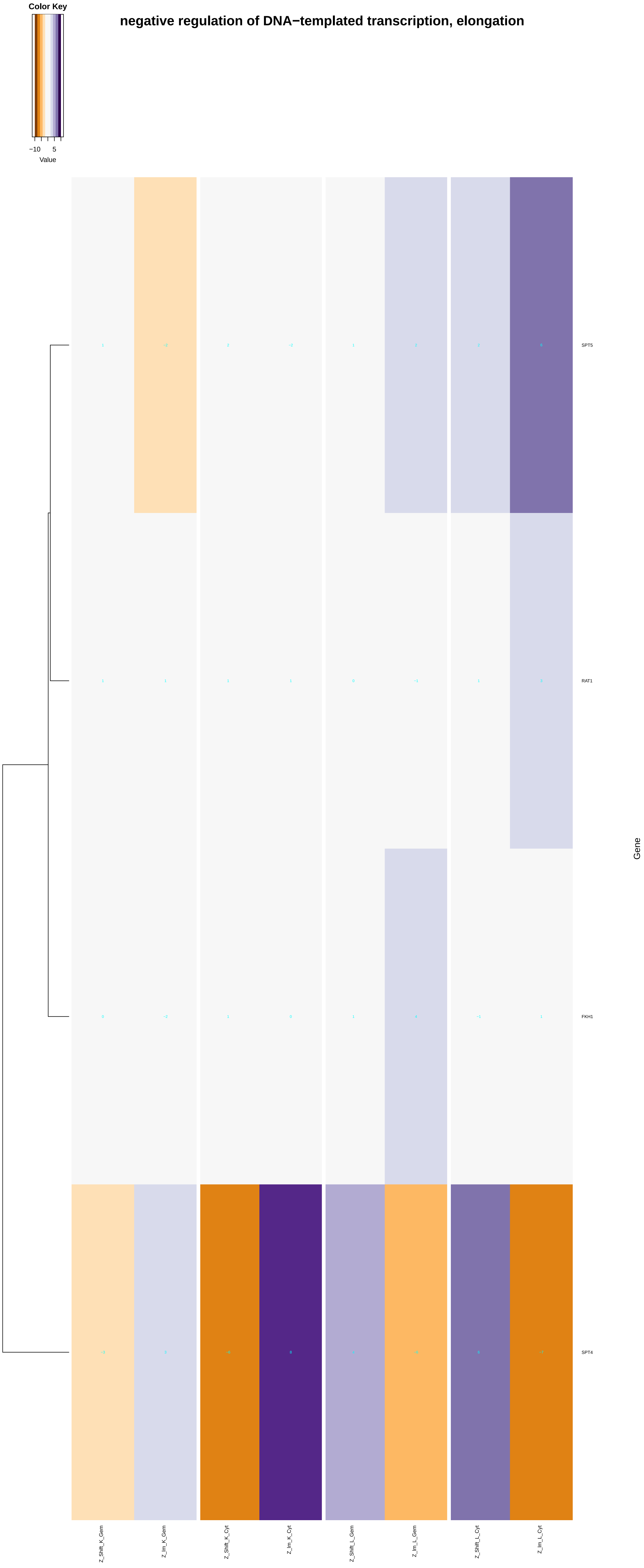

### carbon catabolite repression of transcription

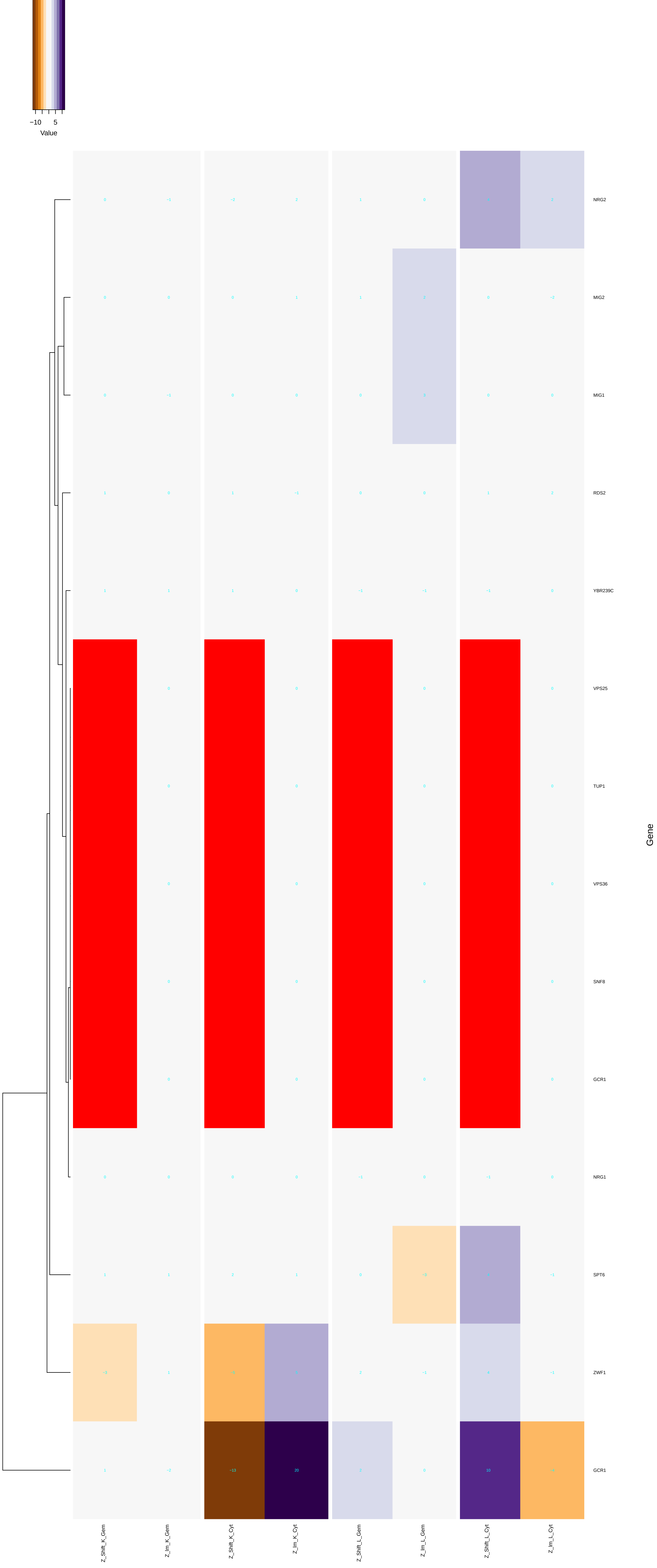

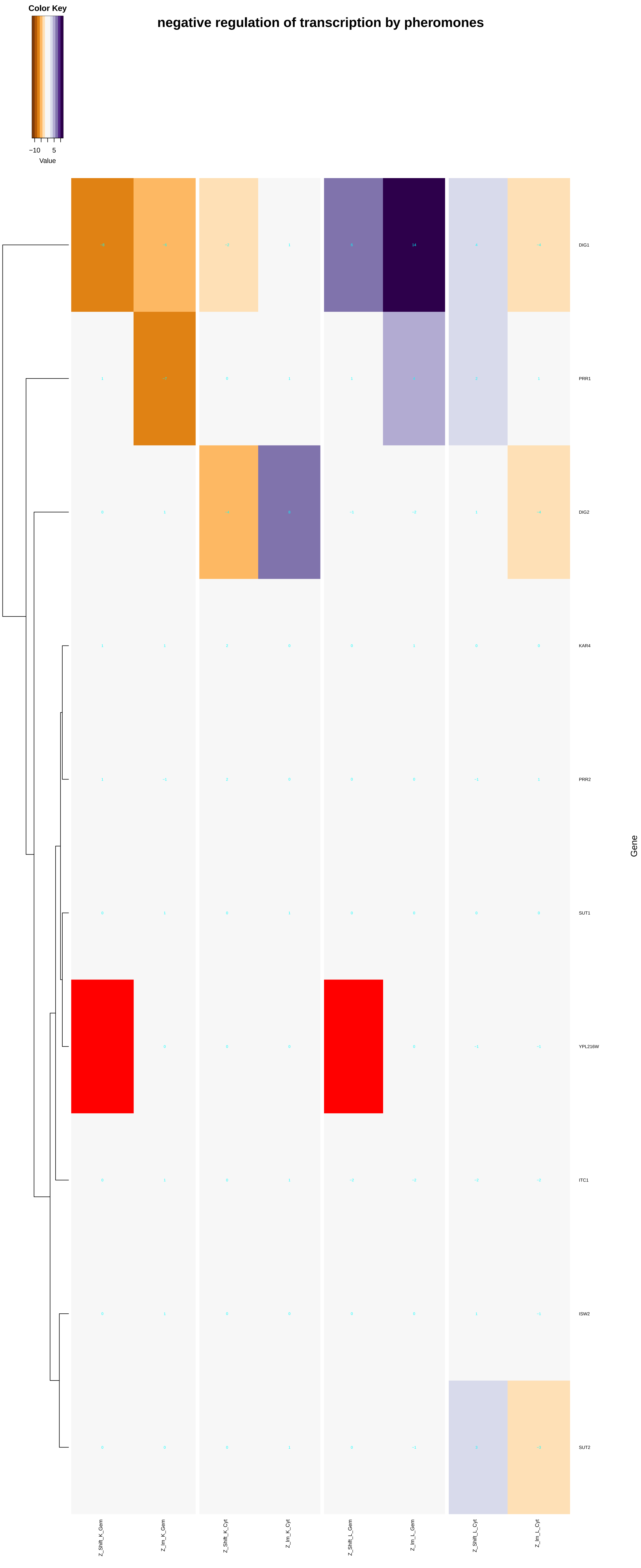

Color Key
−10
5
Value
negative regulation of transcription by pheromones
DIG1
PRR1
DIG2
KAR4
PRR2
SUT1
YPL216W
ITC1
ISW2
SUT2
Gene
Z_Shift_K_Gem
Z_Im_K_Gem
Z_Shift_K_Cyt
Z_Im_K_Cyt
Z_Shift_L_Gem
Z_Im_L_Gem
Z_Shift_L_Cyt
Z_Im_L_Cyt

### negative regulation of transcription involved in meiotic cell cycle

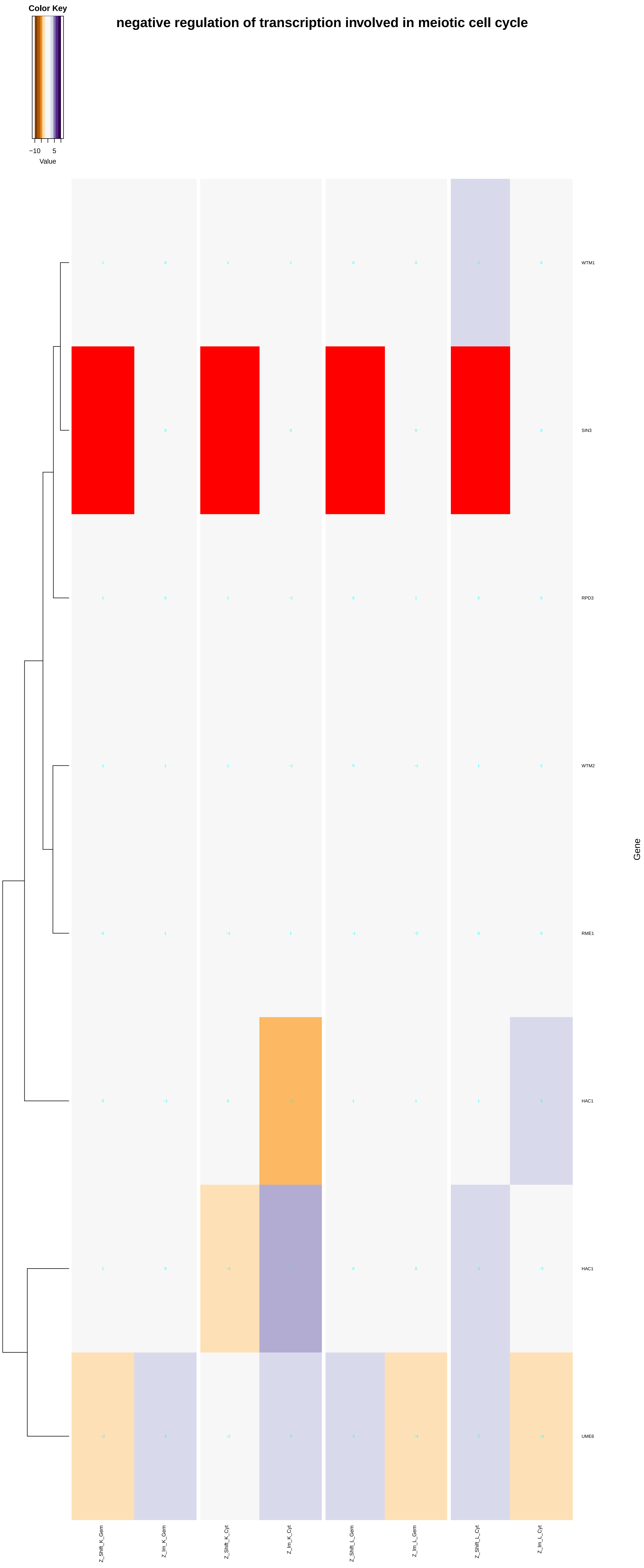

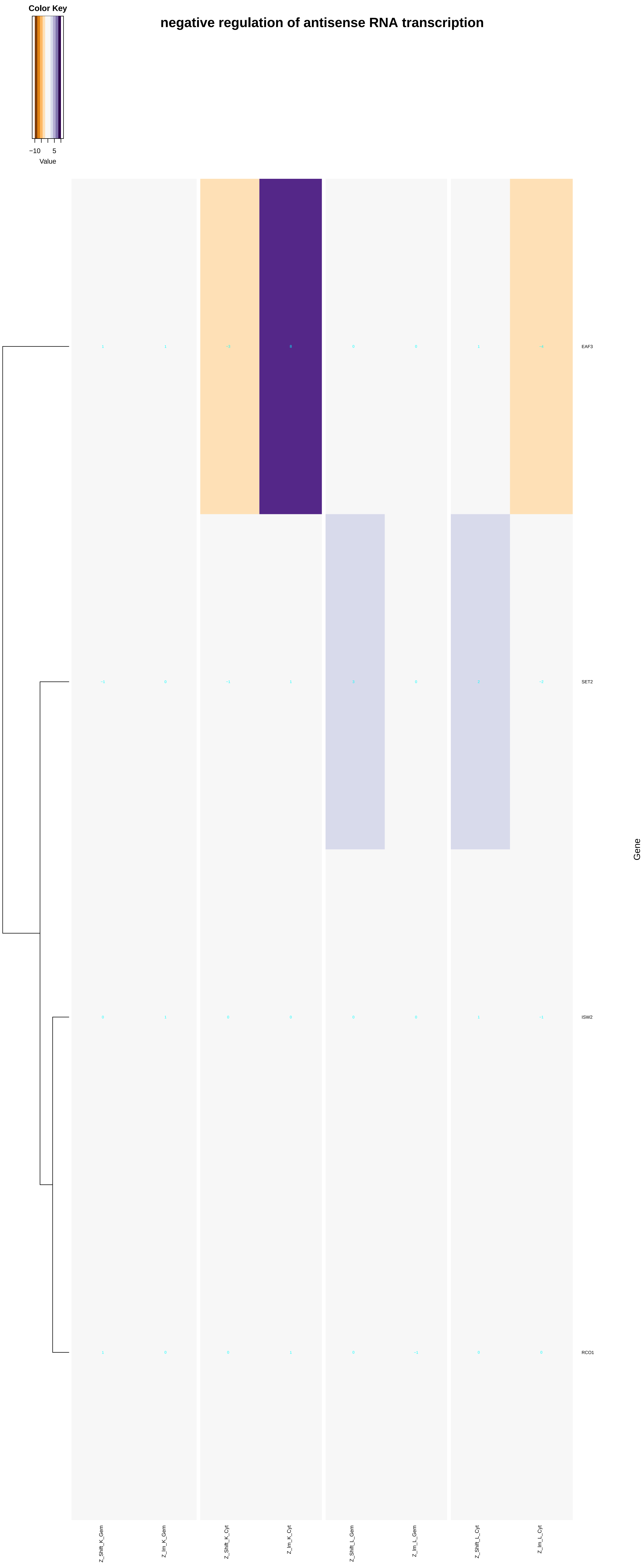

Color Key
−10
5
Value
negative regulation of antisense RNA transcription
EAF3
SET2
ISW2
RCO1
Gene
Z_Shift_K_Gem
Z_Im_K_Gem
Z_Shift_K_Cyt
Z_Im_K_Cyt
Z_Shift_L_Gem
Z_Im_L_Gem
Z_Shift_L_Cyt
Z_Im_L_Cyt

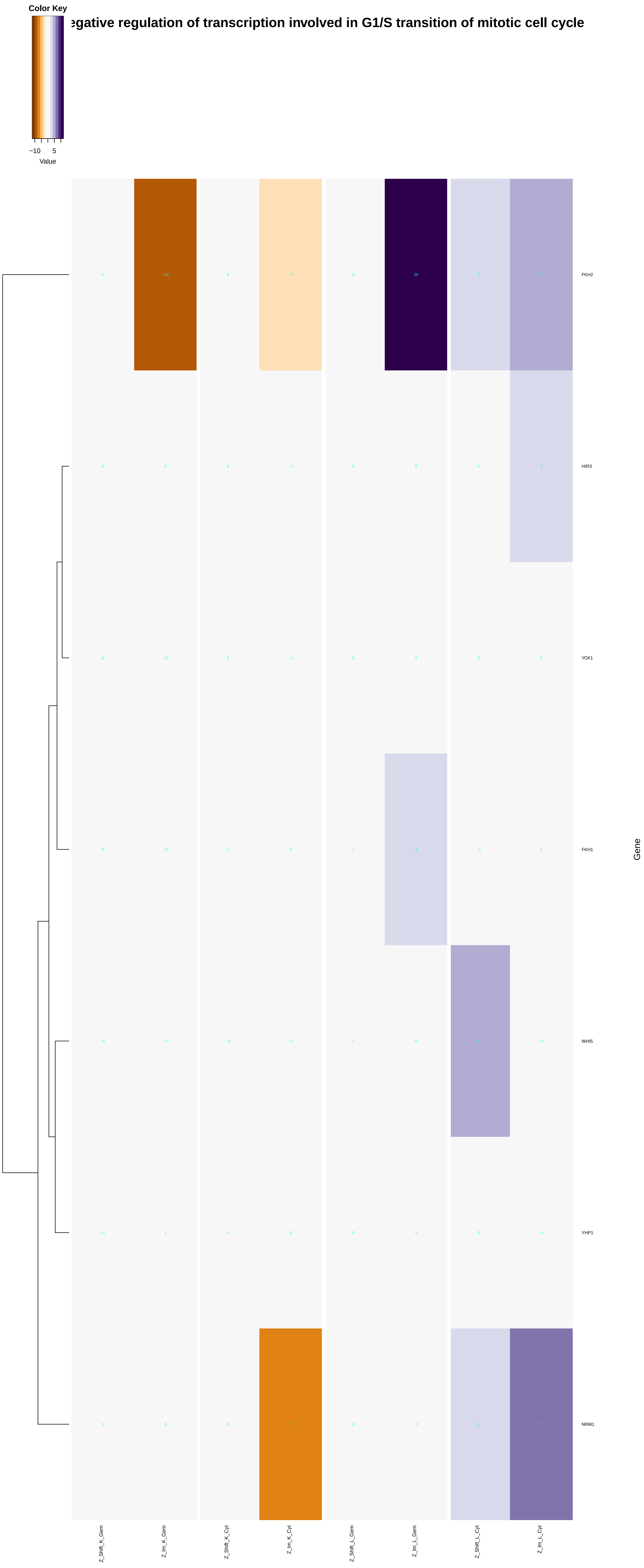
egative regulation of transcription involved in G1/S transition of mitotic cell cycle
Color Key
−10
5
Value
FKH2
HIR3
YOX1
FKH1
WHI5
YHP1
NRM1
Gene
Z_Shift_K_Gem
Z_lm_K_Gem
Z_Shift_K_Cyt
Z_lm_K_Cyt
Z_Shift_L_Gem
Z_lm_L_Gem
Z_Shift_L_Cyt
Z_lm_L_Cyt

### nitrogen catabolite repression of transcription

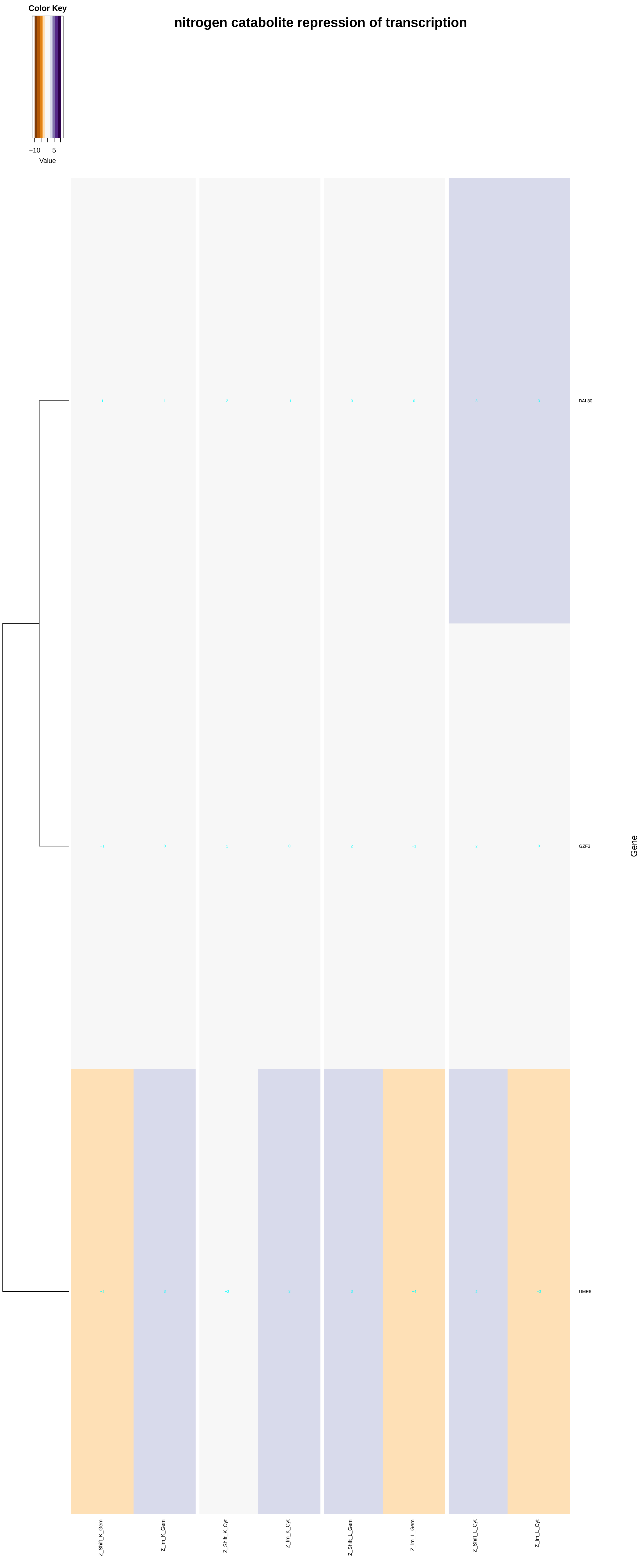

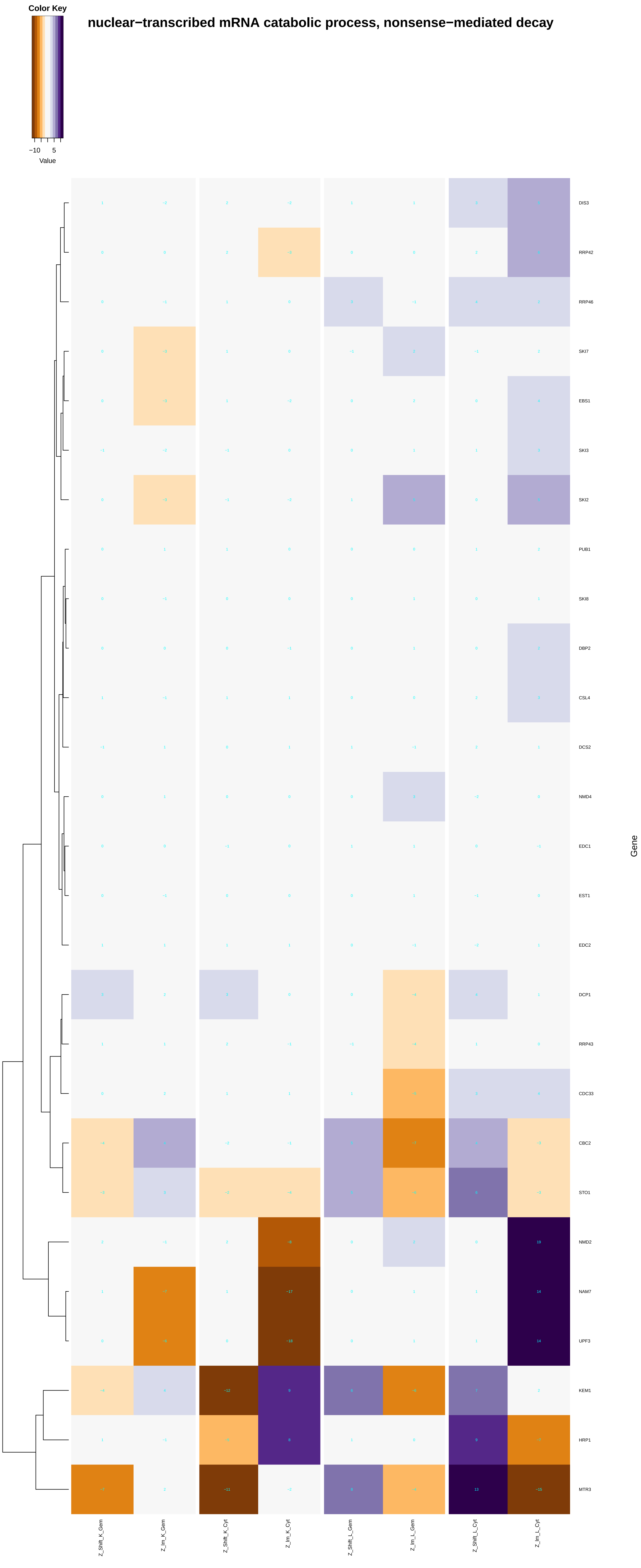
Color Key
−10 5
Value
nuclear−transcribed mRNA catabolic process, nonsense−mediated decay
DIS3
RRP42
RRP46
SKI7
EBS1
SKI3
SKI2
PUB1
SKI8
DBP2
CSL4
DCS2
NMD4
EDC1
EST1
EDC2
DCP1
RRP43
CDC33
CBC2
STO1
NMD2
NAM7
UPF3
KEM1
HRP1
MTR3
Gene
Z_Shift_K_Gem
Z_lm_K_Gem
Z_Shift_K_Cyt
Z_lm_K_Cyt
Z_Shift_L_Gem
Z_lm_L_Gem
Z_Shift_L_Cyt
Z_lm_L_Cyt

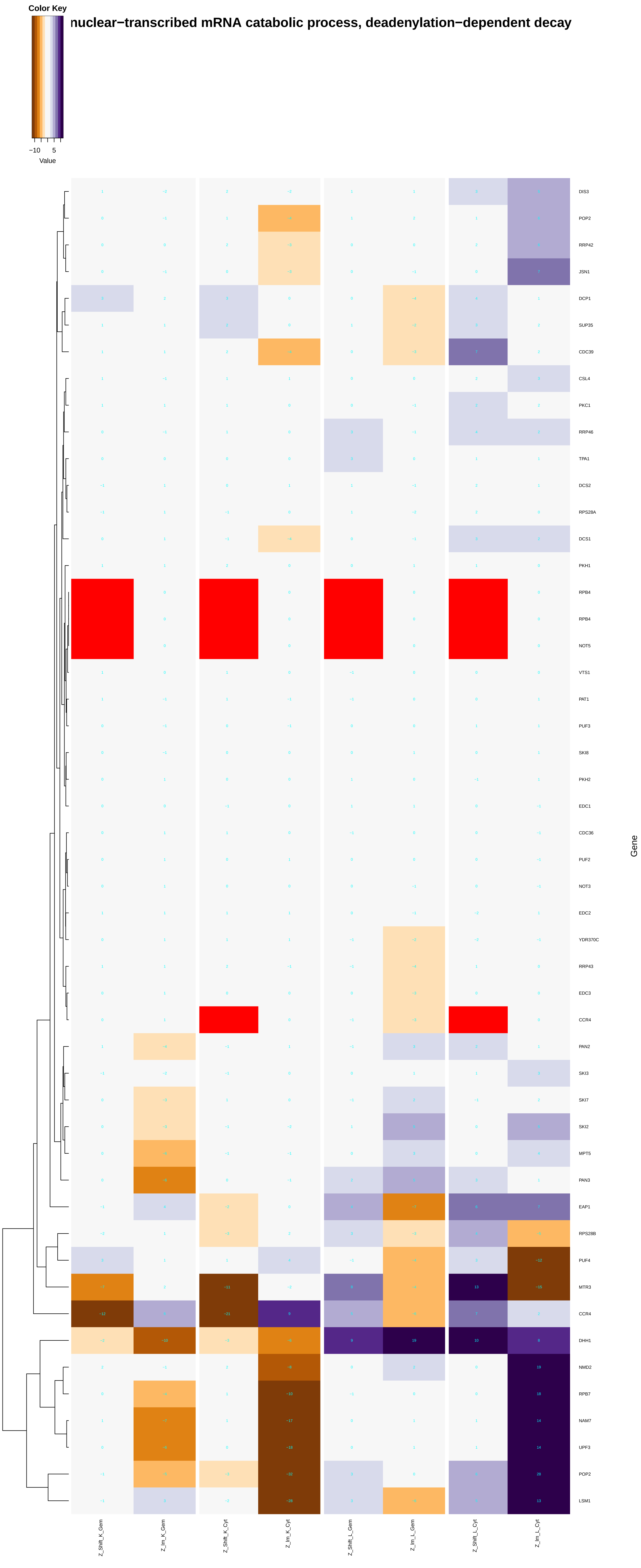
Color Key
−10 5
Value
nuclear−transcribed mRNA catabolic process, deadenylation−dependent decay
DIS3
POP2
RRP42
JSN1
DCP1
SUP35
CDC39
CSL4
PKC1
RRP46
TPA1
DCS2
RPS28A
DCS1
PKH1
RPB4
RPB4
NOT5
VTS1
PAT1
PUF3
SKI8
PKH2
EDC1
CDC36
PUF2
NOT3
EDC2
YDR370C
RRP43
EDC3
CCR4
PAN2
SKI3
SKI7
SKI2
MPT5
PAN3
EAP1
RPS28B
PUF4
MTR3
CCR4
DHH1
NMD2
RPB7
NAM7
UPF3
POP2
LSM1
Gene
Z_Shift_K_Gem
Z_Im_K_Gem
Z_Shift_K_Cyt
Z_Im_K_Cyt
Z_Shift_L_Gem
Z_Im_L_Gem
Z_Shift_L_Cyt
Z_Im_L_Cyt

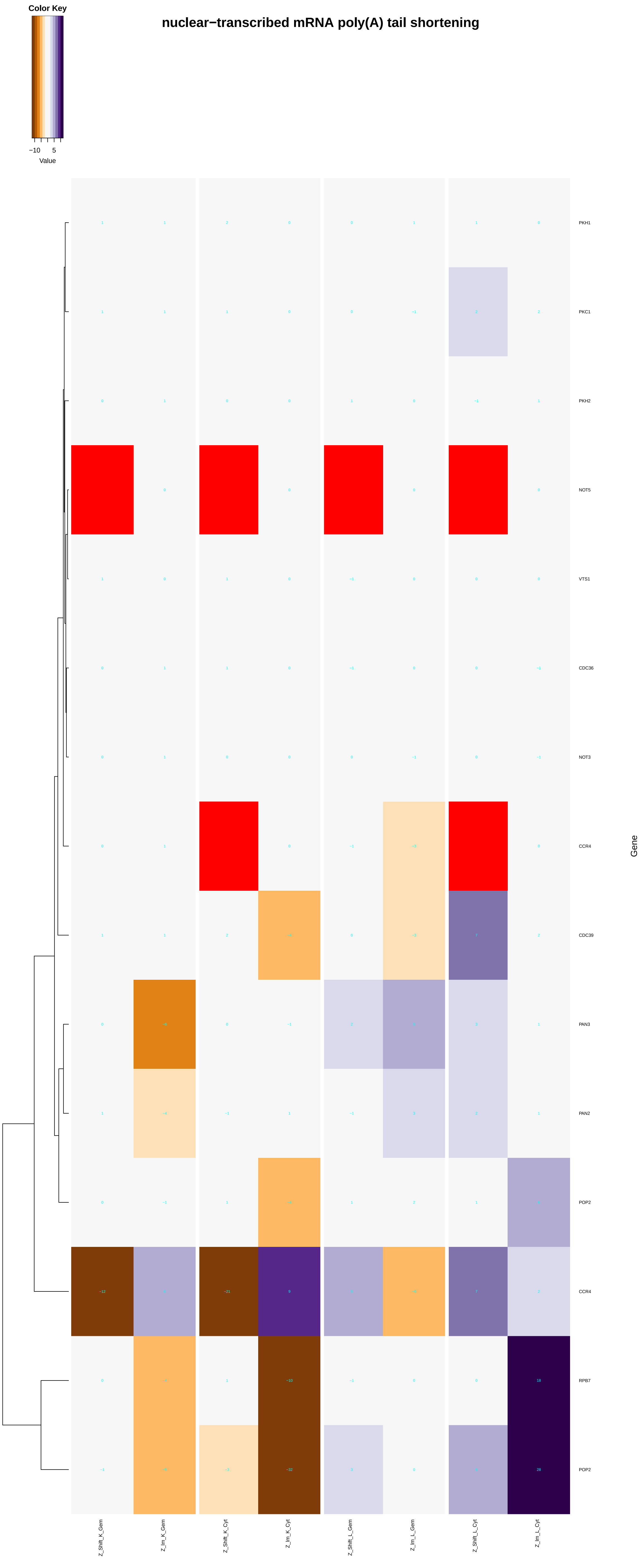

nuclear−transcribed mRNA poly(A) tail shortening
Color Key
−10
5
Value
PKH1
PKC1
PKH2
NOT5
VTS1
CDC36
NOT3
CCR4
CDC39
PAN3
PAN2
POP2
CCR4
RPB7
POP2
Gene
Z_Shift_K_Gem
Z_Im_K_Gem
Z_Shift_K_Cyt
Z_Im_K_Cyt
Z_Shift_L_Gem
Z_Im_L_Gem
Z_Shift_L_Cyt
Z_Im_L_Cyt

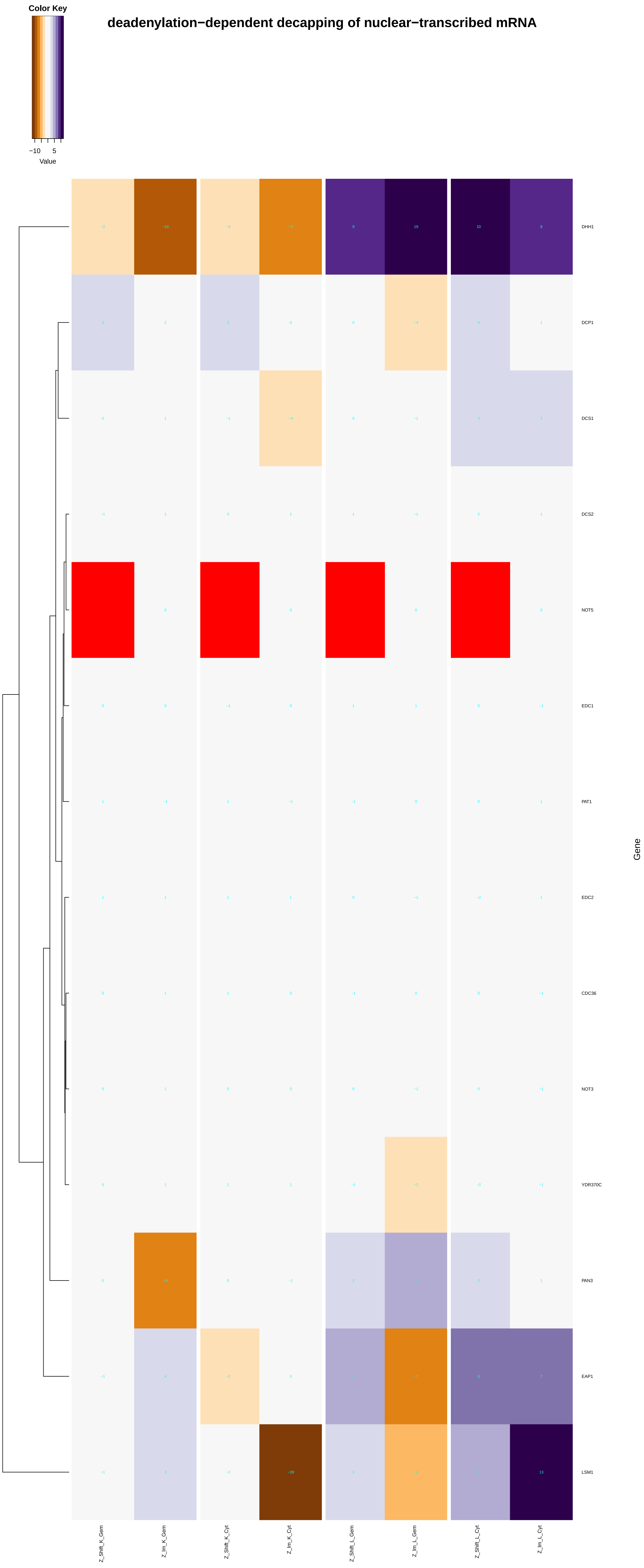

deadenylation−dependent decapping of nuclear−transcribed mRNA
Color Key
−10
5
Value
DHH1
DCP1
DCS1
DCS2
NOT5
EDC1
PAT1
EDC2
CDC36
NOT3
YDR370C
PAN3
EAP1
LSM1
Gene
Z_Shift_K_Gem
Z_Im_K_Gem
Z_Shift_K_Cyt
Z_Im_K_Cyt
Z_Shift_L_Gem
Z_Im_L_Gem
Z_Shift_L_Cyt
Z_Im_L_Cyt

### nuclear−transcribed mRNA catabolic process, exonucleolytic

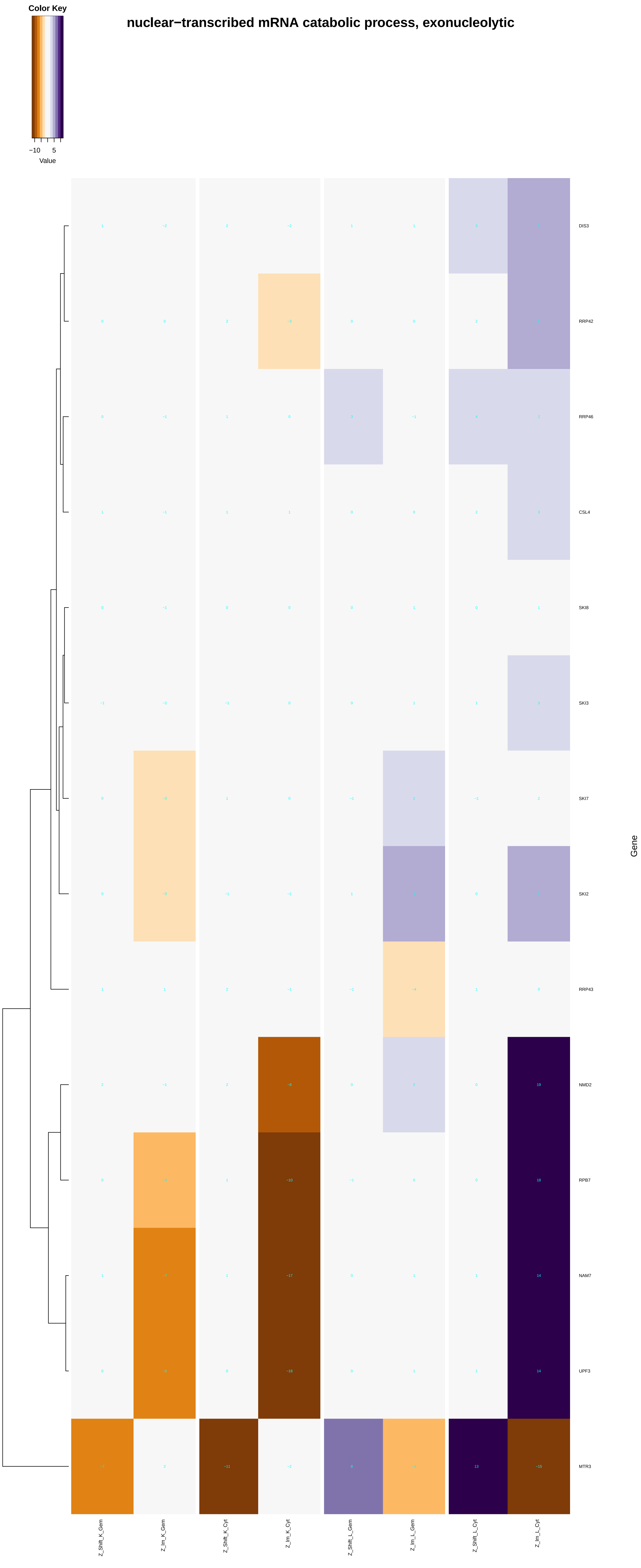

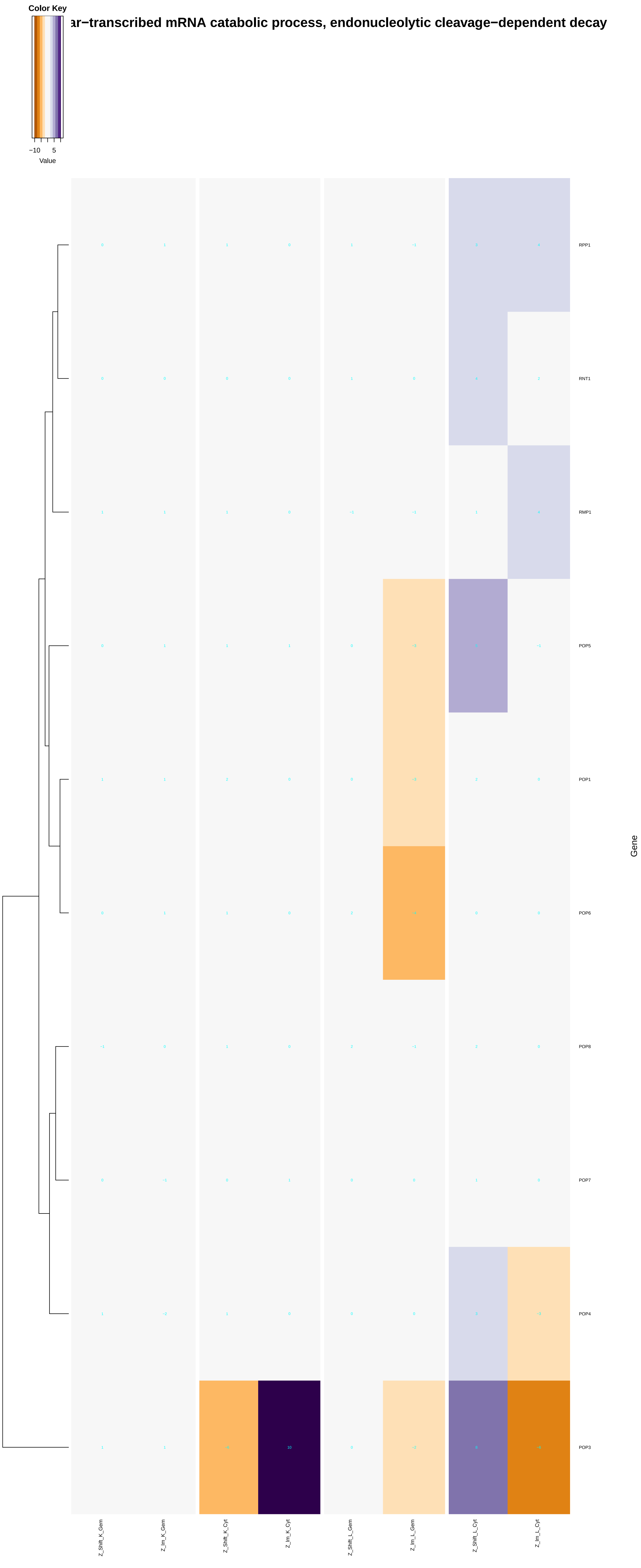

Color Key
−10
5
Value
ar−transcribed mRNA catabolic process, endonucleolytic cleavage−dependent decay
RPP1
RNT1
RMP1
POP5
POP1
POP6
POP8
POP7
POP4
POP3
Gene
Z_Shift_K_Gem
Z_Im_K_Gem
Z_Shift_K_Cyt
Z_Im_K_Cyt
Z_Shift_L_Gem
Z_Im_L_Gem
Z_Shift_L_Cyt
Z_Im_L_Cyt

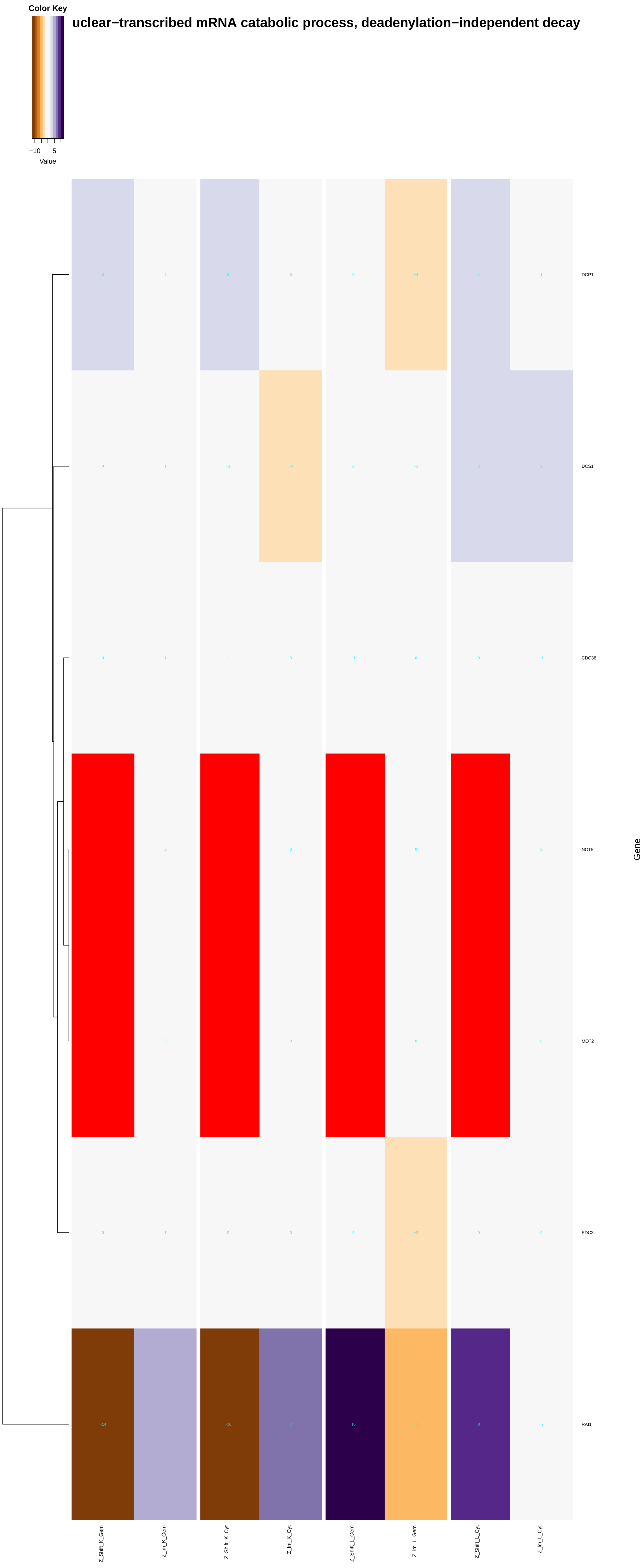
Color Key
−10 5
Value
uclear−transcribed mRNA catabolic process, deadenylation−independent decay
DCP1
DCS1
CDC36
NOT5
MOT2
EDC3
RAI1
Gene
Z_Shift_K_Gem
Z_Im_K_Gem
Z_Shift_K_Cyt
Z_Im_K_Cyt
Z_Shift_L_Gem
Z_Im_L_Gem
Z_Shift_L_Cyt
Z_Im_L_Cyt

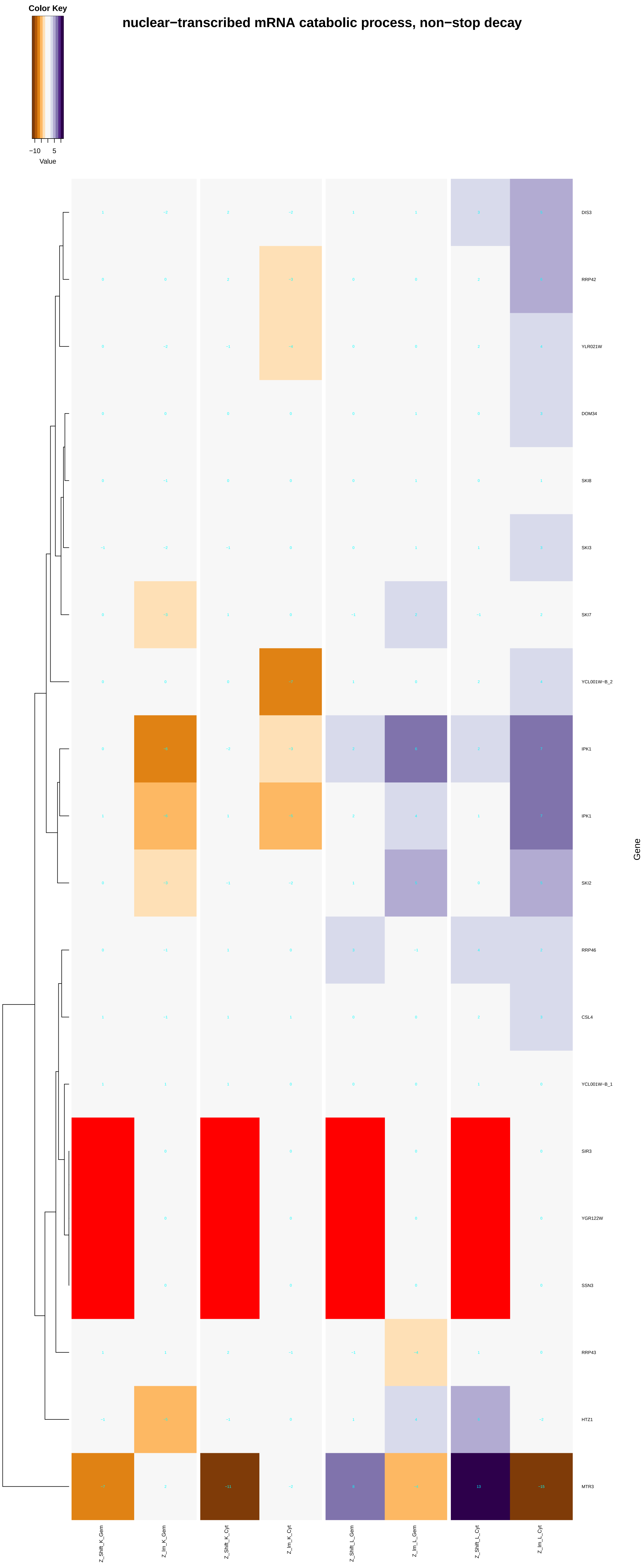
nuclear−transcribed mRNA catabolic process, non−stop decay
Color Key
−10
5
Value
DIS3
RRP42
YLR021W
DOM34
SKI8
SKI3
SKI7
YCL001W−B_2
IPK1
IPK1
SKI2
RRP46
CSL4
YCL001W−B_1
SIR3
YGR122W
SSN3
RRP43
HTZ1
MTR3
Gene
Z_Shift_K_Gem
Z_Im_K_Gem
Z_Shift_K_Cyt
Z_Im_K_Cyt
Z_Shift_L_Gem
Z_Im_L_Gem
Z_Shift_L_Cyt
Z_Im_L_Cyt

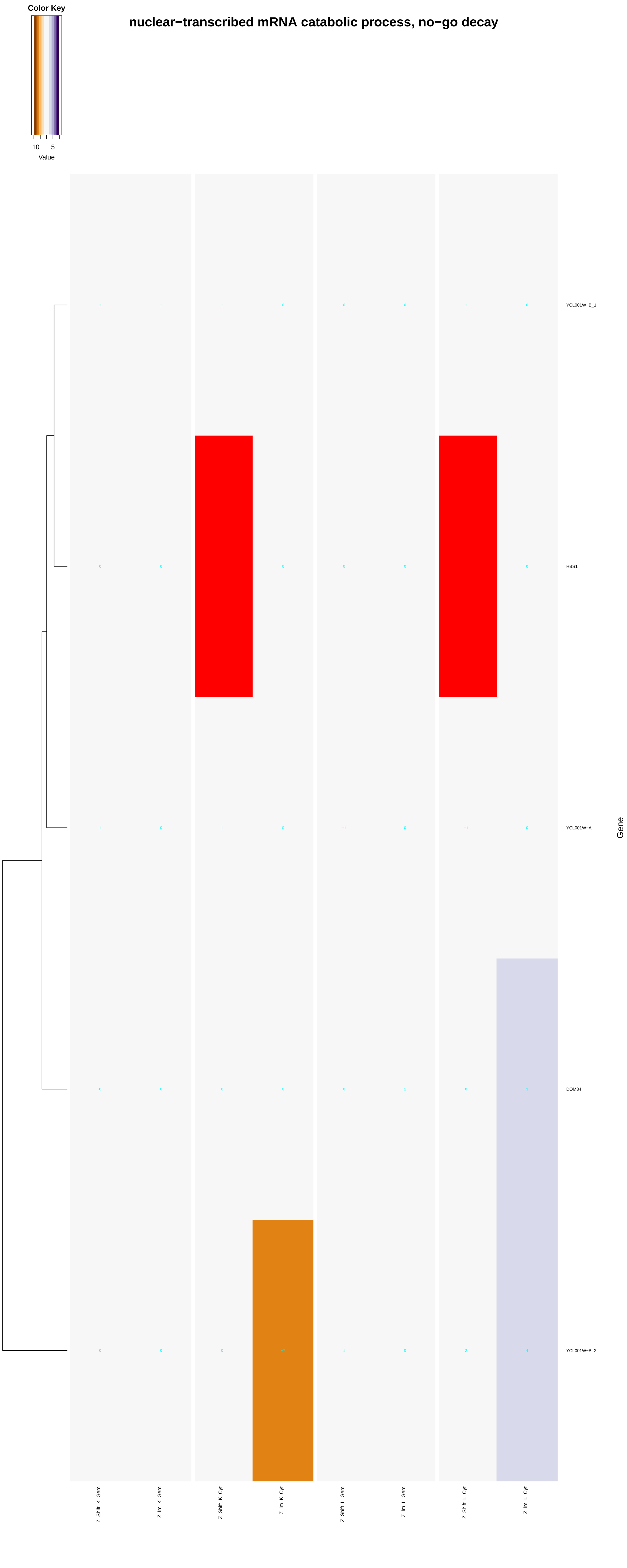
nuclear−transcribed mRNA catabolic process, no−go decay
Color Key
−10
5
Value
YCL001W−B_1
HBS1
YCL001W−A
DOM34
YCL001W−B_2
Gene
Z_Shift_K_Gem
Z_Im_K_Gem
Z_Shift_K_Cyt
Z_Im_K_Cyt
Z_Shift_L_Gem
Z_Im_L_Gem
Z_Shift_L_Cyt
Z_Im_L_Cyt

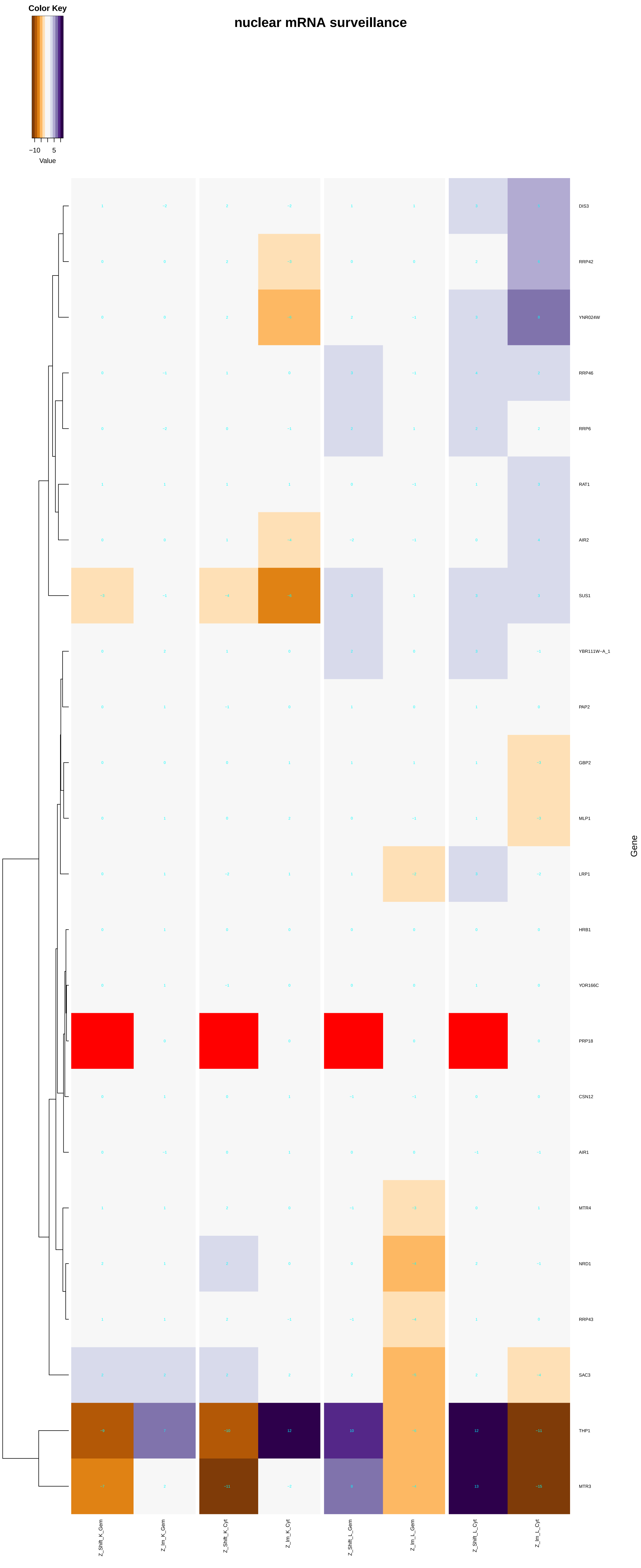
Color Key
−10
5
Value
nuclear mRNA surveillance
DIS3
RRP42
YNR024W
RRP46
RRP6
RAT1
AIR2
SUS1
YBR111W−A_1
PAP2
GBP2
MLP1
LRP1
HRB1
YOR166C
PRP18
CSN12
AIR1
MTR4
NRD1
RRP43
SAC3
THP1
MTR3
Gene
Z_Shift_K_Gem
Z_Im_K_Gem
Z_Shift_K_Cyt
Z_Im_K_Cyt
Z_Shift_L_Gem
Z_Im_L_Gem
Z_Shift_L_Cyt
Z_Im_L_Cyt

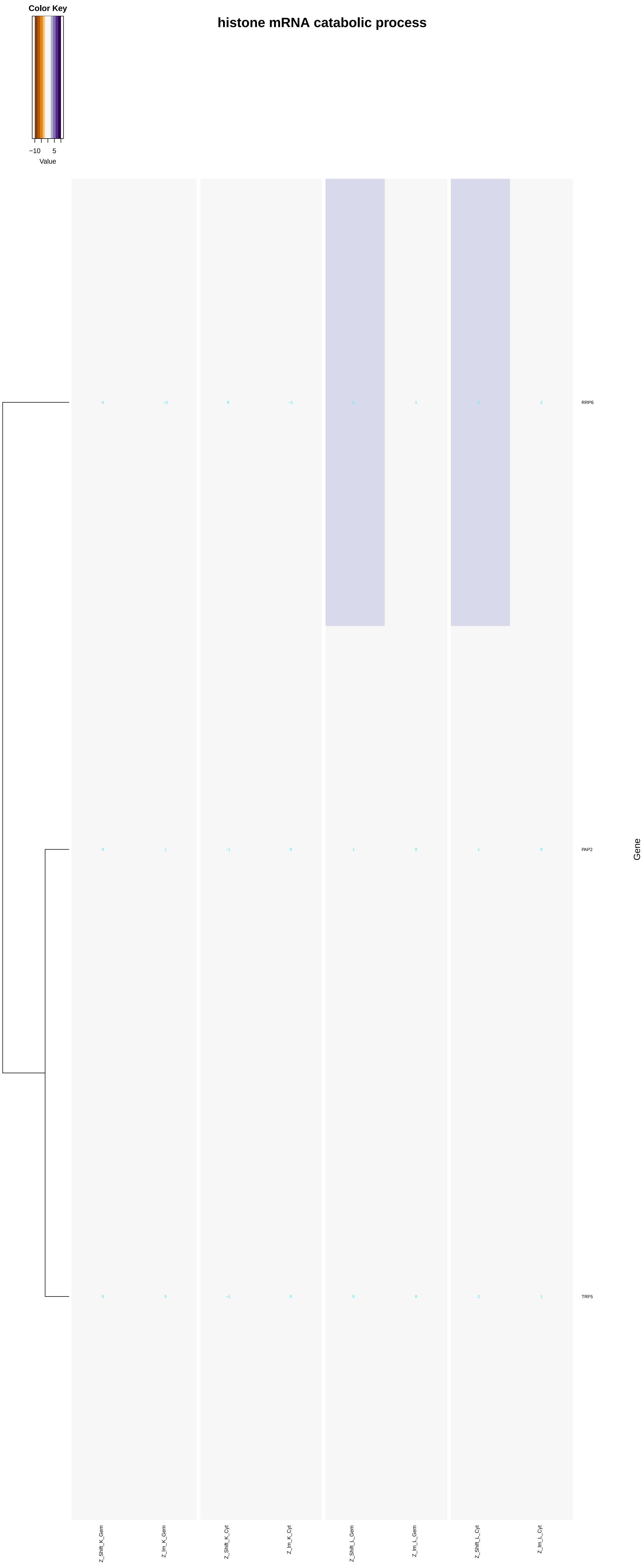
histone mRNA catabolic process
Color Key
−10
5
Value
RRP6
PAP2
TRF5
Gene
Z_Shift_K_Gem
Z_Im_K_Gem
Z_Shift_K_Cyt
Z_Im_K_Cyt
Z_Shift_L_Gem
Z_Im_L_Gem
Z_Shift_L_Cyt
Z_Im_L_Cyt

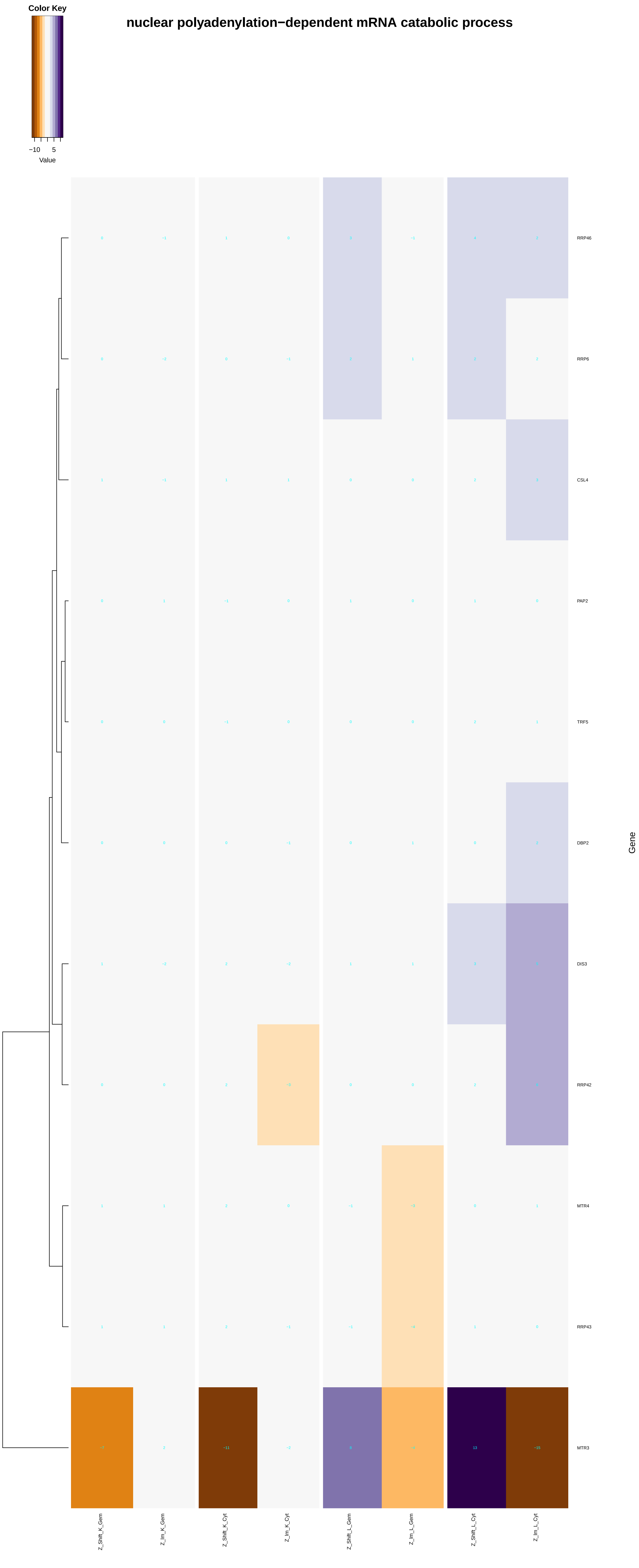
nuclear polyadenylation−dependent mRNA catabolic process
Color Key
−10
5
Value
RRP46
RRP6
CSL4
PAP2
TRF5
DBP2
DIS3
RRP42
MTR4
RRP43
MTR3
Gene
Z_Shift_K_Gem
Z_Im_K_Gem
Z_Shift_K_Cyt
Z_Im_K_Cyt
Z_Shift_L_Gem
Z_Im_L_Gem
Z_Shift_L_Cyt
Z_Im_L_Cyt

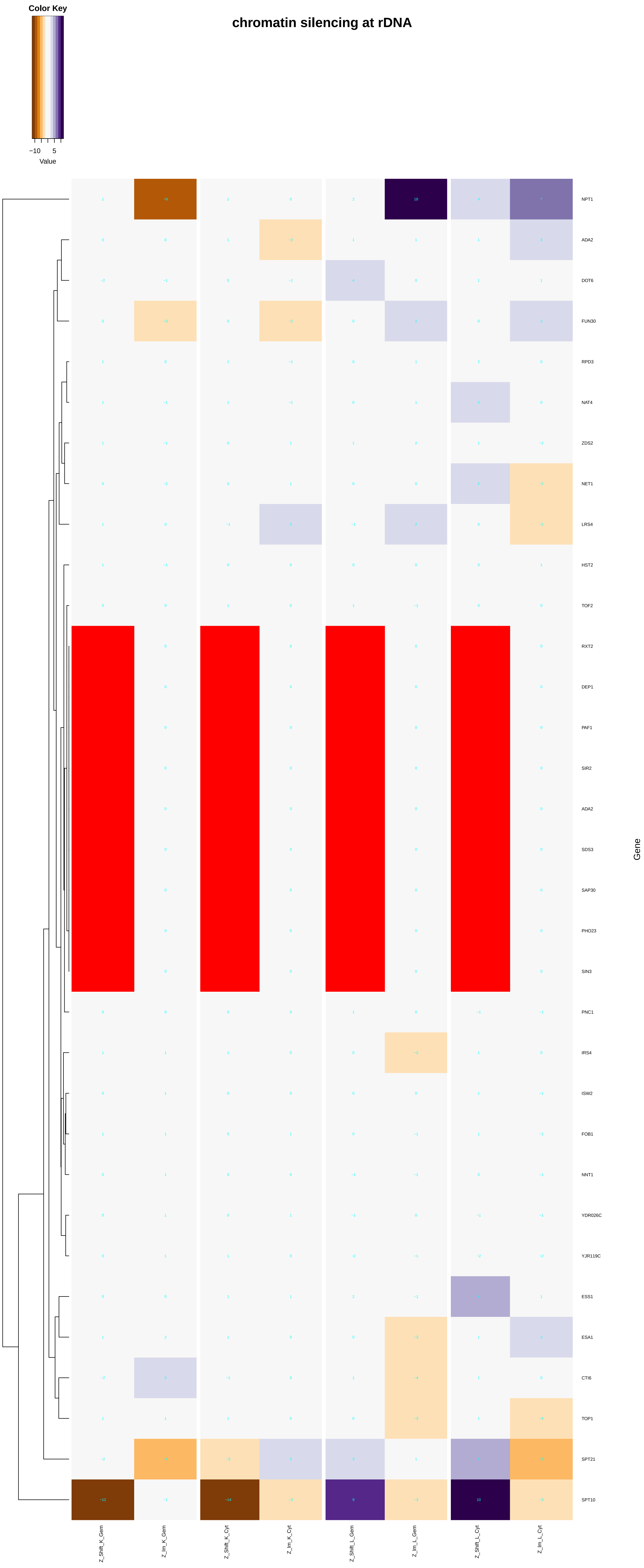

chromatin silencing at rDNA
Color Key
−10
5
Value
NPT1
ADA2
DOT6
FUN30
RPD3
NAT4
ZDS2
NET1
LRS4
HST2
TOF2
RXT2
DEP1
PAF1
SIR2
ADA2
SDS3
SAP30
PHO23
SIN3
PNC1
IRS4
ISW2
FOB1
NNT1
YDR026C
YJR119C
ESS1
ESA1
CTI6
TOP1
SPT21
SPT10
Gene
Z_Shift_K_Gem
Z_lm_K_Gem
Z_Shift_K_Cyt
Z_lm_K_Cyt
Z_Shift_L_Gem
Z_lm_L_Gem
Z_Shift_L_Cyt
Z_lm_L_Cyt

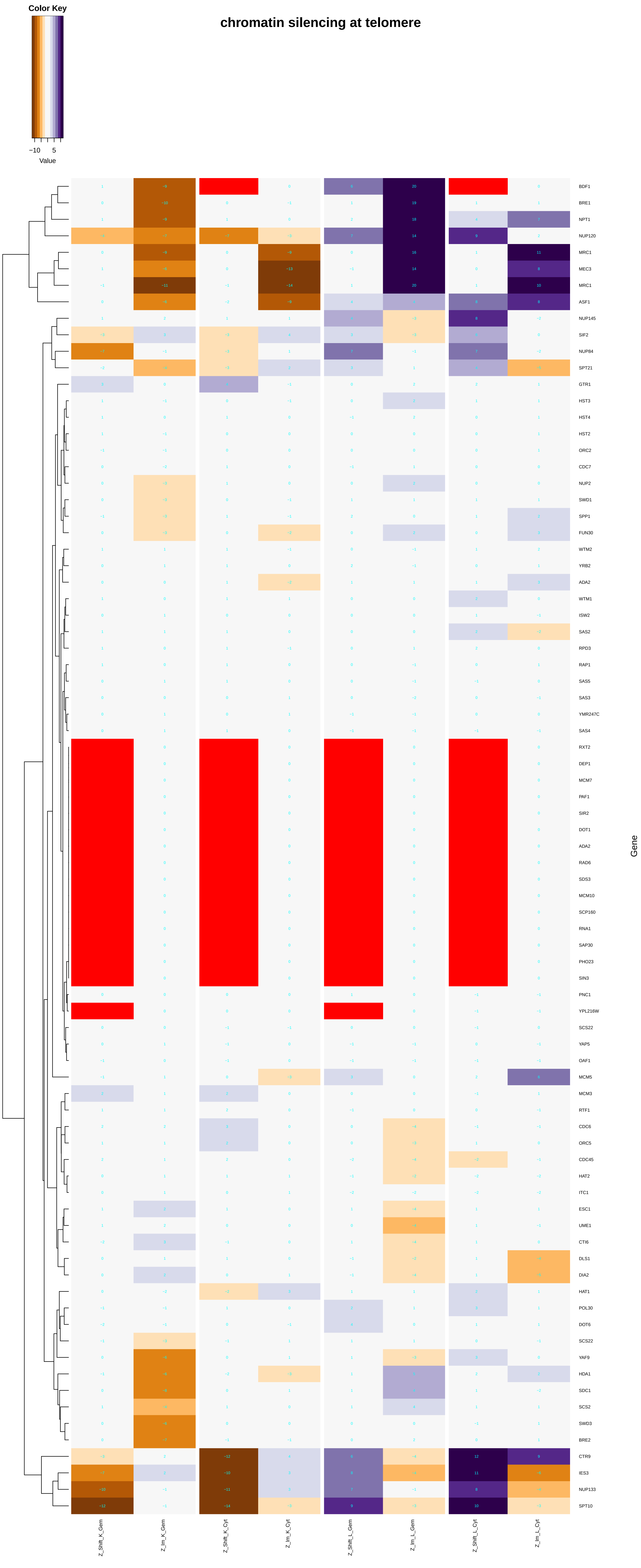

chromatin silencing at telomere
Color Key
−10
5
Value
Gene
BDF1
BRE1
NPT1
NUP120
MRC1
MEC3
MRC1
ASF1
NUP145
SIF2
NUP84
SPT21
GTR1
HST3
HST4
HST2
ORC2
CDC7
NUP2
SWD1
SPP1
FUN30
WTM2
YRB2
ADA2
WTM1
ISW2
SAS2
RPD3
RAP1
SAS5
SAS3
YMR247C
SAS4
RXT2
DEP1
MCM7
PAF1
SIR2
DOT1
ADA2
RAD6
SDS3
MCM10
SCP160
RNA1
SAP30
PHO23
SIN3
PNC1
YPL216W
SCS22
YAP5
OAF1
MCM5
MCM3
RTF1
CDC6
ORC5
CDC45
HAT2
ITC1
ESC1
UME1
CTI6
DLS1
DIA2
HAT1
POL30
DOT6
SCS22
YAF9
HDA1
SDC1
SCS2
SWD3
BRE2
CTR9
IES3
NUP133
SPT10
Z_Shift_K_Gem
Z_Im_K_Gem
Z_Shift_K_Cyt
Z_Im_K_Cyt
Z_Shift_L_Gem
Z_Im_L_Gem
Z_Shift_L_Cyt
Z_Im_L_Cyt

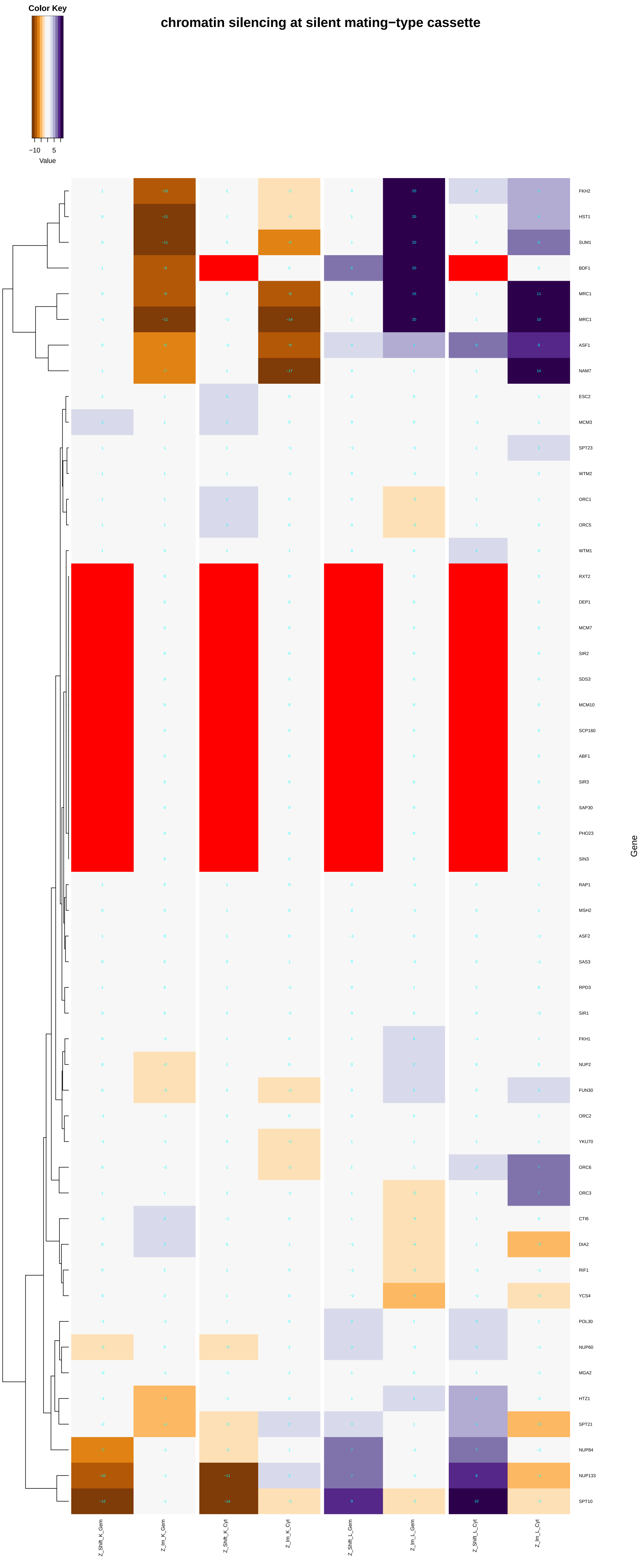
Color Key
−10 5
Value
chromatin silencing at silent mating−type cassette
FKH2
HST1
SUM1
BDF1
MRC1
MRC1
ASF1
NAM7
ESC2
MCM3
SPT23
WTM2
ORC1
ORC5
WTM1
RXT2
DEP1
MCM7
SIR2
SDS3
MCM10
SCP160
ABF1
SIR3
SAP30
PHO23
SIN3
RAP1
MSH2
ASF2
SAS3
RPD3
SIR1
FKH1
NUP2
FUN30
ORC2
YKU70
ORC6
ORC3
CTI6
DIA2
RIF1
YCS4
POL30
NUP60
MGA2
HTZ1
SPT21
NUP84
NUP133
SPT10
Gene
Z_Shift_K_Gem
Z_Im_K_Gem
Z_Shift_K_Cyt
Z_Im_K_Cyt
Z_Shift_L_Gem
Z_Im_L_Gem
Z_Shift_L_Cyt
Z_Im_L_Cyt

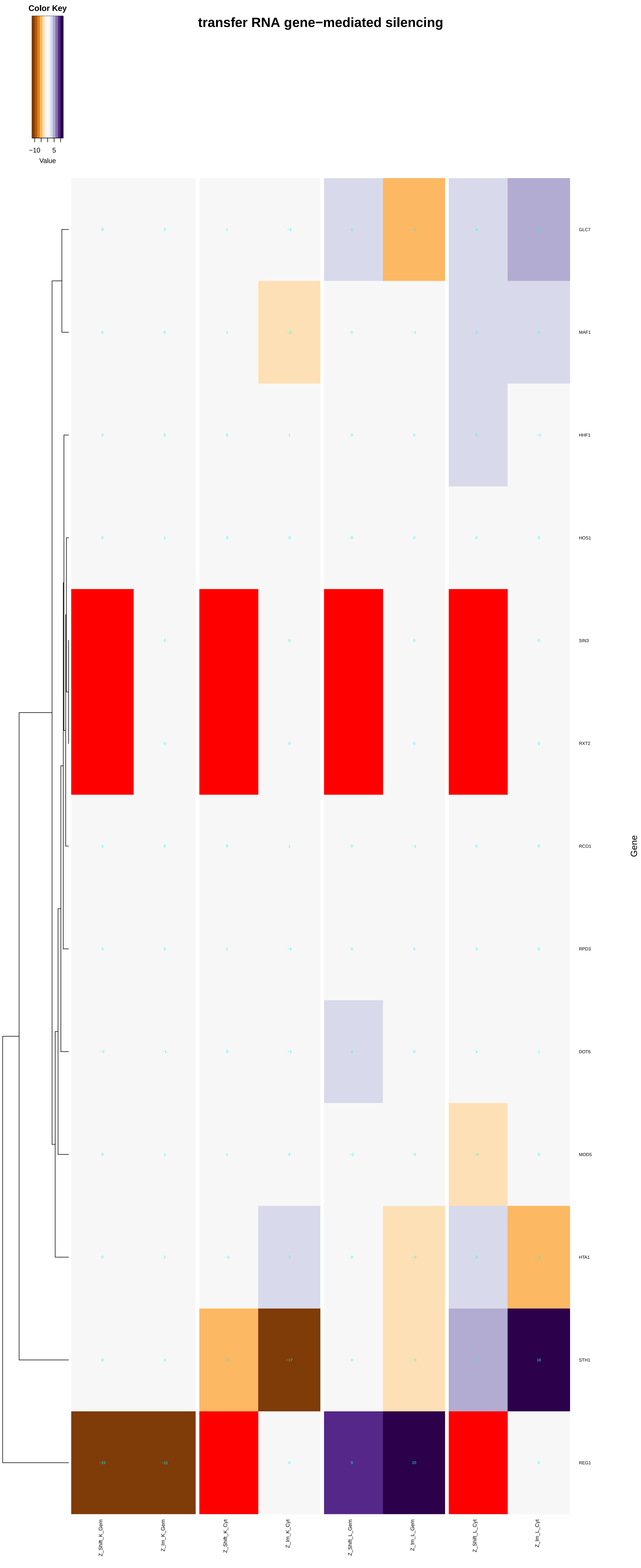

transfer RNA gene−mediated silencing
Color Key
−10
5
Value
GLC7
MAF1
HHF1
HOS1
SIN3
RXT2
RCO1
RPD3
DOT6
MOD5
HTA1
STH1
REG1
Gene
Z_Shift_K_Gem
Z_Im_K_Gem
Z_Shift_K_Cyt
Z_Im_K_Cyt
Z_Shift_L_Gem
Z_Im_L_Gem
Z_Shift_L_Cyt
Z_Im_L_Cyt

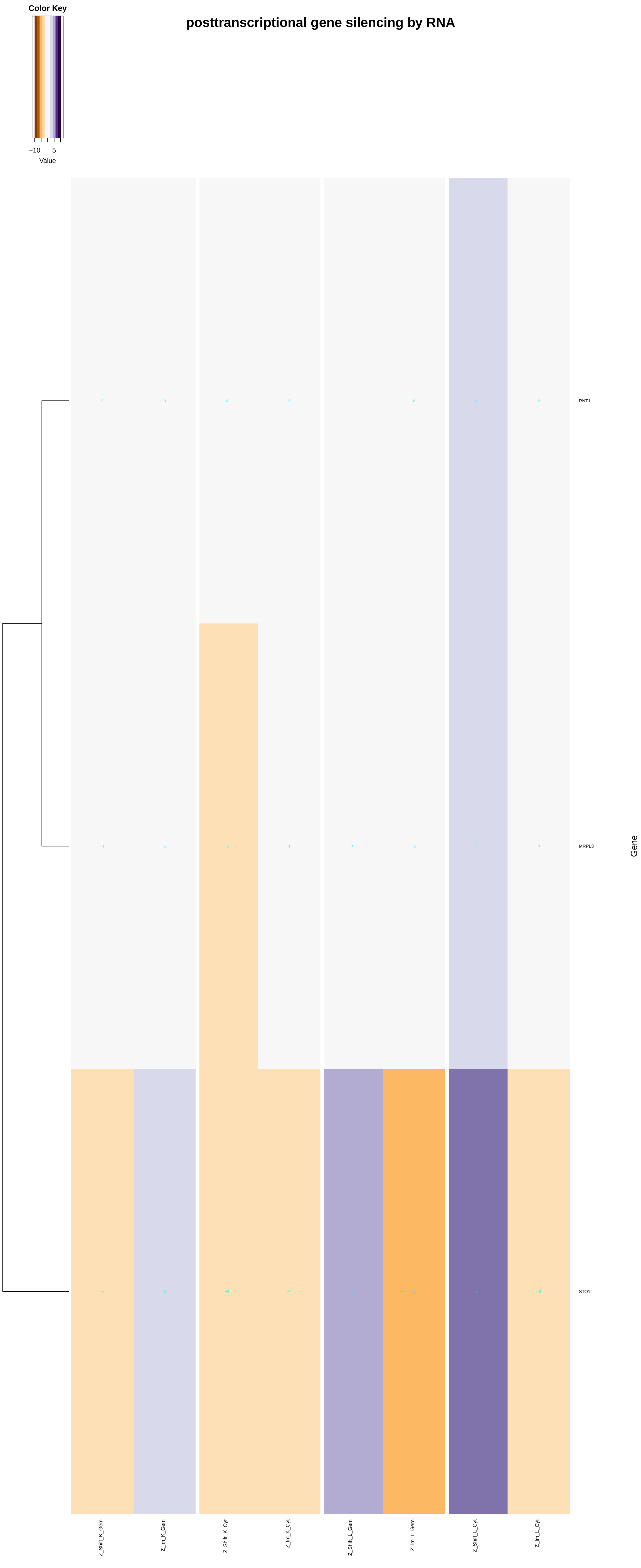
posttranscriptional gene silencing by RNA
Color Key
−10
5
Value
RNT1
MRPL3
STO1
Gene
Z_Shift_K_Gem
Z_Im_K_Gem
Z_Shift_K_Cyt
Z_Im_K_Cyt
Z_Shift_L_Gem
Z_Im_L_Gem
Z_Shift_L_Cyt
Z_Im_L_Cyt

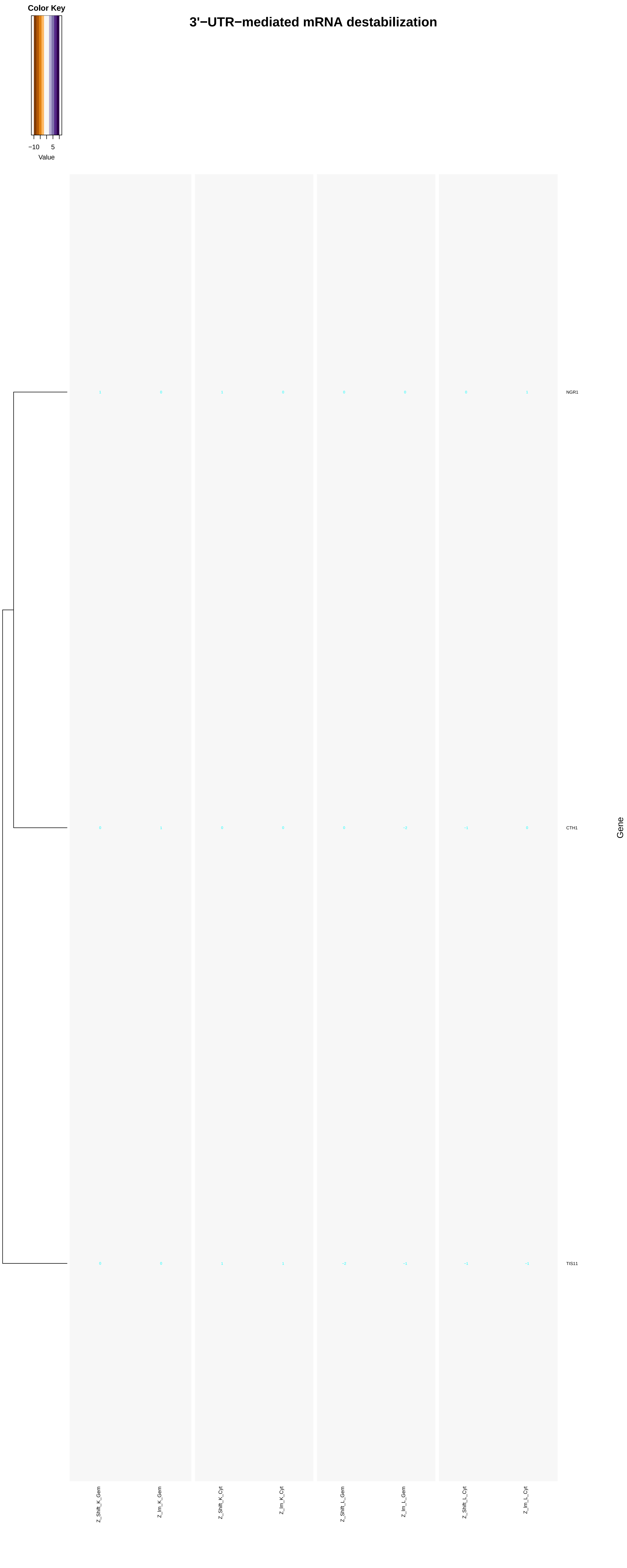

Color Key
−10
5
Value
3'−UTR−mediated mRNA destabilization
1
0
1
0
0
0
0
1
NGR1
0
1
0
0
0
−2
−1
0
CTH1
0
0
1
1
−2
−1
−1
−1
TIS11
Gene
Z_Shift_K_Gem
Z_Im_K_Gem
Z_Shift_K_Cyt
Z_Im_K_Cyt
Z_Shift_L_Gem
Z_Im_L_Gem
Z_Shift_L_Cyt
Z_Im_L_Cyt

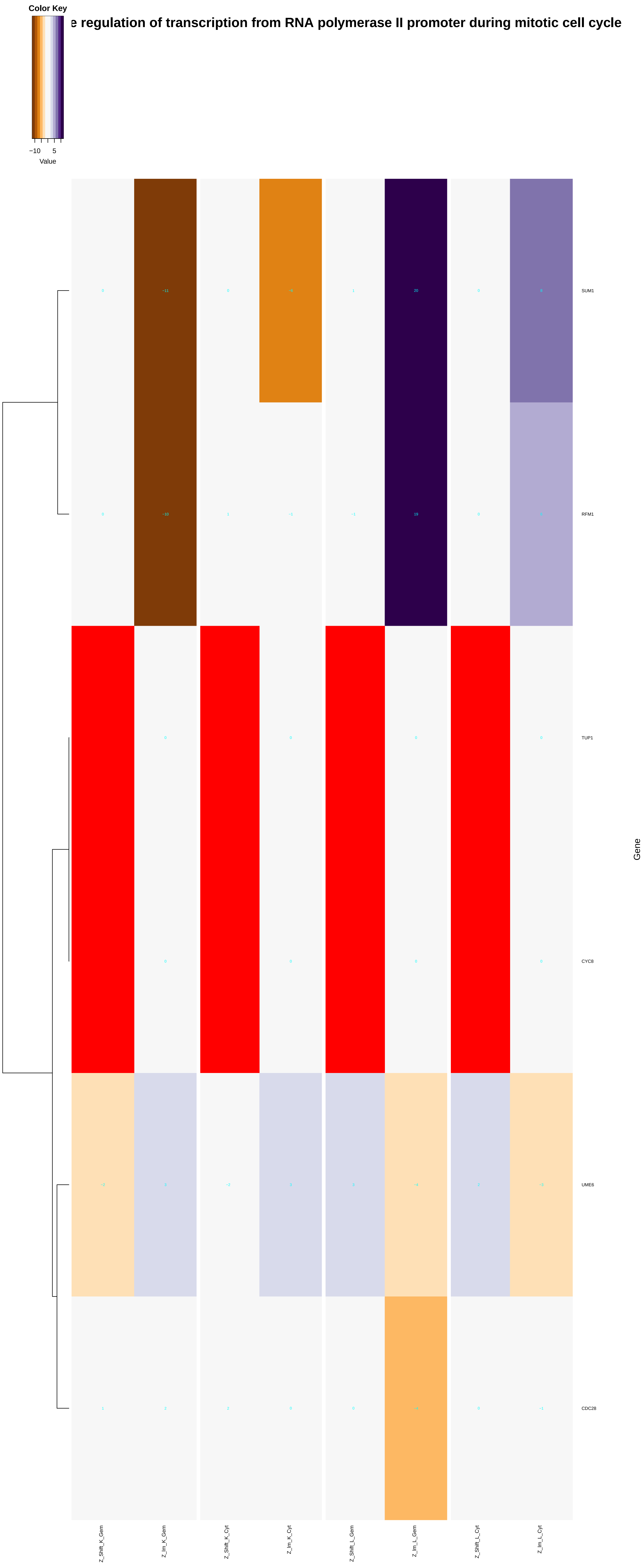
e regulation of transcription from RNA polymerase II promoter during mitotic cell cycle
Color Key
−10 5
Value
SUM1
RFM1
TUP1
CYC8
UME6
CDC28
Gene
Z_Shift_K_Gem
Z_Im_K_Gem
Z_Shift_K_Cyt
Z_Im_K_Cyt
Z_Shift_L_Gem
Z_Im_L_Gem
Z_Shift_L_Cyt
Z_Im_L_Cyt

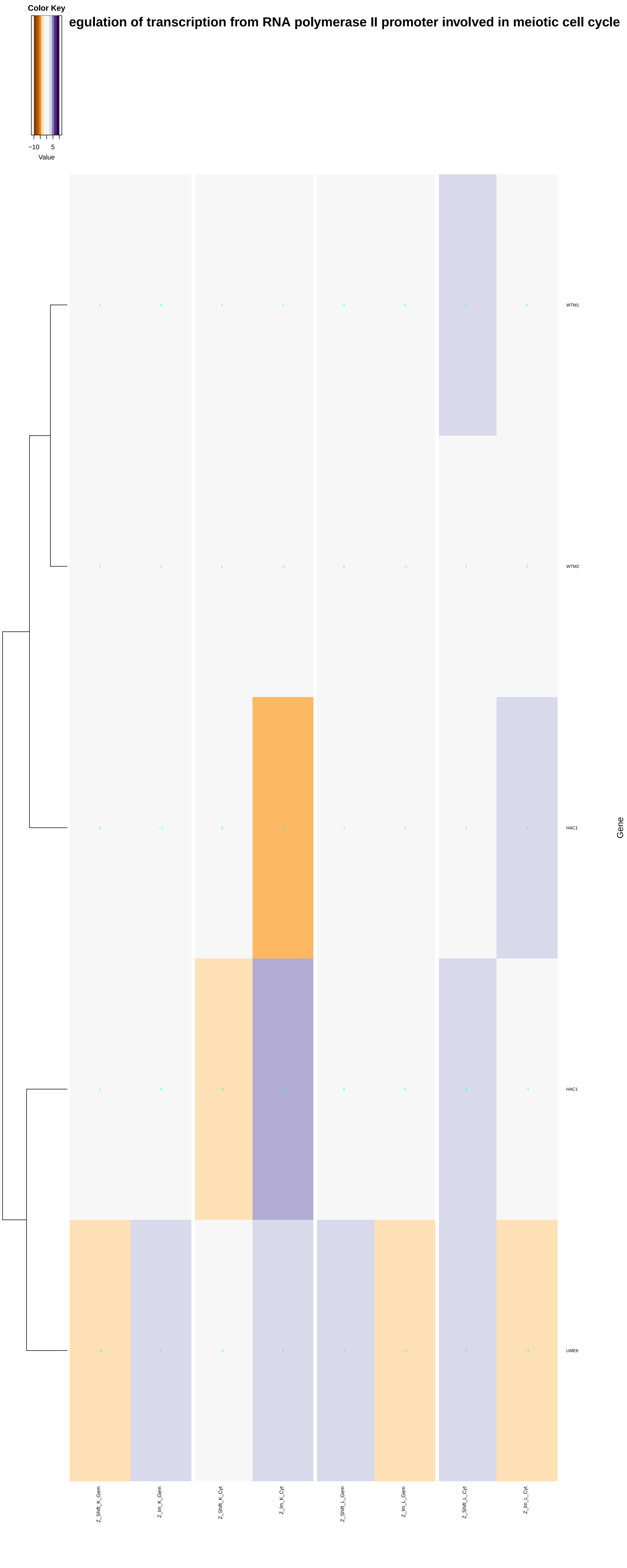
Color Key
−10
5
Value
egulation of transcription from RNA polymerase II promoter involved in meiotic cell cycle
WTM1
WTM2
HAC1
HAC1
UME6
Gene
Z_Shift_K_Gem
Z_lm_K_Gem
Z_Shift_K_Cyt
Z_lm_K_Cyt
Z_Shift_L_Gem
Z_lm_L_Gem
Z_Shift_L_Cyt
Z_lm_L_Cyt

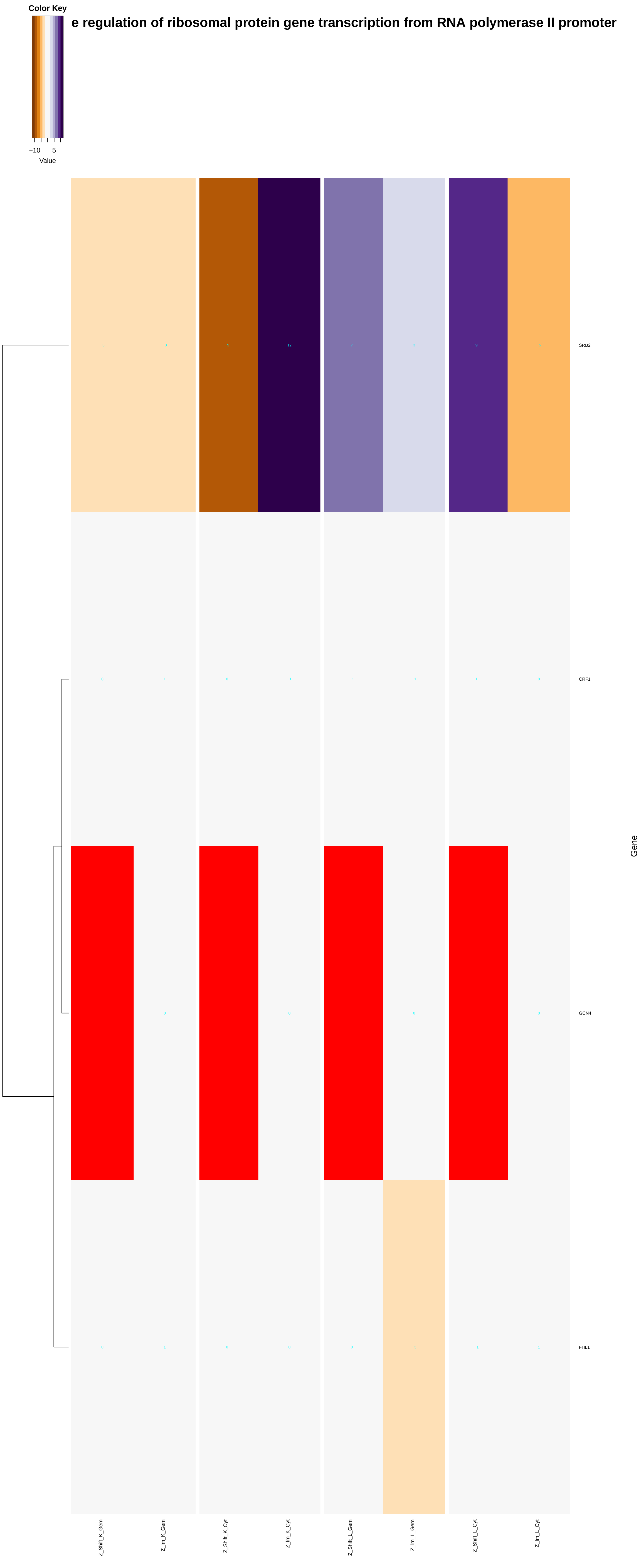
Color Key
−10
5
Value
e regulation of ribosomal protein gene transcription from RNA polymerase II promoter
SRB2
CRF1
GCN4
FHL1
Gene
Z_Shift_K_Gem
Z_Im_K_Gem
Z_Shift_K_Cyt
Z_Im_K_Cyt
Z_Shift_L_Gem
Z_Im_L_Gem
Z_Shift_L_Cyt
Z_Im_L_Cyt

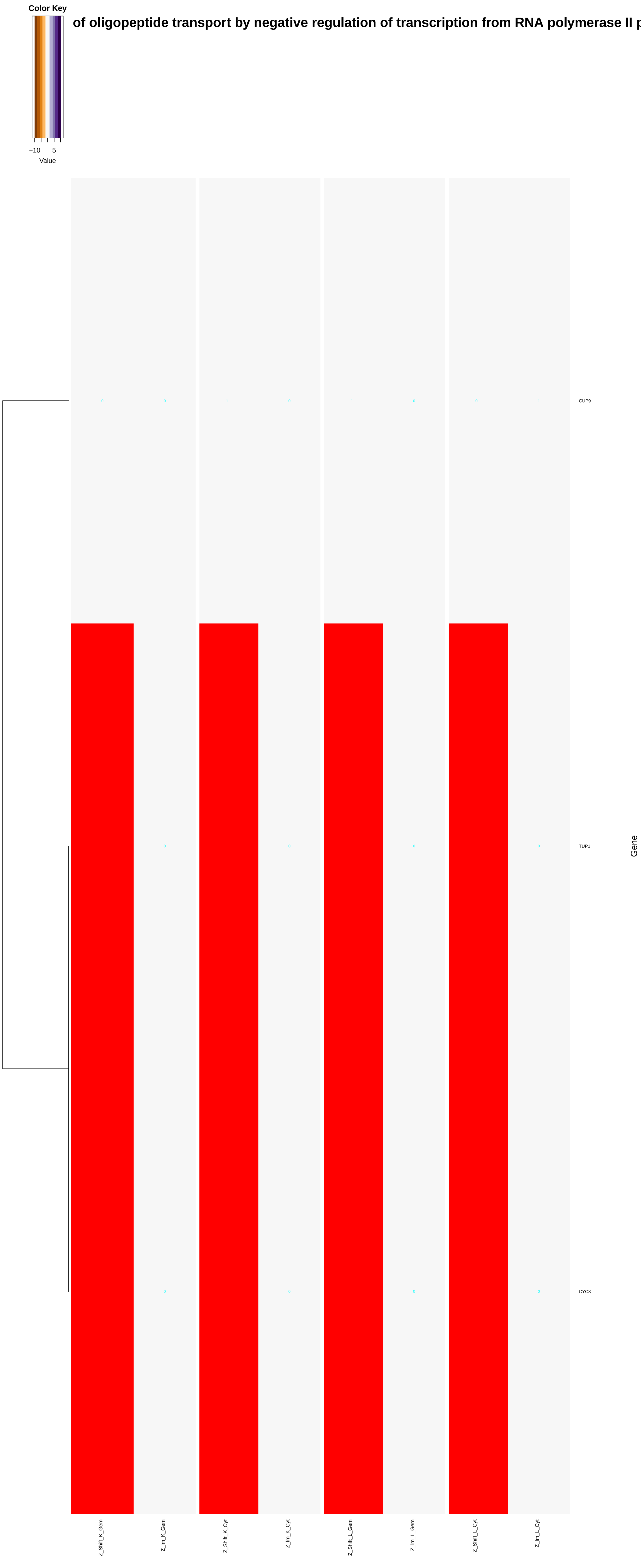
Color Key
−10
5
Value
of oligopeptide transport by negative regulation of transcription from RNA polymerase II p
CUP9
TUP1
CYC8
Gene
Z_Shift_K_Gem
Z_Im_K_Gem
Z_Shift_K_Cyt
Z_Im_K_Cyt
Z_Shift_L_Gem
Z_Im_L_Gem
Z_Shift_L_Cyt
Z_Im_L_Cyt

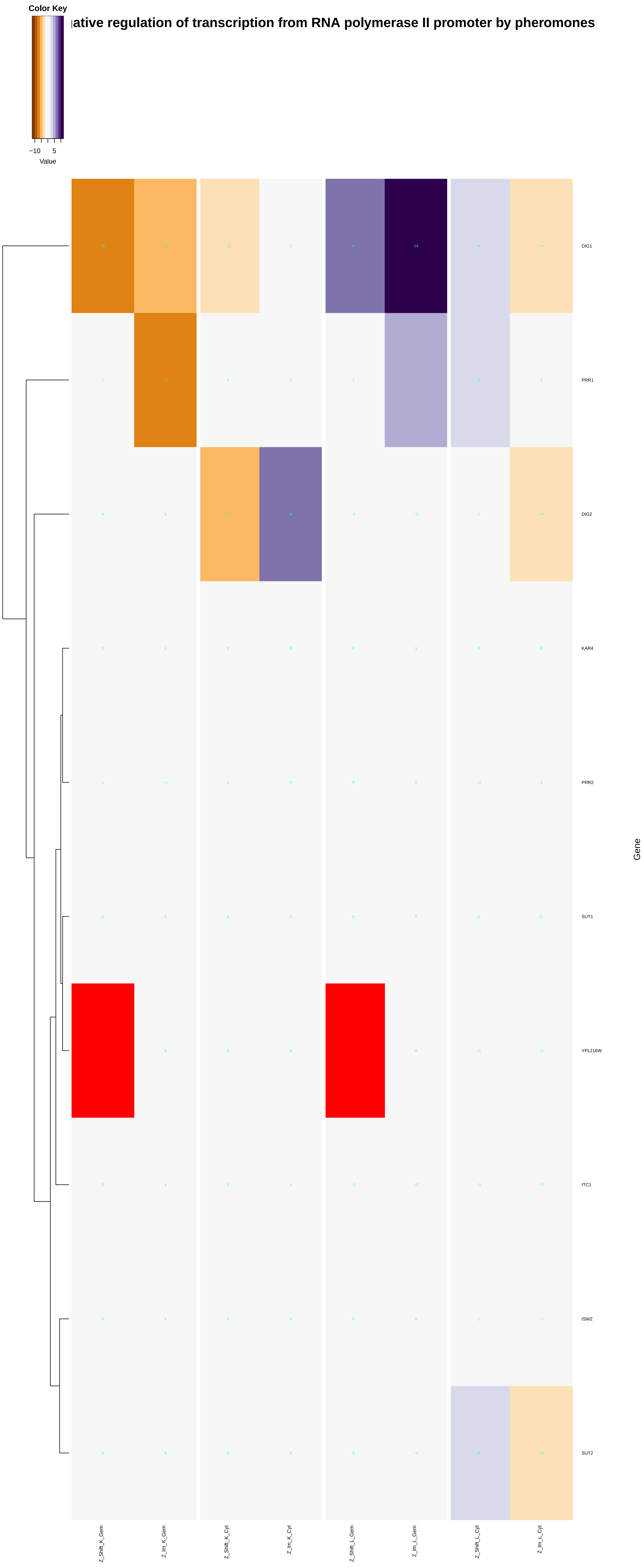

Color Key
−10
5
Value
ative regulation of transcription from RNA polymerase II promoter by pheromones
DIG1
PRR1
DIG2
KAR4
PRR2
SUT1
YPL216W
ITC1
ISW2
SUT2
Gene
Z_Shift_K_Gem
Z_Im_K_Gem
Z_Shift_K_Cyt
Z_Im_K_Cyt
Z_Shift_L_Gem
Z_Im_L_Gem
Z_Shift_L_Cyt
Z_Im_L_Cyt

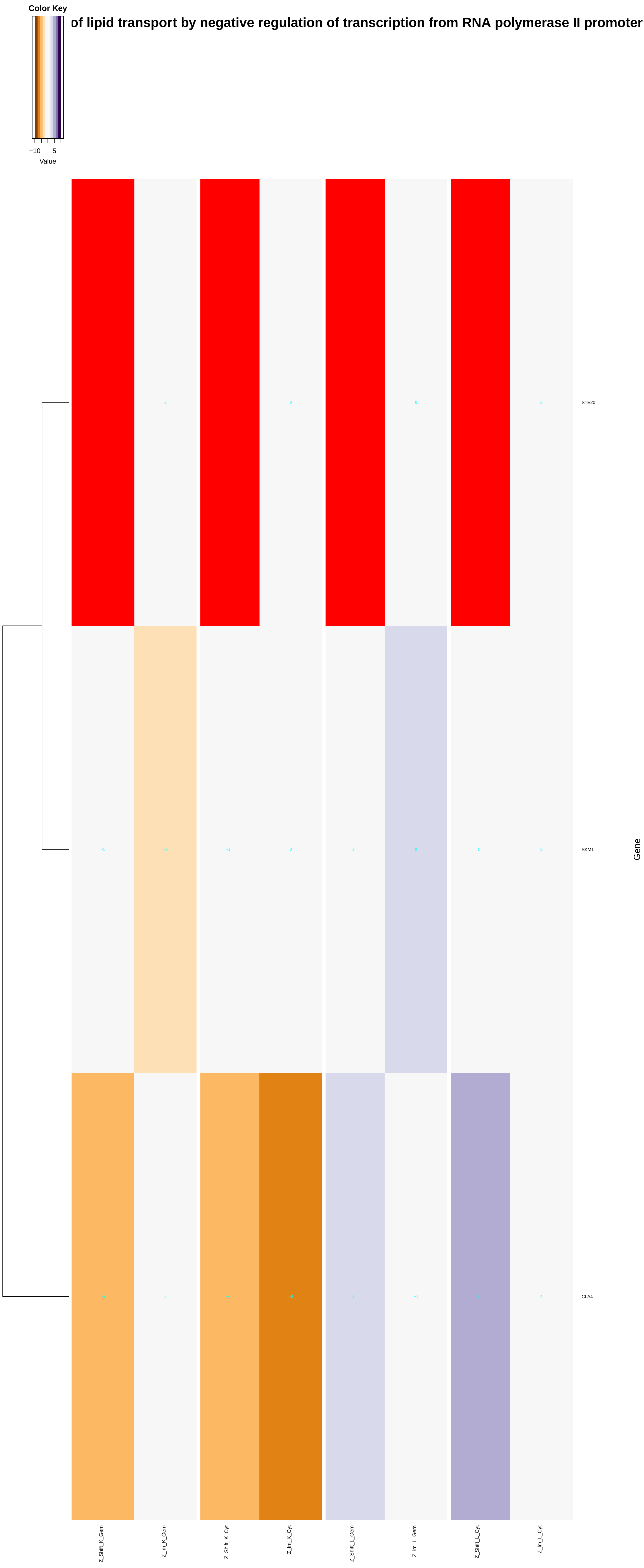

Color Key
−10
5
Value
of lipid transport by negative regulation of transcription from RNA polymerase II promoter
STE20
SKM1
CLA4
Gene
Z_Shift_K_Gem
Z_Im_K_Gem
Z_Shift_K_Cyt
Z_Im_K_Cyt
Z_Shift_L_Gem
Z_Im_L_Gem
Z_Shift_L_Cyt
Z_Im_L_Cyt

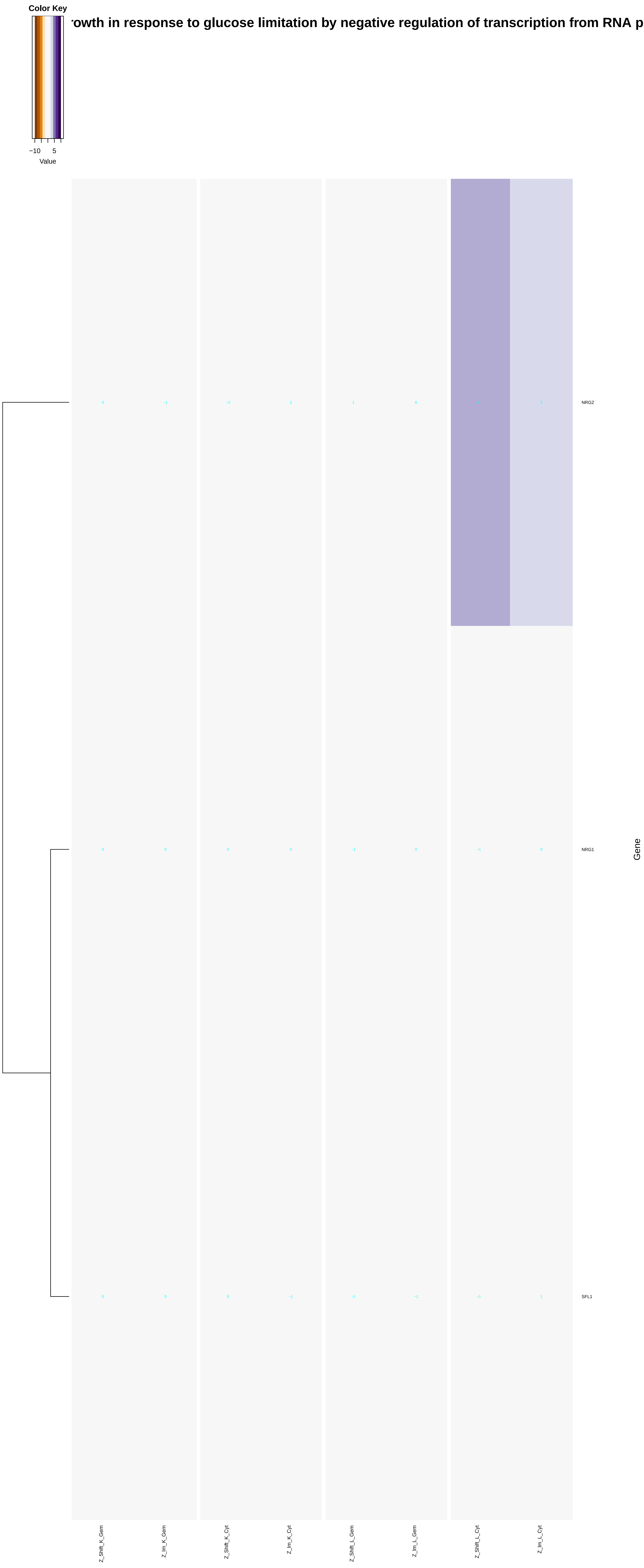

Color Key
−10
5
Value
rowth in response to glucose limitation by negative regulation of transcription from RNA p
NRG2
NRG1
SFL1
Gene
Z_Shift_K_Gem
Z_Im_K_Gem
Z_Shift_K_Cyt
Z_Im_K_Cyt
Z_Shift_L_Gem
Z_Im_L_Gem
Z_Shift_L_Cyt
Z_Im_L_Cyt

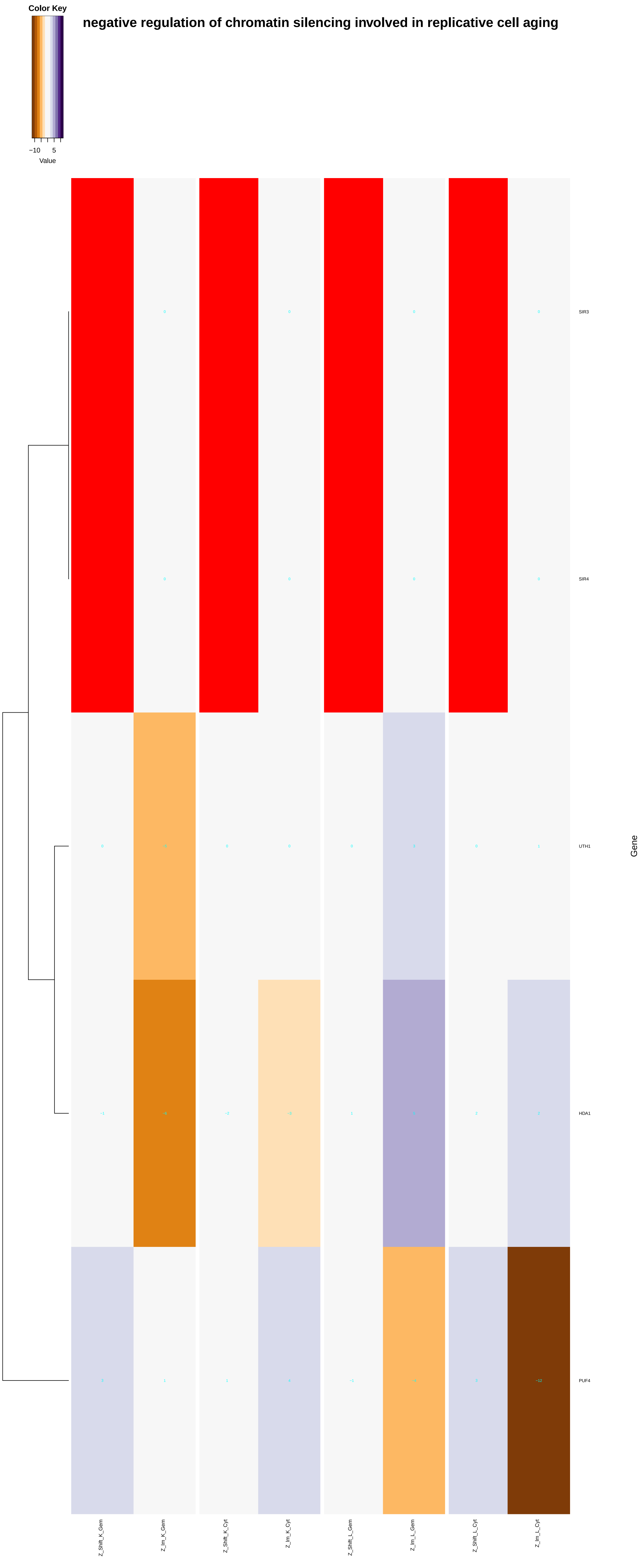
negative regulation of chromatin silencing involved in replicative cell aging
Color Key
−10
5
Value
SIR3
SIR4
UTH1
HDA1
PUF4
Gene
Z_Shift_K_Gem
Z_lm_K_Gem
Z_Shift_K_Cyt
Z_lm_K_Cyt
Z_Shift_L_Gem
Z_lm_L_Gem
Z_Shift_L_Cyt
Z_lm_L_Cyt

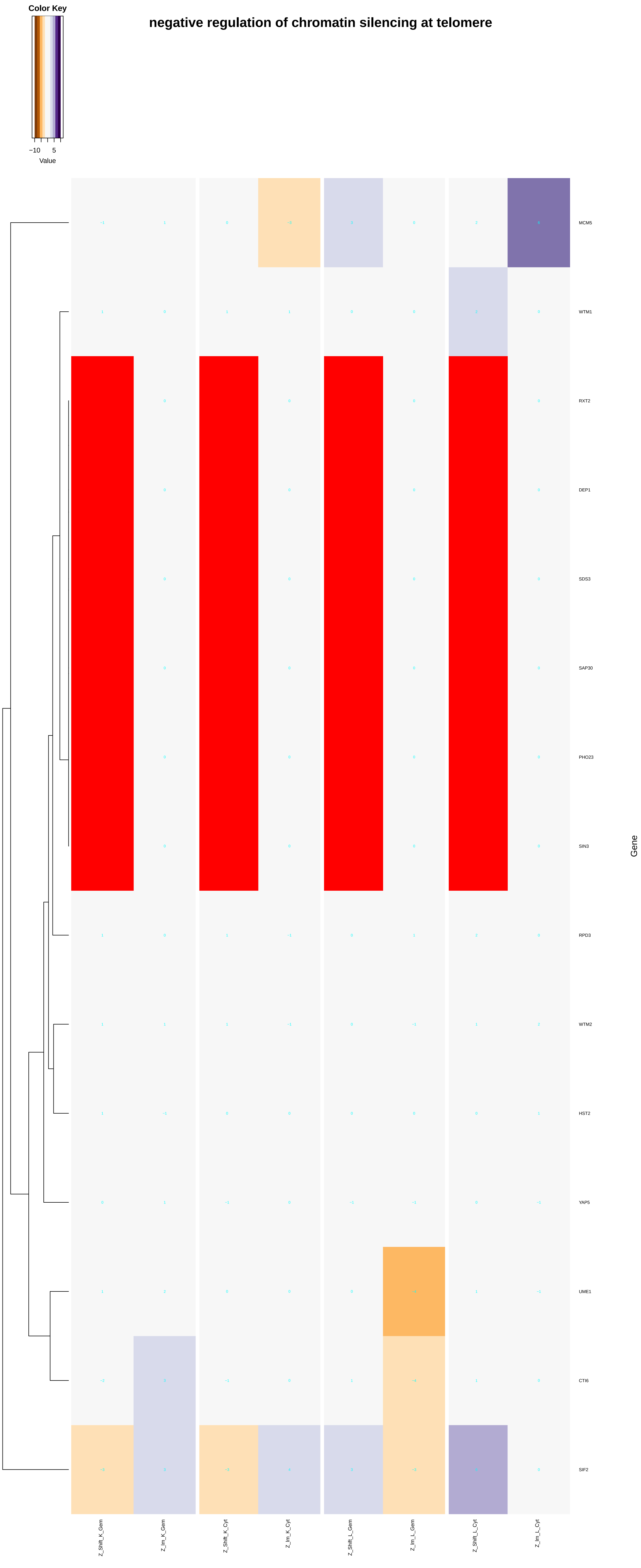

Color Key
−10
5
Value
negative regulation of chromatin silencing at telomere
MCM5
WTM1
RXT2
DEP1
SDS3
SAP30
PHO23
SIN3
RPD3
WTM2
HST2
YAP5
UME1
CTI6
SIF2
Gene
Z_Shift_K_Gem
Z_Im_K_Gem
Z_Shift_K_Cyt
Z_Im_K_Cyt
Z_Shift_L_Gem
Z_Im_L_Gem
Z_Shift_L_Cyt
Z_Im_L_Cyt

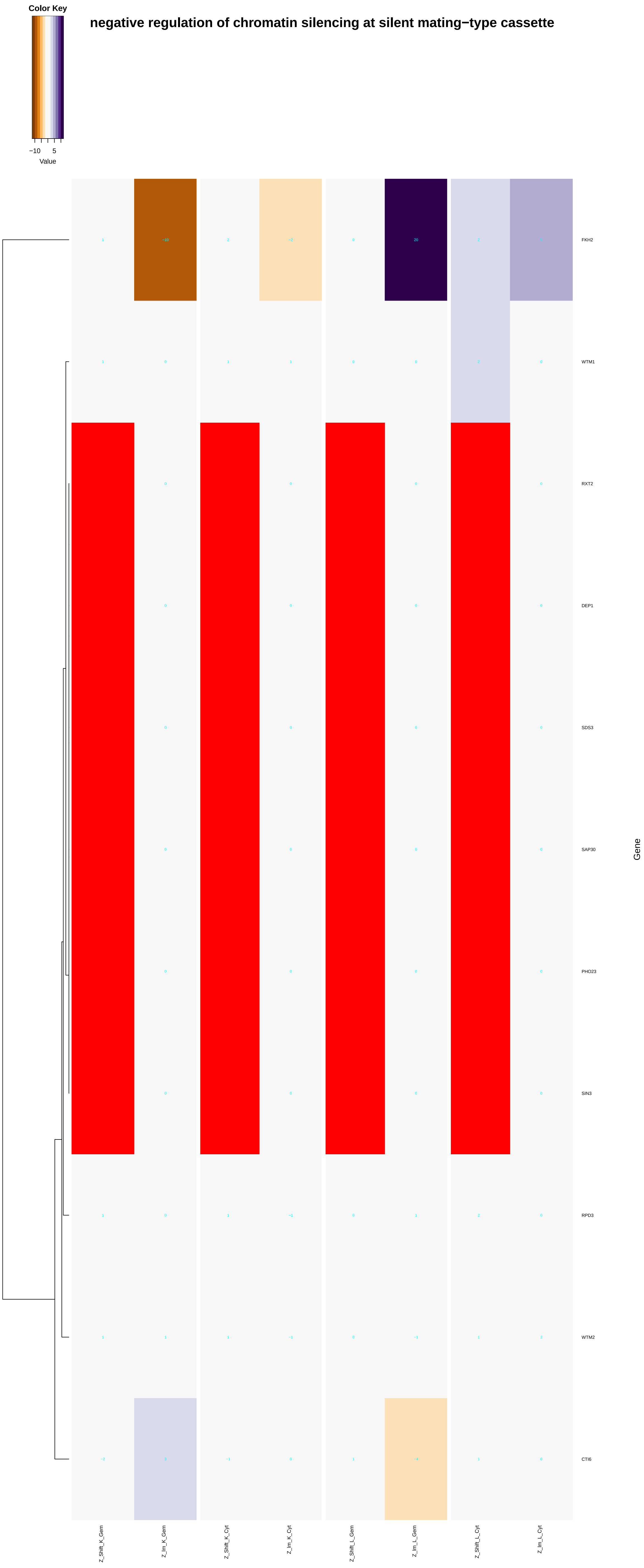
Color Key
−10
5
Value
negative regulation of chromatin silencing at silent mating−type cassette
FKH2
WTM1
RXT2
DEP1
SDS3
SAP30
PHO23
SIN3
RPD3
WTM2
CTI6
Gene
Z_Shift_K_Gem
Z_Im_K_Gem
Z_Shift_K_Cyt
Z_Im_K_Cyt
Z_Shift_L_Gem
Z_Im_L_Gem
Z_Shift_L_Cyt
Z_Im_L_Cyt

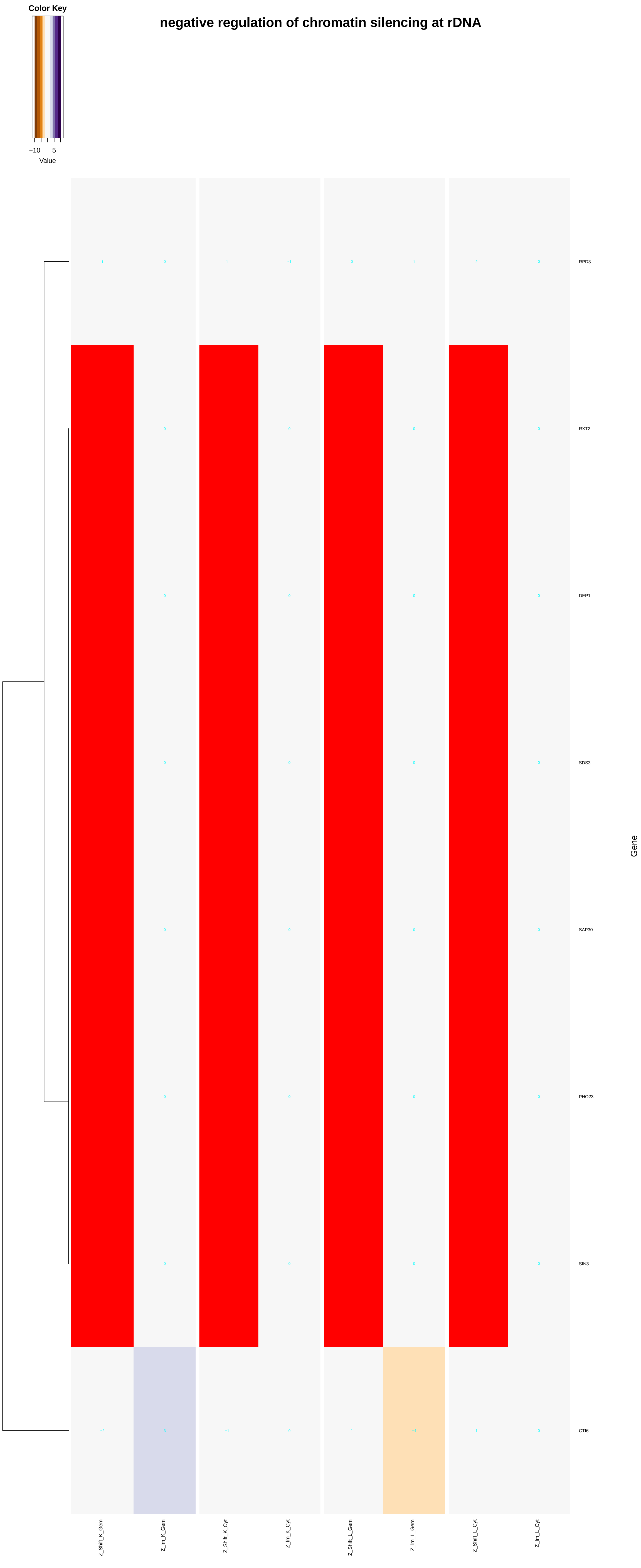
negative regulation of chromatin silencing at rDNA
Color Key
−10
5
Value
RPD3
RXT2
DEP1
SDS3
SAP30
PHO23
SIN3
CTI6
Gene
Z_Shift_K_Gem
Z_Im_K_Gem
Z_Shift_K_Cyt
Z_Im_K_Cyt
Z_Shift_L_Gem
Z_Im_L_Gem
Z_Shift_L_Cyt
Z_Im_L_Cyt

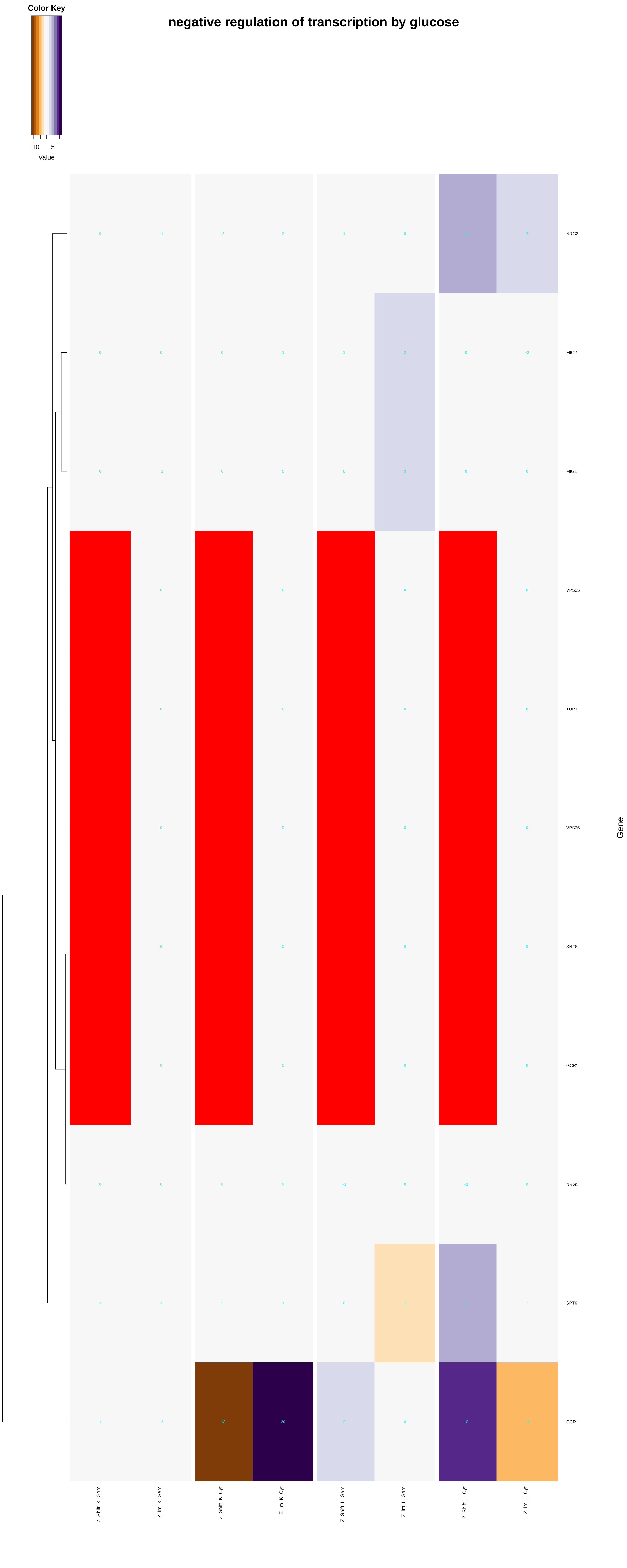
negative regulation of transcription by glucose
Color Key
−10
5
Value
NRG2
MIG2
MIG1
VPS25
TUP1
VPS36
SNF8
GCR1
NRG1
SPT6
GCR1
Gene
Z_Shift_K_Gem
Z_Im_K_Gem
Z_Shift_K_Cyt
Z_Im_K_Cyt
Z_Shift_L_Gem
Z_Im_L_Gem
Z_Shift_L_Cyt
Z_Im_L_Cyt

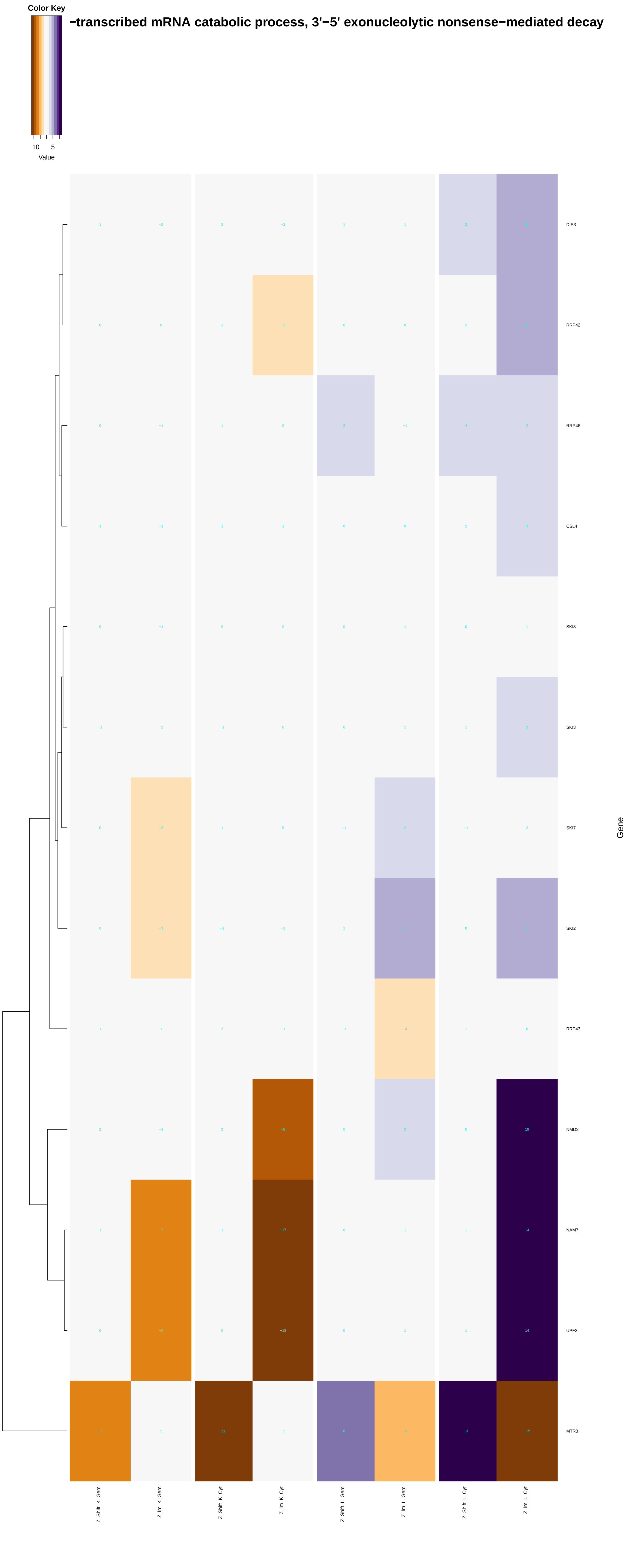

Color Key
−10
5
Value
−transcribed mRNA catabolic process, 3'−5' exonucleolytic nonsense−mediated decay
DIS3
RRP42
RRP46
CSL4
SKI8
SKI3
SKI7
SKI2
RRP43
NMD2
NAM7
UPF3
MTR3
Gene
Z_Shift_K_Gem
Z_Im_K_Gem
Z_Shift_K_Cyt
Z_Im_K_Cyt
Z_Shift_L_Gem
Z_Im_L_Gem
Z_Shift_L_Cyt
Z_Im_L_Cyt

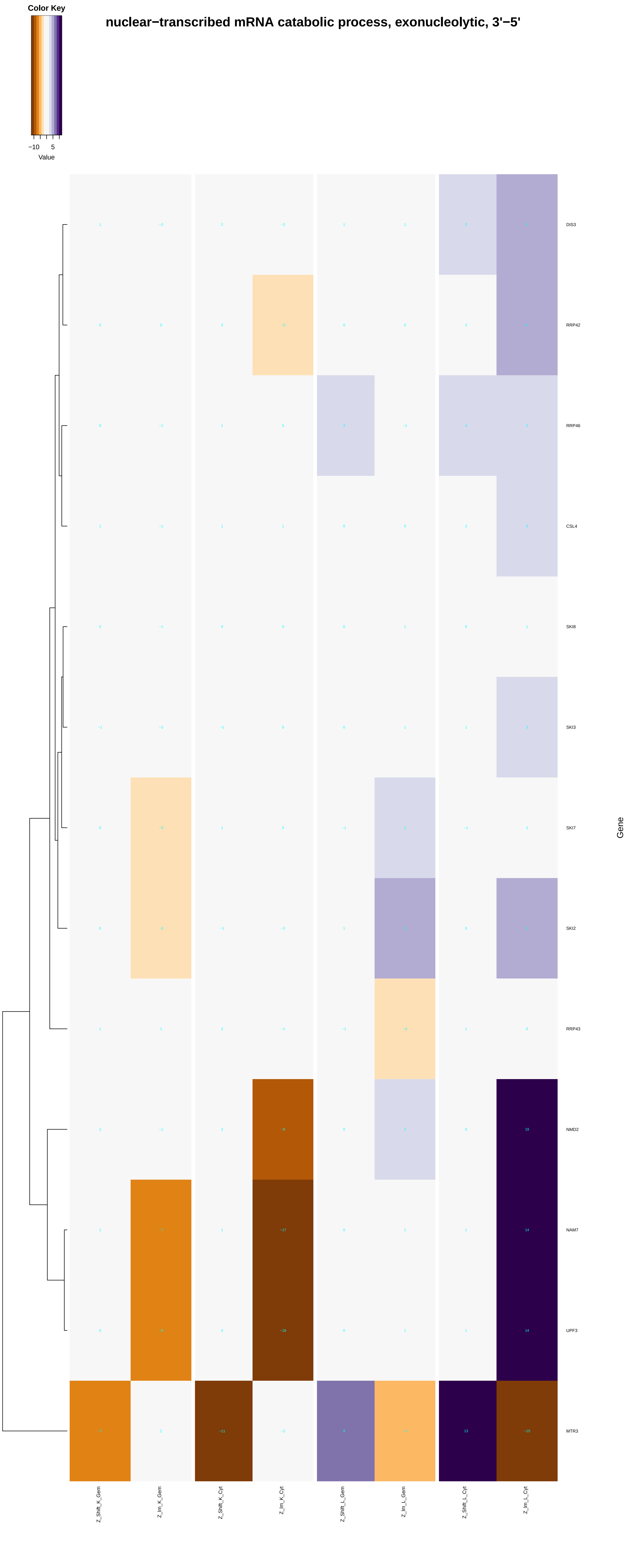

Color Key
−10
5
Value
nuclear−transcribed mRNA catabolic process, exonucleolytic, 3'−5'
DIS3
RRP42
RRP46
CSL4
SKI8
SKI3
SKI7
SKI2
RRP43
NMD2
NAM7
UPF3
MTR3
Gene
Z_Shift_K_Gem
Z_Im_K_Gem
Z_Shift_K_Cyt
Z_Im_K_Cyt
Z_Shift_L_Gem
Z_Im_L_Gem
Z_Shift_L_Cyt
Z_Im_L_Cyt

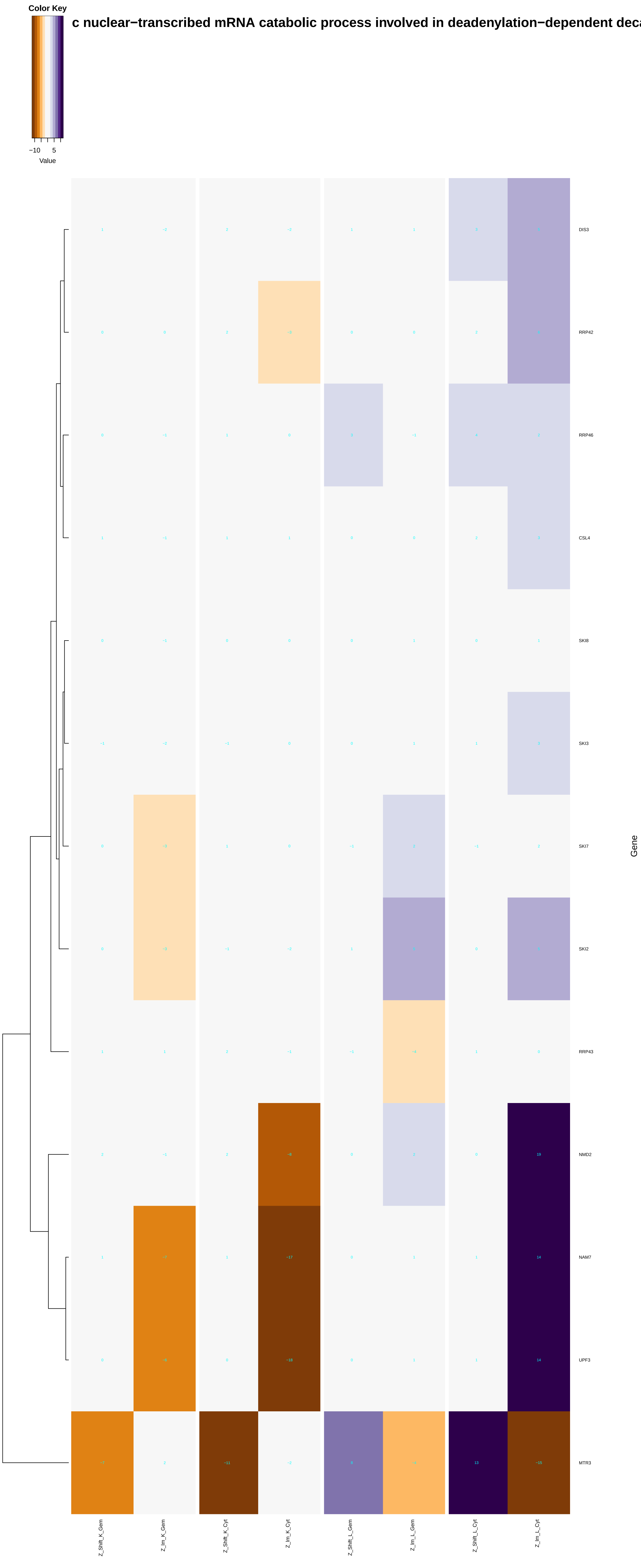
Color Key
−10
5
Value
c nuclear−transcribed mRNA catabolic process involved in deadenylation−dependent deca
DIS3
RRP42
RRP46
CSL4
SKI8
SKI3
SKI7
SKI2
RRP43
NMD2
NAM7
UPF3
MTR3
Gene
Z_Shift_K_Gem
Z_Im_K_Gem
Z_Shift_K_Cyt
Z_Im_K_Cyt
Z_Shift_L_Gem
Z_Im_L_Gem
Z_Shift_L_Cyt
Z_Im_L_Cyt

### nuclear mRNA surveillance of spliceosomal pre−mRNA splicing

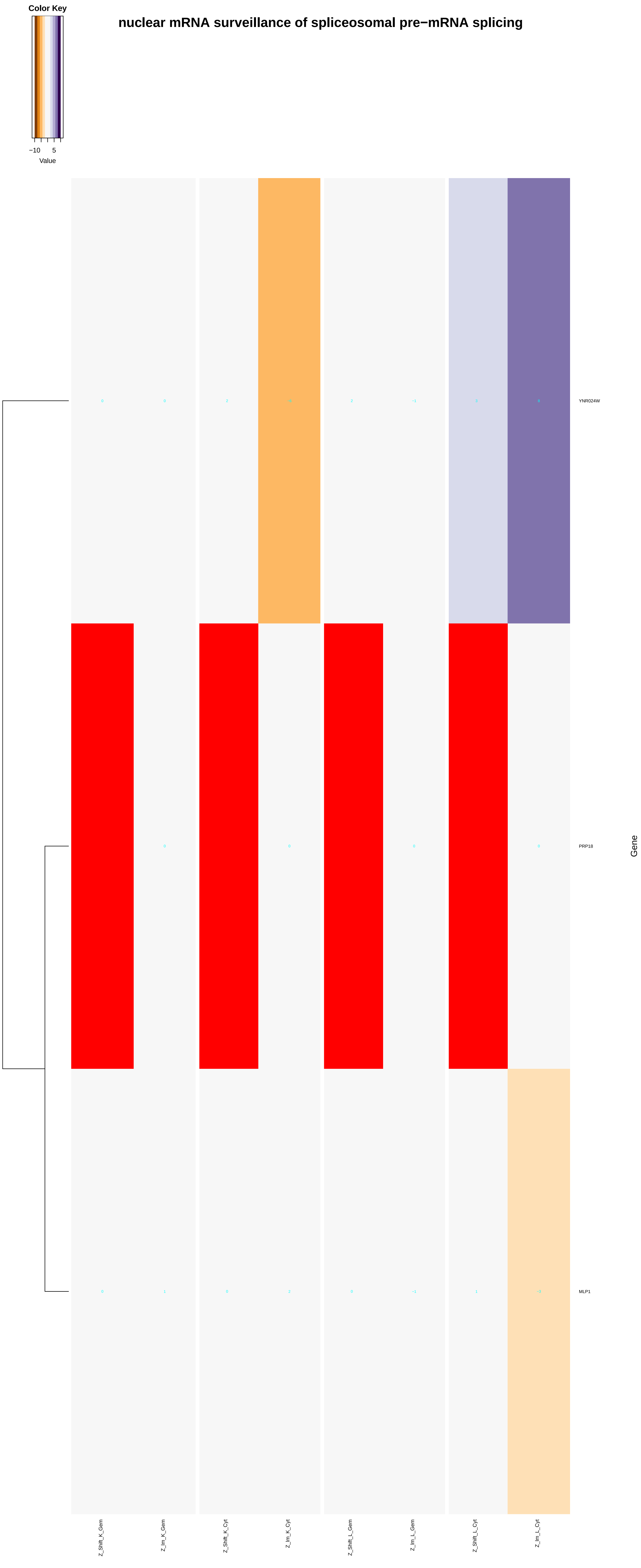

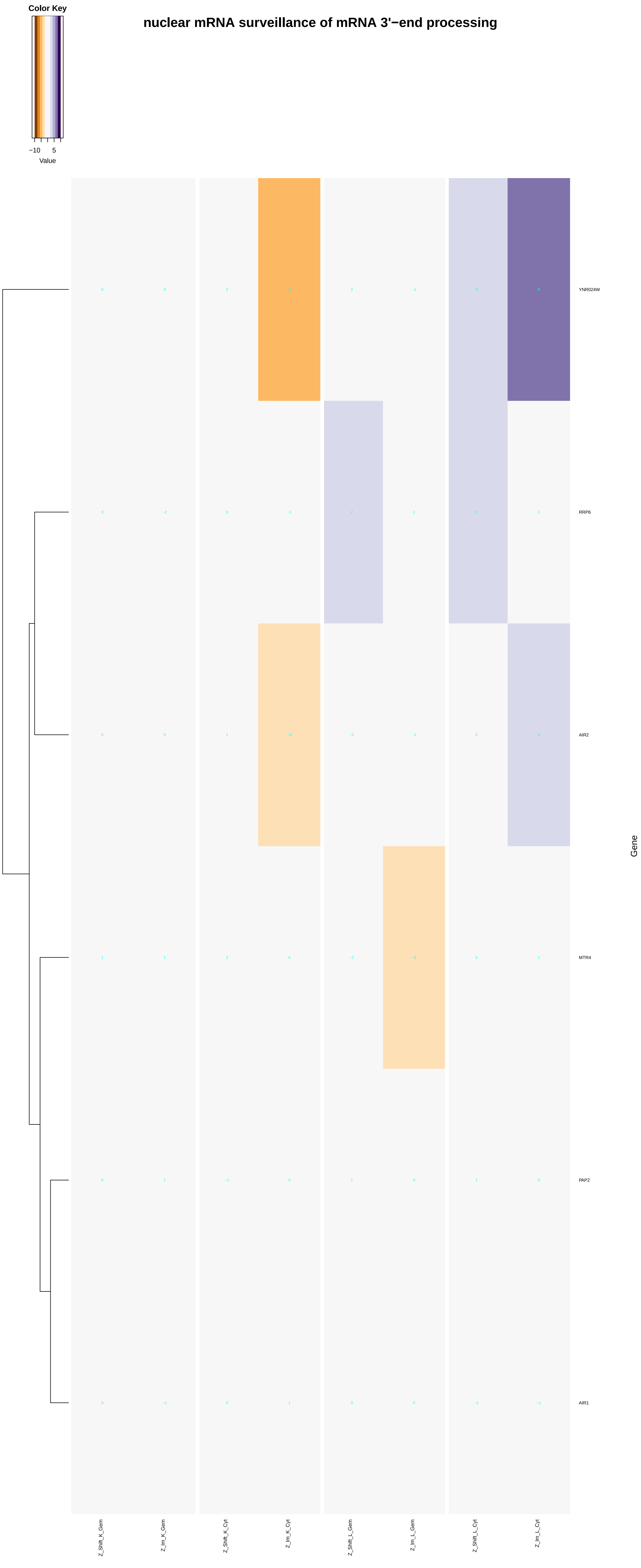
nuclear mRNA surveillance of mRNA 3'−end processing
Color Key
−10
5
Value
YNR024W
RRP6
AIR2
MTR4
PAP2
AIR1
Gene
Z_Shift_K_Gem
Z_Im_K_Gem
Z_Shift_K_Cyt
Z_Im_K_Cyt
Z_Shift_L_Gem
Z_Im_L_Gem
Z_Shift_L_Cyt
Z_Im_L_Cyt

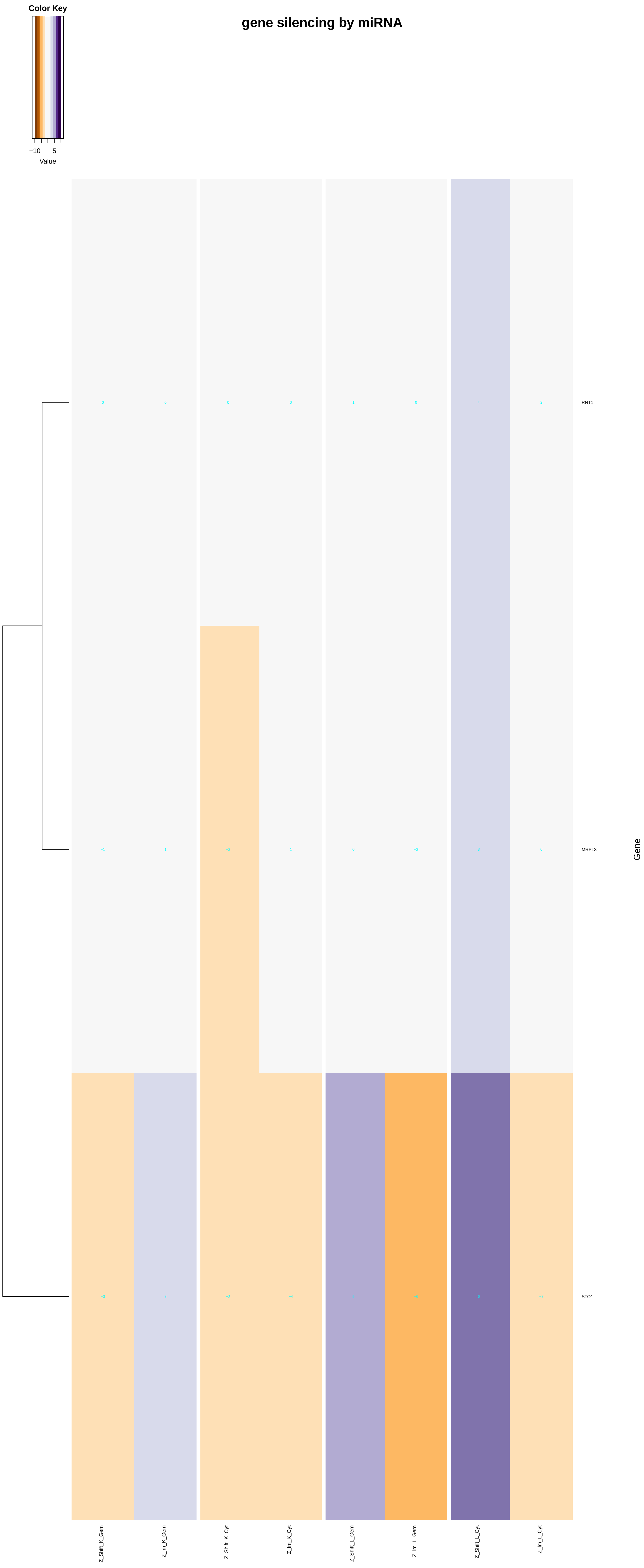

gene silencing by miRNA
Color Key
−10
5
Value
RNT1
MRPL3
STO1
Gene
Z_Shift_K_Gem
Z_Im_K_Gem
Z_Shift_K_Cyt
Z_Im_K_Cyt
Z_Shift_L_Gem
Z_Im_L_Gem
Z_Shift_L_Cyt
Z_Im_L_Cyt

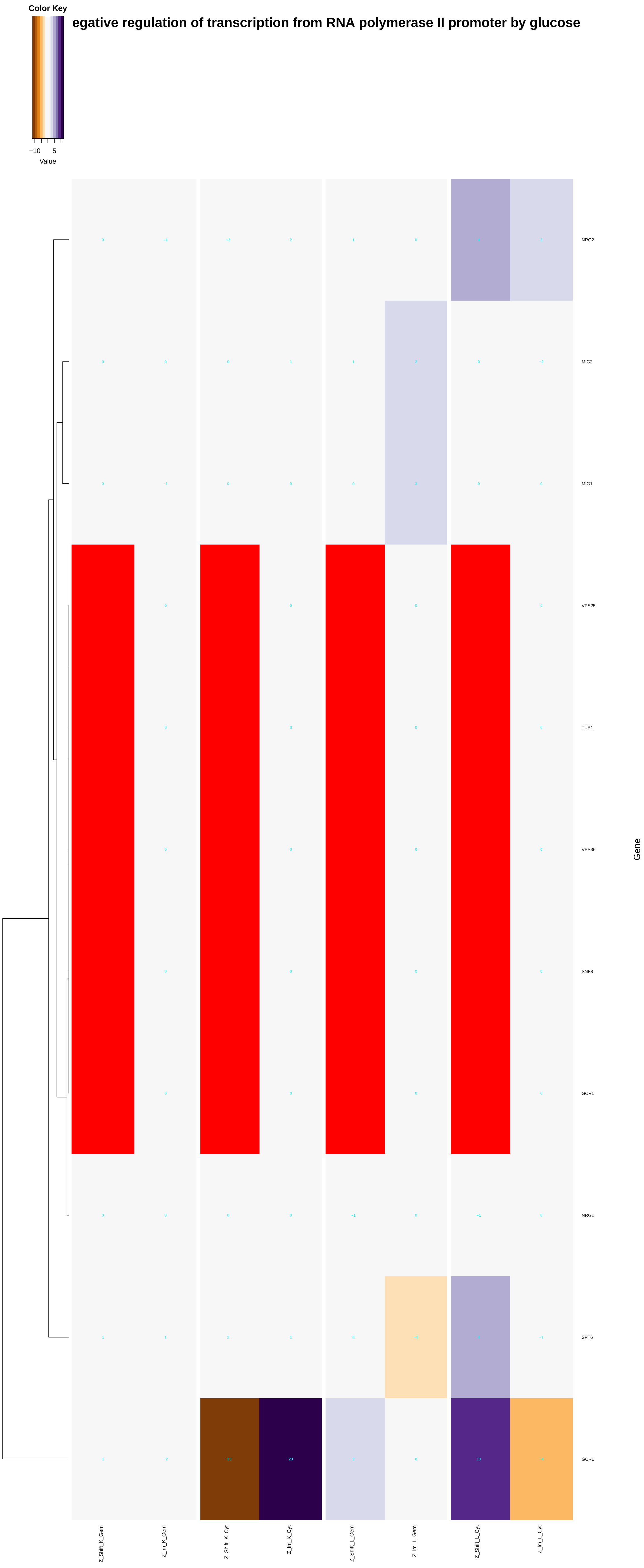

Color Key
−10
5
Value
egative regulation of transcription from RNA polymerase II promoter by glucose
NRG2
MIG2
MIG1
VPS25
TUP1
VPS36
SNF8
GCR1
NRG1
SPT6
GCR1
Gene
Z_Shift_K_Gem
Z_Im_K_Gem
Z_Shift_K_Cyt
Z_Im_K_Cyt
Z_Shift_L_Gem
Z_Im_L_Gem
Z_Shift_L_Cyt
Z_Im_L_Cyt

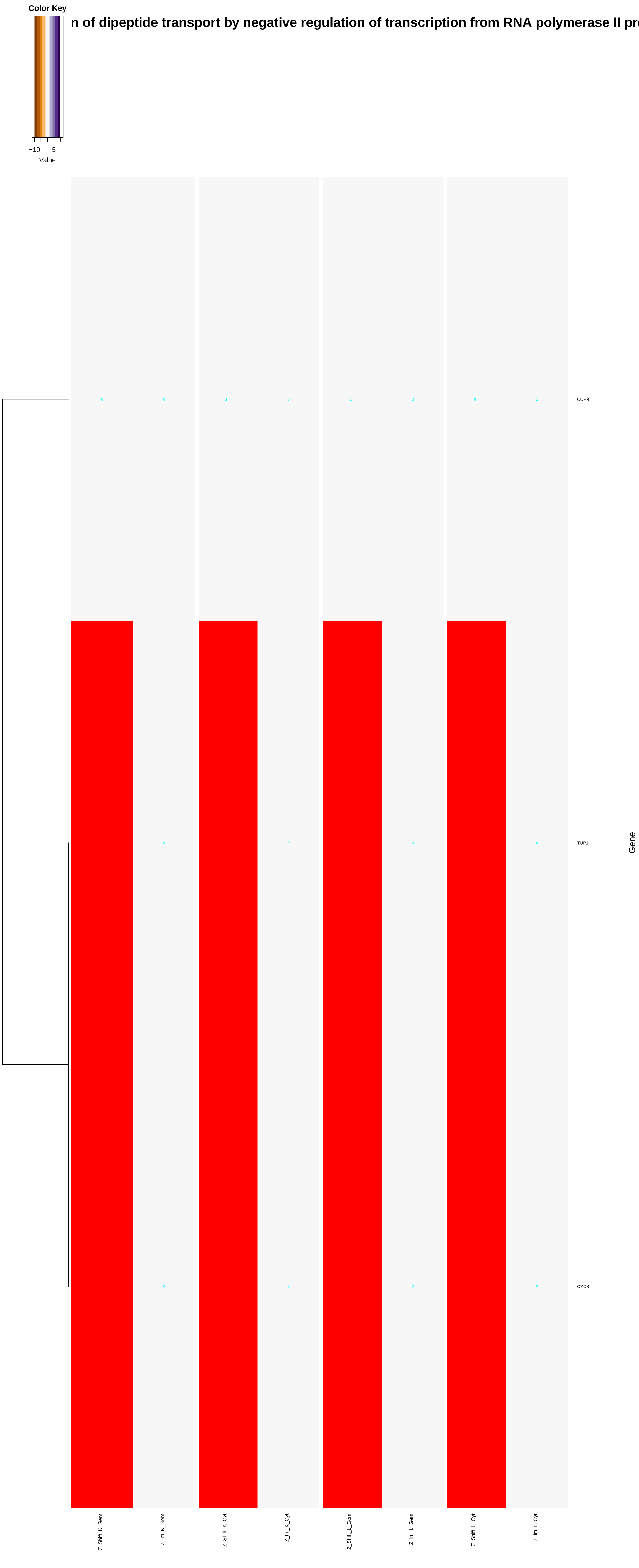

Color Key
−10
5
Value
n of dipeptide transport by negative regulation of transcription from RNA polymerase II pro
CUP9
TUP1
CYC8
Gene
Z_Shift_K_Gem
Z_Im_K_Gem
Z_Shift_K_Cyt
Z_Im_K_Cyt
Z_Shift_L_Gem
Z_Im_L_Gem
Z_Shift_L_Cyt
Z_Im_L_Cyt

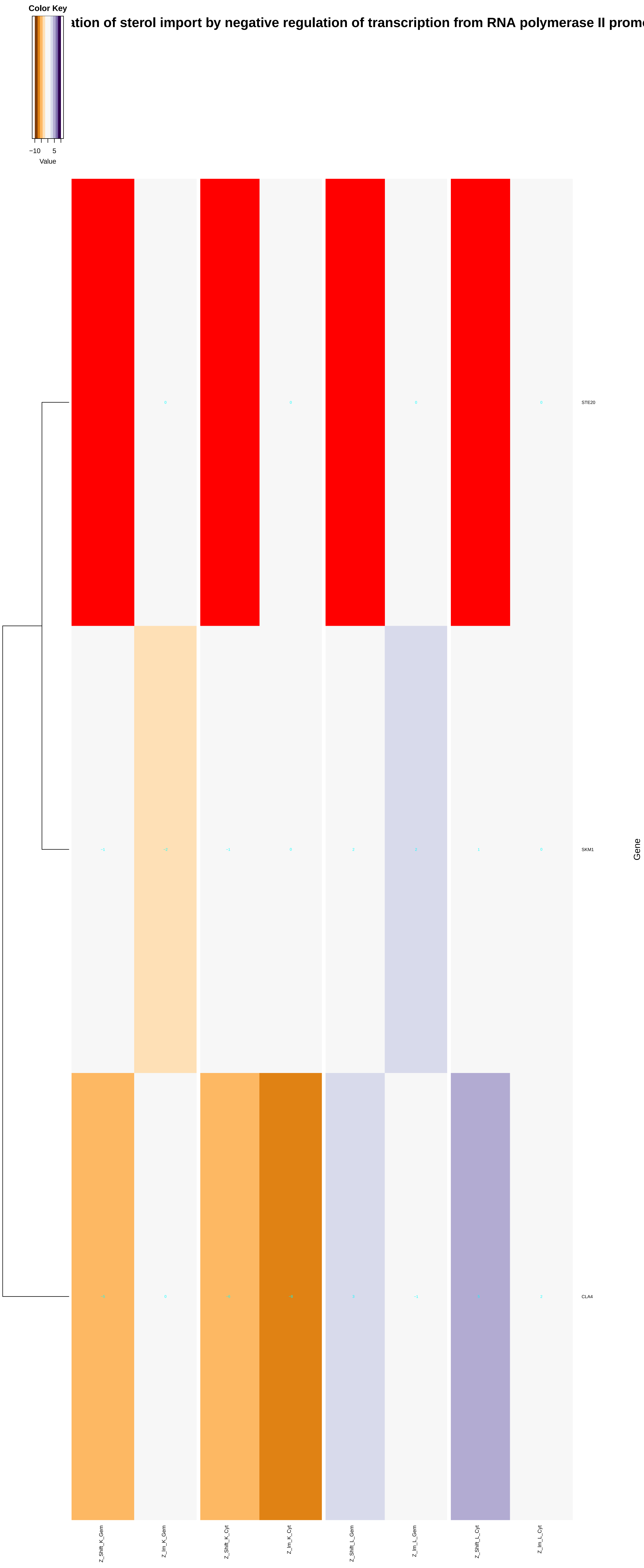

Color Key
−10
5
Value
ation of sterol import by negative regulation of transcription from RNA polymerase II prom
STE20
SKM1
CLA4
Gene
Z_Shift_K_Gem
Z_Im_K_Gem
Z_Shift_K_Cyt
Z_Im_K_Cyt
Z_Shift_L_Gem
Z_Im_L_Gem
Z_Shift_L_Cyt
Z_Im_L_Cyt
