## Additional File 7 for "A humanized yeast phenomic model of deoxycytidine kinase to predict genetic buffering of nucleoside analog cytotoxicity": negative_regulation_of_nucleic_acid-templated_transcription.pdf

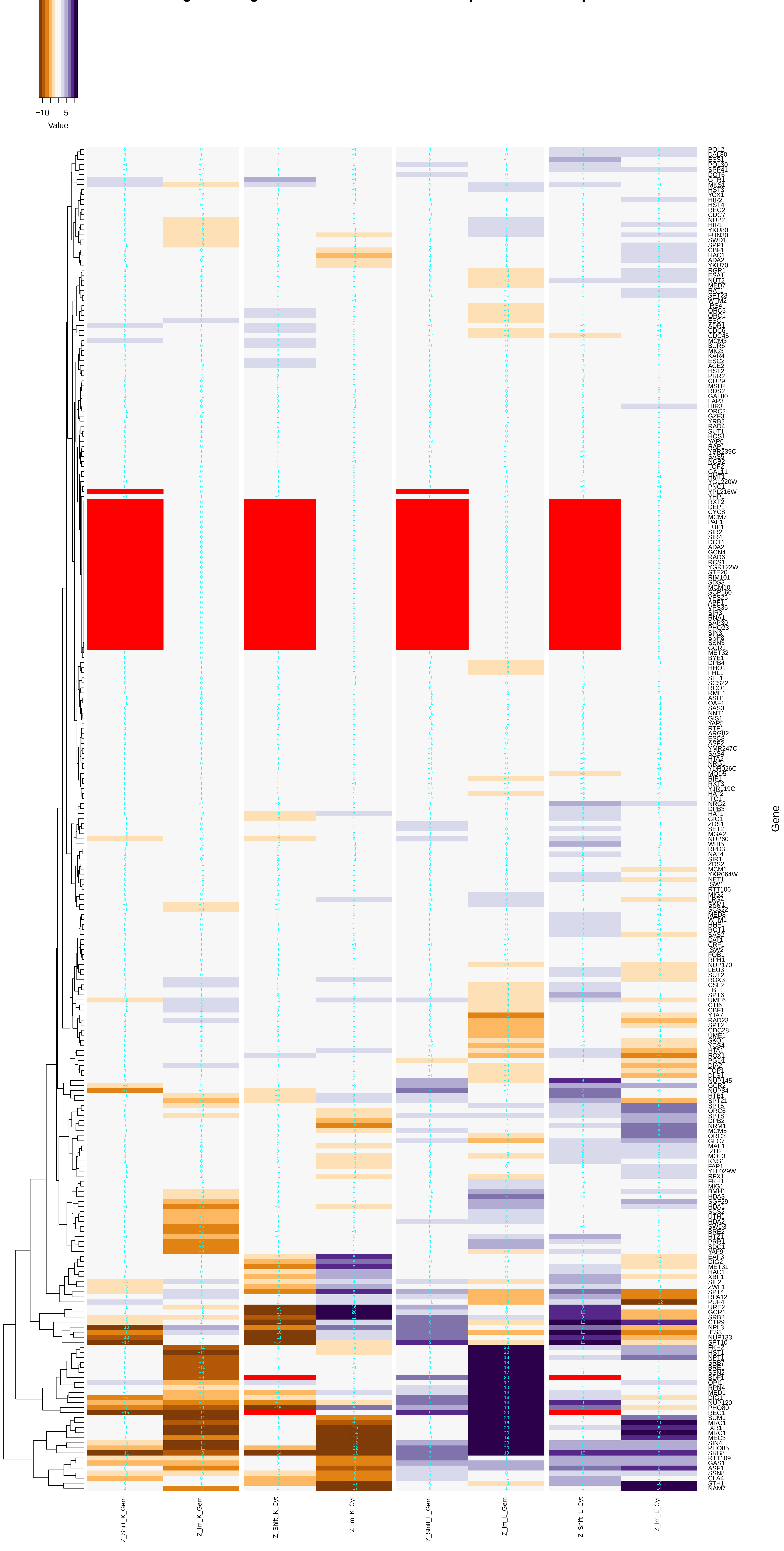

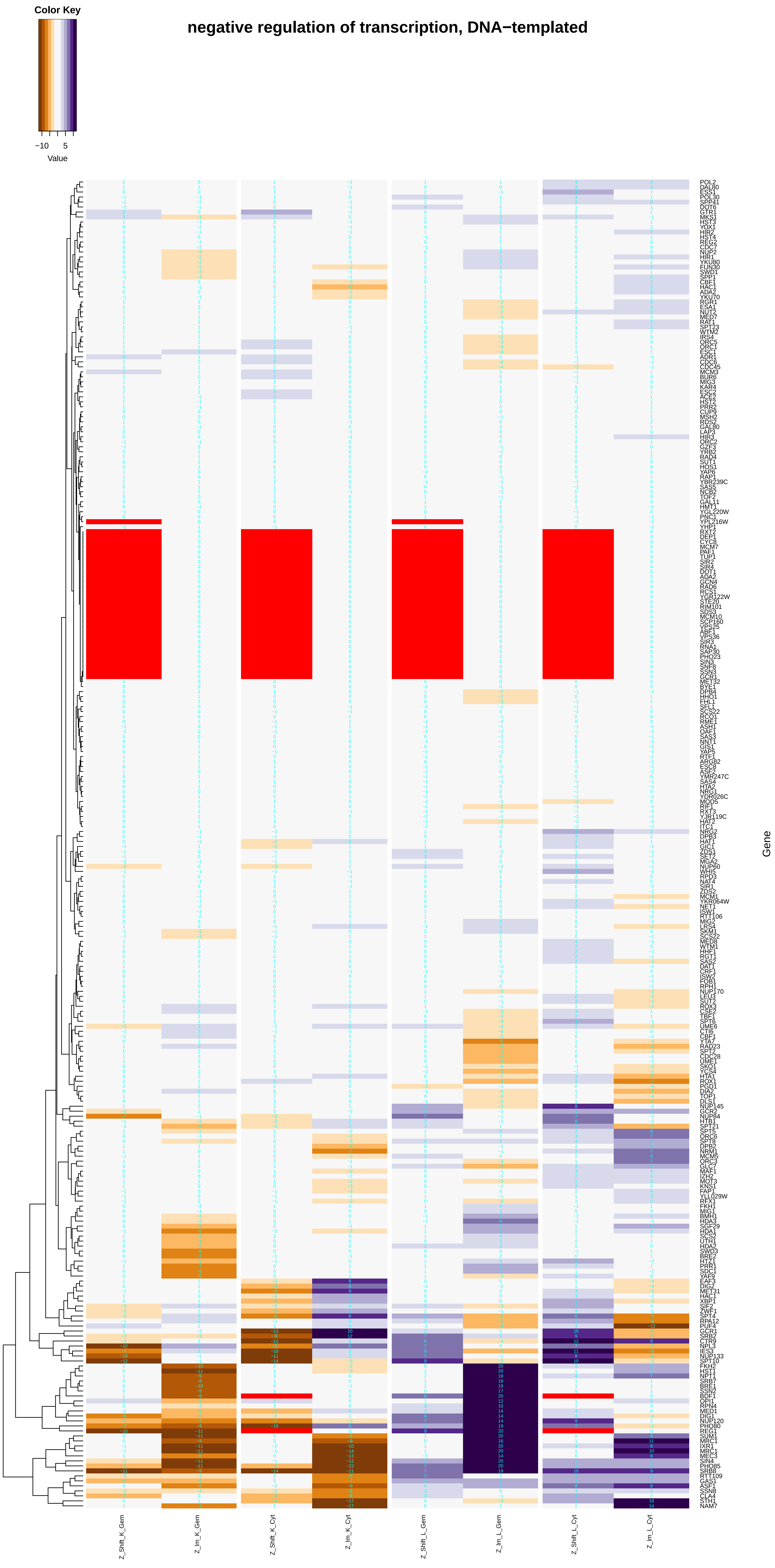
negative regulation of transcription, DNA−templated
Color Key
−10
5
Value
Gene
Z_Shift_K_Gem
Z_Im_K_Gem
Z_Shift_K_Cyt
Z_Im_K_Cyt
Z_Shift_L_Gem
Z_Im_L_Gem
Z_Shift_L_Cyt
Z_Im_L_Cyt

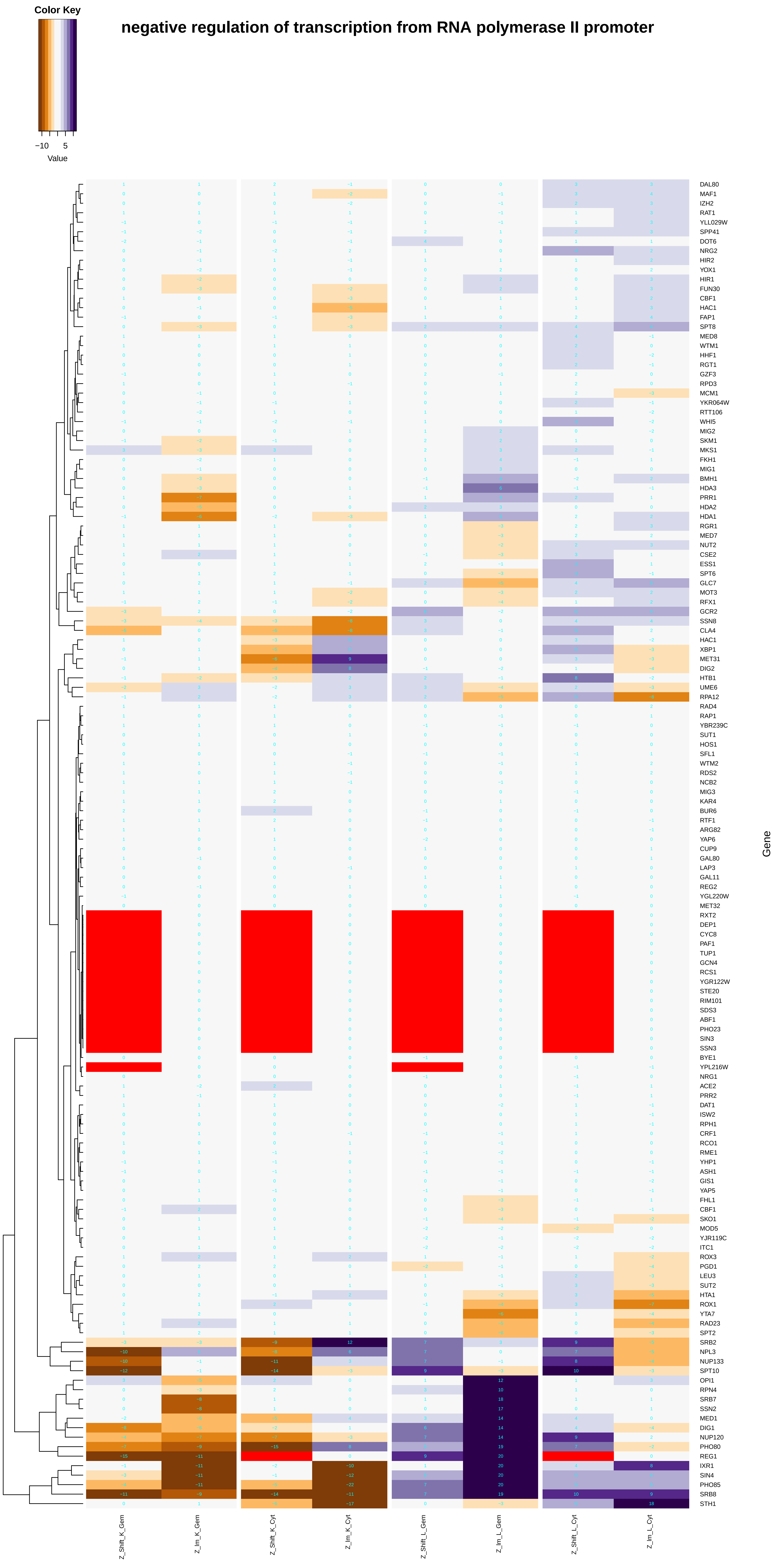
Color Key
−10
5
Value
negative regulation of transcription from RNA polymerase II promoter
DAL80
MAF1
IZH2
RAT1
YLL029W
SPP41
DOT6
NRG2
HIR2
YOX1
HIR1
FUN30
CBF1
HAC1
FAP1
SPT8
MED8
WTM1
HHF1
RGT1
GZF3
RPD3
MCM1
YKR064W
RTT106
WHI5
MIG2
SKM1
MKS1
FKH1
MIG1
BMH1
HDA3
PRR1
HDA2
HDA1
RGR1
MED7
NUT2
CSE2
ESS1
SPT6
GLC7
MOT3
RFX1
GCR2
SSN8
CLA4
HAC1
XBP1
MET31
DIG2
HTB1
UME6
RPA12
RAD4
RAP1
YBR239C
SUT1
HOS1
SFL1
WTM2
RDS2
NCB2
MIG3
KAR4
BUR6
RTF1
ARG82
YAP6
CUP9
GAL80
LAP3
GAL11
REG2
YGL220W
MET32
RXT2
DEP1
CYC8
PAF1
TUP1
GCN4
RCS1
YGR122W
STE20
RIM101
SDS3
ABF1
PHO23
SIN3
SSN3
BYE1
YPL216W
NRG1
ACE2
PRR2
DAT1
ISW2
RPH1
CRF1
RCO1
RME1
YHP1
ASH1
GIS1
YAP5
FHL1
CBF1
SKO1
MOD5
YJR119C
ITC1
ROX3
PGD1
LEU3
SUT2
HTA1
ROX1
YTA7
RAD23
SPT2
SRB2
NPL3
NUP133
SPT10
OPI1
RPN4
SRB7
SSN2
MED1
DIG1
NUP120
PHO80
REG1
IXR1
SIN4
PHO85
SRB8
STH1
Gene
Z_Shift_K_Gem
Z_lm_K_Gem
Z_Shift_K_Cyt
Z_lm_K_Cyt
Z_Shift_L_Gem
Z_lm_L_Gem
Z_Shift_L_Cyt
Z_lm_L_Cyt

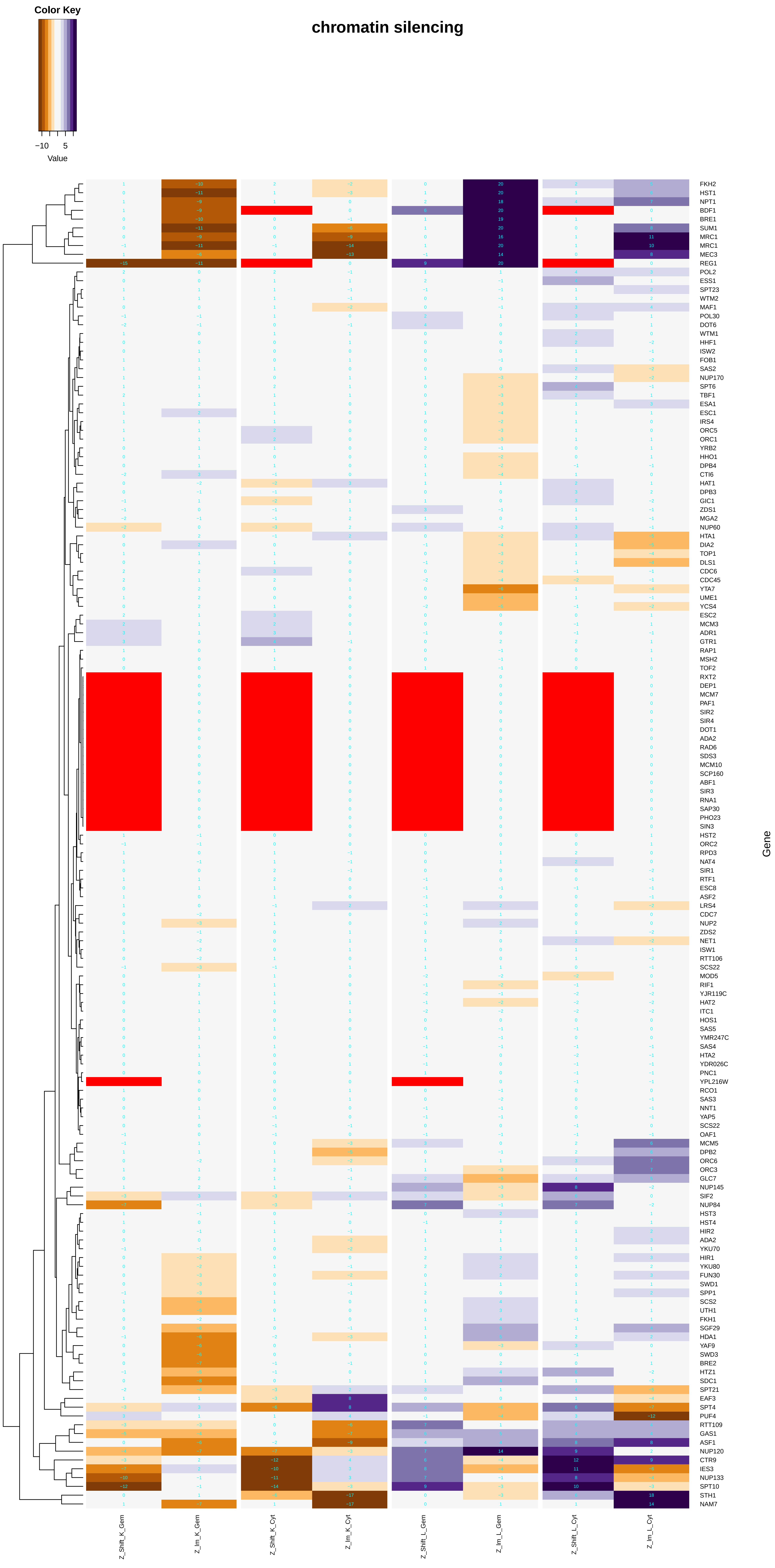
chromatin silencing
Color Key
−10
5
Value
FKH2
HST1
NPT1
BDF1
BRE1
SUM1
MRC1
MRC1
MEC3
REG1
POL2
ESS1
SPT23
WTM2
MAF1
POL30
DOT6
WTM1
HHF1
ISW2
FOB1
SAS2
NUP170
SPT6
TBF1
ESA1
ESC1
IRS4
ORC5
ORC1
YRB2
HHO1
DPB4
CTI6
HAT1
DPB3
GIC1
ZDS1
MGA2
NUP60
HTA1
DIA2
TOP1
DLS1
CDC6
CDC45
YTA7
UME1
YCS4
ESC2
MCM3
ADR1
GTR1
RAP1
MSH2
TOF2
RXT2
DEP1
MCM7
PAF1
SIR2
SIR4
DOT1
ADA2
RAD6
SDS3
MCM10
SCP160
ABF1
SIR3
RNA1
SAP30
PHO23
SIN3
HST2
ORC2
RPD3
NAT4
SIR1
RTF1
ESC8
ASF2
LRS4
CDC7
NUP2
ZDS2
NET1
ISW1
RTT106
SCS22
MOD5
RIF1
YJR119C
HAT2
ITC1
HOS1
SAS5
YMR247C
SAS4
HTA2
YDR026C
PNC1
YPL216W
RCO1
SAS3
NNT1
YAP5
SCS22
OAF1
MCM5
DPB2
ORC6
ORC3
GLC7
NUP145
SIF2
NUP84
HST3
HST4
HIR2
ADA2
YKU70
HIR1
YKU80
FUN30
SWD1
SPP1
SCS2
UTH1
FKH1
SGF29
HDA1
YAF9
SWD3
BRE2
HTZ1
SDC1
SPT21
EAF3
SPT4
PUF4
RTT109
GAS1
ASF1
NUP120
CTR9
IES3
NUP133
SPT10
STH1
NAM7
Gene
Z_Shift_K_Gem
Z_Im_K_Gem
Z_Shift_K_Cyt
Z_Im_K_Cyt
Z_Shift_L_Gem
Z_Im_L_Gem
Z_Shift_L_Cyt
Z_Im_L_Cyt

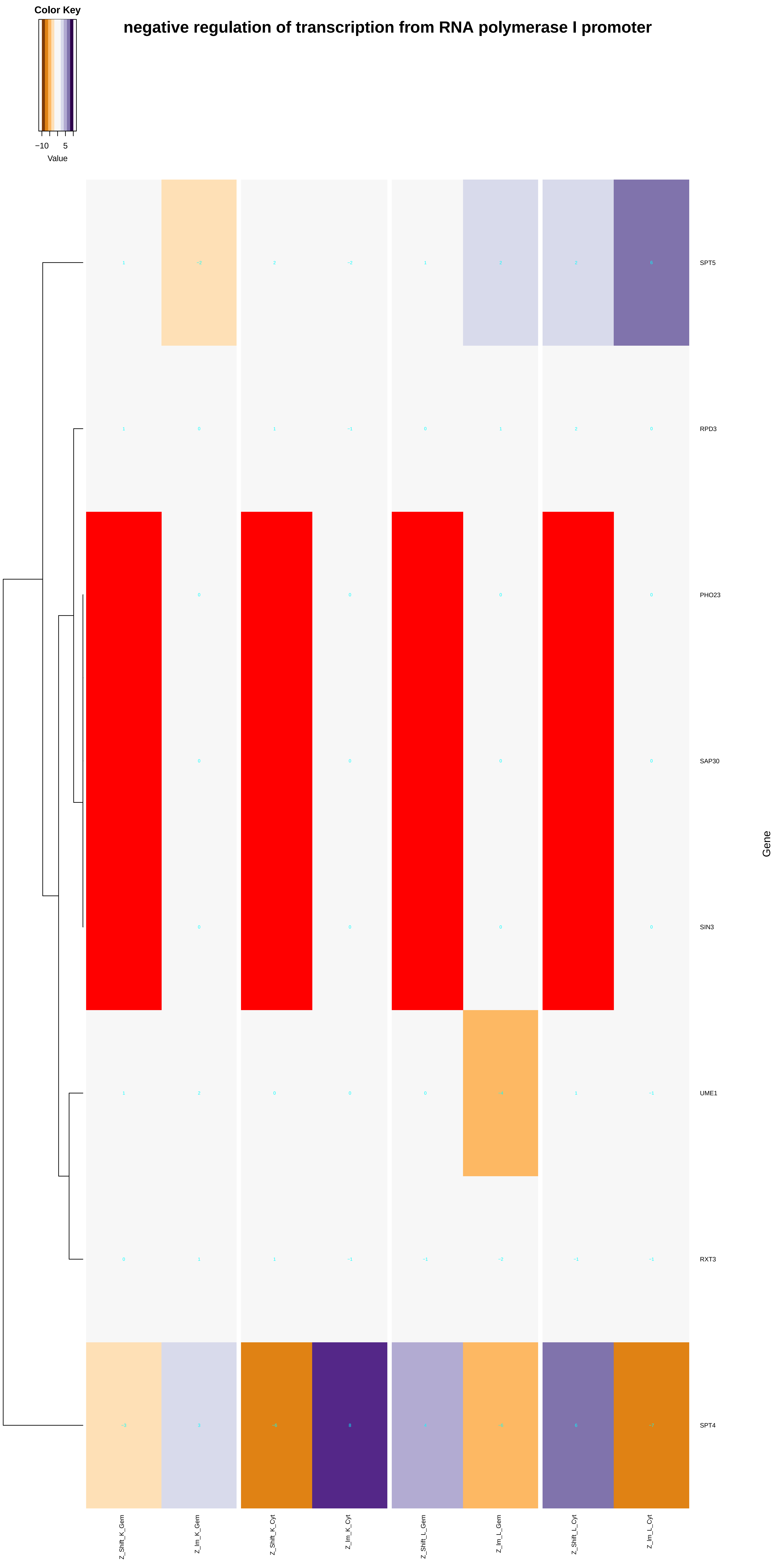

Color Key
−10
5
Value
negative regulation of transcription from RNA polymerase I promoter
SPT5
RPD3
PHO23
SAP30
SIN3
UME1
RXT3
SPT4
Gene
Z_Shift_K_Gem
Z_Im_K_Gem
Z_Shift_K_Cyt
Z_Im_K_Cyt
Z_Shift_L_Gem
Z_Im_L_Gem
Z_Shift_L_Cyt
Z_Im_L_Cyt

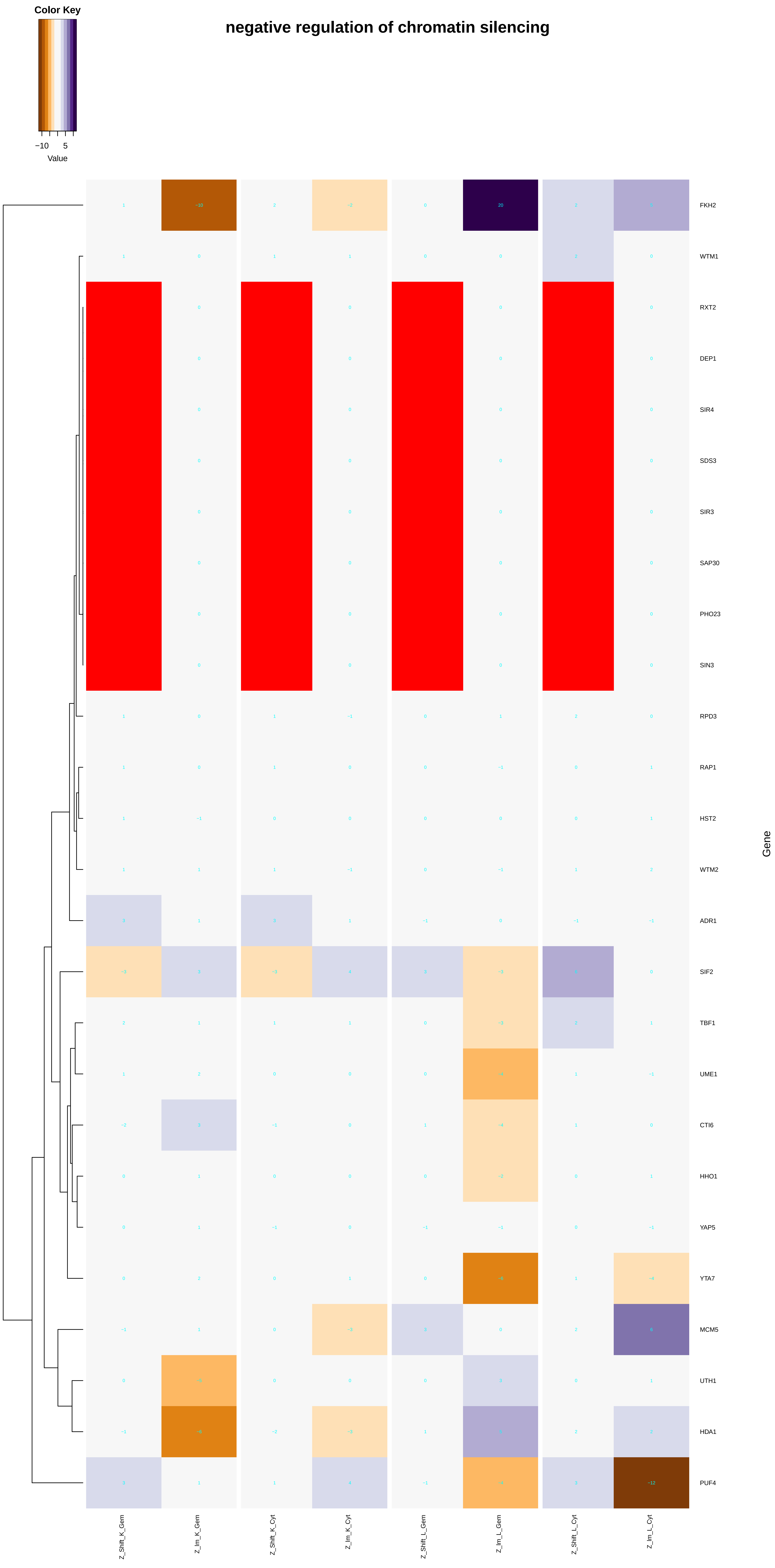
negative regulation of chromatin silencing
Color Key
−10
5
Value
FKH2
WTM1
RXT2
DEP1
SIR4
SDS3
SIR3
SAP30
PHO23
SIN3
RPD3
RAP1
HST2
WTM2
ADR1
SIF2
TBF1
UME1
CTI6
HHO1
YAP5
YTA7
MCM5
UTH1
HDA1
PUF4
Gene
Z_Shift_K_Gem
Z_Im_K_Gem
Z_Shift_K_Cyt
Z_Im_K_Cyt
Z_Shift_L_Gem
Z_Im_L_Gem
Z_Shift_L_Cyt
Z_Im_L_Cyt

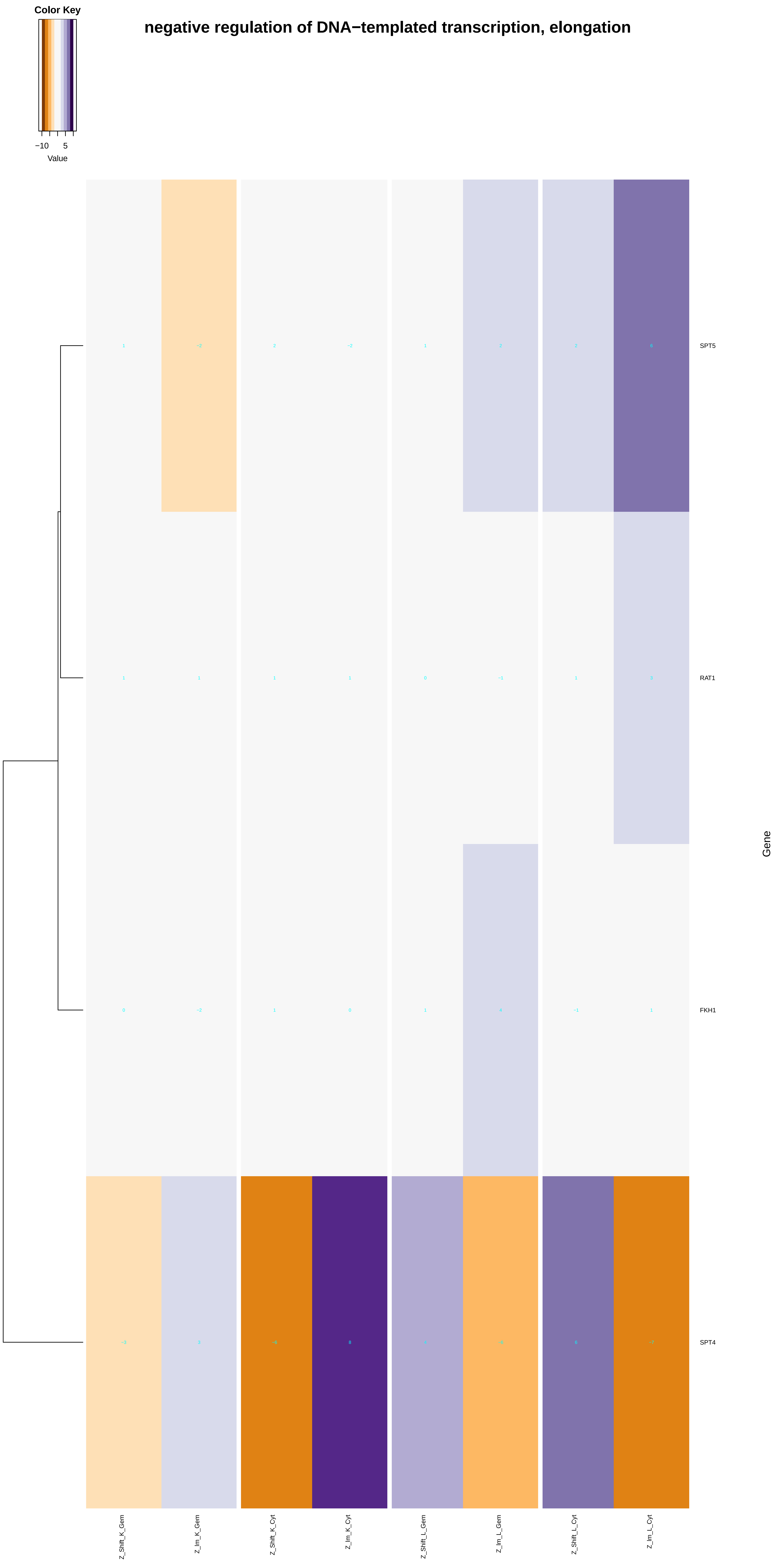

negative regulation of DNA−templated transcription, elongation
Color Key
−10
5
Value
SPT5
RAT1
FKH1
SPT4
Gene
Z_Shift_K_Gem
Z_Im_K_Gem
Z_Shift_K_Cyt
Z_Im_K_Cyt
Z_Shift_L_Gem
Z_Im_L_Gem
Z_Shift_L_Cyt
Z_Im_L_Cyt

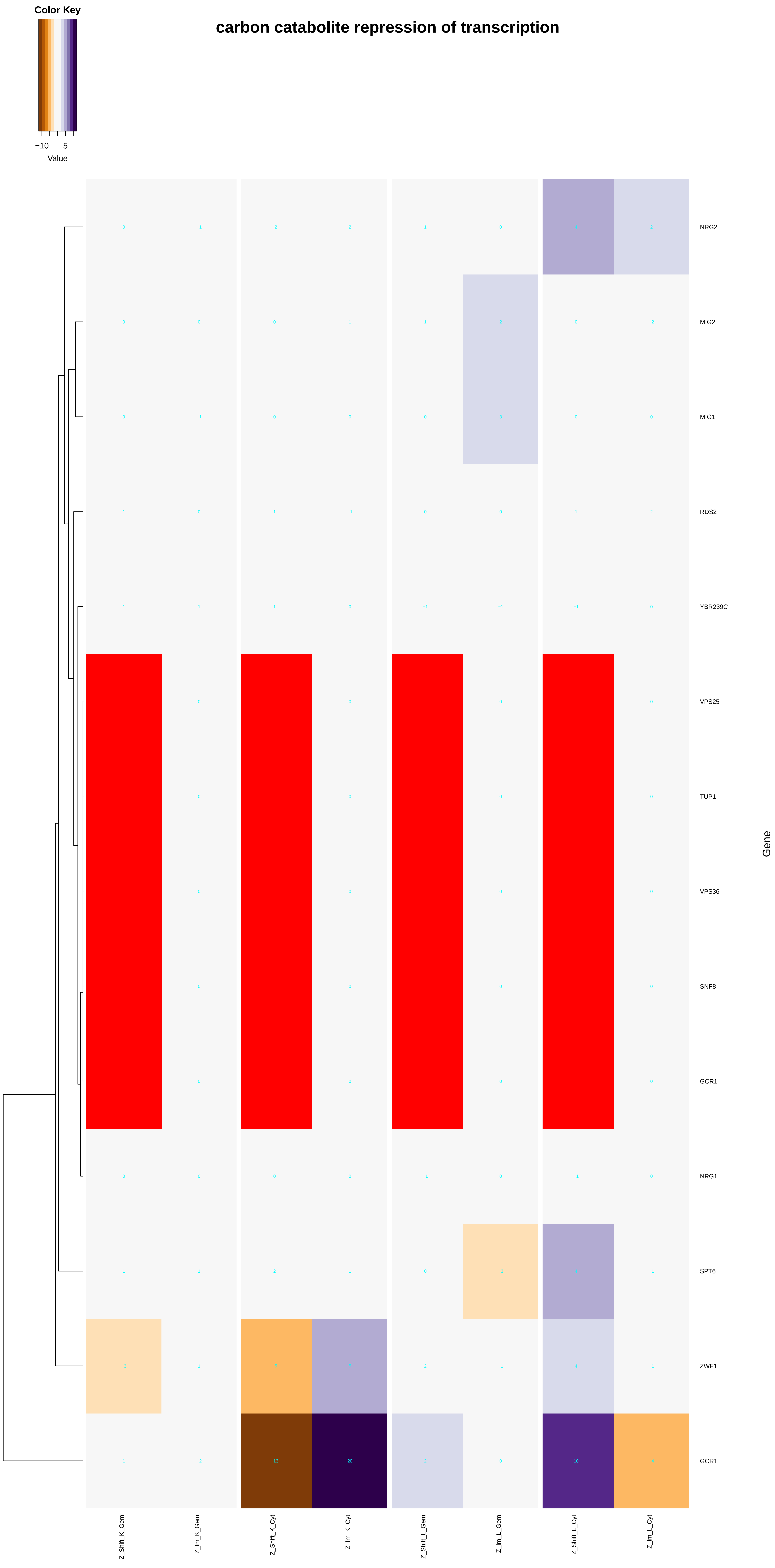

Color Key
−10
5
Value
carbon catabolite repression of transcription
NRG2
MIG2
MIG1
RDS2
YBR239C
VPS25
TUP1
VPS36
SNF8
GCR1
NRG1
SPT6
ZWF1
GCR1
Gene
Z_Shift_K_Gem
Z_Im_K_Gem
Z_Shift_K_Cyt
Z_Im_K_Cyt
Z_Shift_L_Gem
Z_Im_L_Gem
Z_Shift_L_Cyt
Z_Im_L_Cyt

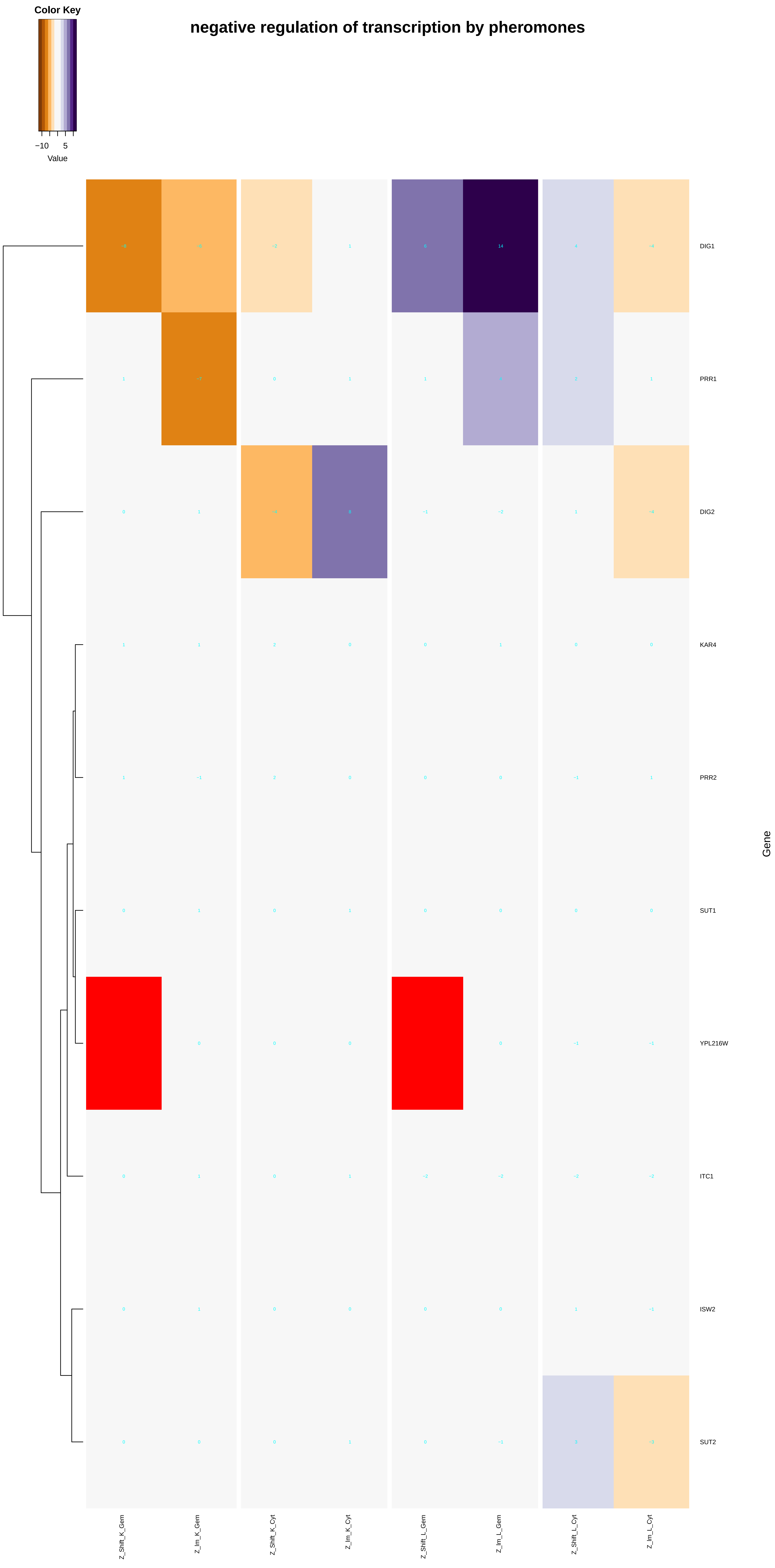

negative regulation of transcription by pheromones
Color Key
−10
5
Value
DIG1
PRR1
DIG2
KAR4
PRR2
SUT1
YPL216W
ITC1
ISW2
SUT2
Gene
Z_Shift_K_Gem
Z_lm_K_Gem
Z_Shift_K_Cyt
Z_lm_K_Cyt
Z_Shift_L_Gem
Z_lm_L_Gem
Z_Shift_L_Cyt
Z_lm_L_Cyt

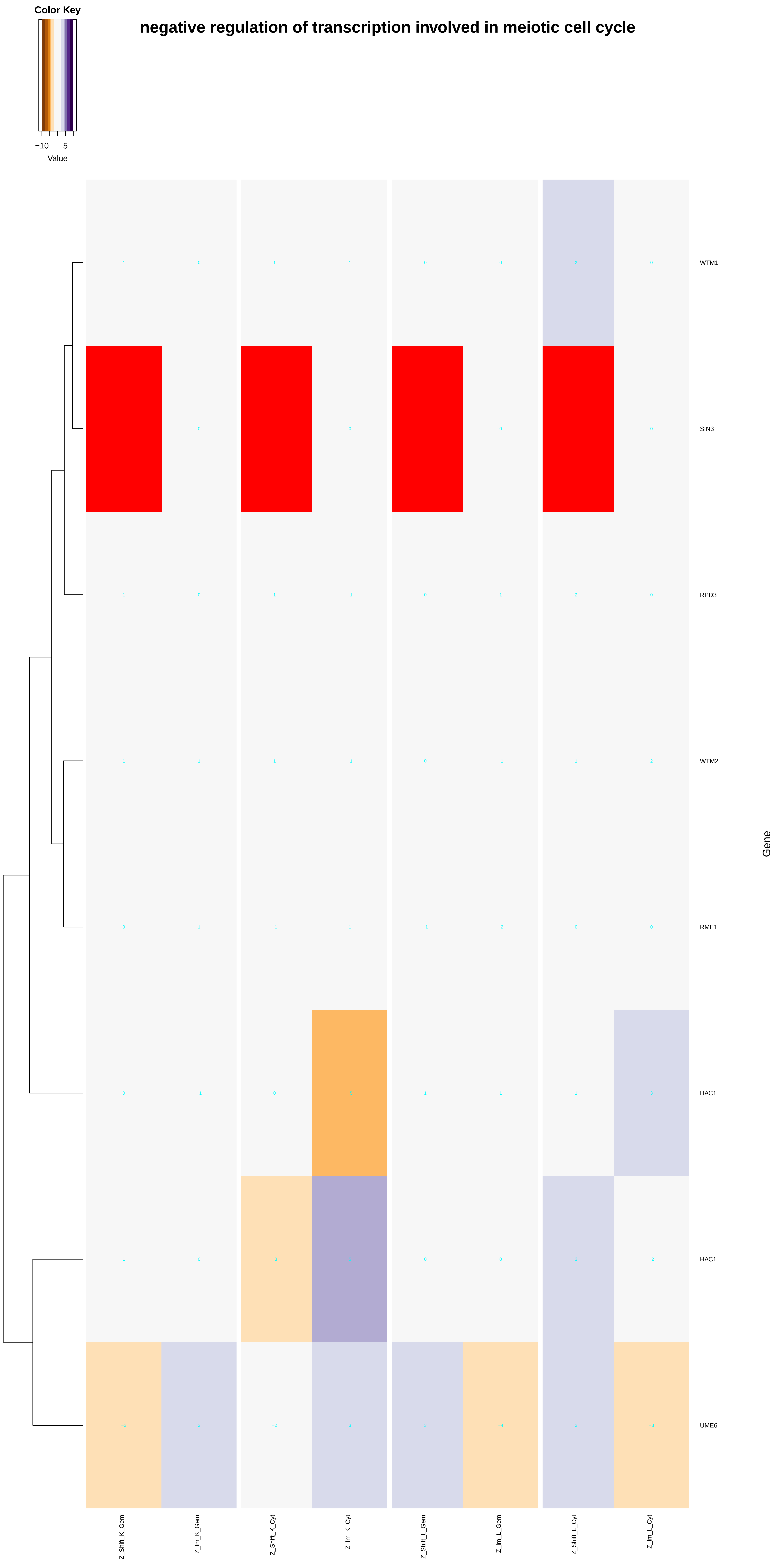
negative regulation of transcription involved in meiotic cell cycle
Color Key
−10
5
Value
WTM1
SIN3
RPD3
WTM2
RME1
HAC1
HAC1
UME6
Gene
Z_Shift_K_Gem
Z_Im_K_Gem
Z_Shift_K_Cyt
Z_Im_K_Cyt
Z_Shift_L_Gem
Z_Im_L_Gem
Z_Shift_L_Cyt
Z_Im_L_Cyt

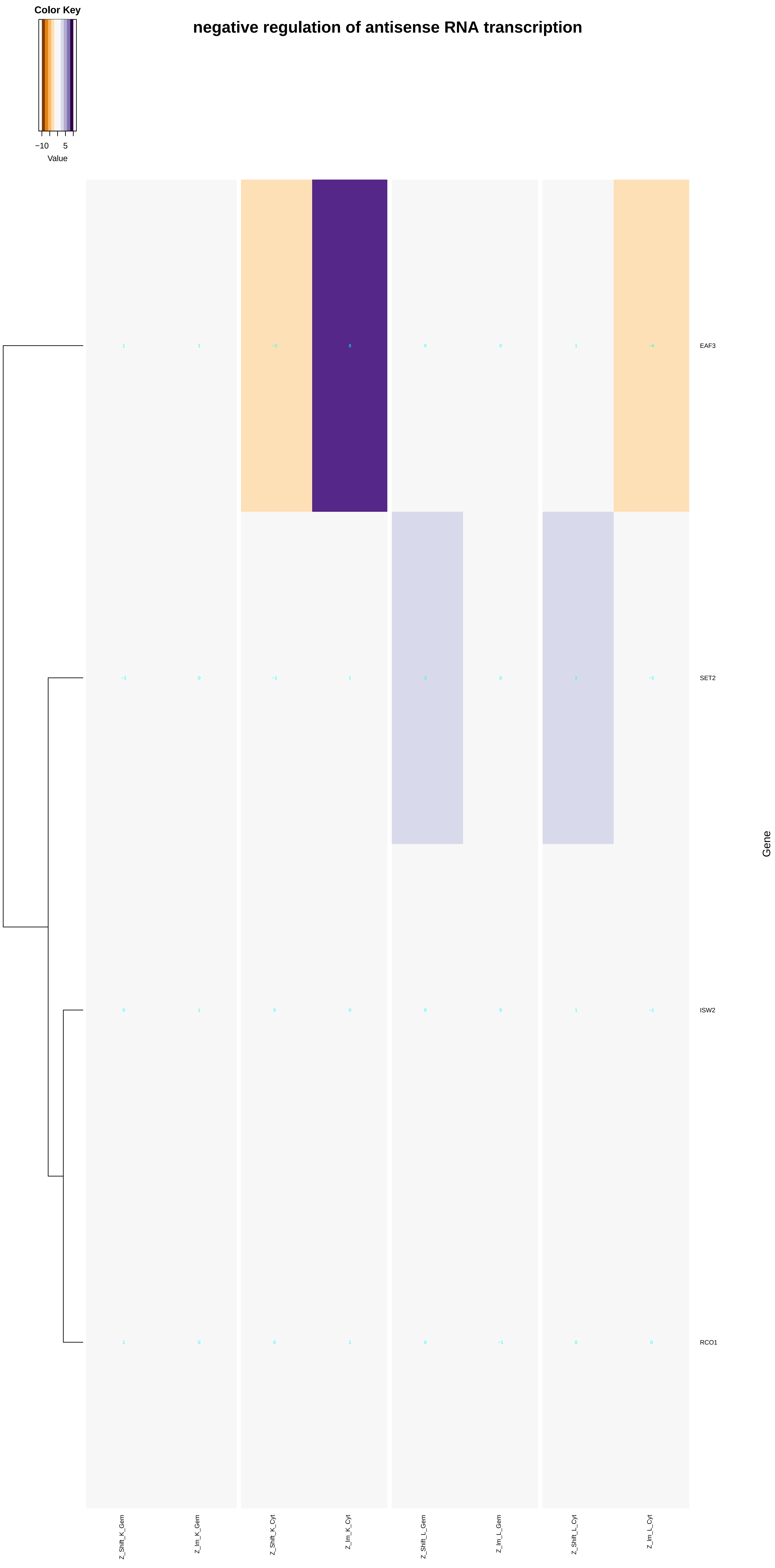
Color Key
−10
5
Value
negative regulation of antisense RNA transcription
EAF3
SET2
ISW2
RCO1
Gene
Z_Shift_K_Gem
Z_Im_K_Gem
Z_Shift_K_Cyt
Z_Im_K_Cyt
Z_Shift_L_Gem
Z_Im_L_Gem
Z_Shift_L_Cyt
Z_Im_L_Cyt

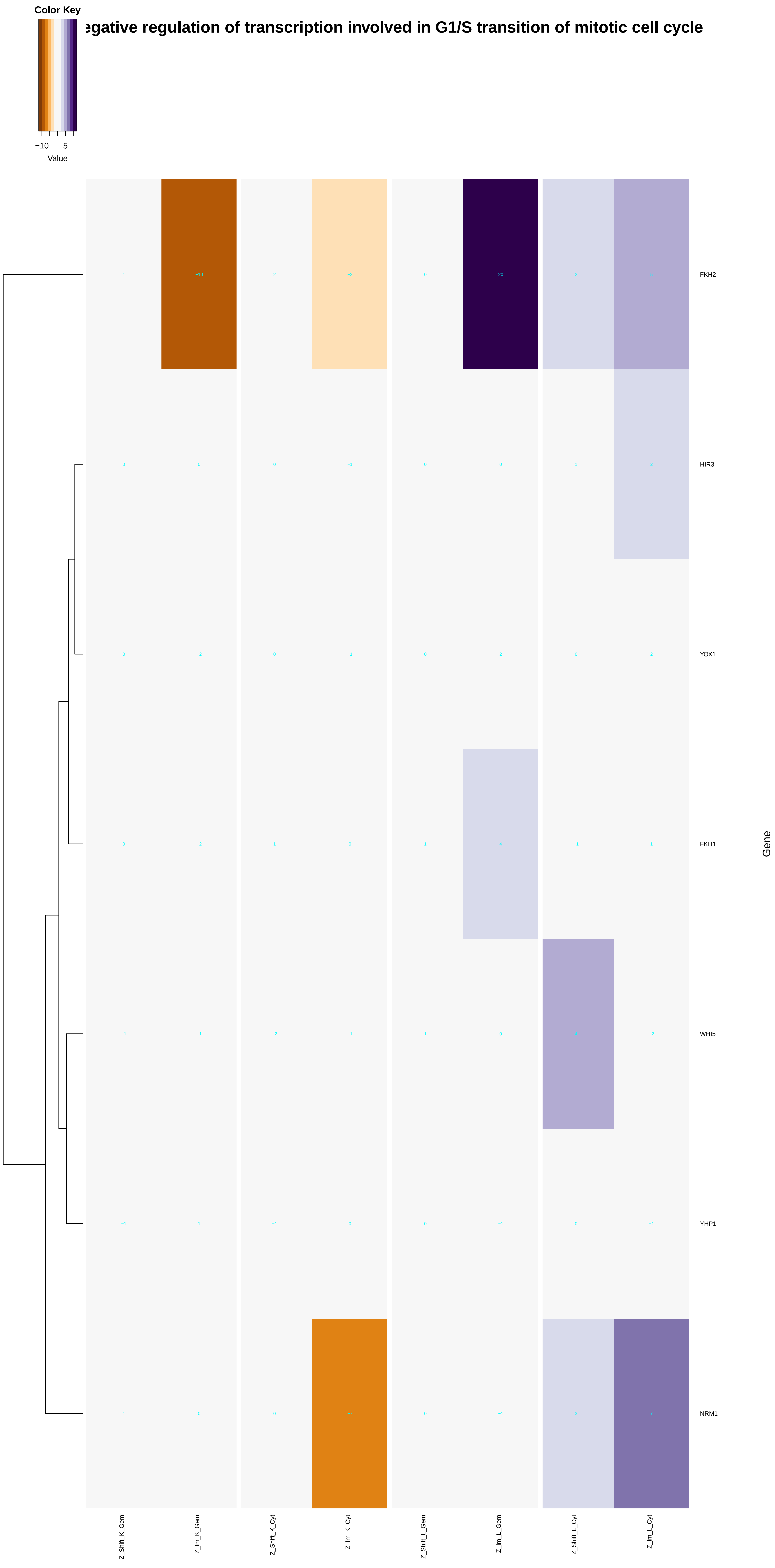

Color Key
−10
5
Value
egative regulation of transcription involved in G1/S transition of mitotic cell cycle
FKH2
HIR3
YOX1
FKH1
WHI5
YHP1
NRM1
Gene
Z_Shift_K_Gem
Z_Im_K_Gem
Z_Shift_K_Cyt
Z_Im_K_Cyt
Z_Shift_L_Gem
Z_Im_L_Gem
Z_Shift_L_Cyt
Z_Im_L_Cyt

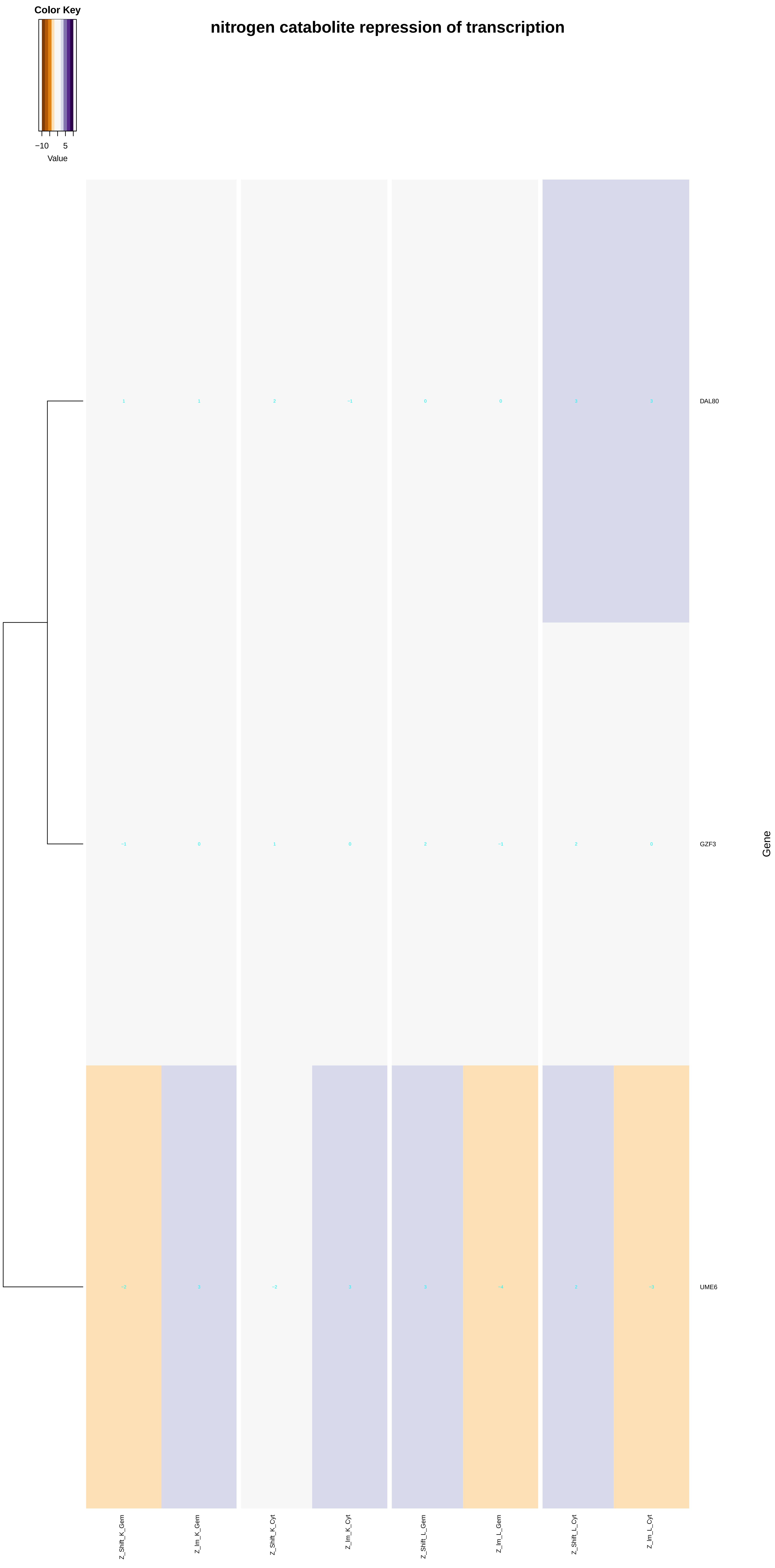
nitrogen catabolite repression of transcription
Color Key
−10
5
Value
DAL80
GZF3
UME6
Gene
Z_Shift_K_Gem
Z_Im_K_Gem
Z_Shift_K_Cyt
Z_Im_K_Cyt
Z_Shift_L_Gem
Z_Im_L_Gem
Z_Shift_L_Cyt
Z_Im_L_Cyt

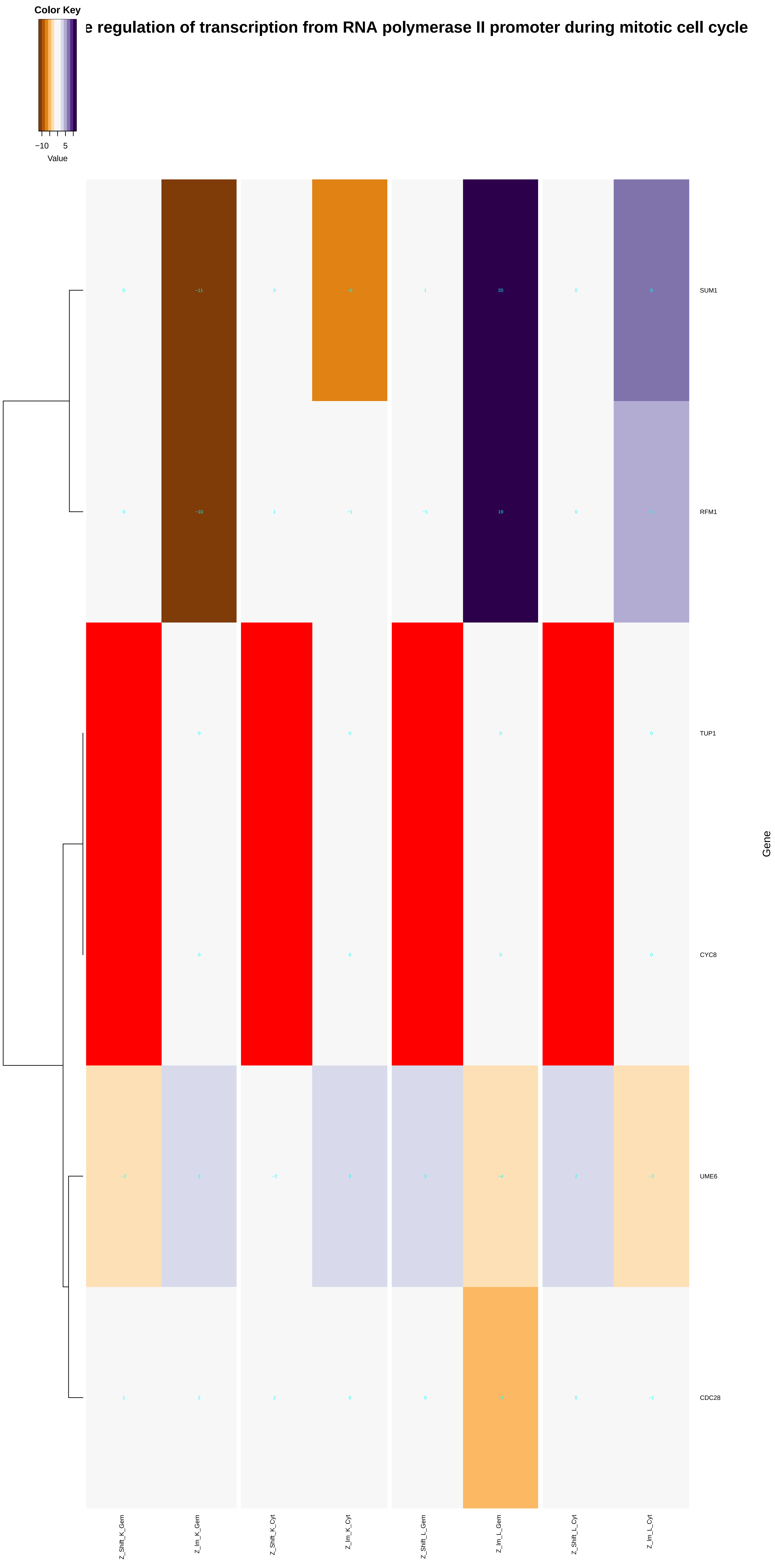
Color Key
−10 5
Value
e regulation of transcription from RNA polymerase II promoter during mitotic cell cycle
SUM1
RFM1
TUP1
CYC8
UME6
CDC28
Gene
Z_Shift_K_Gem
Z_Im_K_Gem
Z_Shift_K_Cyt
Z_Im_K_Cyt
Z_Shift_L_Gem
Z_Im_L_Gem
Z_Shift_L_Cyt
Z_Im_L_Cyt

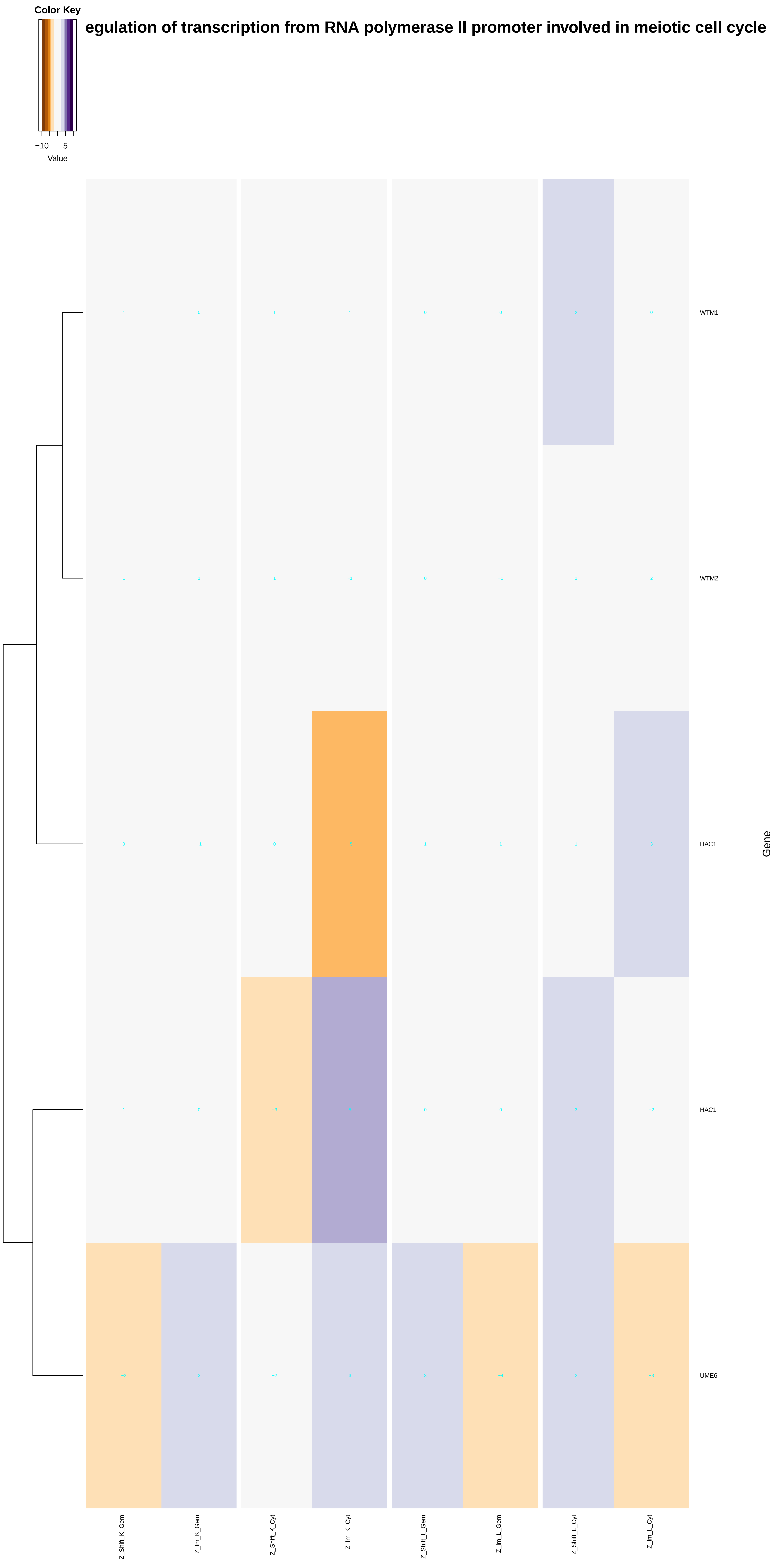

Color Key
egulation of transcription from RNA polymerase II promoter involved in meiotic cell cycle
−10
5
Value
WTM1
WTM2
HAC1
HAC1
UME6
Gene
Z_Shift_K_Gem
Z_Im_K_Gem
Z_Shift_K_Cyt
Z_Im_K_Cyt
Z_Shift_L_Gem
Z_Im_L_Gem
Z_Shift_L_Cyt
Z_Im_L_Cyt

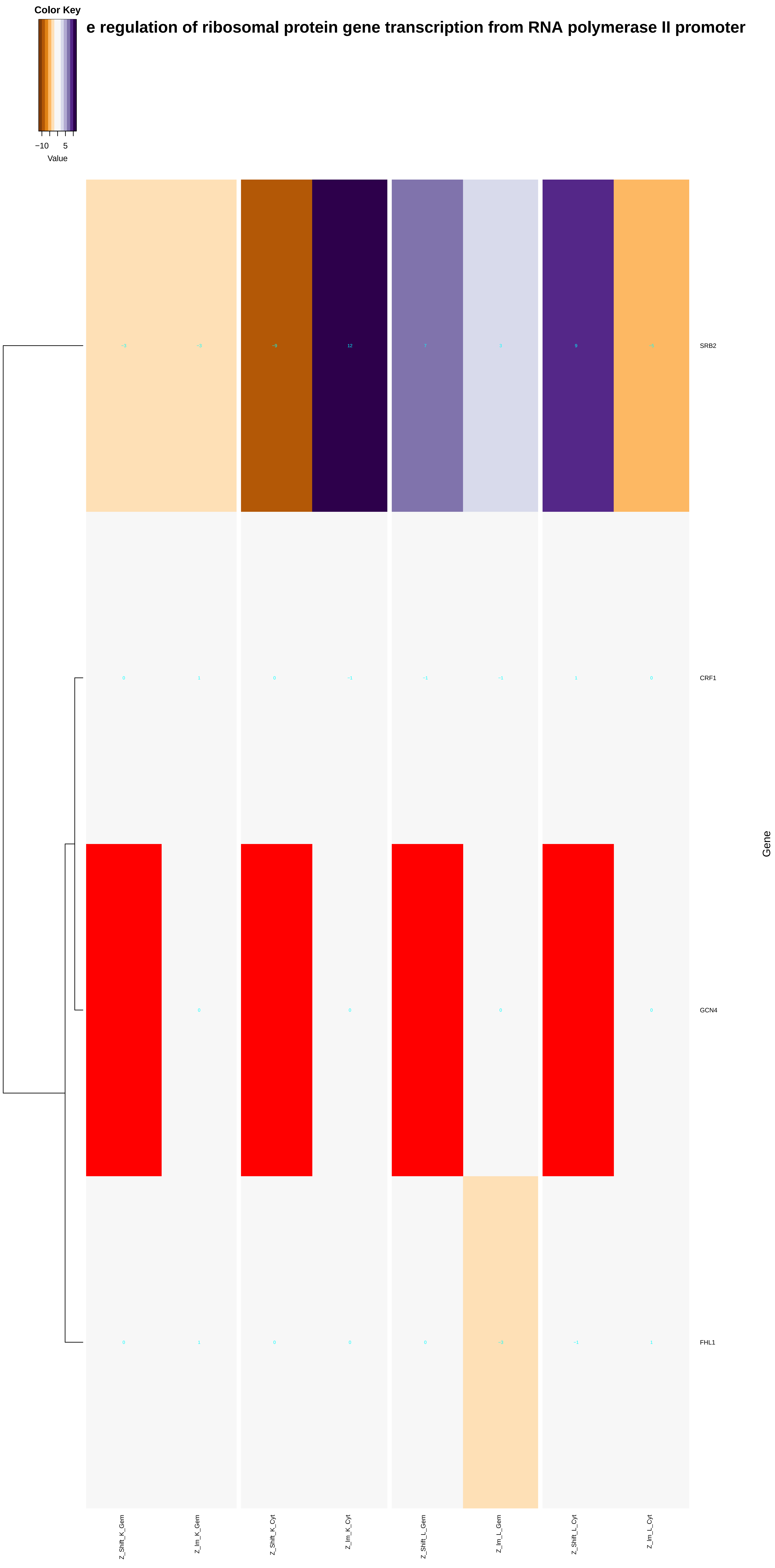
Color Key
−10
5
Value
e regulation of ribosomal protein gene transcription from RNA polymerase II promoter
SRB2
CRF1
GCN4
FHL1
Gene
Z_Shift_K_Gem
Z_Im_K_Gem
Z_Shift_K_Cyt
Z_Im_K_Cyt
Z_Shift_L_Gem
Z_Im_L_Gem
Z_Shift_L_Cyt
Z_Im_L_Cyt

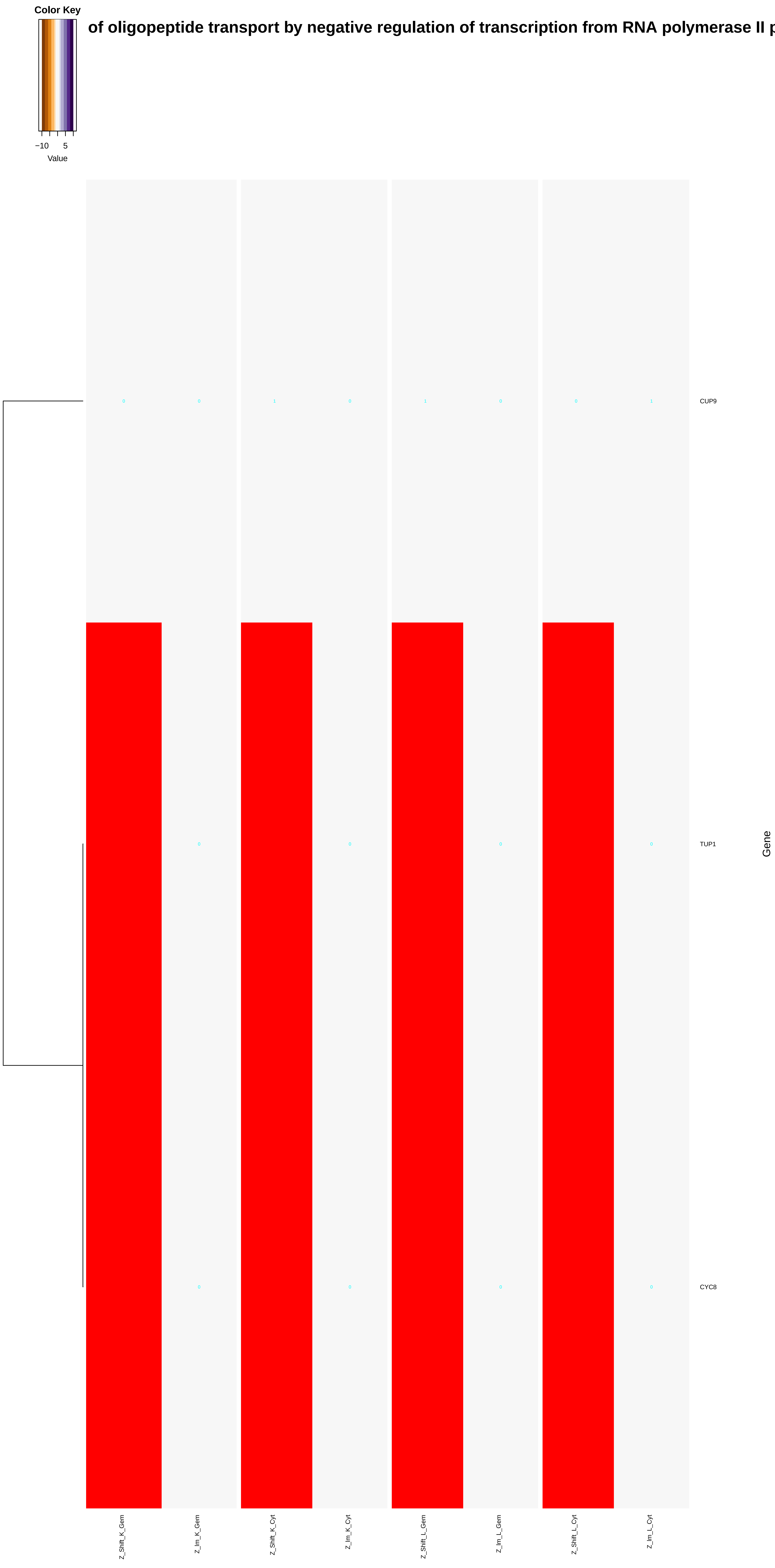
Color Key
−10
5
Value
of oligopeptide transport by negative regulation of transcription from RNA polymerase II p
CUP9
TUP1
CYC8
Gene
Z_Shift_K_Gem
Z_Im_K_Gem
Z_Shift_K_Cyt
Z_Im_K_Cyt
Z_Shift_L_Gem
Z_Im_L_Gem
Z_Shift_L_Cyt
Z_Im_L_Cyt

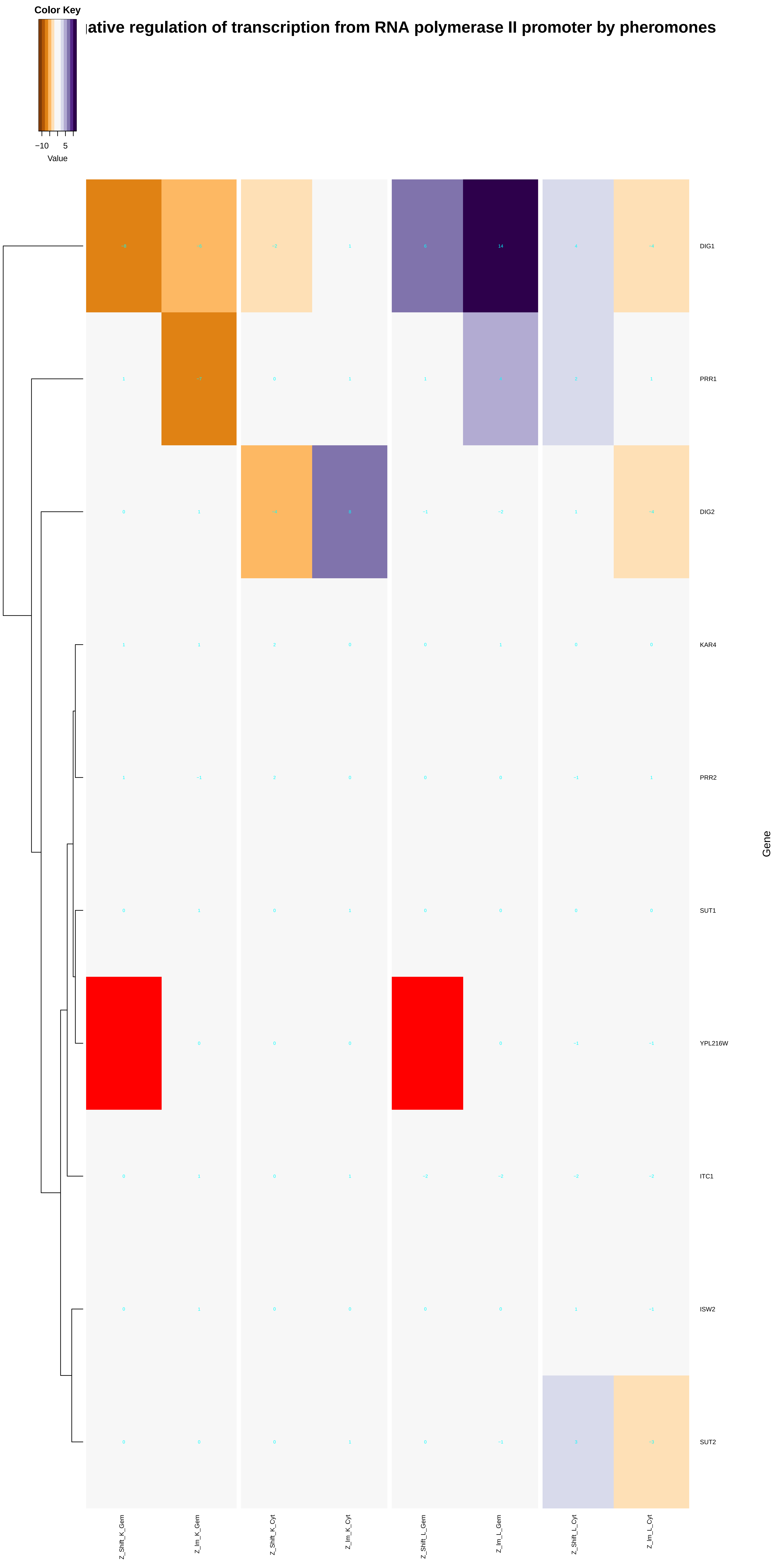

ative regulation of transcription from RNA polymerase II promoter by pheromones
Color Key
−10
5
Value
DIG1
PRR1
DIG2
KAR4
PRR2
SUT1
YPL216W
ITC1
ISW2
SUT2
Gene
Z_Shift_K_Gem
Z_Im_K_Gem
Z_Shift_K_Cyt
Z_Im_K_Cyt
Z_Shift_L_Gem
Z_Im_L_Gem
Z_Shift_L_Cyt
Z_Im_L_Cyt

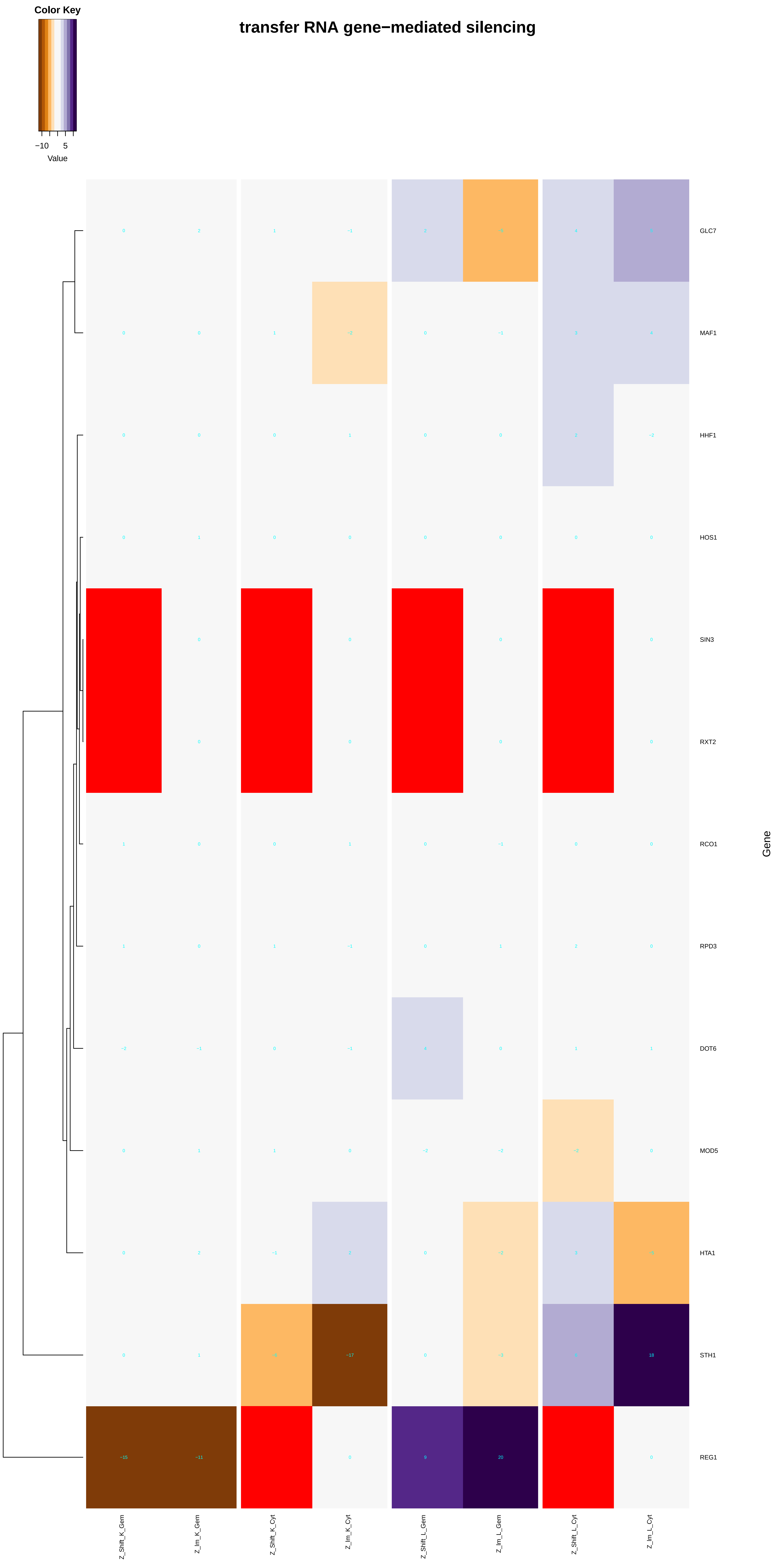

Color Key
−10 5
Value
transfer RNA gene−mediated silencing
GLC7
MAF1
HHF1
HOS1
SIN3
RXT2
RCO1
RPD3
DOT6
MOD5
HTA1
STH1
REG1
Gene
Z_Shift_K_Gem
Z_lm_K_Gem
Z_Shift_K_Cyt
Z_lm_K_Cyt
Z_Shift_L_Gem
Z_lm_L_Gem
Z_Shift_L_Cyt
Z_lm_L_Cyt

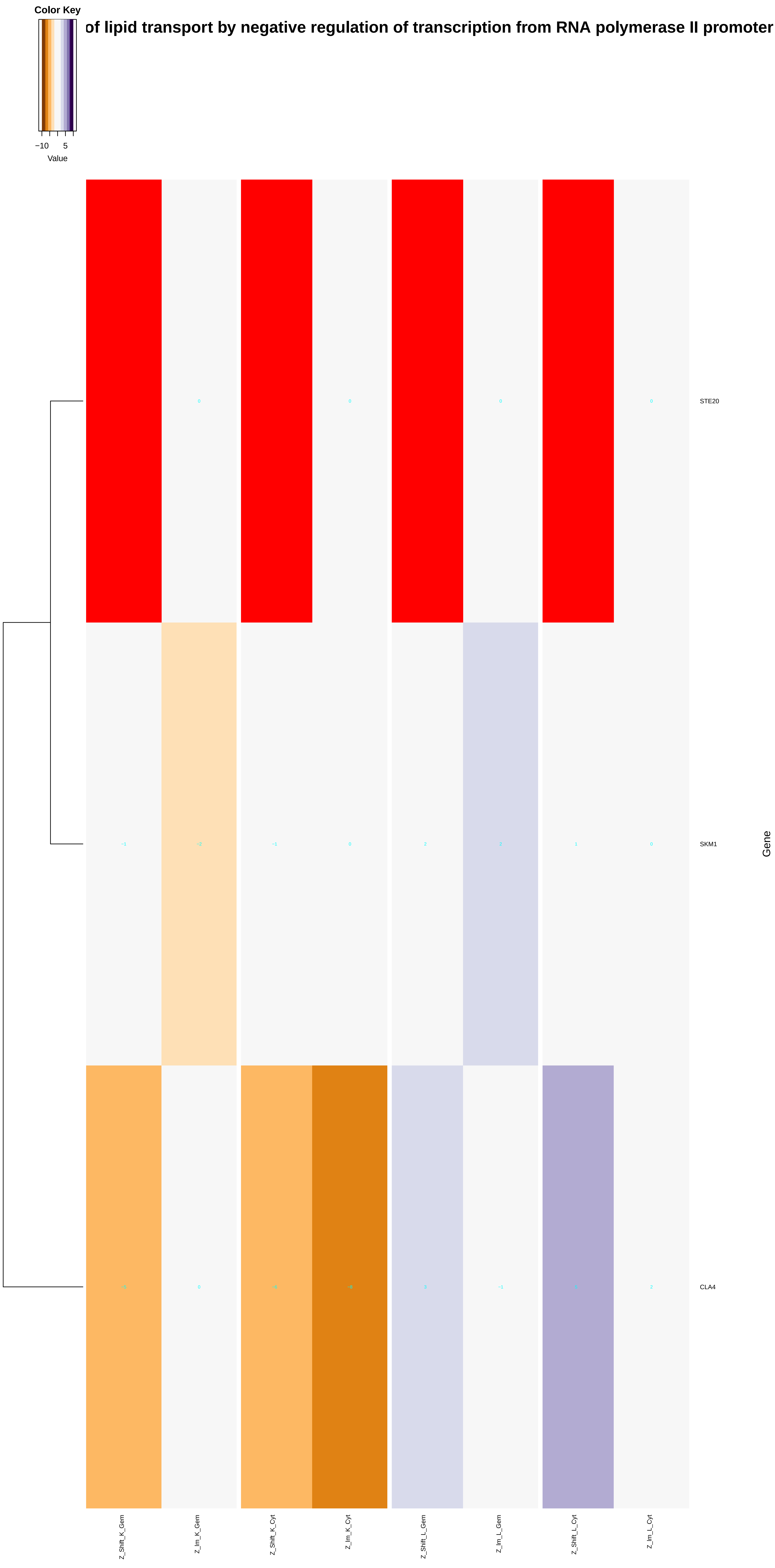
of lipid transport by negative regulation of transcription from RNA polymerase II promoter
Color Key
−10
5
Value
STE20
SKM1
CLA4
Gene
Z_Shift_K_Gem
Z_Im_K_Gem
Z_Shift_K_Cyt
Z_Im_K_Cyt
Z_Shift_L_Gem
Z_Im_L_Gem
Z_Shift_L_Cyt
Z_Im_L_Cyt

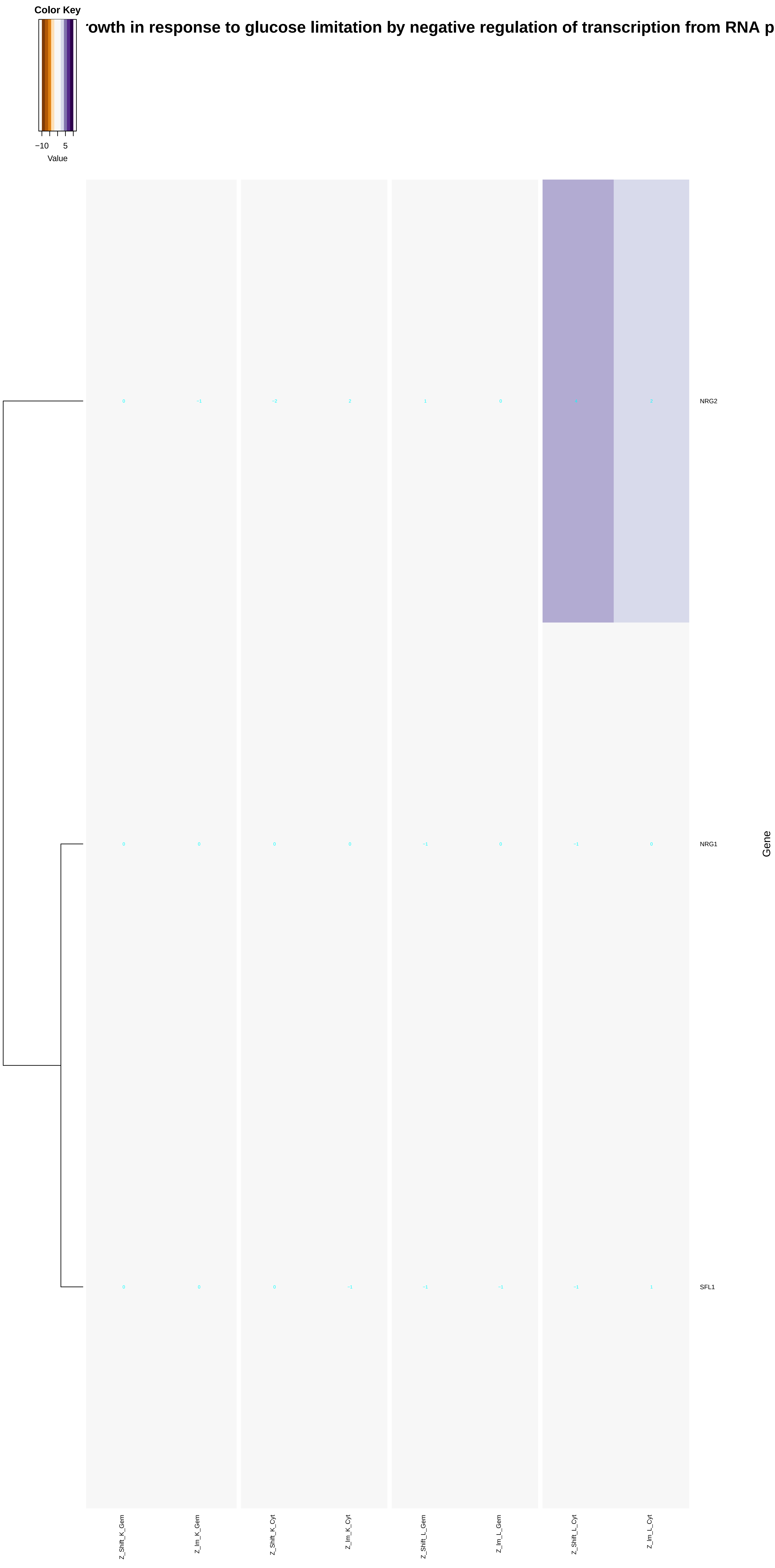
Color Key
−10
5
Value
rowth in response to glucose limitation by negative regulation of transcription from RNA p
NRG2
NRG1
SFL1
Gene
Z_Shift_K_Gem
Z_lm_K_Gem
Z_Shift_K_Cyt
Z_lm_K_Cyt
Z_Shift_L_Gem
Z_lm_L_Gem
Z_Shift_L_Cyt
Z_lm_L_Cyt

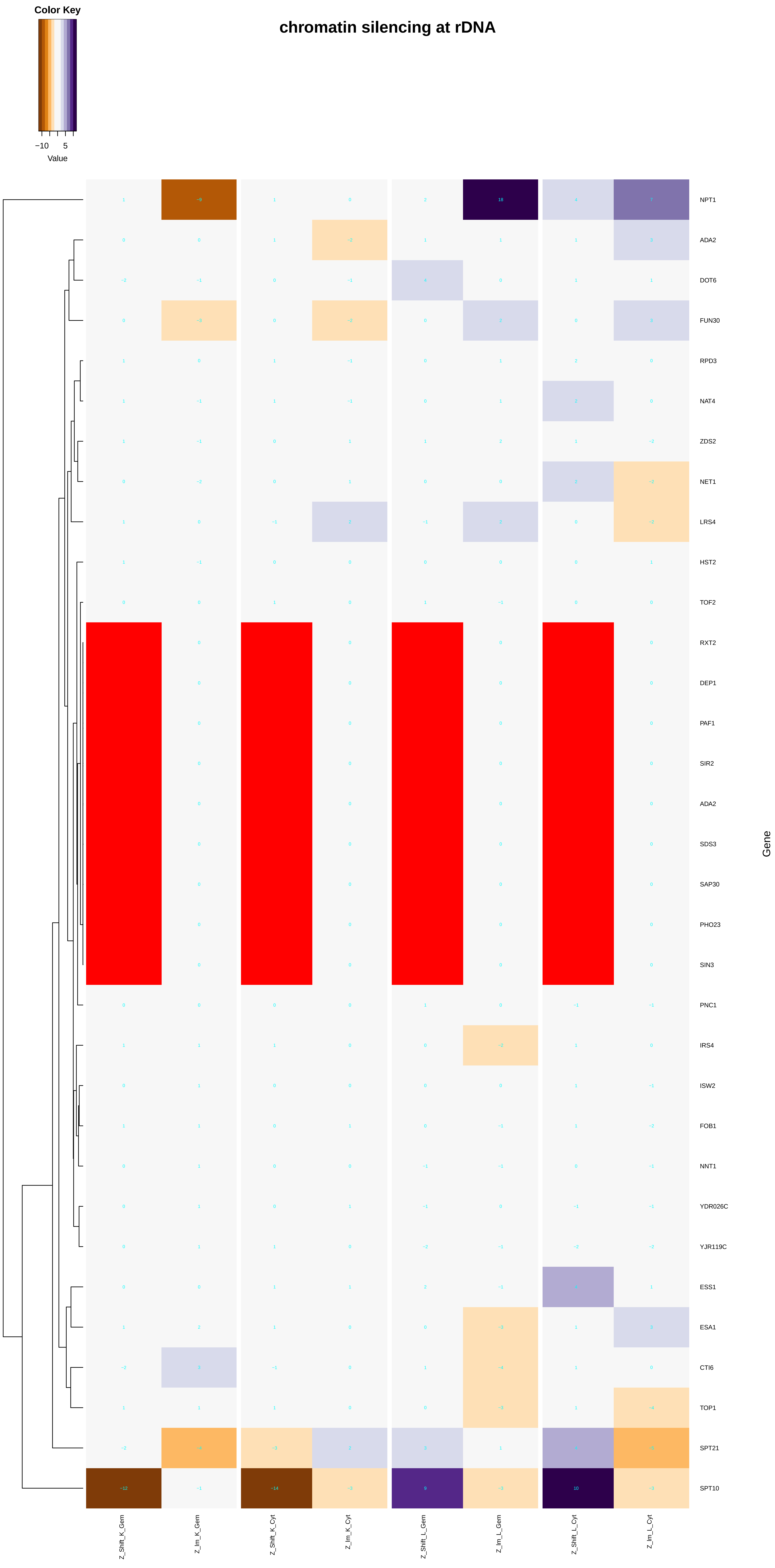
Color Key
−10
5
Value
chromatin silencing at rDNA
NPT1
ADA2
DOT6
FUN30
RPD3
NAT4
ZDS2
NET1
LRS4
HST2
TOF2
RXT2
DEP1
PAF1
SIR2
ADA2
SDS3
SAP30
PHO23
SIN3
PNC1
IRS4
ISW2
FOB1
NNT1
YDR026C
YJR119C
ESS1
ESA1
CTI6
TOP1
SPT21
SPT10
Gene
Z_Shift_K_Gem
Z_Im_K_Gem
Z_Shift_K_Cyt
Z_Im_K_Cyt
Z_Shift_L_Gem
Z_Im_L_Gem
Z_Shift_L_Cyt
Z_Im_L_Cyt

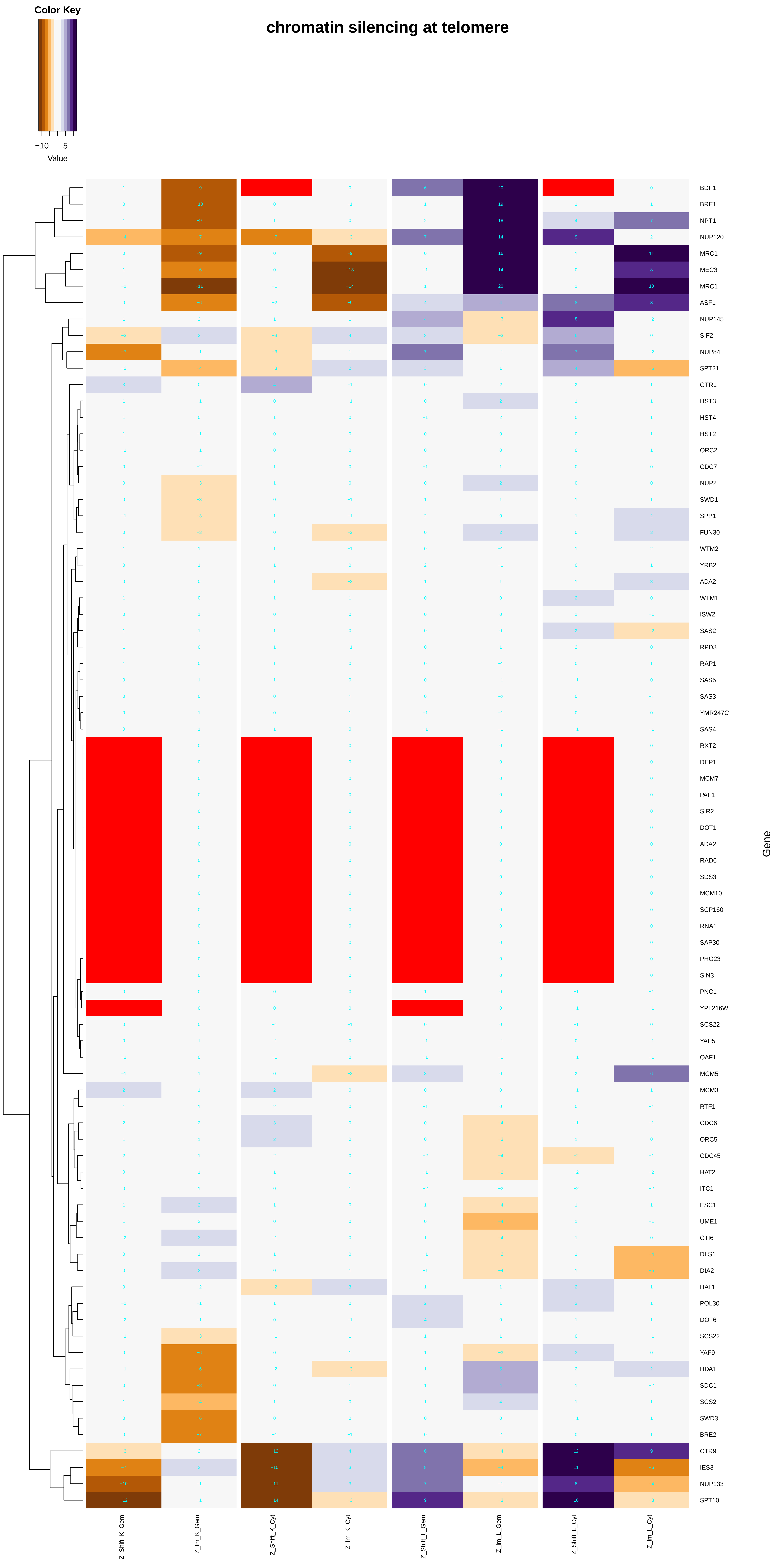

Color Key
−10 5
Value
chromatin silencing at telomere
BDF1
BRE1
NPT1
NUP120
MRC1
MEC3
MRC1
ASF1
NUP145
SIF2
NUP84
SPT21
GTR1
HST3
HST4
HST2
ORC2
CDC7
NUP2
SWD1
SPP1
FUN30
WTM2
YRB2
ADA2
WTM1
ISW2
SAS2
RPD3
RAP1
SAS5
SAS3
YMR247C
SAS4
RXT2
DEP1
MCM7
PAF1
SIR2
DOT1
ADA2
RAD6
SDS3
MCM10
SCP160
RNA1
SAP30
PHO23
SIN3
PNC1
YPL216W
SCS22
YAP5
OAF1
MCM5
MCM3
RTF1
CDC6
ORC5
CDC45
HAT2
ITC1
ESC1
UME1
CTI6
DLS1
DIA2
HAT1
POL30
DOT6
SCS22
YAF9
HDA1
SDC1
SCS2
SWD3
BRE2
CTR9
IES3
NUP133
SPT10
Gene
Z_Shift_K_Gem
Z_Im_K_Gem
Z_Shift_K_Cyt
Z_Im_K_Cyt
Z_Shift_L_Gem
Z_Im_L_Gem
Z_Shift_L_Cyt
Z_Im_L_Cyt

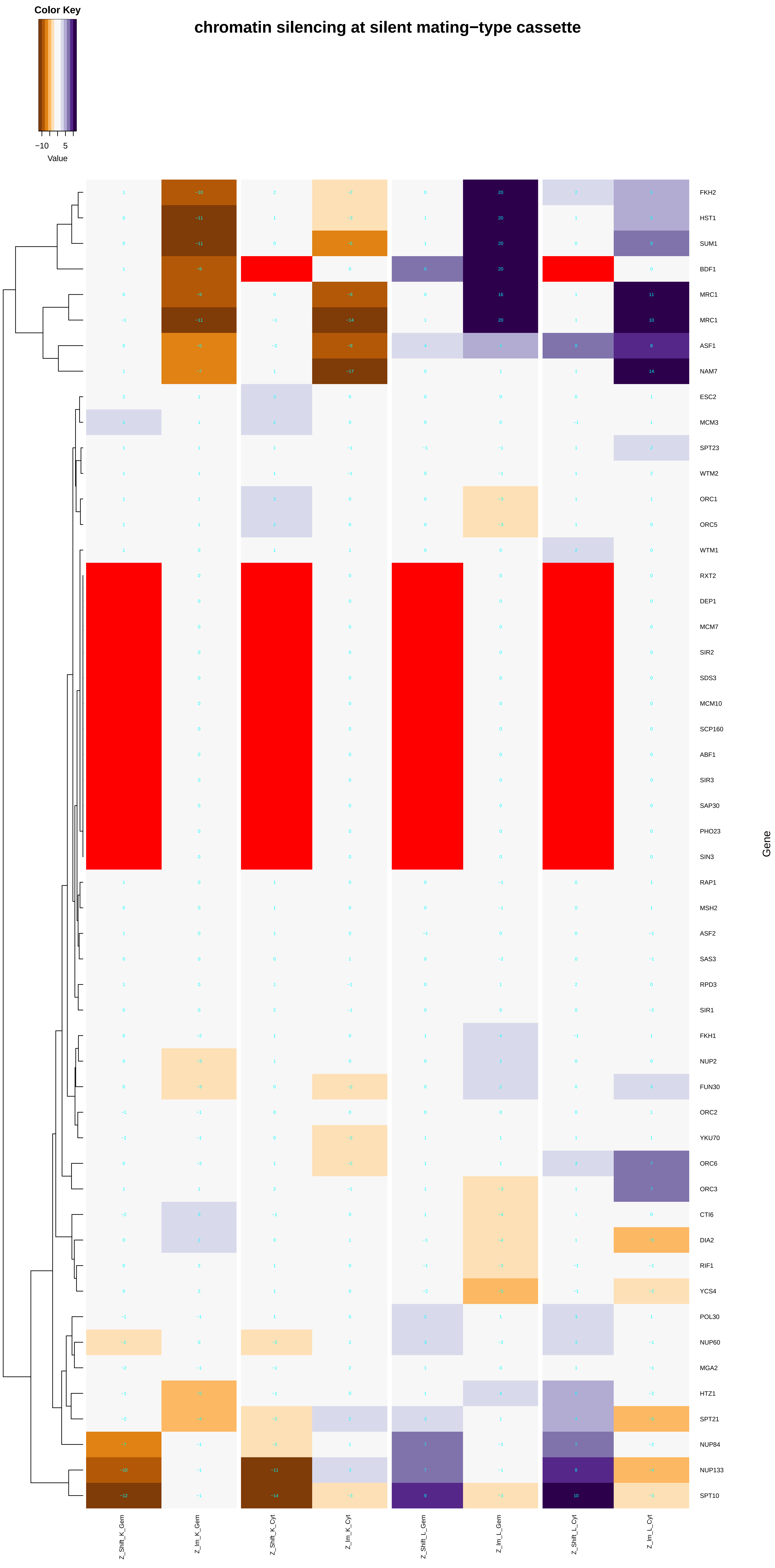

Color Key
−10 5
Value
chromatin silencing at silent mating−type cassette
FKH2
HST1
SUM1
BDF1
MRC1
MRC1
ASF1
NAM7
ESC2
MCM3
SPT23
WTM2
ORC1
ORC5
WTM1
RXT2
DEP1
MCM7
SIR2
SDS3
MCM10
SCP160
ABF1
SIR3
SAP30
PHO23
SIN3
RAP1
MSH2
ASF2
SAS3
RPD3
SIR1
FKH1
NUP2
FUN30
ORC2
YKU70
ORC6
ORC3
CTI6
DIA2
RIF1
YCS4
POL30
NUP60
MGA2
HTZ1
SPT21
NUP84
NUP133
SPT10
Gene
Z_Shift_K_Gem
Z_Im_K_Gem
Z_Shift_K_Cyt
Z_Im_K_Cyt
Z_Shift_L_Gem
Z_Im_L_Gem
Z_Shift_L_Cyt
Z_Im_L_Cyt

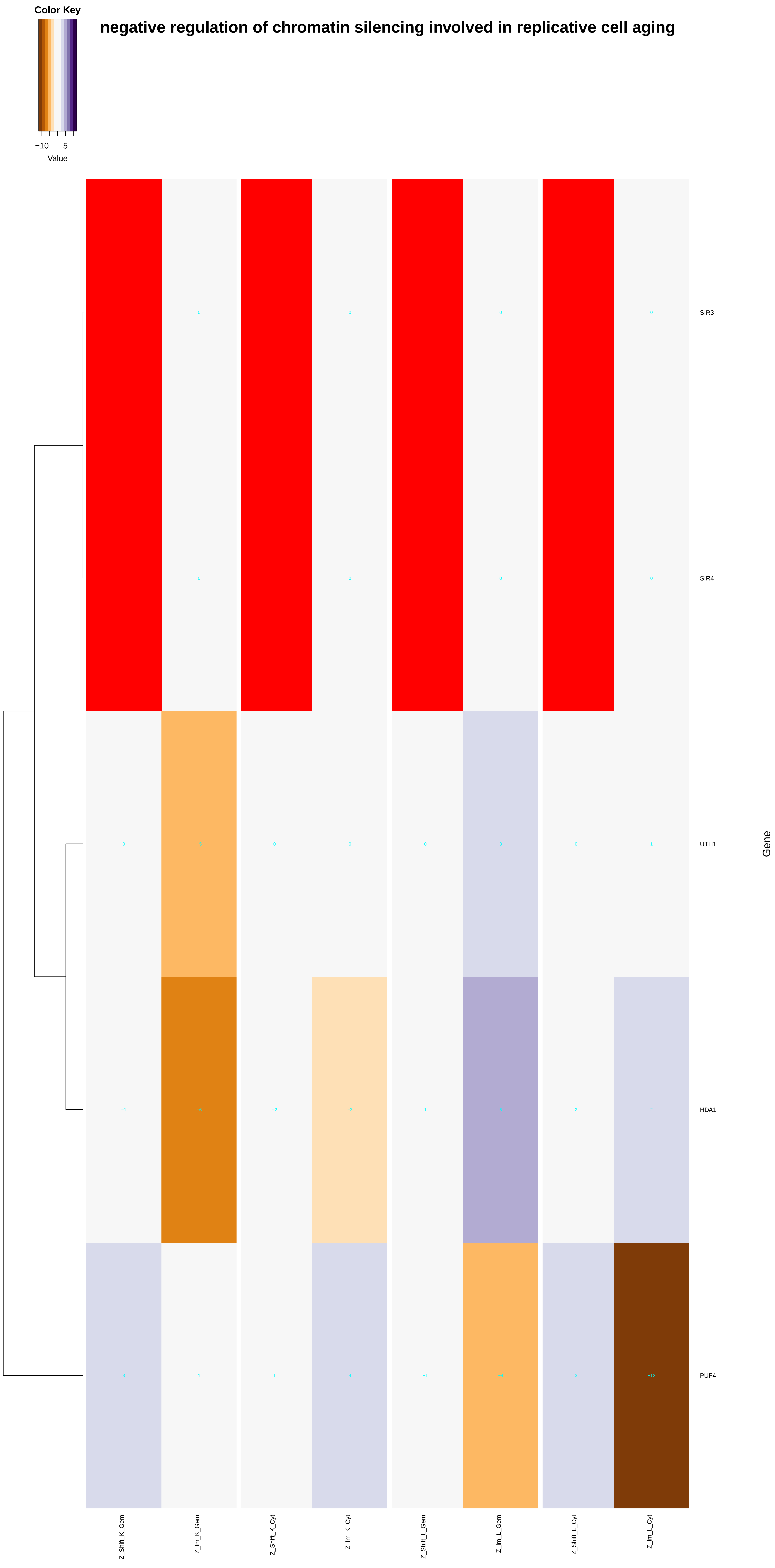
Color Key
−10
5
Value
negative regulation of chromatin silencing involved in replicative cell aging
SIR3
SIR4
UTH1
HDA1
PUF4
Gene
Z_Shift_K_Gem
Z_Im_K_Gem
Z_Shift_K_Cyt
Z_Im_K_Cyt
Z_Shift_L_Gem
Z_Im_L_Gem
Z_Shift_L_Cyt
Z_Im_L_Cyt

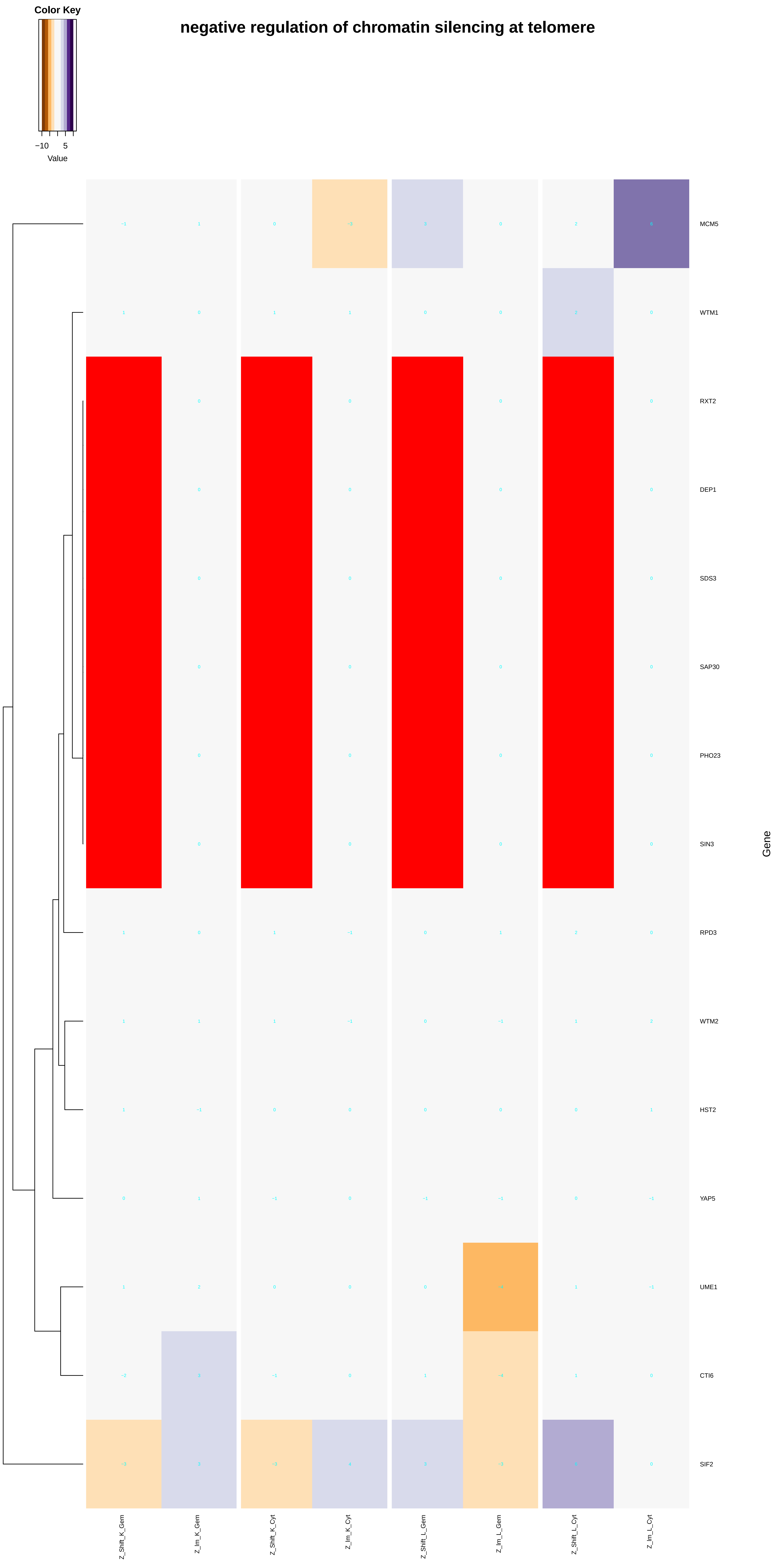

Color Key
−10
5
Value
negative regulation of chromatin silencing at telomere
MCM5
WTM1
RXT2
DEP1
SDS3
SAP30
PHO23
SIN3
RPD3
WTM2
HST2
YAP5
UME1
CTI6
SIF2
Gene
Z_Shift_K_Gem
Z_Im_K_Gem
Z_Shift_K_Cyt
Z_Im_K_Cyt
Z_Shift_L_Gem
Z_Im_L_Gem
Z_Shift_L_Cyt
Z_Im_L_Cyt

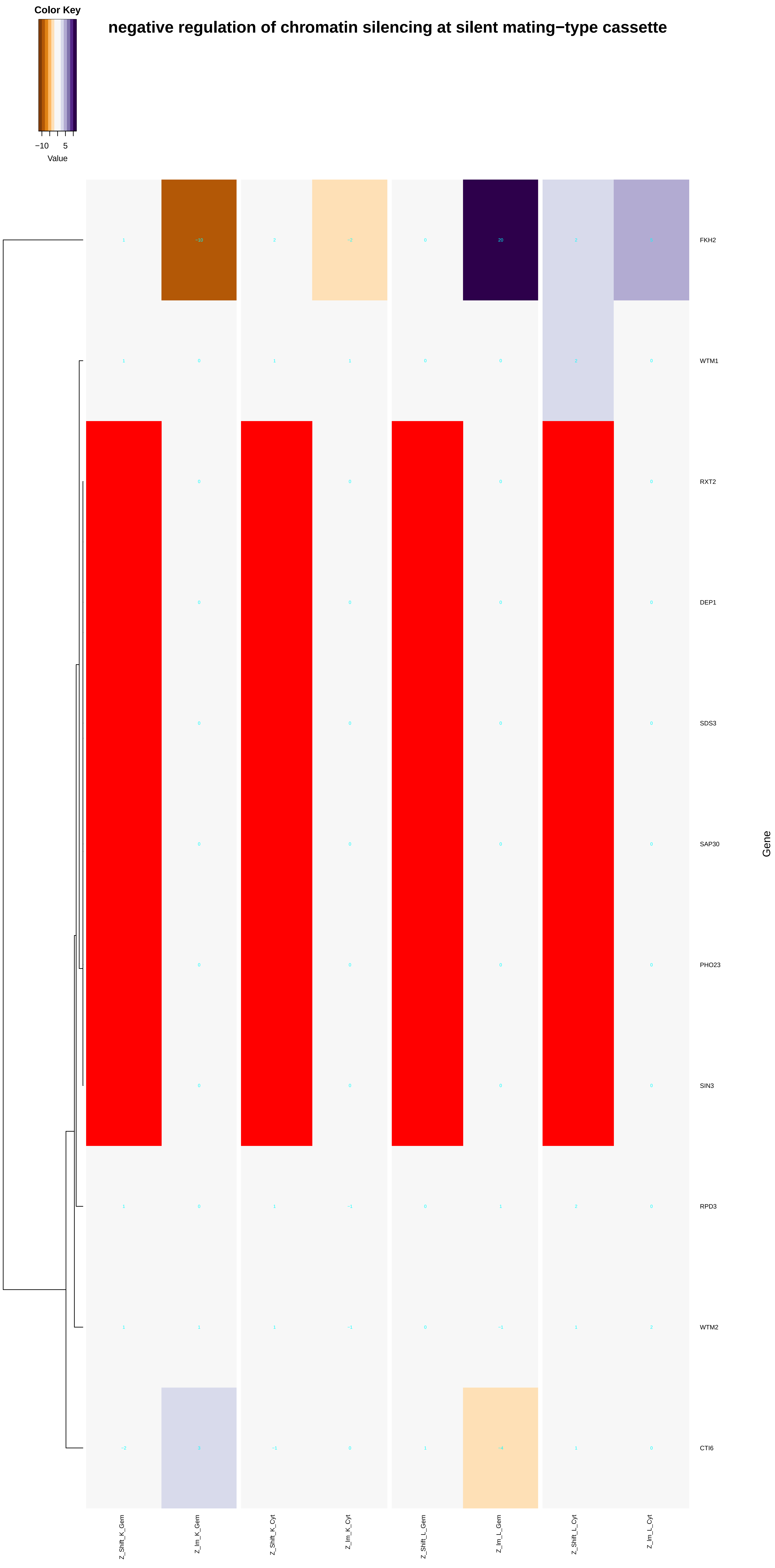
Color Key
−10
5
Value
negative regulation of chromatin silencing at silent mating−type cassette
FKH2
WTM1
RXT2
DEP1
SDS3
SAP30
PHO23
SIN3
RPD3
WTM2
CTI6
Gene
Z_Shift_K_Gem
Z_lm_K_Gem
Z_Shift_K_Cyt
Z_lm_K_Cyt
Z_Shift_L_Gem
Z_lm_L_Gem
Z_Shift_L_Cyt
Z_lm_L_Cyt

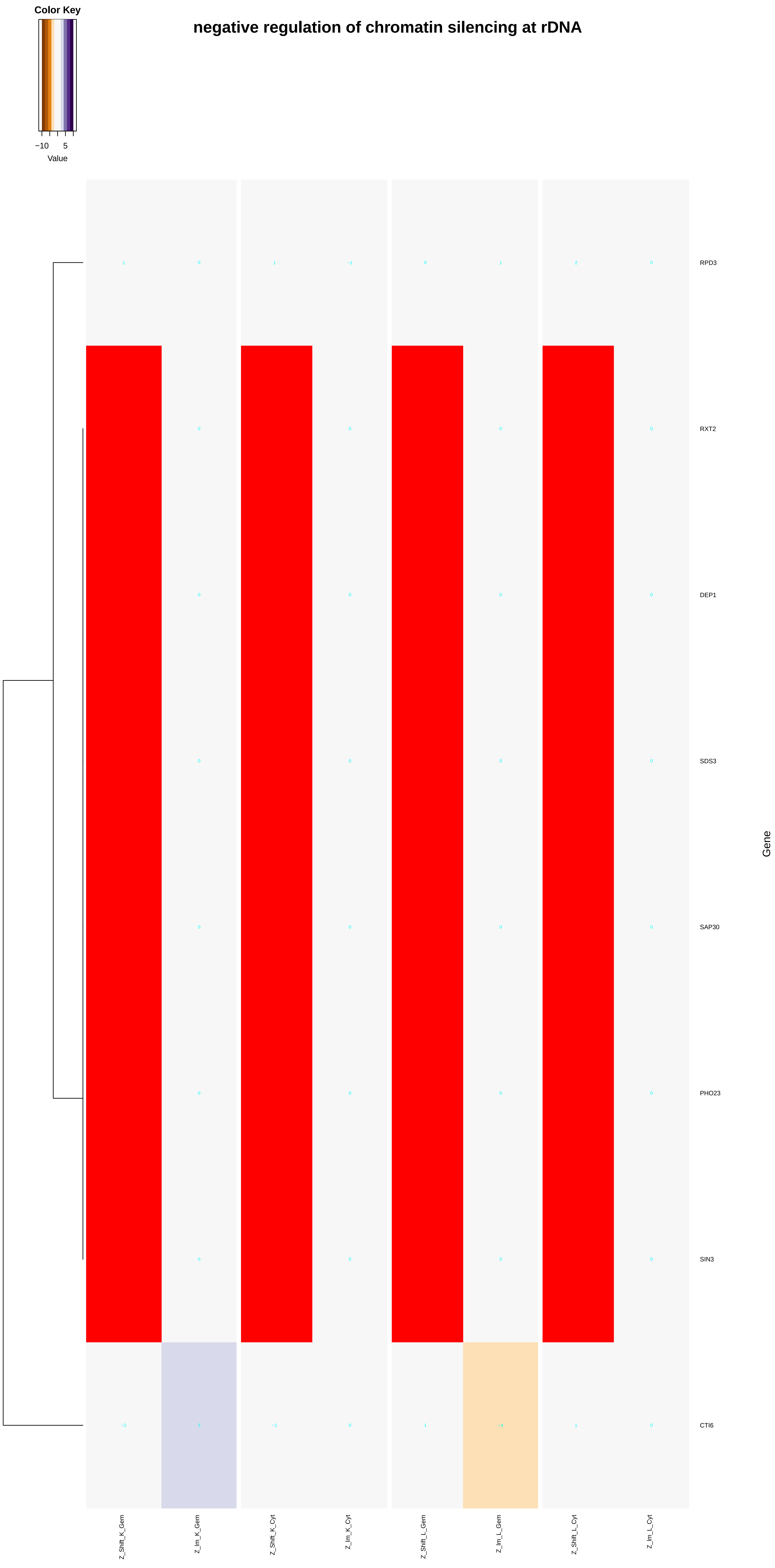
Color Key
−10
5
Value
negative regulation of chromatin silencing at rDNA
RPD3
RXT2
DEP1
SDS3
SAP30
PHO23
SIN3
CTI6
Gene
Z_Shift_K_Gem
Z_Im_K_Gem
Z_Shift_K_Cyt
Z_Im_K_Cyt
Z_Shift_L_Gem
Z_Im_L_Gem
Z_Shift_L_Cyt
Z_Im_L_Cyt

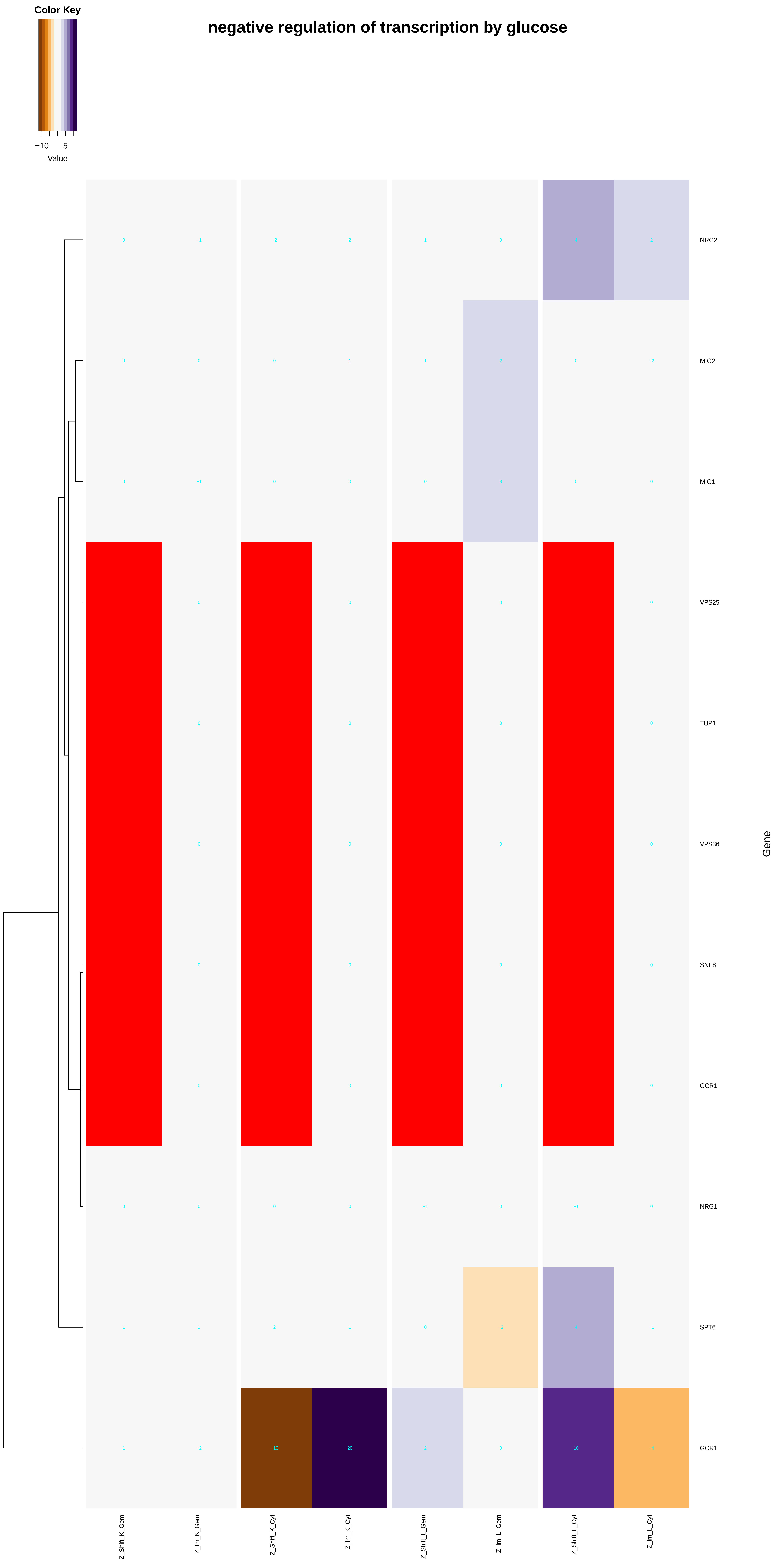
Color Key
−10
5
Value
negative regulation of transcription by glucose
NRG2
MIG2
MIG1
VPS25
TUP1
VPS36
SNF8
GCR1
NRG1
SPT6
GCR1
Gene
Z_Shift_K_Gem
Z_Im_K_Gem
Z_Shift_K_Cyt
Z_Im_K_Cyt
Z_Shift_L_Gem
Z_Im_L_Gem
Z_Shift_L_Cyt
Z_Im_L_Cyt

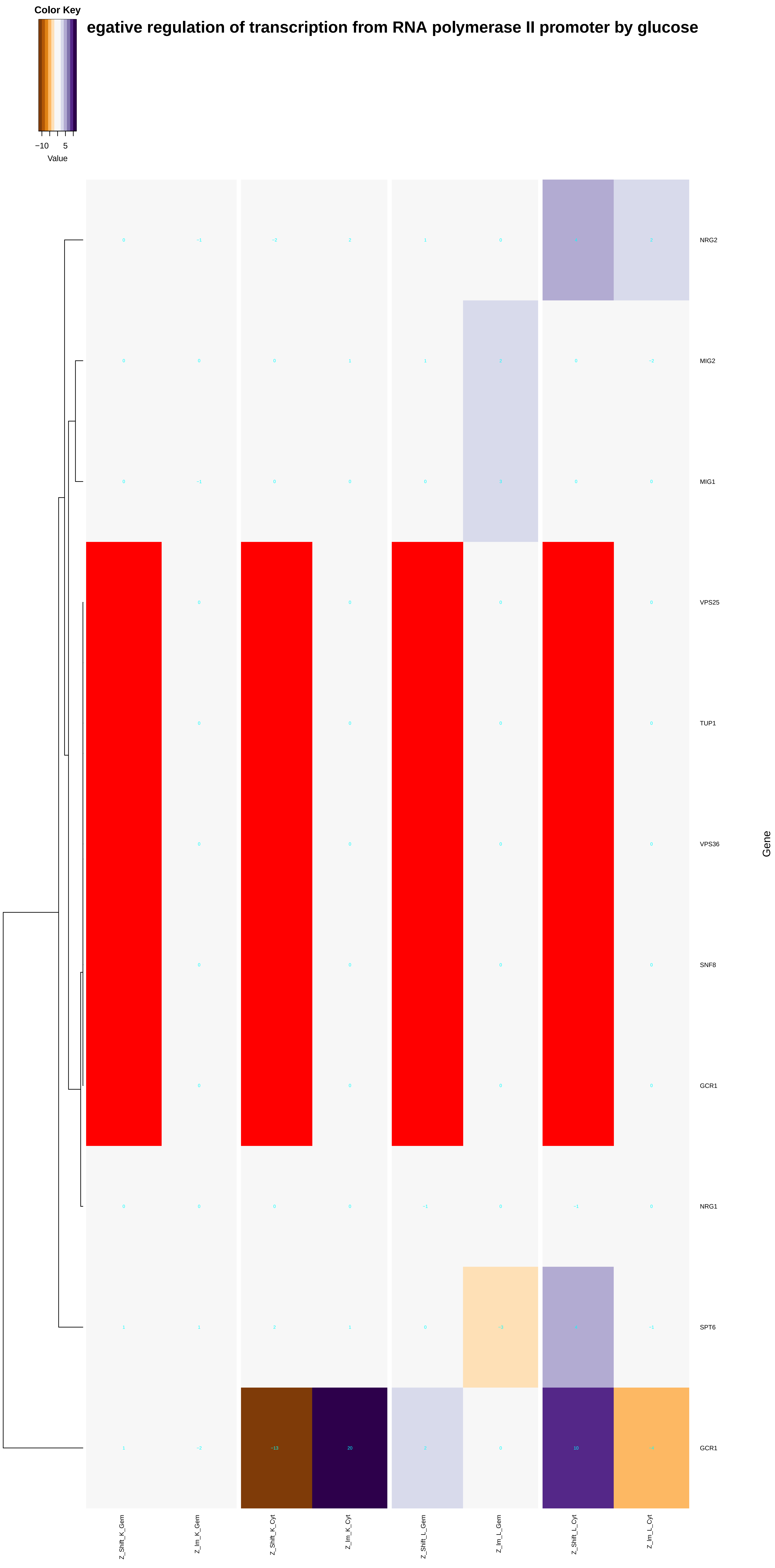

Color Key
−10
5
Value
egative regulation of transcription from RNA polymerase II promoter by glucose
NRG2
MIG2
MIG1
VPS25
TUP1
VPS36
SNF8
GCR1
NRG1
SPT6
GCR1
Gene
Z_Shift_K_Gem
Z_Im_K_Gem
Z_Shift_K_Cyt
Z_Im_K_Cyt
Z_Shift_L_Gem
Z_Im_L_Gem
Z_Shift_L_Cyt
Z_Im_L_Cyt

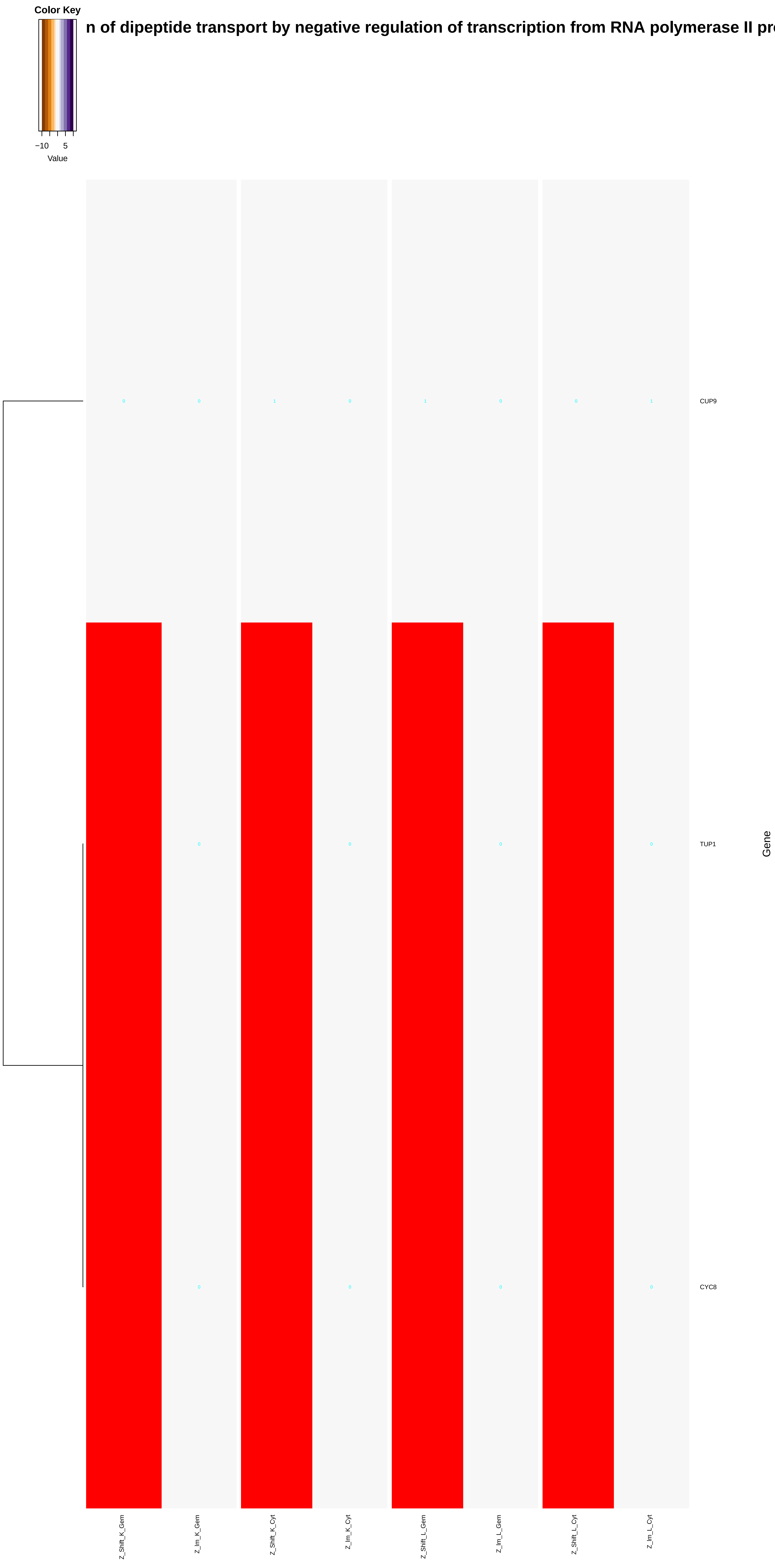

Color Key
−10
5
Value
n of dipeptide transport by negative regulation of transcription from RNA polymerase II pro
CUP9
TUP1
CYC8
Gene
Z_Shift_K_Gem
Z_Im_K_Gem
Z_Shift_K_Cyt
Z_Im_K_Cyt
Z_Shift_L_Gem
Z_Im_L_Gem
Z_Shift_L_Cyt
Z_Im_L_Cyt

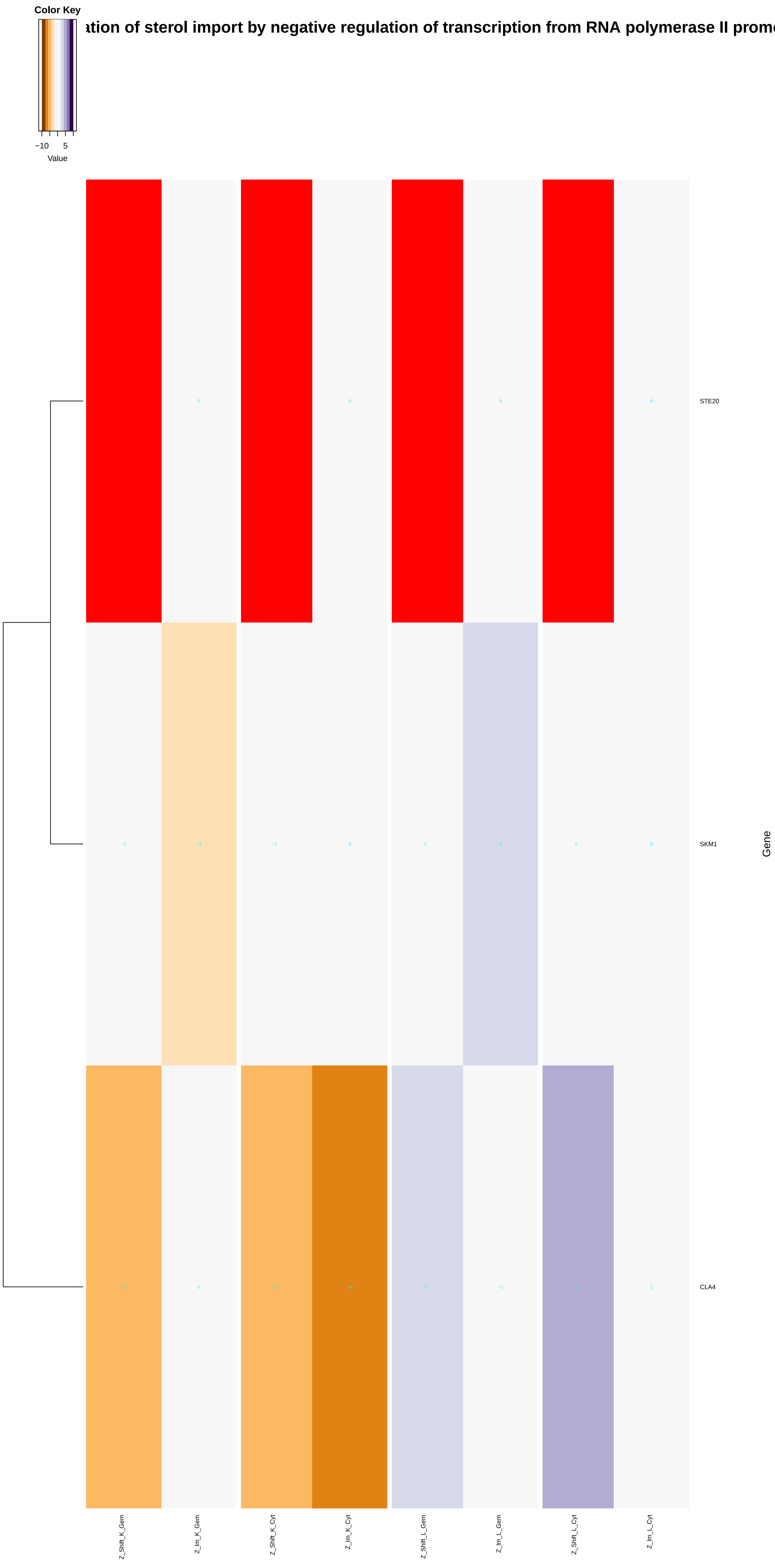

Color Key
−10
5
Value
ation of sterol import by negative regulation of transcription from RNA polymerase II promo
STE20
SKM1
CLA4
Gene
Z_Shift_K_Gem
Z_Im_K_Gem
Z_Shift_K_Cyt
Z_Im_K_Cyt
Z_Shift_L_Gem
Z_Im_L_Gem
Z_Shift_L_Cyt
Z_Im_L_Cyt
