## Additional File 7 for "A humanized yeast phenomic model of deoxycytidine kinase to predict genetic buffering of nucleoside analog cytotoxicity": negative_regulation_of_transcription_from_RNA_polymerase_II_promoter.pdf

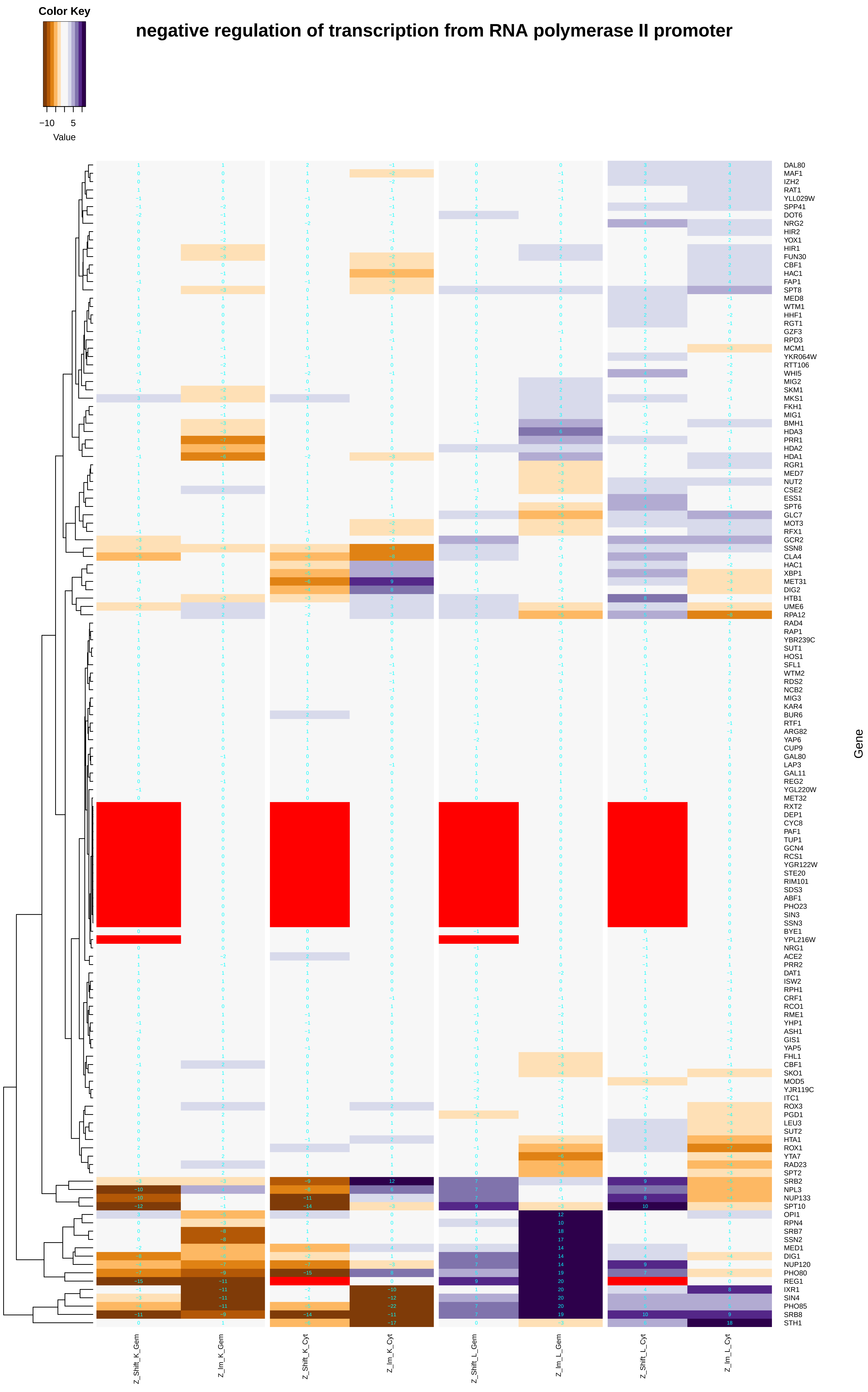

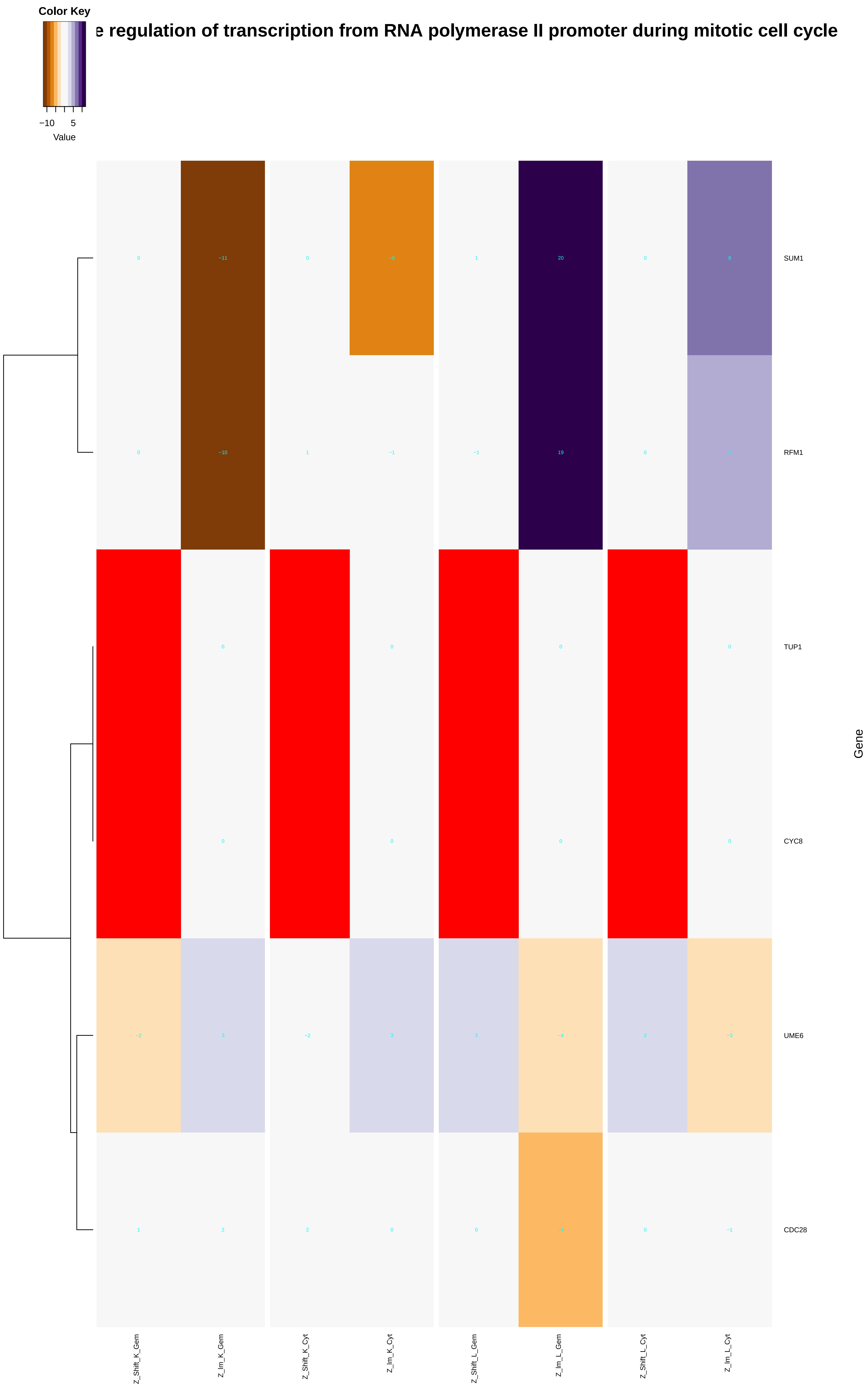

Color Key
−10
5
Value
e regulation of transcription from RNA polymerase II promoter during mitotic cell cycle
SUM1
RFM1
TUP1
CYC8
UME6
CDC28
Gene
Z_Shift_K_Gem
Z_Im_K_Gem
Z_Shift_K_Cyt
Z_Im_K_Cyt
Z_Shift_L_Gem
Z_Im_L_Gem
Z_Shift_L_Cyt
Z_Im_L_Cyt

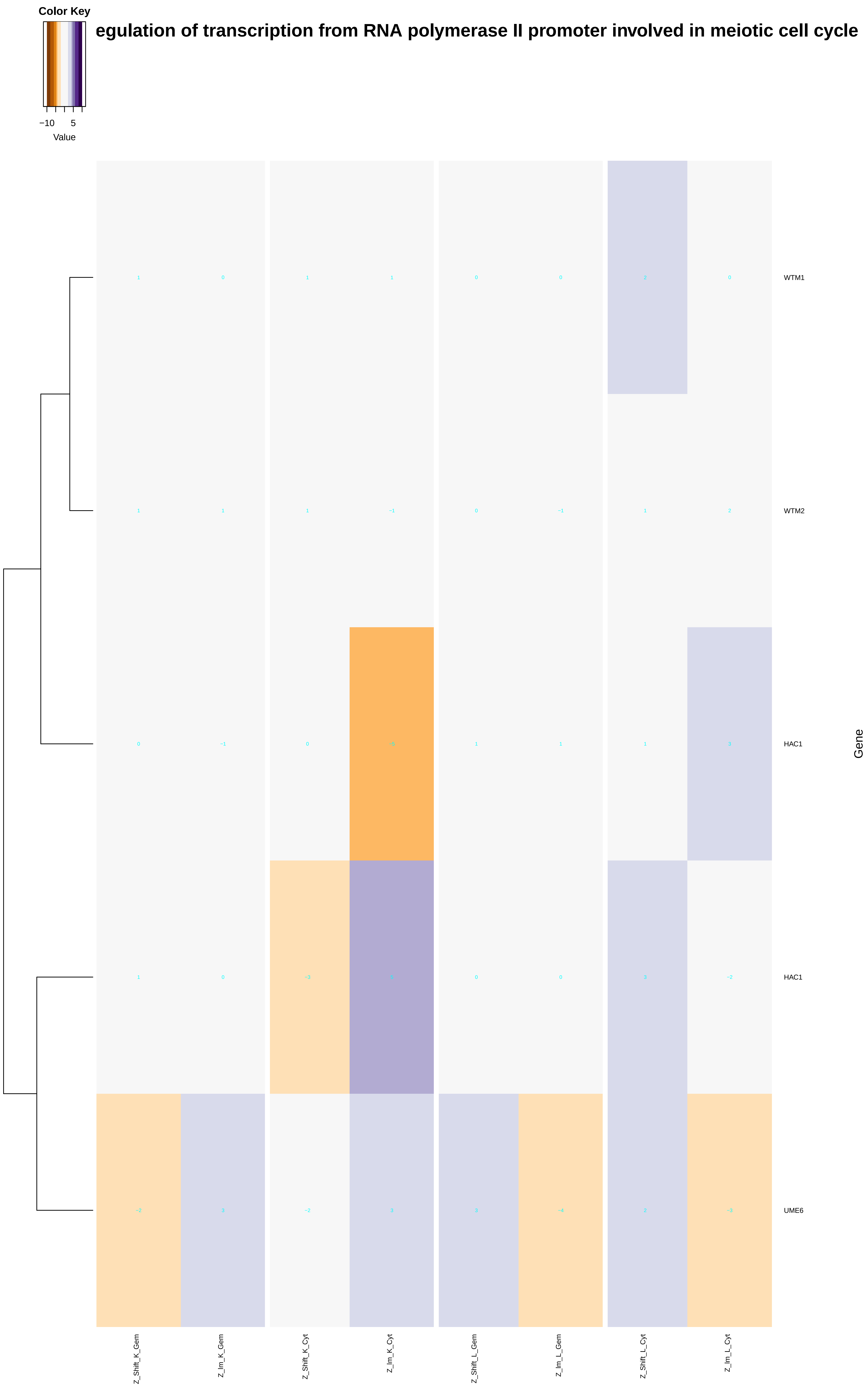
Color Key
−10
5
Value
egulation of transcription from RNA polymerase II promoter involved in meiotic cell cycle
WTM1
WTM2
HAC1
HAC1
UME6
Gene
Z_Shift_K_Gem
Z_Im_K_Gem
Z_Shift_K_Cyt
Z_Im_K_Cyt
Z_Shift_L_Gem
Z_Im_L_Gem
Z_Shift_L_Cyt
Z_Im_L_Cyt

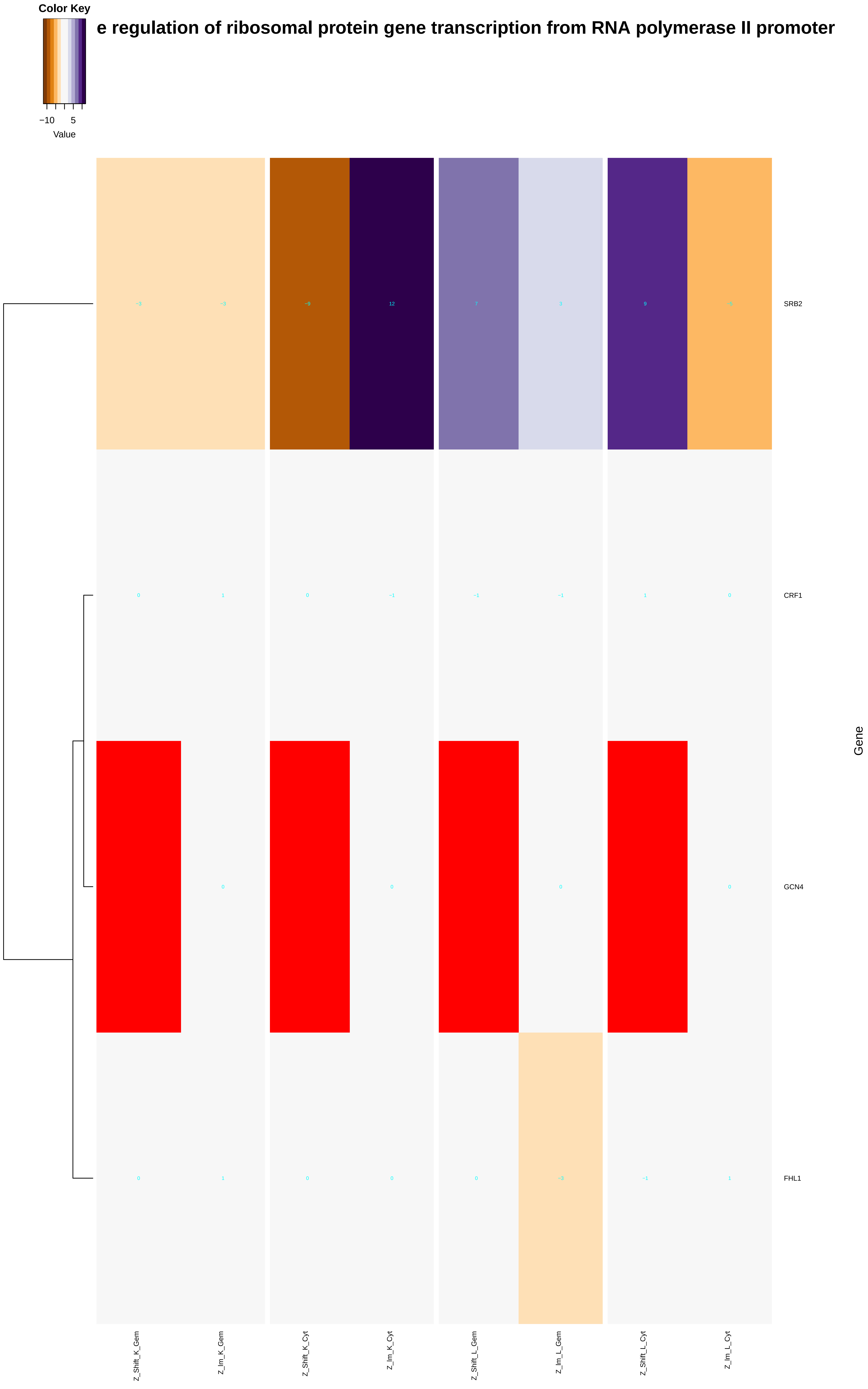

e regulation of ribosomal protein gene transcription from RNA polymerase II promoter
Color Key
−10
5
Value
SRB2
CRF1
GCN4
FHL1
Gene
Z_Shift_K_Gem
Z_Im_K_Gem
Z_Shift_K_Cyt
Z_Im_K_Cyt
Z_Shift_L_Gem
Z_Im_L_Gem
Z_Shift_L_Cyt
Z_Im_L_Cyt

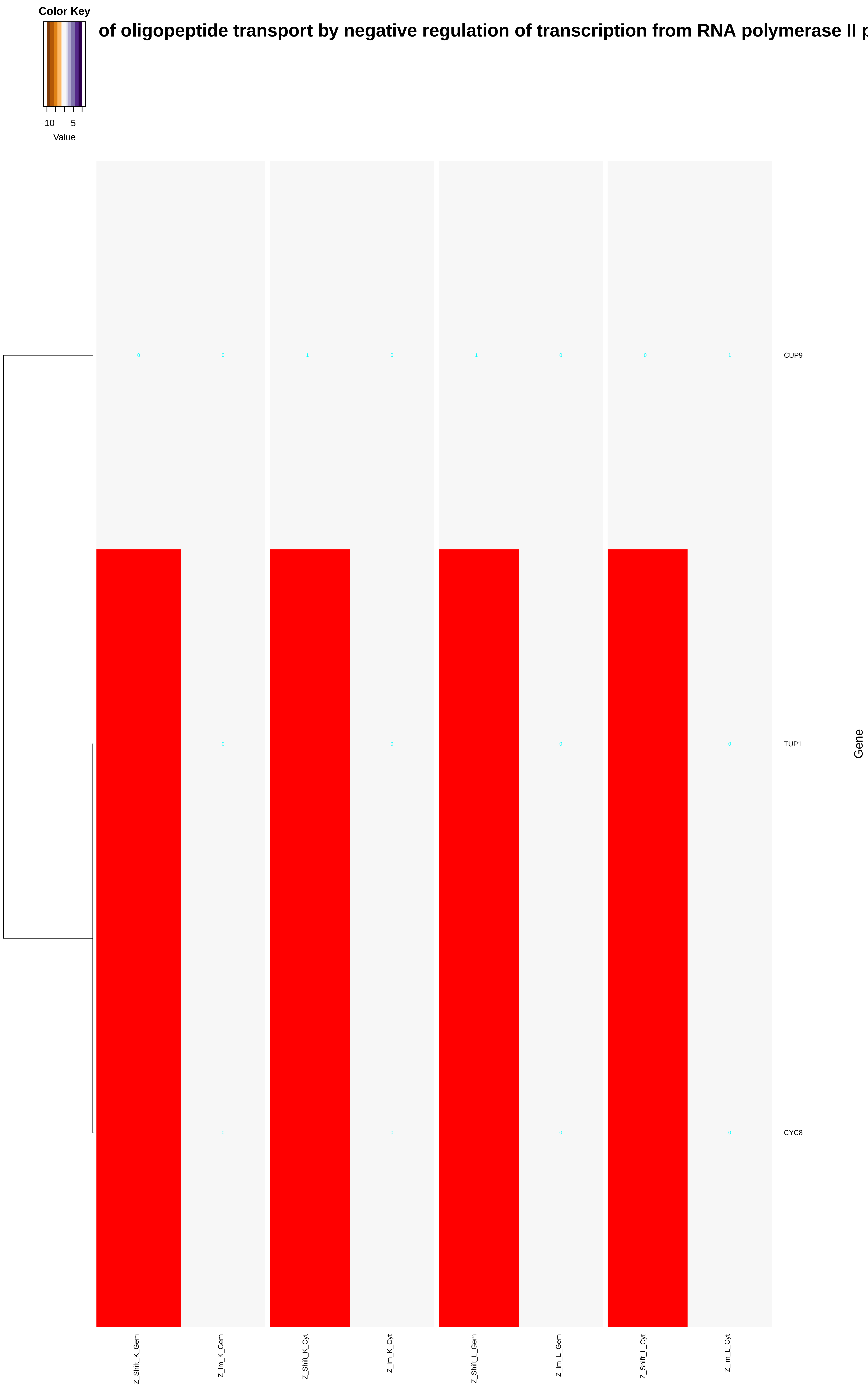

Color Key
−10
5
Value
of oligopeptide transport by negative regulation of transcription from RNA polymerase II p
0
0
1
0
1
0
0
1
CUP9
0
0
0
0
TUP1
0
0
0
0
CYC8
Gene
Z_Shift_K_Gem
Z_Im_K_Gem
Z_Shift_K_Cyt
Z_Im_K_Cyt
Z_Shift_L_Gem
Z_Im_L_Gem
Z_Shift_L_Cyt
Z_Im_L_Cyt

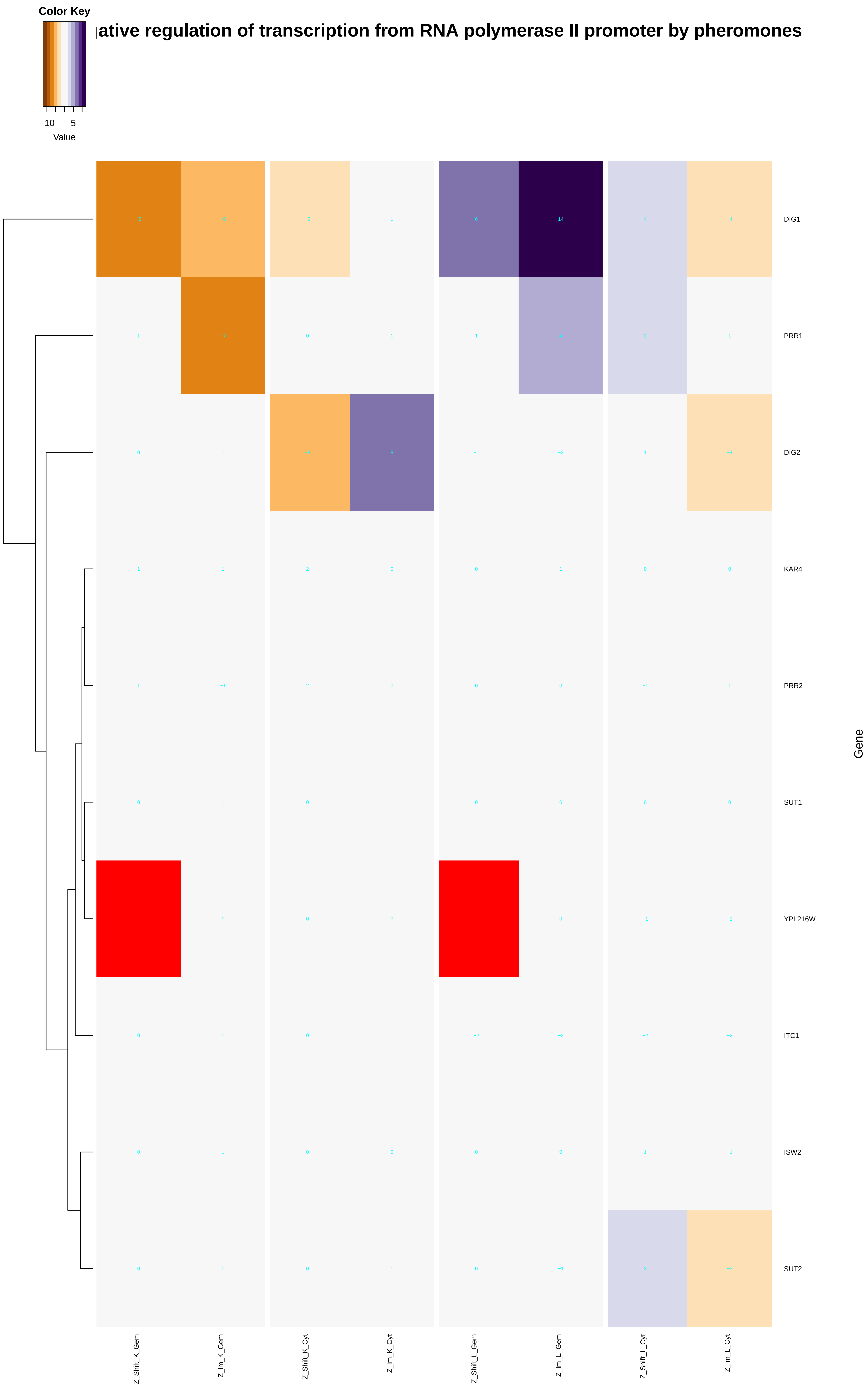
Color Key
−10
5
Value
ative regulation of transcription from RNA polymerase II promoter by pheromones
DIG1
PRR1
DIG2
KAR4
PRR2
SUT1
YPL216W
ITC1
ISW2
SUT2
Gene
Z_Shift_K_Gem
Z_Im_K_Gem
Z_Shift_K_Cyt
Z_Im_K_Cyt
Z_Shift_L_Gem
Z_Im_L_Gem
Z_Shift_L_Cyt
Z_Im_L_Cyt

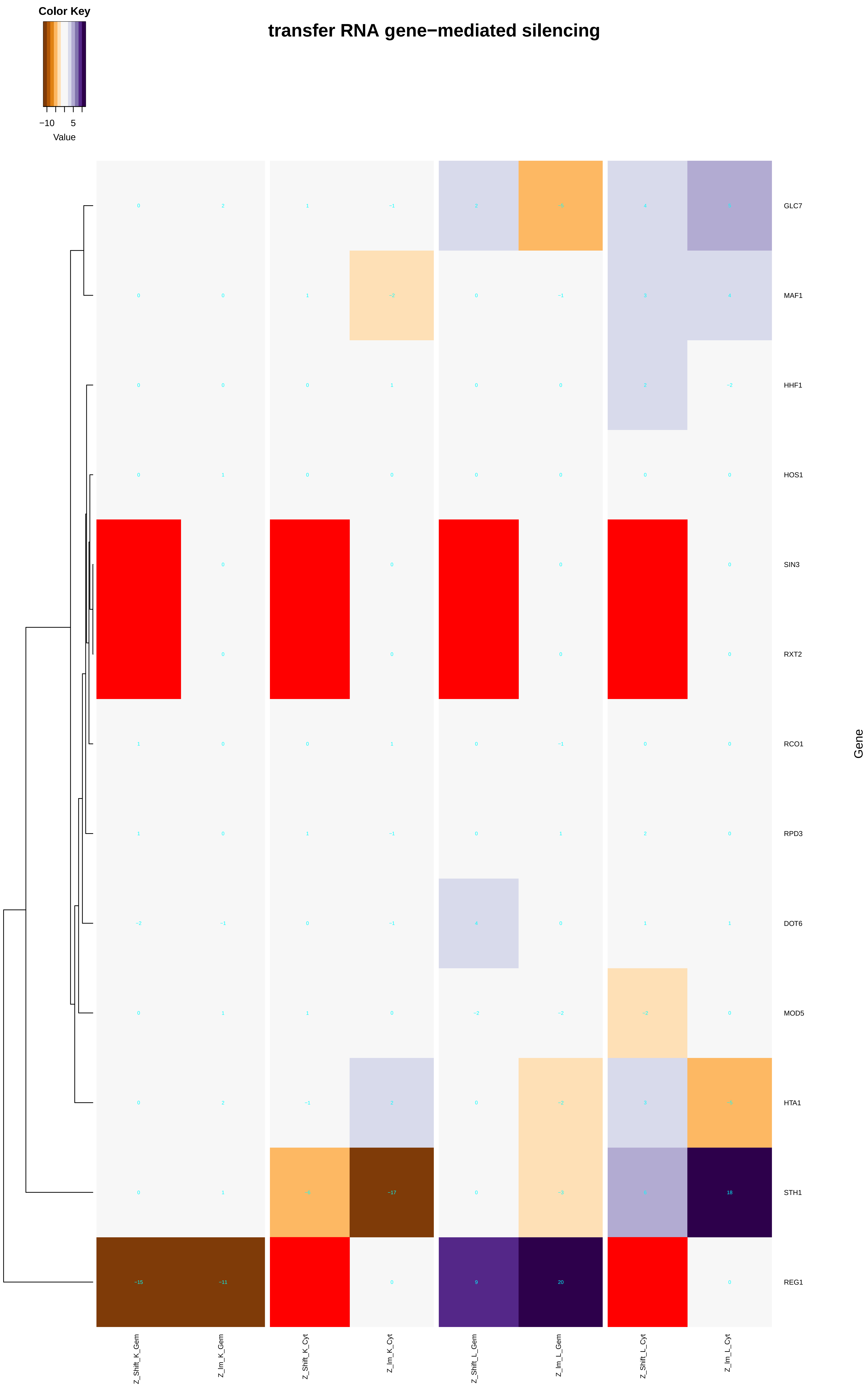

Color Key
−10
5
Value
transfer RNA gene−mediated silencing
GLC7
MAF1
HHF1
HOS1
SIN3
RXT2
RCO1
RPD3
DOT6
MOD5
HTA1
STH1
REG1
Gene
Z_Shift_K_Gem
Z_lm_K_Gem
Z_Shift_K_Cyt
Z_lm_K_Cyt
Z_Shift_L_Gem
Z_lm_L_Gem
Z_Shift_L_Cyt
Z_lm_L_Cyt

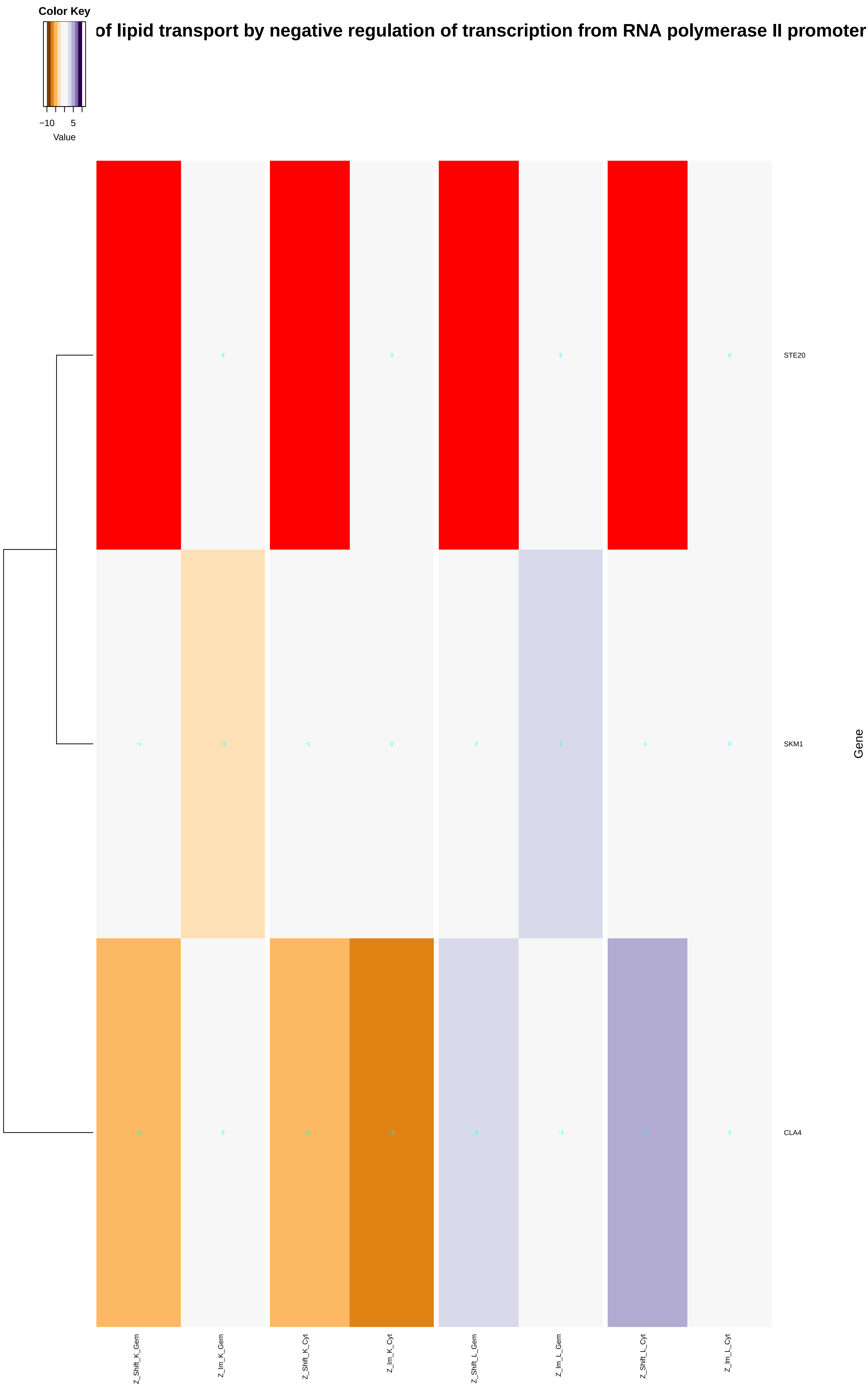
Color Key
−10
5
Value
of lipid transport by negative regulation of transcription from RNA polymerase II promoter
0
0
0
0
STE20
−1
−2
−1
0
2
2
1
0
SKM1
Gene
−5
0
−6
−8
3
−1
5
2
CLA4
Z_Shift_K_Gem
Z_Im_K_Gem
Z_Shift_K_Cyt
Z_Im_K_Cyt
Z_Shift_L_Gem
Z_Im_L_Gem
Z_Shift_L_Cyt
Z_Im_L_Cyt

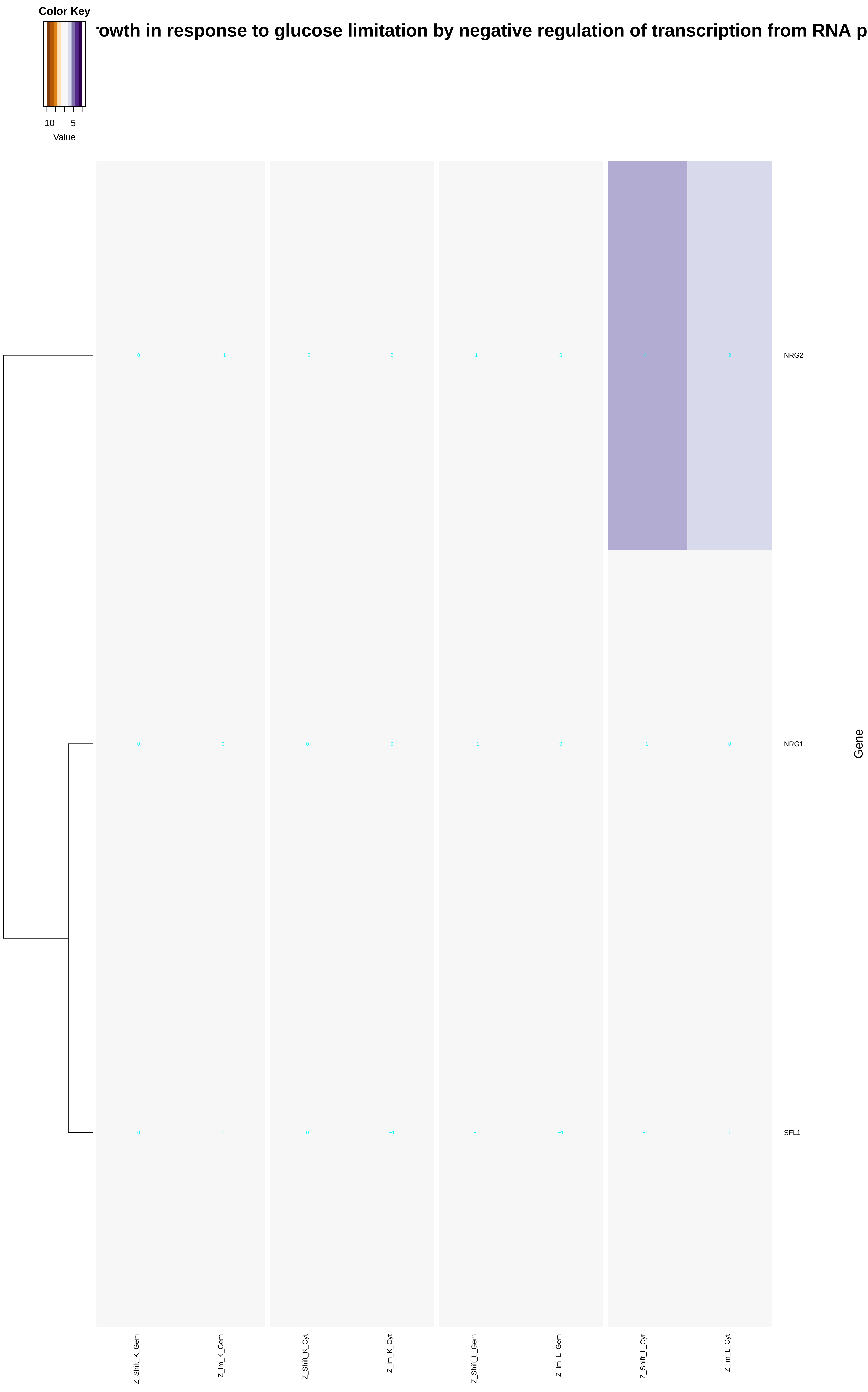

rowth in response to glucose limitation by negative regulation of transcription from RNA p
Color Key
−10
5
Value
NRG2
0 −1 −2 2 1 0 2
NRG1
0 0 0 0 −1 0 −1 0
SFL1
0 0 0 −1 −1 −1 −1 1
Gene
Z_Shift_K_Gem
Z_Im_K_Gem
Z_Shift_K_Cyt
Z_Im_K_Cyt
Z_Shift_L_Gem
Z_Im_L_Gem
Z_Shift_L_Cyt
Z_Im_L_Cyt

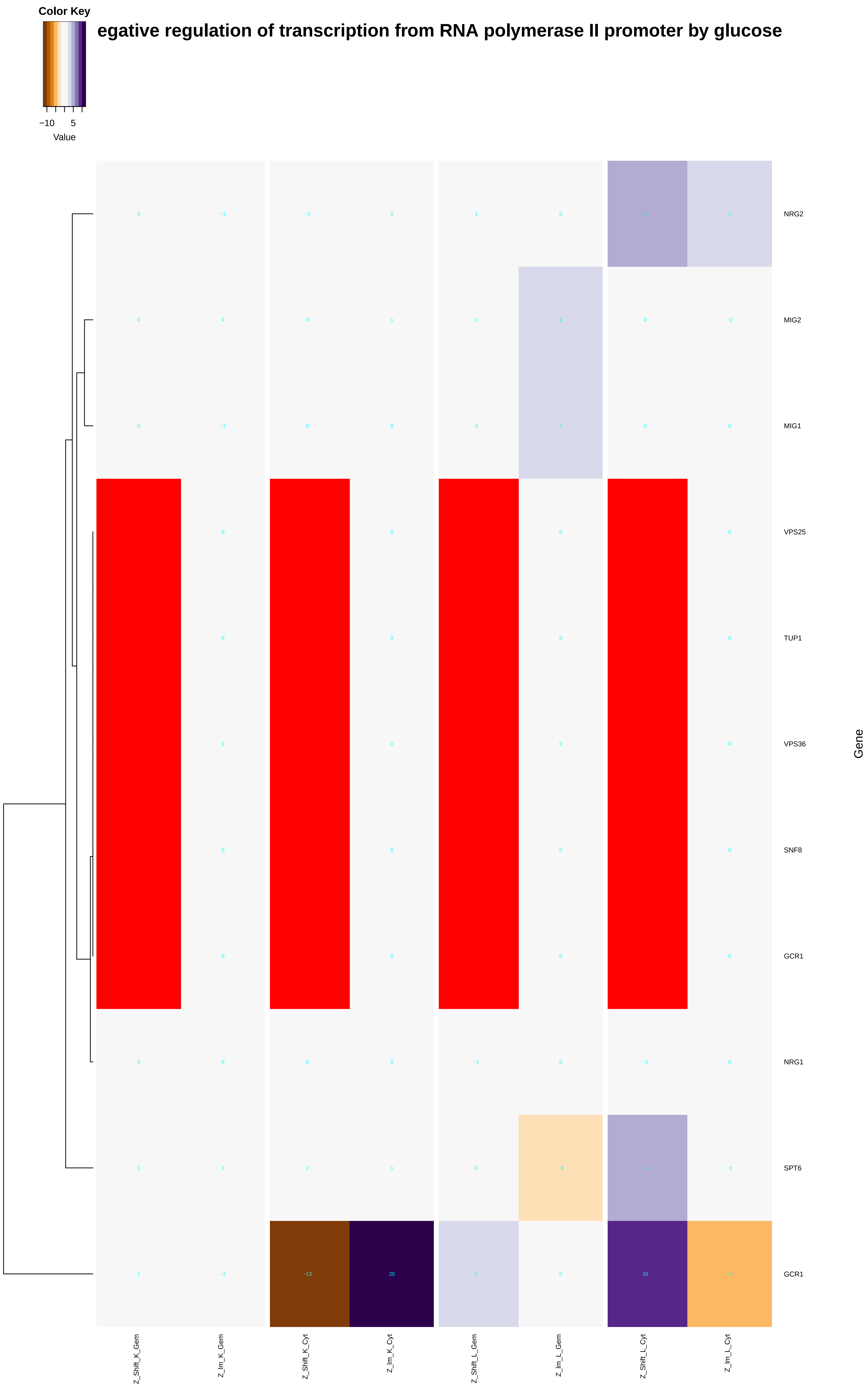

Color Key
−10
5
Value
egative regulation of transcription from RNA polymerase II promoter by glucose
NRG2
MIG2
MIG1
VPS25
TUP1
VPS36
SNF8
GCR1
NRG1
SPT6
GCR1
Gene
Z_Shift_K_Gem
Z_Im_K_Gem
Z_Shift_K_Cyt
Z_Im_K_Cyt
Z_Shift_L_Gem
Z_Im_L_Gem
Z_Shift_L_Cyt
Z_Im_L_Cyt

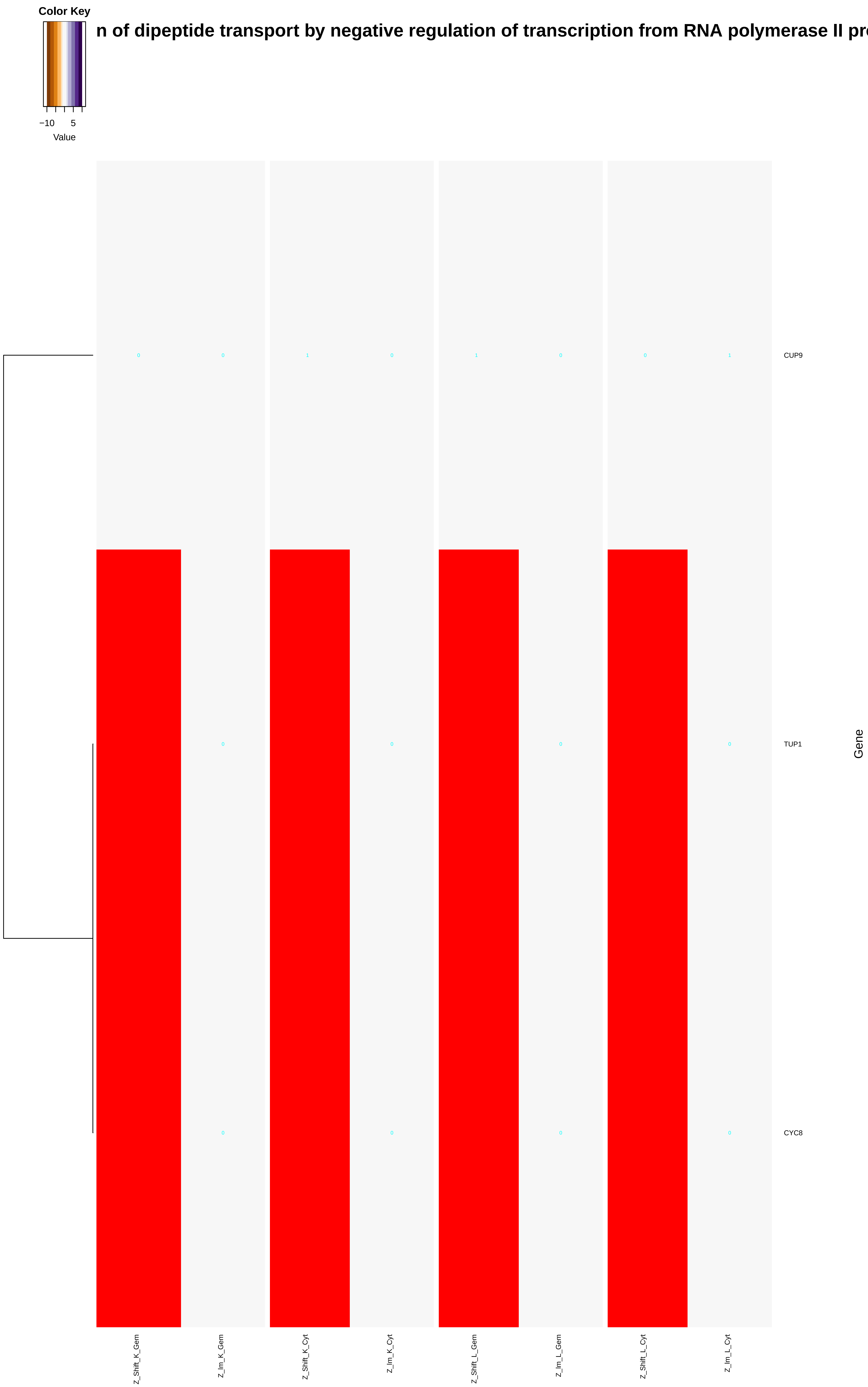
Color Key
−10
5
Value
n of dipeptide transport by negative regulation of transcription from RNA polymerase II pr
CUP9
TUP1
CYC8
Gene
Z_Shift_K_Gem
Z_Im_K_Gem
Z_Shift_K_Cyt
Z_Im_K_Cyt
Z_Shift_L_Gem
Z_Im_L_Gem
Z_Shift_L_Cyt
Z_Im_L_Cyt

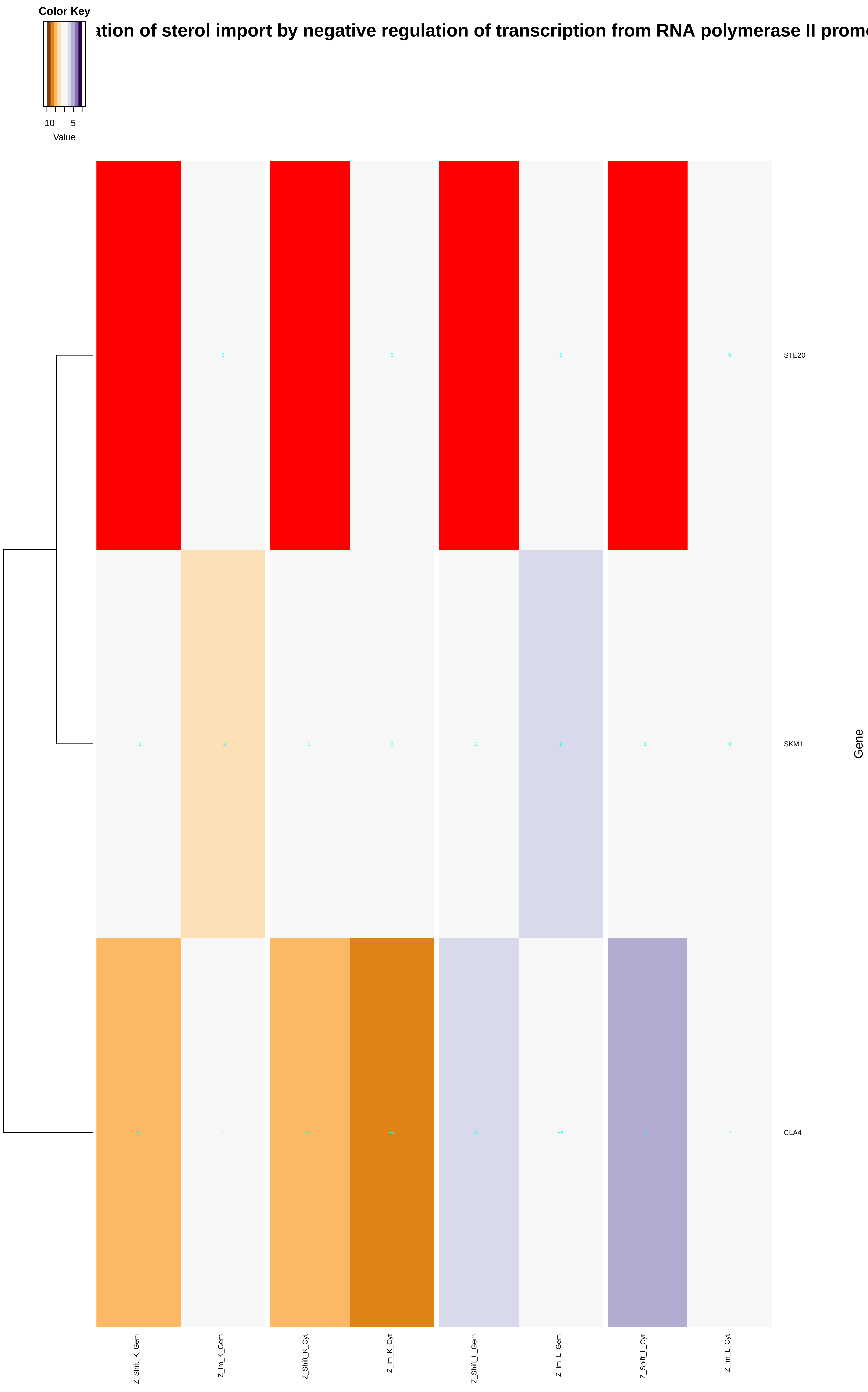

ation of sterol import by negative regulation of transcription from RNA polymerase II prom
Color Key
−10
5
Value
STE20
SKM1
CLA4
Gene
Z_Shift_K_Gem
Z_Im_K_Gem
Z_Shift_K_Cyt
Z_Im_K_Cyt
Z_Shift_L_Gem
Z_Im_L_Gem
Z_Shift_L_Cyt
Z_Im_L_Cyt
