## Supplementary figures and images for "A humanized yeast phenomic model of deoxycytidine kinase to predict genetic buffering of nucleoside analog cytotoxicity"

### histone-lysine_N-methyltransferase_activity.pdf

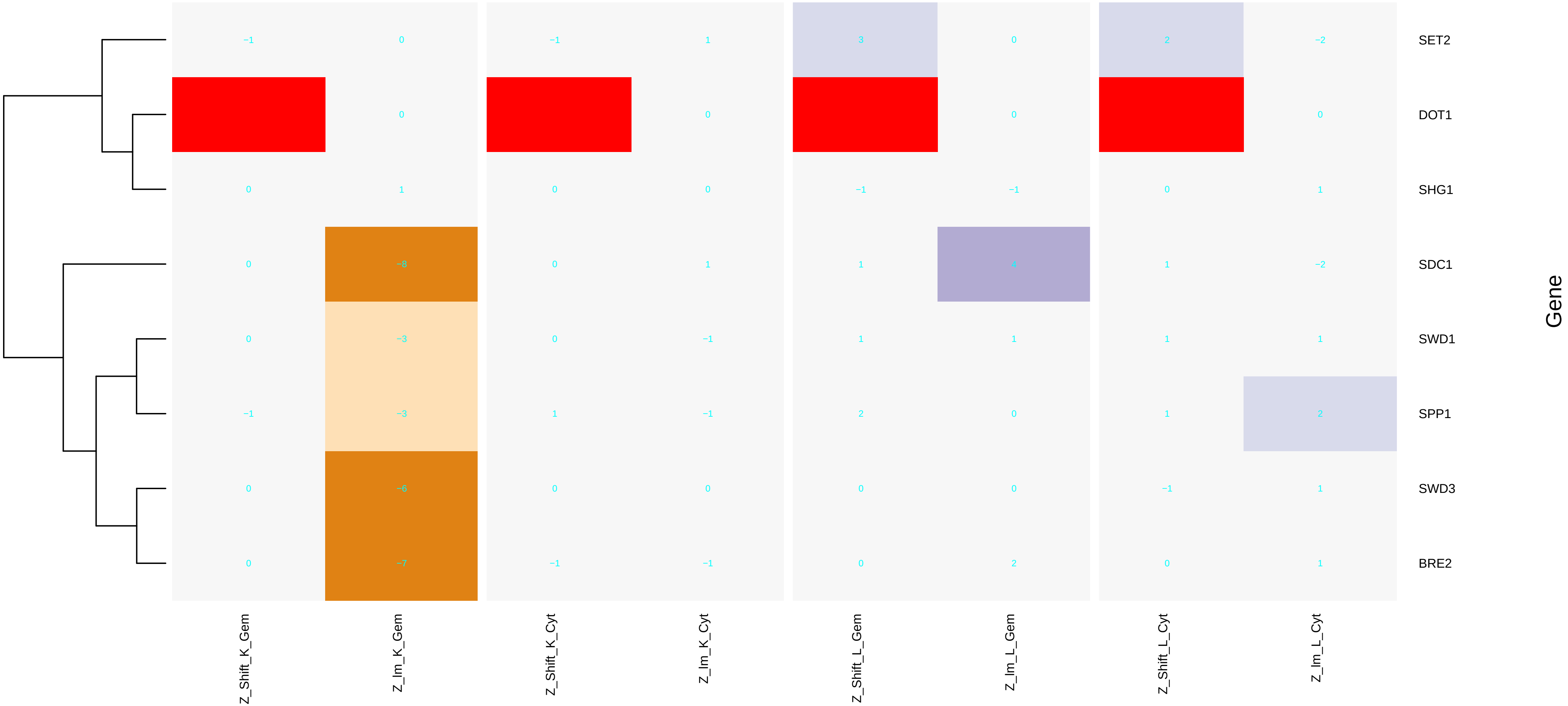

### histone_ubiquitination.pdf

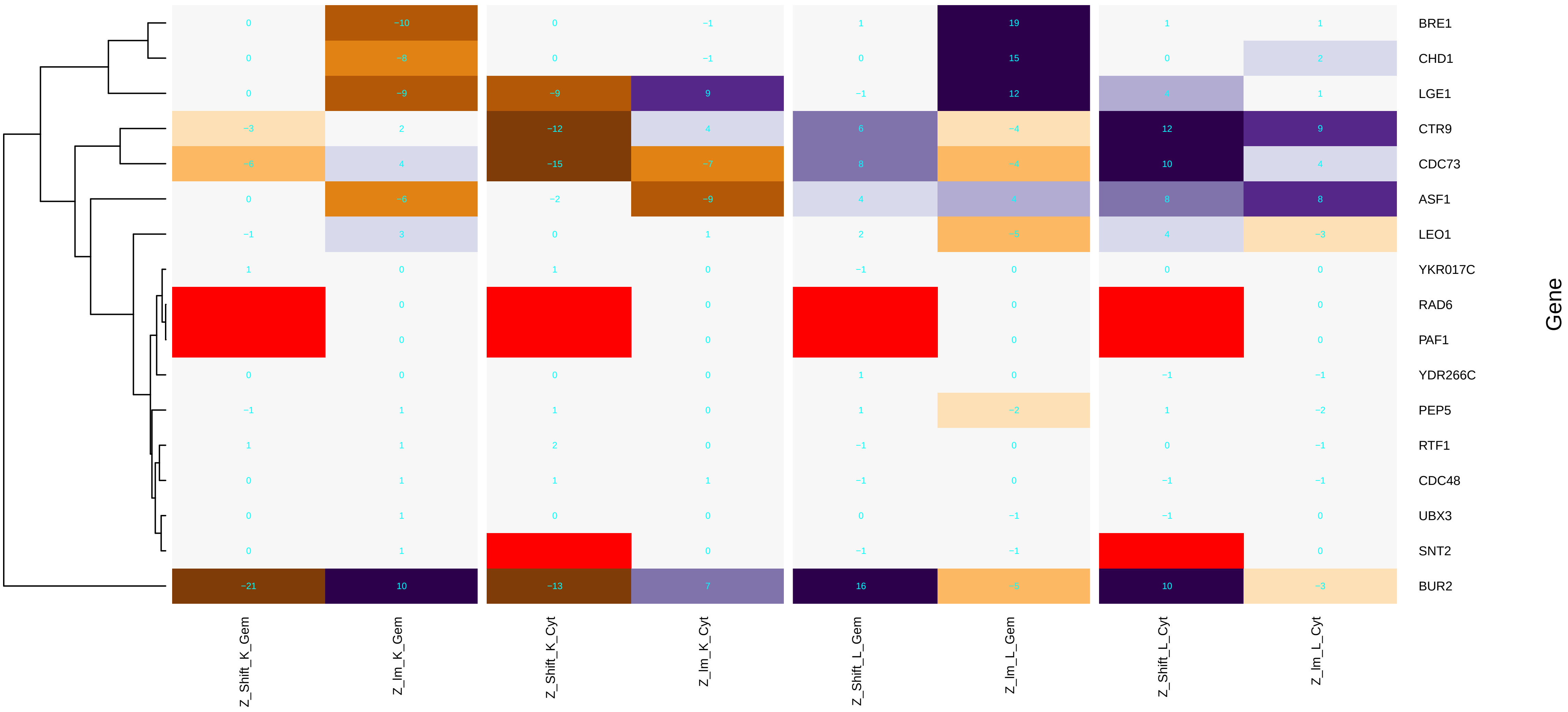

### holo_TFIIH_complex.pdf

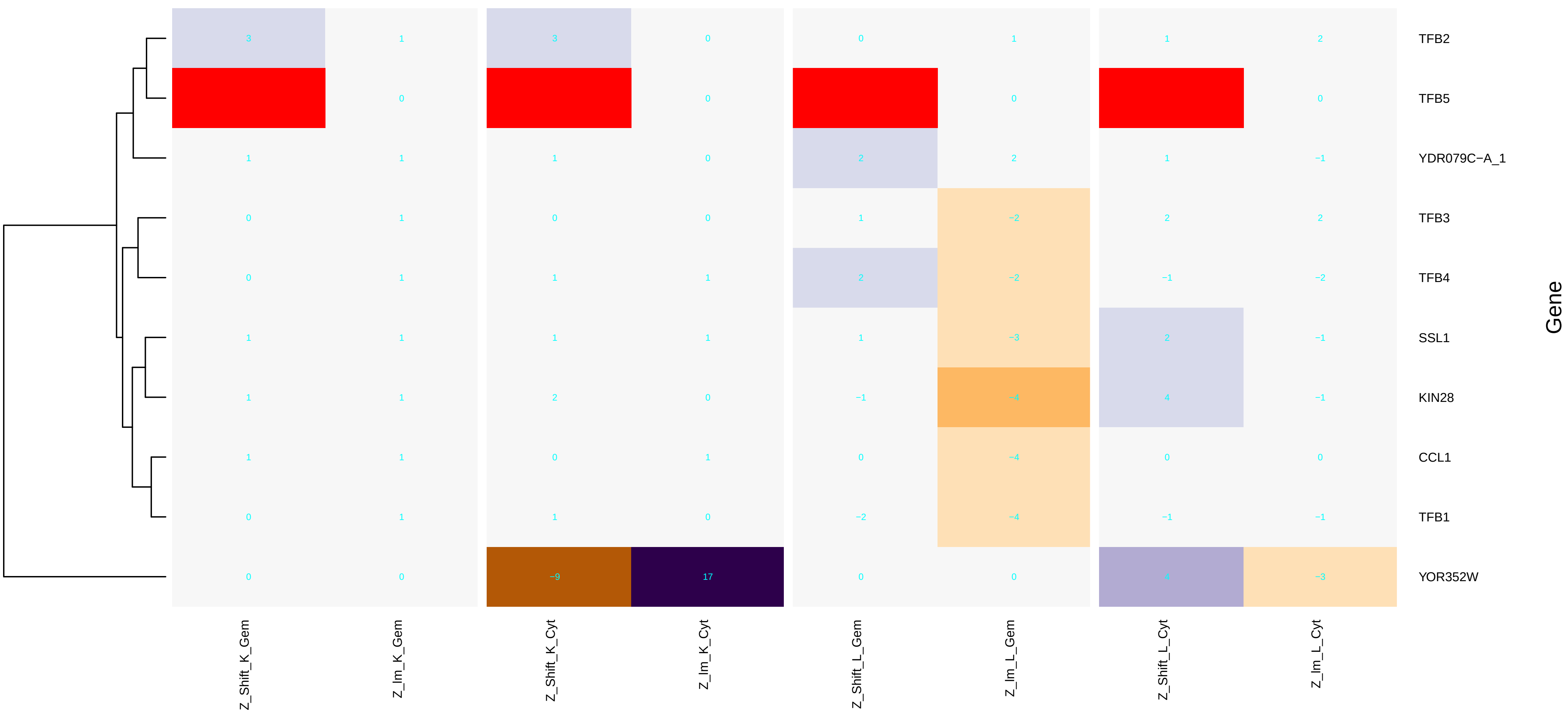

### HOPS_complex.pdf

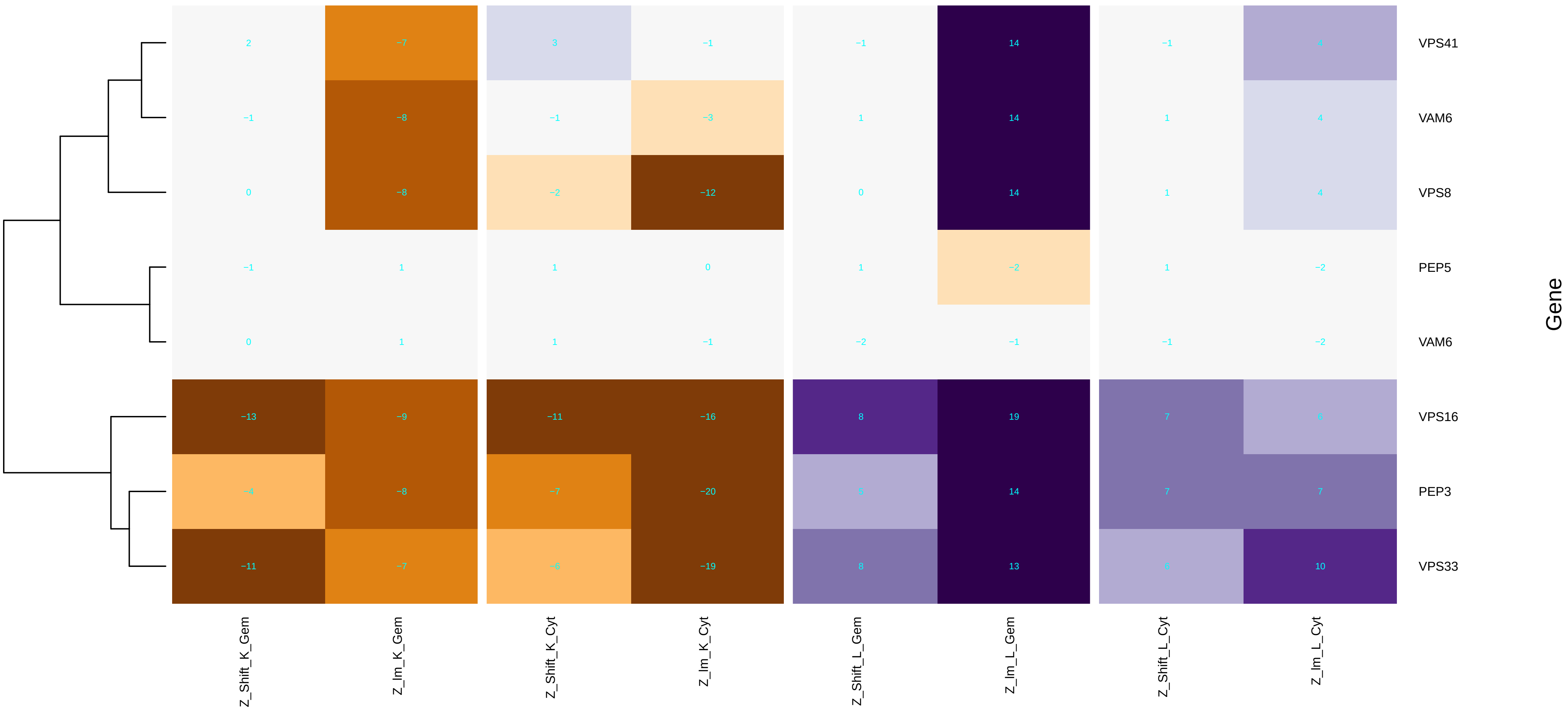

### hydrogen_ion_transmembrane_transporter_activity.pdf

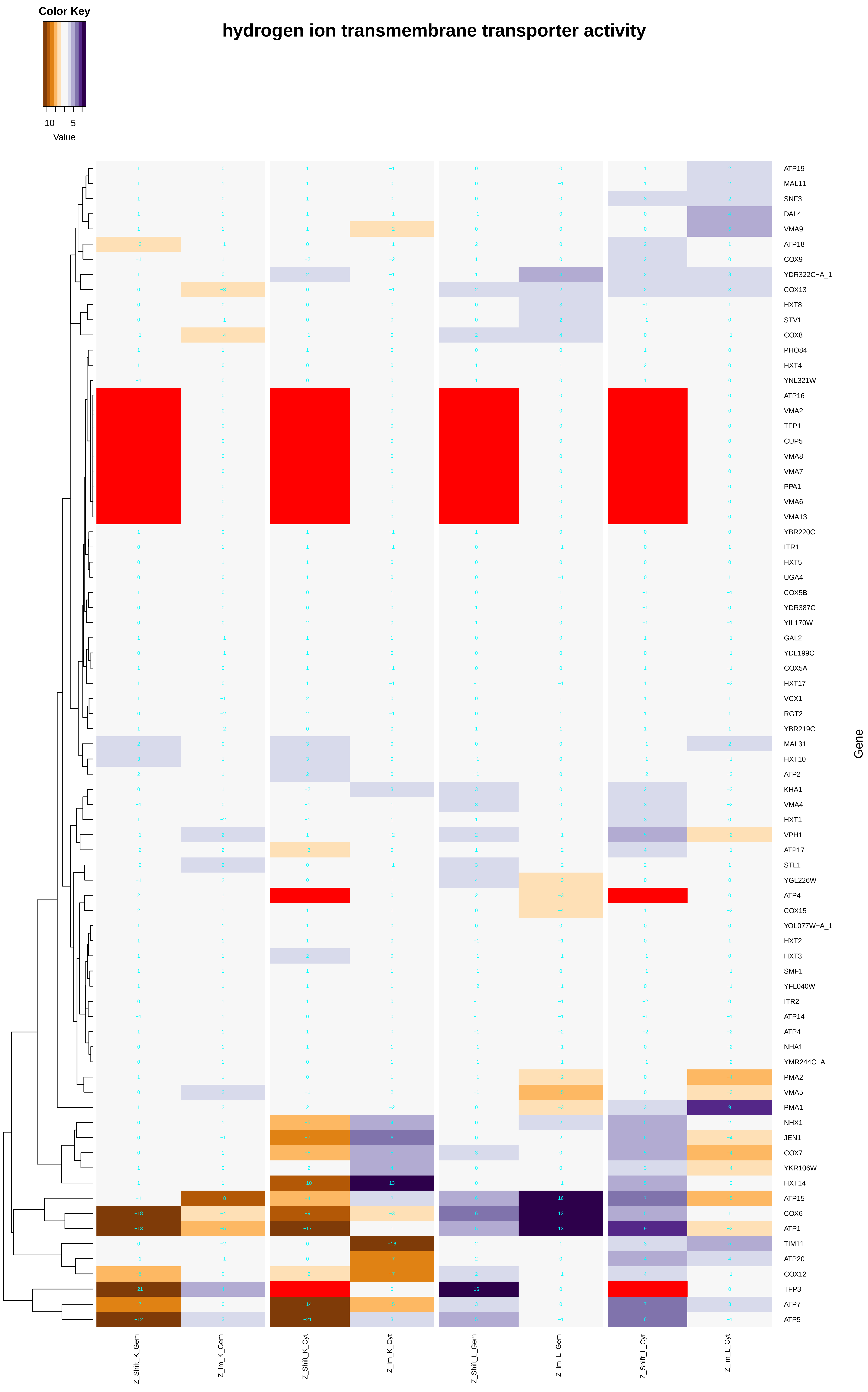

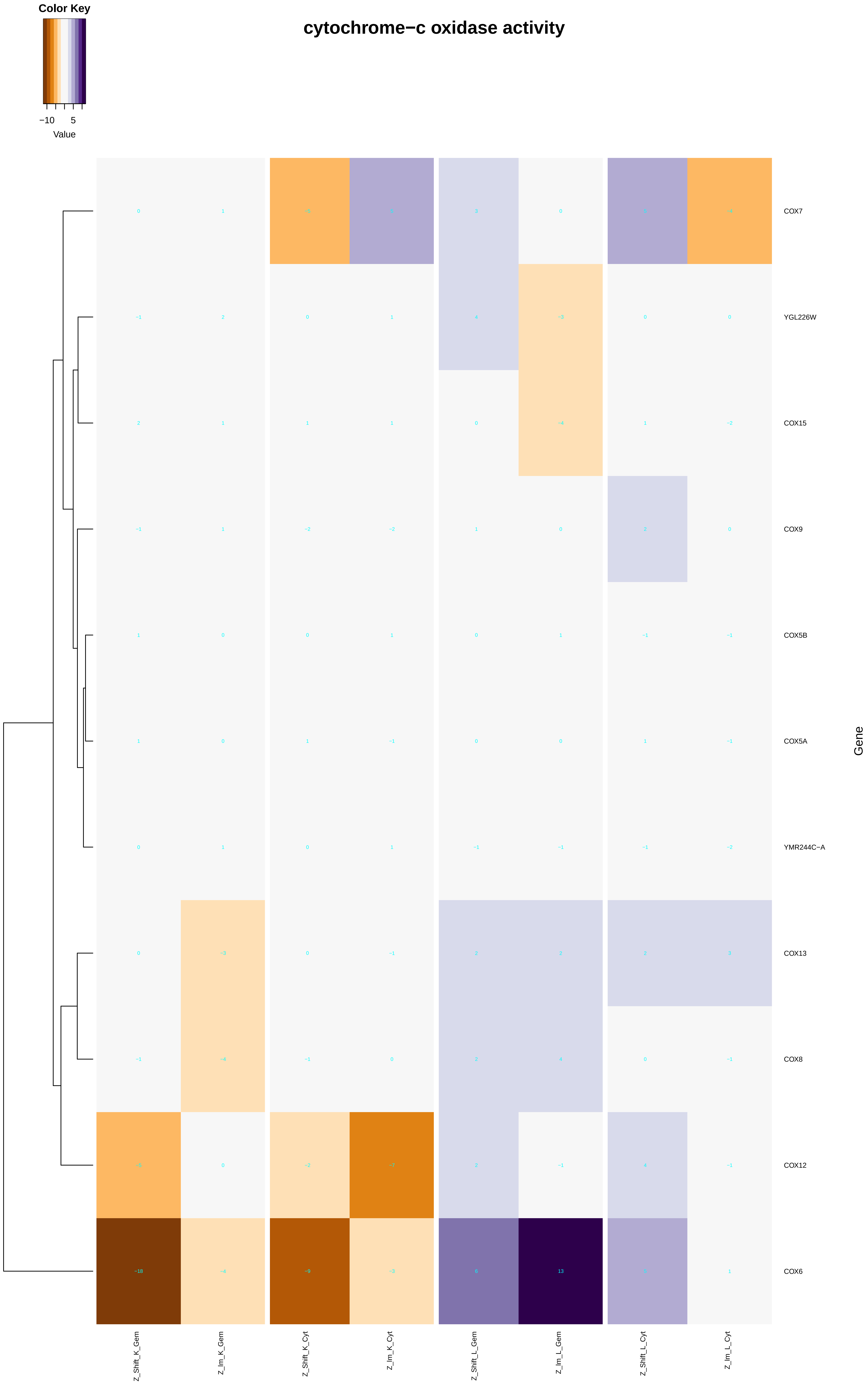

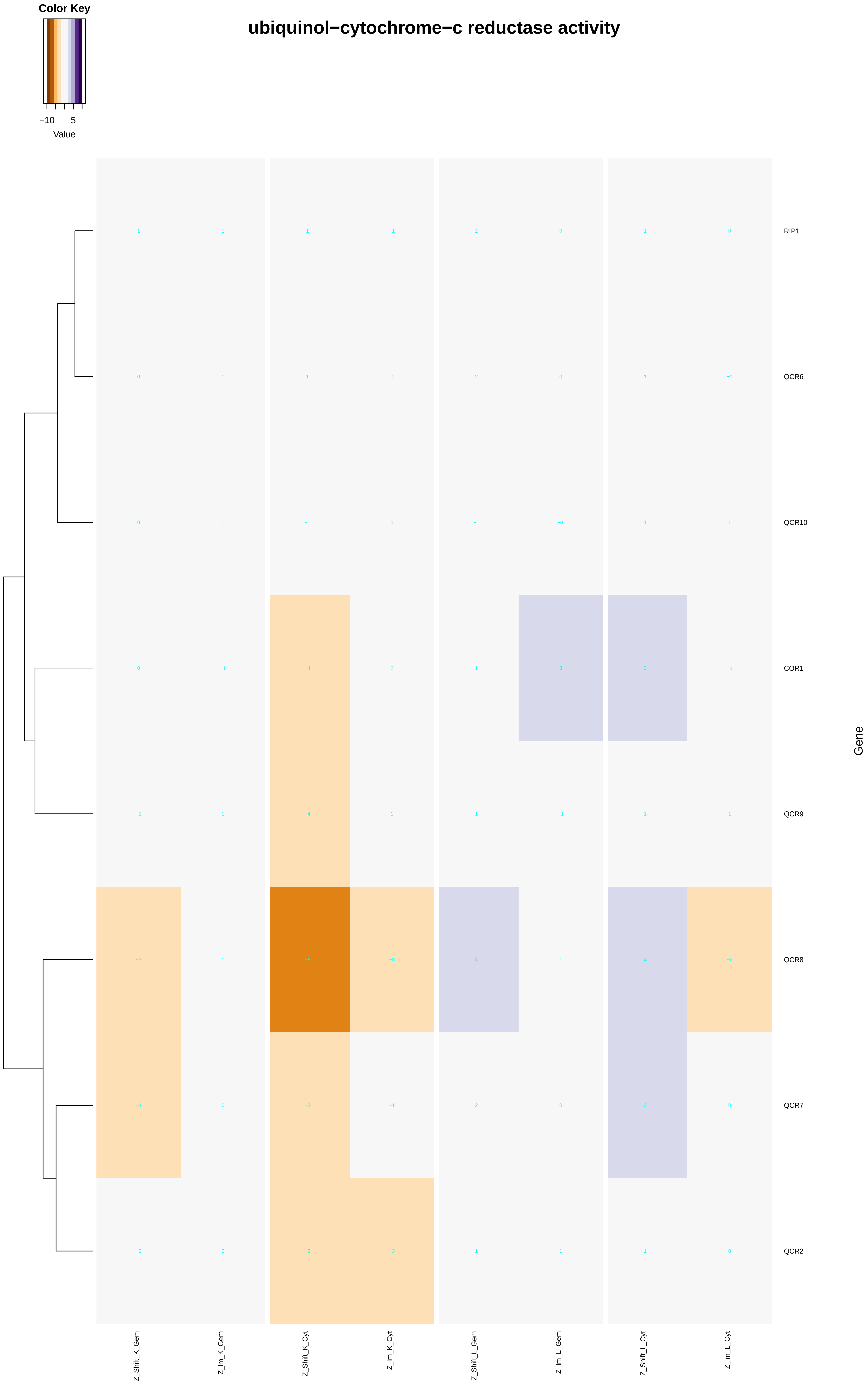

##### hydrogen−exporting ATPase activity, phosphorylative mechanism

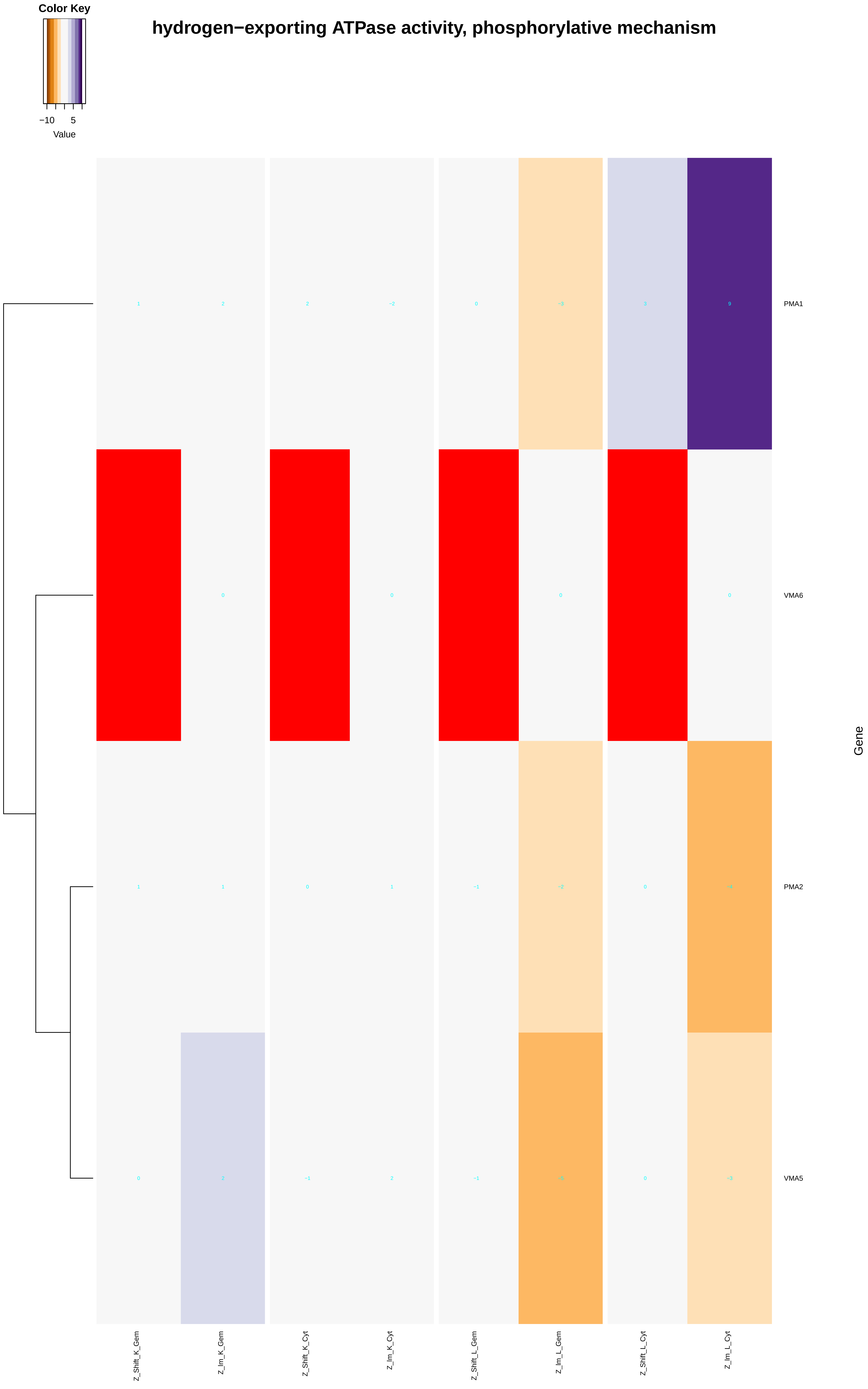

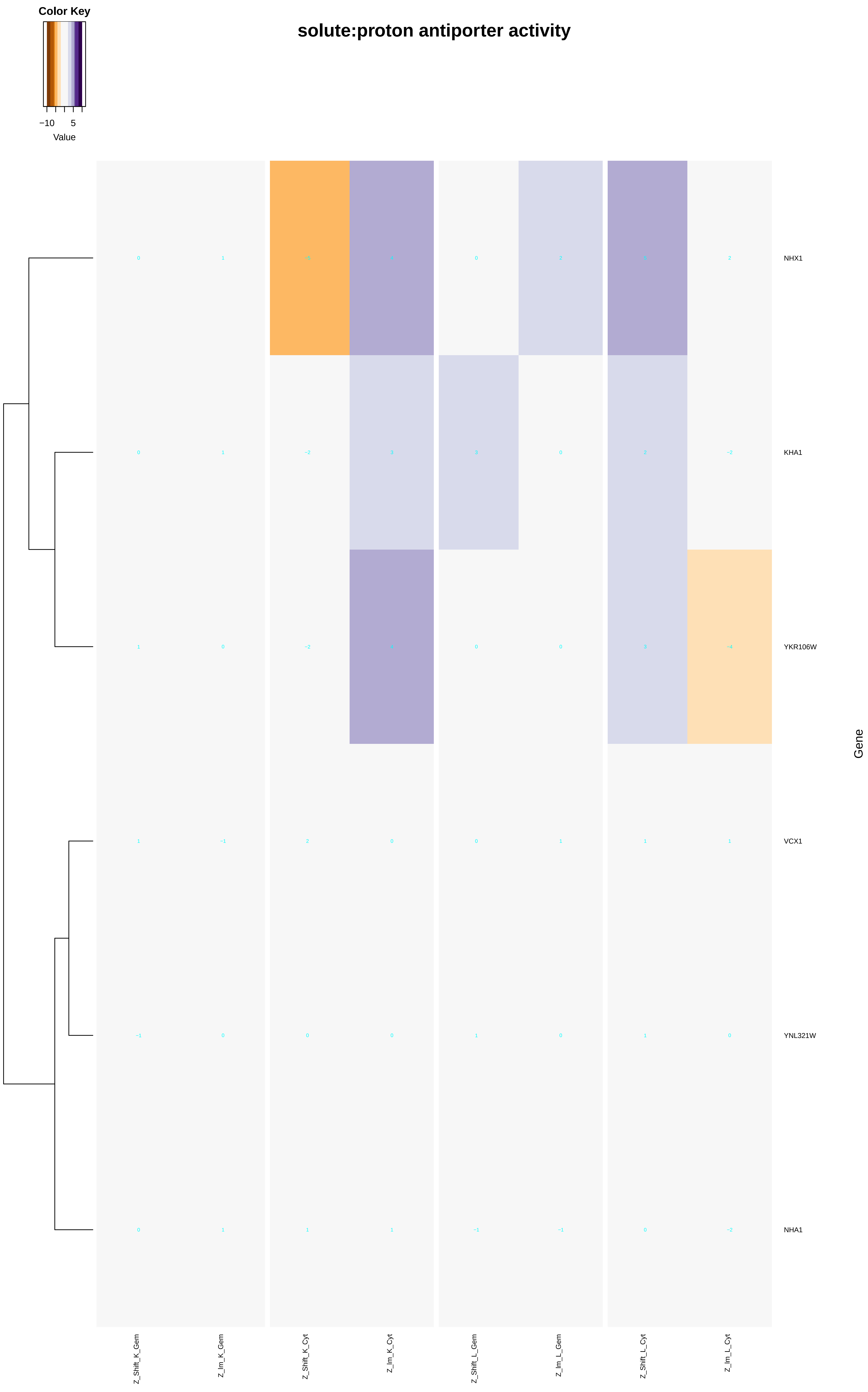

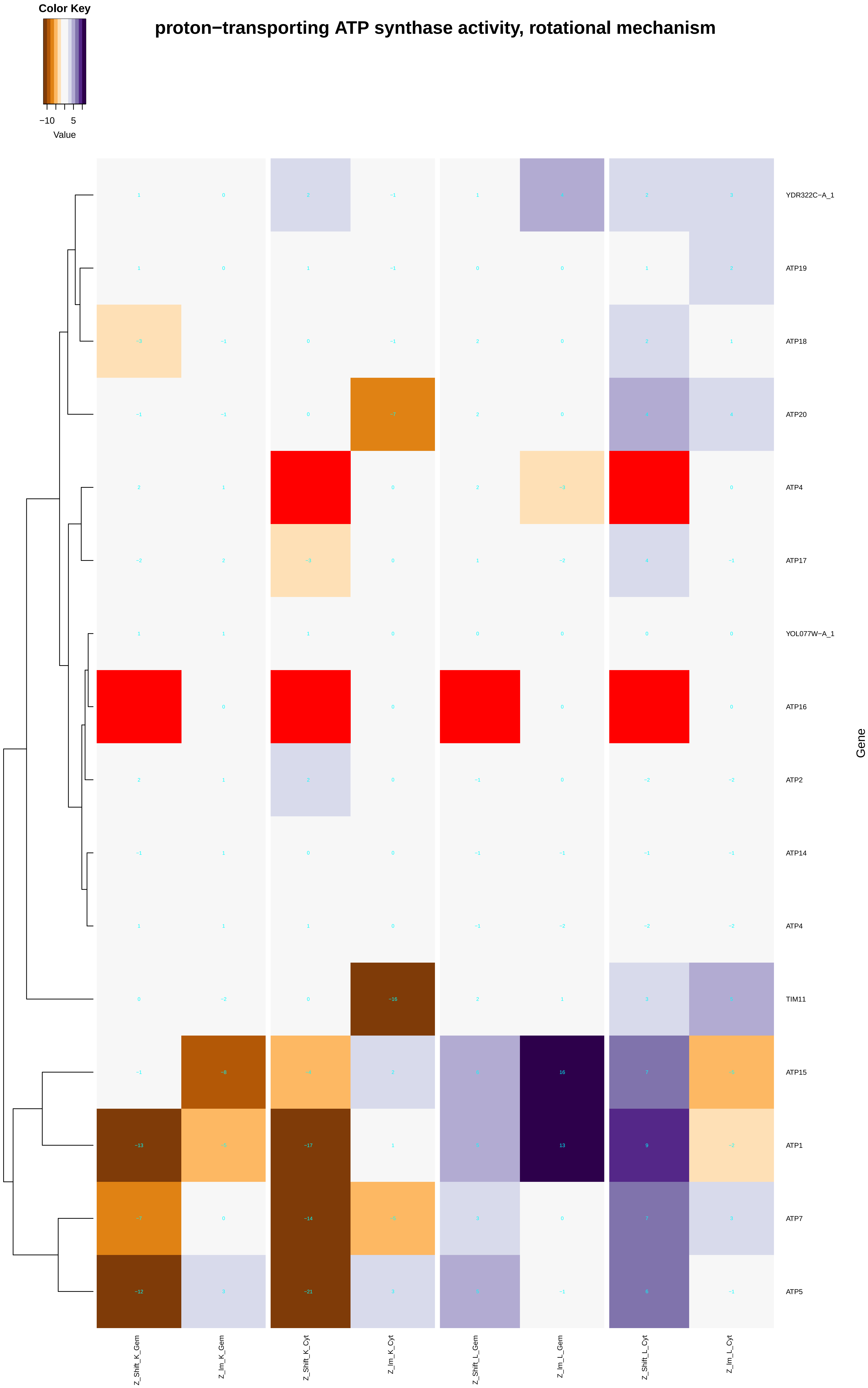

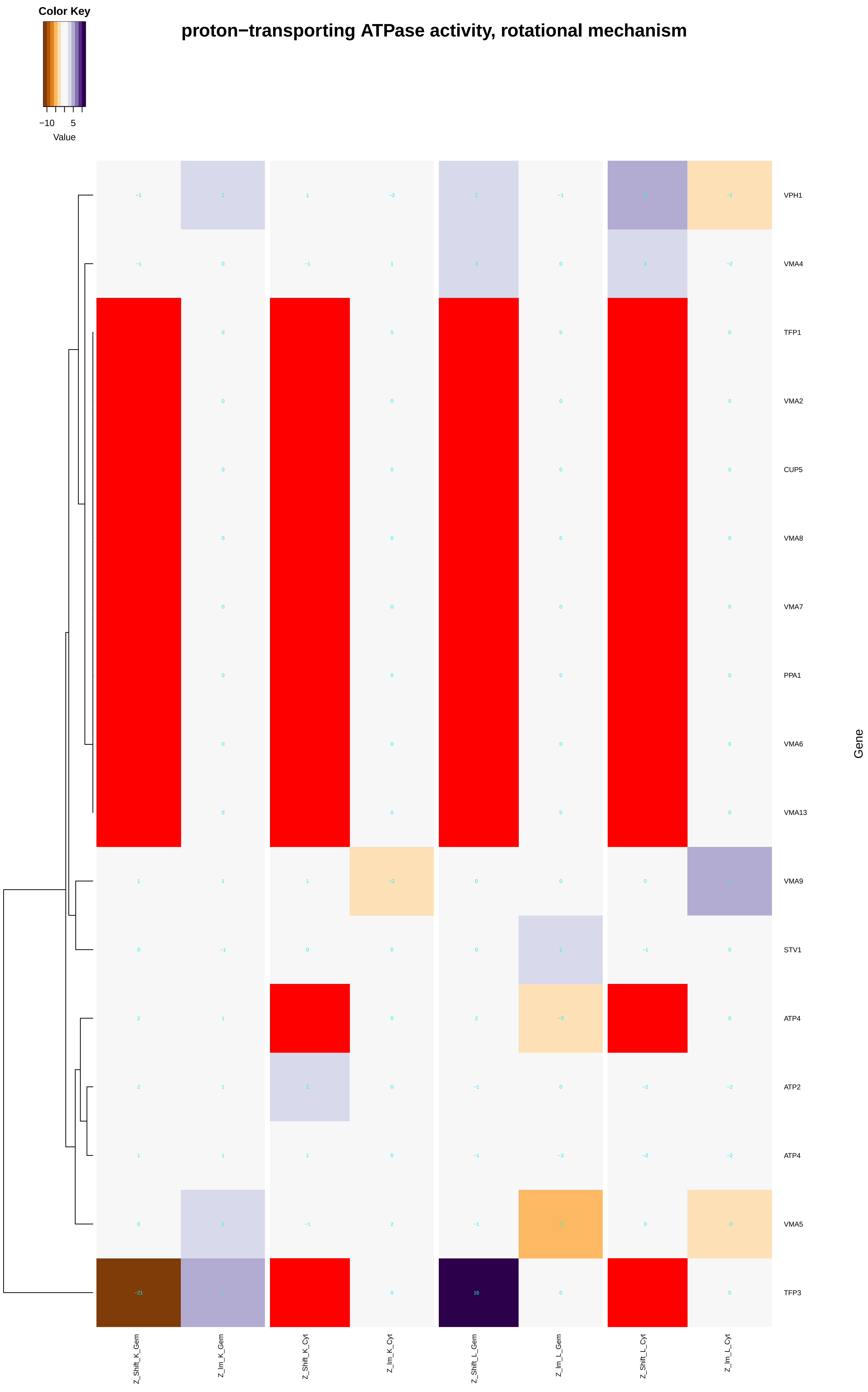

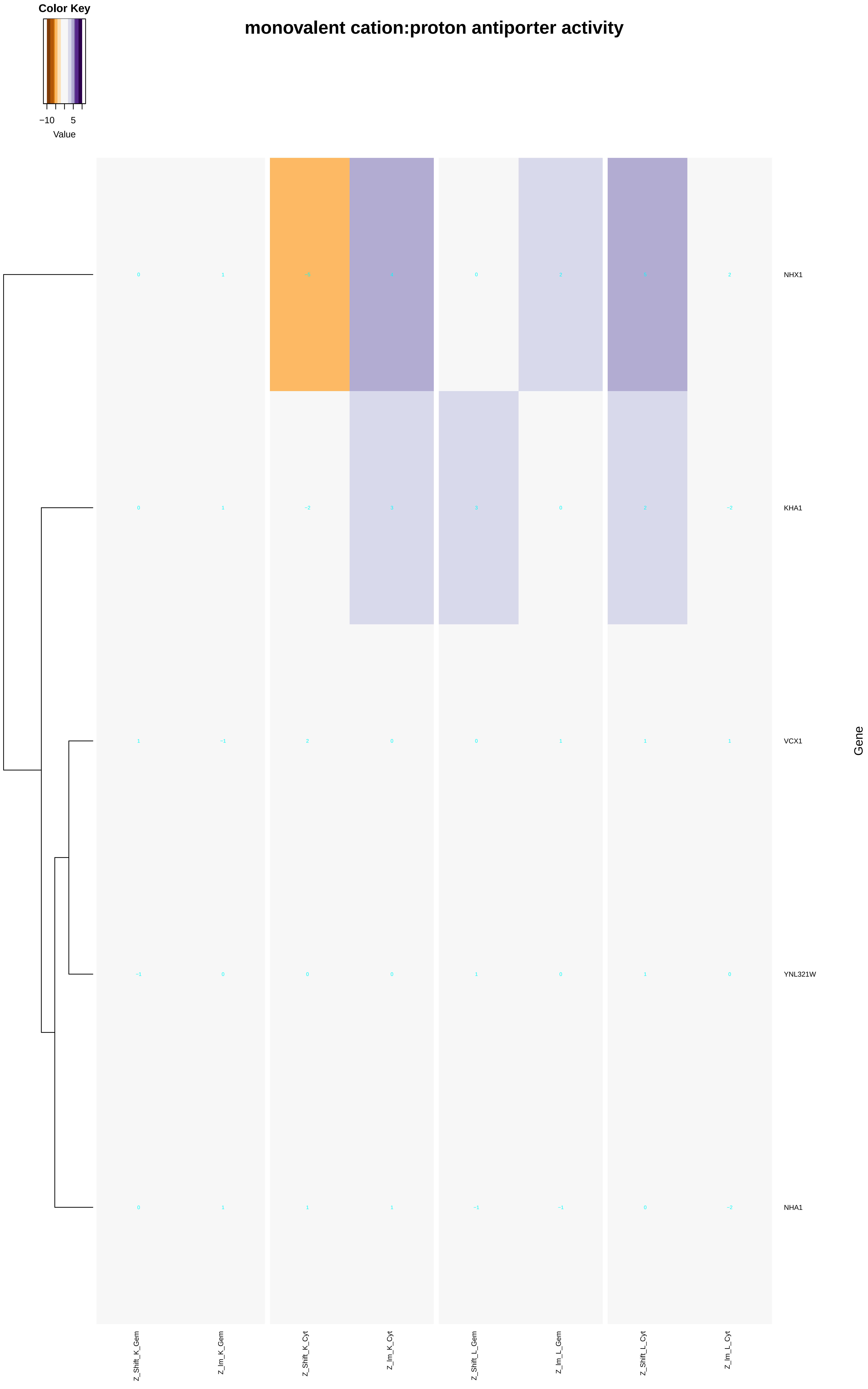

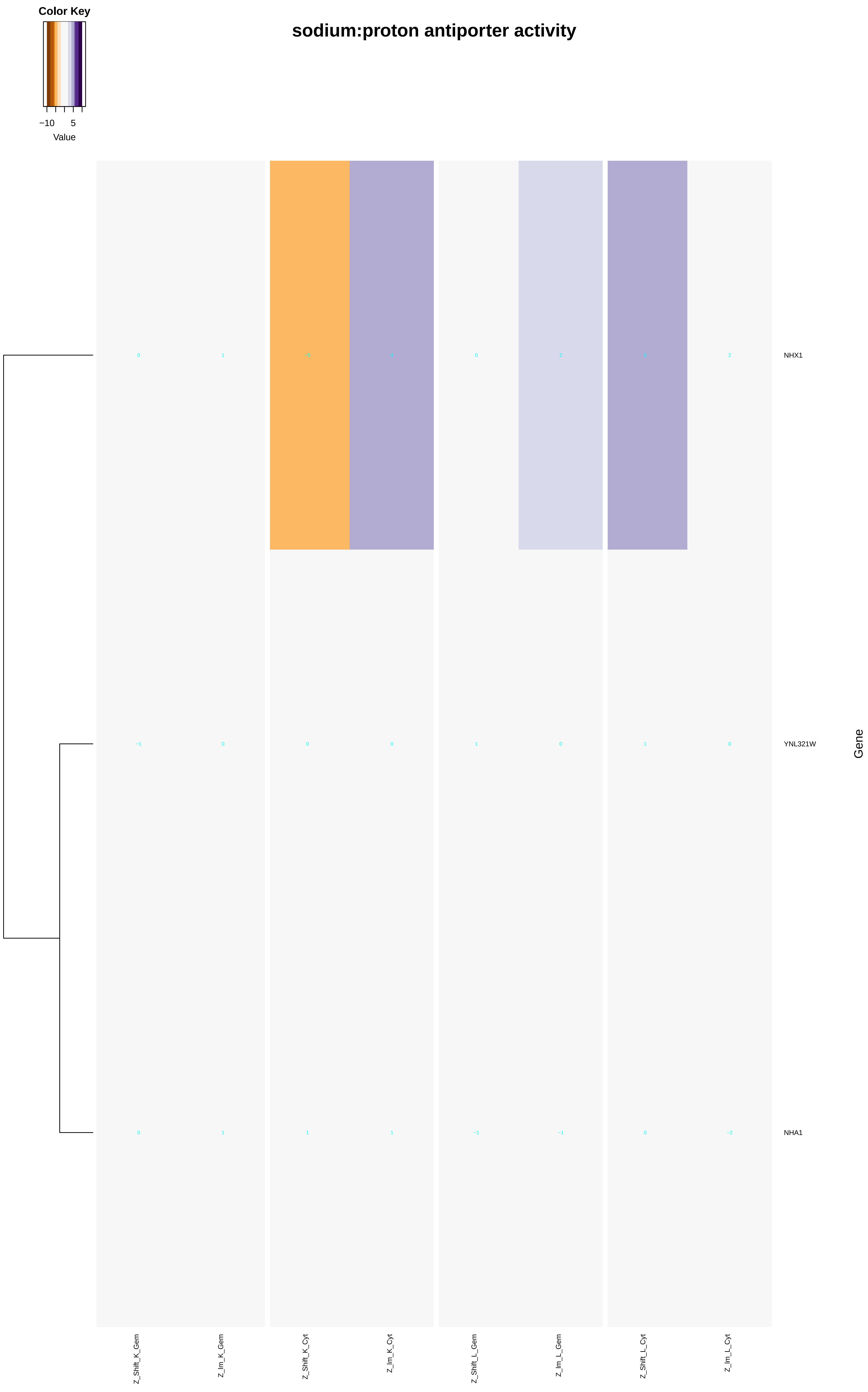

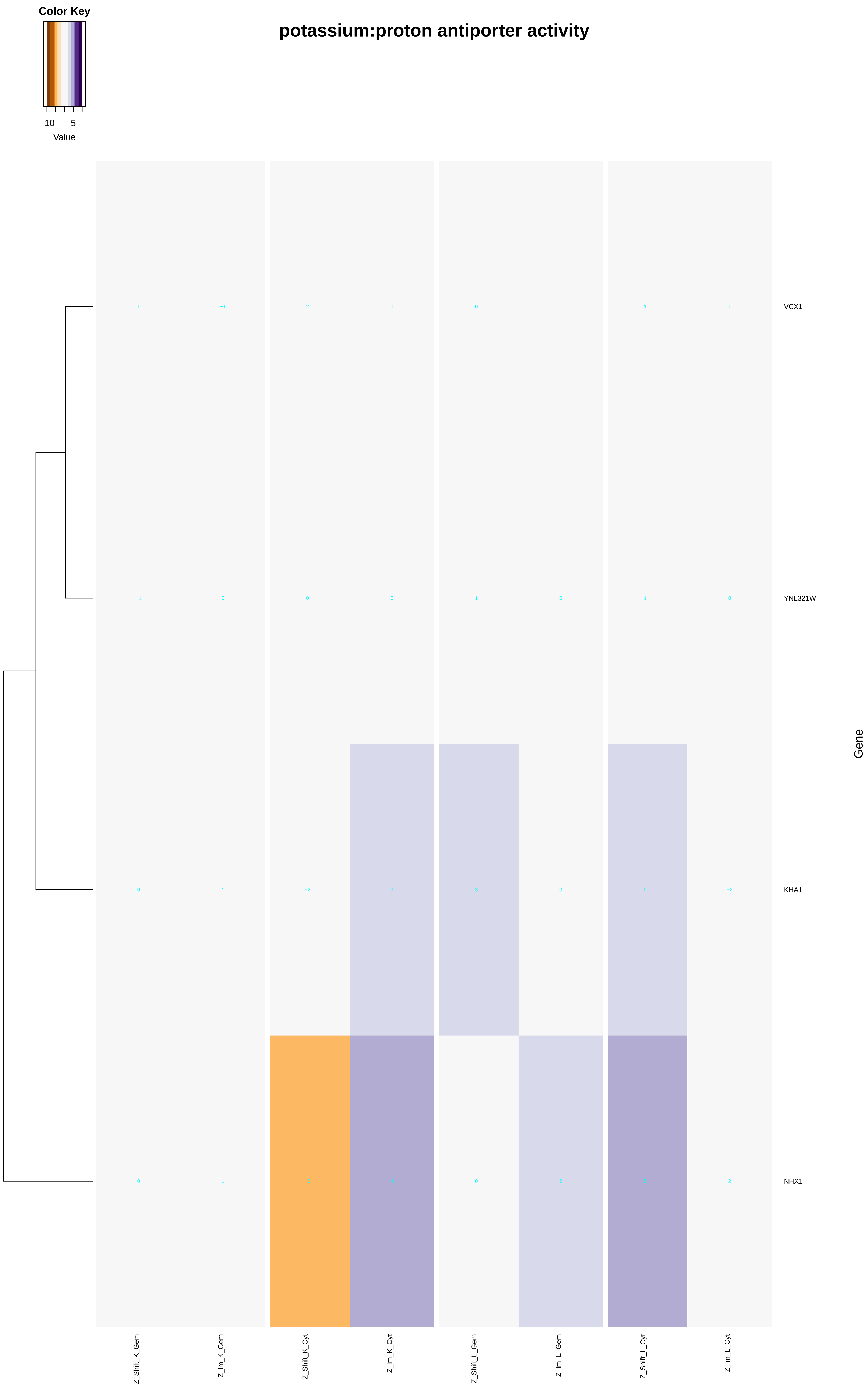

### integral_component_of_nuclear_inner_membrane.pdf

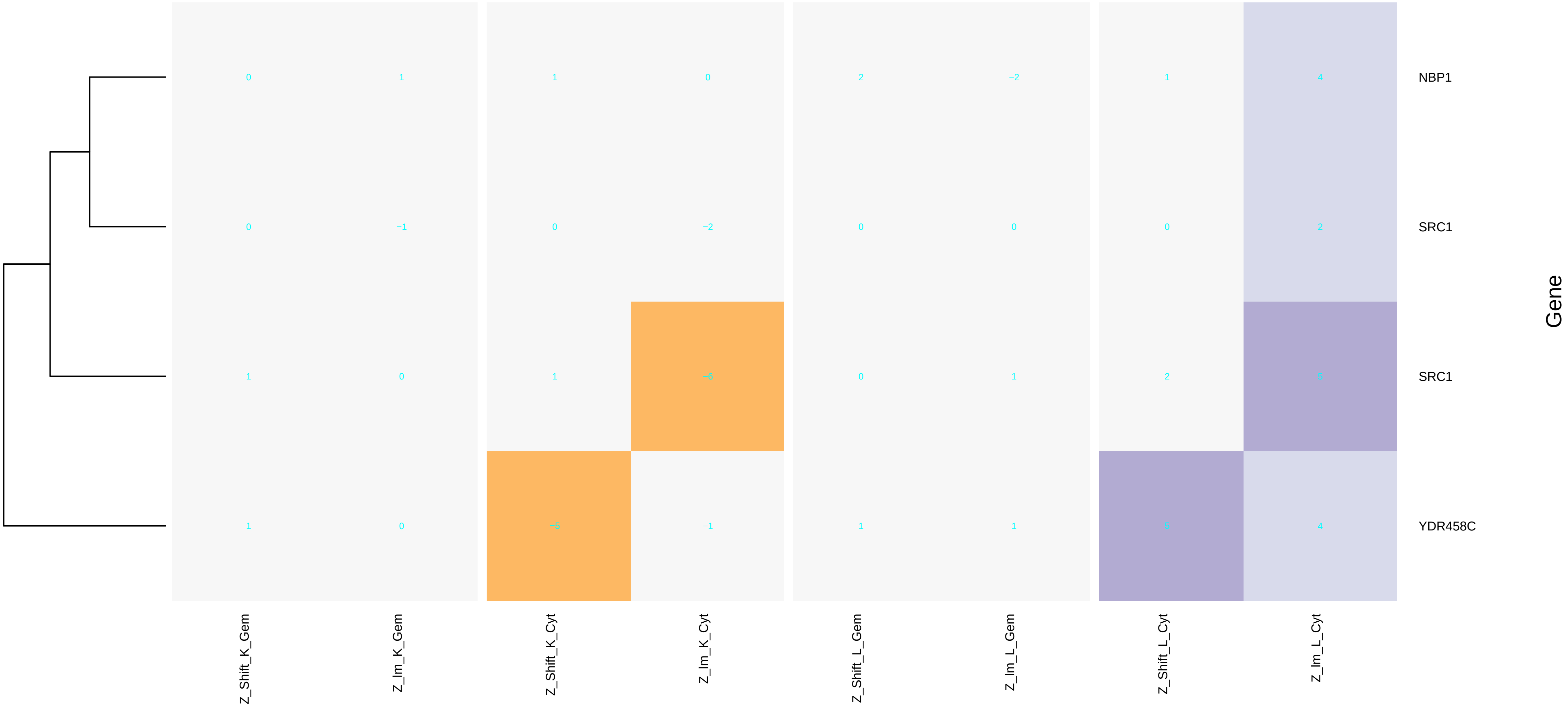

### intra-S_DNA_damage_checkpoint.pdf

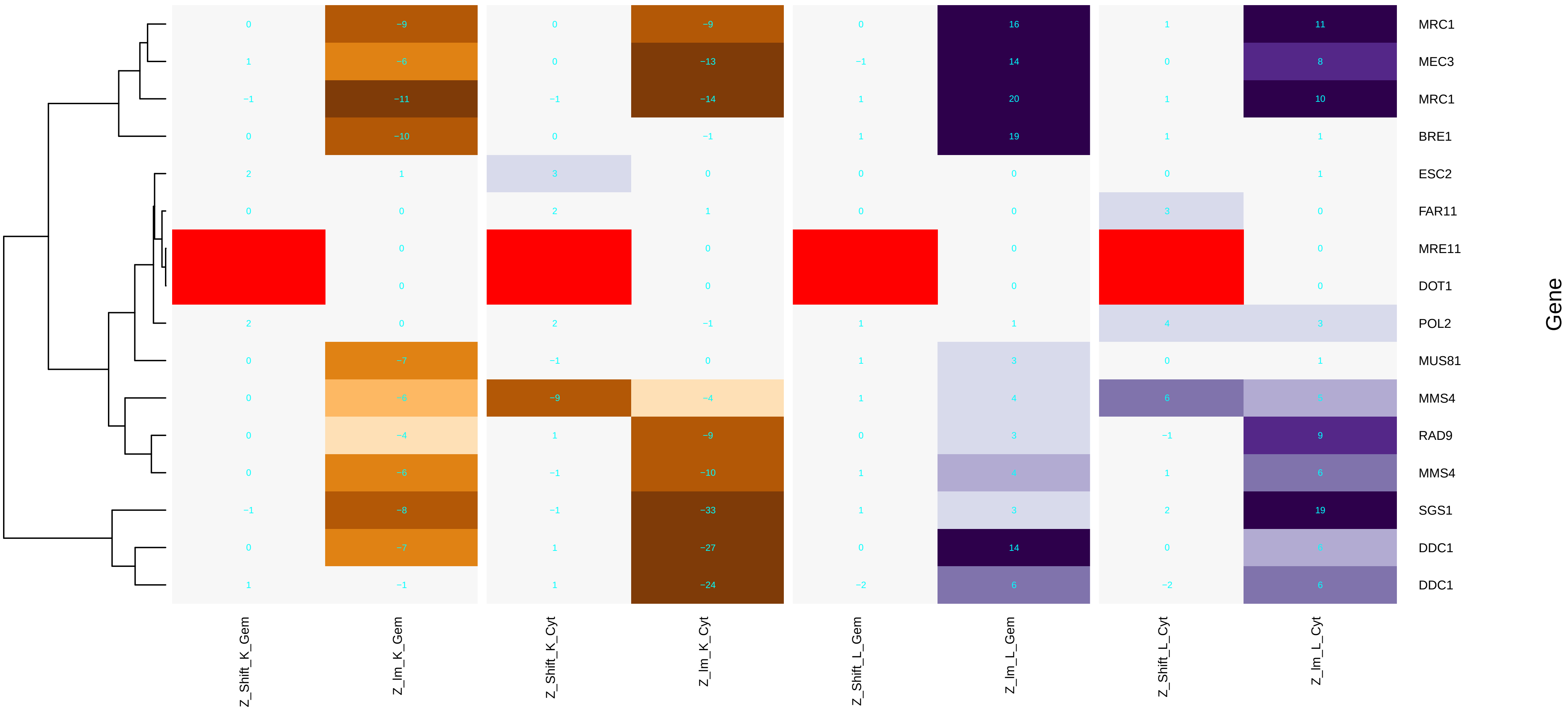

### intralumenal_vesicle_formation.pdf

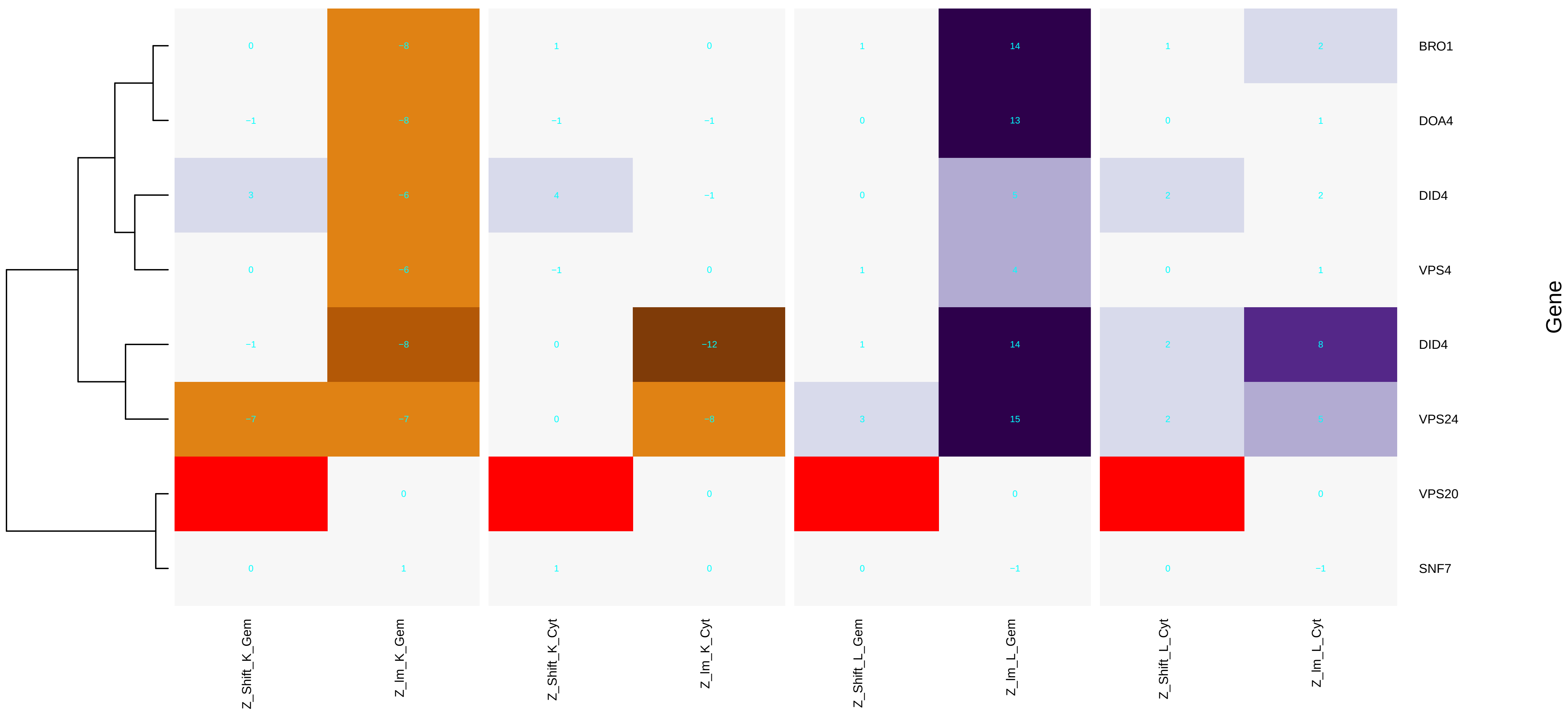

### large_ribosomal_subunit.pdf

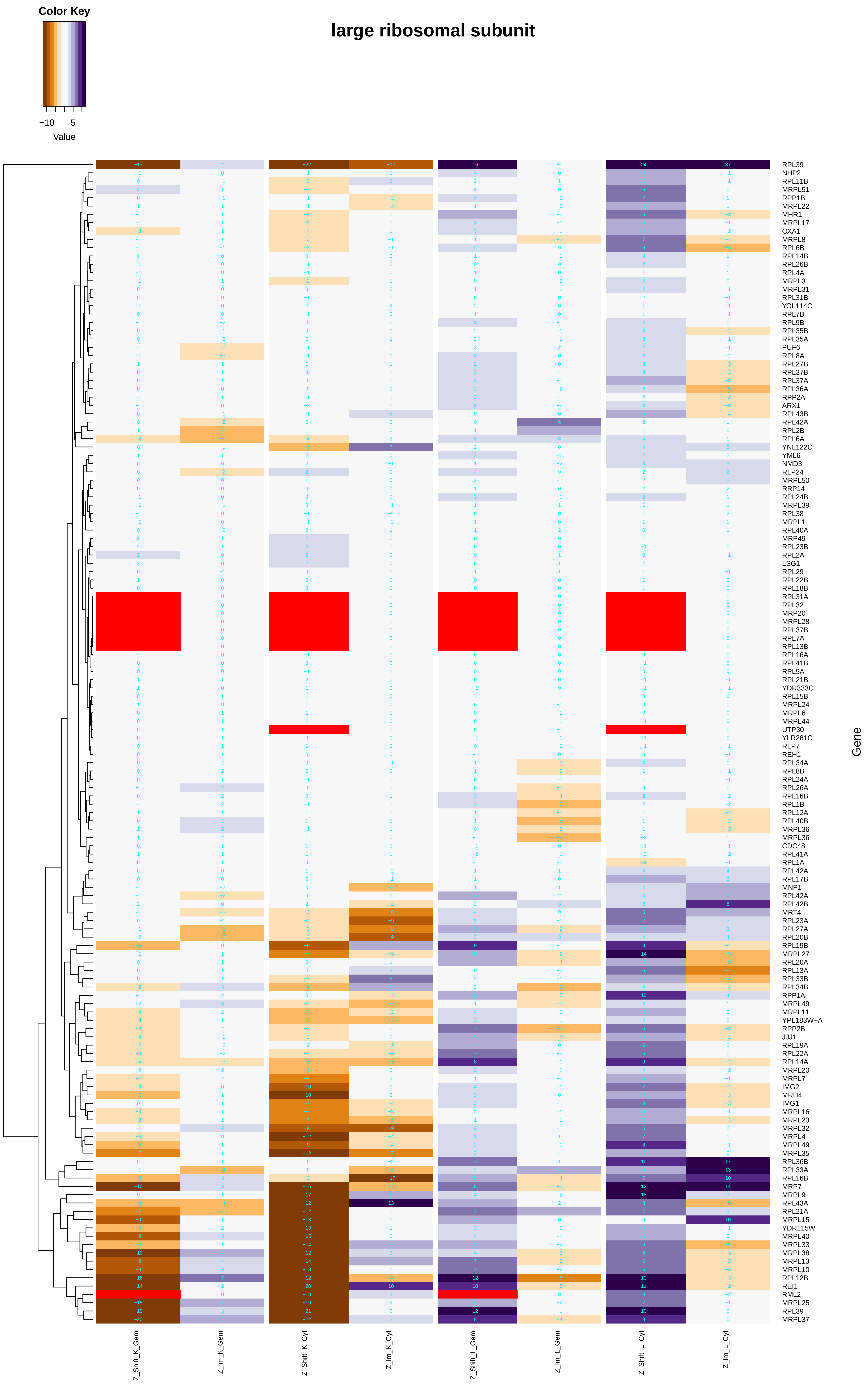

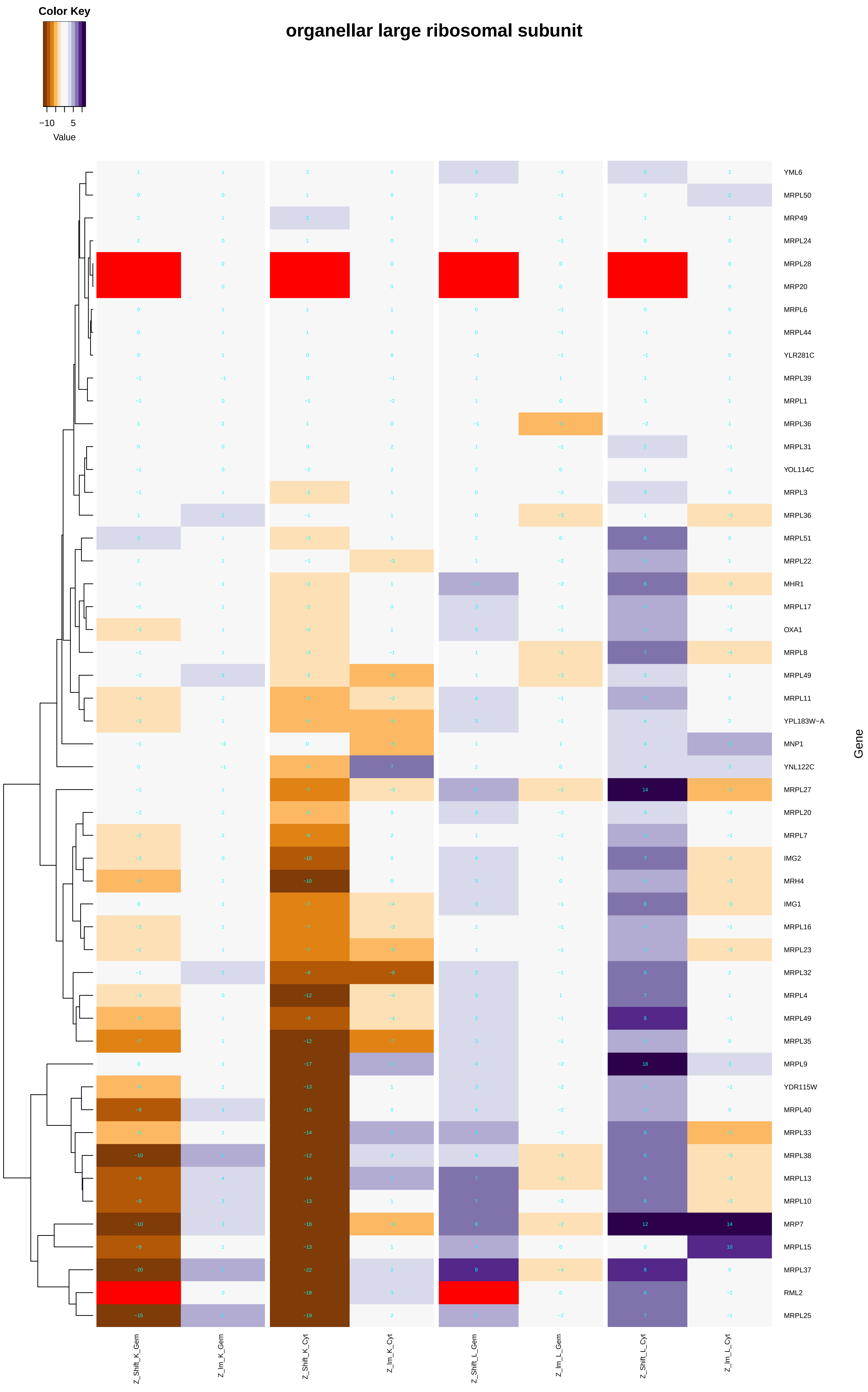

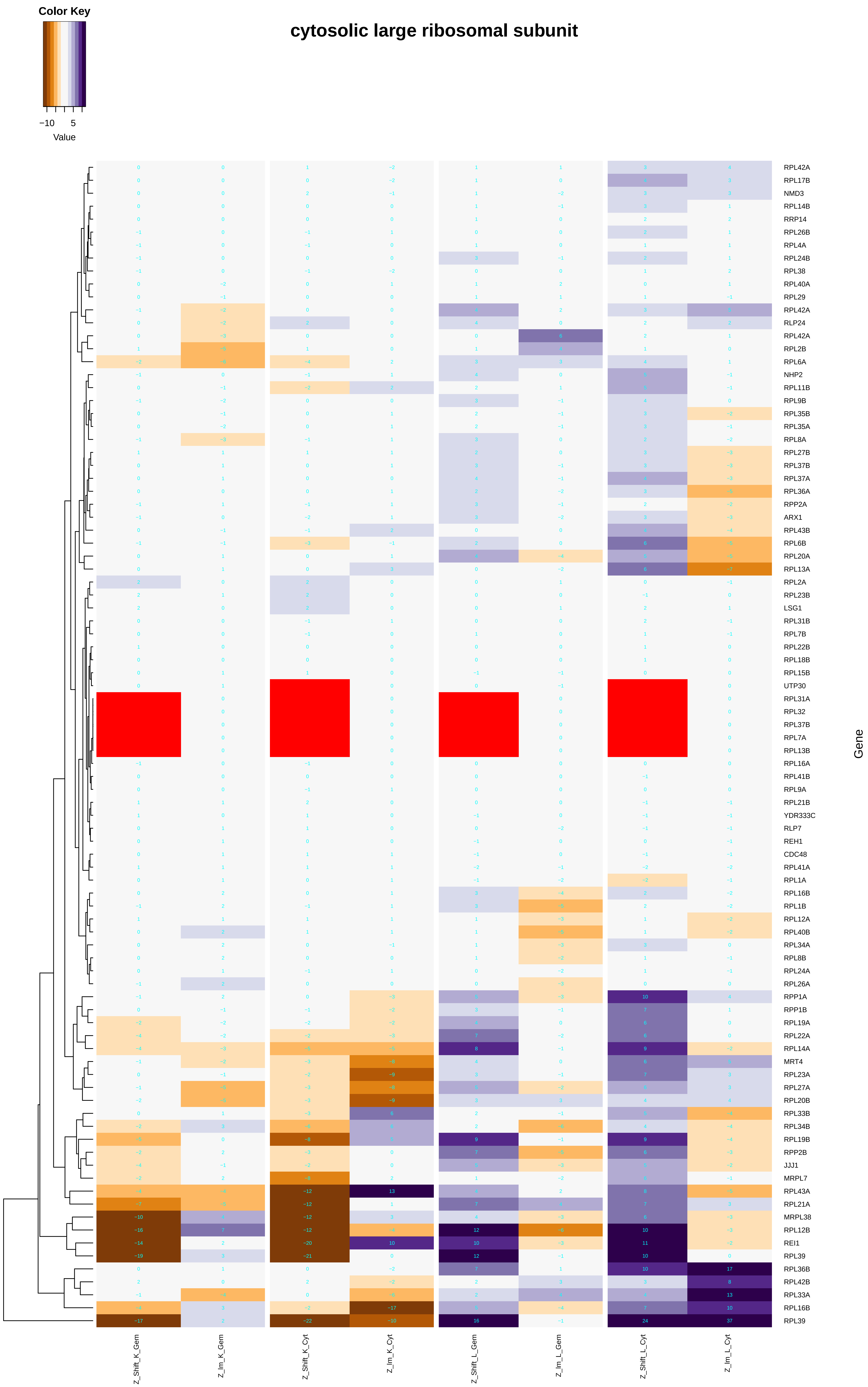

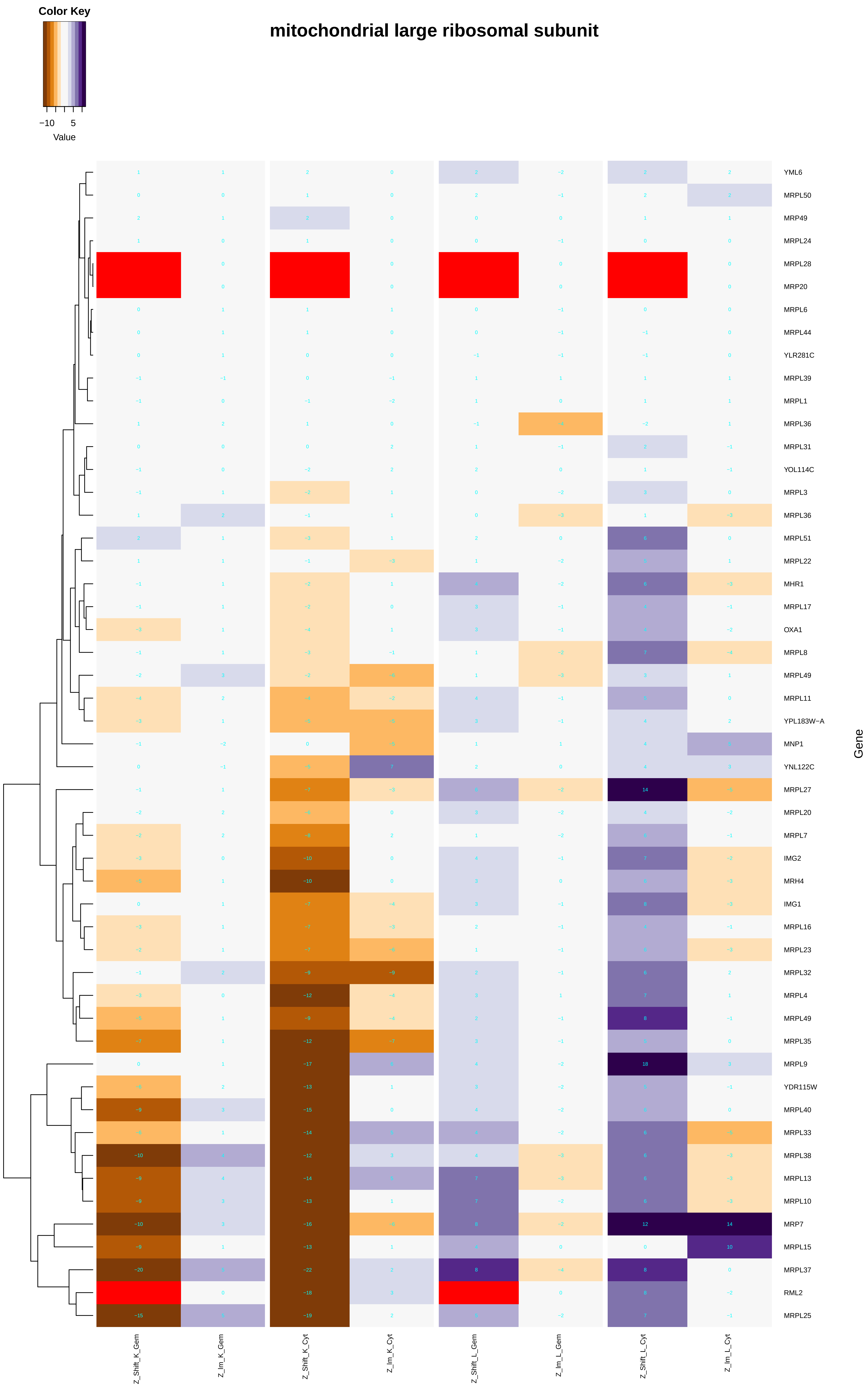

### late_endosome_to_vacuole_transport.pdf

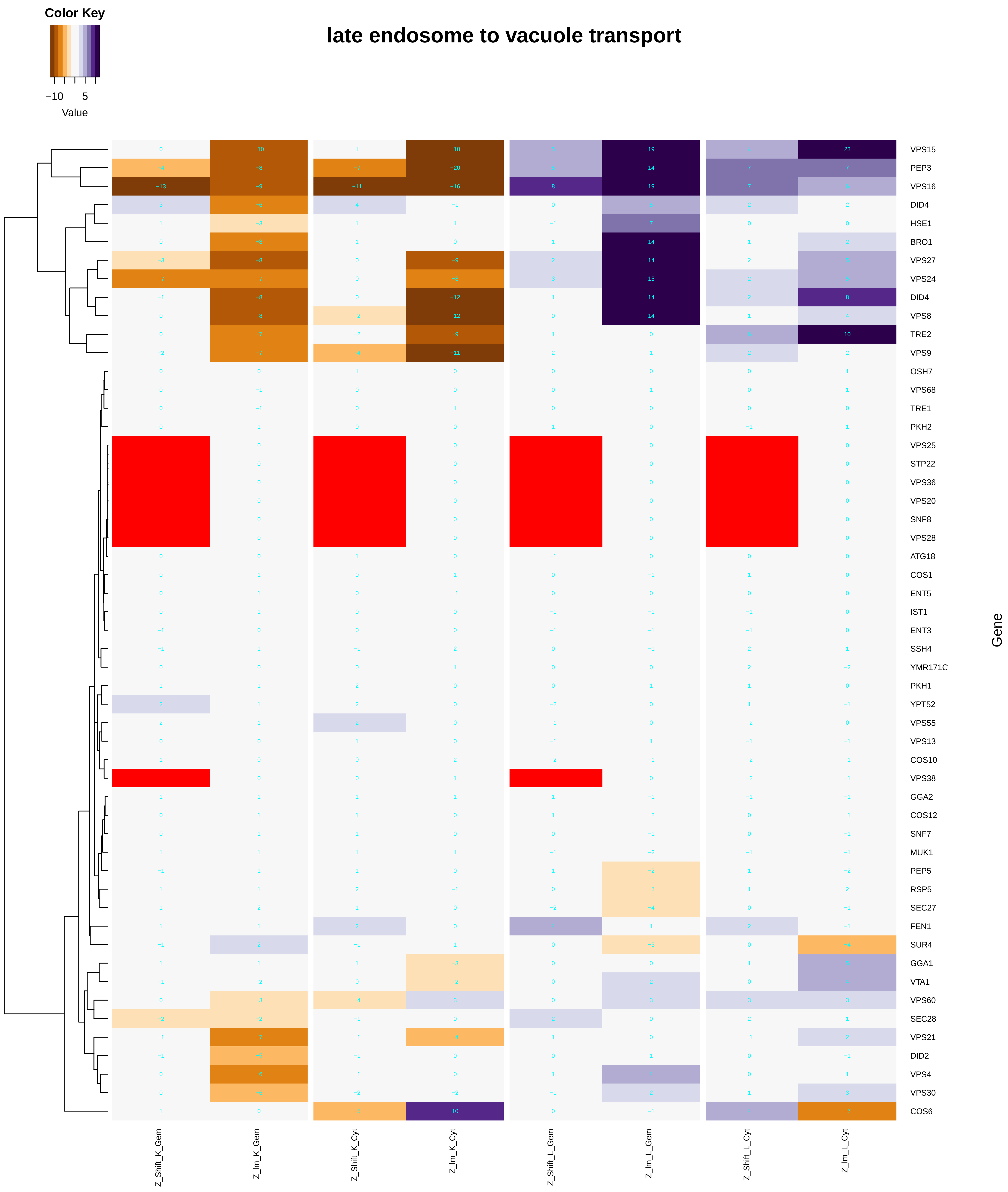

#### late endosome to vacuole transport via multivesicular body sorting pathway

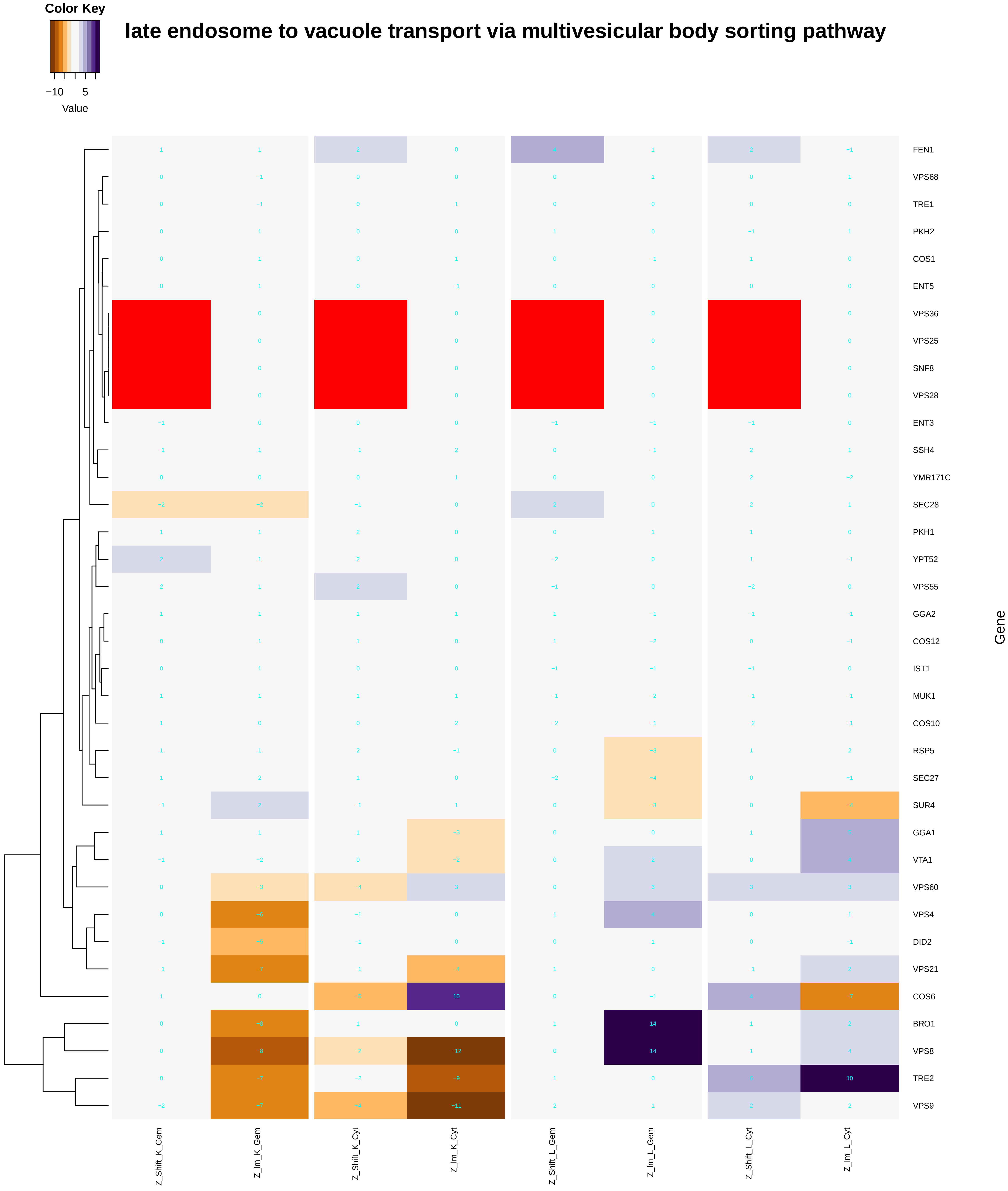

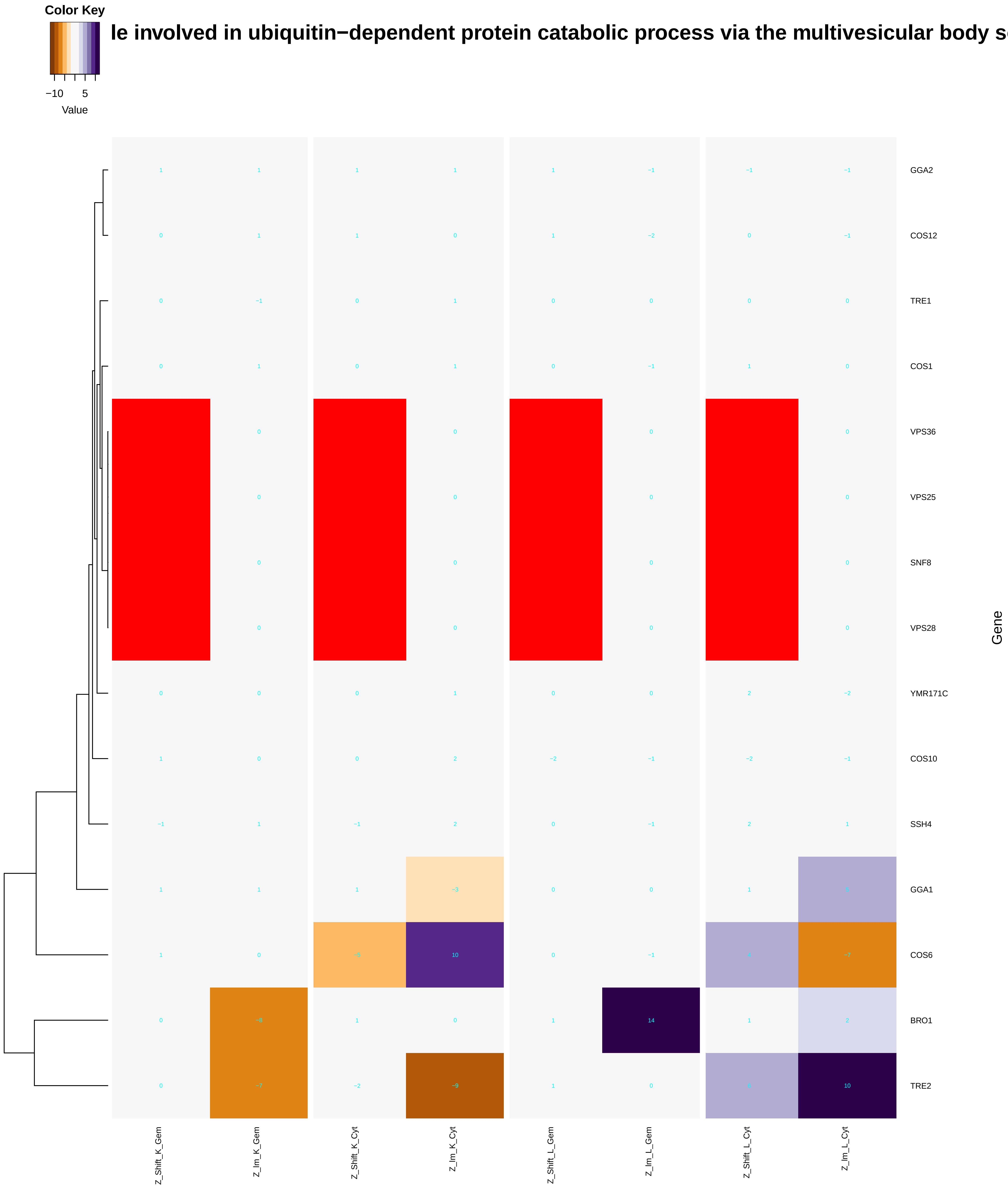

### Lsm1-7-Pat1_complex.pdf

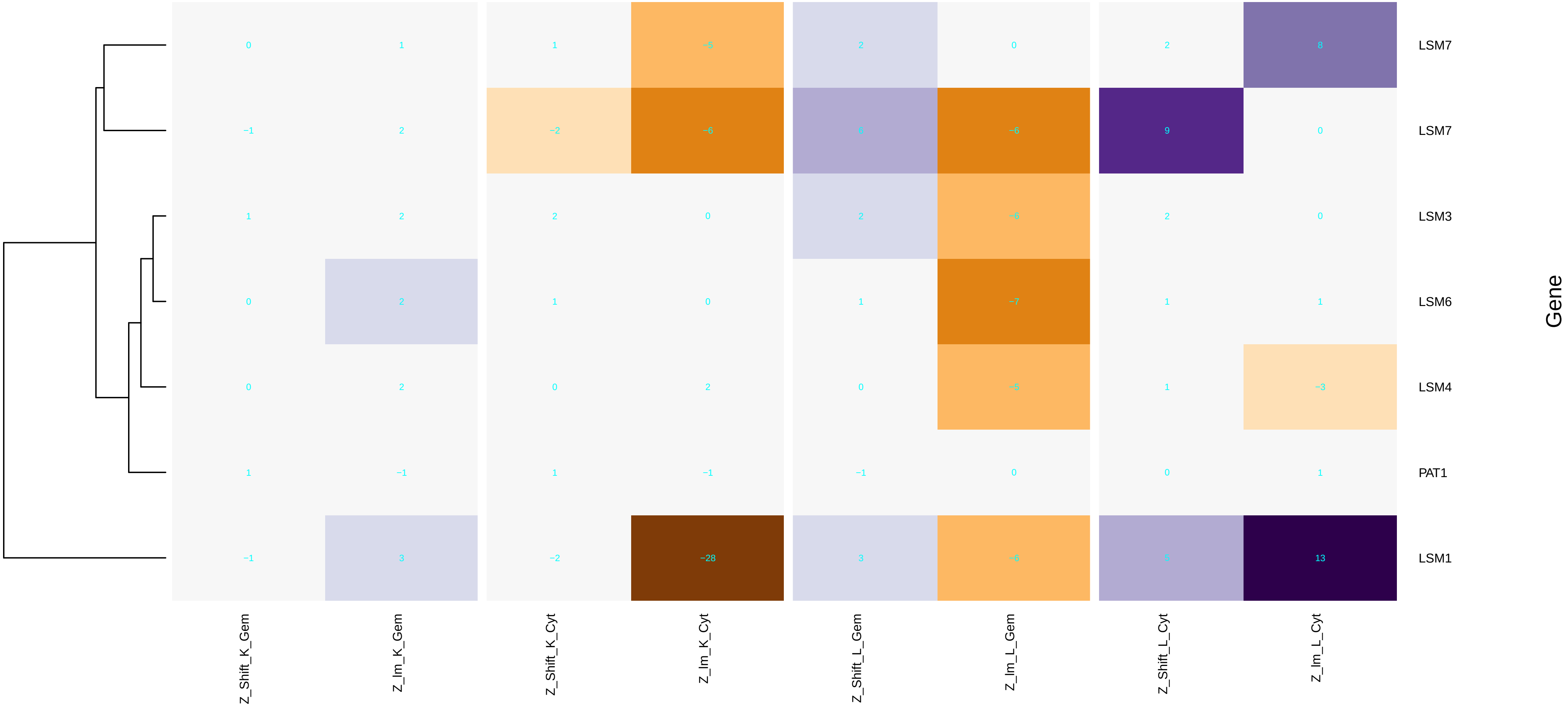

### macroautophagy.pdf

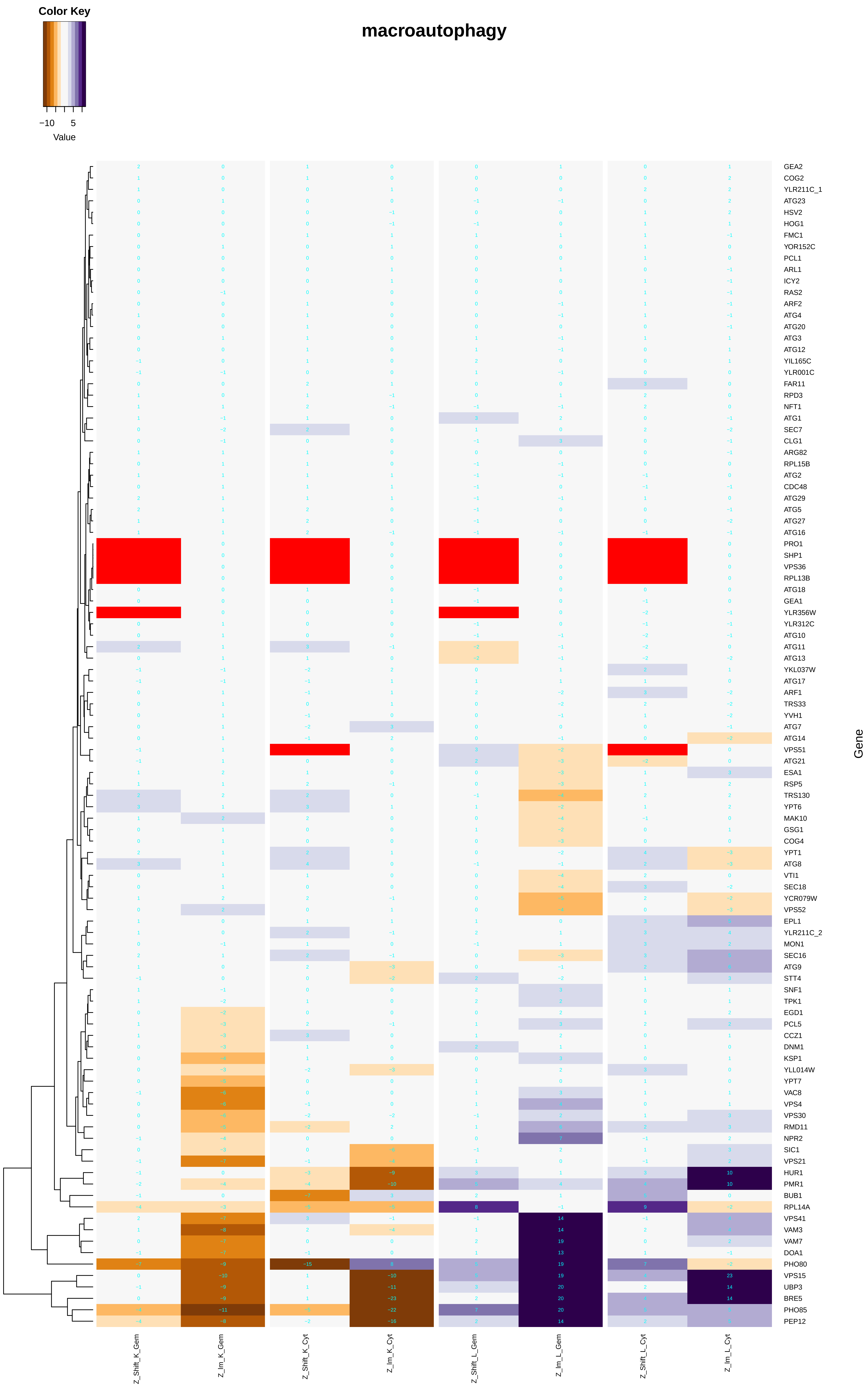

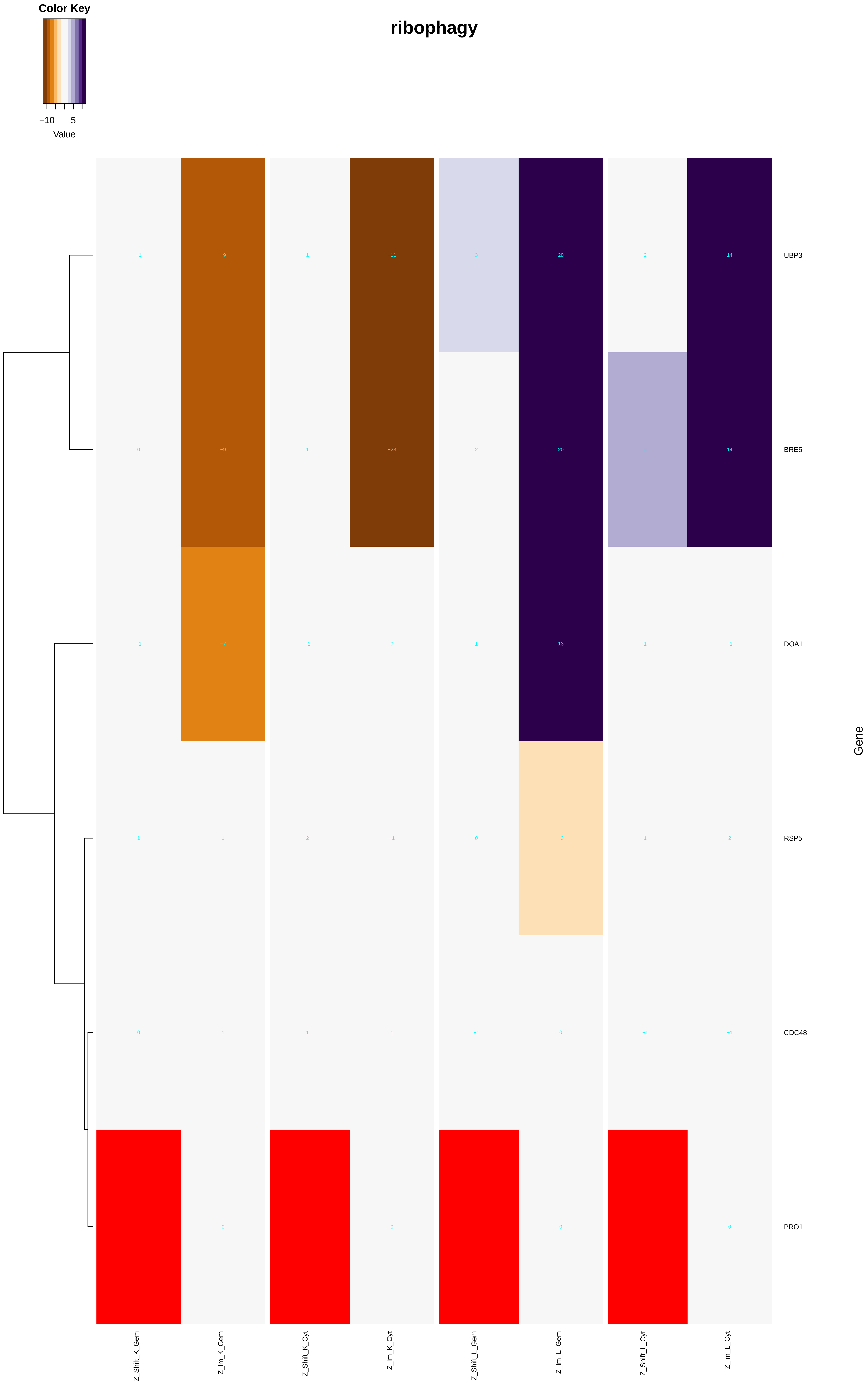

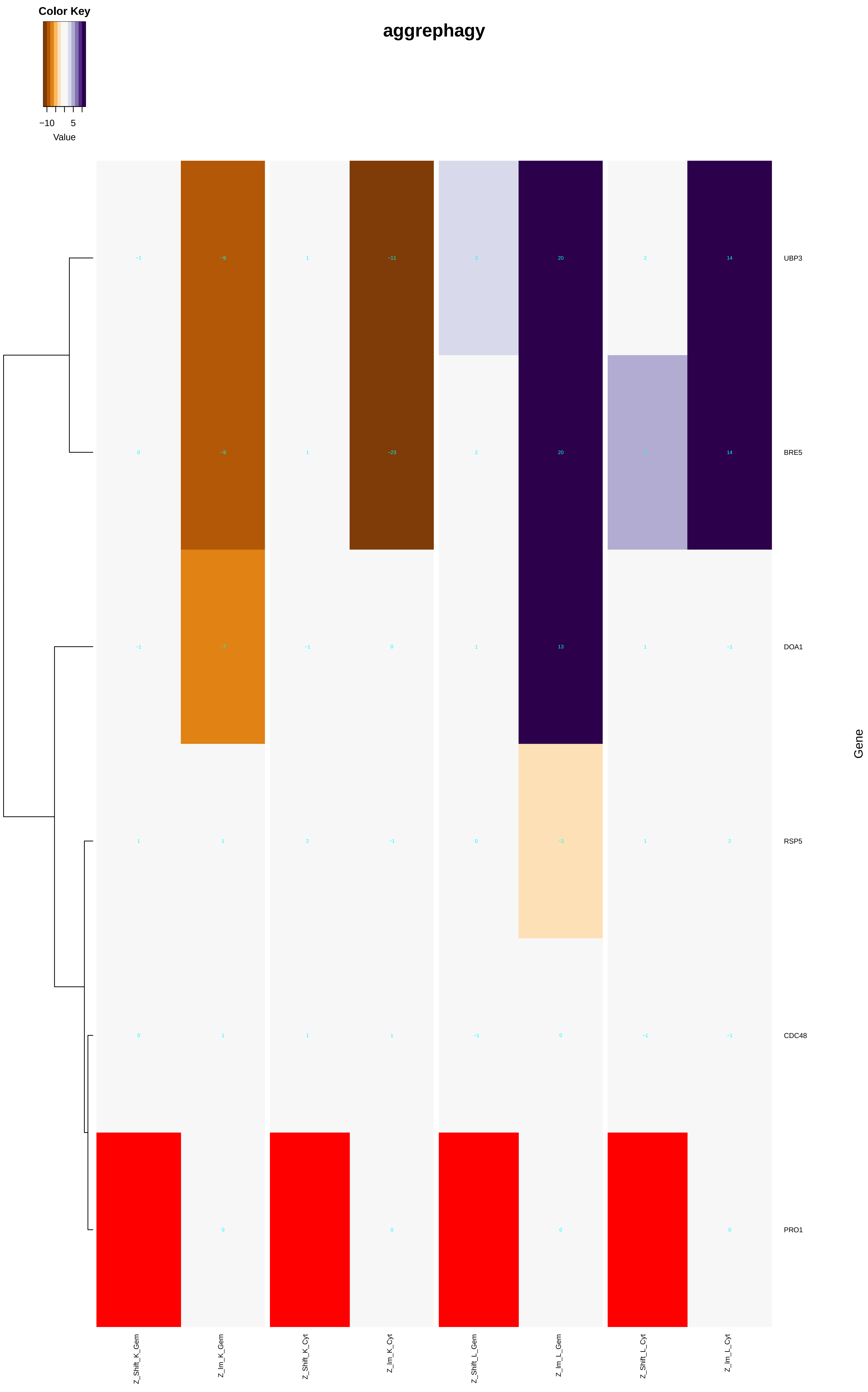

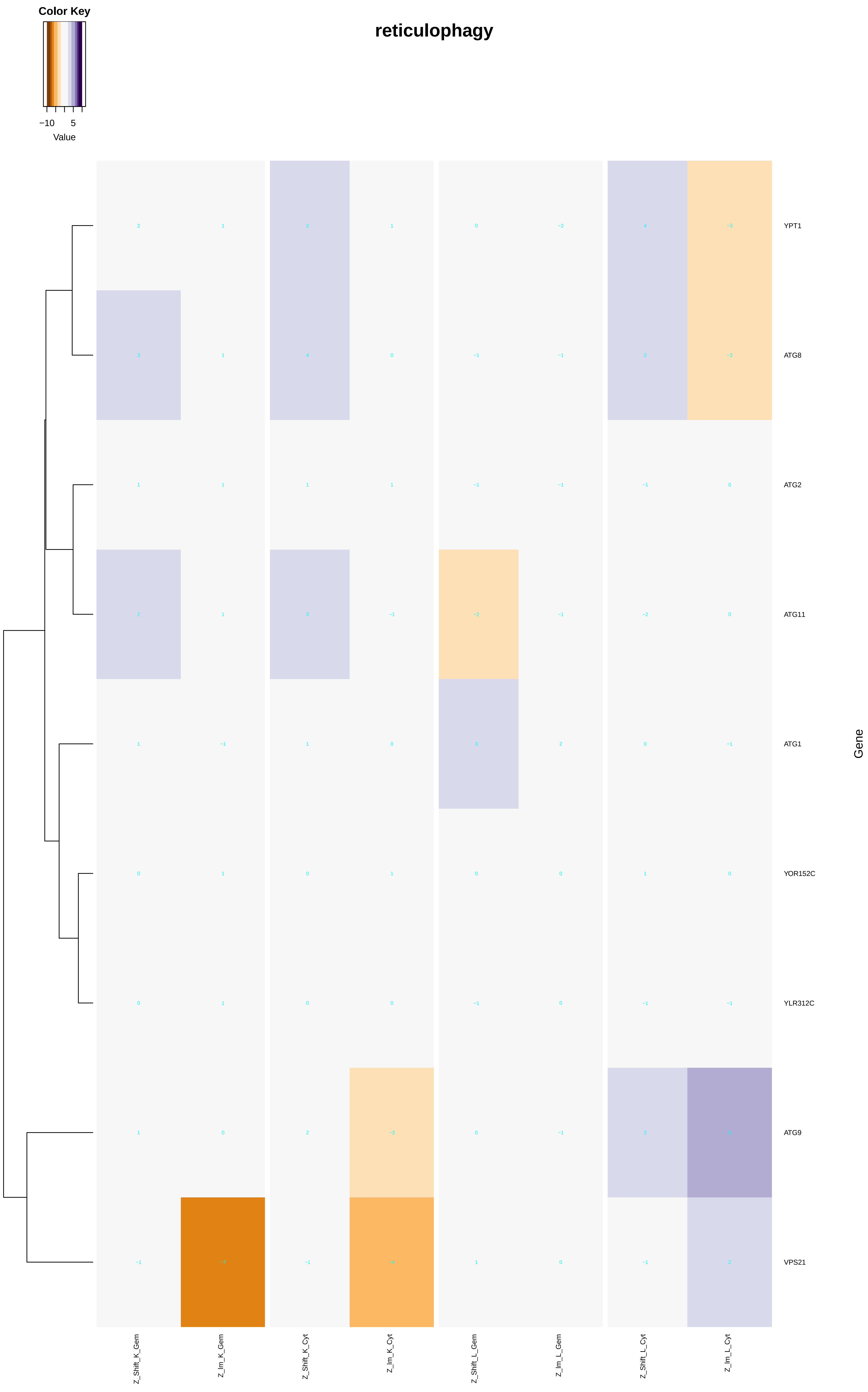

### maintenance_of_DNA_trinucleotide_repeats.pdf

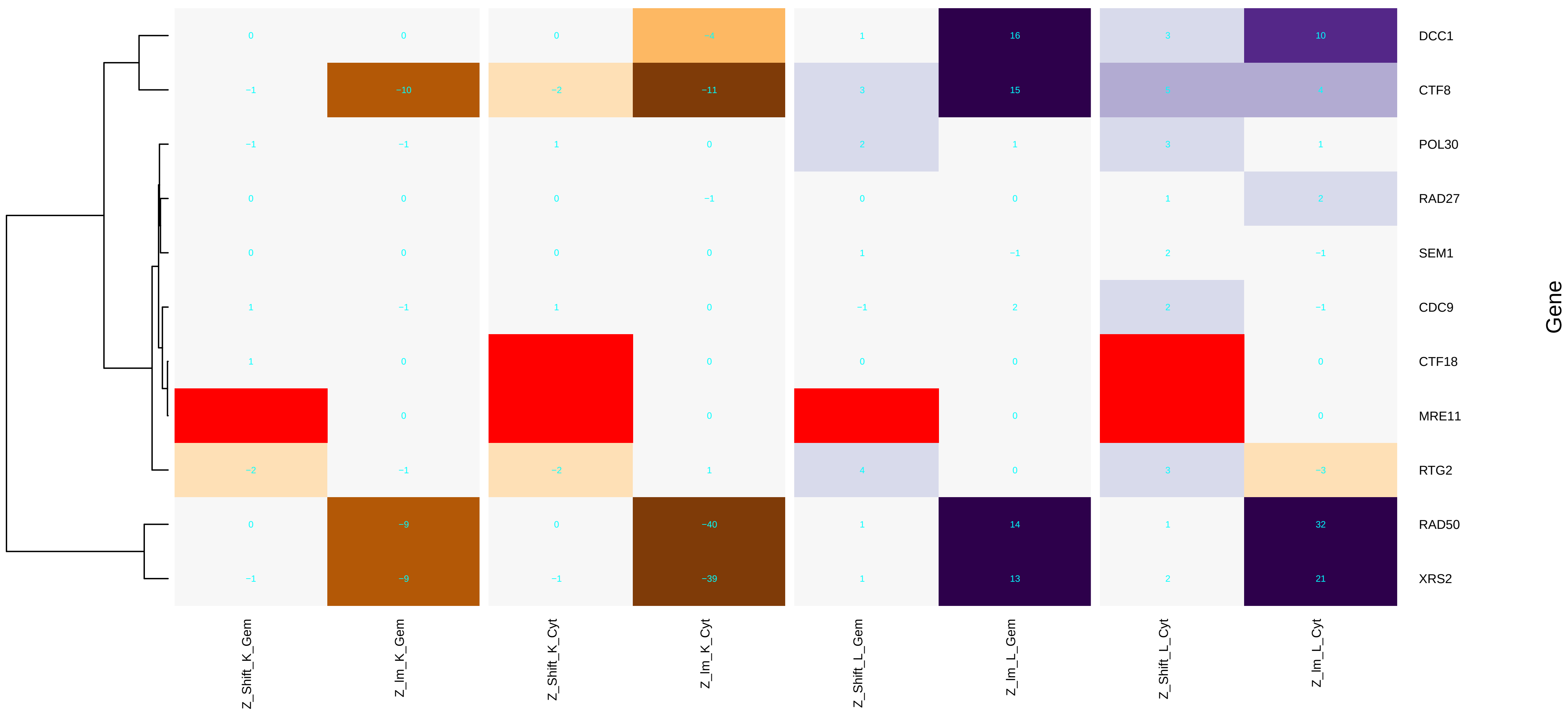

### maturation_of_5.8S_rRNA.pdf

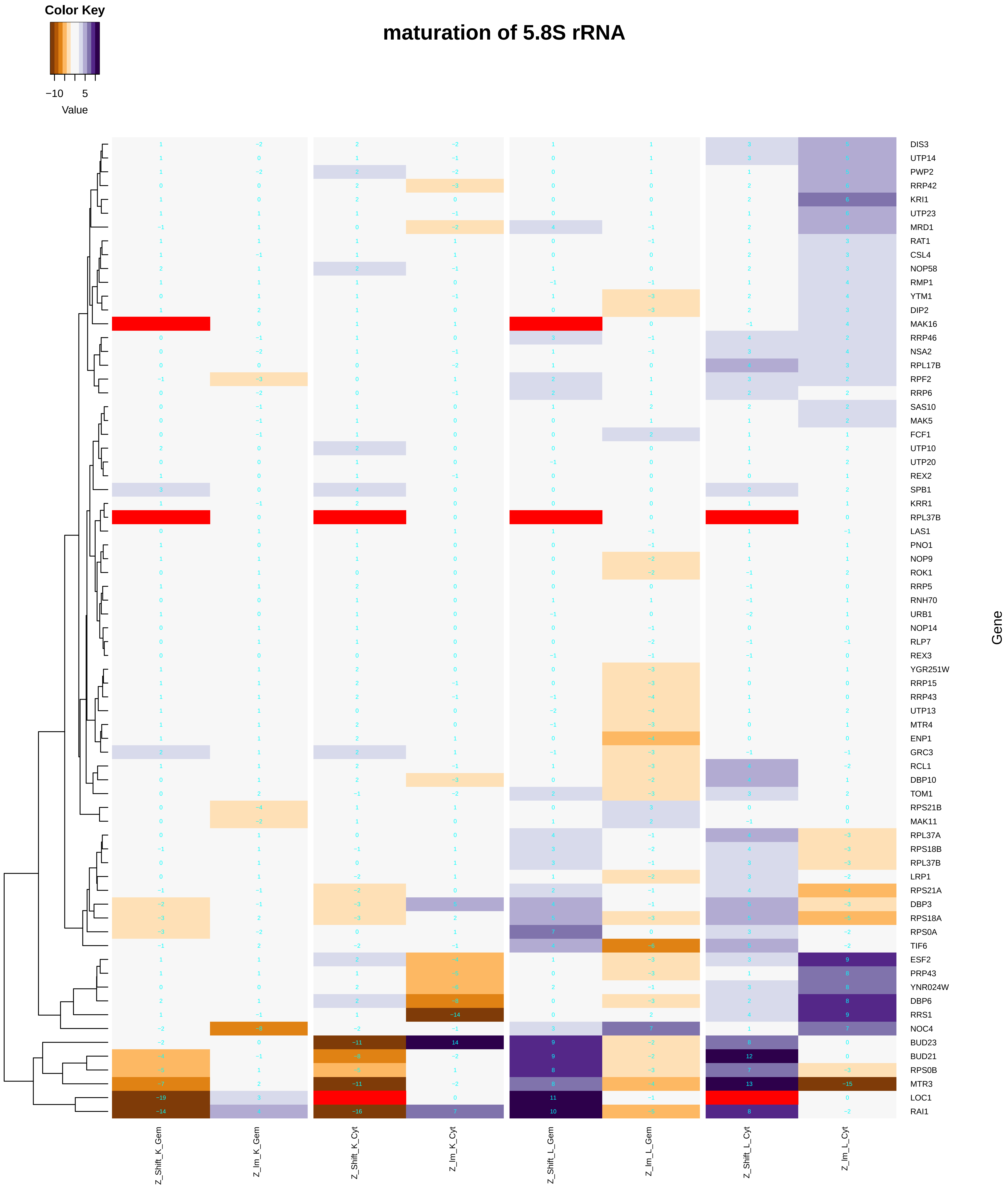

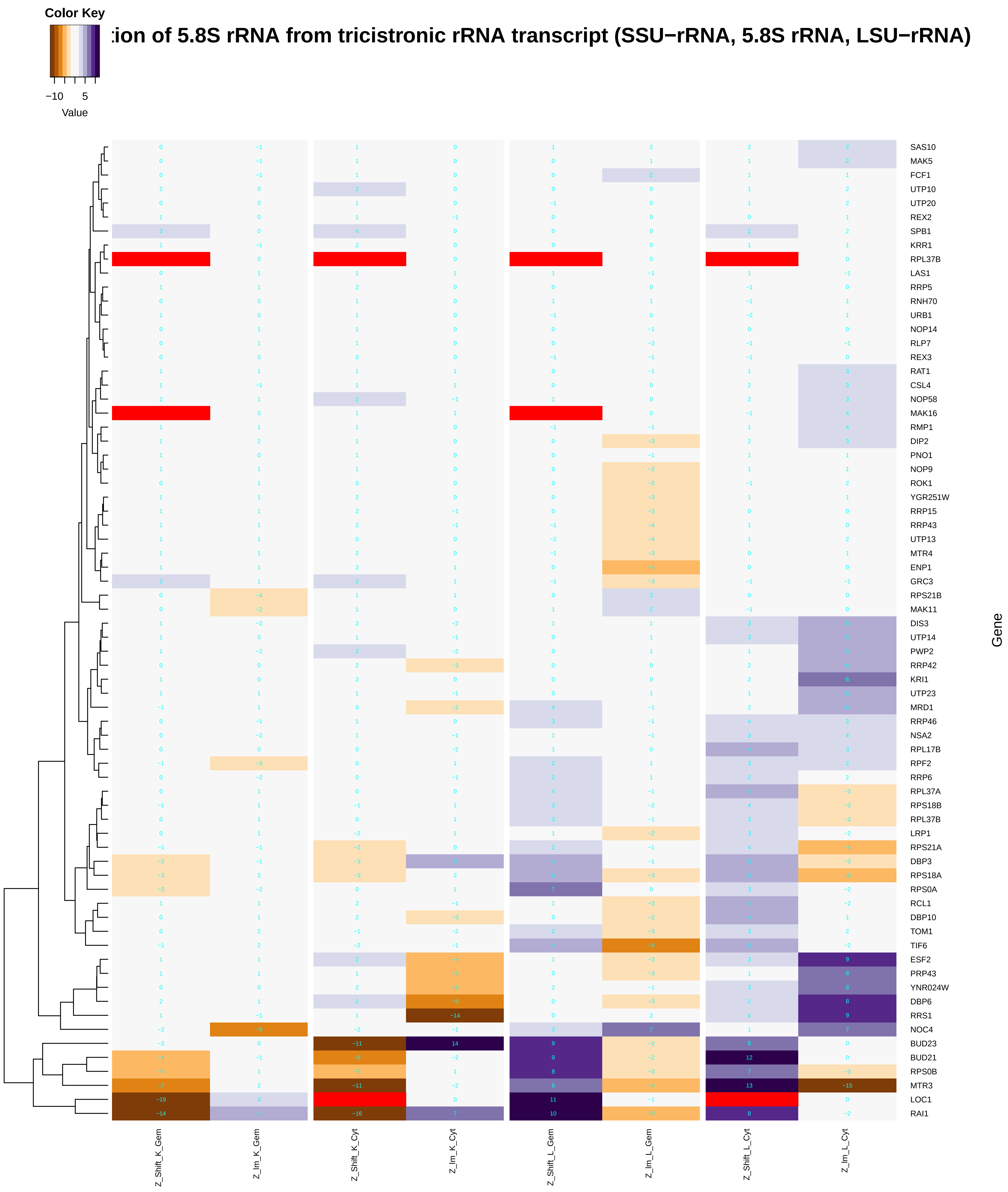

### maturation_of_5.8S_rRNA_from_tricistronic_rRNA_transcript_(SSU-rRNA,_5.8S_rRNA,_LSU-rRNA).pdf

##### :ion of 5.8S rRNA from tricistronic rRNA transcript (SSU−rRNA, 5.8S rRNA, LSU−rRNA)

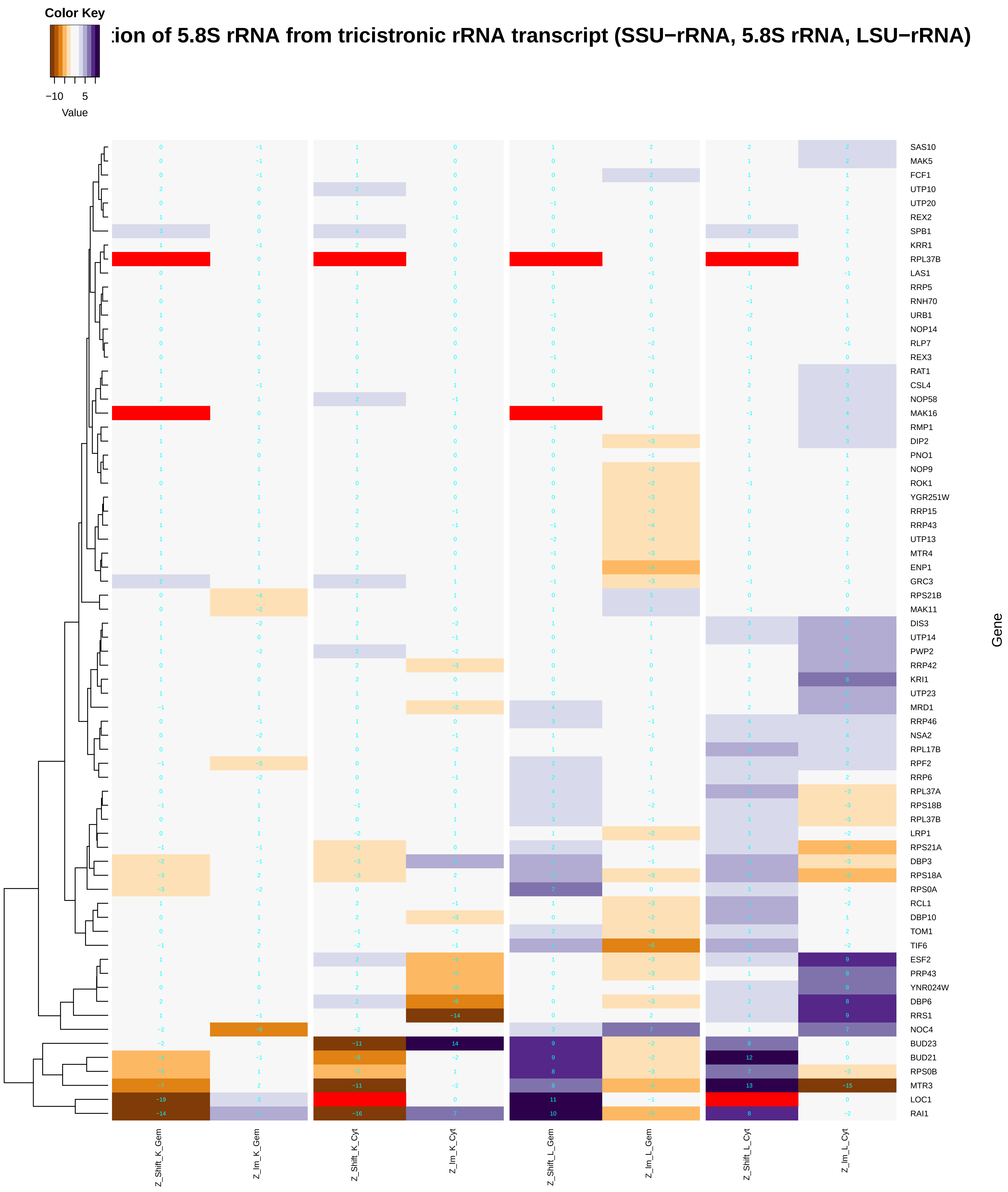

### maturation_of_LSU-rRNA.pdf

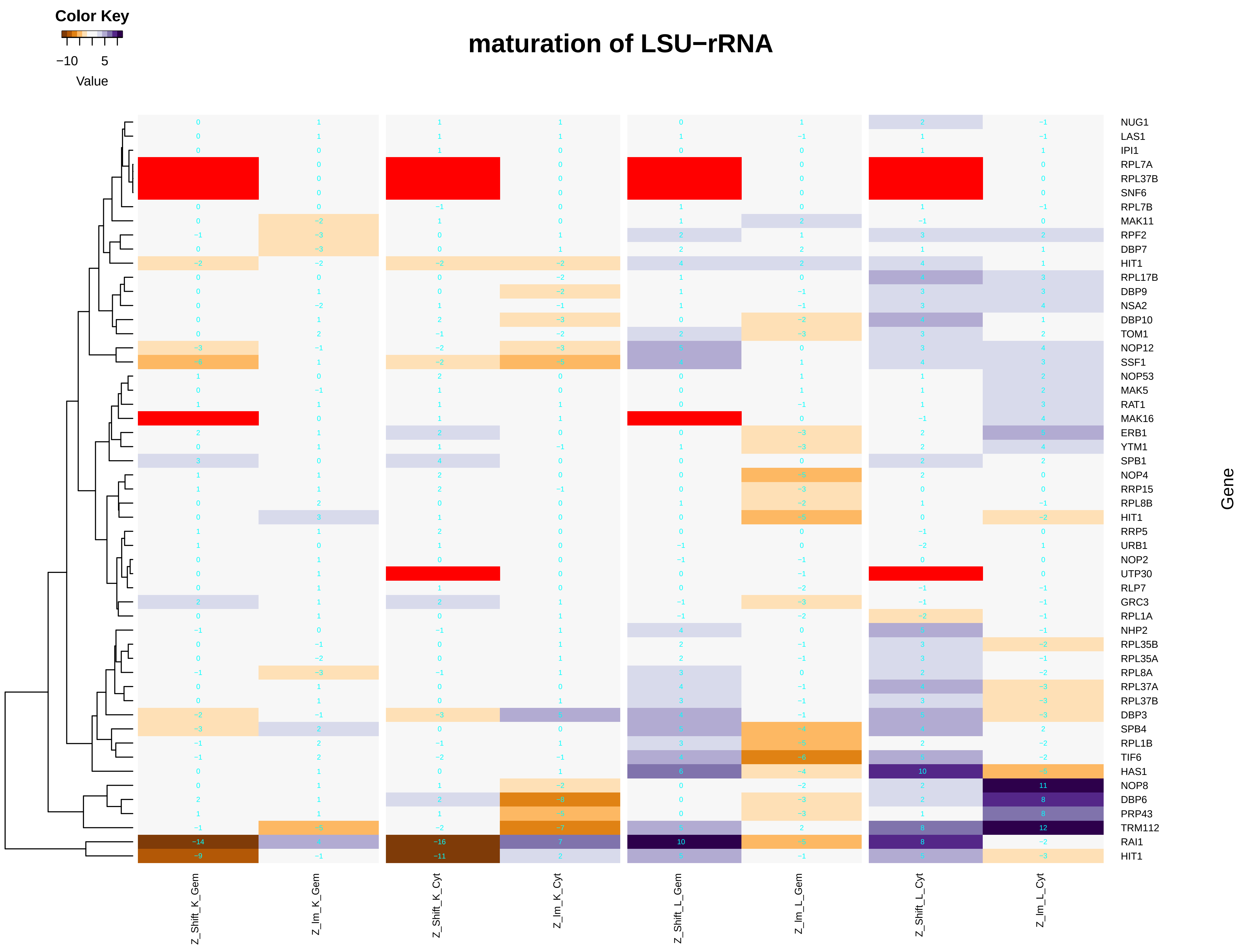

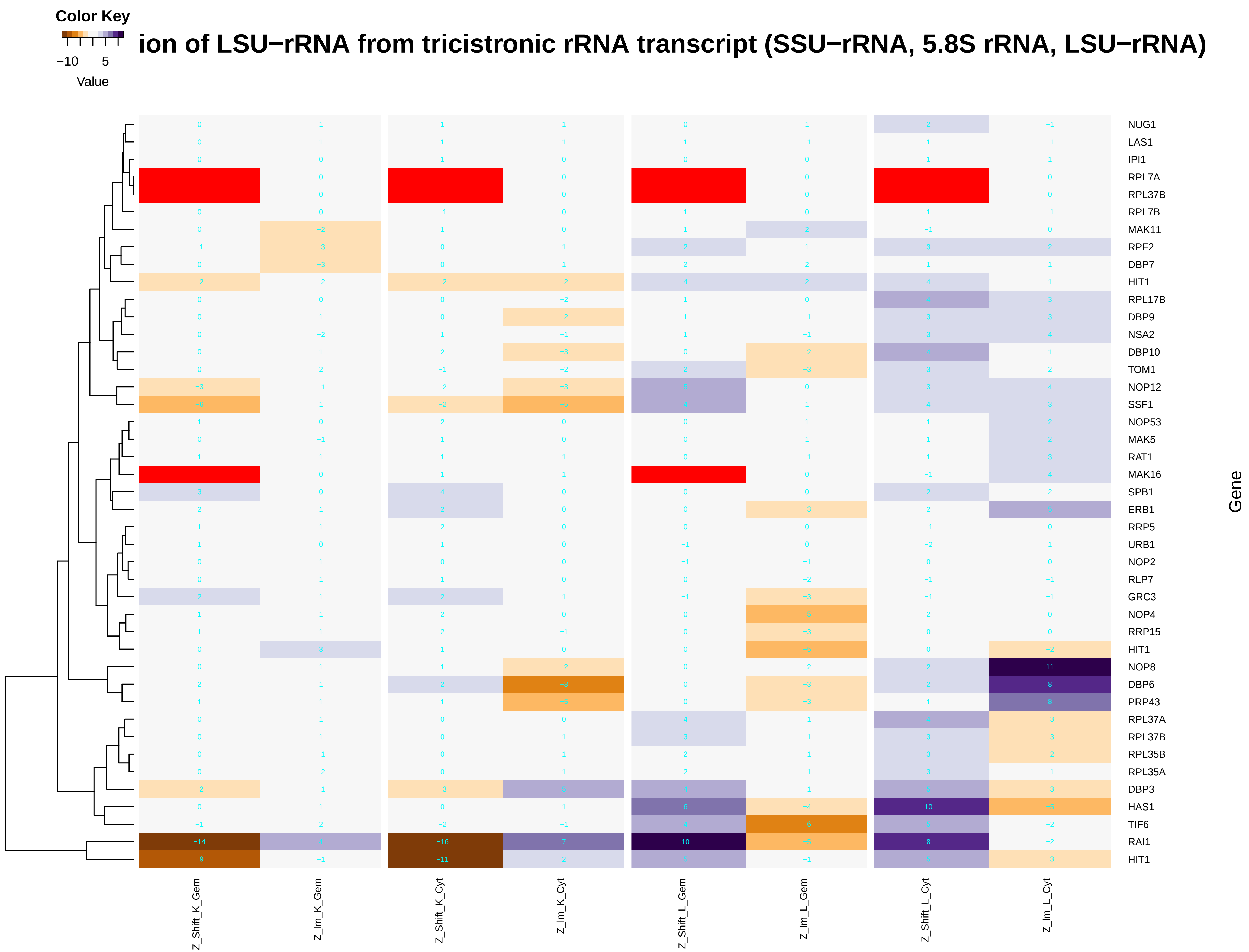

### maturation_of_LSU-rRNA_from_tricistronic_rRNA_transcript_(SSU-rRNA,_5.8S_rRNA,_LSU-rRNA).pdf

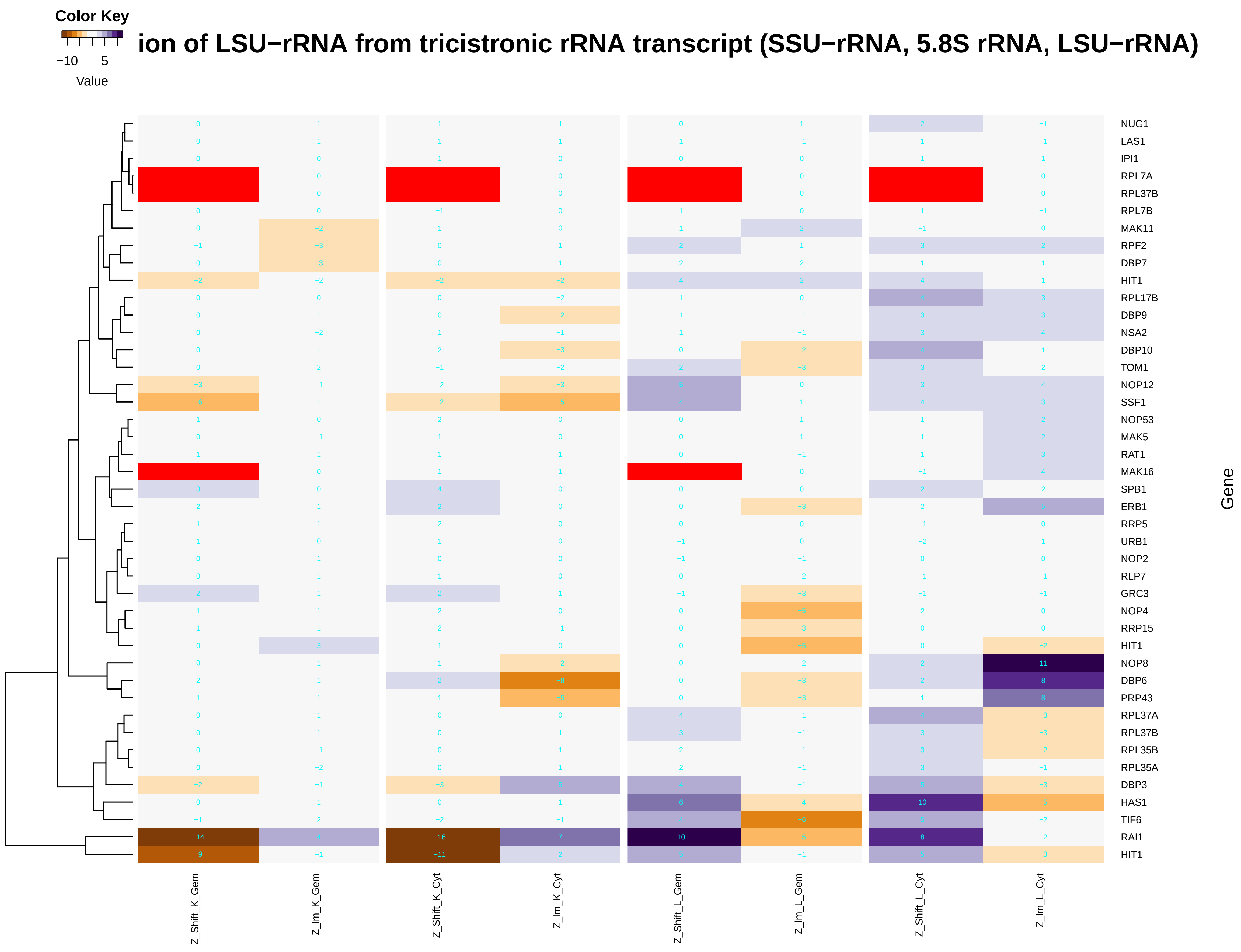

### maturation_of_SSU-rRNA.pdf

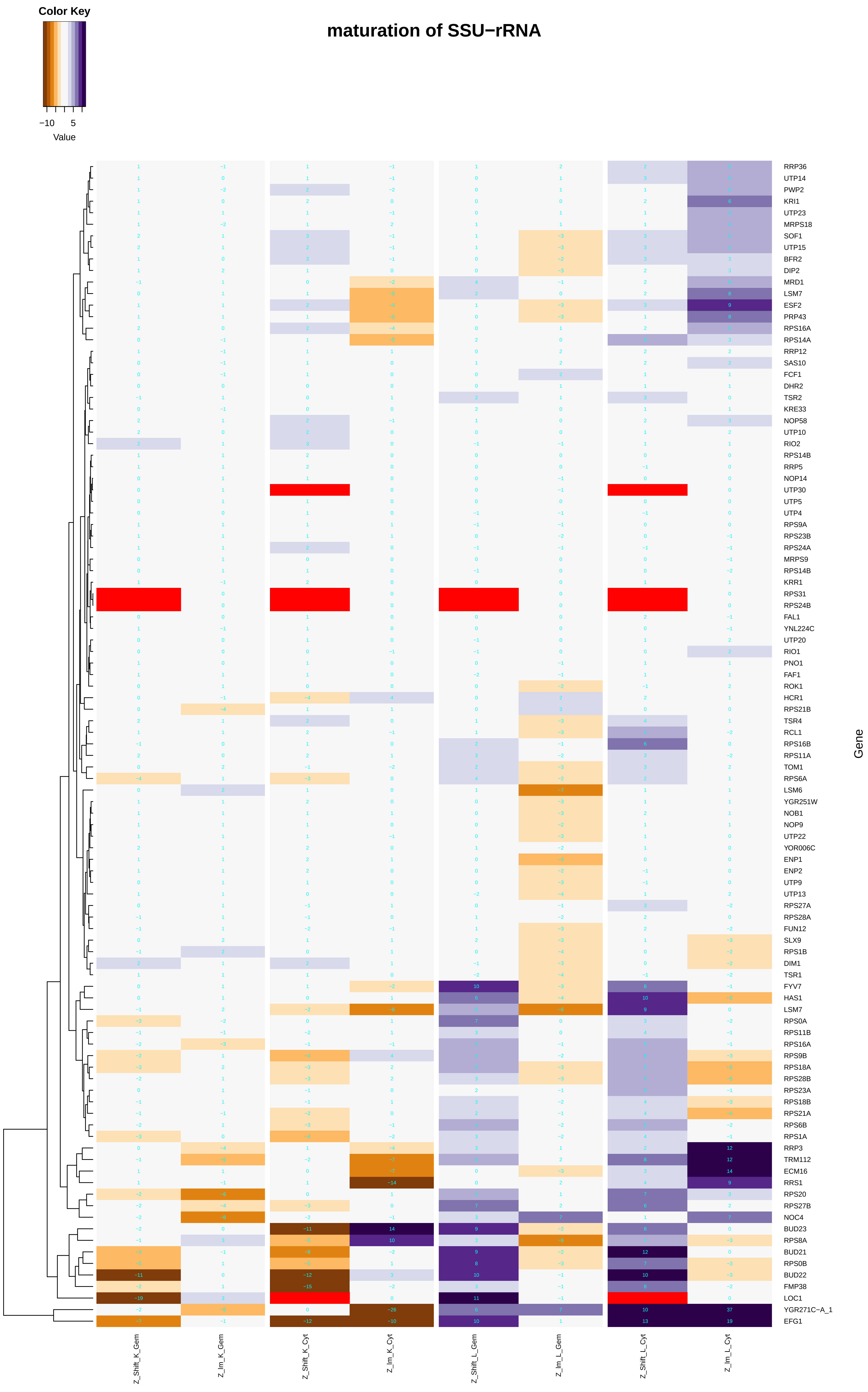

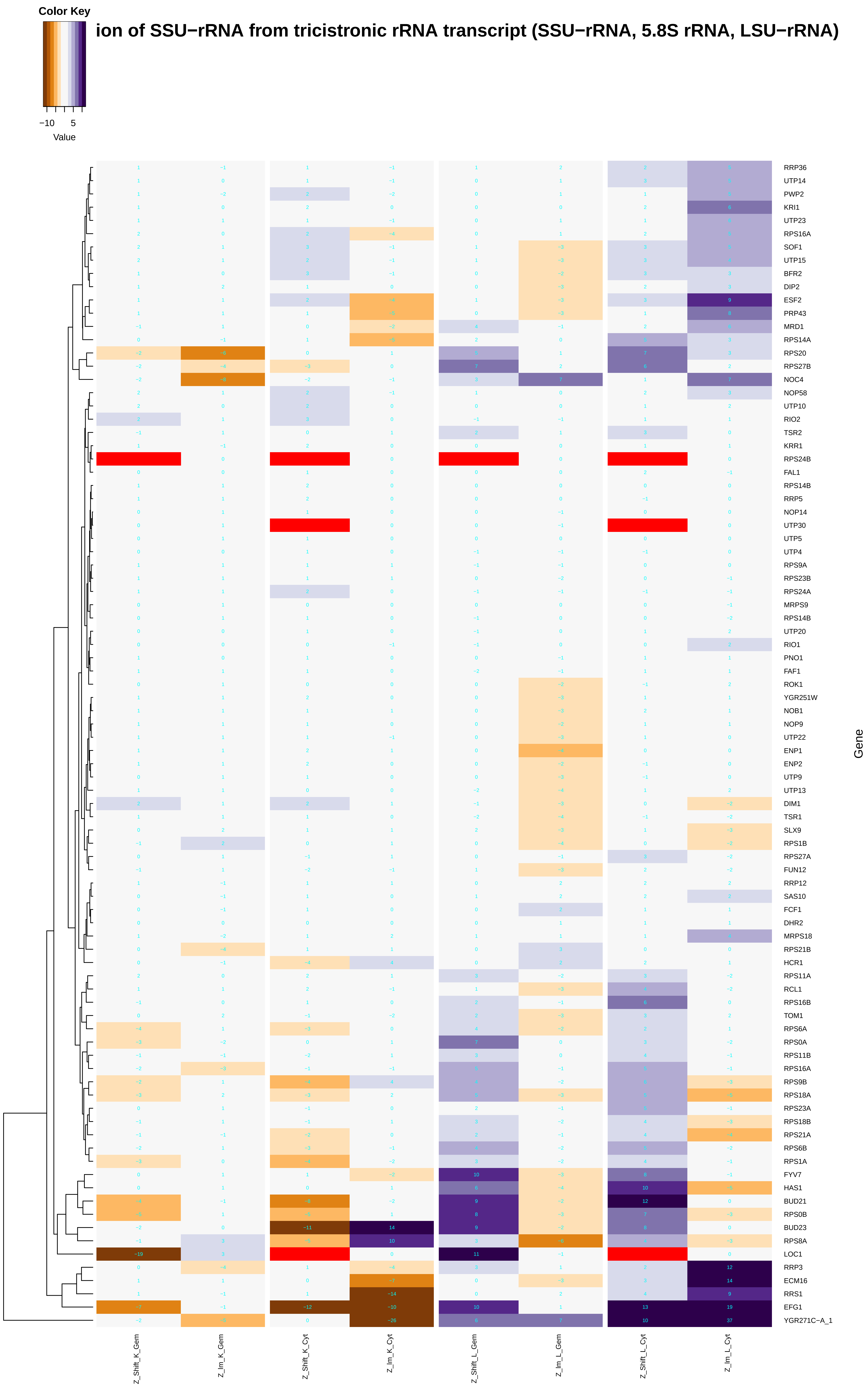

### maturation_of_SSU-rRNA_from_tricistronic_rRNA_transcript_(SSU-rRNA,_5.8S_rRNA,_LSU-rRNA).pdf

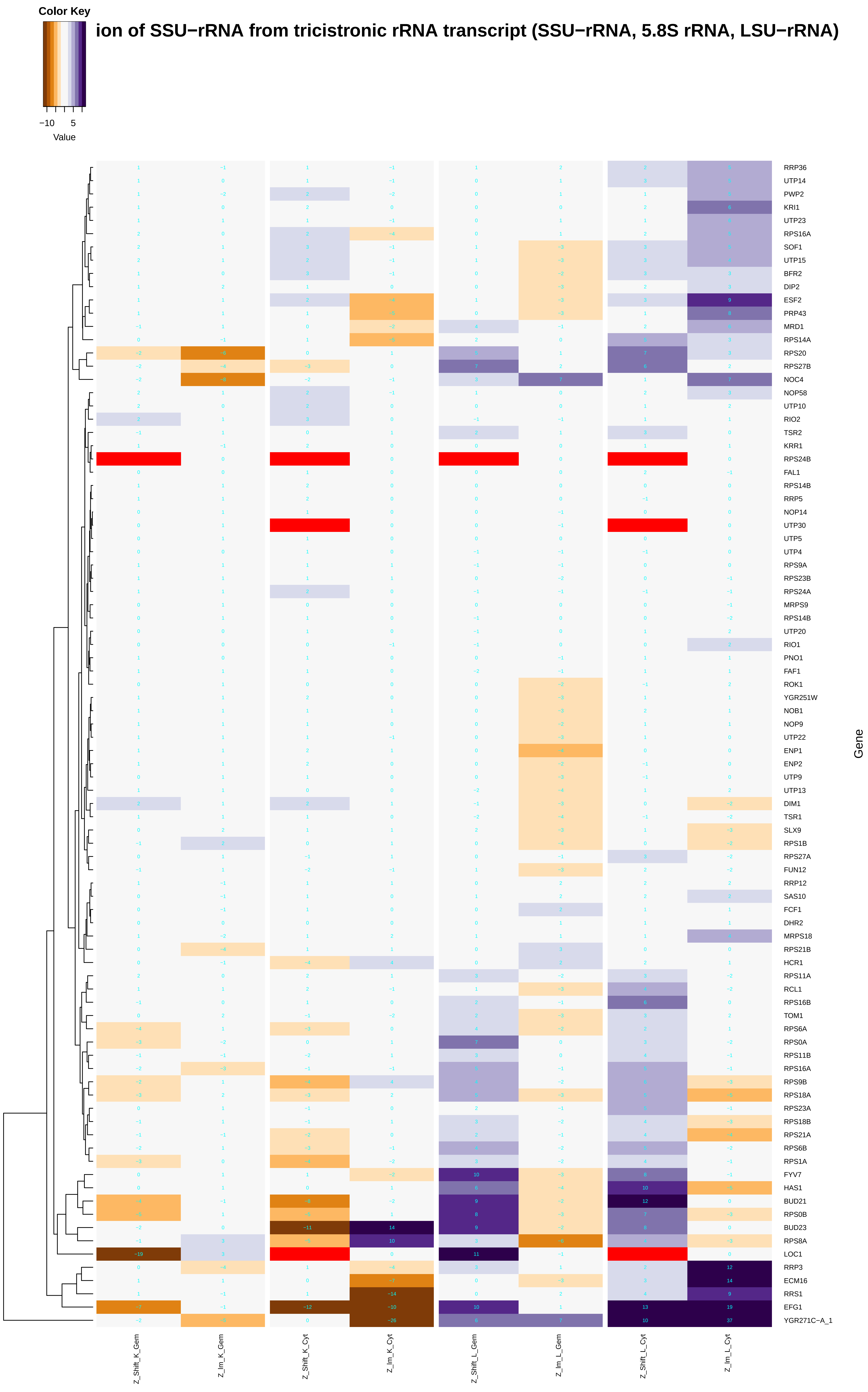

### mediator_complex.pdf

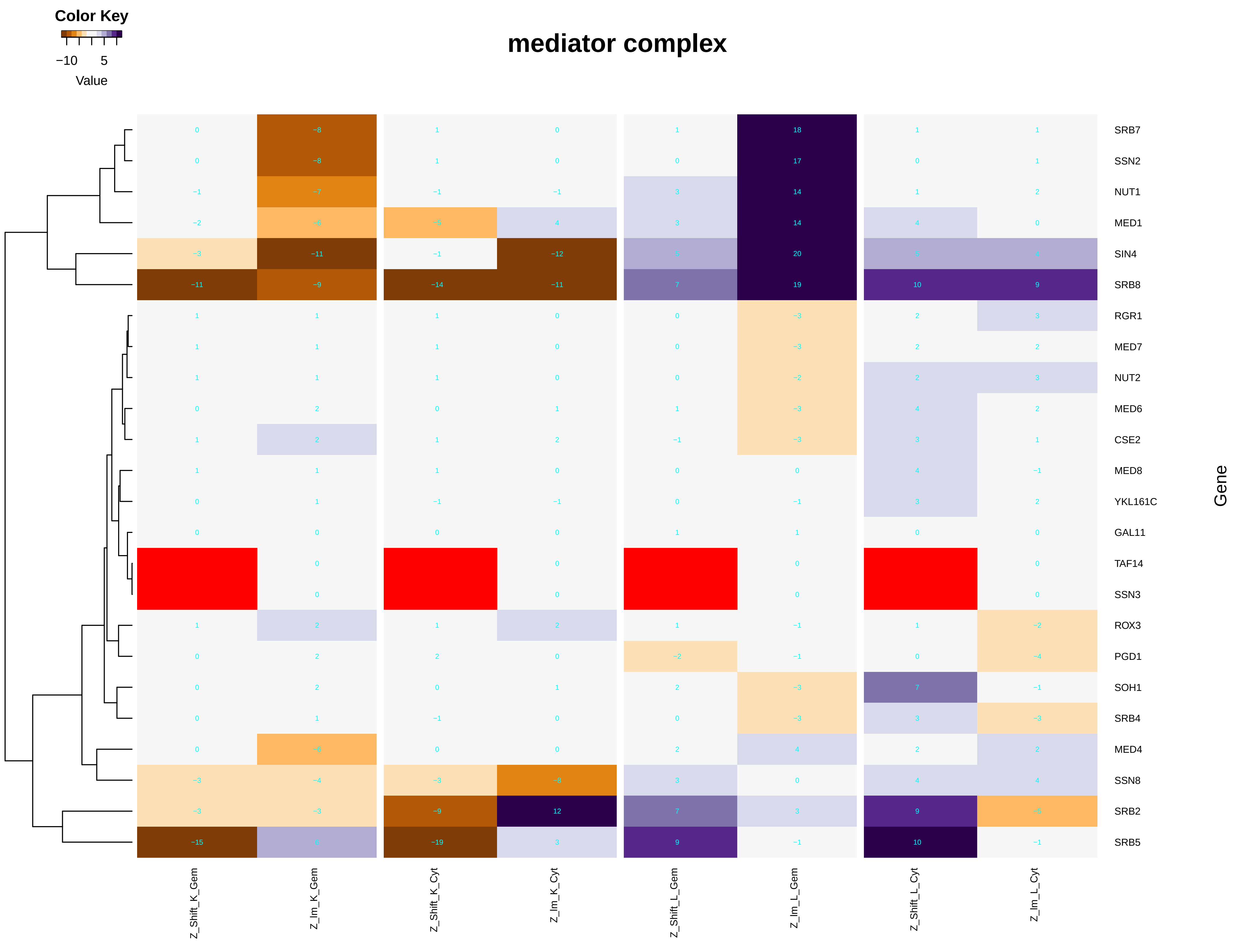

### meiosis_I.pdf

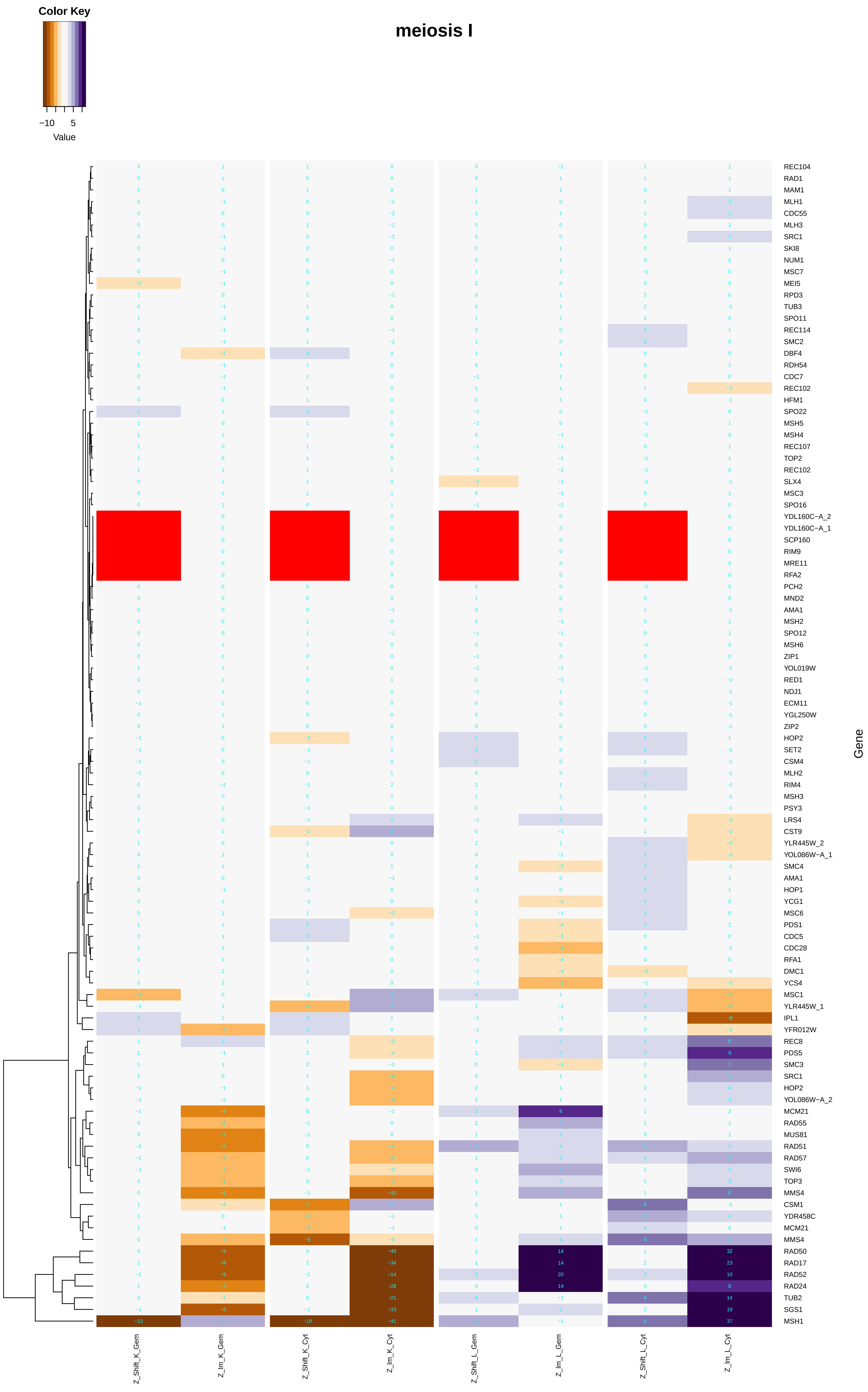

### meiotic_cell_cycle.pdf

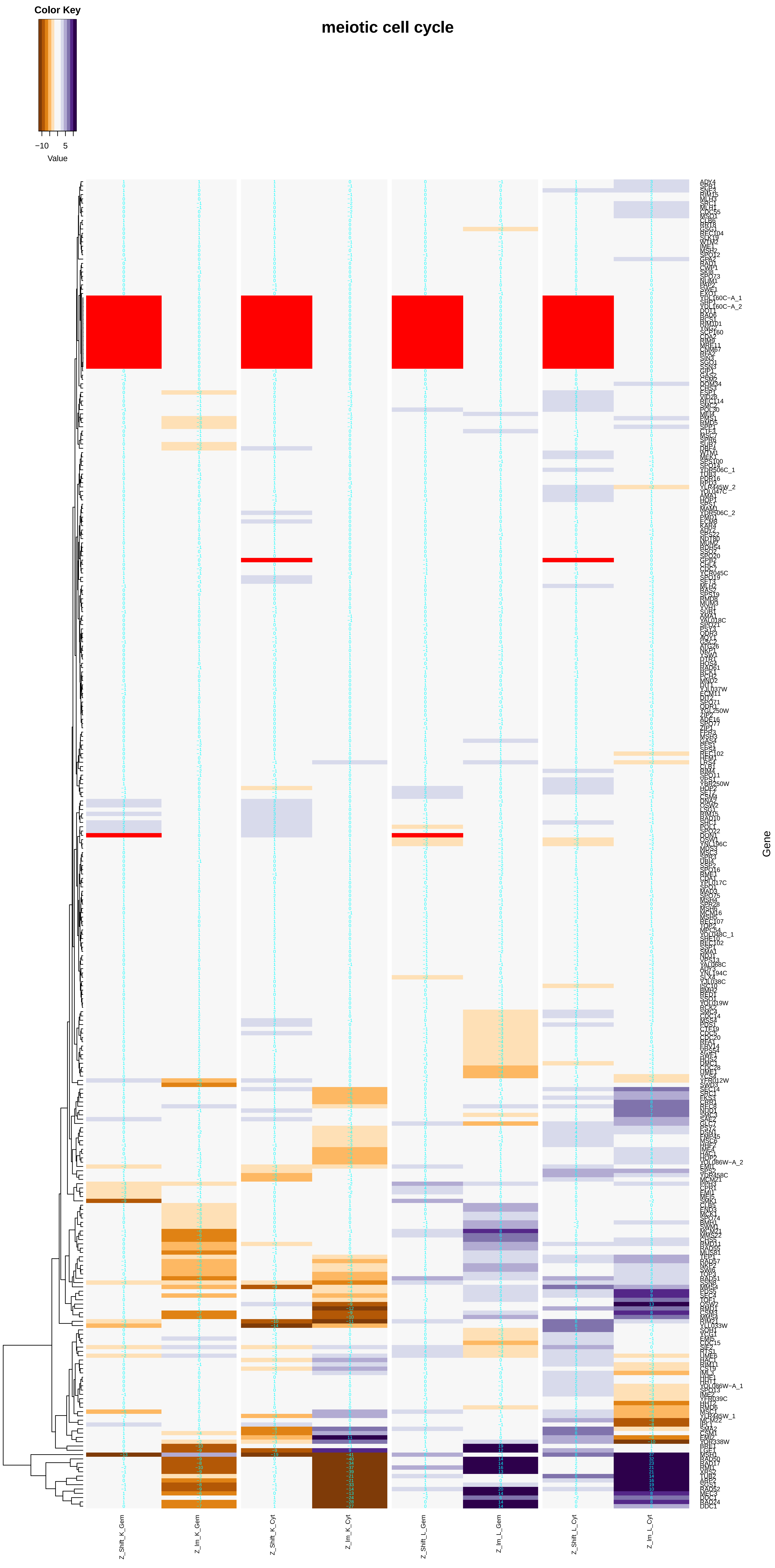

### meiotic_DNA_double-strand_break_formation.pdf

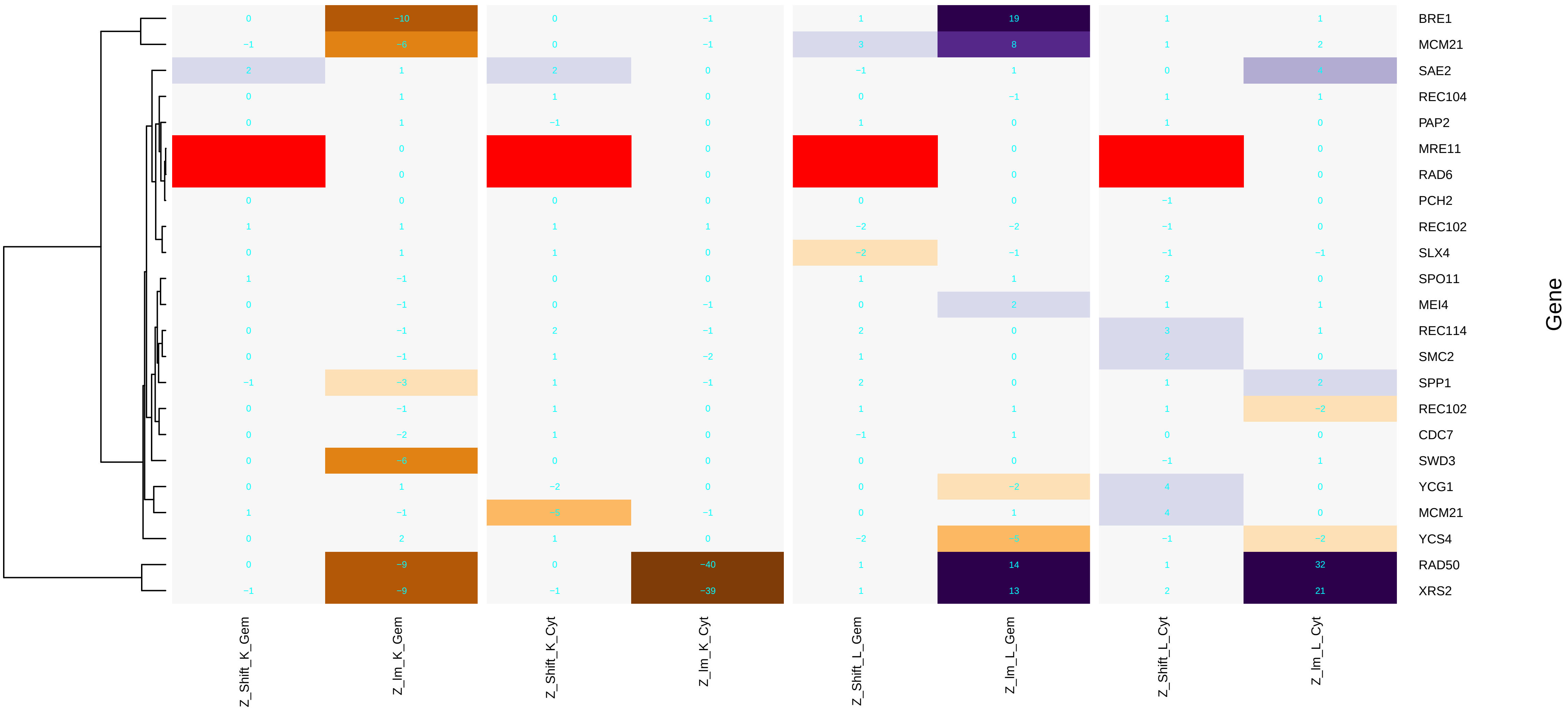

### meiotic_DNA_double-strand_break_processing.pdf

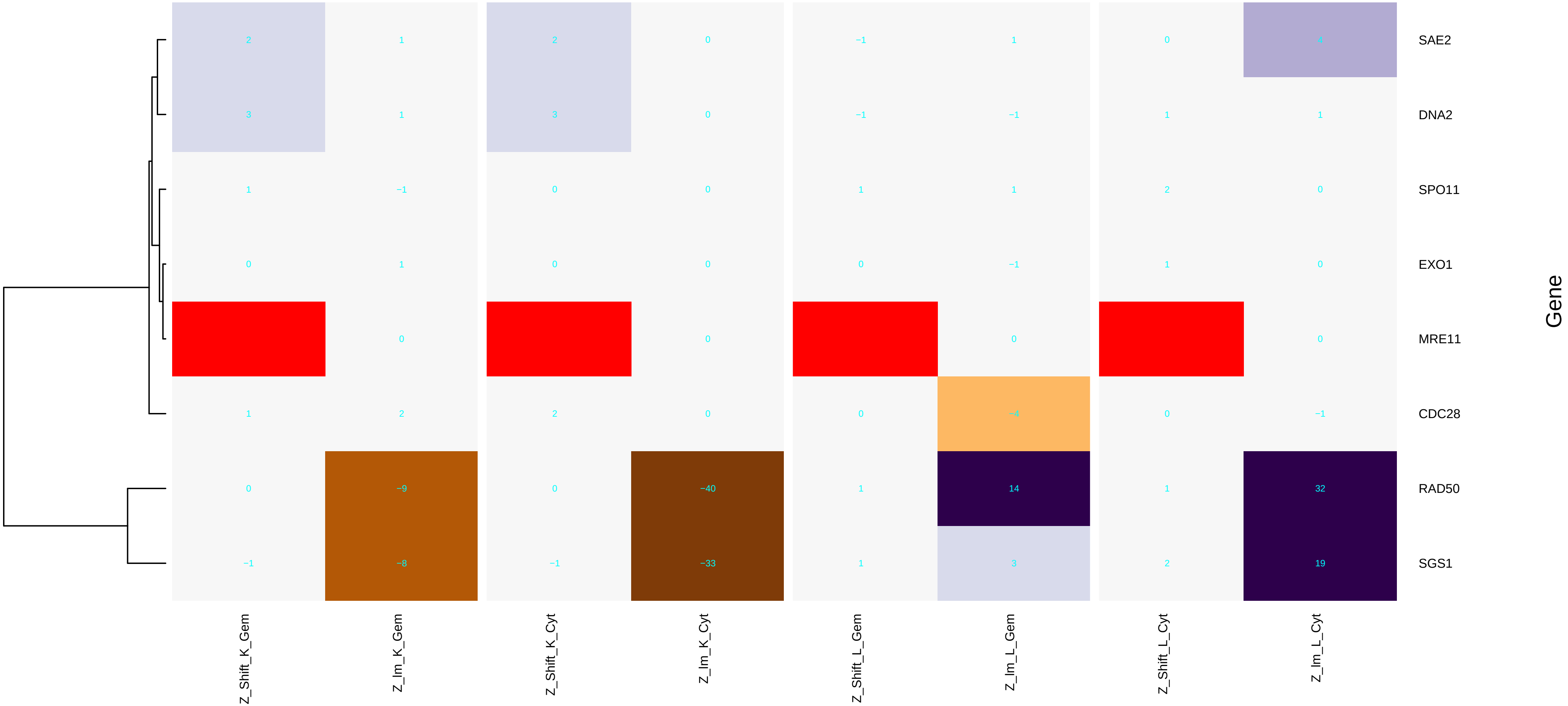

### membrane_fusion.pdf

#### membrane fusion

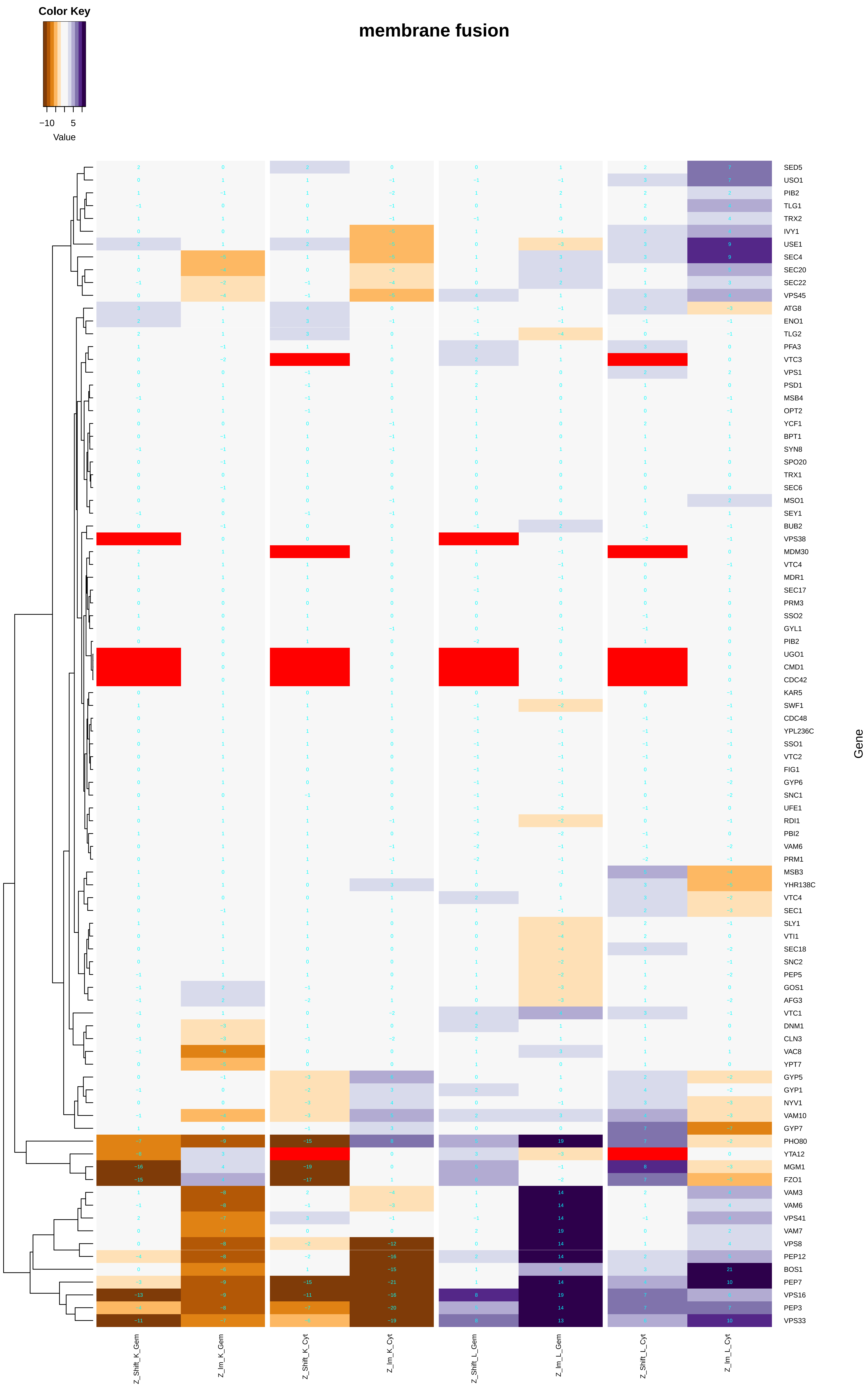

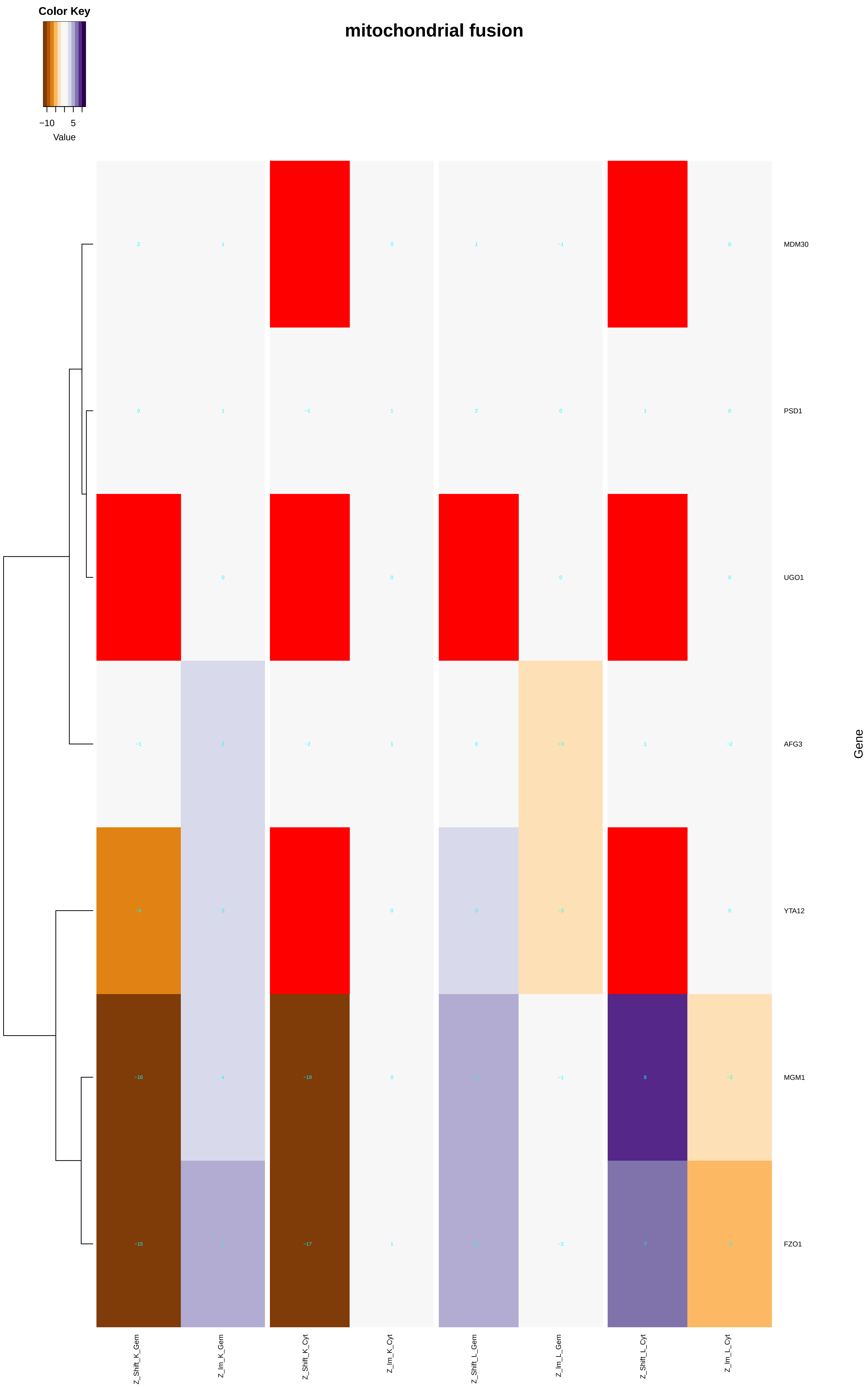

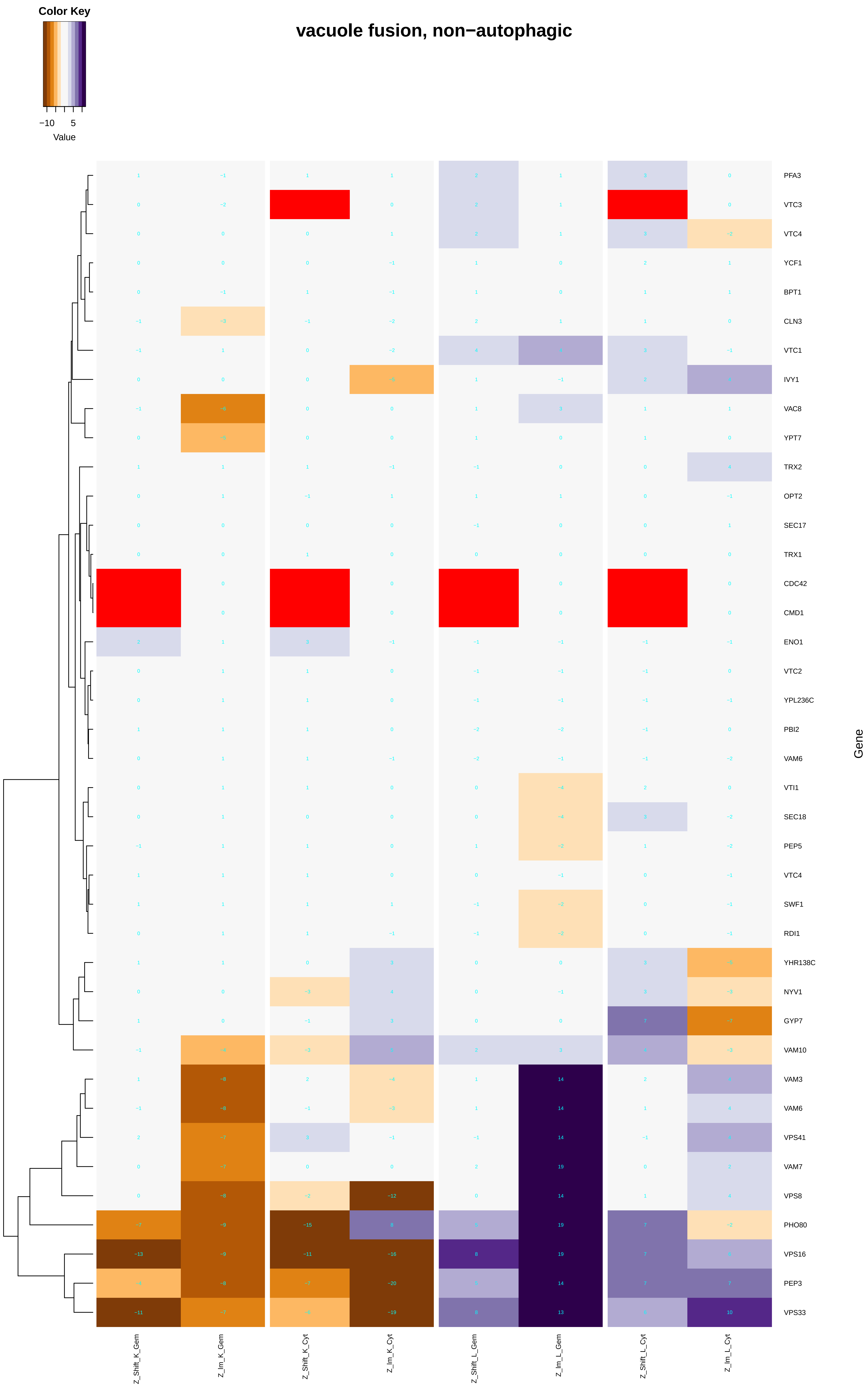

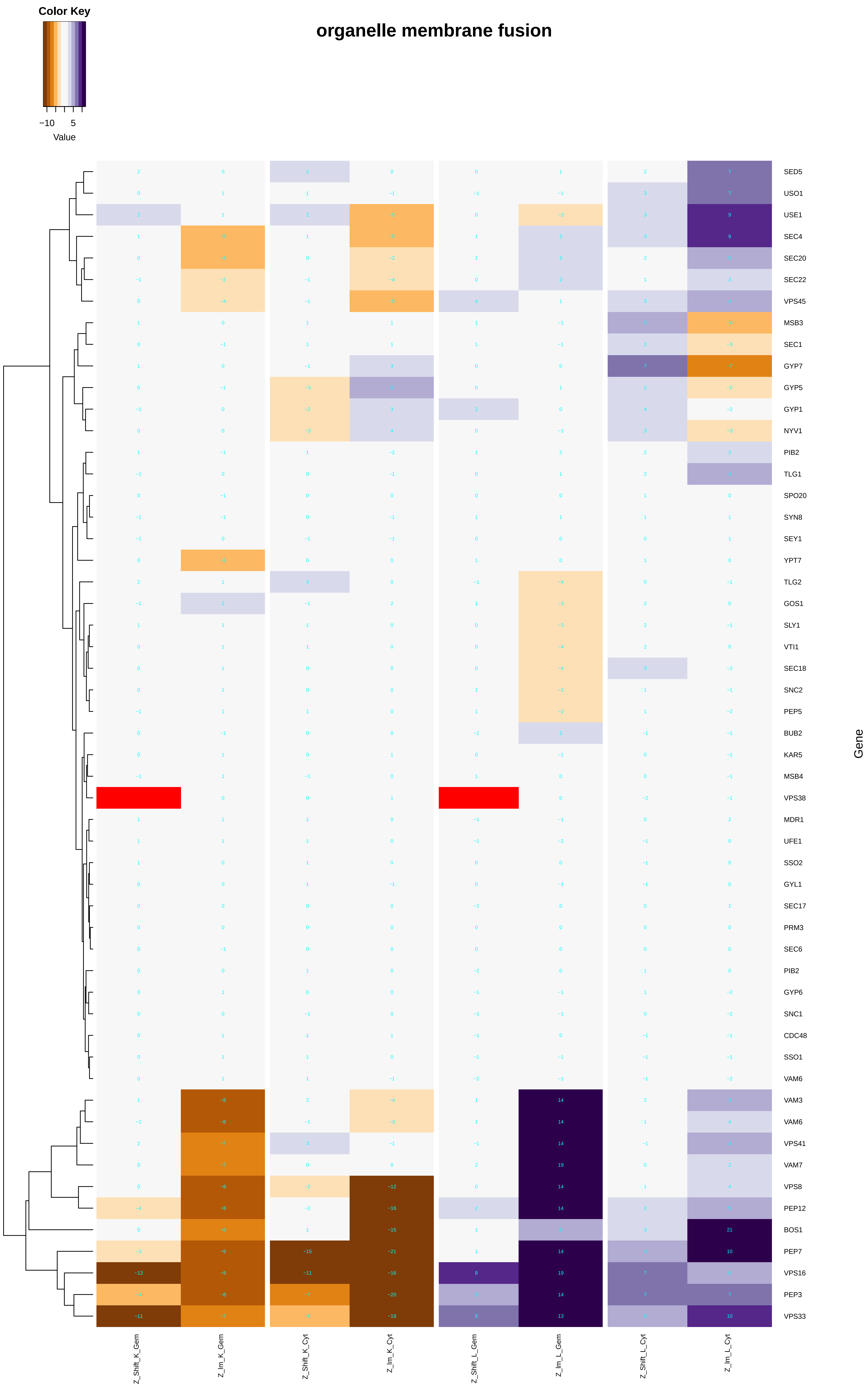

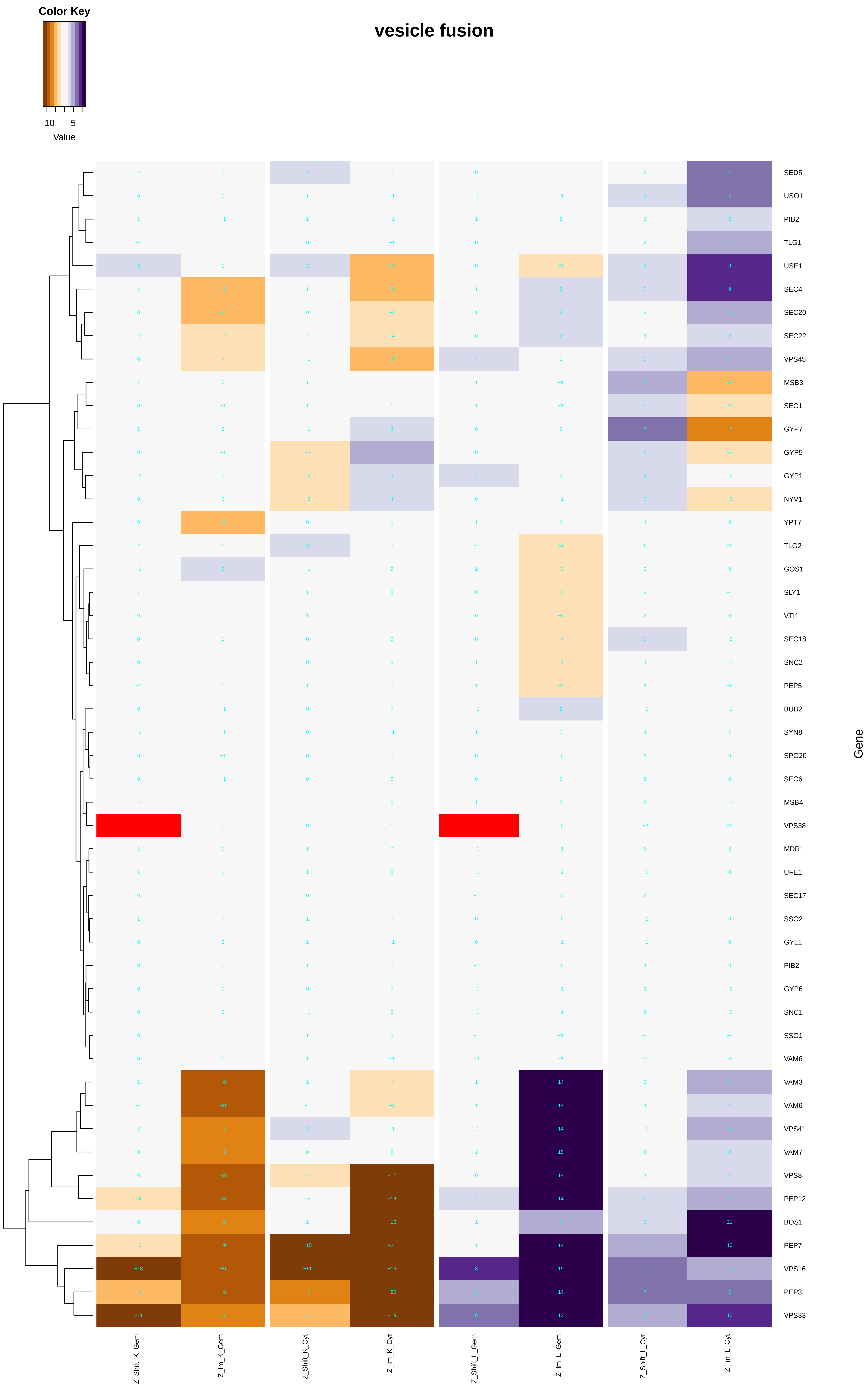

#### endoplasmic reticulum membrane fusion

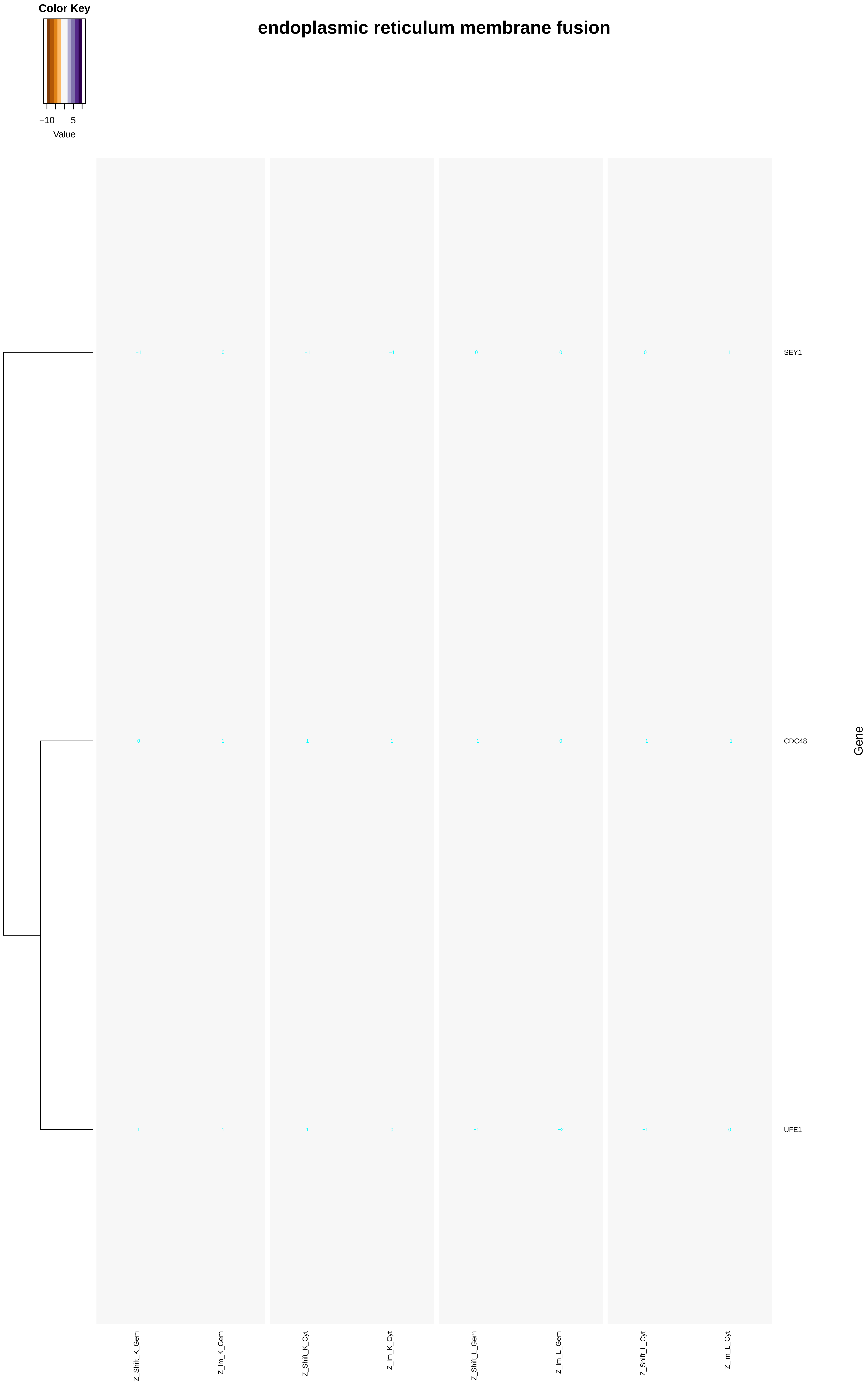

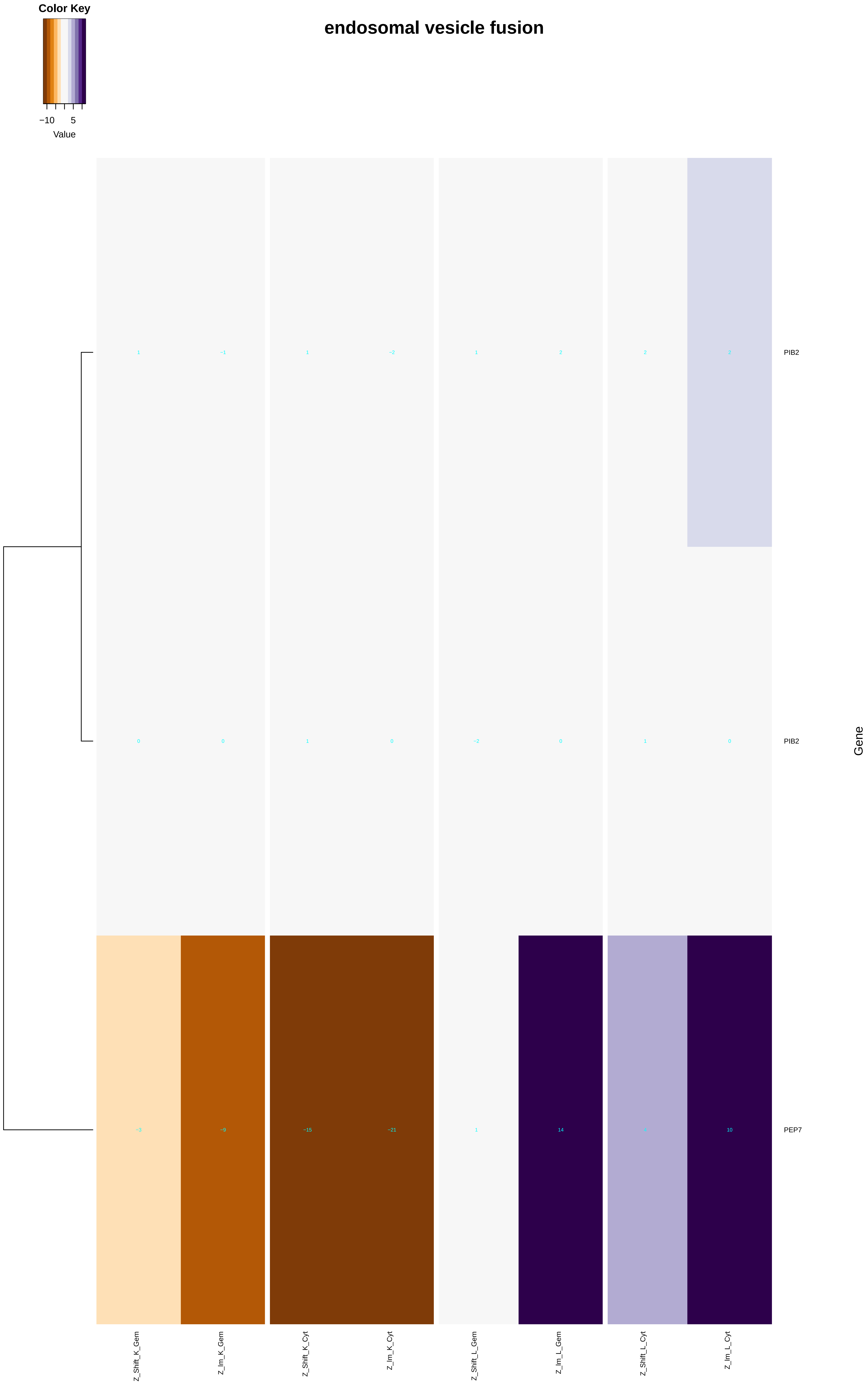

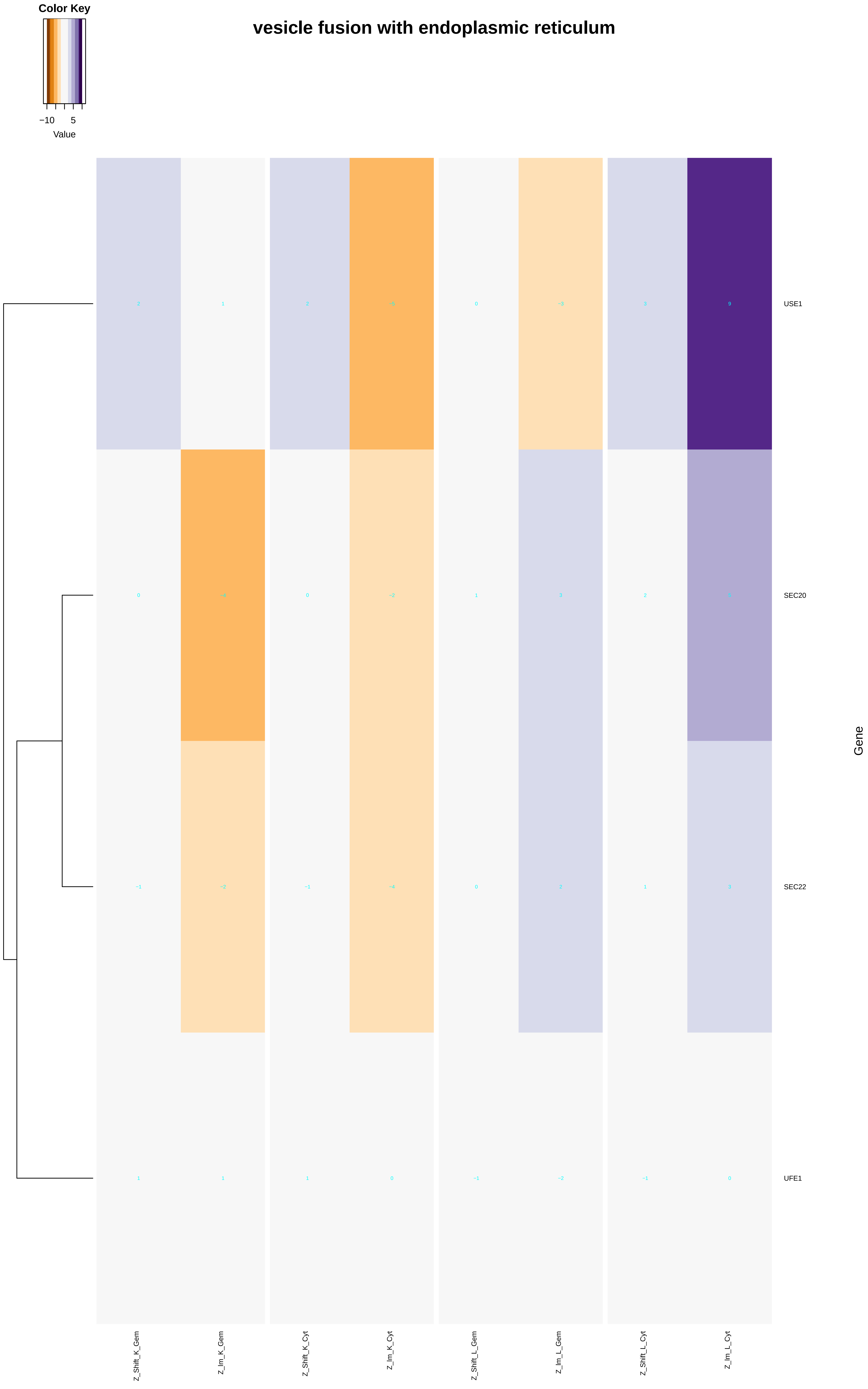

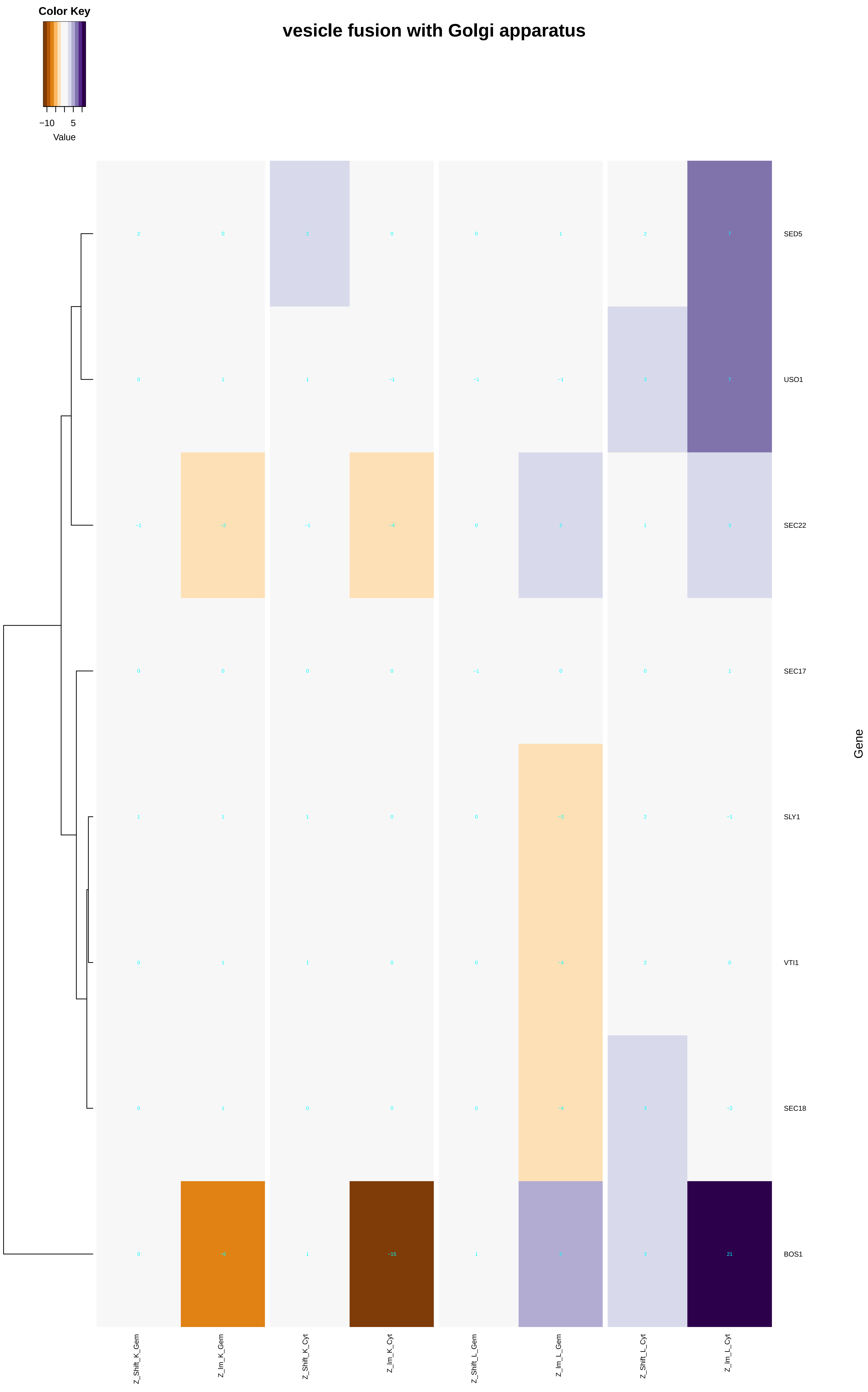

### mitochondrial_double-strand_break_repair_via_homologous_recombination.pdf

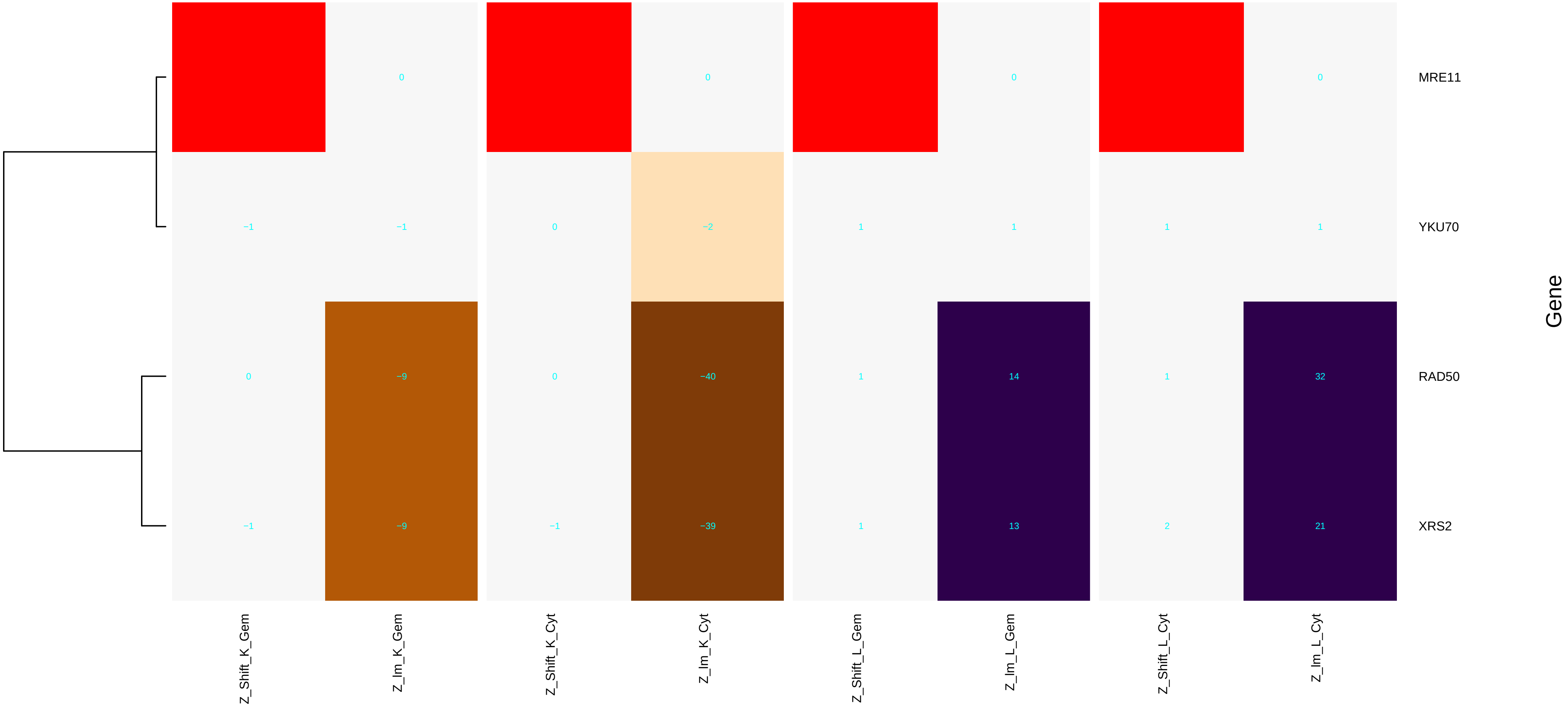

### mitochondrial_fusion.pdf

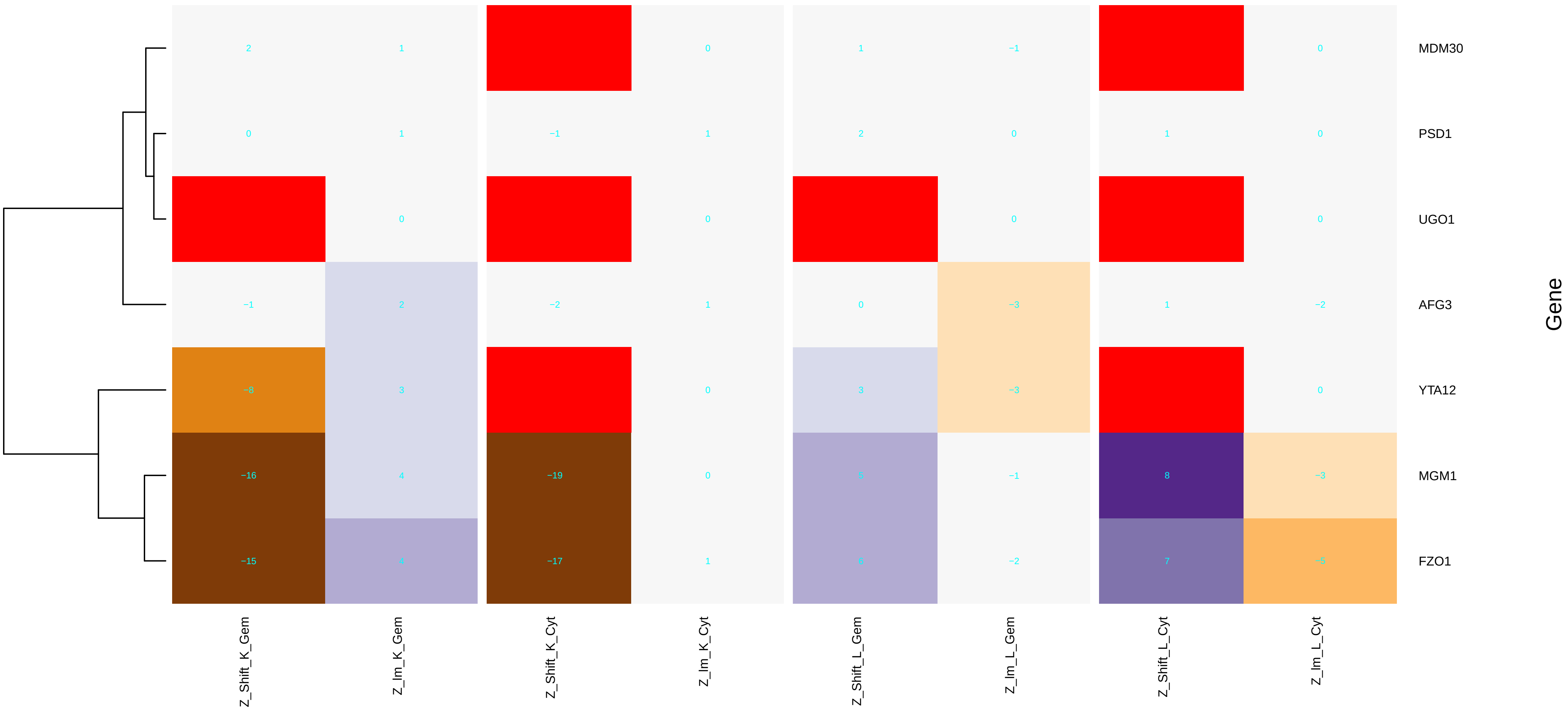

### mitochondrial_inner_boundary_membrane.pdf

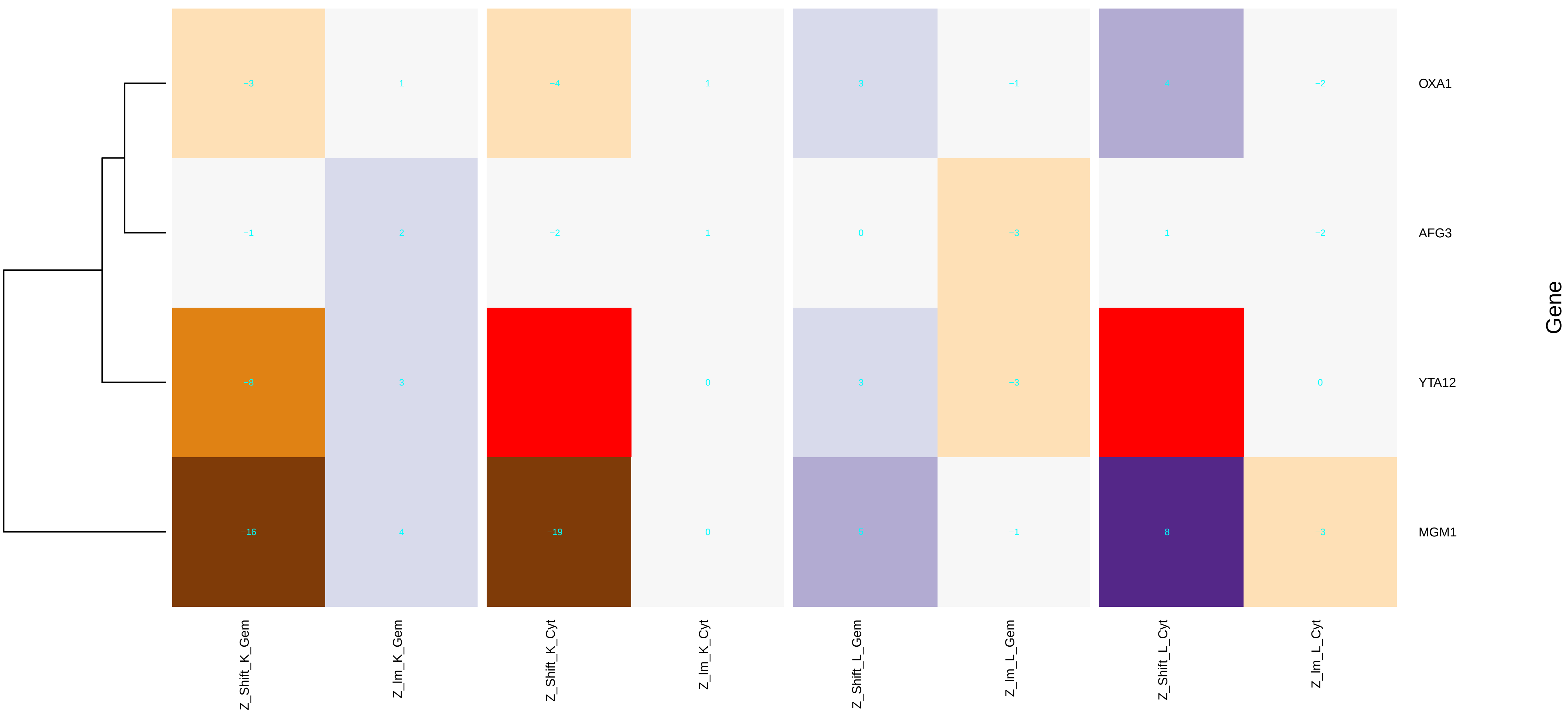

### mitochondrial_inner_membrane.pdf

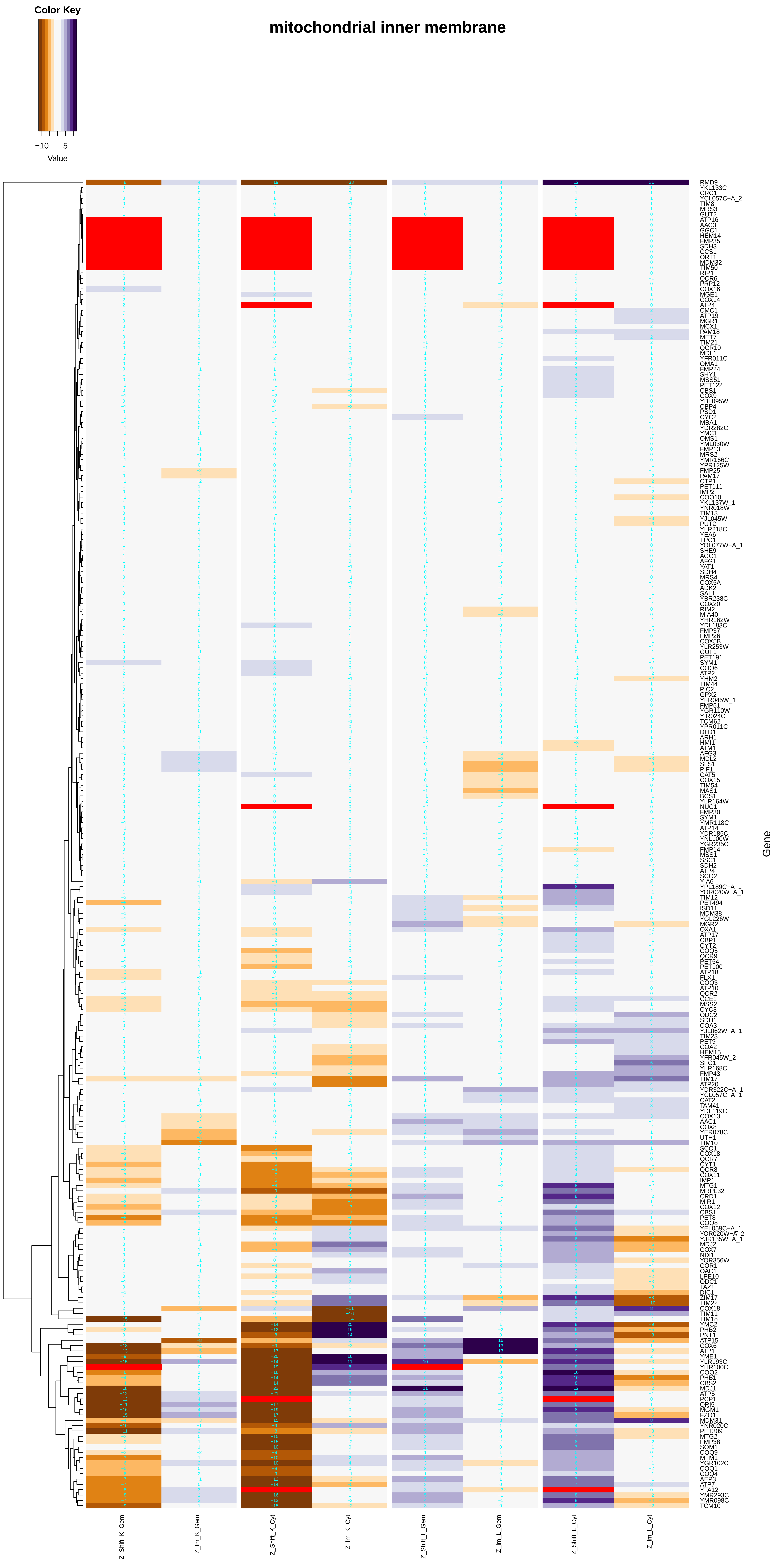

### mitochondrial_large_ribosomal_subunit.pdf

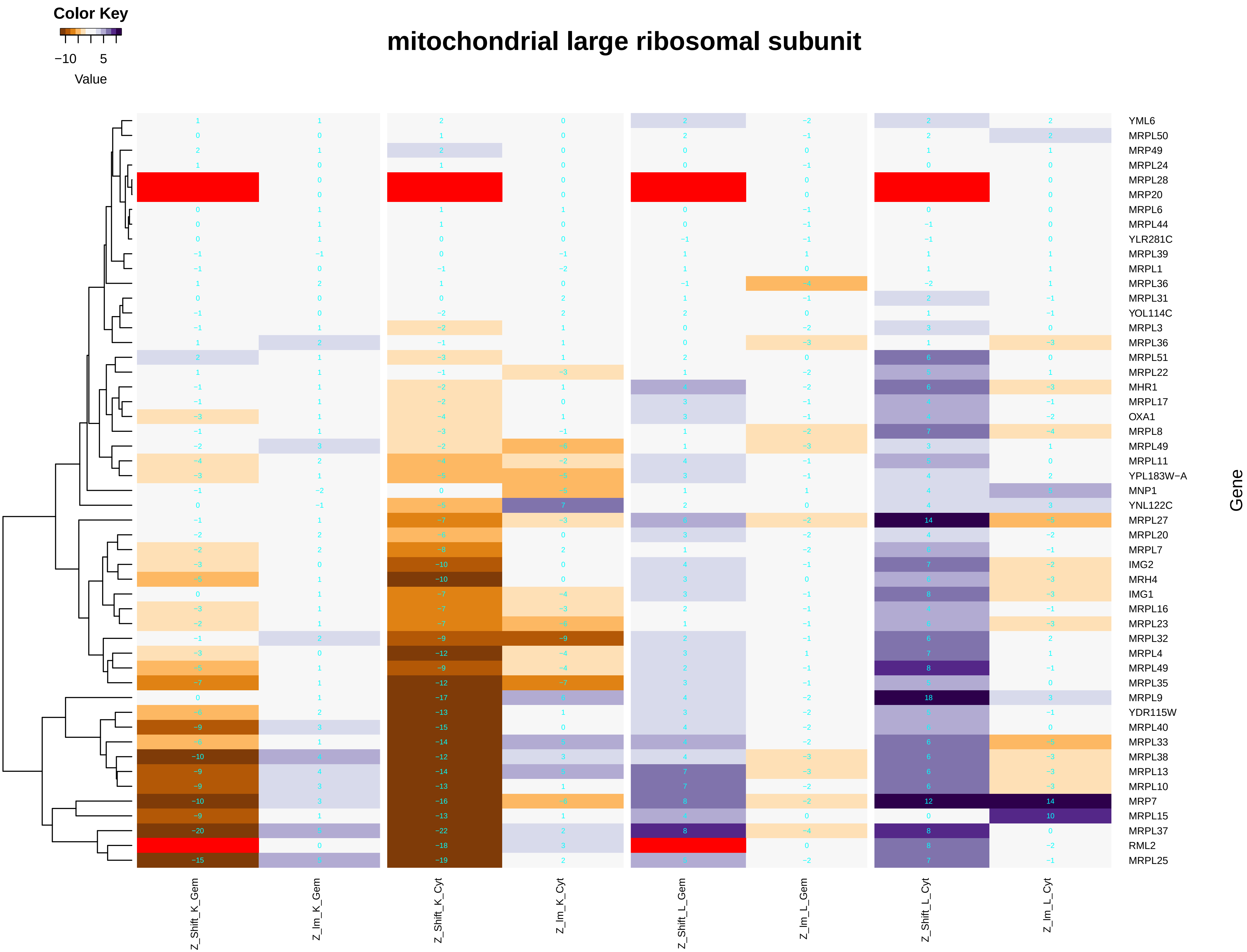

### mitochondrial_matrix.pdf

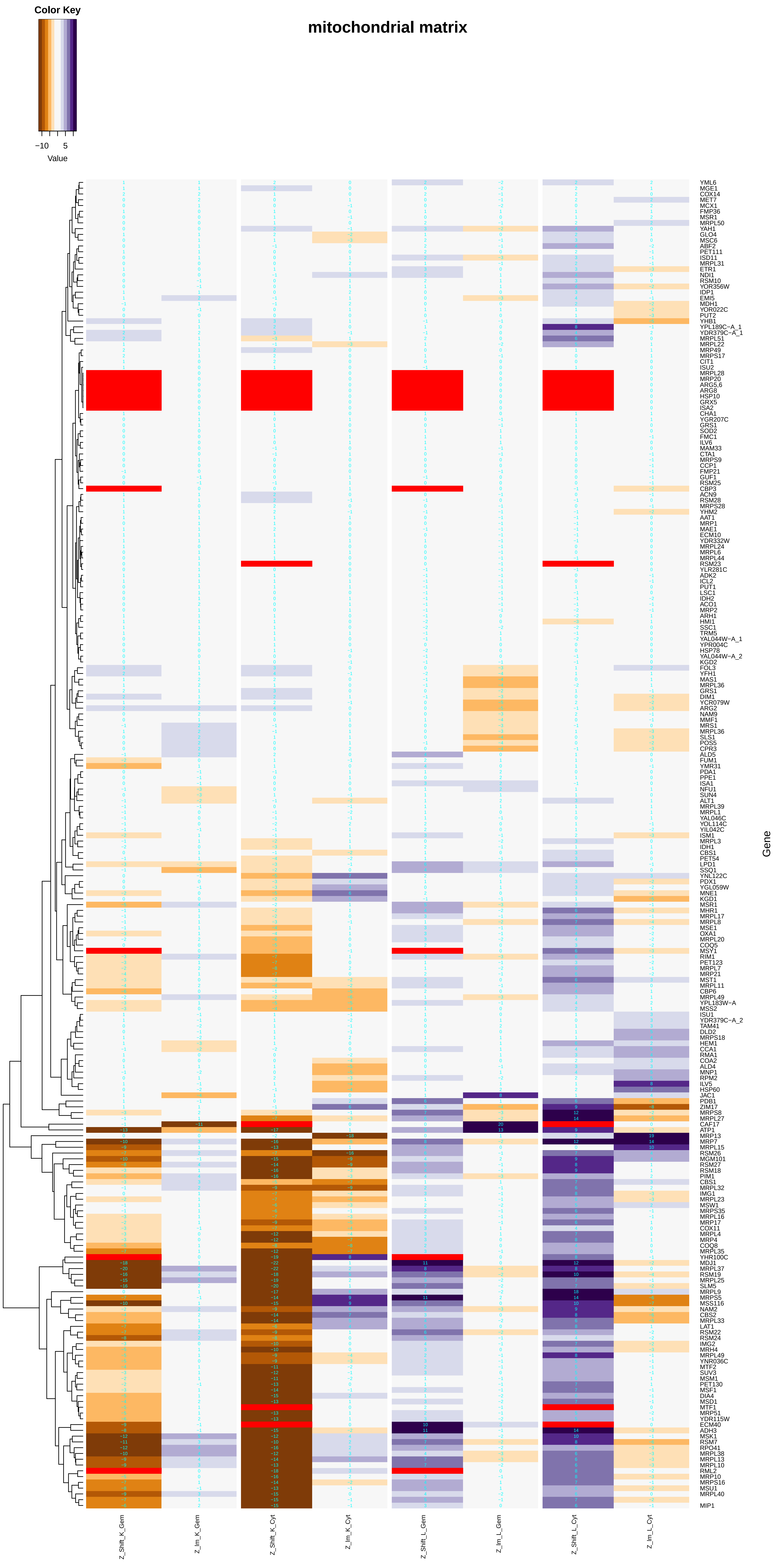

### mitochondrial_proton-transporting_ATP_synthase_complex,_catalytic_core_F(1).pdf

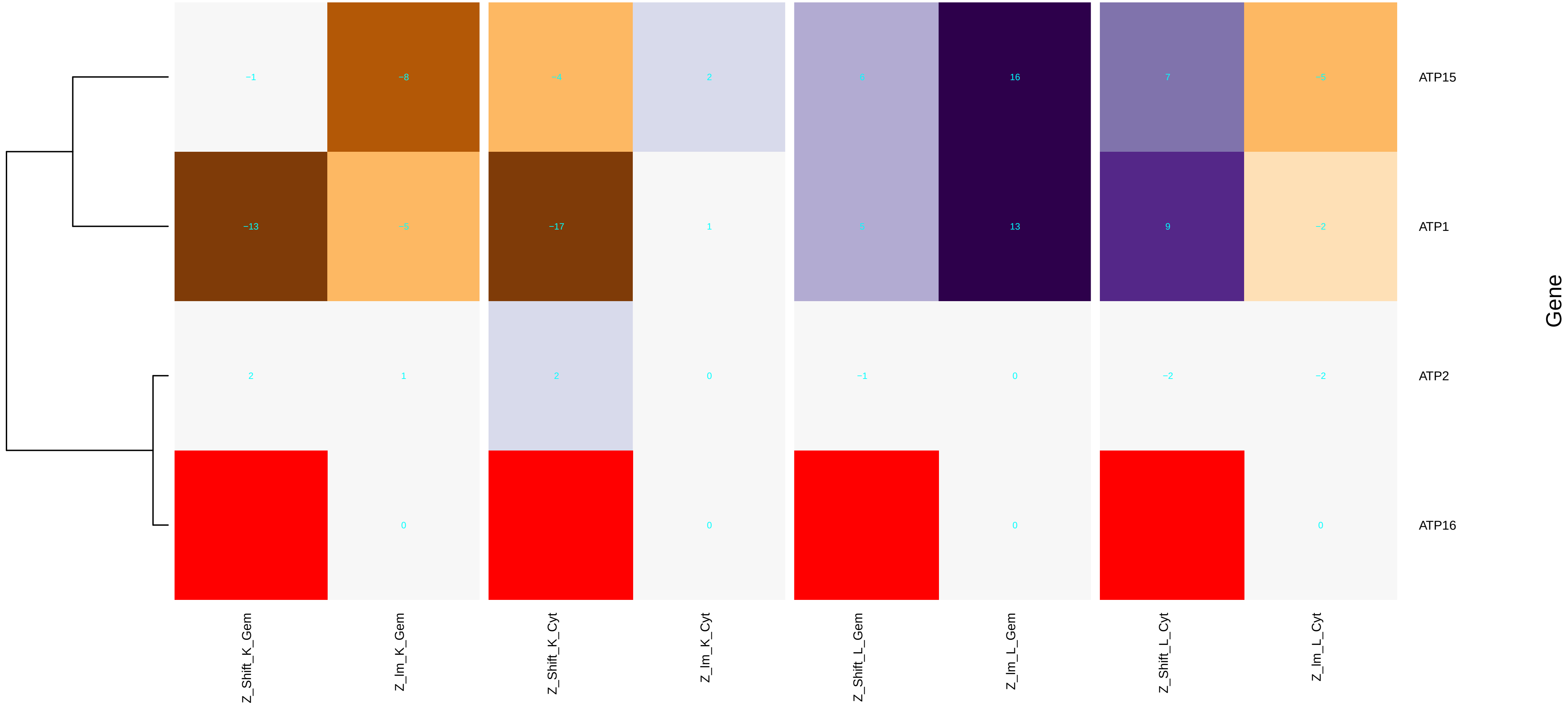

### mitochondrial_proton-transporting_ATP_synthase_complex.pdf

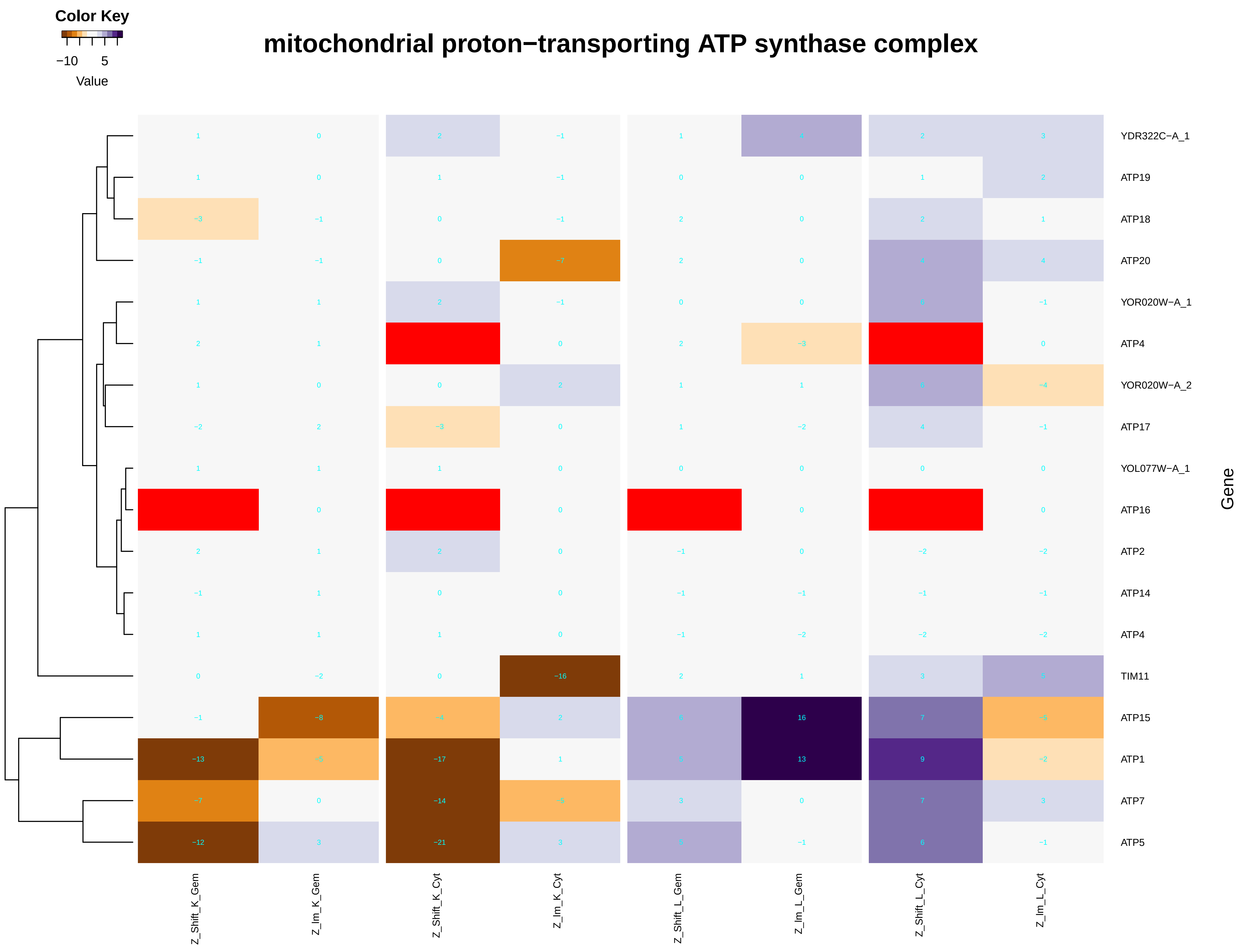

### mitochondrial_ribosome.pdf

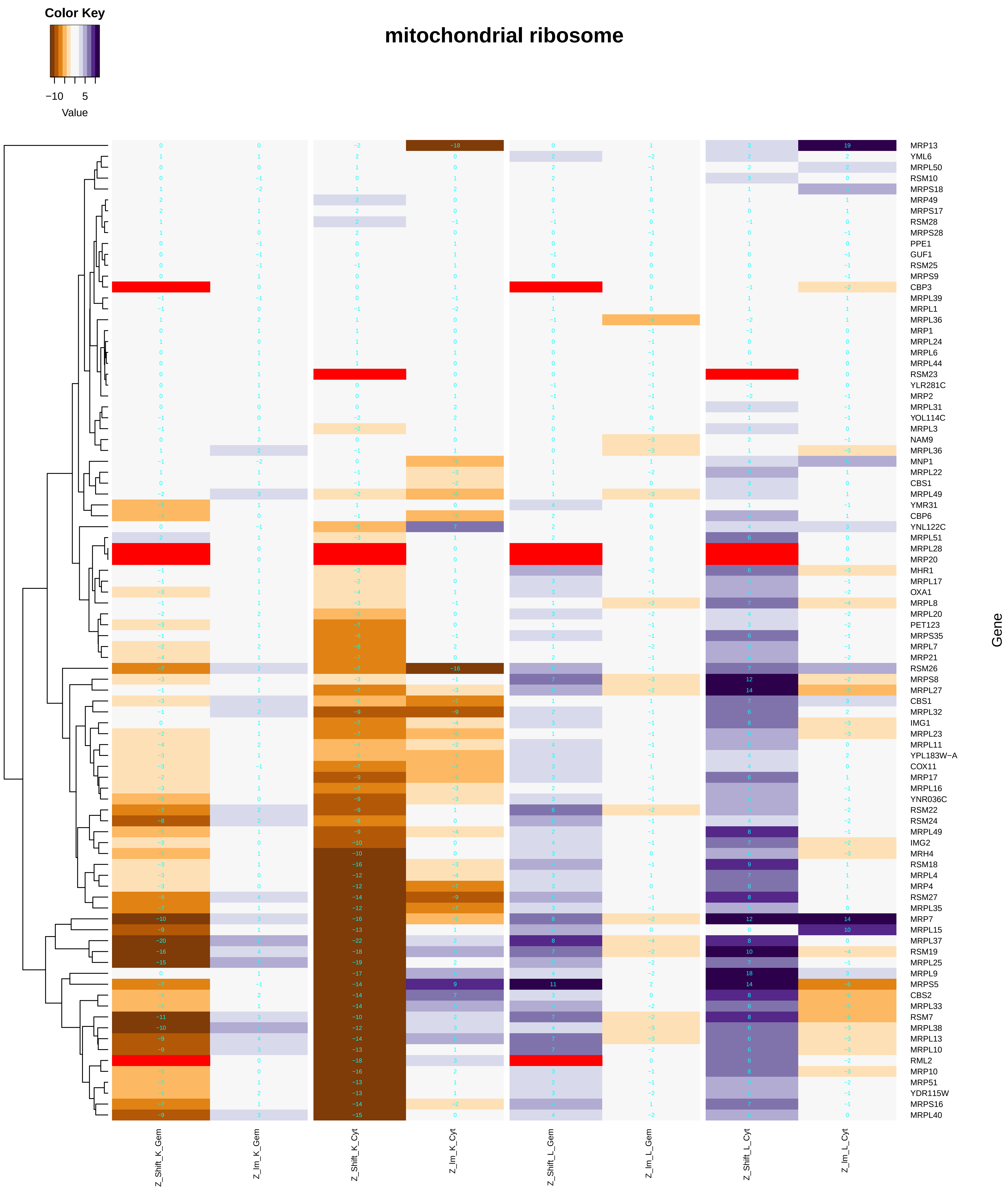

### mitochondrial_small_ribosomal_subunit.pdf

#### mitochondrial small ribosomal subunit

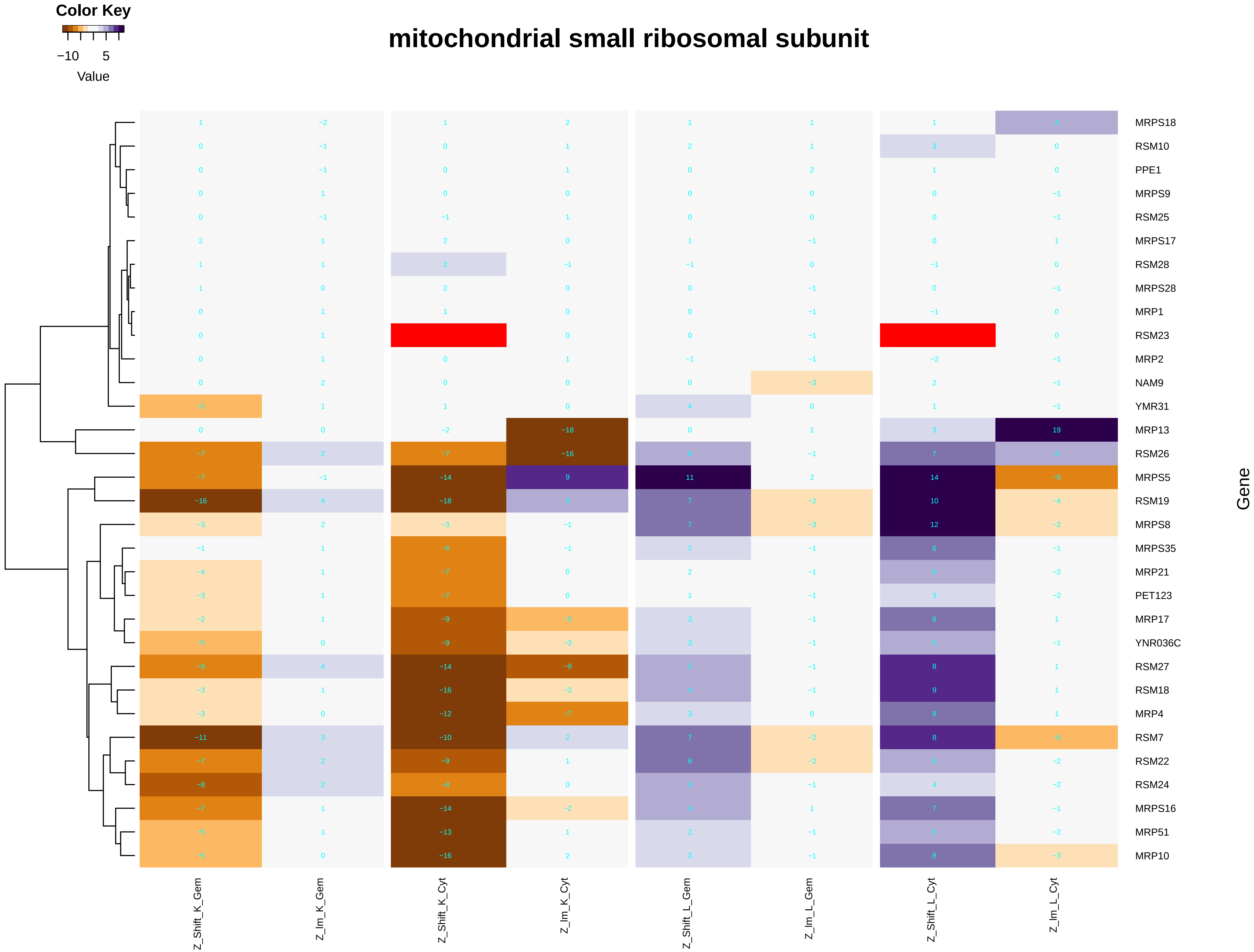

### mitochondrial_translation.pdf

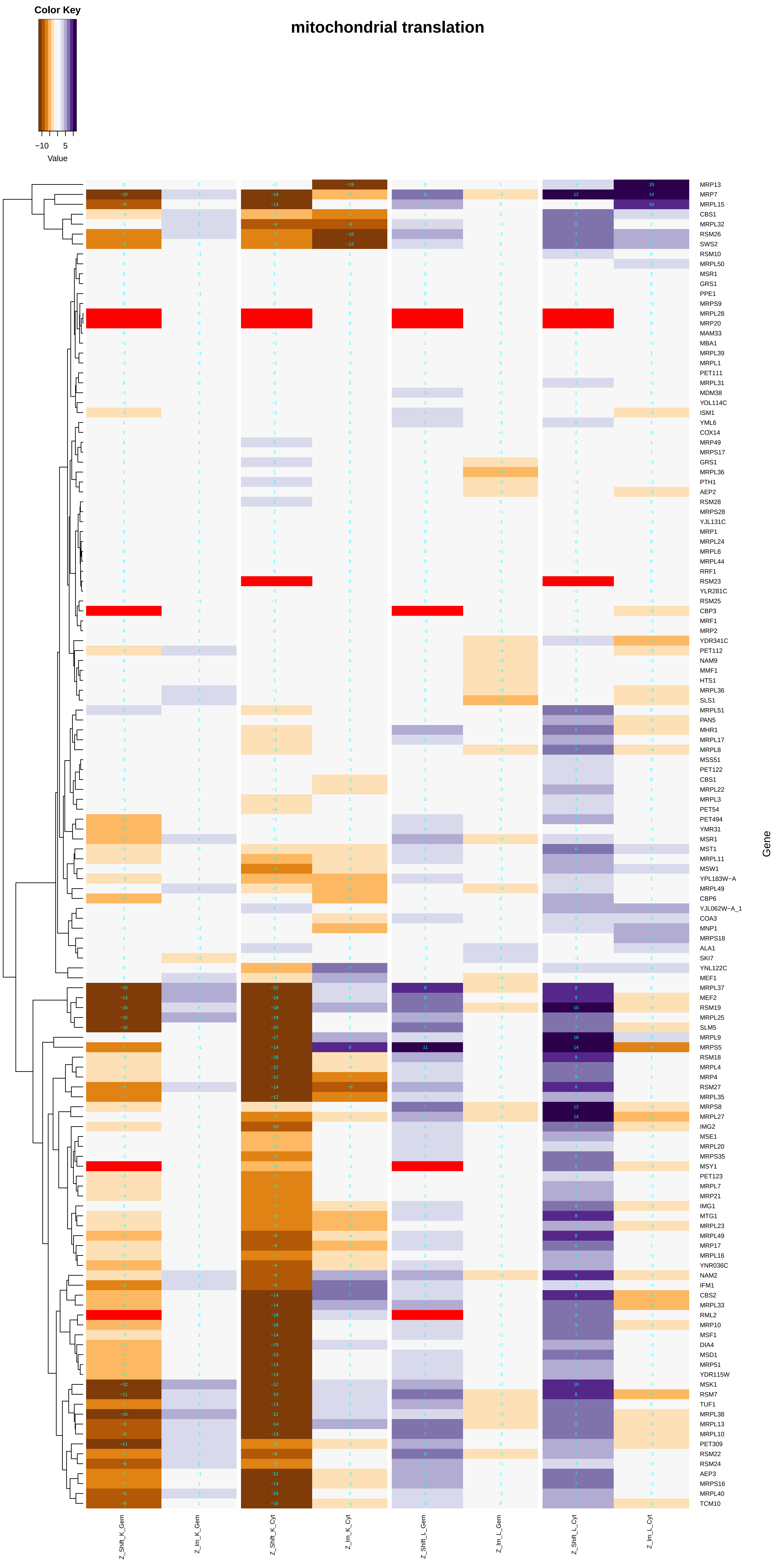

### mitochondrion.pdf

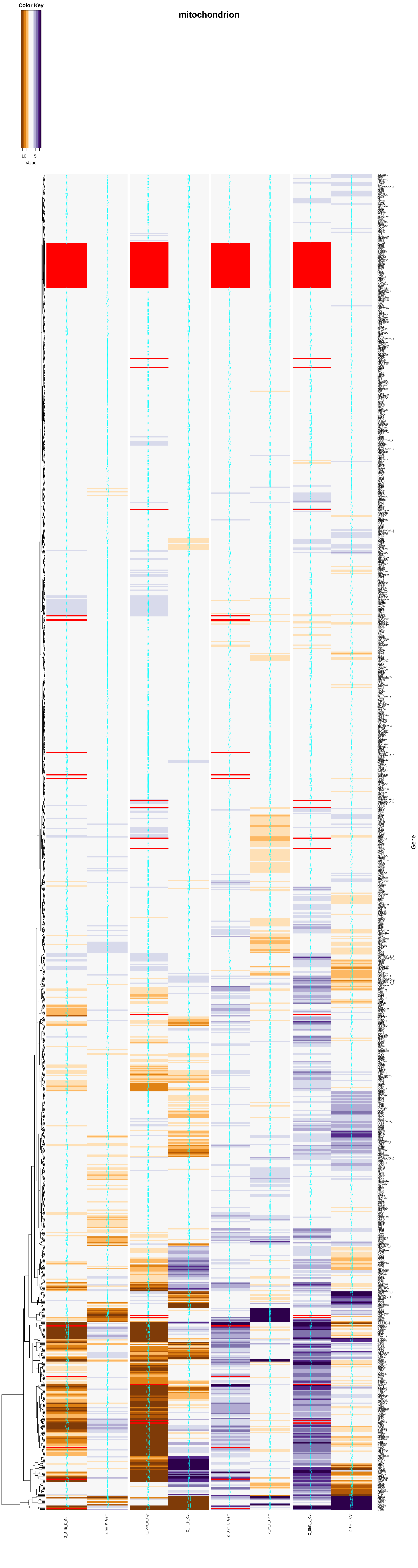

### mitotic_cell_cycle.pdf

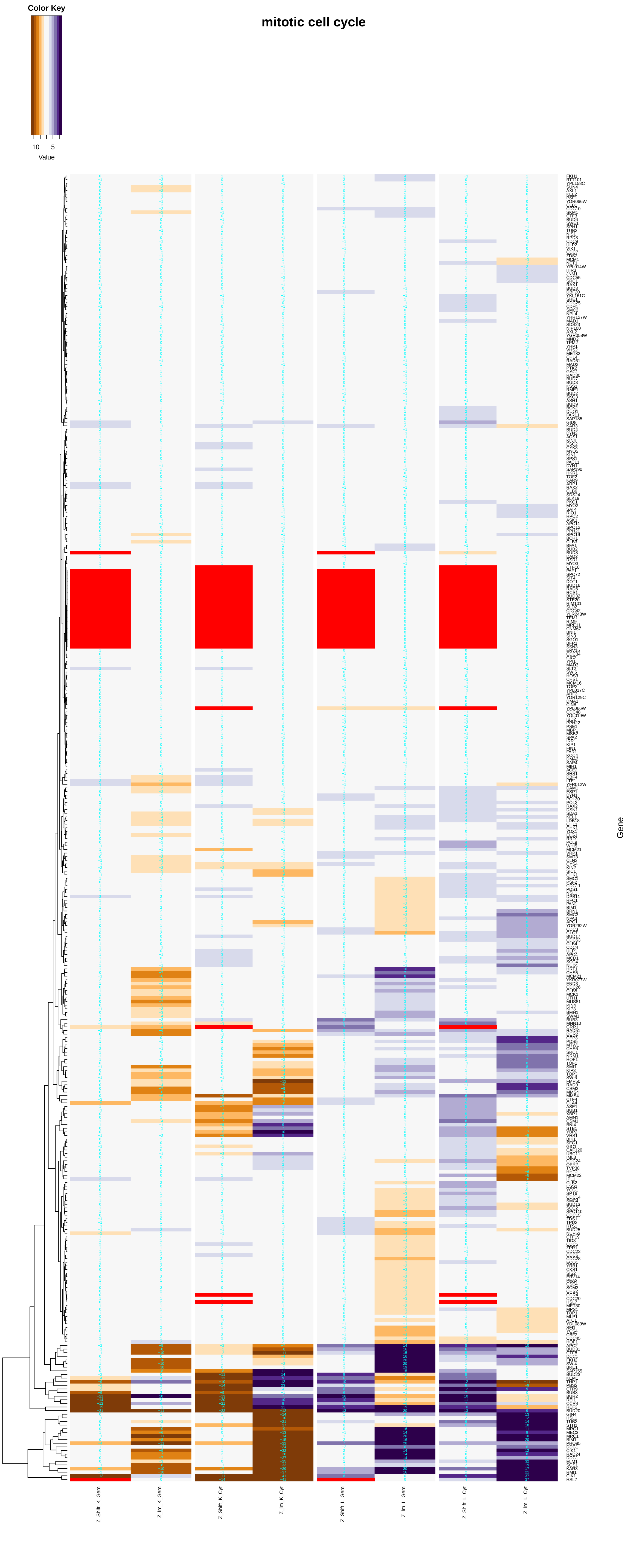

### mitotic_DNA_replication_checkpoint.pdf

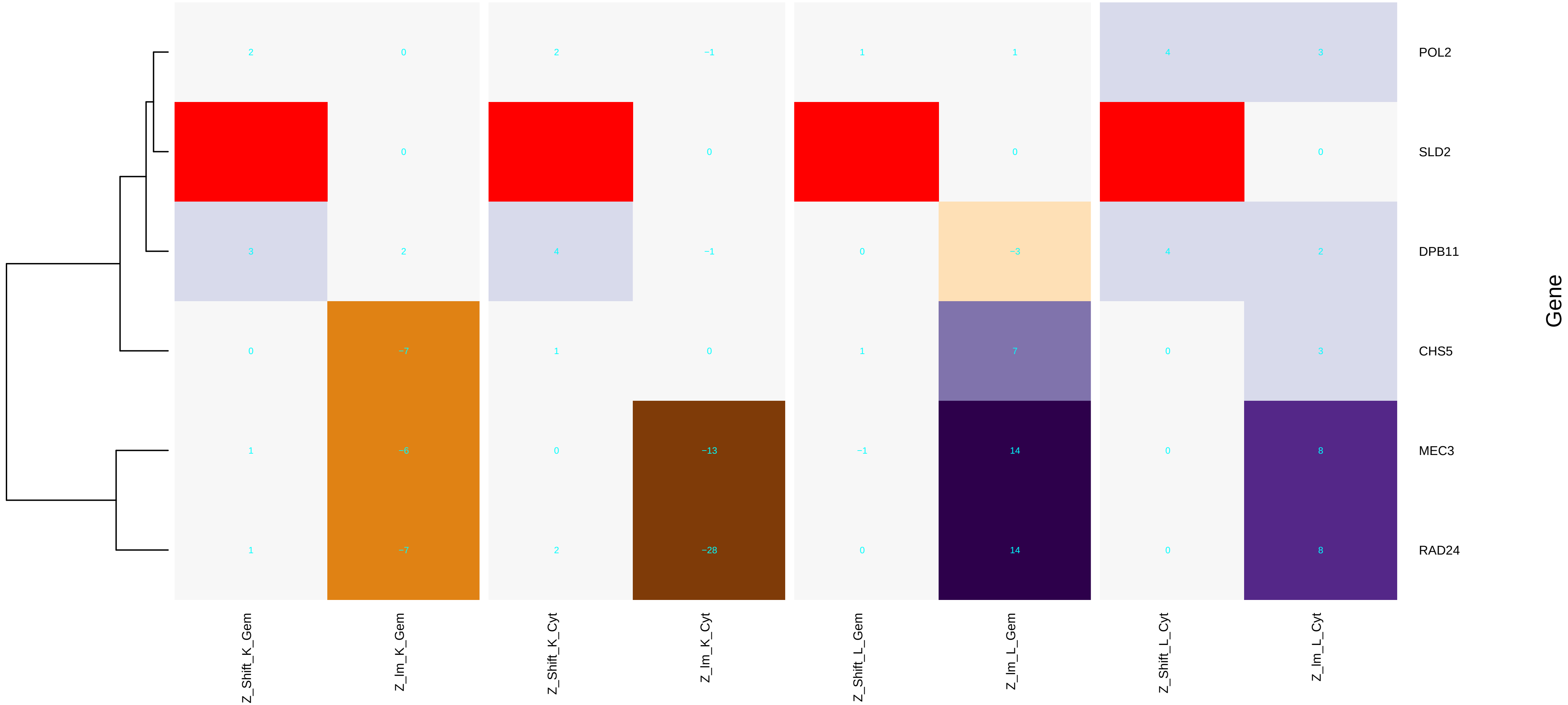

### mitotic_G1_DNA_damage_checkpoint.pdf

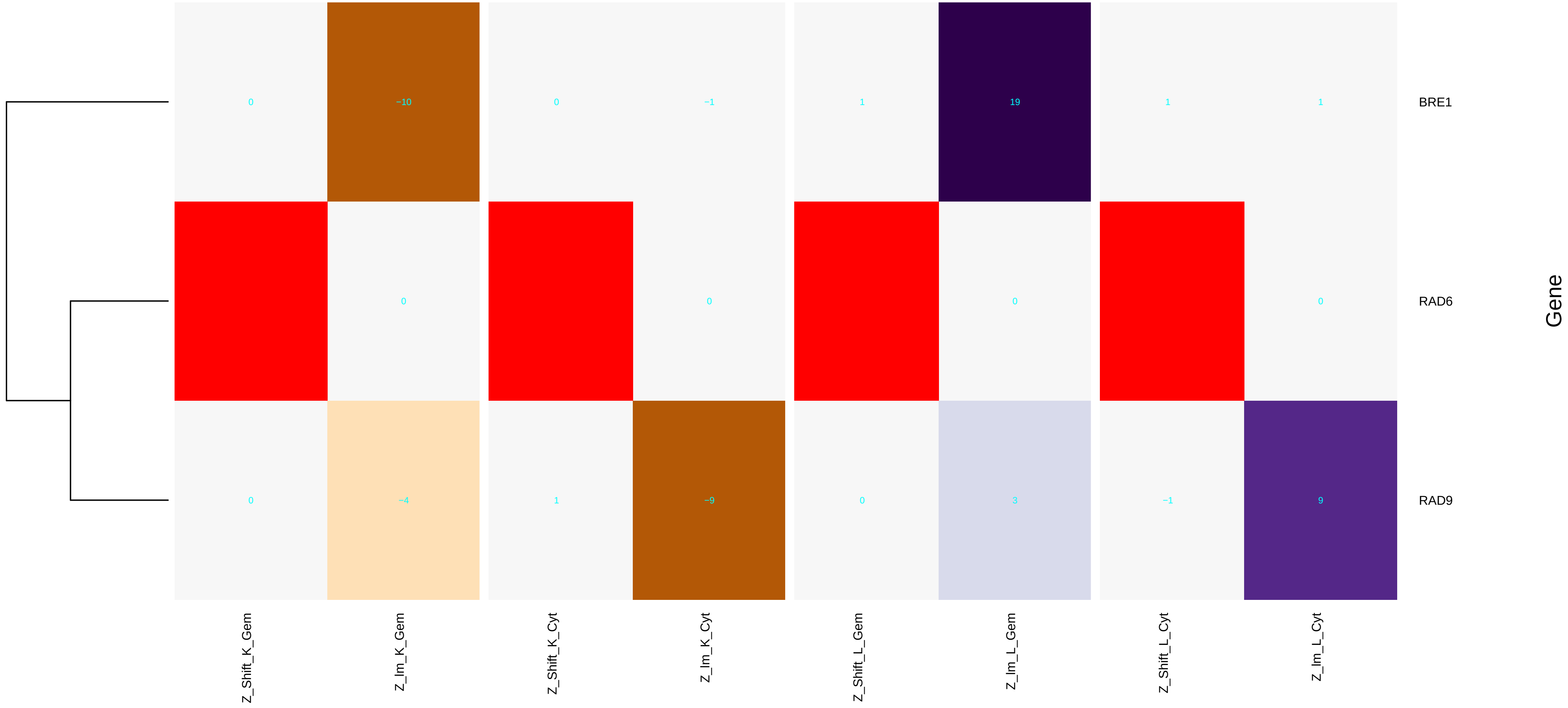

### mitotic_G2_M_transition_checkpoint.pdf

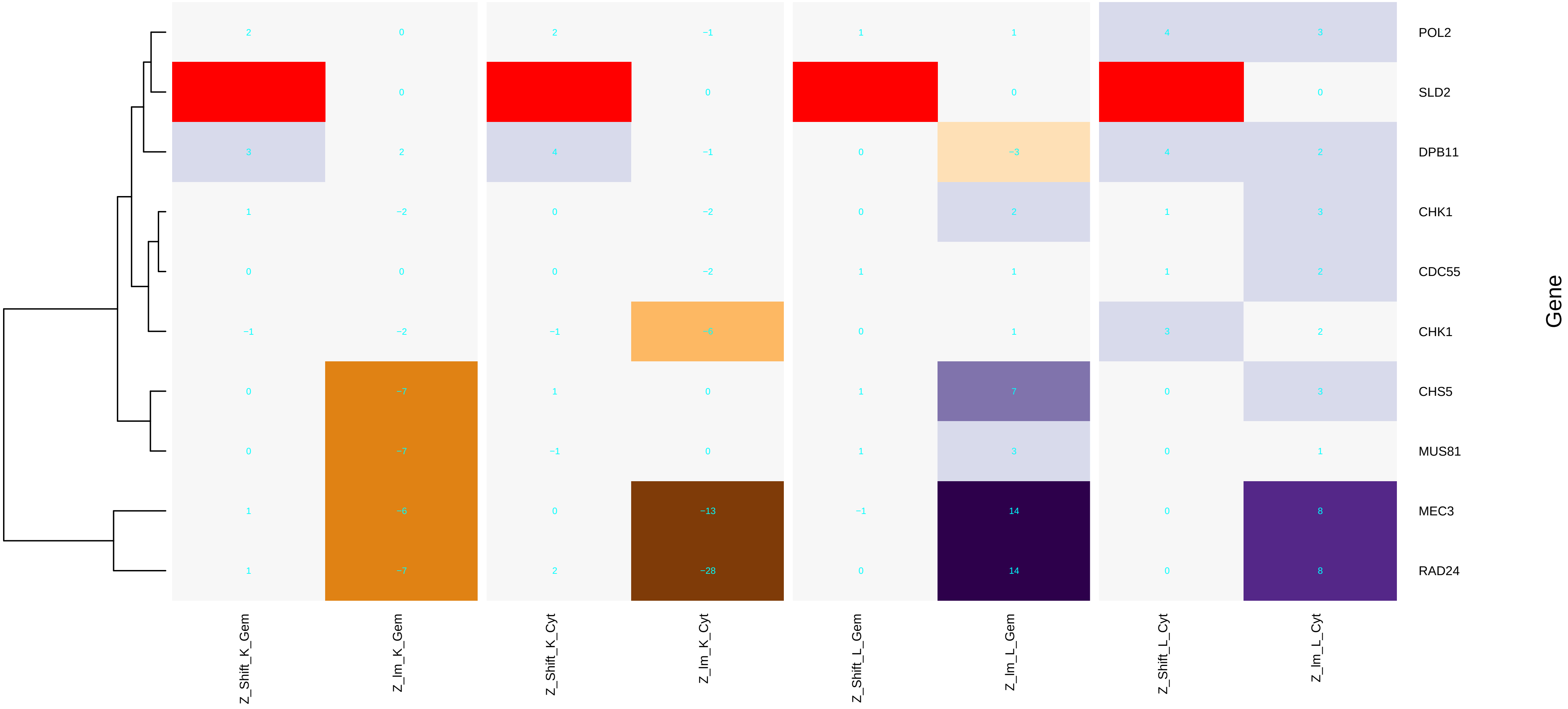

### mitotic_spindle_pole_body.pdf

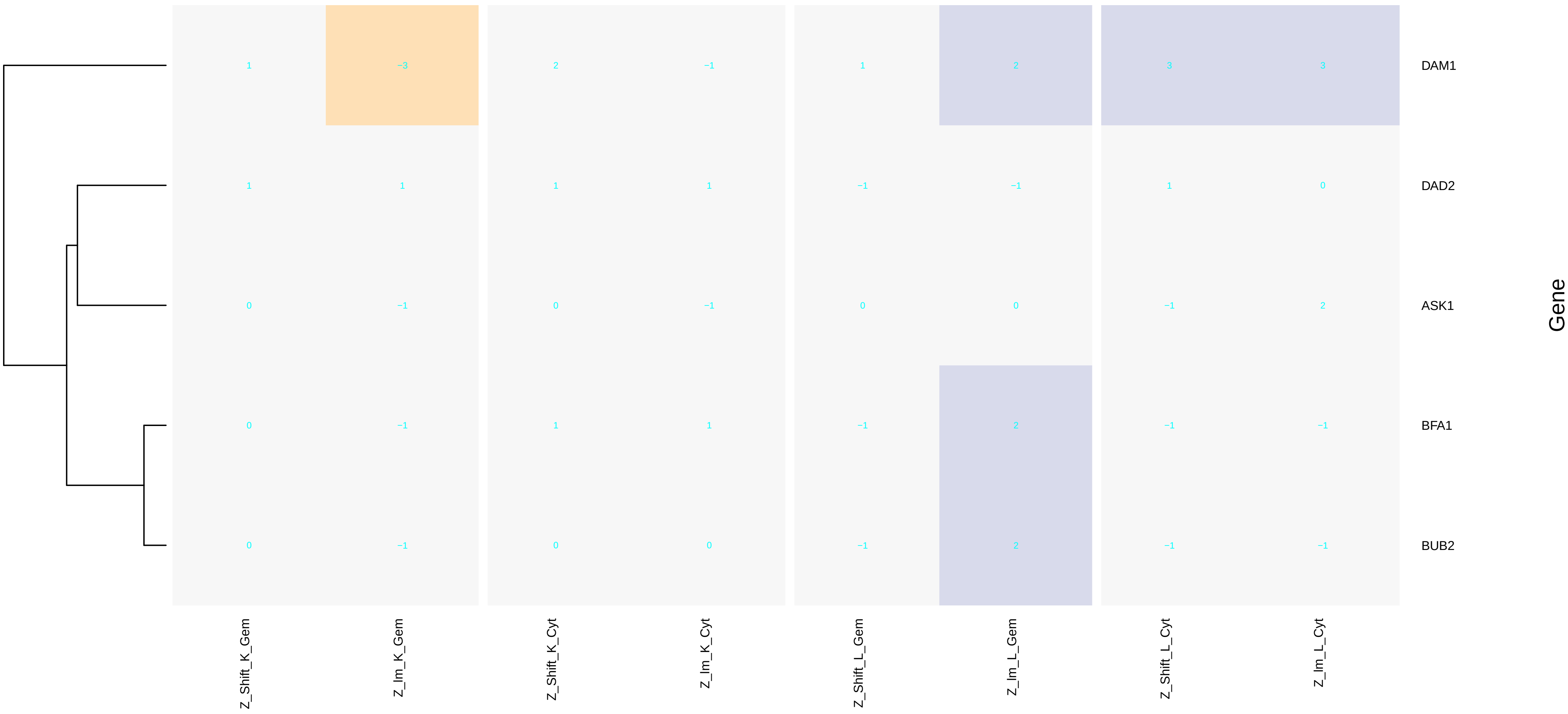

### Mre11_complex.pdf

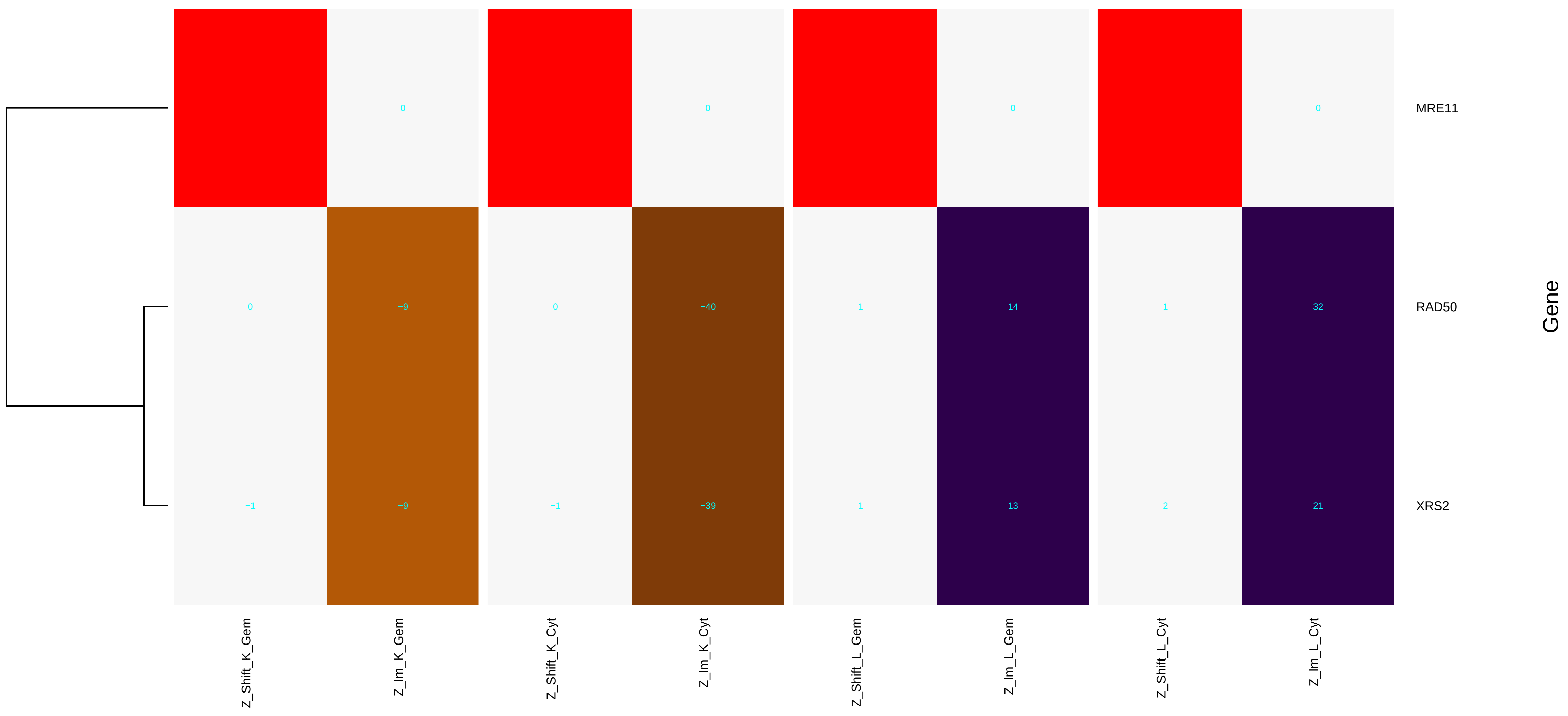

### mRNA_binding.pdf

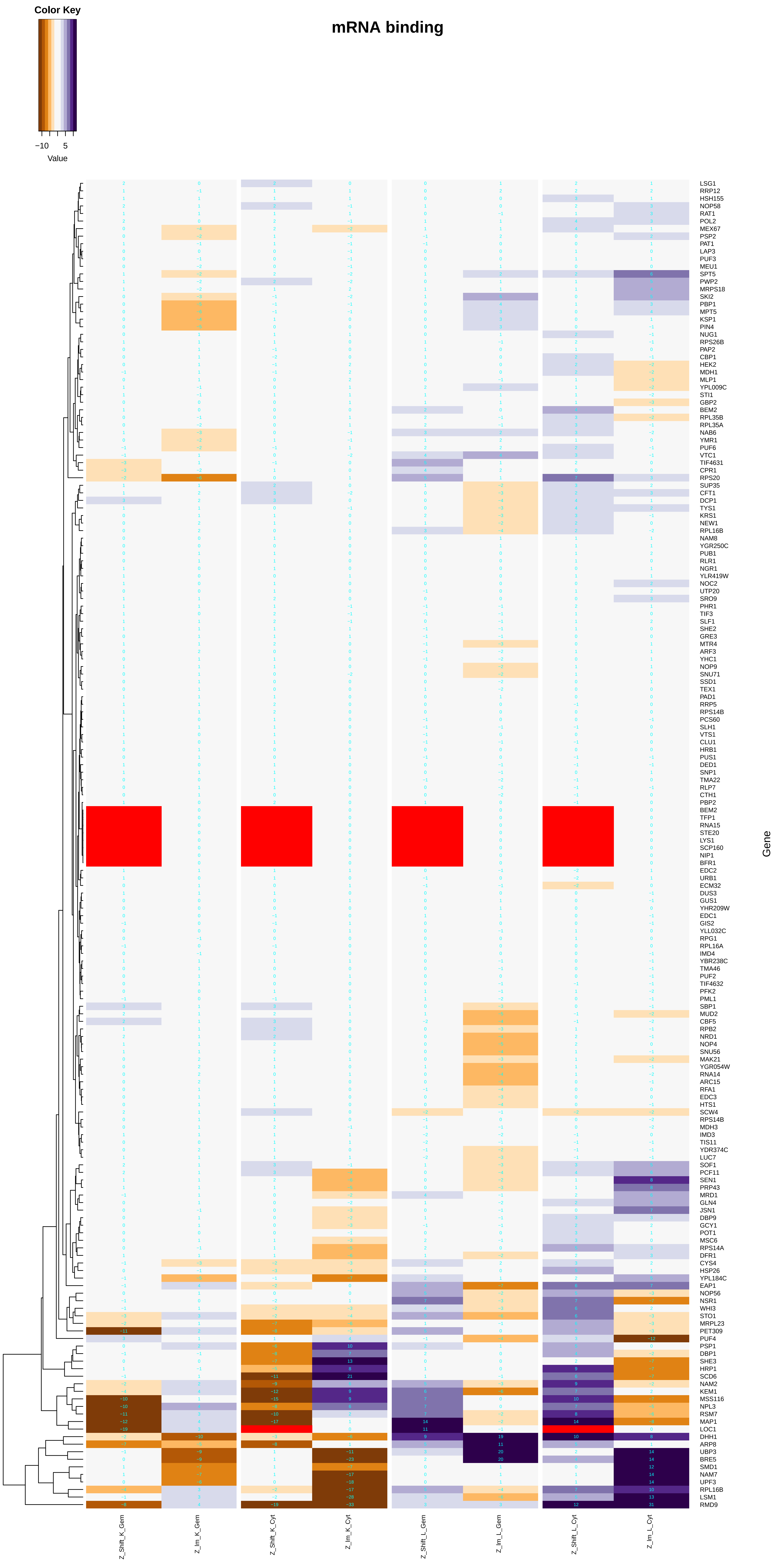

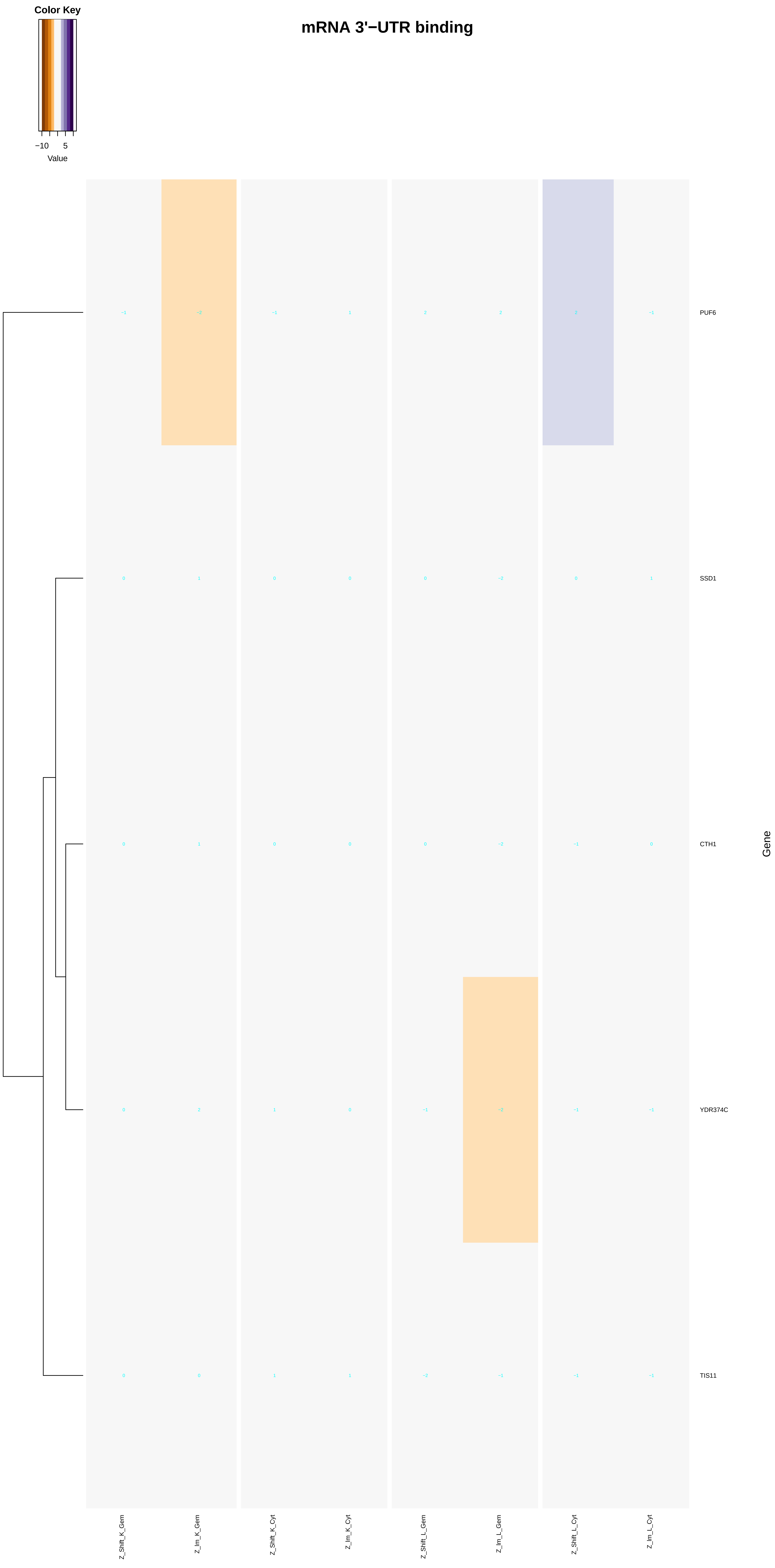

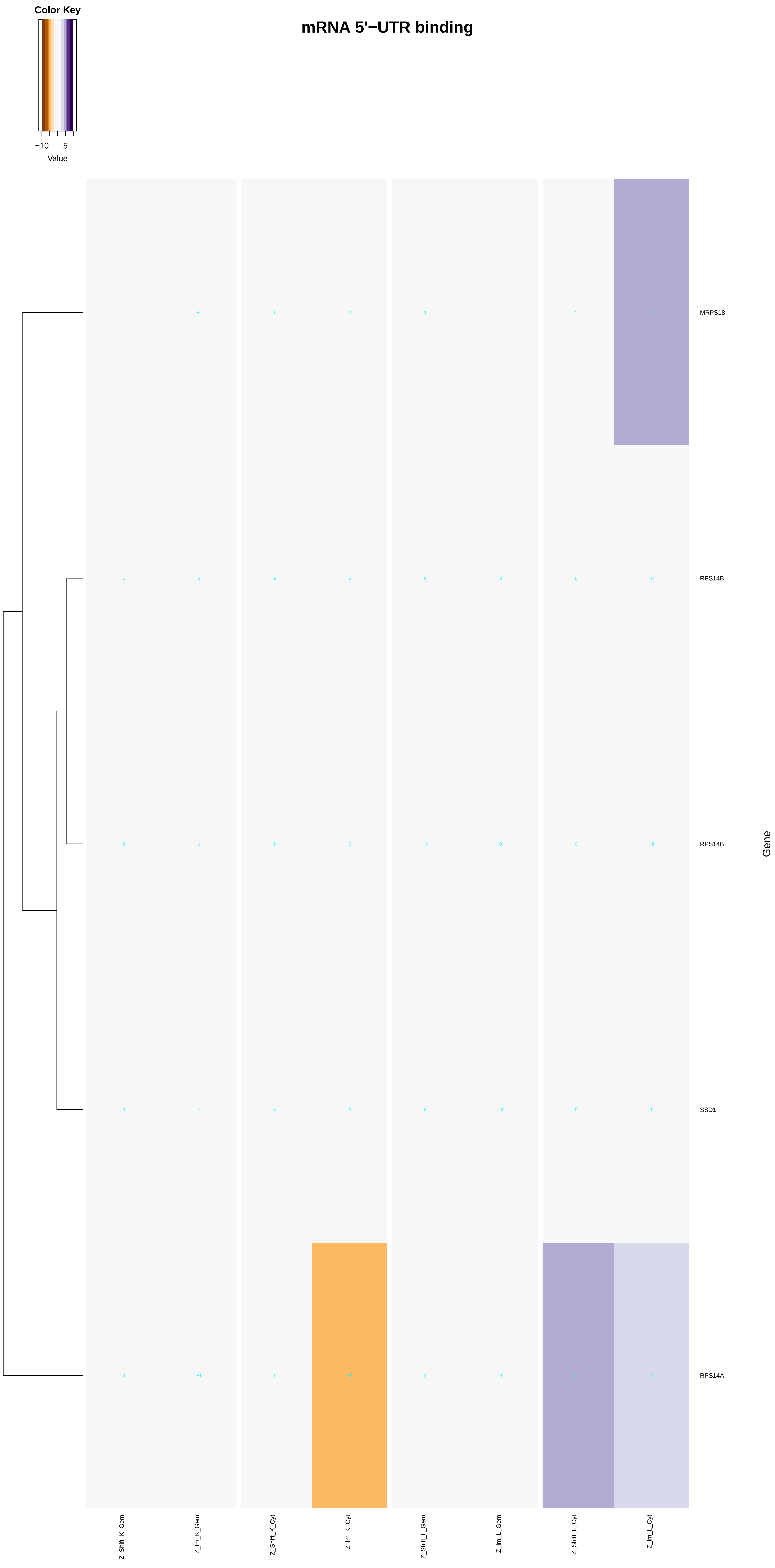

### mRNA_cap_binding_complex.pdf

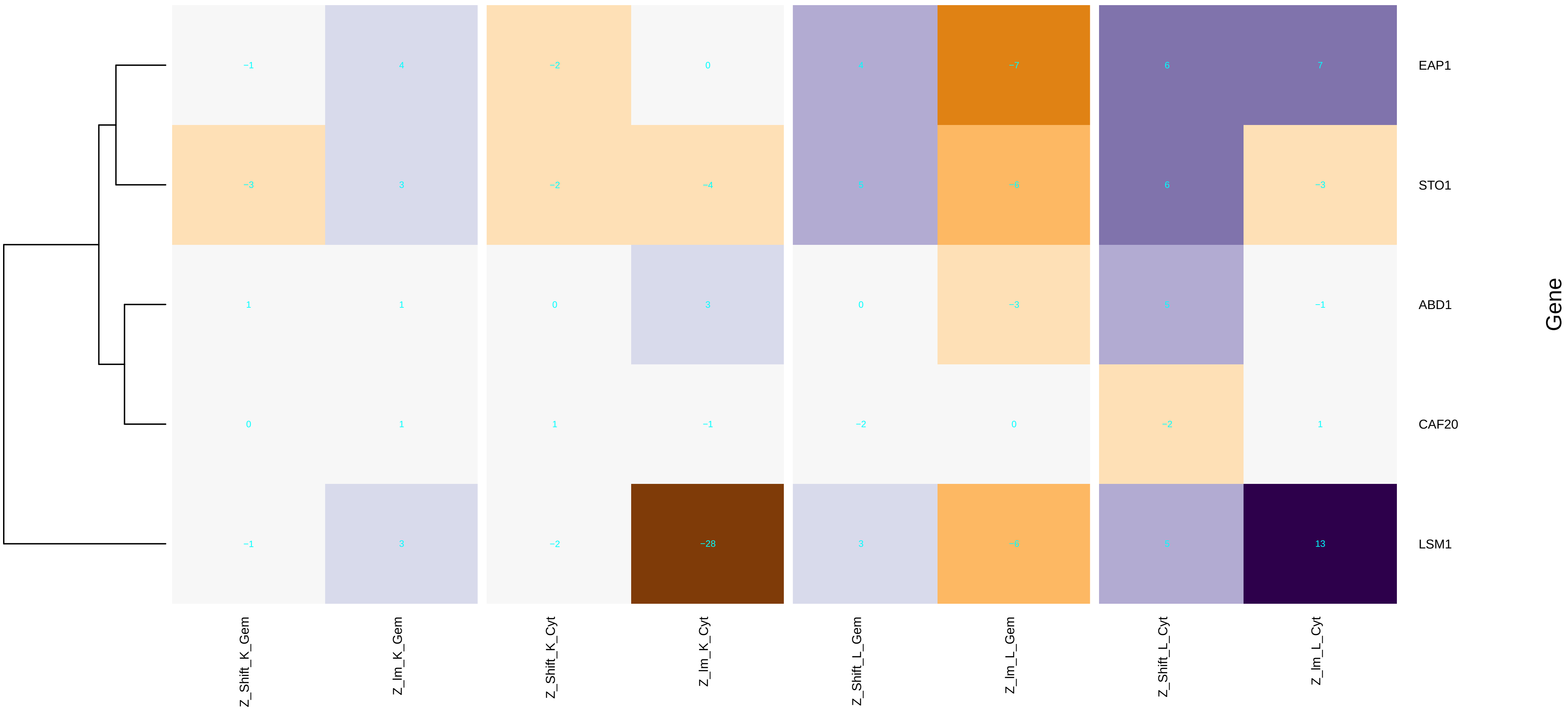

### mRNA_cleavage_factor_complex.pdf

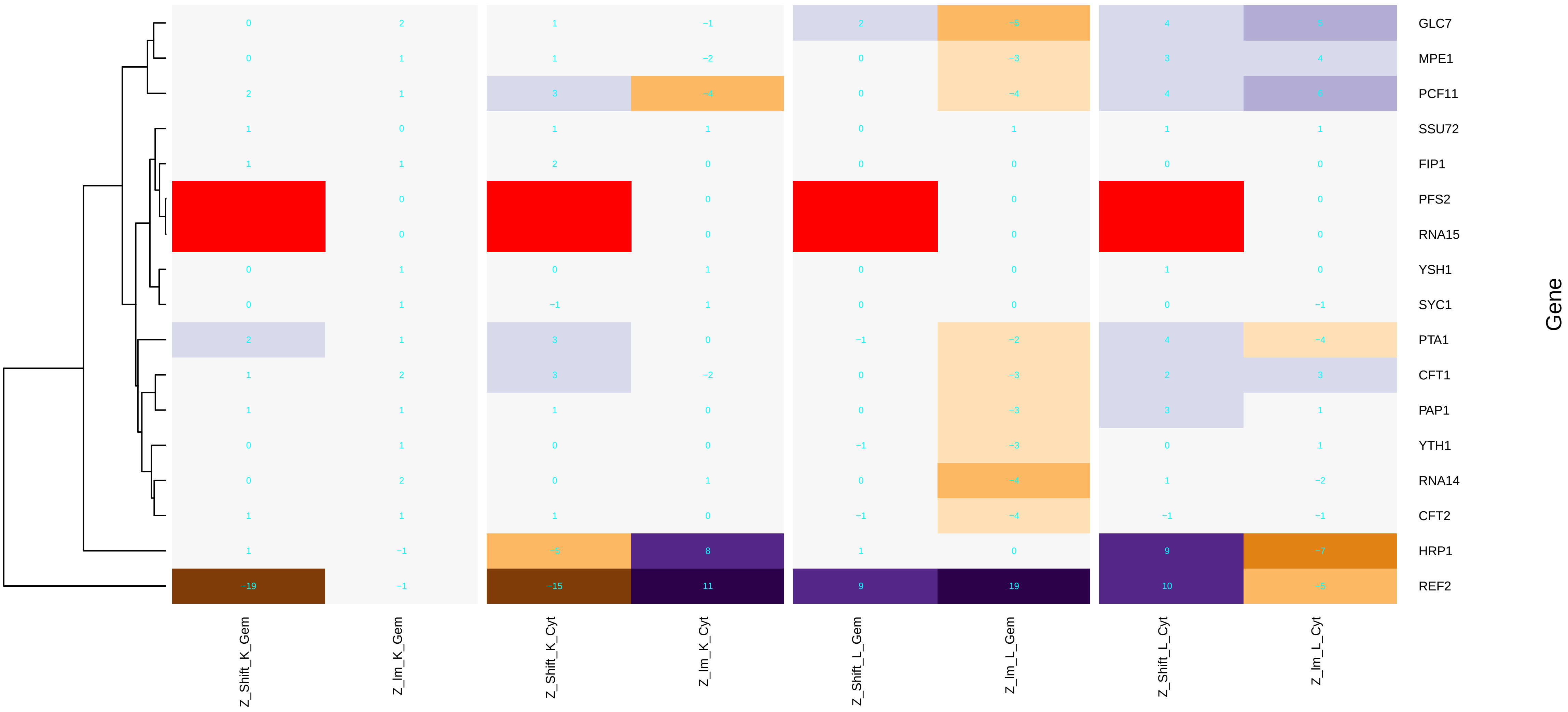

### mRNA_processing.pdf

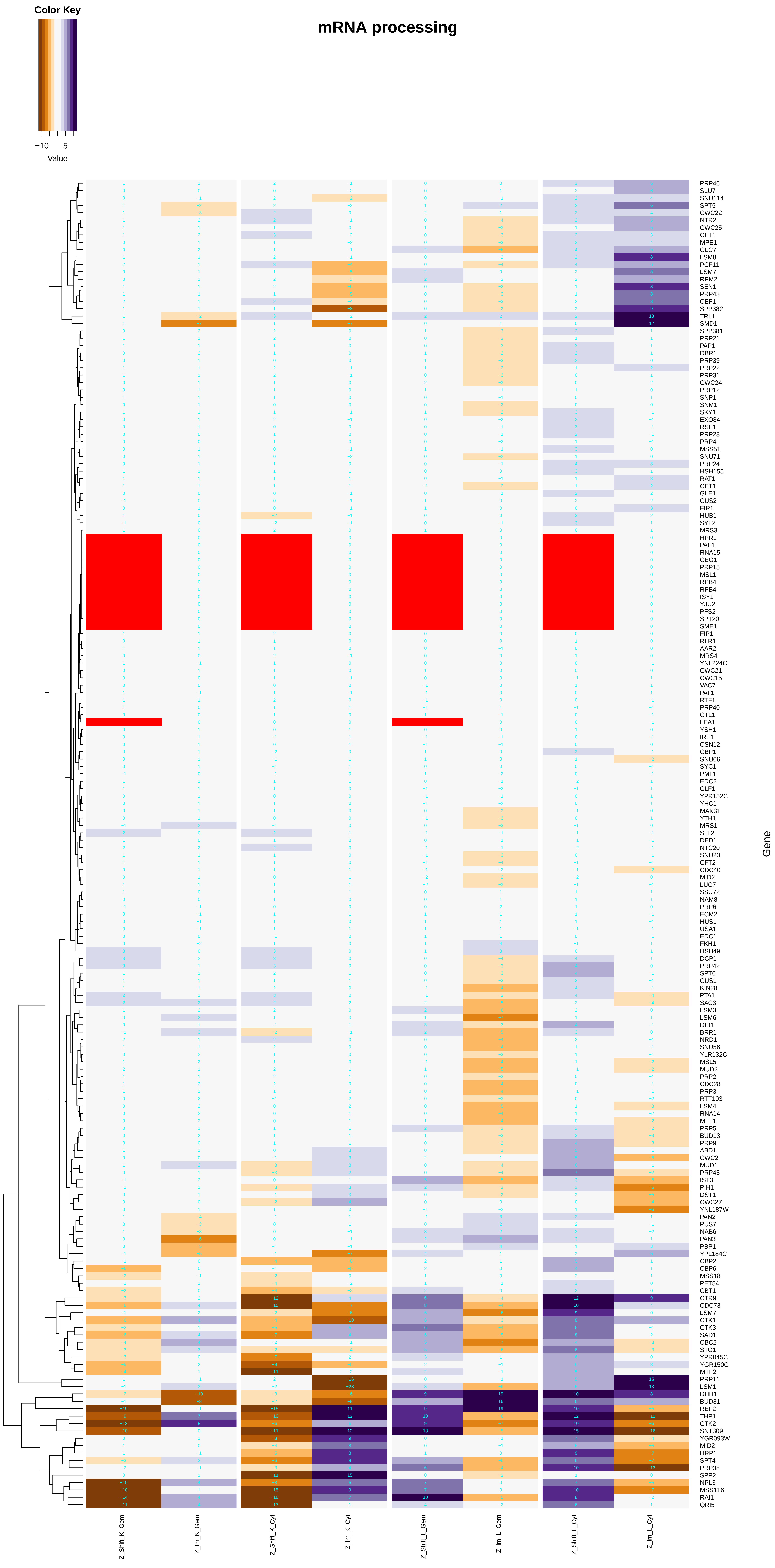

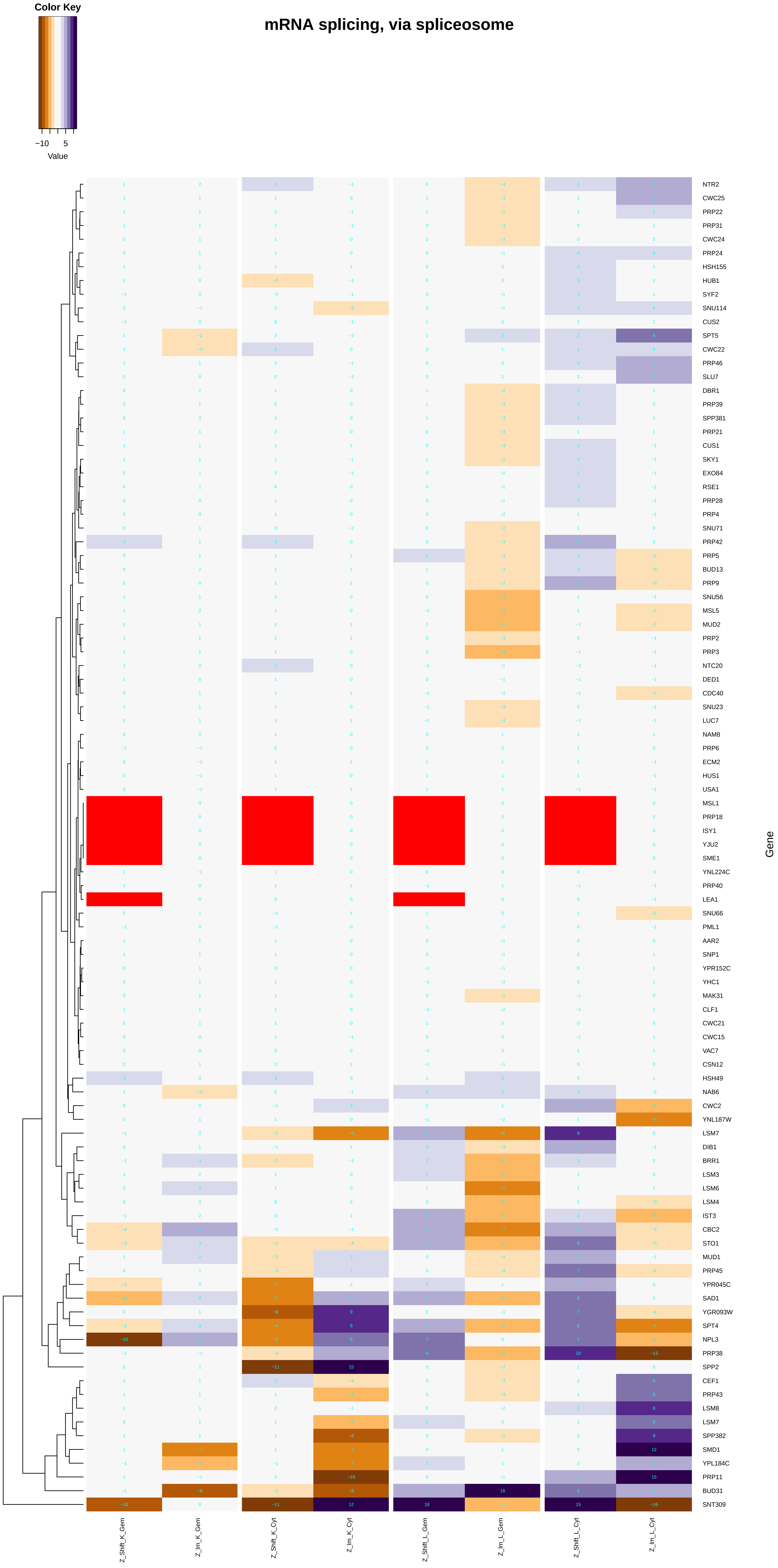

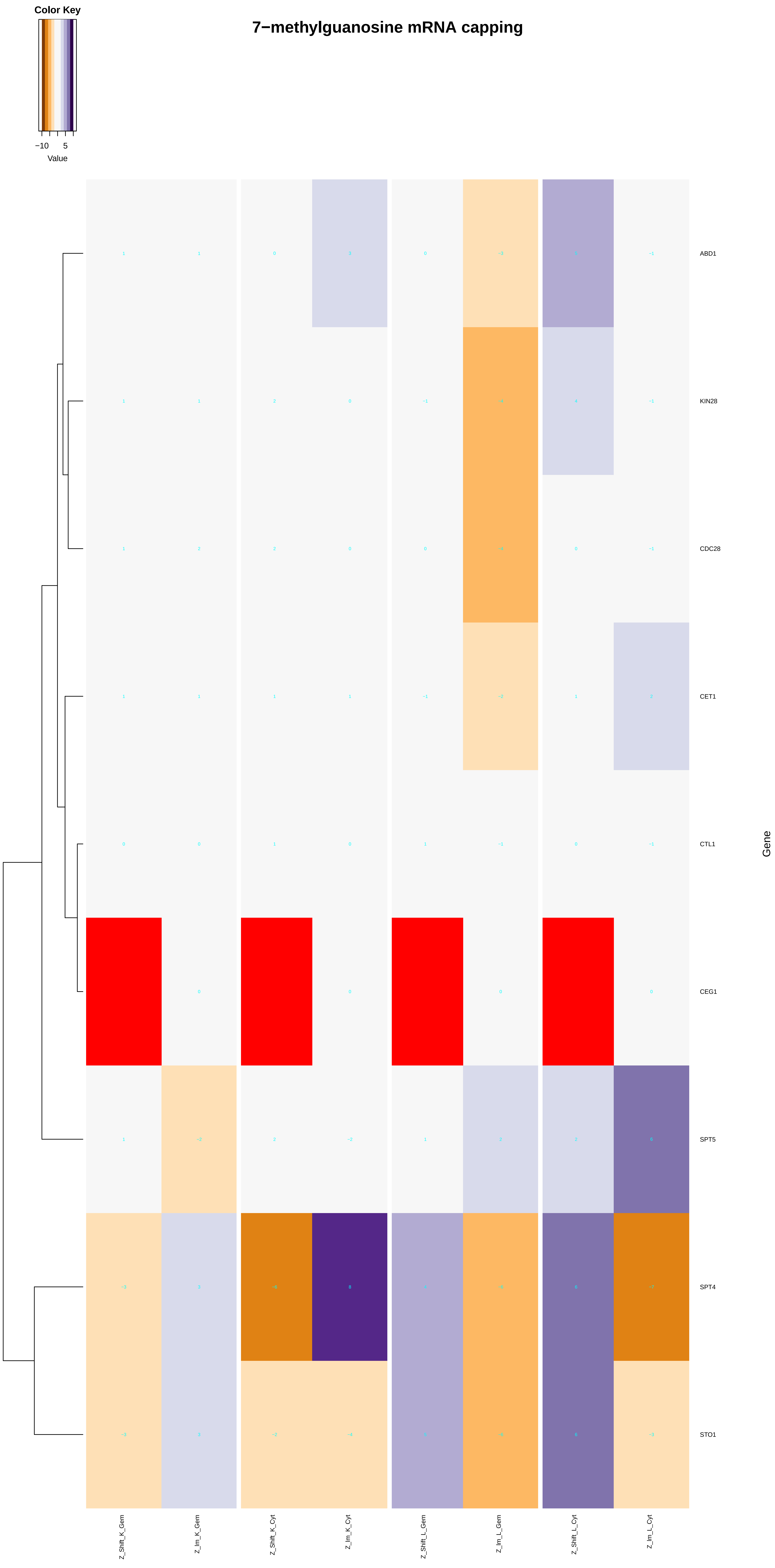

#### mRNA 3'−end processing

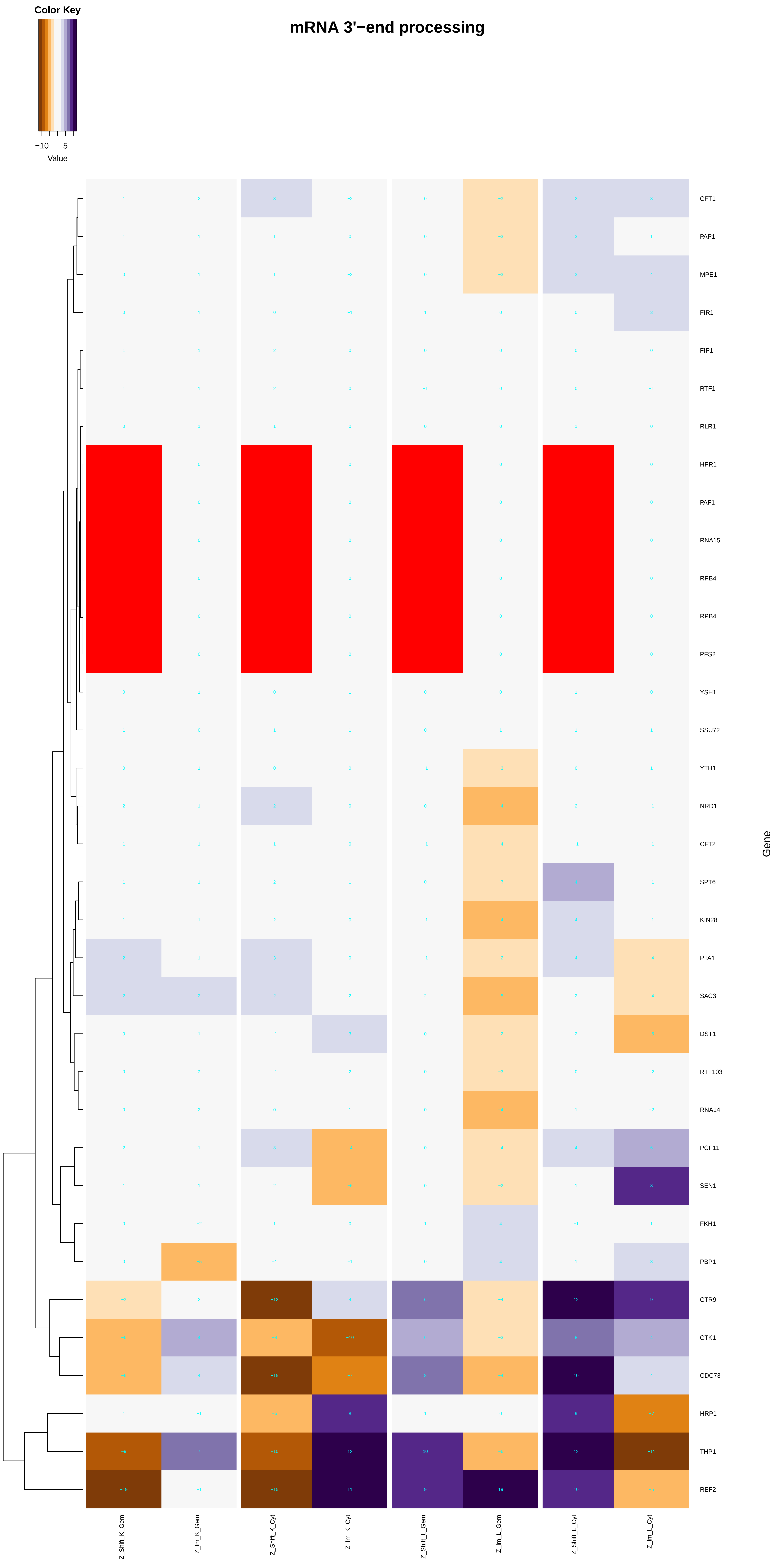

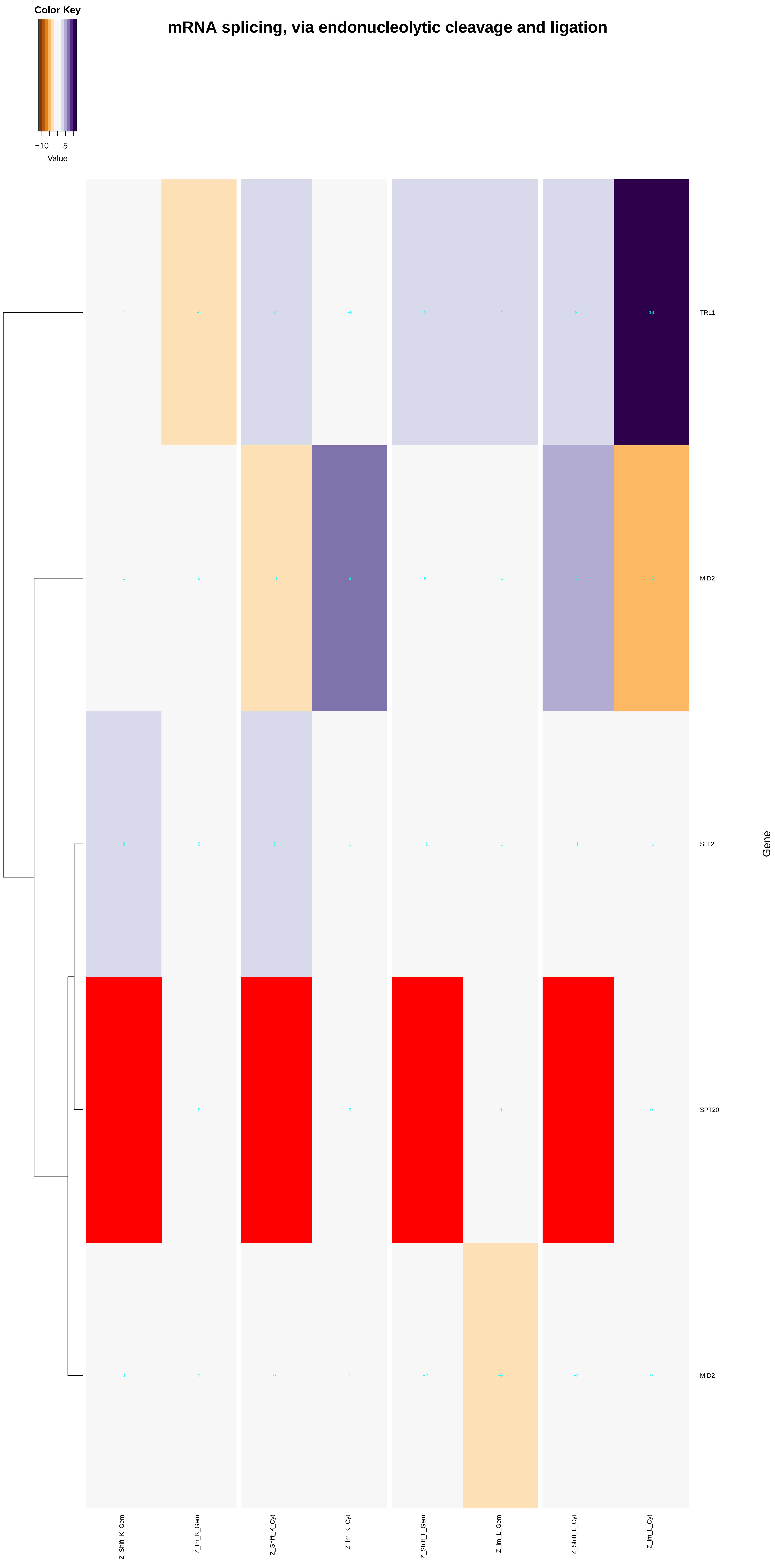

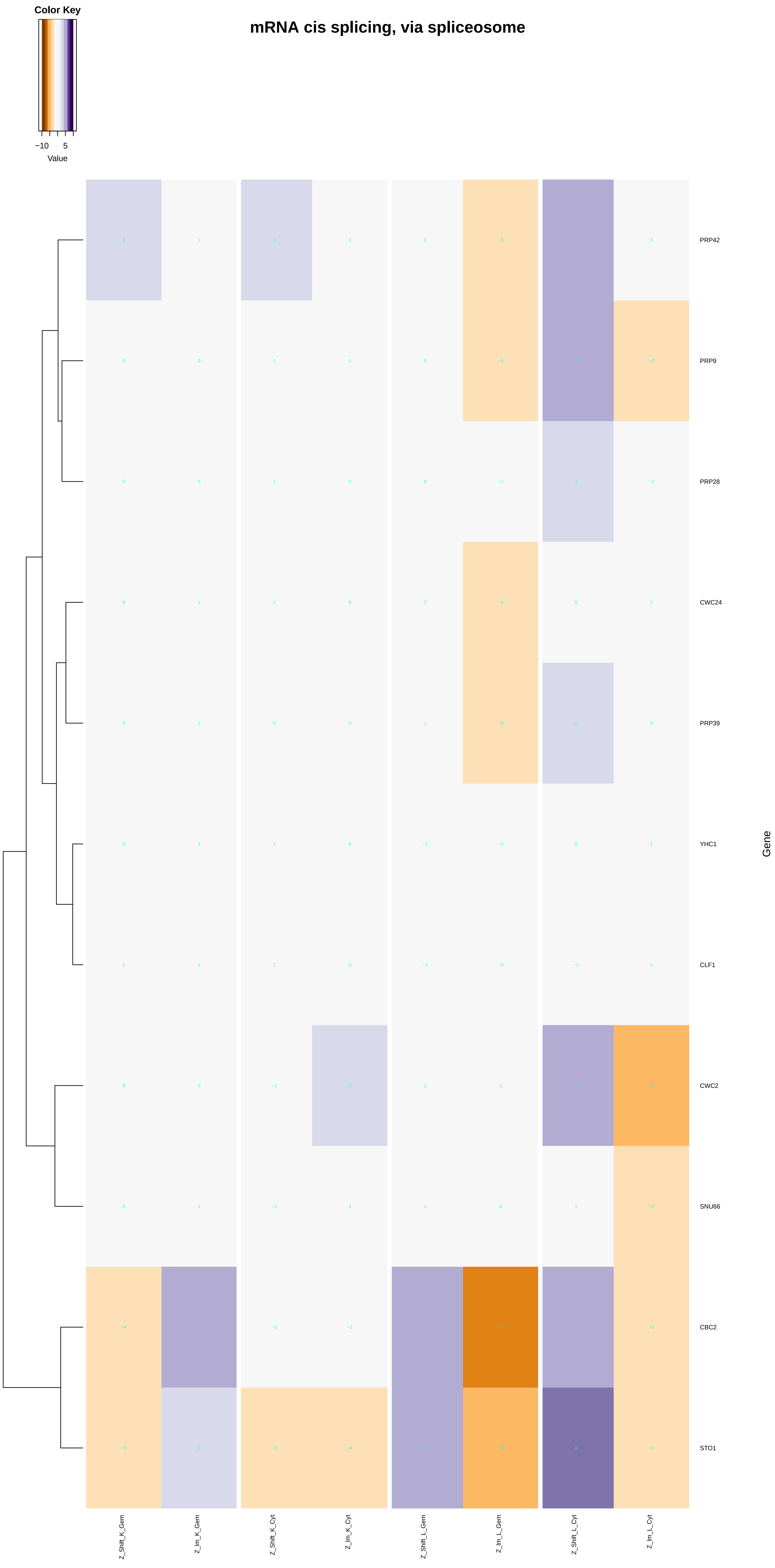

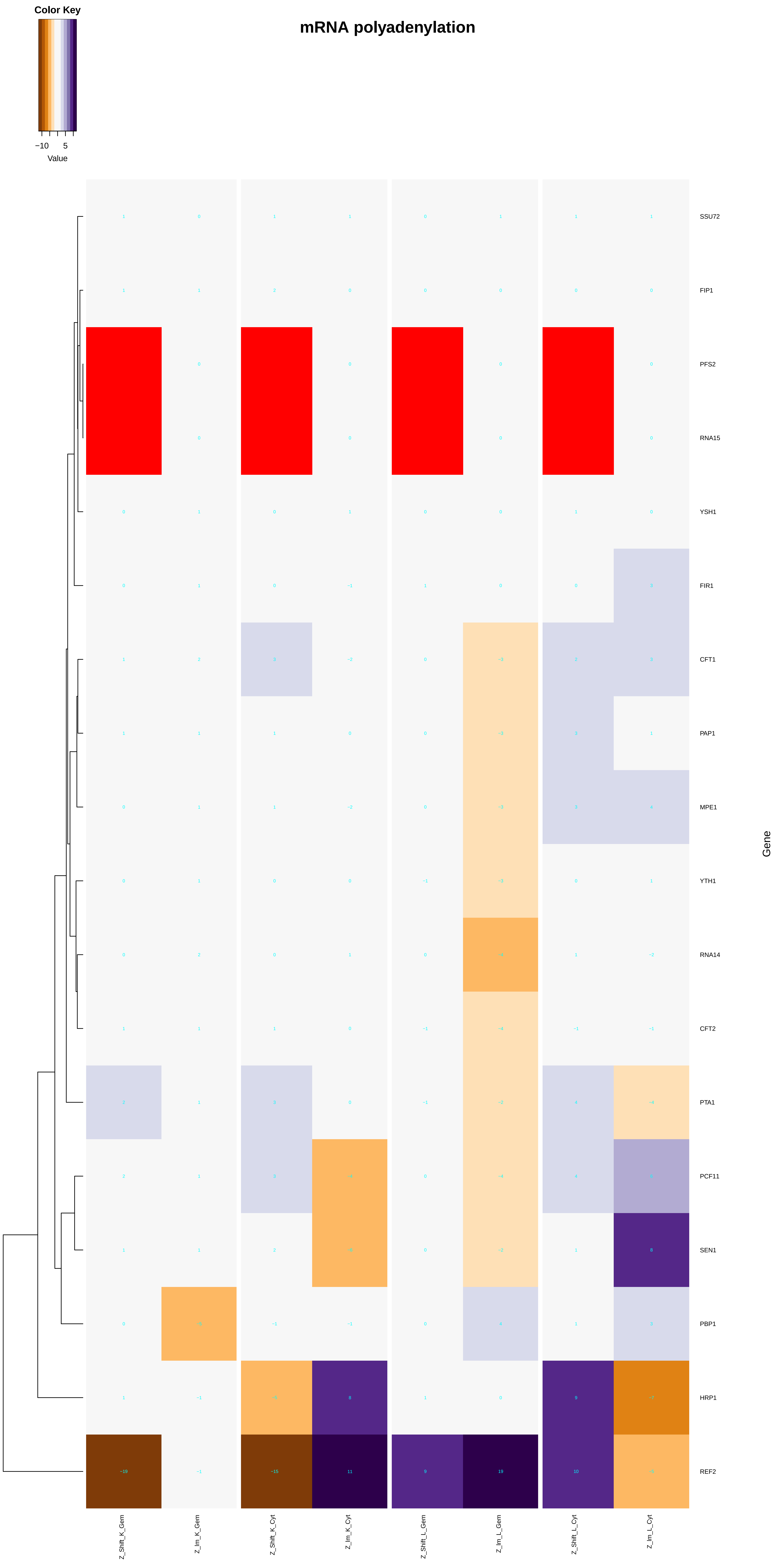

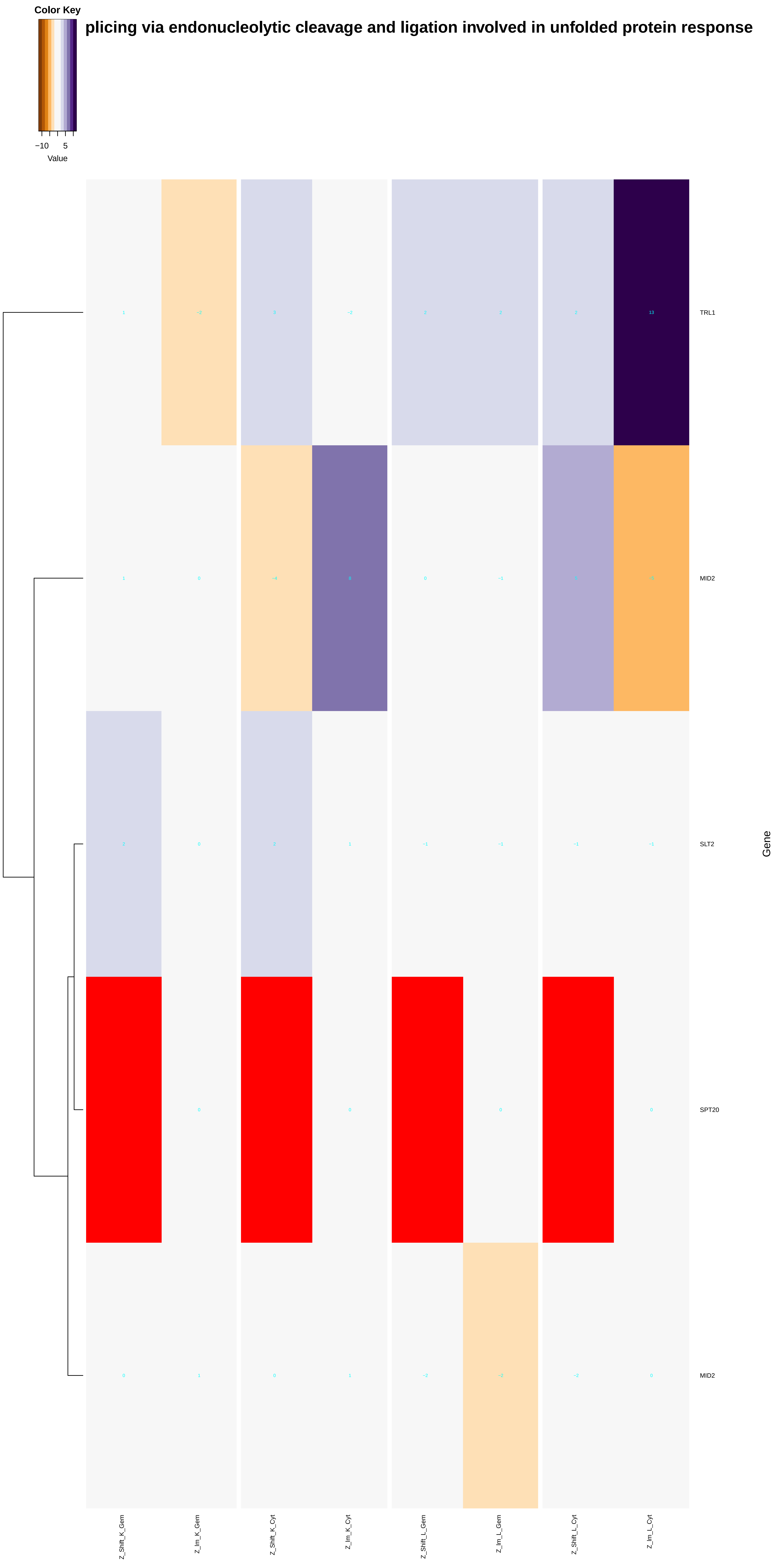

### mRNA_splicing,_via_spliceosome.pdf

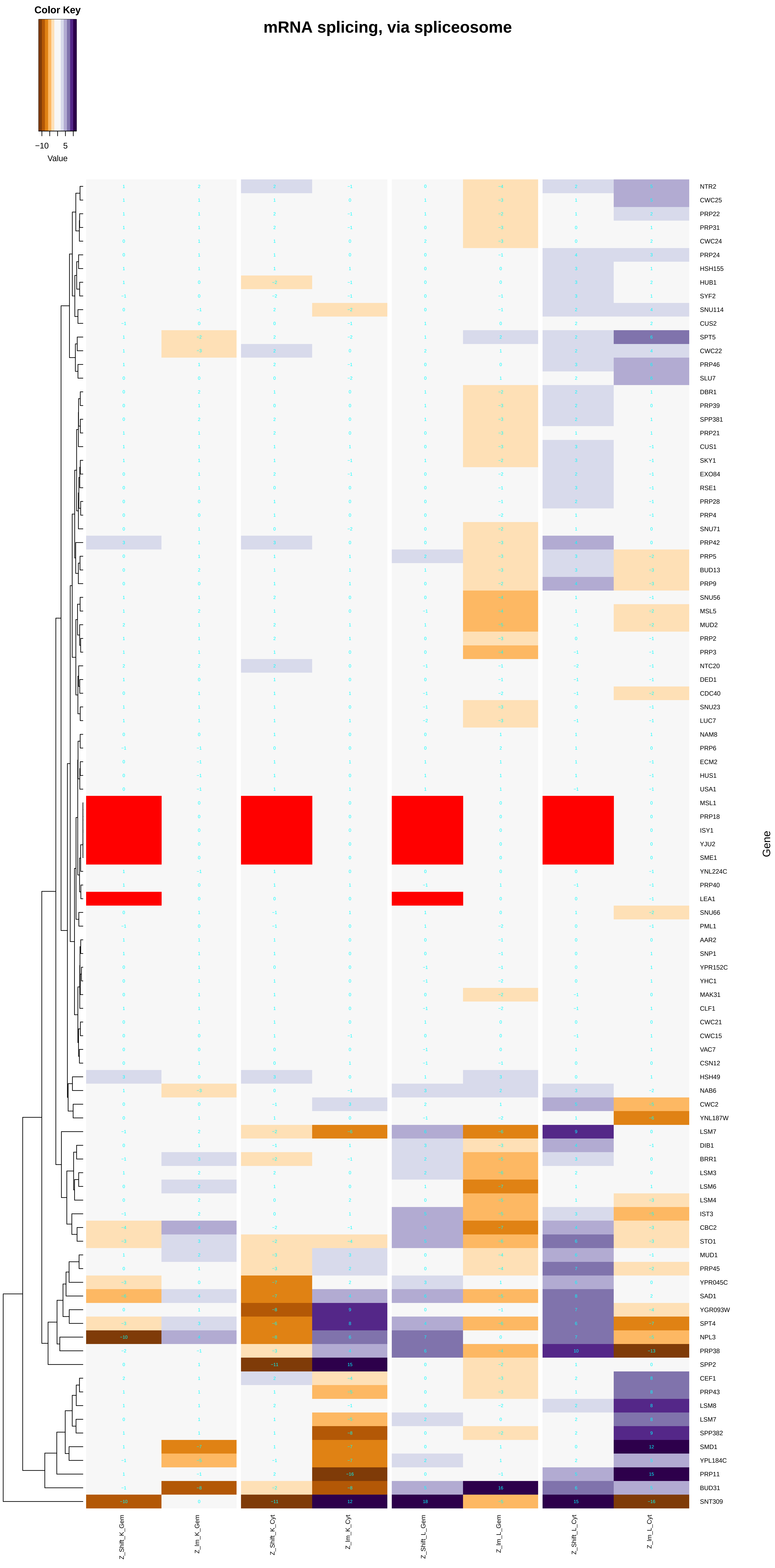

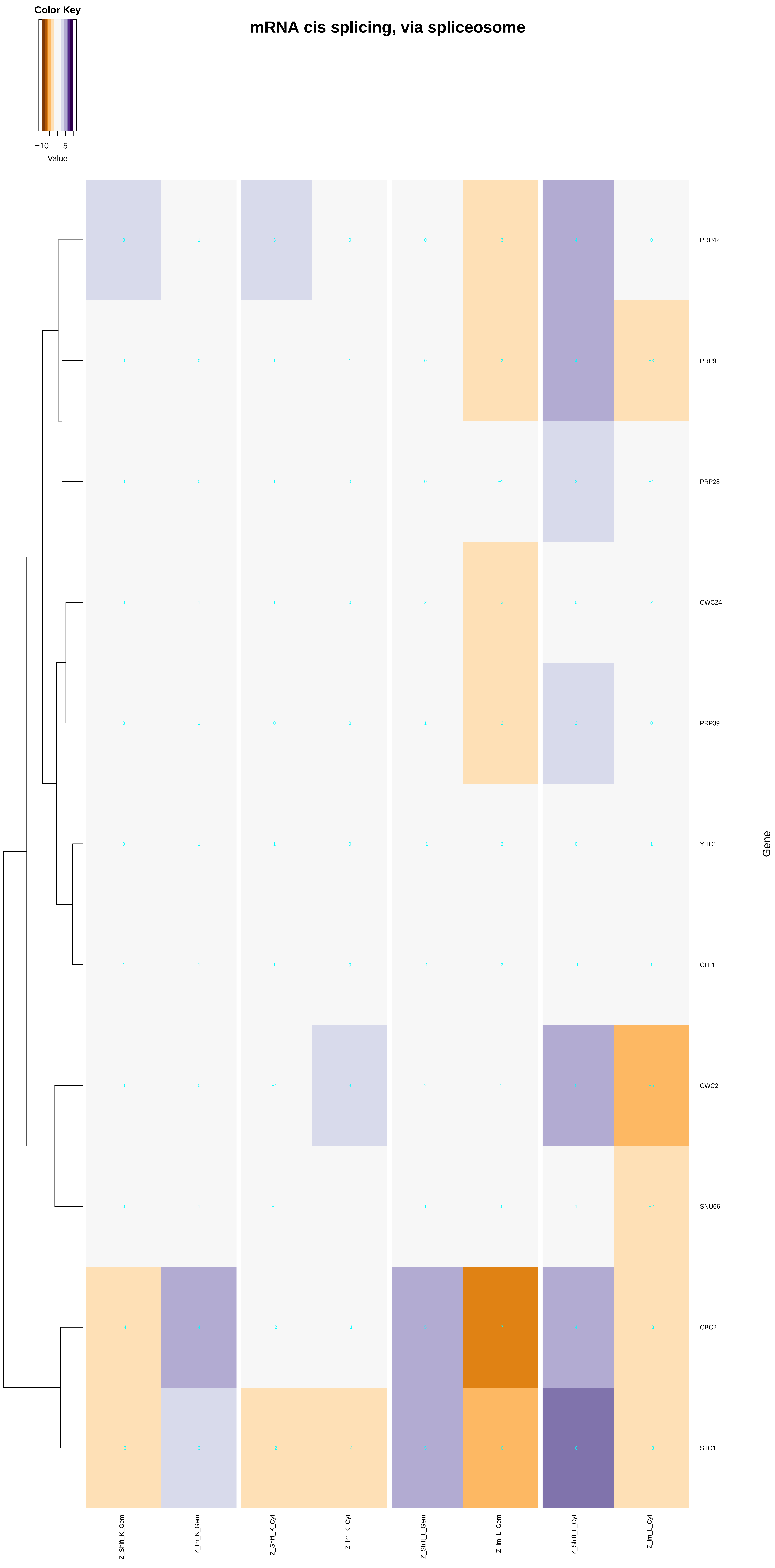

### multi-eIF_complex.pdf

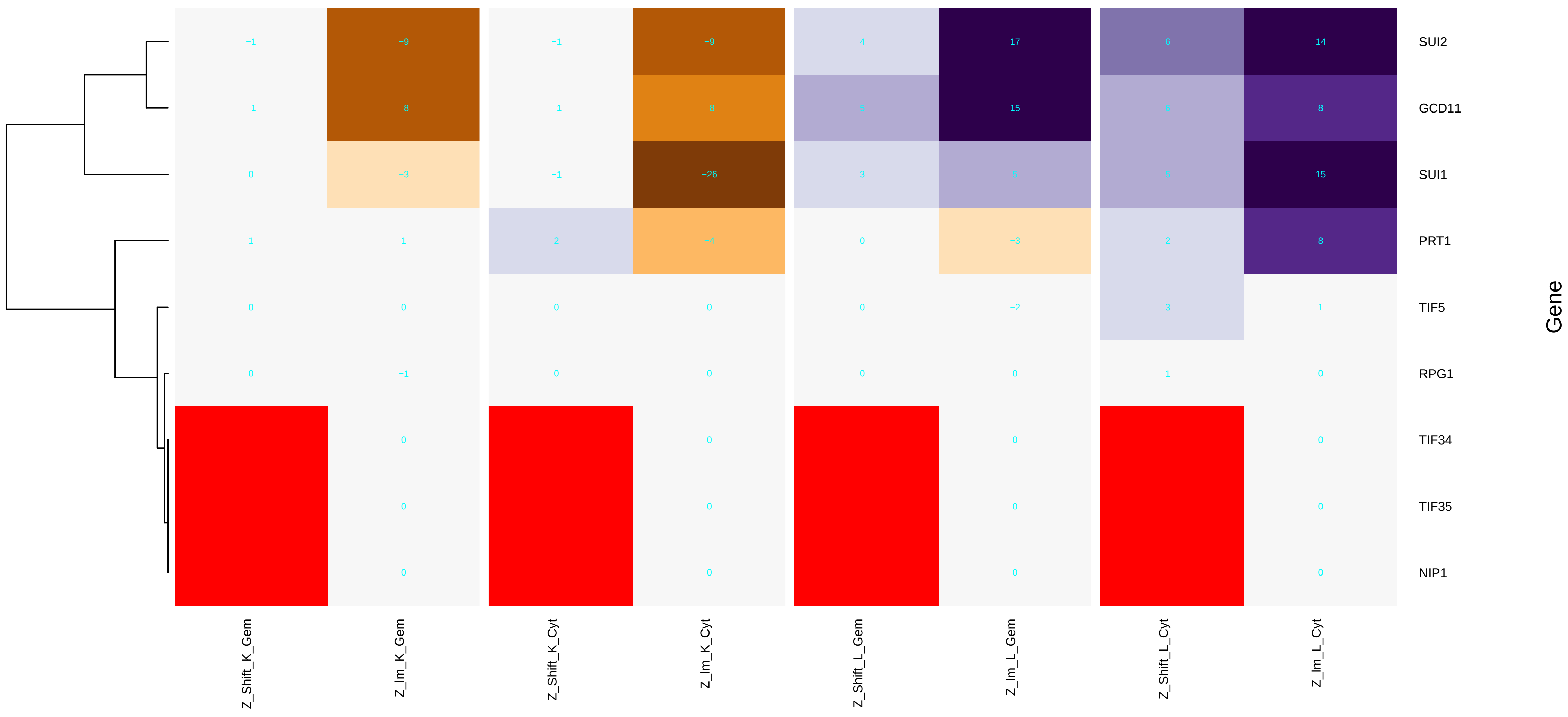

### N-terminal_peptidyl-methionine_acetylation.pdf

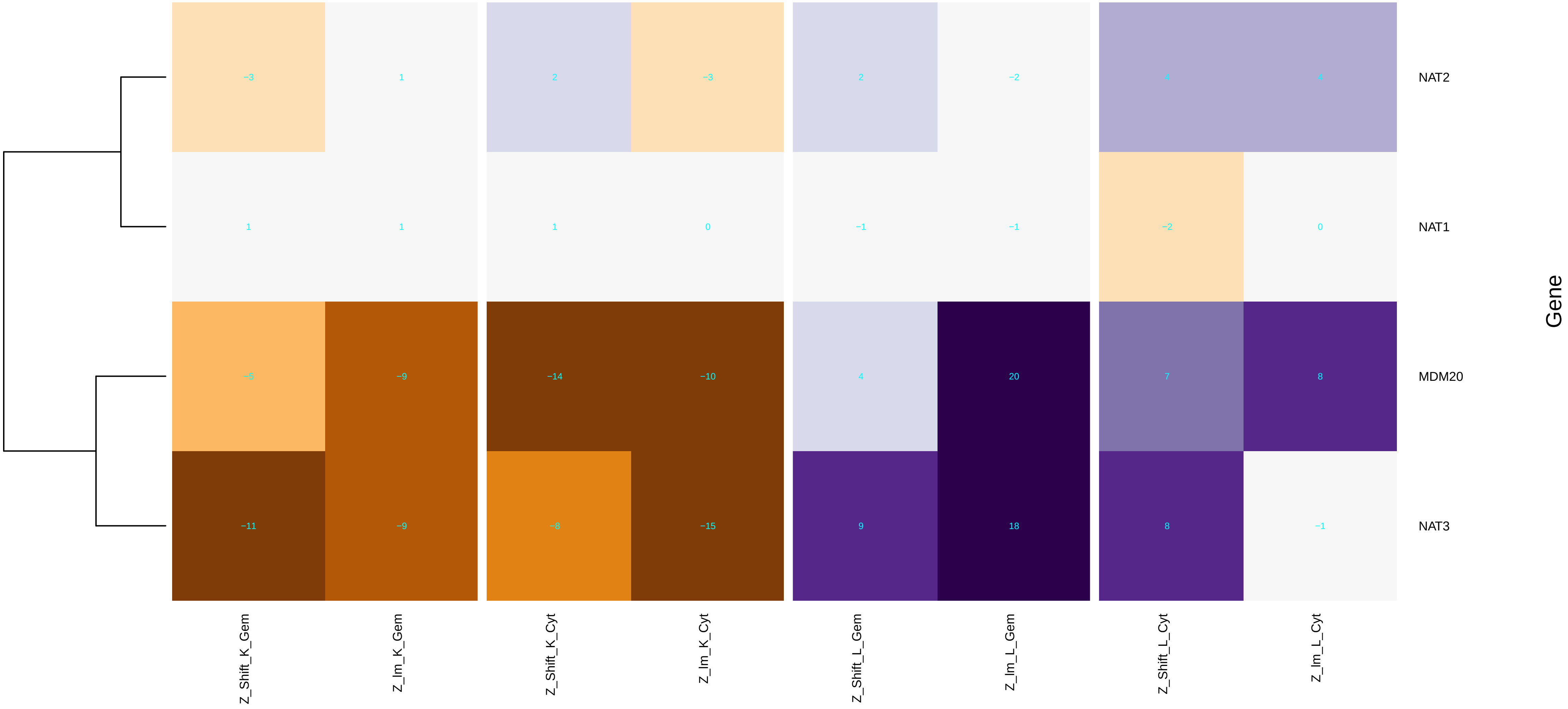

### N-terminal_protein_amino_acid_acetylation.pdf

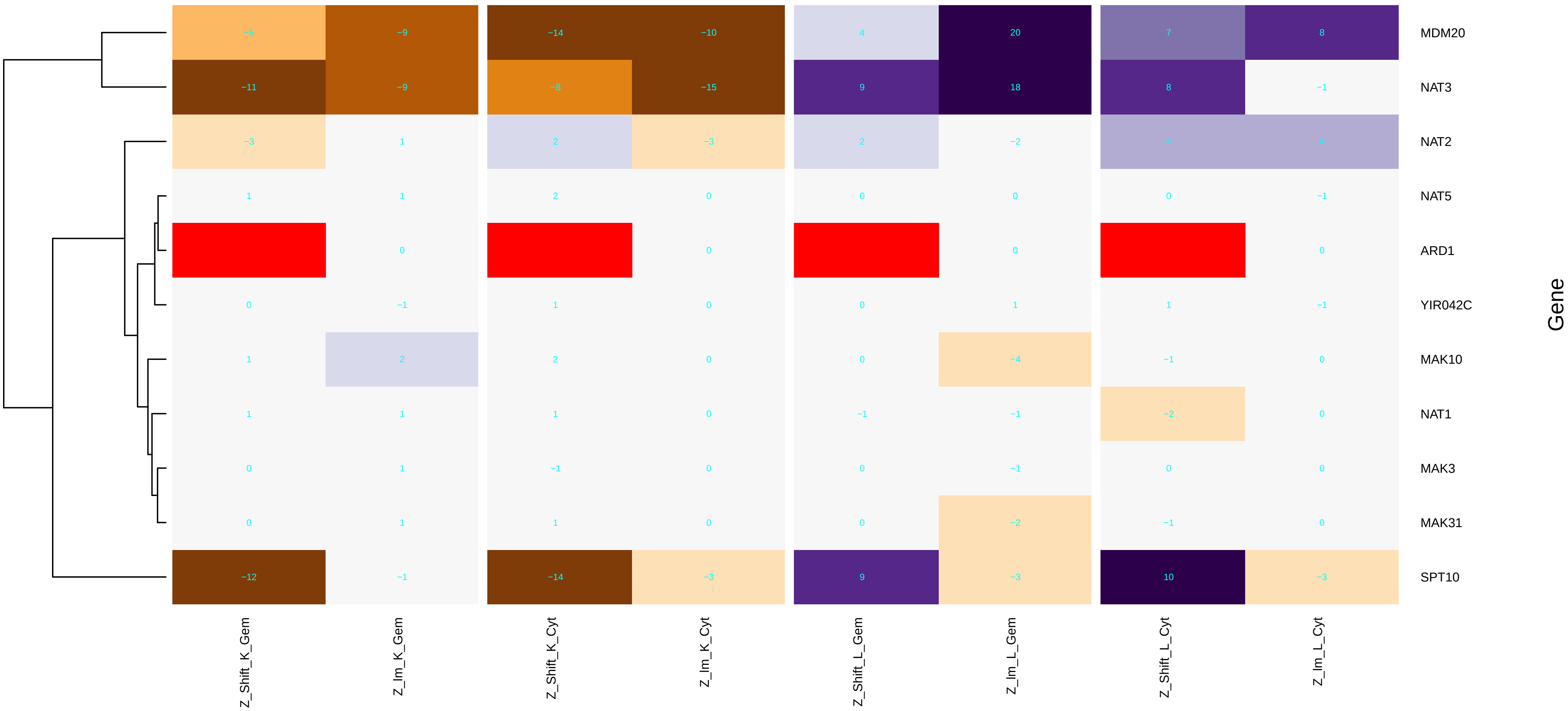

### NatB_complex.pdf

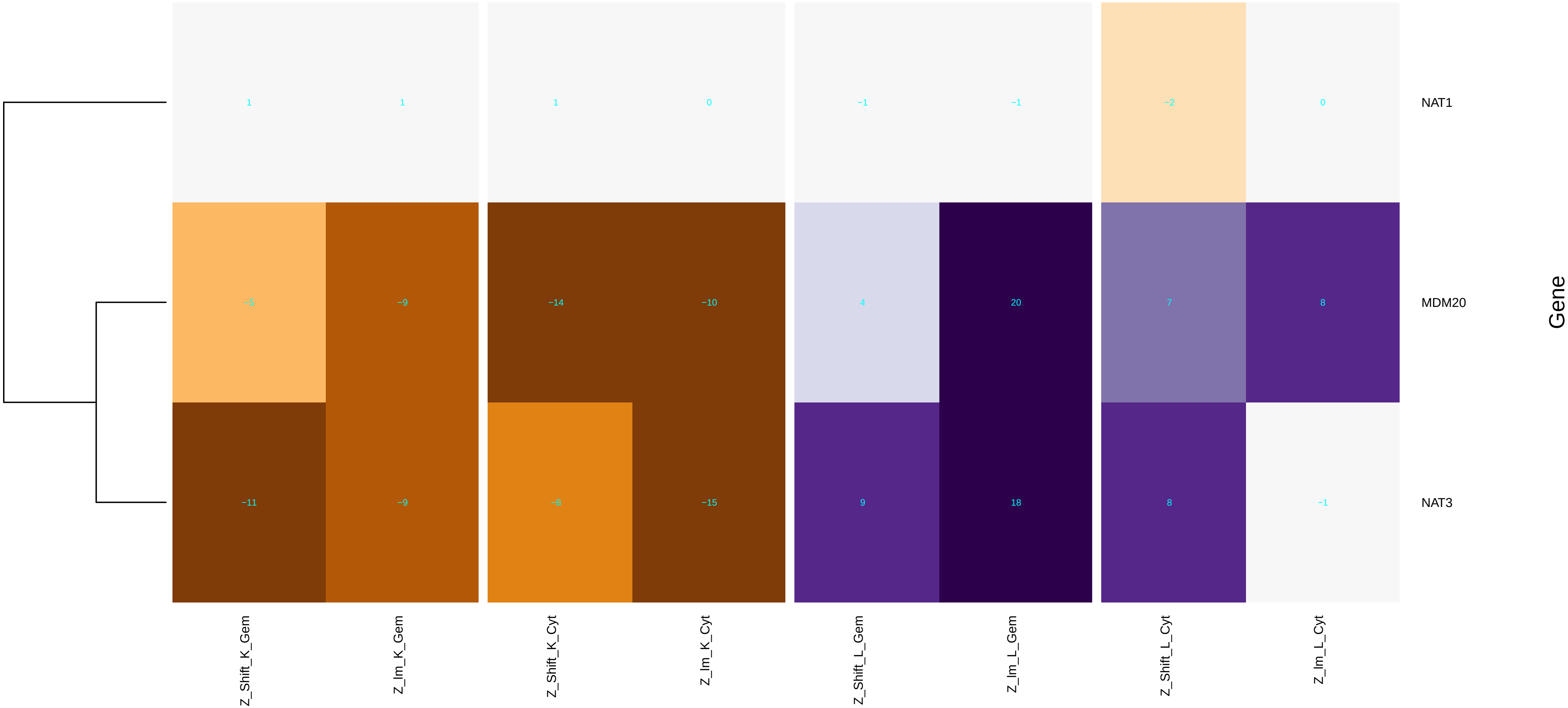

### negative_regulation_of_macroautophagy.pdf

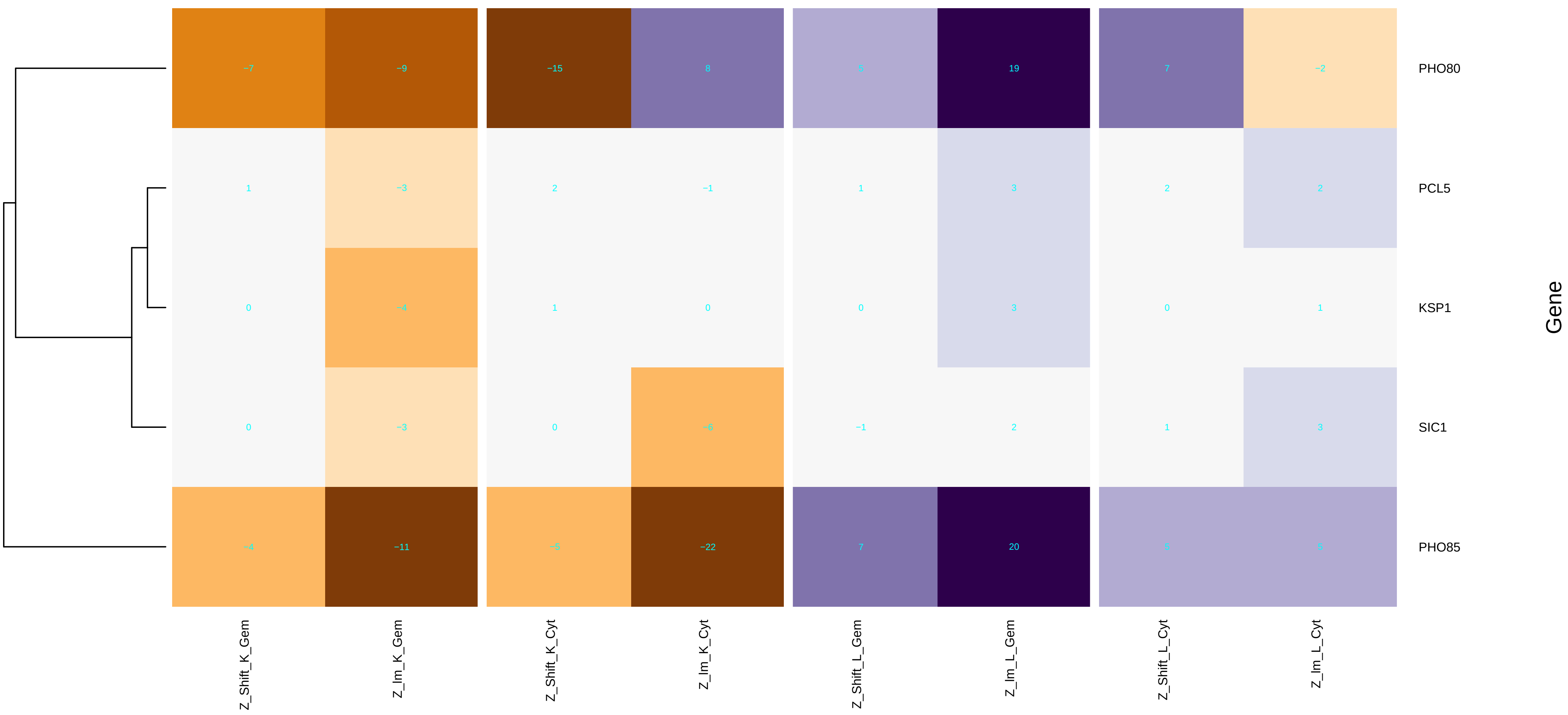

### negative_regulation_of_phosphatidylcholine_biosynthetic_process.pdf

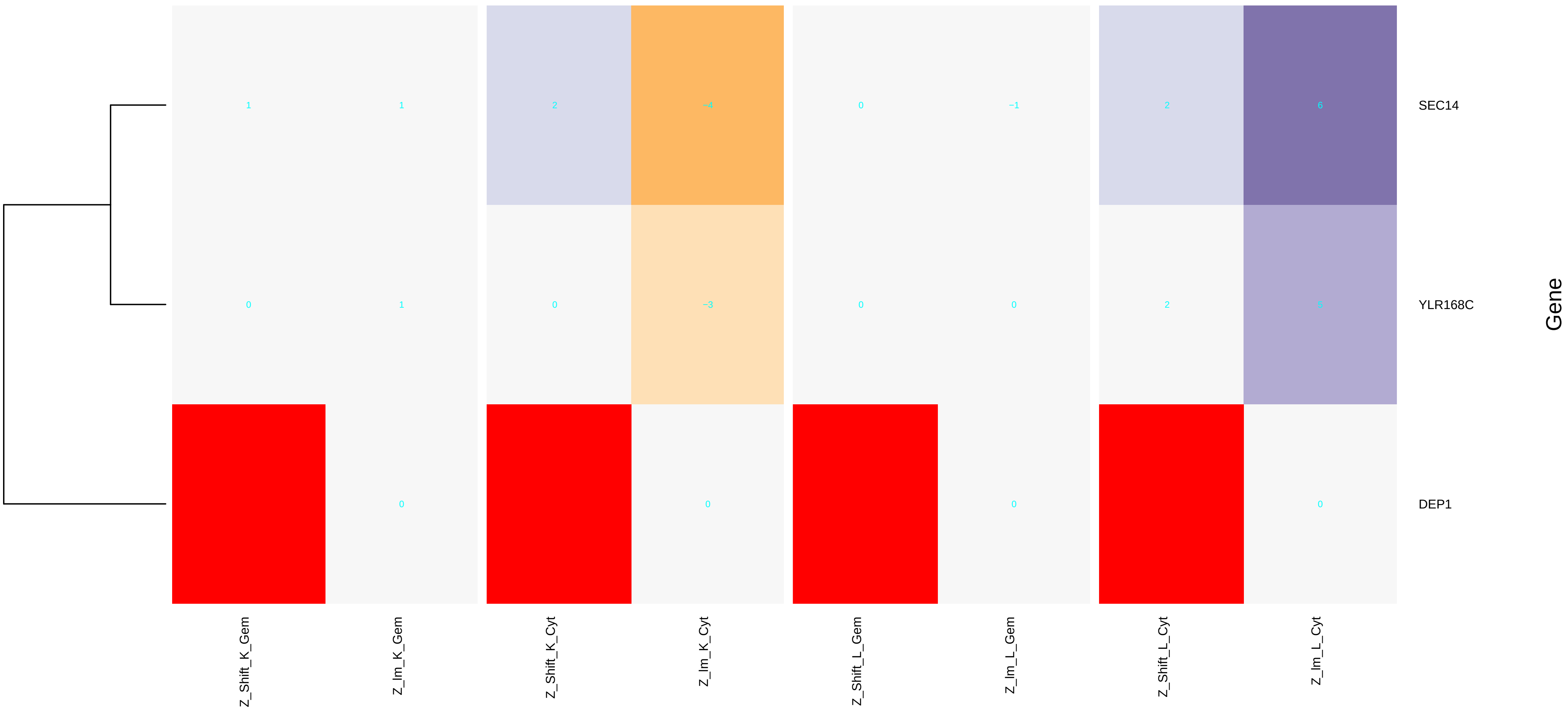

### nuclear_chromatin.pdf

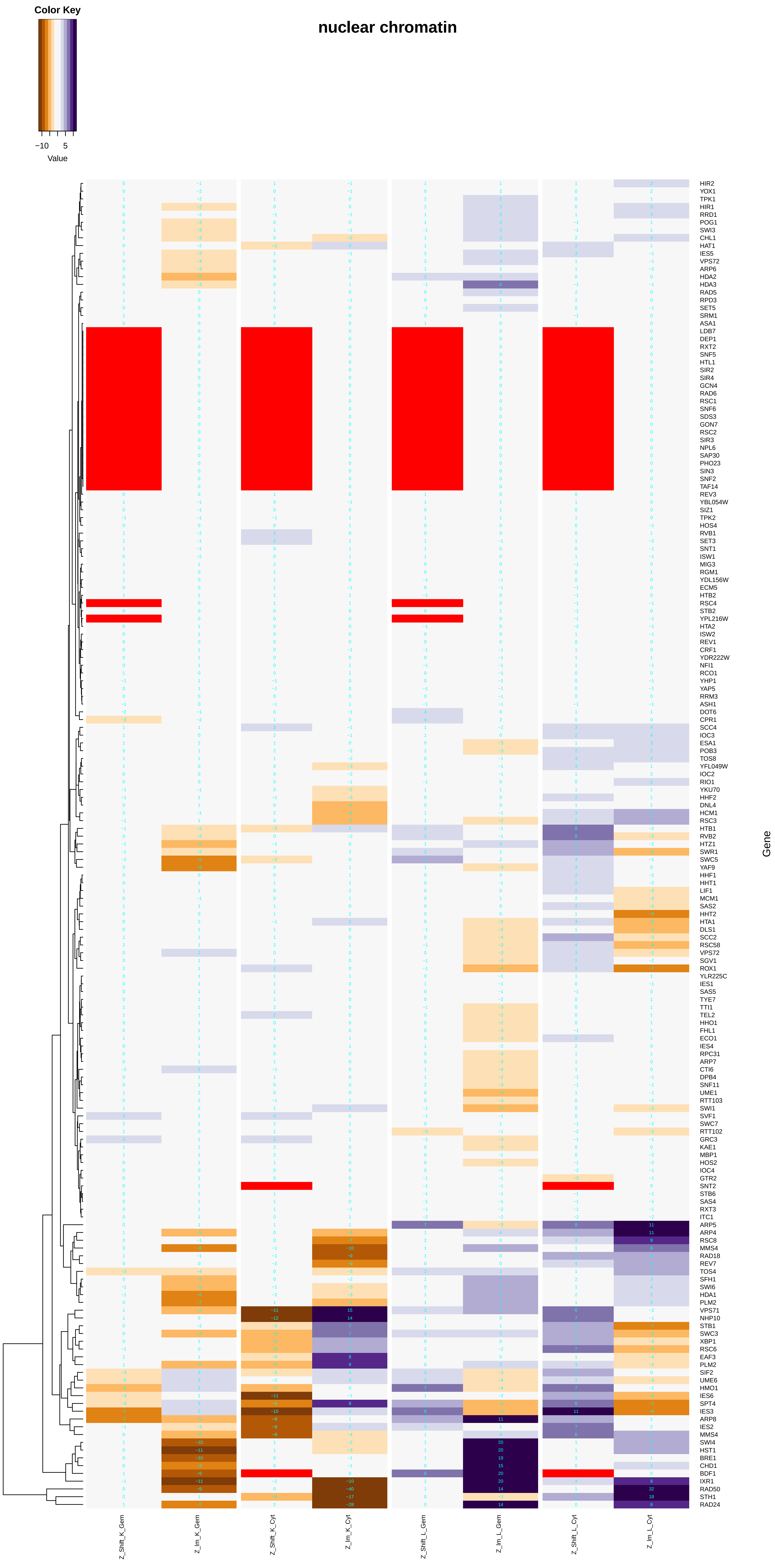

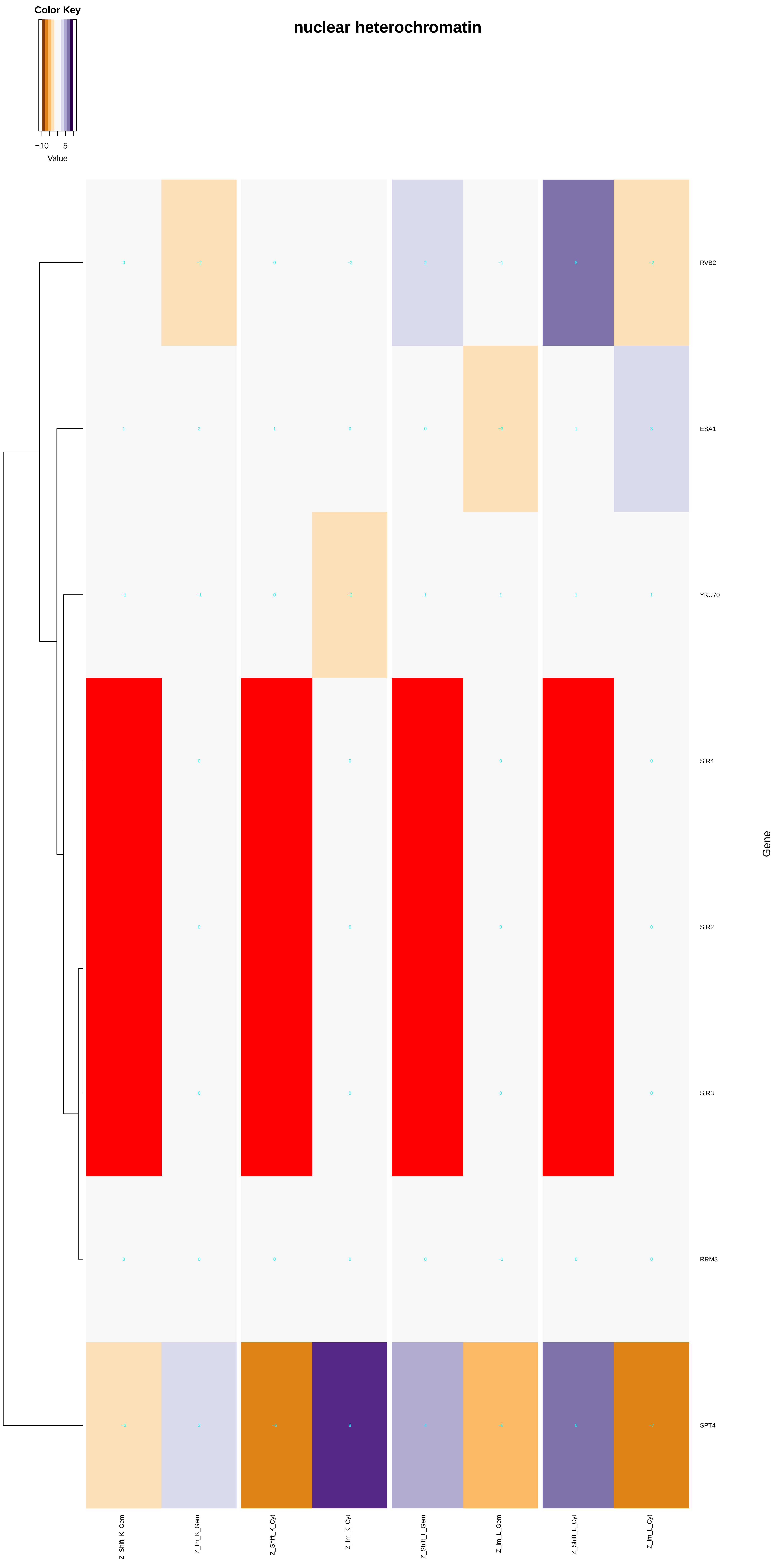

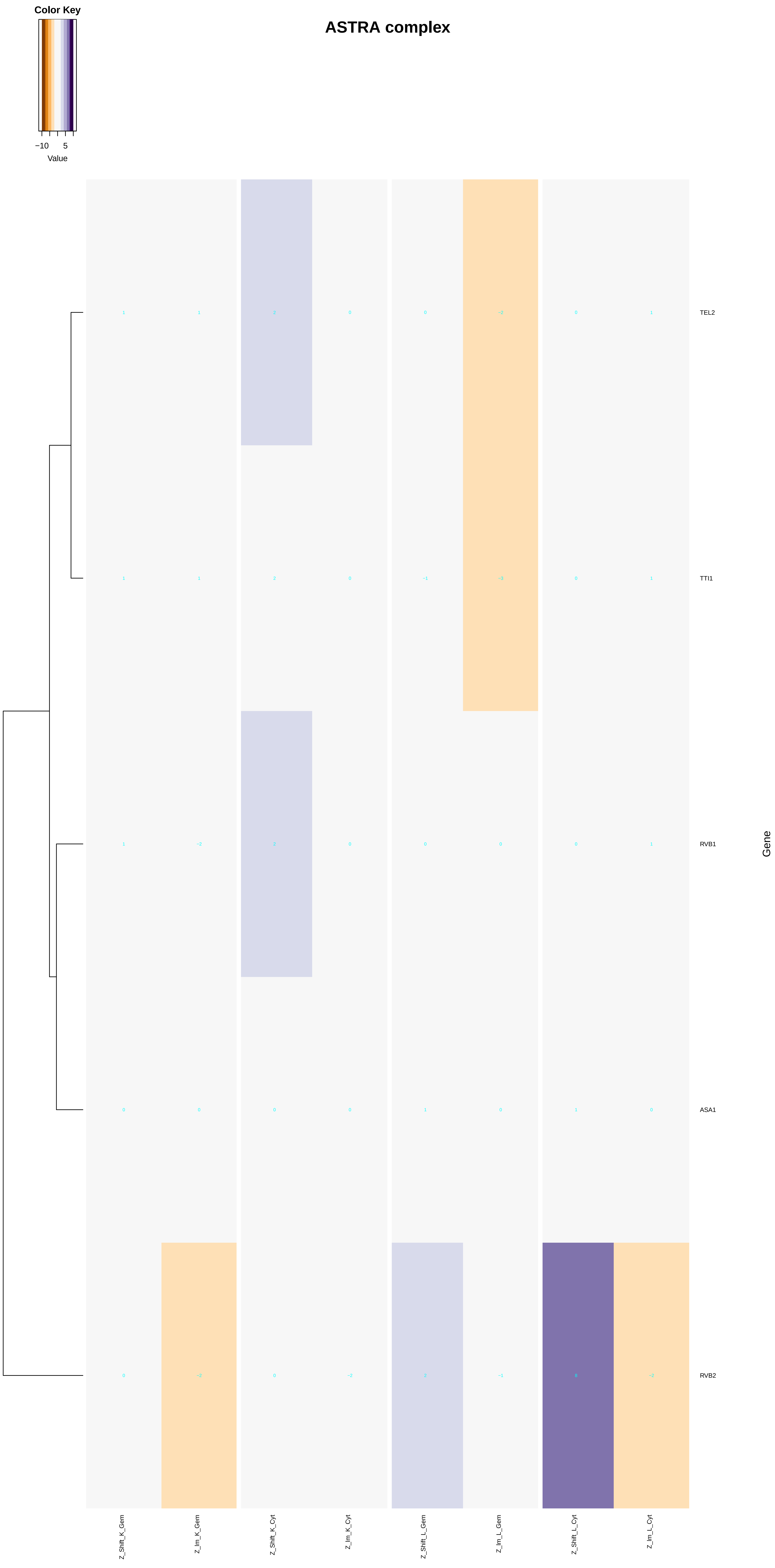

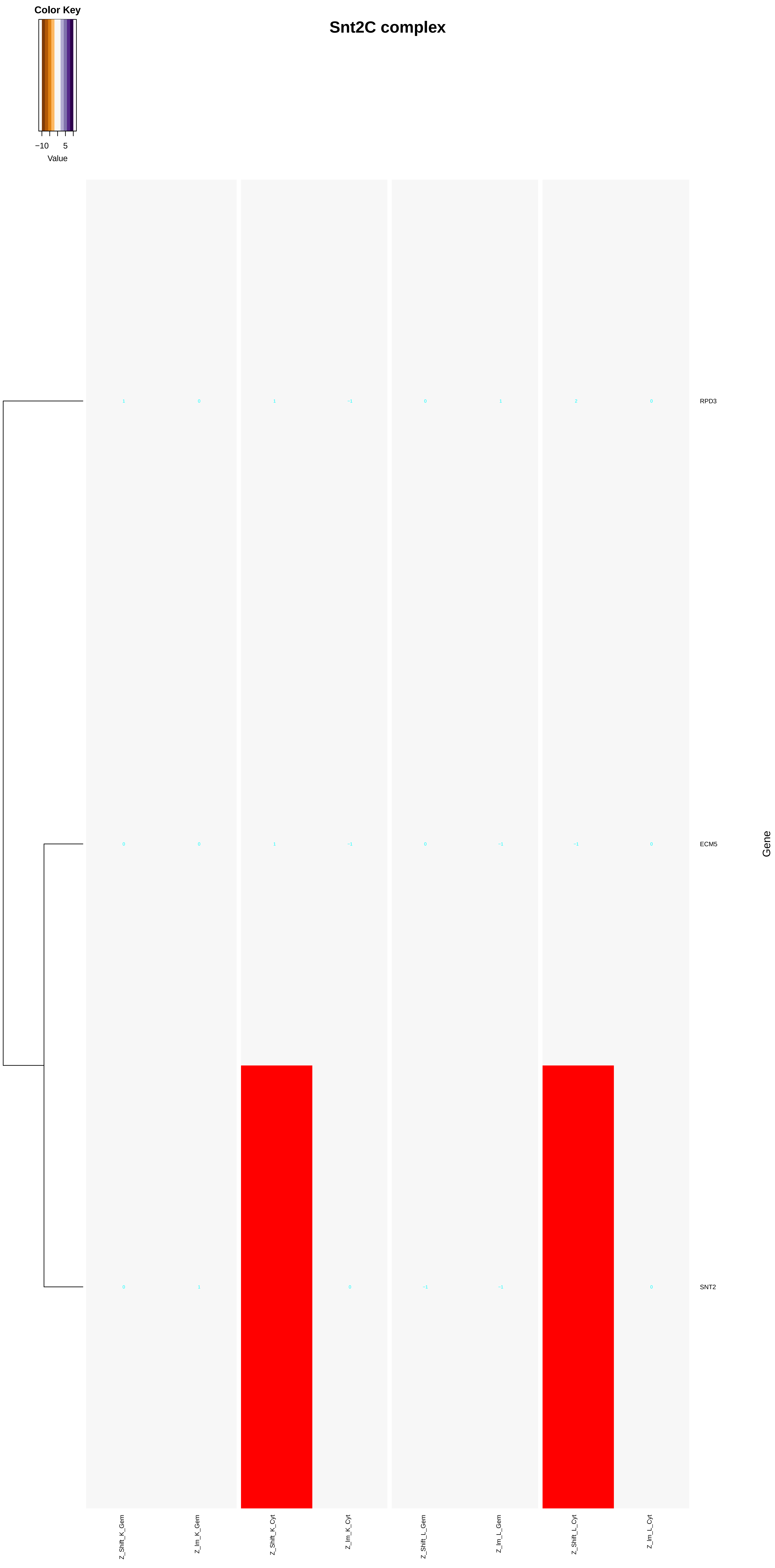

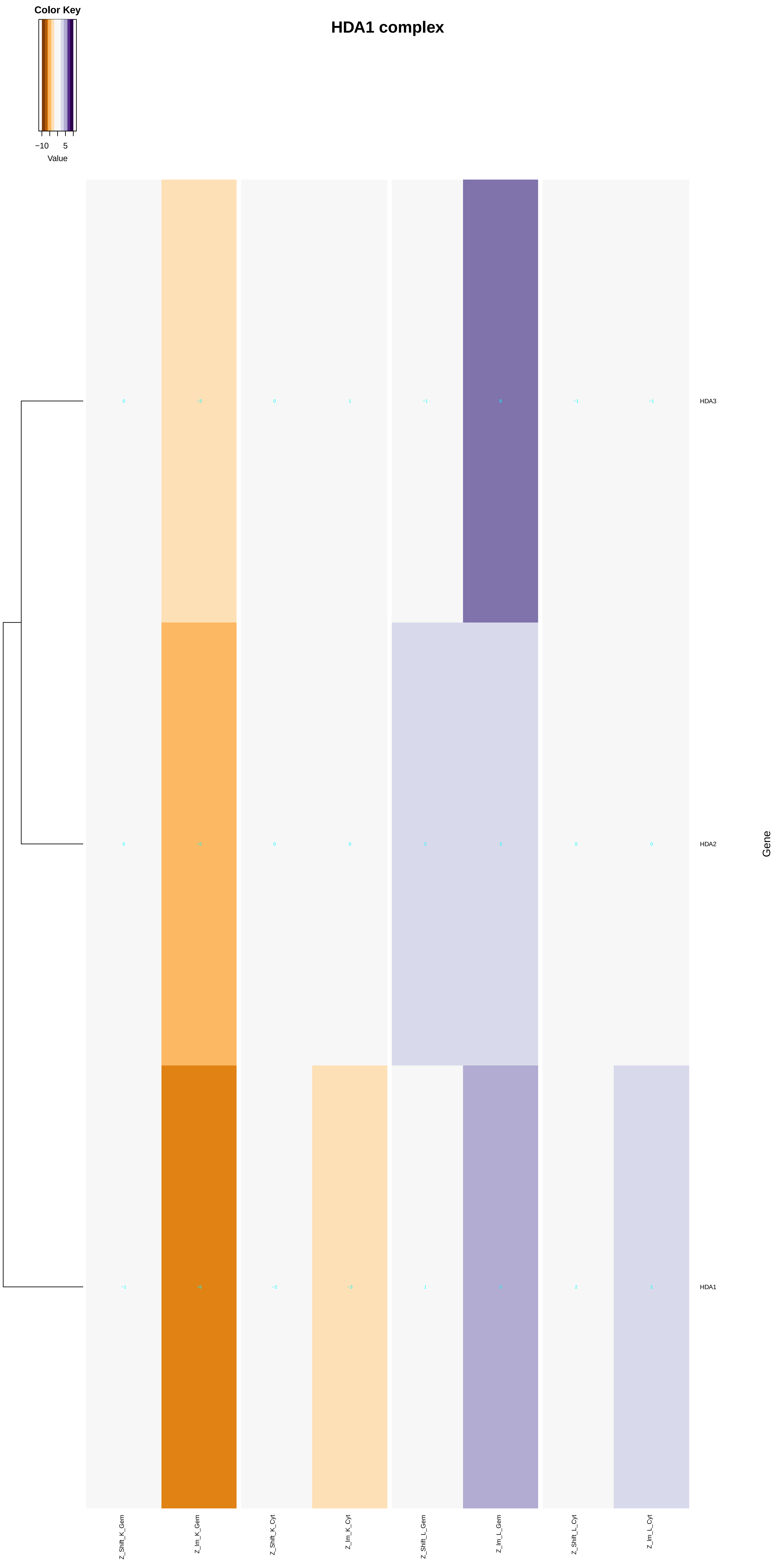

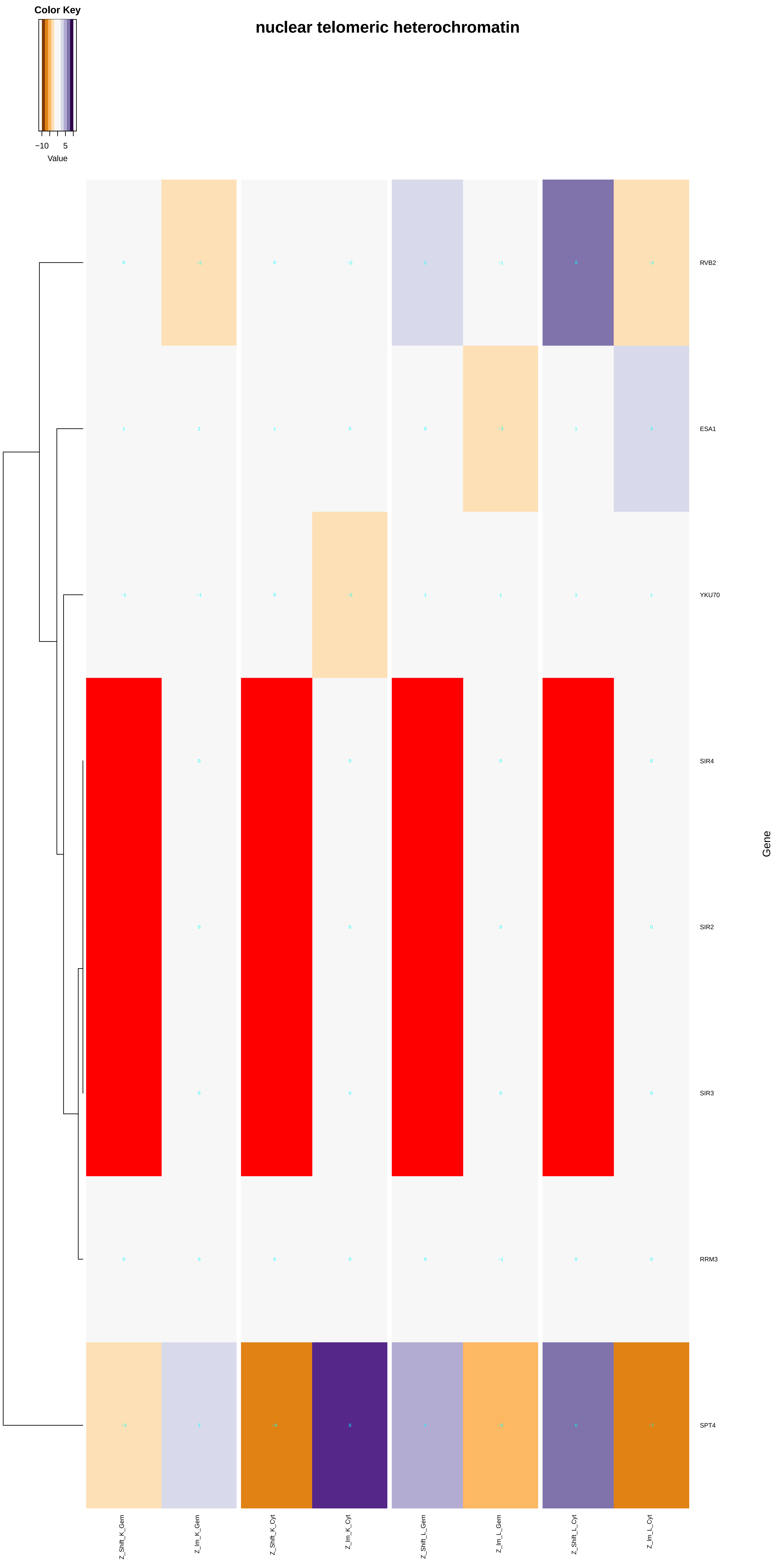

### nuclear_chromosome.pdf

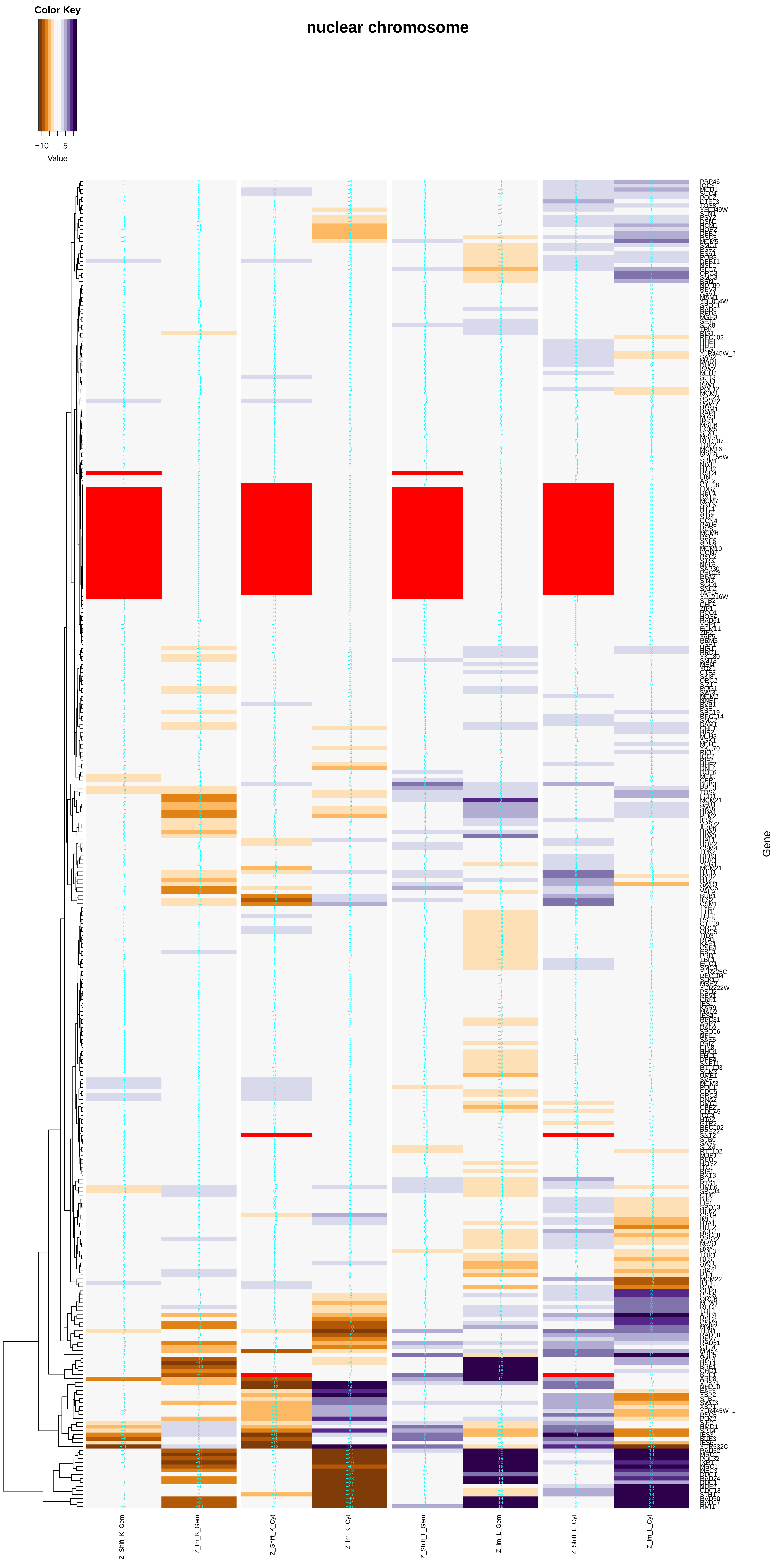

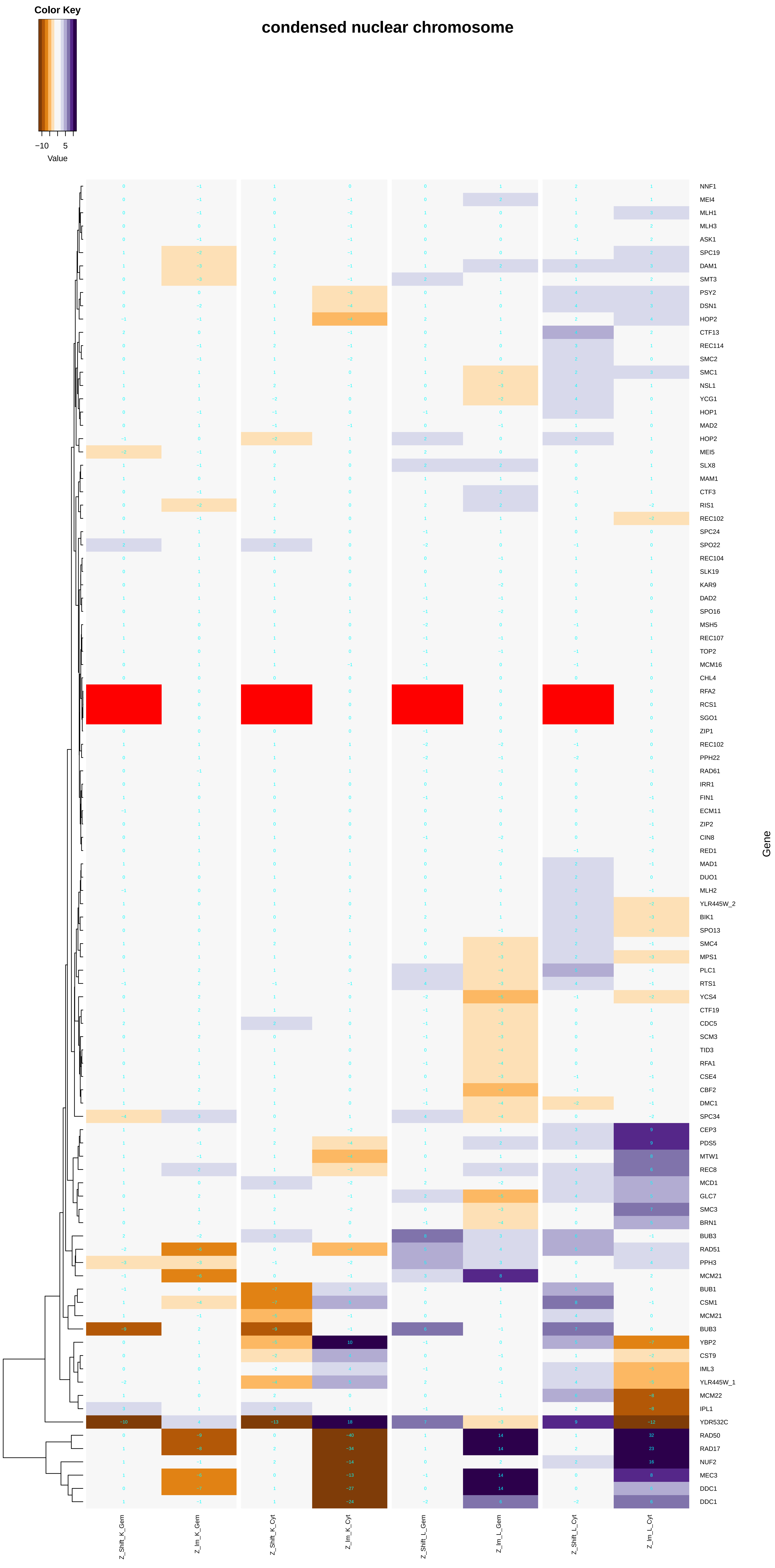

### nuclear_cohesin_complex.pdf

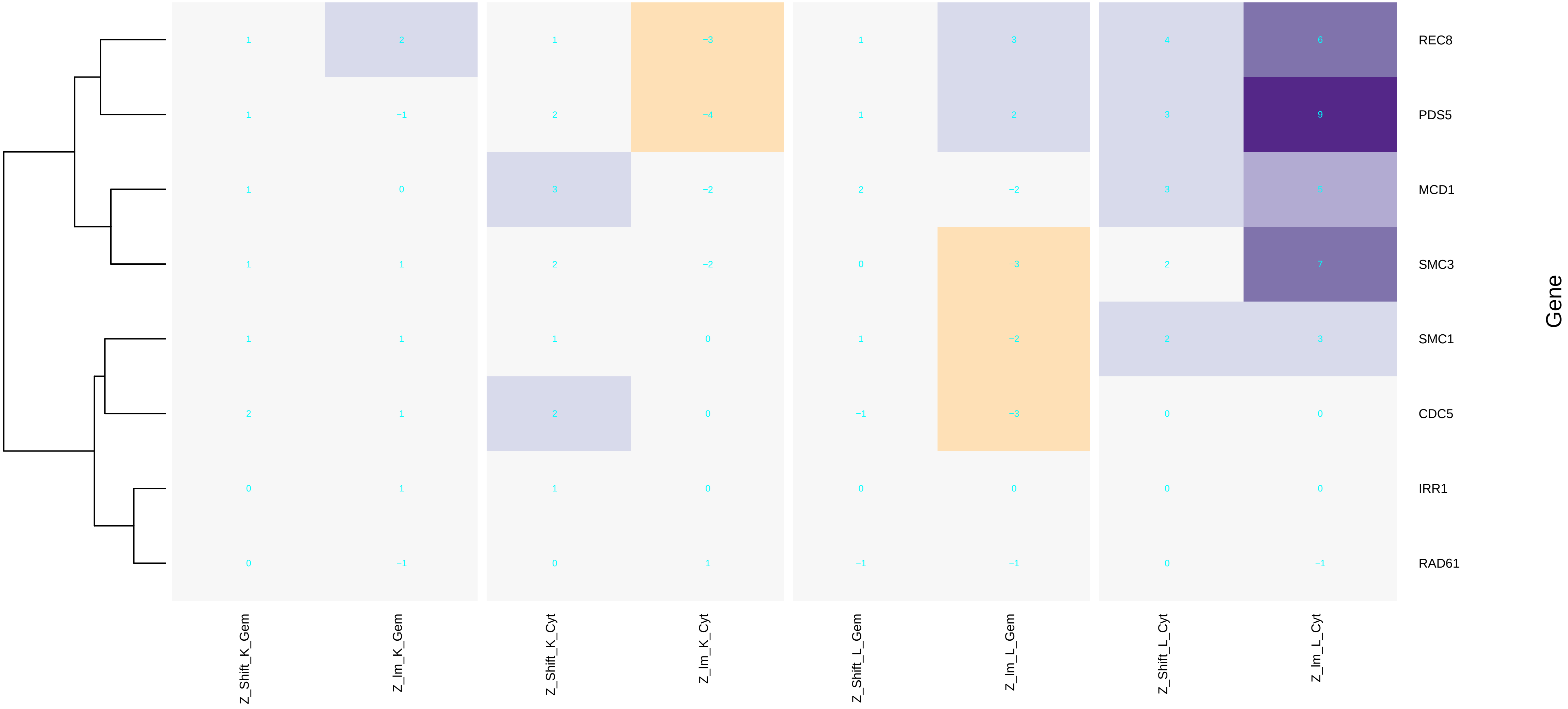

### nuclear_division.pdf

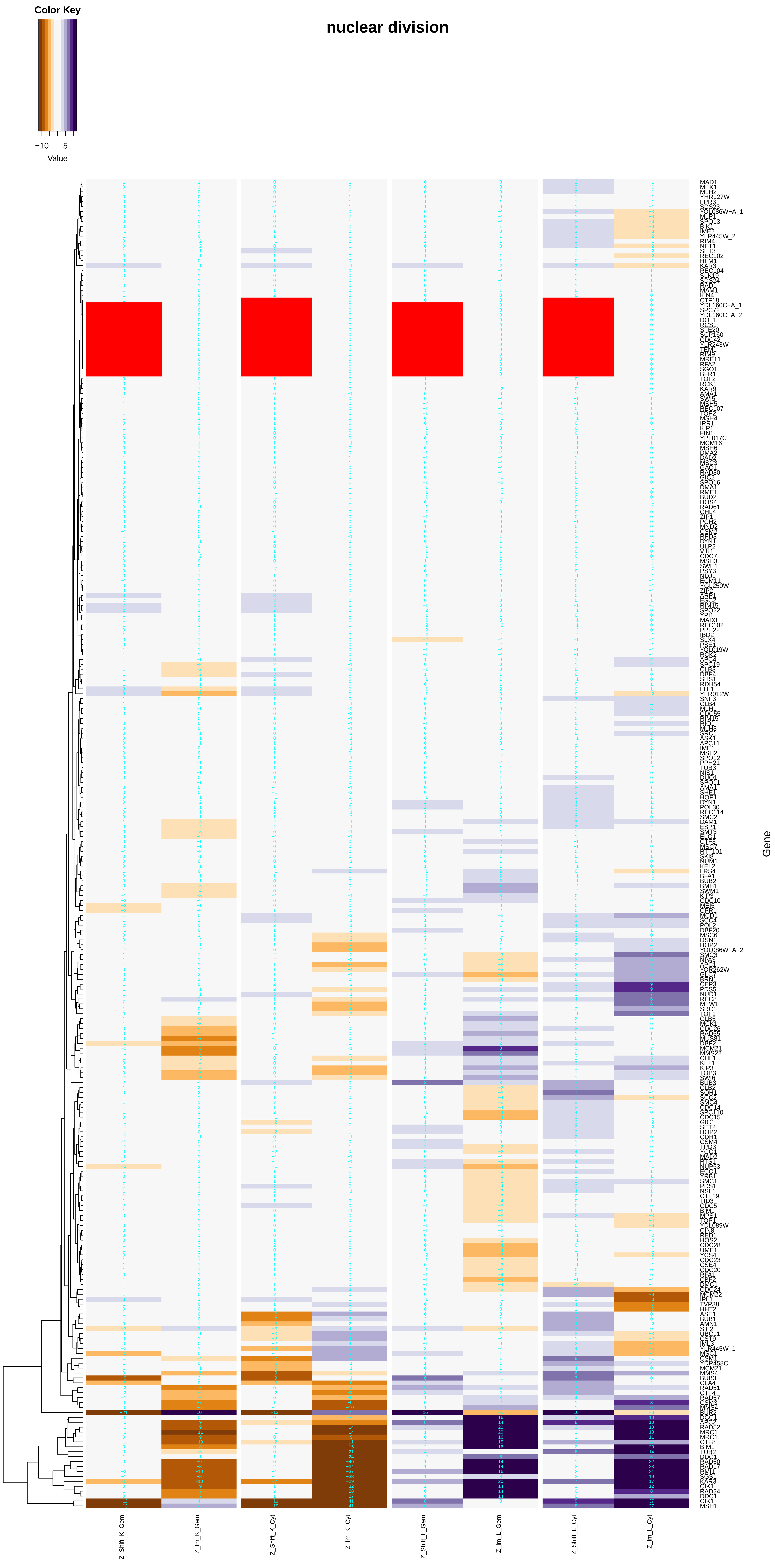

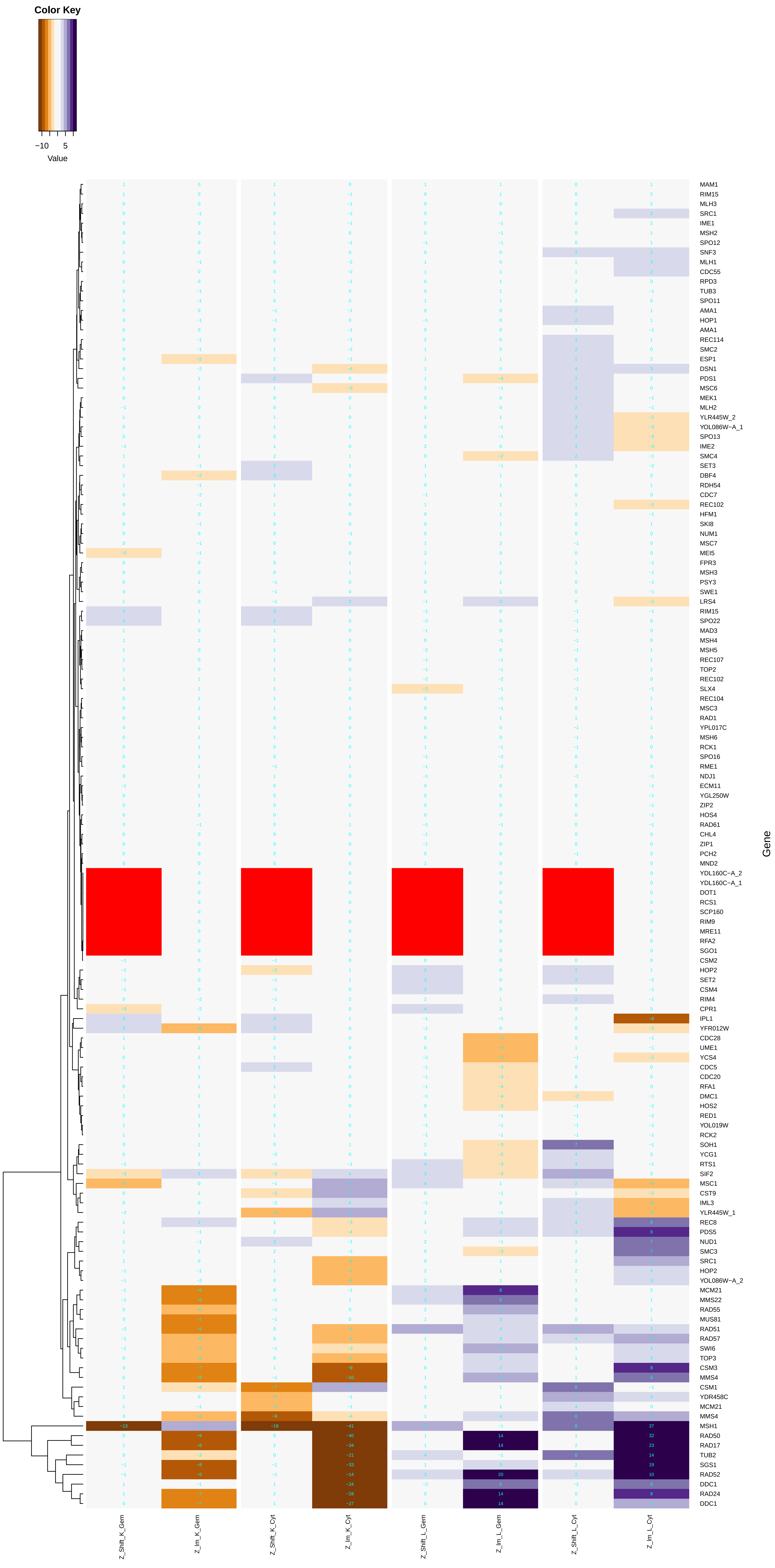

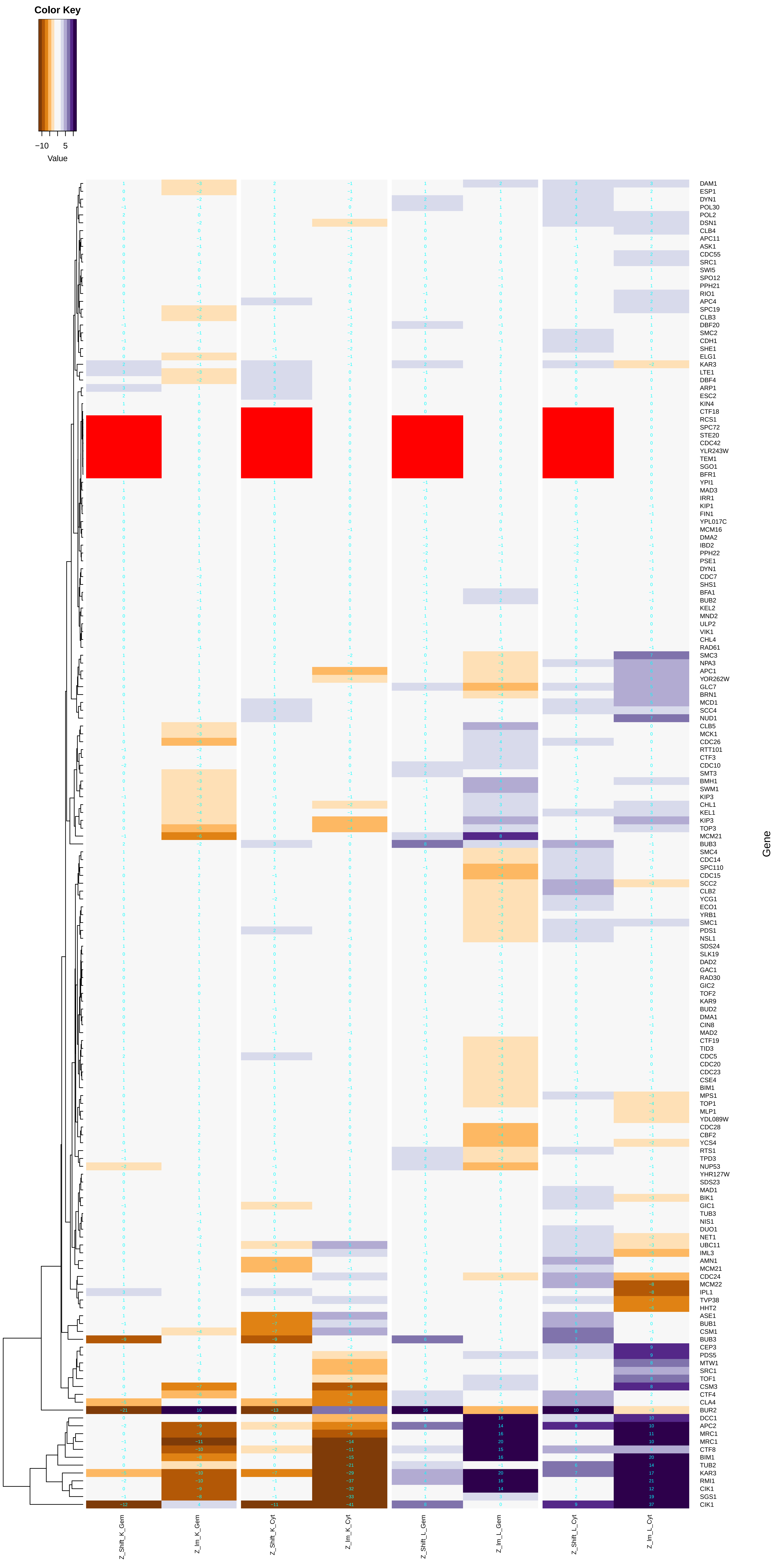

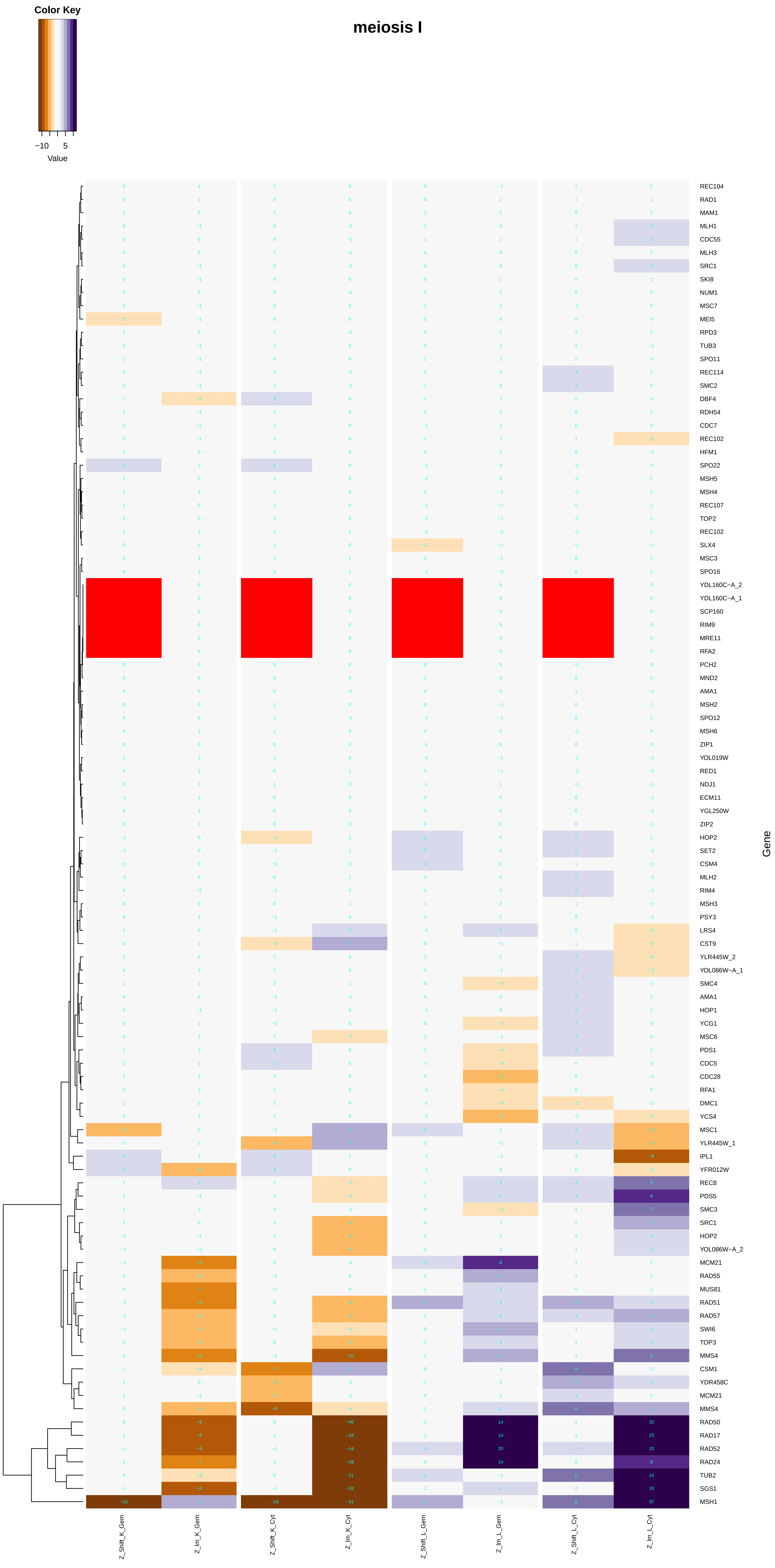

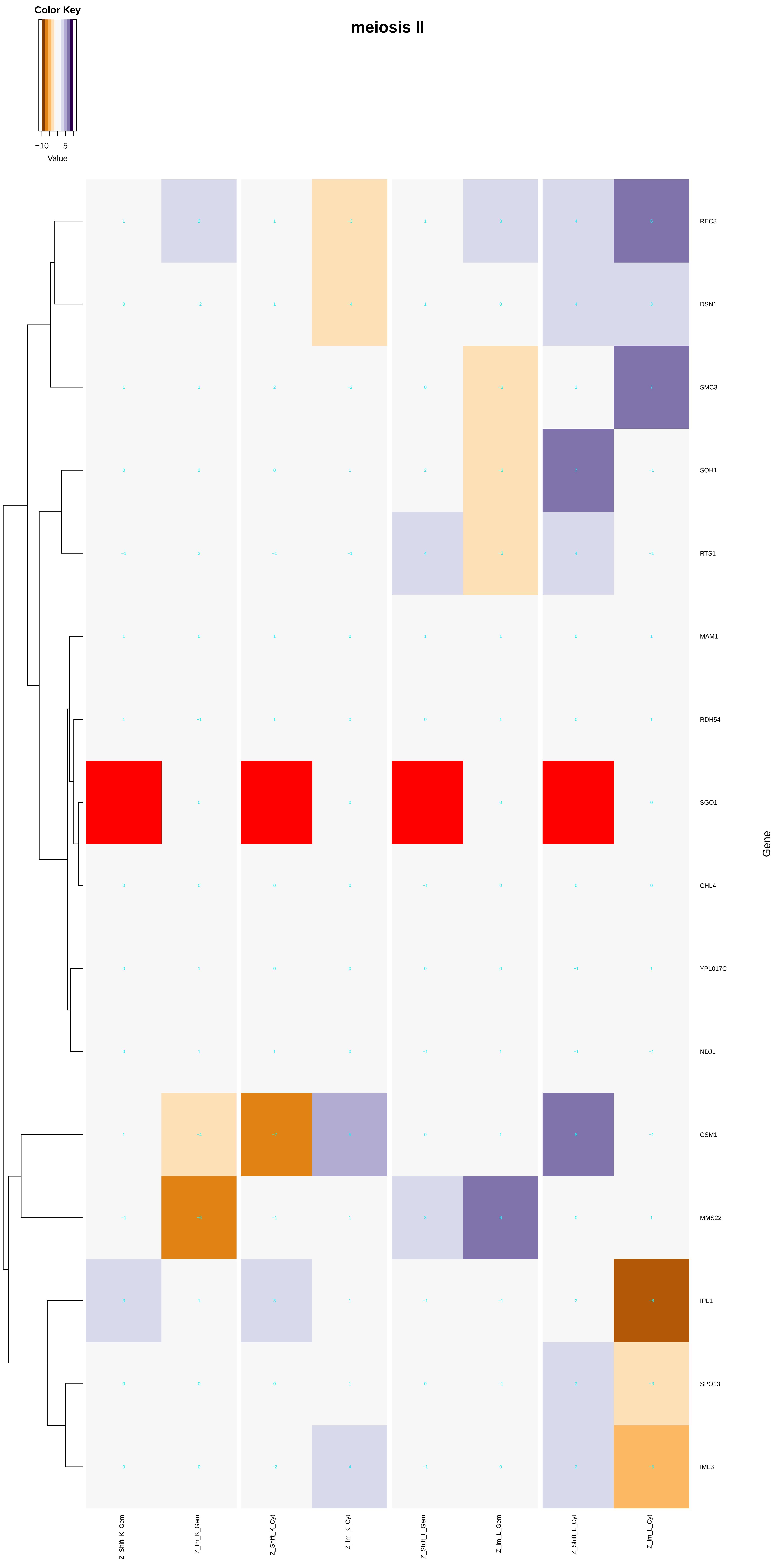

### nuclear_envelope_lumen.pdf

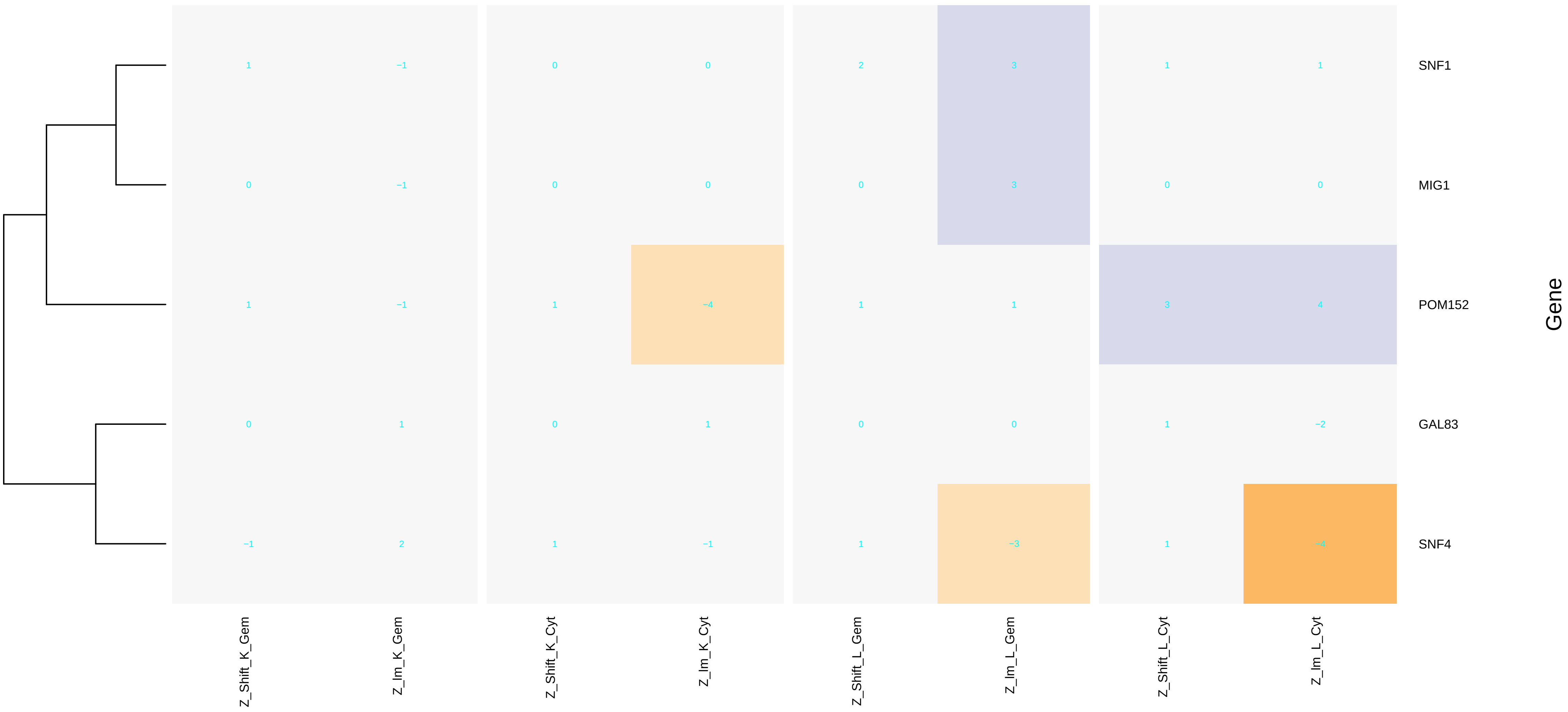
