## Supplementary figures and images for "A humanized yeast phenomic model of deoxycytidine kinase to predict genetic buffering of nucleoside analog cytotoxicity"

### 5S_class_rRNA_transcription_from_RNA_polymerase_III_type_1_promoter.pdf

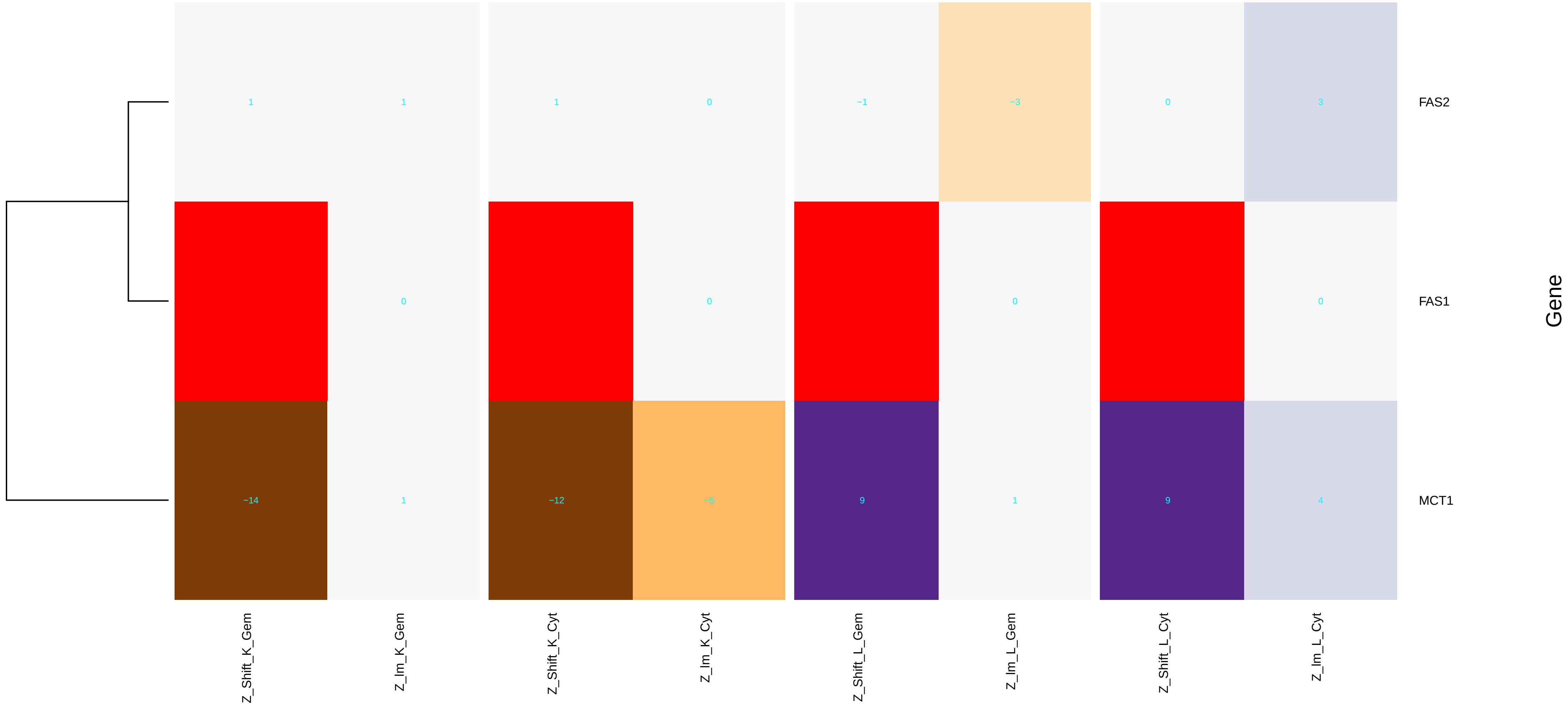

Supplement: Additional File 7 [file 700153_file09.zip › Additional_File7_GOTermHeatmaps/[acyl-carrier-protein]_S-malonyltransferase_activity.pdf]

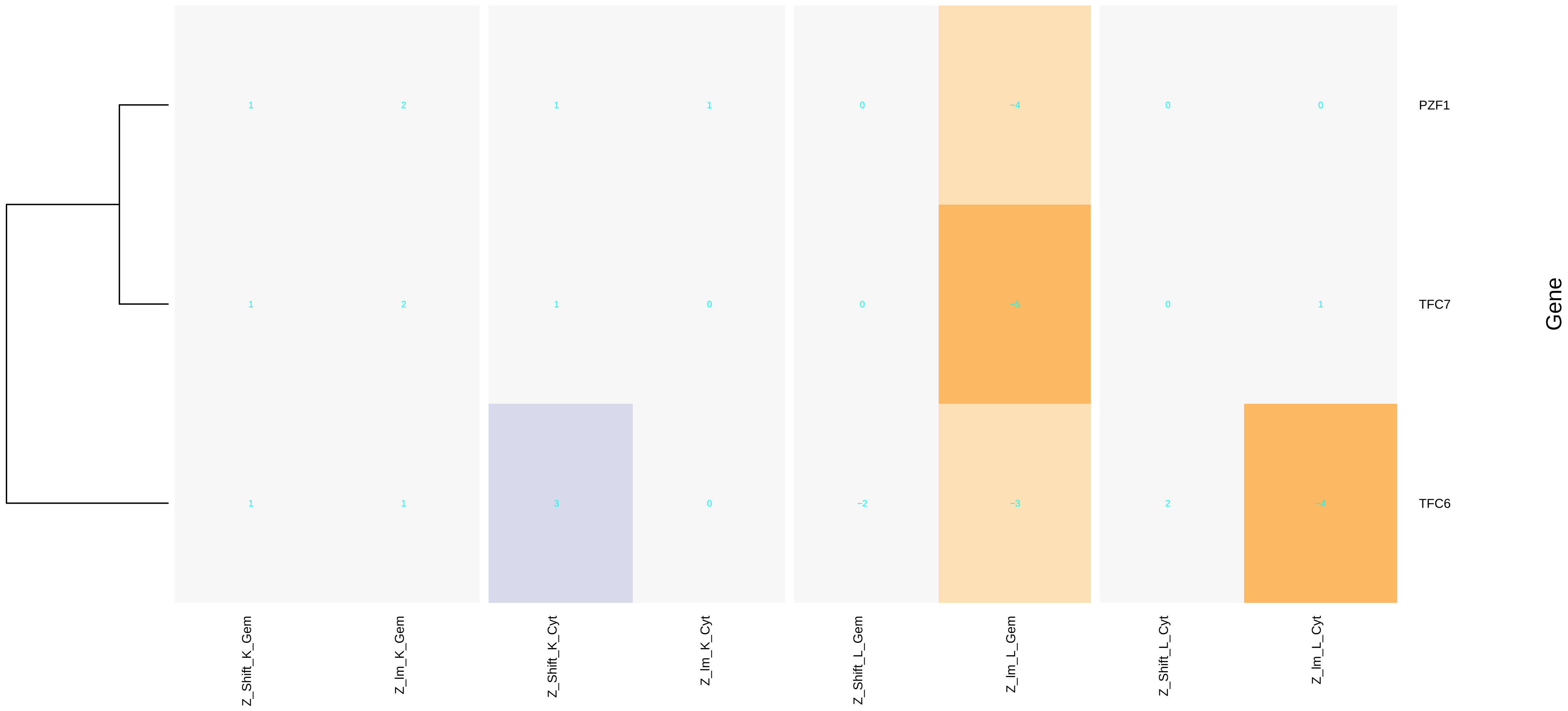

### 7-methylguanosine_RNA_capping.pdf

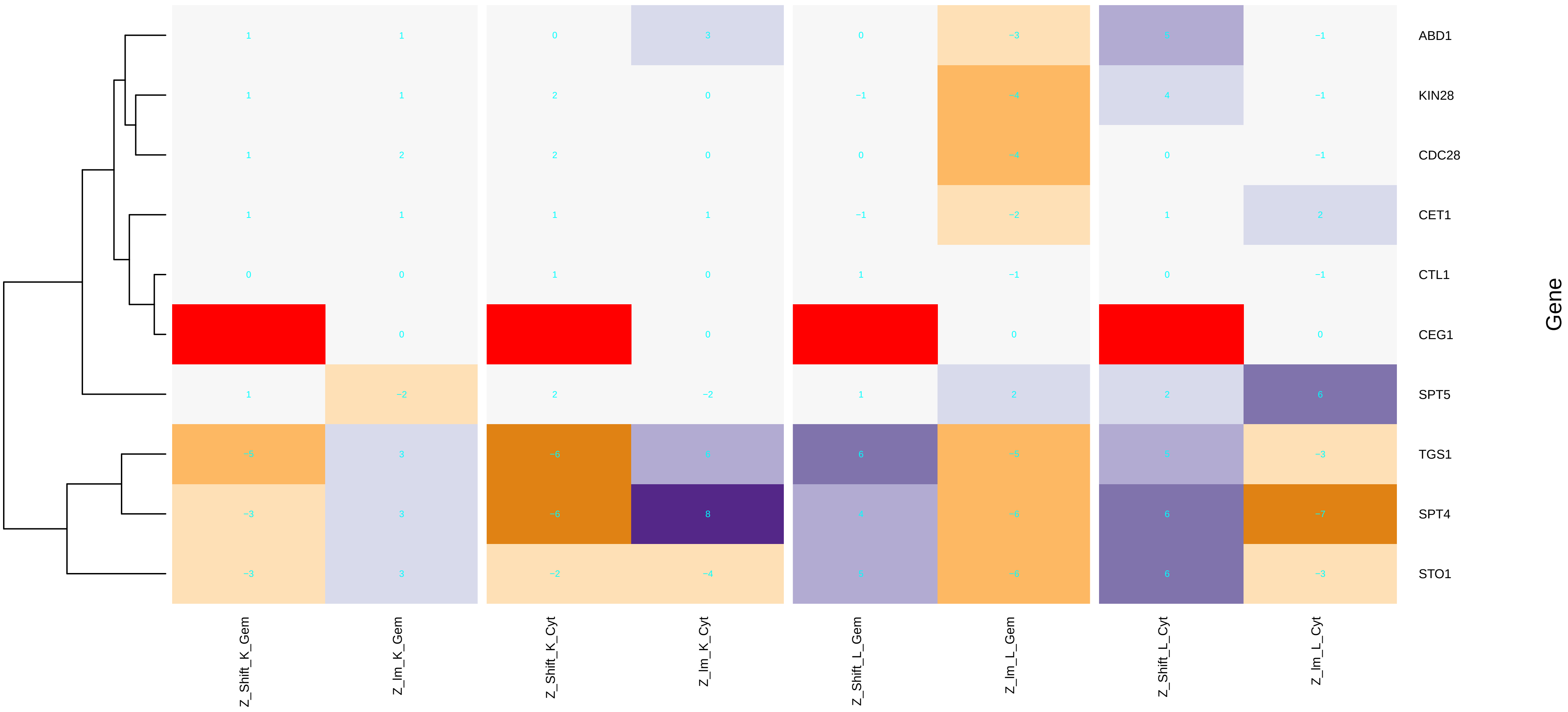

### 90S_preribosome.pdf

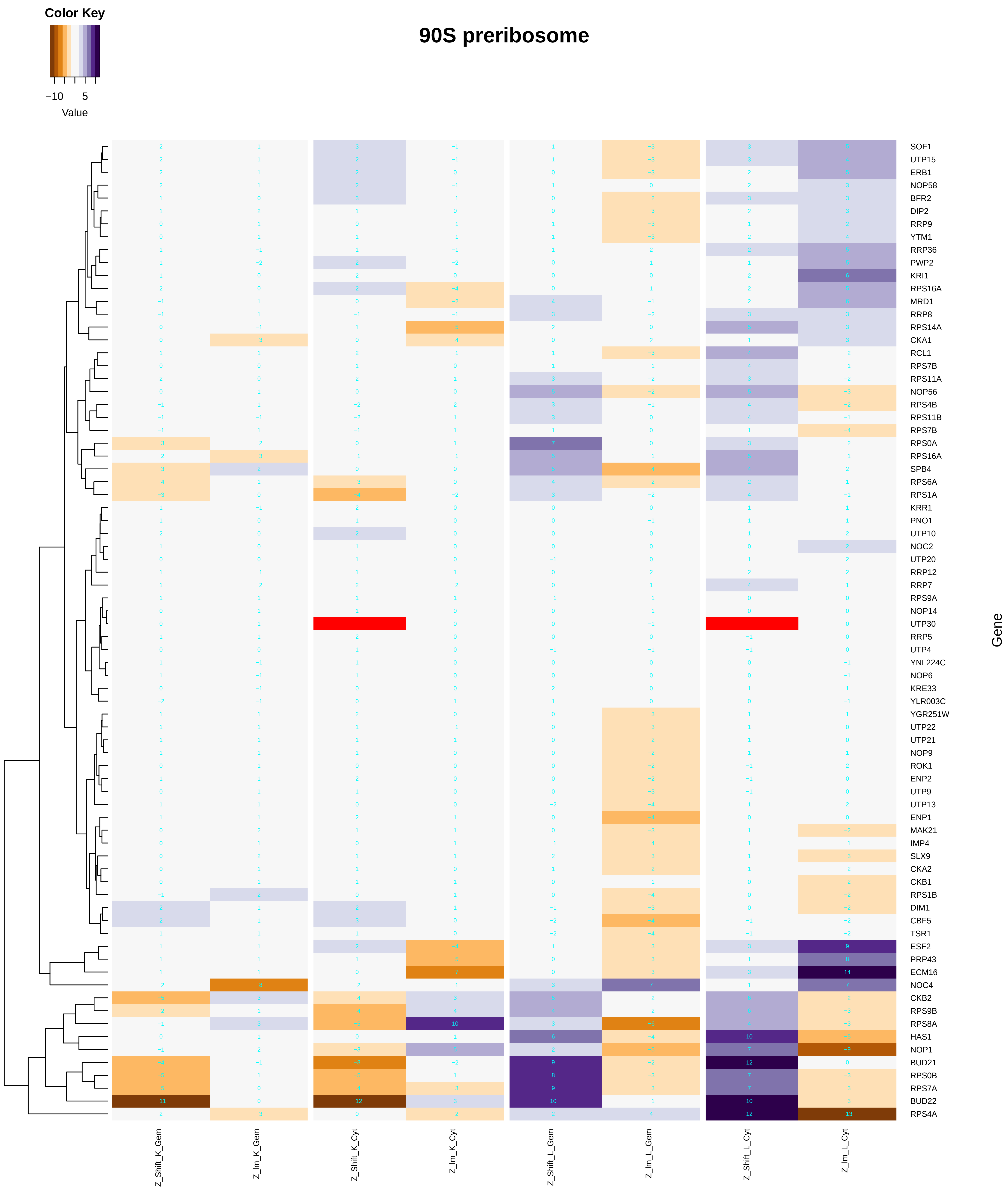

### adenyl-nucleotide_exchange_factor_activity.pdf

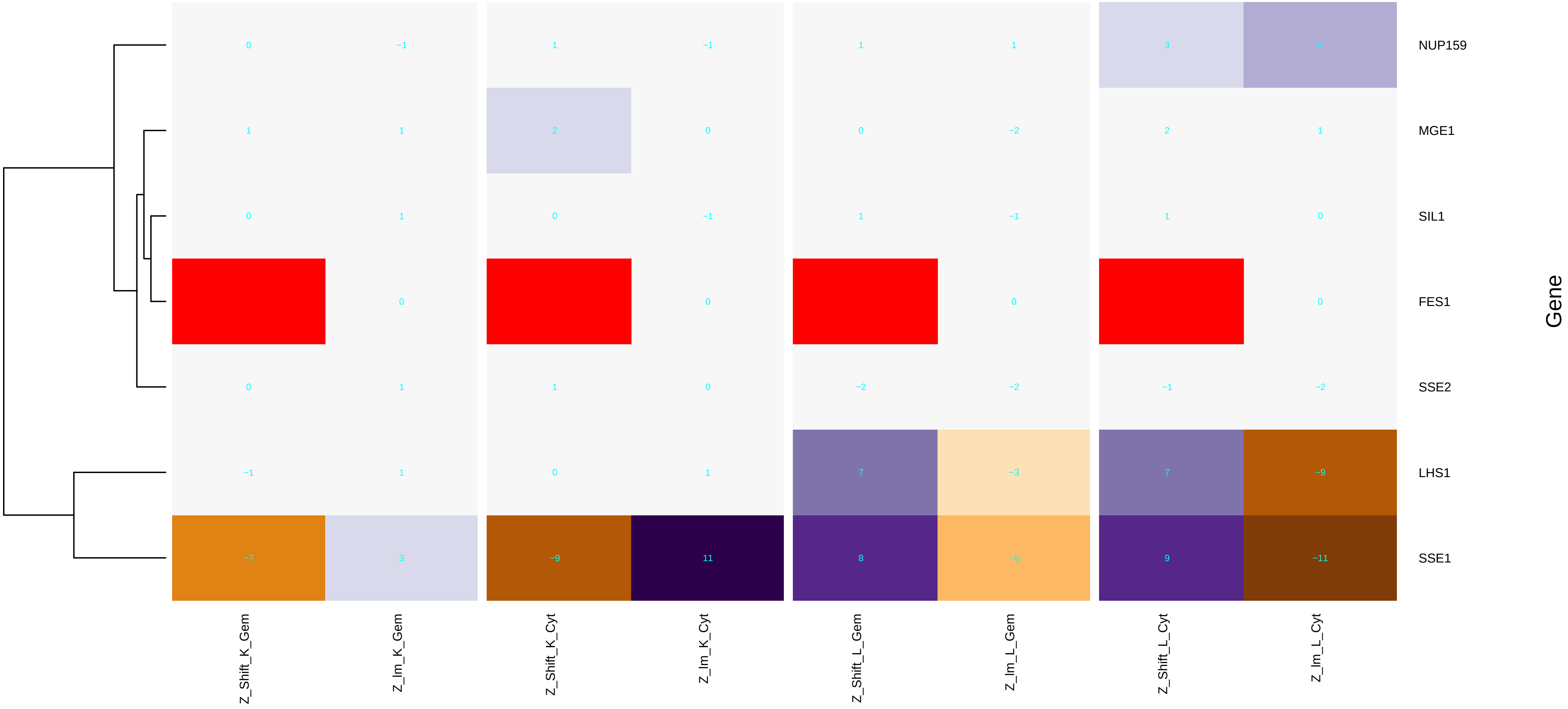

### adenyl_ribonucleotide_binding.pdf

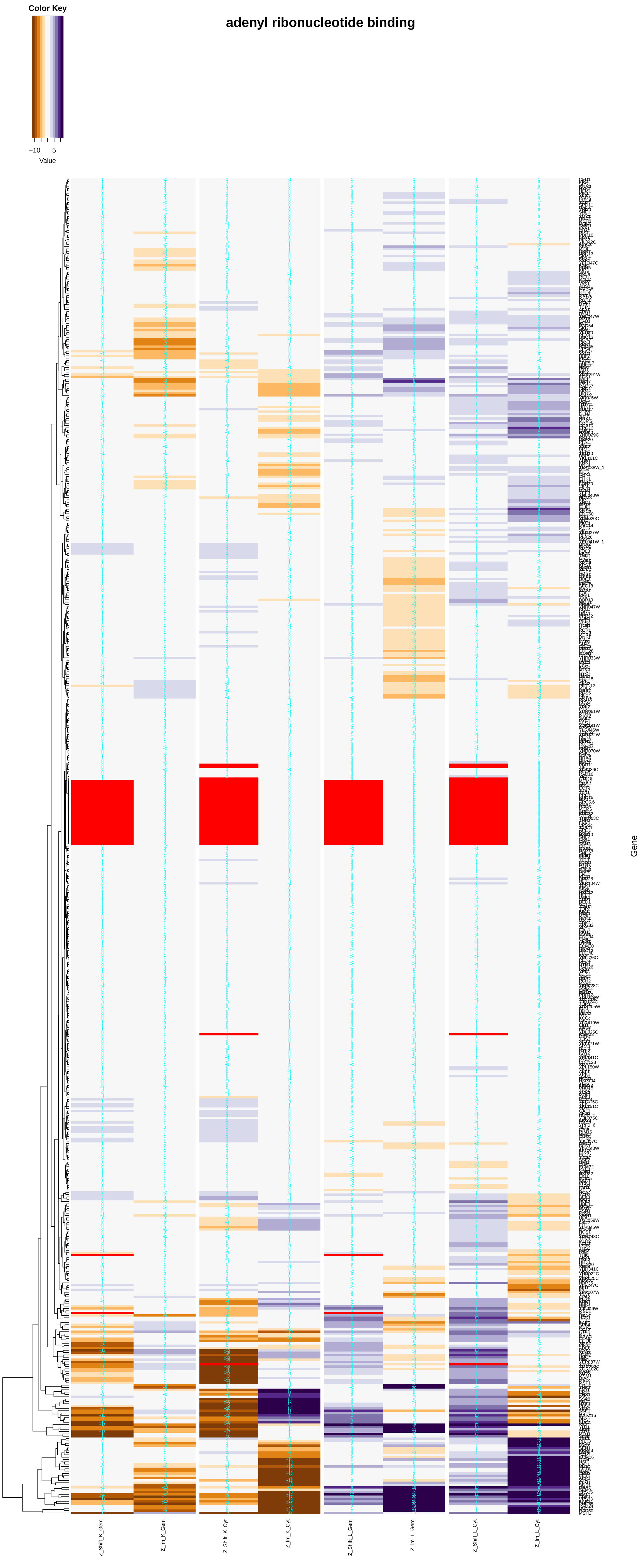

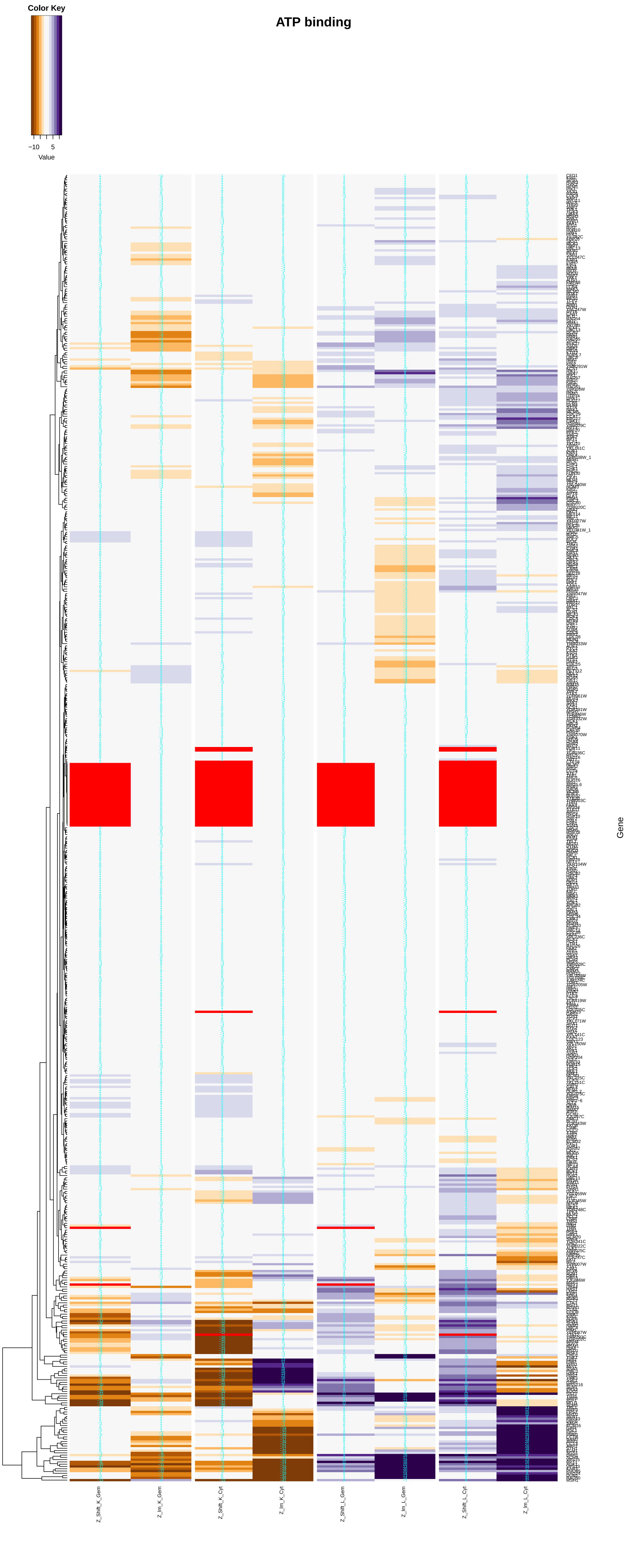

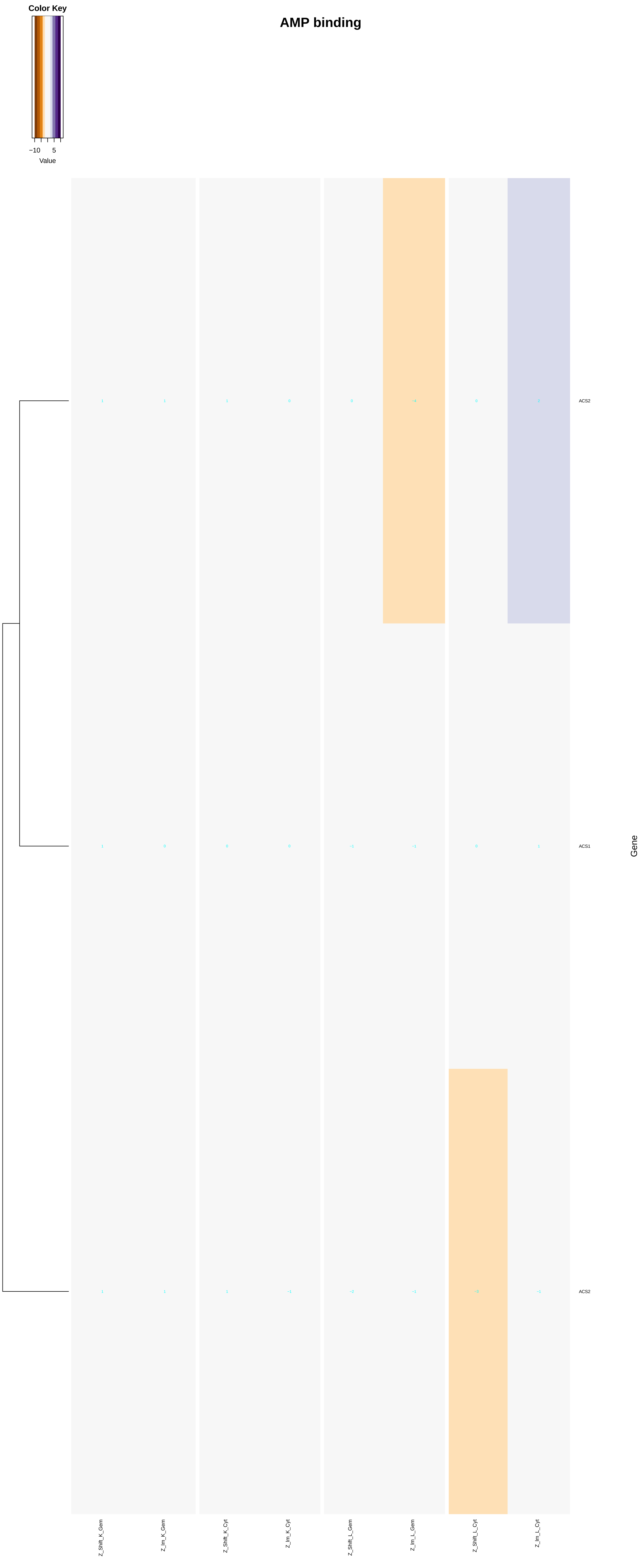

### aminoacyl-tRNA_editing_activity.pdf

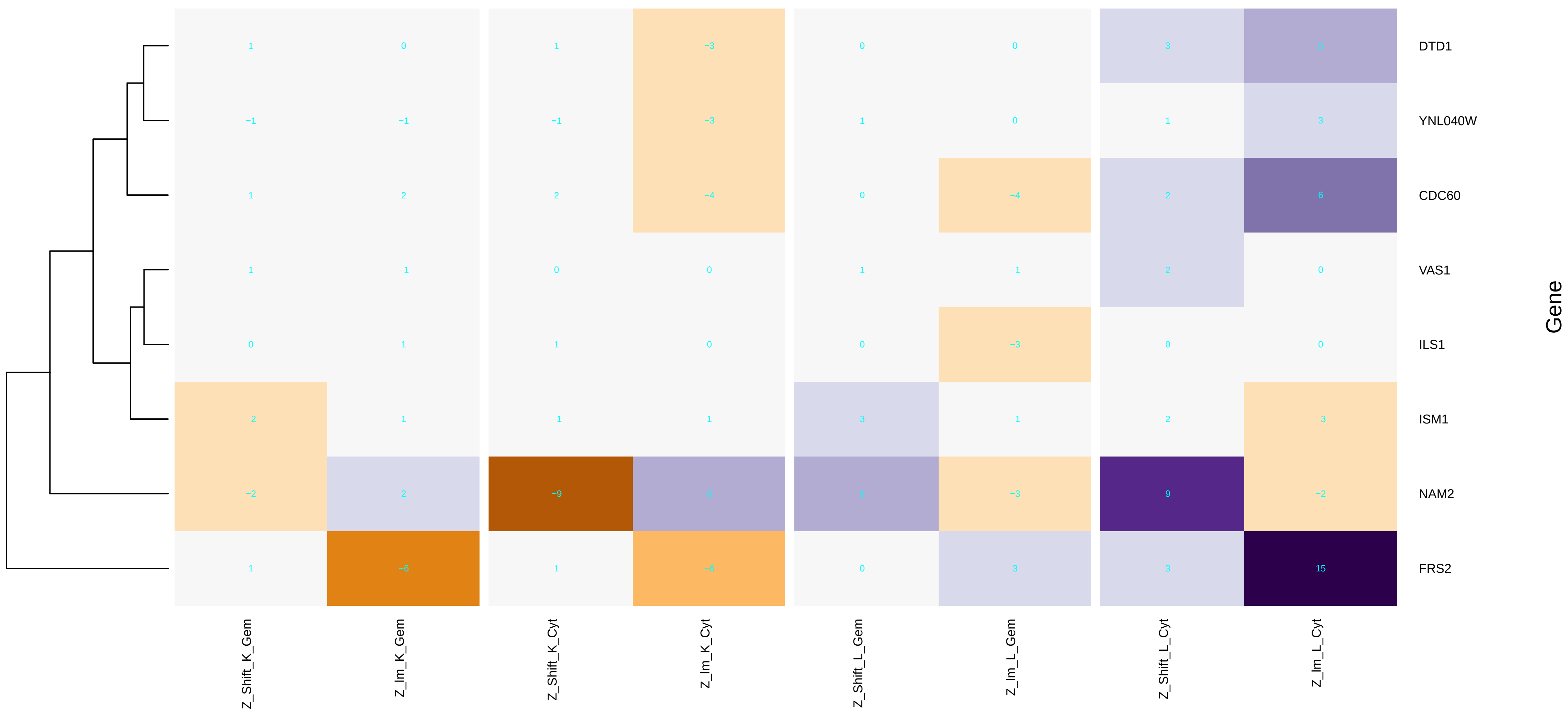

### AMP_salvage.pdf

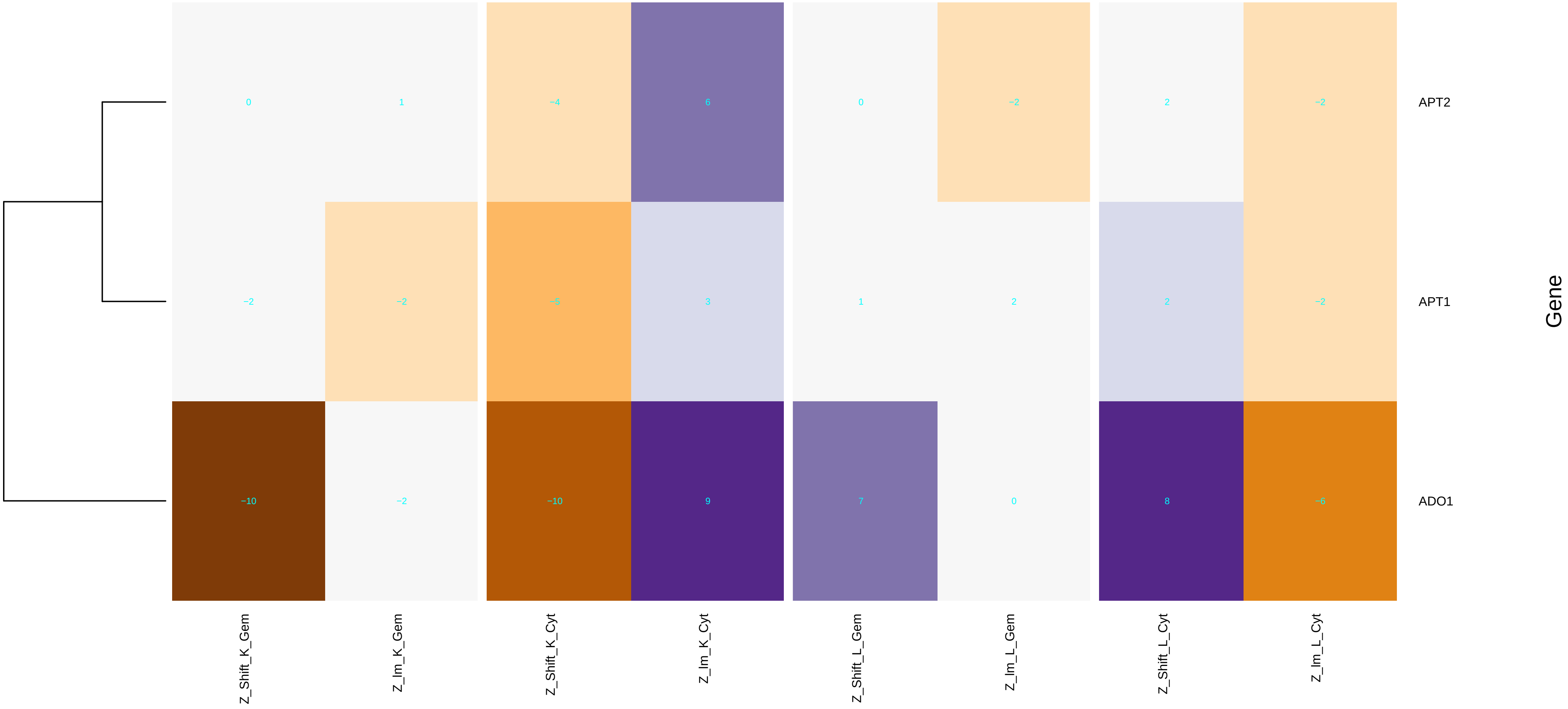

### Arp2_3_complex-mediated_actin_nucleation.pdf

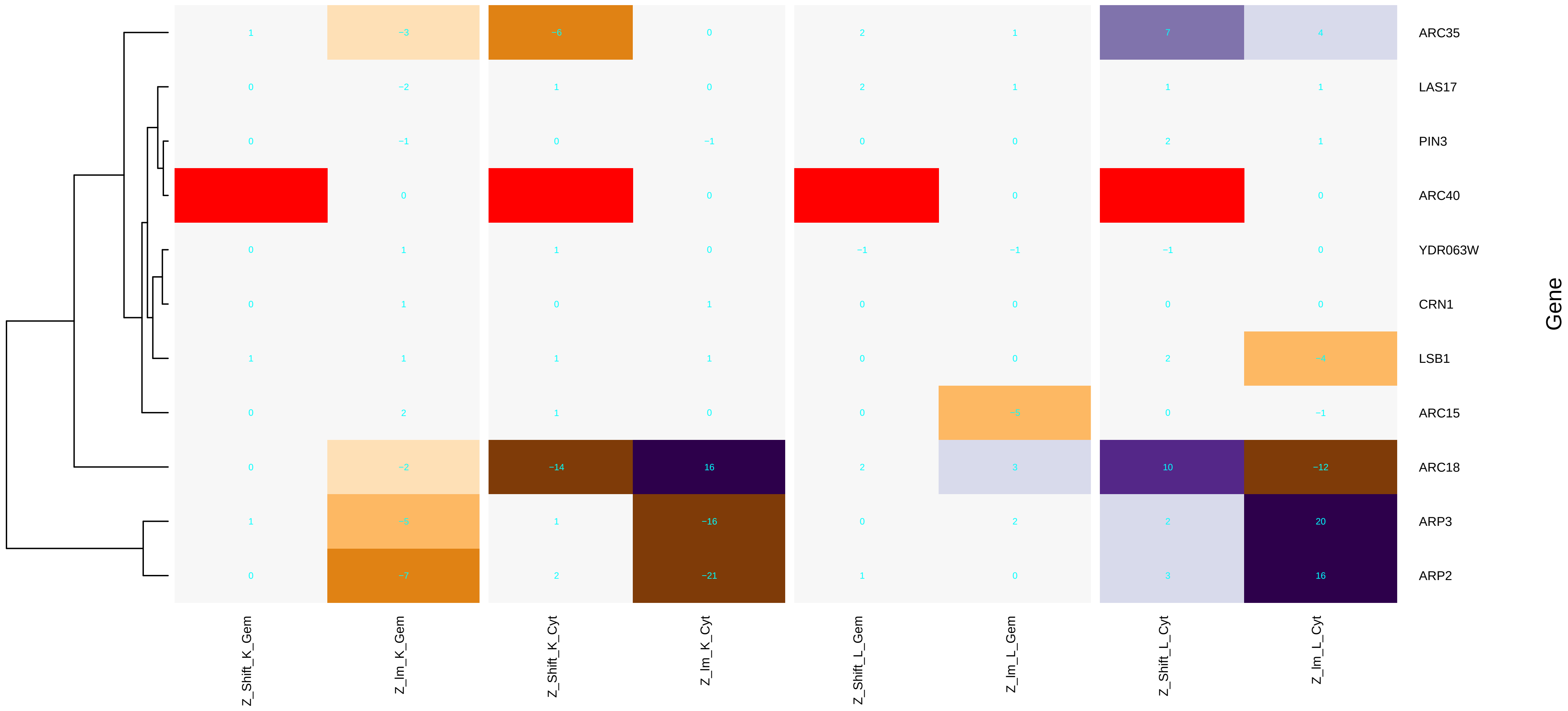

### Arp2_3_protein_complex.pdf

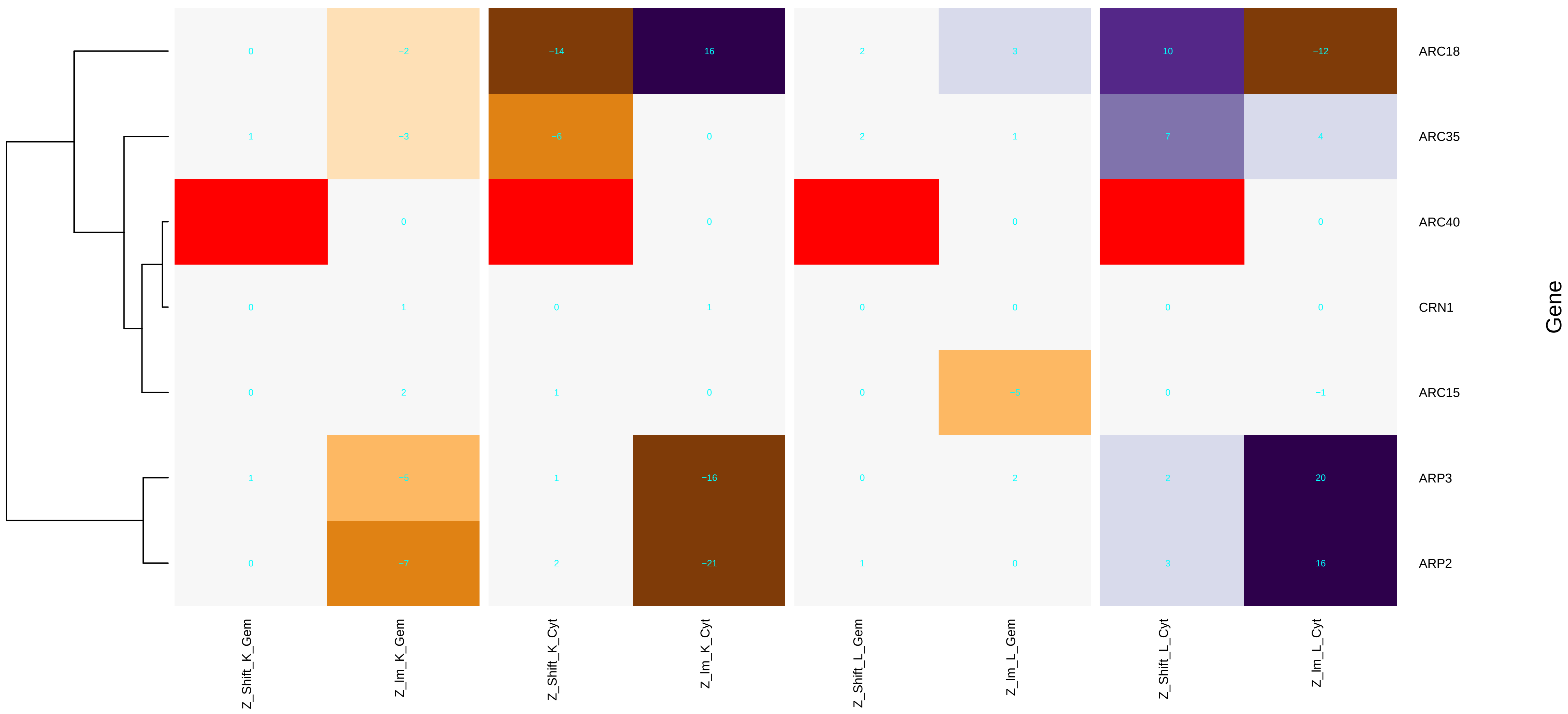

### ATP-dependent_DNA_helicase_activity.pdf

#### ATP−dependent DNA helicase activity

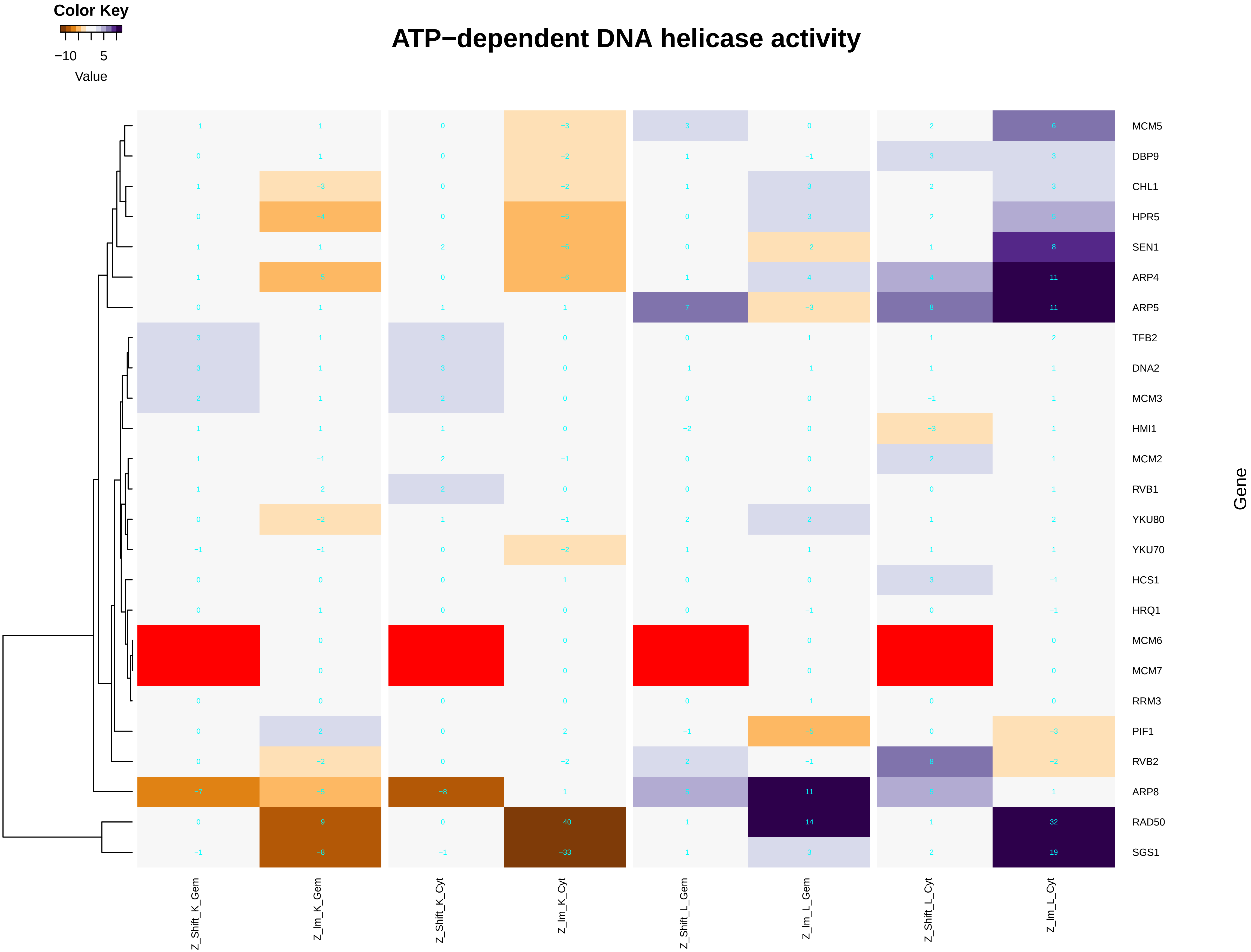

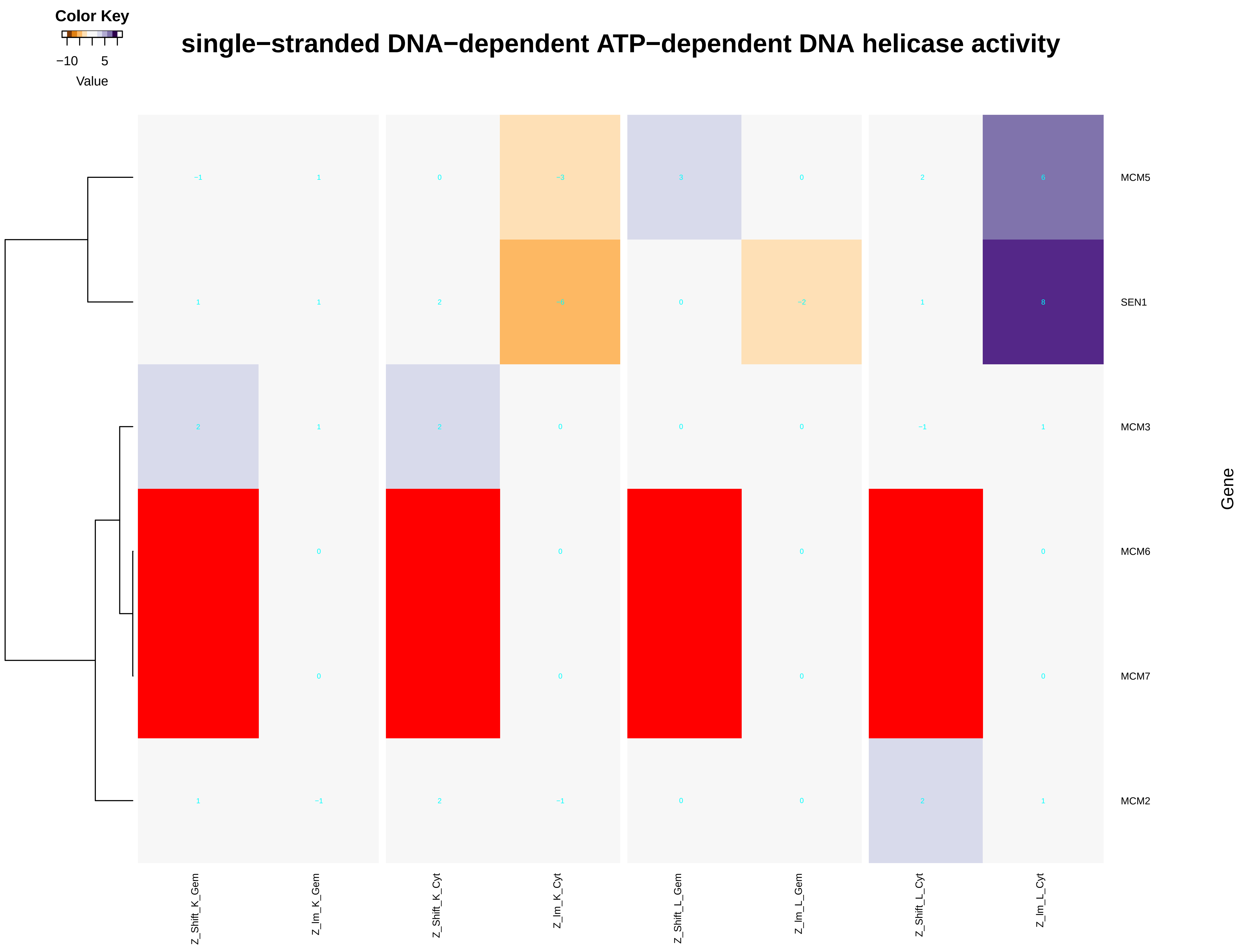

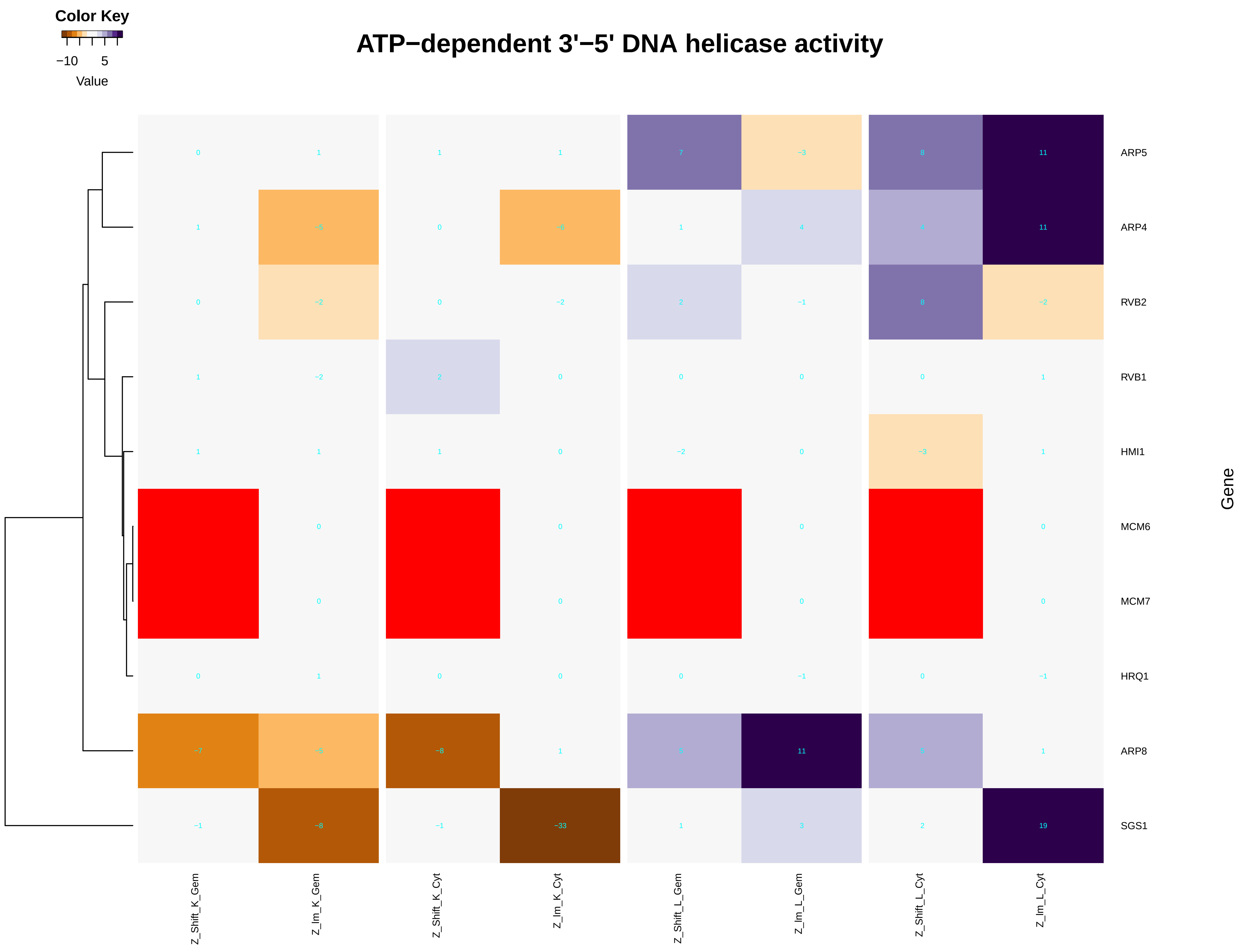

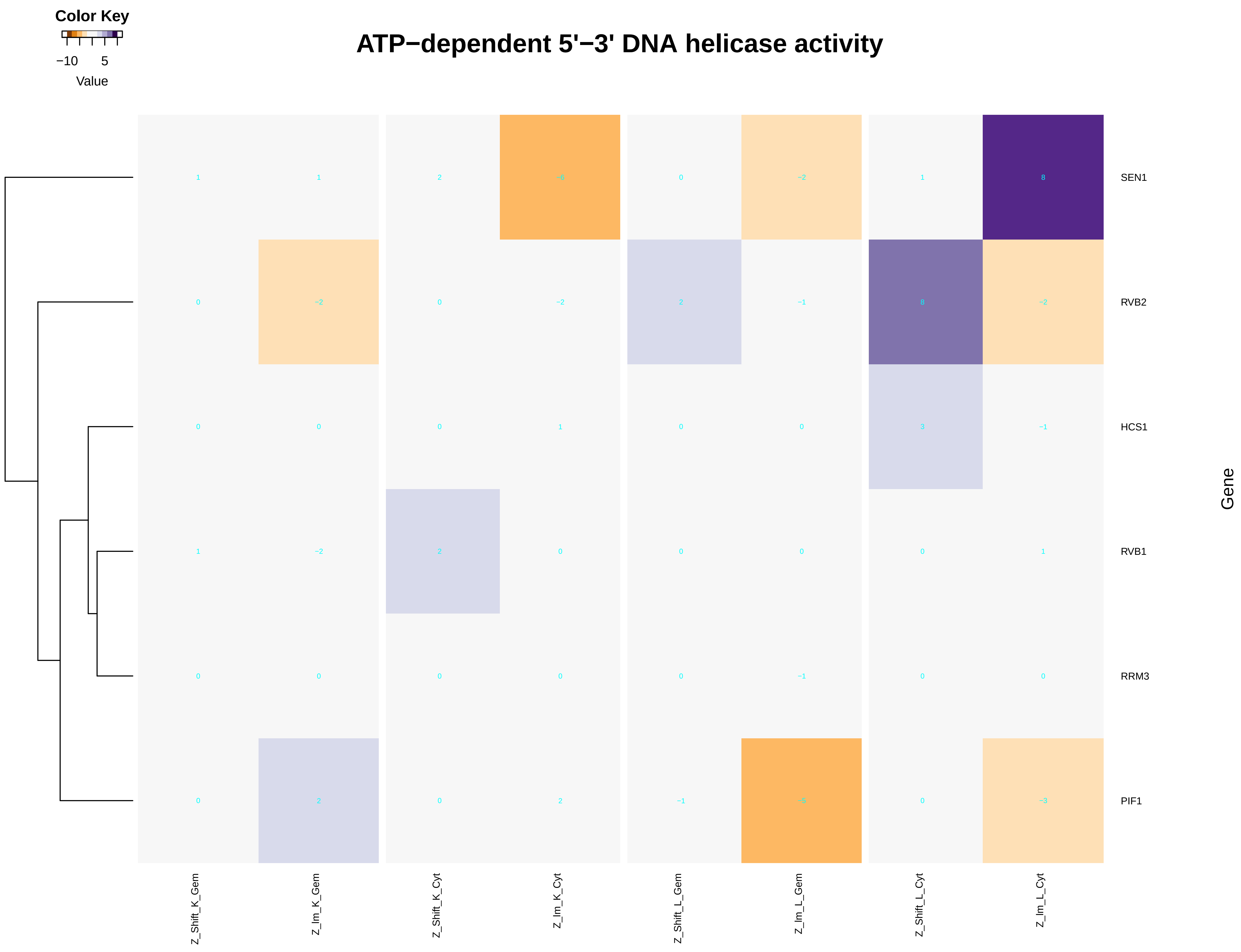

### ATP-dependent_helicase_activity.pdf

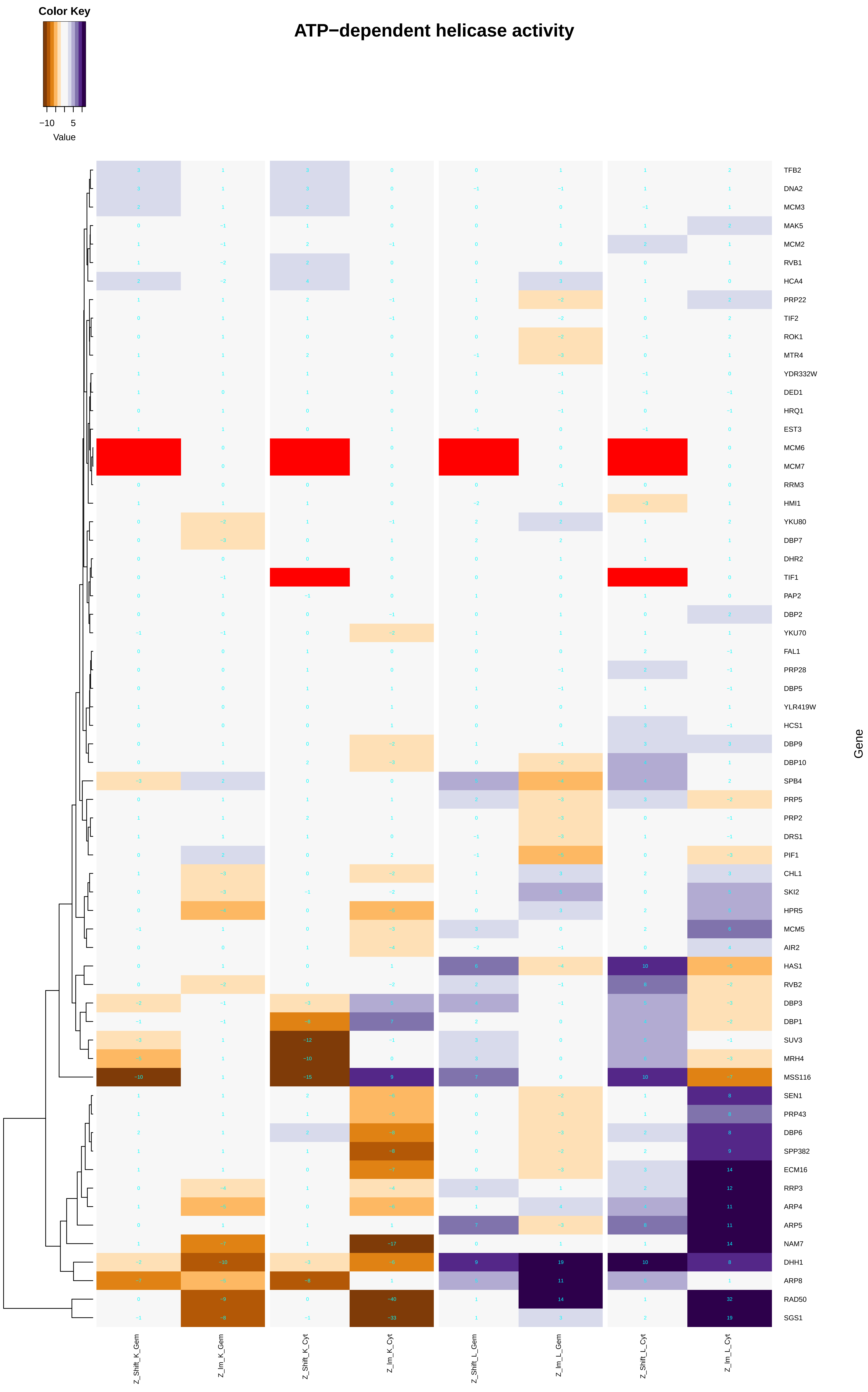

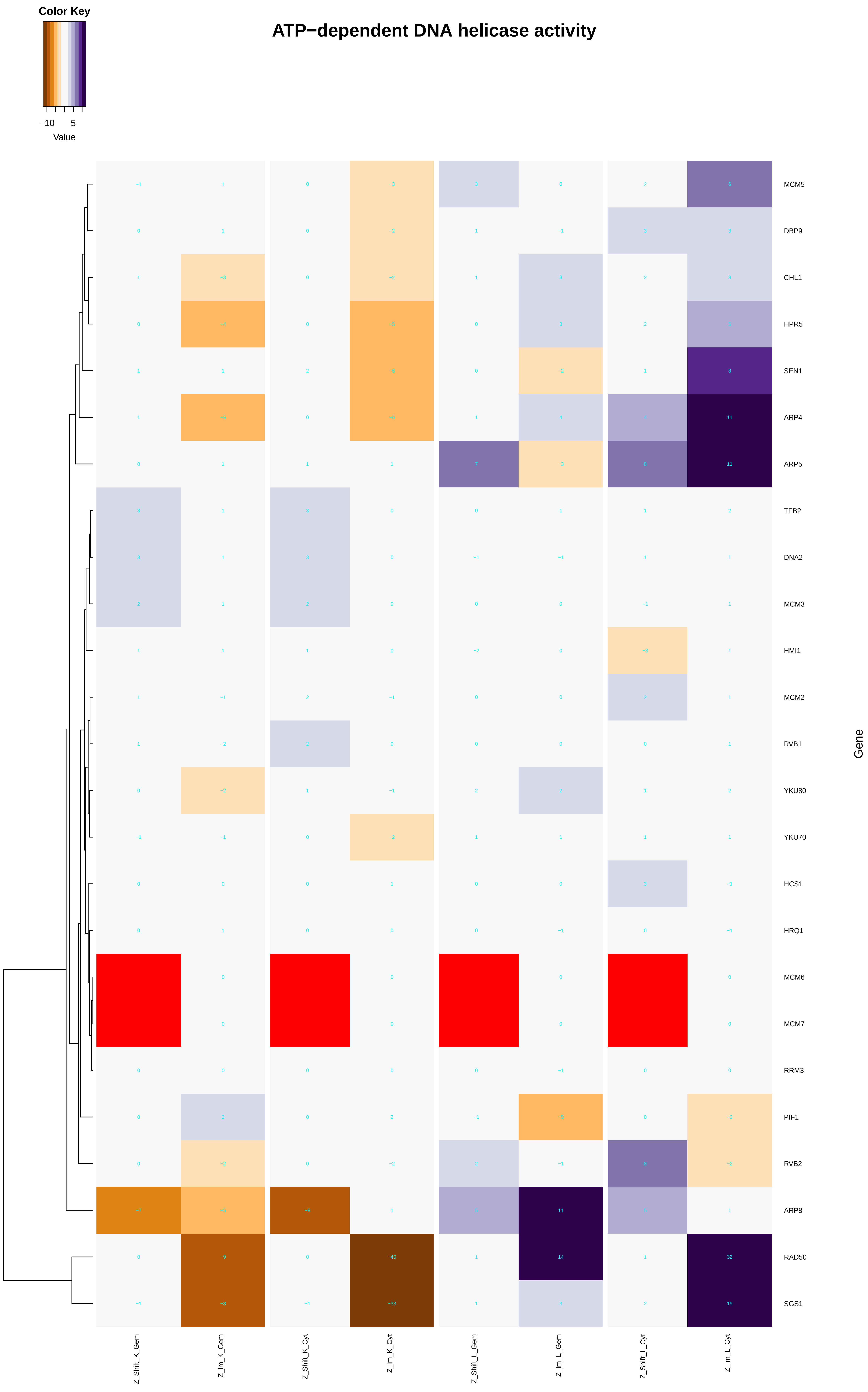

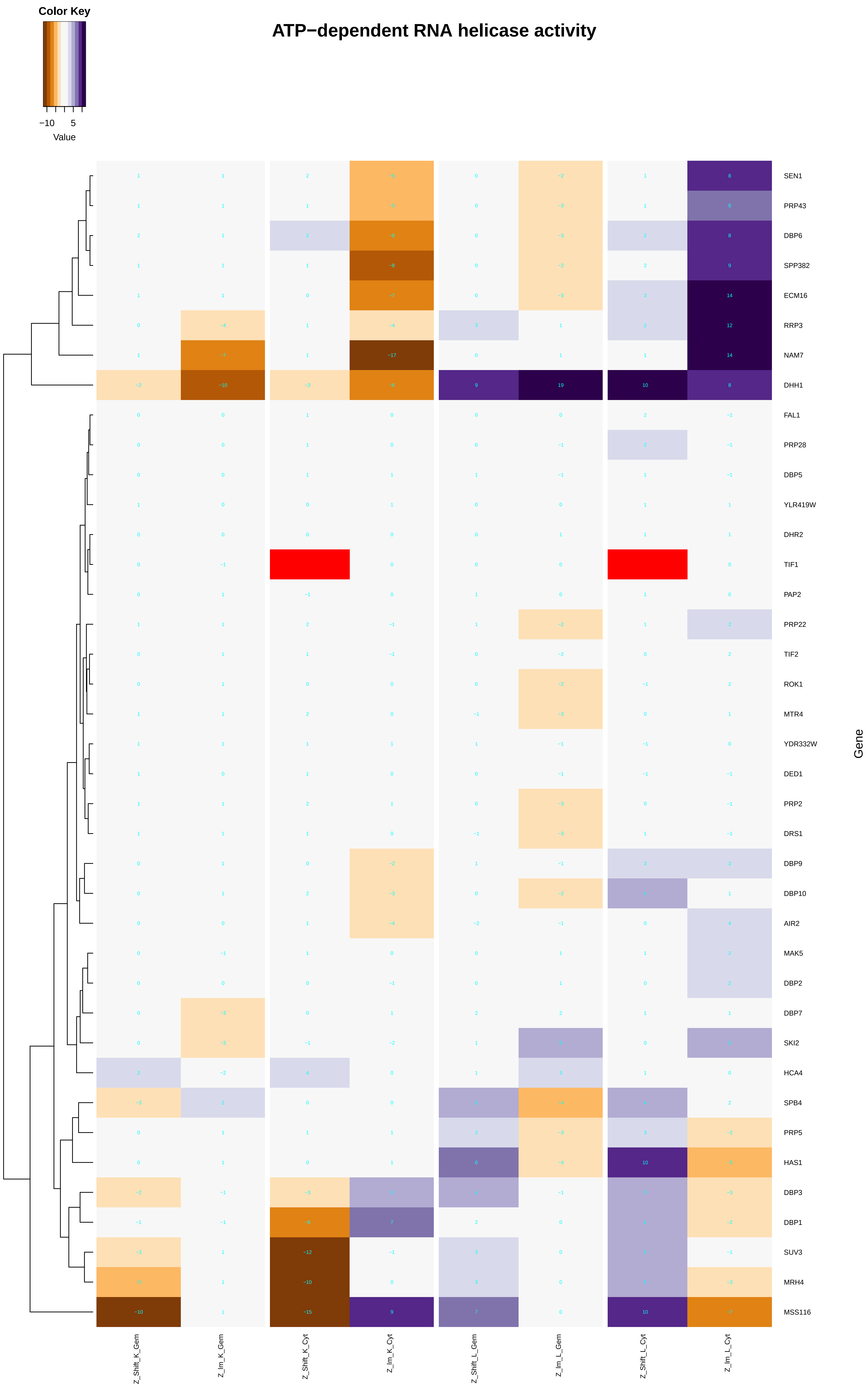

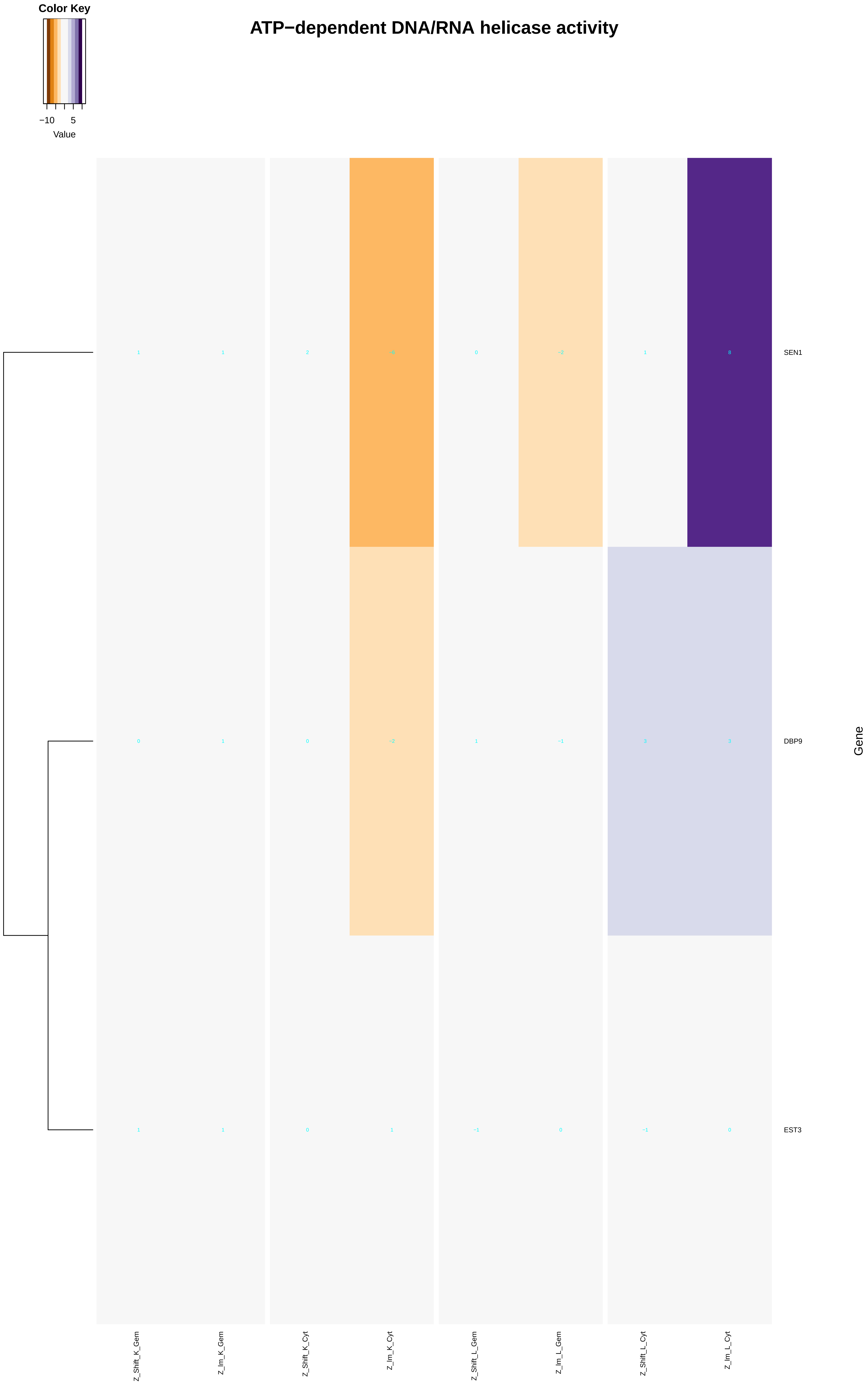

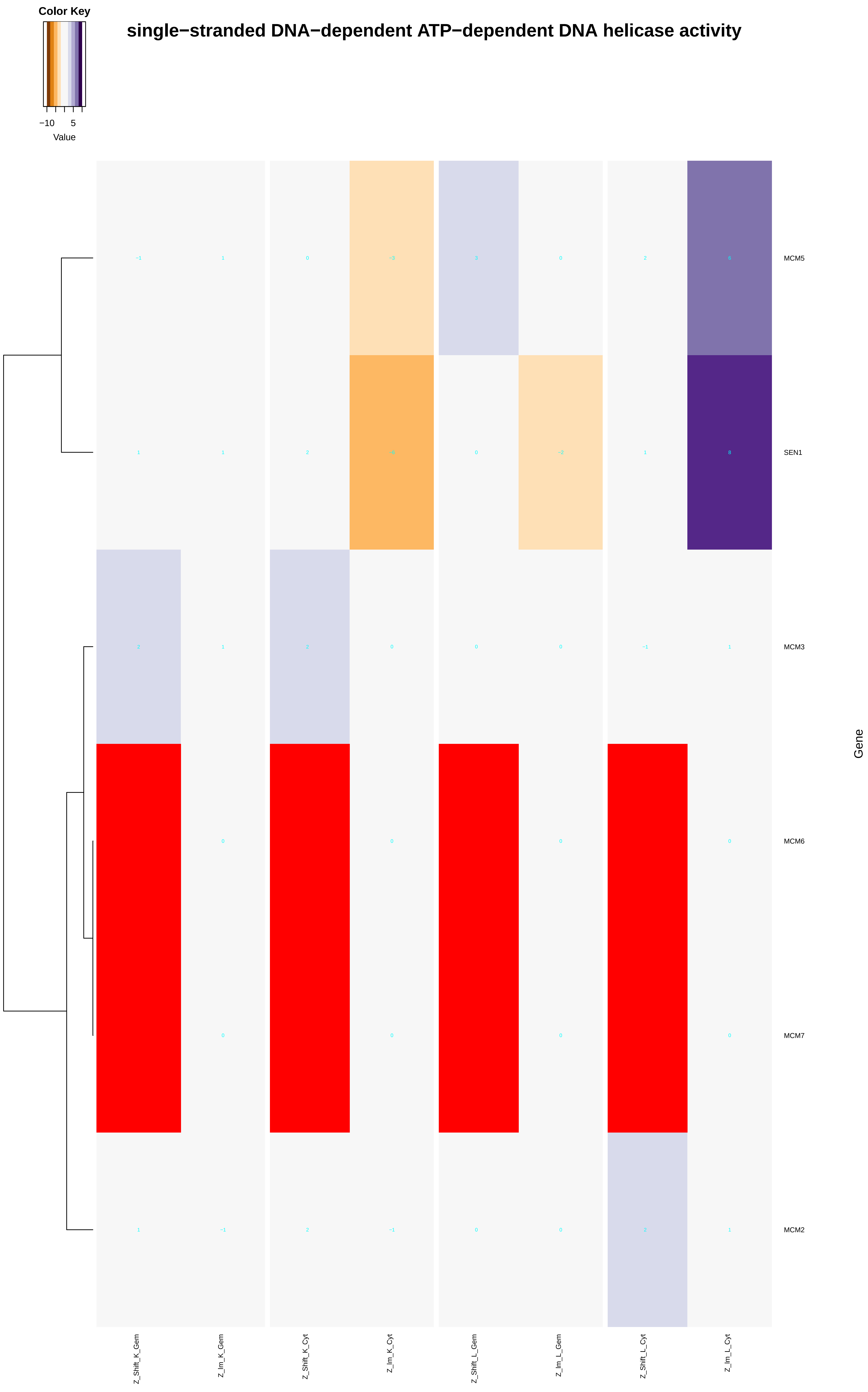

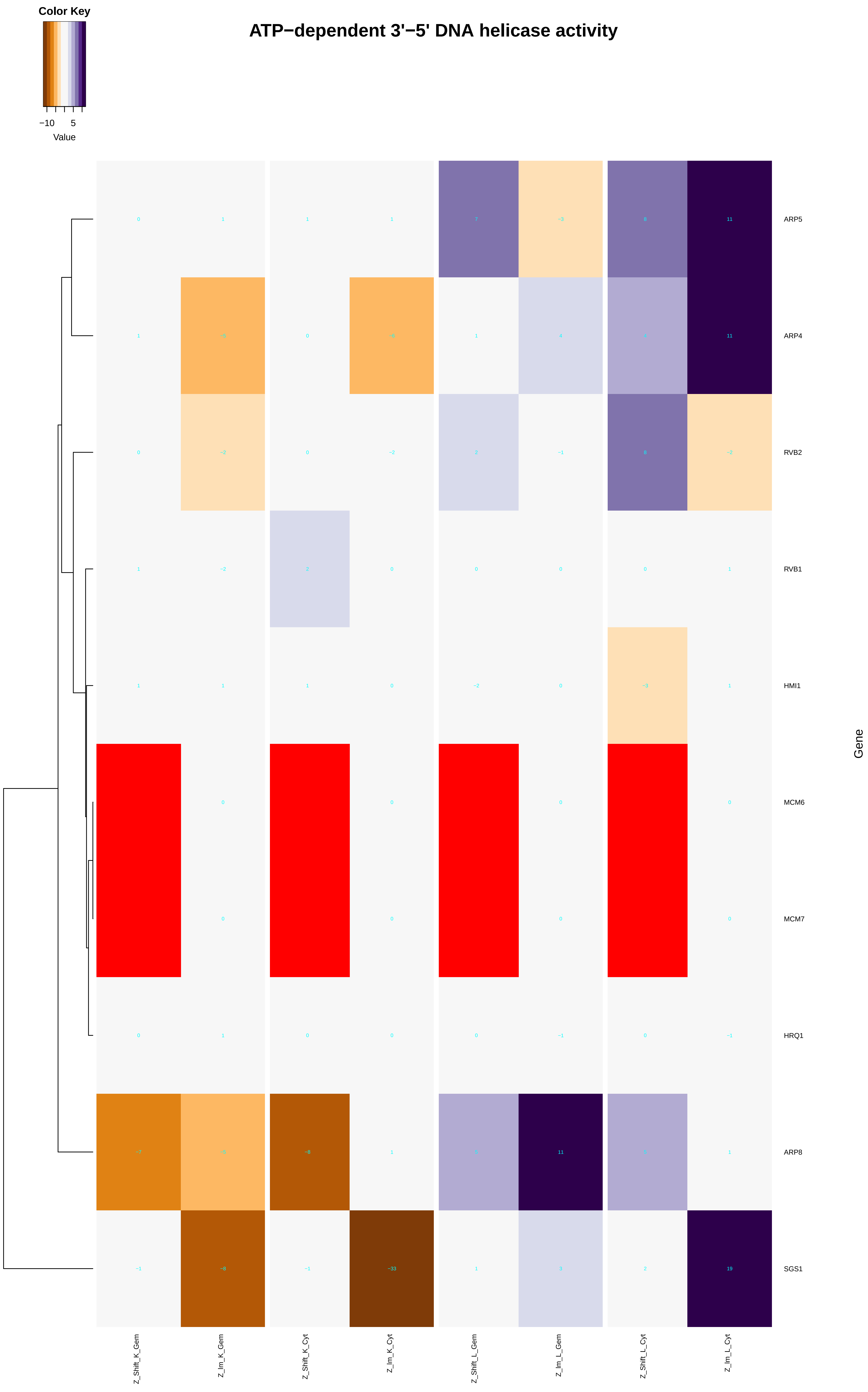

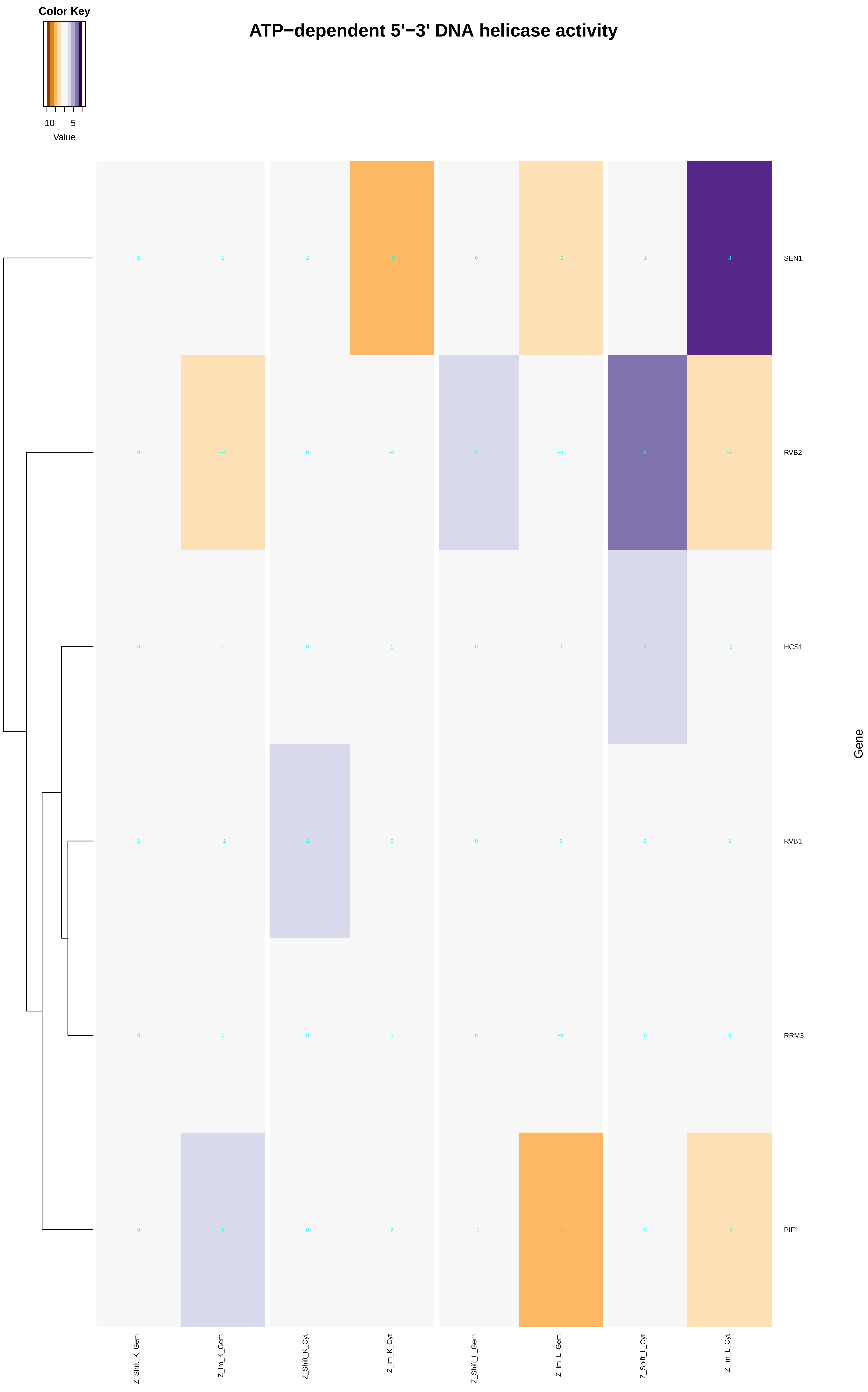

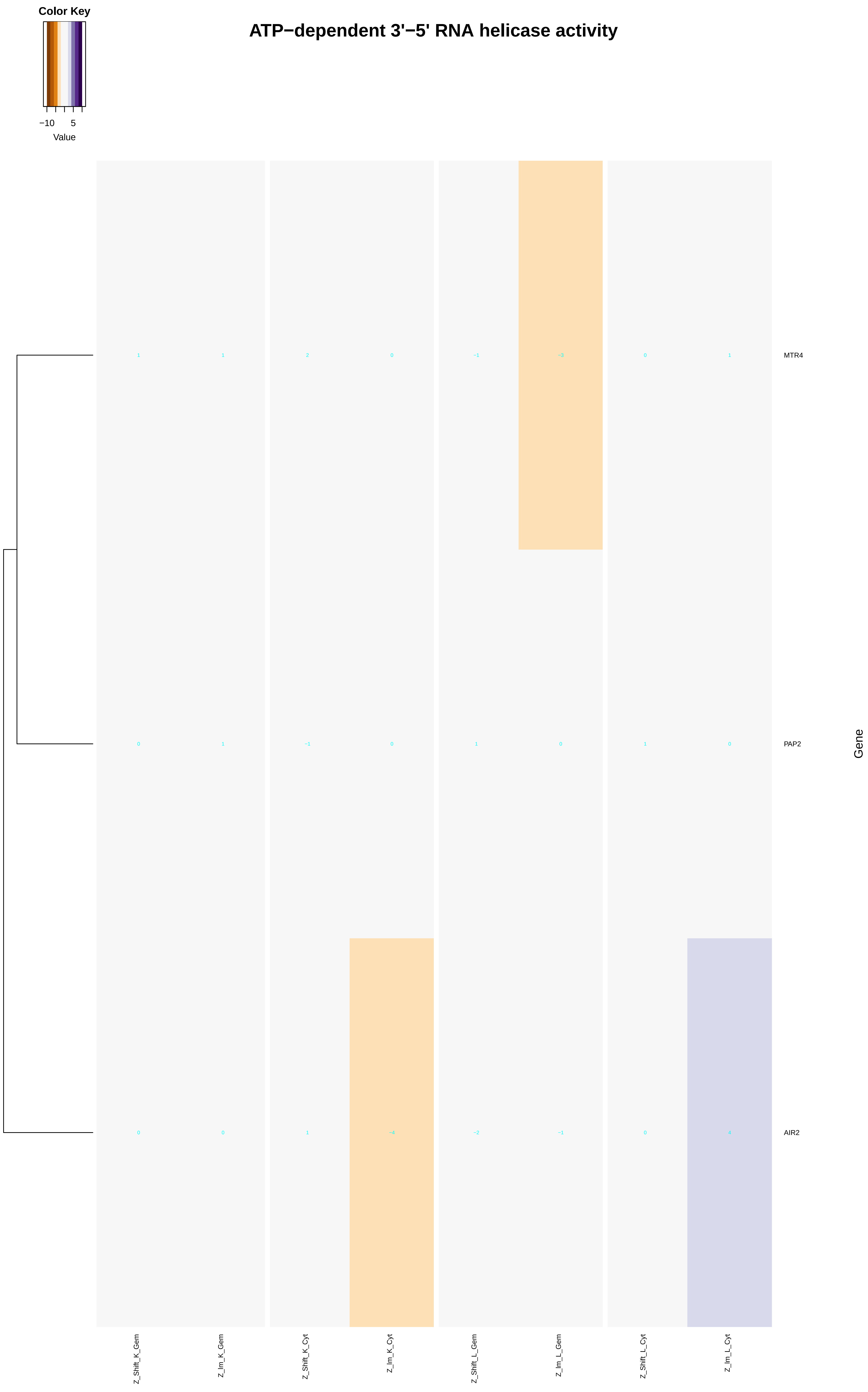

### ATP-dependent_RNA_helicase_activity.pdf

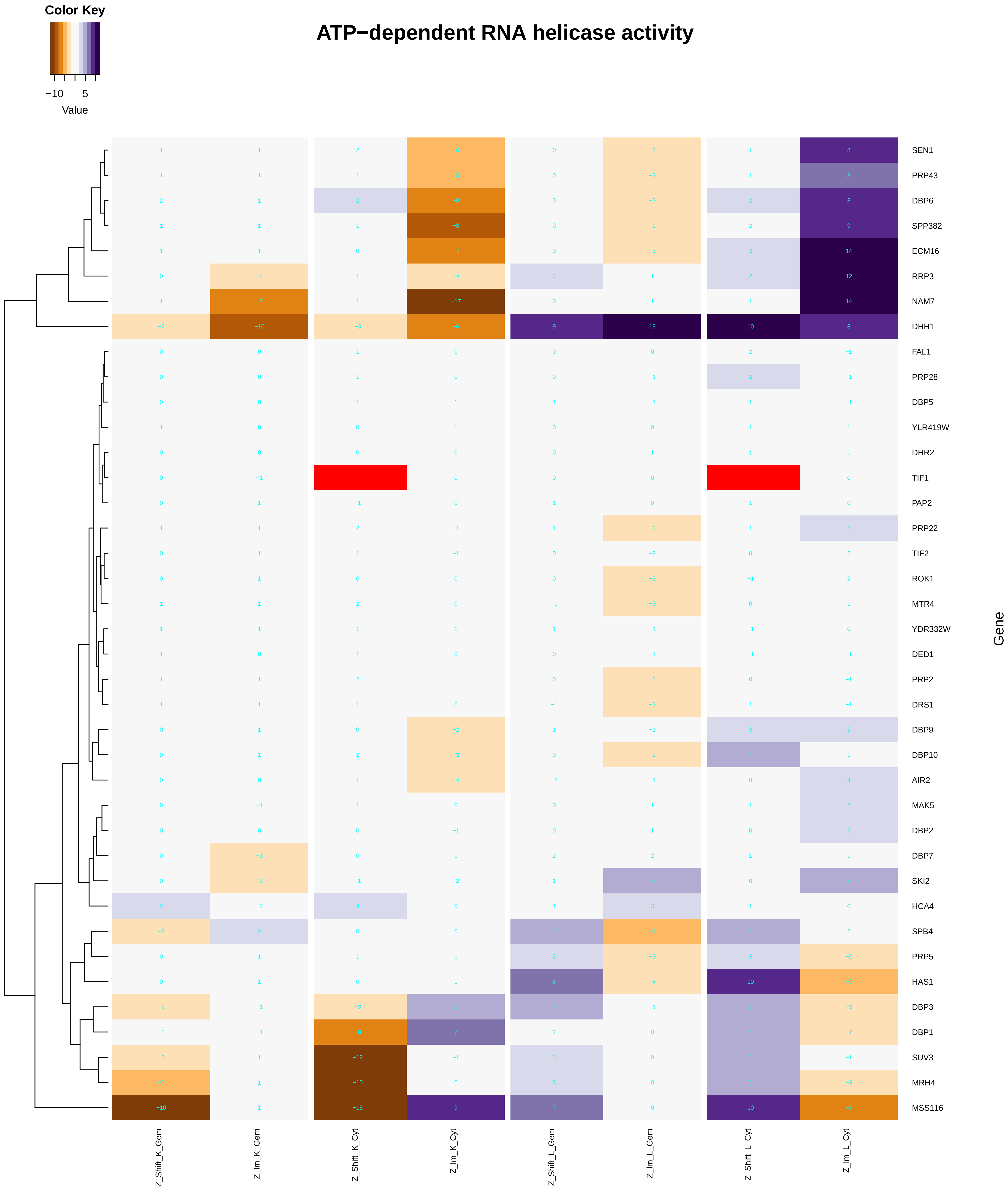

#### ATP−dependent 3'−5' RNA helicase activity

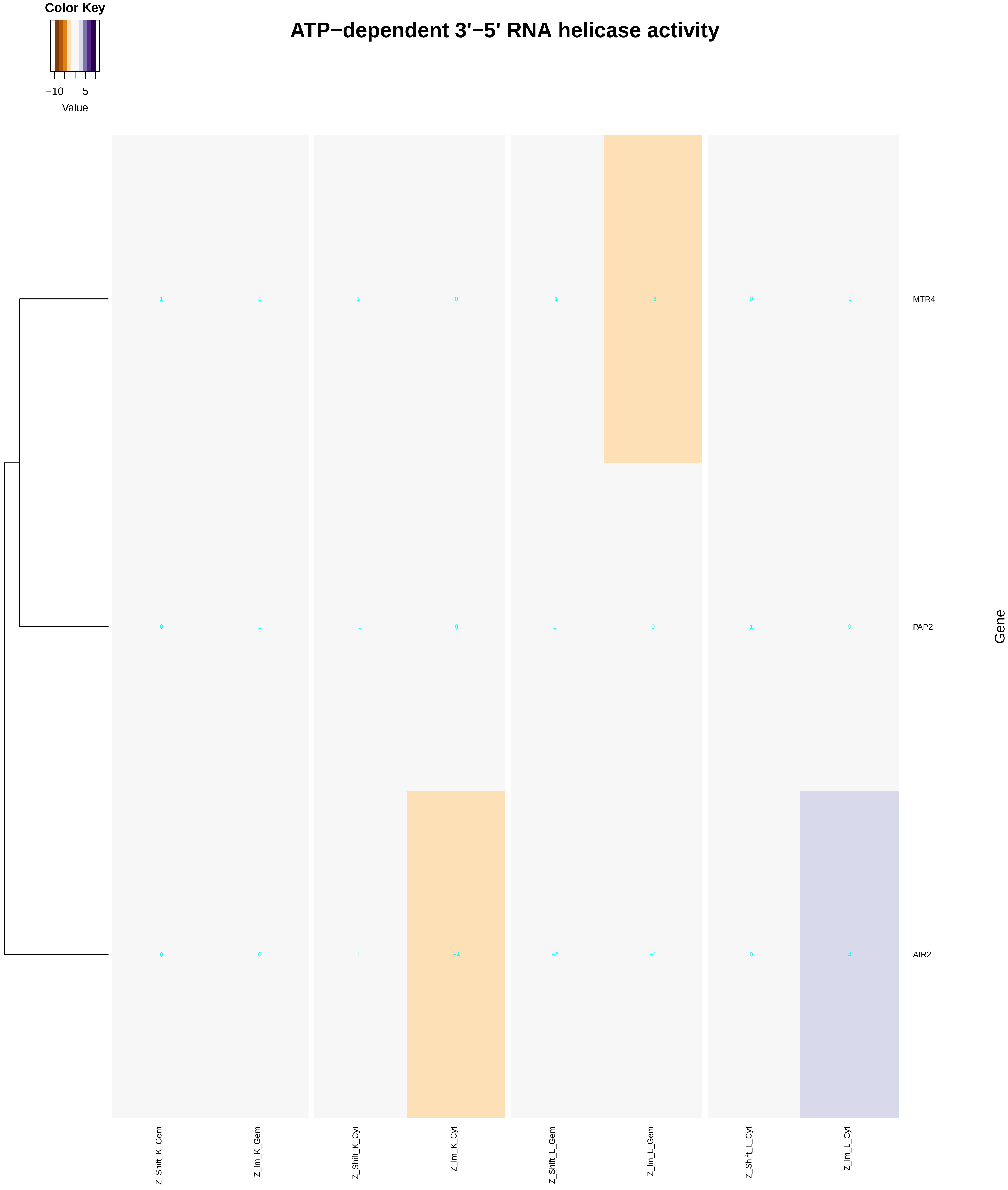

### ATP_binding.pdf

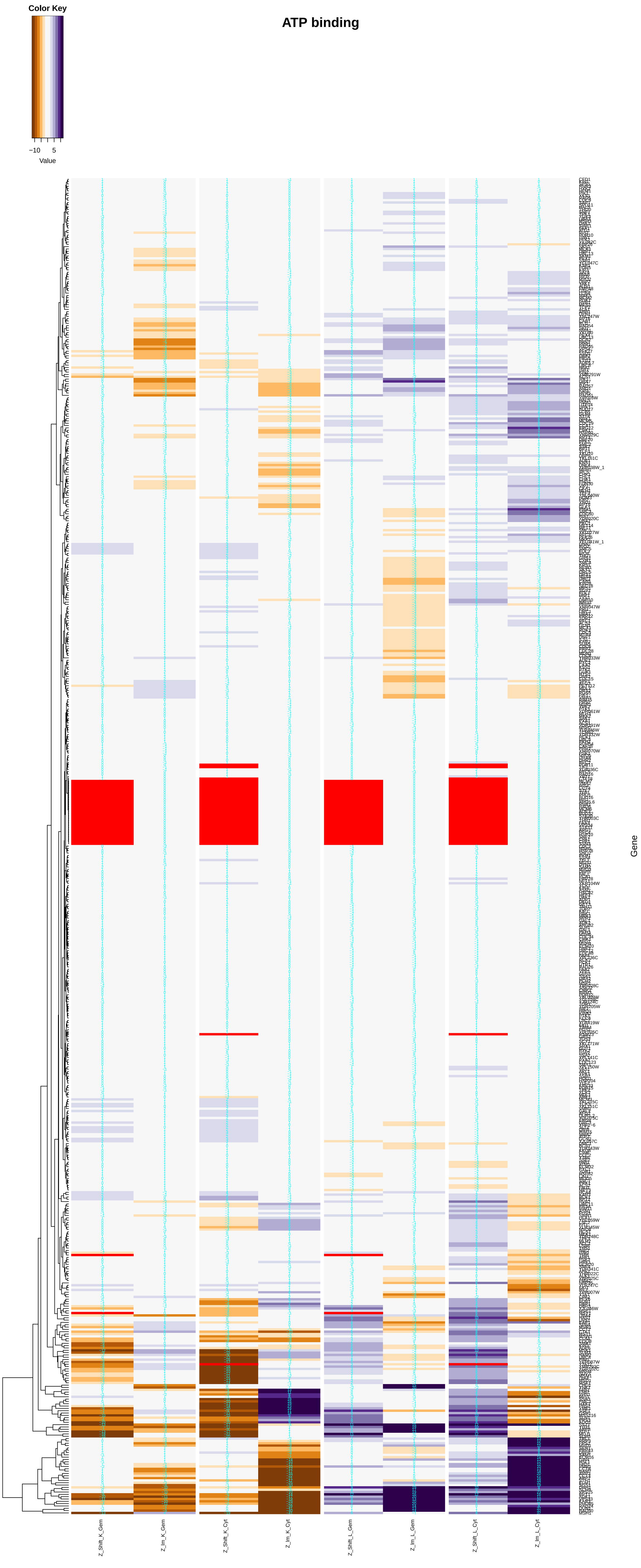

### ATP_export.pdf

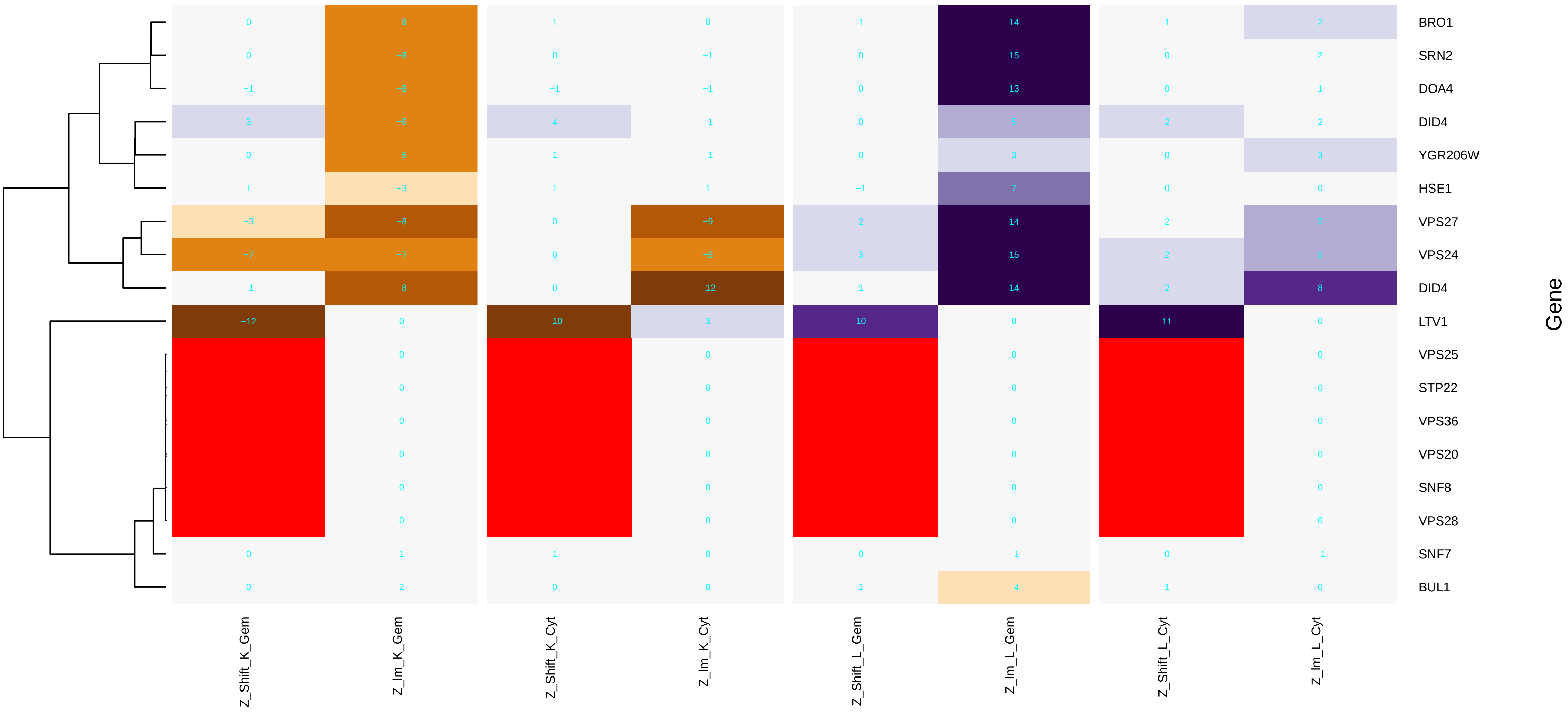

### ATP_synthesis_coupled_proton_transport.pdf

#### ATP synthesis coupled proton transport

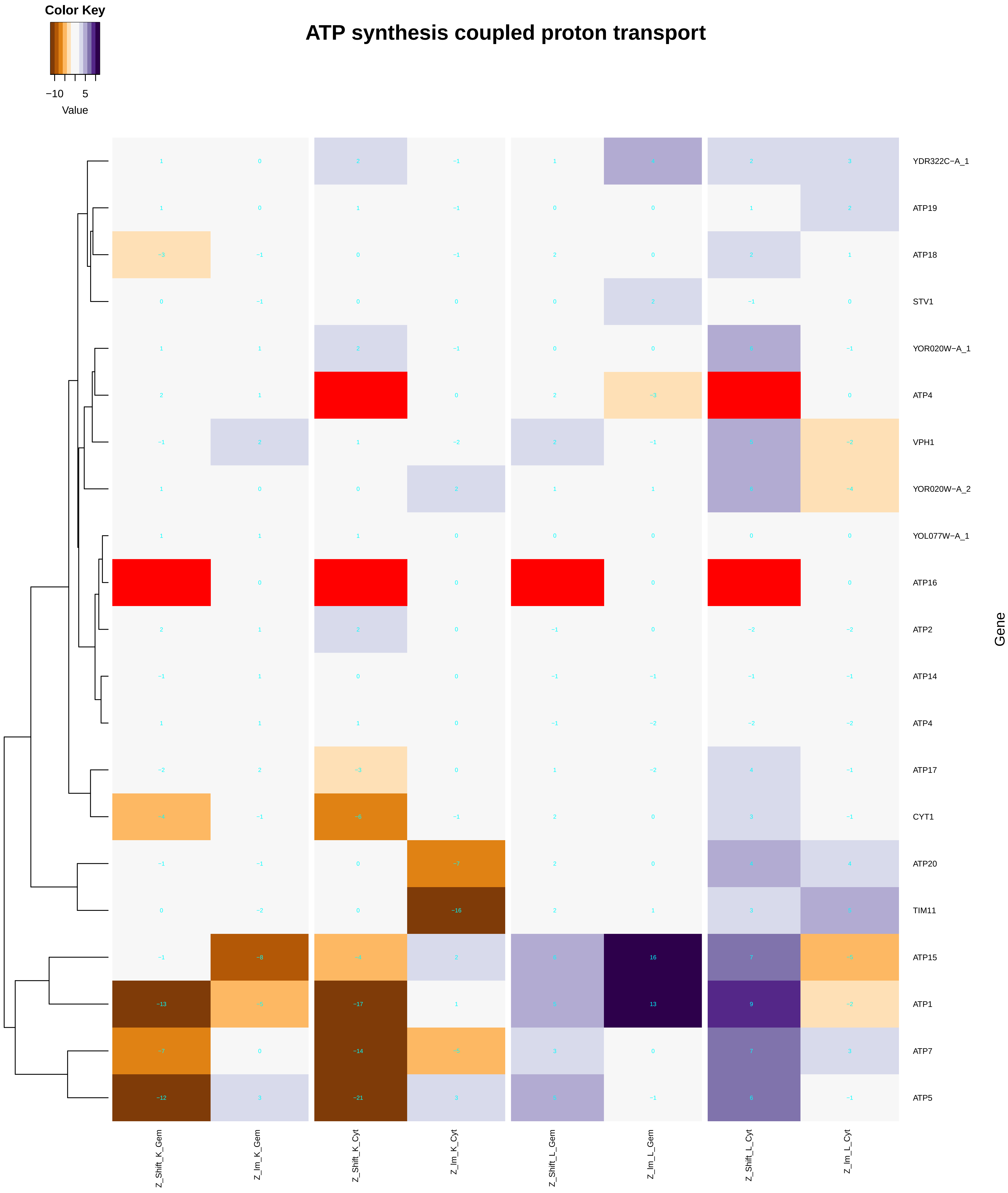

### ATP_transport.pdf

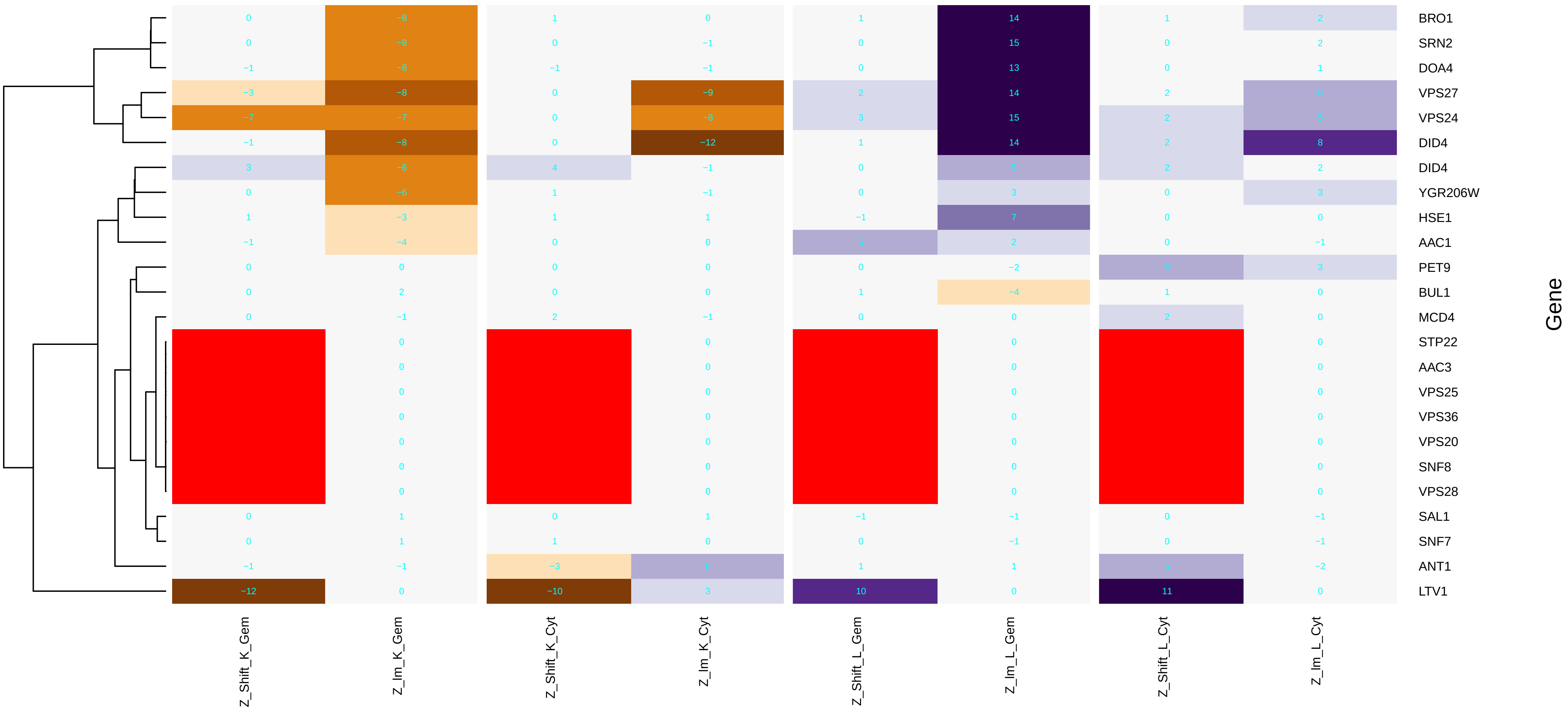

### base-excision_repair.pdf

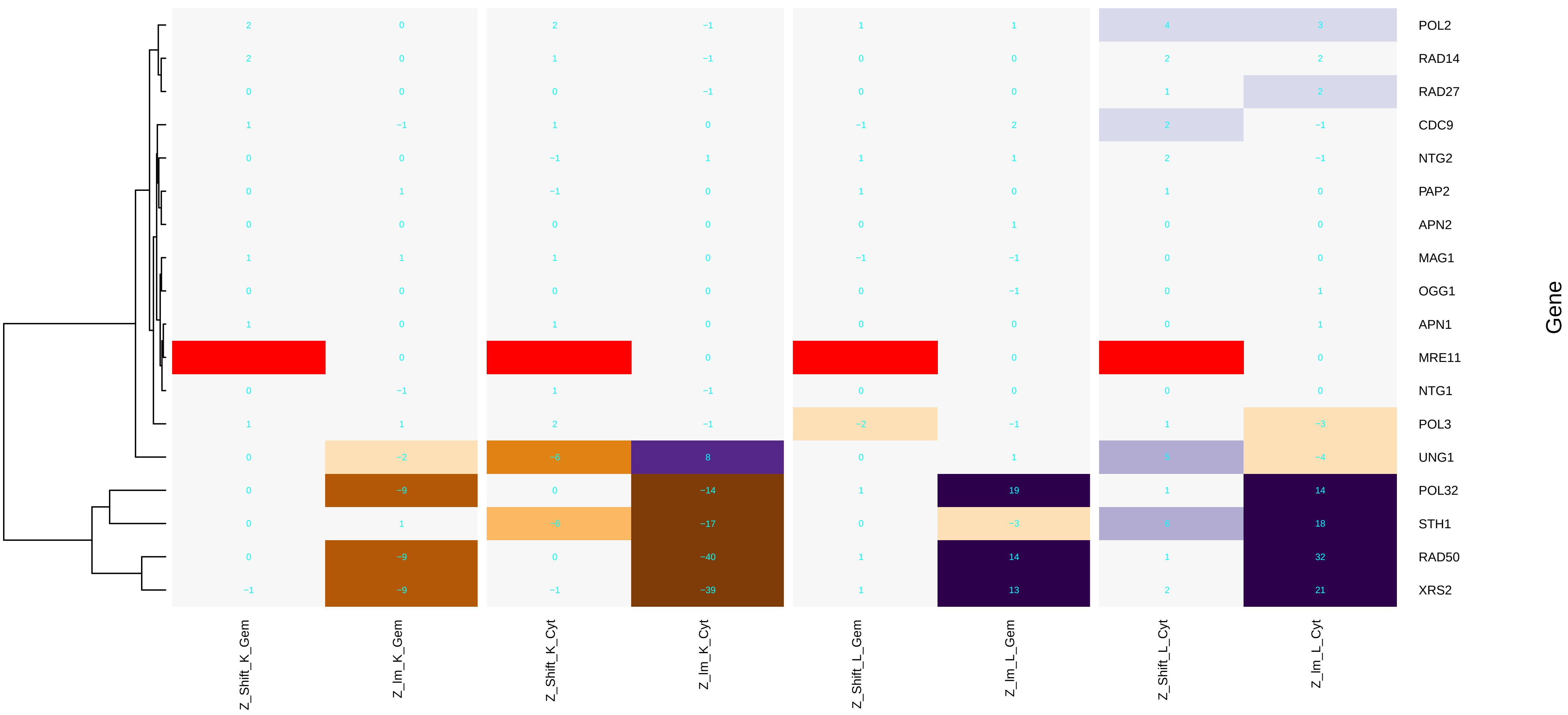

### Bfa1-Bub2_complex.pdf

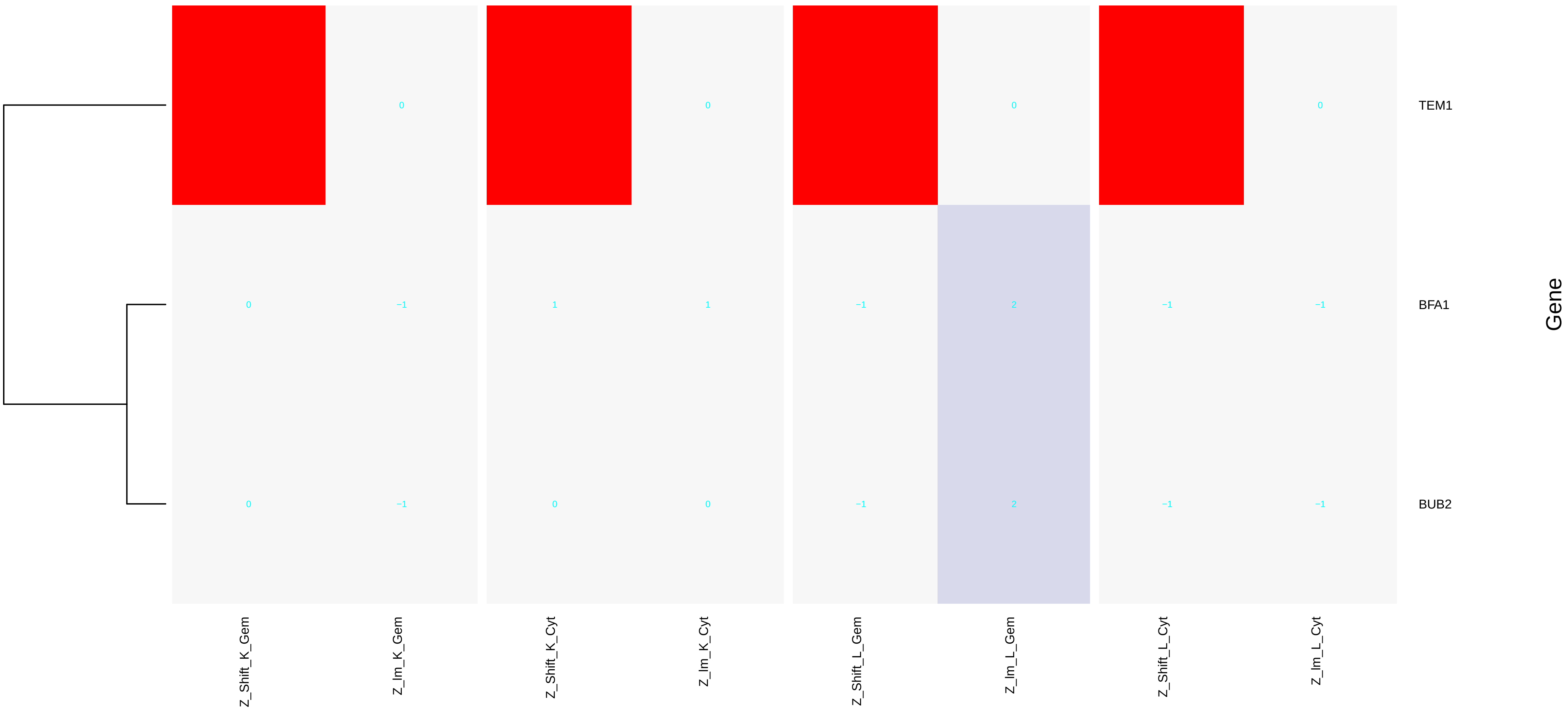

### carbon-nitrogen_ligase_activity,_with_glutamine_as_amido-N-donor.pdf

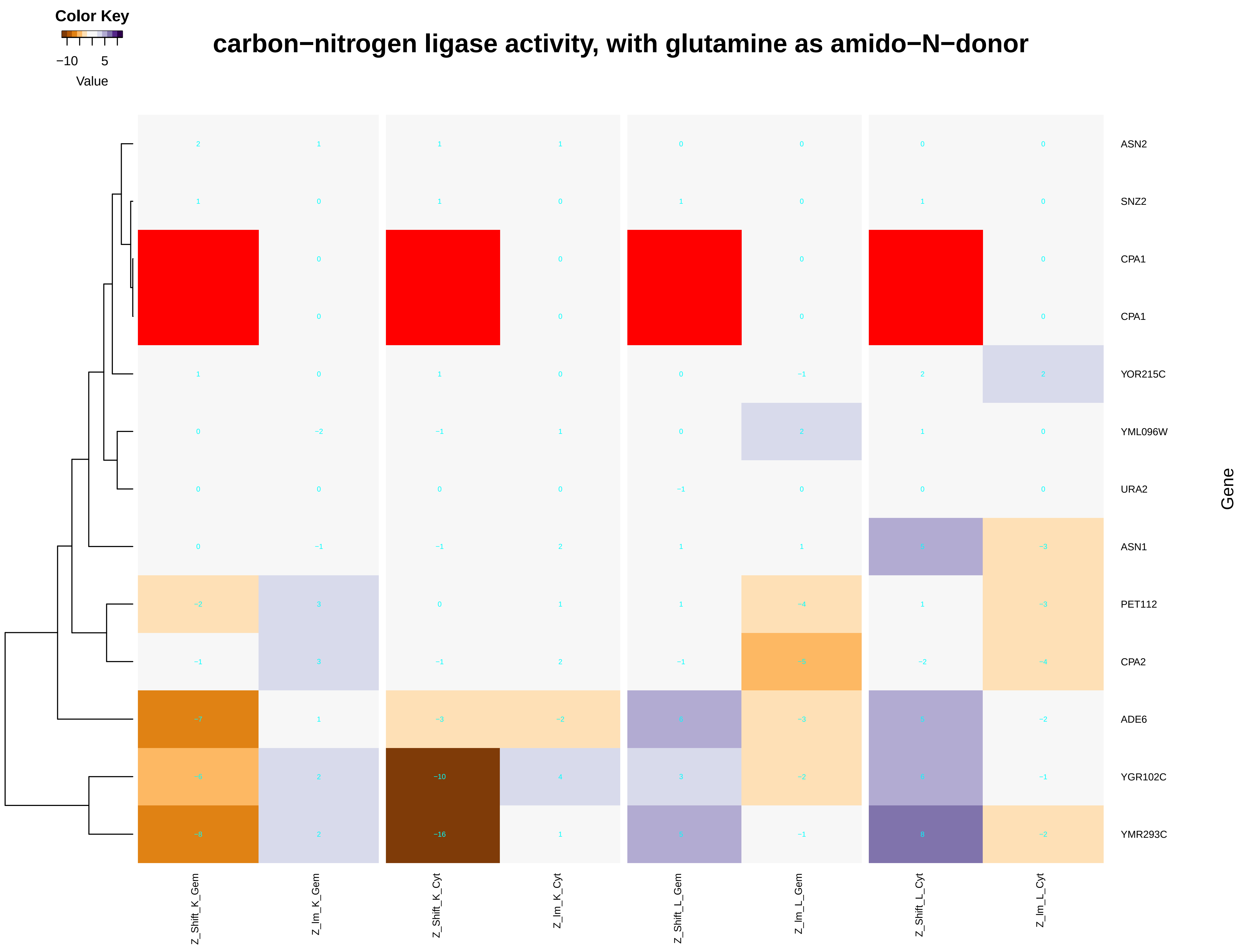

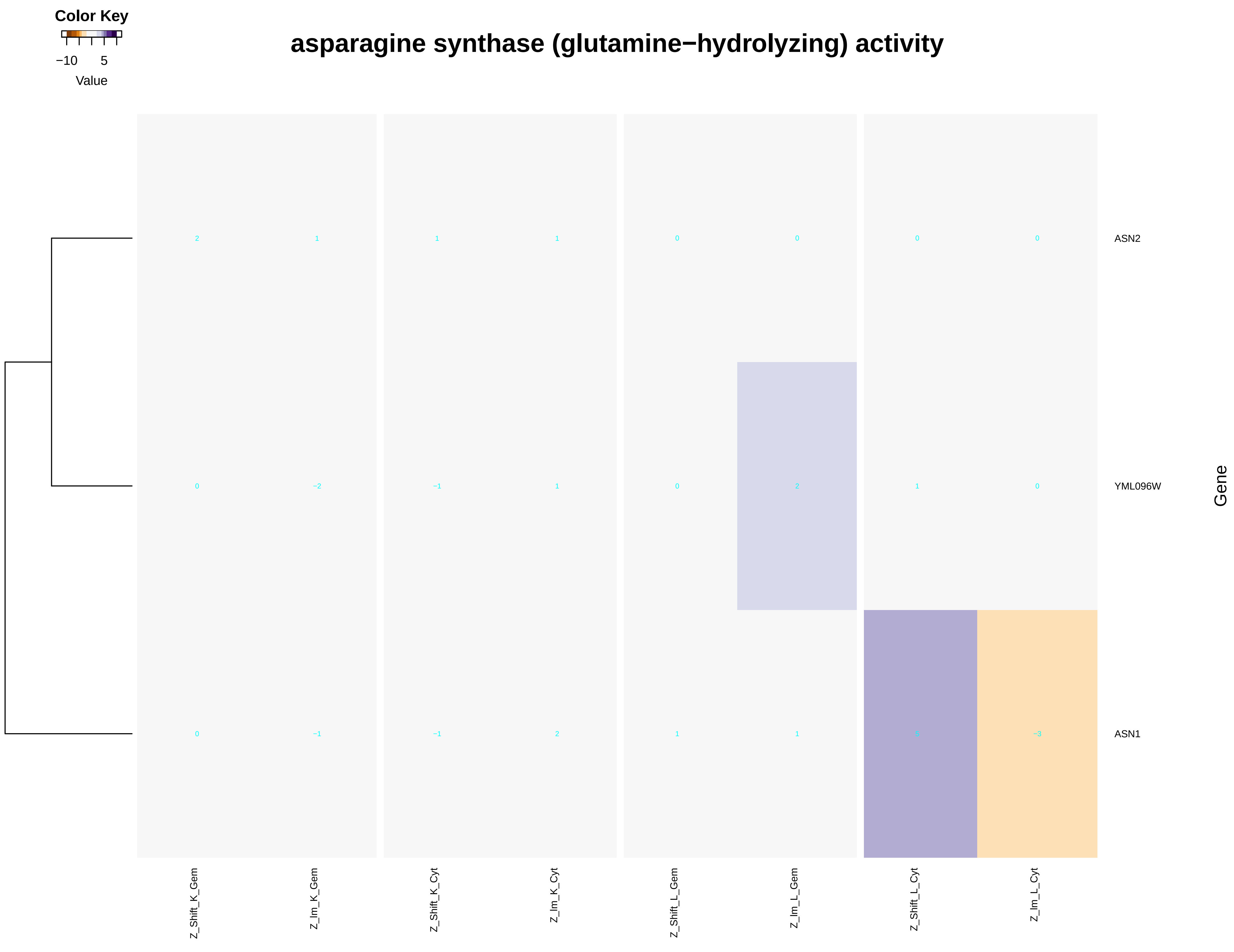

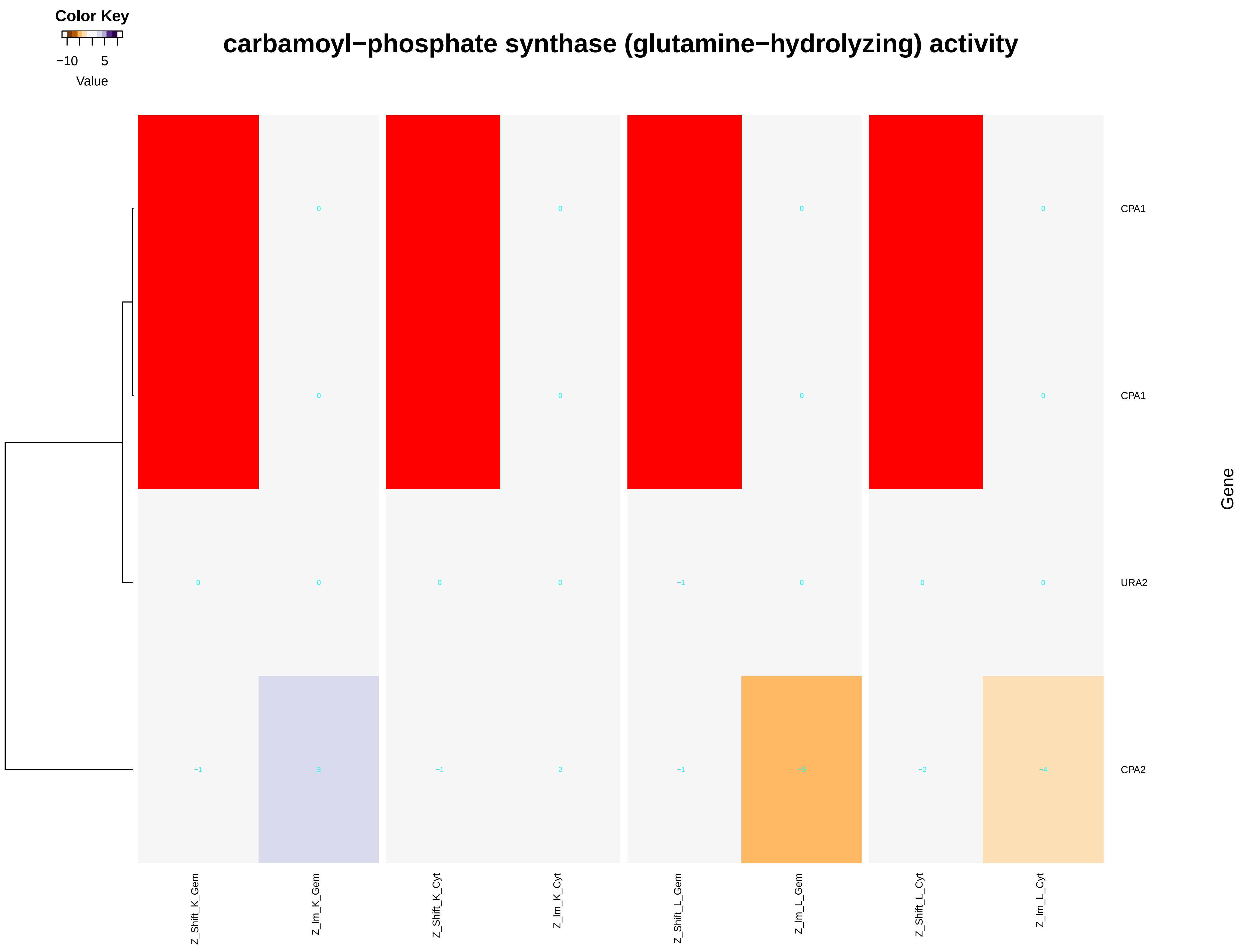

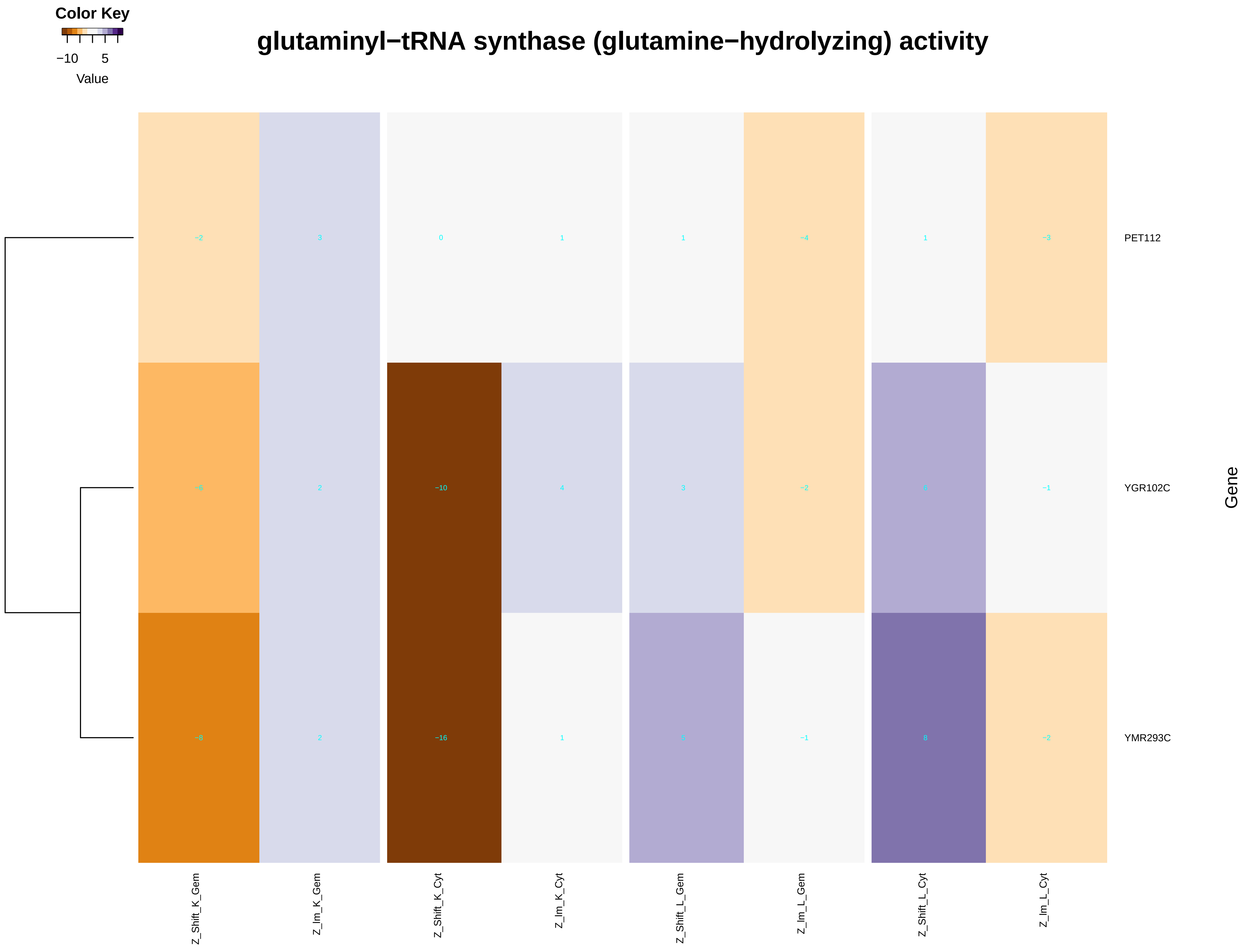

### cell_cycle.pdf

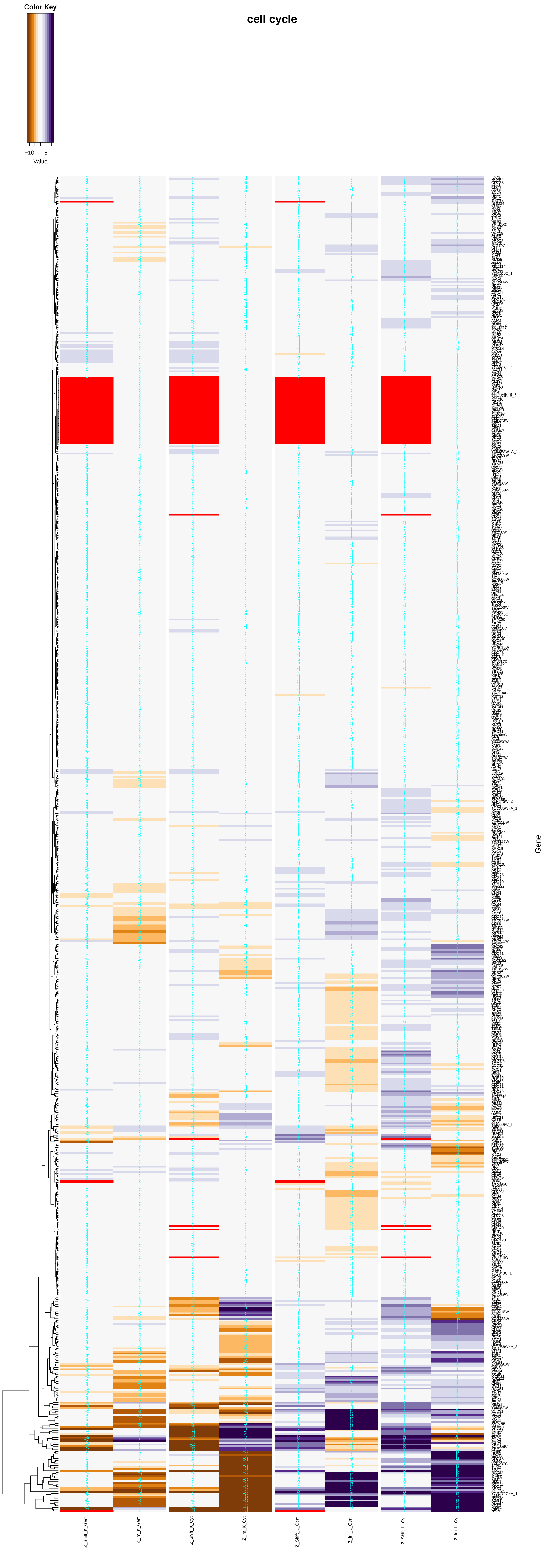

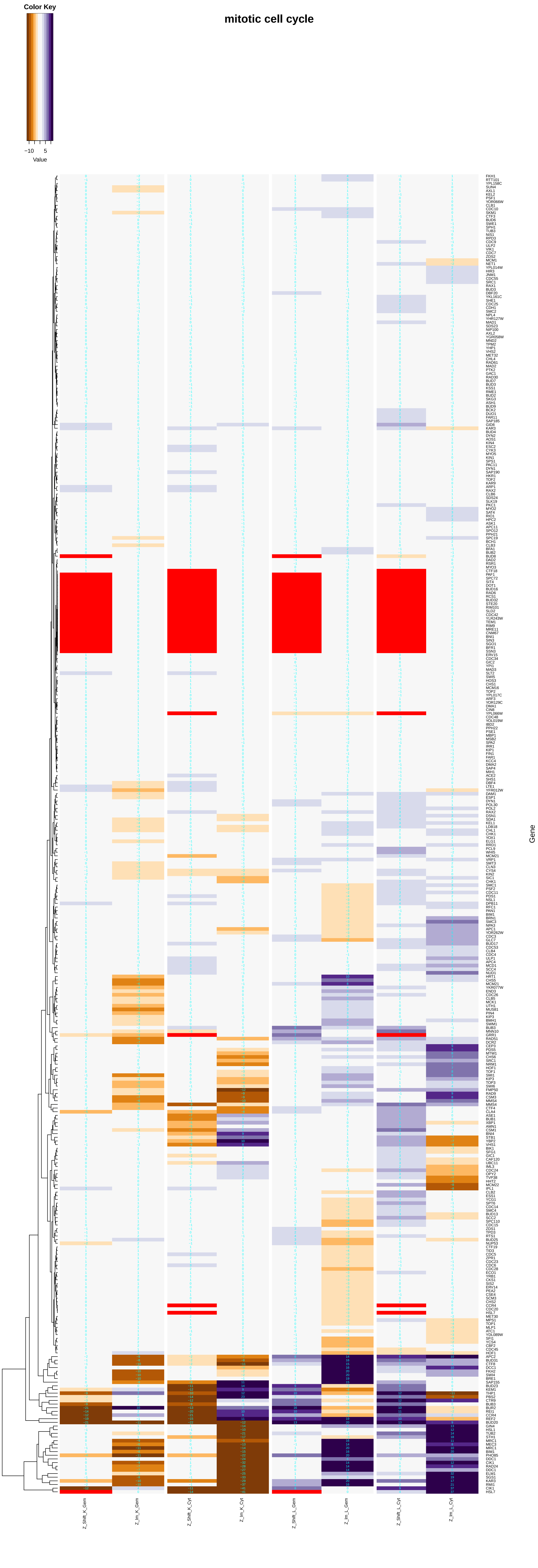

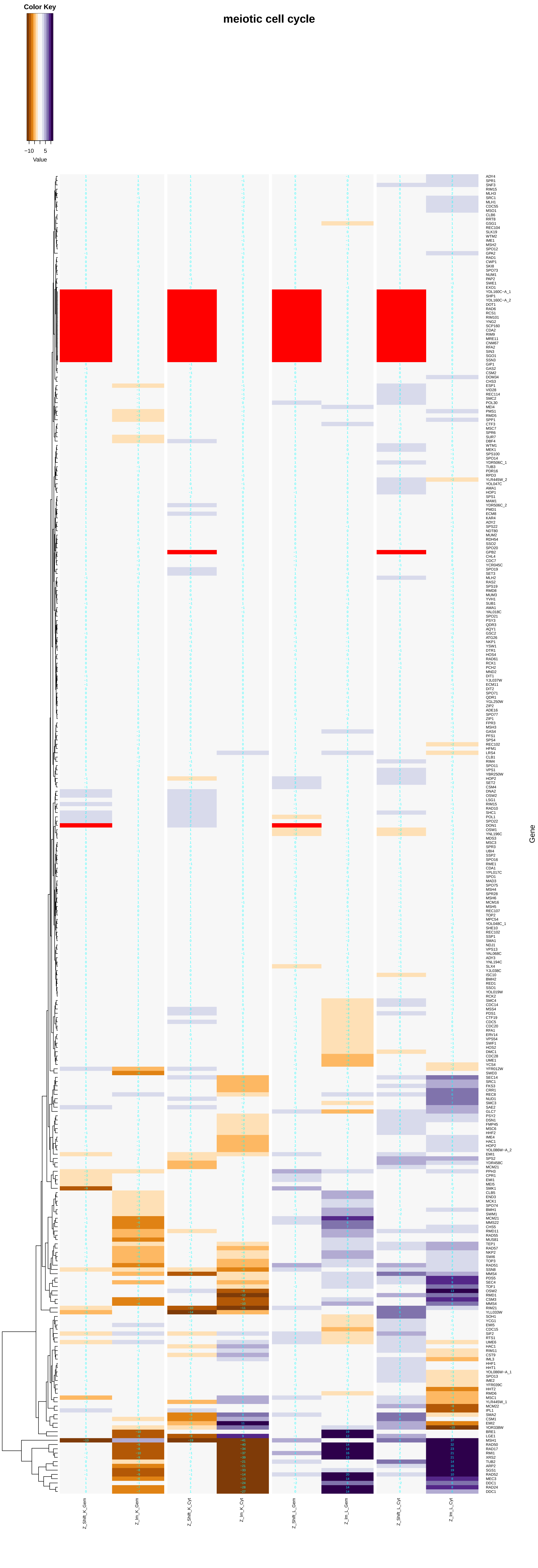

### cell_division.pdf

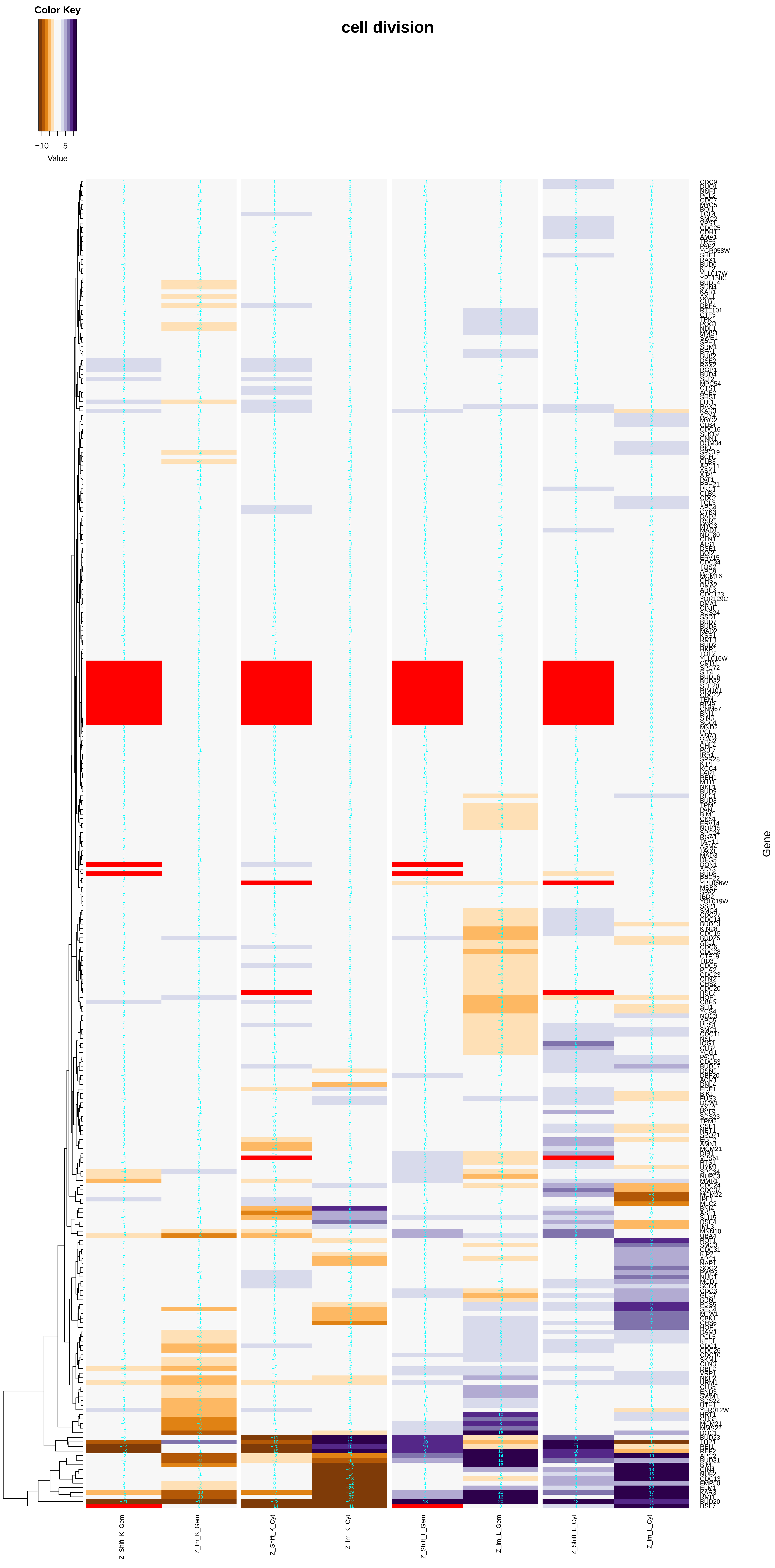

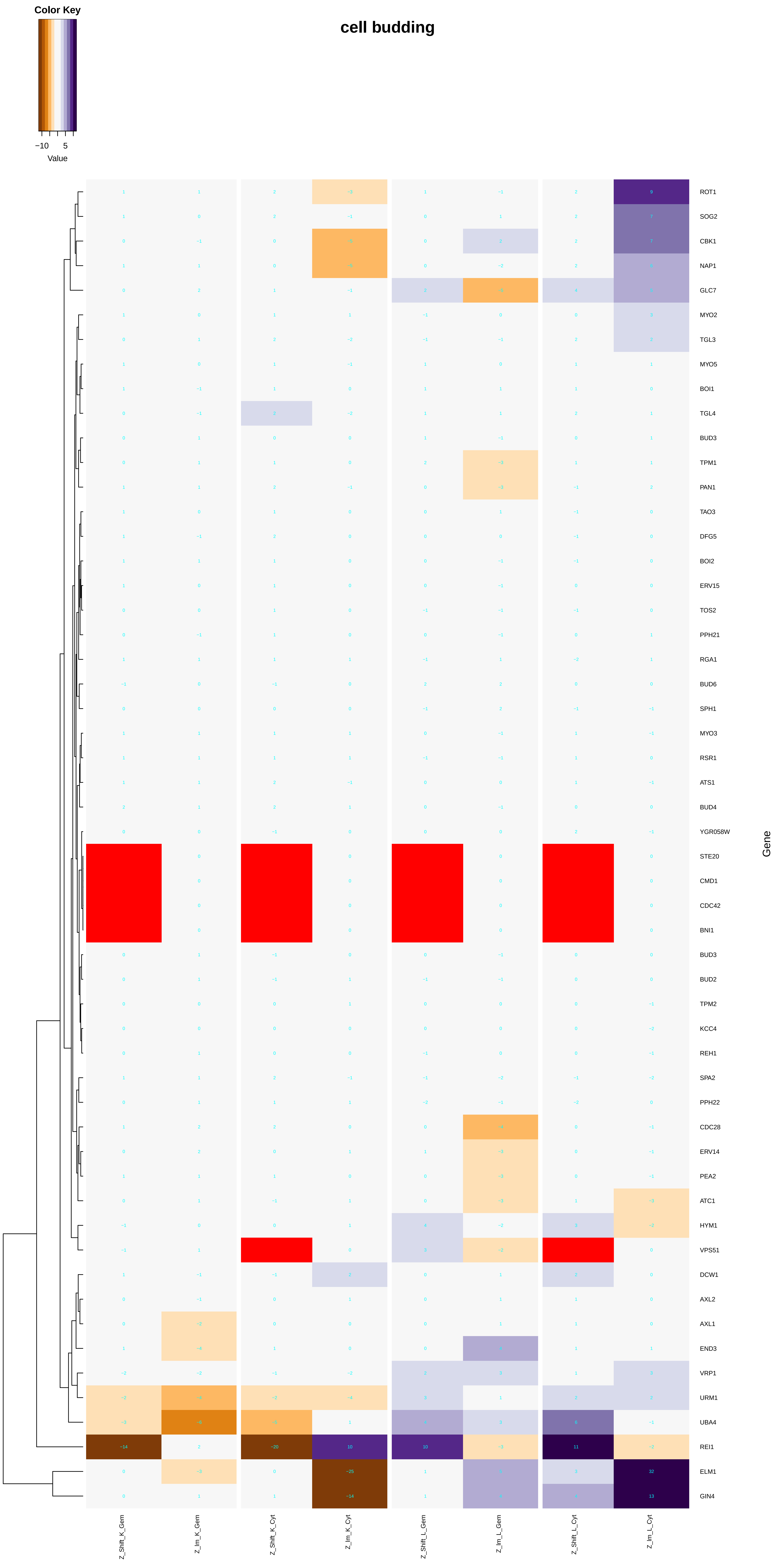

#### axial cellular bud site selection

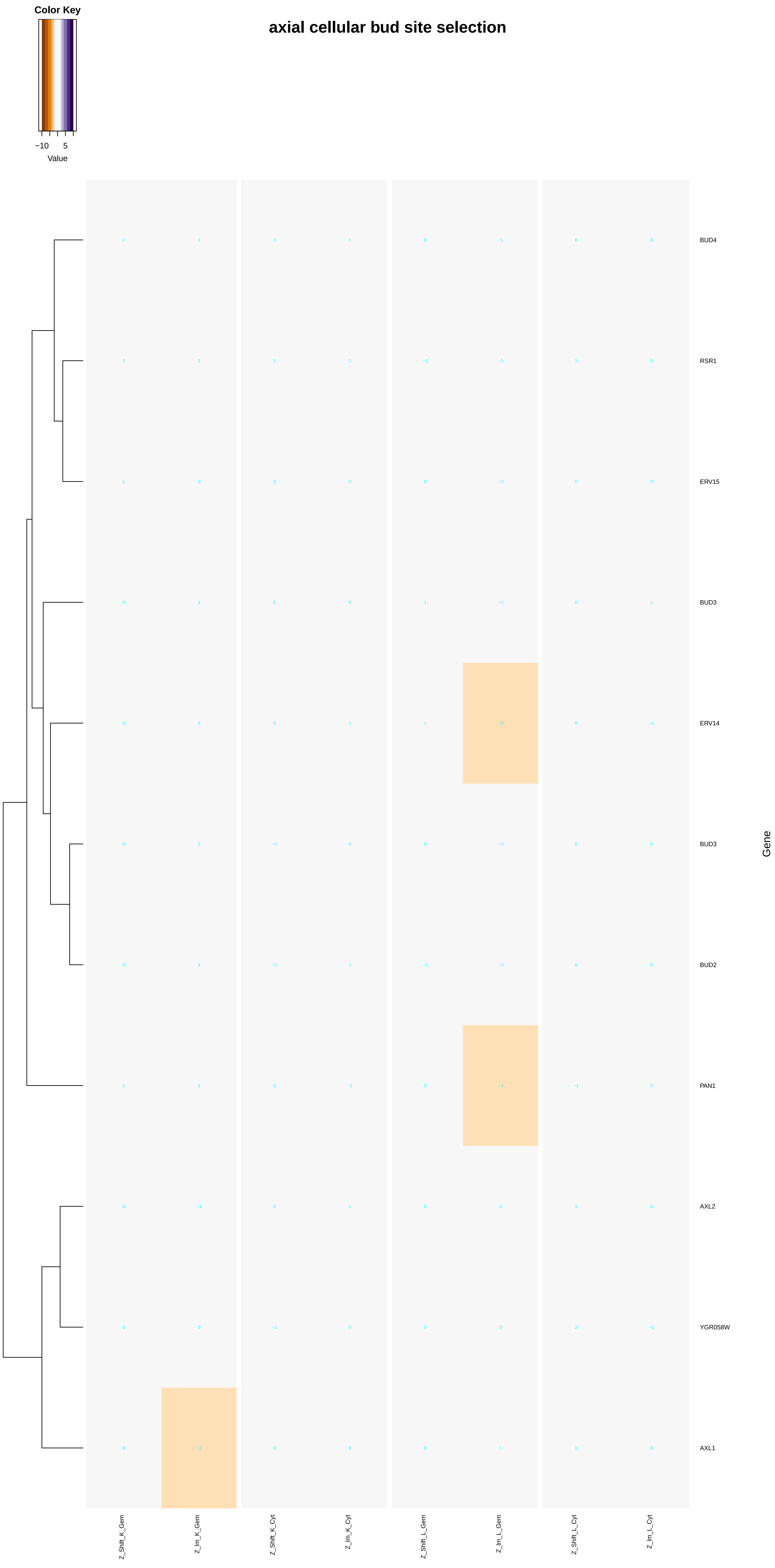

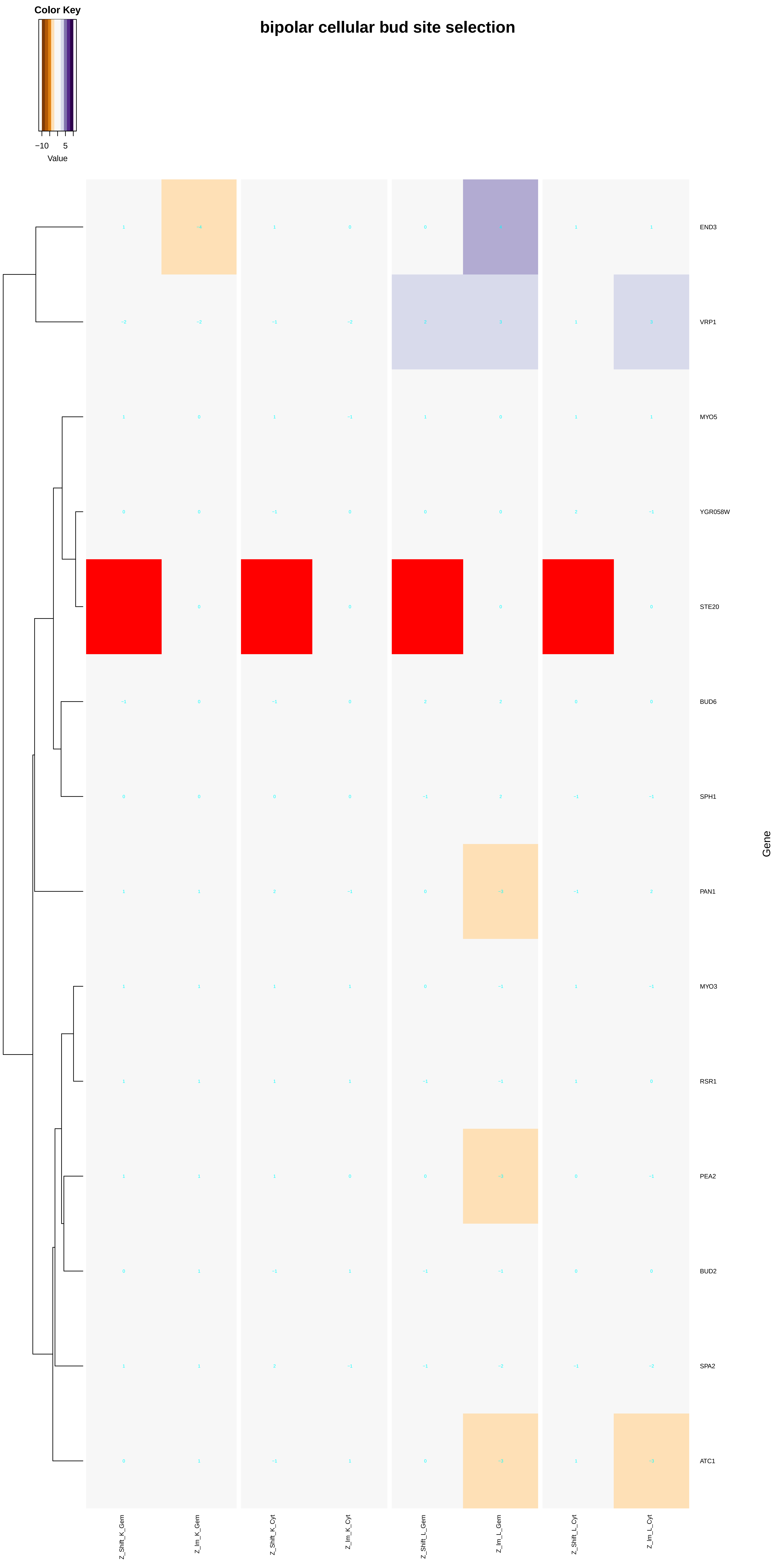

### chaperonin-containing_T-complex.pdf

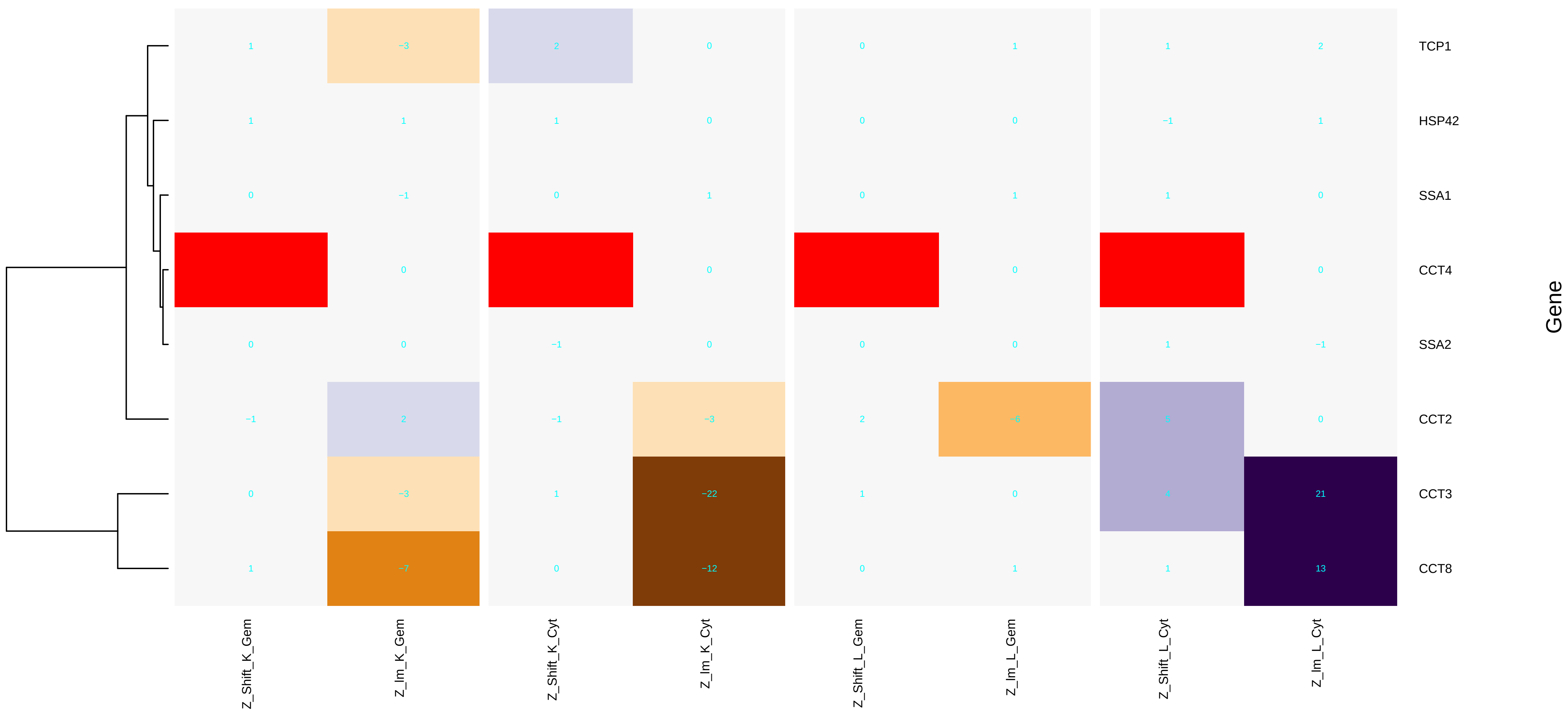

### checkpoint_clamp_complex.pdf

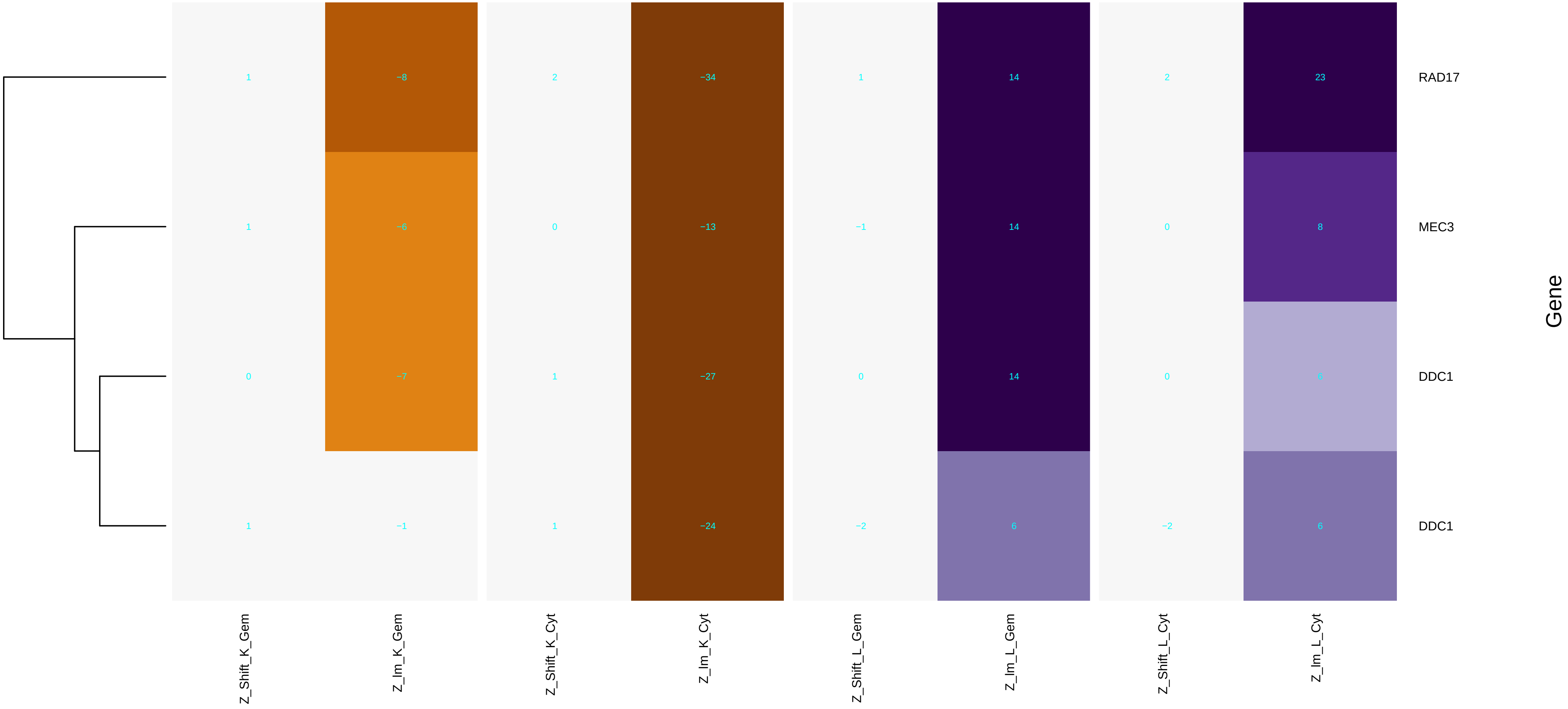

### chromatin_binding.pdf

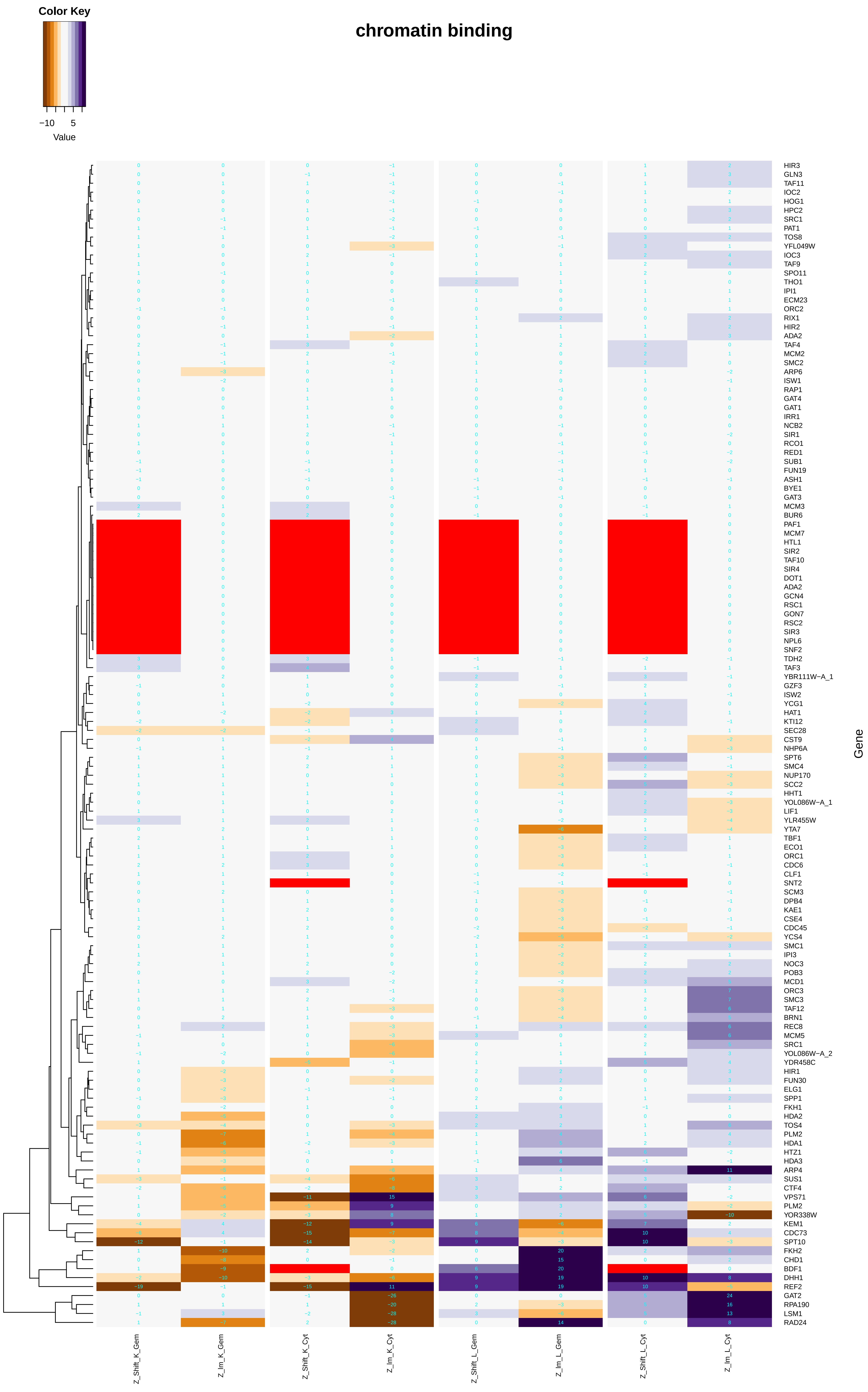

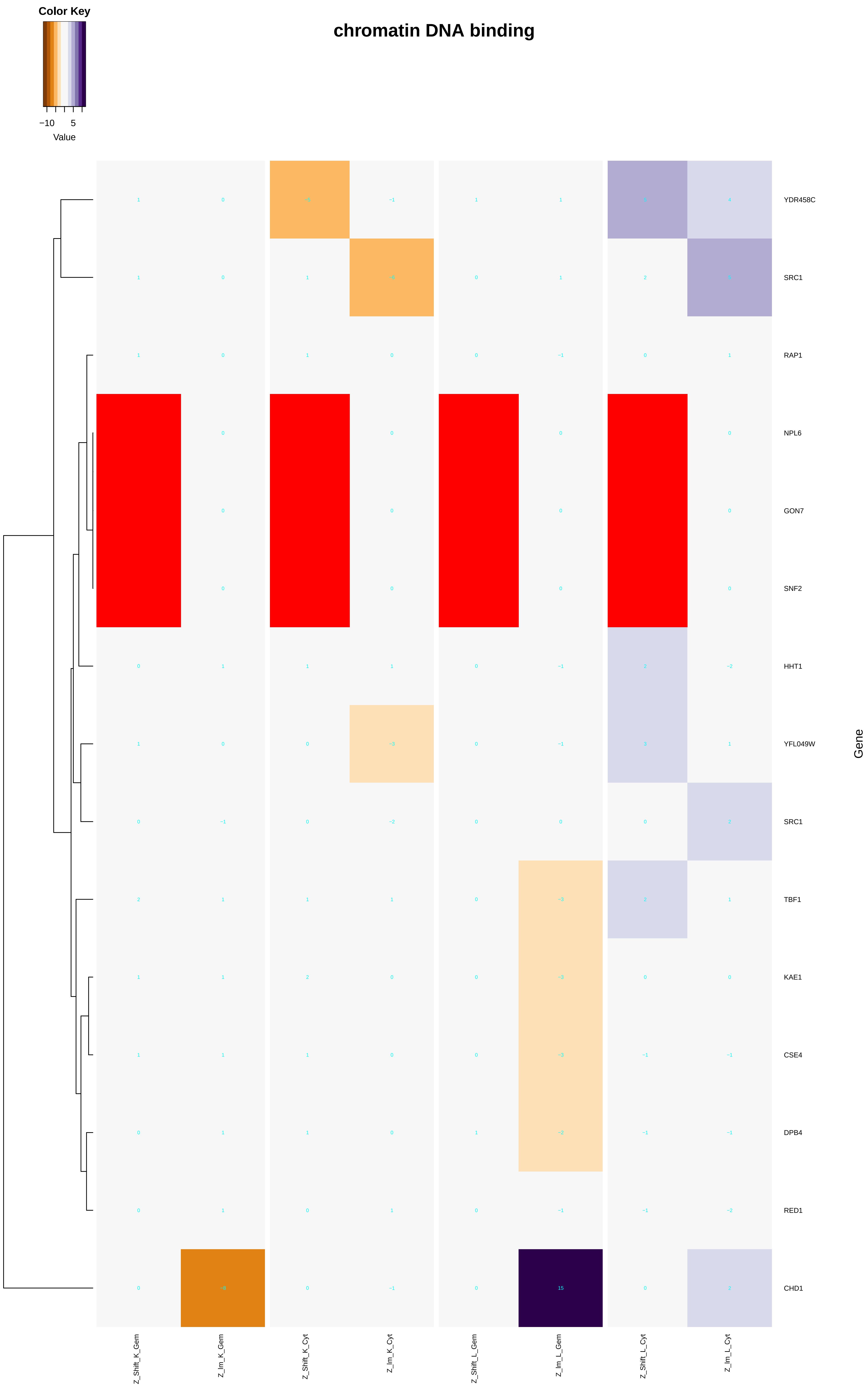

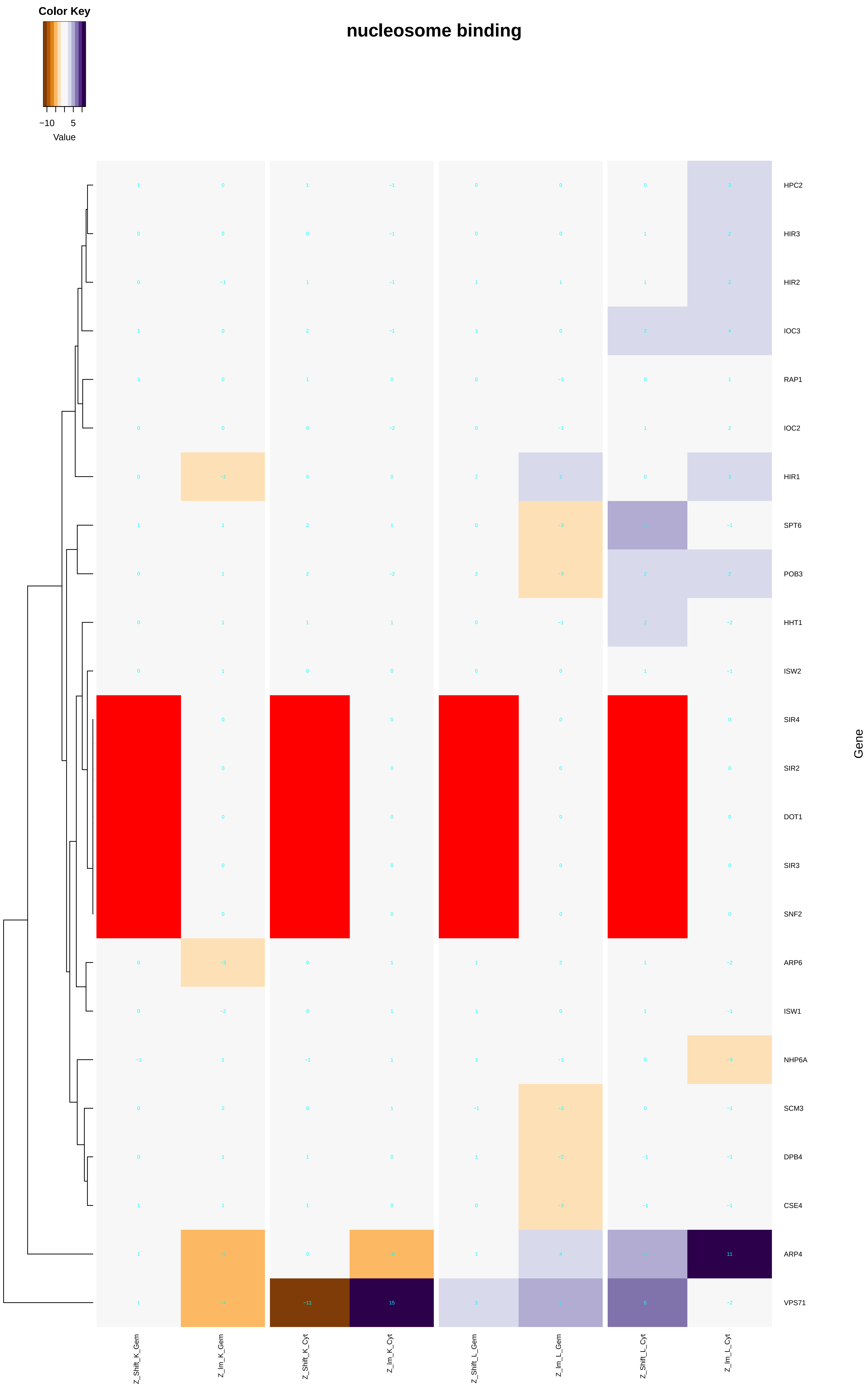

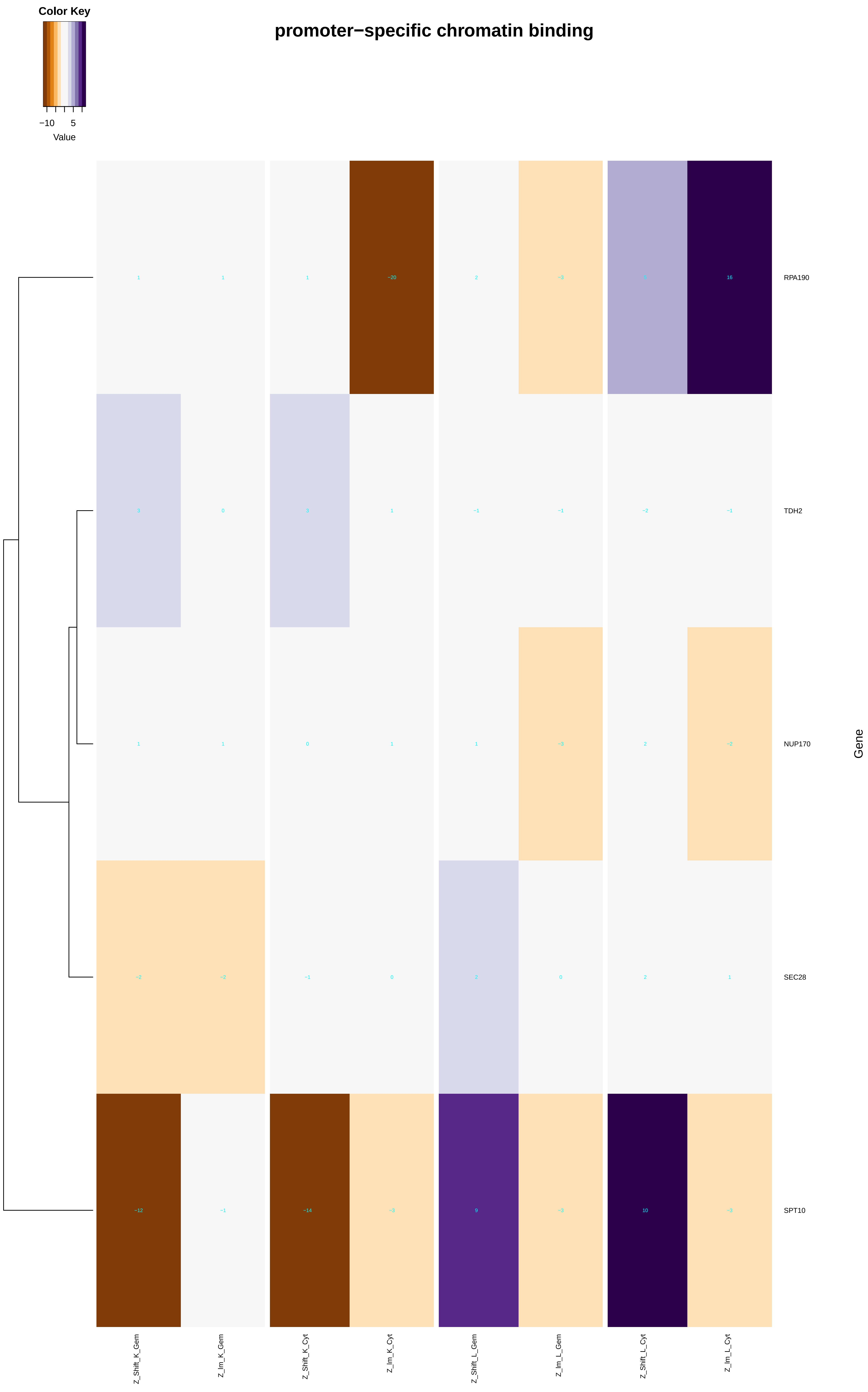

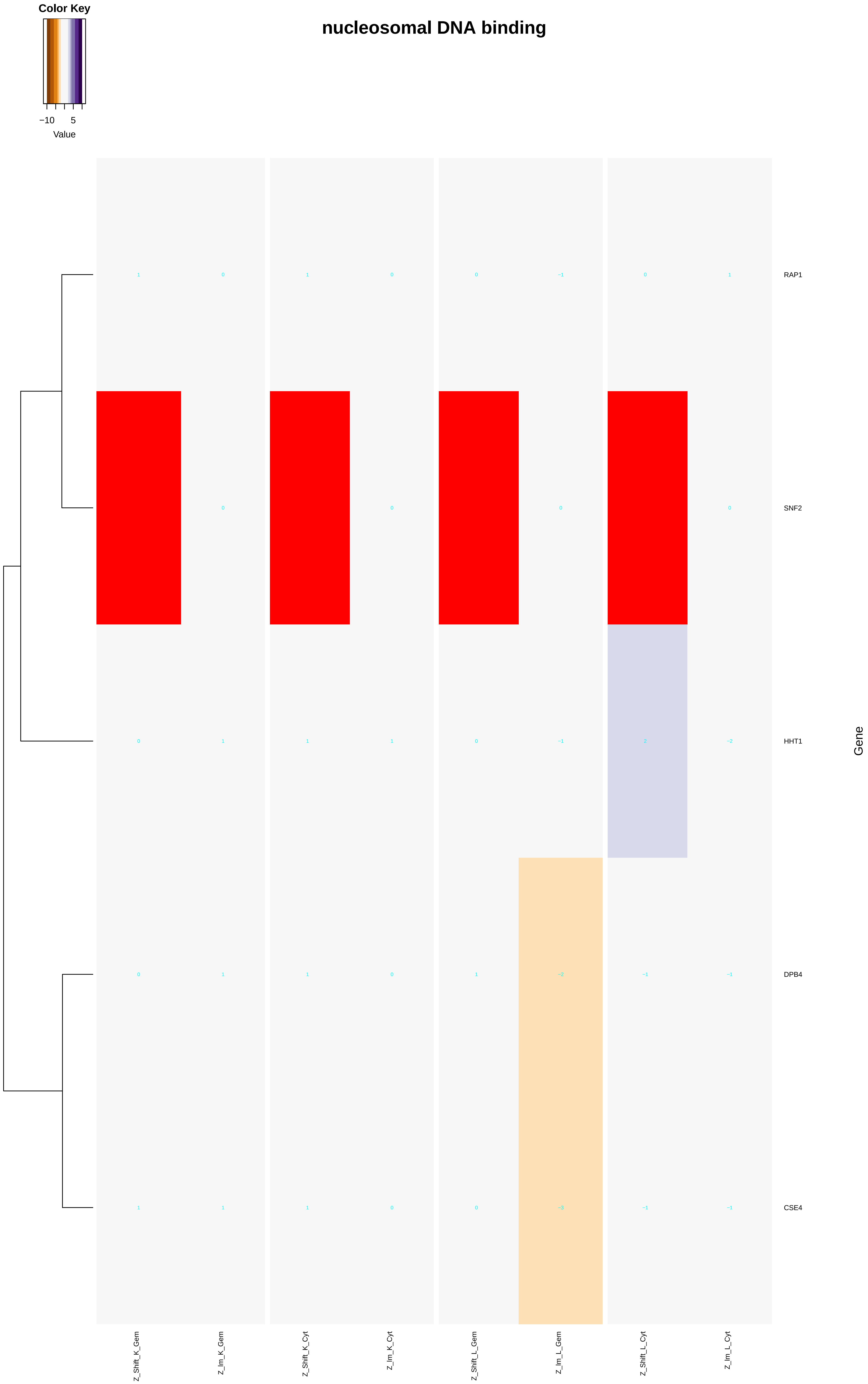

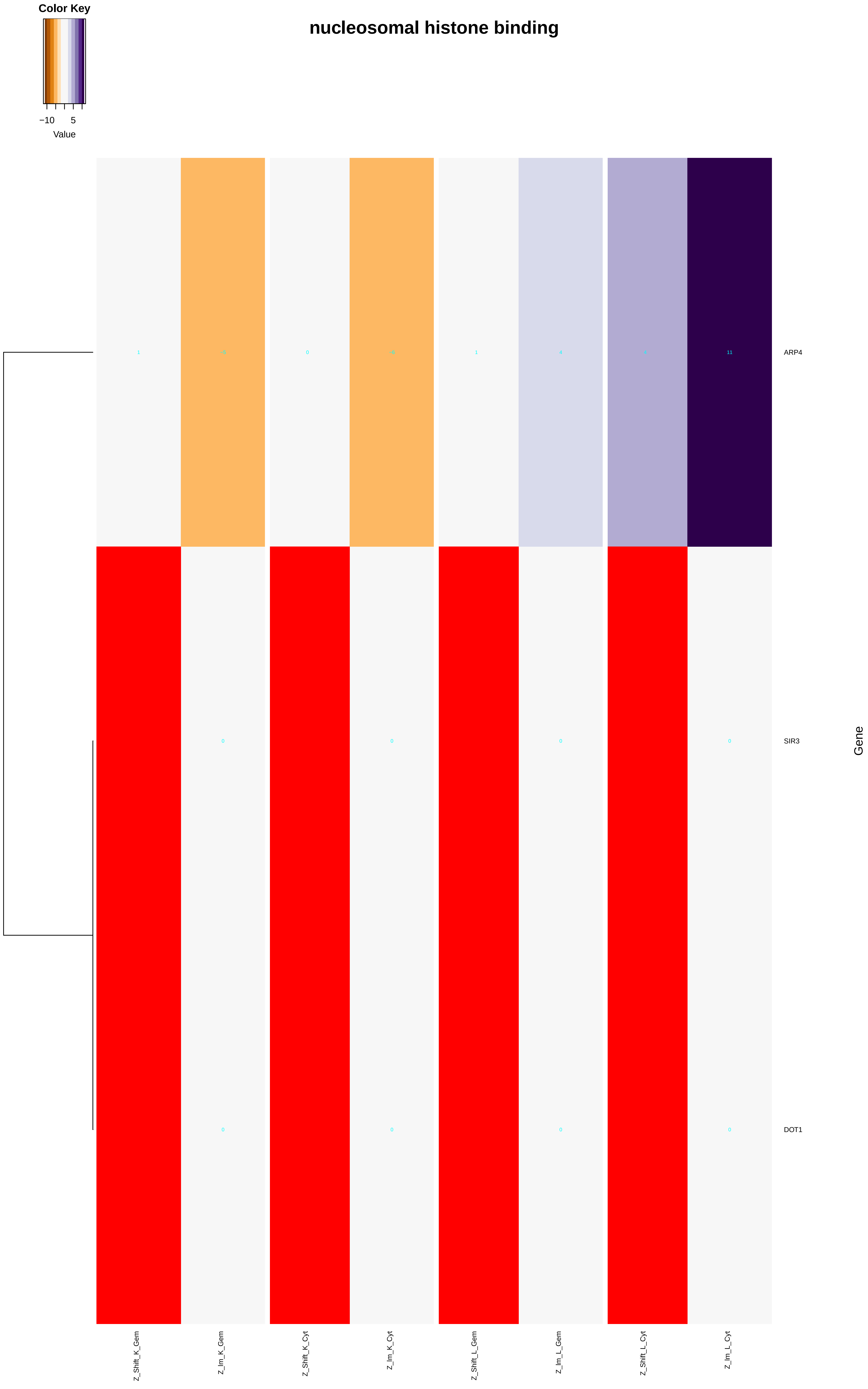

### chromatin_silencing.pdf

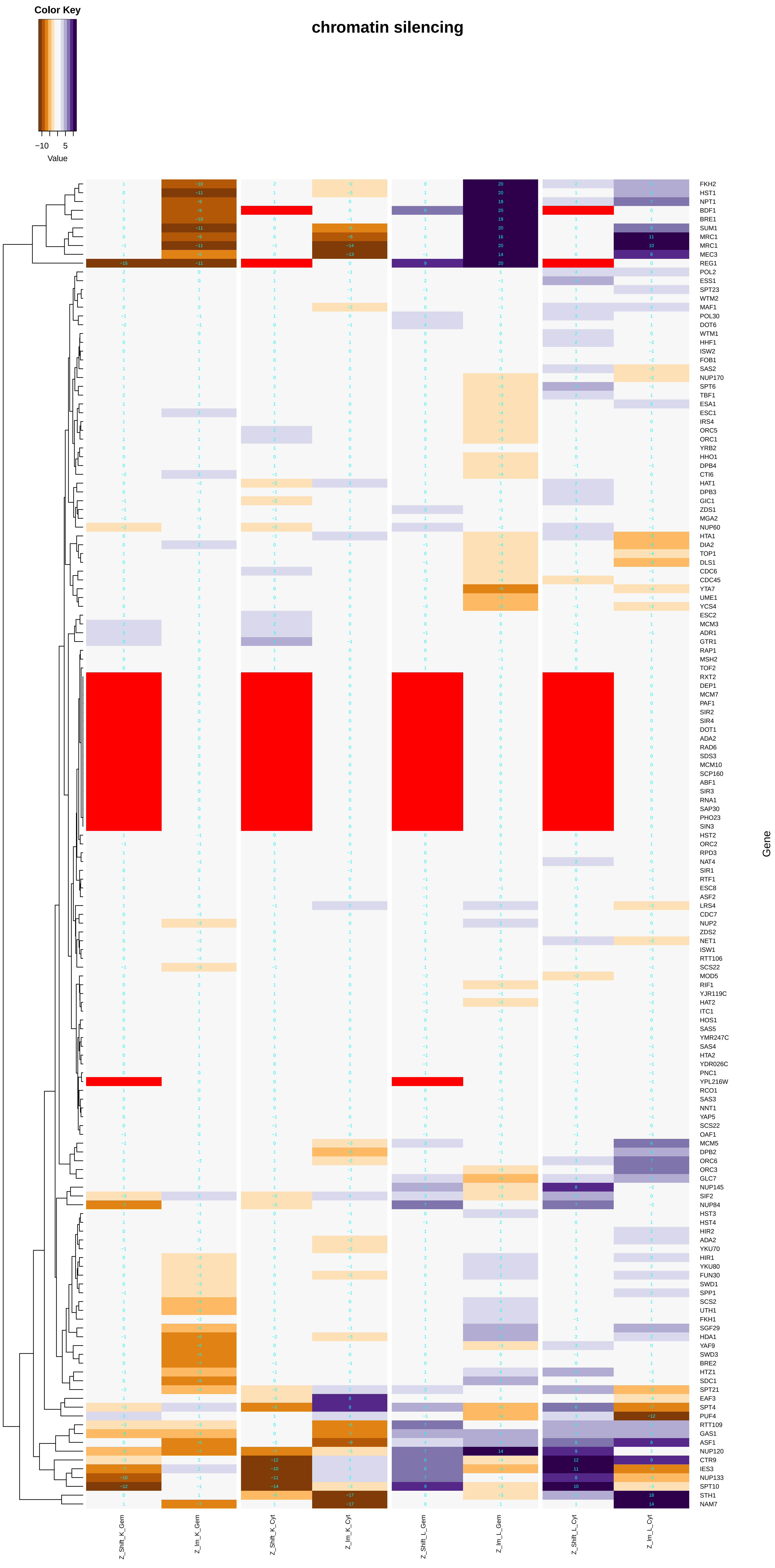

#### chromatin silencing at rDNA

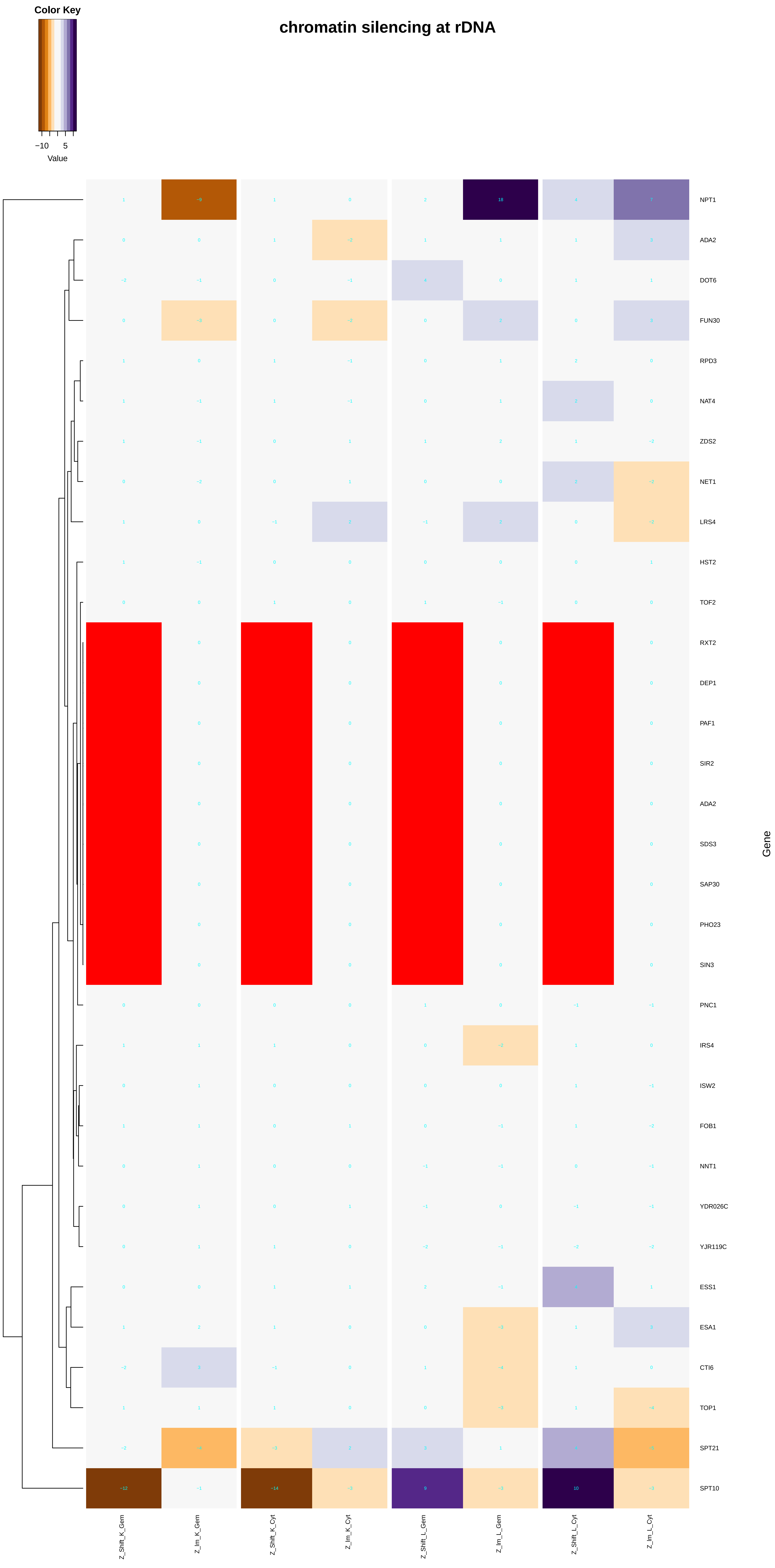

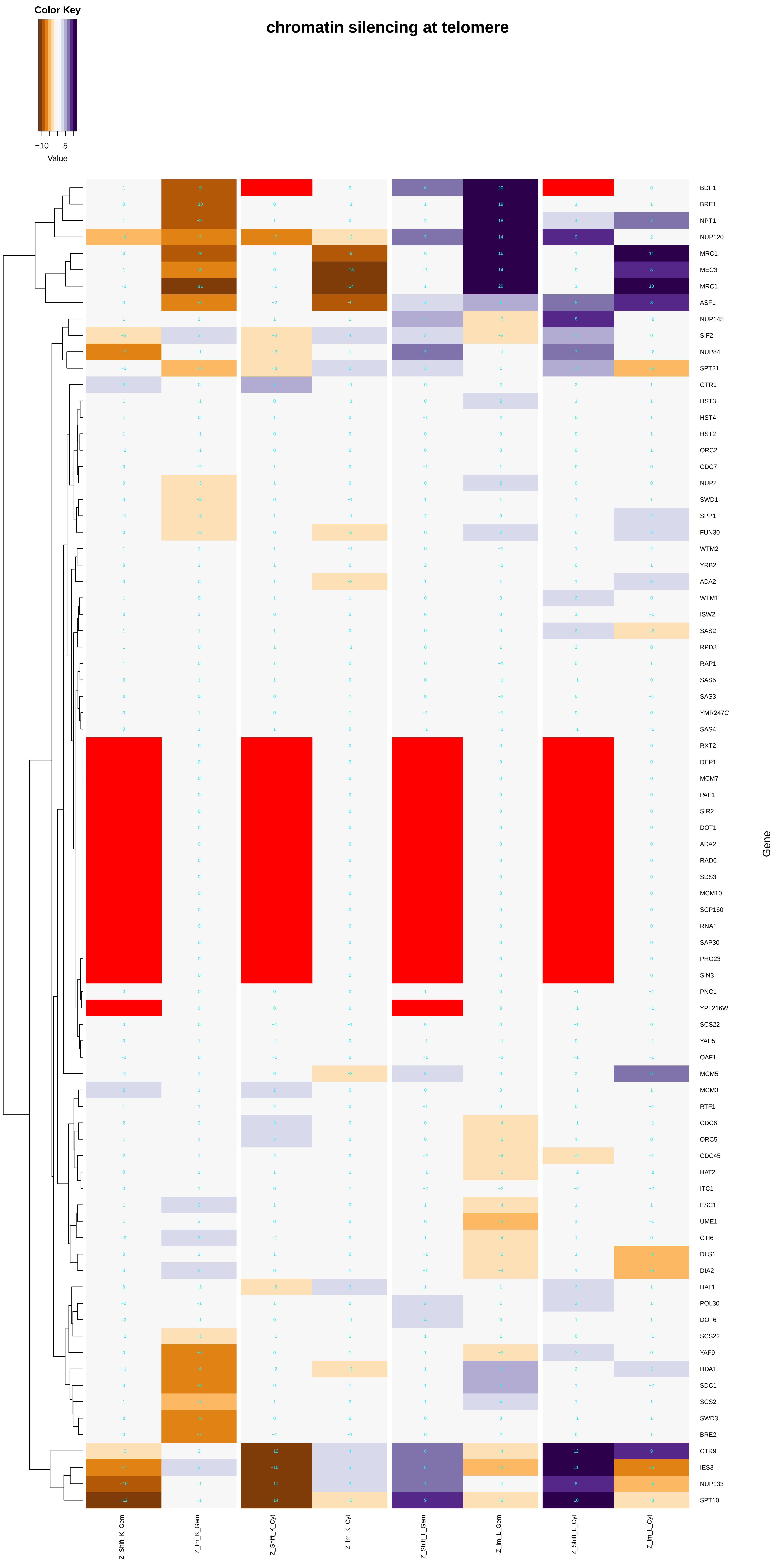

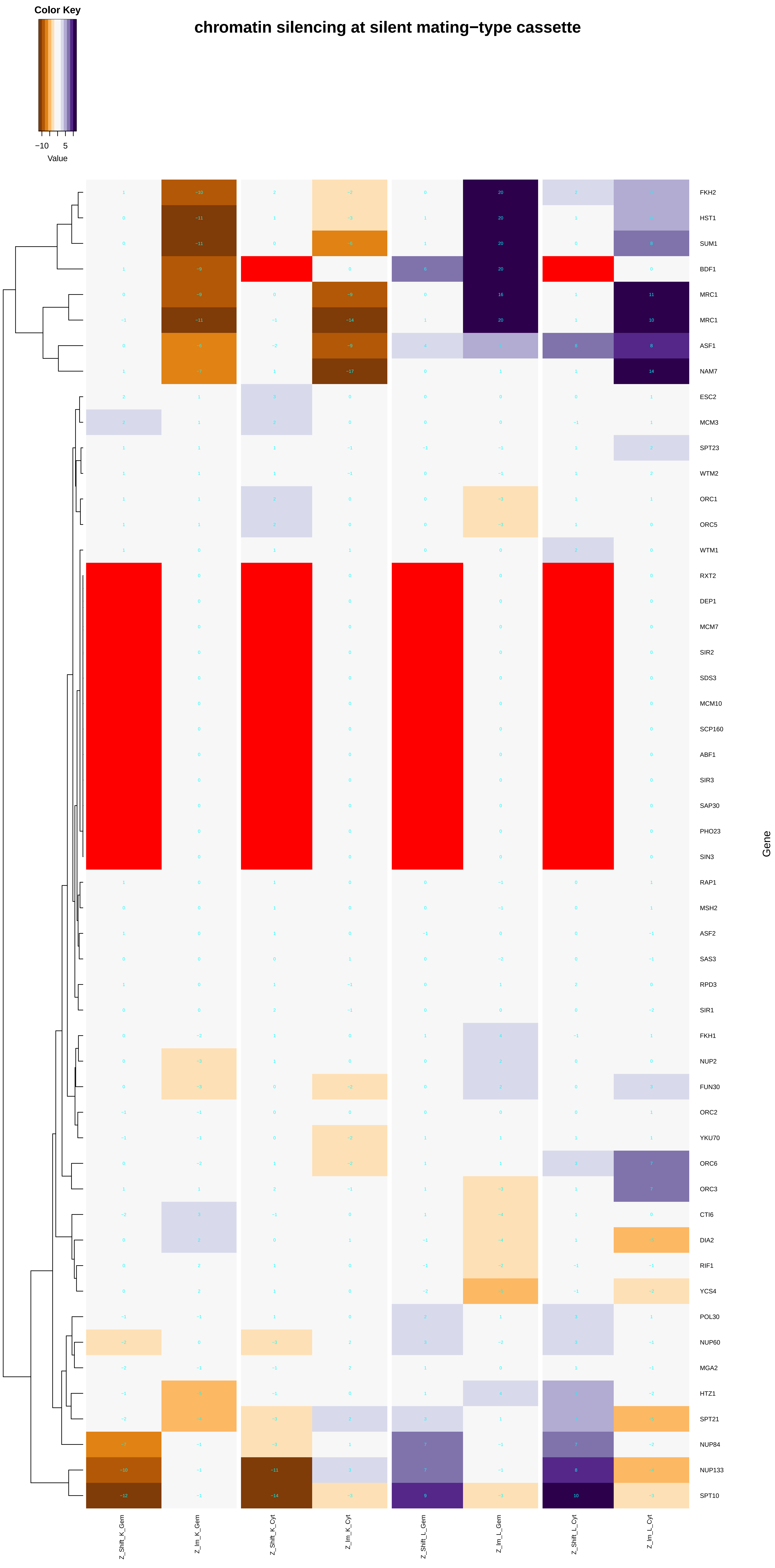

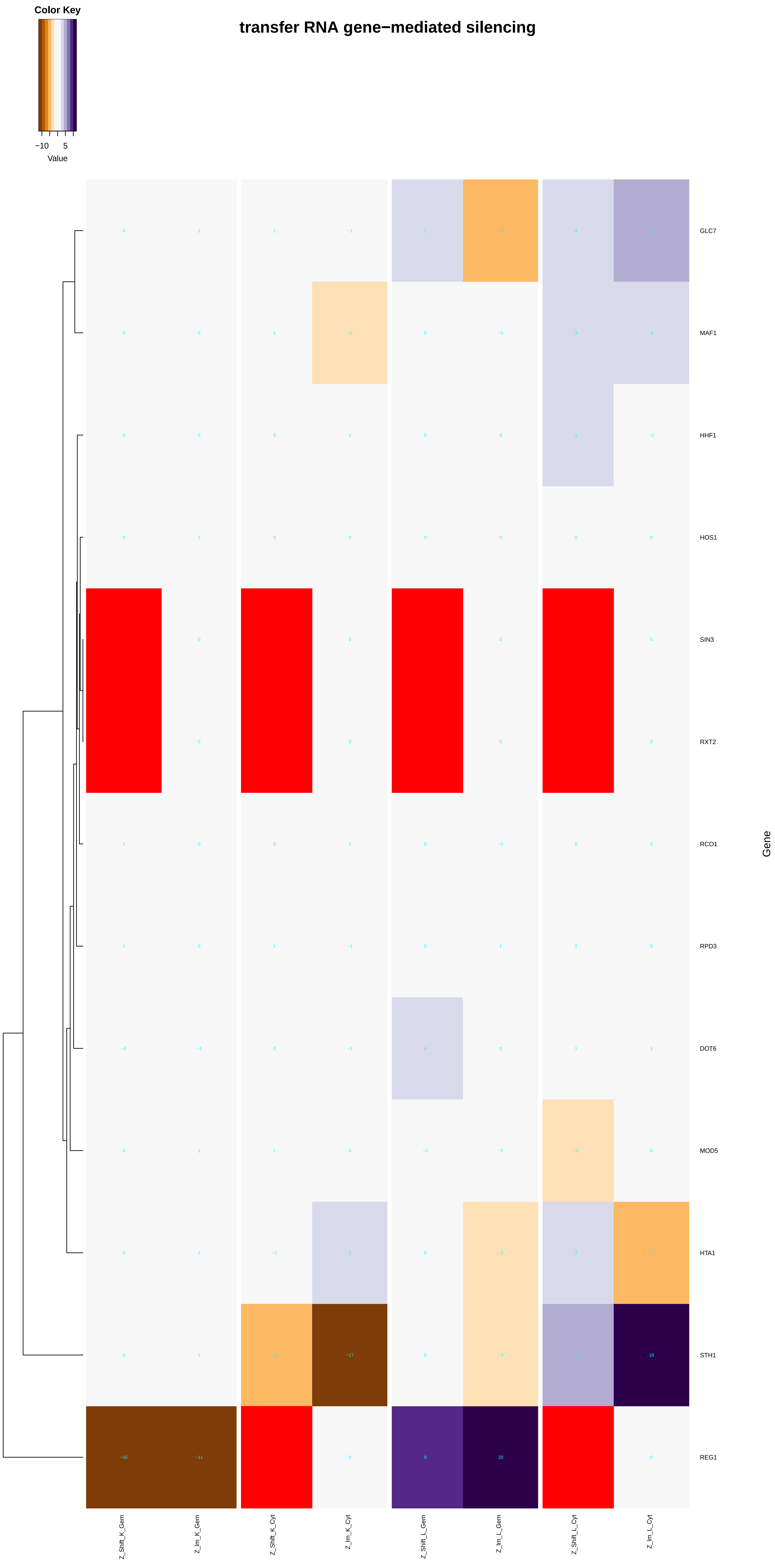

### chromosome.pdf

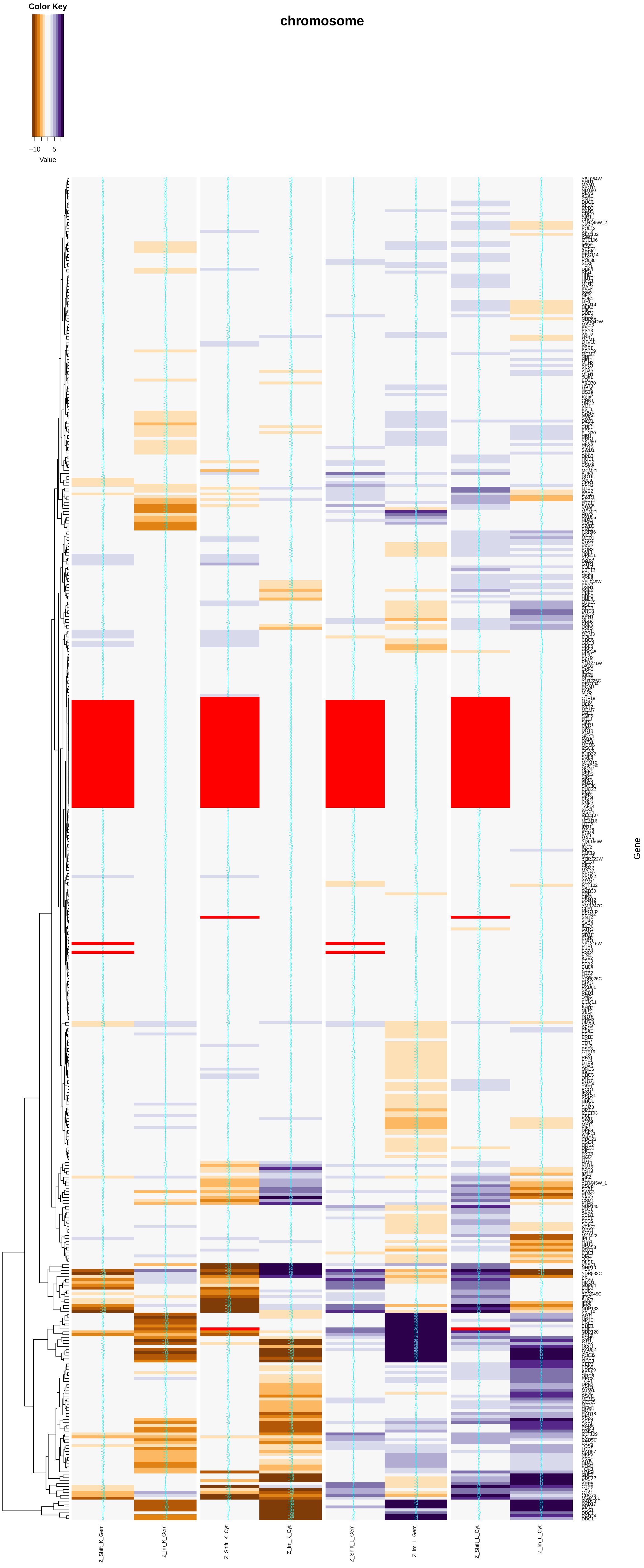

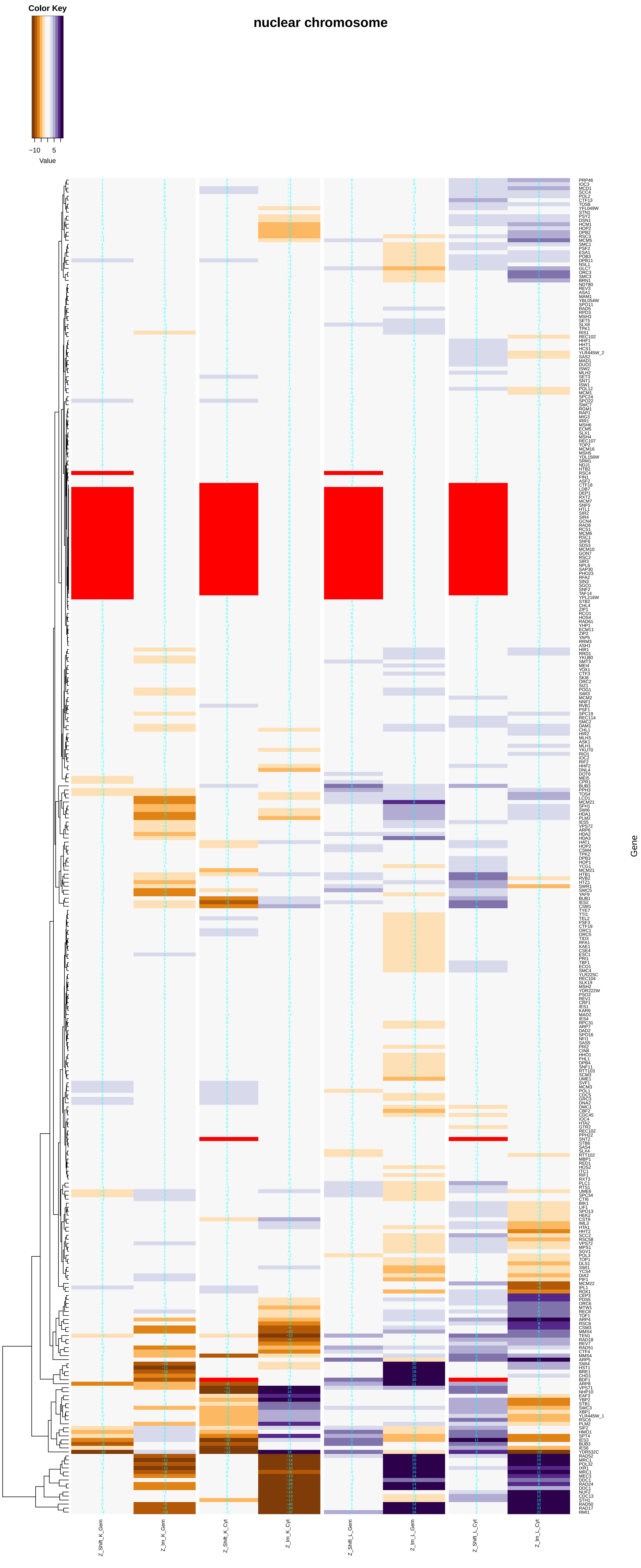

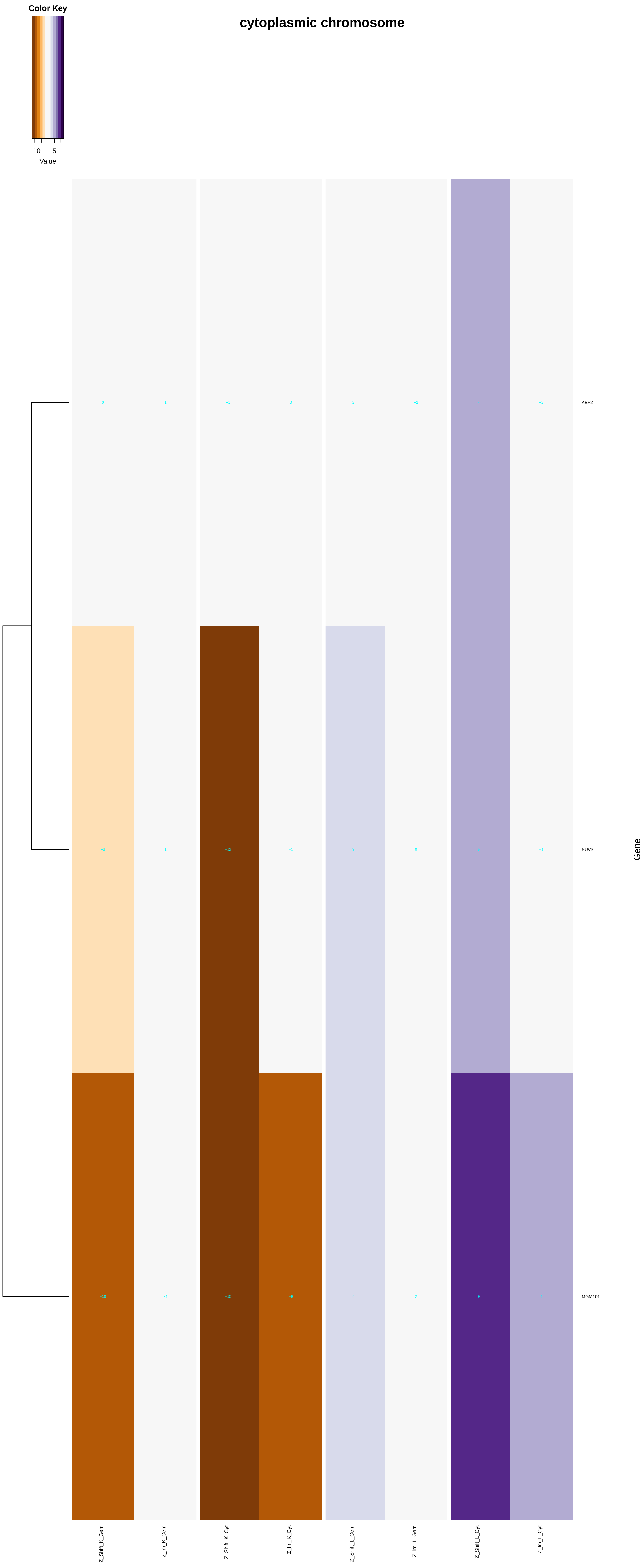

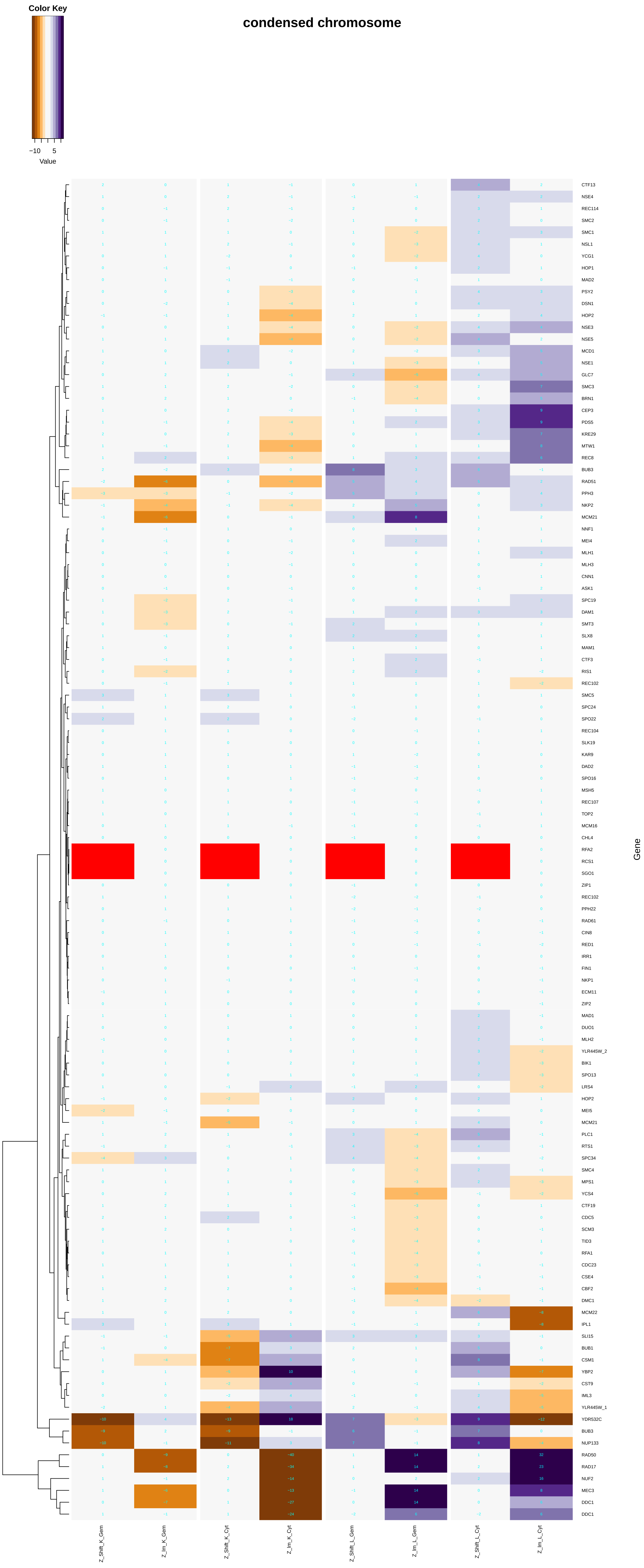

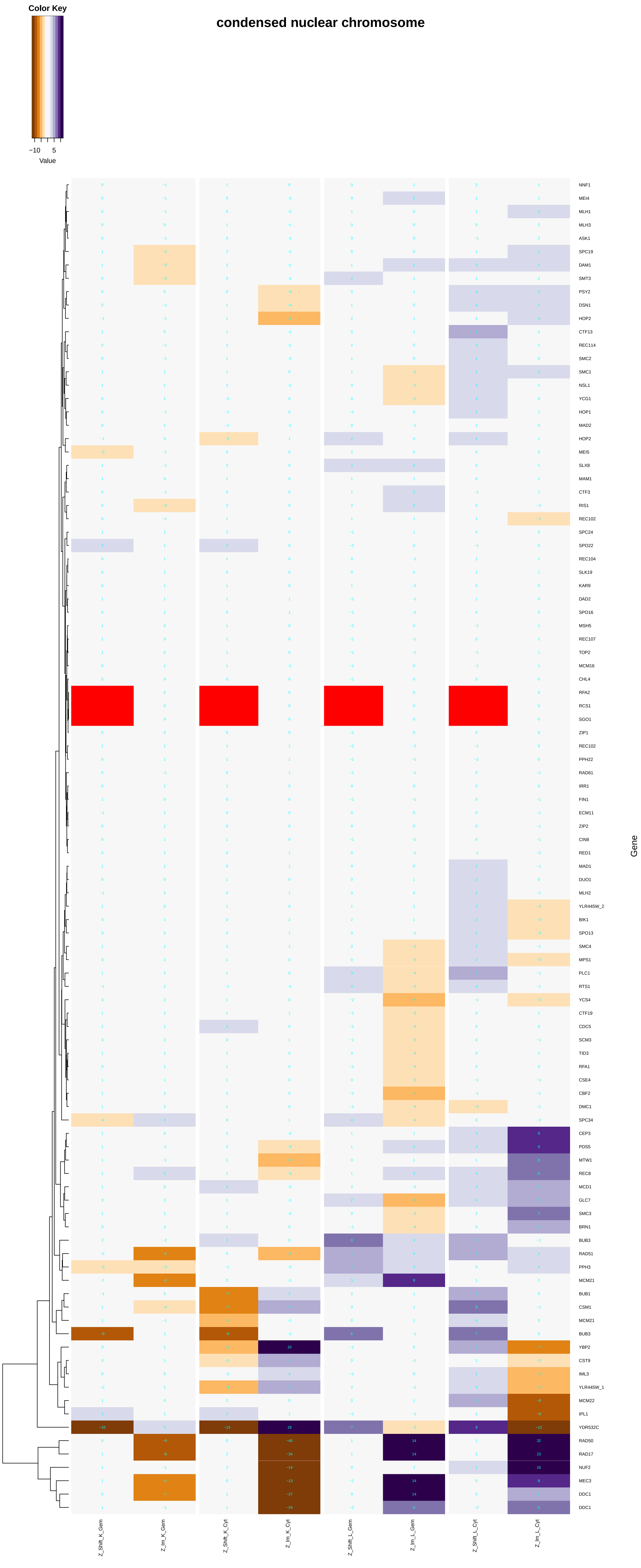

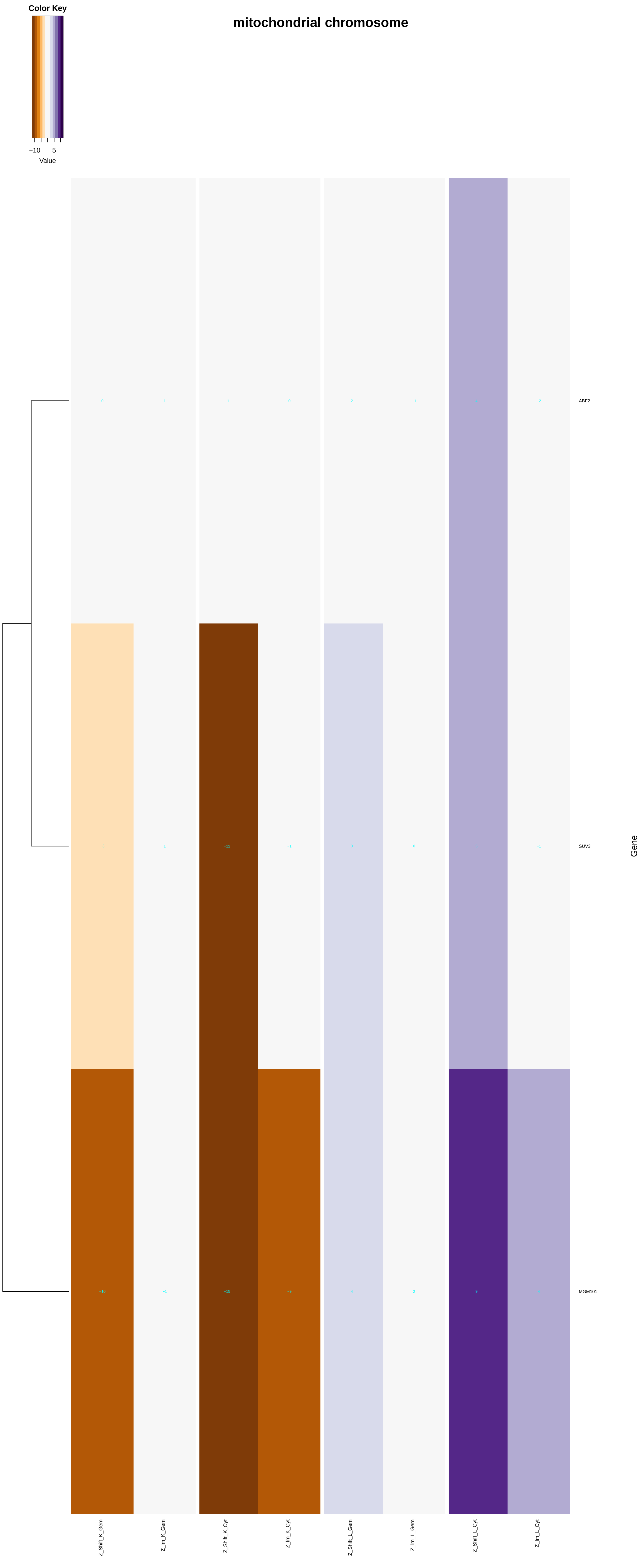

### cleavage_involved_in_rRNA_processing.pdf

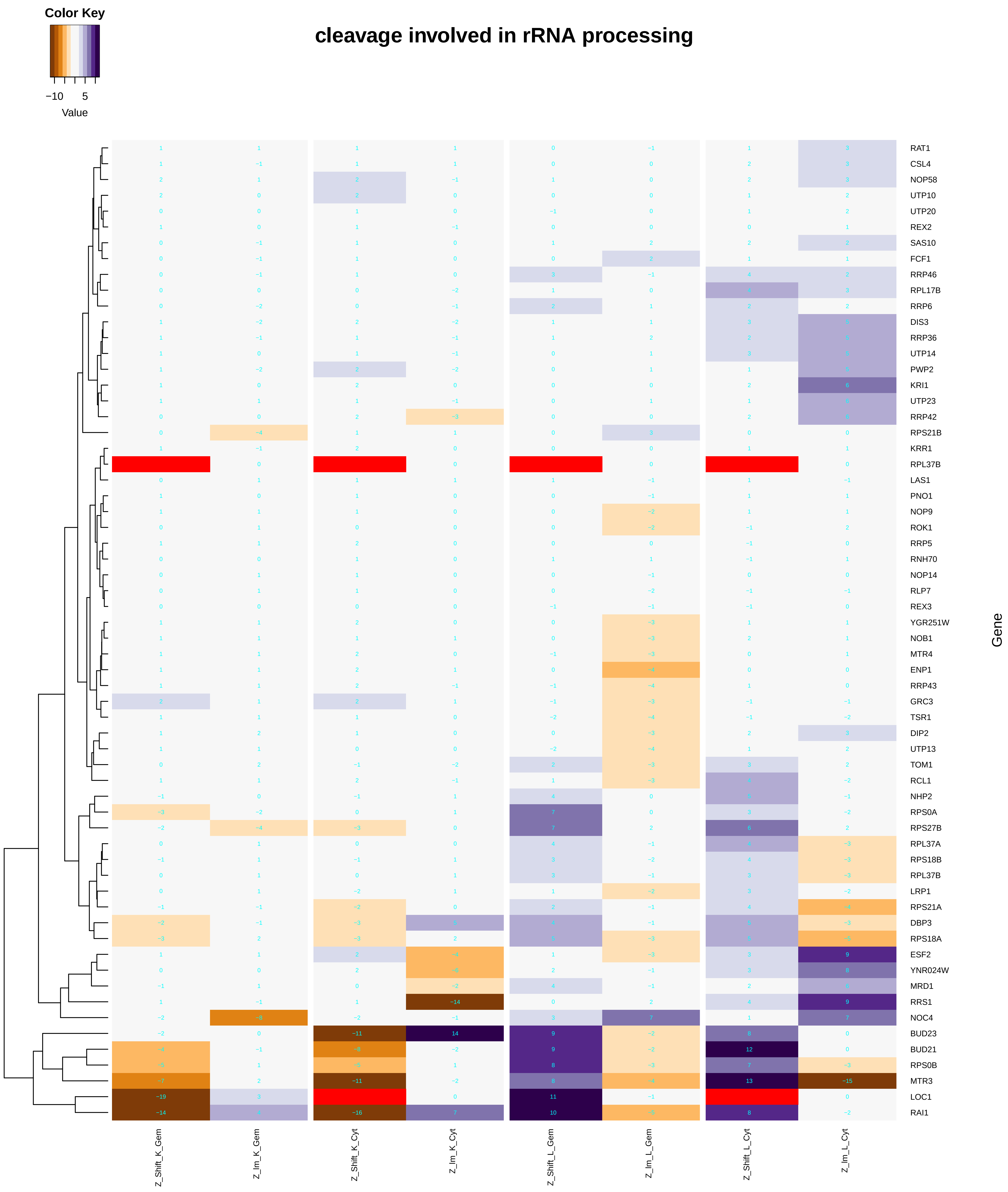

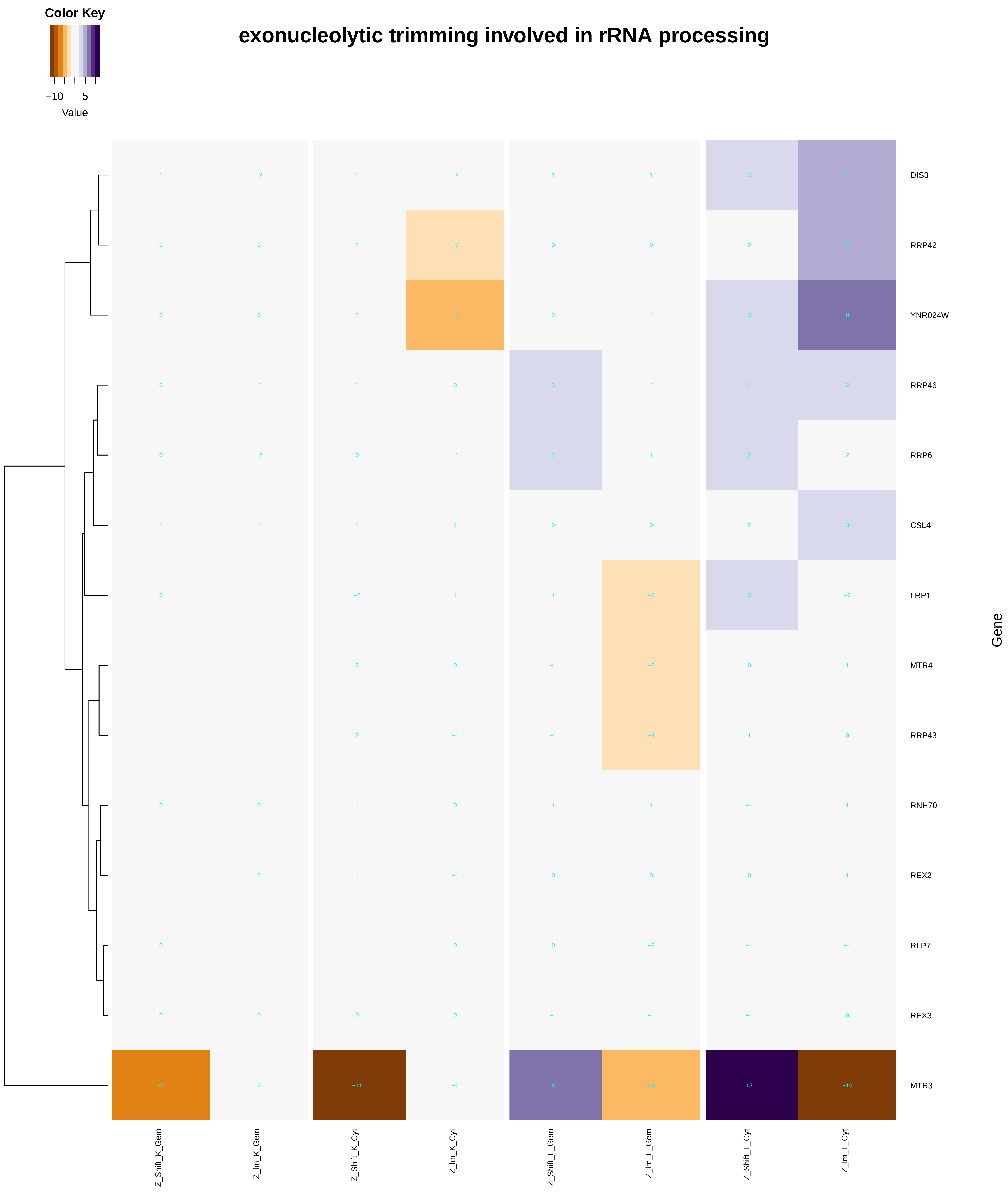

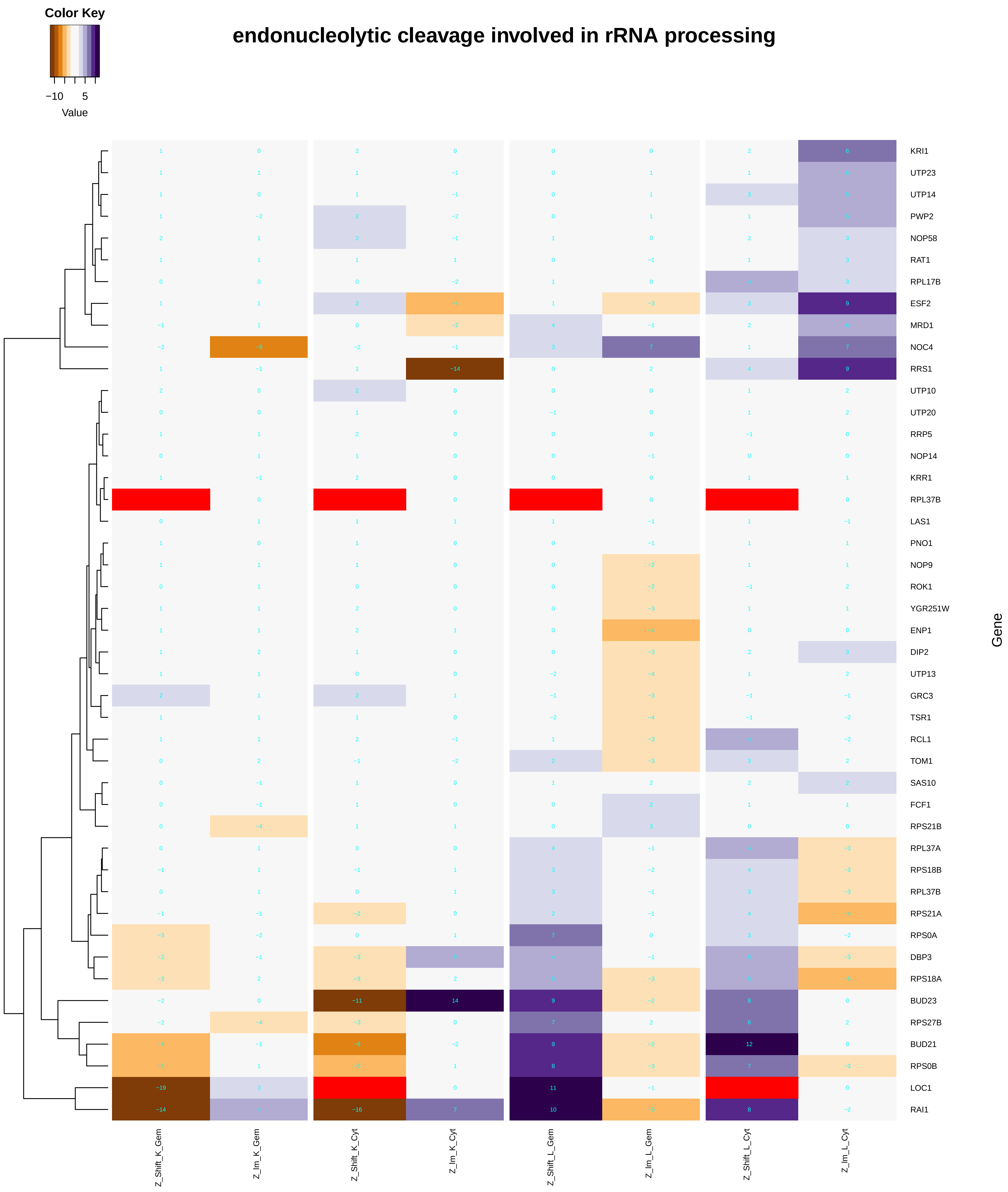

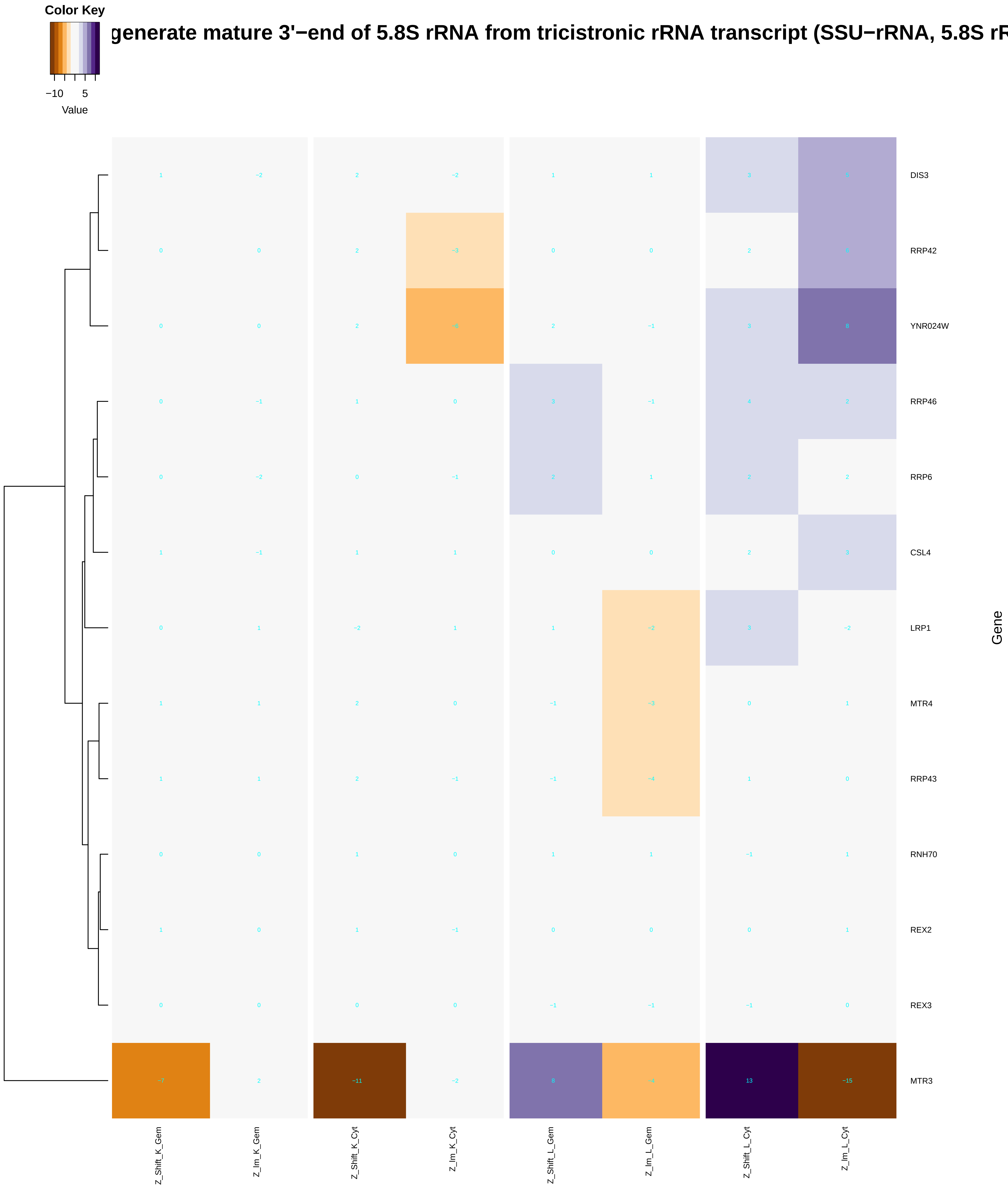

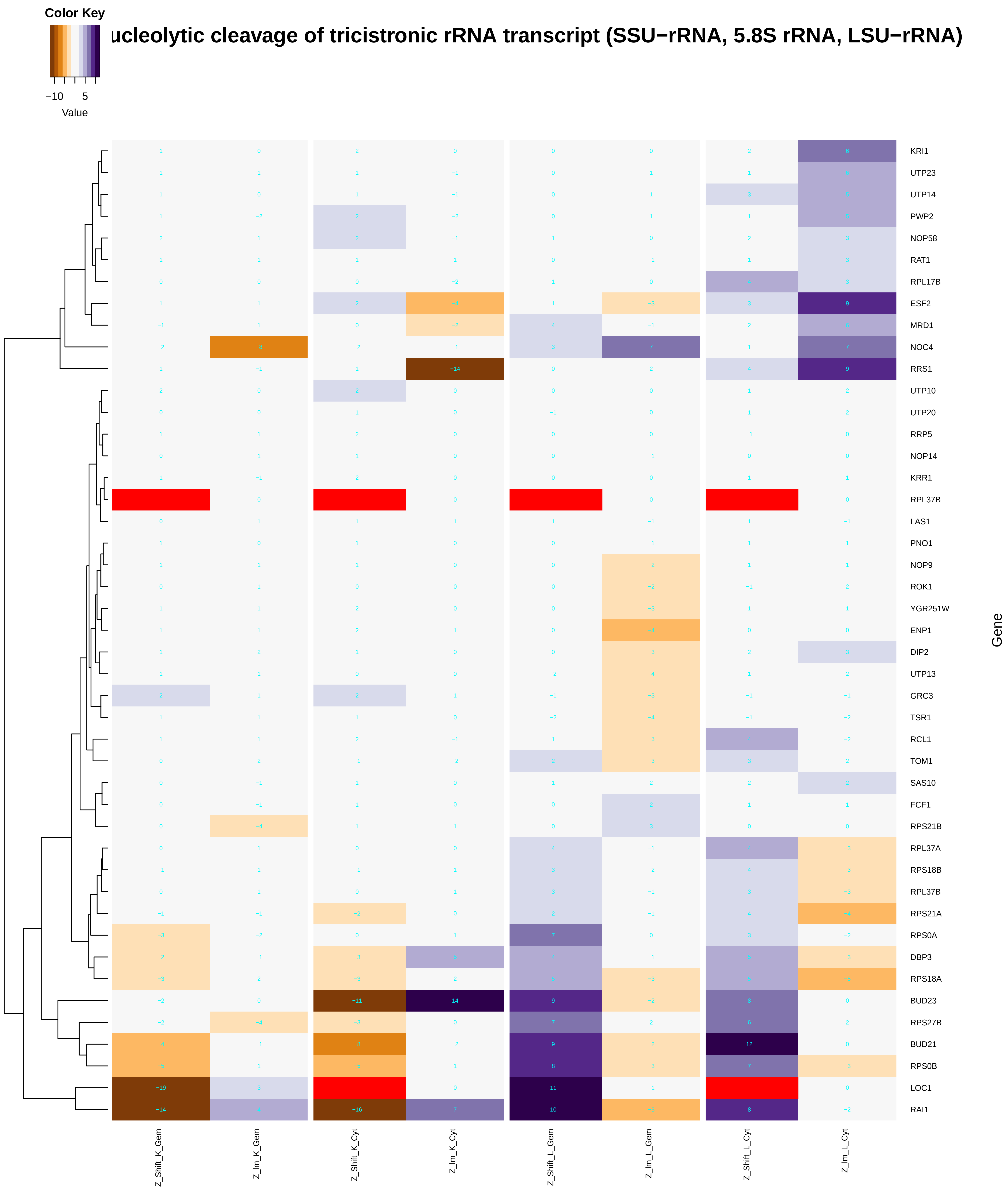

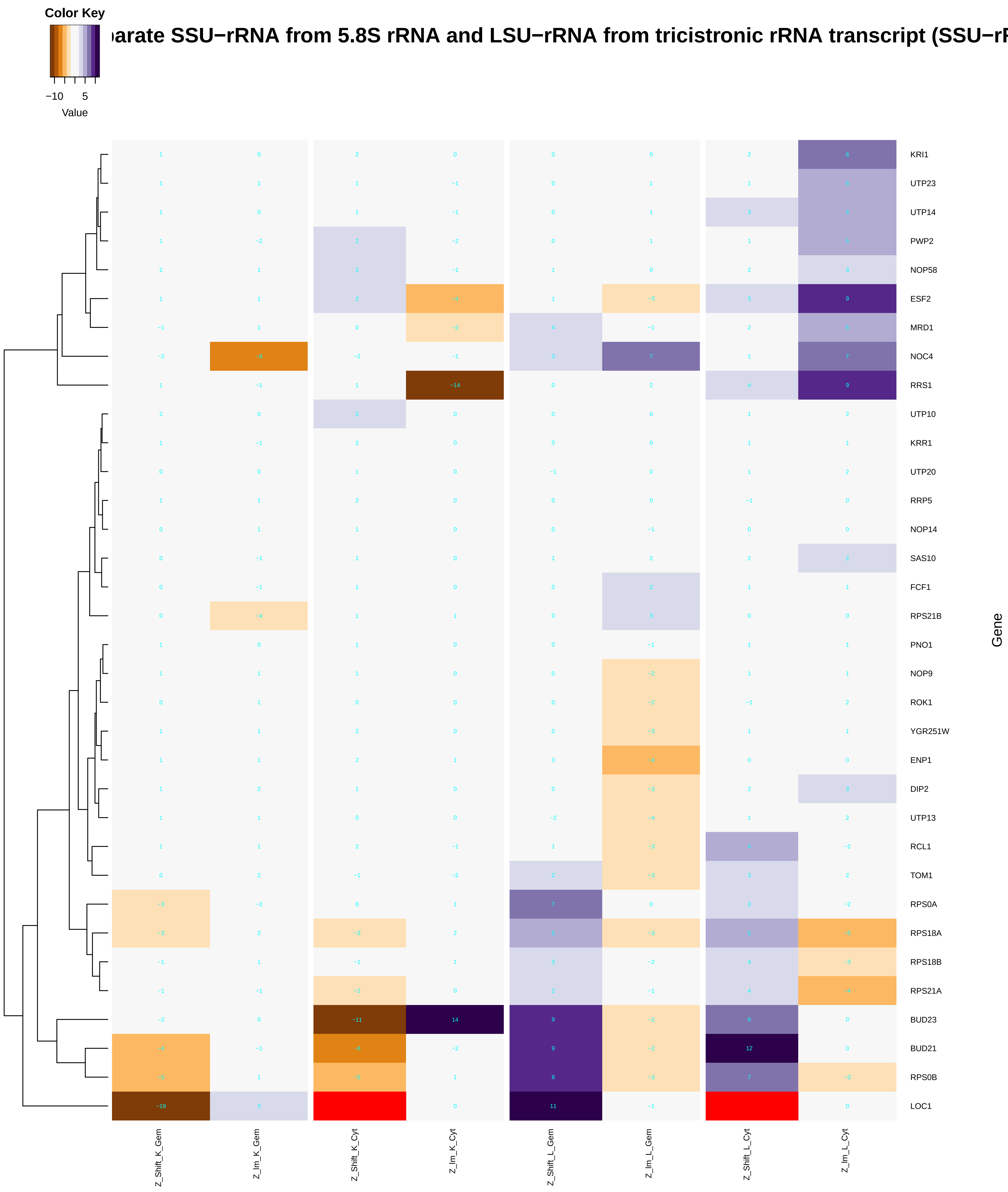

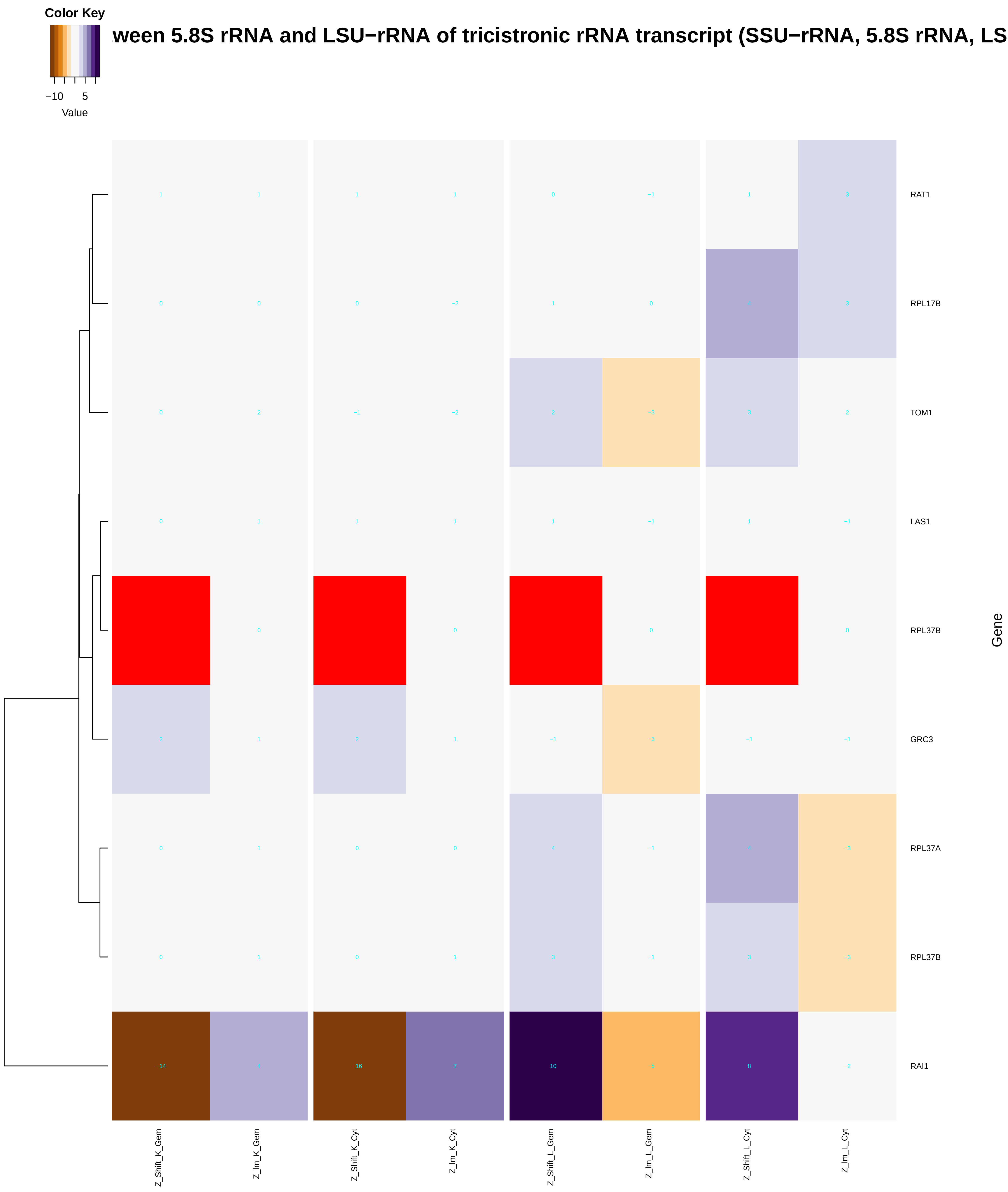

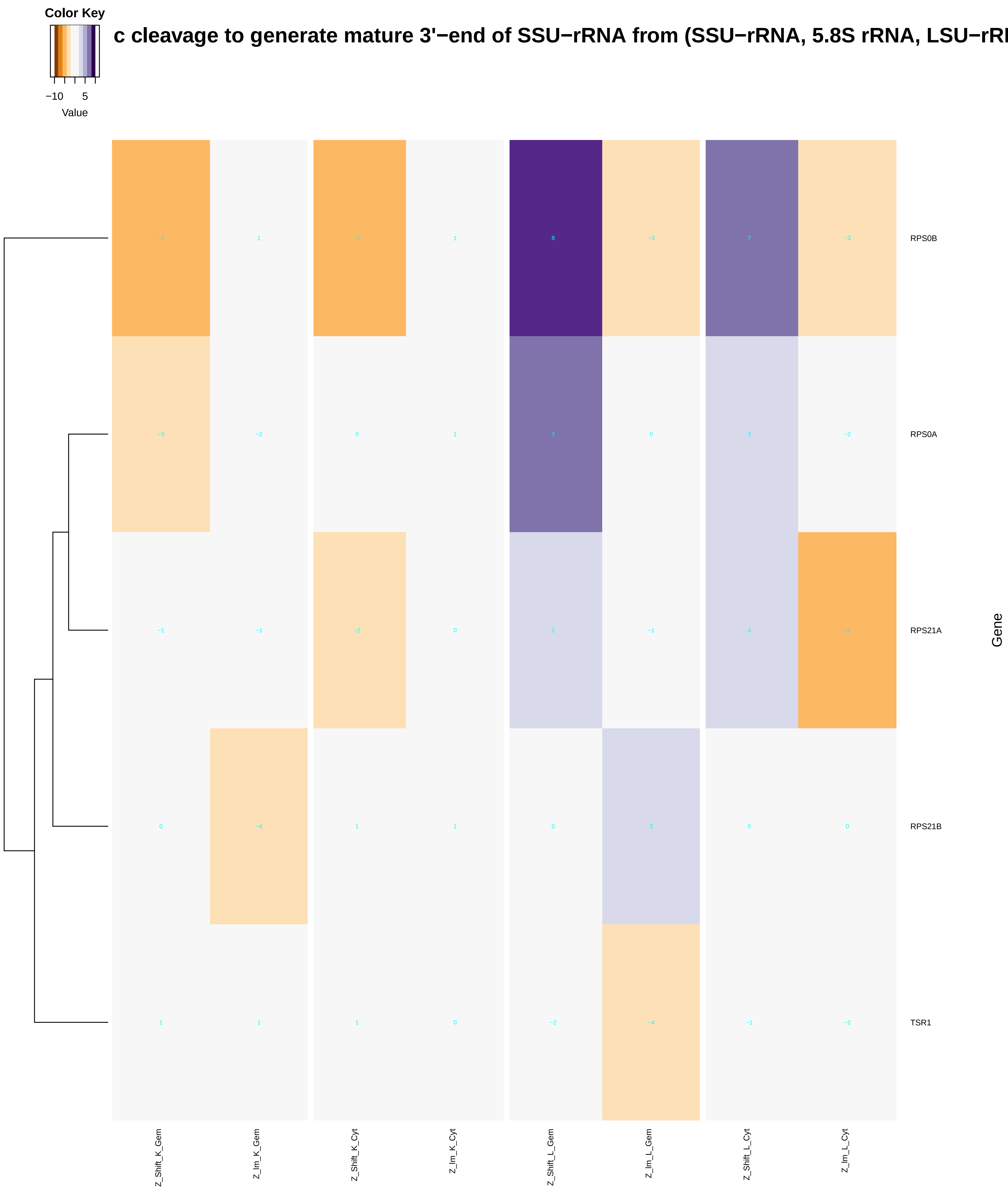

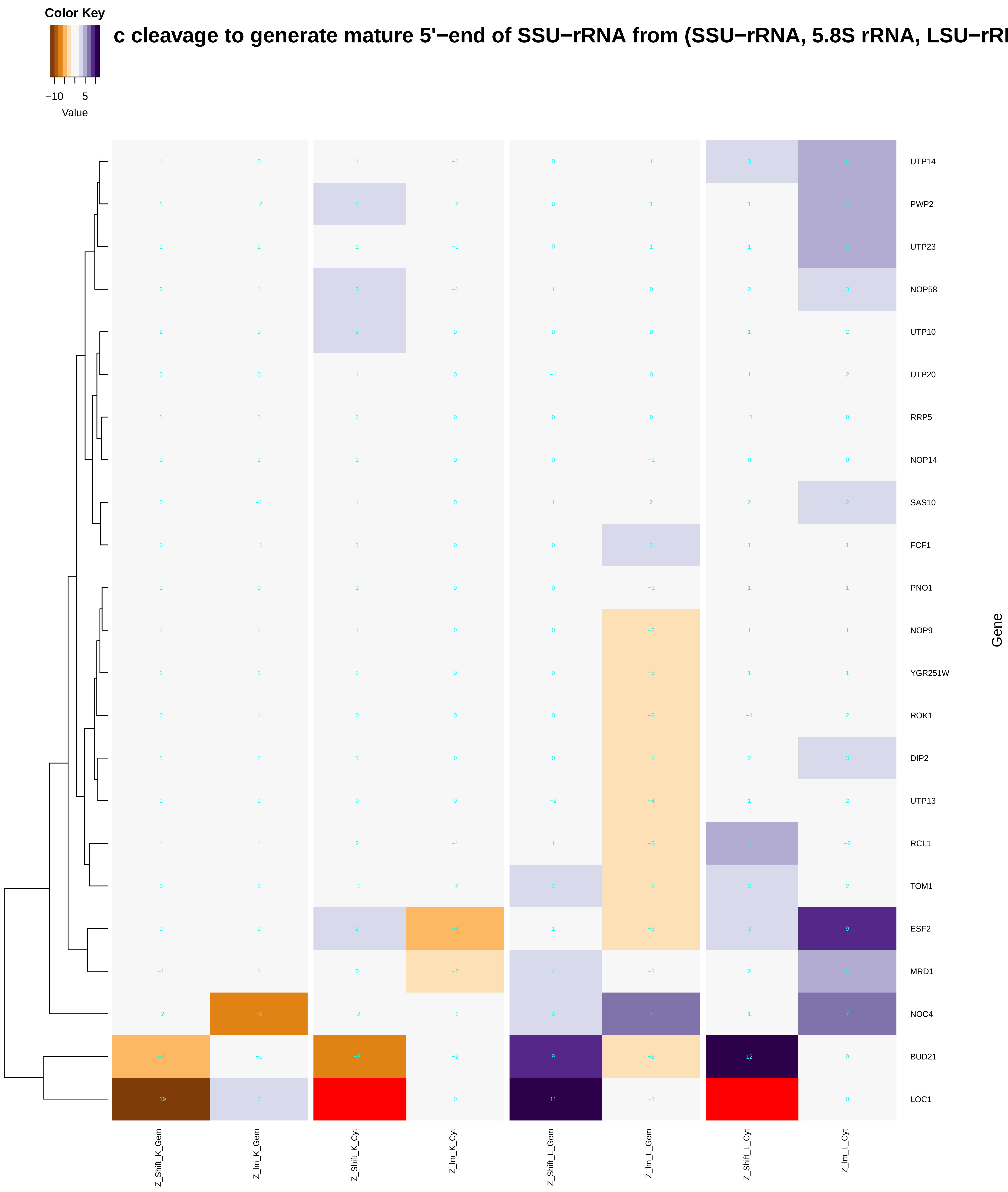

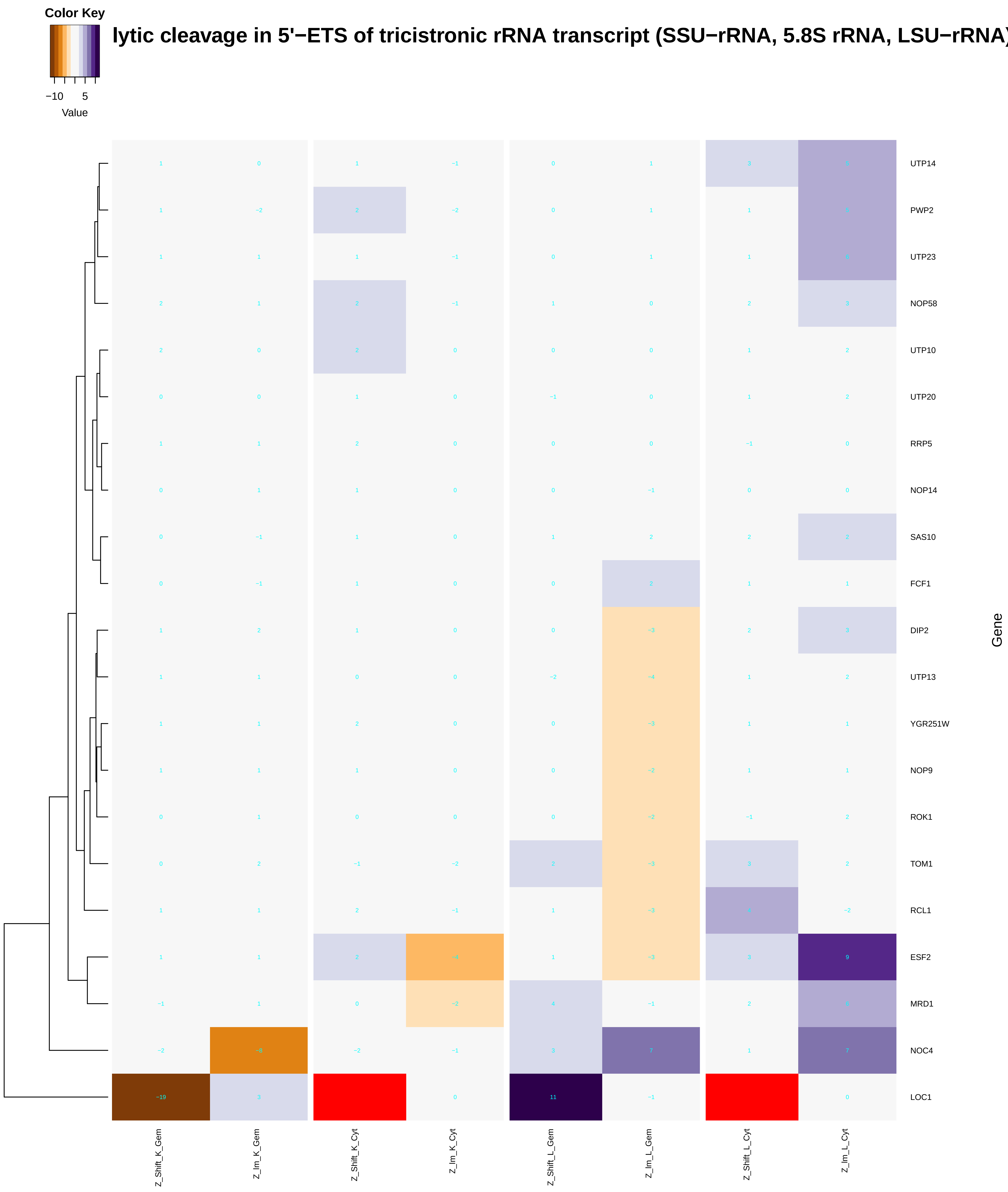

### cohesin_complex.pdf

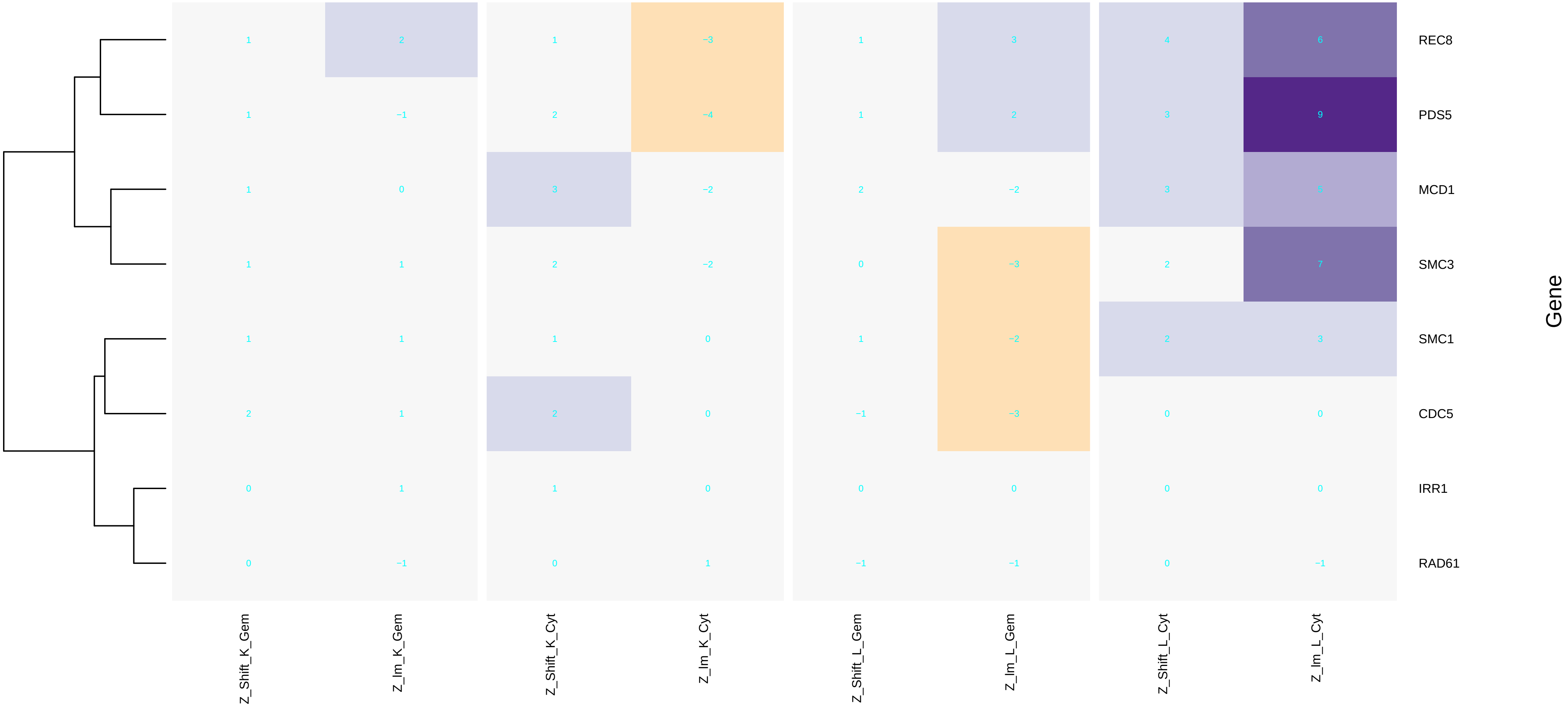

### commitment_complex.pdf

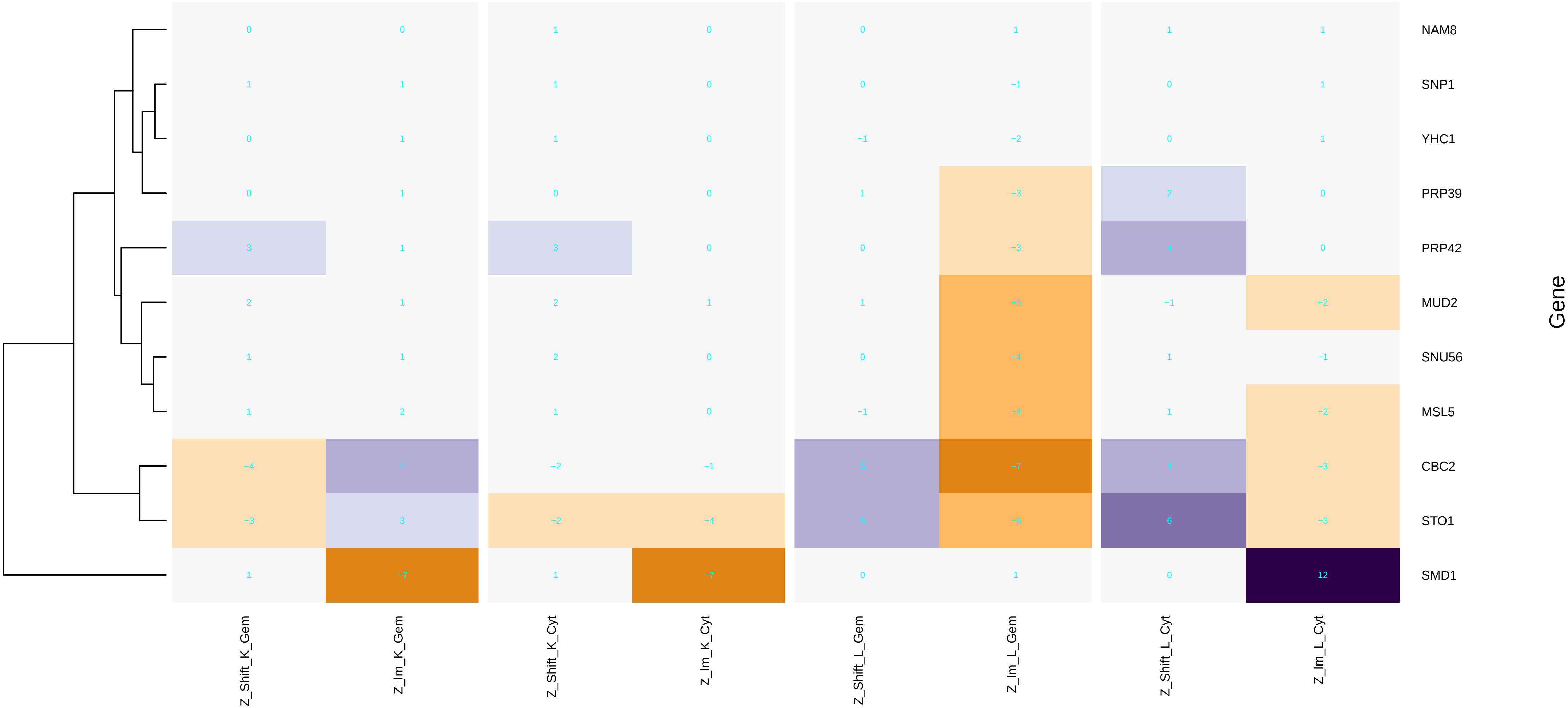

### core_mediator_complex.pdf

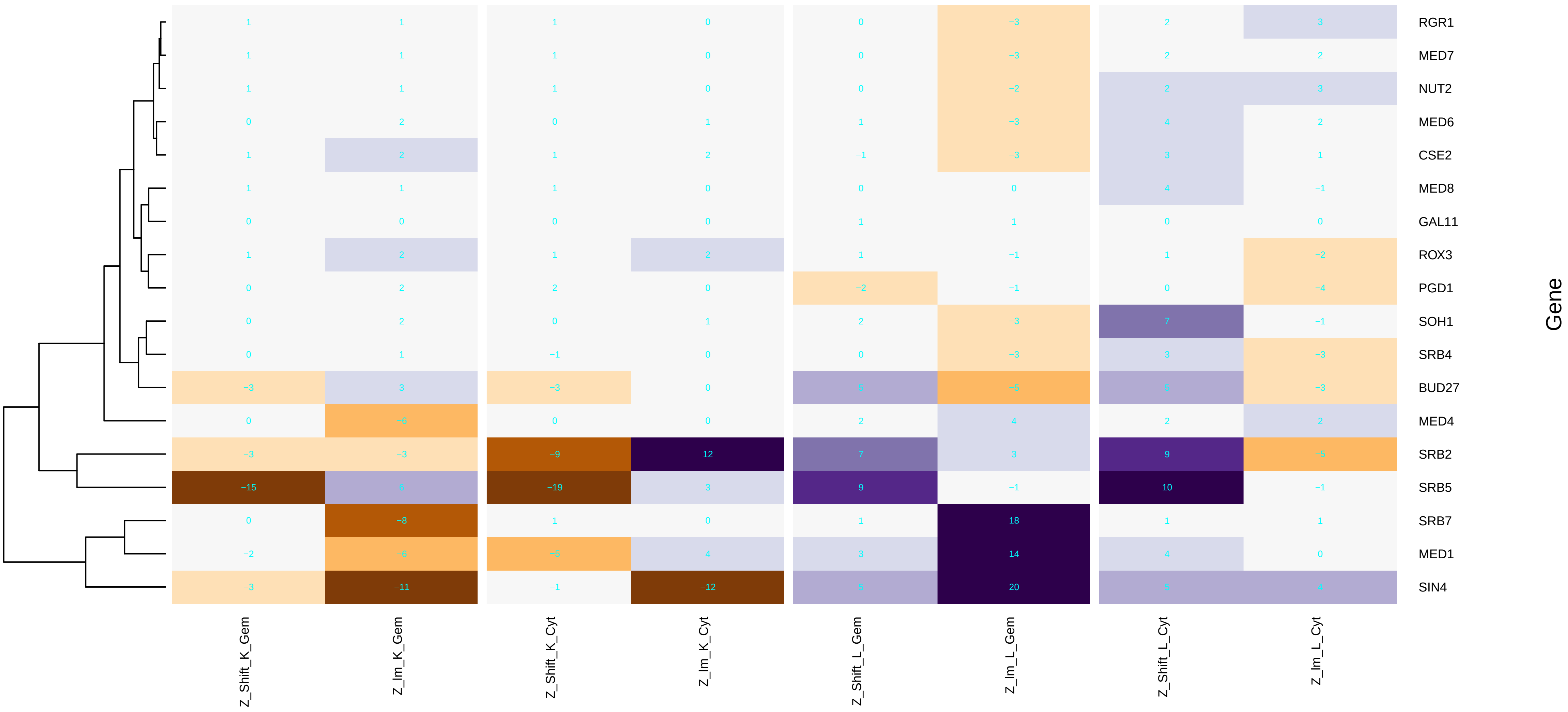

### CORVET_complex.pdf

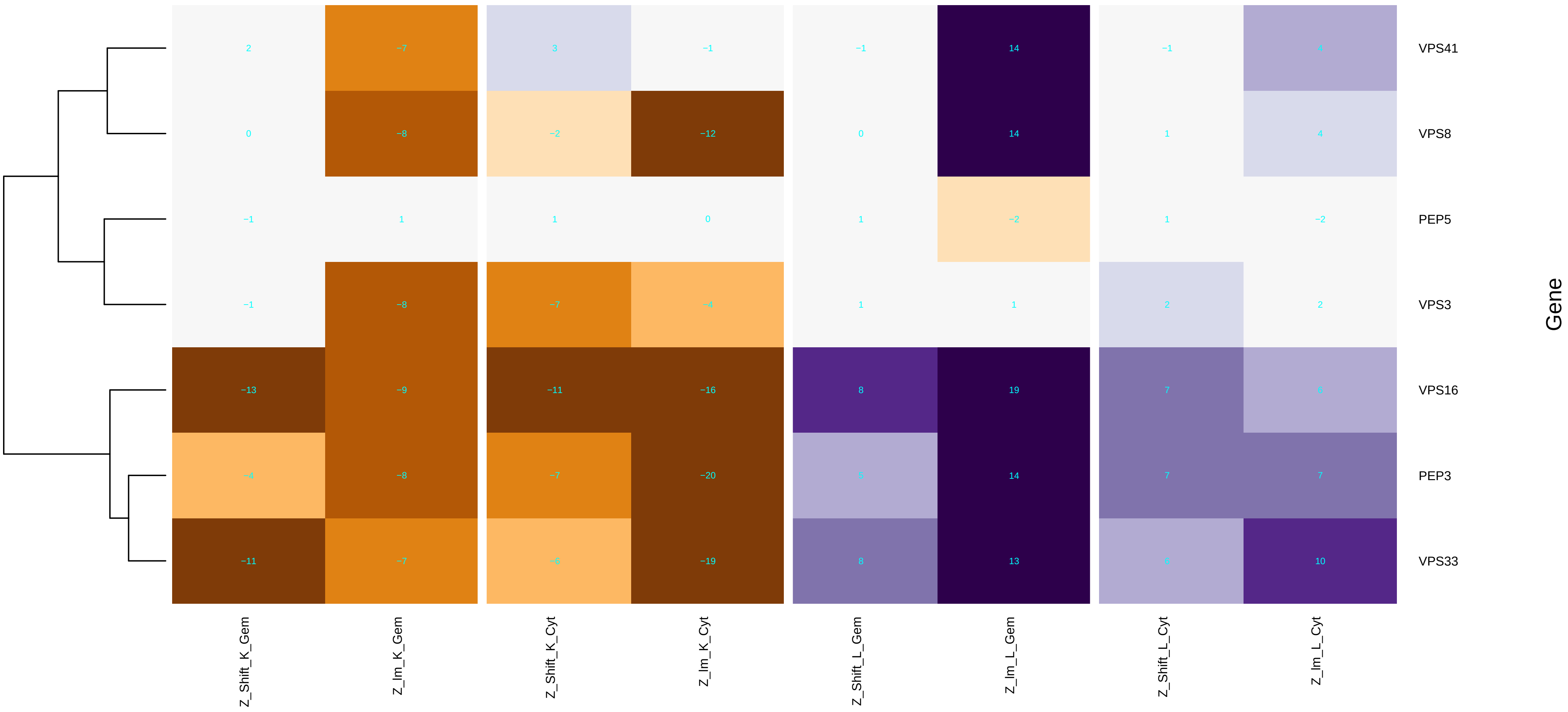

### CTDK-1_complex.pdf

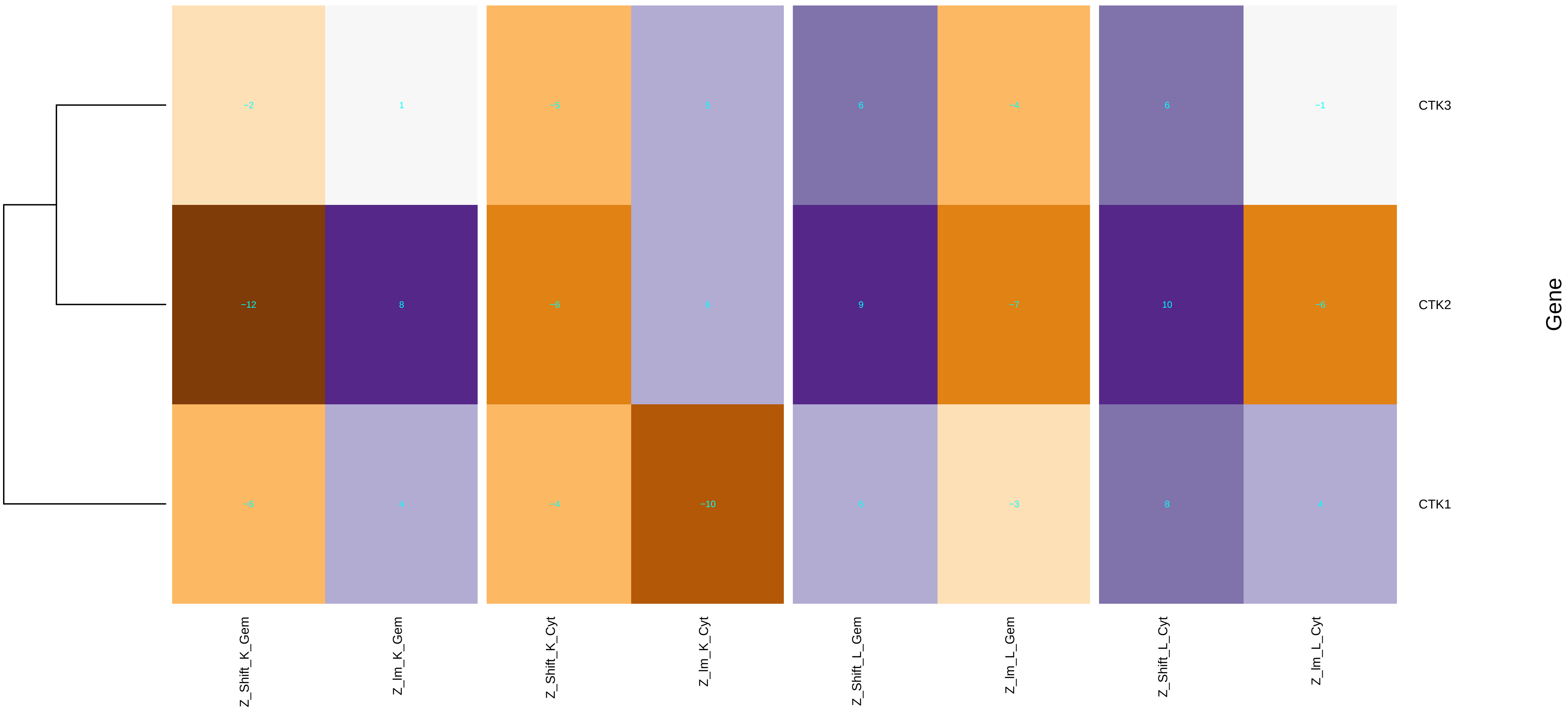

### Cul4-RING_E3_ubiquitin_ligase_complex.pdf

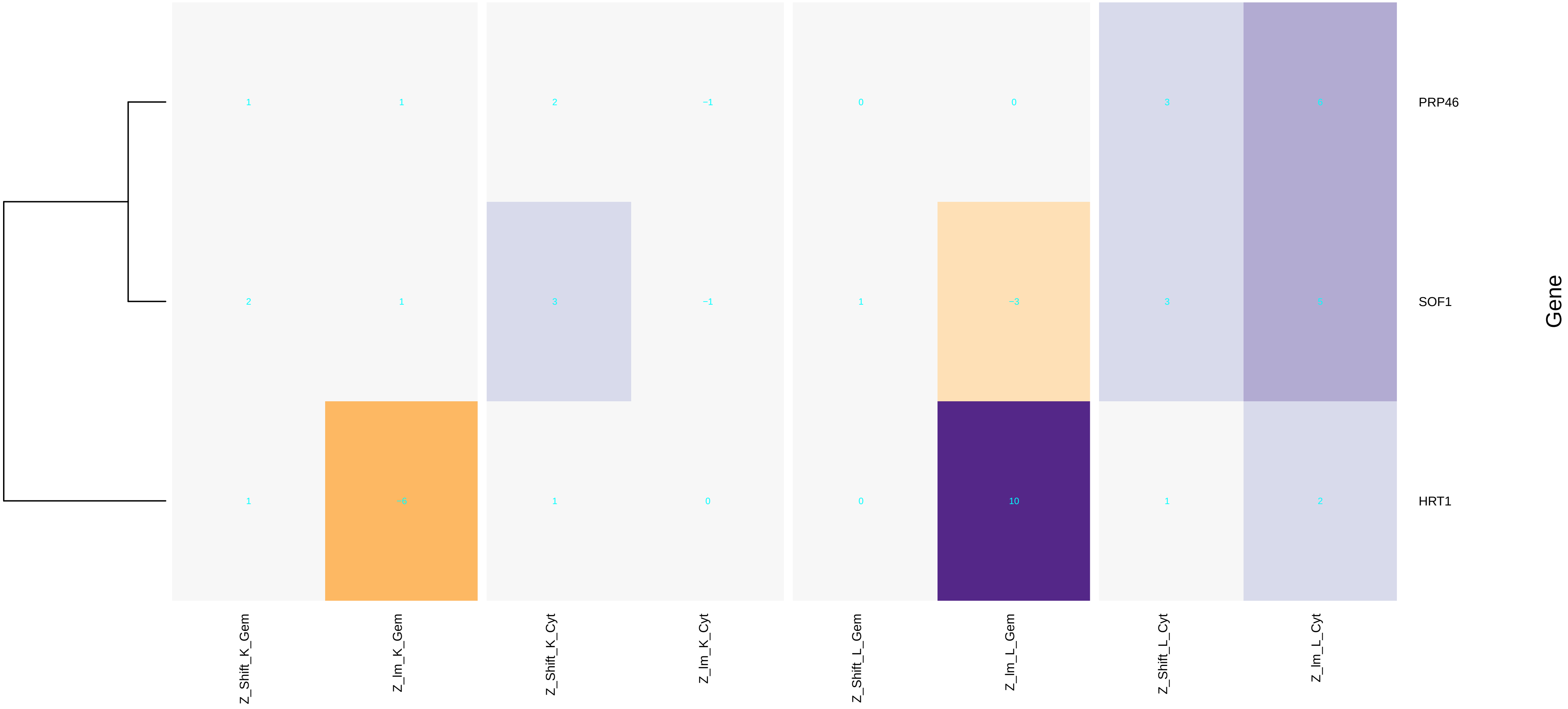

### cytosol.pdf

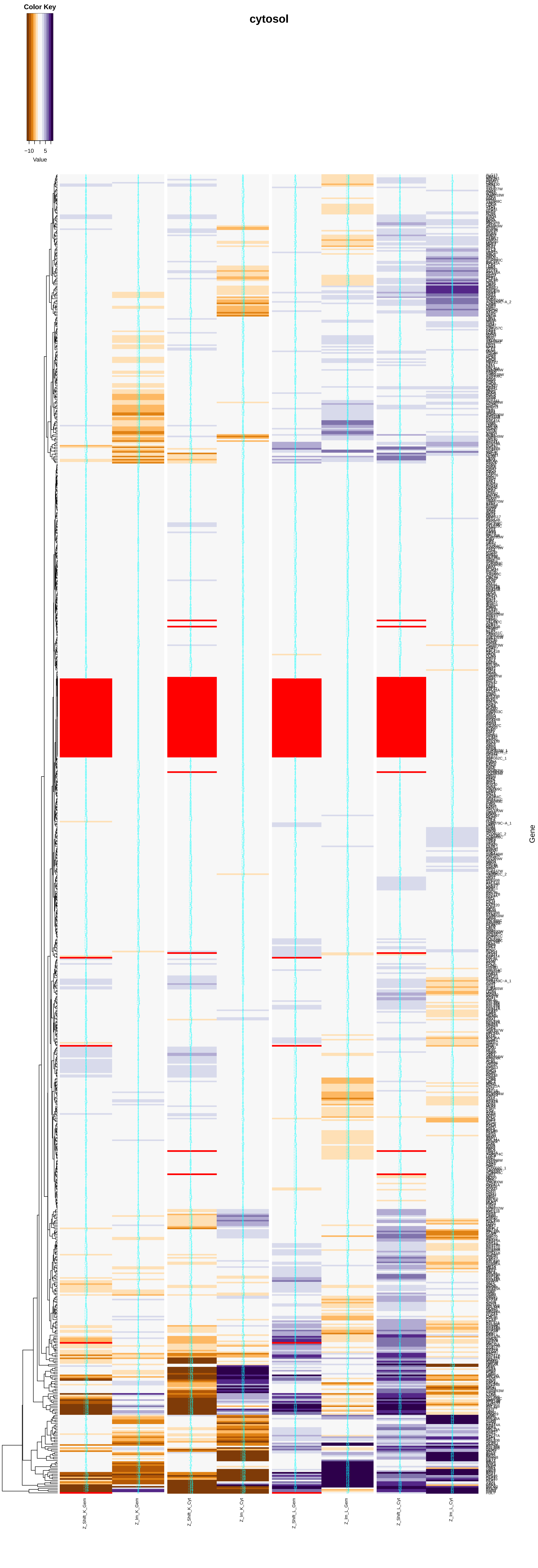

### cytosolic_proteasome_complex.pdf

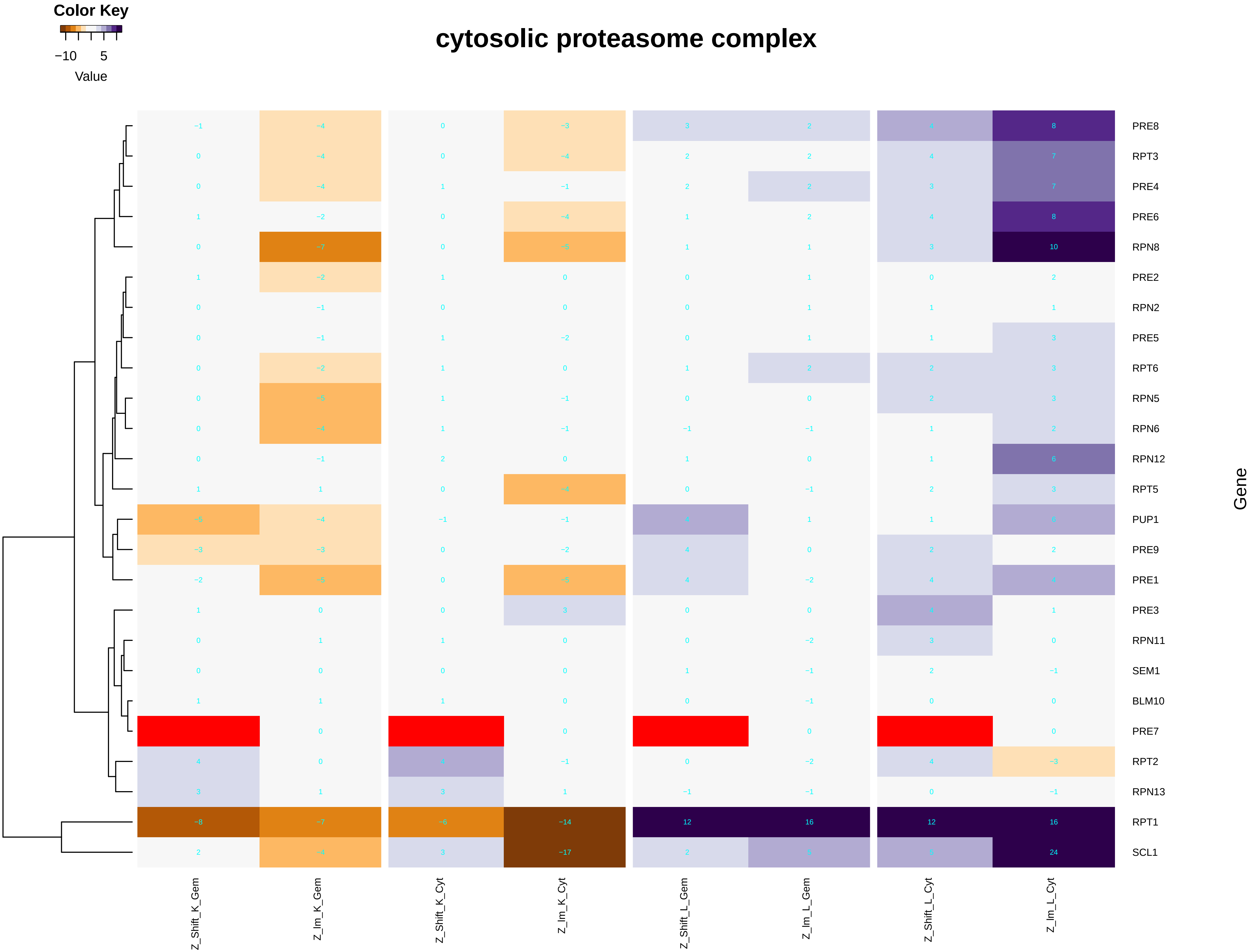

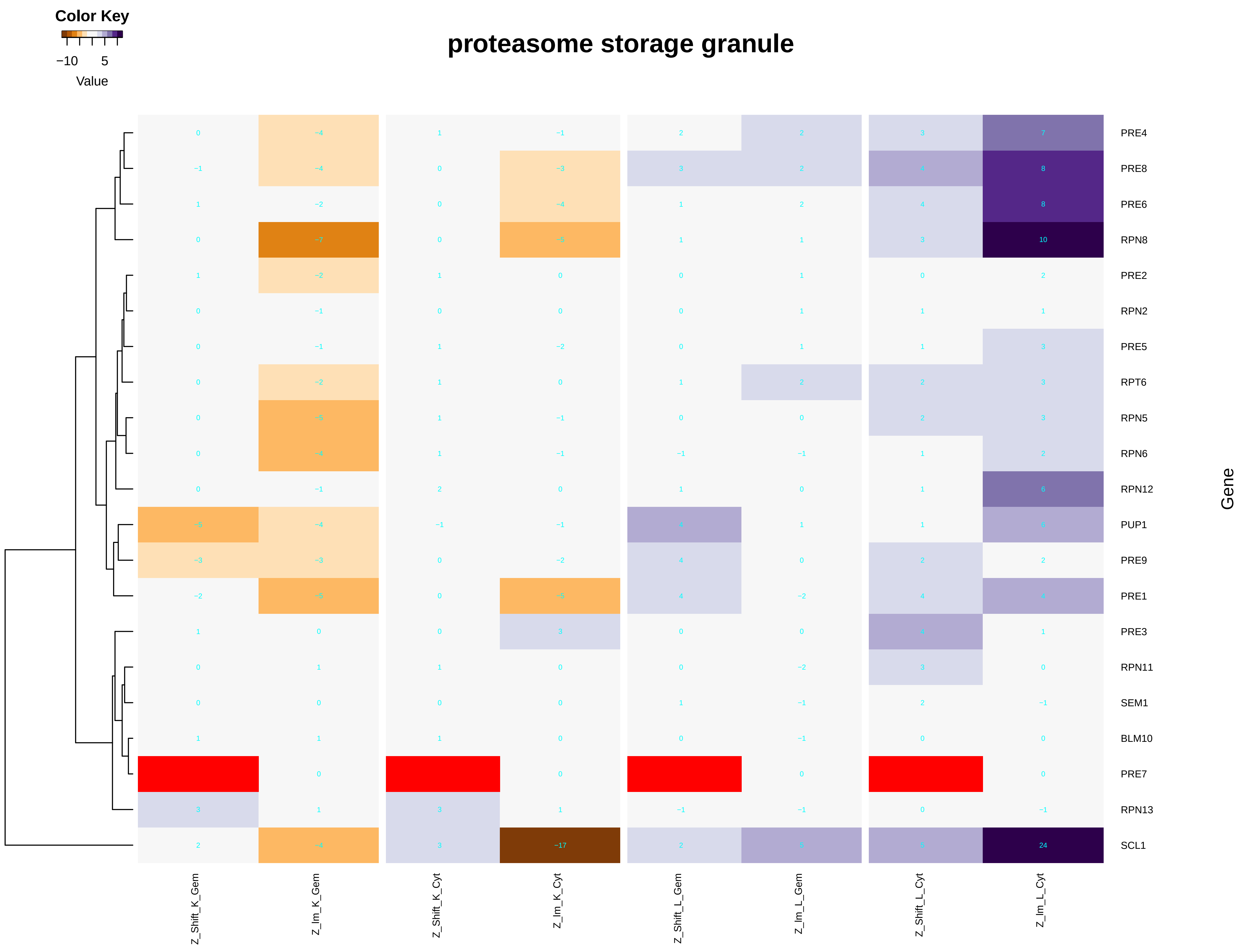

### cytosolic_ribosome.pdf

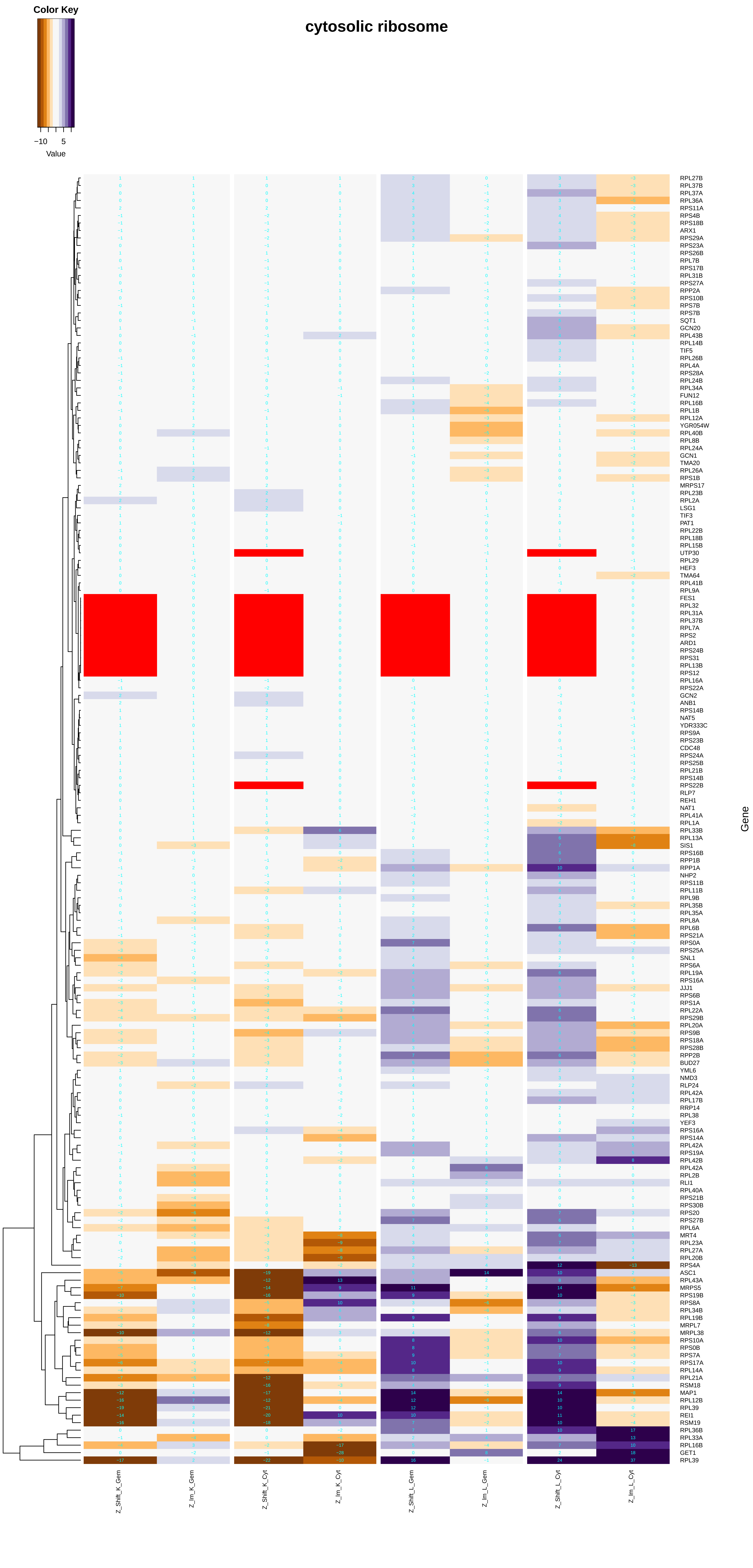

### cytosolic_small_ribosomal_subunit.pdf

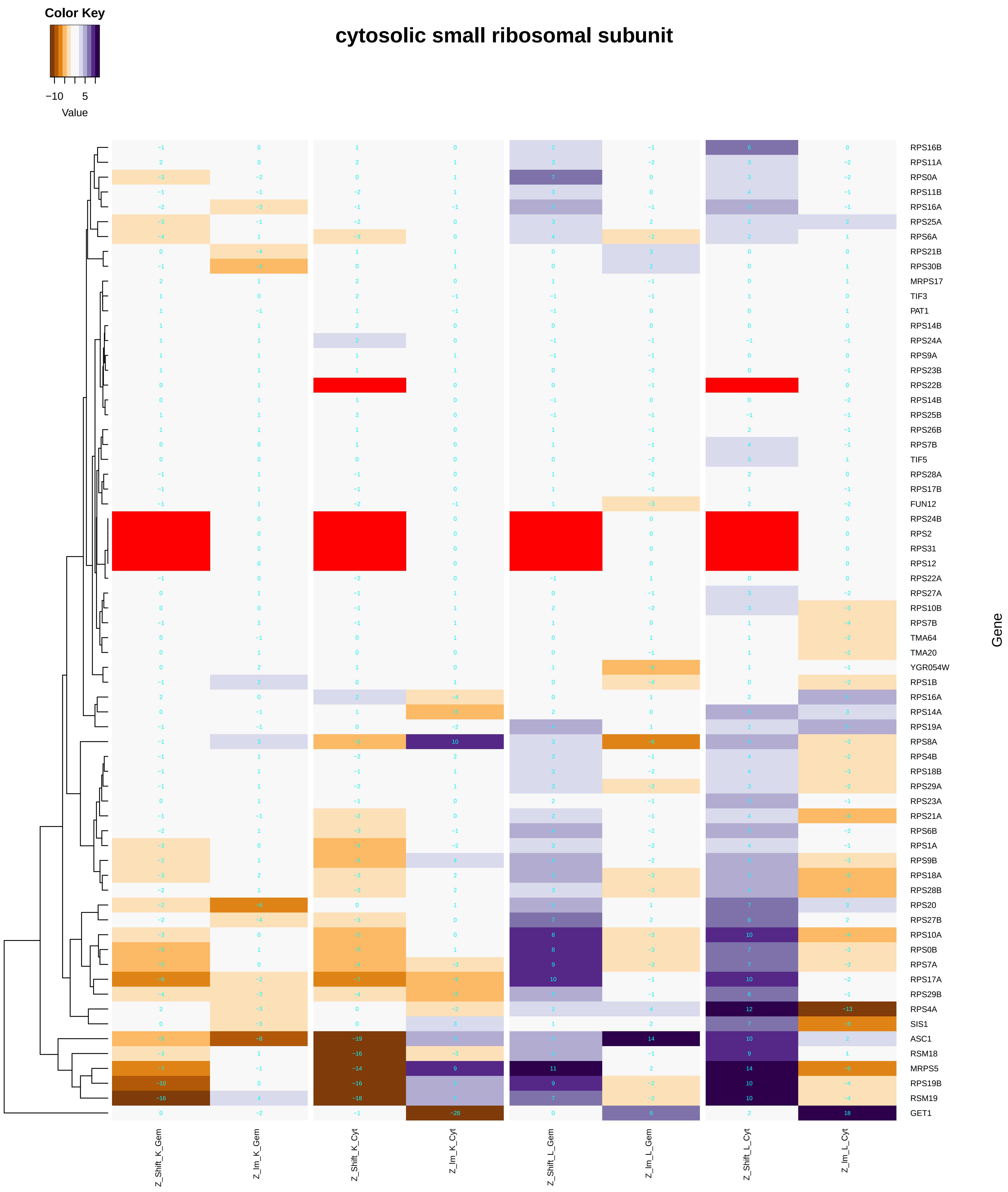

### dihydrofolate_metabolic_process.pdf

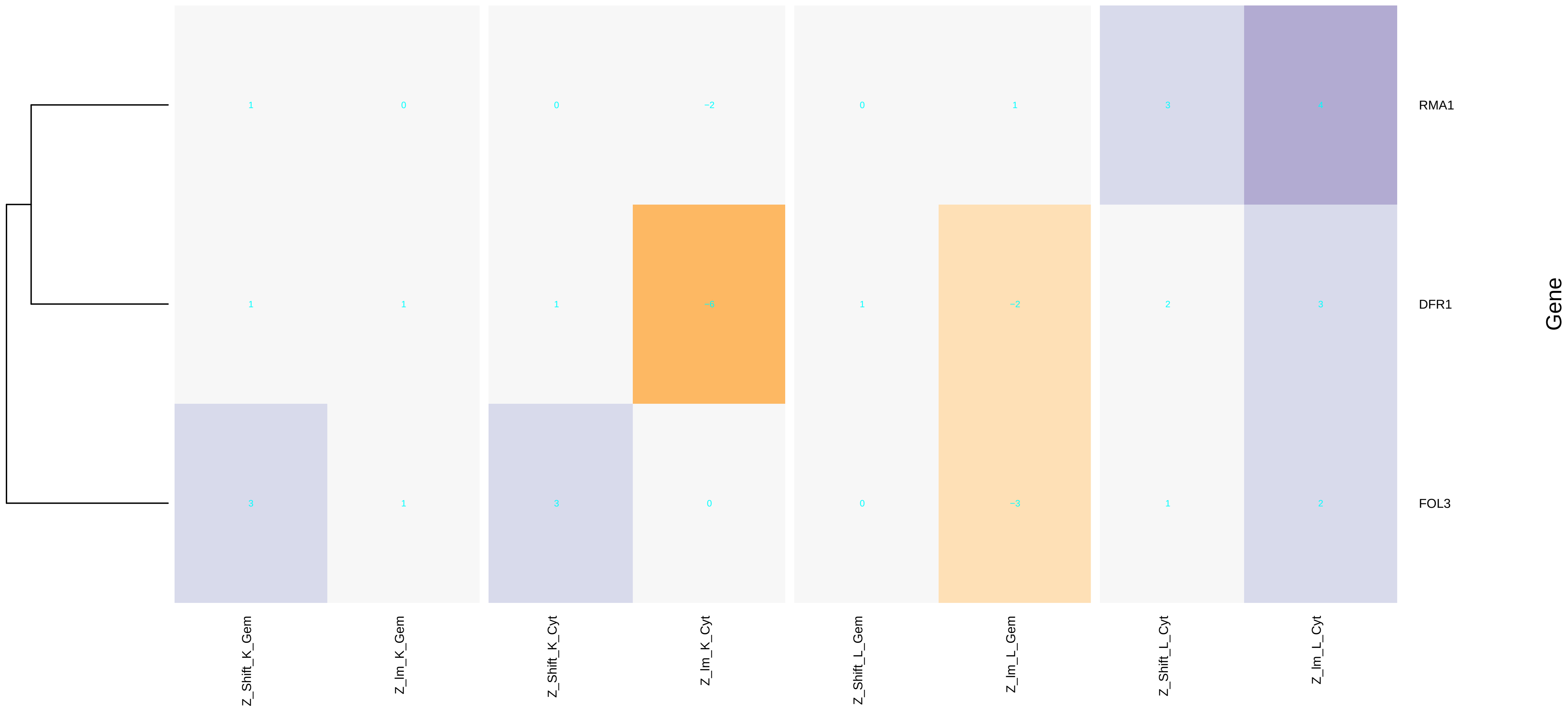

### DNA-dependent_ATPase_activity.pdf

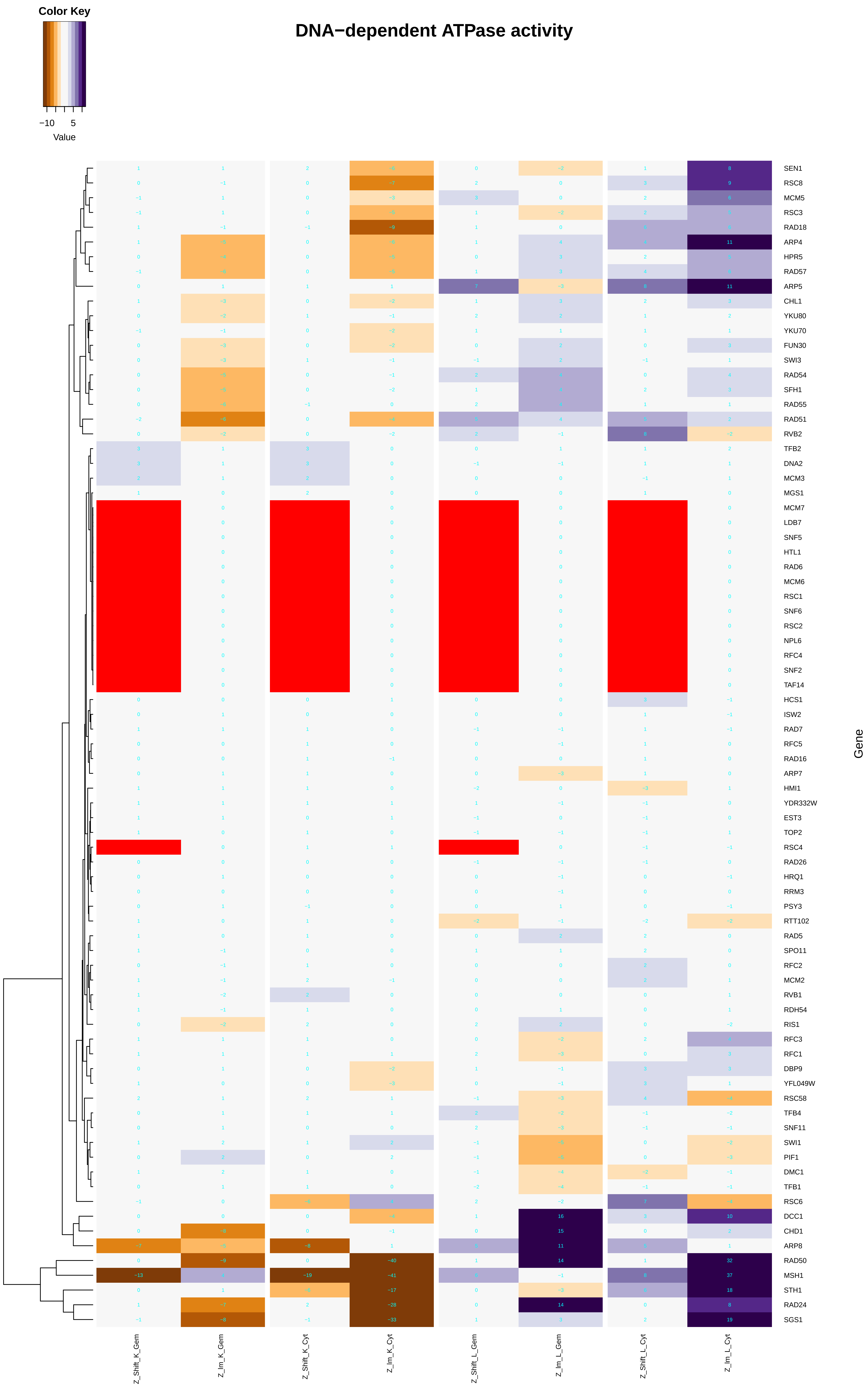

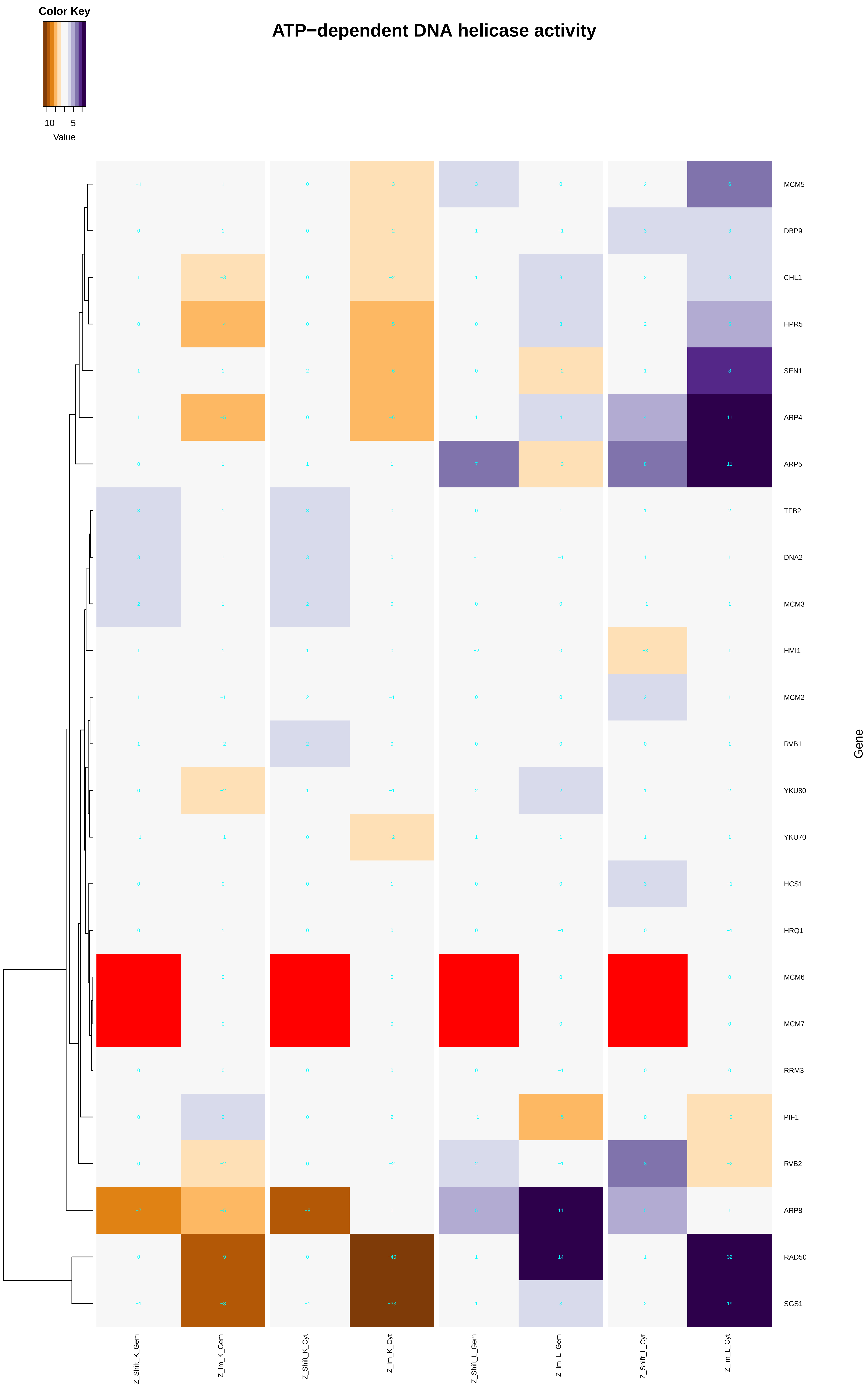

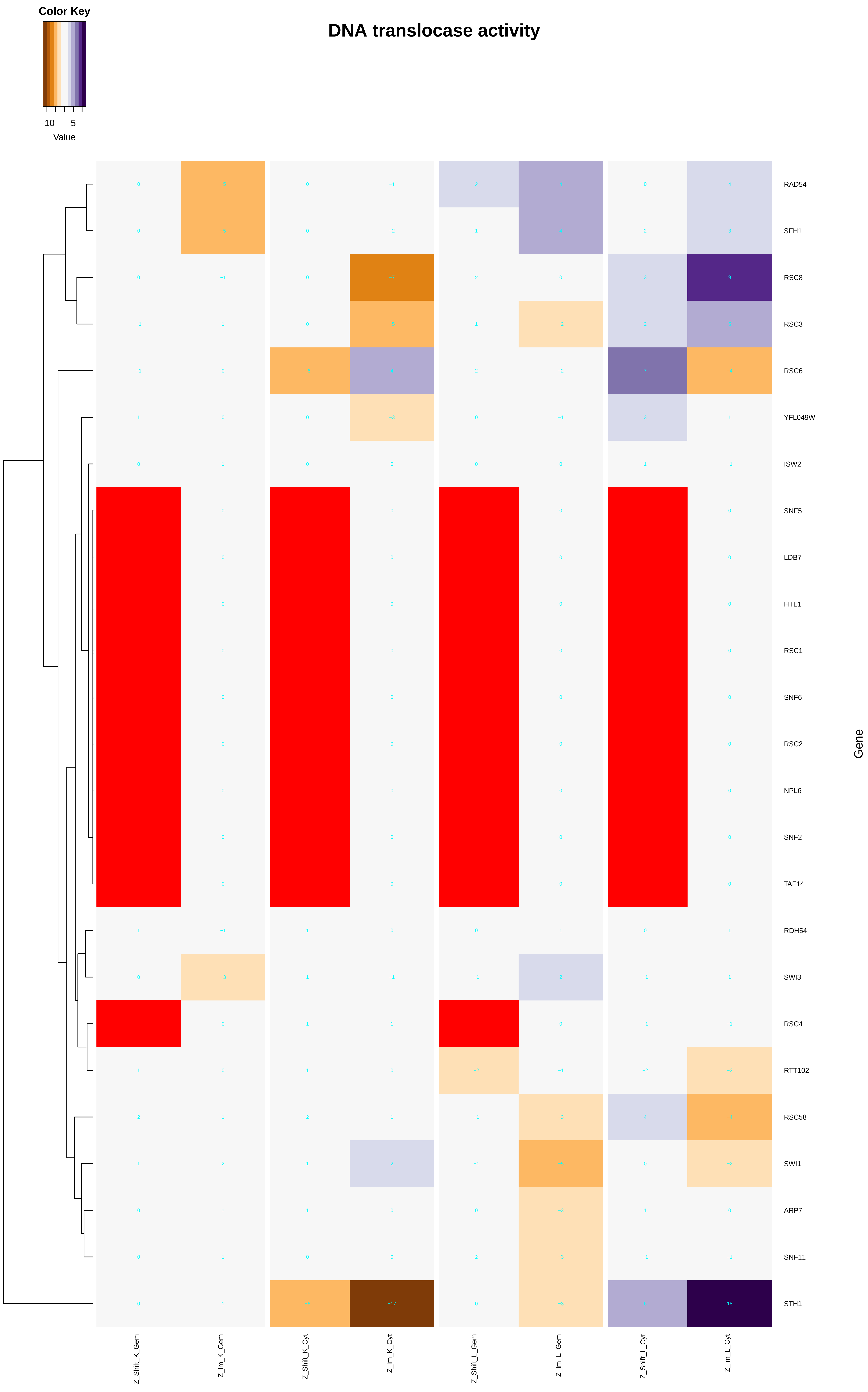

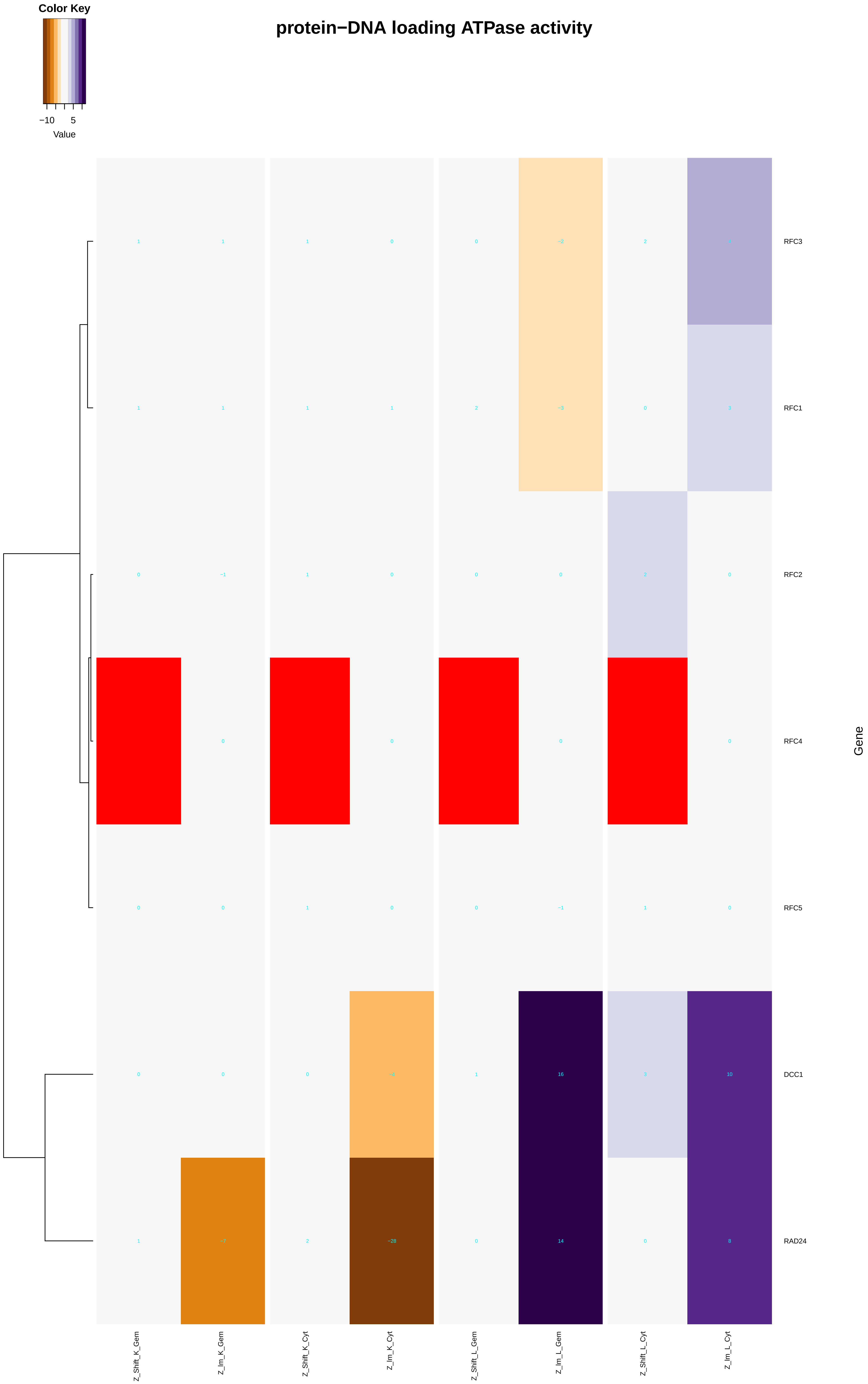

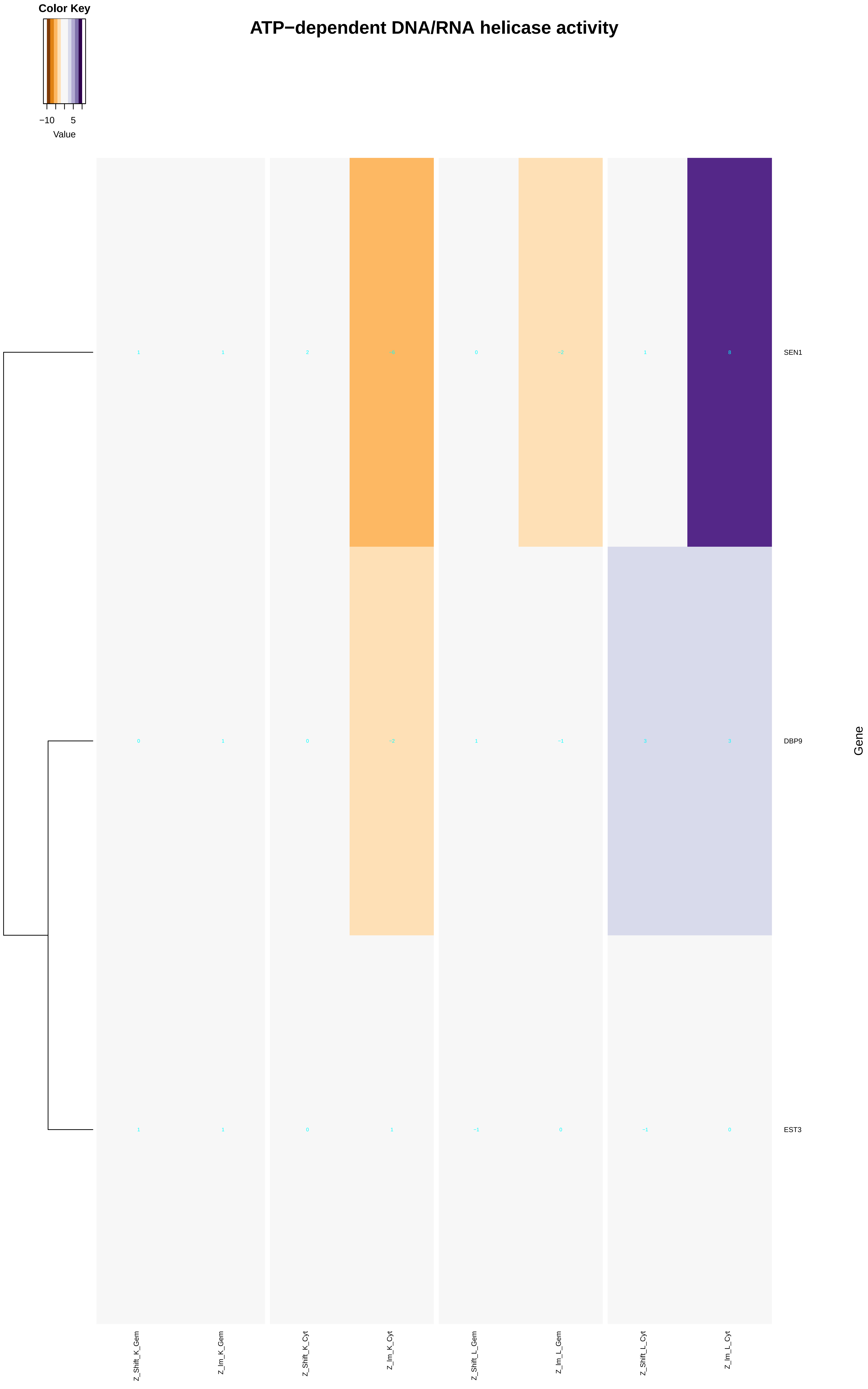

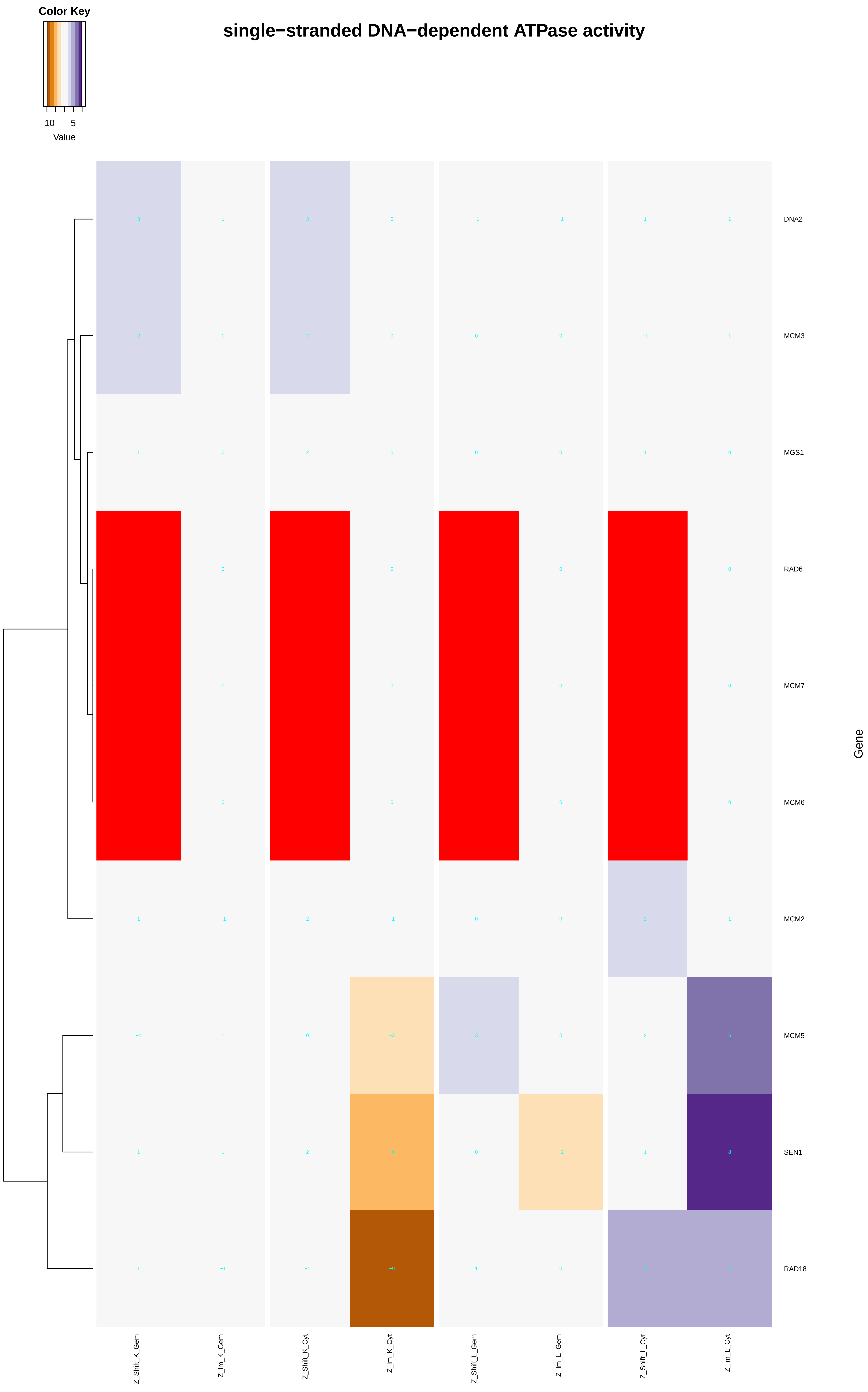

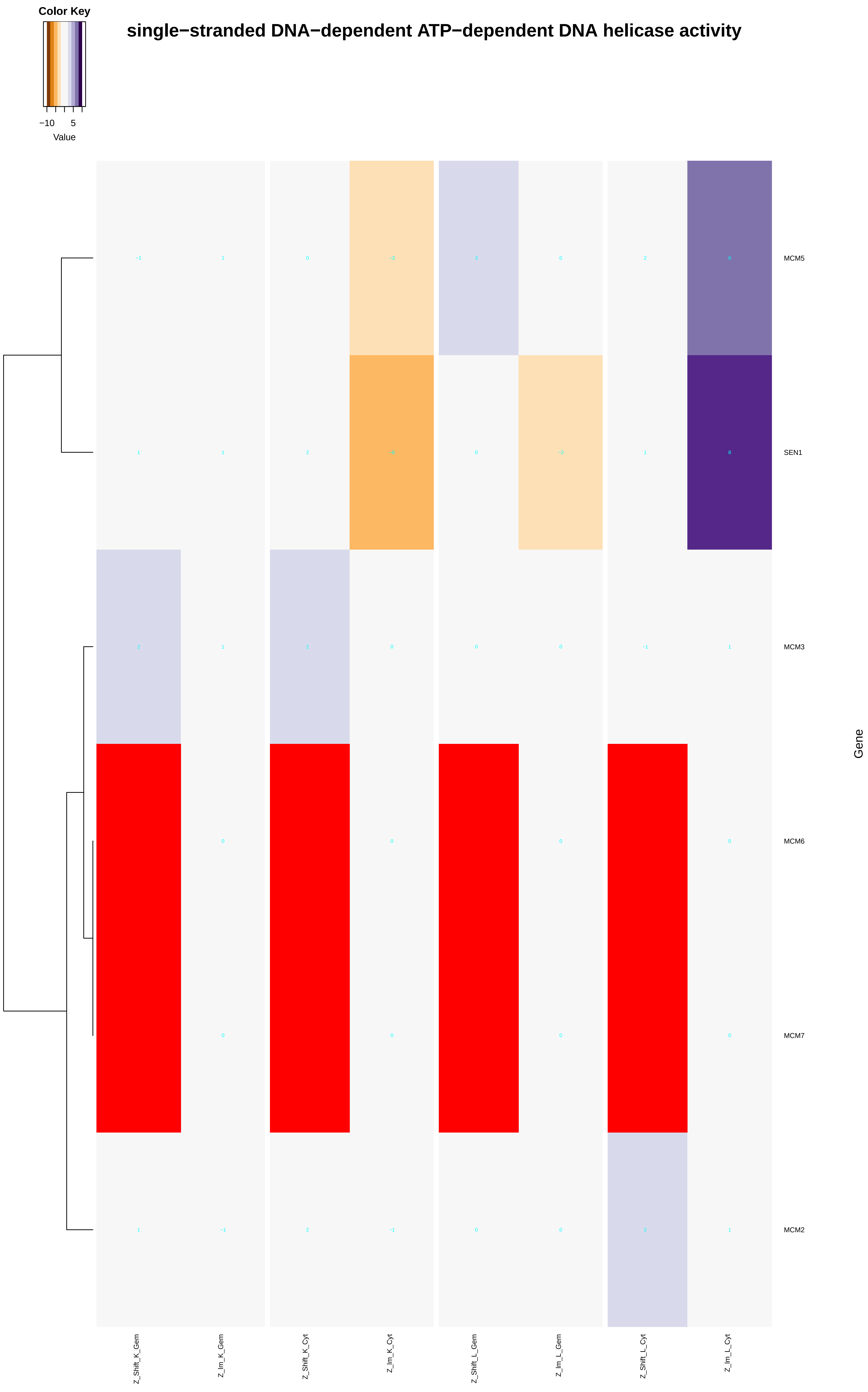

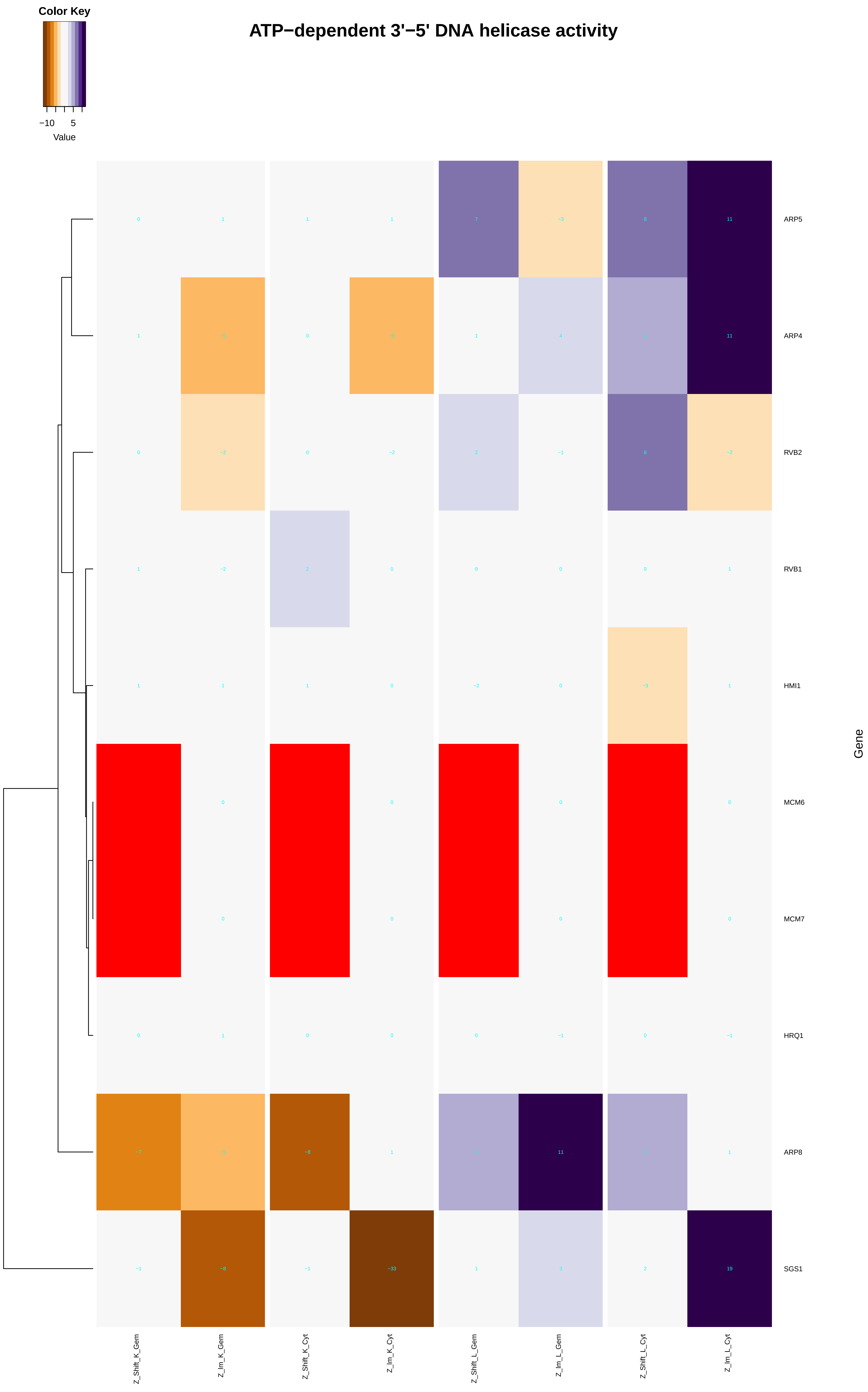

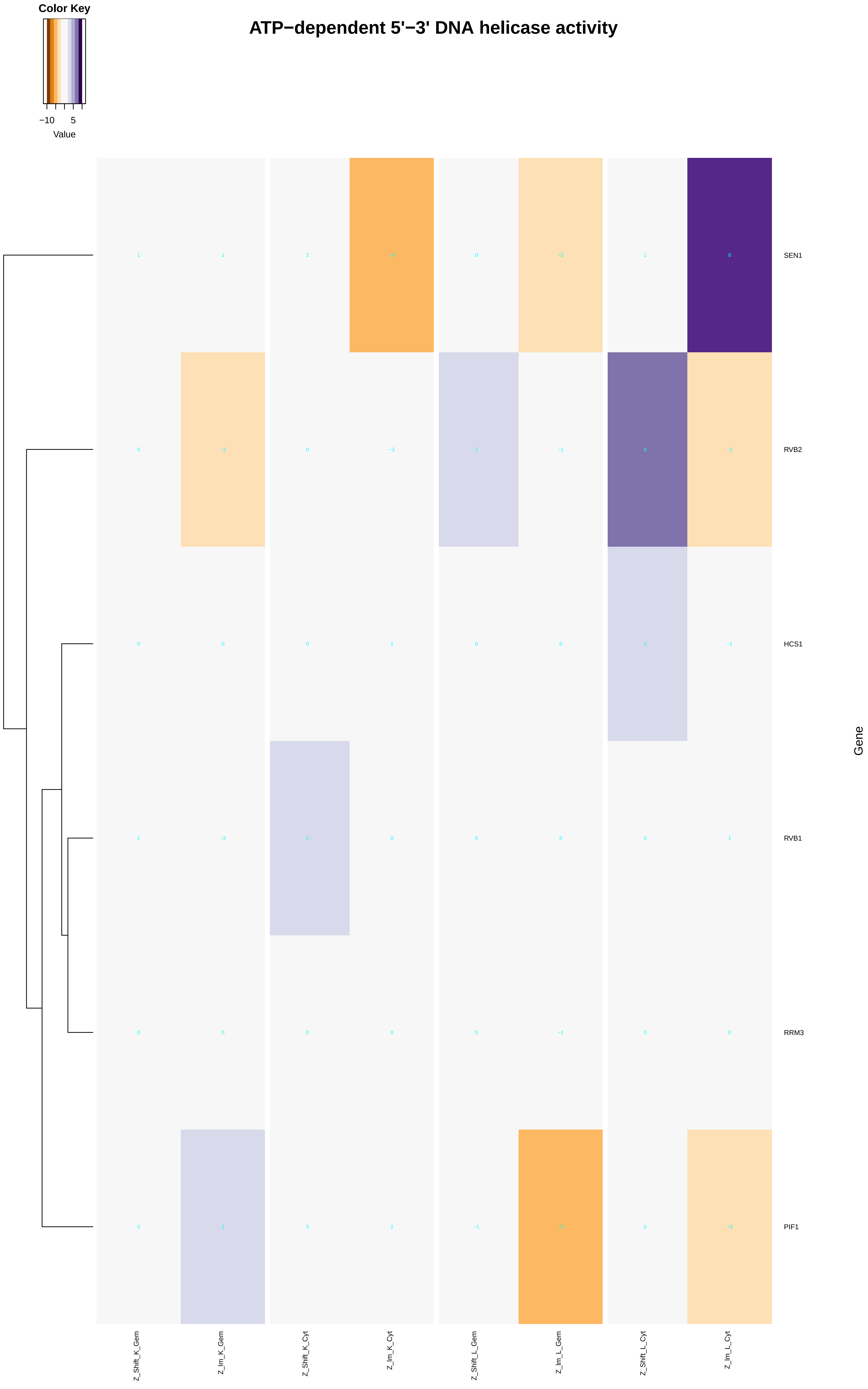

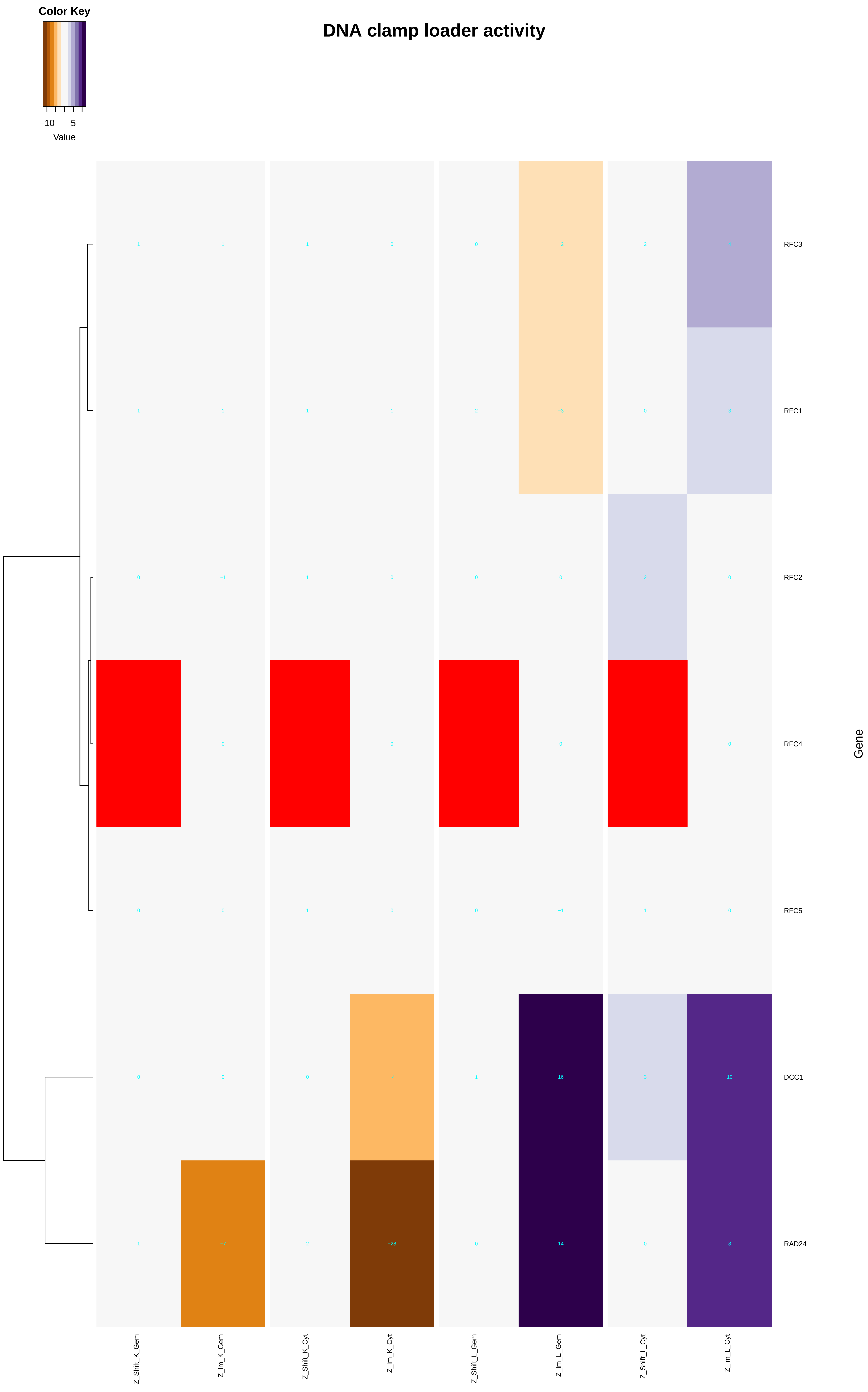

### DNA-dependent_DNA_replication.pdf

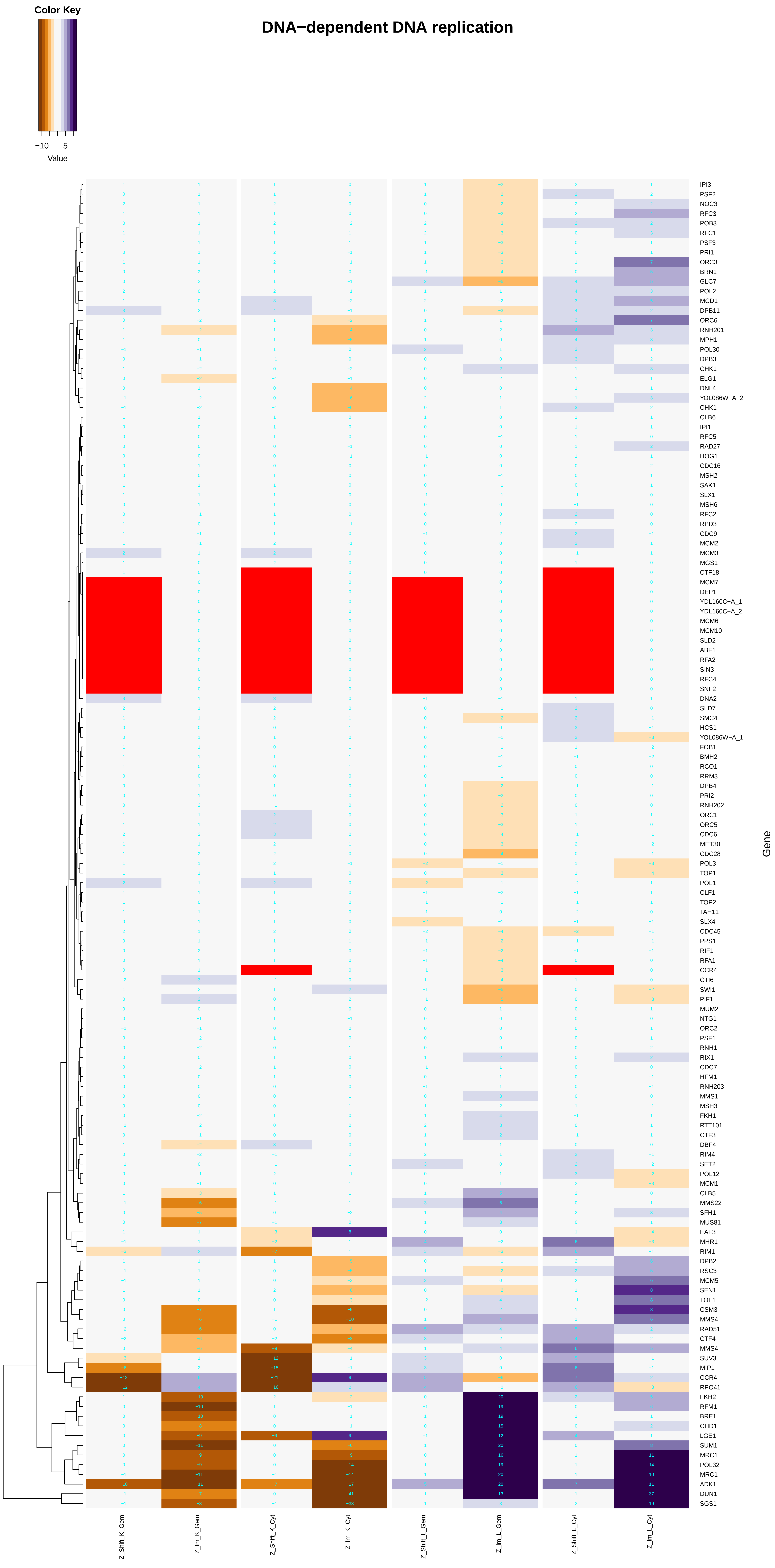

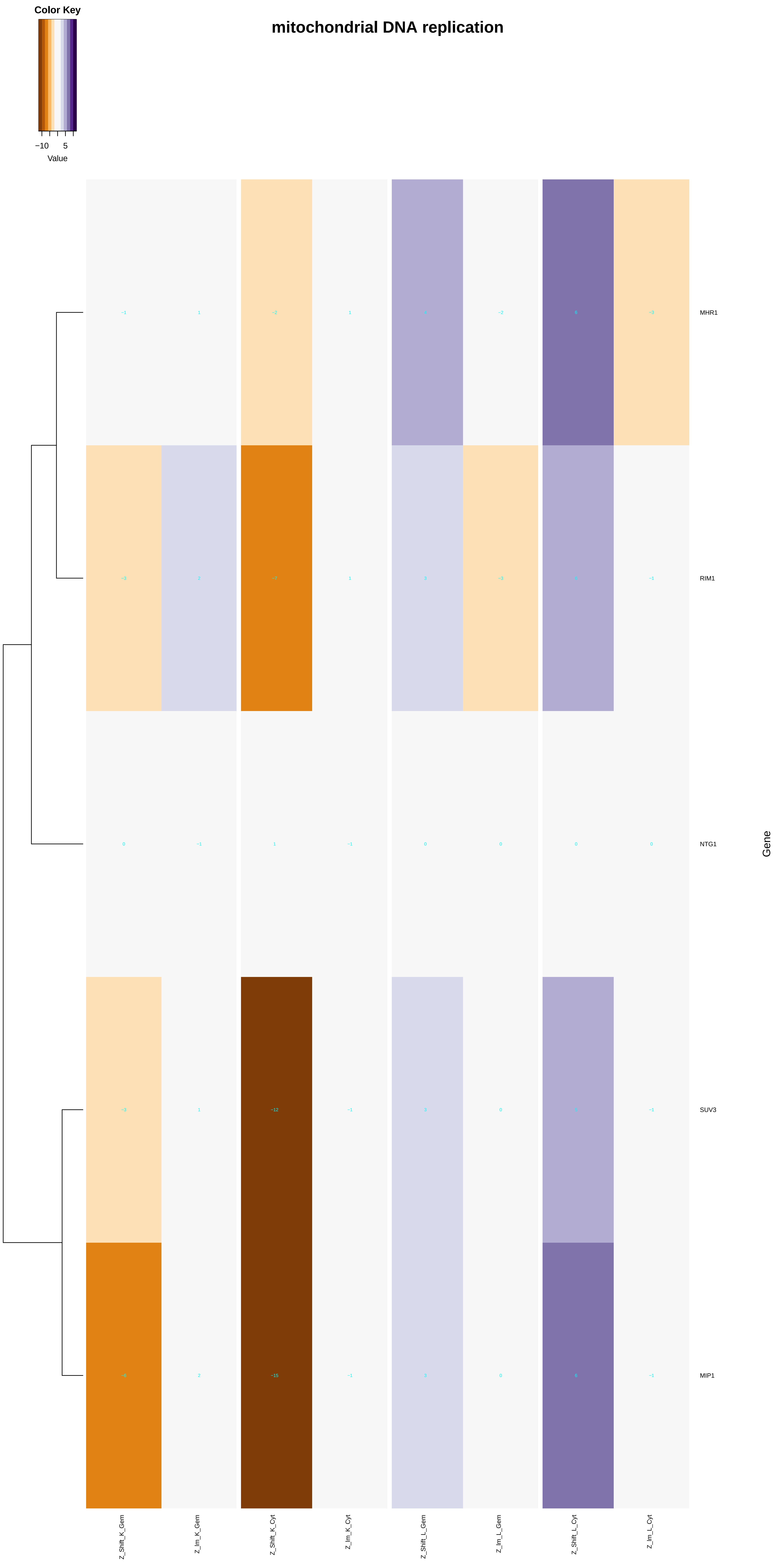

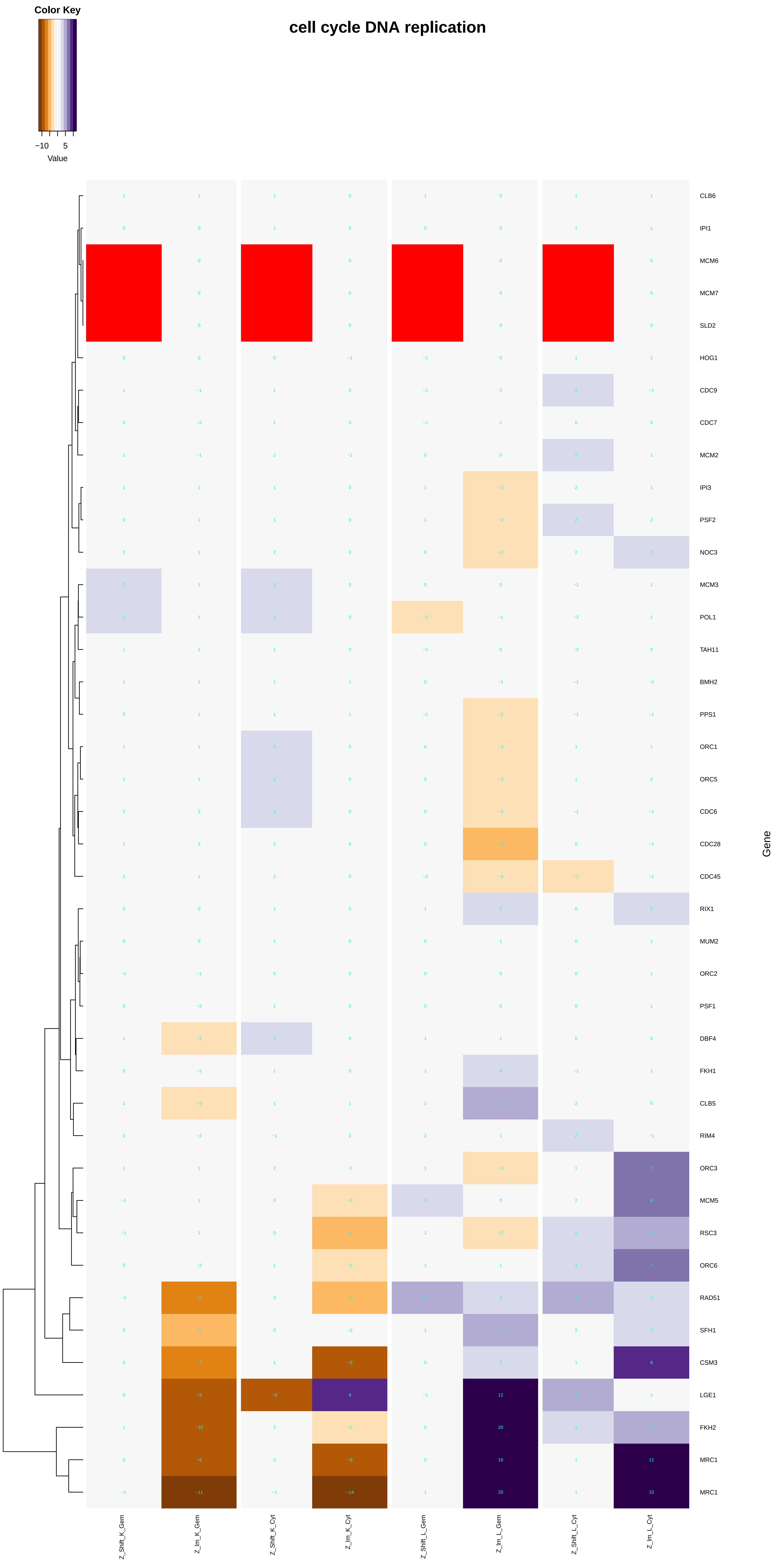

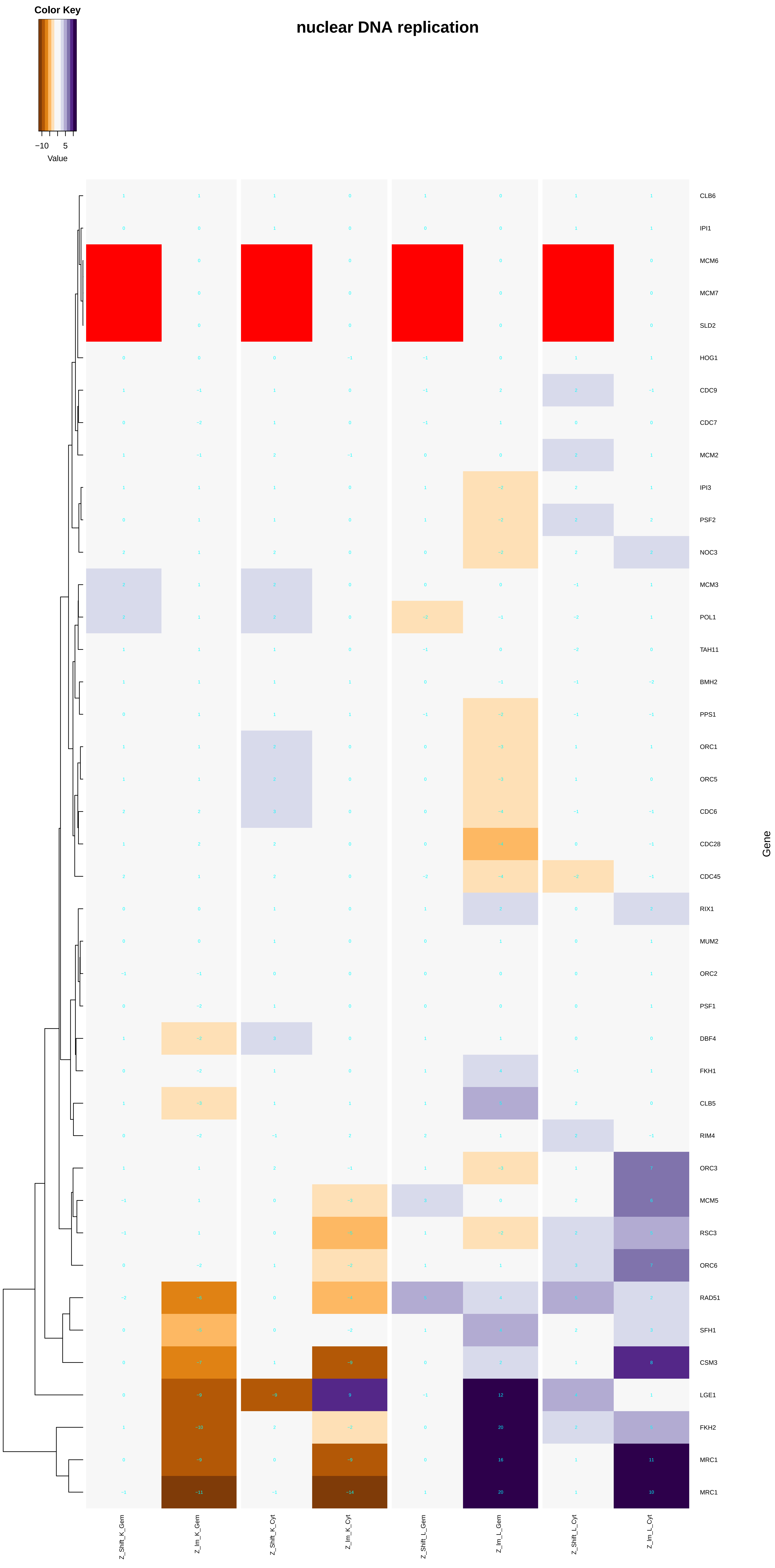

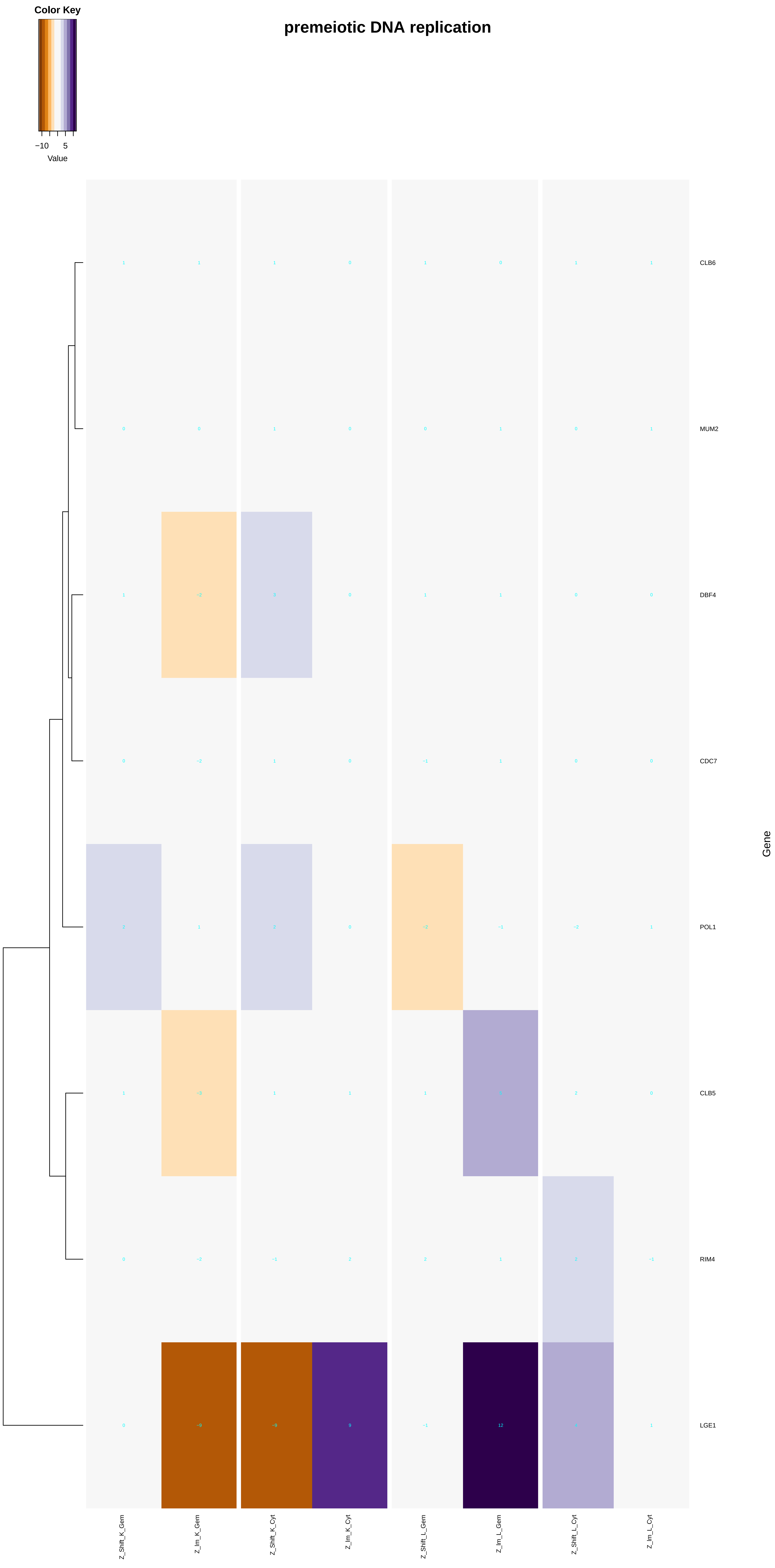

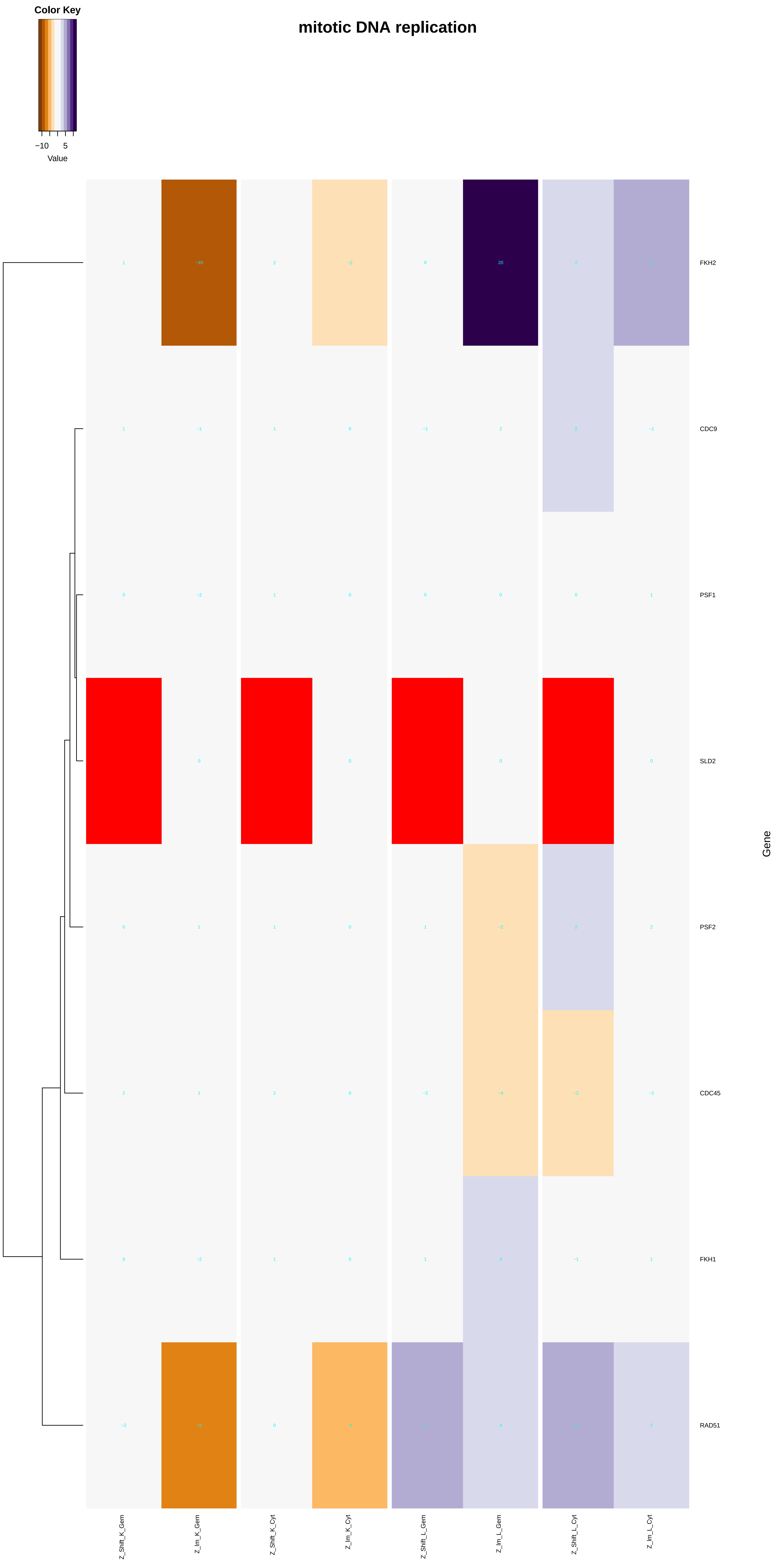

### DNA-directed_5'-3'_RNA_polymerase_activity.pdf

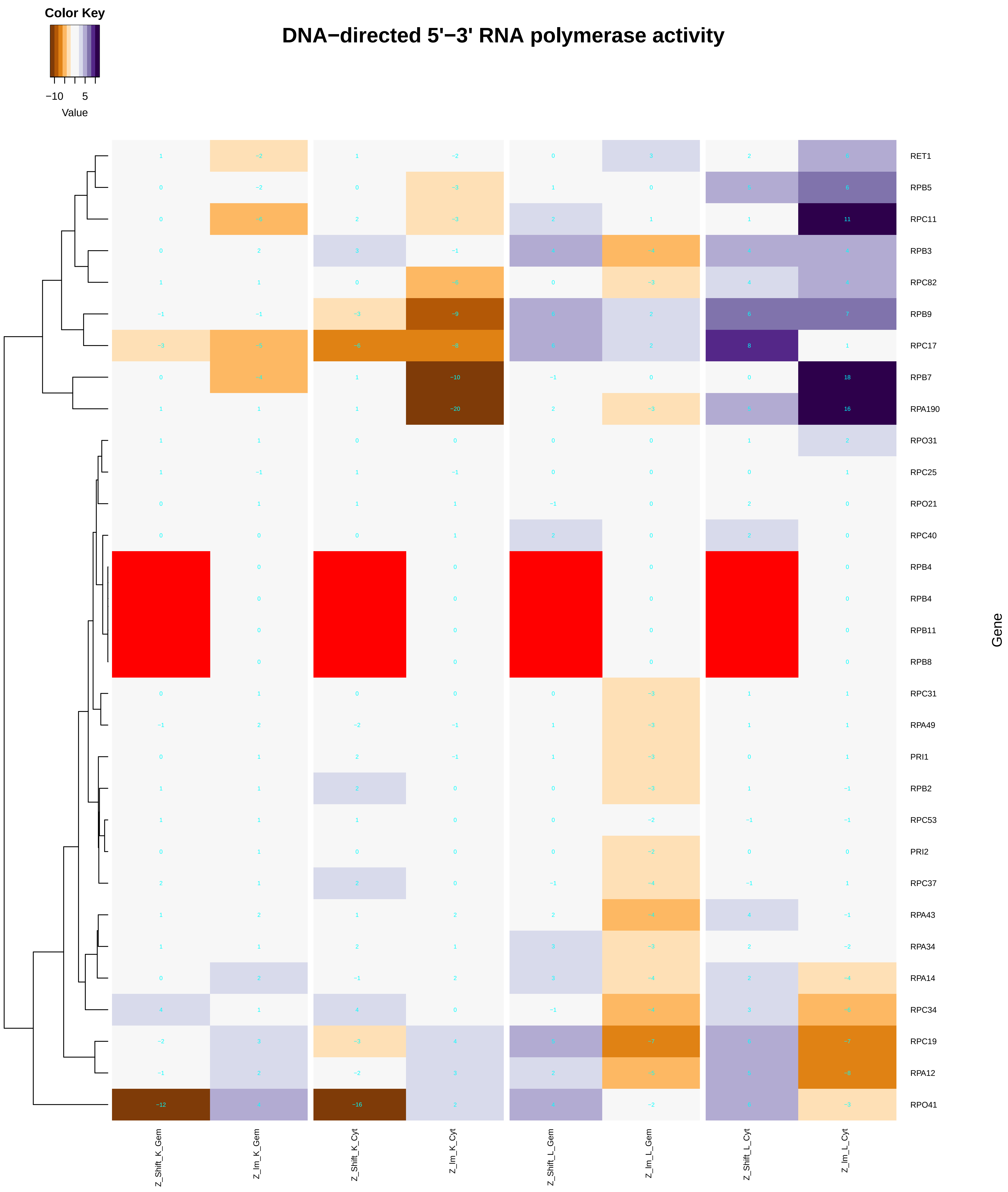

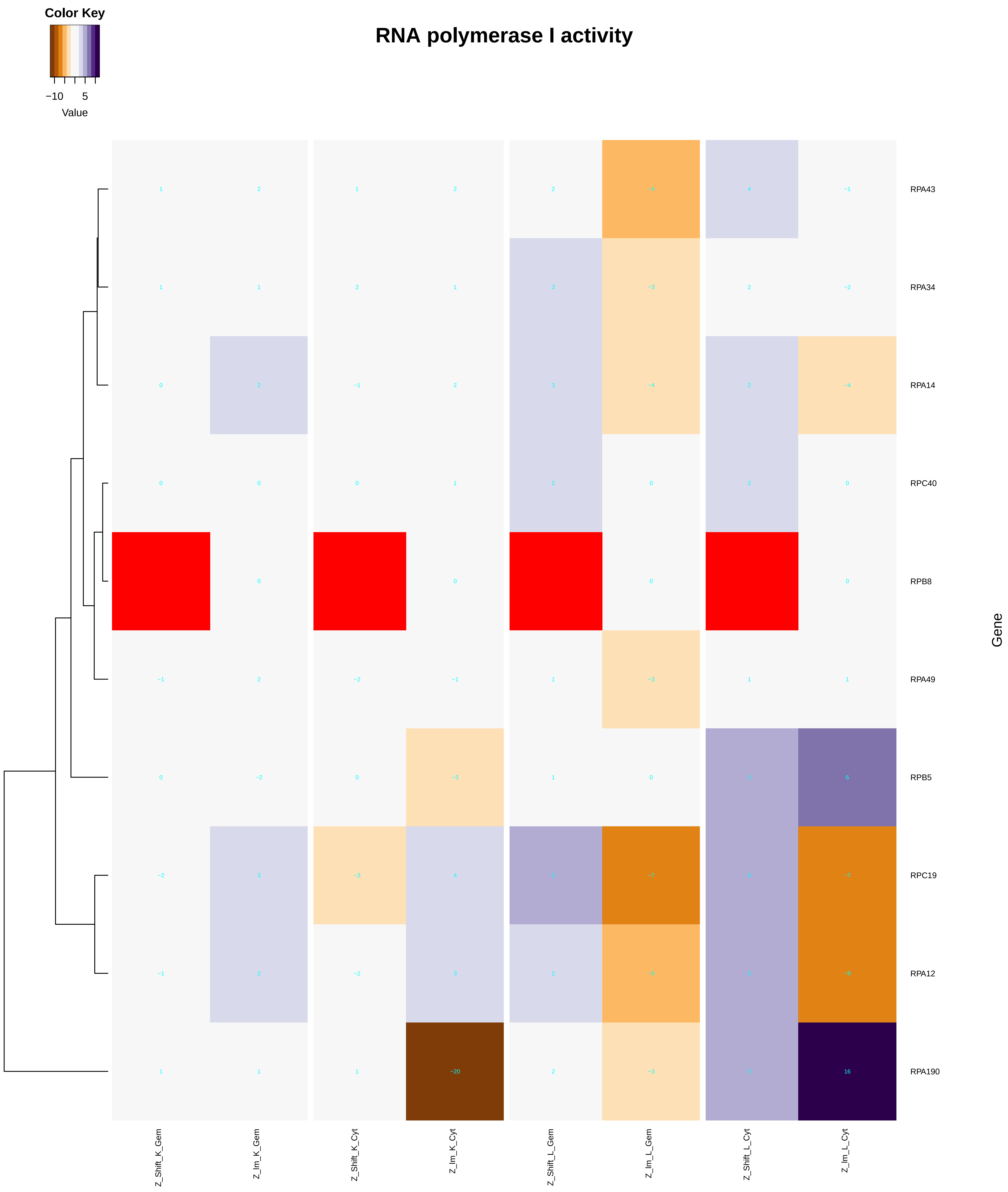

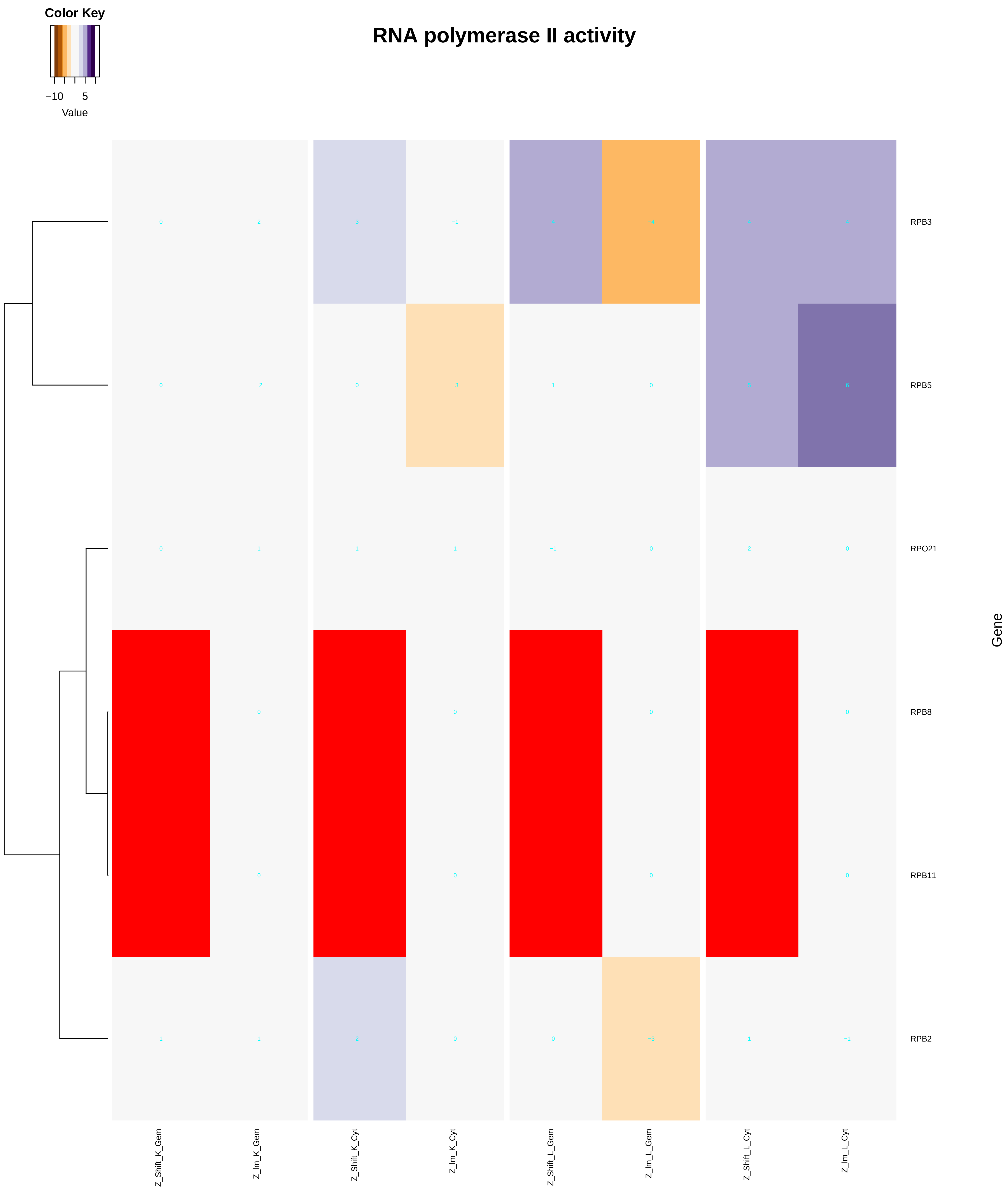

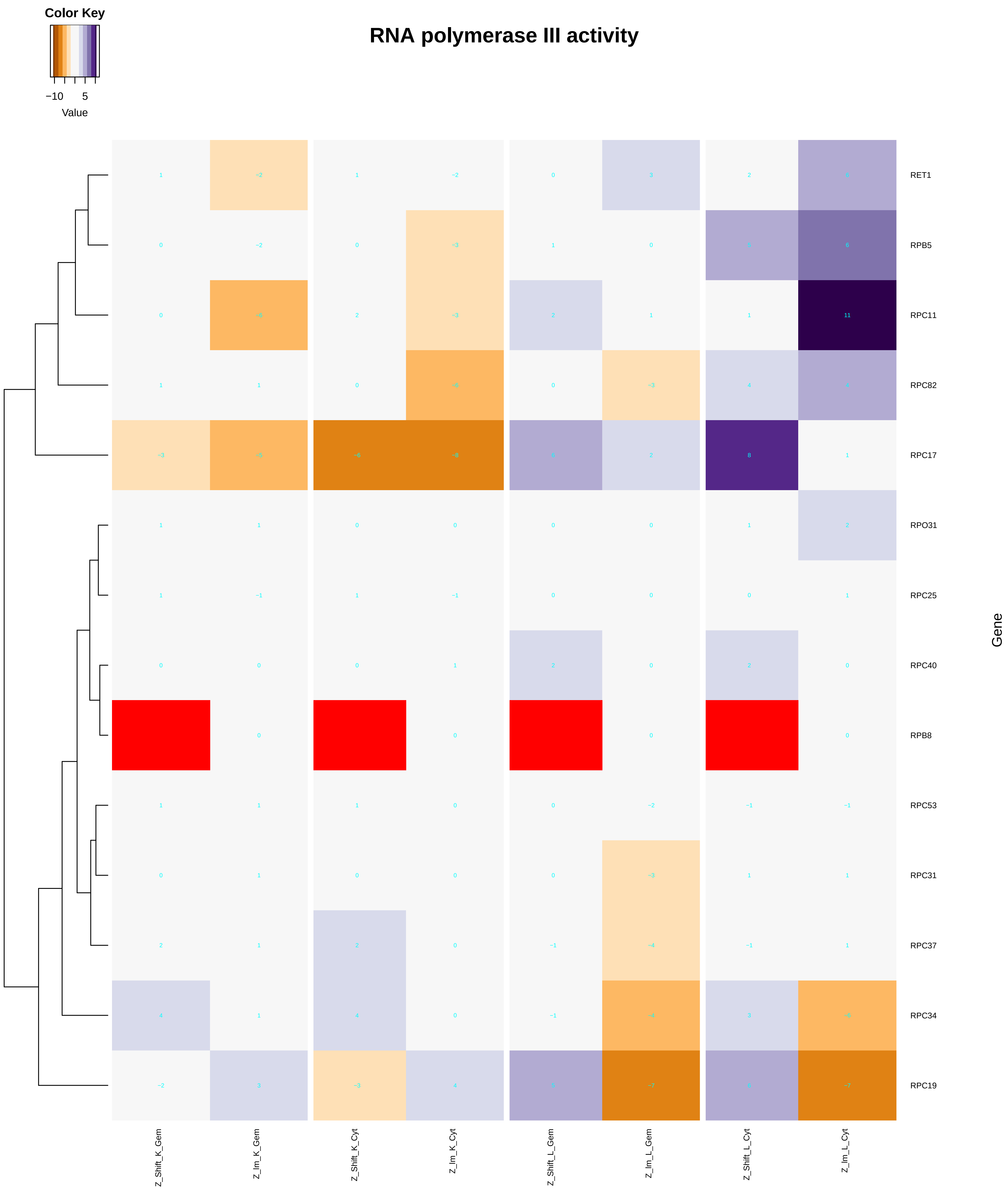

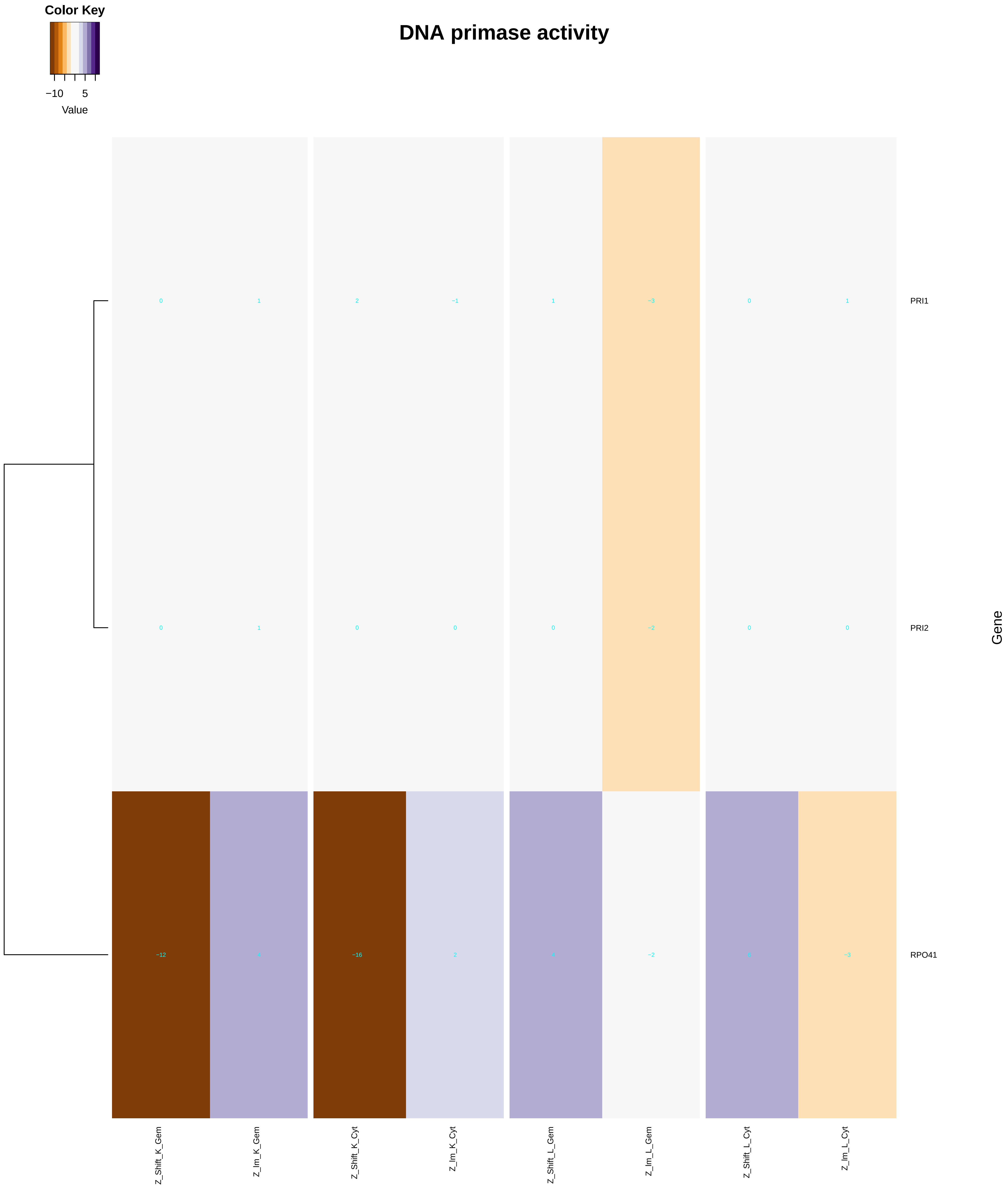

### DNA-directed_RNA_polymerase_II,_holoenzyme.pdf

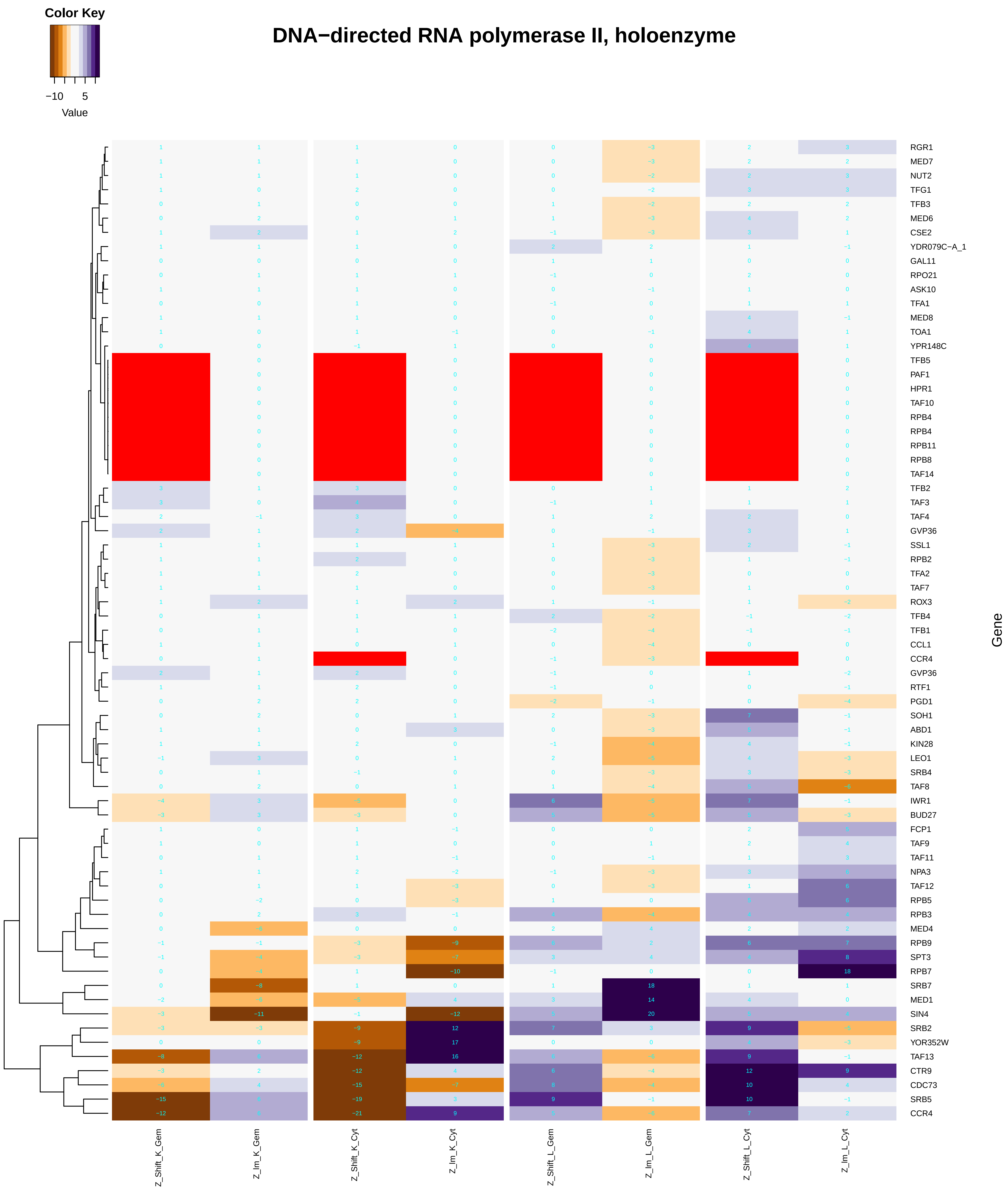

### DNA-templated_transcription,_termination.pdf

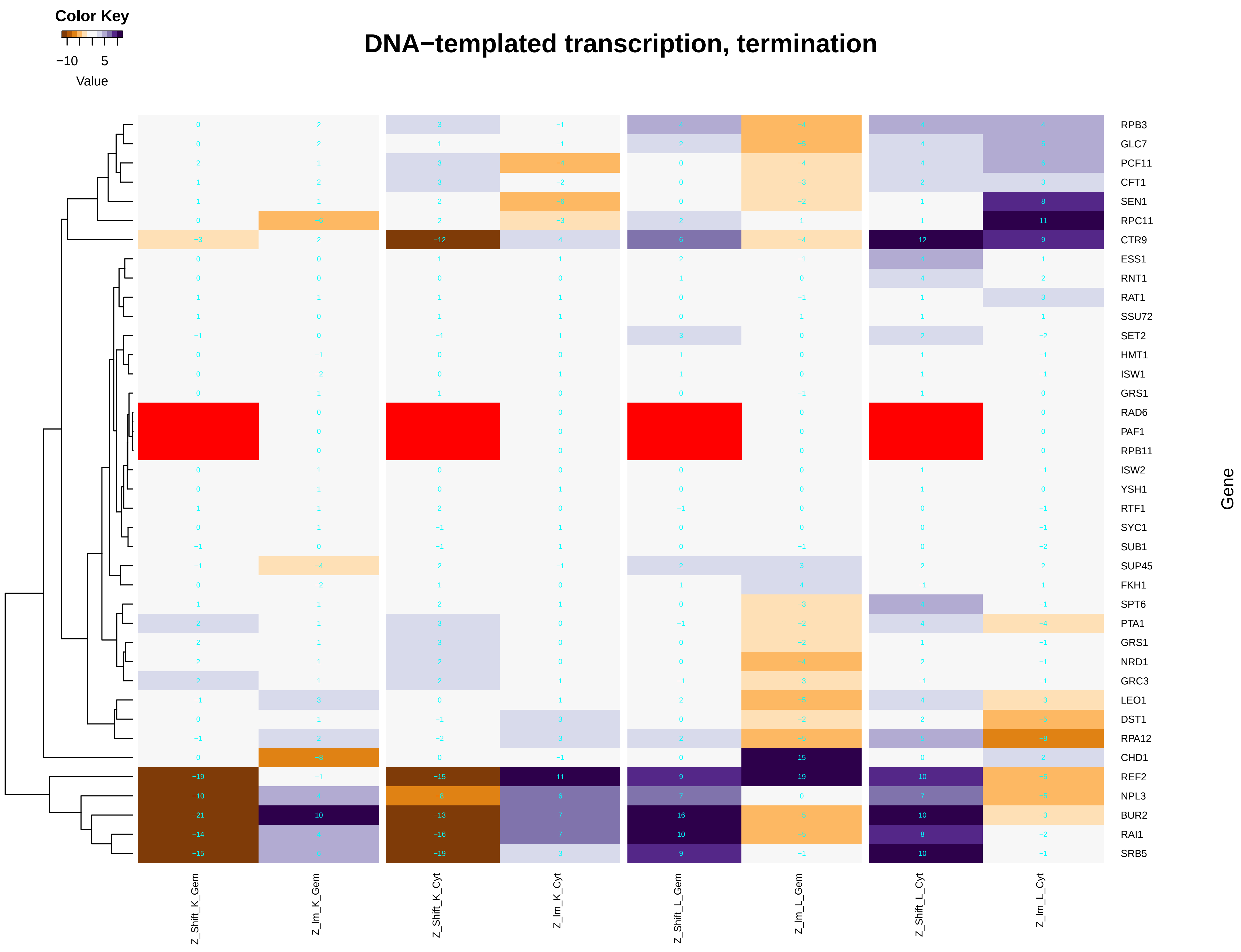

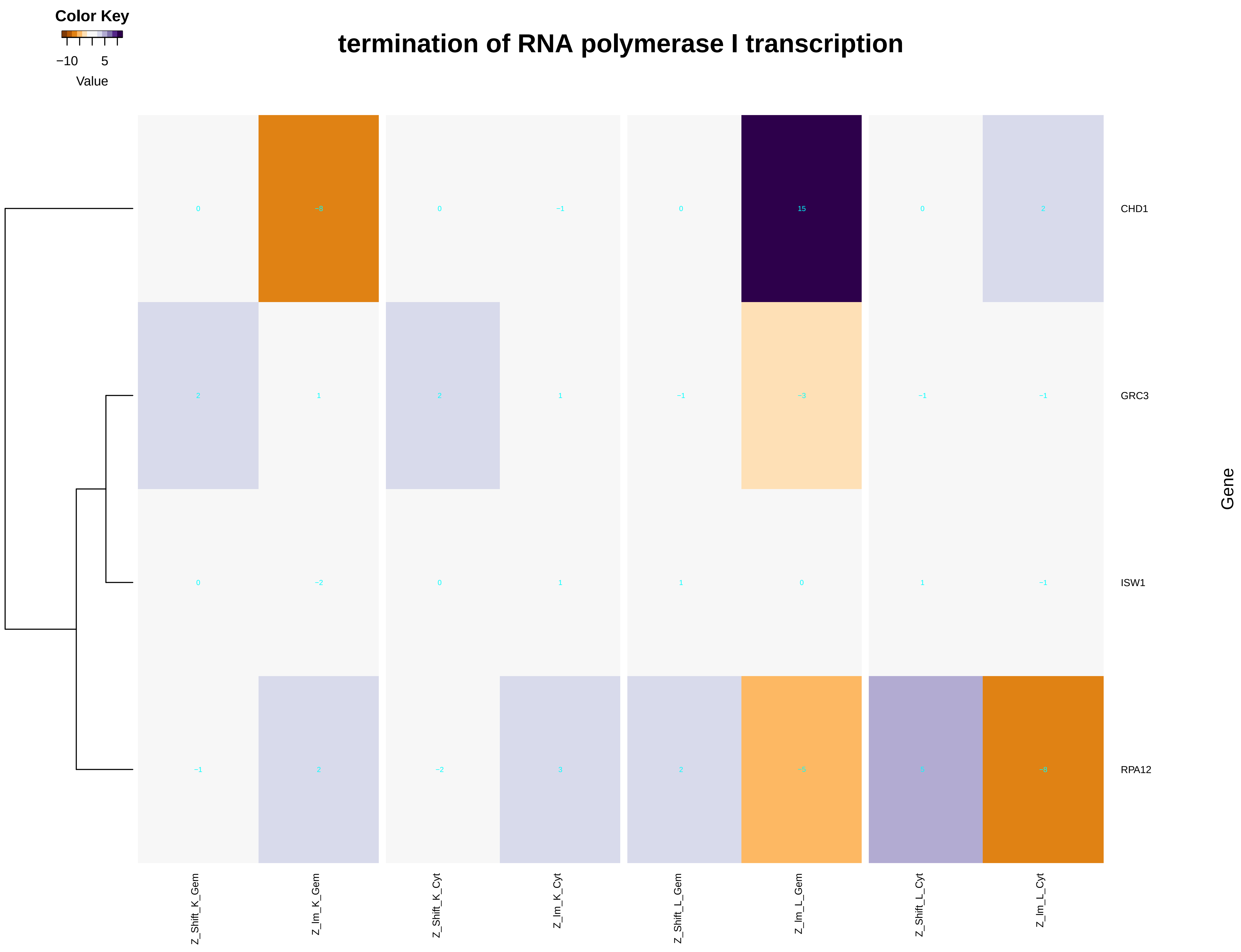

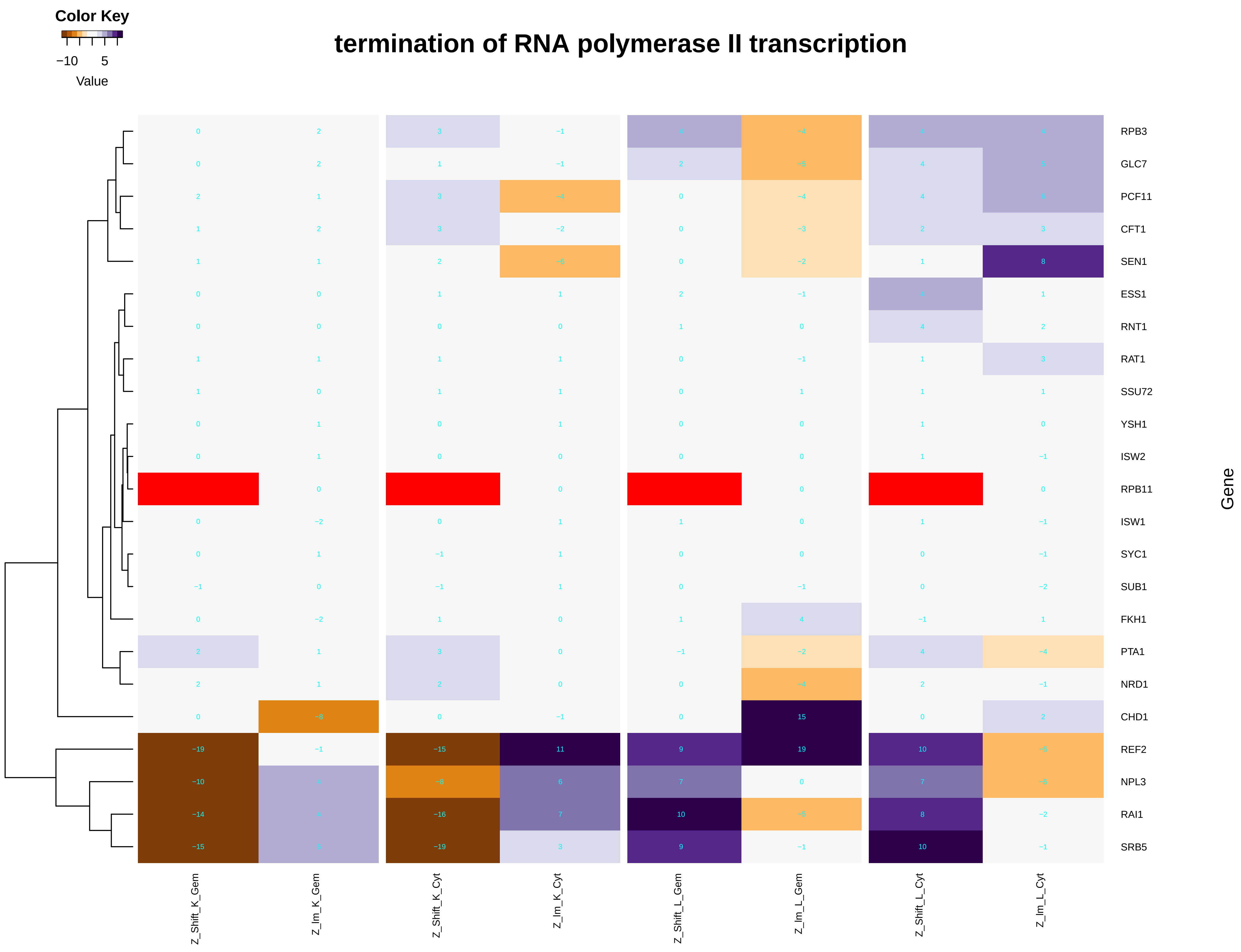

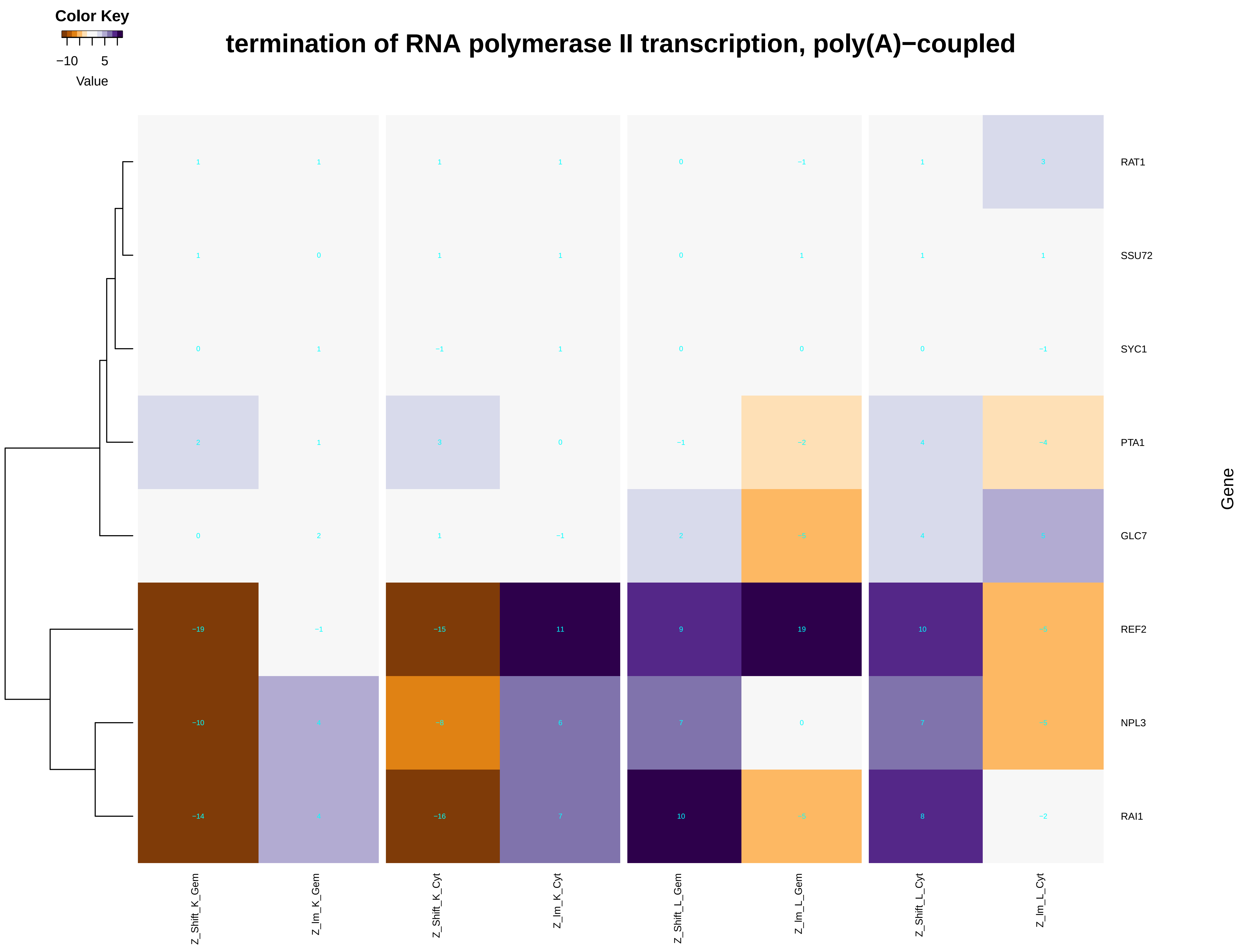

#### termination of RNA polymerase II transcription, exosome−dependent

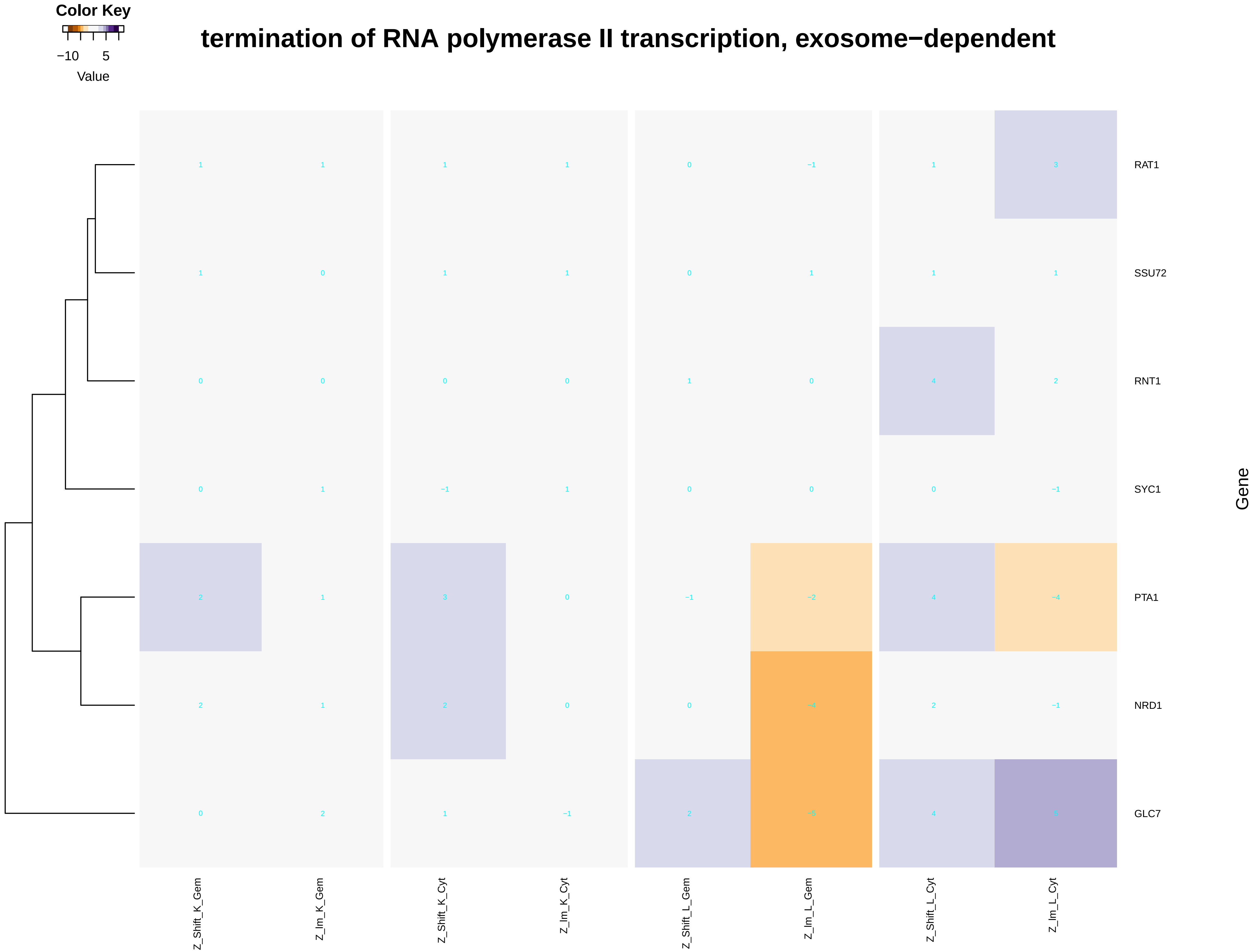

### DNA_amplification.pdf

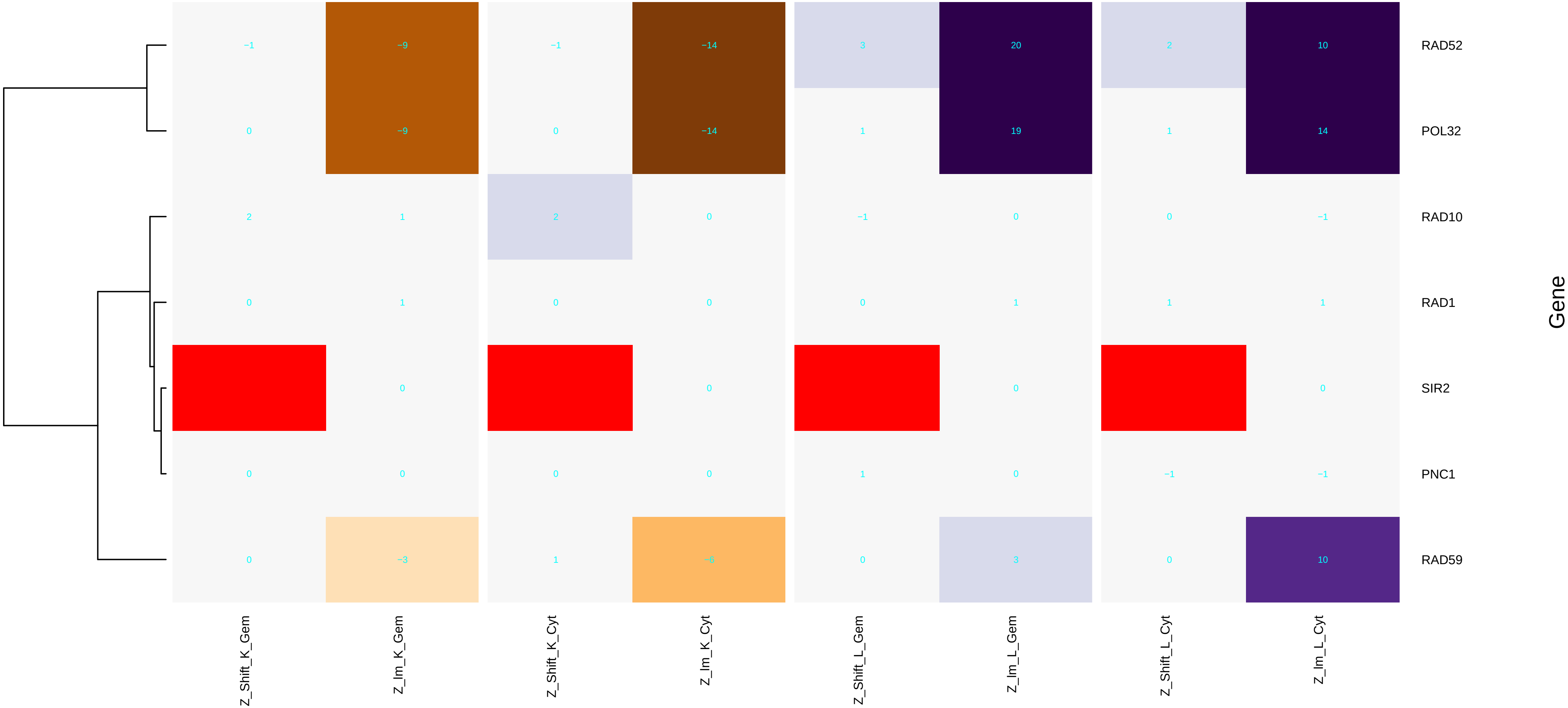

### DNA_biosynthetic_process.pdf

#### DNA biosynthetic process

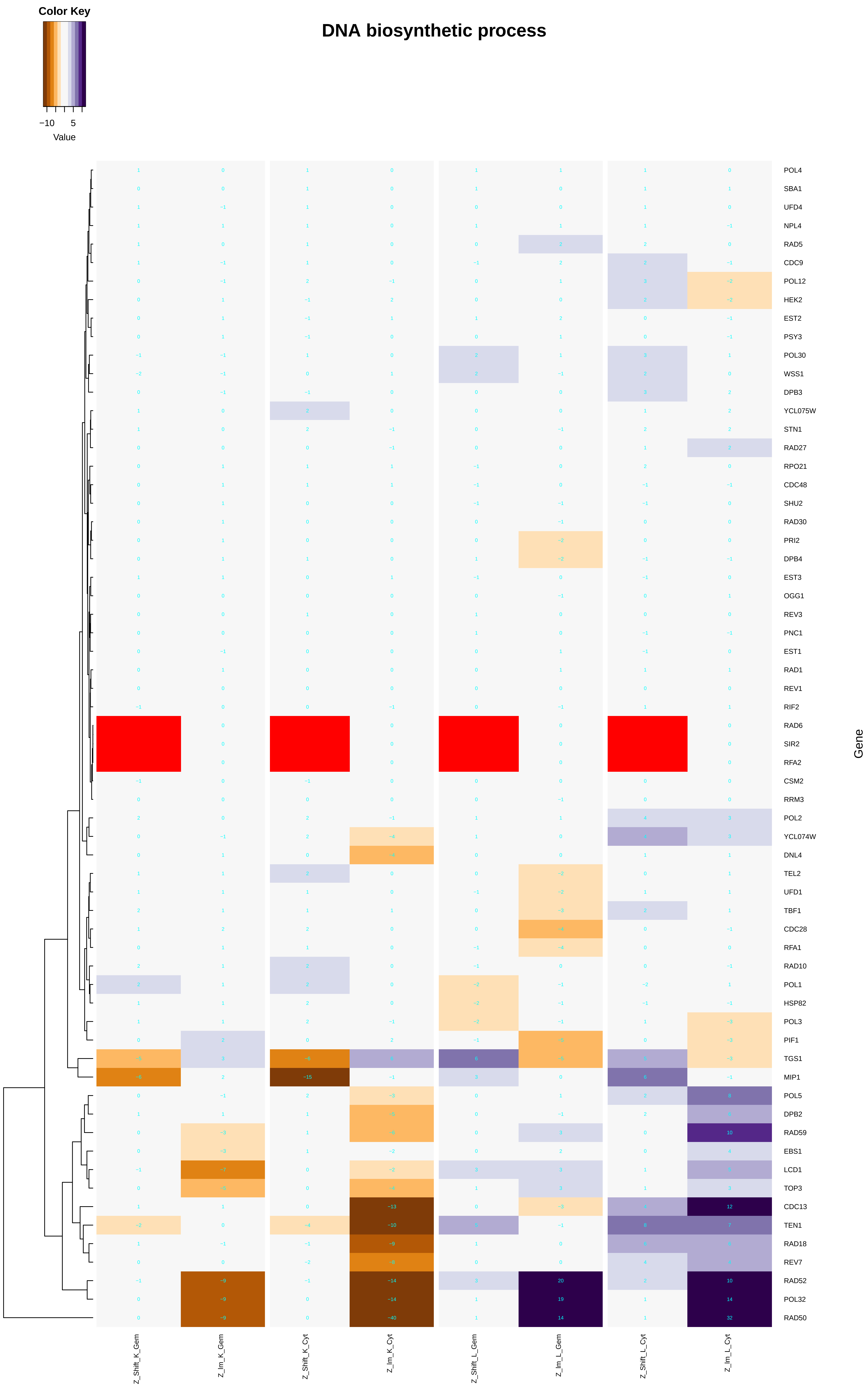

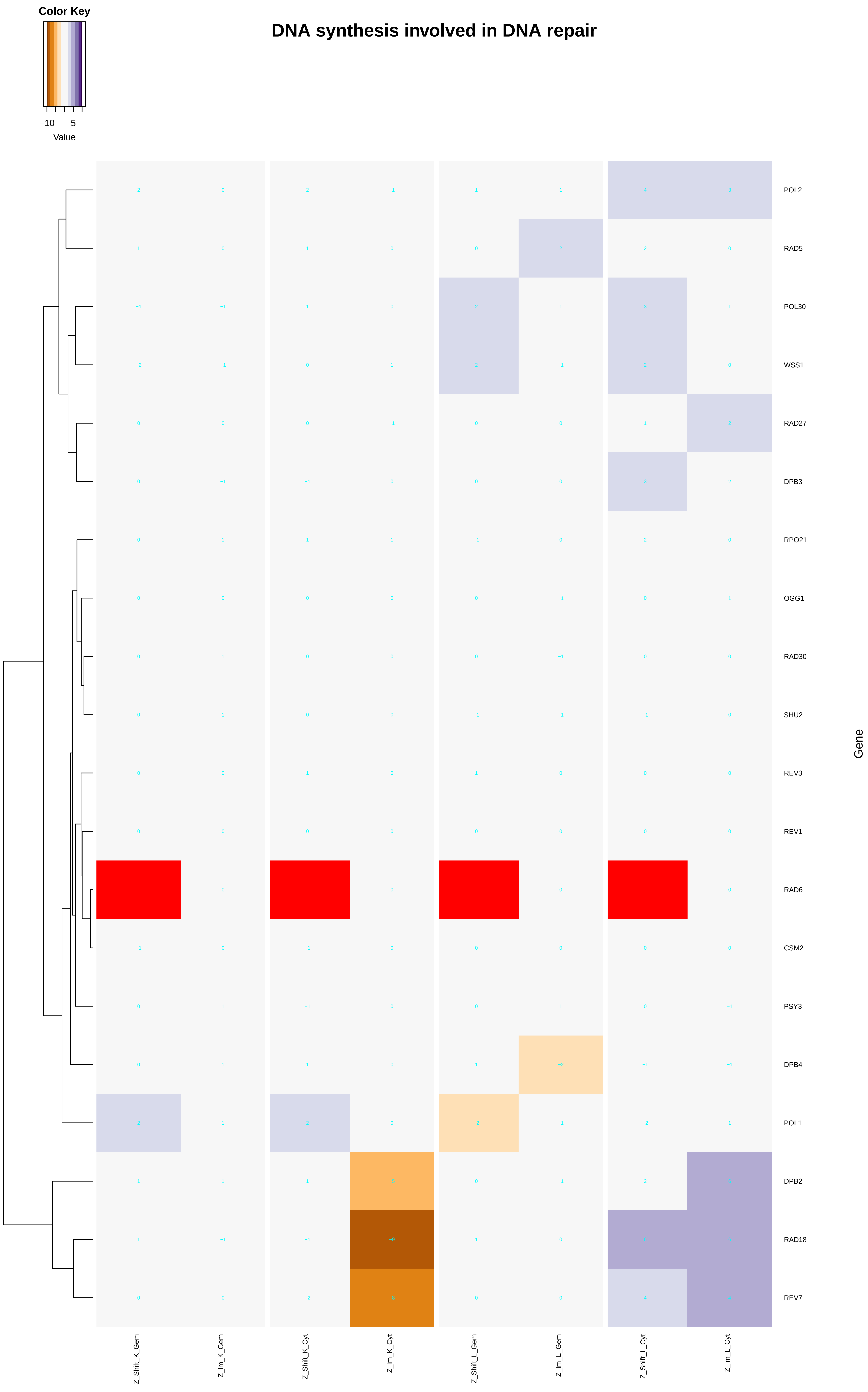

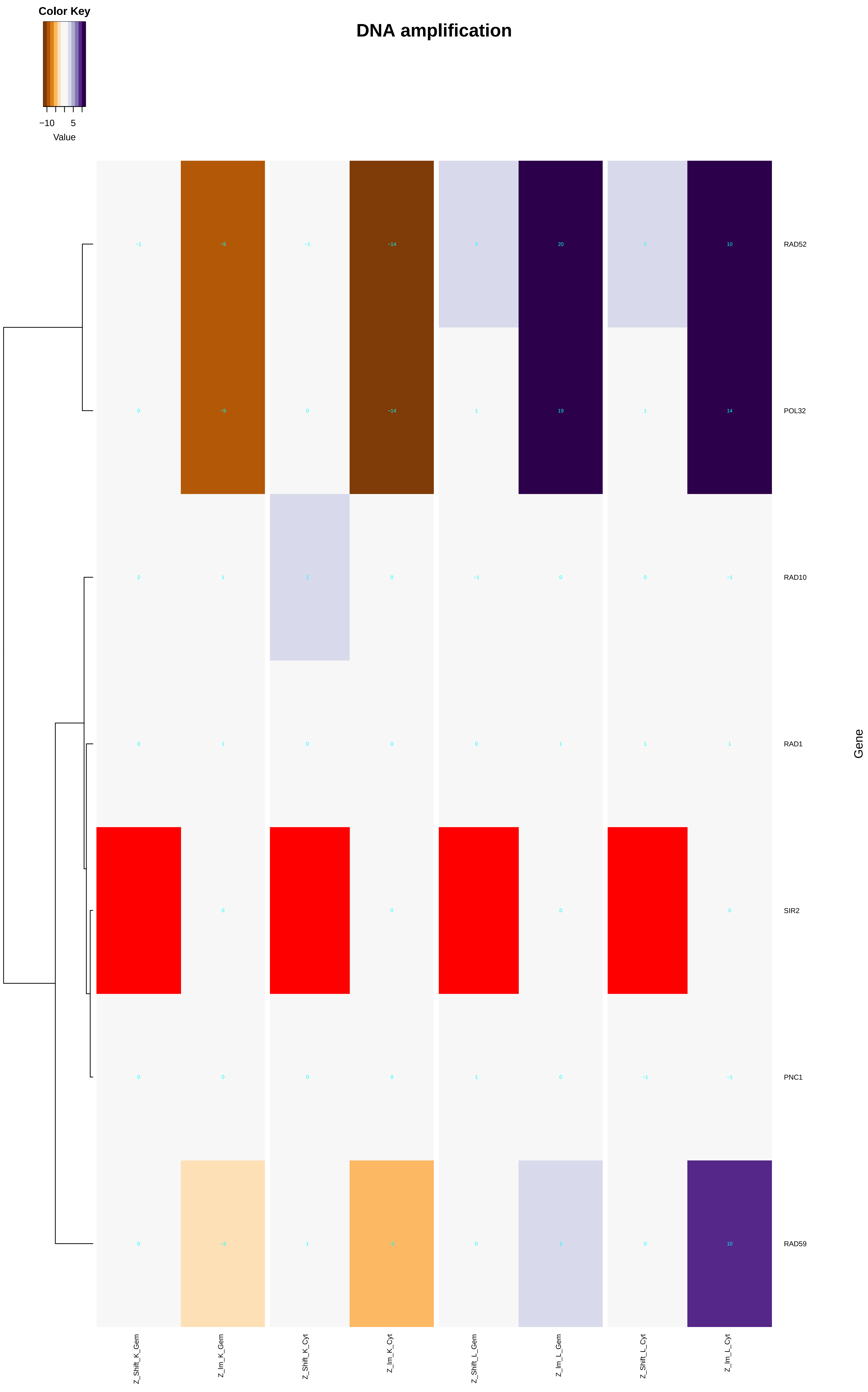

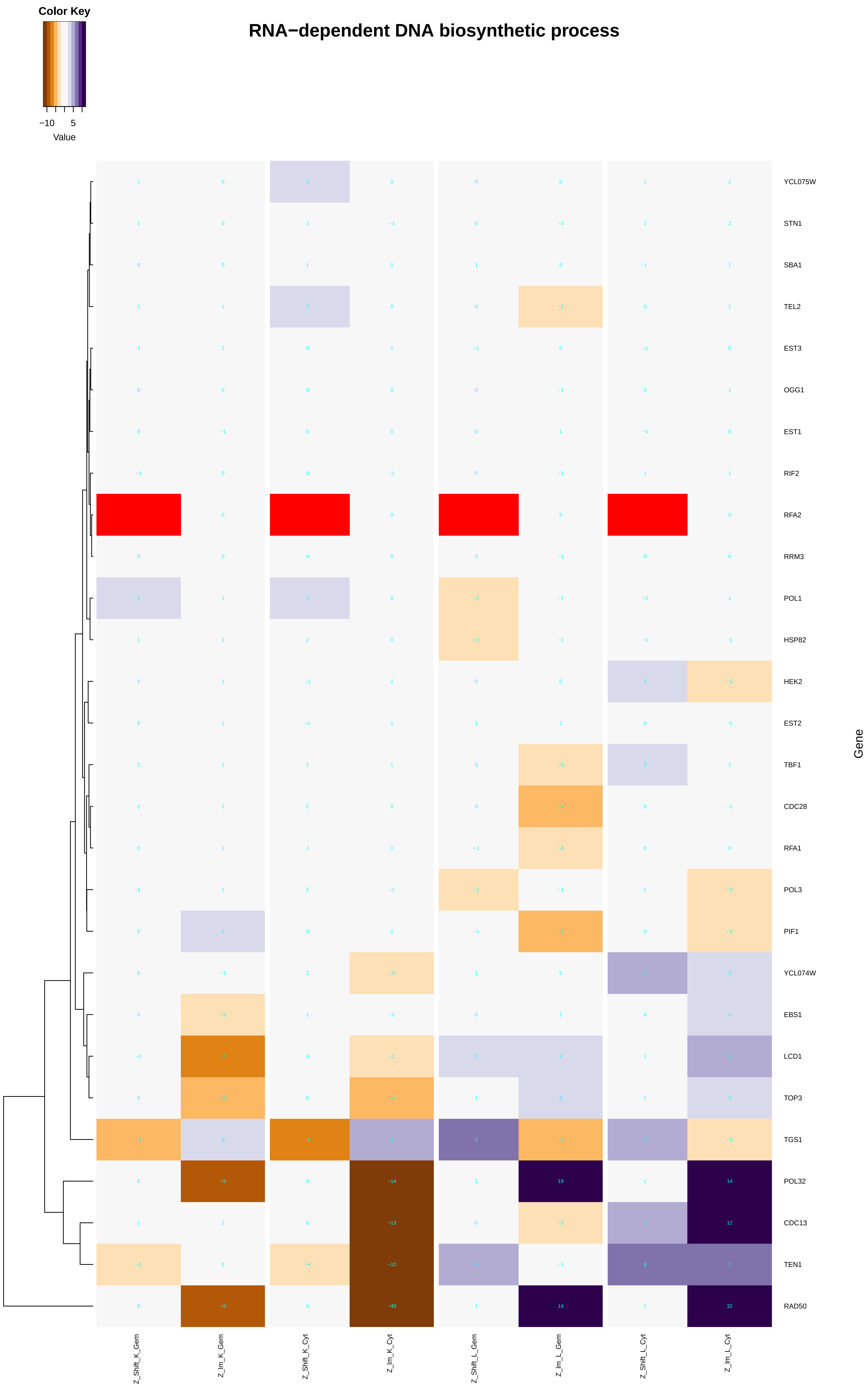

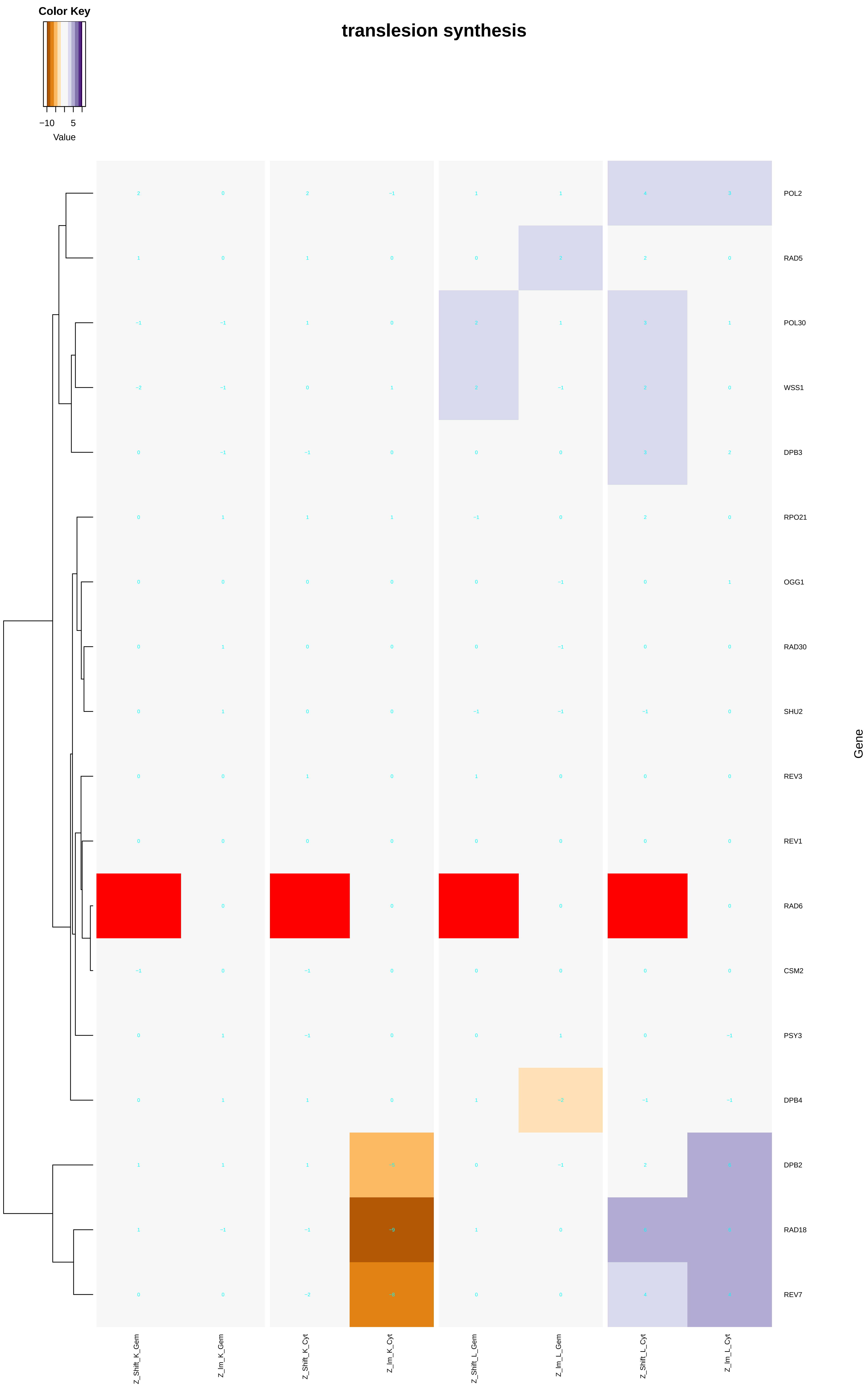

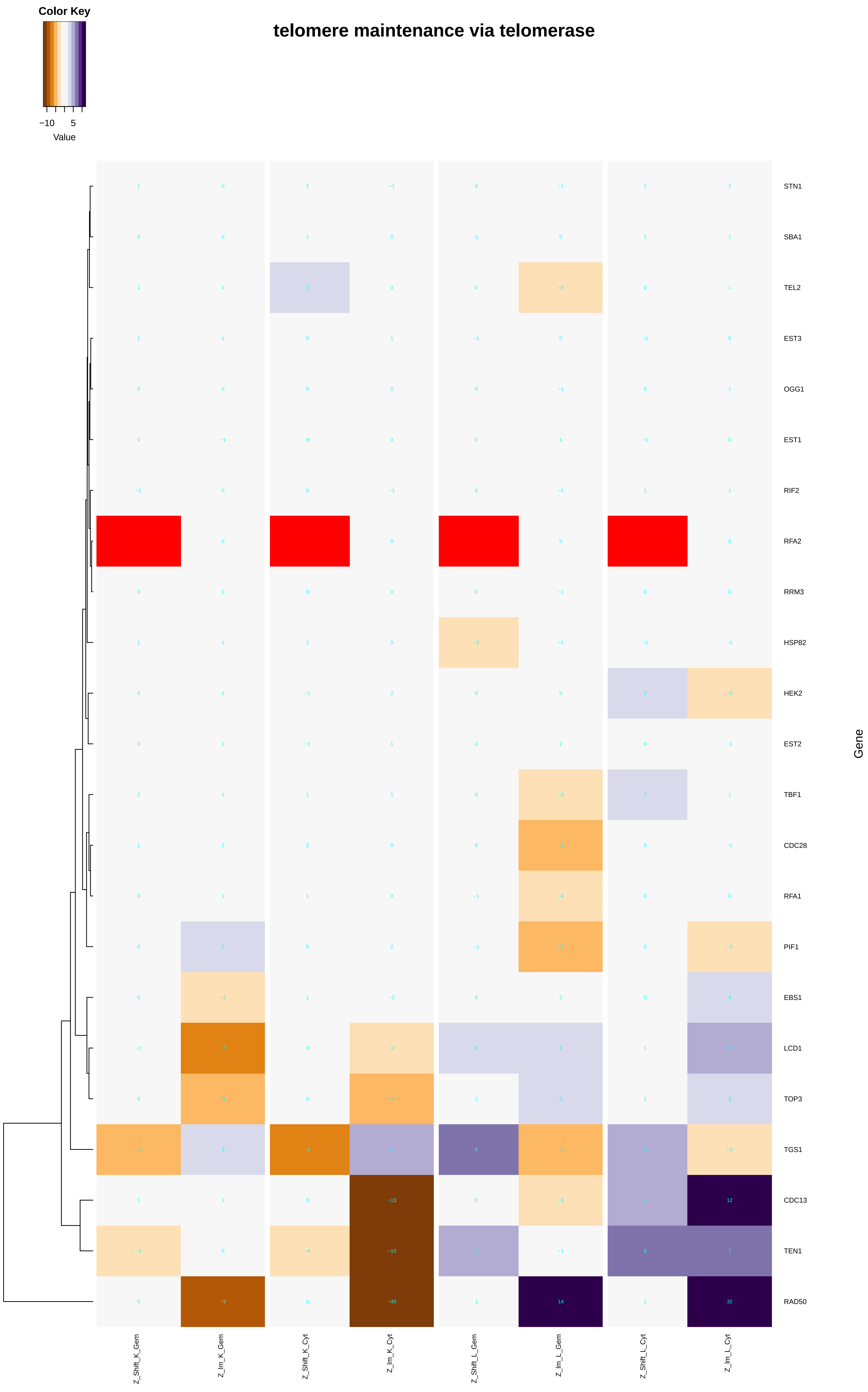

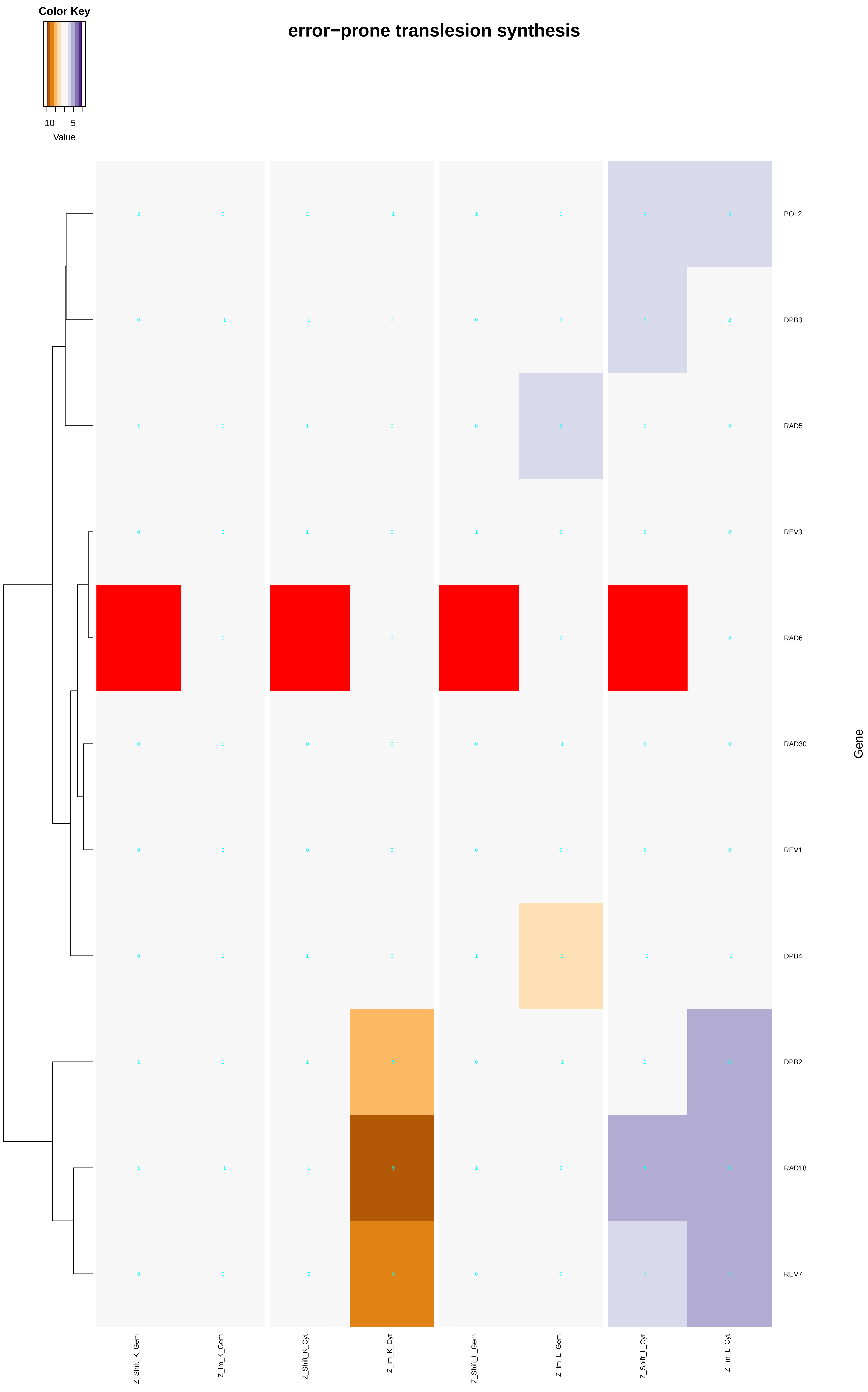

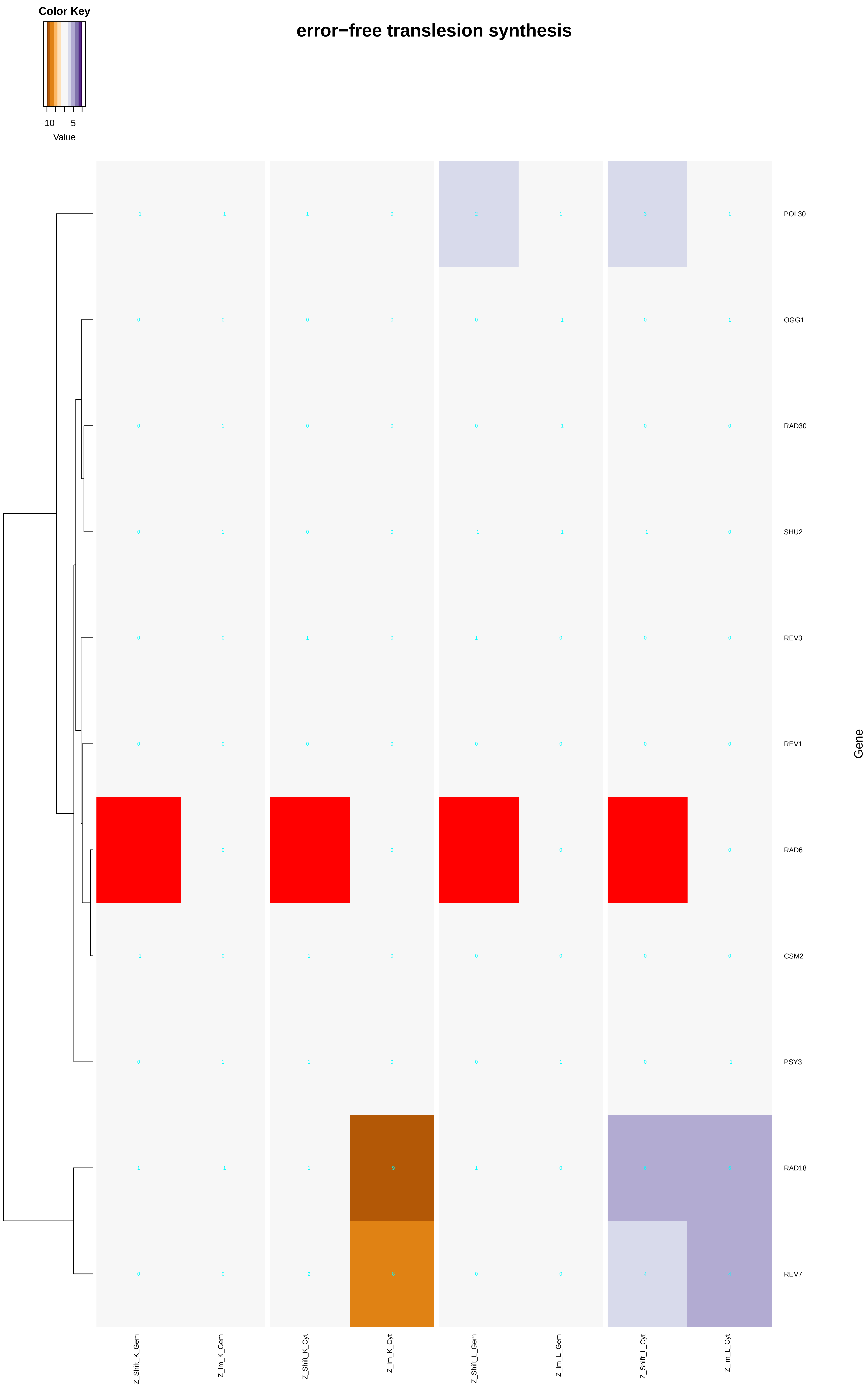

### DNA_damage_checkpoint.pdf

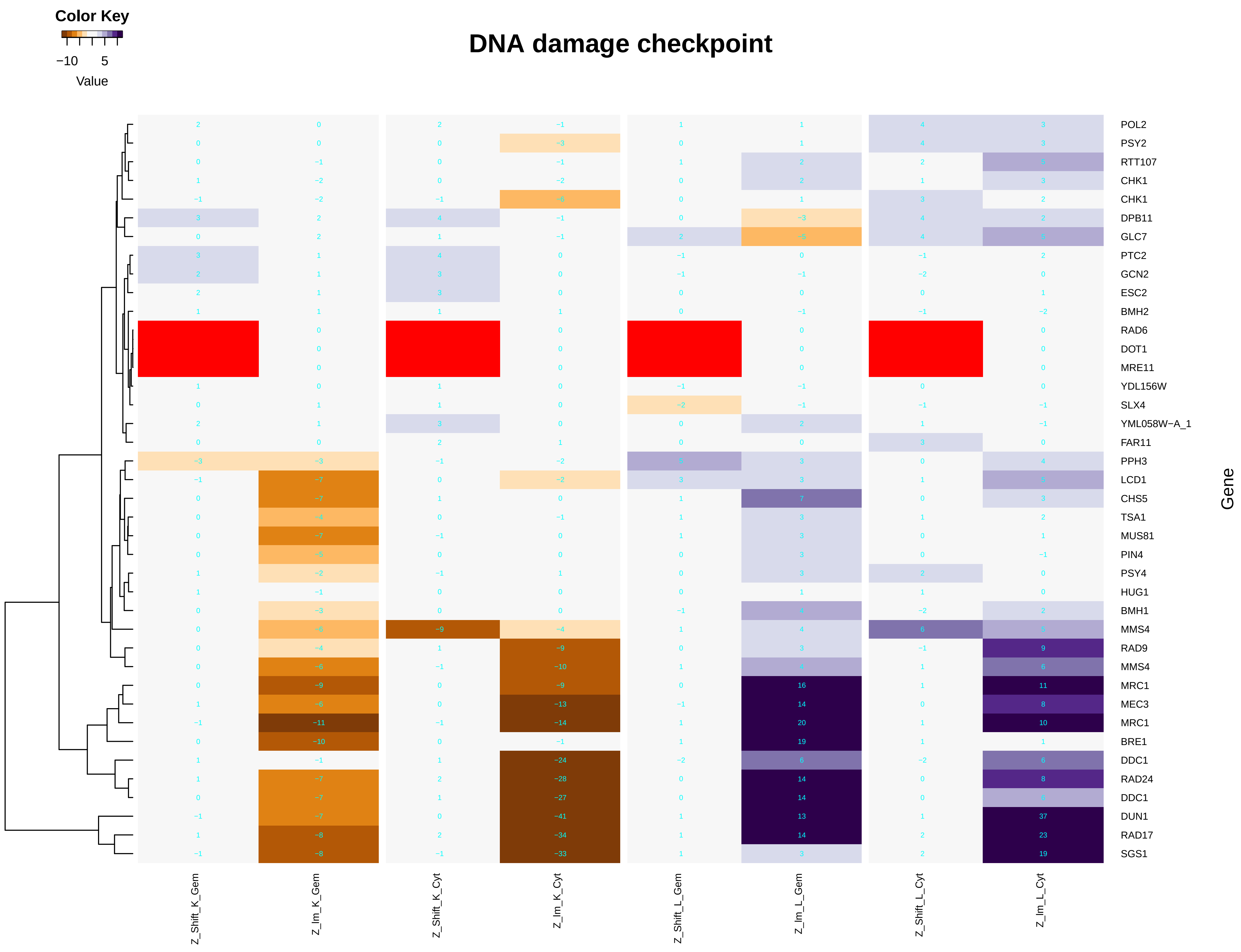

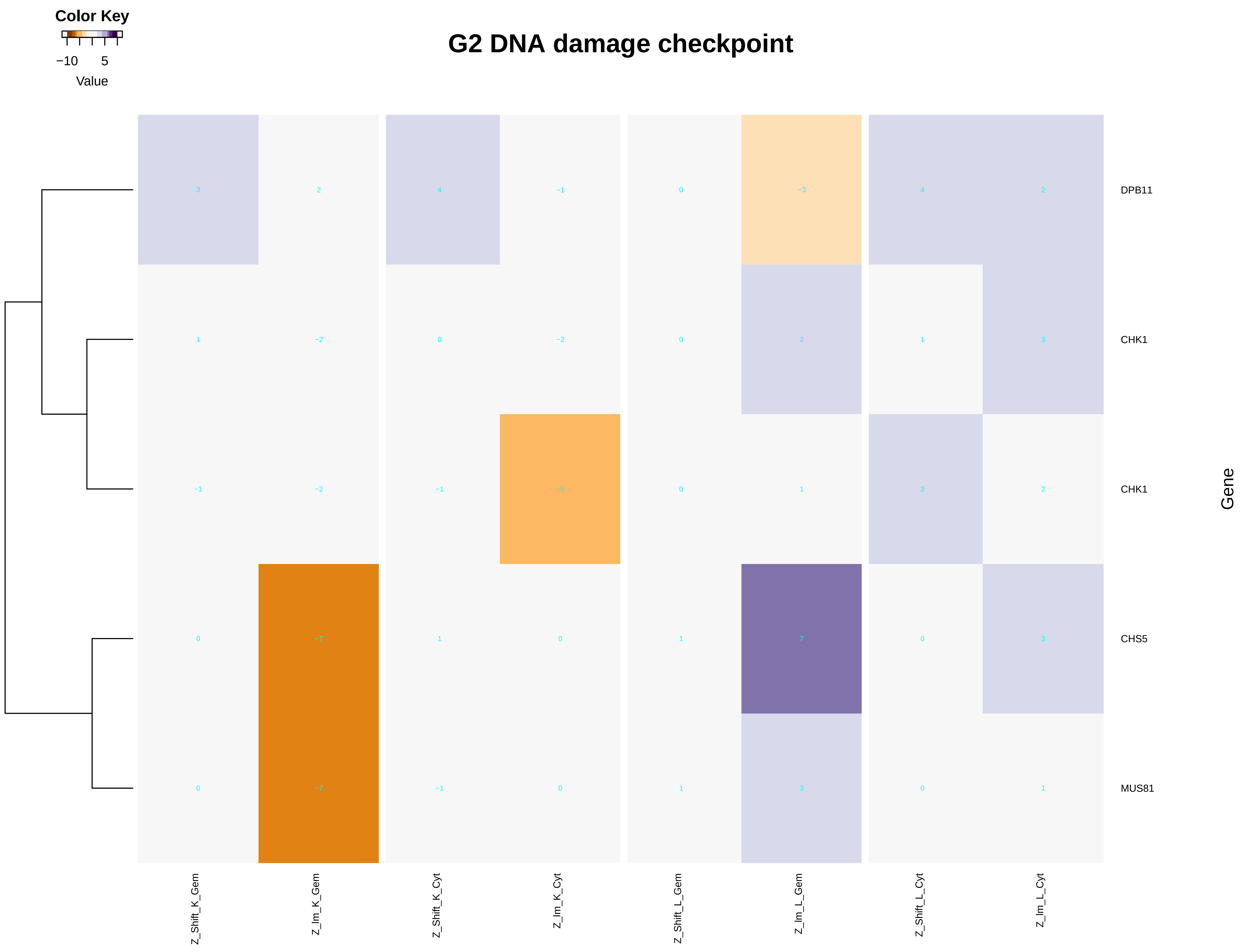

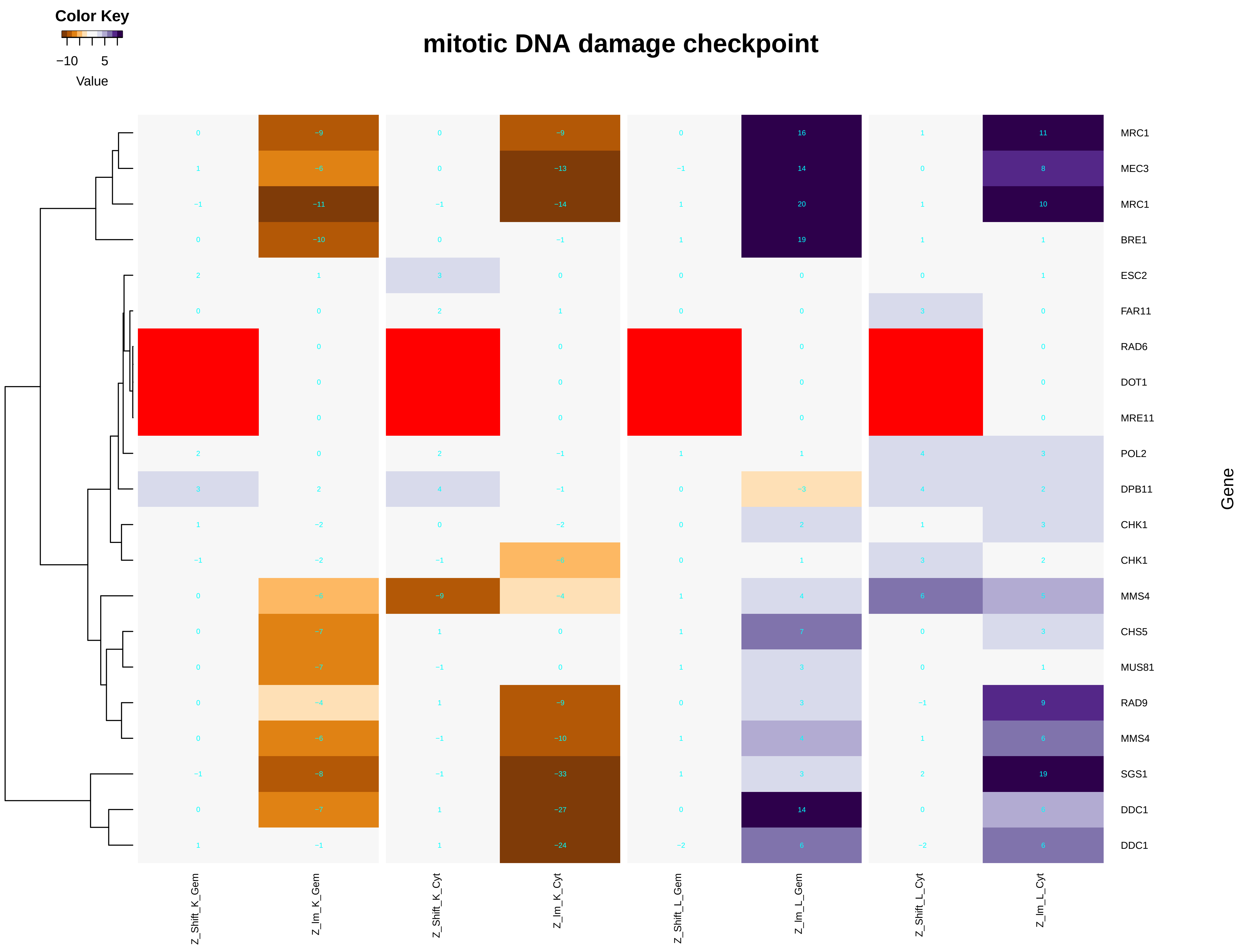

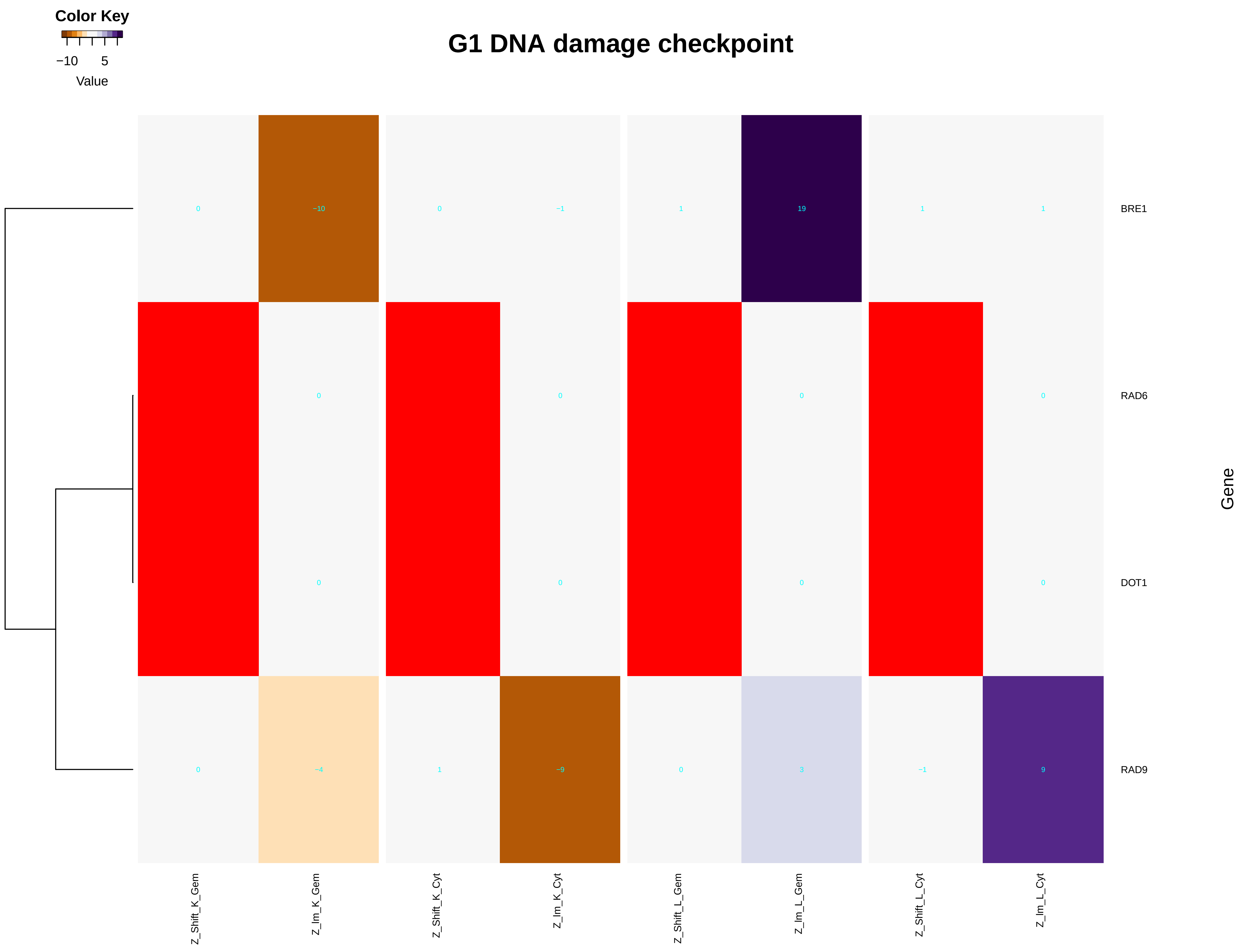

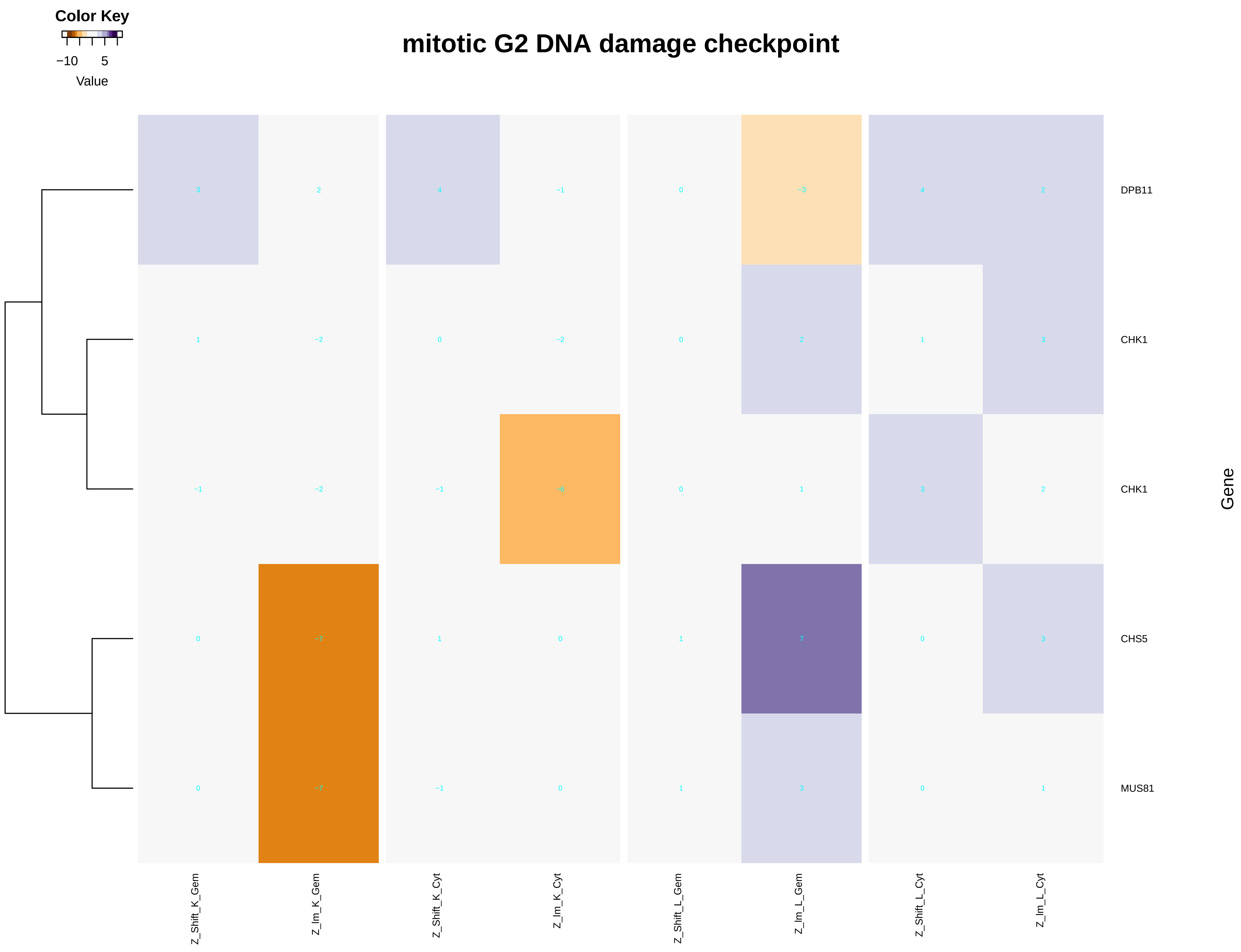

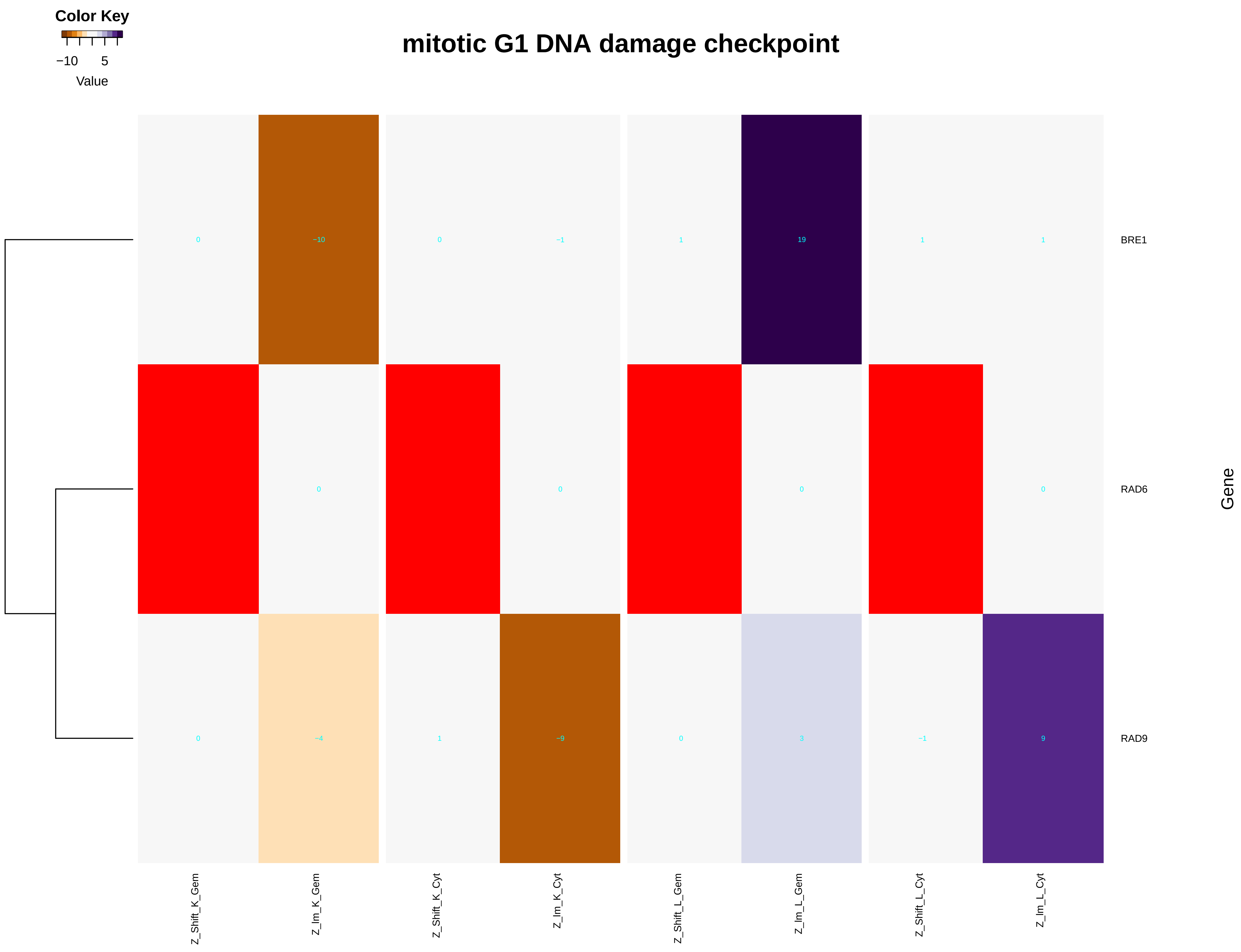

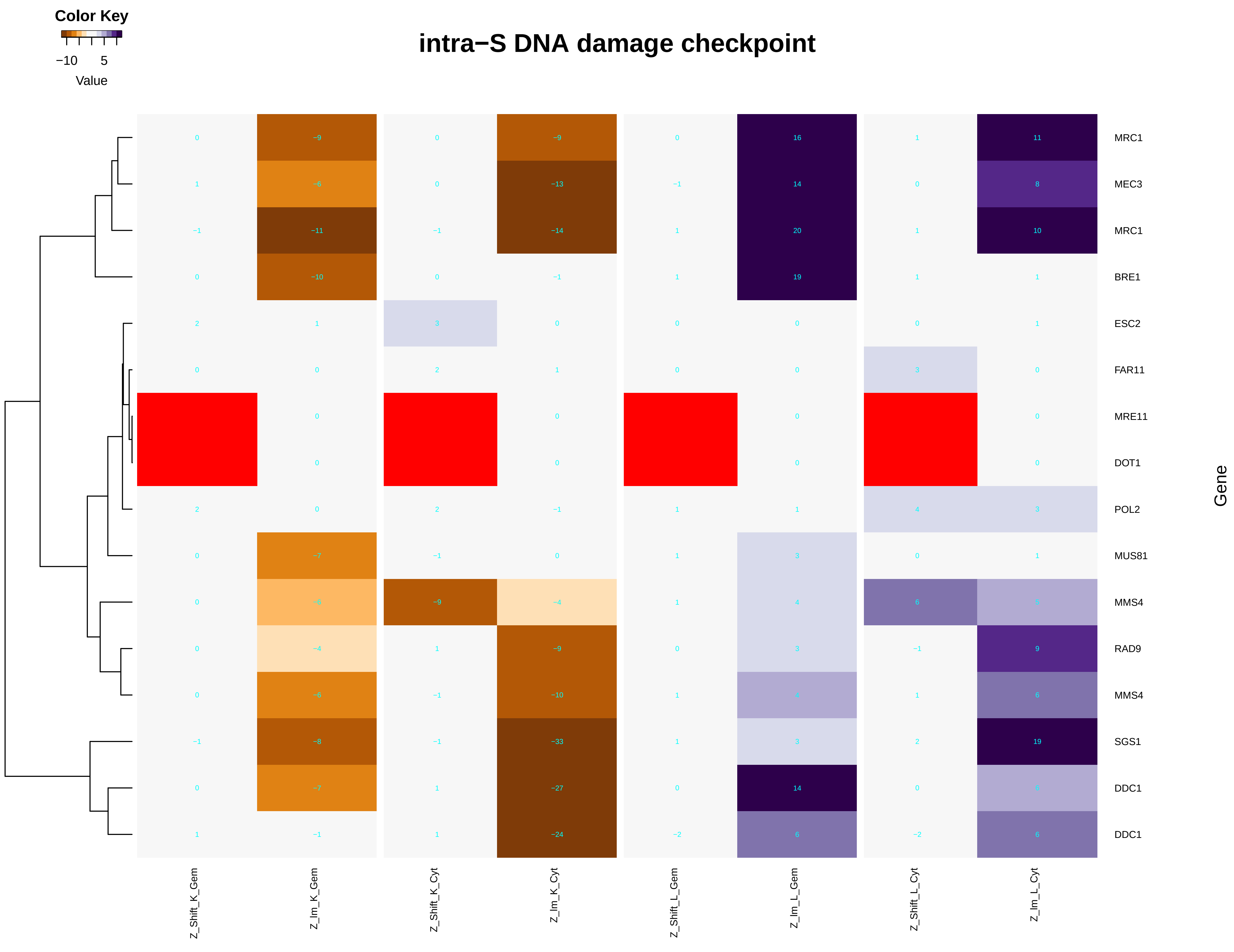

### DNA_double-strand_break_processing.pdf

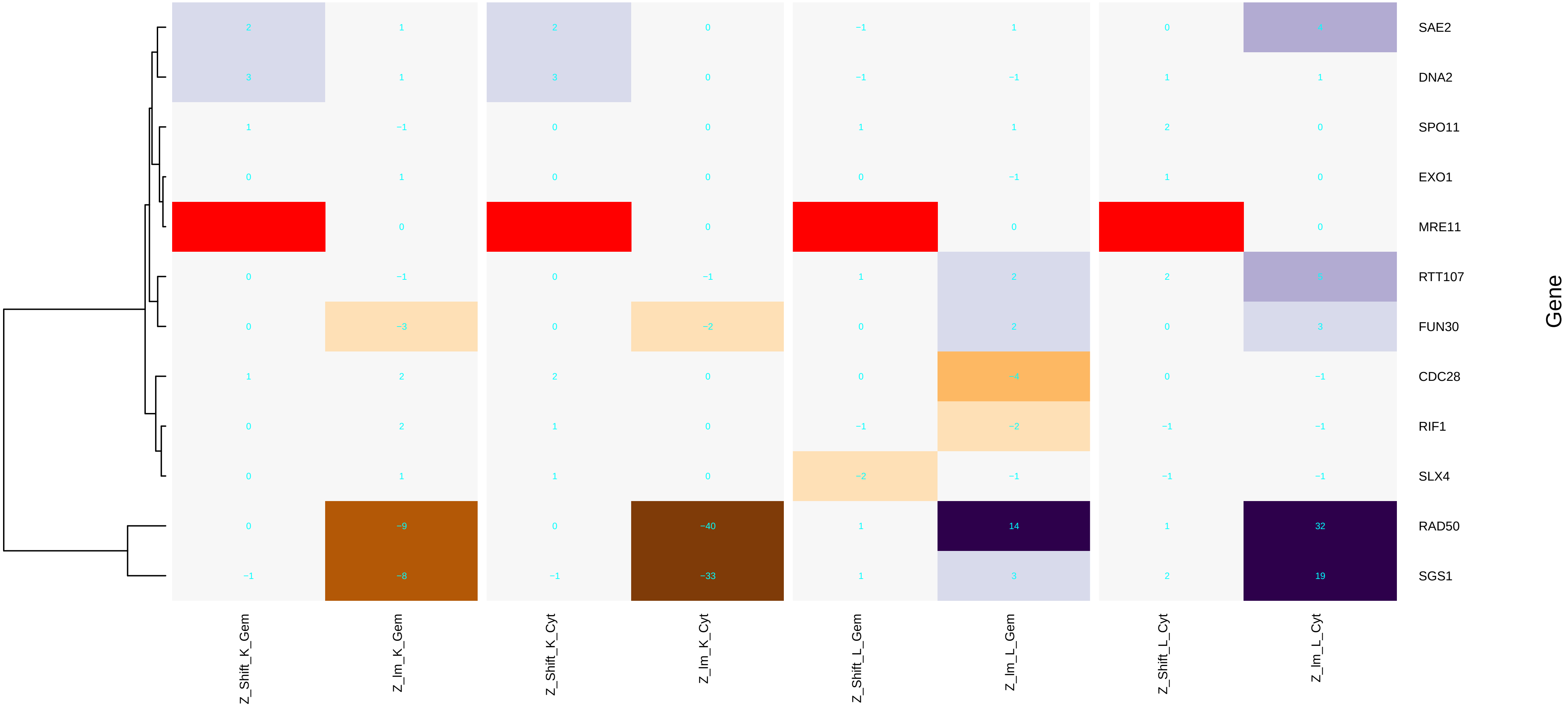

### DNA_helicase_activity.pdf

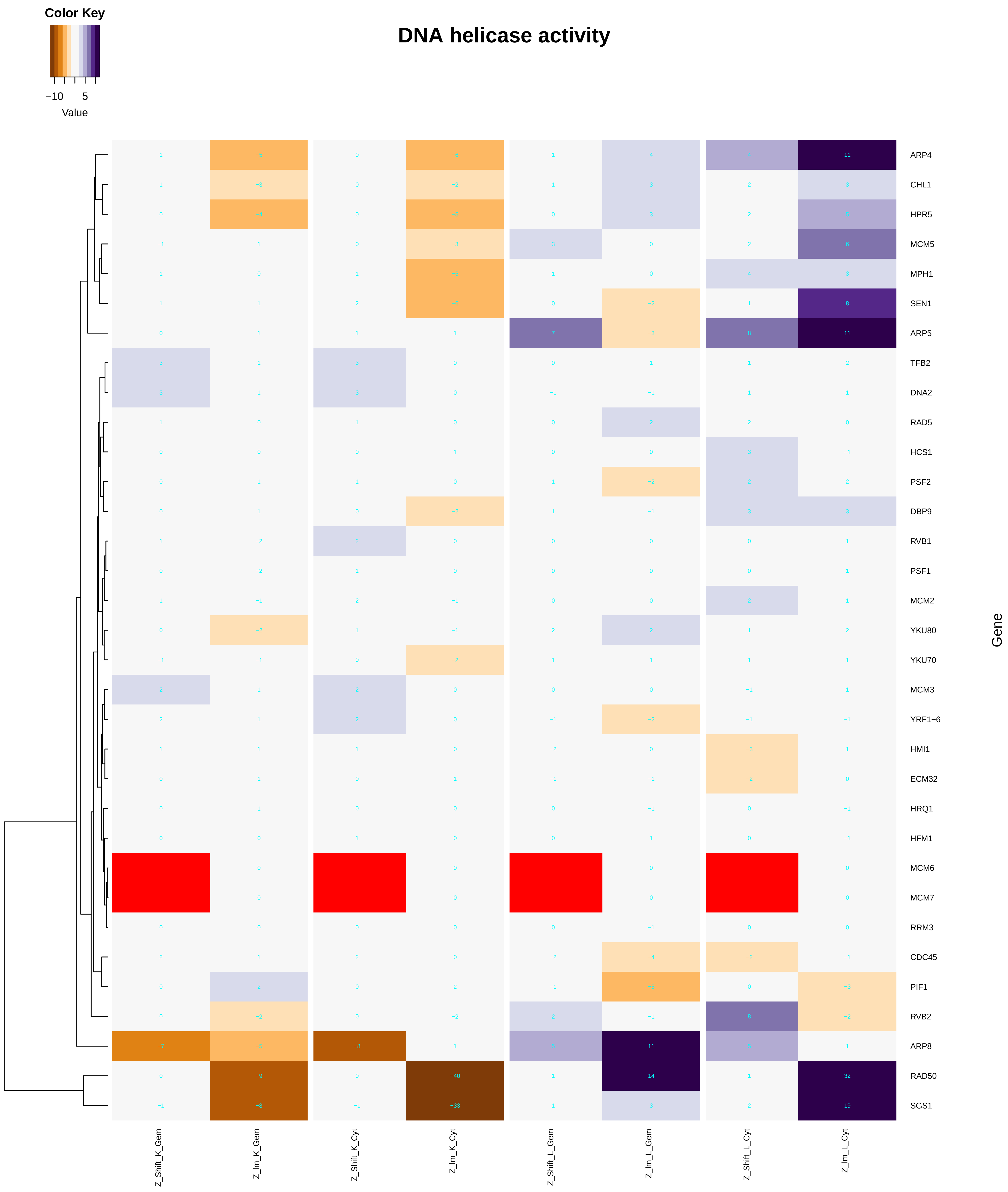

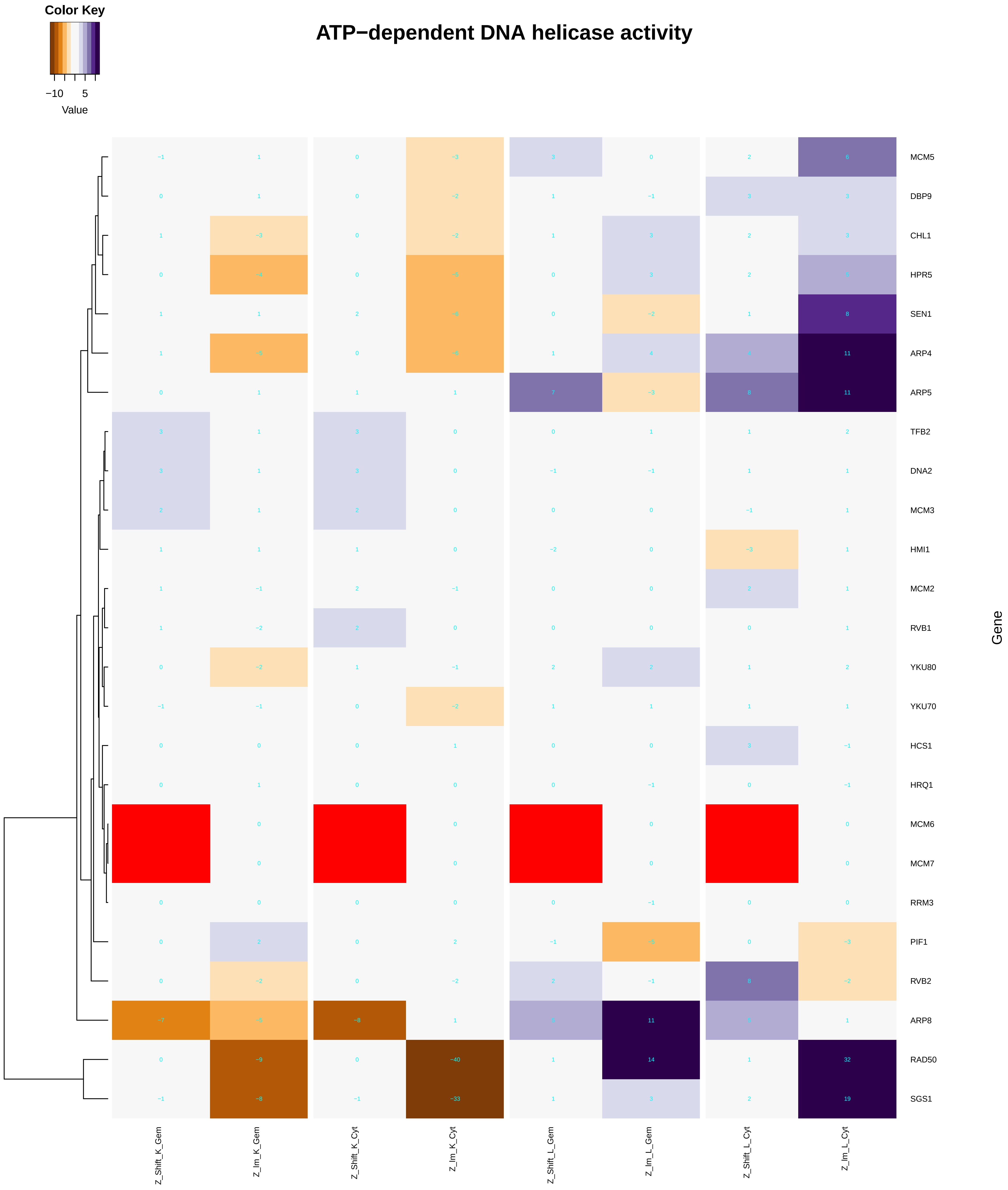

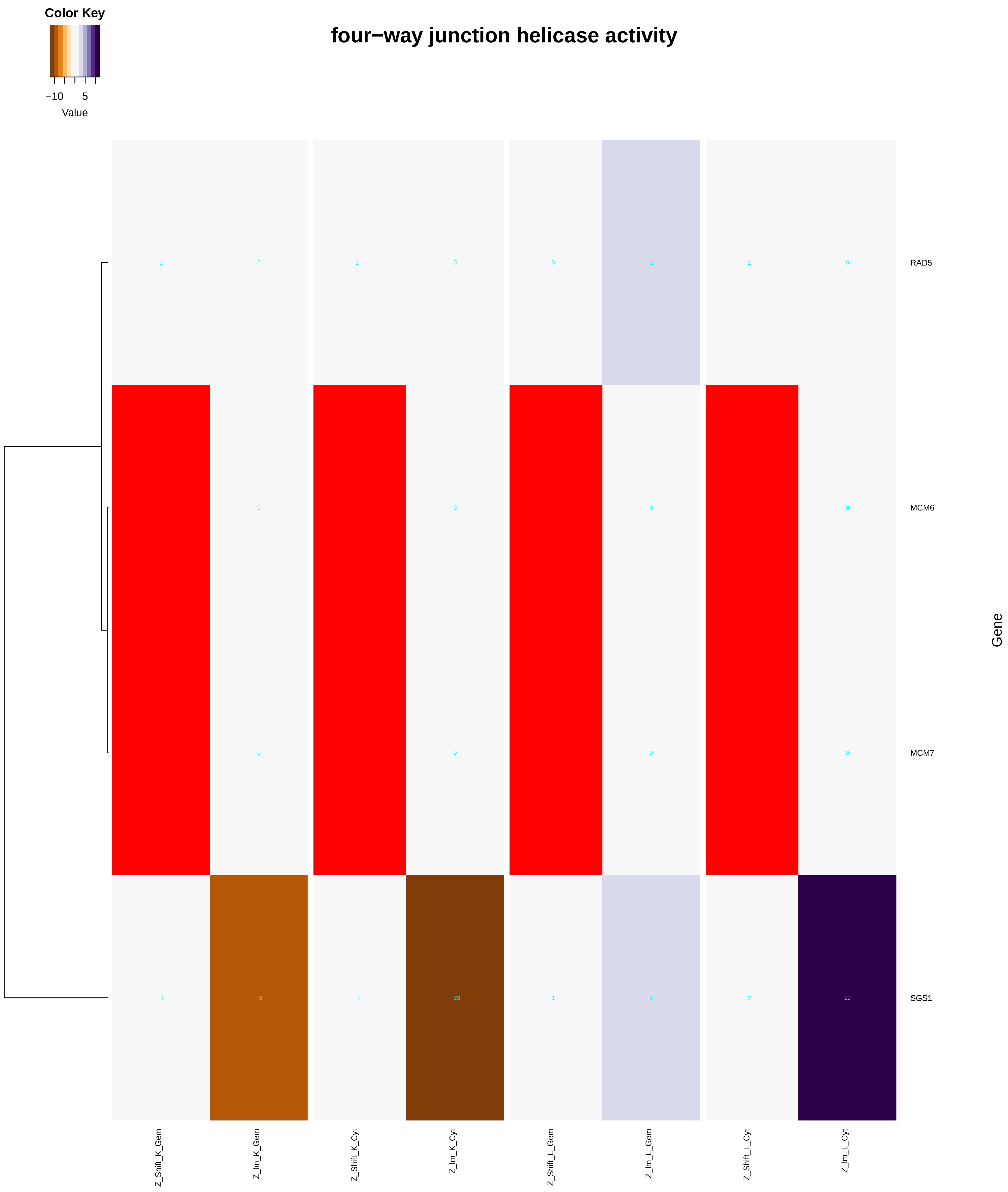

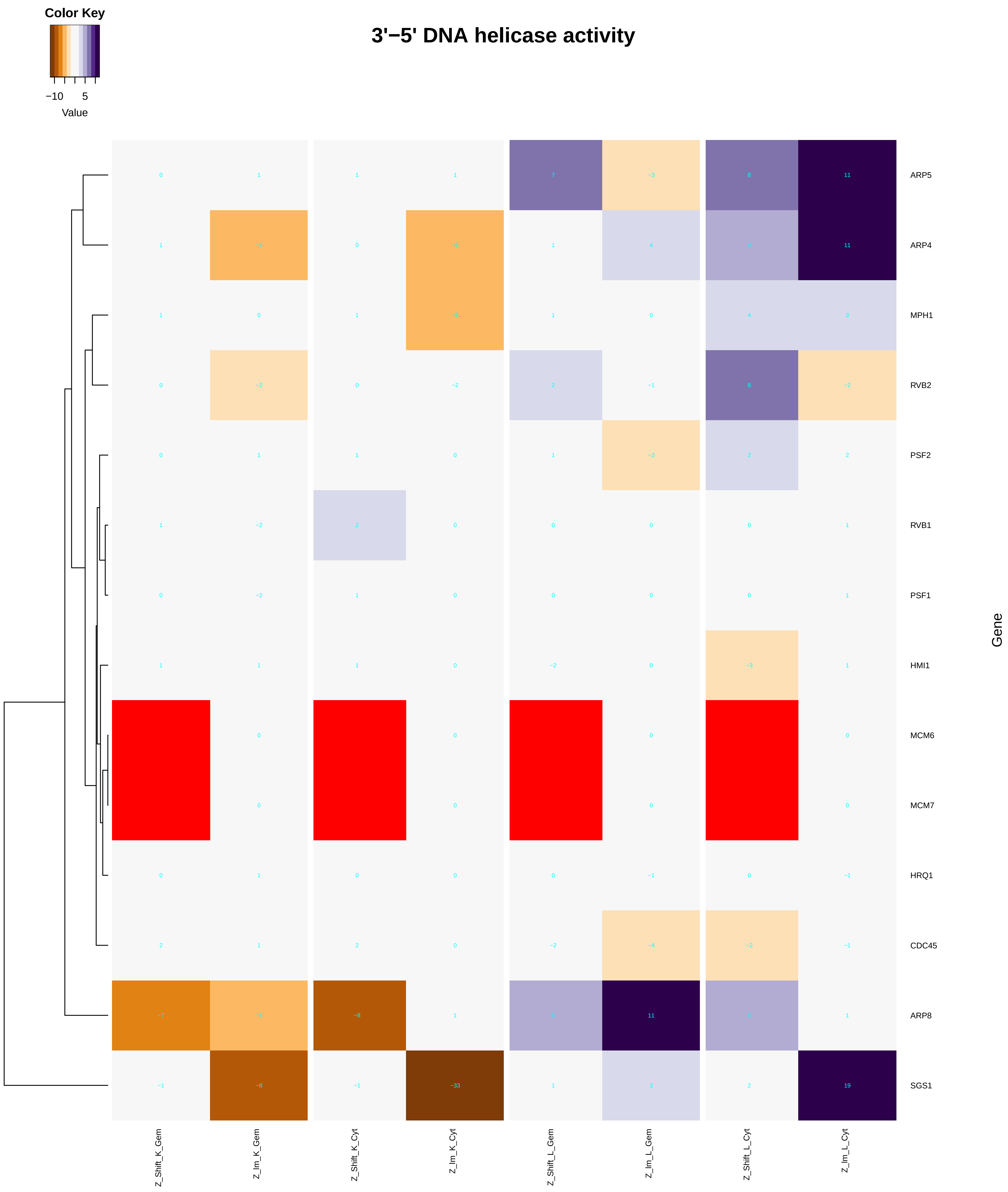

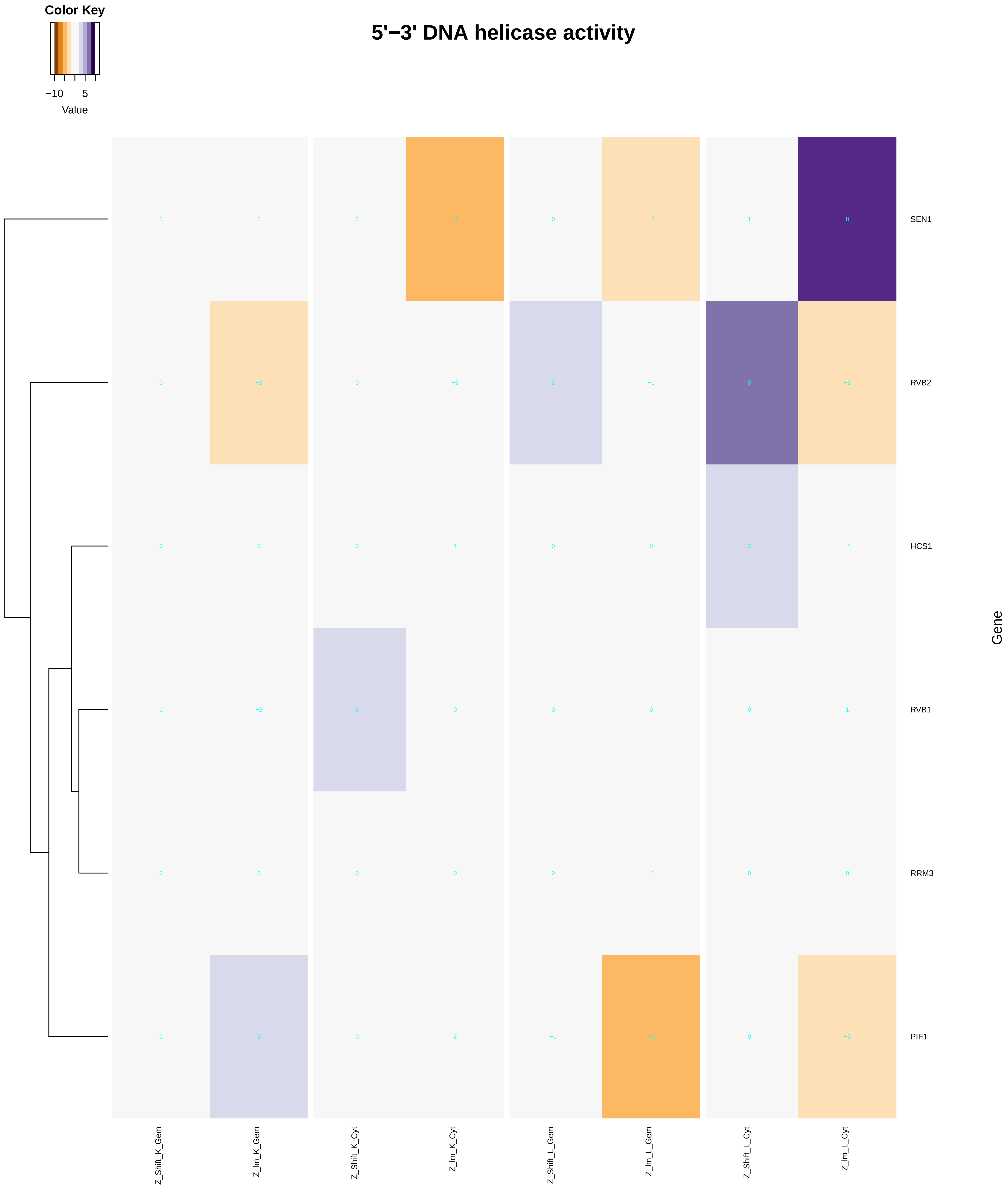

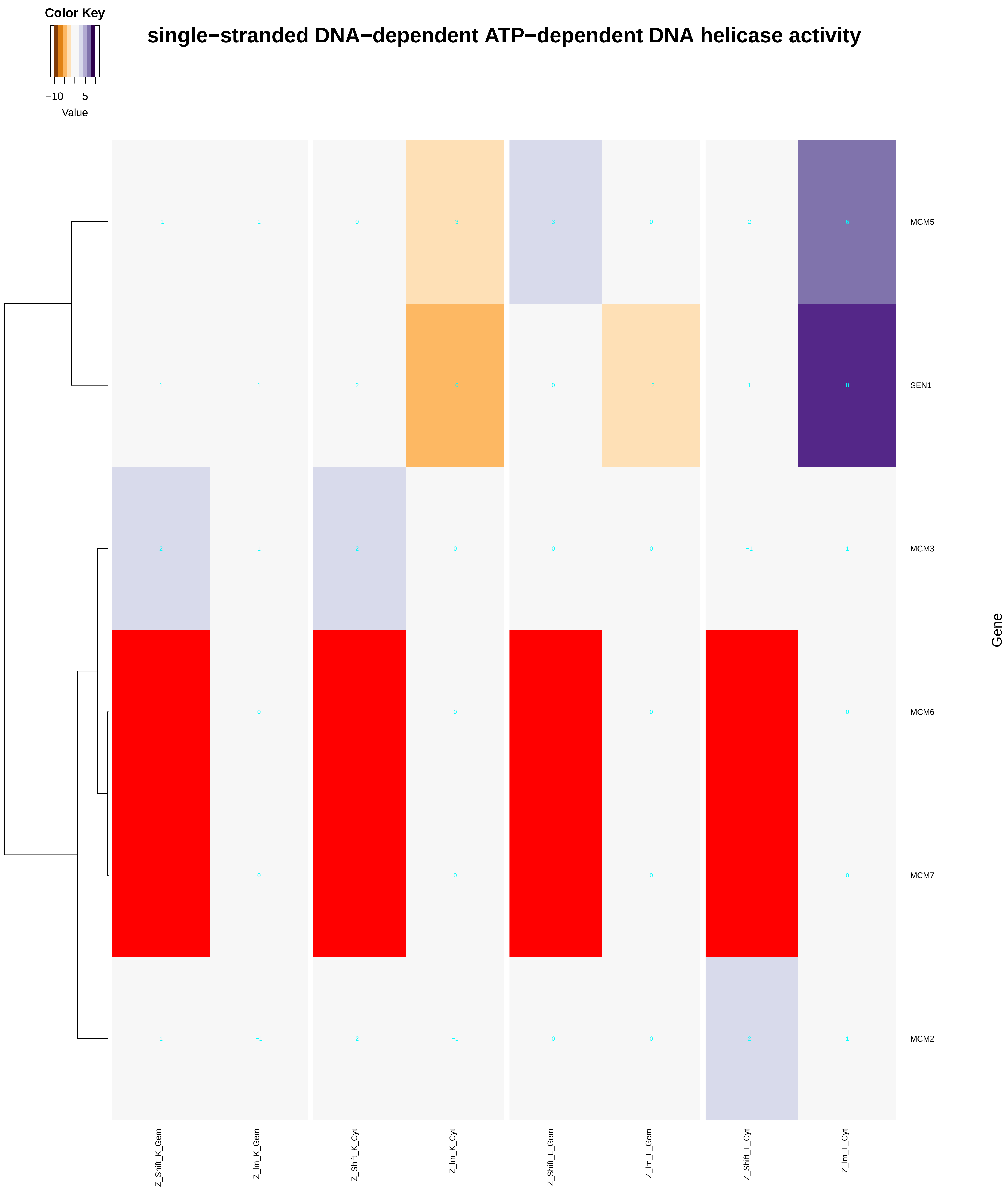

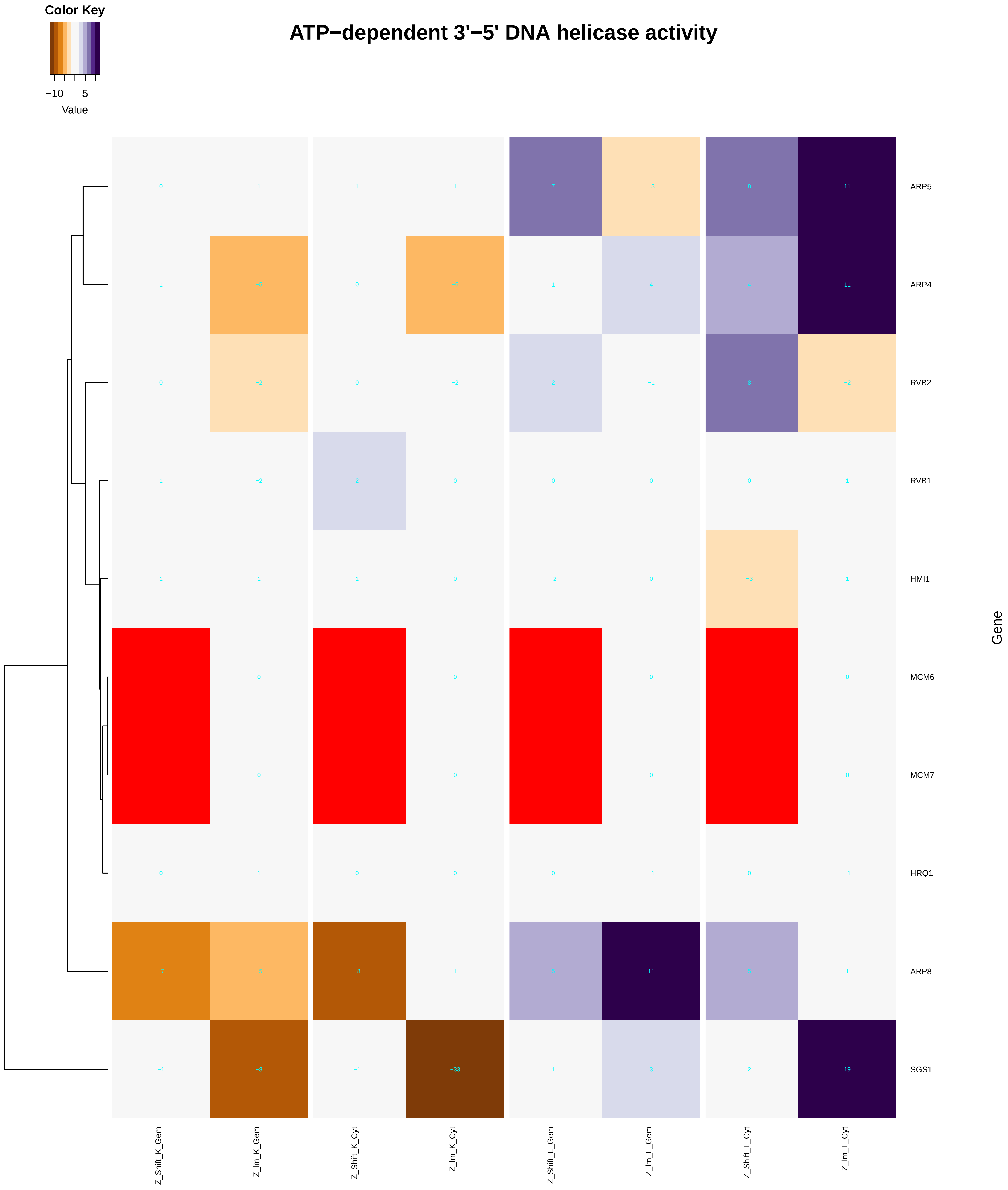

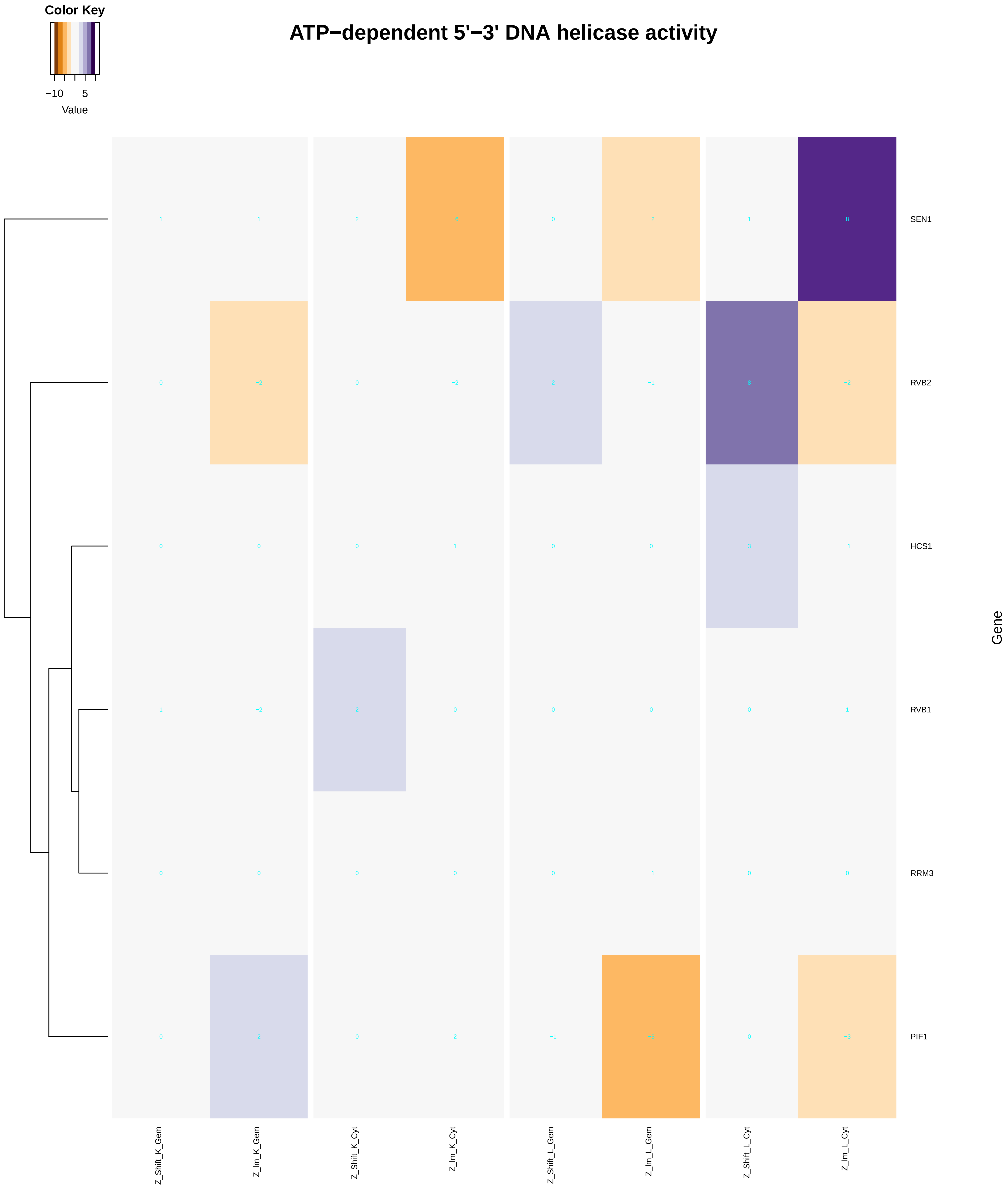

### DNA_packaging.pdf

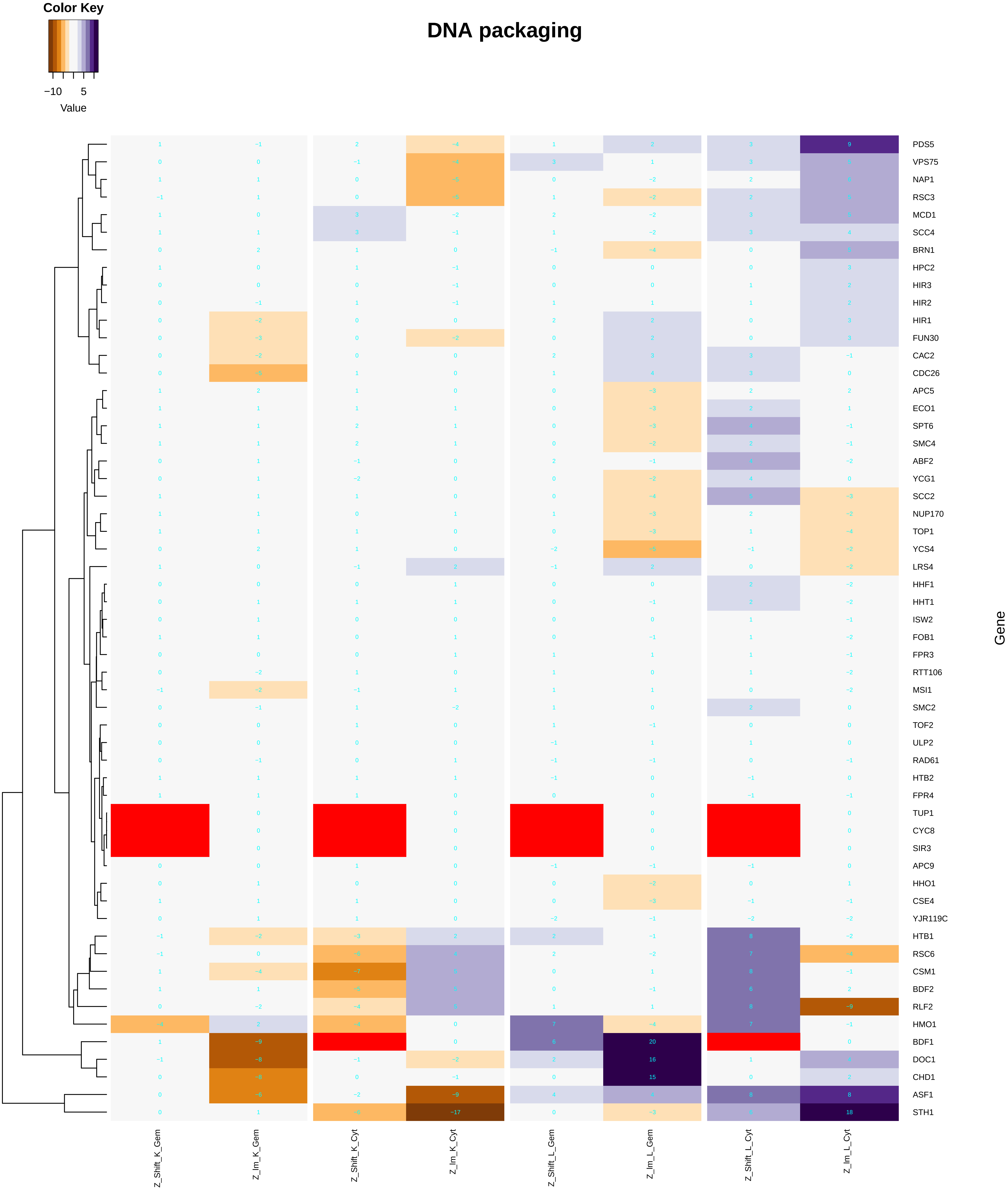

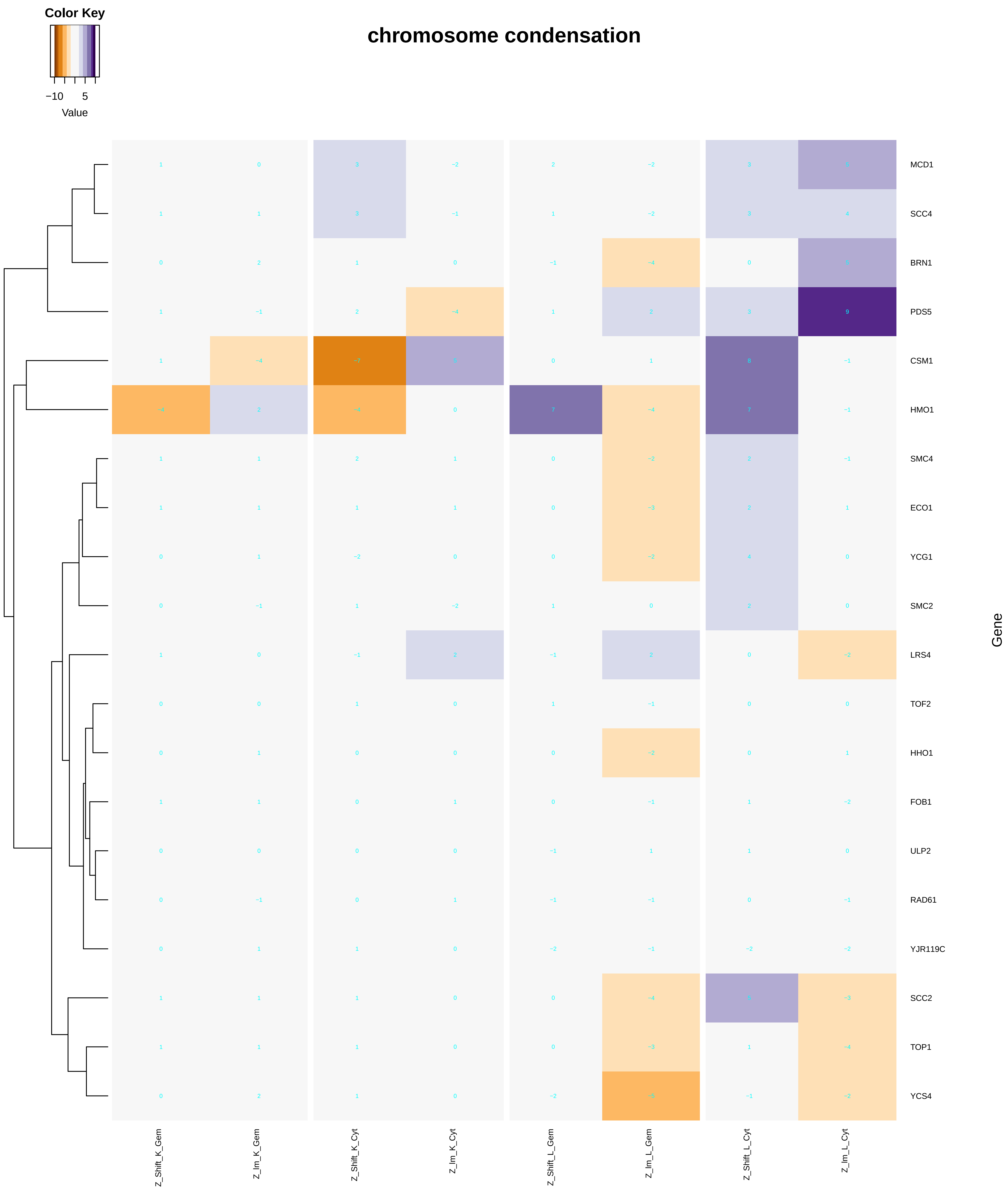

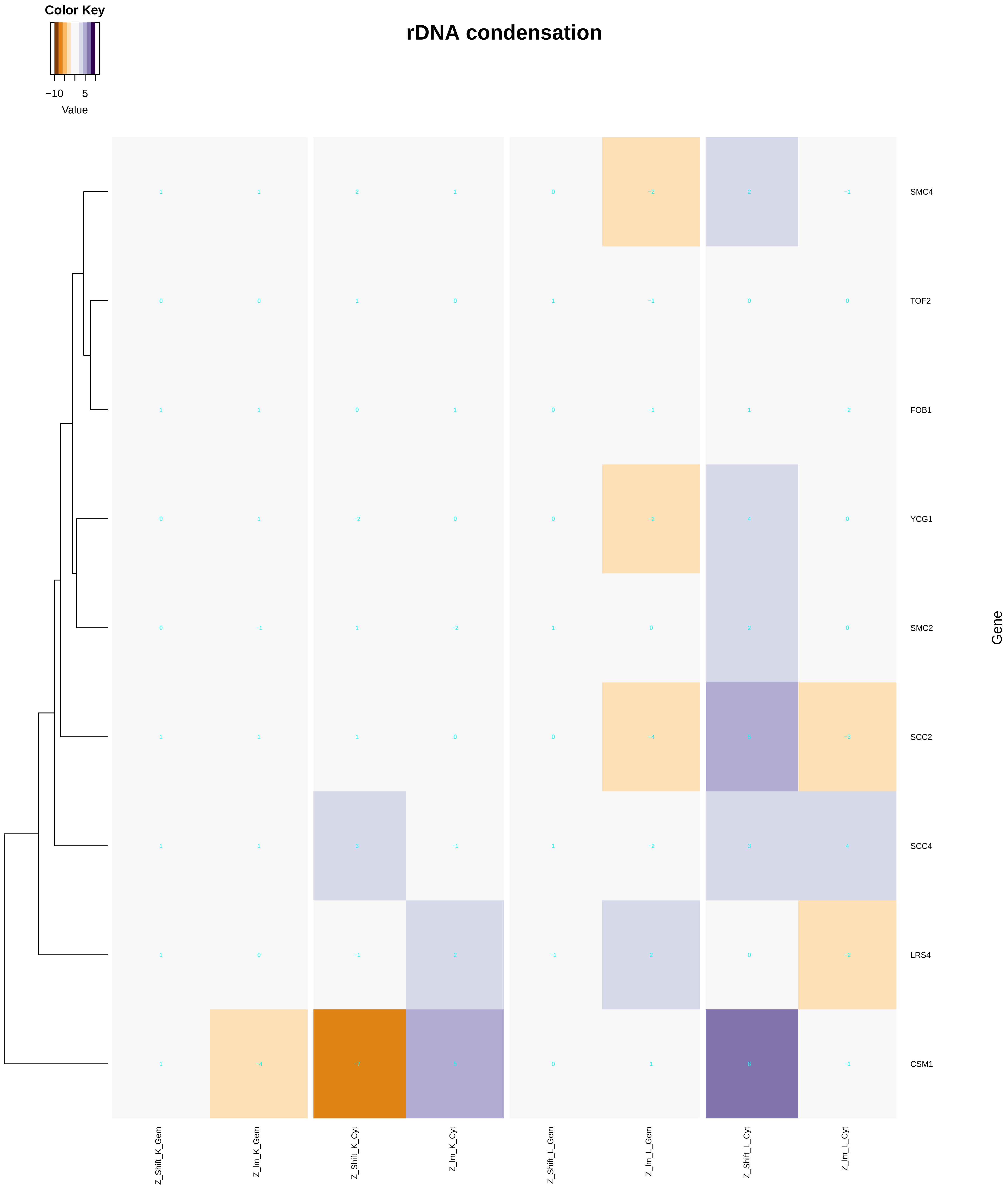

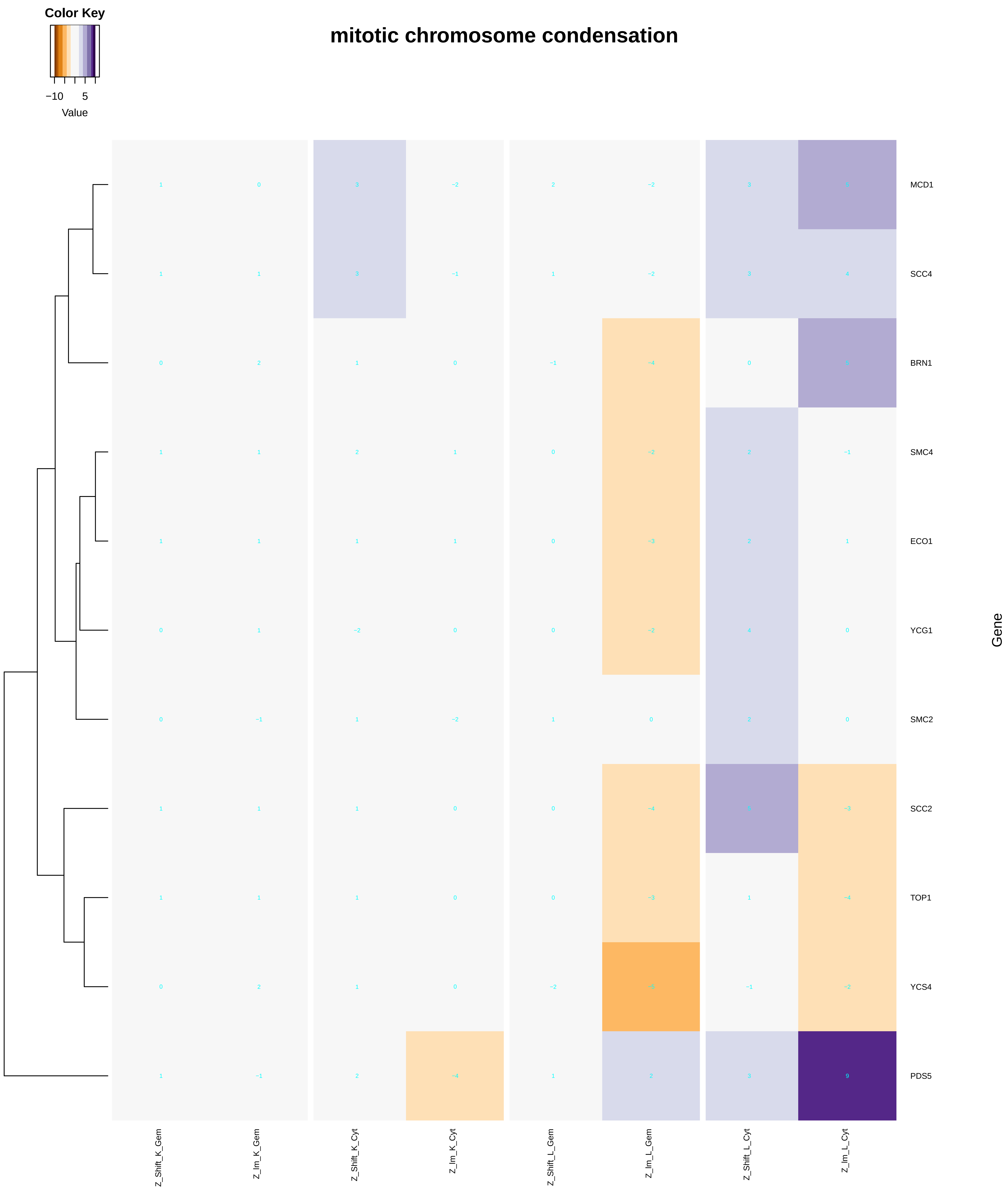

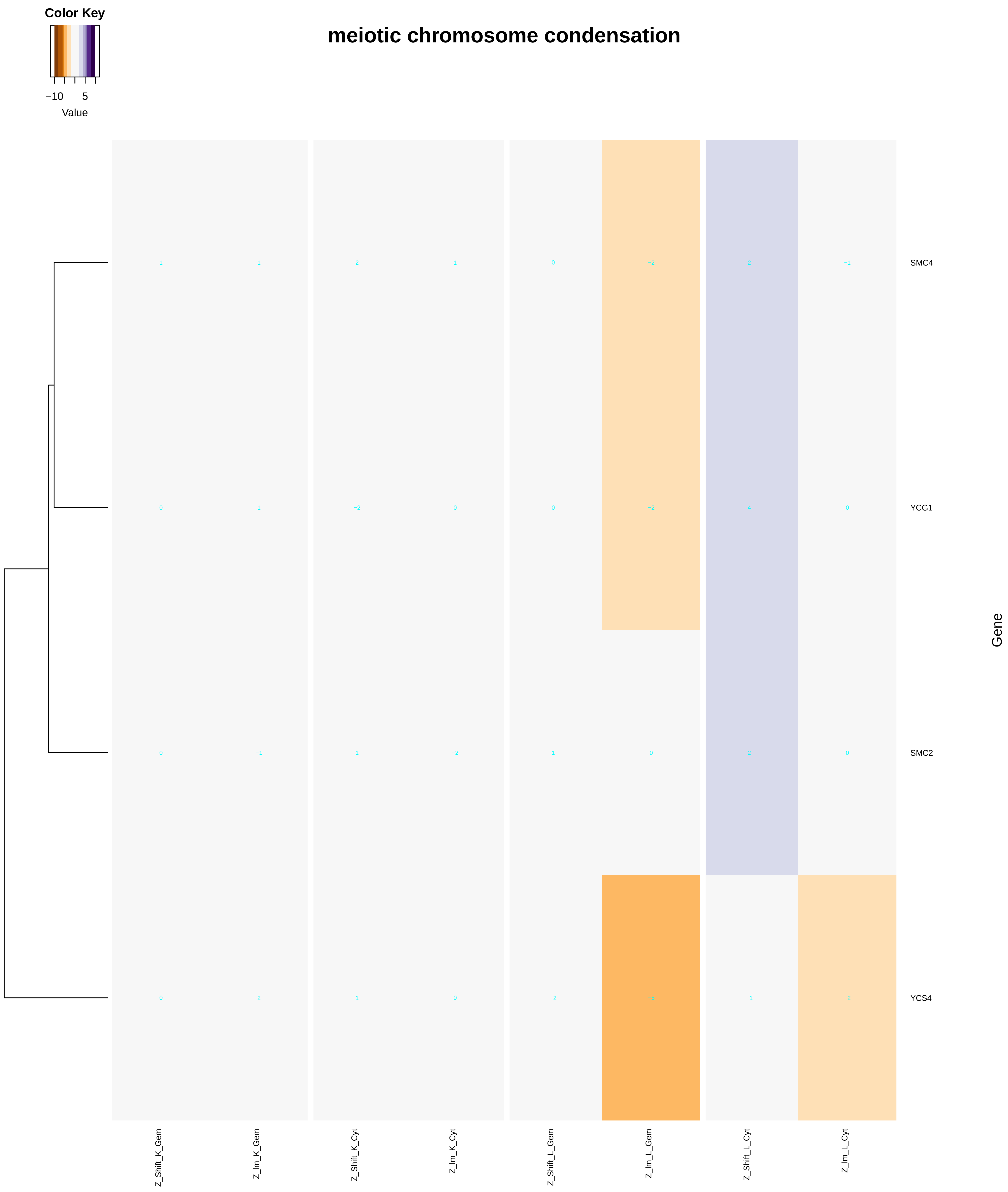

### DNA_replication.pdf

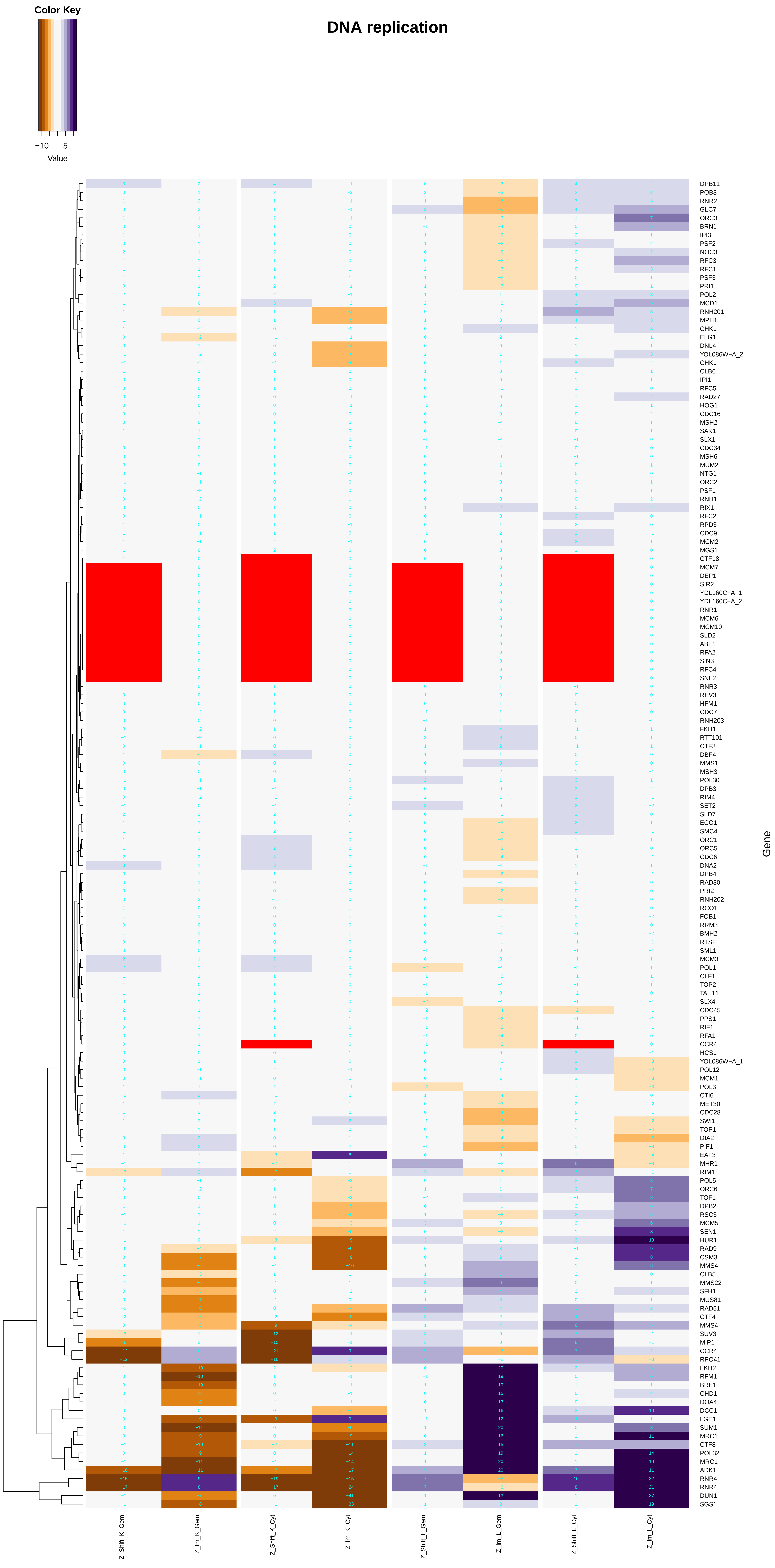

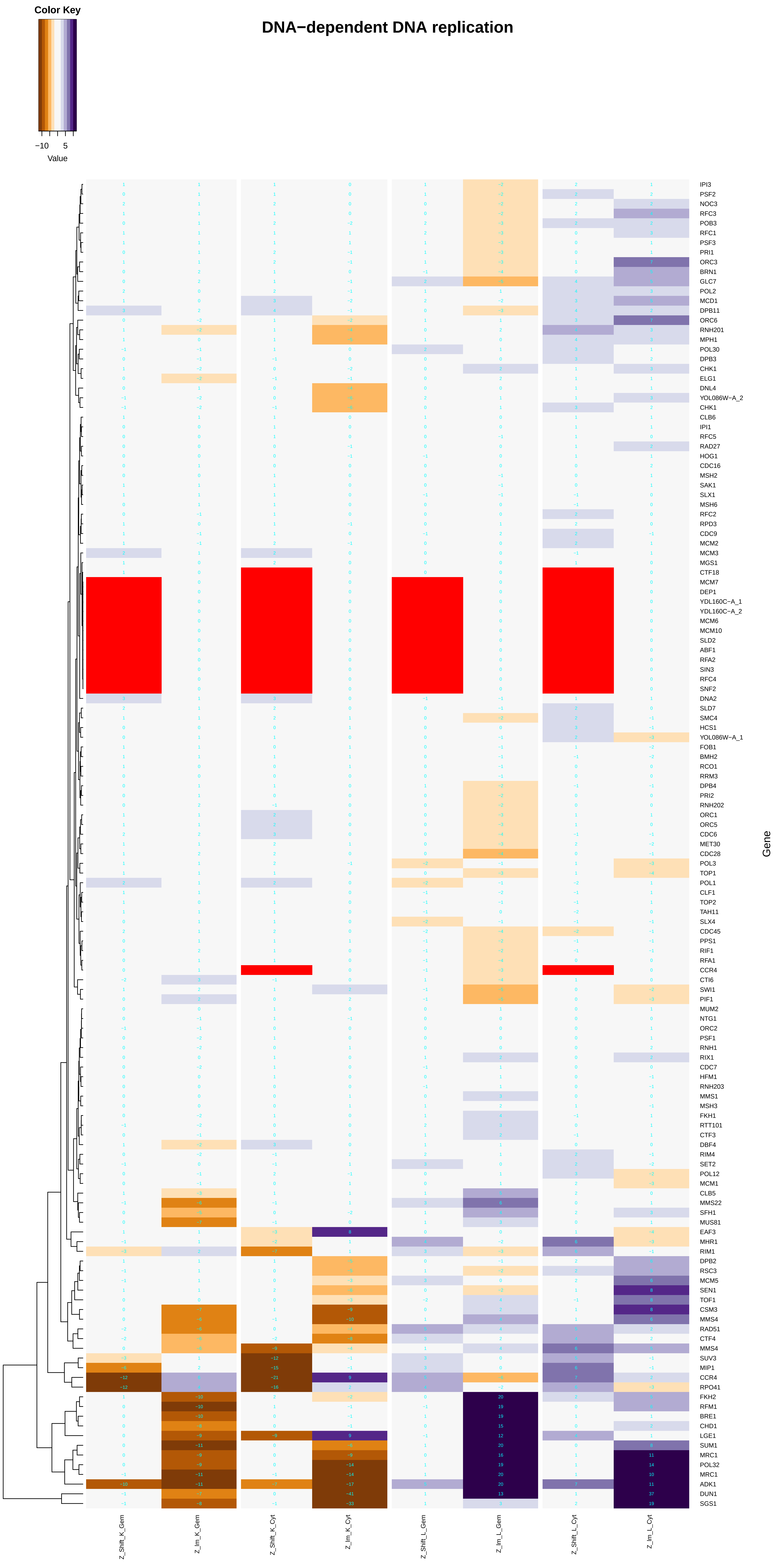

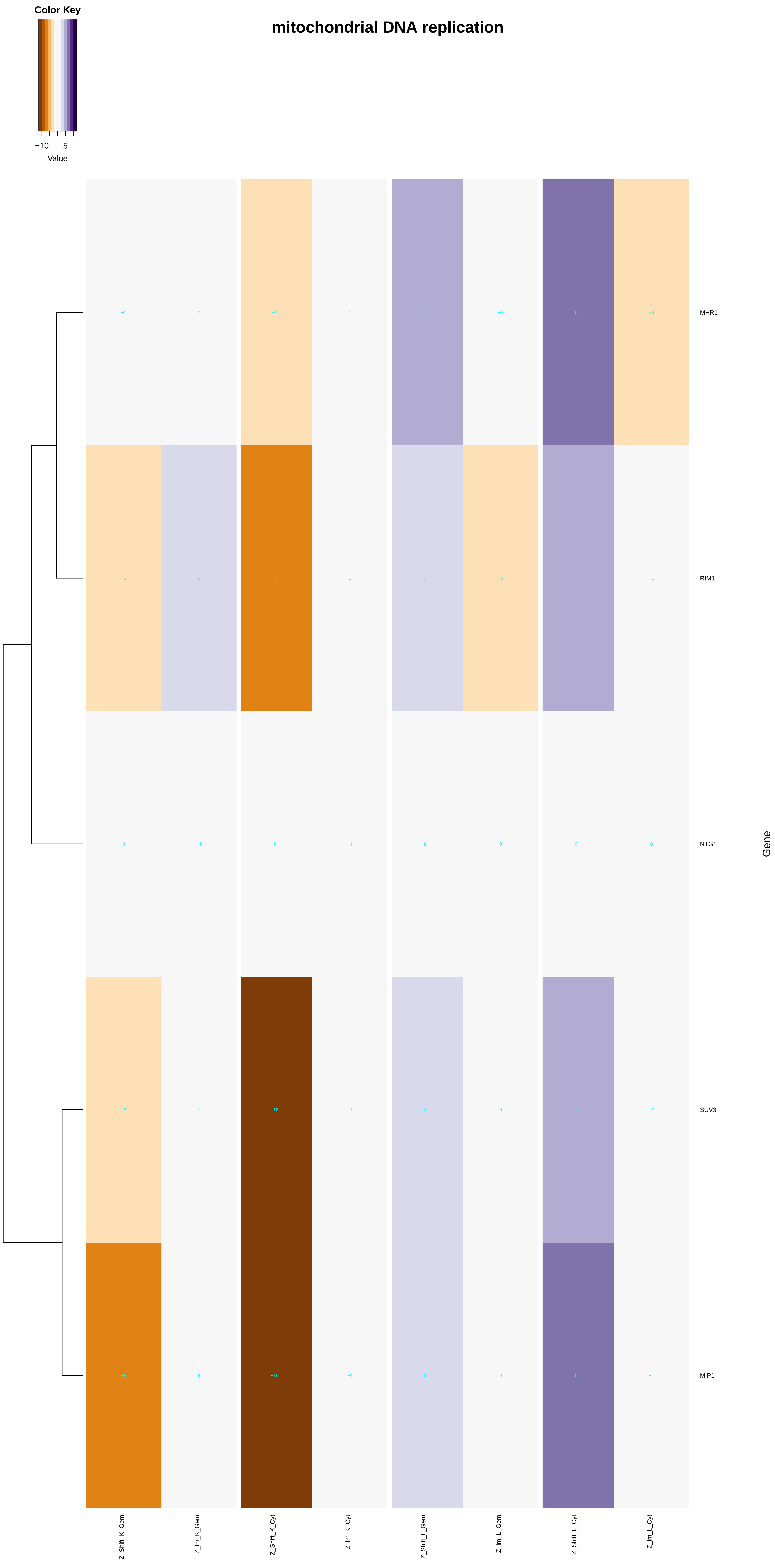

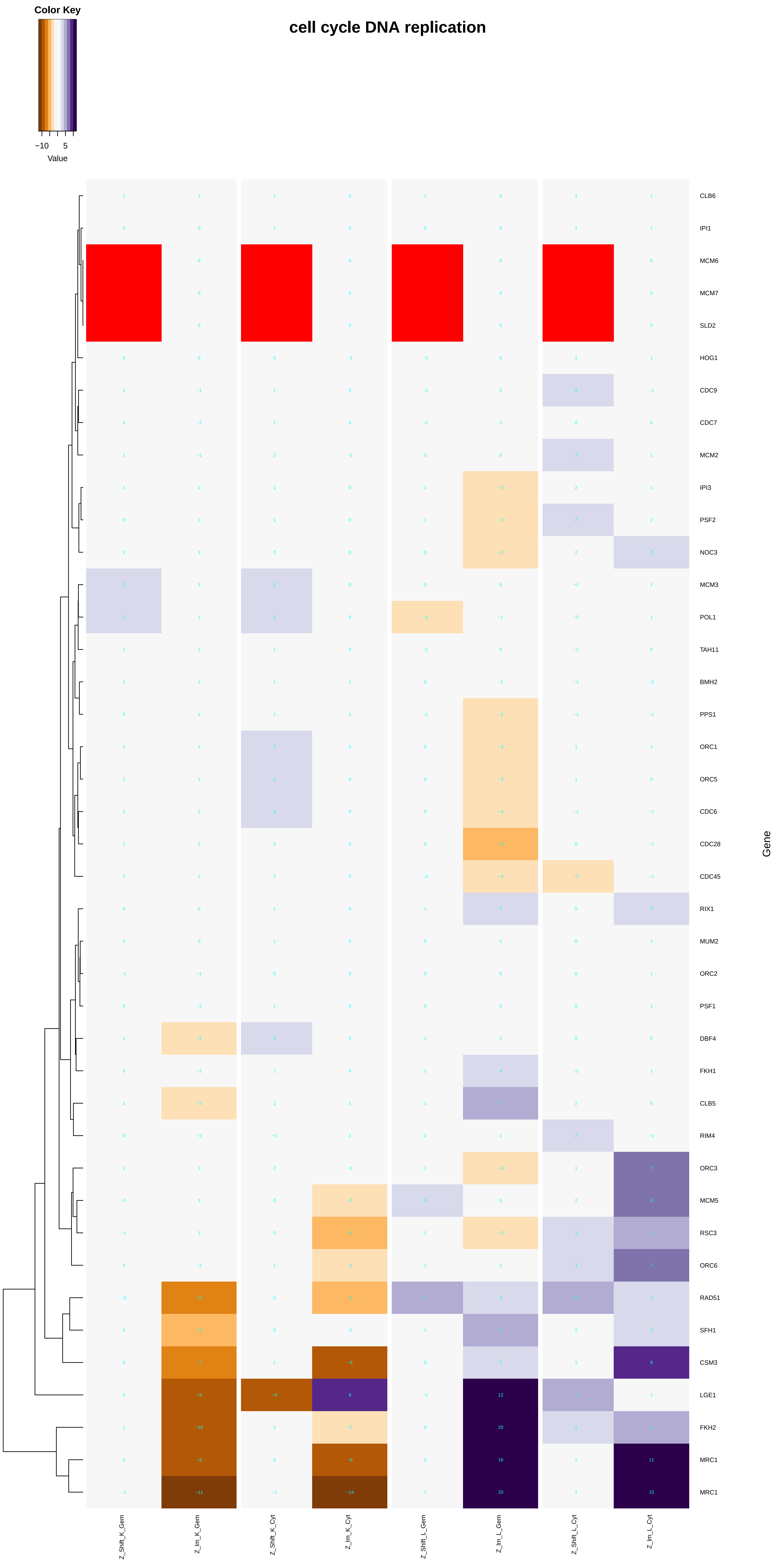

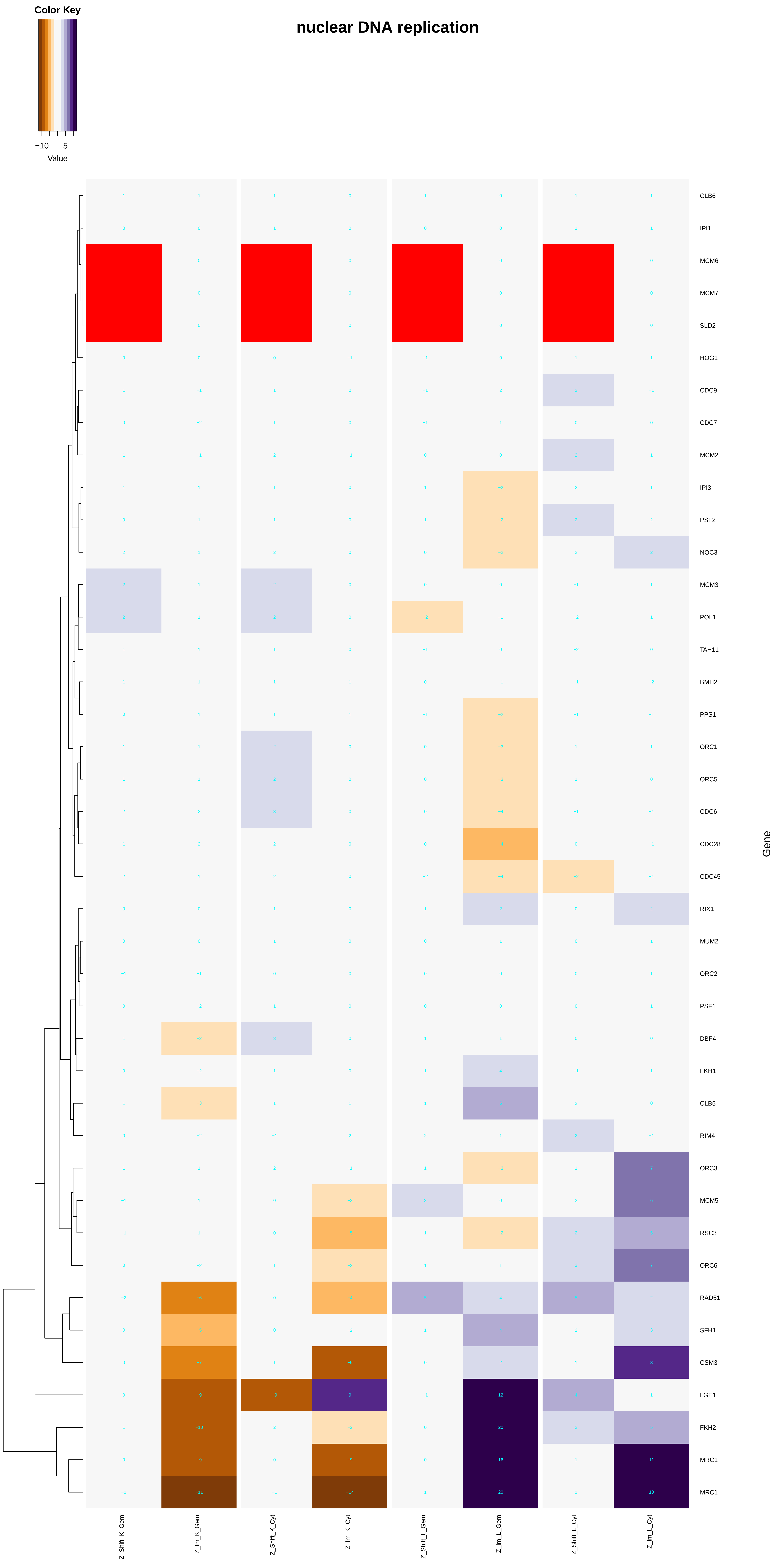

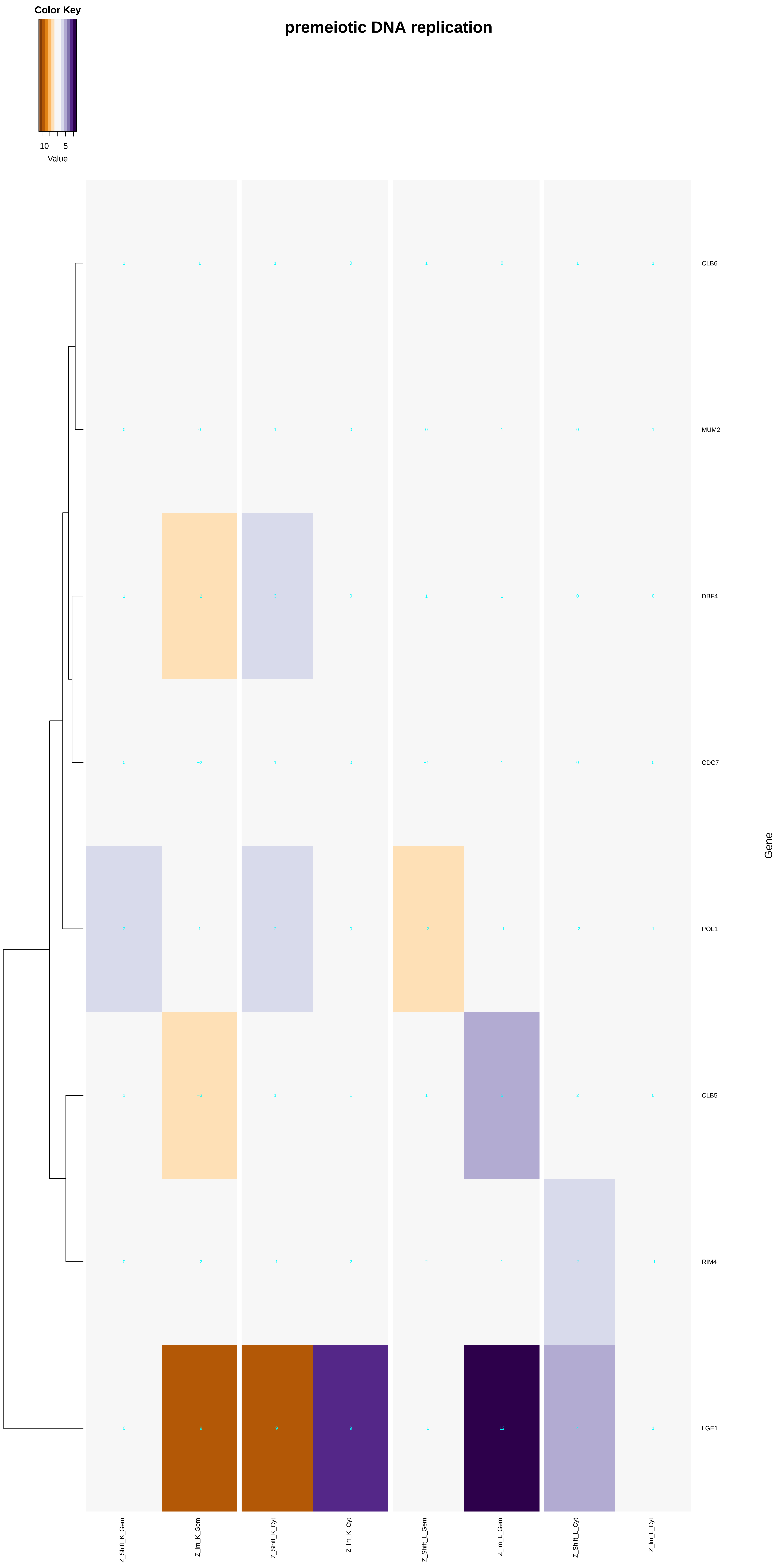

#### mitotic DNA replication

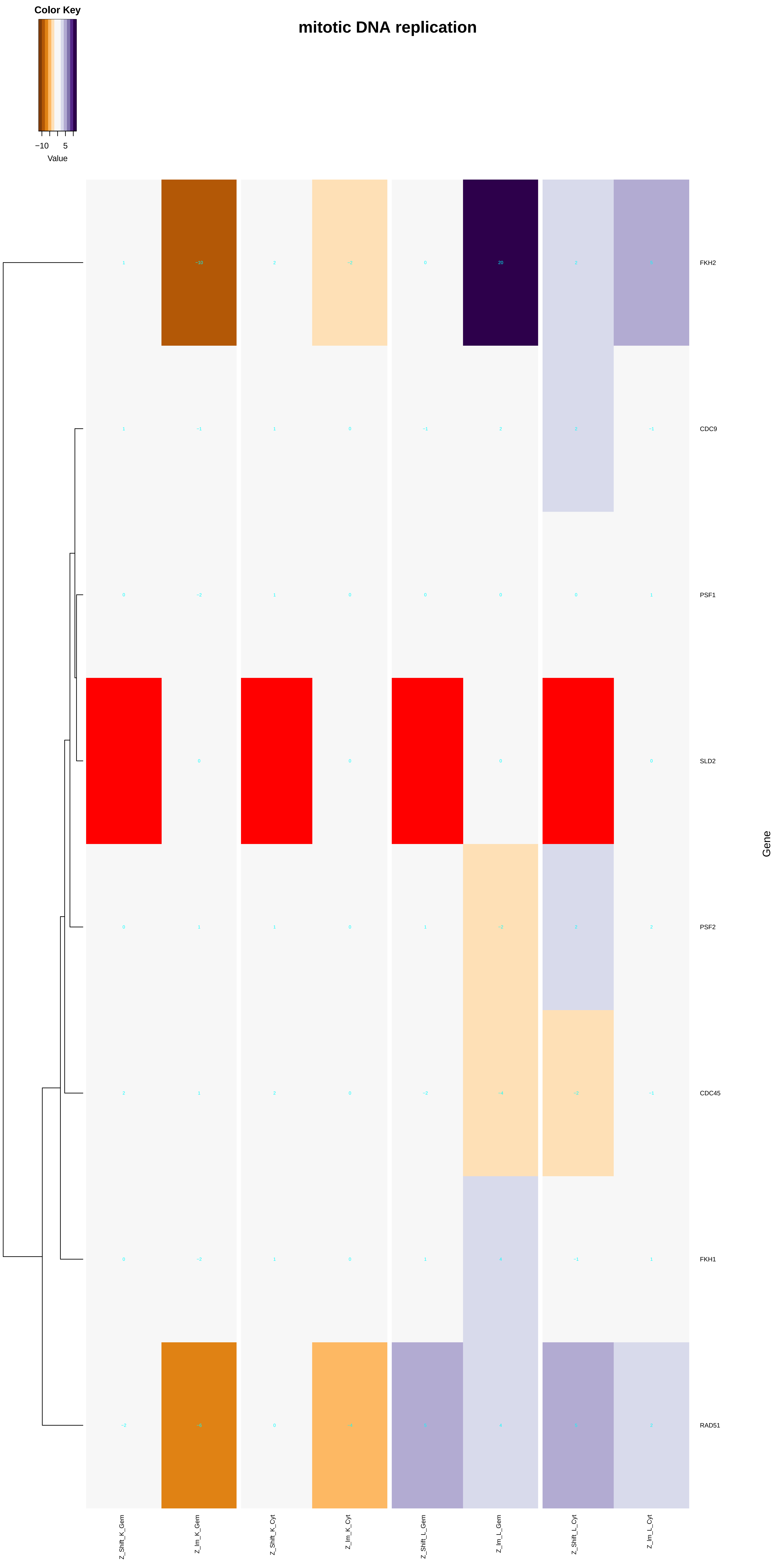

### DNA_replication_initiation.pdf

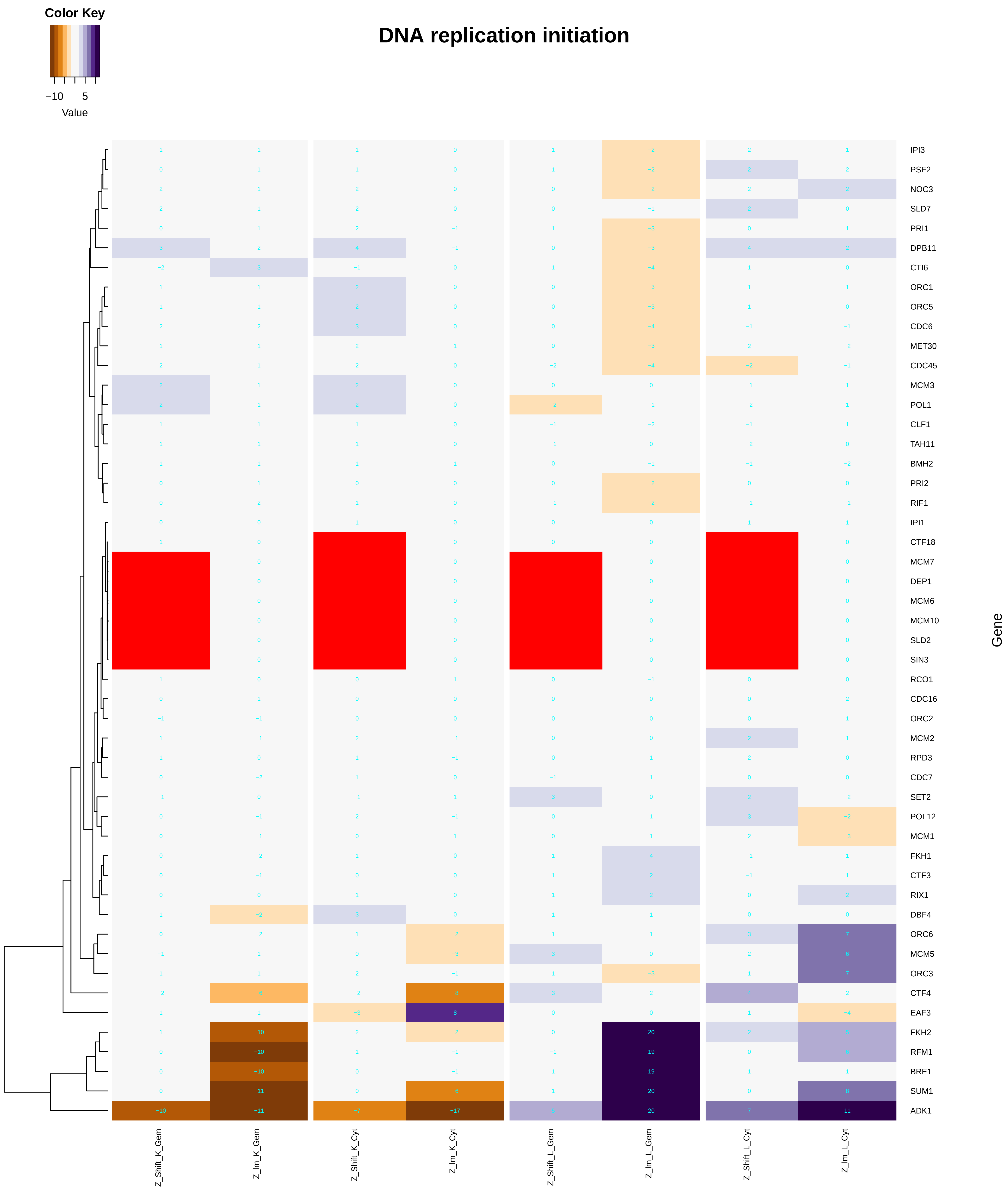

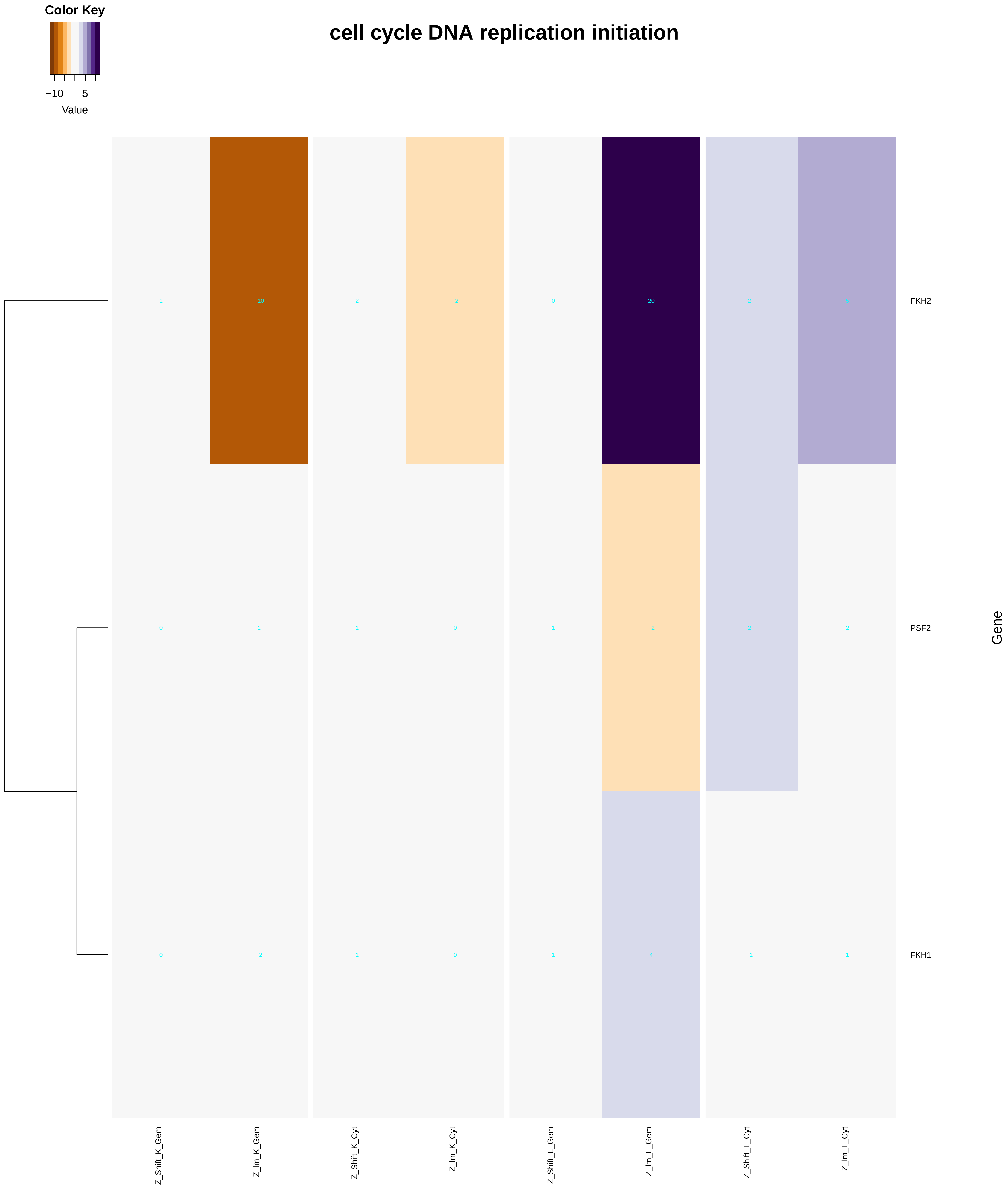

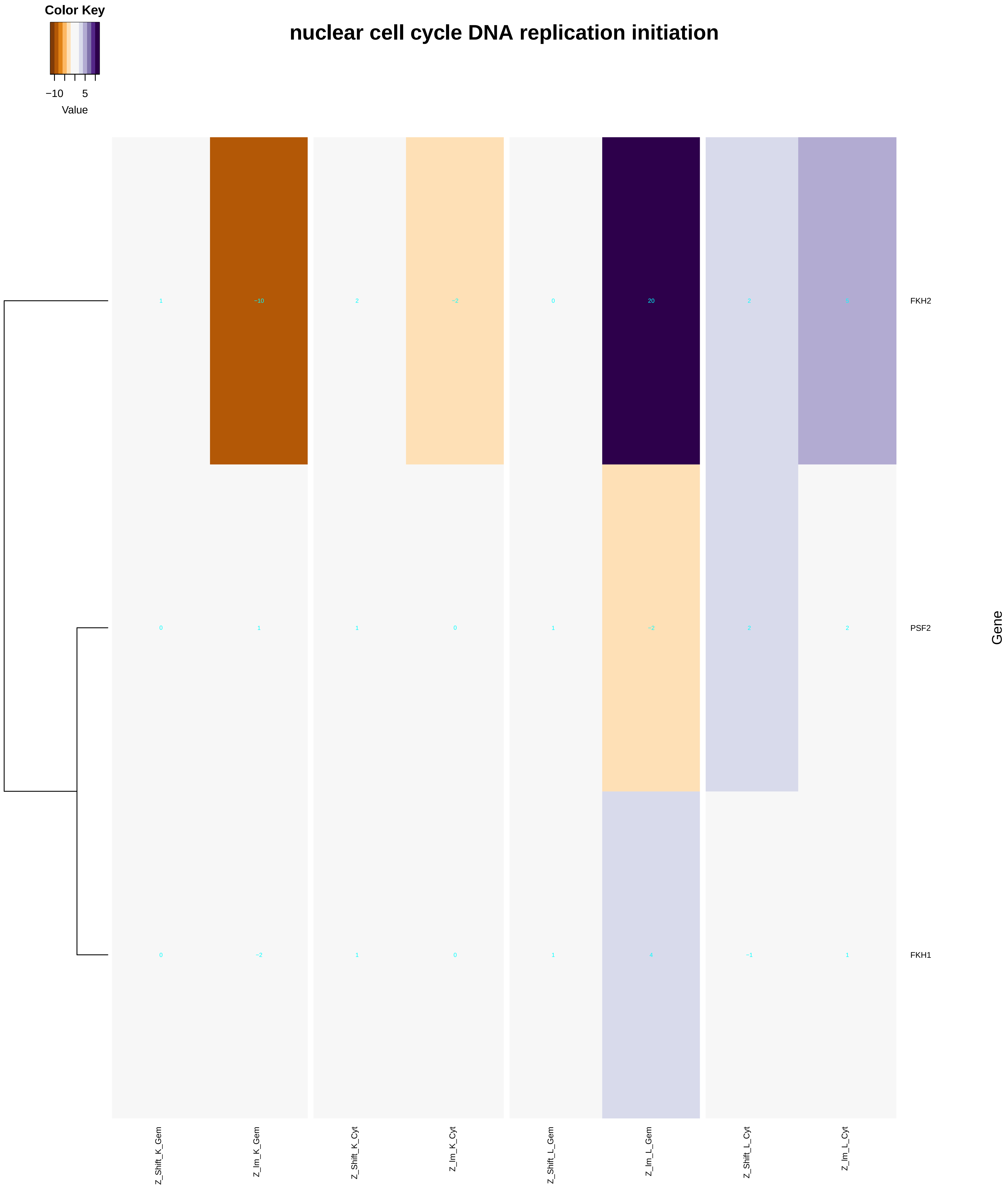

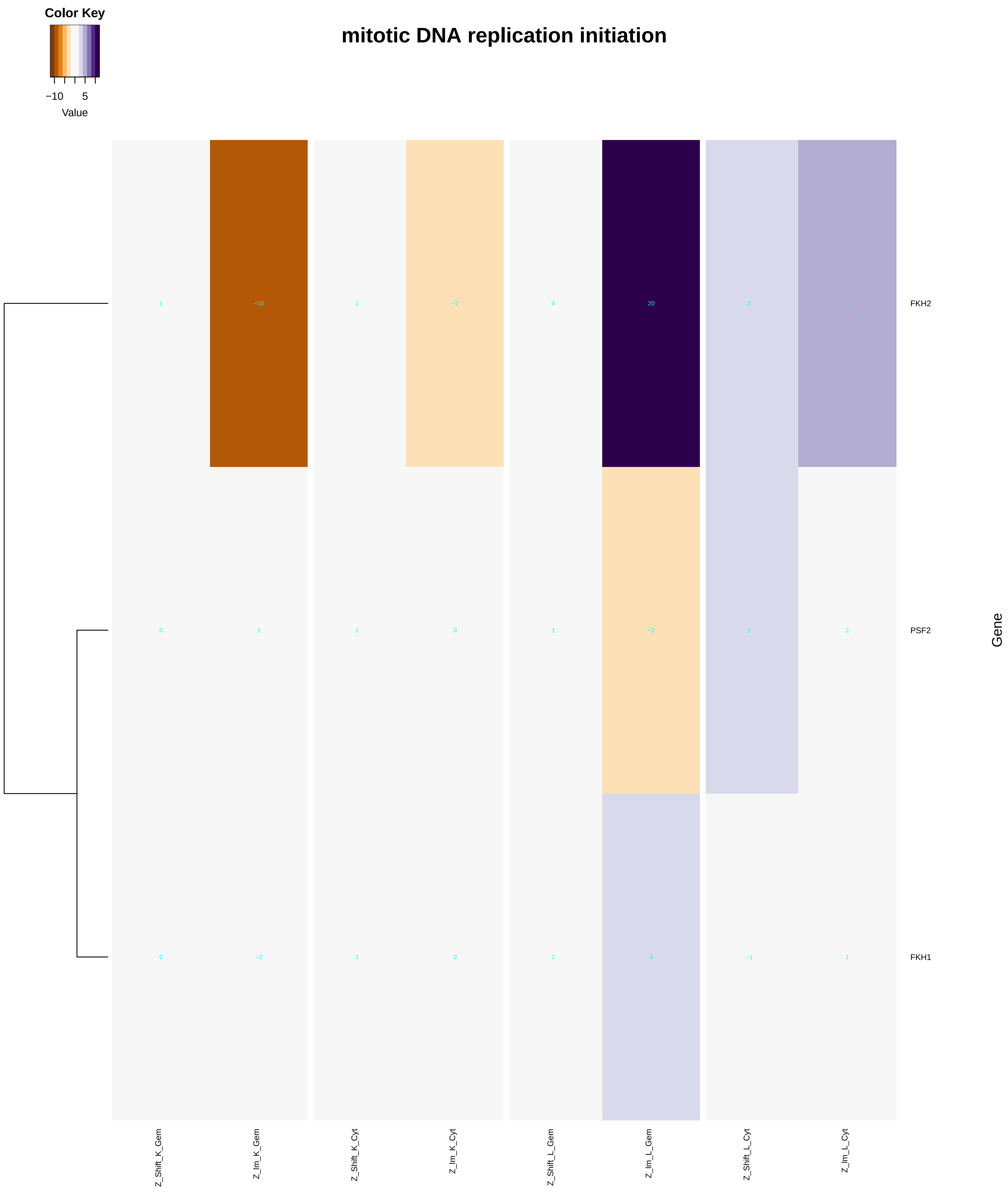

### DNA_replication_origin_binding.pdf

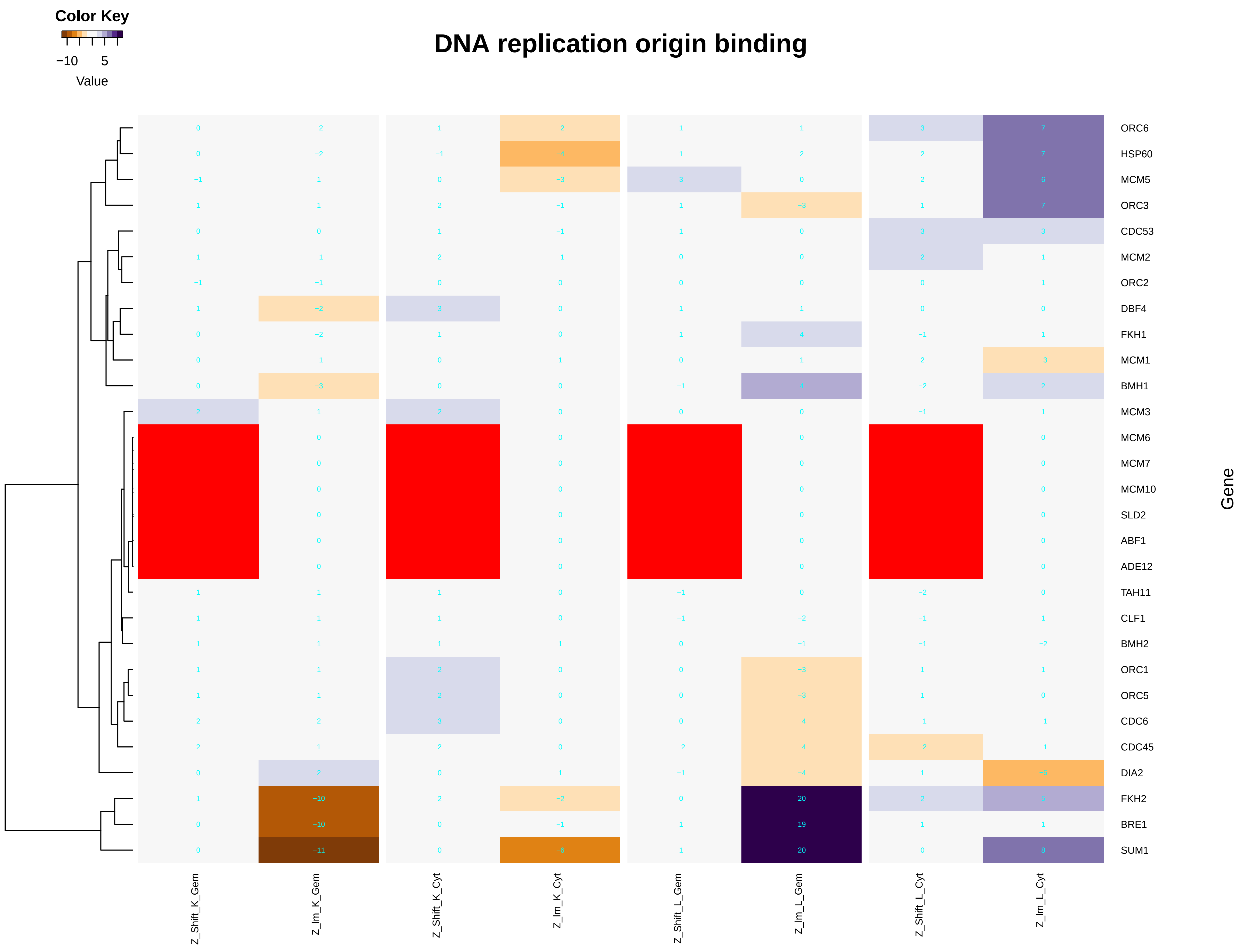

### DNA_replication_preinitiation_complex.pdf

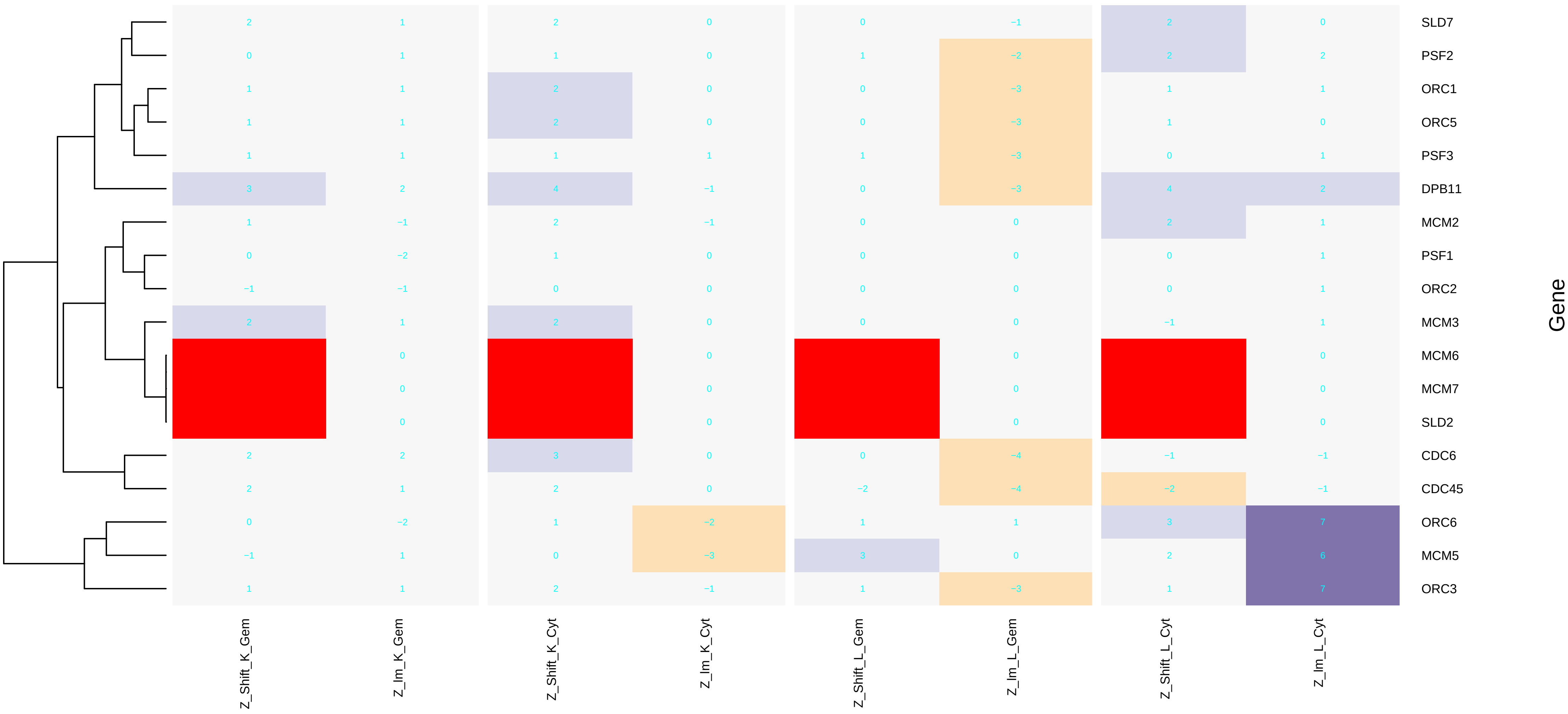

### DNA_topological_change.pdf

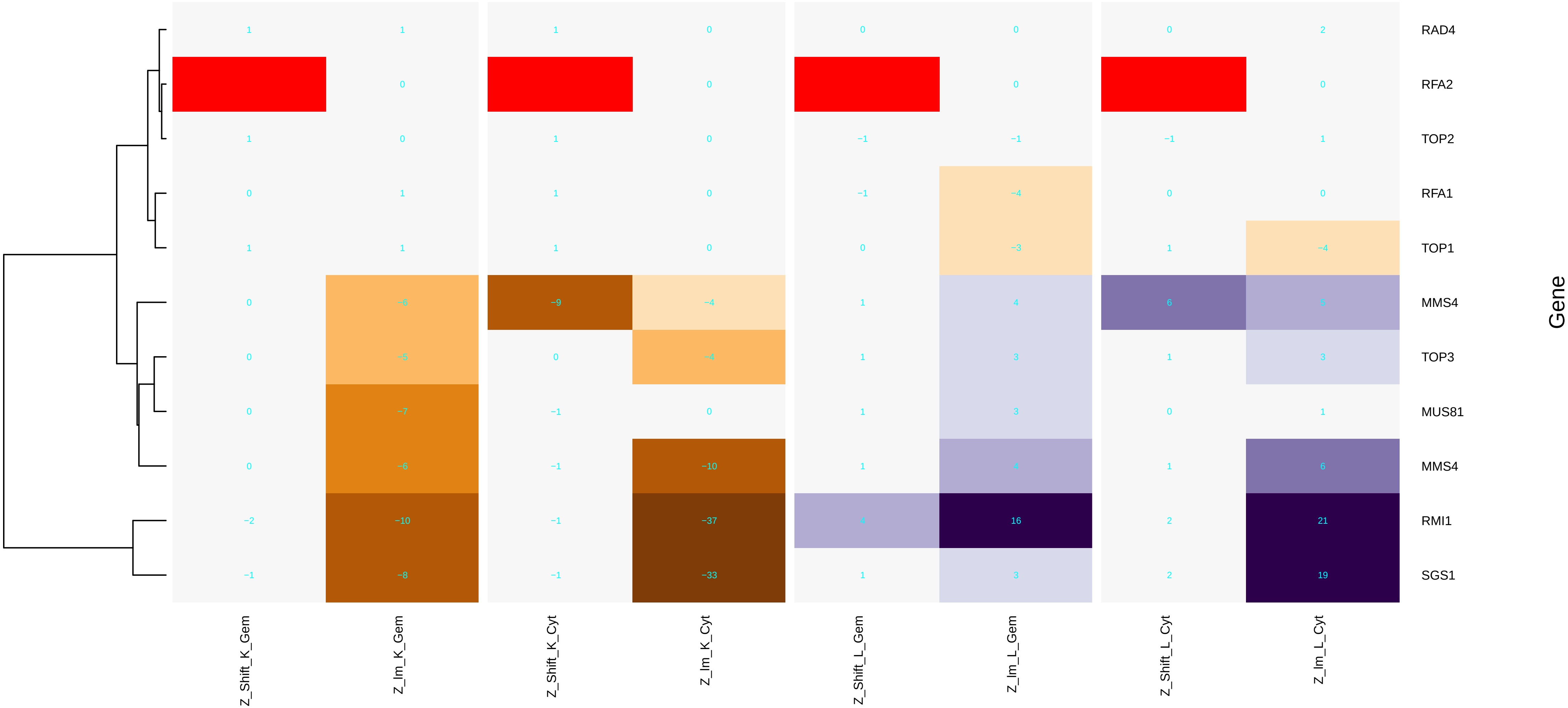

### double-strand_break_repair.pdf

#### double−strand break repair

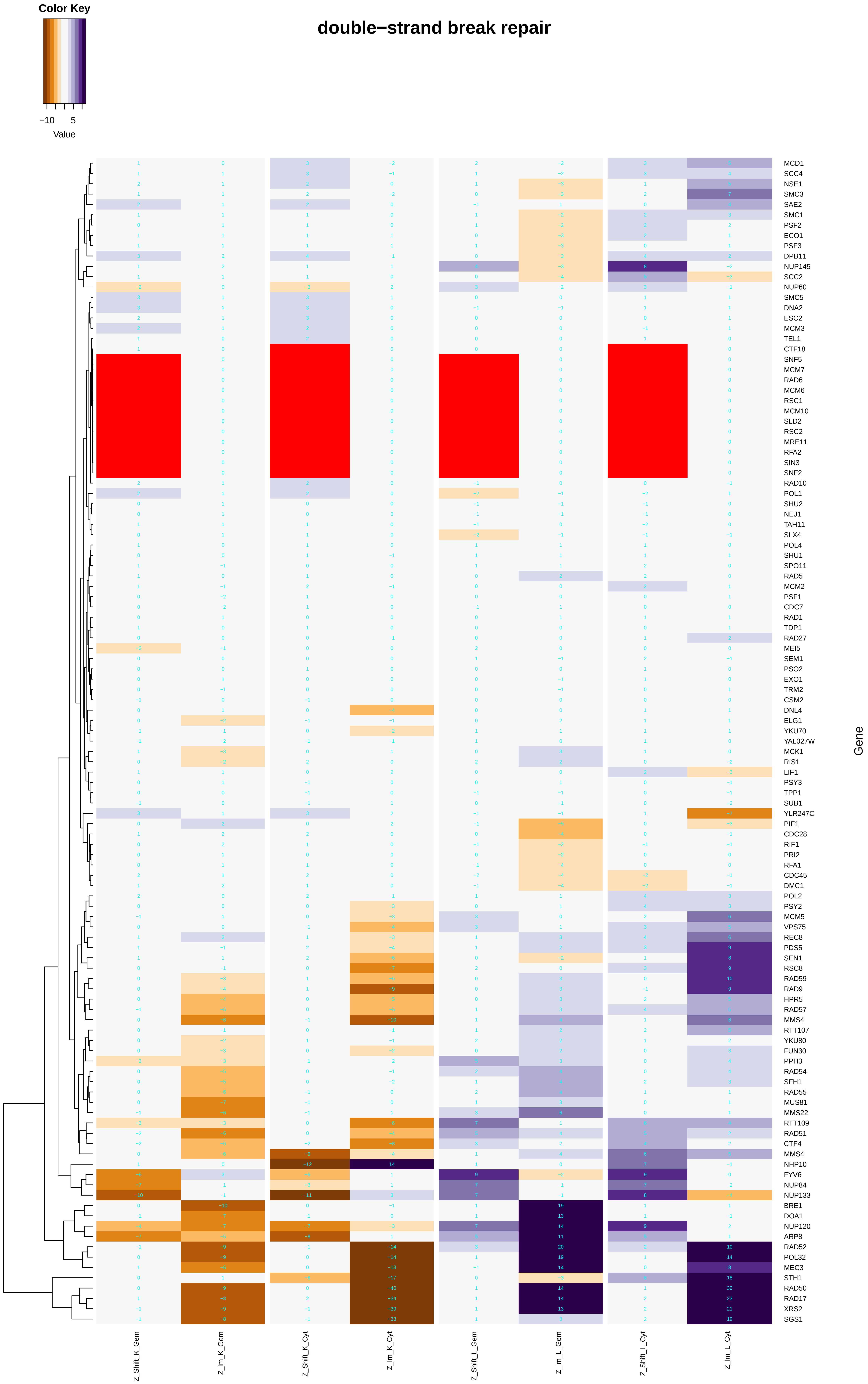

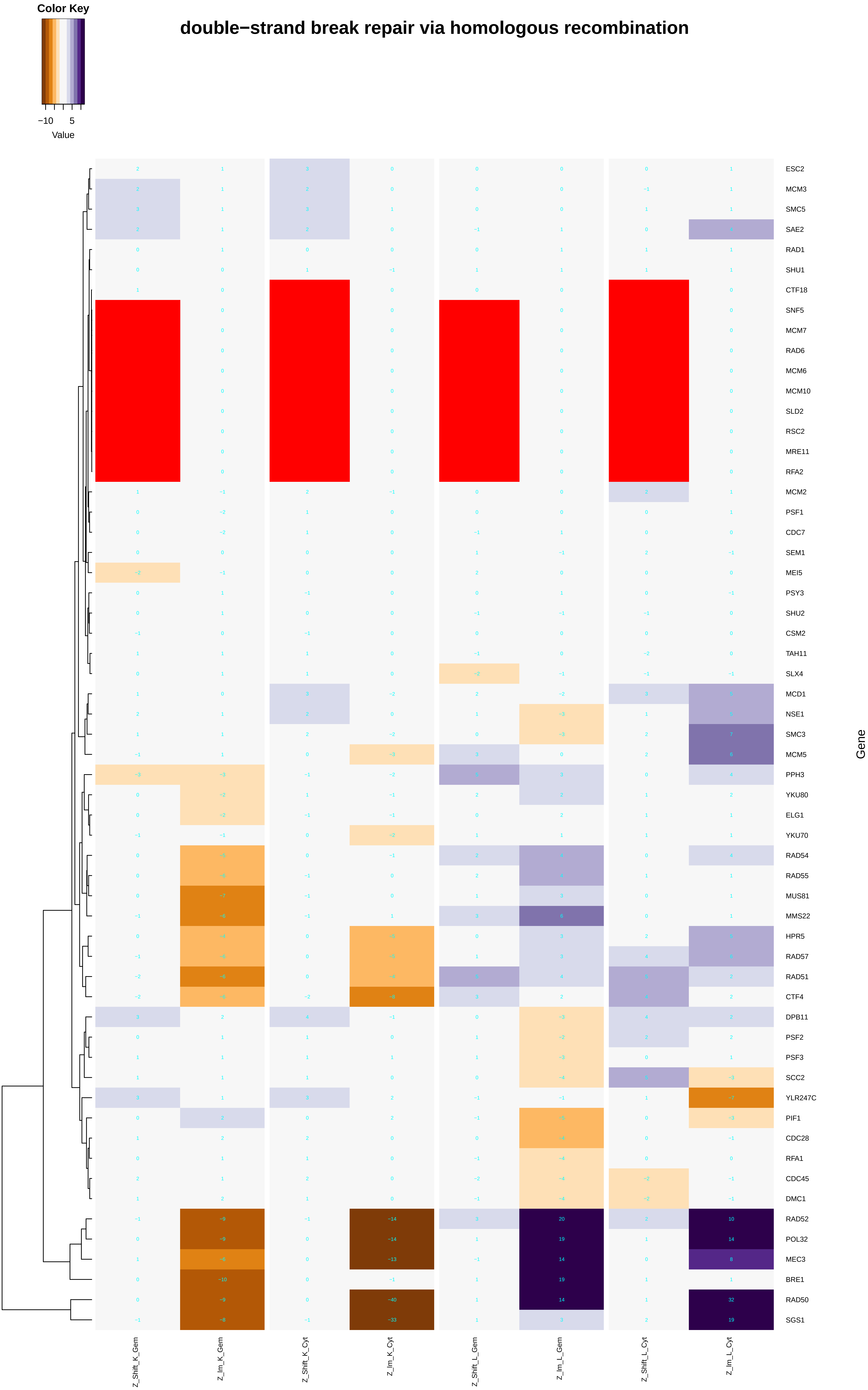

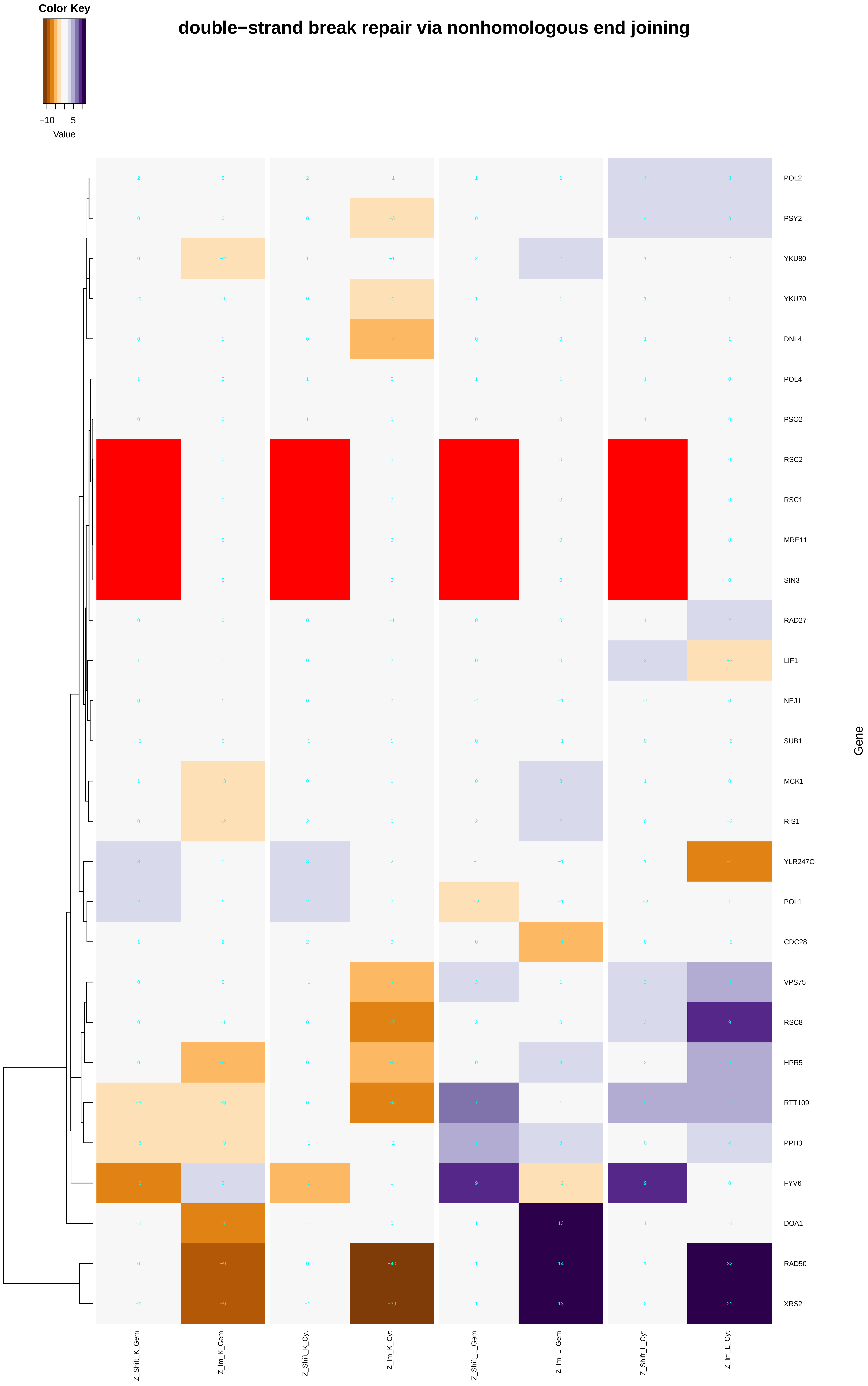

#### double−strand break repair via single−strand annealing

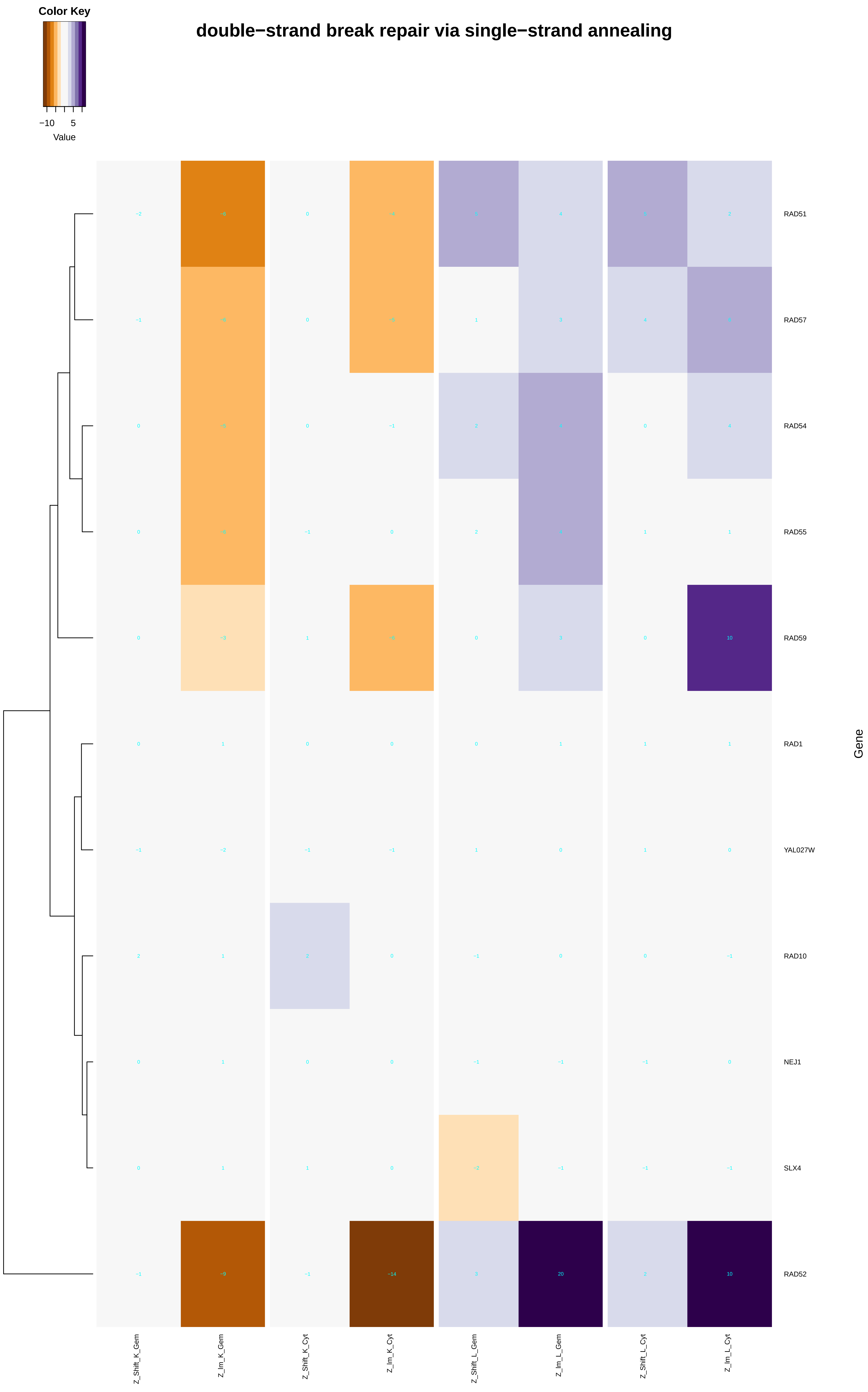

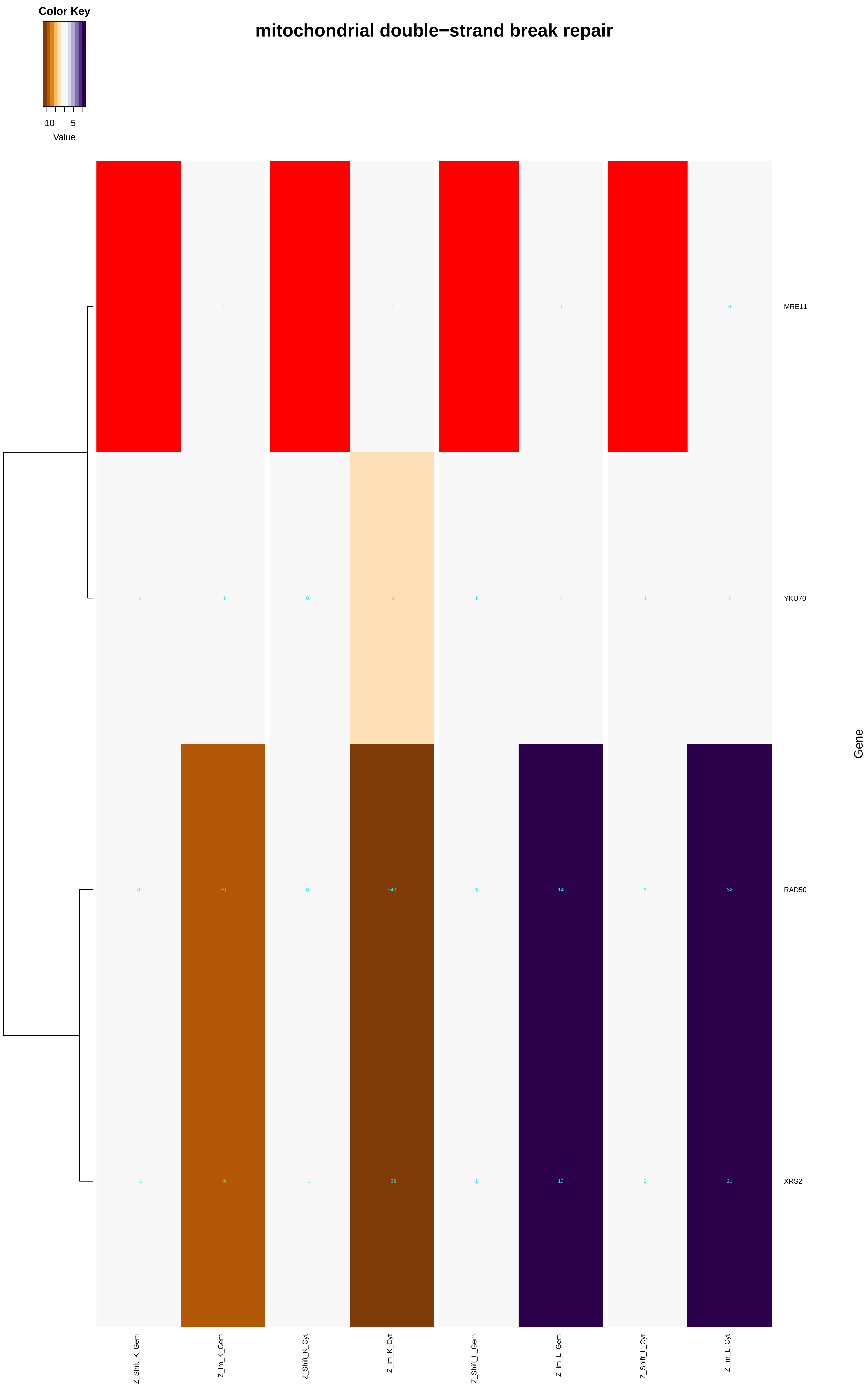

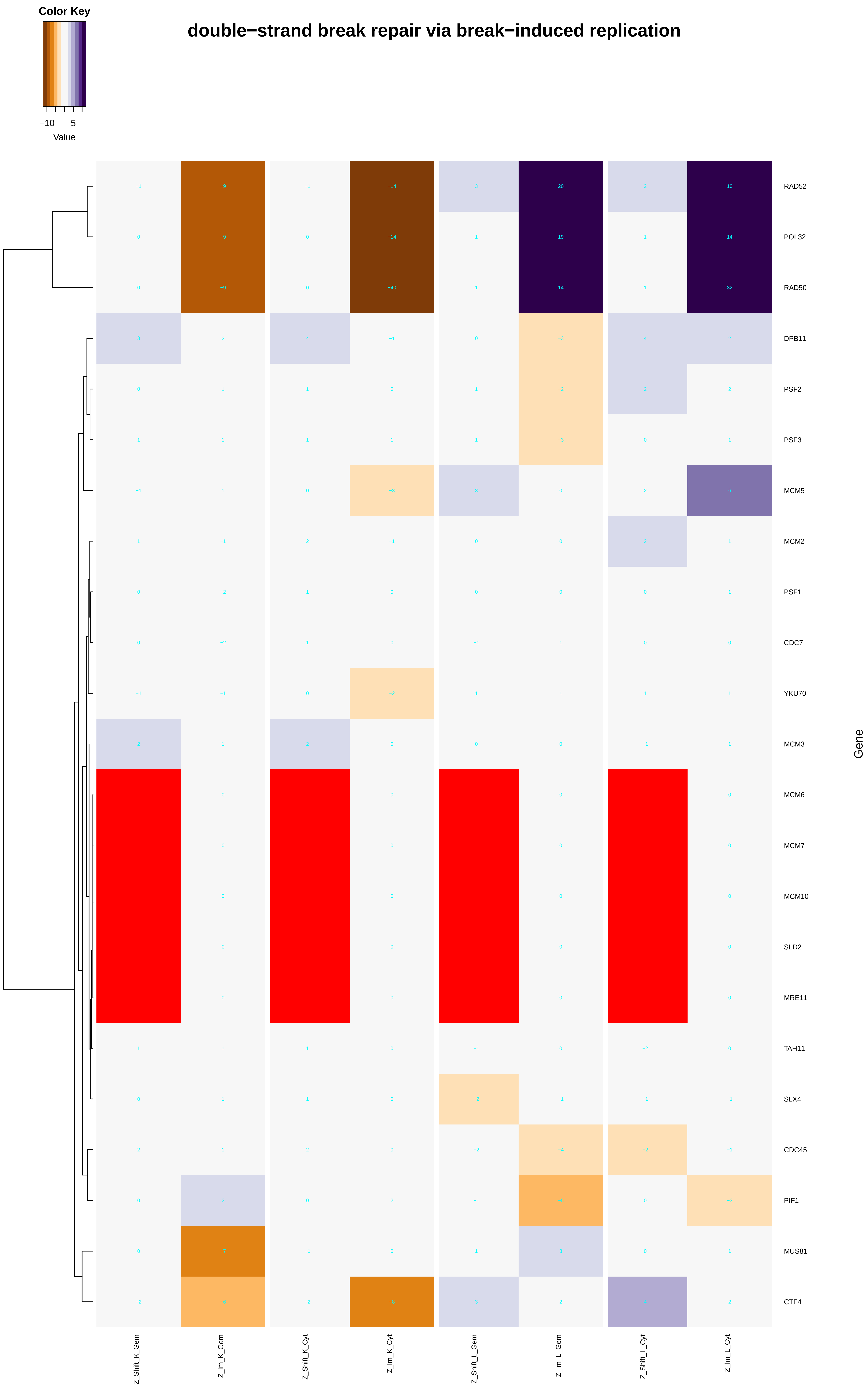

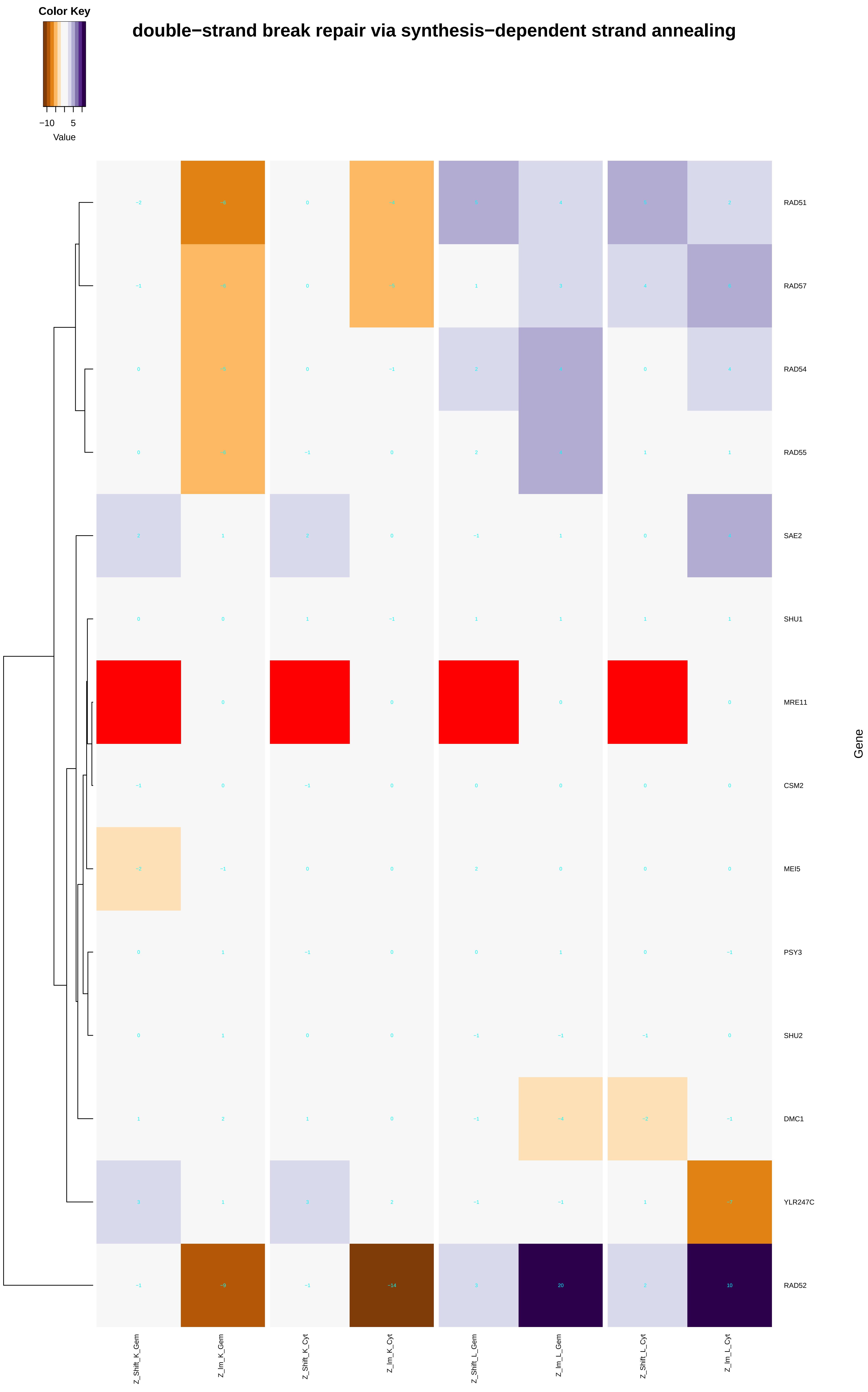

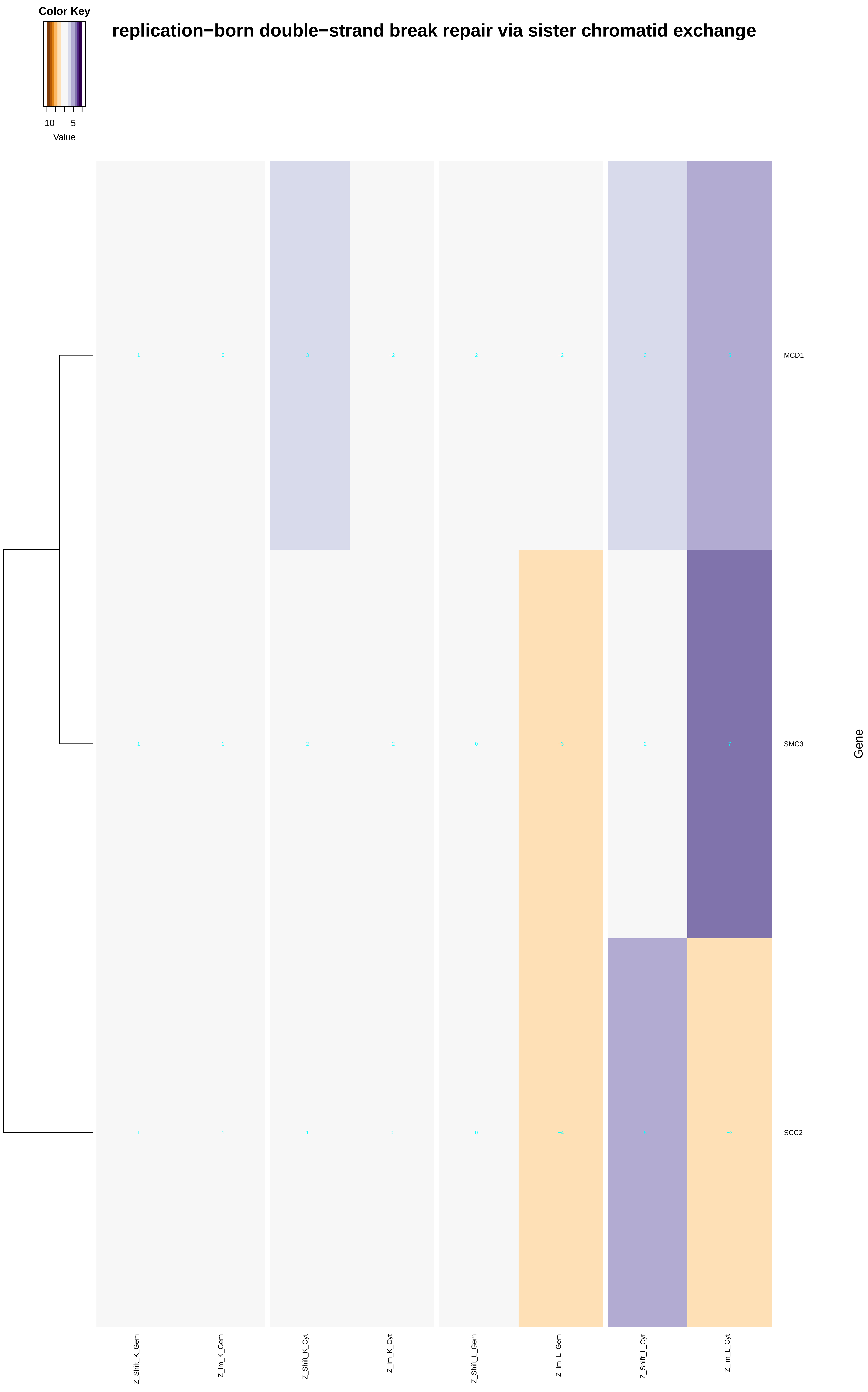

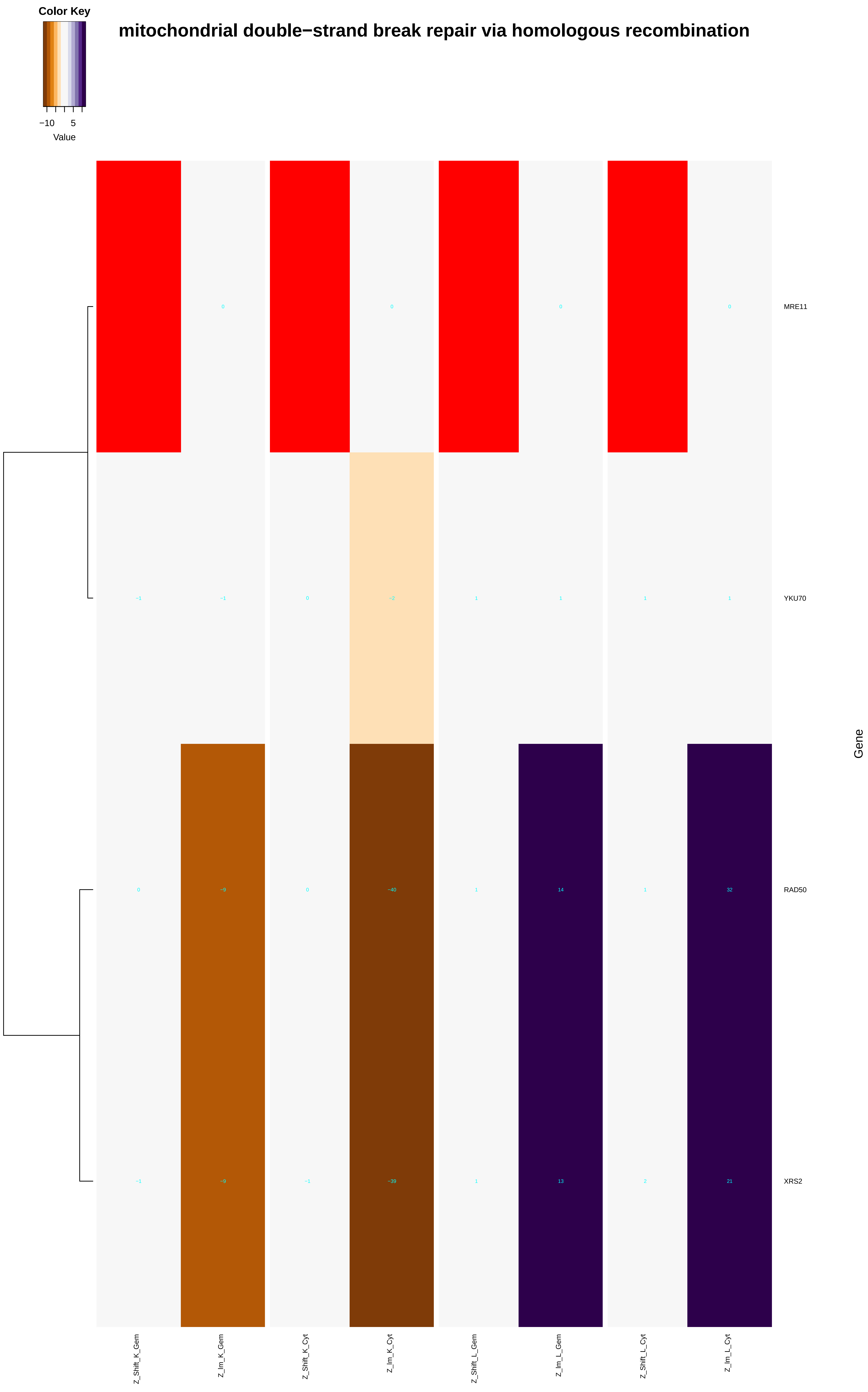

### double-strand_break_repair_via_break-induced_replication.pdf

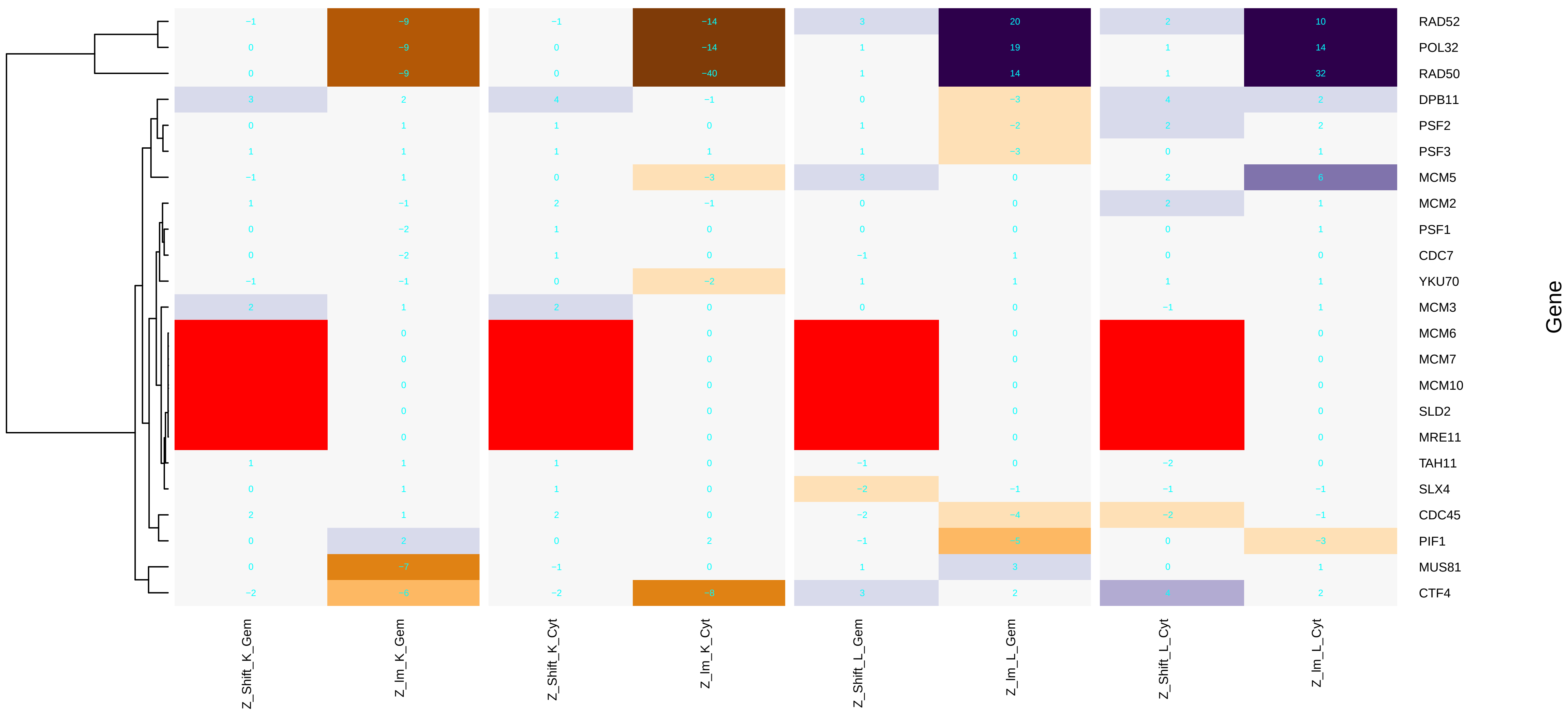

### double-strand_break_repair_via_homologous_recombination.pdf

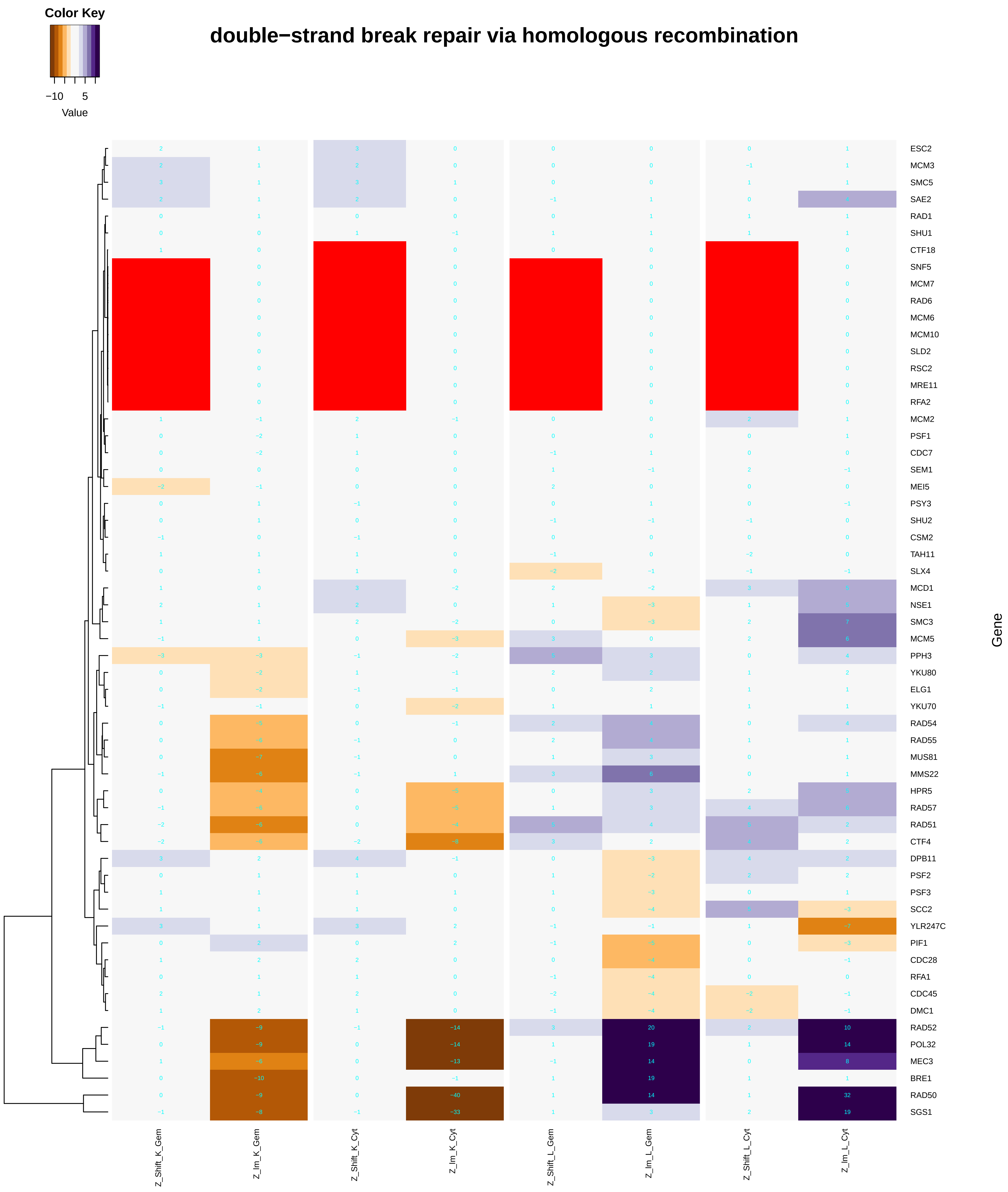

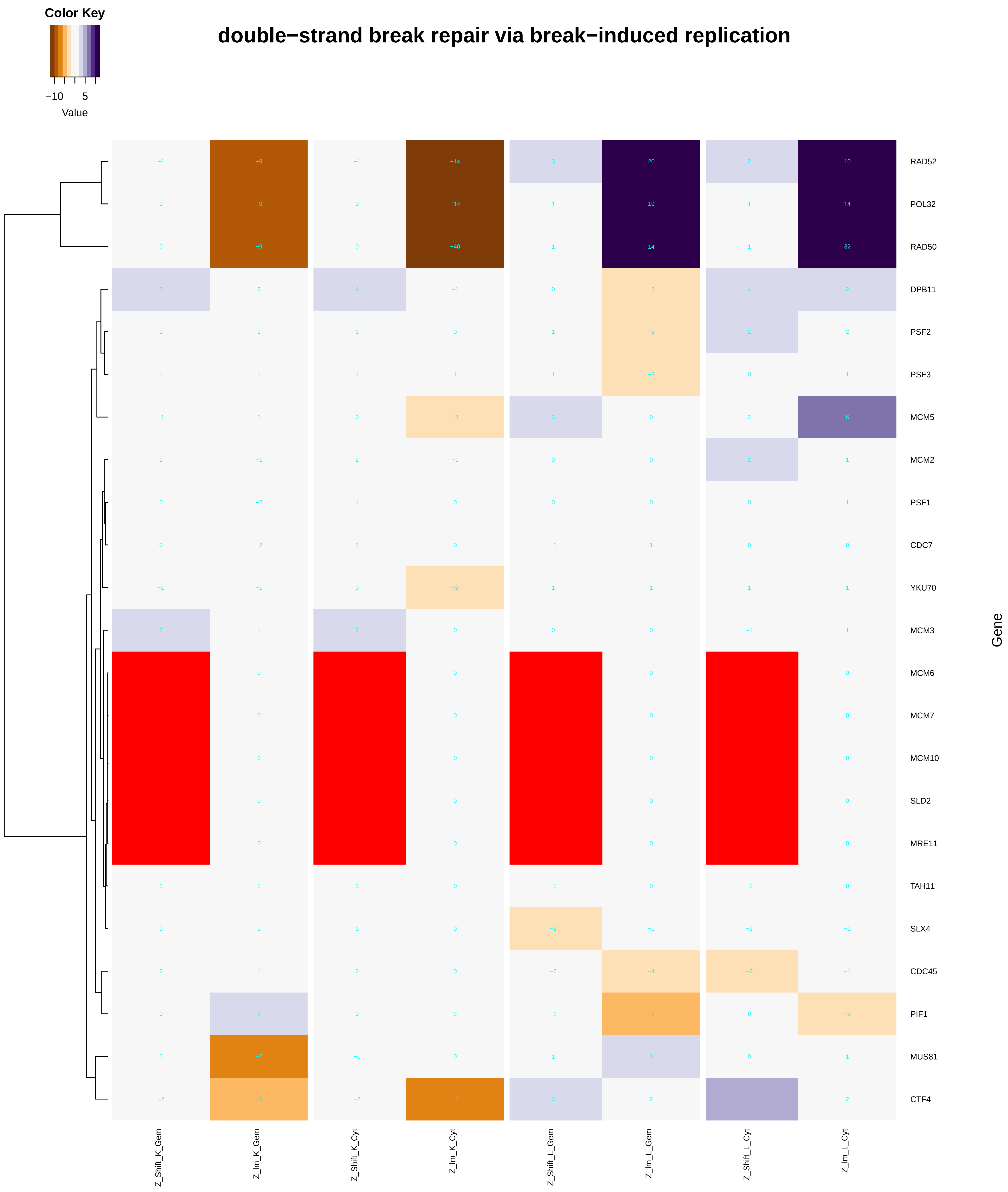

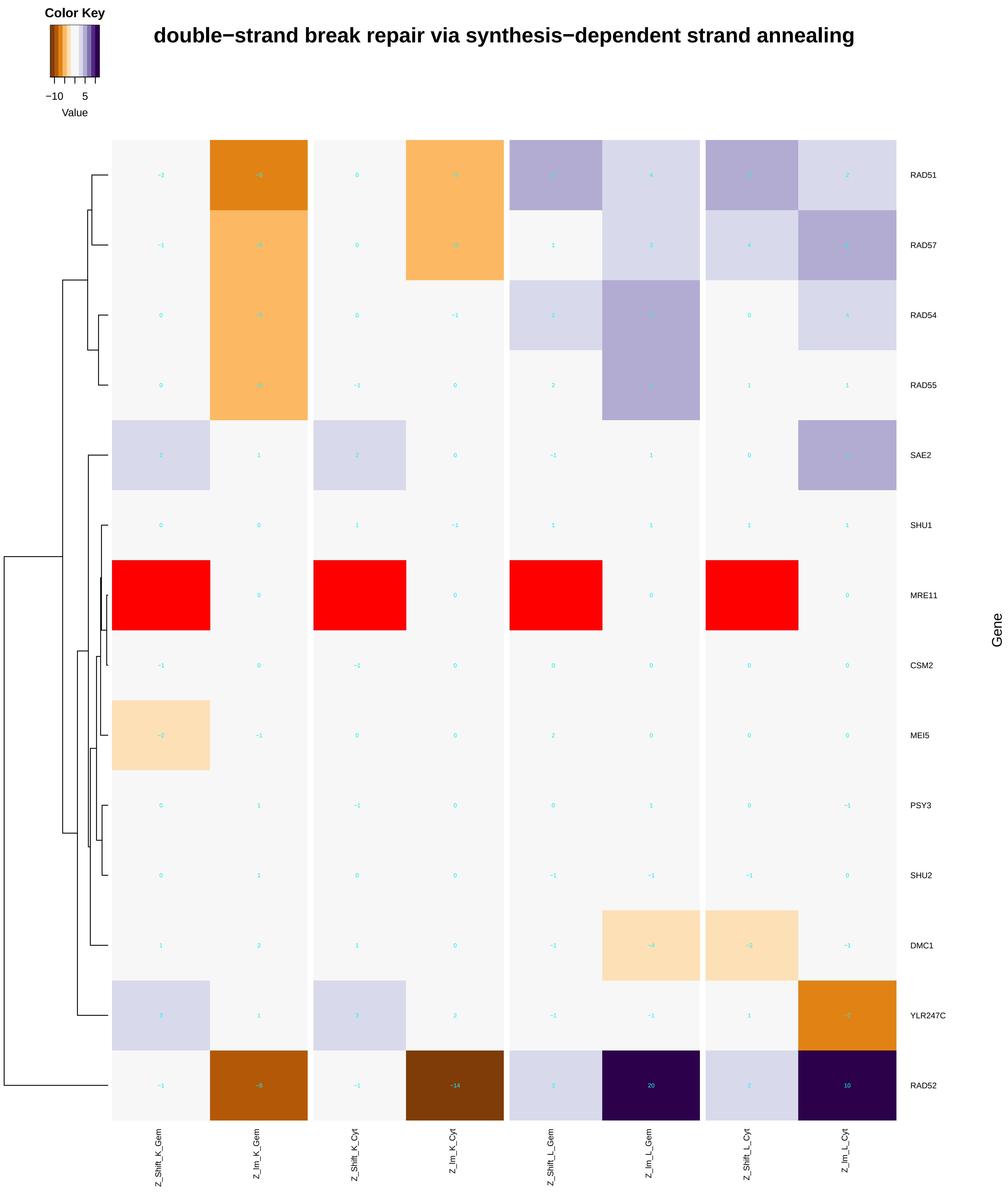

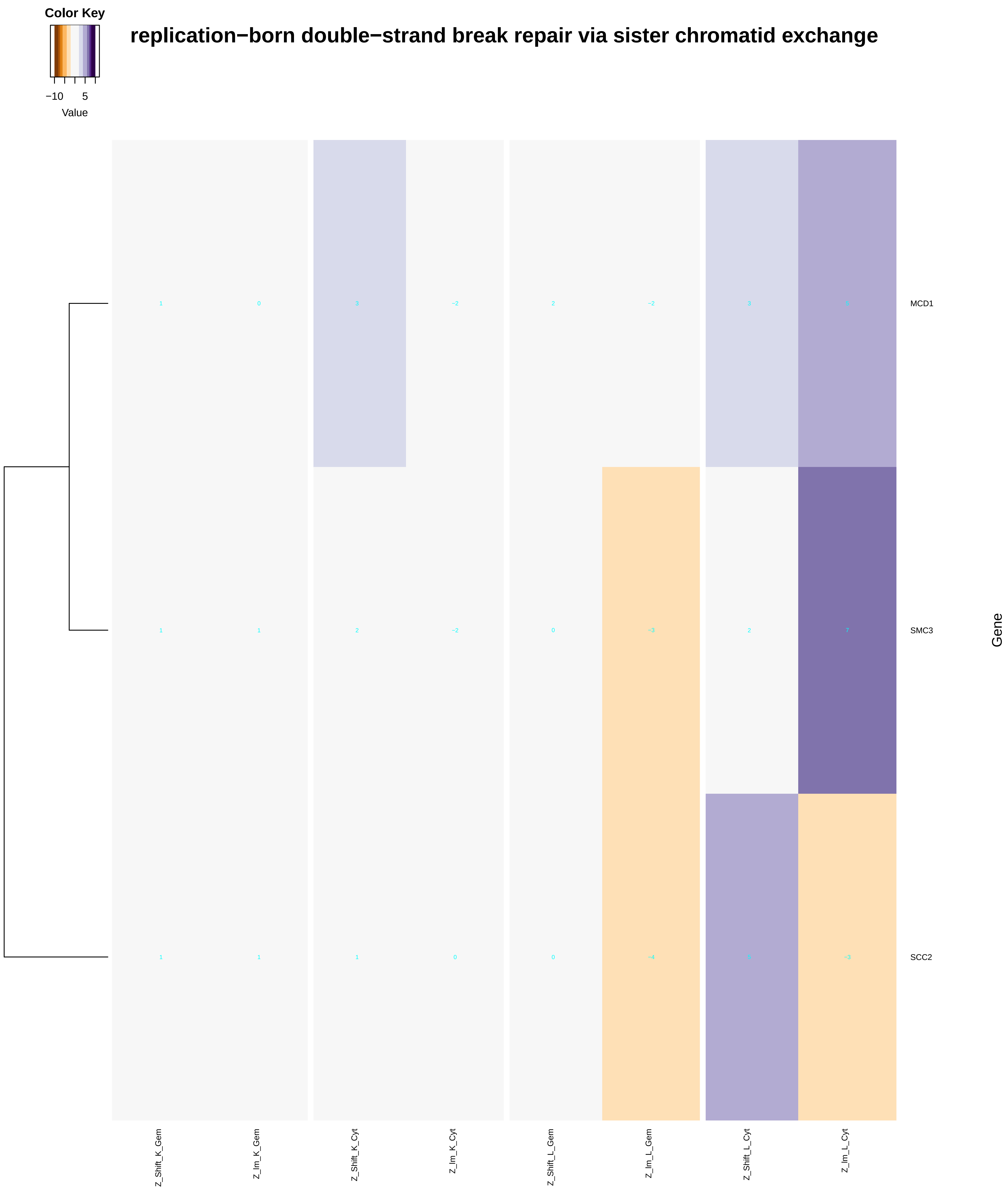

### double-strand_break_repair_via_synthesis-dependent_strand_annealing.pdf

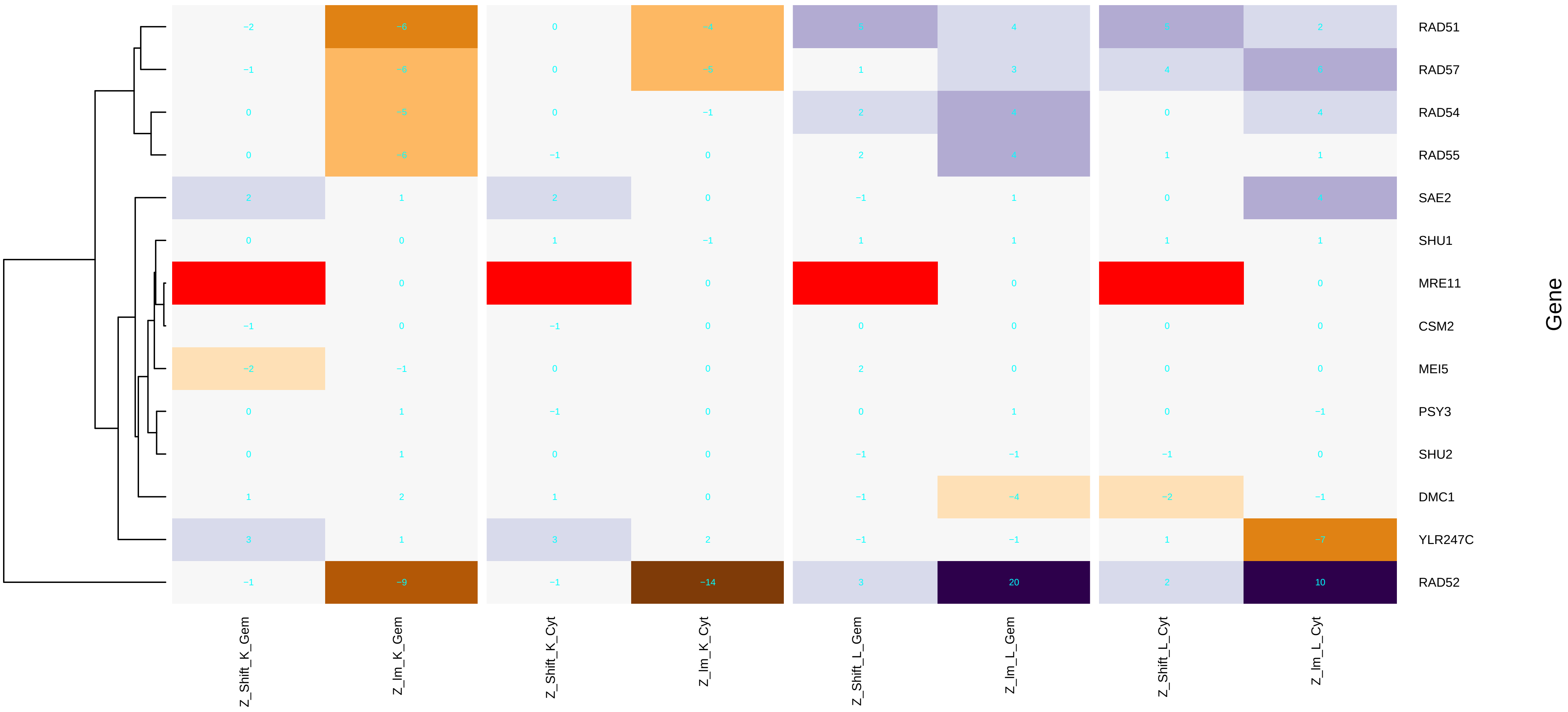

### double-stranded_telomeric_DNA_binding.pdf

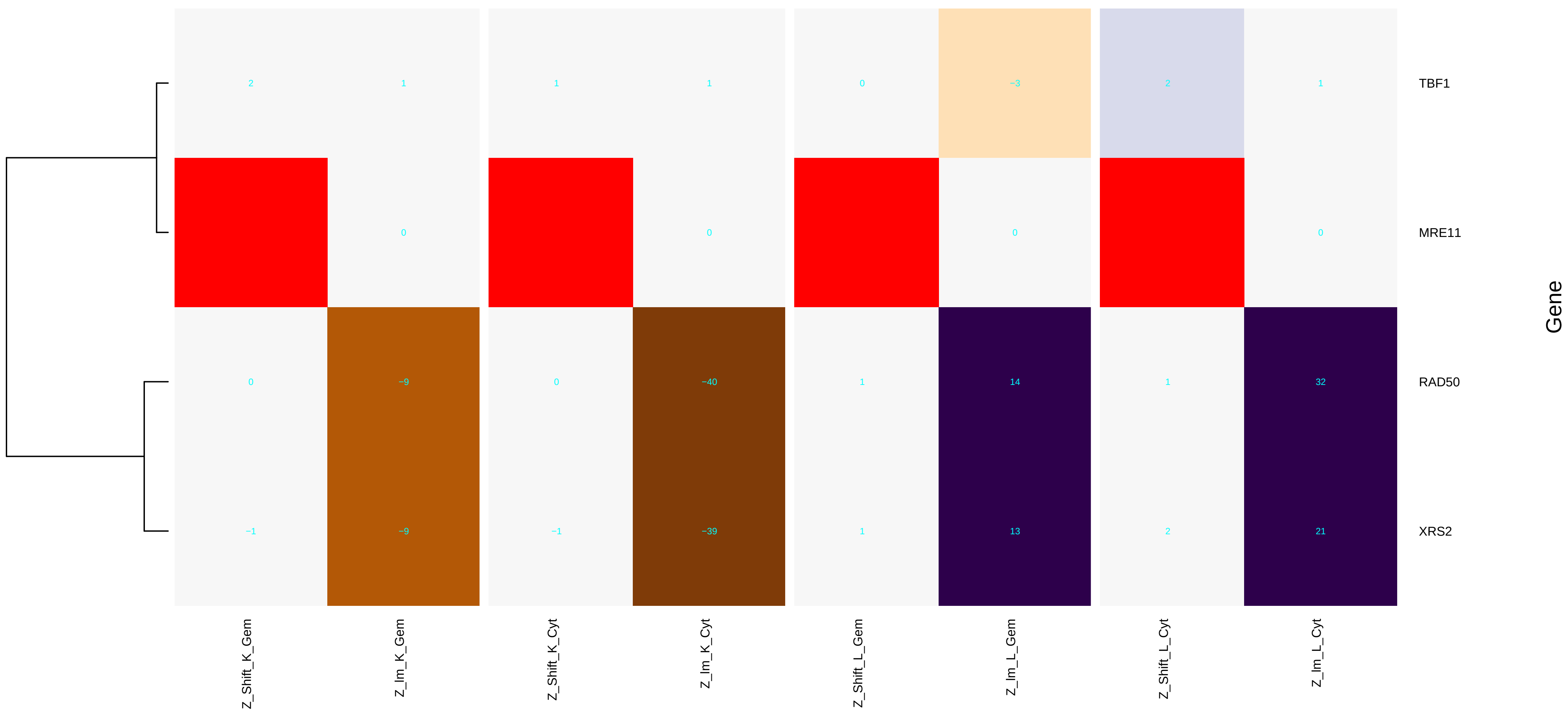

### Elongator_holoenzyme_complex.pdf

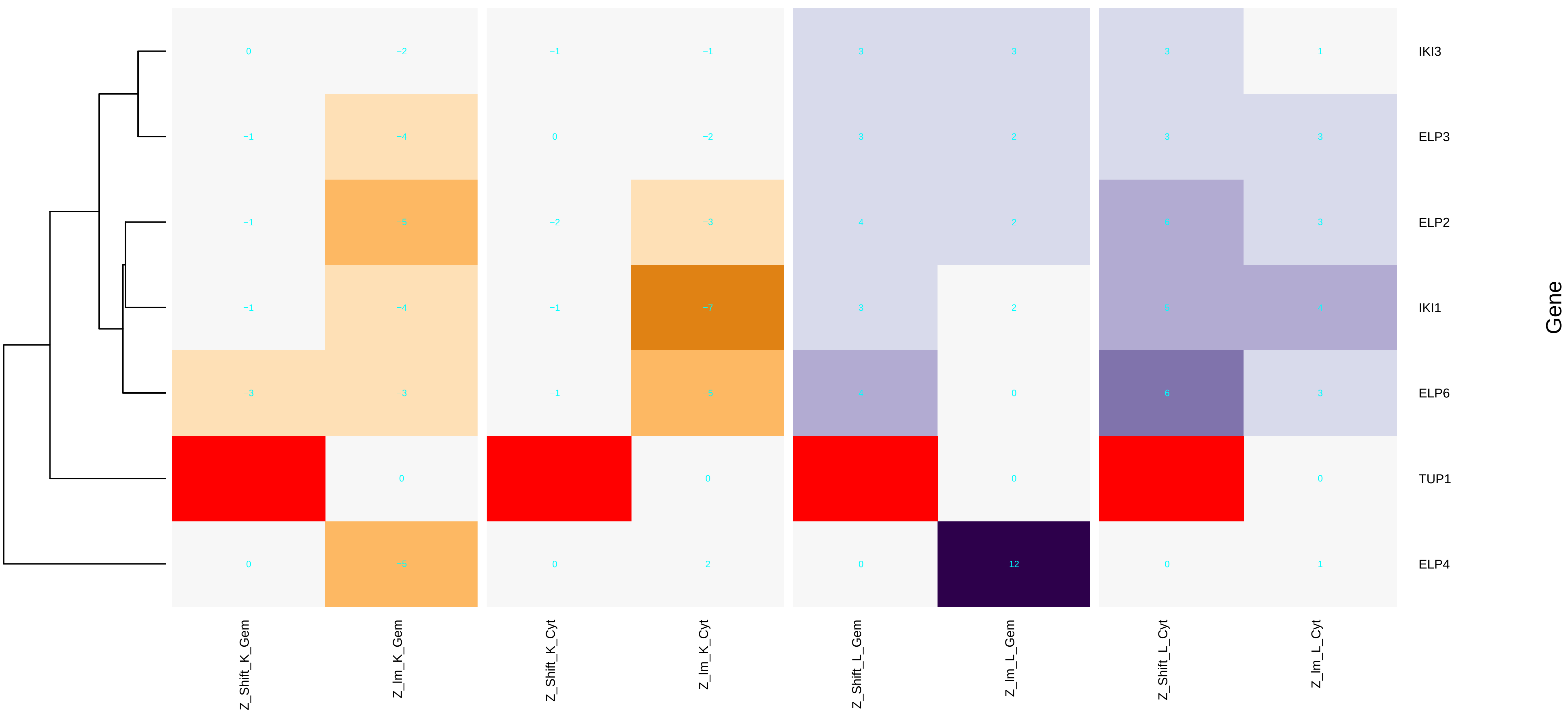

### endomembrane_system.pdf

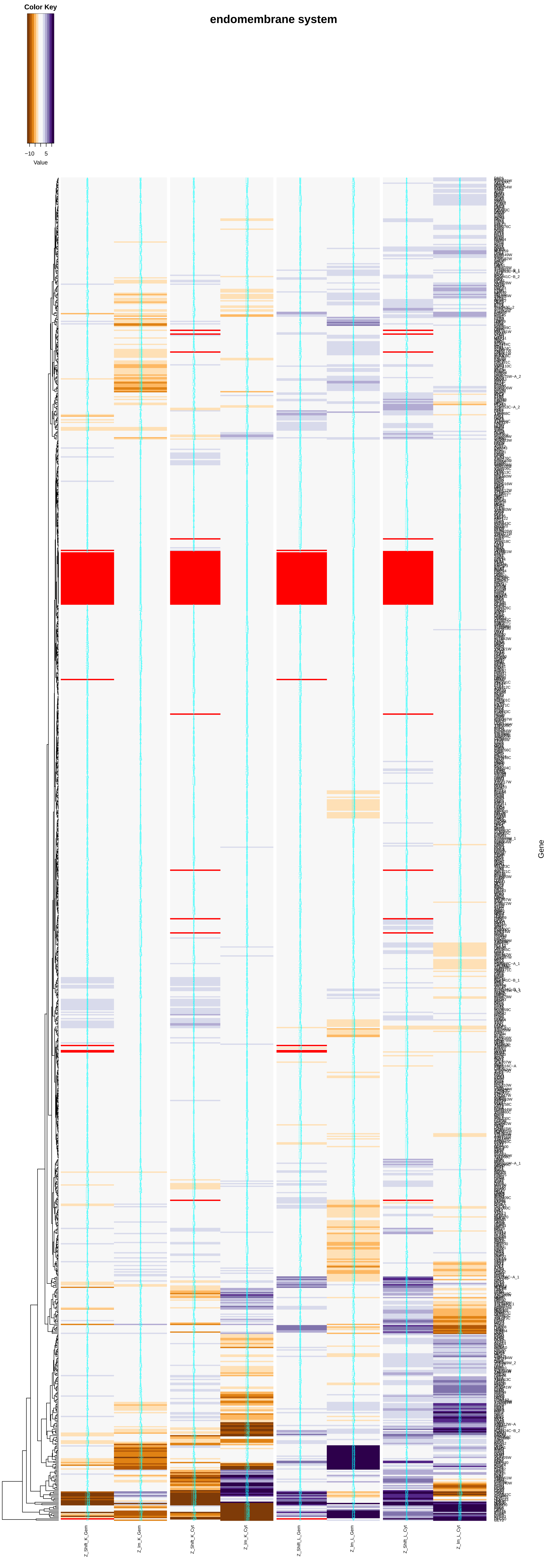

### endonucleolytic_cleavage_in_ITS1_to_separate_SSU-rRNA_from_5.8S_rRNA_and_LSU-rRNA_from_tricistronic_rRNA_transcript_(SSU-rRNA,_5.8S_rRNA,_LSU-rRNA).pdf

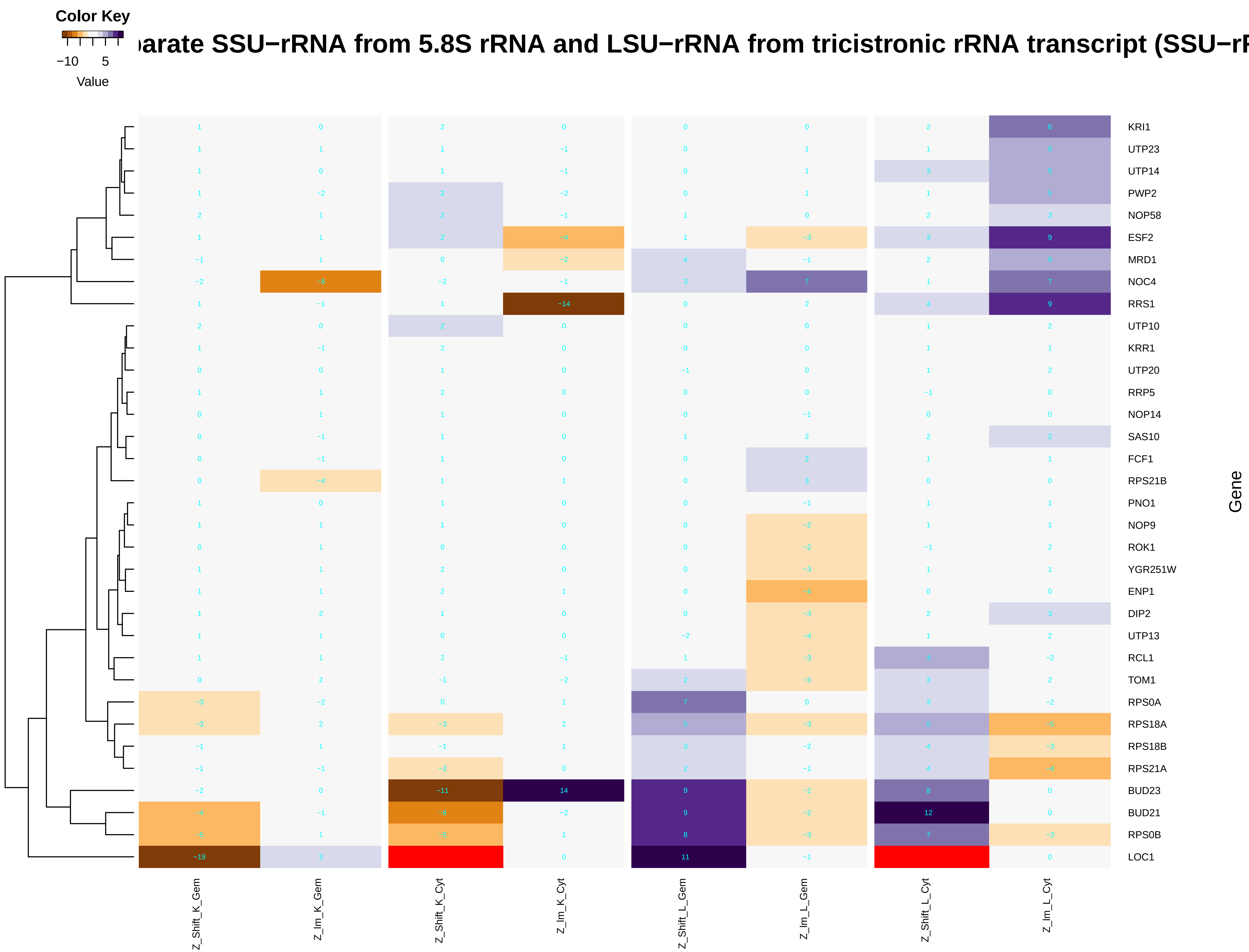

### endonucleolytic_cleavage_of_tricistronic_rRNA_transcript_(SSU-rRNA,_5.8S_rRNA,_LSU-rRNA).pdf

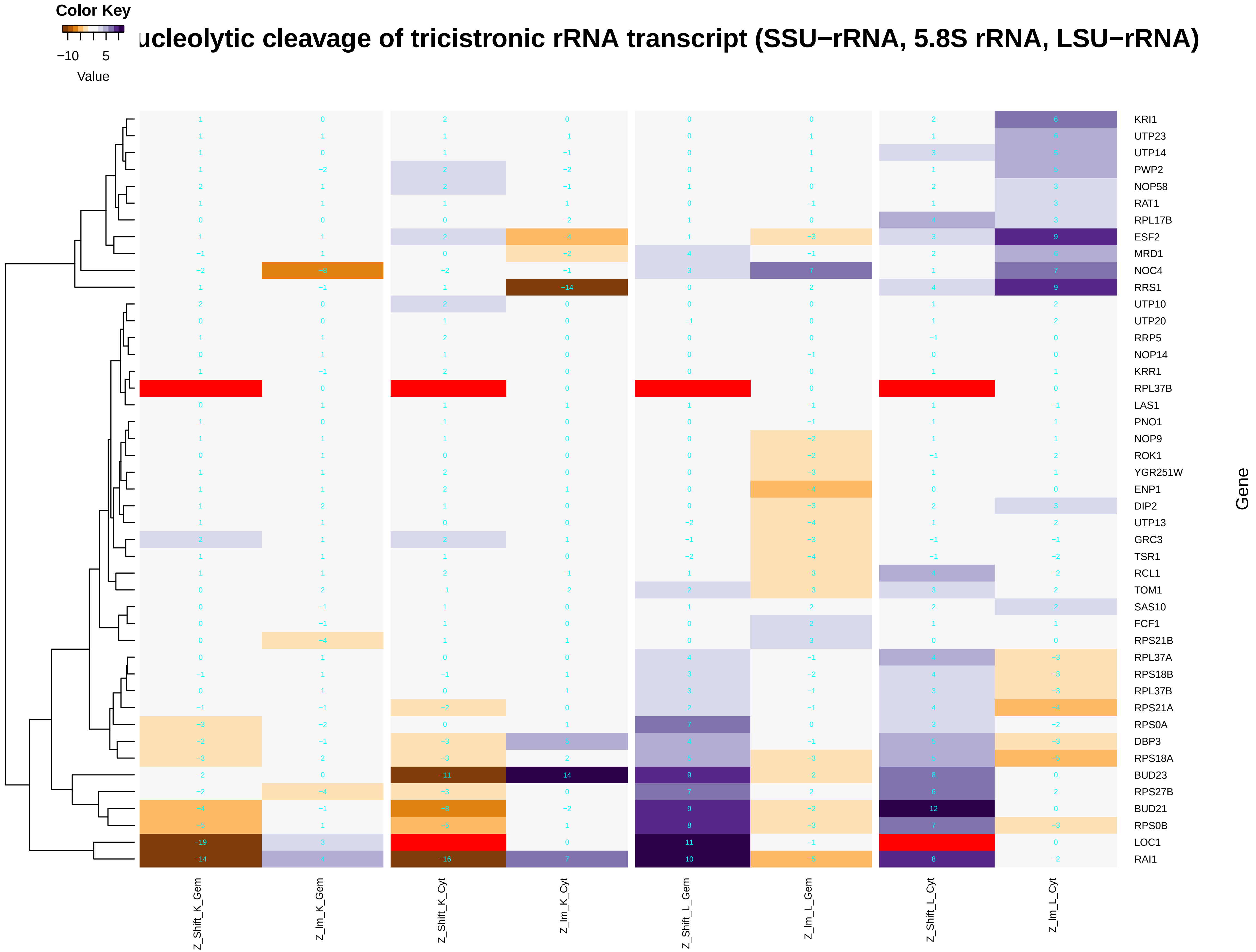

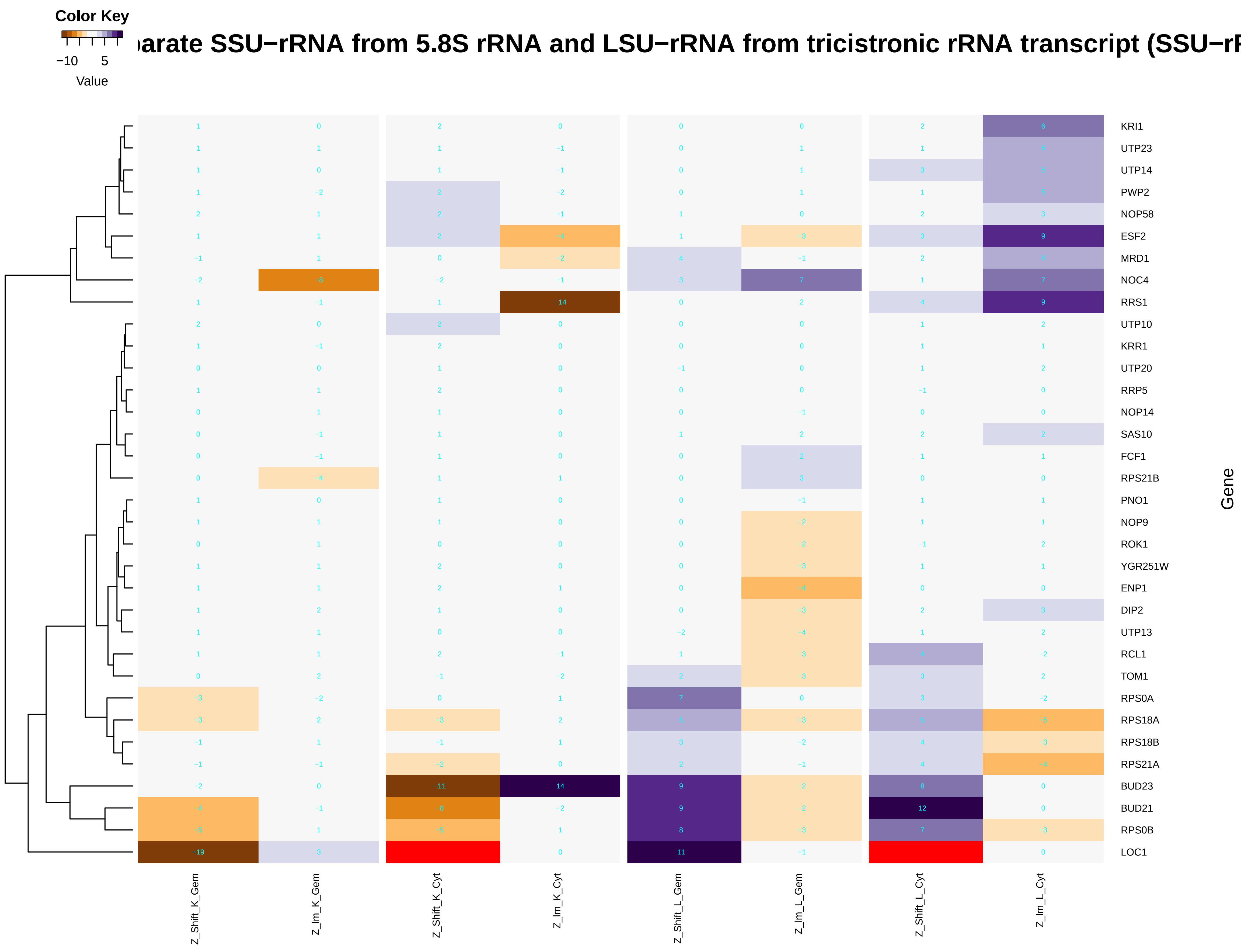

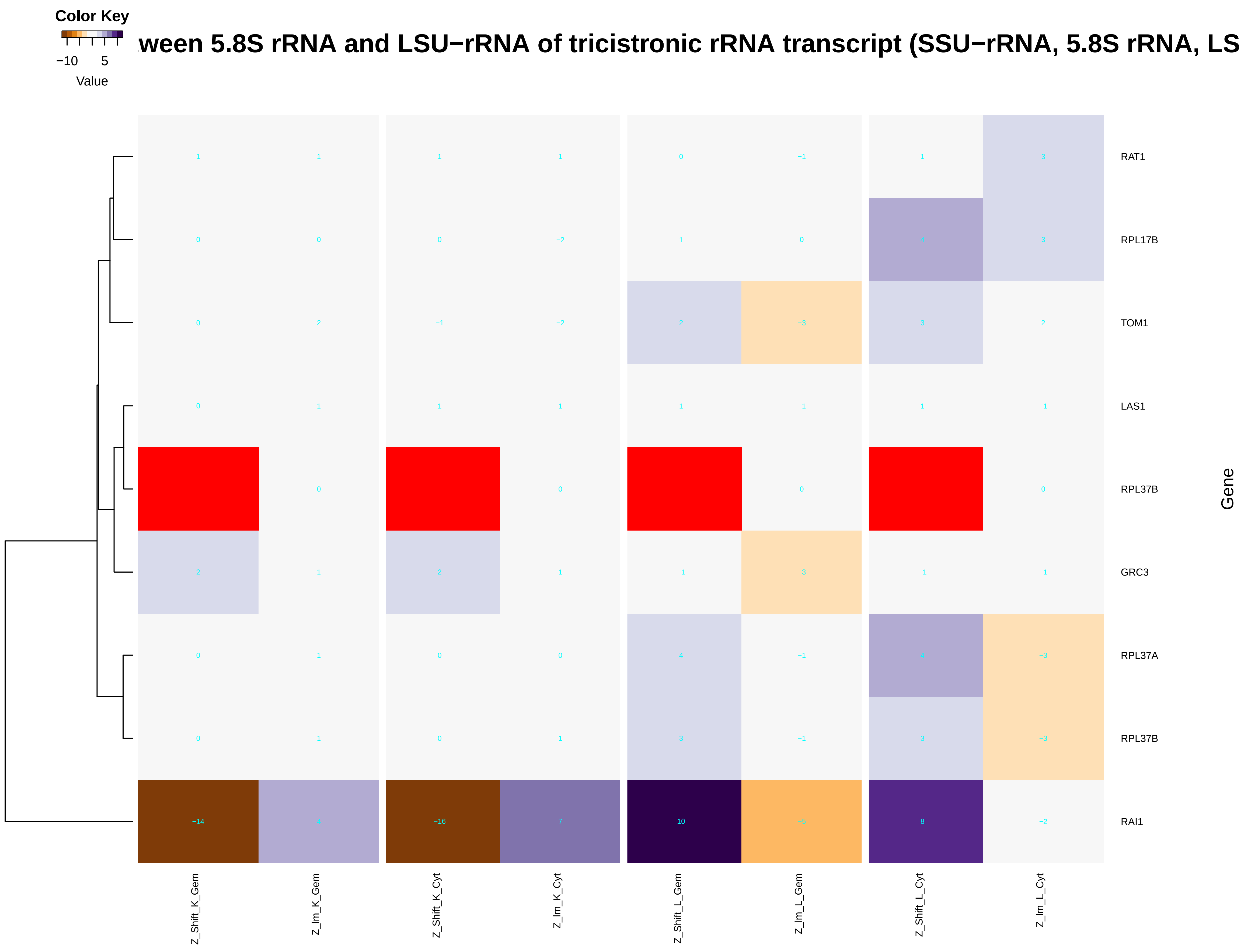

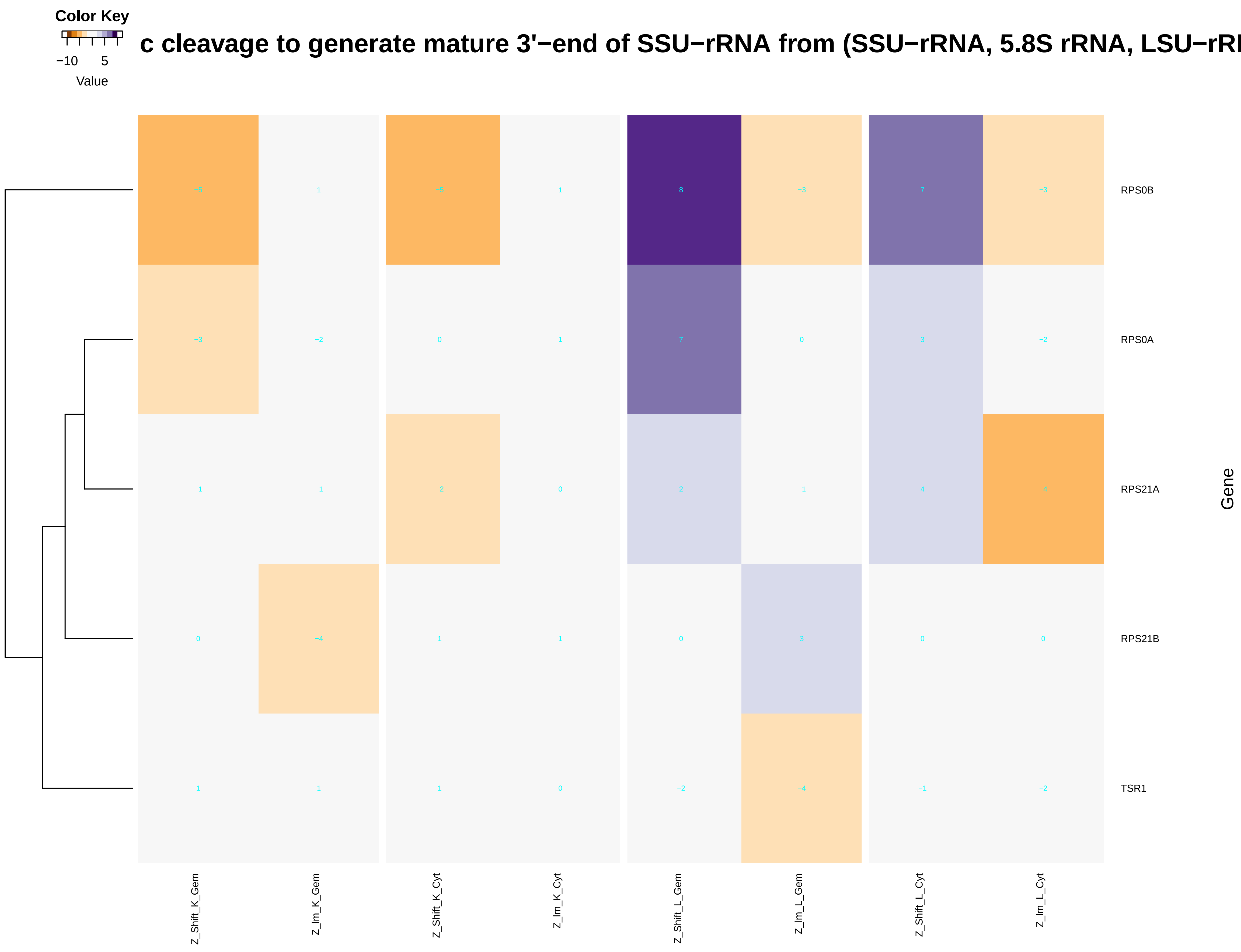

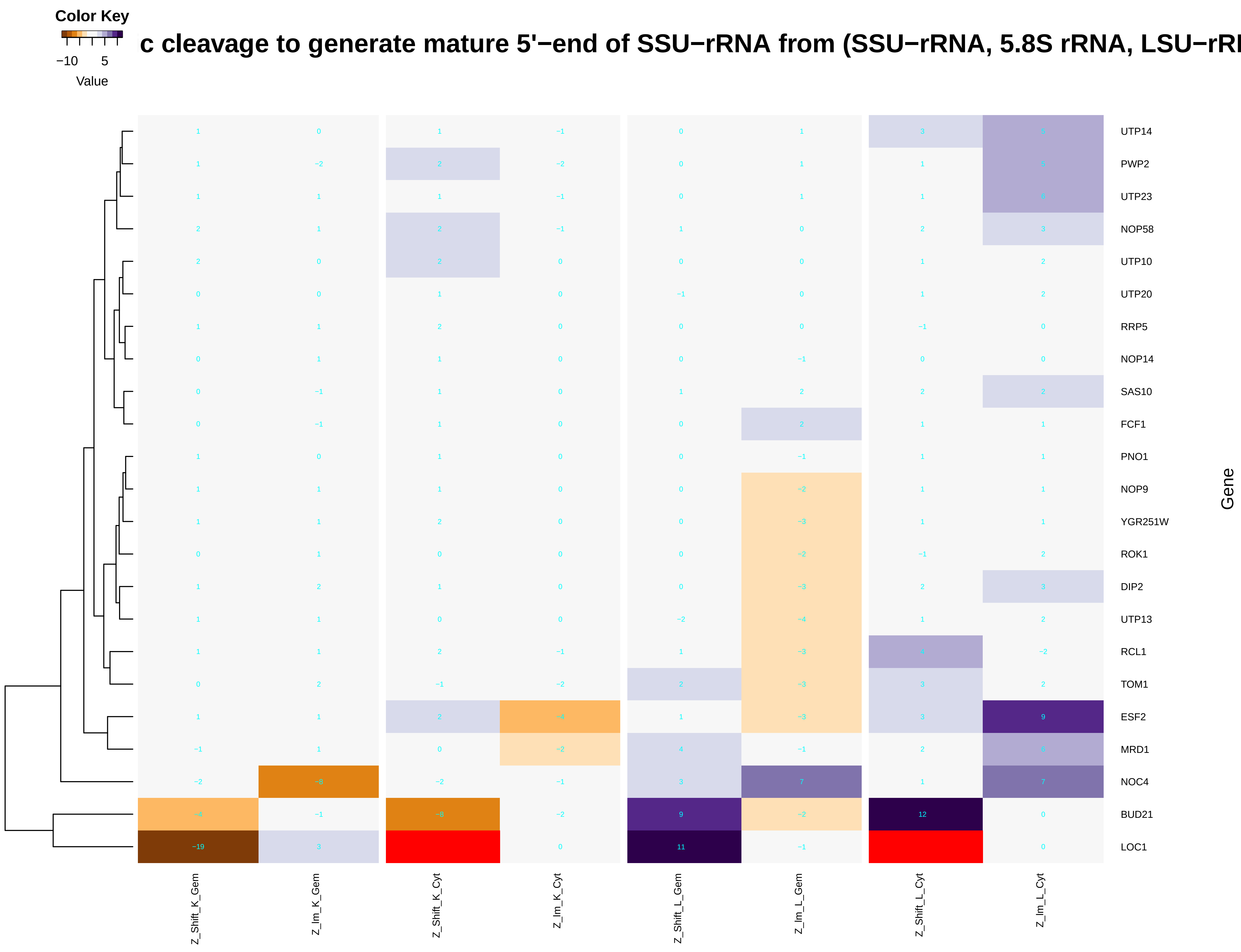

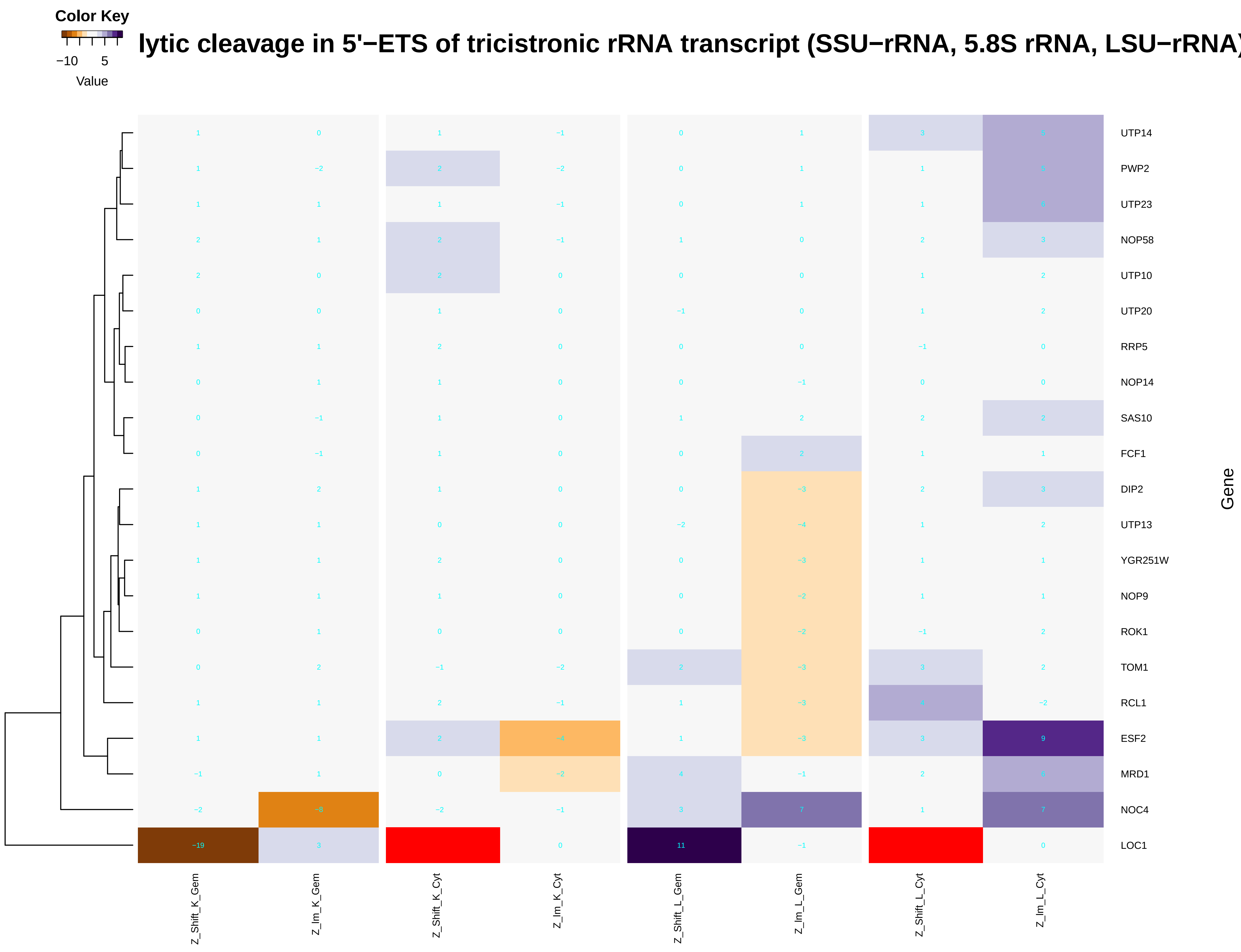

### endoribonuclease_activity.pdf

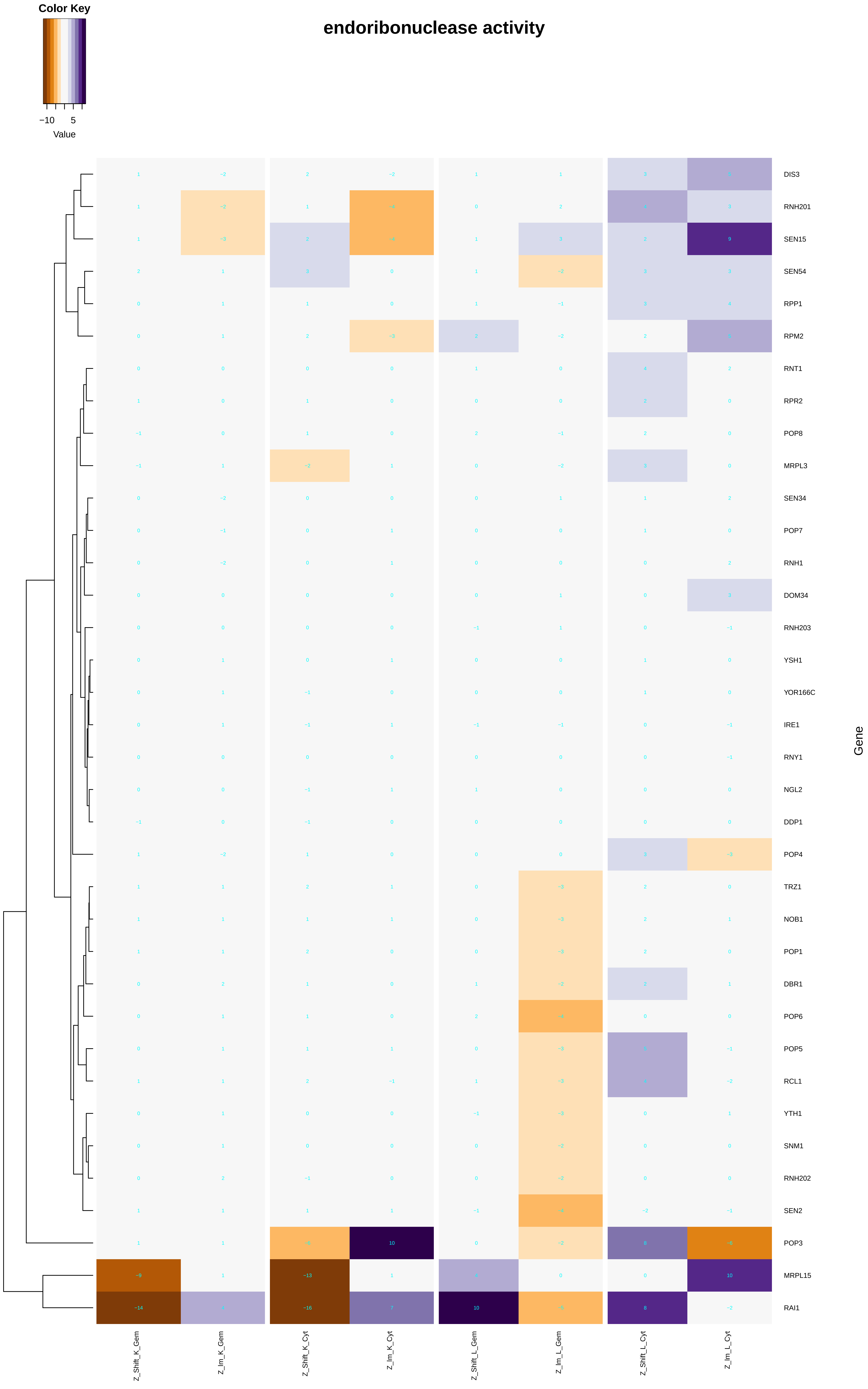

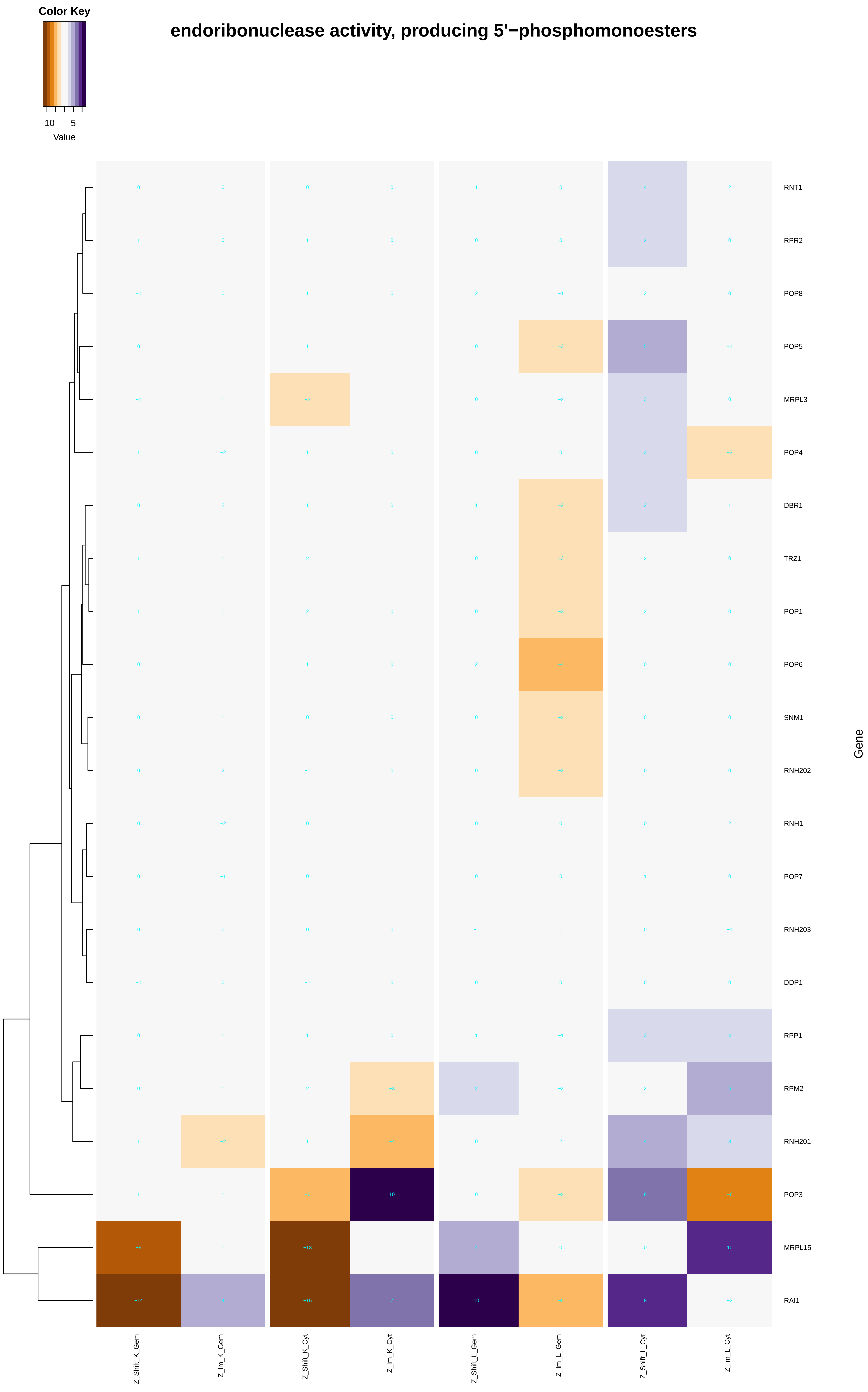

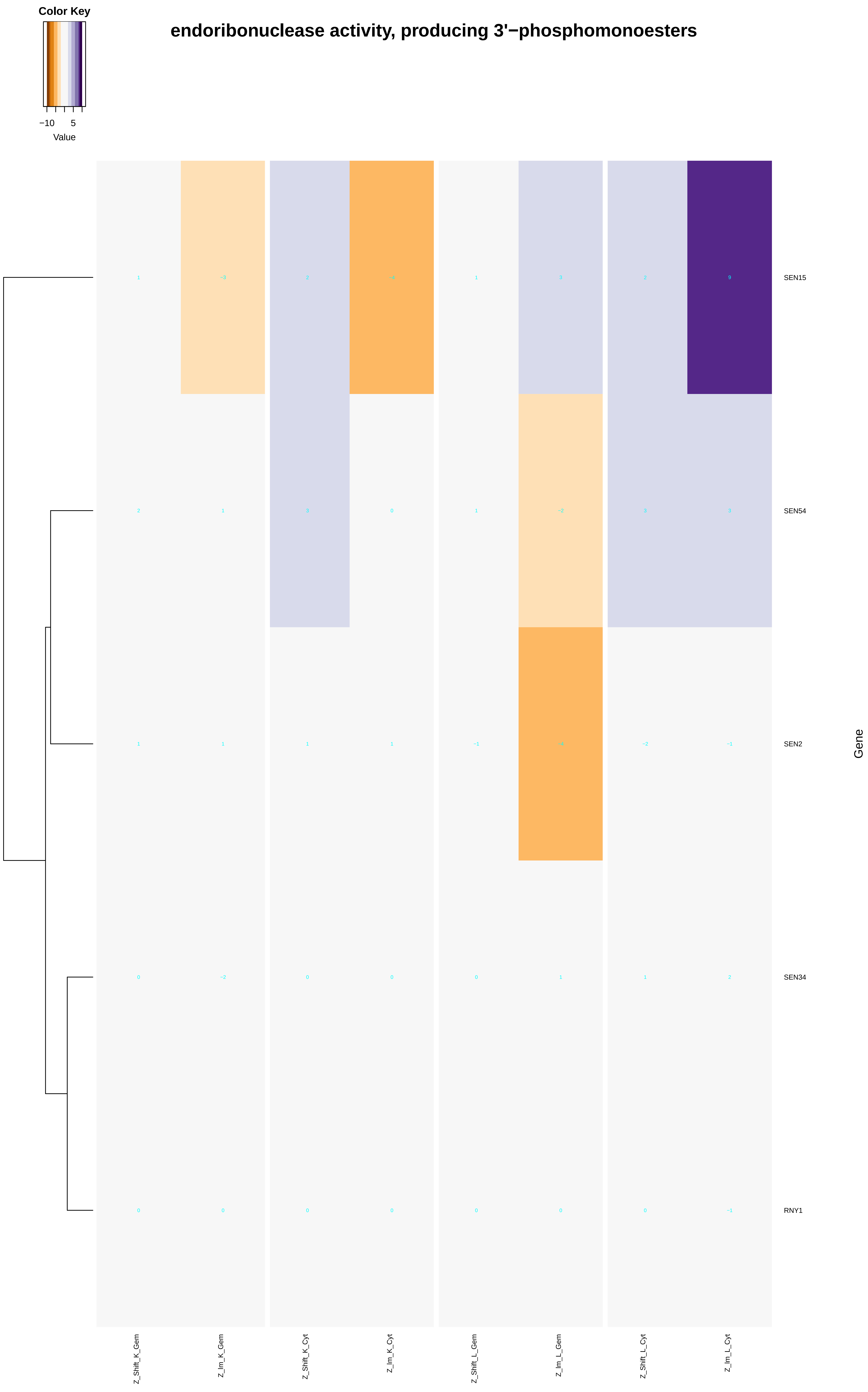

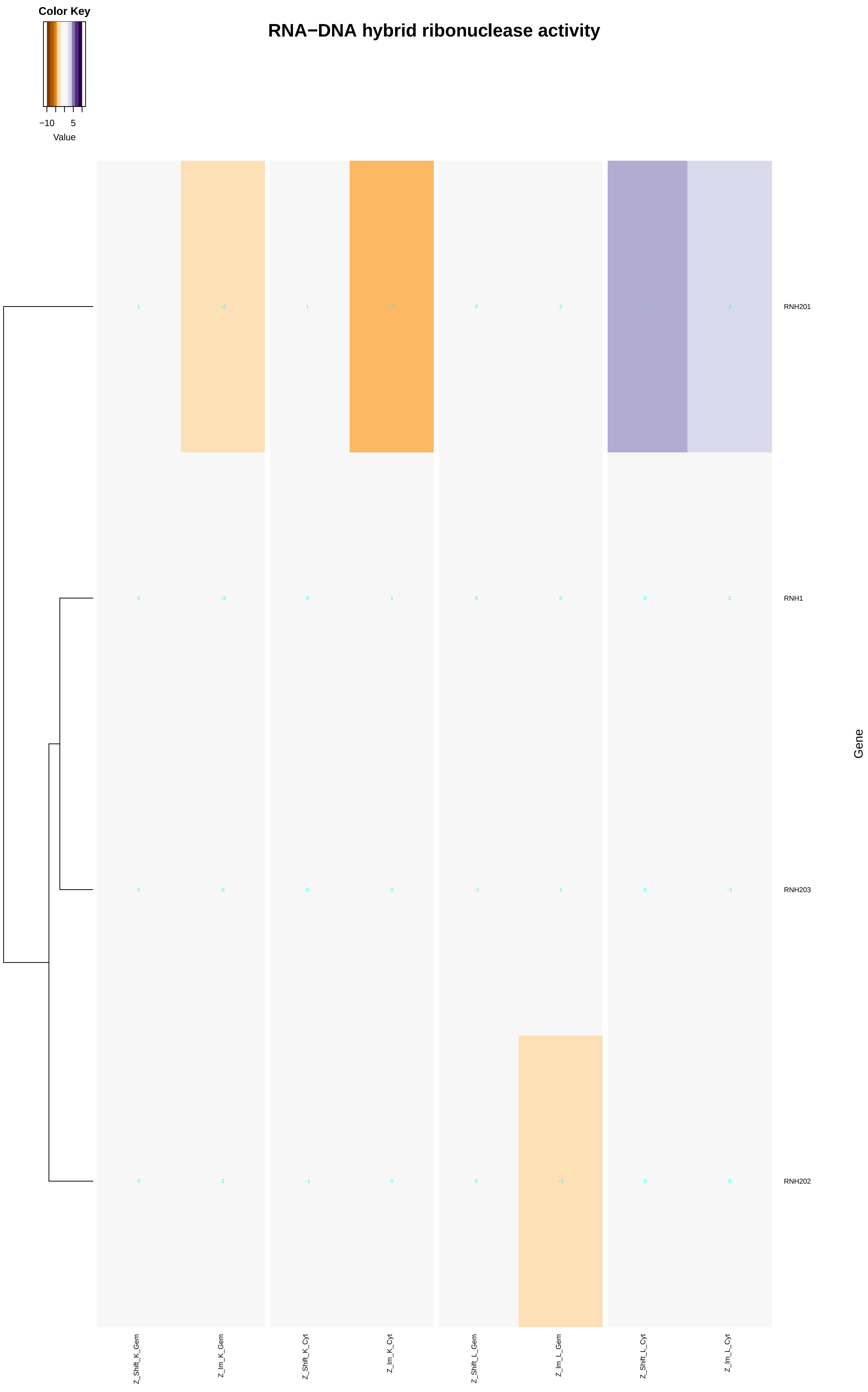

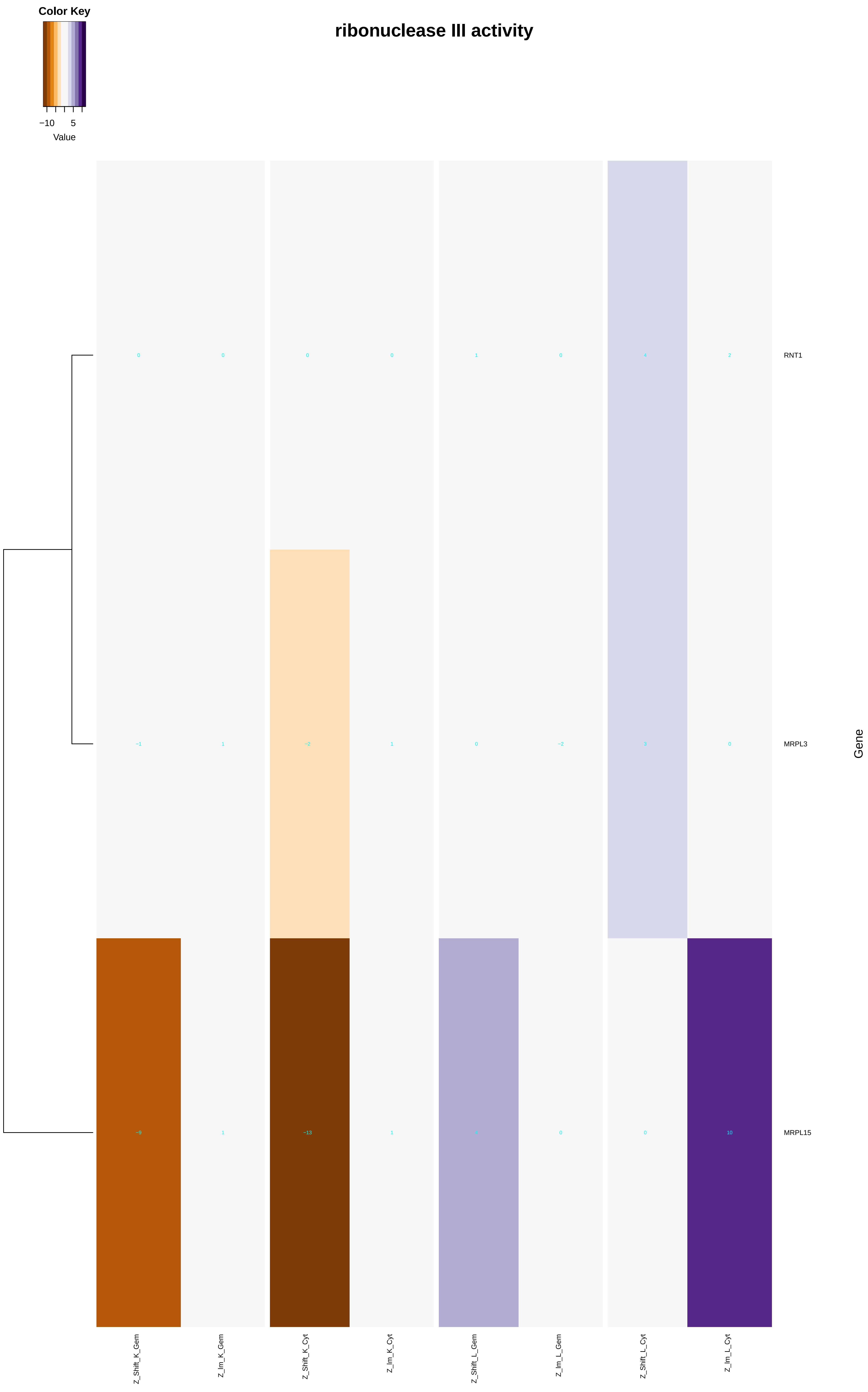

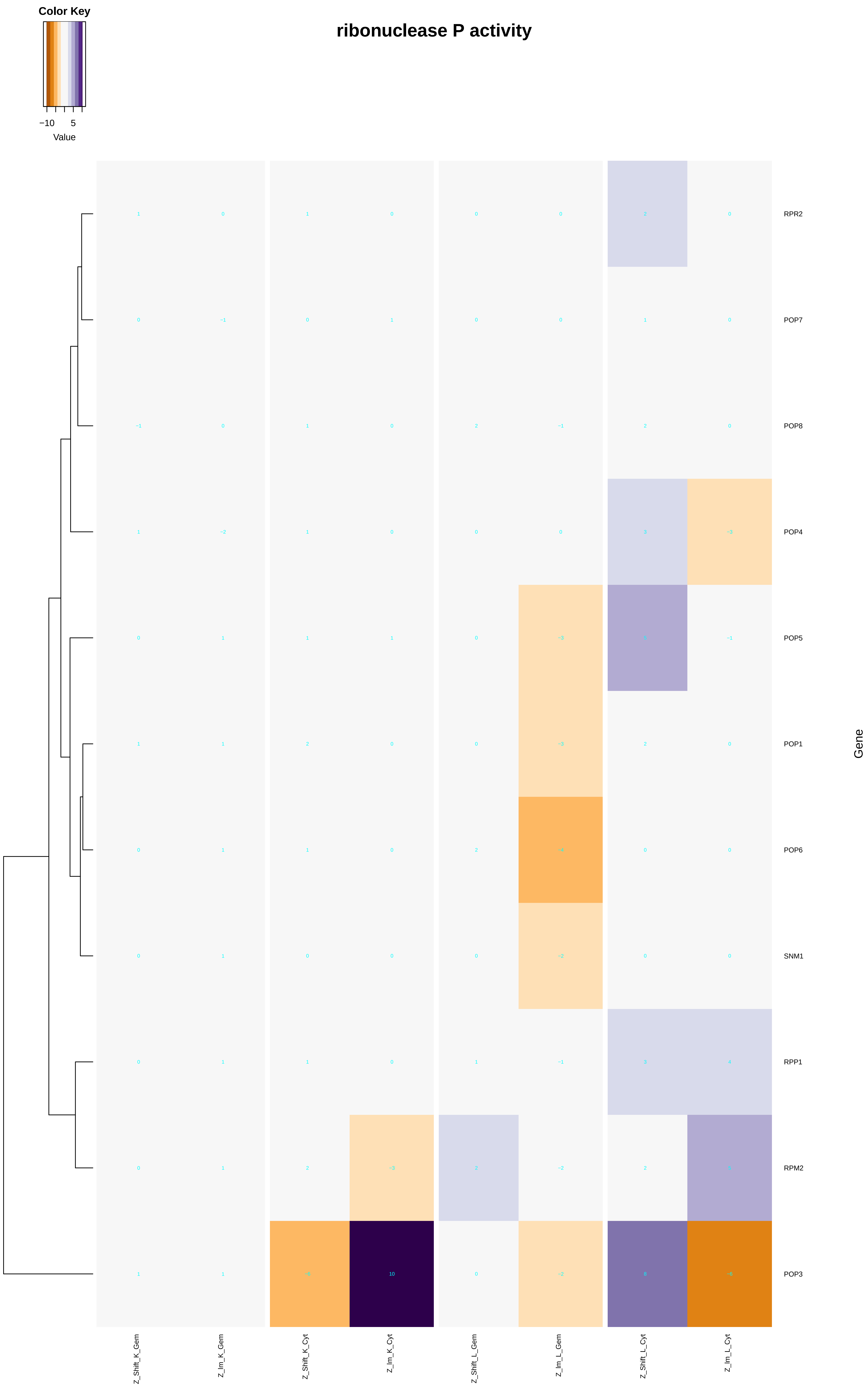

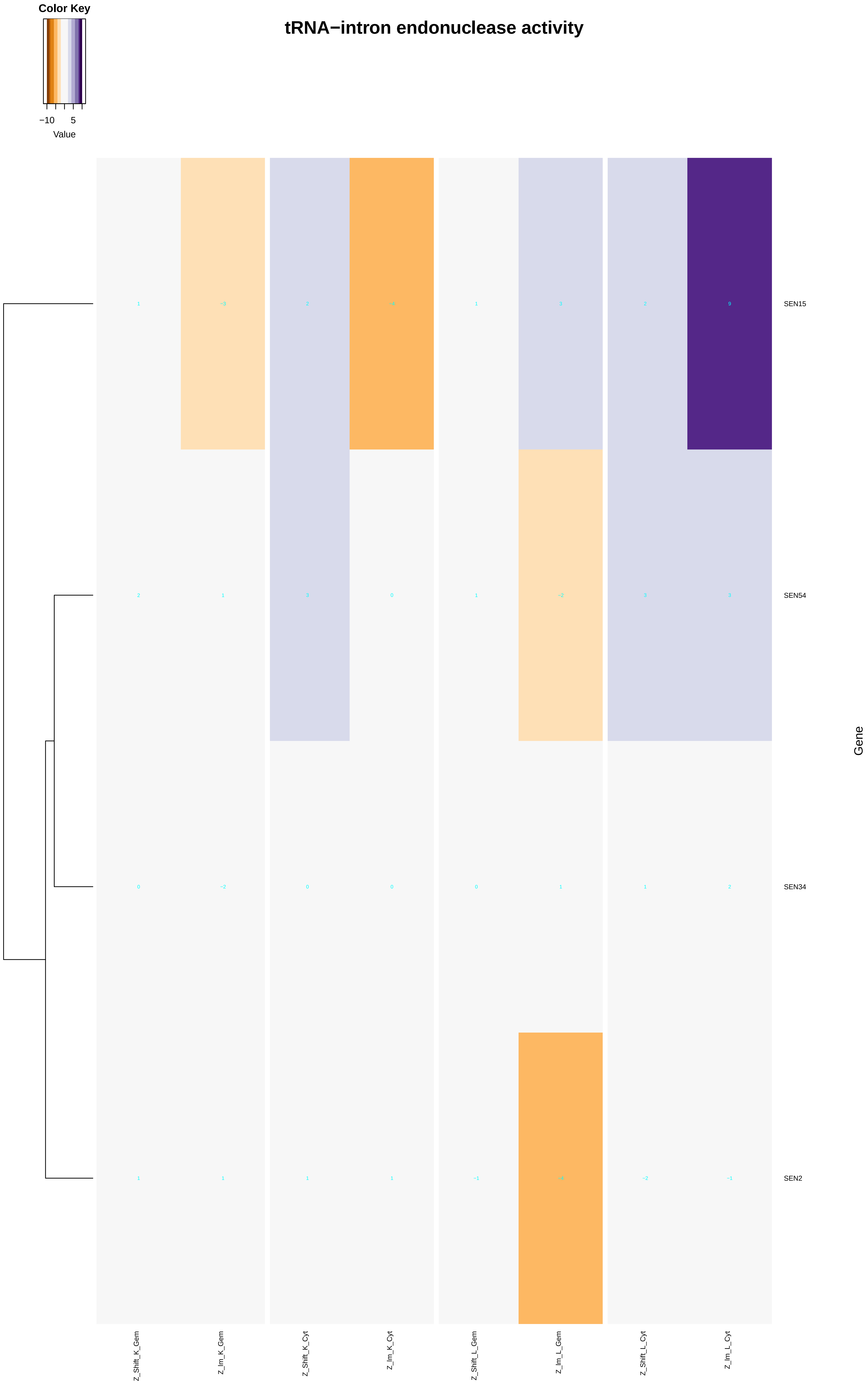

### endosomal_transport.pdf

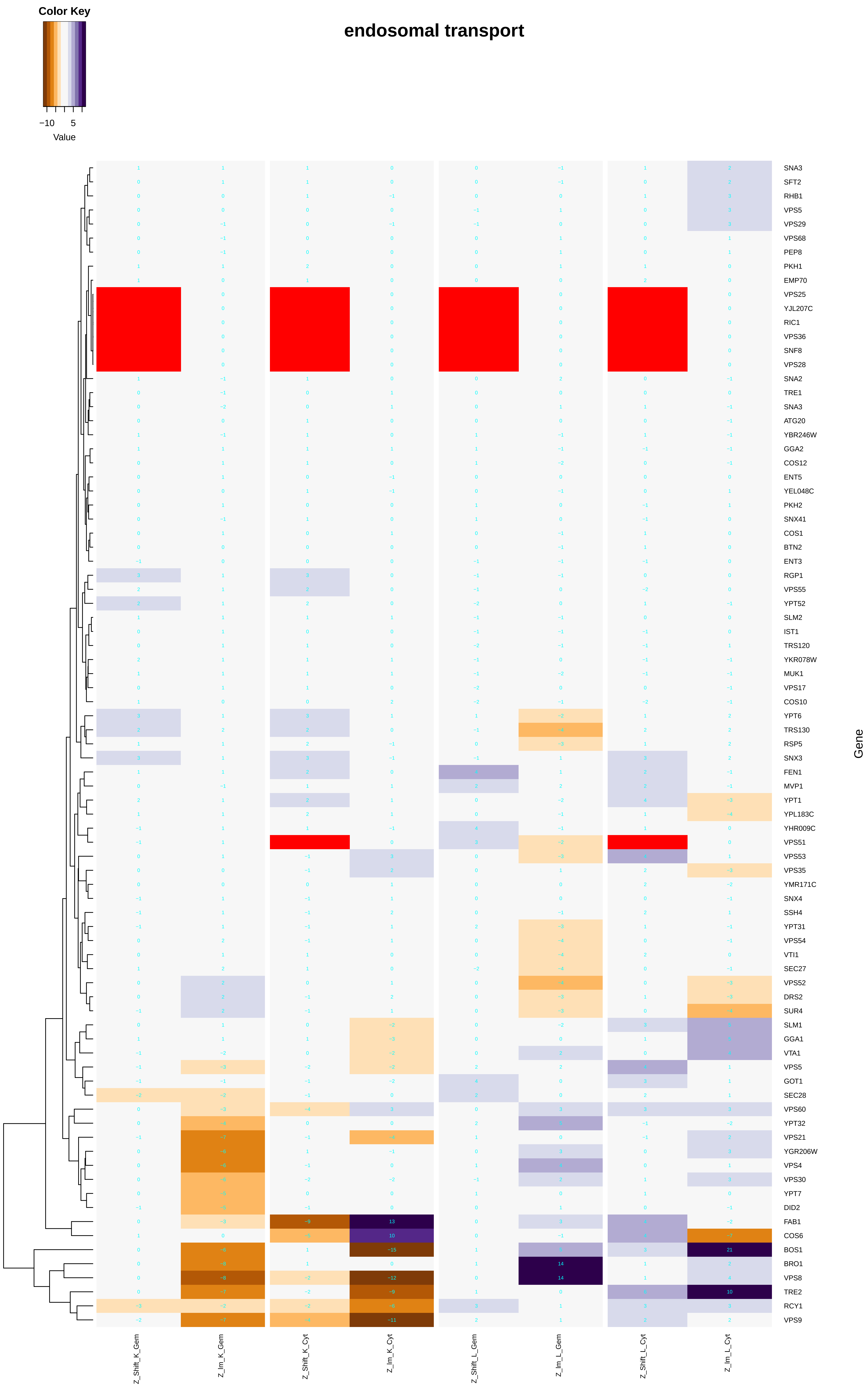

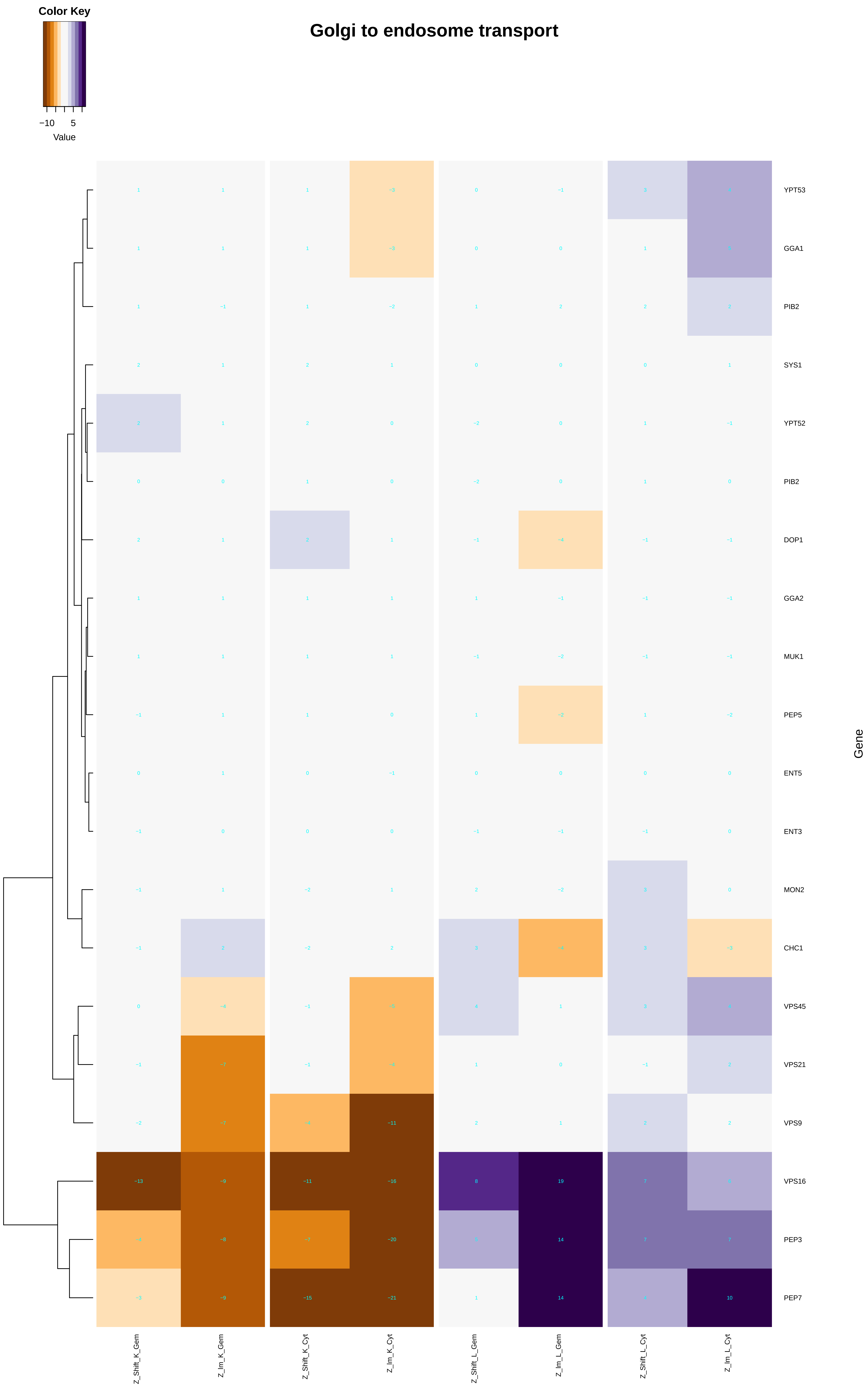

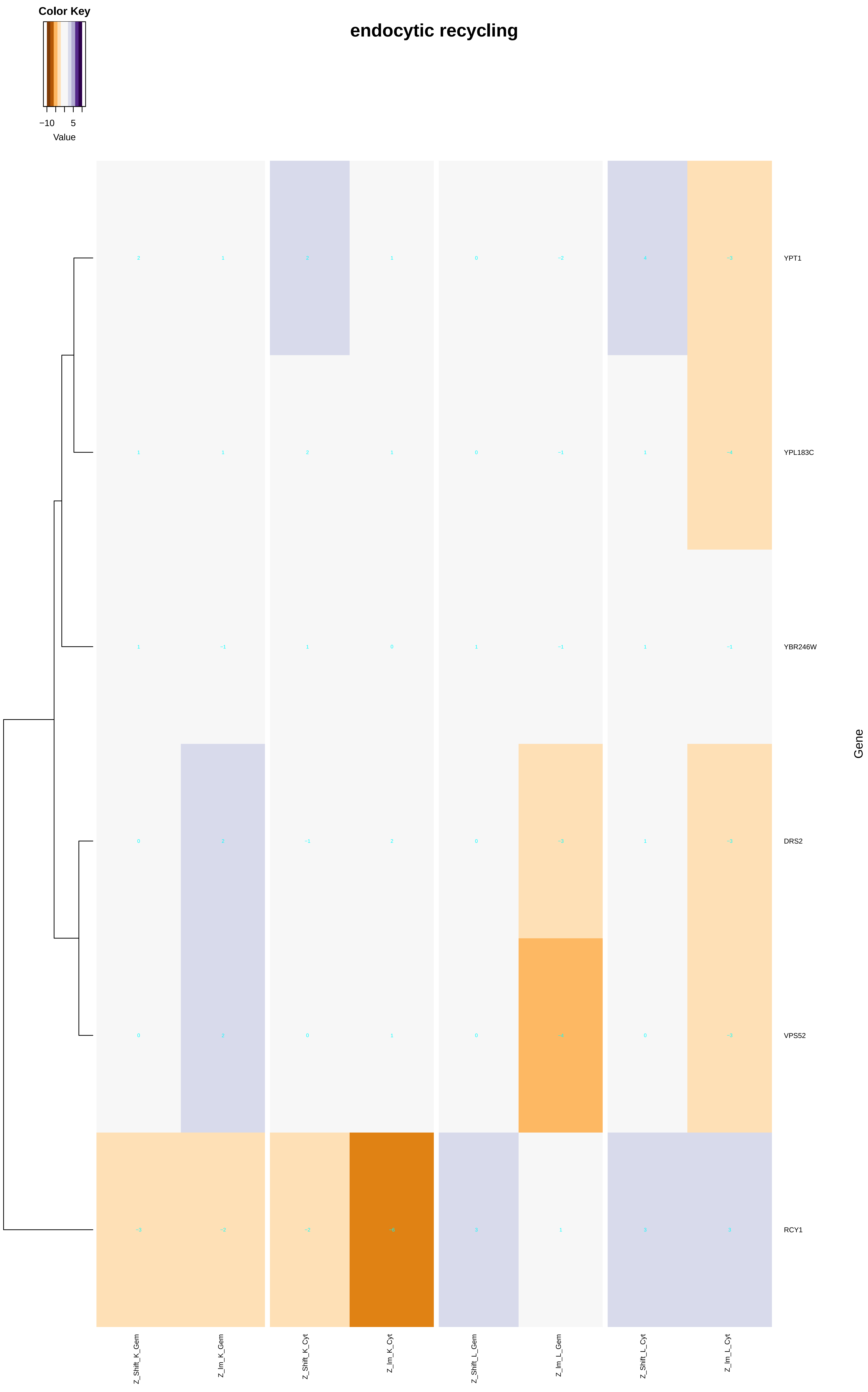

#### endosome transport via multivesicular body sorting pathway

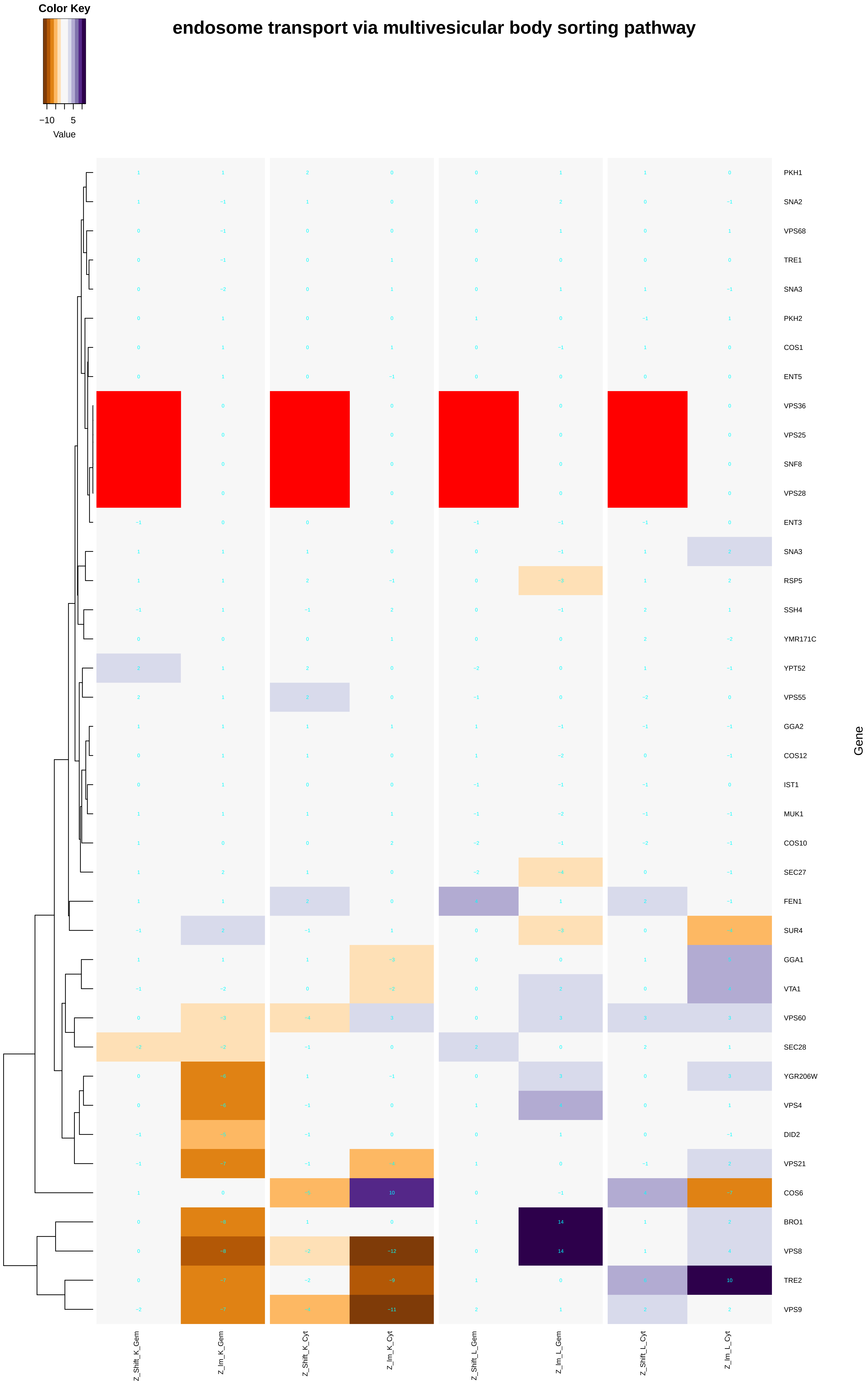

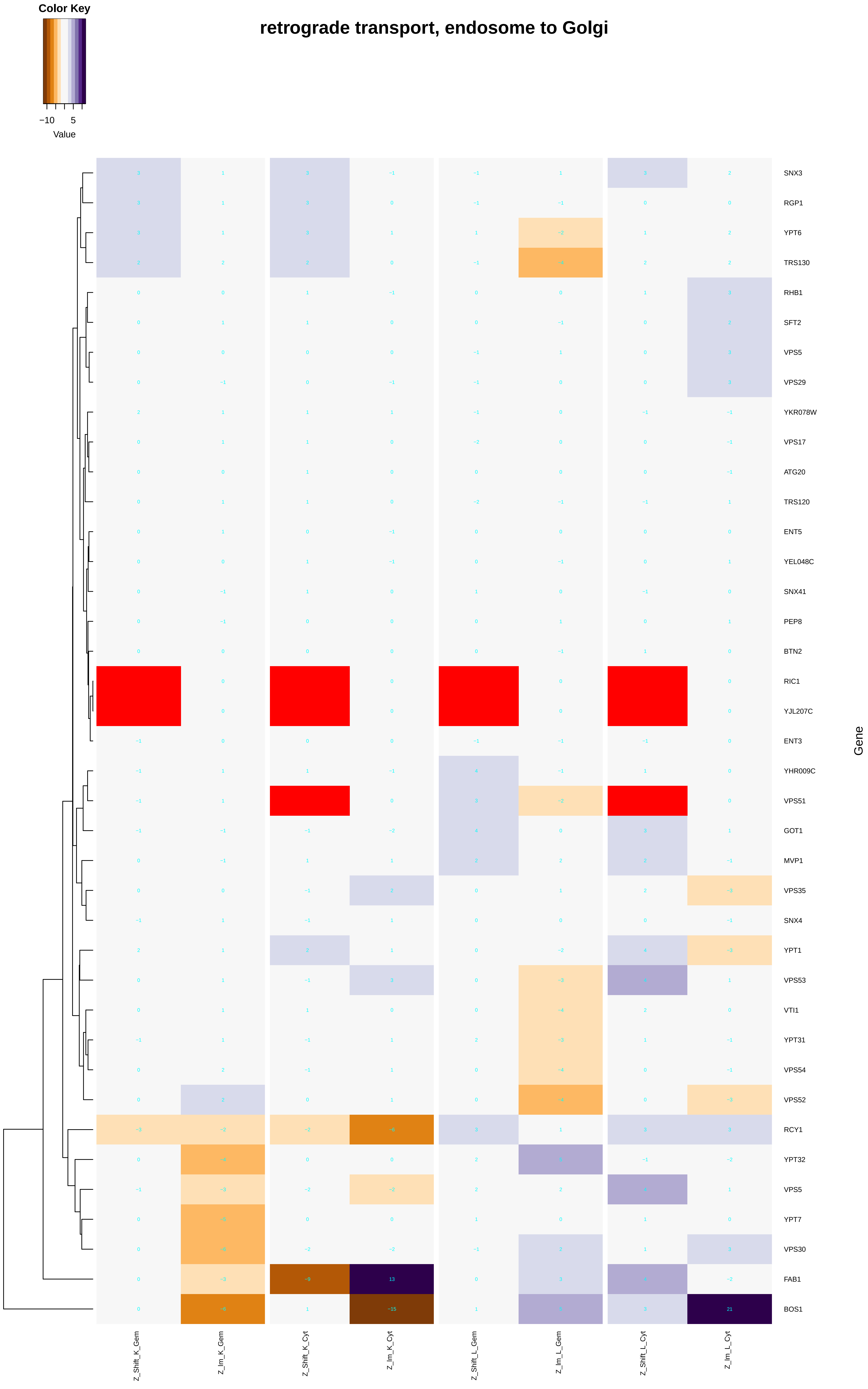

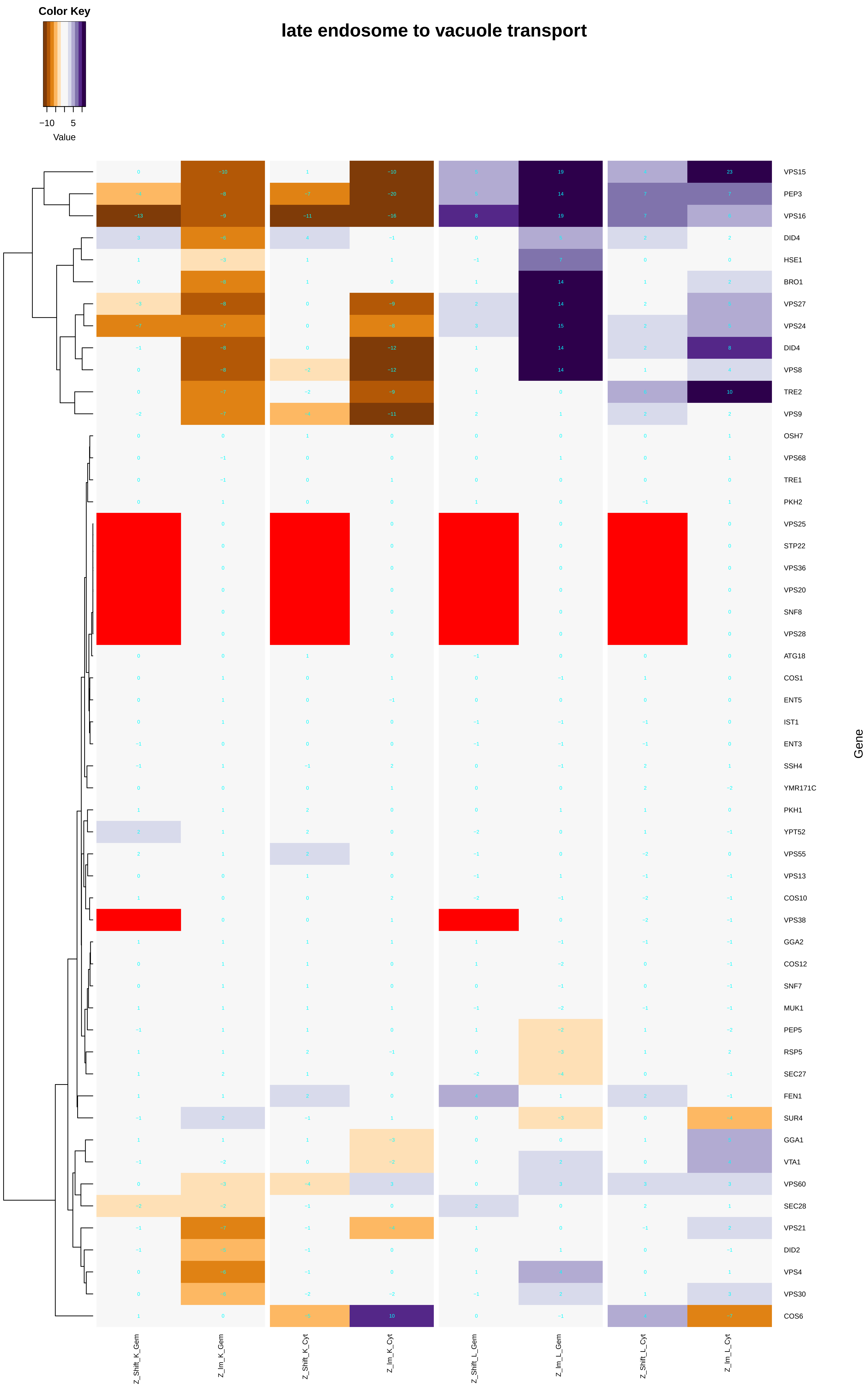

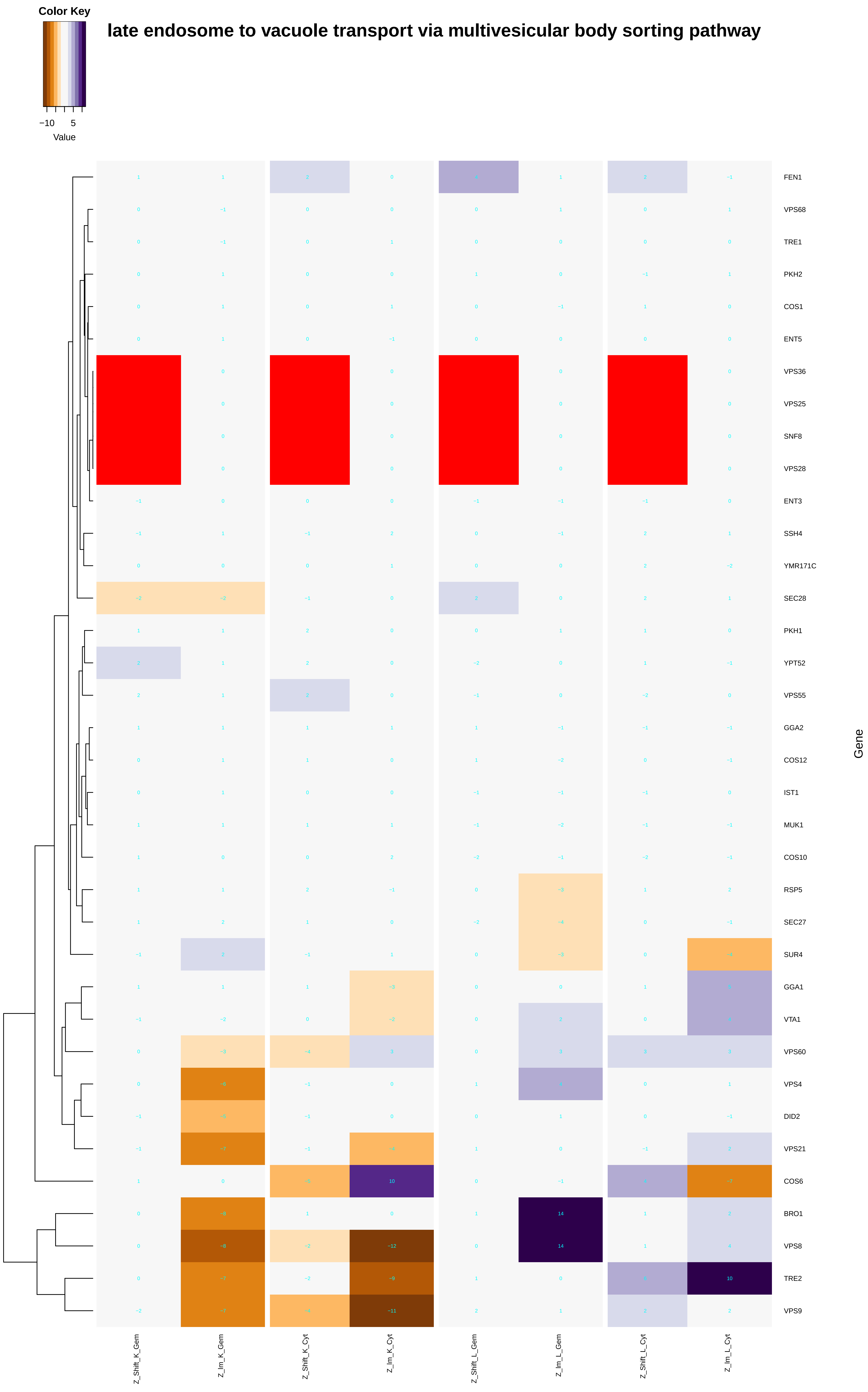

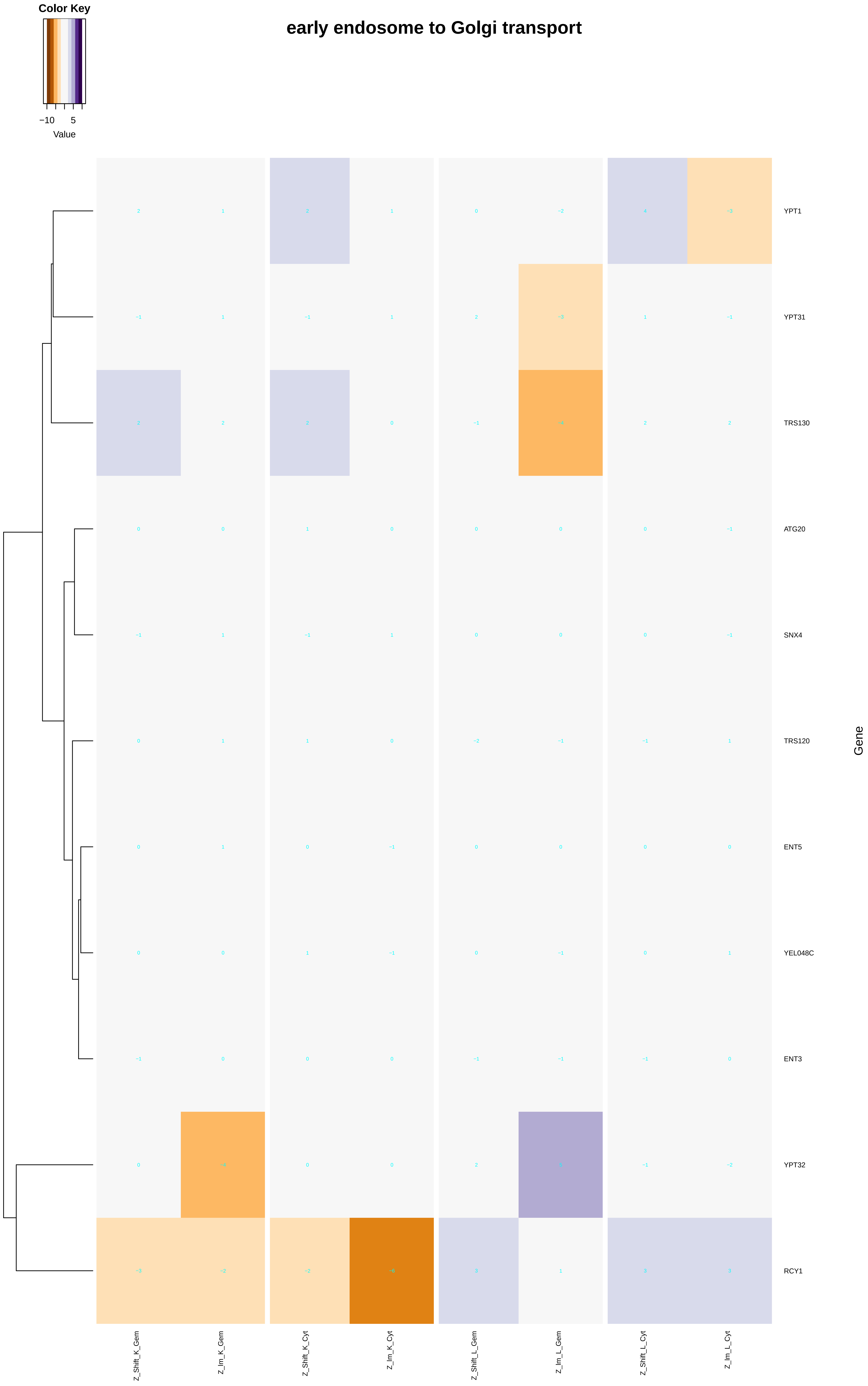

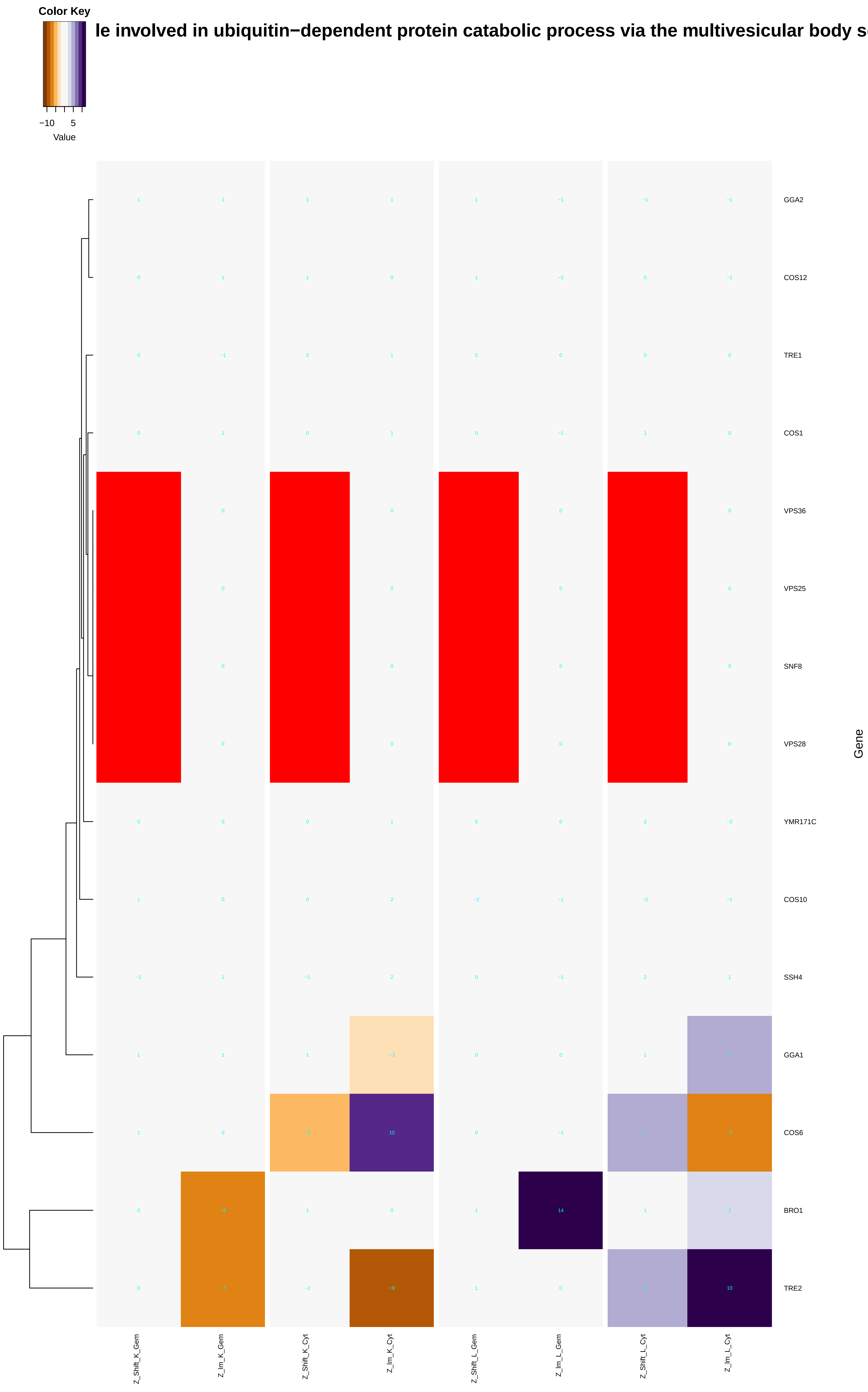

### endosome.pdf

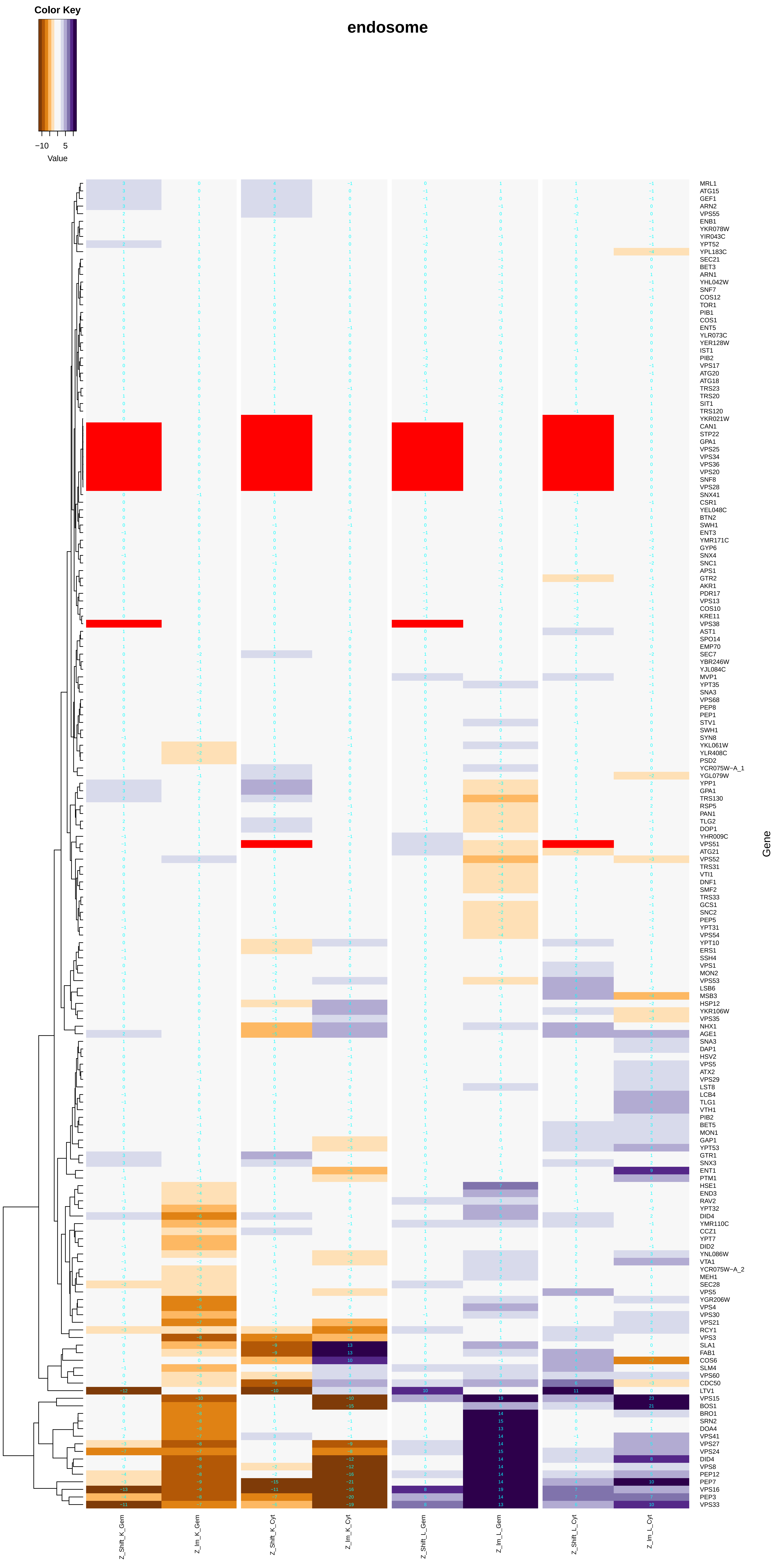

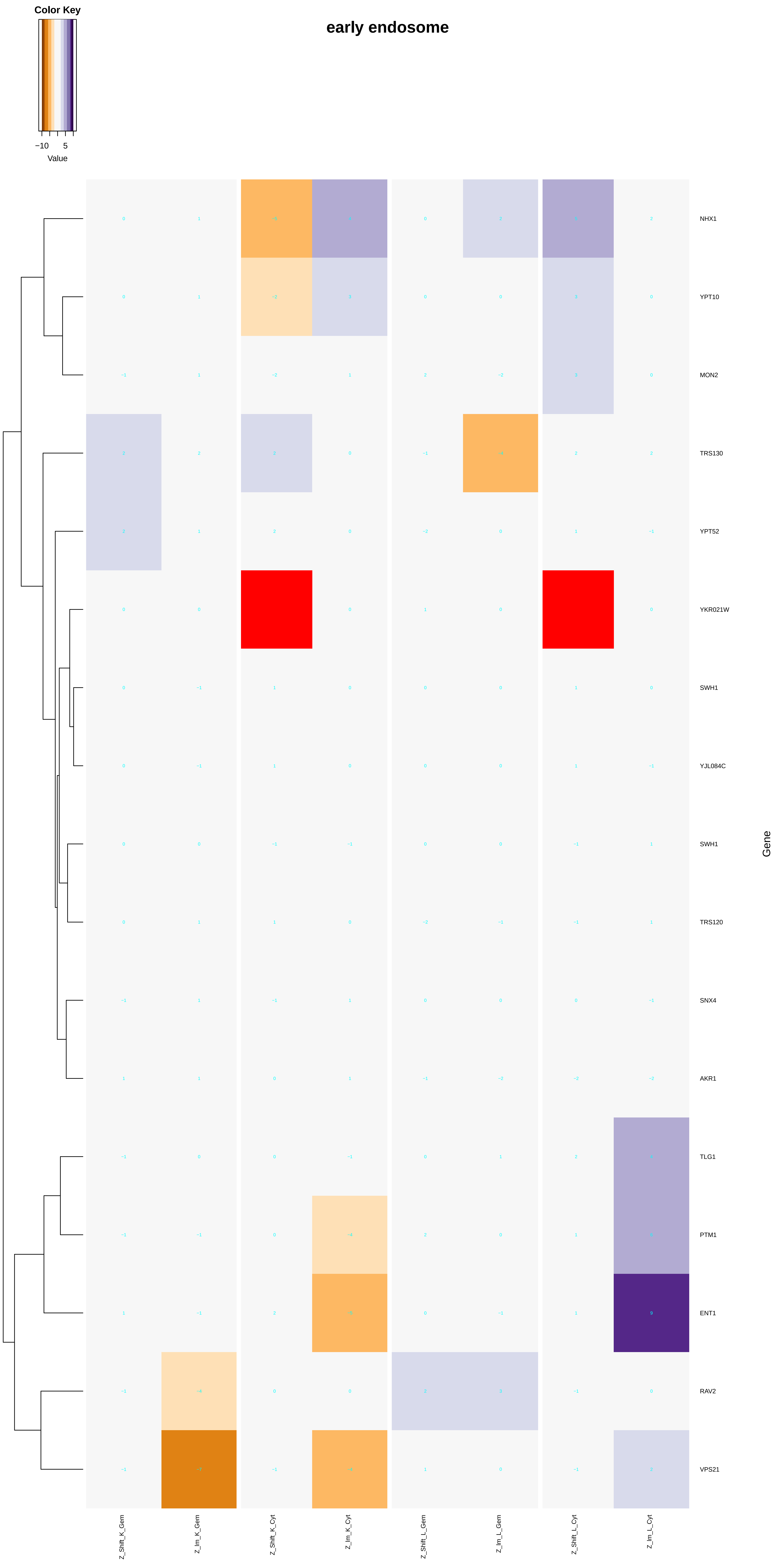

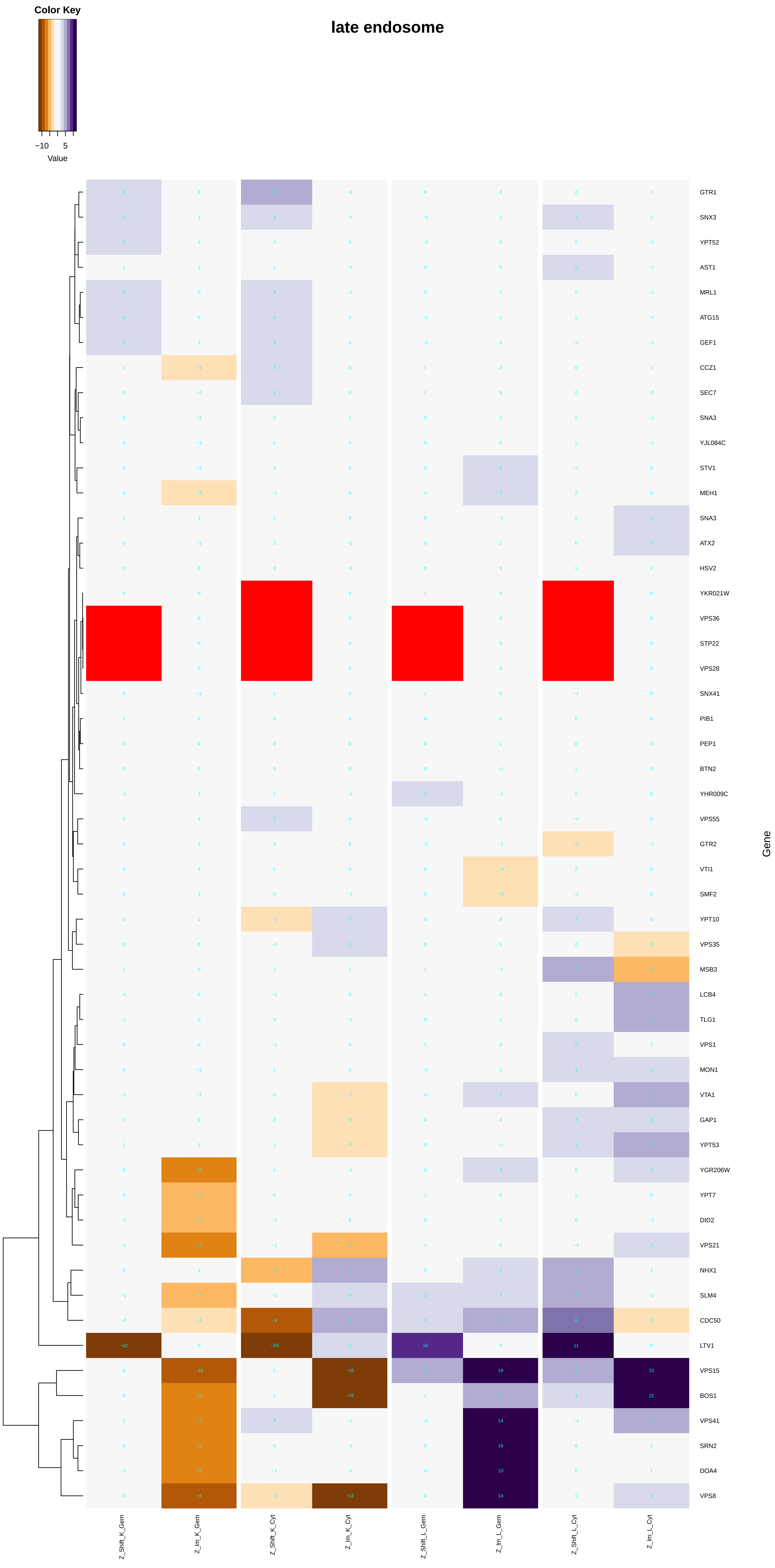

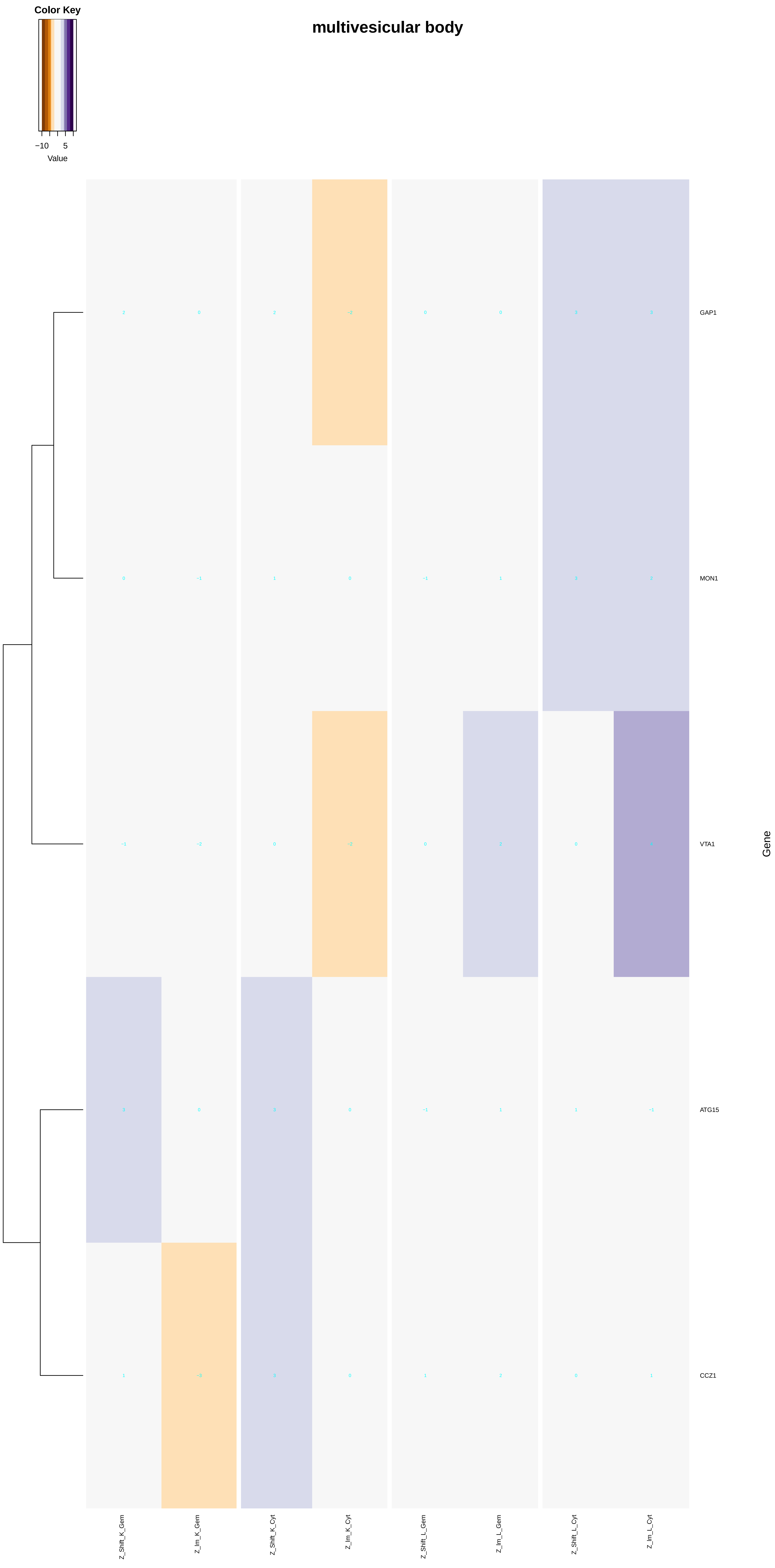

### endosome_membrane.pdf

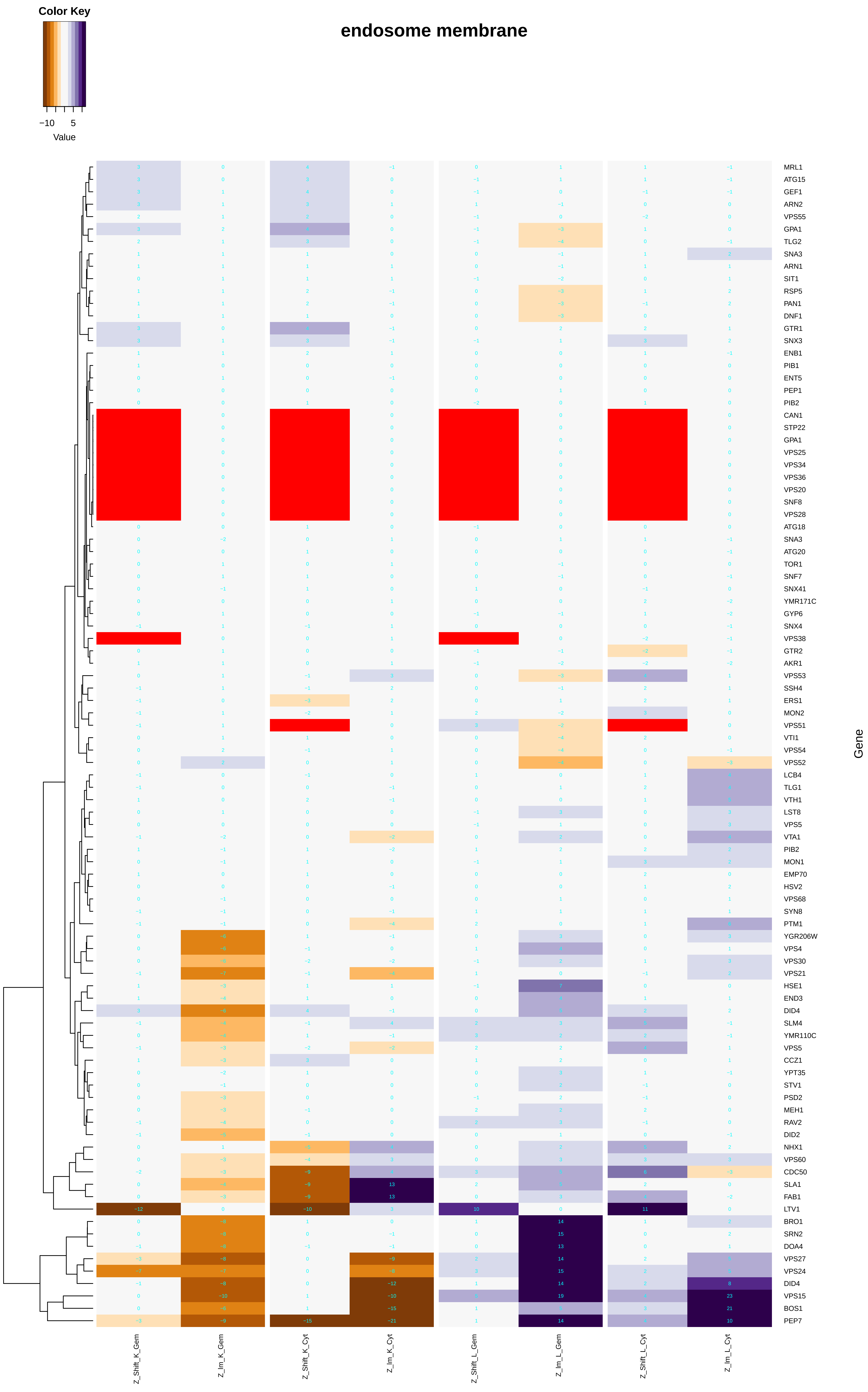

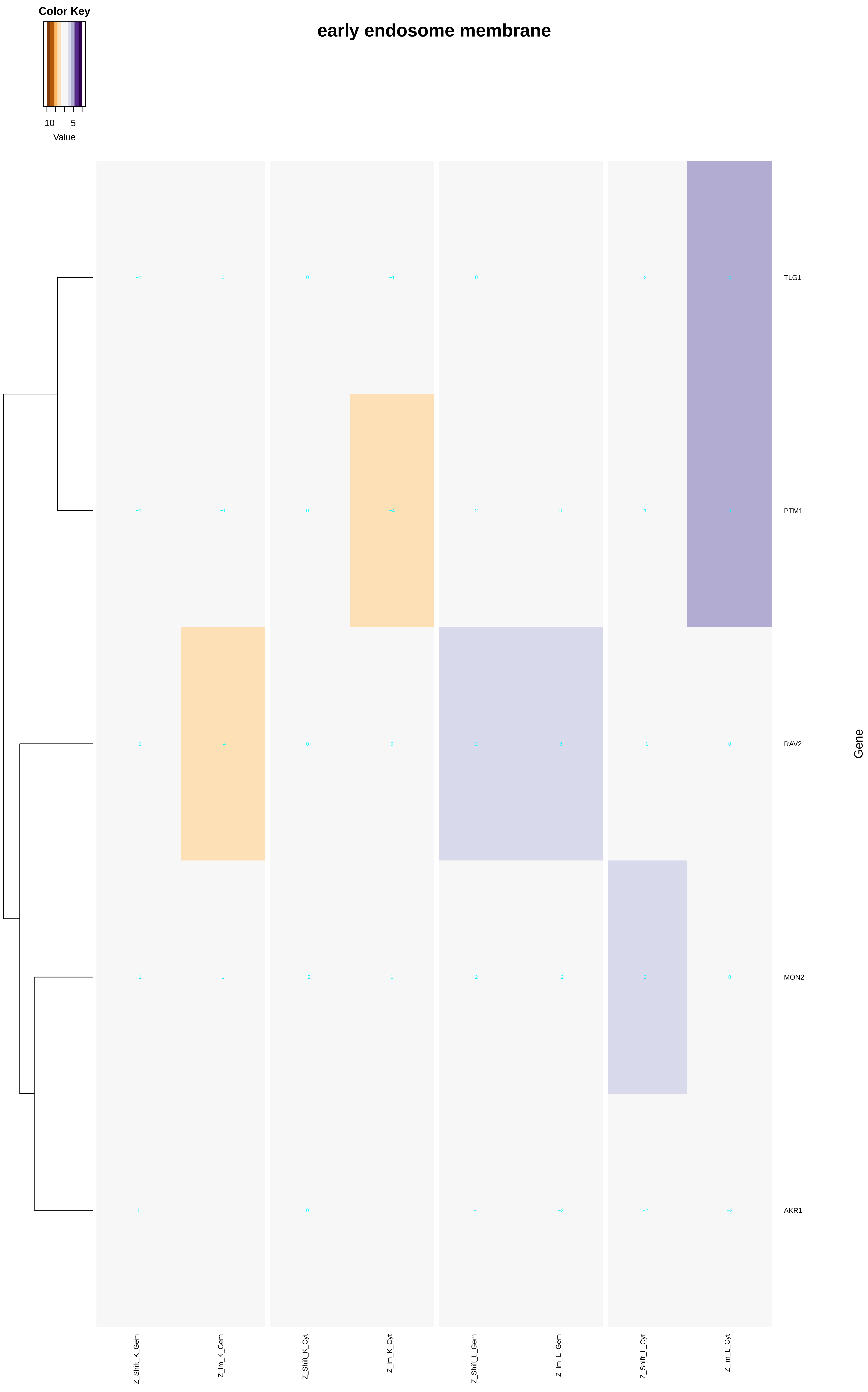

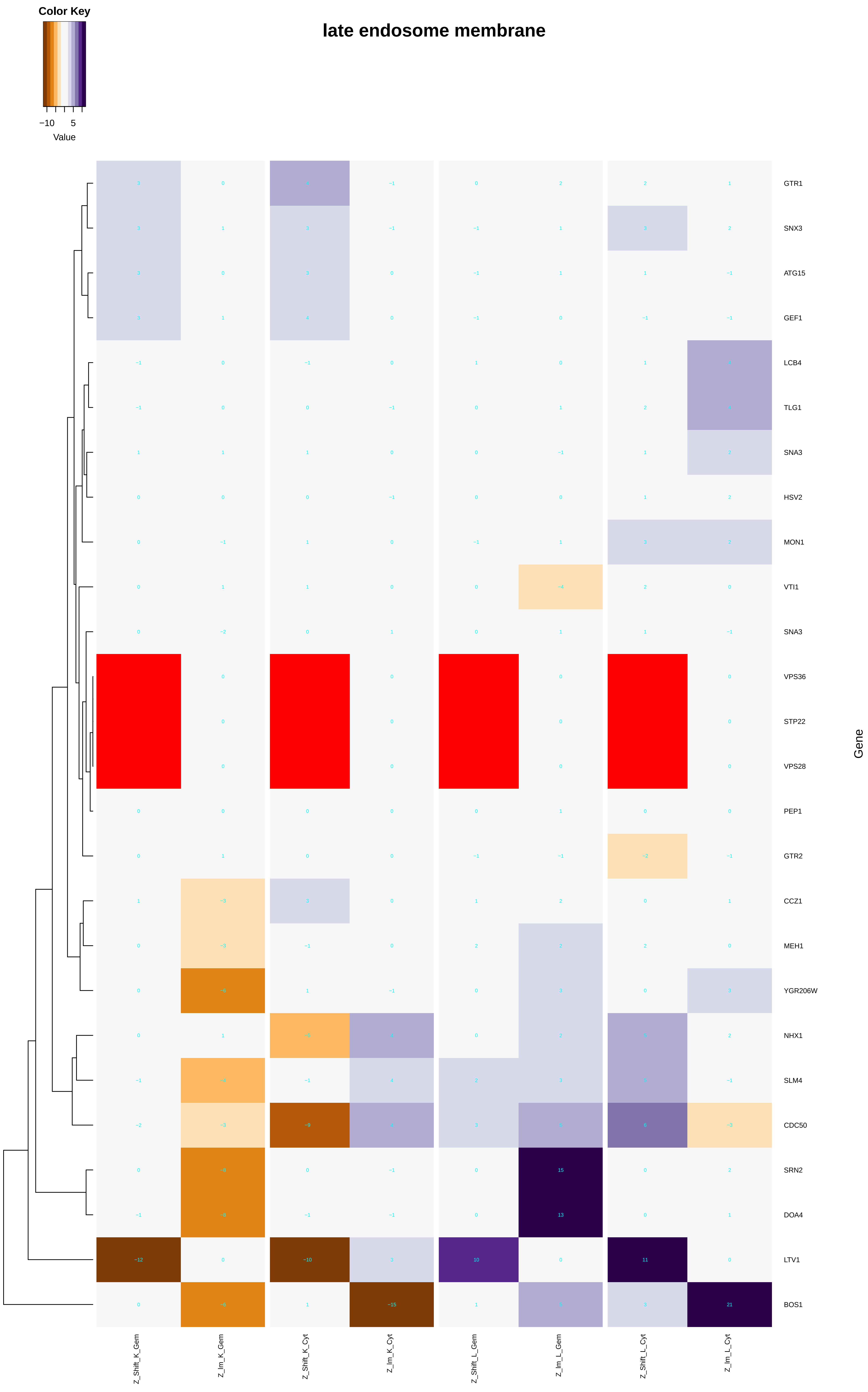

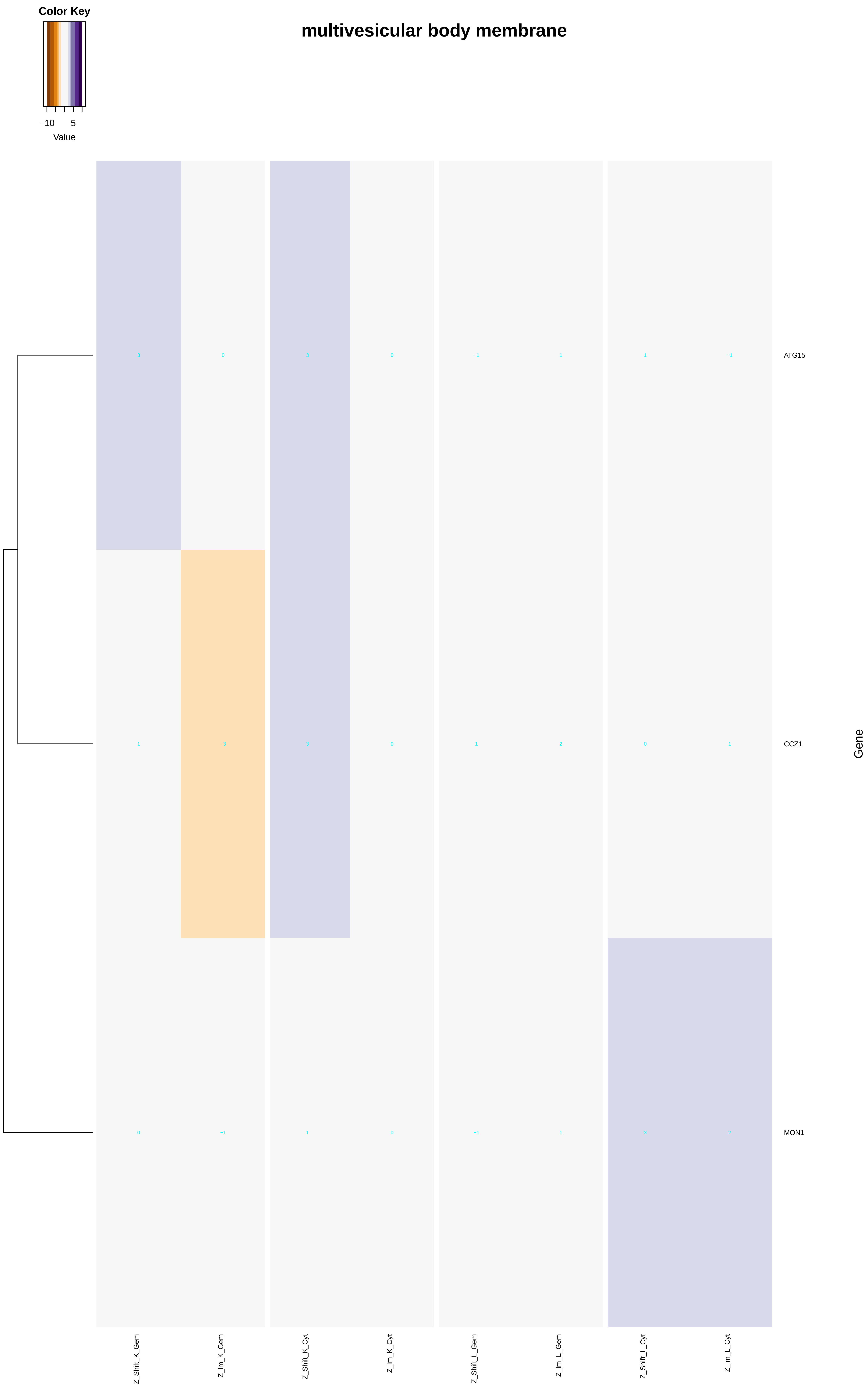

### enzyme_activator_activity.pdf

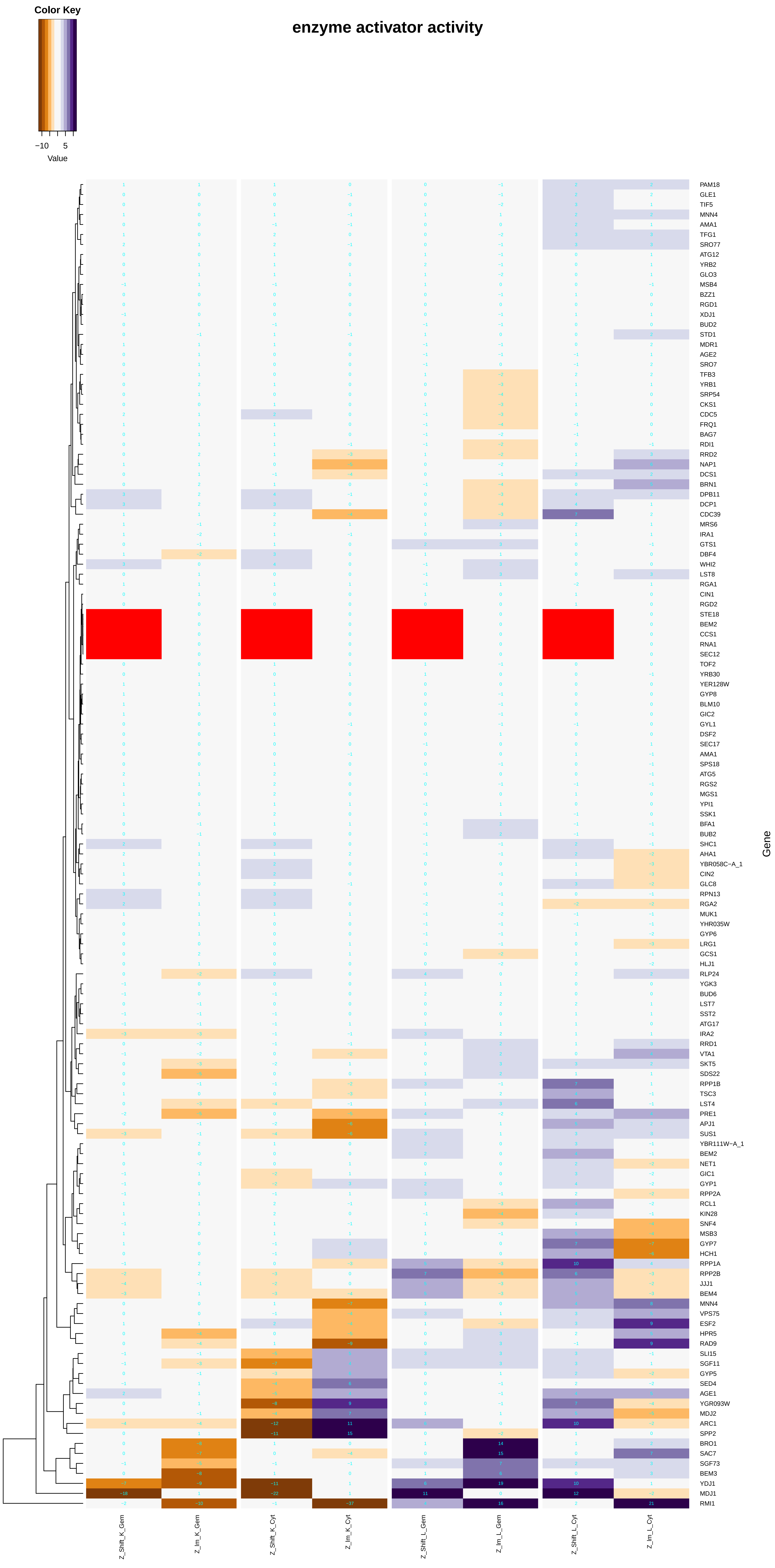

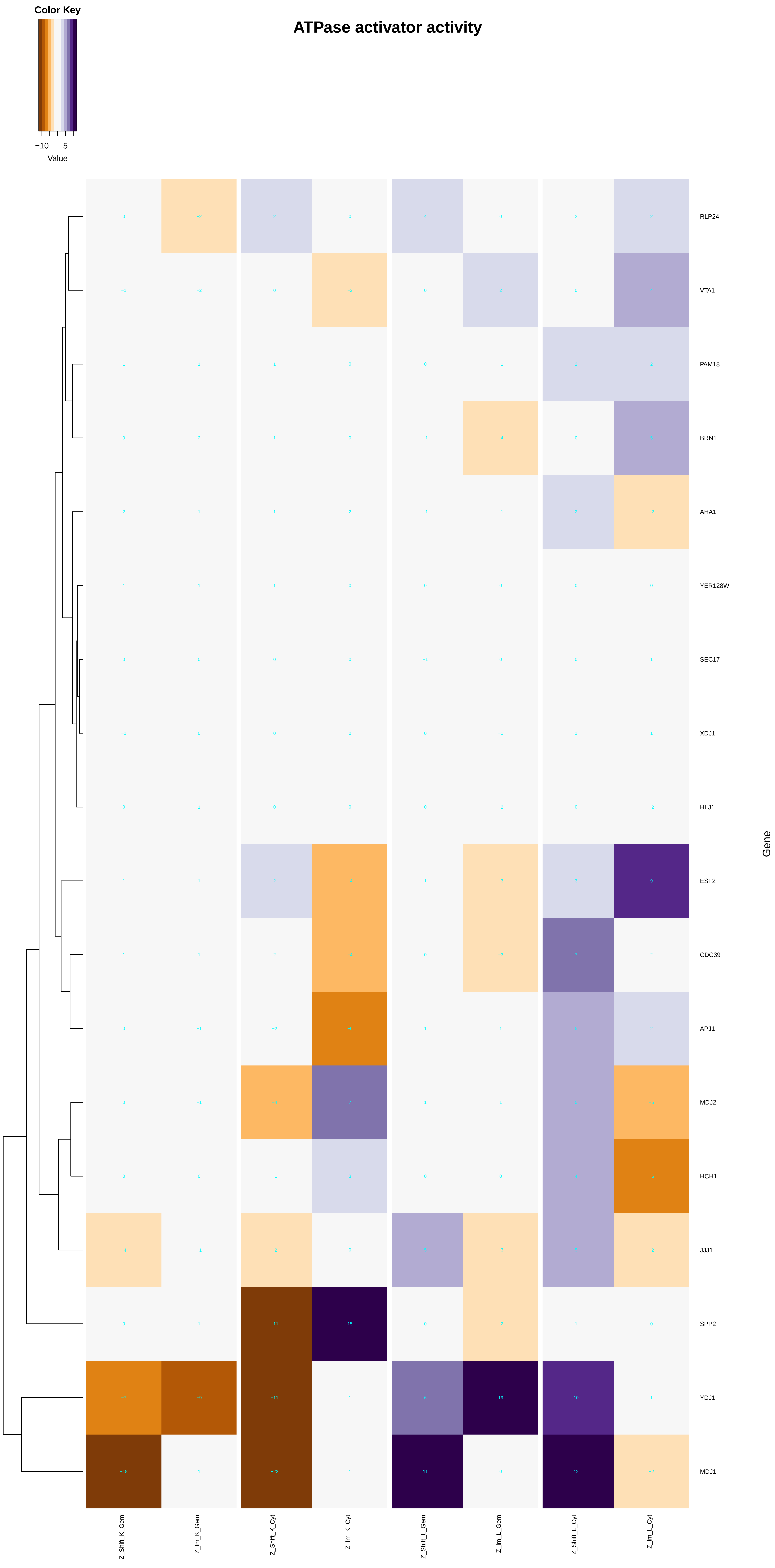

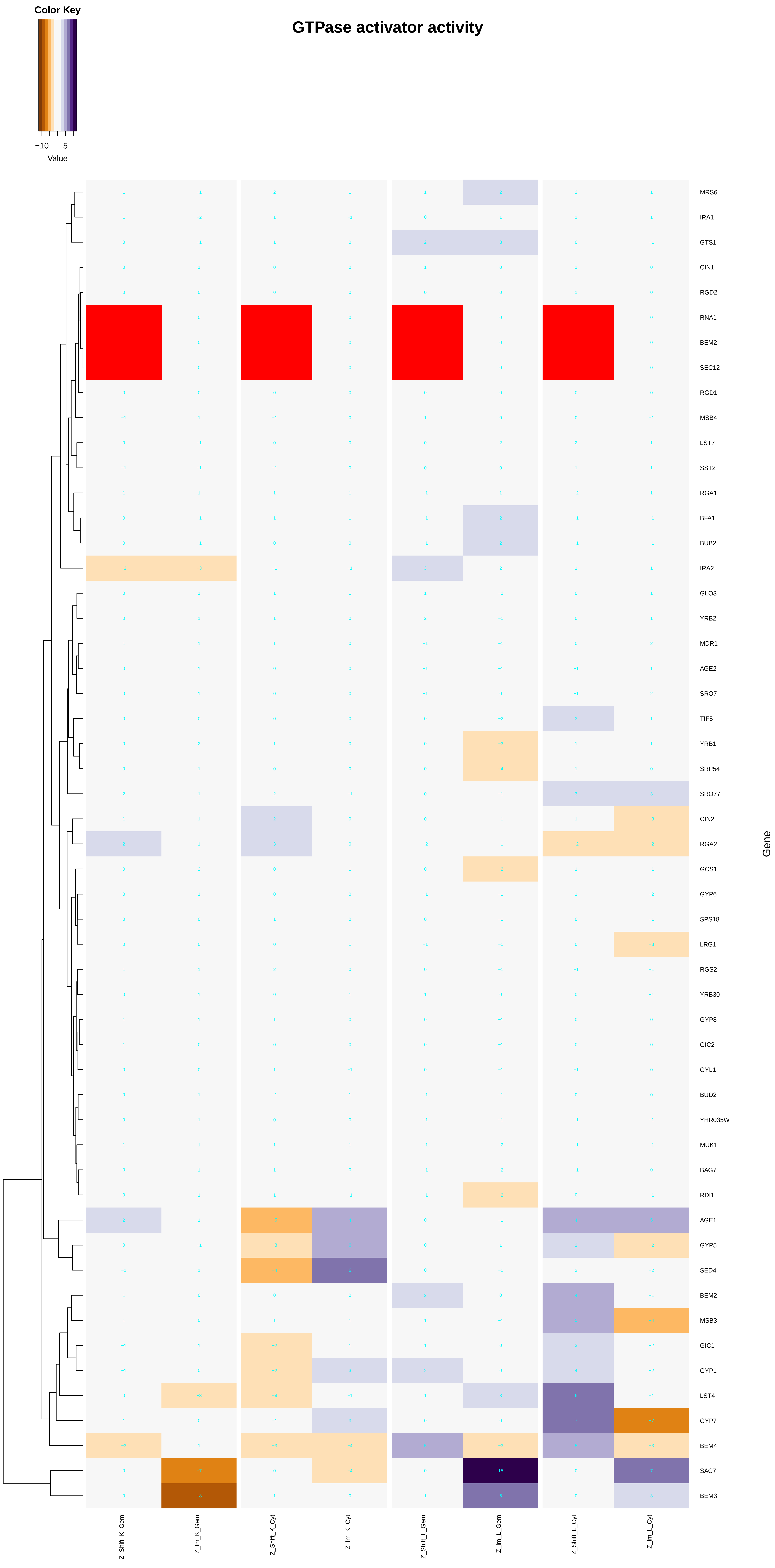

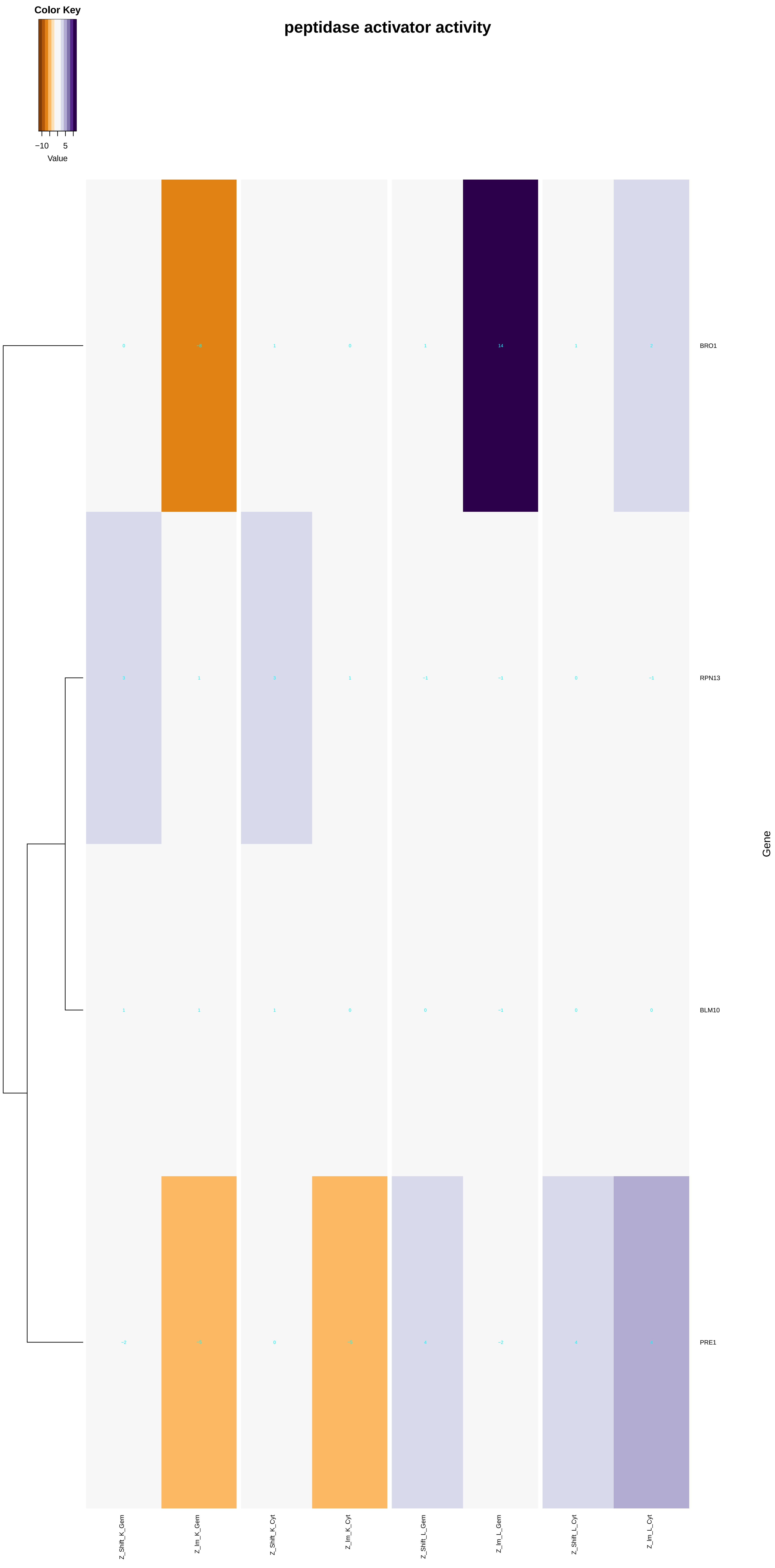

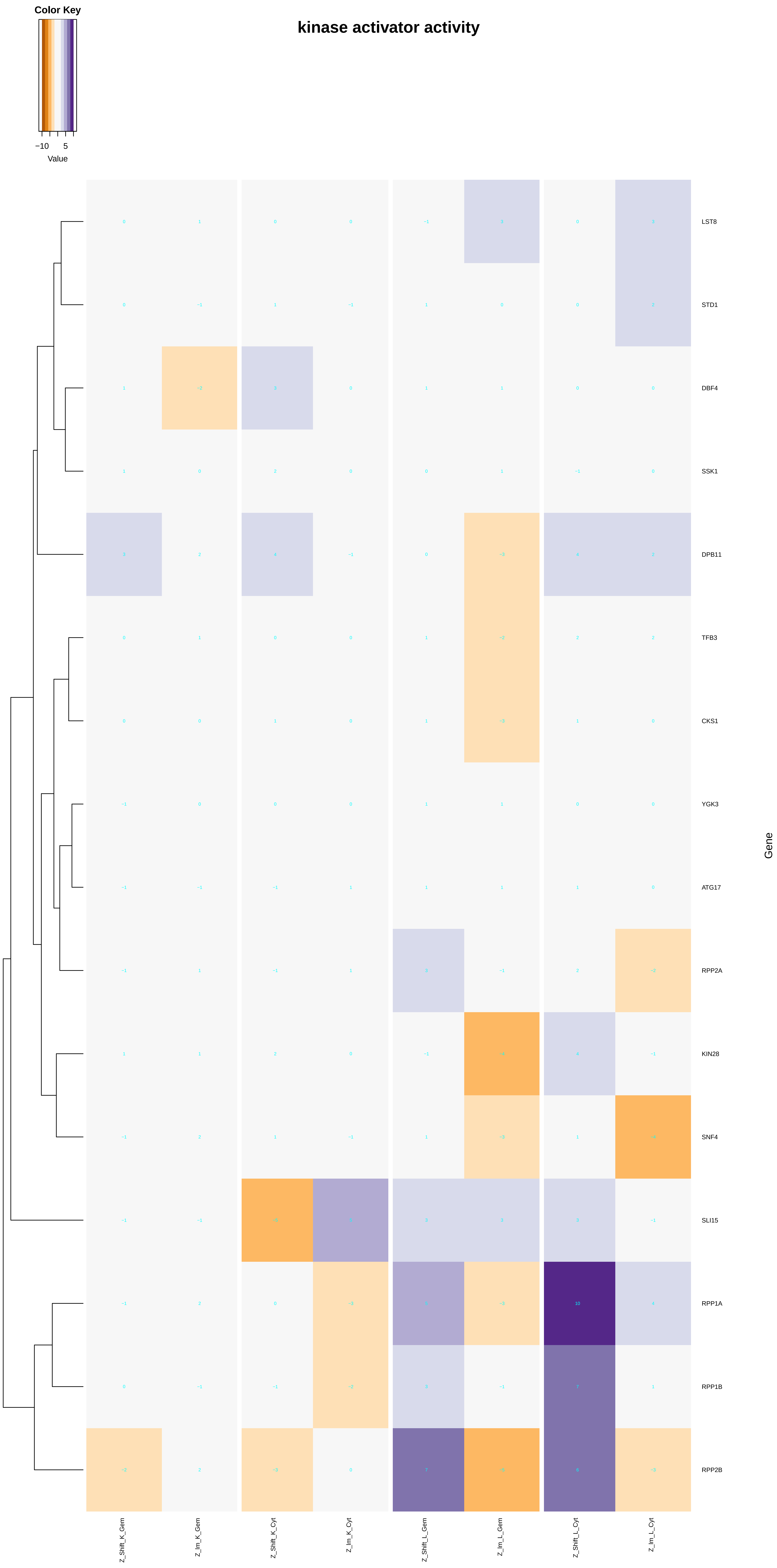

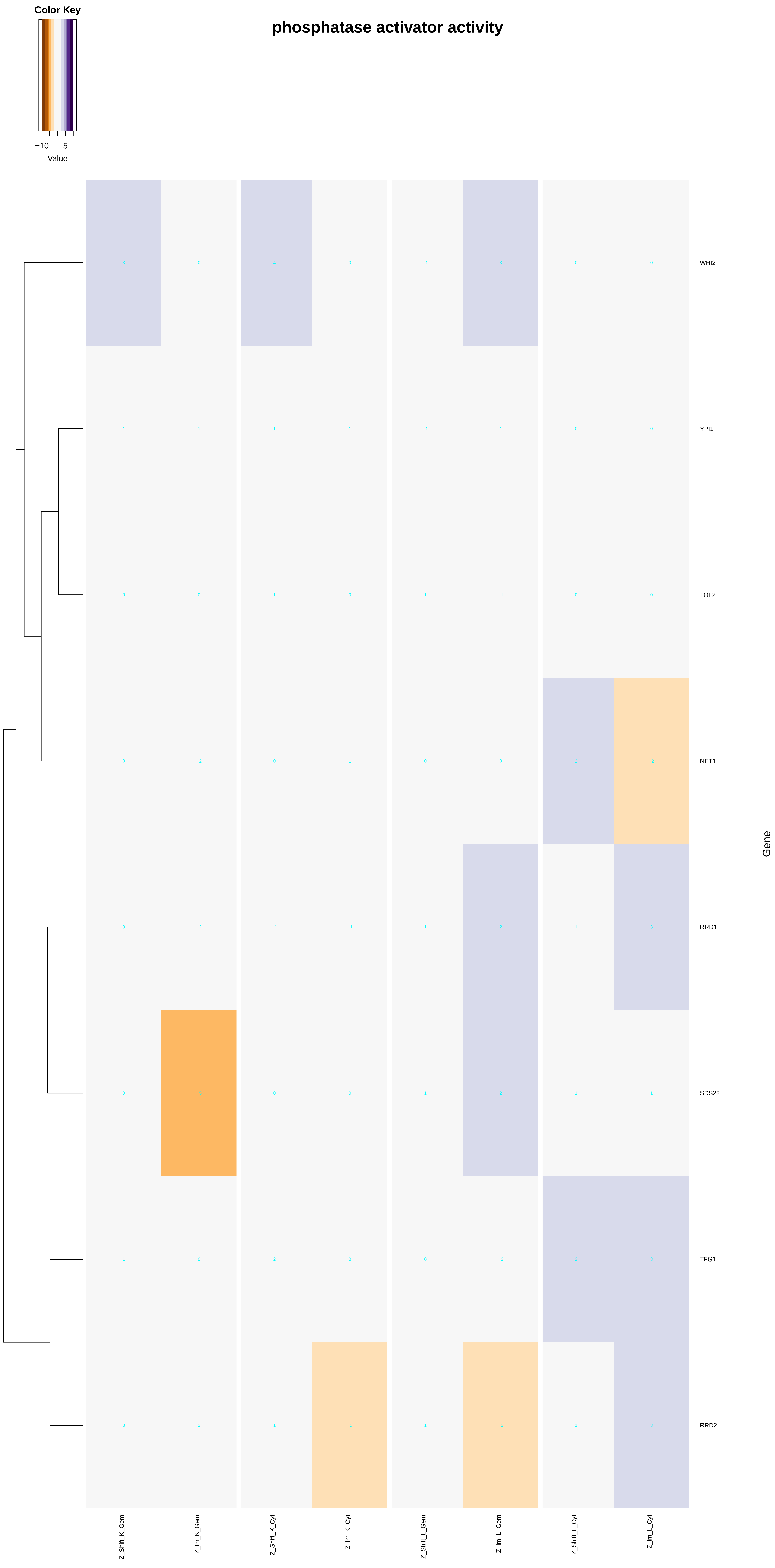

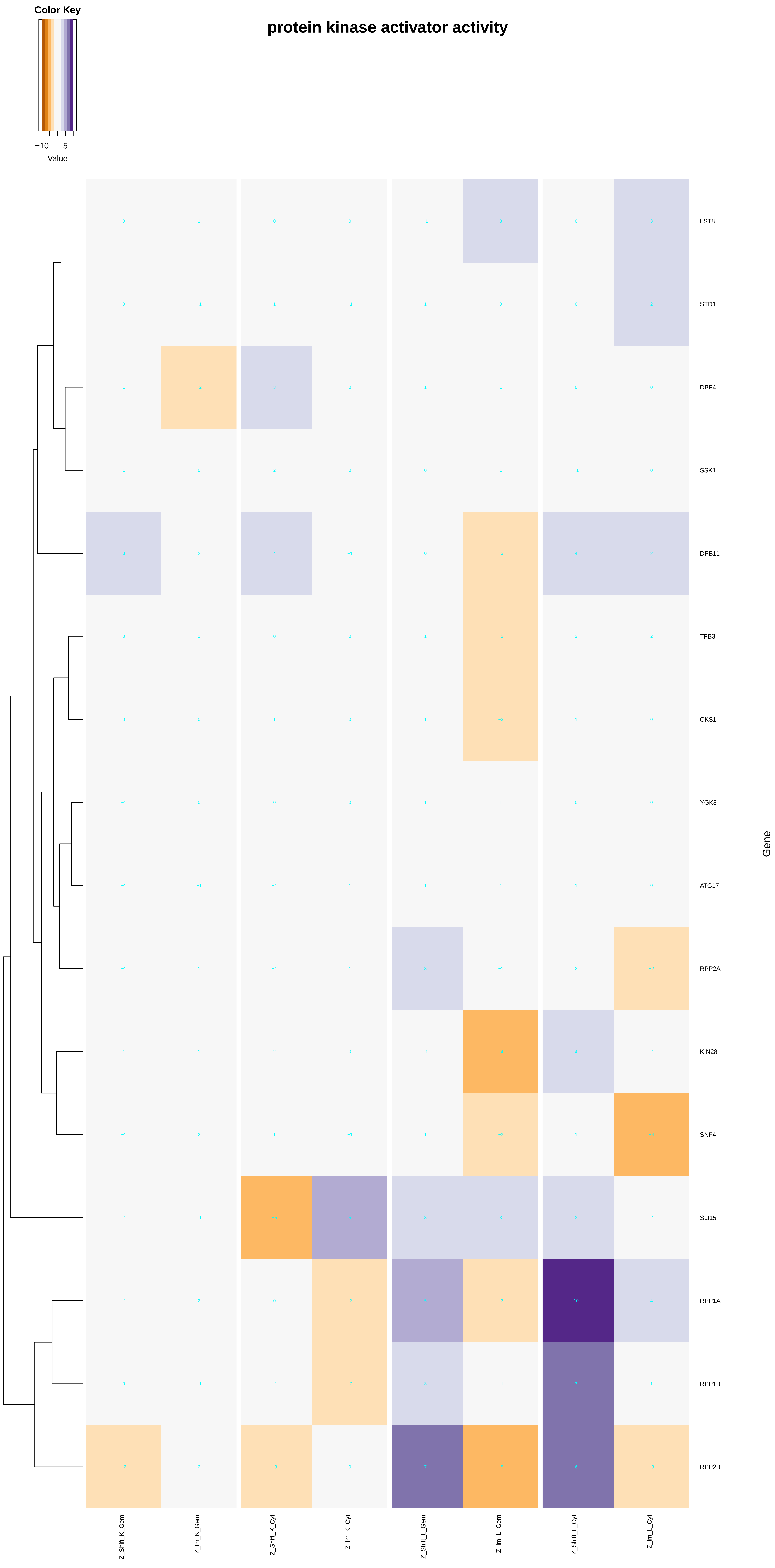

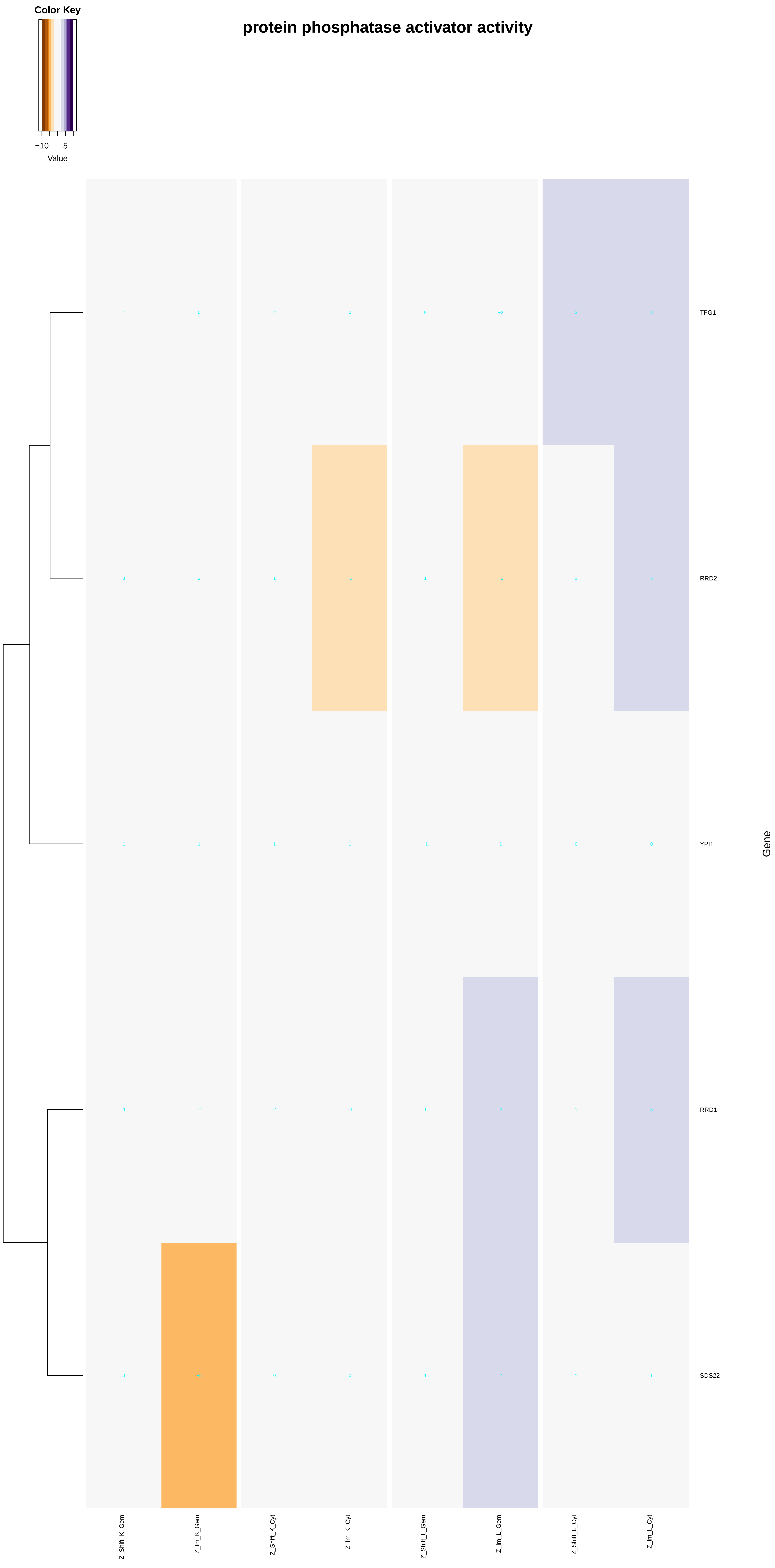

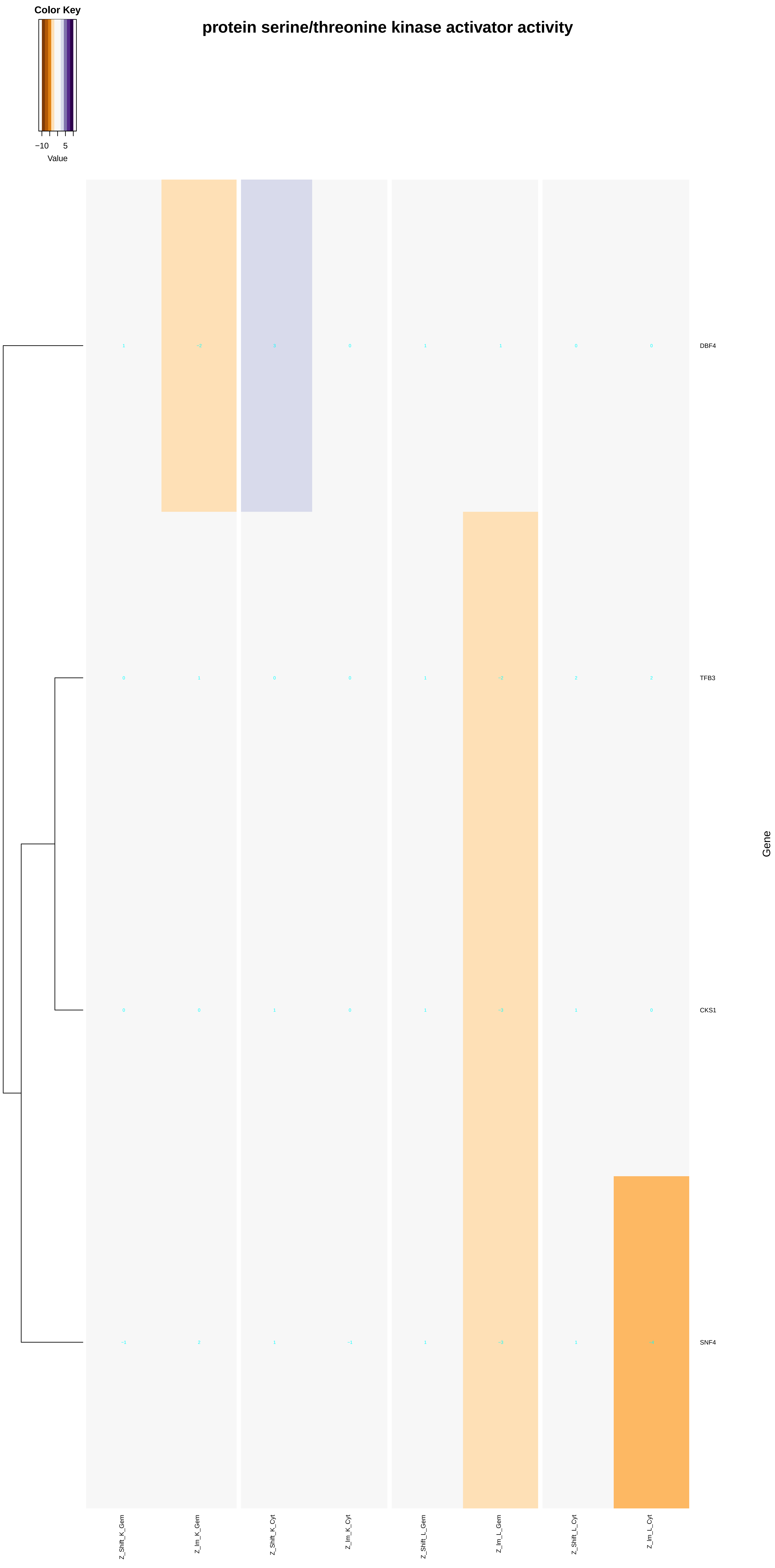

### ER_to_Golgi_transport_vesicle.pdf

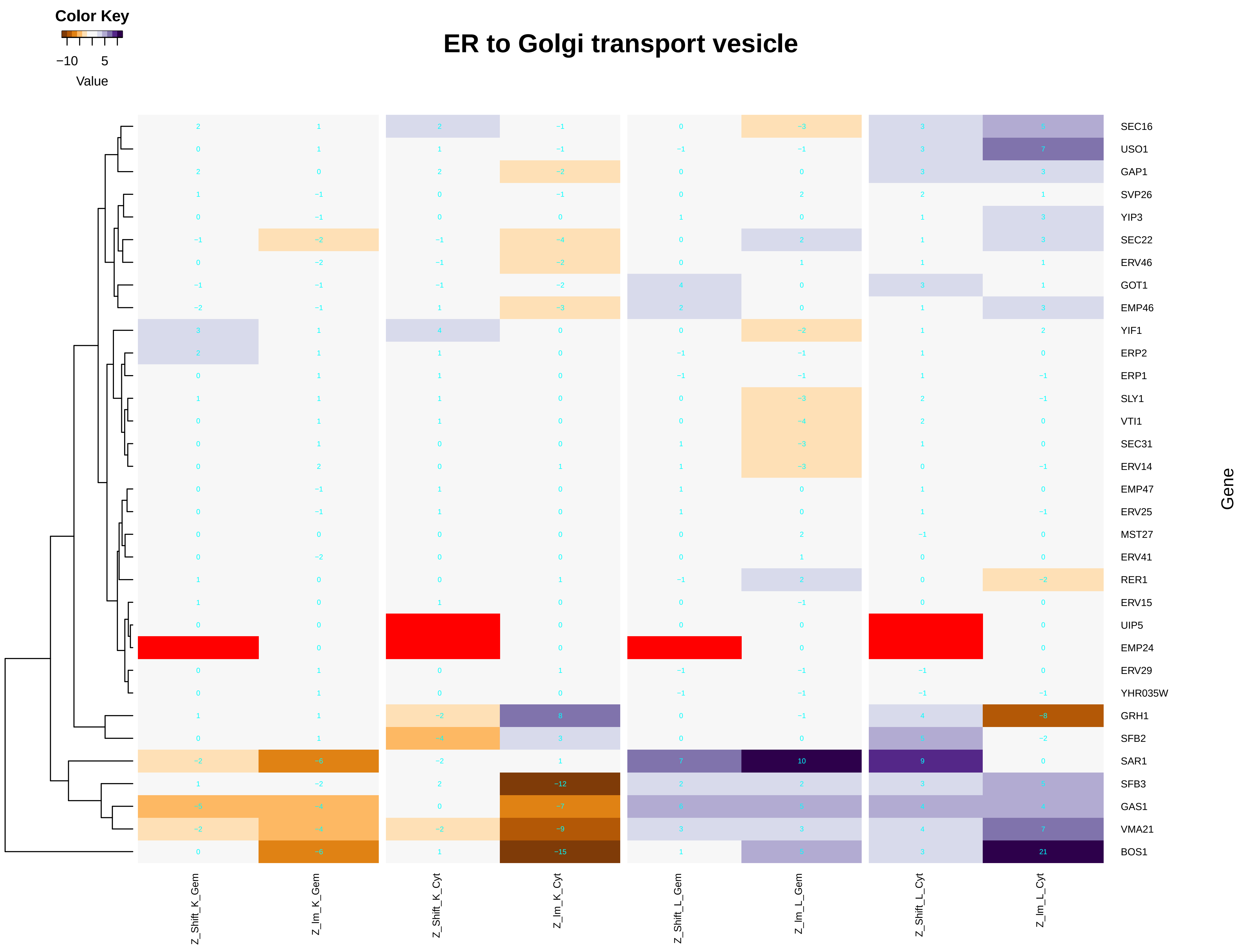

### ER_to_Golgi_transport_vesicle_membrane.pdf

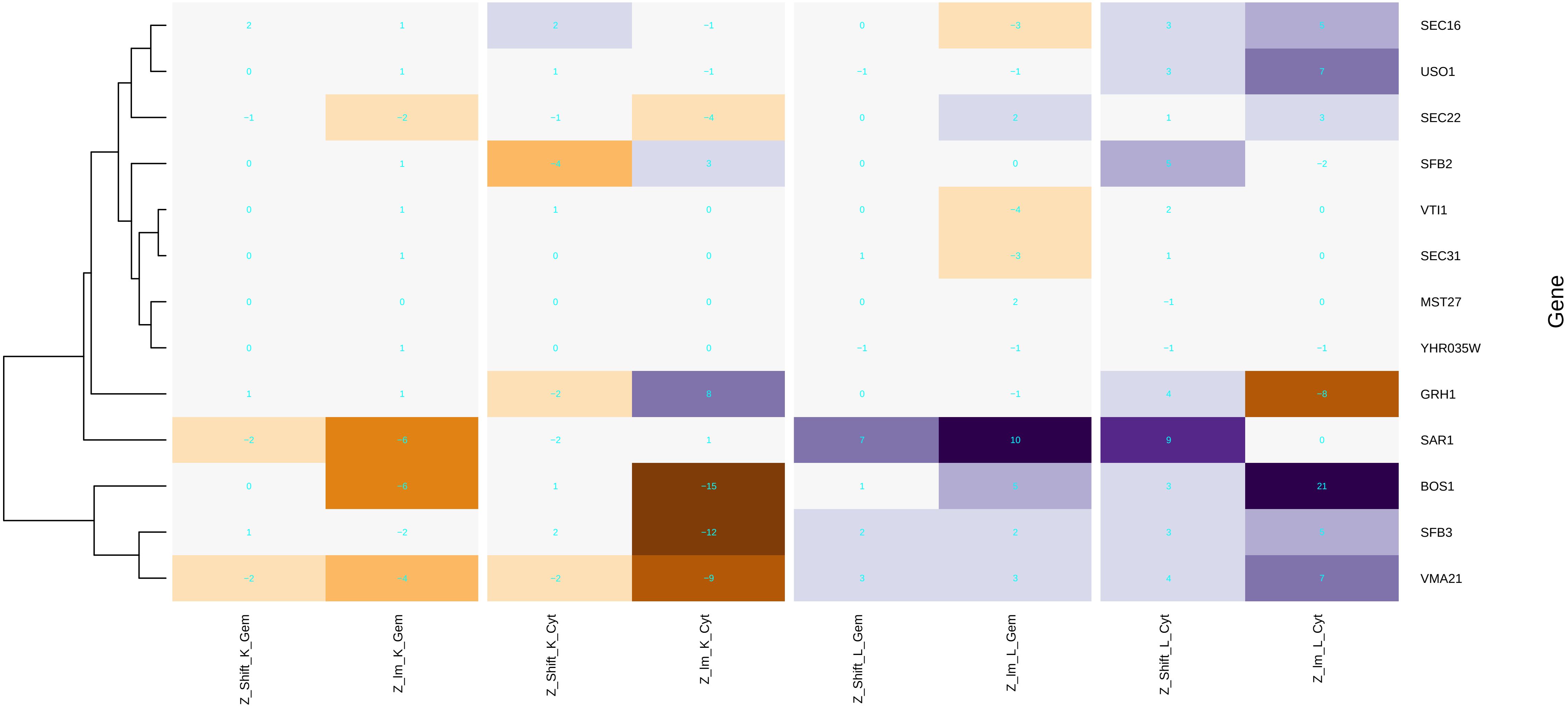

### ER_to_Golgi_vesicle-mediated_transport.pdf

#### ER to Golgi vesicle−mediated transport

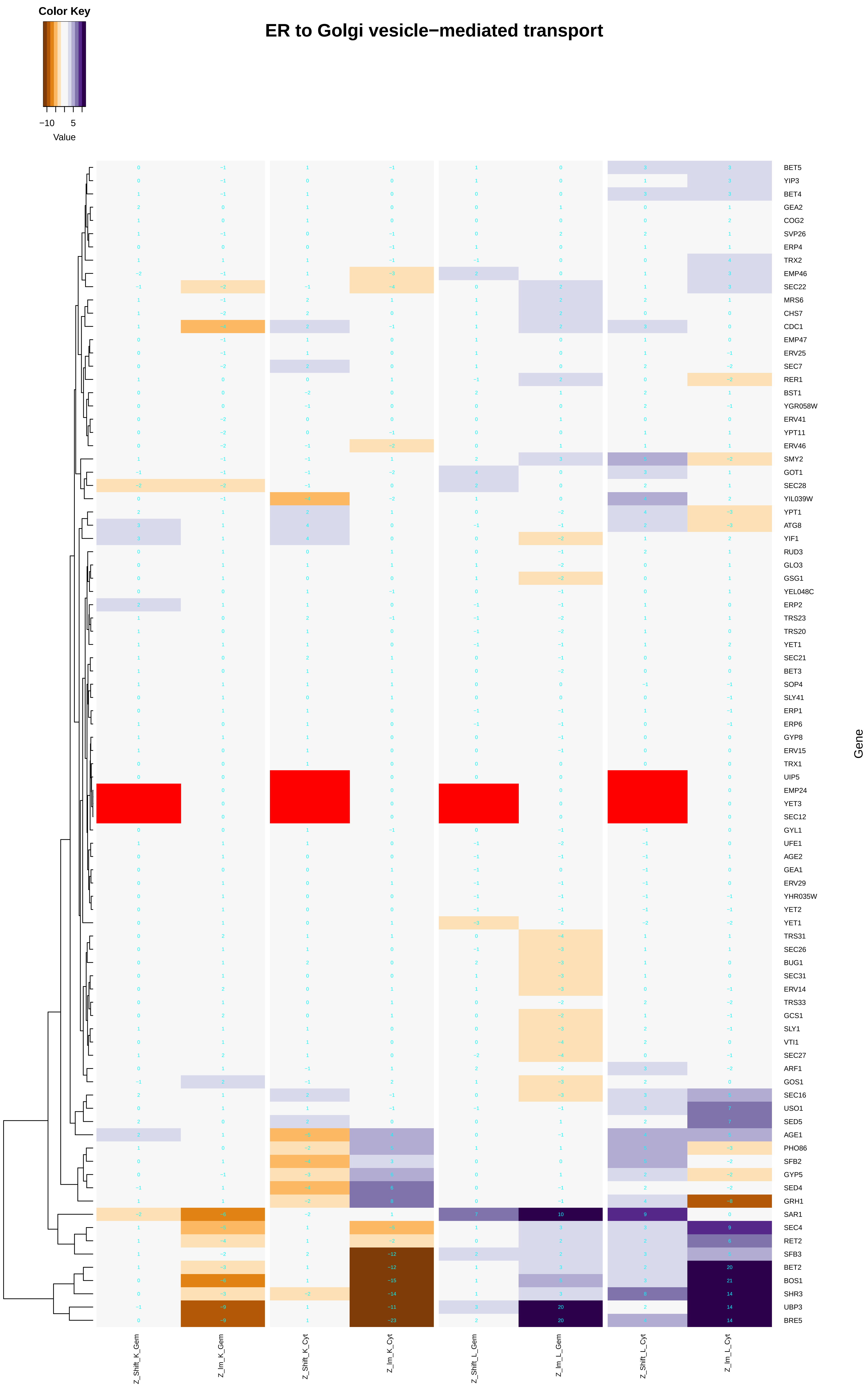

### ESCRT_I_complex.pdf

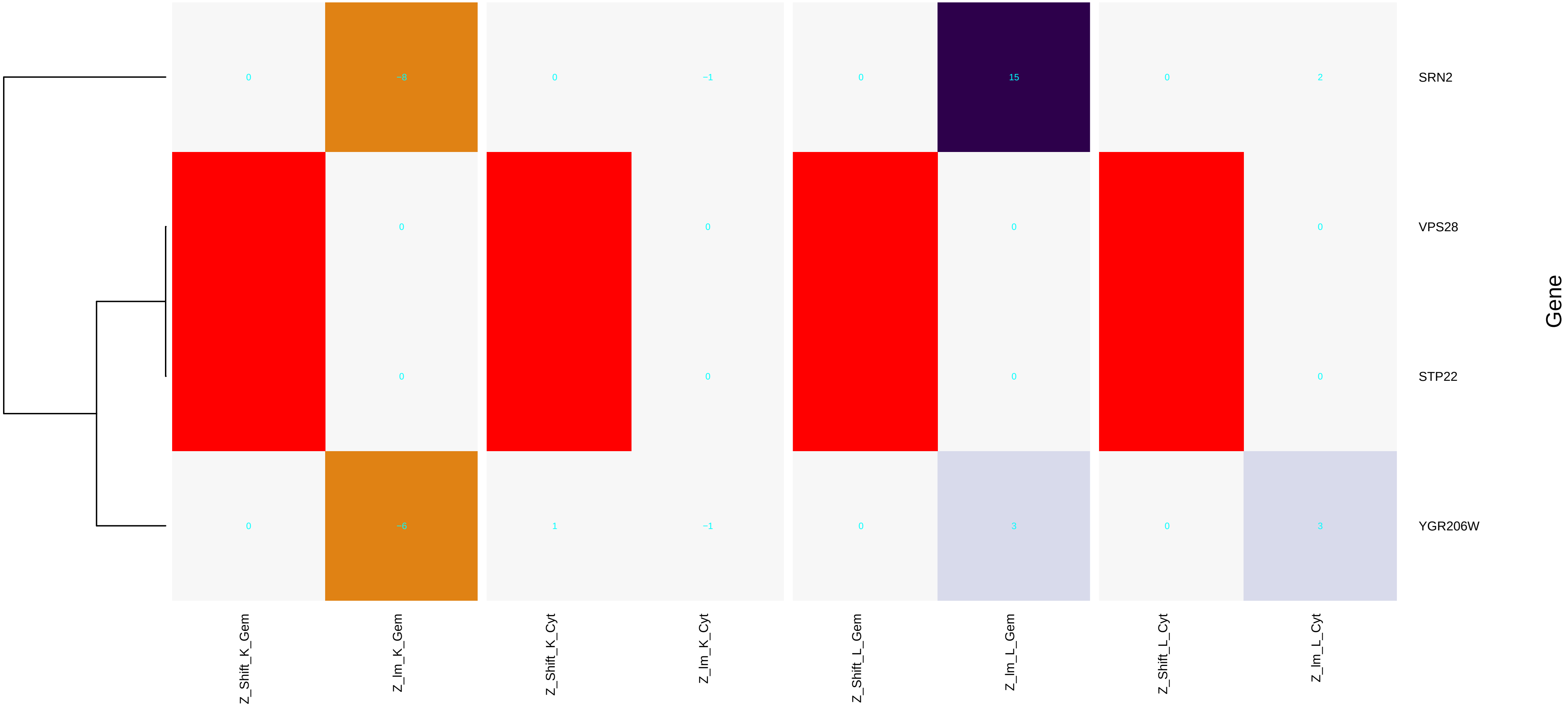

### eukaryotic_43S_preinitiation_complex.pdf

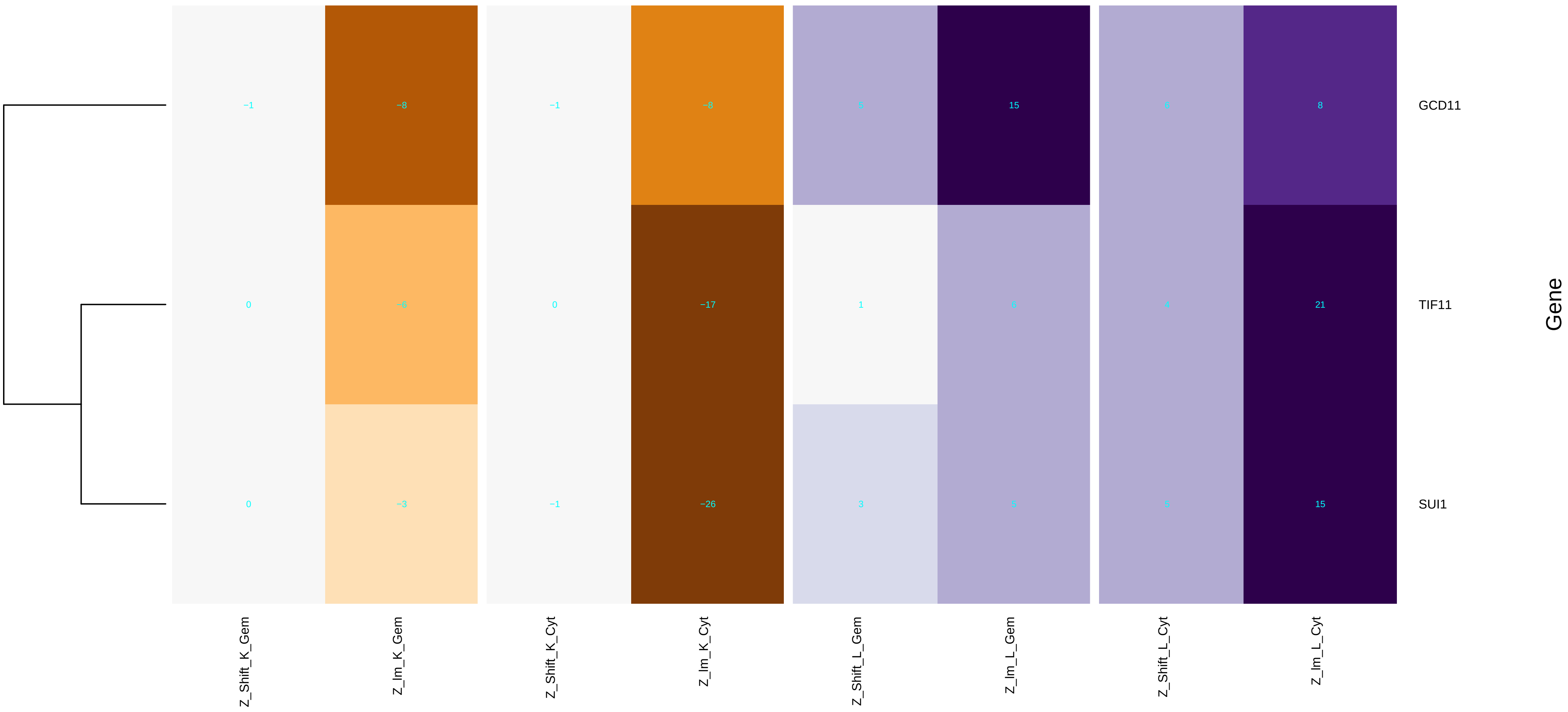

### eukaryotic_48S_preinitiation_complex.pdf

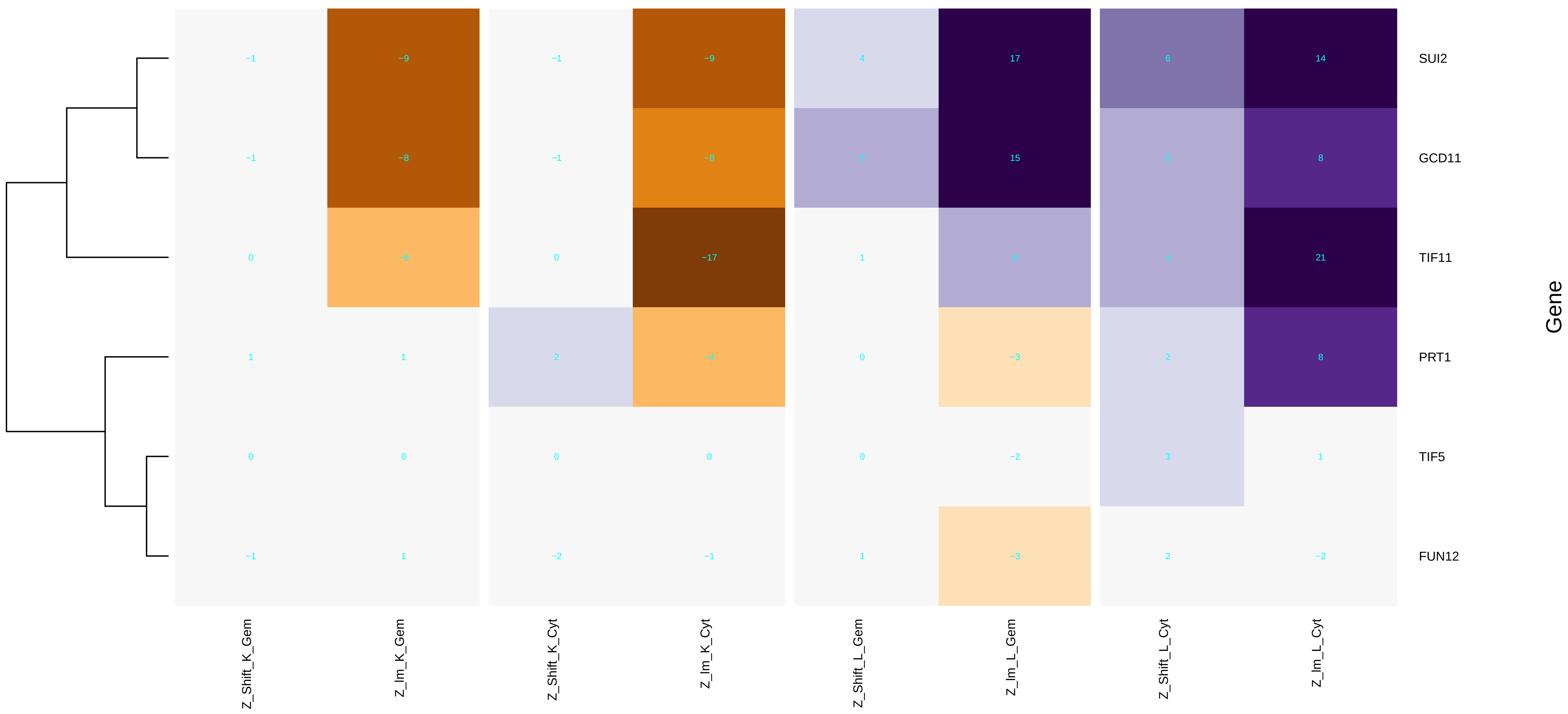

### exonucleolytic_nuclear-transcribed_mRNA_catabolic_process_involved_in_deadenylation-dependent_decay.pdf

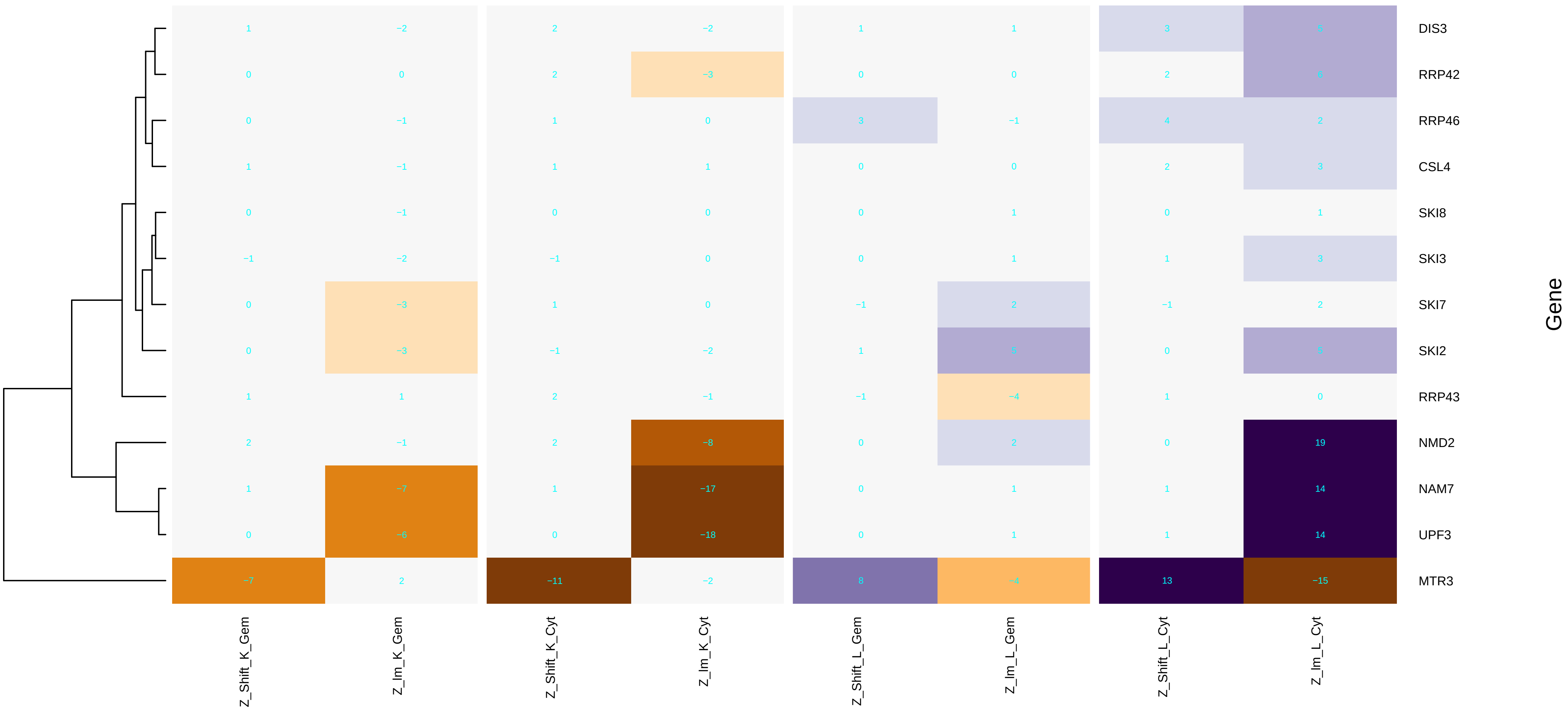

### four-way_junction_helicase_activity.pdf

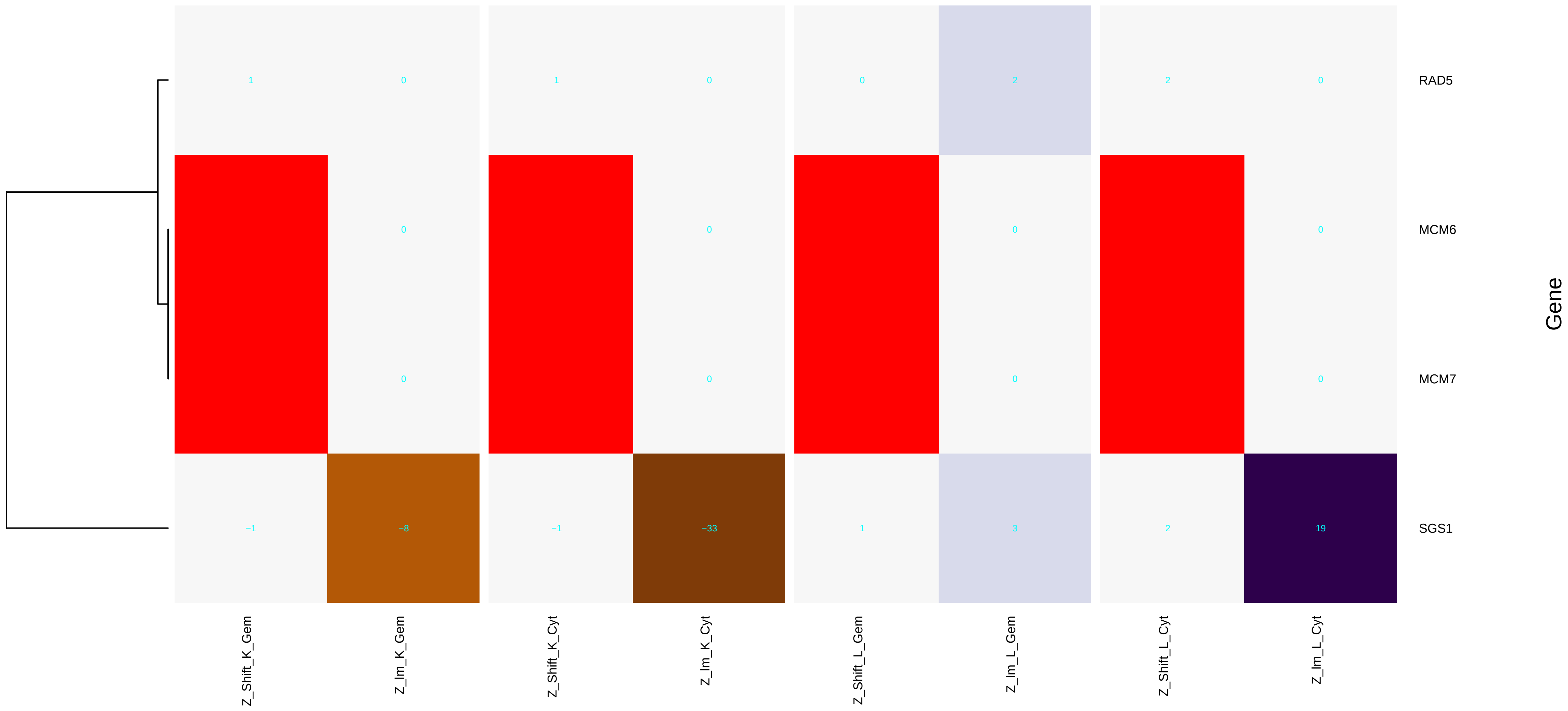

### fungal-type_vacuole_lumen.pdf

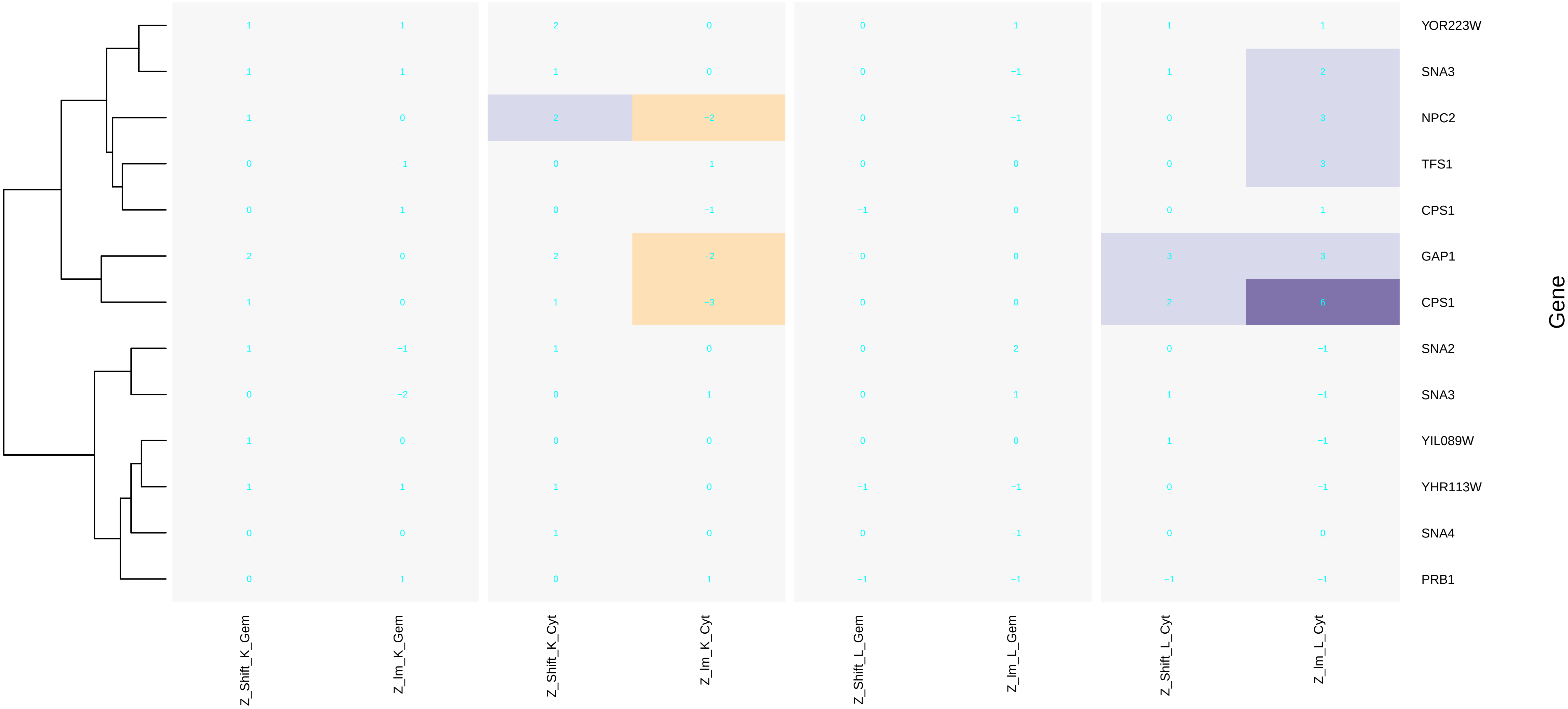

### G1_DNA_damage_checkpoint.pdf

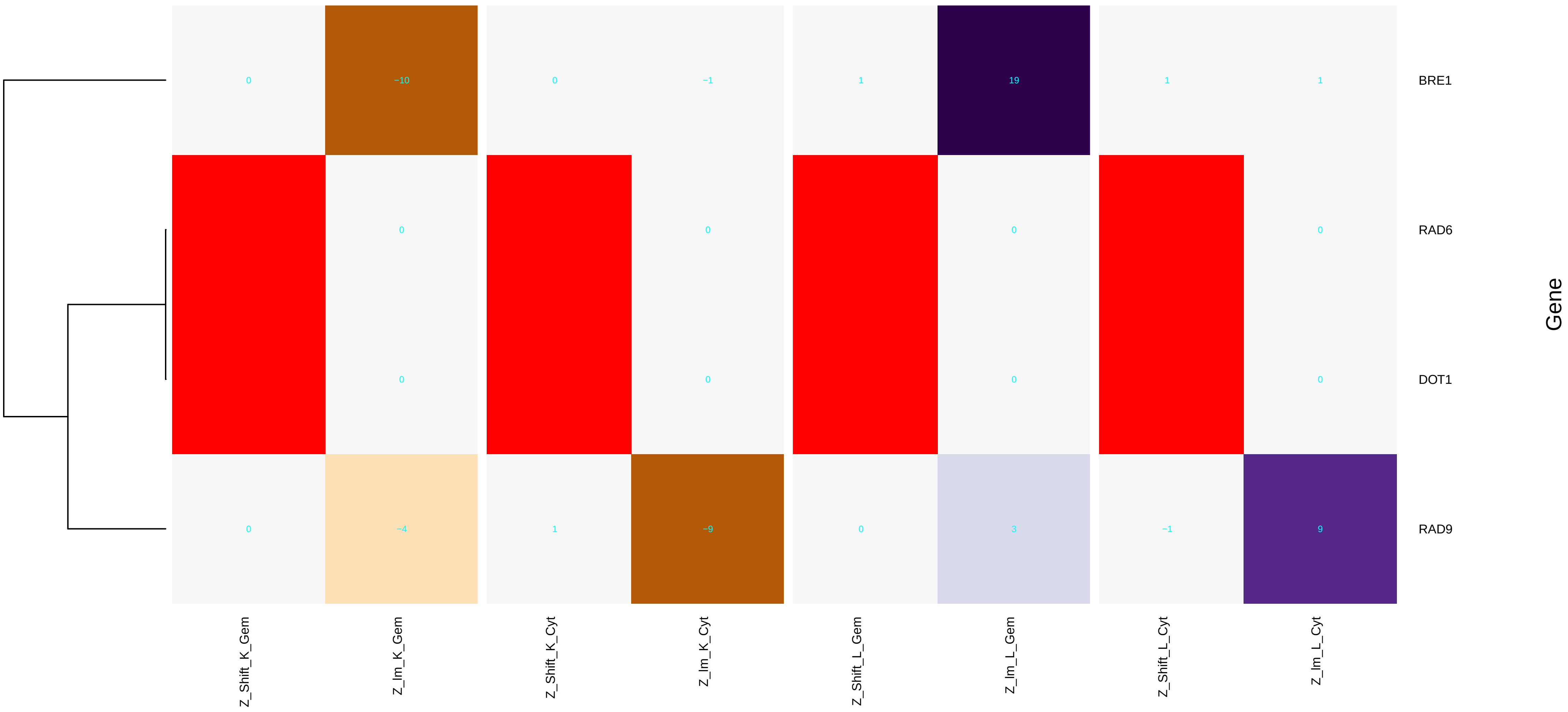

### G-quadruplex_DNA_binding.pdf

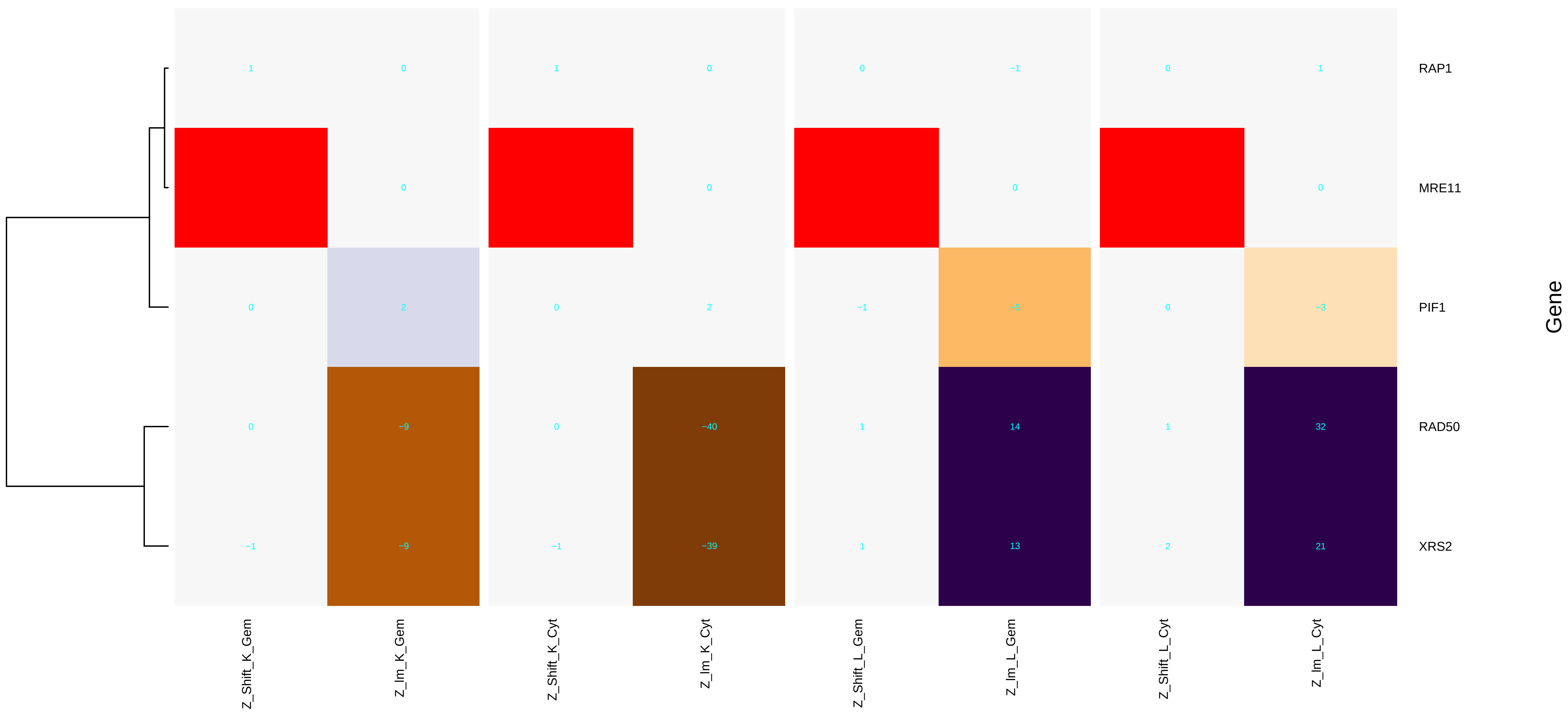

### GARP_complex.pdf

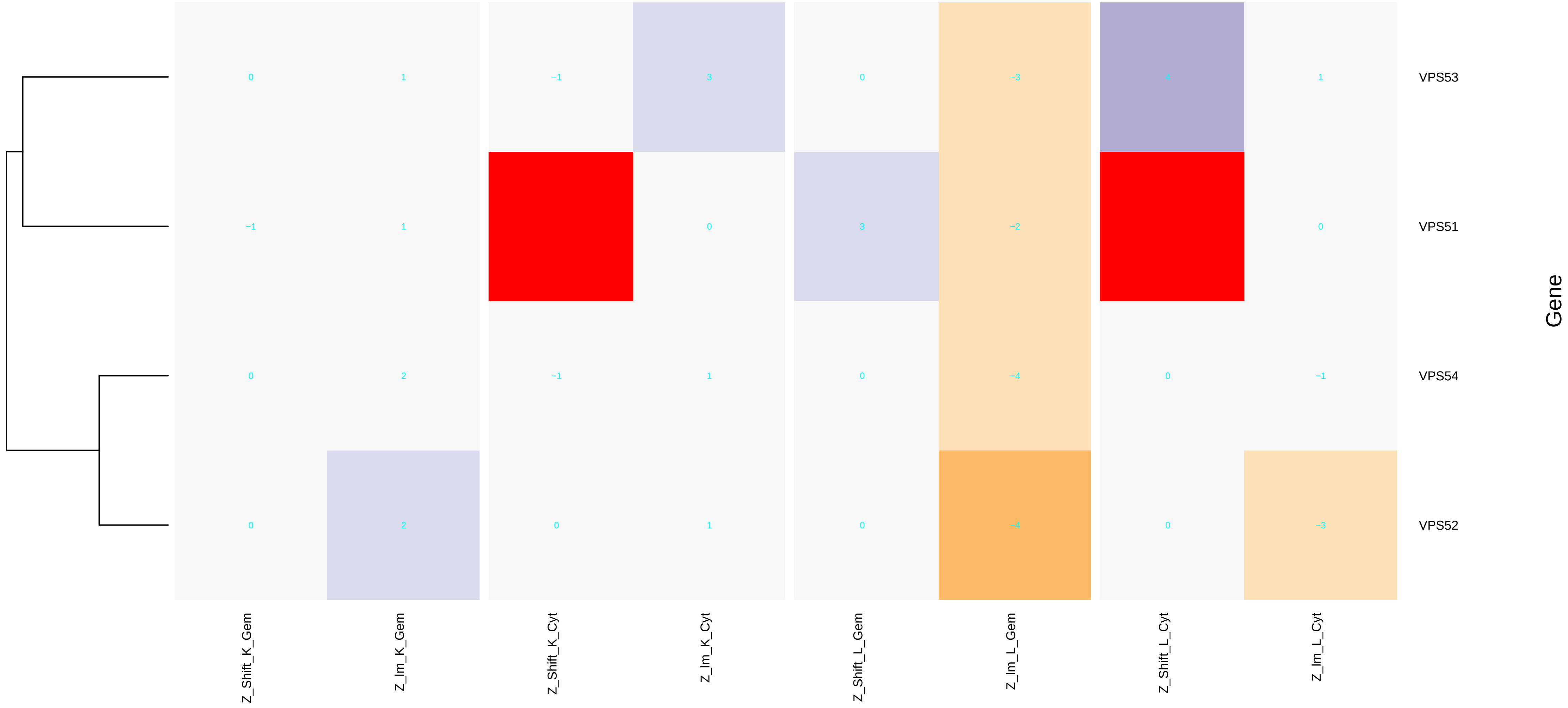

### GDP-dissociation_inhibitor_activity.pdf

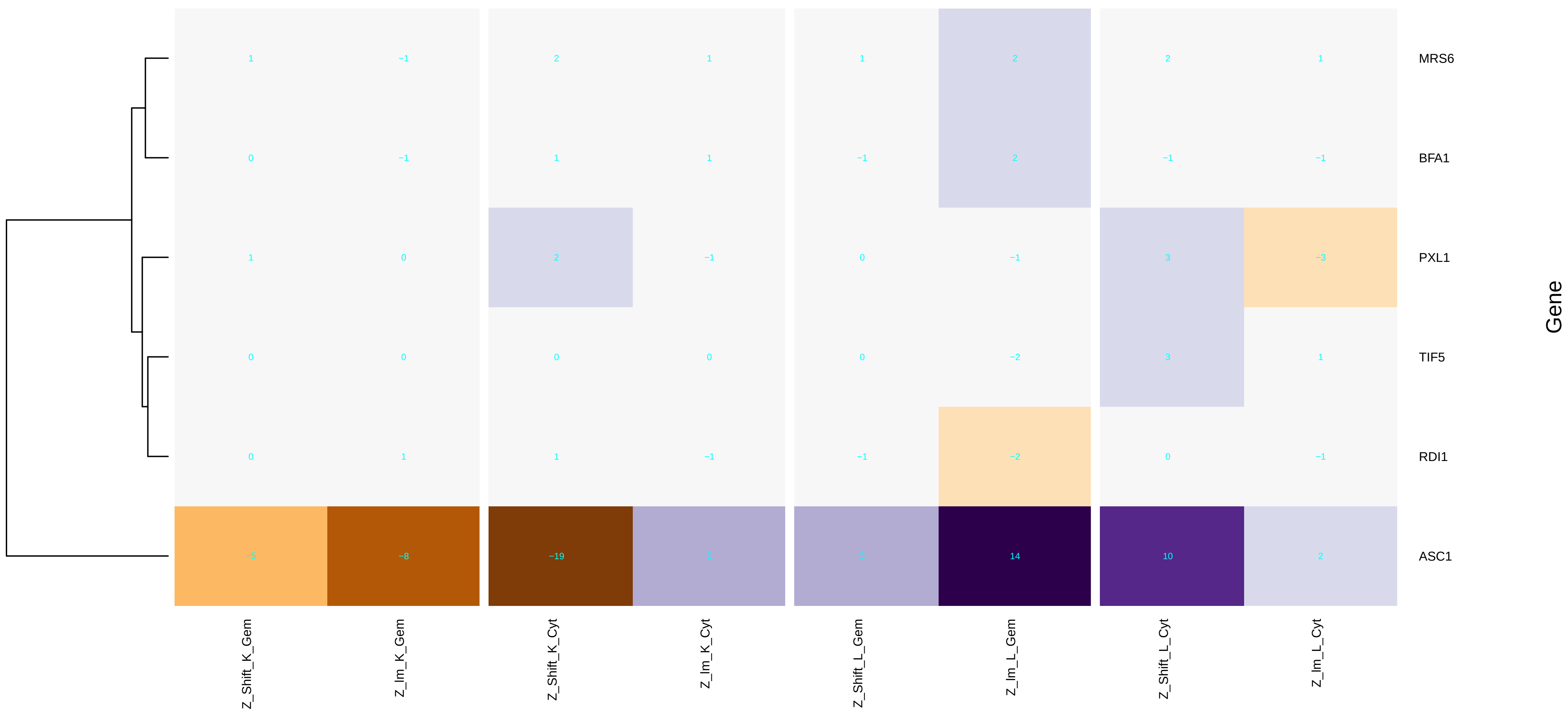

### gene_silencing_involved_in_chronological_cell_aging.pdf

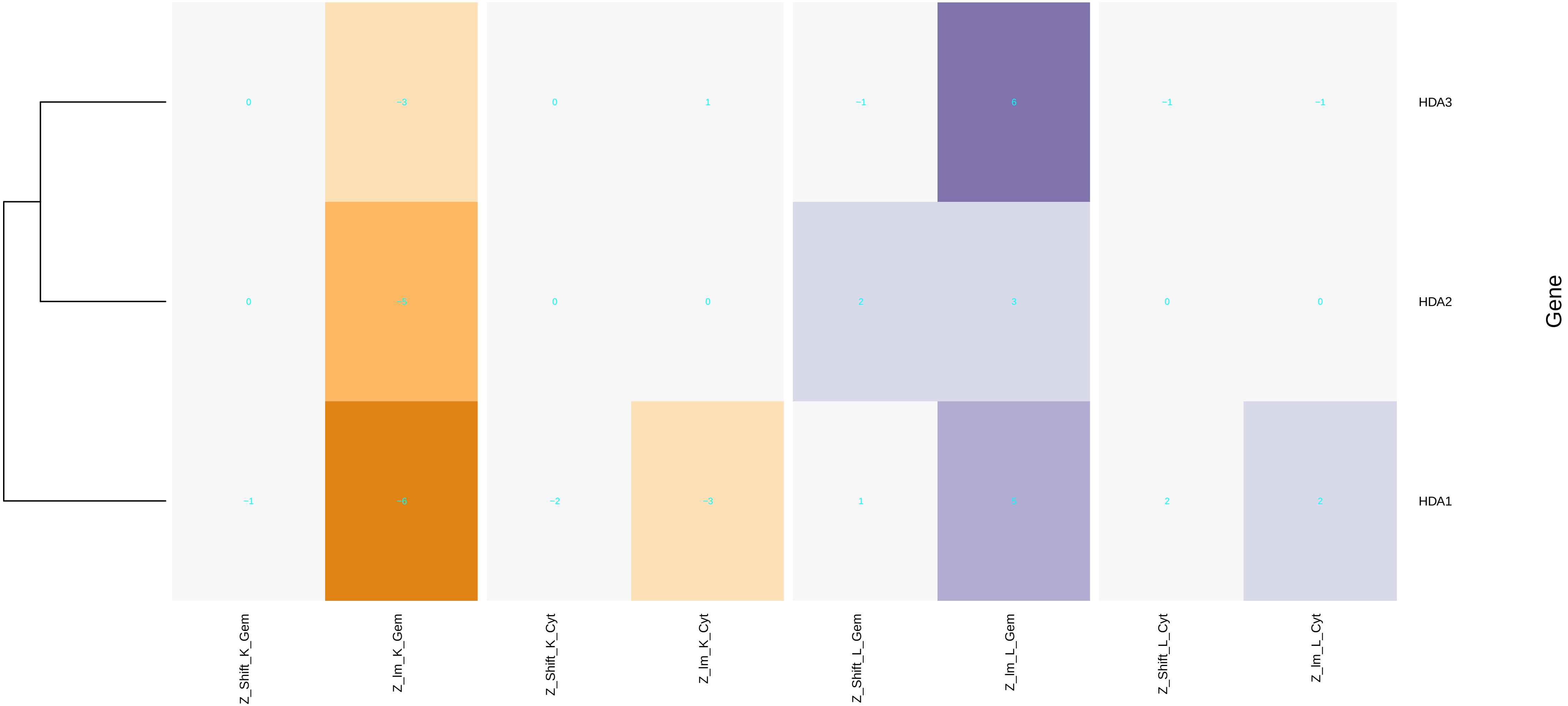

### glutaminyl-tRNA_synthase_(glutamine-hydrolyzing)_activity.pdf

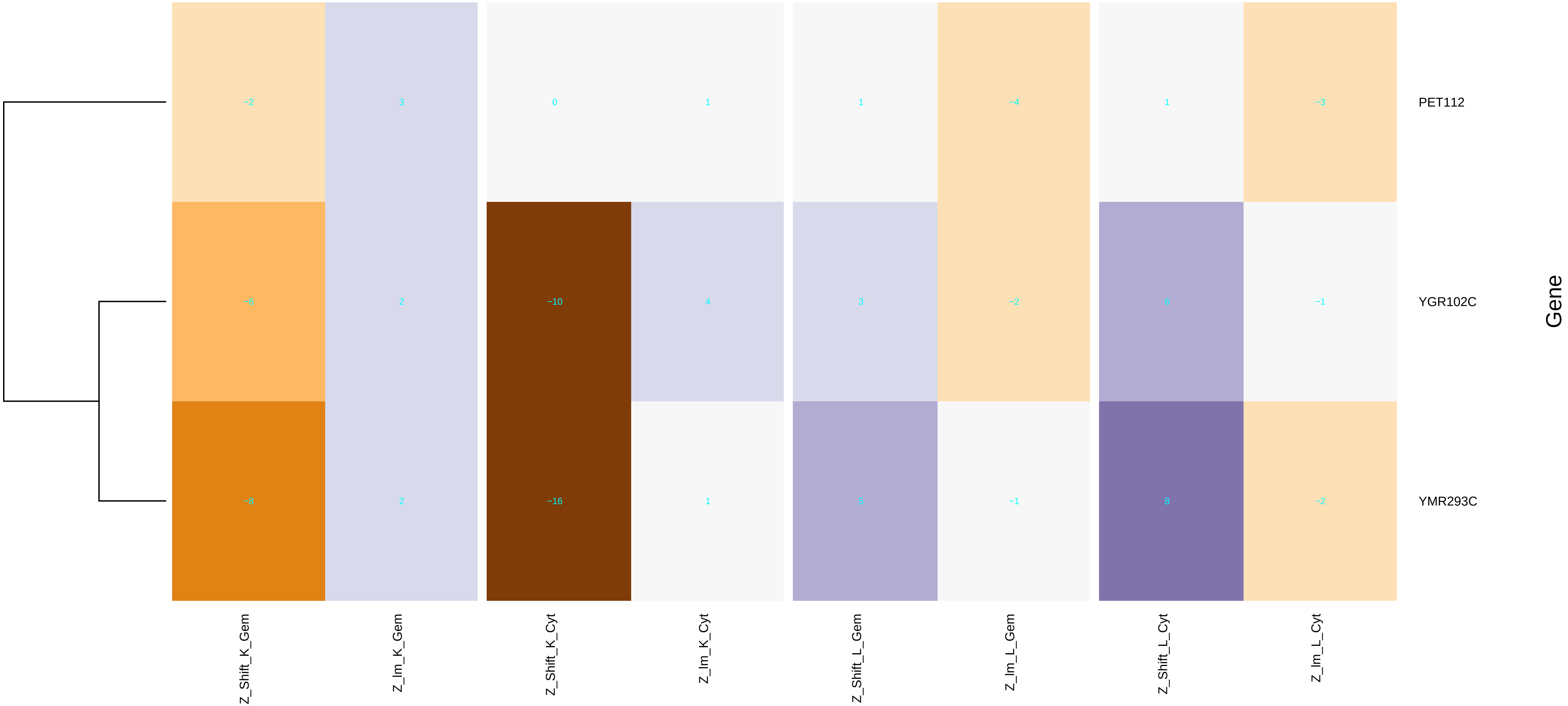

### glycine_biosynthetic_process.pdf

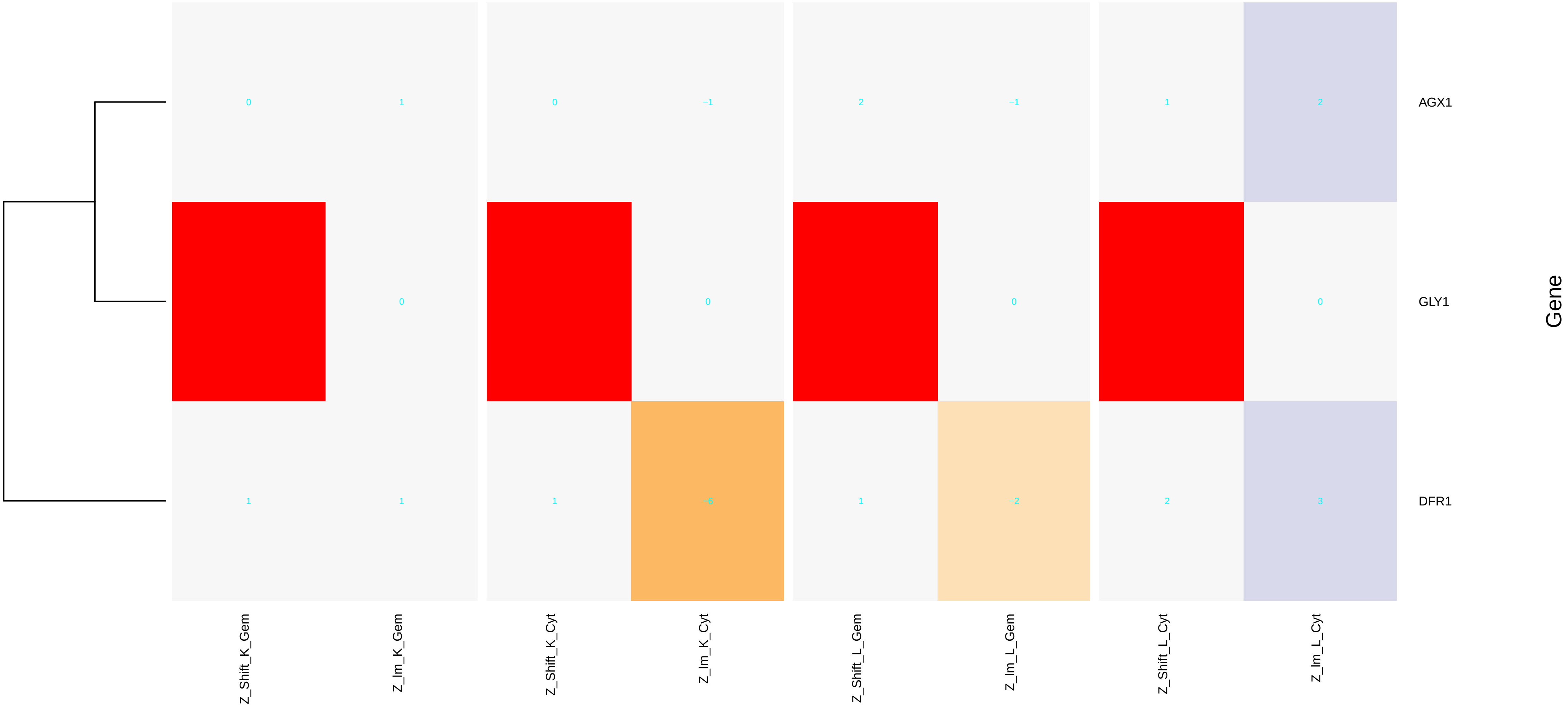

### glycosphingolipid_biosynthetic_process.pdf

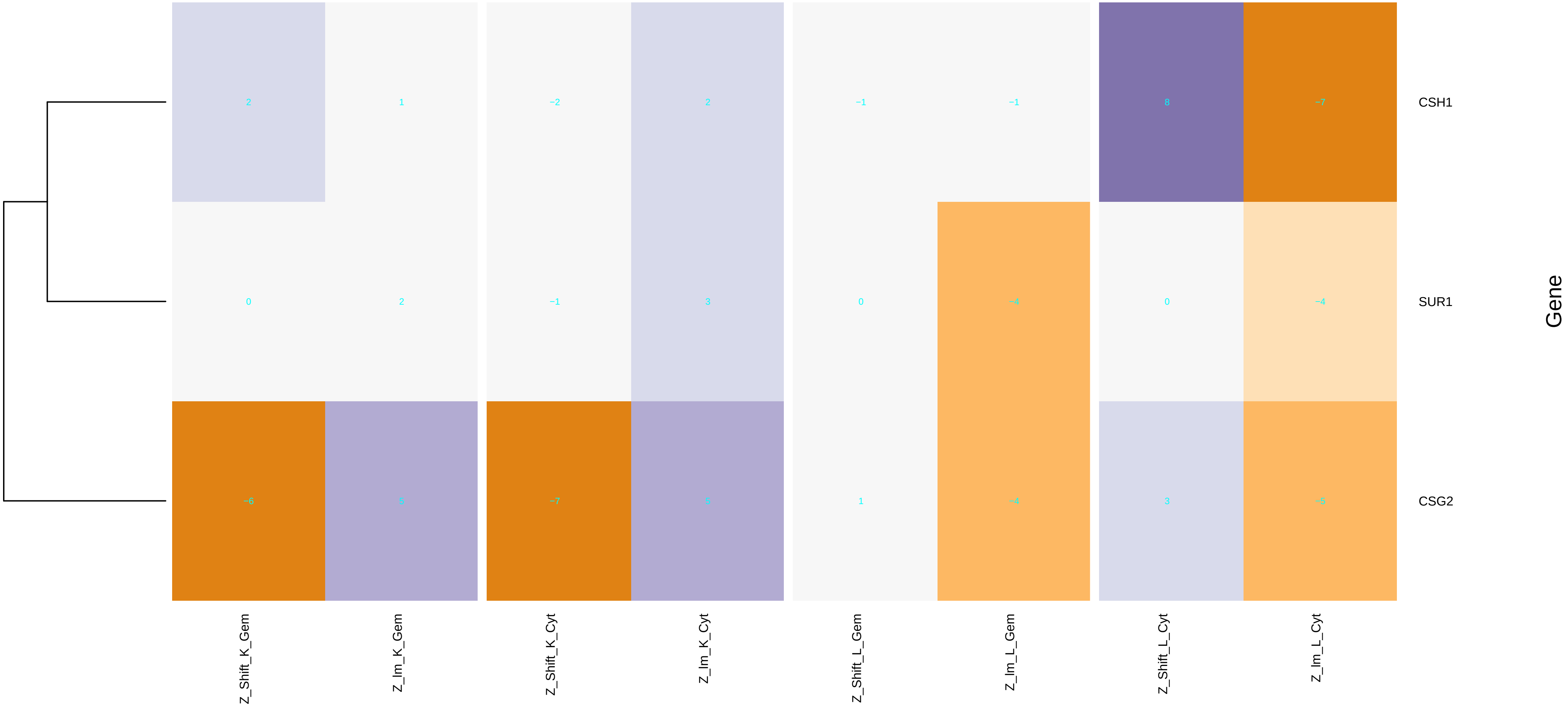

### Golgi_membrane.pdf

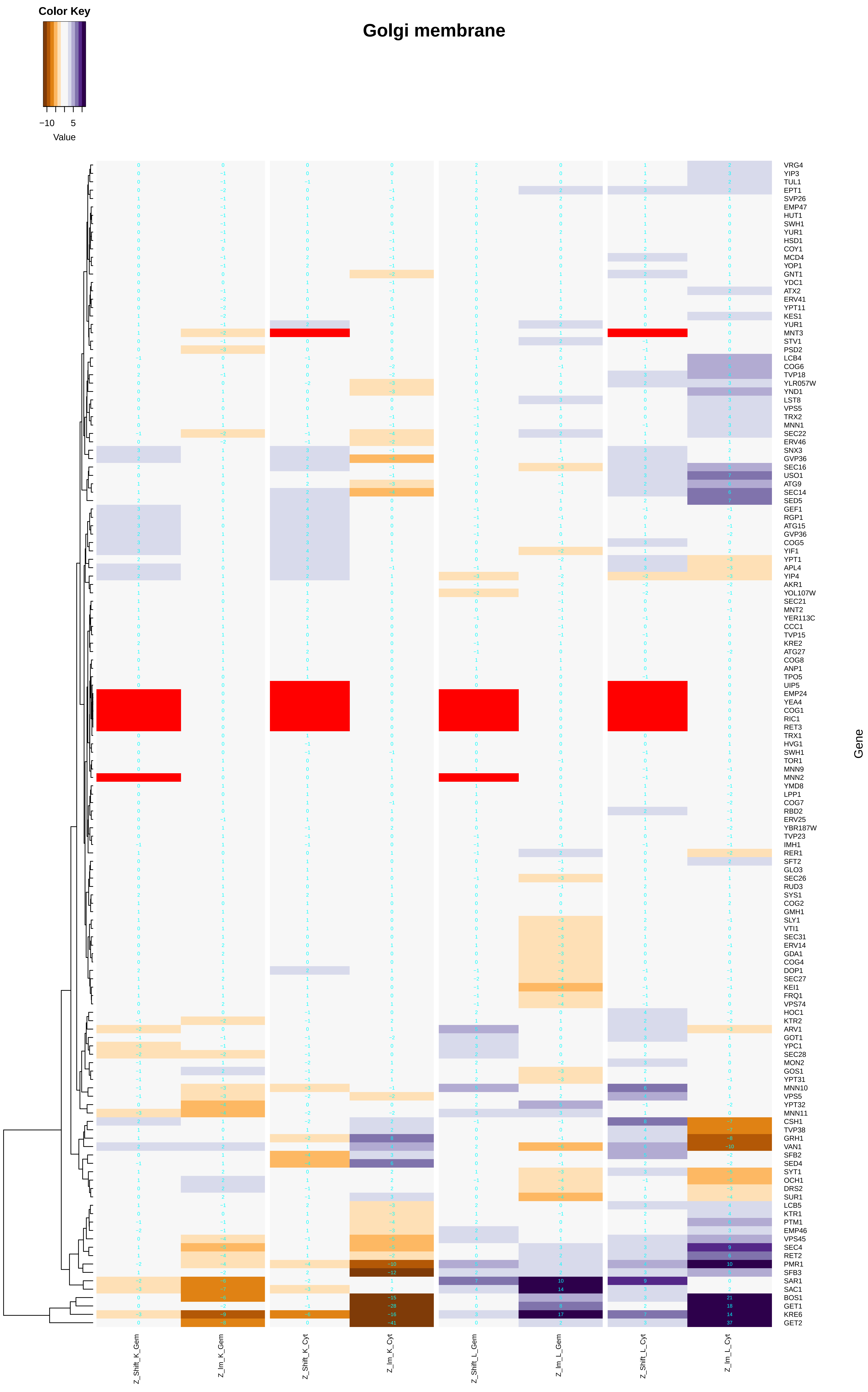

### Golgi_to_endosome_transport.pdf

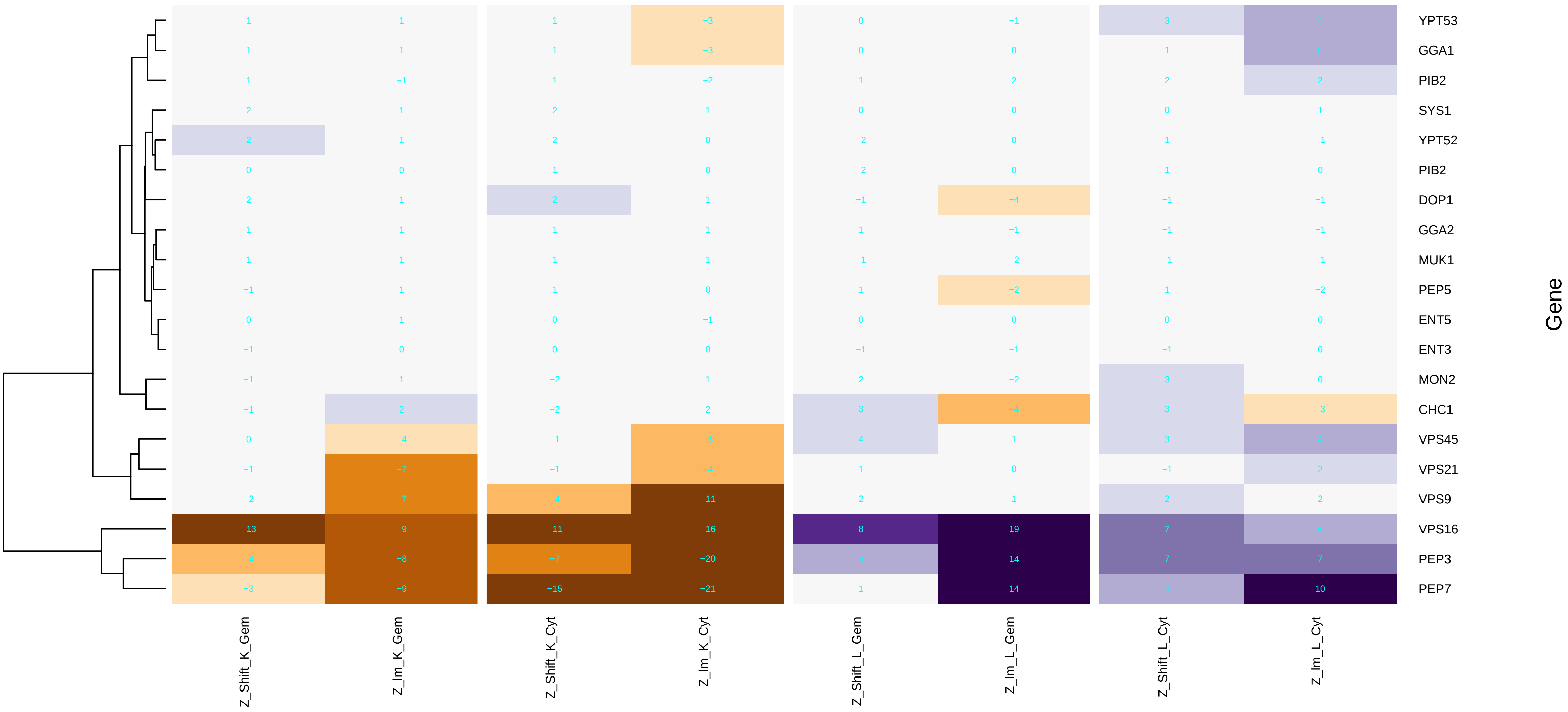

### Golgi_to_vacuole_transport.pdf

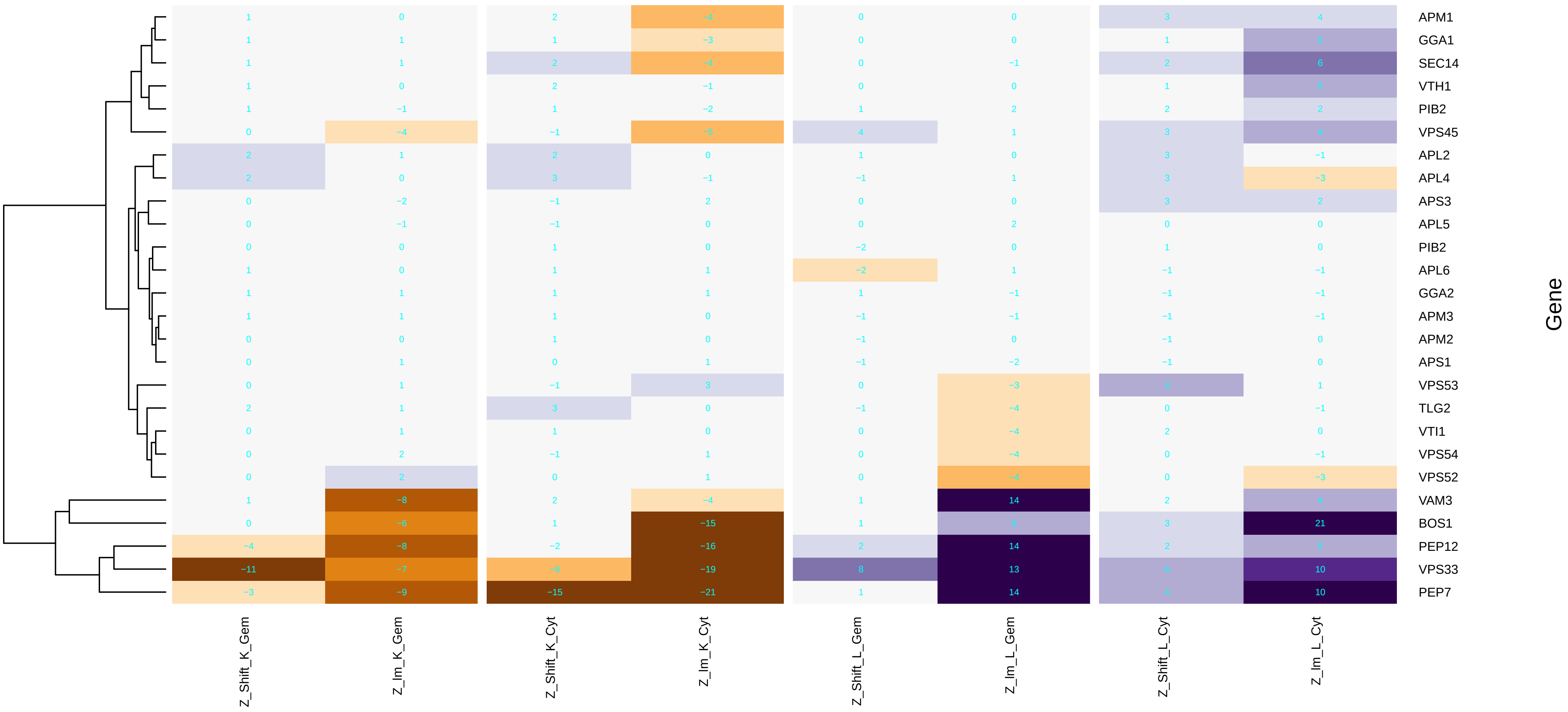

### GTP_binding.pdf

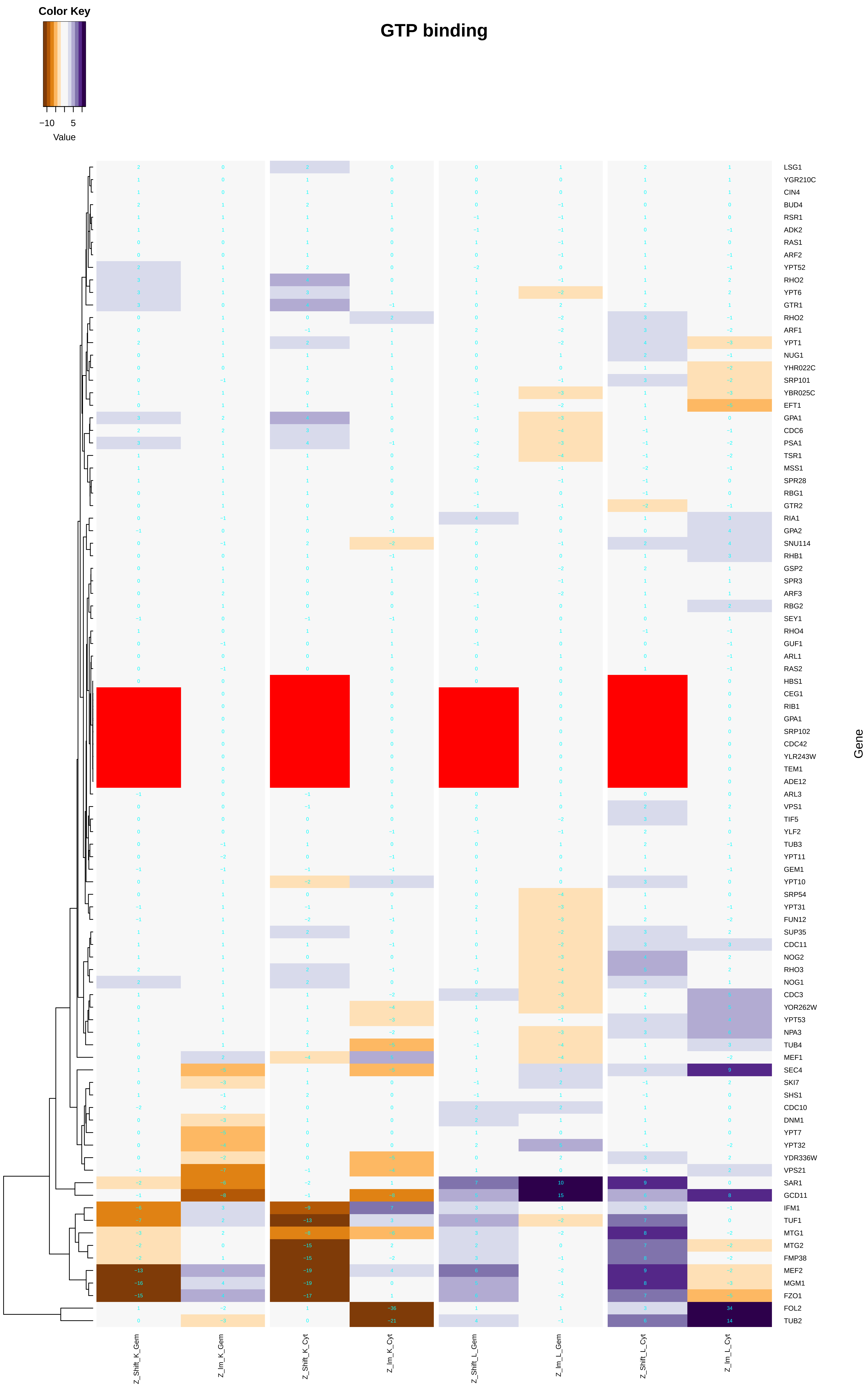

### GTPase_activity.pdf

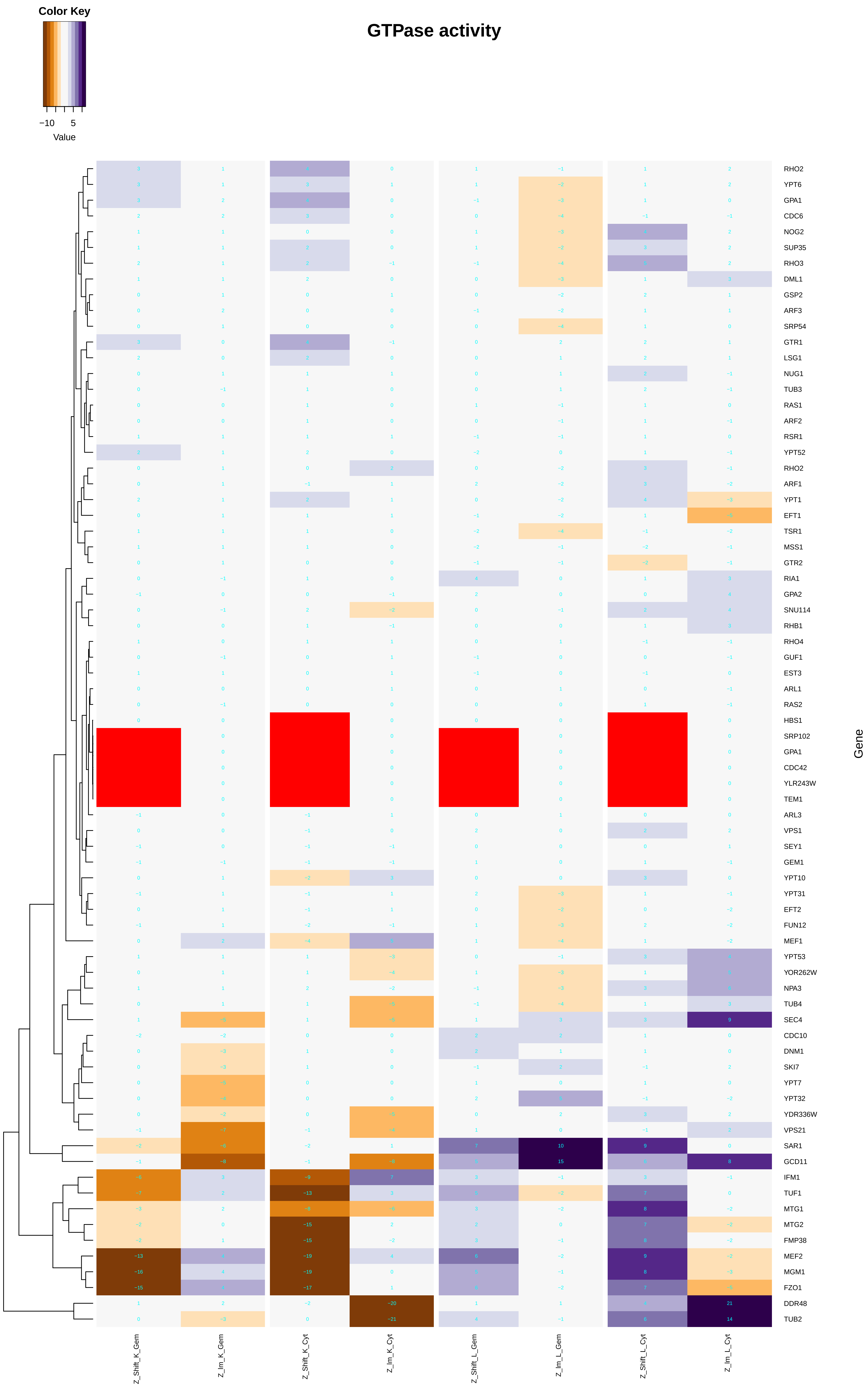

### guanyl_nucleotide_binding.pdf

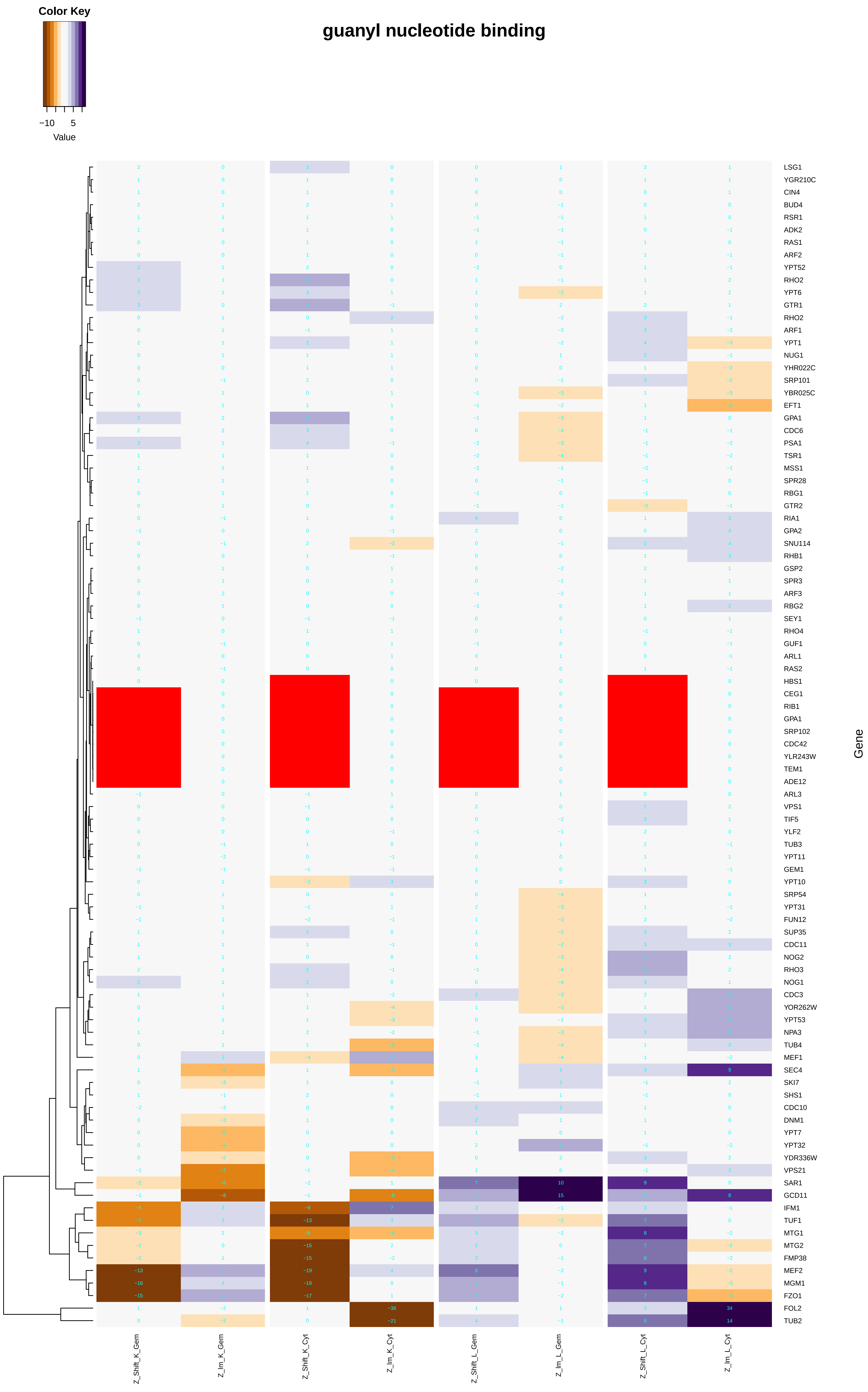

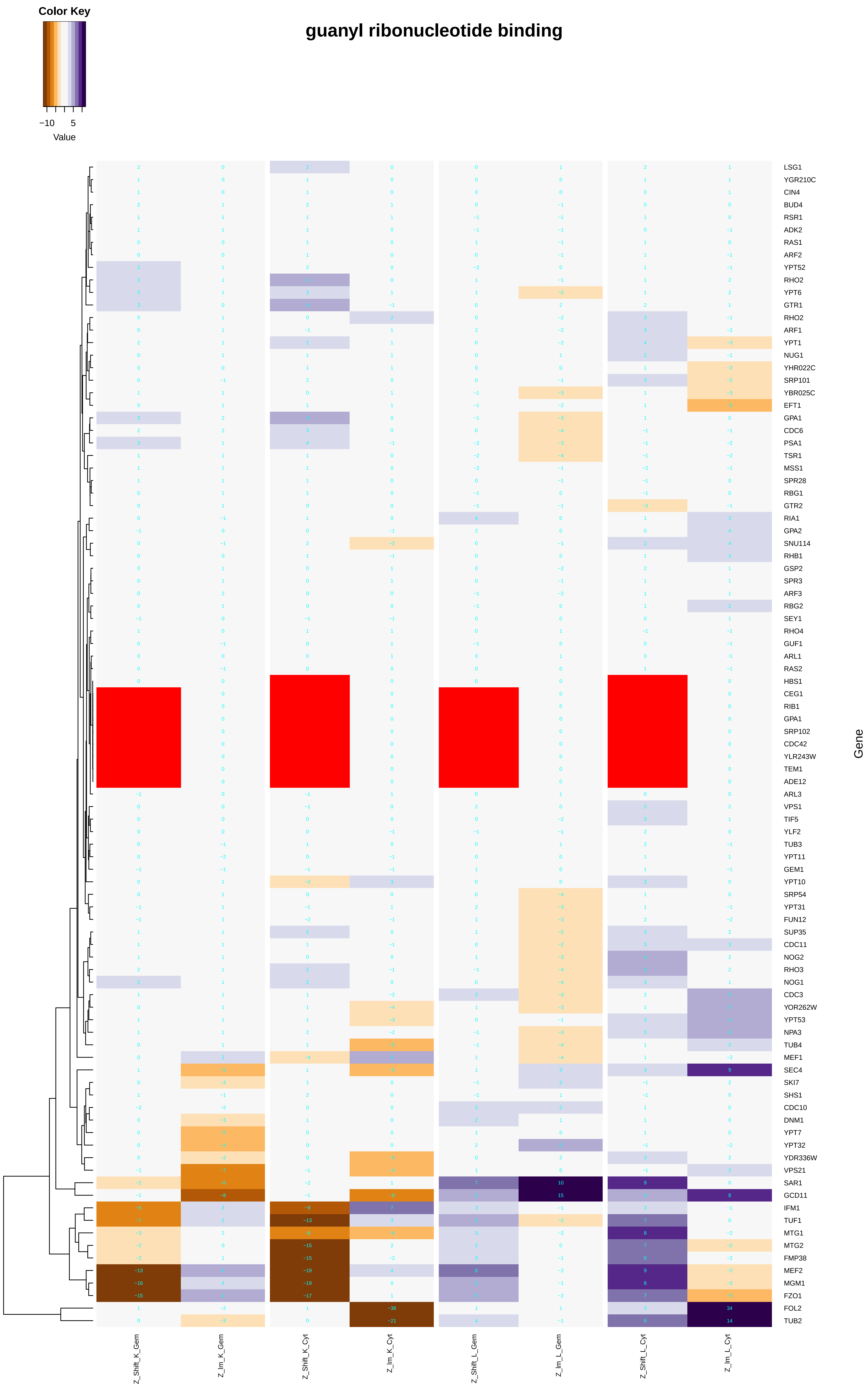

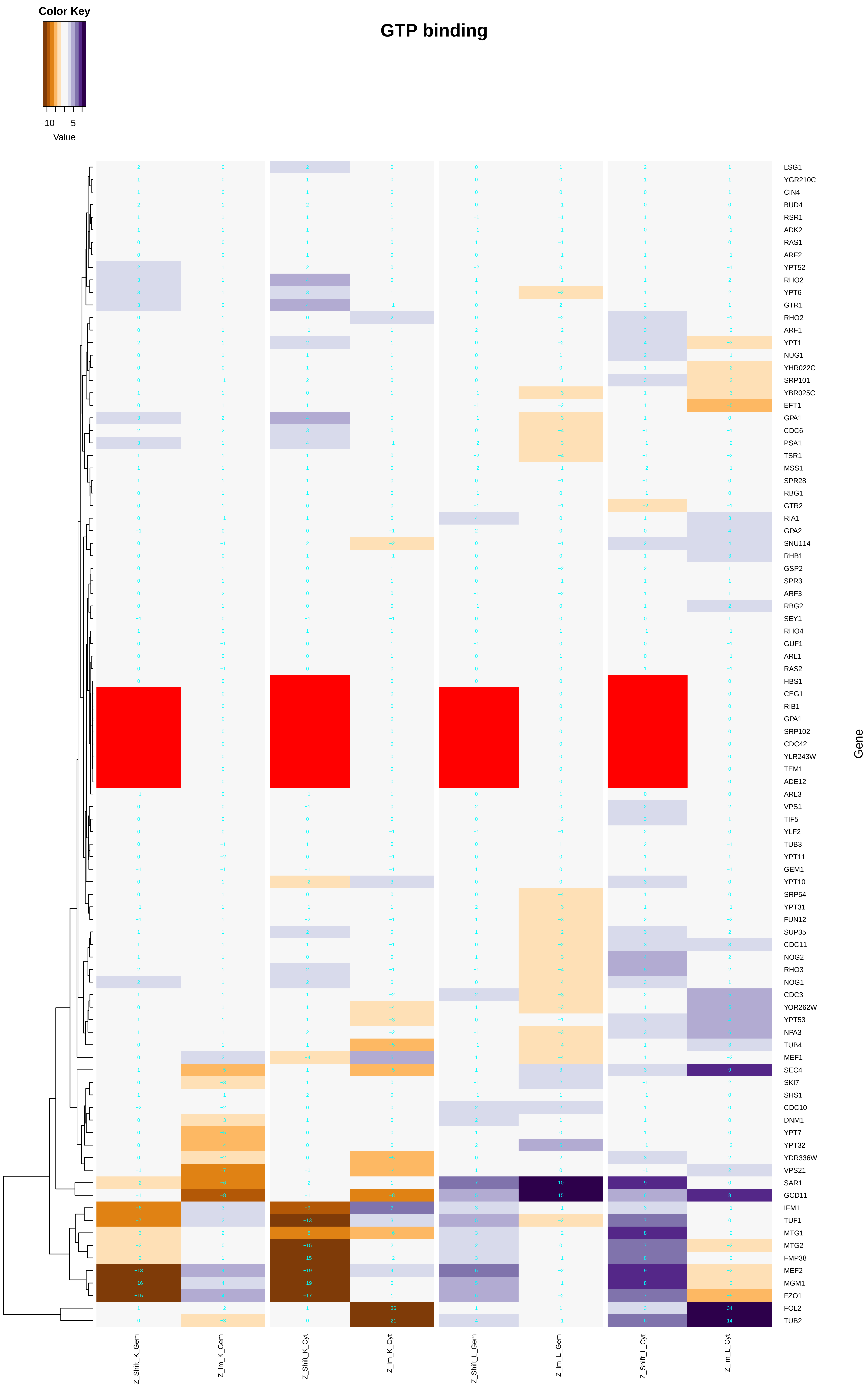

### HDA1_complex.pdf

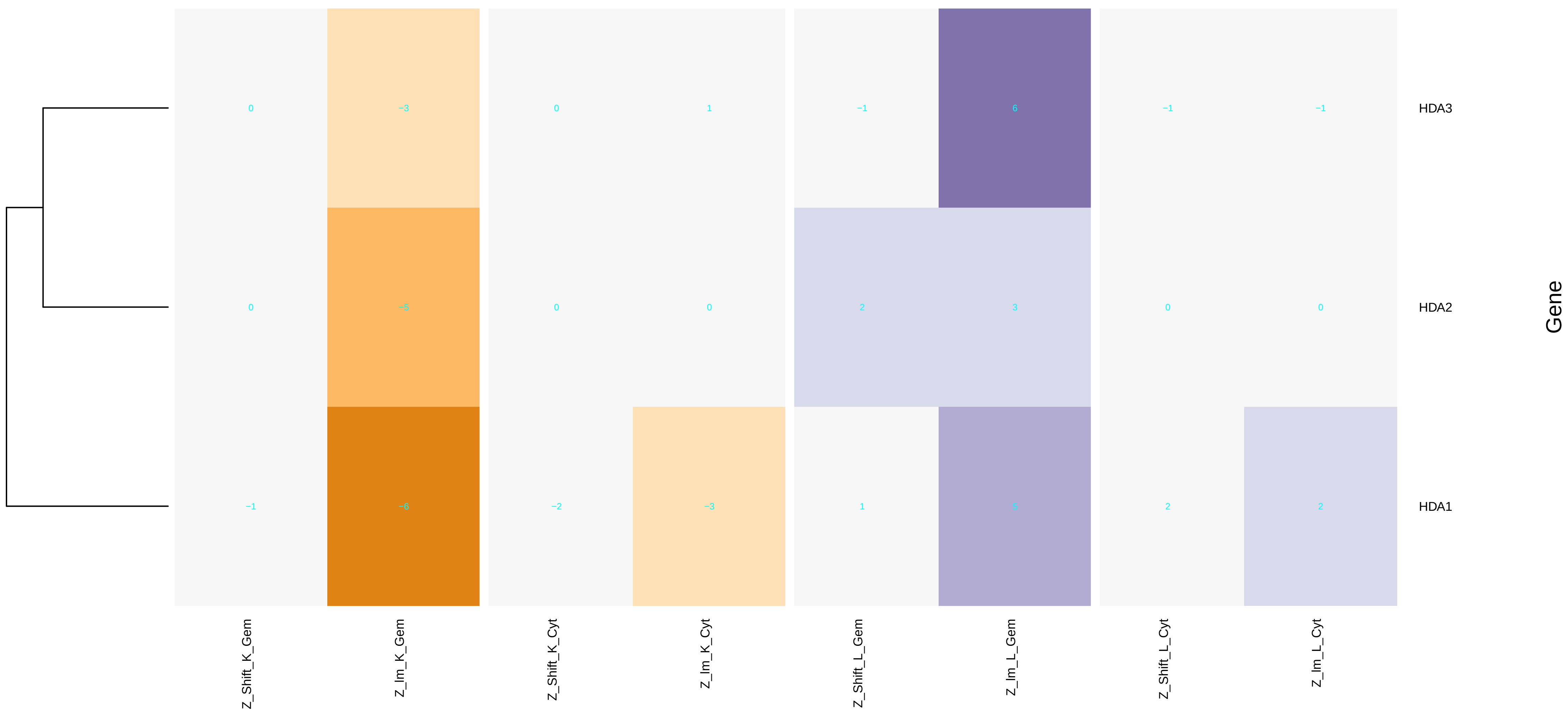

### heteroduplex_formation.pdf

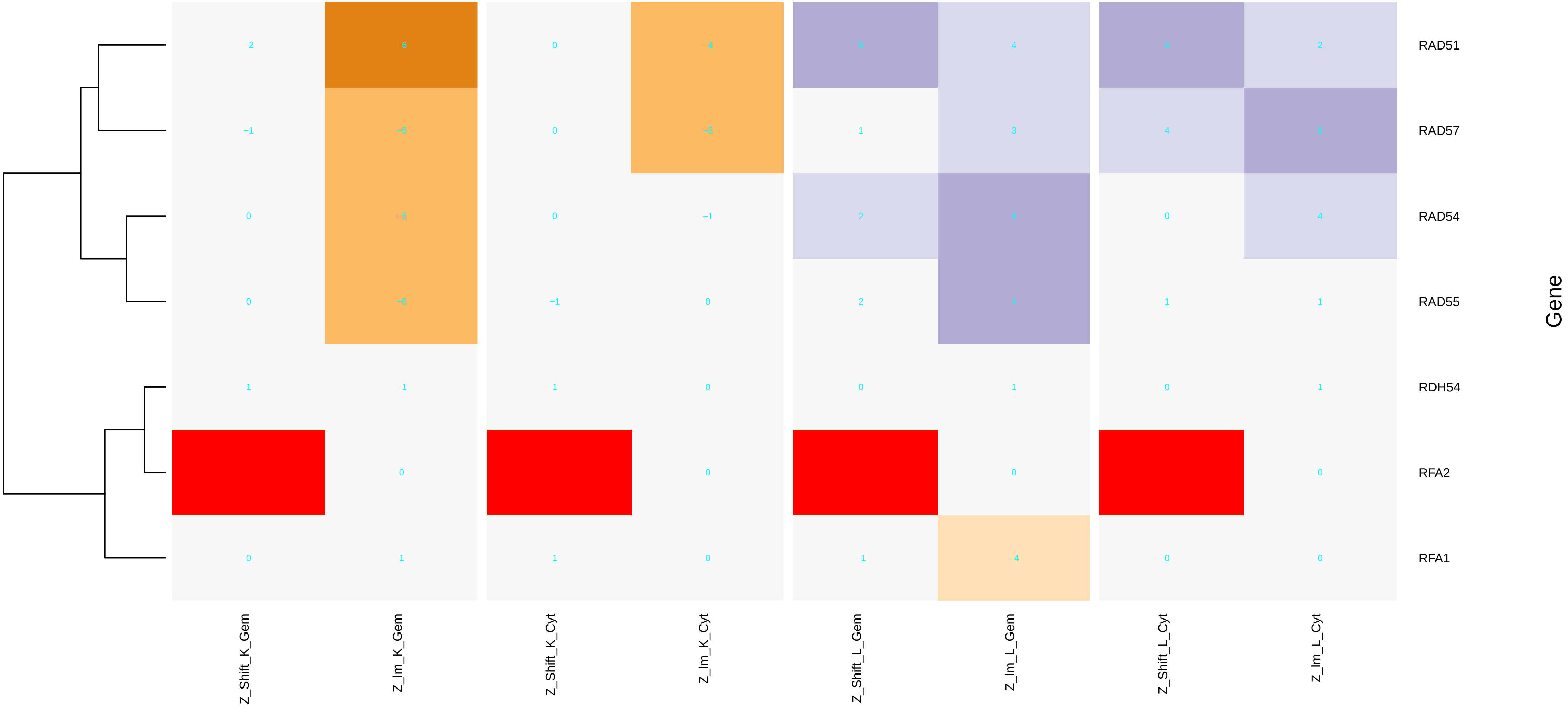

### HIR_complex.pdf

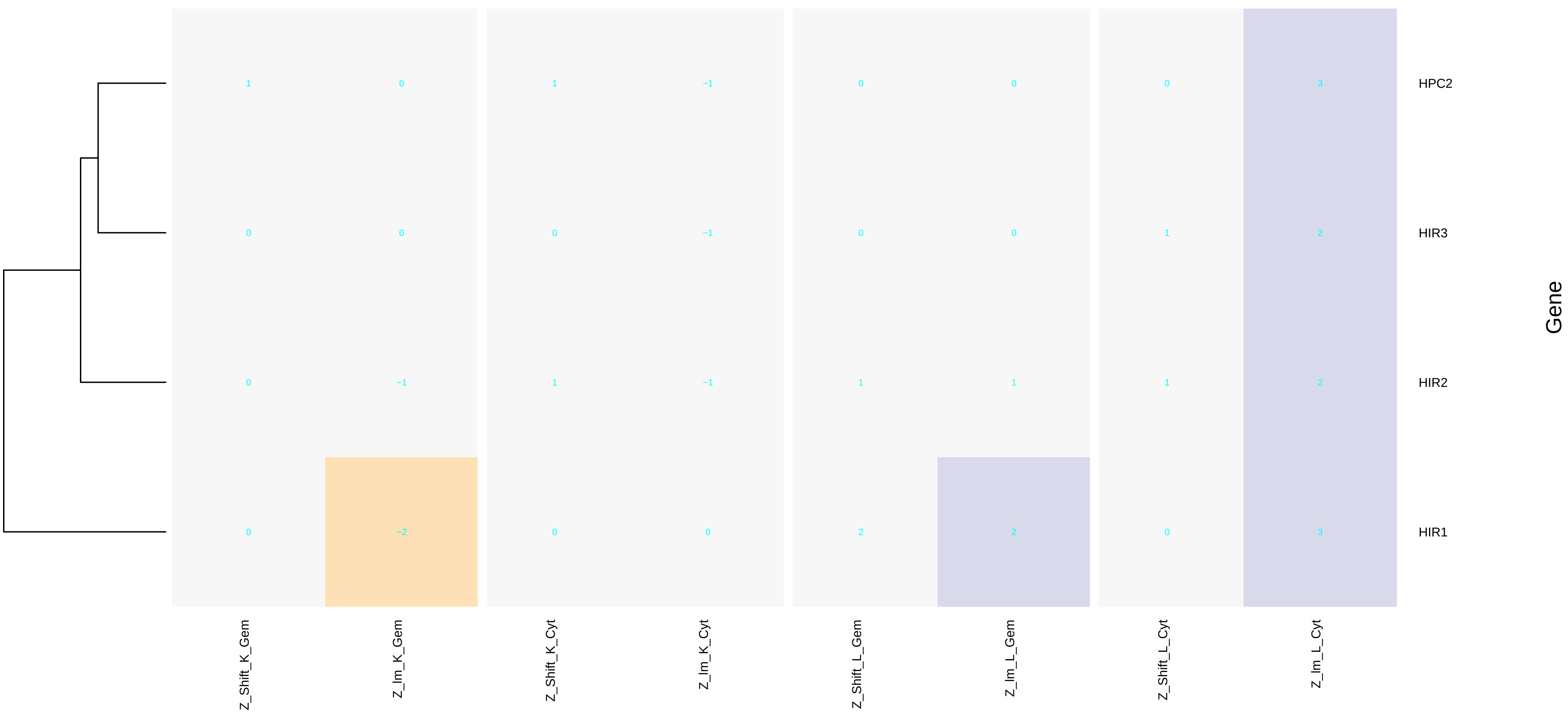

### histone_deacetylase_complex.pdf

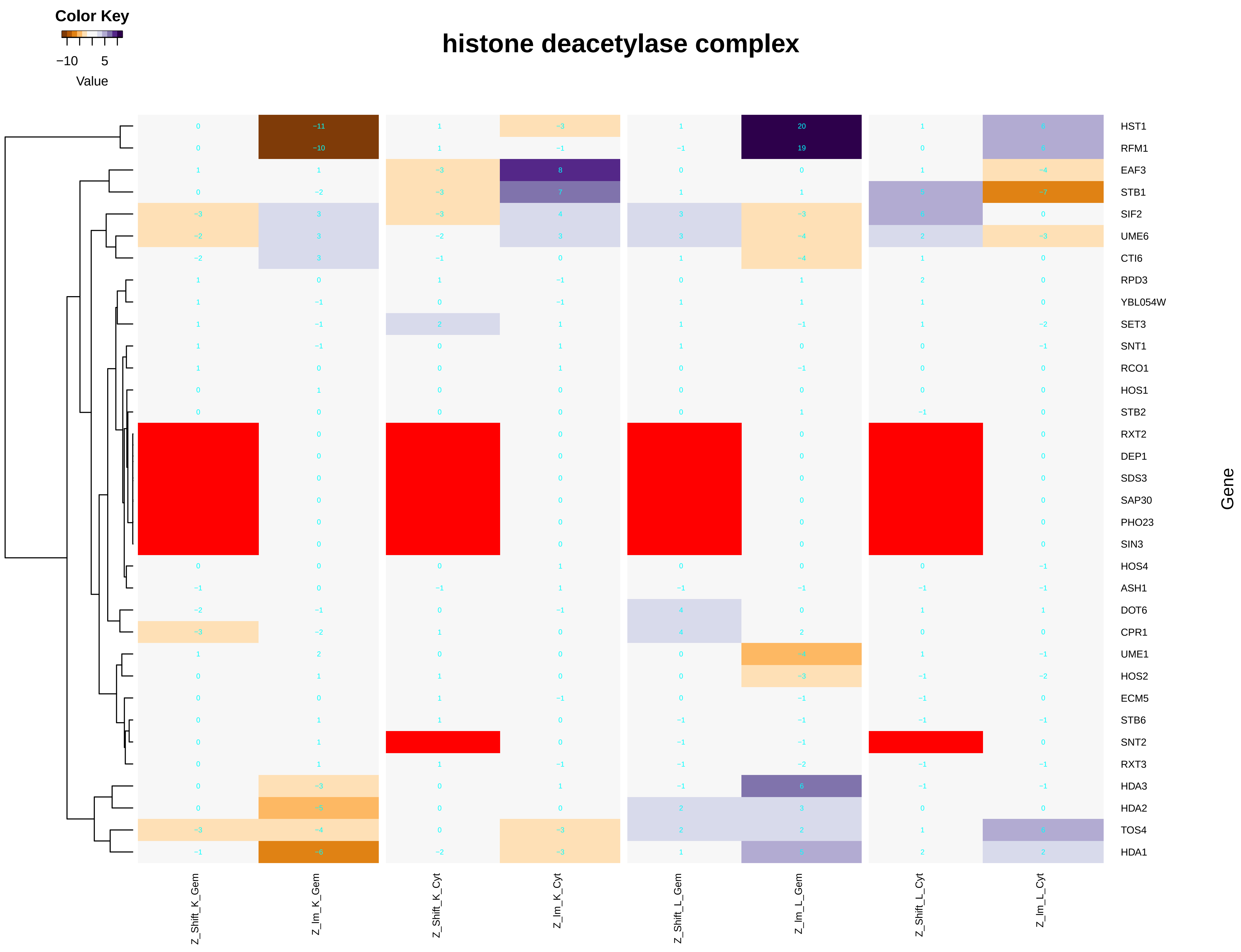

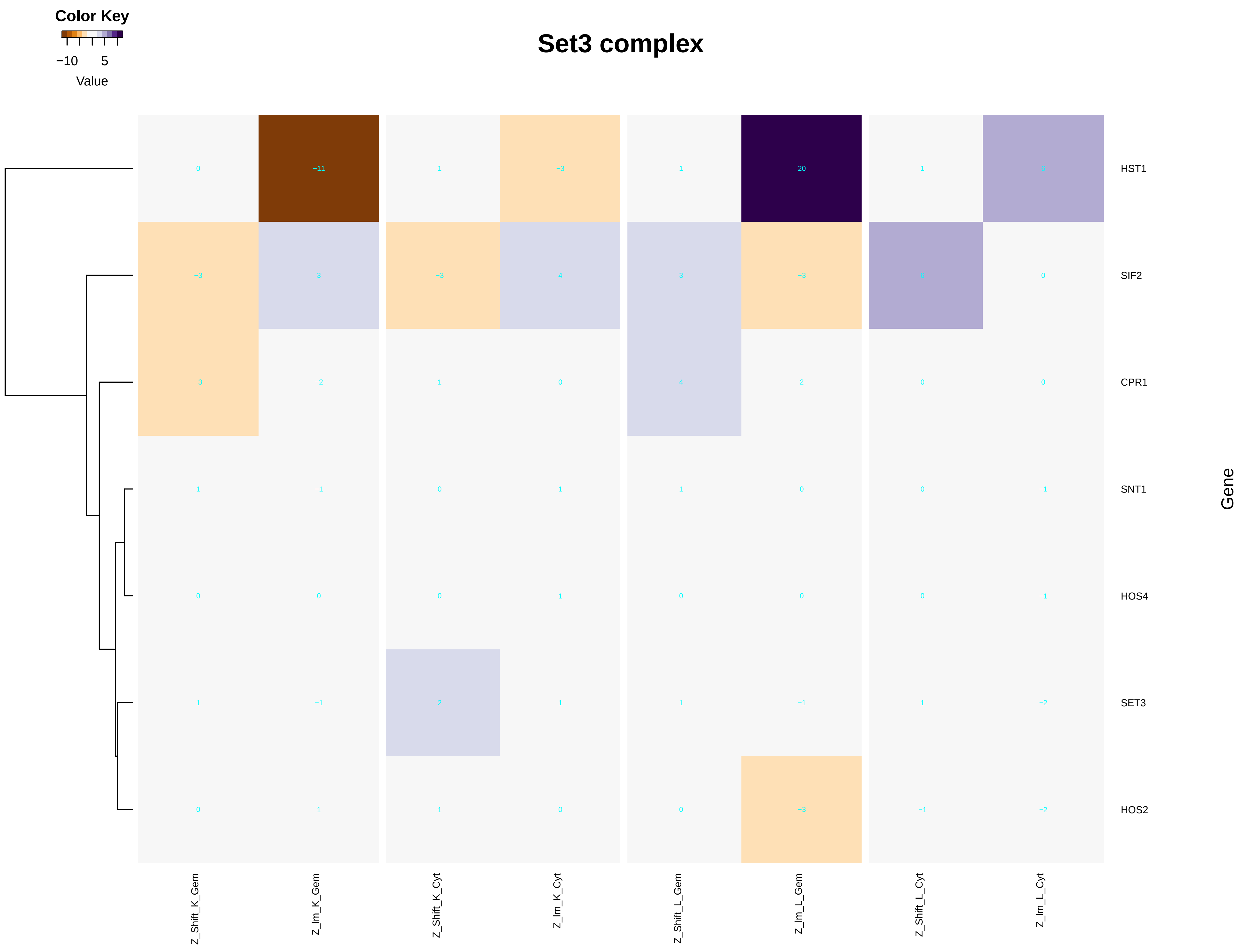

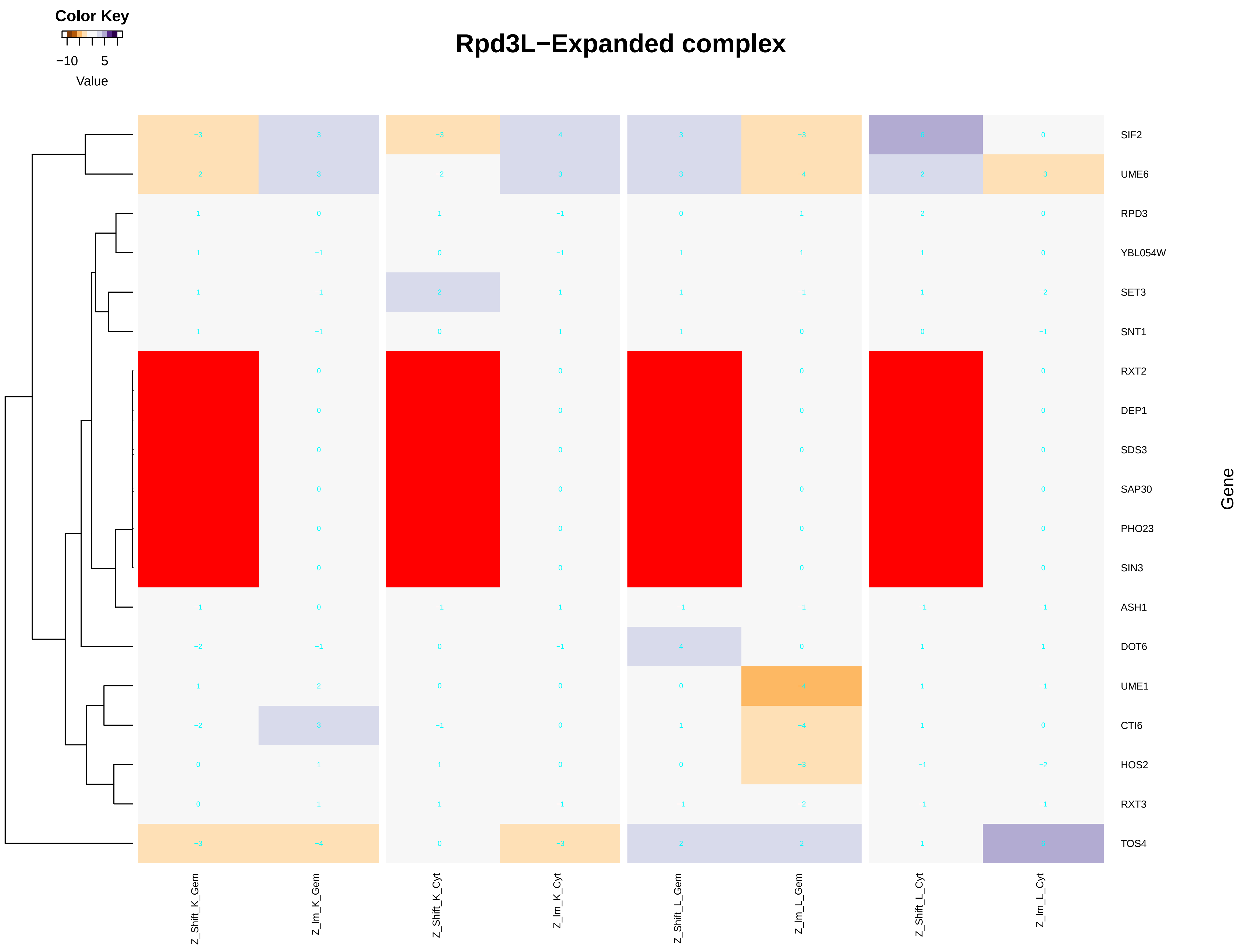

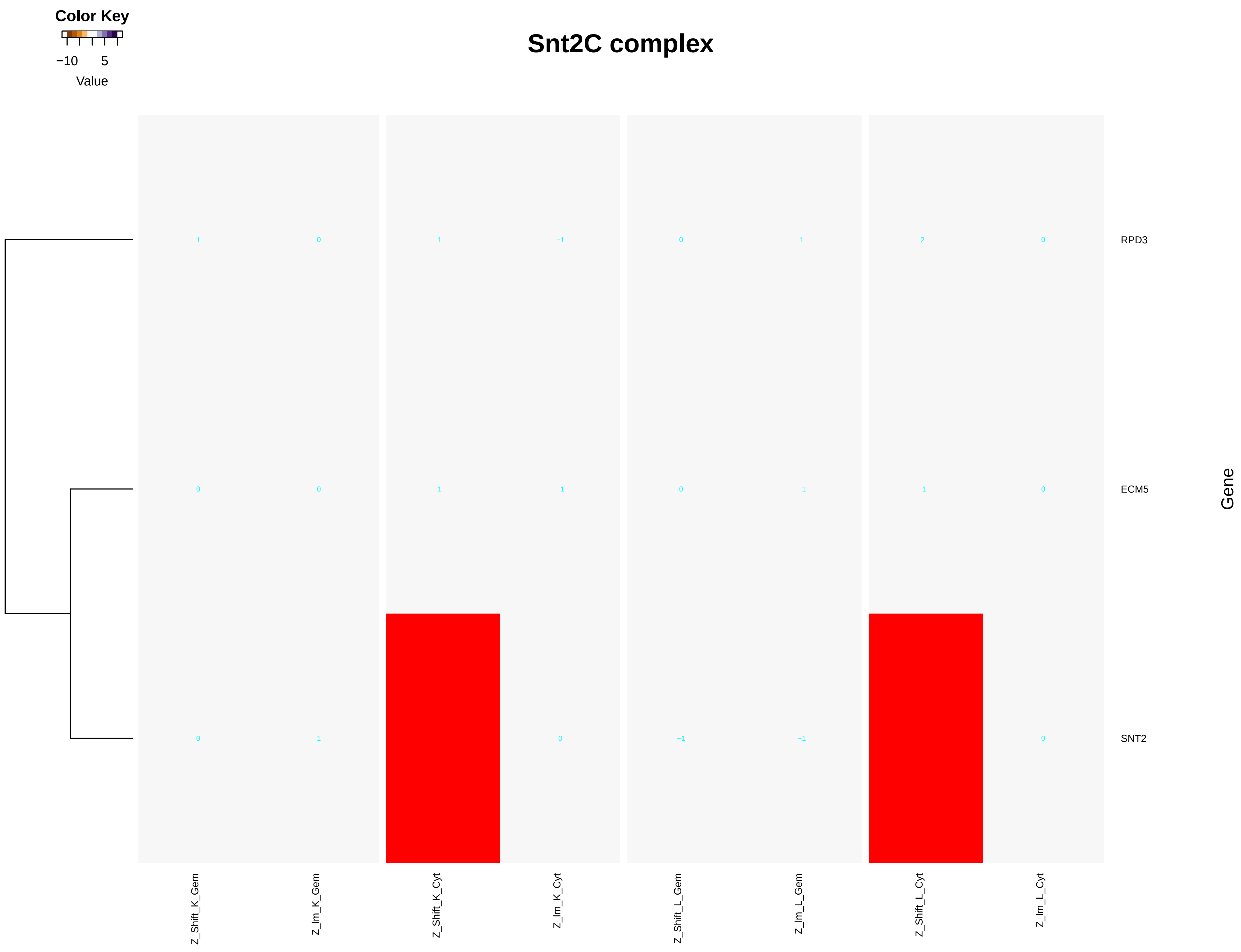

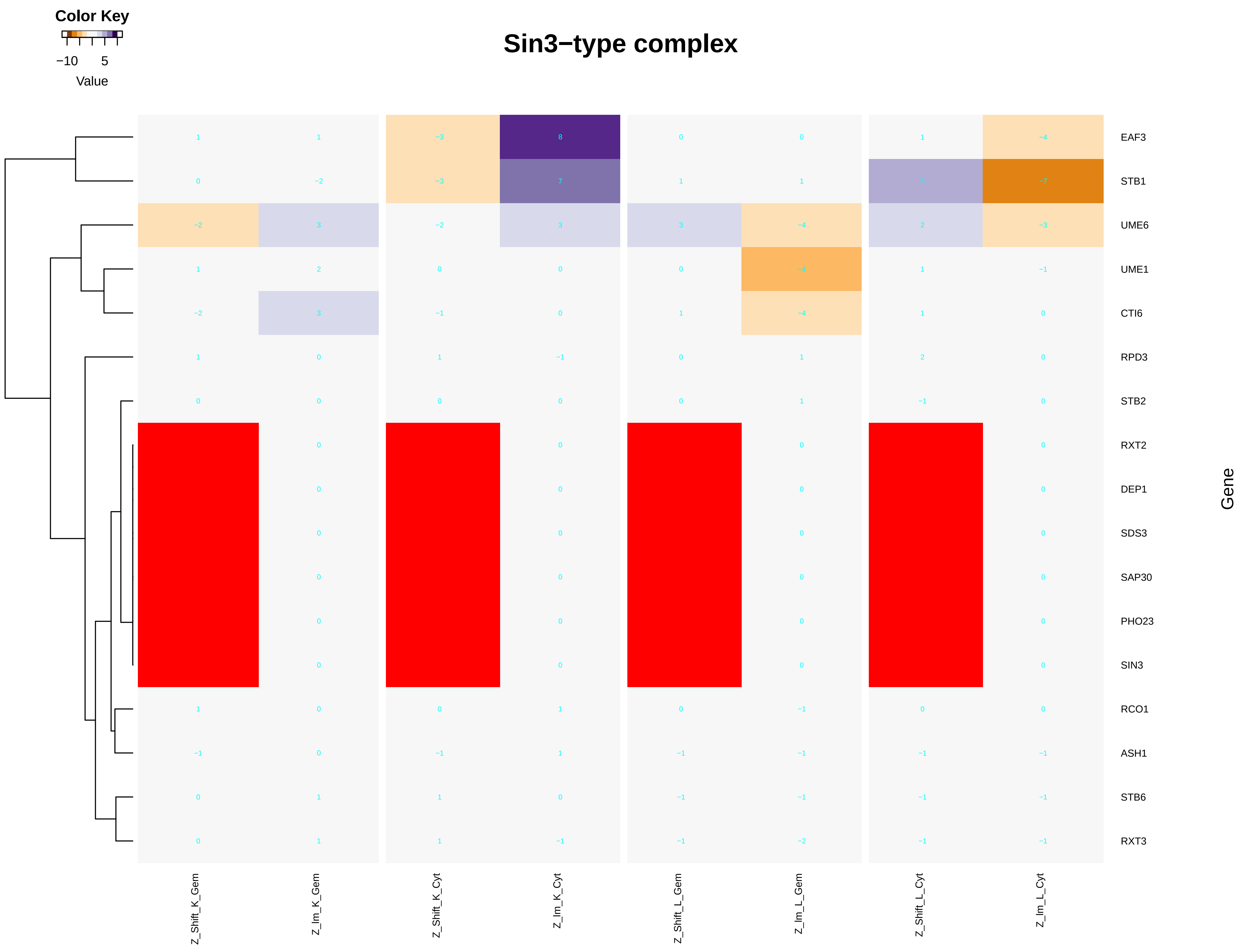

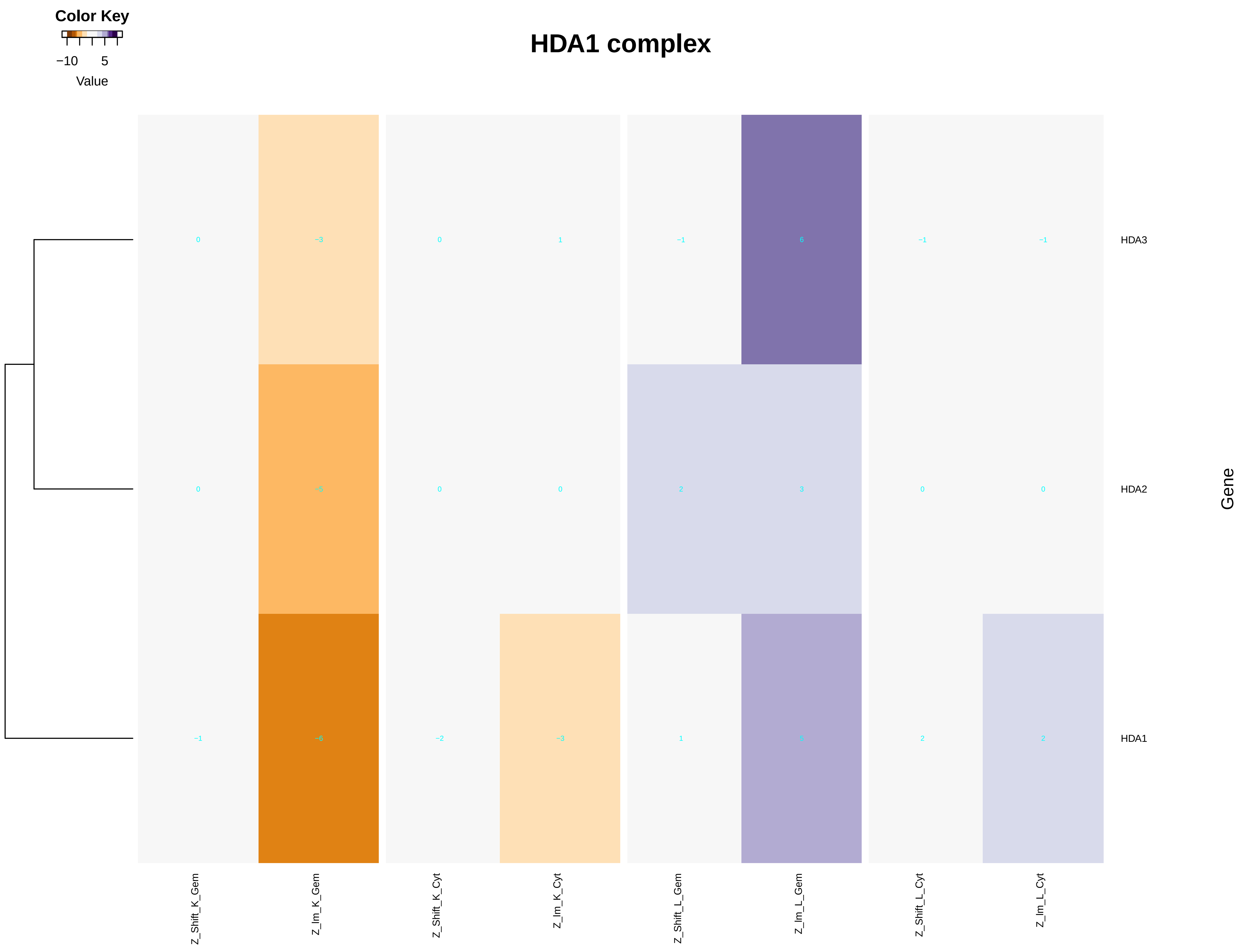

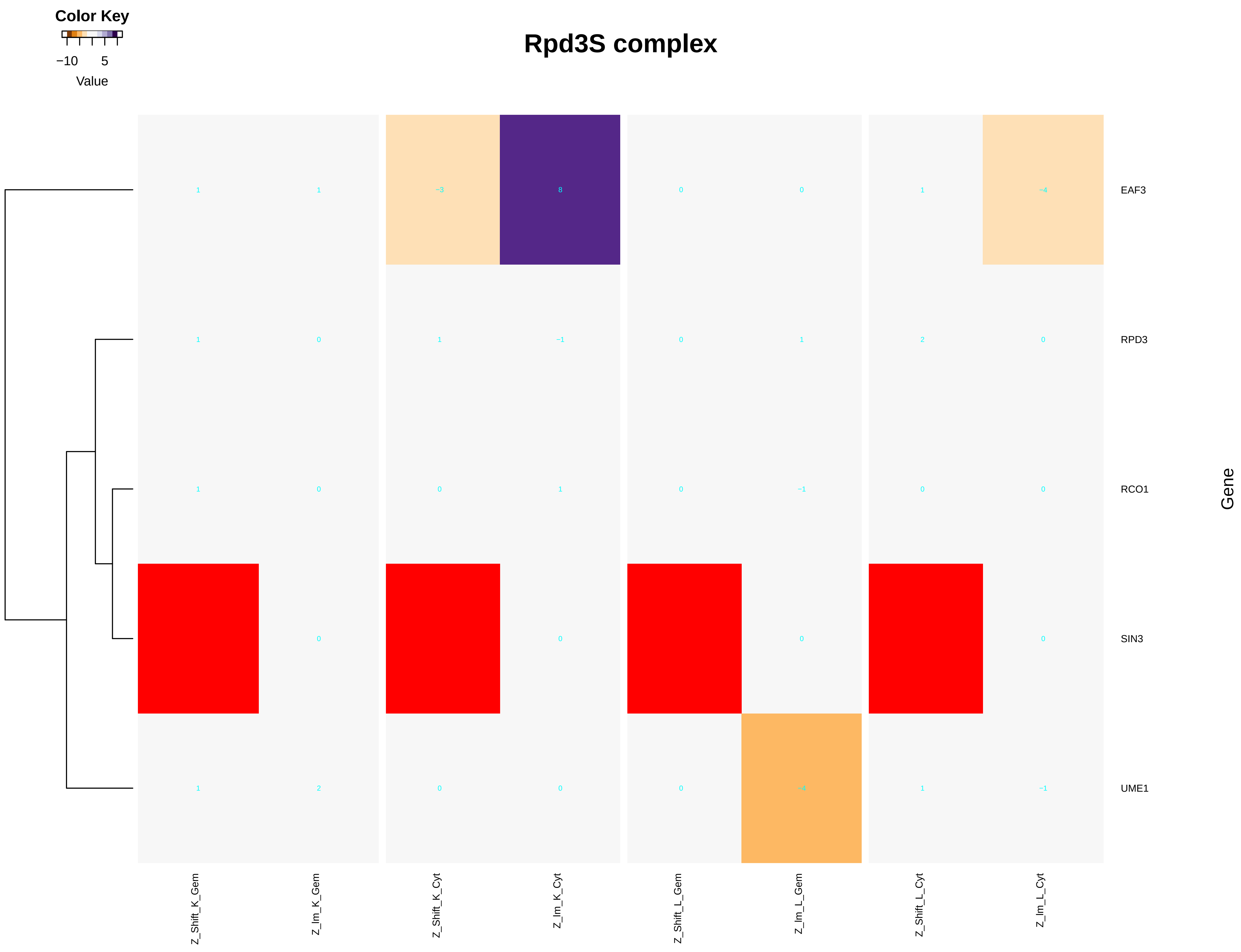

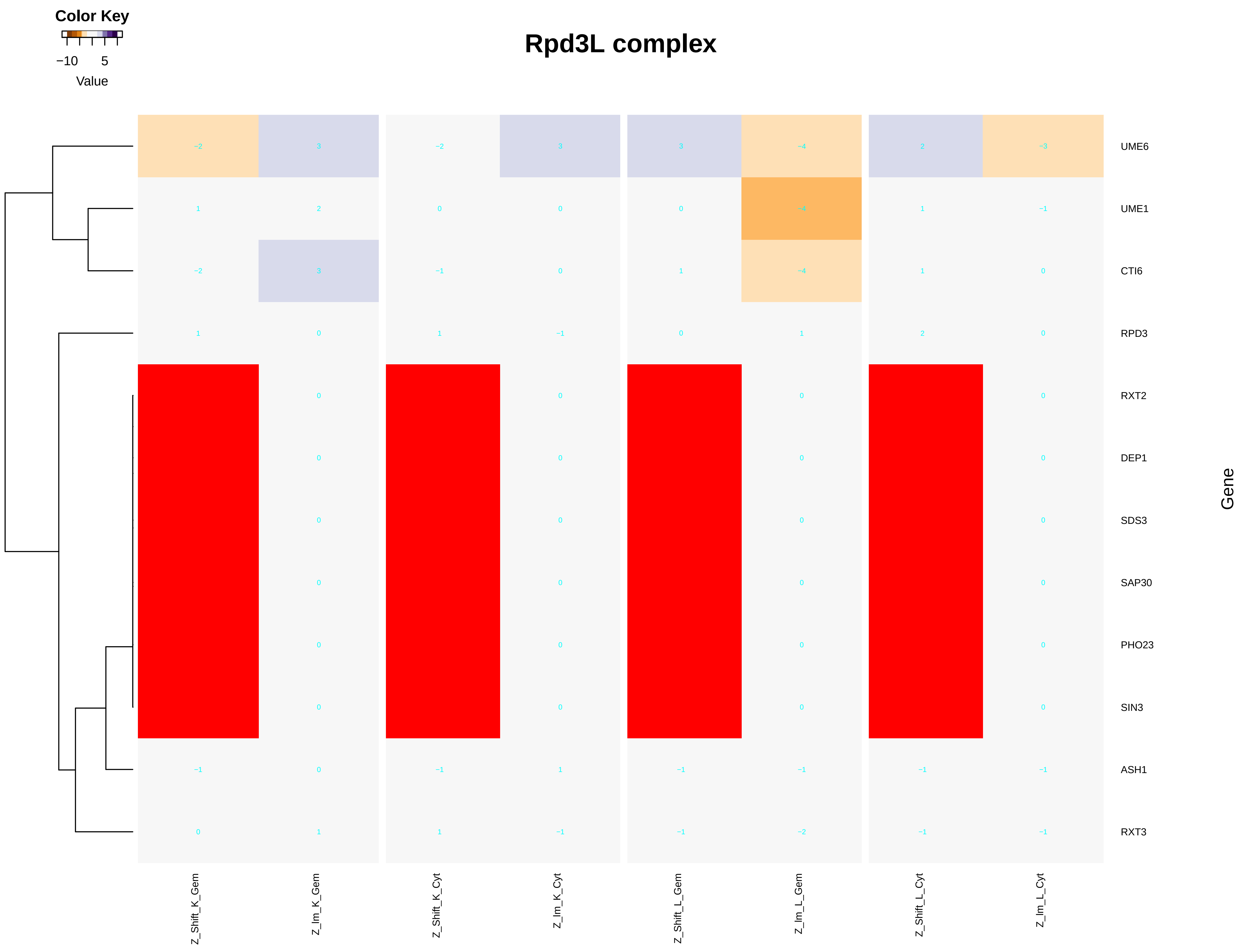

### histone_exchange.pdf

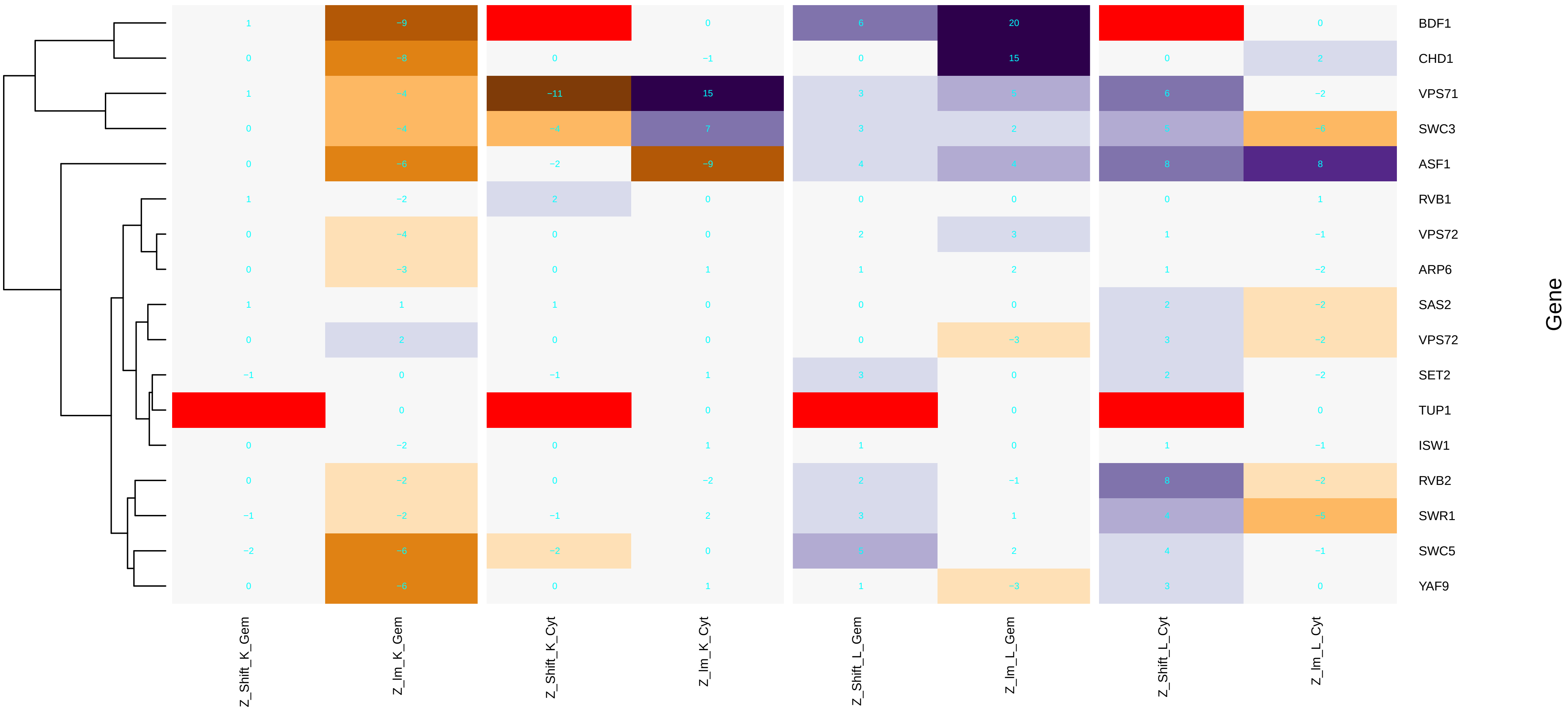

### histone_H3-K4_methylation.pdf

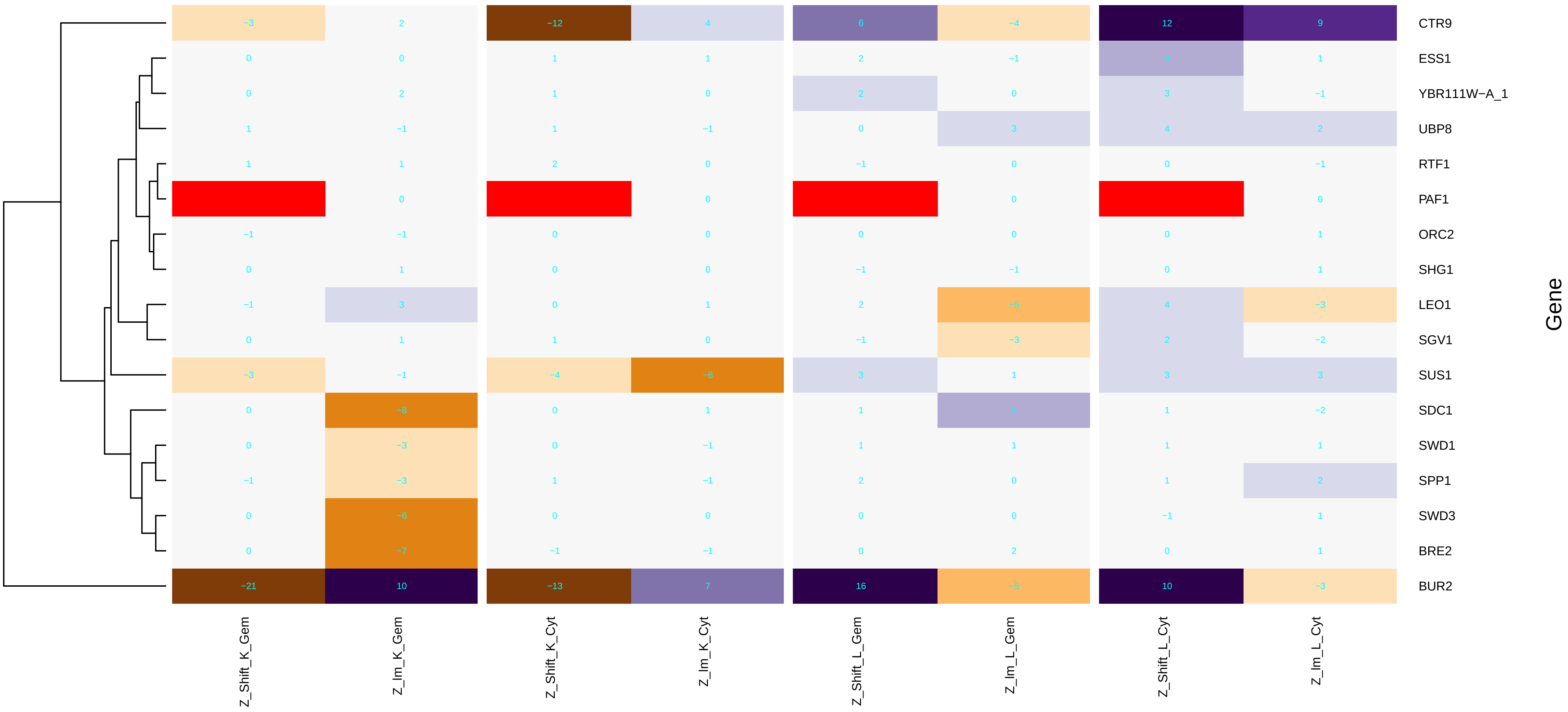

### histone_methyltransferase_activity.pdf

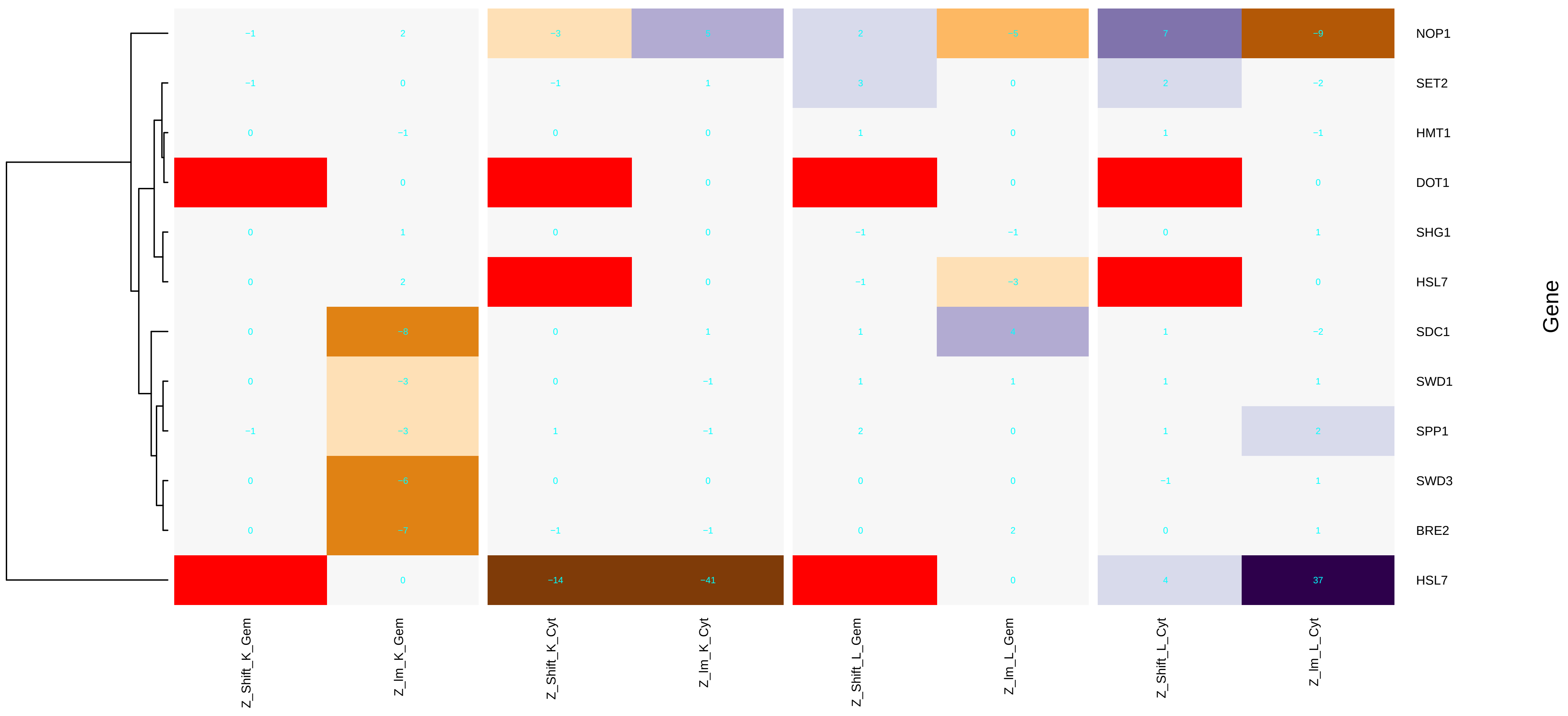

### histone_methyltransferase_activity_(H3-K4_specific).pdf

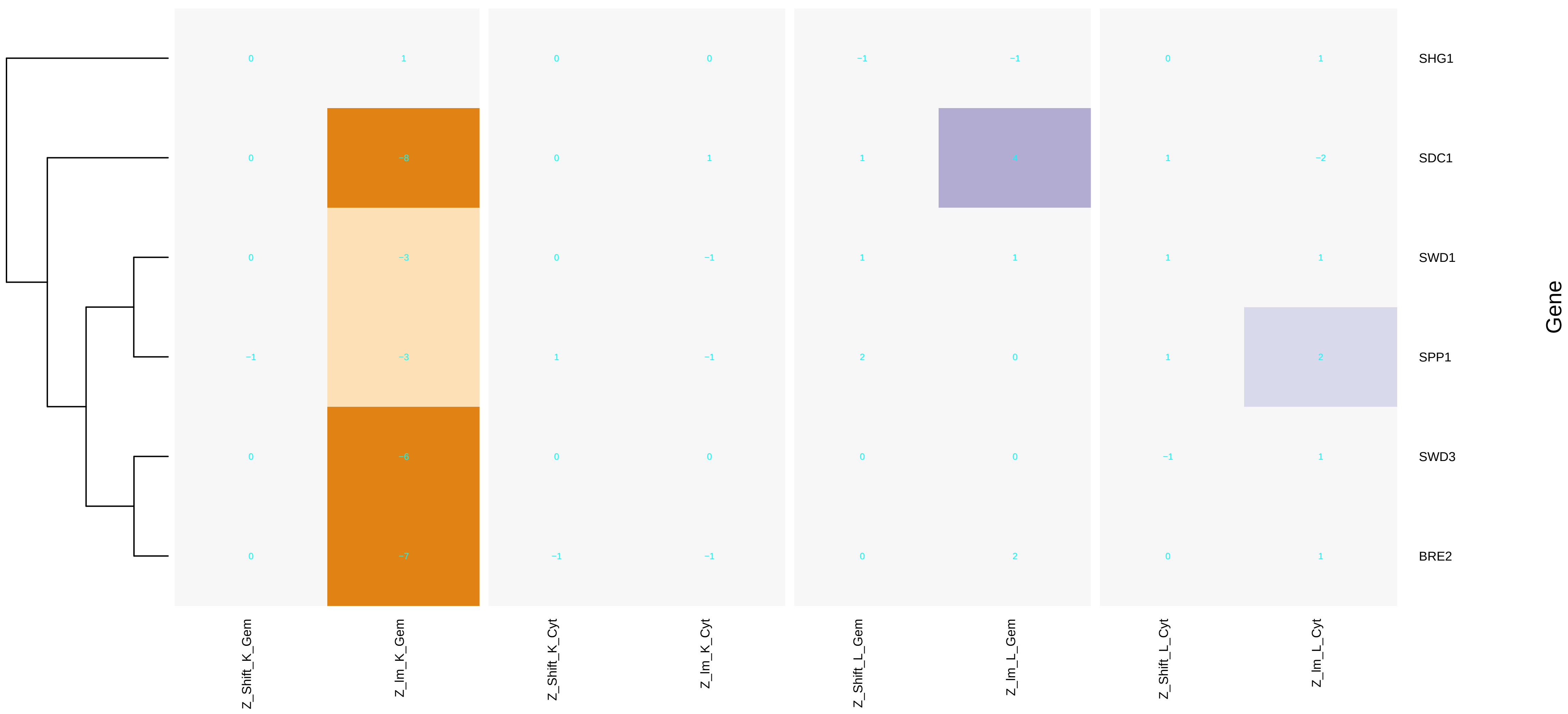
