## Additional File 8 for "A humanized yeast phenomic model of deoxycytidine kinase to predict genetic buffering of nucleoside analog cytotoxicity": 19_0314_AcrossAll_to_orig_hmap_Heatmaps.pdf

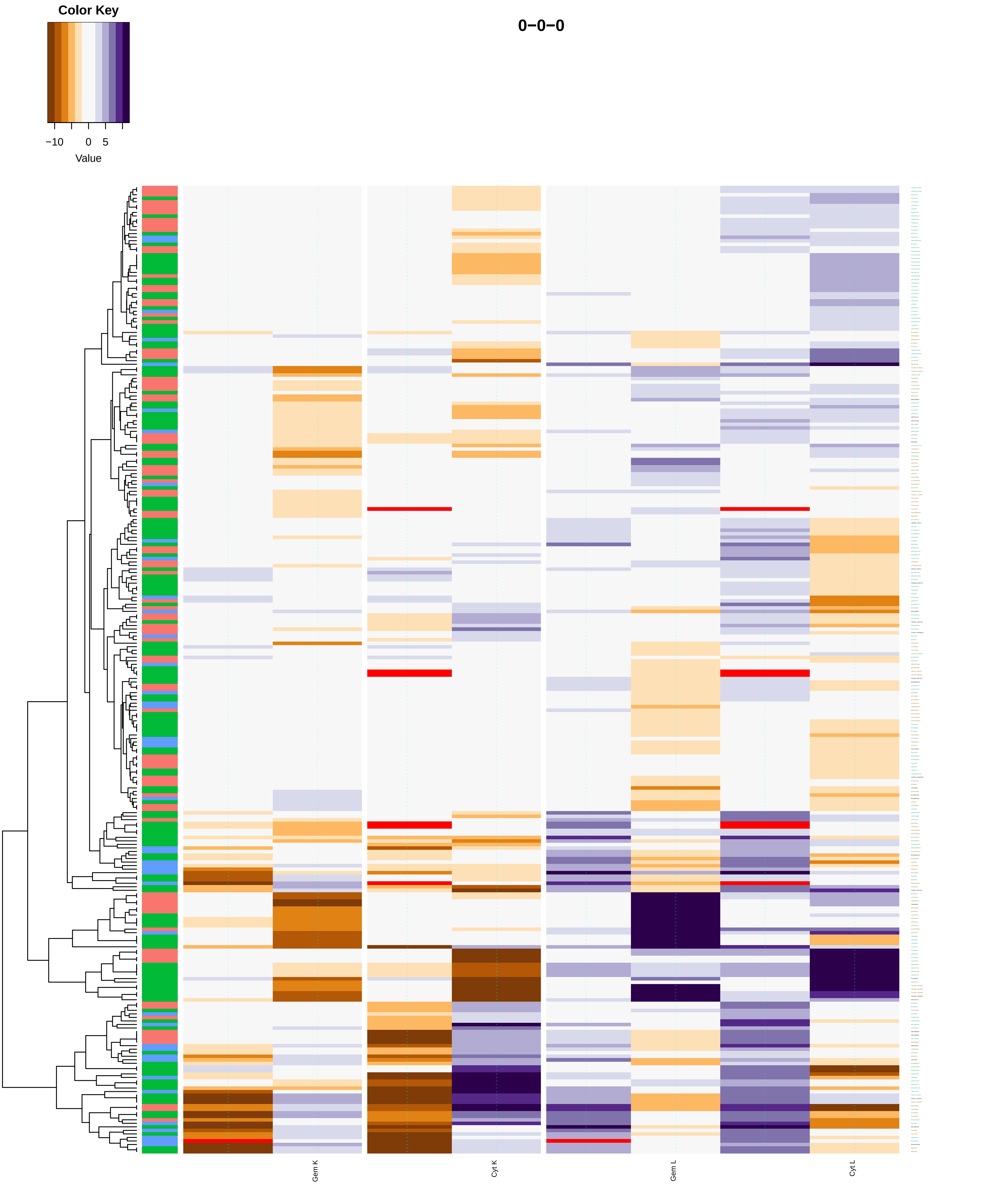

Color Key
−10
0
5
Value
0−0−0
Gem K
Cyt K
Gem L
Cyt L

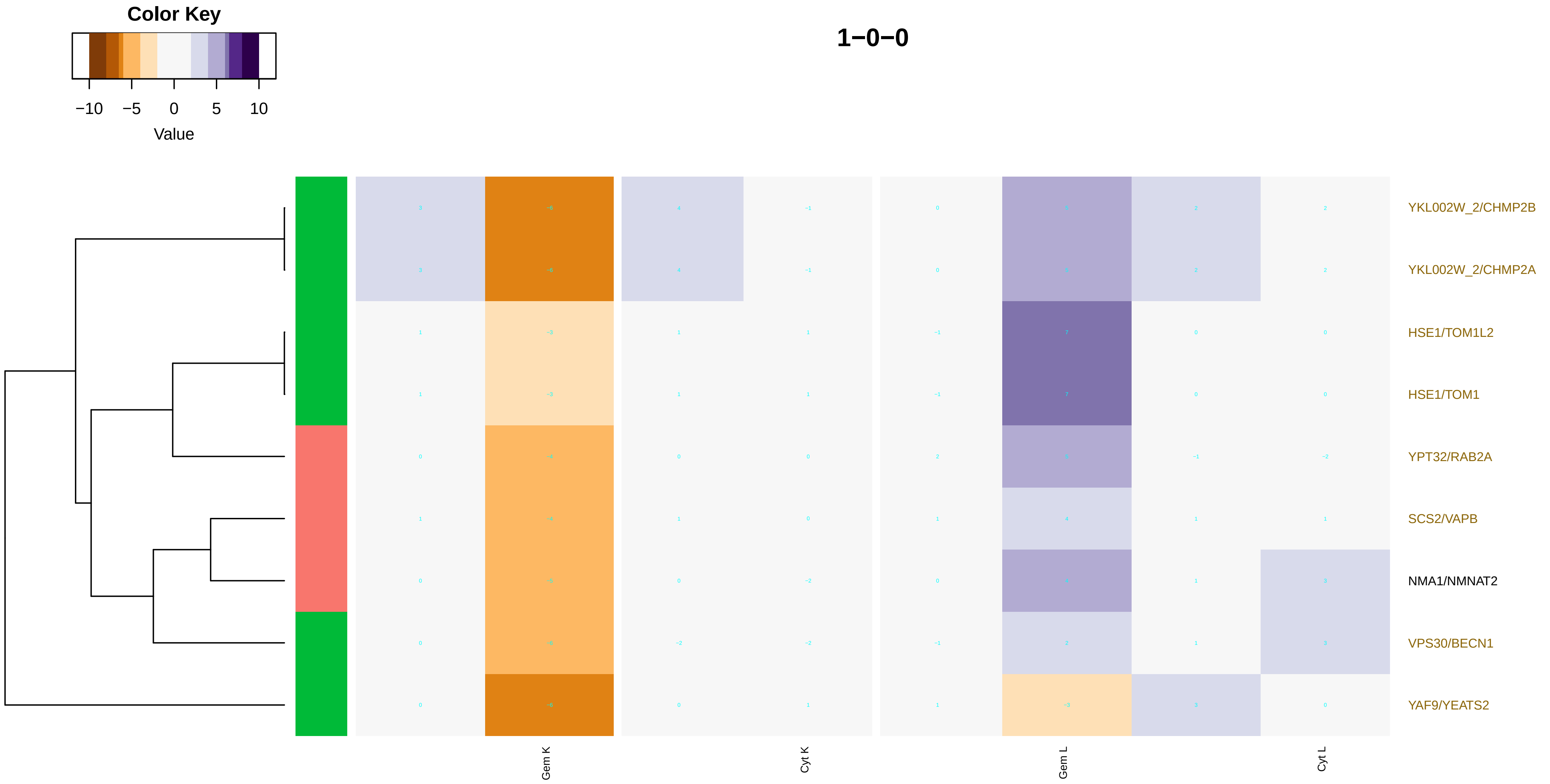

Color Key
−10
−5
0
5
10
Value
1−0−0
YKL002W_2/CHMP2B
YKL002W_2/CHMP2A
HSE1/TOM1L2
HSE1/TOM1
YPT32/RAB2A
SCS2/VAPB
NMA1/NMNAT2
VPS30/BECN1
YAF9/YEATS2
Gem K
Cyt K
Gem L
Cyt L

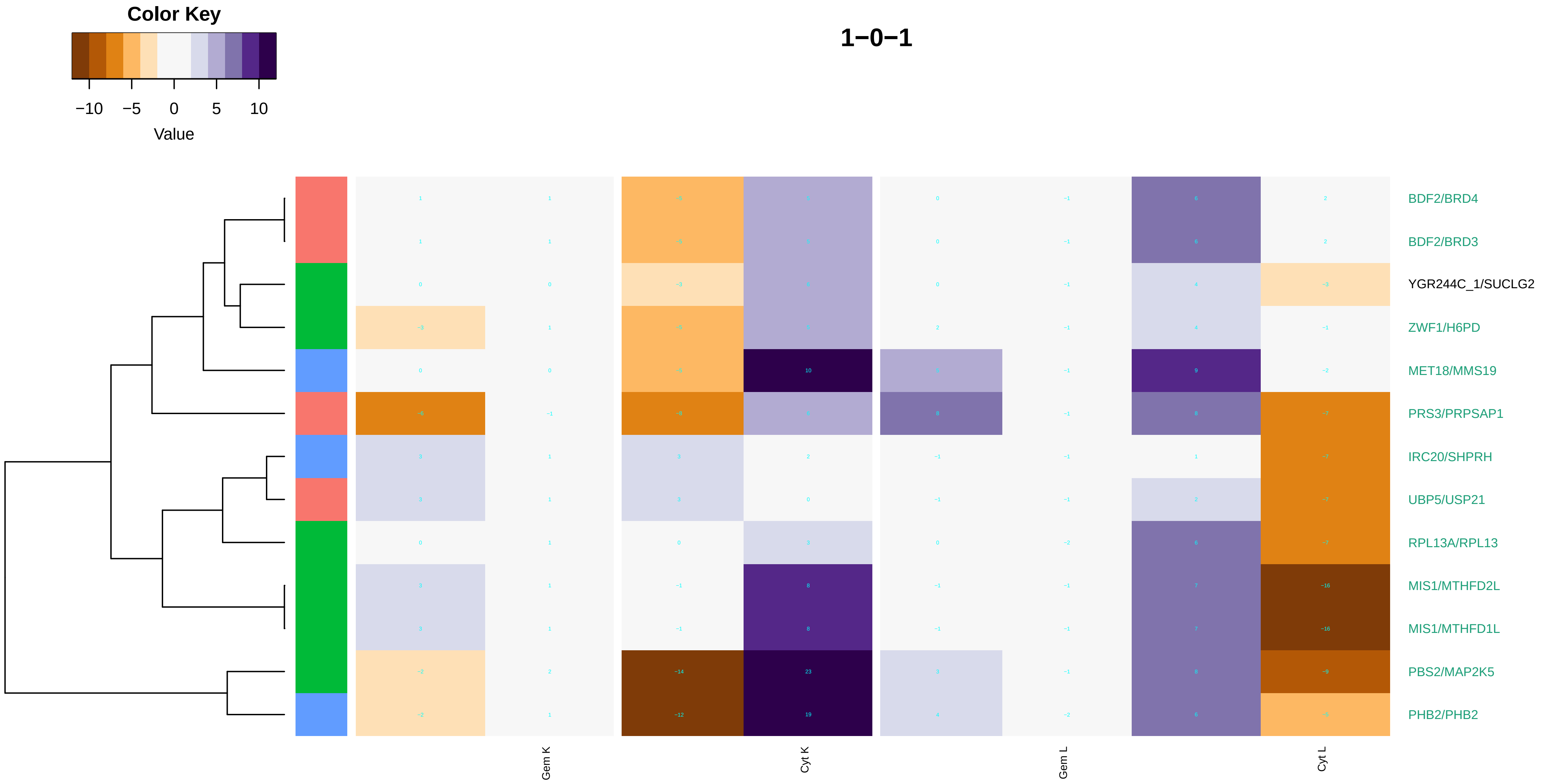

Color Key
−10
−5
0
5
10
Value
1−0−1
BDF2/BRD4
BDF2/BRD3
YGR244C_1/SUCLG2
ZWF1/H6PD
MET18/MMS19
PRS3/PRPSAP1
IRC20/SHPRH
UBP5/USP21
RPL13A/RPL13
MIS1/MTHFD2L
MIS1/MTHFD1L
PBS2/MAP2K5
PHB2/PHB2
Gem K
Cyt K
Gem L
Cyt L

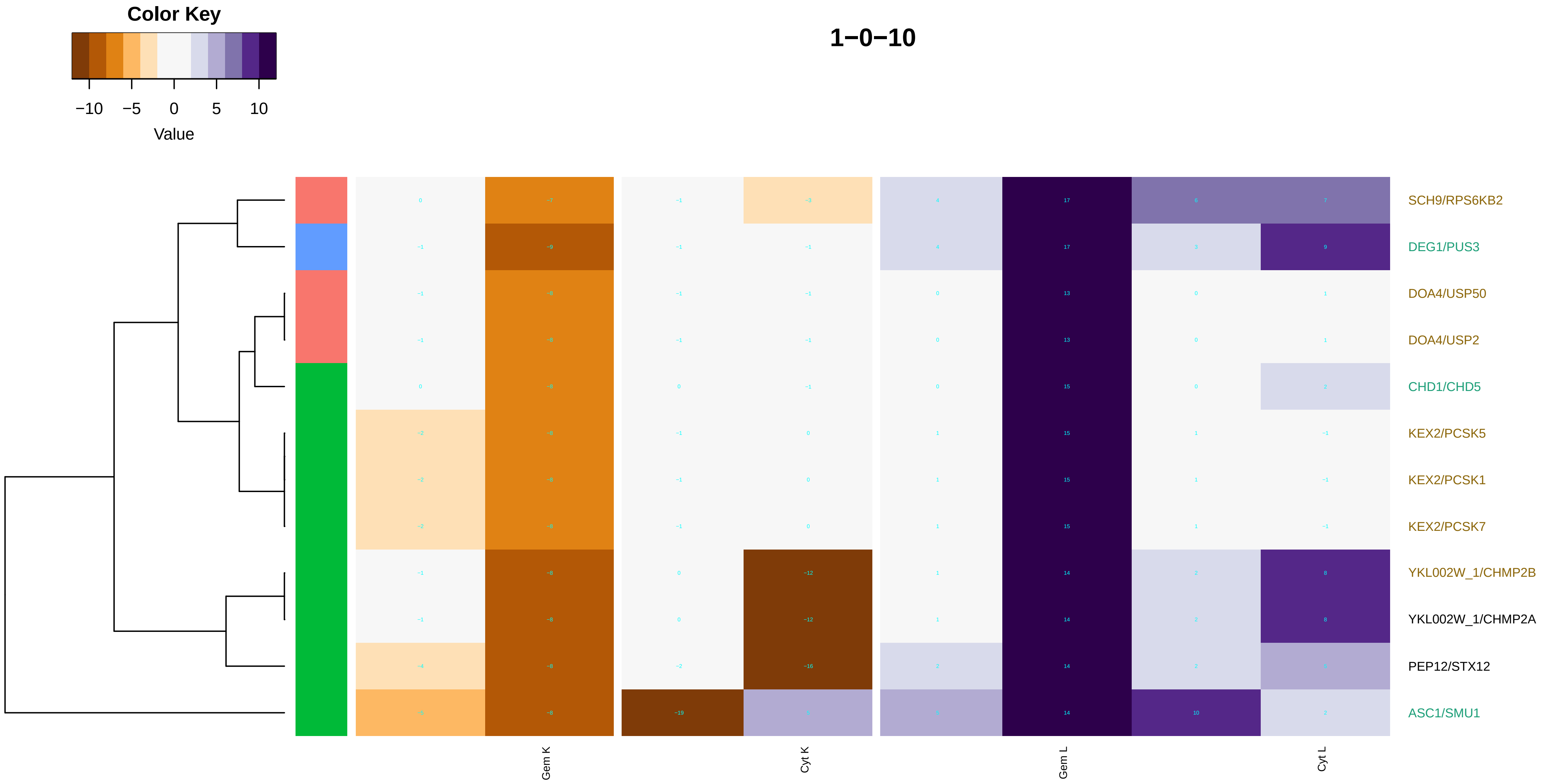
Color Key
−10 −5 0 5 10
Value
1−0−10
SCH9/RPS6KB2
DEG1/PUS3
DOA4/USP50
DOA4/USP2
CHD1/CHD5
KEX2/PCSK5
KEX2/PCSK1
KEX2/PCSK7
YKL002W_1/CHMP2B
YKL002W_1/CHMP2A
PEP12/STX12
ASC1/SMU1
Gem K
Cyt K
Gem L
Cyt L

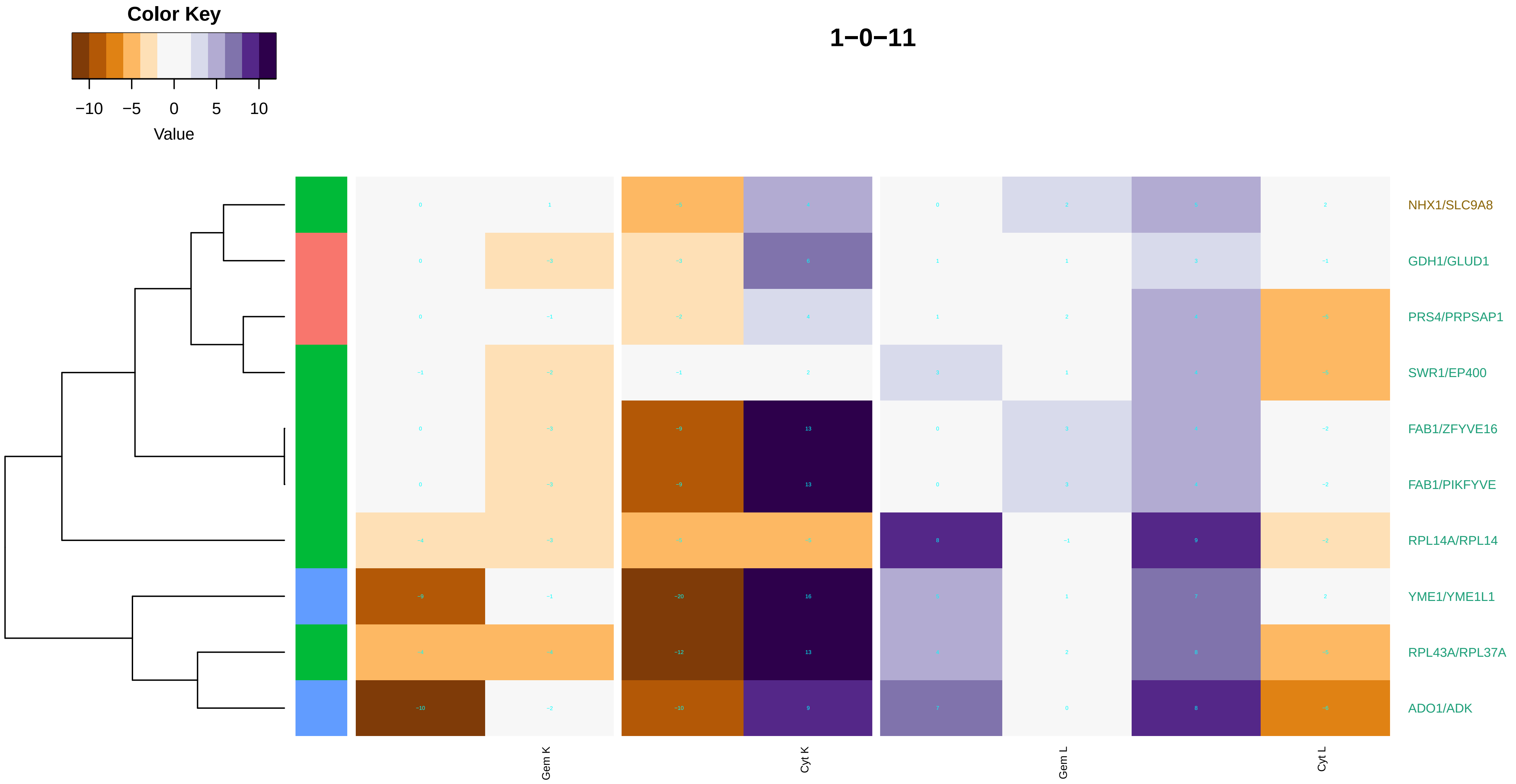
Color Key
−10 −5 0 5 10
Value
1−0−11
NHX1/SLC9A8
GDH1/GLUD1
PRS4/PRPSAP1
SWR1/EP400
FAB1/ZFYVE16
FAB1/PIKFYVE
RPL14A/RPL14
YME1/YME1L1
RPL43A/RPL37A
ADO1/ADK
Gem K
Cyt K
Gem L
Cyt L

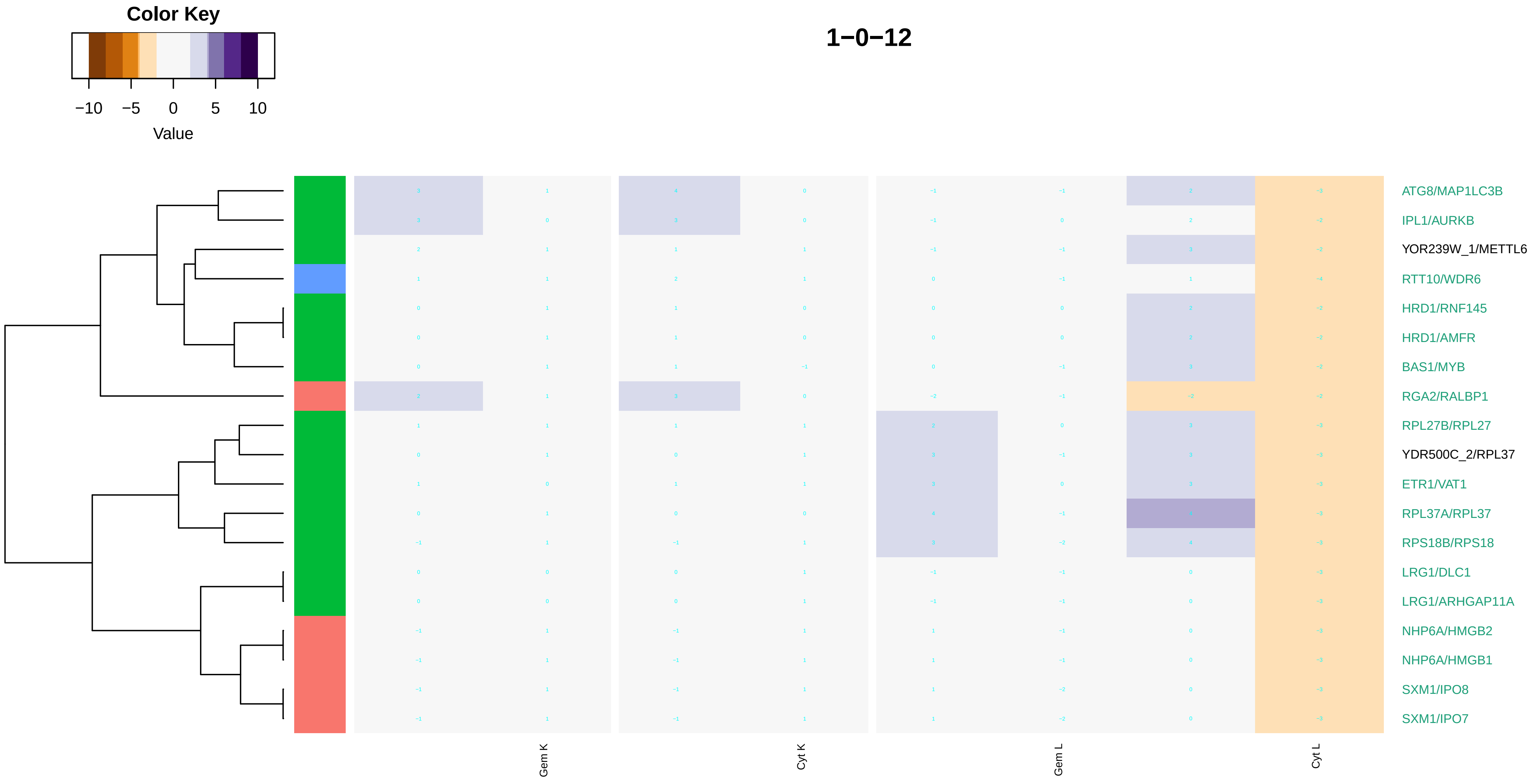

Color Key
−10 −5 0 5 10
Value
1−0−12
ATG8/MAP1LC3B
IPL1/AURKB
YOR239W_1/METTL6
RTT10/WDR6
HRD1/RNF145
HRD1/AMFR
BAS1/MYB
RGA2/RALBP1
RPL27B/RPL27
YDR500C_2/RPL37
ETR1/VAT1
RPL37A/RPL37
RPS18B/RPS18
LRG1/DLC1
LRG1/ARHGAP11A
NHP6A/HMGB2
NHP6A/HMGB1
SXM1/IPO8
SXM1/IPO7
Gem K
Cyt K
Gem L
Cyt L

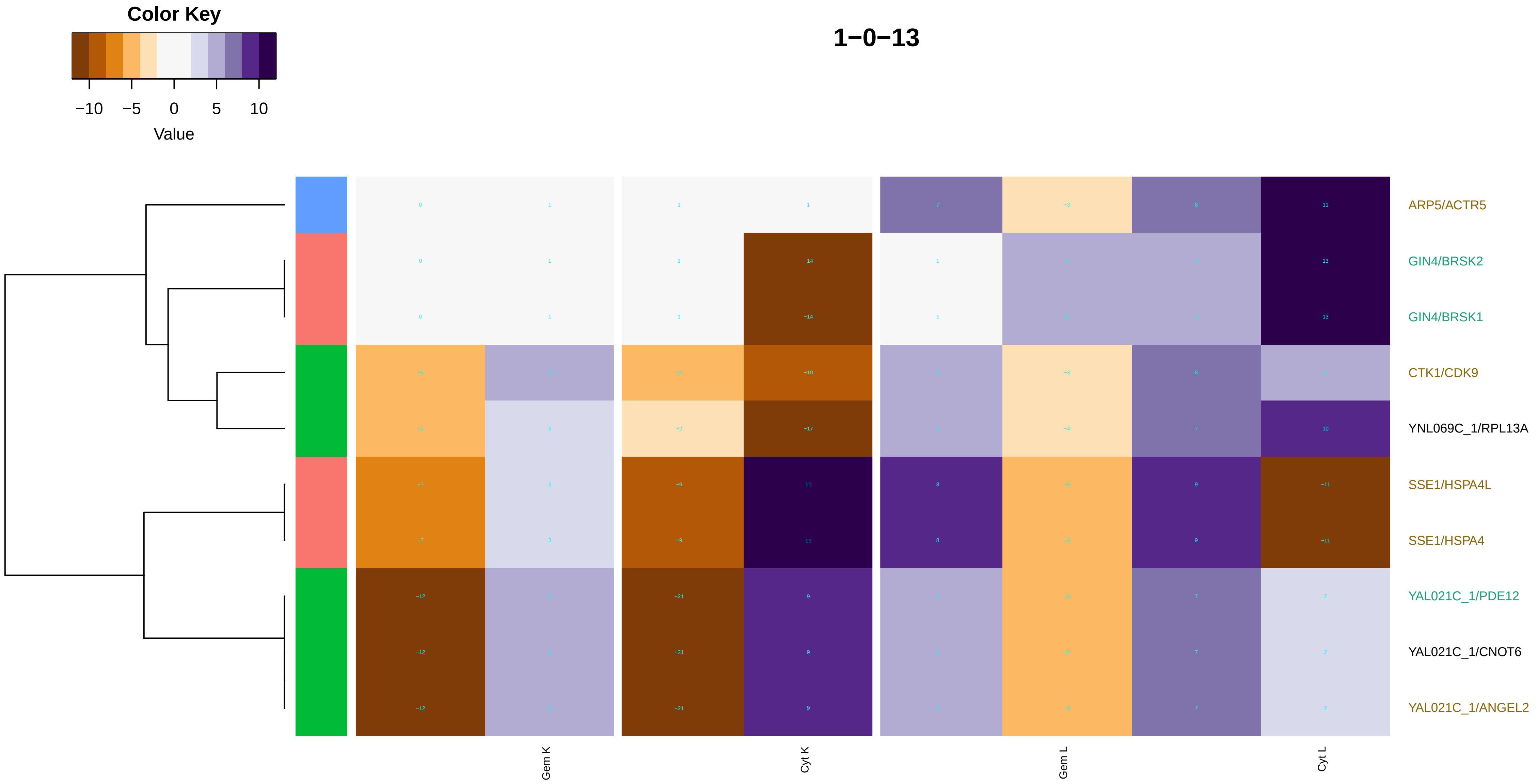
Color Key
−10
−5
0
5
10
Value
1−0−13
ARP5/ACTR5
GIN4/BRSK2
GIN4/BRSK1
CTK1/CDK9
YNL069C_1/RPL13A
SSE1/HSPA4L
SSE1/HSPA4
YAL021C_1/PDE12
YAL021C_1/CNOT6
YAL021C_1/ANGEL2
Gem K
Cyt K
Gem L
Cyt L

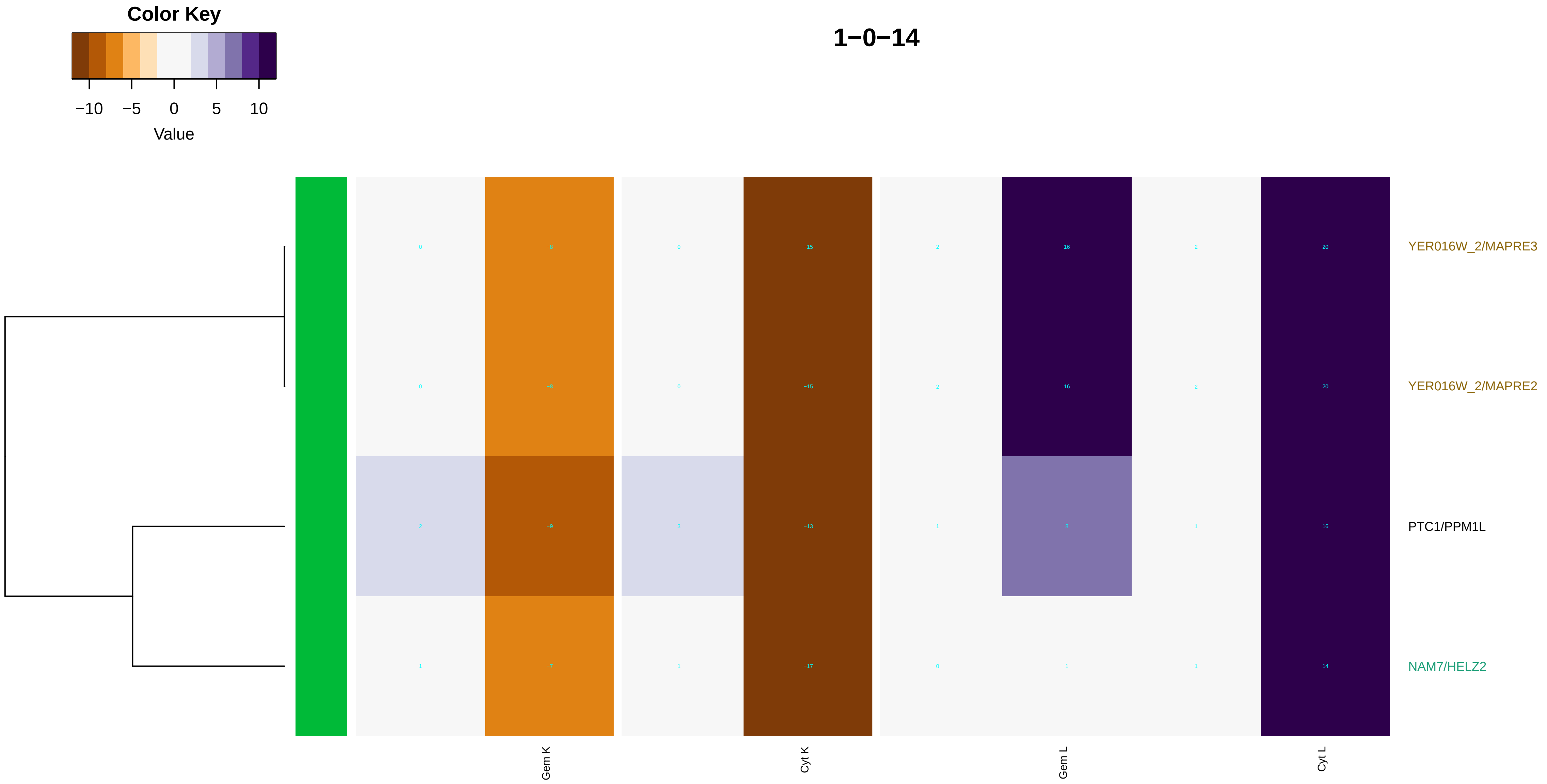

1−0−14
Color Key
−10
−5
0
5
10
Value
YER016W_2/MAPRE3
YER016W_2/MAPRE2
PTC1/PPM1L
NAM7/HELZ2
Gem K
Cyt K
Gem L
Cyt L

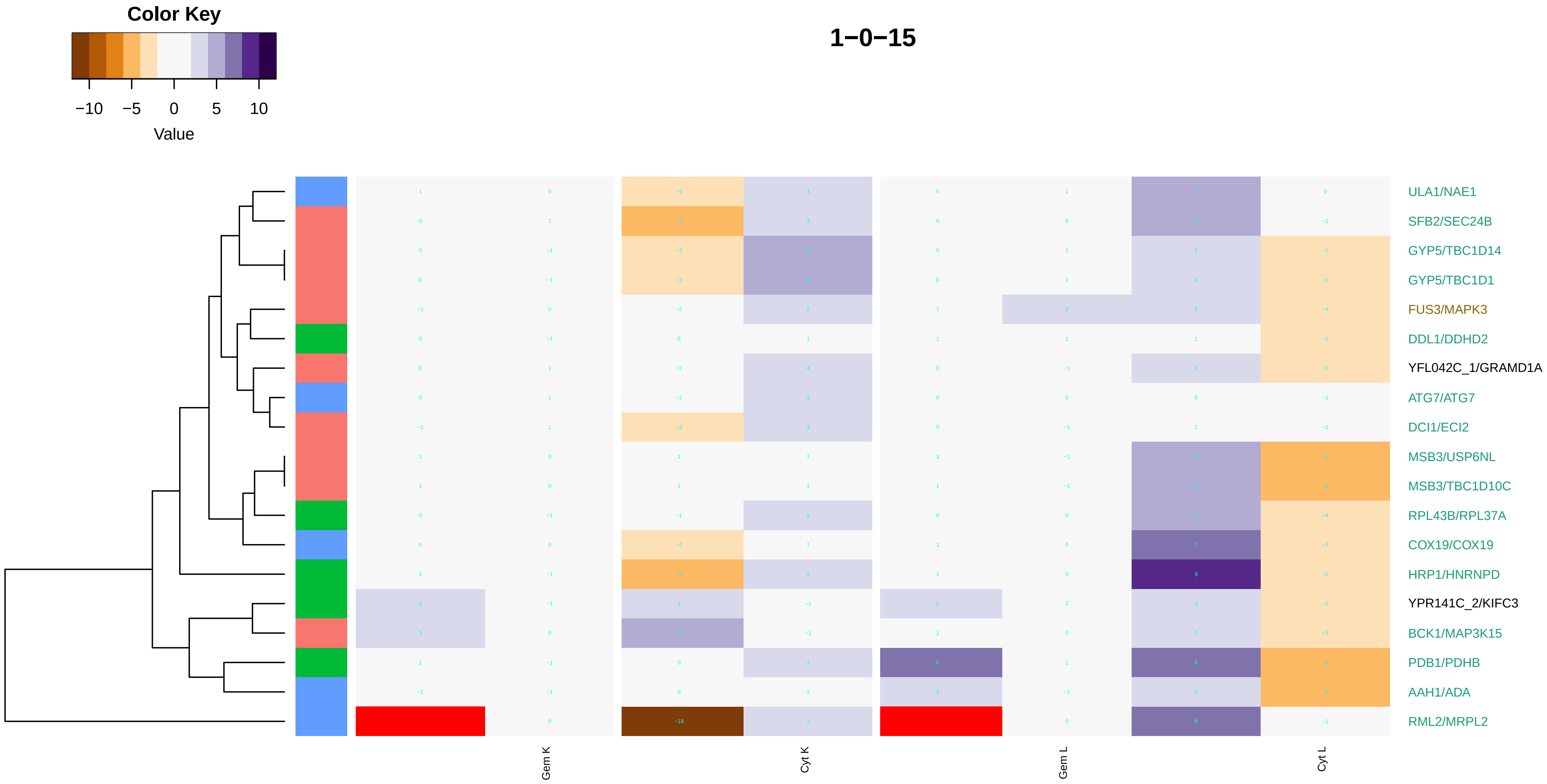

Color Key
−10
−5
0
5
10
Value
1−0−15
ULA1/NAE1
SFB2/SEC24B
GYP5/TBC1D14
GYP5/TBC1D1
FUS3/MAPK3
DDL1/DDHD2
YFL042C_1/GRAMD1A
ATG7/ATG7
DCI1/ECI2
MSB3/USP6NL
MSB3/TBC1D10C
RPL43B/RPL37A
COX19/COX19
HRP1/HNRNPD
YPR141C_2/KIFC3
BCK1/MAP3K15
PDB1/PDHB
AAH1/ADA
RML2/MRPL2
Gem K
Cyt K
Gem L
Cyt L

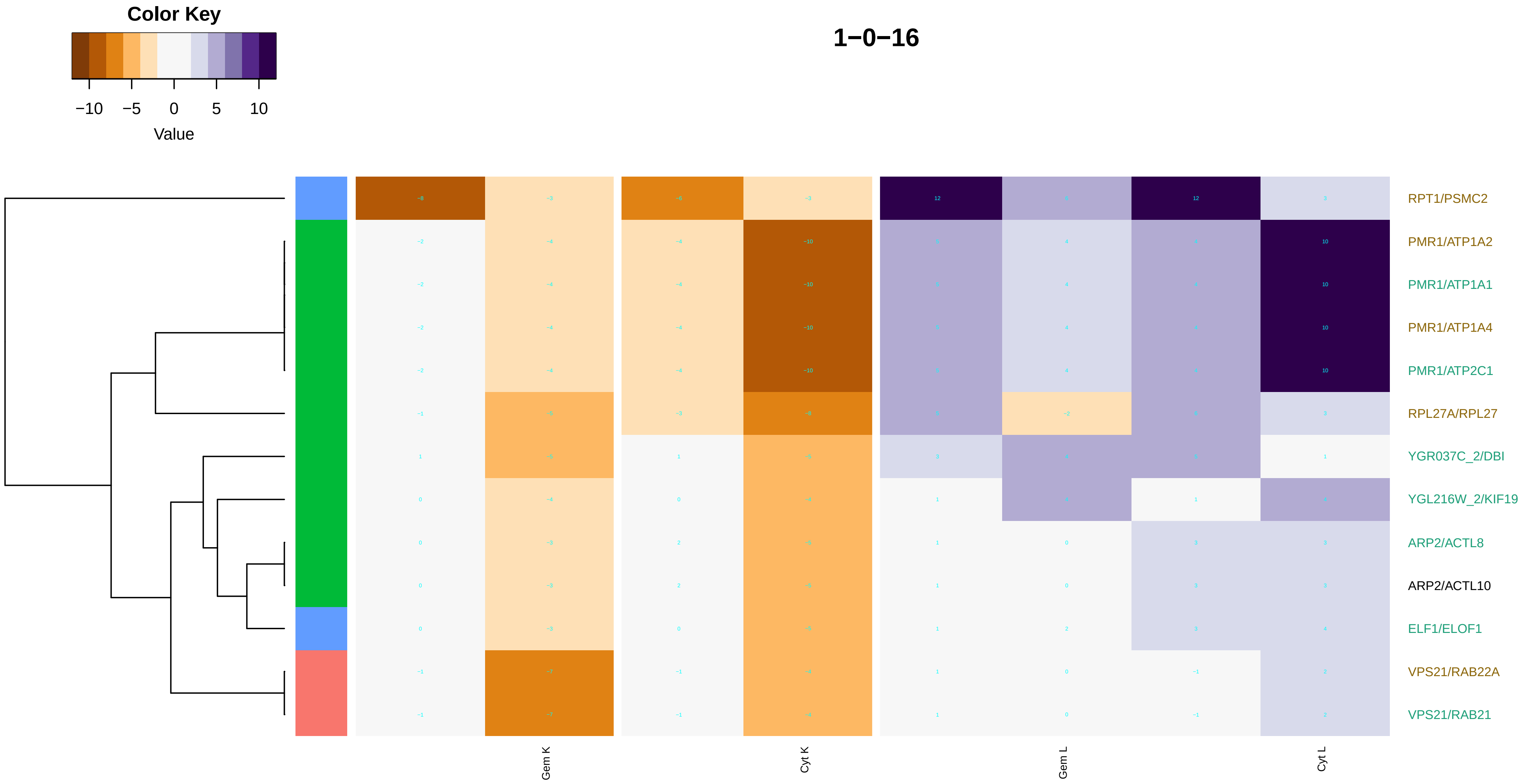
Color Key
−10
−5
0
5
10
Value
1−0−16
RPT1/PSMC2
PMR1/ATP1A2
PMR1/ATP1A1
PMR1/ATP1A4
PMR1/ATP2C1
RPL27A/RPL27
YGR037C_2/DBI
YGL216W_2/KIF19
ARP2/ACTL8
ARP2/ACTL10
ELF1/ELOF1
VPS21/RAB22A
VPS21/RAB21
Gem K
Cyt K
Gem L
Cyt L

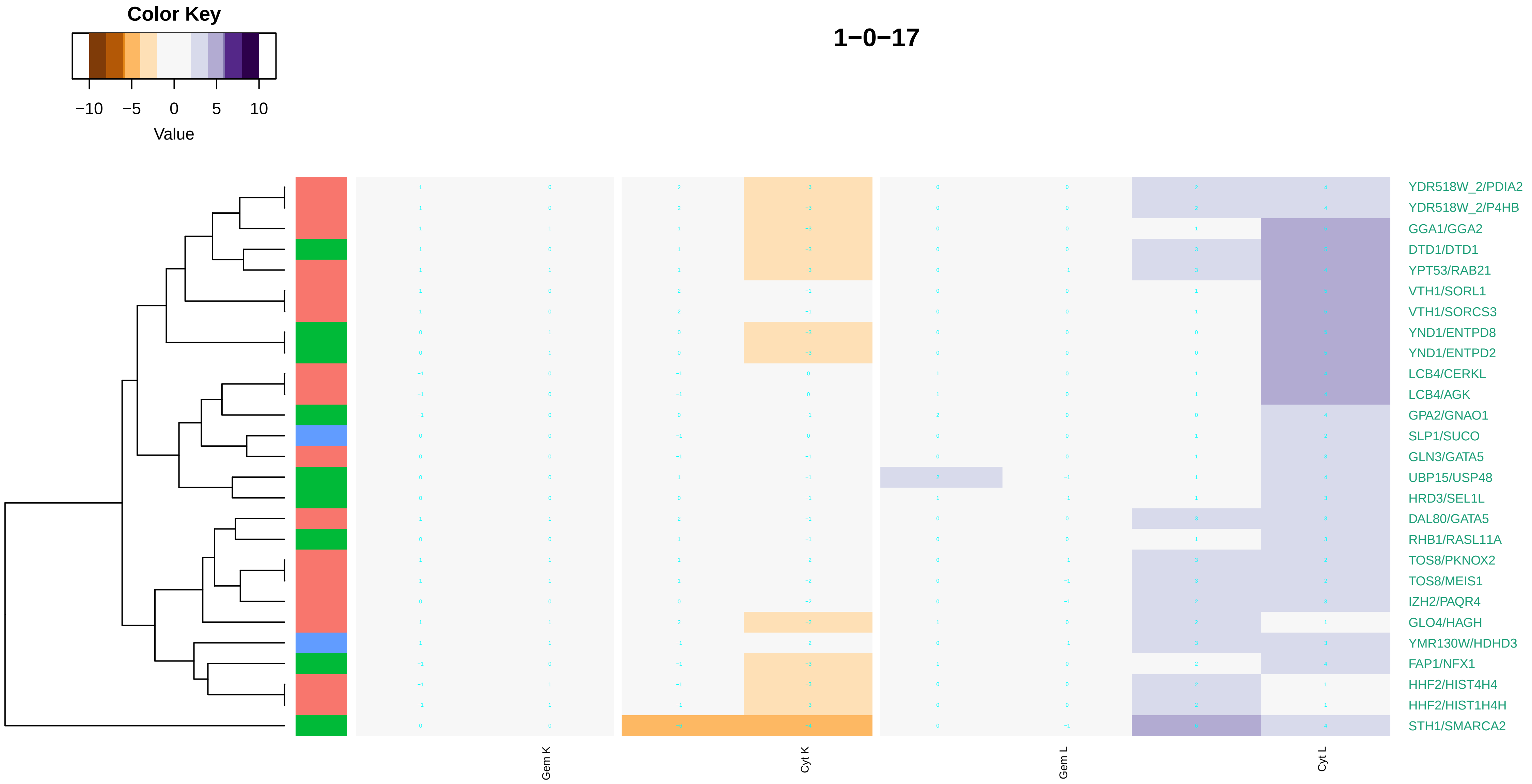
Color Key
−10
−5
0
5
10
Value
1−0−17
Gem K
Cyt K
Gem L
Cyt L
YDR518W_2/PDIA2
YDR518W_2/P4HB
GGA1/GGA2
DTD1/DTD1
YPT53/RAB21
VTH1/SORL1
VTH1/SORCS3
YND1/ENTPD8
YND1/ENTPD2
LCB4/CERKL
LCB4/AGK
GPA2/GNAO1
SLP1/SUCO
GLN3/GATA5
UBP15/USP48
HRD3/SEL1L
DAL80/GATA5
RHB1/RASL11A
TOS8/PKNOX2
TOS8/MEIS1
IZH2/PAQR4
GLO4/HAGH
YMR130W/HDHD3
FAP1/NFX1
HHF2/HIST4H4
HHF2/HIST1H4H
STH1/SMARCA2

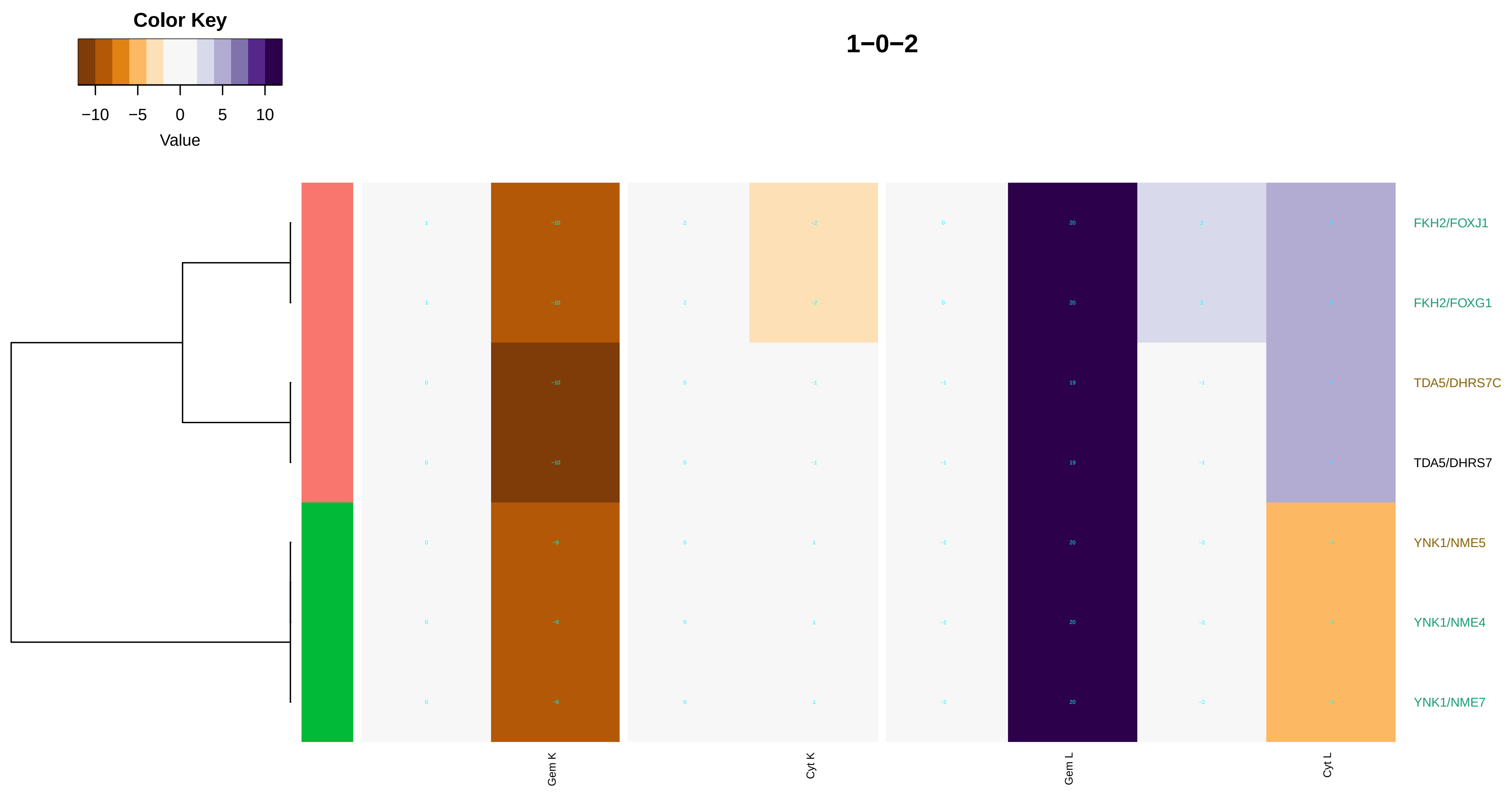
Color Key
−10
−5
0
5
10
Value
1−0−2
FKH2/FOXJ1
FKH2/FOXG1
TDA5/DHRS7C
TDA5/DHRS7
YNK1/NME5
YNK1/NME4
YNK1/NME7
Gem K
Cyt K
Gem L
Cyt L

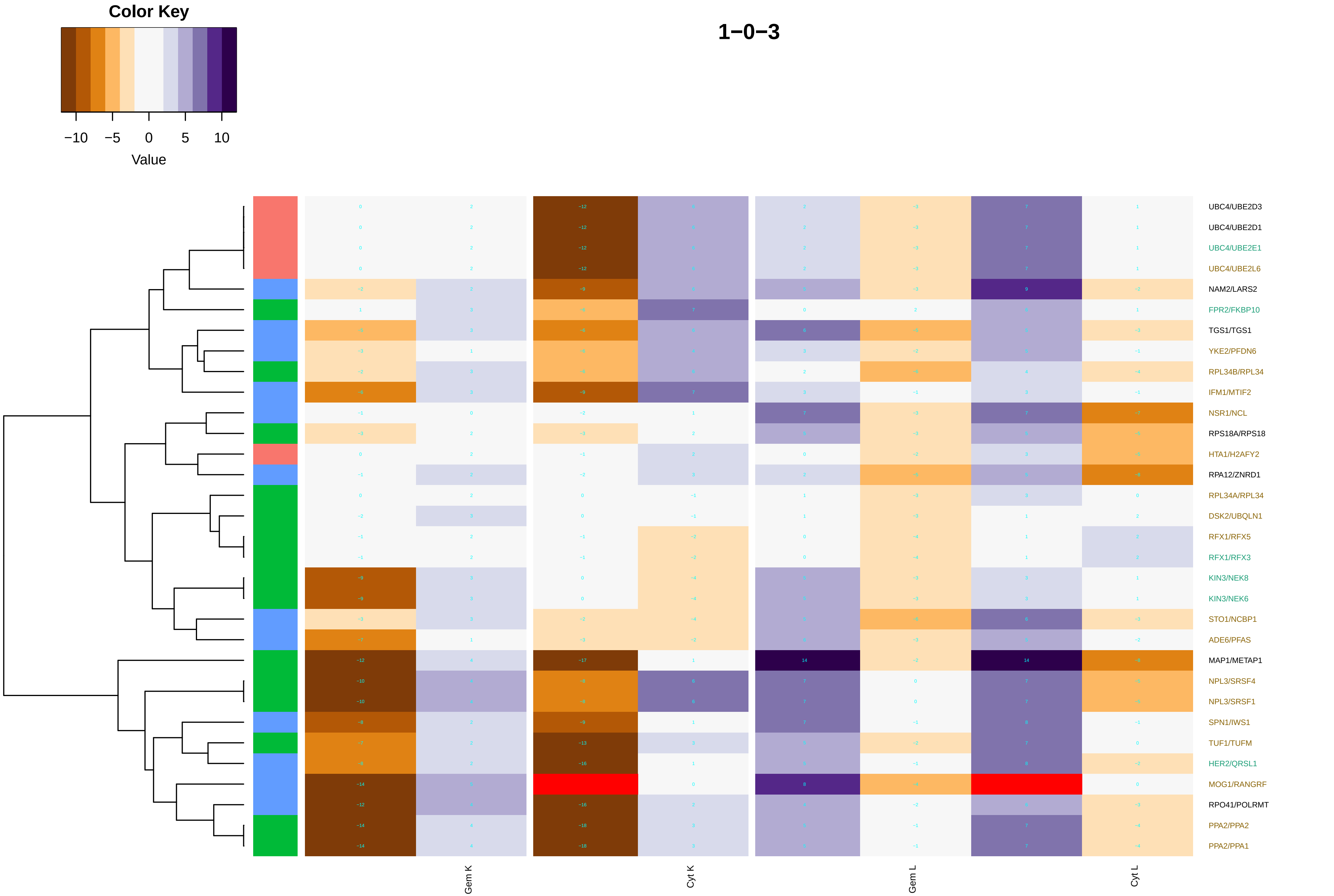
Color Key
−10
−5
0
5
10
Value
1−0−3
UBC4/UBE2D3
UBC4/UBE2D1
UBC4/UBE2E1
UBC4/UBE2L6
NAM2/LARS2
FPR2/FKBP10
TGS1/TGS1
YKE2/PFDN6
RPL34B/RPL34
IFM1/MTIF2
NSR1/NCL
RPS18A/RPS18
HTA1/H2AFY2
RPA12/ZNRD1
RPL34A/RPL34
DSK2/UBQLN1
RFX1/RFX5
RFX1/RFX3
KIN3/NEK8
KIN3/NEK6
STO1/NCBP1
ADE6/PFAS
MAP1/METAP1
NPL3/SRSF4
NPL3/SRSF1
SPN1/IWS1
TUF1/TUFM
HER2/QRSL1
MOG1/RANGRF
RPO41/POLRMT
PPA2/PPA2
PPA2/PPA1
Gem K
Cyt K
Gem L
Cyt L

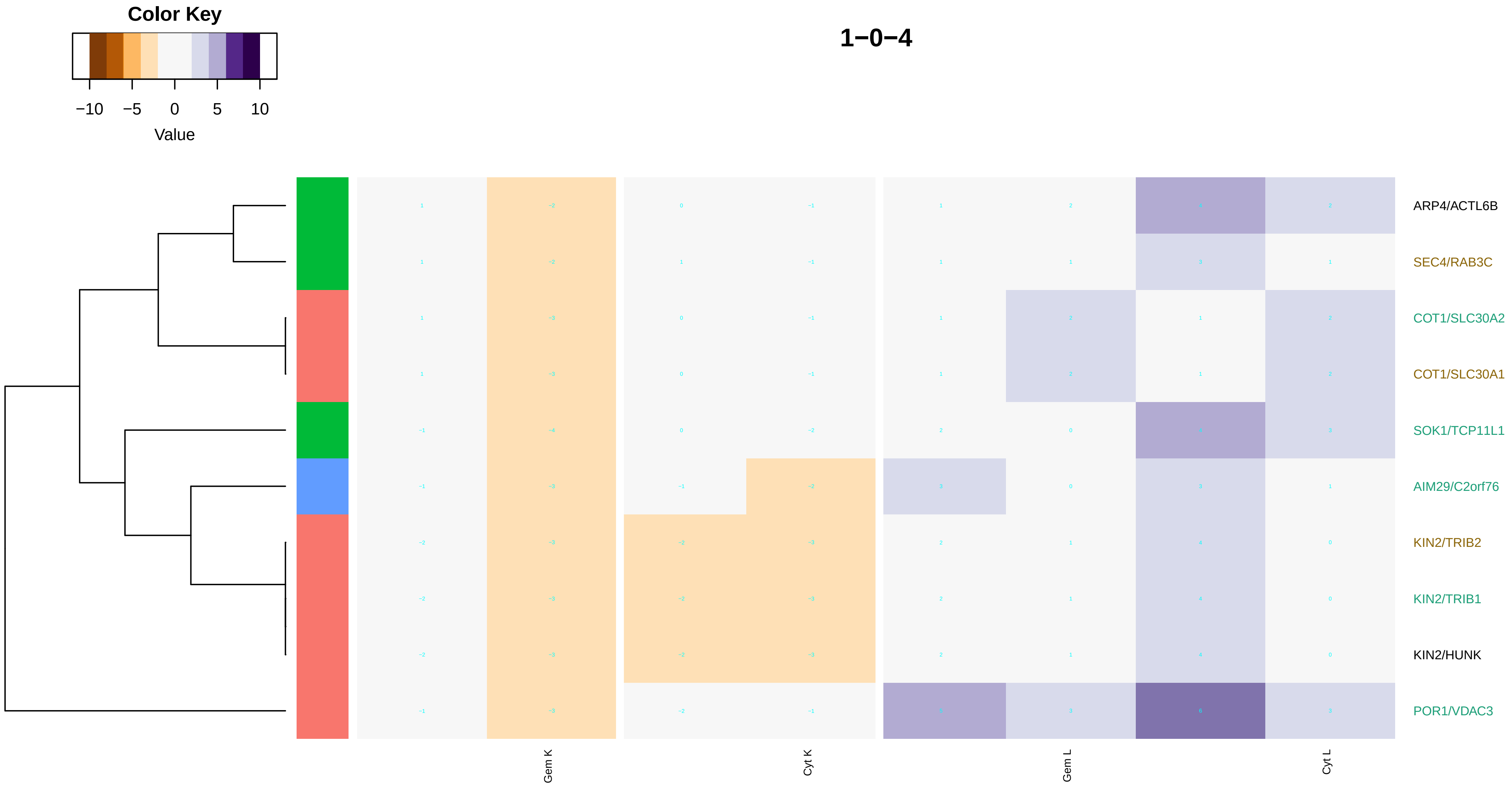

Color Key
−10
−5
0
5
10
Value
1−0−4
ARP4/ACTL6B
SEC4/RAB3C
COT1/SLC30A2
COT1/SLC30A1
SOK1/TCP11L1
AIM29/C2orf76
KIN2/TRIB2
KIN2/TRIB1
KIN2/HUNK
POR1/VDAC3
Gem K
Cyt K
Gem L
Cyt L

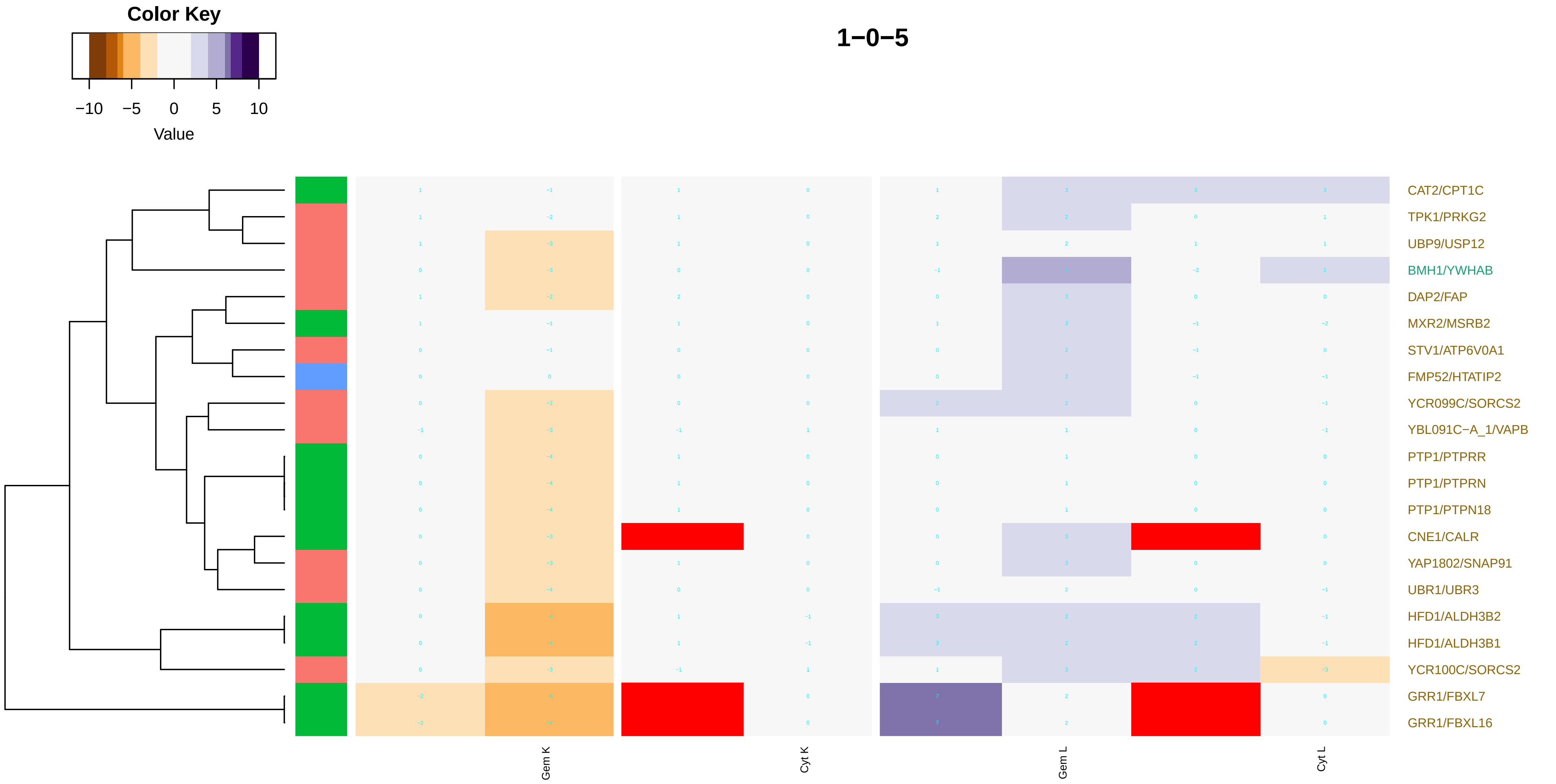

Color Key
−10
−5
0
5
10
Value
1−0−5
Gem K
Cyt K
Gem L
Cyt L
CAT2/CPT1C
TPK1/PRKG2
UBP9/USP12
BMH1/YWHAB
DAP2/FAP
MXR2/MSRB2
STV1/ATP6V0A1
FMP52/HTATIP2
YCR099C/SORCS2
YBL091C−A_1/VAPB
PTP1/PTPRR
PTP1/PTPRN
PTP1/PTPN18
CNE1/CALR
YAP1802/SNAP91
UBR1/UBR3
HFD1/ALDH3B2
HFD1/ALDH3B1
YCR100C/SORCS2
GRR1/FBXL7
GRR1/FBXL16

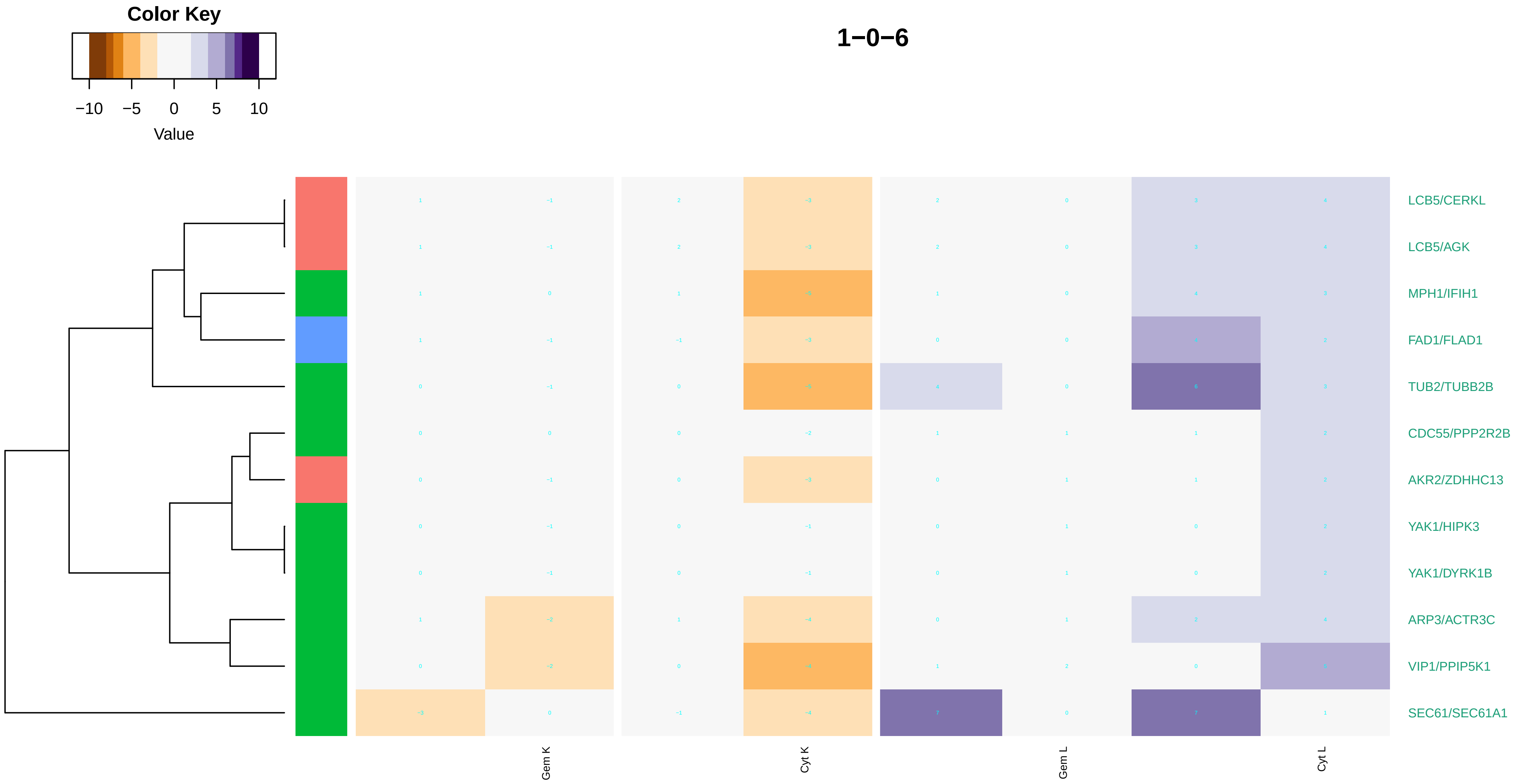

Color Key
−10
−5
0
5
10
Value
1−0−6
LCB5/CERKL
LCB5/AGK
MPH1/IFIH1
FAD1/FLAD1
TUB2/TUBB2B
CDC55/PPP2R2B
AKR2/ZDHHC13
YAK1/HIPK3
YAK1/DYRK1B
ARP3/ACTR3C
VIP1/PPIP5K1
SEC61/SEC61A1
Gem K
Cyt K
Gem L
Cyt L

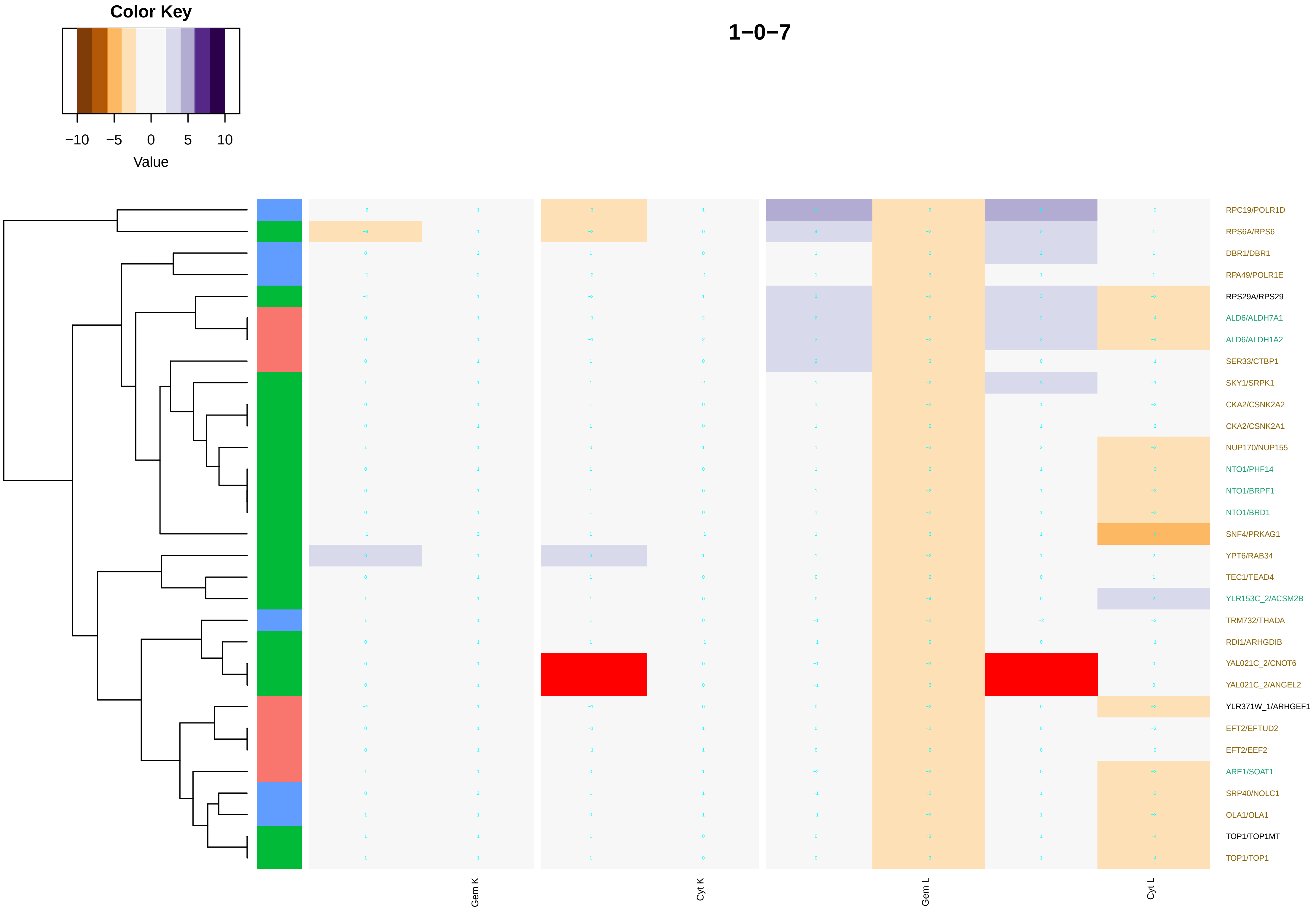
Color Key
−10
−5
0
5
10
Value
1−0−7
RPC19/POLR1D
RPS6A/RPS6
DBR1/DBR1
RPA49/POLR1E
RPS29A/RPS29
ALD6/ALDH7A1
ALD6/ALDH1A2
SER33/CTBP1
SKY1/SRPK1
CKA2/CSNK2A2
CKA2/CSNK2A1
NUP170/NUP155
NTO1/PHF14
NTO1/BRPF1
NTO1/BRD1
SNF4/PRKAG1
YPT6/RAB34
TEC1/TEAD4
YLR153C_2/ACSM2B
TRM732/THADA
RDI1/ARHGDIB
YAL021C_2/CNOT6
YAL021C_2/ANGEL2
YLR371W_1/ARHGEF1
EFT2/EFTUD2
EFT2/EEF2
ARE1/SOAT1
SRP40/NOLC1
OLA1/OLA1
TOP1/TOP1MT
TOP1/TOP1
Gem K
Cyt K
Gem L
Cyt L

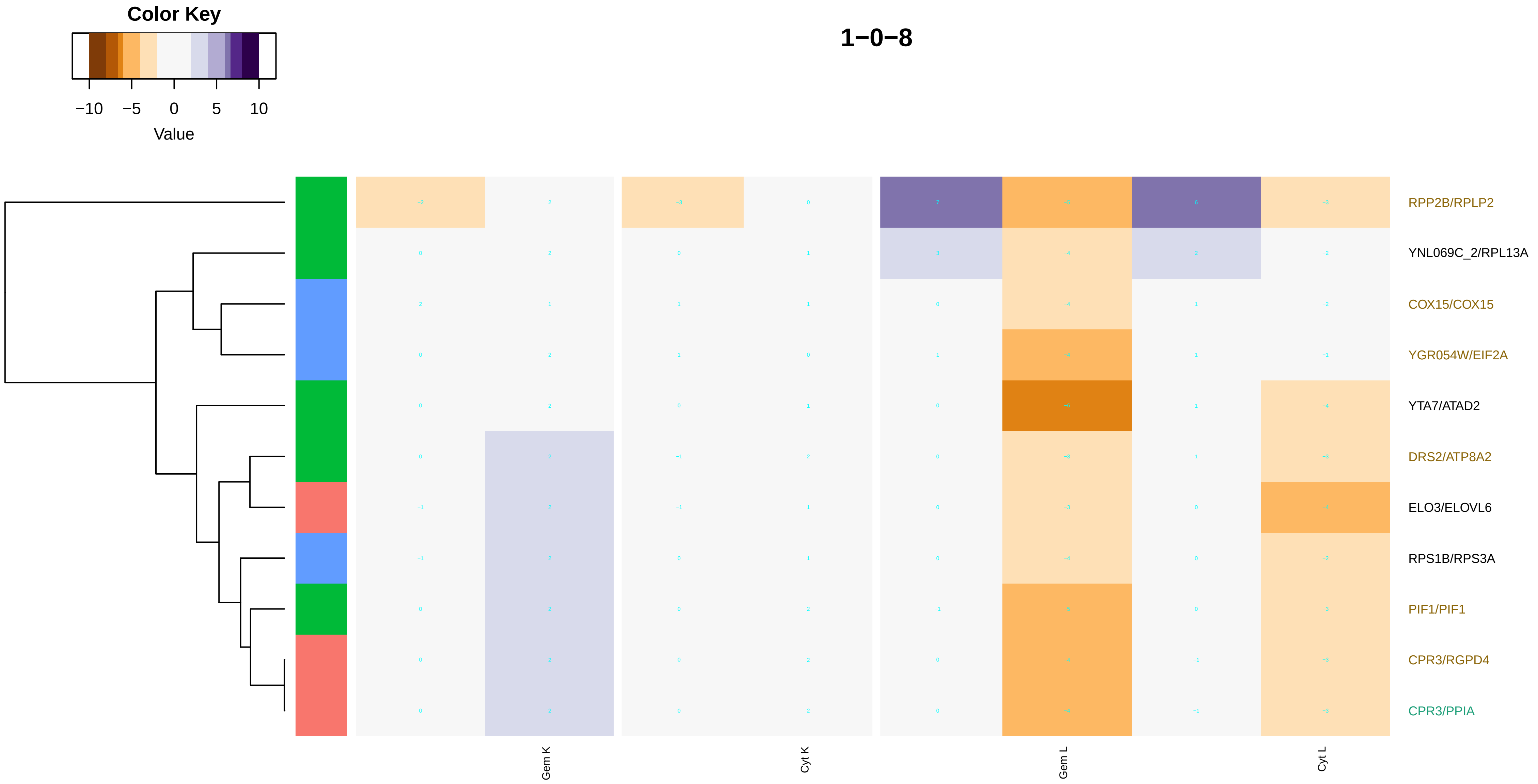

Color Key
−10
−5
0
5
10
Value
1−0−8
RPP2B/RPLP2
YNL069C_2/RPL13A
COX15/COX15
YGR054W/EIF2A
YTA7/ATAD2
DRS2/ATP8A2
ELO3/ELOVL6
RPS1B/RPS3A
PIF1/PIF1
CPR3/RGPD4
CPR3/PPIA
Gem K
Cyt K
Gem L
Cyt L

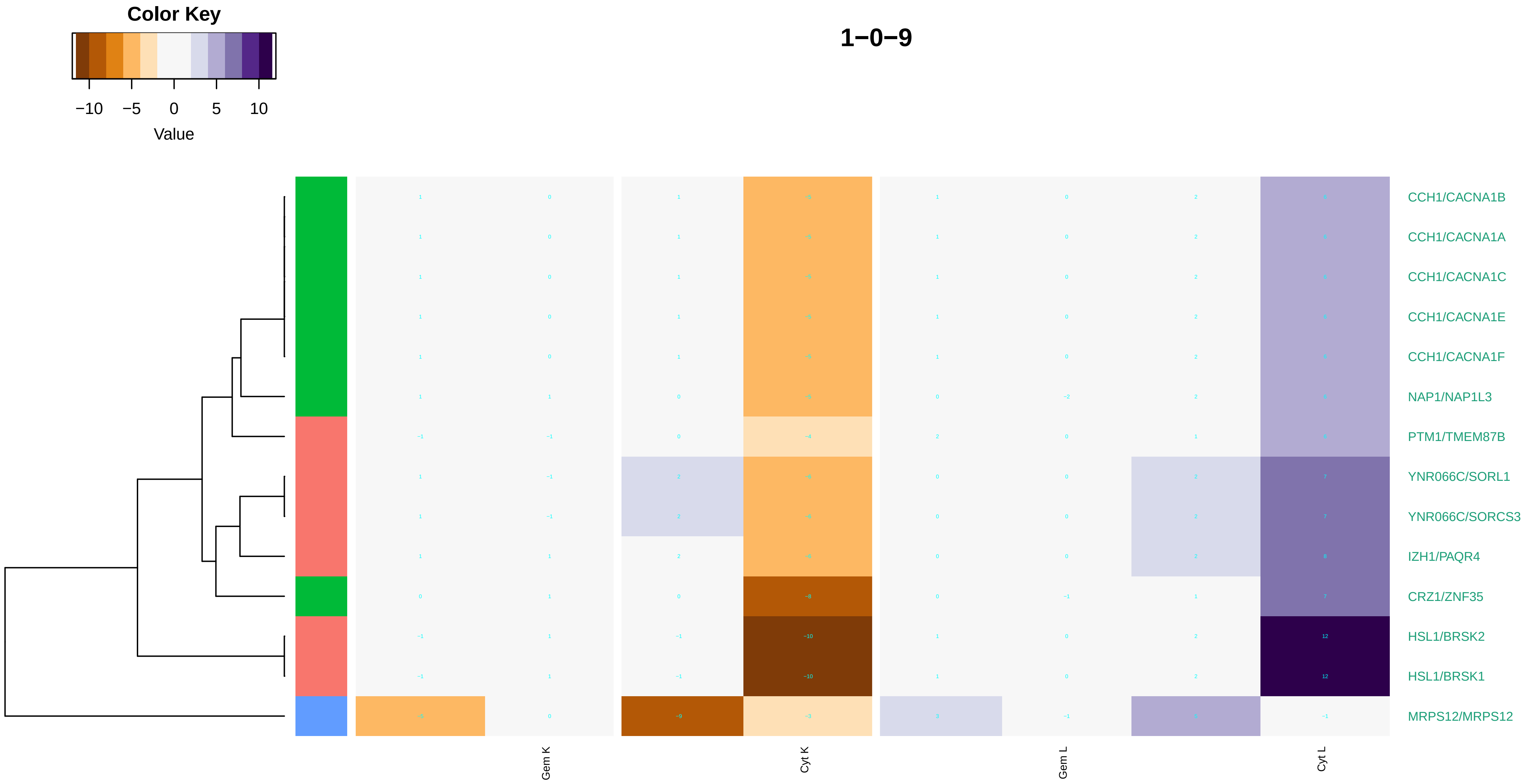

Color Key
−10
−5
0
5
10
Value
1−0−9
CCH1/CACNA1B
CCH1/CACNA1A
CCH1/CACNA1C
CCH1/CACNA1E
CCH1/CACNA1F
NAP1/NAP1L3
PTM1/TMEM87B
YNR066C/SORL1
YNR066C/SORCS3
IZH1/PAQR4
CRZ1/ZNF35
HSL1/BRSK2
HSL1/BRSK1
MRPS12/MRPS12
Gem K
Cyt K
Gem L
Cyt L

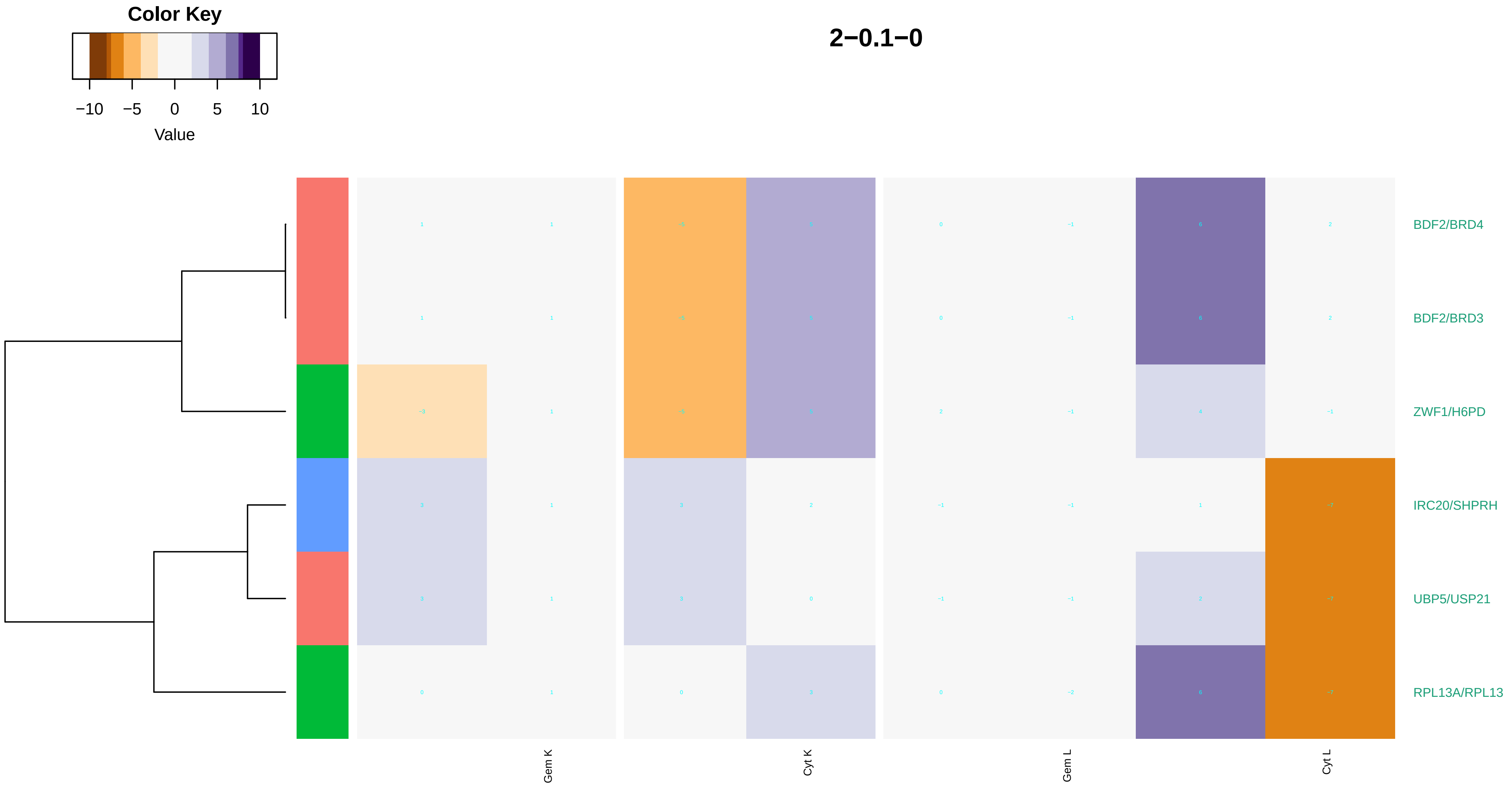

Color Key
−10
−5
0
5
10
Value
2−0.1−0
BDF2/BRD4
BDF2/BRD3
ZWF1/H6PD
IRC20/SHPRH
UBP5/USP21
RPL13A/RPL13
Gem K
Cyt K
Gem L
Cyt L

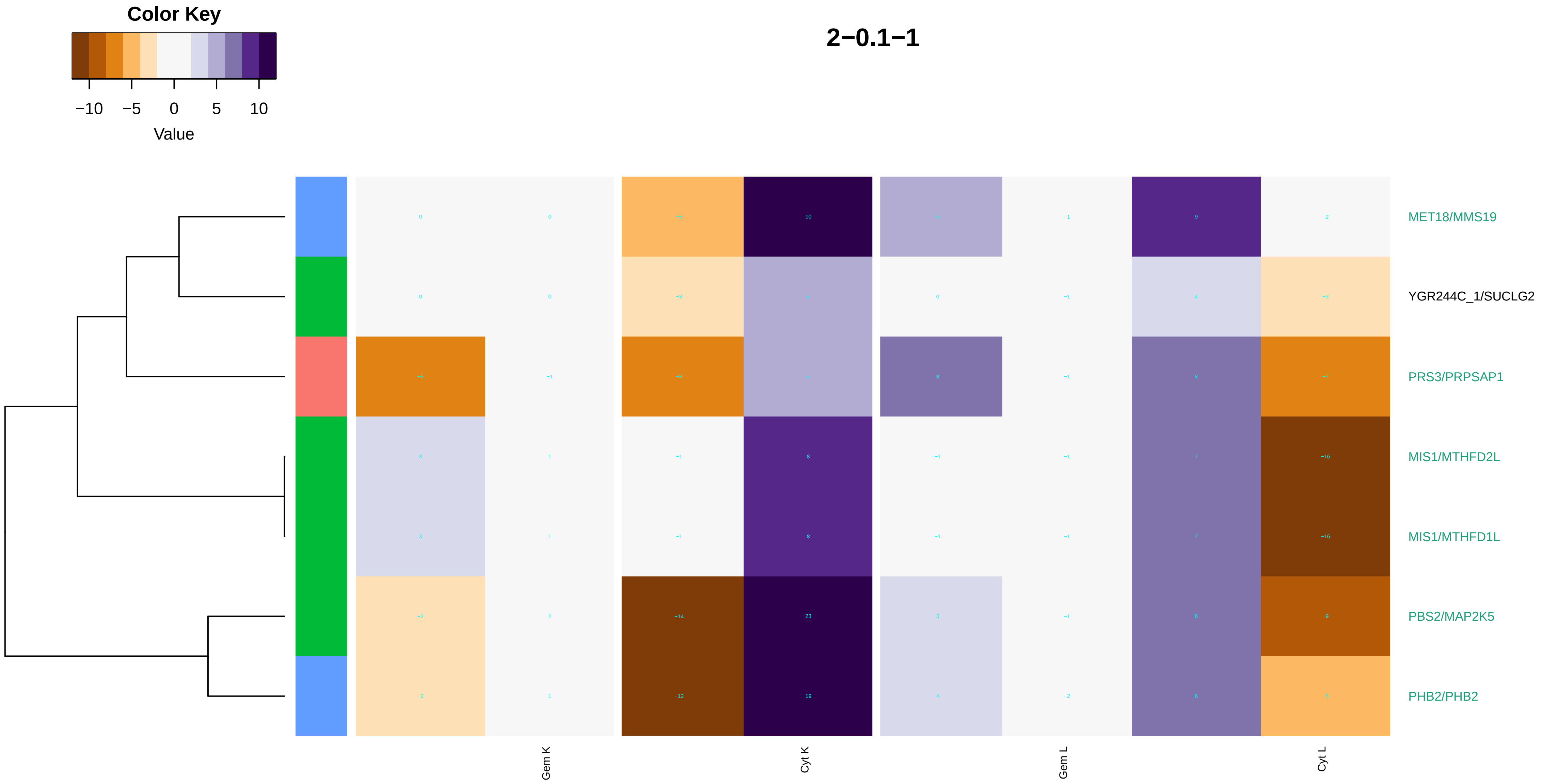

Color Key
−10 −5 0 5 10
Value
2−0.1−1
MET18/MMS19
YGR244C_1/SUCLG2
PRS3/PRPSAP1
MIS1/MTHFD2L
MIS1/MTHFD1L
PBS2/MAP2K5
PHB2/PHB2
Gem K
Cyt K
Gem L
Cyt L

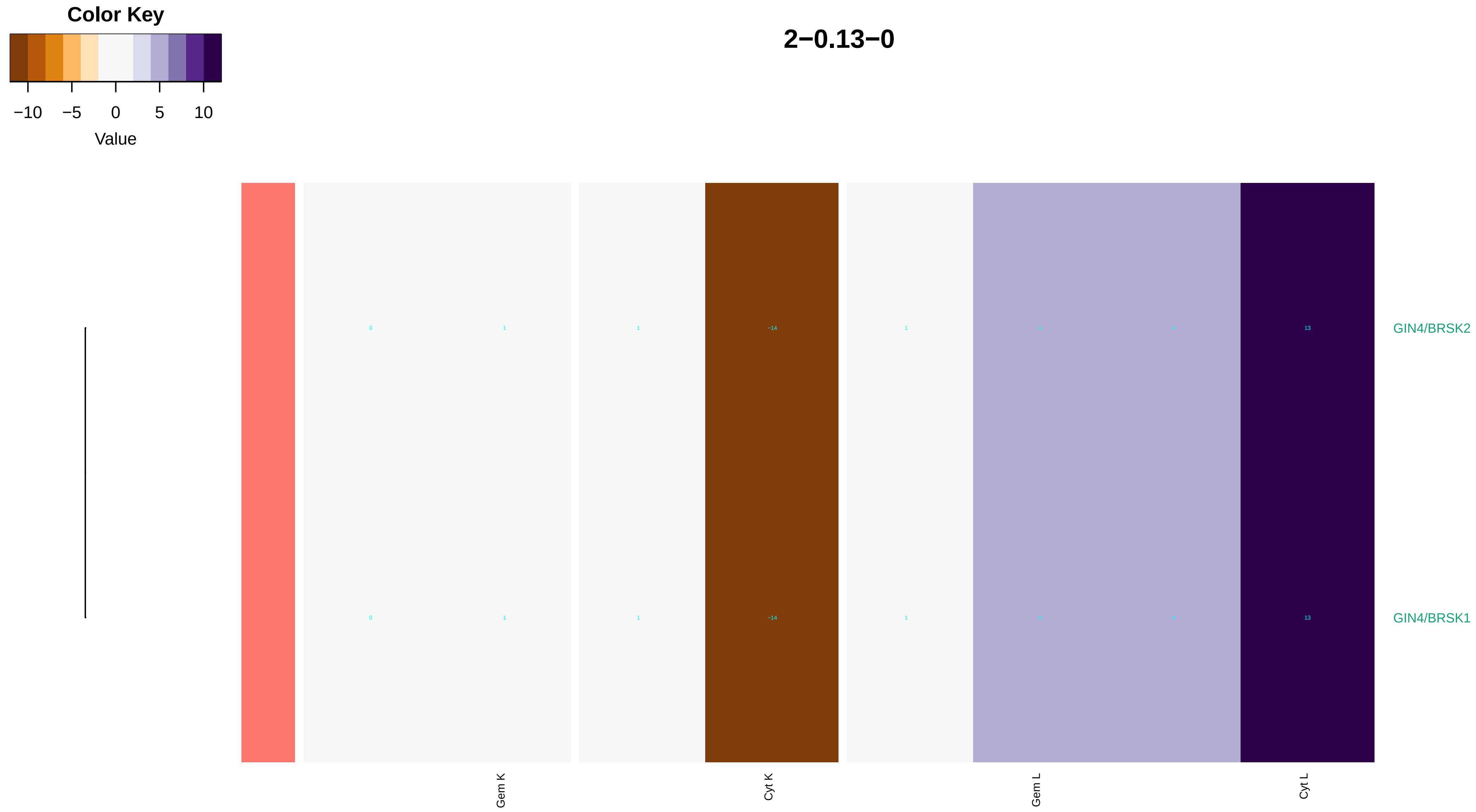
Color Key
−10
−5
0
5
10
Value
2−0.13−0
0
1
1
−14
1
4
4
13
GIN4/BRSK2
0
1
1
−14
1
4
4
13
GIN4/BRSK1
Gem K
Cyt K
Gem L
Cyt L

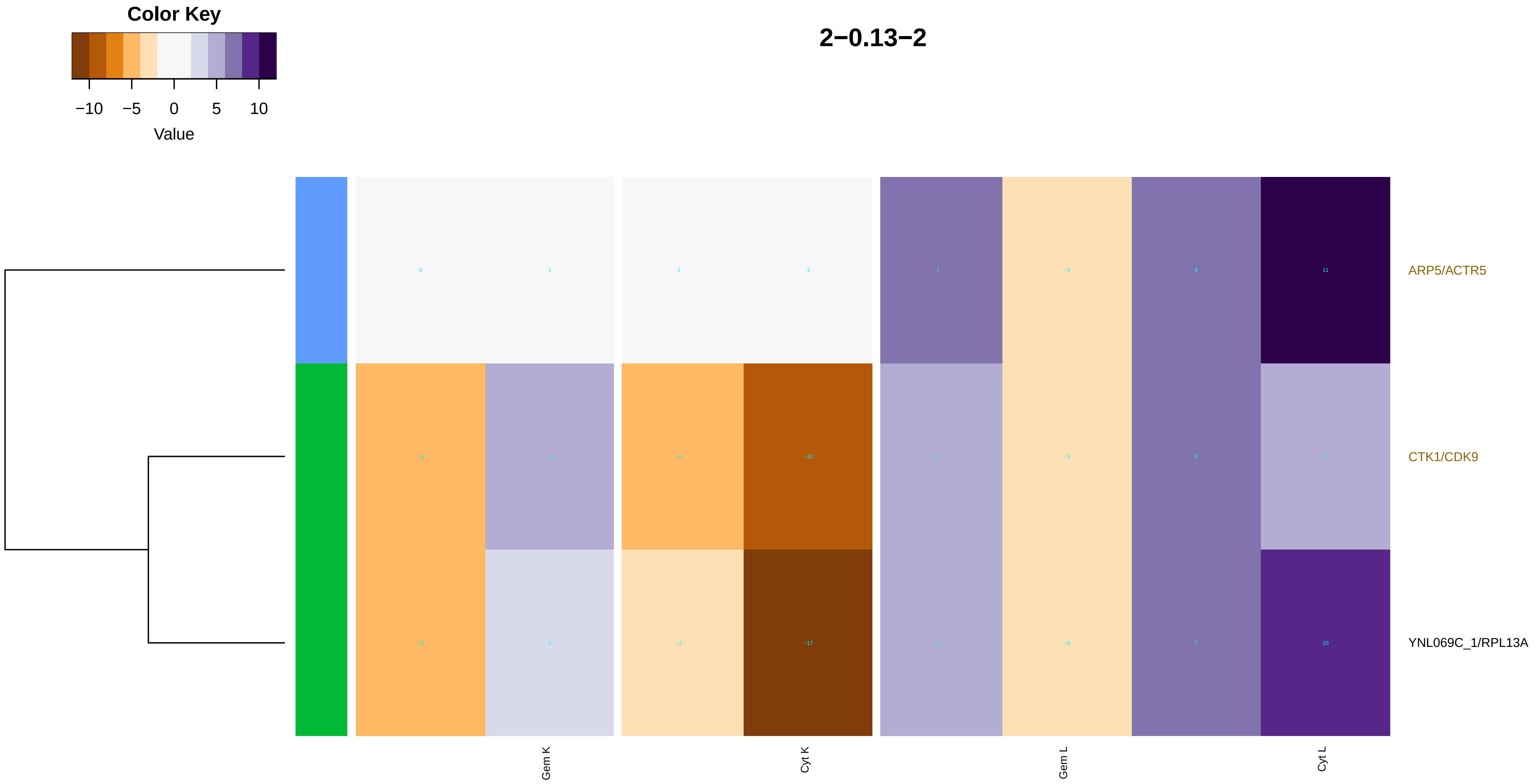
Color Key
−10
−5
0
5
10
Value
2−0.13−2
ARP5/ACTR5
CTK1/CDK9
YNL069C_1/RPL13A
Gem K
Cyt K
Gem L
Cyt L

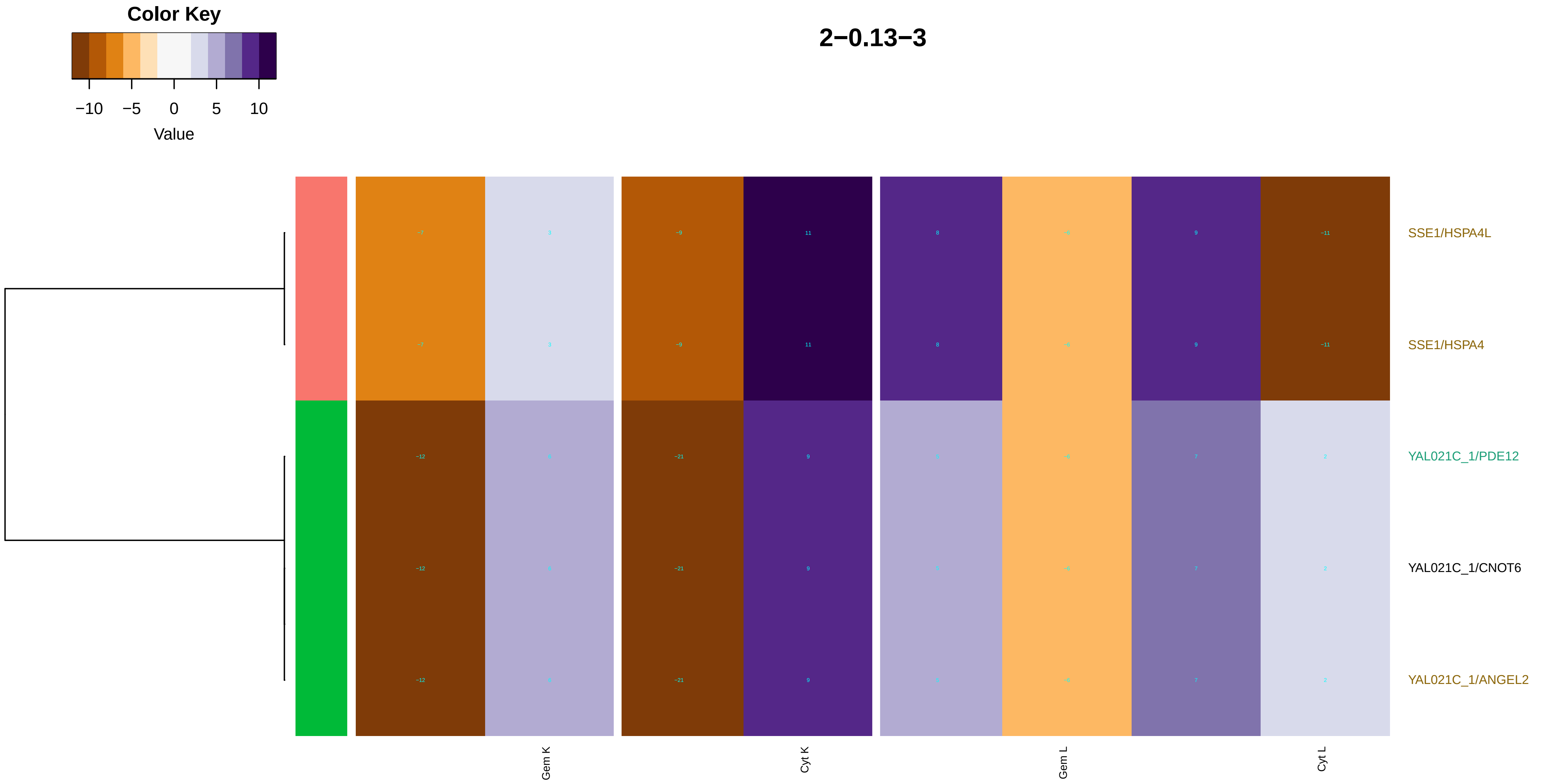
2−0.13−3
Color Key
−10 −5 0 5 10
Value
SSE1/HSPA4L
SSE1/HSPA4
YAL021C_1/PDE12
YAL021C_1/CNOT6
YAL021C_1/ANGEL2
Gem K
Cyt K
Gem L
Cyt L

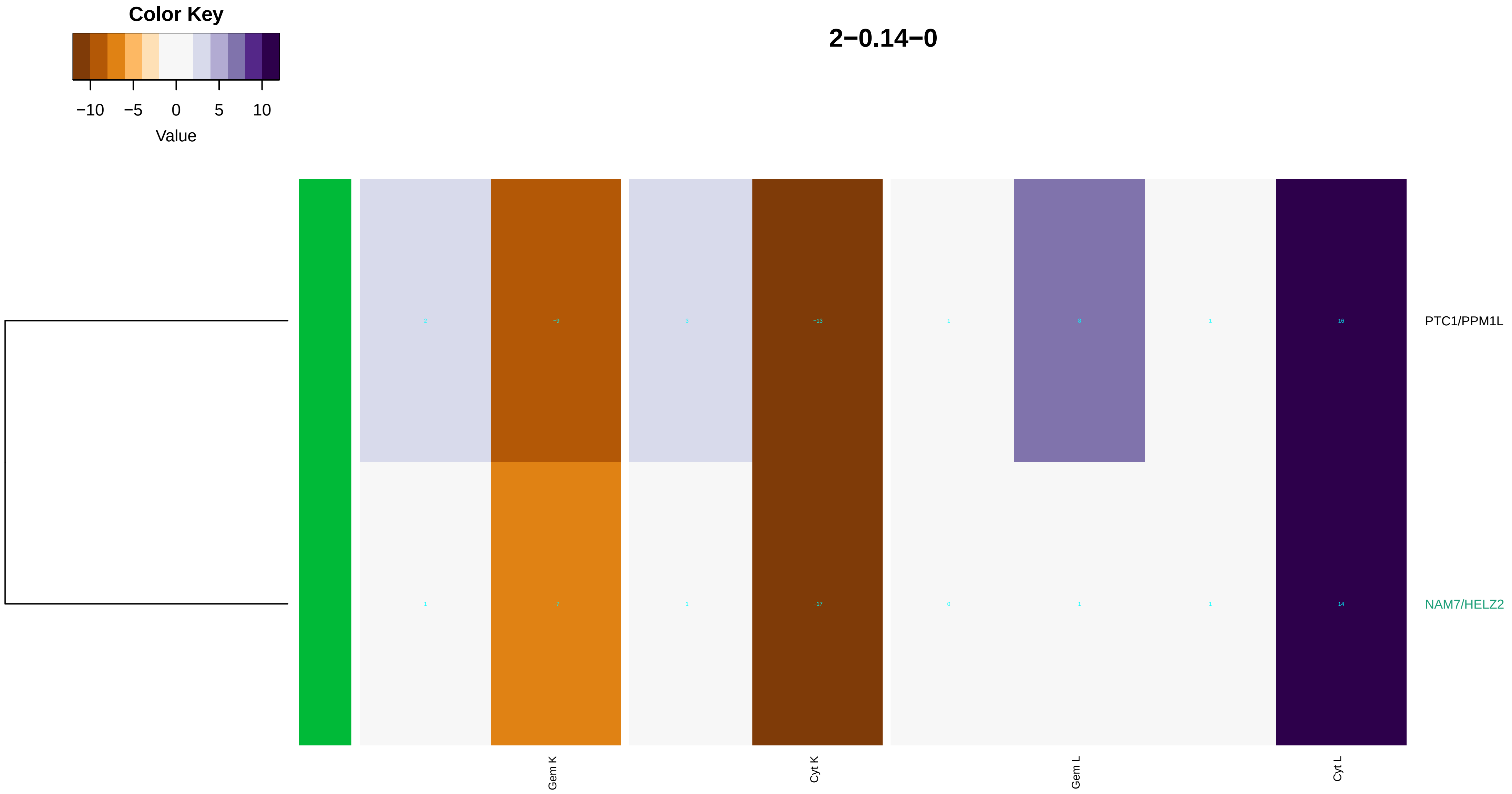
Color Key
−10
−5
0
5
10
Value
2−0.14−0
PTC1/PPM1L
NAM7/HELZ2
Gem K
Cyt K
Gem L
Cyt L

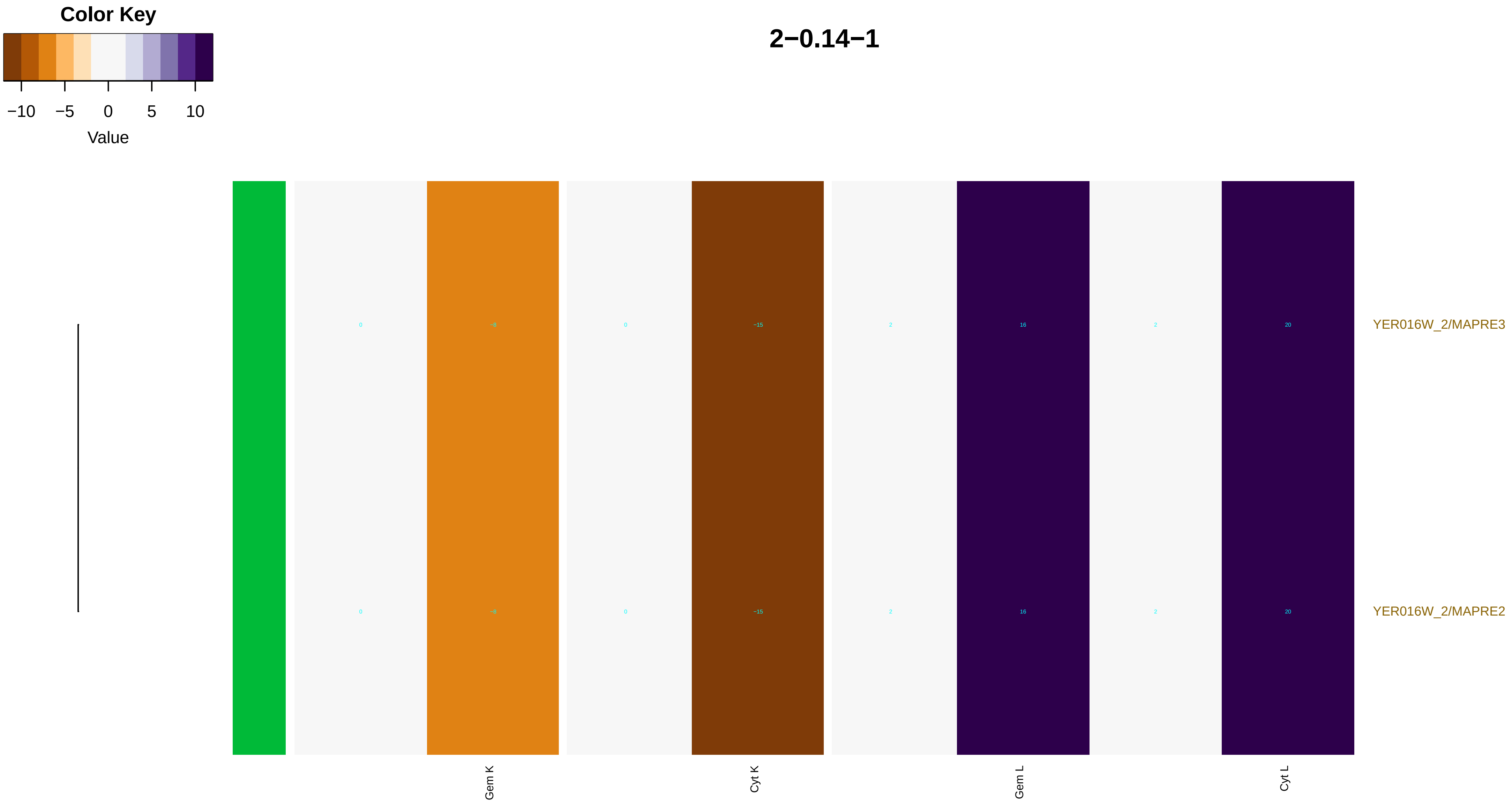

Color Key
−10
−5
0
5
10
Value
2−0.14−1
0
−8
0
−15
2
16
2
20
YER016W_2/MAPRE3
0
−8
0
−15
2
16
2
20
YER016W_2/MAPRE2
Gem K
Cyt K
Gem L
Cyt L

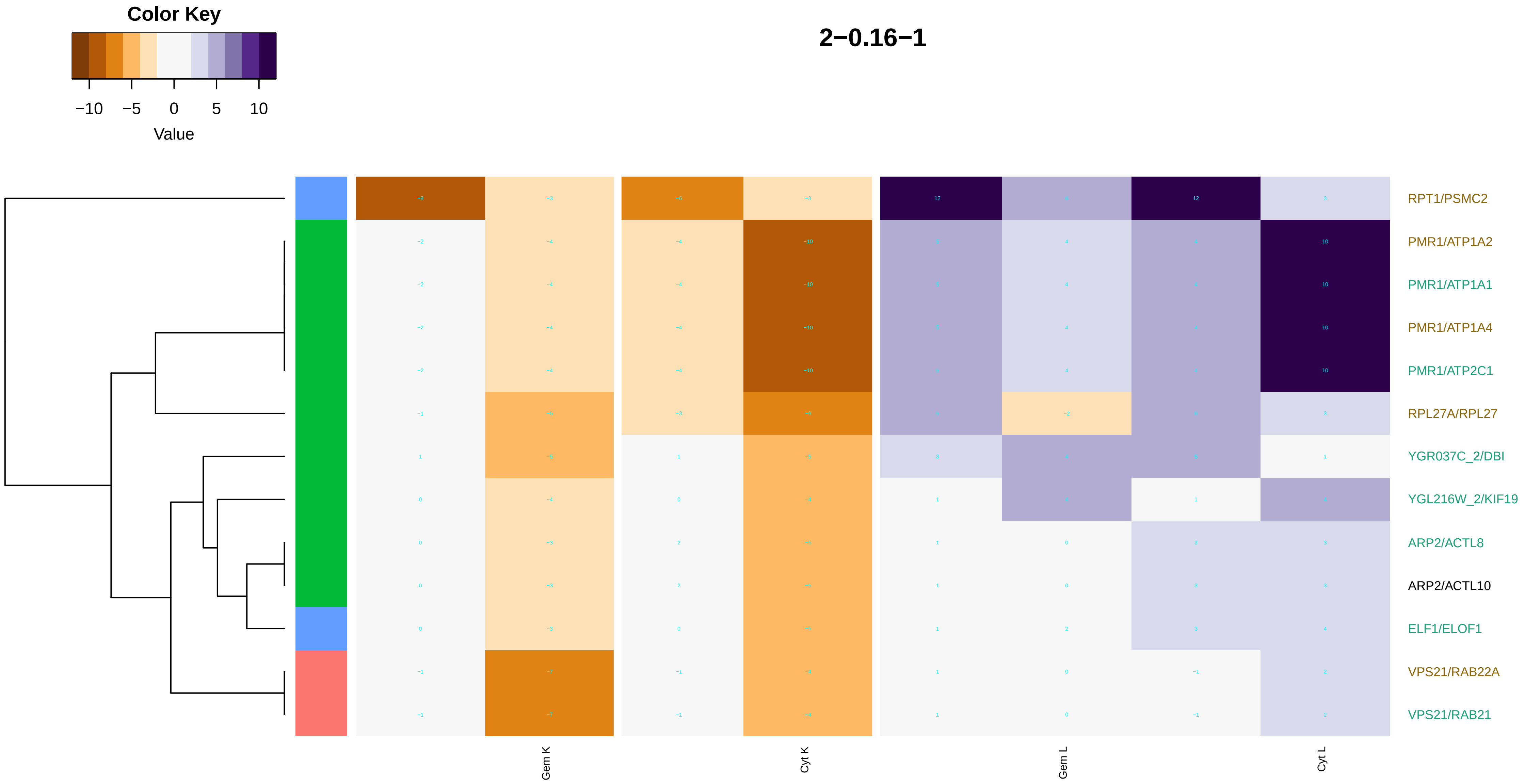

Color Key
−10
−5
0
5
10
Value
2−0.16−1
RPT1/PSMC2
PMR1/ATP1A2
PMR1/ATP1A1
PMR1/ATP1A4
PMR1/ATP2C1
RPL27A/RPL27
YGR037C_2/DBI
YGL216W_2/KIF19
ARP2/ACTL8
ARP2/ACTL10
ELF1/ELOF1
VPS21/RAB22A
VPS21/RAB21
Gem K
Cyt K
Gem L
Cyt L

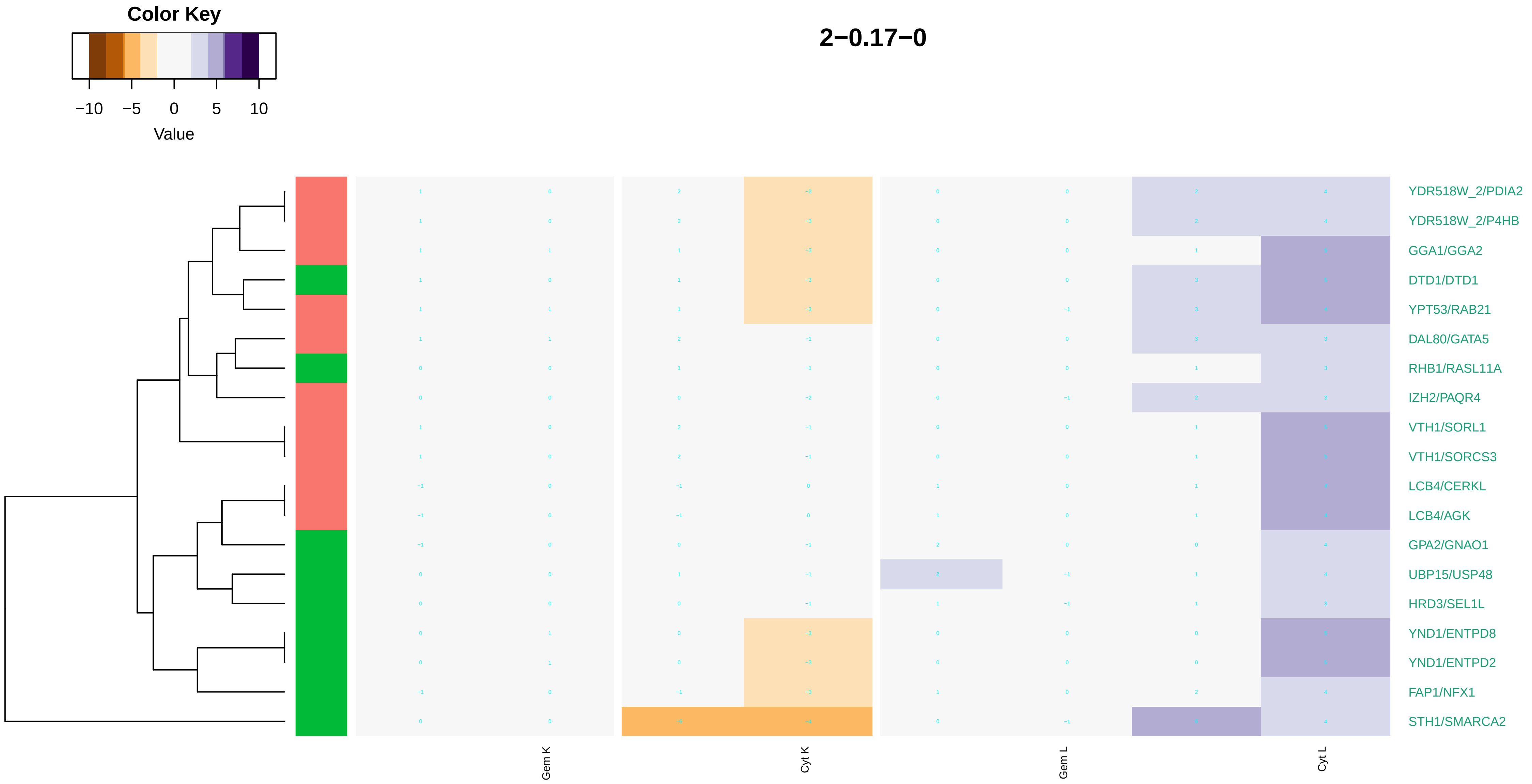

Color Key
−10 −5 0 5 10
Value
2−0.17−0
Gem K
Cyt K
Gem L
Cyt L
YDR518W_2/PDIA2
YDR518W_2/P4HB
GGA1/GGA2
DTD1/DTD1
YPT53/RAB21
DAL80/GATA5
RHB1/RASL11A
IZH2/PAQR4
VTH1/SORL1
VTH1/SORCS3
LCB4/CERKL
LCB4/AGK
GPA2/GNAO1
UBP15/USP48
HRD3/SEL1L
YND1/ENTPD8
YND1/ENTPD2
FAP1/NFX1
STH1/SMARCA2

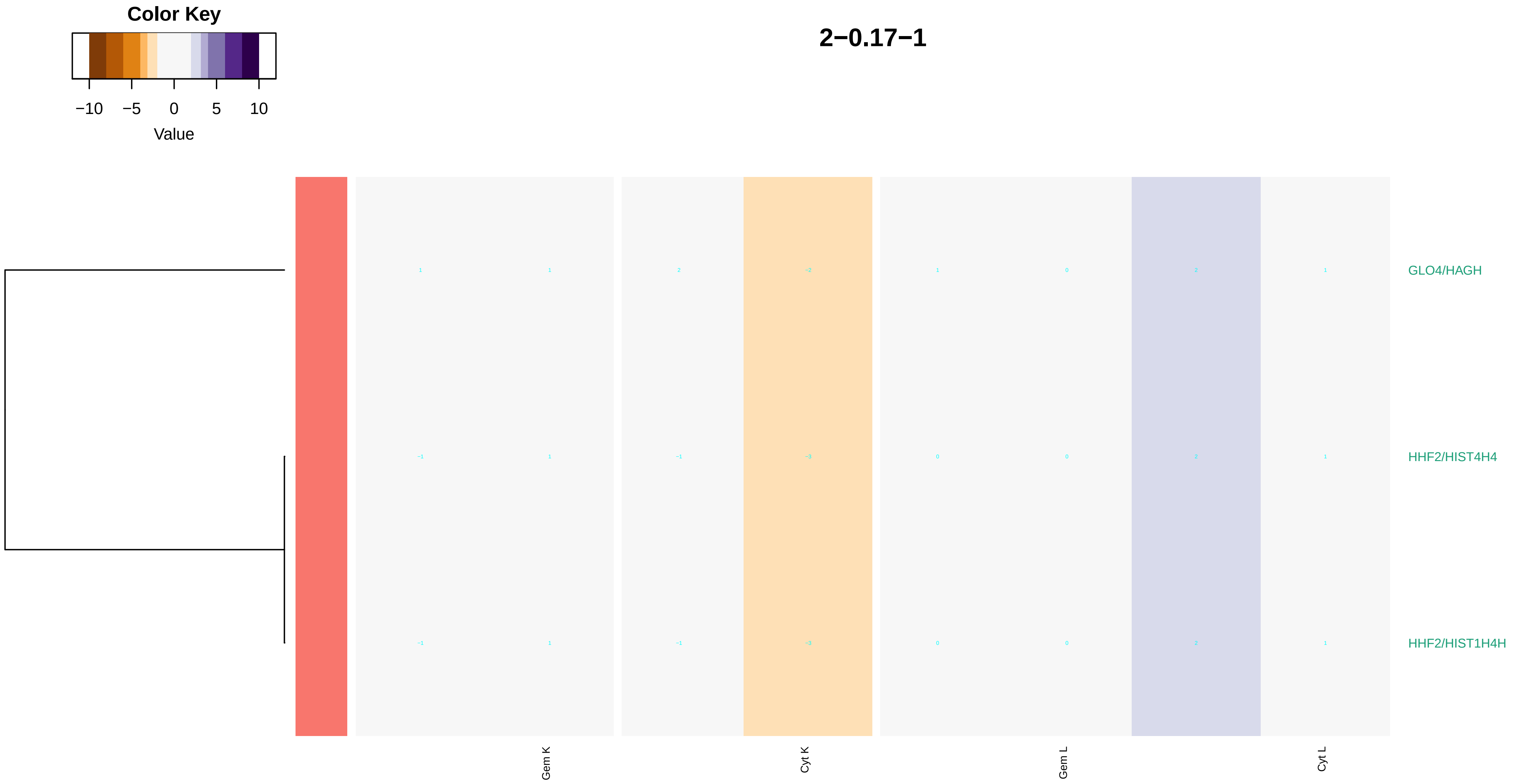
Color Key
−10 −5 0 5 10
Value
2−0.17−1
GLO4/HAGH
HHF2/HIST4H4
HHF2/HIST1H4H
Gem K
Cyt K
Gem L
Cyt L

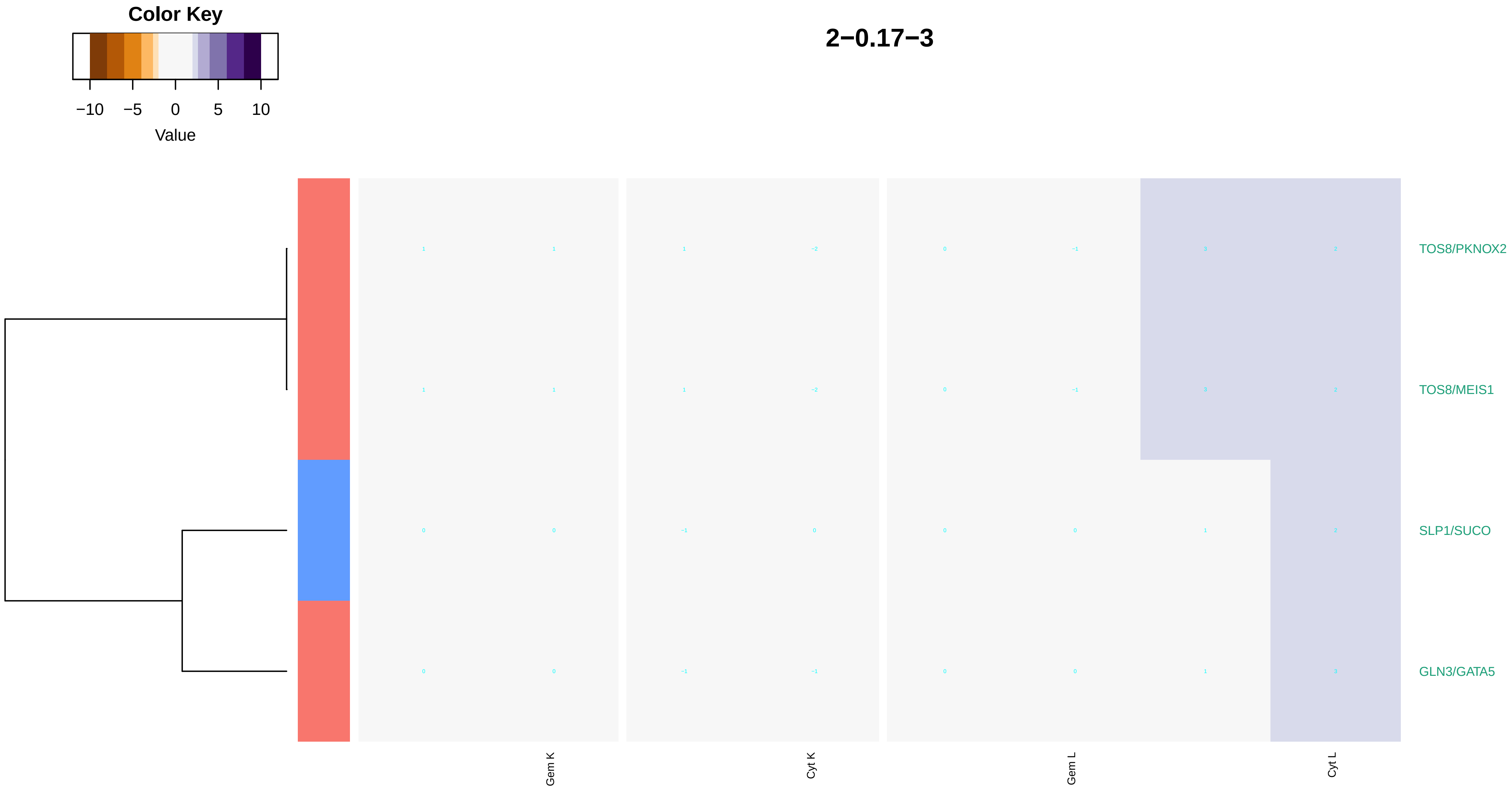
Color Key
−10
−5
0
5
10
Value
2−0.17−3
TOS8/PKNOX2
TOS8/MEIS1
SLP1/SUCO
GLN3/GATA5
Gem K
Cyt K
Gem L
Cyt L

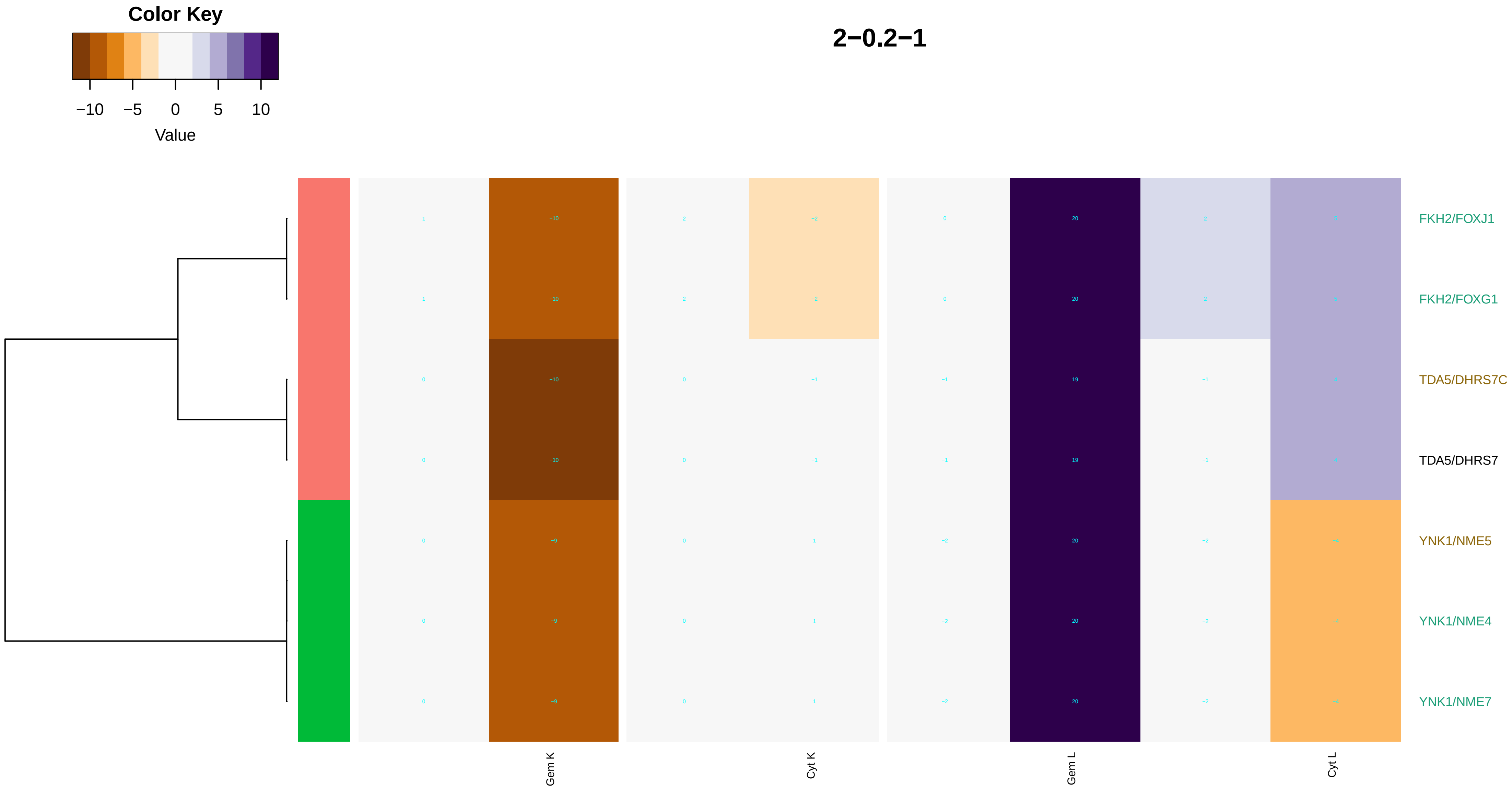
Color Key
−10
−5
0
5
10
Value
2−0.2−1
FKH2/FOXJ1
FKH2/FOXG1
TDA5/DHRS7C
TDA5/DHRS7
YNK1/NME5
YNK1/NME4
YNK1/NME7
Gem K
Cyt K
Gem L
Cyt L

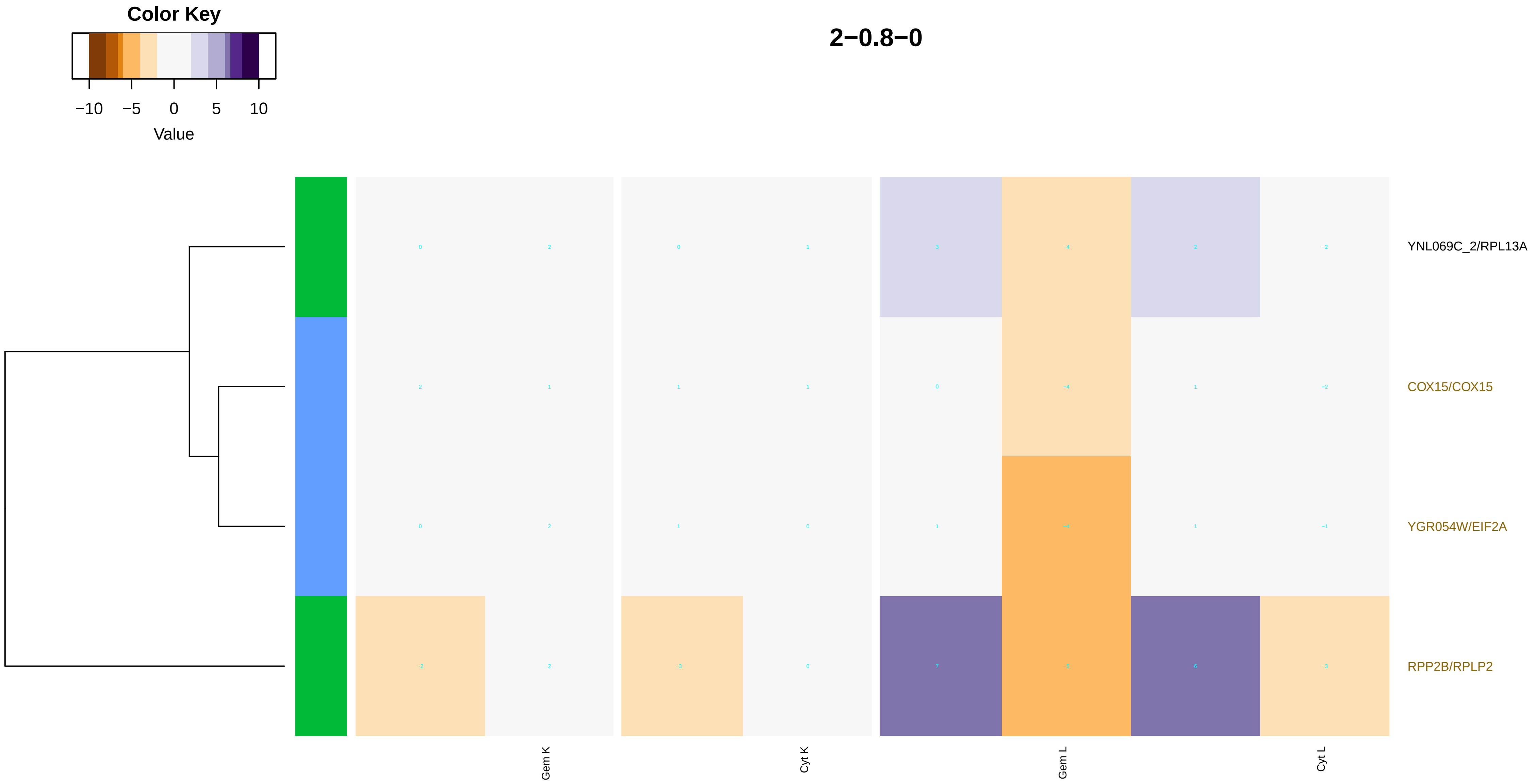
Color Key
−10
−5
0
5
10
Value
2−0.8−0
YNL069C_2/RPL13A
COX15/COX15
YGR054W/EIF2A
RPP2B/RPLP2
Gem K
Cyt K
Gem L
Cyt L

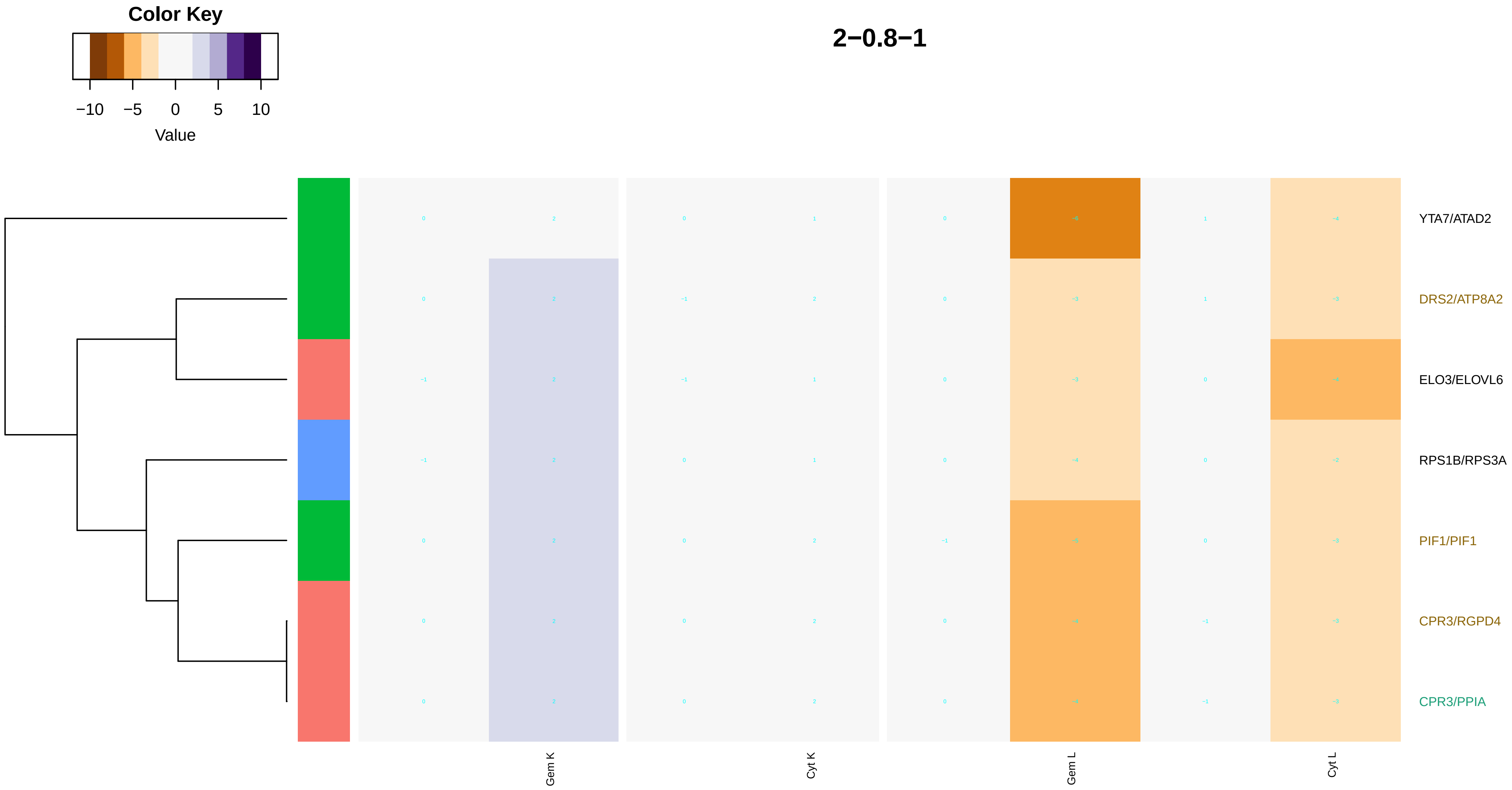

Color Key
−10
−5
0
5
10
Value
2−0.8−1
YTA7/ATAD2
DRS2/ATP8A2
ELO3/ELOVL6
RPS1B/RPS3A
PIF1/PIF1
CPR3/RGPD4
CPR3/PPIA
Gem K
Cyt K
Gem L
Cyt L

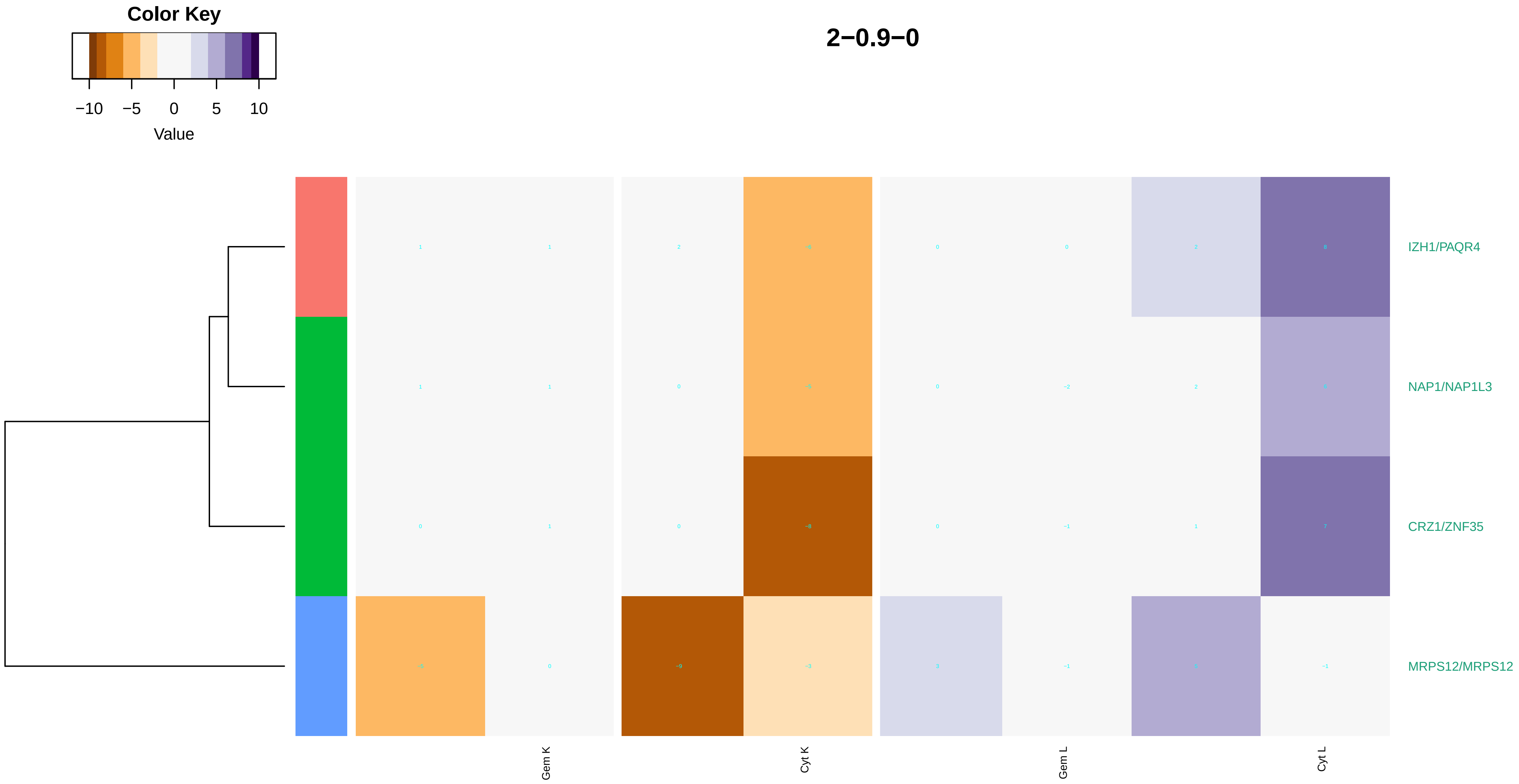
Color Key
−10
−5
0
5
10
Value
2−0.9−0
IZH1/PAQR4
NAP1/NAP1L3
CRZ1/ZNF35
MRPS12/MRPS12
Gem K
Cyt K
Gem L
Cyt L

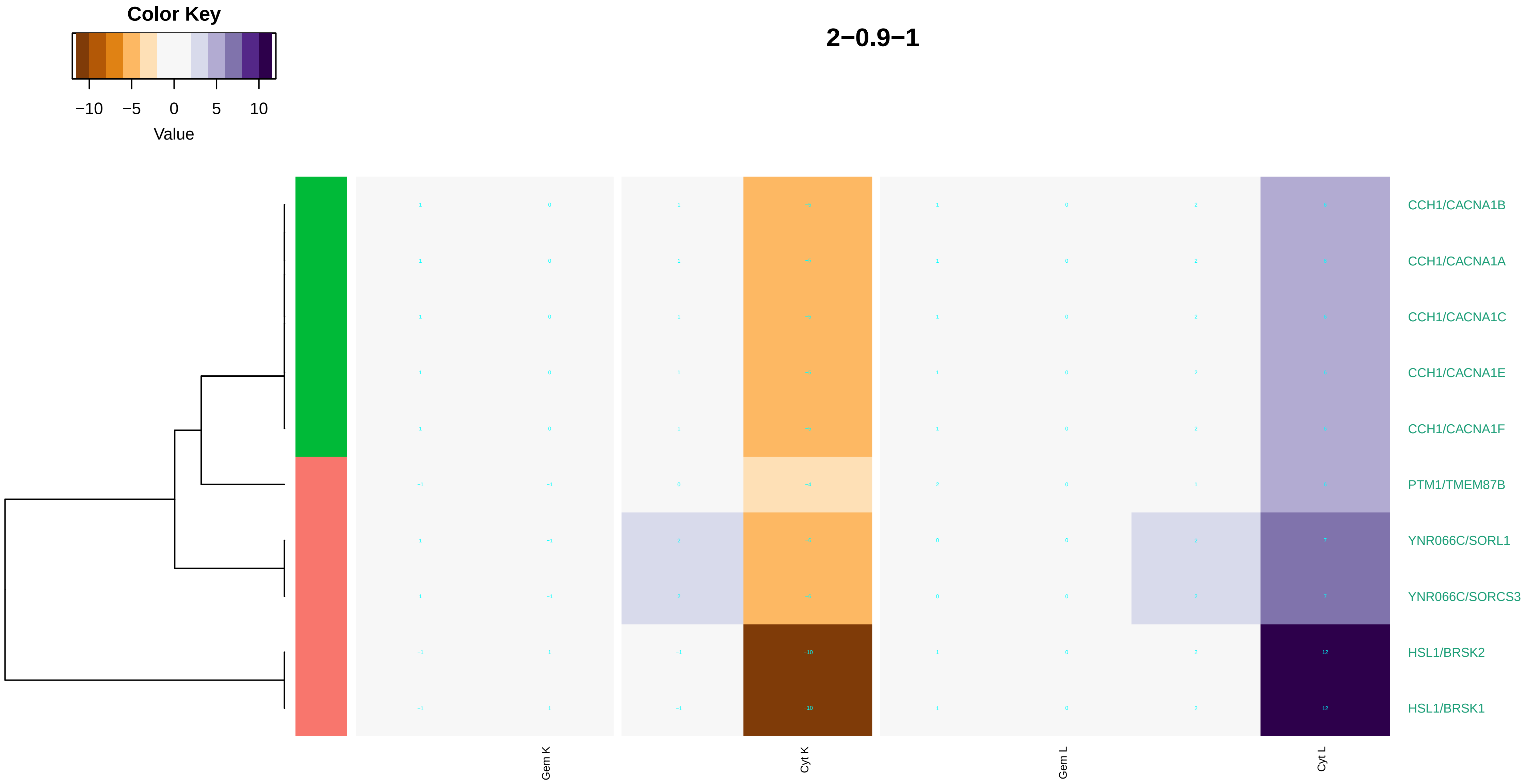

Color Key
−10
−5
0
5
10
Value
2−0.9−1
CCH1/CACNA1B
CCH1/CACNA1A
CCH1/CACNA1C
CCH1/CACNA1E
CCH1/CACNA1F
PTM1/TMEM87B
YNR066C/SORL1
YNR066C/SORCS3
HSL1/BRSK2
HSL1/BRSK1
Gem K
Cyt K
Gem L
Cyt L

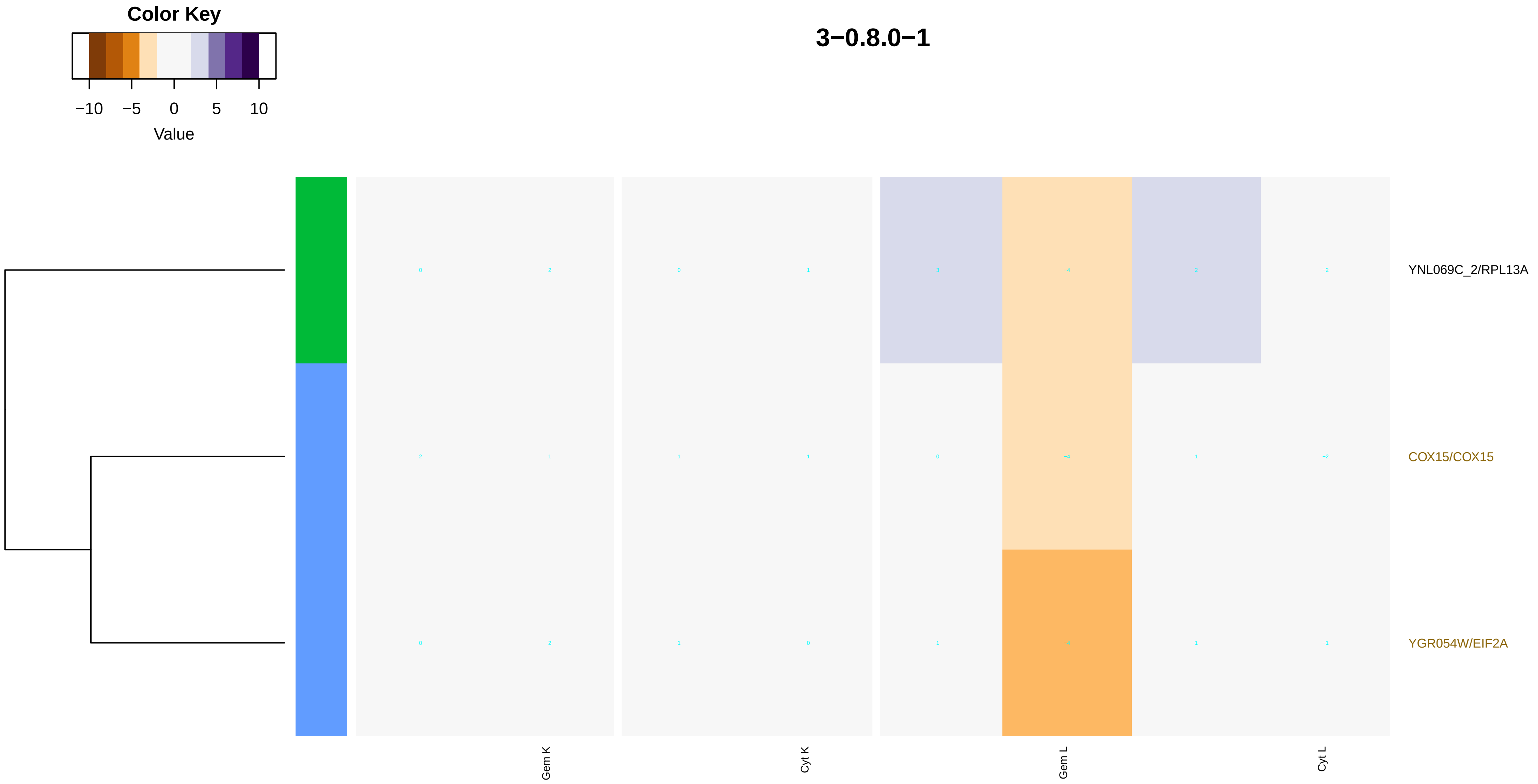

Color Key
−10
−5
0
5
10
Value
3−0.8.0−1
YNL069C_2/RPL13A
COX15/COX15
YGR054W/EIF2A
Gem K
Cyt K
Gem L
Cyt L
