## Additional File 8 for "A humanized yeast phenomic model of deoxycytidine kinase to predict genetic buffering of nucleoside analog cytotoxicity": 19_0327_Lung_to_orig_hmap_Heatmaps.pdf

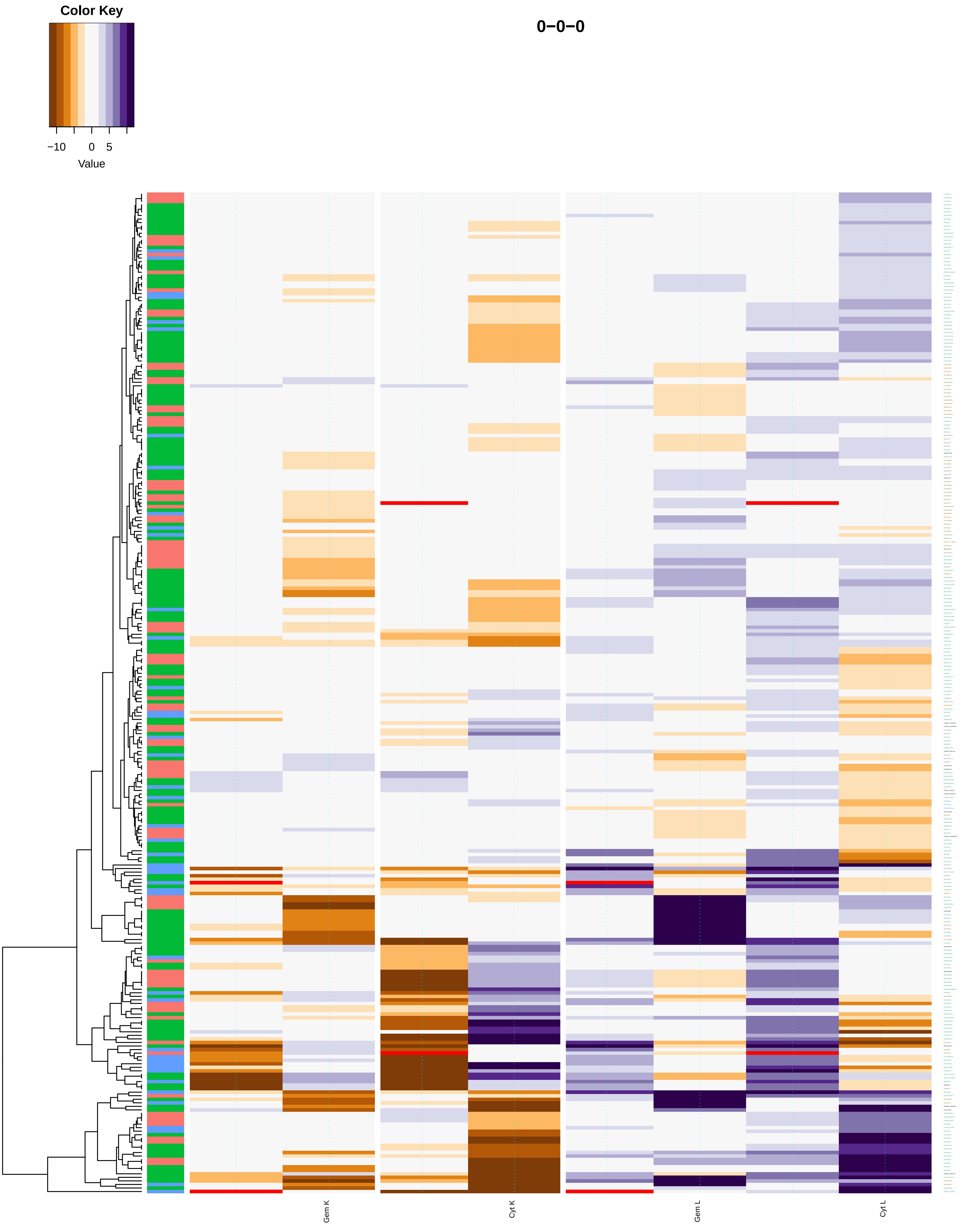

Color Key
−10
0
5
Value
0−0−0
Gem K
Cyt K
Gem L
Cyt L

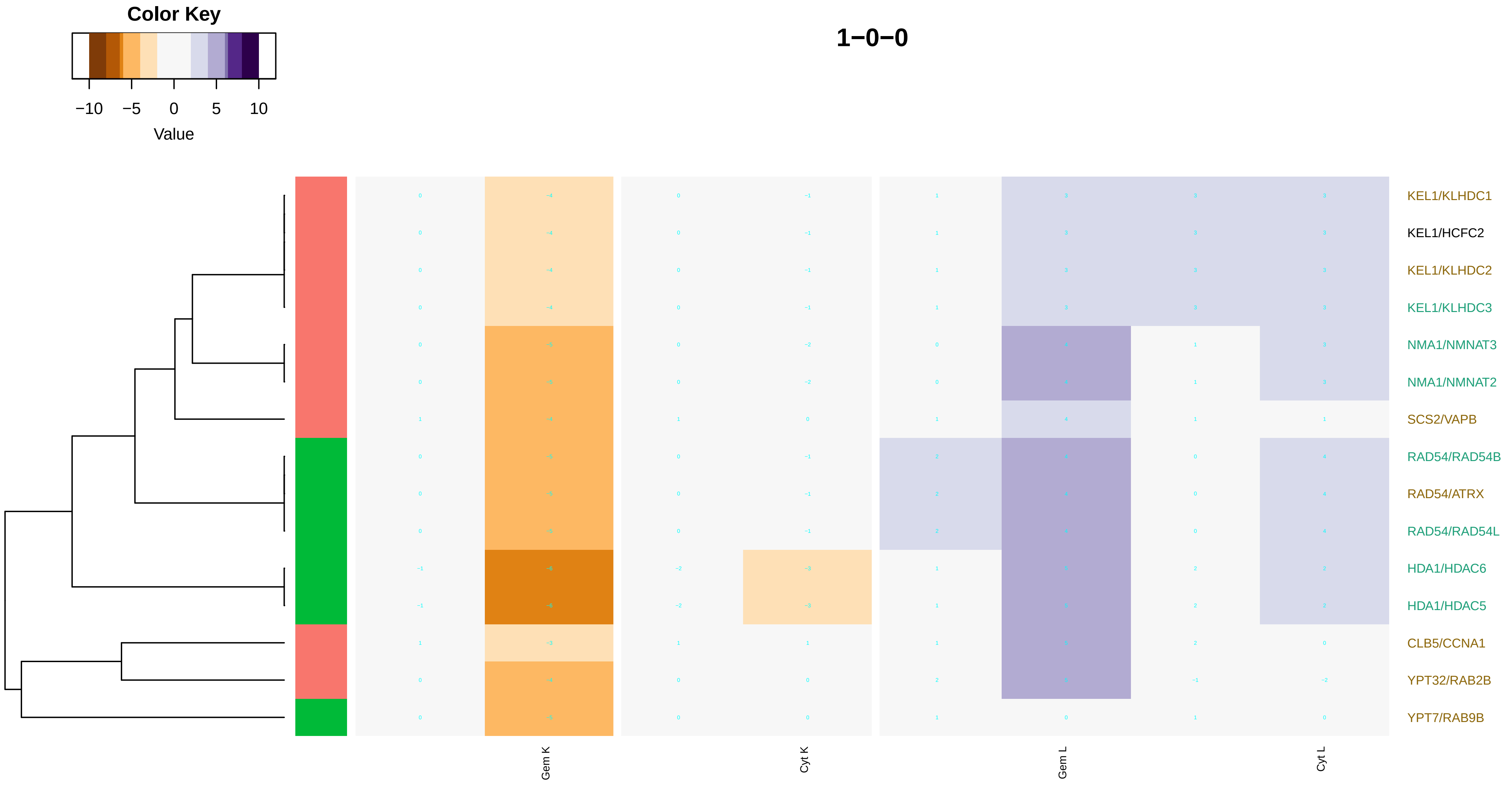

Color Key
−10
−5
0
5
10
Value
1−0−0
KEL1/KLHDC1
KEL1/HCFC2
KEL1/KLHDC2
KEL1/KLHDC3
NMA1/NMNAT3
NMA1/NMNAT2
SCS2/VAPB
RAD54/RAD54B
RAD54/ATRX
RAD54/RAD54L
HDA1/HDAC6
HDA1/HDAC5
CLB5/CCNA1
YPT32/RAB2B
YPT7/RAB9B
Gem K
Cyt K
Gem L
Cyt L

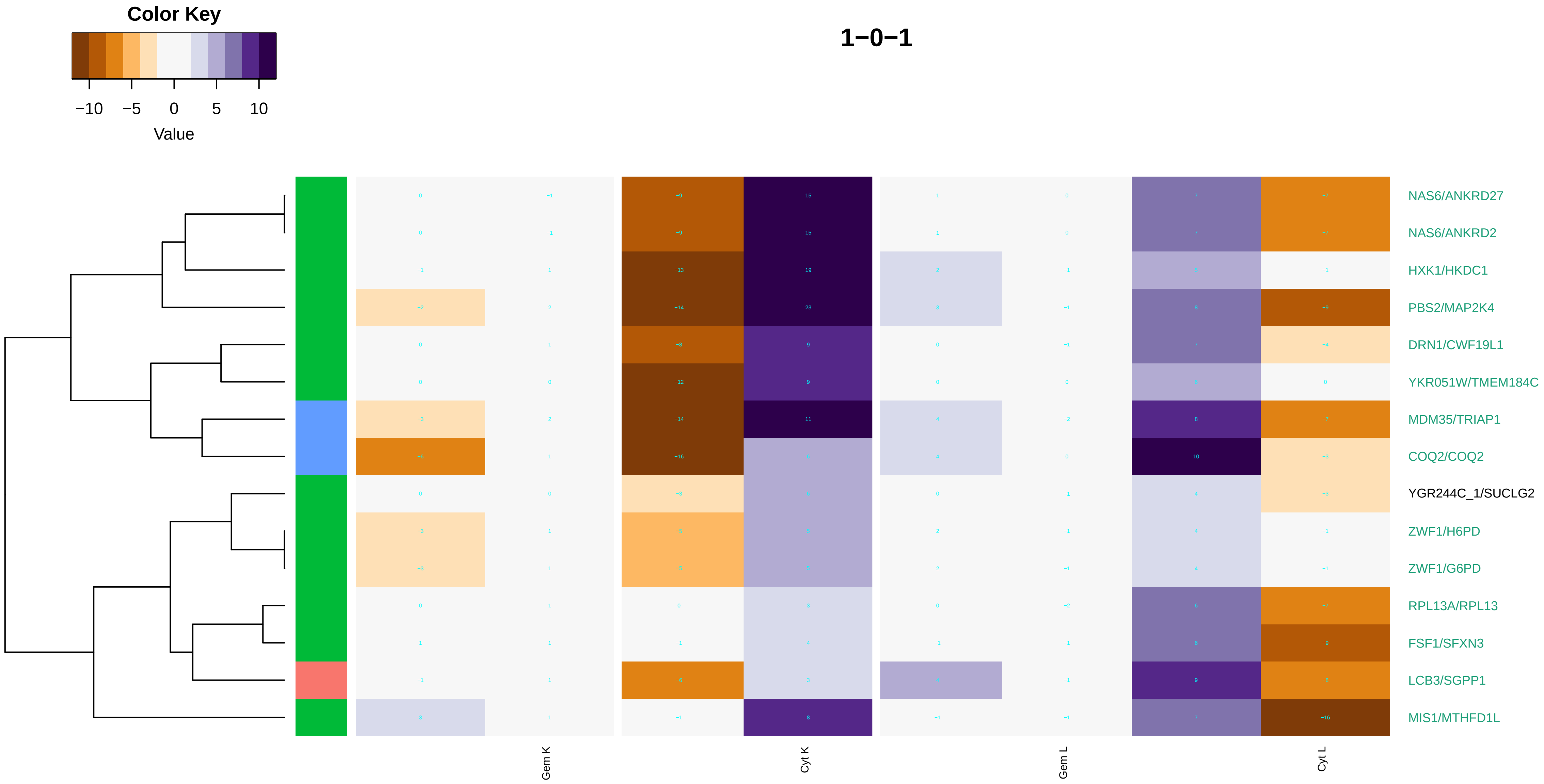

Color Key
−10
−5
0
5
10
Value
1−0−1
NAS6/ANKRD27
NAS6/ANKRD2
HXK1/HKDC1
PBS2/MAP2K4
DRN1/CWF19L1
YKR051W/TMEM184C
MDM35/TRIAP1
COQ2/COQ2
YGR244C_1/SUCLG2
ZWF1/H6PD
ZWF1/G6PD
RPL13A/RPL13
FSF1/SFXN3
LCB3/SGPP1
MIS1/MTHFD1L
Gem K
Cyt K
Gem L
Cyt L

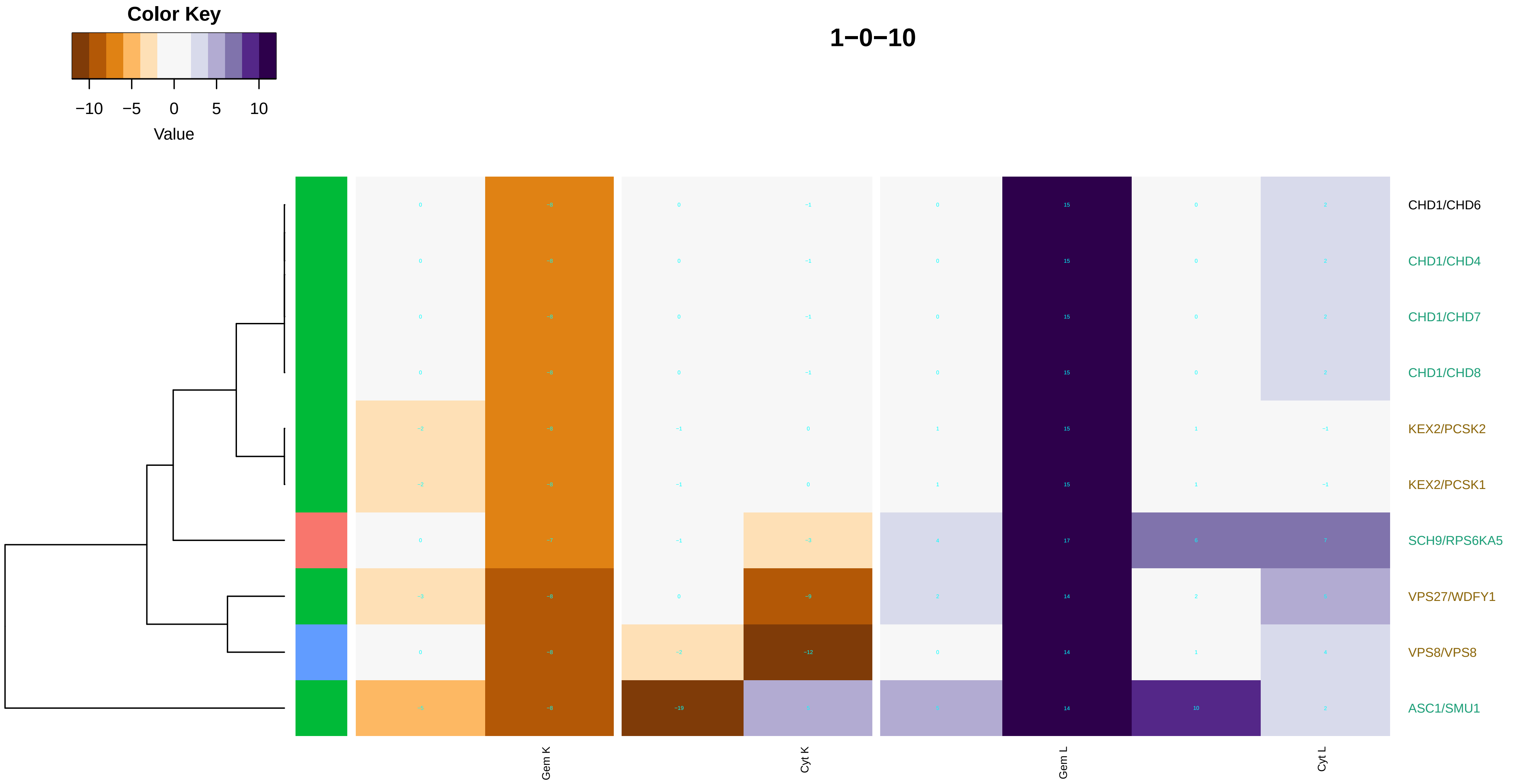
Color Key
−10
−5
0
5
10
Value
1−0−10
CHD1/CHD6
CHD1/CHD4
CHD1/CHD7
CHD1/CHD8
KEX2/PCSK2
KEX2/PCSK1
SCH9/RPS6KA5
VPS27/WDFY1
VPS8/VPS8
ASC1/SMU1
Gem K
Cyt K
Gem L
Cyt L

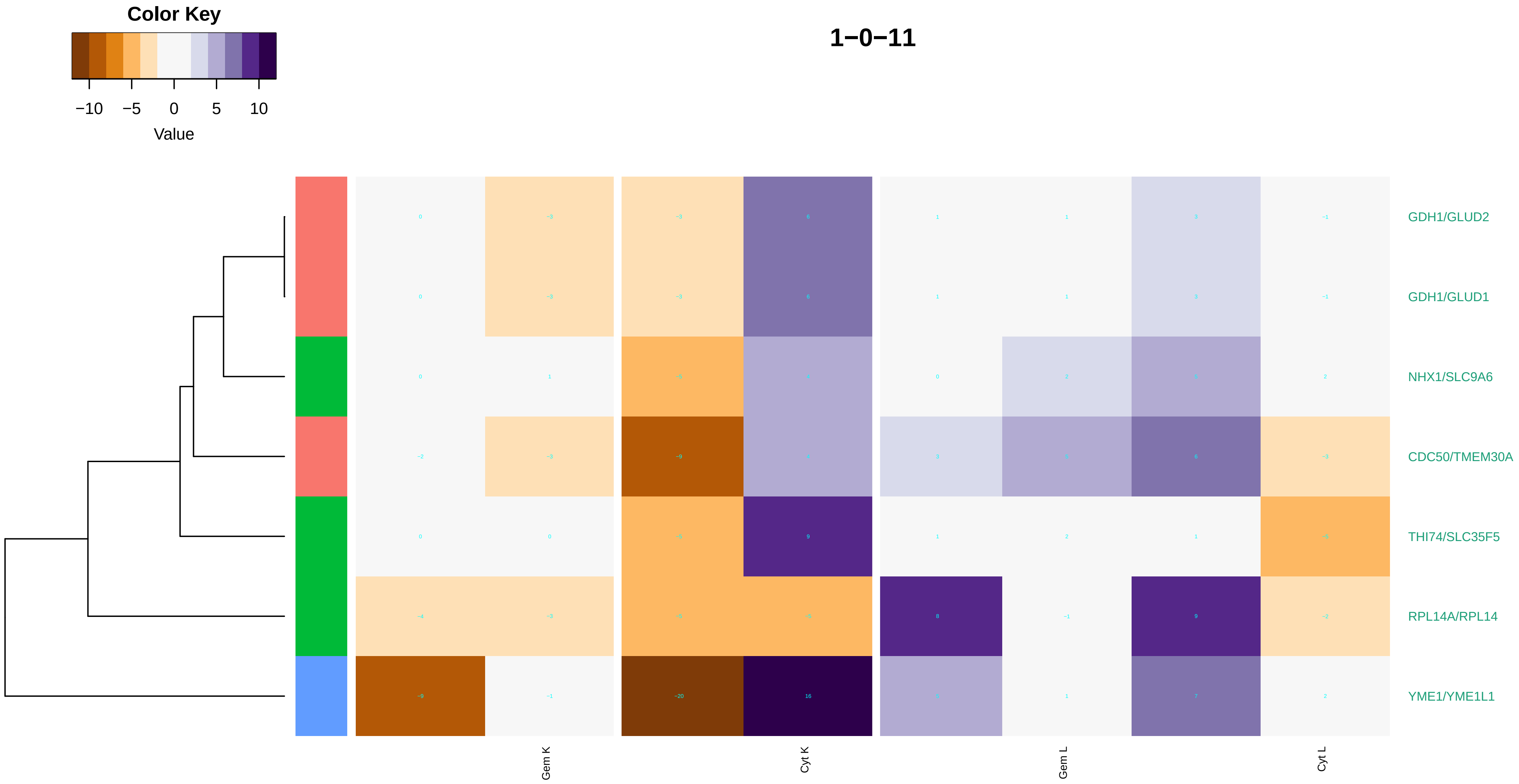

Color Key
−10 −5 0 5 10
Value
1−0−11
GDH1/GLUD2
GDH1/GLUD1
NHX1/SLC9A6
CDC50/TMEM30A
THI74/SLC35F5
RPL14A/RPL14
YME1/YME1L1
Gem K
Cyt K
Gem L
Cyt L

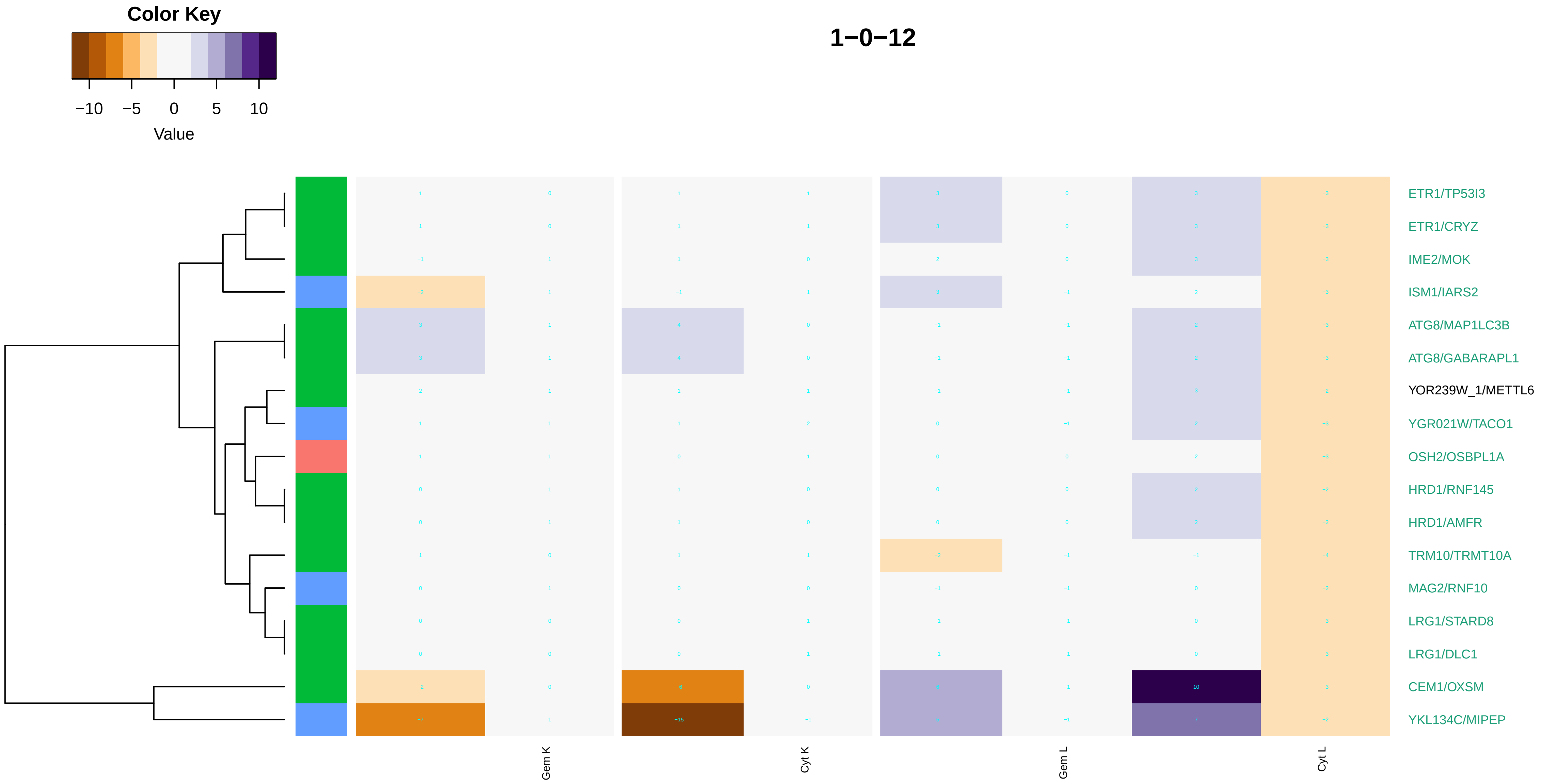
Color Key
−10
−5
0
5
10
Value
1−0−12
ETR1/TP53I3
ETR1/CRYZ
IME2/MOK
ISM1/IARS2
ATG8/MAP1LC3B
ATG8/GABARAPL1
YOR239W_1/METTL6
YGR021W/TACO1
OSH2/OSBPL1A
HRD1/RNF145
HRD1/AMFR
TRM10/TRMT10A
MAG2/RNF10
LRG1/STARD8
LRG1/DLC1
CEM1/OXSM
YKL134C/MIPEP
Gem K
Cyt K
Gem L
Cyt L

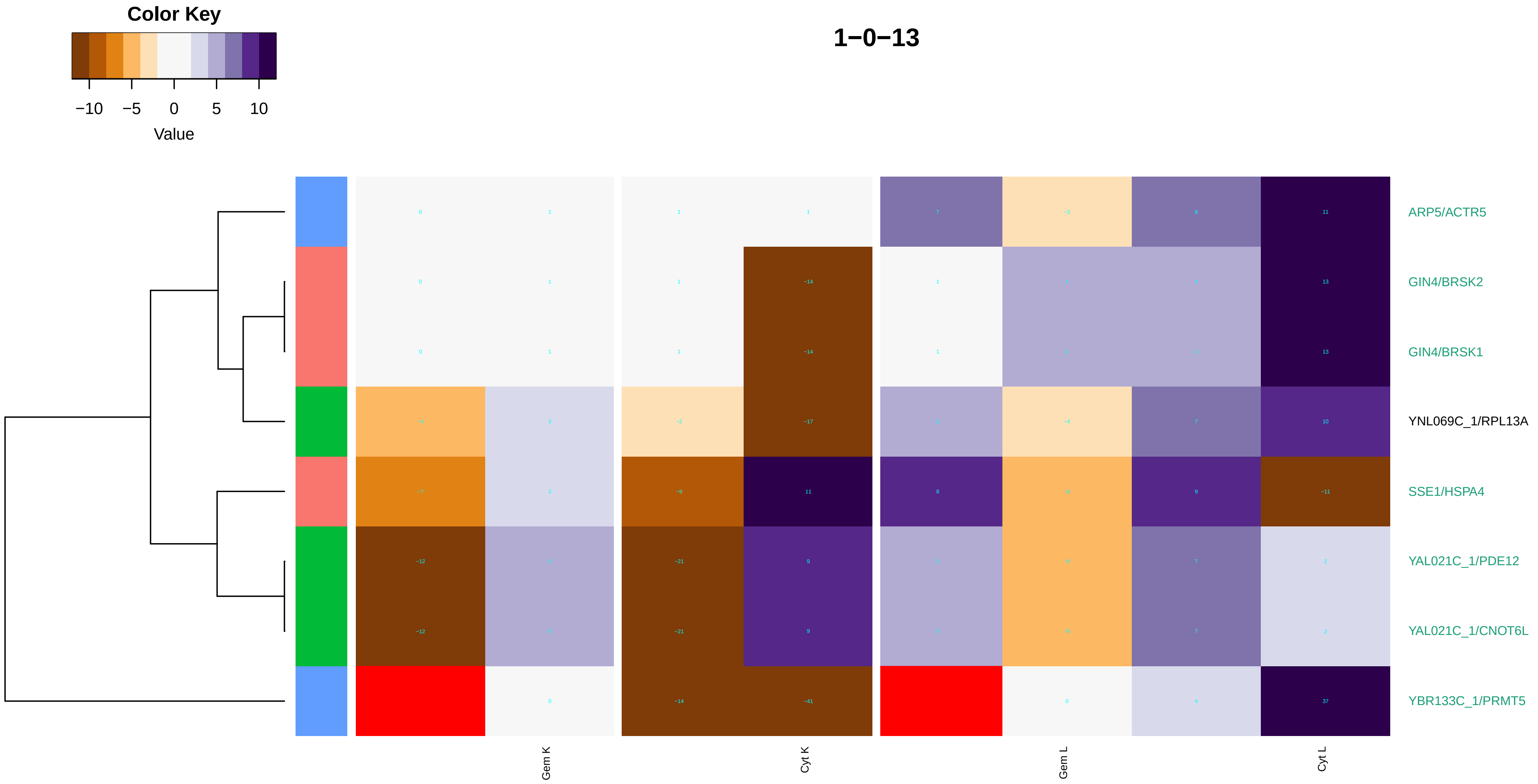
Color Key
−10 −5 0 5 10
Value
1−0−13
ARP5/ACTR5
GIN4/BRSK2
GIN4/BRSK1
YNL069C_1/RPL13A
SSE1/HSPA4
YAL021C_1/PDE12
YAL021C_1/CNOT6L
YBR133C_1/PRMT5
Gem K
Cyt K
Gem L
Cyt L

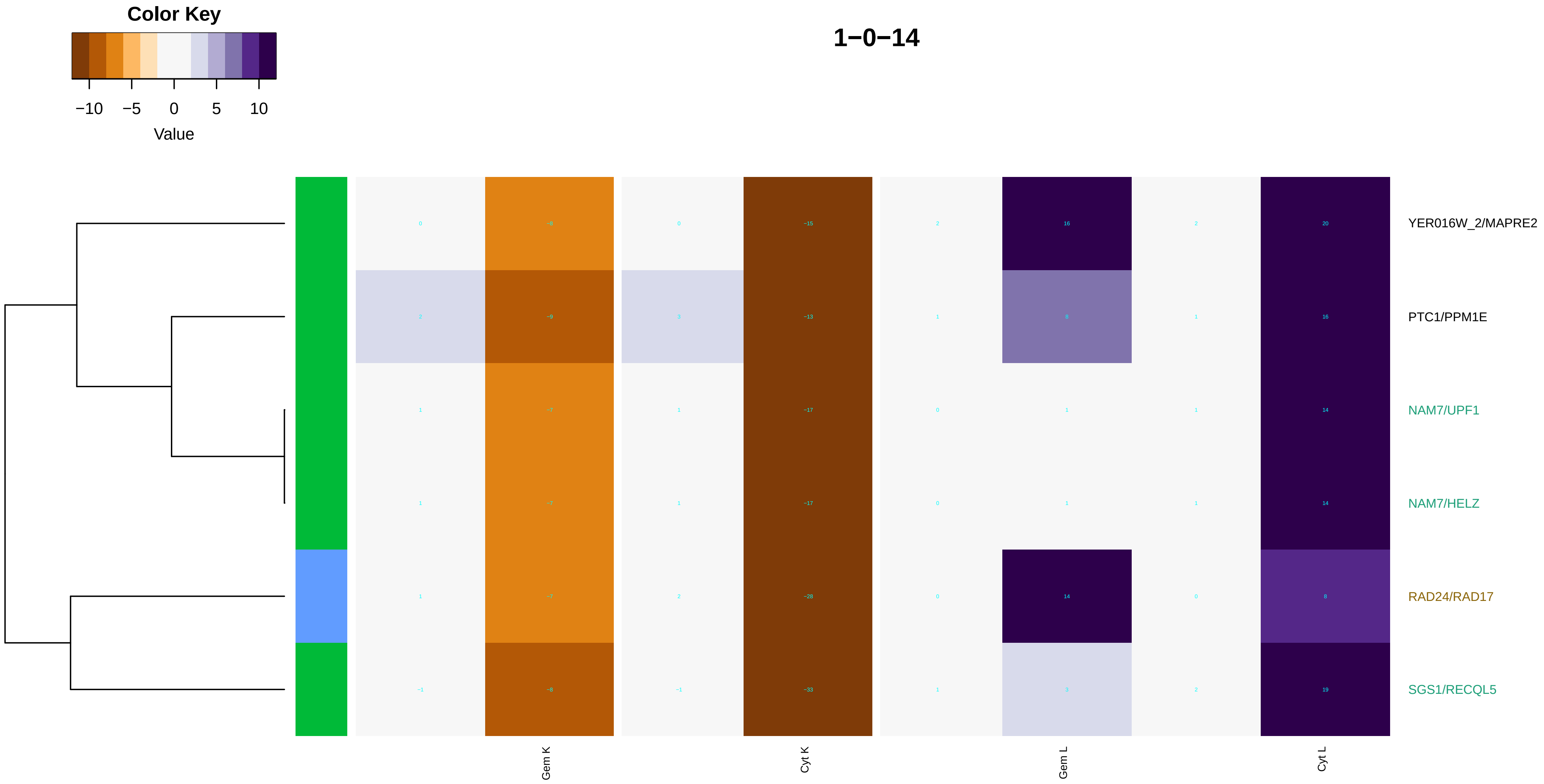
Color Key
−10
−5
0
5
10
Value
1−0−14
YER016W_2/MAPRE2
PTC1/PPM1E
NAM7/UPF1
NAM7/HELZ
RAD24/RAD17
SGS1/RECQL5
Gem K
Cyt K
Gem L
Cyt L

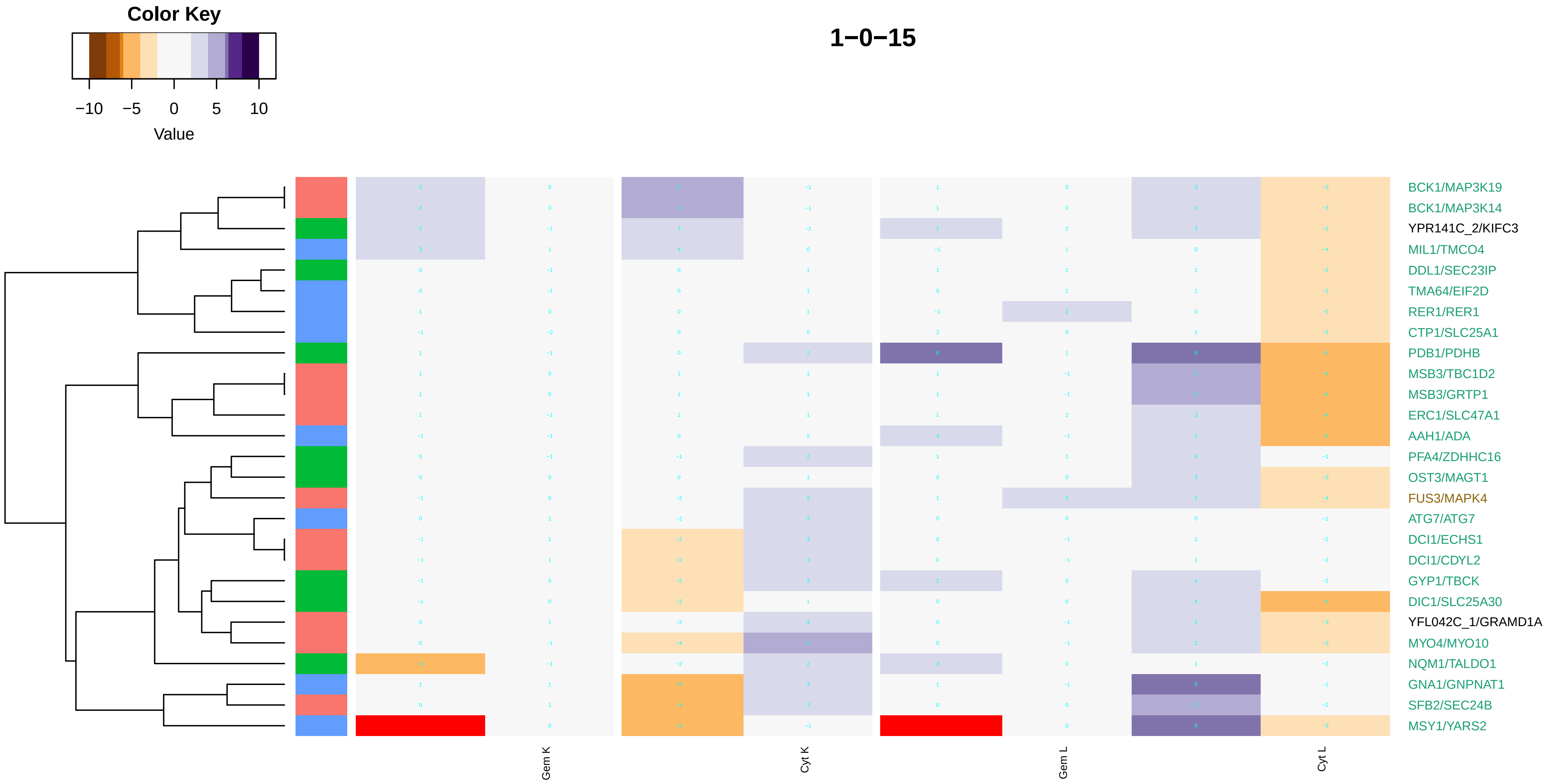

Color Key
−10 −5 0 5 10
Value
1−0−15
BCK1/MAP3K19
BCK1/MAP3K14
YPR141C_2/KIFC3
MIL1/TMCO4
DDL1/SEC23IP
TMA64/EIF2D
RER1/RER1
CTP1/SLC25A1
PDB1/PDHB
MSB3/TBC1D2
MSB3/GRTP1
ERC1/SLC47A1
AAH1/ADA
PFA4/ZDHHC16
OST3/MAGT1
FUS3/MAPK4
ATG7/ATG7
DCI1/ECHS1
DCI1/CDYL2
GYP1/TBCK
DIC1/SLC25A30
YFL042C_1/GRAMD1A
MYO4/MYO10
NQM1/TALDO1
GNA1/GNPNAT1
SFB2/SEC24B
MSY1/YARS2
Gem K
Cyt K
Gem L
Cyt L

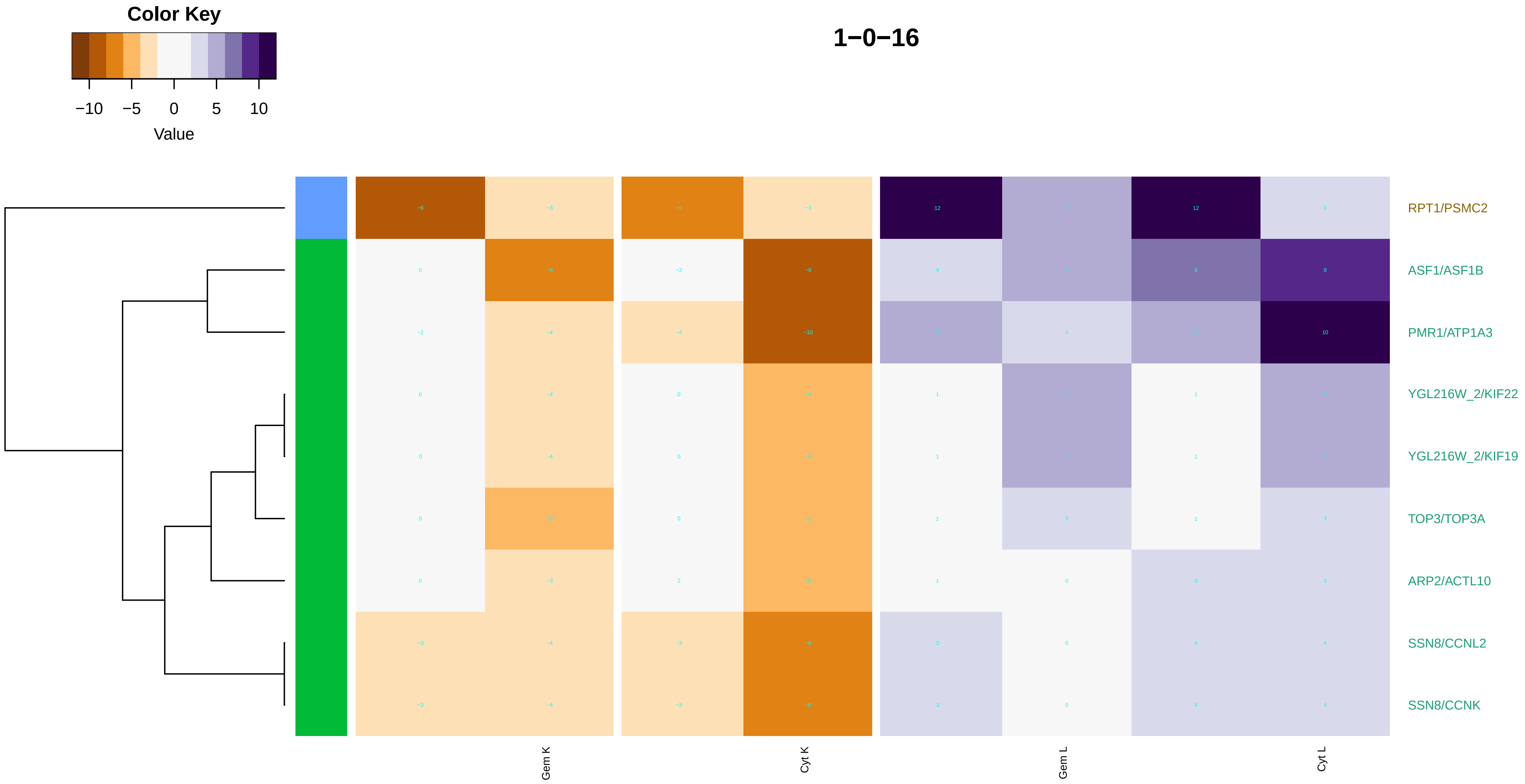
Color Key
−10 −5 0 5 10
Value
1−0−16
RPT1/PSMC2
ASF1/ASF1B
PMR1/ATP1A3
YGL216W_2/KIF22
YGL216W_2/KIF19
TOP3/TOP3A
ARP2/ACTL10
SSN8/CCNL2
SSN8/CCNK
Gem K
Cyt K
Gem L
Cyt L

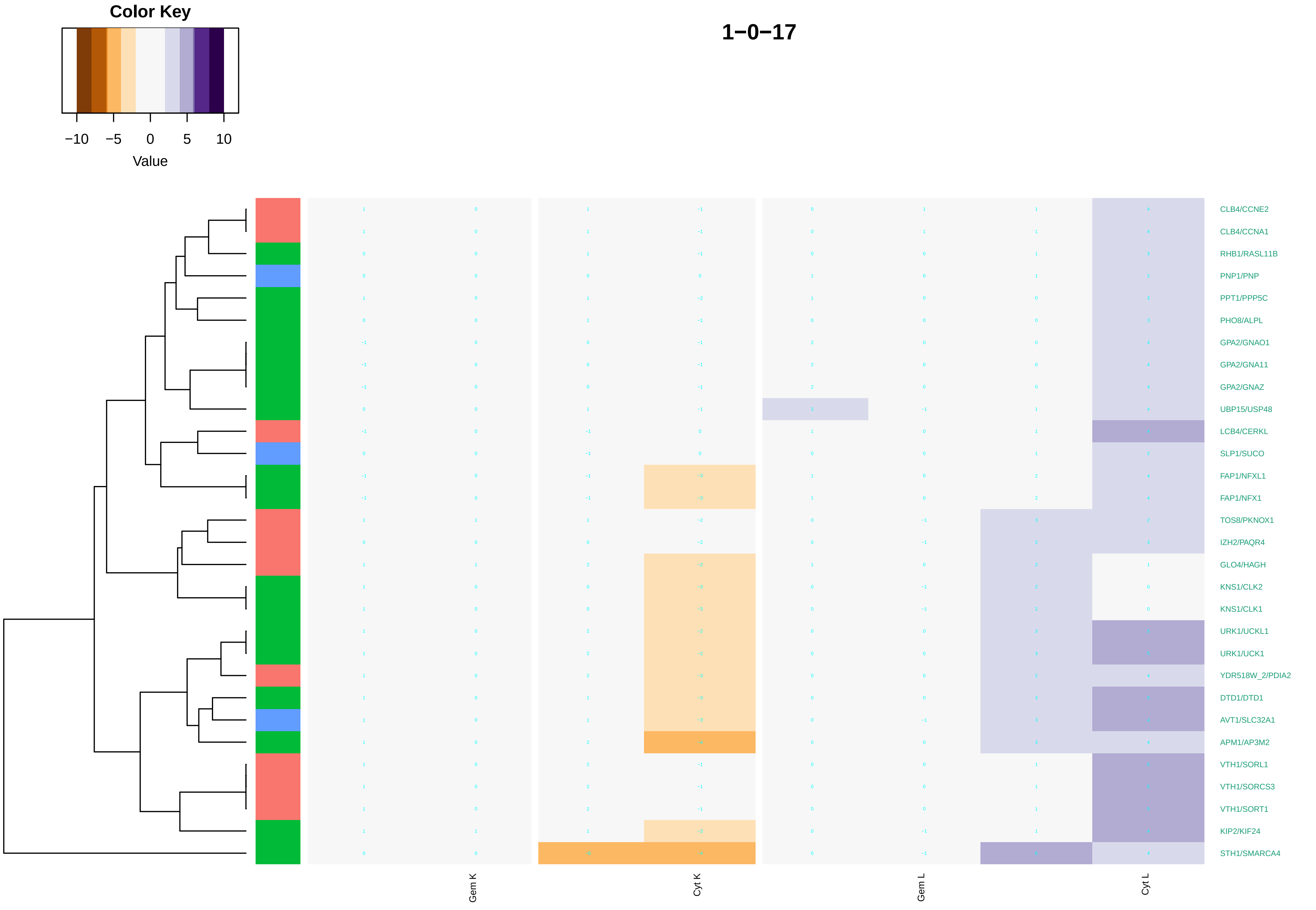

Color Key
−10
−5
0
5
10
Value
1−0−17
CLB4/CCNE2
CLB4/CCNA1
RHB1/RASL11B
PNP1/PNP
PPT1/PPP5C
PHO8/ALPL
GPA2/GNAO1
GPA2/GNA11
GPA2/GNAZ
UBP15/USP48
LCB4/CERKL
SLP1/SUCO
FAP1/NFXL1
FAP1/NFX1
TOS8/PKNOX1
IZH2/PAQR4
GLO4/HAGH
KNS1/CLK2
KNS1/CLK1
URK1/UCKL1
URK1/UCK1
YDR518W_2/PDIA2
DTD1/DTD1
AVT1/SLC32A1
APM1/AP3M2
VTH1/SORL1
VTH1/SORCS3
VTH1/SORT1
KIP2/KIF24
STH1/SMARCA4
Gem K
Cyt K
Gem L
Cyt L

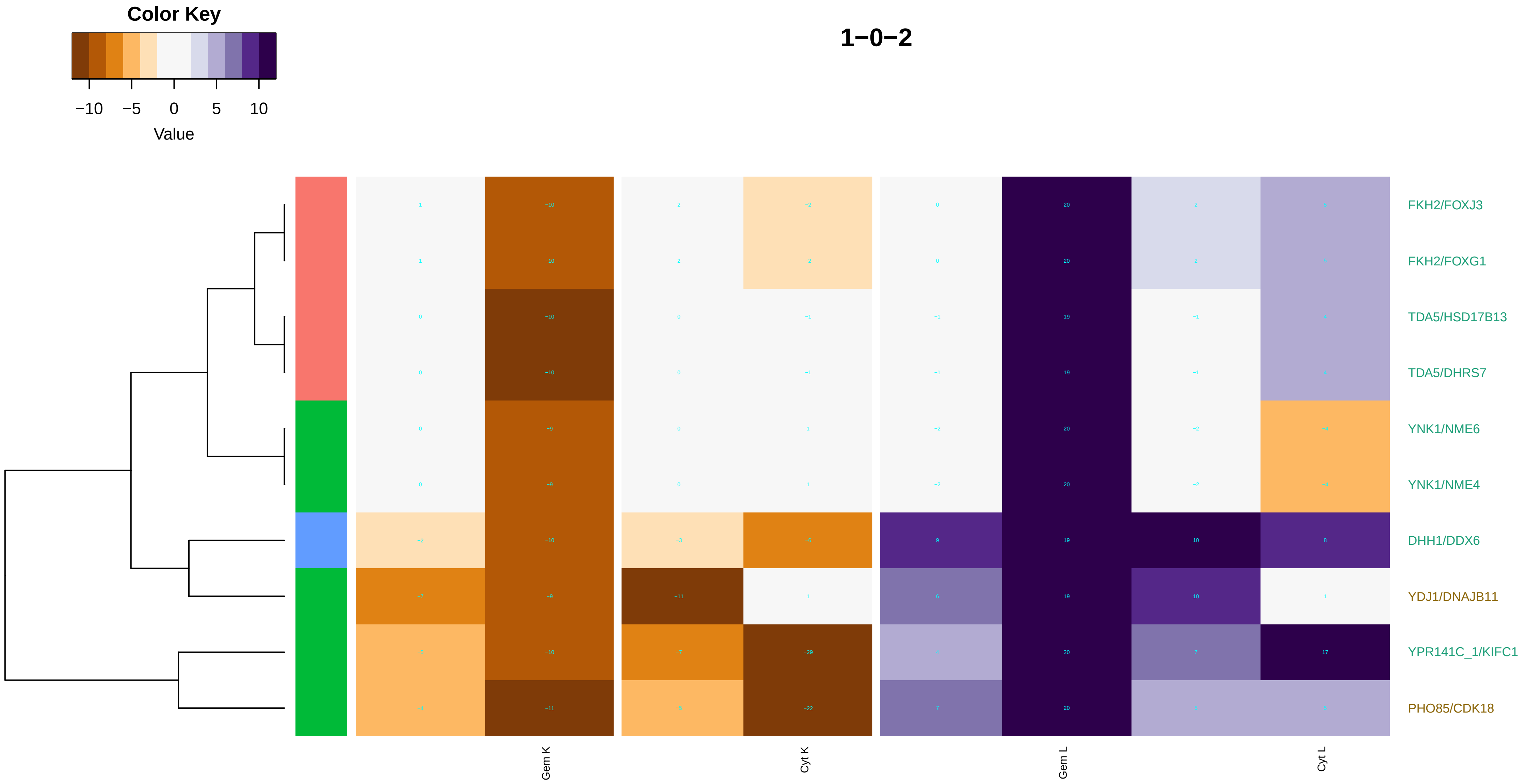

Color Key
−10 −5 0 5 10
Value
1−0−2
Gem K
Cyt K
Gem L
Cyt L
FKH2/FOXJ3
FKH2/FOXG1
TDA5/HSD17B13
TDA5/DHRS7
YNK1/NME6
YNK1/NME4
DHH1/DDX6
YDJ1/DNAJB11
YPR141C_1/KIFC1
PHO85/CDK18

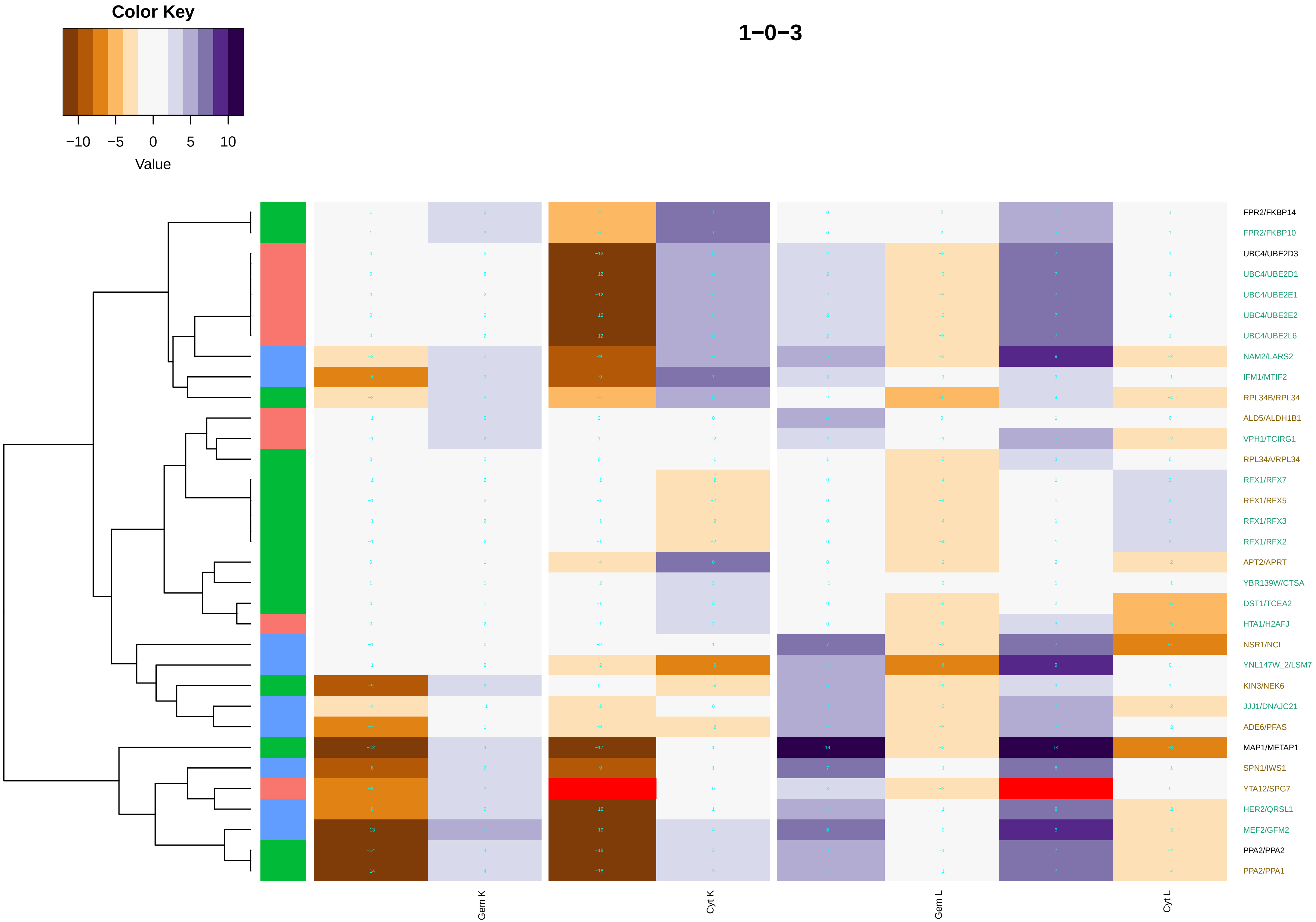

Color Key
−10
−5
0
5
10
Value
1−0−3
Gem K
Cyt K
Gem L
Cyt L
FPR2/FKBP14
FPR2/FKBP10
UBC4/UBE2D3
UBC4/UBE2D1
UBC4/UBE2E1
UBC4/UBE2E2
UBC4/UBE2L6
NAM2/LARS2
IFM1/MTIF2
RPL34B/RPL34
ALD5/ALDH1B1
VPH1/TCIRG1
RPL34A/RPL34
RFX1/RFX7
RFX1/RFX5
RFX1/RFX3
RFX1/RFX2
APT2/APRT
YBR139W/CTSA
DST1/TCEA2
HTA1/H2AFJ
NSR1/NCL
YNL147W_2/LSM7
KIN3/NEK6
JJJ1/DNAJC21
ADE6/PFAS
MAP1/METAP1
SPN1/IWS1
YTA12/SPG7
HER2/QRSL1
MEF2/GFM2
PPA2/PPA2
PPA2/PPA1

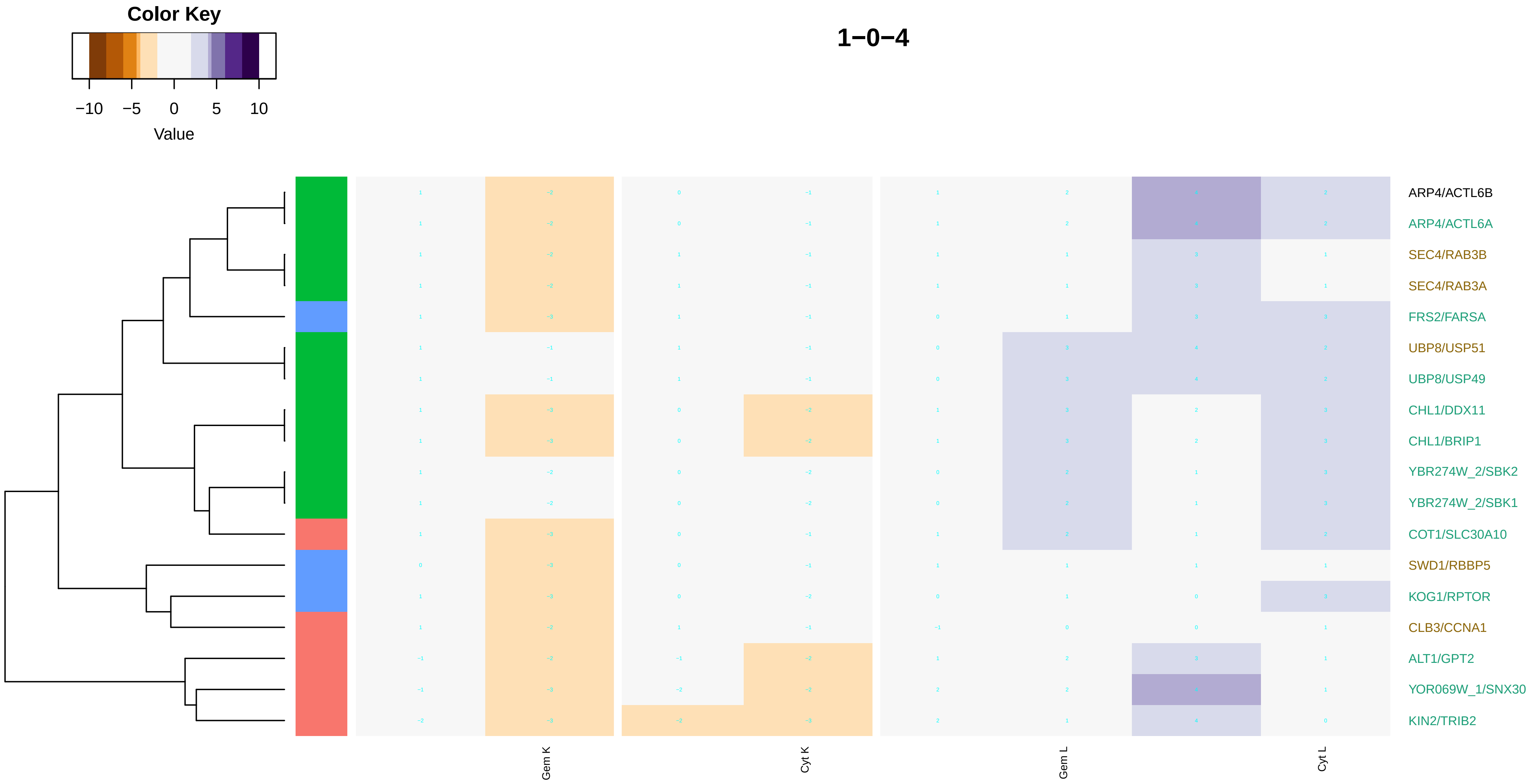
Color Key
−10 −5 0 5 10
Value
1−0−4
ARP4/ACTL6B
ARP4/ACTL6A
SEC4/RAB3B
SEC4/RAB3A
FRS2/FARSA
UBP8/USP51
UBP8/USP49
CHL1/DDX11
CHL1/BRIP1
YBR274W_2/SBK2
YBR274W_2/SBK1
COT1/SLC30A10
SWD1/RBBP5
KOG1/RPTOR
CLB3/CCNA1
ALT1/GPT2
YOR069W_1/SNX30
KIN2/TRIB2
Gem K
Cyt K
Gem L
Cyt L

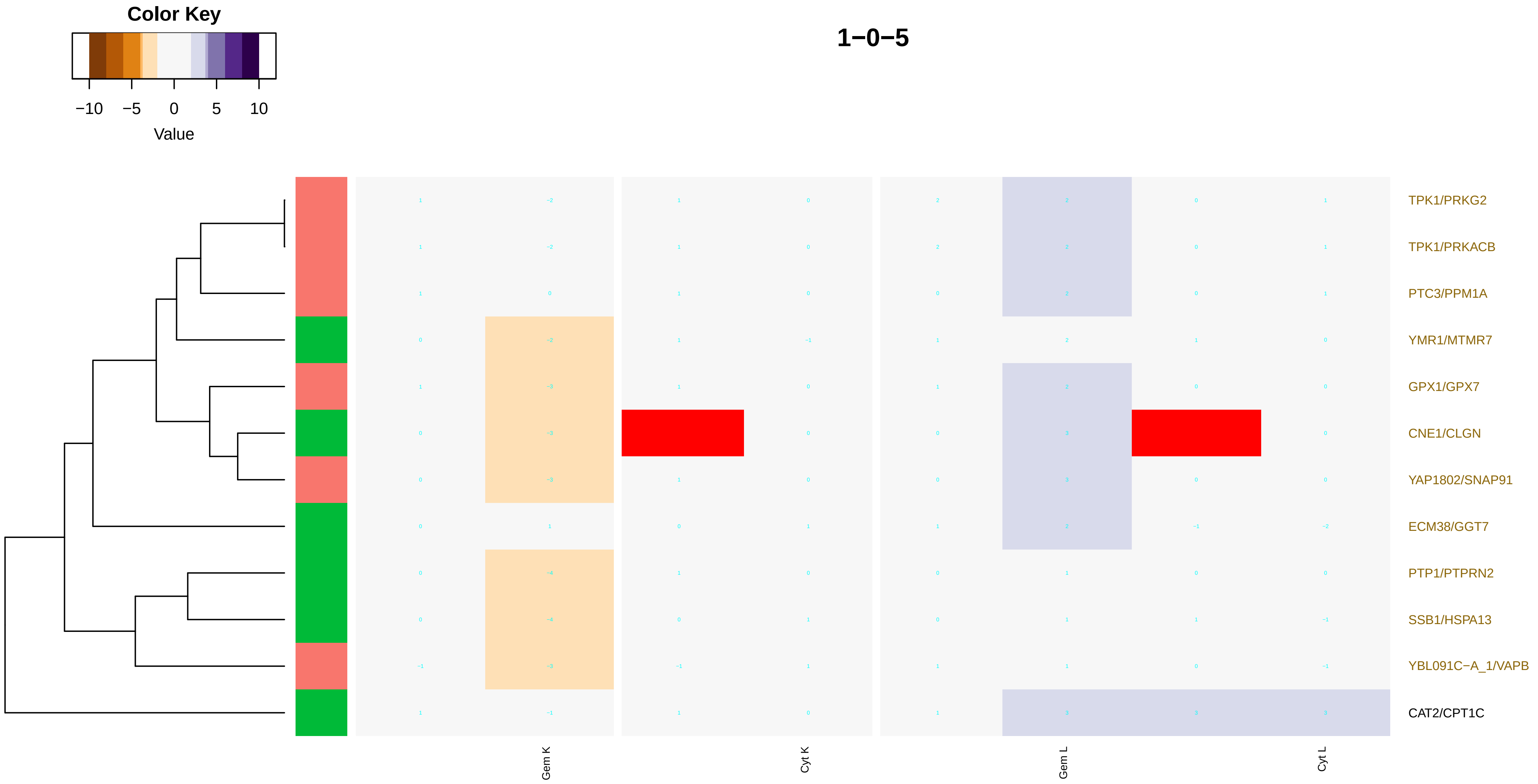

Color Key
−10 −5 0 5 10
Value
1−0−5
TPK1/PRKG2
TPK1/PRKACB
PTC3/PPM1A
YMR1/MTMR7
GPX1/GPX7
CNE1/CLGN
YAP1802/SNAP91
ECM38/GGT7
PTP1/PTPRN2
SSB1/HSPA13
YBL091C−A_1/VAPB
CAT2/CPT1C
Gem K
Cyt K
Gem L
Cyt L

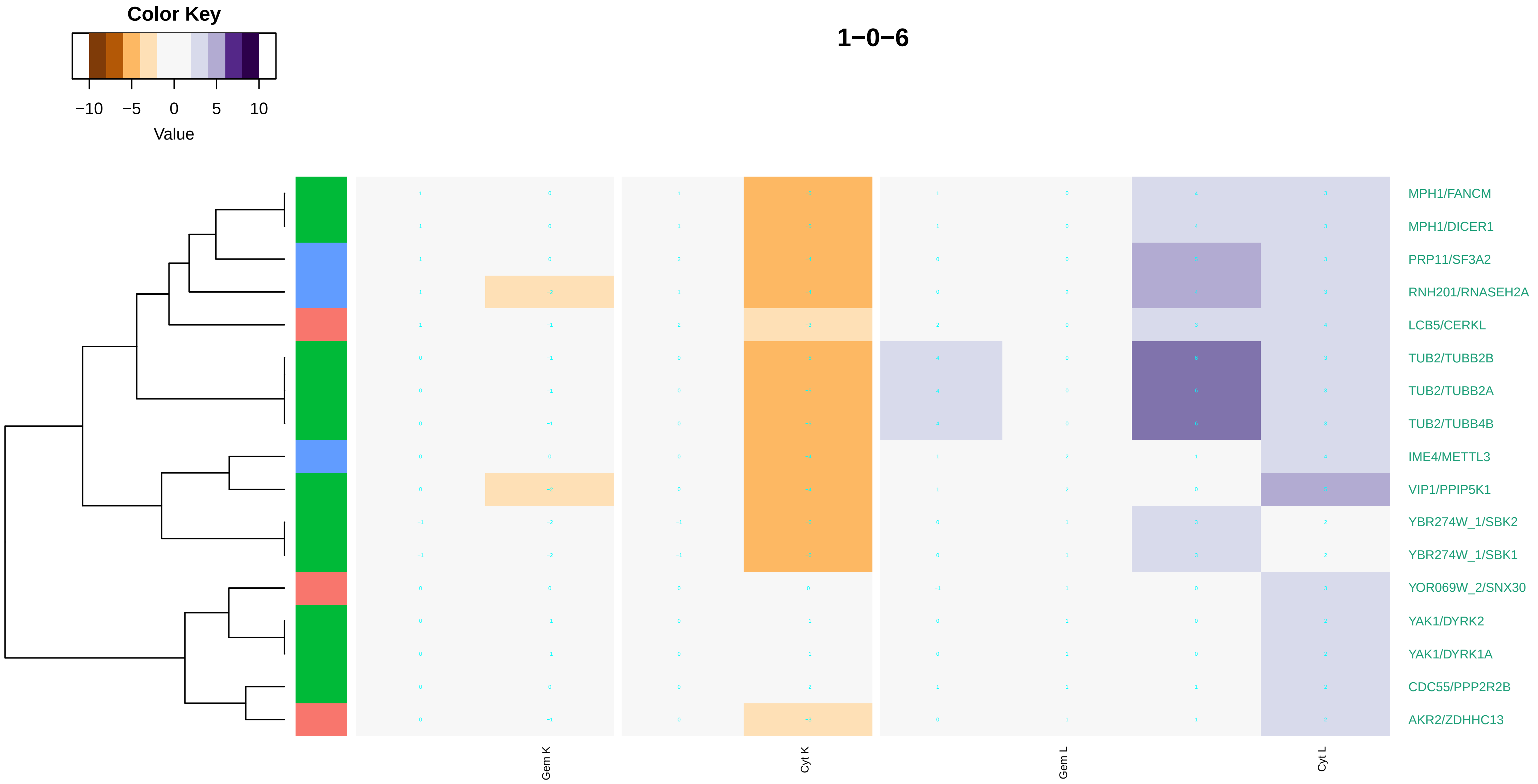
Color Key
−10
−5
0
5
10
Value
1−0−6
MPH1/FANCM
MPH1/DICER1
PRP11/SF3A2
RNH201/RNASEH2A
LCB5/CERKL
TUB2/TUBB2B
TUB2/TUBB2A
TUB2/TUBB4B
IME4/METTL3
VIP1/PPIP5K1
YBR274W_1/SBK2
YBR274W_1/SBK1
YOR069W_2/SNX30
YAK1/DYRK2
YAK1/DYRK1A
CDC55/PPP2R2B
AKR2/ZDHHC13
Gem K
Cyt K
Gem L
Cyt L

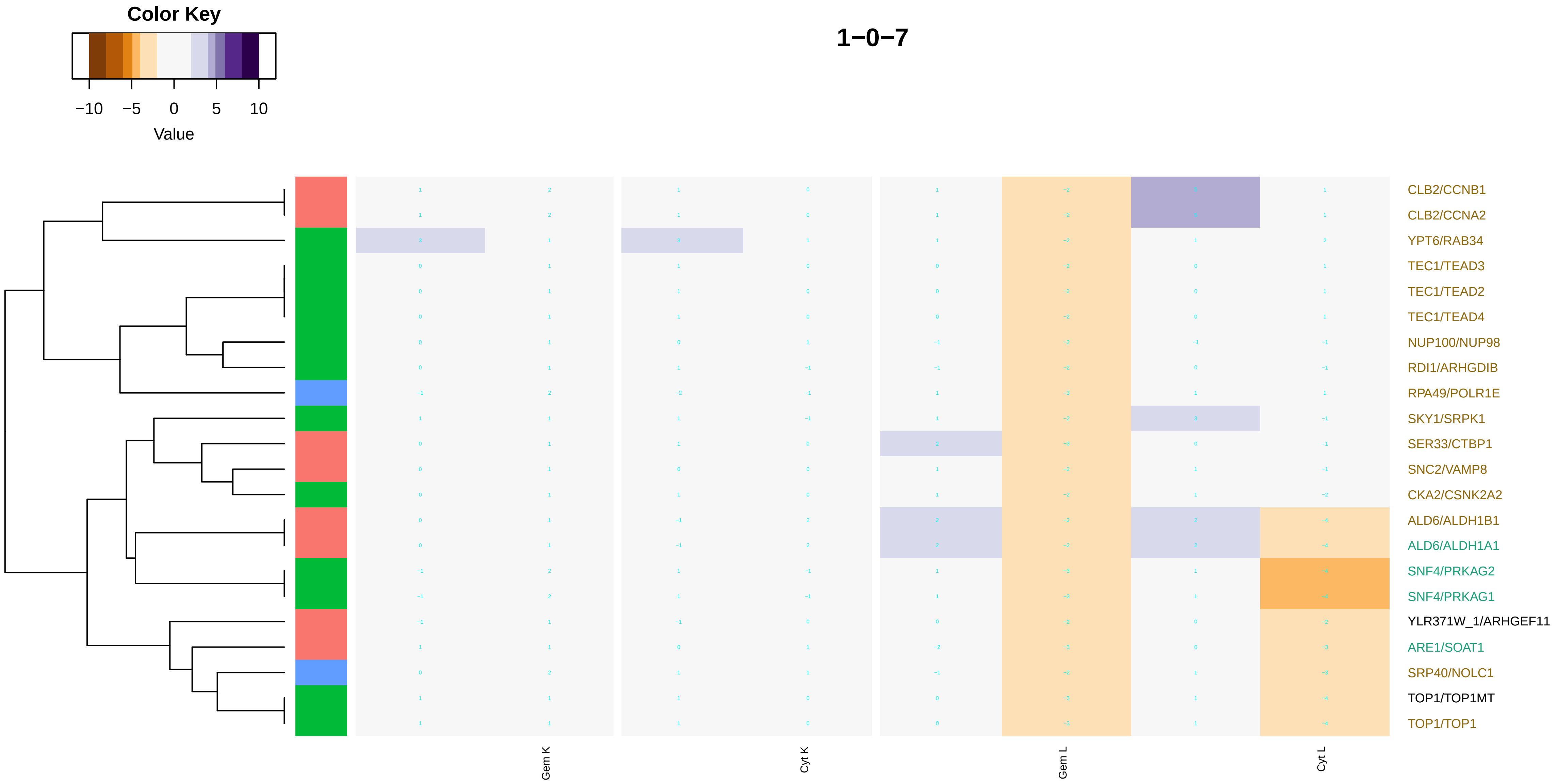

Color Key
−10
−5
0
5
10
Value
1−0−7
CLB2/CCNB1
CLB2/CCNA2
YPT6/RAB34
TEC1/TEAD3
TEC1/TEAD2
TEC1/TEAD4
NUP100/NUP98
RDI1/ARHGDIB
RPA49/POLR1E
SKY1/SRPK1
SER33/CTBP1
SNC2/VAMP8
CKA2/CSNK2A2
ALD6/ALDH1B1
ALD6/ALDH1A1
SNF4/PRKAG2
SNF4/PRKAG1
YLR371W_1/ARHGEF11
ARE1/SOAT1
SRP40/NOLC1
TOP1/TOP1MT
TOP1/TOP1
Gem K
Cyt K
Gem L
Cyt L

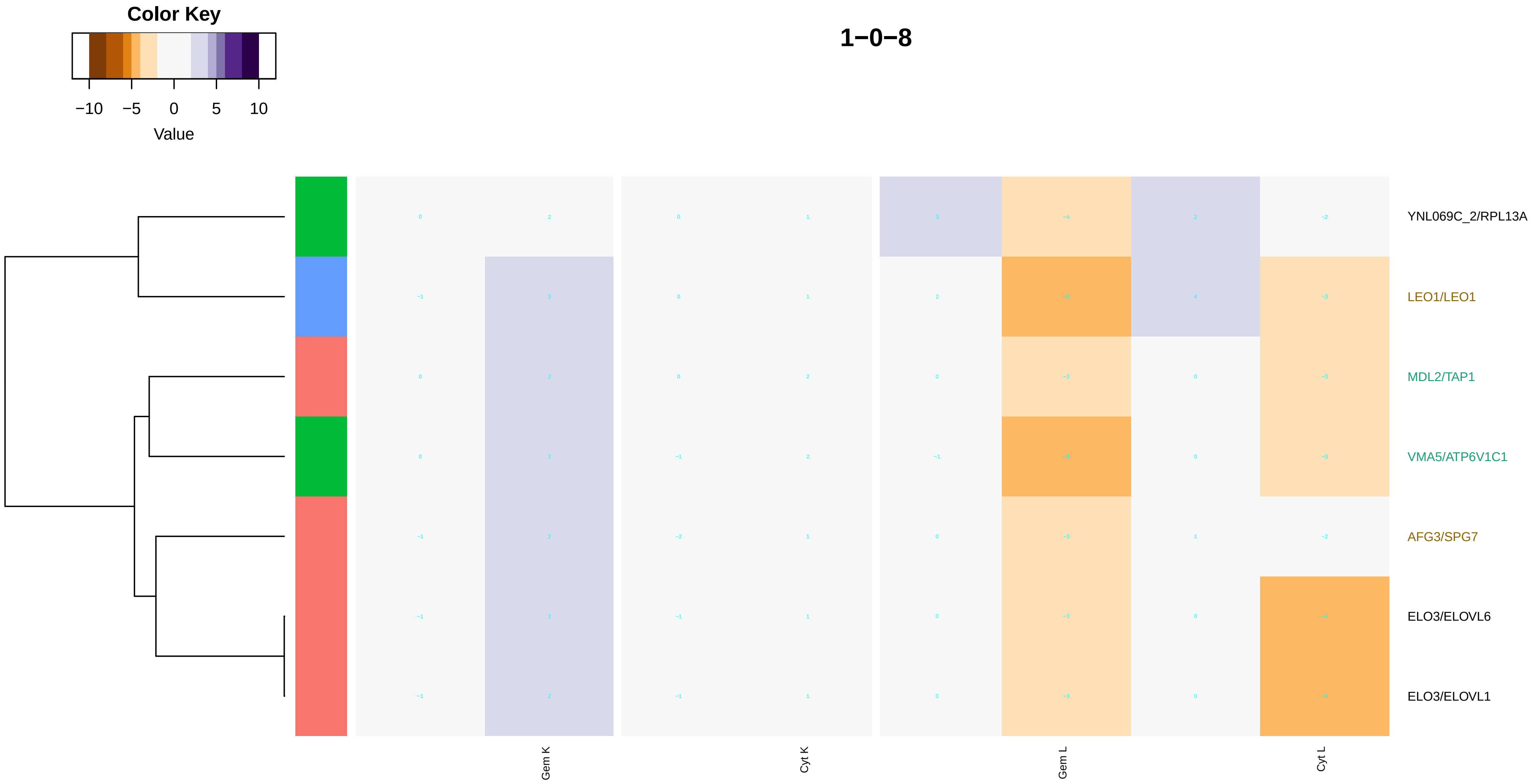
Color Key
−10 −5 0 5 10
Value
1−0−8
YNL069C_2/RPL13A
LEO1/LEO1
MDL2/TAP1
VMA5/ATP6V1C1
AFG3/SPG7
ELO3/ELOVL6
ELO3/ELOVL1
Gem K
Cyt K
Gem L
Cyt L

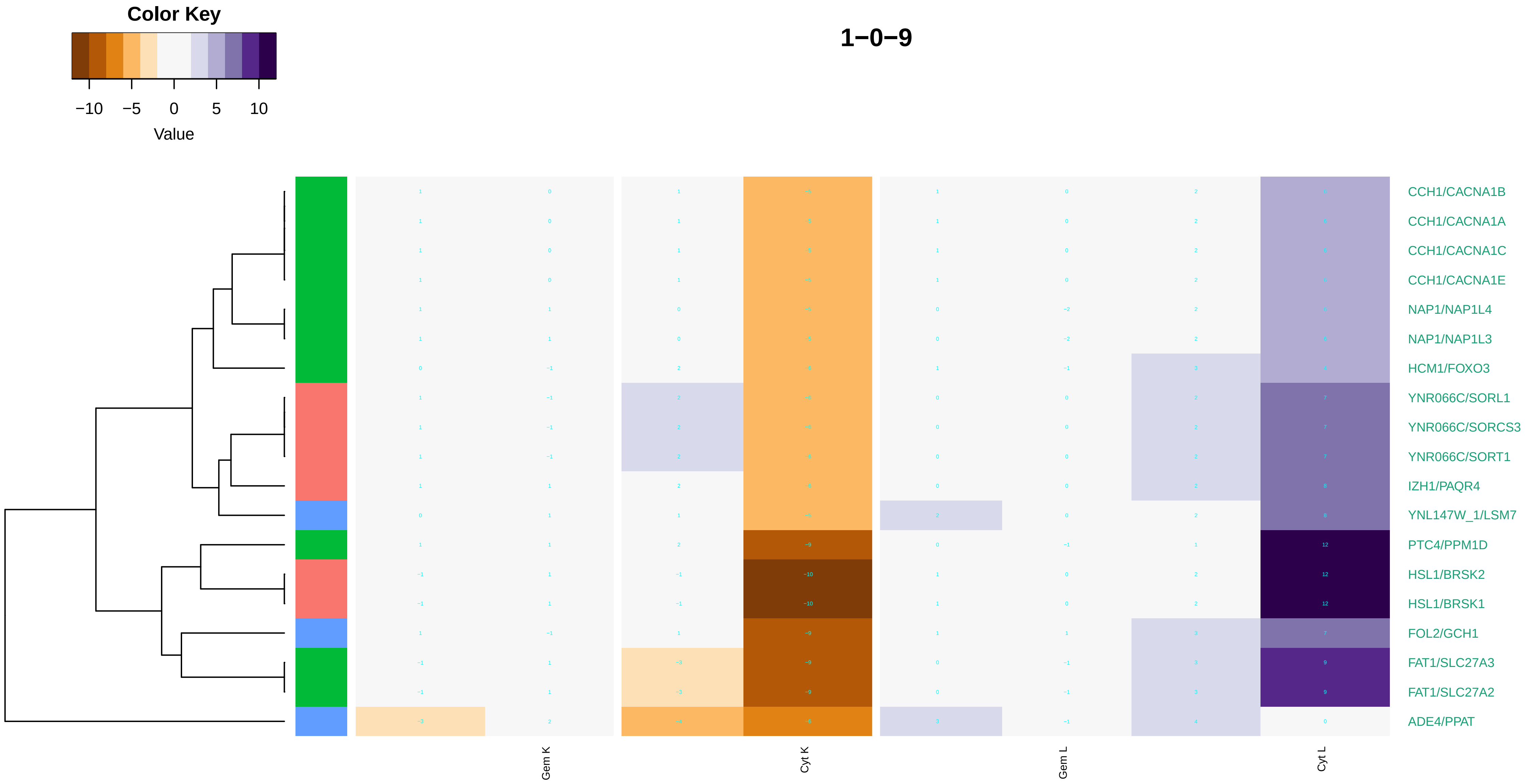
Color Key
−10
−5
0
5
10
Value
1−0−9
CCH1/CACNA1B
CCH1/CACNA1A
CCH1/CACNA1C
CCH1/CACNA1E
NAP1/NAP1L4
NAP1/NAP1L3
HCM1/FOXO3
YNR066C/SORL1
YNR066C/SORCS3
YNR066C/SORT1
IZH1/PAQR4
YNL147W_1/LSM7
PTC4/PPM1D
HSL1/BRSK2
HSL1/BRSK1
FOL2/GCH1
FAT1/SLC27A3
FAT1/SLC27A2
ADE4/PPAT
Gem K
Cyt K
Gem L
Cyt L

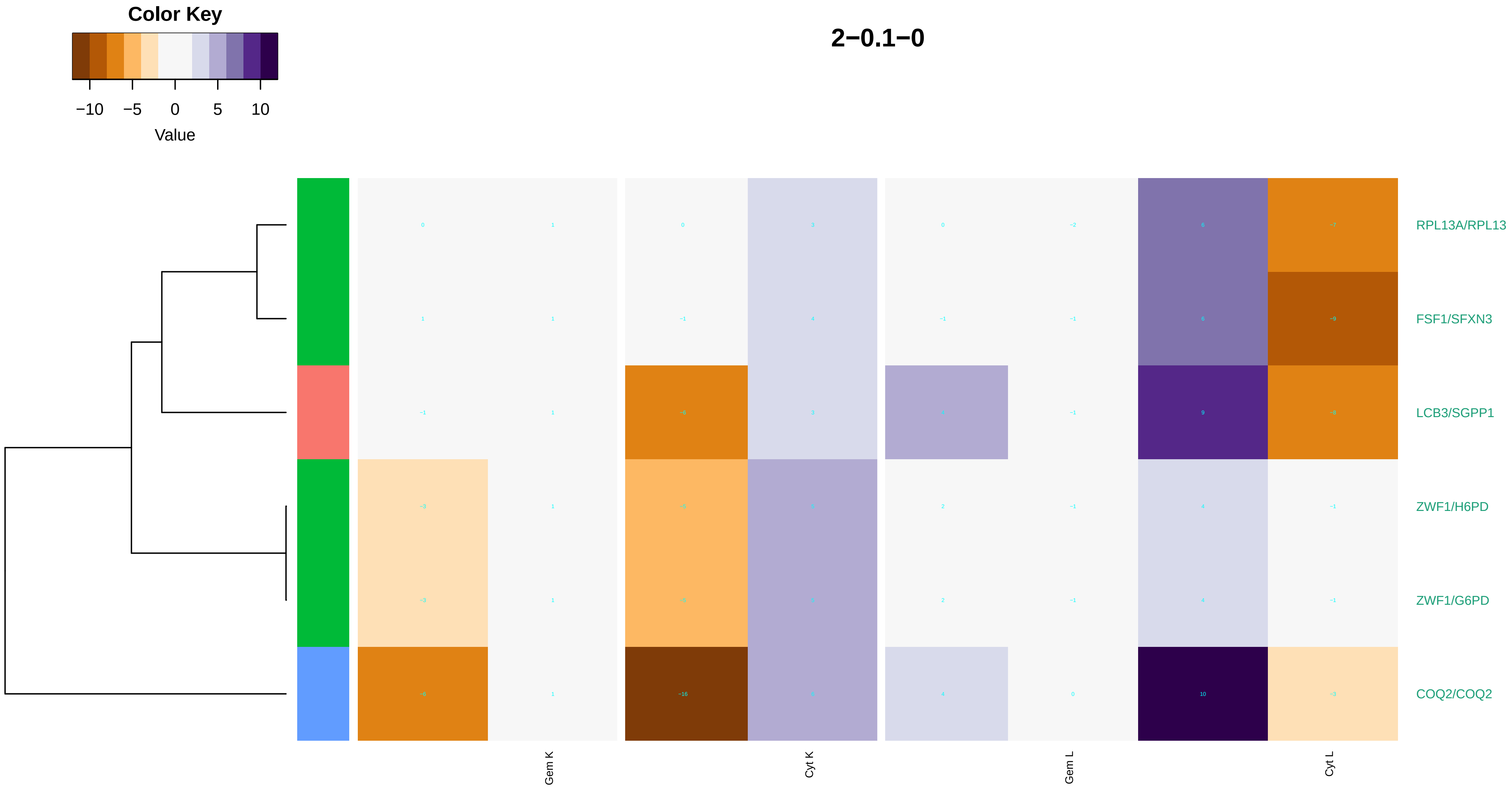
Color Key
−10
−5
0
5
10
Value
2−0.1−0
RPL13A/RPL13
FSF1/SFXN3
LCB3/SGPP1
ZWF1/H6PD
ZWF1/G6PD
COQ2/COQ2
Gem K
Cyt K
Gem L
Cyt L

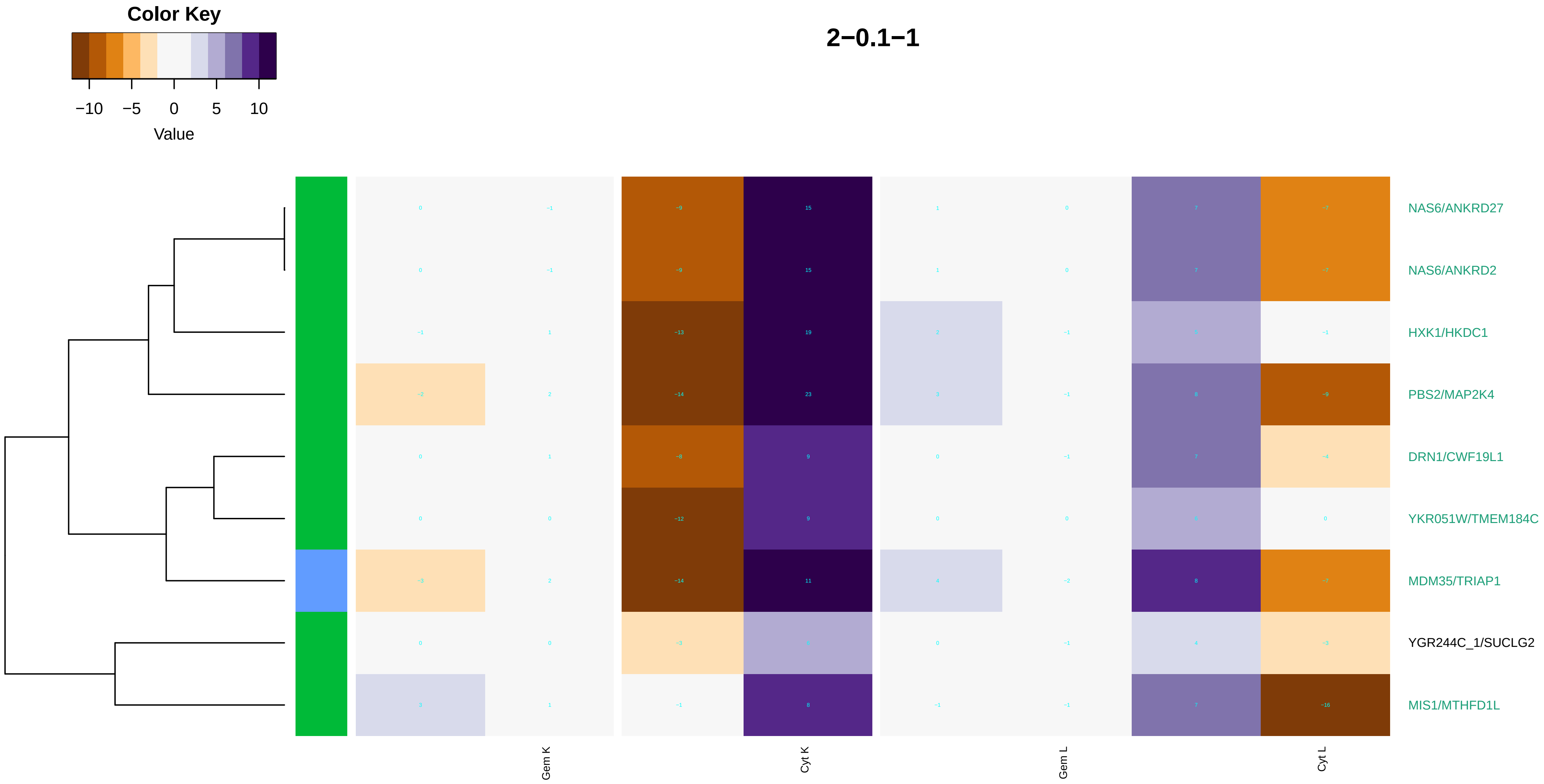

Color Key
−10
−5
0
5
10
Value
2−0.1−1
NAS6/ANKRD27
NAS6/ANKRD2
HXK1/HKDC1
PBS2/MAP2K4
DRN1/CWF19L1
YKR051W/TMEM184C
MDM35/TRIAP1
YGR244C_1/SUCLG2
MIS1/MTHFD1L
Gem K
Cyt K
Gem L
Cyt L

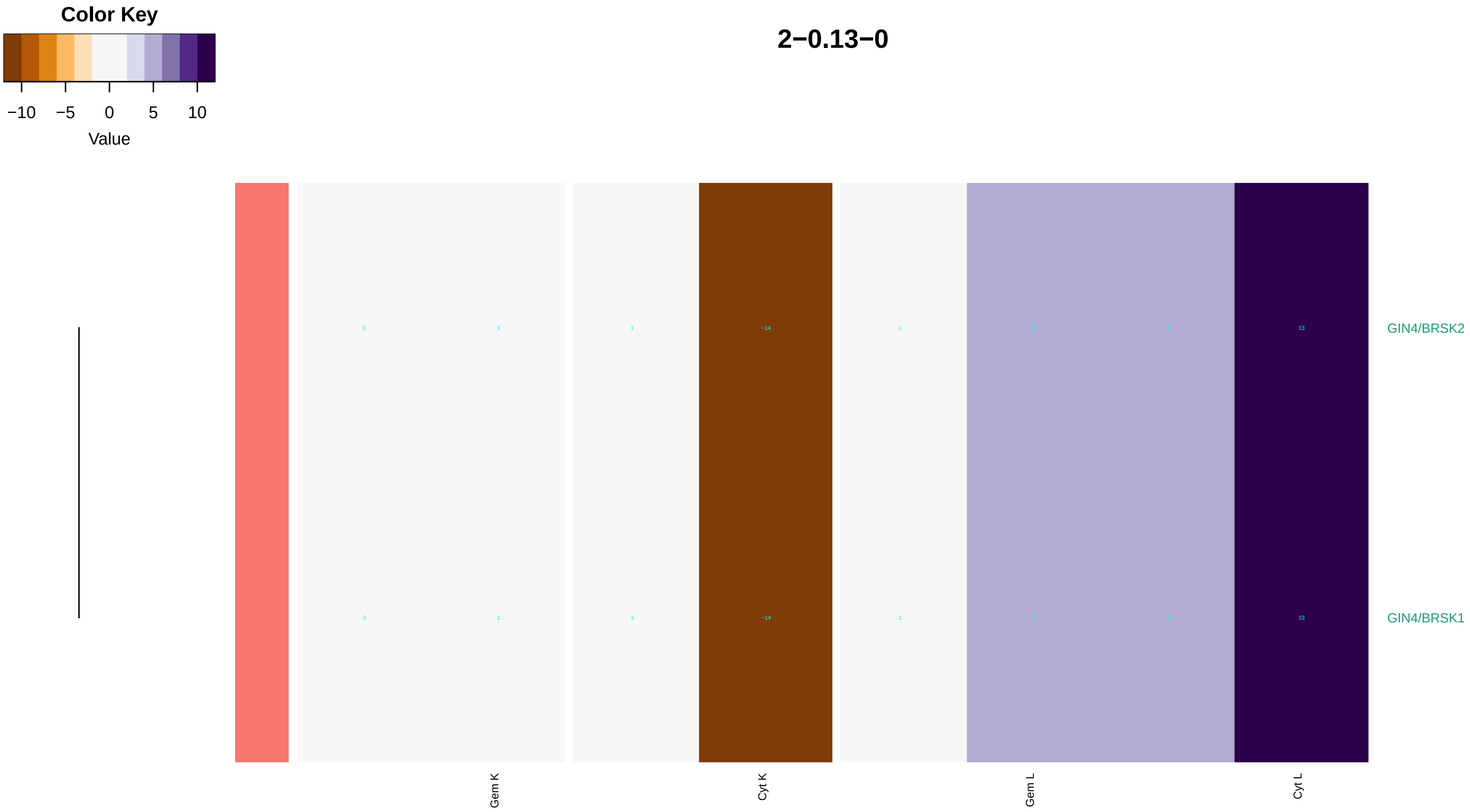
Color Key
−10
−5
0
5
10
Value
2−0.13−0
GIN4/BRSK2
GIN4/BRSK1
Gem K
Cyt K
Gem L
Cyt L

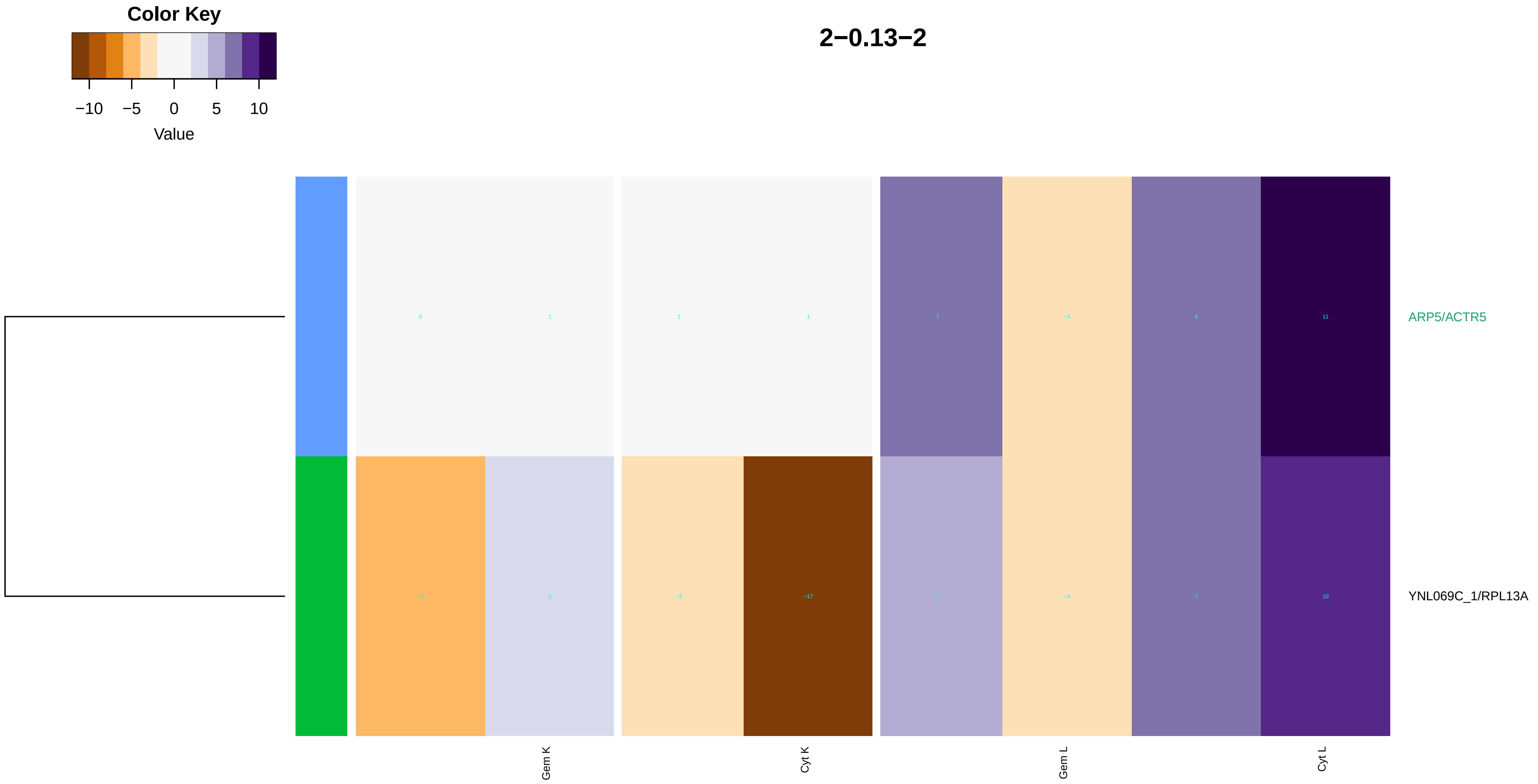

Color Key
−10
−5
0
5
10
Value
2−0.13−2
ARP5/ACTR5
YNL069C_1/RPL13A
Gem K
Cyt K
Gem L
Cyt L

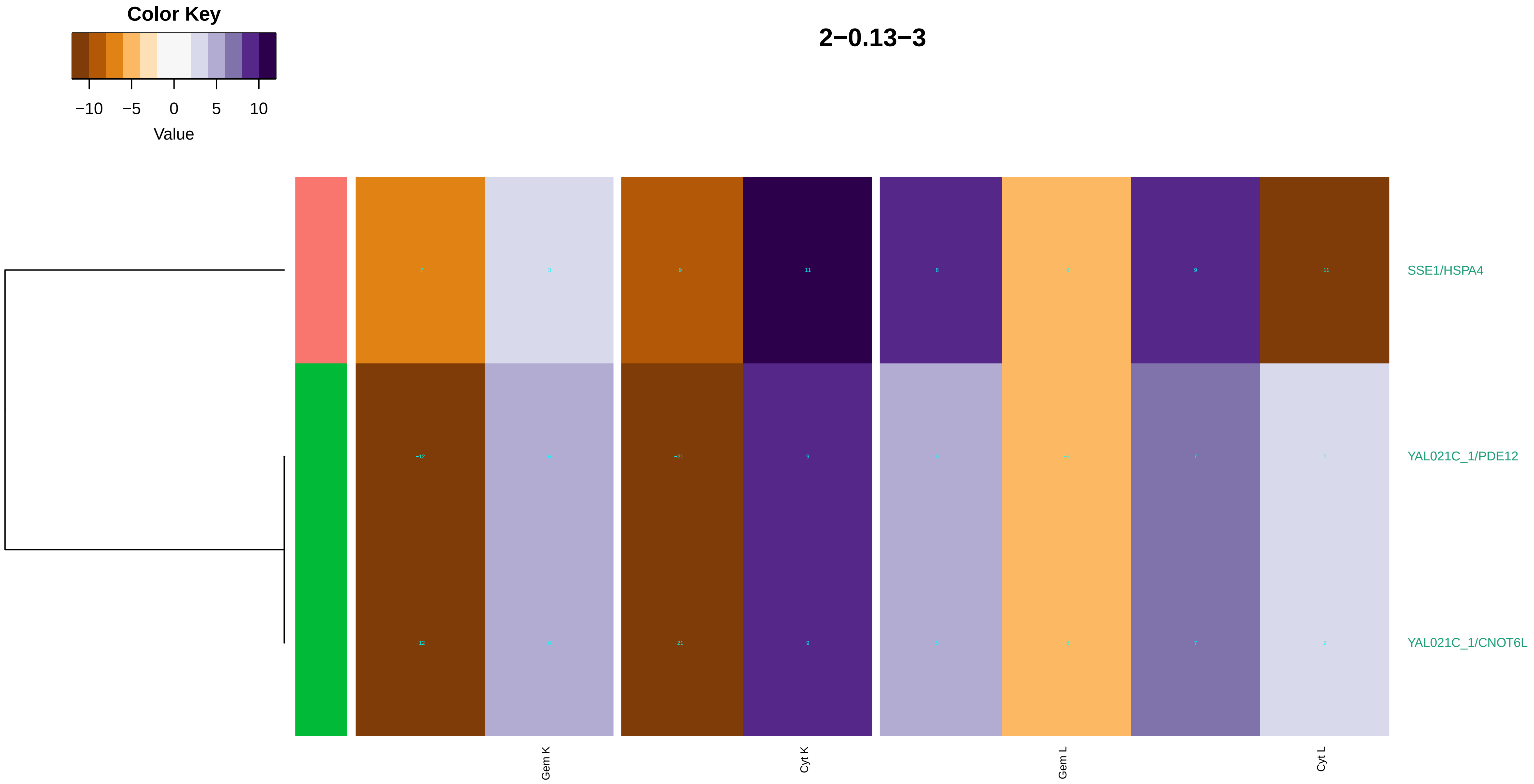
Color Key
−10
−5
0
5
10
Value
2−0.13−3
SSE1/HSPA4
YAL021C_1/PDE12
YAL021C_1/CNOT6L
Gem K
Cyt K
Gem L
Cyt L

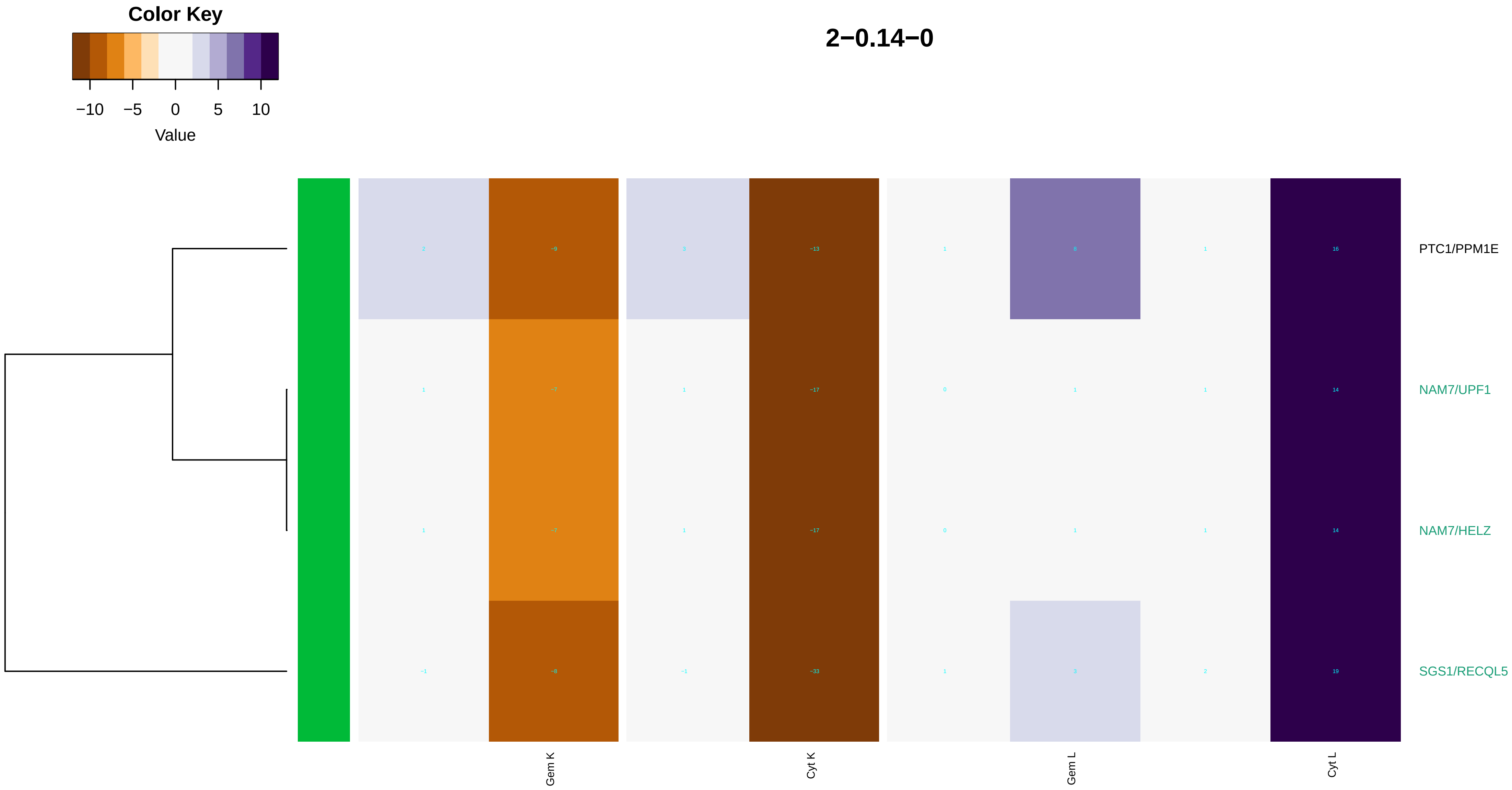
Color Key
−10 −5 0 5 10
Value
2−0.14−0
PTC1/PPM1E
NAM7/UPF1
NAM7/HELZ
SGS1/RECQL5
Gem K
Cyt K
Gem L
Cyt L

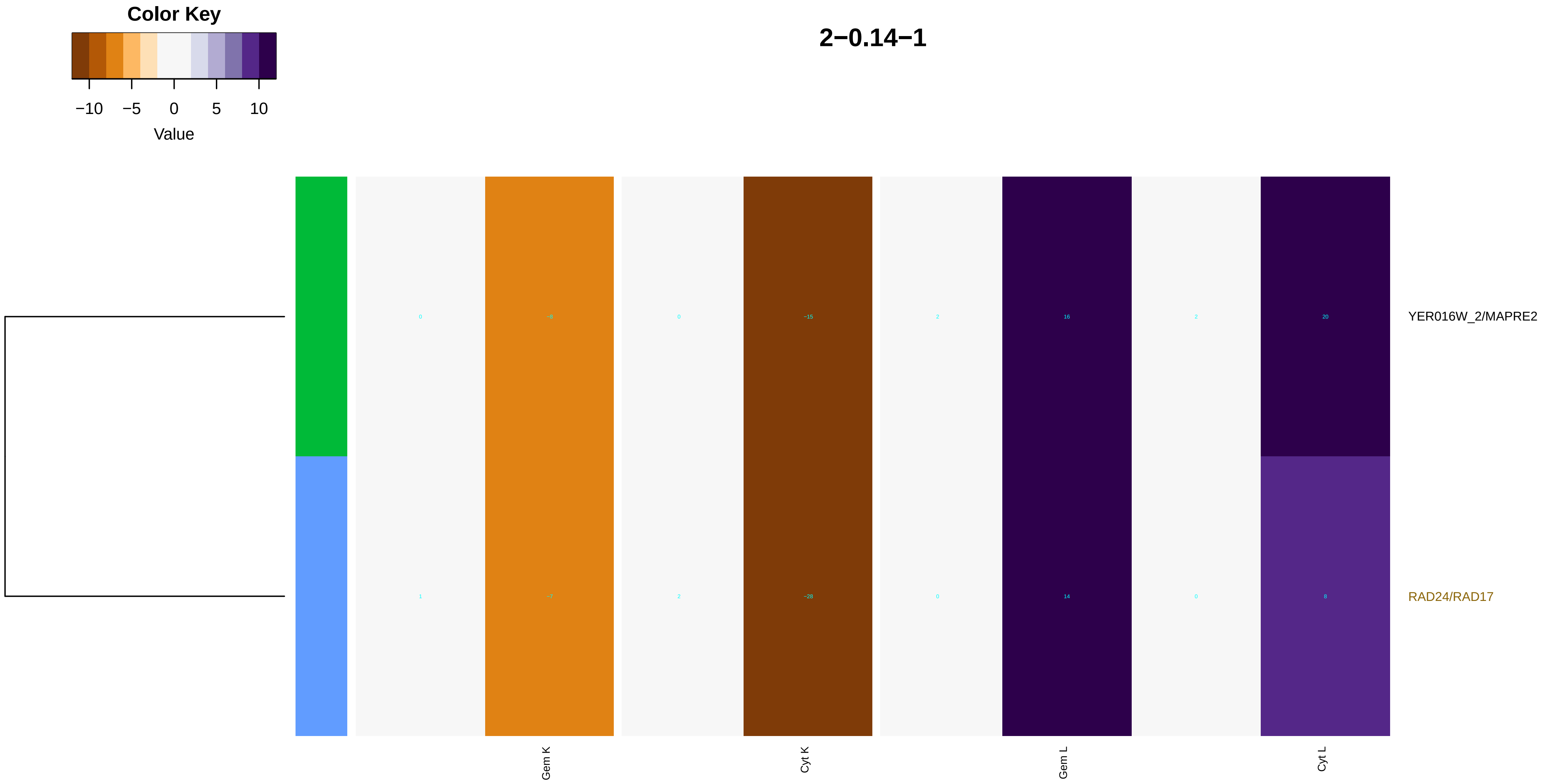
Color Key
−10
−5
0
5
10
Value
2−0.14−1
YER016W_2/MAPRE2
RAD24/RAD17
Gem K
Cyt K
Gem L
Cyt L

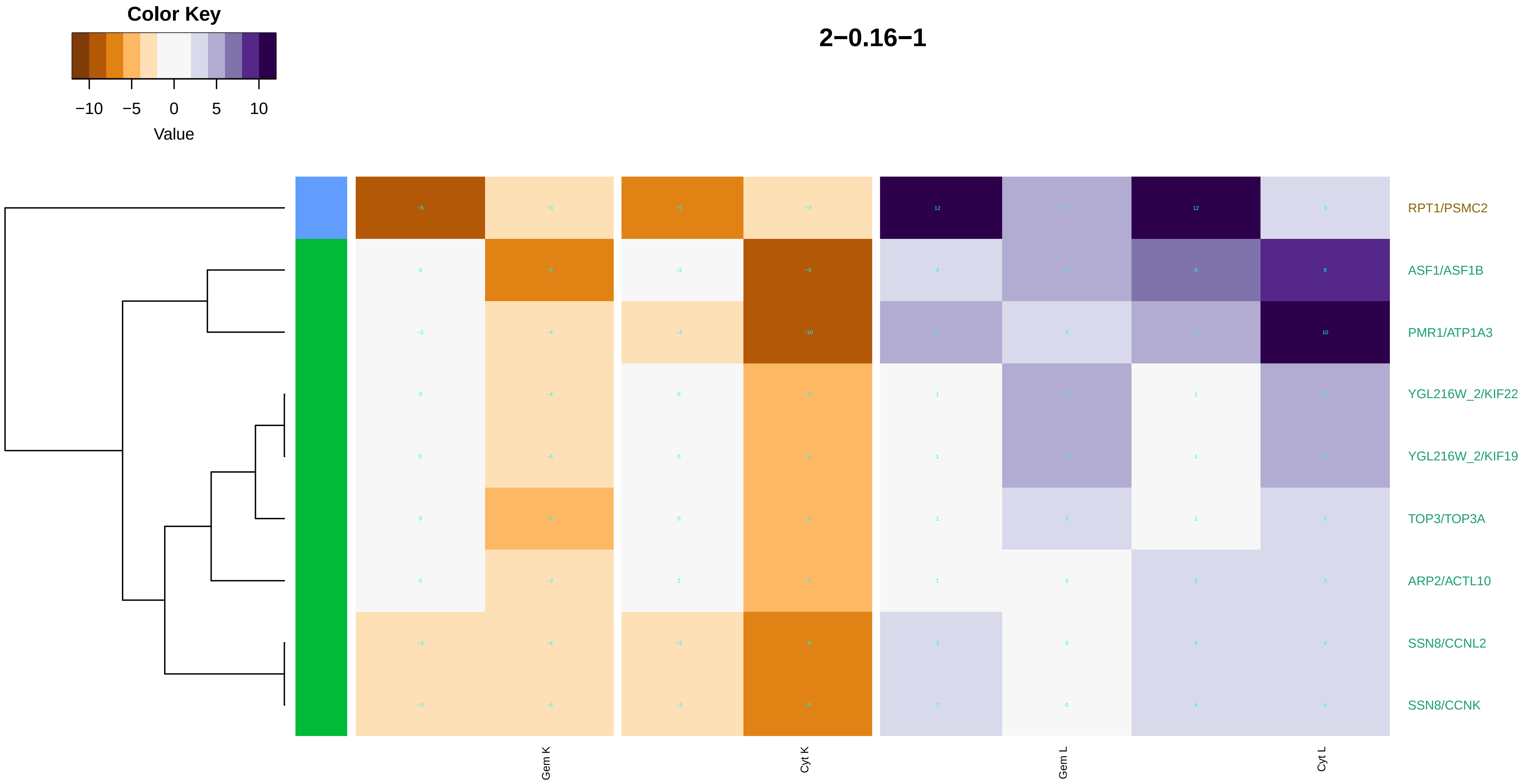
Color Key
−10 −5 0 5 10
Value
2−0.16−1
RPT1/PSMC2
ASF1/ASF1B
PMR1/ATP1A3
YGL216W_2/KIF22
YGL216W_2/KIF19
TOP3/TOP3A
ARP2/ACTL10
SSN8/CCNL2
SSN8/CCNK
Gem K
Cyt K
Gem L
Cyt L

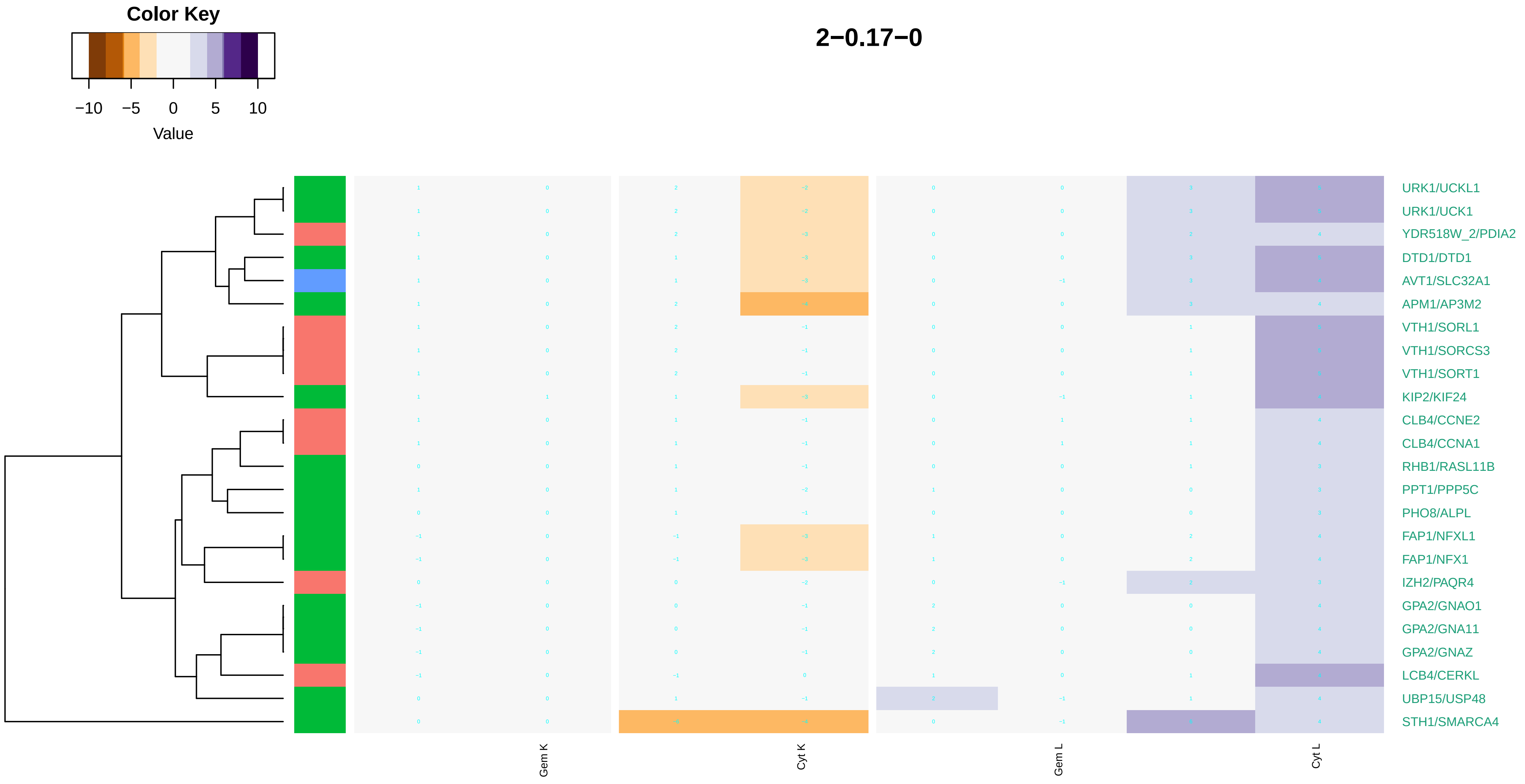
Color Key
−10
−5
0
5
10
Value
2−0.17−0
Gem K
Cyt K
Gem L
Cyt L
URK1/UCKL1
URK1/UCK1
YDR518W_2/PDIA2
DTD1/DTD1
AVT1/SLC32A1
APM1/AP3M2
VTH1/SORL1
VTH1/SORCS3
VTH1/SORT1
KIP2/KIF24
CLB4/CCNE2
CLB4/CCNA1
RHB1/RASL11B
PPT1/PPP5C
PHO8/ALPL
FAP1/NFXL1
FAP1/NFX1
IZH2/PAQR4
GPA2/GNAO1
GPA2/GNA11
GPA2/GNAZ
LCB4/CERKL
UBP15/USP48
STH1/SMARCA4

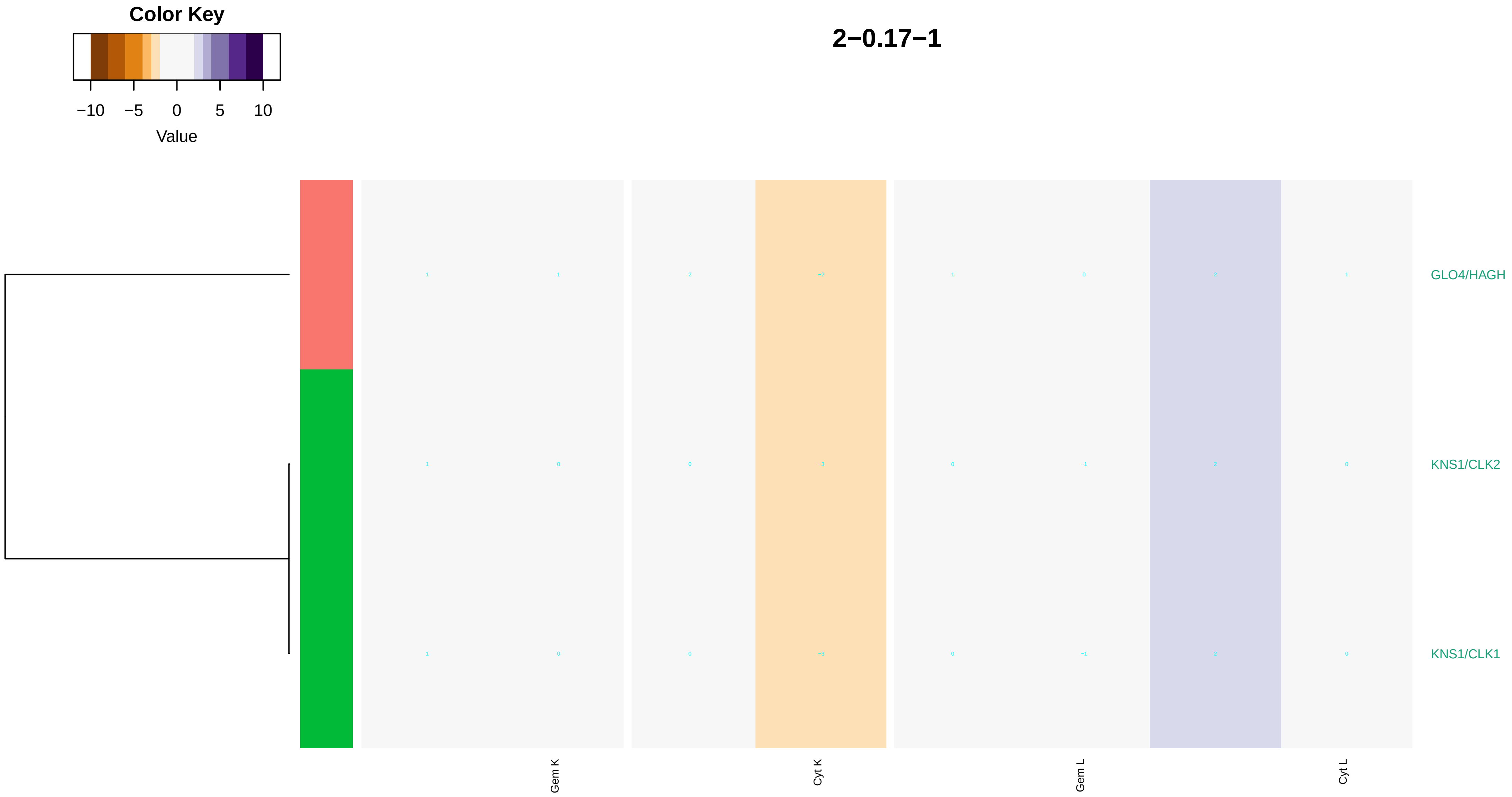
Color Key
−10
−5
0
5
10
Value
2−0.17−1
GLO4/HAGH
KNS1/CLK2
KNS1/CLK1
Gem K
Cyt K
Gem L
Cyt L

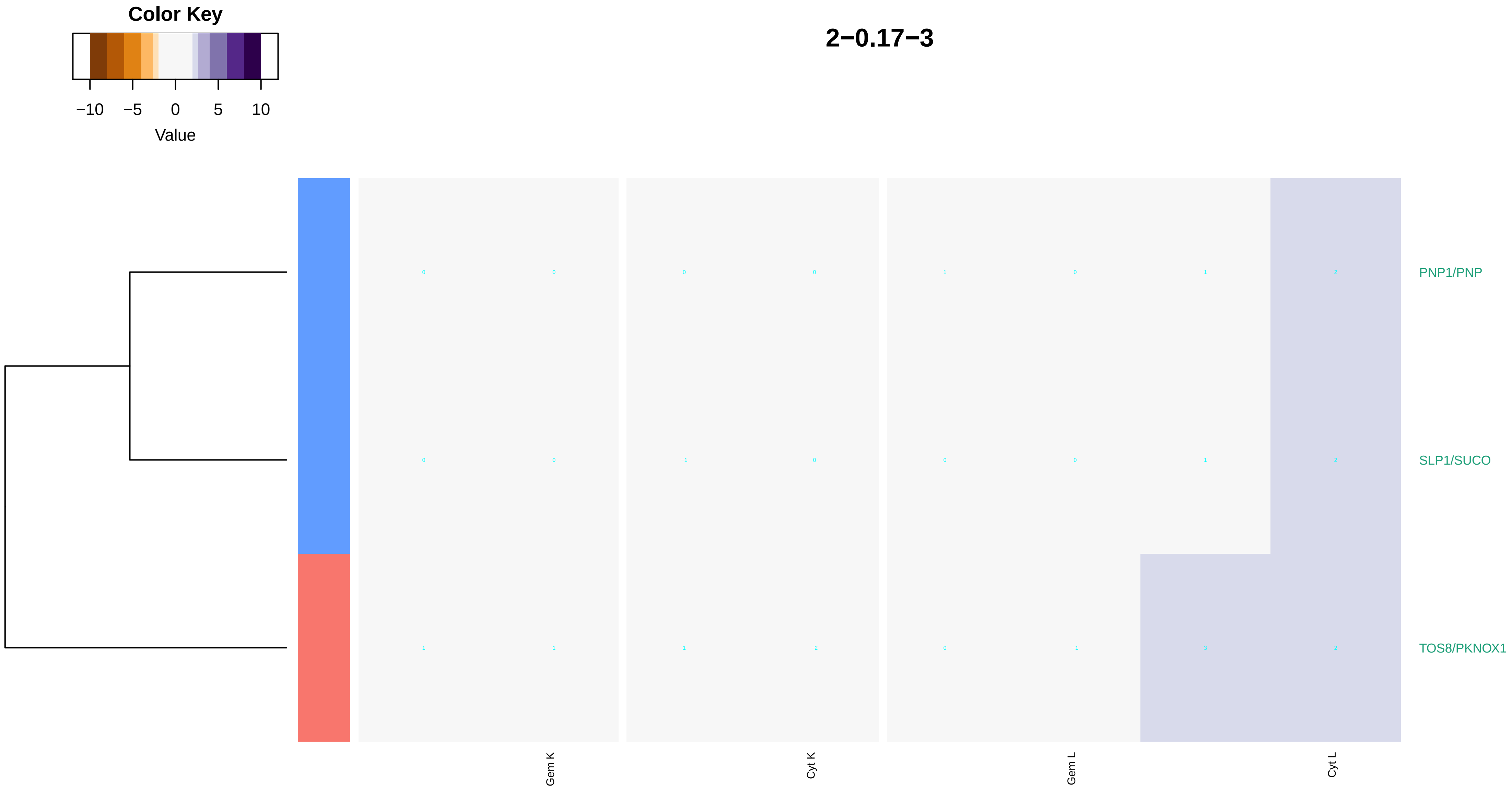
Color Key
−10
−5
0
5
10
Value
2−0.17−3
PNP1/PNP
SLP1/SUCO
TOS8/PKNOX1
Gem K
Cyt K
Gem L
Cyt L

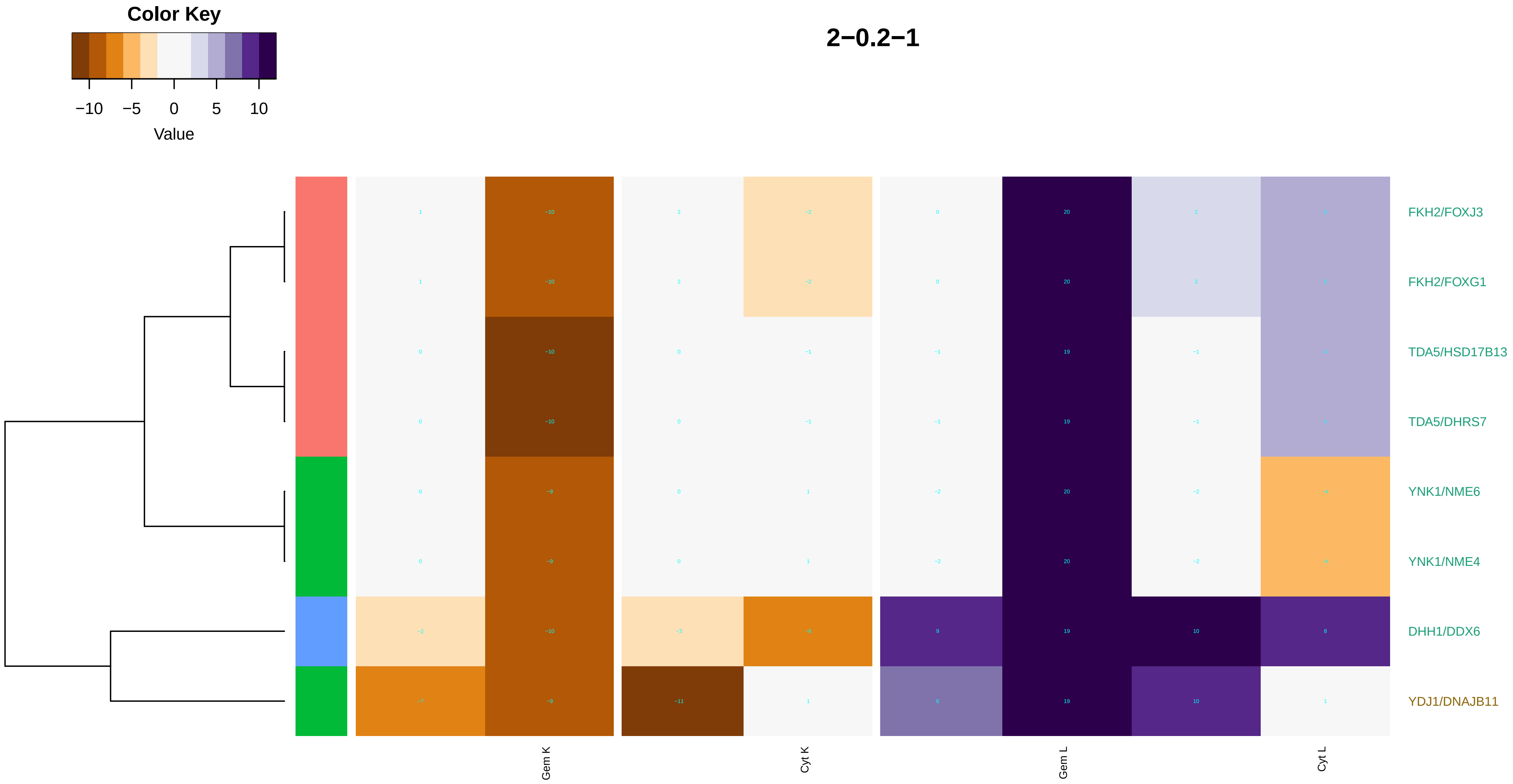
Color Key
−10 −5 0 5 10
Value
2−0.2−1
FKH2/FOXJ3
FKH2/FOXG1
TDA5/HSD17B13
TDA5/DHRS7
YNK1/NME6
YNK1/NME4
DHH1/DDX6
YDJ1/DNAJB11
Gem K
Cyt K
Gem L
Cyt L

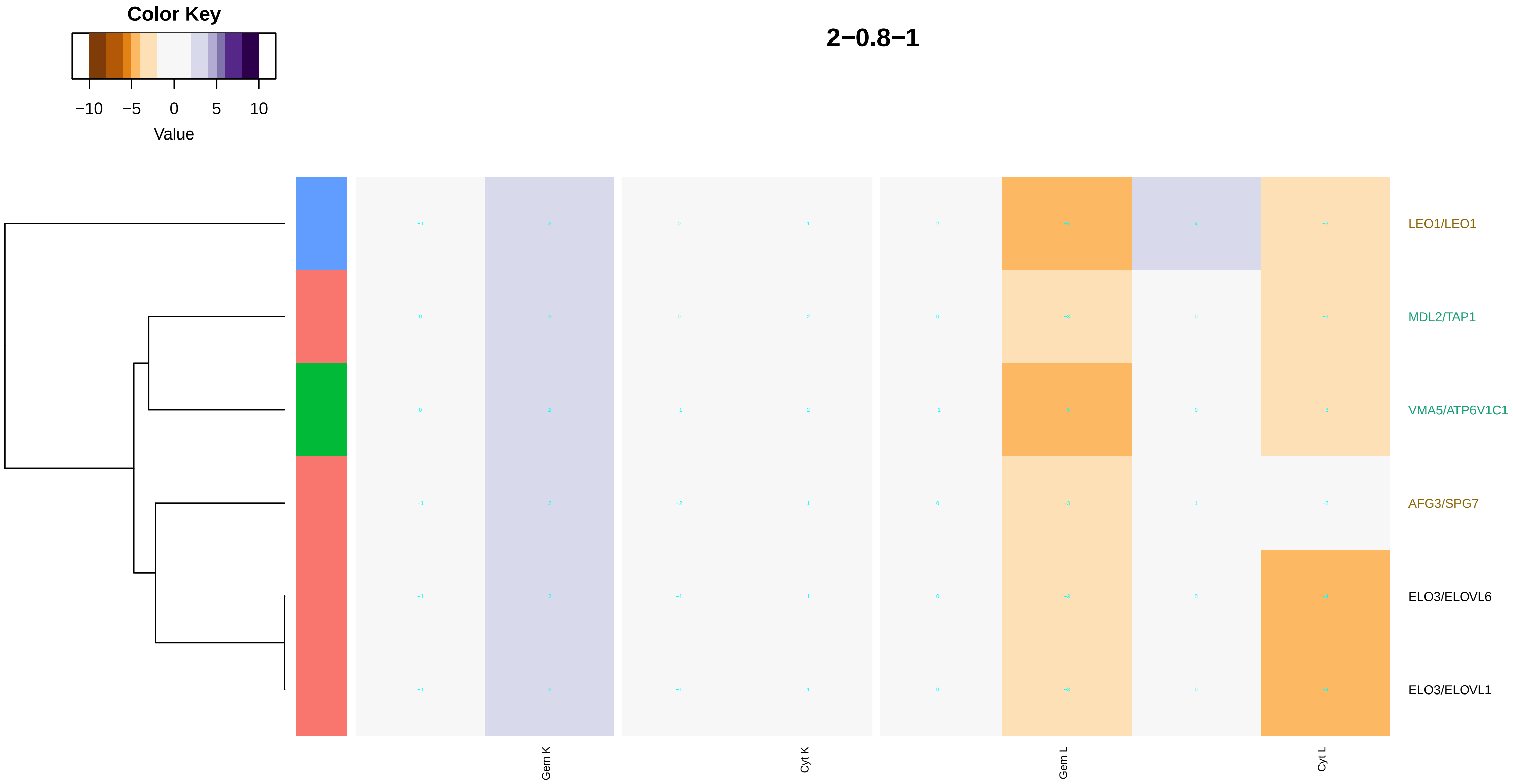
Color Key
−10
−5
0
5
10
Value
2−0.8−1
LEO1/LEO1
MDL2/TAP1
VMA5/ATP6V1C1
AFG3/SPG7
ELO3/ELOVL6
ELO3/ELOVL1
Gem K
Cyt K
Gem L
Cyt L

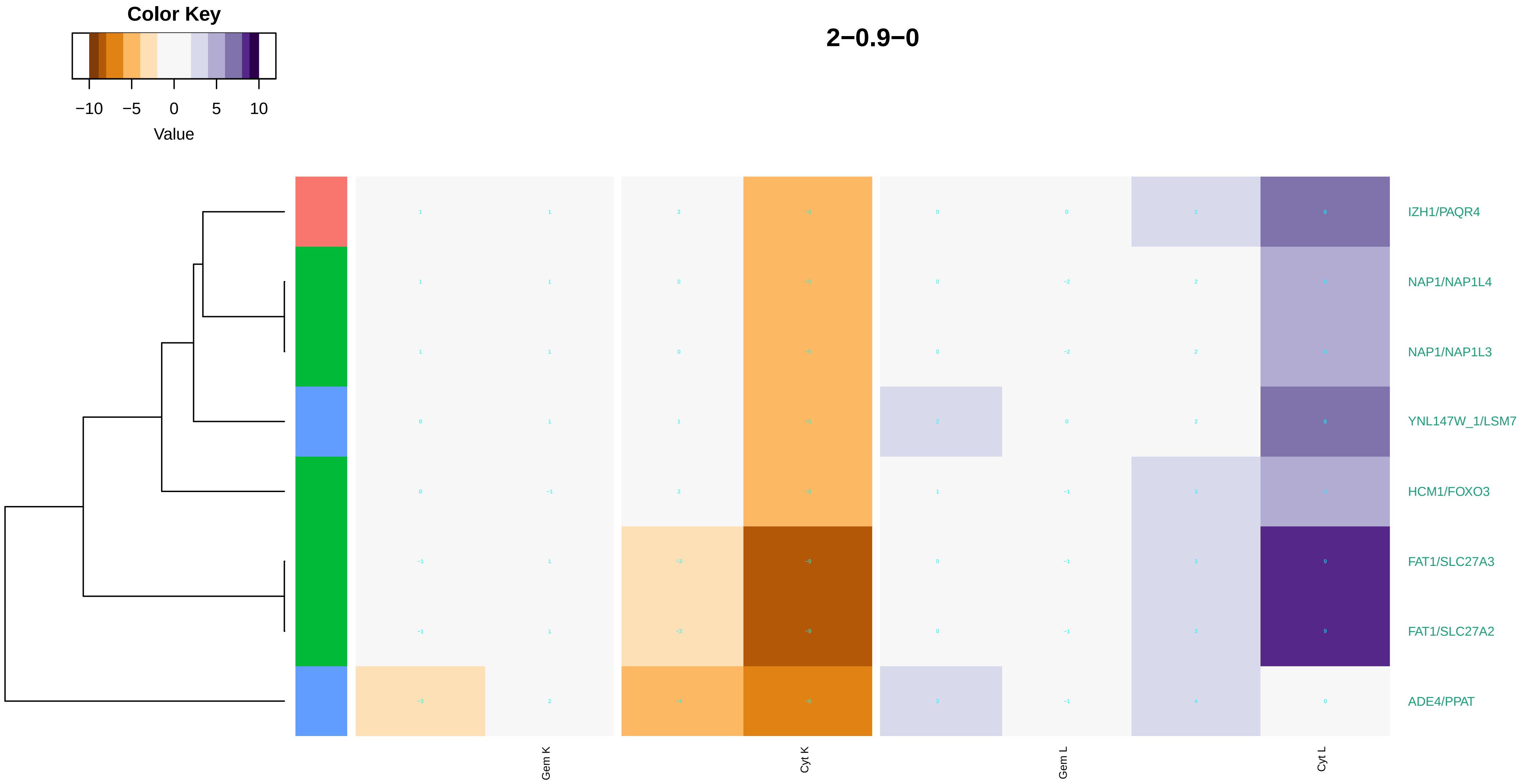

Color Key
−10
−5
0
5
10
Value
2−0.9−0
IZH1/PAQR4
NAP1/NAP1L4
NAP1/NAP1L3
YNL147W_1/LSM7
HCM1/FOXO3
FAT1/SLC27A3
FAT1/SLC27A2
ADE4/PPAT
Gem K
Cyt K
Gem L
Cyt L

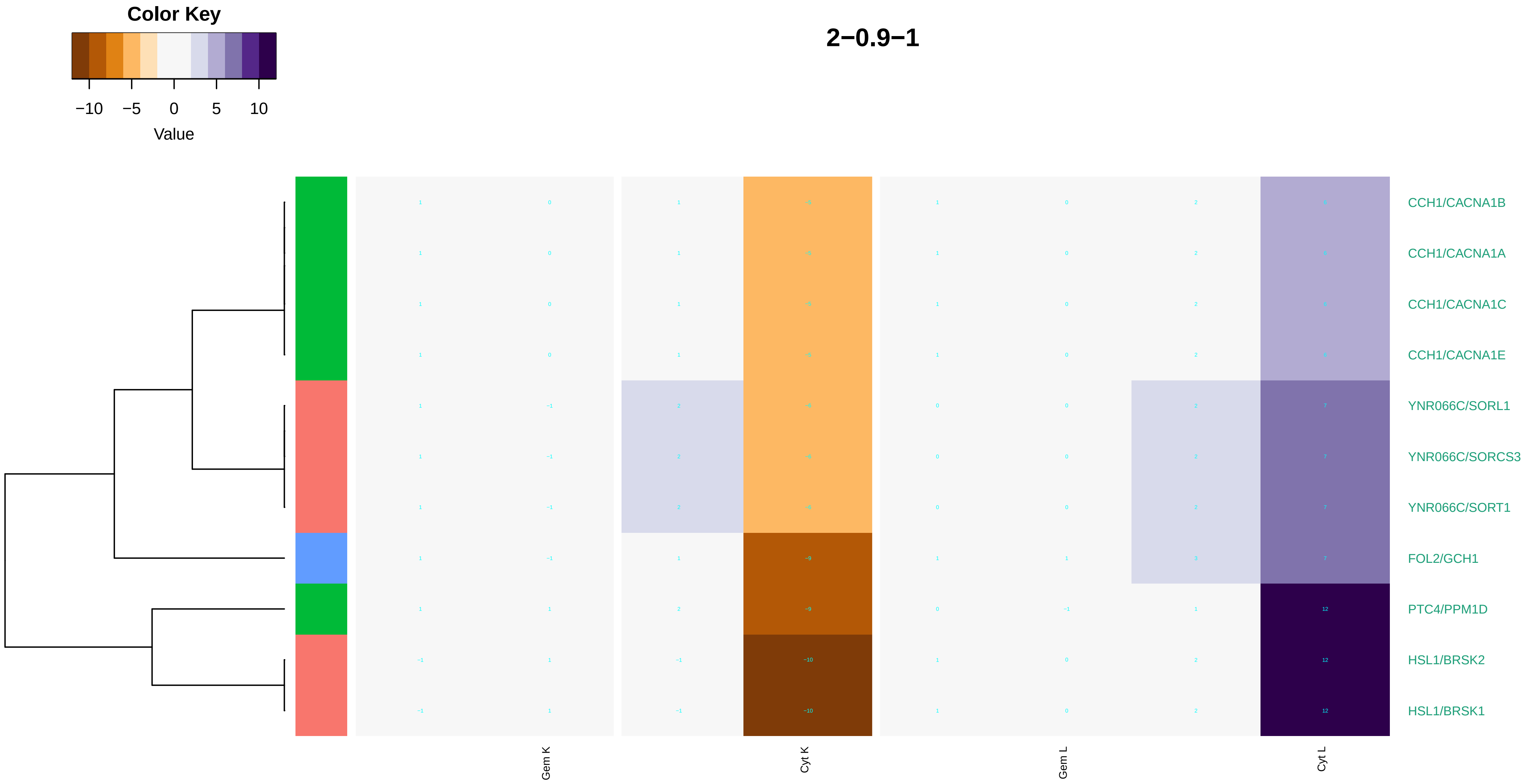

Color Key
−10
−5
0
5
10
Value
2−0.9−1
CCH1/CACNA1B
CCH1/CACNA1A
CCH1/CACNA1C
CCH1/CACNA1E
YNR066C/SORL1
YNR066C/SORCS3
YNR066C/SORT1
FOL2/GCH1
PTC4/PPM1D
HSL1/BRSK2
HSL1/BRSK1
Gem K
Cyt K
Gem L
Cyt L
