## Additional File 8 for "A humanized yeast phenomic model of deoxycytidine kinase to predict genetic buffering of nucleoside analog cytotoxicity": 19_0327_HaL_to_orig_hmap_Heatmaps.pdf

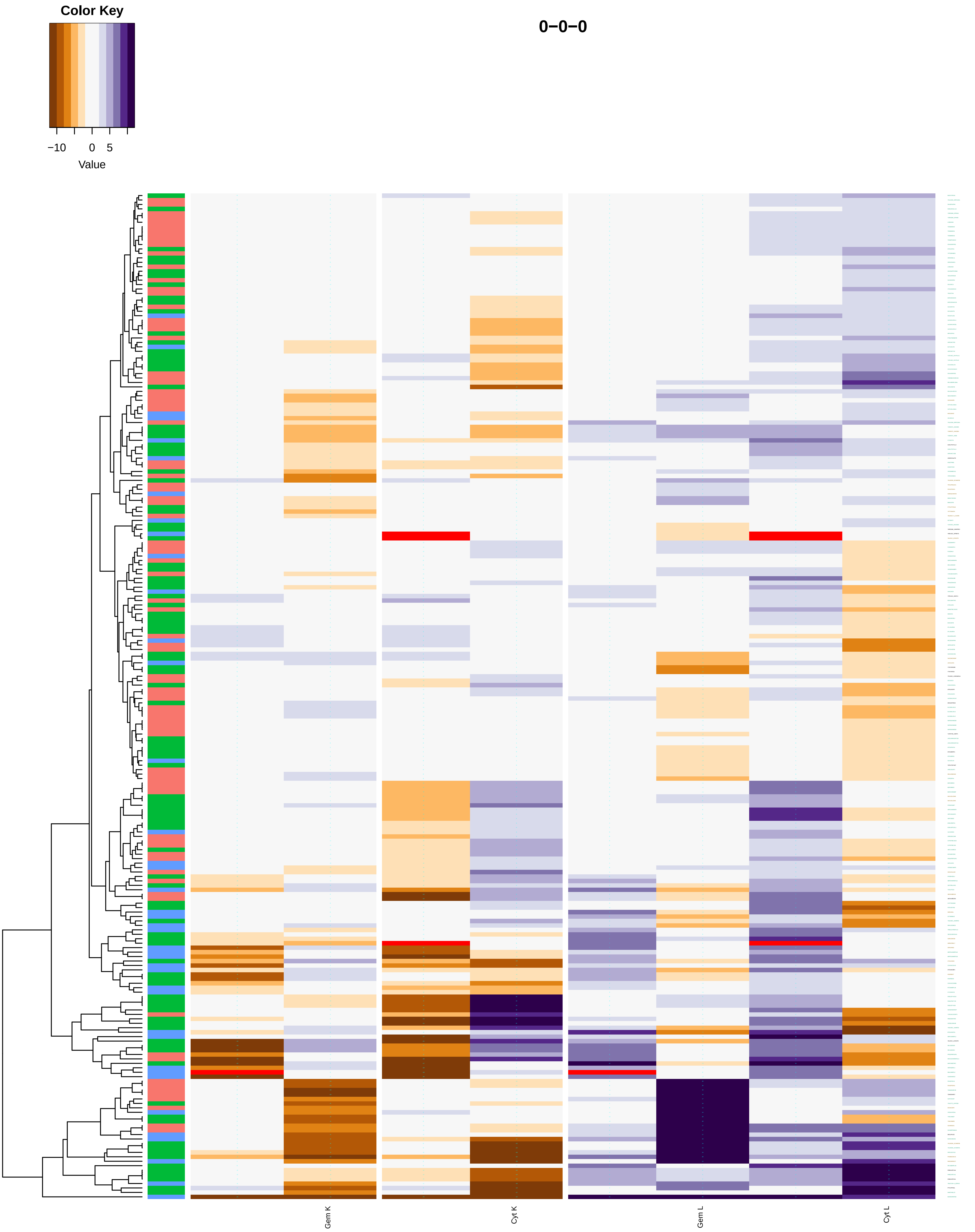

Color Key
−10
0
5
Value
0−0−0
Gem K
Cyt K
Gem L
Cyt L

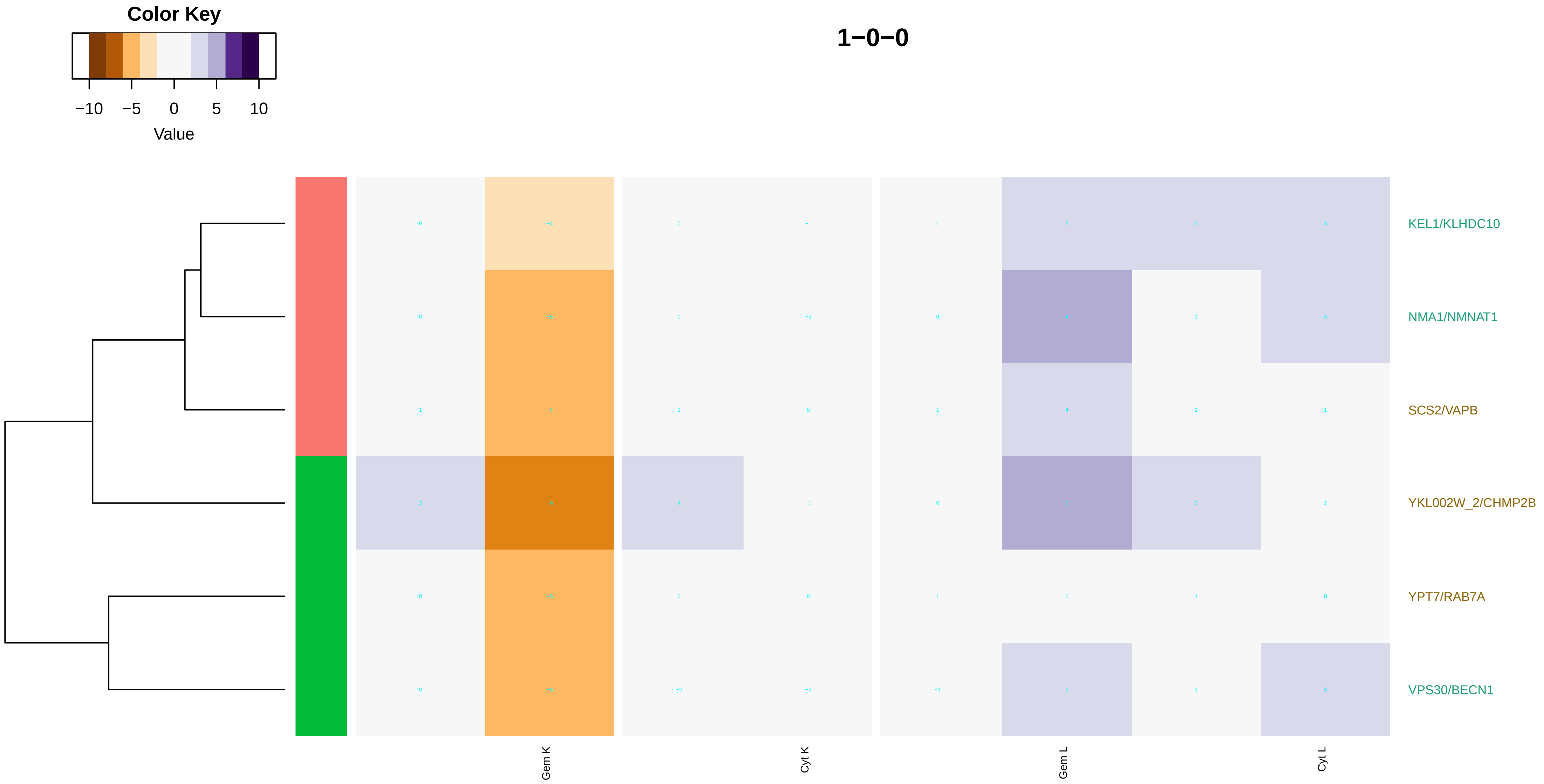
Color Key
−10 −5 0 5 10
Value
1−0−0
KEL1/KLHDC10
NMA1/NMNAT1
SCS2/VAPB
YKL002W_2/CHMP2B
YPT7/RAB7A
VPS30/BECN1
Gem K
Cyt K
Gem L
Cyt L

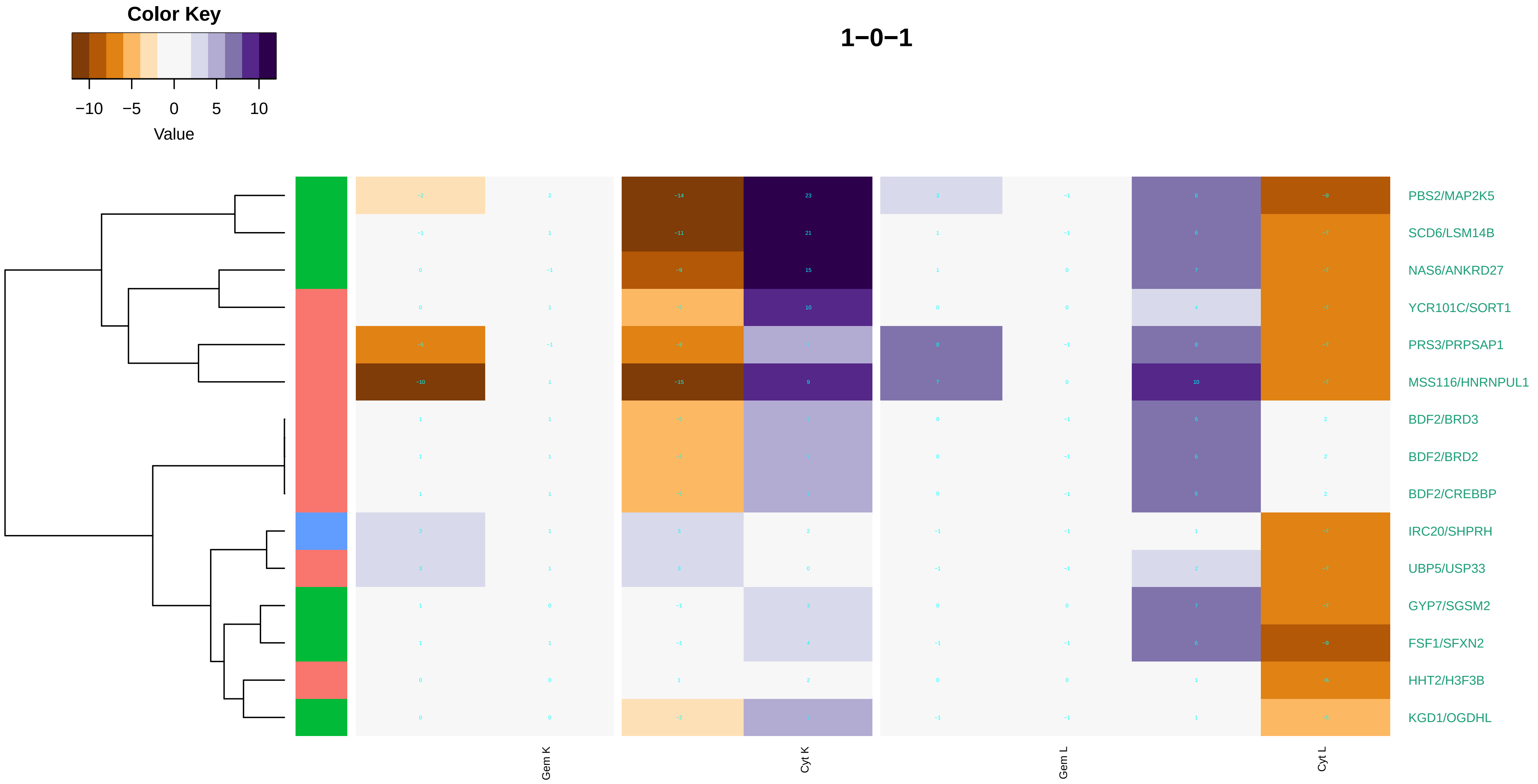
Color Key
−10 −5 0 5 10
Value
1−0−1
PBS2/MAP2K5
SCD6/LSM14B
NAS6/ANKRD27
YCR101C/SORT1
PRS3/PRPSAP1
MSS116/HNRNPUL1
BDF2/BRD3
BDF2/BRD2
BDF2/CREBBP
IRC20/SHPRH
UBP5/USP33
GYP7/SGSM2
FSF1/SFXN2
HHT2/H3F3B
KGD1/OGDHL
Gem K
Cyt K
Gem L
Cyt L

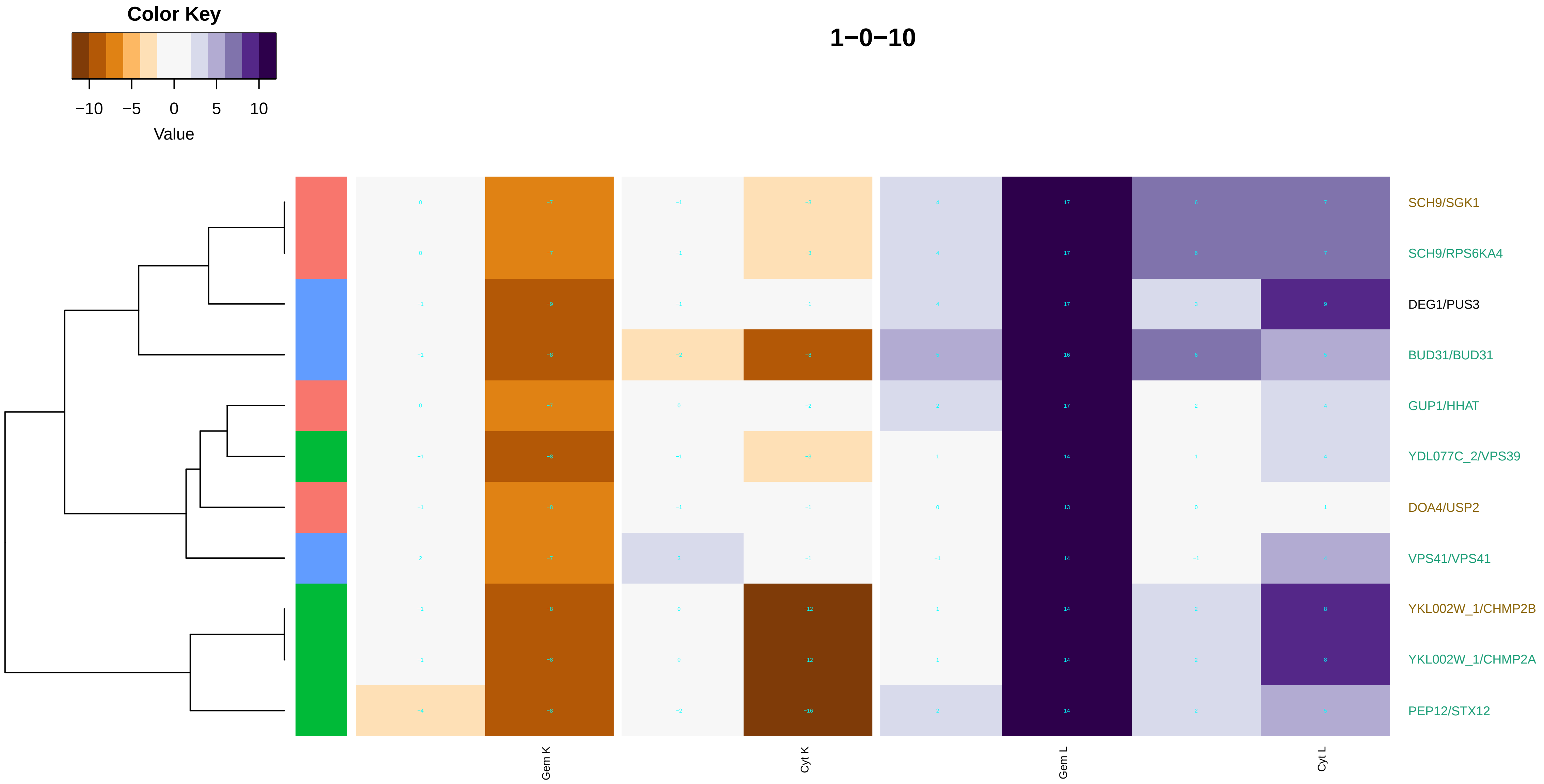
Color Key
−10
−5
0
5
10
Value
1−0−10
Gem K
Cyt K
Gem L
Cyt L
SCH9/SGK1
SCH9/RPS6KA4
DEG1/PUS3
BUD31/BUD31
GUP1/HHAT
YDL077C_2/VPS39
DOA4/USP2
VPS41/VPS41
YKL002W_1/CHMP2B
YKL002W_1/CHMP2A
PEP12/STX12

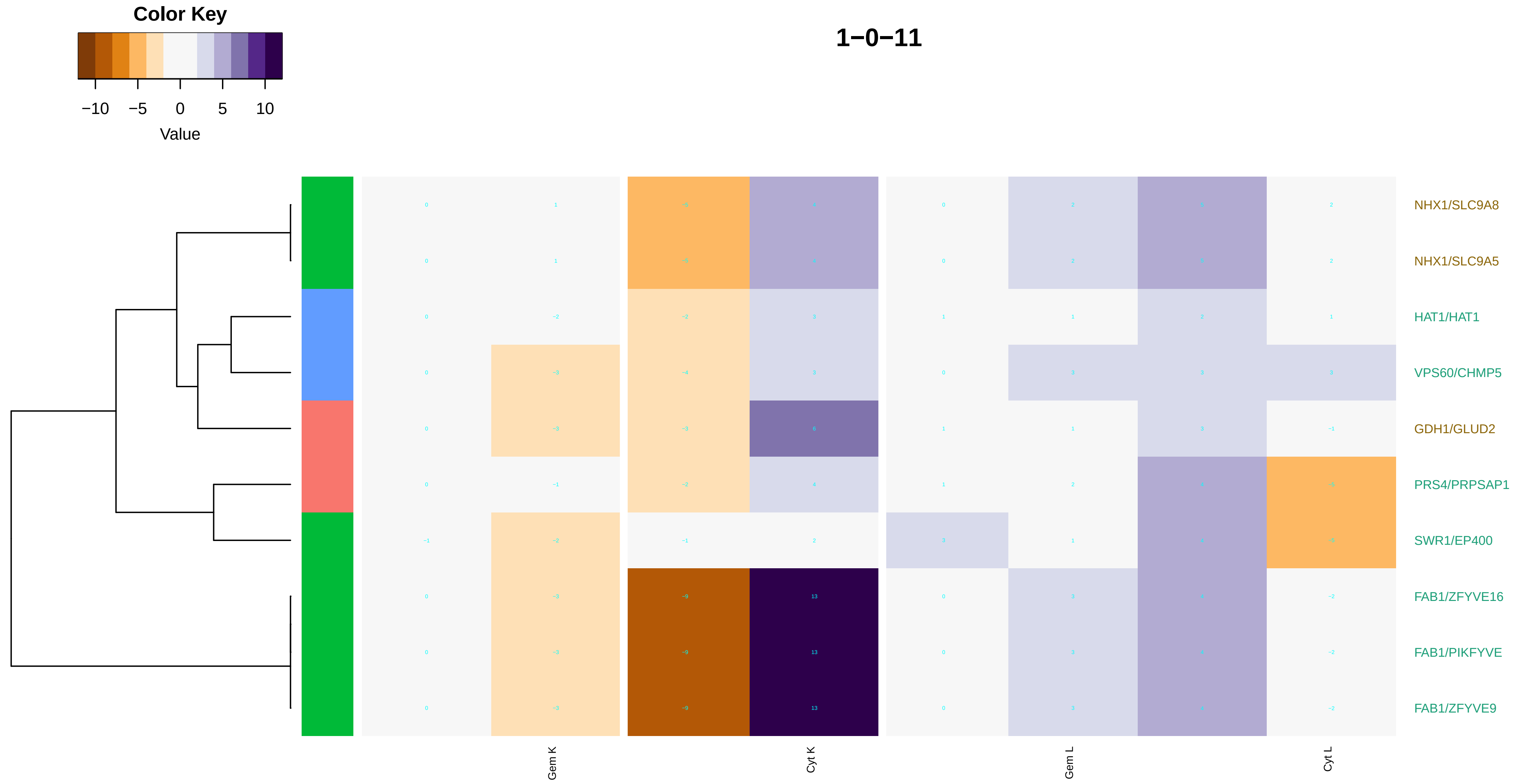

Color Key
−10
−5
0
5
10
Value
1−0−11
NHX1/SLC9A8
NHX1/SLC9A5
HAT1/HAT1
VPS60/CHMP5
GDH1/GLUD2
PRS4/PRPSAP1
SWR1/EP400
FAB1/ZFYVE16
FAB1/PIKFYVE
FAB1/ZFYVE9
Gem K
Cyt K
Gem L
Cyt L

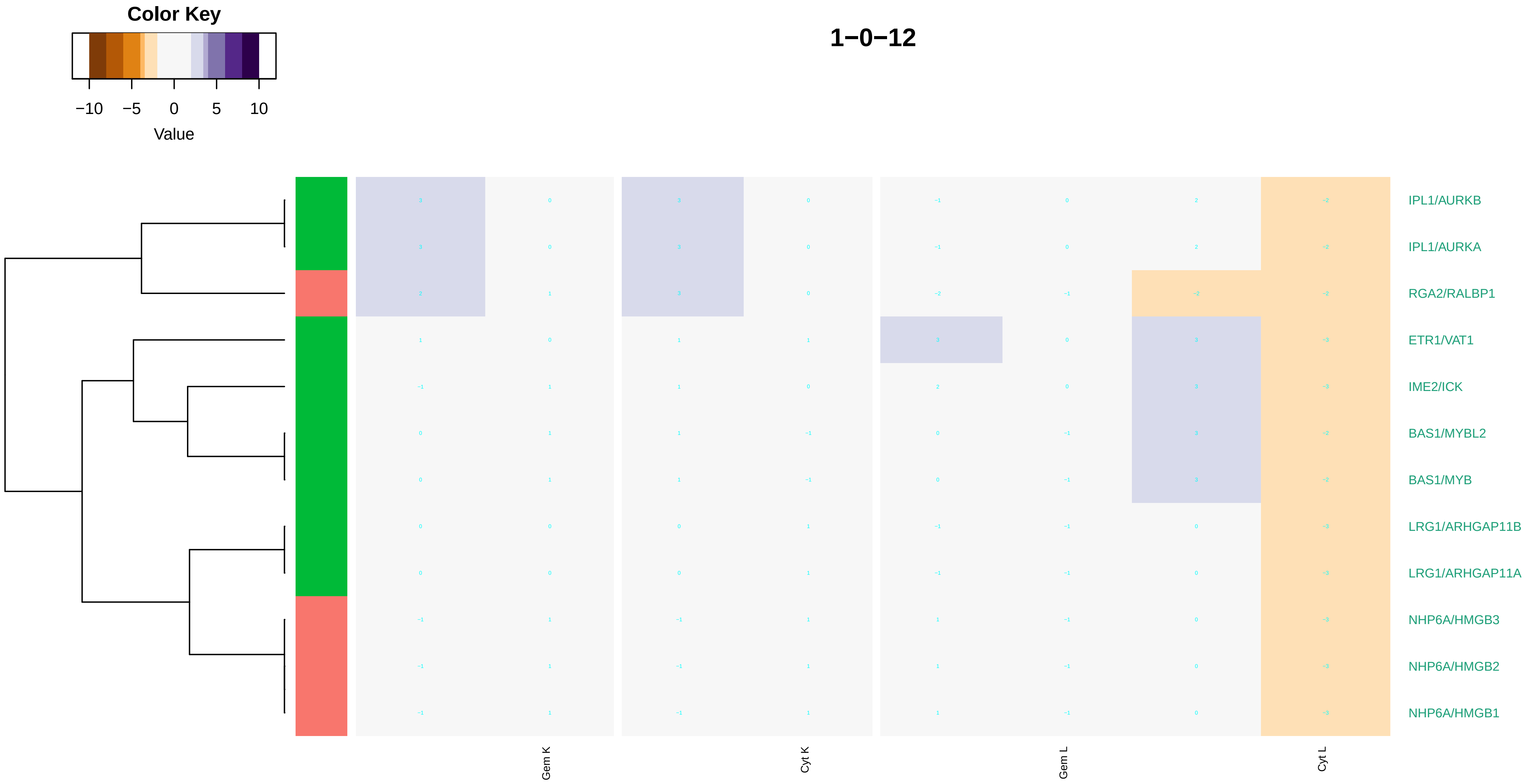
Color Key
−10
−5
0
5
10
Value
1−0−12
IPL1/AURKB
IPL1/AURKA
RGA2/RALBP1
ETR1/VAT1
IME2/ICK
BAS1/MYBL2
BAS1/MYB
LRG1/ARHGAP11B
LRG1/ARHGAP11A
NHP6A/HMGB3
NHP6A/HMGB2
NHP6A/HMGB1
Gem K
Cyt K
Gem L
Cyt L

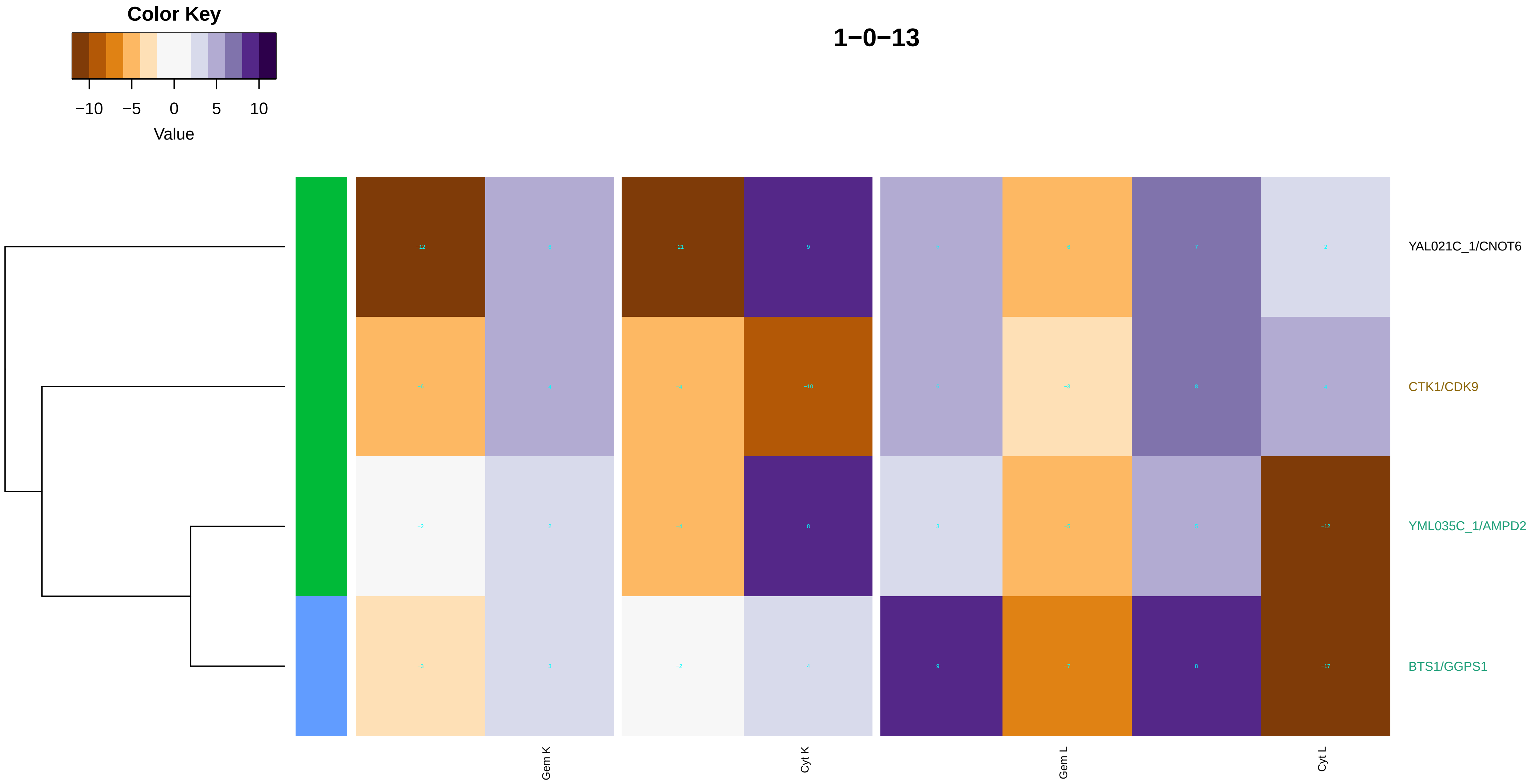
Color Key
−10
−5
0
5
10
Value
1−0−13
YAL021C_1/CNOT6
CTK1/CDK9
YML035C_1/AMPD2
BTS1/GGPS1
Gem K
Cyt K
Gem L
Cyt L

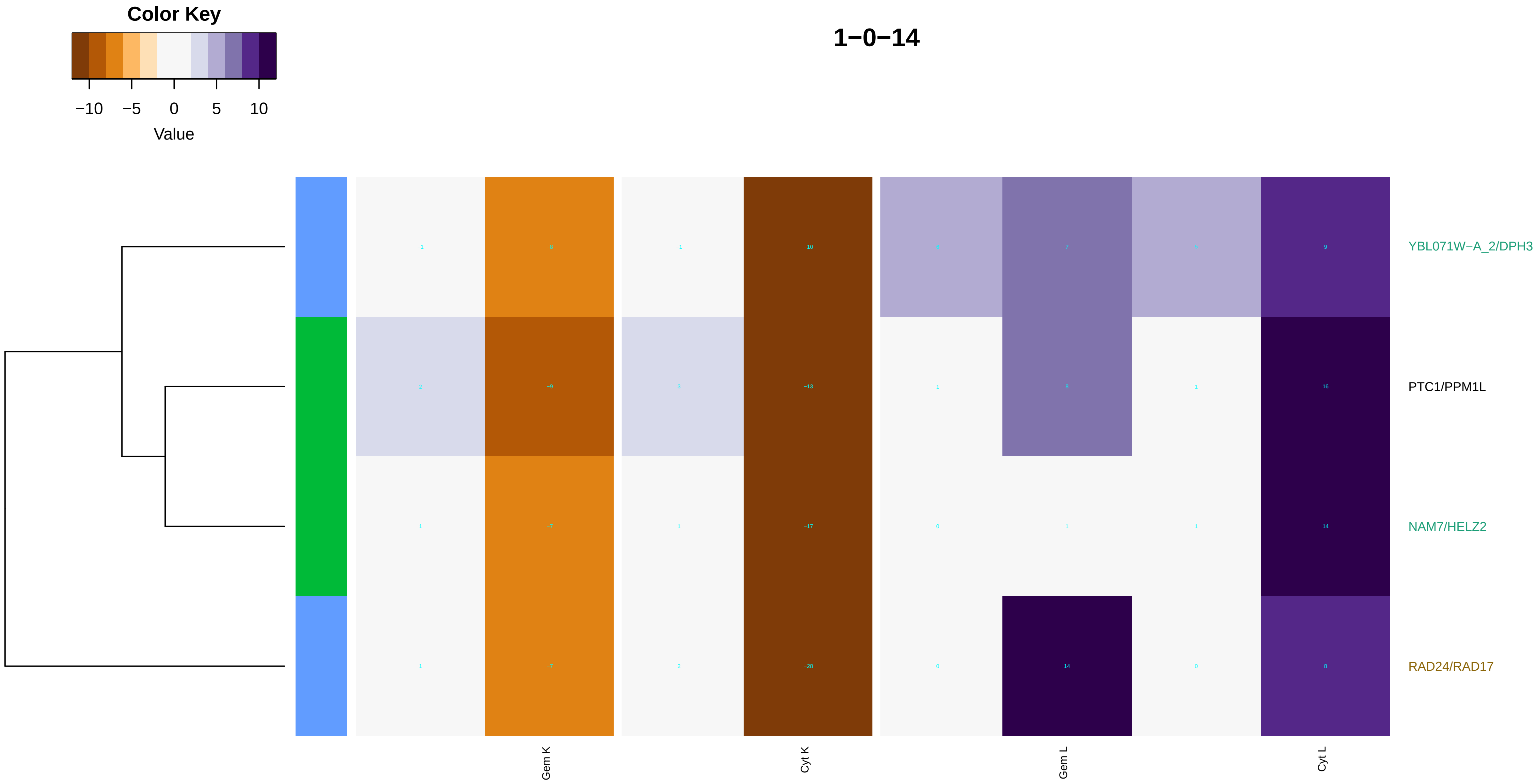
Color Key
−10
−5
0
5
10
Value
1−0−14
YBL071W−A_2/DPH3
PTC1/PPM1L
NAM7/HELZ2
RAD24/RAD17
Gem K
Cyt K
Gem L
Cyt L

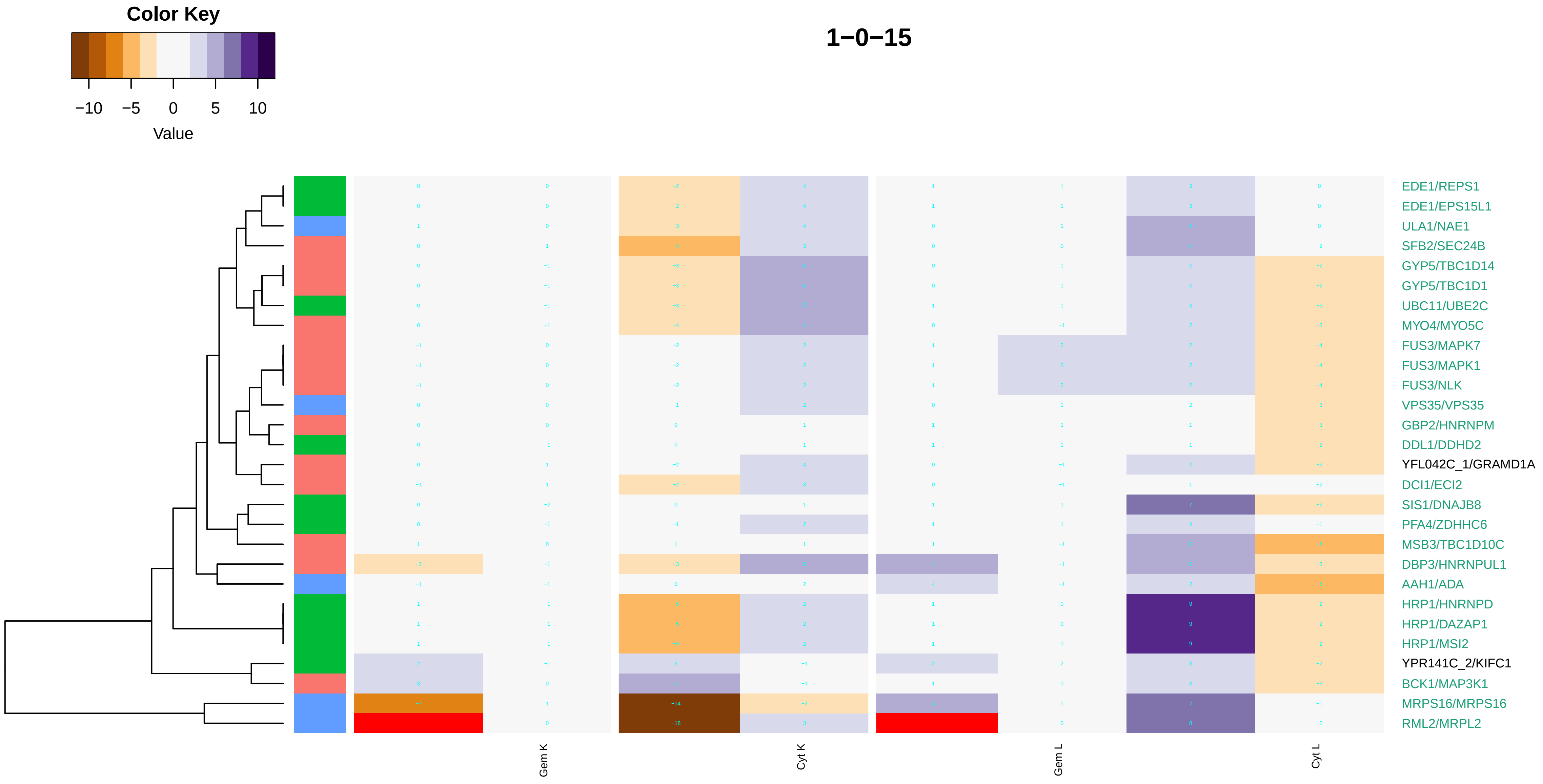
Color Key
−10 −5 0 5 10
Value
1−0−15
EDE1/REPS1
EDE1/EPS15L1
ULA1/NAE1
SFB2/SEC24B
GYP5/TBC1D14
GYP5/TBC1D1
UBC11/UBE2C
MYO4/MYO5C
FUS3/MAPK7
FUS3/MAPK1
FUS3/NLK
VPS35/VPS35
GBP2/HNRNPM
DDL1/DDHD2
YFL042C_1/GRAMD1A
DCI1/ECI2
SIS1/DNAJB8
PFA4/ZDHHC6
MSB3/TBC1D10C
DBP3/HNRNPUL1
AAH1/ADA
HRP1/HNRNPD
HRP1/DAZAP1
HRP1/MSI2
YPR141C_2/KIFC1
BCK1/MAP3K1
MRPS16/MRPS16
RML2/MRPL2
Gem K
Cyt K
Gem L
Cyt L

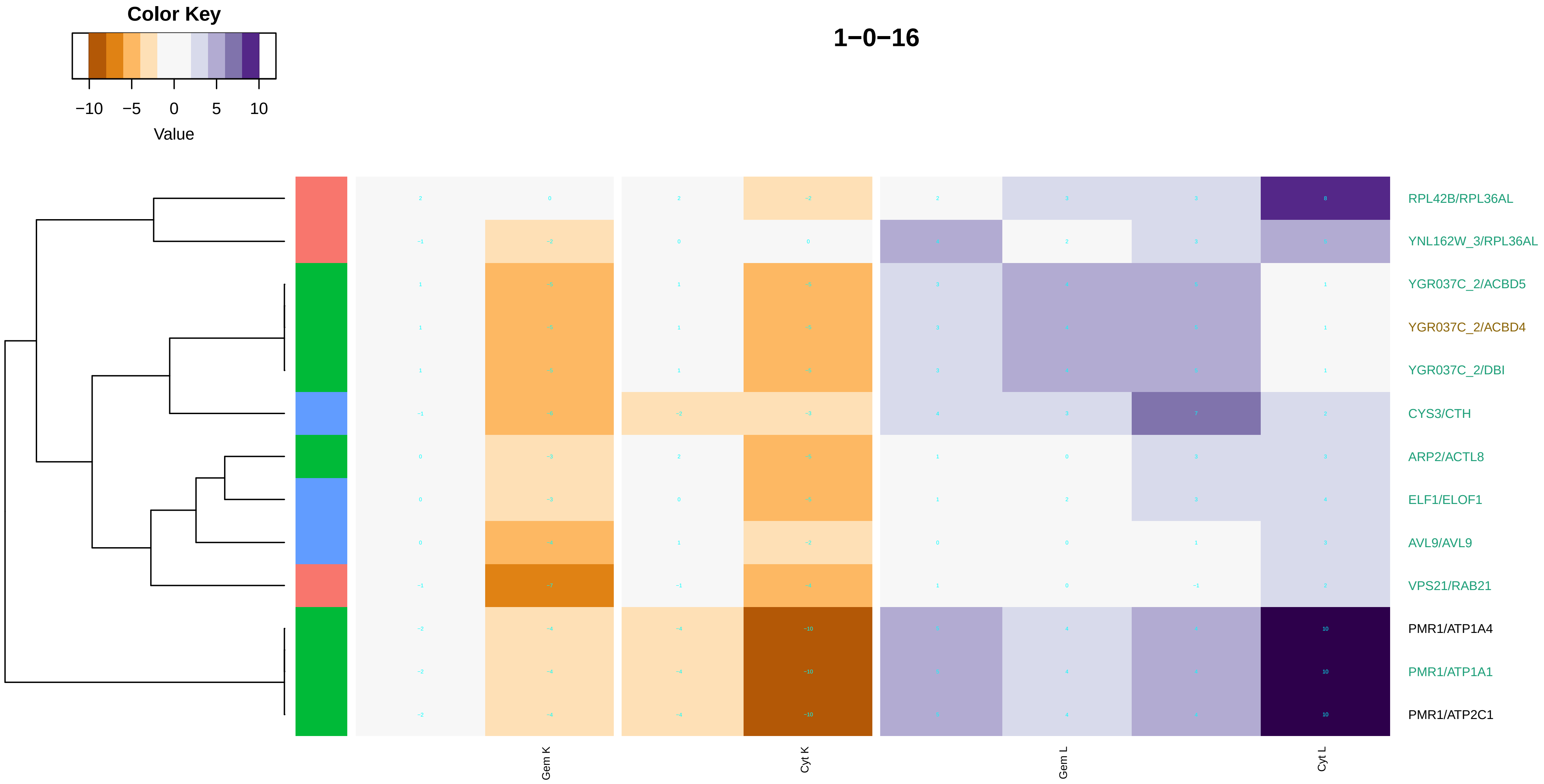
Color Key
−10
−5
0
5
10
Value
1−0−16
RPL42B/RPL36AL
YNL162W_3/RPL36AL
YGR037C_2/ACBD5
YGR037C_2/ACBD4
YGR037C_2/DBI
CYS3/CTH
ARP2/ACTL8
ELF1/ELOF1
AVL9/AVL9
VPS21/RAB21
PMR1/ATP1A4
PMR1/ATP1A1
PMR1/ATP2C1
Gem K
Cyt K
Gem L
Cyt L

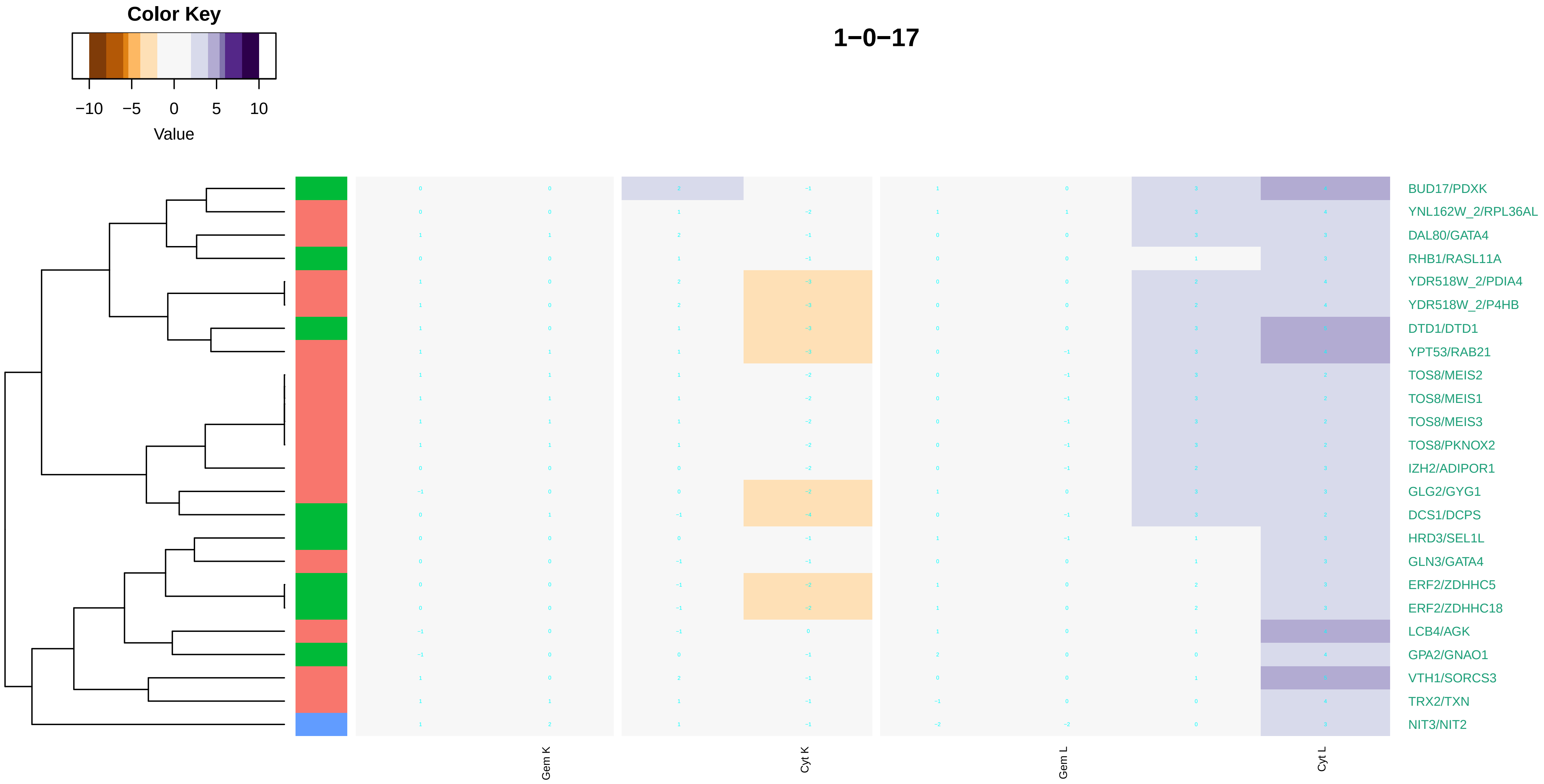

Color Key
−10 −5 0 5 10
Value
1−0−17
Gem K
Cyt K
Gem L
Cyt L
BUD17/PDXK
YNL162W_2/RPL36AL
DAL80/GATA4
RHB1/RASL11A
YDR518W_2/PDIA4
YDR518W_2/P4HB
DTD1/DTD1
YPT53/RAB21
TOS8/MEIS2
TOS8/MEIS1
TOS8/MEIS3
TOS8/PKNOX2
IZH2/ADIPOR1
GLG2/GYG1
DCS1/DCPS
HRD3/SEL1L
GLN3/GATA4
ERF2/ZDHHC5
ERF2/ZDHHC18
LCB4/AGK
GPA2/GNAO1
VTH1/SORCS3
TRX2/TXN
NIT3/NIT2

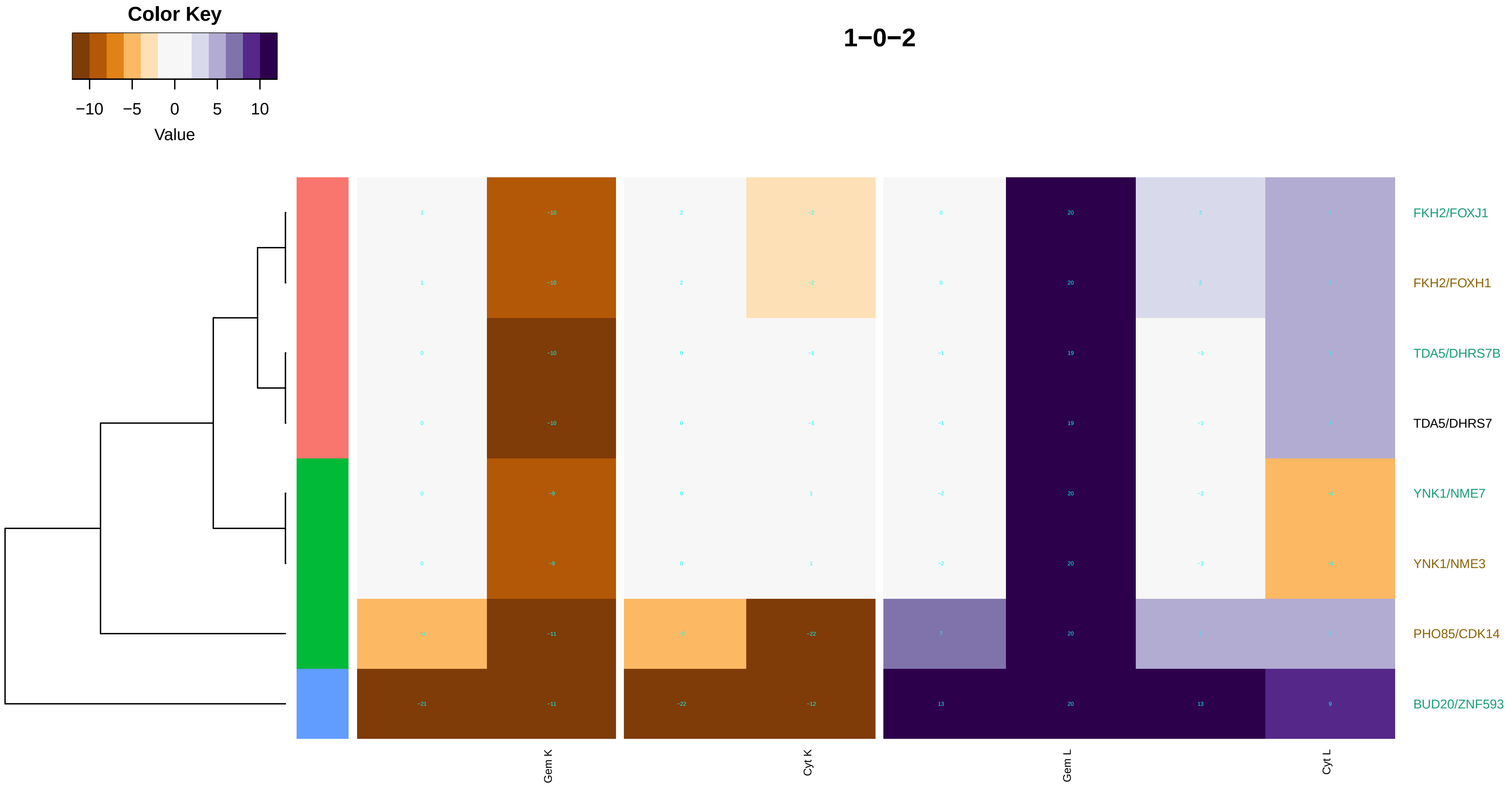
Color Key
−10
−5
0
5
10
Value
1−0−2
FKH2/FOXJ1
FKH2/FOXH1
TDA5/DHRS7B
TDA5/DHRS7
YNK1/NME7
YNK1/NME3
PHO85/CDK14
BUD20/ZNF593
Gem K
Cyt K
Gem L
Cyt L

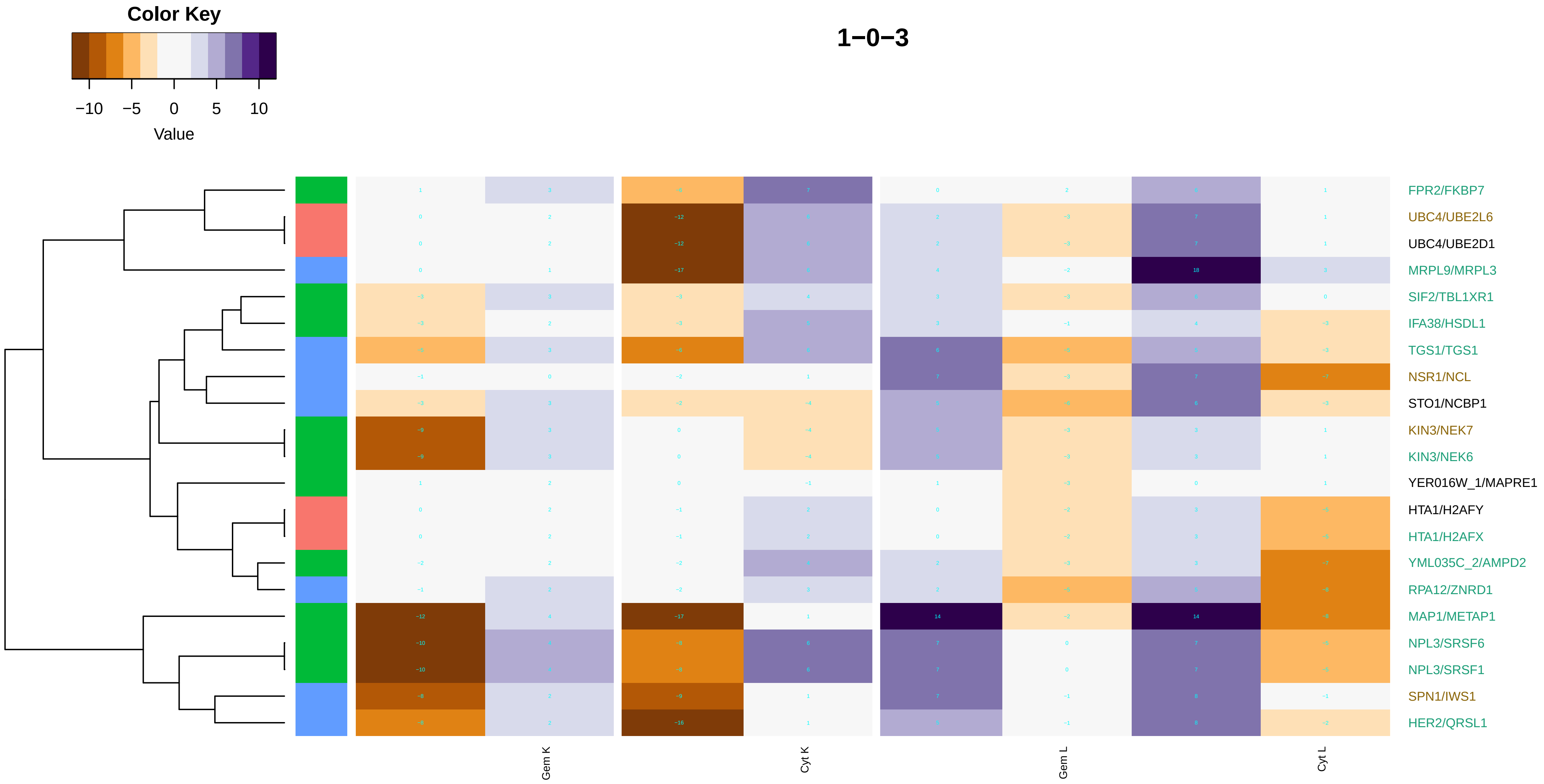

Color Key
−10 −5 0 5 10
Value
1−0−3
FPR2/FKBP7
UBC4/UBE2L6
UBC4/UBE2D1
MRPL9/MRPL3
SIF2/TBL1XR1
IFA38/HSDL1
TGS1/TGS1
NSR1/NCL
STO1/NCBP1
KIN3/NEK7
KIN3/NEK6
YER016W_1/MAPRE1
HTA1/H2AFY
HTA1/H2AFX
YML035C_2/AMPD2
RPA12/ZNRD1
MAP1/METAP1
NPL3/SRSF6
NPL3/SRSF1
SPN1/IWS1
HER2/QRSL1
Gem K
Cyt K
Gem L
Cyt L

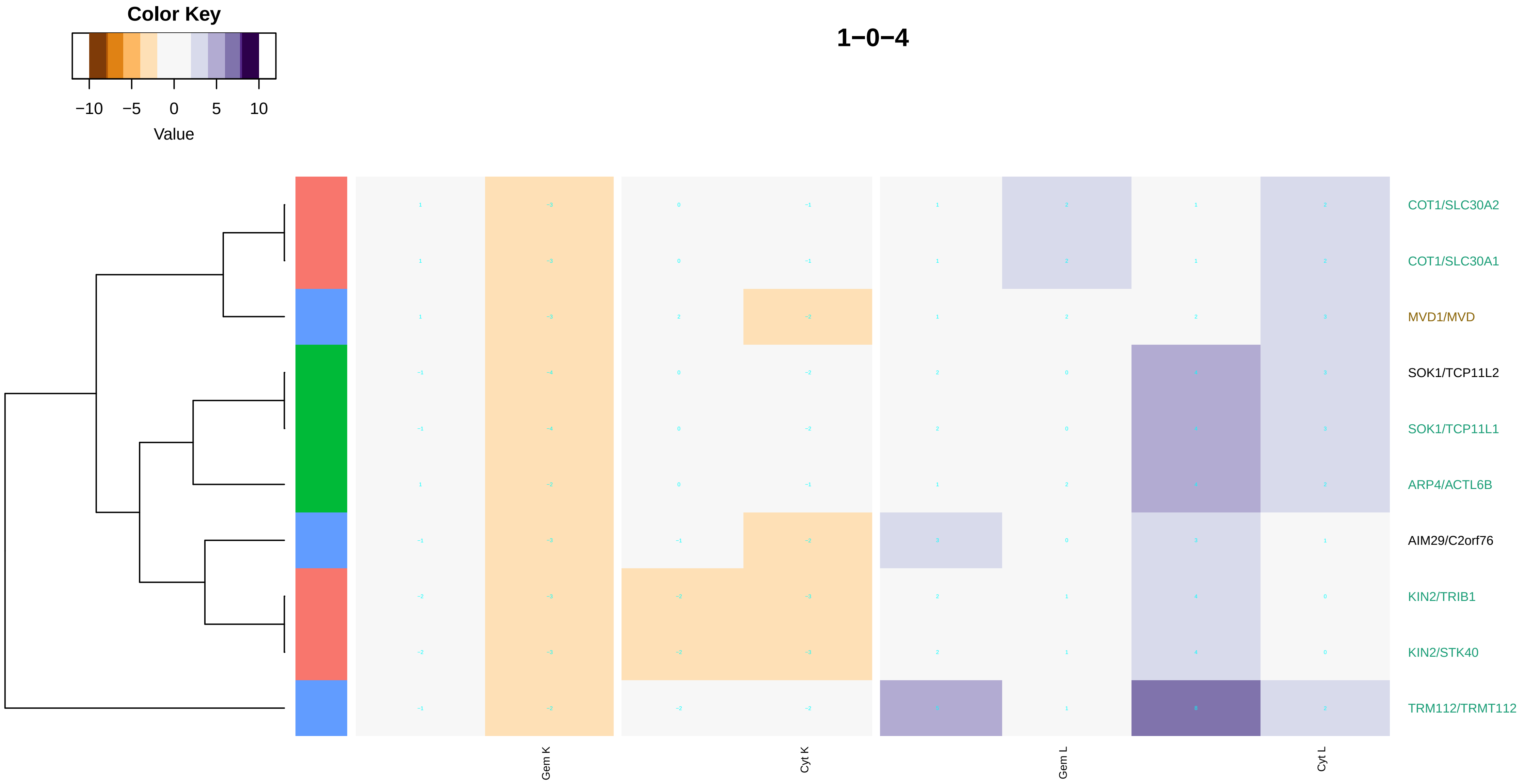
Color Key
−10 −5 0 5 10
Value
1−0−4
COT1/SLC30A2
COT1/SLC30A1
MVD1/MVD
SOK1/TCP11L2
SOK1/TCP11L1
ARP4/ACTL6B
AIM29/C2orf76
KIN2/TRIB1
KIN2/STK40
TRM112/TRMT112
Gem K
Cyt K
Gem L
Cyt L

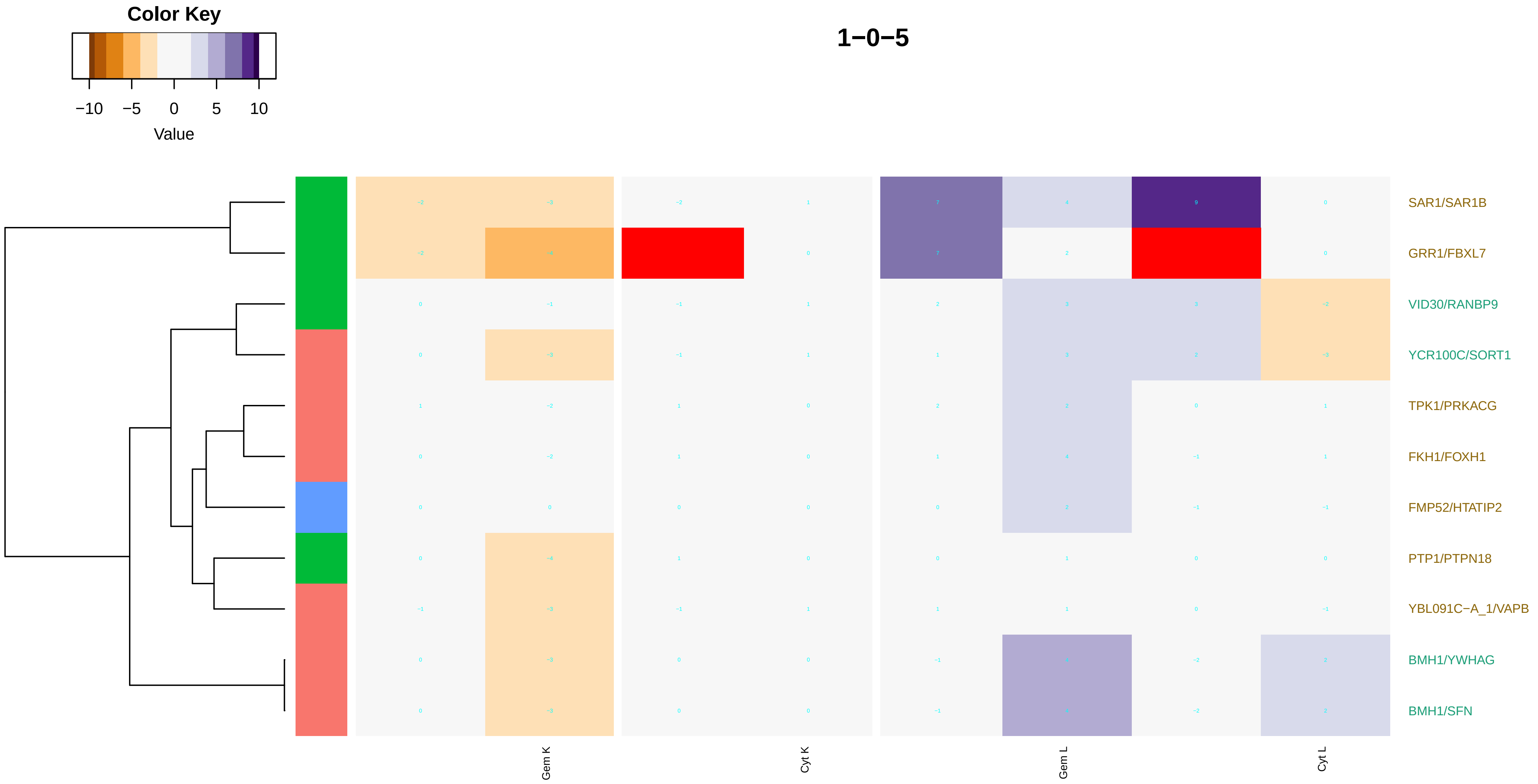
Color Key
−10 −5 0 5 10
Value
1−0−5
SAR1/SAR1B
GRR1/FBXL7
VID30/RANBP9
YCR100C/SORT1
TPK1/PRKACG
FKH1/FOXH1
FMP52/HTATIP2
PTP1/PTPN18
YBL091C−A_1/VAPB
BMH1/YWHAG
BMH1/SFN
Gem K
Cyt K
Gem L
Cyt L

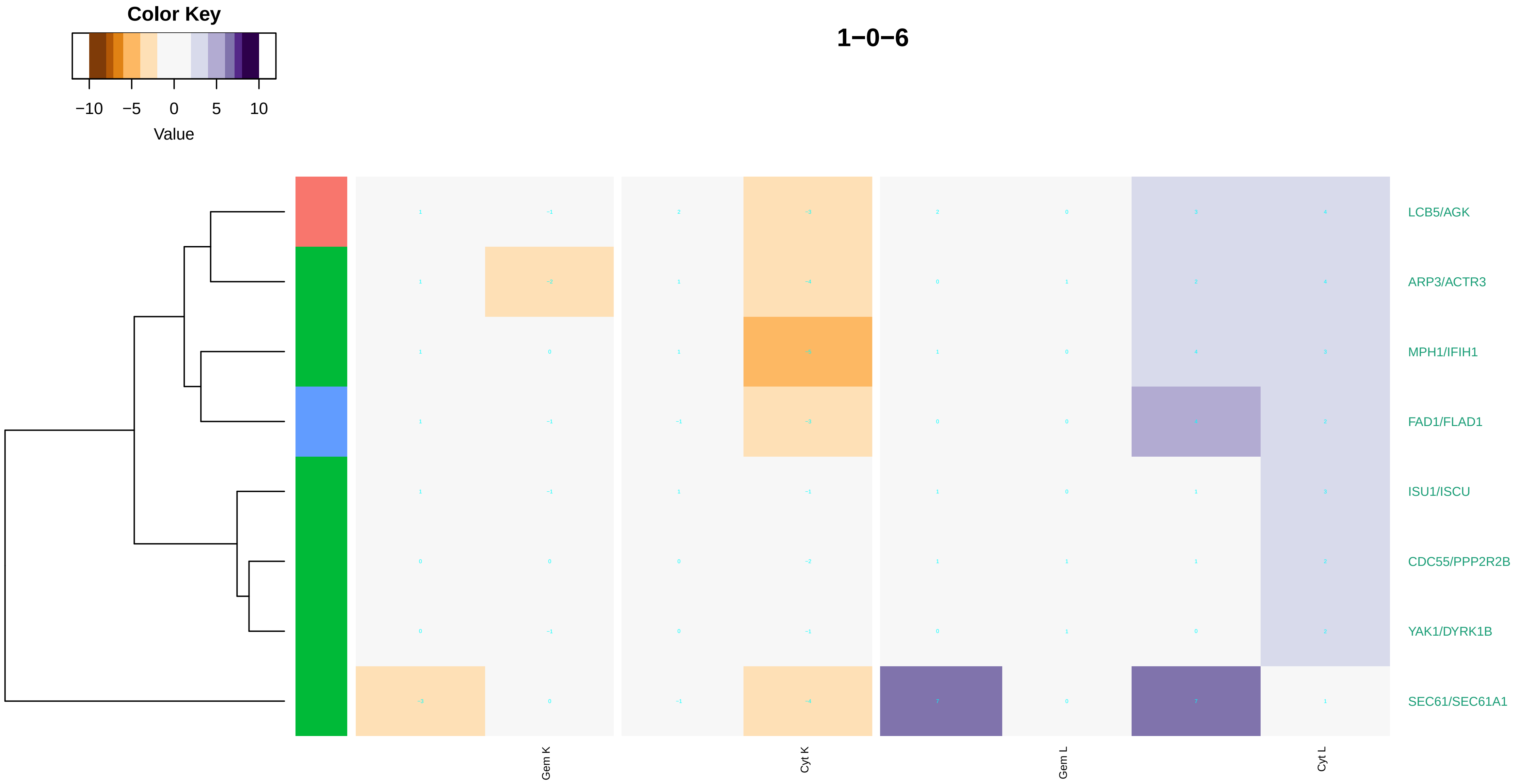
Color Key
−10
−5
0
5
10
Value
1−0−6
LCB5/AGK
ARP3/ACTR3
MPH1/IFIH1
FAD1/FLAD1
ISU1/ISCU
CDC55/PPP2R2B
YAK1/DYRK1B
SEC61/SEC61A1
Gem K
Cyt K
Gem L
Cyt L

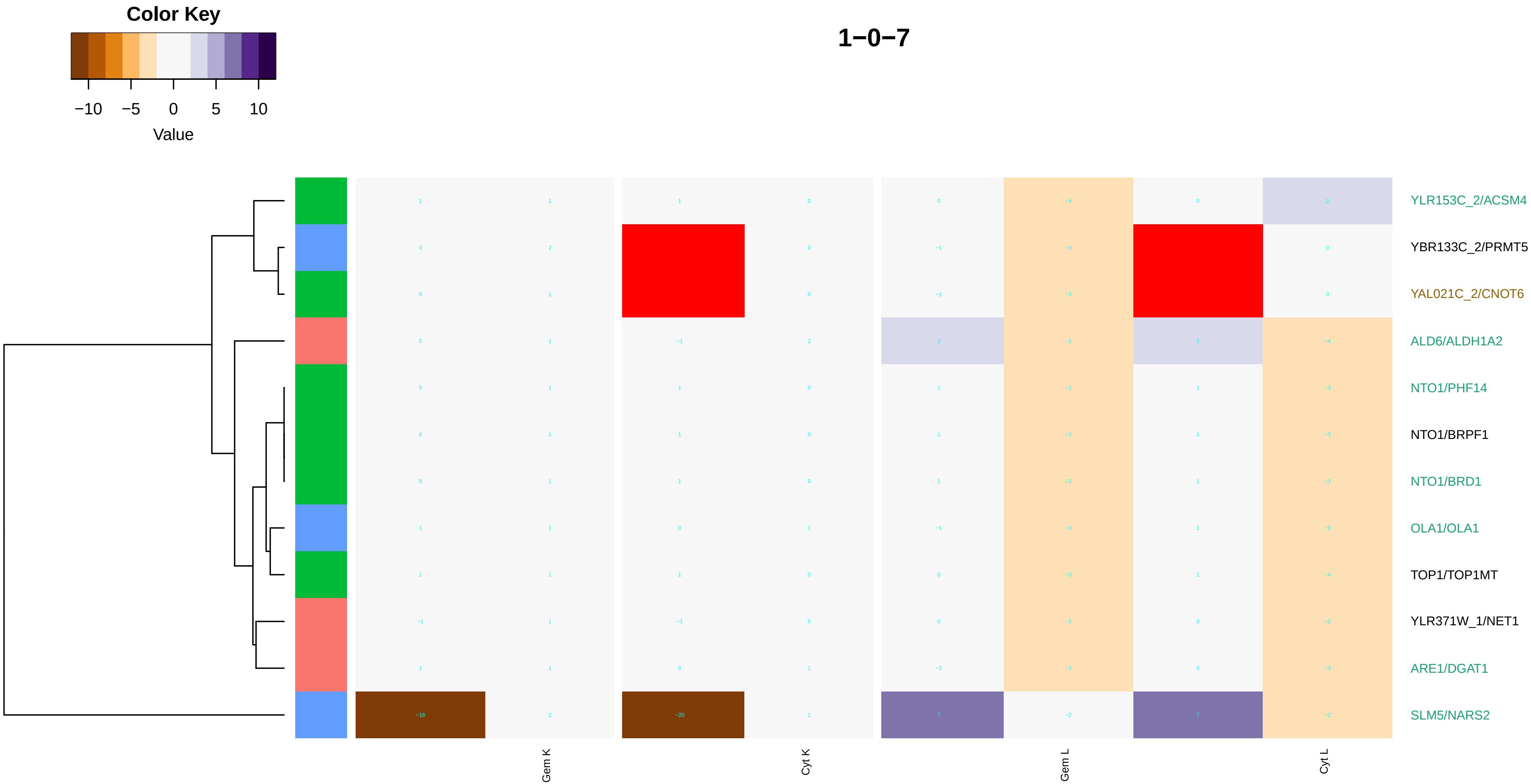
Color Key
−10
−5
0
5
10
Value
1−0−7
YLR153C_2/ACSM4
YBR133C_2/PRMT5
YAL021C_2/CNOT6
ALD6/ALDH1A2
NTO1/PHF14
NTO1/BRPF1
NTO1/BRD1
OLA1/OLA1
TOP1/TOP1MT
YLR371W_1/NET1
ARE1/DGAT1
SLM5/NARS2
Gem K
Cyt K
Gem L
Cyt L

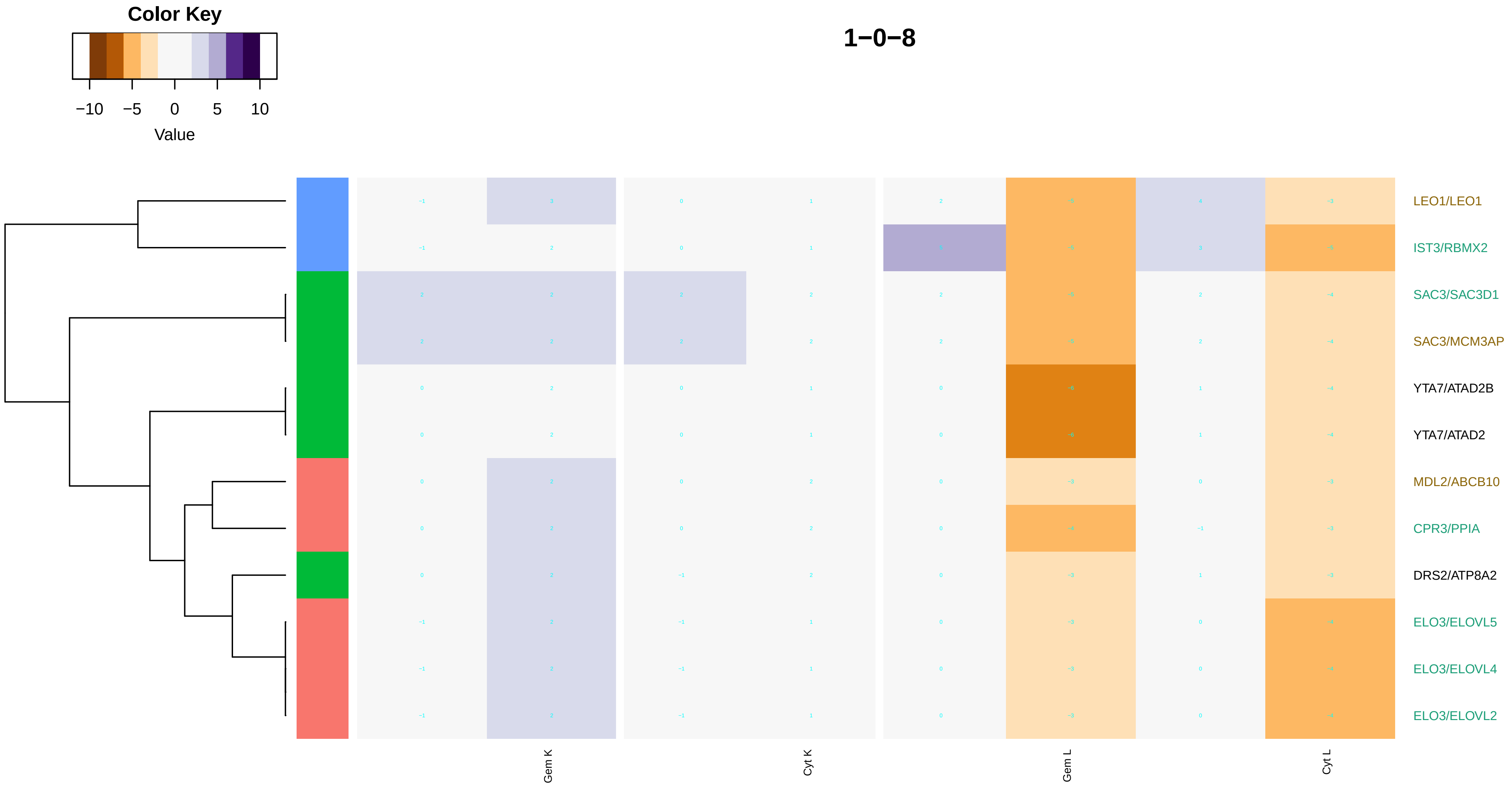
Color Key
−10
−5
0
5
10
Value
1−0−8
LEO1/LEO1
IST3/RBMX2
SAC3/SAC3D1
SAC3/MCM3AP
YTA7/ATAD2B
YTA7/ATAD2
MDL2/ABCB10
CPR3/PPIA
DRS2/ATP8A2
ELO3/ELOVL5
ELO3/ELOVL4
ELO3/ELOVL2
Gem K
Cyt K
Gem L
Cyt L

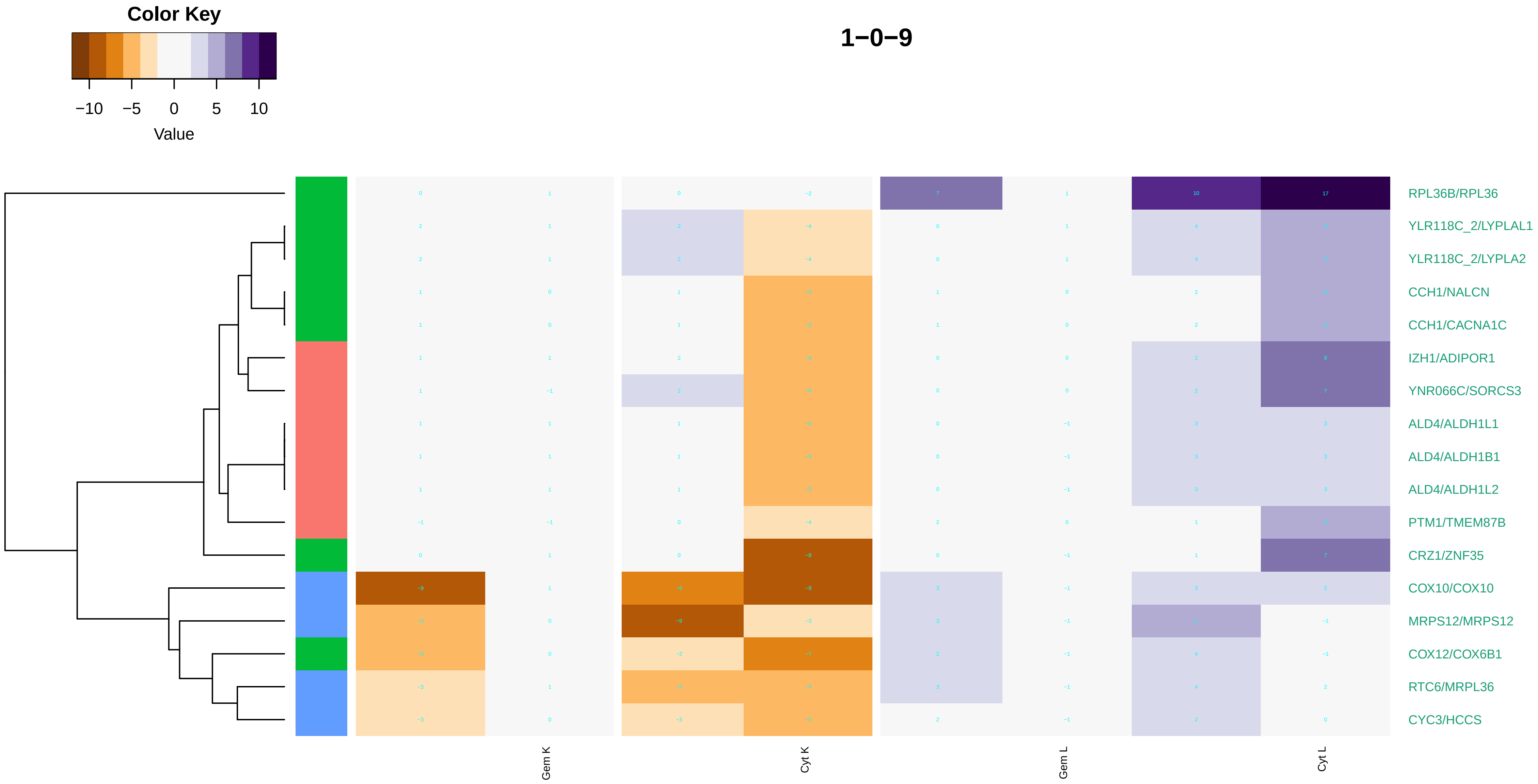
Color Key
−10 −5 0 5 10
Value
1−0−9
RPL36B/RPL36
YLR118C_2/LYPLAL1
YLR118C_2/LYPLA2
CCH1/NALCN
CCH1/CACNA1C
IZH1/ADIPOR1
YNR066C/SORCS3
ALD4/ALDH1L1
ALD4/ALDH1B1
ALD4/ALDH1L2
PTM1/TMEM87B
CRZ1/ZNF35
COX10/COX10
MRPS12/MRPS12
COX12/COX6B1
RTC6/MRPL36
CYC3/HCCS
Gem K
Cyt K
Gem L
Cyt L

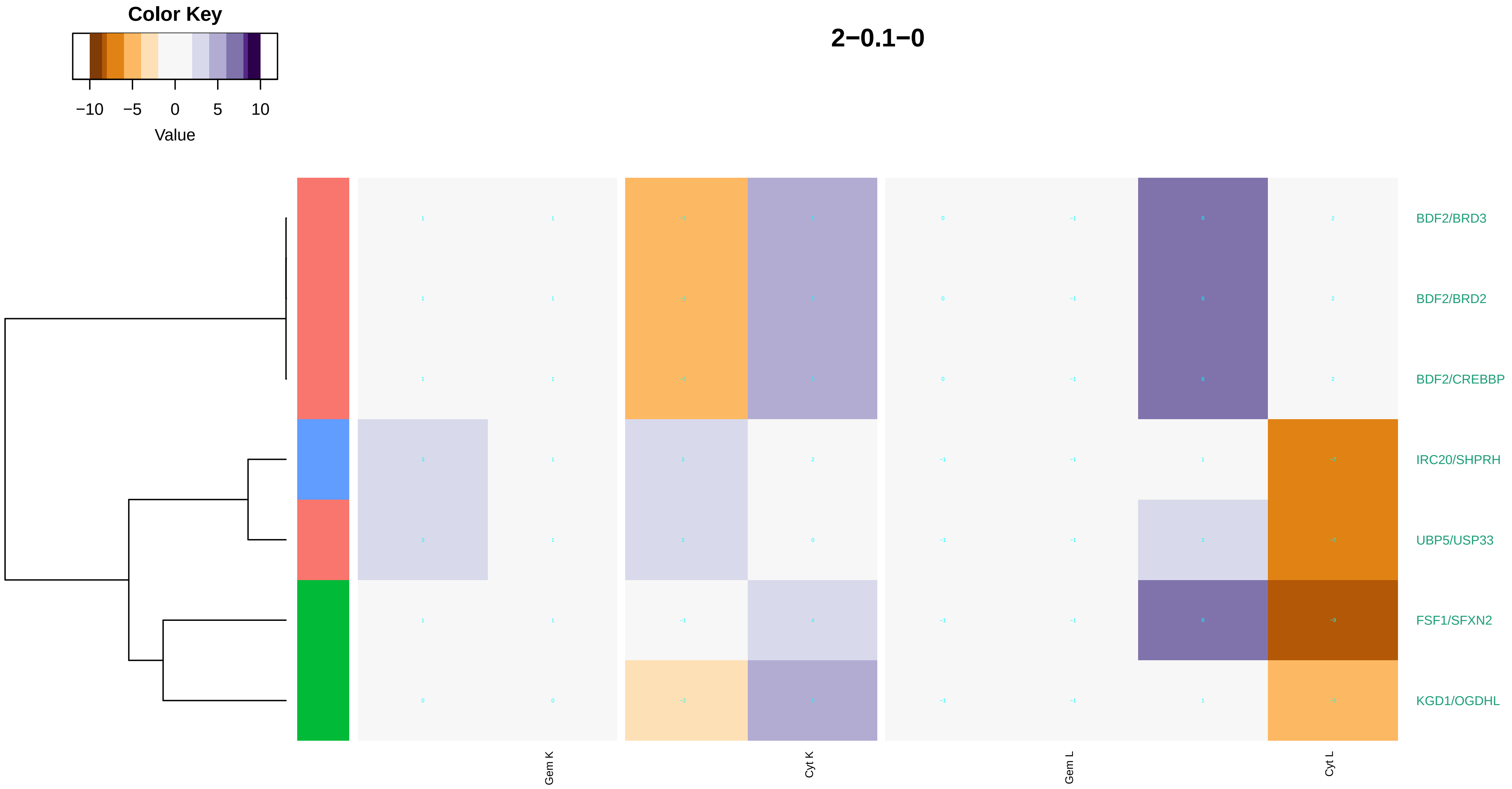

Color Key
−10
−5
0
5
10
Value
2−0.1−0
BDF2/BRD3
BDF2/BRD2
BDF2/CREBBP
IRC20/SHPRH
UBP5/USP33
FSF1/SFXN2
KGD1/OGDHL
Gem K
Cyt K
Gem L
Cyt L

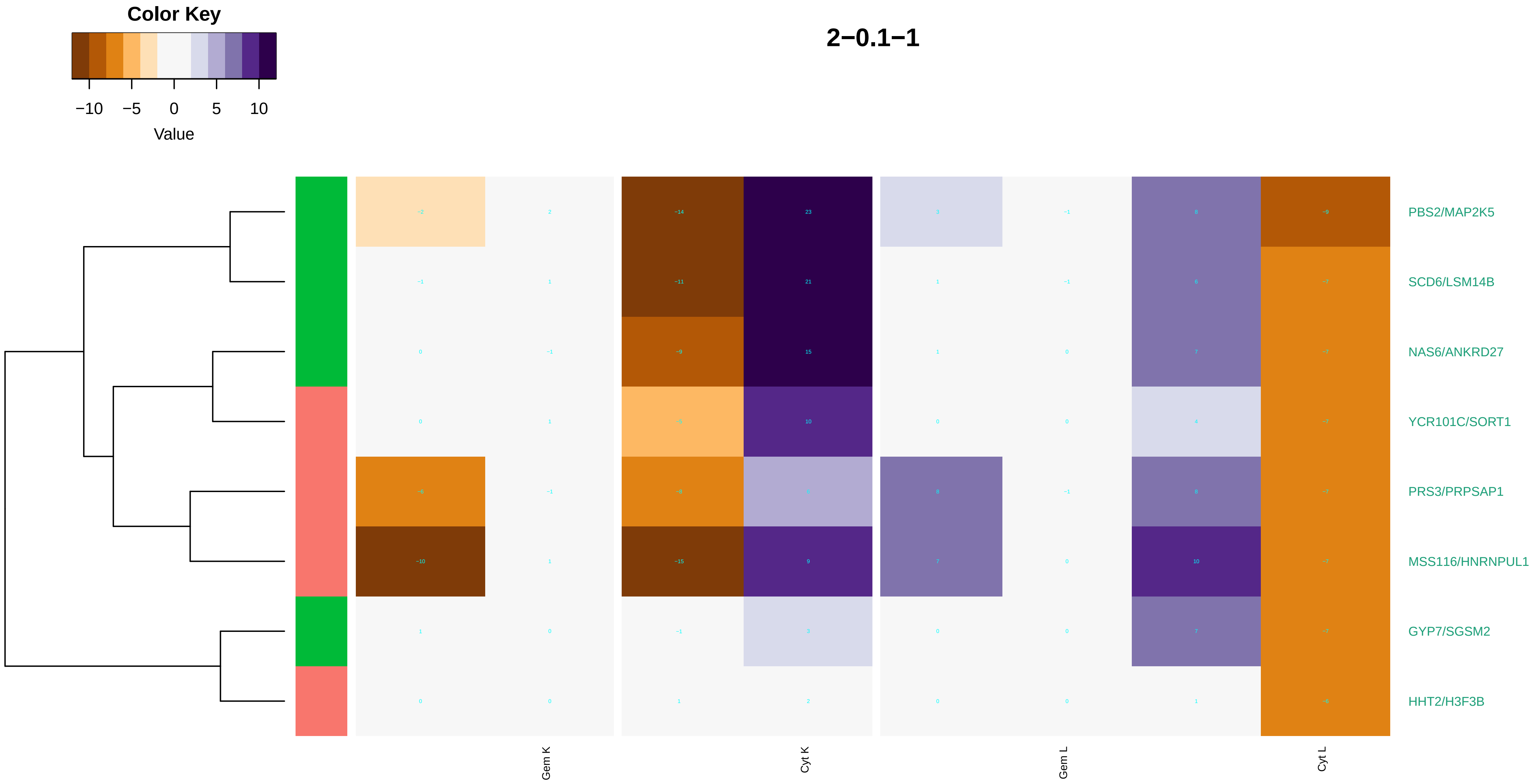
Color Key
−10
−5
0
5
10
Value
2−0.1−1
PBS2/MAP2K5
SCD6/LSM14B
NAS6/ANKRD27
YCR101C/SORT1
PRS3/PRPSAP1
MSS116/HNRNPUL1
GYP7/SGSM2
HHT2/H3F3B
Gem K
Cyt K
Gem L
Cyt L

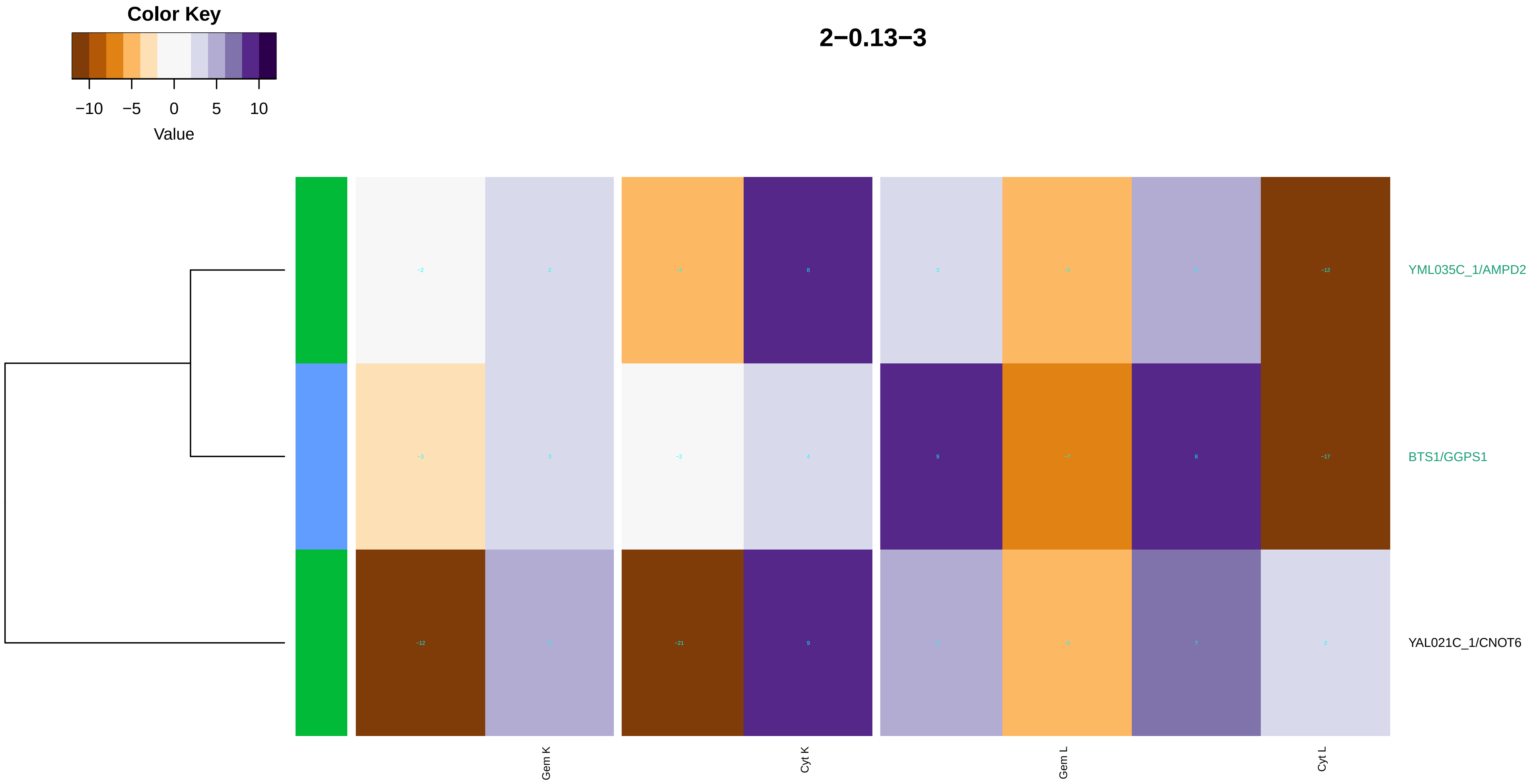
Color Key
−10
−5
0
5
10
Value
2−0.13−3
YML035C_1/AMPD2
BTS1/GGPS1
YAL021C_1/CNOT6
Gem K
Cyt K
Gem L
Cyt L

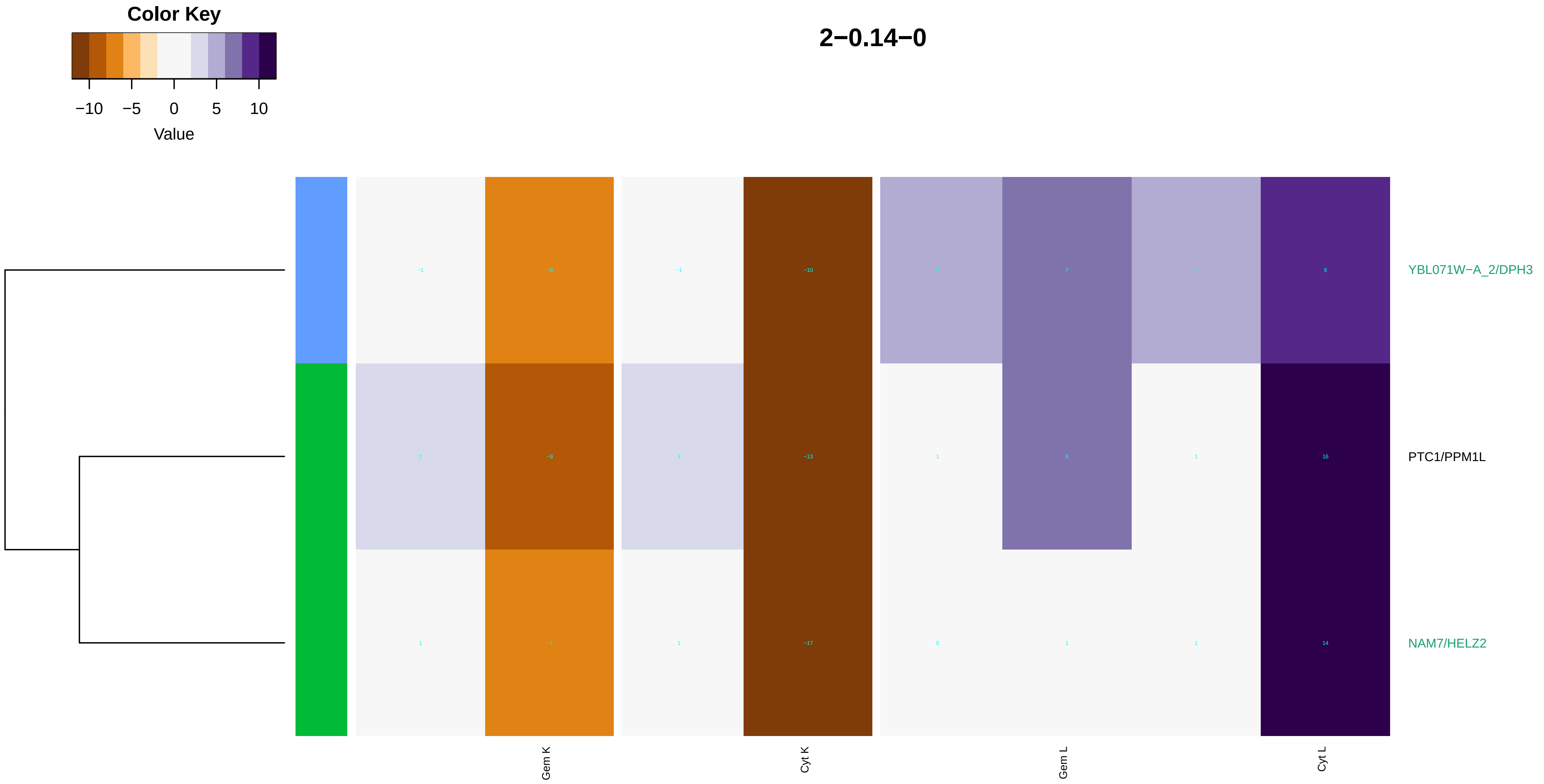
Color Key
−10 −5 0 5 10
Value
2−0.14−0
YBL071W−A_2/DPH3
PTC1/PPM1L
NAM7/HELZ2
Gem K
Cyt K
Gem L
Cyt L

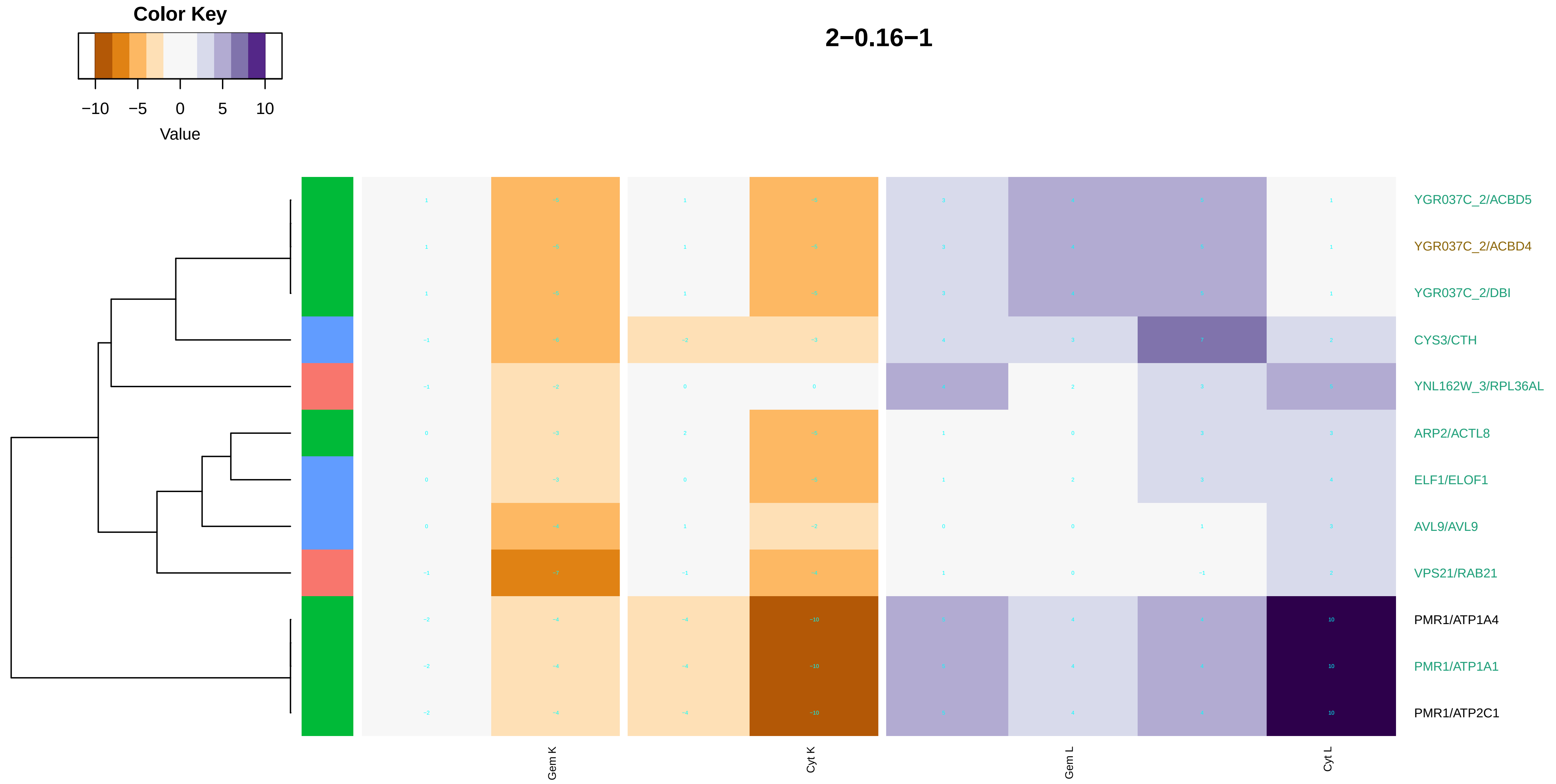

Color Key
−10
−5
0
5
10
Value
2−0.16−1
YGR037C_2/ACBD5
YGR037C_2/ACBD4
YGR037C_2/DBI
CYS3/CTH
YNL162W_3/RPL36AL
ARP2/ACTL8
ELF1/ELOF1
AVL9/AVL9
VPS21/RAB21
PMR1/ATP1A4
PMR1/ATP1A1
PMR1/ATP2C1
Gem K
Cyt K
Gem L
Cyt L

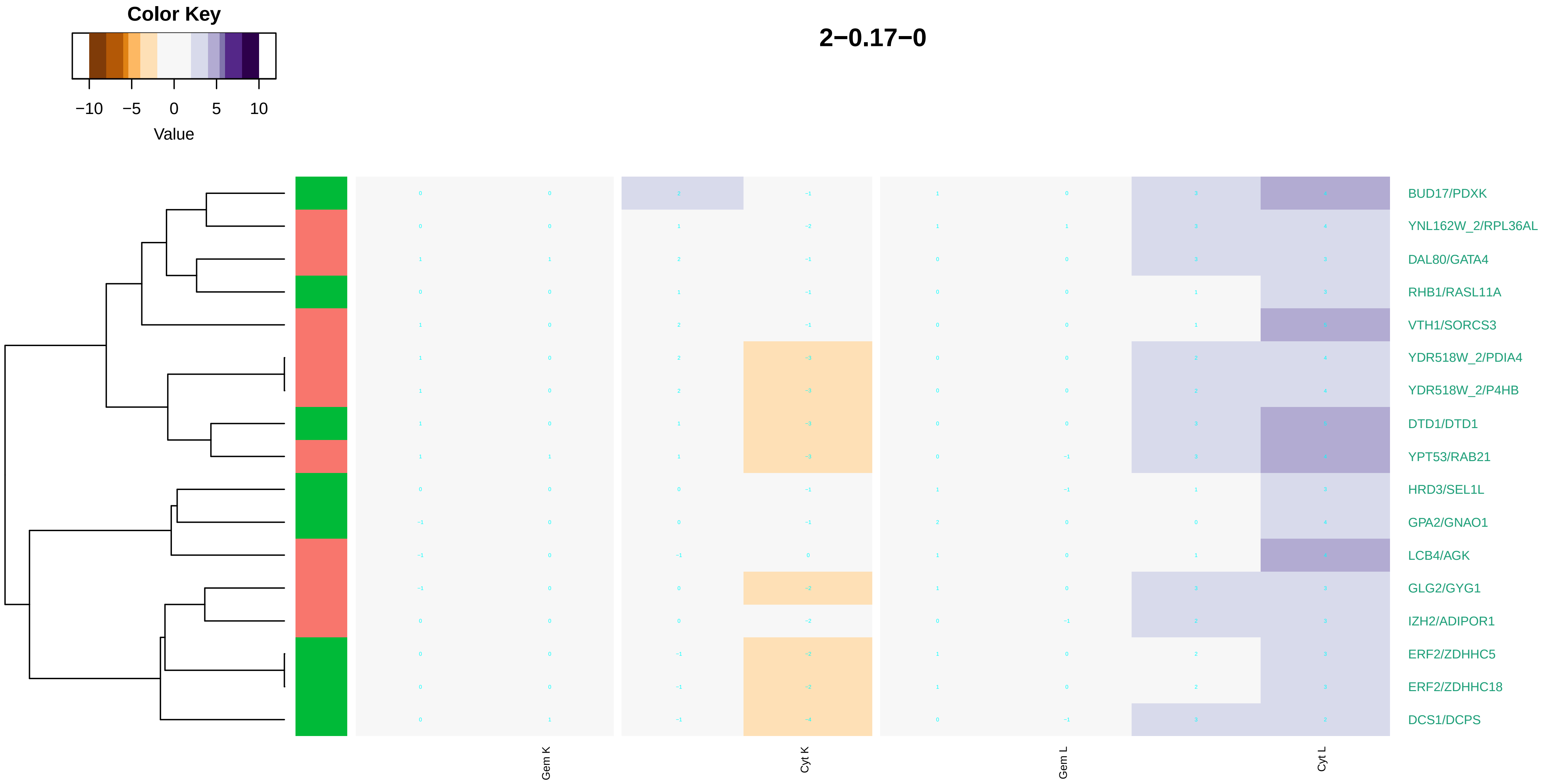
Color Key
−10
−5
0
5
10
Value
2−0.17−0
BUD17/PDXK
YNL162W_2/RPL36AL
DAL80/GATA4
RHB1/RASL11A
VTH1/SORCS3
YDR518W_2/PDIA4
YDR518W_2/P4HB
DTD1/DTD1
YPT53/RAB21
HRD3/SEL1L
GPA2/GNAO1
LCB4/AGK
GLG2/GYG1
IZH2/ADIPOR1
ERF2/ZDHHC5
ERF2/ZDHHC18
DCS1/DCPS
Gem K
Cyt K
Gem L
Cyt L

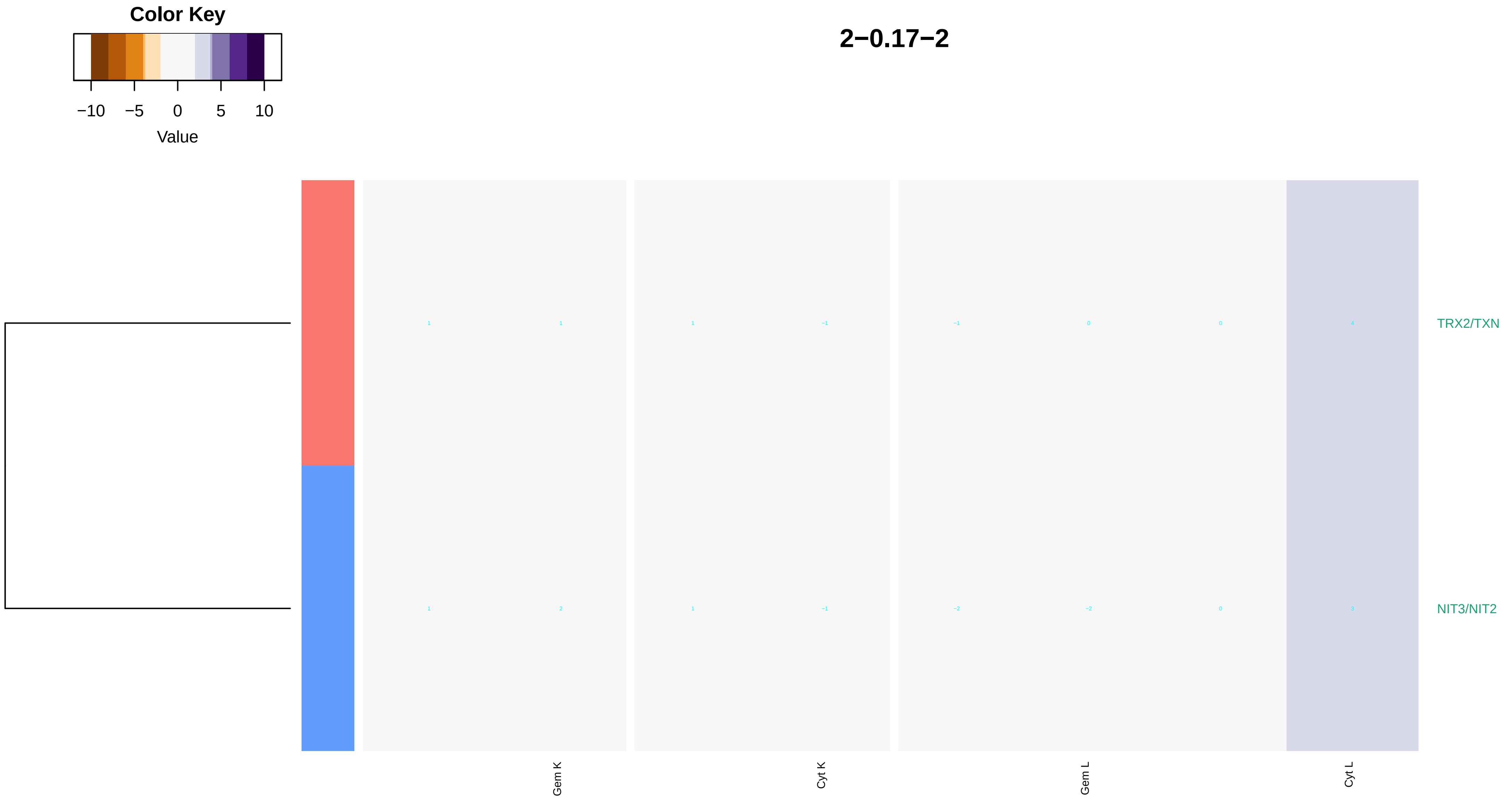

Color Key
−10
−5
0
5
10
Value
2−0.17−2
TRX2/TXN
NIT3/NIT2
Gem K
Cyt K
Gem L
Cyt L

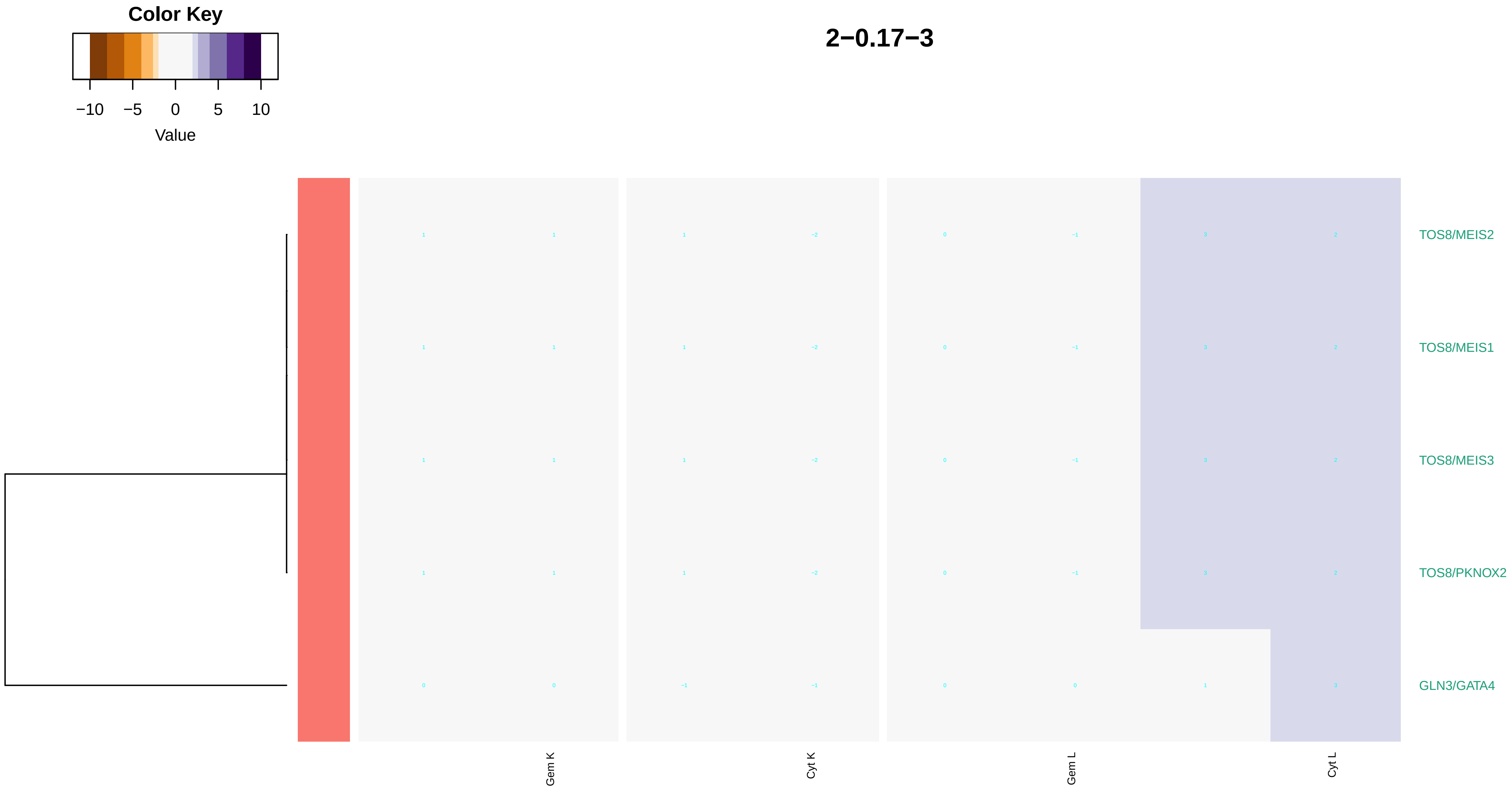
Color Key
−10 −5 0 5 10
Value
2−0.17−3
TOS8/MEIS2
TOS8/MEIS1
TOS8/MEIS3
TOS8/PKNOX2
GLN3/GATA4
Gem K
Cyt K
Gem L
Cyt L

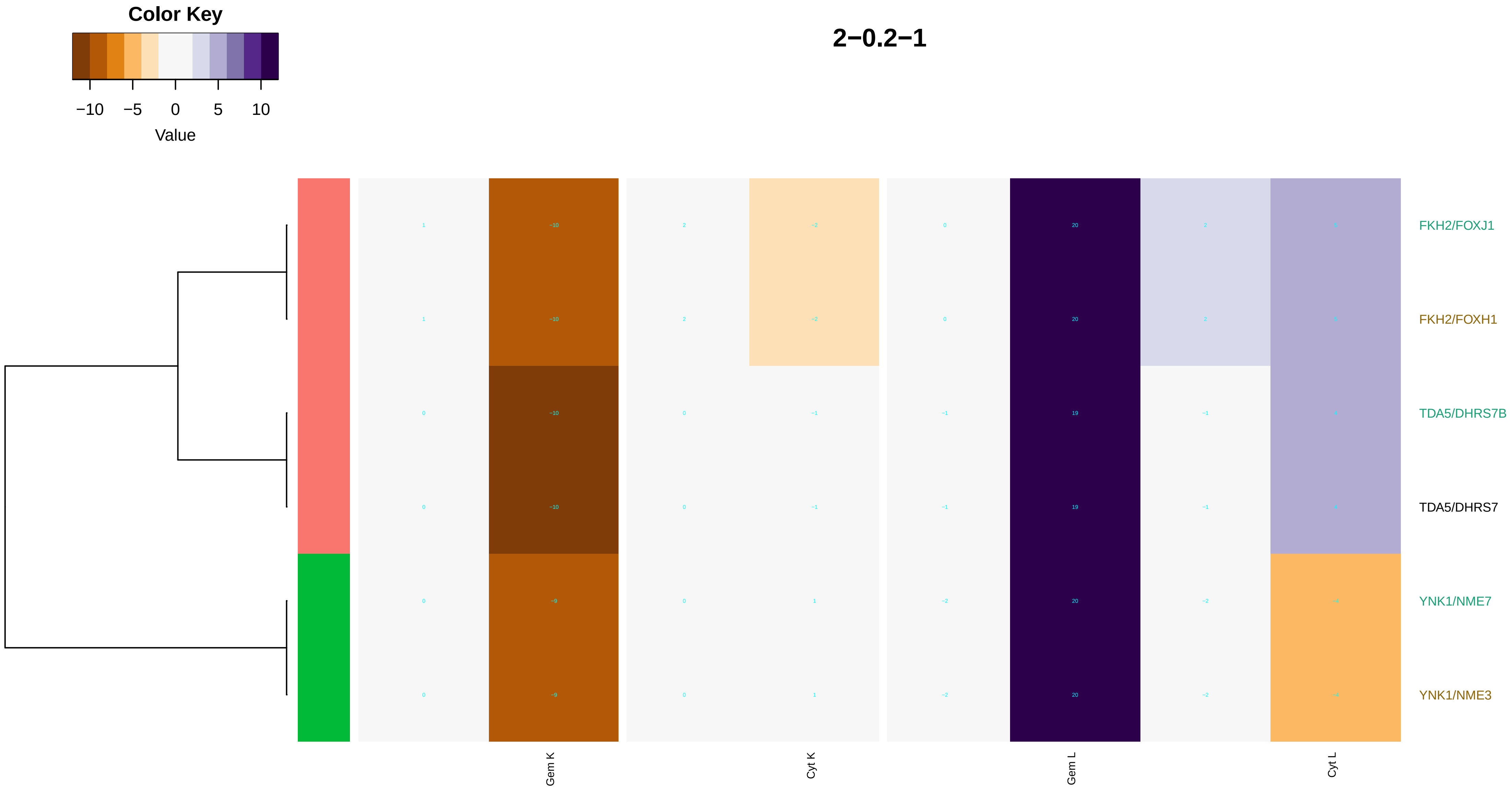

Color Key
−10
−5
0
5
10
Value
2−0.2−1
FKH2/FOXJ1
FKH2/FOXH1
TDA5/DHRS7B
TDA5/DHRS7
YNK1/NME7
YNK1/NME3
Gem K
Cyt K
Gem L
Cyt L

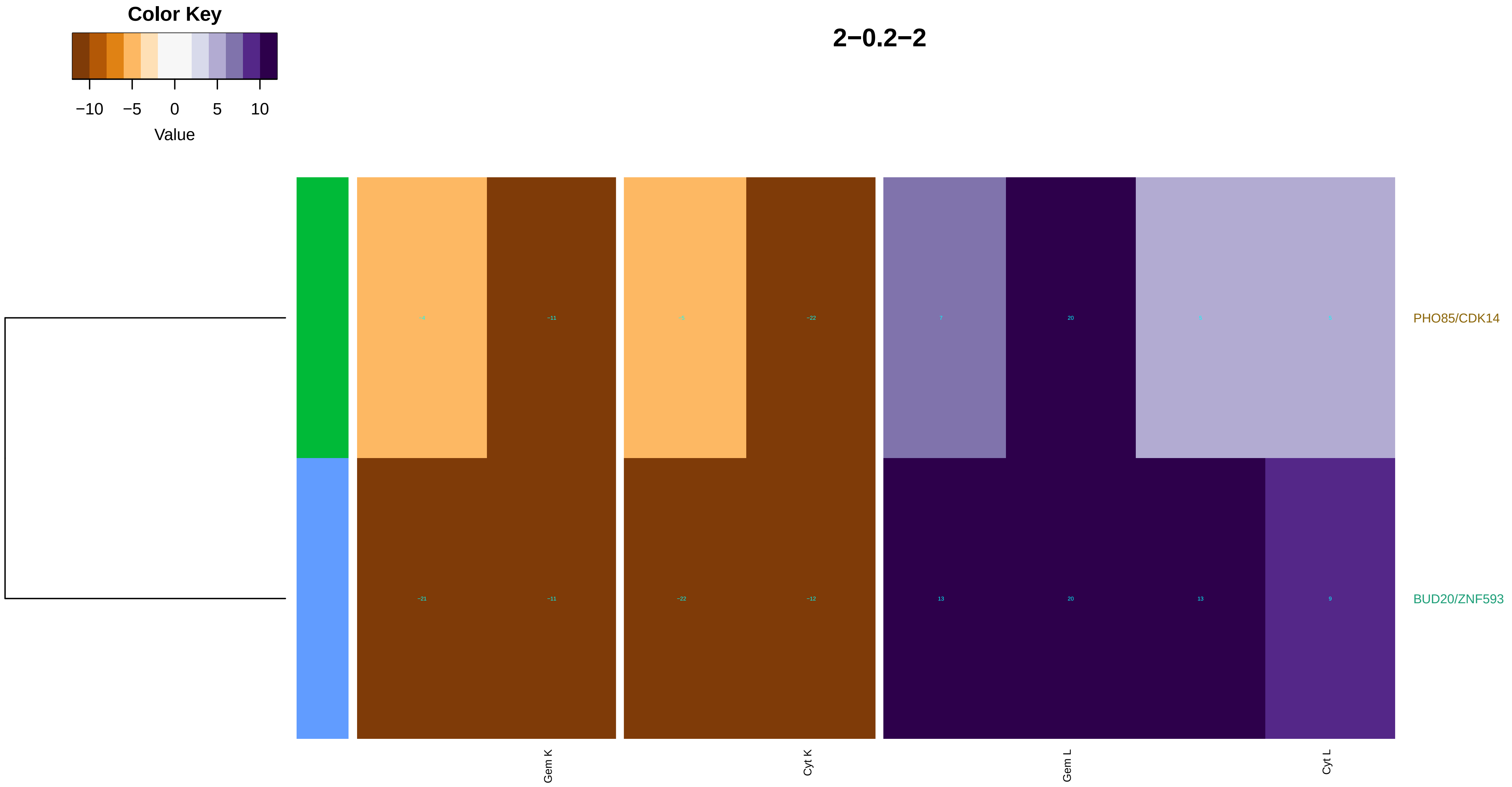
Color Key
−10
−5
0
5
10
Value
2−0.2−2
−4
−11
−5
−22
7
20
5
5
PHO85/CDK14
−21
−11
−22
−12
13
20
13
9
BUD20/ZNF593
Gem K
Cyt K
Gem L
Cyt L

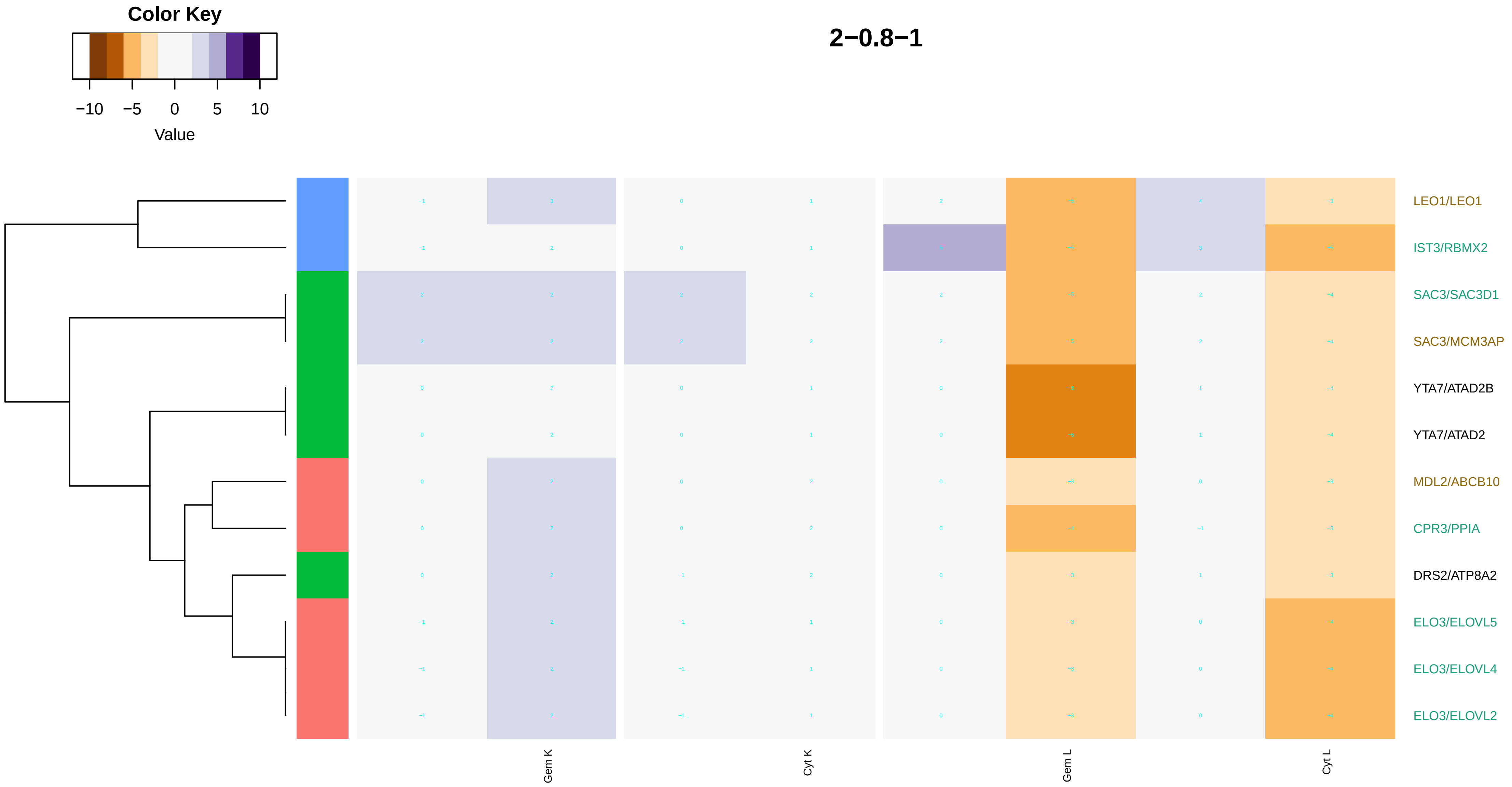
Color Key
−10
−5
0
5
10
Value
2−0.8−1
LEO1/LEO1
IST3/RBMX2
SAC3/SAC3D1
SAC3/MCM3AP
YTA7/ATAD2B
YTA7/ATAD2
MDL2/ABCB10
CPR3/PPIA
DRS2/ATP8A2
ELO3/ELOVL5
ELO3/ELOVL4
ELO3/ELOVL2
Gem K
Cyt K
Gem L
Cyt L

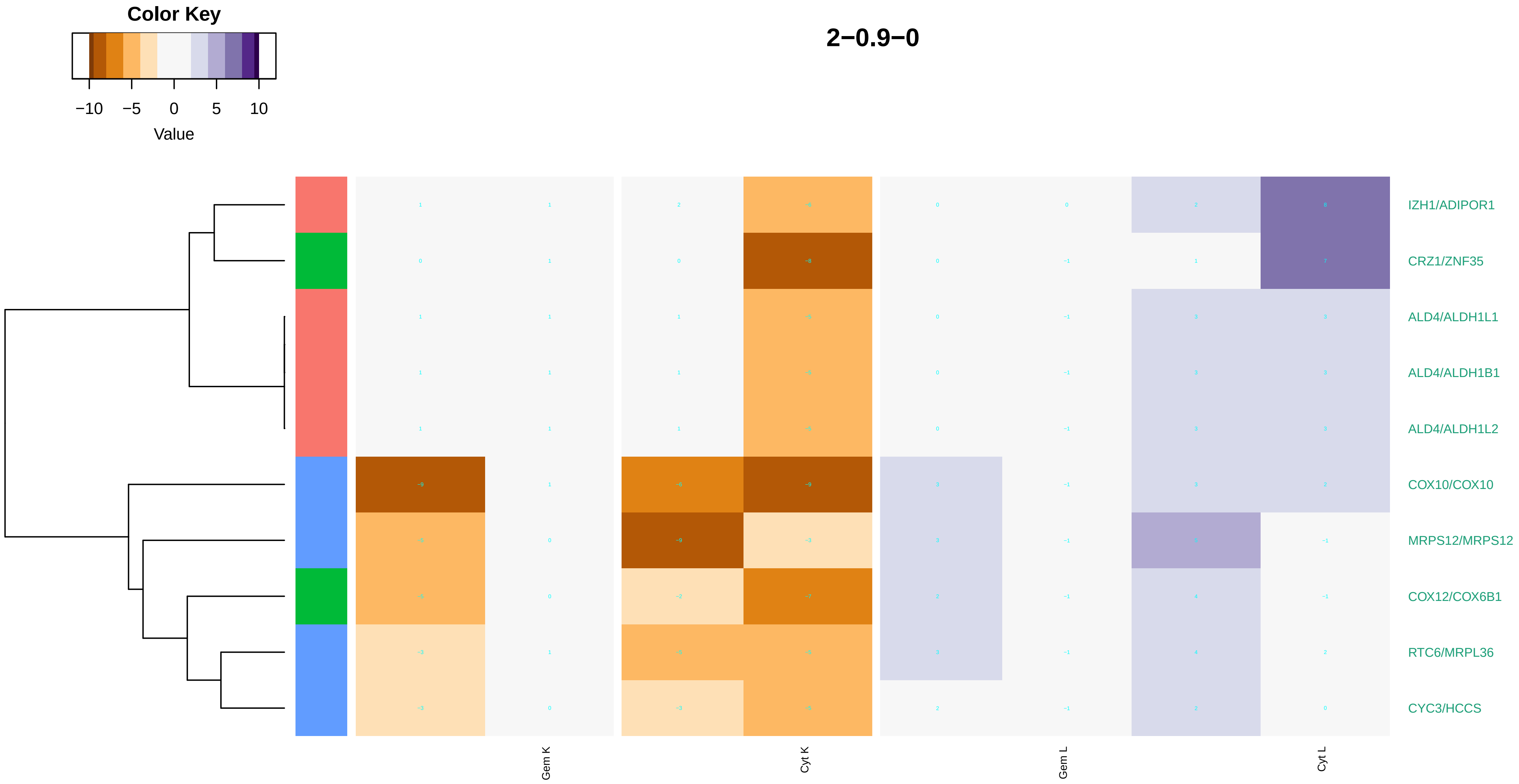
Color Key
−10
−5
0
5
10
Value
2−0.9−0
IZH1/ADIPOR1
CRZ1/ZNF35
ALD4/ALDH1L1
ALD4/ALDH1B1
ALD4/ALDH1L2
COX10/COX10
MRPS12/MRPS12
COX12/COX6B1
RTC6/MRPL36
CYC3/HCCS
Gem K
Cyt K
Gem L
Cyt L

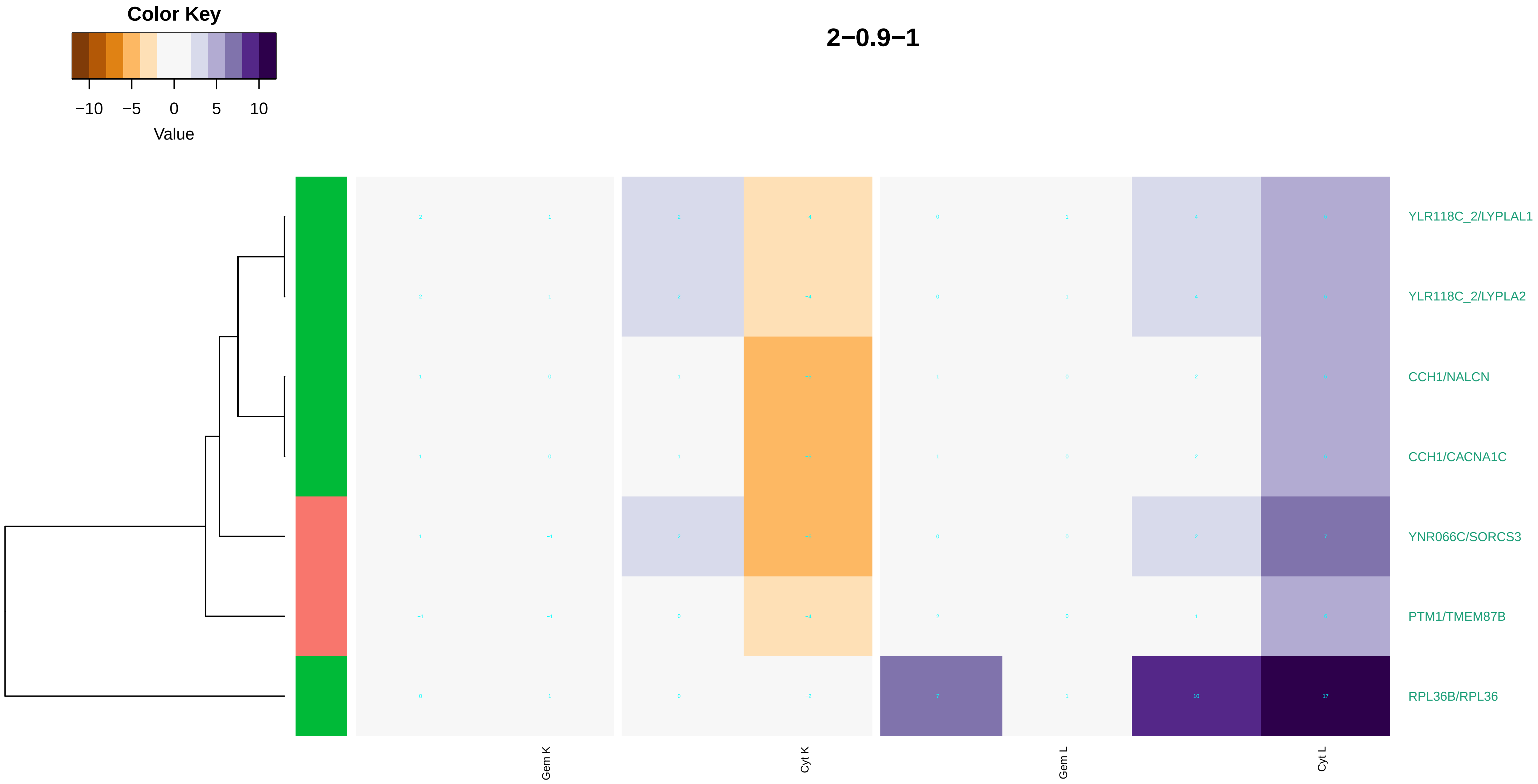
Color Key
−10
−5
0
5
10
Value
2−0.9−1
YLR118C_2/LYPLAL1
YLR118C_2/LYPLA2
CCH1/NALCN
CCH1/CACNA1C
YNR066C/SORCS3
PTM1/TMEM87B
RPL36B/RPL36
Gem K
Cyt K
Gem L
Cyt L

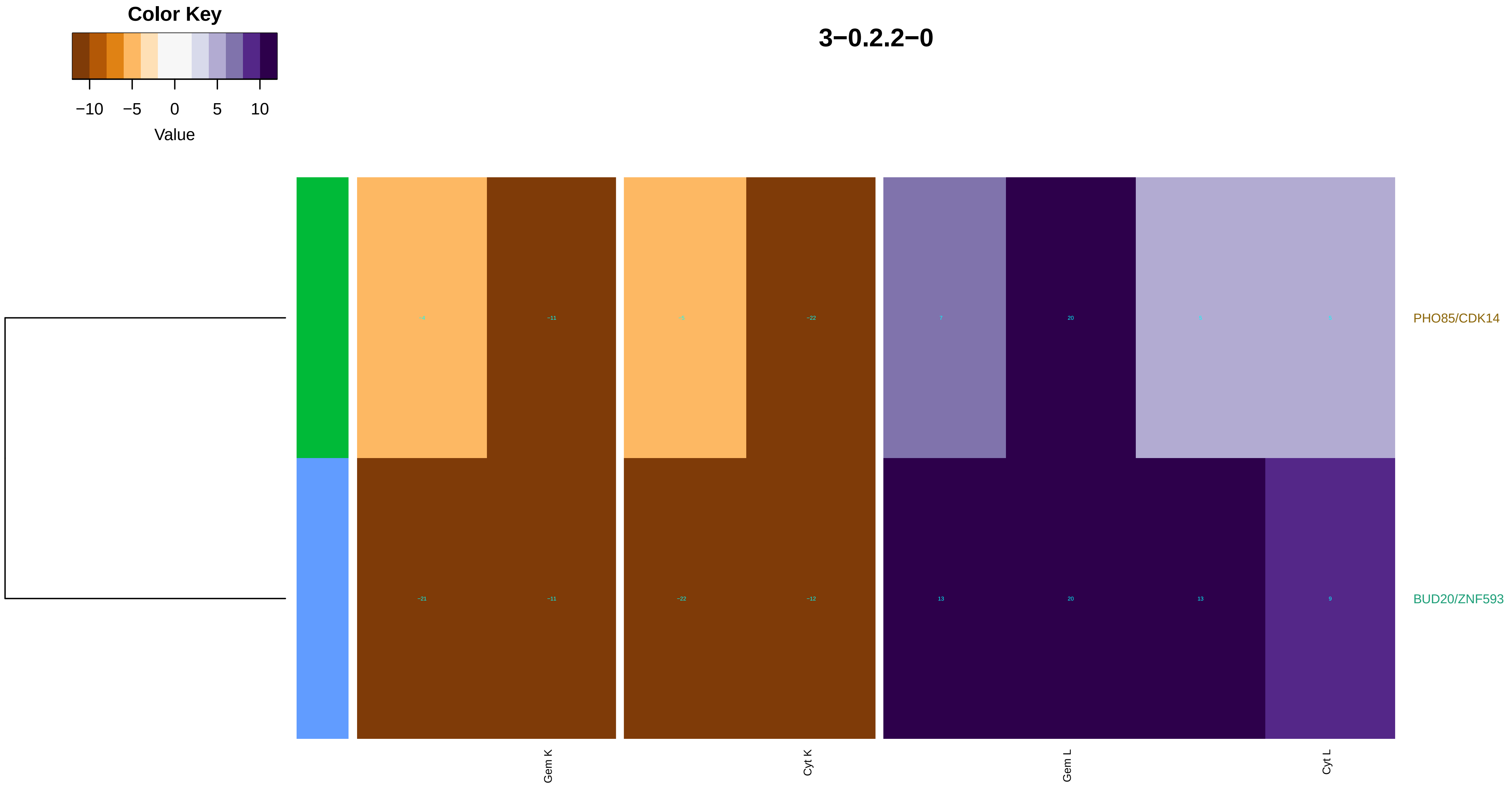
Color Key
−10
−5
0
5
10
Value
3−0.2.2−0
−4
−11
−5
−22
7
20
5
5
PHO85/CDK14
−21
−11
−22
−12
13
20
13
9
BUD20/ZNF593
Gem K
Cyt K
Gem L
Cyt L
